# Phosphatases modified by LH signaling in ovarian follicles: testing their role in regulating the NPR2 guanylyl cyclase

**DOI:** 10.1101/2023.06.12.544636

**Authors:** Jeremy R. Egbert, Ivan Silbern, Tracy F. Uliasz, Katie M. Lowther, Siu-Pok Yee, Henning Urlaub, Laurinda A. Jaffe

## Abstract

In response to luteinizing hormone, multiple proteins in rat and mouse granulosa cells are rapidly dephosphorylated, but the responsible phosphatases remain to be identified. Because the phosphorylation state of phosphatases can regulate their interaction with substrates, we searched for phosphatases that might function in LH signaling by using quantitative mass spectrometry. We identified all proteins in rat ovarian follicles whose phosphorylation state changed detectably in response to a 30-minute exposure to LH, and within this list, identified protein phosphatases or phosphatase regulatory subunits that showed changes in phosphorylation. Phosphatases in the PPP family were of particular interest because of their requirement for dephosphorylating the natriuretic peptide receptor 2 (NPR2) guanylyl cyclase in the granulosa cells, which triggers oocyte meiotic resumption. Among the PPP family regulatory subunits, PPP1R12A and PPP2R5D showed the largest increases in phosphorylation, with 4-10 fold increases in signal intensity on several sites. Although follicles from mice in which these phosphorylations were prevented by serine-to-alanine mutations in either *Ppp1r12a* or *Ppp2r5d* showed normal LH-induced NPR2 dephosphorylation, these regulatory subunits and others could act redundantly to dephosphorylate NPR2. Our identification of phosphatases and other proteins whose phosphorylation state is rapidly modified by LH provides clues about multiple signaling pathways in ovarian follicles.

**Graphical Abstract:** 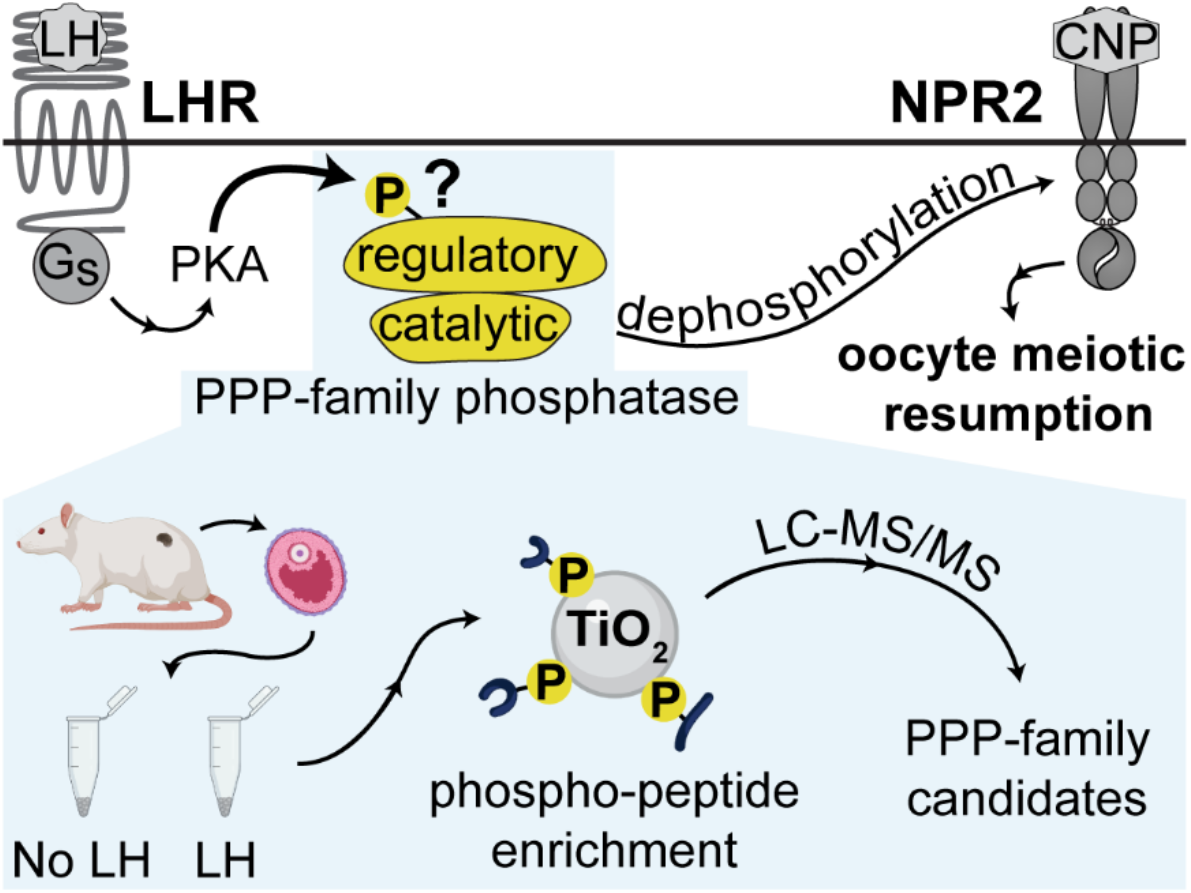

**Summary sentence:** Quantitative mass spectrometric analysis of phosphatases whose phosphorylation state is rapidly modified by luteinizing hormone provides clues about how LH signaling dephosphorylates NPR2 as well as a resource for future studies.

## Introduction

Luteinizing hormone (LH) acts on its receptor in the mural granulosa cells of mammalian preovulatory follicles to initiate a complex signaling cascade that leads to resumption of meiosis in the oocyte, ovulation, and formation of the corpus luteum. The LH receptor activates G_s_ and other G-proteins, which activate protein kinase A (PKA) and other kinases (1–4). LH signaling also results in the rapid dephosphorylation of multiple regulatory proteins in the granulosa cells, including the guanylyl cyclase NPR2 (5), the actin regulator cofilin (6), and the multifunctional protein MAP2D (7), but little is known about which phosphatases are responsible. The goal of this study was to identify protein phosphatases that are modified by LH signaling, and then to investigate if the identified phosphatases function to dephosphorylate the NPR2 guanylyl cyclase. Because the phosphorylation state of phosphatases can regulate their interaction with substrates (8–11), we used quantitative mass spectrometry (MS) to identify phosphatases in rat ovarian follicles that showed rapid LH-induced changes in phosphorylation.

One centrally important LH-induced response that requires phosphatase activity in the granulosa cells is the resumption of meiosis in the prophase-arrested oocyte. Mammalian preovulatory follicles send a signal to the oocyte that maintains meiotic arrest, and then in response to LH, reverses the arrest (1–4). In mice (12) and other mammalian species (reviewed in ref 2), prophase arrest is maintained by the activity of the natriuretic peptide receptor (NPR2) guanylyl cyclase in the granulosa cells around the oocyte. NPR2 generates cGMP that diffuses through gap junctions into the oocyte in the center of the follicle, inhibiting meiotic progression (13). LH-induced dephosphorylation of NPR2 decreases its activity, causing meiosis to resume (5, 14, 15). If dephosphorylation of NPR2 is prevented by changing its juxtamembrane serines and threonines to the phosphomimetic amino acid glutamate, meiotic resumption is delayed by several hours (14). The LH-induced dephosphorylation of NPR2 is prevented by small molecule inhibitors of phosphatases, cantharidin and okadaic acid (5), which inhibit most PPP family phosphatases (16), but not PPM family phosphatases (17, 18). However, the particular PPP holoenzymes that are responsible for the dephosphorylation and inactivation of NPR2 or other related guanylyl cyclases are unknown.

Each PPP holoenzyme is comprised of a catalytic subunit belonging to 1 of 7 subfamilies (PPP1c-PPP7c), in complex with 1 or 2 other subunits. Over 250 proteins are known to form complexes with PPP catalytic subunits (19, 20), and these have been characterized with varying degrees of certainty to be regulators of PPP family phosphatase activity. ∼60 of the best characterized are given names including “PPP” (19, 20). We analyzed proteins in rat preovulatory follicles using quantitative mass spectrometry to obtain clues about which among this large number of candidates might regulate NPR2 dephosphorylation and other responses to LH signaling in ovarian follicles.

## Materials and Methods

### Animals

Rats (CD-Sprague-Dawley, Charles River Laboratories, Kingston, NY) and mice (C57BL/6J, Jackson Laboratory, Bar Harbor, ME) were used for these studies. Where indicated, mice were genetically modified using CRISPR-Cas9 to insert a hemagglutinin (HA) tag on *Npr2* (21), or to insert point mutations in *Ppp1r12a* (S507A, Figure S1) or *Ppp2r5d* (S53A/S81A/S82A/S566A, Figure S2). Mice with these mutations in phosphatase regulatory subunits also had an HA tag on *Npr2*. All procedures were approved by the animal care committee of the University of Connecticut Health Center. The genetically modified mice described here are available from Dr. Siu-Pok Yee upon request. The HA-*Npr2* mice are also available from the MMRRC Repository at The Jackson Laboratory (Bar Harbor, ME; RRID:MMRRC_071304-JAX).

### Ovarian follicles: isolation, culture and sample preparation

Preovulatory rat follicles, 700 - 900 µm in diameter, were dissected from the ovaries of 25- day old prepubertal animals that had been injected 48 hours previously with equine chorionic gonadotropin (National Hormone and Peptide Program, 12 I.U.) to stimulate follicle growth and LH receptor expression (5). The follicles were cultured for 1-2 hours on 30 mm Millicell cell culture inserts (PICM0RG50, MilliporeSigma, St. Louis, MO) as previously described (5). LH (ovine LH-26, National Hormone and Peptide Program) or a control solution of PBS was applied to the medium under the culture membrane, and a small drop was also added to the top of the membrane around each follicle, to ensure rapid exposure of the follicles. LH was used at 350 nM (10 µg/ml), a concentration that results in a maximal percentage of nuclear envelope breakdown in rat follicles under the conditions used here (22), and maximal dephosphorylation of NPR2 at 30 min (5). At 30 min after applying LH or control vehicle (PBS), follicles were washed in PBS, then frozen in liquid nitrogen and stored at −80°C, for mass spectrometry. Each rat follicle contains ∼20 µg of protein as determined by a BCA assay (22), and 17-27 follicles were obtained per rat. Each sample for mass spectrometry contained 1.0-1.6 mg protein (50-81 follicles).

Fully grown mouse follicles, 300 - 400 µm in diameter, were dissected from the ovaries of 24- 26 day old prepubertal animals that had not been injected with hormones. The follicles were then cultured for 23-26 hours on 30 mm Millicell cell culture inserts in the presence of 1 nM follicle stimulating hormone (FSH; 30 ng/ml, ovine FSH, National Hormone and Peptide Program) to stimulate synthesis of LH receptors and progression to the preovulatory stage (23). For some experiments, the FSH-cultured follicles were then incubated with or without the PPP family phosphatase inhibitor cantharidin (C7632, Sigma-Aldrich) or the PKA inhibitor Rp-8-CPT-cAMPS (C011-05, BioLog, Bremen, Germany). LH or control vehicle (PBS) was then applied as described for rat follicles. LH was used at 10 nM (∼0.3 µg/ml), the lowest concentration needed to achieve maximal nuclear envelope breakdown in mouse follicle-enclosed oocytes (23). For experiments with or without Rp-8-CPT-cAMPS, the volume required was minimized by transferring follicles to 12 mm Millicells (PICM01250) in a 4-well plate containing 250 µl medium. For western blotting, follicles that had been treated with or without LH for 30 minutes were washed in PBS, and then lysed by probe sonication in Laemmli sample buffer with 75 mM dithiothreitol and protease and phosphatase inhibitors as previously described (24). Each mouse follicle contains ∼3.5 µg of protein as determined by a BCA assay (25).

### Sample preparation for LC-MS/MS

For liquid chromatography-tandem mass spectrometry (LC-MS/MS) sample preparation (Figure 1), LC/MS-grade water, methanol, and acetonitrile (ACN) (Merck, Darmstadt, Germany) were used. Frozen rat ovarian follicles were homogenized in 400 µL lysis buffer (100 mM HEPES, pH 8, 4% SDS, 1 mM EDTA, and 1× Halt protease/phosphatase inhibitor cocktail (ThermoFisher Scientific) using Zirconia/glass beads in a bead beating grinder (FastPrep24, MP Biomedicals). The lysates were sonicated in a BioRuptor (Diagenode) for 5 min (30 s on/30 s off cycle). After clearing by centrifugation, protein concentration in the supernatant was determined using a BCA assay (ThermoFisher Scientific). Equal protein amounts were reduced/alkylated by incubating with 10 mM TCEP/ 40 mM CAA for 30 min at 37°C, and proteins were precipitated with methanol-chloroform (26). Precipitated proteins were resuspended in 1% RapiGest (Waters) in 100 mM TEAB, and digested as previously described (27) with the following modification: Equal amounts of proteins were digested overnight using either trypsin (1:40 trypsin-to-protein ratio) or LysC (1:100 LysC-to-protein ratio), and then LysC- or trypsin-digested aliquots of the same sample were respectively combined before phosphopeptide enrichment. To enrich for phosphopeptides, peptides were incubated with TiO_2_ beads (GL Sciences, Tokyo, Japan) (27). Unbound (non-phosphorylated) peptides were collected, then bound (phosphorylated) peptides were eluted from the beads. Isobaric tandem mass tag (TMT)6 labeling, sample desalting, and basic-reversed phase (bRP) fractionation were performed as described previously (27). Aliquots of protein digests containing mostly non-phosphorylated peptides were separately labeled using TMT reagents processed within the same pipeline omitting the enrichment step. Following bRP fractionation, 20 concatenated fractions were collected and analyzed.

**Figure 1.**
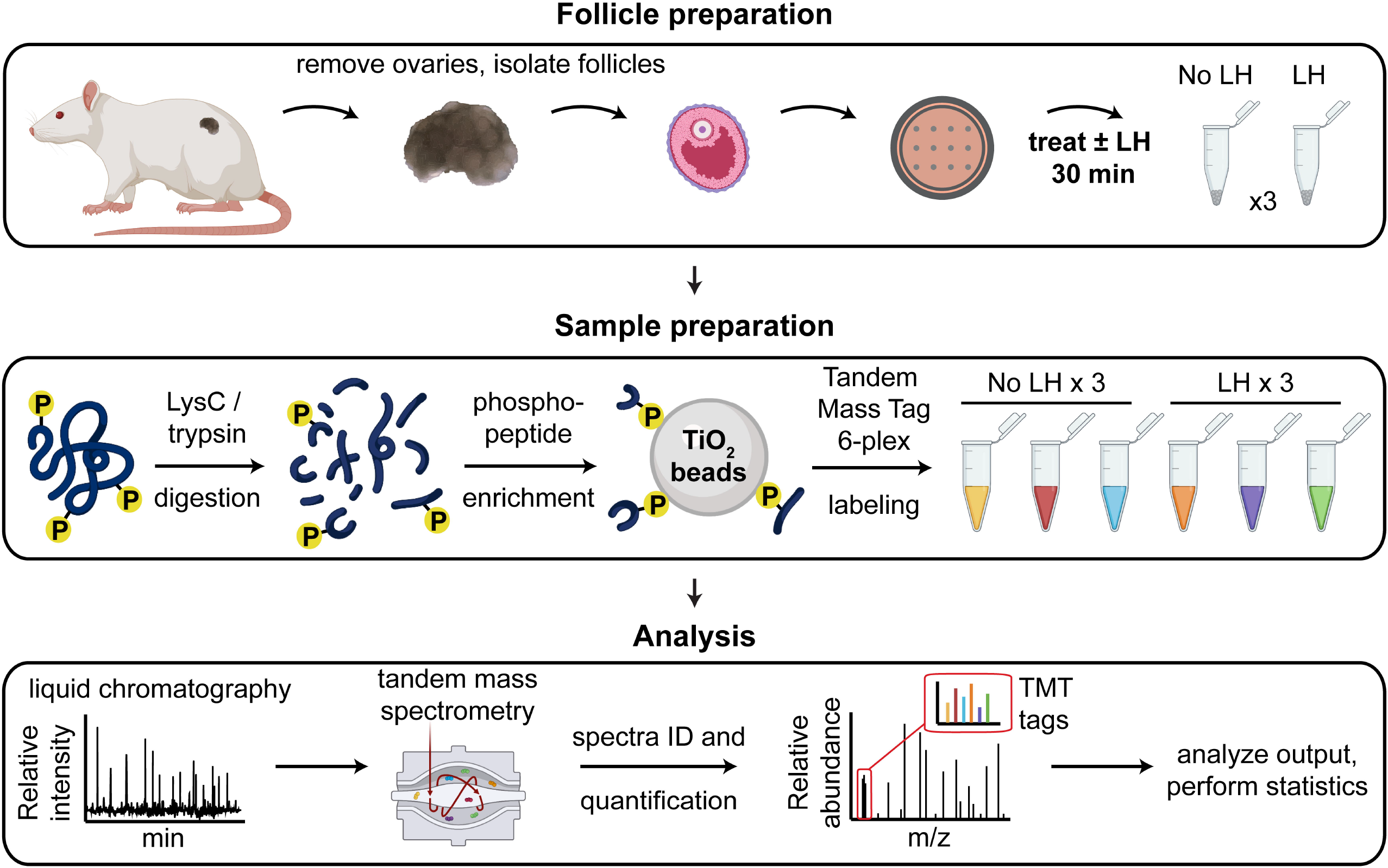
Workflow for the phosphoproteome analyses of ovarian follicles with or without LH stimulation.

### LC-MS/MS

Dried bRP fractions of peptides prior to and after phosphopeptide enrichment were dissolved in 2% (v/v) ACN 0.1% (v/v) TFA in water and subjected to LC-MS/MS analysis using a setup as previously described (27, Figure 1). Phosphorylated peptides were injected in technical duplicates and separated using an 88 min gradient consisting of loading at 98% buffer A (0.1% (v/v) FA in water) 2% buffer B (80% ACN (v/v) 0.1% FA (v/v) in water) for 4 min; linear gradient ranging from 7% to 32% B over 56 min; followed by linear increase to 50% B over 18 min, washing step at 90% B for 5 min and re-equilibration step at 2% B for 5 min. For the separation of non-phosphorylated peptides, the first two steps were adjusted accordingly: 1) loading at 5% B for 5 min; 2) linear gradient from 8 to 34% B over 56 min.

An Orbitrap/Ion Trap mass spectrometer (Tribrid Fusion, ThermoFisher Scientific) was operated in data-dependent acquisition mode. Survey MS1 scans were acquired in Orbitrap at the resolution of 120,000, scan range 350-2000 *m/z,* normalized automatic gain control (AGC) target of 250%, and maximum injection time (maxIT) of 50 ms. The most abundant precursors of charge state 2-7 were subjected to fragmentation using HCD and normalized collision energy (NCE) of 33%. The length of the duty cycle was fixed to 3 s. Precursor ions were excluded from repetitive fragmentation for 30 s. MS2 spectra were acquired at the resolution of 30,000, normalized AGC target of 1000%, and 200 ms maxIT. The top 10 fragment ions were isolated using synchronous precursor selection (SPS; 28) and fragmented in HCD at NCE of 50%. Resulting SPS-MS3 spectra were acquired in Orbitrap at 50,000 resolution, normalized AGC target of 500% and maxIT of 200ms. When analyzing non-phosphorylated peptides, the resolution and maxIT of MS2 scans were reduced to 15,000 and 54 ms, respectively. Accordingly, maxIT of SPS-MS3 scans was set to 86 ms.

### LC-MS/MS data analysis

Raw LC-MS/MS data were processed using MaxQuant software (version 1.6.2.10; 29) using default settings if not stated otherwise. MS-data from phosphorylated and non-phosphorylated peptides were processed simultaneously after defining “Phospho (S,T,Y)” modification as variable modification in the group parameters for the samples after the enrichment step. Other default modifications included carbamidomethylation of cysteine residues (fixed) and oxidation of methionine and acetylation of protein N-terms (both variable modifications). Up to five variable modifications were allowed per peptide. Trypsin was selected as the protease. “Reporter ion MS3” was selected as quantification parameter. Default mass tolerances were preserved at 4.5 ppm and 20 ppm for precursor and fragment masses, respectively. Canonical protein sequences of *Rattus norvegicus* were downloaded from UniProt (February 2019, 29951 entries; 30). Further steps of statistical analysis were conducted in R programming language using custom scripts as previously described (27). In brief, impurity-corrected reporter ion intensities were used to assess differential abundance of phosphorylated peptides at the level of phosphorylation sites (“Phospho(S,T,Y).txt” output table of MaxQuant). Only phosphorylation sites with localization probability >0.75 were considered for the analysis. Reporter ion intensities were log_2_-transformed, normalized by median polishing, and subjected to differential expression testing using *limma* (31)*. Limma*-moderated p-values were corrected for multiple comparisons using a q-value approach (32). Phosphorylation events were considered as significantly altered (“candidates”) if satisfying the following criteria: 1) absolute log_2_ (Treated/Control) > log_2_(1.5) and 2) *q-*value < 0.01, corresponding to a false discovery rate (FDR) of <1%. Impurity-corrected protein group intensities (“proteinGroups.txt” output table) were analyzed in a similar way in order to confirm that treatment had no significant impact on the proteome level.

Because of the similarity in amino acid sequences for PPP1R12A and R12B, peptides from these 2 proteins were indistinguishable. The assignment of these peptides to PPP1R12A was determined by ddPCR analysis of the relative abundances of PPP1R12A and R12B mRNAs (Figure S3).

### Bioinformatics analysis

Lists of ovarian follicle proteins that had at least one site with one statistically significant difference in phosphorylation intensity following LH stimulation were subjected to gene set enrichment analysis using DAVID (2021 Update; 33). Gene ontology (GO) terms were considered significant if the Benjamini-Hochberg adjusted p < 0.01. Follicle GO terms with at least 5 gene products were downloaded March 18, 2022.

### Western blotting

For analysis of LH-induced NPR2 dephosphorylation (Figures 2, 5, 6), we used follicles from mice with HA-tagged NPR2 (see above). Proteins were separated by SDS-polyacrylamide gel electrophoresis (PAGE) using a gel containing the phosphate binding molecule Phos-tag, which retards the migration of phosphorylated proteins (34). Methods were as previously described (5, 35), except that follicle lysates (30-60 µg protein) were directly loaded onto the gels without immunoprecipitation. Phos-tag reagent and compatible molecular weight markers were obtained from Fujifilm Wako Chemicals USA (Richmond, VA; cat. #AAL-107 and #230-02461, respectively). The antibody used to detect the HA tag on NPR2, as well as other antibodies used for western blotting, are listed in Table S1. Signals were detected using a horseradish peroxidase-conjugated secondary antibody (Table S1) and a Western Bright Sirius detection kit (Advansta, San Jose, CA; cat. #K-12043-D20). Blots were imaged using a charge-coupled device camera (G:Box Chemi XX6; Syngene, Frederick, MD). Relative levels of HA-NPR2 phosphorylation were compared by ratioing the signal intensity in the upper vs lower bands. Phos-tag gel electrophoresis was also used to detect LH-induced phosphorylation of PPP2R5D (Figure 4), using an antibody against total PPP2 (Table S1).

**Figure 2.**
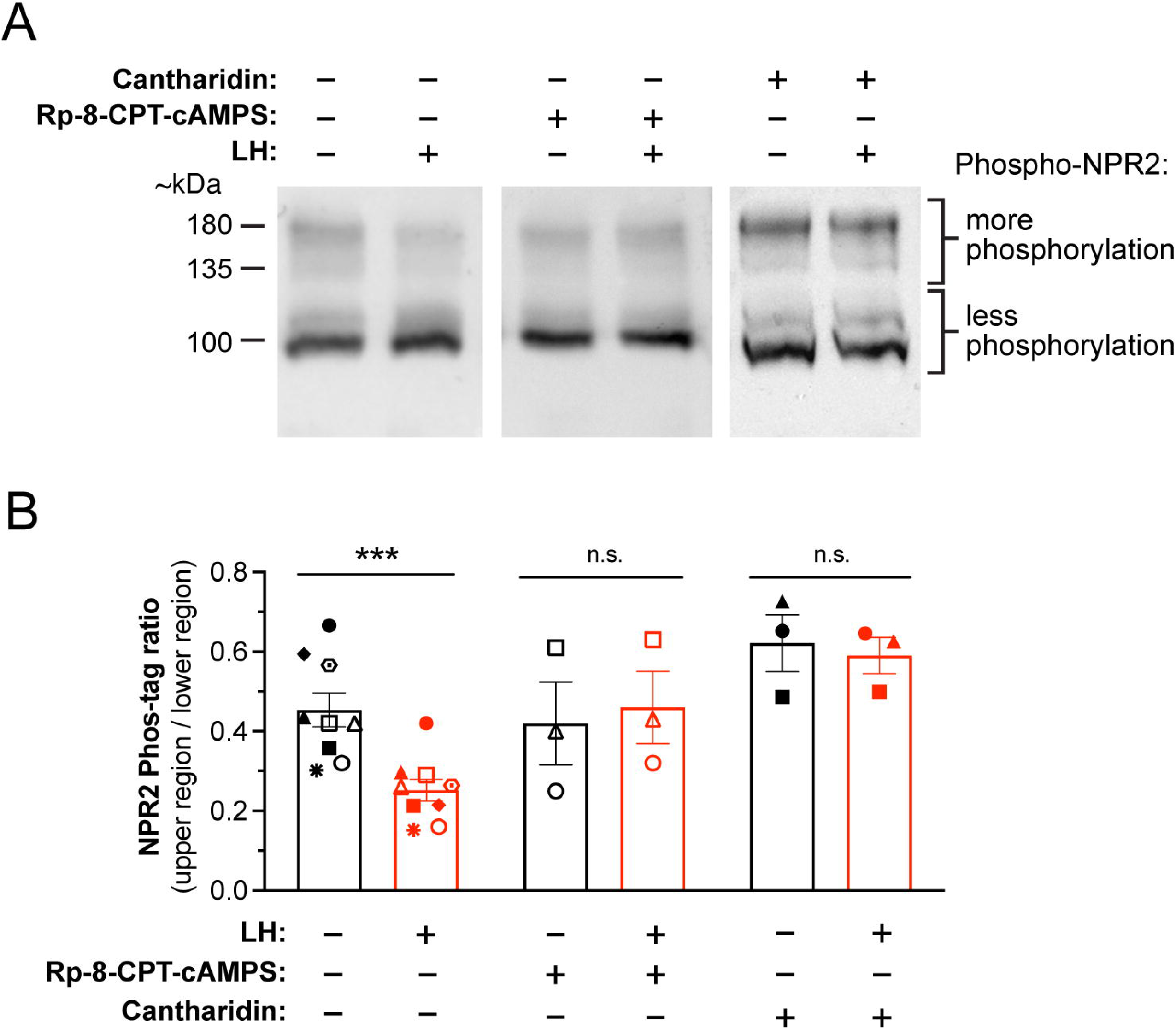
LH-induced NPR2 dephosphorylation in mouse preovulatory follicles requires PKA and PPP family phosphatase activities. **(A)** Representative western blots of Phos-tag-containing SDS-PAGE gels, showing that the PKA inhibitor Rp-8-CPT-cAMPS or the PPP family phosphatase inhibitor cantharidin inhibits LH-induced HA-NPR2 dephosphorylation. Isolated preovulatory follicles from mice in which endogenous NPR2 was tagged with the HA epitope were treated with or without Rp-8-CPT-cAMPS (10 mM, 4 hr) or cantharidin (10 μM, 4 hr), then stimulated with 10 nM LH (30 min). Data shown in Figure S4 confirmed that under these conditions, Rp-8-CPT-cAMPS effectively inhibits PKA. Follicle proteins were separated on a Phos-tag gel, and western blots were probed with an antibody recognizing HA (see methods). Upper bands indicate more phosphorylated NPR2, and lower bands indicate less phosphorylated NPR2. Note that molecular weight markers are not exact on a Phos-tag gel, and primarily correspond to the least phosphorylated form of a protein. (**B**) Summary of the ratios of signal intensities for upper/ lower NPR2 bands, for multiple blots like those shown in A. Each symbol represents a set of samples that were treated as indicated and then analyzed together on the same blot. Bars show mean ± SEM. Data were analyzed by paired t-tests with correction for multiple comparisons. *** p < 0.001.

To quantify LH-induced changes in PPP1R12A phosphorylation (Figure 3), proteins from follicles treated with or without LH were separated on an SDS-PAGE gel (no Phostag), and blots were probed with phosphospecific antibodies (Table S1). To provide signal linearity, fluorescently labeled secondary antibodies (Table S1) were detected with an Odyssey imager (LI-COR, Lincoln, NE). To ensure that equal amounts of protein were loaded in each lane and transferred equally to the blot, blots were stained with Ponceau S (Sigma Chemical P3504), which allows comparison of total protein amounts in each lane (36). LH-induced changes were calculated by first normalizing each phospho-PPP1R12A band intensity to the Ponceau S signal in the corresponding lane, and then normalizing values for LH-treated lanes to those for control lanes. The effect of Rp-8-CPT-cAMPS on PPP1R12A phosphorylation was quantified similarly.

**Figure 3.**
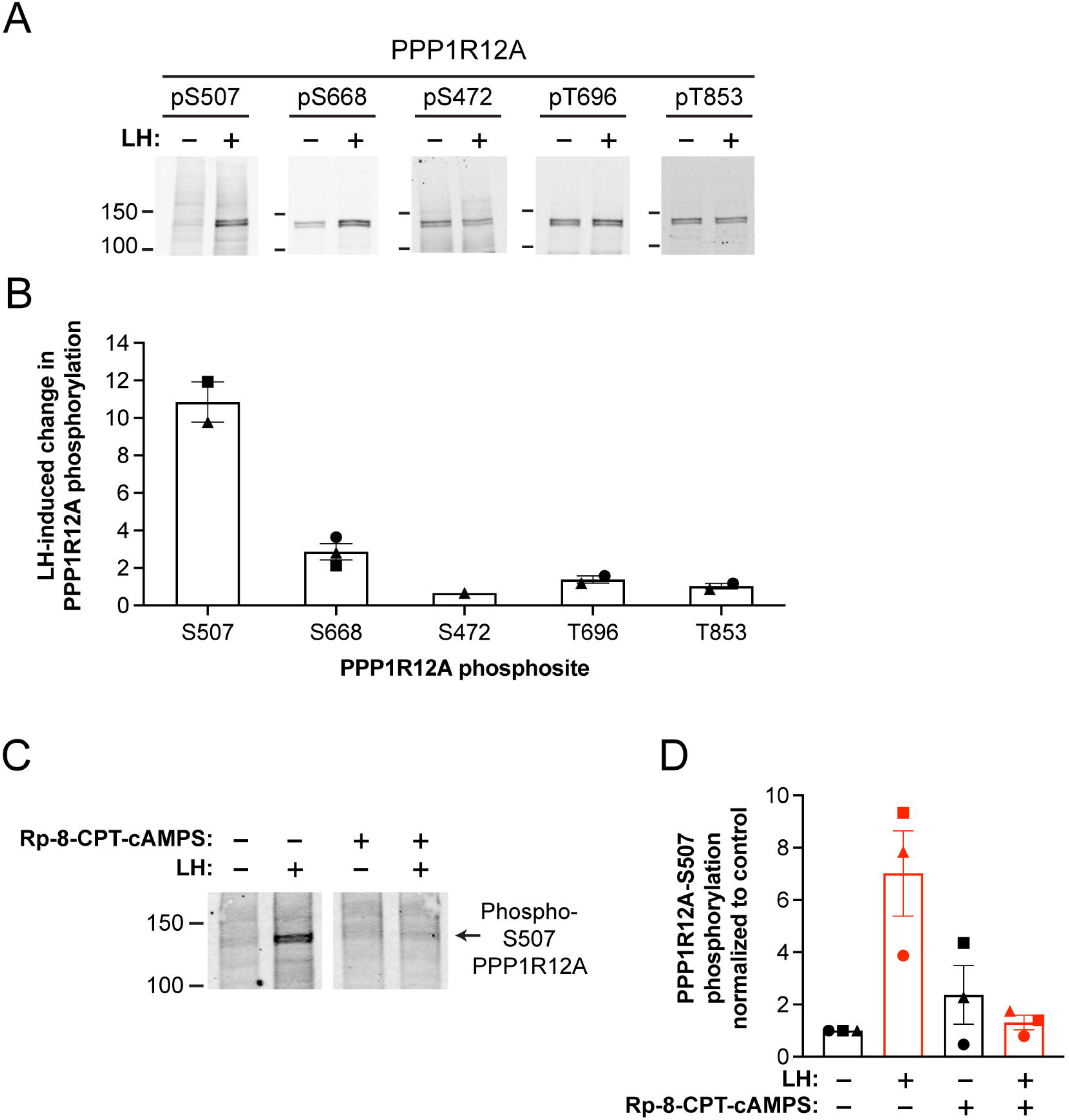
Quantitative western blot analysis of PPP1R12A phosphosite changes in LH-treated mouse preovulatory follicles. Binding of phosphospecific antibodies was quantified using a fluorescent secondary antibody and a LI-COR Odyssey imager (see methods). (**A**) Representative images of blots of PPP1R12A phosphosites in ovarian follicles treated ± 10 nM LH (30 min). (**B**) Summary of experiments like that in (A). To quantify, each phosphospecific antibody signal was normalized to the total protein signal in that lane, as determined by Ponceau-S staining (see methods). The normalized signal with LH was then divided by the normalized signal without LH to determine the LH-induced change in PPP1R12A phosphorylation at each site. (**C**) PKA-dependence of PPP1R12A-S507 phosphorylation. Follicles were treated with or without the PKA inhibitor Rp-8-CPT-cAMPS (Rp; 10 mM, 8 hr), then stimulated with 10 nM LH (30 min). (**D**) Summary (mean ± SEM) of 3 experiments like that in (C), with Rp-8-CPT-cAMPS applied for either 4 or 8 hr. Values for the graph were generated by normalizing to the Ponceau stain intensity for the lane, and then normalizing to the value with no Rp-8-CPT-cAMPS or LH treatment. For B and D, each symbol (circle, square, triangle) represents a set of samples that were treated as indicated and then analyzed together on the same blot. For each experiment, changes in phospho-S507 intensity of the different treatment groups are plotted relative to the control with no Rp-8-CPT-cAMPS and no LH.

LH-induced changes in CREB-S133 phosphorylation (Figure S4) were detected using a phosphospecific mouse antibody and a fluorescently labelled goat-anti-mouse secondary antibody (Table S1). Signal intensities were measured using an Odyssey imager and were normalized to those for total CREB, which was measured by simultaneously probing the same blot with a rabbit antibody against total CREB and a fluorescently labelled goat-anti-rabbit secondary antibody (Table S1). Phospho and total CREB signals were detected simultaneously with 800 and 680 nm laser lines.

### Statistics

Data were analyzed using Prism 9 (GraphPad Software, Boston, MA) as described in the figure legends.

## Results and Discussion

### Mass spectrometric analysis of LH-induced changes in the phosphoproteome of rat preovulatory follicles

Using a mass spectrometry (MS)-based quantification strategy relying on tandem mass tag (TMT) labelling (27) (Figure 1), we first analyzed all proteins in rat ovarian follicles that showed detectable changes in phosphorylation in response to LH. Rats rather than mice were used in order to obtain sufficient protein. Isolated preovulatory ovarian follicles were treated with or without LH for 30 min, then lysates were prepared for TMT-based quantitative MS. The 30 min time point was chosen because maximal dephosphorylation of NPR2 is reached at this time point (5). Proteins extracted from LH-stimulated and non-stimulated ovarian follicles were digested with endoproteinase and enriched for phosphopeptides in parallel; the phosphopeptides were then labeled with isotopically labeled TMT6 reagents. Peptides were pooled, prefractionated by bRP chromatography, and finally analyzed by LC-MS/MS (Figure 1) (27).

Our analysis identified 4097 phosphorylated proteins in ovarian follicles, of which 866 proteins had at least one site with a statistically significant difference in phosphorylation intensity following LH stimulation (Supplementary Data Files S1-S3). Most but not all previously described LH-induced protein phosphorylation changes were identified by this analysis. The absence of some known phosphorylation sites within proteins that have been detected in other studies could result because of technical limitations of the applied experimental protocol, which utilized trypsin and LysC proteases for peptide generation, leading to difficulties in quantifying phosphorylation sites in lysine or arginine rich regions, for example for NPR2 and CREB1. Furthermore, despite the presence of protease/phosphatase inhibitors and strong denaturing agent such as SDS in the sample lysis buffer, it still cannot be completely excluded that particular peptides were degraded by the proteases used in generating the samples used for MS, or were not recovered during the initial fraction steps (37).

Strikingly, of the 866 proteins that showed changes in phosphorylation in response to LH stimulation, 347 proteins had sites that showed significant *decreases* in phosphorylation intensity in response to LH stimulation. Though inhibitory phosphorylations of some kinases could explain some of these changes, this finding suggests that signaling by protein kinase A and other kinases that are activated by the G_q_ G-protein (1, 38, 39) massively and rapidly increases phosphatase activity targeting multiple substrates. Identification of such phosphatases could contribute to understanding of multiple aspects of LH signaling.

### Mass spectrometric analysis of LH-induced changes in phosphorylation of PPP family phosphatase regulatory subunits in rat preovulatory follicles

Phosphatases in the PPP family are estimated to be responsible for ∼90% of dephosphorylation events in eukaryotic cells (40). These phosphatases are of particular interest because their activity is required for LH-stimulated and PKA-dependent dephosphorylation of the NPR2 guanylyl cyclase in the granulosa cells, which triggers meiotic resumption the oocyte (Figures 2A,B,S4). We confirmed that, as for rat follicles (5), LH-induced dephosphorylation of NPR2 in mouse follicles is inhibited by the PPP-family phosphatase inhibitor, cantharidin (Figure 2A,B).

None of the PPP family phosphatase catalytic subunits showed changes in phosphorylation in response to LH (Supplementary Data File S1). However, among the ∼60 interacting proteins that have been best characterized as regulatory subunits of PPP family phosphatases, as indicated by their assignment of official names including “PPP” (19, 20), 6 showed statistically significant changes in phosphorylation in response to LH (Table 1). The 2 largest phosphorylation increases were in PPP2R5D (also known as PP2A-B56δ) and PPP1R12A (also known as MYPT1) (Table 1, Figure S4). Among the over 200 other proteins that are known to interact with PPP family phosphatase catalytic subunits (with non-“PPP” gene names), those that showed changes in phosphorylation in response to LH are listed in Table S2.

**Table 1.** Statistically significant LH-induced changes in phosphopeptide intensity of well- characterized PPP-family phosphatase regulatory subunits in rat ovarian follicles. This list includes all proteins in the Supplementary Data File 1 that had official names including “PPP”, and for which LH stimulation resulted in a significant change in phosphopeptide intensity. Multiplicity refers to either singly phosphorylated (1) or doubly phosphorylated (2) peptides. Significant increases in phosphopeptide intensity following LH treatment are above the dashed line; significant decreases in phosphopeptide intensity with LH treatment are below the dashed line.

| Gene name | UniProt accession | Amino acid | Site # | Multiplicity | Phosphopeptide intensity<br>(log<sub>2</sub> mean ± SEM) |  | Fold change |
| --- | --- | --- | --- | --- | --- | --- | --- |
|  |  |  |  |  | Control | 350 nM LH |  |
| PPP2R5D | F1MAA3 | S | 53 | 1 | 16.20 ± 0.03 | 19.49 ± 0.27 | 10.4 |
| PPP1R12A* | A0A0G2JWJ0 | S | 507* | 1 | 16.42 ± 0.06 | 18.57 ± 0.59 | 4.7 |
| PPP2R5D | F1MAA3 | S | 566 | 1 | 19.94 ± 0.32 | 21.90 ± 0.35 | 4.1 |
| PPP2R5A | D3ZDI7 | S | 38 | 1 | 16.45 ± 0.13 | 17.92 ± 0.07 | 2.9 |
| PPP1R9A | O35867 | S | 185 | 1 | 14.20 ± 0.07 | 15.36 ± 0.14 | 2.4 |
| PPP2R5C | D4A1A5 | S | 497 | 1 | 19.03 ± 0.18 | 20.08 ± 0.31 | 2.2 |
| PPP2R5D | F1MAA3 | S | 81<sup>#</sup> | 2 | 22.39 ± 0.12 | 23.15 ± 0.15 | 1.8 |
| PPP2R5D | F1MAA3 | S | 82<sup>#</sup> | 2 | 22.12 ± 0.11 | 22.84 ± 0.17 | 1.7 |
| PPP1R14A | Q99MC0 | S | 26 | 1 | 22.38 ± 0.02 | 19.48 ± 0.44 | -7.0 |
| PPP1R14A | Q99MC0 | S | 136 | 1 | 18.75 ± 0.24 | 16.95 ± 0.63 | -3.3 |
| PPP1R14A | Q99MC0 | S | 16 | 2 | 17.80 ± 0.05 | 16.11 ± 0.23 | -3.1 |
| PPP1R14A | Q99MC0 | S | 26 | 2 | 17.80 ± 0.05 | 16.11 ± 0.23 | -3.1 |

\* The fragment ion sequence containing this phosphosites is almost identical for PPP1R12A-S507 and PPP1R12B-S502, such that its assignment to PPP1R12A or PPP1R12B is ambiguous. We assigned it to PPP1R12A because ddPCR analysis showed that PPP1R12A mRNA is present in mouse ovarian follicles at a concentration ~5X higher than PPP1R12B mRNA (Figure S3).
<sup>#</sup> q-values for these sites (0.041 and 0.046, respectively) were just above the significance threshold of 0.01.

PPP2R5D showed an ∼10x increase in phosphorylation signal intensity on serine 53, and a 4x increase on serine 566. PPP2R5D-S566 phosphorylation in response to LH receptor stimulation in rat ovarian granulosa cells has been reported previously (7), and phosphorylation of serine 566 is critical for activation of PPP2 by PKA in other cells (8). A marginally significant increase in phosphorylation intensity on serine 81 and serine 82 of PPP2R5D was also detected. PPP1R12A showed an ∼4x increase in the signal intensity of phosphorylation of on serine 507, known to be a regulatory site in other cells (11). Another PPP1 regulatory subunit, PPP1R14A (also known as CPI-17), showed a 3-7 X *decrease* in phosphorylation signal intensity on serines 16 and 26 (Table 1).

These LH-dependent changes in phosphorylation of PPP family phosphatase regulatory subunits identified candidates for phosphatase complexes that might mediate dephosphorylation of NPR2 in response to LH signaling. Because LH-induced NPR2 dephosphorylation depends on the activation of protein kinase A, as demonstrated by inhibition by the PKA-specific inhibitor Rp-8-CPT-cAMPS (41) (Figure 2A,B), we focused on investigating regulatory subunits whose phosphorylation increased rather than decreased in response to LH.

### Western blot confirmation of PPP1R12A and PPP2R5D phosphorylation increases in LH-treated mouse preovulatory follicles

Consistent with the MS analysis, quantitative western blotting of mouse ovarian follicles using a phosphospecific antibody showed that LH caused an 11x increase in phosphorylation of PPP1R12A on serine 507 (Figure 3A,B). A 3x increase in phosphorylation on serine 668 was also detected (Figure 3A,B), as previously seen with LH receptor stimulation of granulosa cells from rat preovulatory follicles (42). 3 other sites on PPP1R12A showed no change in phosphorylation in response to LH (Figure 3A,B). Inhibition of serine 507 phosphorylation by the PKA-specific inhibitor Rp-8-CPT-cAMPS (41) showed that the LH response depended on protein kinase A (Figure 3C,D), like the LH-induced dephosphorylation of NPR2 (Figure 2A,B).

Phosphospecific antibodies were not available for PPP2R5D, so as an alternative, we used Phos-tag gel electrophoresis (34) followed by western blotting with antibodies against the total proteins. Using this approach, we confirmed that LH increased phosphorylation of PPP2R5D in mouse ovarian follicles, and determined that the phosphorylation was PKA-dependent (Figure 4A,B).

**Figure 4.**
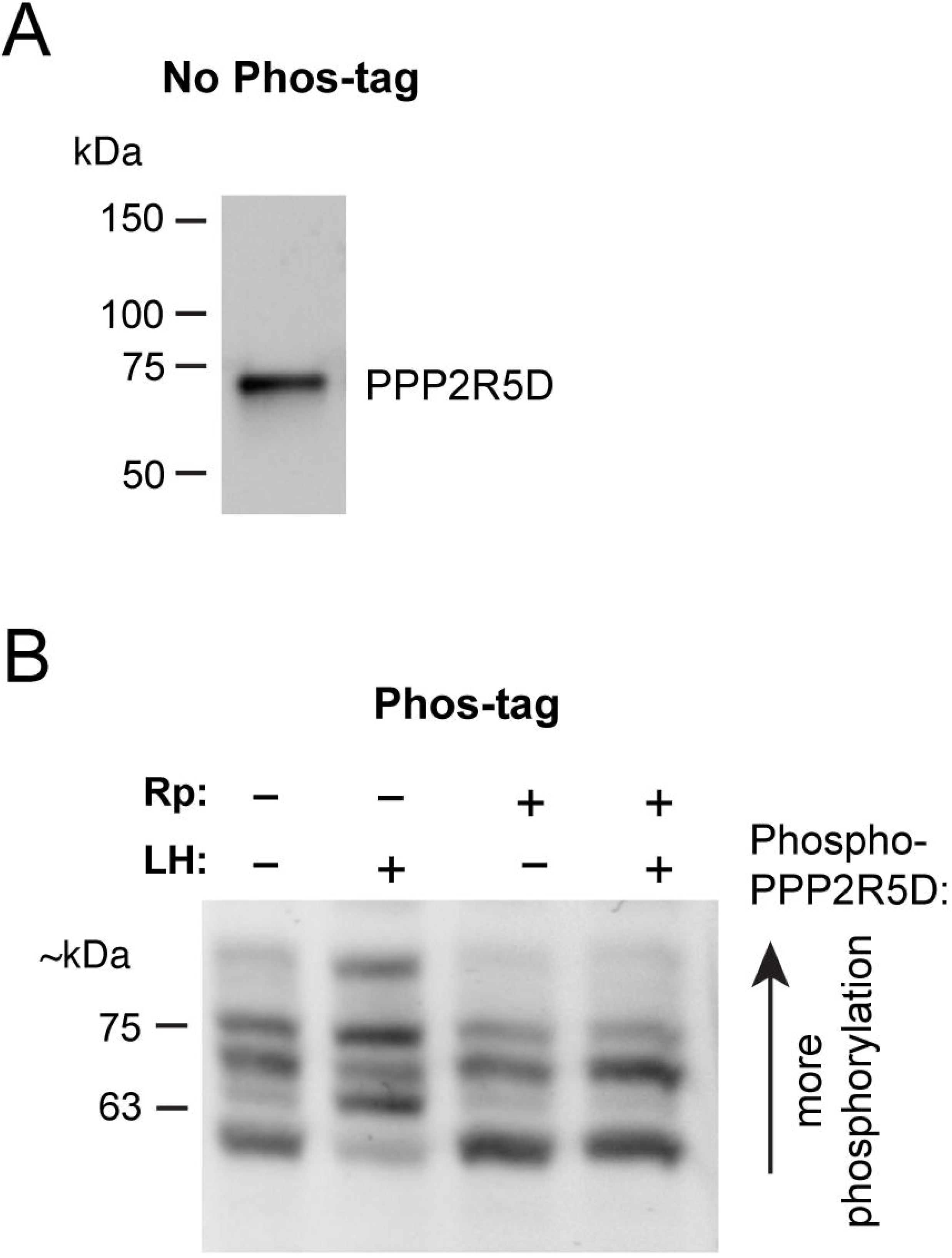
Phos-tag gel/ western blot analysis of LH-induced phosphosite changes in PPP2R5D in mouse ovarian follicles. (**A**) Standard non-Phos-tag SDS-PAGE and western blot, showing that PPP2R5D runs as a single band at the expected molecular weight (69 kDa). (**B**) Phos-tag SDS-PAGE and western blot, showing LH-induced PPP2R5D phosphorylation and its PKA-dependence. Follicles were treated with or without the PKA inhibitor Rp-8-CPT-cAMPS (Rp; 10 mM, 4 hr) then stimulated with 10 nM LH (30 min). Similar results were obtained in 3 other experiments.

### Investigation of the function of LH-induced phosphorylation of PPP family phosphatase regulatory subunits in pathways leading to NPR2 dephosphorylation, using genetically modified mice with serine-to-alanine mutations

Our MS and western blotting results suggested PKA-dependent phosphorylation of PPP1R12A on serine 507, and of PPP2R5D on serines 53 and 566, as possible mediators of LH-induced NPR2 dephosphorylation. Phosphorylations of these sites could potentially increase the interaction of the regulatory subunits with the NPR2 protein, bringing the associated catalytic subunits close to NPR2. The known substrate interaction motif for PPP2R5D, LxxIxE within an intrinsically disordered region (40), is not found in NPR2, and although PPP1R12A contains a motif that binds its catalytic subunit, a general motif that serves to recruit PPP1R12A to substrates is thus far unknown (43). However, other as yet uncharacterized interaction sites could be present in the NPR2 protein that mediate either direct or indirect binding to PPP1R12A or PPP2R5D.

Due to the low abundance of NPR2 in follicles, and the limited amount of follicle protein that could be obtained, searching for such interaction sites by coimmunoprecipitation was not attempted. Instead, we tested the function of the phosphorylated sites by making serine-to-alanine substitutions, an approach that was used effectively to analyze phosphorylation changes detected in a previous mass spectrometry study (27). The serine-to-alanine modifications were made in vivo rather than in isolated granulosa cells because, like some other responses to LH that fail to occur in isolated cells (44), LH-induced NPR2 dephosphorylation is not preserved in isolated granulosa cells (our unpublished results). We generated 2 mouse lines in which the serines in PPP1R12A and PPP2R5D that were phosphorylated in response to LH were changed to alanines (Figures S1, S2), to test whether these LH-induced phosphorylations are required for NPR2 dephosphorylation. The mouse lines that we made with global serine-to-alanine modifications were viable and did not show obvious morphological or physiological defects. Homozygous mice were fertile, although fertility was not investigated quantitatively.

In the first of these mouse lines, in which serine 507 of *Ppp1r12a* was changed to alanine, western blots confirmed that ovarian follicles did not show the LH-induced increase in PPP1R12A-S507 phosphorylation that was seen in wildtype follicles (Figure 5A). We then investigated whether the S507A mutation inhibited the LH-stimulated dephosphorylation of NPR2, and found that it did not (Figure 5B,C).

**Figure 5.**
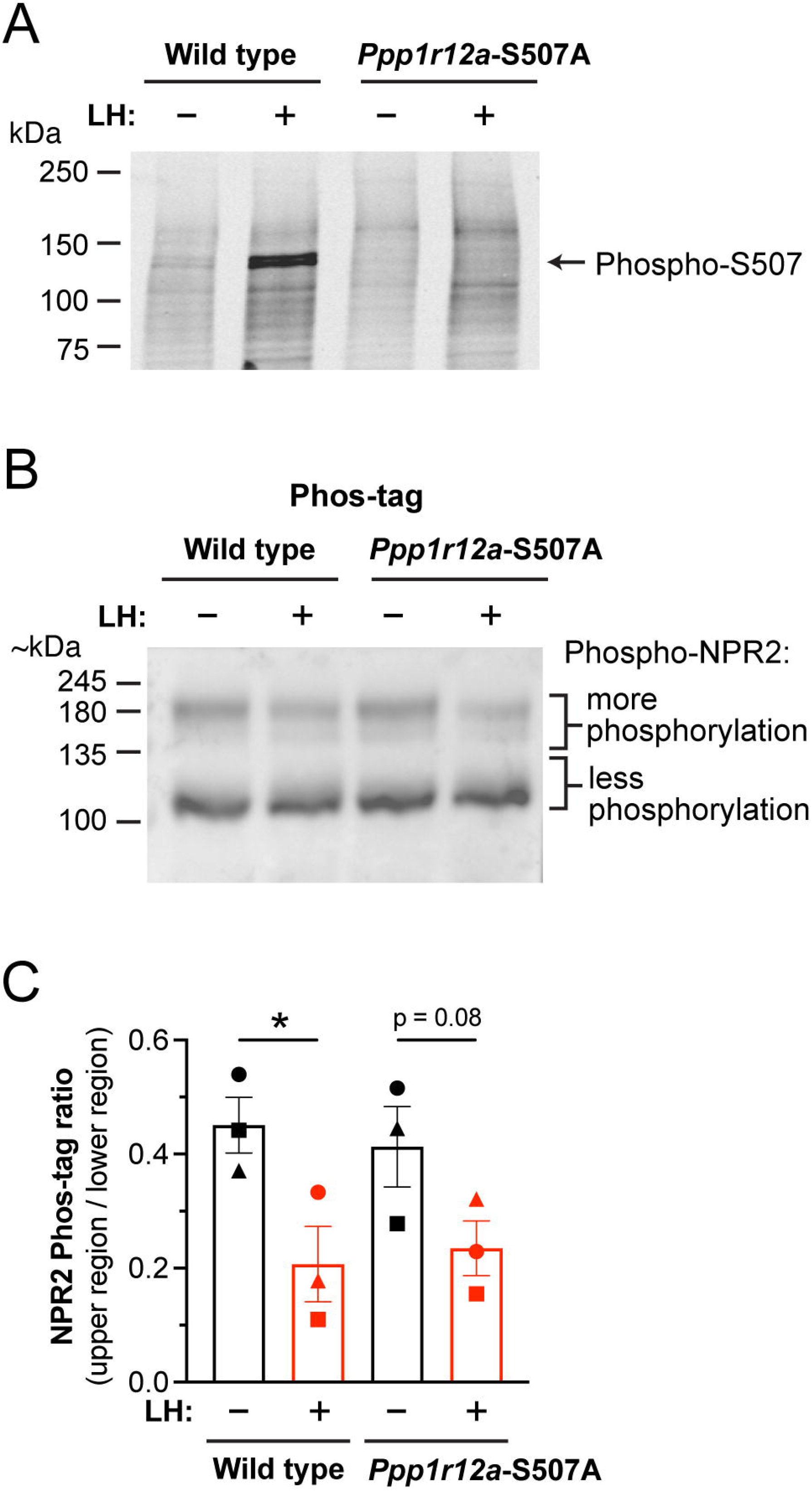
No effect of preventing PPP1R12A-S507 phosphorylation on LH-induced NPR2 dephosphorylation. (**A**) Western blot confirming inhibition of LH-stimulated PPP1R12A-S507 phosphorylation in ovarian follicles from *Ppp1r12a*-S507A mice. Similar results were obtained with 2 other sets of samples. (**B**) LH-induced NPR2 dephosphorylation is not inhibited in ovarian follicles from *Ppp1r12a*-S507A mice. (**C**) Summary (mean ± SEM) of 3 experiments like that shown in B. Each symbol (circle, square, triangle) represents the results from an independently prepared set of samples, each analyzed on a separate blot. Data were analyzed by two-tailed paired t-tests. * p < 0.05.

In a second mouse line, serines 53 and 566 of *Ppp2r5d* were changed to alanines, corresponding to the 2 sites that showed significantly increased phosphorylation in response to LH (Table 1). Because serines 81 and 82 also showed close to significant increases in phosphorylation (Table 1), these sites were also changed to alanines. The resulting mouse line with the 4 serine-to-alanine mutations was called *Ppp2r5d*-S53A/S81A/S82A/S566A or *Ppp2r5d*-4A for short. Phos-tag gel electrophoresis of protein in follicles from mice with these 4 mutations showed much reduced basal phosphorylation of the PPP2R5D protein, and no increase in phosphorylation in response to LH (Figure 6A). However, as with the *Ppp1r12a*- S507A mice, the *Ppp2r5d*-4A mice showed no inhibition of LH-induced dephosphorylation of NPR2 (Figure 6B,C). These results indicated that neither of these individual regulatory subunits of PPP1 and PPP2 is by itself responsible for LH-induced dephosphorylation of NPR2.

**Figure 6.**
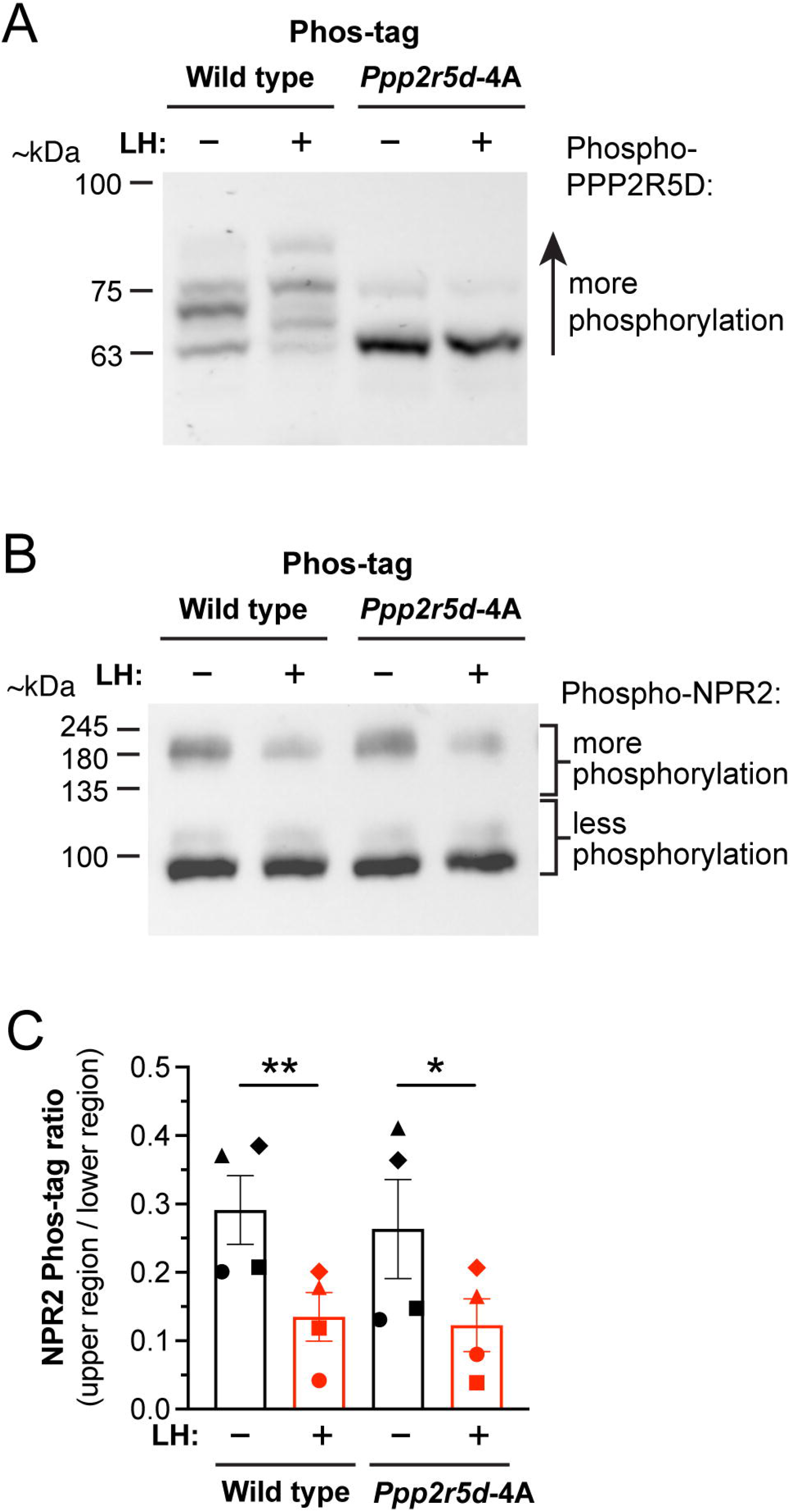
No effect of preventing PPP2R5D-S53/S81/S82/S566 phosphorylation on LH-induced NPR2 dephosphorylation in mouse ovarian follicles. (**A**) Western blot confirming inhibition of LH- stimulated PPP2R5D phosphorylation in ovarian follicles from *Ppp2r5d*-4A mice. Similar results were obtained with 3 other sets of samples. (**B**) LH-induced NPR2 dephosphorylation is not inhibited in ovarian follicles from *Ppp2r5d*-4A mice. (**C**) Summary (mean ± SEM) of 4 experiments like that shown in (B). Each symbol (circle, square, triangle, diamond) represents the results from an independently prepared set of samples, each analyzed on a separate blot. Bars show mean ± SEM. Data were analyzed by two-tailed paired t-tests. *p < 0.05, **p < 0.01.

Whether mice with these mutations in *both Ppp1r12a* and *Ppp2r5d* would show inhibition of LH-induced dephosphorylation of NPR2 will be useful to examine, as different PPP family proteins can have overlapping functions. In a recent study, mice lacking *Ppp2r5d* or *Ppp2r5c* had no overt phenotypes, but mice lacking both of these genes were arrested at embryonic day 12 due to a heart defect (45). LH also induced smaller (2-3 fold) but significant increases in phosphorylation signal intensity of serines in PPP2R5A, PPP1R9A, and PPP2R5C (Table 1). Smaller fold changes may well be important, particularly if they reflect larger changes that occur in particular cellular compartments (37). In addition, significant increases in phosphorylation occur for several other less well characterized PPP interacting proteins (Table S2), raising the possibility of even more complex redundancy. The large LH-induced *decrease* in phosphorylation of several sites of the R14A regulatory subunit of PPP1, also known as CPI-17, (Table 1) suggests PPP1R14A as another possible regulator of LH-induced dephosphorylation of NPR2, although the decrease in R14A phosphorylation could not be a direct action of PKA.

Alternatively, LH signaling could cause NPR2 dephosphorylation by decreasing kinase activity, although the kinases responsible for phosphorylating the juxtamembrane serines of NPR2 are unknown (46). Stimulation of the LH receptor increases phosphorylation of glycogen synthase kinase-3β (GSK3B) on serine 9 (7) and this PKA-dependent phosphorylation inhibits GSK3B activity (47), suggesting a possible candidate. Our mass spectrometric analysis identified multiple serine/threonine kinases that are phosphorylated in response to LH (Supplementary Data Files S1 and S3), suggesting other candidates that could be considered.

Identification of the PPP family phosphatases responsible for NPR2 dephosphorylation, or of kinases whose phosphorylation of NPR2 is decreased by LHR activation, will be useful not only for understanding of signaling in the ovary, but also in other systems in which the phosphorylation state of NPR2, or of the related guanylyl cyclase NPR1, controls function (46, 48). For example, inhibition of bone growth by fibroblast growth factor depends at least in part on NPR2 dephosphorylation by a PPP family phosphatase (35, 49–51).

### LH-induced changes in phosphorylation of protein phosphatases that regulate the cytoskeleton and cell motility

Although our studies have been primarily directed at identifying LH-modified phosphatase regulatory proteins in relation to the question of how LH signaling dephosphorylates the NPR2 guanylyl cyclase, the phosphatases identified by our screen could regulate multiple other pathways in the preovulatory follicle. Strikingly, among both the PPP family phosphatases (Tables 1 and S2) and the non-PPP family phosphatases (Table 2) whose phosphorylation changes in response to LH, many have known functions in regulating the cytoskeleton and cell motility. Among the PPP family phosphatase regulatory proteins showing large LH-induced changes in phosphorylation intensity (Table 1), PPP1R12A (MYPT1) and PPP1R14A (CPI-17) are regulators of myosin light chain phosphorylation (52), which controls non-muscle myosin assembly and contractility (53). Other PPP family regulatory subunits that show large LH-induced changes in phosphorylation intensity (Table S2) include HDAC6, which regulates tubulin (54), and FARP1, PHACTR4, and PHACTR2, which regulate actin (55, 56).

**Table 2.**
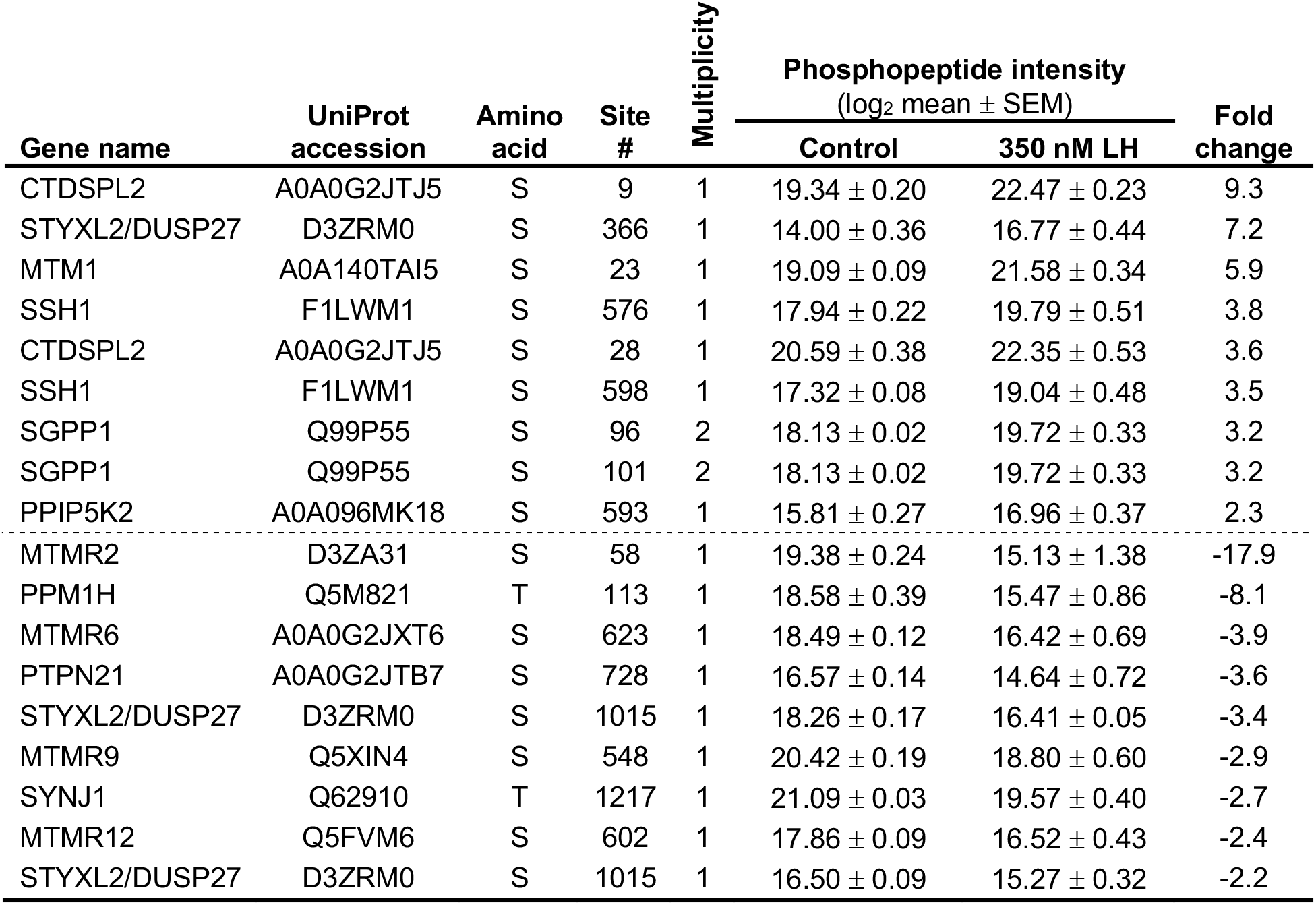
Statistically significant LH-induced changes in phosphopeptide intensity of non-PPP- family phosphatases in rat ovarian follicles. Previously published lists of phosphatase genes (64, 65) were used to query the phosphopeptide database (Supplementary Data File 1) for significant differences in response to LH. Multiplicity refers to either singly phosphorylated (1) or doubly phosphorylated (2) peptides. Significant increases in phosphopeptide intensity following LH treatment are above the dashed line; significant decreases in phosphopeptide intensity with LH treatment are below the dashed line.

In addition to the PPP family phosphatases described above, our quantitative mass spectrometry analysis indicated that LH causes large changes in phosphorylation intensity of multiple protein phosphatases outside of the PPP family (Table 2). These include the Slingshot protein phosphatase 1 (SSH1), for which LH causes an ∼4x increase in phosphorylation intensity on serines 576 and 598 (Figure 7). SSH1 dephosphorylates and activates proteins in the cofilin/actin depolymerizing factor family (CFL1, CFL2, DSTN) (57), and is of high interest because LH signaling decreases cofilin phosphorylation in isolated granulosa cells, causing an increase in motility (6). Granulosa cell motility increases in intact ovarian follicles after the preovulatory surge of LH, and may contribute to the ovulatory process (58).

**Figure 7.**
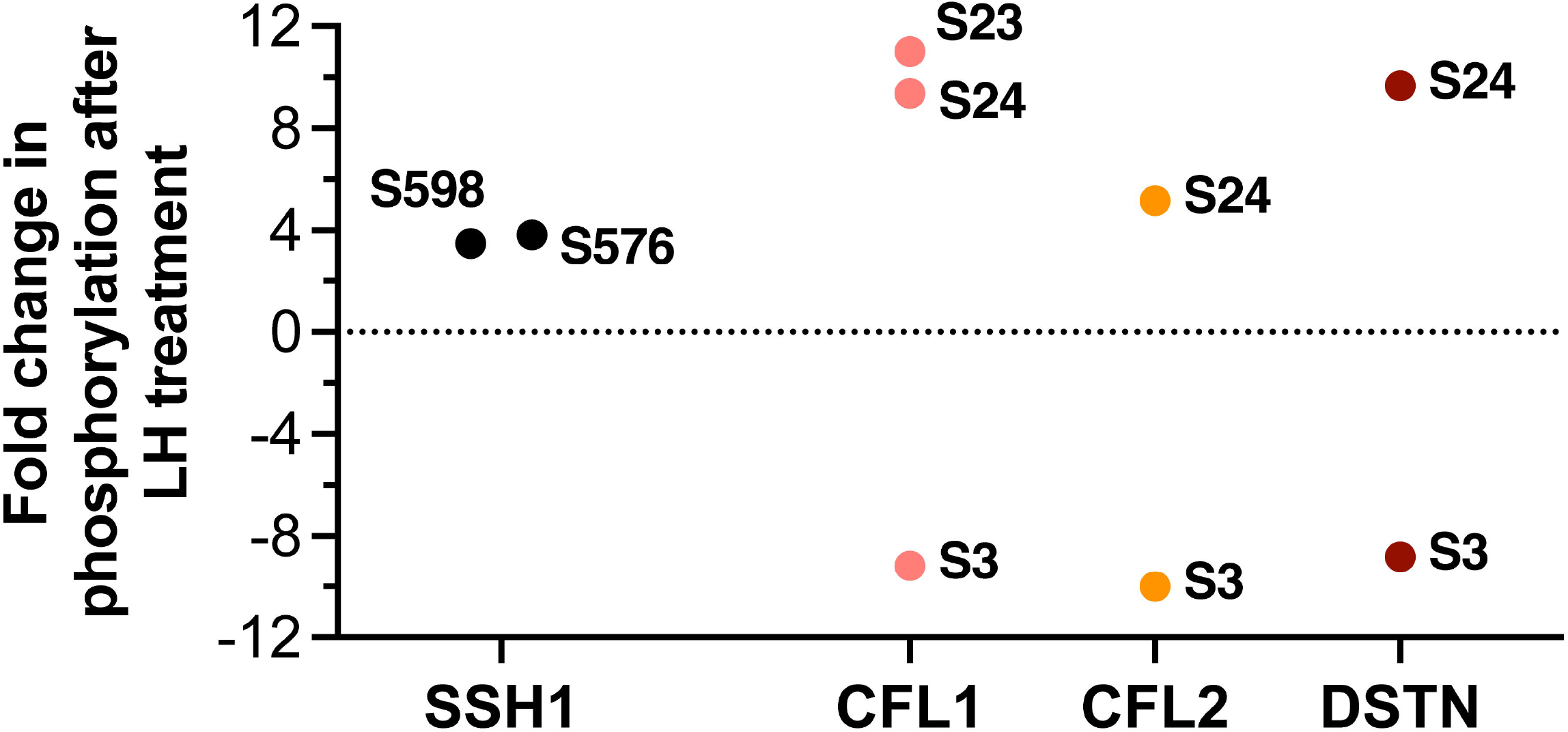
LH-induced phosphorylation changes in the Slingshot phosphatase SSH1 and in cofilin/actin depolymerizing factor family proteins CFL1, CFL2, and DSTN that are dephosphorylated by SH1. SSH1 showed ∼4x increases in phosphorylation intensity at S576 and S598. CFL1, CFL2, and DSTN all showed a decrease in phosphorylation intensity of serine 3; dephosphorylation at this conserved site activates actin-depolymerizing activity of these proteins (57). LH-induced changes in phosphorylation intensity of other sites (S23, S24) of CFL1, CFL2, and DSTN were also detected. Primary data for SSH1 is listed in Table 2 and Supplementary Data File 1. Primary data for CFL1, CFL2, and DSTN is listed in Supplementary Data File 1. All of the indicated changes are statistically significant (see Methods).

Although it is unknown if the modifications of SSH1 on serines 576 and 598 increase its phosphatase activity, our MS screen revealed LH-induced decreases in phosphorylation intensity of the serine 3 regulatory sites of CFL1, CFL2, and DSTN to ∼10% of baseline (Figure 7). Dephosphorylation of serine 3 on these proteins increases their activity (57). The LH-induced phosphorylation of serines 576 and 598 of SSH1 suggests that LH might dephosphorylate and activate cofilin family members by way of SSH1.

### A database of LH-induced protein phosphorylation changes as a resource for future studies

In the present study, we examined changes in the global phosphoproteome in response to LH signaling in rat preovulatory follicles. We then used this dataset to identify protein phosphatases that are modified by LH signaling, and to investigate the function of phosphatases in the PPP family in dephosphorylating the NPR2 guanylyl cyclase, which triggers oocyte meiotic resumption (5, 14, 15). As discussed above, the mass spec data also suggested that LH-induced changes in phosphorylation of protein phosphatases may regulate LH-induced changes in the cytoskeleton and cell motility. In addition, inhibitor studies in rat preovulatory follicles have indicated that PPP-family phosphatase activity is also required for LH stimulation of synthesis of progesterone (59). Whether LH-induced changes in protein phosphatase phosphorylation affect their subcellular localization is unknown, but studies in other cells have shown that PPP family phosphatases can shuttle between the cytoplasm and nucleus, potentially regulating transcription (60). Thus, our identification of phosphatase proteins whose phosphorylation state is modified by LH (Tables 1,2, S2) may provide clues about multiple signaling pathways in preovulatory follicles that are mediated by phosphatases.

More generally, the data from our phospho-mass spectrometric analysis (Supplementary Data Files S1-S3) will be broadly useful for future studies of the many aspects of LH signaling that are mediated by changes in protein phosphorylation. Many of the LH-induced phosphorylation changes identified by our analysis have not been previously described, and the data provide a general resource for future studies, extending beyond our particular interest in phosphatases. Because changes in protein phosphorylation are dynamic (37), and because additional changes could occur as a consequence of newly synthesized kinase and phosphatase enzymes and their ligands, further insights could be gained from similar analyses of samples prepared at different time points after applying LH.

To understand the broader implications of the phosphoproteome changes induced by LH, we applied gene ontology (GO) enrichment analysis to the proteins showing at least one significantly regulated phosphosite (Figure 8; Supplementary Data Files S4-S6). This analysis revealed that terms related to signaling, the cytoskeleton, and DNA were consistently enriched across all three GO domains in this dataset. The robust and rapid effects of LH signaling on the cytoskeleton of follicle cells, likely primarily represented by granulosa cells, are particularly striking. For instance, half of the 20 most significant GO “Biological Process” terms are directly related to the cytoskeleton and its reorganization (Figure 8A; Supplementary Data File S4). Similarly, among the 10 most significant GO “Cellular Compartment” terms, 5 are related to aspects of the cytoskeleton or motile processes (Figure 8B; Supplementary Data File S5). Additionally, many of the significant GO “Molecular Function” terms reflect binding to elements of the cytoskeleton, as well as binding or activity of cytoskeletal regulators (Figure 8C; Supplementary Data File S6).

**Figure 8.**
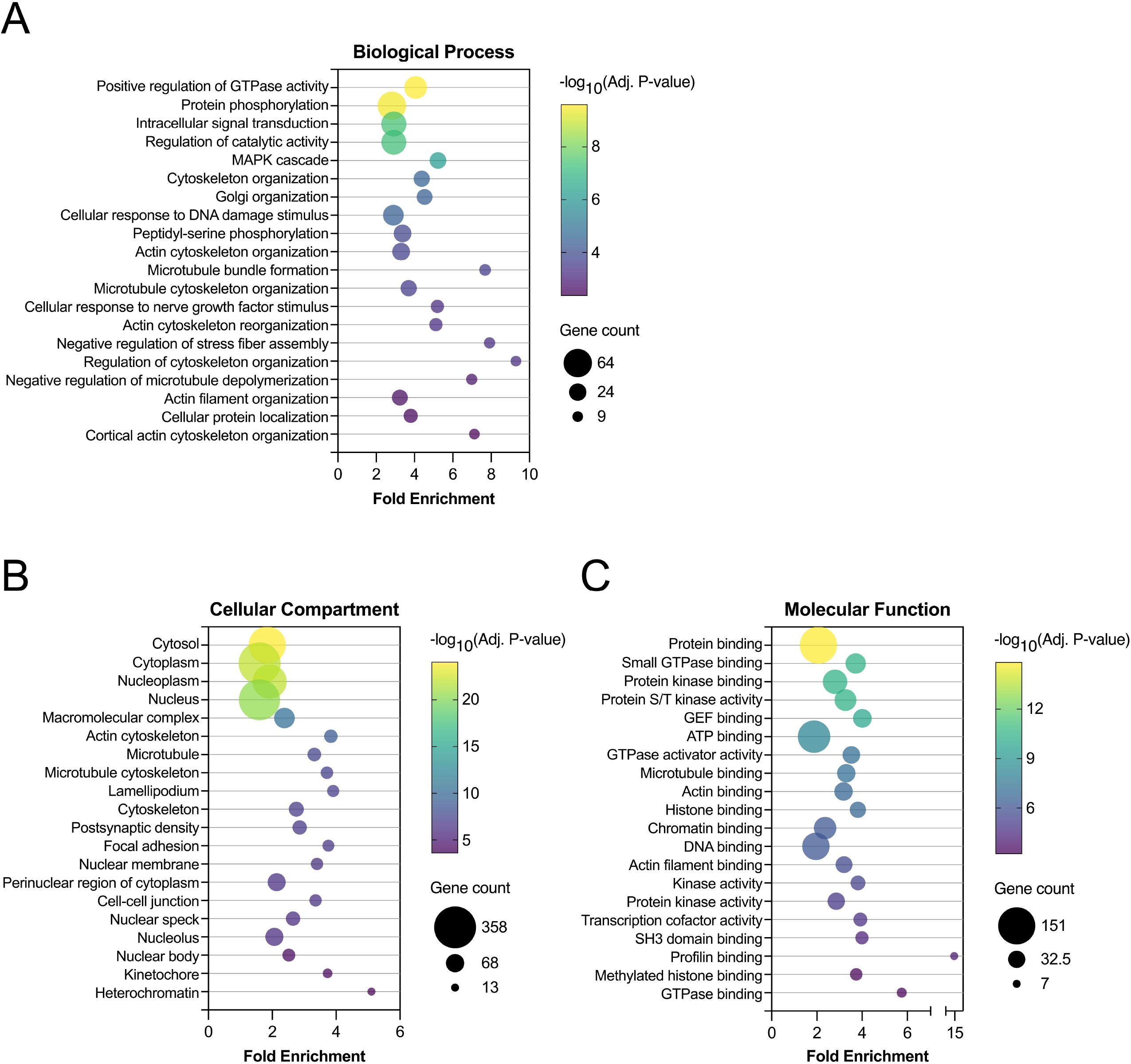
Gene ontology (GO) summary of rat preovulatory follicle phosphoproteome dataset. Genes encoding proteins with at least one phosphosite that was significantly regulated by LH treatment were subjected to GO enrichment analysis using DAVID 2021 (see Methods). For each of the three GO domains, (**A**) Biological Process, (**B**) Cellular Compartment, and (**C**) Molecular Function, the 20 terms with the smallest Benjamini-Hochberg-adjusted p-values are plotted. Fold enrichment refers to the enrichment of the GO term in the list of genes encoding proteins with a significantly regulated phosphosite, compared to the background population of genes. Bubble size reflects the number of genes in the GO term that encode proteins with at least one significantly regulated phosphosite. The largest and smallest bubbles reflect the terms with the most and fewest genes, respectively; the middle bubble reflects the geometric mean. Bubble color reflects the inverse-log10 of the adjusted p-values, as indicated on each color bar scale. Complete GO enrichment analysis results are provided in Supplementary Data Sets 4-6.

Another topic on which our dataset may be illuminating is steroidogenesis. Phosphoproteomic studies identifying proteins phosphorylated in response to cAMP elevation have provided clues for understanding of LH regulation of steroidogenesis in Leydig cells of the testis (37, 61). Our phosphoproteomic data revealed similar regulation of many of the same proteins (Supplementary Data File S1), and could provide insights into how LH rapidly stimulates steroid production in the follicle (62). Our data also identified >300 nuclear proteins whose phosphorylation state changes by 30 minutes after LH stimulation (Supplementary Data File S5), which could provide clues about the transcriptional regulation of events leading to ovulation. In summary, phosphoproteomic data described here could benefit studies of many aspects of LH signaling in preovulatory follicles.

## Supplementary material

This article contains supplementary material (Figures S1-S4, Tables S1 and S2, Supplementary Data Files S1-S6).

## Data availability

The mass spectrometry data have been deposited to the ProteomeXchange Consortium via the PRIDE (63) partner repository with the data set identifier PXD033878.

## Author contributions

JRE, IS, HU, and LAJ designed the study. IS and HU carried out the mass spectrometry studies. JRE, TFU, and LAJ carried out the western blotting studies. KML and SPY generated the genetically modified mice. JRE, IS, SPY, HU, and LAJ analyzed the data and wrote the manuscript; all authors approved the manuscript.

## Acknowledgments

We thank Monika Raabe for assistance in the sample preparation for LC-MS/MS, and Deborah Kaback and Iris Nakashima for assistance in generating the genetically modified mice. We also thank Lincoln Potter, Rebecca Page, Joe Beavo, Melina Schuh, Corie Owen, Leia Shuhaibar, and Rachael Norris for helpful discussions and critical reading of the manuscript. The graphical abstract and Figure 1 were created with BioRender.com. We dedicate this paper to the memory of Alexei Evsikov, who inspired our studies of the bioinformatics of reproduction.

## Abbreviations

bRP: basic reverse-phase
FSH: follicle stimulating hormone
GO: gene ontology
HA: hemagglutinin
LC-MS/MS: liquid chromatography-tandem mass spectrometry
LH: luteinizing hormone
NPR2: natriuretic peptide receptor 2
PKA: protein kinase A
PPP: phosphoprotein phosphatase
TMT: tandem mass tag

## References

1. Hunzicker-Dunn M, Mayo K. Gonadotropin signaling in the ovary. In Knobil and Neill’s Physiology of Reproduction, 4th Edition, 2015. TM Plant, AJ Zeleznik, editors. Academic Press/San Diego, pp. 895–945.

2. Jaffe LA, Egbert JR. Regulation of mammalian oocyte meiosis by intercellular communication within the ovarian follicle. Annu Rev Physiol 2017; 79:237–260.

3. Clarke HJ. Regulation of germ cell development by intercellular signaling in the mammalian ovarian follicle. WIRE’s Dev Biol 2018; 7:e294.

4. Richards JS, Ascoli M. Endocrine, paracrine, and autocrine signaling pathways that regulate ovulation. Trends Endocrinol Metab 2018; 29:313–325.

5. Egbert JR, Shuhaibar LC, Edmund AB, Van Helden DA, Robinson JW, Uliasz TF, Baena V, Geerts A, Wunder F, Potter LR, Jaffe LA. Dephosphorylation and inactivation of the NPR2 guanylyl cyclase in the granulosa cells contributes to the LH-induced cGMP decrease that causes resumption of meiosis in rat oocytes. Development 2014; 141:3594–3604.

6. Karlsson AB, Maizels ET, Flynn MP, Jones JC, Shelden EA, Bamburg JR, Hunzicker-Dunn M. Luteinizing hormone receptor-stimulated progesterone production by preovulatory granulosa cells requires protein kinase A-dependent activation/dephosphorylation of the actin dynamizing protein cofilin. Mol Endocrinol 2010; 24:1765–1781.

7. Flynn MP, Maizels ET, Karlsson AB, McAvoy T, Ahn JH, Nairn AC, Hunzicker-Dunn M. Luteinizing hormone receptor activation in ovarian granulosa cells promotes protein kinase A-dependent dephosphorylation of microtubule-associated protein 2D. Mol Endocrinol 2008; 22:1695–1710.

8. Ahn J-H, McAvoy T, Rakhilin, SV, Nishi A, Greengard P, Nairn AC. Protein kinase A activates protein phosphatase 2A by phosphorylation of the B56δ subunit. Proc Natl Acad Sci USA 2007; 104:2979–2984.

9. Yamashiro S, Yamakita Y, Totsukawa G, Goto H, Kaibuchi K, Ito M, Hartshorne DJ, Matsumura F. (2008) Myosin phosphatase-targeting subunit 1 regulates mitosis by antagonizing polo-like kinase 1. Develop Cell 2008; 14:787–797.

10. Brautigan DL, Shenolikar S. Protein serine/threonine phosphatases: keys to unlocking regulators and substrates. Ann Rev Biochem 2018; 87:921–964.

11. Samson SC, Elliott A, Mueller BD, Kim Y, Carney KR, Bergman JP, Blenis J, Mendoza MC. p90 ribosomal S6 kinase (RSK) phosphorylates myosin phosphatase and thereby controls edge dynamics during cell migration. J Biol Chem 2019; 294:10846–10862.

12. Zhang M, Su Y-Q, Sugiura K, Xia G, Eppig JJ. Granulosa cell ligand NPPC and its receptor NPR2 maintain meiotic arrest in mouse oocytes. Science 2010; 330:366–369.

13. Shuhaibar LC, Egbert JR, Norris RP, Lampe PD, Nikolaev VO, Thunemann M, Wen L, Feil R, Jaffe LA. Intercellular signaling via cyclic GMP diffusion through gap junctions in the mouse ovarian follicle. Proc Natl Acad Sci USA 2015; 112:5527–5532.

14. Shuhaibar LC, Egbert JR, Edmund AB, Uliasz TF, Dickey DM, Yee S-P, Potter LR, Jaffe LA. Dephosphorylation of juxtamembrane serines and threonines of the NPR2 guanylyl cyclase is required for rapid resumption of oocyte meiosis in response to luteinizing hormone. Dev Biol 2016; 409:194–201.

15. Egbert JR, Robinson JW, Uliasz TF, Potter LR, Jaffe LA. Cyclic AMP links luteinizing hormone signaling to dephosphorylation and inactivation of the NPR2 guanylyl cyclase in ovarian follicles. Biol Reprod 2021; 104:939–941.

16. Swingle M, Ni L, Honkanen RE. Small molecule inhibitors of ser/thr protein phosphatases: specificity, use and common forms of abuse. Methods Mol Biol 2007; 365:23–38.

17. Li Y-M, MacKintosh C, Casida JE. Protein phosphatase 2A and its [^3^H]cantharidin/[^3^H]endothall thioanhydride binding site. Inhibitor specificity of cantharidin and ATP analogues. Biochem Pharmacol 1993; 46:1435–1443.

18. Wang Y, Santini F, Qin K, Huang CY. A Mg^2+-^dependent, Ca^2+^-inhibitable serine/threonine protein phosphatase from bovine brain. J Biol Chem 1995; 270:25607–25612.

19. Bollen M, Peti W, Ragusa MJ, Beullens M. The extended PP1 toolkit: designed to create specificity. Trends Biochem Sci 2010; 35:450–458.

20. Brauer BL, Wiredu K, Mitchell S, Moorhead GB, Gerber SA, Kettenbach AN. Affinity-based profiling of endogenous phosphoprotein phosphatases by mass spectrometry. Nat Protoc 2021; 16:4919–4943.

21. Baena V, Owen CM, Uliasz TF, Lowther KM, Yee S-P, Terasaki M, Egbert JR, Jaffe LA. Cellular heterogeneity of the LH receptor and its significance for cyclic GMP signaling in mouse preovulatory follicles. Endocrinology 2020; 161:bqaa074.

22. Egbert JR, Uliasz TF, Shuhaibar LC, Geerts A, Wunder F, Kleiman RJ, Humphrey JM, Lampe PD, Artemyev NO, Rybalkin SD, Beavo JA, Movsesian MA, Jaffe LA. Luteinizing hormone causes phosphorylation and activation of the cyclic GMP phosphodiesterase PDE5 in rat ovarian follicles, contributing, together with PDE1 activity, to the resumption of meiosis. Biol Reprod 2016; 94(5):110.

23. Vigone G, Shuhaibar LC, Egbert JR, Uliasz TF, Movsesian MA, Jaffe LA. Multiple cAMP phosphodiesterases act together to prevent premature oocyte meiosis and ovulation. Endocrinology 2018; 159:2142–2152.

24. Egbert JR, Yee S-P, Jaffe LA. Luteinizing hormone signaling phosphorylates and activates the cyclic GMP phosphodiesterase PDE5 in mouse ovarian follicles, contributing an additional component to the hormonally induced decrease in cyclic GMP that reinitiates meiosis. Dev Biol 2018; 435:6–14.

25. Norris RP, Freudzon M, Mehlmann LM, Cowan AE, Simon AM, Paul DL, Lampe PD, Jaffe LA. Luteinizing hormone causes MAP kinase-dependent phosphorylation and closure of connexin 43 gap junctions in mouse ovarian follicles: one of two paths to meiotic resumption. Development 2008; 135:3229–3238.

26. Wessel D, Fluegge UI. A method for the recovery of protein in dilute solution in the presence of detergents and lipids. Anal Biochem 1984; 138:141–143.

27. Silbern I, Pan K-T, Fiosins M, Bonn S, Rizzoli SO, Fornasiero EF, Urlaub H, Jahn R. Protein phosphorylation in depolarized synaptosomes: dissecting primary effects of calcium from synaptic vesicle cycling. Mol Cell Proteom 2021; 20:100061.

28. McAlister GC, Nusinow DP, Jedrychowski MP, Wühr M, Huttlin EL, Erickson BK, Rad R, Haas W, Gygi SP. MultiNotch MS3 enables accurate, sensitive, and multiplexed detection of differential expression across cancer cell proteomes. Anal Chem 2014; 86:7150–7158.

29. Tyanova S, Temu T, Cox J. The MaxQuant computational platform for mass spectrometry-based shotgun approaches. Nat Protoc 2016; 11:2301–2319.

30. UniProt Consortium. UniProt: a worldwide hub of protein knowledge. Nucleic Acids Res 2019; 47:D506–D515.

31. Smyth GK. limma: Linear models for microarray data. 2005; In Bioinformatics and Computational Biology Solutions Using R and Bioconductor. R Gentleman, VJ Carey, W Huber, RA Irizarry, S Dudoit, (eds) Springer, New York, pp. 397–420.

32. Storey JD, Tibshirani R. Statistical significance for genomewide studies. Proc Natl Acad Sci USA 2003; 100:9440–9445.

33. Sherman BT, Hao M, Qiu J, Jiao X, Baseler MW, Lane HC, Imamichi T, Chang W. DAVID: a web server for functional enrichment analysis and functional annotation of gene lists (2021 update). Nucleic Acids Res 2022; 50:W216–W221.

34. Kinoshita E, Kinoshita-Kikuta E, Takiyama K, Koike T. Phosphate-binding tag, a new tool to visualize phosphorylated proteins. Mol Cell Proteom 2006; 5:749–757.

35. Shuhaibar LC, Kaci N, Egbert JR, Horville T, Loisay L, Vigone G, Uliasz TF, Dambroise E, Swingle MR, Honkanen RE, Biosse Duplan M, Jaffe LA, Legeai-Mallet L. Phosphatase inhibition by LB-100 enhances BMN-111 stimulation of bone growth. JCI Insight 2021; 6:e141426.

36. Sander H, Wallace S, Plouse R, Tiwari S, Gomes AV. Ponceau S waste: Ponceau S staining for total protein normalization. Anal Biochem 2019; 575:44–53.

37. Beavo JA, Golkowski M, Shimizu-Albergine M, Beltejar M-C, Bornfeldt KE, Ong S-E. Phosphoproteomic analysis as an approach for understanding molecular mechanisms of cAMP-dependent actions. Mol Pharmacol 2021; 99:342–357.

38. Breen SM, Andric N, Ping T, Xie F, Offermans S, Gossen JA, Ascoli M. Ovulation involves the luteinizing hormone-dependent activation of G_q/11_ in granulosa cells. Mol Endocrinol 2013; 27:1483–1491.

39. Egbert JR, Fahey PG, Reimer J, Owen CM, Evsikov AV, Nikolaev VO, Griesbeck O, Ray RS, Tolias AS, Jaffe LA. Follicle-stimulating hormone and luteinizing hormone increase Ca^2+^ in the granulosa cells of mouse ovarian follicles. Biol Reprod 2019; 101:433–444.

40. Wang X, Garvanska DH, Nasa I, Ueki Y, Zhang G, Kettenbach AN, Peti W, Nilsson J, Page R. A dynamic charge-charge interaction modulates PP2A:B56 substrate recruitment. eLife 2020; 9:e55966.

41. Schwede F, Maronde E, Genieser H-G, Jastorff B. Cyclic nucleotide analogs as biochemical tools and prospective drugs. Pharmacol Ther 2000; 87:199–226.

42. Law NC, White MF, Hunzicker-Dunn ME. G protein-coupled receptors (GPCRs) that signal via protein kinase A (PKA) cross-talk at insulin receptor substrate 1 (IRS1) to activate the phosphatidylinositol 3-kinase (PI3K)/AKT pathway. J Biol Chem 2016; 291:27160–27169.

43. Kiss A, Erdodi F, Lontay B. Myosin phosphatase: unexpected functions of a long-known enzyme. BBA – Mol Cell Res 2019; 1866:2–15.

44. Lee K-B, Zhang M, Sugiura K, Wigglesworth K, Uliasz T, Jaffe LA, Eppig JJ. Hormonal coordination of natriuretic peptide type C and natriuretic peptide receptor 3 expression in mouse granulosa cells. Biol Reprod 2013; 88(2):42.

45. Dyson JJ, Abbasi F, Varadkar P, McCright B. Growth arrest of PPP2R5C and PPP2R5D double knockout mice indicates a genetic interaction and conserved function for these PP2A B subunits. FASEB BioAdvances 2022; 4:273–282.

46. Kuhn M. Molecular physiology of membrane guanylyl cyclase receptors. Physiol Rev 2016; 96:751–804.

47. Fang X, Yu SX, Lu Y, Bast RC, Woodgett JR, Mills GB. Phosphorylation and inactivation of glycogen synthase kinase 3 by protein kinase A. Proc Natl Acad Sci 2000; 97:11960–11965.

48. Egbert JR, Uliasz TF, Lowther KM, Kaback D, Wagner BM, Healy CL, O’Connell TD, Potter LR, Jaffe LA, Yee S-P. Epitope-tagged and phosphomimetic mouse models for investigating natriuretic peptide-stimulated receptor guanylyl cyclases. Front Mol Neurosci 2022; 15:1007026.

49. Shuhaibar LC, Robinson JW, Vigone G, Shuhaibar NP, Egbert JR, Baena V, Uliasz TF, Kaback D, Yee S-P, Feil R, Fisher MC, Dealy CN, Potter LR, Jaffe LA. Dephosphorylation of the NPR2 guanylyl cyclase contributes to inhibition of bone growth by fibroblast growth factor. eLife 2017; 6:e31343.

50. Robinson JW, Blixt NC, Norton A, Mansky KC, Ye Z, Aparicio C, Wagner BM, Benton AM, Warren GL, Khosla S, Gaddy D, Suva LJ, Potter LR. Male mice with elevated C-type natriuretic peptide-dependent guanylyl cyclase-B activity have increased osteoblasts, bone mass and bone strength. Bone 2020; 135:115320.

51. Wagner BM, Robinson JR, Lin Y-W, Li Y-C, Kaci N, Legeai-Mallet L, Potter LR. Prevention of guanylyl cyclase-B dephosphorylation rescues achondroplastic dwarfism. JCI Insight 2021; 6:e147832.

52. Ito M, Okamoto R, Ito H, Zhe Y, Dohi K. Regulation of myosin light-chain phosphorylation and its roles in cardiovascular physiology and pathophysiology. Hypertension Res 2022; 45:40–52.

53. Vincente-Manzanares M, Ma X, Adelstein RS, Horwitz AR. Non-muscle myosin II takes centre stage in cell adhesion and migration. Nat Rev Mol Cell Biol 2009; 10:778–790.

54. Hubbert C, Guardiola A, Shao R, Kawaguchi Y, Ito A, Nixon A, Yoshida M, Wang X-F, Yao T-P. HDAC6 is a microtubule-associated deacetylase. Nature 2002; 417:455–458.

55. Cheadle L, Biederer T. The novel synaptogenic protein Farp1 links postsynaptic cytoskeletal dynamics and transsynaptic organization. J Cell Biol 2012; 199:985–1001.

56. Huet G, Rajakylä EK, Viita T, Skarp K-P, Crivaro M, Dopie J, Vartiainen, MK. Actin-regulated feedback loop based on Phactr4, PP1 and cofilin maintains the actin monomer pool. J Cell Sci 2013; 126:497–507.

57. Huang TY, DerMardirossian C, Bokoch GM. Cofilin phosphatases and regulation of actin dynamics. Curr Opin Cell Biol 2006; 18:26–31.

58. Owen CM, Jaffe LA. Luteinizing hormone stimulates ingression of granulosa cells within the mouse preovulatory follicle. BioRxiv 2023; 04.21.537855; doi: 10.1101/2023.04.21.537855.

59. Yu C-C, Chen W-Y, Li PS. Protein phosphatase inhibitor cantharidin inhibits steroidogenesis and steroidogenic acute regulatory protein expression in cultured rat preovulatory follicles. Life Sci 2001; 70:57–72.

60. Nakatsumi H, Oka T, Higa T, Shirane M, Nakayama KI. Nuclear-cytoplasmic shuttling protein PP2A^B56^ contributes to mTORC1-dependent dephosphorylation of FOXK1. Genes Cells 2018; 23:599–605.

61. Golkowski M, Shimizu-Albergine M, Suh HW, Beavo JA, Ong S-E. Studying mechanisms of cAMP and cyclic nucleotide phosphodiesterase signaling in Leydig cell function with phosphoproteomics. Cell Signalling 2016; 28:764–778.

62. Light A, Hammes SR. LH-induced steroidogenesis in the mouse ovary, but not testis, requires matrix metalloproteinase 2- and 9-mediated cleavage of upregulated EGF receptor ligands. Biol Reprod 2015; 93(3):65,1–13.

63. Perez-Riverol Y, Csordas A, Bai J, Bernal-Llinares M, Hewapathirana S, Kundu DJ, Inuganti A, Griss J, Mayer G, Eisenacher M, Perez E, Uszkoreit J, Pfeuffer J, Sachsenberg T, Yilmaz S, Tiwary S, Cox J, Audain E, Walzer M, Jarnuczak AF, Ternent T, Brazma A, Vizcaíno JA. The PRIDE database and related tools and resources in 2019: improving support for quantification data. Nucleic Acids Res 2019; 47:D442–D450.

64. Sacco F, Perfetto L, Castagnoli L, Cesareni G. The human phosphatase interactome: An intricate family portrait. FEBS Lett 2012; 586:2732–2739.

65. Chen MJ, Dixon JE, Manning G. Genomics and evolution of protein phosphatases. Sci Signal 2017; 10:eaag1796.

