## Supplemental Figures and Tables for "Phosphatases modified by LH signaling in ovarian follicles: testing their role in regulating the NPR2 guanylyl cyclase"

**Figure S1.** Generation of *Ppp1r12a*-S507A mice.

**Figure S2.** Generation of *Ppp2r5d*-S53A/S81A/S82A/S566A mice.

**Figure S3.** PPP1R12A mRNA is ~5 times more abundant than PPP1R12B mRNA in mouse ovarian follicles.

**Figure S4.** Rp-8-CPT-cAMPS effectively inhibits LH-induced PKA activation in mouse preovulatory follicles.

**Table S1.** Antibodies used for western blotting.

**Table S2.** Statistically significant LH-induced changes in phosphopeptide intensity of proteins in rat ovarian follicles that have been reported to form complexes with PPP family phosphatase catalytic subunits, other than those listed in Table 1.

**Supplementary Data File 1.** Full table of phosphorylation changes in rat follicles ± LH (Excel)

**Supplementary Data File 2.** Full table of quantified protein intensities in rat follicles ± LH (Excel)

**Supplementary Data File 3.** Full list of plots visualizing phosphorylation changes in rat follicles (PDF)

**Supplementary Data File 4.** Gene Ontology—Biological Process enrichment analysis of rat follicle proteins that are significantly differentially phosphorylated in response to LH (Excel)

**Supplementary Data File 5.** Gene Ontology—Cellular Compartment enrichment analysis of rat follicle proteins that are significantly differentially phosphorylated in response to LH (Excel)

**Supplementary Data File 6.** Gene Ontology—Molecular Function enrichment analysis of rat follicle proteins that are significantly differentially phosphorylated in response to LH (Excel)

**Figure S1**. Generation of *Ppp1r12a*-S507A mice. To introduce the S507A mutation into *Ppp1r12a* (indicated in bold pink type), we employed CRISPOR (http://crispor.tefor.net) and identified a CRISPR cleavage site (5'- CTC TTC AAT GTC AGA TGT AC) adjacent to S507 in exon 11 of *Ppp1r12a*. The PAM sequence is underlined. CRISPR/Cas9 ribonucleoprotein together with single-strand DNA donor were electroporated into one-cell embryos from C57BL/6J mice that had been previously modified to insert an HA tag on the N-terminus of *Npr2* (21). The embryos were then transferred into foster females for subsequent development. PCR genotyping was performed to screen for potential founders using the primer pair PS507AF and P1r12a I12R, which amplifies a fragment of 135 bp specific to the S507A knockin mutation. Founders were further confirmed by PCR using primer pair P1r12a I11F and P1r12a I12R to amplify a fragment of 258 bp containing the S507A mutation followed by sequencing of the PCR product. Founders were then bred with C57BL/6J mice (with HA-tagged *Npr2*) to establish the *Ppp1r12a*-S507A mouse line.

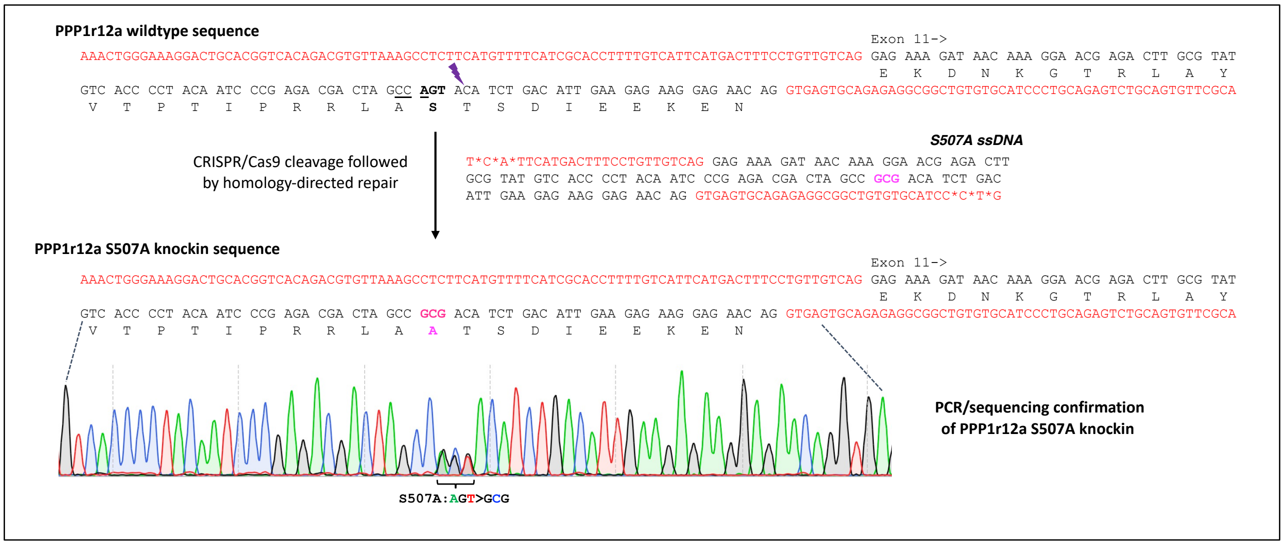


DNA sequences for generation and genotyping of PPP1R12A-S507A mice.
(* represents phosphorothioate linkages to prevent degradation from exonuclease)

| Type of sequence | Sequence |
|---|---|
| sgRNA | 5'- CTCTTCAATGTCAGATGTAC |
| S507A ssDNA donor (sense) | 5'- T*C*A*TTCATGACTTTCCTGTTGTCAG GAG AAA GAT AAC AAA GGA ACG AGA CTT GCG TAT GTC ACC CCT ACA ATC CCG AGA CGA CTA GCC **GCG** ACA TCT GAC ATT GAA GAG AAG GAG AAC AG GTGAGTGCAGAGAGGCGGCTGTGTGCATCC*C*T*G |
| PS507AF (Fwd genotyping primer) | 5'- CAATCCCGAGACGACTAGCCGCG |
| P1r12a I12R (Rev genotyping) | 5'- GCTCGCATCAGTCCTCACATC |
| P1r12a I11F (PCR/sequencing) | 5'- GACTGCACGGTCACAGACGTG |
| P1r12a I12R (PCR/sequencing) | 5'- GCTCGCATCAGTCCTCACATC |

**Figure S2**. Generation of *Ppp2r5d*-S53A/S81A/S82A/S566A mice, nicknamed *Ppp2r5d*-4A. These mice were generated in a two-step process. *Ppp2r5d*-S566A mice were generated first, then bred to homozygosity to provide embryos for generation of the final *Ppp2r5d*-4A mice. (**A**) We identified a CRISPR site cleavage site (5'- GAG GAA GTC AGA GCT GCC AC) close to S566 in exon 15 using CRISPOR (http://crispor.tefor.net). The PAM site is underlined. A G>A silent mutation in Q560 was included in the S566A ssDNA donor to eliminate the PAM in order to prevent re-cleavage of the S566A knockin allele. S566A ssDNA and CRISPR/Cas9 ribonucleoprotein were electroporated into one-cell embryos from C57BL/6J mice that had been previously modified to insert an HA tag on the N-terminus of *Npr2* (21). The embryos were then transferred into a foster mother for subsequent development. Founder pups were identified by PCR genotyping using the primer pair PS566AF and PppE15R1, to amplify a 157 bp fragment specific to the S566A knockin. Their genotypes were confirmed by PCR using primer pair Ppp I14F1 and Ppp E15R1 to amplify a fragment of 255 bp containing the S566A knockin mutation followed by sequencing of the PCR product. (**B**) Two CRISPR sites adjacent to S53 (sgRNA1) and S81/S82 (sgRNA2) were identified. CRISPR/Cas9 ribonucleoproteins and 3A ssDNA were electroporated into homozygous *Ppp2r5d*-S566A one-cell embryos. Founders containing the S53A/S81A/S82A knockin mutations were identified using the primer pair S53.E3F1 and S81.82E3R1 to specifically amplify a fragment of 120 bp, and their identities were further confirmed by PCR using primer pair P2r5d.SeqF and P2r5d.SeqR to amplify a fragment of 383 bp containing the S53A/S81A/S82A mutations followed by sequencing of the PCR product. Founders were bred with C57BL/6J mice (with HA-tagged *Npr2*) to establish the *Ppp2r5d*-4A mouse line.

A

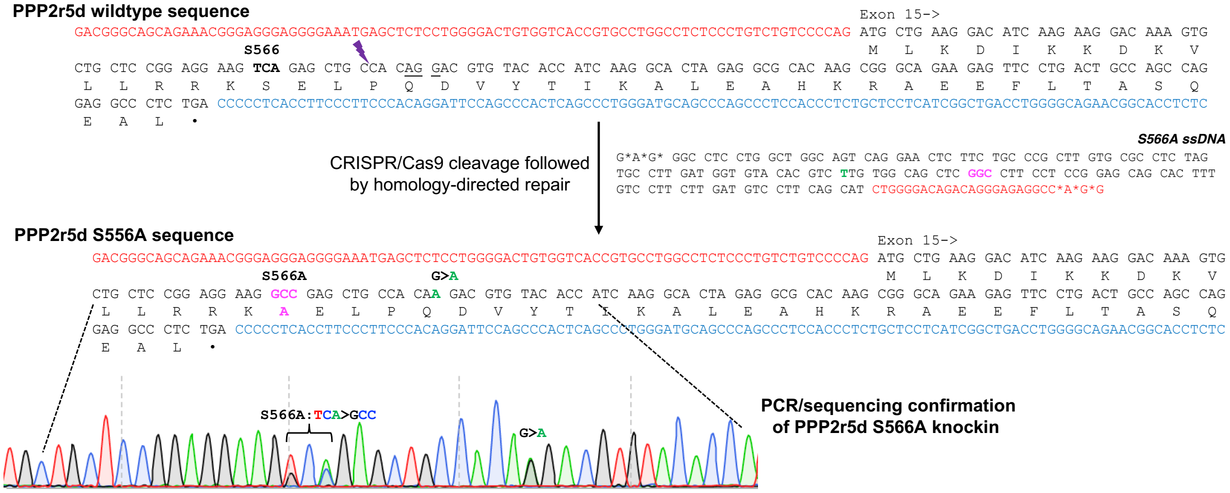

DNA sequences for generation and genotyping of PPP2R5D-S566A mice.
(* represents phosphorothioate linkages to prevent degradation from exonuclease)

| Type of sequence | Sequence |
|---|---|
| sgRNA | 5’- GAGGAAGTCAGAGCTGCCAC |
| S566A ssDNA donor (antisense) | 5’- G*A*G*GGCCTCCTGGCTGGCAGTCAGGAACTCTTCTGCCCGCTTGTGCGCCTCTAGTGCCTTGATGGTGTACACGTCTTGTGGCAGCTC**GGC**CTTCCTCCGGAGCAGCACTTTGTCCTTCTTGATGTCCTTCAGCATCTGGGGACAGACAGGGAGAGGCC*A*G*G |
| PppS566AF (Forward genotyping primer) | 5’- AAGTGCTGCTCCGGAGGAAGGCC |
| PppE15R1 (Reverse genotyping primer) | 5’- GGGCTGAGTGGGCTGGAATCC |
| Ppp I14F1 (PCR/sequencing) | 5’ - ACGGGAGGGAGGGGAAATGAGC |
| Ppp E15R1 (PCR/sequencing) | 5’ - GGGCTGAGTGGGCTGGAATCC |

B

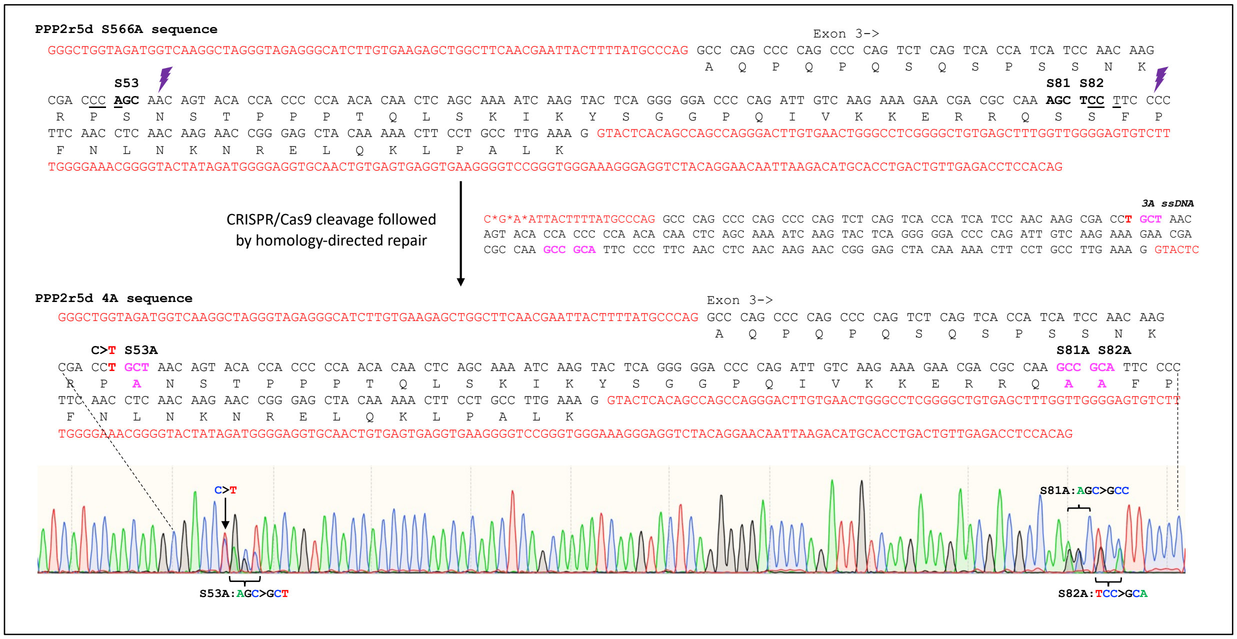

DNA sequences for generation and genotyping of PPP2R5D-4A mice.
(* represents phosphorothioate linkages to prevent degradation from exonuclease)

| Type of sequence | Sequence |
|---|---|
| sgRNA1 | 5'- TGGGGGTGGTGTACTGTTGC |
| sgRNA2 | 5'- CTTGTTGAGGTTGAAGGGGA |
| 3A ssDNA donor (sense) | 5'- C*G*A*ATTACTTTTATGCCCAGGCCCAGCCCCAGCCCCAGTCTCAGTCACCATCATCCAACAAGCGACC**TGCT**AACAGTACACCACCCCCAACACAACTCAGCAAAATCAAGTACTCAGGGGGACCCCAGATTGTCAAGAAAGAACGACGCCAA**GCCGCA**TTCCCCTTCAACCTCAACAAGAACCGGGAGCTACAAAAACTTCCTGCCTTGAAAGGTACTC |
| S53.E3F1 (Forward genotyping primer) | 5'- CATCCAACAAGCGACCTGCT |
| S81.82E3R1 (Reverse genotyping primer) | 5'- GAGGTTGAAGGGGAATGCGGC |
| P2r5d.SeqR (PCR/sequencing) | 5' - GAGGGCTGGTAGATGGTCAAG |
| P2r5d.SeqR (PCR/sequencing) | 5' - CTCACAGTTGCACCTCCCCAT |

**Figure S3.** PPP1R12A mRNA is ~5 times more abundant than PPP1R12B mRNA in mouse ovarian follicles. The bars show the mean ± SEM for analysis of 3 independent preparations of RNA.

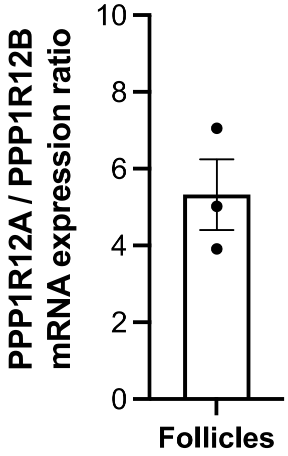


<u>Methods:</u> RNA from mouse preovulatory follicles was extracted with Trizol and treated with DNAse. ddPCR was performed using a Bio-Rad QX-200 Droplet Digital system and primers as listed below. We thank Andrew Collins and Angela Ross (Bio-Rad) for performing the ddPCR assay.

Primer sequences used for ddPCR to compare amounts of PPP1R12A and PPP2R12B mRNA.

| Gene name | Forward | Reverse | Amplicon |
|---|---|---|---|
| *Ppp1r12a* | 5'-CTCTATGCCTCAAGTCAGCTC | 5'-GCTGTGACTTATCTTCCCCTTC | 139 bp |
| *Ppp1r12b* | 5'-AGAAGCTTGAAGATCCTGGTG | 5'-TGGGTTTGTCTGGTTGAGTTG | 150 bp |

**Figure S4.** Rp-8-CPT-cAMPS effectively inhibits LH-induced PKA activation in mouse preovulatory follicles. (**A**) Follicles were treated with or without Rp-8-CPT-cAMPS (10 mM, 4 hr) then stimulated with 10 nM LH (30 min). Follicle proteins were separated on a standard SDS-PAGE gel (no Phostag) and western blots were probed with an antibody recognizing phosphorylated serine 133 of the PKA substrate CREB, and with an antibody against total CREB. Signals were detected with fluorescent secondary antibodies to ensure linearity (see methods). (**B**) Summary of ratios of signal intensities for pS133 CREB/total CREB, for 2 blots like those shown in A. Each symbol represents a set of samples that were treated as indicated and then analyzed together on the same blot. Bars show mean ± SEM. Values were normalized to the control with no Rp-8-CPT-cAMPS and no LH.

A

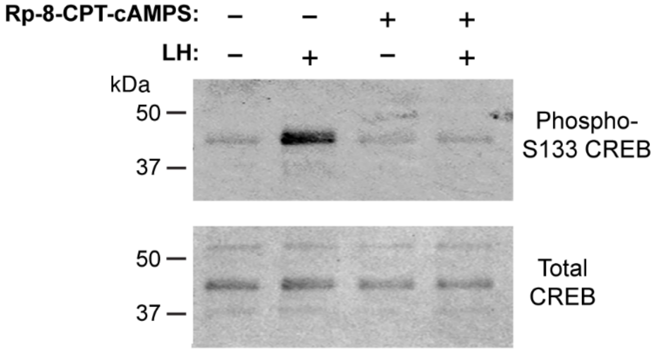


B

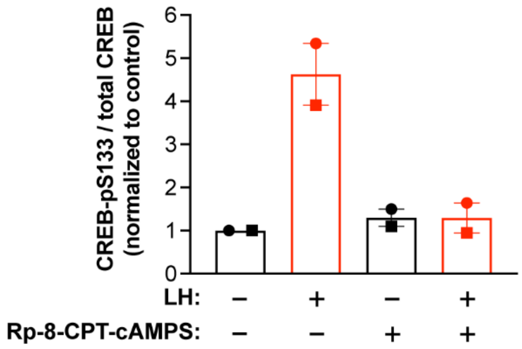

**Table S1.** Antibodies used for western blotting

| Target | Supplier | Host species | Catalog # | Dilution | RRID |
|---|---|---|---|---|---|
| HA epitope tag (6E2) | Cell Signaling Technology | mouse | 2367 | 1:1000 | AB_10691311 |
| Phospho-S507 PPP1R12A (P-MYPT1)[a] | Cell Signaling Technology | rabbit | 3040 | 1:1000 | AB_2168424 |
| Phospho-S668 PPP1R12A (P-MYPT1) | Cell Signaling Technology | rabbit | 3048 | 1:1000 | AB_2168418 |
| Phospho-S472 PPP1R12A (P-MYPT1) | Invitrogen | rabbit | PA5-114608 | 1:1000 | AB_2899244 |
| Phospho-T696 PPP1R12A (P-MYPT1) | Cell Signaling Technology | rabbit | 5163 | 1:1000 | AB_10691830 |
| Phospho-T853 PPP1R12A (P-MYPT1) | Cell Signaling Technology | rabbit | 4563 | 1:1000 | AB_1031185 |
| Total PPP2R5D | Abcam | rabbit | ab188323 | 1:10000 | -- |
| Phospho-S133 CREB | EMD Millipore | mouse | 05-667 | 1:500 | AB_309889 |
| Total CREB | Cell Signaling Technology | rabbit | 9197 | 1:1000 | AB_331277 |
| Goat-anti-mouse IgG (H+L), HRP conjugate | Advansta | goat | R-05071-500 | 1:20000 | -- |
| IRDye® 800CW Goat anti-Rabbit IgG Secondary Antibody | LI-COR | goat | 925-32211 | 1:15000 | AB_10956166 |
| IRDye® 680RD Goat anti-Rabbit IgG Secondary Antibody | LI-COR | goat | 926-68071 | 1:15000 | AB_621842 |
| IRDye® 800CW Goat anti-Mouse IgG Secondary Antibody | LI-COR | goat | 926-32210 | 1:15000 | AB_10718209 |

[a]The antigen used to make this antibody is 100% identical to the mouse PPP1R12A sequence, and 85% identical to the mouse PPP1R12B sequence. Therefore, this antibody would also recognize Phospho-S502 of PPP1R12B.

**Table S2.** Statistically significant LH-induced changes in phosphopeptide intensity of proteins in rat ovarian follicles that have been reported to form complexes with PPP family phosphatase catalytic subunits, other than those listed in Table 1. Previously published lists of validated PPP family-interacting proteins (19, 20) were used to query the phosphopeptide database for significant differences in response to LH. Multiplicity refers to either singly phosphorylated (1) or doubly phosphorylated (2) peptides. Significant increases in phosphopeptide intensity following LH treatment are above the dashed line; significant decreases in phosphopeptide intensity with LH treatment are below the dashed line.

| Gene name | UniProt accession | Amino acid | Site # | Multiplicity | Phosphopeptide intensity ($log_2$ mean ± SEM) Control | 350 nM LH | Fold change |
|---|---|---|---|---|---|---|---|
| HDAC6* | A0A0G2QC41 | S | 59 | 1 | 14.56 ± 0.61 | 18.64 ± 1.07 | 18.0 |
| STRN3* | E9PT82 | S | 239 | 2 | 15.24 ± 0.08 | 19.02 ± 0.18 | 14.6 |
| STRN3* | E9PT82 | S | 255 | 2 | 15.24 ± 0.08 | 19.02 ± 0.18 | 14.6 |
| PLCL1 | F1LP62 | S | 19 | 2 | 17.04 ± 0.13 | 20.14 ± 0.06 | 9.1 |
| EIF2AK2 | M0RDJ3 | S | 212 | 1 | 18.95 ± 0.18 | 21.94 ± 0.26 | 8.5 |
| FARP1* | F1LYQ8 | S | 513 | 2 | 13.80 ± 0.21 | 16.70 ± 0.18 | 7.9 |
| FARP1* | F1LYQ8 | S | 517 | 2 | 13.80 ± 0.21 | 16.70 ± 0.18 | 7.9 |
| MAP1B* | P15205 | S | 561 | 1 | 22.90 ± 0.12 | 25.52 ± 0.31 | 6.5 |
| ZFYVE16 | D4ADF6 | S | 903 | 2 | 17.57 ± 0.24 | 20.17 ± 0.09 | 6.5 |
| ZFYVE16 | D4ADF6 | S | 907 | 2 | 17.57 ± 0.24 | 20.17 ± 0.09 | 6.5 |
| PHACTR4* | M0R7T1 | S | 159 | 2 | 18.97 ± 0.16 | 21.52 ± 0.12 | 6.2 |
| APC* | G3V8Q9 | S | 2793 | 1 | 18.43 ± 0.11 | 20.87 ± 0.19 | 5.8 |
| MKI67 | D4A0Y6 | S | 124 | 1 | 17.98 ± 0.02 | 20.41 ± 0.24 | 5.7 |
| PHACTR4* | M0R7T1 | S | 162 | 2 | 19.29 ± 0.20 | 21.68 ± 0.13 | 5.5 |
| AKAP1 | D4A9M6 | S | 425 | 1 | 17.01 ± 0.06 | 29.39 ± 0.11 | 5.5 |
| APC* | G3V8Q9 | S | 2817 | 2 | 15.38 ± 0.33 | 17.67 ± 0.21 | 5.2 |
| APC* | G3V8Q9 | S | 2829 | 2 | 15.67 ± 0.27 | 17.77 ± 0.18 | 4.6 |
| APC* | G3V8Q9 | T | 2819 | 2 | 14.37 ± 0.25 | 16.23 ± 0.10 | 3.8 |
| PLCL1 | F1LP62 | T | 15 | 2 | 19.78 ± 0.11 | 21.58 ± 0.07 | 3.7 |
| SH2D4A | Q6AYC8 | S | 51 | 1 | 18.21 ± 0.21 | 20.01 ± 0.39 | 3.7 |
| PCIF1 | D4A417 | T | 135 | 3 | 14.49 ± 0.25 | 16.29 ± 0.14 | 3.7 |
| SLC9A1 | P26431 | S | 707 | 2 | 19.21 ± 0.14 | 20.98 ± 0.09 | 3.6 |
| AKAP11 | A0A0G2JZI9 | S | 21 | 1 | 17.30 ± 0.37 | 19.05 ± 0.37 | 3.6 |
| PLCL1 | F1LP62 | S | 491 | 1 | 11.53 ± 0.35 | 13.23 ± 0.24 | 3.4 |
| PCIF1 | D4A417 | T | 150 | 2 | 18.36 ± 0.37 | 20.05 ± 0.09 | 3.4 |
| PCIF1 | D4A417 | T | 135 | 2 | 16.05 ± 0.44 | 17.73 ± 0.28 | 3.4 |
| PCIF1 | D4A417 | S | 140 | 3 | 16.06 ± 0.23 | 17.66 ± 0.16 | 3.2 |
| PLCL1 | F1LP62 | S | 17 | 2 | 19.58 ± 0.13 | 21.12 ± 0.07 | 3.1 |
| SLC9A1 | P26431 | S | 697 | 2 | 20.56 ± 0.11 | 22.09 ± 0.14 | 3.1 |
| SLC9A1 | P26431 | S | 776 | 2 | 16.38 ± 0.10 | 17.86 ± 0.17 | 3.0 |
| MAP1B* | P15205 | S | 985 | 2 | 23.10 ± 0.09 | 24.56 ± 0.16 | 2.9 |

| | | | | | | | |
|---|---|---|---|---|---|---|---|
| MAP1B* | P15205 | S | 988 | 2 | 23.10 ± 0.09 | 24.56 ± 0.16 | 2.9 |
| MAP1B* | P15205 | S | 1454 | 2 | 17.42 ± 0.41 | 18.86 ± 0.14 | 2.9 |
| MAP1B* | P15205 | S | 1465 | 2 | 17.42 ± 0.41 | 18.86 ± 0.14 | 2.9 |
| MAP1B* | P15205 | T | 1262 | 3 | 19.12 ± 0.26 | 20.51 ± 0.06 | 2.8 |
| SLC9A1 | P26431 | S | 606 | 2 | 17.61 ± 0.02 | 19.00 ± 0.15 | 2.8 |
| MKI67 | D4A0Y6 | S | 336 | 1 | 19.80 ± 0.08 | 21.09 ± 0.22 | 2.6 |
| SLC9A1 | P26431 | S | 790 | 2 | 19.02 ± 0.07 | 20.22 ± 0.12 | 2.4 |
| SET* | Q63945 | S | 28 | 2 | 18.89 ± 0.06 | 20.09 ± 0.11 | 2.4 |
| SET* | Q63945 | S | 30 | 2 | 18.89 ± 0.06 | 20.09 ± 0.11 | 2.4 |
| MAP1B* | P15205 | S | 1471 | 1 | 17.76 ± 0.11 | 18.90 ± 0.32 | 2.3 |
| CASP9 | Q9JHK1 | S | 348 | 2 | 14.65 ± 0.09 | 15.70 ± 0.26 | 2.2 |
| CASP9 | Q9JHK1 | S | 350 | 2 | 14.65 ± 0.09 | 15.70 ± 0.26 | 2.2 |
| AKAP1 | D4A9M6 | S | 101 | 1 | 18.57 ± 0.07 | 19.60 ± 0.31 | 2.2 |
| HDAC6* | A0A0G2QC41 | S | 43 | 1 | 22.25 ± 0.11 | 19.52 ± 0.37 | -6.2 |
| HDAC6* | A0A0G2QC41 | S | 859 | 1 | 21.50 ± 0.12 | 19.27 ± 0.61 | -4.4 |
| MAP1B* | P15205 | T | 527 | 1 | 21.25 ± 0.23 | 19.10 ± 0.73 | -4.2 |
| AKAP1 | D4A9M6 | S | 103 | 1 | 19.38 ± 0.19 | 17.48 ± 0.51 | -3.5 |
| PHACTR2* | A0A0G2K8Q4 | T | 22 | 1 | 20.29 ± 0.23 | 18.48 ± 0.73 | -3.3 |
| FARP1* | F1LYQ8 | S | 427 | 2 | 20.51 ± 0.31 | 18.85 ± 0.13 | -3.0 |
| FARP1* | F1LYQ8 | S | 435 | 2 | 20.51 ± 0.31 | 18.85 ± 0.13 | -3.0 |
| PHACTR2* | A0A0G2K8Q4 | S | 154 | 1 | 18.30 ± 0.08 | 16.65 ± 0.40 | -3.0 |
| MAP1B* | P15205 | S | 541 | 1 | 24.80 ± 0.03 | 23.33 ± 0.40 | -2.6 |
| APC* | G3V8Q9 | S | 2556 | 2 | 17.80 ± 0.18 | 16.35 ± 0.19 | -2.6 |
| MAP1B* | P15205 | T | 1959 | 1 | 17.86 ± 0.12 | 16.45 ± 0.35 | -2.5 |
| SLC9A1 | P26431 | S | 790 | 1 | 22.26 ± 0.11 | 20.91 ± 0.47 | -2.4 |
| APC* | G3V8Q9 | S | 2556 | 2 | 19.43 ± 0.08 | 18.22 ± 0.36 | -2.2 |

* Proteins known to regulate the cytoskeleton and/or cell motility according to the National Center for Biotechnology Information (NCBI) gene summaries.
