## Supplemental Data File 3 for "Phosphatases modified by LH signaling in ovarian follicles: testing their role in regulating the NPR2 guanylyl cyclase"

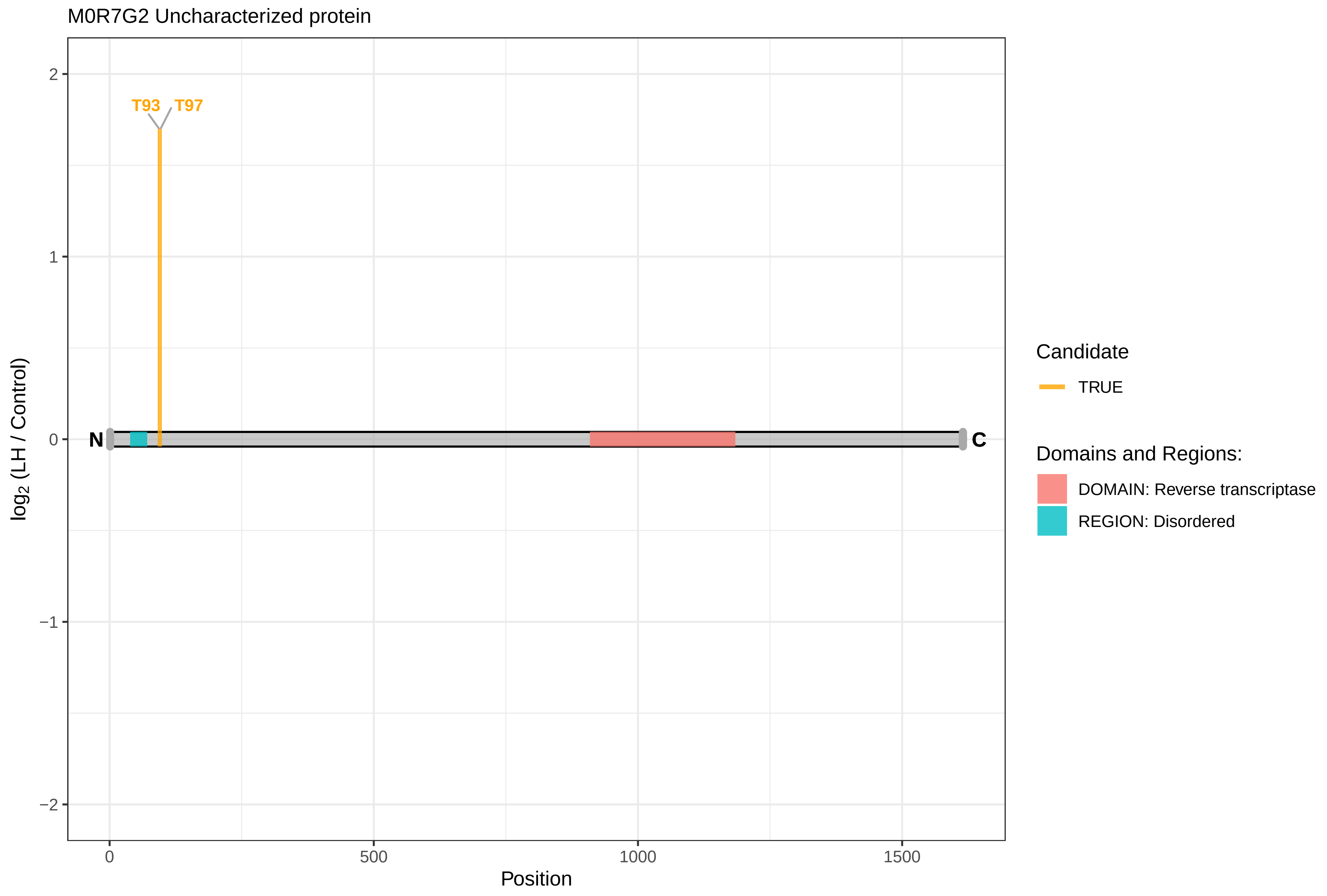

M0R7G2 Uncharacterized protein
T93
T97
N
C
2
1
0
−1
−2
log2 (LH / Control)
0
500
1000
1500
Position
Candidate
TRUE
Domains and Regions:
DOMAIN: Reverse transcriptase
REGION: Disordered

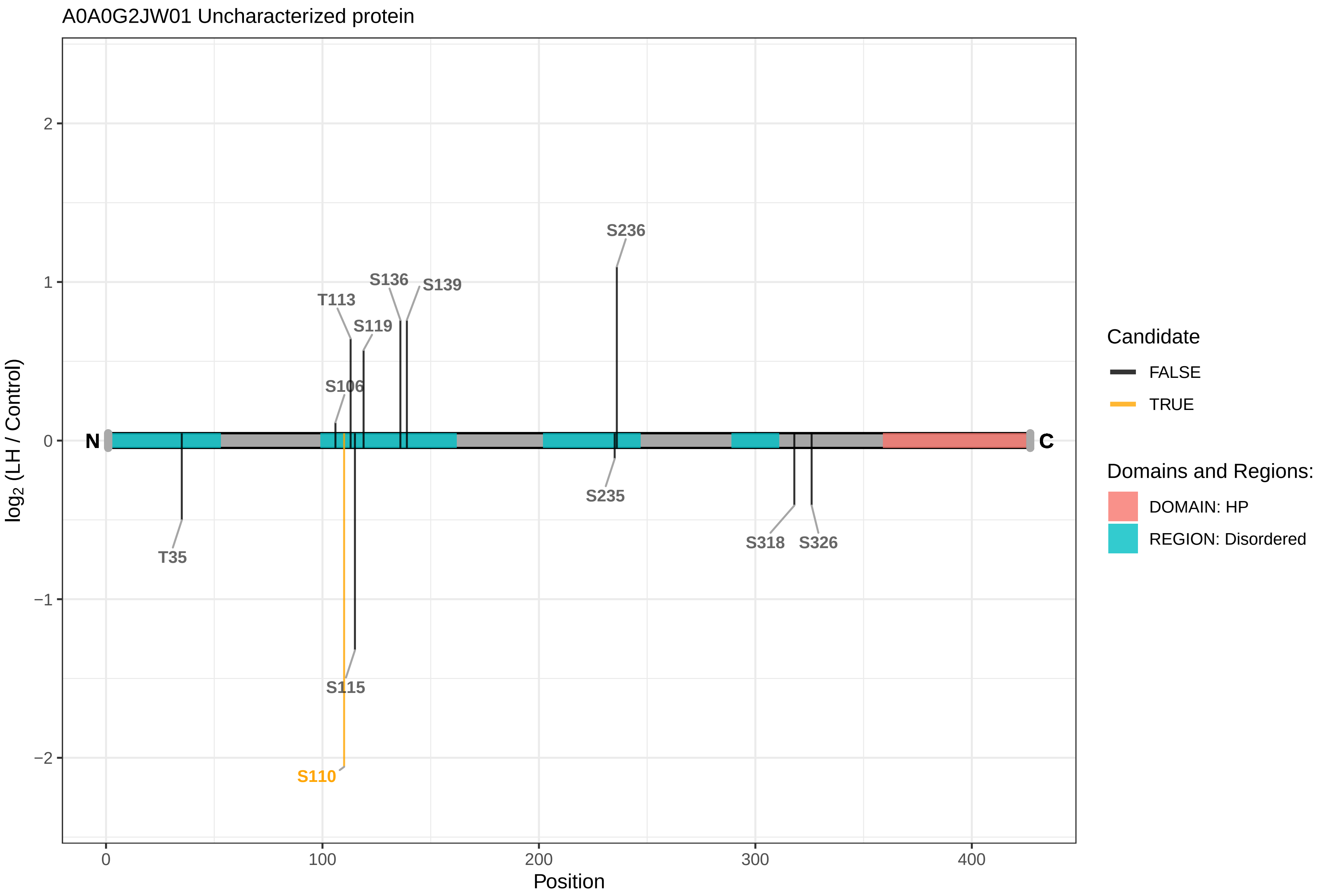
A0A0G2JW01 Uncharacterized protein
log2 (LH / Control)
Position
2
1
0
−1
−2
0
100
200
300
400
N
C
S236
S136
S139
T113
S119
S106
T35
S115
S110
S235
S318
S326
Candidate
FALSE
TRUE
Domains and Regions:
DOMAIN: HP
REGION: Disordered

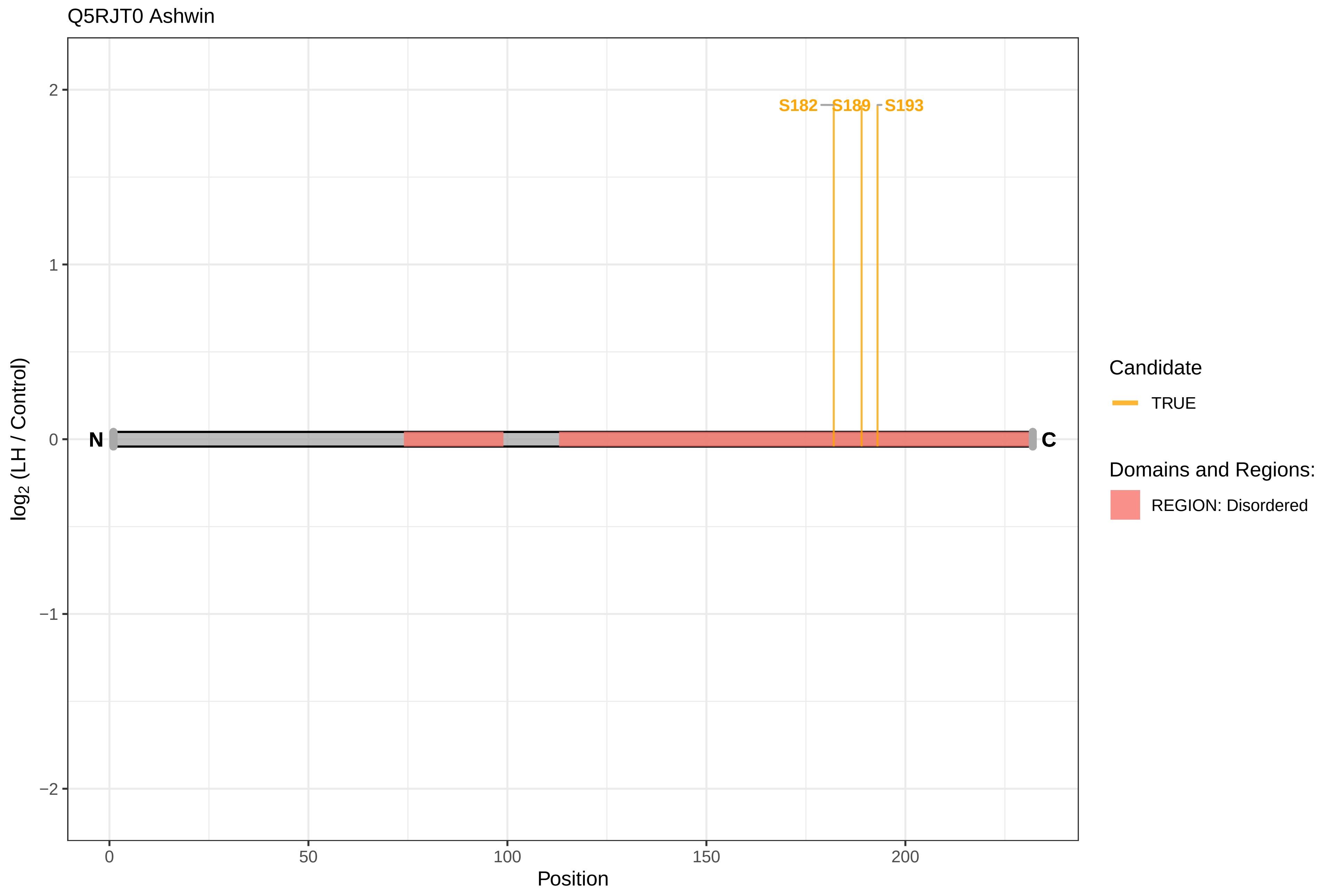

Q5RJT0 Ashwin
S182
S189
S193
N
C
2
1
0
−1
−2
log2 (LH / Control)
0
50
100
150
200
Position
Candidate
TRUE
Domains and Regions:
REGION: Disordered

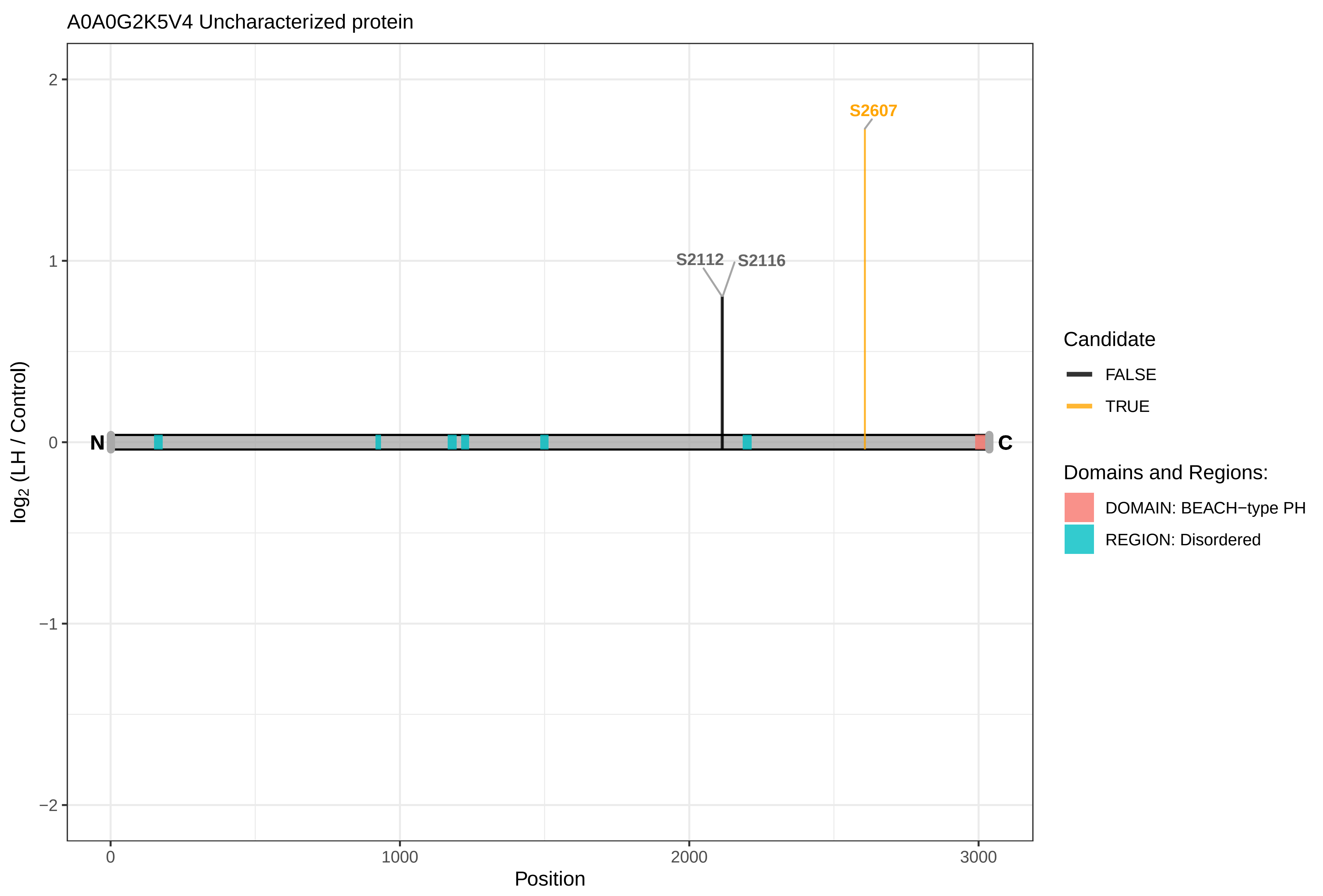

A0A0G2K5V4 Uncharacterized protein
S2607
S2112
S2116
N
C
2
1
0
−1
−2
log2 (LH / Control)
0
1000
2000
3000
Position
Candidate
FALSE
TRUE
Domains and Regions:
DOMAIN: BEACH−type PH
REGION: Disordered

AAK1

F1LRI7 AP2−associated protein kinase 1

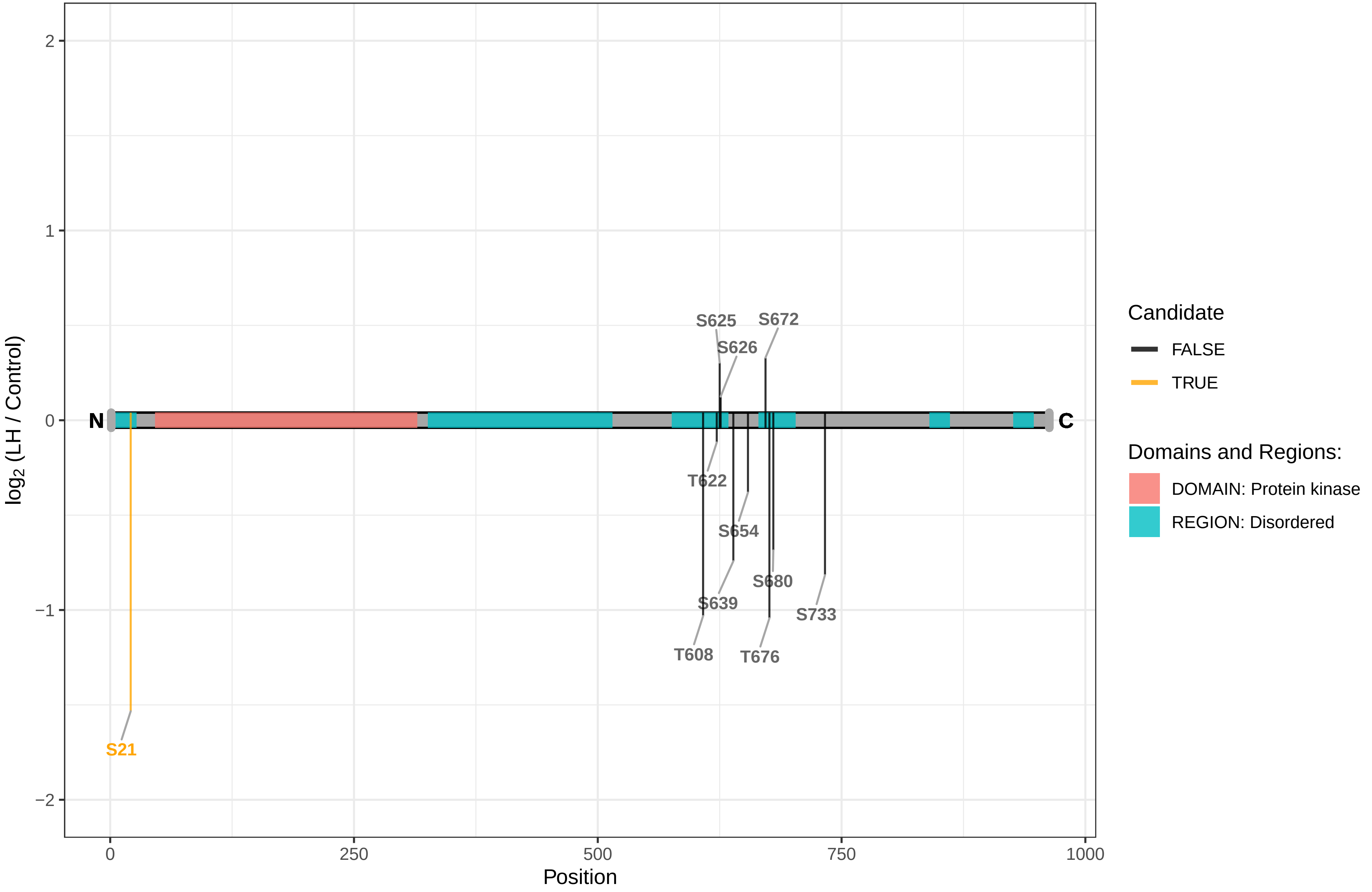

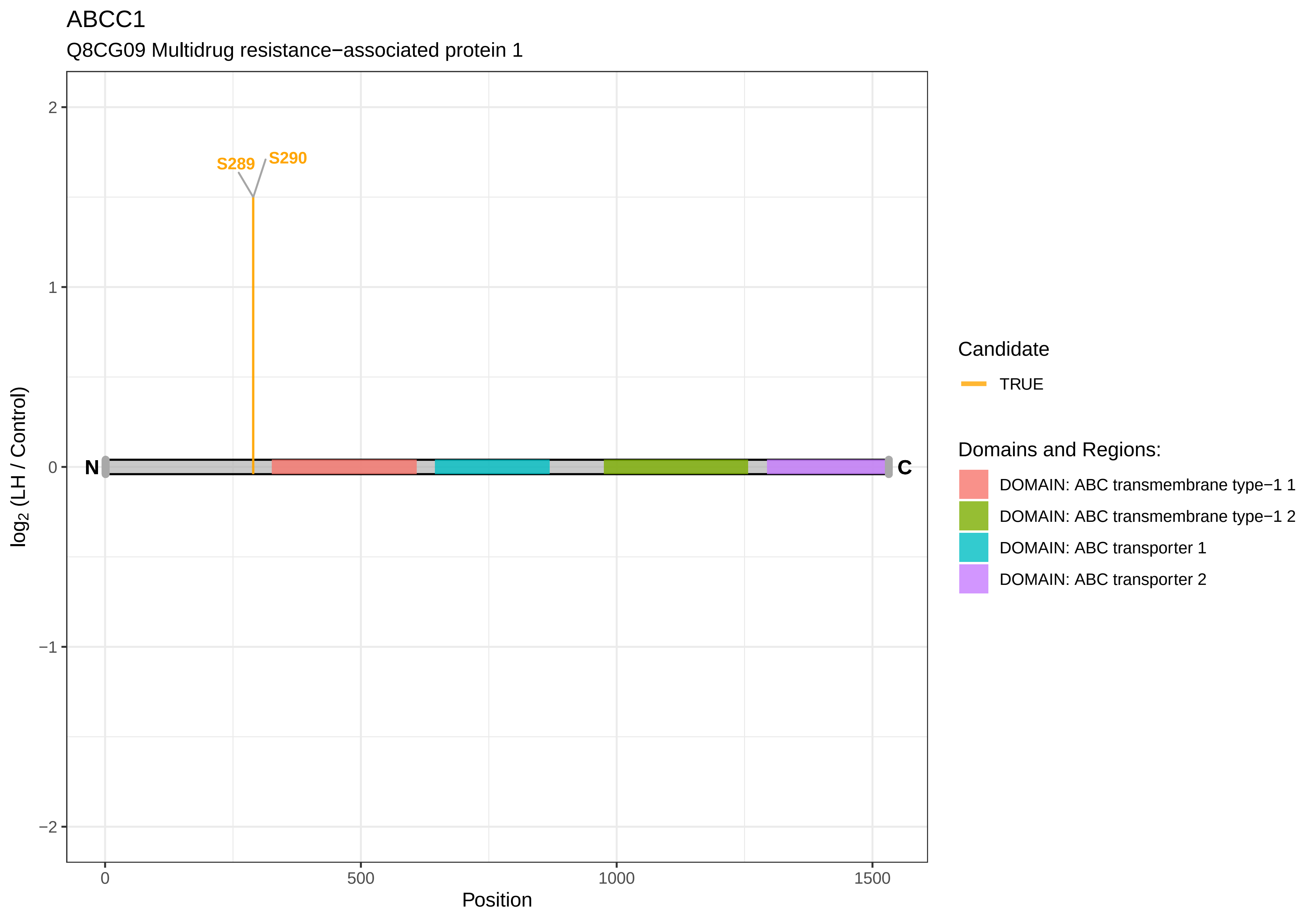
ABCC1
Q8CG09 Multidrug resistance−associated protein 1
S289
S290
N
C
2
1
0
−1
−2
log2 (LH / Control)
0
500
1000
1500
Position
Candidate
TRUE
Domains and Regions:
DOMAIN: ABC transmembrane type−1 1
DOMAIN: ABC transmembrane type−1 2
DOMAIN: ABC transporter 1
DOMAIN: ABC transporter 2

ABI1

Q9QZM5 Abl interactor 1

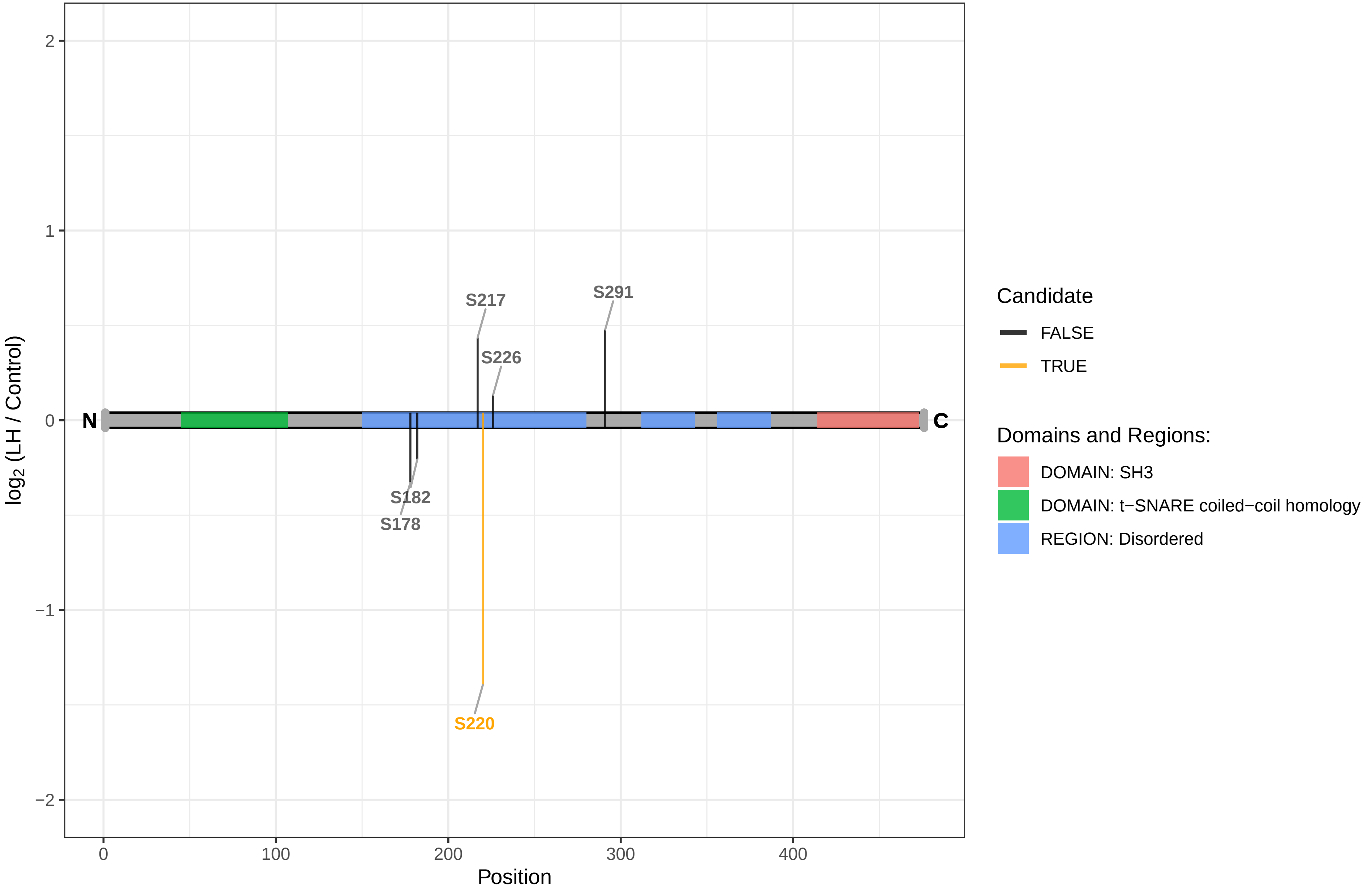

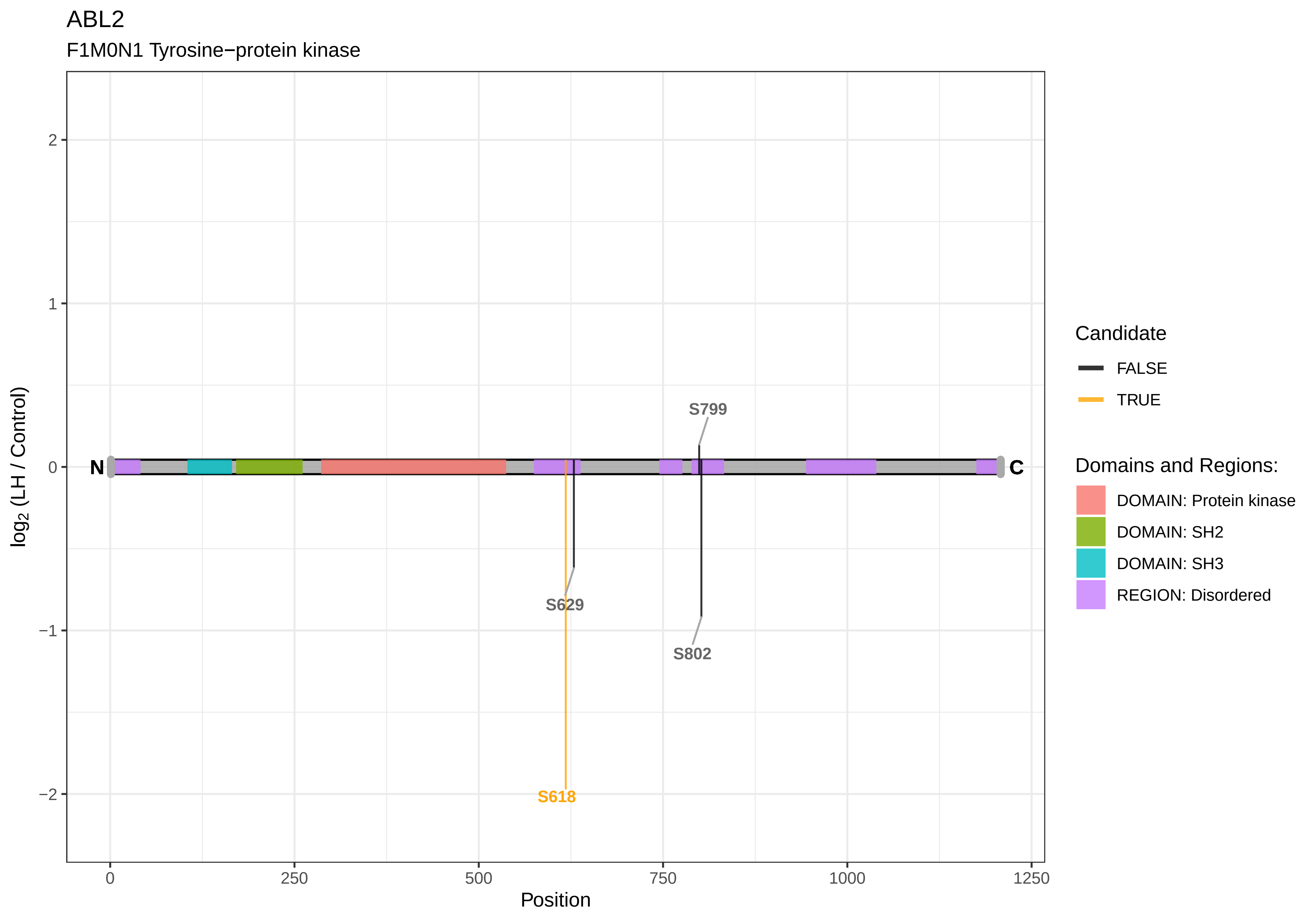

ABL2
F1M0N1 Tyrosine−protein kinase
log2 (LH / Control)
Position
2
1
0
−1
−2
0
250
500
750
1000
1250
N
C
S799
S629
S802
S618
Candidate
FALSE
TRUE
Domains and Regions:
DOMAIN: Protein kinase
DOMAIN: SH2
DOMAIN: SH3
REGION: Disordered

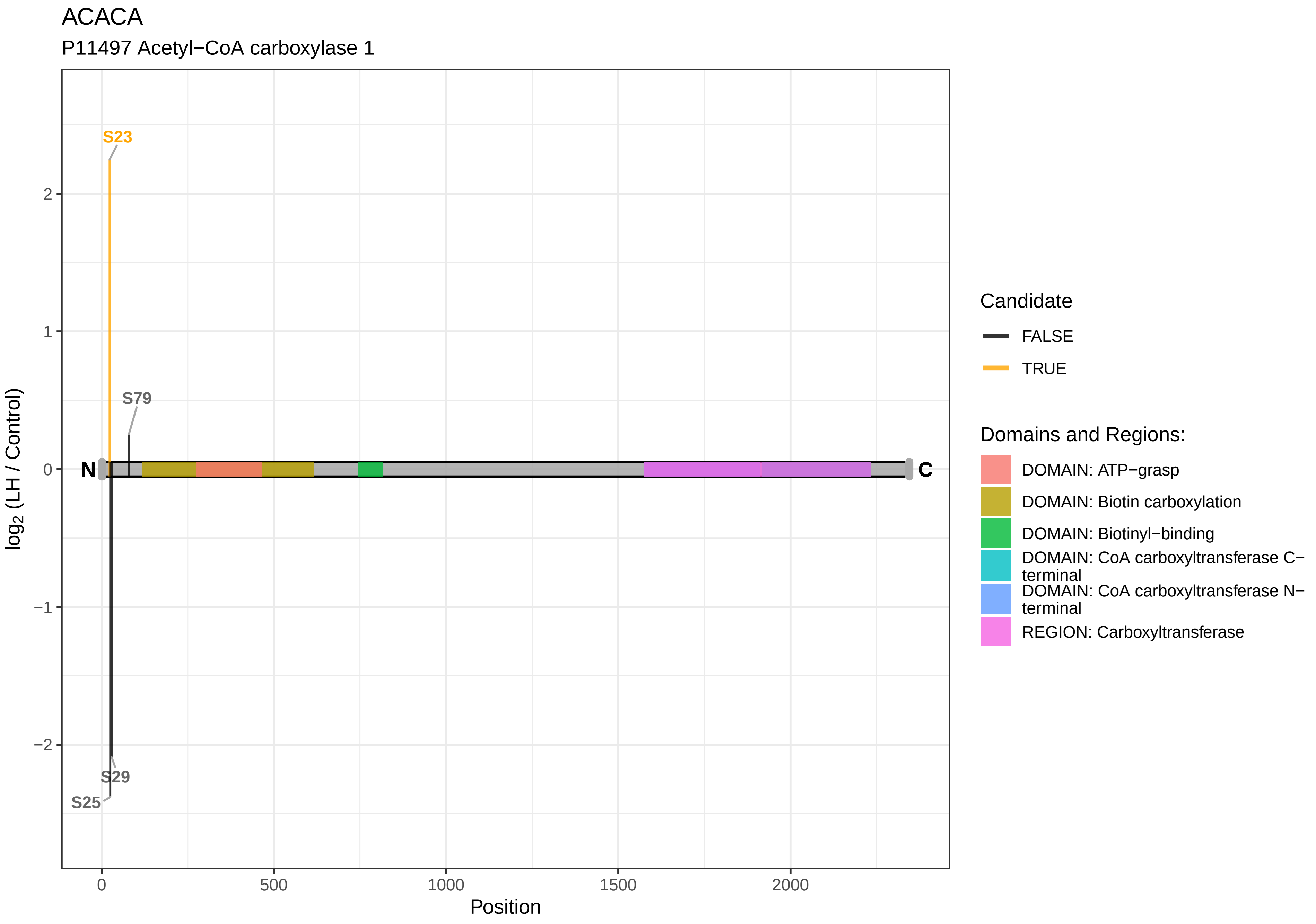

ACACA
P11497 Acetyl−CoA carboxylase 1
S23
S79
N
C
S29
S25
2
1
0
−1
−2
log2 (LH / Control)
0
500
1000
1500
2000
Position
Candidate
FALSE
TRUE
Domains and Regions:
DOMAIN: ATP−grasp
DOMAIN: Biotin carboxylation
DOMAIN: Biotinyl−binding
DOMAIN: CoA carboxyltransferase C−terminal
DOMAIN: CoA carboxyltransferase N−terminal
REGION: Carboxyltransferase

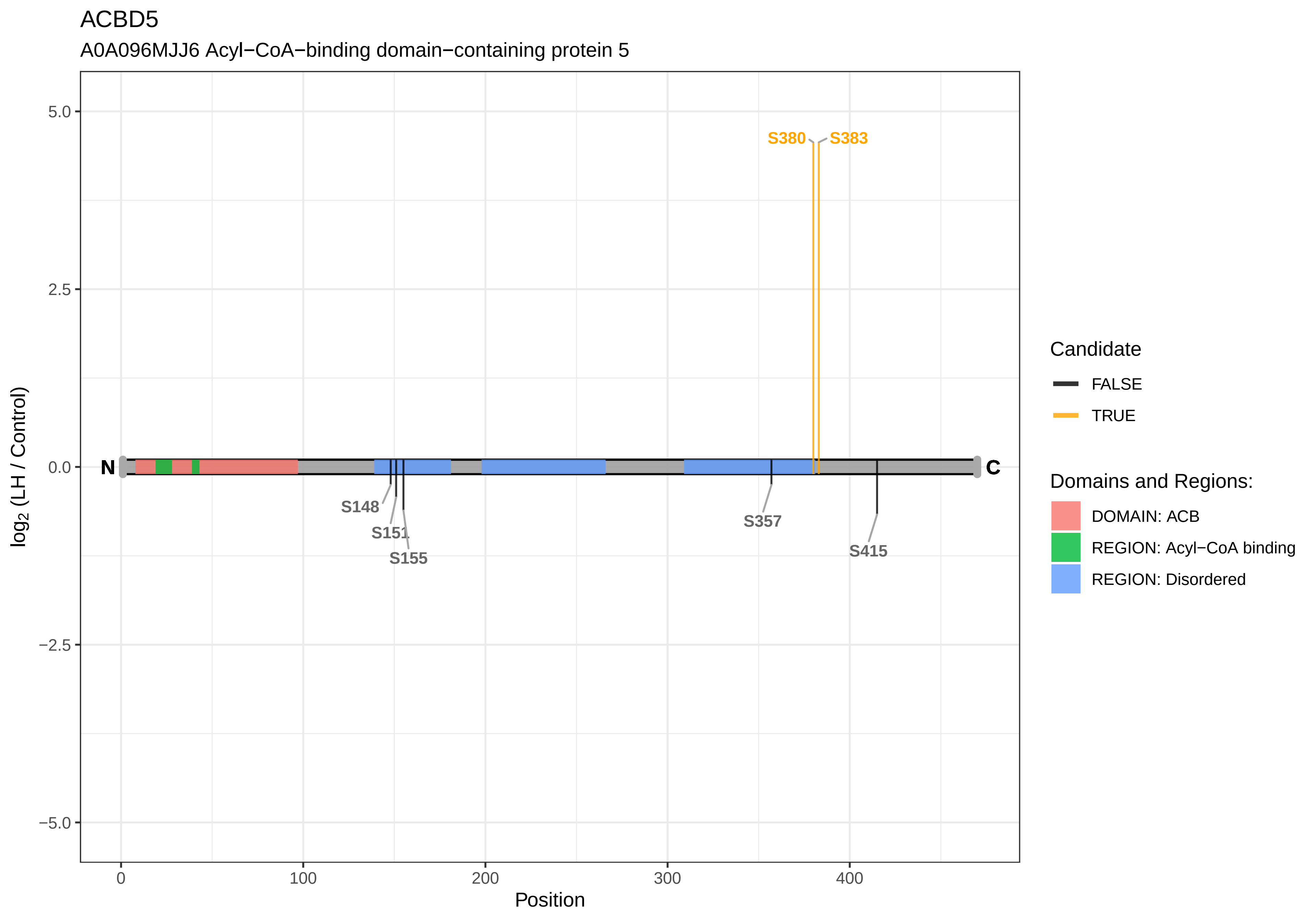
ACBD5
A0A096MJJ6 Acyl−CoA−binding domain−containing protein 5
S380
S383
N
C
S148
S151
S155
S357
S415
5.0
2.5
0.0
−2.5
−5.0
log2 (LH / Control)
0
100
200
300
400
Position
Candidate
FALSE
TRUE
Domains and Regions:
DOMAIN: ACB
REGION: Acyl−CoA binding
REGION: Disordered

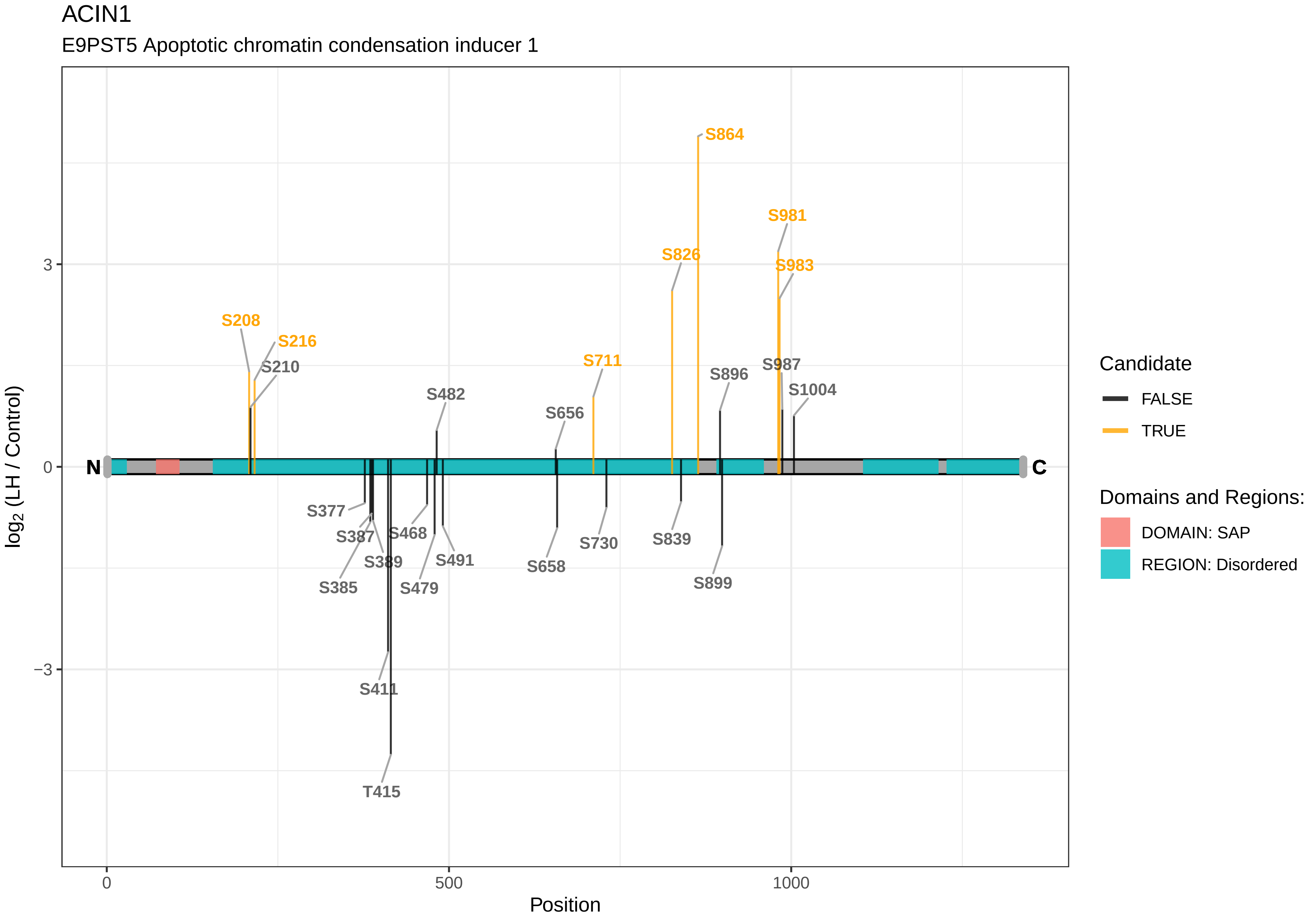
ACIN1
E9PST5 Apoptotic chromatin condensation inducer 1
S864
S981
S826
S983
S208
S216
S210
S711
S896
S987
S1004
S482
S656
N
C
S377
S387
S468
S389
S491
S385
S479
S658
S730
S839
S899
S411
T415
3
0
−3
0
500
1000
Position
log2 (LH / Control)
Candidate
FALSE
TRUE
Domains and Regions:
DOMAIN: SAP
REGION: Disordered

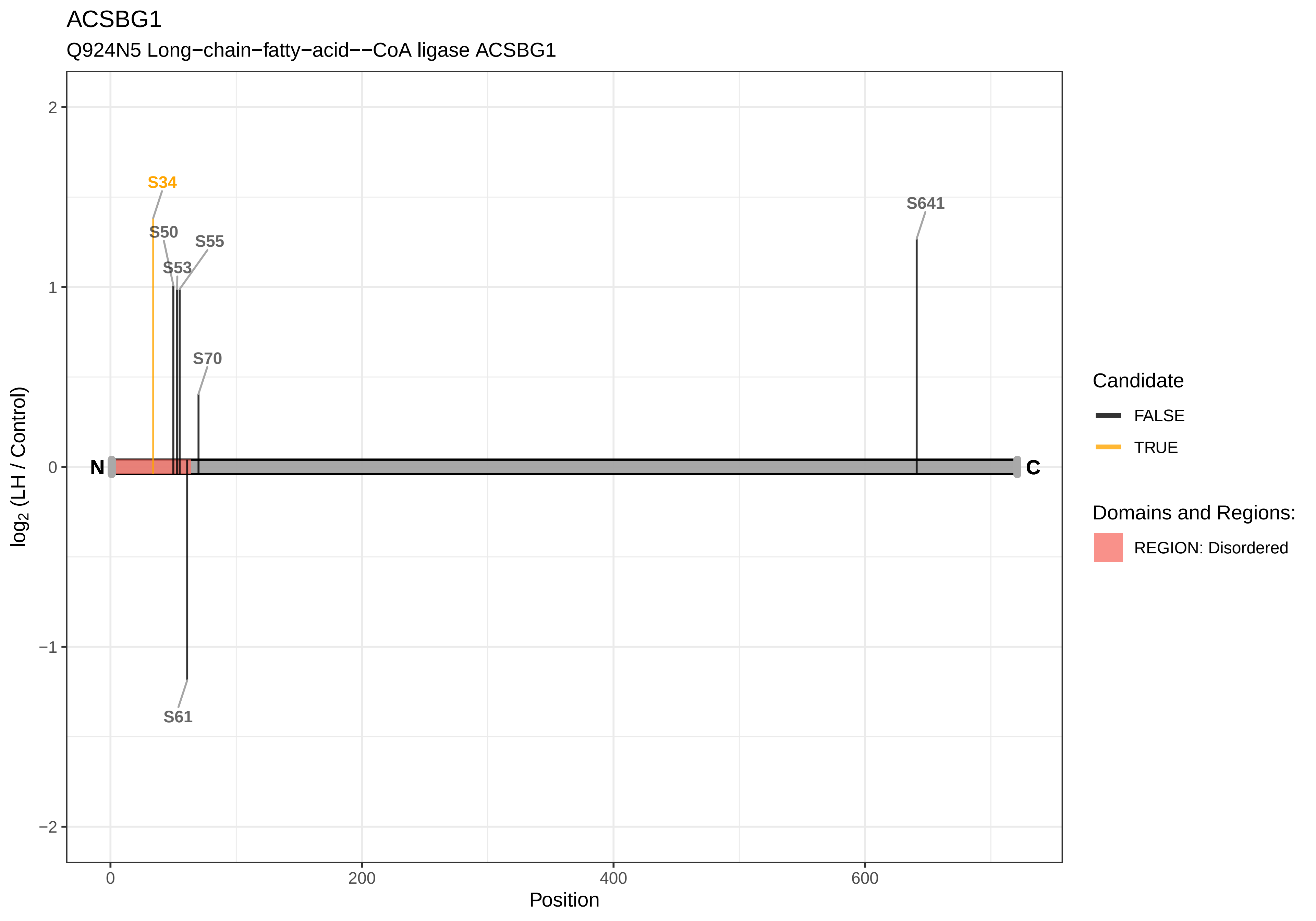
ACSBG1
Q924N5 Long−chain−fatty−acid−−CoA ligase ACSBG1
S34
S50
S55
S53
S70
S641
S61
N
C
2
1
0
−1
−2
0
200
400
600
Position
log2 (LH / Control)
Candidate
FALSE
TRUE
Domains and Regions:
REGION: Disordered

### ADCY5

G3V9G1 Adenylate cyclase

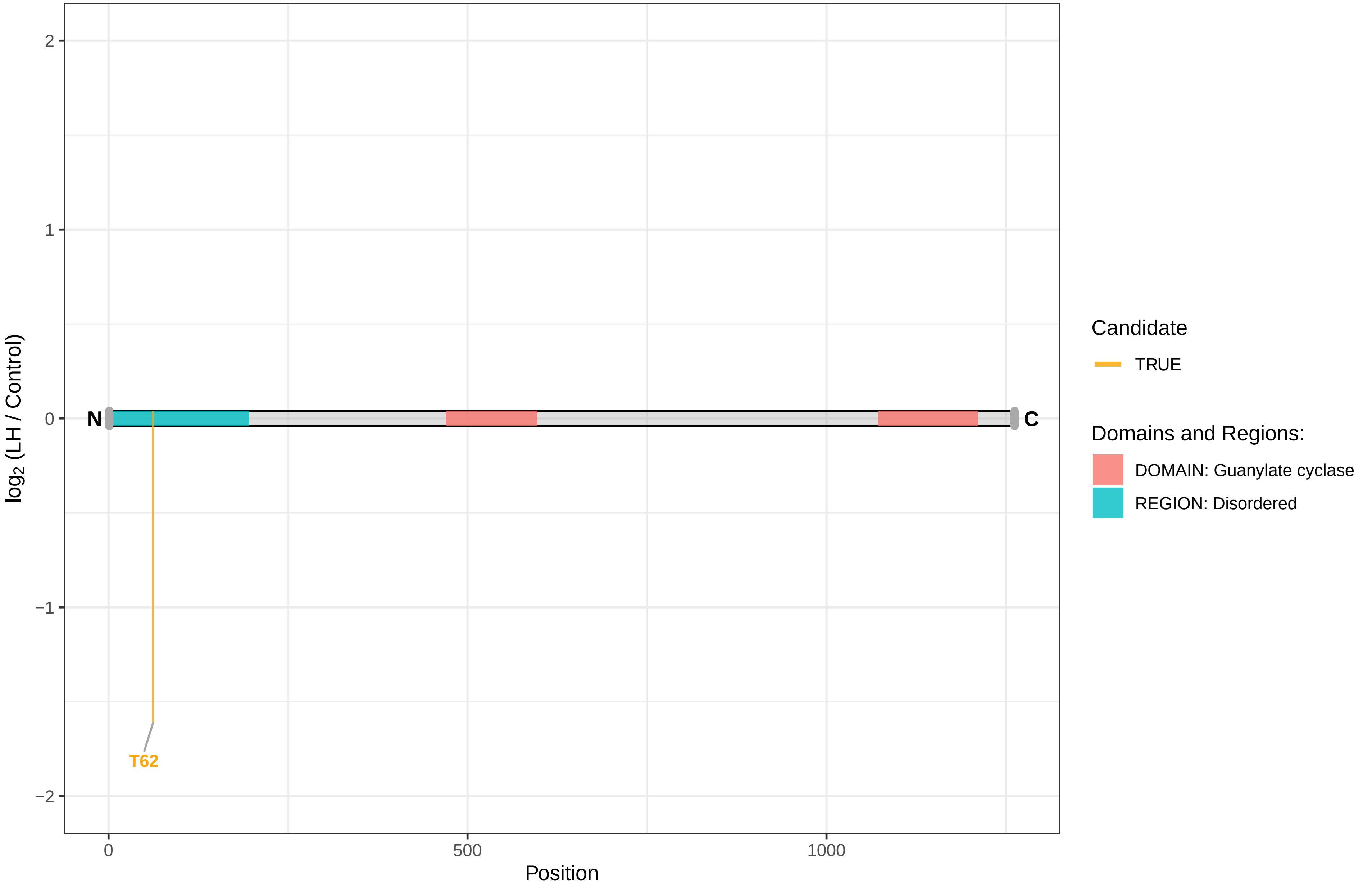

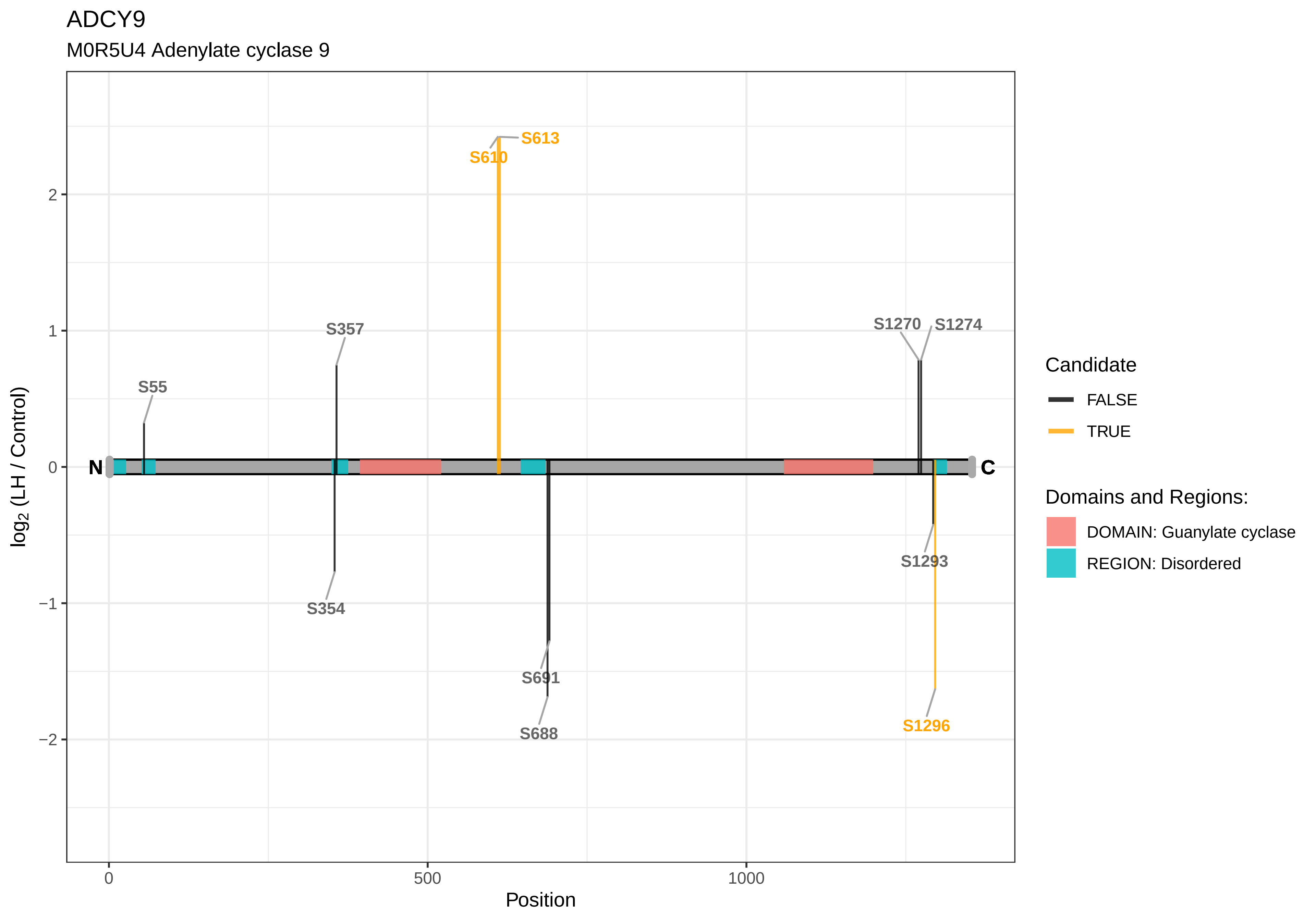

ADCY9
M0R5U4 Adenylate cyclase 9
S613
S610
S357
S55
S1270
S1274
S354
S691
S688
S1293
S1296
N
C
log2 (LH / Control)
Position
0
500
1000
2
1
0
−1
−2
Candidate
FALSE
TRUE
Domains and Regions:
DOMAIN: Guanylate cyclase
REGION: Disordered

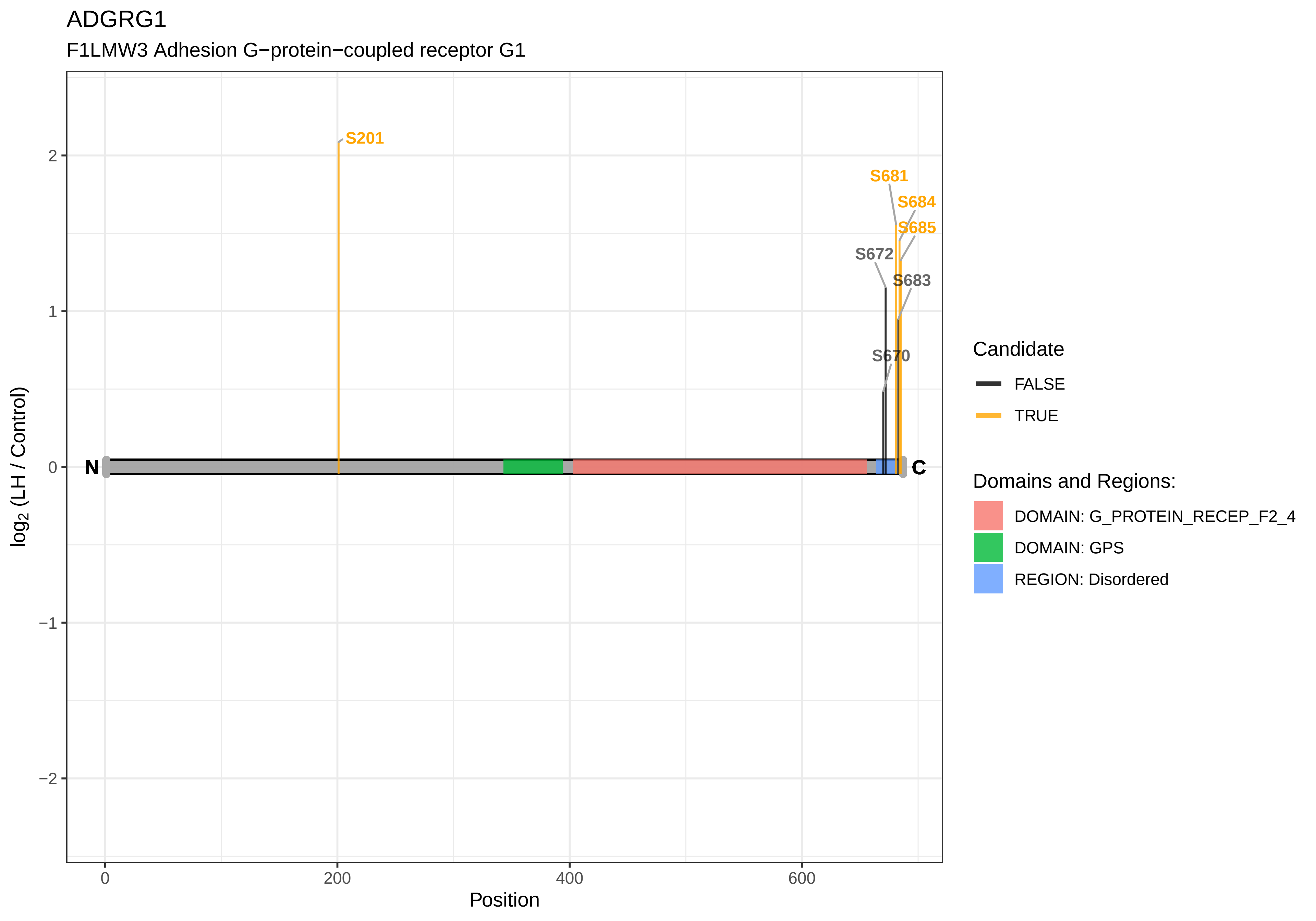
ADGRG1
F1LMW3 Adhesion G−protein−coupled receptor G1
S201
S681
S684
S685
S672
S683
S670
N
C
log2 (LH / Control)
Position
2
1
0
−1
−2
0
200
400
600
Candidate
FALSE
TRUE
Domains and Regions:
DOMAIN: G_PROTEIN_RECEP_F2_4
DOMAIN: GPS
REGION: Disordered

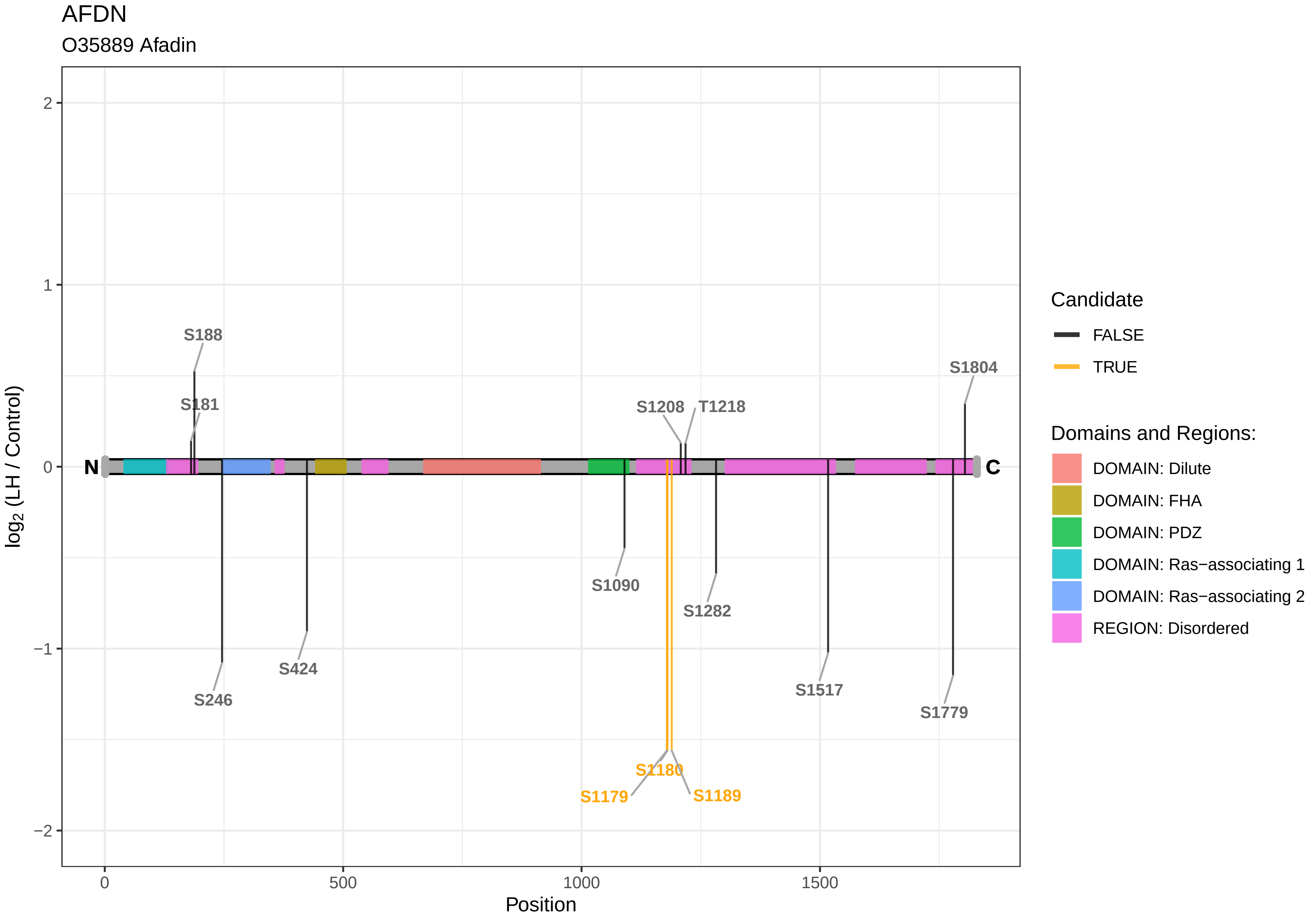

AFDN
O35889 Afadin
log2 (LH / Control)
Position
0
500
1000
1500
2
1
0
−1
−2
N
C
S188
S181
S246
S424
S1090
S1208
T1218
S1282
S1517
S1779
S1804
S1180
S1179
S1189
Candidate
FALSE
TRUE
Domains and Regions:
DOMAIN: Dilute
DOMAIN: FHA
DOMAIN: PDZ
DOMAIN: Ras−associating 1
DOMAIN: Ras−associating 2
REGION: Disordered

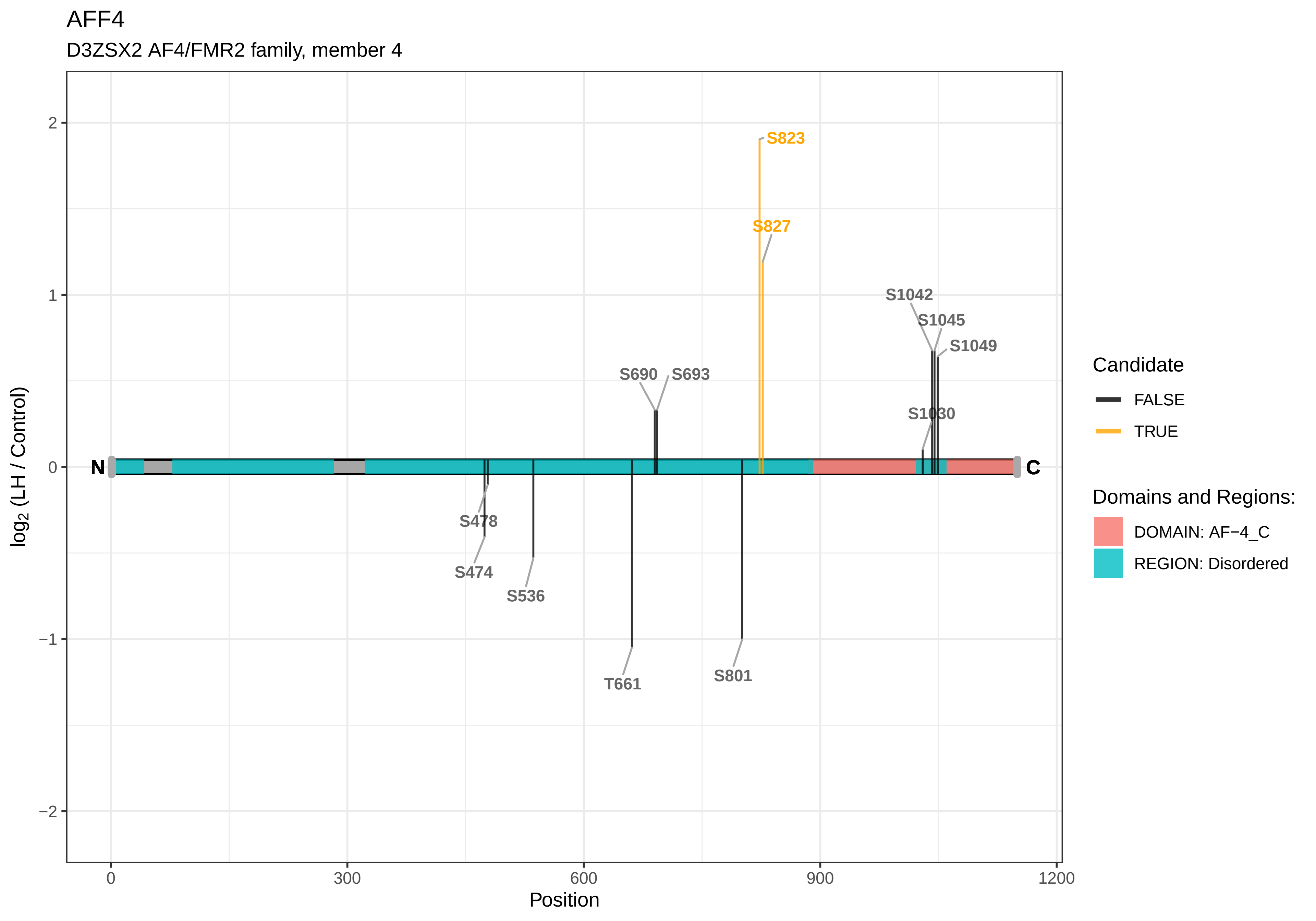
AFF4
D3ZSX2 AF4/FMR2 family, member 4
log2 (LH / Control)
Position
0
300
600
900
1200
2
1
0
−1
−2
N
C
S823
S827
S1042
S1045
S1049
S690
S693
S1030
S478
S474
S536
T661
S801
Candidate
FALSE
TRUE
Domains and Regions:
DOMAIN: AF−4_C
REGION: Disordered

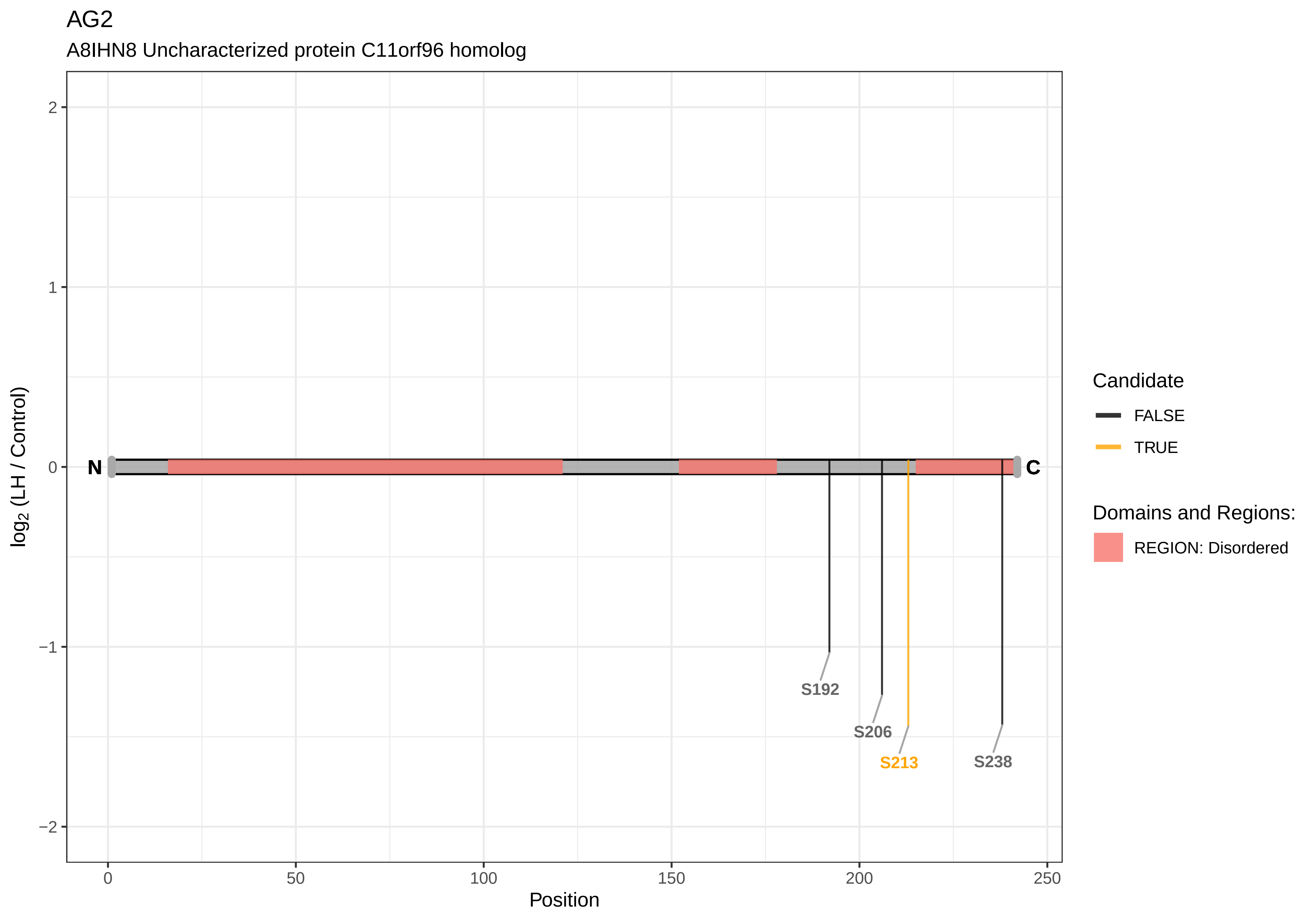

AG2
A8IHN8 Uncharacterized protein C11orf96 homolog
log2 (LH / Control)
Position
N
C
S192
S206
S213
S238
Candidate
FALSE
TRUE
Domains and Regions:
REGION: Disordered
2
1
0
−1
−2
0
50
100
150
200
250

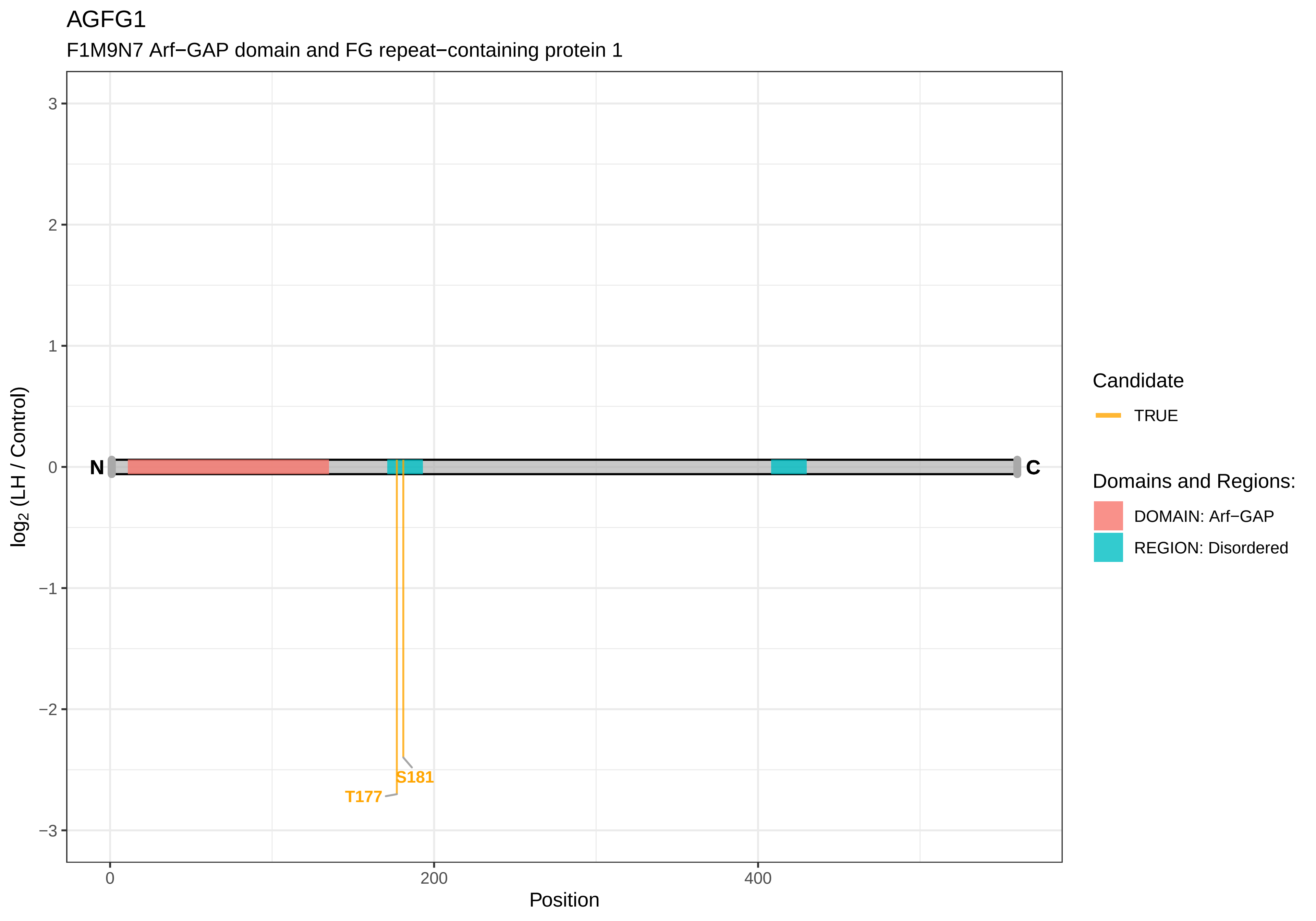

AGFG1
F1M9N7 Arf−GAP domain and FG repeat−containing protein 1
log2 (LH / Control)
3
2
1
0
−1
−2
−3
N
C
S181
T177
0
200
400
Position
Candidate
TRUE
Domains and Regions:
DOMAIN: Arf−GAP
REGION: Disordered

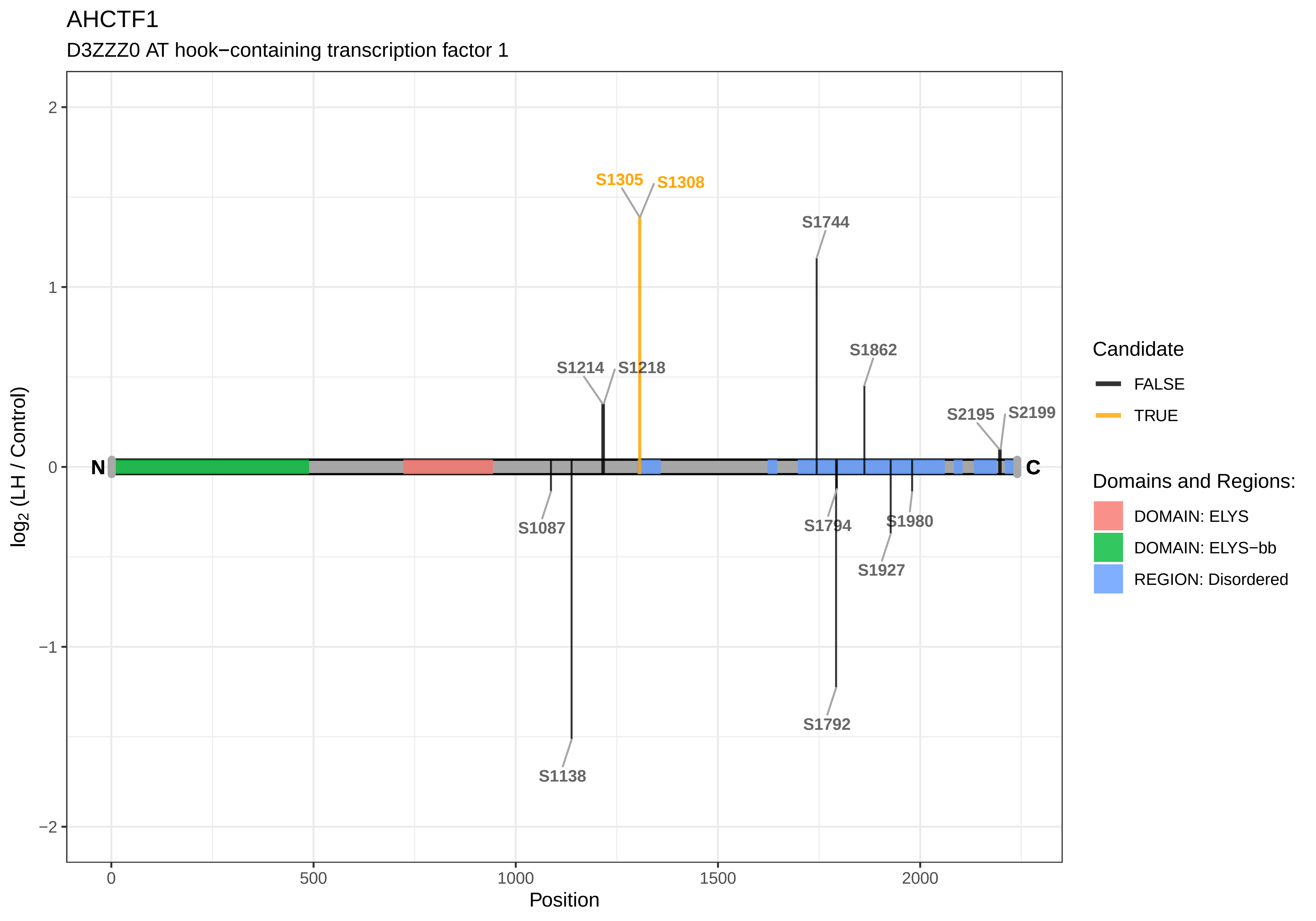

AHCTF1
D3ZZZ0 AT hook−containing transcription factor 1
S1305
S1308
S1744
S1862
S1214
S1218
S2195
S2199
N
C
S1087
S1794
S1980
S1927
S1792
S1138
2
1
0
−1
−2
0
500
1000
1500
2000
Position
log2 (LH / Control)
Candidate
FALSE
TRUE
Domains and Regions:
DOMAIN: ELYS
DOMAIN: ELYS−bb
REGION: Disordered

AHDC1

D4A5R3 AT hook, DNA−binding motif,−containing 1

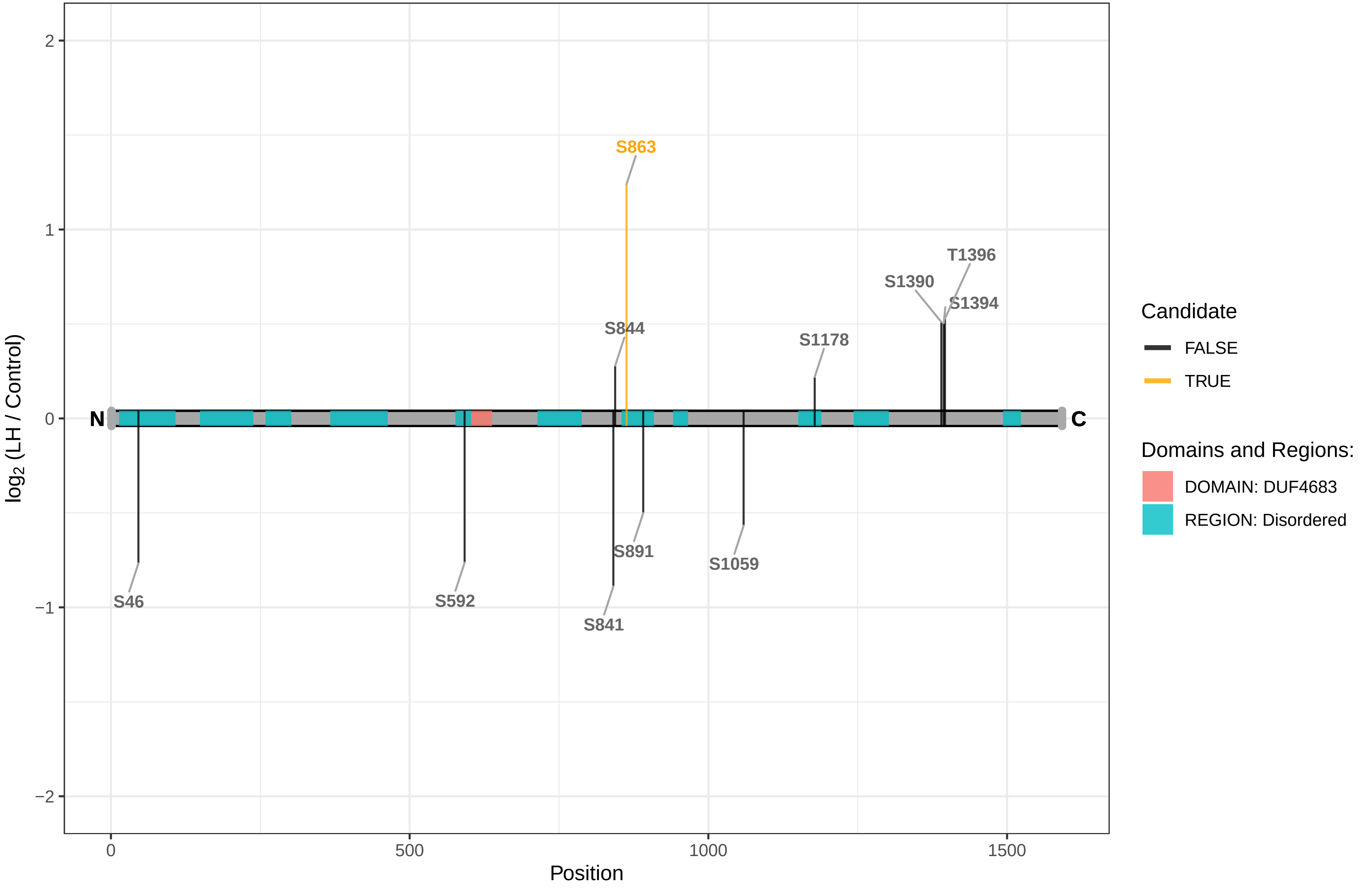

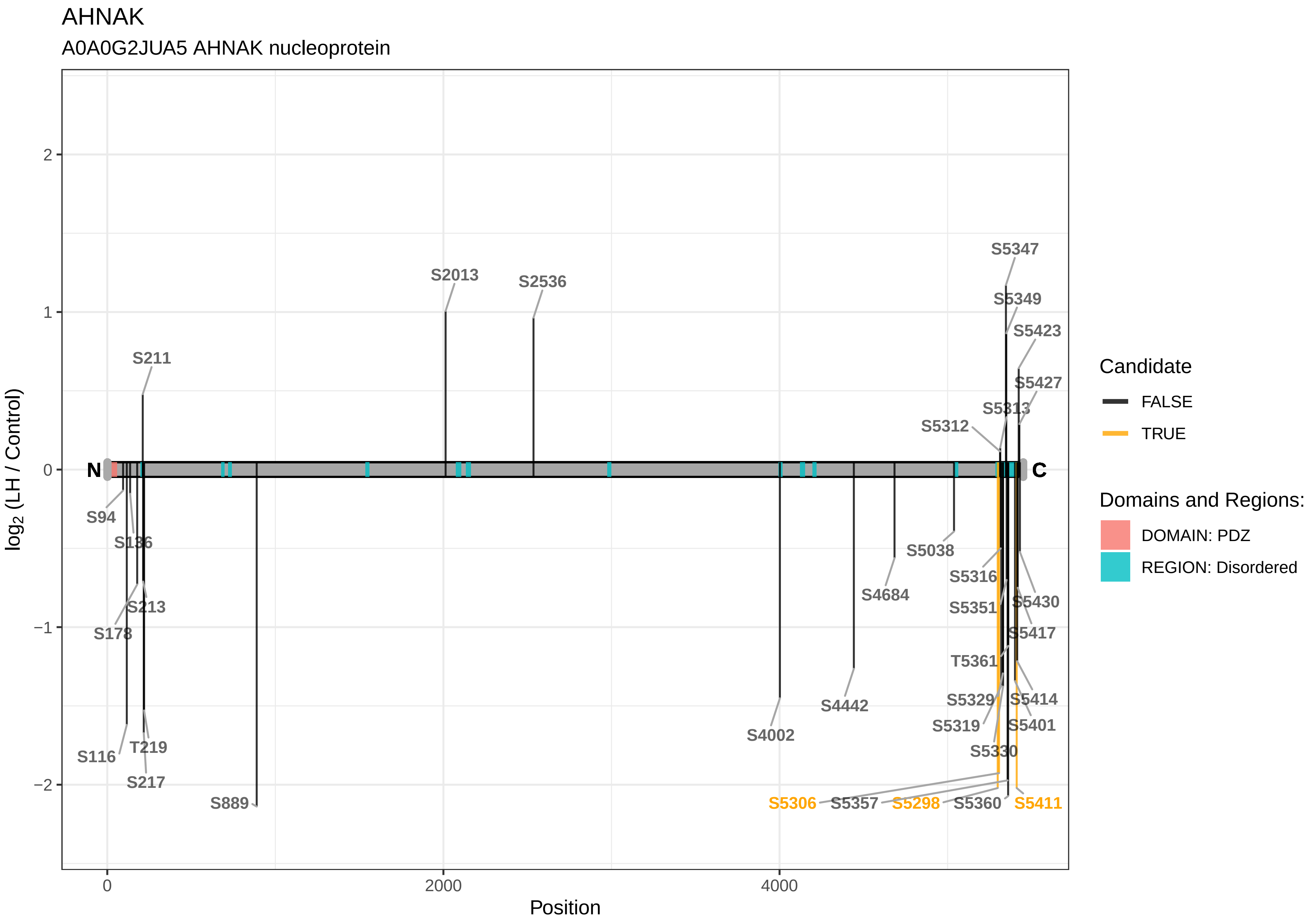

AHNAK
A0A0G2JUA5 AHNAK nucleoprotein
log2 (LH / Control)
Position
S2013
S2536
S211
S5347
S5349
S5423
S5427
S5313
S5312
S94
S136
S213
S178
S116
T219
S217
S889
S4002
S4442
S4684
S5038
S5316
S5351
S5430
S5417
T5361
S5329
S5414
S5319
S5401
S5330
S5306
S5357
S5298
S5360
S5411
N
C
Candidate
FALSE
TRUE
Domains and Regions:
DOMAIN: PDZ
REGION: Disordered

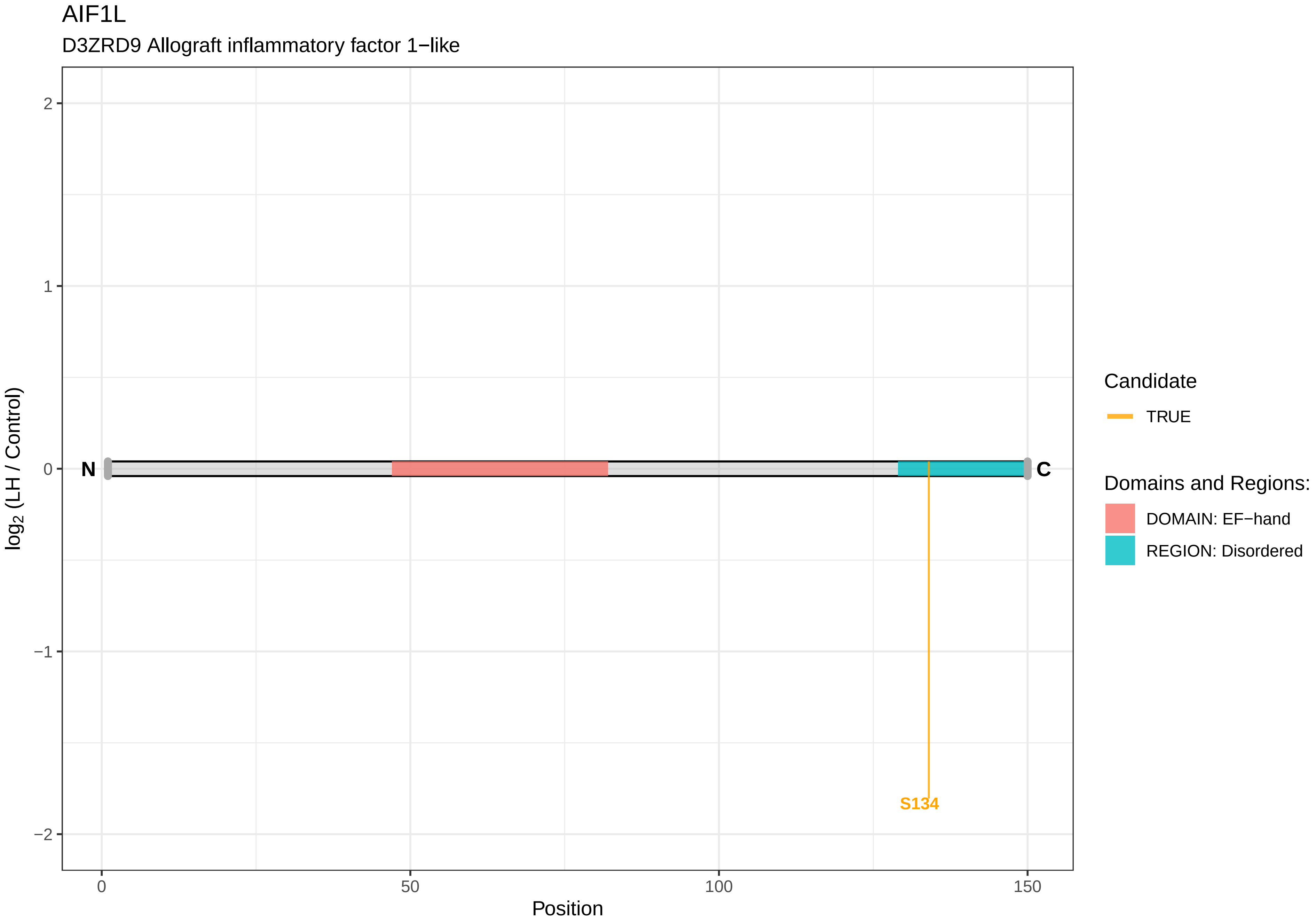
AIF1L
D3ZRD9 Allograft inflammatory factor 1−like
log2 (LH / Control)
2
1
0
−1
−2
N
C
S134
0
50
100
150
Position
Candidate
TRUE
Domains and Regions:
DOMAIN: EF−hand
REGION: Disordered

AIMP1

Q4G079 Aminoacyl tRNA synthetase complex−interacting multifunctional protein 1

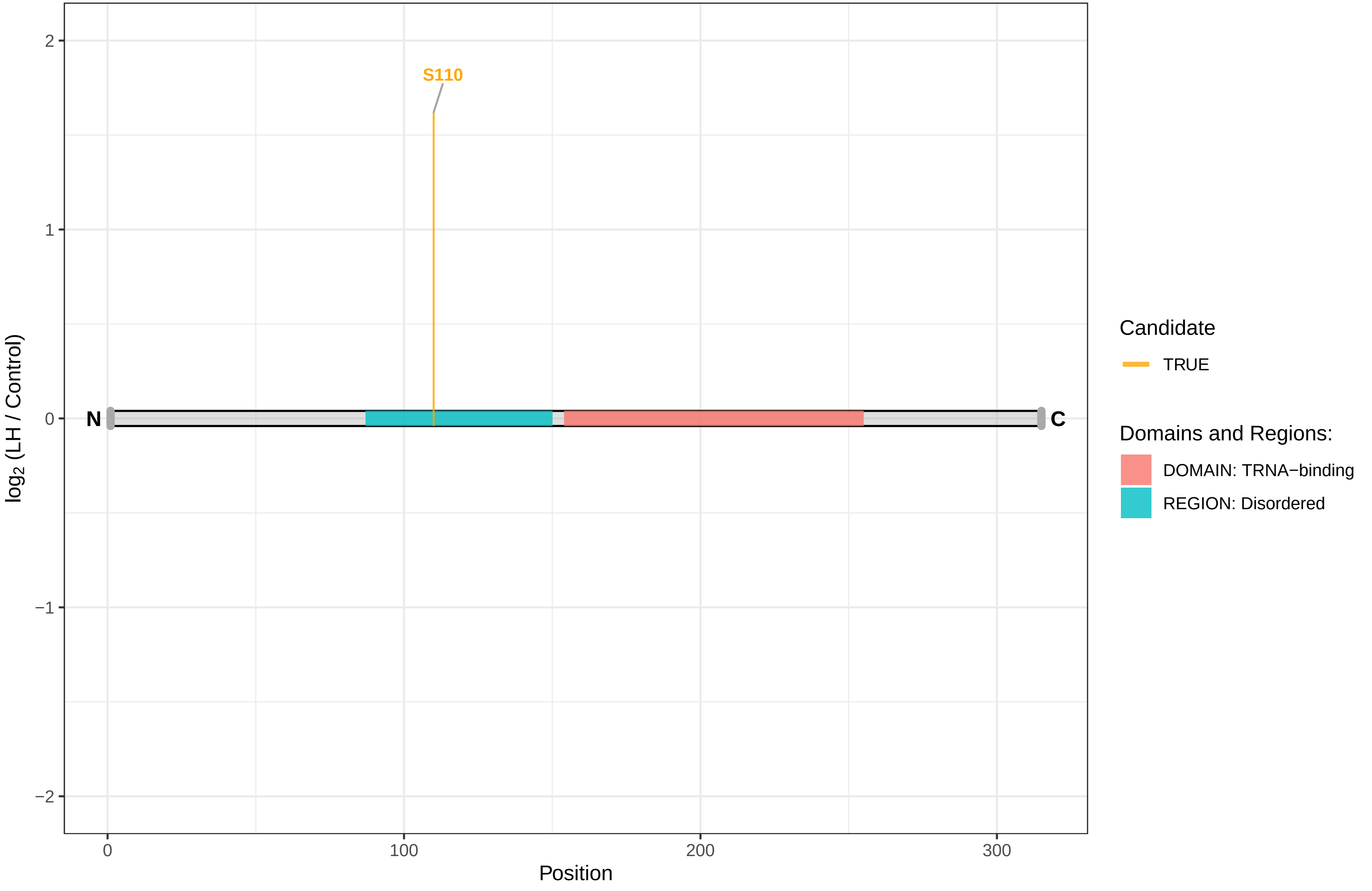

AKAP1

D4A9M6 A kinase (PRKA) anchor protein 1, isoform CRA_b

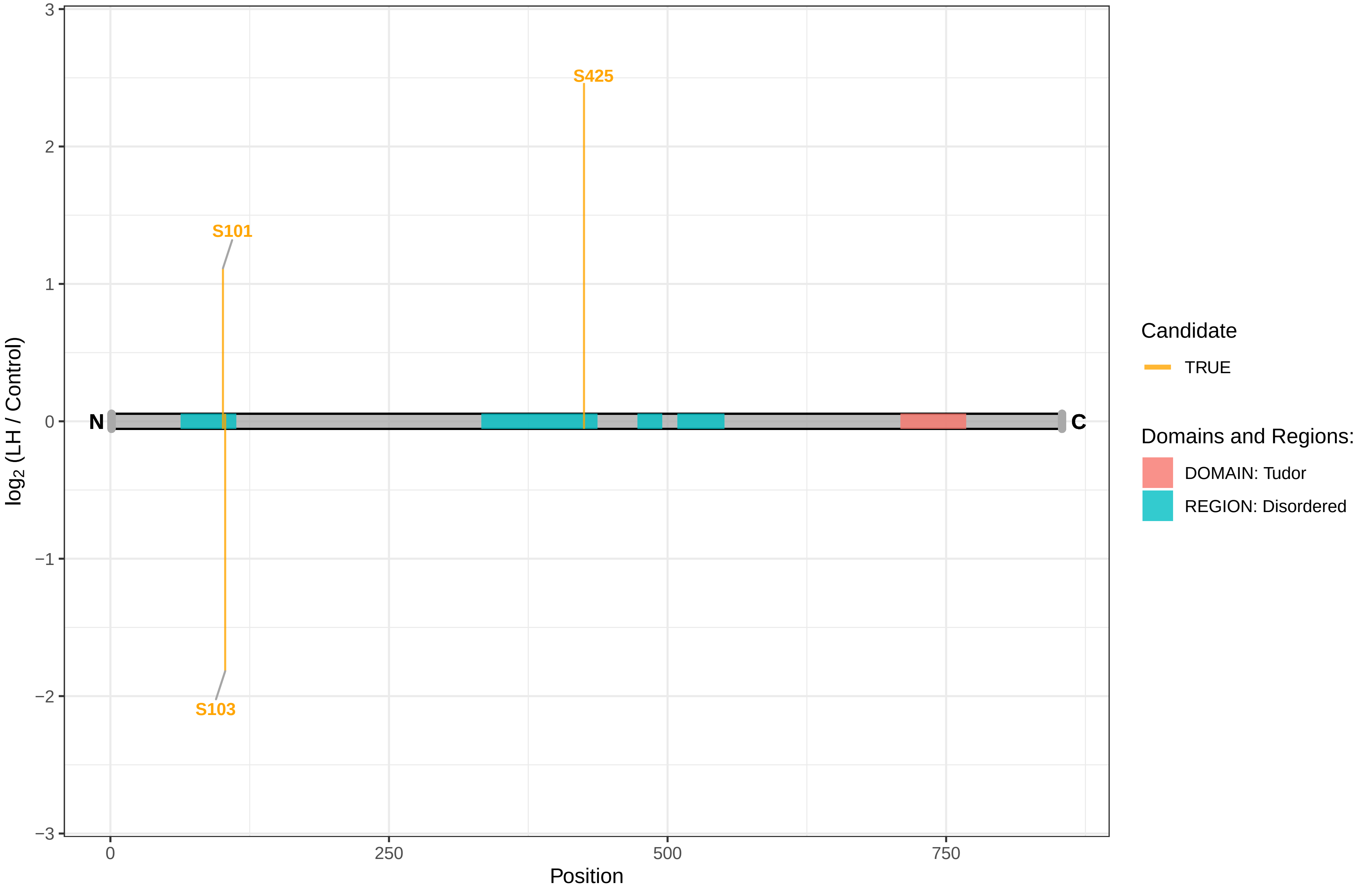

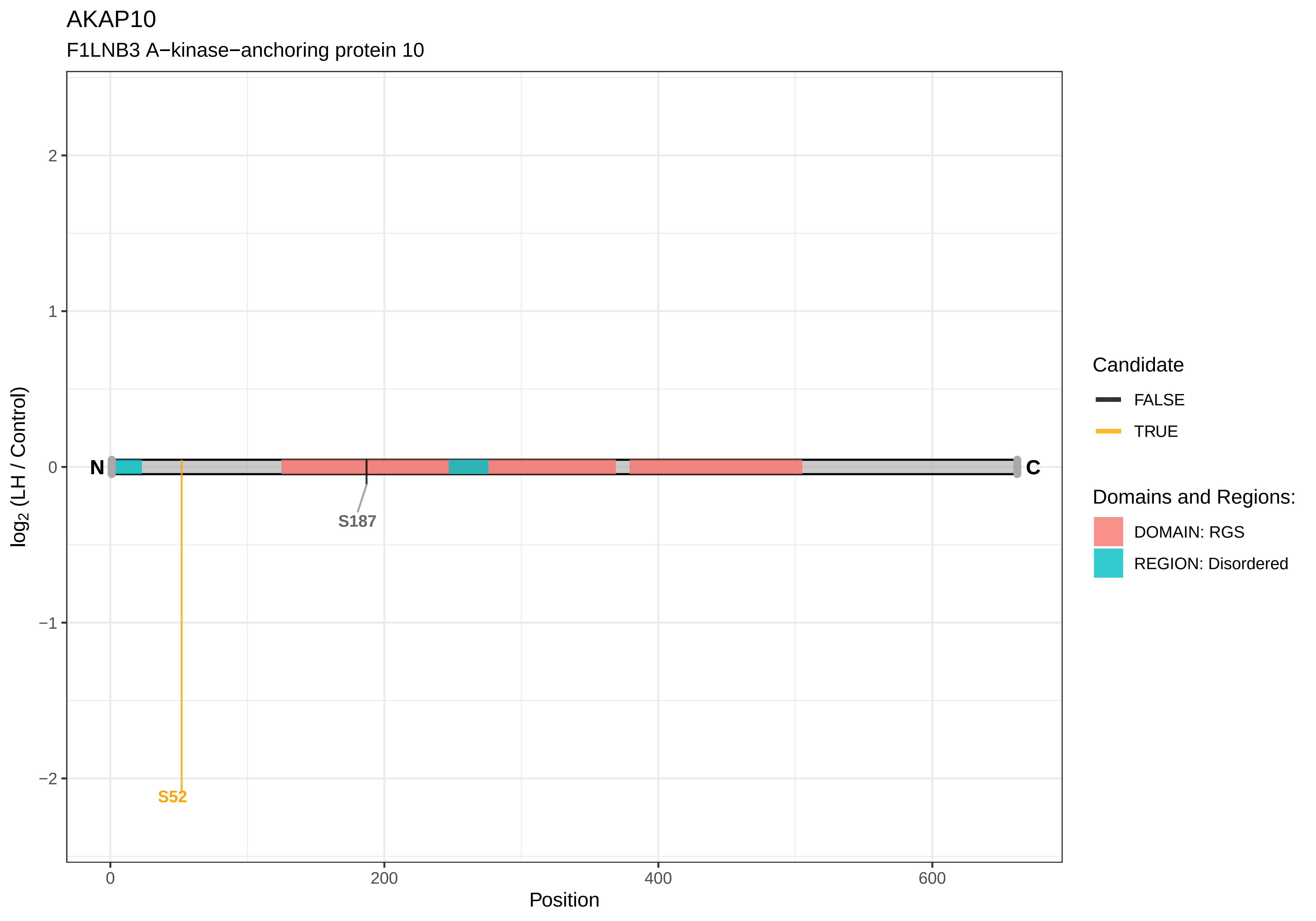

AKAP10
F1LNB3 A−kinase−anchoring protein 10
log2 (LH / Control)
2
1
0
−1
−2
N
C
S187
S52
0
200
400
600
Position
Candidate
FALSE
TRUE
Domains and Regions:
DOMAIN: RGS
REGION: Disordered

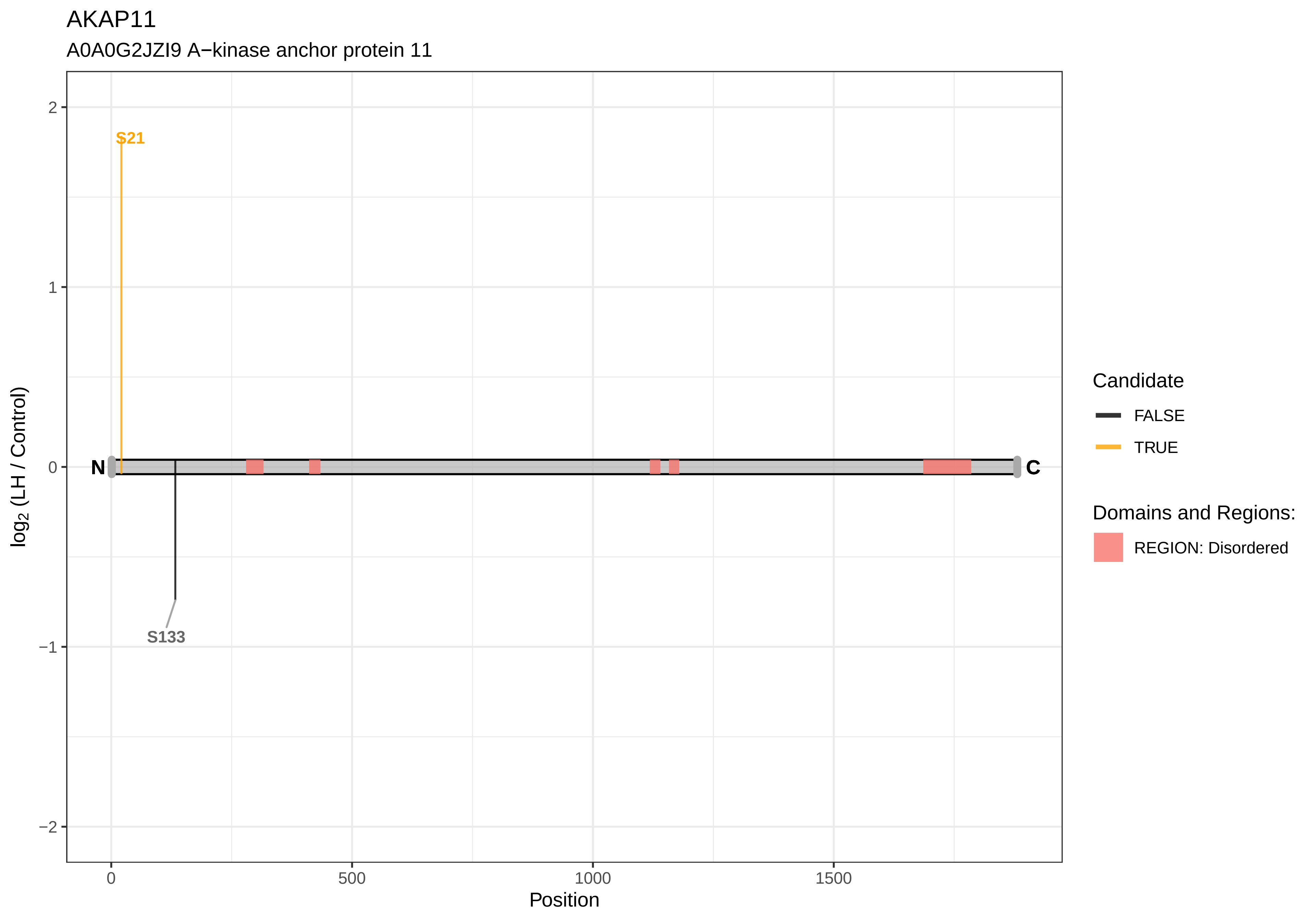

AKAP11
A0A0G2JZI9 A−kinase anchor protein 11
S21
S133
N
C
2
1
0
−1
−2
log2 (LH / Control)
0
500
1000
1500
Position
Candidate
FALSE
TRUE
Domains and Regions:
REGION: Disordered

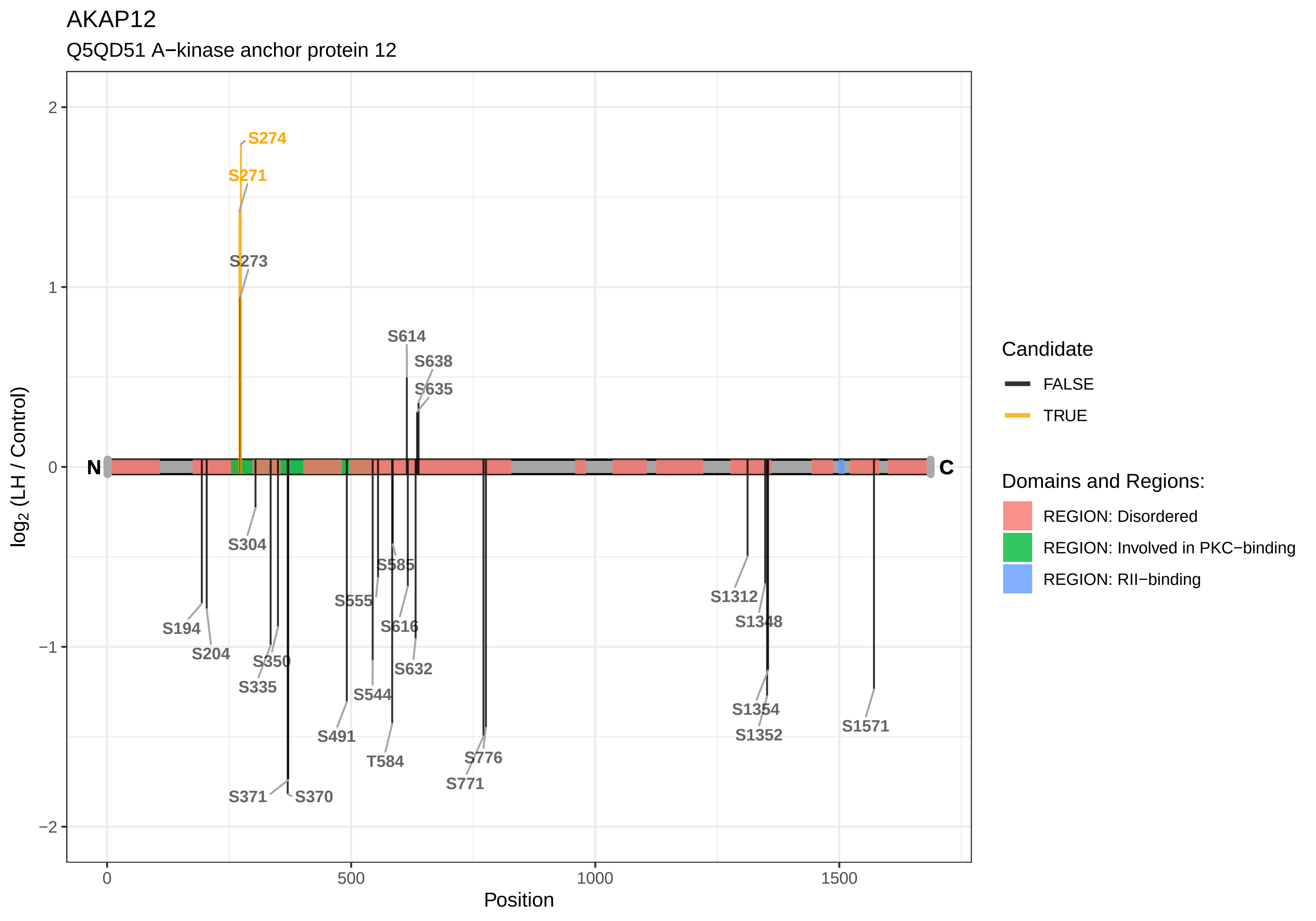
AKAP12
Q5QD51 A−kinase anchor protein 12
S274
S271
S273
S614
S638
S635
S304
S585
S555
S616
S194
S204
S350
S335
S632
S544
S491
T584
S776
S771
S371
S370
S1312
S1348
S1354
S1352
S1571
N
C
log2 (LH / Control)
Position
Candidate
FALSE
TRUE
Domains and Regions:
REGION: Disordered
REGION: Involved in PKC−binding
REGION: RII−binding

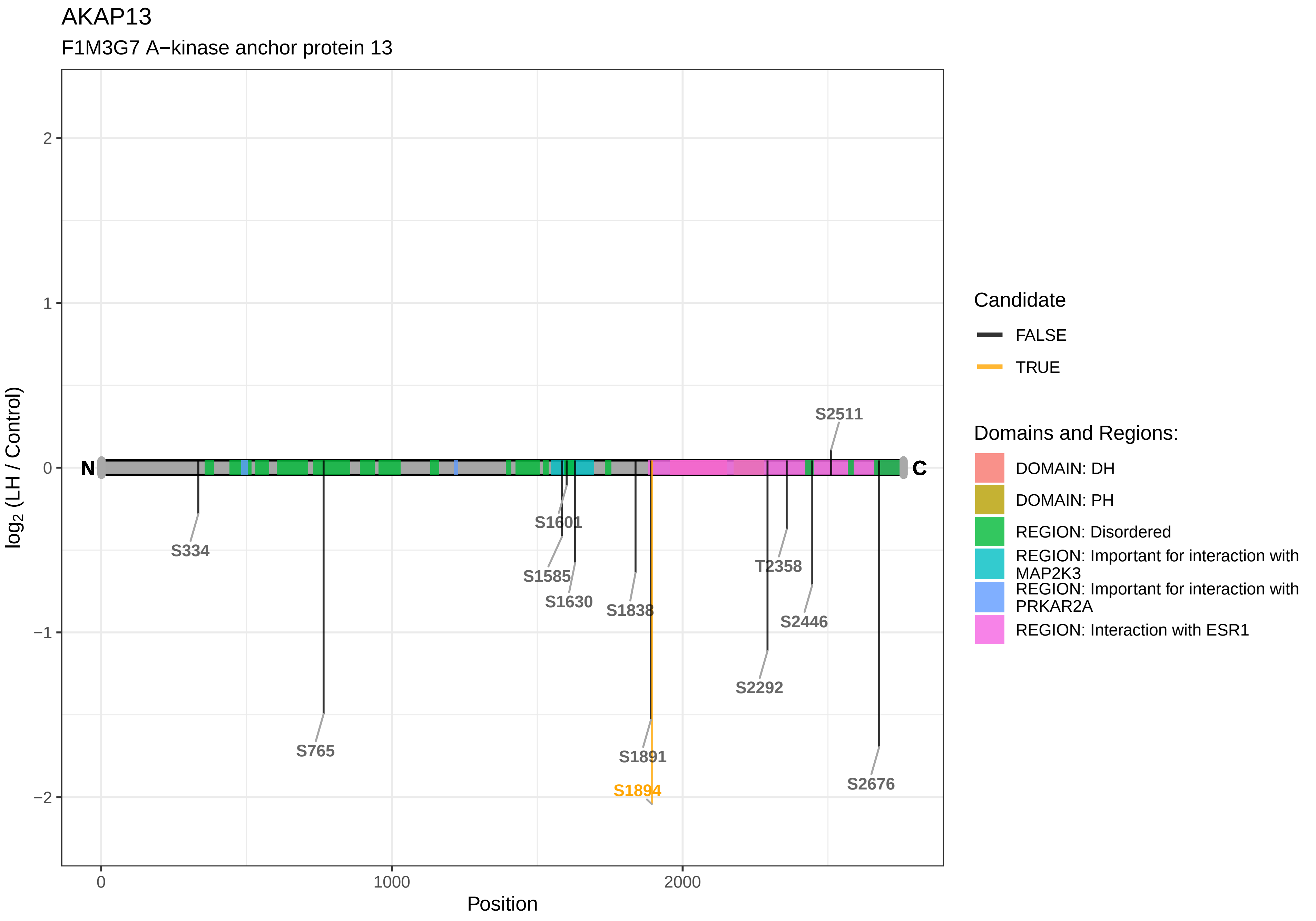
AKAP13
F1M3G7 A−kinase anchor protein 13
log2 (LH / Control)
Position
N
C
S334
S765
S1601
S1585
S1630
S1838
S1891
S1894
T2358
S2446
S2292
S2511
S2676
Candidate
FALSE
TRUE
Domains and Regions:
DOMAIN: DH
DOMAIN: PH
REGION: Disordered
REGION: Important for interaction with MAP2K3
REGION: Important for interaction with PRKAR2A
REGION: Interaction with ESR1

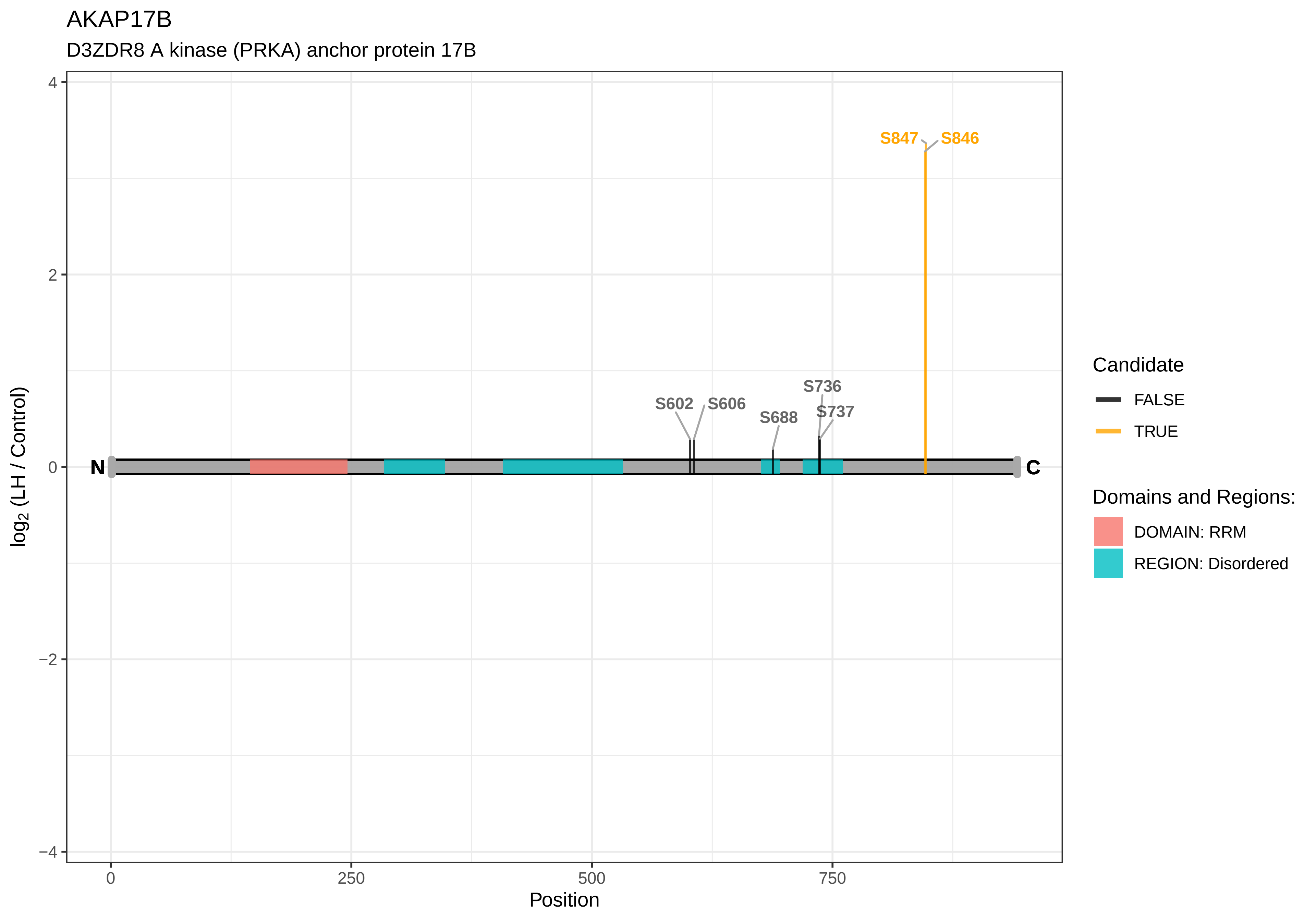
AKAP17B
D3ZDR8 A kinase (PRKA) anchor protein 17B
S847
S846
S602
S606
S736
S688
S737
N
C
4
2
0
−2
−4
log2 (LH / Control)
0
250
500
750
Position
Candidate
FALSE
TRUE
Domains and Regions:
DOMAIN: RRM
REGION: Disordered

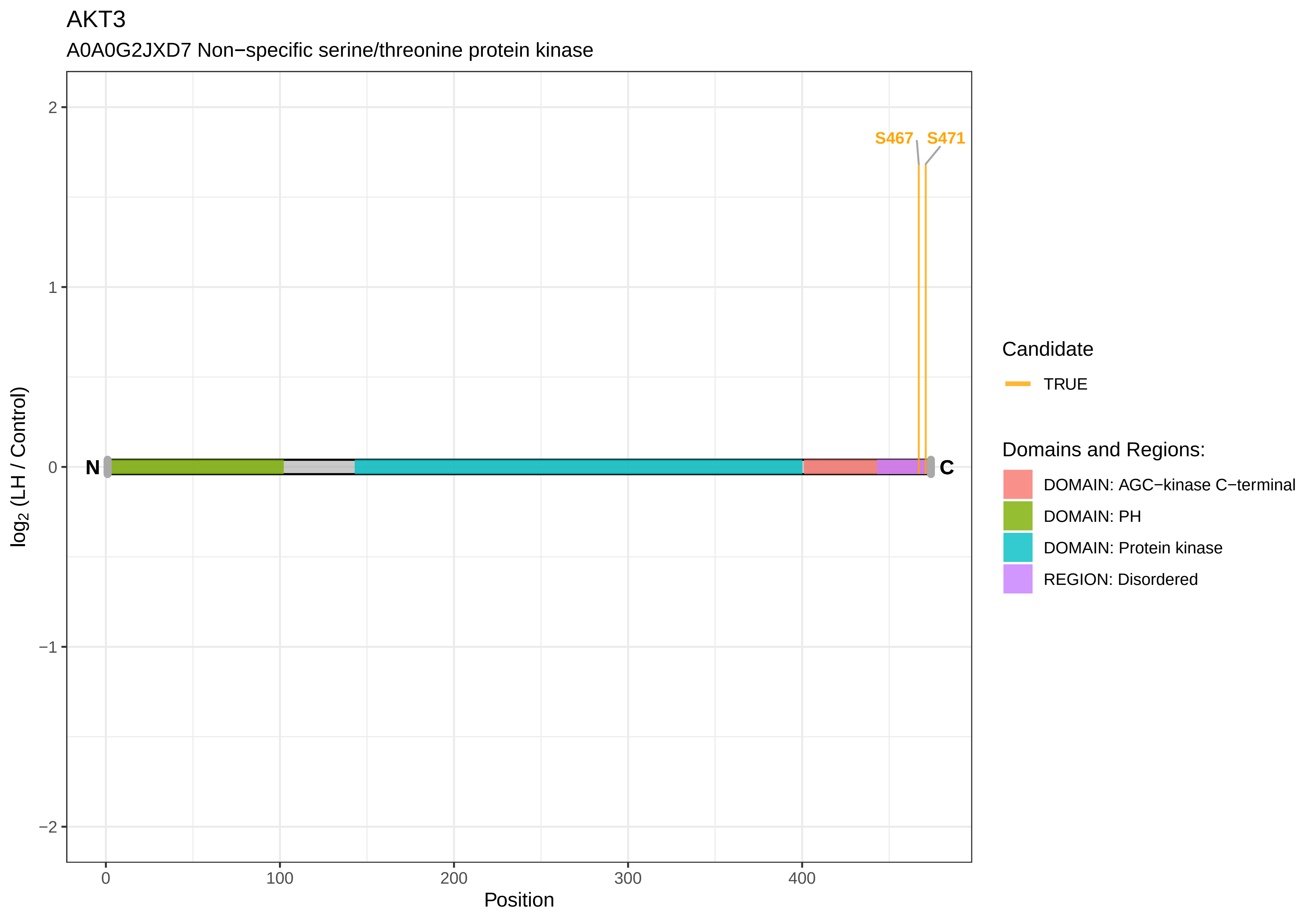

AKT3
A0A0G2JXD7 Non−specific serine/threonine protein kinase
S467
S471
N
C
2
1
0
−1
−2
log2 (LH / Control)
0
100
200
300
400
Position
Candidate
TRUE
Domains and Regions:
DOMAIN: AGC−kinase C−terminal
DOMAIN: PH
DOMAIN: Protein kinase
REGION: Disordered

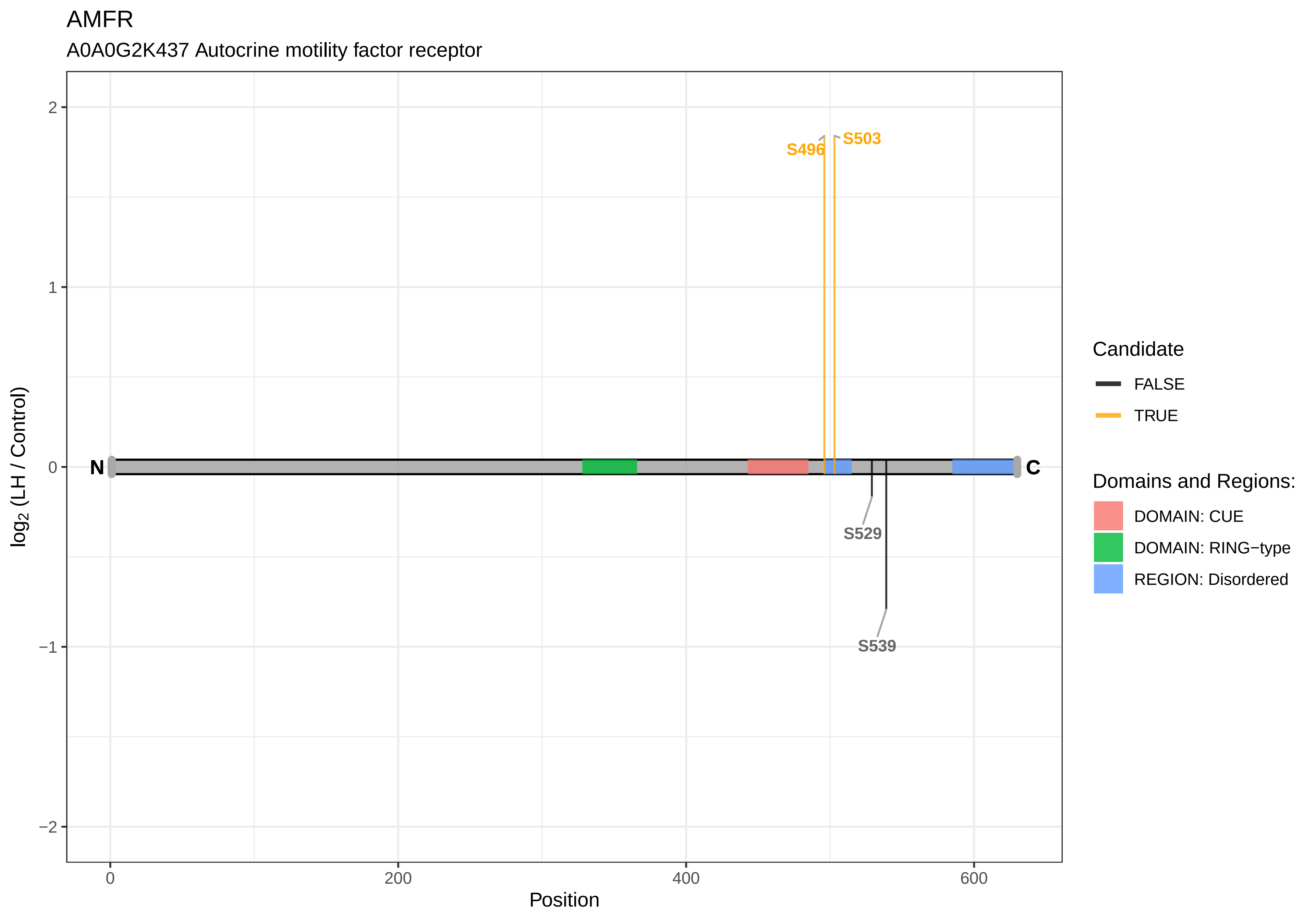
AMFR
A0A0G2K437 Autocrine motility factor receptor
S496
S503
S529
S539
N
C
2
1
0
−1
−2
0
200
400
600
Position
log2 (LH / Control)
Candidate
FALSE
TRUE
Domains and Regions:
DOMAIN: CUE
DOMAIN: RING−type
REGION: Disordered

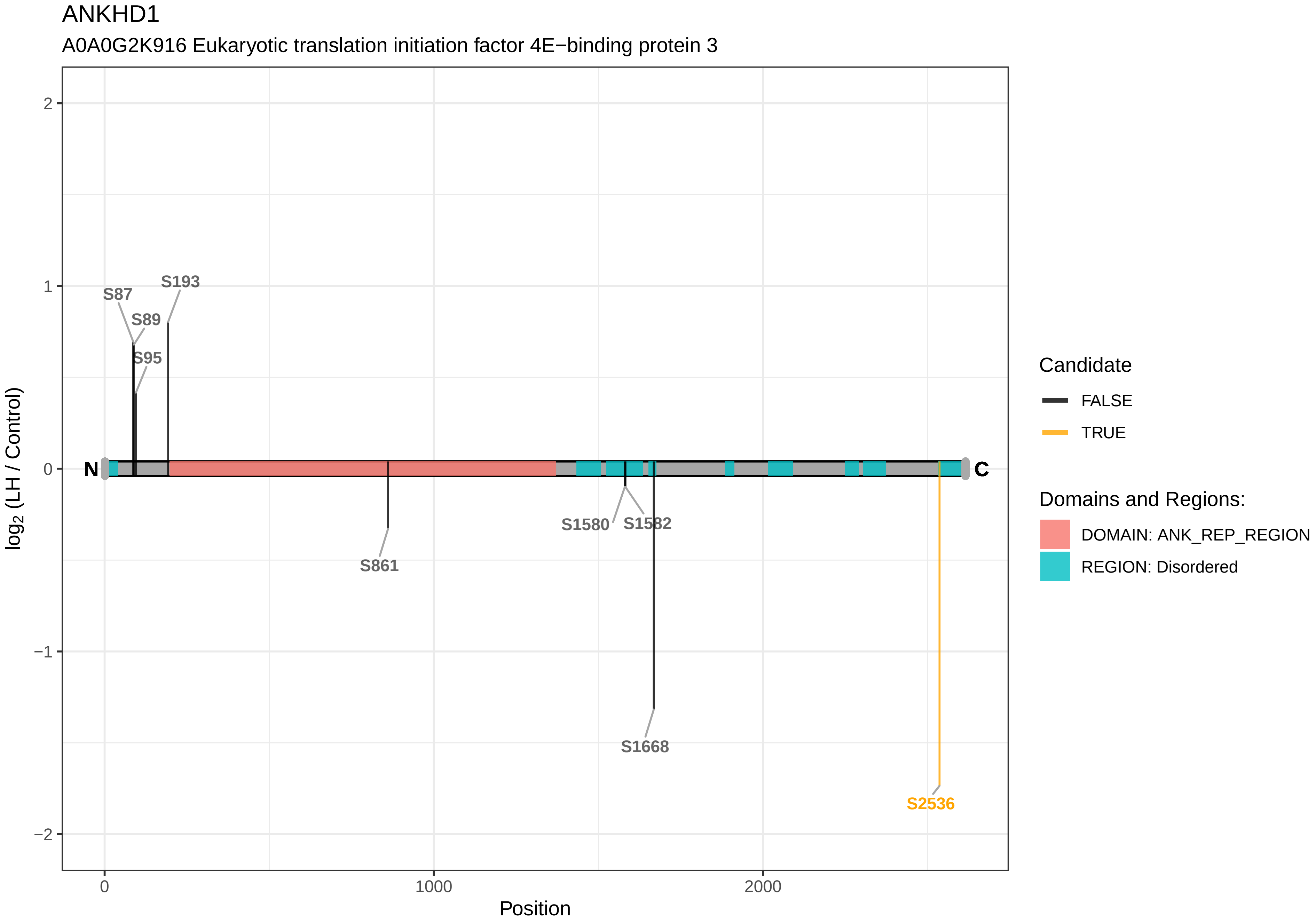
ANKHD1
A0A0G2K916 Eukaryotic translation initiation factor 4E−binding protein 3
log2 (LH / Control)
Position
0
1000
2000
2
1
0
−1
−2
N
C
S87
S89
S95
S193
S861
S1580
S1582
S1668
S2536
Candidate
FALSE
TRUE
Domains and Regions:
DOMAIN: ANK_REP_REGION
REGION: Disordered

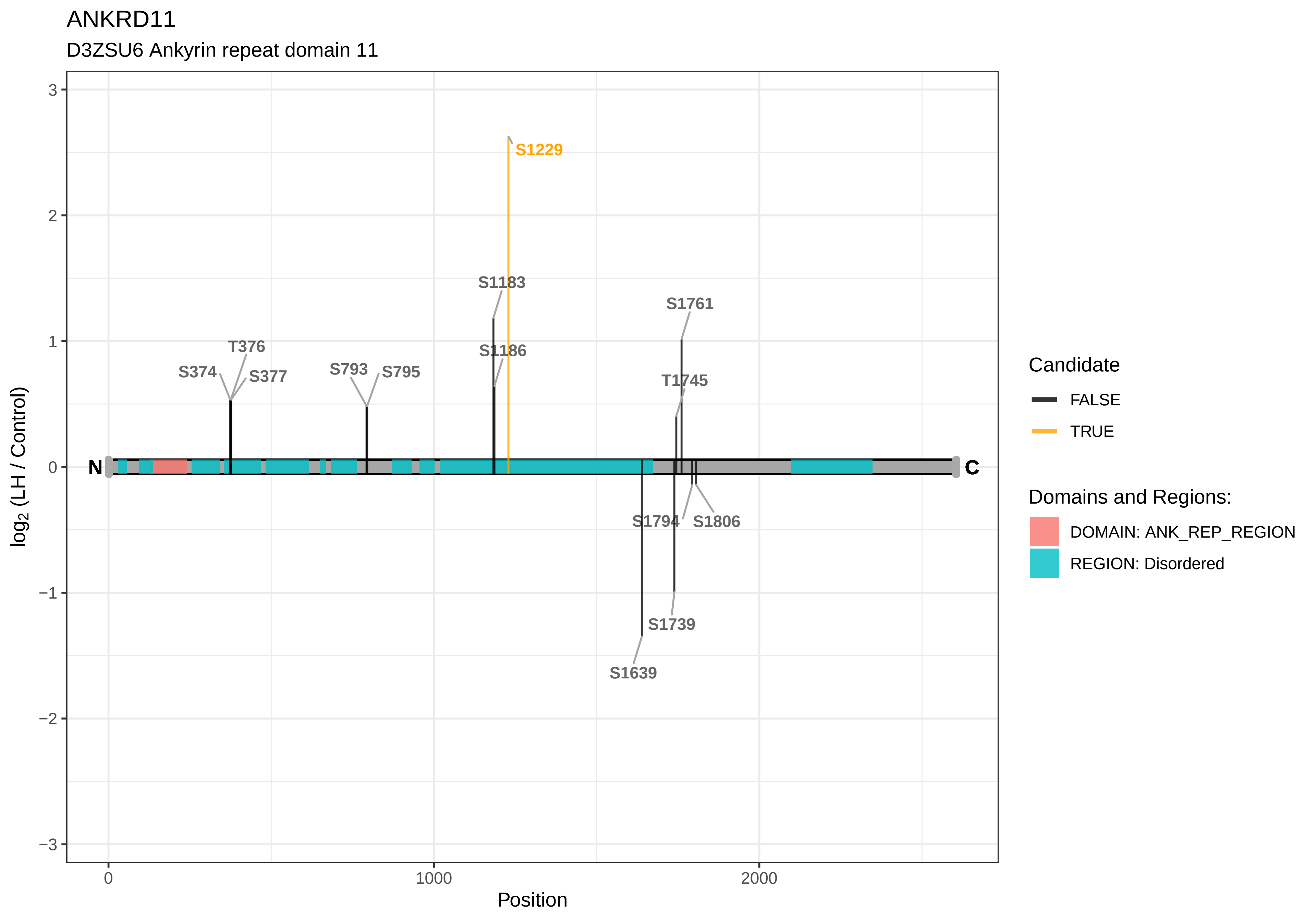

ANKRD11
D3ZSU6 Ankyrin repeat domain 11
S1229
S1183
S1761
T376
S1186
S374
S377
S793
S795
T1745
N
C
S1794
S1806
S1739
S1639
log2 (LH / Control)
Position
0
1000
2000
3
2
1
0
−1
−2
−3
Candidate
FALSE
TRUE
Domains and Regions:
DOMAIN: ANK_REP_REGION
REGION: Disordered

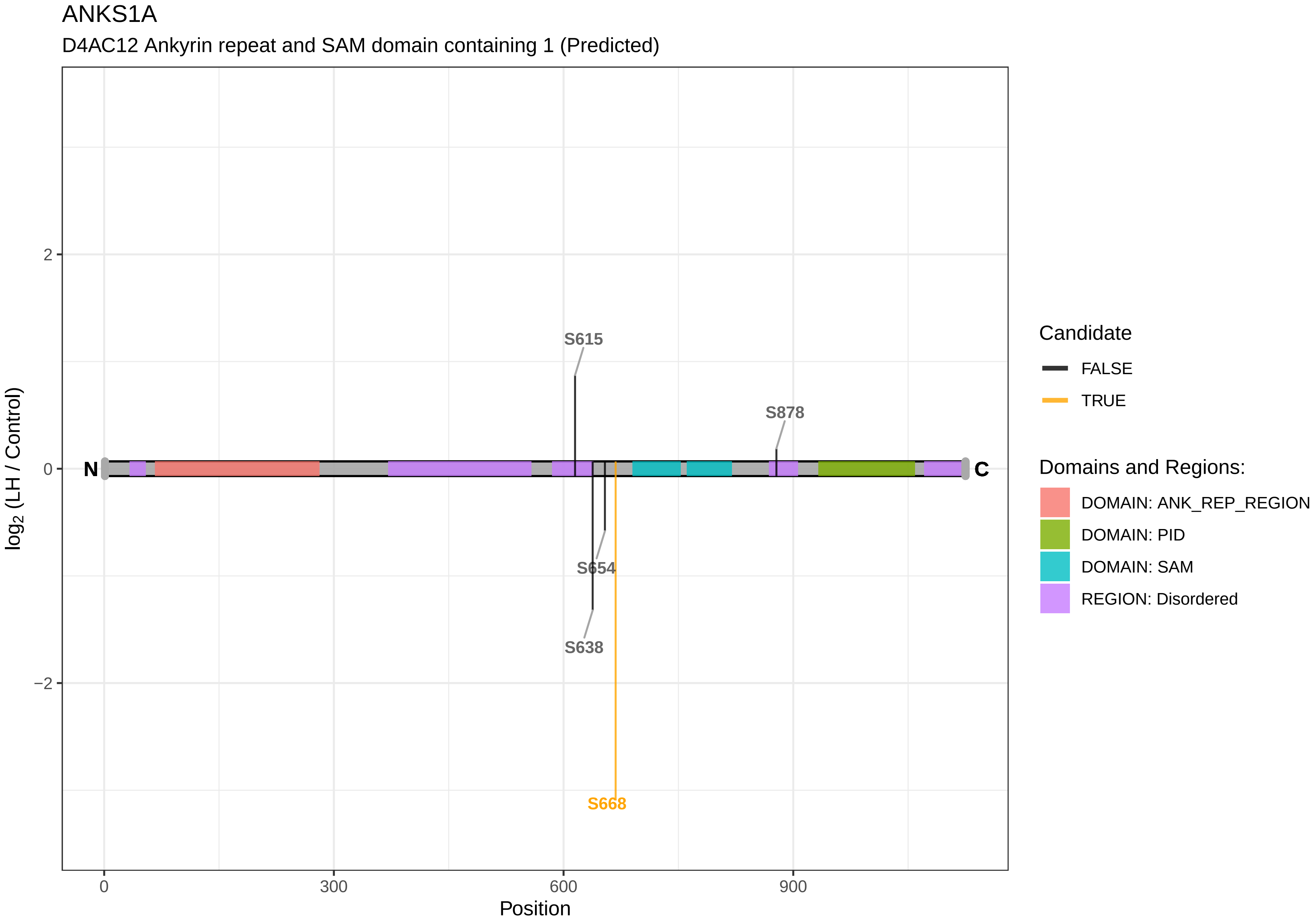
ANKS1A
D4AC12 Ankyrin repeat and SAM domain containing 1 (Predicted)
S615
S878
S654
S638
S668
N
C
2
0
−2
0
300
600
900
Position
log2 (LH / Control)
Candidate
FALSE
TRUE
Domains and Regions:
DOMAIN: ANK_REP_REGION
DOMAIN: PID
DOMAIN: SAM
REGION: Disordered

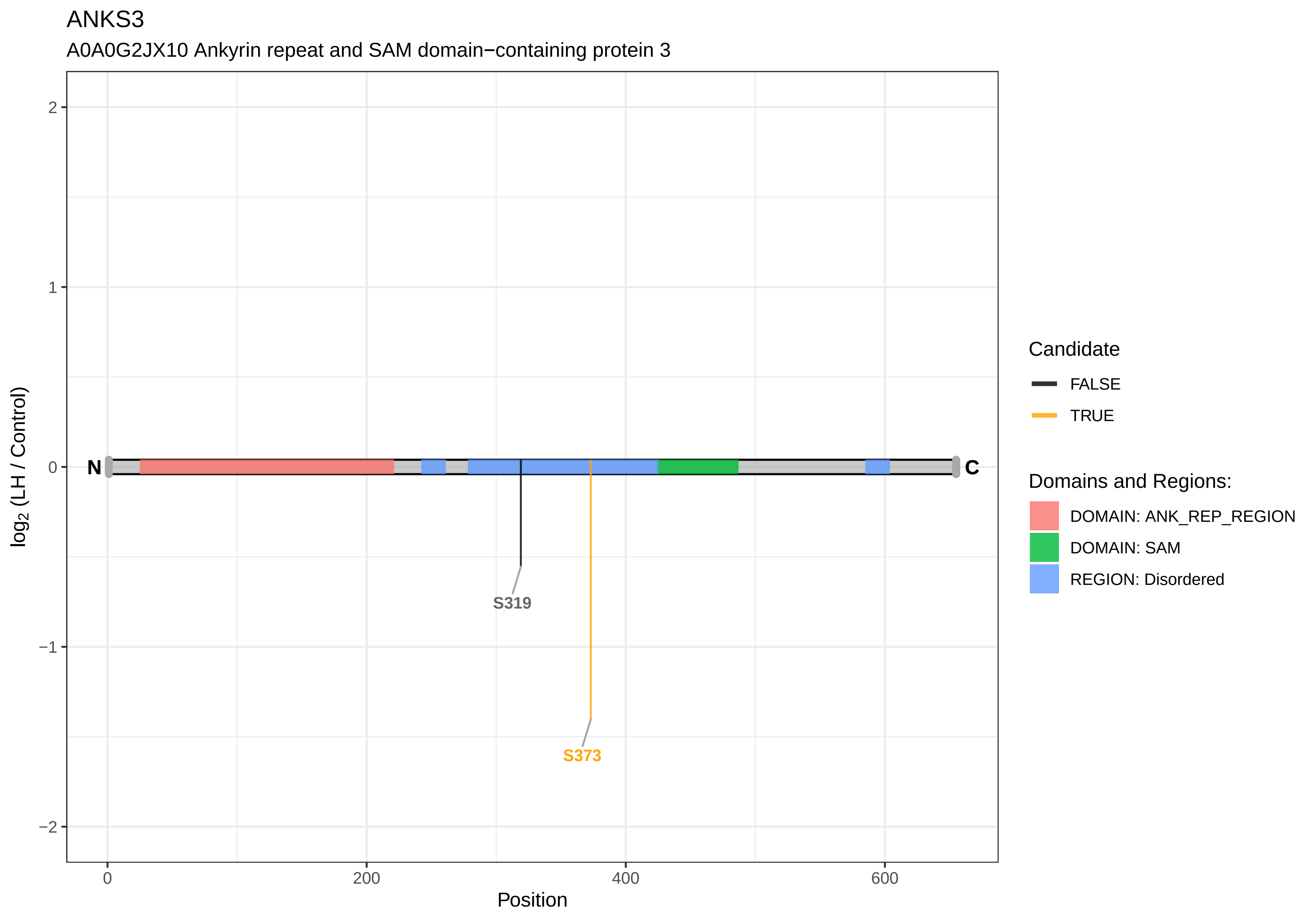

ANKS3
A0A0G2JX10 Ankyrin repeat and SAM domain−containing protein 3
log2 (LH / Control)
2
1
0
−1
−2
N
C
S319
S373
0
200
400
600
Position
Candidate
FALSE
TRUE
Domains and Regions:
DOMAIN: ANK_REP_REGION
DOMAIN: SAM
REGION: Disordered

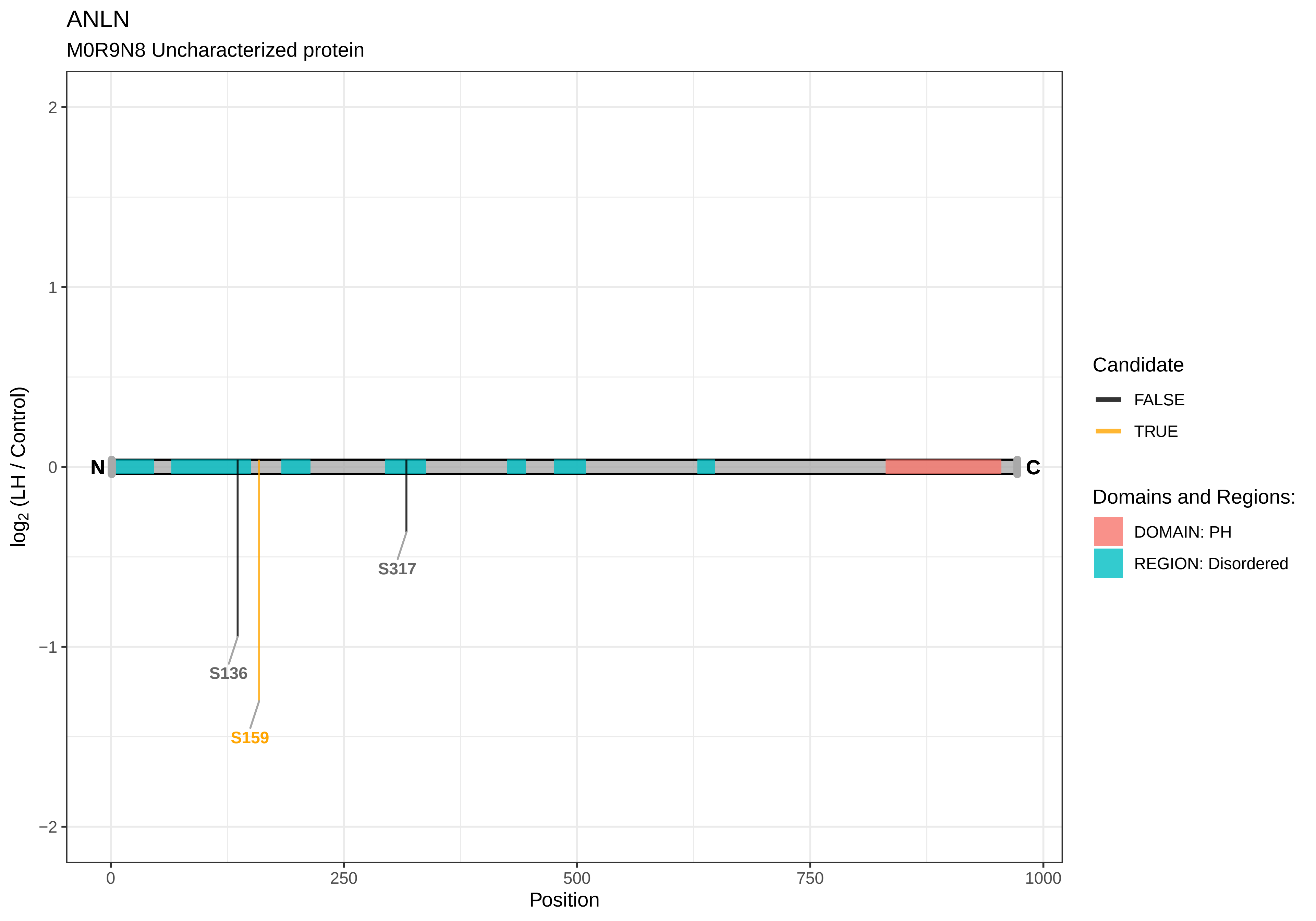
ANLN
M0R9N8 Uncharacterized protein
2
1
0
−1
−2
log2 (LH / Control)
N
C
S317
S136
S159
0
250
500
750
1000
Position
Candidate
FALSE
TRUE
Domains and Regions:
DOMAIN: PH
REGION: Disordered

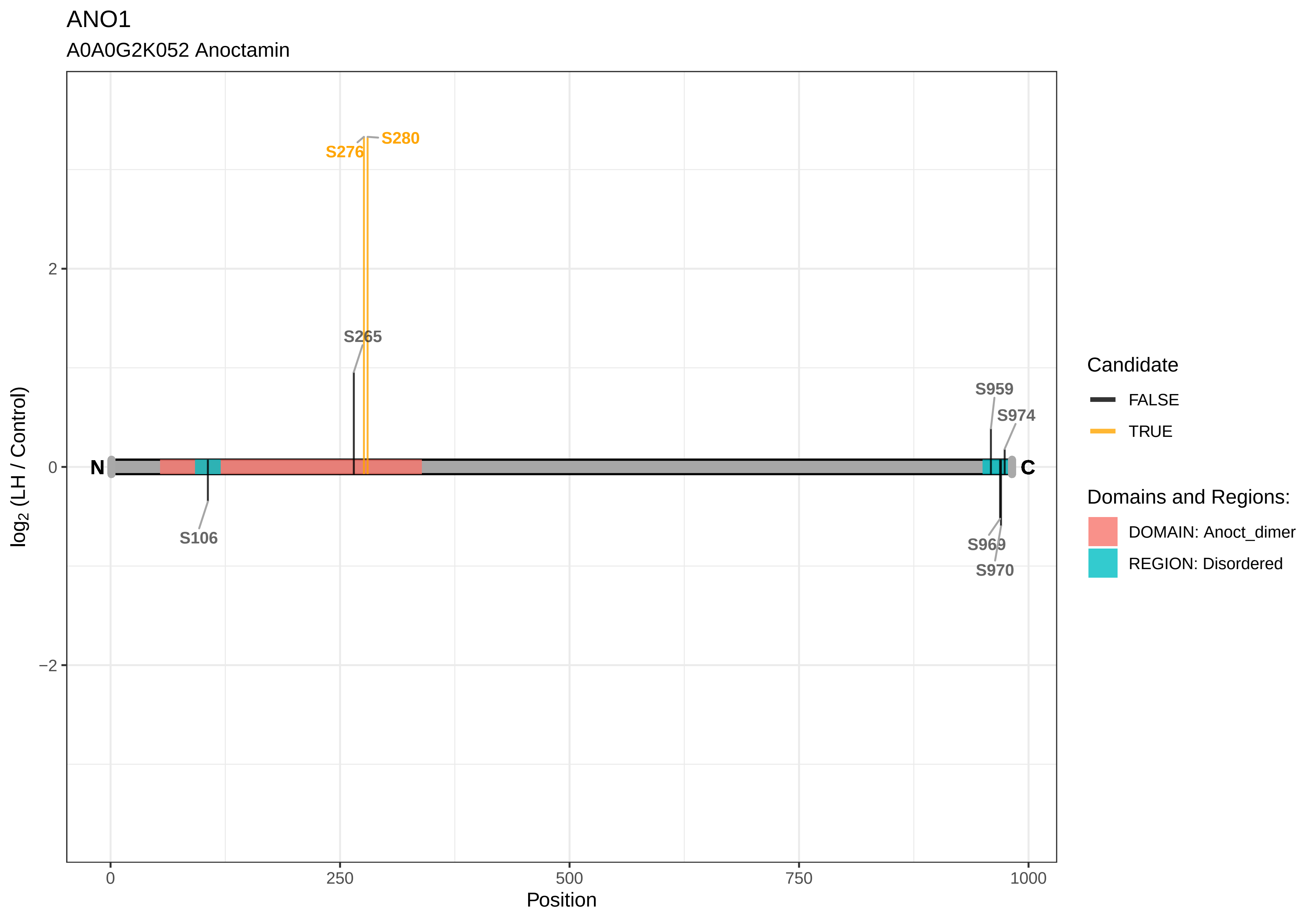

ANO1
A0A0G2K052 Anoctamin
S280
S276
S265
S106
S959
S974
S969
S970
N
C
2
0
−2
0
250
500
750
1000
Position
log2 (LH / Control)
Candidate
FALSE
TRUE
Domains and Regions:
DOMAIN: Anoct_dimer
REGION: Disordered

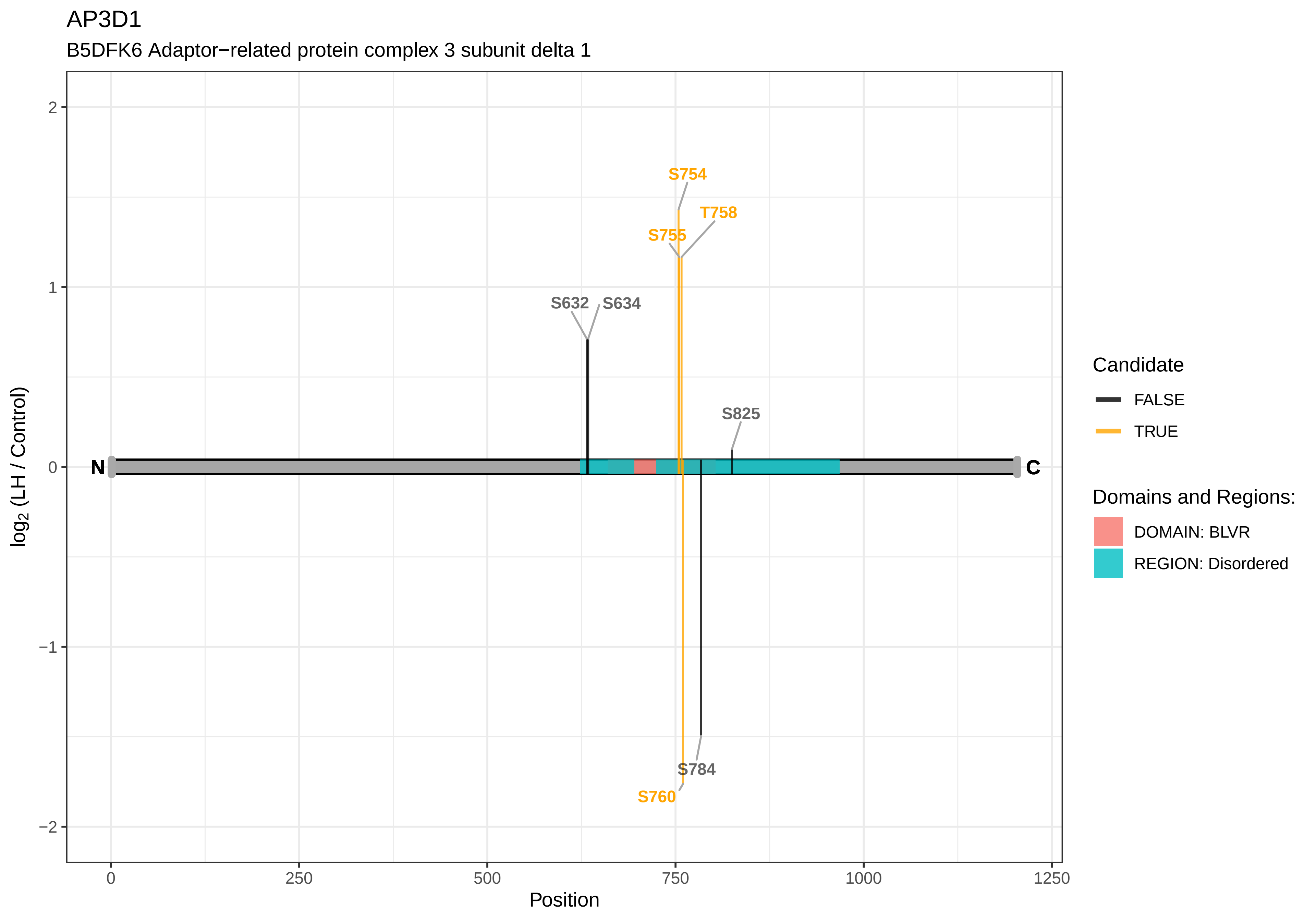

AP3D1
B5DFK6 Adaptor−related protein complex 3 subunit delta 1
S754
T758
S755
S632
S634
S825
S784
S760
N
C
log2 (LH / Control)
Position
2
1
0
−1
−2
0
250
500
750
1000
1250
Candidate
FALSE
TRUE
Domains and Regions:
DOMAIN: BLVR
REGION: Disordered

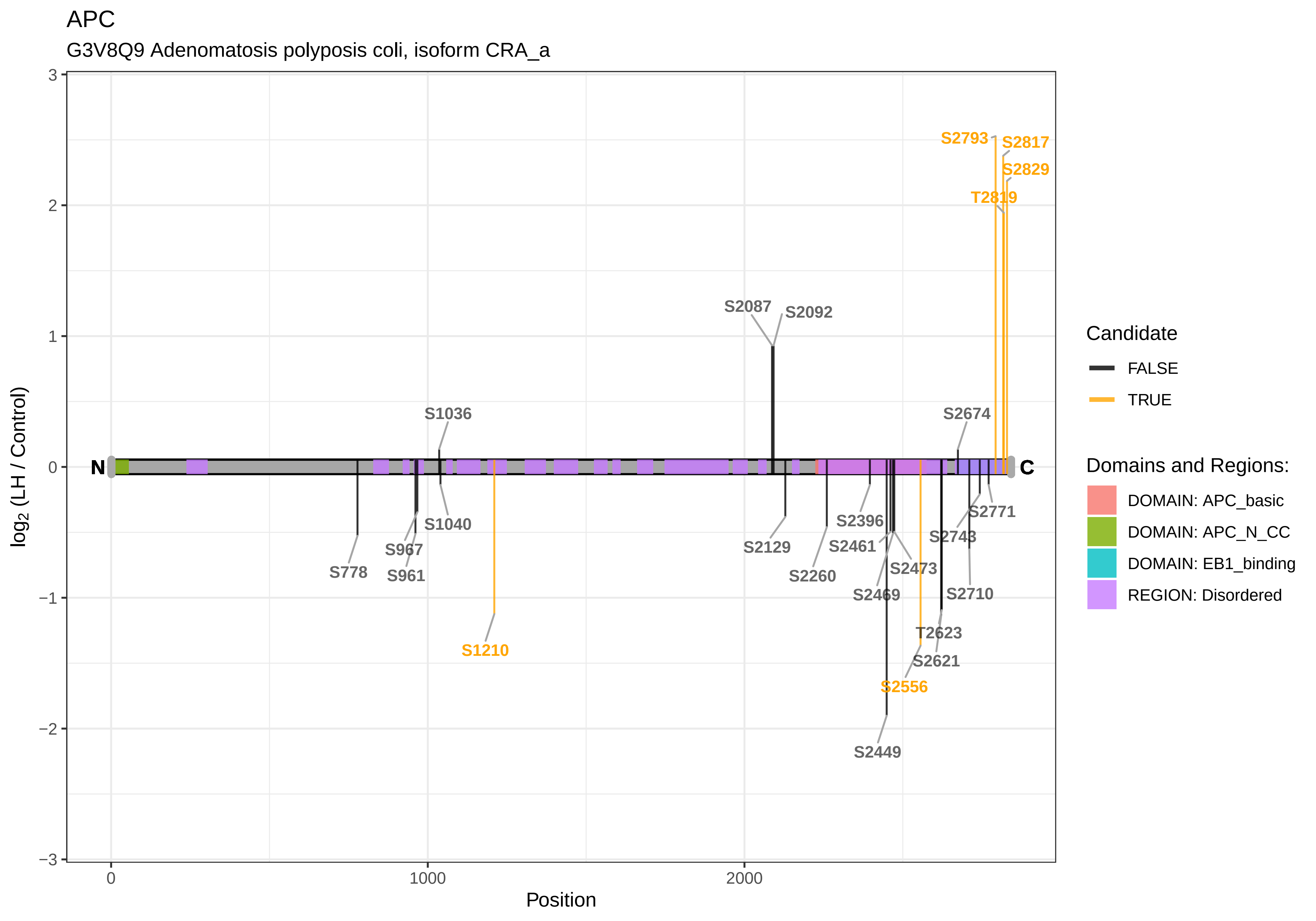

APC
G3V8Q9 Adenomatosis polyposis coli, isoform CRA_a
log2 (LH / Control)
Position
N
C
S2793
S2817
S2829
T2819
S2087
S2092
S1036
S2674
S1040
S967
S961
S778
S1210
S2129
S2260
S2396
S2461
S2473
S2469
S2771
S2743
S2710
T2623
S2621
S2556
S2449
Candidate
FALSE
TRUE
Domains and Regions:
DOMAIN: APC_basic
DOMAIN: APC_N_CC
DOMAIN: EB1_binding
REGION: Disordered

API5

B1WC49 Api5 protein

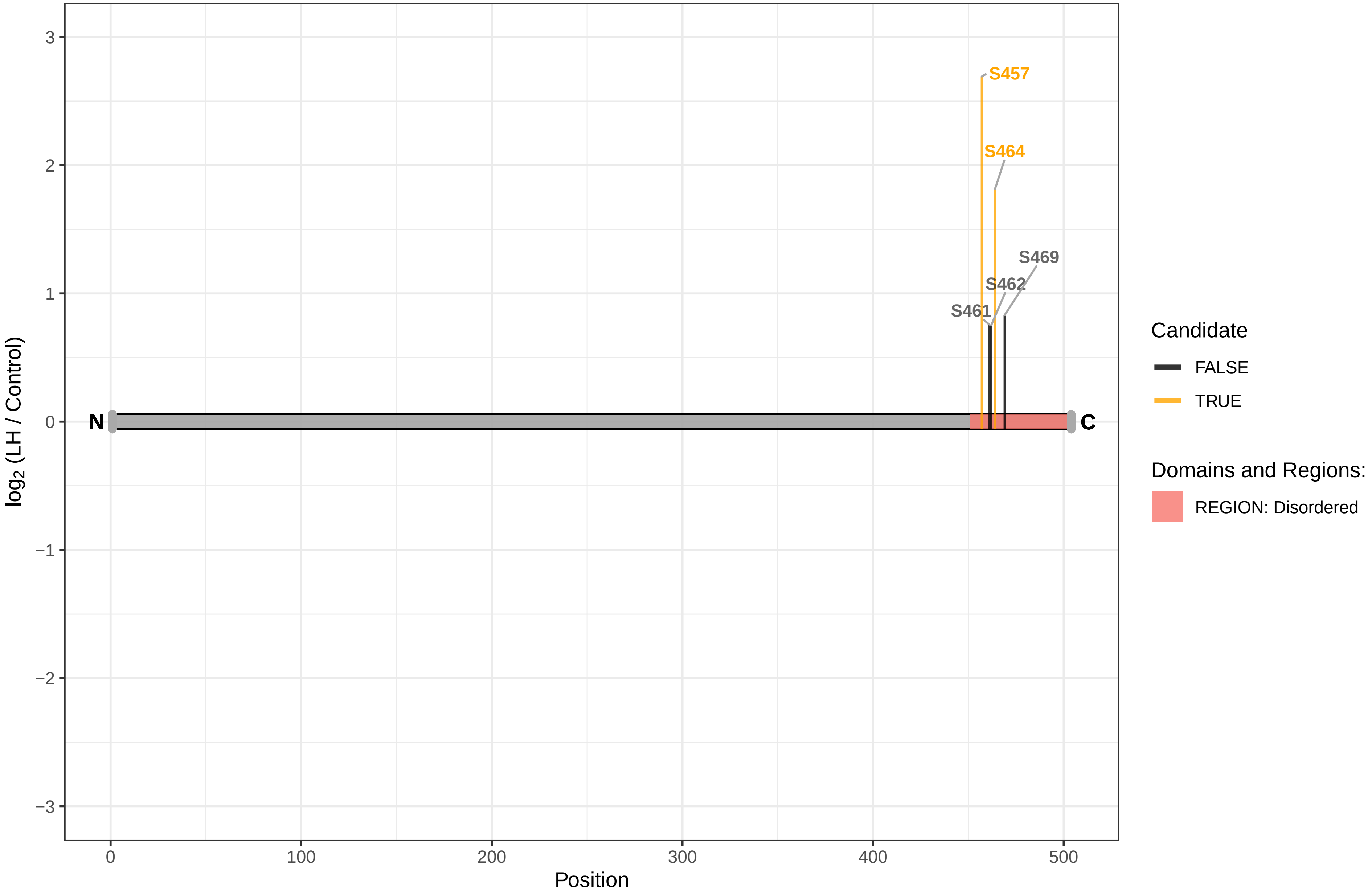

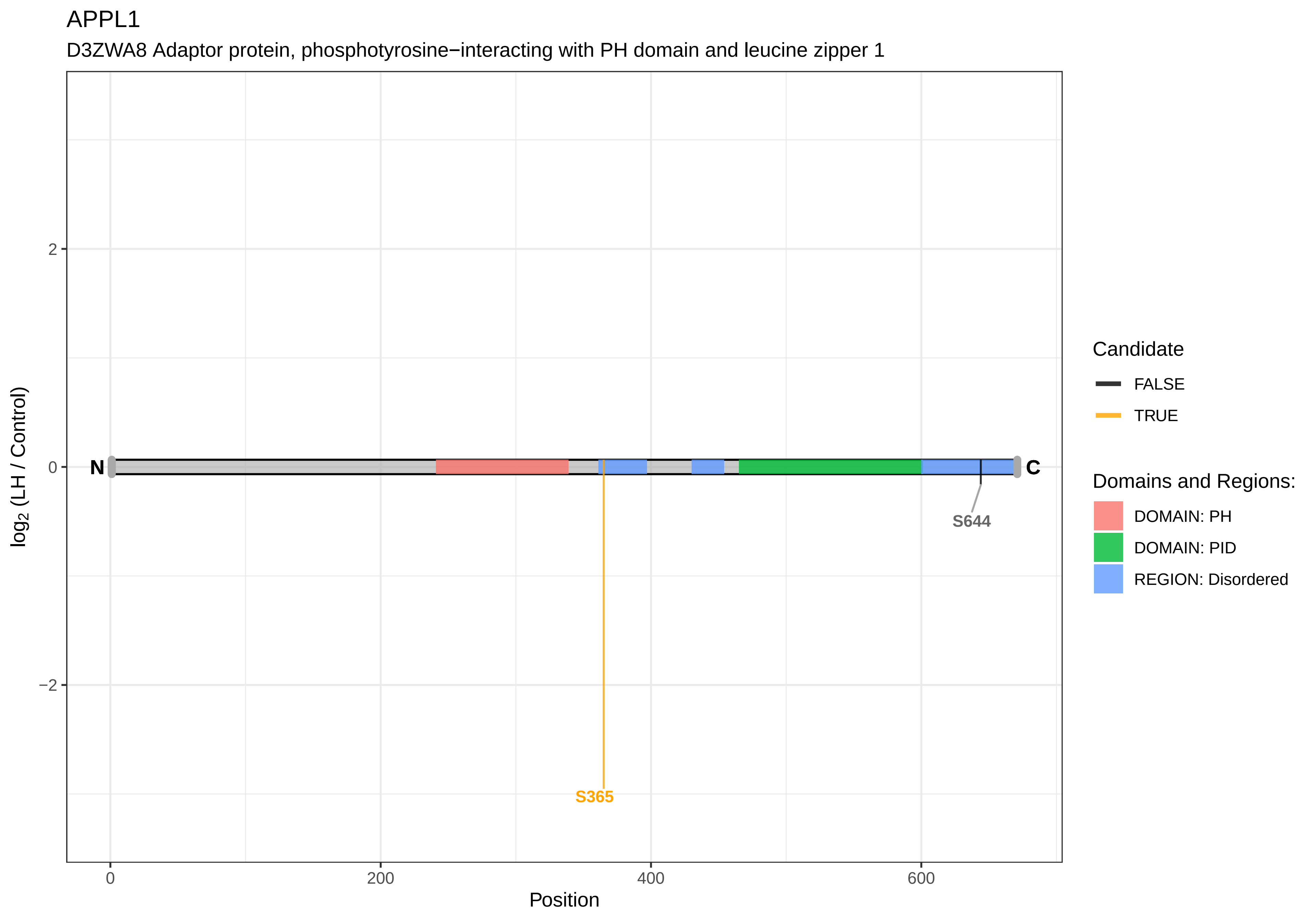
APPL1
D3ZWA8 Adaptor protein, phosphotyrosine−interacting with PH domain and leucine zipper 1
2
0
−2
log2 (LH / Control)
N
C
S644
S365
0
200
400
600
Position
Candidate
FALSE
TRUE
Domains and Regions:
DOMAIN: PH
DOMAIN: PID
REGION: Disordered

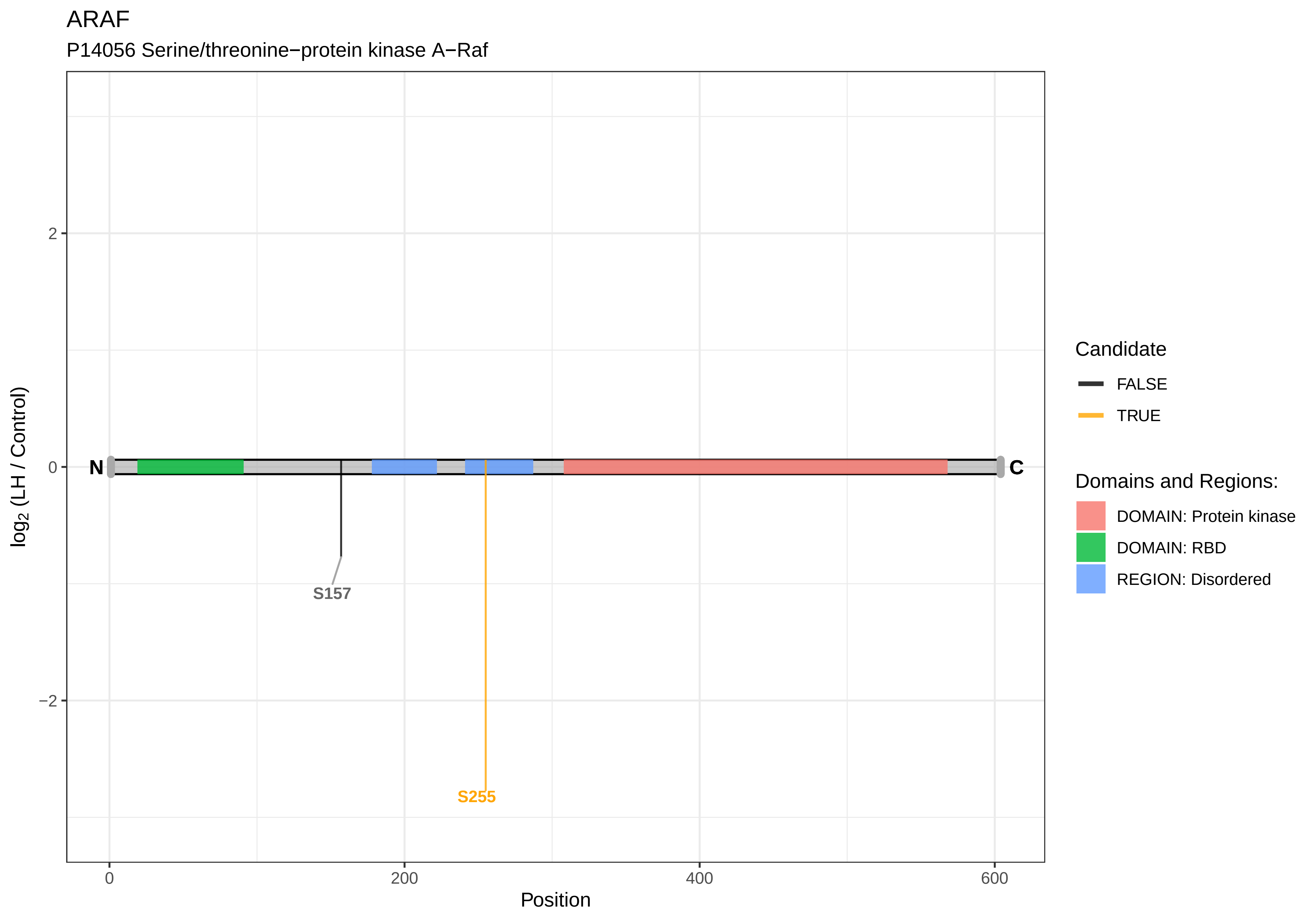
ARAF
P14056 Serine/threonine−protein kinase A−Raf
log2 (LH / Control)
2
0
−2
N
C
S157
S255
0
200
400
600
Position
Candidate
FALSE
TRUE
Domains and Regions:
DOMAIN: Protein kinase
DOMAIN: RBD
REGION: Disordered

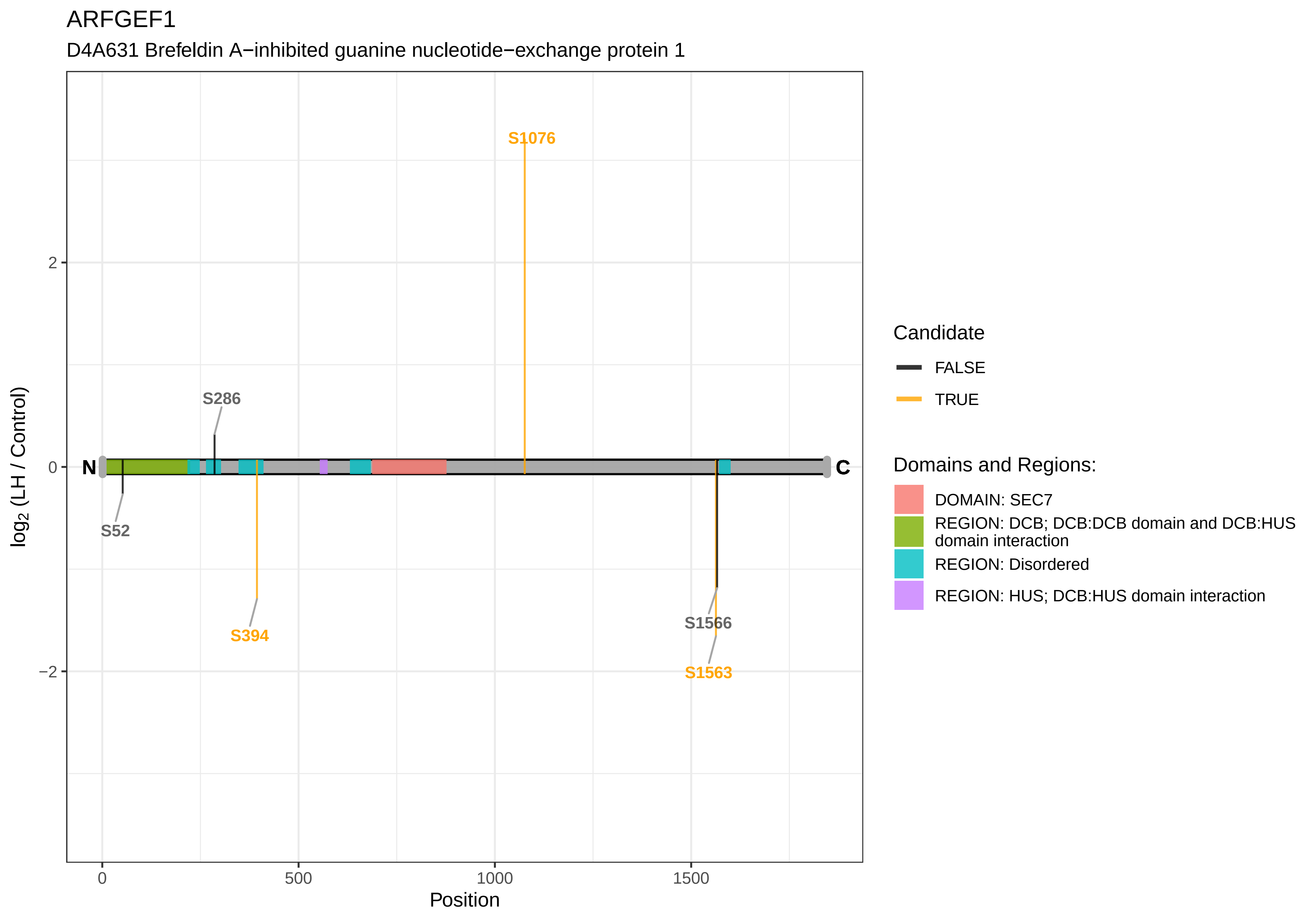

ARFGEF1
D4A631 Brefeldin A−inhibited guanine nucleotide−exchange protein 1
S1076
S286
S52
S394
S1566
S1563
N
C
2
0
−2
0
500
1000
1500
Position
log2 (LH / Control)
Candidate
FALSE
TRUE
Domains and Regions:
DOMAIN: SEC7
REGION: DCB; DCB:DCB domain and DCB:HUS domain interaction
REGION: Disordered
REGION: HUS; DCB:HUS domain interaction

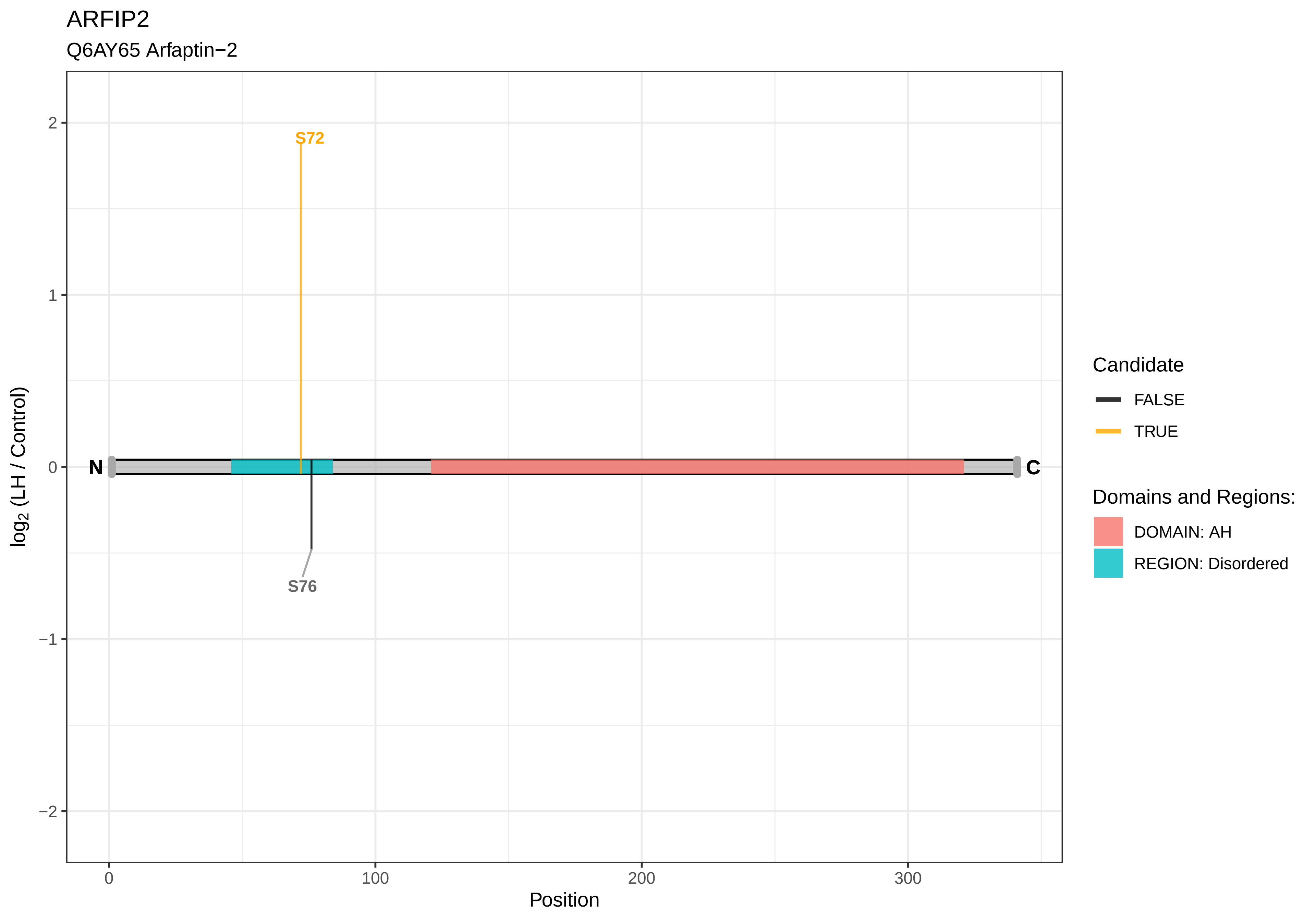
ARFIP2
Q6AY65 Arfaptin−2
S72
S76
N
C
2
1
0
−1
−2
log2 (LH / Control)
0
100
200
300
Position
Candidate
FALSE
TRUE
Domains and Regions:
DOMAIN: AH
REGION: Disordered

### ARHGAP10

D4AB50 RCG51408

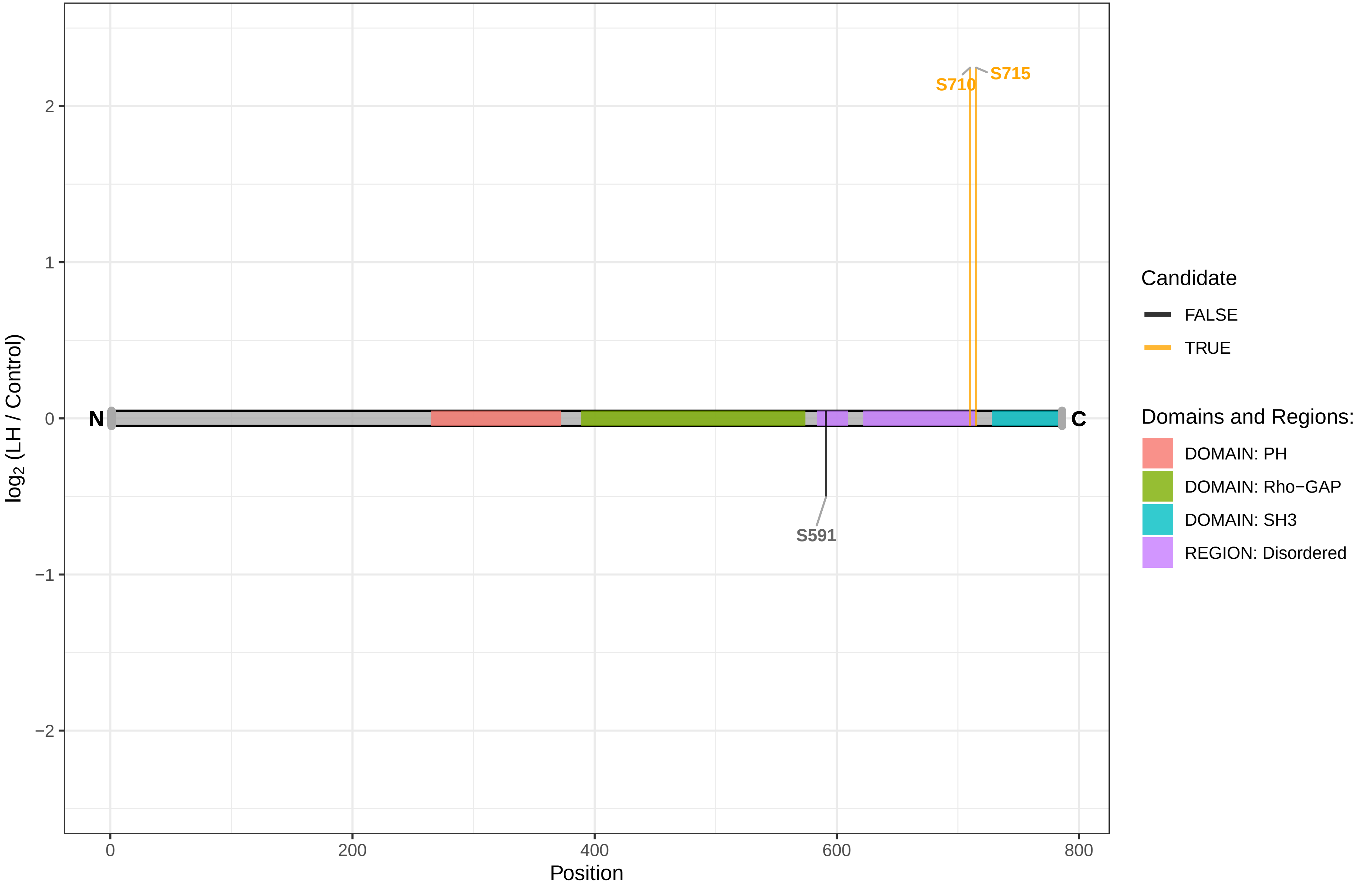

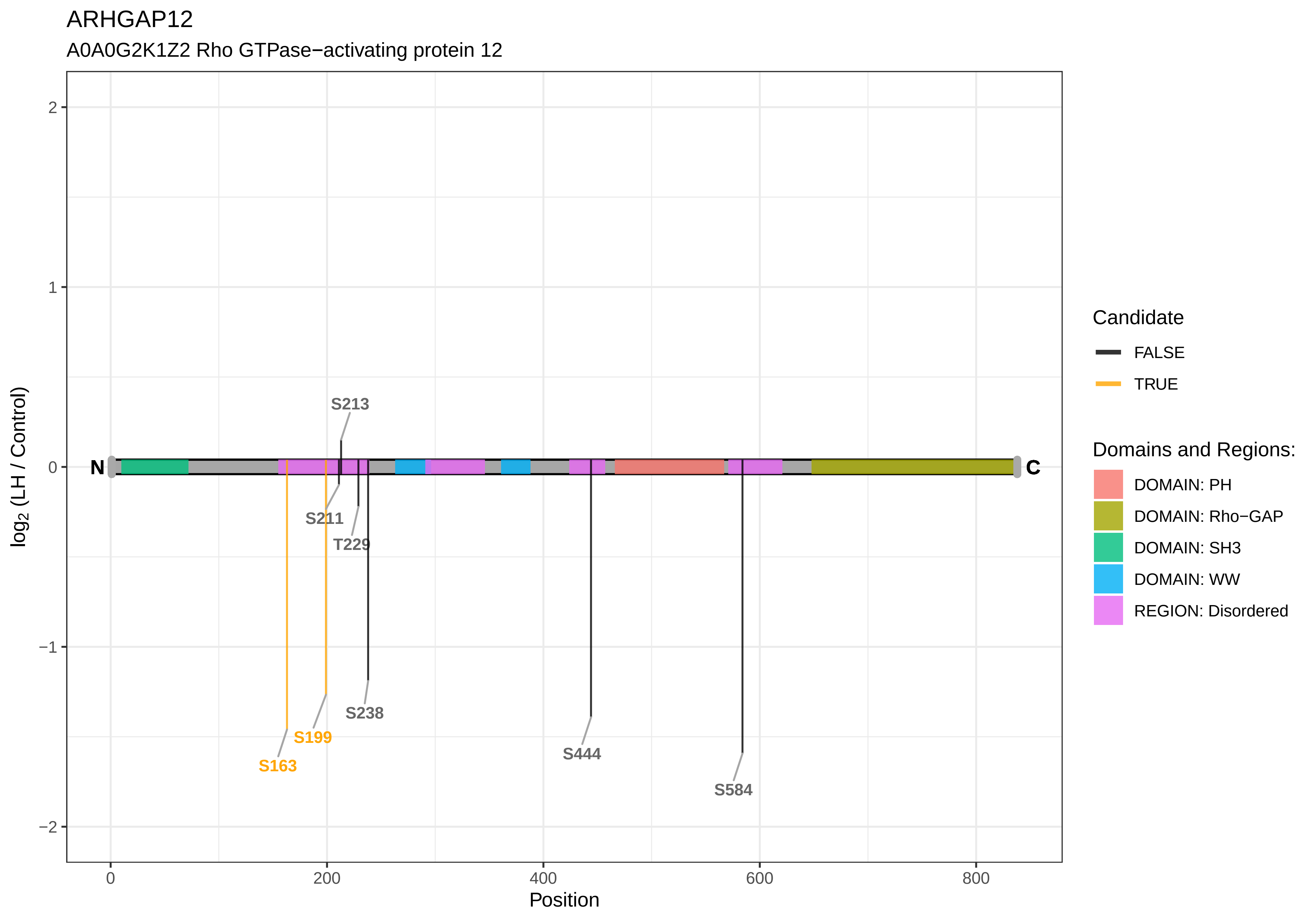
ARHGAP12
A0A0G2K1Z2 Rho GTPase−activating protein 12
log2 (LH / Control)
Position
N
C
S213
S211
T229
S238
S199
S163
S444
S584
2
1
0
−1
−2
0
200
400
600
800
Candidate
FALSE
TRUE
Domains and Regions:
DOMAIN: PH
DOMAIN: Rho−GAP
DOMAIN: SH3
DOMAIN: WW
REGION: Disordered

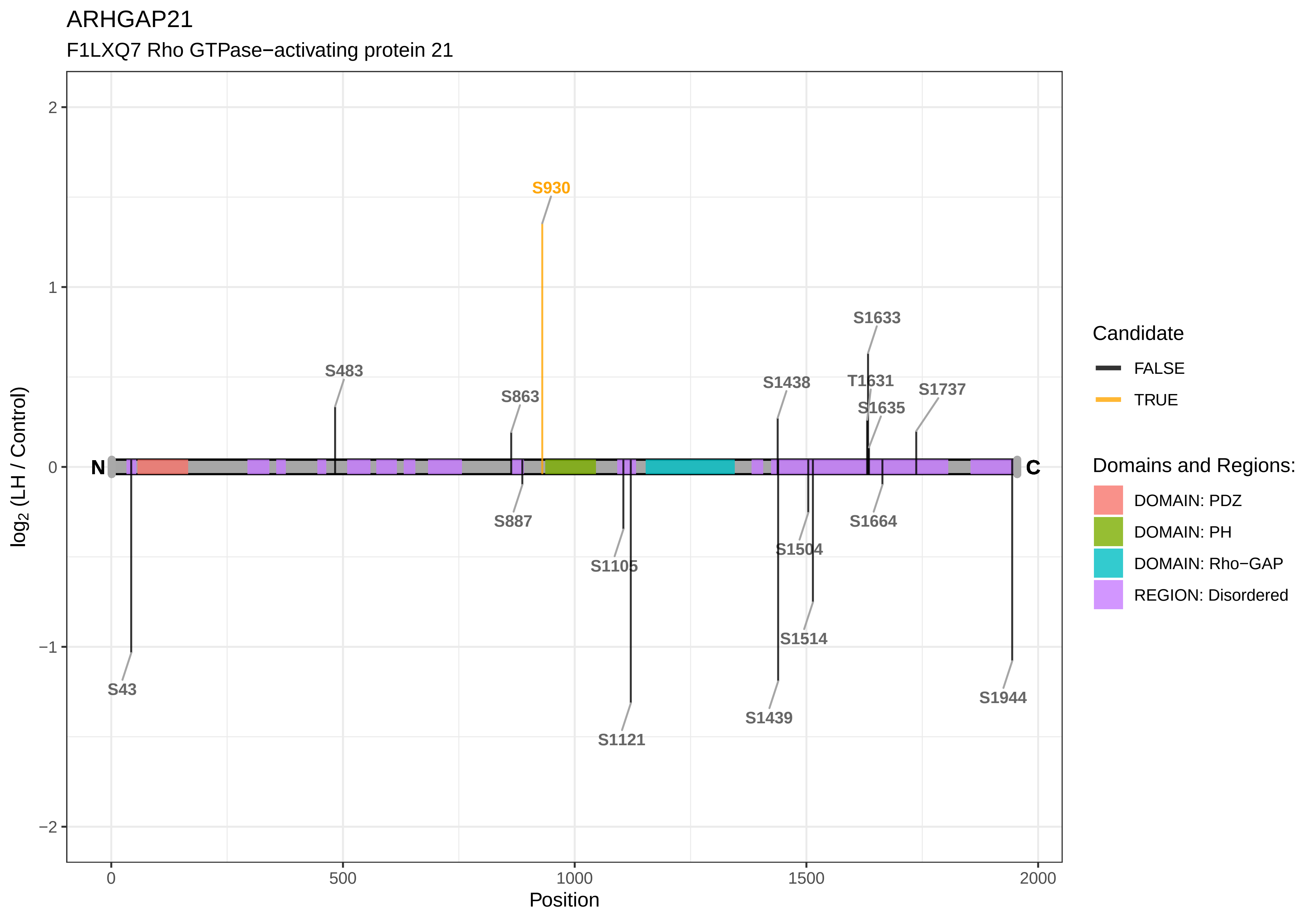

ARHGAP21
F1LXQ7 Rho GTPase−activating protein 21
S930
S1633
S483
S1438
T1631
S1737
S863
S1635
N
C
S887
S1664
S1504
S1105
S1514
S43
S1944
S1439
S1121
log2 (LH / Control)
Position
Candidate
FALSE
TRUE
Domains and Regions:
DOMAIN: PDZ
DOMAIN: PH
DOMAIN: Rho−GAP
REGION: Disordered

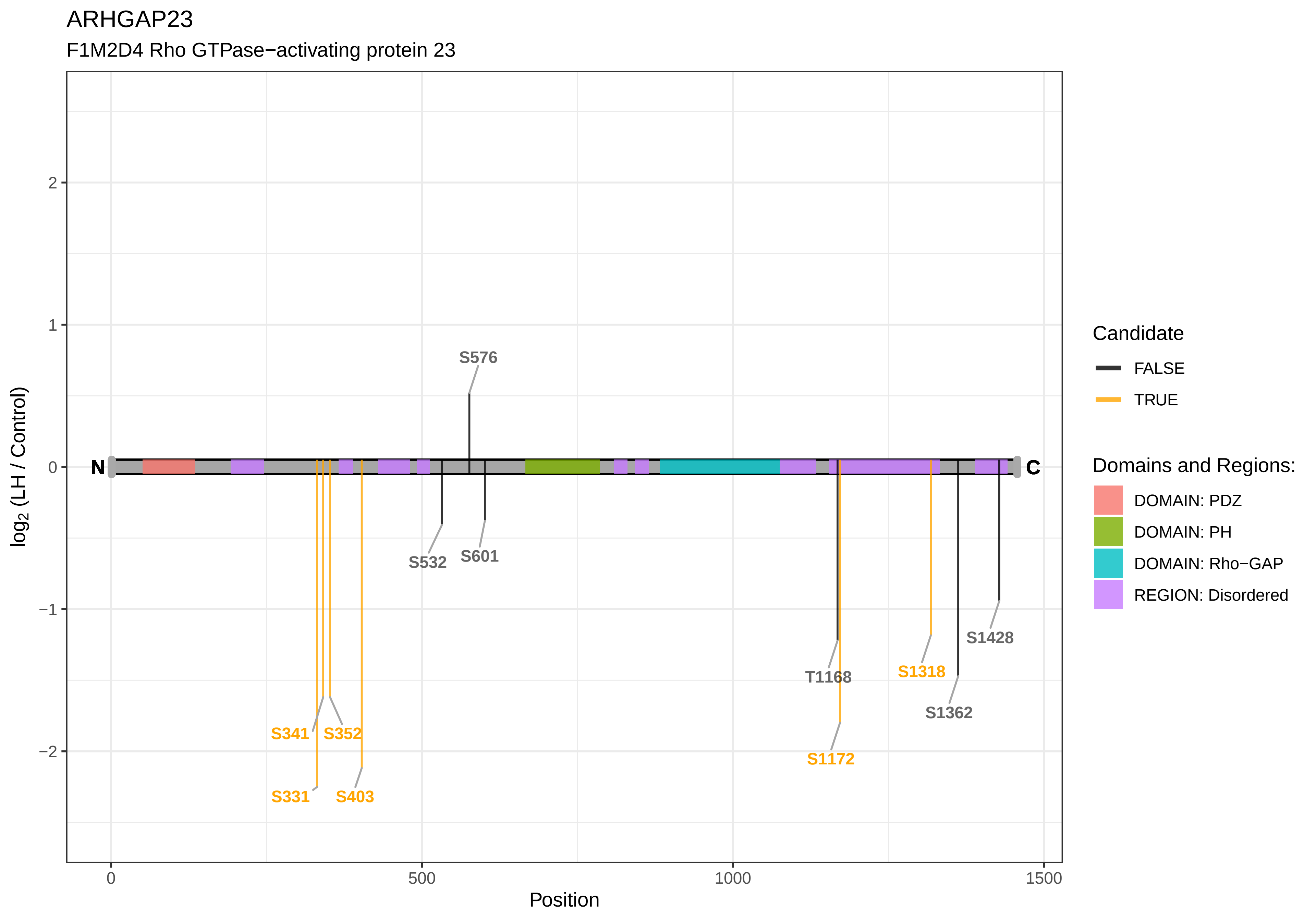
ARHGAP23
F1M2D4 Rho GTPase−activating protein 23
log2 (LH / Control)
Position
N
C
S576
S532
S601
S341
S352
S331
S403
T1168
S1172
S1318
S1362
S1428
Candidate
FALSE
TRUE
Domains and Regions:
DOMAIN: PDZ
DOMAIN: PH
DOMAIN: Rho−GAP
REGION: Disordered

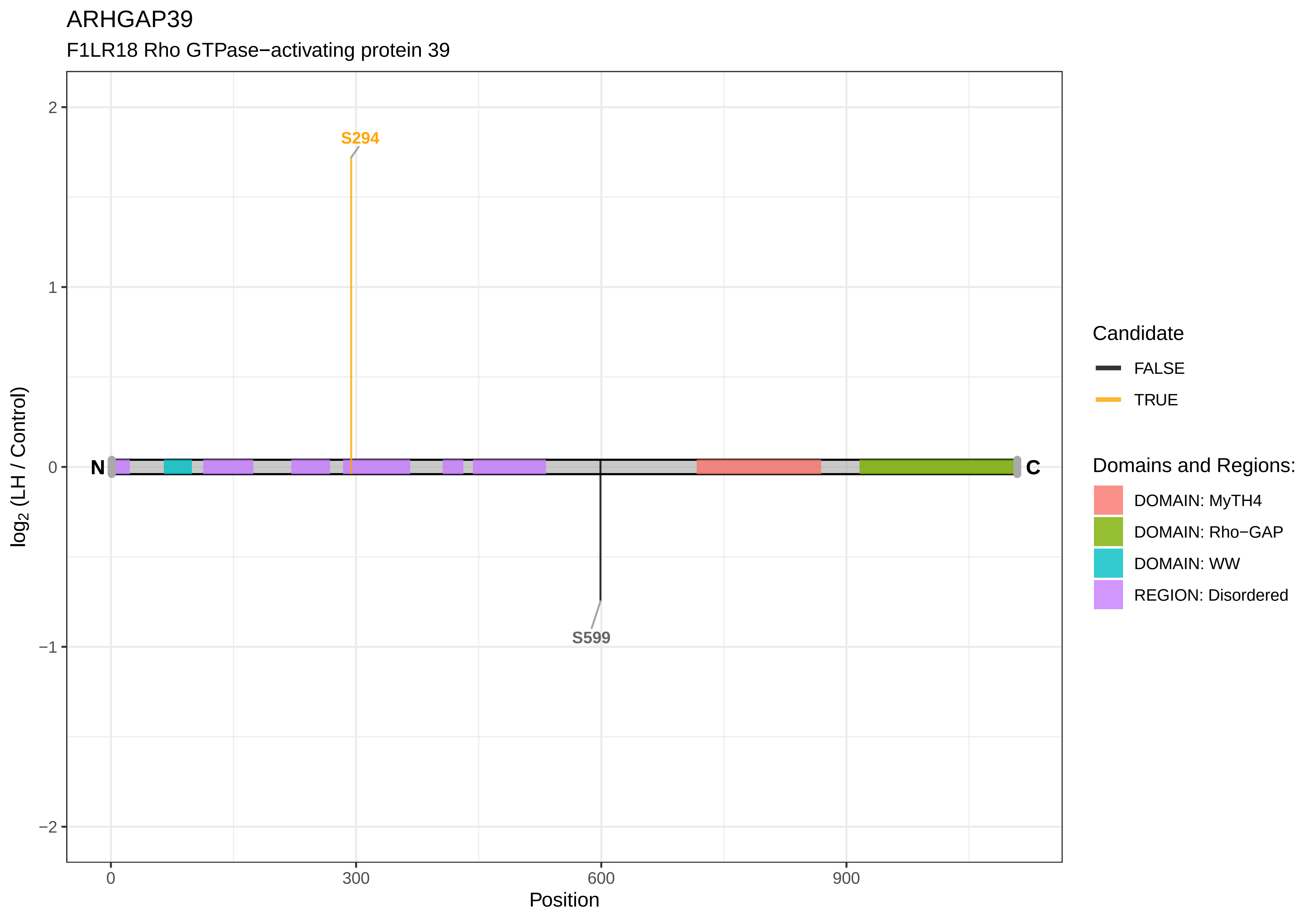

ARHGAP39
F1LR18 Rho GTPase−activating protein 39
S294
S599
N
C
2
1
0
−1
−2
log2 (LH / Control)
0
300
600
900
Position
Candidate
FALSE
TRUE
Domains and Regions:
DOMAIN: MyTH4
DOMAIN: Rho−GAP
DOMAIN: WW
REGION: Disordered

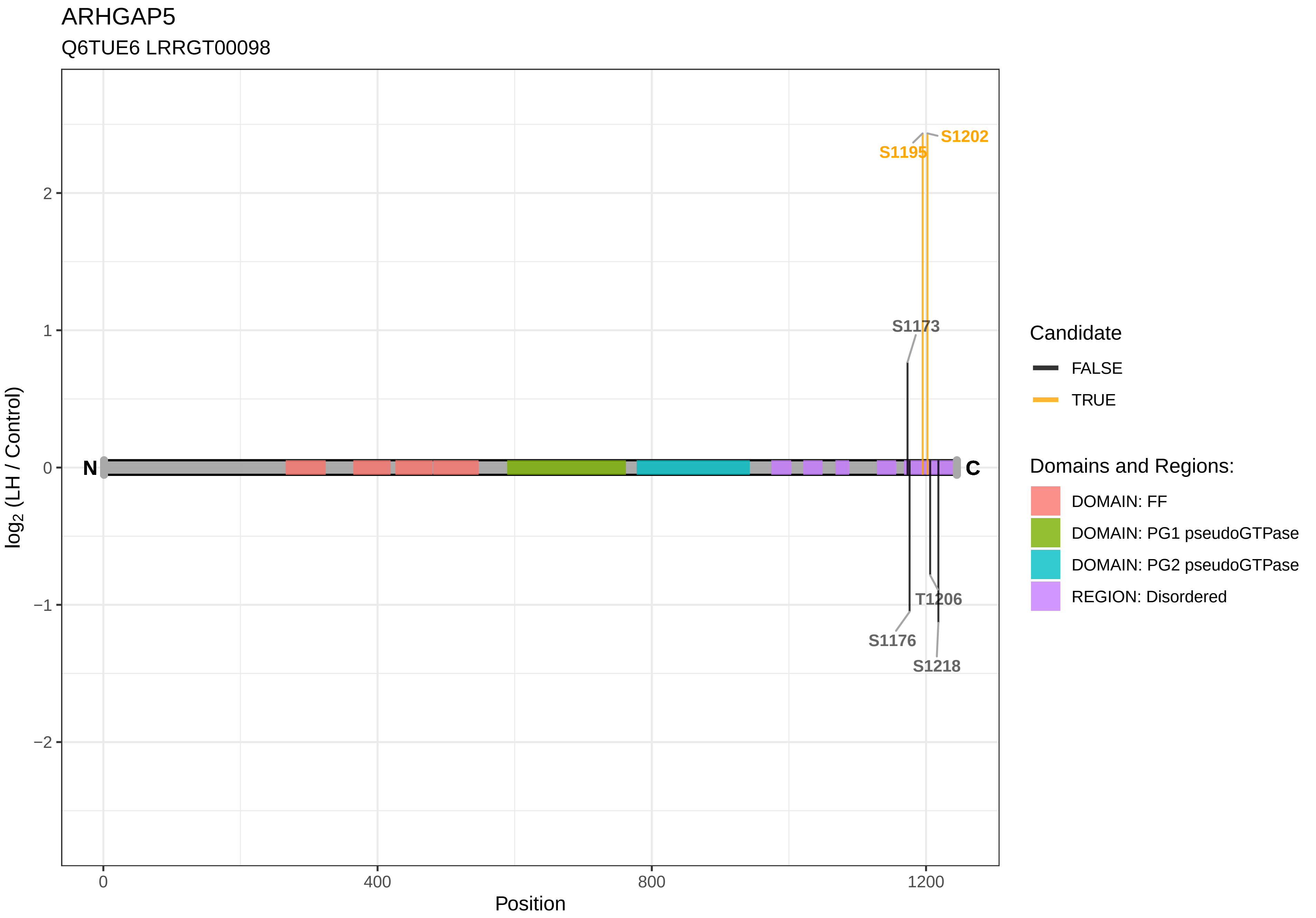
ARHGAP5
Q6TUE6 LRRGT00098
S1202
S1195
S1173
T1206
S1176
S1218
N
C
log2 (LH / Control)
2
1
0
−1
−2
0
400
800
1200
Position
Candidate
FALSE
TRUE
Domains and Regions:
DOMAIN: FF
DOMAIN: PG1 pseudoGTPase
DOMAIN: PG2 pseudoGTPase
REGION: Disordered

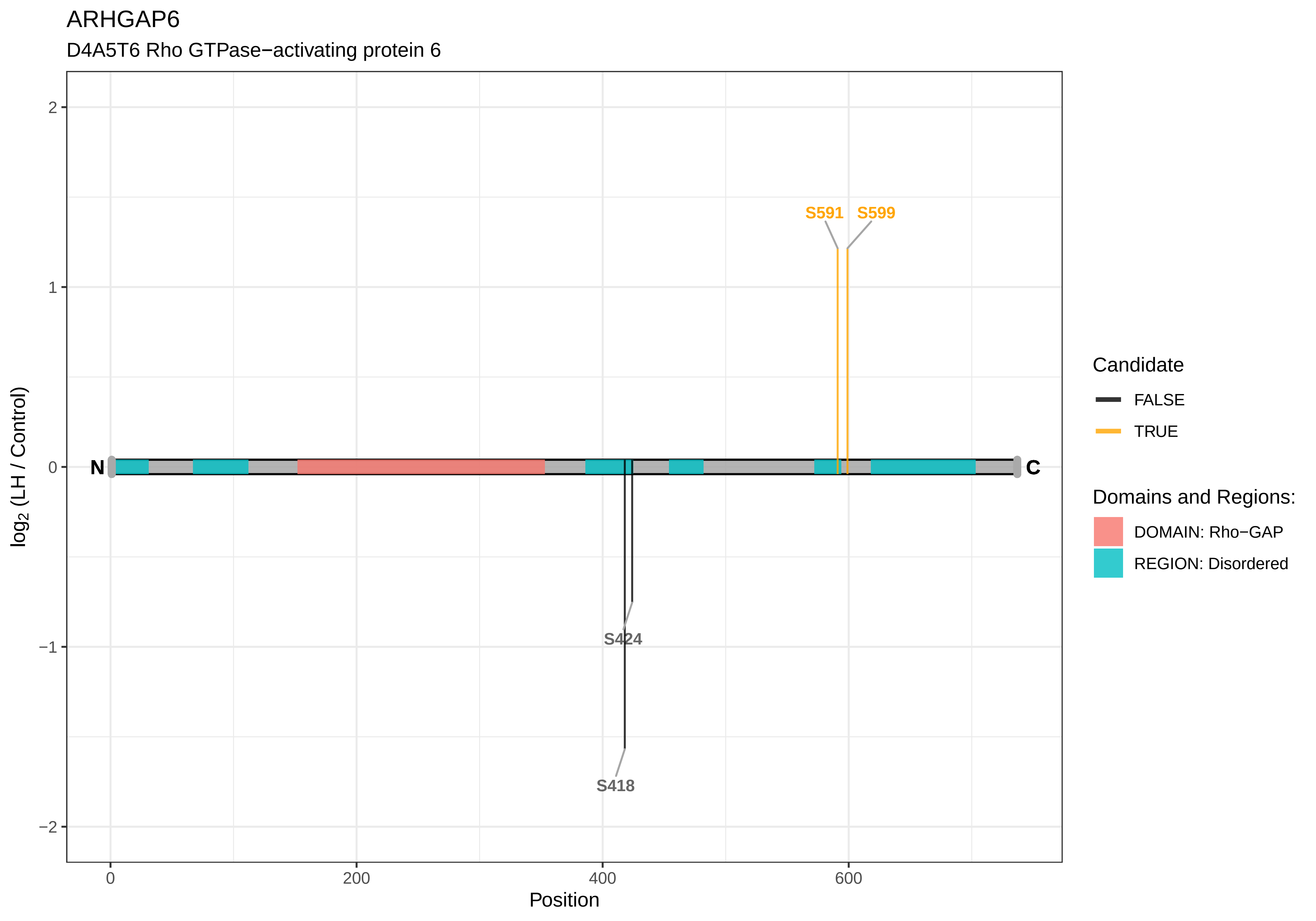

ARHGAP6
D4A5T6 Rho GTPase−activating protein 6
S591
S599
S424
S418
N
C
2
1
0
−1
−2
log2 (LH / Control)
0
200
400
600
Position
Candidate
FALSE
TRUE
Domains and Regions:
DOMAIN: Rho−GAP
REGION: Disordered

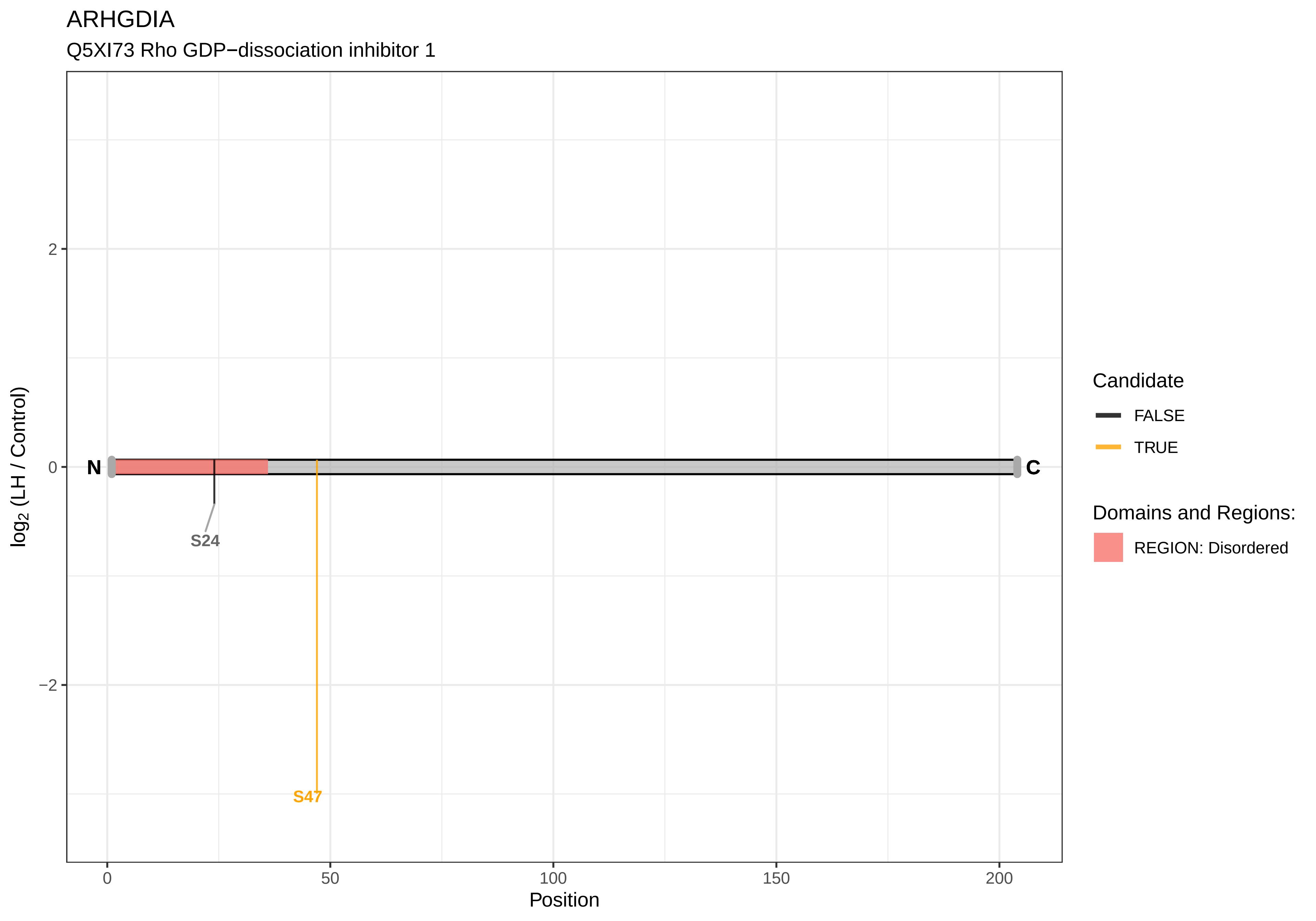

ARHGDIA
Q5XI73 Rho GDP−dissociation inhibitor 1
log2 (LH / Control)
Position
2
0
−2
0
50
100
150
200
N
C
S24
S47
Candidate
FALSE
TRUE
Domains and Regions:
REGION: Disordered

### ARHGEF1

A0A0G2K4L2 Rho guanine nucleotide exchange factor 1

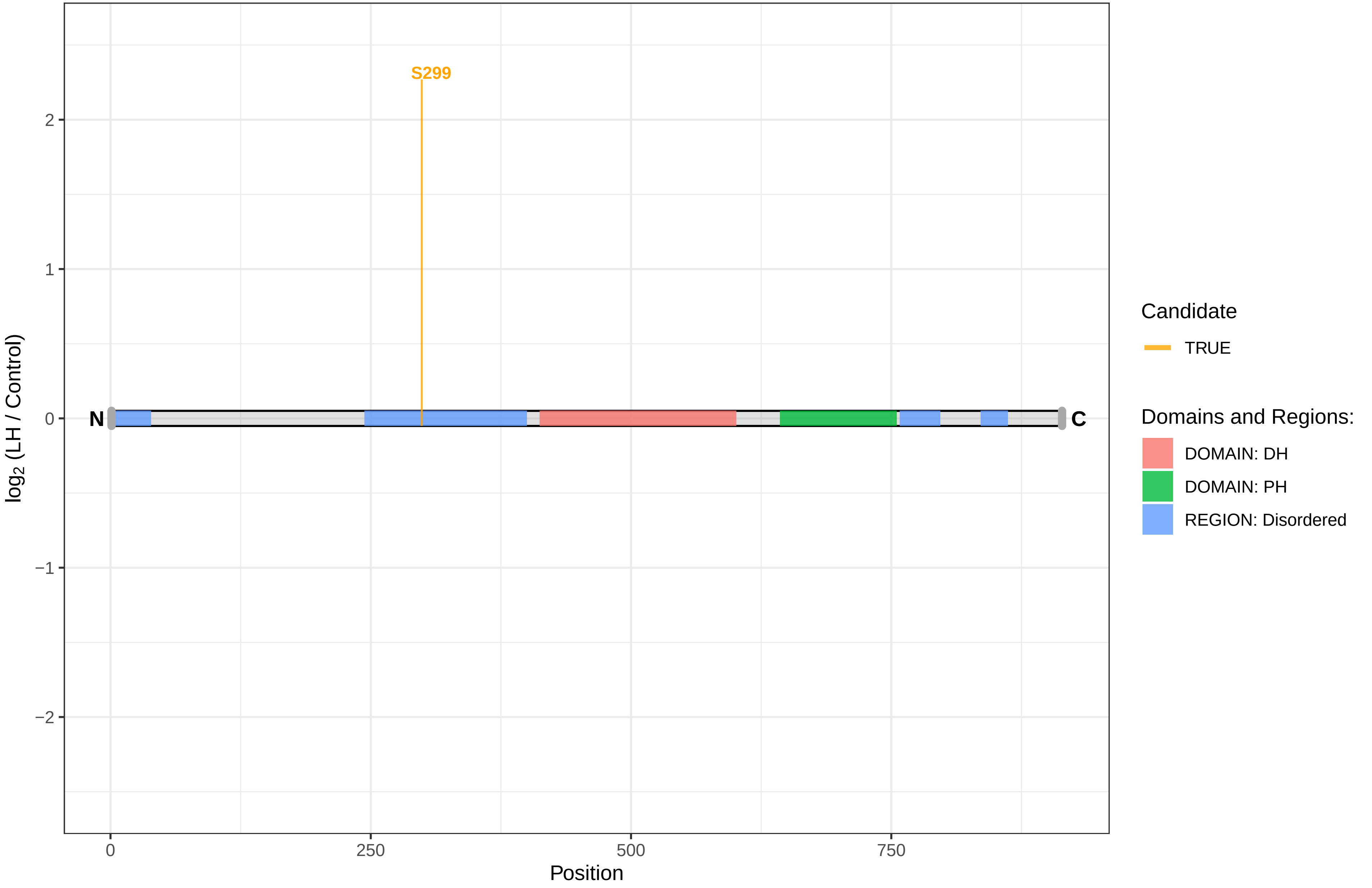

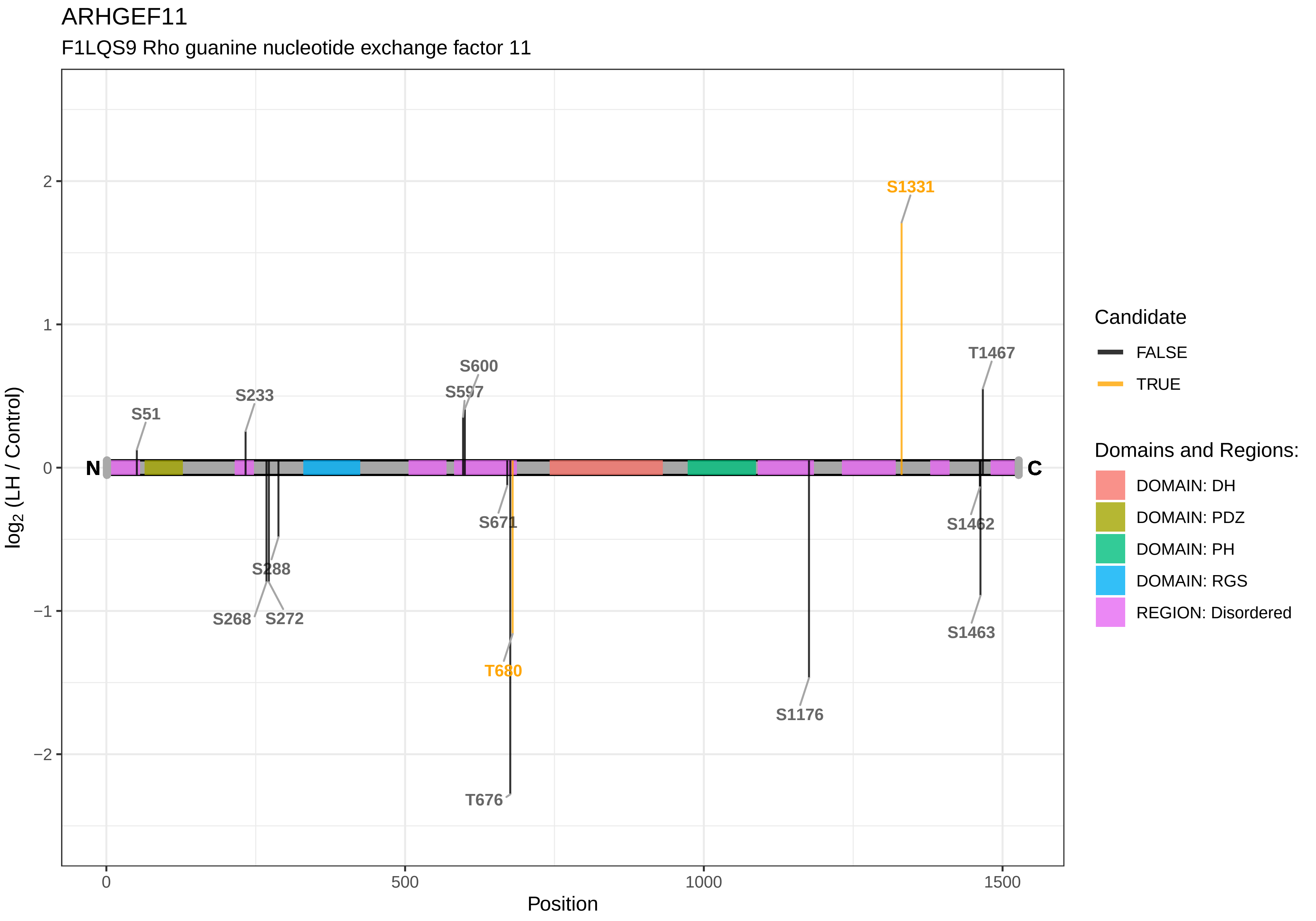
ARHGEF11
F1LQS9 Rho guanine nucleotide exchange factor 11
S1331
T1467
S600
S597
S233
S51
N
C
S671
S1462
S288
S268
S272
T680
S1463
S1176
T676
2
1
0
−1
−2
0
500
1000
1500
Position
log2 (LH / Control)
Candidate
FALSE
TRUE
Domains and Regions:
DOMAIN: DH
DOMAIN: PDZ
DOMAIN: PH
DOMAIN: RGS
REGION: Disordered

ARHGEF17

A0A0G2JXT9 Rho guanine nucleotide exchange factor (GEF) 17

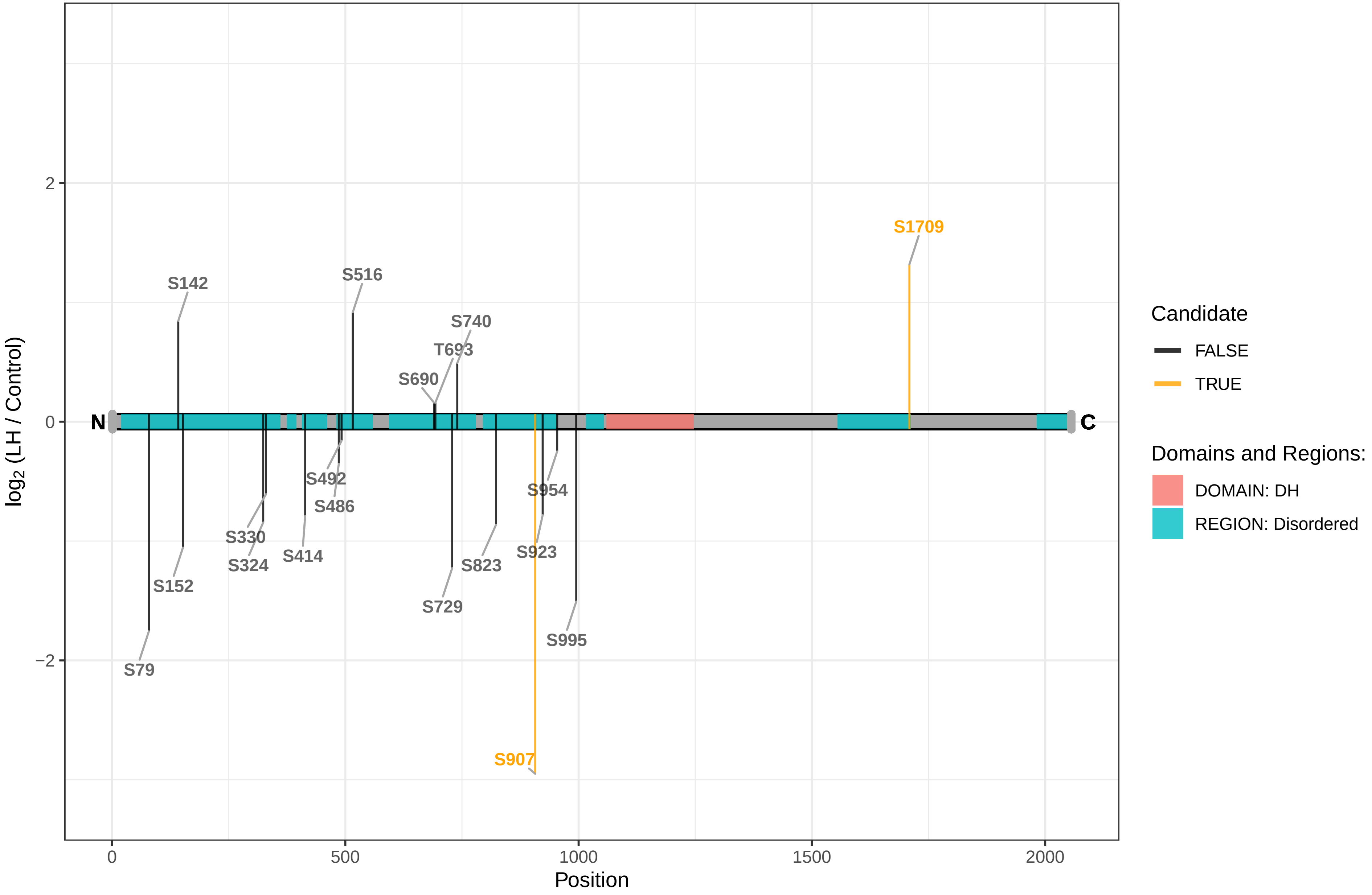

ARHGEF18

F1LT94 Rho/Rac guanine nucleotide exchange factor 18

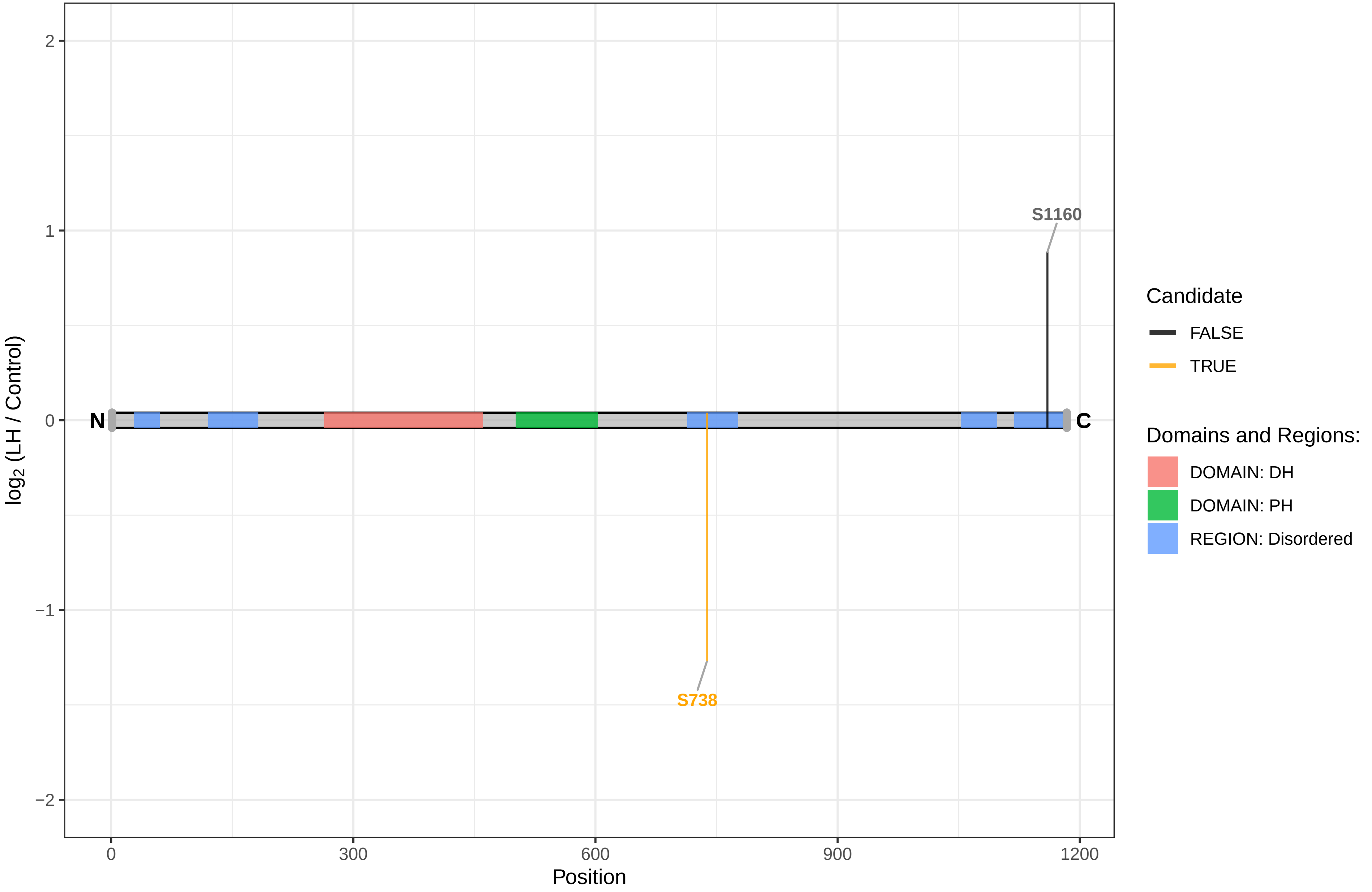

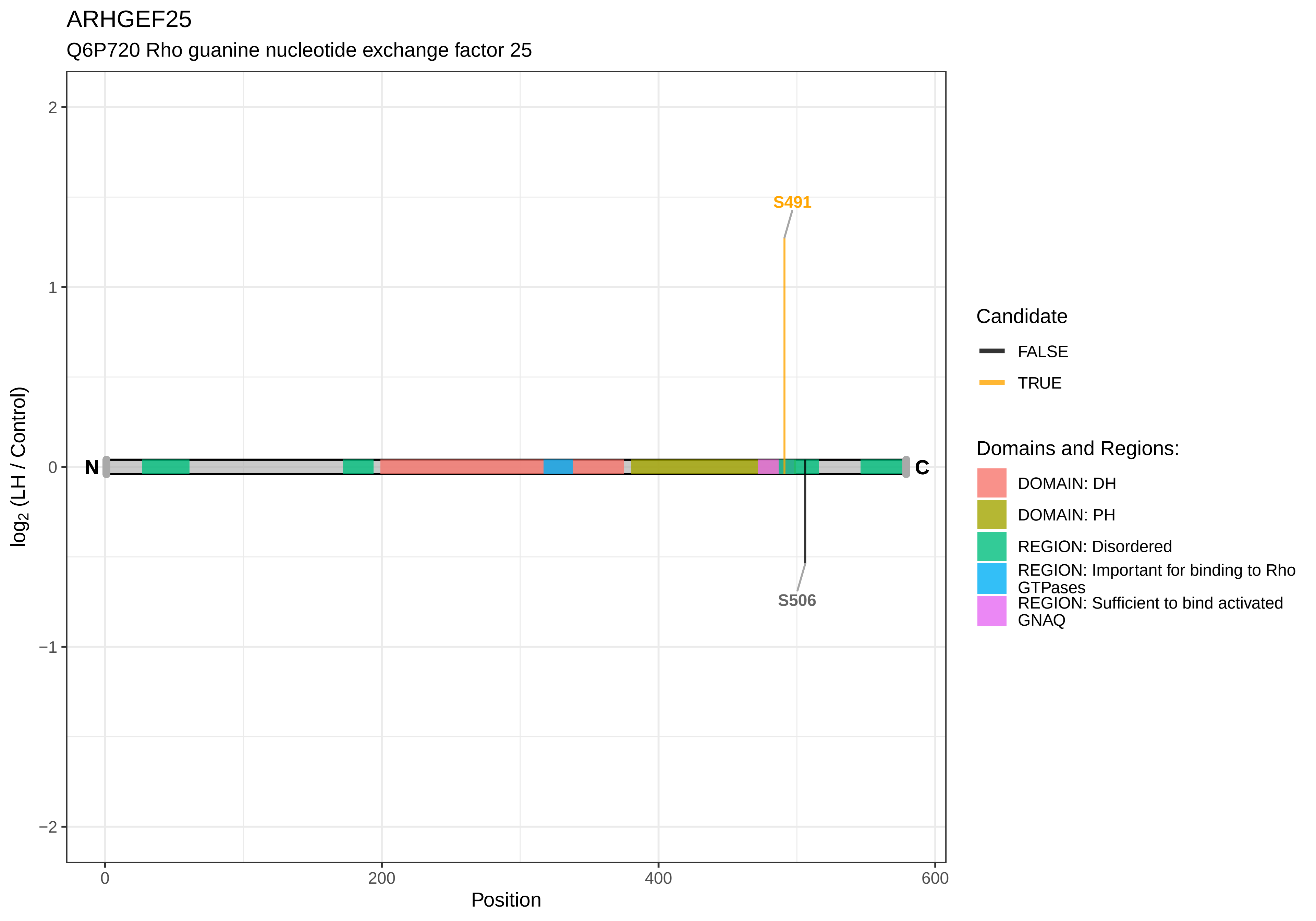
ARHGEF25
Q6P720 Rho guanine nucleotide exchange factor 25
S491
S506
N
C
2
1
0
−1
−2
log2 (LH / Control)
0
200
400
600
Position
Candidate
FALSE
TRUE
Domains and Regions:
DOMAIN: DH
DOMAIN: PH
REGION: Disordered
REGION: Important for binding to Rho GTPases
REGION: Sufficient to bind activated GNAQ

ARHGEF40

A0A0G2JZE7 Rho guanine nucleotide exchange factor 40

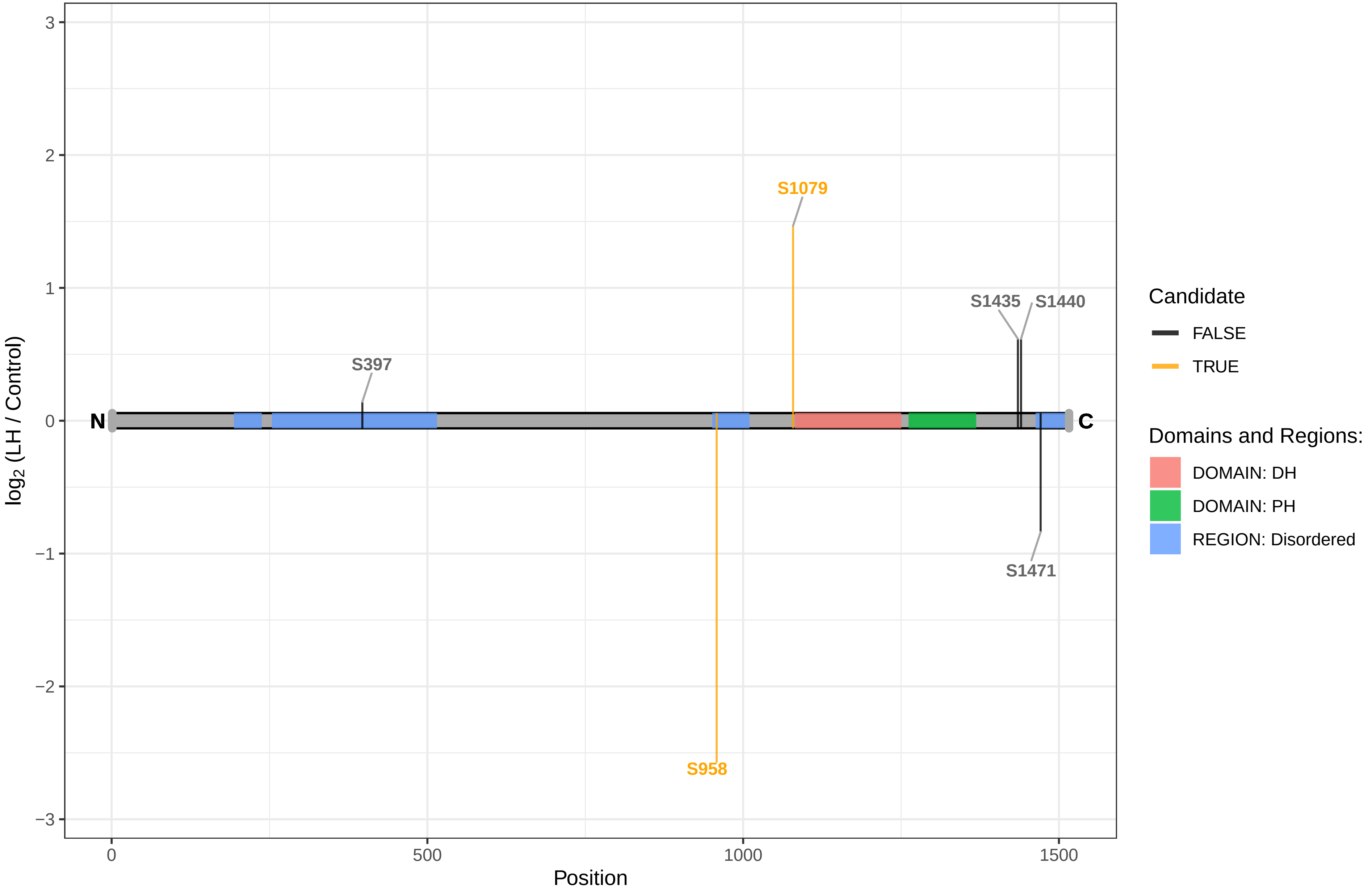

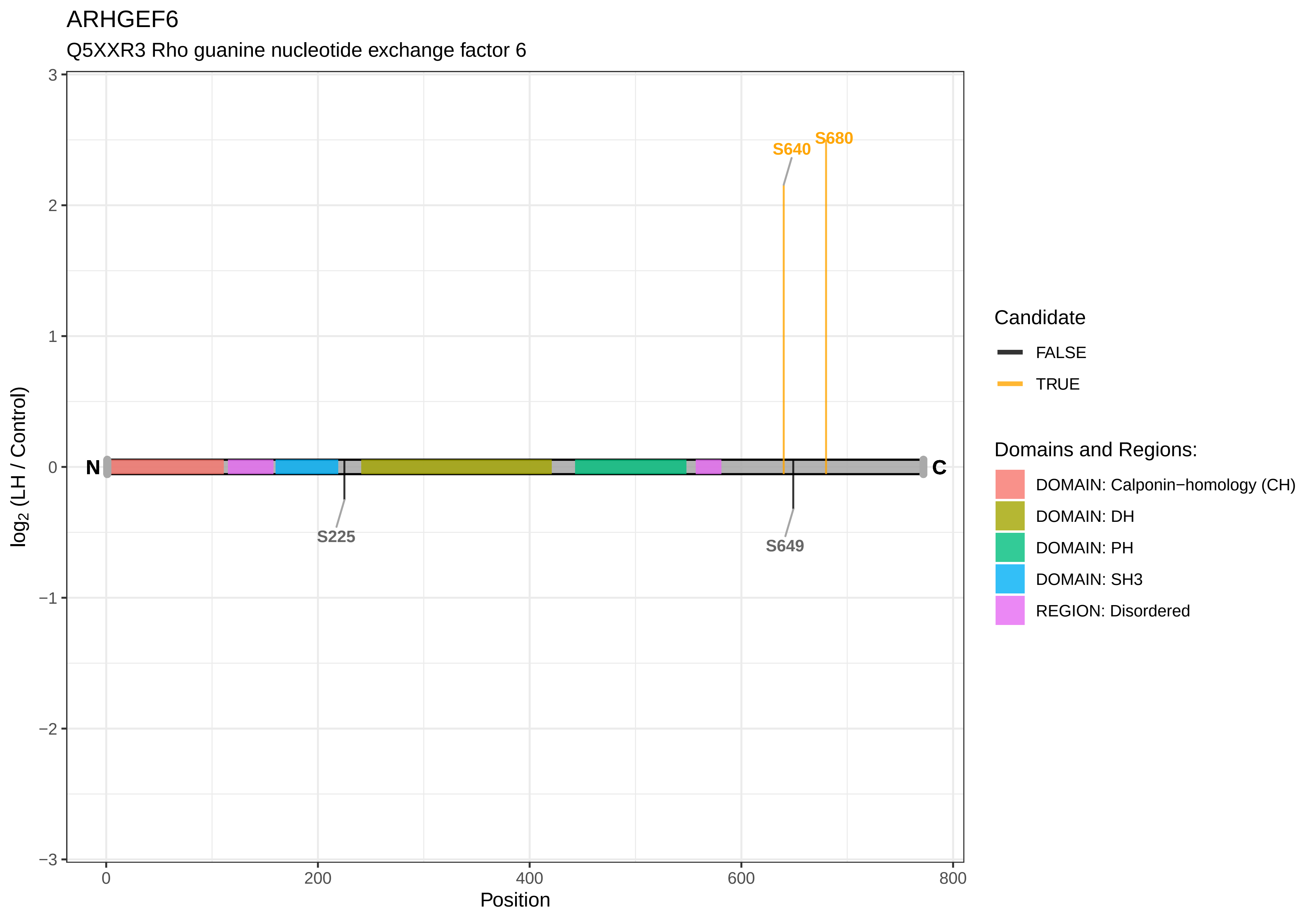

ARHGEF6
Q5XXR3 Rho guanine nucleotide exchange factor 6
S640
S680
S225
S649
N
C
log2 (LH / Control)
Position
3
2
1
0
−1
−2
−3
0
200
400
600
800
Candidate
FALSE
TRUE
Domains and Regions:
DOMAIN: Calponin−homology (CH)
DOMAIN: DH
DOMAIN: PH
DOMAIN: SH3
REGION: Disordered

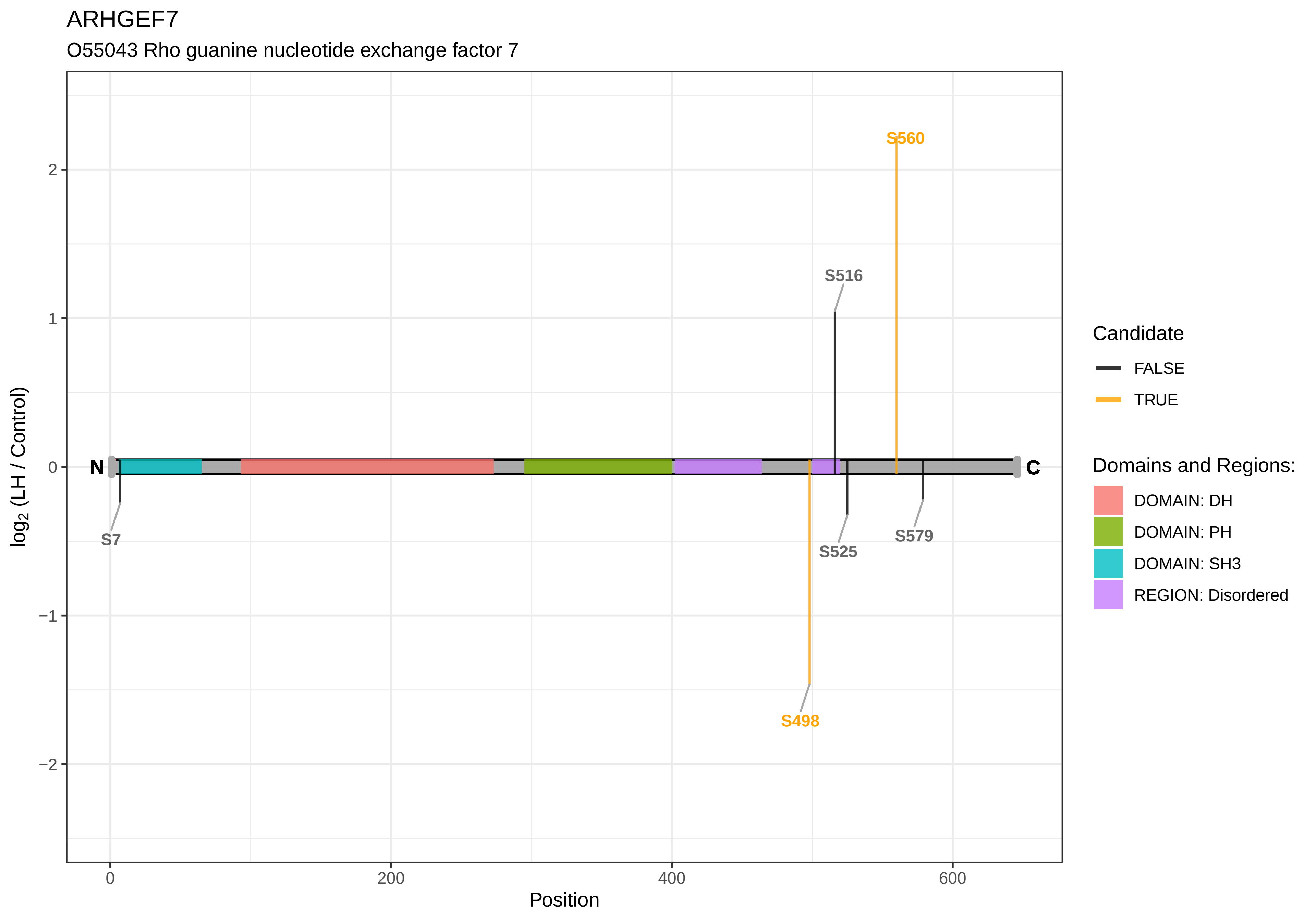
ARHGEF7
O55043 Rho guanine nucleotide exchange factor 7
log2 (LH / Control)
2
1
0
−1
−2
N
C
S560
S516
S7
S525
S579
S498
0
200
400
600
Position
Candidate
FALSE
TRUE
Domains and Regions:
DOMAIN: DH
DOMAIN: PH
DOMAIN: SH3
REGION: Disordered

ARMC10

B1WBW4 Armadillo repeat−containing protein 10

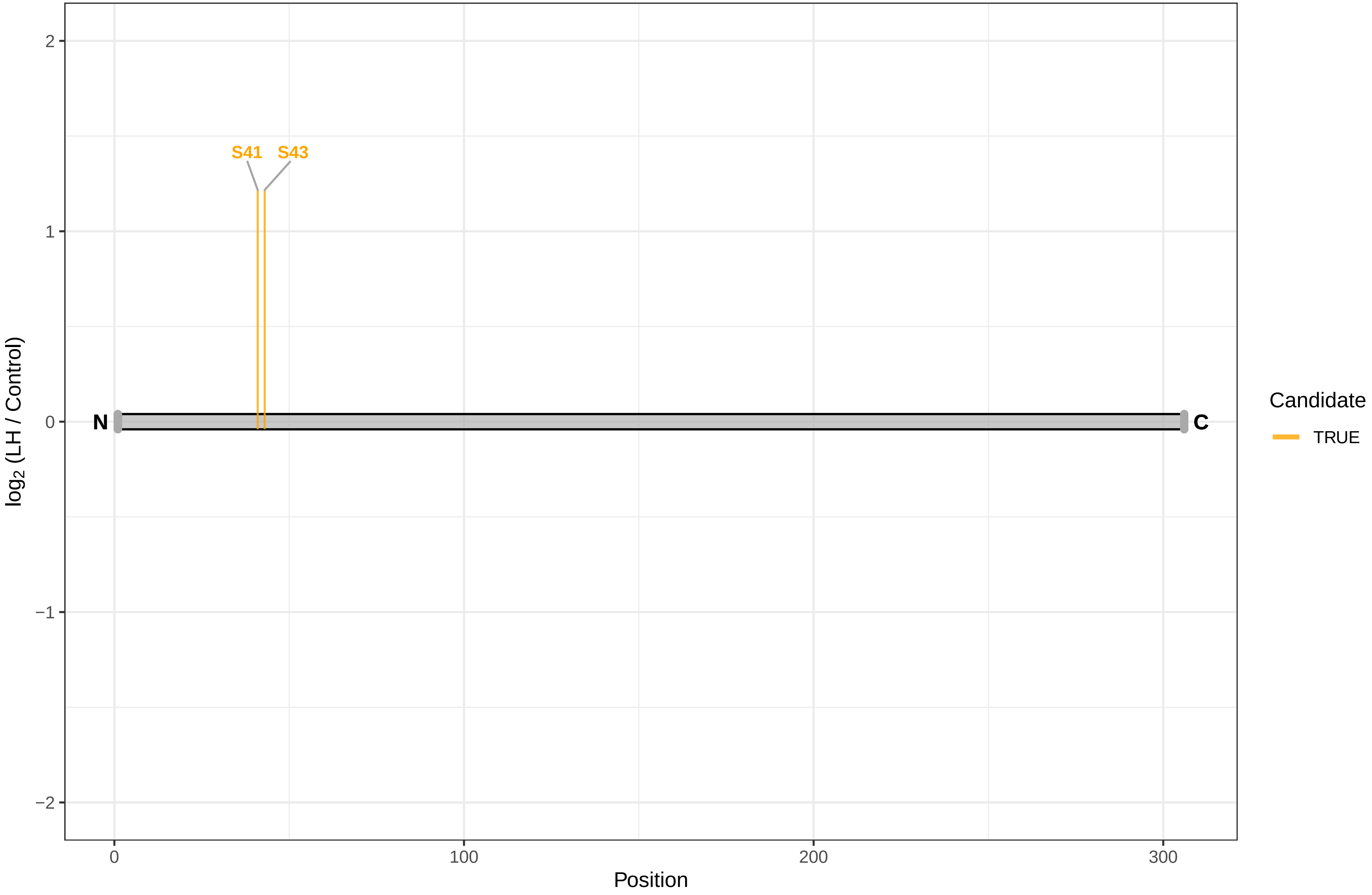

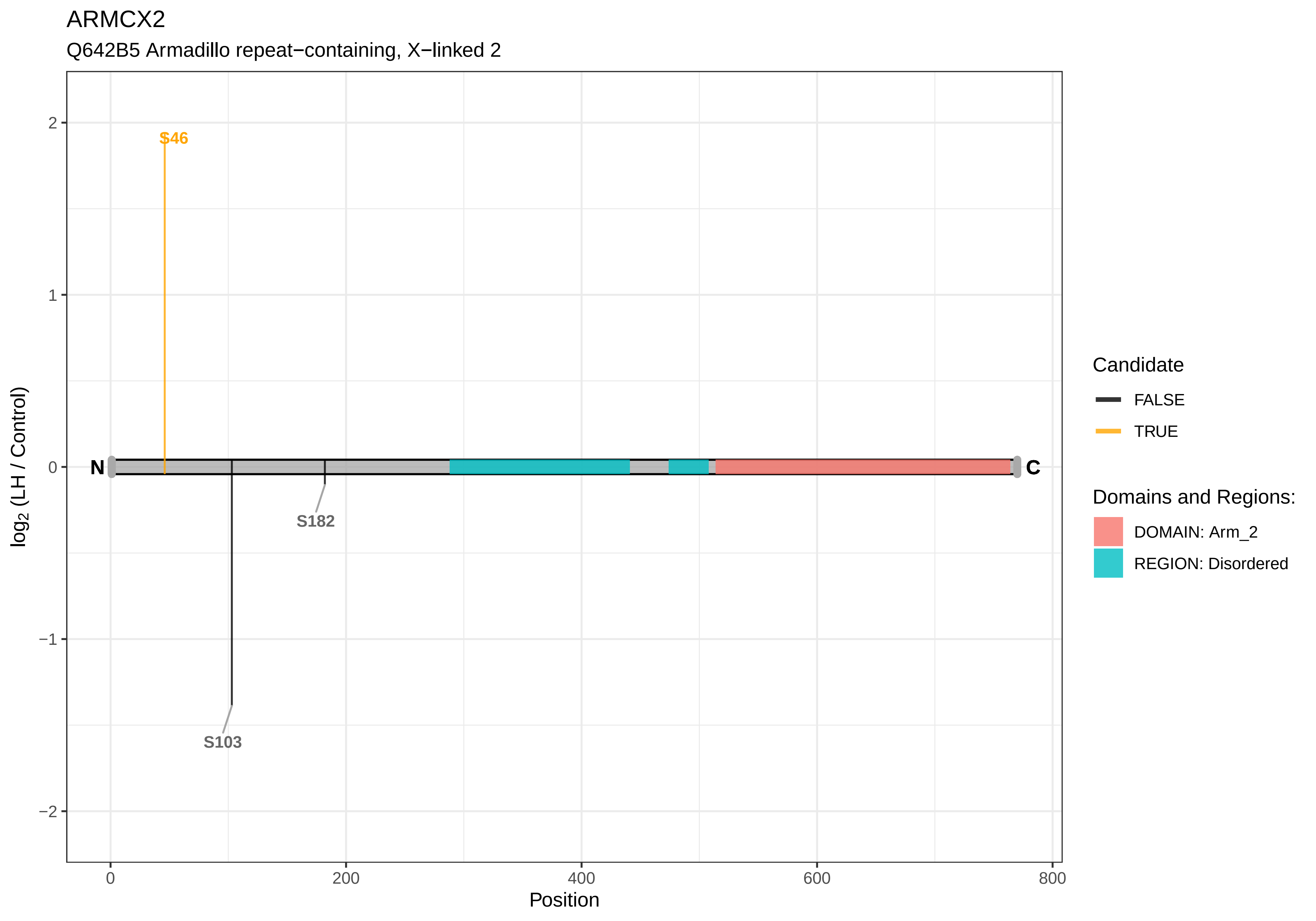

ARMCX2
Q642B5 Armadillo repeat−containing, X−linked 2
S46
S182
S103
N
C
2
1
0
−1
−2
0
200
400
600
800
Position
log2 (LH / Control)
Candidate
FALSE
TRUE
Domains and Regions:
DOMAIN: Arm_2
REGION: Disordered

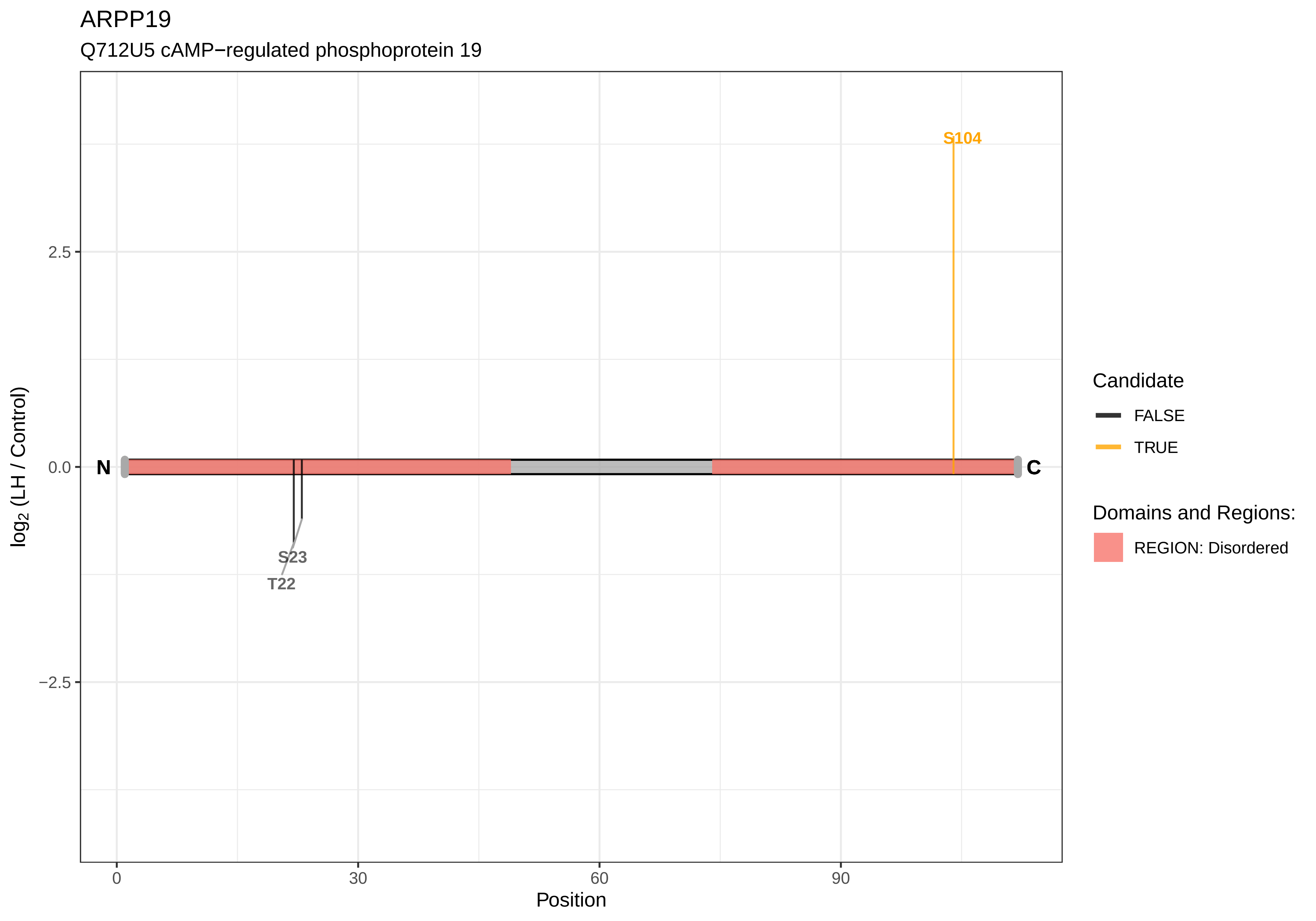
ARPP19
Q712U5 cAMP−regulated phosphoprotein 19
S104
2.5
0.0
−2.5
N
C
S23
T22
log2 (LH / Control)
0
30
60
90
Position
Candidate
FALSE
TRUE
Domains and Regions:
REGION: Disordered

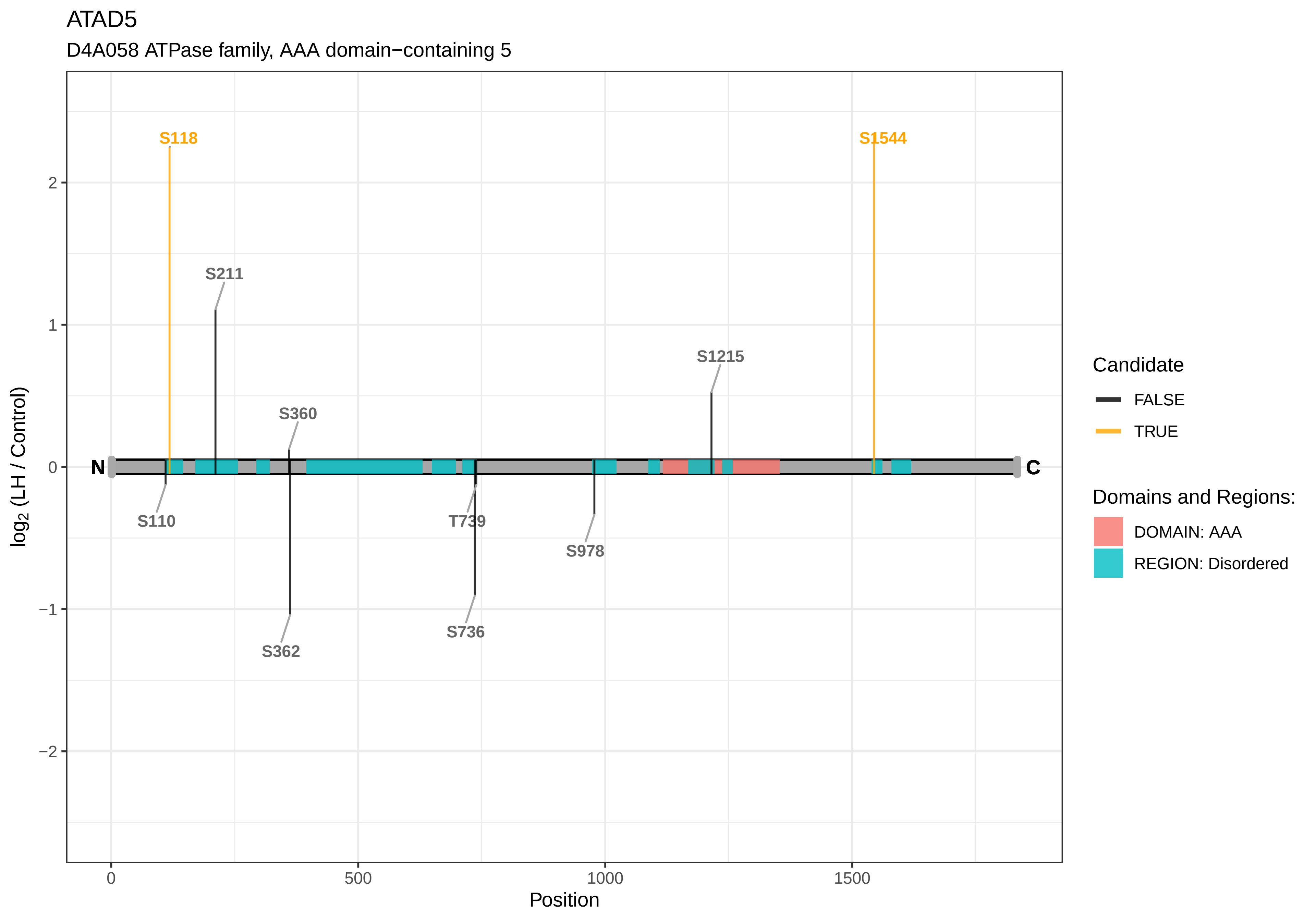
ATAD5
D4A058 ATPase family, AAA domain−containing 5
S118
S1544
S211
S1215
S360
N
C
S110
T739
S978
S362
S736
log2 (LH / Control)
Position
2
1
0
−1
−2
0
500
1000
1500
Candidate
FALSE
TRUE
Domains and Regions:
DOMAIN: AAA
REGION: Disordered

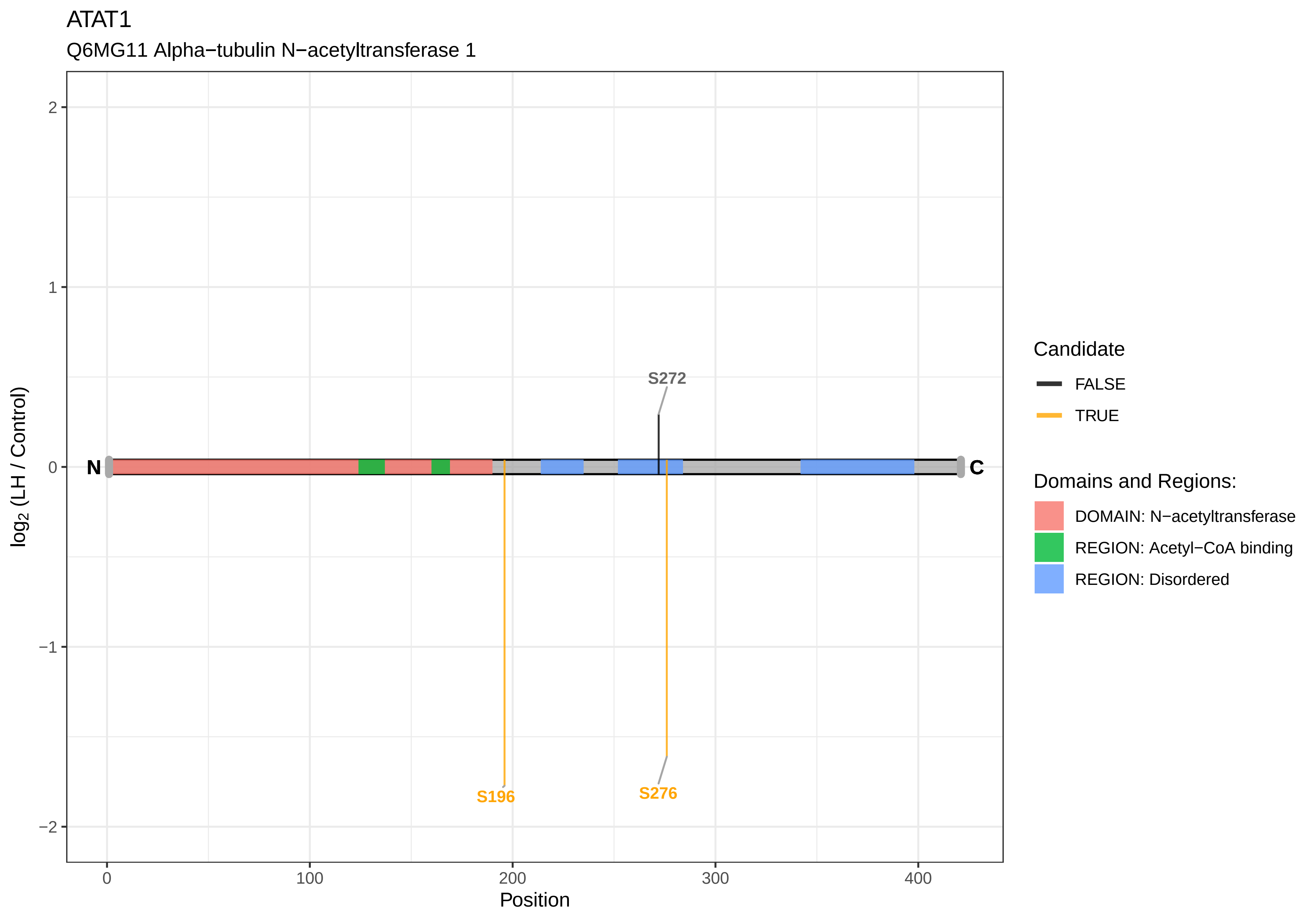

ATAT1
Q6MG11 Alpha−tubulin N−acetyltransferase 1
log2 (LH / Control)
Position
N
C
S272
S196
S276
2
1
0
−1
−2
0
100
200
300
400
Candidate
FALSE
TRUE
Domains and Regions:
DOMAIN: N−acetyltransferase
REGION: Acetyl−CoA binding
REGION: Disordered

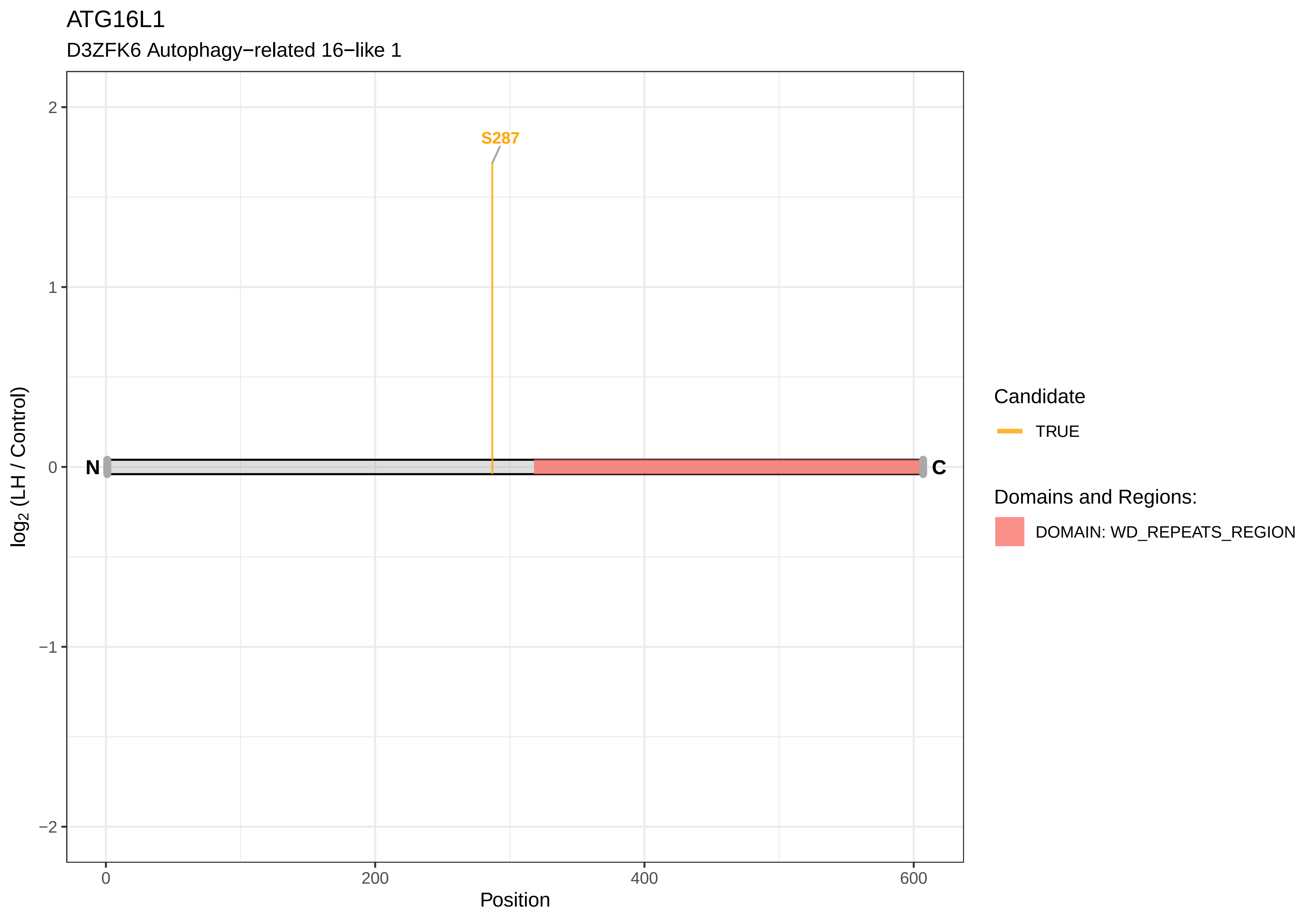

ATG16L1
D3ZFK6 Autophagy−related 16−like 1
S287
N
C
2
1
0
−1
−2
log2 (LH / Control)
0
200
400
600
Position
Candidate
TRUE
Domains and Regions:
DOMAIN: WD_REPEATS_REGION

ATG2B

F1MAF8 Autophagy related 2B

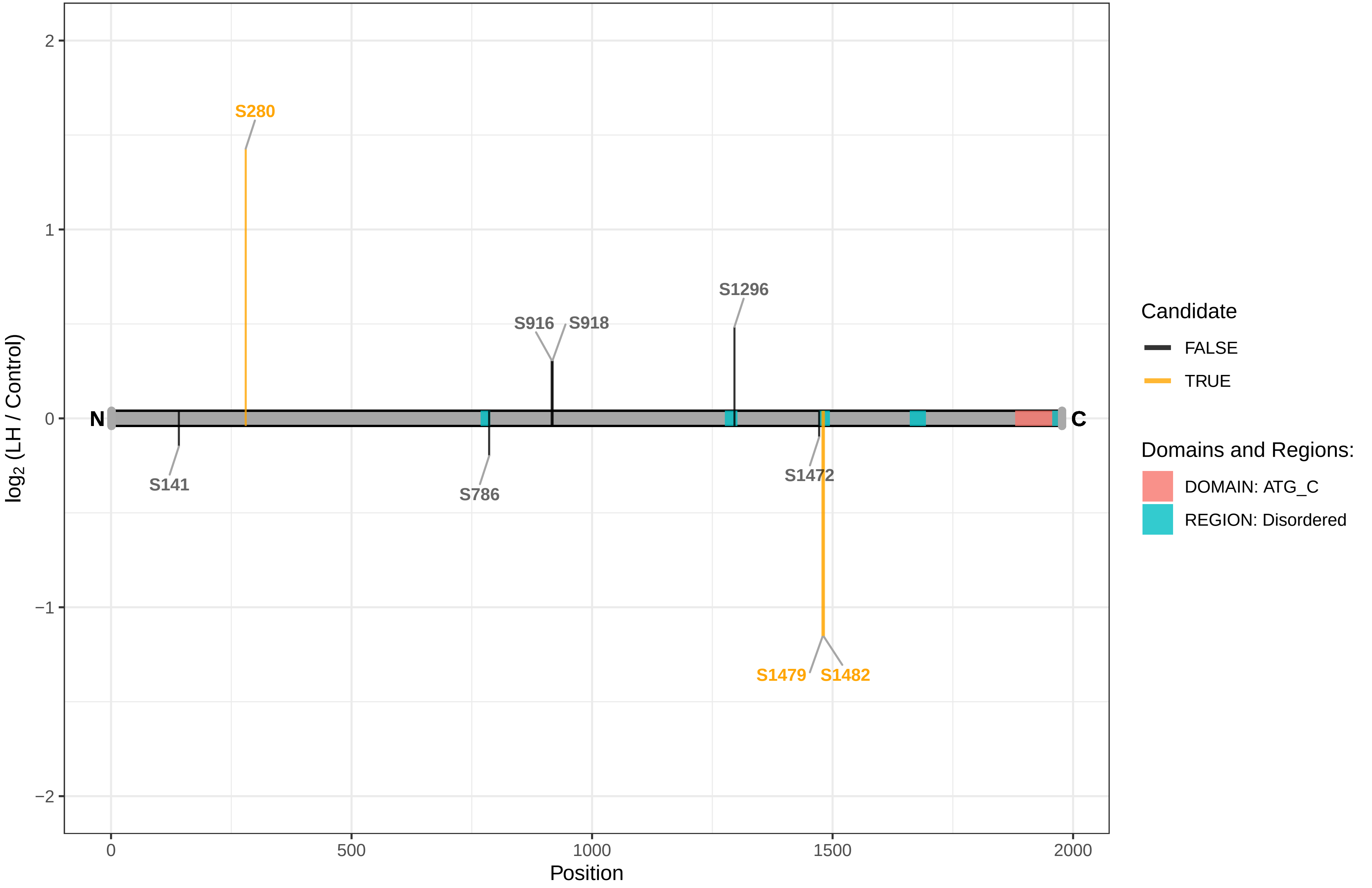

### ATG9A

Q5FWU3 Autophagy−related protein 9A

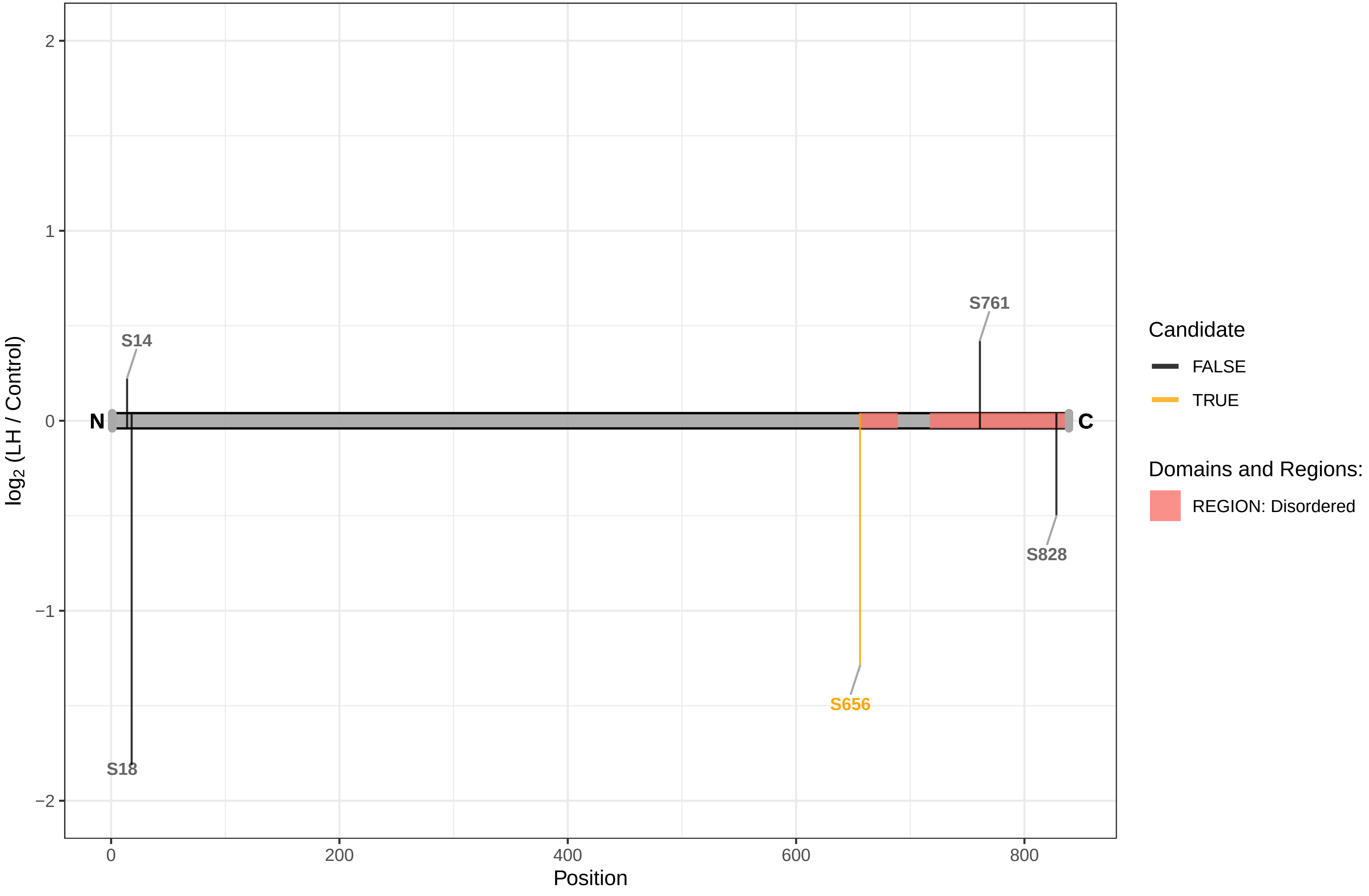

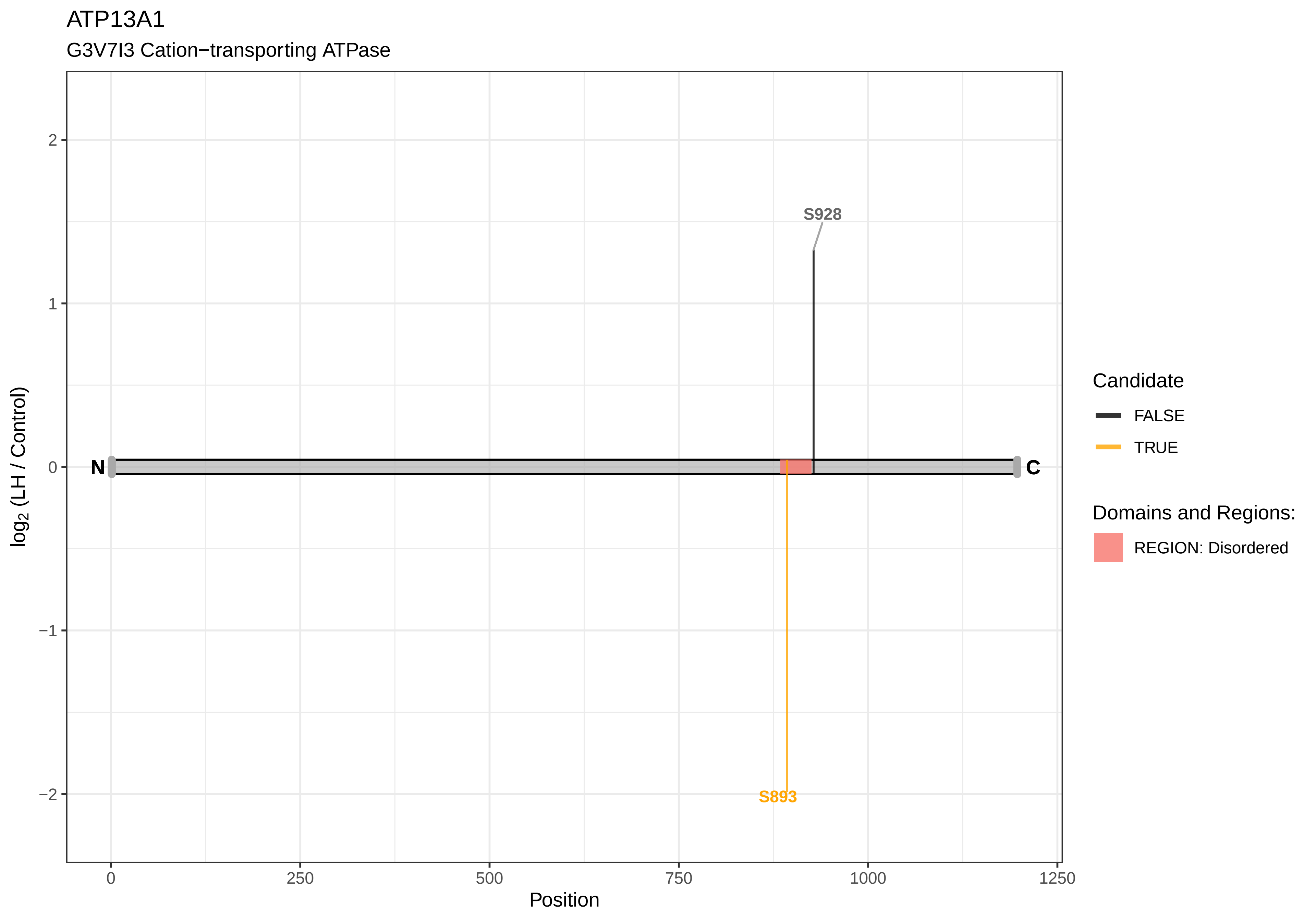
ATP13A1
G3V7I3 Cation−transporting ATPase
S928
S893
N
C
2
1
0
−1
−2
log2 (LH / Control)
0
250
500
750
1000
1250
Position
Candidate
FALSE
TRUE
Domains and Regions:
REGION: Disordered

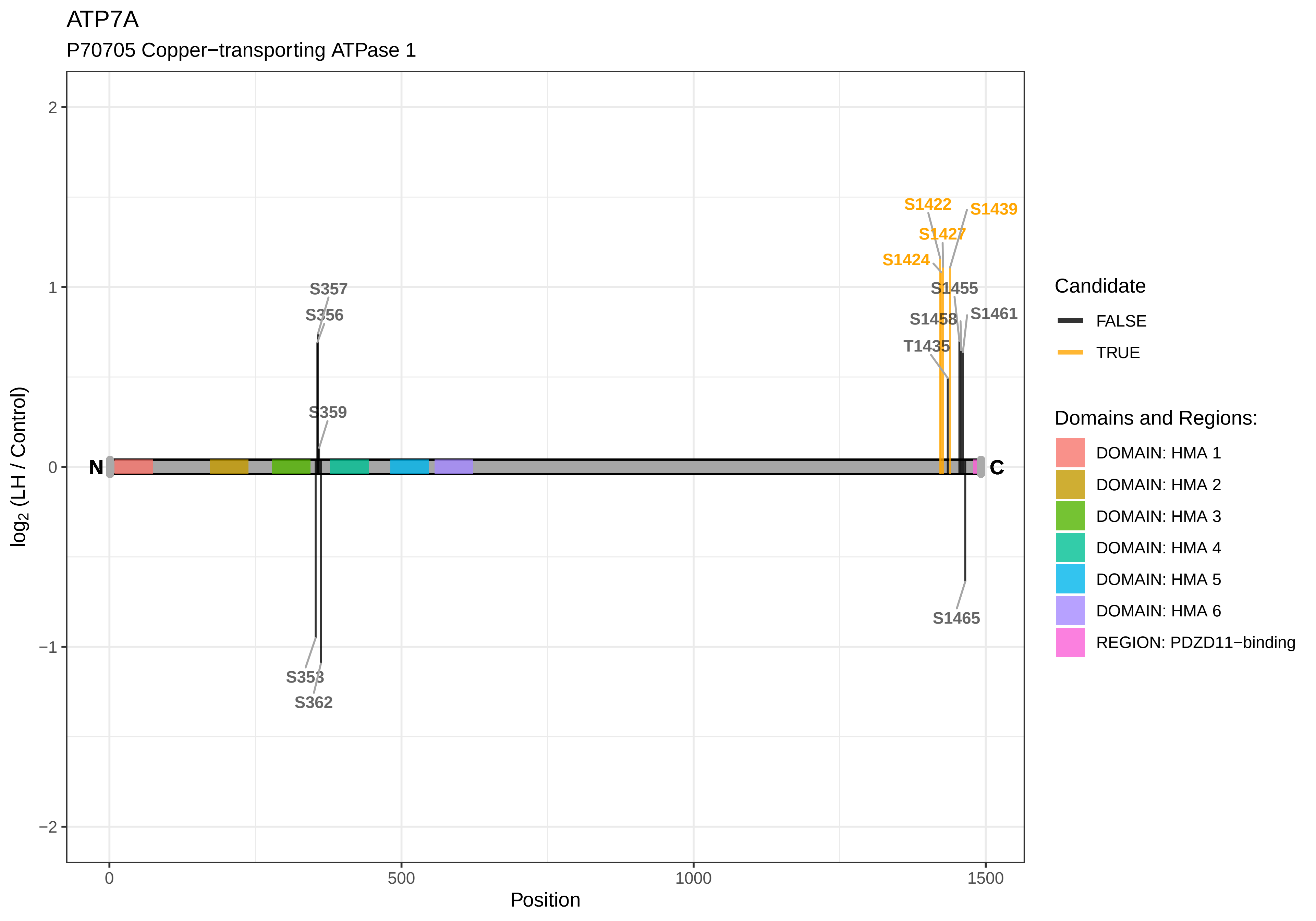

ATP7A
P70705 Copper−transporting ATPase 1
S1422
S1439
S1427
S1424
S1455
S1461
S1458
T1435
S357
S356
S359
S353
S362
S1465
N
C
log2 (LH / Control)
Position
Candidate
FALSE
TRUE
Domains and Regions:
DOMAIN: HMA 1
DOMAIN: HMA 2
DOMAIN: HMA 3
DOMAIN: HMA 4
DOMAIN: HMA 5
DOMAIN: HMA 6
REGION: PDZD11−binding

ATP8B2

D4A509 Phospholipid−transporting ATPase

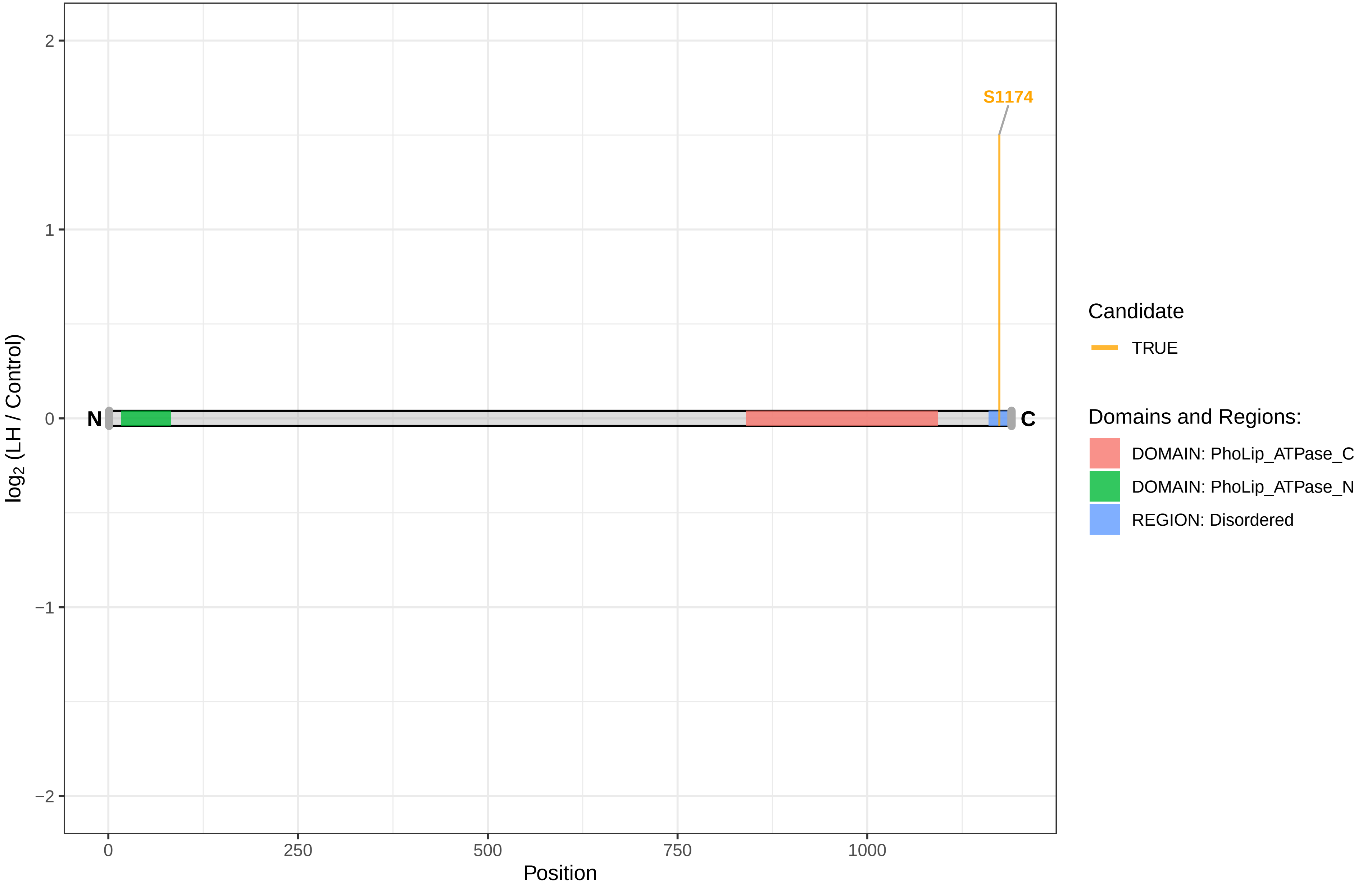

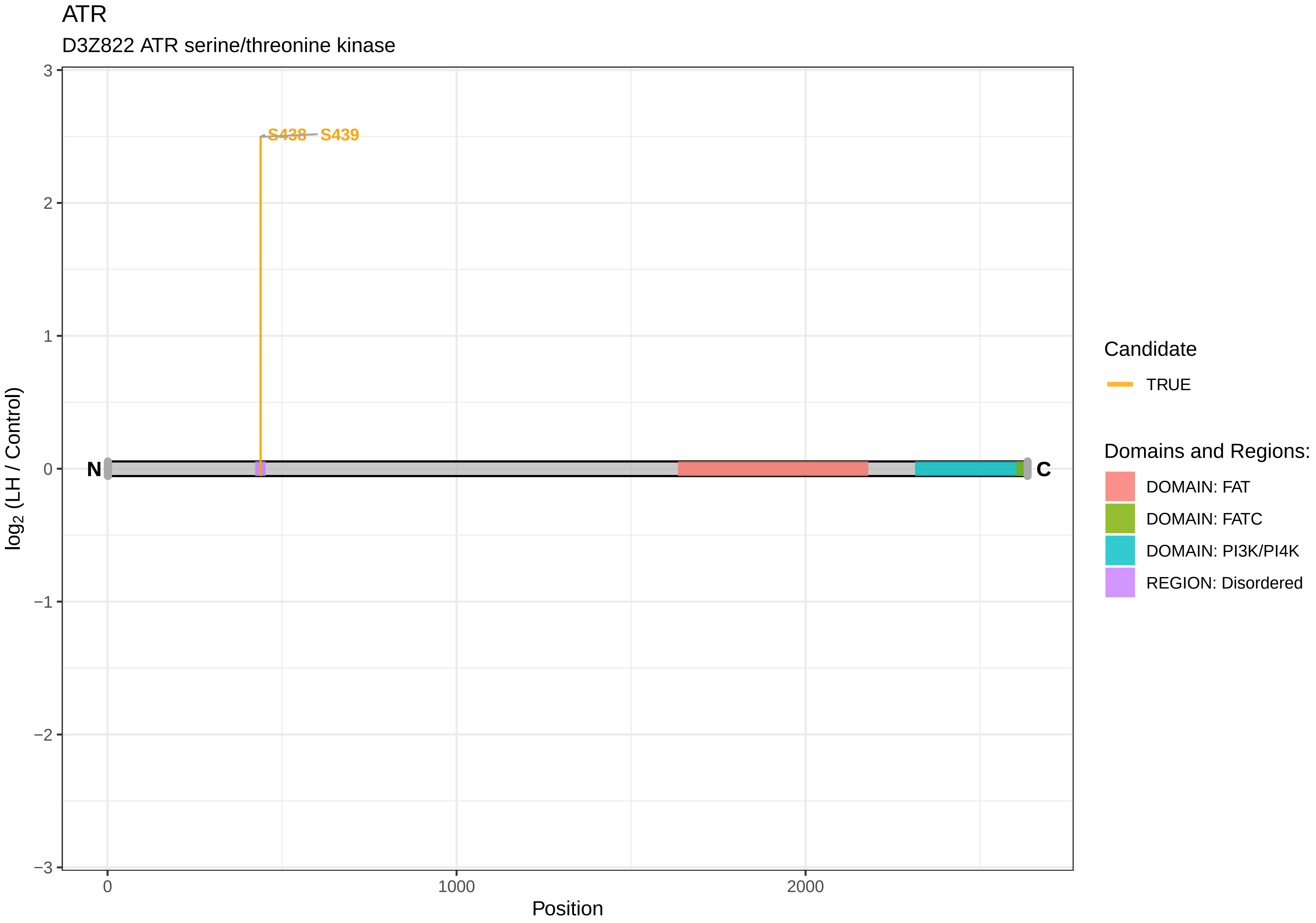

ATR
D3Z822 ATR serine/threonine kinase
S438
S439
N
C
log2 (LH / Control)
3
2
1
0
−1
−2
−3
0
1000
2000
Position
Candidate
TRUE
Domains and Regions:
DOMAIN: FAT
DOMAIN: FATC
DOMAIN: PI3K/PI4K
REGION: Disordered

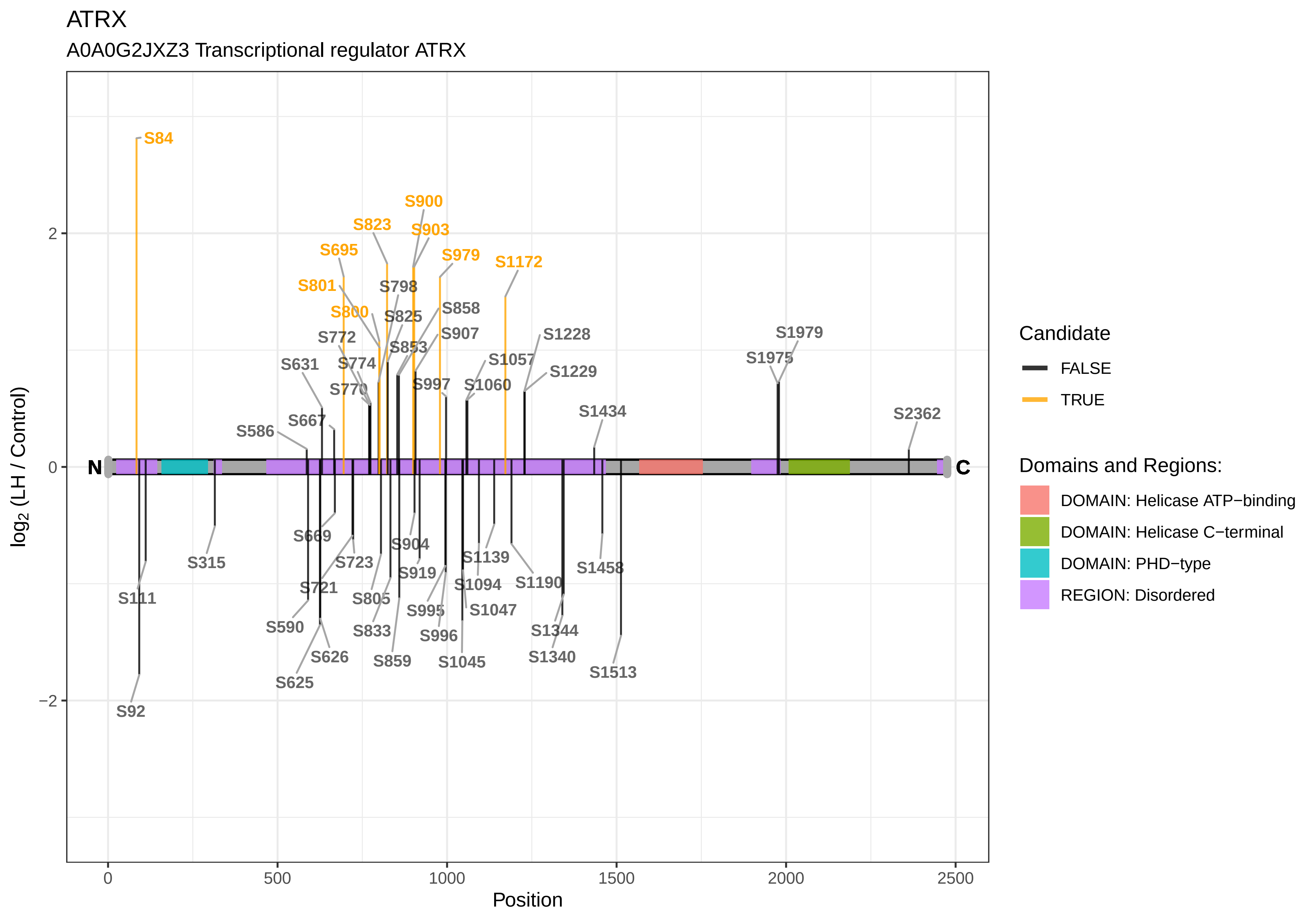

ATRX
A0A0G2JXZ3 Transcriptional regulator ATRX
log2 (LH / Control)
Position
0
500
1000
1500
2000
2500
2
0
−2
N
C
S84
S900
S823
S903
S695
S979
S1172
S801
S798
S800
S825
S858
S772
S907
S853
S1228
S631
S774
S1057
S1229
S770
S997
S1060
S1979
S1975
S667
S586
S1434
S2362
S669
S315
S904
S723
S1139
S111
S721
S919
S1094
S1190
S1458
S805
S590
S995
S1047
S833
S996
S1344
S626
S859
S1045
S1340
S1513
S625
S92
Candidate
FALSE
TRUE
Domains and Regions:
DOMAIN: Helicase ATP−binding
DOMAIN: Helicase C−terminal
DOMAIN: PHD−type
REGION: Disordered

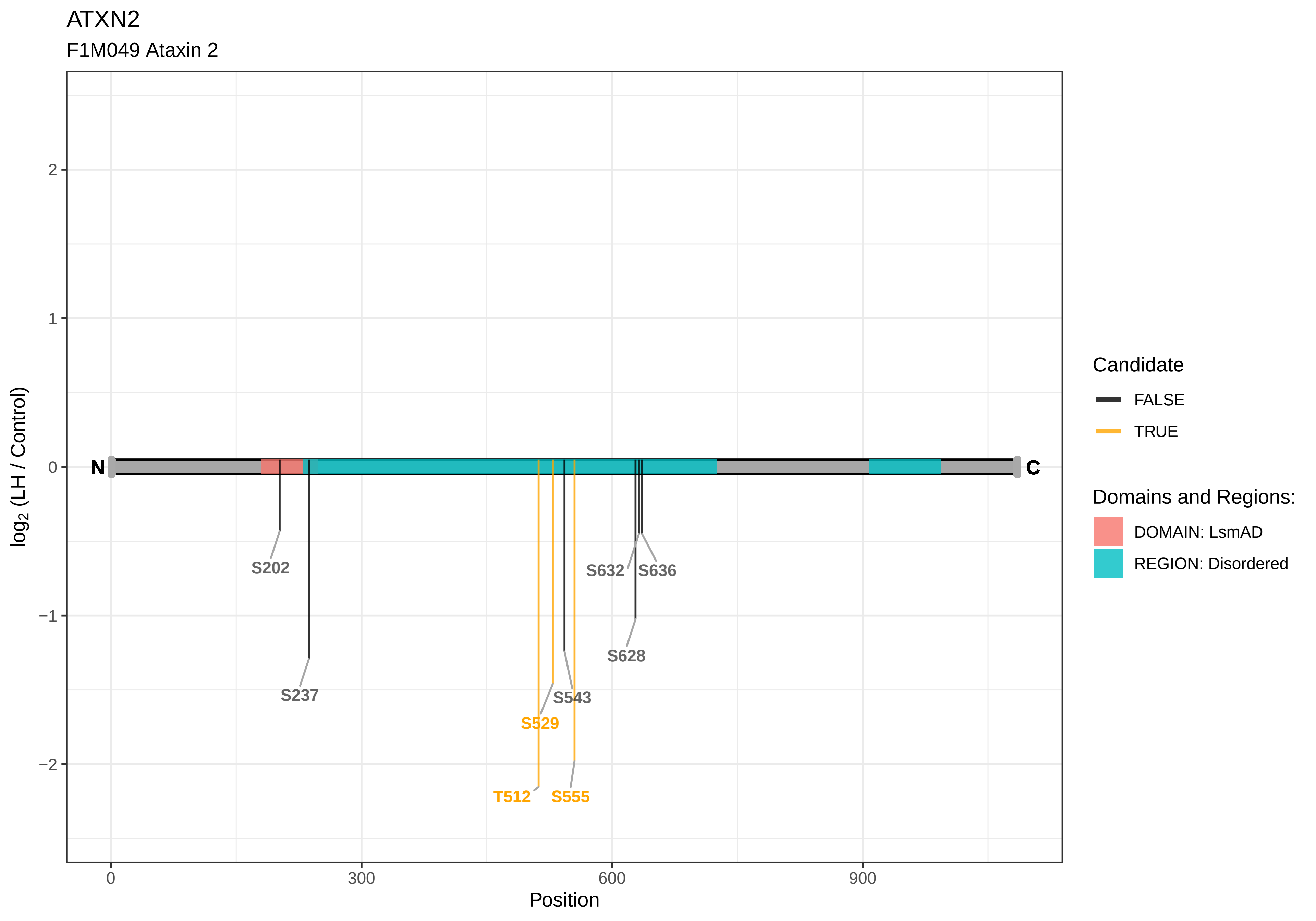

ATXN2
F1M049 Ataxin 2
log2 (LH / Control)
Position
N
C
S202
S237
S529
S543
T512
S555
S632
S636
S628
Candidate
FALSE
TRUE
Domains and Regions:
DOMAIN: LsmAD
REGION: Disordered

ATXN2L

B3DMA1 Ataxin 2−like

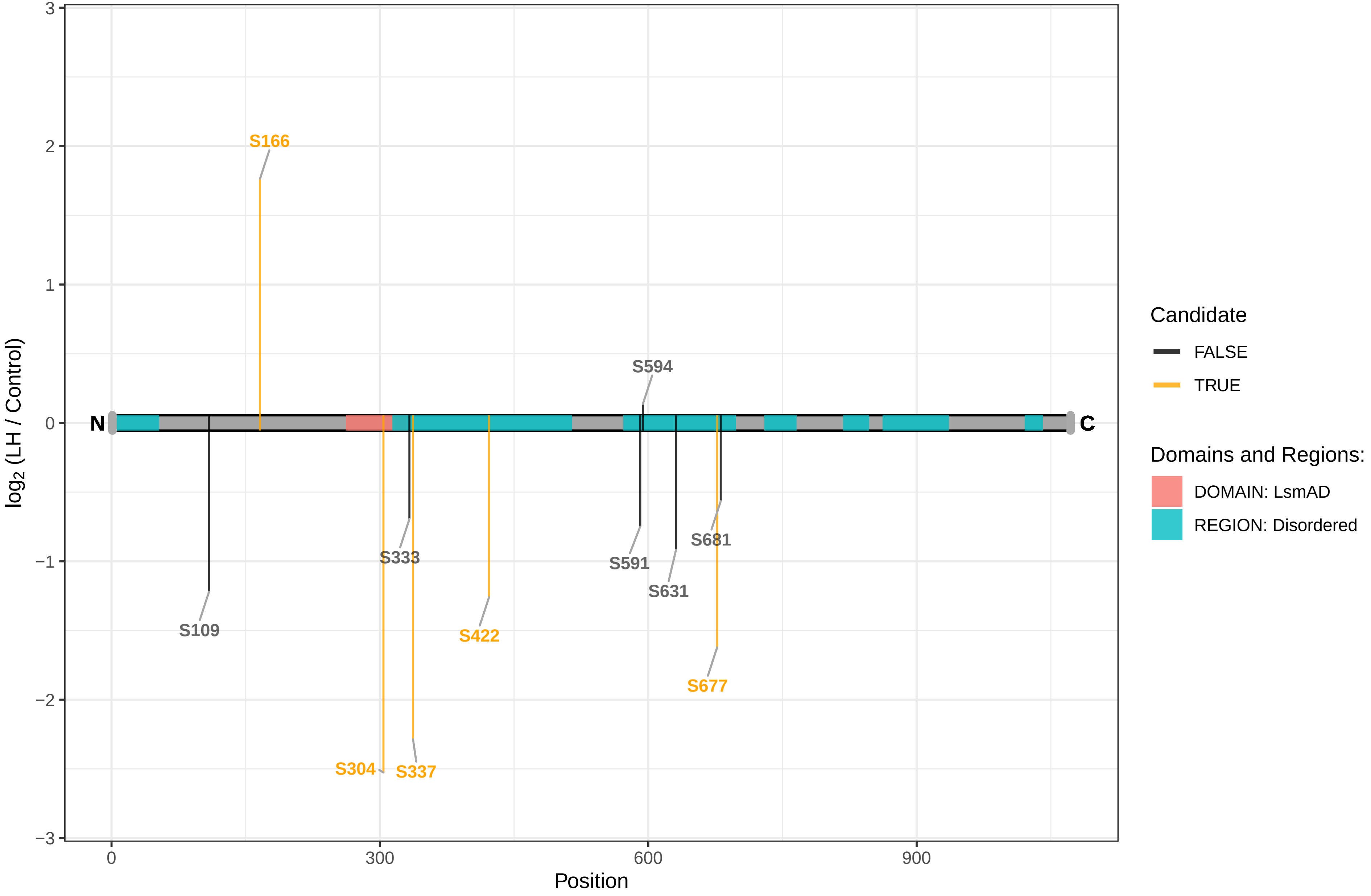

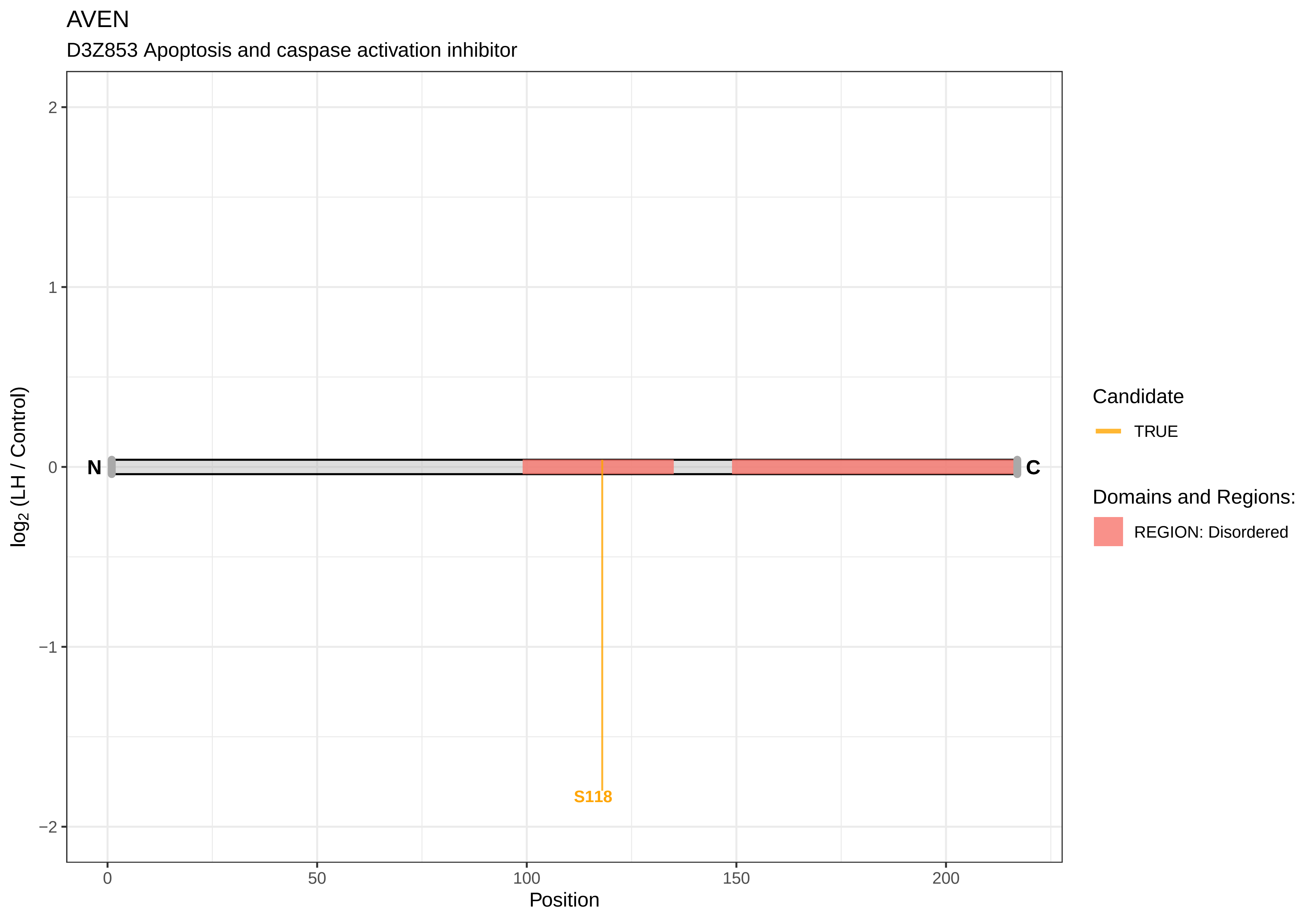
AVEN
D3Z853 Apoptosis and caspase activation inhibitor
2
1
0
−1
−2
log2 (LH / Control)
N
C
S118
0
50
100
150
200
Position
Candidate
TRUE
Domains and Regions:
REGION: Disordered

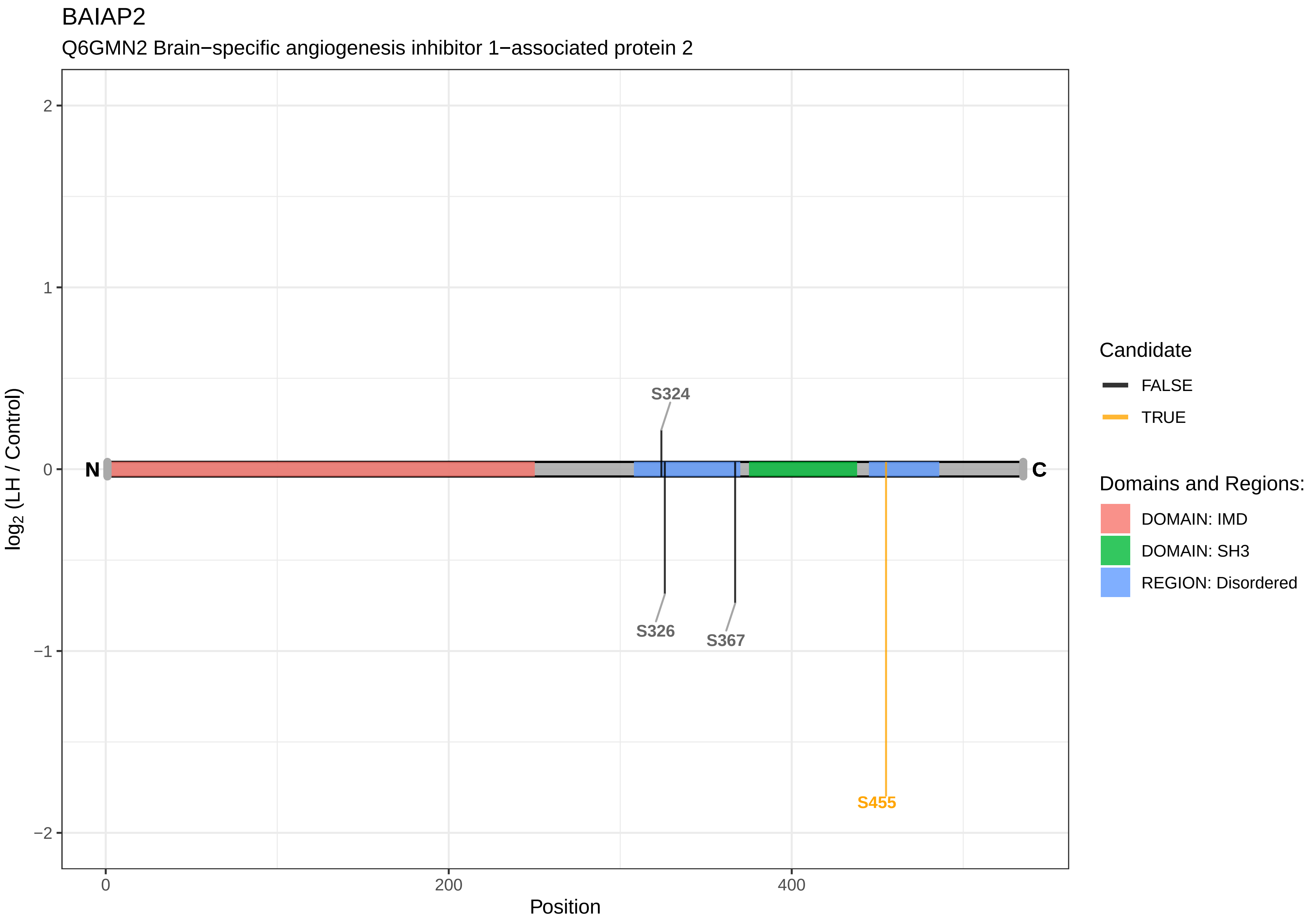
BAIAP2
Q6GMN2 Brain−specific angiogenesis inhibitor 1−associated protein 2
S324
S326
S367
S455
N
C
2
1
0
−1
−2
0
200
400
Position
log2 (LH / Control)
Candidate
FALSE
TRUE
Domains and Regions:
DOMAIN: IMD
DOMAIN: SH3
REGION: Disordered

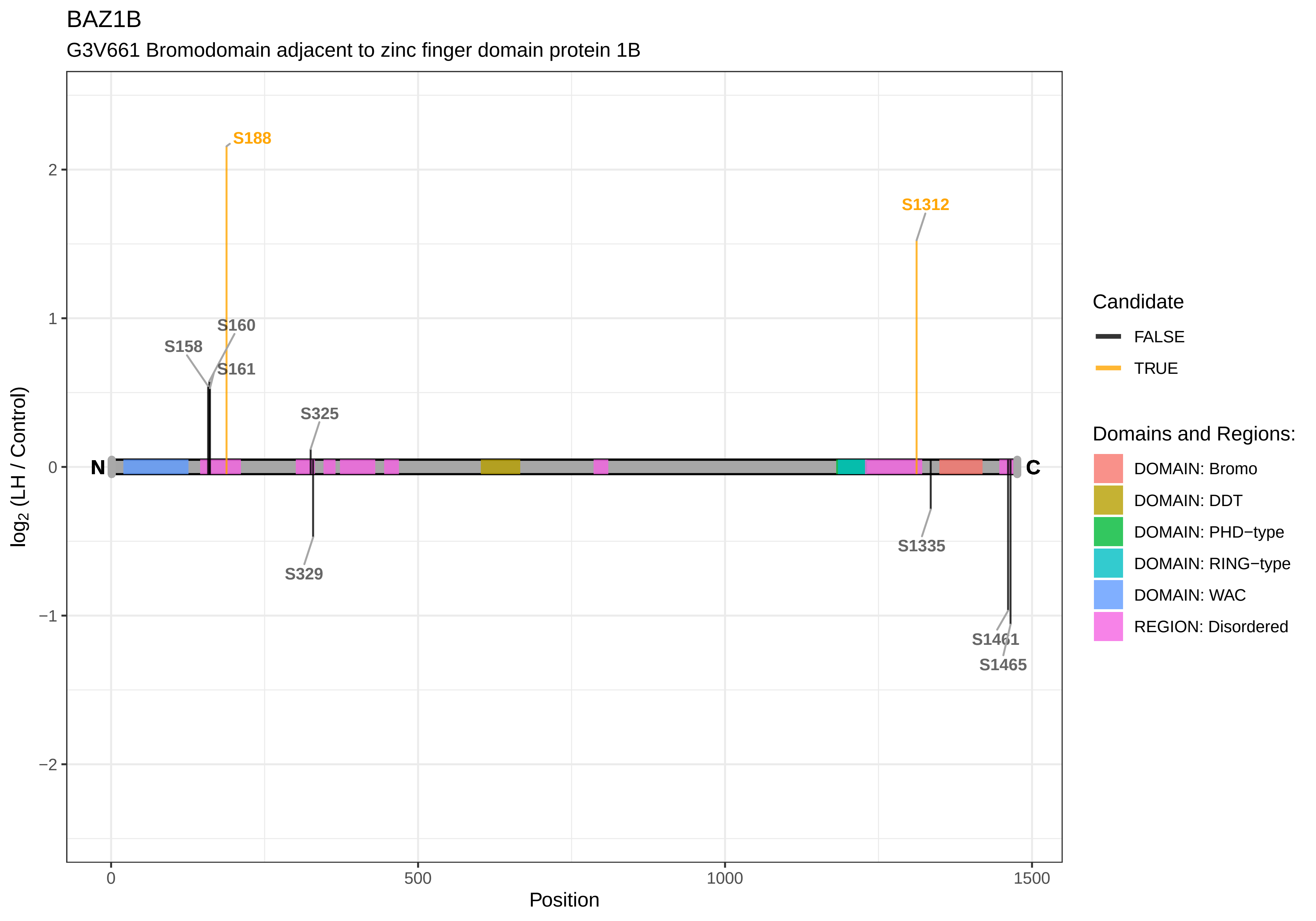
BAZ1B
G3V661 Bromodomain adjacent to zinc finger domain protein 1B
S188
S1312
S160
S158
S161
S325
S329
S1335
S1461
S1465
N
C
2
1
0
−1
−2
log2 (LH / Control)
0
500
1000
1500
Position
Candidate
FALSE
TRUE
Domains and Regions:
DOMAIN: Bromo
DOMAIN: DDT
DOMAIN: PHD−type
DOMAIN: RING−type
DOMAIN: WAC
REGION: Disordered

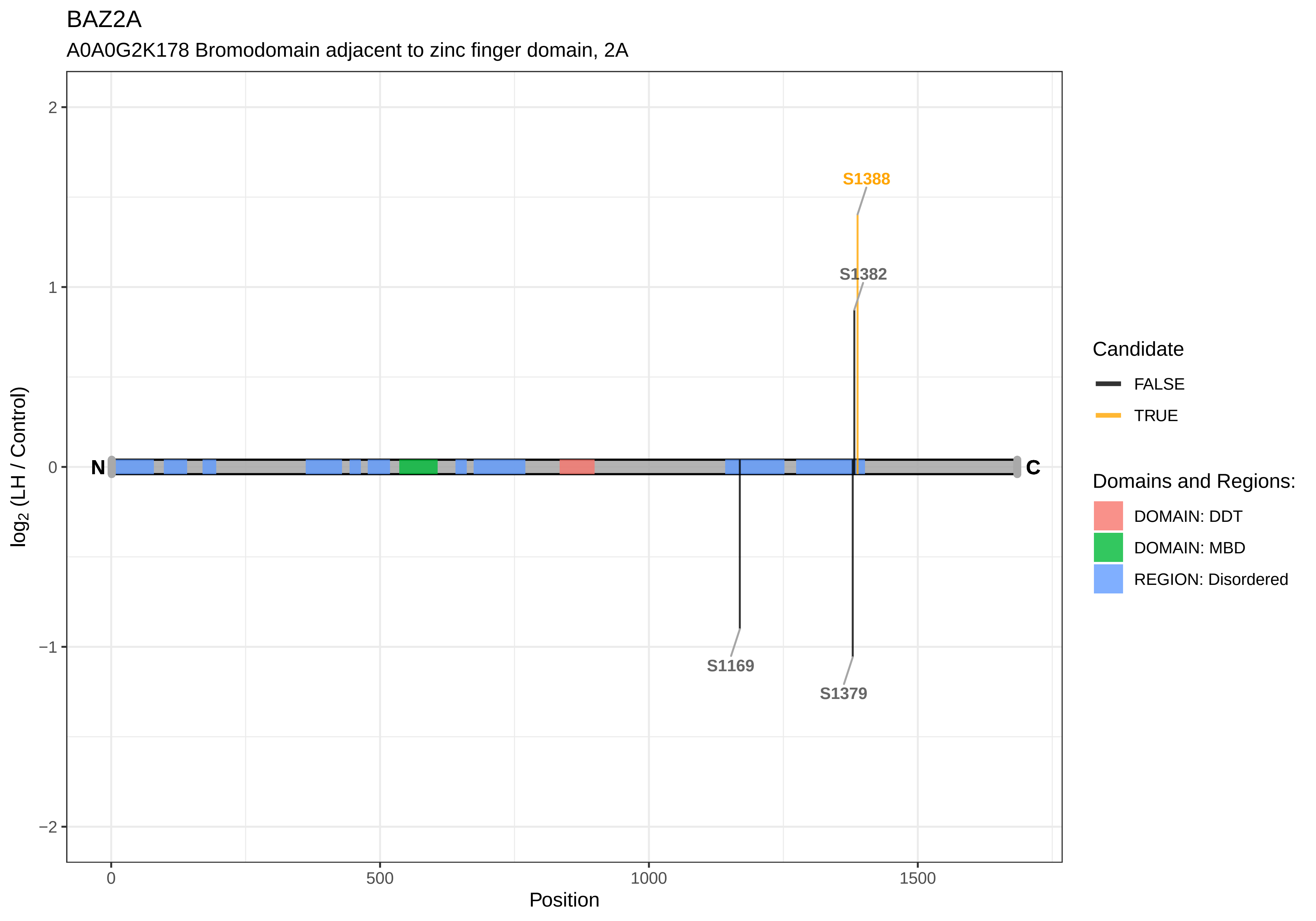

BAZ2A
A0A0G2K178 Bromodomain adjacent to zinc finger domain, 2A
S1388
S1382
S1169
S1379
N
C
log2 (LH / Control)
Position
2
1
0
−1
−2
0
500
1000
1500
Candidate
FALSE
TRUE
Domains and Regions:
DOMAIN: DDT
DOMAIN: MBD
REGION: Disordered

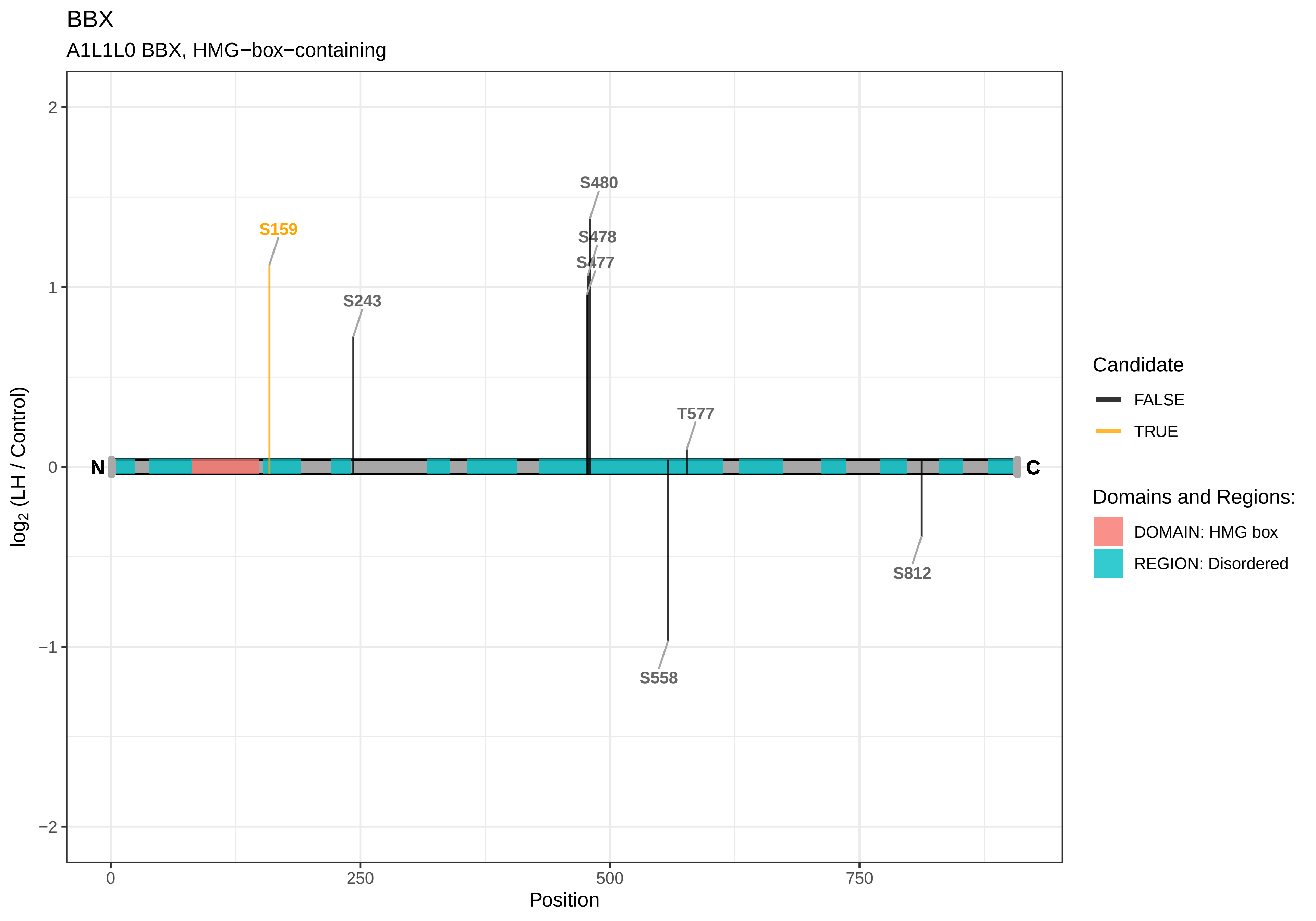
BBX
A1L1L0 BBX, HMG−box−containing
S480
S478
S477
S159
S243
T577
S558
S812
N
C
2
1
0
−1
−2
log2 (LH / Control)
0
250
500
750
Position
Candidate
FALSE
TRUE
Domains and Regions:
DOMAIN: HMG box
REGION: Disordered

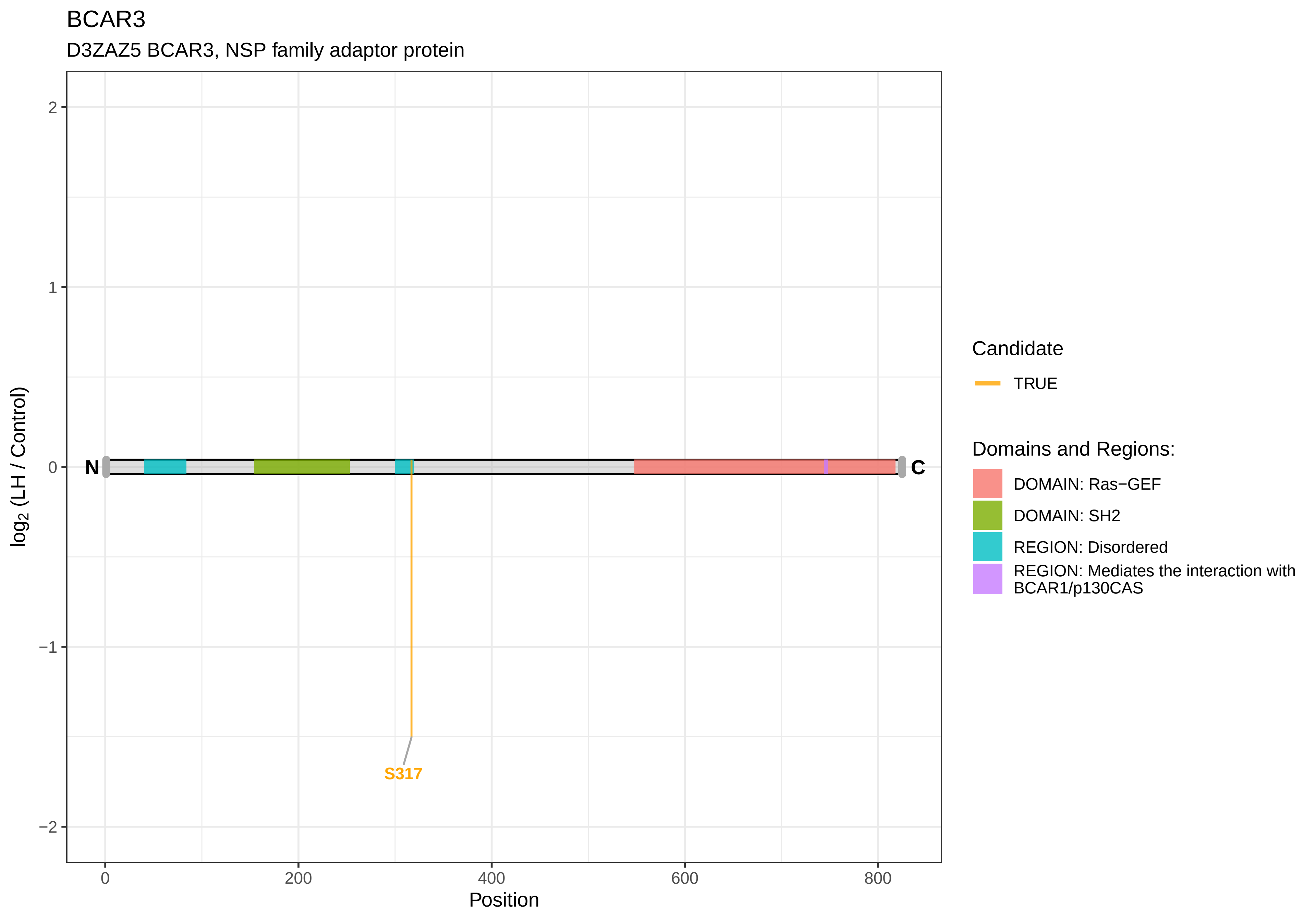
BCAR3
D3ZAZ5 BCAR3, NSP family adaptor protein
log2 (LH / Control)
2
1
0
−1
−2
N
C
S317
0
200
400
600
800
Position
Candidate
TRUE
Domains and Regions:
DOMAIN: Ras−GEF
DOMAIN: SH2
REGION: Disordered
REGION: Mediates the interaction with BCAR1/p130CAS

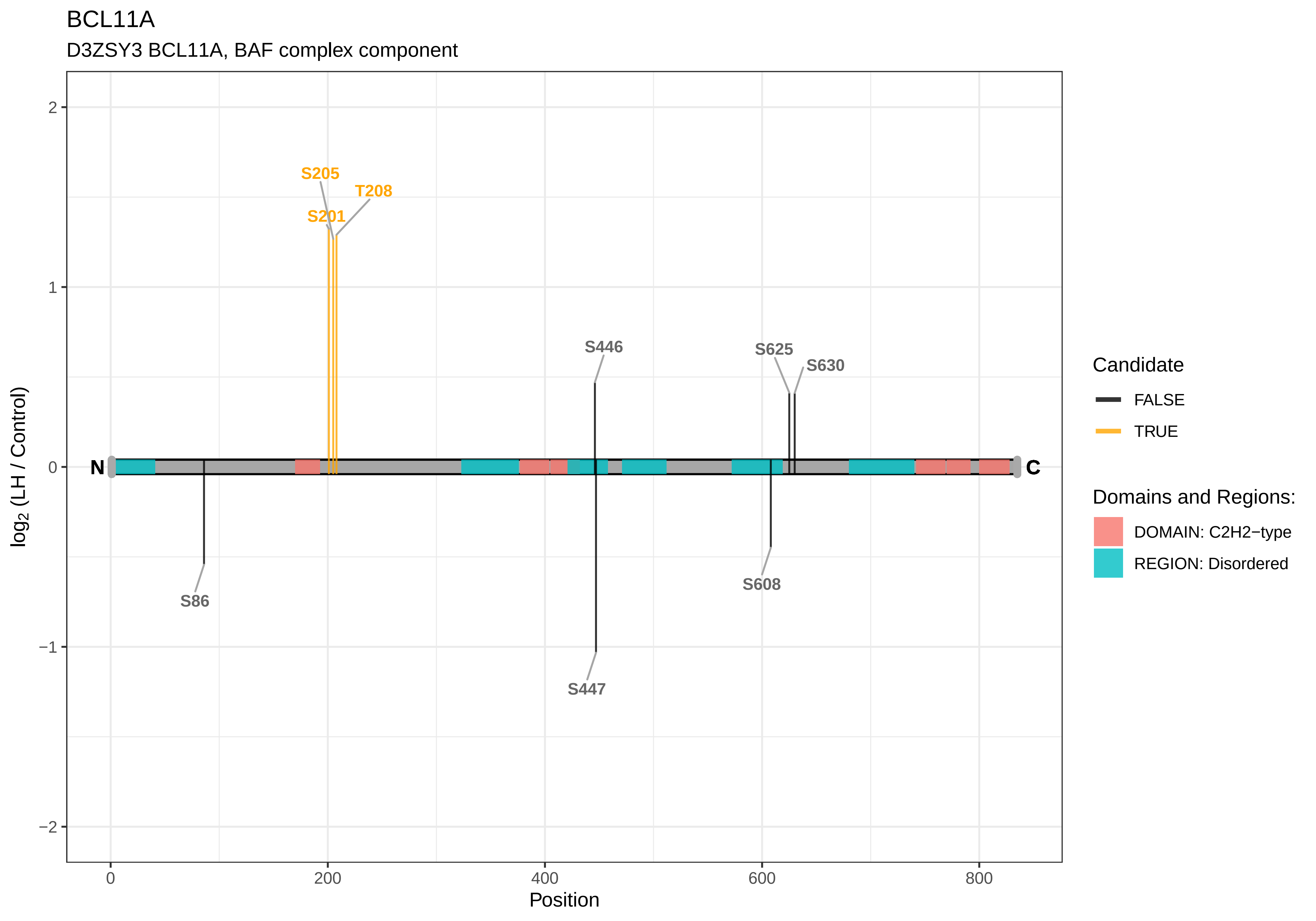
BCL11A
D3ZSY3 BCL11A, BAF complex component
log2 (LH / Control)
Position
S205
T208
S201
S446
S625
S630
S86
S447
S608
N
C
2
1
0
−1
−2
0
200
400
600
800
Candidate
FALSE
TRUE
Domains and Regions:
DOMAIN: C2H2−type
REGION: Disordered

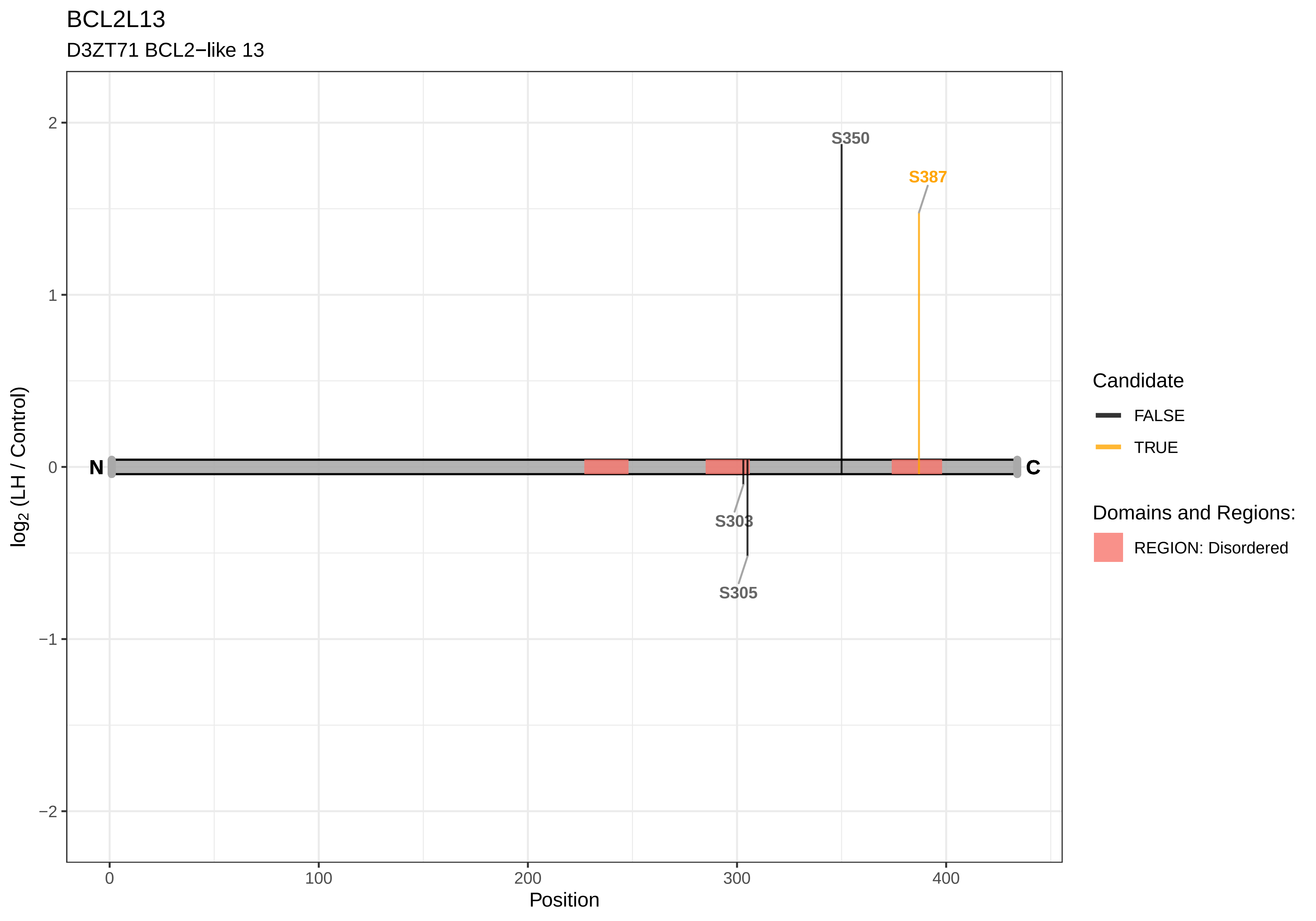

BCL2L13
D3ZT71 BCL2−like 13
S350
S387
S303
S305
N
C
log2 (LH / Control)
Position
2
1
0
−1
−2
0
100
200
300
400
Candidate
FALSE
TRUE
Domains and Regions:
REGION: Disordered

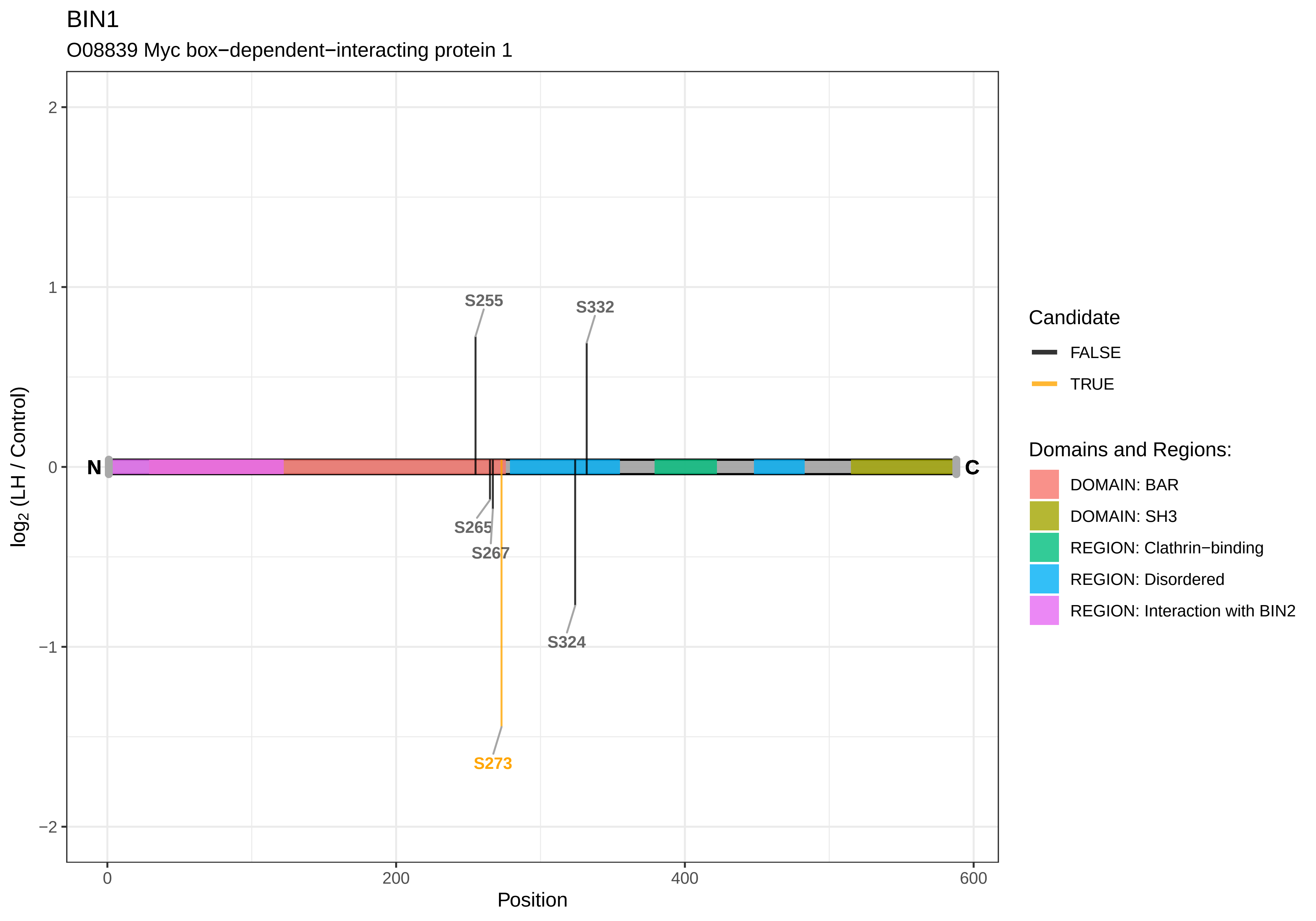

BIN1
O08839 Myc box−dependent−interacting protein 1
log2 (LH / Control)
Position
0
200
400
600
2
1
0
−1
−2
N
C
S255
S332
S265
S267
S324
S273
Candidate
FALSE
TRUE
Domains and Regions:
DOMAIN: BAR
DOMAIN: SH3
REGION: Clathrin−binding
REGION: Disordered
REGION: Interaction with BIN2

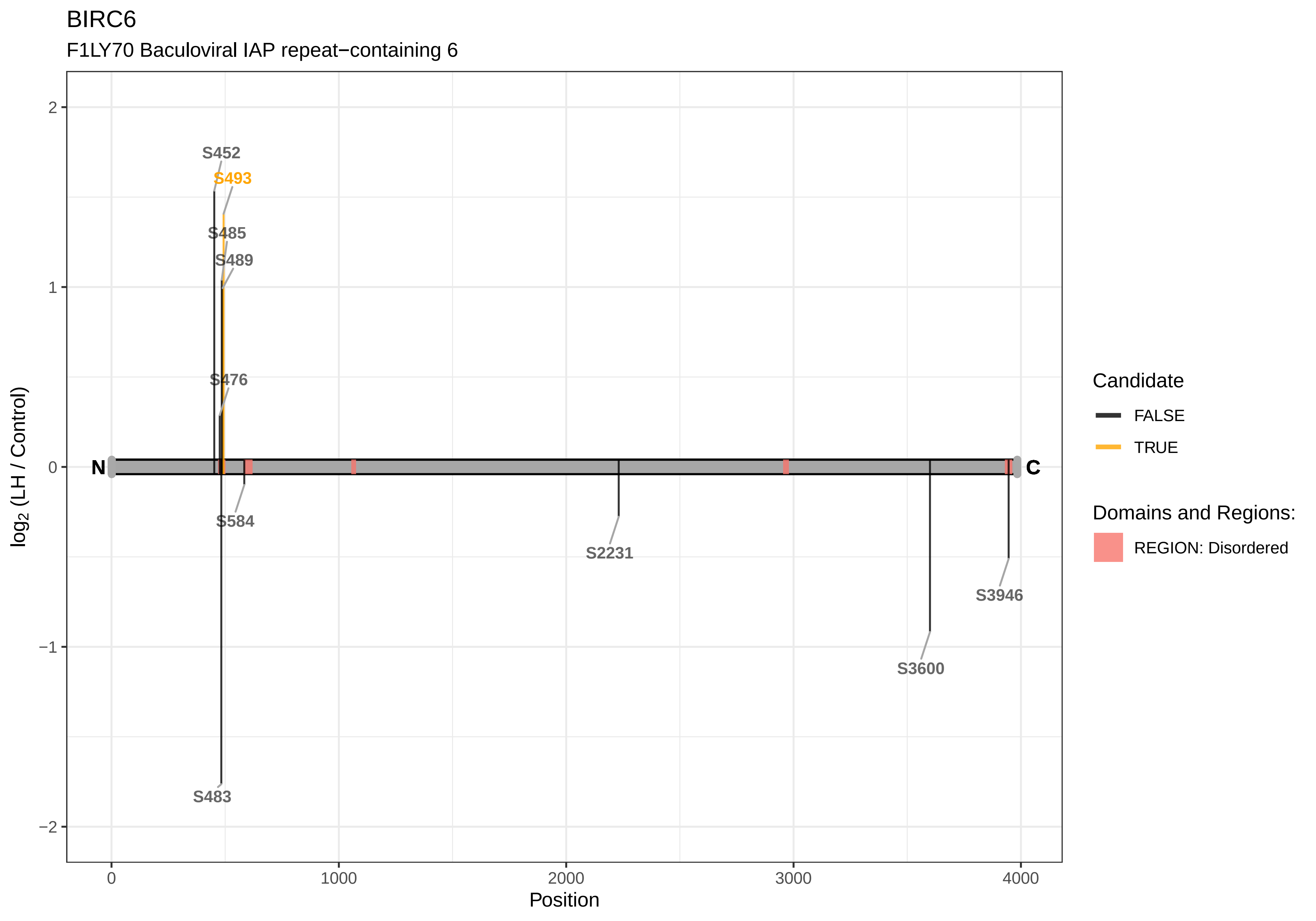
BIRC6
F1LY70 Baculoviral IAP repeat−containing 6
log2 (LH / Control)
Position
2
1
0
−1
−2
0
1000
2000
3000
4000
N
C
S452
S493
S485
S489
S476
S584
S2231
S3946
S3600
S483
Candidate
FALSE
TRUE
Domains and Regions:
REGION: Disordered

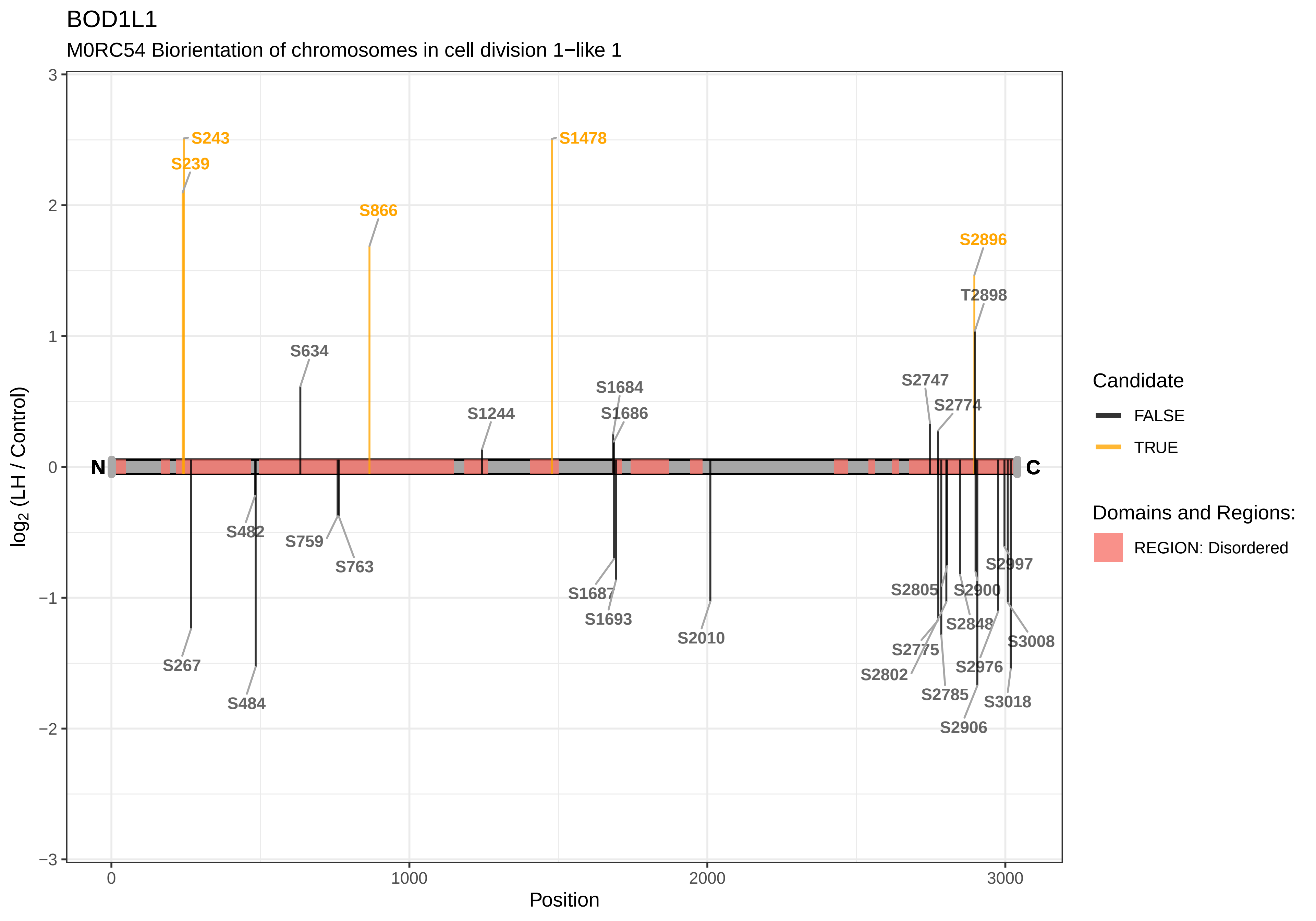
BOD1L1
M0RC54 Biorientation of chromosomes in cell division 1−like 1
S243
S239
S866
S1478
S2896
T2898
S634
S1244
S1684
S1686
S2747
S2774
N
C
S482
S759
S763
S267
S484
S1687
S1693
S2010
S2997
S2805
S2900
S2848
S3008
S2775
S2802
S2976
S2785
S3018
S2906
log2 (LH / Control)
Position
Candidate
FALSE
TRUE
Domains and Regions:
REGION: Disordered

### BOLA1

Q06C60 BolA family member 1

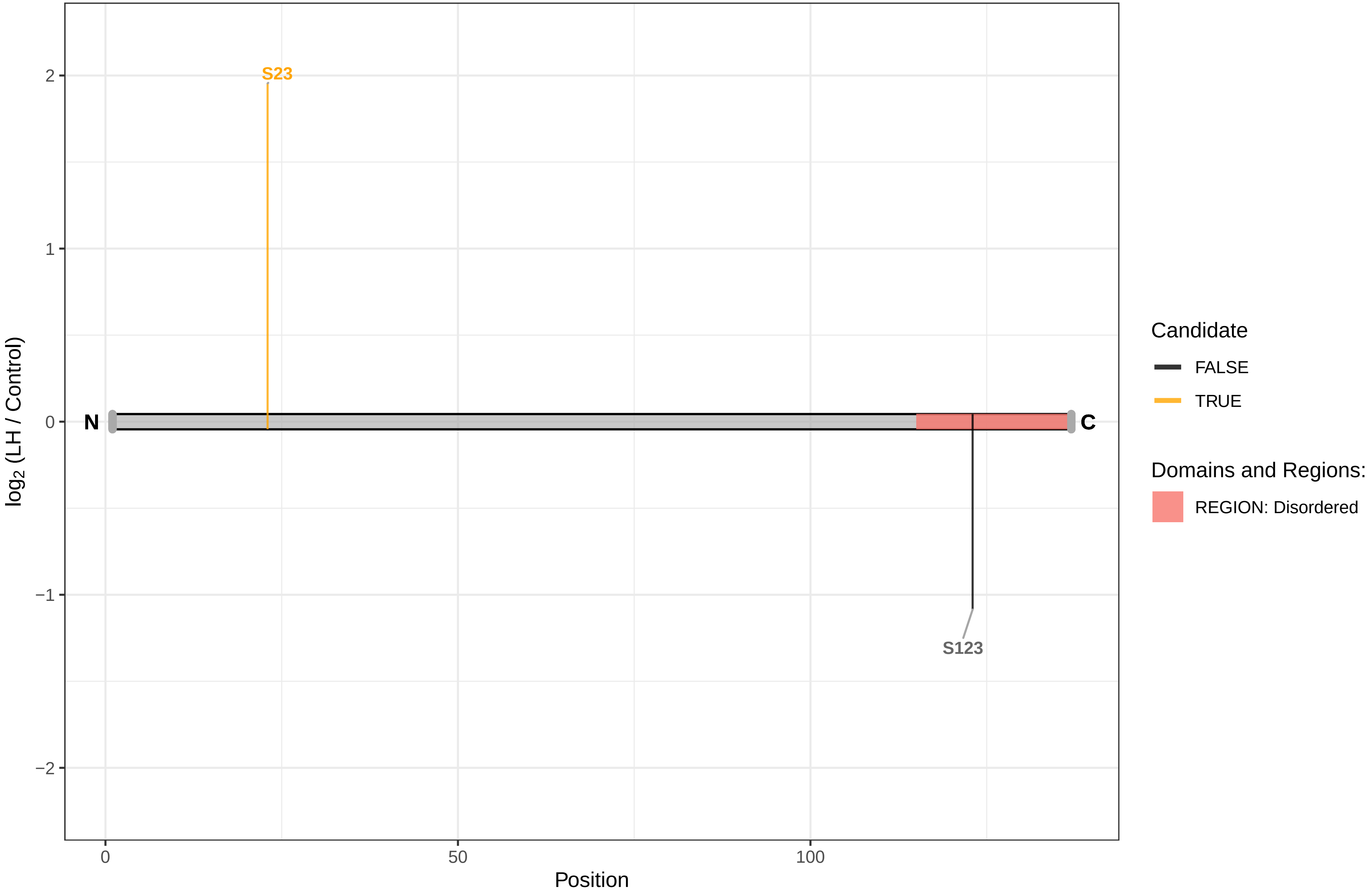

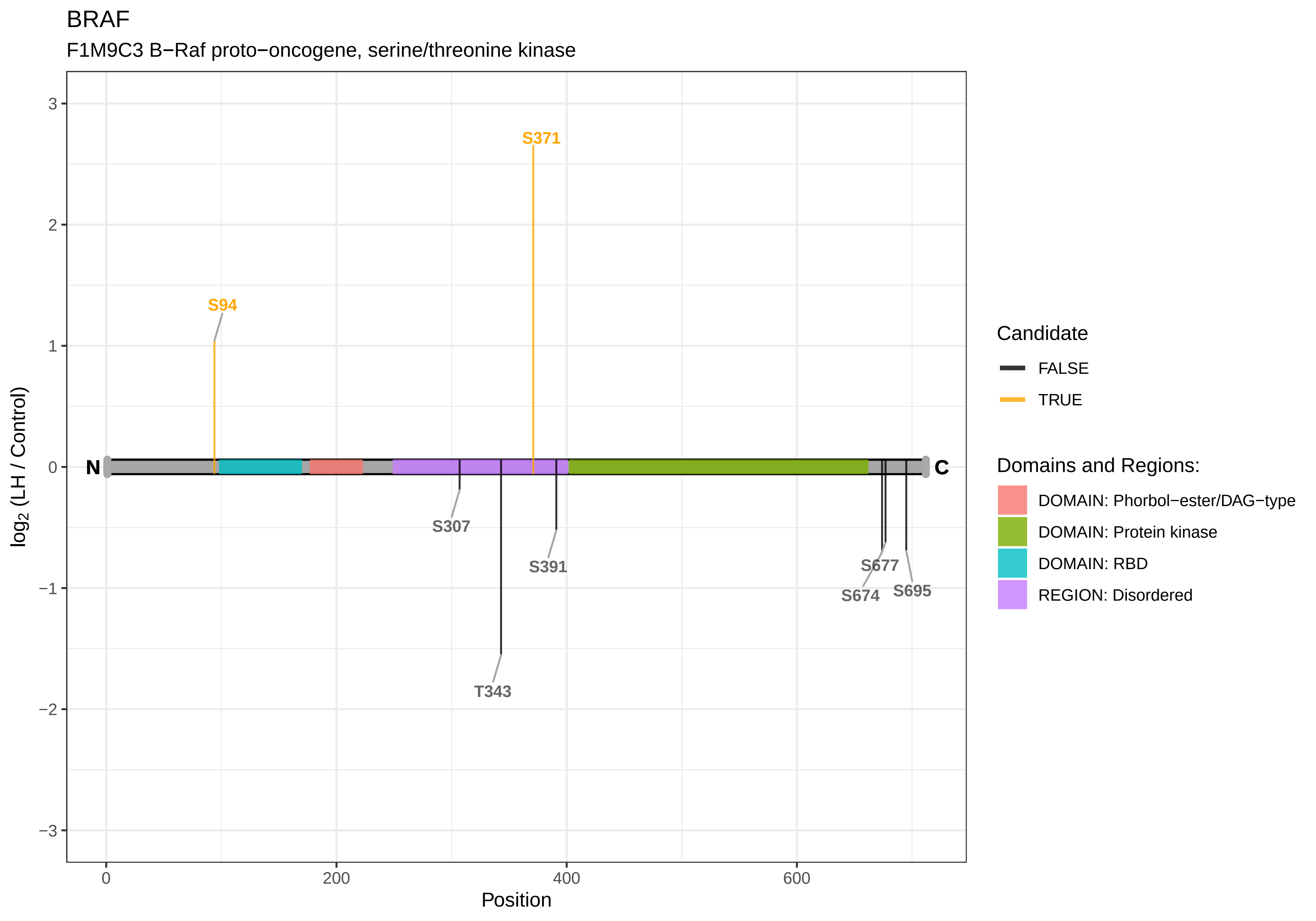
BRAF
F1M9C3 B−Raf proto−oncogene, serine/threonine kinase
S371
S94
N
C
S307
S391
T343
S677
S674
S695
log2 (LH / Control)
Position
3
2
1
0
−1
−2
−3
0
200
400
600
Candidate
FALSE
TRUE
Domains and Regions:
DOMAIN: Phorbol−ester/DAG−type
DOMAIN: Protein kinase
DOMAIN: RBD
REGION: Disordered

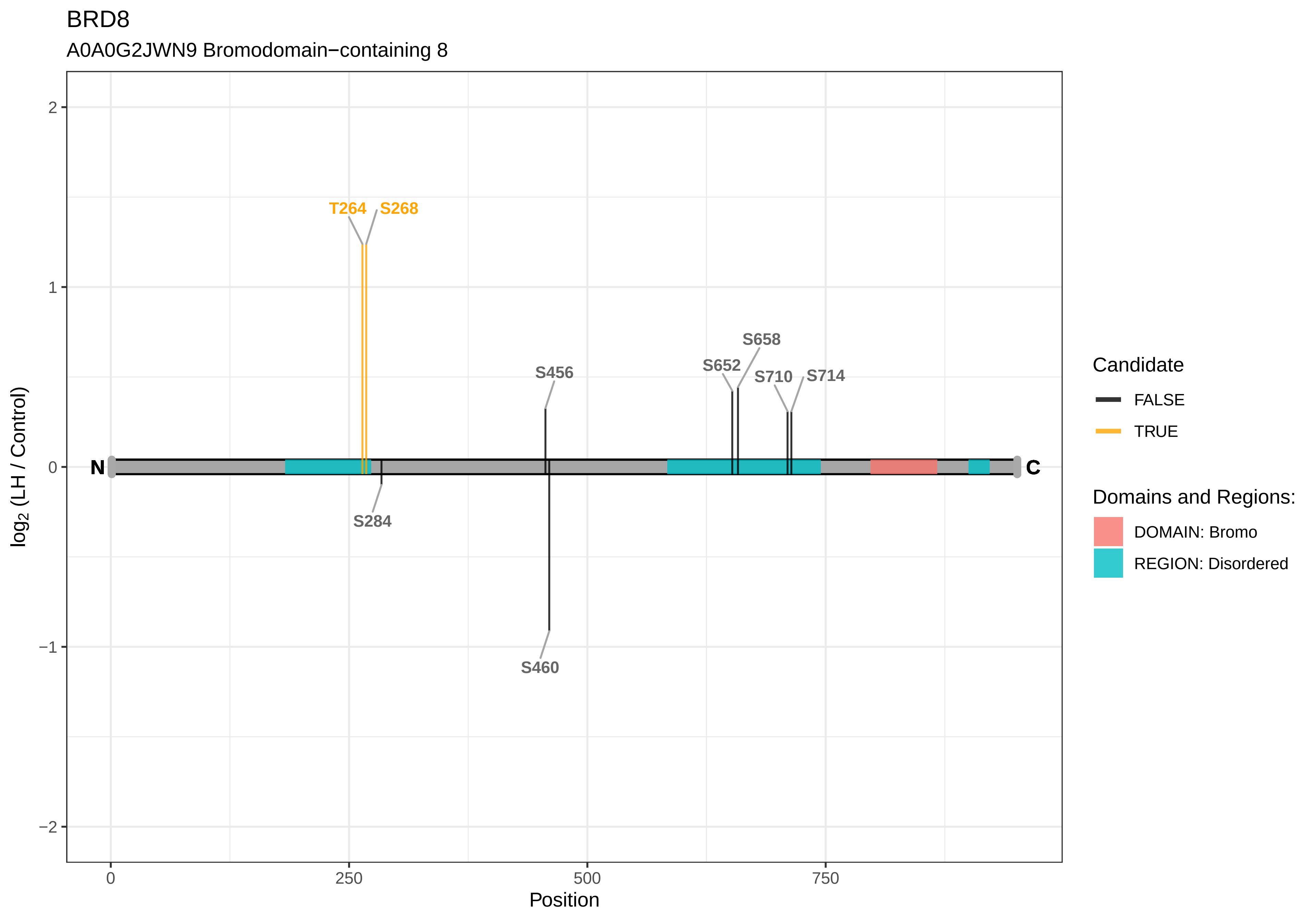
BRD8
A0A0G2JWN9 Bromodomain−containing 8
T264
S268
S456
S652
S658
S710
S714
S284
S460
N
C
2
1
0
−1
−2
0
250
500
750
Position
log2 (LH / Control)
Candidate
FALSE
TRUE
Domains and Regions:
DOMAIN: Bromo
REGION: Disordered

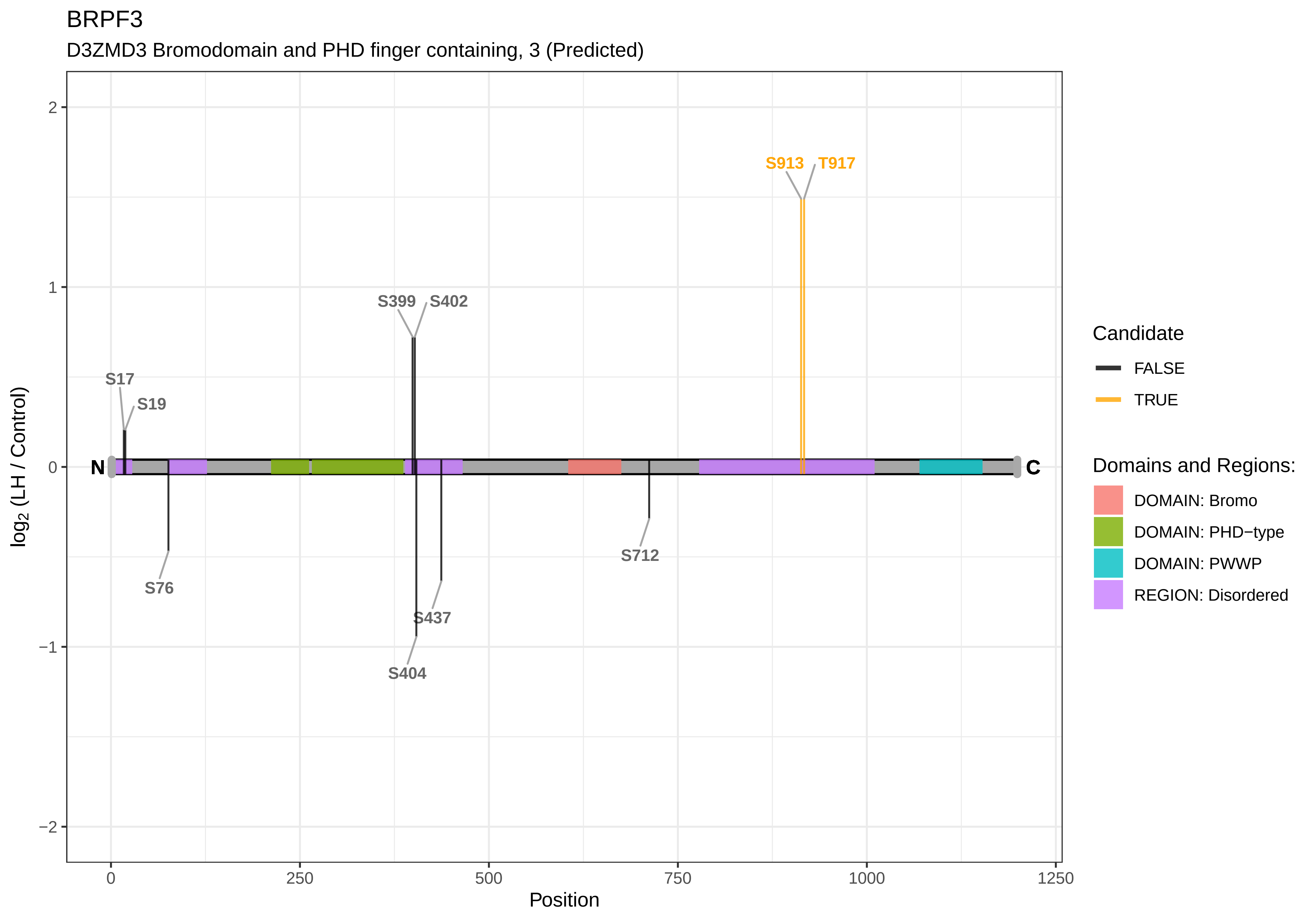

BRPF3
D3ZMD3 Bromodomain and PHD finger containing, 3 (Predicted)
log2 (LH / Control)
Position
2
1
0
−1
−2
0
250
500
750
1000
1250
N
C
S17
S19
S76
S399
S402
S404
S437
S712
S913
T917
Candidate
FALSE
TRUE
Domains and Regions:
DOMAIN: Bromo
DOMAIN: PHD−type
DOMAIN: PWWP
REGION: Disordered

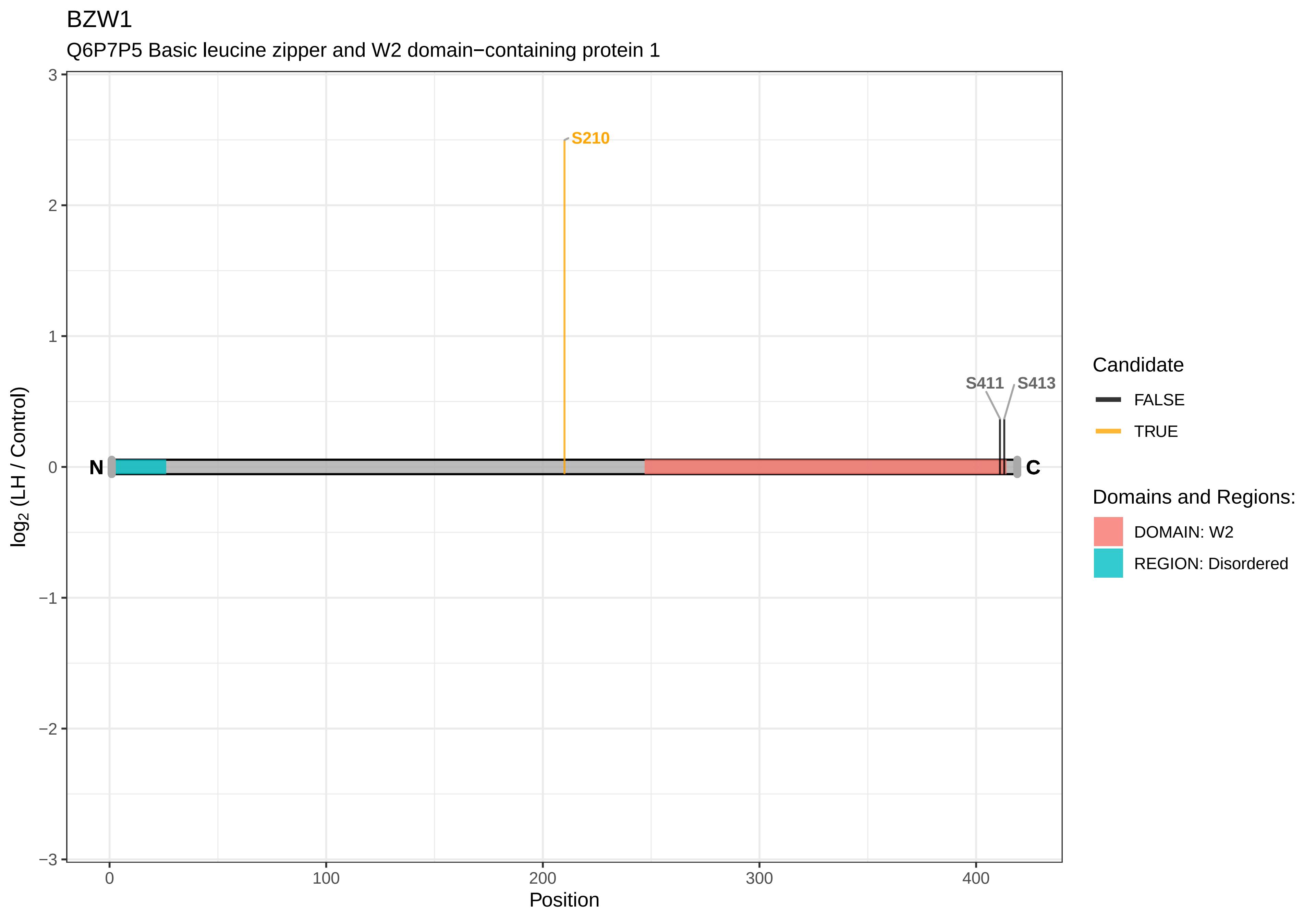
BZW1
Q6P7P5 Basic leucine zipper and W2 domain−containing protein 1
S210
S411
S413
N
C
log2 (LH / Control)
Position
Candidate
FALSE
TRUE
Domains and Regions:
DOMAIN: W2
REGION: Disordered

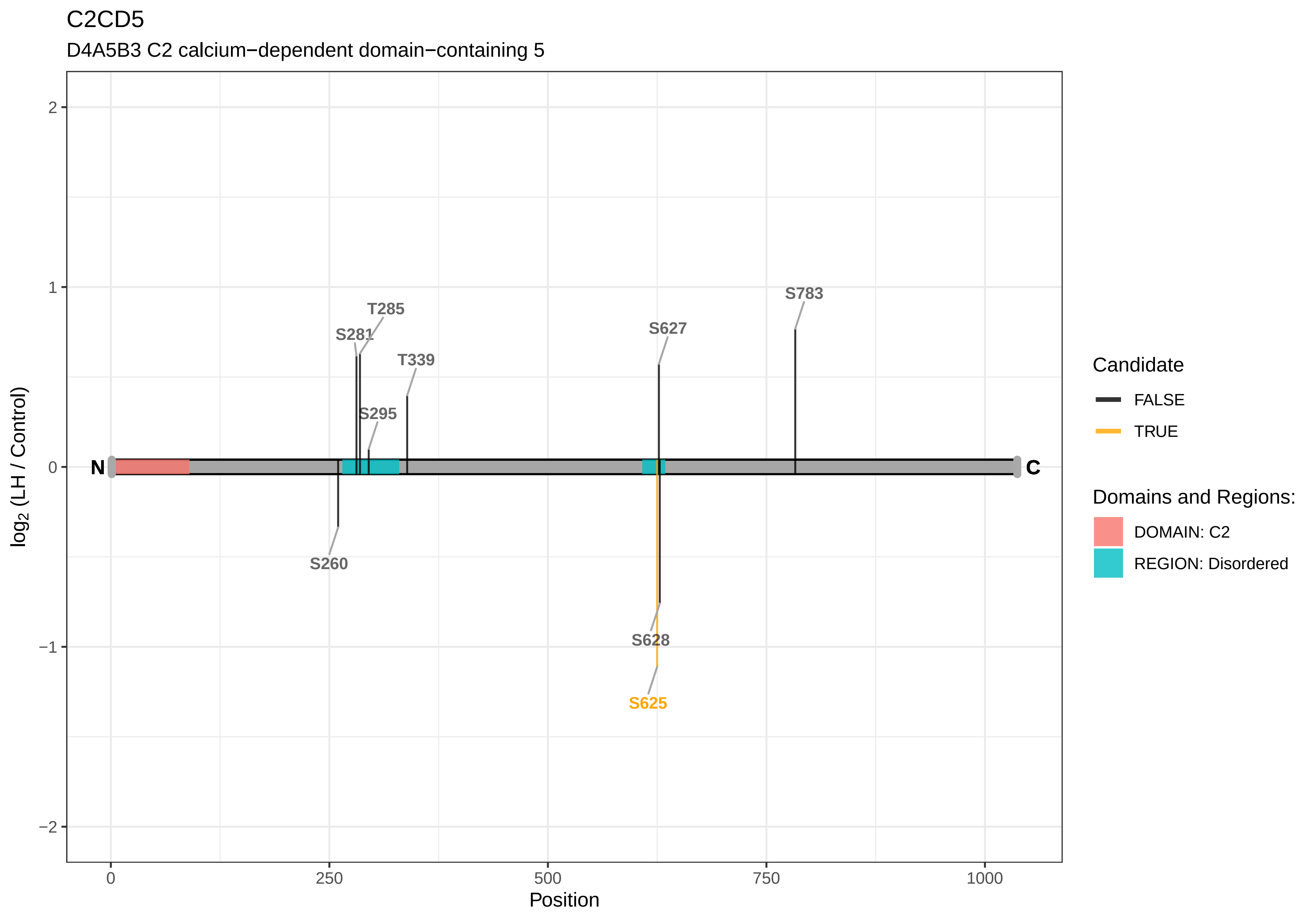

C2CD5
D4A5B3 C2 calcium−dependent domain−containing 5
log2 (LH / Control)
Position
N
C
S260
S281
T285
S295
T339
S627
S628
S625
S783
2
1
0
−1
−2
0
250
500
750
1000
Candidate
FALSE
TRUE
Domains and Regions:
DOMAIN: C2
REGION: Disordered

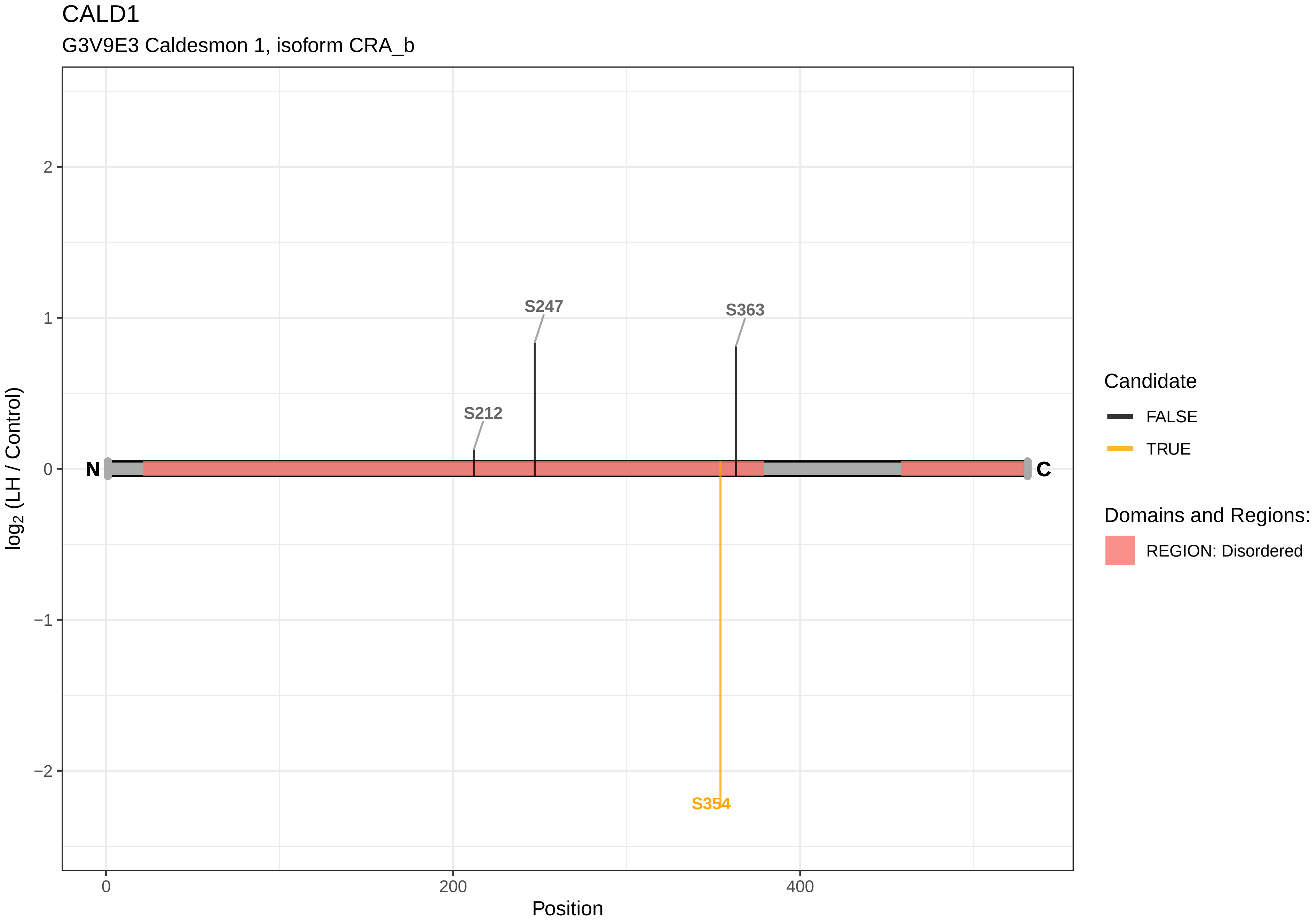

CALD1
G3V9E3 Caldesmon 1, isoform CRA_b
S247
S363
S212
N
C
S354
2
1
0
−1
−2
log2 (LH / Control)
0
200
400
Position
Candidate
FALSE
TRUE
Domains and Regions:
REGION: Disordered

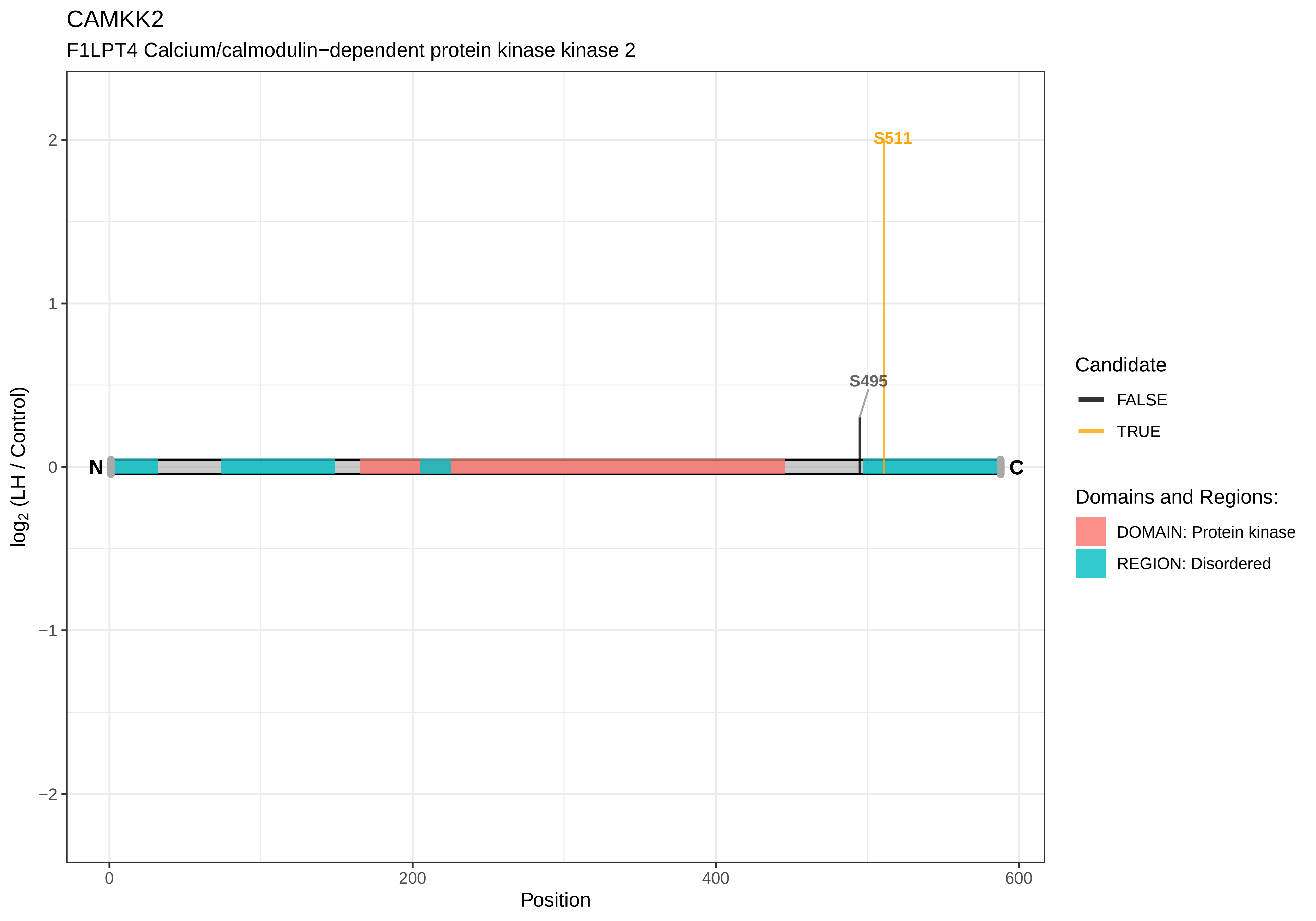
CAMKK2
F1LPT4 Calcium/calmodulin−dependent protein kinase kinase 2
S511
S495
N
C
2
1
0
−1
−2
log2 (LH / Control)
0
200
400
600
Position
Candidate
FALSE
TRUE
Domains and Regions:
DOMAIN: Protein kinase
REGION: Disordered

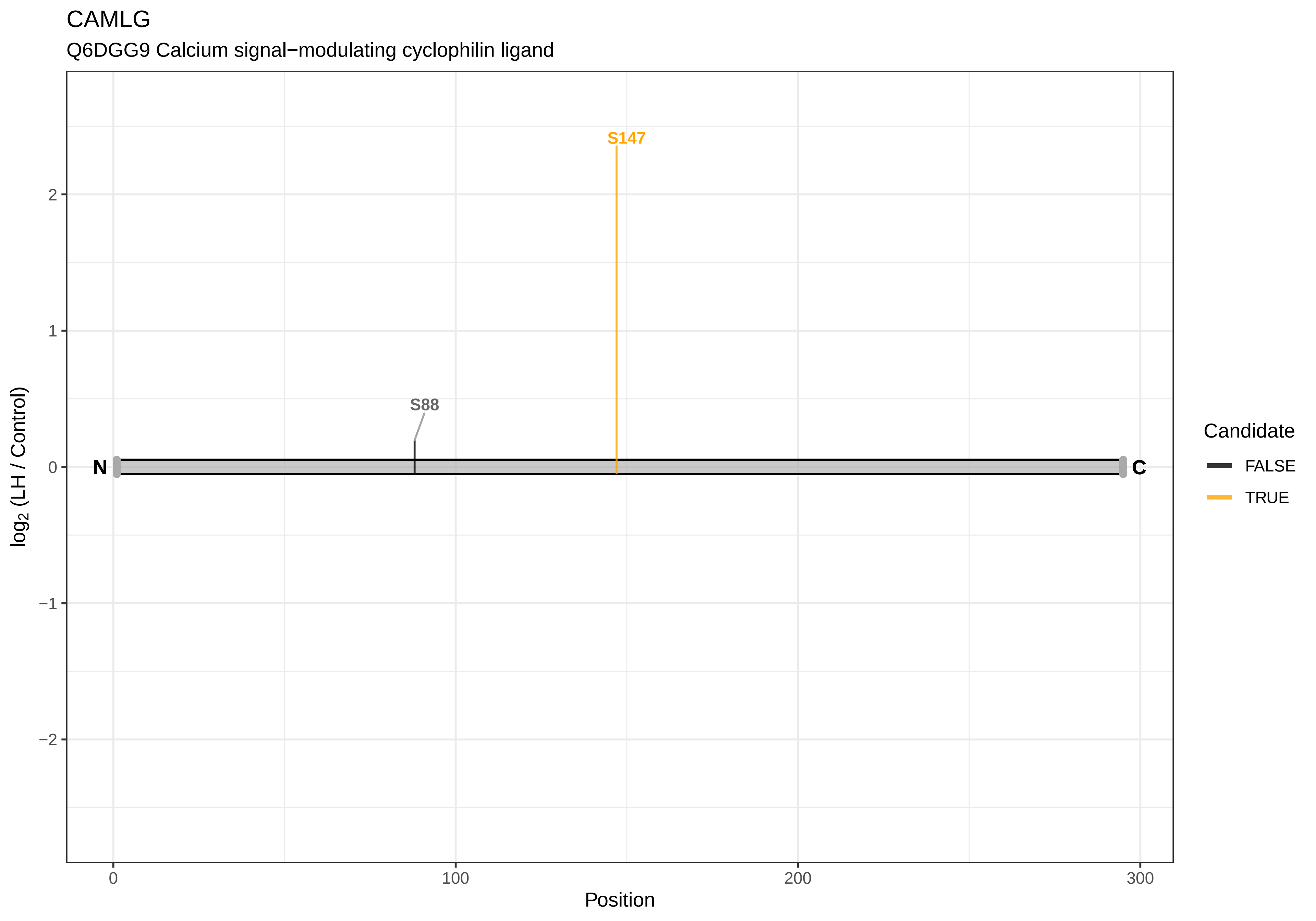
CAMLG
Q6DGG9 Calcium signal−modulating cyclophilin ligand
S147
S88
N
C
log2 (LH / Control)
2
1
0
−1
−2
0
100
200
300
Position
Candidate
FALSE
TRUE

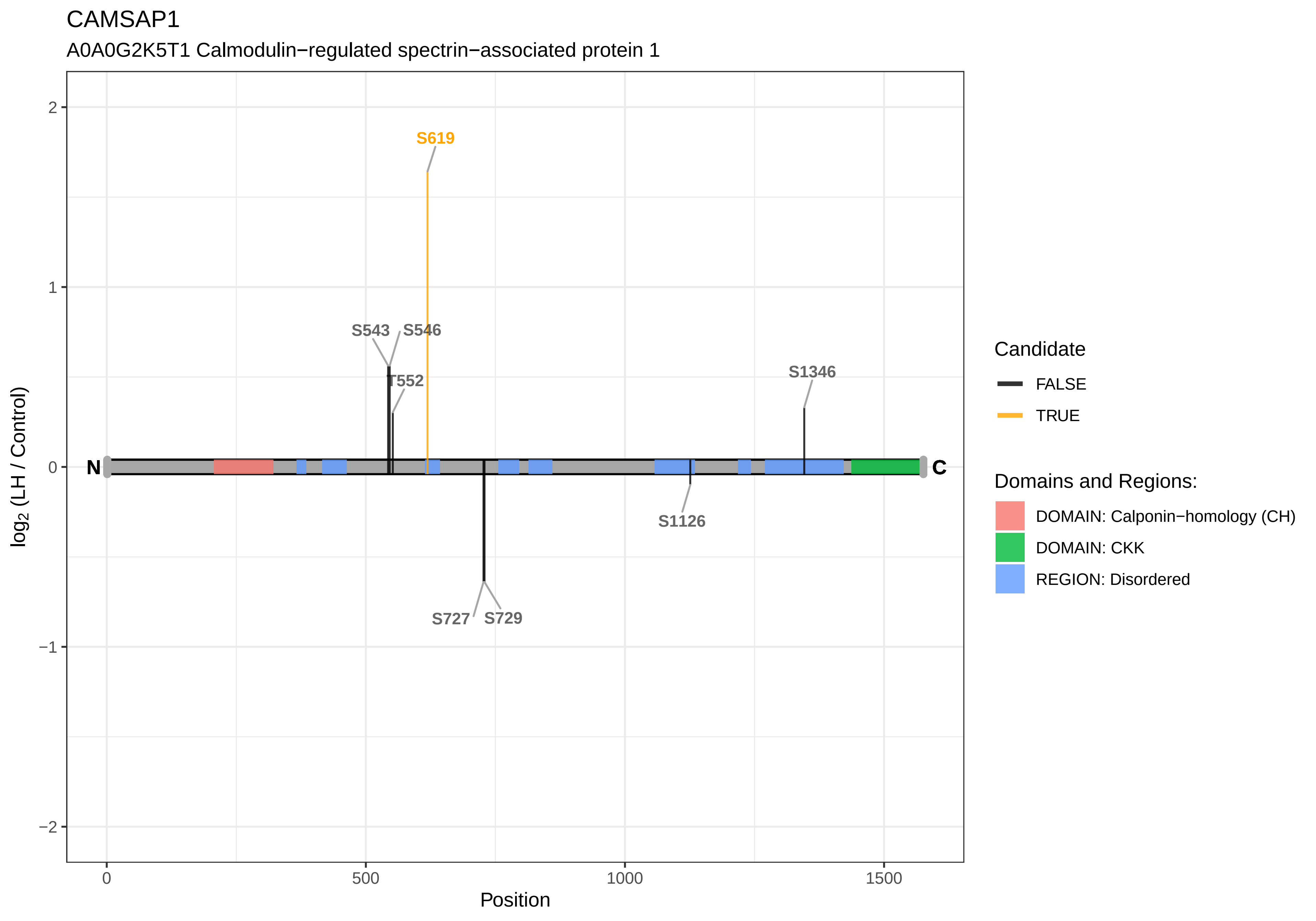
CAMSAP1
A0A0G2K5T1 Calmodulin−regulated spectrin−associated protein 1
S619
S543
S546
T552
S1346
N
C
S1126
S727
S729
2
1
0
−1
−2
0
500
1000
1500
Position
log2 (LH / Control)
Candidate
FALSE
TRUE
Domains and Regions:
DOMAIN: Calponin−homology (CH)
DOMAIN: CKK
REGION: Disordered

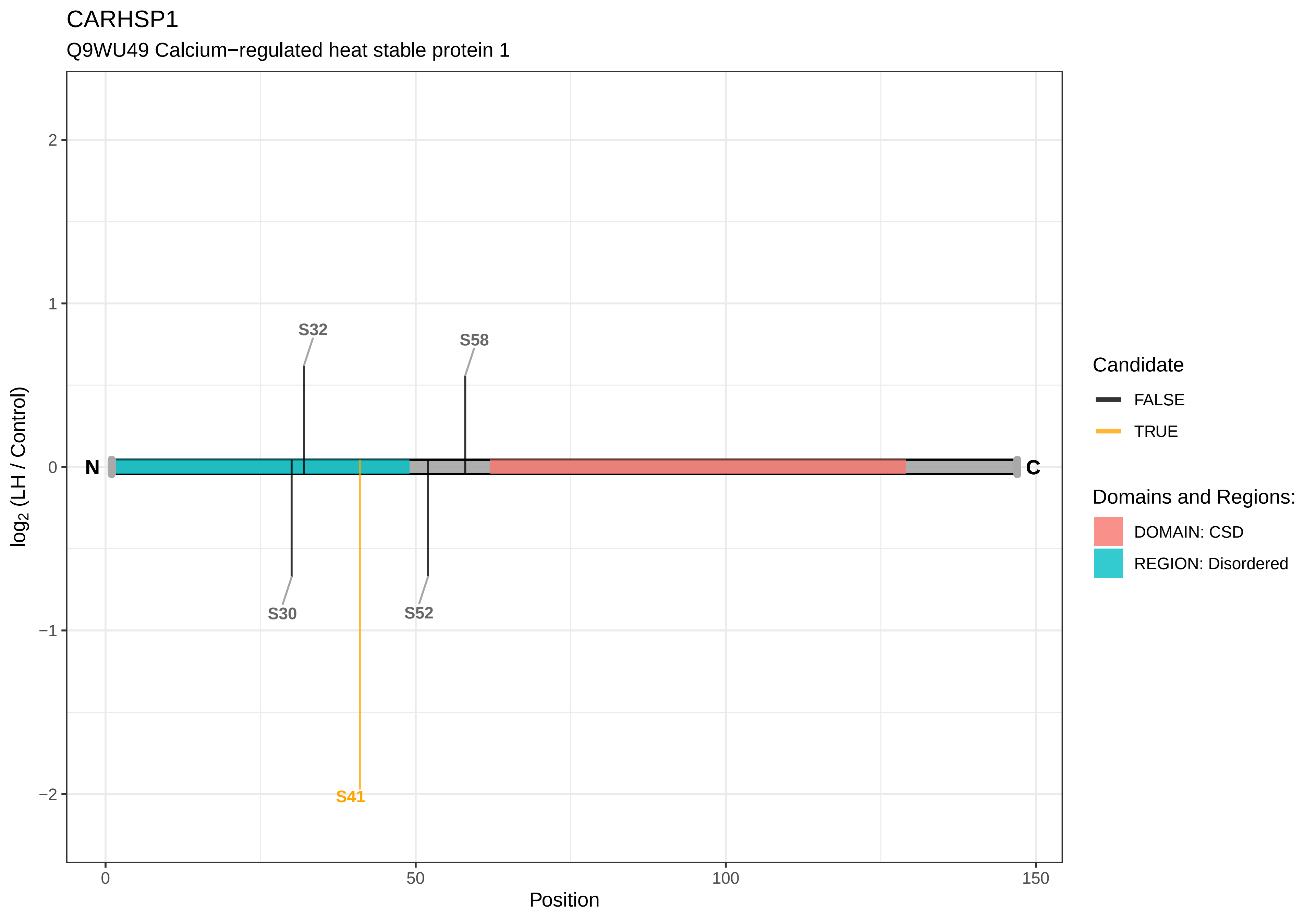
CARHSP1
Q9WU49 Calcium−regulated heat stable protein 1
S32
S58
S30
S52
S41
N
C
2
1
0
−1
−2
log2 (LH / Control)
0
50
100
150
Position
Candidate
FALSE
TRUE
Domains and Regions:
DOMAIN: CSD
REGION: Disordered

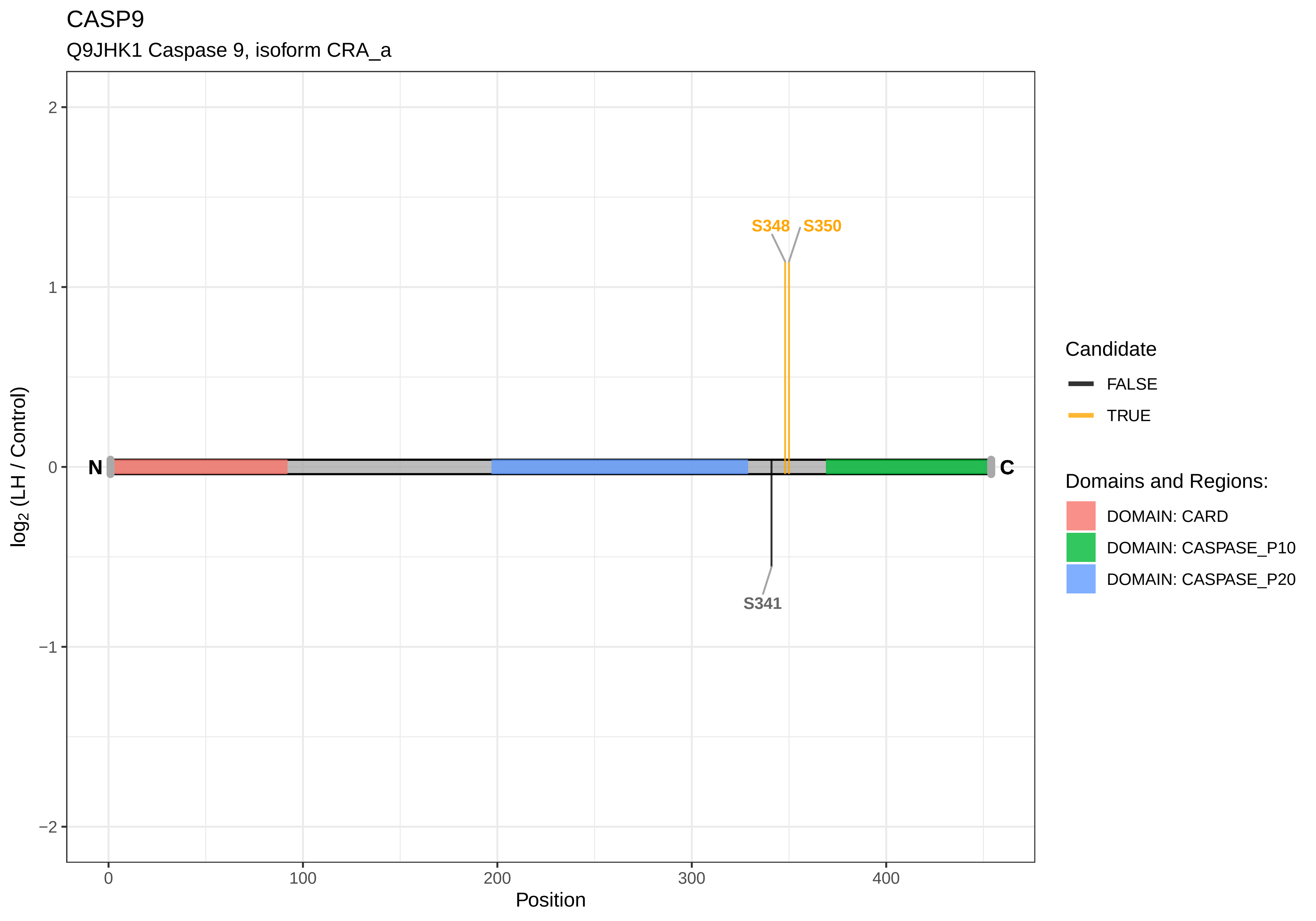
CASP9
Q9JHK1 Caspase 9, isoform CRA_a
S348
S350
S341
N
C
log2 (LH / Control)
Position
0
100
200
300
400
2
1
0
−1
−2
Candidate
FALSE
TRUE
Domains and Regions:
DOMAIN: CARD
DOMAIN: CASPASE_P10
DOMAIN: CASPASE_P20

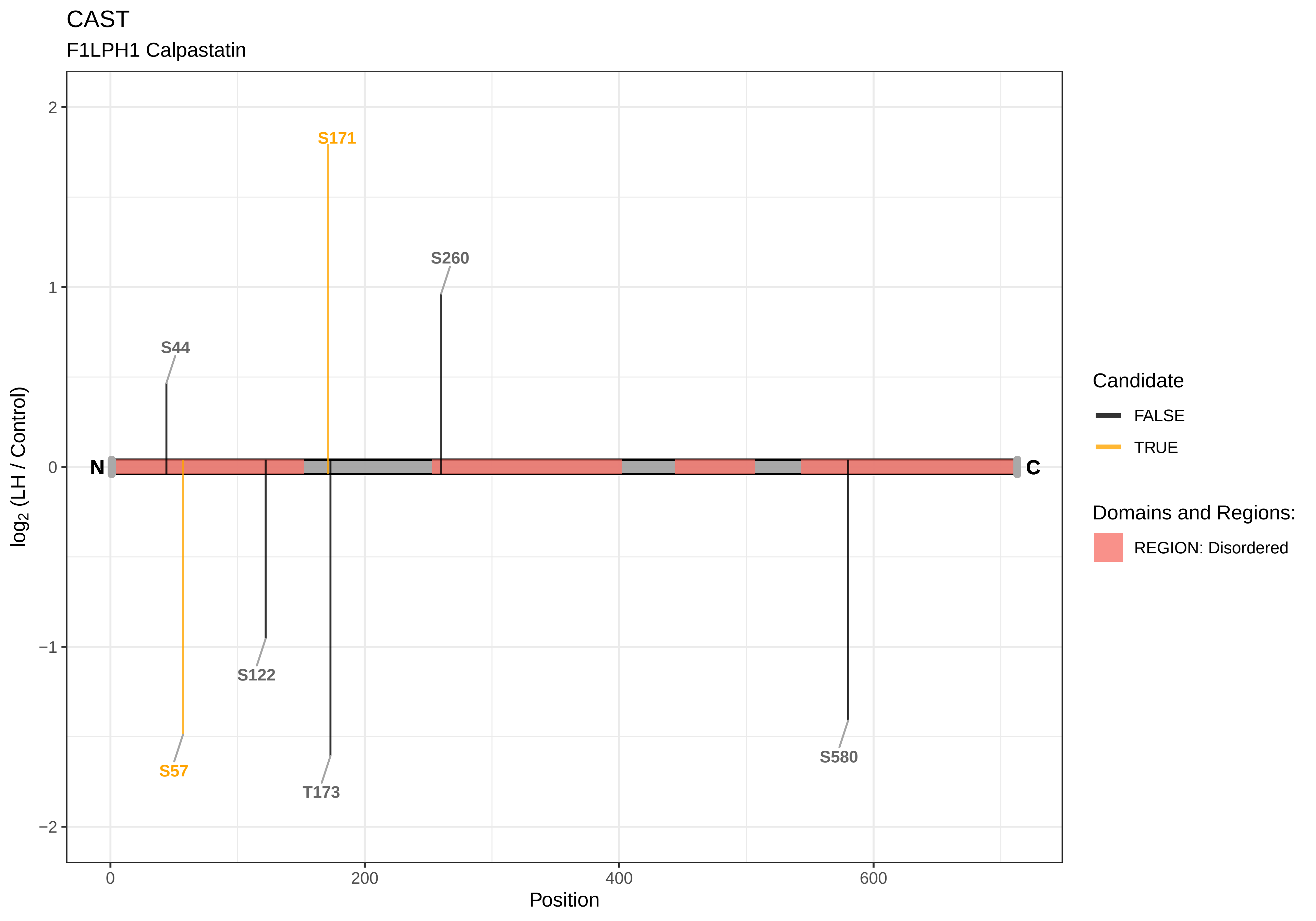
CAST
F1LPH1 Calpastatin
S171
S260
S44
N
C
S122
S57
T173
S580
log2 (LH / Control)
2
1
0
−1
−2
0
200
400
600
Position
Candidate
FALSE
TRUE
Domains and Regions:
REGION: Disordered

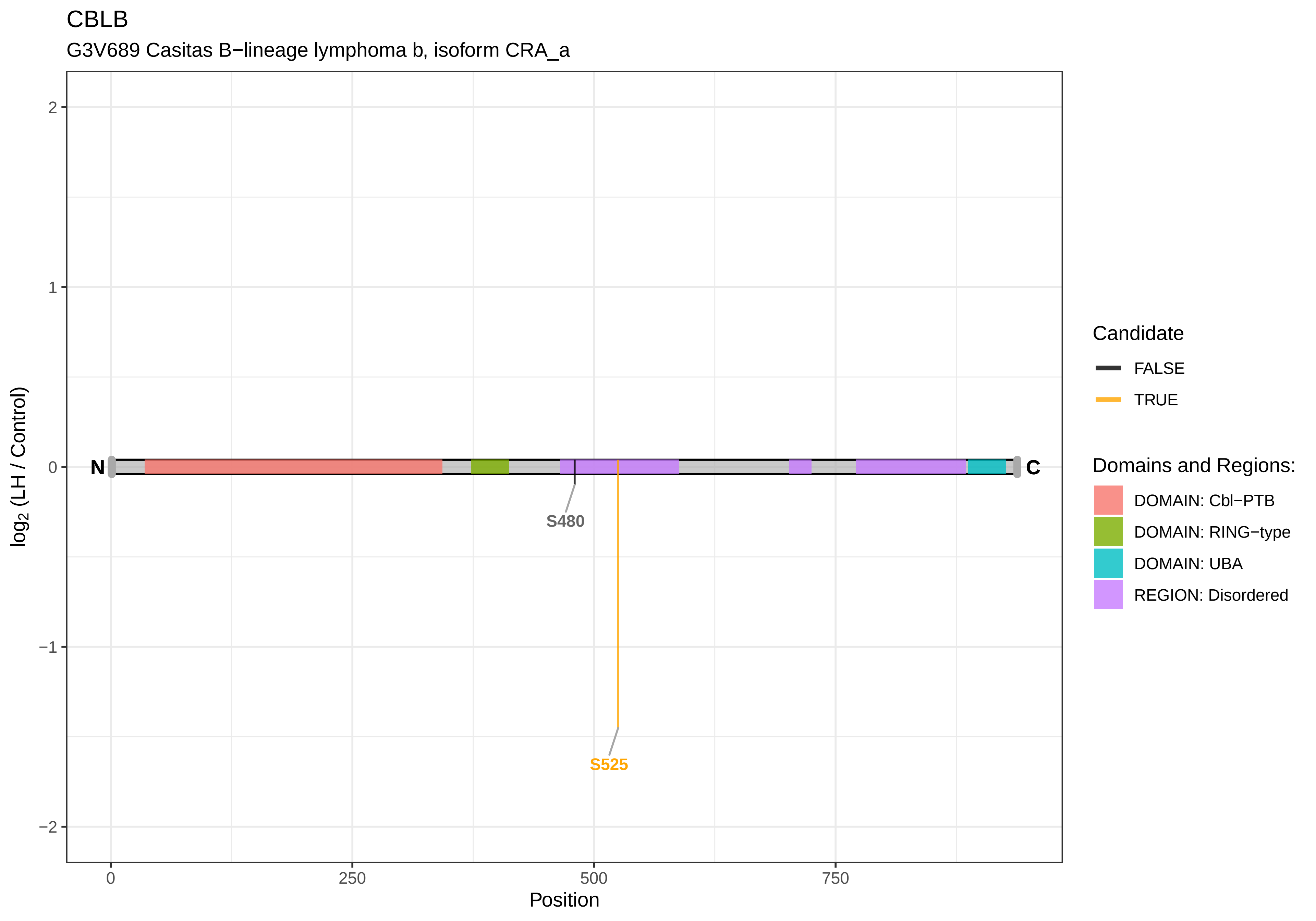
CBLB
G3V689 Casitas B−lineage lymphoma b, isoform CRA_a
log2 (LH / Control)
2
1
0
−1
−2
N
C
S480
S525
0
250
500
750
Position
Candidate
FALSE
TRUE
Domains and Regions:
DOMAIN: Cbl−PTB
DOMAIN: RING−type
DOMAIN: UBA
REGION: Disordered

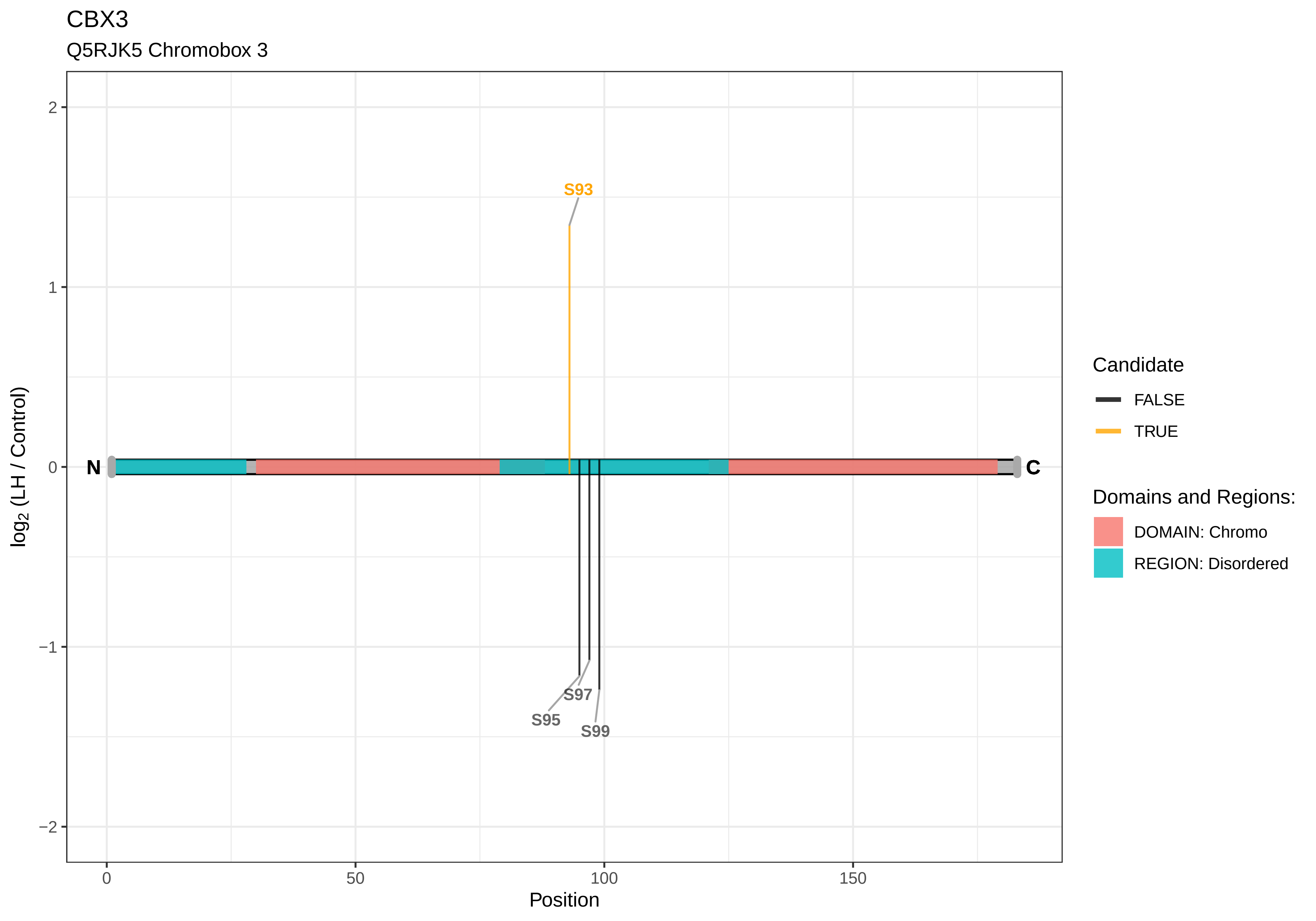
CBX3
Q5RJK5 Chromobox 3
S93
S95
S97
S99
N
C
2
1
0
−1
−2
0
50
100
150
Position
log2 (LH / Control)
Candidate
FALSE
TRUE
Domains and Regions:
DOMAIN: Chromo
REGION: Disordered

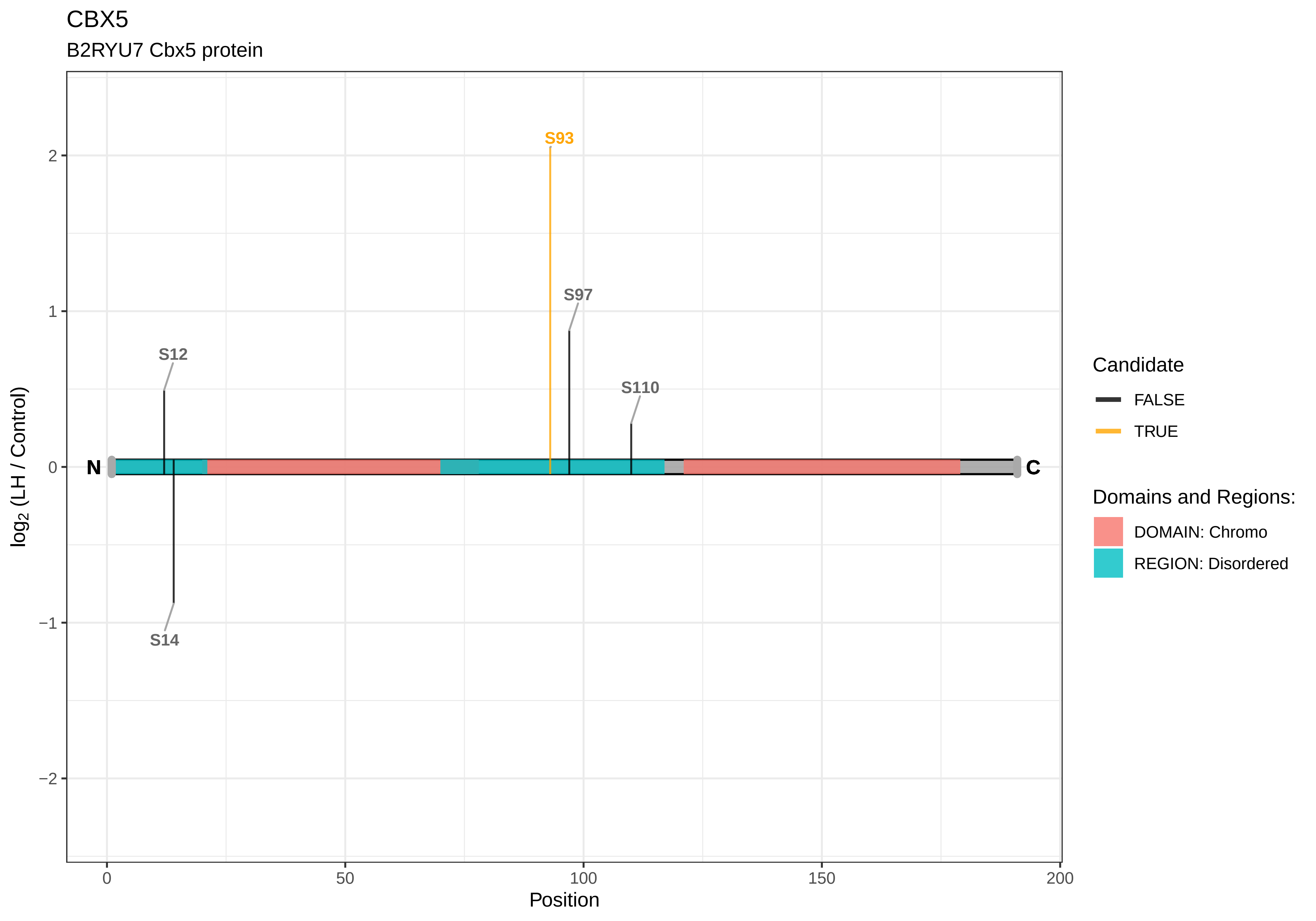
CBX5
B2RYU7 Cbx5 protein
S93
S97
S12
S110
S14
N
C
log2 (LH / Control)
Position
0
50
100
150
200
2
1
0
−1
−2
Candidate
FALSE
TRUE
Domains and Regions:
DOMAIN: Chromo
REGION: Disordered

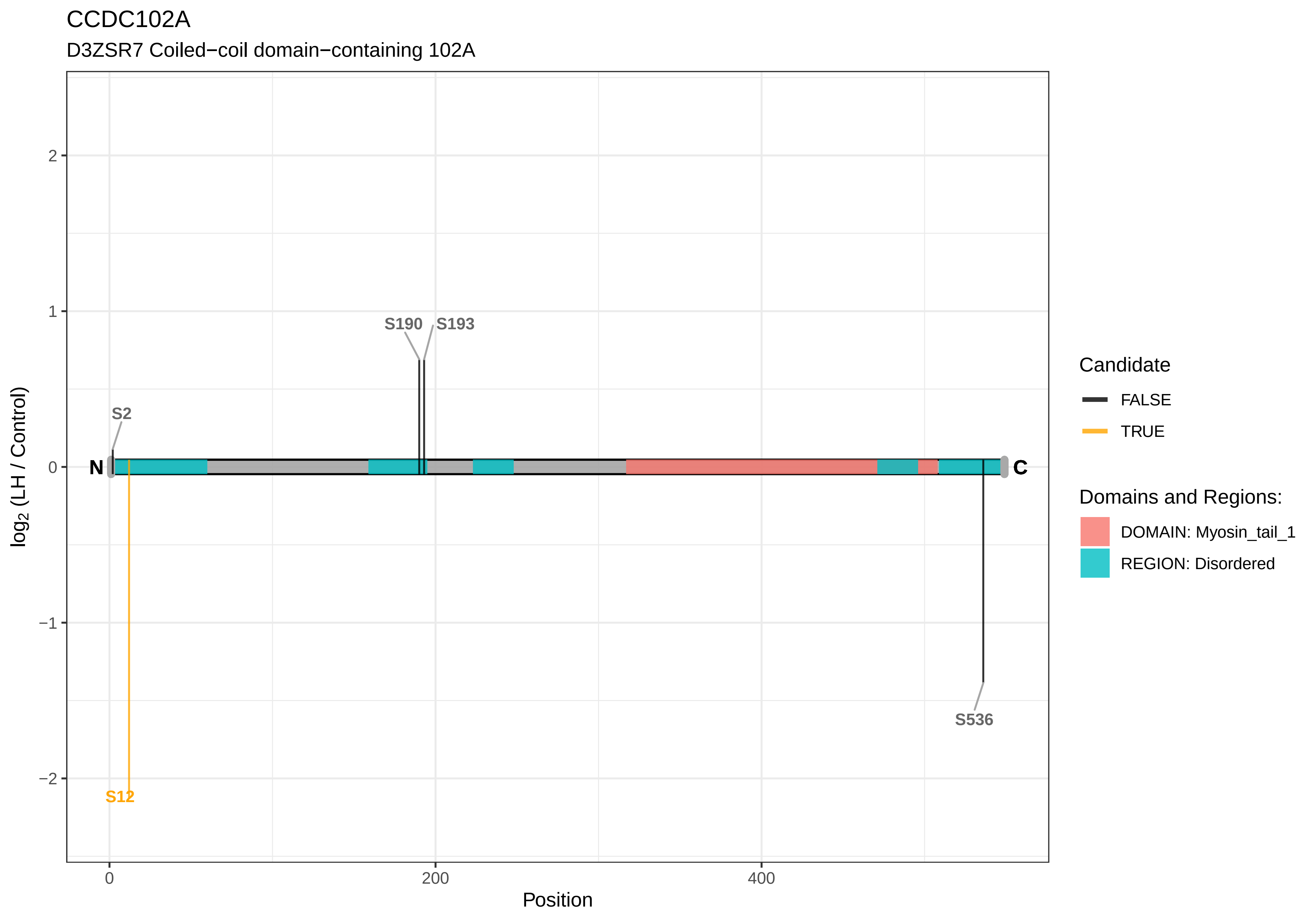
CCDC102A
D3ZSR7 Coiled−coil domain−containing 102A
S190
S193
S2
N
C
S536
S12
2
1
0
−1
−2
log2 (LH / Control)
0
200
400
Position
Candidate
FALSE
TRUE
Domains and Regions:
DOMAIN: Myosin_tail_1
REGION: Disordered

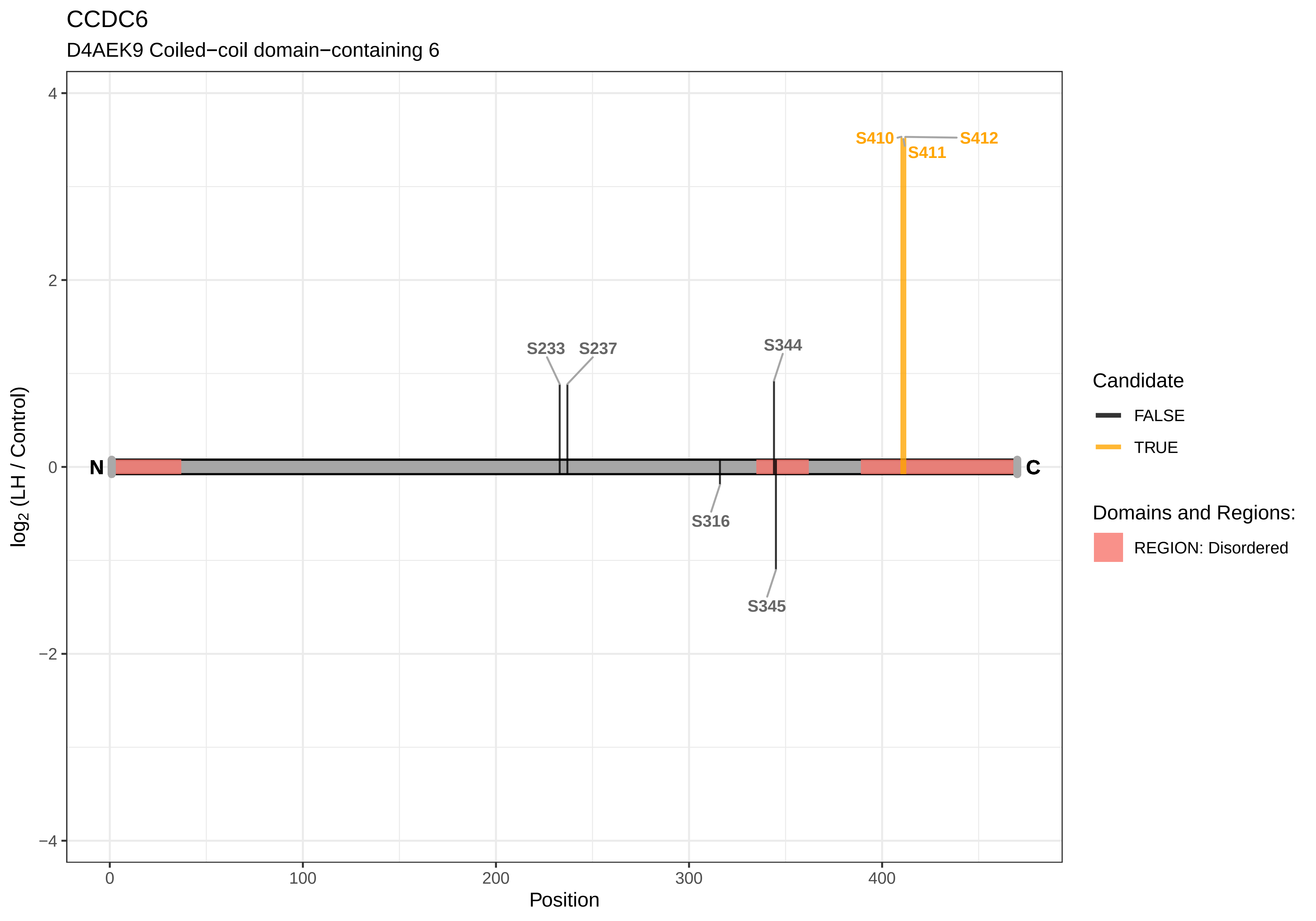
CCDC6
D4AEK9 Coiled−coil domain−containing 6
log2 (LH / Control)
Position
N
C
S233
S237
S316
S344
S345
S410
S411
S412
4
2
0
−2
−4
0
100
200
300
400
Candidate
FALSE
TRUE
Domains and Regions:
REGION: Disordered

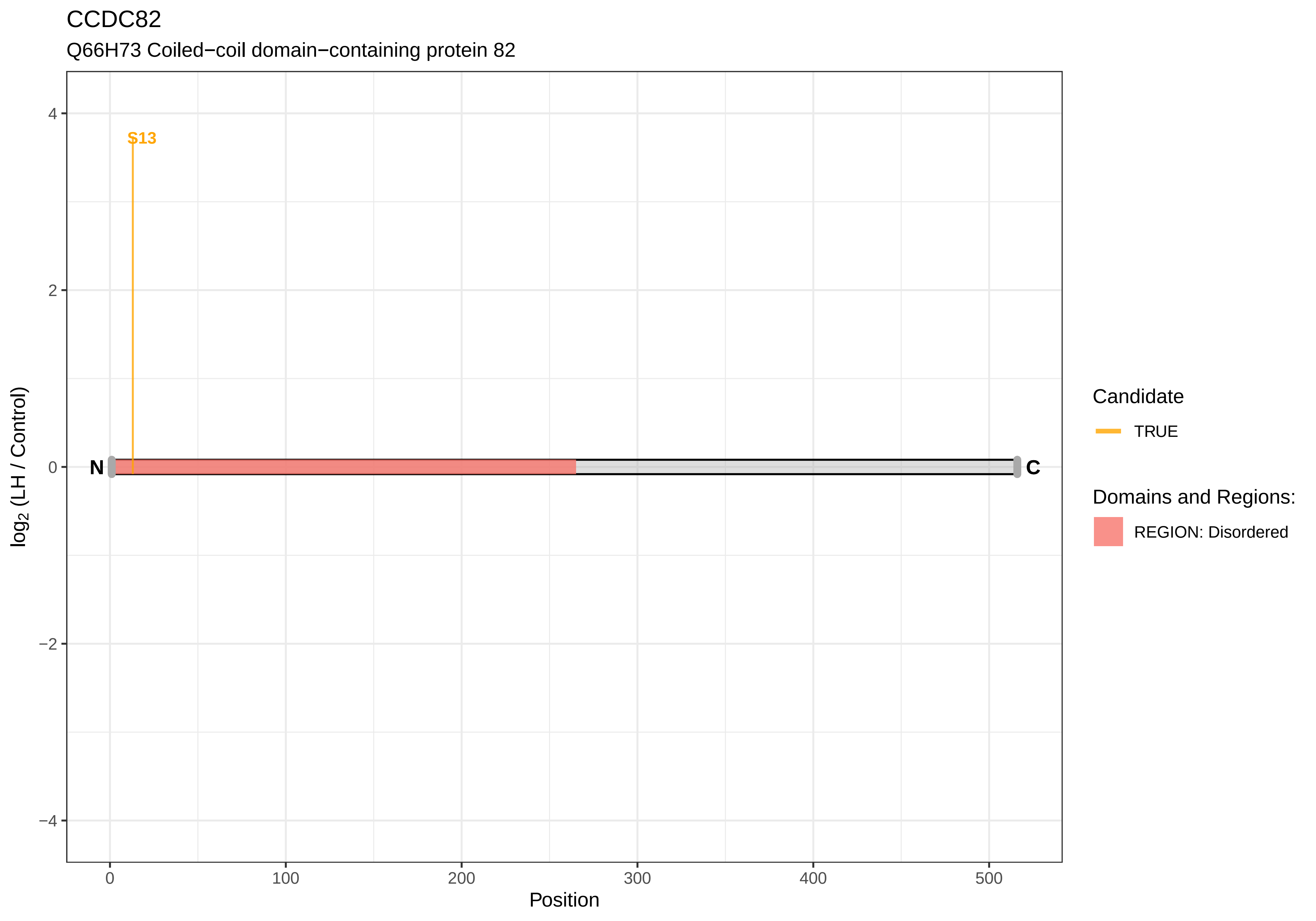
CCDC82
Q66H73 Coiled−coil domain−containing protein 82
S13
N
C
4
2
0
−2
−4
log2 (LH / Control)
0
100
200
300
400
500
Position
Candidate
TRUE
Domains and Regions:
REGION: Disordered

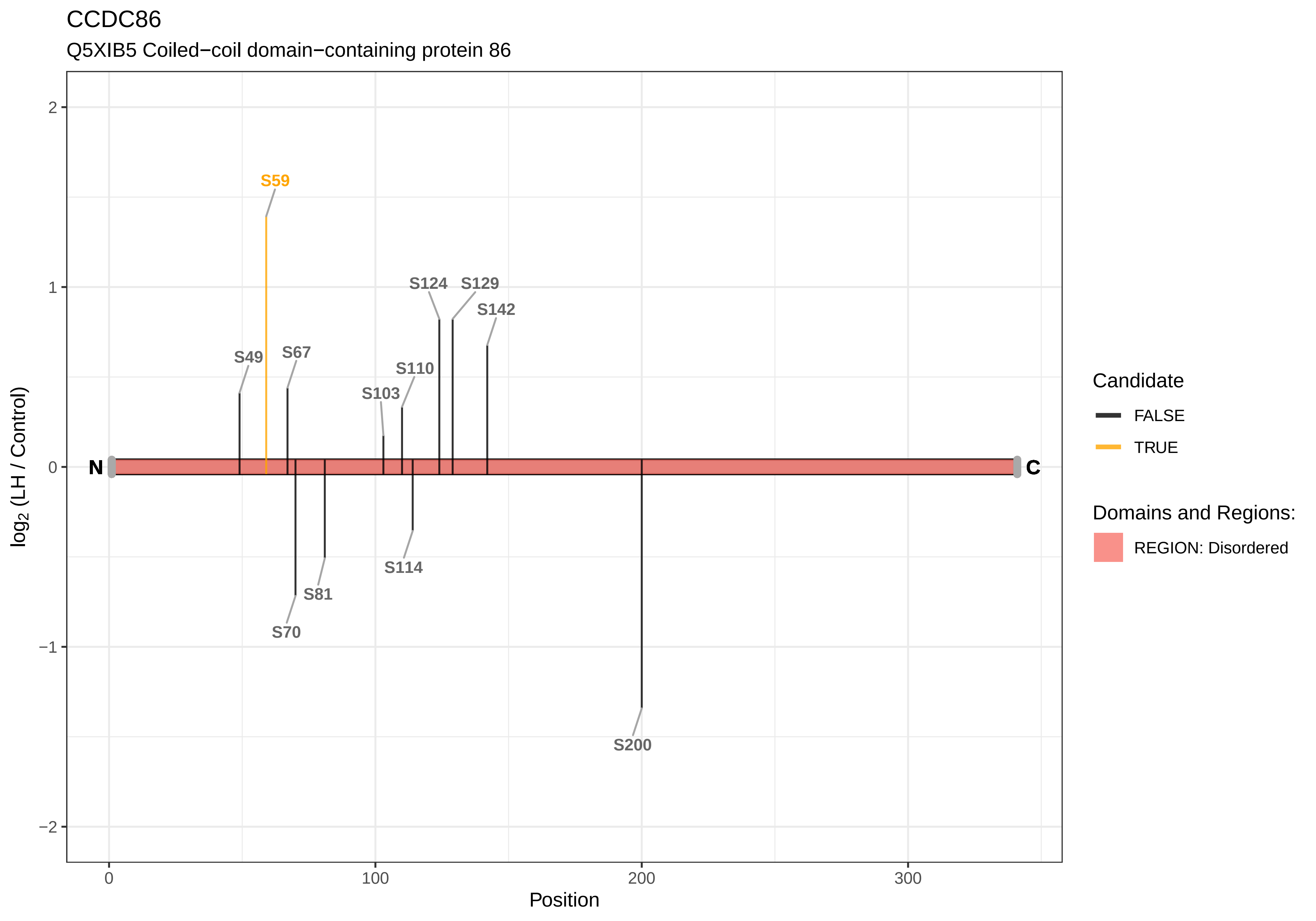

CCDC86
Q5XIB5 Coiled−coil domain−containing protein 86
S59
S124
S129
S142
S49
S67
S110
S103
S114
S81
S70
S200
N
C
2
1
0
−1
−2
0
100
200
300
Position
log2 (LH / Control)
Candidate
FALSE
TRUE
Domains and Regions:
REGION: Disordered

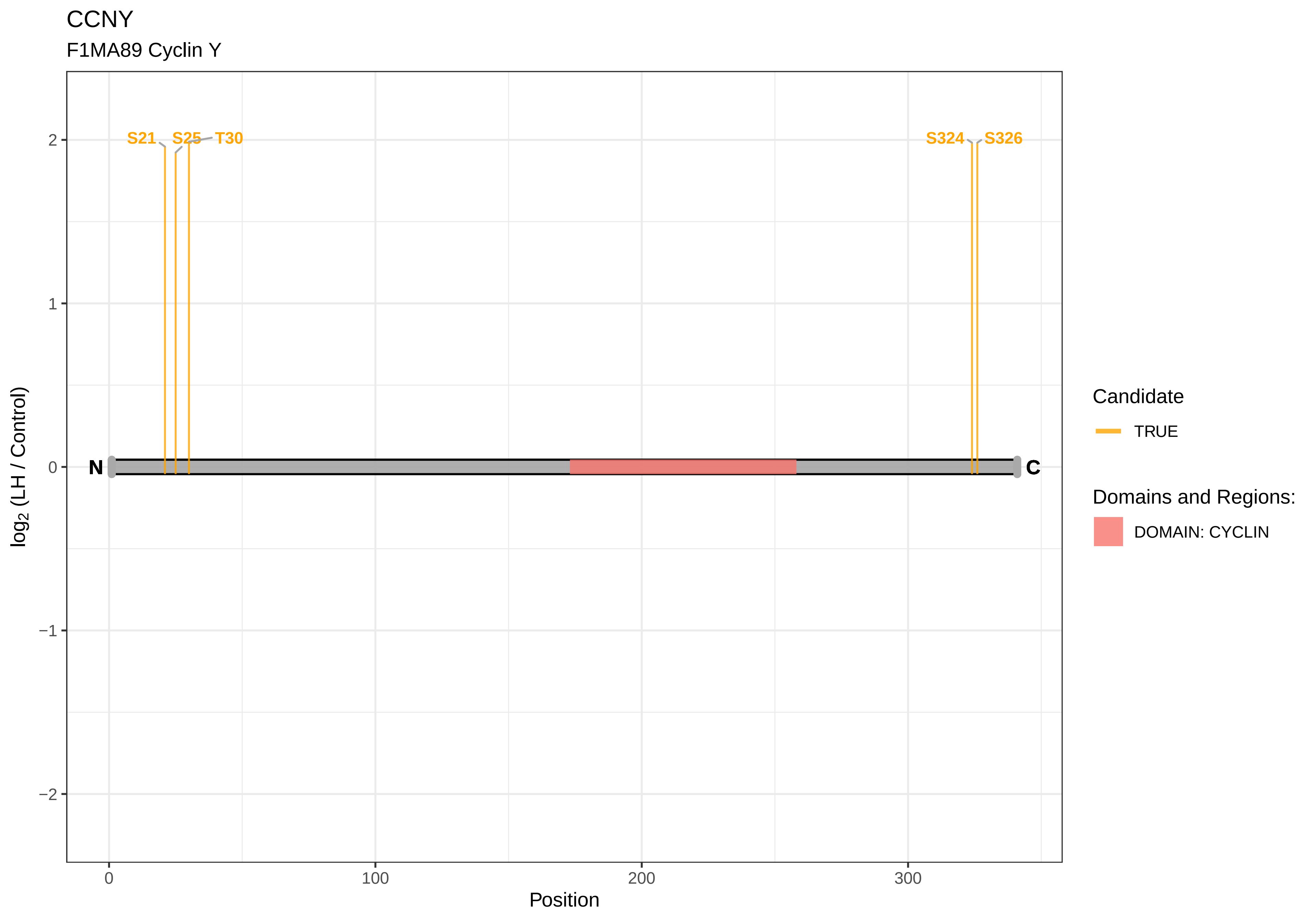

CCNY
F1MA89 Cyclin Y
S21
S25
T30
S324
S326
2
1
0
−1
−2
N
C
log2 (LH / Control)
0
100
200
300
Position
Candidate
TRUE
Domains and Regions:
DOMAIN: CYCLIN

CCS

A0A0G2JSZ9 Copper chaperone for superoxide dismutase

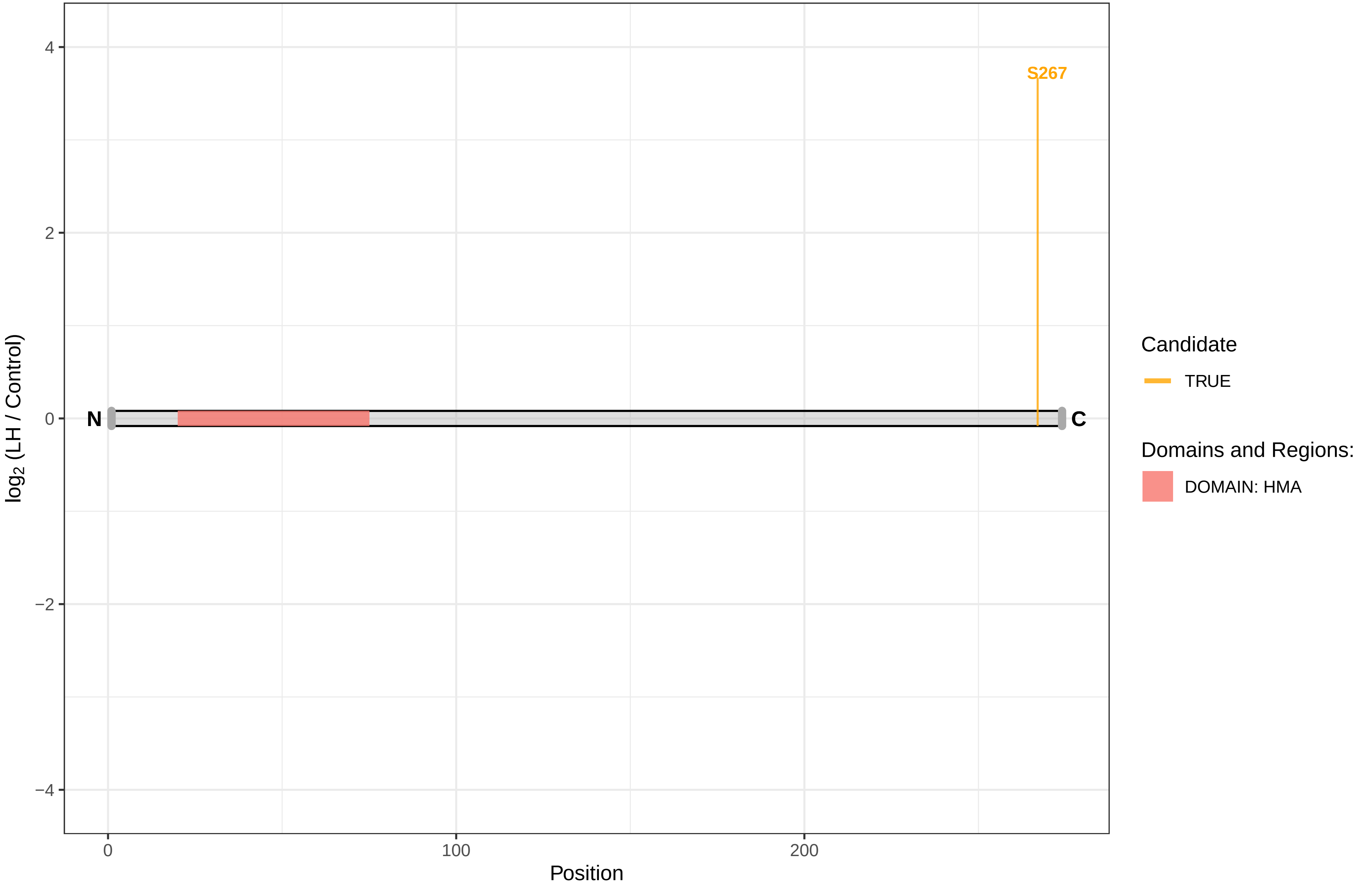

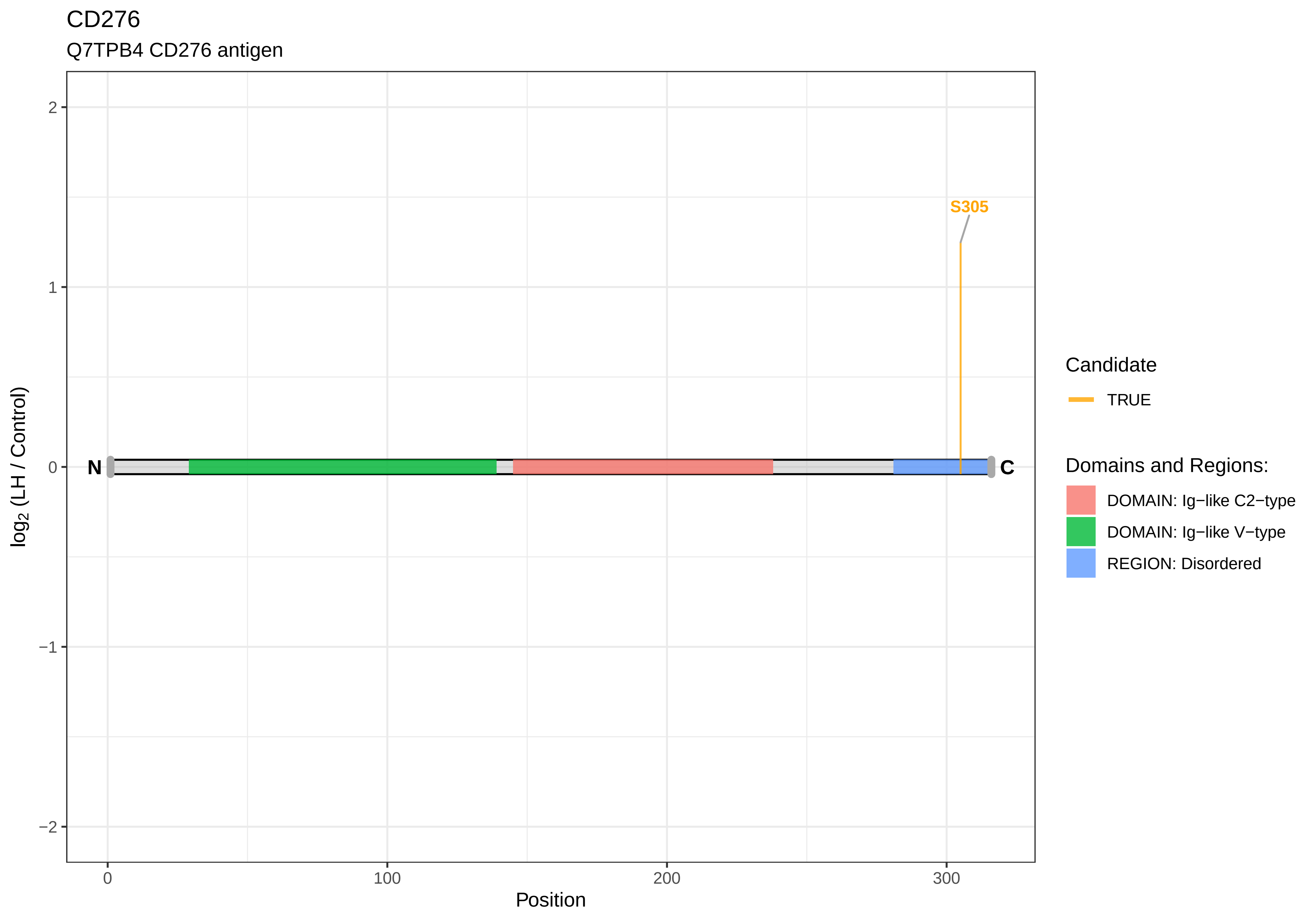
CD276
Q7TPB4 CD276 antigen
S305
N
C
2
1
0
−1
−2
0
100
200
300
Position
log2 (LH / Control)
Candidate
TRUE
Domains and Regions:
DOMAIN: Ig−like C2−type
DOMAIN: Ig−like V−type
REGION: Disordered

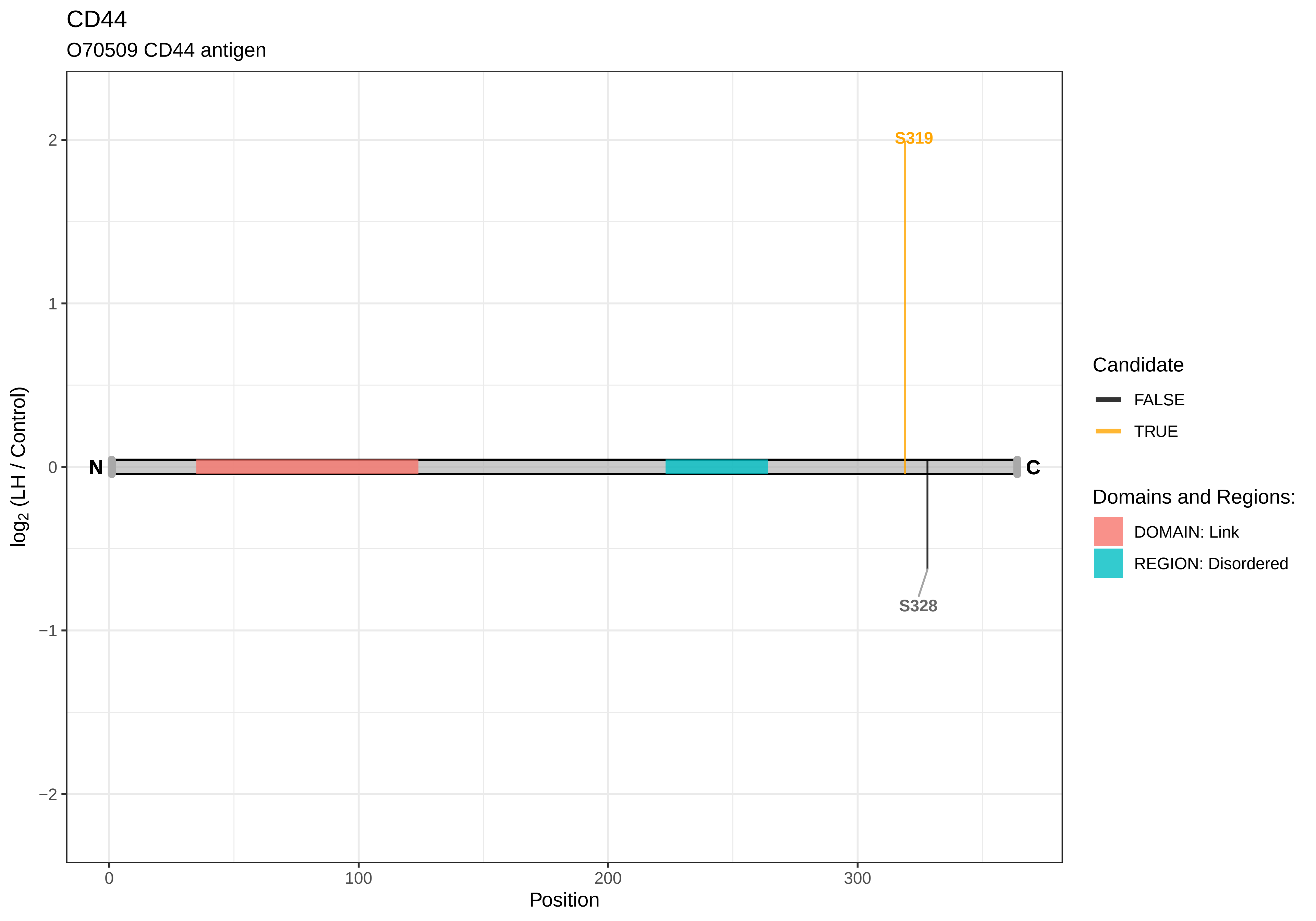
CD44
O70509 CD44 antigen
S319
S328
N
C
2
1
0
−1
−2
0
100
200
300
Position
log2 (LH / Control)
Candidate
FALSE
TRUE
Domains and Regions:
DOMAIN: Link
REGION: Disordered

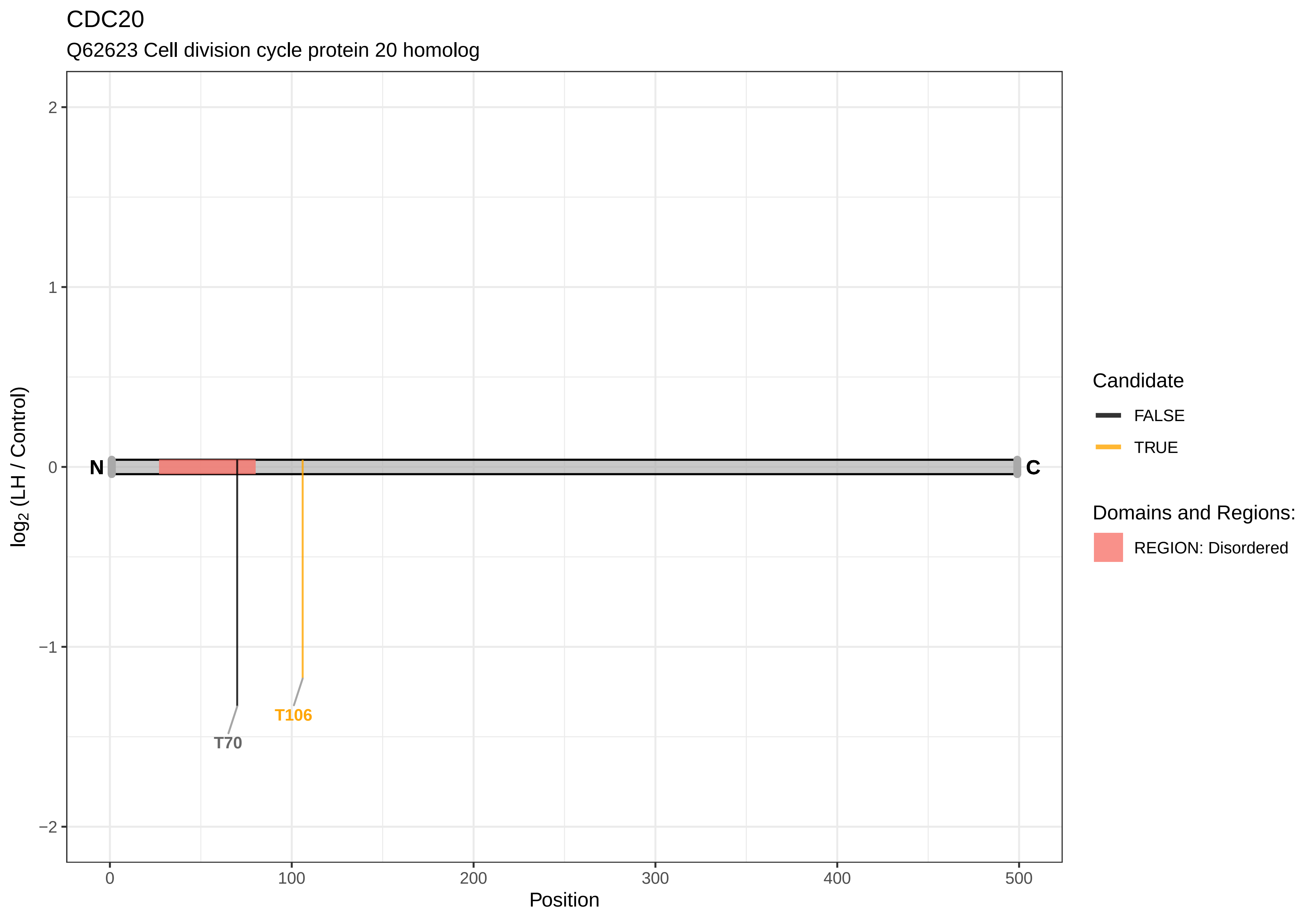

CDC20
Q62623 Cell division cycle protein 20 homolog
log2 (LH / Control)
2
1
0
−1
−2
N
C
T70
T106
0
100
200
300
400
500
Position
Candidate
FALSE
TRUE
Domains and Regions:
REGION: Disordered

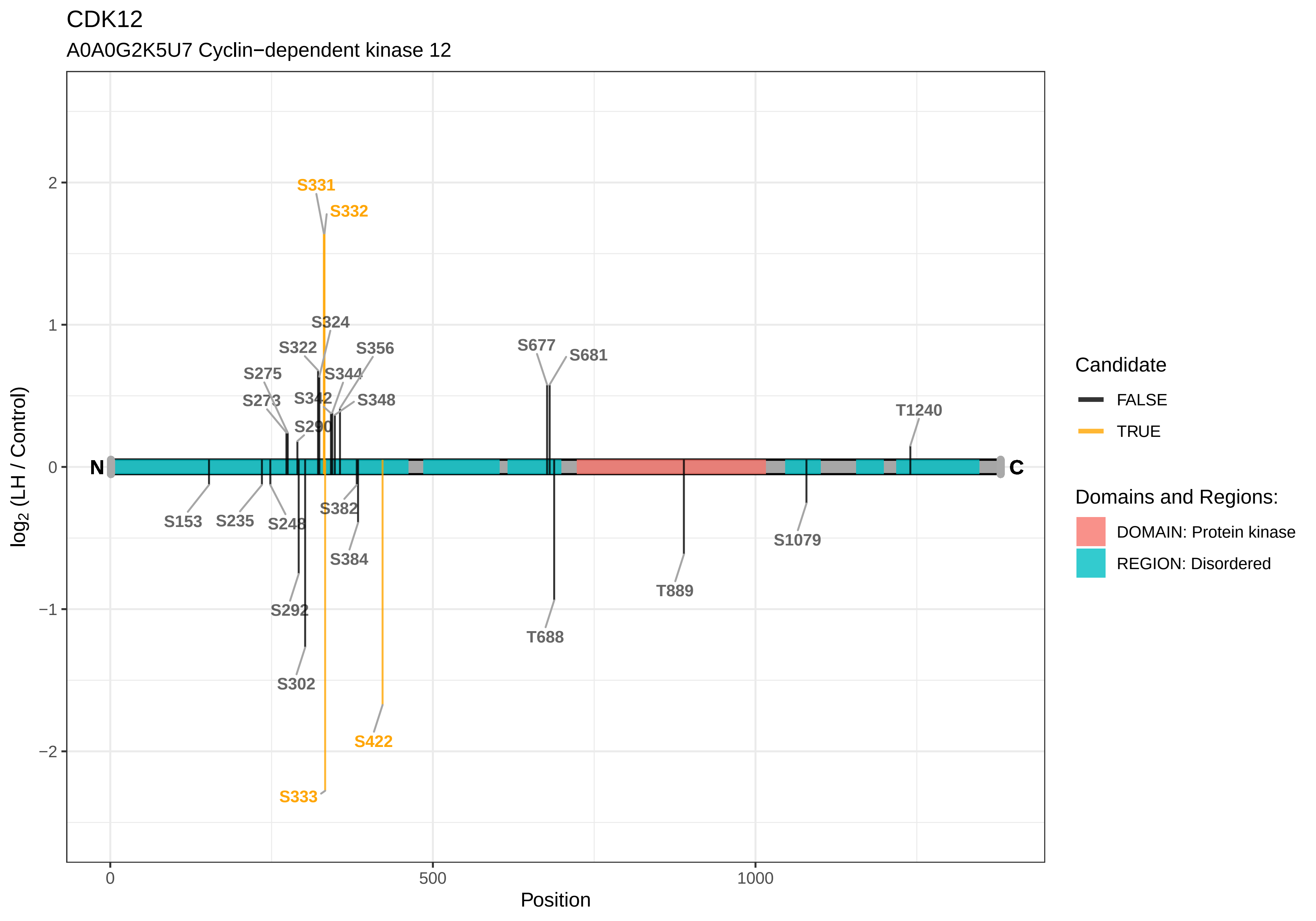

CDK12
A0A0G2K5U7 Cyclin−dependent kinase 12
S331
S332
S324
S322
S356
S677
S681
S275
S344
S273
S342
S348
T1240
S290
N
C
S153
S235
S248
S382
S384
S1079
T889
S292
T688
S302
S422
S333
2
1
0
−1
−2
log2 (LH / Control)
0
500
1000
Position
Candidate
FALSE
TRUE
Domains and Regions:
DOMAIN: Protein kinase
REGION: Disordered

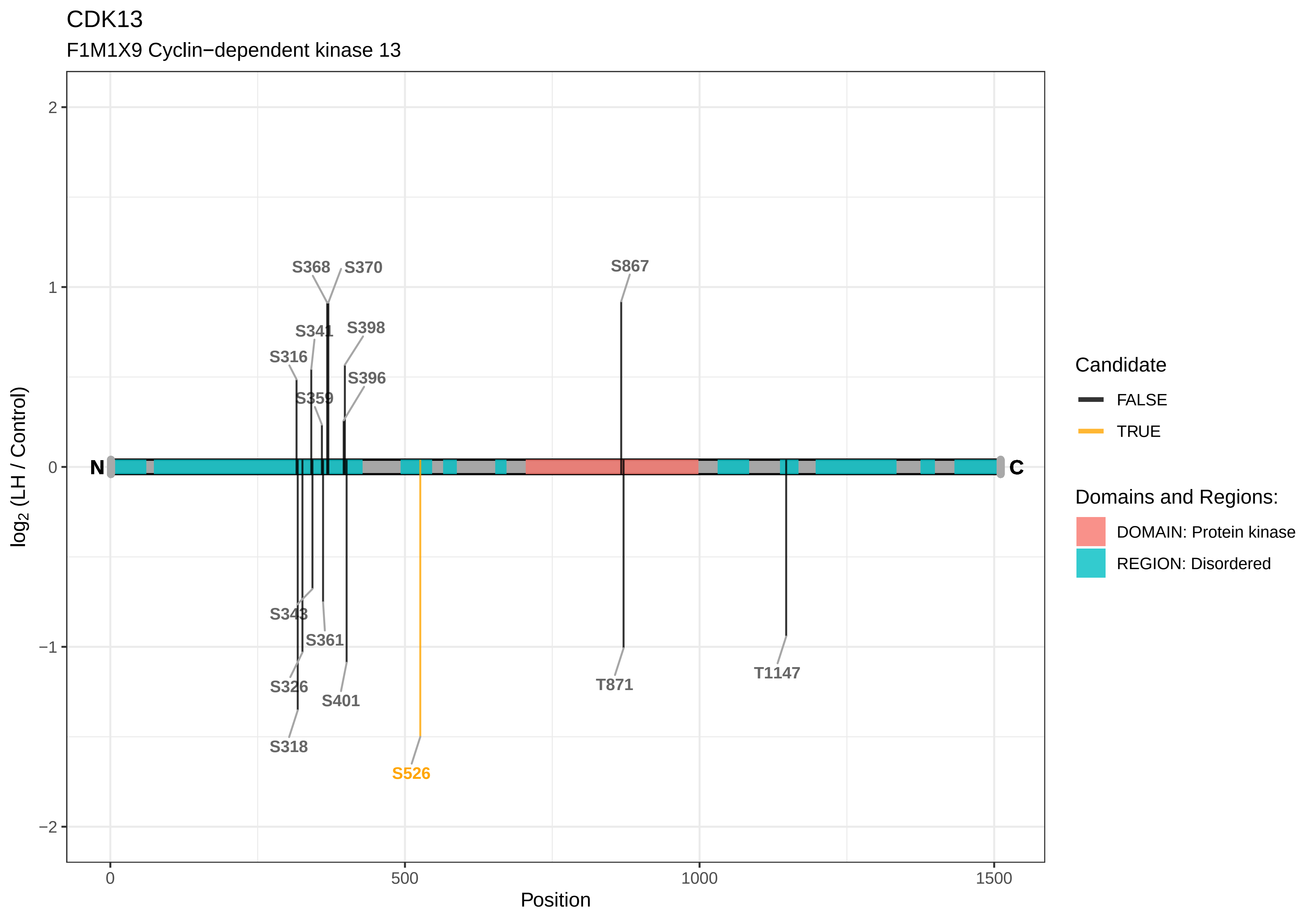

CDK13
F1M1X9 Cyclin−dependent kinase 13
S368
S370
S867
S341
S398
S316
S396
S359
N
C
S343
S361
S326
S401
T871
T1147
S318
S526
log2 (LH / Control)
2
1
0
−1
−2
0
500
1000
1500
Position
Candidate
FALSE
TRUE
Domains and Regions:
DOMAIN: Protein kinase
REGION: Disordered

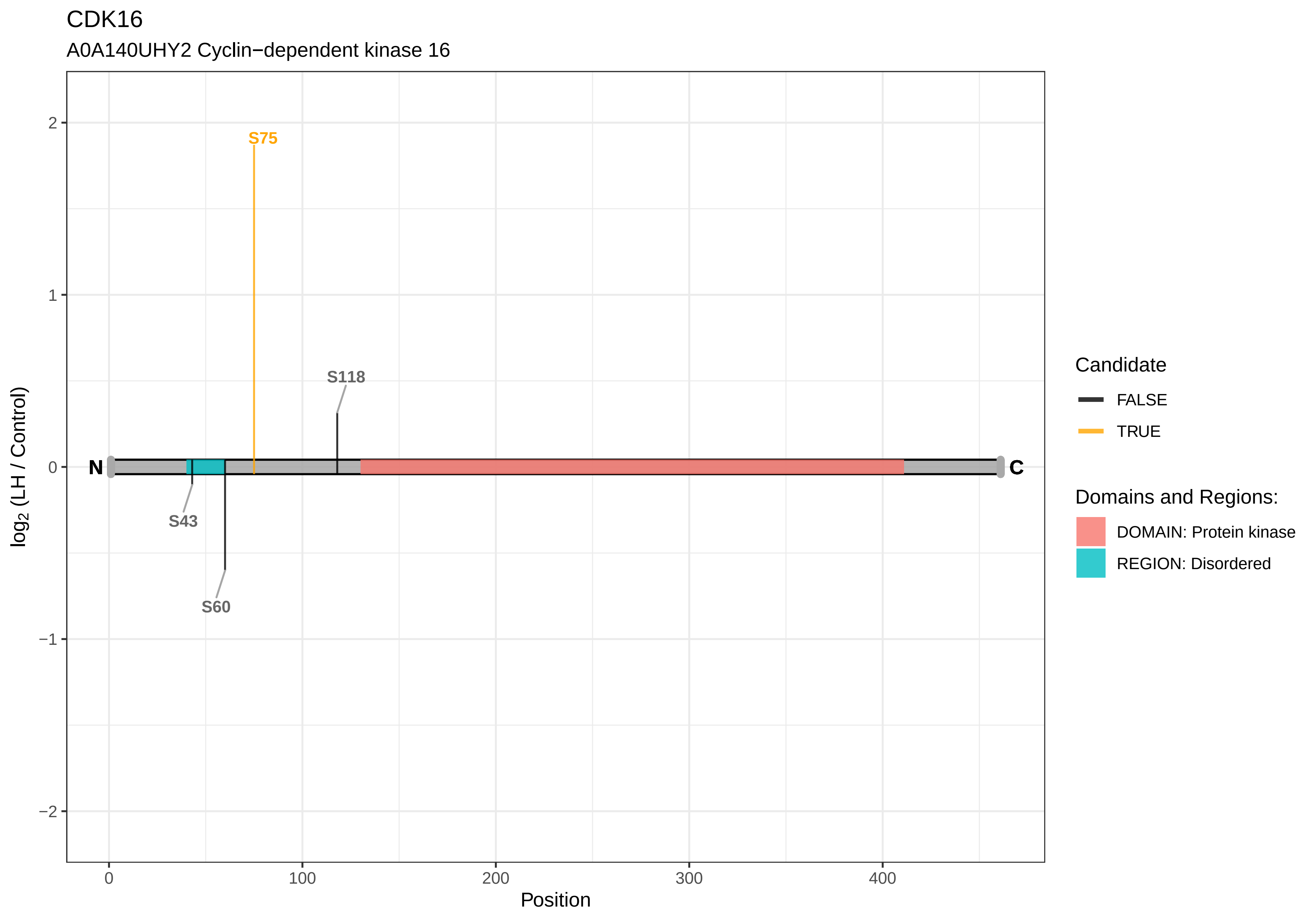

CDK16
A0A140UHY2 Cyclin−dependent kinase 16
S75
S118
S43
S60
N
C
2
1
0
−1
−2
0
100
200
300
400
Position
log2 (LH / Control)
Candidate
FALSE
TRUE
Domains and Regions:
DOMAIN: Protein kinase
REGION: Disordered

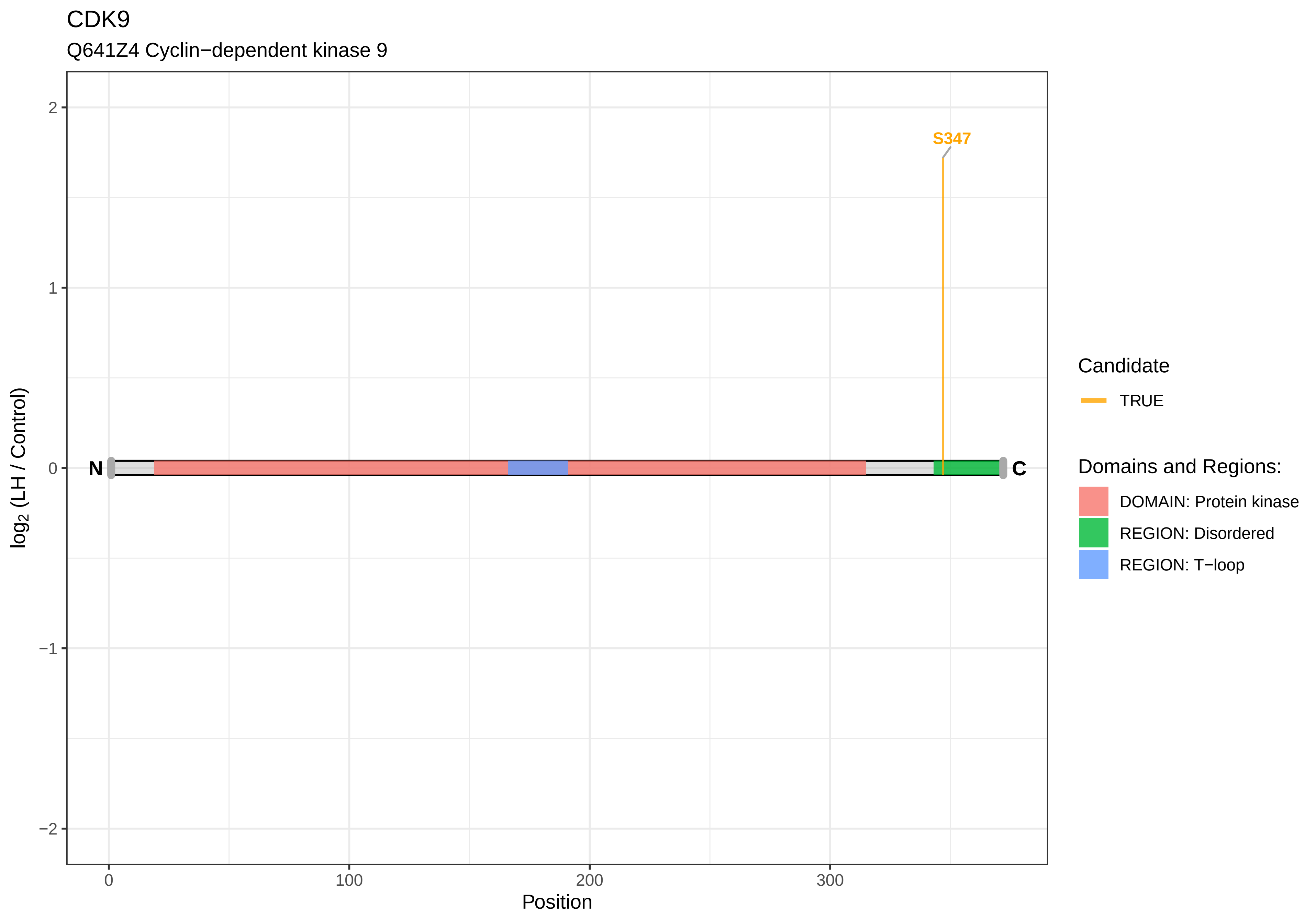

CDK9
Q641Z4 Cyclin−dependent kinase 9
S347
N
C
2
1
0
−1
−2
log2 (LH / Control)
0
100
200
300
Position
Candidate
TRUE
Domains and Regions:
DOMAIN: Protein kinase
REGION: Disordered
REGION: T−loop

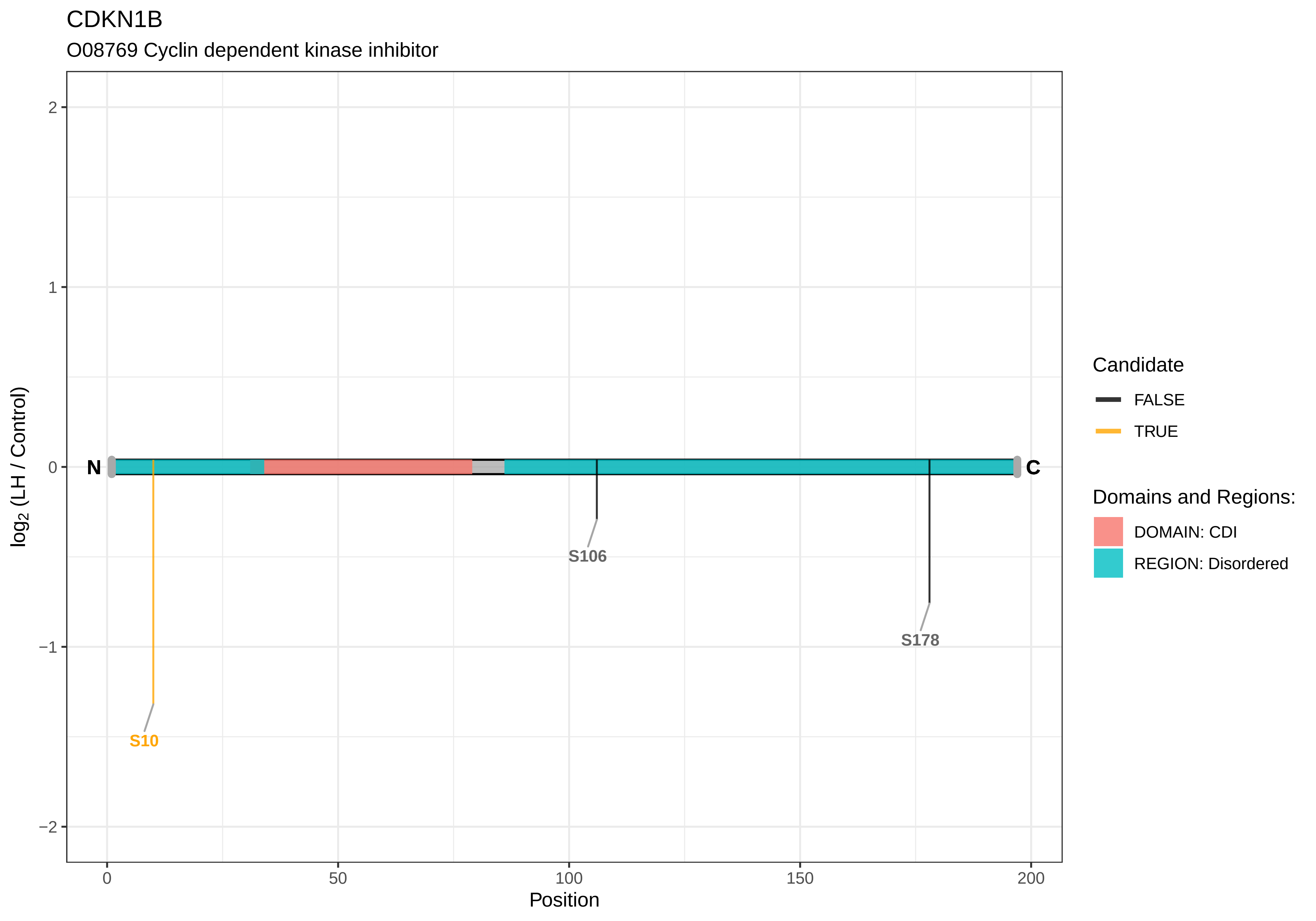
CDKN1B
O08769 Cyclin dependent kinase inhibitor
log2 (LH / Control)
2
1
0
−1
−2
N
C
S10
S106
S178
0
50
100
150
200
Position
Candidate
FALSE
TRUE
Domains and Regions:
DOMAIN: CDI
REGION: Disordered

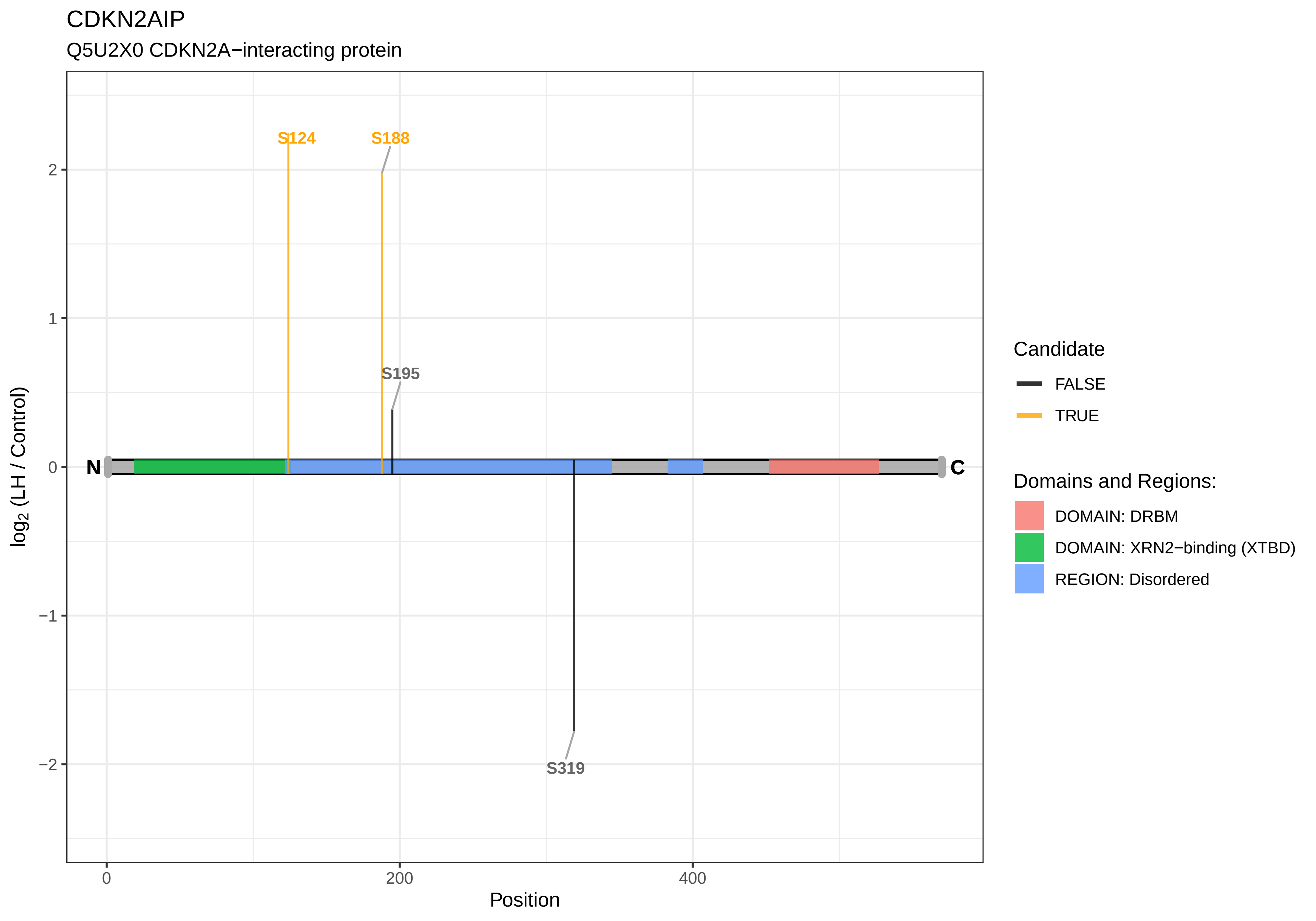

CDKN2AIP
Q5U2X0 CDKN2A−interacting protein
S124
S188
S195
S319
N
C
2
1
0
−1
−2
log2 (LH / Control)
0
200
400
Position
Candidate
FALSE
TRUE
Domains and Regions:
DOMAIN: DRBM
DOMAIN: XRN2−binding (XTBD)
REGION: Disordered

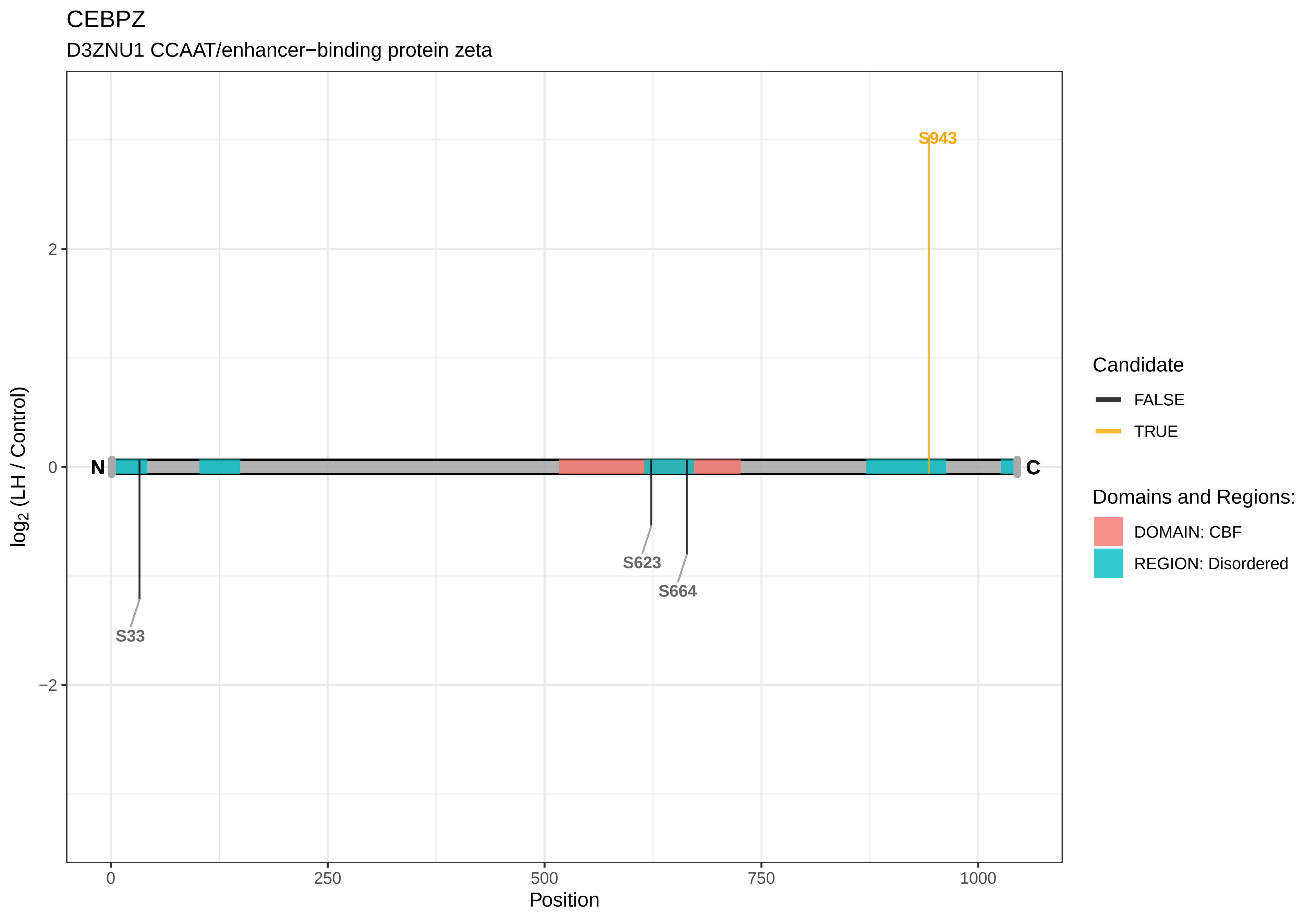
CEBPZ
D3ZNU1 CCAAT/enhancer−binding protein zeta
S943
S623
S664
S33
N
C
2
0
−2
0
250
500
750
1000
Position
log2 (LH / Control)
Candidate
FALSE
TRUE
Domains and Regions:
DOMAIN: CBF
REGION: Disordered

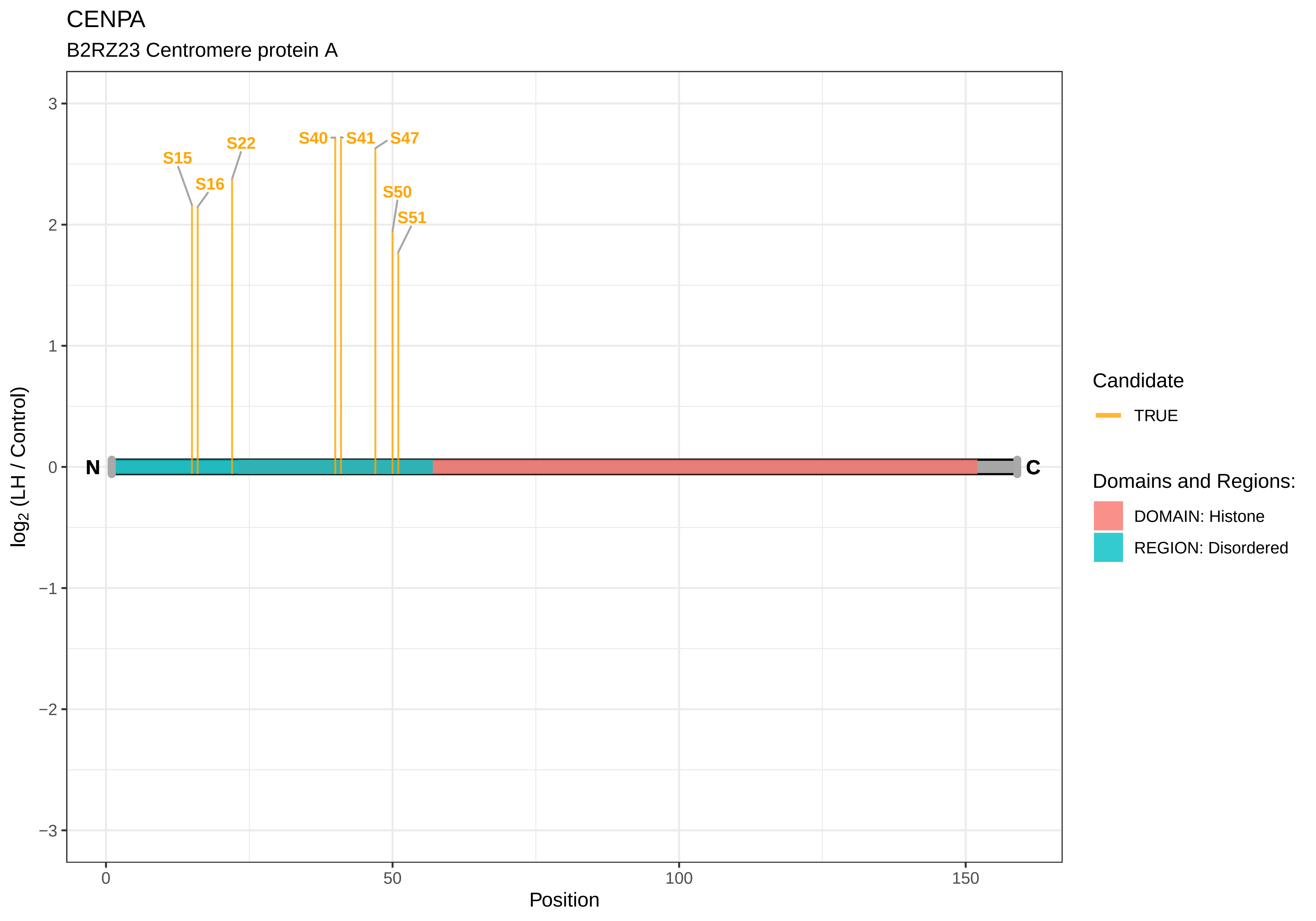

CENPA
B2RZ23 Centromere protein A
S15
S16
S22
S40
S41
S47
S50
S51
N
C
3
2
1
0
−1
−2
−3
0
50
100
150
Position
log2 (LH / Control)
Candidate
TRUE
Domains and Regions:
DOMAIN: Histone
REGION: Disordered

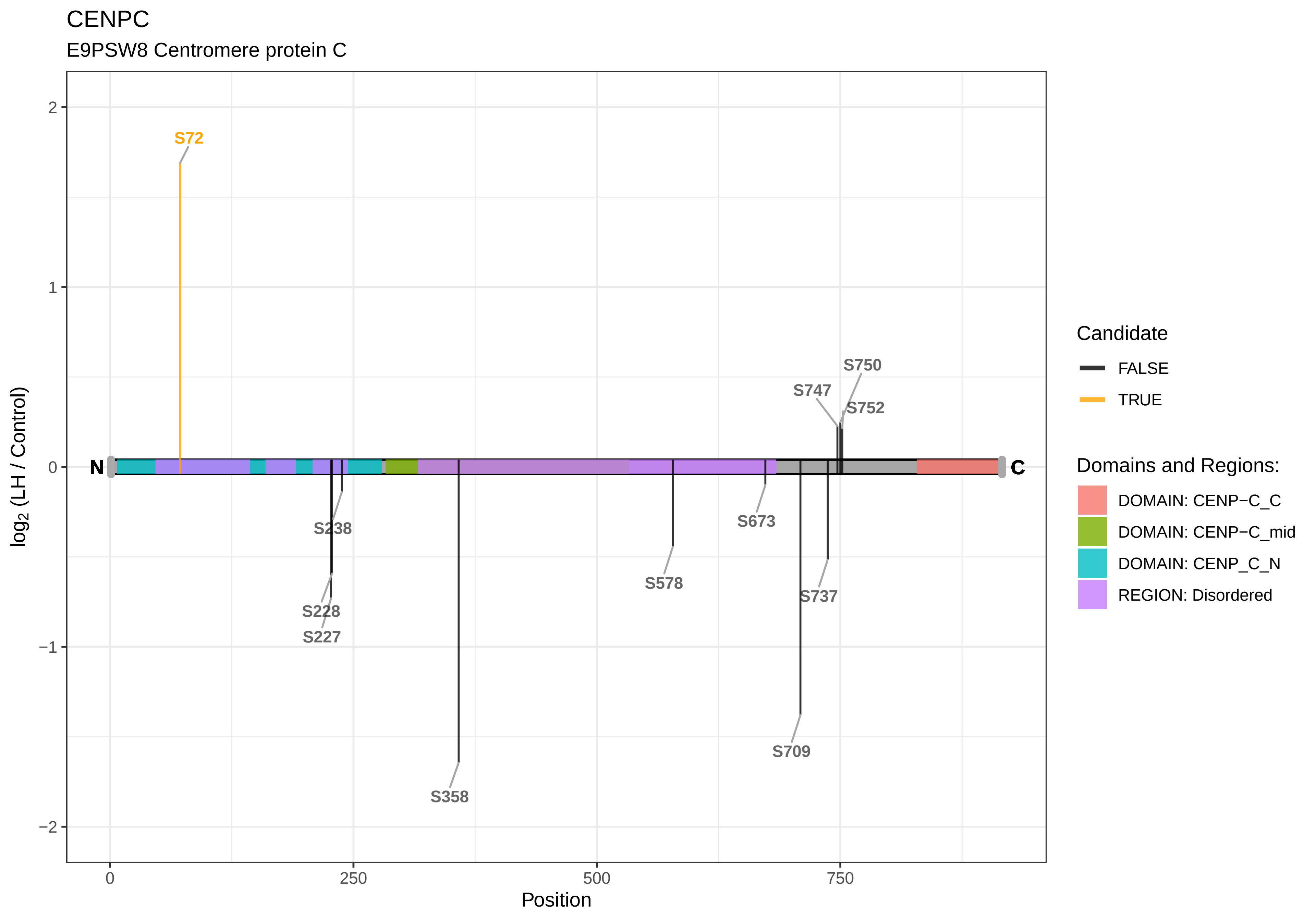
CENPC
E9PSW8 Centromere protein C
log2 (LH / Control)
Position
N
C
S72
S238
S228
S227
S358
S578
S673
S709
S737
S747
S750
S752
2
1
0
−1
−2
0
250
500
750
Candidate
FALSE
TRUE
Domains and Regions:
DOMAIN: CENP−C_C
DOMAIN: CENP−C_mid
DOMAIN: CENP_C_N
REGION: Disordered

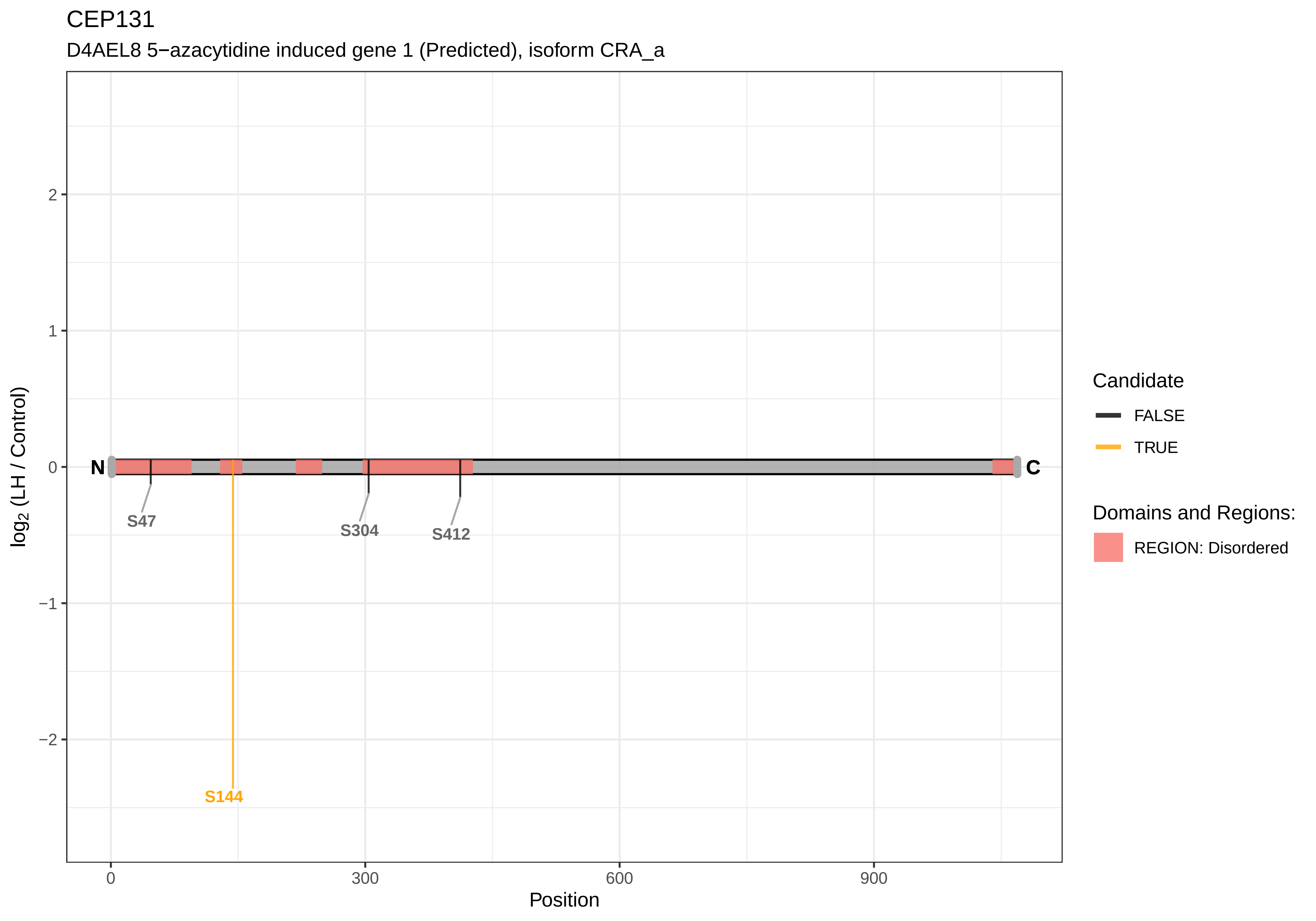
CEP131
D4AEL8 5−azacytidine induced gene 1 (Predicted), isoform CRA_a
log2 (LH / Control)
2
1
0
−1
−2
N
C
S47
S144
S304
S412
0
300
600
900
Position
Candidate
FALSE
TRUE
Domains and Regions:
REGION: Disordered

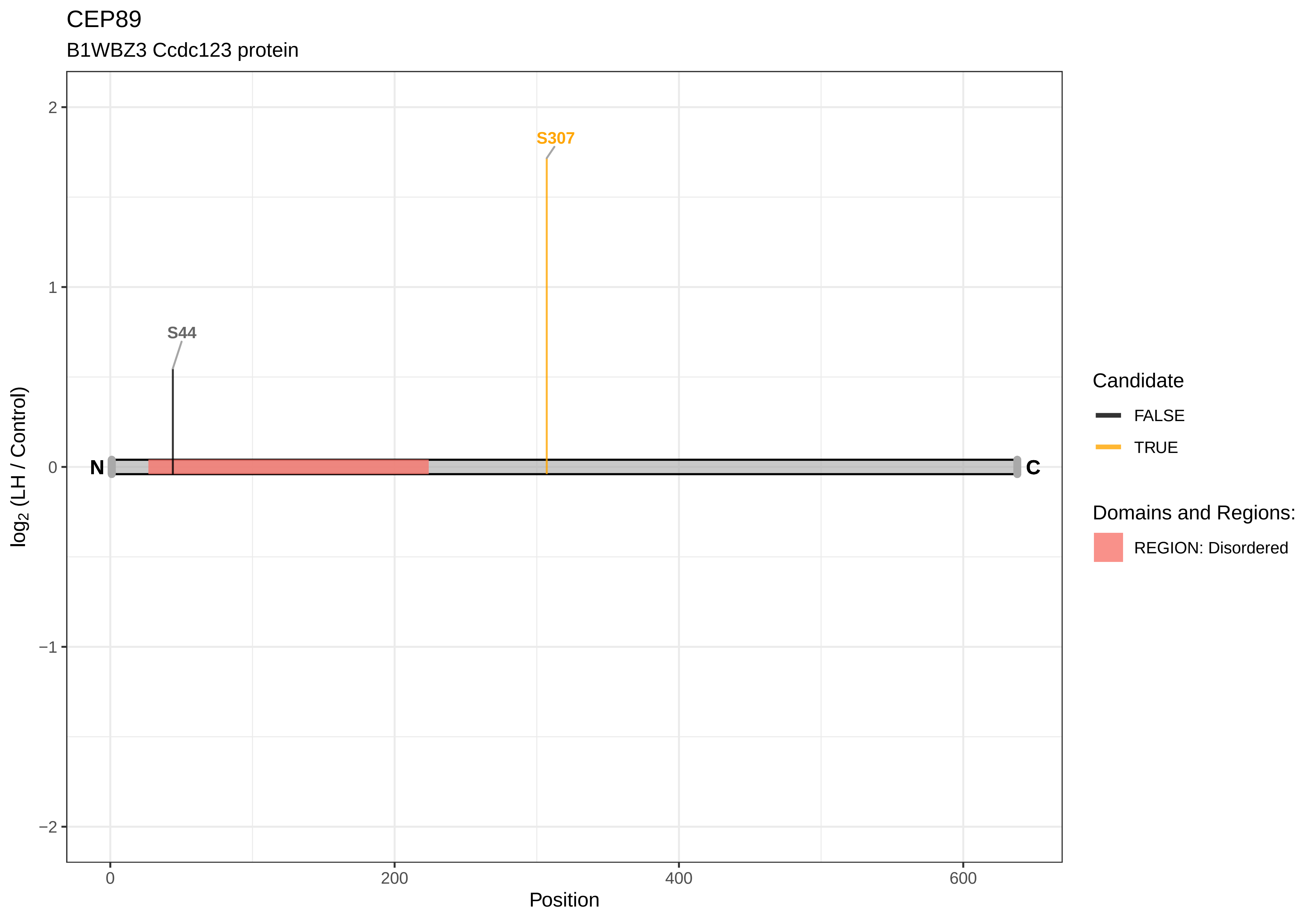

CEP89
B1WBZ3 Ccdc123 protein
S307
S44
N
C
2
1
0
−1
−2
log2 (LH / Control)
0
200
400
600
Position
Candidate
FALSE
TRUE
Domains and Regions:
REGION: Disordered

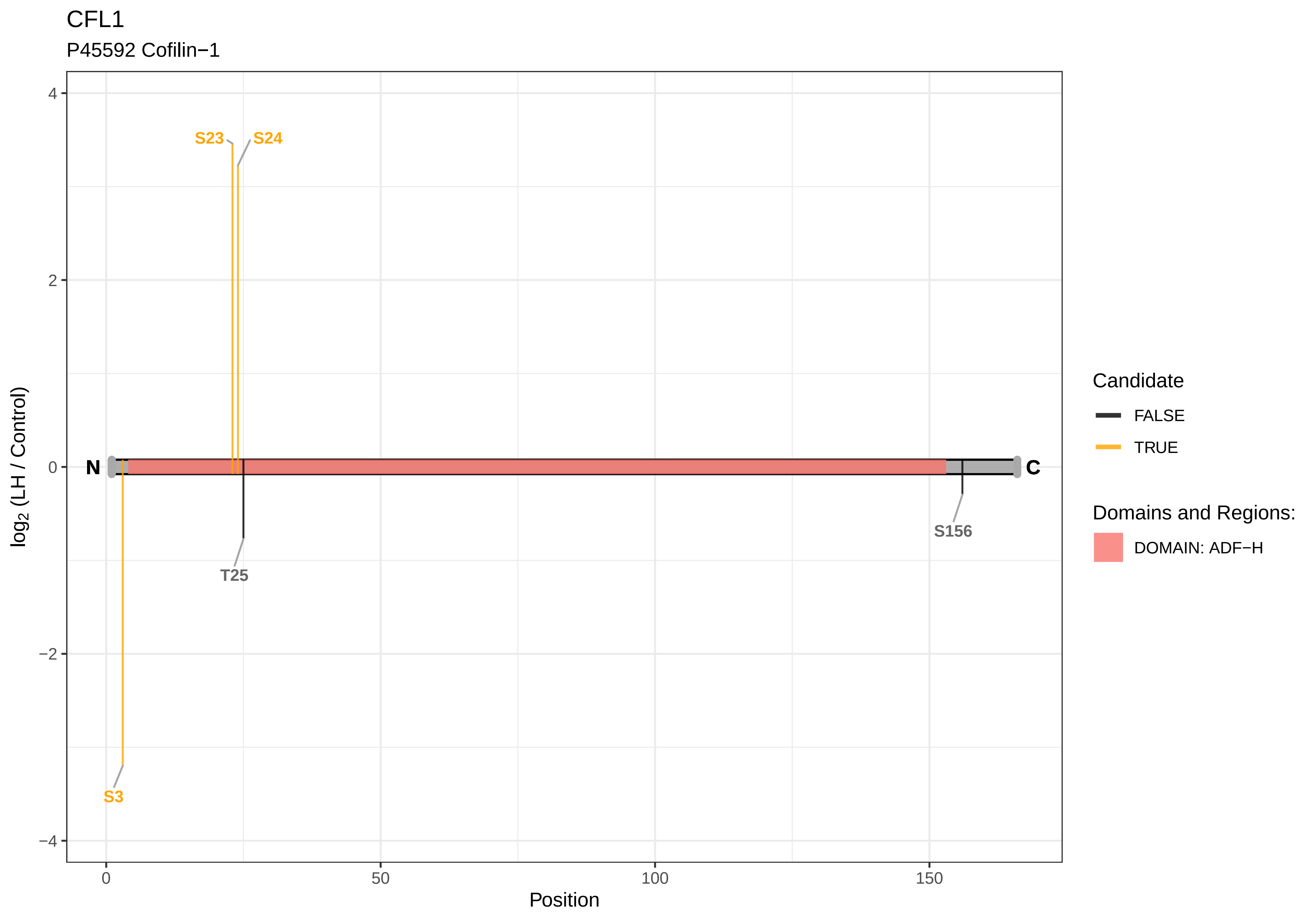

CFL1
P45592 Cofilin−1
S23
S24
N
C
T25
S156
S3
4
2
0
−2
−4
log2 (LH / Control)
0
50
100
150
Position
Candidate
FALSE
TRUE
Domains and Regions:
DOMAIN: ADF−H

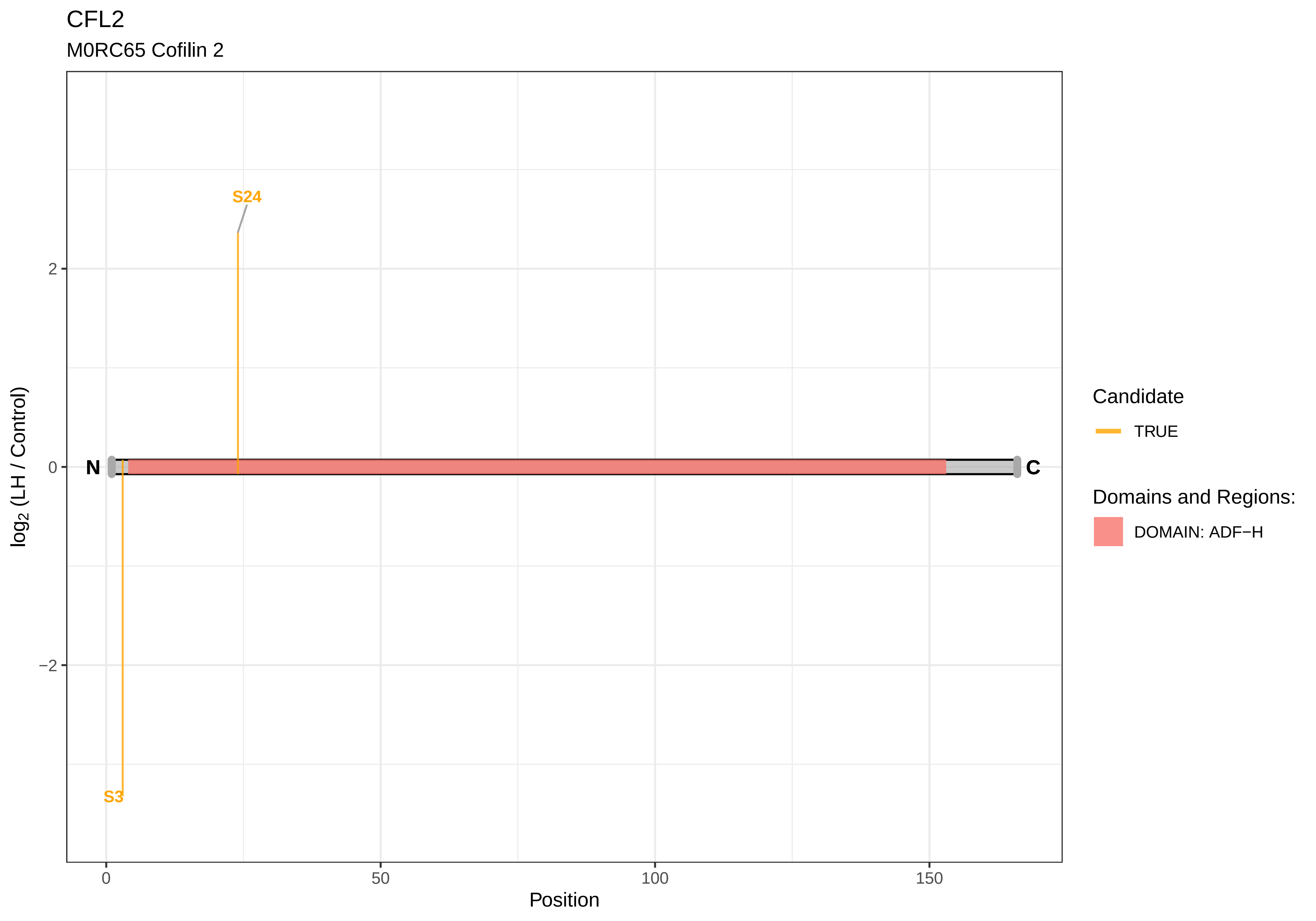
CFL2
M0RC65 Cofilin 2
S24
S3
N
C
2
0
−2
0
50
100
150
Position
log2 (LH / Control)
Candidate
TRUE
Domains and Regions:
DOMAIN: ADF−H

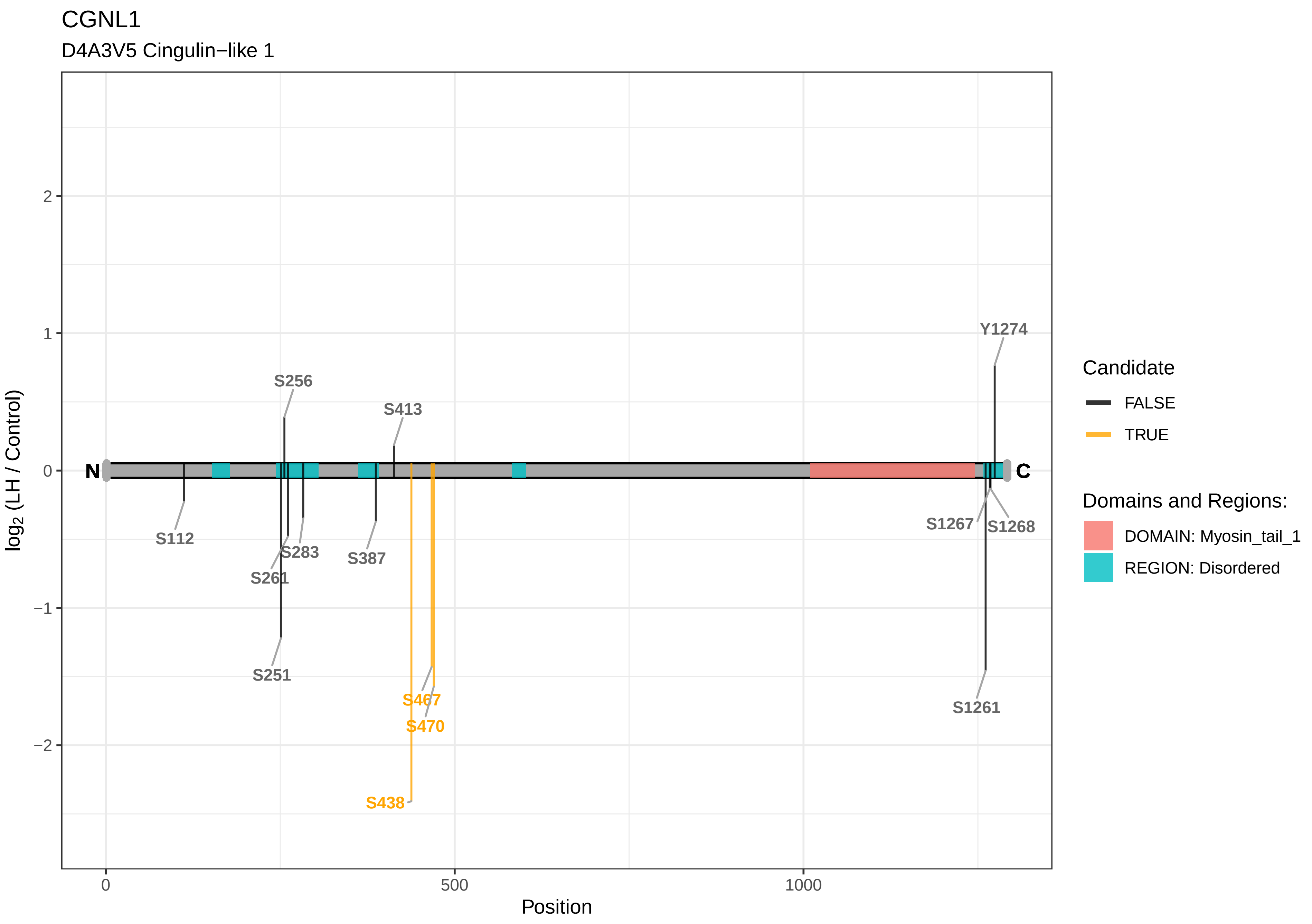

CGNL1
D4A3V5 Cingulin−like 1
log2 (LH / Control)
Position
0
500
1000
2
1
0
−1
−2
N
C
S256
S413
Y1274
S112
S283
S261
S387
S251
S467
S470
S438
S1267
S1268
S1261
Candidate
FALSE
TRUE
Domains and Regions:
DOMAIN: Myosin_tail_1
REGION: Disordered

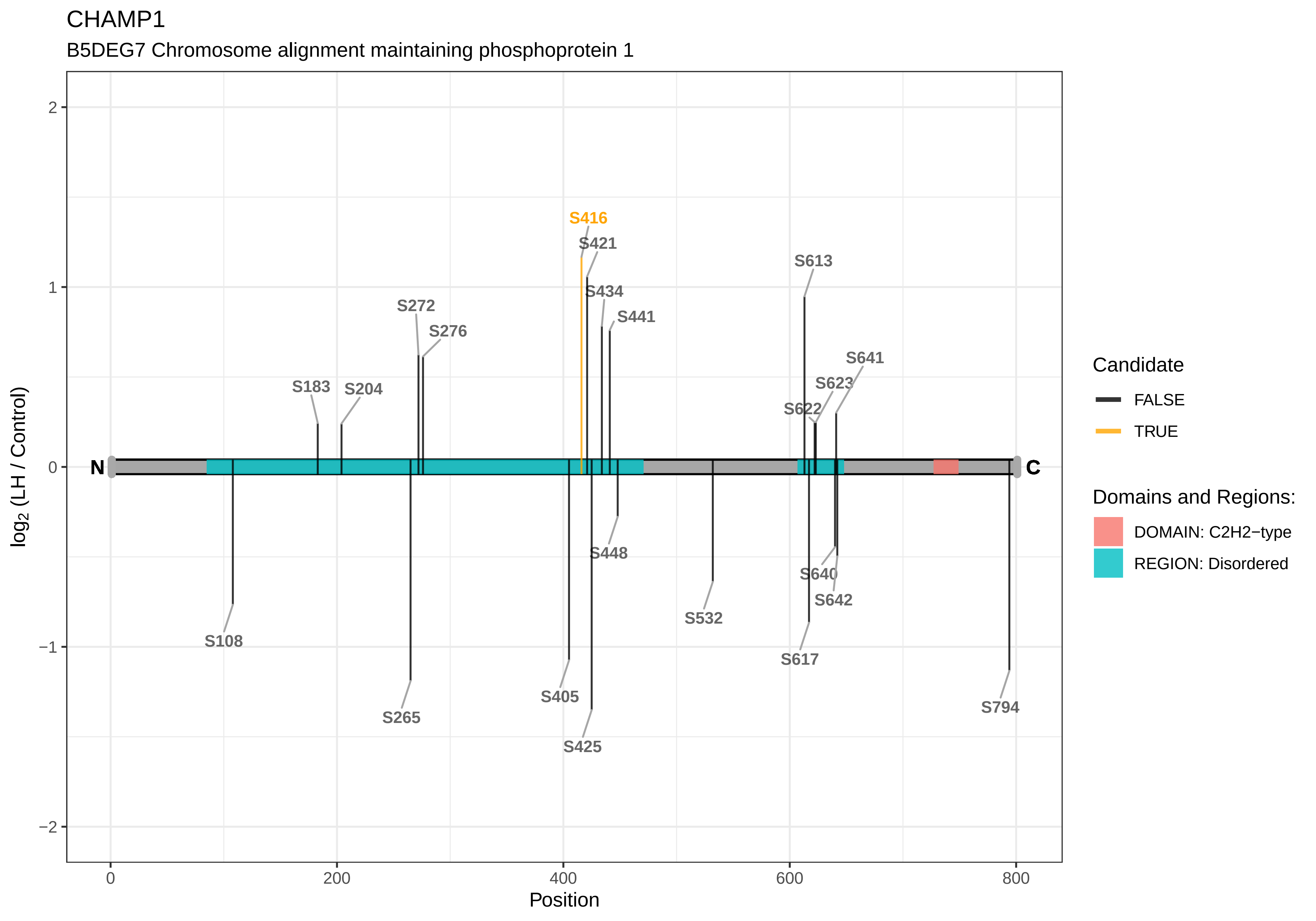
CHAMP1
B5DEG7 Chromosome alignment maintaining phosphoprotein 1
S416
S421
S613
S434
S441
S272
S276
S641
S623
S183
S204
S622
N
C
S448
S640
S642
S532
S108
S617
S405
S265
S794
S425
2
1
0
−1
−2
0
200
400
600
800
Position
log2 (LH / Control)
Candidate
FALSE
TRUE
Domains and Regions:
DOMAIN: C2H2−type
REGION: Disordered

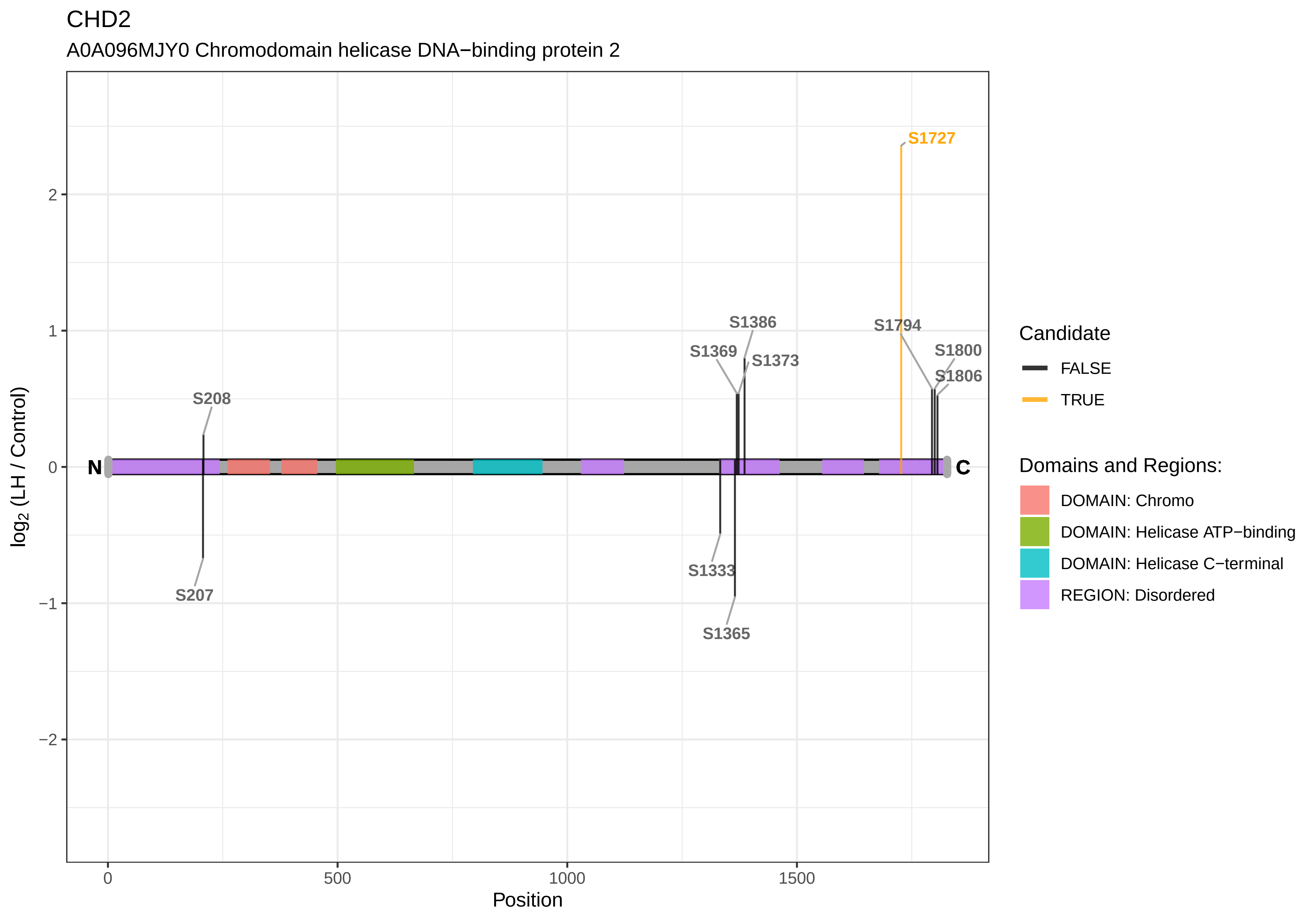

CHD2
A0A096MJY0 Chromodomain helicase DNA−binding protein 2
S1727
S1386
S1794
S1369
S1373
S1800
S1806
S208
N
C
S207
S1333
S1365
2
1
0
−1
−2
0
500
1000
1500
Position
log2 (LH / Control)
Candidate
FALSE
TRUE
Domains and Regions:
DOMAIN: Chromo
DOMAIN: Helicase ATP−binding
DOMAIN: Helicase C−terminal
REGION: Disordered

CIAPIN1

Q5XID1 Anamorsin

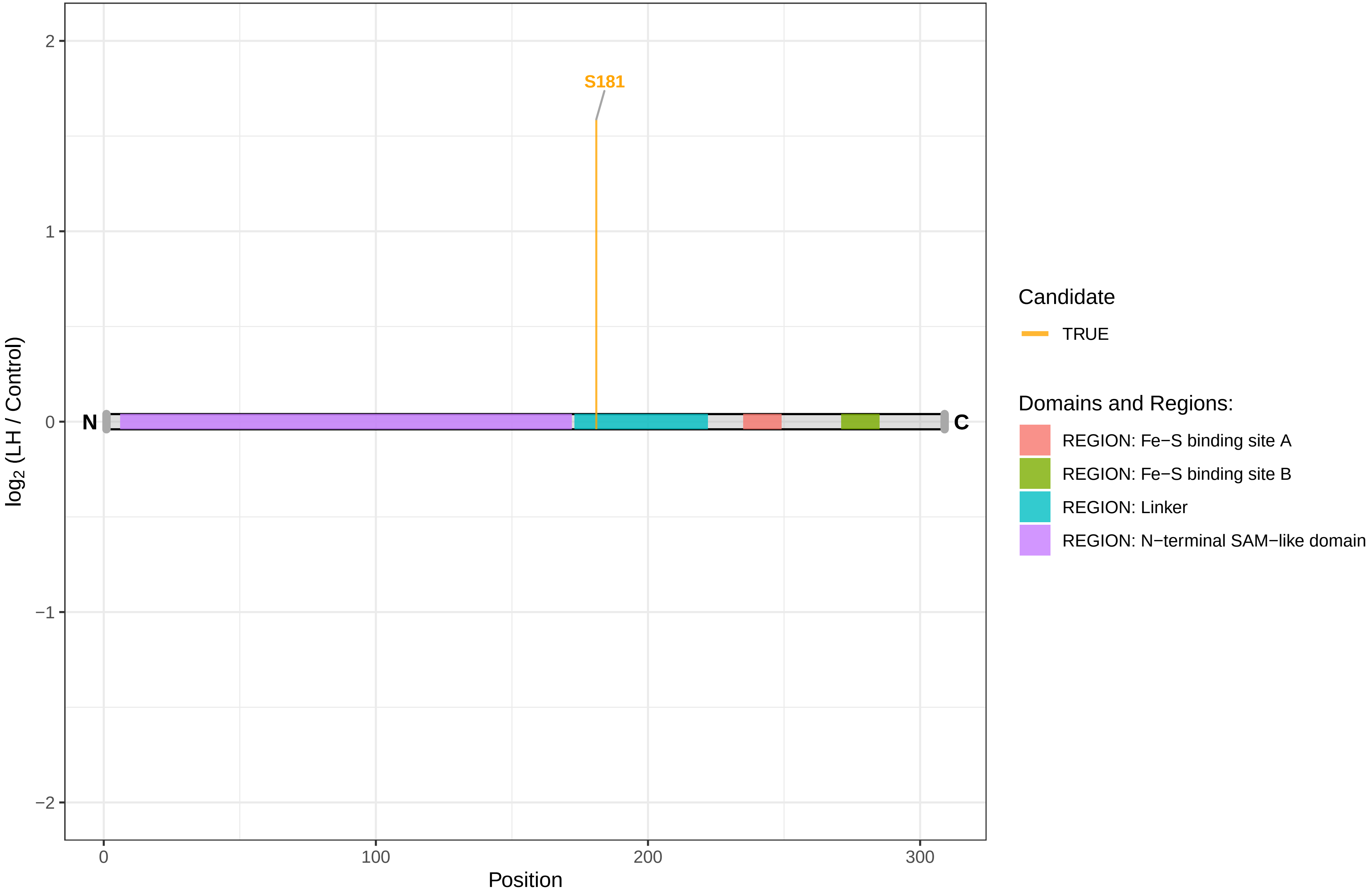

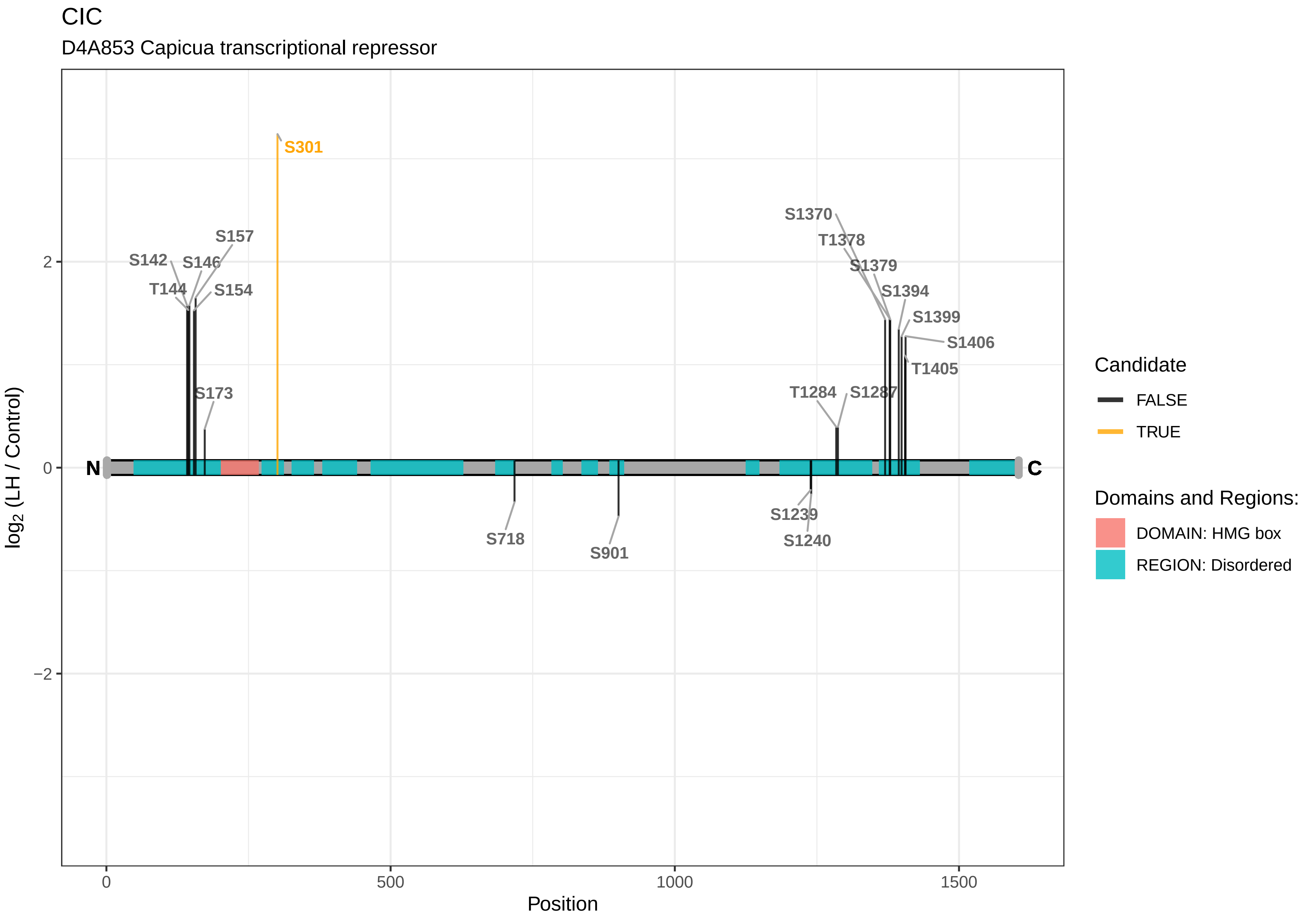

CIC
D4A853 Capicua transcriptional repressor
S301
S157
S142
S146
T144
S154
S173
S1370
T1378
S1379
S1394
S1399
S1406
T1405
T1284
S1287
N
C
S718
S901
S1239
S1240
2
0
−2
0
500
1000
1500
Position
log2 (LH / Control)
Candidate
FALSE
TRUE
Domains and Regions:
DOMAIN: HMG box
REGION: Disordered

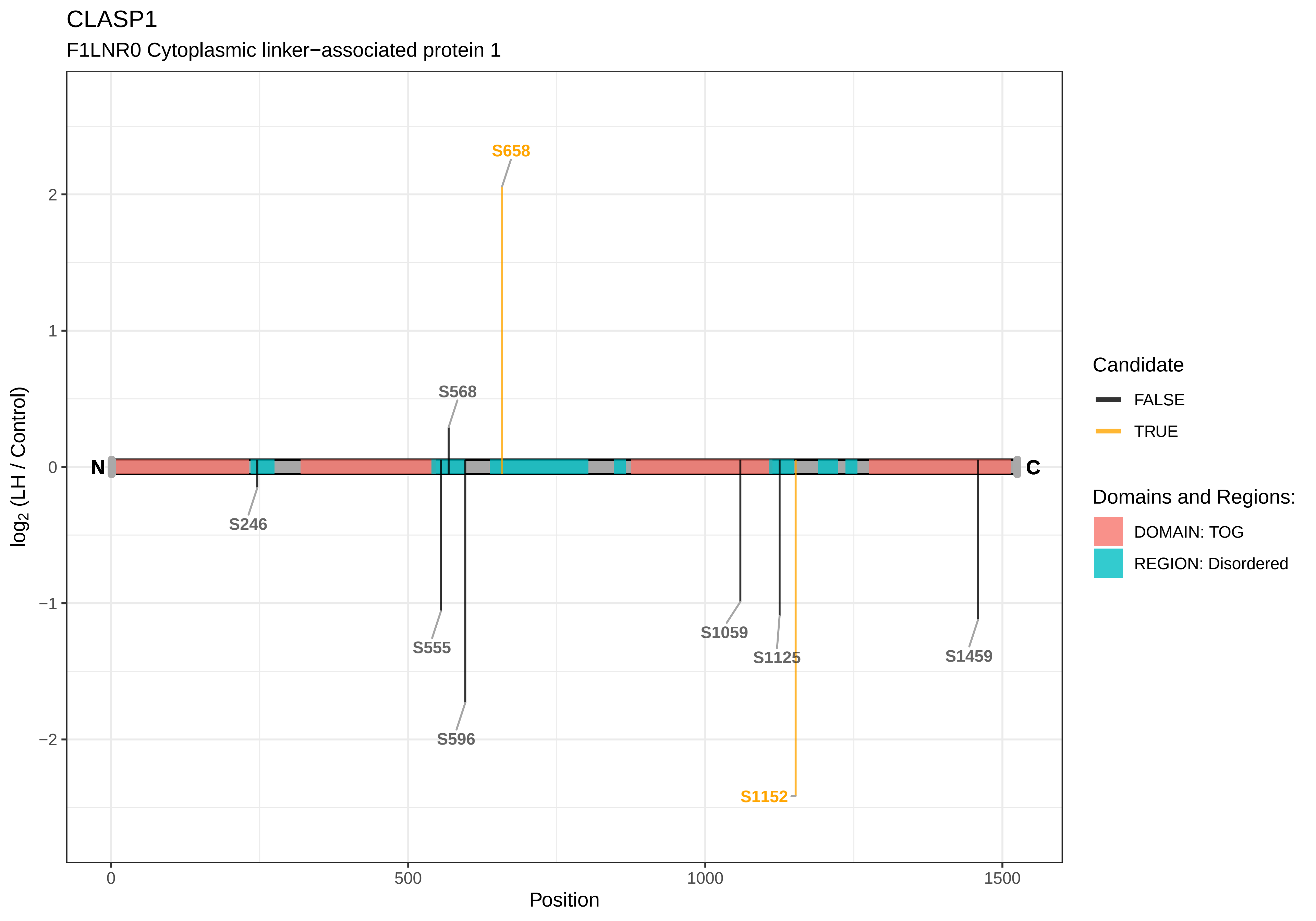

CLASP1
F1LNR0 Cytoplasmic linker−associated protein 1
S658
S568
S246
S555
S596
S1059
S1125
S1152
S1459
N
C
2
1
0
−1
−2
log2 (LH / Control)
0
500
1000
1500
Position
Candidate
FALSE
TRUE
Domains and Regions:
DOMAIN: TOG
REGION: Disordered

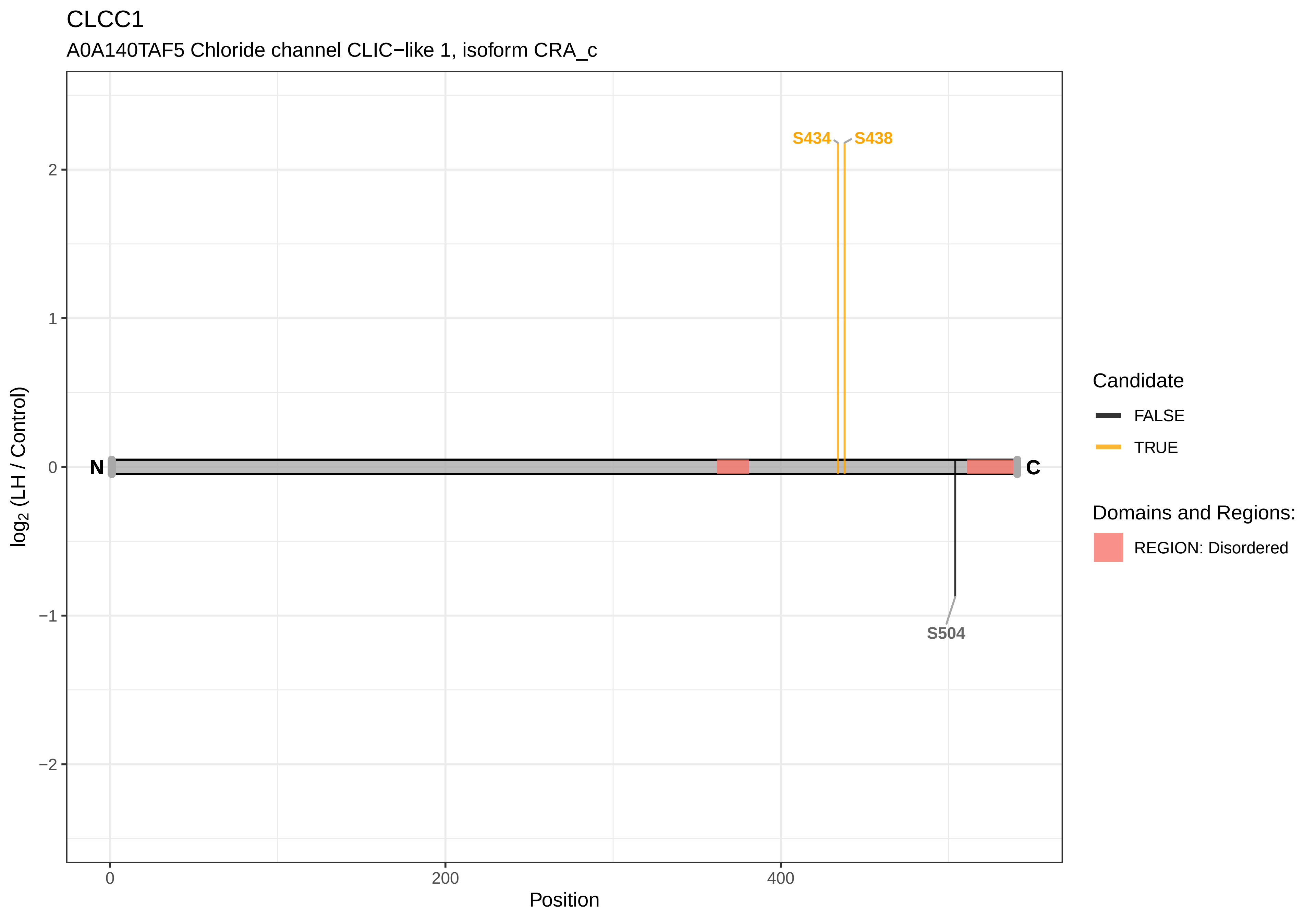

CLCC1
A0A140TAF5 Chloride channel CLIC−like 1, isoform CRA_c
S434
S438
S504
N
C
2
1
0
−1
−2
log2 (LH / Control)
0
200
400
Position
Candidate
FALSE
TRUE
Domains and Regions:
REGION: Disordered

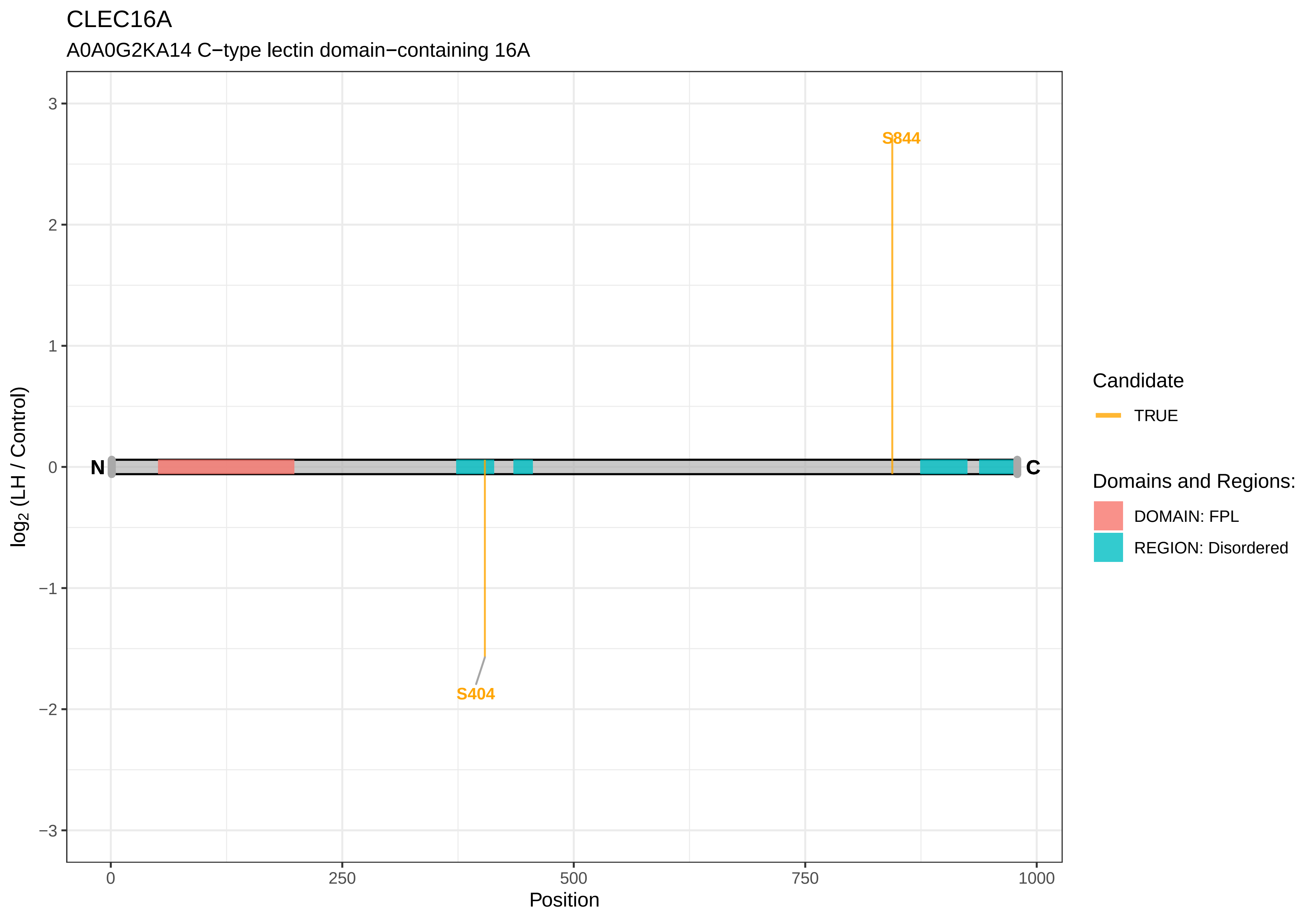

CLEC16A
A0A0G2KA14 C−type lectin domain−containing 16A
S844
S404
N
C
log2 (LH / Control)
Position
Candidate
TRUE
Domains and Regions:
DOMAIN: FPL
REGION: Disordered

CLIP1

A0A0G2K681 CAP−Gly domain−containing linker protein 1

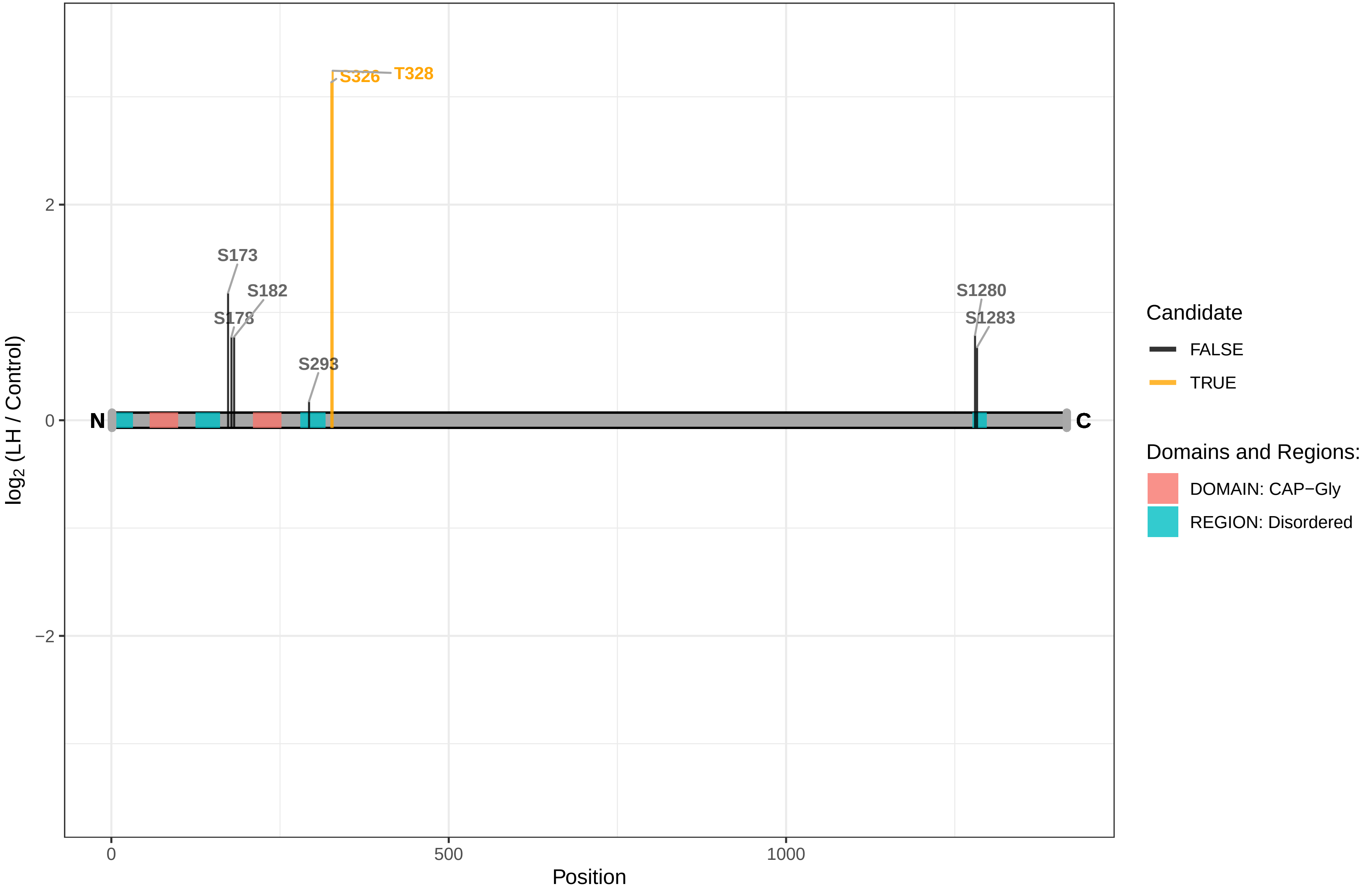

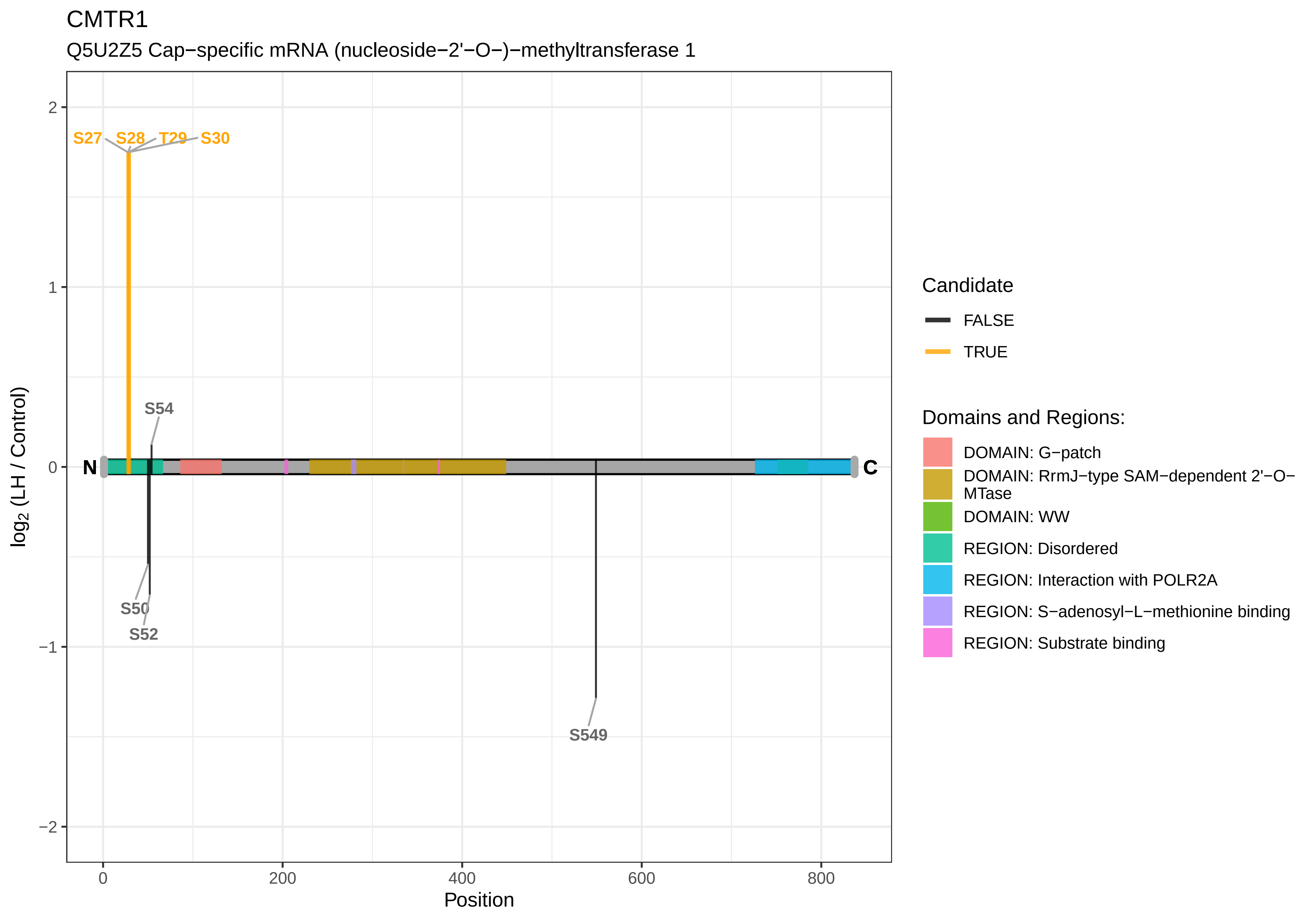
CMTR1
Q5U2Z5 Cap−specific mRNA (nucleoside−2'−O−)−methyltransferase 1
S27
S28
T29
S30
S54
N
C
S50
S52
S549
log2 (LH / Control)
2
1
0
−1
−2
0
200
400
600
800
Position
Candidate
FALSE
TRUE
Domains and Regions:
DOMAIN: G−patch
DOMAIN: RrmJ−type SAM−dependent 2'−O−MTase
DOMAIN: WW
REGION: Disordered
REGION: Interaction with POLR2A
REGION: S−adenosyl−L−methionine binding
REGION: Substrate binding

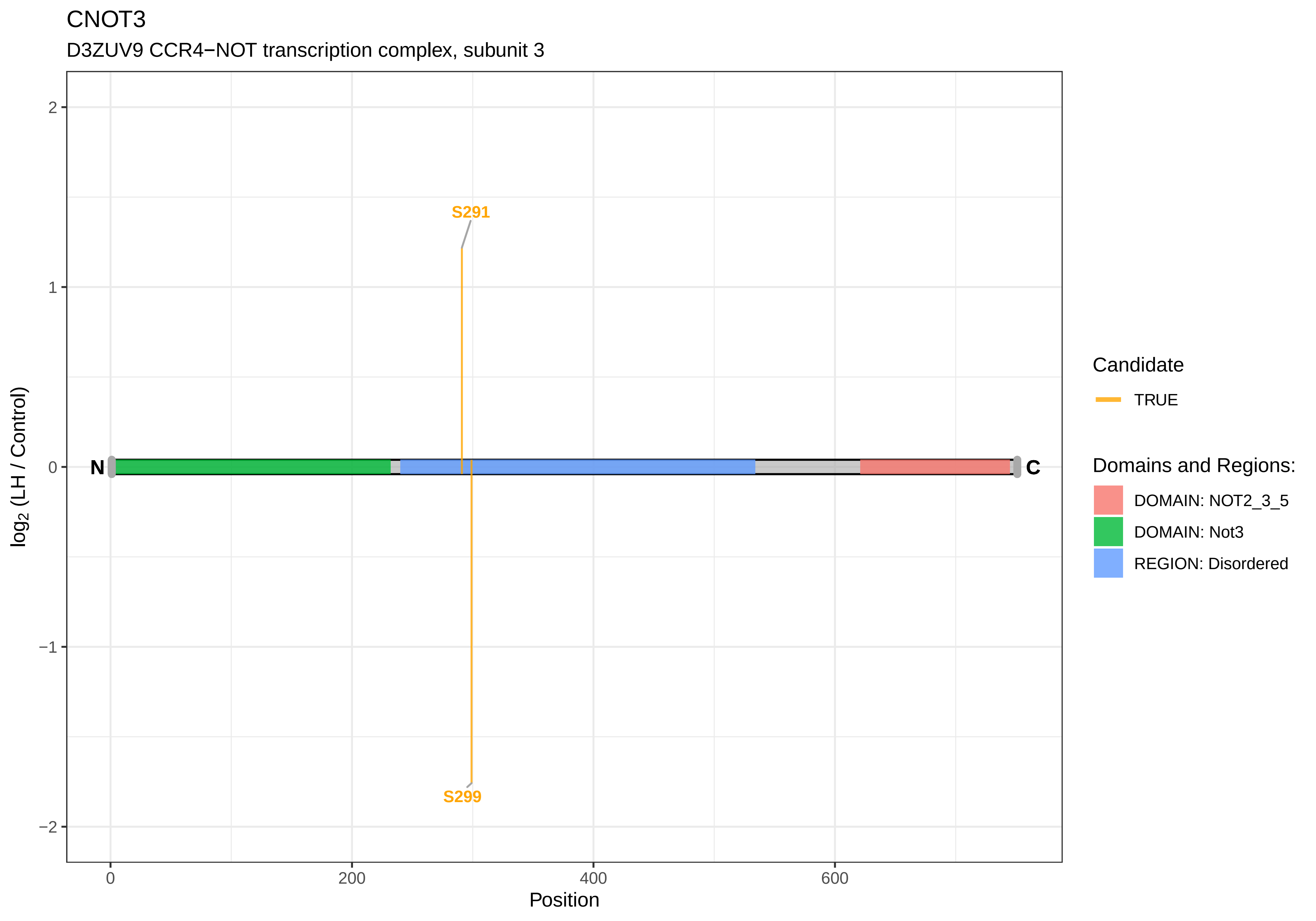
CNOT3
D3ZUV9 CCR4−NOT transcription complex, subunit 3
S291
S299
N
C
log2 (LH / Control)
2
1
0
−1
−2
0
200
400
600
Position
Candidate
TRUE
Domains and Regions:
DOMAIN: NOT2_3_5
DOMAIN: Not3
REGION: Disordered

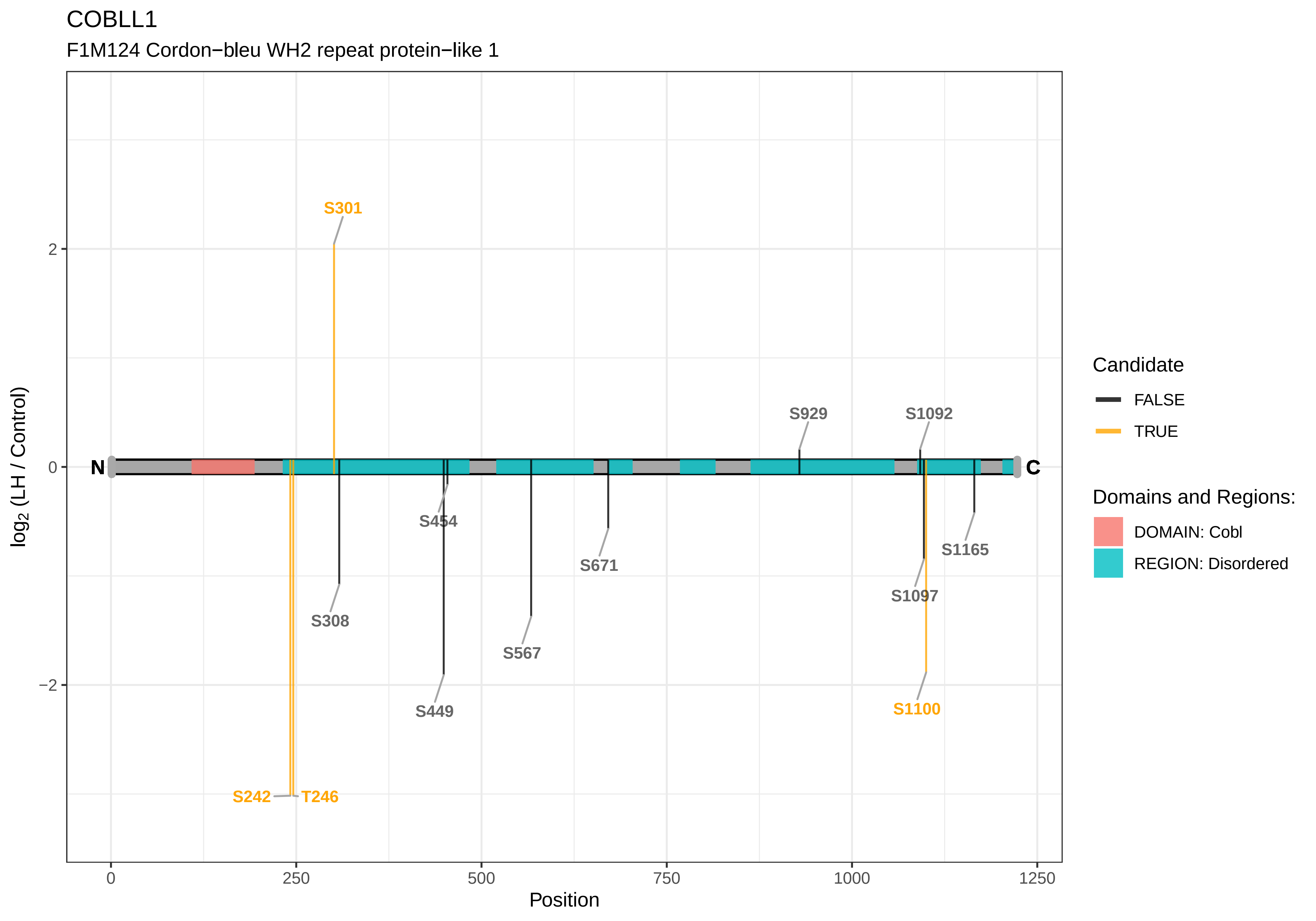
COBLL1
F1M124 Cordon−bleu WH2 repeat protein−like 1
S301
S929
S1092
N
C
S454
S671
S1165
S1097
S308
S567
S449
S1100
S242
T246
2
0
−2
0
250
500
750
1000
1250
Position
log2 (LH / Control)
Candidate
FALSE
TRUE
Domains and Regions:
DOMAIN: Cobl
REGION: Disordered

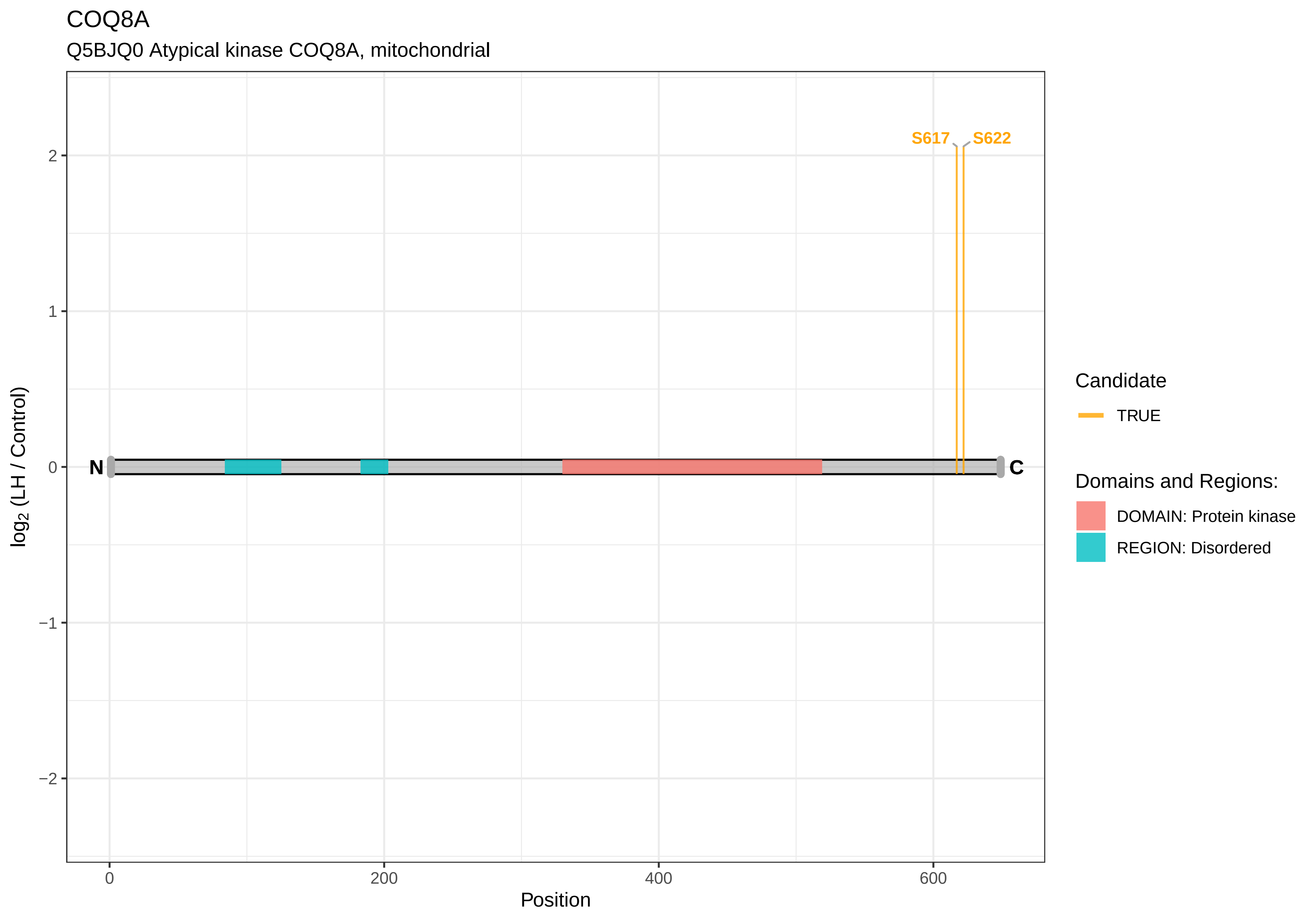

COQ8A
Q5BJQ0 Atypical kinase COQ8A, mitochondrial
S617
S622
2
1
0
−1
−2
N
C
log2 (LH / Control)
0
200
400
600
Position
Candidate
TRUE
Domains and Regions:
DOMAIN: Protein kinase
REGION: Disordered

CPD

F1LPC6 Carboxypeptidase D

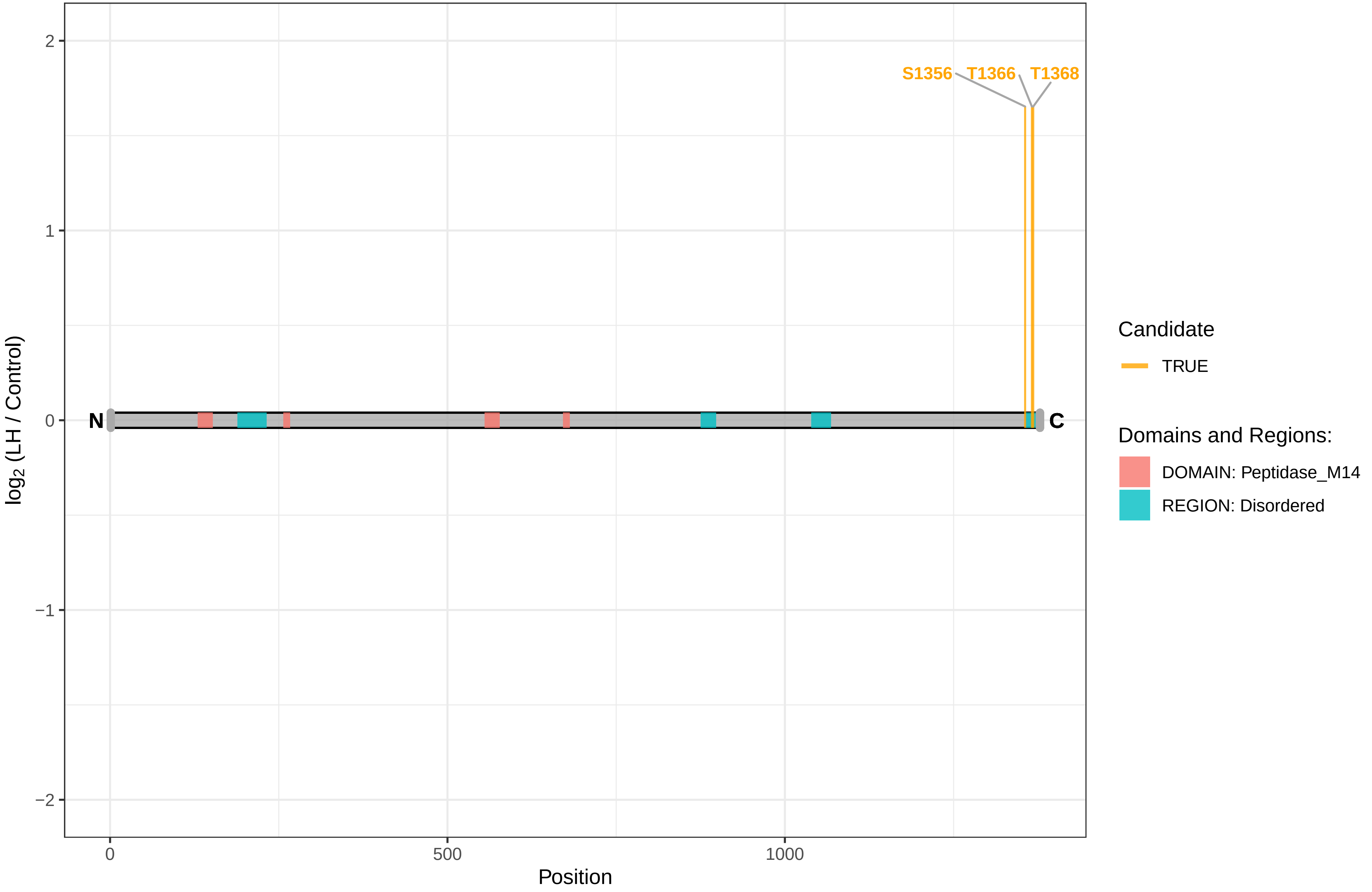

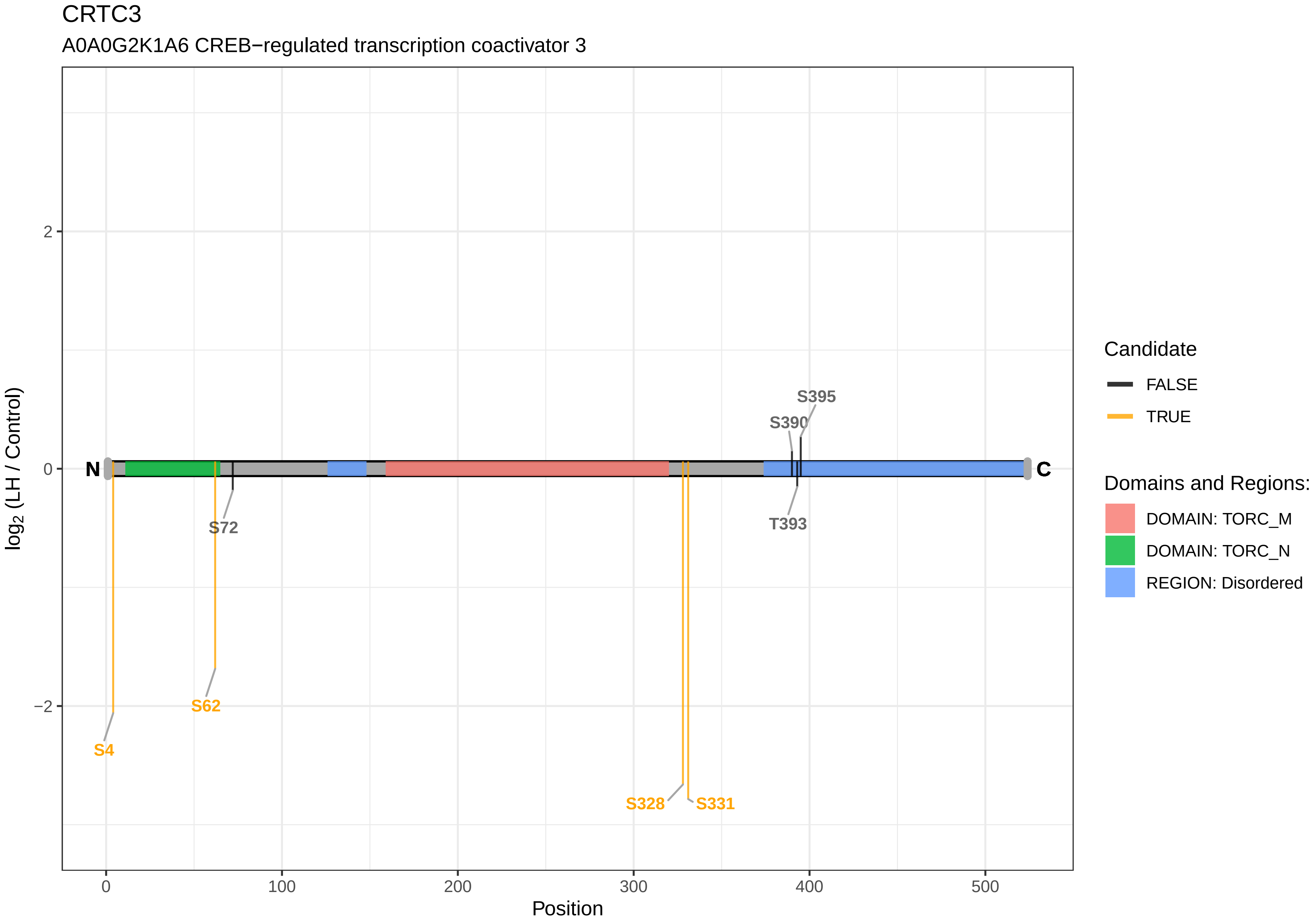

CRTC3
A0A0G2K1A6 CREB−regulated transcription coactivator 3
log2 (LH / Control)
Position
2
0
−2
0
100
200
300
400
500
N
C
S395
S390
S72
T393
S62
S4
S328
S331
Candidate
FALSE
TRUE
Domains and Regions:
DOMAIN: TORC_M
DOMAIN: TORC_N
REGION: Disordered

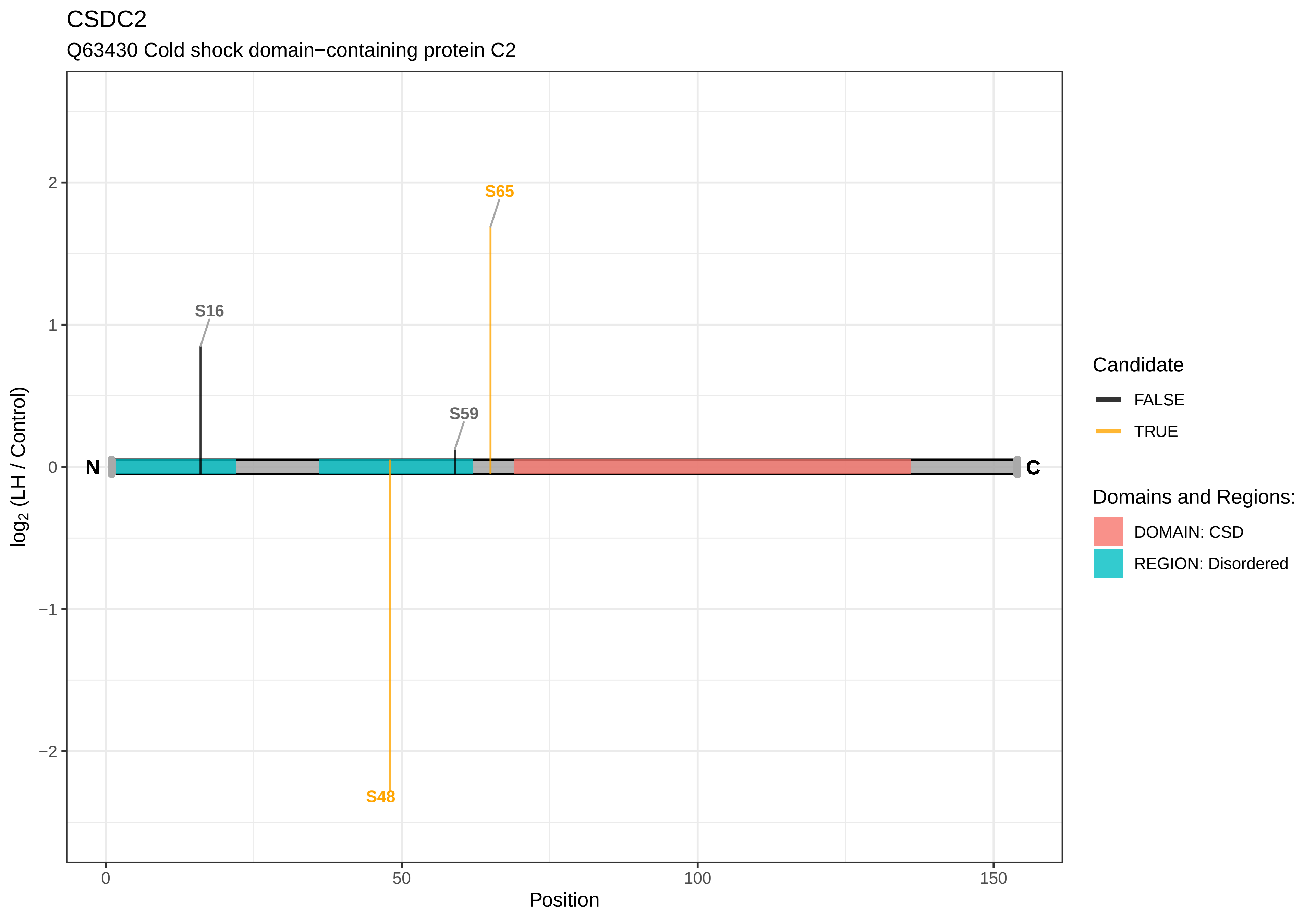
CSDC2
Q63430 Cold shock domain−containing protein C2
S65
S16
S59
S48
N
C
log2 (LH / Control)
2
1
0
−1
−2
0
50
100
150
Position
Candidate
FALSE
TRUE
Domains and Regions:
DOMAIN: CSD
REGION: Disordered

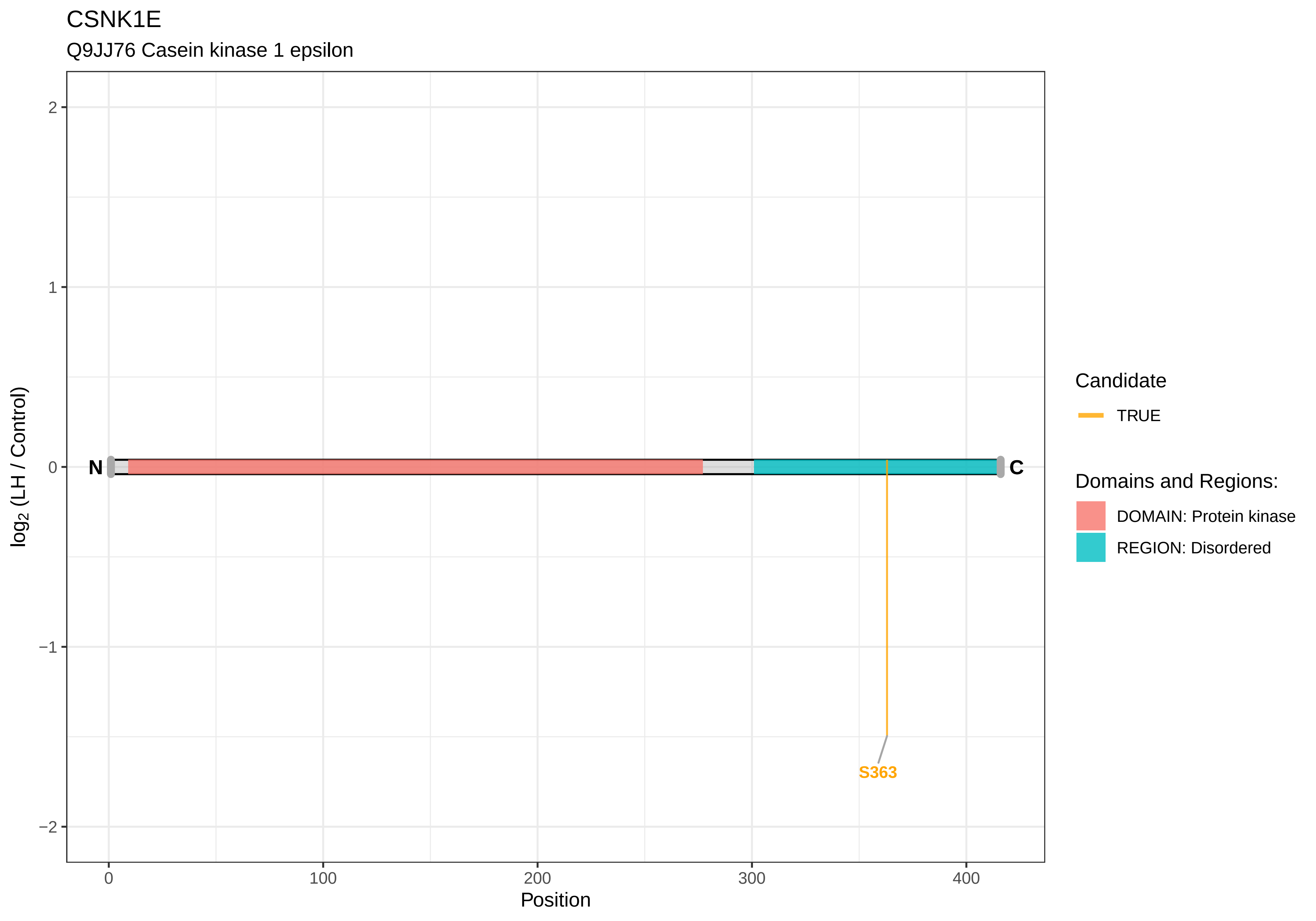
CSNK1E
Q9JJ76 Casein kinase 1 epsilon
log2 (LH / Control)
2
1
0
−1
−2
N
C
S363
0
100
200
300
400
Position
Candidate
TRUE
Domains and Regions:
DOMAIN: Protein kinase
REGION: Disordered

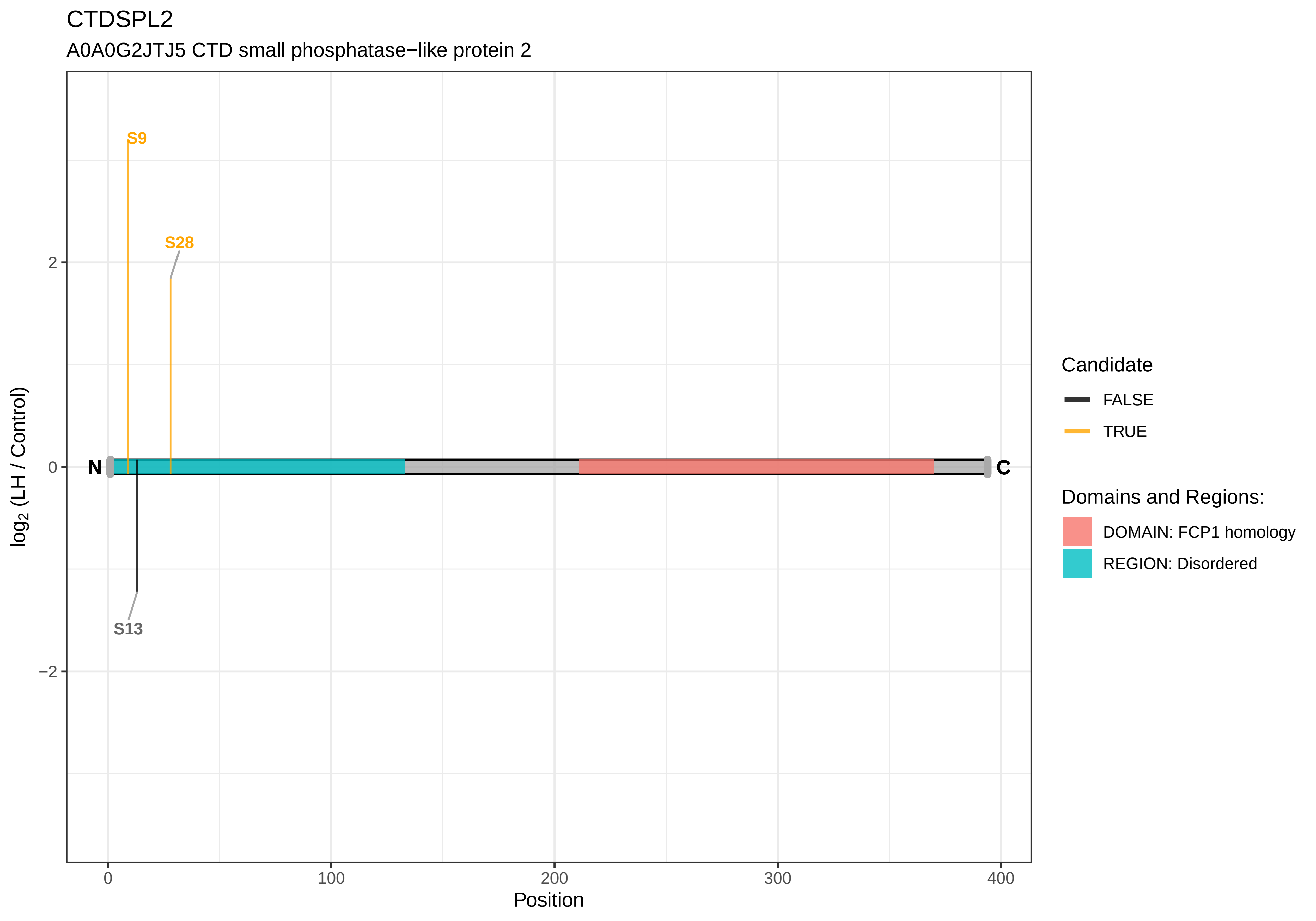
CTDSPL2
A0A0G2JTJ5 CTD small phosphatase−like protein 2
S9
S28
S13
N
C
2
0
−2
0
100
200
300
400
Position
log2 (LH / Control)
Candidate
FALSE
TRUE
Domains and Regions:
DOMAIN: FCP1 homology
REGION: Disordered

### CTNNB1

A0A0G2JT93 Catenin (Cadherin associated protein), beta 1, isoform CRA_a

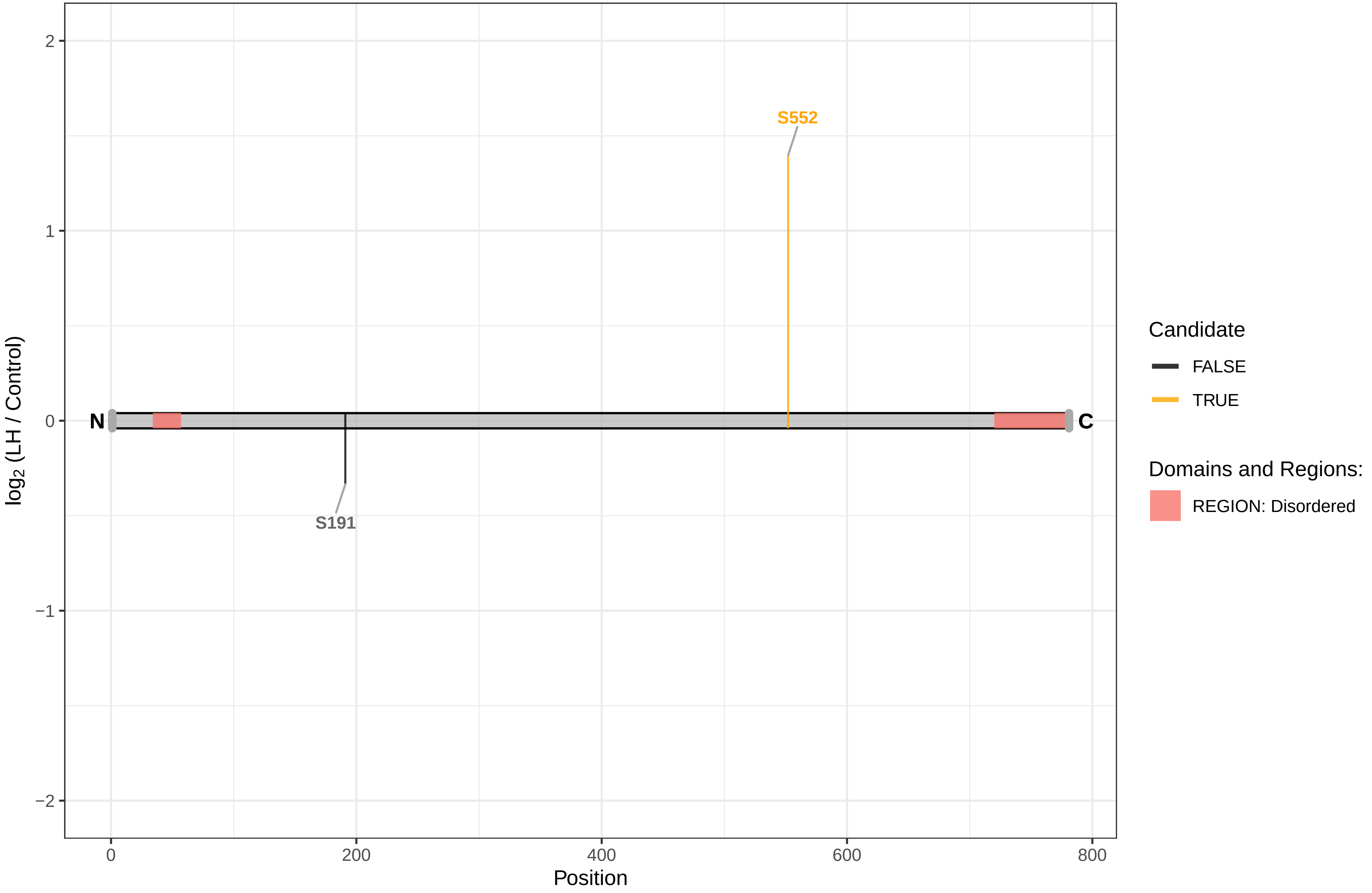

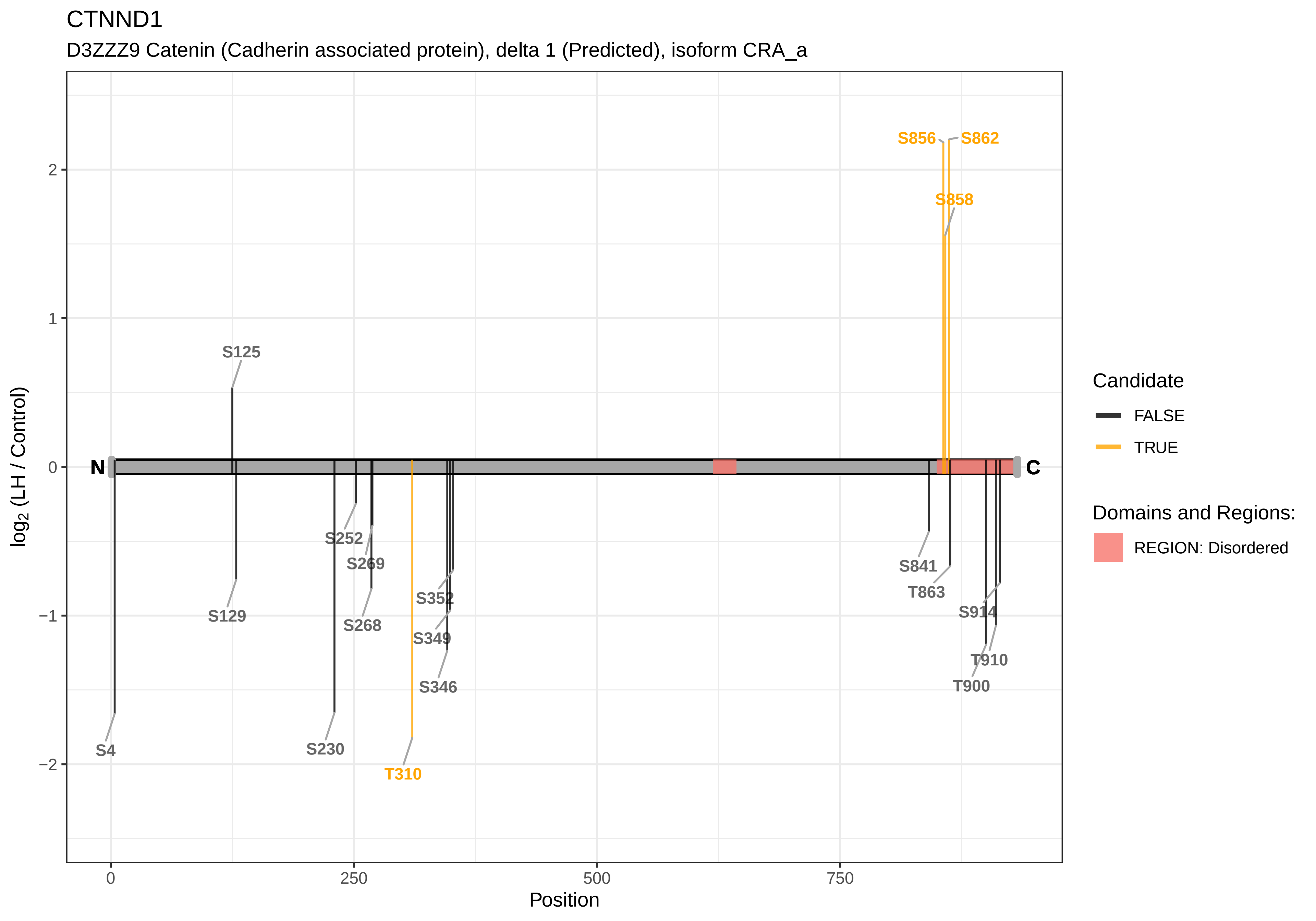
CTNND1
D3ZZZ9 Catenin (Cadherin associated protein), delta 1 (Predicted), isoform CRA_a
log2 (LH / Control)
Position
N
C
S856
S862
S858
S125
S252
S269
S129
S268
S352
S349
S346
S4
S230
T310
S841
T863
S914
T910
T900
Candidate
FALSE
TRUE
Domains and Regions:
REGION: Disordered

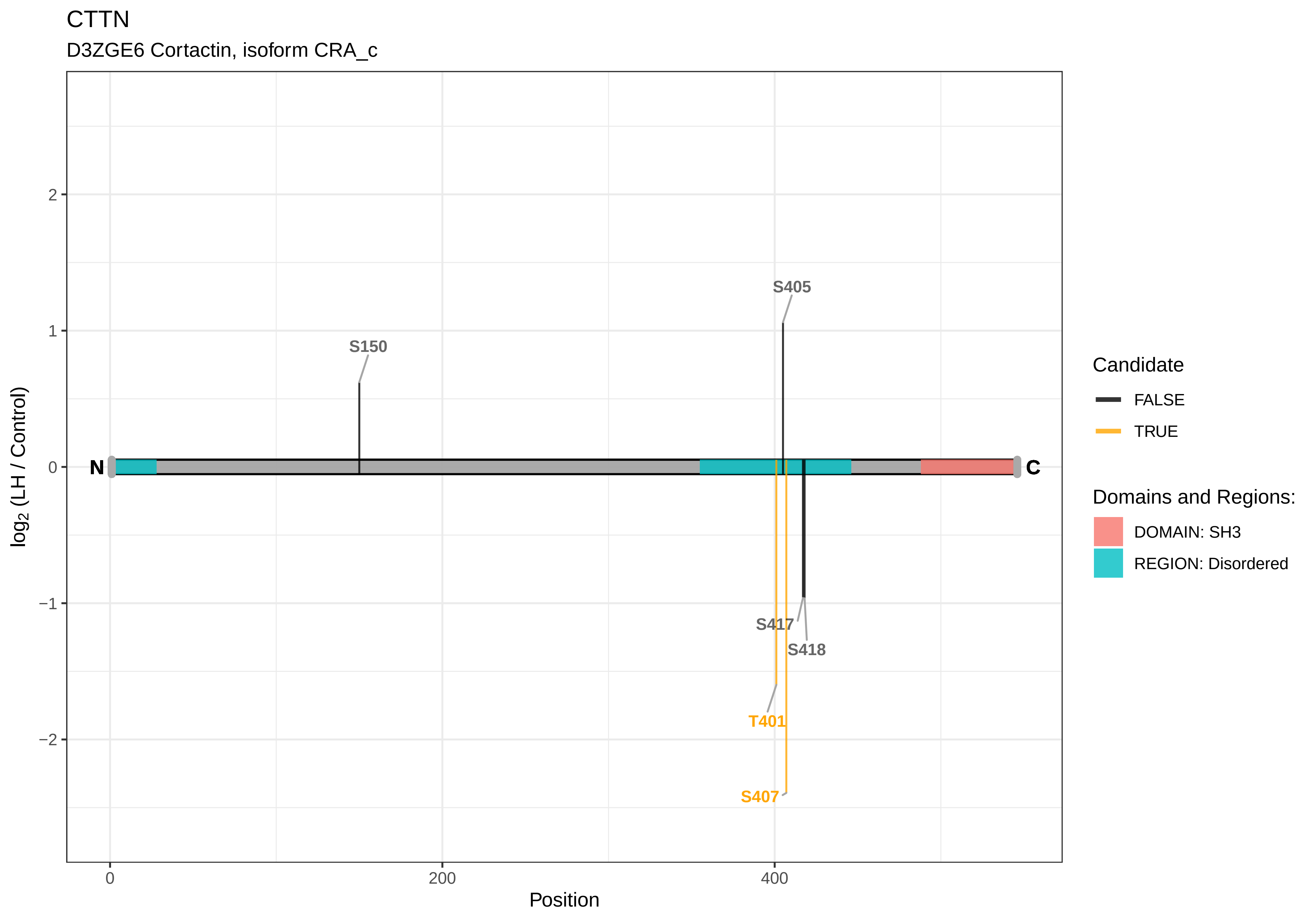
CTTN
D3ZGE6 Cortactin, isoform CRA_c
S405
S150
S417
S418
T401
S407
N
C
2
1
0
−1
−2
0
200
400
log2 (LH / Control)
Position
Candidate
FALSE
TRUE
Domains and Regions:
DOMAIN: SH3
REGION: Disordered

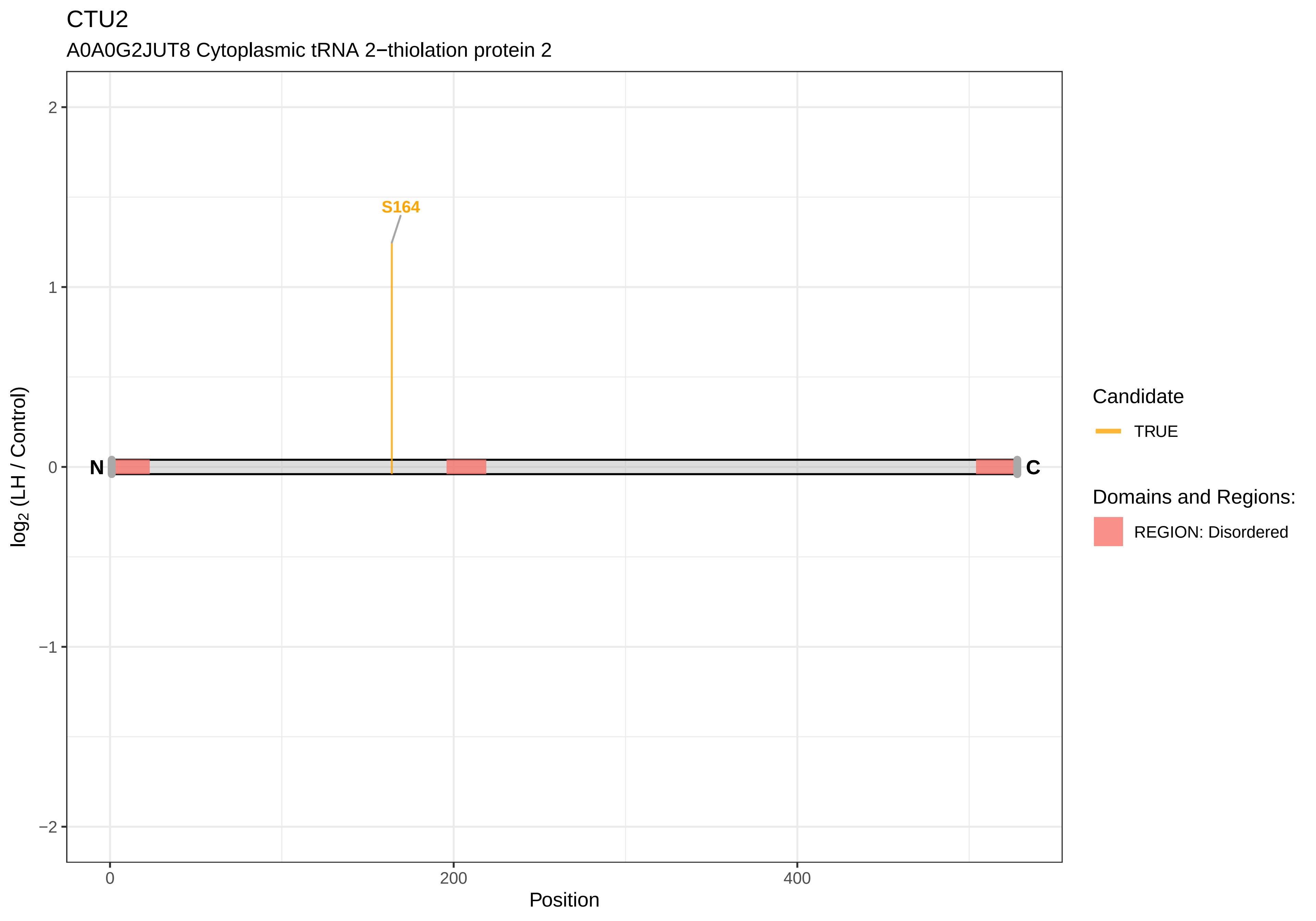
CTU2
A0A0G2JUT8 Cytoplasmic tRNA 2−thiolation protein 2
S164
N
C
2
1
0
−1
−2
0
200
400
Position
log2 (LH / Control)
Candidate
TRUE
Domains and Regions:
REGION: Disordered

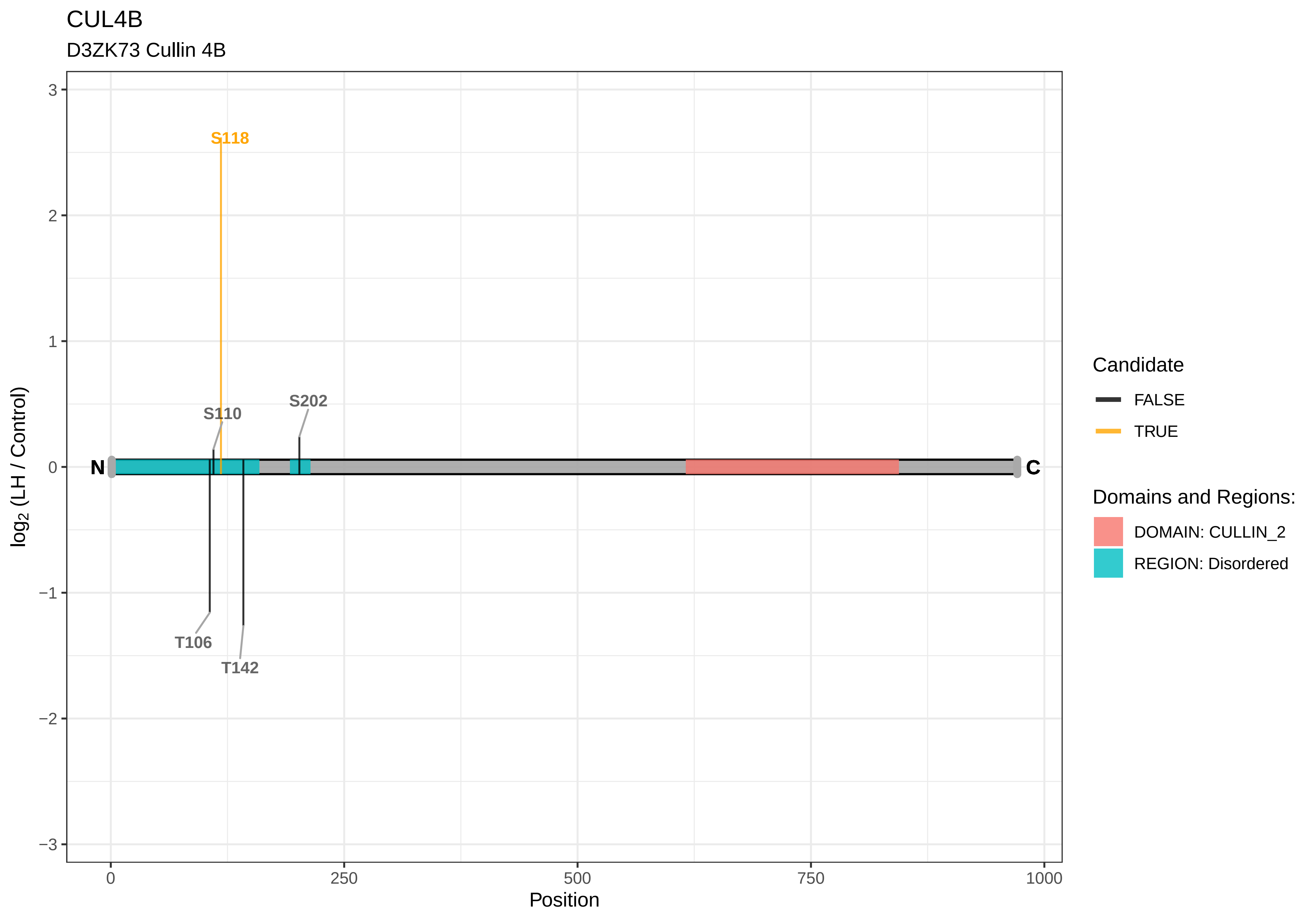
CUL4B
D3ZK73 Cullin 4B
S118
S110
S202
T106
T142
N
C
log2 (LH / Control)
Position
3
2
1
0
−1
−2
−3
0
250
500
750
1000
Candidate
FALSE
TRUE
Domains and Regions:
DOMAIN: CULLIN_2
REGION: Disordered

CUX1

P53565 Homeobox protein cut−like 1

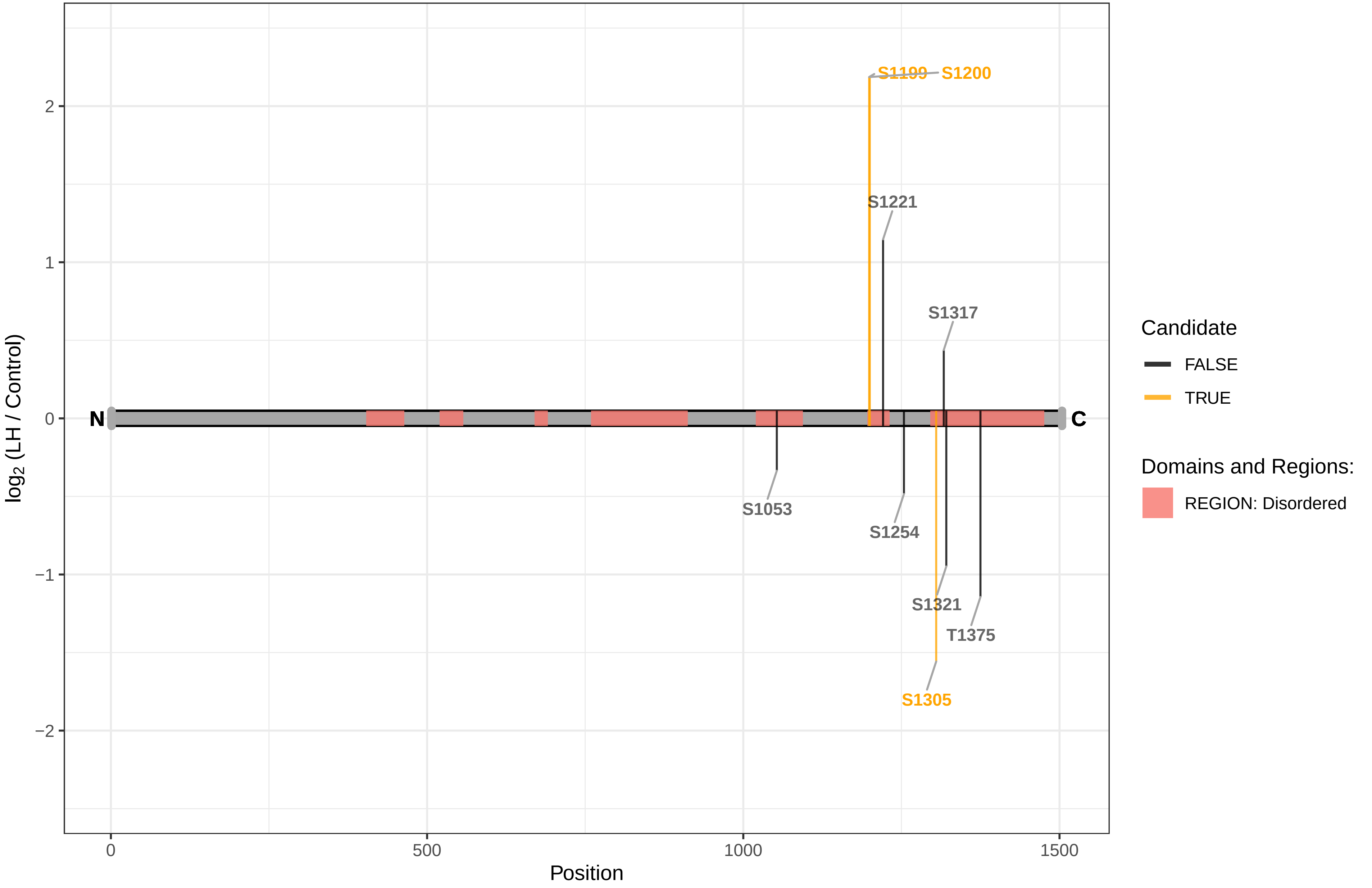

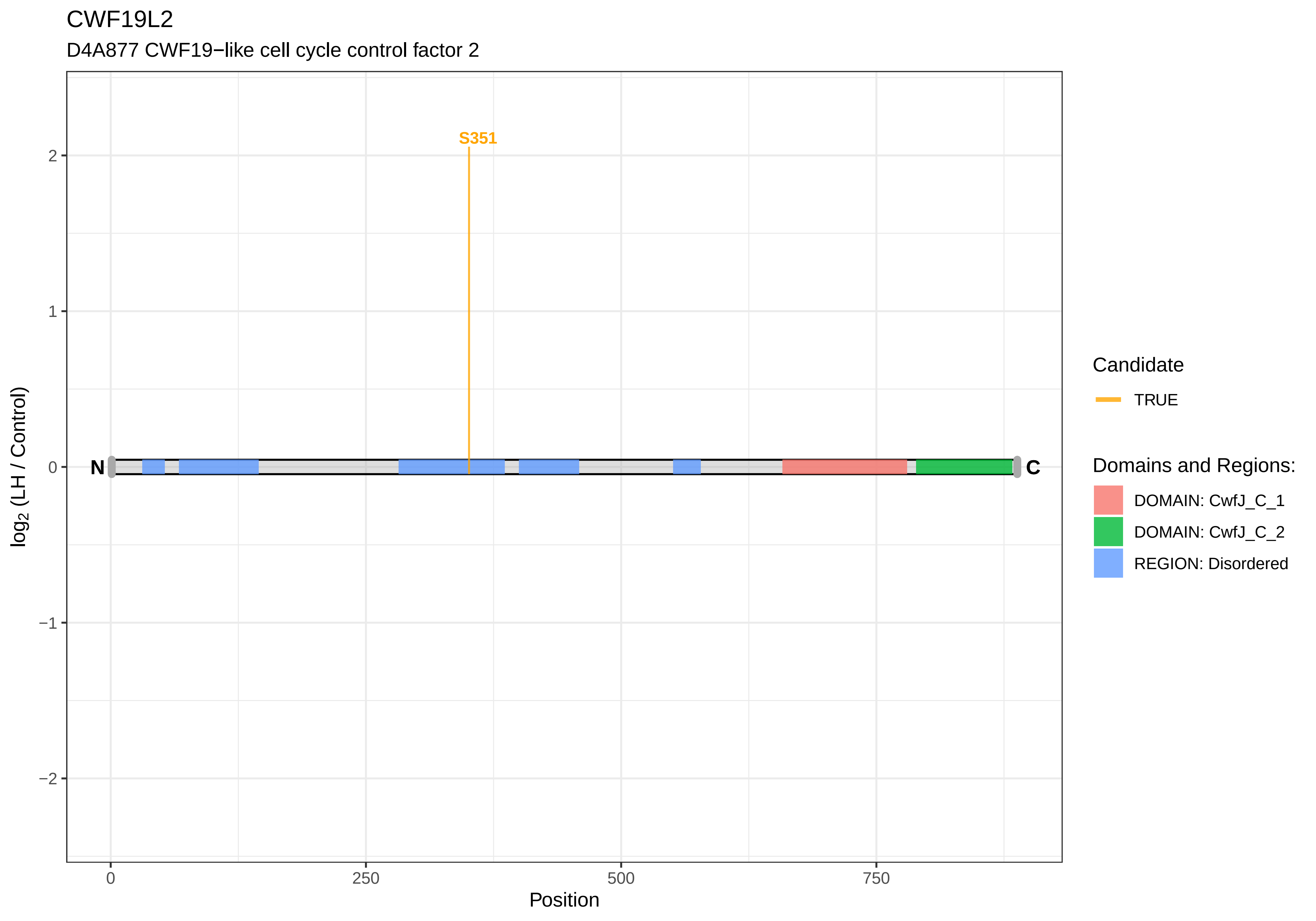
CWF19L2
D4A877 CWF19−like cell cycle control factor 2
S351
N
C
2
1
0
−1
−2
log2 (LH / Control)
0
250
500
750
Position
Candidate
TRUE
Domains and Regions:
DOMAIN: CwfJ_C_1
DOMAIN: CwfJ_C_2
REGION: Disordered

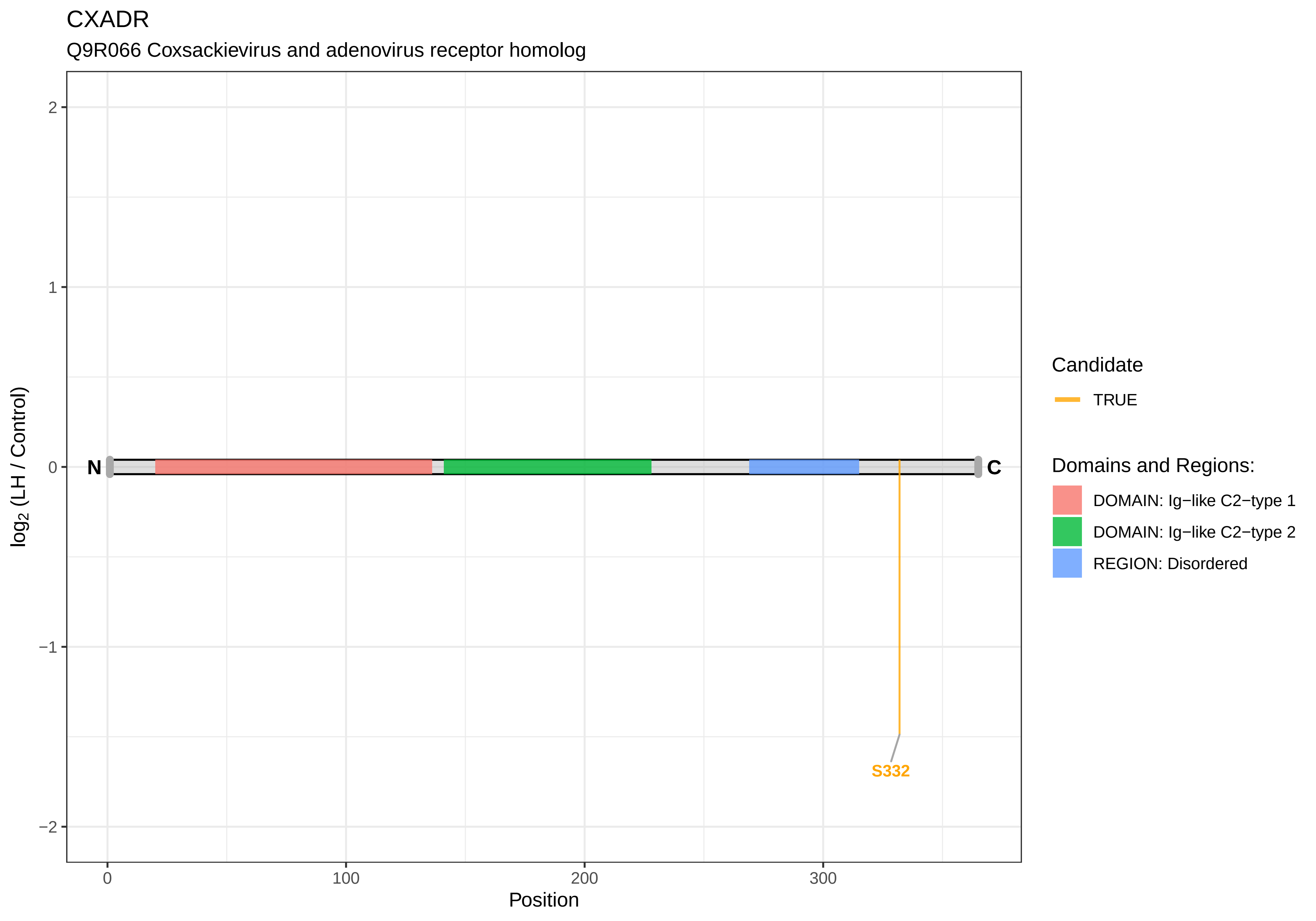
CXADR
Q9R066 Coxsackievirus and adenovirus receptor homolog
log2 (LH / Control)
2
1
0
−1
−2
N
C
S332
0
100
200
300
Position
Candidate
TRUE
Domains and Regions:
DOMAIN: Ig−like C2−type 1
DOMAIN: Ig−like C2−type 2
REGION: Disordered

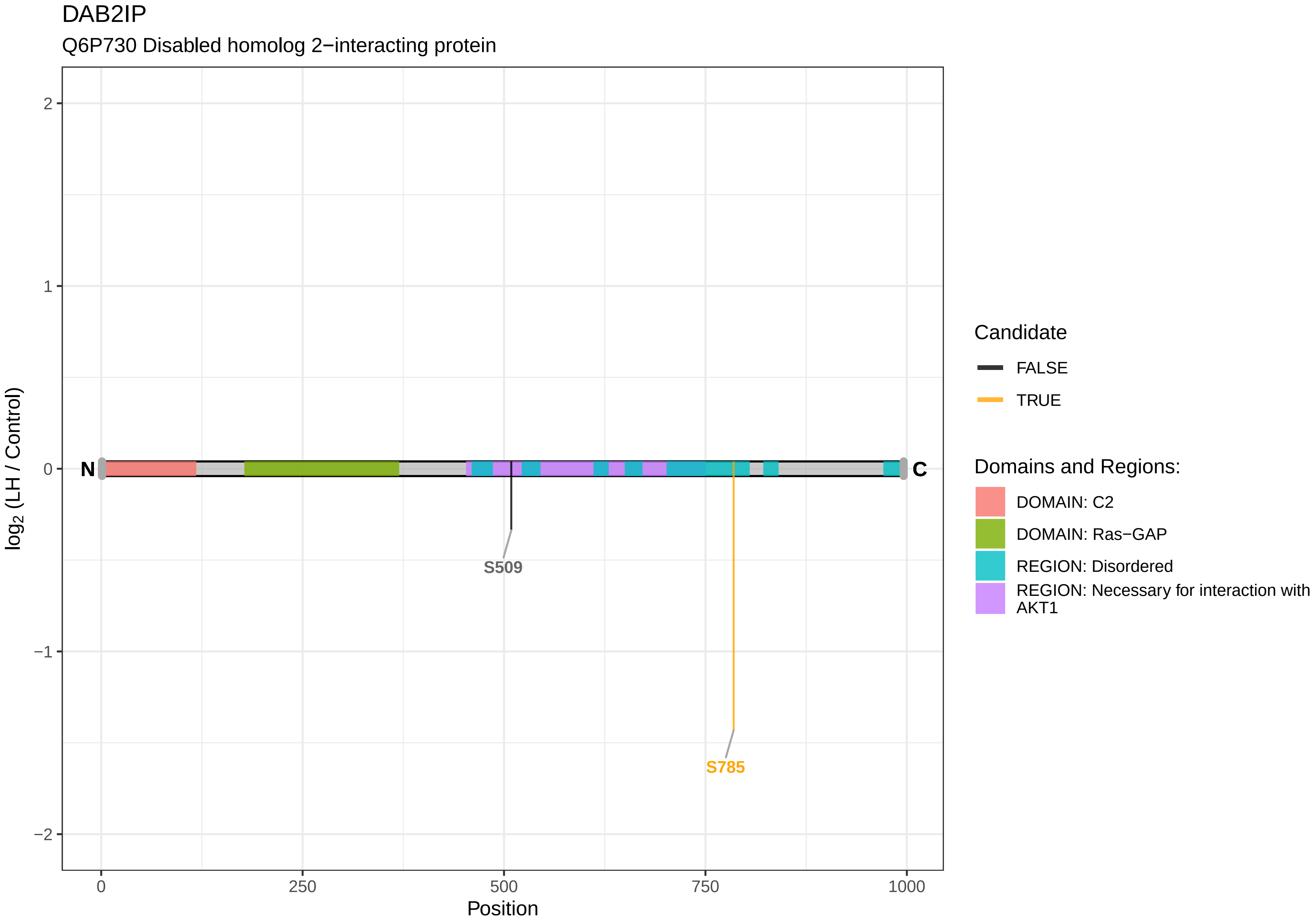

DAB2IP
Q6P730 Disabled homolog 2−interacting protein
2
1
0
−1
−2
log2 (LH / Control)
N
C
S509
S785
0
250
500
750
1000
Position
Candidate
FALSE
TRUE
Domains and Regions:
DOMAIN: C2
DOMAIN: Ras−GAP
REGION: Disordered
REGION: Necessary for interaction with AKT1

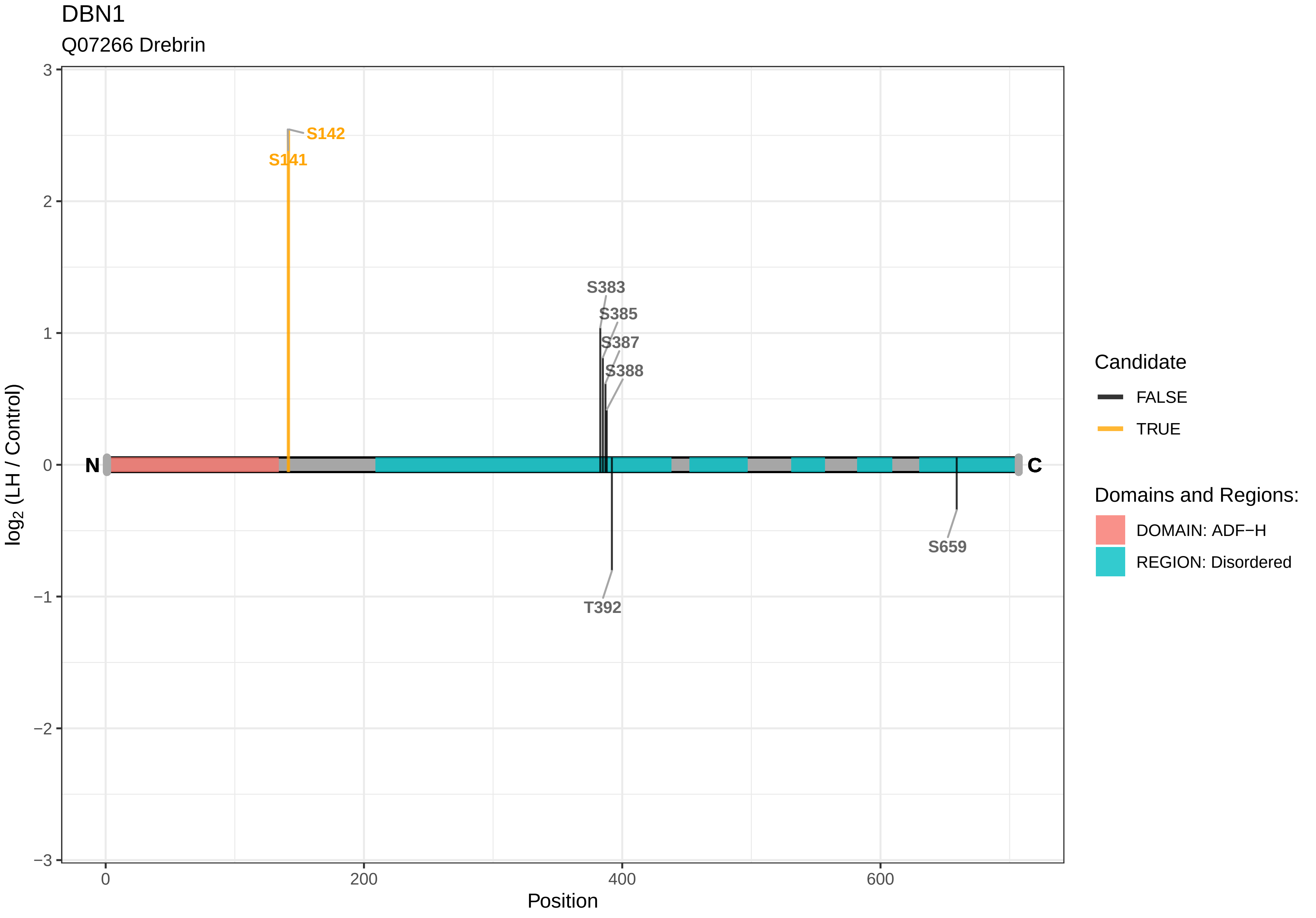
DBN1
Q07266 Drebrin
S142
S141
S383
S385
S387
S388
T392
S659
N
C
log2 (LH / Control)
Position
3
2
1
0
−1
−2
−3
0
200
400
600
Candidate
FALSE
TRUE
Domains and Regions:
DOMAIN: ADF−H
REGION: Disordered

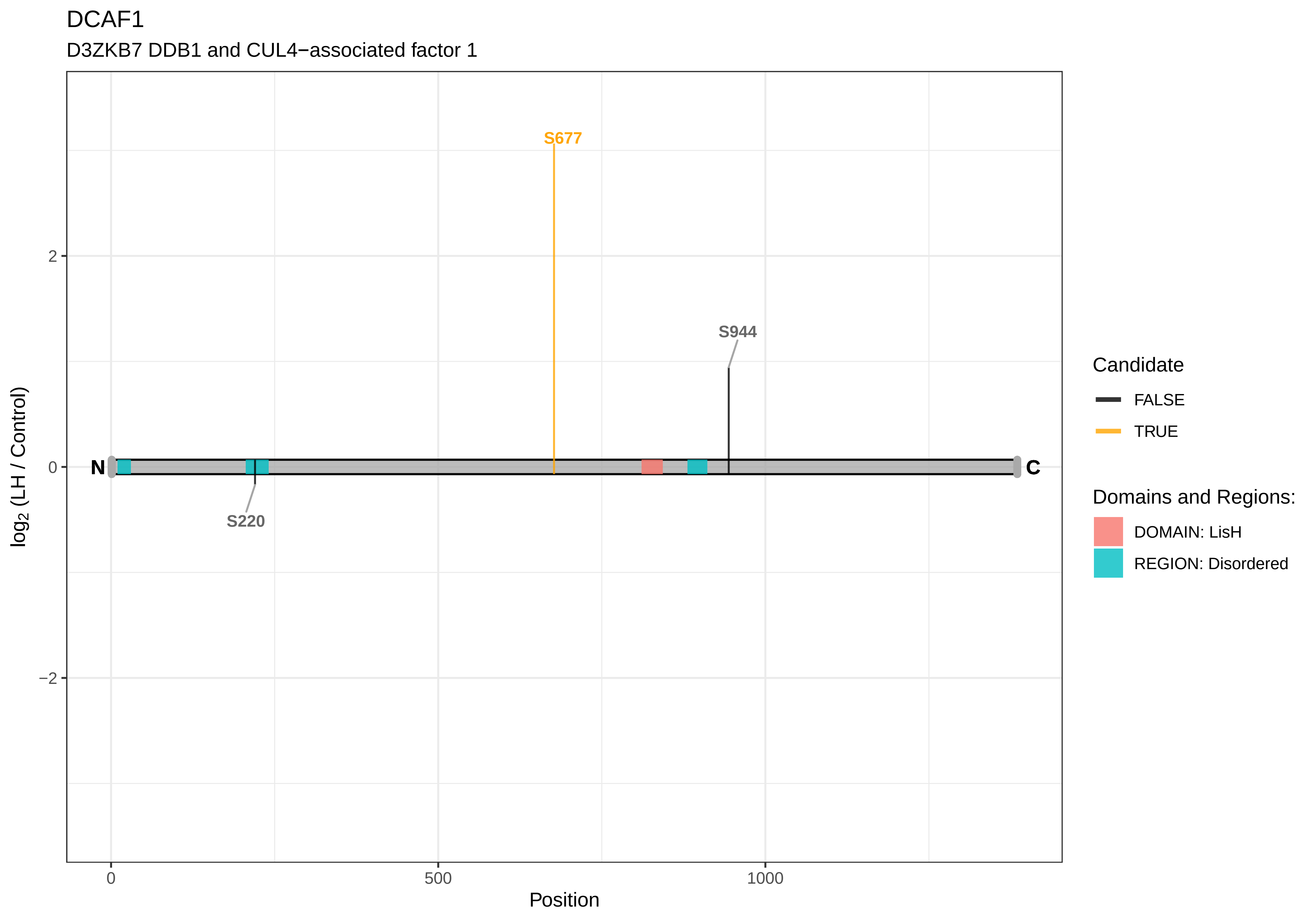

DCAF1
D3ZKB7 DDB1 and CUL4−associated factor 1
S677
S944
S220
N
C
2
0
−2
0
500
1000
Position
log2 (LH / Control)
Candidate
FALSE
TRUE
Domains and Regions:
DOMAIN: LisH
REGION: Disordered

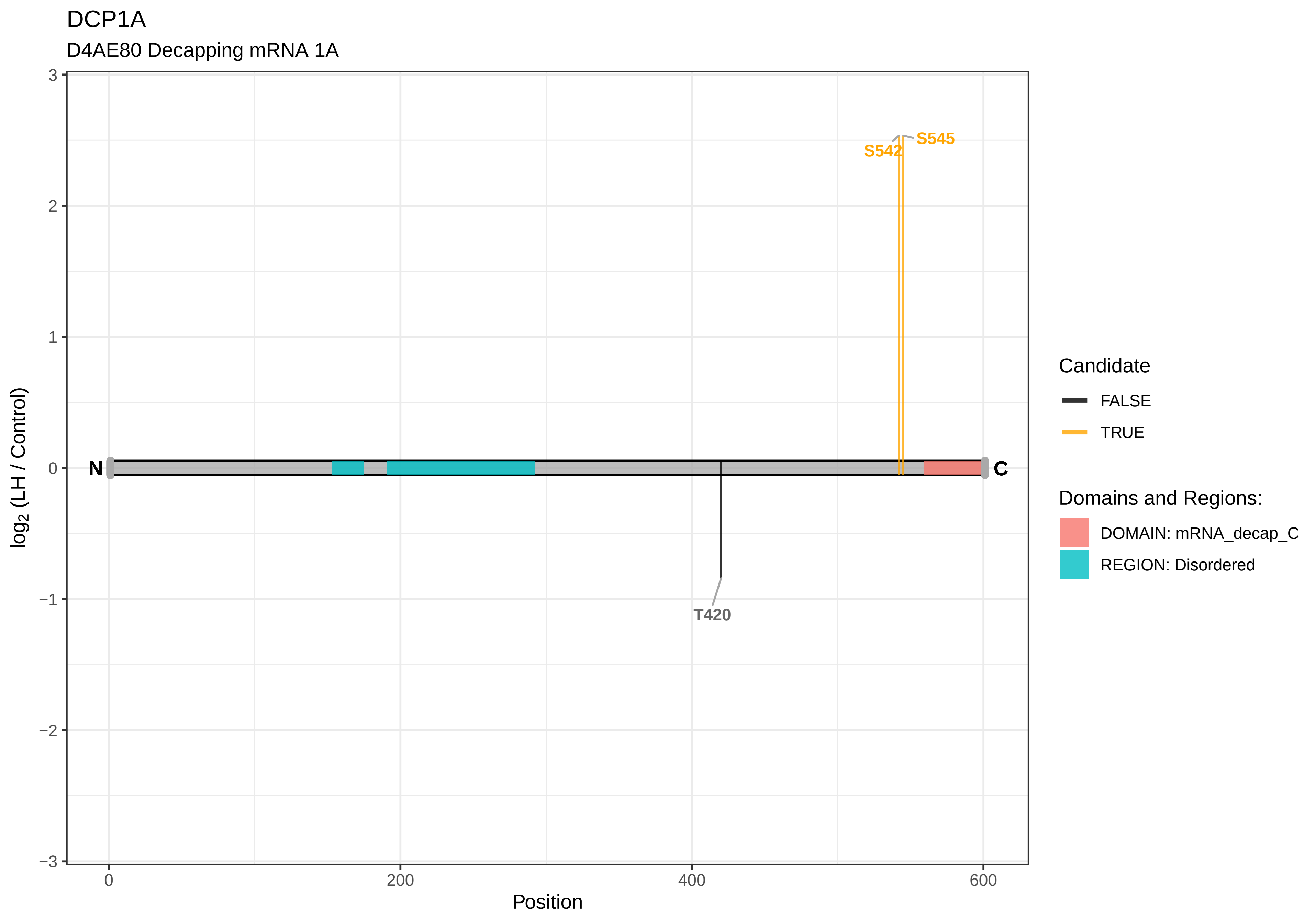
DCP1A
D4AE80 Decapping mRNA 1A
S545
S542
T420
N
C
log2 (LH / Control)
Position
Candidate
FALSE
TRUE
Domains and Regions:
DOMAIN: mRNA_decap_C
REGION: Disordered

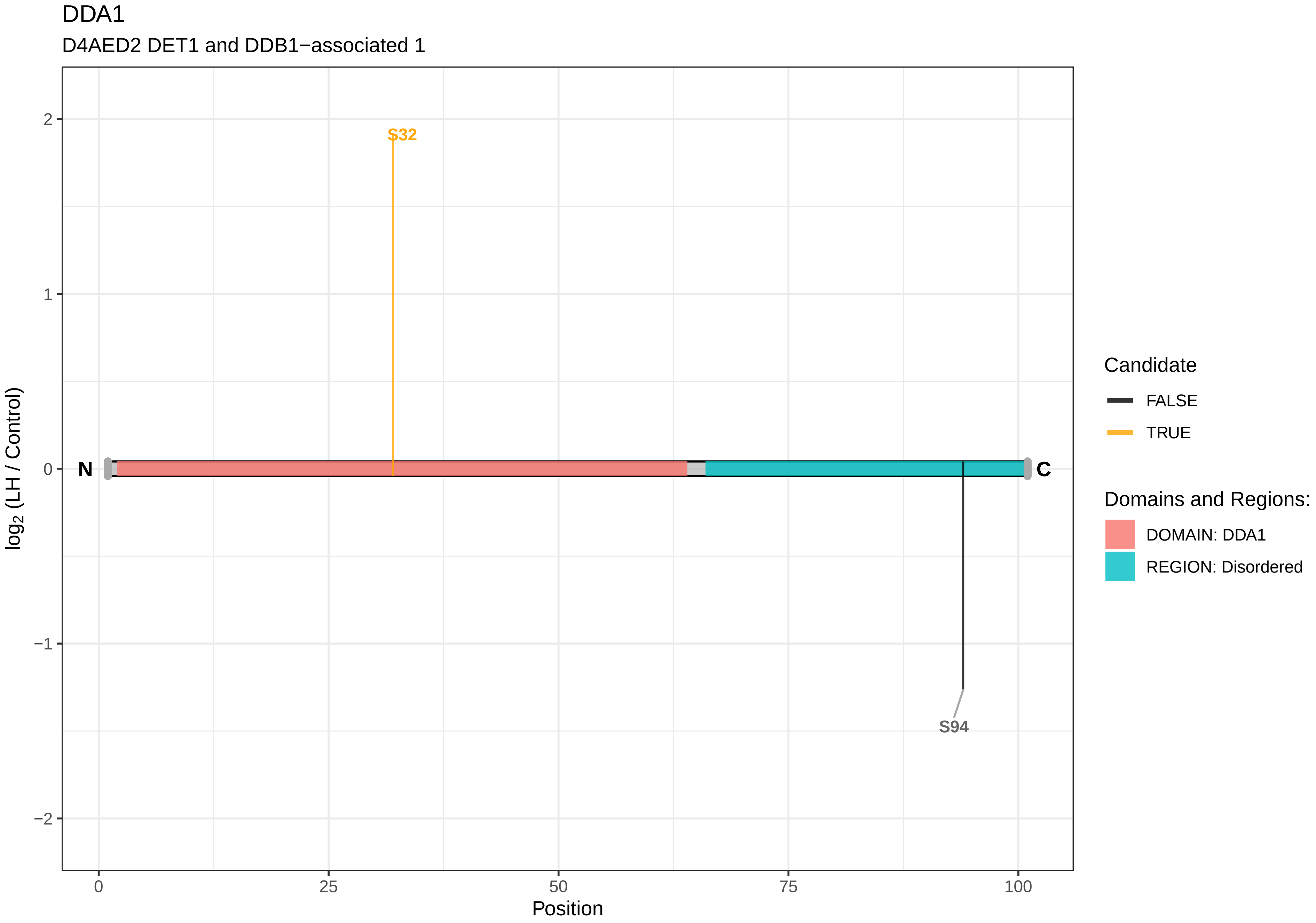

DDA1
D4AED2 DET1 and DDB1−associated 1
S32
S94
N
C
log2 (LH / Control)
2
1
0
−1
−2
0
25
50
75
100
Position
Candidate
FALSE
TRUE
Domains and Regions:
DOMAIN: DDA1
REGION: Disordered

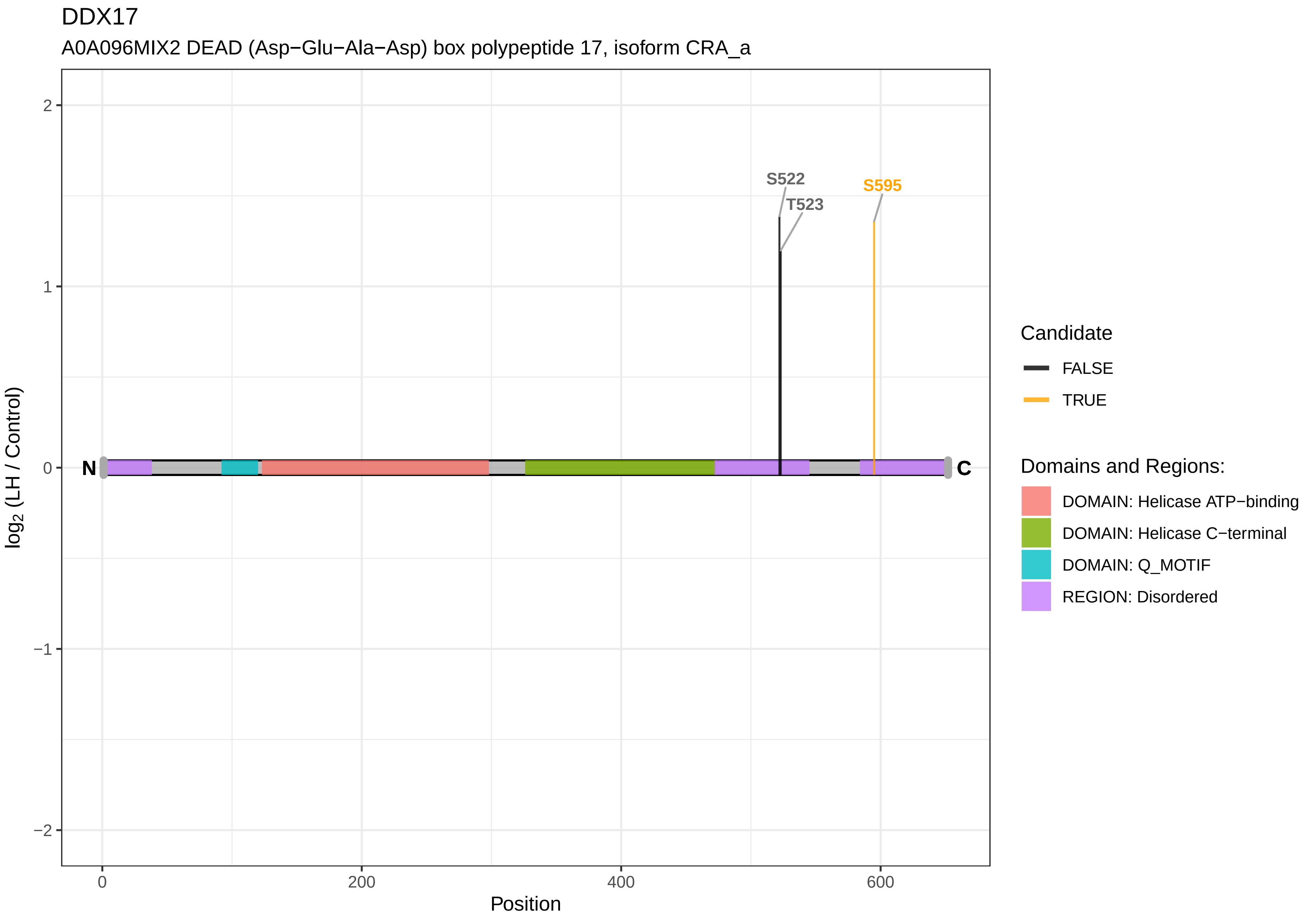

DDX17
A0A096MIX2 DEAD (Asp−Glu−Ala−Asp) box polypeptide 17, isoform CRA_a
S522
T523
S595
N
C
2
1
0
−1
−2
log2 (LH / Control)
0
200
400
600
Position
Candidate
FALSE
TRUE
Domains and Regions:
DOMAIN: Helicase ATP−binding
DOMAIN: Helicase C−terminal
DOMAIN: Q_MOTIF
REGION: Disordered

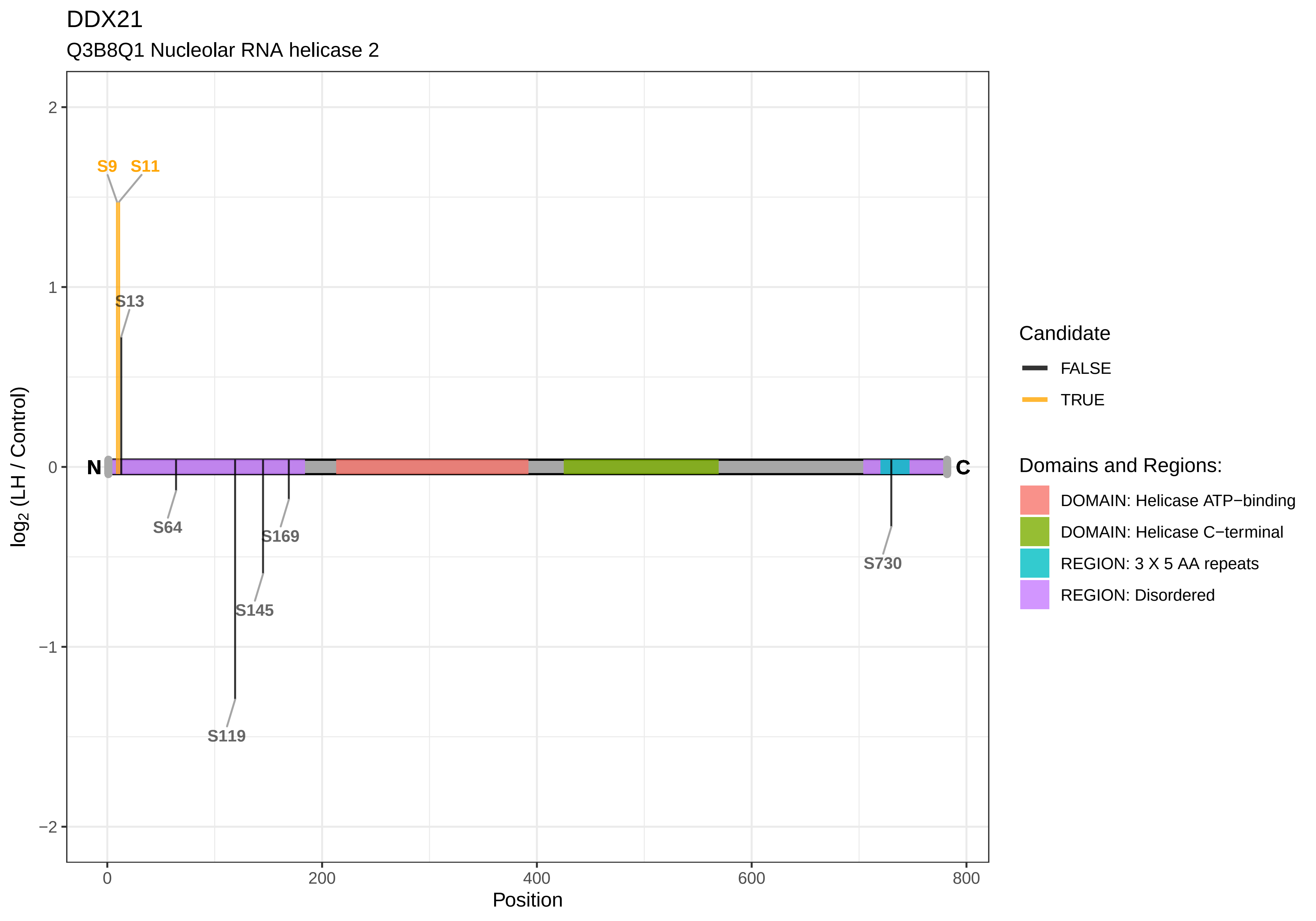
DDX21
Q3B8Q1 Nucleolar RNA helicase 2
S9
S11
S13
S64
S169
S145
S119
S730
N
C
2
1
0
−1
−2
0
200
400
600
800
Position
log2 (LH / Control)
Candidate
FALSE
TRUE
Domains and Regions:
DOMAIN: Helicase ATP−binding
DOMAIN: Helicase C−terminal
REGION: 3 X 5 AA repeats
REGION: Disordered

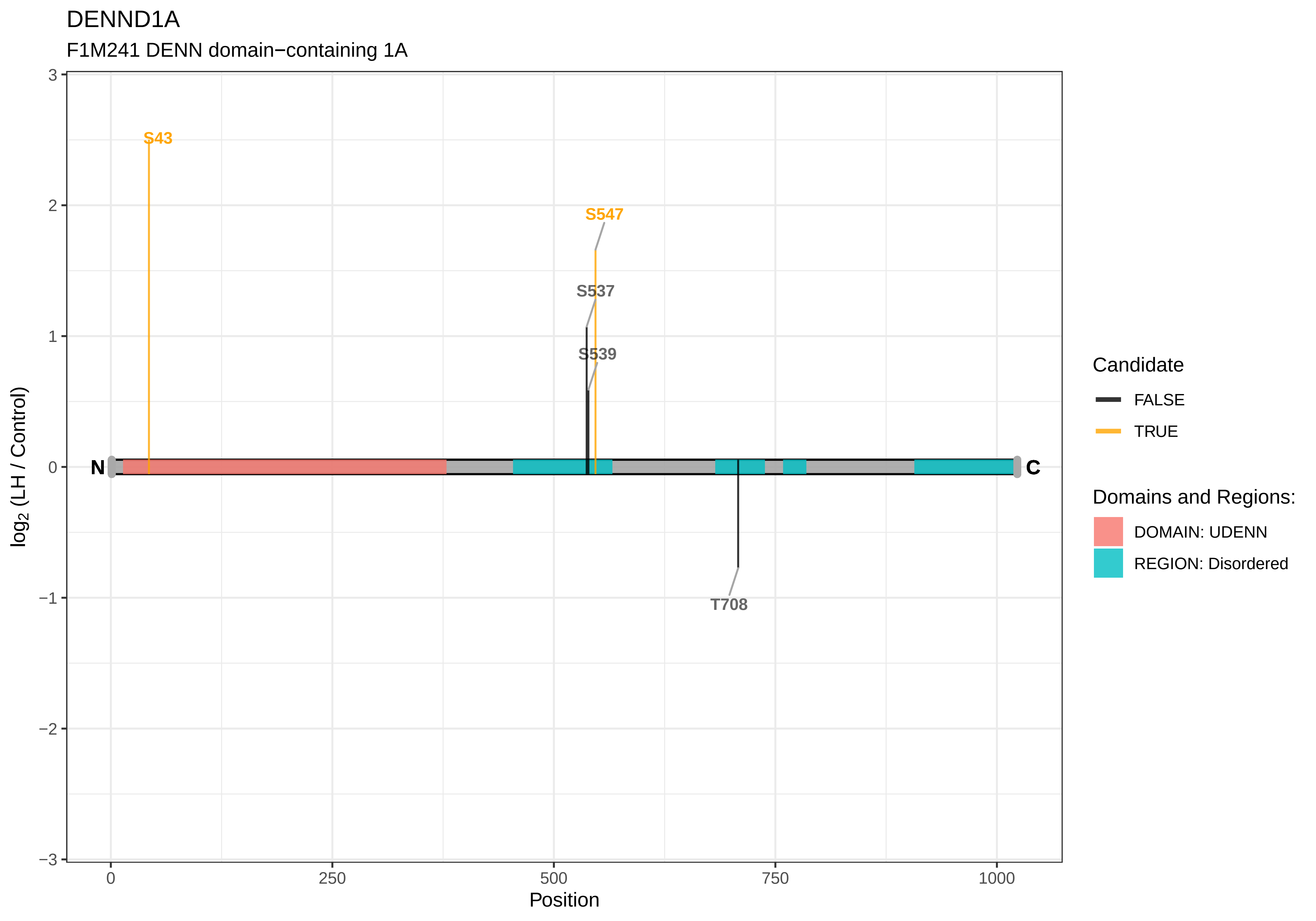

DENND1A
F1M241 DENN domain−containing 1A
S43
S547
S537
S539
T708
N
C
3
2
1
0
−1
−2
−3
log2 (LH / Control)
0
250
500
750
1000
Position
Candidate
FALSE
TRUE
Domains and Regions:
DOMAIN: UDENN
REGION: Disordered

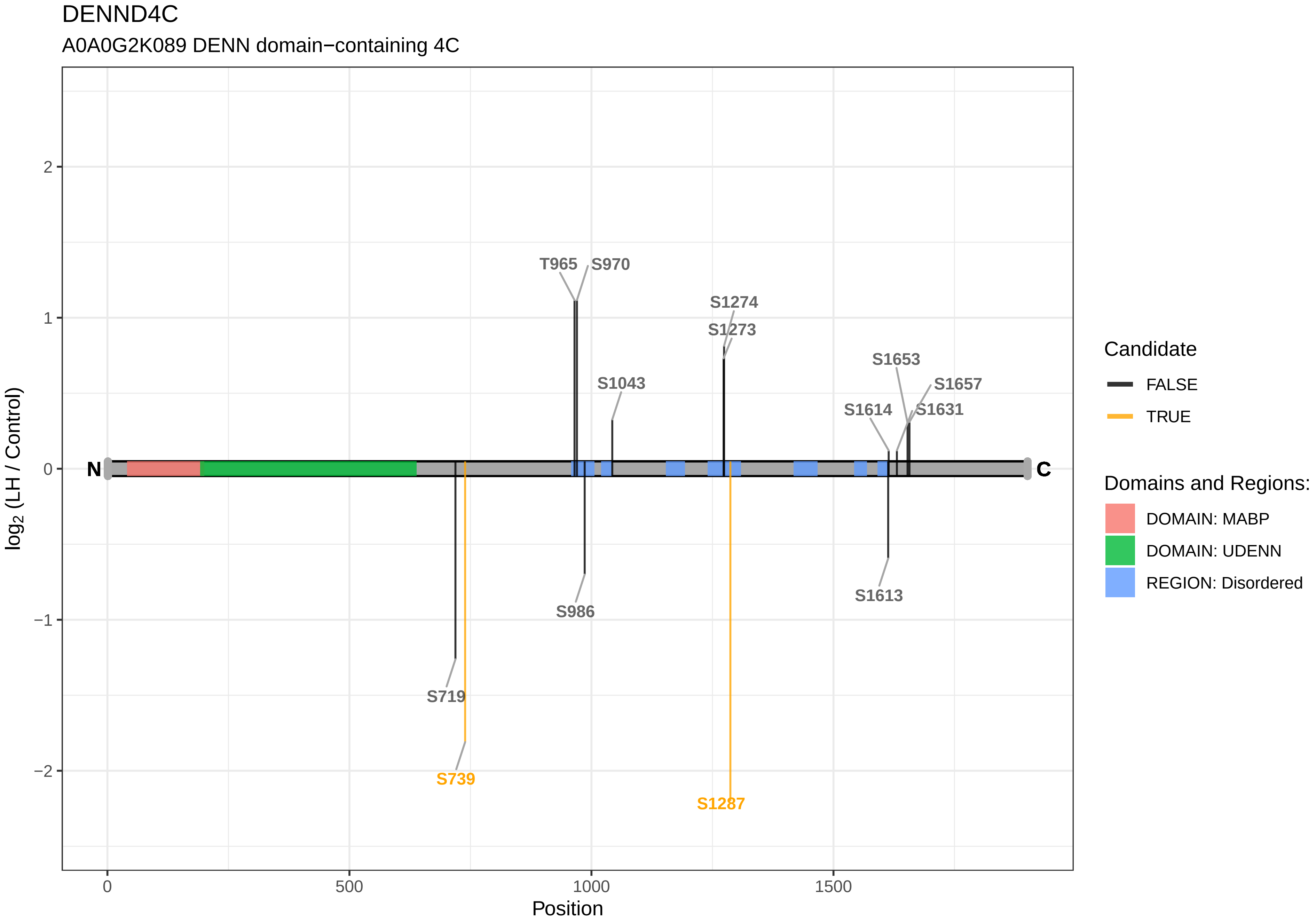

DENND4C
A0A0G2K089 DENN domain−containing 4C
log2 (LH / Control)
Position
T965
S970
S1274
S1273
S1653
S1657
S1043
S1614
S1631
S986
S1613
S719
S739
S1287
N
C
2
1
0
−1
−2
0
500
1000
1500
Candidate
FALSE
TRUE
Domains and Regions:
DOMAIN: MABP
DOMAIN: UDENN
REGION: Disordered

### DENND5A

G3V7Q0 DENN domain−containing protein 5A

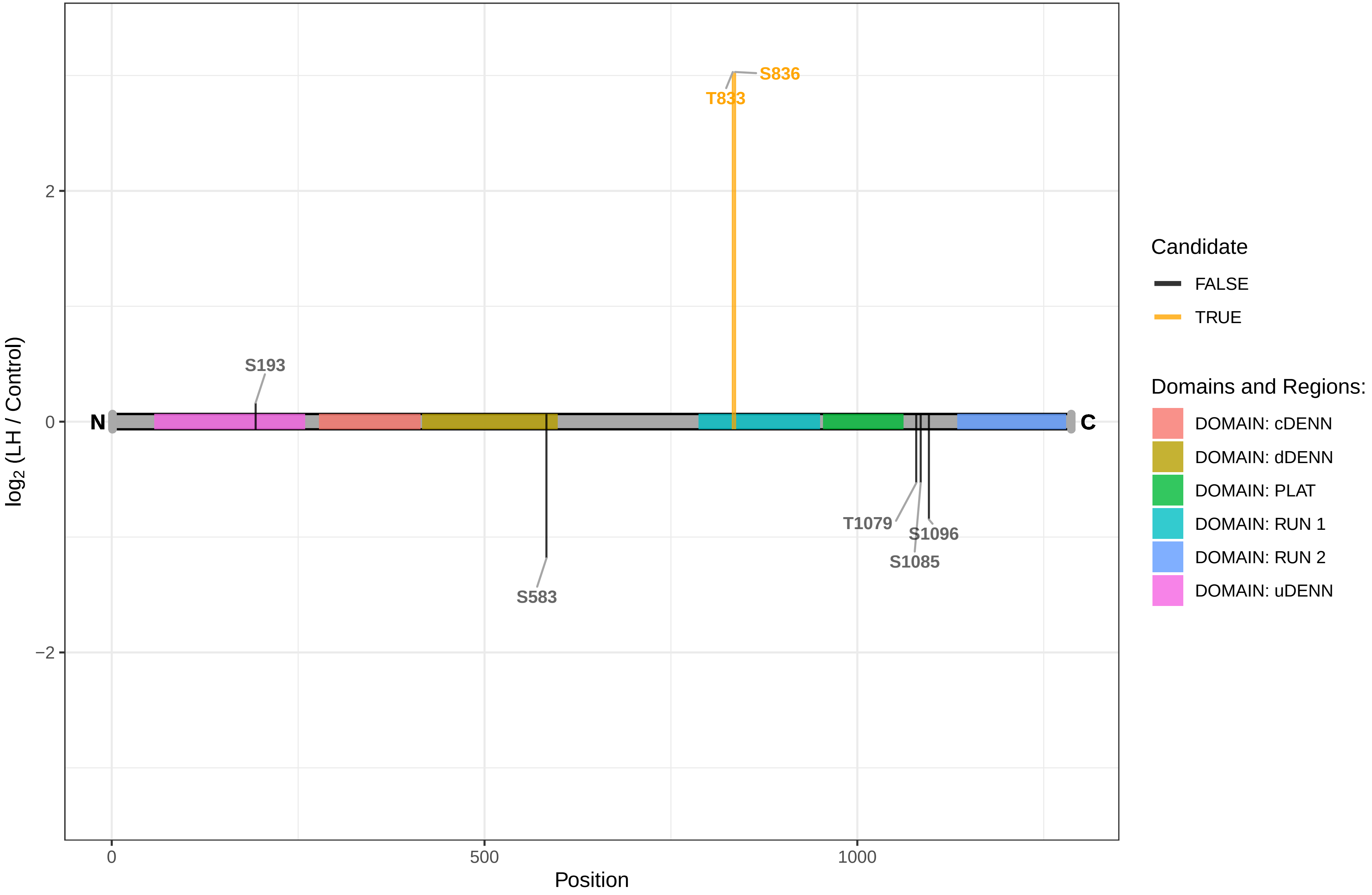

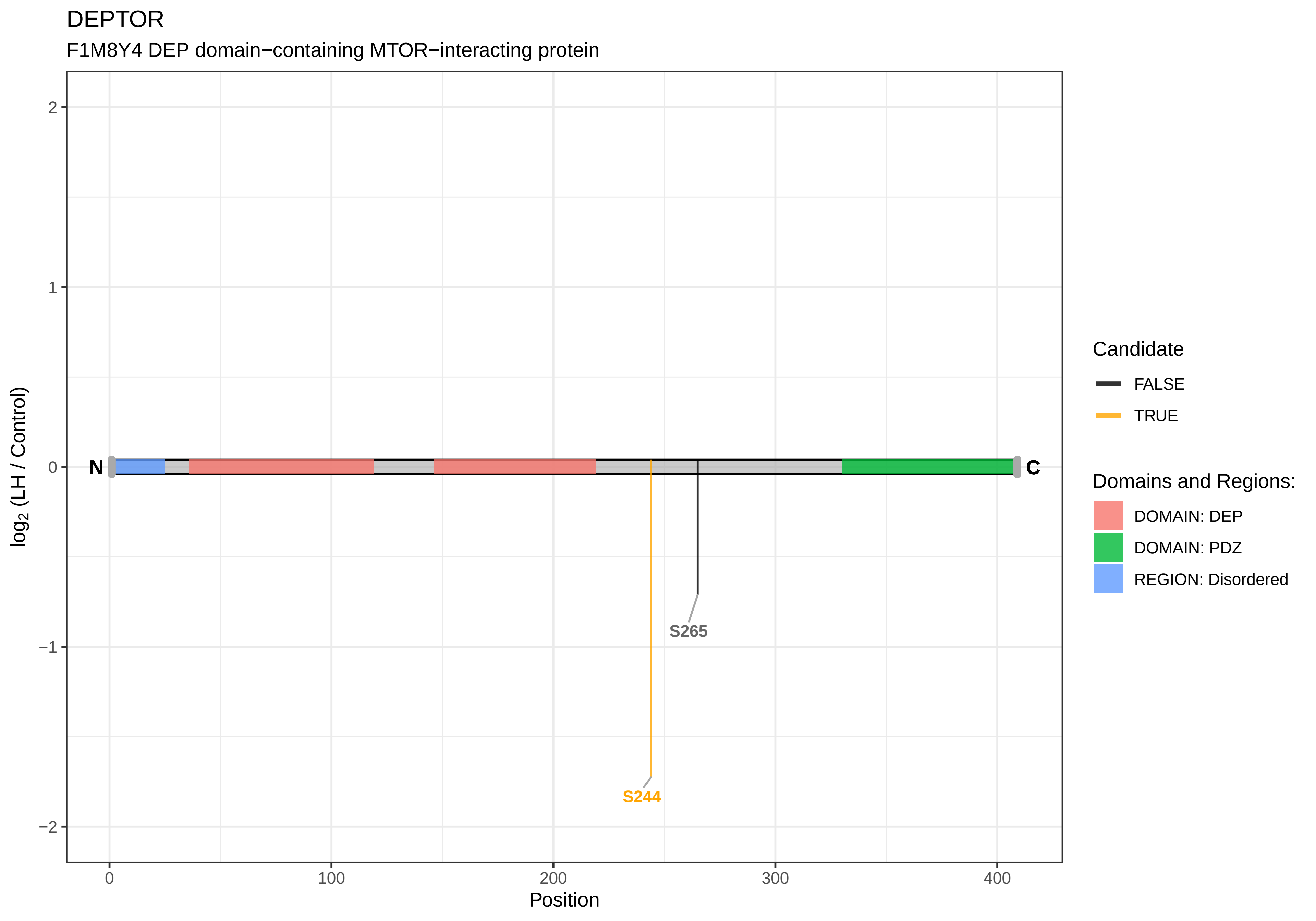
DEPTOR
F1M8Y4 DEP domain−containing MTOR−interacting protein
log2 (LH / Control)
2
1
0
−1
−2
N
C
S265
S244
0
100
200
300
400
Position
Candidate
FALSE
TRUE
Domains and Regions:
DOMAIN: DEP
DOMAIN: PDZ
REGION: Disordered

DGAT1

Q8BHI5 O−acyltransferase

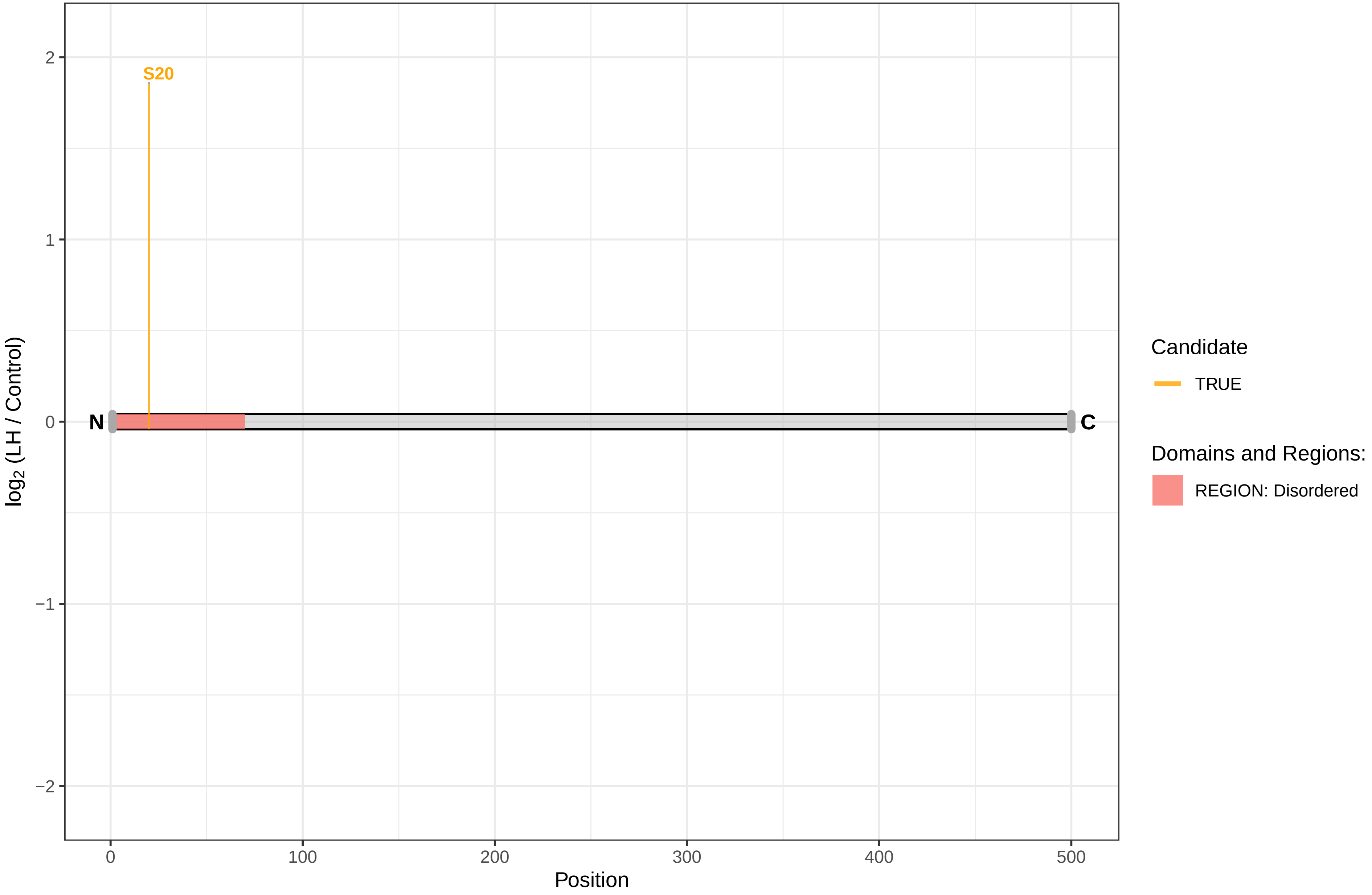

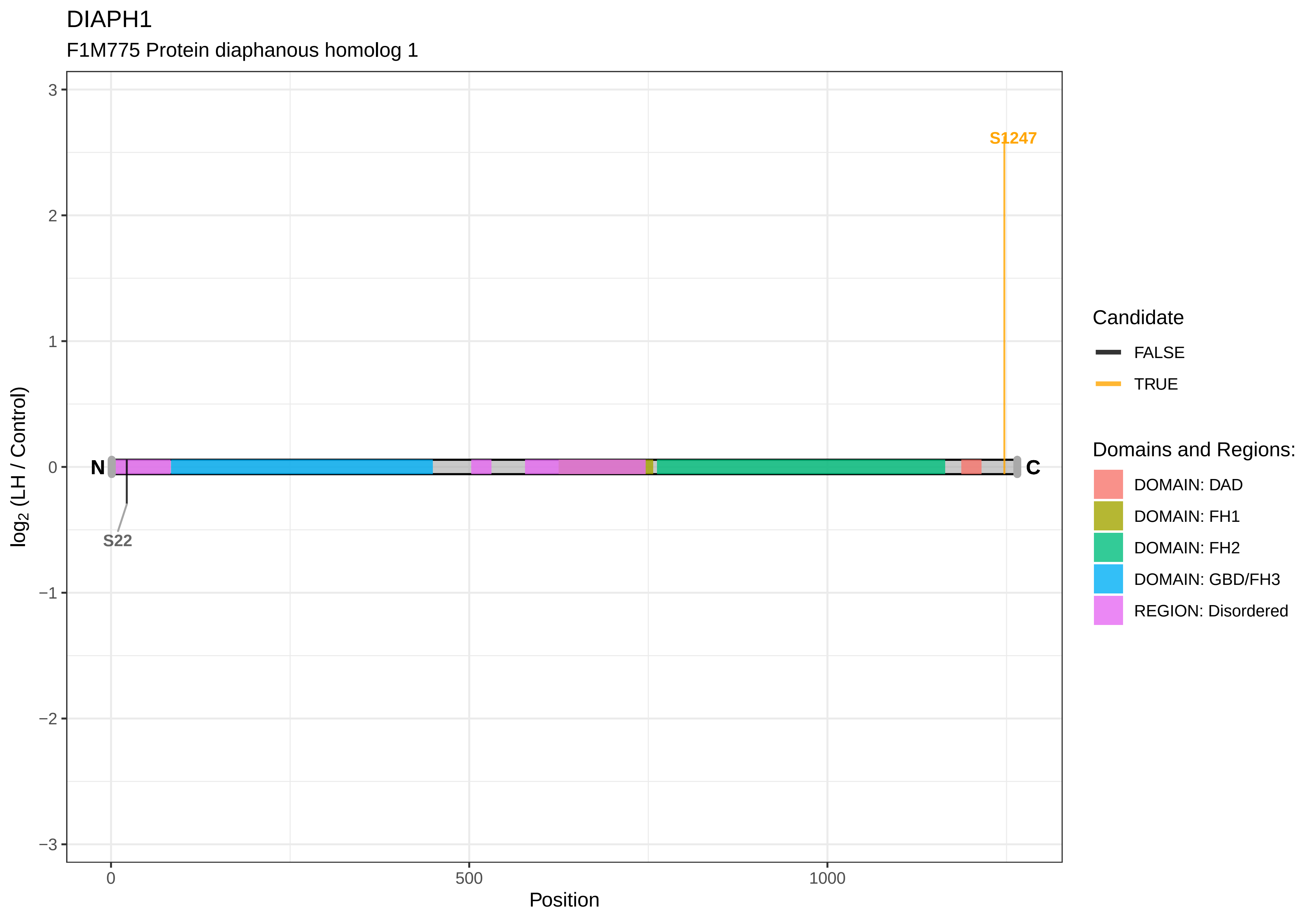

DIAPH1
F1M775 Protein diaphanous homolog 1
S1247
S22
N
C
3
2
1
0
−1
−2
−3
log2 (LH / Control)
0
500
1000
Position
Candidate
FALSE
TRUE
Domains and Regions:
DOMAIN: DAD
DOMAIN: FH1
DOMAIN: FH2
DOMAIN: GBD/FH3
REGION: Disordered

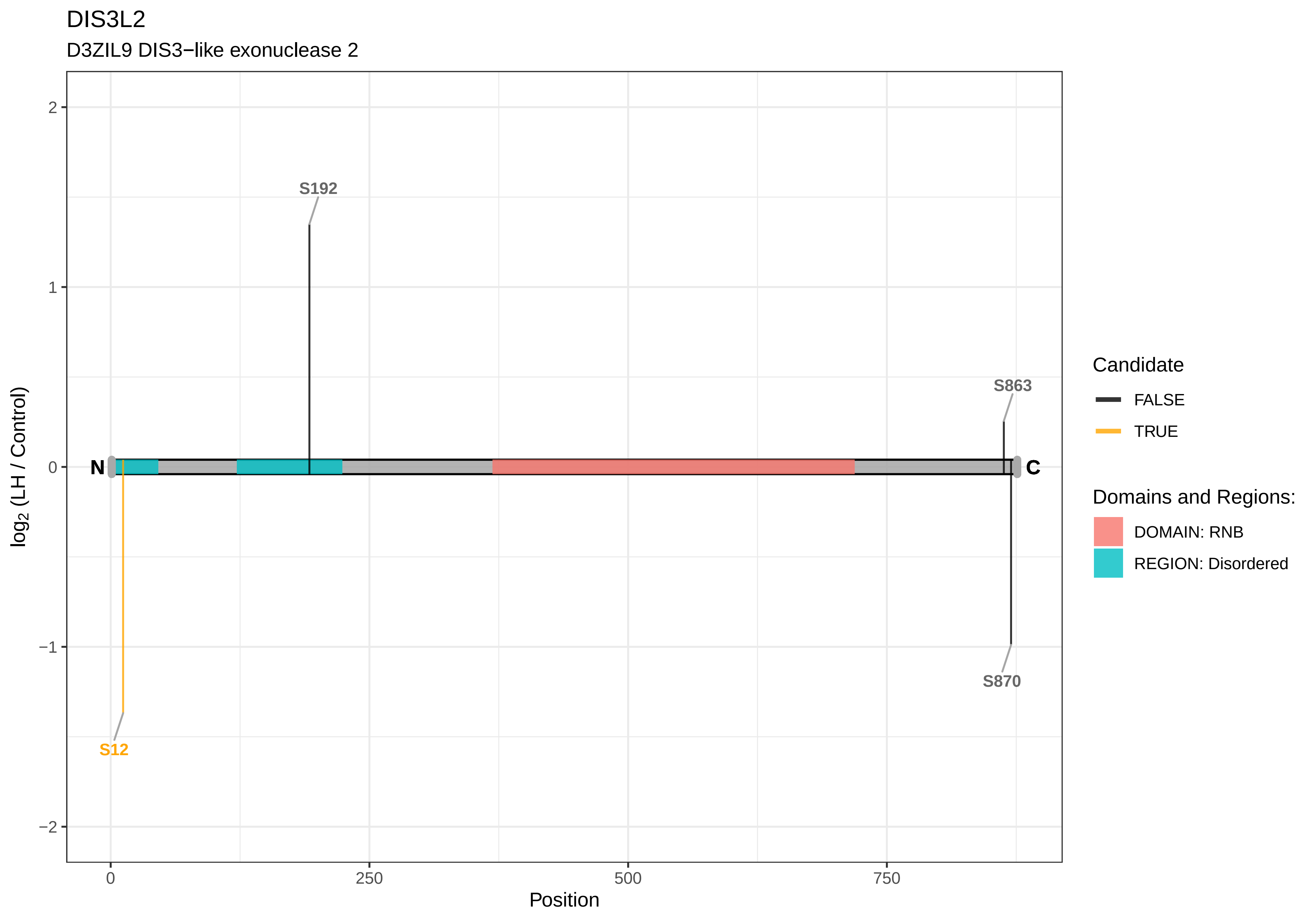
DIS3L2
D3ZIL9 DIS3−like exonuclease 2
S192
S863
S870
S12
N
C
2
1
0
−1
−2
log2 (LH / Control)
0
250
500
750
Position
Candidate
FALSE
TRUE
Domains and Regions:
DOMAIN: RNB
REGION: Disordered

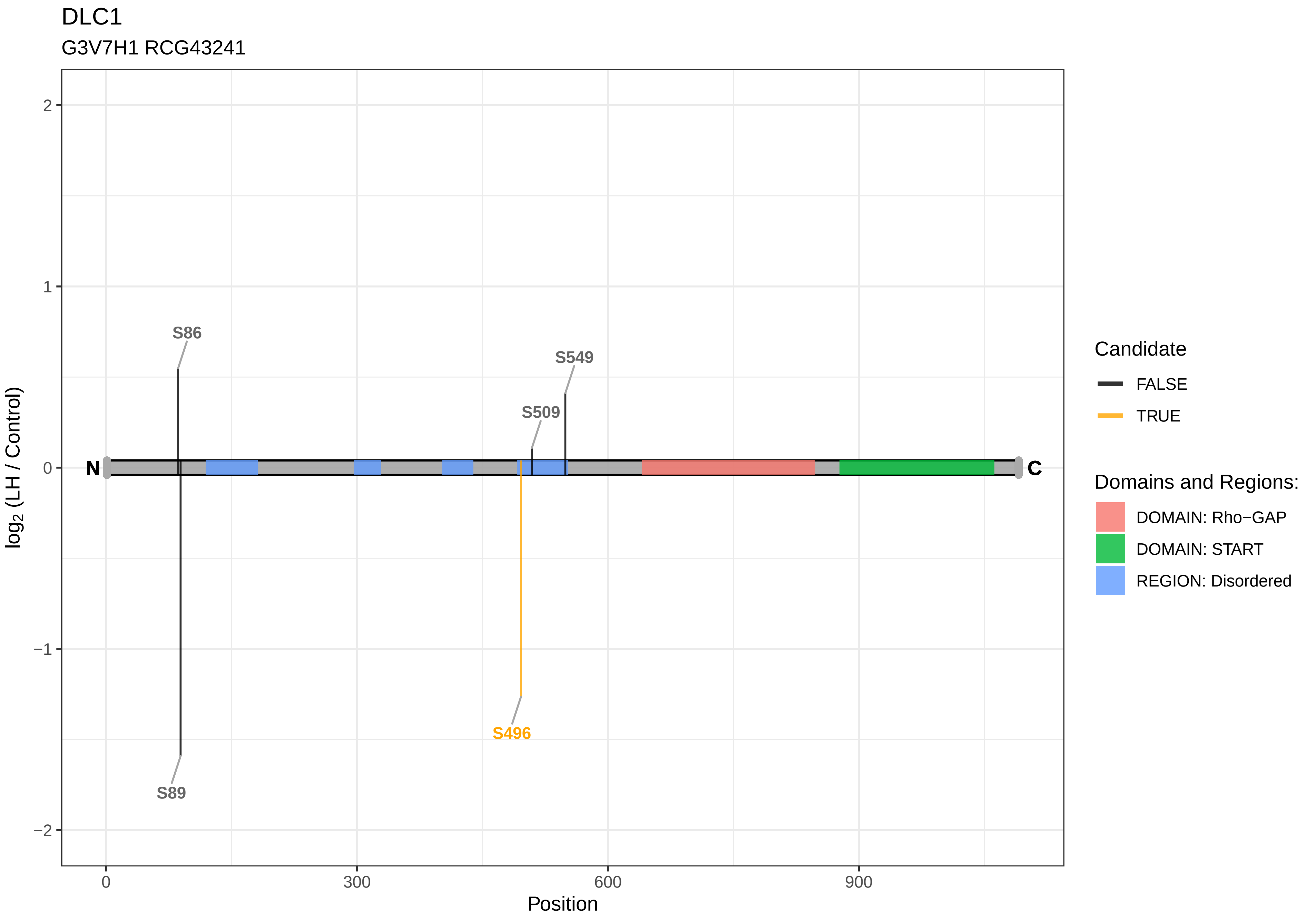
DLC1
G3V7H1 RCG43241
S86
S549
S509
S496
S89
N
C
2
1
0
−1
−2
log2 (LH / Control)
0
300
600
900
Position
Candidate
FALSE
TRUE
Domains and Regions:
DOMAIN: Rho−GAP
DOMAIN: START
REGION: Disordered

### DMAP1

Q568Y6 DNA methyltransferase 1−associated protein 1

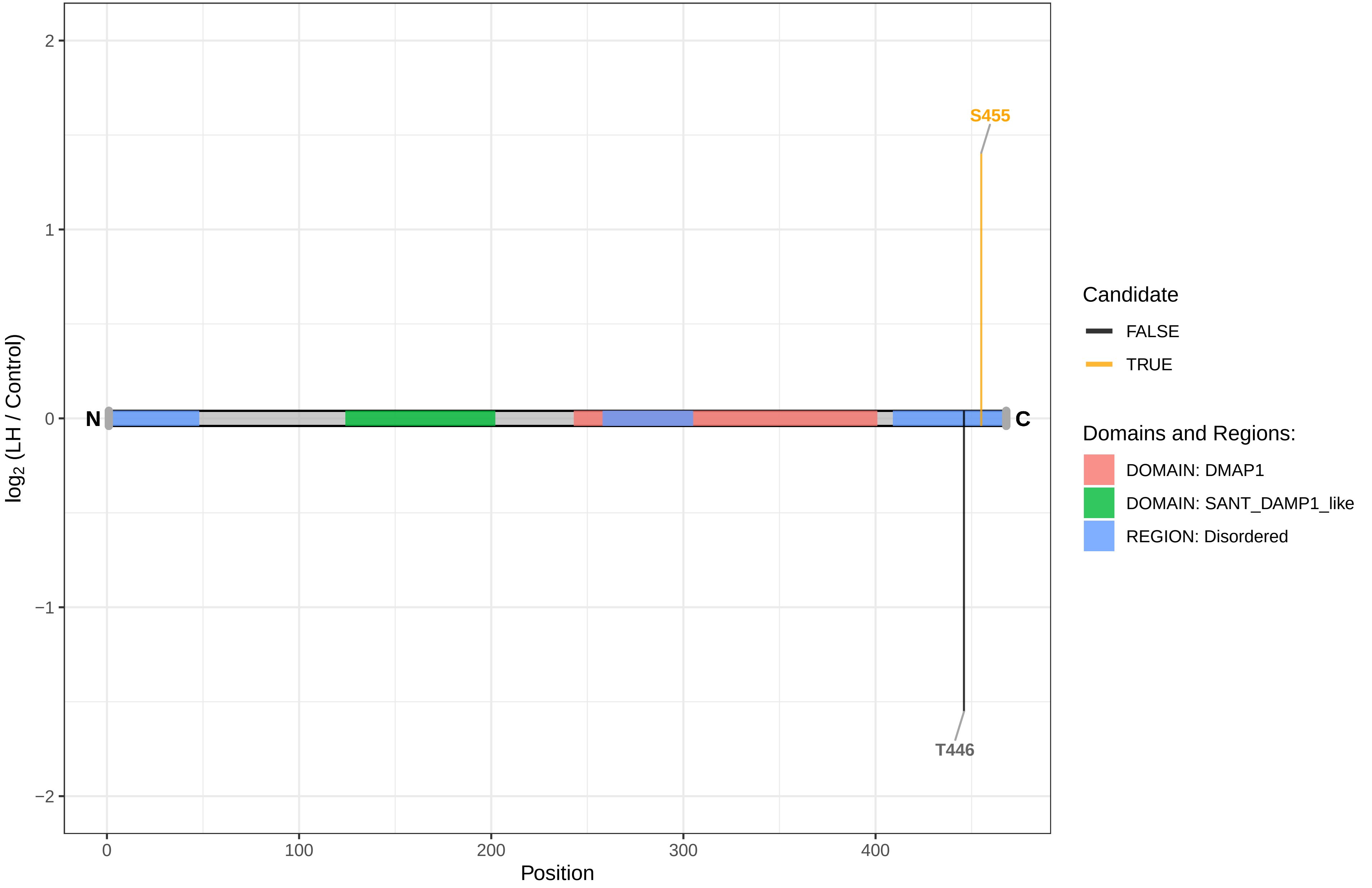

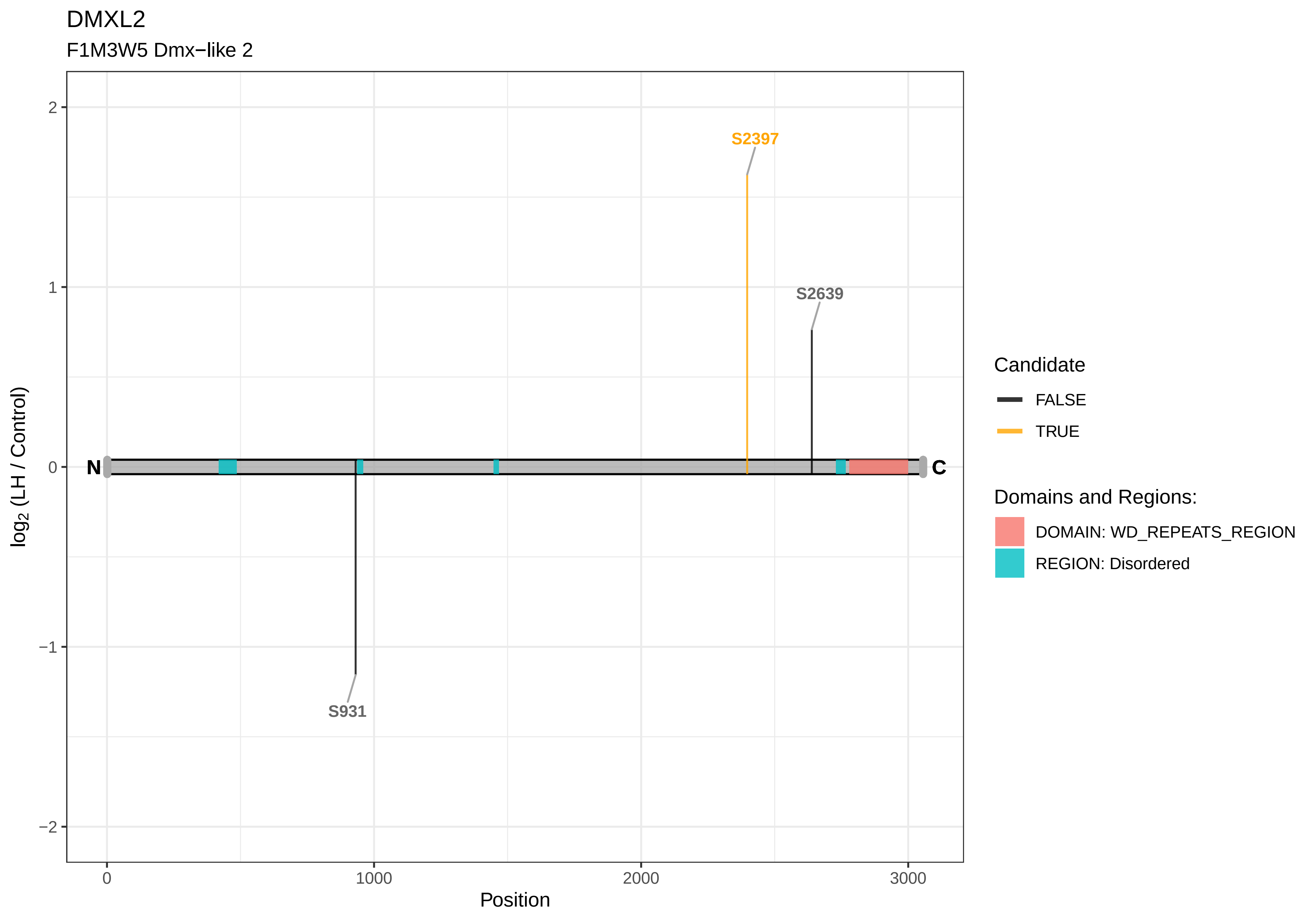

DMXL2
F1M3W5 Dmx−like 2
S2397
S2639
S931
N
C
2
1
0
−1
−2
log2 (LH / Control)
0
1000
2000
3000
Position
Candidate
FALSE
TRUE
Domains and Regions:
DOMAIN: WD_REPEATS_REGION
REGION: Disordered

DNAJB12

Q5FVC4 DnaJ (Hsp40) homolog, subfamily B, member 12

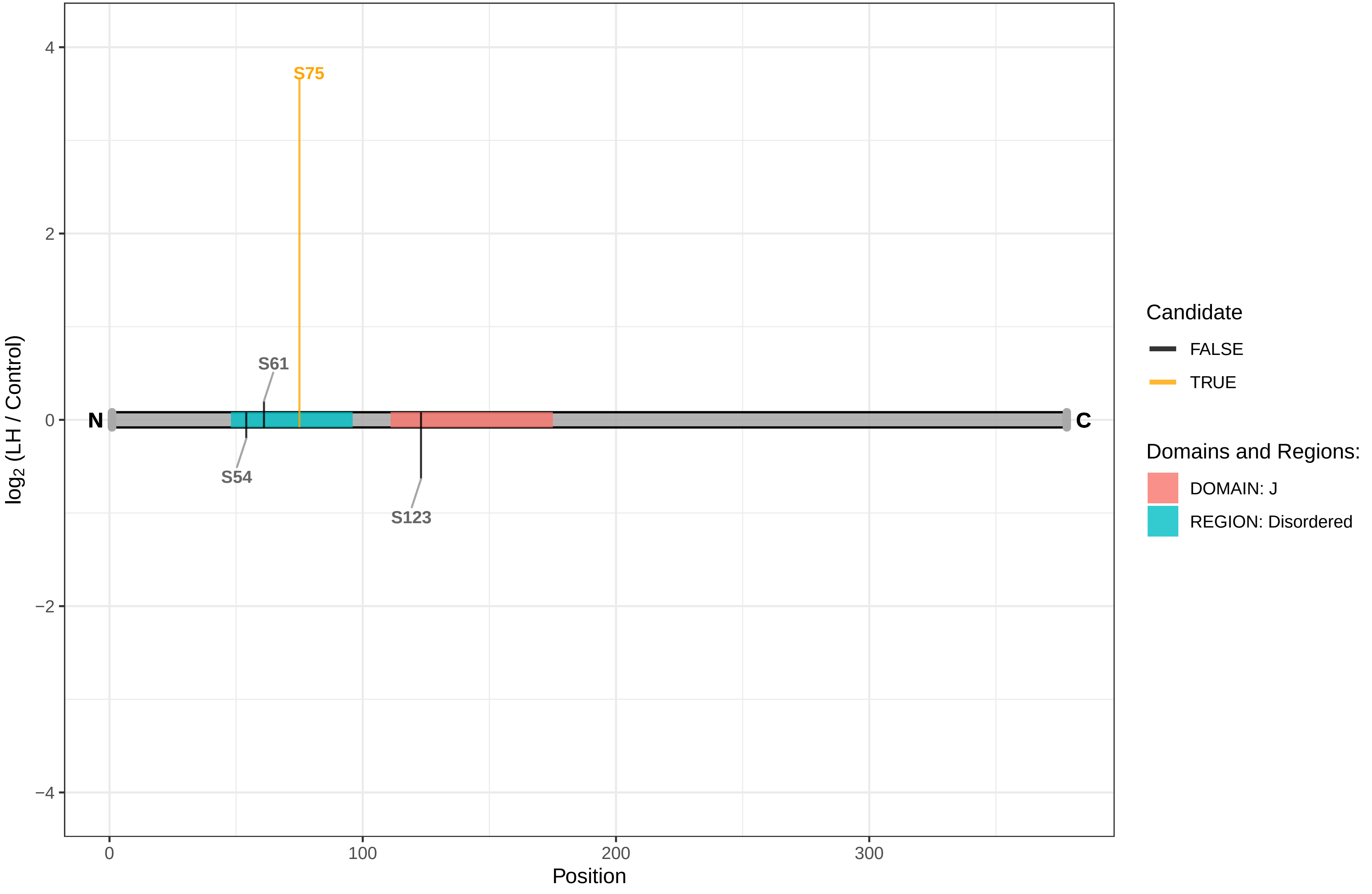

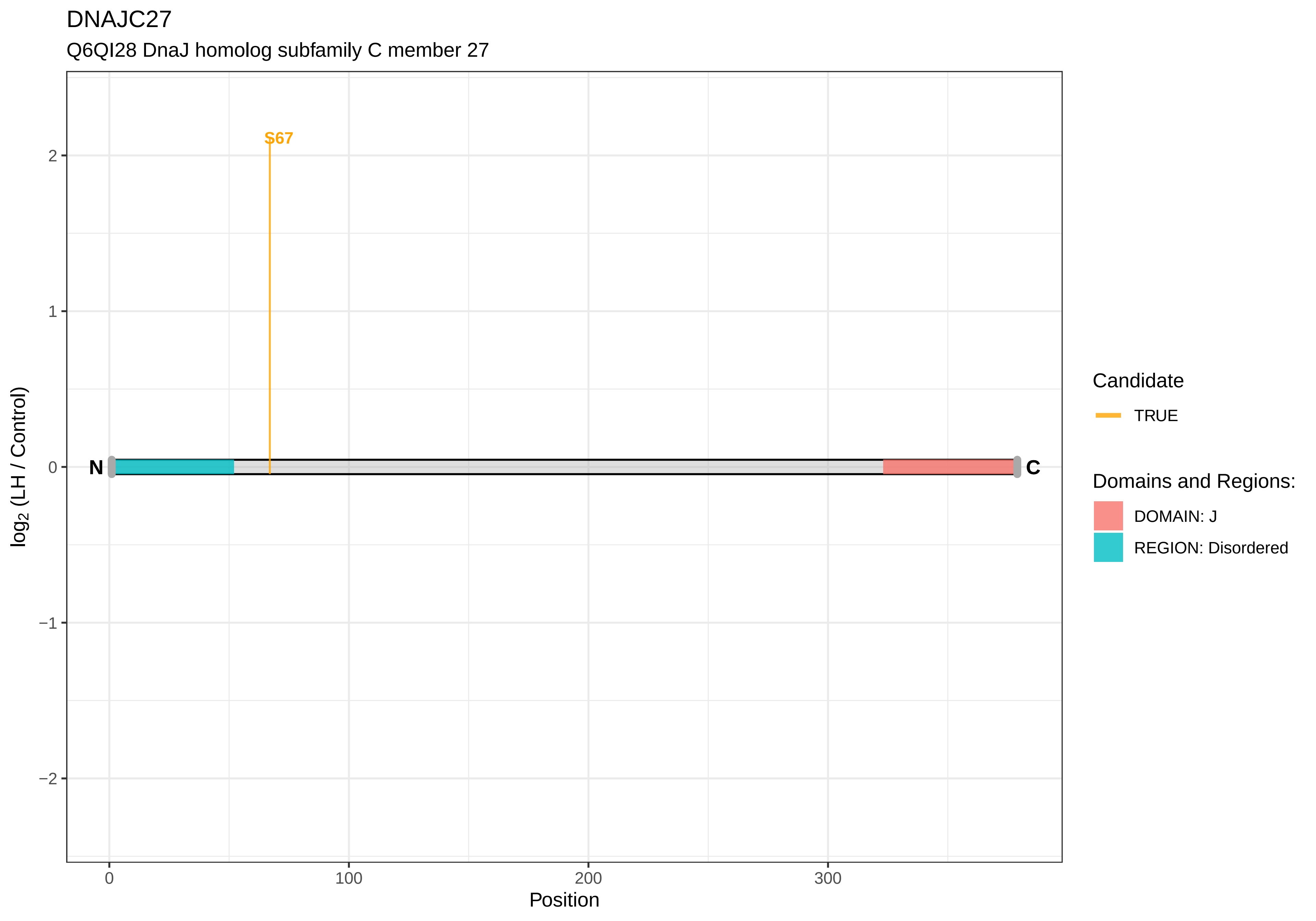

DNAJC27
Q6QI28 DnaJ homolog subfamily C member 27
S67
N
C
2
1
0
−1
−2
0
100
200
300
Position
log2 (LH / Control)
Candidate
TRUE
Domains and Regions:
DOMAIN: J
REGION: Disordered

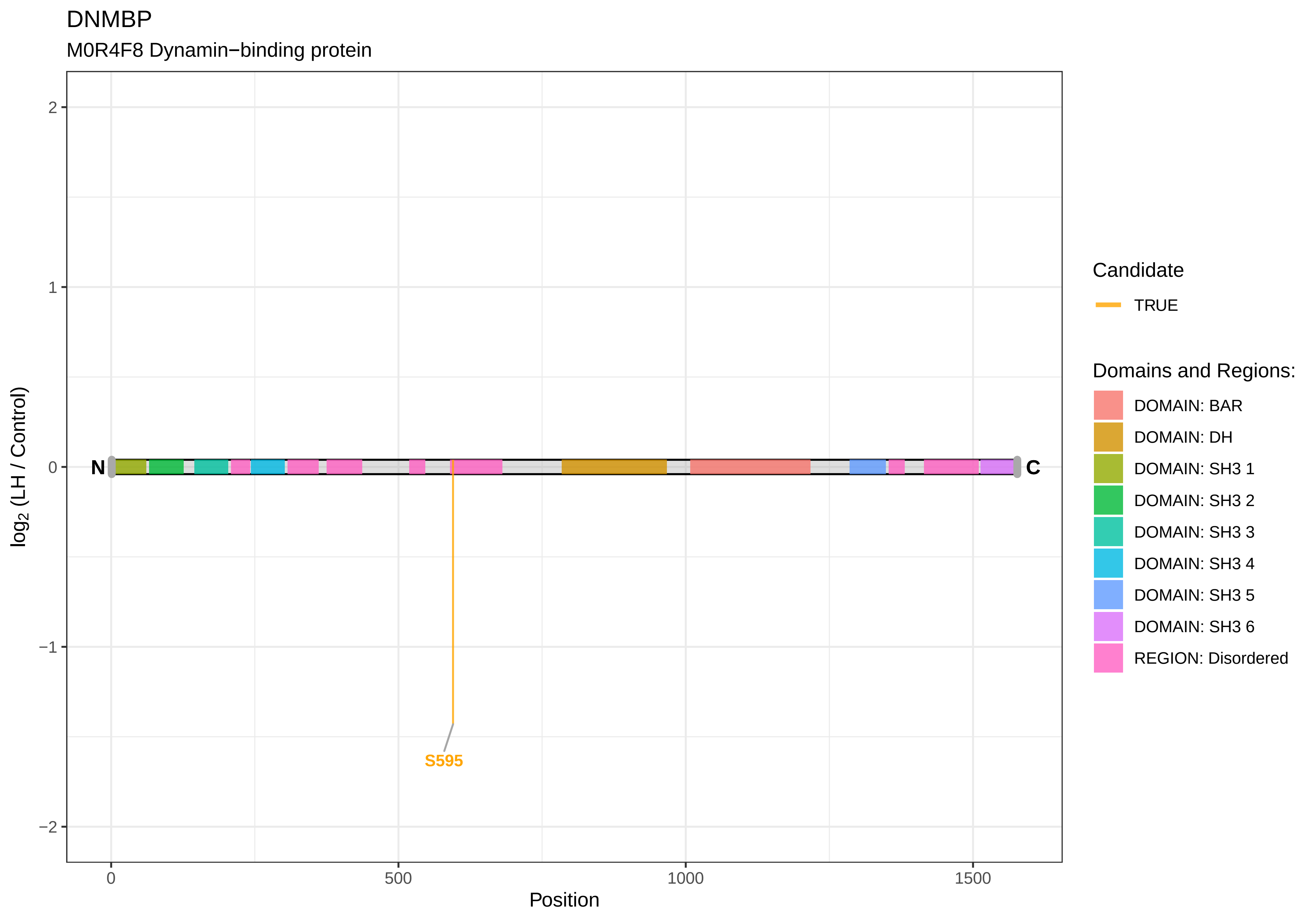
DNMBP
M0R4F8 Dynamin−binding protein
log2 (LH / Control)
2
1
0
−1
−2
N
C
S595
0
500
1000
1500
Position
Candidate
TRUE
Domains and Regions:
DOMAIN: BAR
DOMAIN: DH
DOMAIN: SH3 1
DOMAIN: SH3 2
DOMAIN: SH3 3
DOMAIN: SH3 4
DOMAIN: SH3 5
DOMAIN: SH3 6
REGION: Disordered

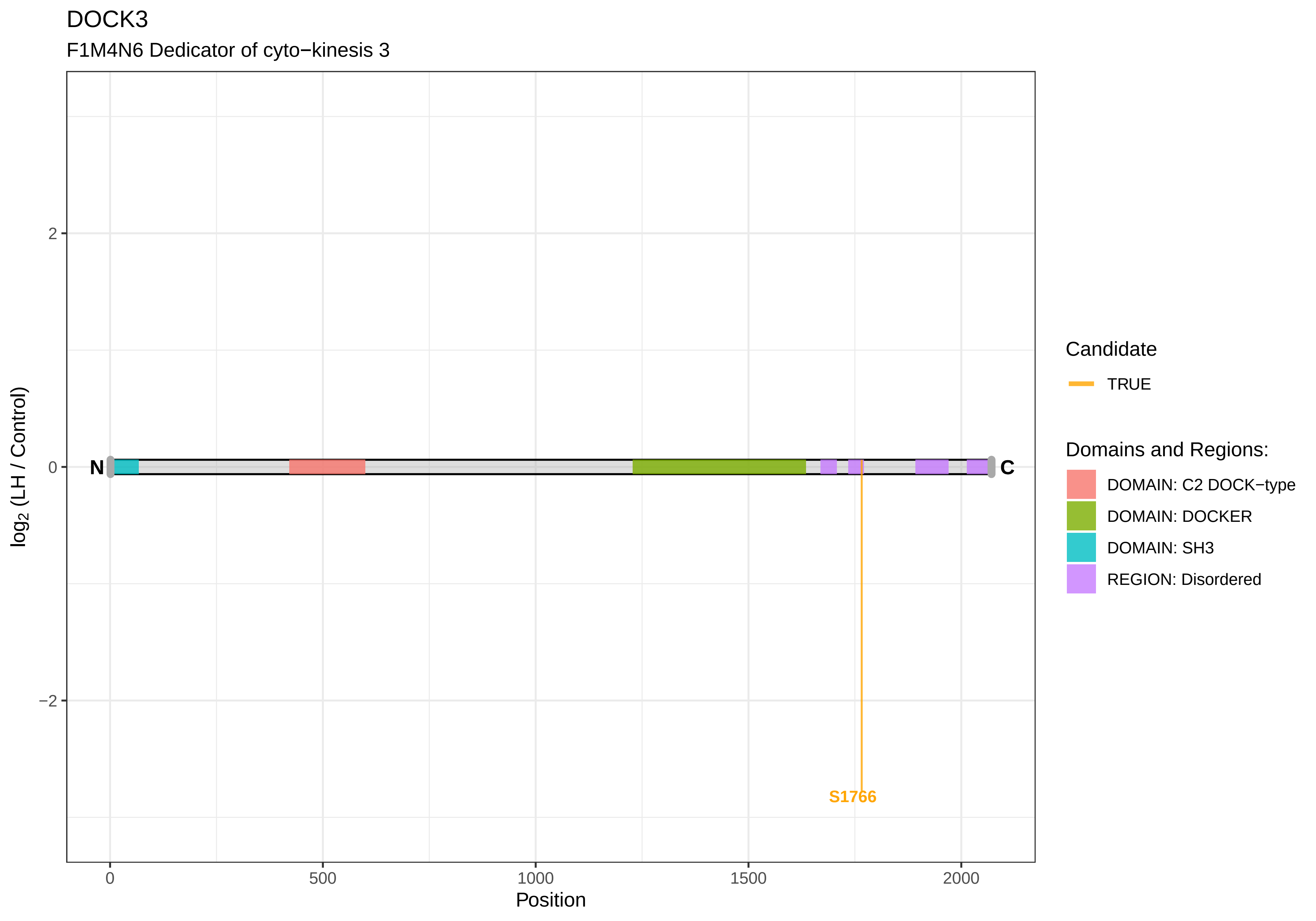

DOCK3
F1M4N6 Dedicator of cyto−kinesis 3
log2 (LH / Control)
2
0
−2
N
C
S1766
0
500
1000
1500
2000
Position
Candidate
TRUE
Domains and Regions:
DOMAIN: C2 DOCK−type
DOMAIN: DOCKER
DOMAIN: SH3
REGION: Disordered

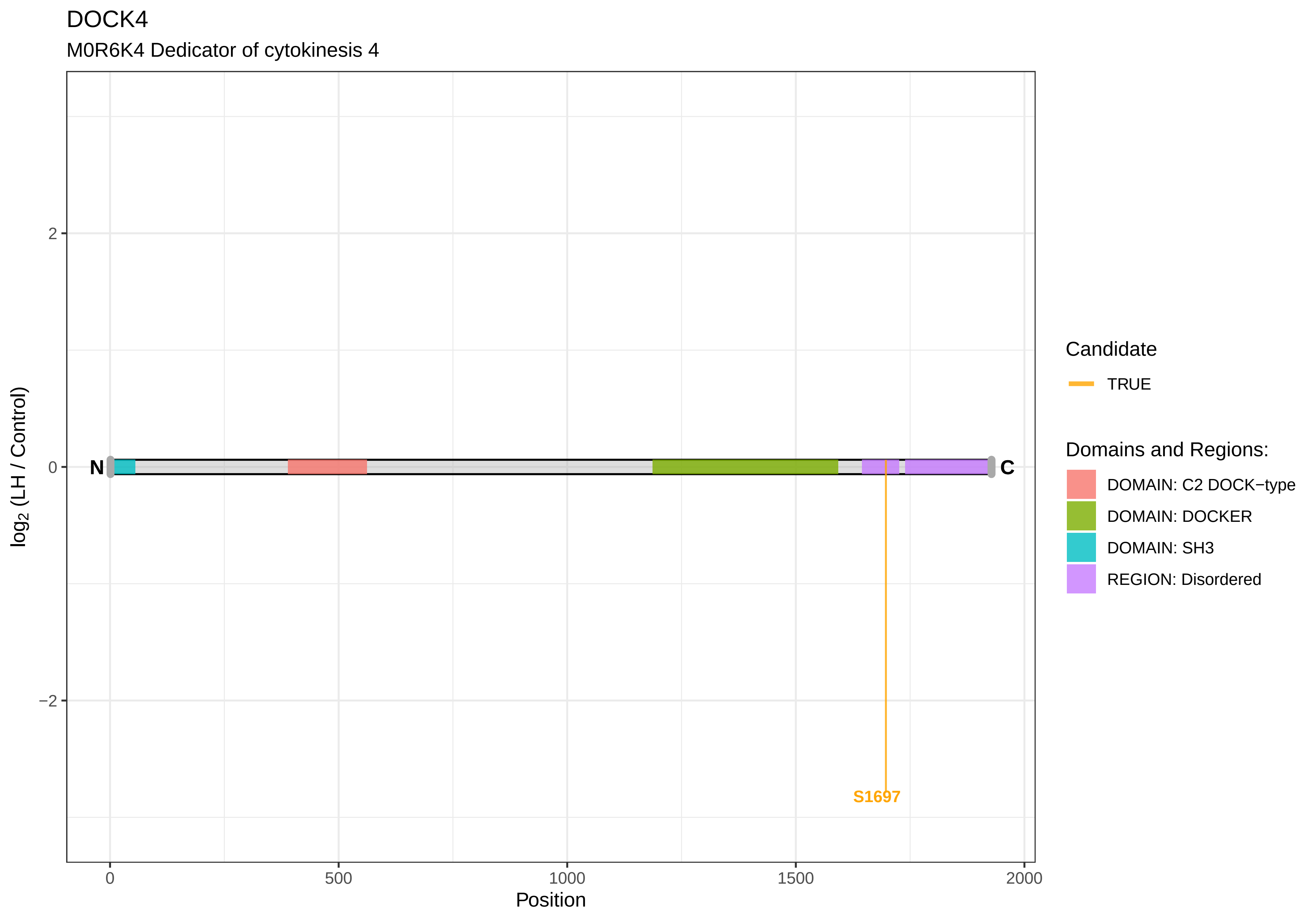
DOCK4
M0R6K4 Dedicator of cytokinesis 4
log2 (LH / Control)
2
0
−2
N
C
S1697
0
500
1000
1500
2000
Position
Candidate
TRUE
Domains and Regions:
DOMAIN: C2 DOCK−type
DOMAIN: DOCKER
DOMAIN: SH3
REGION: Disordered

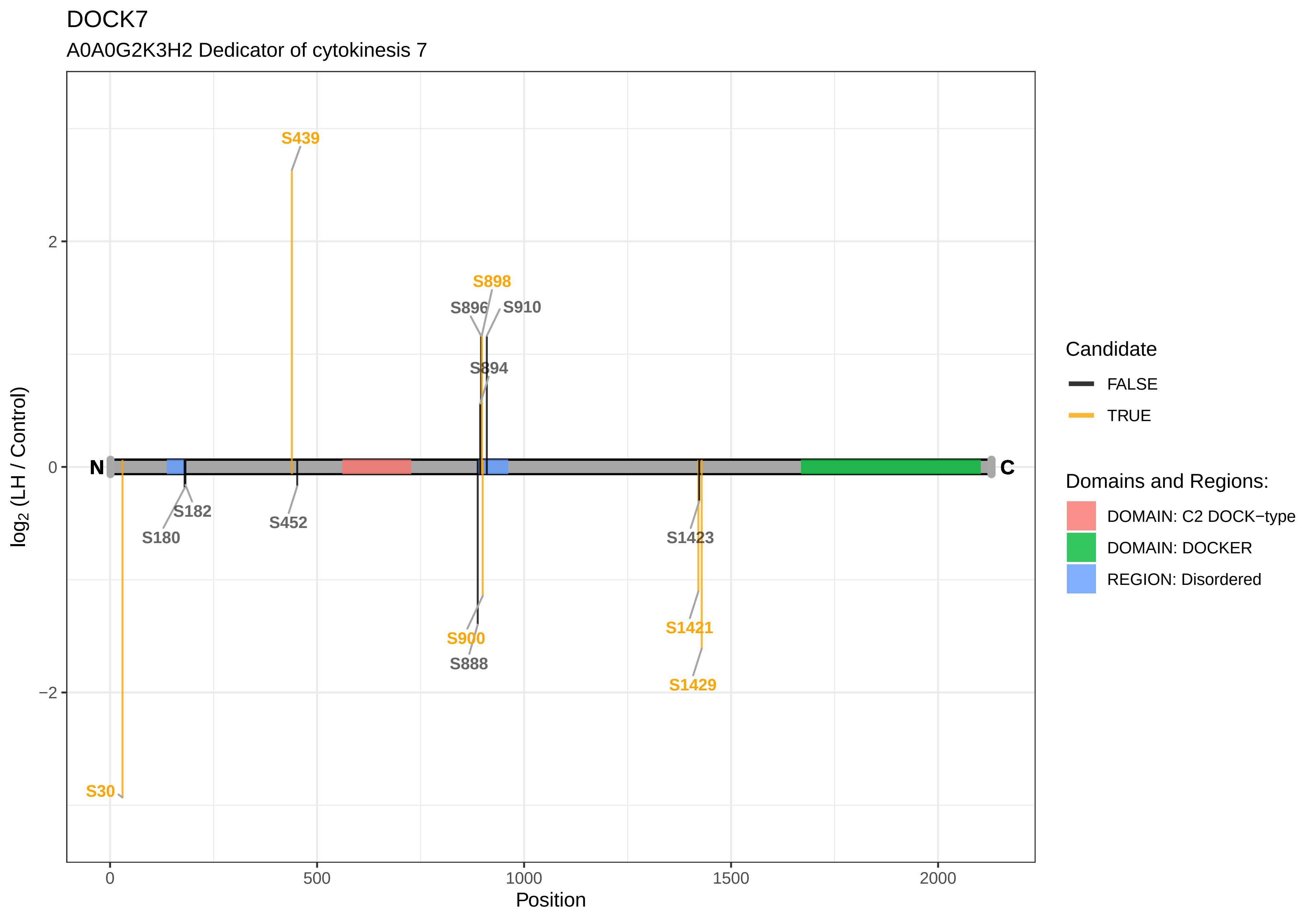

DOCK7
A0A0G2K3H2 Dedicator of cytokinesis 7
S439
S898
S896
S910
S894
N
C
S182
S180
S452
S1423
S900
S888
S1421
S1429
S30
2
0
−2
0
500
1000
1500
2000
Position
log2 (LH / Control)
Candidate
FALSE
TRUE
Domains and Regions:
DOMAIN: C2 DOCK−type
DOMAIN: DOCKER
REGION: Disordered

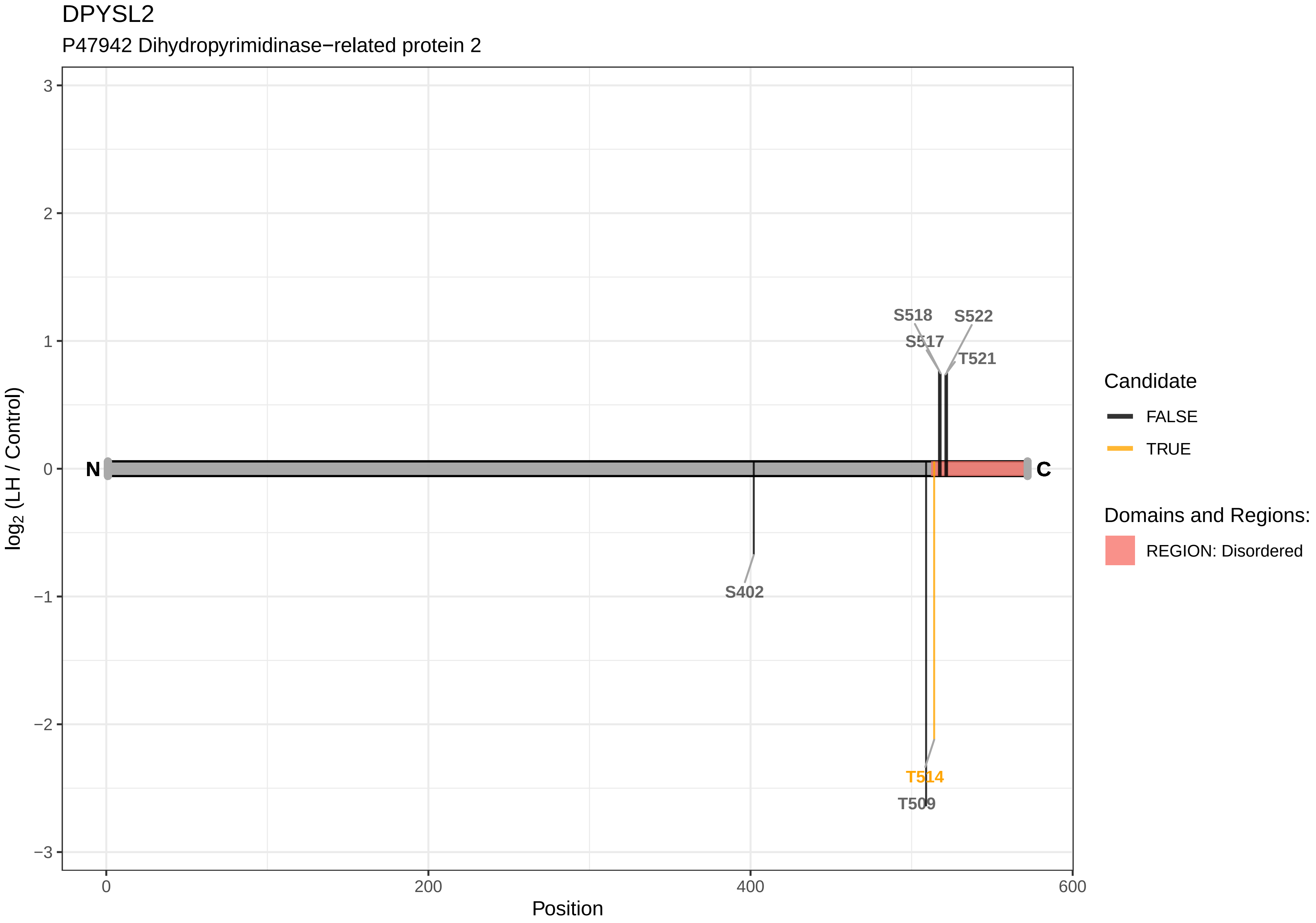
DPYSL2
P47942 Dihydropyrimidinase−related protein 2
3
2
1
0
−1
−2
−3
log2 (LH / Control)
0
200
400
600
Position
N
C
S518
S522
S517
T521
S402
T514
T509
Candidate
FALSE
TRUE
Domains and Regions:
REGION: Disordered

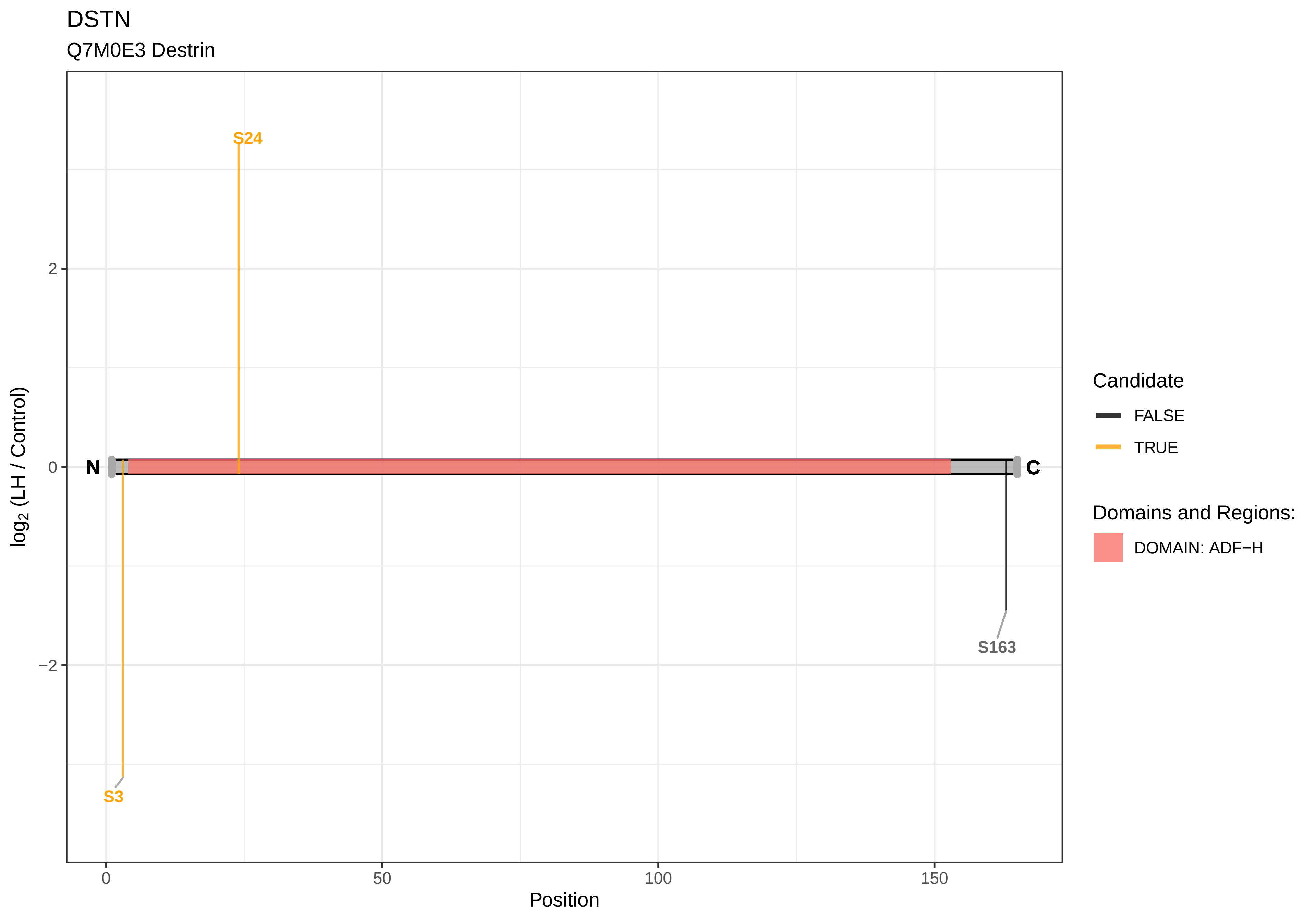
DSTN
Q7M0E3 Destrin
S24
S3
S163
N
C
2
0
−2
0
50
100
150
Position
log2 (LH / Control)
Candidate
FALSE
TRUE
Domains and Regions:
DOMAIN: ADF−H

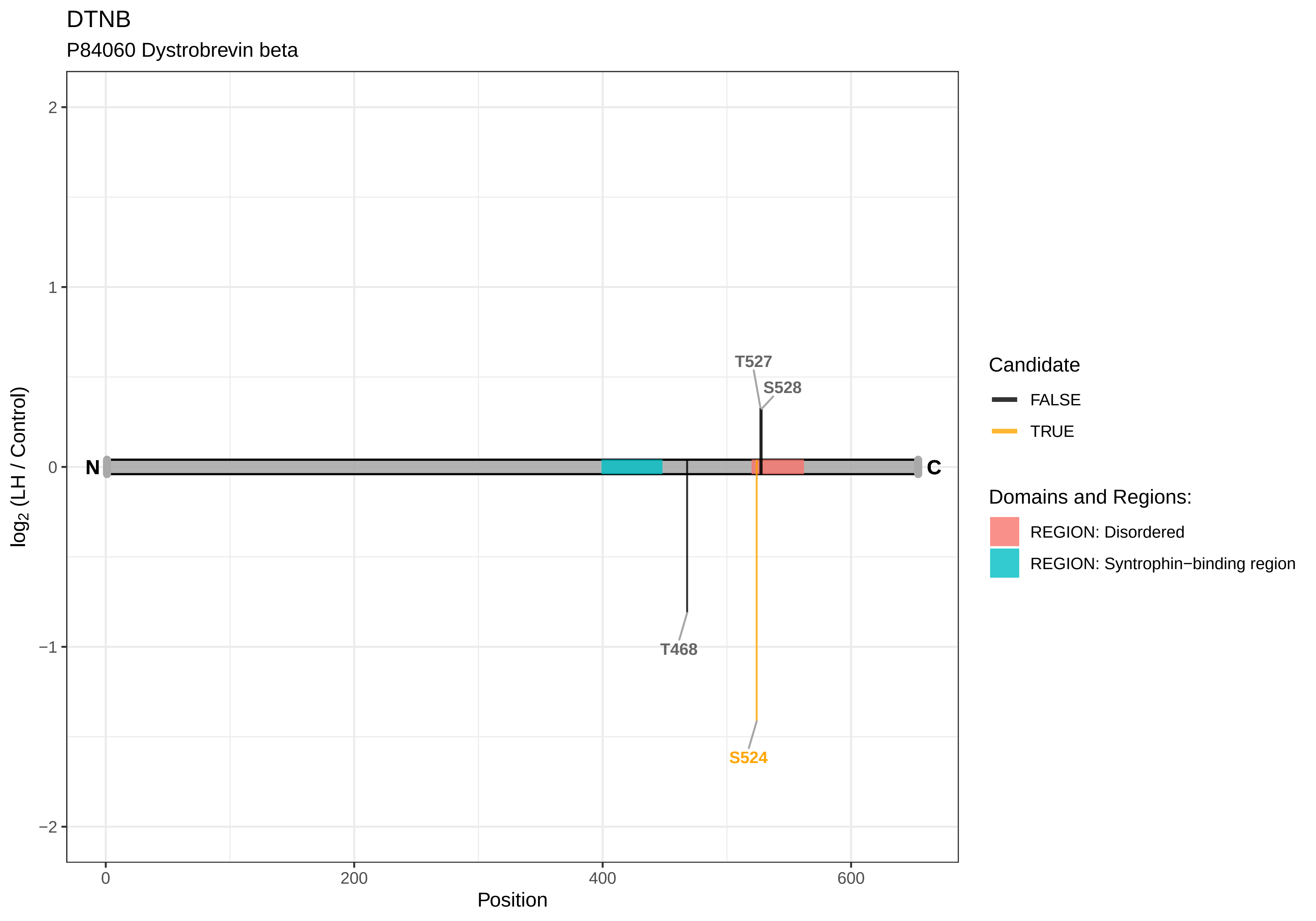

DTNB
P84060 Dystrobrevin beta
2
1
0
−1
−2
log2 (LH / Control)
N
C
T527
S528
T468
S524
0
200
400
600
Position
Candidate
FALSE
TRUE
Domains and Regions:
REGION: Disordered
REGION: Syntrophin−binding region

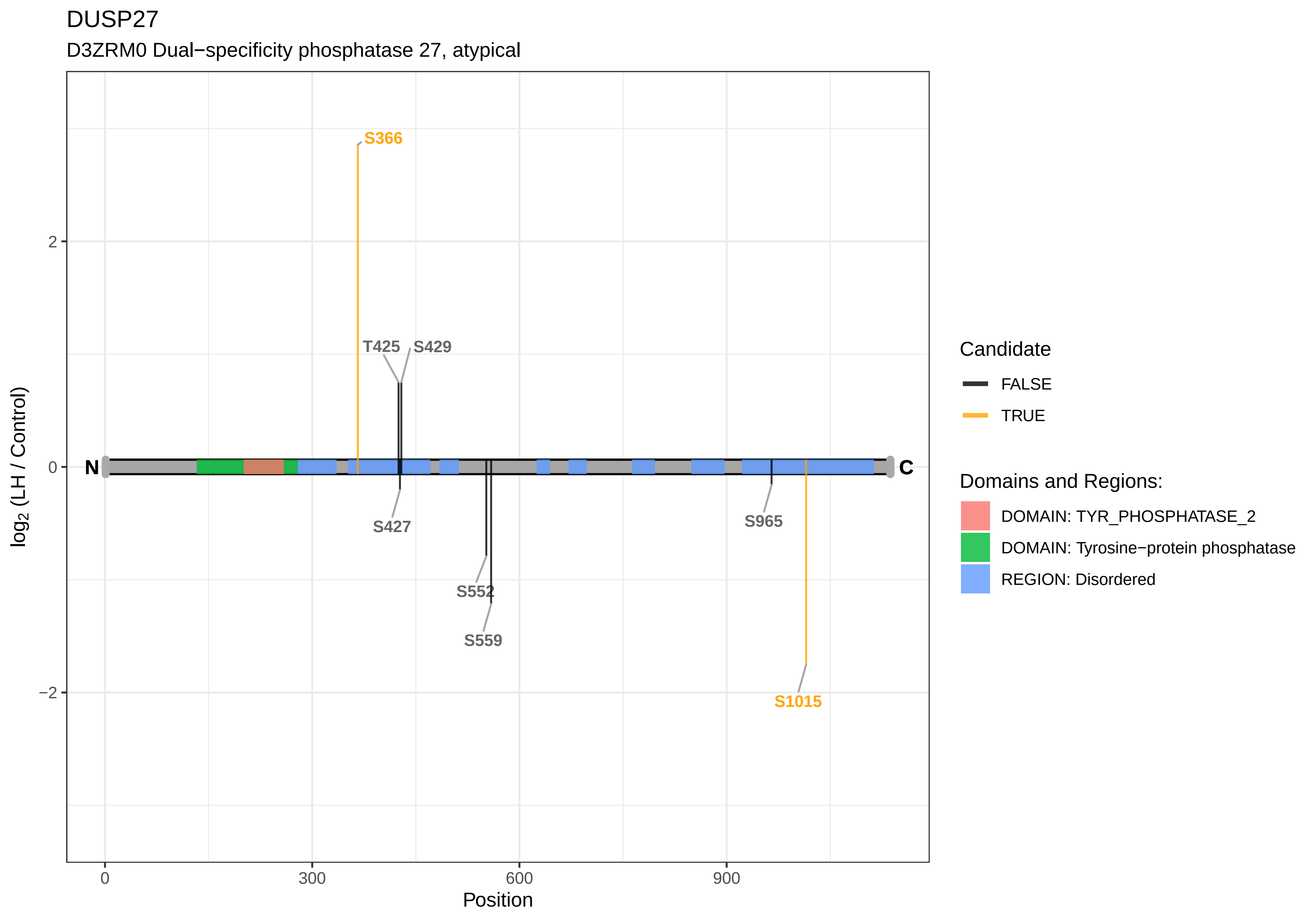
DUSP27
D3ZRM0 Dual−specificity phosphatase 27, atypical
S366
T425
S429
S427
S552
S559
S965
S1015
N
C
2
0
−2
0
300
600
900
Position
log2 (LH / Control)
Candidate
FALSE
TRUE
Domains and Regions:
DOMAIN: TYR_PHOSPHATASE_2
DOMAIN: Tyrosine−protein phosphatase
REGION: Disordered

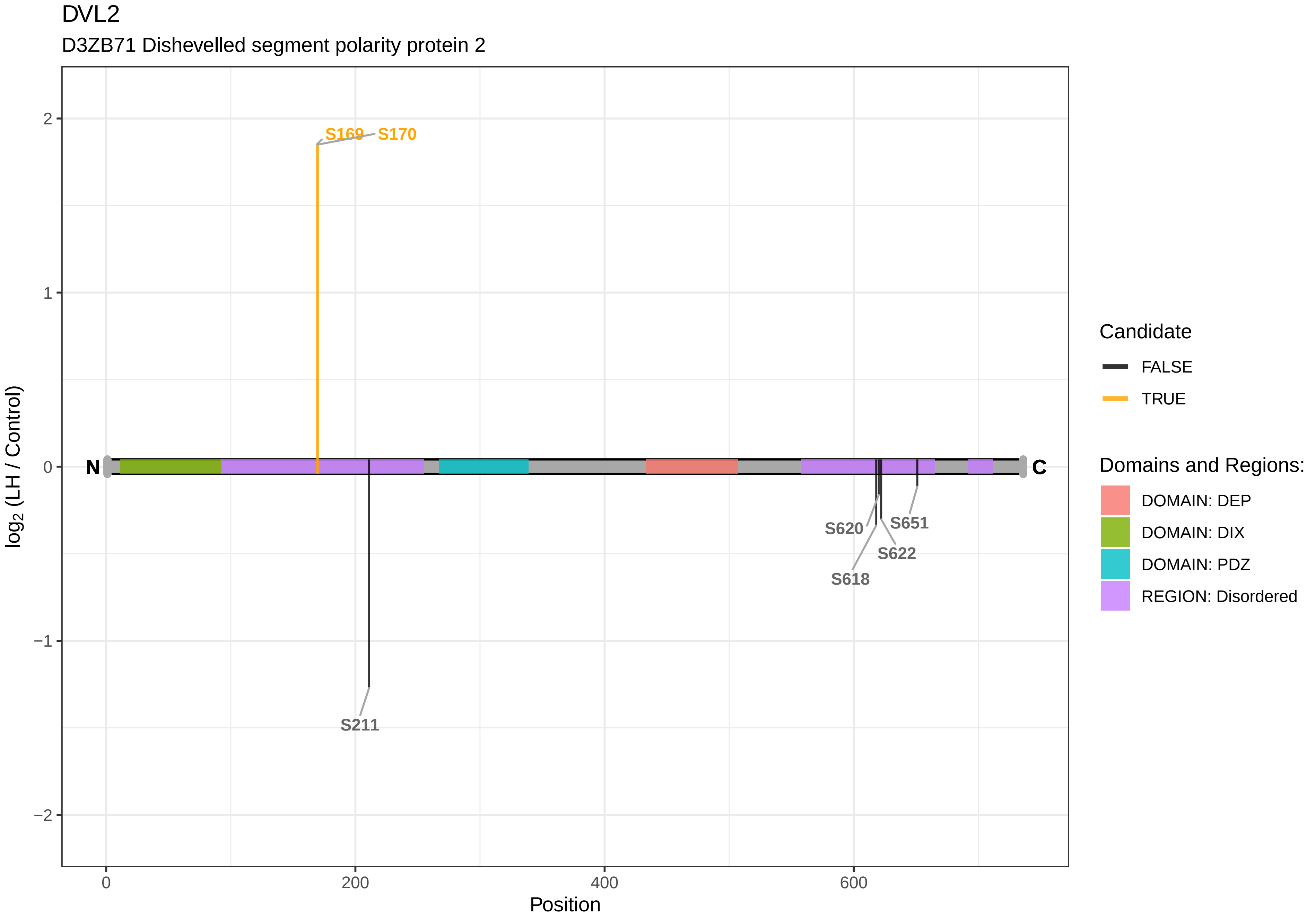

DVL2
D3ZB71 Dishevelled segment polarity protein 2
S169
S170
S211
S618
S620
S622
S651
N
C
log2 (LH / Control)
Position
2
1
0
−1
−2
0
200
400
600
Candidate
FALSE
TRUE
Domains and Regions:
DOMAIN: DEP
DOMAIN: DIX
DOMAIN: PDZ
REGION: Disordered

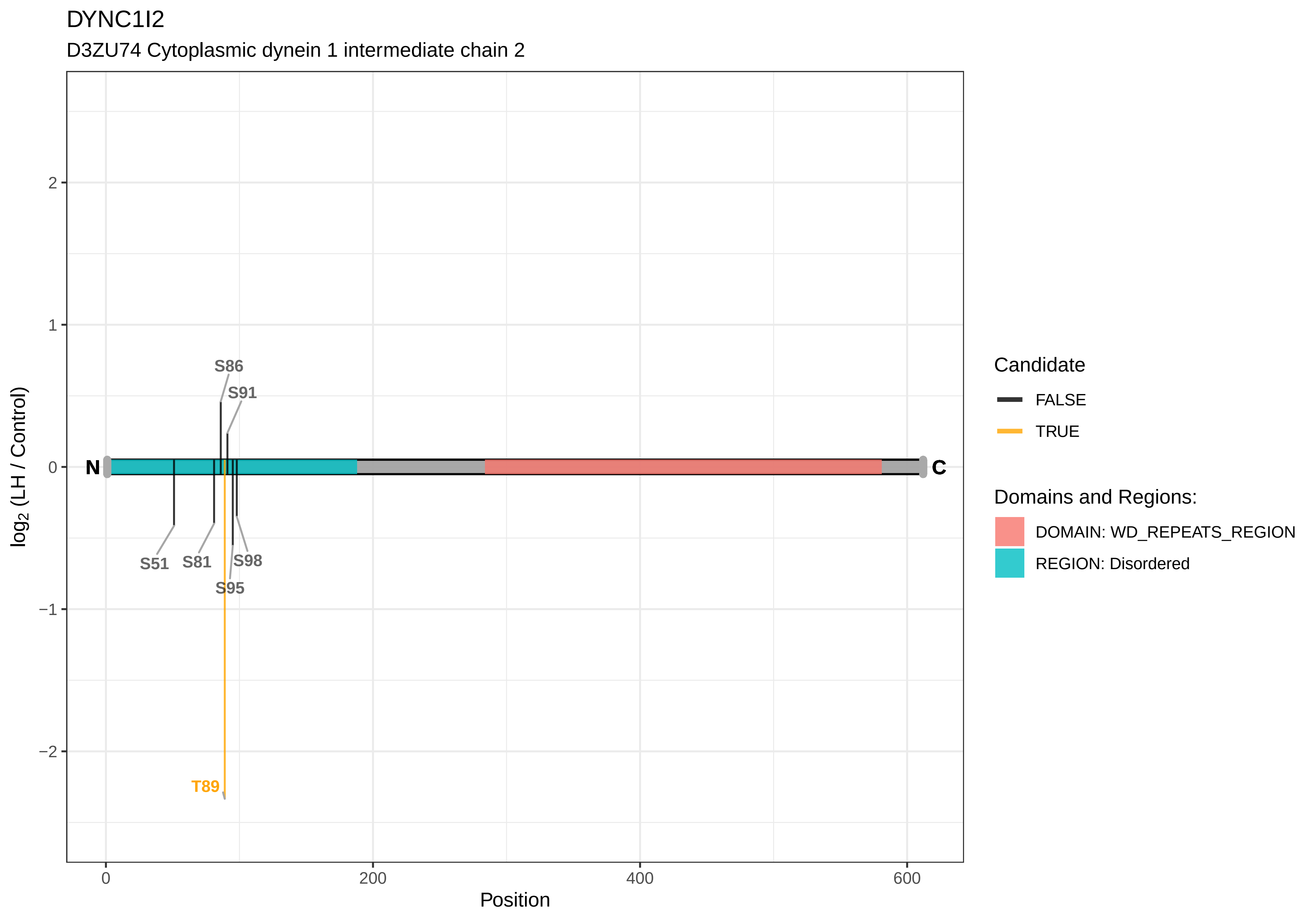

DYNC1I2
D3ZU74 Cytoplasmic dynein 1 intermediate chain 2
log2 (LH / Control)
Position
N
C
S86
S91
S51
S81
S98
S95
T89
Candidate
FALSE
TRUE
Domains and Regions:
DOMAIN: WD_REPEATS_REGION
REGION: Disordered

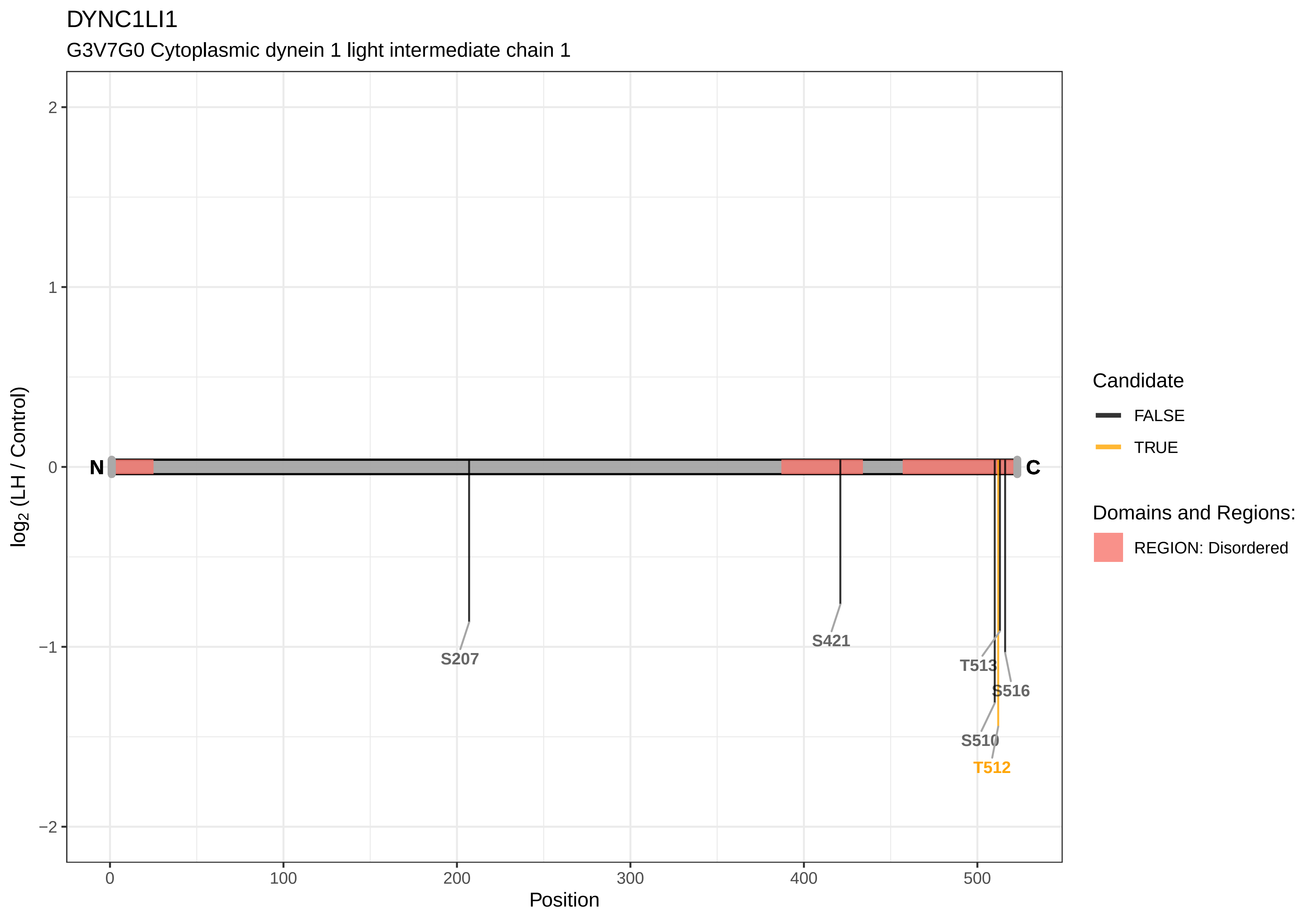

DYNC1LI1
G3V7G0 Cytoplasmic dynein 1 light intermediate chain 1
log2 (LH / Control)
Position
N
C
S207
S421
T513
S516
S510
T512
2
1
0
−1
−2
0
100
200
300
400
500
Candidate
FALSE
TRUE
Domains and Regions:
REGION: Disordered

DYNC1LI2

Q5D023 Cytoplasmic dynein 1 light intermediate chain 2

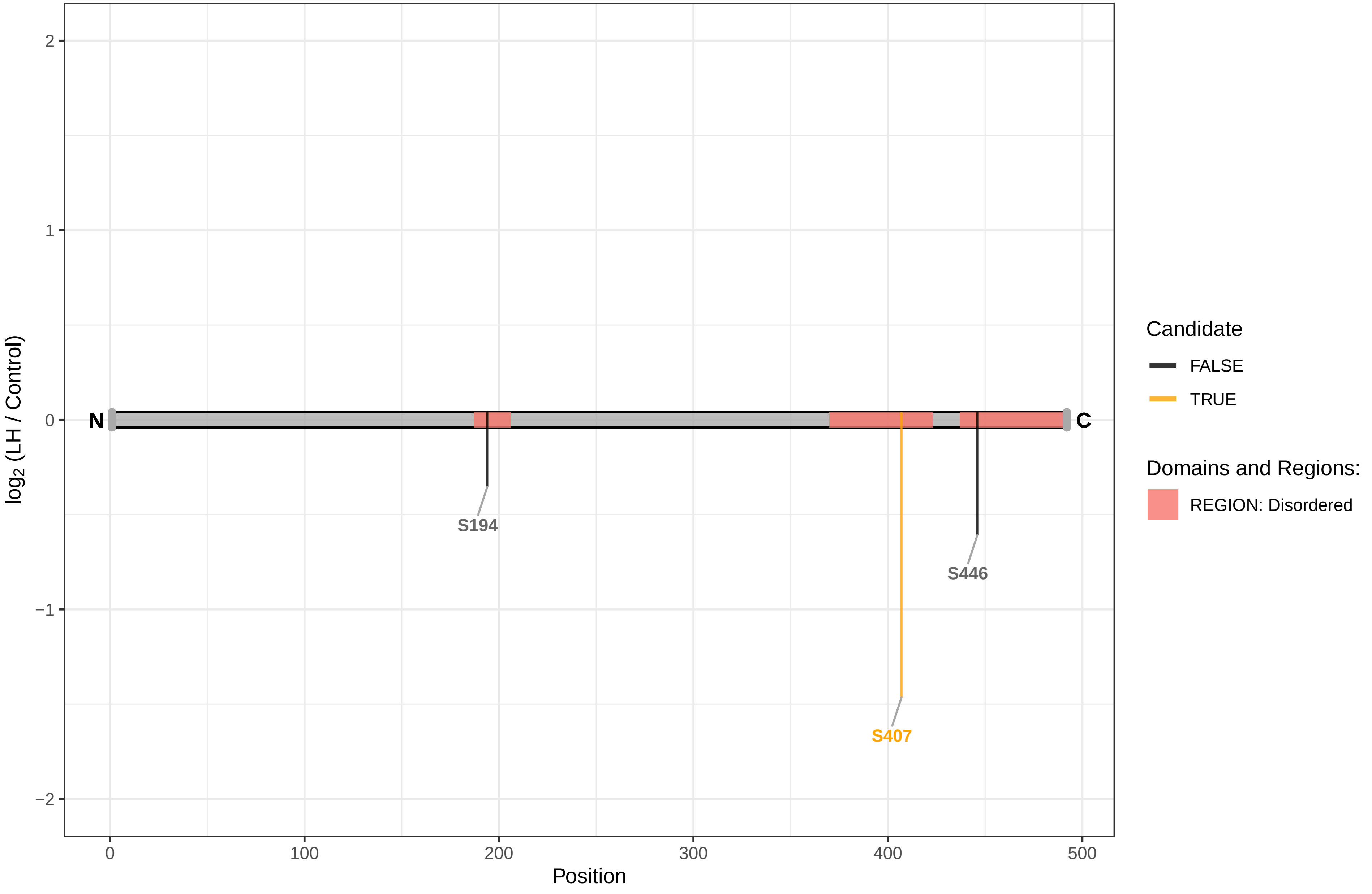

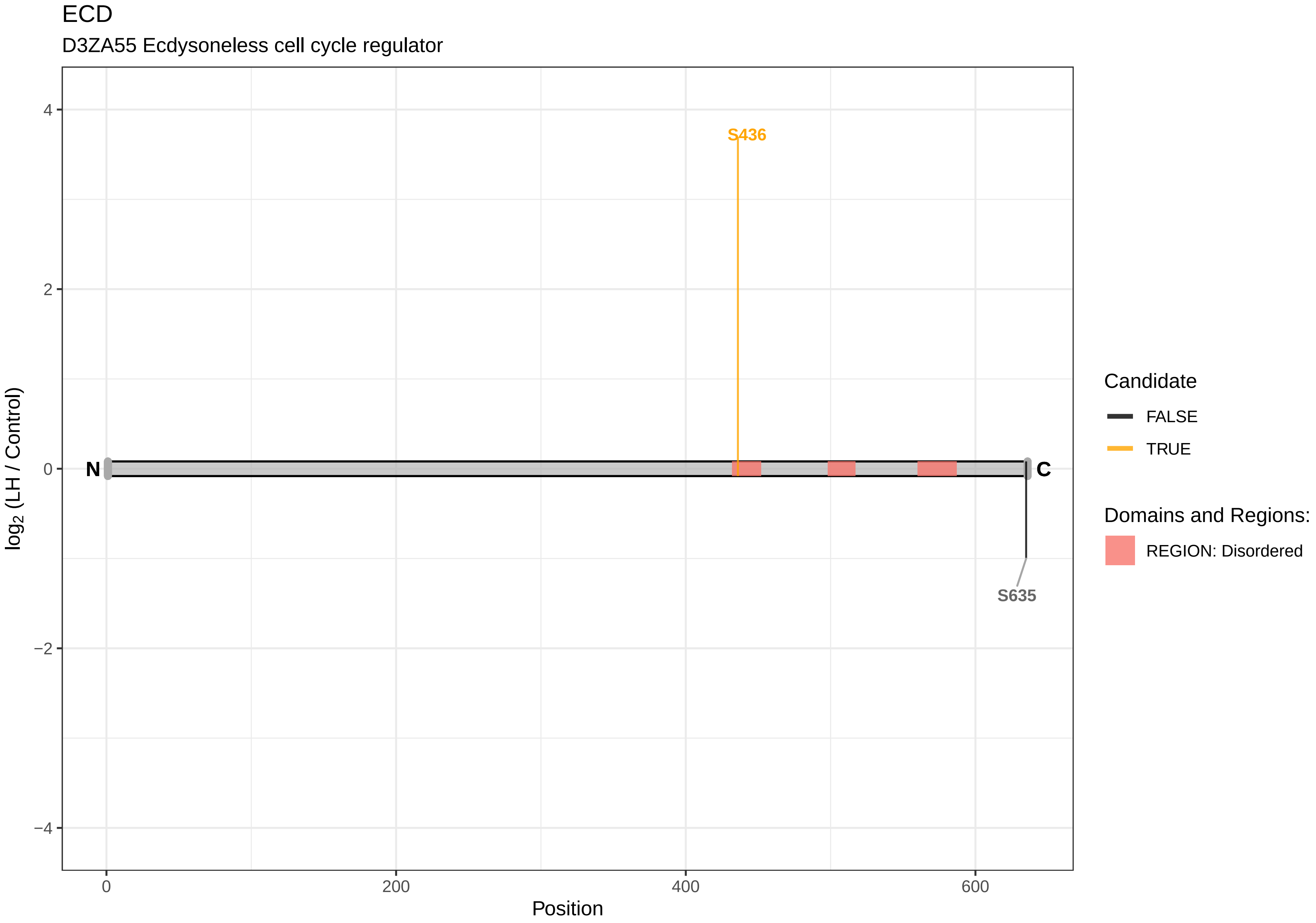

ECD
D3ZA55 Ecdysoneless cell cycle regulator
S436
S635
N
C
4
2
0
−2
−4
log2 (LH / Control)
0
200
400
600
Position
Candidate
FALSE
TRUE
Domains and Regions:
REGION: Disordered

EEPD1

Q5XI74 Endonuclease/exonuclease/phosphatase family domain−containing protein 1

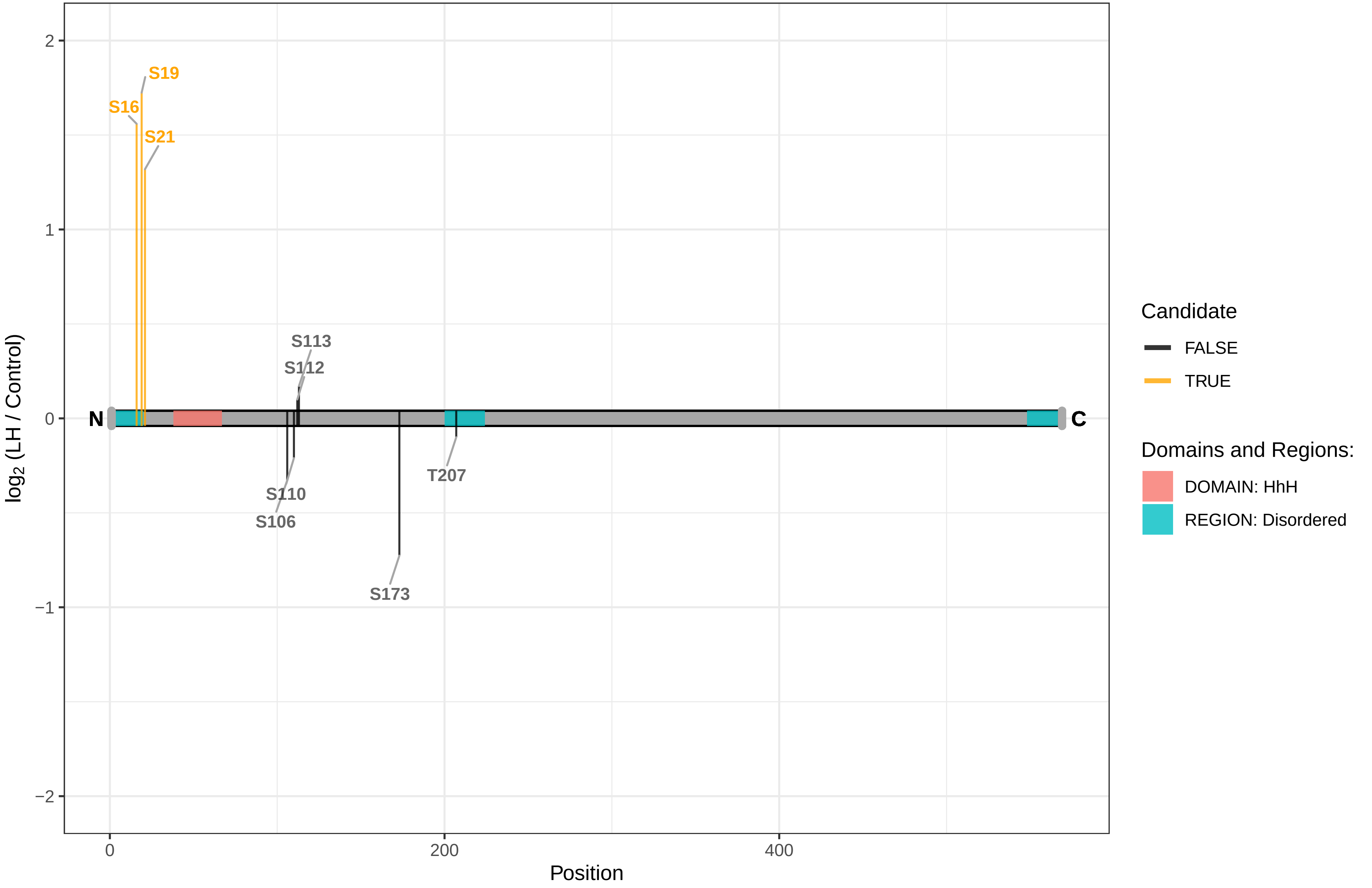

### EFCAB7

D3ZZK0 EF−hand calcium−binding domain 7

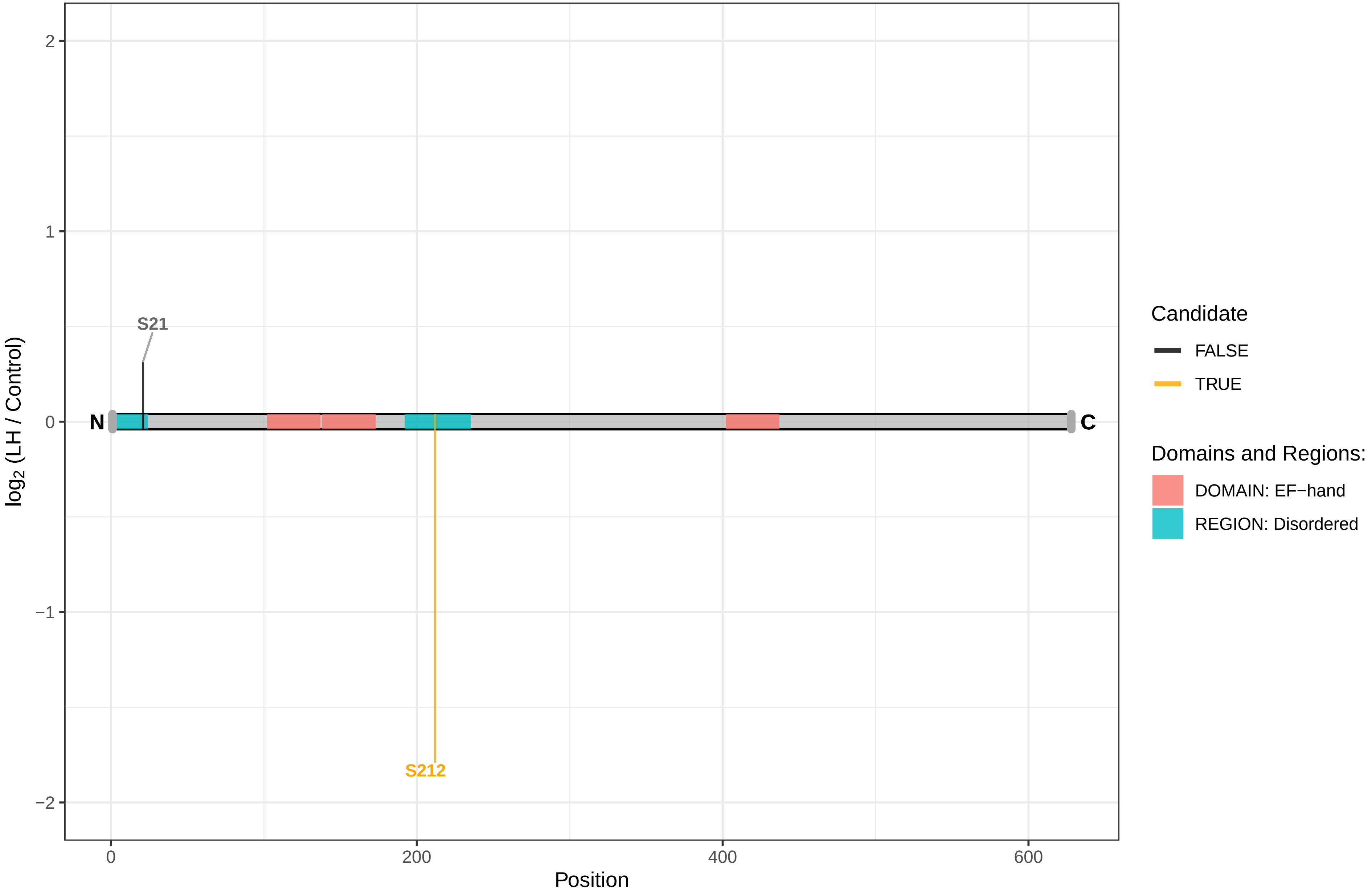

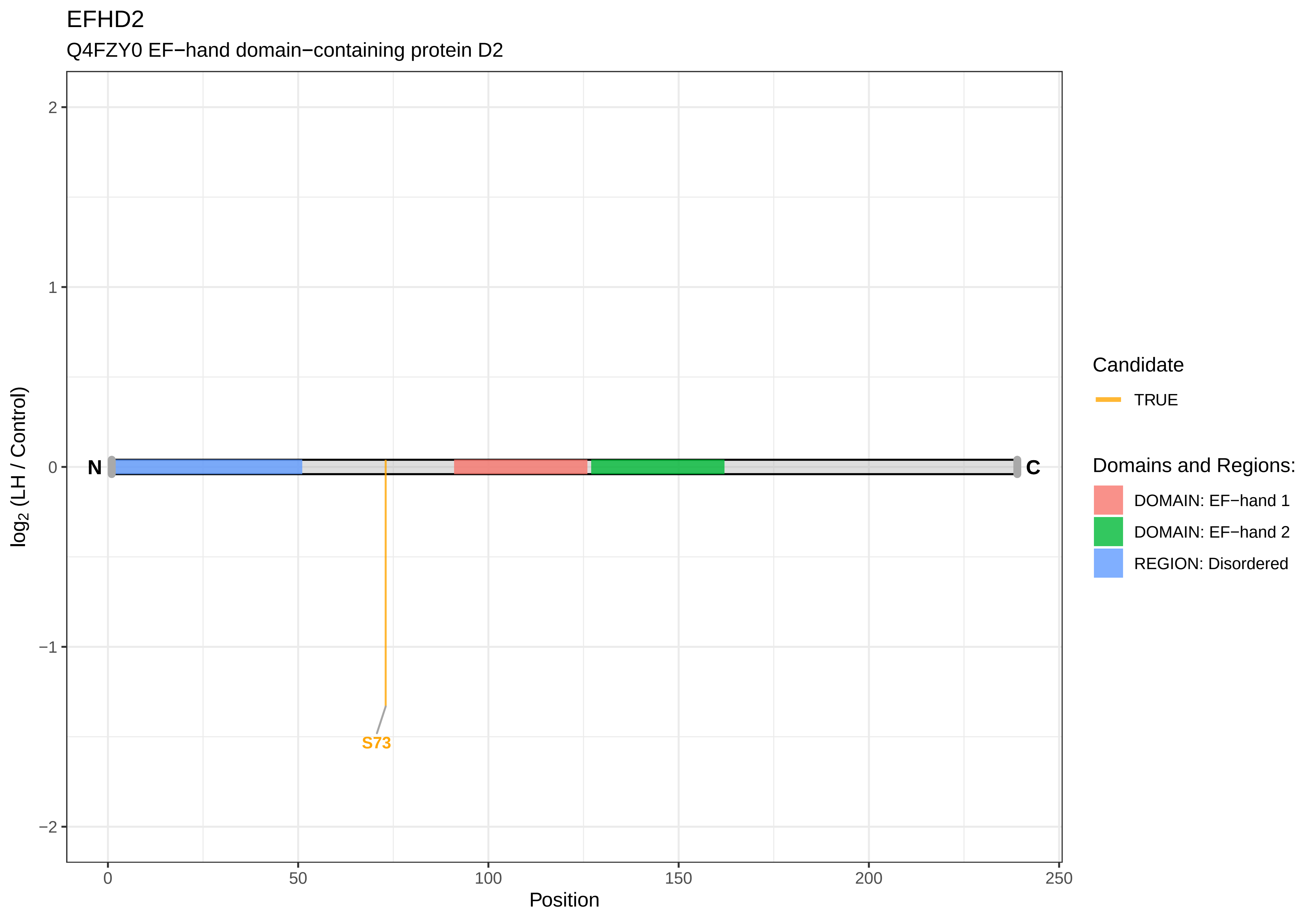
EFHD2
Q4FZY0 EF−hand domain−containing protein D2
log2 (LH / Control)
Position
N
C
S73
Candidate
TRUE
Domains and Regions:
DOMAIN: EF−hand 1
DOMAIN: EF−hand 2
REGION: Disordered
2
1
0
−1
−2
0
50
100
150
200
250

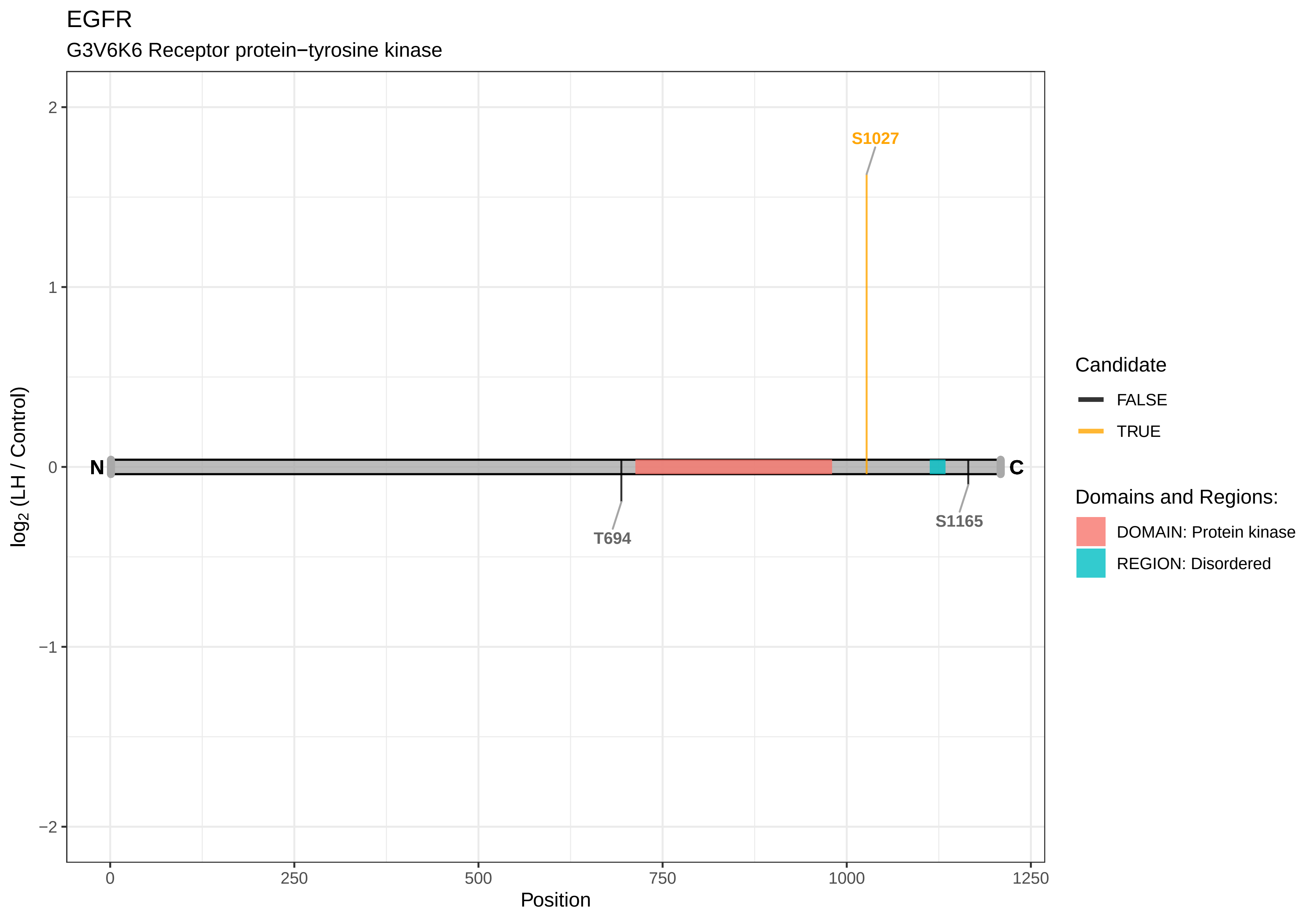

EGFR
G3V6K6 Receptor protein−tyrosine kinase
S1027
N
C
T694
S1165
2
1
0
−1
−2
log2 (LH / Control)
0
250
500
750
1000
1250
Position
Candidate
FALSE
TRUE
Domains and Regions:
DOMAIN: Protein kinase
REGION: Disordered

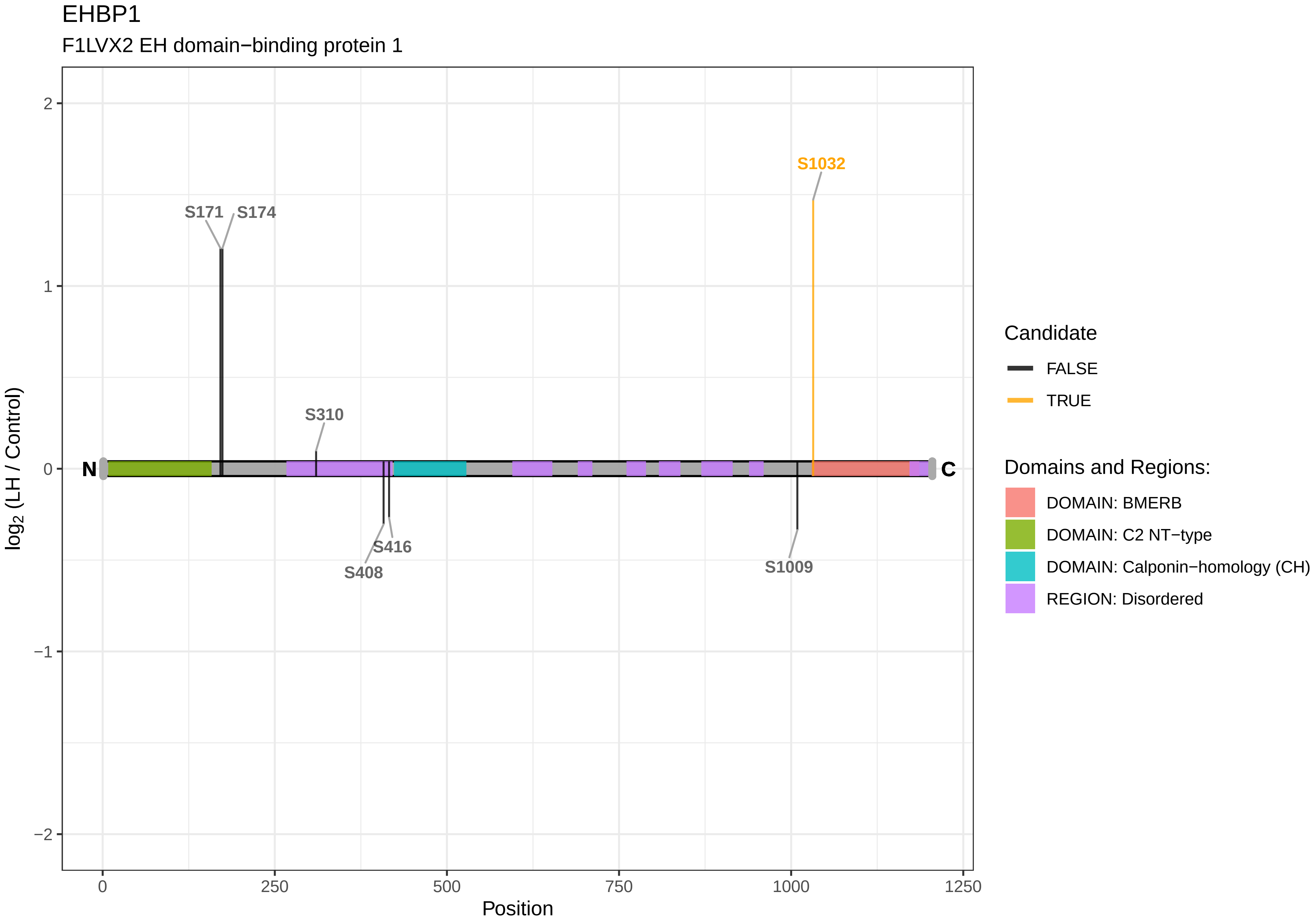

EHBP1
F1LVX2 EH domain−binding protein 1
S1032
S171
S174
S310
S416
S408
S1009
N
C
2
1
0
−1
−2
0
250
500
750
1000
1250
Position
log2 (LH / Control)
Candidate
FALSE
TRUE
Domains and Regions:
DOMAIN: BMERB
DOMAIN: C2 NT−type
DOMAIN: Calponin−homology (CH)
REGION: Disordered

EHMT2

A0A0G2JTA5 Euchromatic histone lysine methyltransferase 2

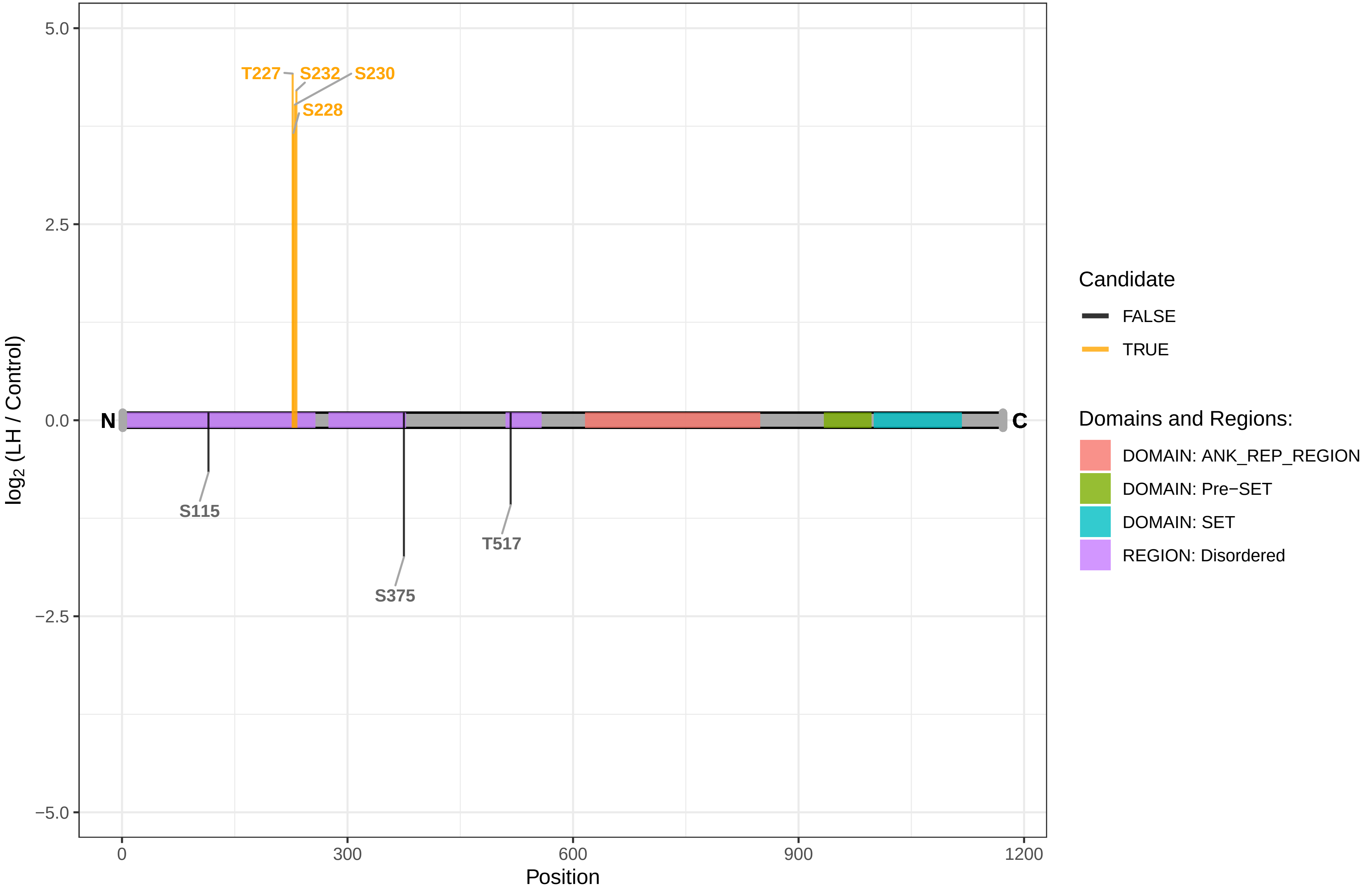

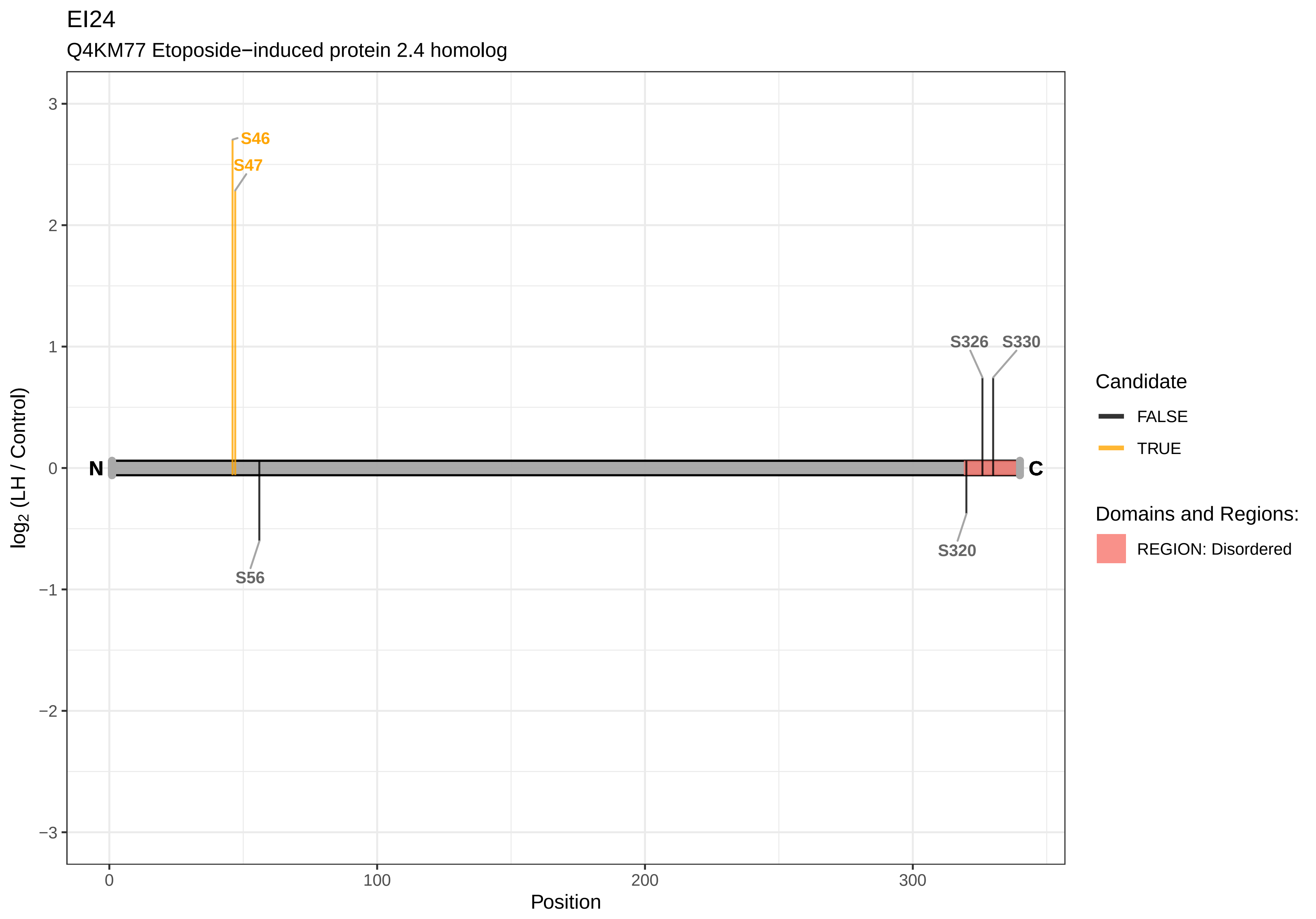

EI24
Q4KM77 Etoposide−induced protein 2.4 homolog
S46
S47
S326
S330
S56
S320
N
C
3
2
1
0
−1
−2
−3
0
100
200
300
Position
log2 (LH / Control)
Candidate
FALSE
TRUE
Domains and Regions:
REGION: Disordered

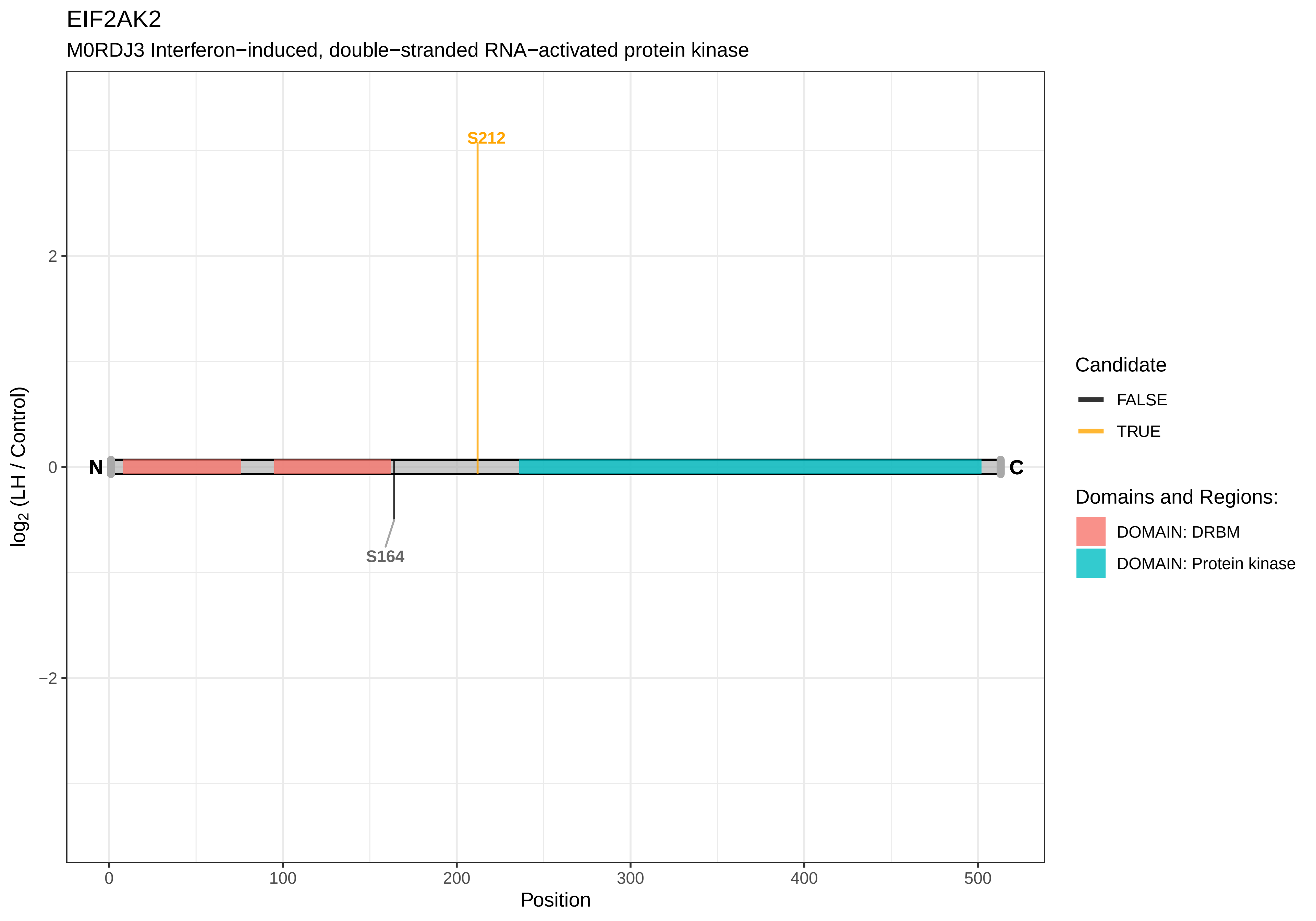
EIF2AK2
M0RDJ3 Interferon−induced, double−stranded RNA−activated protein kinase
S212
S164
N
C
log2 (LH / Control)
2
0
−2
0
100
200
300
400
500
Position
Candidate
FALSE
TRUE
Domains and Regions:
DOMAIN: DRBM
DOMAIN: Protein kinase

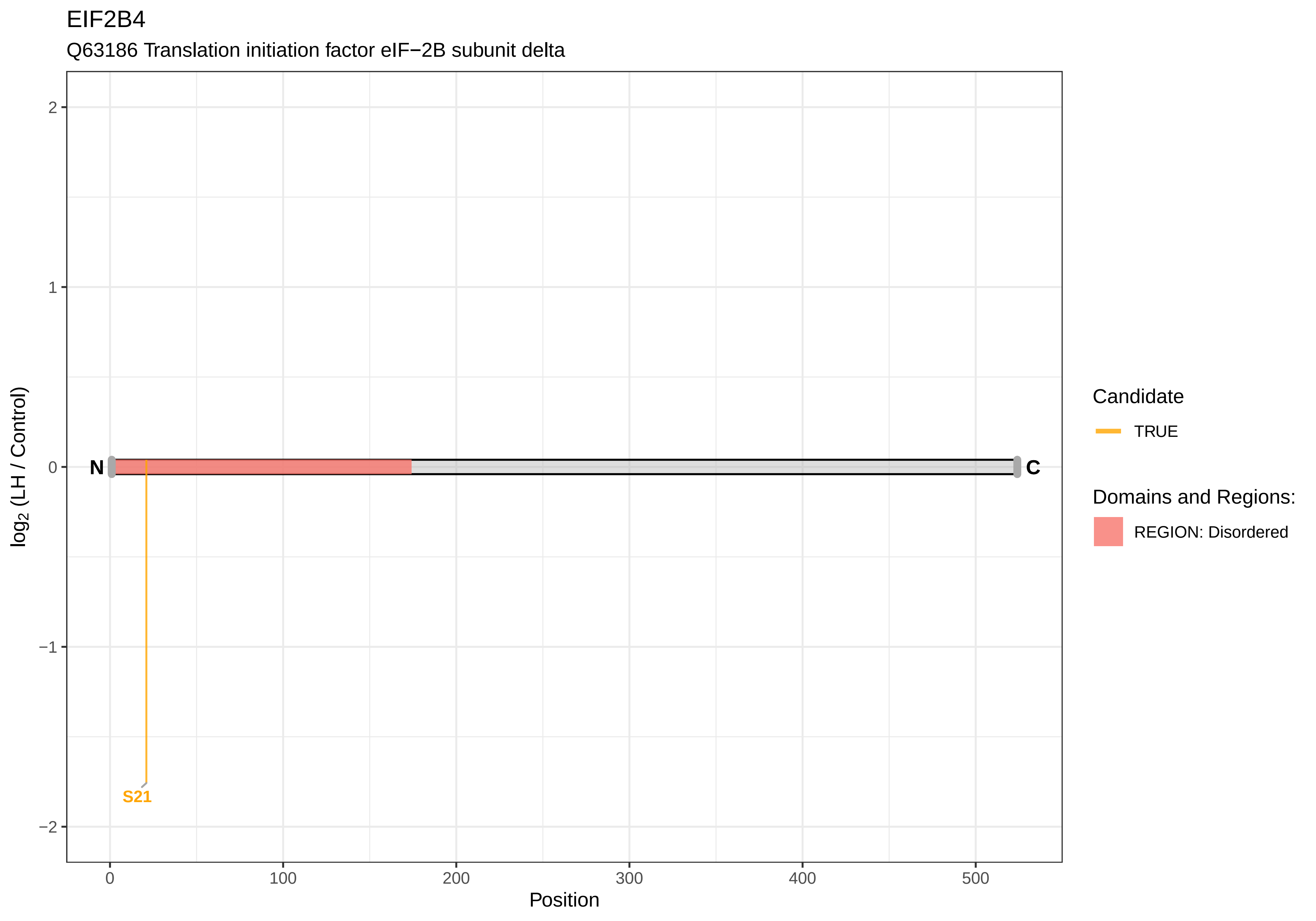
EIF2B4
Q63186 Translation initiation factor eIF−2B subunit delta
log2 (LH / Control)
2
1
0
−1
−2
N
C
S21
0
100
200
300
400
500
Position
Candidate
TRUE
Domains and Regions:
REGION: Disordered

### EIF3B

Q4G061 Eukaryotic translation initiation factor 3 subunit B

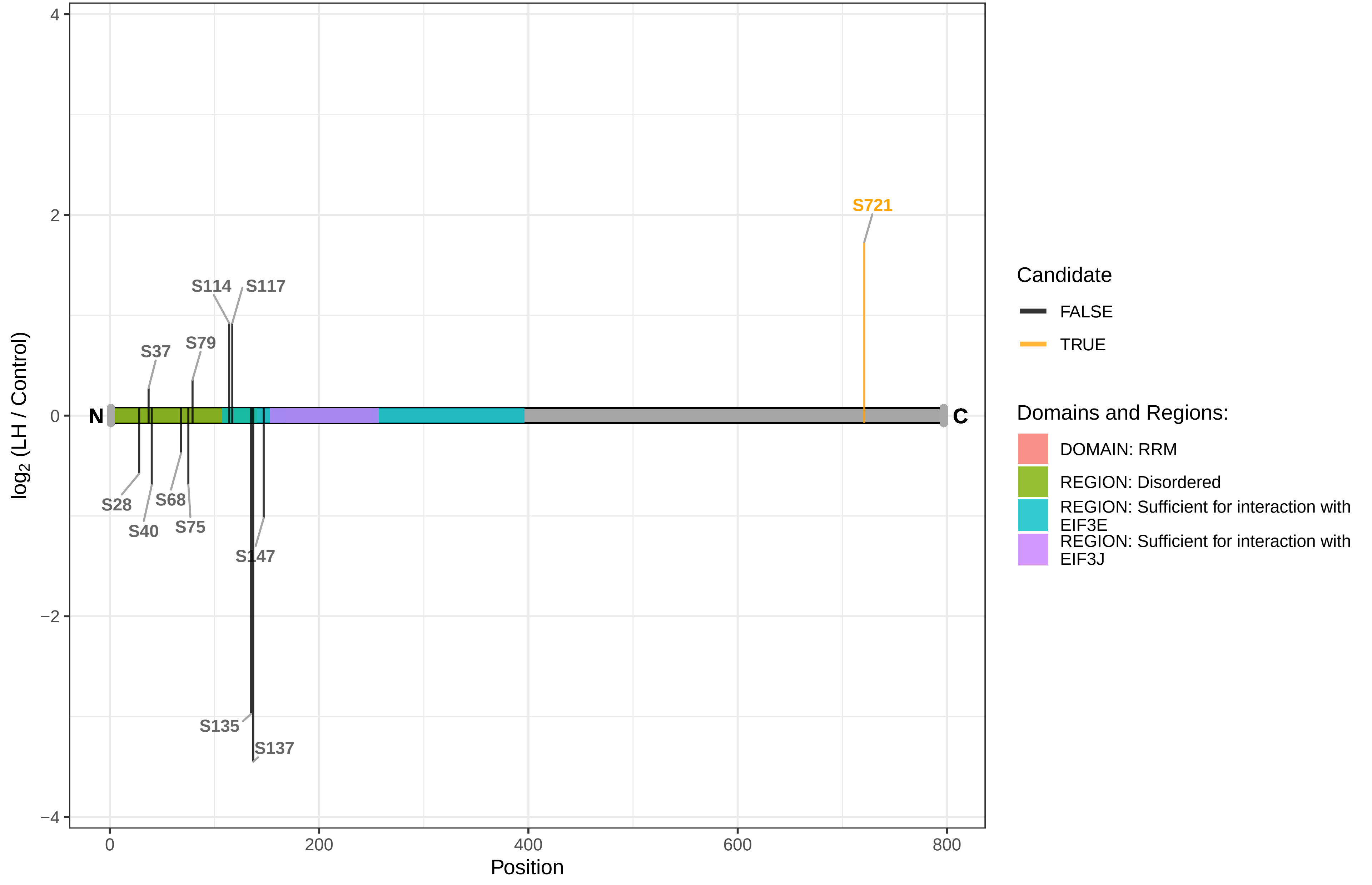

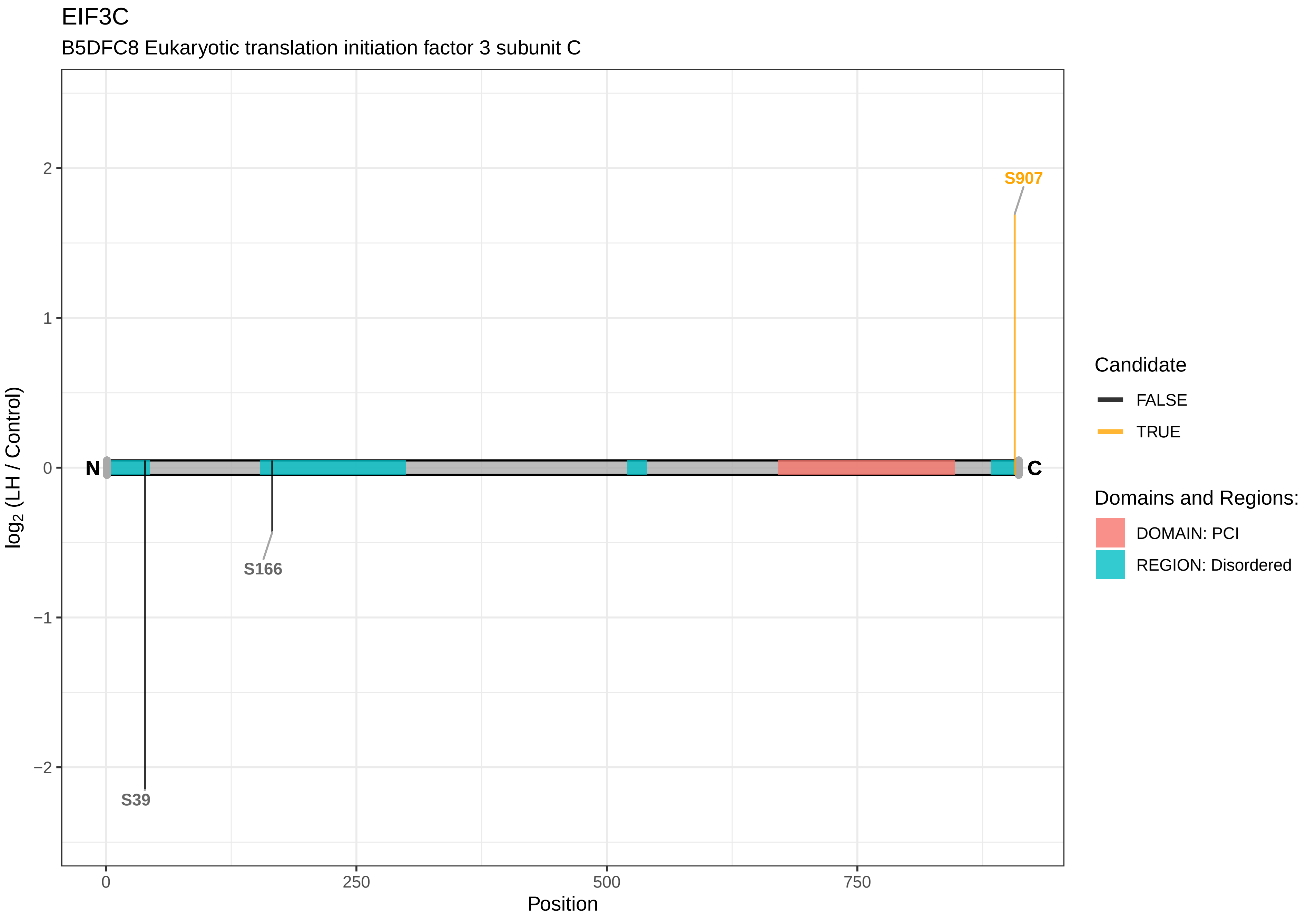
EIF3C
B5DFC8 Eukaryotic translation initiation factor 3 subunit C
S907
S166
S39
N
C
2
1
0
−1
−2
log2 (LH / Control)
0
250
500
750
Position
Candidate
FALSE
TRUE
Domains and Regions:
DOMAIN: PCI
REGION: Disordered

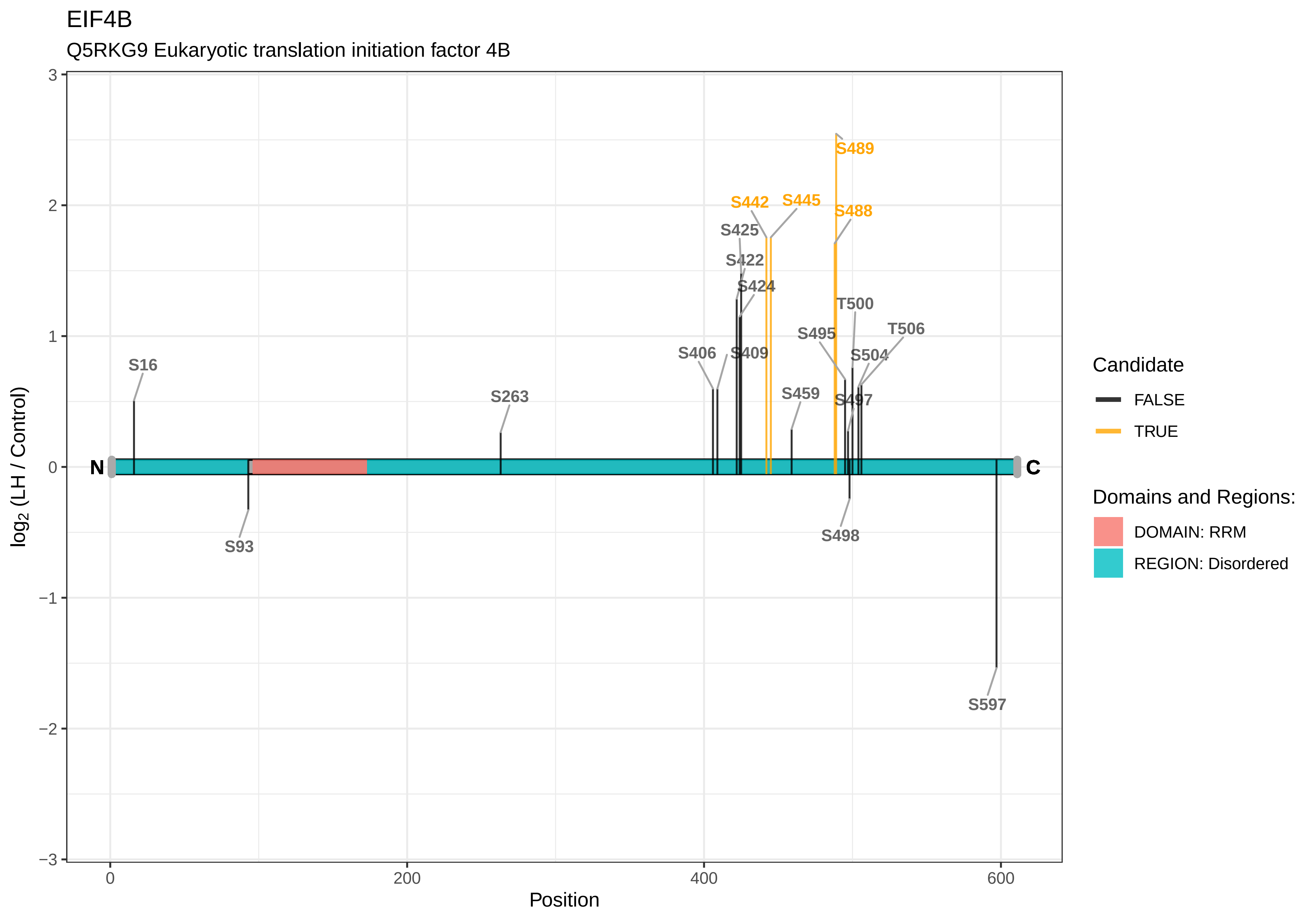
EIF4B
Q5RKG9 Eukaryotic translation initiation factor 4B
log2 (LH / Control)
Position
N
C
S16
S93
S263
S406
S409
S422
S424
S425
S442
S445
S459
S488
S489
S495
S497
S498
T500
S504
T506
S597
3
2
1
0
−1
−2
−3
0
200
400
600
Candidate
FALSE
TRUE
Domains and Regions:
DOMAIN: RRM
REGION: Disordered

EIF4G2

F1LN59 Eukaryotic translation initiation factor 4, gamma 2

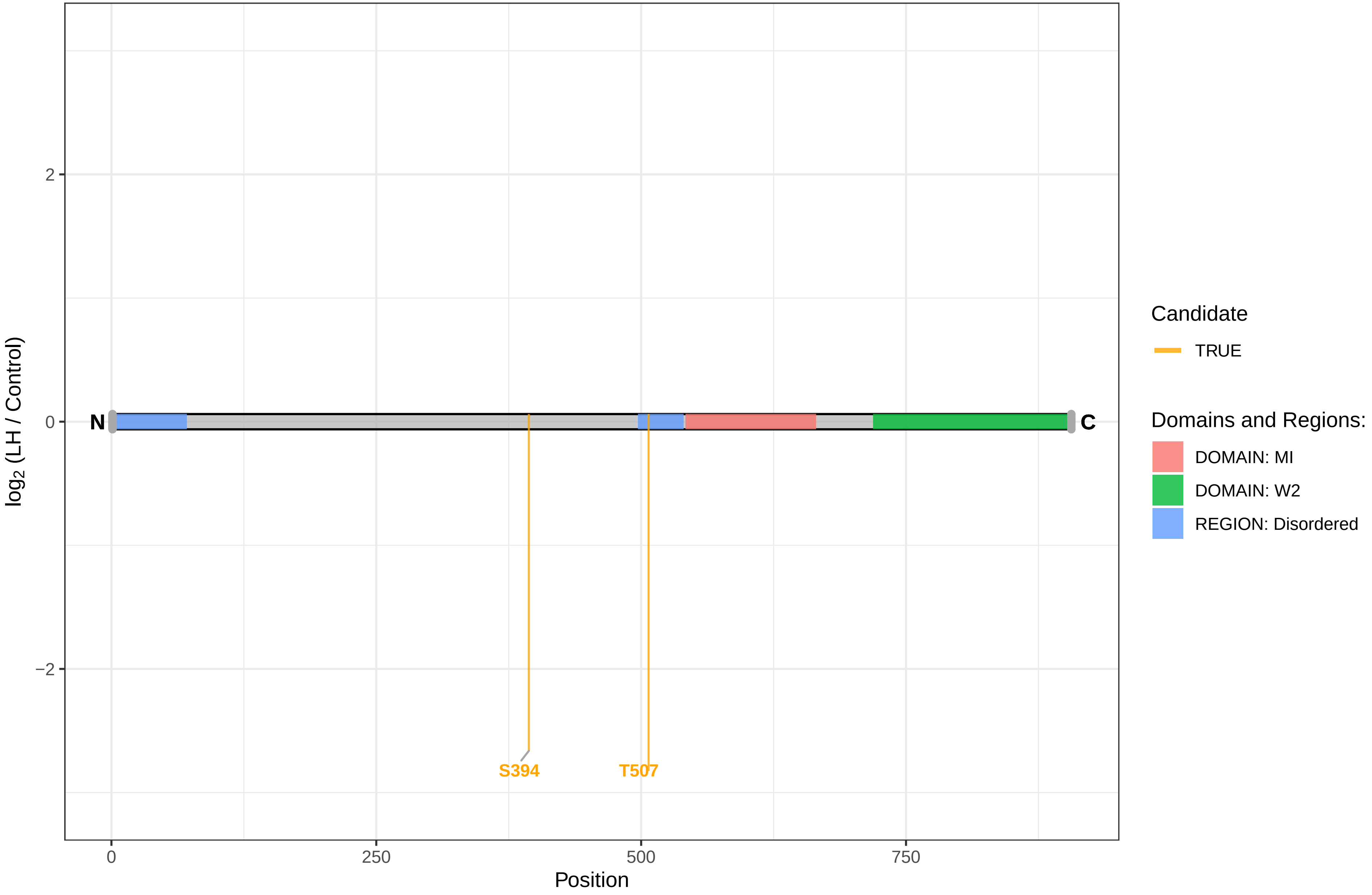

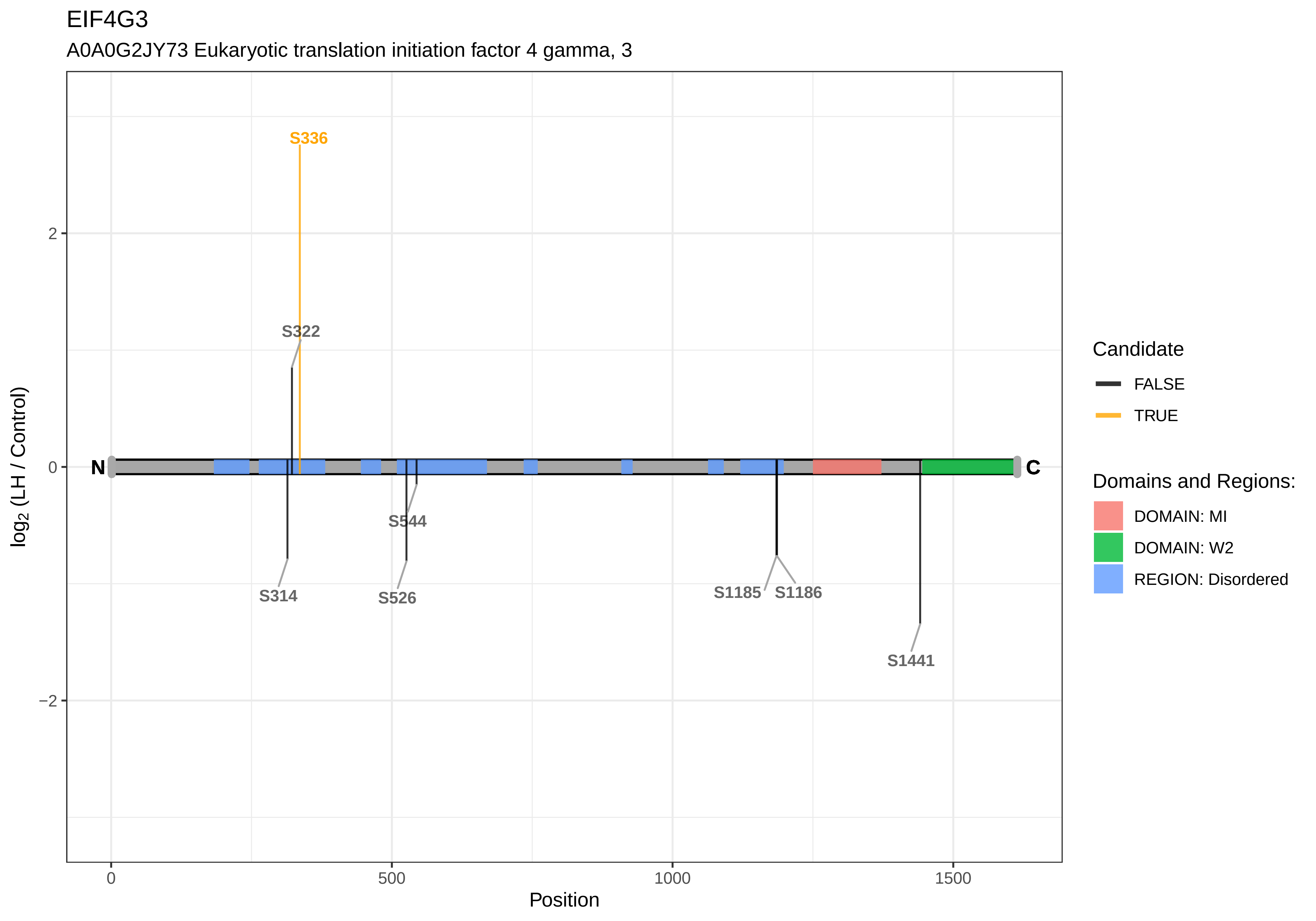
EIF4G3
A0A0G2JY73 Eukaryotic translation initiation factor 4 gamma, 3
S336
S322
S544
S314
S526
S1185
S1186
S1441
N
C
2
0
−2
0
500
1000
1500
Position
log2 (LH / Control)
Candidate
FALSE
TRUE
Domains and Regions:
DOMAIN: MI
DOMAIN: W2
REGION: Disordered

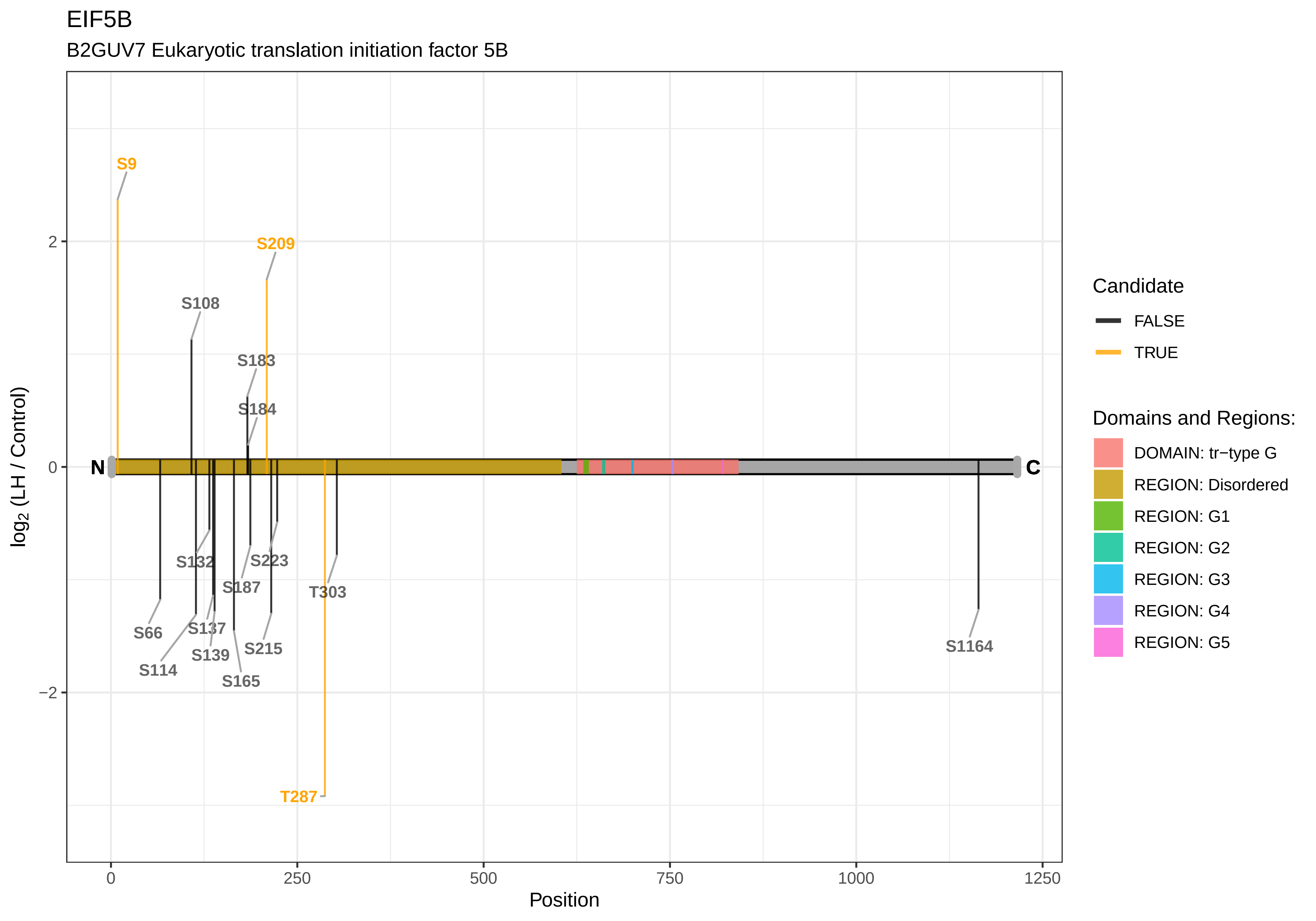
EIF5B
B2GUV7 Eukaryotic translation initiation factor 5B
S9
S209
S108
S183
S184
S132
S223
S187
T303
S66
S137
S139
S215
S114
S165
T287
S1164
N
C
log2 (LH / Control)
2
0
−2
0
250
500
750
1000
1250
Position
Candidate
FALSE
TRUE
Domains and Regions:
DOMAIN: tr−type G
REGION: Disordered
REGION: G1
REGION: G2
REGION: G3
REGION: G4
REGION: G5

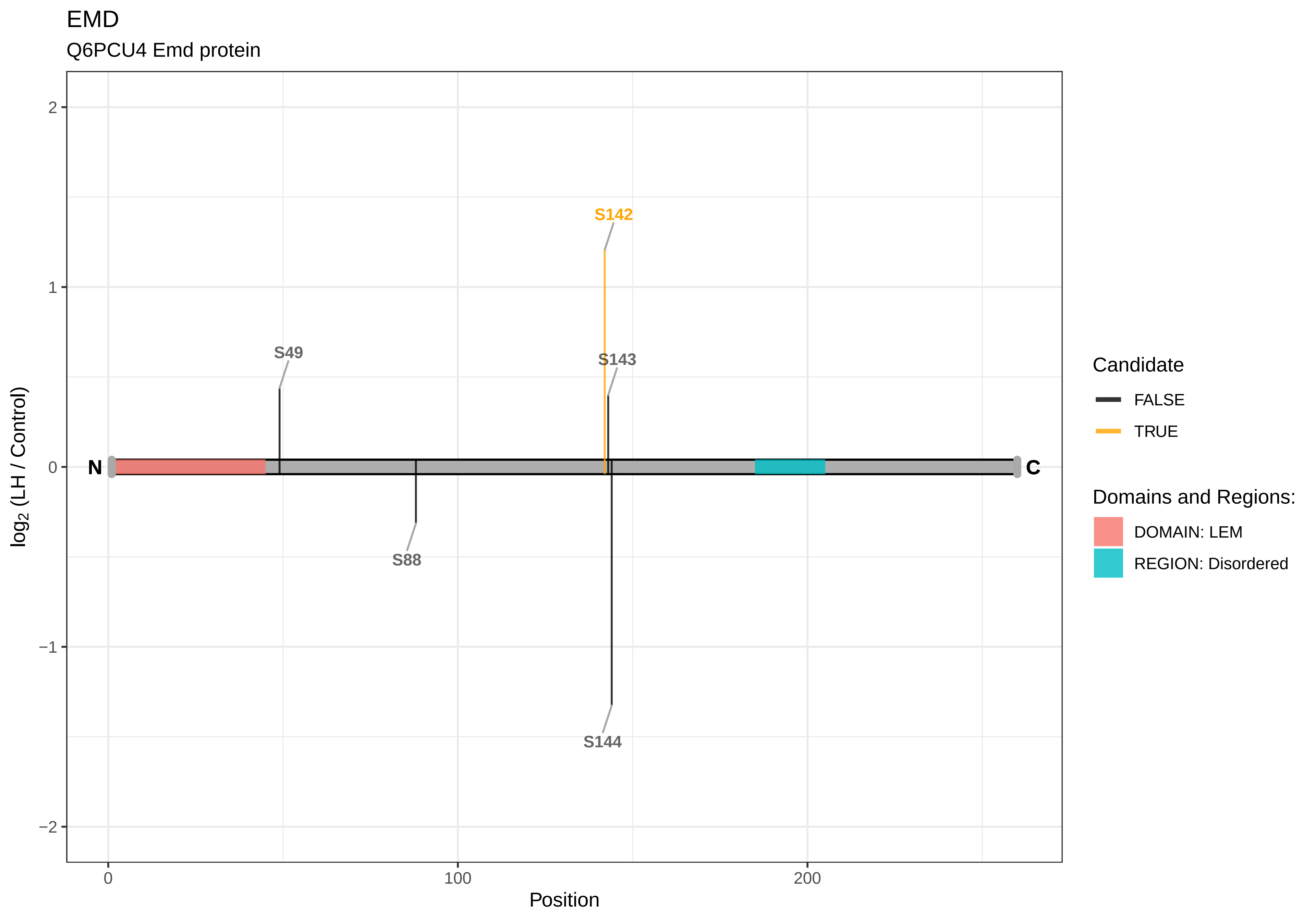
EMD
Q6PCU4 Emd protein
S142
S49
S143
S88
S144
N
C
log2 (LH / Control)
Position
Candidate
FALSE
TRUE
Domains and Regions:
DOMAIN: LEM
REGION: Disordered
2
1
0
−1
−2
0
100
200

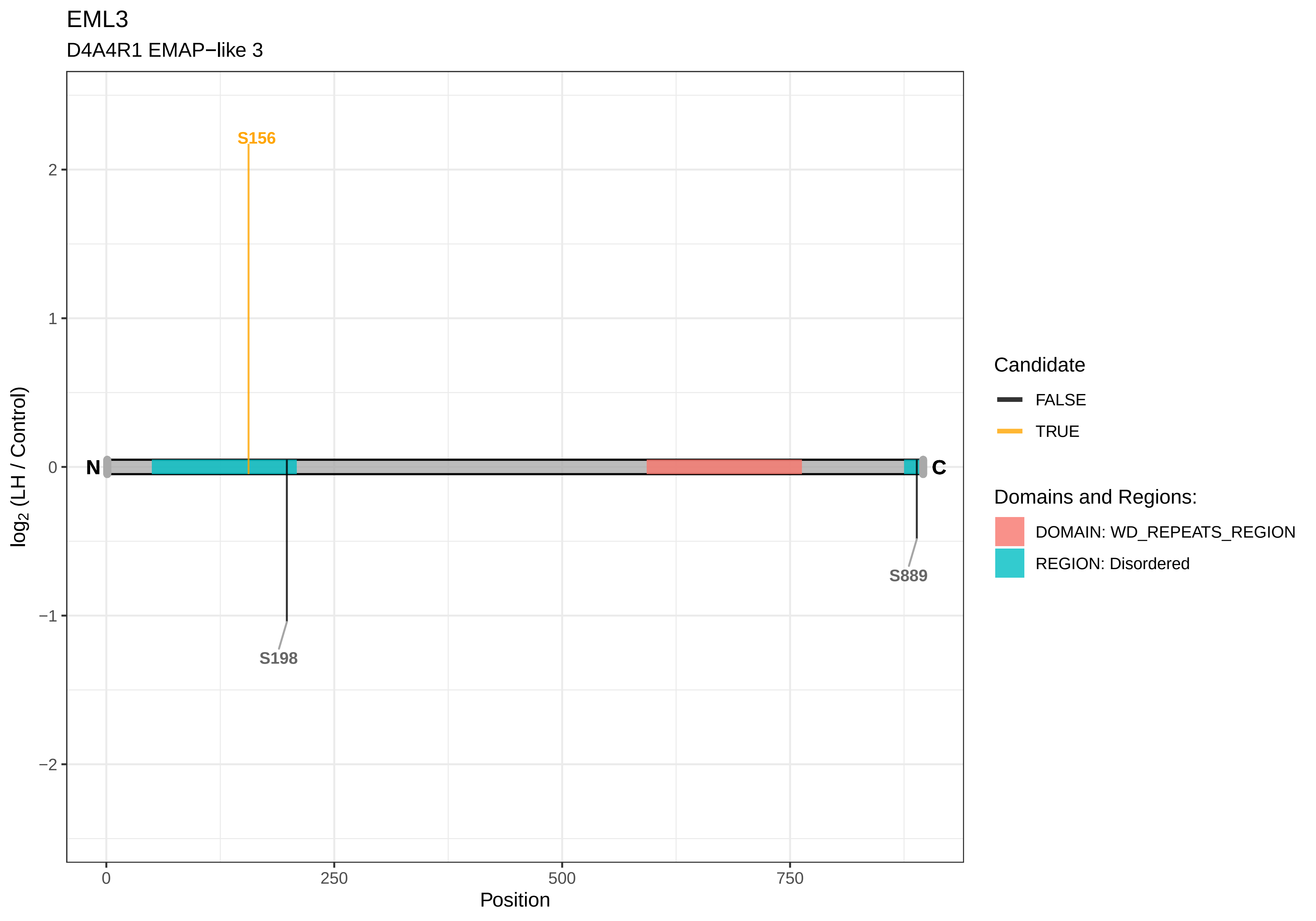

EML3
D4A4R1 EMAP−like 3
S156
S198
S889
N
C
2
1
0
−1
−2
log2 (LH / Control)
0
250
500
750
Position
Candidate
FALSE
TRUE
Domains and Regions:
DOMAIN: WD_REPEATS_REGION
REGION: Disordered

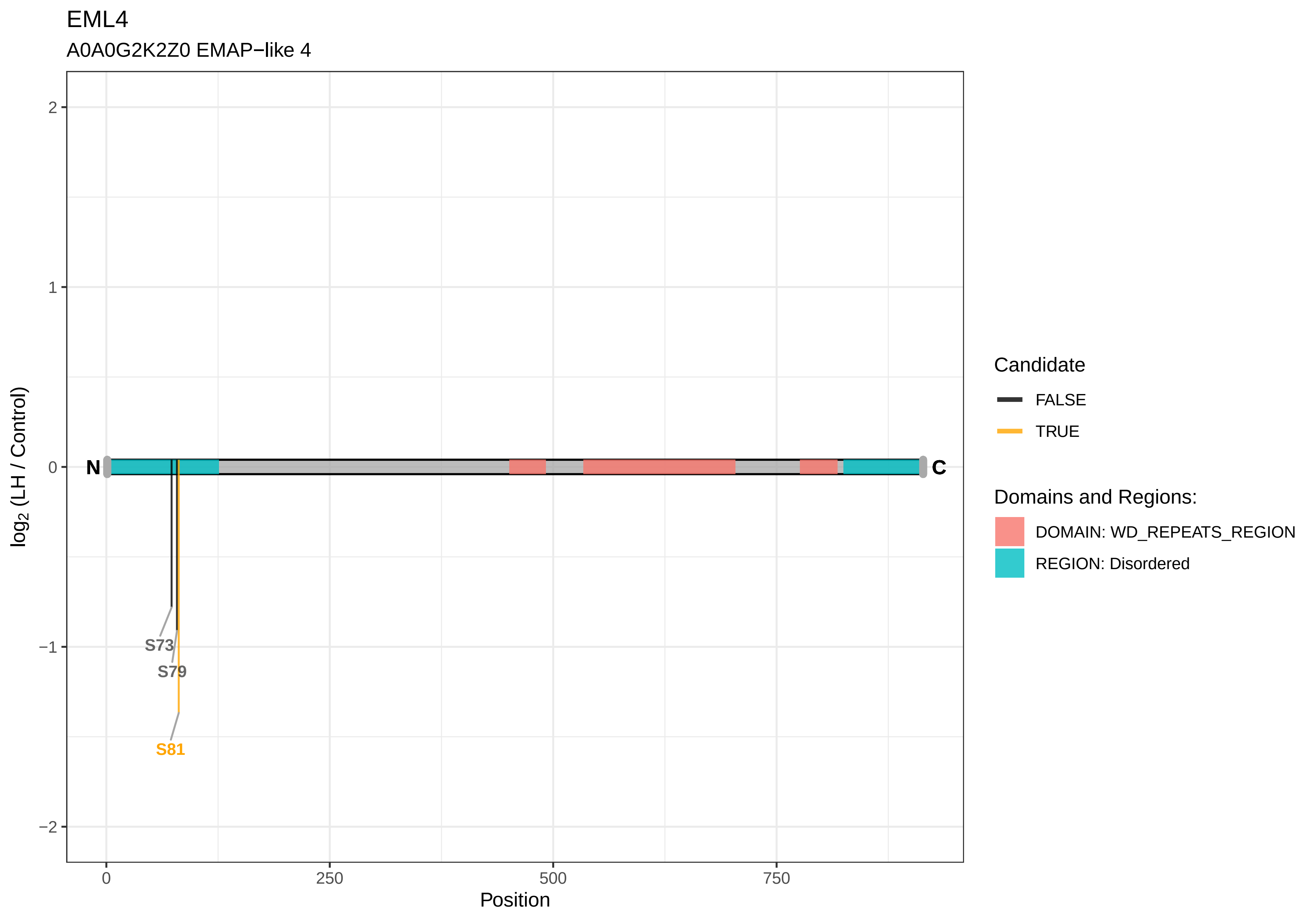

EML4
A0A0G2K2Z0 EMAP−like 4
log2 (LH / Control)
2
1
0
−1
−2
N
C
S73
S79
S81
0
250
500
750
Position
Candidate
FALSE
TRUE
Domains and Regions:
DOMAIN: WD_REPEATS_REGION
REGION: Disordered

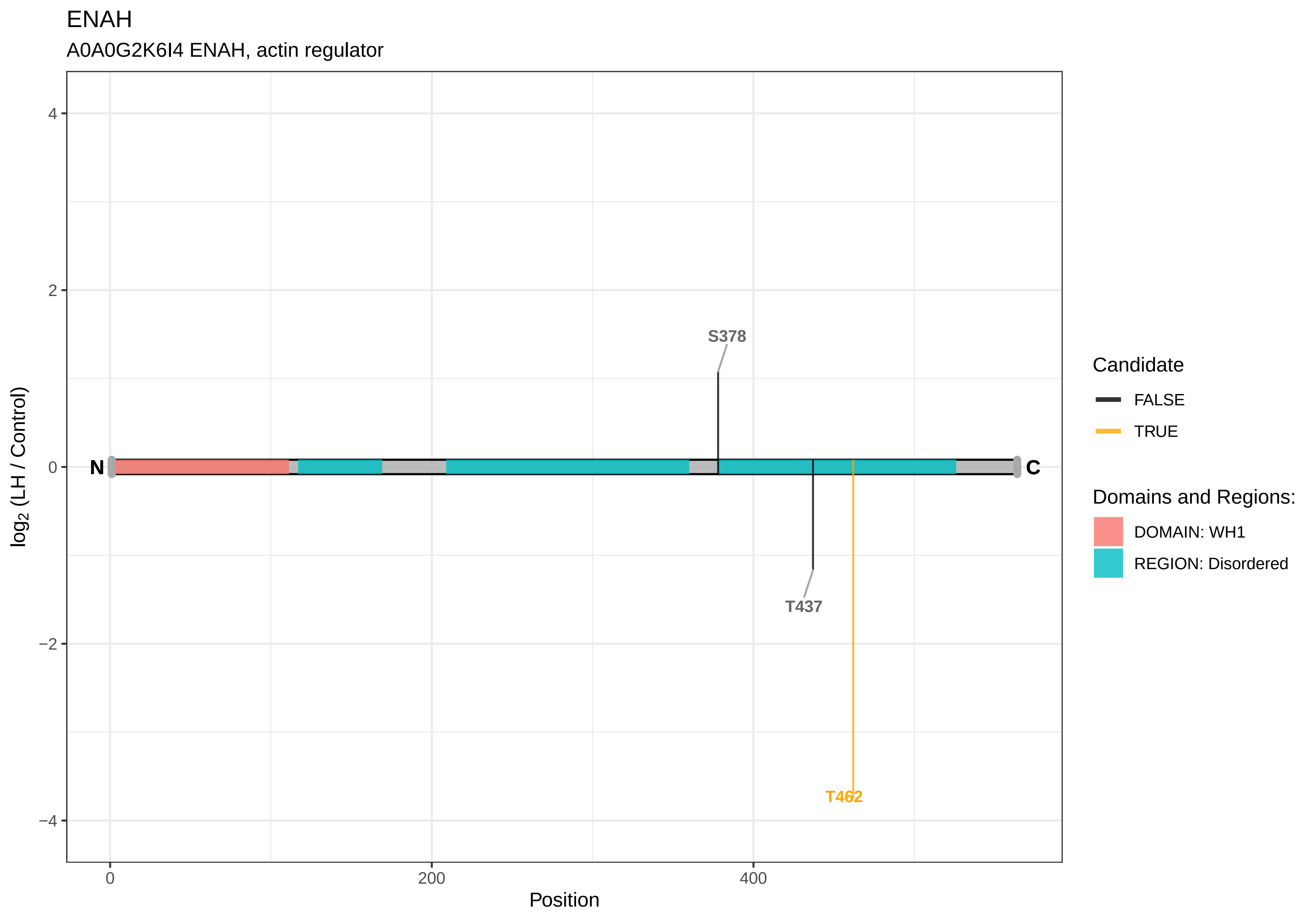

ENAH
A0A0G2K6I4 ENAH, actin regulator
4
2
0
−2
−4
log2 (LH / Control)
N
C
S378
T437
T462
0
200
400
Position
Candidate
FALSE
TRUE
Domains and Regions:
DOMAIN: WH1
REGION: Disordered

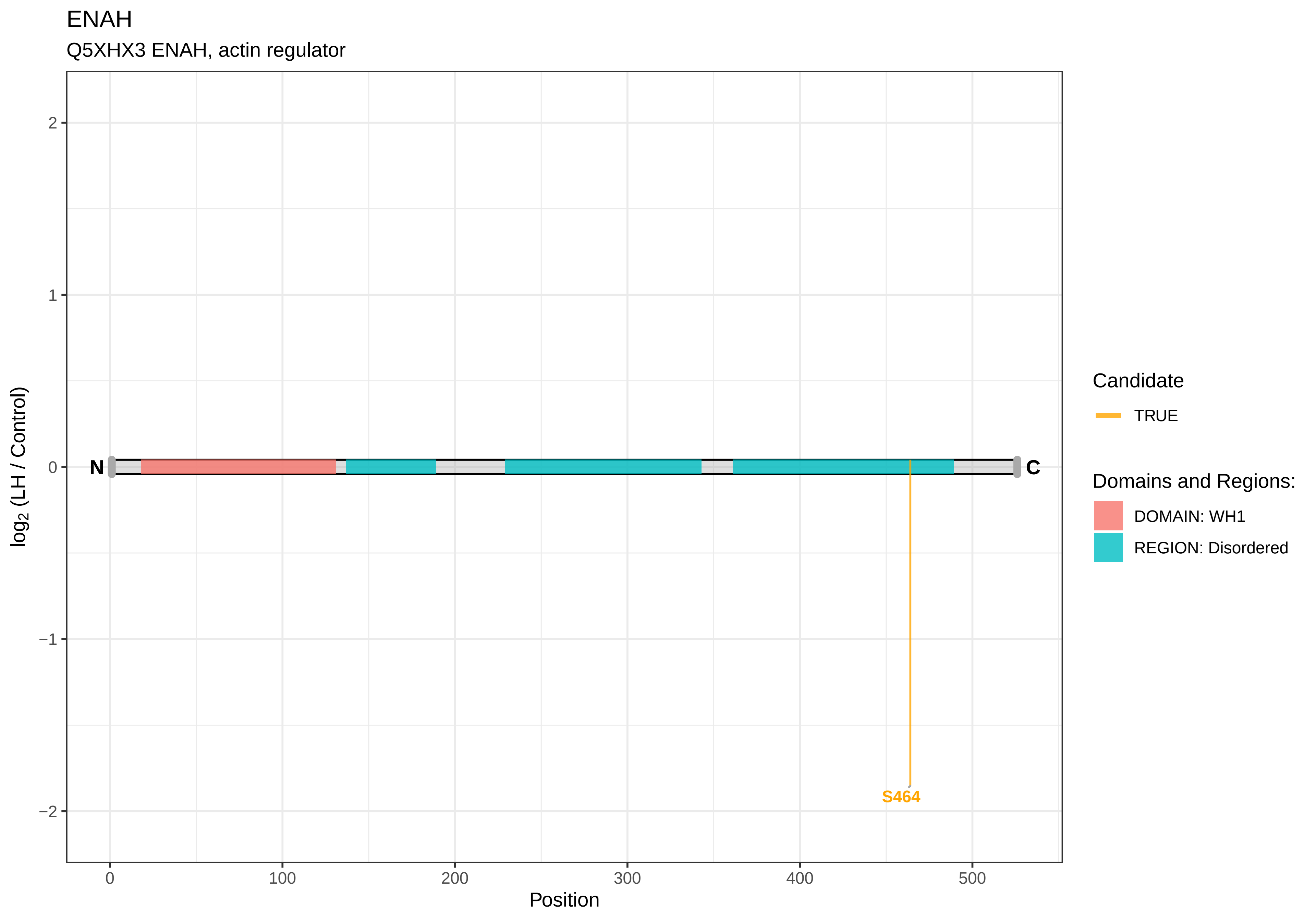
ENAH
Q5XHX3 ENAH, actin regulator
log2 (LH / Control)
2
1
0
−1
−2
N
C
S464
0
100
200
300
400
500
Position
Candidate
TRUE
Domains and Regions:
DOMAIN: WH1
REGION: Disordered

ENSA

P60841 Alpha−endosulfine

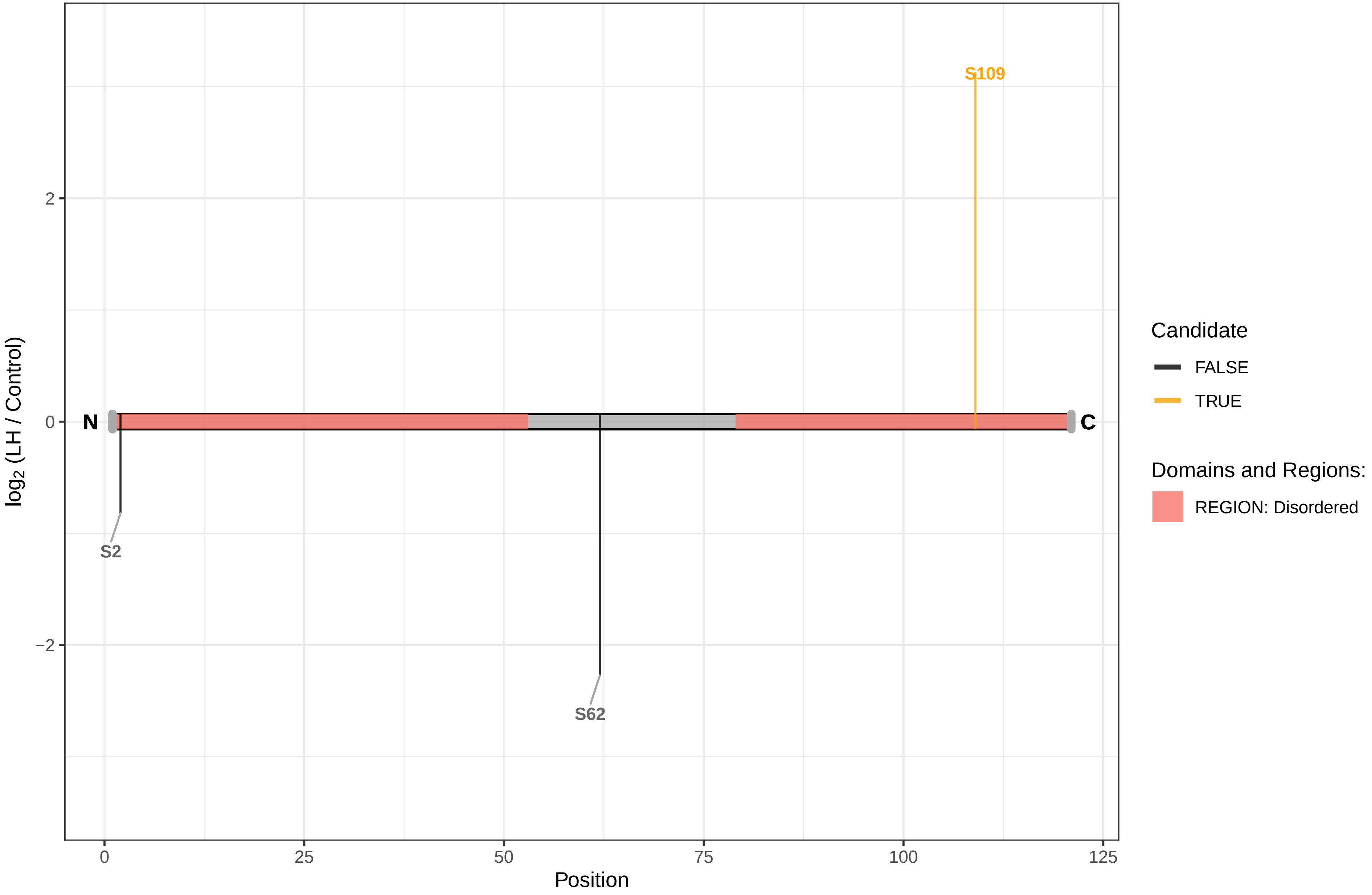

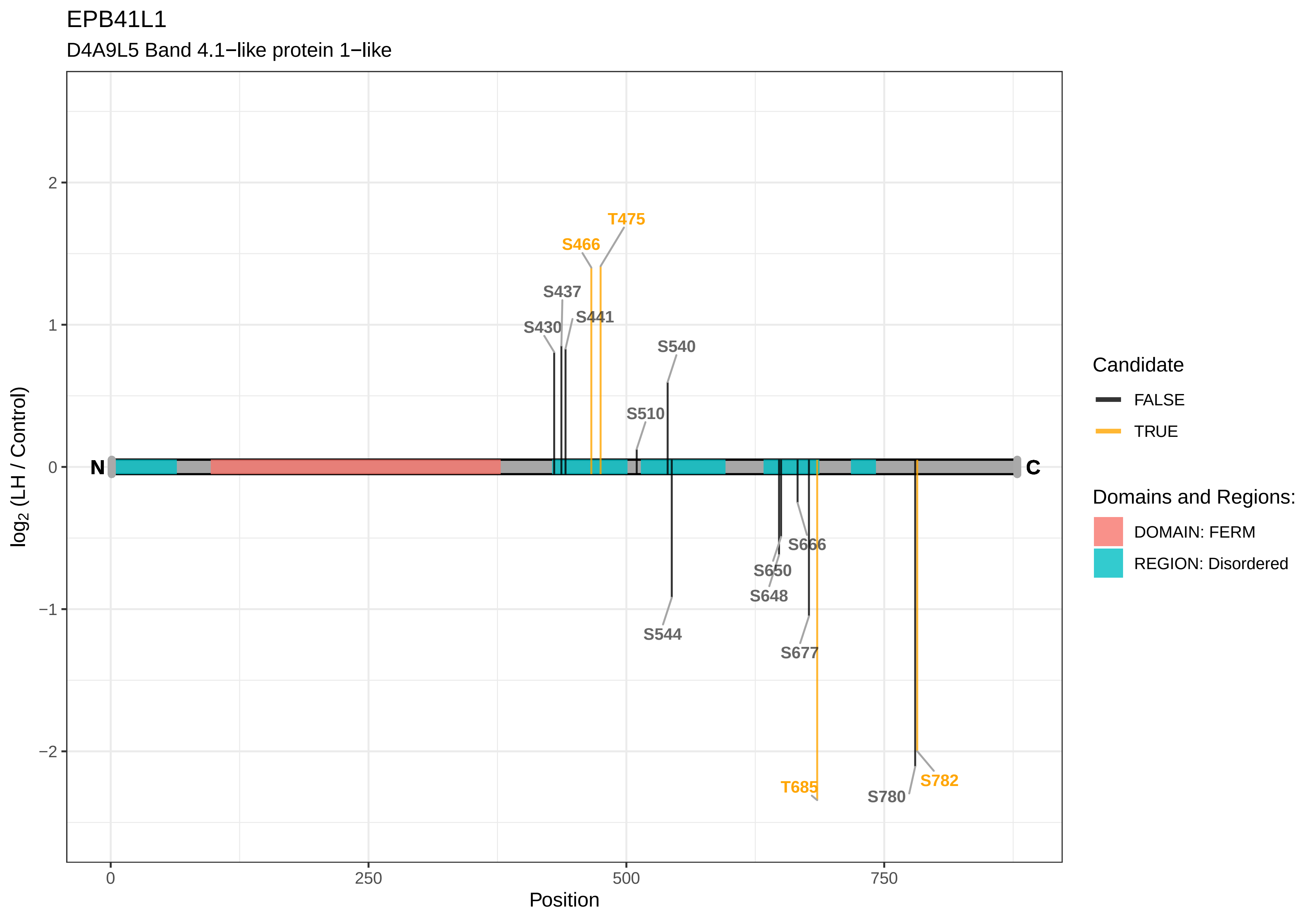

EPB41L1
D4A9L5 Band 4.1−like protein 1−like
T475
S466
S437
S441
S430
S540
S510
N
C
S666
S650
S648
S544
S677
T685
S782
S780
2
1
0
−1
−2
0
250
500
750
Position
log2 (LH / Control)
Candidate
FALSE
TRUE
Domains and Regions:
DOMAIN: FERM
REGION: Disordered

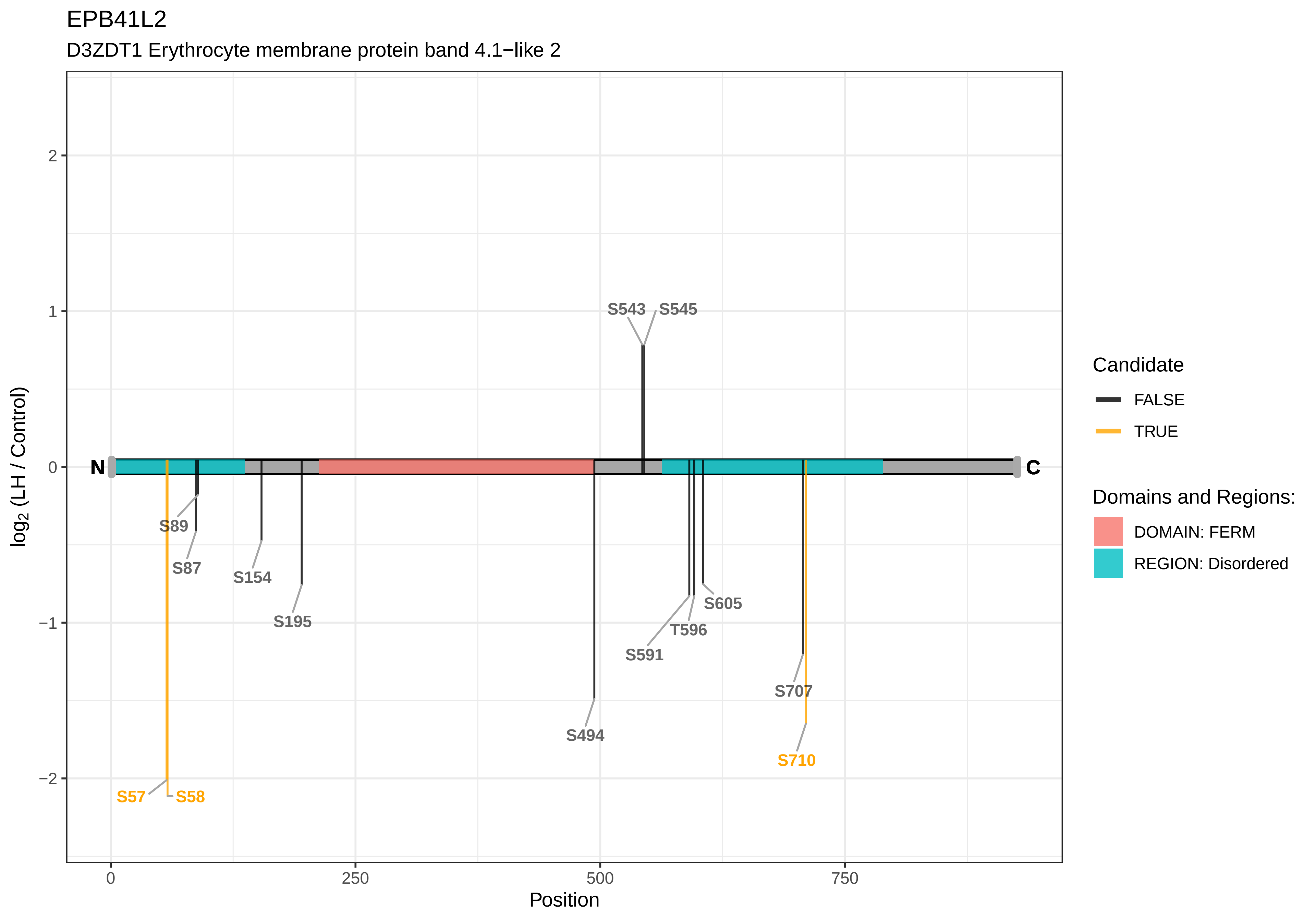

EPB41L2
D3ZDT1 Erythrocyte membrane protein band 4.1−like 2
log2 (LH / Control)
Position
0
250
500
750
2
1
0
−1
−2
N
C
S543
S545
S89
S87
S154
S195
S494
S591
T596
S605
S707
S710
S57
S58
Candidate
FALSE
TRUE
Domains and Regions:
DOMAIN: FERM
REGION: Disordered

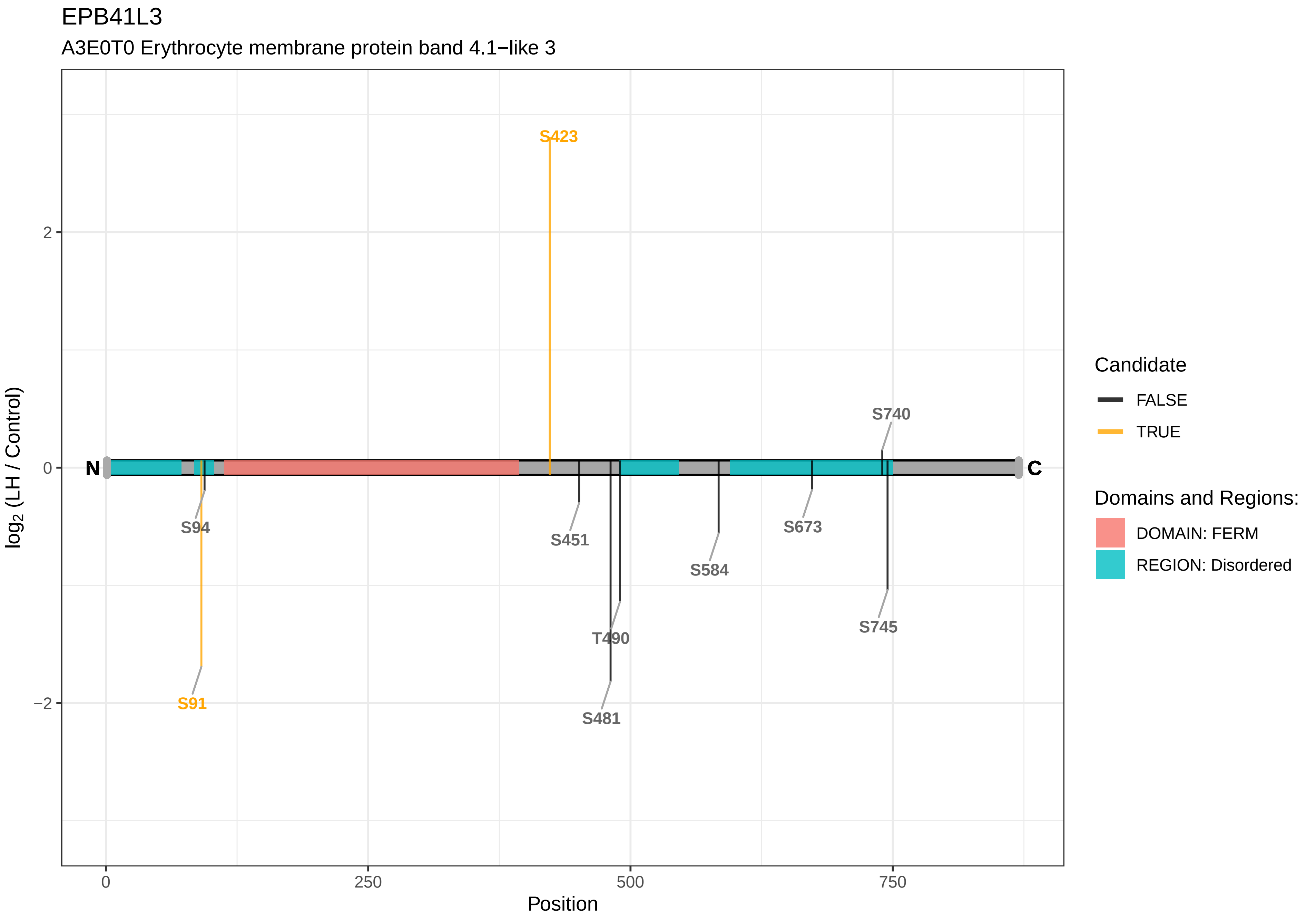

EPB41L3
A3E0T0 Erythrocyte membrane protein band 4.1−like 3
S423
S740
N
C
S94
S451
S673
S584
T490
S745
S91
S481
2
0
−2
log2 (LH / Control)
0
250
500
750
Position
Candidate
FALSE
TRUE
Domains and Regions:
DOMAIN: FERM
REGION: Disordered

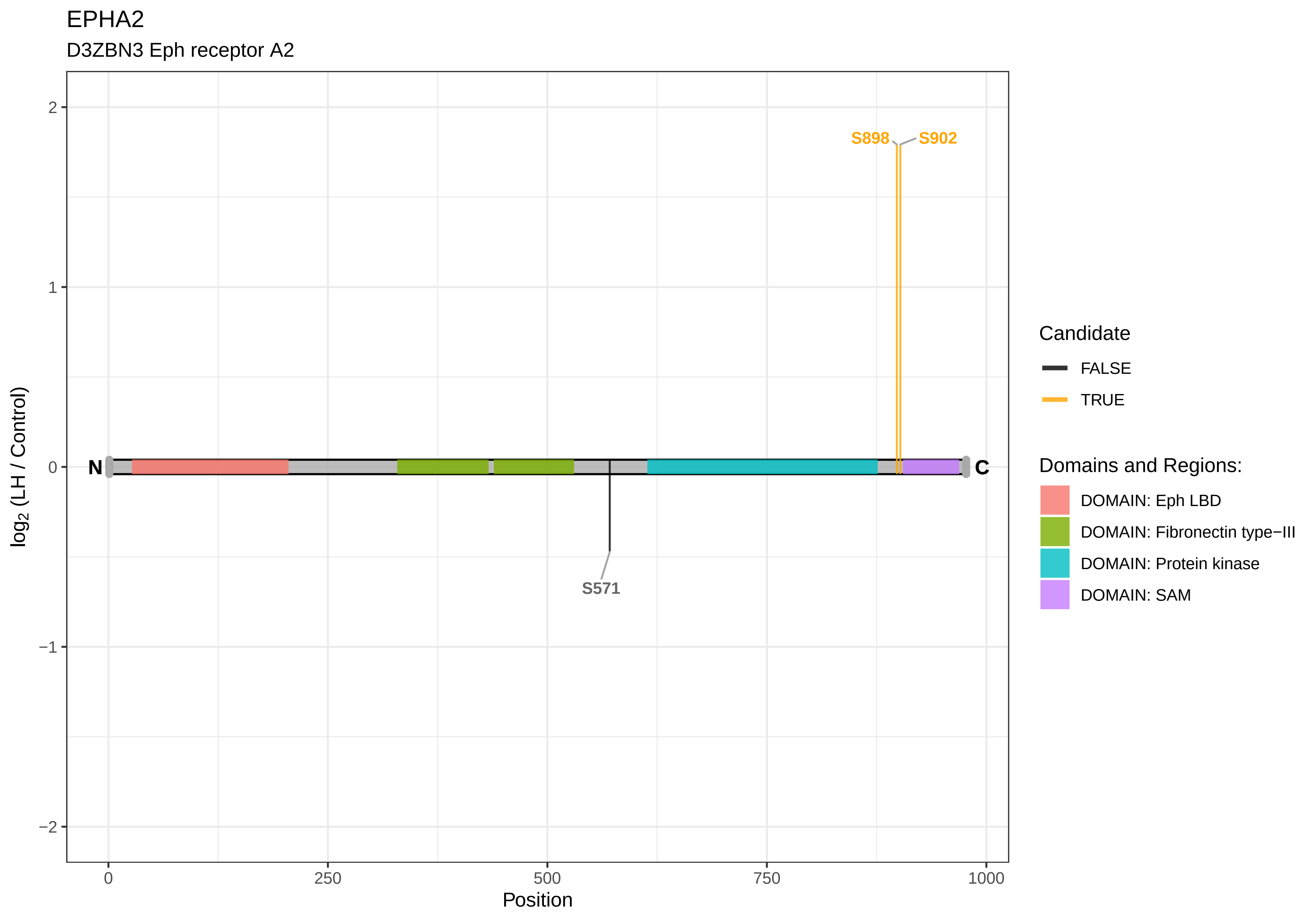

EPHA2
D3ZBN3 Eph receptor A2
S898
S902
S571
N
C
2
1
0
−1
−2
log2 (LH / Control)
0
250
500
750
1000
Position
Candidate
FALSE
TRUE
Domains and Regions:
DOMAIN: Eph LBD
DOMAIN: Fibronectin type−III
DOMAIN: Protein kinase
DOMAIN: SAM

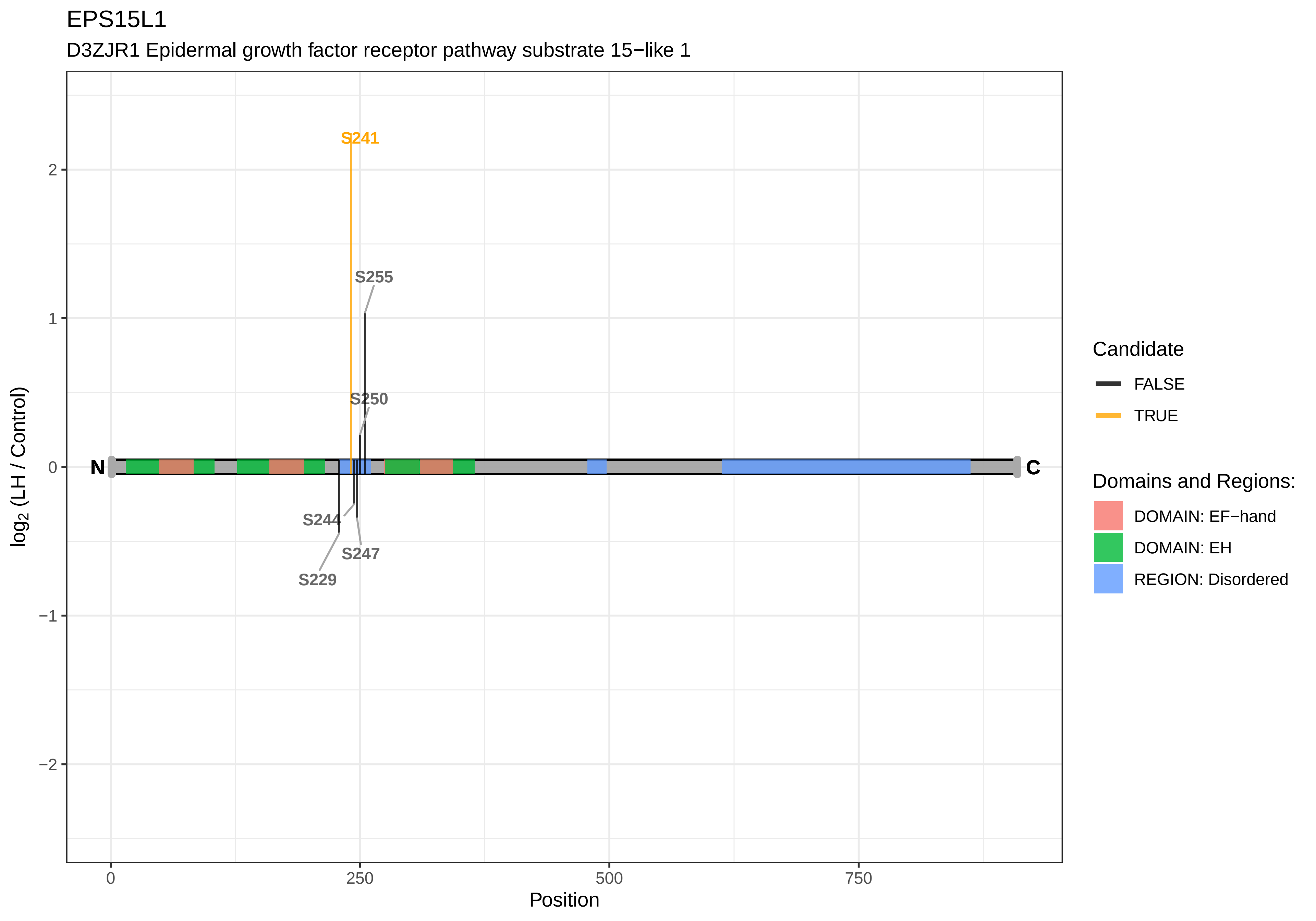

EPS15L1
D3ZJR1 Epidermal growth factor receptor pathway substrate 15−like 1
S241
S255
S250
S244
S247
S229
N
C
2
1
0
−1
−2
log2 (LH / Control)
0
250
500
750
Position
Candidate
FALSE
TRUE
Domains and Regions:
DOMAIN: EF−hand
DOMAIN: EH
REGION: Disordered

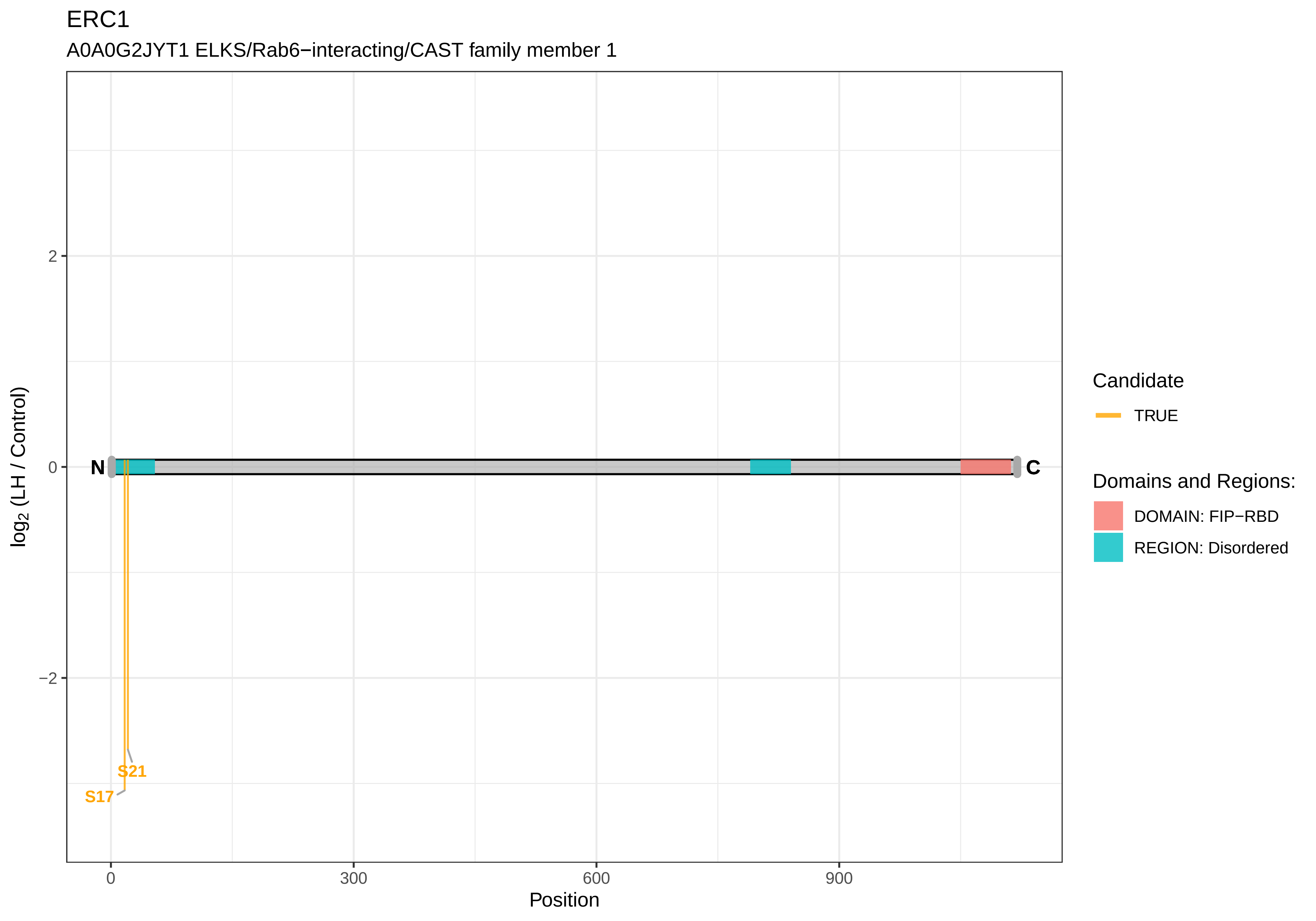

ERC1
A0A0G2JYT1 ELKS/Rab6−interacting/CAST family member 1
log2 (LH / Control)
2
0
−2
N
C
S21
S17
0
300
600
900
Position
Candidate
TRUE
Domains and Regions:
DOMAIN: FIP−RBD
REGION: Disordered

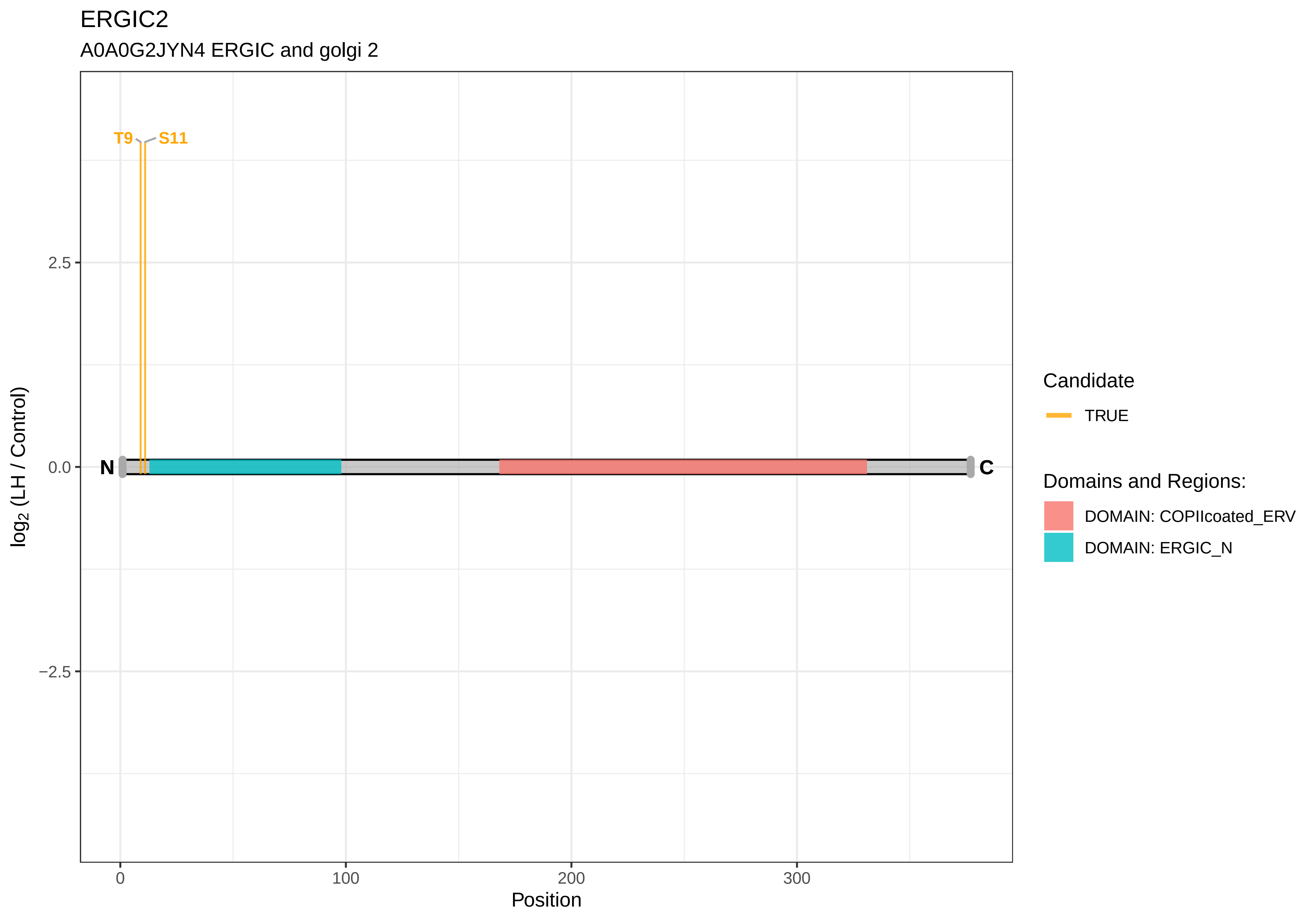

ERGIC2
A0A0G2JYN4 ERGIC and golgi 2
T9
S11
N
C
2.5
0.0
−2.5
0
100
200
300
Position
log2 (LH / Control)
Candidate
TRUE
Domains and Regions:
DOMAIN: COPIIcoated_ERV
DOMAIN: ERGIC_N

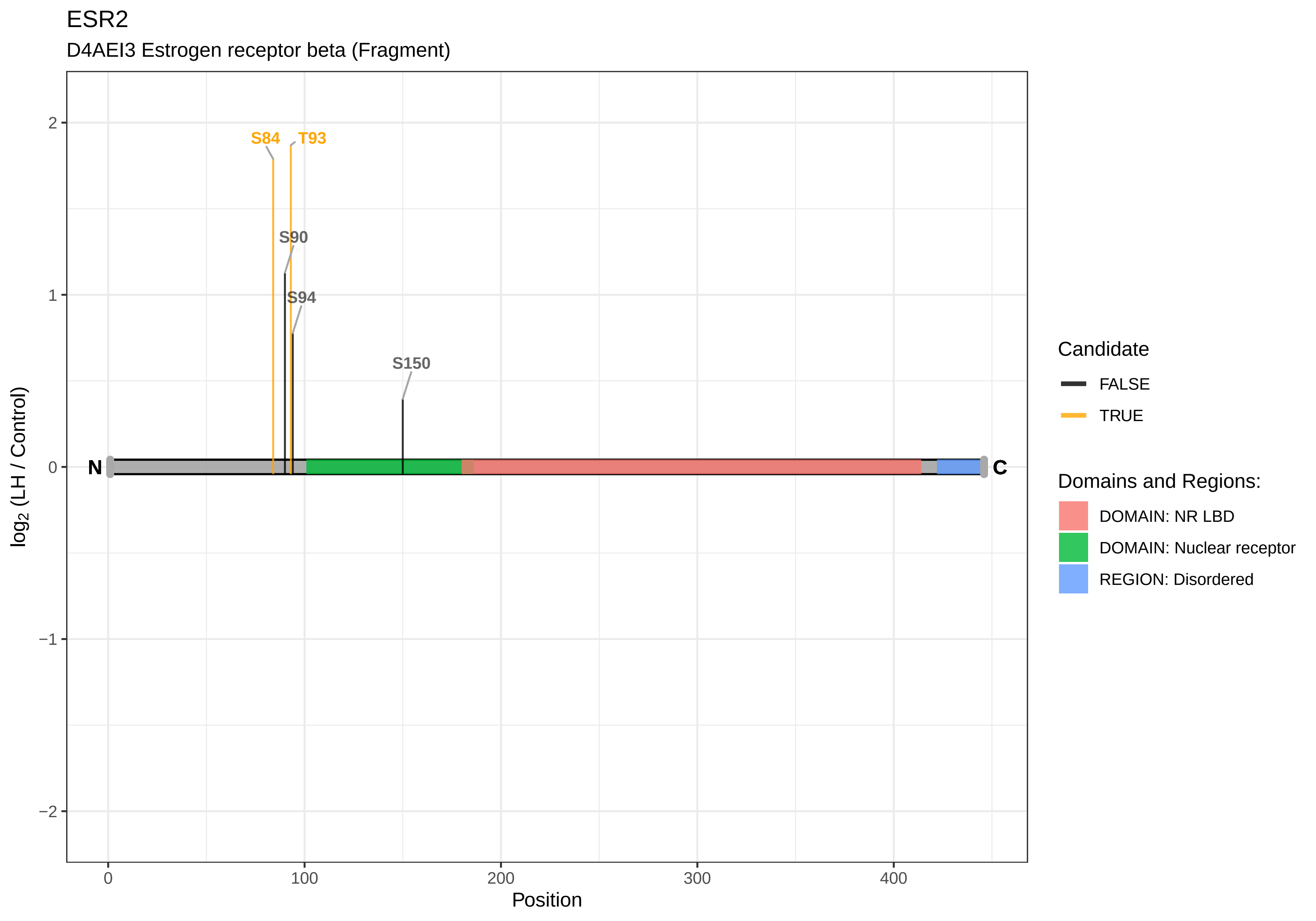

ESR2
D4AEI3 Estrogen receptor beta (Fragment)
S84
T93
S90
S94
S150
N
C
2
1
0
−1
−2
0
100
200
300
400
Position
log2 (LH / Control)
Candidate
FALSE
TRUE
Domains and Regions:
DOMAIN: NR LBD
DOMAIN: Nuclear receptor
REGION: Disordered

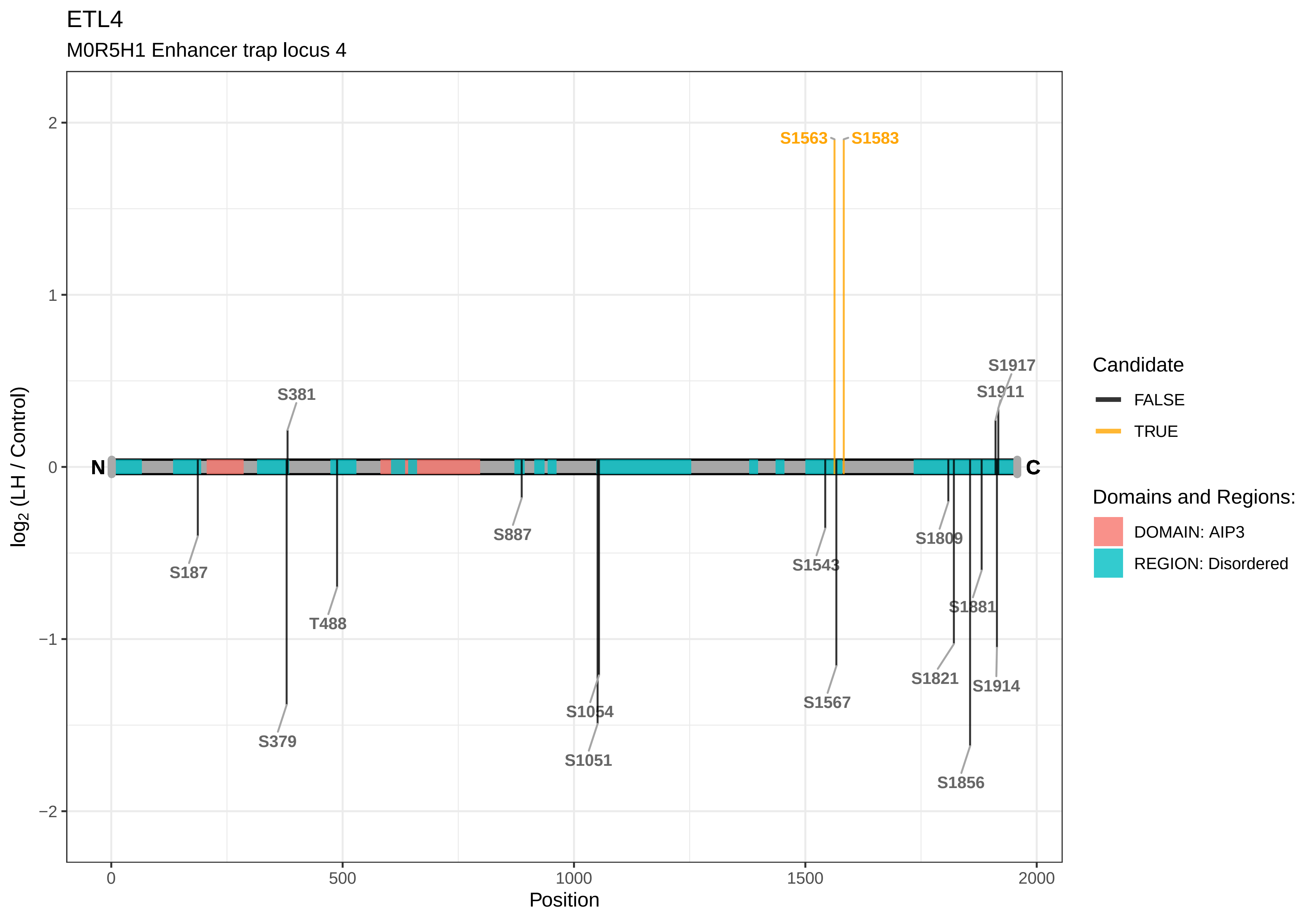
ETL4
M0R5H1 Enhancer trap locus 4
log2 (LH / Control)
Position
N
C
S1563
S1583
S1917
S1911
S381
S187
T488
S379
S887
S1054
S1051
S1543
S1567
S1809
S1881
S1821
S1914
S1856
2
1
0
−1
−2
0
500
1000
1500
2000
Candidate
FALSE
TRUE
Domains and Regions:
DOMAIN: AIP3
REGION: Disordered

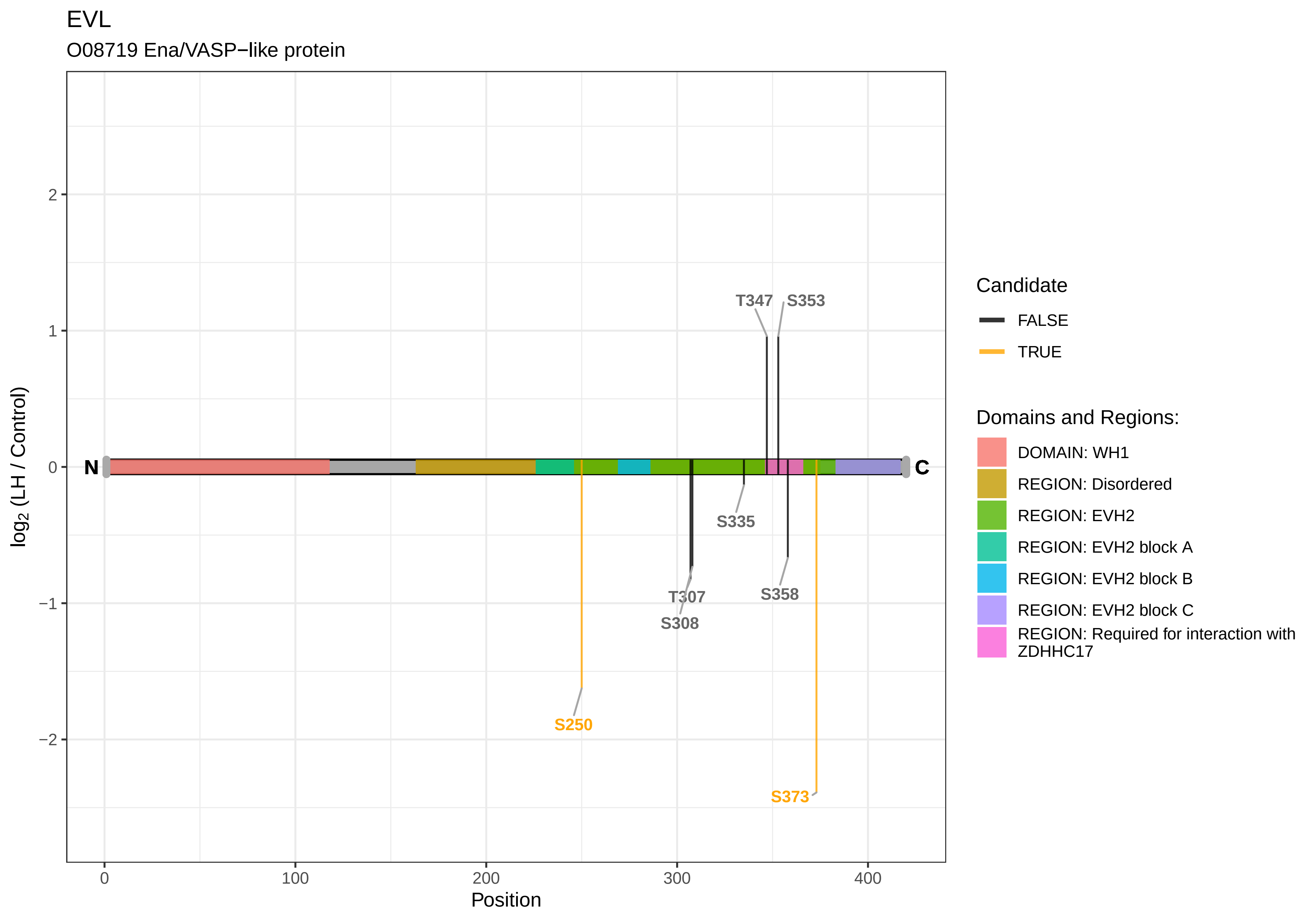
EVL
O08719 Ena/VASP−like protein
log2 (LH / Control)
Position
N
C
T347
S353
S335
T307
S308
S358
S250
S373
0
100
200
300
400
2
1
0
−1
−2
Candidate
FALSE
TRUE
Domains and Regions:
DOMAIN: WH1
REGION: Disordered
REGION: EVH2
REGION: EVH2 block A
REGION: EVH2 block B
REGION: EVH2 block C
REGION: Required for interaction with ZDHHC17

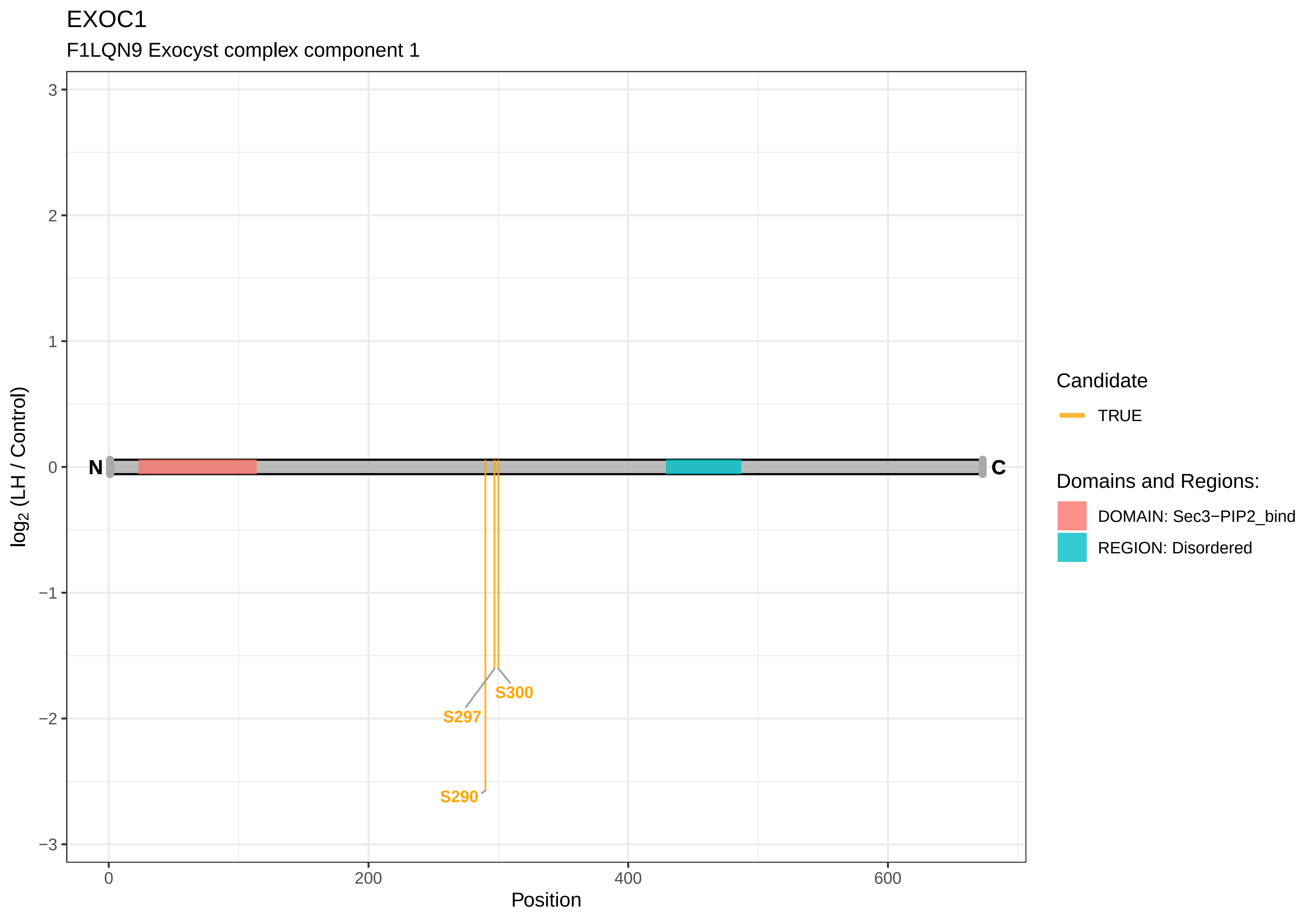
EXOC1
F1LQN9 Exocyst complex component 1
log2 (LH / Control)
3
2
1
0
−1
−2
−3
N
C
S300
S297
S290
0
200
400
600
Position
Candidate
TRUE
Domains and Regions:
DOMAIN: Sec3−PIP2_bind
REGION: Disordered

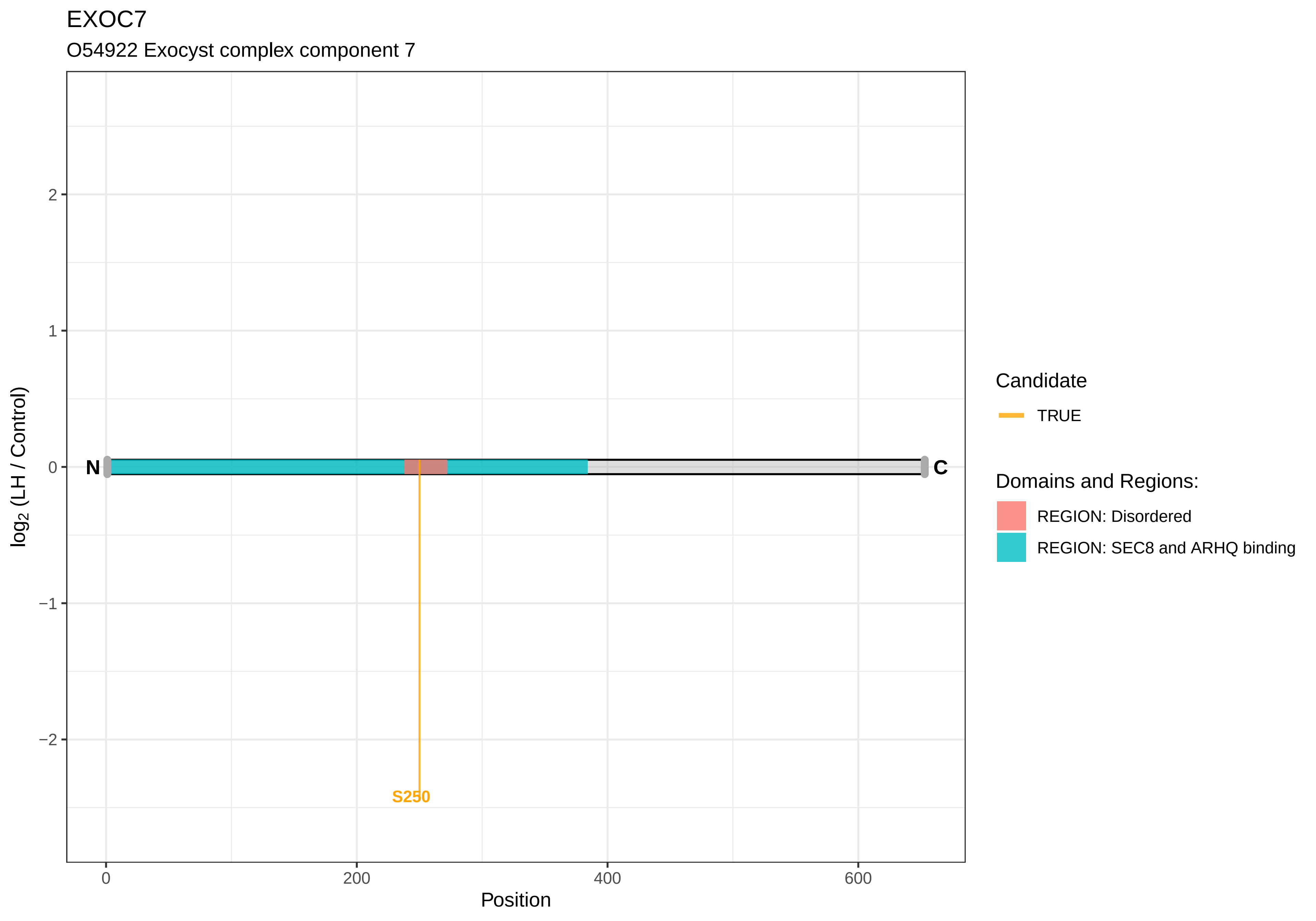

EXOC7
O54922 Exocyst complex component 7
log2 (LH / Control)
2
1
0
−1
−2
N
C
S250
0
200
400
600
Position
Candidate
TRUE
Domains and Regions:
REGION: Disordered
REGION: SEC8 and ARHQ binding

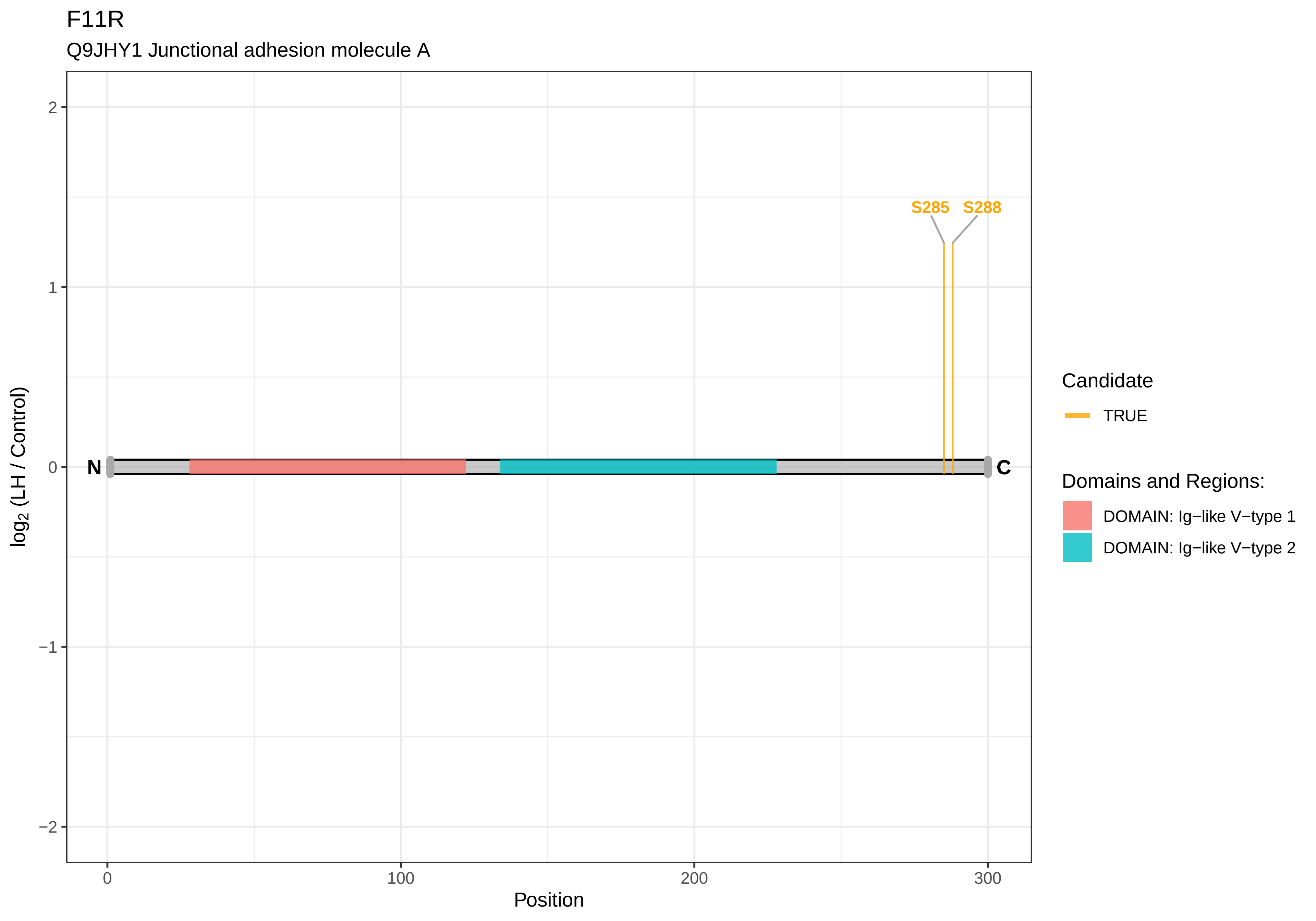

F11R
Q9JHY1 Junctional adhesion molecule A
S285
S288
N
C
2
1
0
−1
−2
0
100
200
300
Position
log2 (LH / Control)
Candidate
TRUE
Domains and Regions:
DOMAIN: Ig−like V−type 1
DOMAIN: Ig−like V−type 2

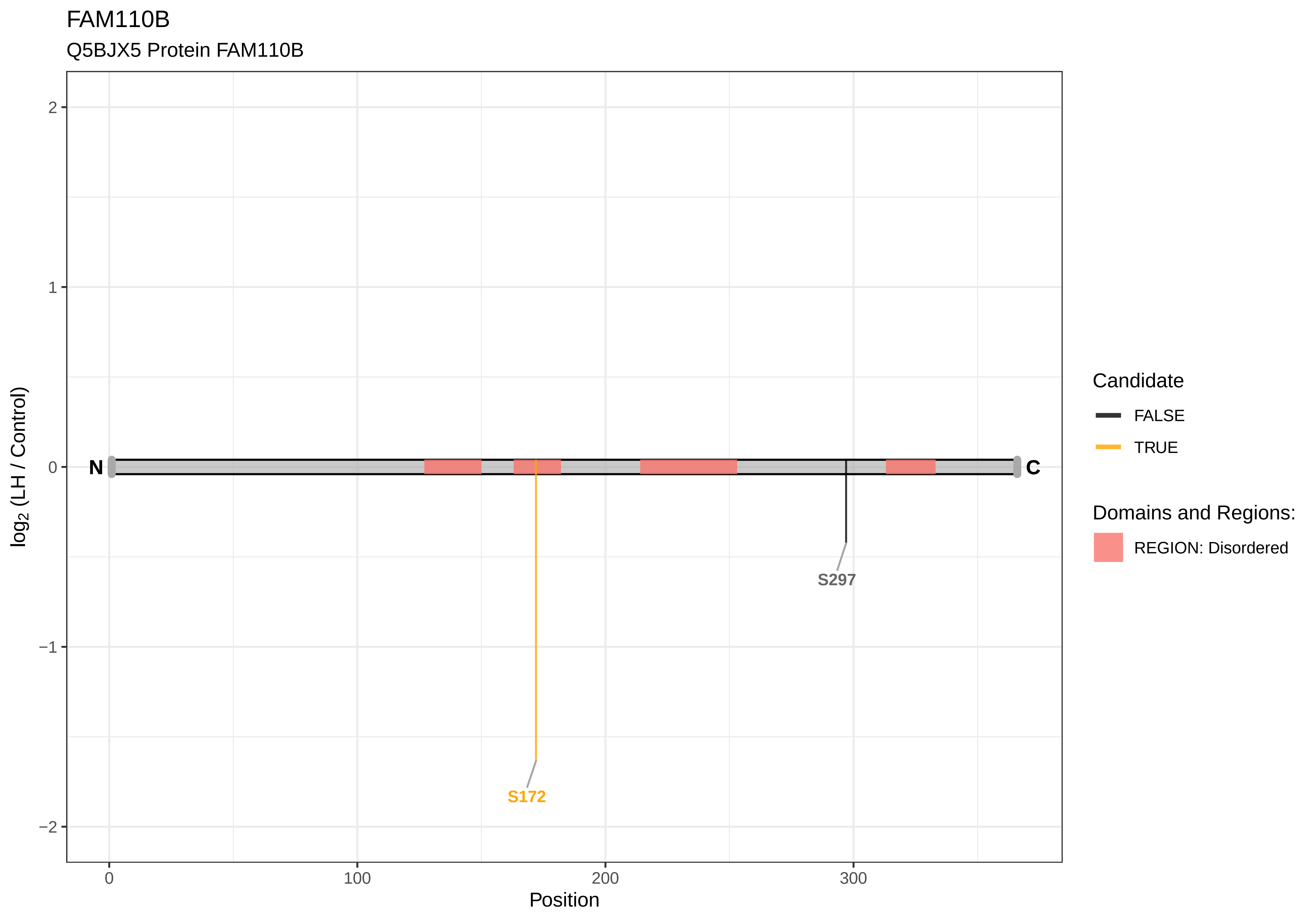

FAM110B
Q5BJX5 Protein FAM110B
log2 (LH / Control)
2
1
0
−1
−2
N
C
S172
S297
0
100
200
300
Position
Candidate
FALSE
TRUE
Domains and Regions:
REGION: Disordered

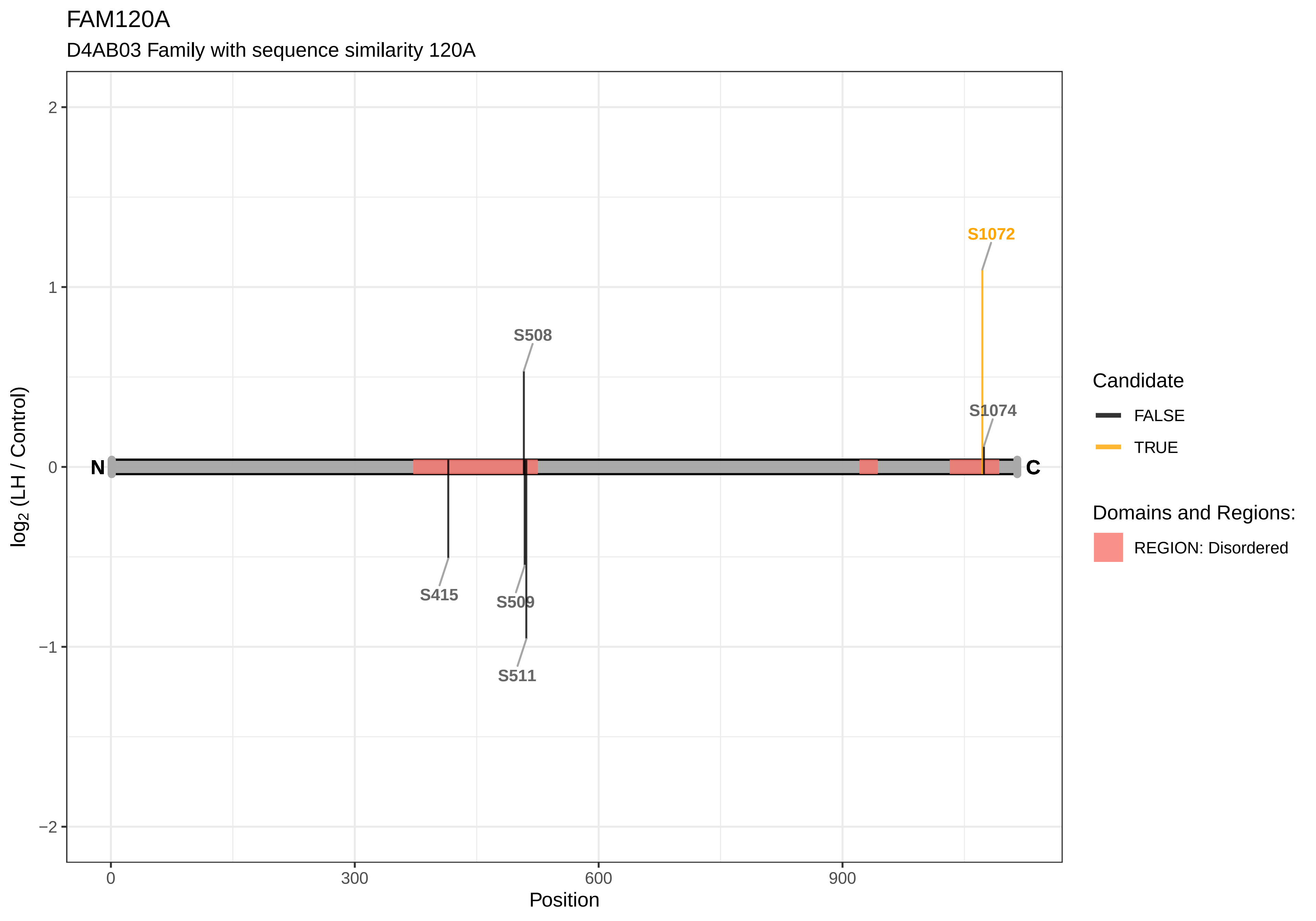

FAM120A
D4AB03 Family with sequence similarity 120A
S1072
S508
S1074
S415
S509
S511
N
C
log2 (LH / Control)
Position
2
1
0
−1
−2
0
300
600
900
Candidate
FALSE
TRUE
Domains and Regions:
REGION: Disordered

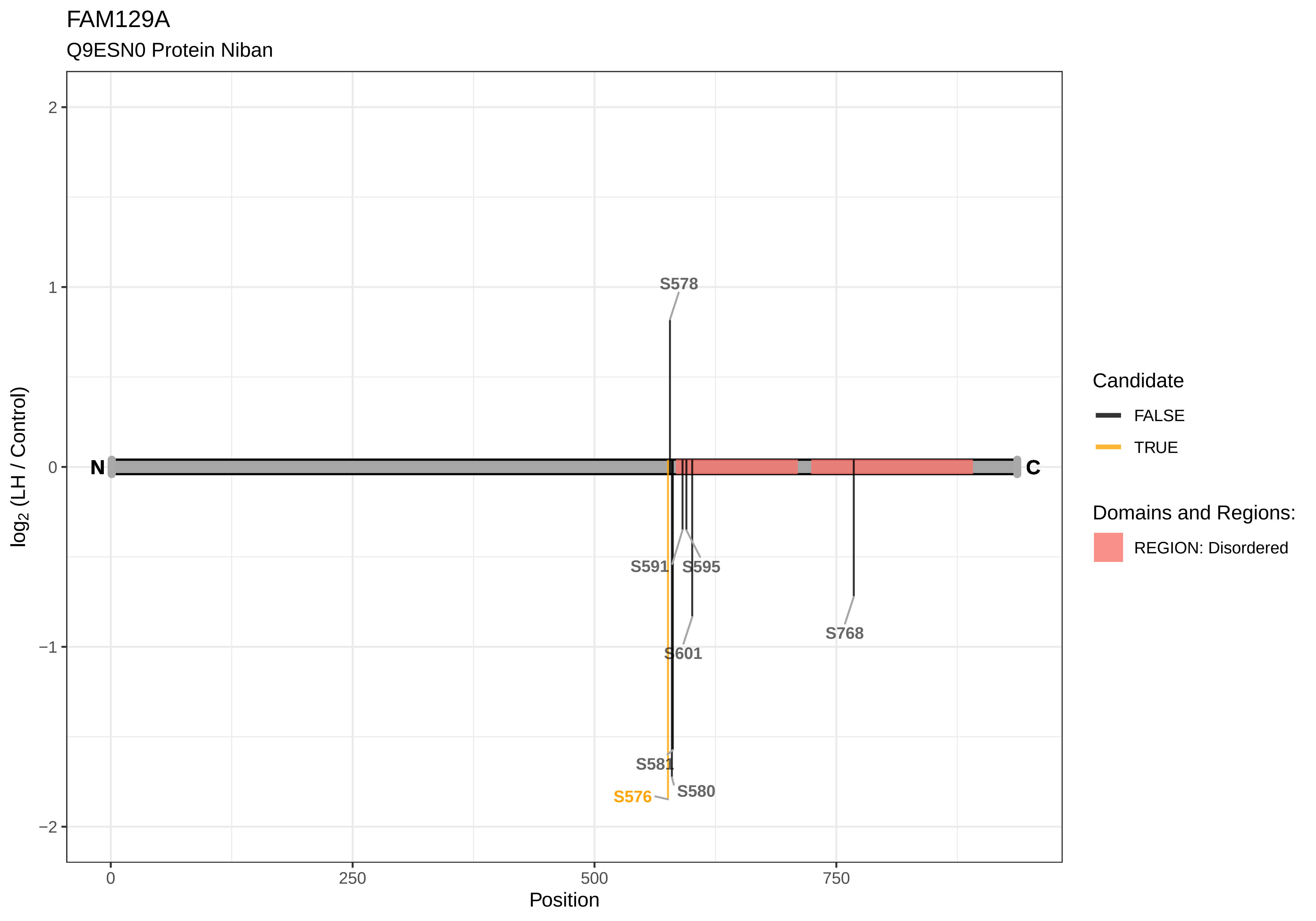
FAM129A
Q9ESN0 Protein Niban
S578
S591
S595
S601
S768
S581
S576
S580
N
C
2
1
0
−1
−2
0
250
500
750
Position
log2 (LH / Control)
Candidate
FALSE
TRUE
Domains and Regions:
REGION: Disordered

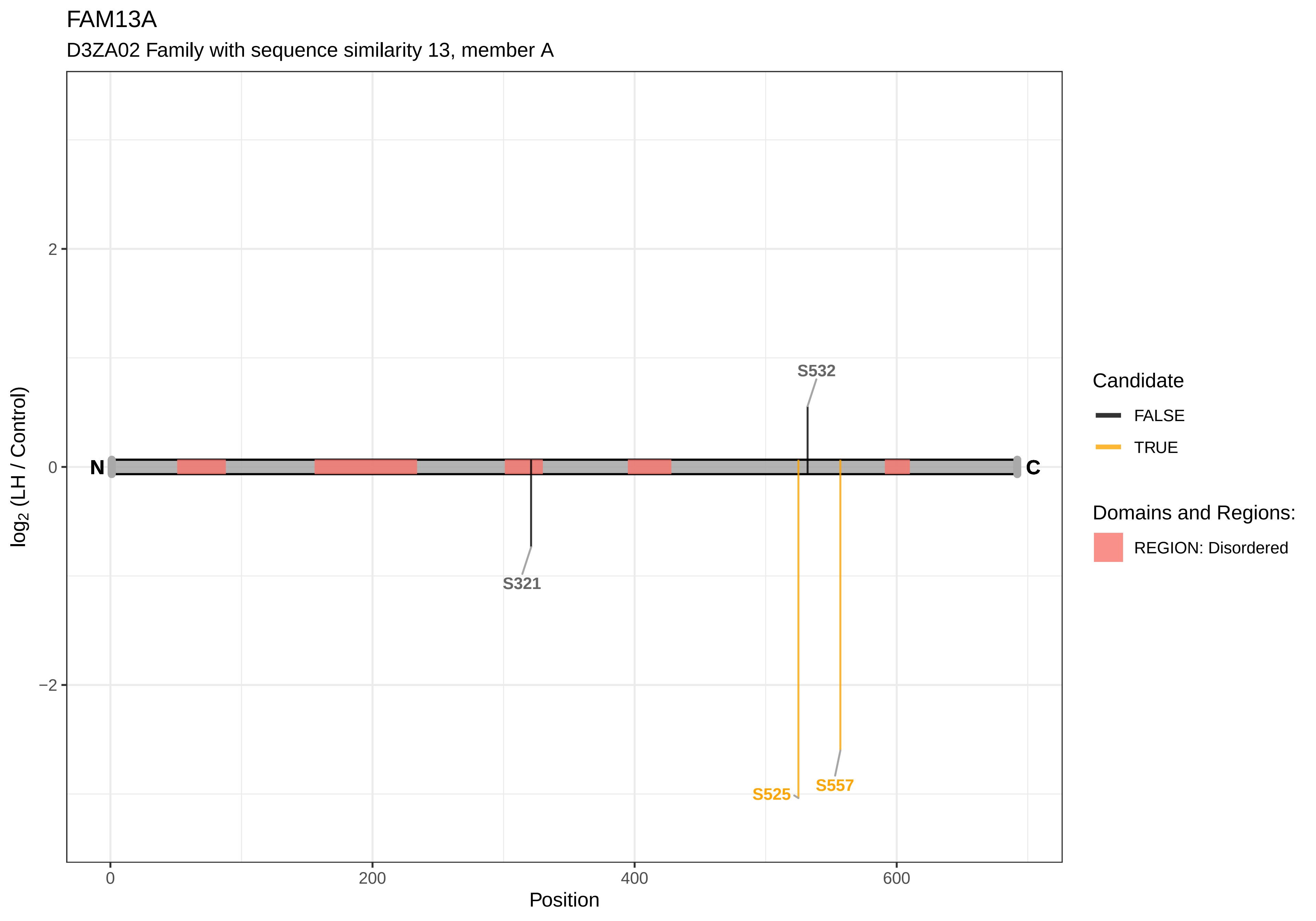

FAM13A
D3ZA02 Family with sequence similarity 13, member A
2
0
−2
0
200
400
600
Position
log2 (LH / Control)
N
C
S532
S321
S525
S557
Candidate
FALSE
TRUE
Domains and Regions:
REGION: Disordered

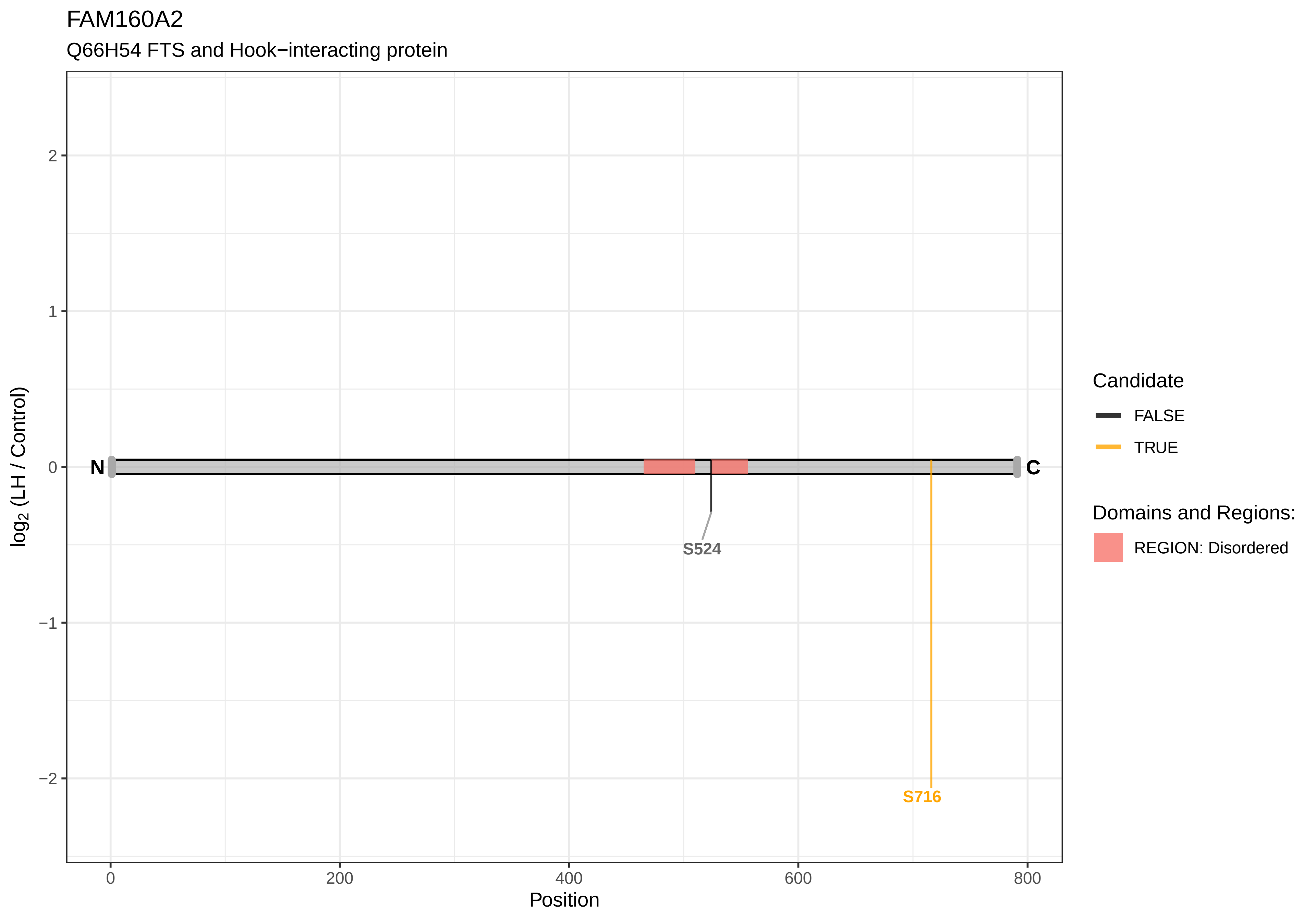

FAM160A2
Q66H54 FTS and Hook−interacting protein
log2 (LH / Control)
2
1
0
−1
−2
N
C
S524
S716
0
200
400
600
800
Position
Candidate
FALSE
TRUE
Domains and Regions:
REGION: Disordered

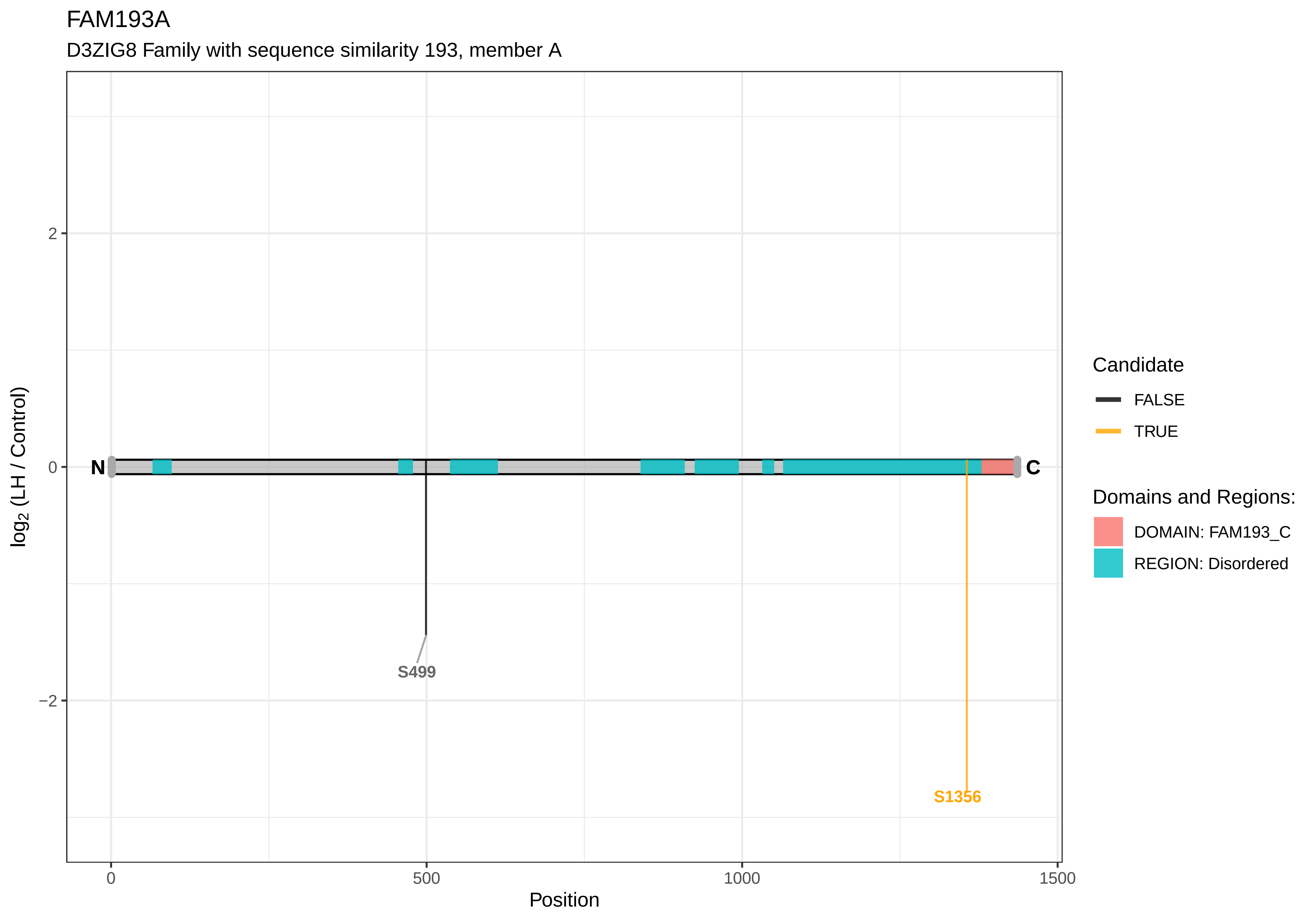

FAM193A
D3ZIG8 Family with sequence similarity 193, member A
log2 (LH / Control)
2
0
−2
N
C
S499
S1356
0
500
1000
1500
Position
Candidate
FALSE
TRUE
Domains and Regions:
DOMAIN: FAM193_C
REGION: Disordered

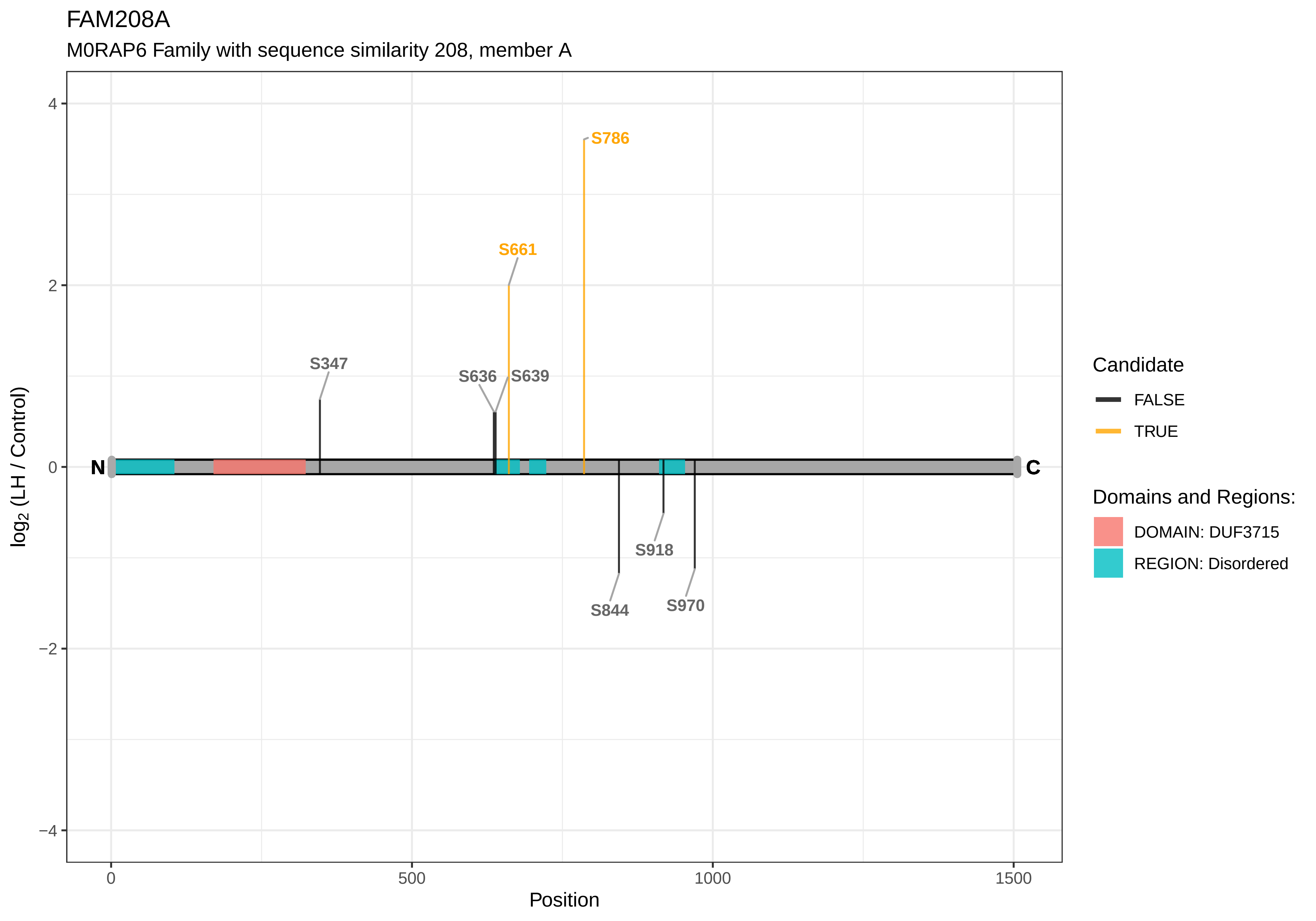
FAM208A
M0RAP6 Family with sequence similarity 208, member A
S786
S661
S347
S636
S639
S918
S844
S970
N
C
4
2
0
−2
−4
log2 (LH / Control)
0
500
1000
1500
Position
Candidate
FALSE
TRUE
Domains and Regions:
DOMAIN: DUF3715
REGION: Disordered

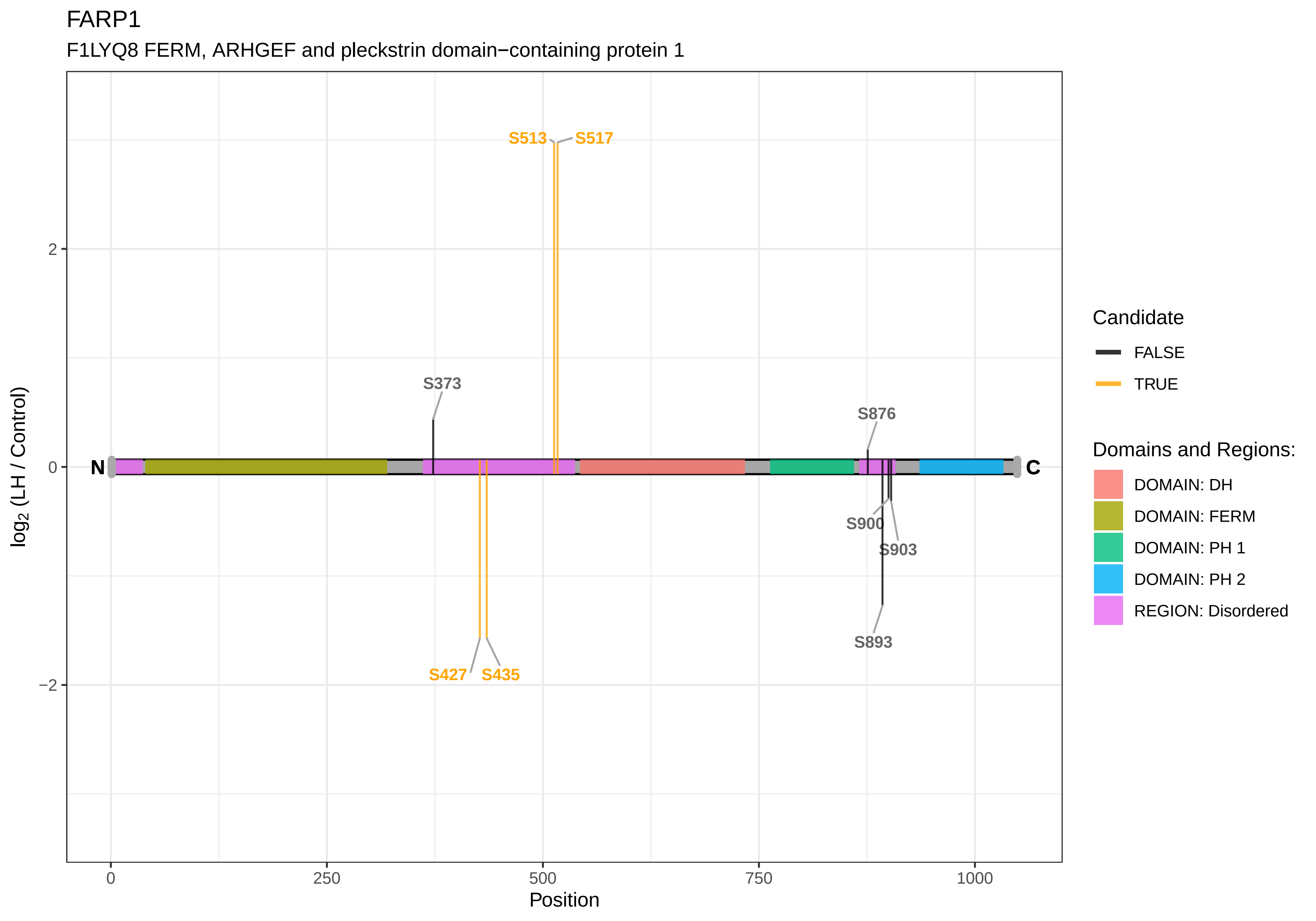
FARP1
F1LYQ8 FERM, ARHGEF and pleckstrin domain−containing protein 1
S513
S517
S373
S876
N
C
S900
S903
S893
S427
S435
2
0
−2
0
250
500
750
1000
Position
log2 (LH / Control)
Candidate
FALSE
TRUE
Domains and Regions:
DOMAIN: DH
DOMAIN: FERM
DOMAIN: PH 1
DOMAIN: PH 2
REGION: Disordered

FARP2

D3ZFK8 FERM, ARH/RhoGEF and pleckstrin domain protein 2

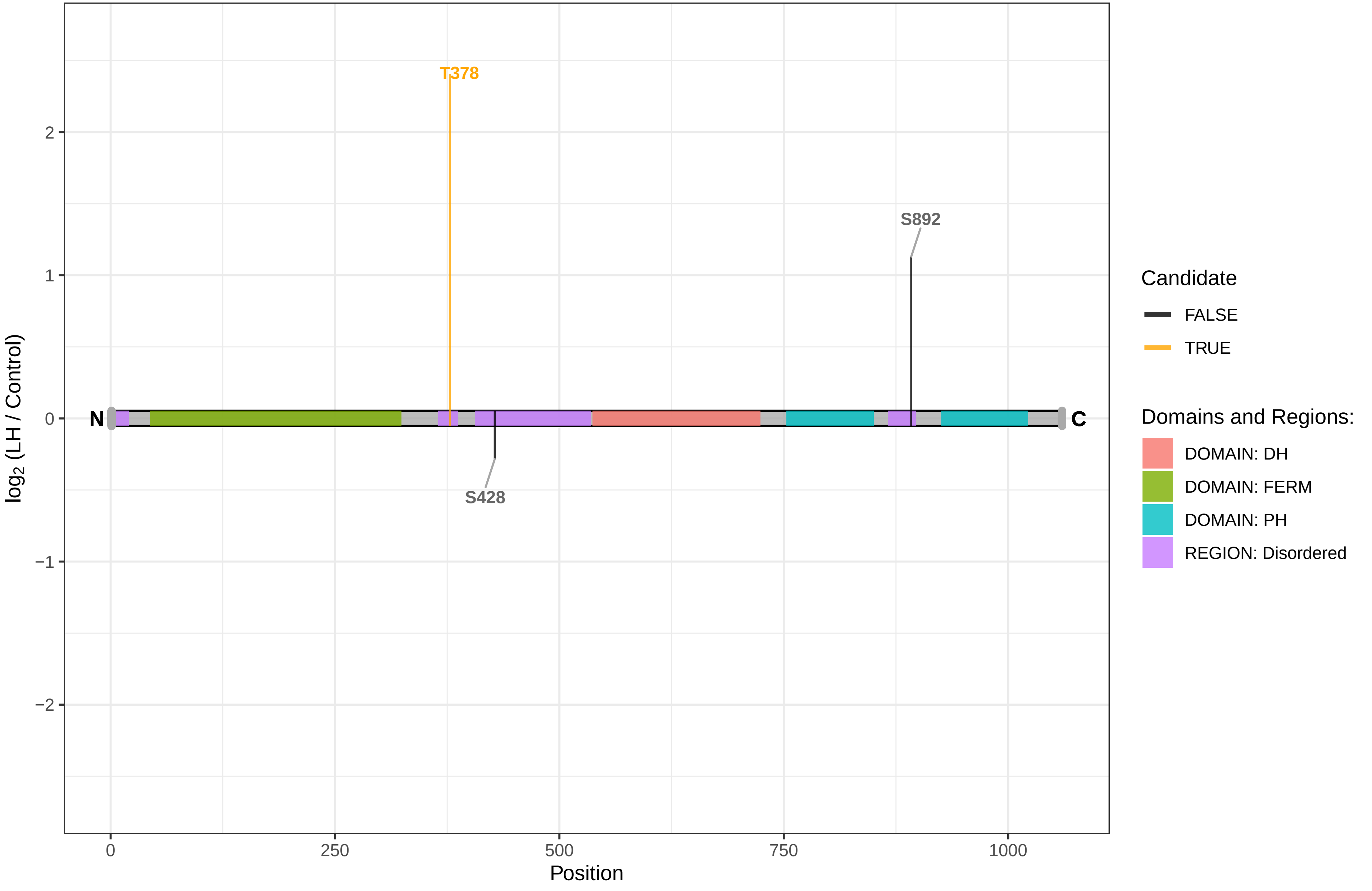

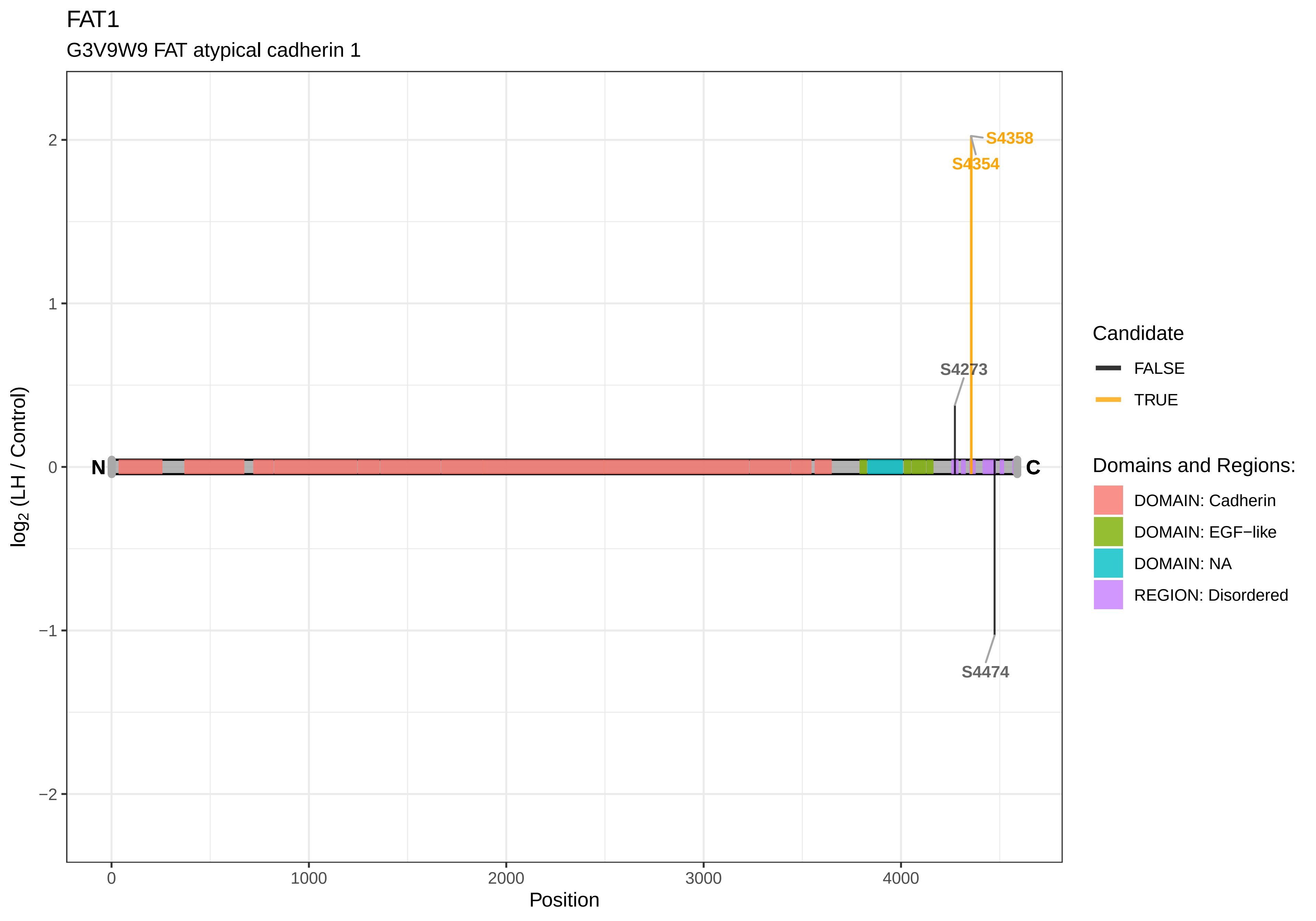

FAT1
G3V9W9 FAT atypical cadherin 1
log2 (LH / Control)
Position
N
C
S4358
S4354
S4273
S4474
2
1
0
−1
−2
0
1000
2000
3000
4000
Candidate
FALSE
TRUE
Domains and Regions:
DOMAIN: Cadherin
DOMAIN: EGF−like
DOMAIN: NA
REGION: Disordered

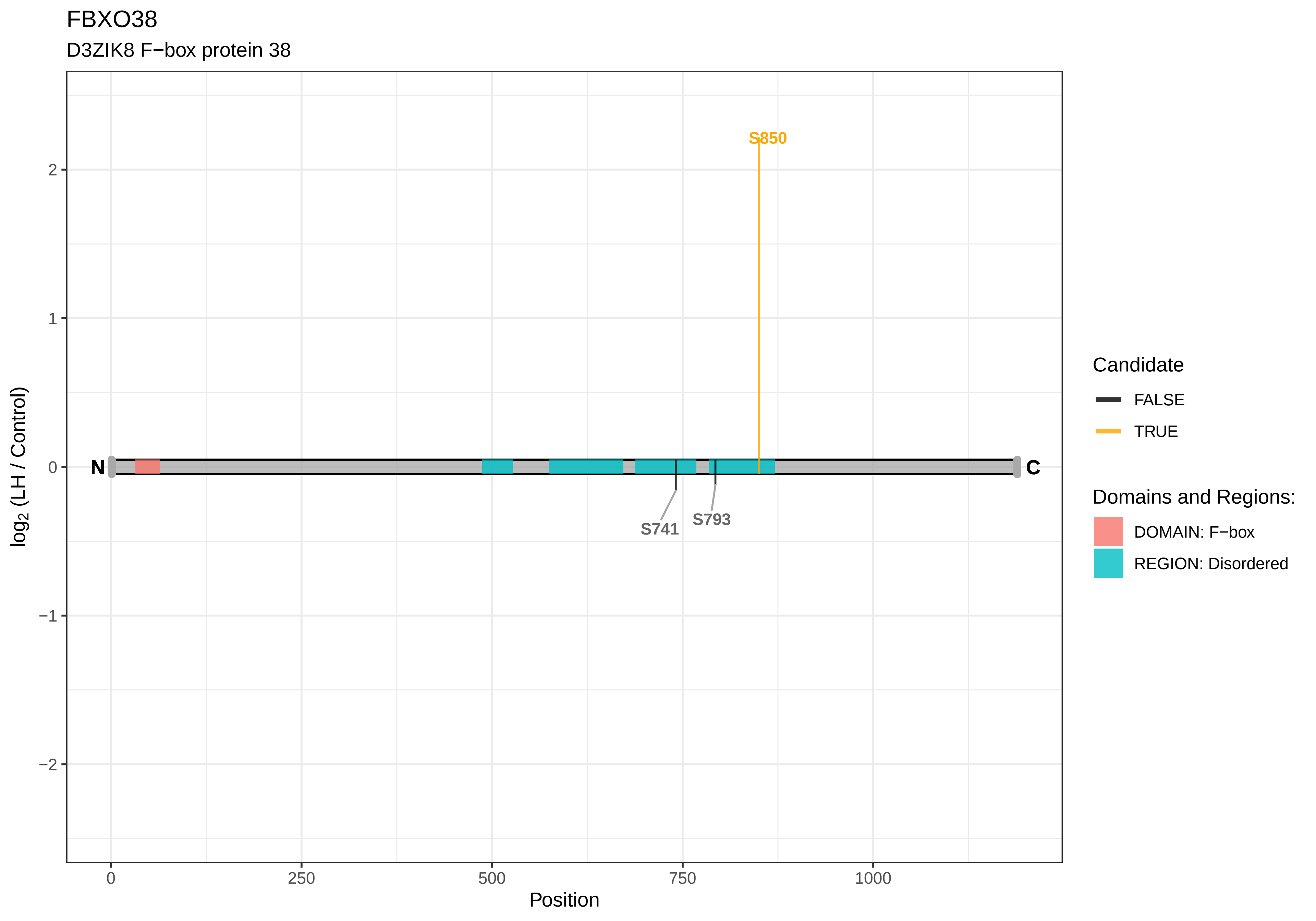

FBXO38
D3ZIK8 F−box protein 38
S850
S741
S793
N
C
2
1
0
−1
−2
log2 (LH / Control)
0
250
500
750
1000
Position
Candidate
FALSE
TRUE
Domains and Regions:
DOMAIN: F−box
REGION: Disordered

FBXO42

A0A0G2JTS9 F−box protein 42

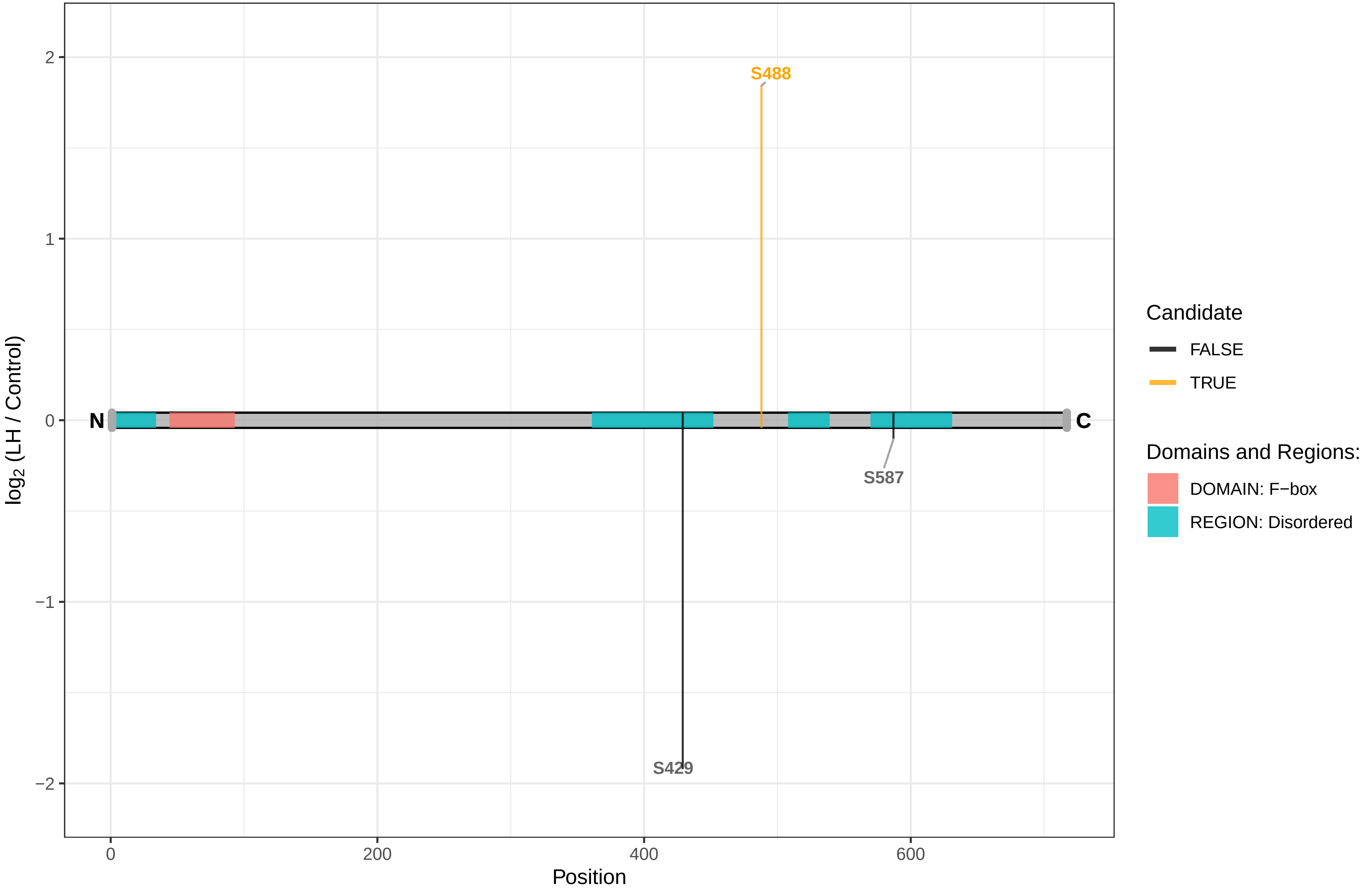

### FEN1

Q5XIP6 Flap endonuclease 1

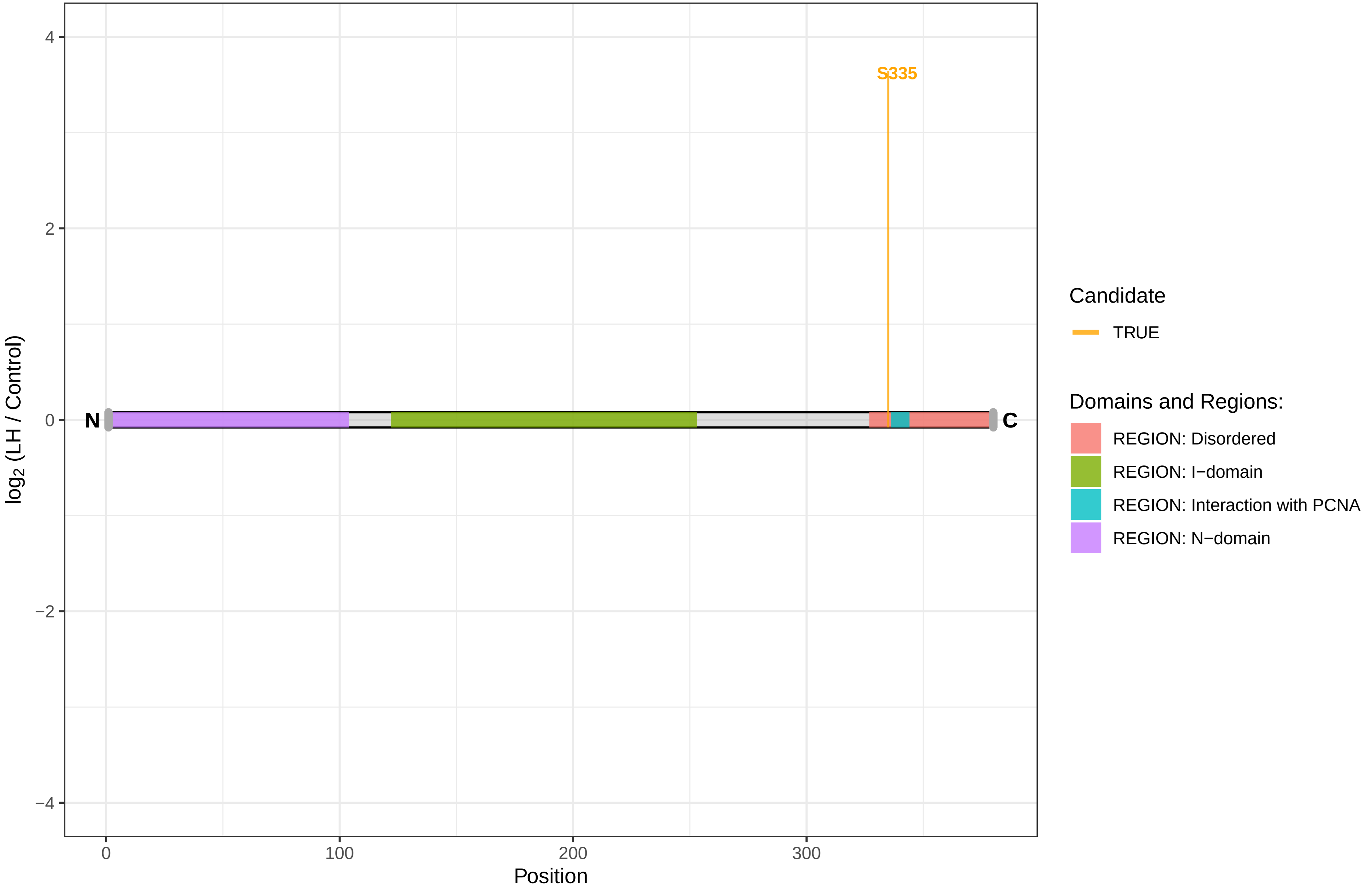

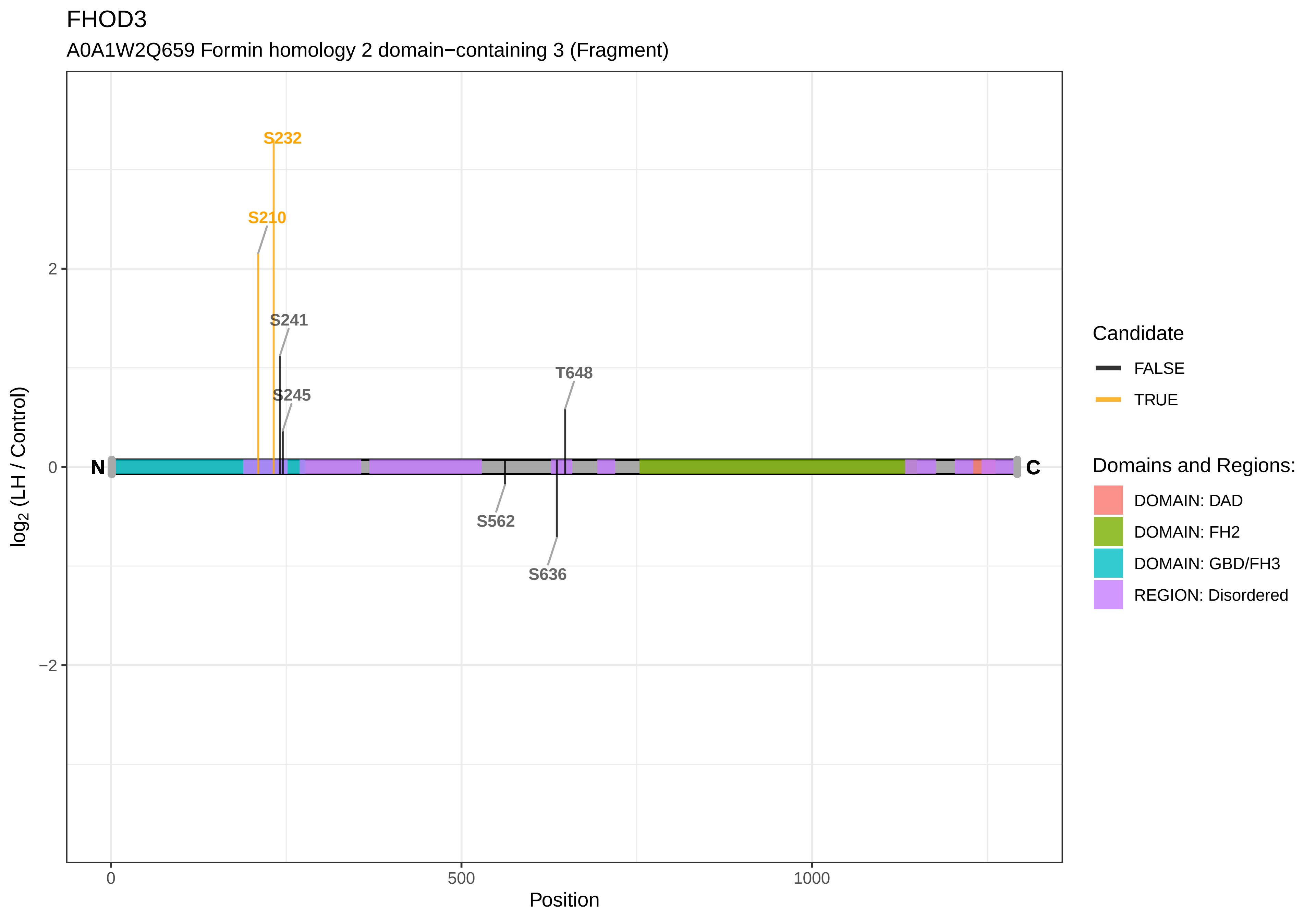
FHOD3
A0A1W2Q659 Formin homology 2 domain−containing 3 (Fragment)
S232
S210
S241
S245
T648
S562
S636
N
C
2
0
−2
0
500
1000
Position
log2 (LH / Control)
Candidate
FALSE
TRUE
Domains and Regions:
DOMAIN: DAD
DOMAIN: FH2
DOMAIN: GBD/FH3
REGION: Disordered

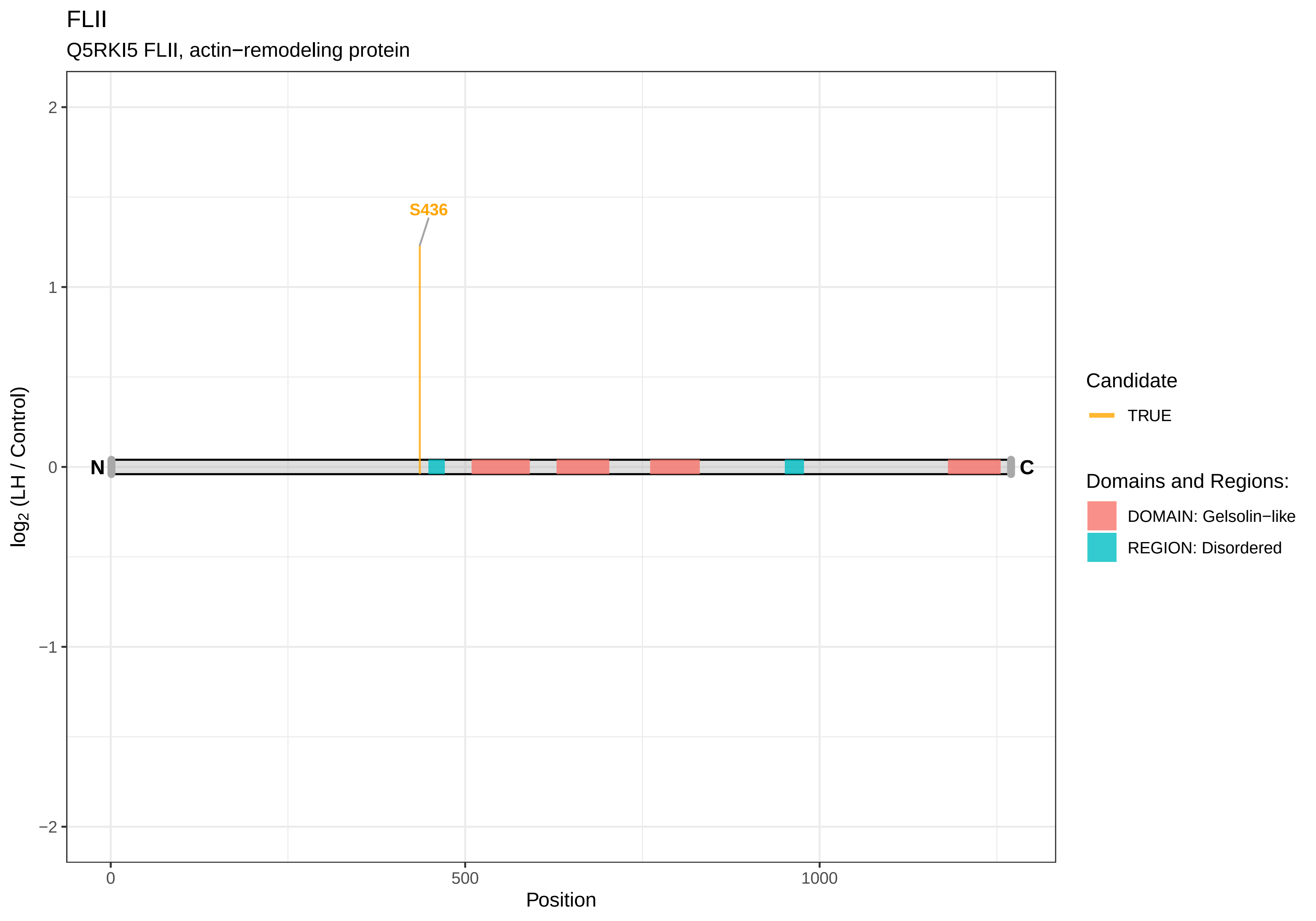

FLII
Q5RKI5 FLII, actin−remodeling protein
S436
N
C
2
1
0
−1
−2
log2 (LH / Control)
0
500
1000
Position
Candidate
TRUE
Domains and Regions:
DOMAIN: Gelsolin−like
REGION: Disordered

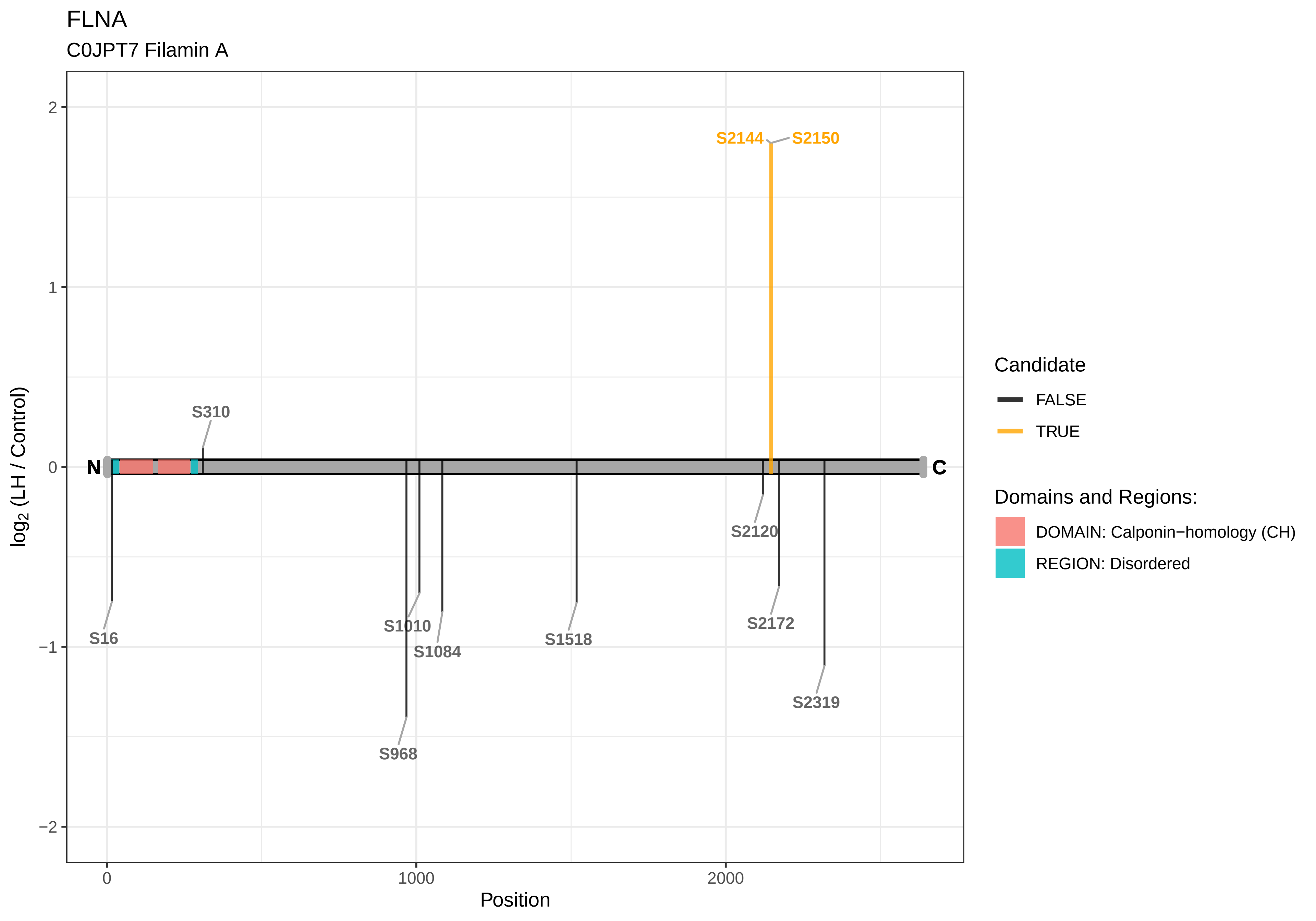

FLNA
C0JPT7 Filamin A
S2144
S2150
S310
N
C
S2120
S16
S1010
S1084
S1518
S2172
S2319
S968
2
1
0
−1
−2
log2 (LH / Control)
0
1000
2000
Position
Candidate
FALSE
TRUE
Domains and Regions:
DOMAIN: Calponin−homology (CH)
REGION: Disordered

FLNB

D4A8D5 Filamin B

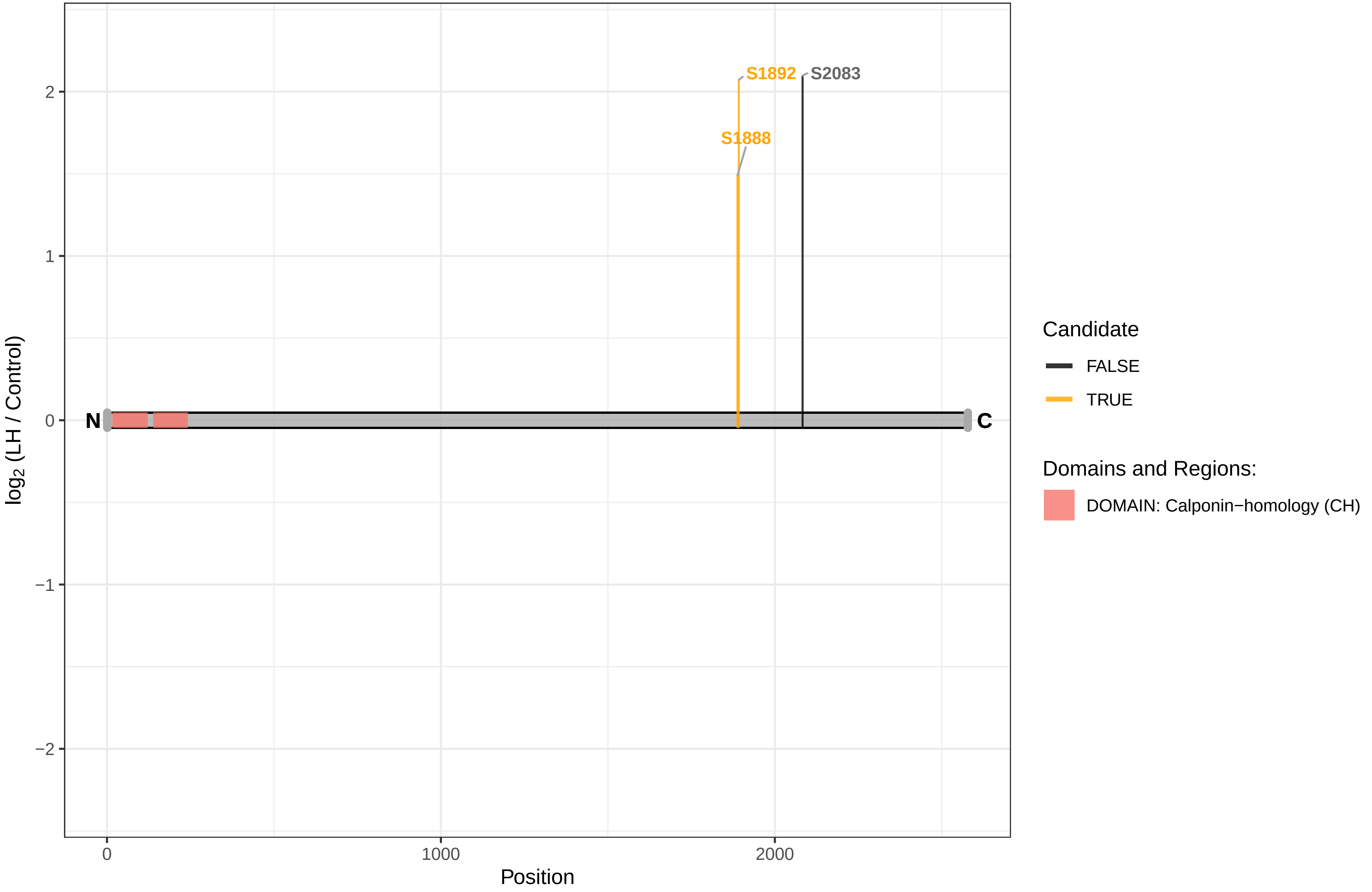

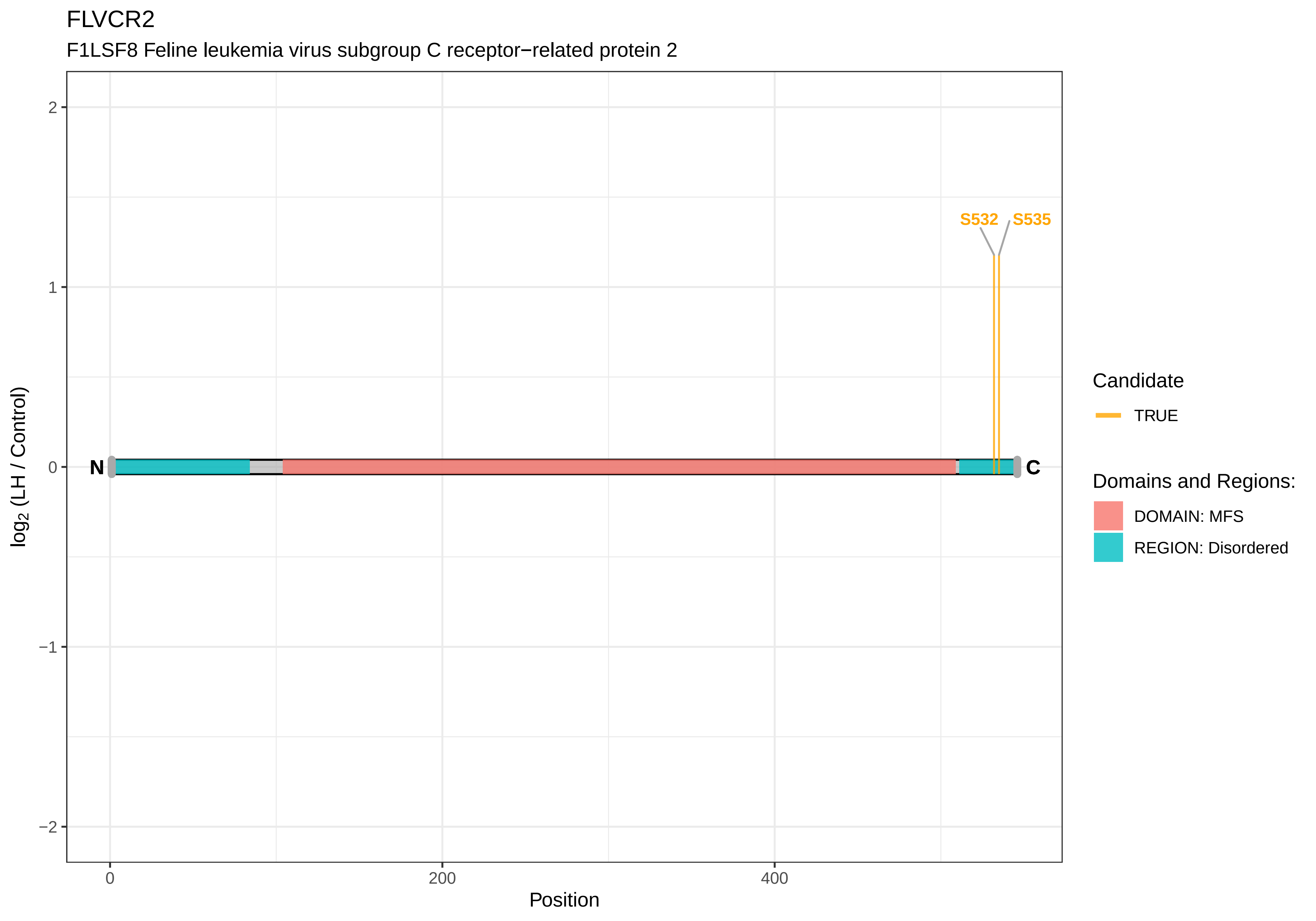
FLVCR2
F1LSF8 Feline leukemia virus subgroup C receptor−related protein 2
S532
S535
N
C
2
1
0
−1
−2
log2 (LH / Control)
0
200
400
Position
Candidate
TRUE
Domains and Regions:
DOMAIN: MFS
REGION: Disordered

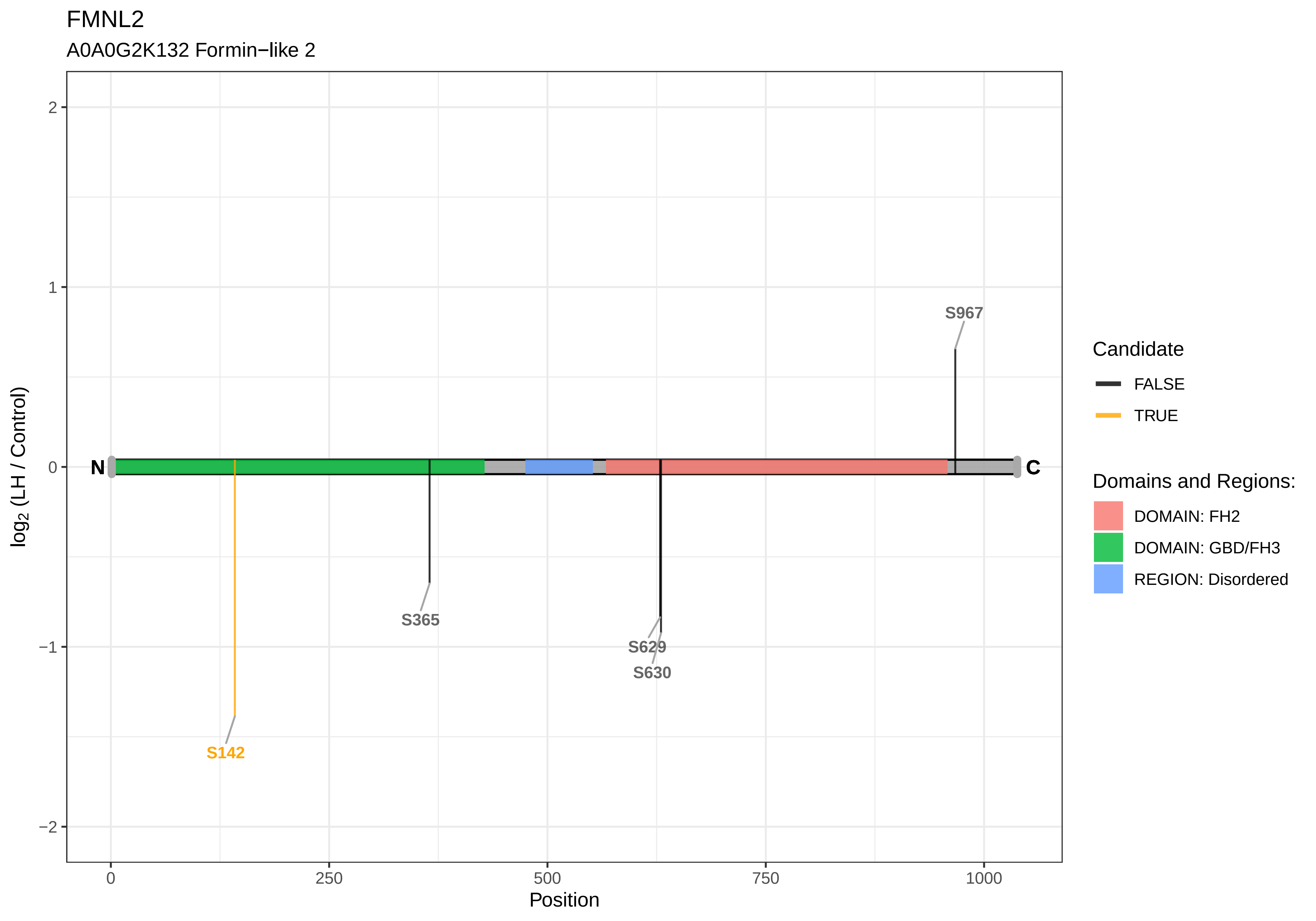
FMNL2
A0A0G2K132 Formin−like 2
2
1
0
−1
−2
log2 (LH / Control)
N
C
S967
S365
S629
S630
S142
0
250
500
750
1000
Position
Candidate
FALSE
TRUE
Domains and Regions:
DOMAIN: FH2
DOMAIN: GBD/FH3
REGION: Disordered

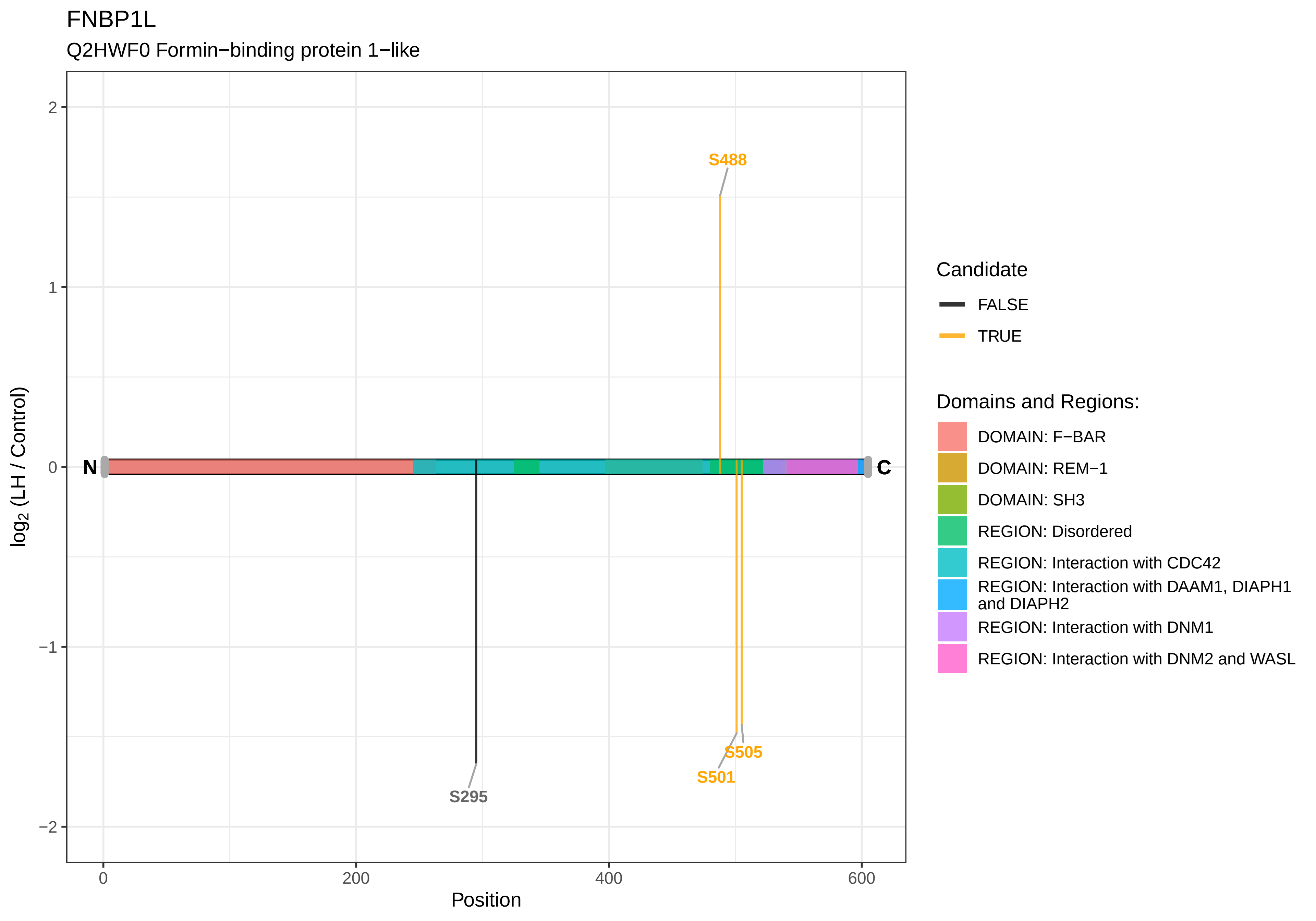

FNBP1L
Q2HWF0 Formin−binding protein 1−like
S488
S295
S505
S501
N
C
2
1
0
−1
−2
log2 (LH / Control)
0
200
400
600
Position
Candidate
FALSE
TRUE
Domains and Regions:
DOMAIN: F−BAR
DOMAIN: REM−1
DOMAIN: SH3
REGION: Disordered
REGION: Interaction with CDC42
REGION: Interaction with DAAM1, DIAPH1 and DIAPH2
REGION: Interaction with DNM1
REGION: Interaction with DNM2 and WASL

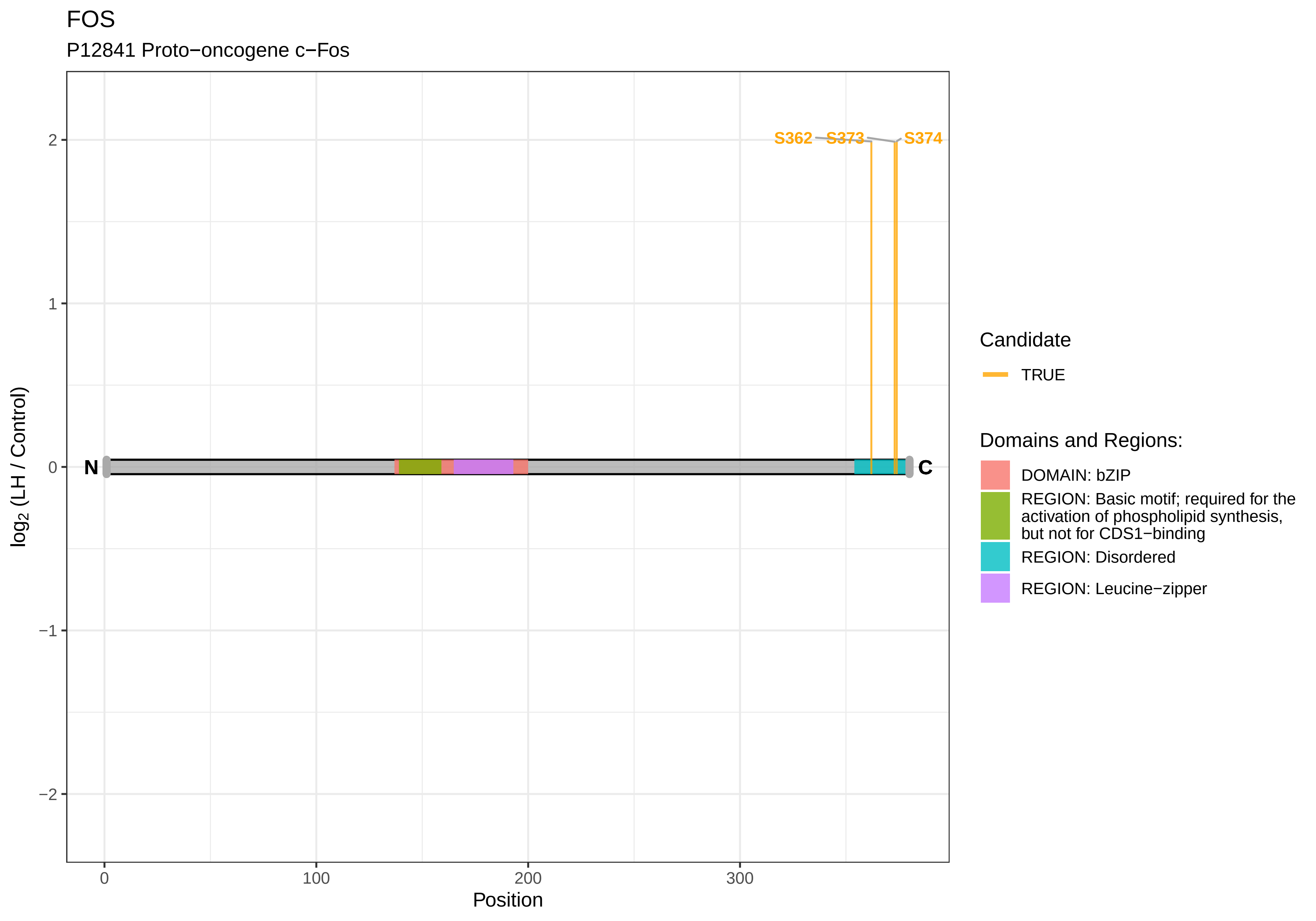
FOS
P12841 Proto−oncogene c−Fos
S362
S373
S374
N
C
2
1
0
−1
−2
log2 (LH / Control)
0
100
200
300
Position
Candidate
TRUE
Domains and Regions:
DOMAIN: bZIP
REGION: Basic motif; required for the activation of phospholipid synthesis, but not for CDS1−binding
REGION: Disordered
REGION: Leucine−zipper

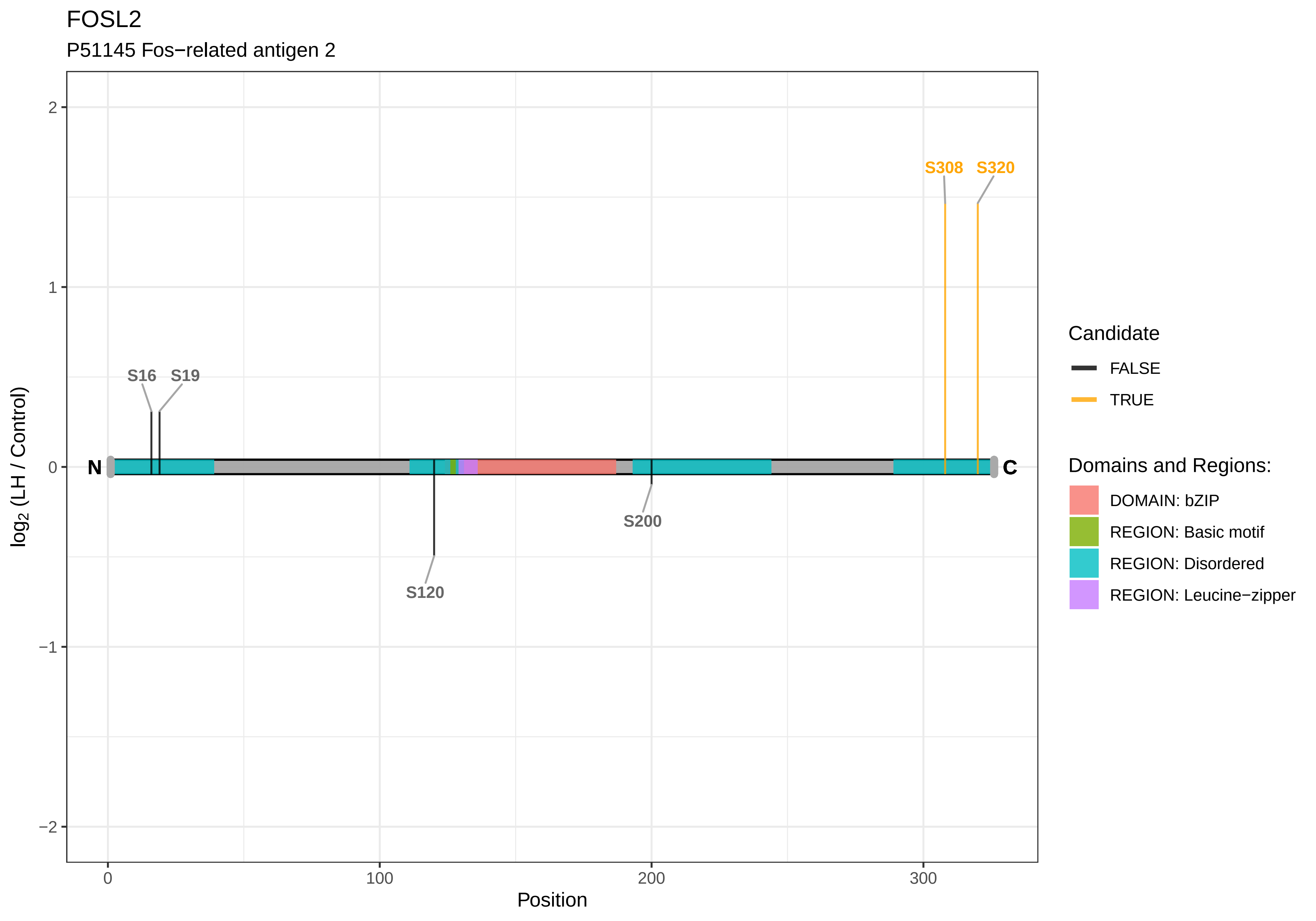

FOSL2
P51145 Fos−related antigen 2
S308
S320
S16
S19
S120
S200
N
C
2
1
0
−1
−2
0
100
200
300
Position
log2 (LH / Control)
Candidate
FALSE
TRUE
Domains and Regions:
DOMAIN: bZIP
REGION: Basic motif
REGION: Disordered
REGION: Leucine−zipper

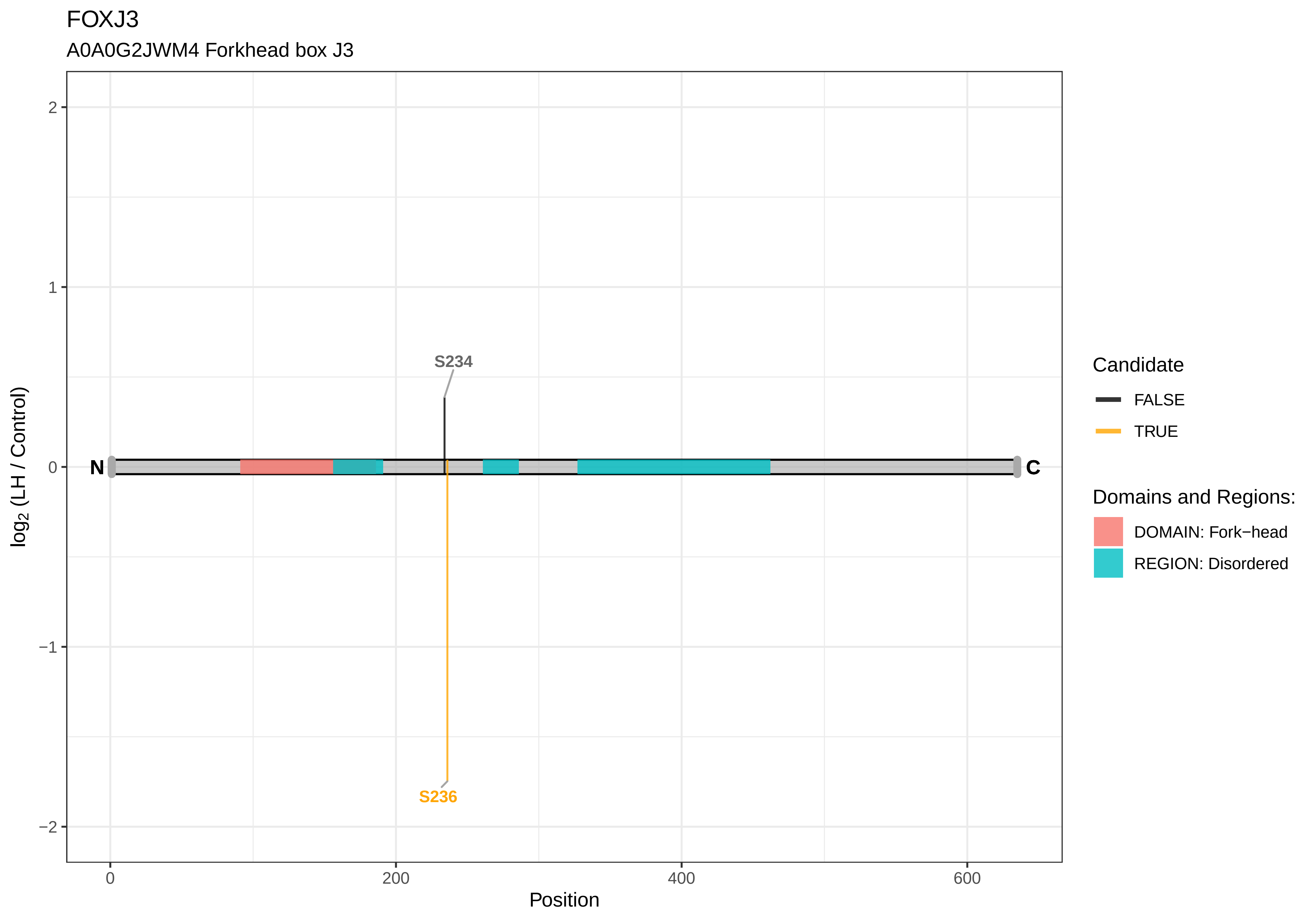

FOXJ3
A0A0G2JWM4 Forkhead box J3
S234
S236
N
C
2
1
0
−1
−2
0
200
400
600
Position
log2 (LH / Control)
Candidate
FALSE
TRUE
Domains and Regions:
DOMAIN: Fork−head
REGION: Disordered

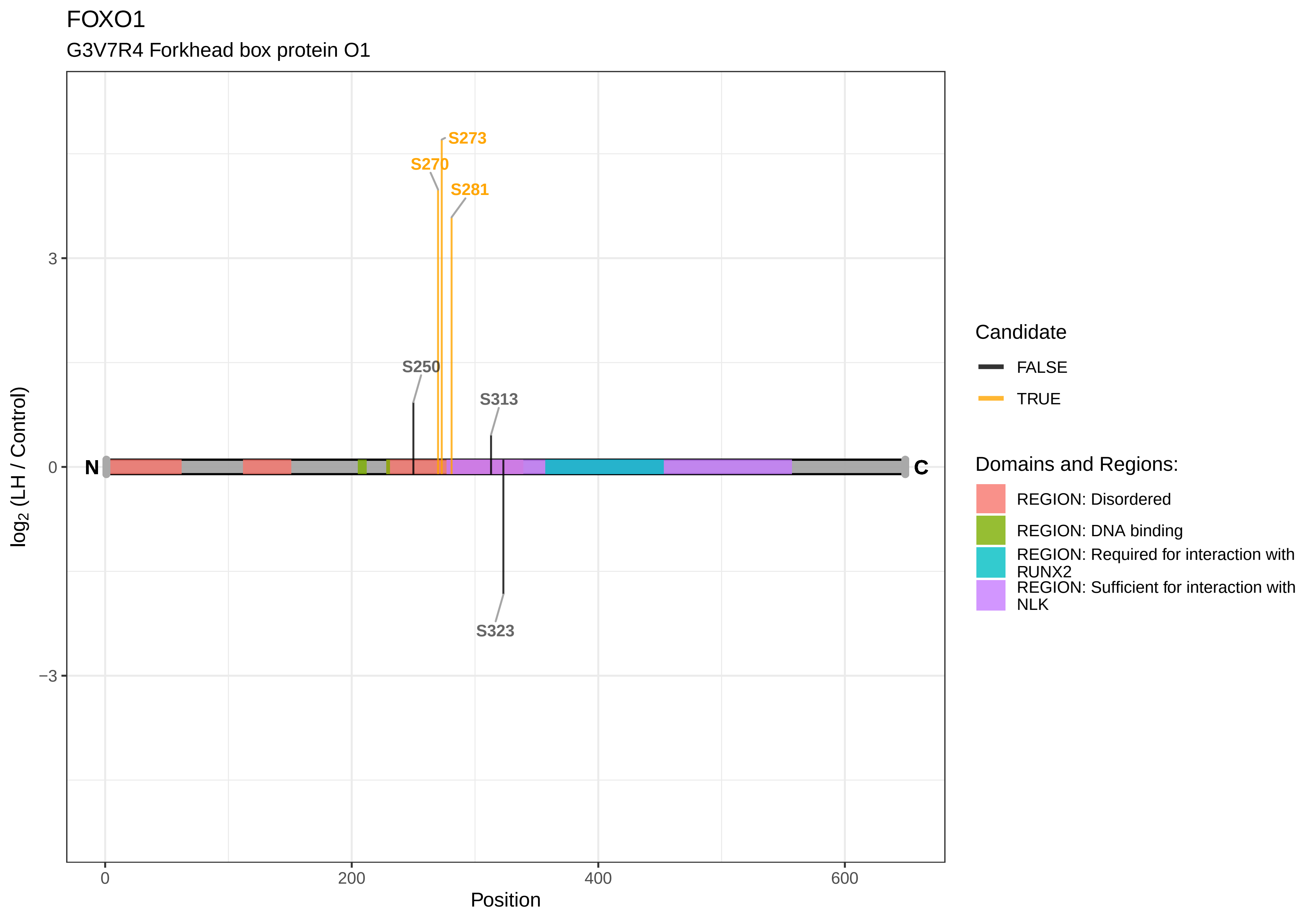
FOXO1
G3V7R4 Forkhead box protein O1
S273
S270
S281
S250
S313
S323
N
C
3
0
−3
0
200
400
600
Position
log2 (LH / Control)
Candidate
FALSE
TRUE
Domains and Regions:
REGION: Disordered
REGION: DNA binding
REGION: Required for interaction with RUNX2
REGION: Sufficient for interaction with NLK

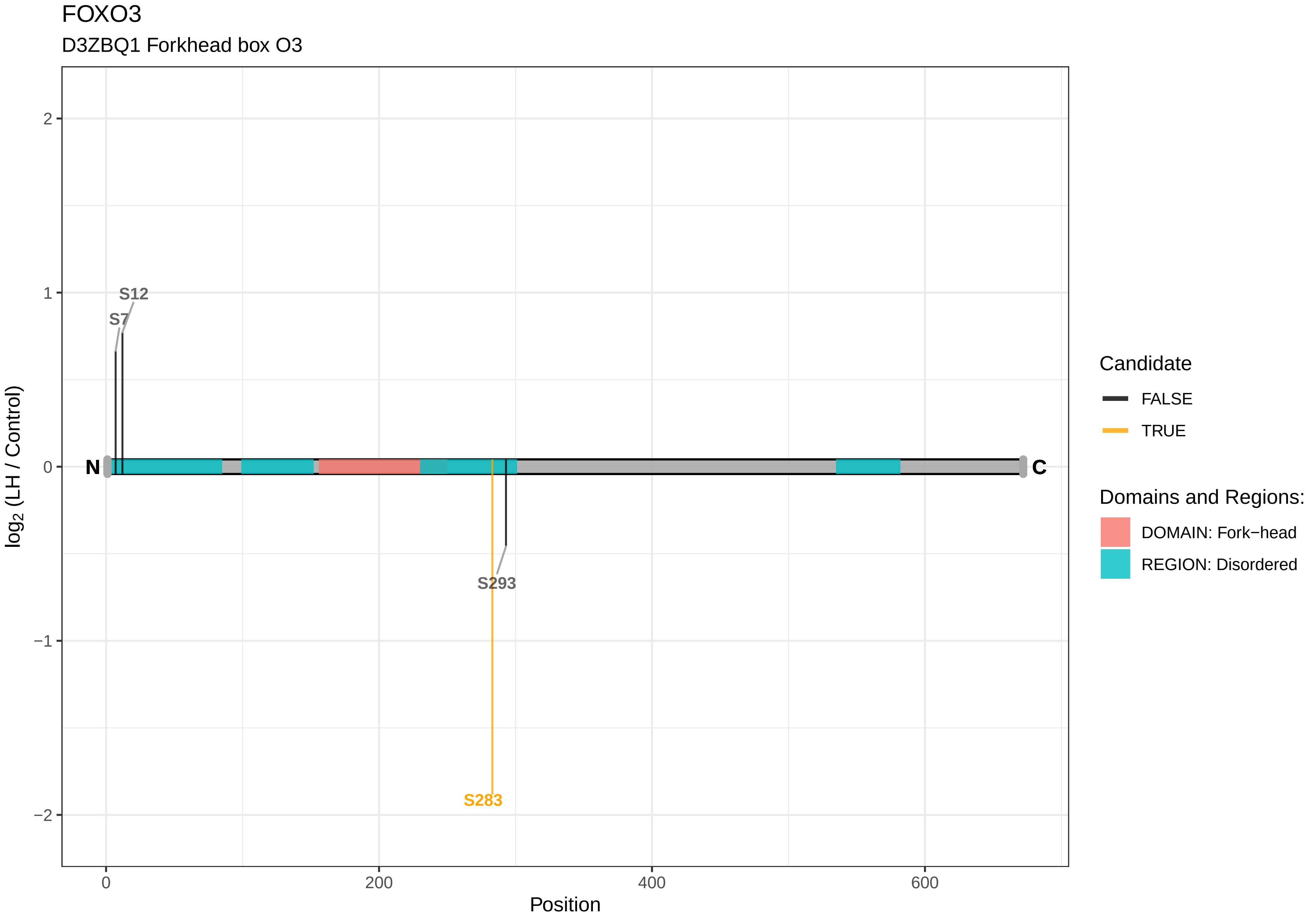
FOXO3
D3ZBQ1 Forkhead box O3
S12
S7
S293
S283
N
C
2
1
0
−1
−2
log2 (LH / Control)
0
200
400
600
Position
Candidate
FALSE
TRUE
Domains and Regions:
DOMAIN: Fork−head
REGION: Disordered

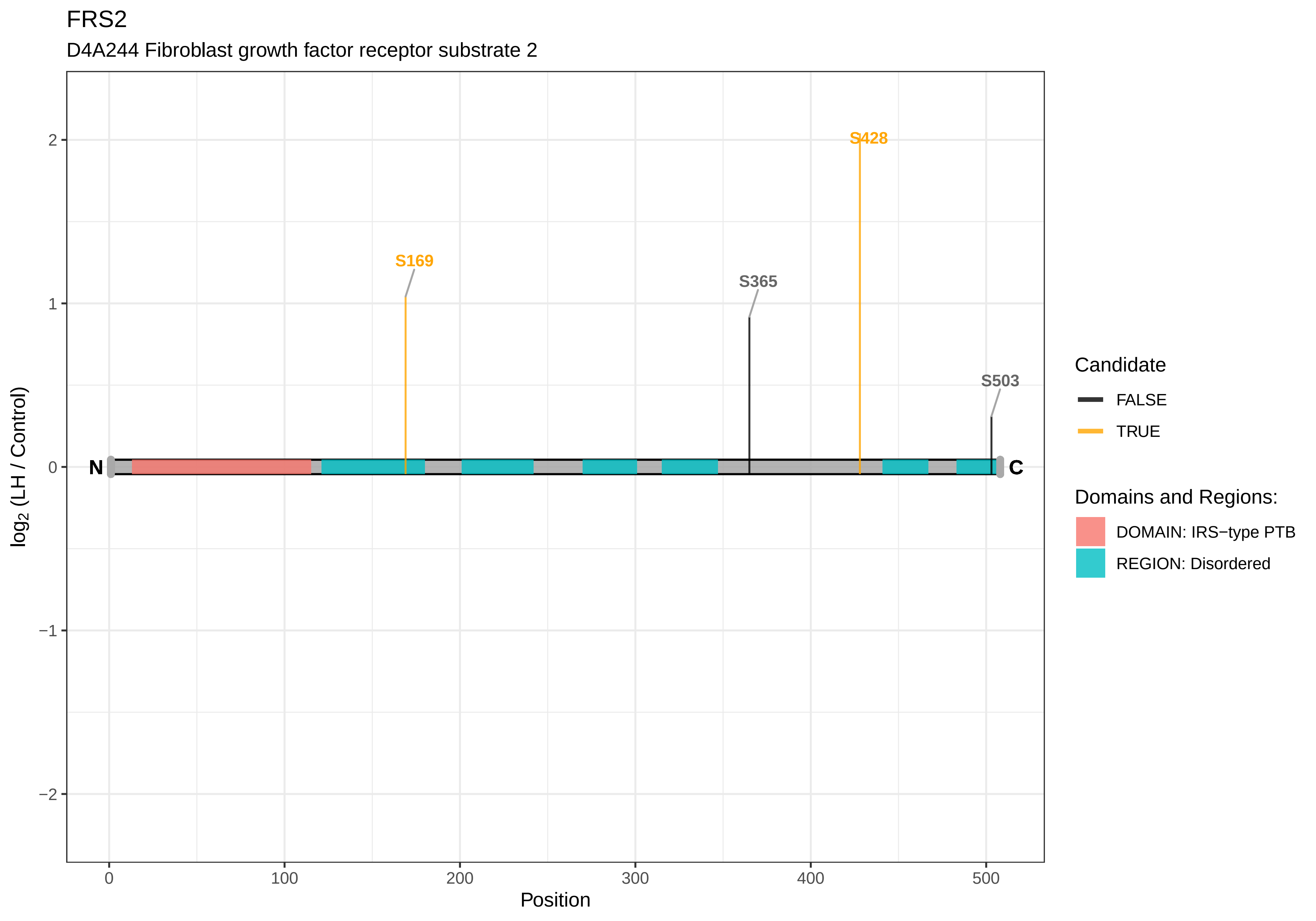

FRS2
D4A244 Fibroblast growth factor receptor substrate 2
S428
S169
S365
S503
N
C
2
1
0
−1
−2
log2 (LH / Control)
0
100
200
300
400
500
Position
Candidate
FALSE
TRUE
Domains and Regions:
DOMAIN: IRS−type PTB
REGION: Disordered

### FSHR

P20395 Follicle−stimulating hormone receptor

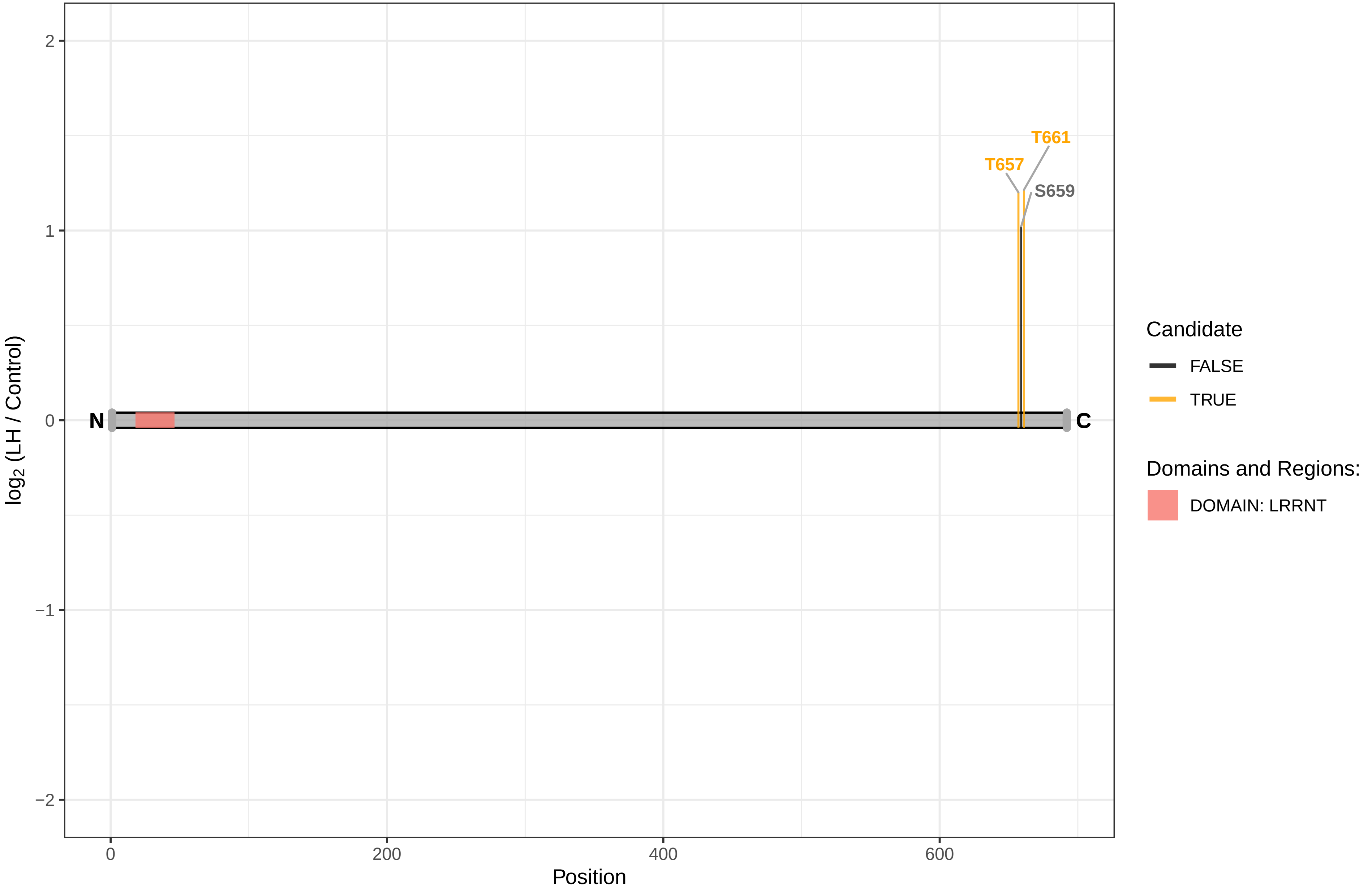

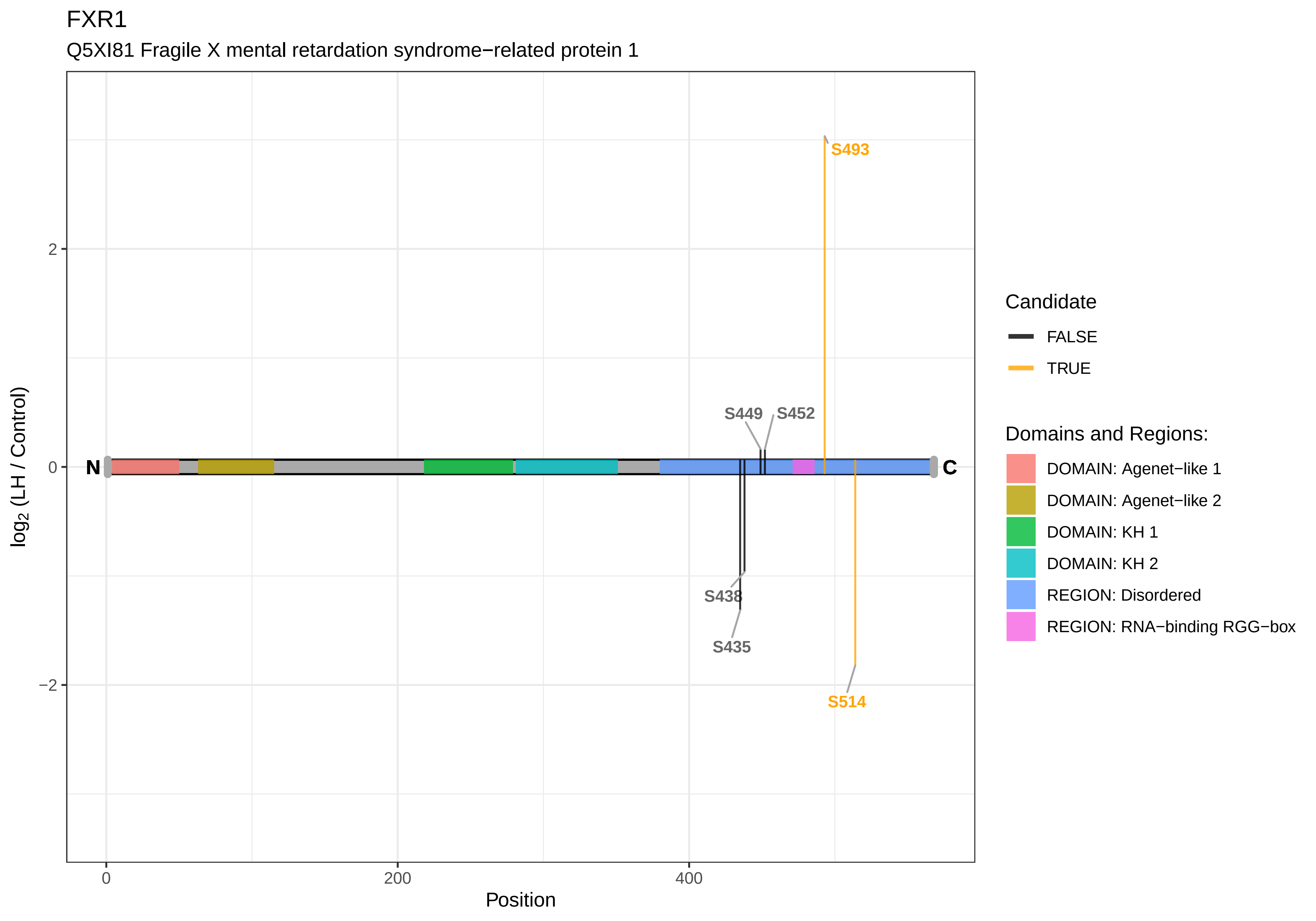

FXR1
Q5XI81 Fragile X mental retardation syndrome−related protein 1
S493
S449
S452
S438
S435
S514
N
C
2
0
−2
0
200
400
Position
log2 (LH / Control)
Candidate
FALSE
TRUE
Domains and Regions:
DOMAIN: Agenet−like 1
DOMAIN: Agenet−like 2
DOMAIN: KH 1
DOMAIN: KH 2
REGION: Disordered
REGION: RNA−binding RGG−box

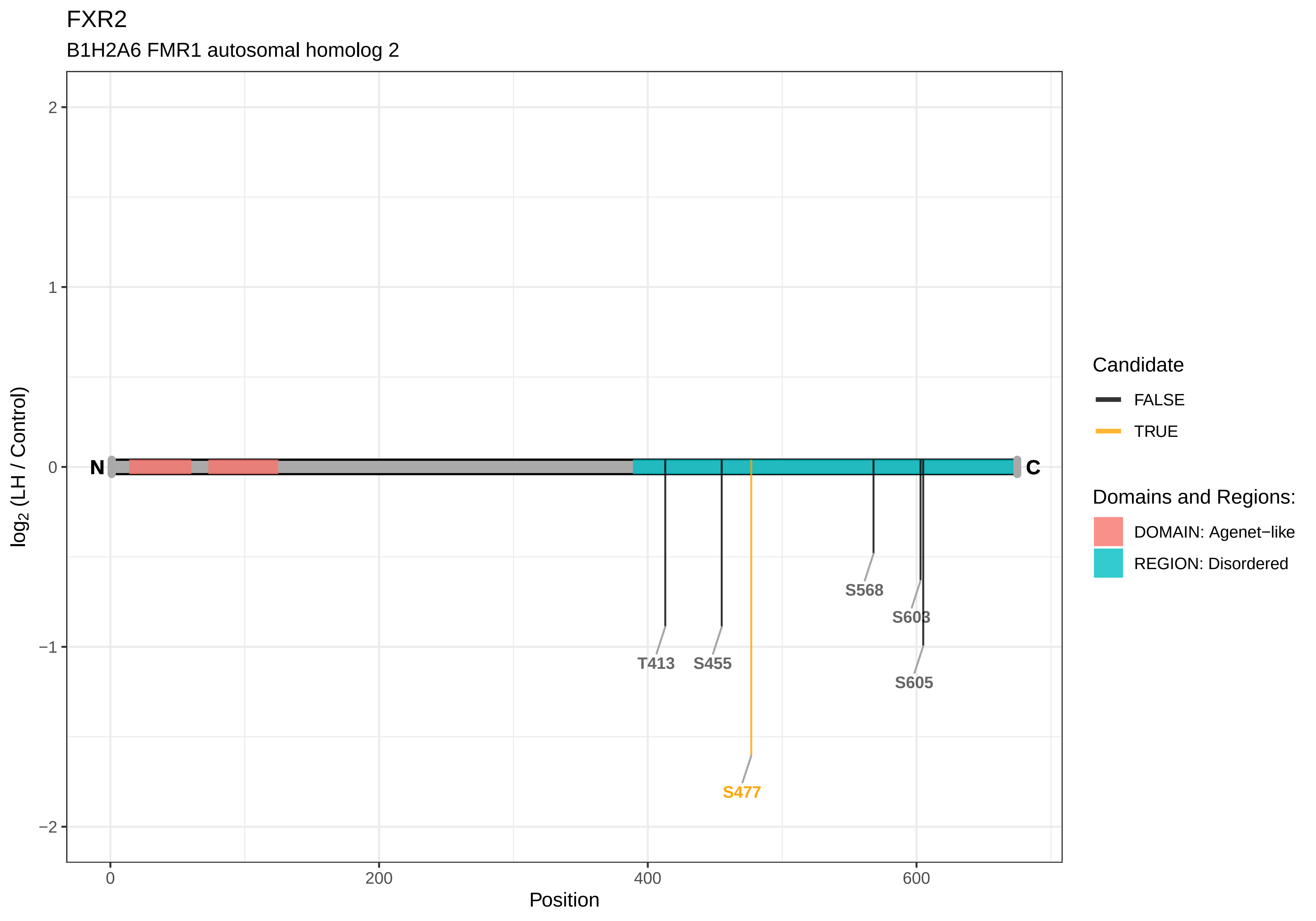

FXR2
B1H2A6 FMR1 autosomal homolog 2
log2 (LH / Control)
Position
N
C
T413
S455
S477
S568
S603
S605
2
1
0
−1
−2
0
200
400
600
Candidate
FALSE
TRUE
Domains and Regions:
DOMAIN: Agenet−like
REGION: Disordered

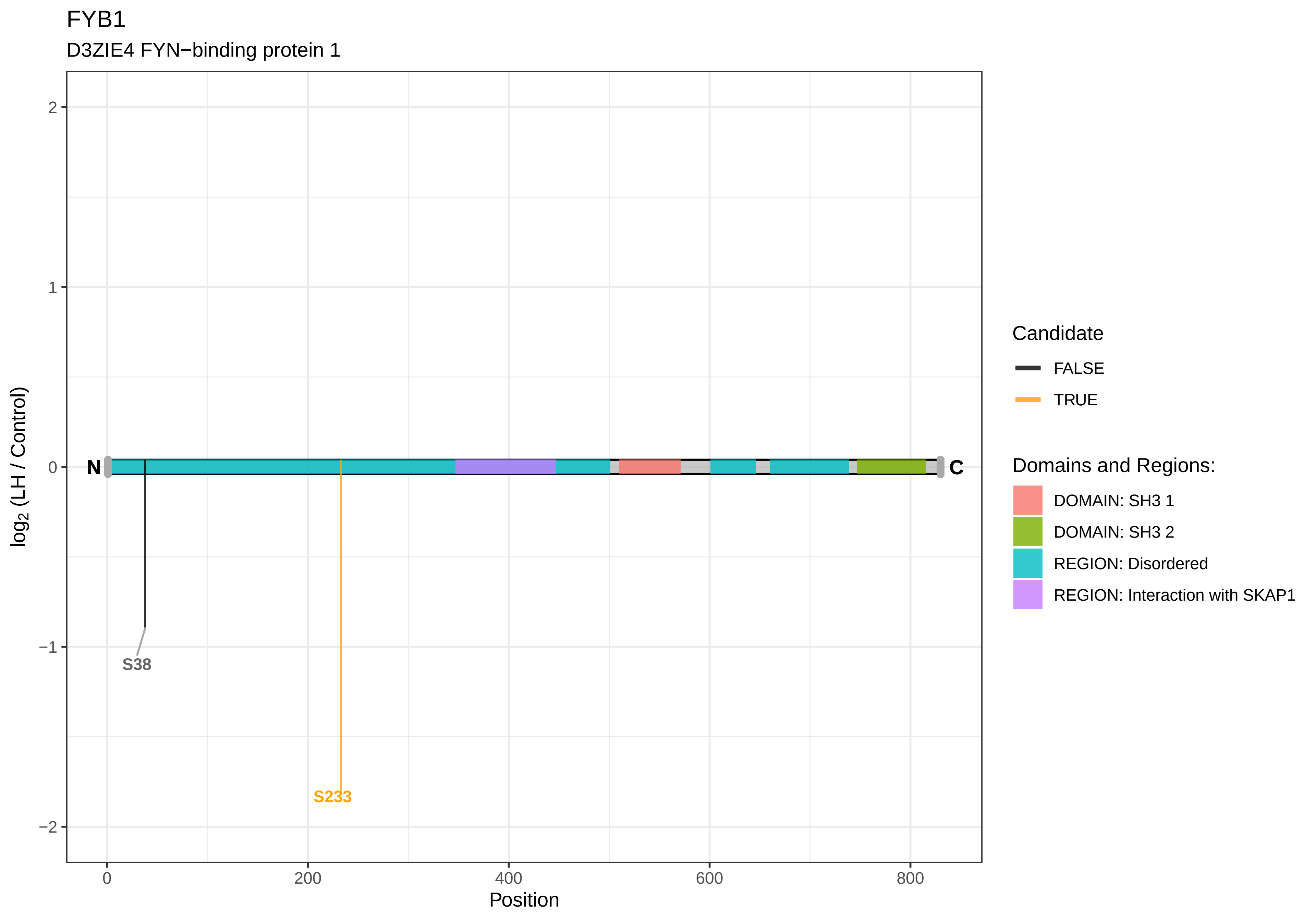

FYB1
D3ZIE4 FYN−binding protein 1
2
1
0
−1
−2
log2 (LH / Control)
N
C
S38
S233
0
200
400
600
800
Position
Candidate
FALSE
TRUE
Domains and Regions:
DOMAIN: SH3 1
DOMAIN: SH3 2
REGION: Disordered
REGION: Interaction with SKAP1

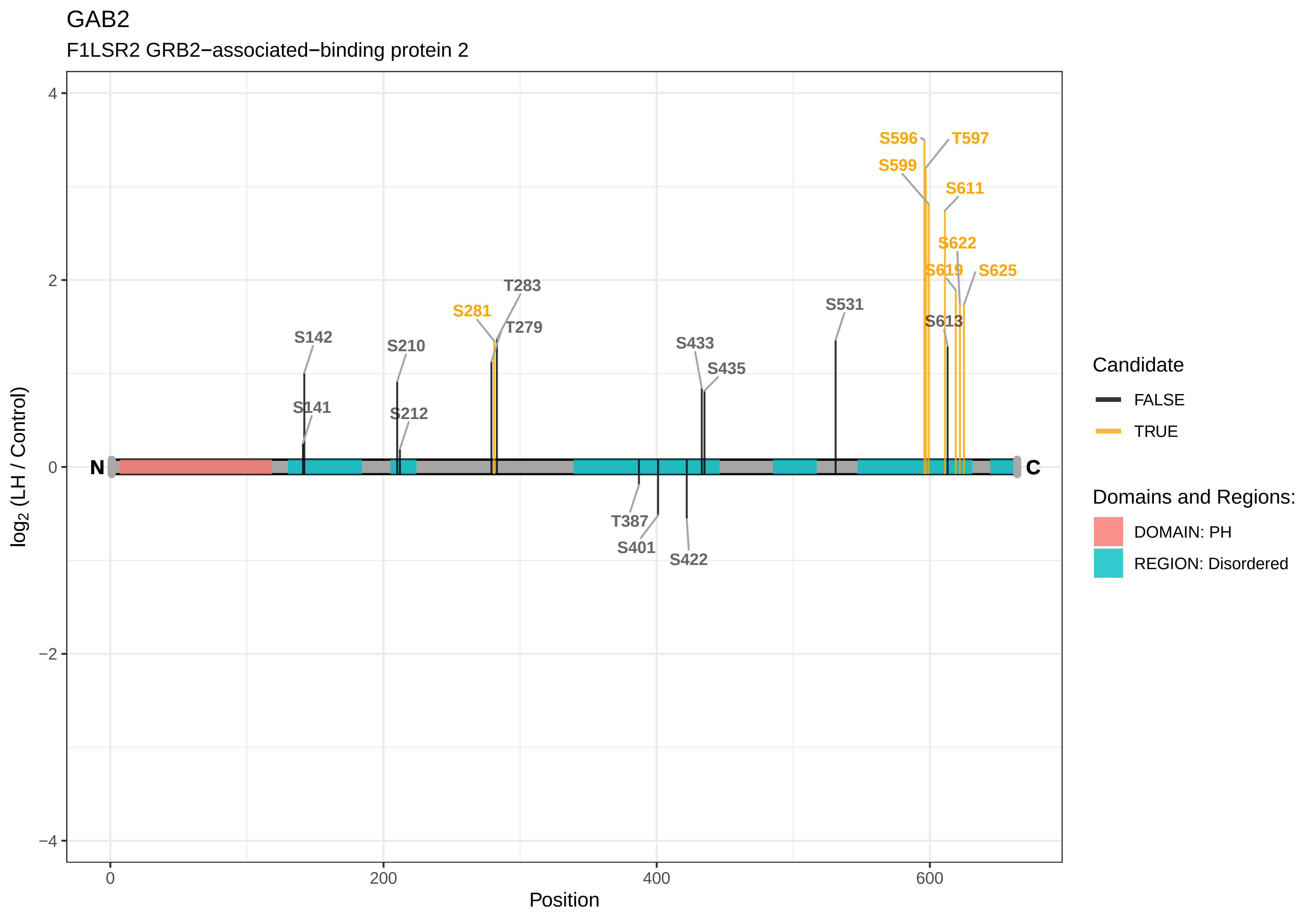

GAB2
F1LSR2 GRB2−associated−binding protein 2
log2 (LH / Control)
Position
N
C
S596
T597
S599
S611
S622
S619
S625
S531
S613
T283
S281
T279
S142
S210
S433
S435
S141
S212
T387
S401
S422
4
2
0
−2
−4
0
200
400
600
Candidate
FALSE
TRUE
Domains and Regions:
DOMAIN: PH
REGION: Disordered

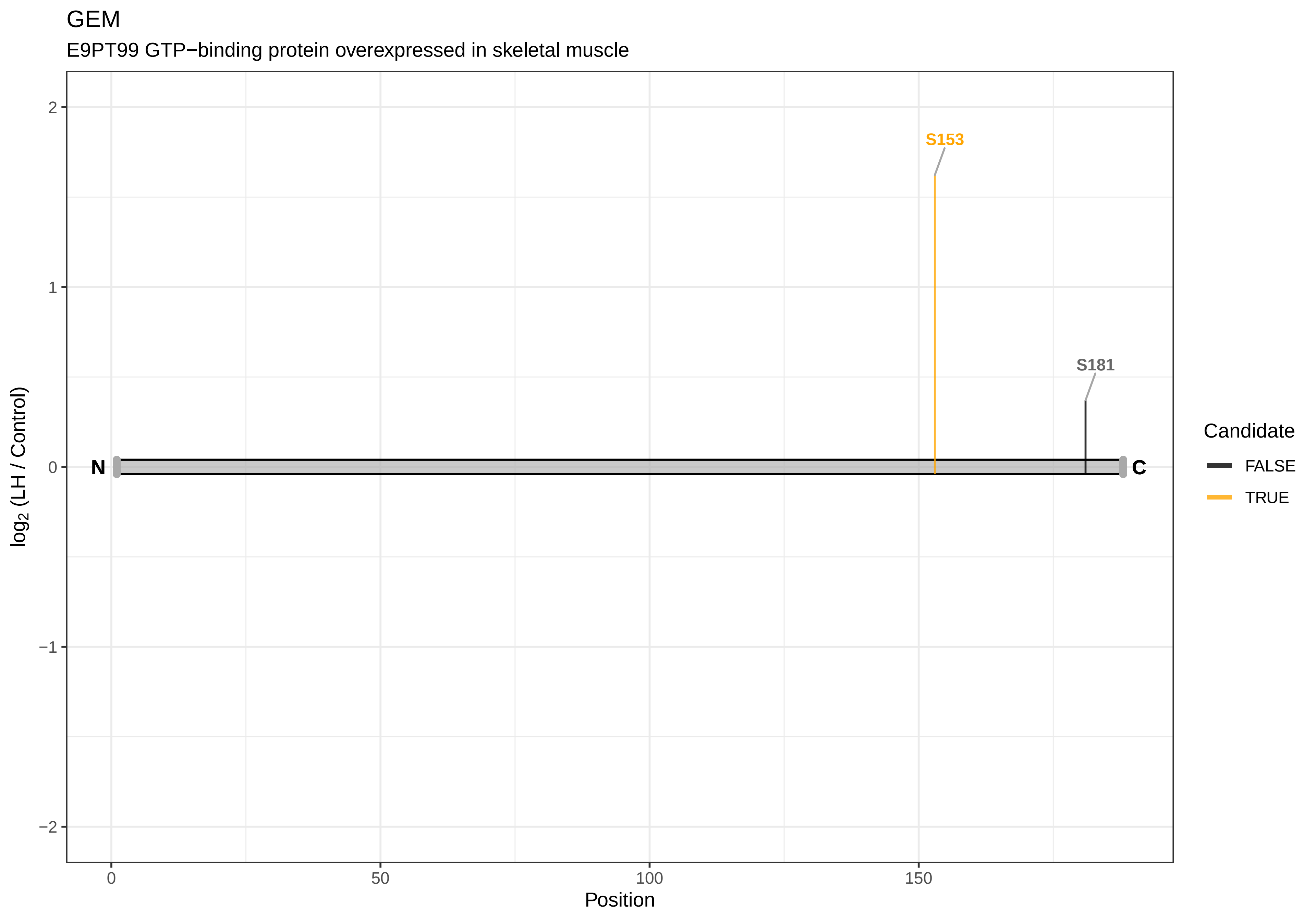
GEM
E9PT99 GTP−binding protein overexpressed in skeletal muscle
S153
S181
N
C
2
1
0
−1
−2
0
50
100
150
Position
log2 (LH / Control)
Candidate
FALSE
TRUE

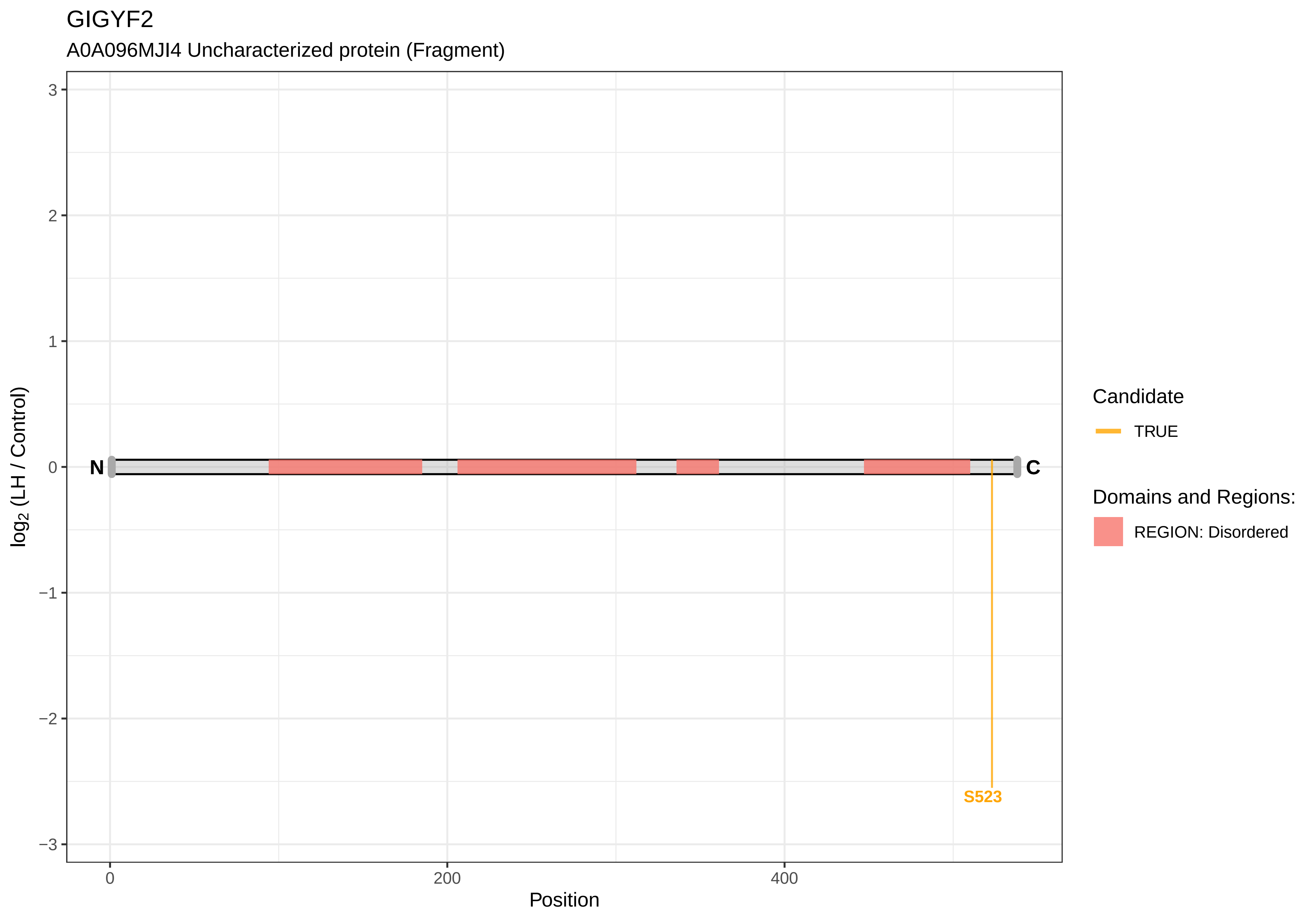
GIGYF2
A0A096MJI4 Uncharacterized protein (Fragment)
log2 (LH / Control)
3
2
1
0
−1
−2
−3
N
C
S523
0
200
400
Position
Candidate
TRUE
Domains and Regions:
REGION: Disordered

GKAP1

Q5XIG5 G kinase−anchoring protein 1

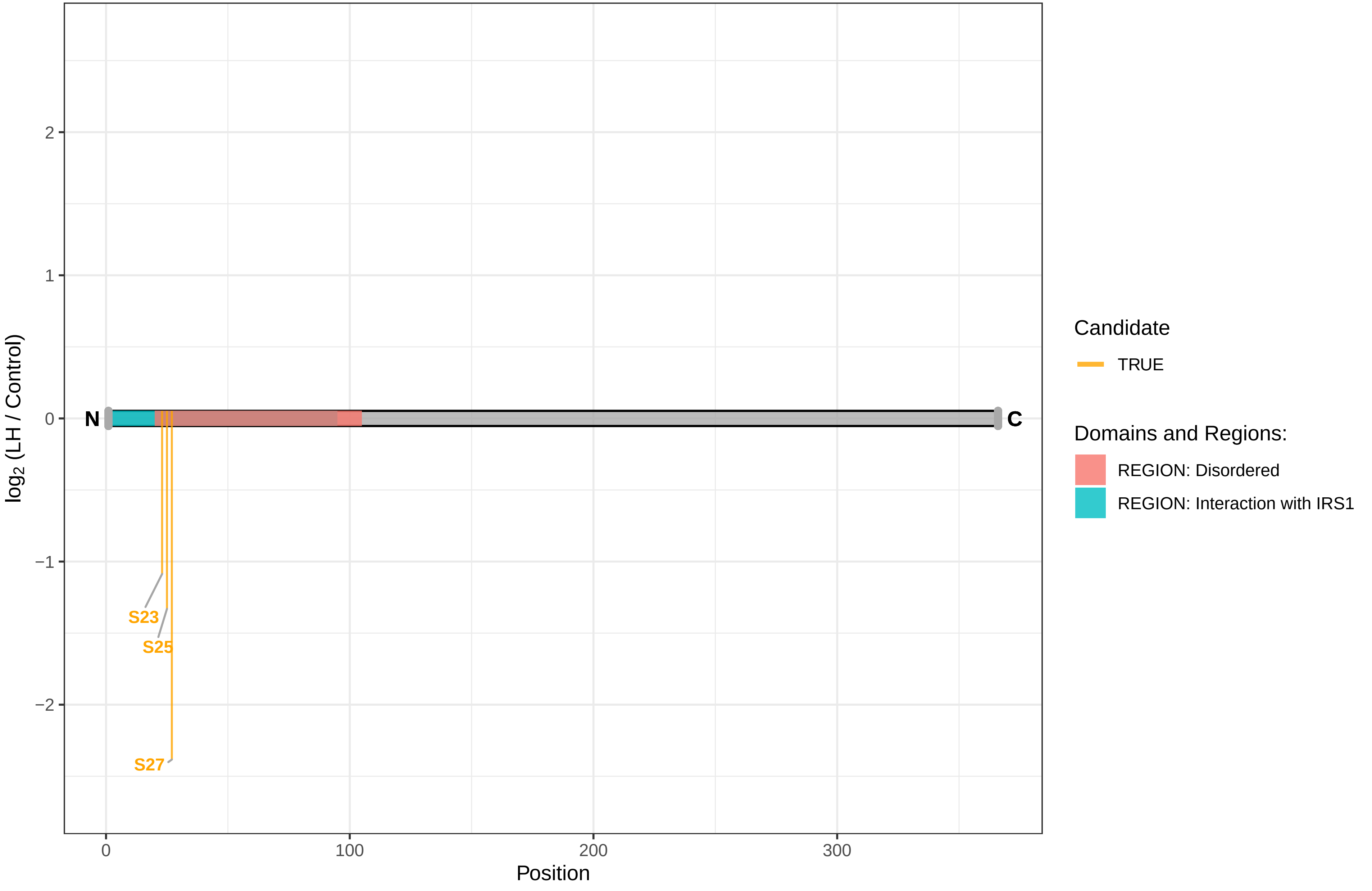

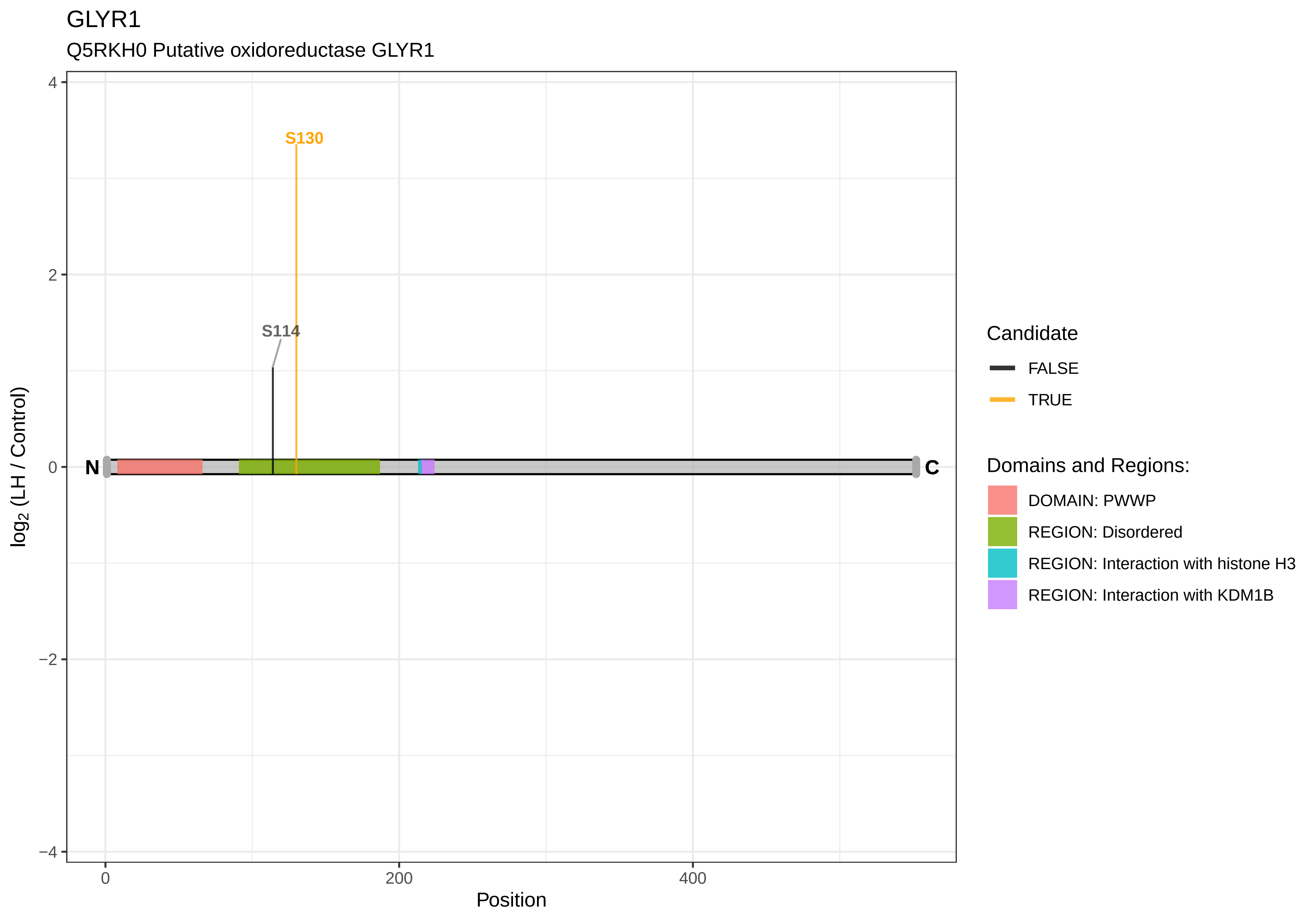
GLYR1
Q5RKH0 Putative oxidoreductase GLYR1
S130
S114
N
C
4
2
0
−2
−4
0
200
400
Position
log2 (LH / Control)
Candidate
FALSE
TRUE
Domains and Regions:
DOMAIN: PWWP
REGION: Disordered
REGION: Interaction with histone H3
REGION: Interaction with KDM1B

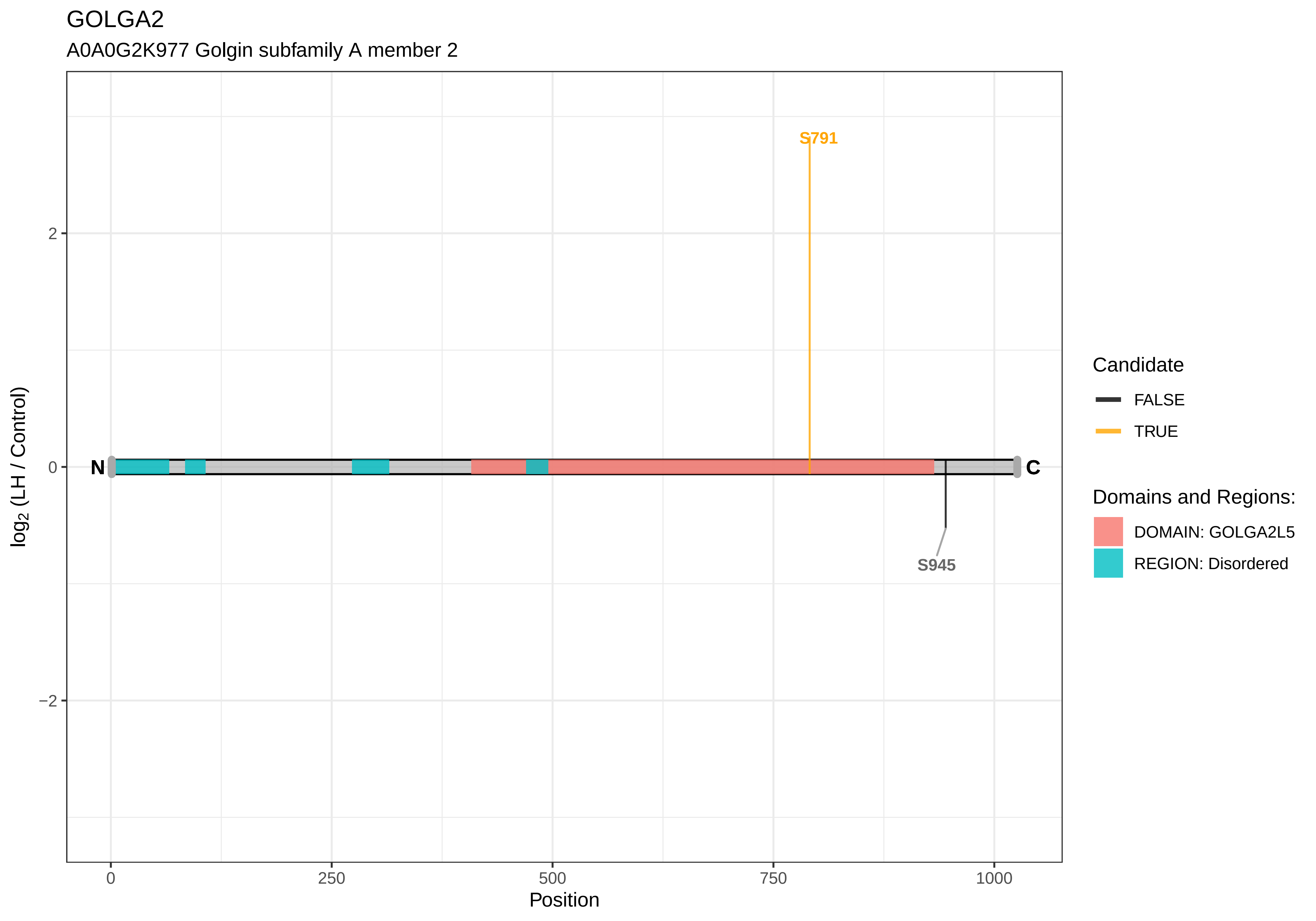

GOLGA2
A0A0G2K977 Golgin subfamily A member 2
S791
S945
N
C
2
0
−2
0
250
500
750
1000
Position
log2 (LH / Control)
Candidate
FALSE
TRUE
Domains and Regions:
DOMAIN: GOLGA2L5
REGION: Disordered

GOLGA3

A0A0G2K5A4 Uncharacterized protein

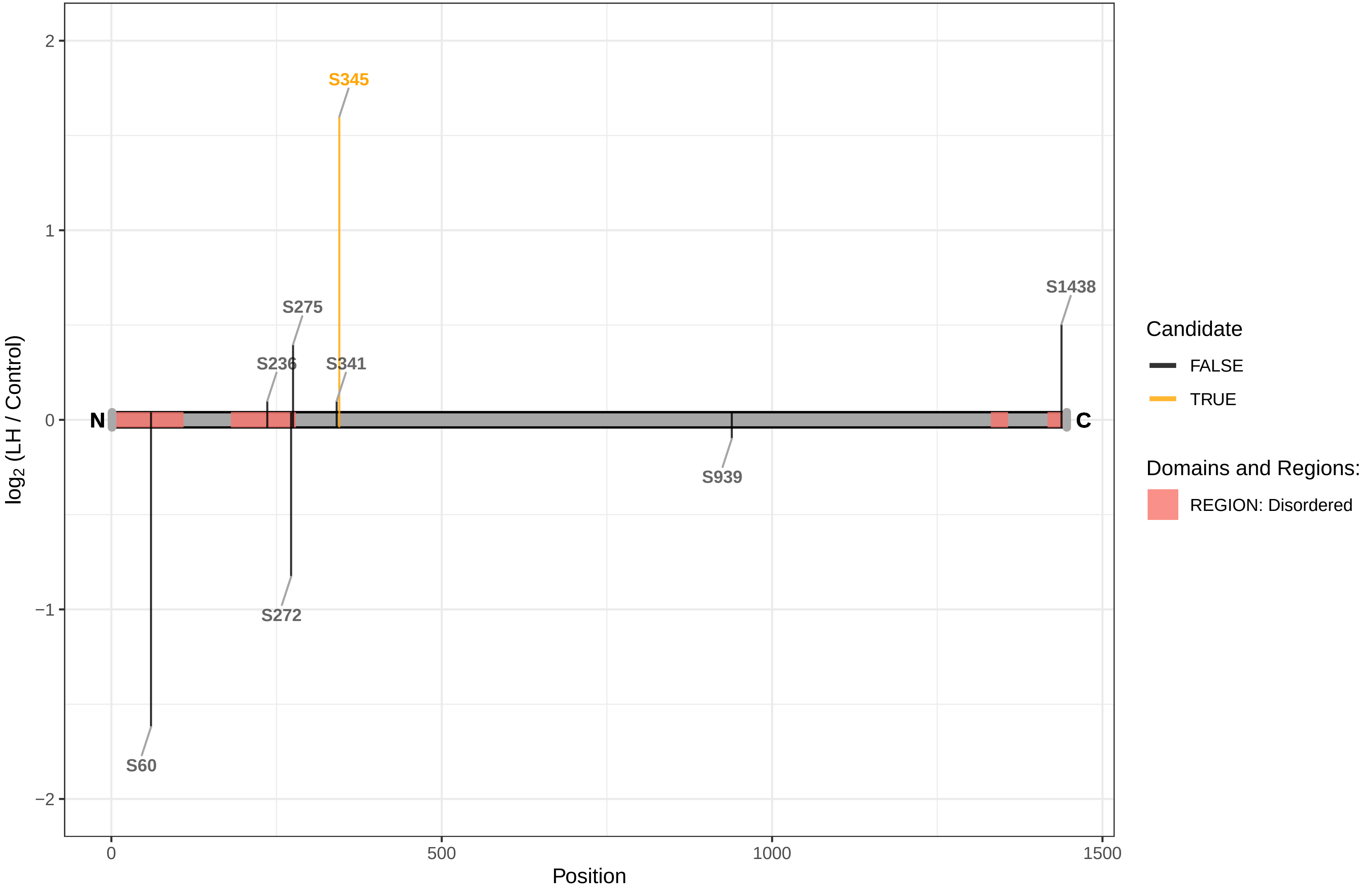

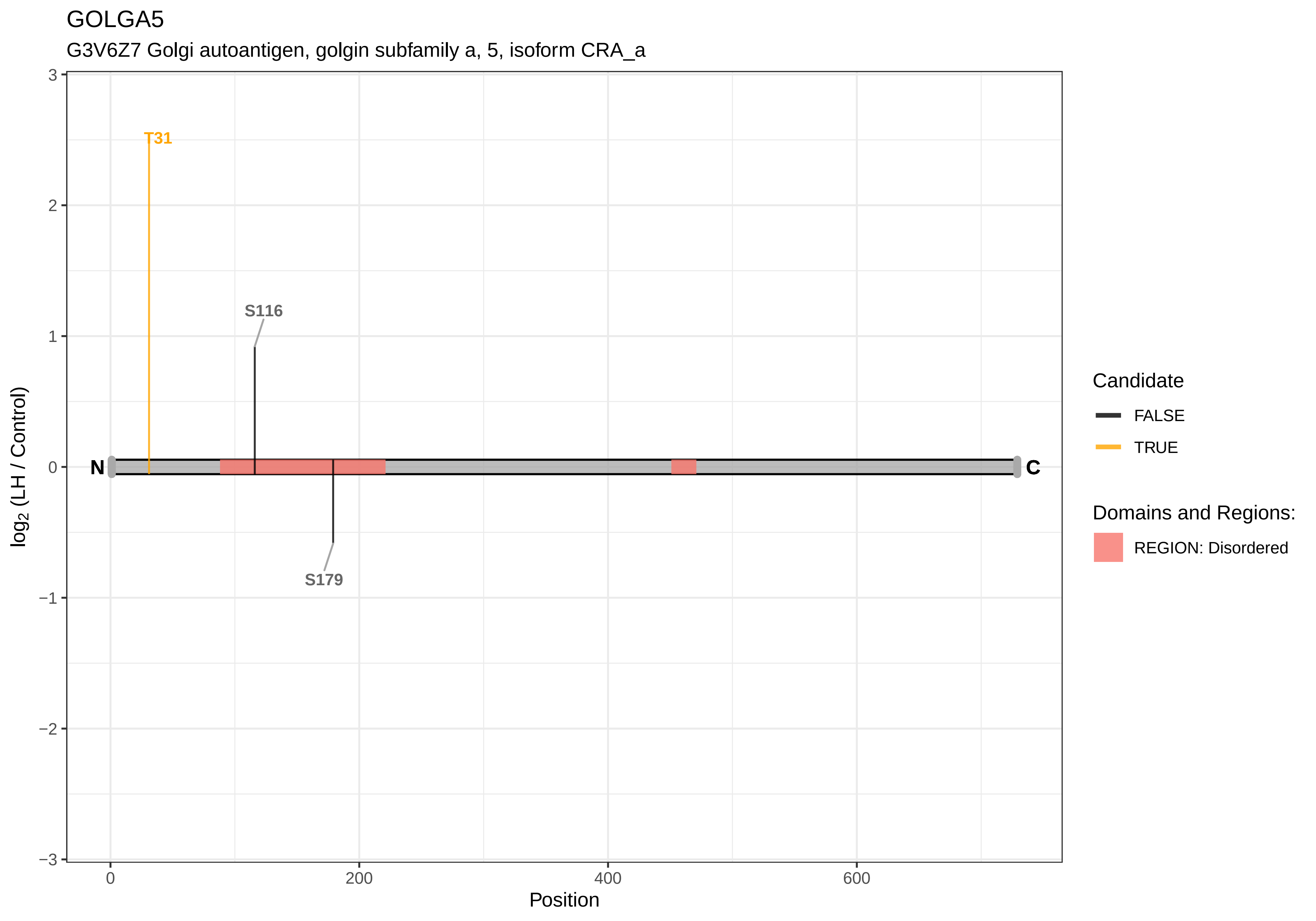
GOLGA5
G3V6Z7 Golgi autoantigen, golgin subfamily a, 5, isoform CRA_a
T31
S116
S179
N
C
log2 (LH / Control)
Position
Candidate
FALSE
TRUE
Domains and Regions:
REGION: Disordered

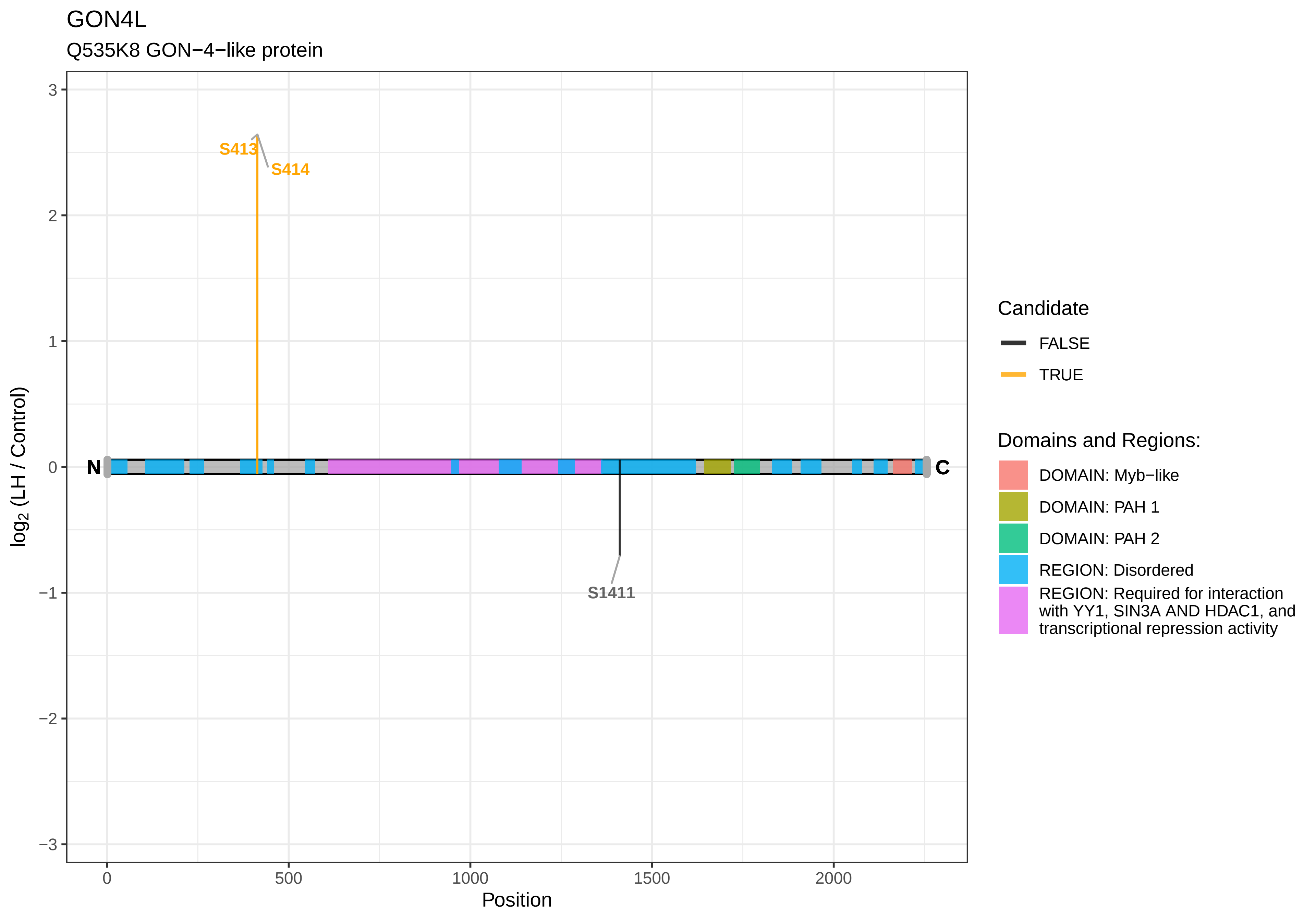
GON4L
Q535K8 GON−4−like protein
log2 (LH / Control)
Position
S413
S414
S1411
N
C
3
2
1
0
−1
−2
−3
0
500
1000
1500
2000
Candidate
FALSE
TRUE
Domains and Regions:
DOMAIN: Myb−like
DOMAIN: PAH 1
DOMAIN: PAH 2
REGION: Disordered
REGION: Required for interaction with YY1, SIN3A AND HDAC1, and transcriptional repression activity

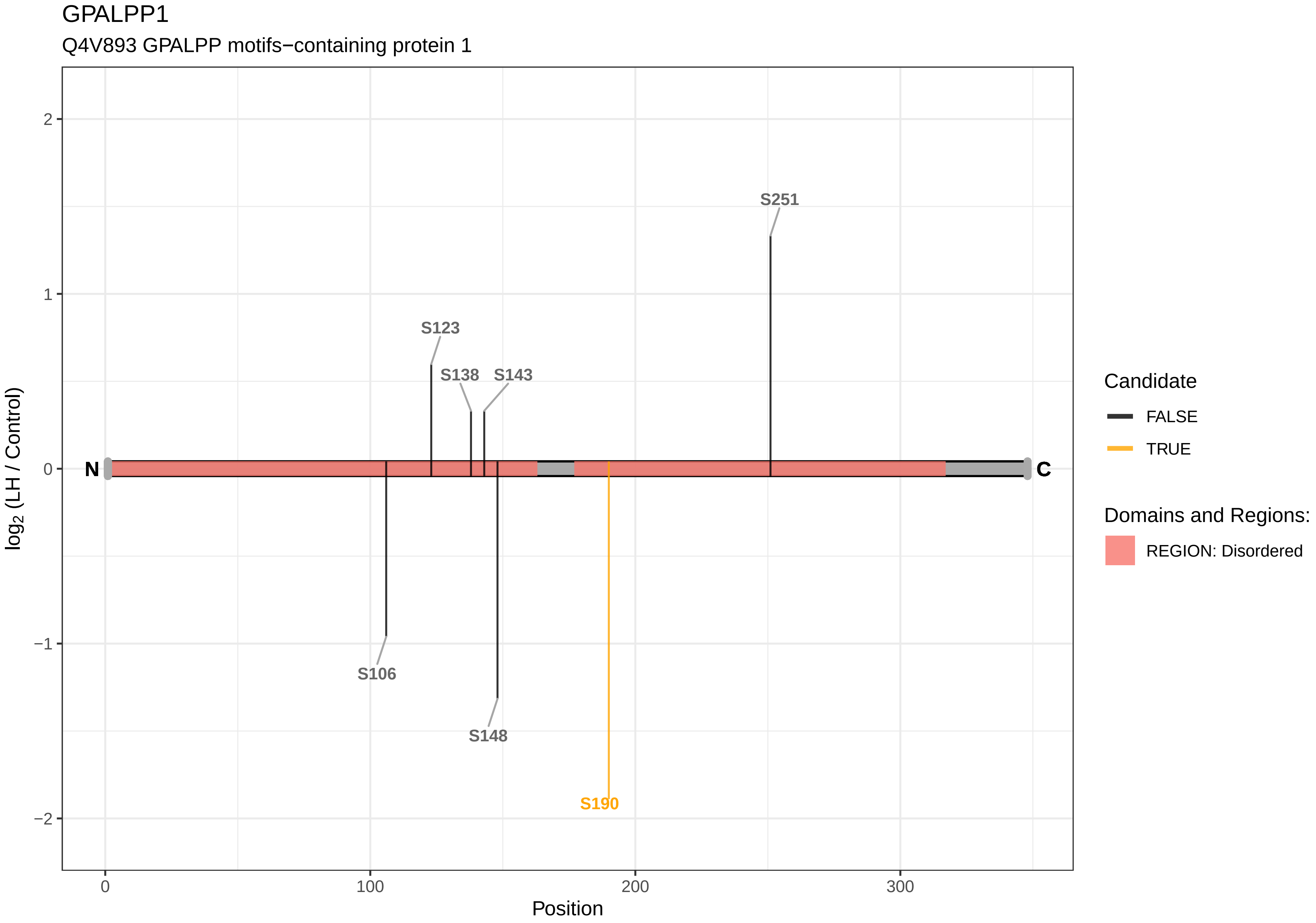
GPALPP1
Q4V893 GPALPP motifs−containing protein 1
log2 (LH / Control)
Position
S251
S123
S138
S143
S106
S148
S190
N
C
2
1
0
−1
−2
0
100
200
300
Candidate
FALSE
TRUE
Domains and Regions:
REGION: Disordered

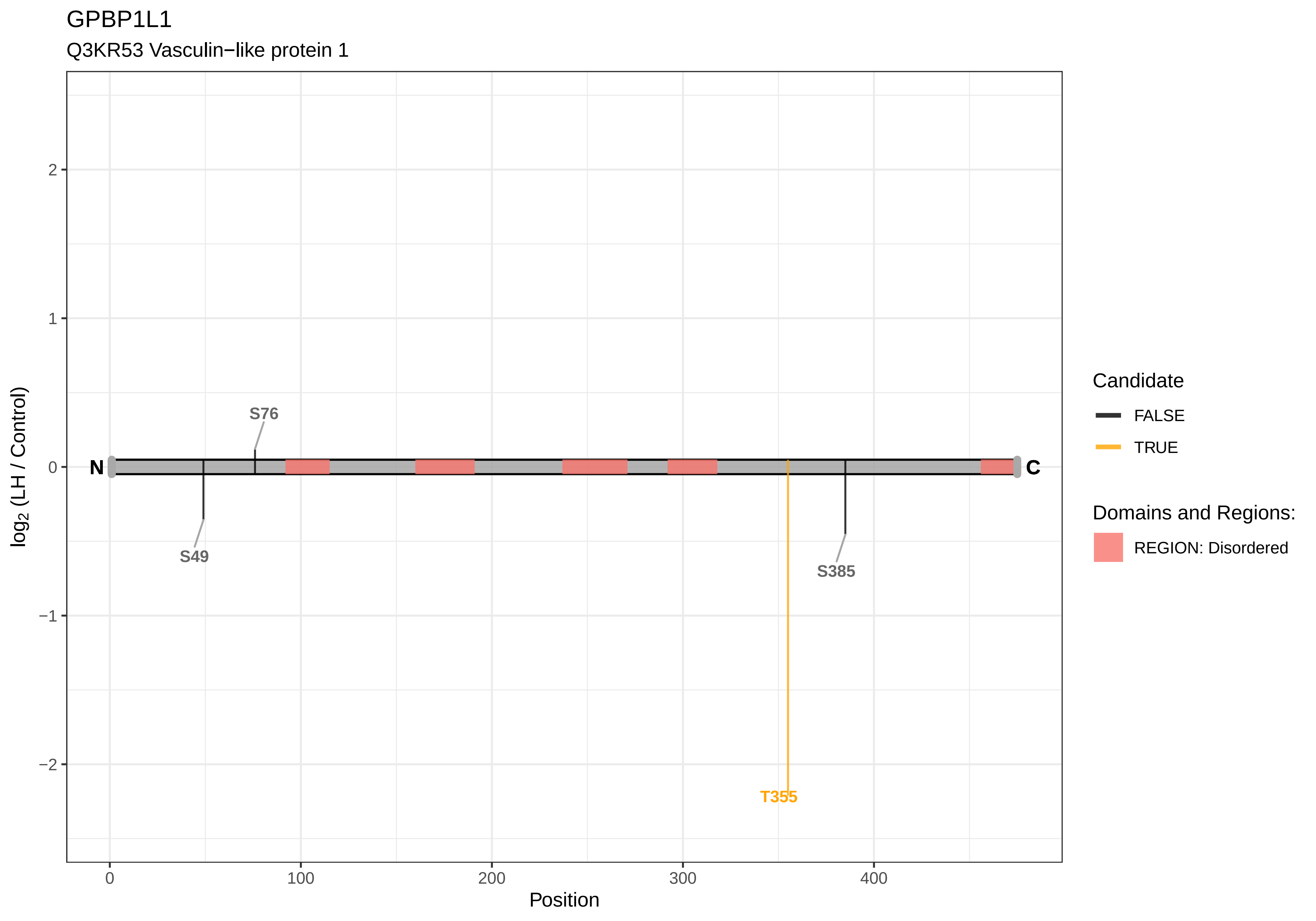

GPBP1L1
Q3KR53 Vasculin−like protein 1
2
1
0
−1
−2
log2 (LH / Control)
N
C
S76
S49
S385
T355
0
100
200
300
400
Position
Candidate
FALSE
TRUE
Domains and Regions:
REGION: Disordered

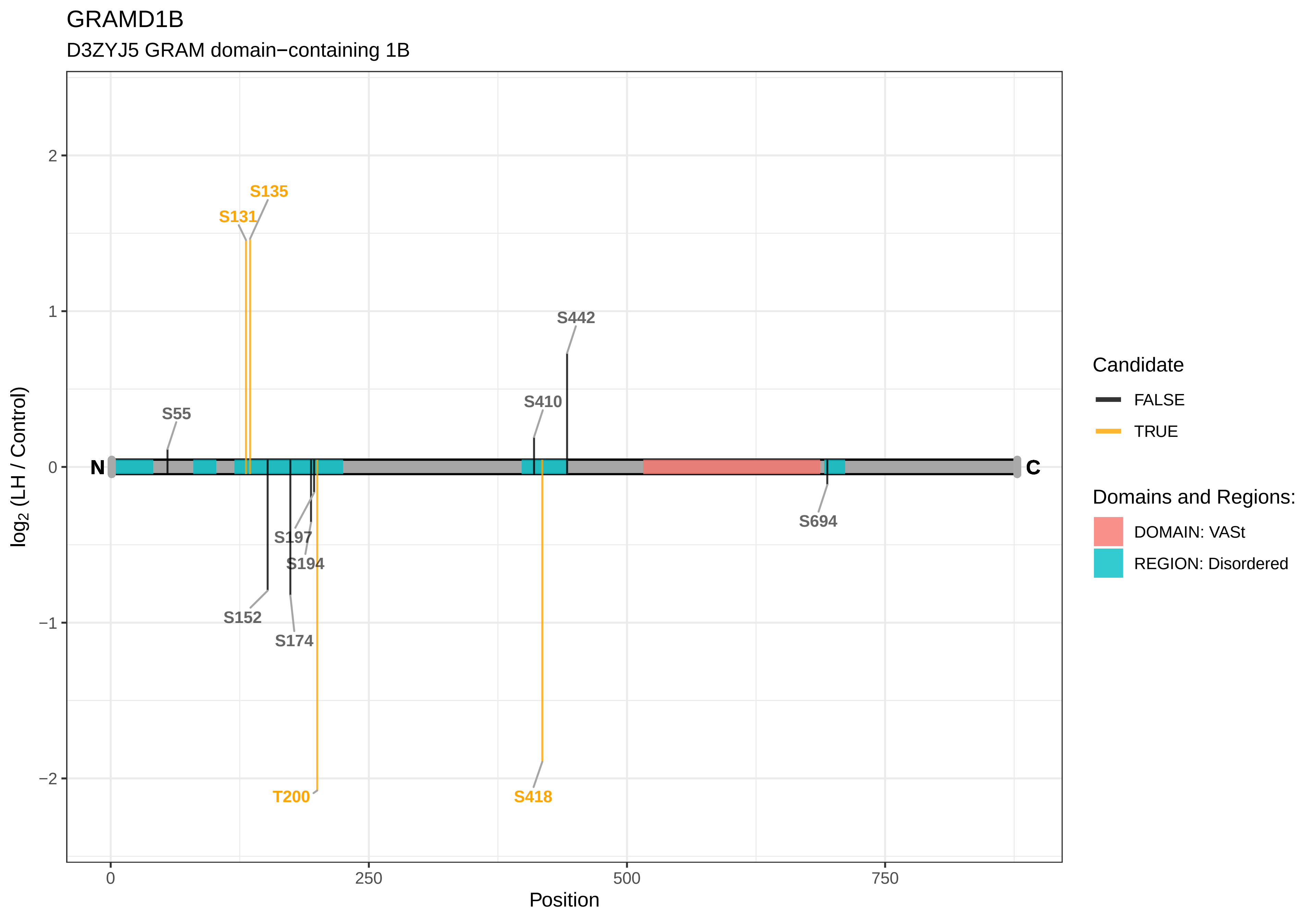
GRAMD1B
D3ZYJ5 GRAM domain−containing 1B
S135
S131
S442
S410
S55
S197
S194
S152
S174
S694
T200
S418
N
C
2
1
0
−1
−2
0
250
500
750
Position
log2 (LH / Control)
Candidate
FALSE
TRUE
Domains and Regions:
DOMAIN: VASt
REGION: Disordered

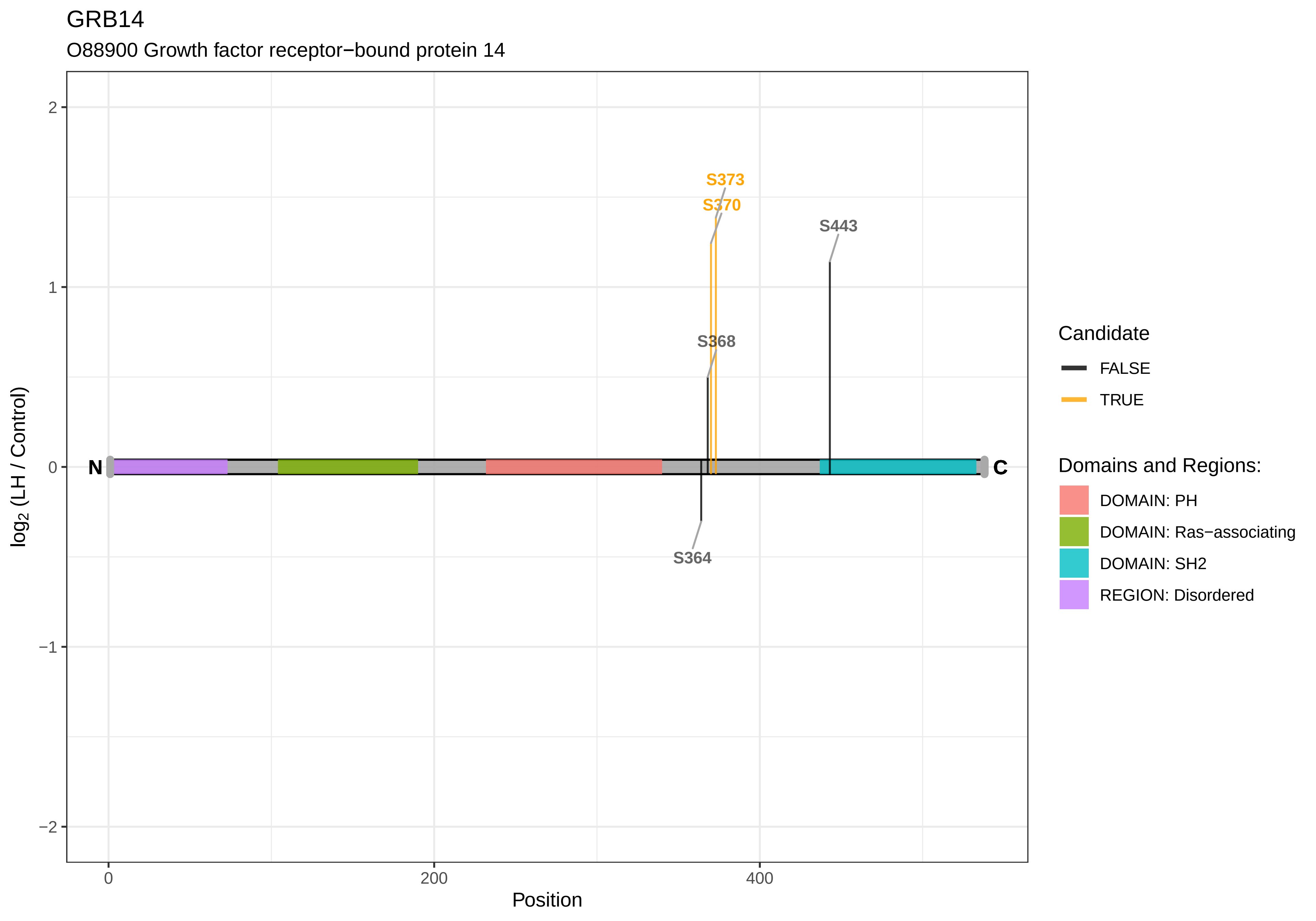
GRB14
O88900 Growth factor receptor−bound protein 14
S373
S370
S443
S368
S364
N
C
2
1
0
−1
−2
log2 (LH / Control)
0
200
400
Position
Candidate
FALSE
TRUE
Domains and Regions:
DOMAIN: PH
DOMAIN: Ras−associating
DOMAIN: SH2
REGION: Disordered

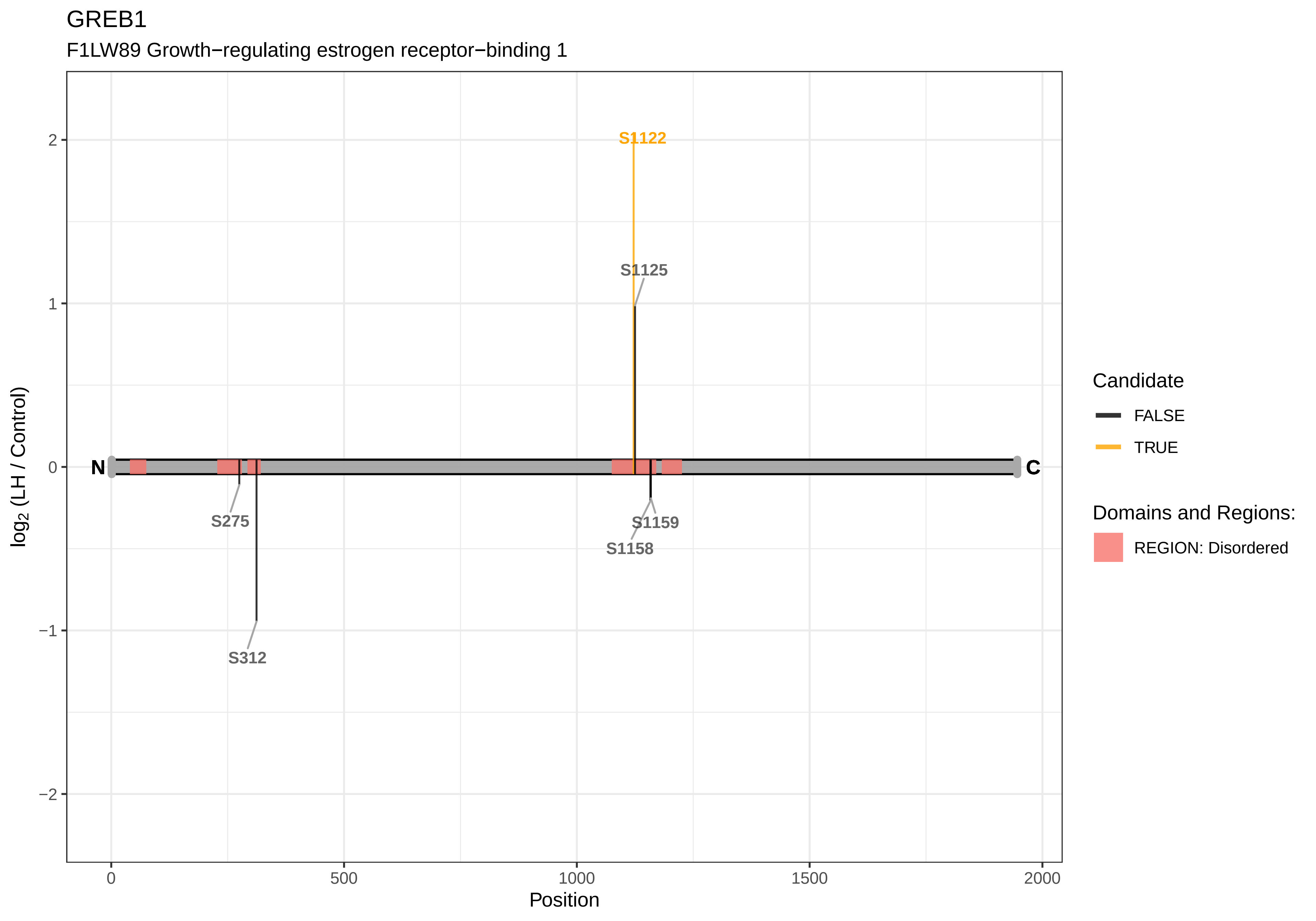

GREB1
F1LW89 Growth−regulating estrogen receptor−binding 1
S1122
S1125
S275
S312
S1159
S1158
N
C
2
1
0
−1
−2
log2 (LH / Control)
0
500
1000
1500
2000
Position
Candidate
FALSE
TRUE
Domains and Regions:
REGION: Disordered

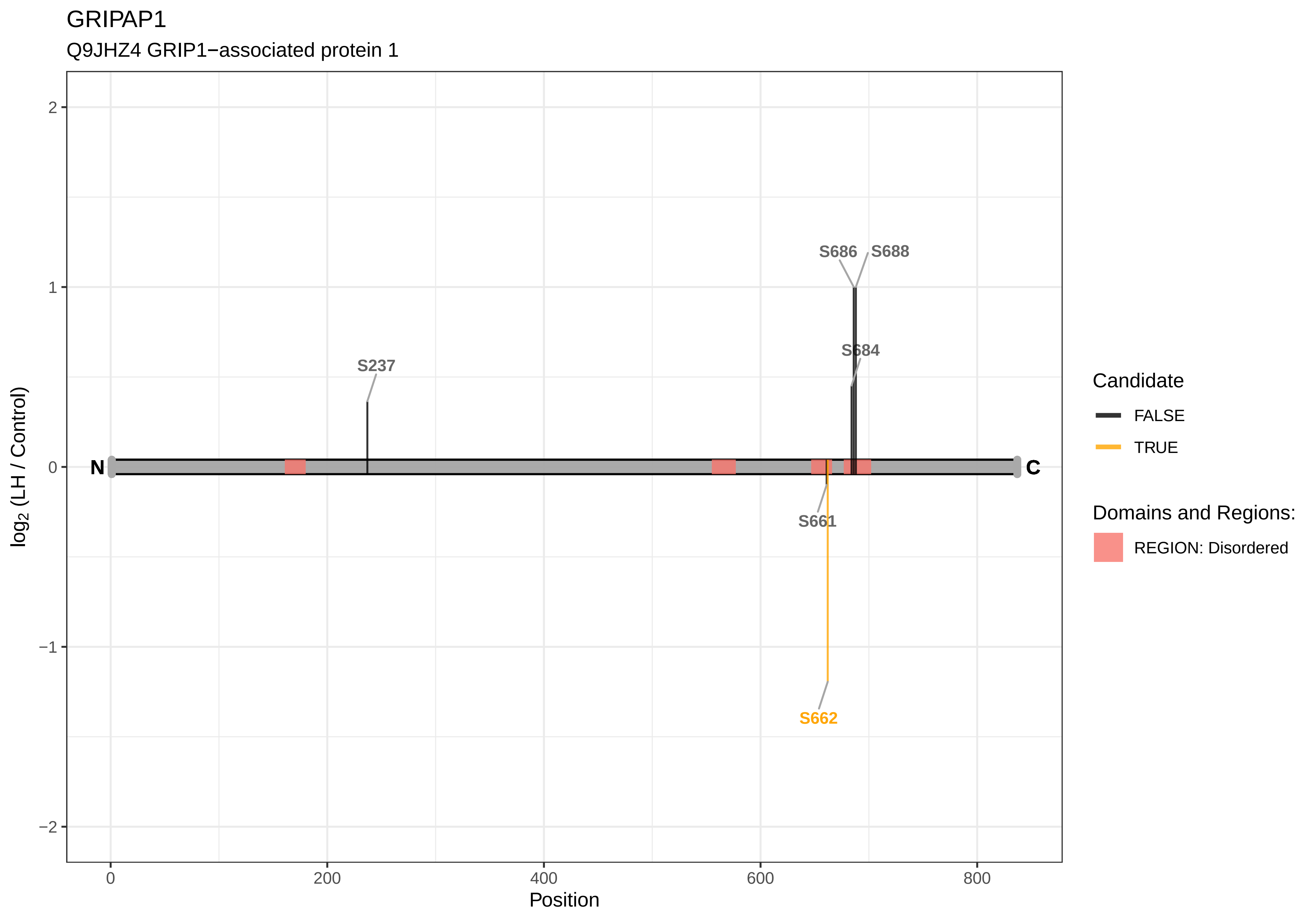
GRIPAP1
Q9JHZ4 GRIP1−associated protein 1
log2 (LH / Control)
Position
N
C
S237
S686
S688
S684
S661
S662
Candidate
FALSE
TRUE
Domains and Regions:
REGION: Disordered
2
1
0
−1
−2
0
200
400
600
800

GTF2F1

Q6AY96 General transcription factor IIF subunit 1

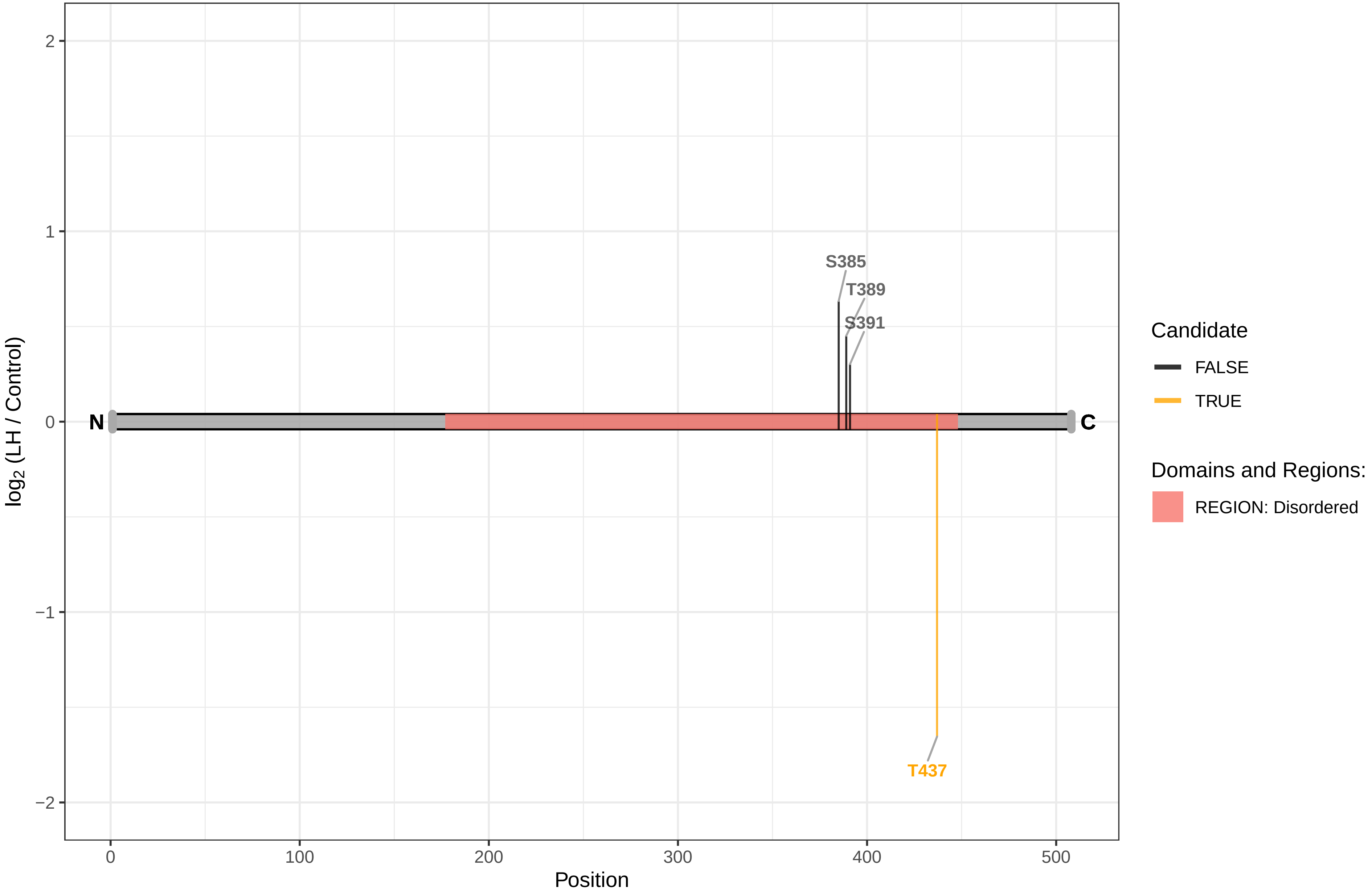

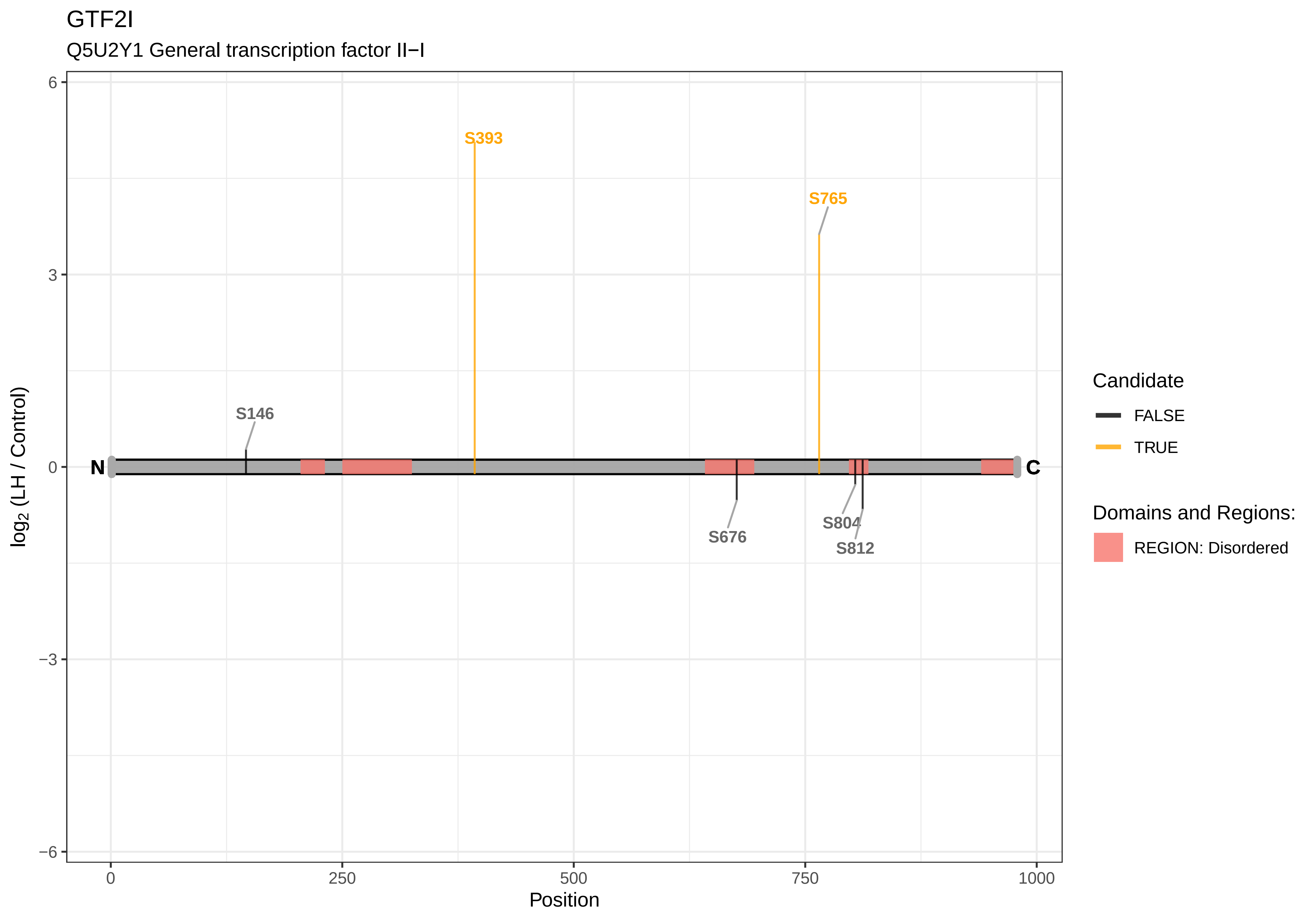
GTF2I
Q5U2Y1 General transcription factor II−I
log2 (LH / Control)
6
3
0
−3
−6
S393
S765
S146
N
C
S676
S804
S812
0
250
500
750
1000
Position
Candidate
FALSE
TRUE
Domains and Regions:
REGION: Disordered

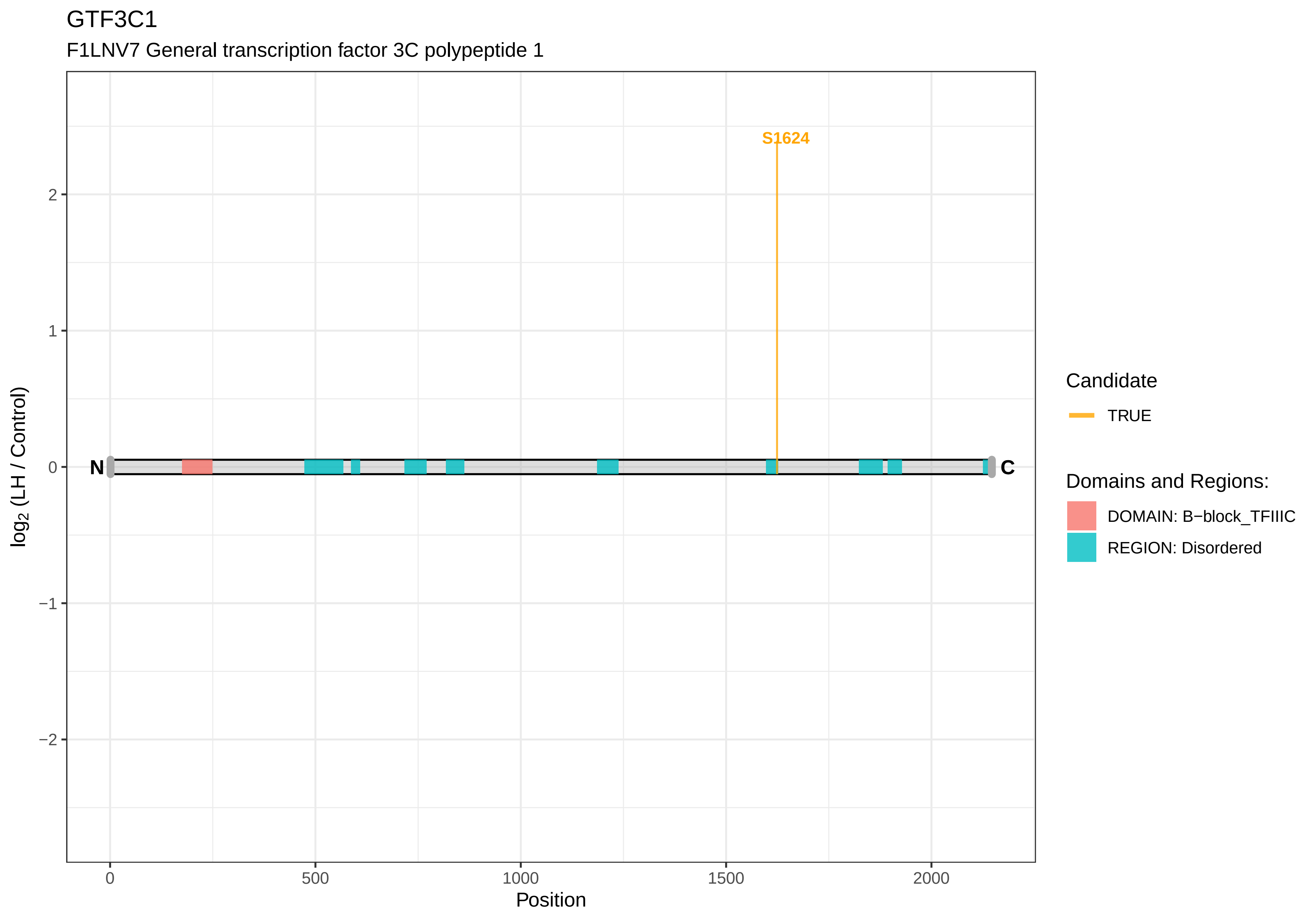
GTF3C1
F1LNV7 General transcription factor 3C polypeptide 1
S1624
N
C
log2 (LH / Control)
Position
0
500
1000
1500
2000
2
1
0
−1
−2
Candidate
TRUE
Domains and Regions:
DOMAIN: B−block_TFIIIC
REGION: Disordered

GTF3C2

A0A0G2JY75 General transcription factor IIIC subunit 2

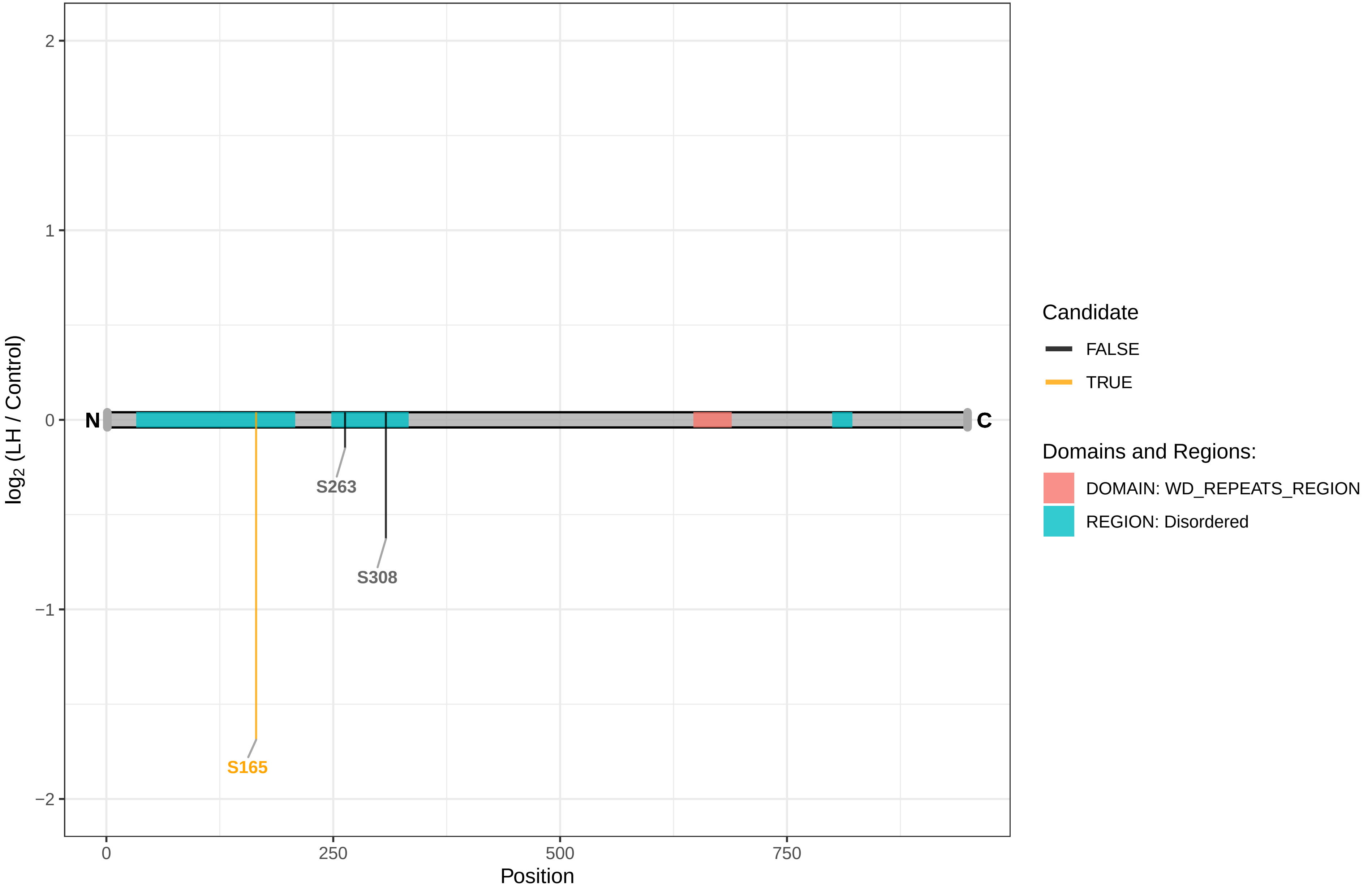

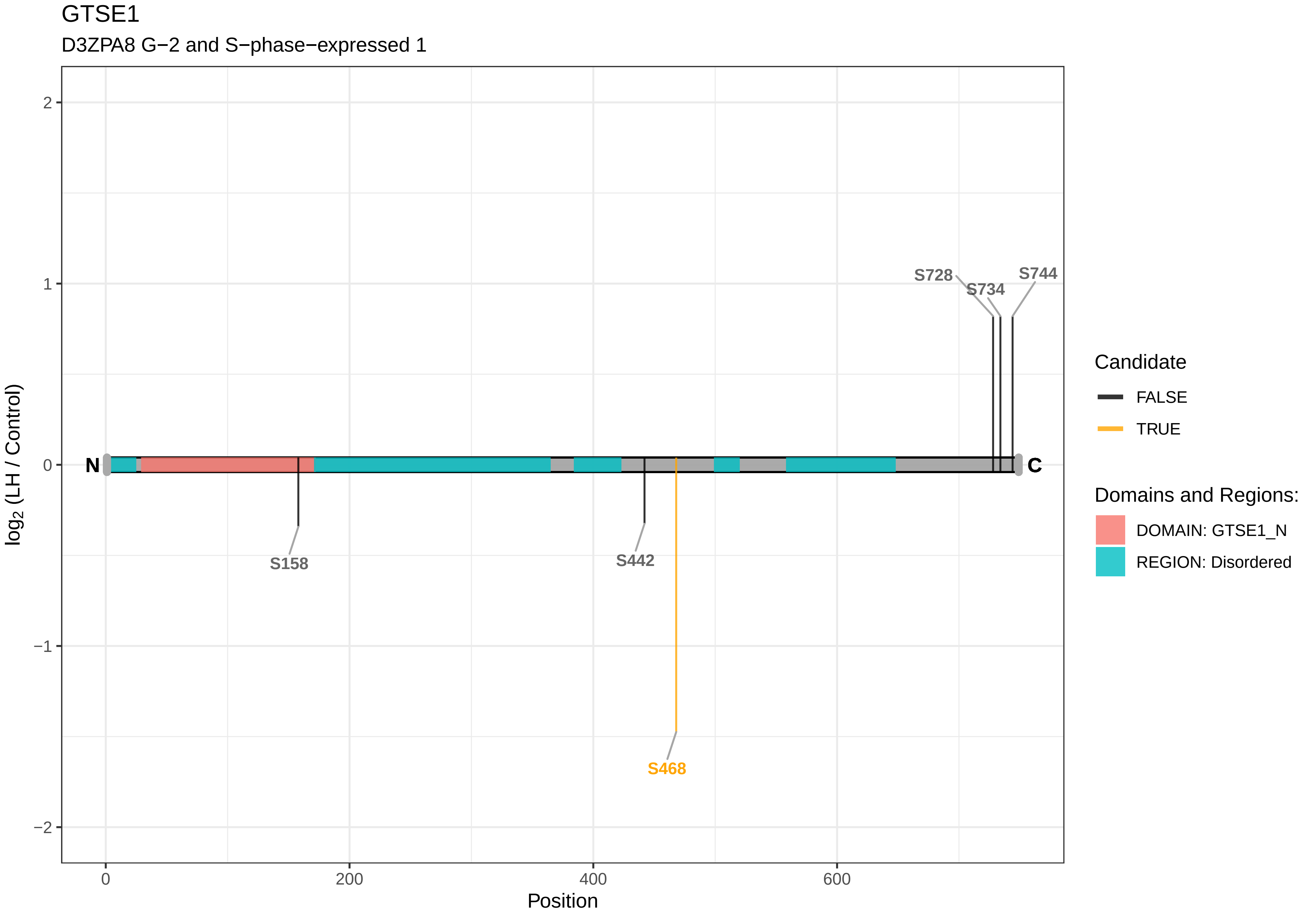
GTSE1
D3ZPA8 G−2 and S−phase−expressed 1
log2 (LH / Control)
Position
N
C
S158
S442
S468
S728
S734
S744
2
1
0
−1
−2
0
200
400
600
Candidate
FALSE
TRUE
Domains and Regions:
DOMAIN: GTSE1_N
REGION: Disordered

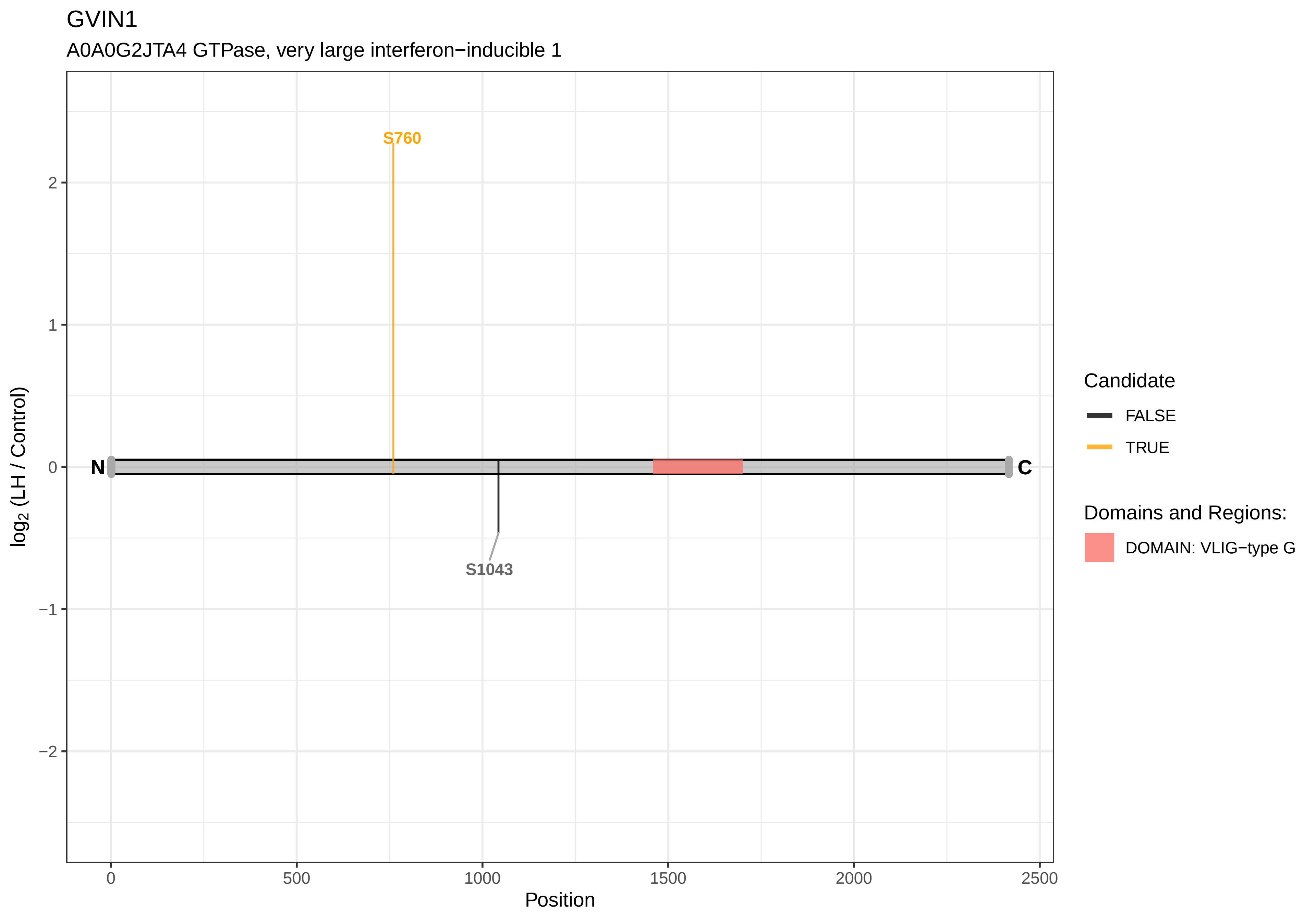

GVIN1
A0A0G2JTA4 GTPase, very large interferon−inducible 1
S760
S1043
N
C
log2 (LH / Control)
Position
0
500
1000
1500
2000
2500
2
1
0
−1
−2
Candidate
FALSE
TRUE
Domains and Regions:
DOMAIN: VLIG−type G

GYS1

A2RRU1 Glycogen [starch] synthase, muscle

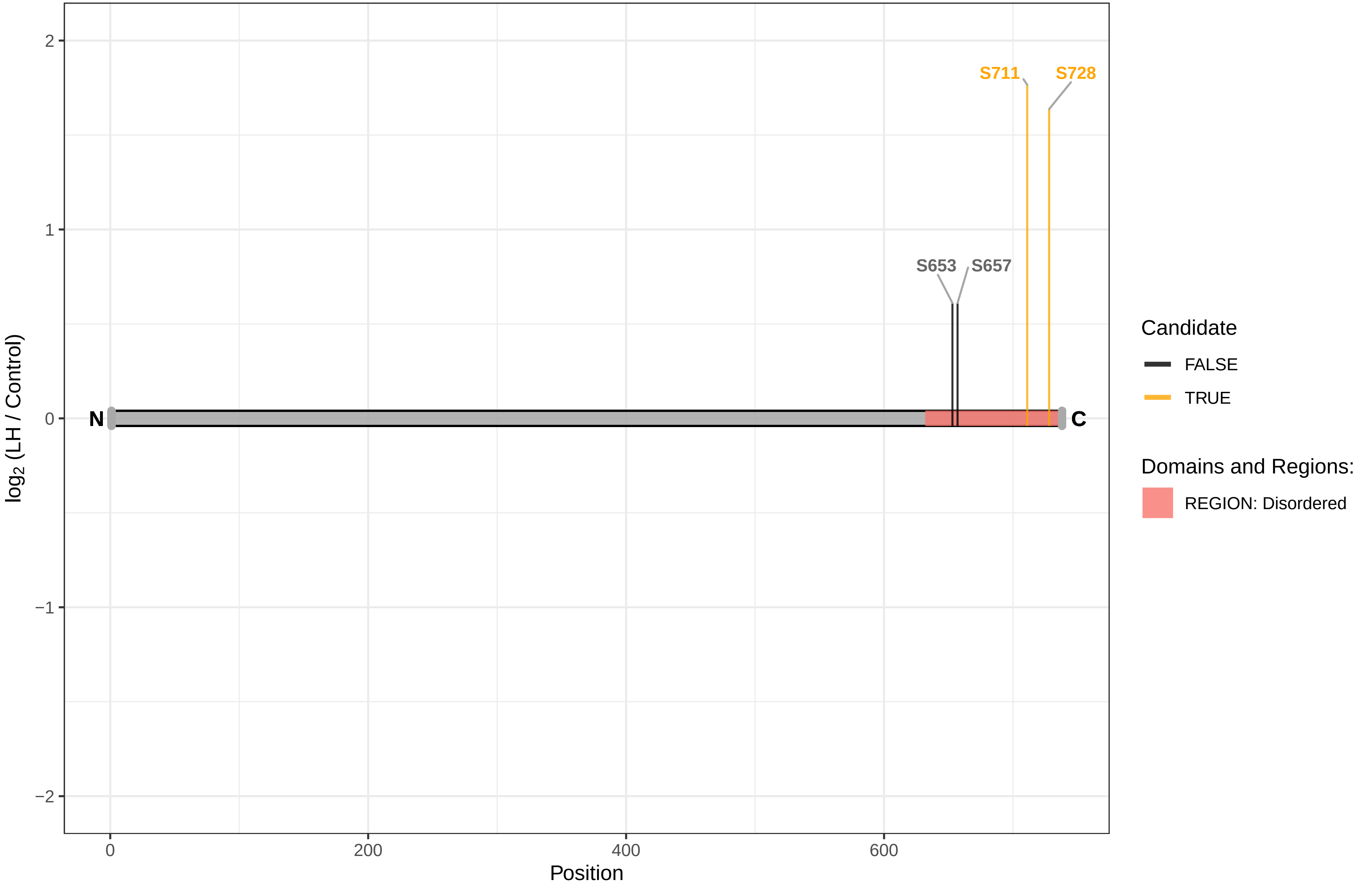

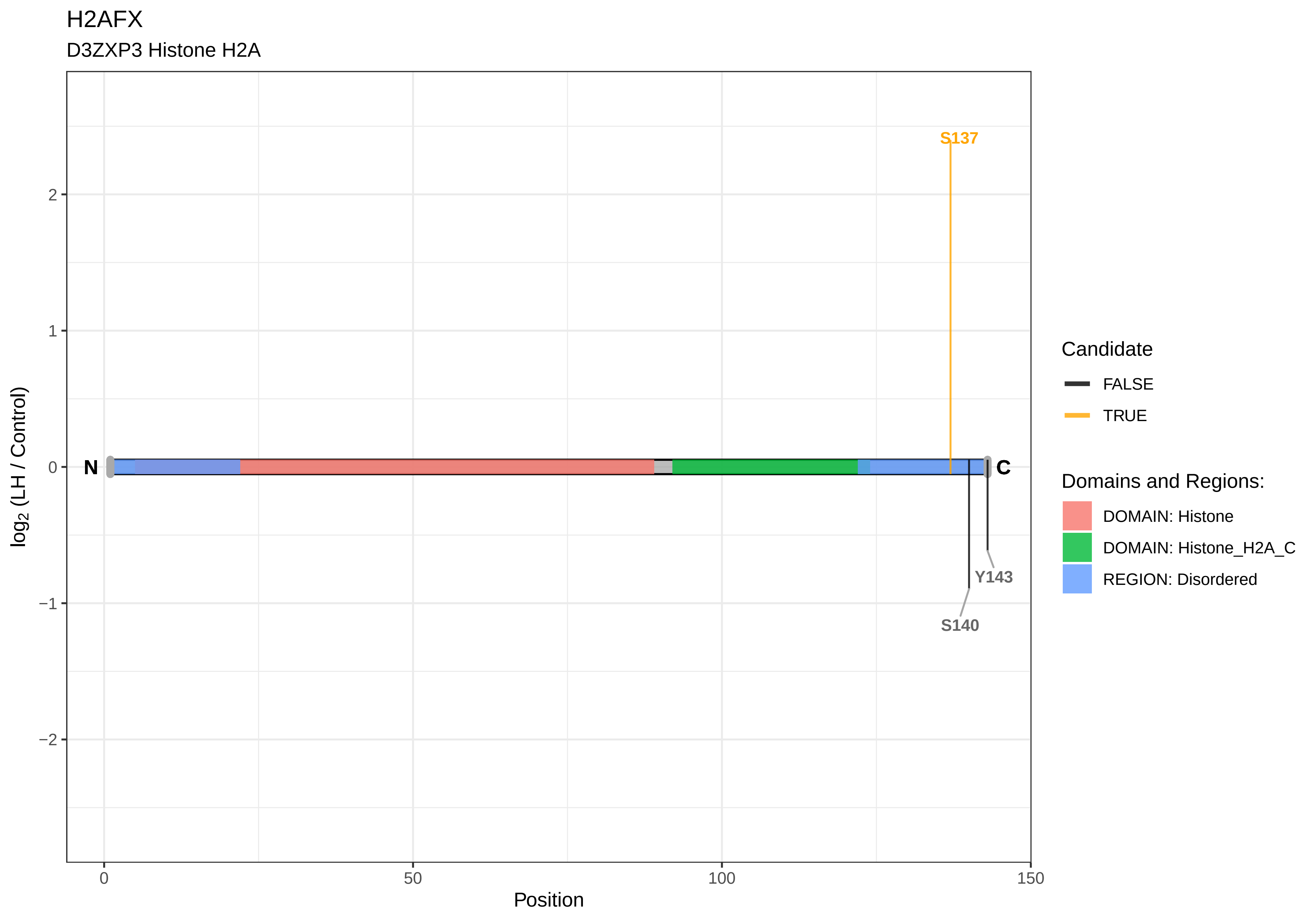

H2AFX
D3ZXP3 Histone H2A
S137
Y143
S140
N
C
2
1
0
−1
−2
0
50
100
150
Position
log2 (LH / Control)
Candidate
FALSE
TRUE
Domains and Regions:
DOMAIN: Histone
DOMAIN: Histone_H2A_C
REGION: Disordered

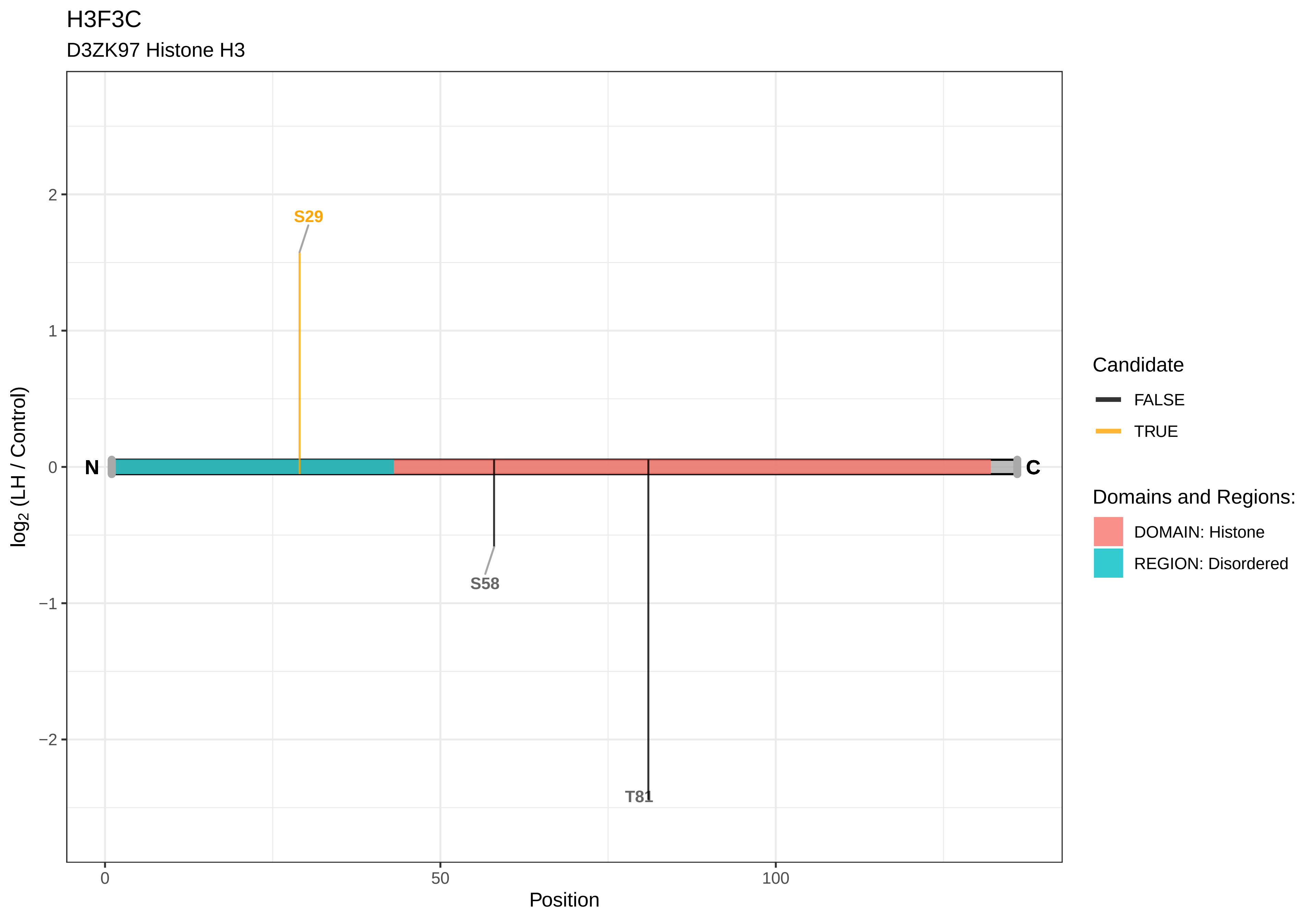

H3F3C
D3ZK97 Histone H3
S29
S58
T81
N
C
2
1
0
−1
−2
0
50
100
Position
log2 (LH / Control)
Candidate
FALSE
TRUE
Domains and Regions:
DOMAIN: Histone
REGION: Disordered

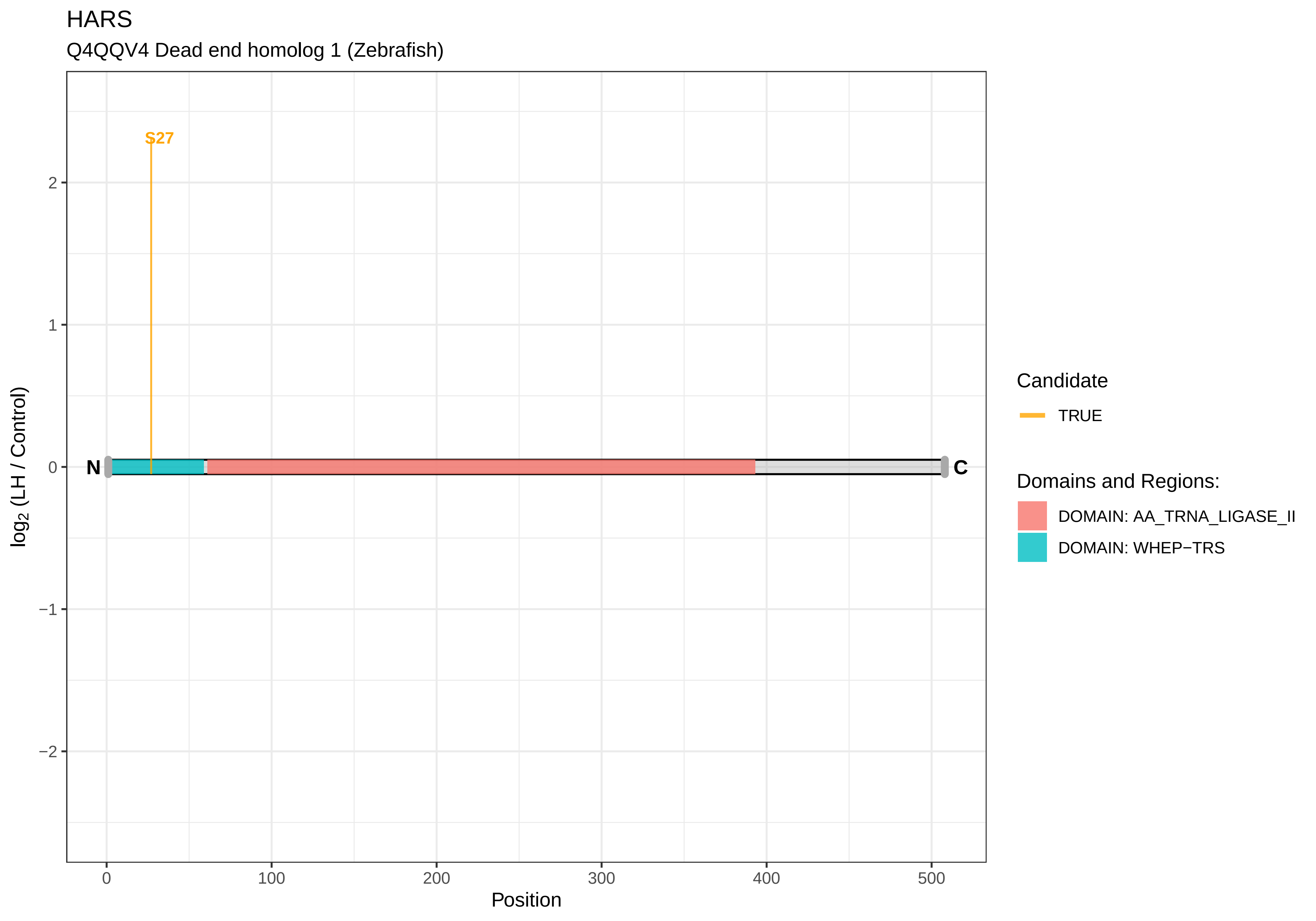

HARS
Q4QQV4 Dead end homolog 1 (Zebrafish)
S27
N
C
2
1
0
−1
−2
log2 (LH / Control)
0
100
200
300
400
500
Position
Candidate
TRUE
Domains and Regions:
DOMAIN: AA_TRNA_LIGASE_II
DOMAIN: WHEP−TRS

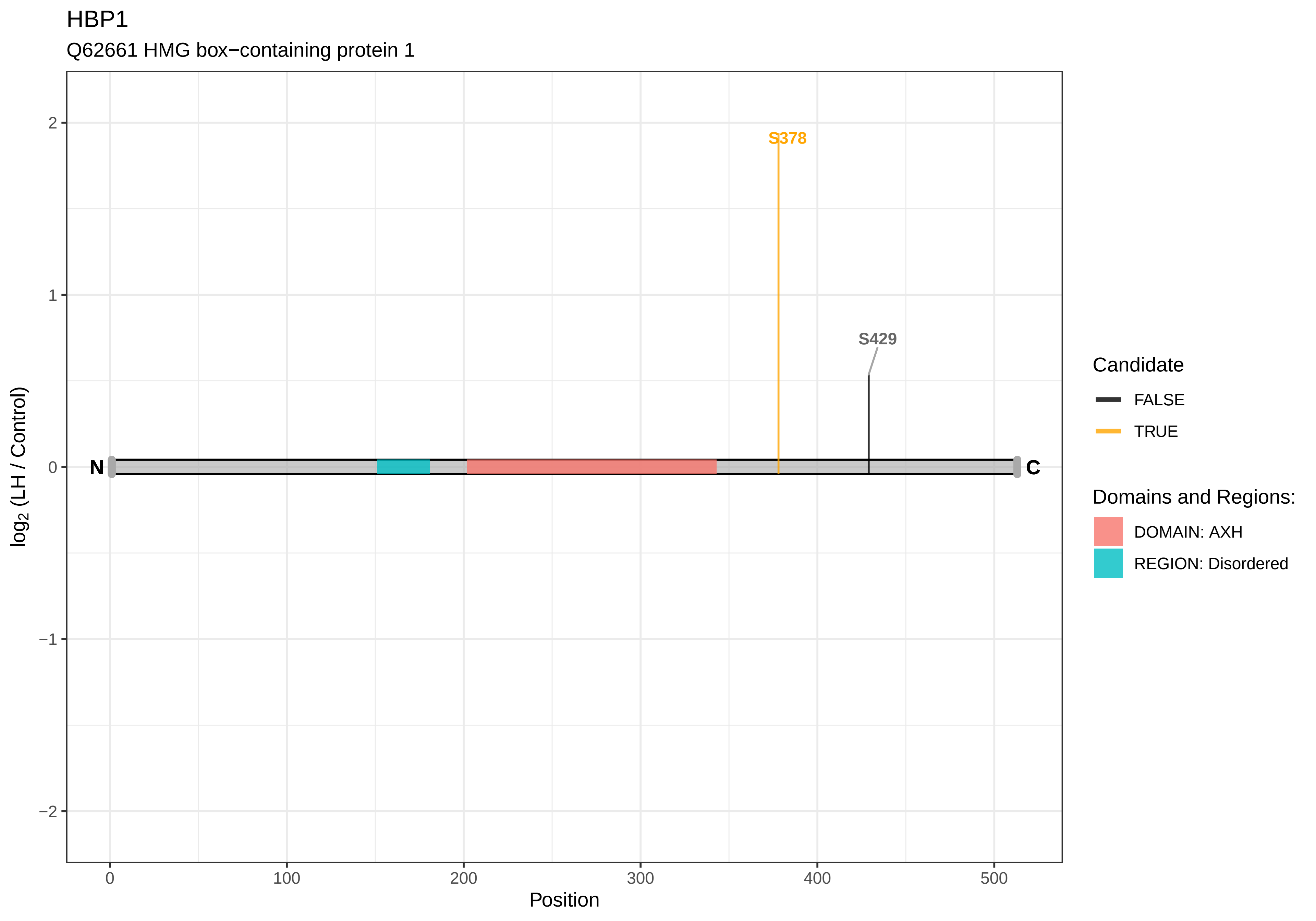

HBP1
Q62661 HMG box−containing protein 1
S378
S429
N
C
log2 (LH / Control)
2
1
0
−1
−2
0
100
200
300
400
500
Position
Candidate
FALSE
TRUE
Domains and Regions:
DOMAIN: AXH
REGION: Disordered

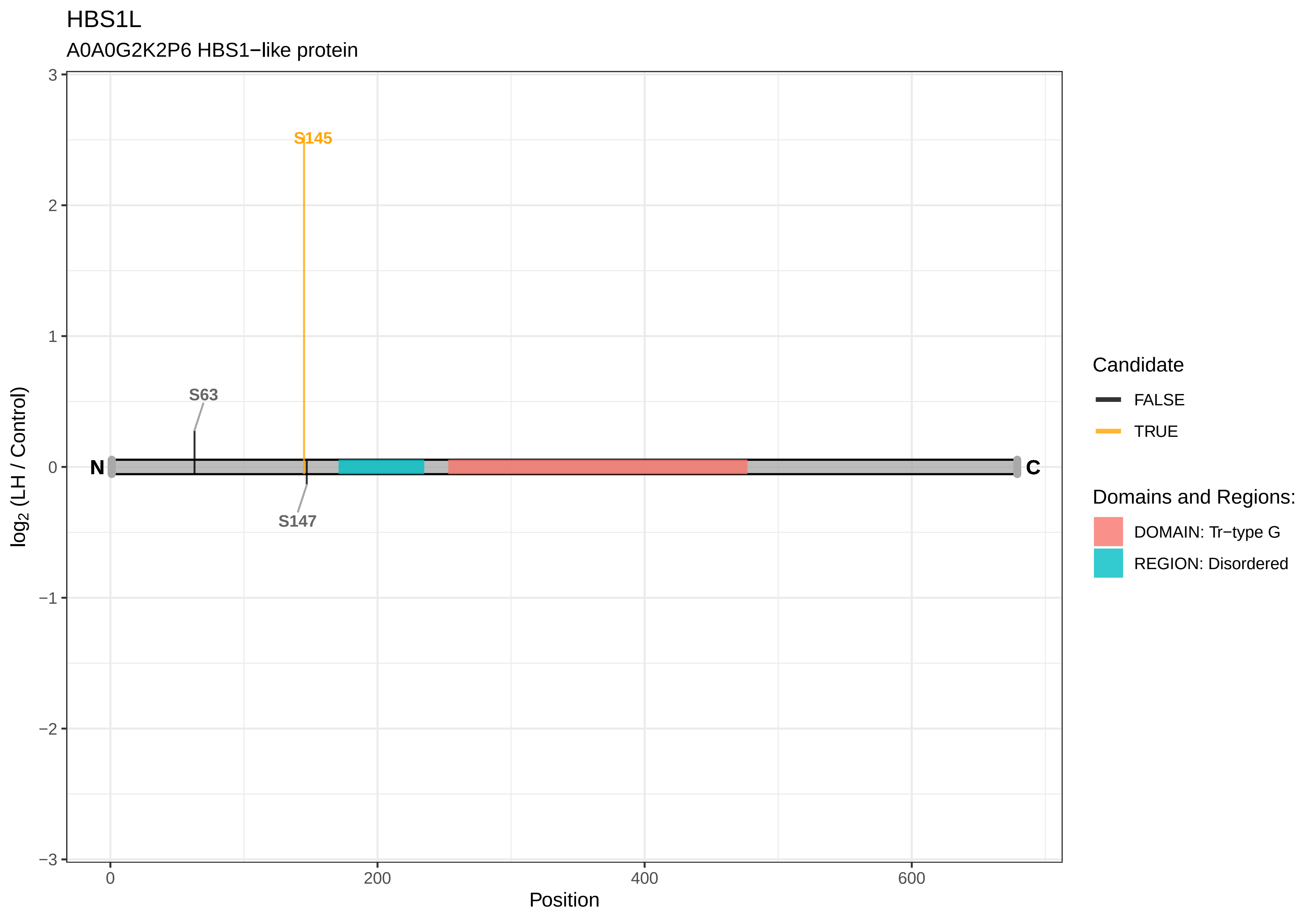
HBS1L
A0A0G2K2P6 HBS1−like protein
S145
S63
S147
N
C
log2 (LH / Control)
Position
3
2
1
0
−1
−2
−3
0
200
400
600
Candidate
FALSE
TRUE
Domains and Regions:
DOMAIN: Tr−type G
REGION: Disordered

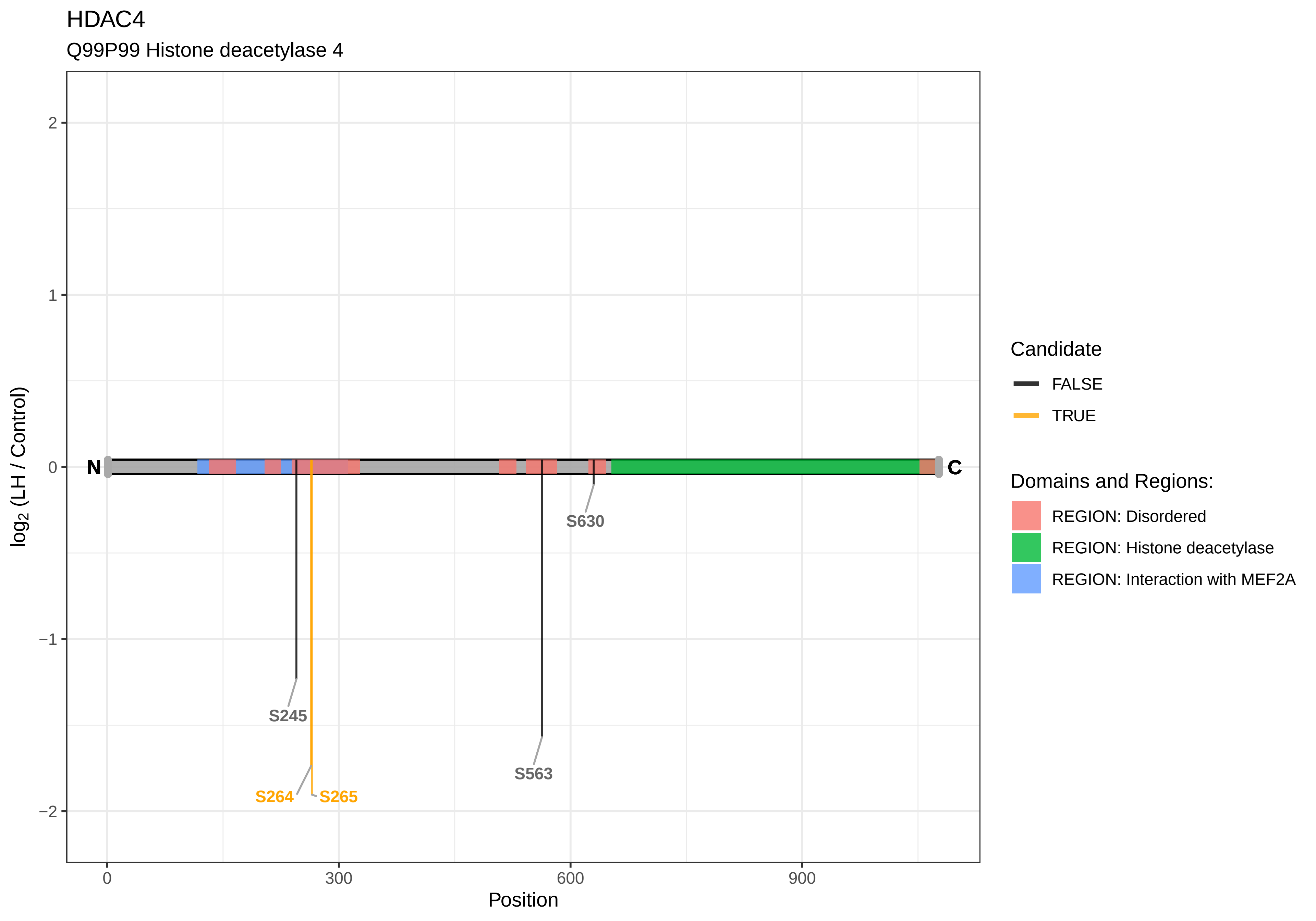

HDAC4
Q99P99 Histone deacetylase 4
log2 (LH / Control)
Position
N
C
S245
S264
S265
S563
S630
2
1
0
−1
−2
0
300
600
900
Candidate
FALSE
TRUE
Domains and Regions:
REGION: Disordered
REGION: Histone deacetylase
REGION: Interaction with MEF2A

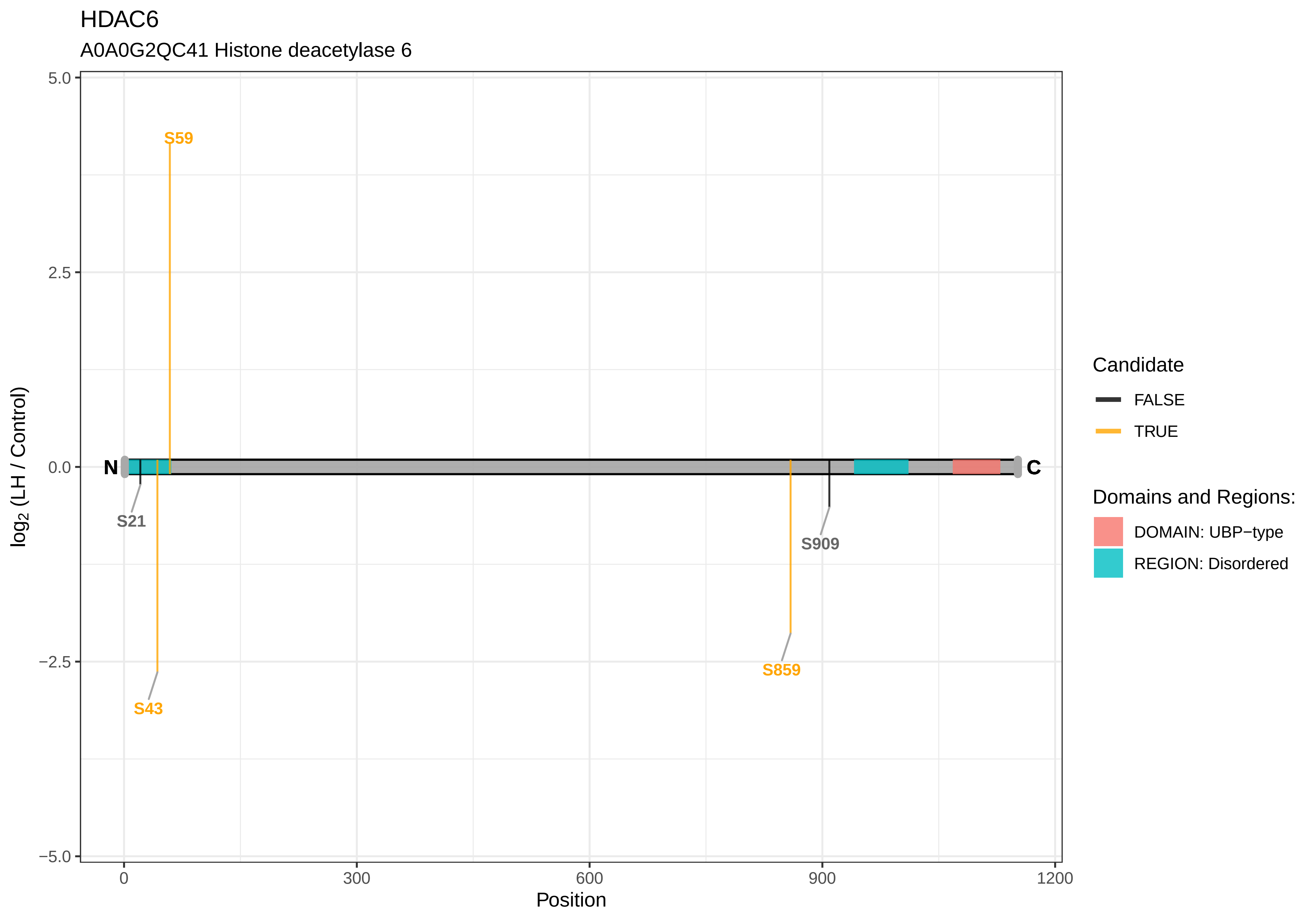
HDAC6
A0A0G2QC41 Histone deacetylase 6
5.0
2.5
0.0
−2.5
−5.0
log2 (LH / Control)
S59
N
C
S21
S43
S859
S909
0
300
600
900
1200
Position
Candidate
FALSE
TRUE
Domains and Regions:
DOMAIN: UBP−type
REGION: Disordered

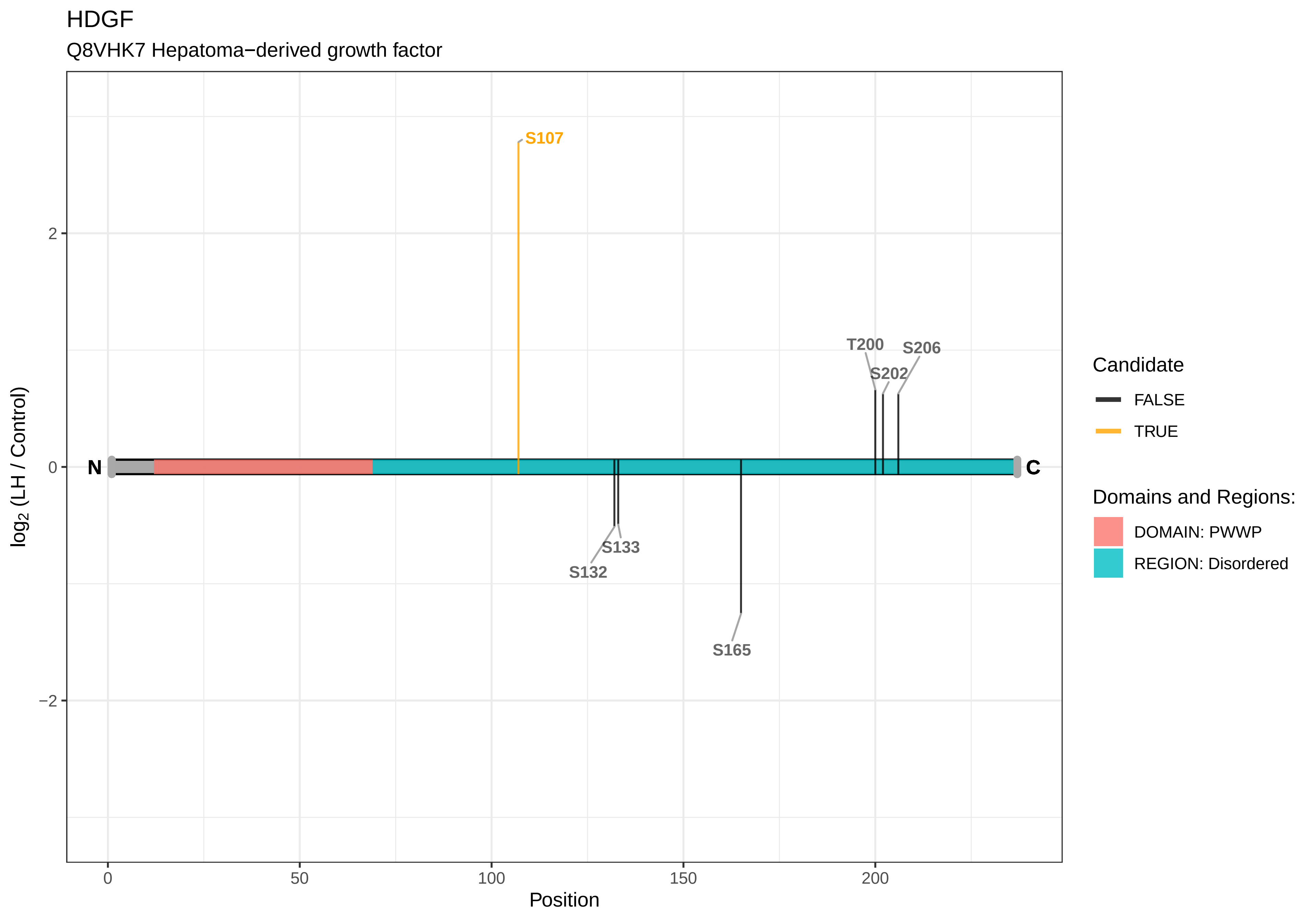

HDGF
Q8VHK7 Hepatoma−derived growth factor
S107
T200
S206
S202
N
C
S133
S132
S165
2
0
−2
log2 (LH / Control)
0
50
100
150
200
Position
Candidate
FALSE
TRUE
Domains and Regions:
DOMAIN: PWWP
REGION: Disordered

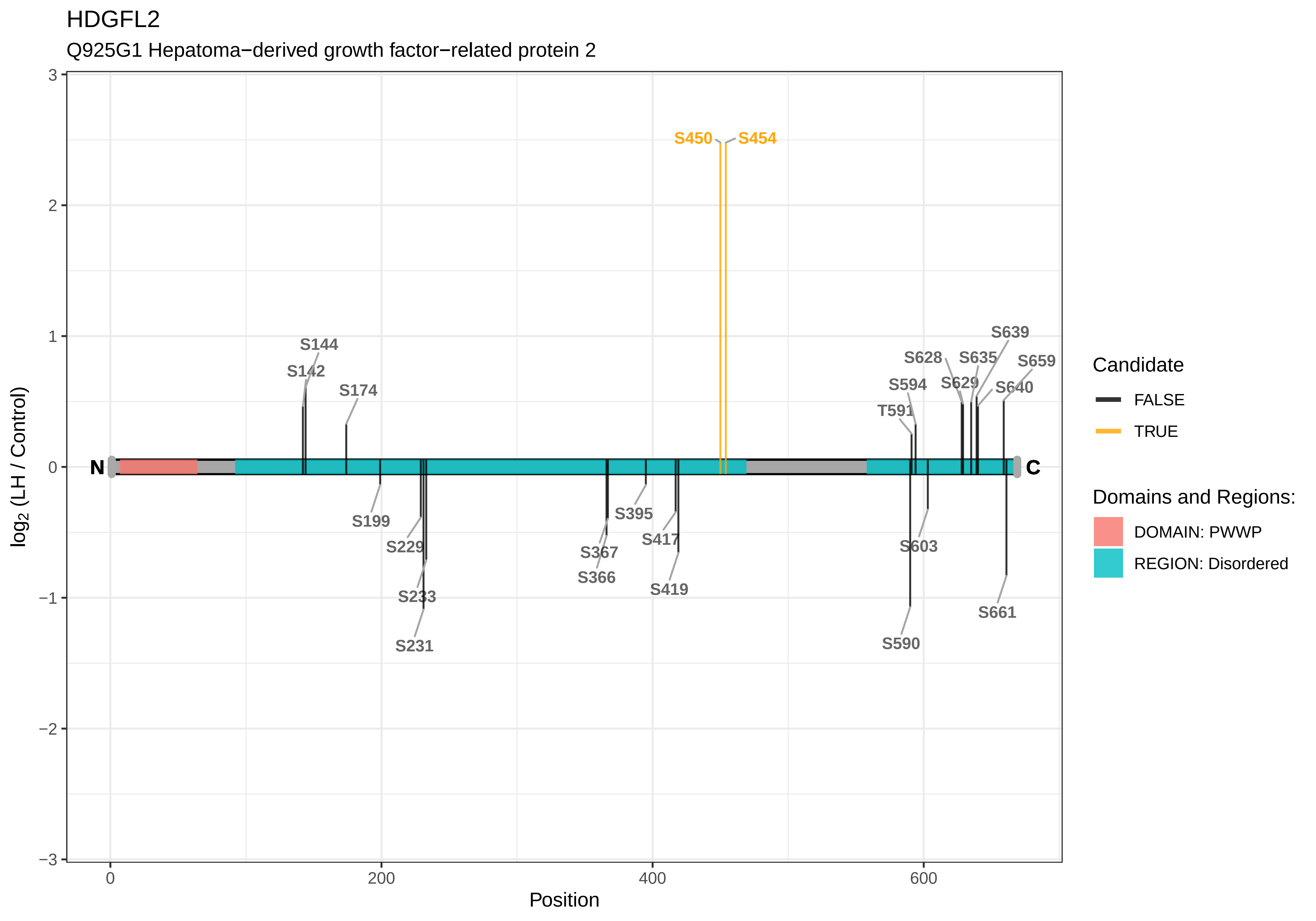
HDGFL2
Q925G1 Hepatoma−derived growth factor−related protein 2
S450
S454
S144
S142
S174
S199
S229
S233
S231
S367
S366
S395
S417
S419
S639
S628
S635
S659
S594
S629
S640
T591
S603
S590
S661
N
C
log2 (LH / Control)
Position
Candidate
FALSE
TRUE
Domains and Regions:
DOMAIN: PWWP
REGION: Disordered

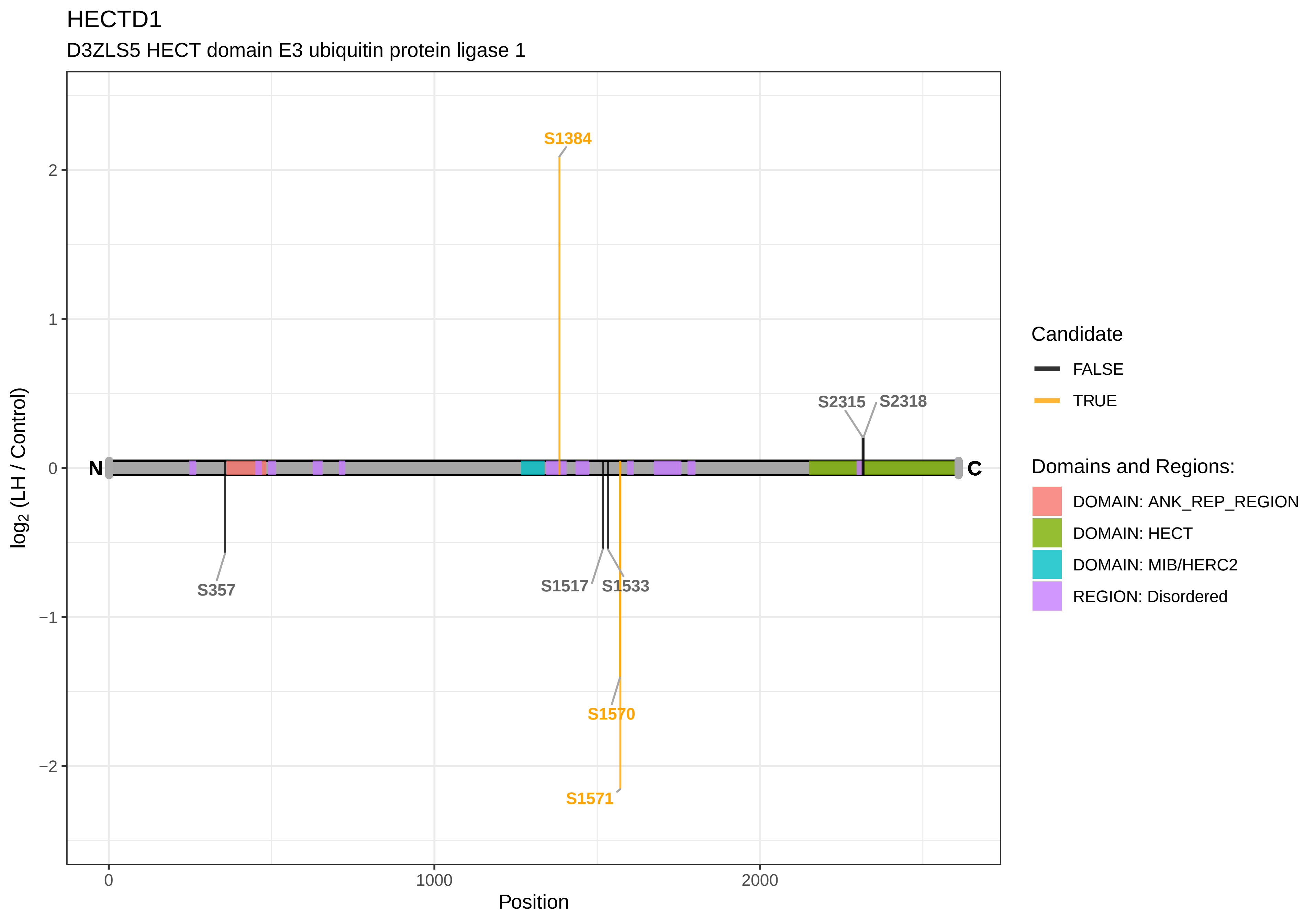

HECTD1
D3ZLS5 HECT domain E3 ubiquitin protein ligase 1
S1384
S2315
S2318
S357
S1517
S1533
S1570
S1571
N
C
2
1
0
−1
−2
log2 (LH / Control)
0
1000
2000
Position
Candidate
FALSE
TRUE
Domains and Regions:
DOMAIN: ANK_REP_REGION
DOMAIN: HECT
DOMAIN: MIB/HERC2
REGION: Disordered

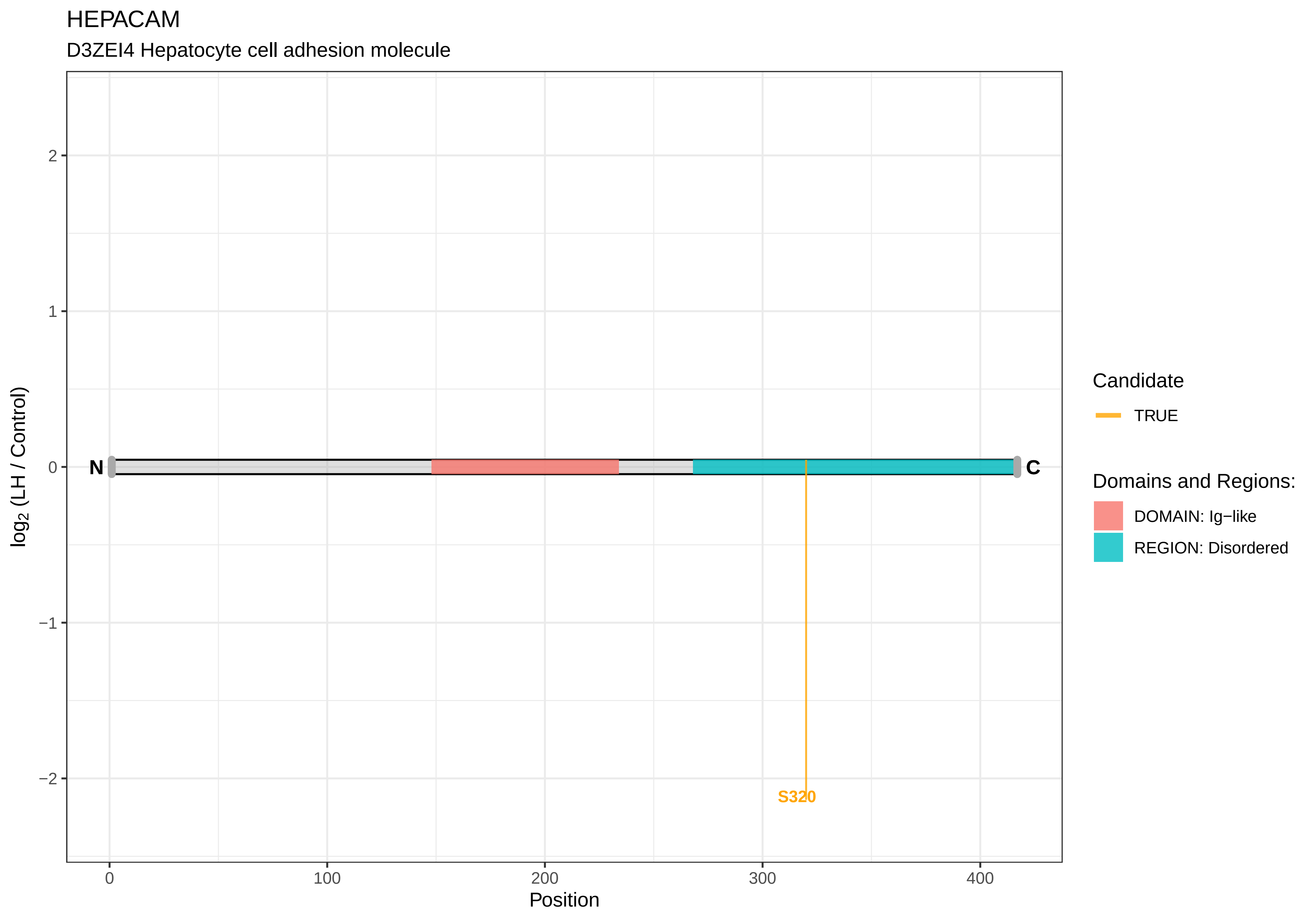

HEPACAM
D3ZEI4 Hepatocyte cell adhesion molecule
log2 (LH / Control)
2
1
0
−1
−2
N
C
S320
0
100
200
300
400
Position
Candidate
TRUE
Domains and Regions:
DOMAIN: Ig−like
REGION: Disordered

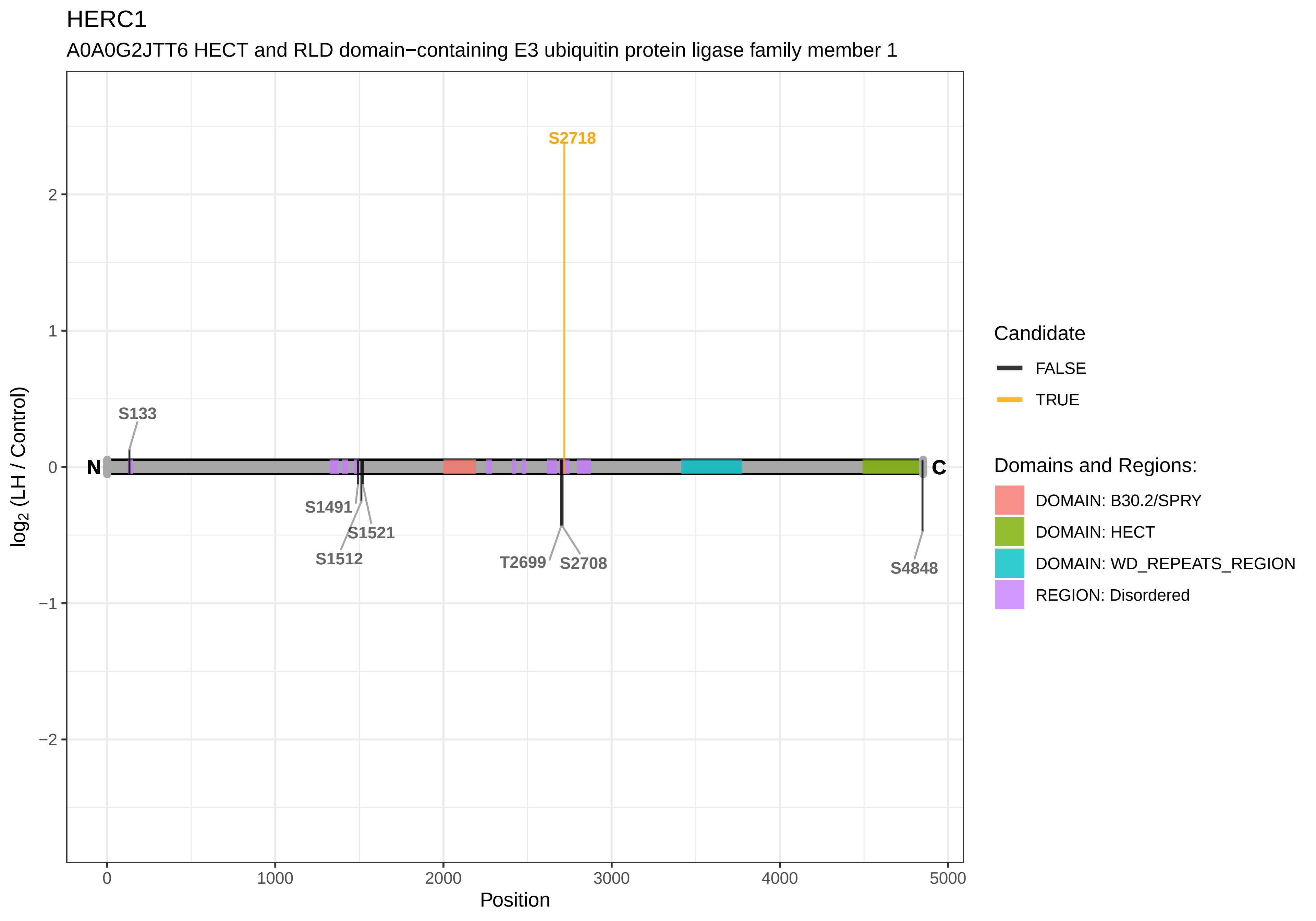

HERC1
A0A0G2JTT6 HECT and RLD domain−containing E3 ubiquitin protein ligase family member 1
S2718
S133
N
C
S1491
S1521
S1512
T2699
S2708
S4848
log2 (LH / Control)
2
1
0
−1
−2
0
1000
2000
3000
4000
5000
Position
Candidate
FALSE
TRUE
Domains and Regions:
DOMAIN: B30.2/SPRY
DOMAIN: HECT
DOMAIN: WD_REPEATS_REGION
REGION: Disordered

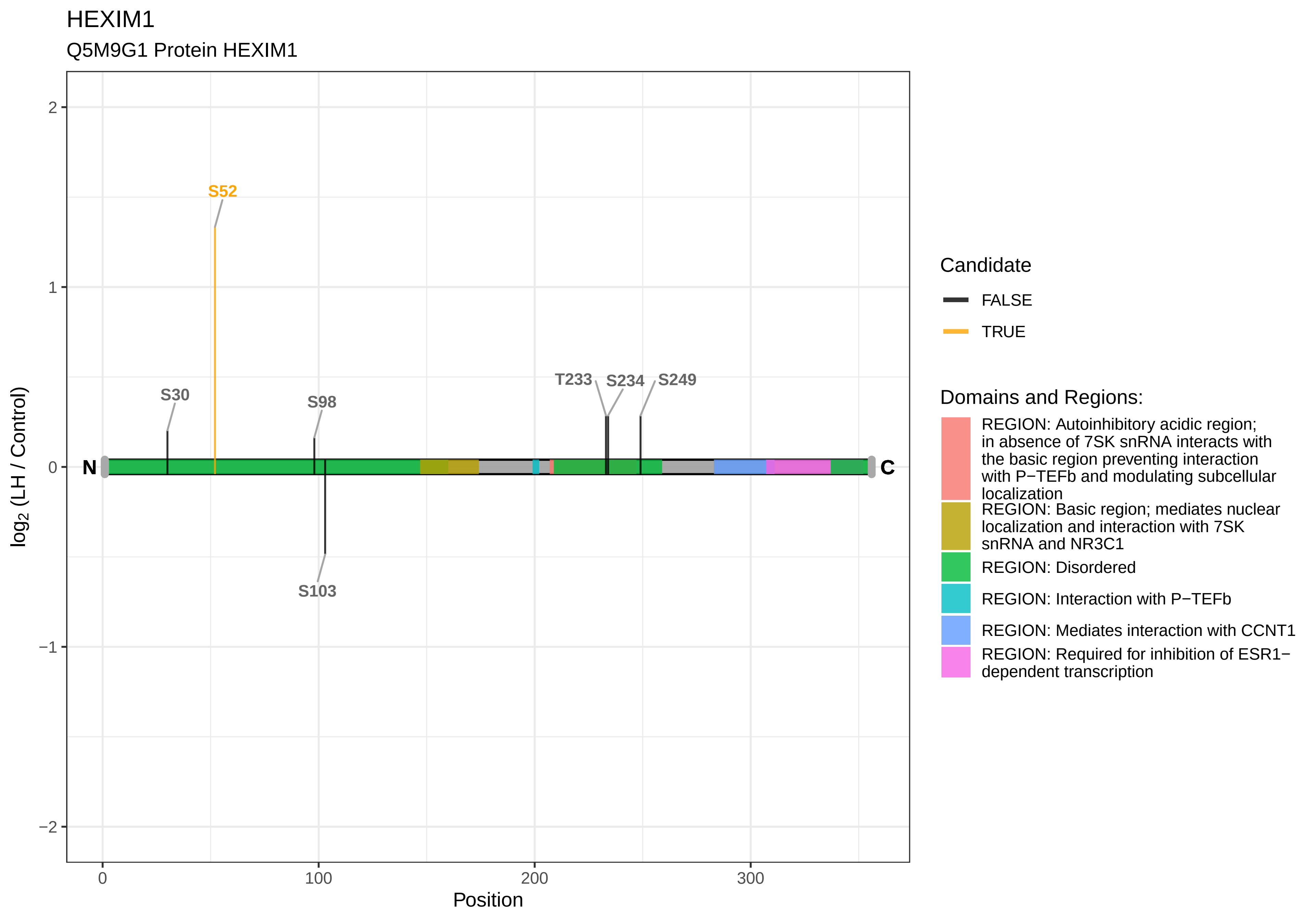

HEXIM1
Q5M9G1 Protein HEXIM1
S52
S30
S98
S103
T233
S234
S249
N
C
2
1
0
−1
−2
0
100
200
300
Position
log2 (LH / Control)
Candidate
FALSE
TRUE
Domains and Regions:
REGION: Autoinhibitory acidic region; in absence of 7SK snRNA interacts with the basic region preventing interaction with P−TEFb and modulating subcellular localization
REGION: Basic region; mediates nuclear localization and interaction with 7SK snRNA and NR3C1
REGION: Disordered
REGION: Interaction with P−TEFb
REGION: Mediates interaction with CCNT1
REGION: Required for inhibition of ESR1−dependent transcription

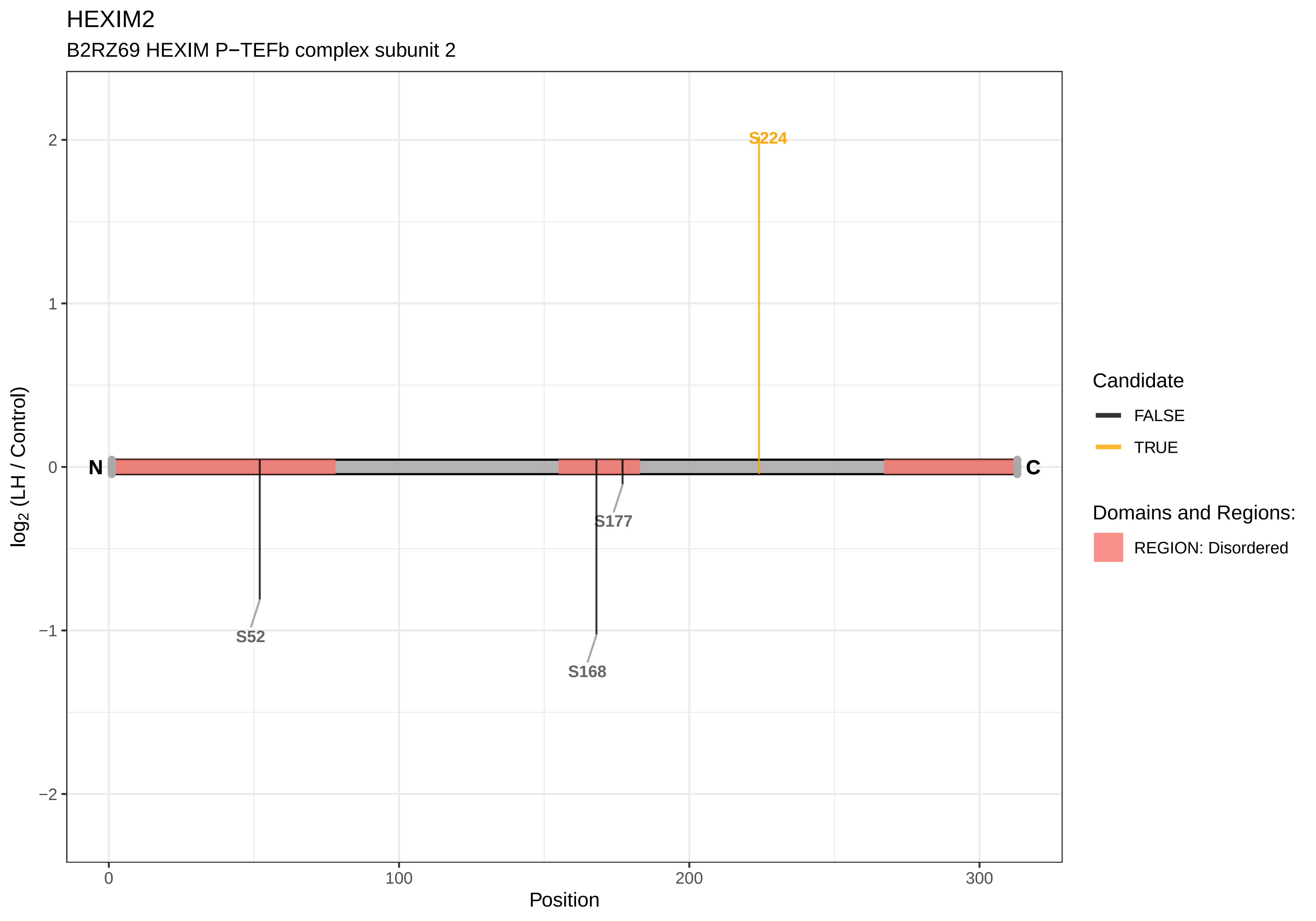
HEXIM2
B2RZ69 HEXIM P−TEFb complex subunit 2
S224
S177
S52
S168
N
C
2
1
0
−1
−2
0
100
200
300
Position
log2 (LH / Control)
Candidate
FALSE
TRUE
Domains and Regions:
REGION: Disordered

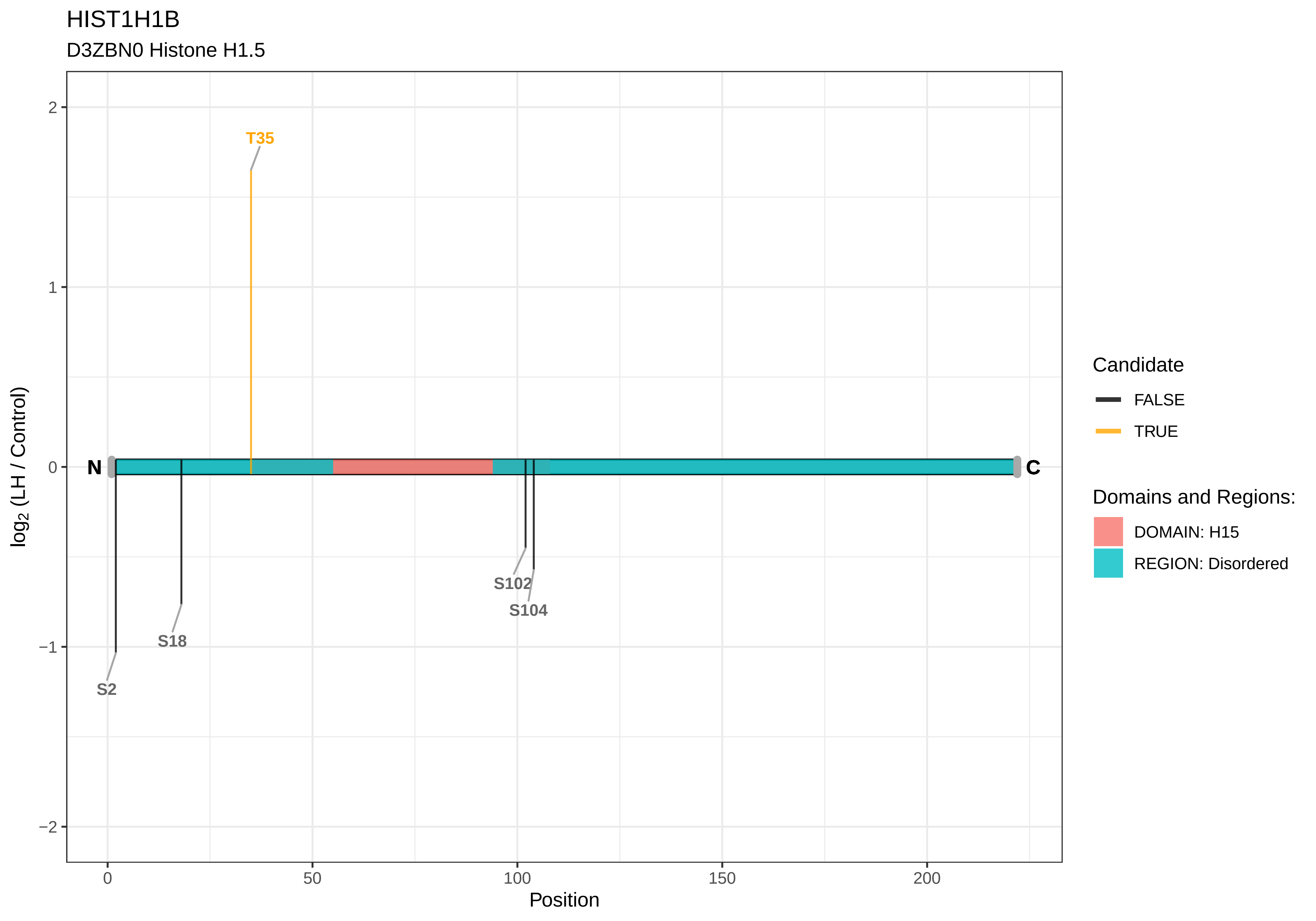

HIST1H1B
D3ZBN0 Histone H1.5
T35
S2
S18
S102
S104
N
C
log2 (LH / Control)
Position
Candidate
FALSE
TRUE
Domains and Regions:
DOMAIN: H15
REGION: Disordered

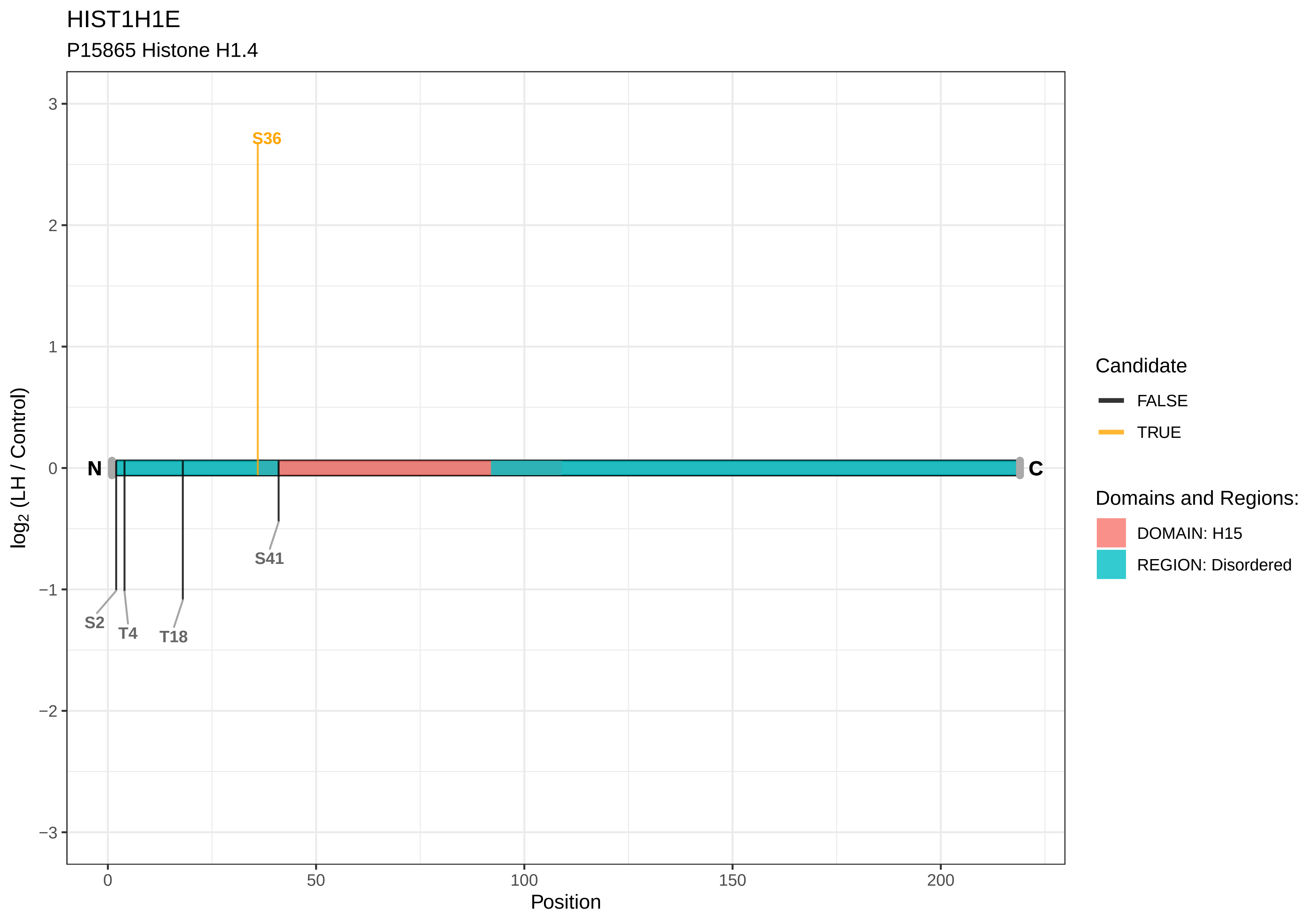
HIST1H1E
P15865 Histone H1.4
S36
S41
S2
T4
T18
N
C
3
2
1
0
−1
−2
−3
0
50
100
150
200
Position
log2 (LH / Control)
Candidate
FALSE
TRUE
Domains and Regions:
DOMAIN: H15
REGION: Disordered

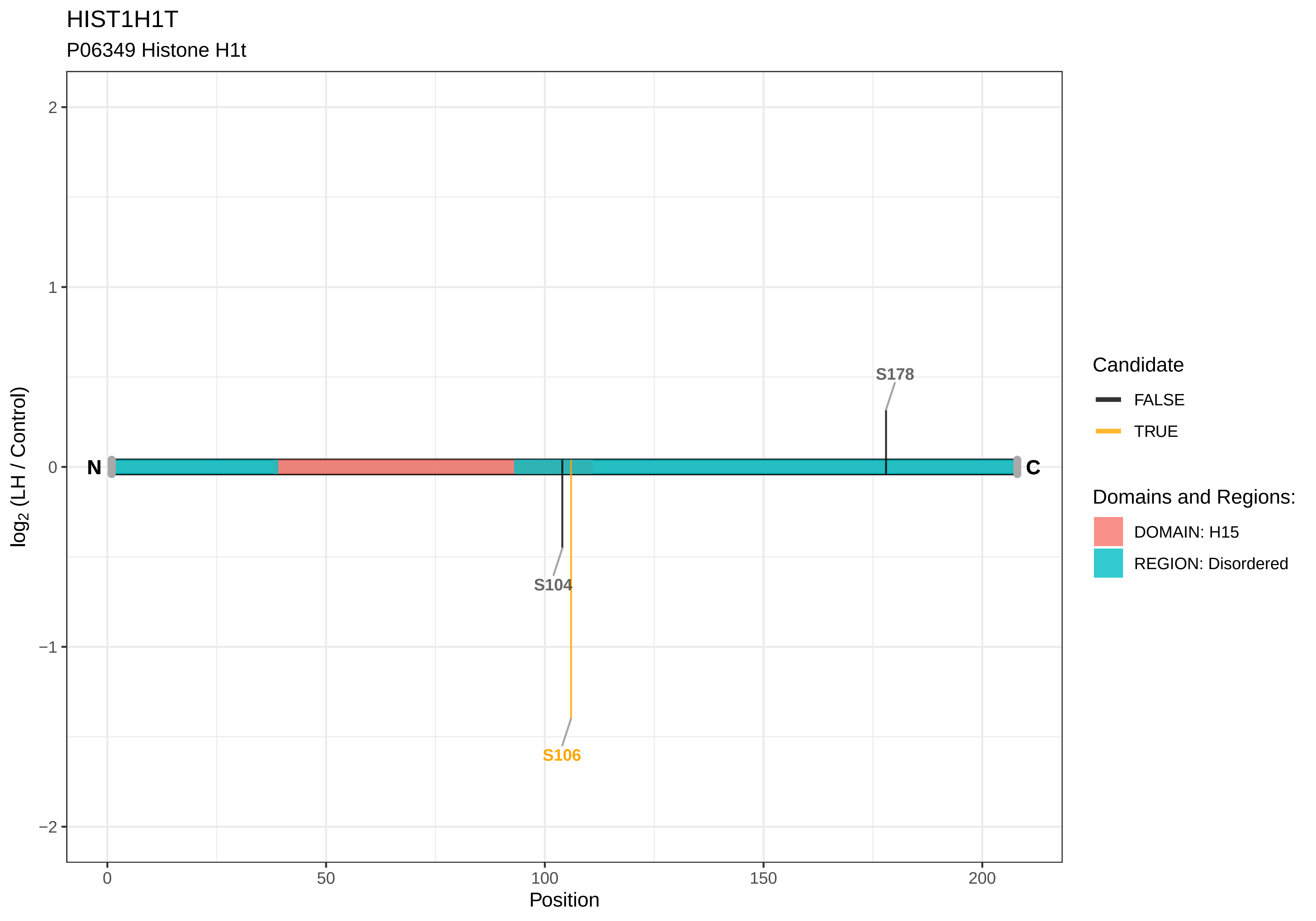
HIST1H1T
P06349 Histone H1t
log2 (LH / Control)
Position
N
C
S178
S104
S106
Candidate
FALSE
TRUE
Domains and Regions:
DOMAIN: H15
REGION: Disordered
2
1
0
−1
−2
0
50
100
150
200

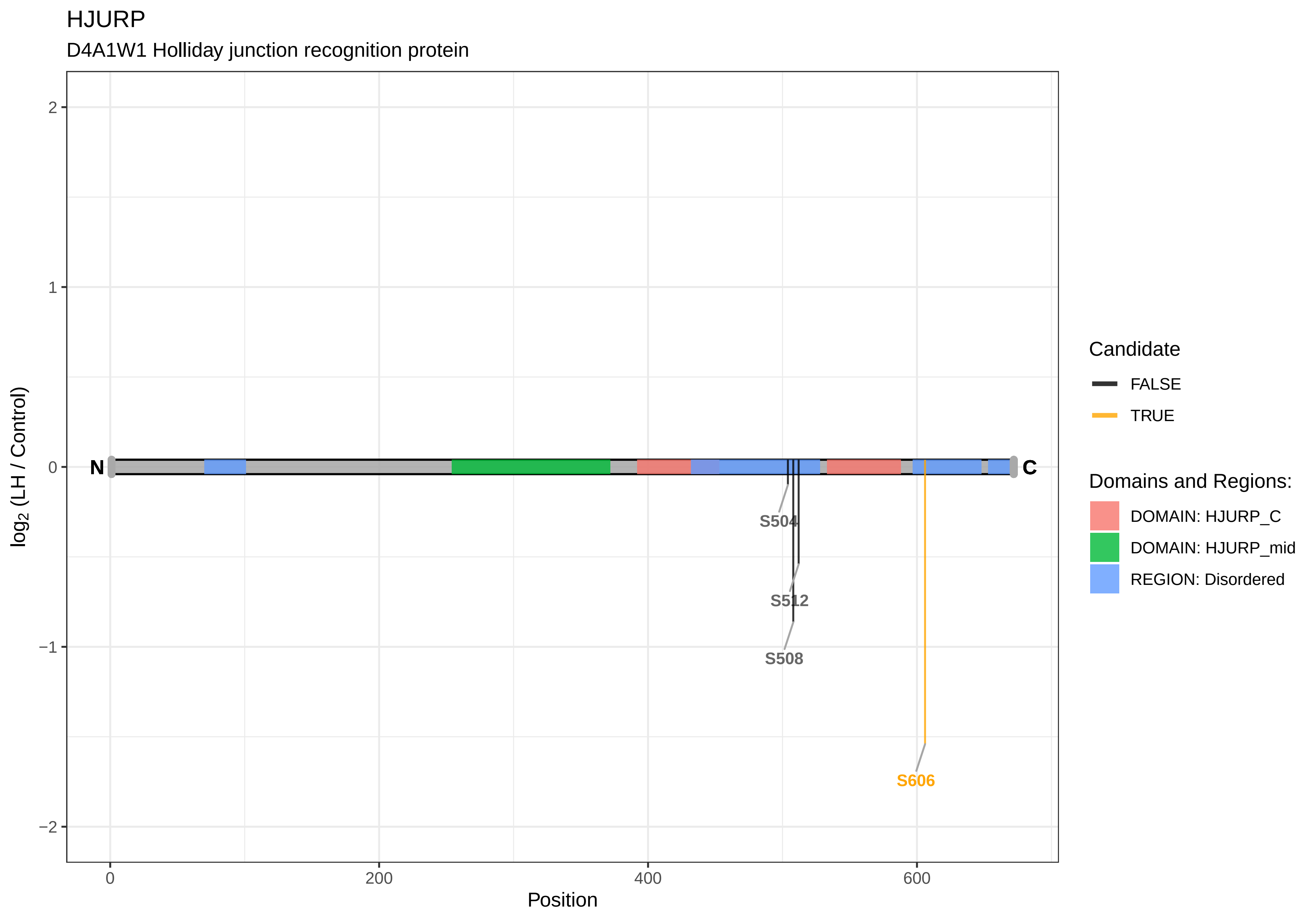
HJURP
D4A1W1 Holliday junction recognition protein
2
1
0
−1
−2
log2 (LH / Control)
N
C
S504
S512
S508
S606
0
200
400
600
Position
Candidate
FALSE
TRUE
Domains and Regions:
DOMAIN: HJURP_C
DOMAIN: HJURP_mid
REGION: Disordered

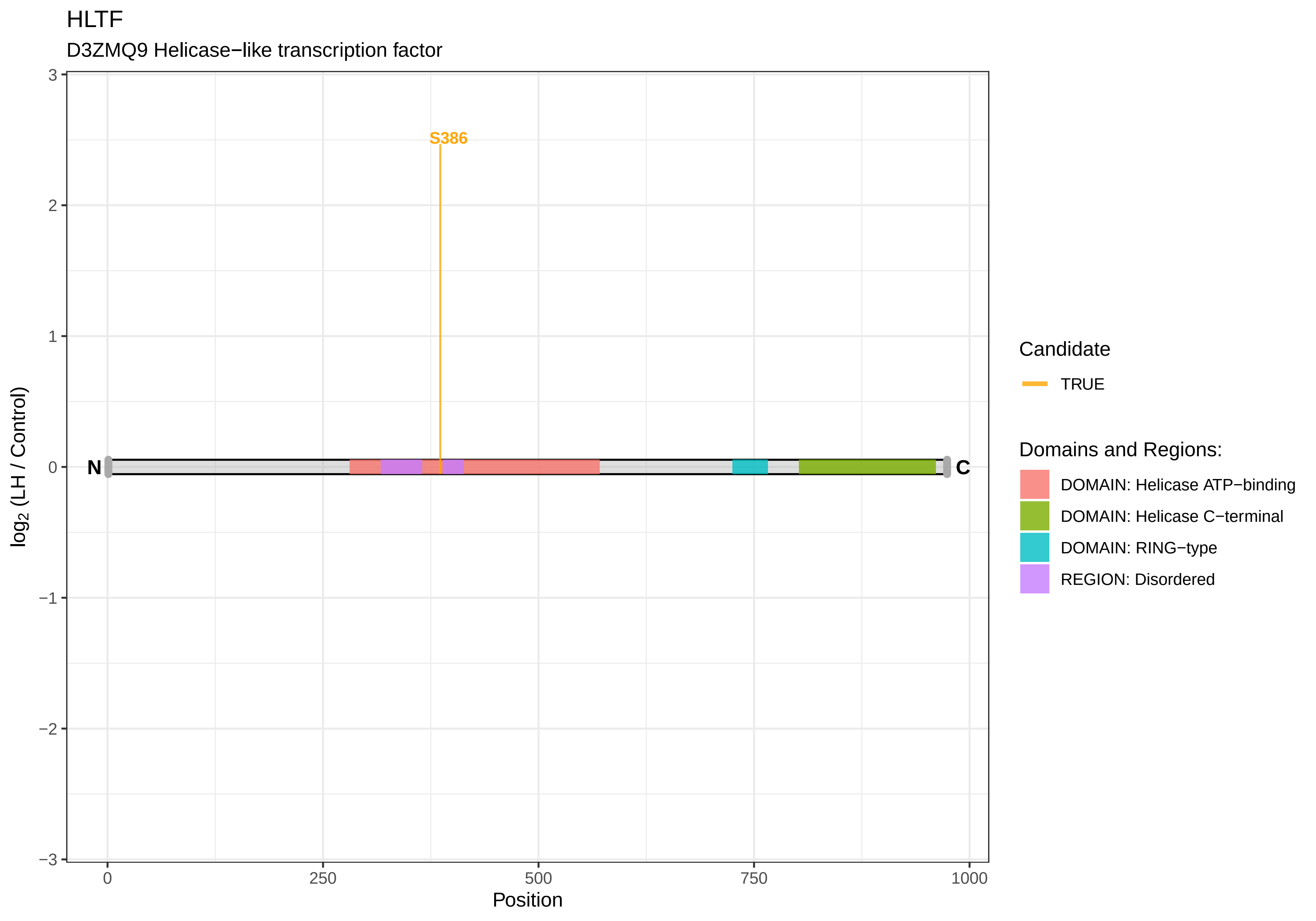

HLTF
D3ZMQ9 Helicase−like transcription factor
S386
N
C
3
2
1
0
−1
−2
−3
0
250
500
750
1000
Position
log2 (LH / Control)
Candidate
TRUE
Domains and Regions:
DOMAIN: Helicase ATP−binding
DOMAIN: Helicase C−terminal
DOMAIN: RING−type
REGION: Disordered

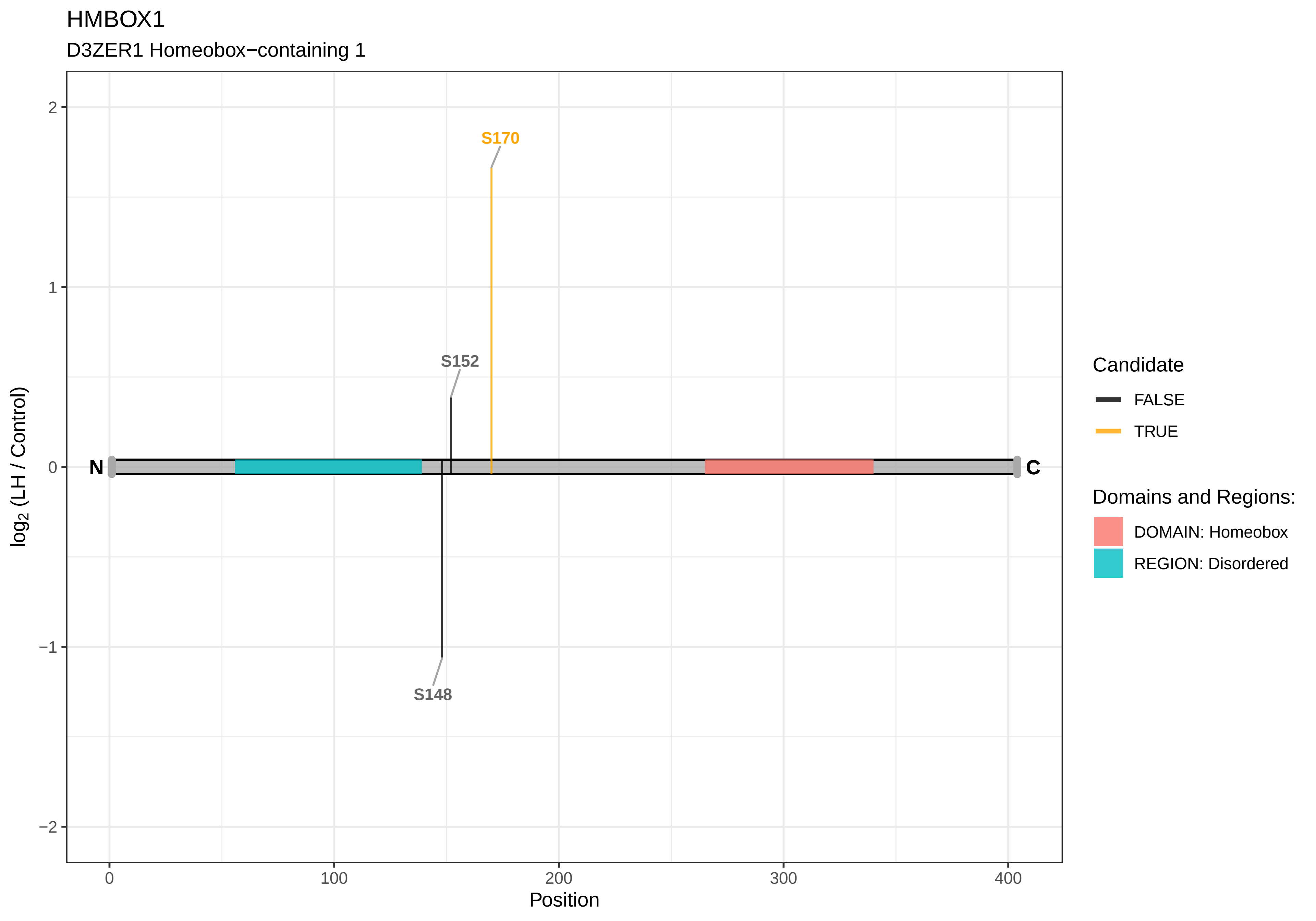
HMBOX1
D3ZER1 Homeobox−containing 1
S170
S152
S148
N
C
2
1
0
−1
−2
log2 (LH / Control)
0
100
200
300
400
Position
Candidate
FALSE
TRUE
Domains and Regions:
DOMAIN: Homeobox
REGION: Disordered

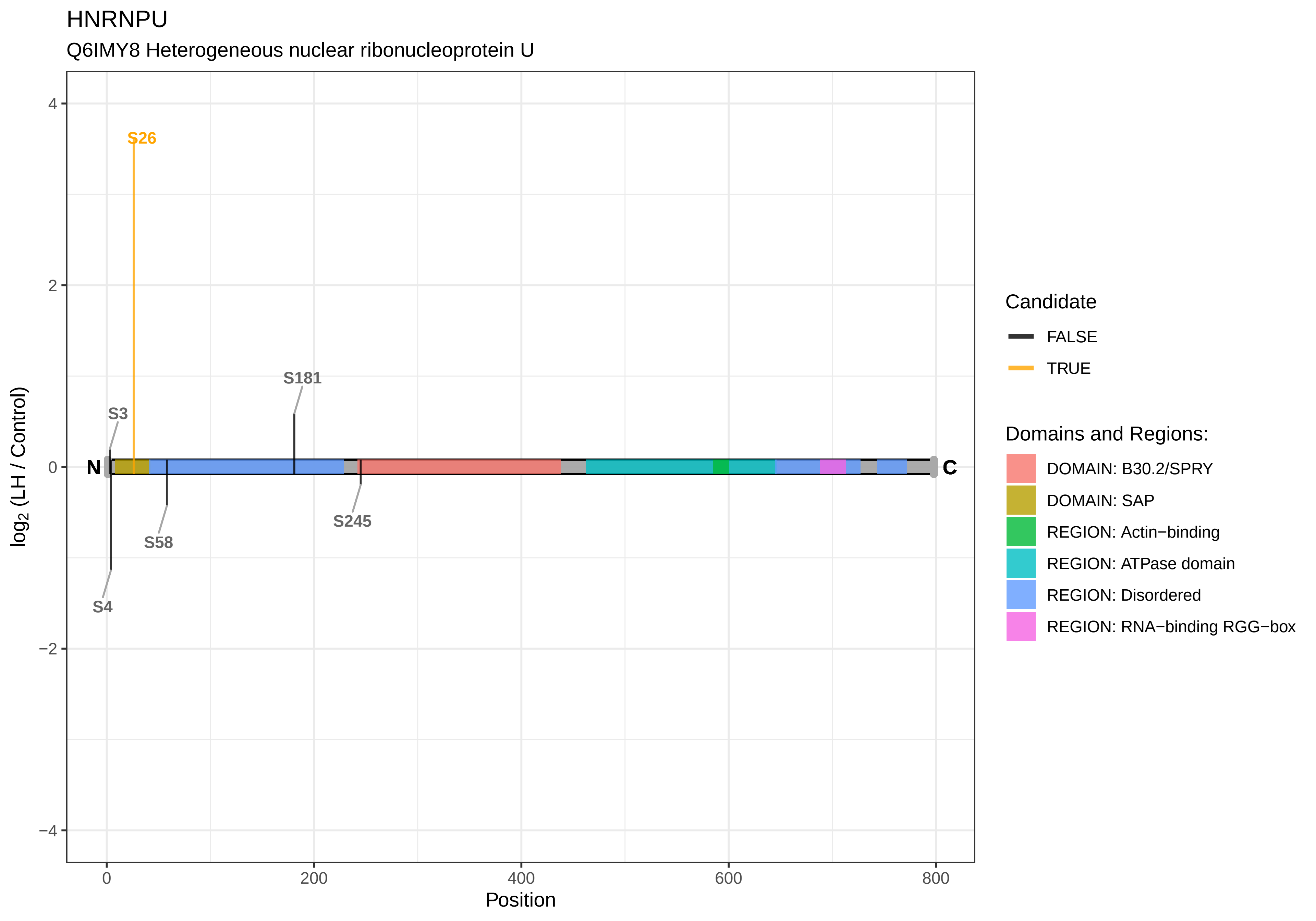
HNRNPU
Q6IMY8 Heterogeneous nuclear ribonucleoprotein U
S26
S181
S3
S58
S245
S4
N
C
4
2
0
−2
−4
0
200
400
600
800
Position
log2 (LH / Control)
Candidate
FALSE
TRUE
Domains and Regions:
DOMAIN: B30.2/SPRY
DOMAIN: SAP
REGION: Actin−binding
REGION: ATPase domain
REGION: Disordered
REGION: RNA−binding RGG−box

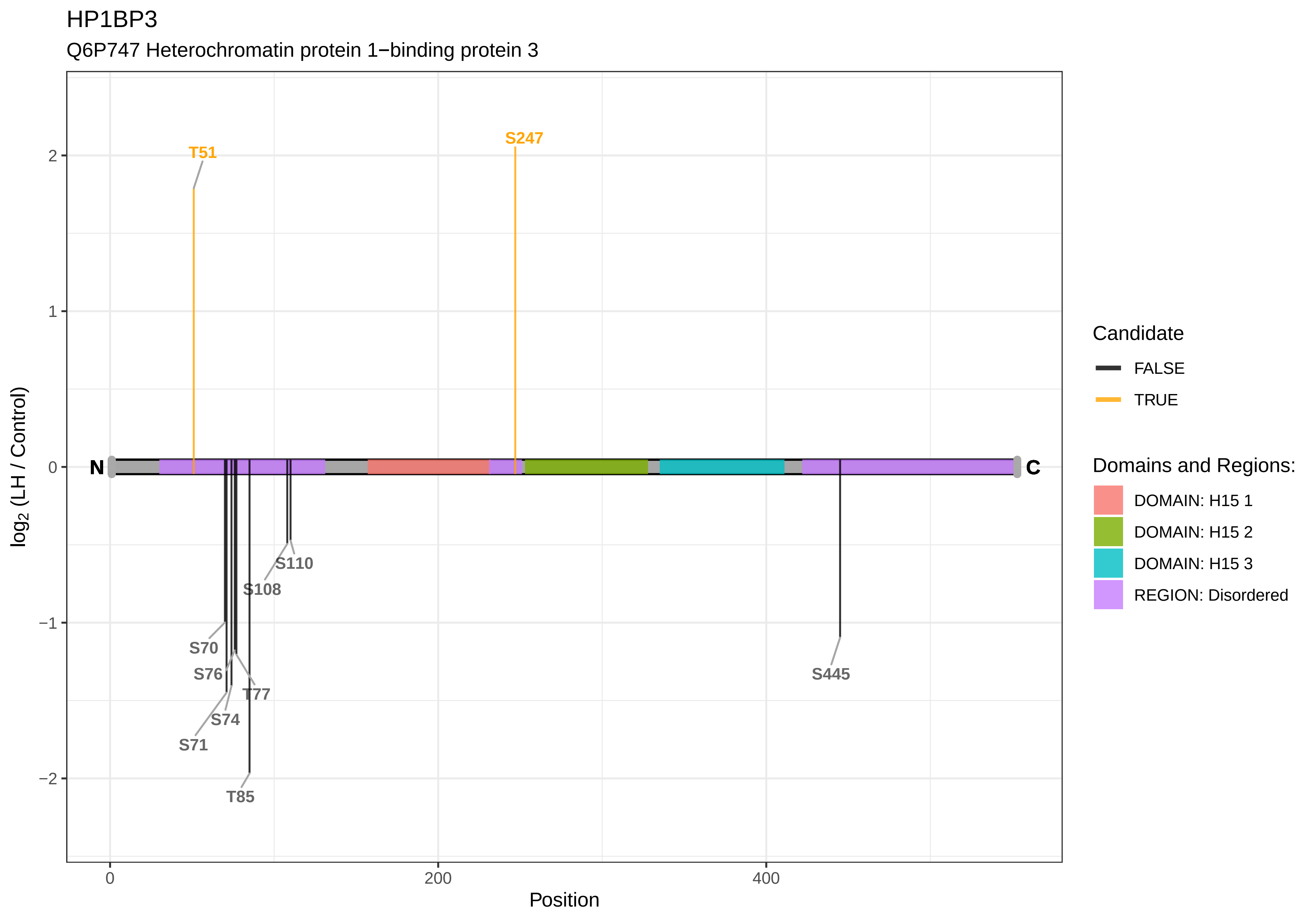

HP1BP3
Q6P747 Heterochromatin protein 1−binding protein 3
T51
S247
S110
S108
S70
S76
T77
S74
S71
T85
S445
N
C
2
1
0
−1
−2
log2 (LH / Control)
0
200
400
Position
Candidate
FALSE
TRUE
Domains and Regions:
DOMAIN: H15 1
DOMAIN: H15 2
DOMAIN: H15 3
REGION: Disordered

HSF1

F1MAF1 Heat shock transcription factor 1

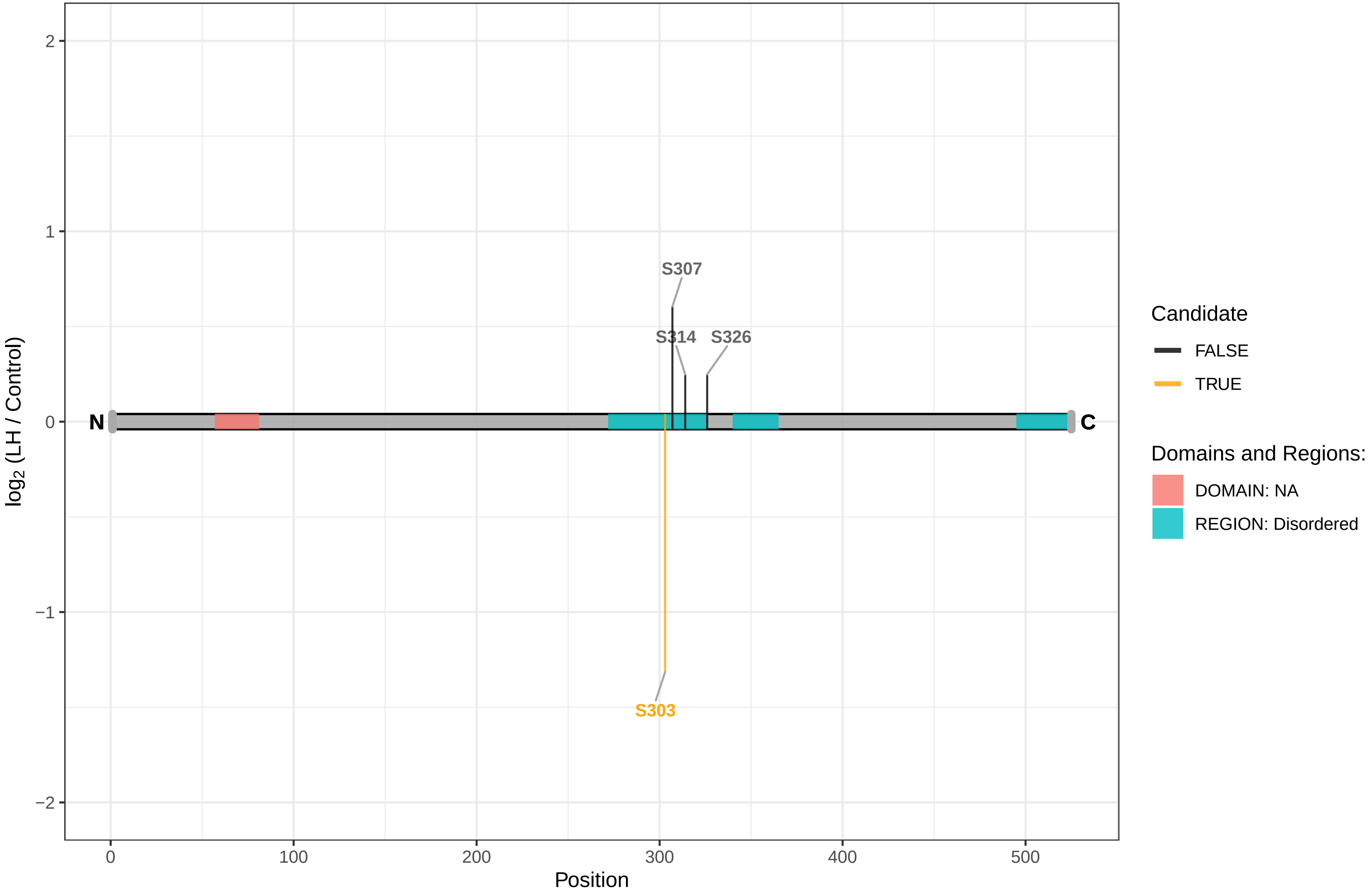

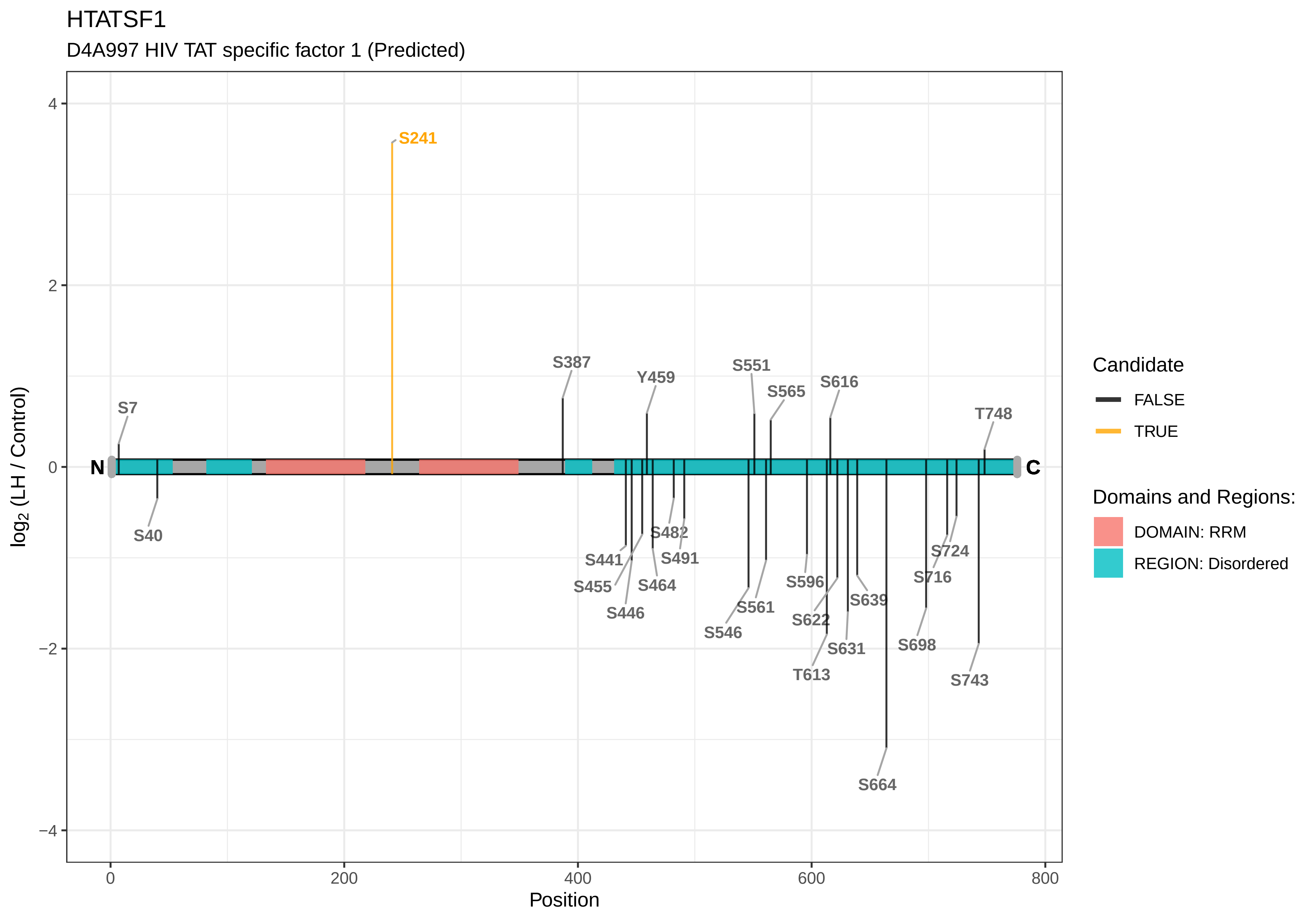

HTATSF1
D4A997 HIV TAT specific factor 1 (Predicted)
log2 (LH / Control)
Position
S241
S387
Y459
S551
S565
S616
T748
S7
S40
S482
S491
S441
S455
S464
S446
S596
S561
S546
S622
S639
S631
T613
S724
S716
S698
S743
S664
N
C
Candidate
FALSE
TRUE
Domains and Regions:
DOMAIN: RRM
REGION: Disordered
4
2
0
−2
−4
0
200
400
600
800

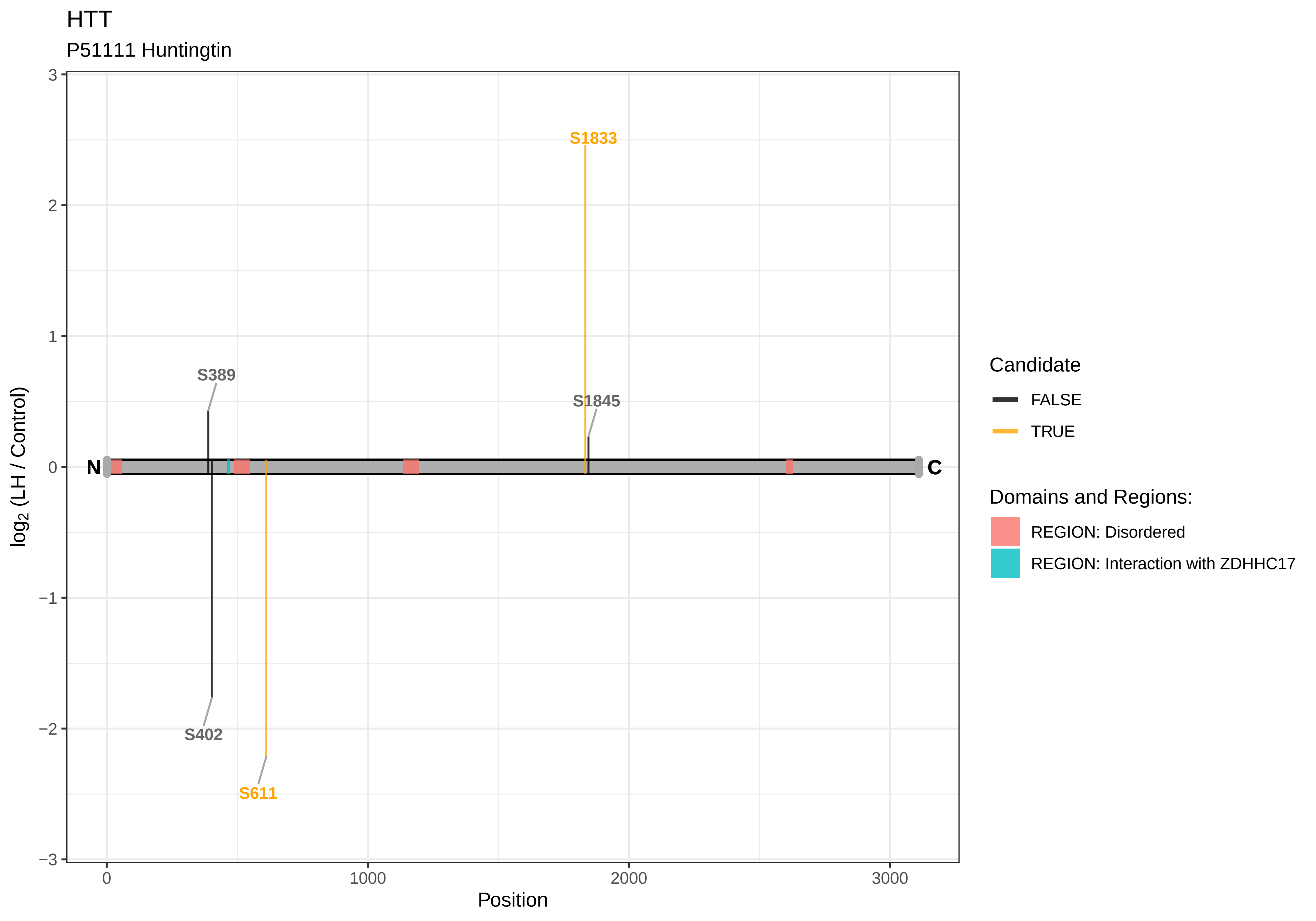

HTT
P51111 Huntingtin
S1833
S389
S1845
S402
S611
N
C
3
2
1
0
−1
−2
−3
0
1000
2000
3000
Position
log2 (LH / Control)
Candidate
FALSE
TRUE
Domains and Regions:
REGION: Disordered
REGION: Interaction with ZDHHC17

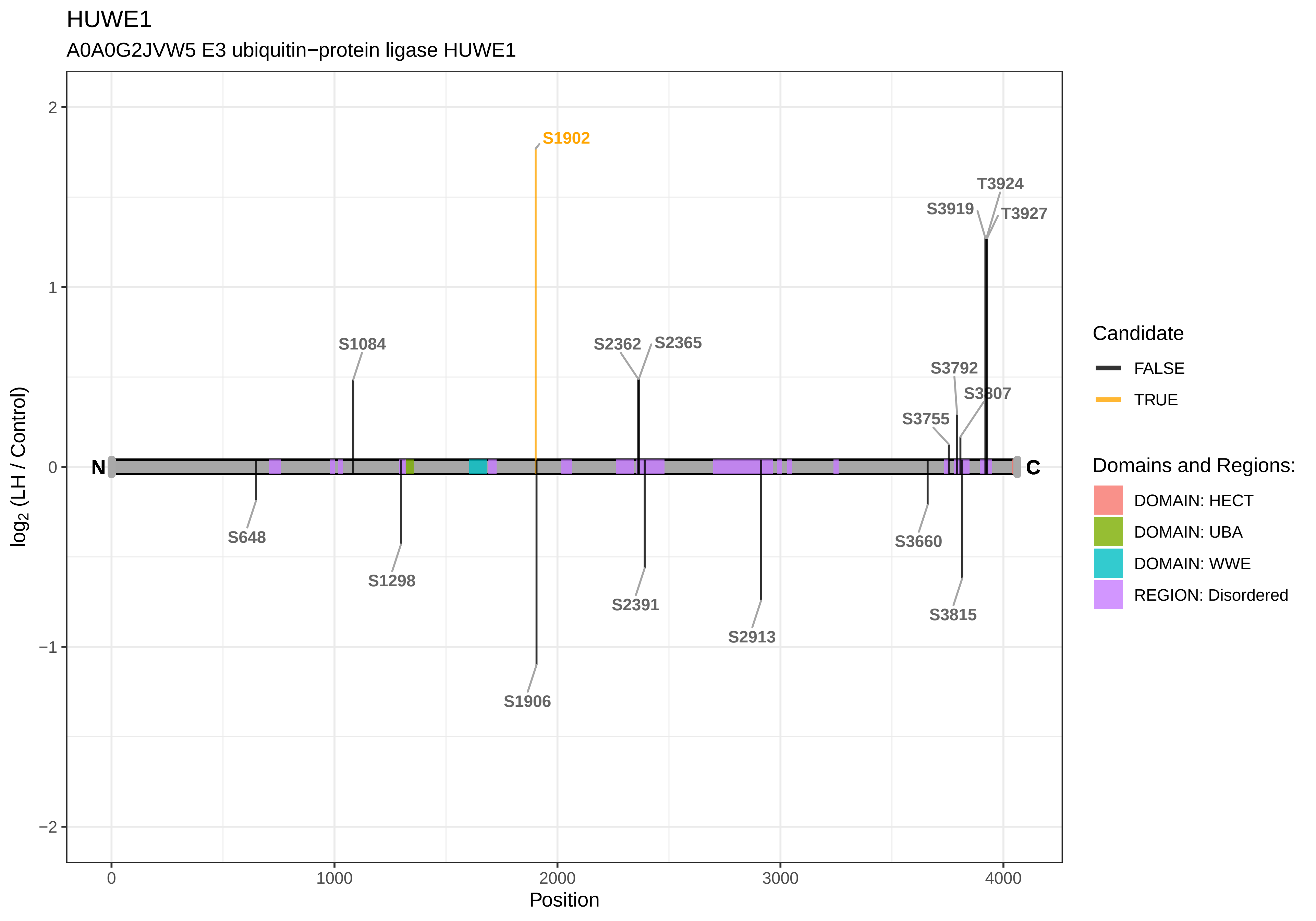

HUWE1
A0A0G2JVW5 E3 ubiquitin−protein ligase HUWE1
S1902
T3924
S3919
T3927
S1084
S2362
S2365
S3792
S3807
S3755
N
C
S648
S1298
S1906
S2391
S2913
S3660
S3815
log2 (LH / Control)
Position
Candidate
FALSE
TRUE
Domains and Regions:
DOMAIN: HECT
DOMAIN: UBA
DOMAIN: WWE
REGION: Disordered

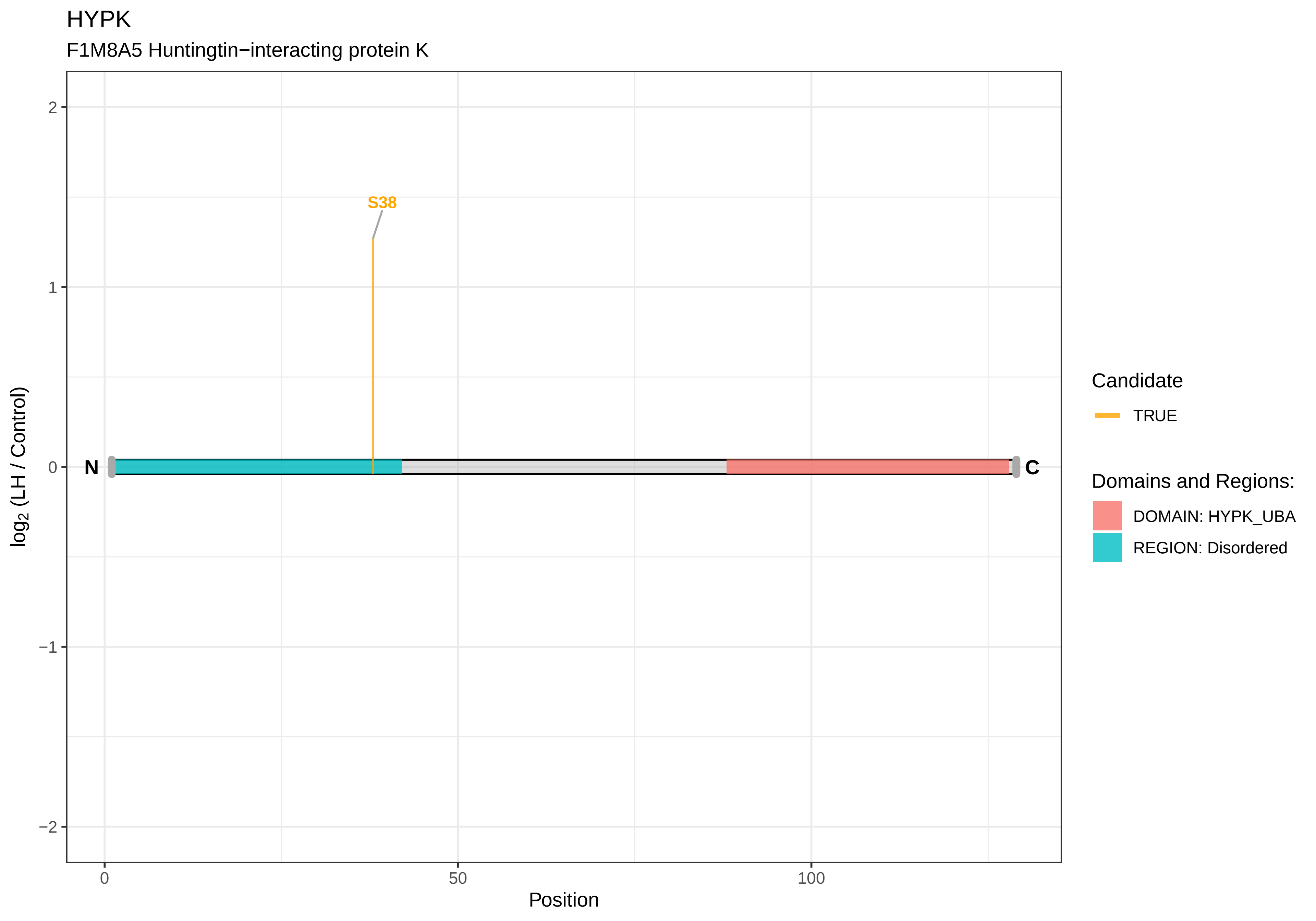

HYPK
F1M8A5 Huntingtin−interacting protein K
S38
N
C
2
1
0
−1
−2
0
50
100
Position
log2 (LH / Control)
Candidate
TRUE
Domains and Regions:
DOMAIN: HYPK_UBA
REGION: Disordered

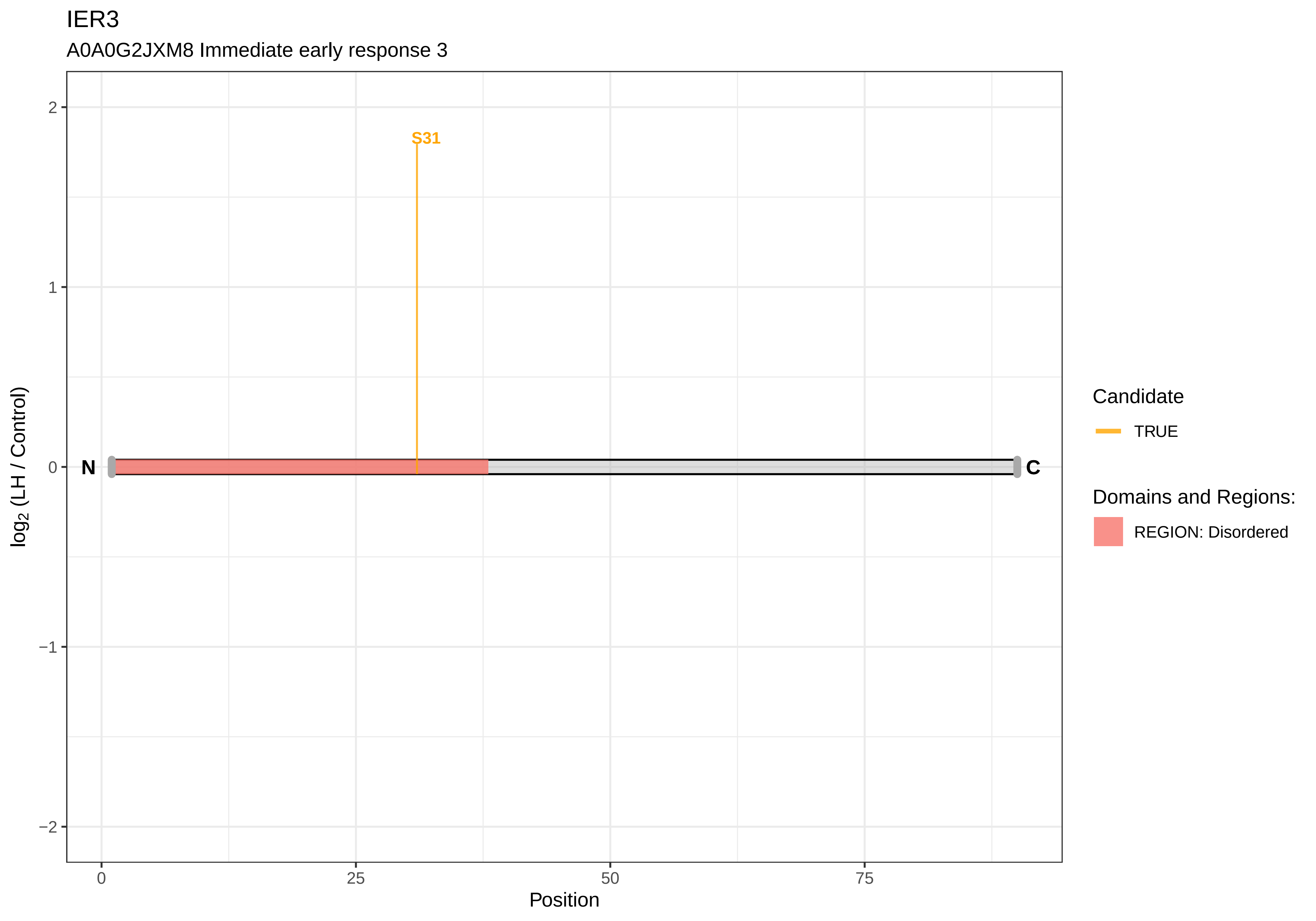

IER3
A0A0G2JXM8 Immediate early response 3
S31
N
C
2
1
0
−1
−2
log2 (LH / Control)
0
25
50
75
Position
Candidate
TRUE
Domains and Regions:
REGION: Disordered

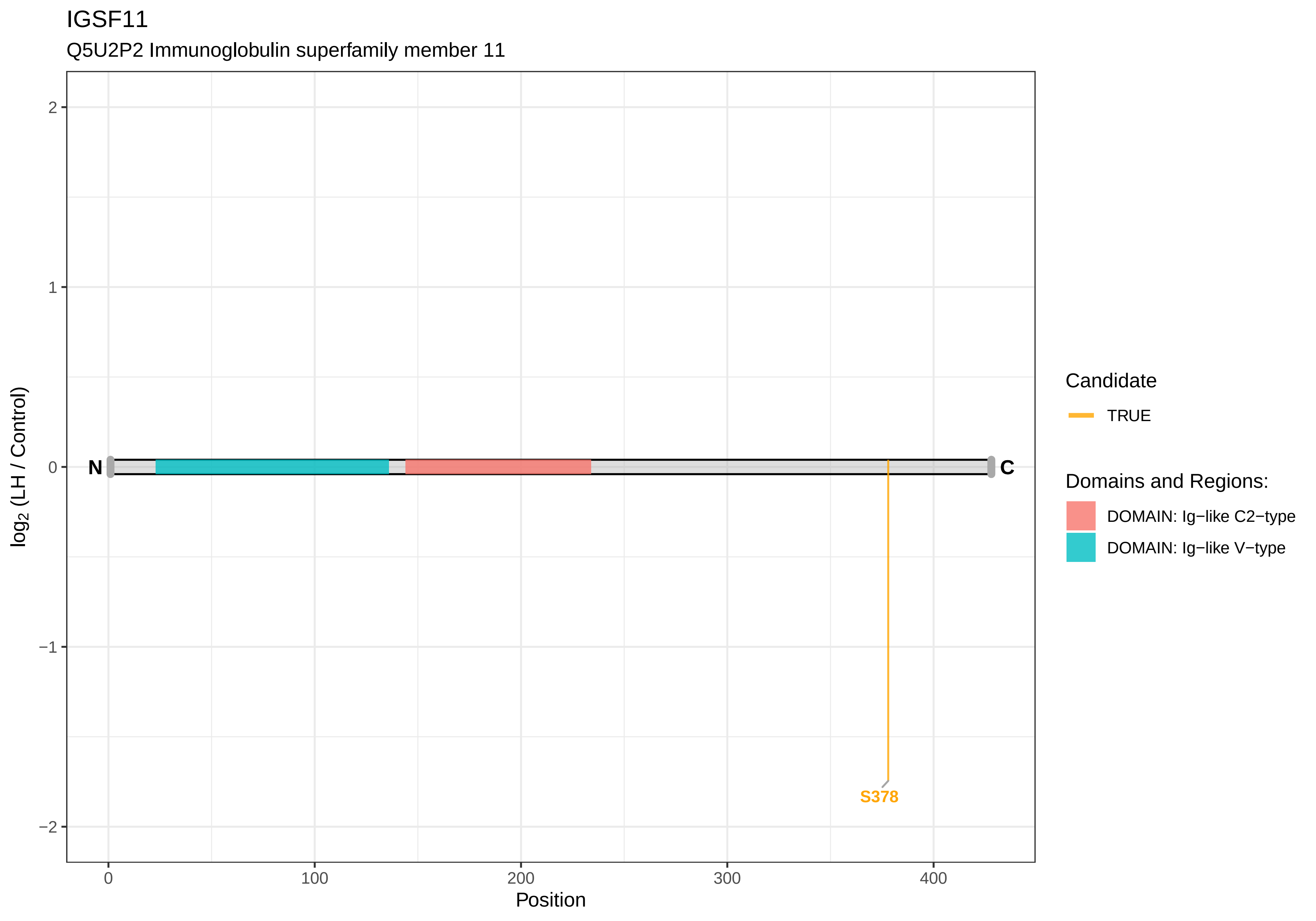

IGSF11
Q5U2P2 Immunoglobulin superfamily member 11
log2 (LH / Control)
2
1
0
−1
−2
N
C
S378
0
100
200
300
400
Position
Candidate
TRUE
Domains and Regions:
DOMAIN: Ig−like C2−type
DOMAIN: Ig−like V−type

IK

Q66HG8 Protein Red

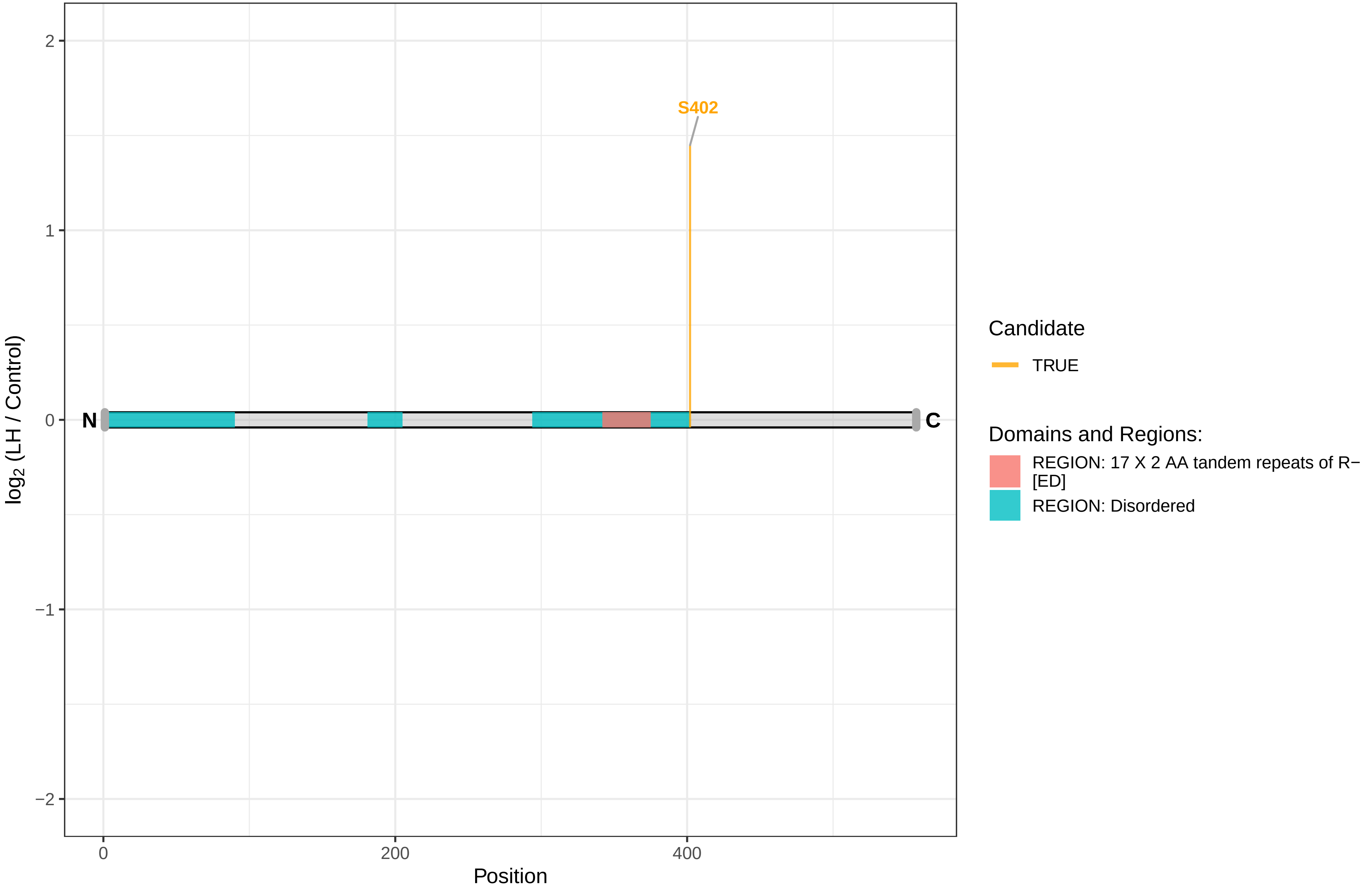

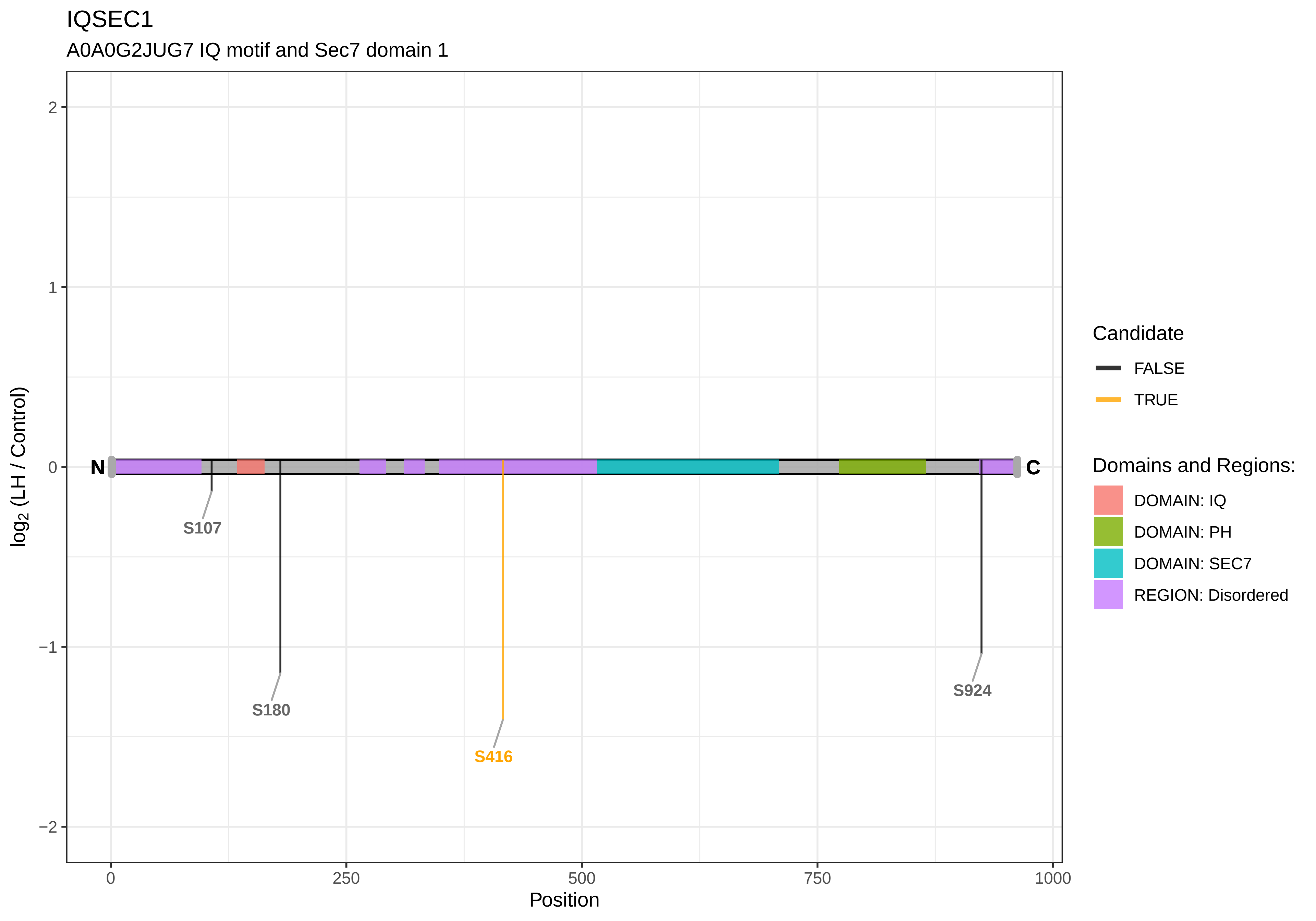

IQSEC1
A0A0G2JUG7 IQ motif and Sec7 domain 1
2
1
0
−1
−2
log2 (LH / Control)
N
C
S107
S180
S416
S924
0
250
500
750
1000
Position
Candidate
FALSE
TRUE
Domains and Regions:
DOMAIN: IQ
DOMAIN: PH
DOMAIN: SEC7
REGION: Disordered

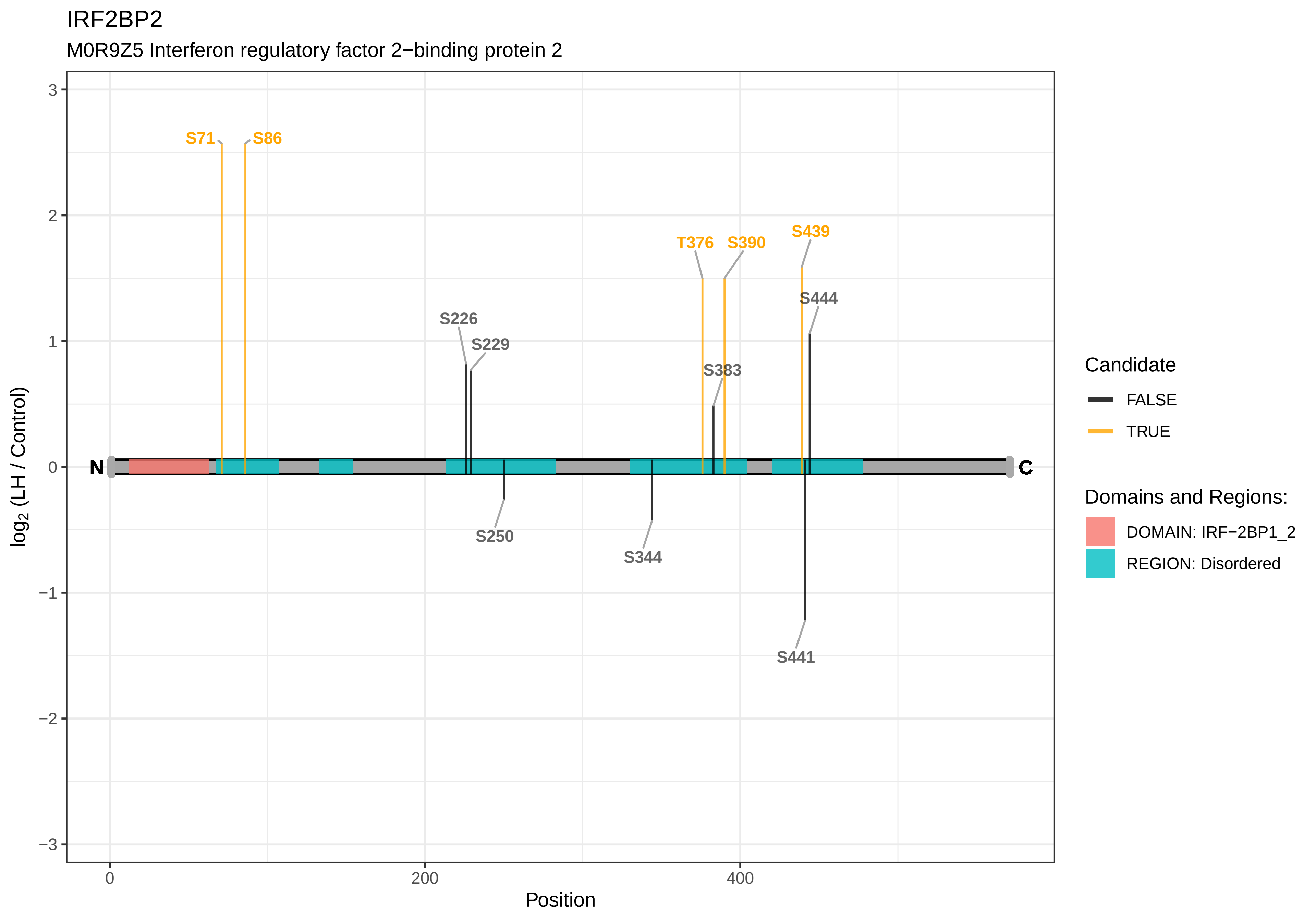
IRF2BP2
M0R9Z5 Interferon regulatory factor 2−binding protein 2
S71
S86
T376
S390
S439
S444
S226
S229
S383
N
C
S250
S344
S441
log2 (LH / Control)
Position
3
2
1
0
−1
−2
−3
0
200
400
Candidate
FALSE
TRUE
Domains and Regions:
DOMAIN: IRF−2BP1_2
REGION: Disordered

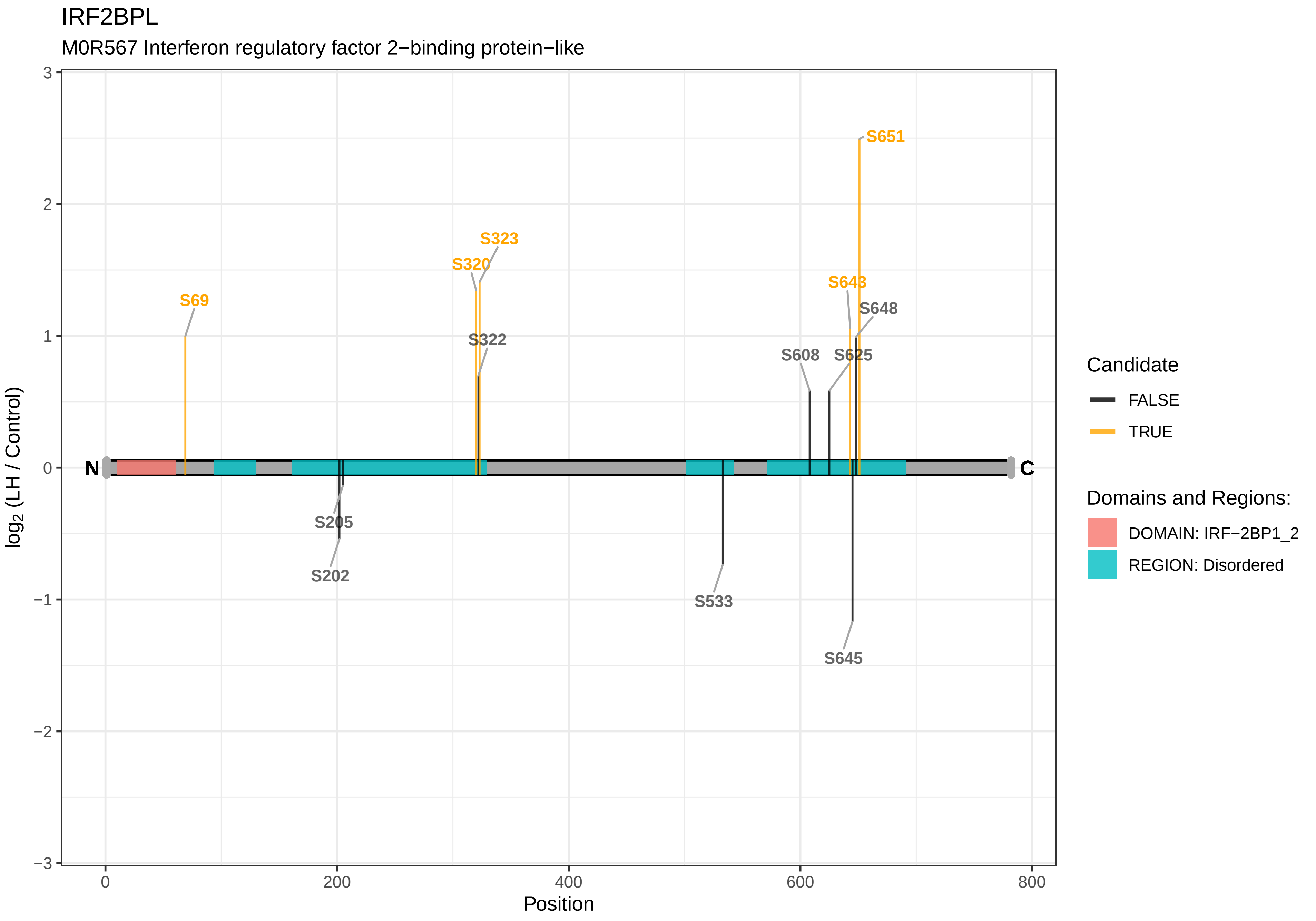

IRF2BPL
M0R567 Interferon regulatory factor 2−binding protein−like
S651
S323
S320
S643
S69
S648
S322
S608
S625
N
C
S205
S202
S533
S645
3
2
1
0
−1
−2
−3
log2 (LH / Control)
0
200
400
600
800
Position
Candidate
FALSE
TRUE
Domains and Regions:
DOMAIN: IRF−2BP1_2
REGION: Disordered

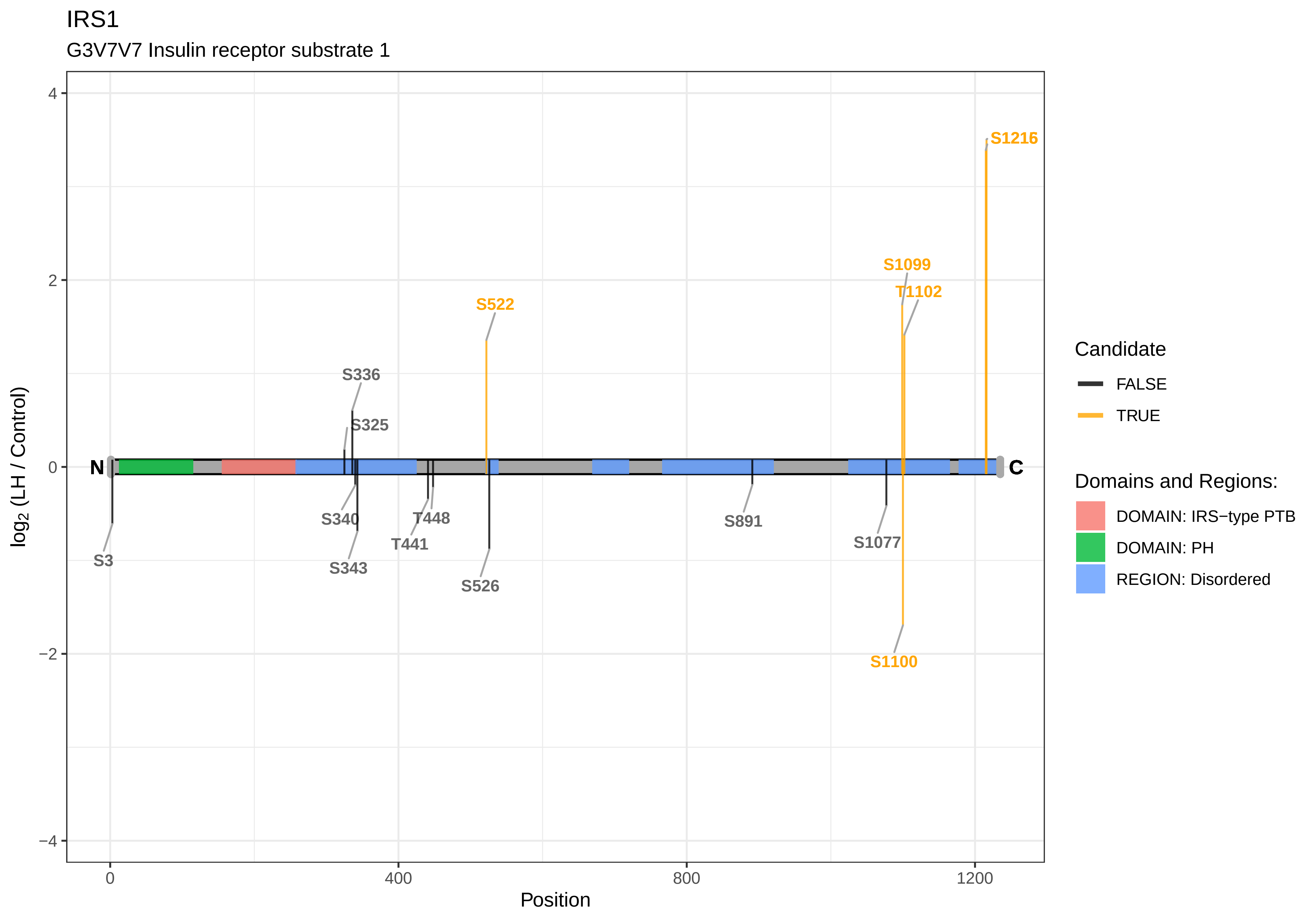
IRS1
G3V7V7 Insulin receptor substrate 1
S1216
S1099
T1102
S522
S336
S325
N
C
S340
T448
S891
S1077
T441
S3
S343
S526
S1100
log2 (LH / Control)
Position
4
2
0
−2
−4
0
400
800
1200
Candidate
FALSE
TRUE
Domains and Regions:
DOMAIN: IRS−type PTB
DOMAIN: PH
REGION: Disordered

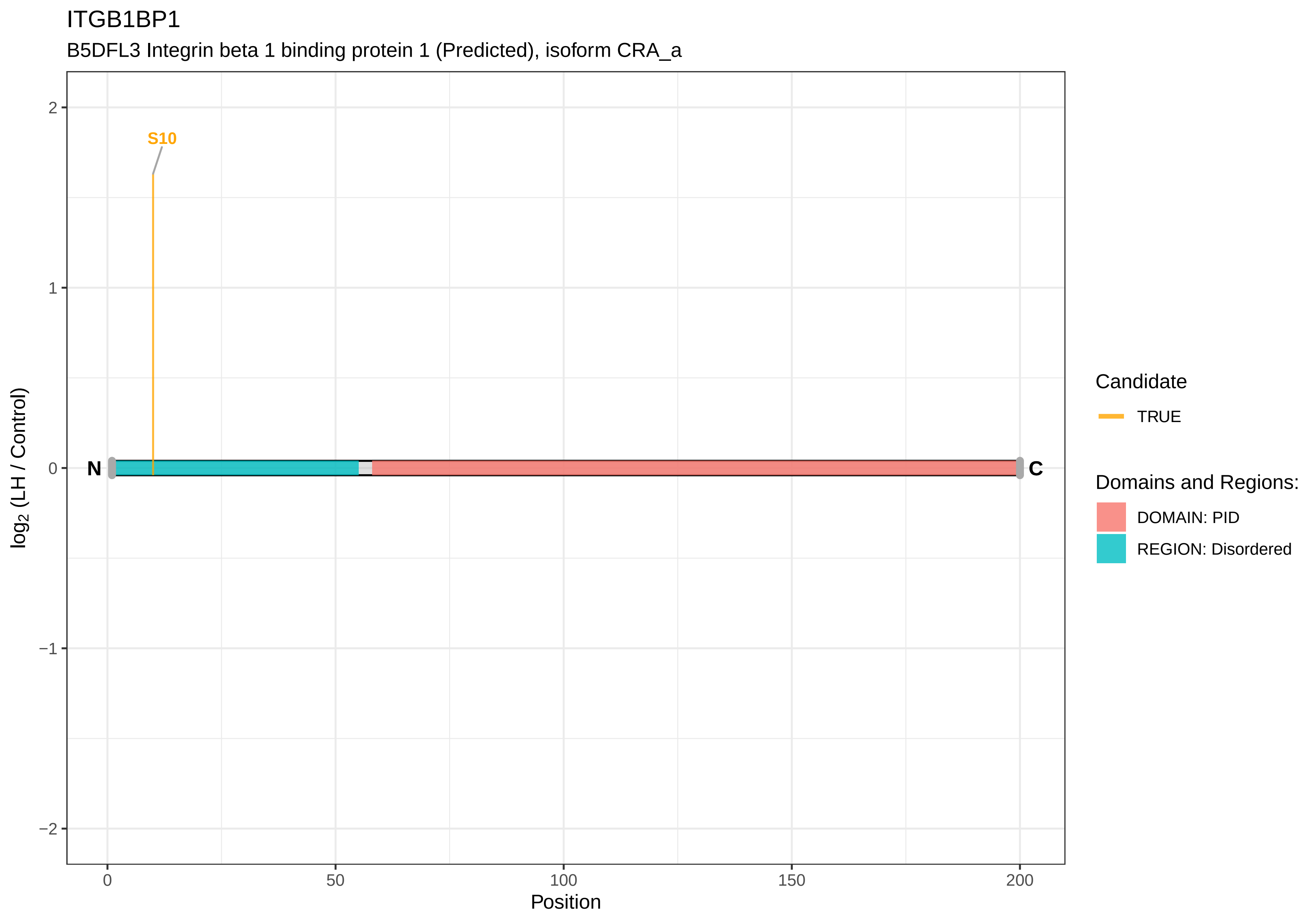
ITGB1BP1
B5DFL3 Integrin beta 1 binding protein 1 (Predicted), isoform CRA_a
S10
N
C
2
1
0
−1
−2
0
50
100
150
200
Position
log2 (LH / Control)
Candidate
TRUE
Domains and Regions:
DOMAIN: PID
REGION: Disordered

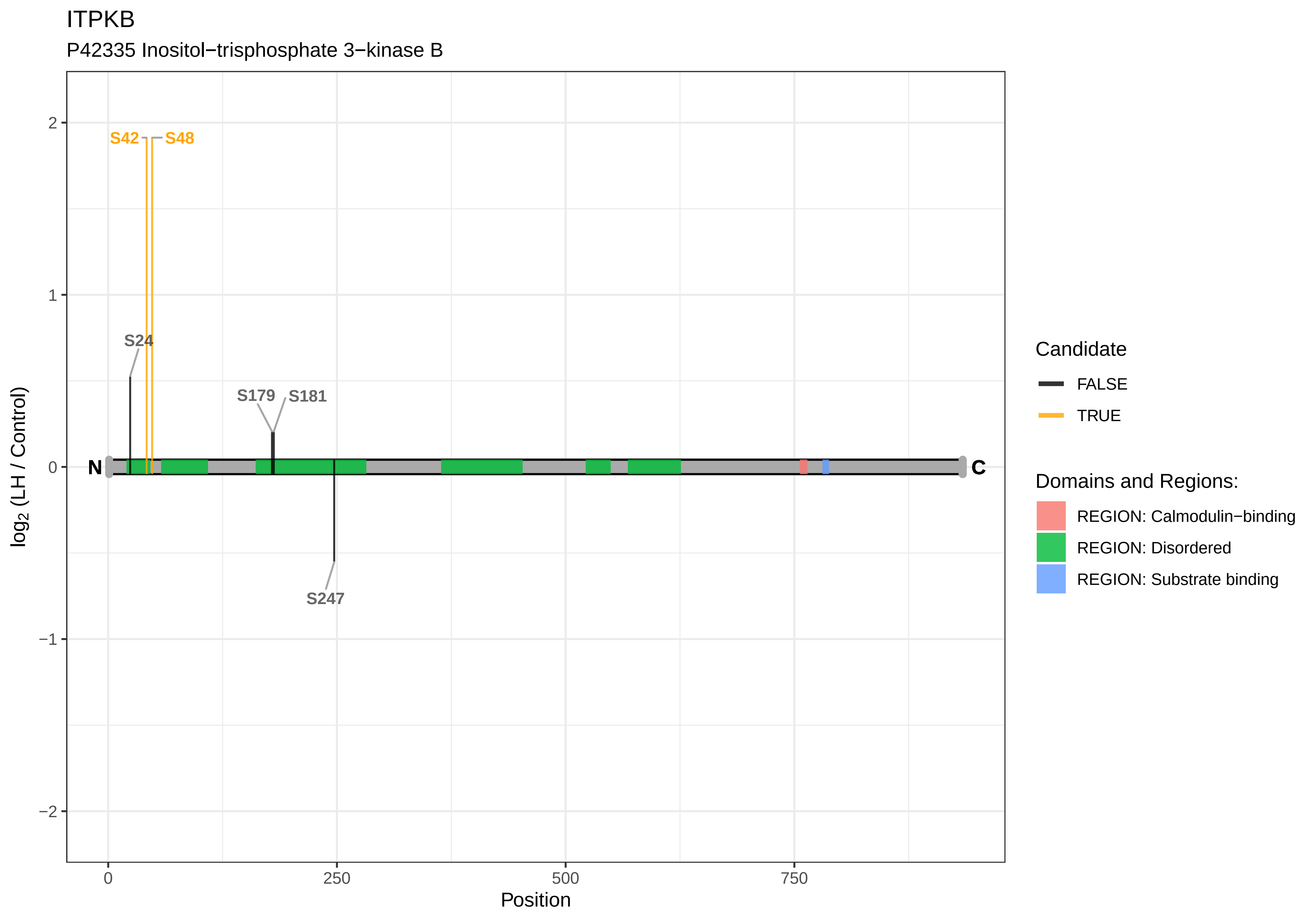
ITPKB
P42335 Inositol−trisphosphate 3−kinase B
S42
S48
S24
S179
S181
S247
N
C
2
1
0
−1
−2
log2 (LH / Control)
0
250
500
750
Position
Candidate
FALSE
TRUE
Domains and Regions:
REGION: Calmodulin−binding
REGION: Disordered
REGION: Substrate binding

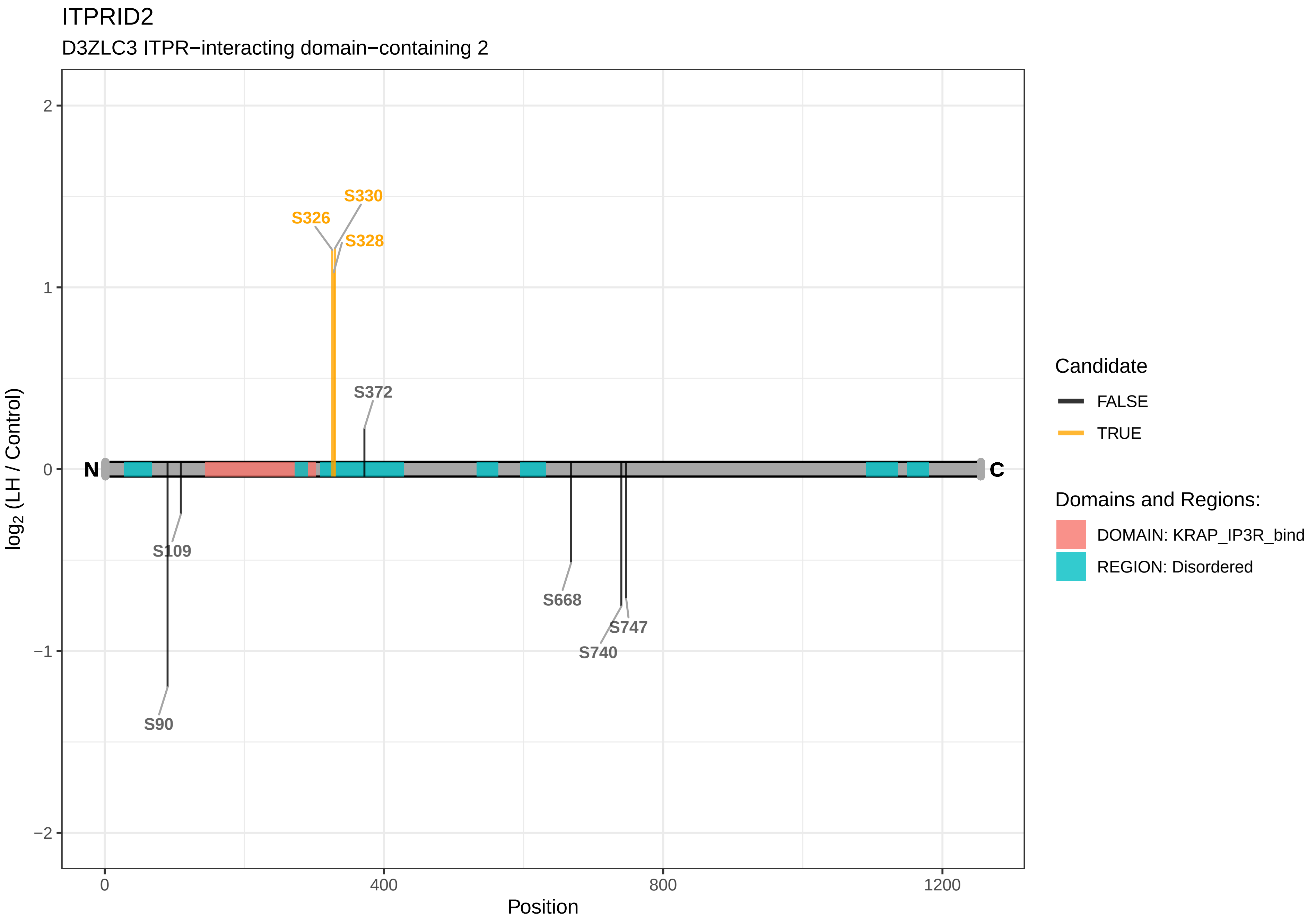
ITPRID2
D3ZLC3 ITPR−interacting domain−containing 2
S330
S326
S328
S372
S109
S668
S747
S740
S90
N
C
2
1
0
−1
−2
log2 (LH / Control)
0
400
800
1200
Position
Candidate
FALSE
TRUE
Domains and Regions:
DOMAIN: KRAP_IP3R_bind
REGION: Disordered

IWS1

Q3SWT4 Protein IWS1 homolog

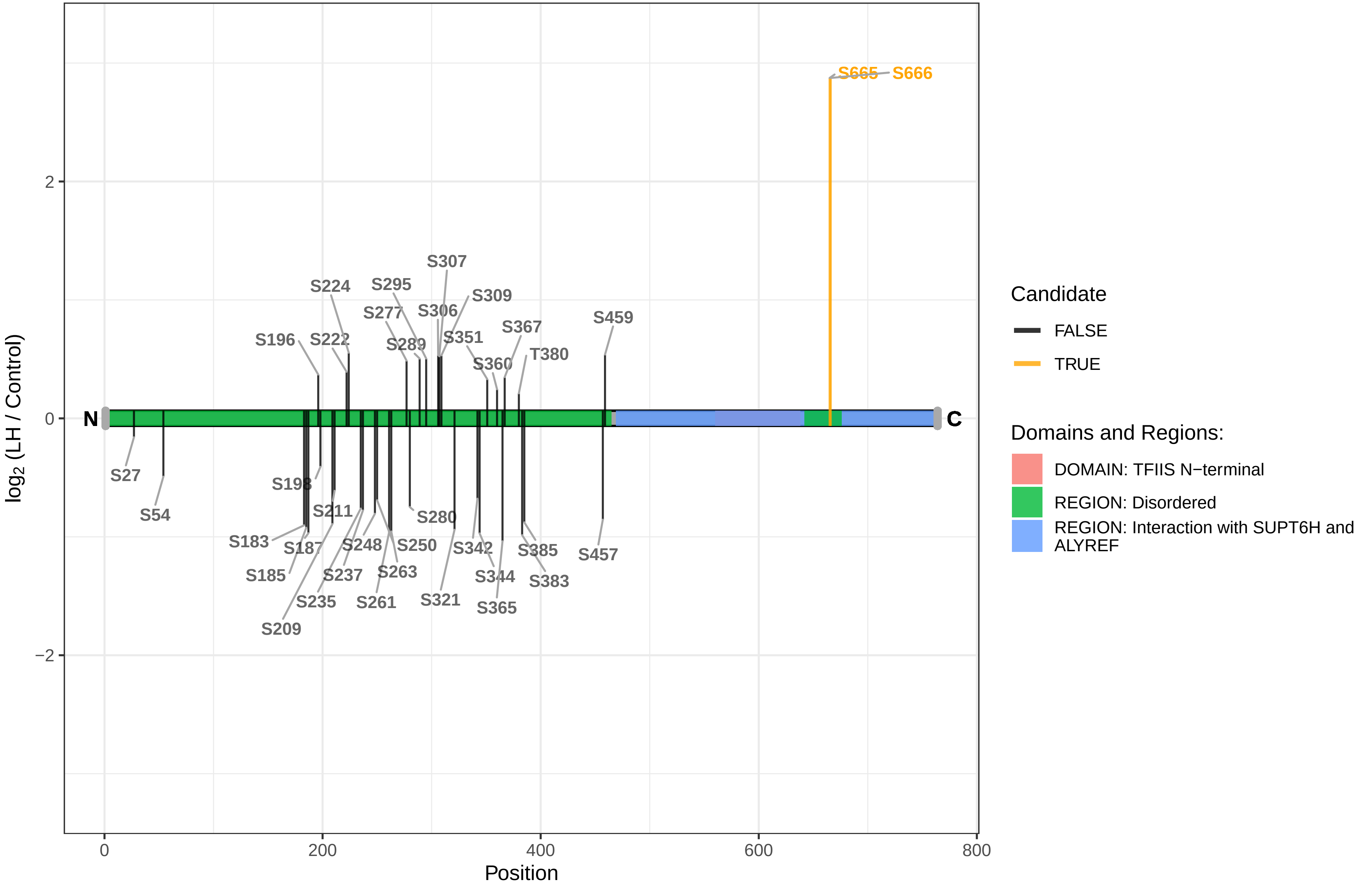

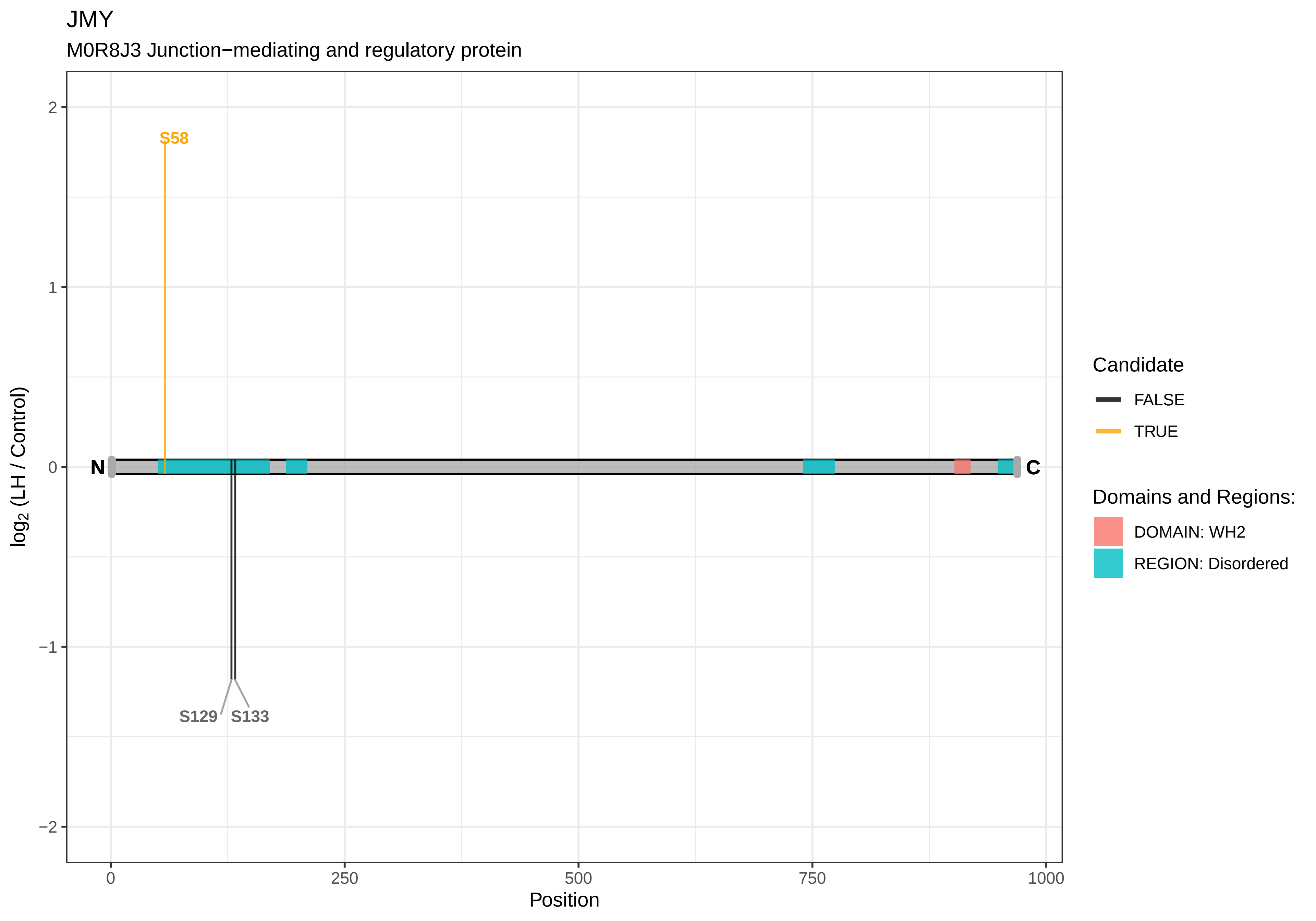

JMY
M0R8J3 Junction−mediating and regulatory protein
S58
S129
S133
N
C
2
1
0
−1
−2
log2 (LH / Control)
0
250
500
750
1000
Position
Candidate
FALSE
TRUE
Domains and Regions:
DOMAIN: WH2
REGION: Disordered

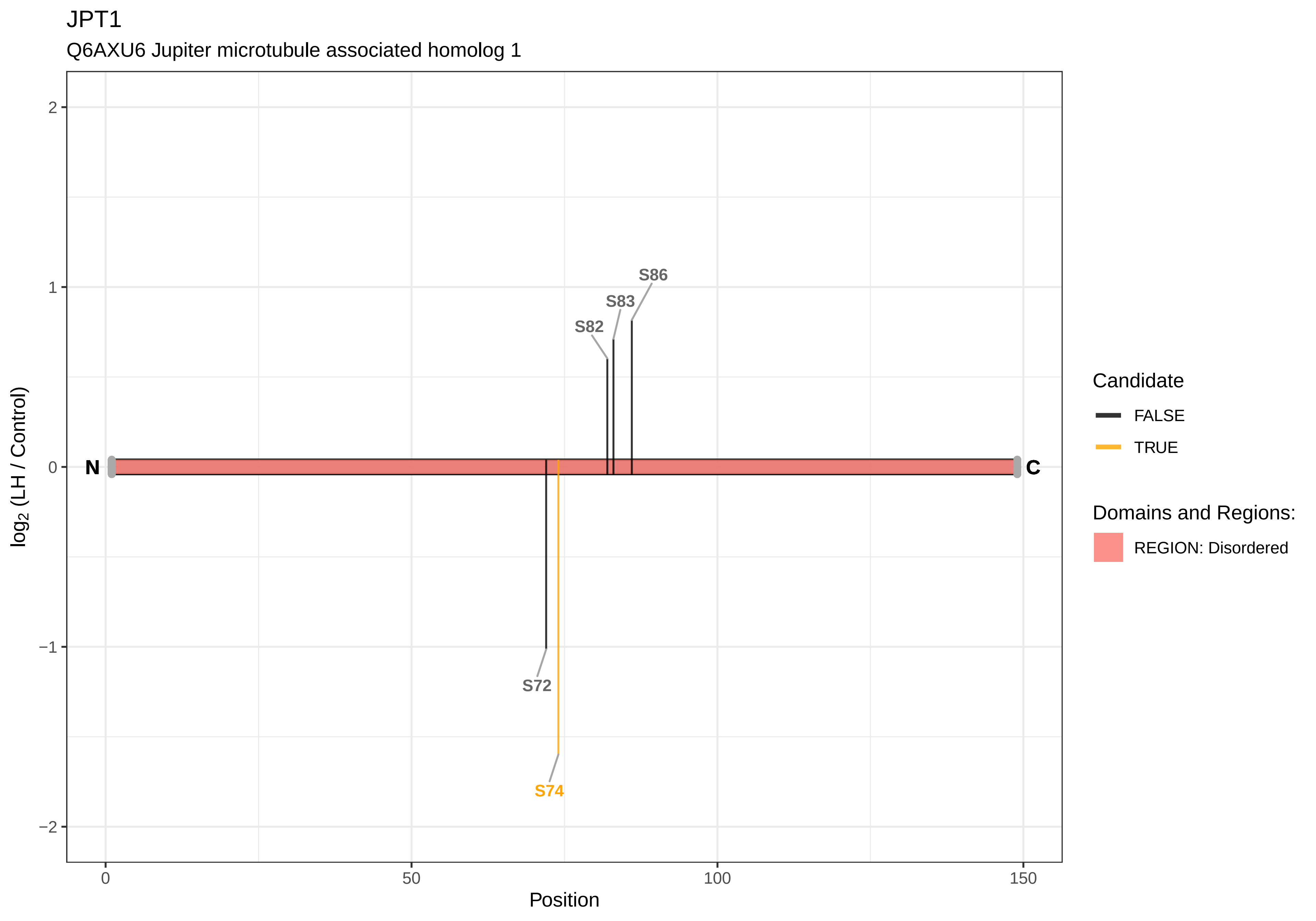
JPT1
Q6AXU6 Jupiter microtubule associated homolog 1
log2 (LH / Control)
Position
2
1
0
−1
−2
0
50
100
150
N
C
S86
S83
S82
S72
S74
Candidate
FALSE
TRUE
Domains and Regions:
REGION: Disordered

KALRN

P97924 Kalirin

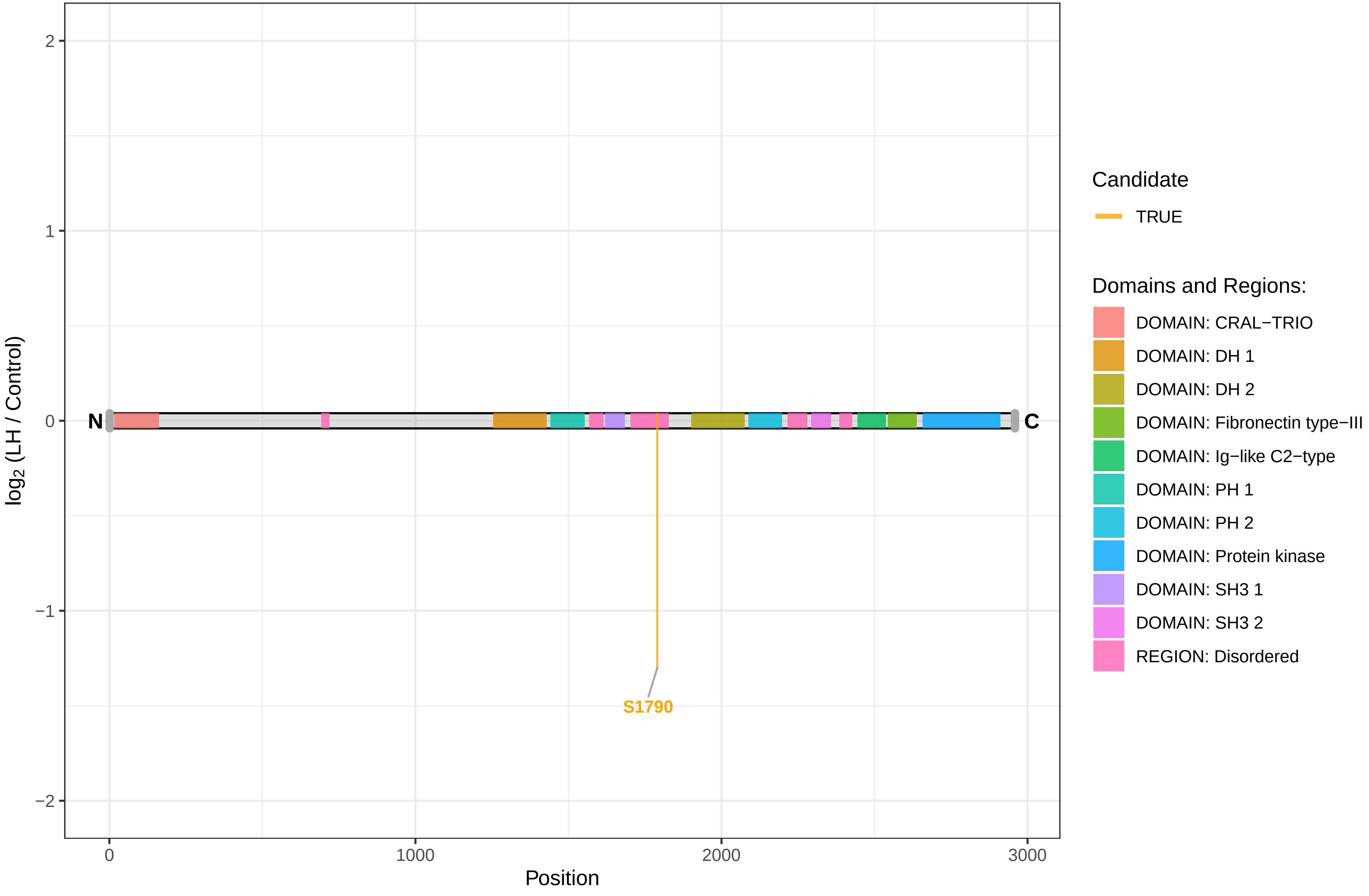

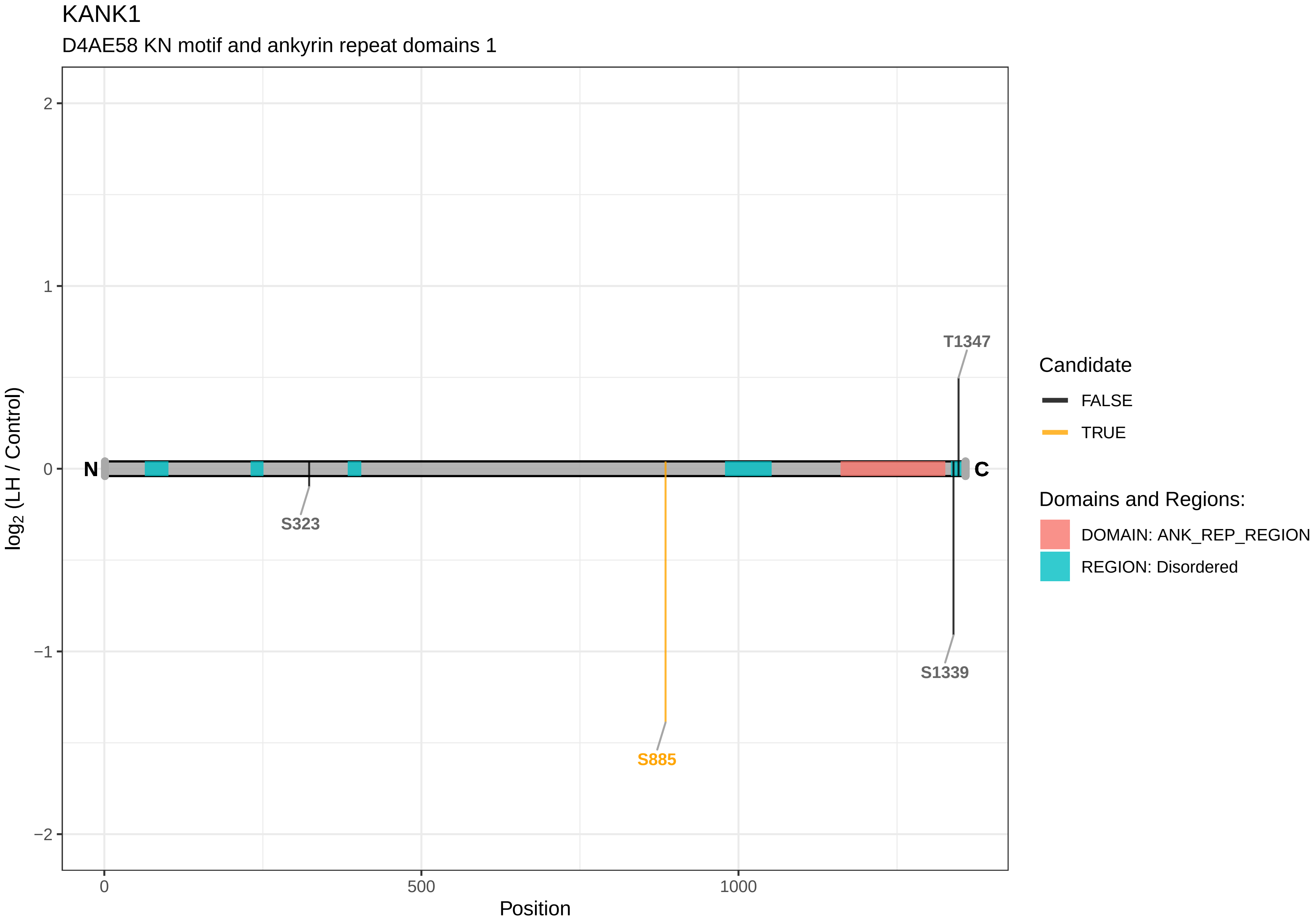
KANK1
D4AE58 KN motif and ankyrin repeat domains 1
log2 (LH / Control)
2
1
0
−1
−2
N
C
T1347
S323
S885
S1339
0
500
1000
Position
Candidate
FALSE
TRUE
Domains and Regions:
DOMAIN: ANK_REP_REGION
REGION: Disordered

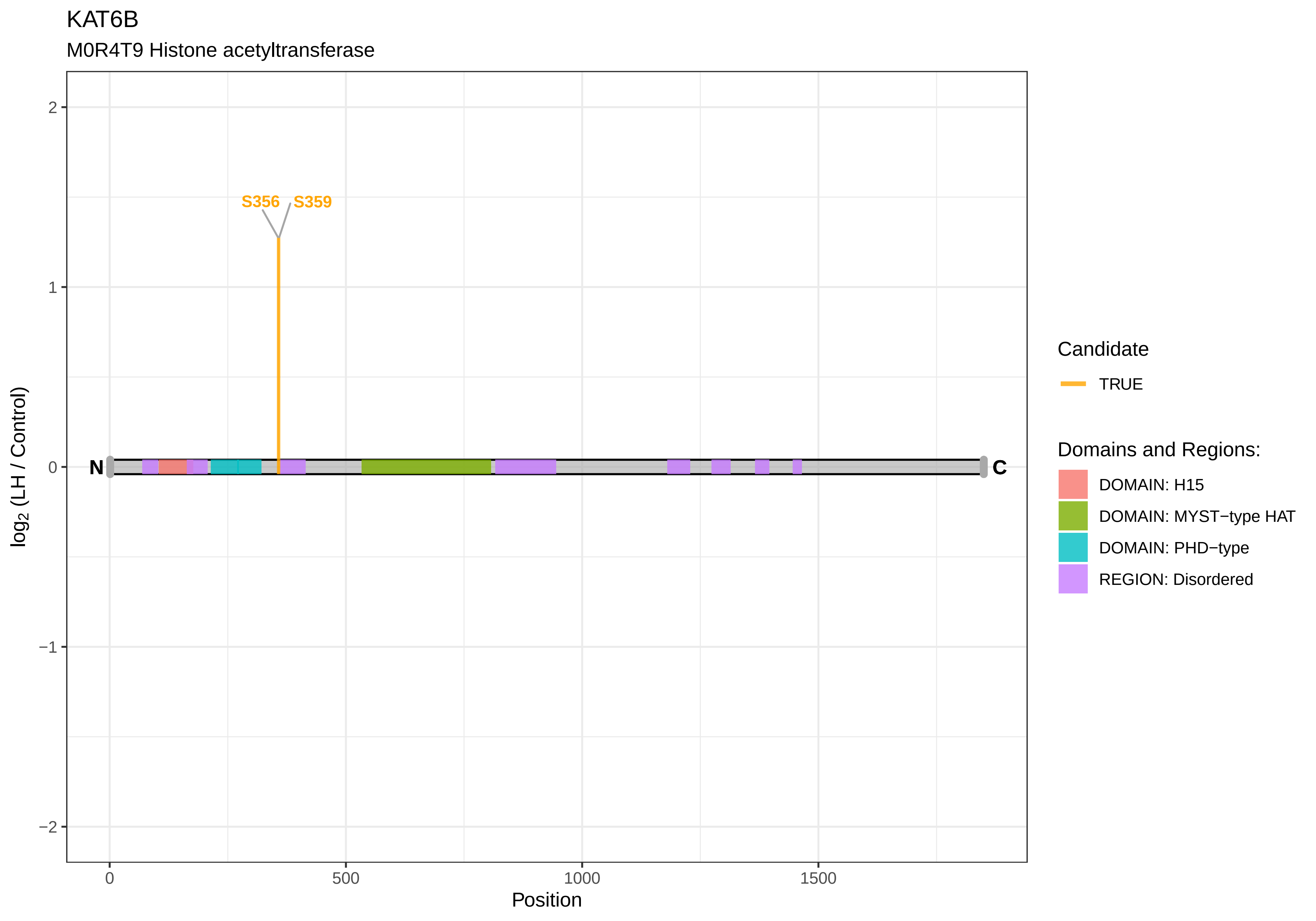
KAT6B
M0R4T9 Histone acetyltransferase
S356
S359
N
C
2
1
0
−1
−2
log2 (LH / Control)
0
500
1000
1500
Position
Candidate
TRUE
Domains and Regions:
DOMAIN: H15
DOMAIN: MYST−type HAT
DOMAIN: PHD−type
REGION: Disordered

### KAT7

Q810T5 Histone acetyltransferase KAT7

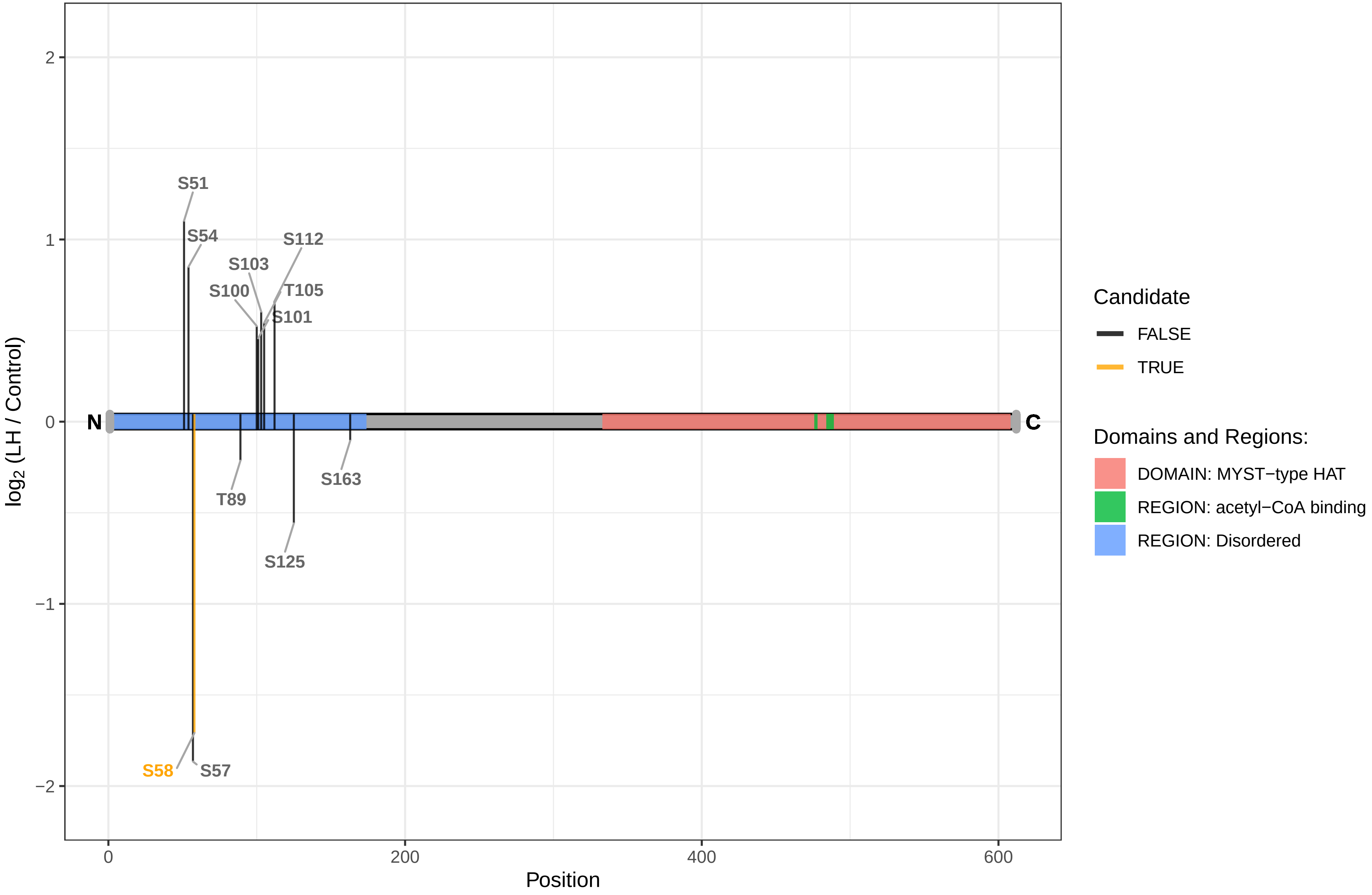

KDM2B

F1LLW9 Lysine demethylase 2B

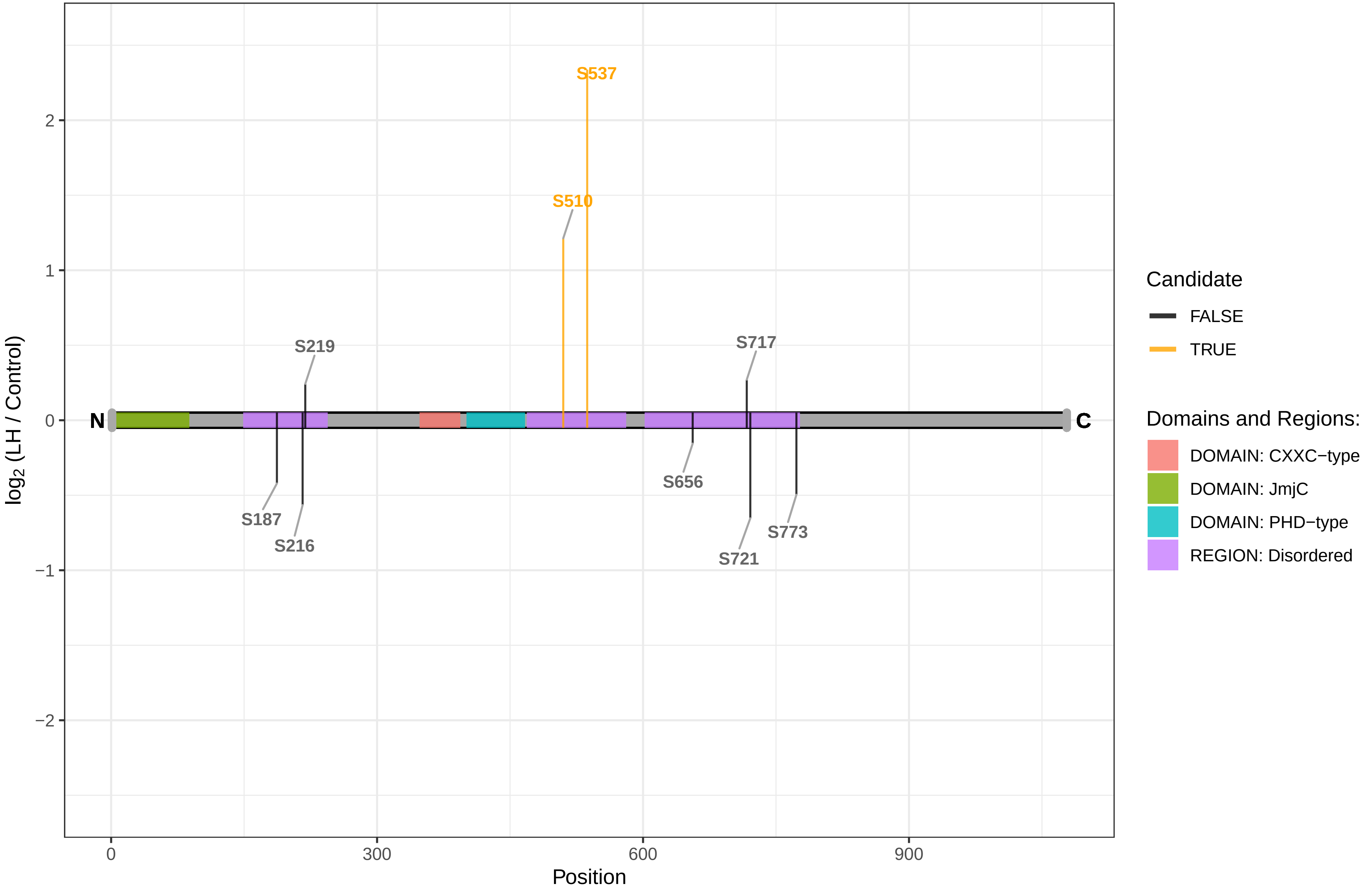

KDM3B

M0RDF1 Lysine demethylase 3B

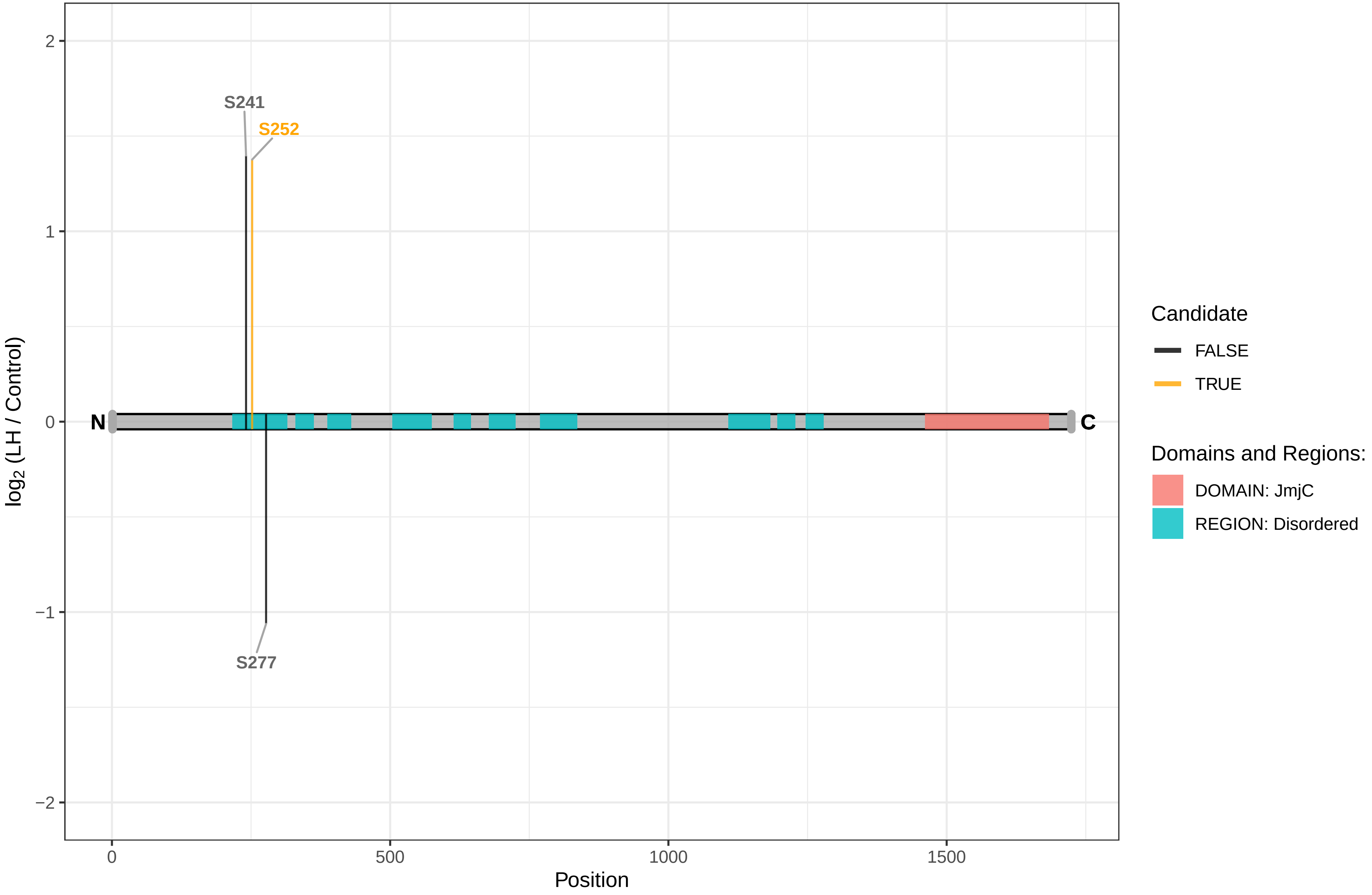

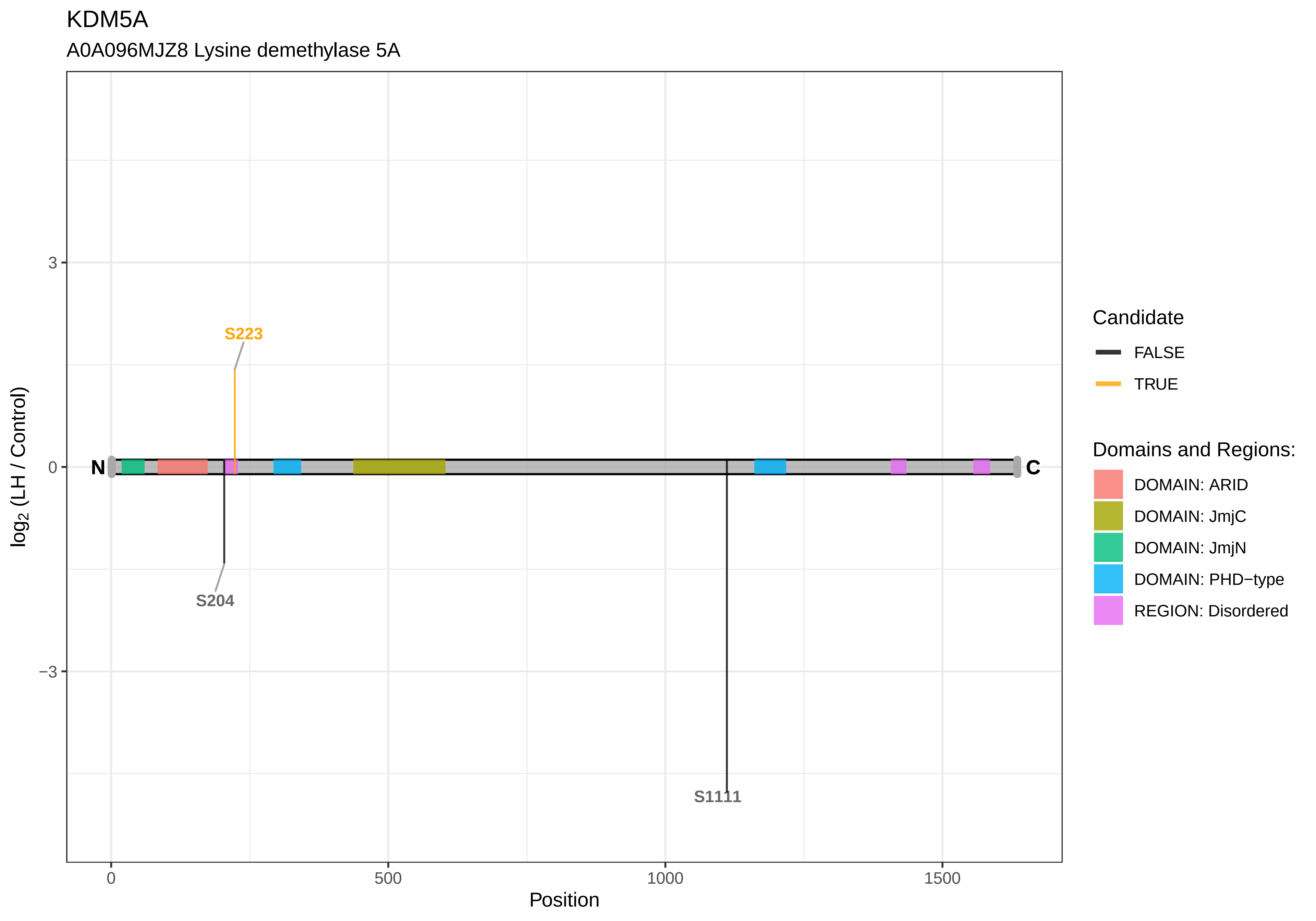
KDM5A
A0A096MJZ8 Lysine demethylase 5A
S223
S204
S1111
N
C
3
0
−3
0
500
1000
1500
Position
log2 (LH / Control)
Candidate
FALSE
TRUE
Domains and Regions:
DOMAIN: ARID
DOMAIN: JmjC
DOMAIN: JmjN
DOMAIN: PHD−type
REGION: Disordered

KIDINS220

D3ZWB2 Kinase D−interacting substrate of 220 kDa

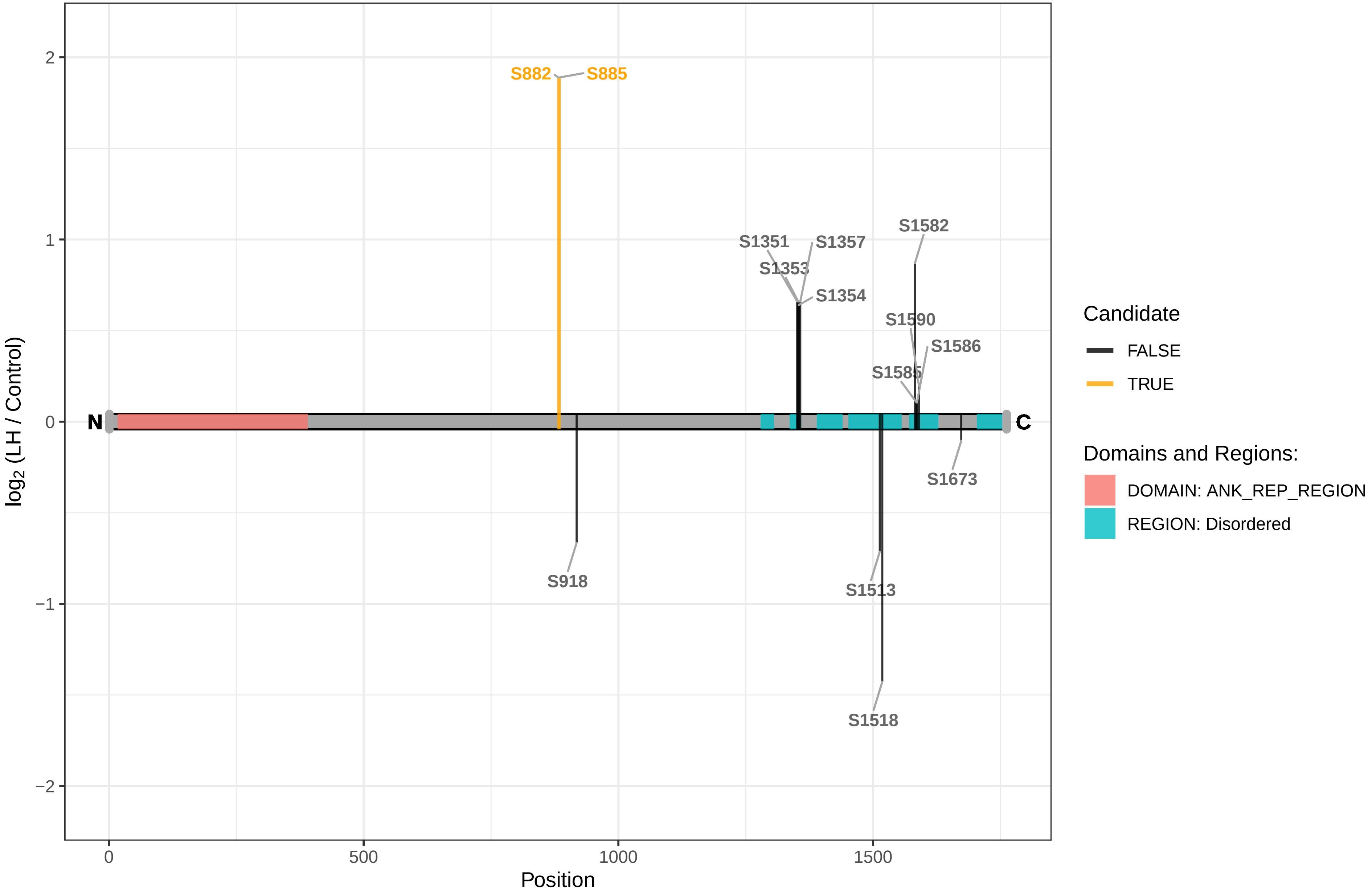

KIF13B

A0A0G2K8Z9 Kinesin family member 13B

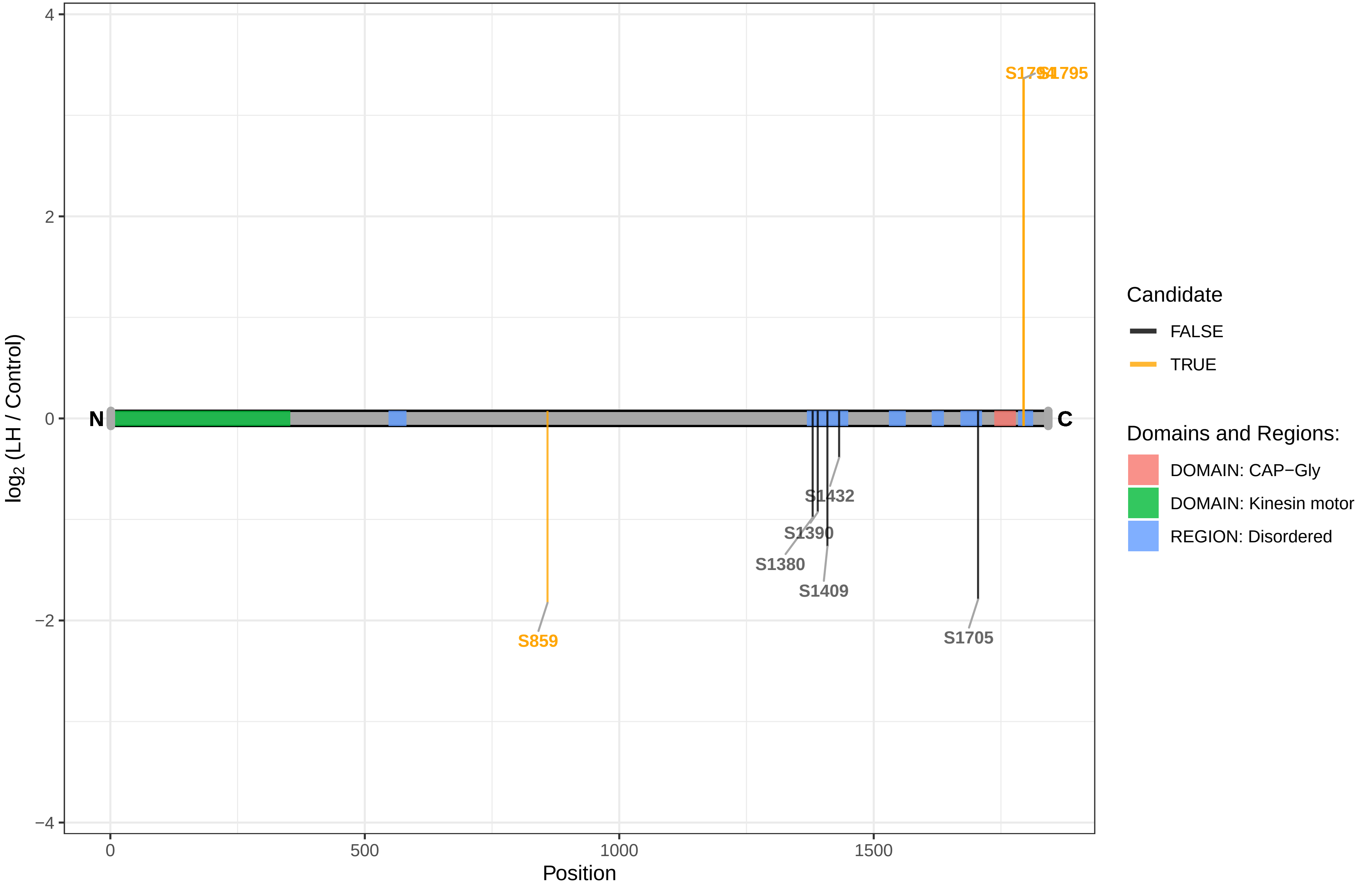

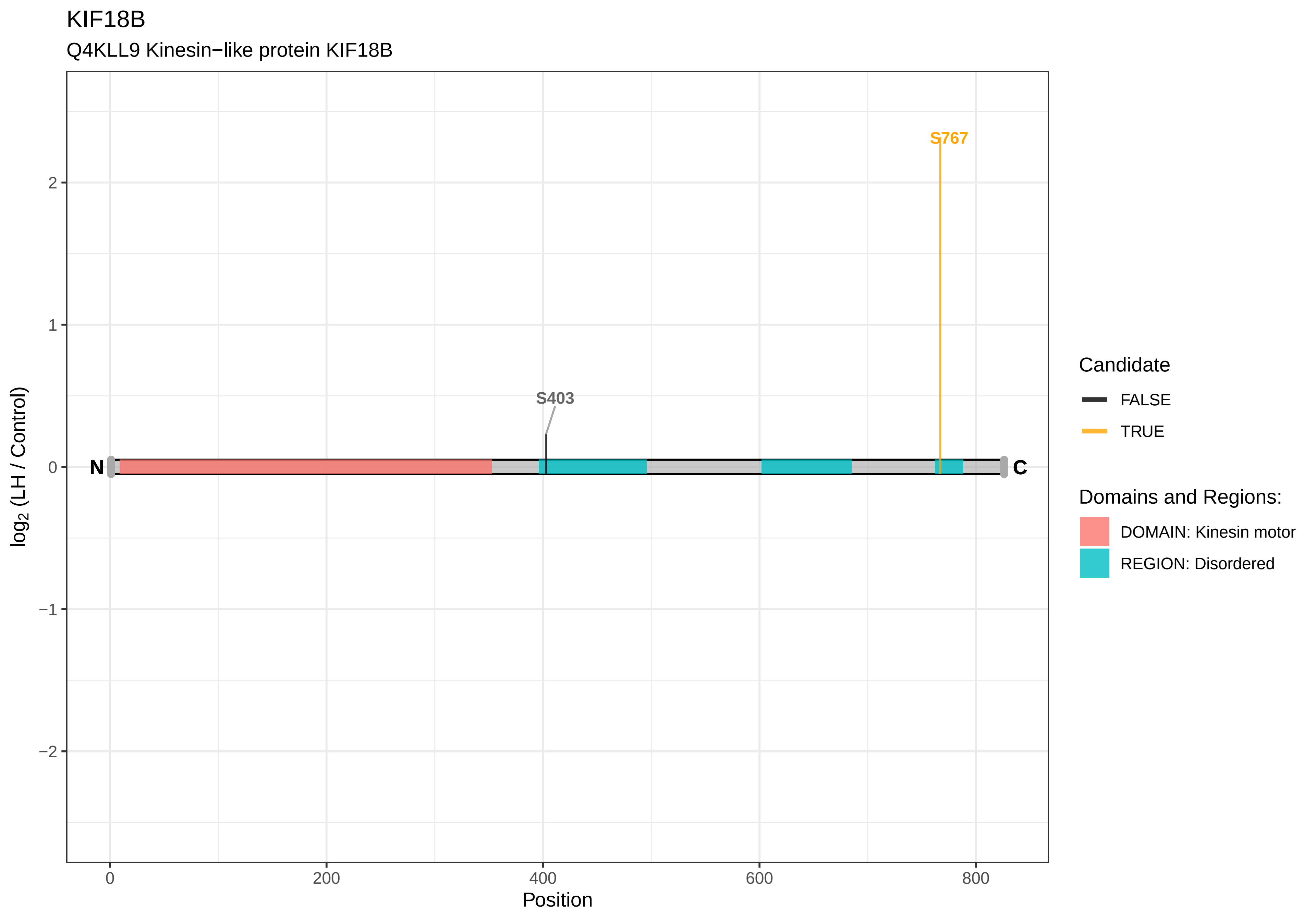
KIF18B
Q4KLL9 Kinesin−like protein KIF18B
S767
S403
N
C
2
1
0
−1
−2
log2 (LH / Control)
0
200
400
600
800
Position
Candidate
FALSE
TRUE
Domains and Regions:
DOMAIN: Kinesin motor
REGION: Disordered

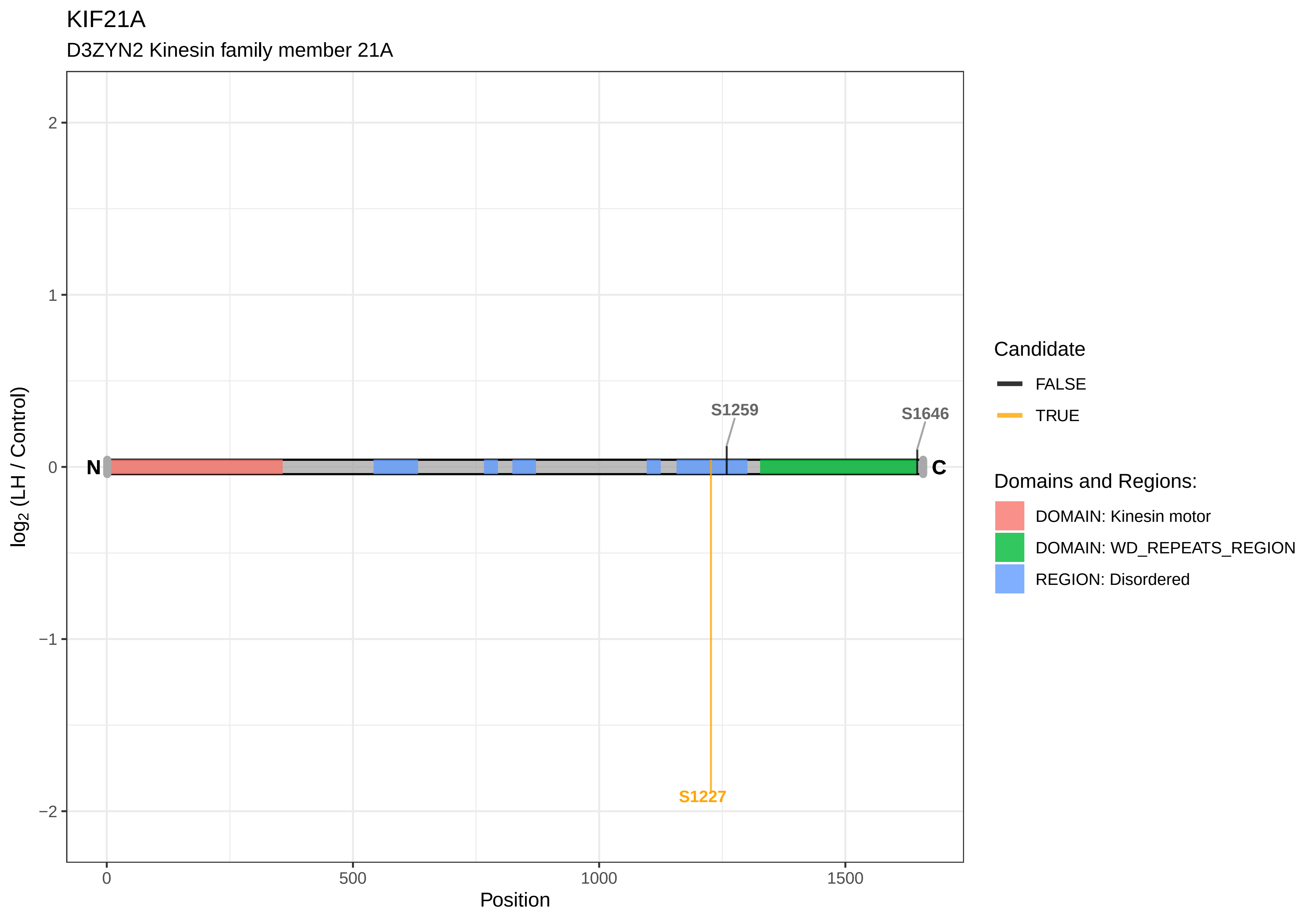

KIF21A
D3ZYN2 Kinesin family member 21A
S1259
S1646
N
C
S1227
2
1
0
−1
−2
log2 (LH / Control)
0
500
1000
1500
Position
Candidate
FALSE
TRUE
Domains and Regions:
DOMAIN: Kinesin motor
DOMAIN: WD_REPEATS_REGION
REGION: Disordered

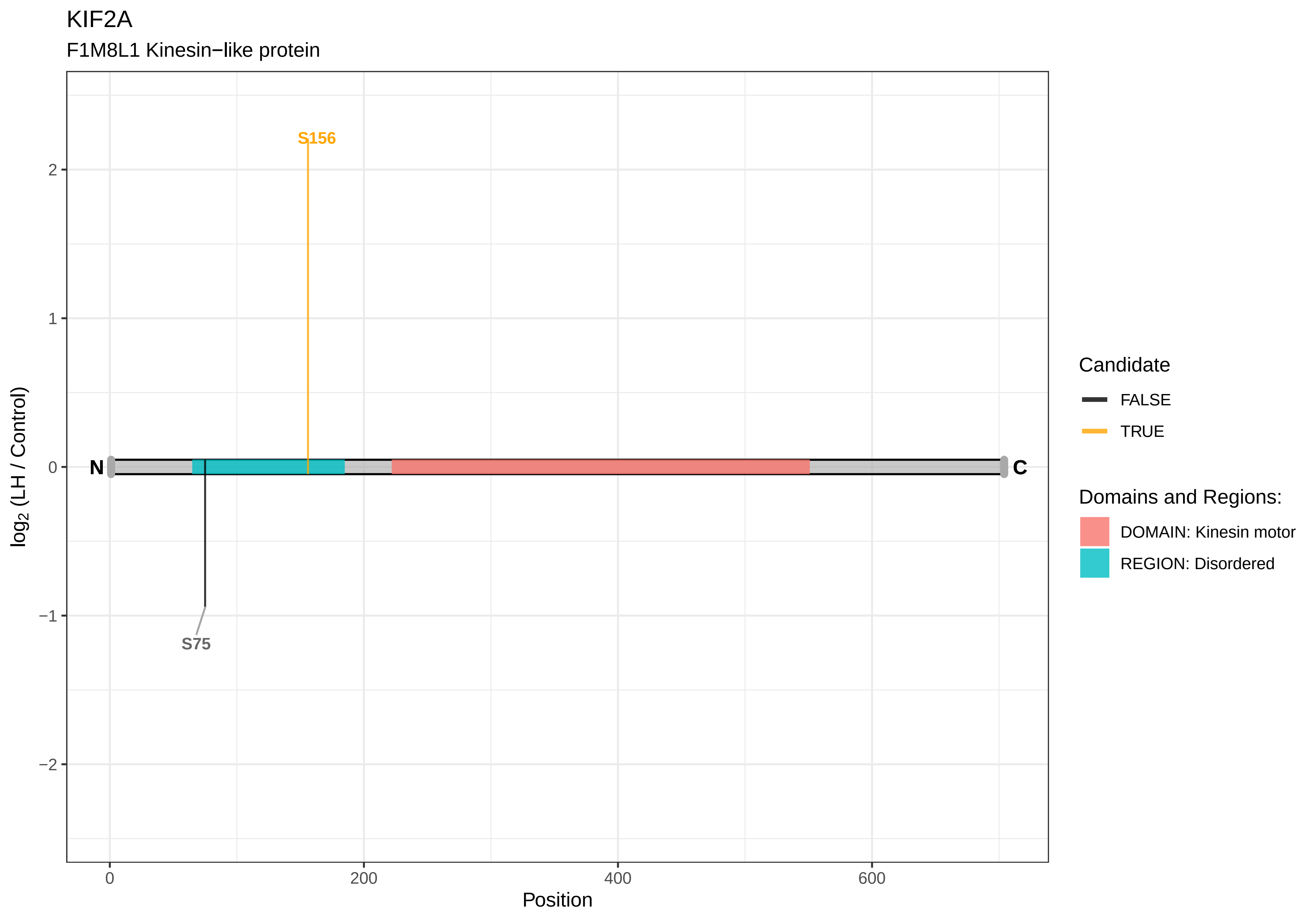
KIF2A
F1M8L1 Kinesin−like protein
S156
S75
N
C
log2 (LH / Control)
Position
2
1
0
−1
−2
0
200
400
600
Candidate
FALSE
TRUE
Domains and Regions:
DOMAIN: Kinesin motor
REGION: Disordered

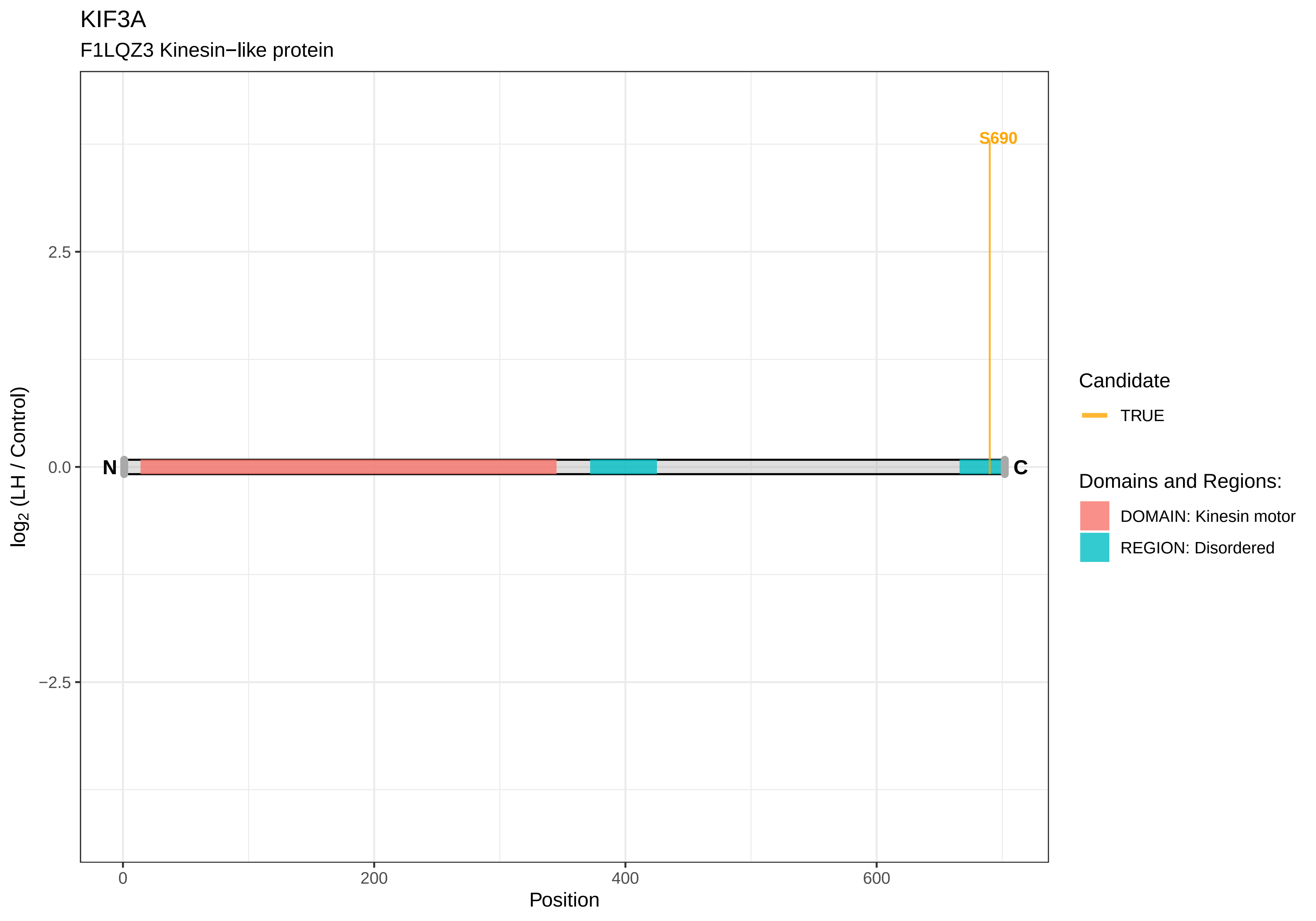
KIF3A
F1LQZ3 Kinesin−like protein
S690
N
C
2.5
0.0
−2.5
log2 (LH / Control)
0
200
400
600
Position
Candidate
TRUE
Domains and Regions:
DOMAIN: Kinesin motor
REGION: Disordered

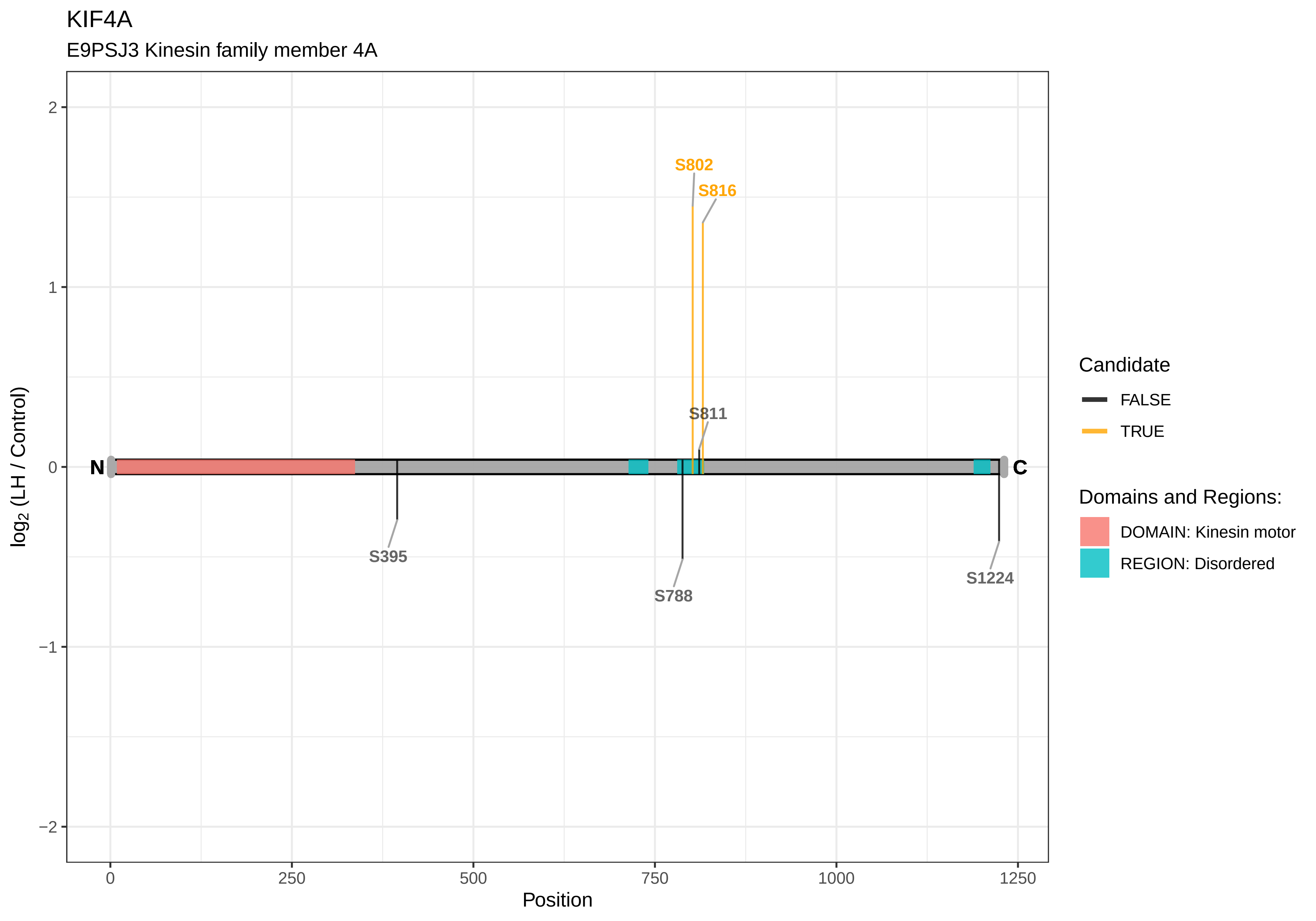

KIF4A
E9PSJ3 Kinesin family member 4A
S802
S816
S811
S395
S788
S1224
N
C
2
1
0
−1
−2
0
250
500
750
1000
1250
Position
log2 (LH / Control)
Candidate
FALSE
TRUE
Domains and Regions:
DOMAIN: Kinesin motor
REGION: Disordered

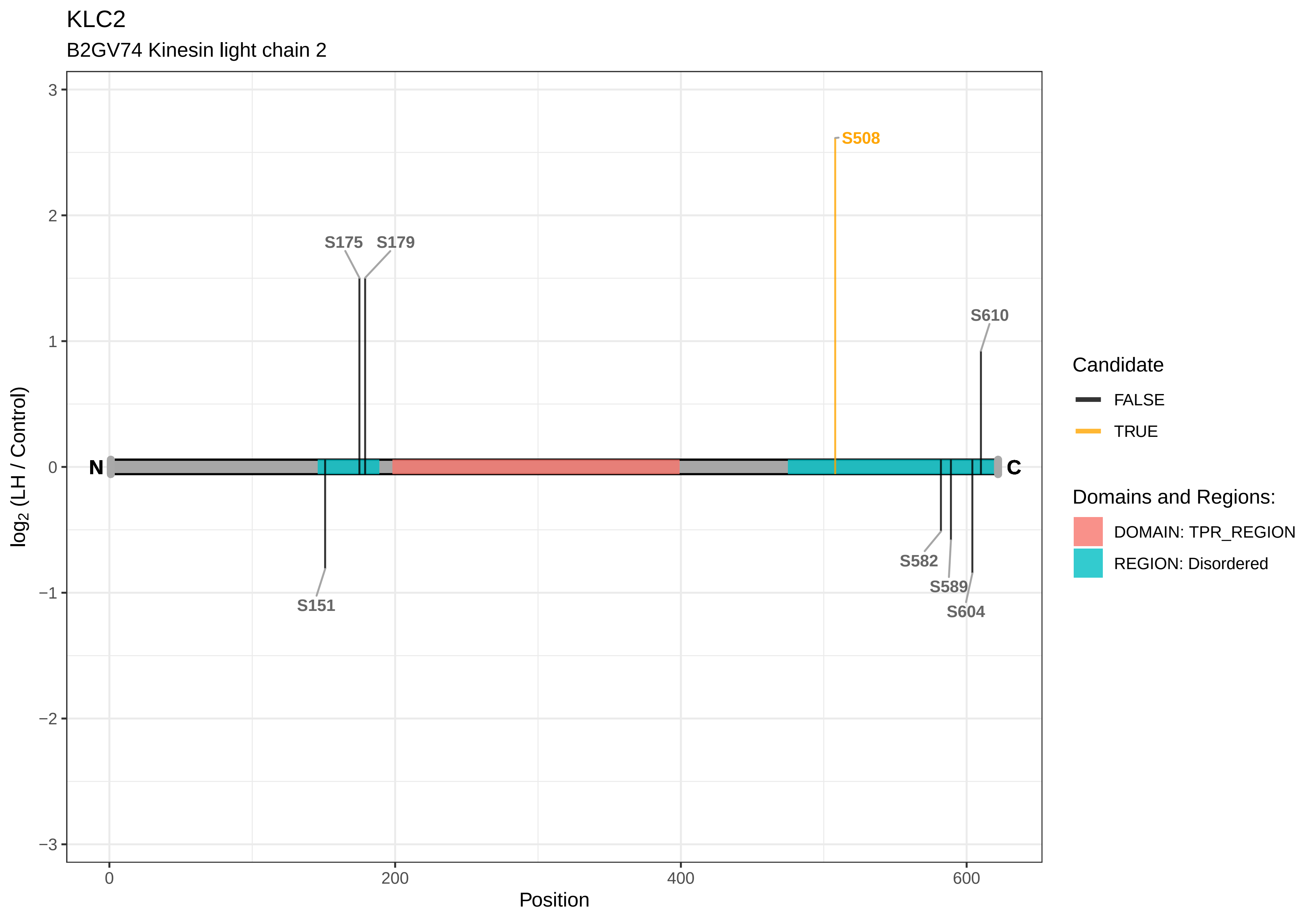
KLC2
B2GV74 Kinesin light chain 2
S508
S175
S179
S610
S151
S582
S589
S604
N
C
log2 (LH / Control)
Position
3
2
1
0
−1
−2
−3
0
200
400
600
Candidate
FALSE
TRUE
Domains and Regions:
DOMAIN: TPR_REGION
REGION: Disordered

KLC4

Q5PQM2 Kinesin light chain 4

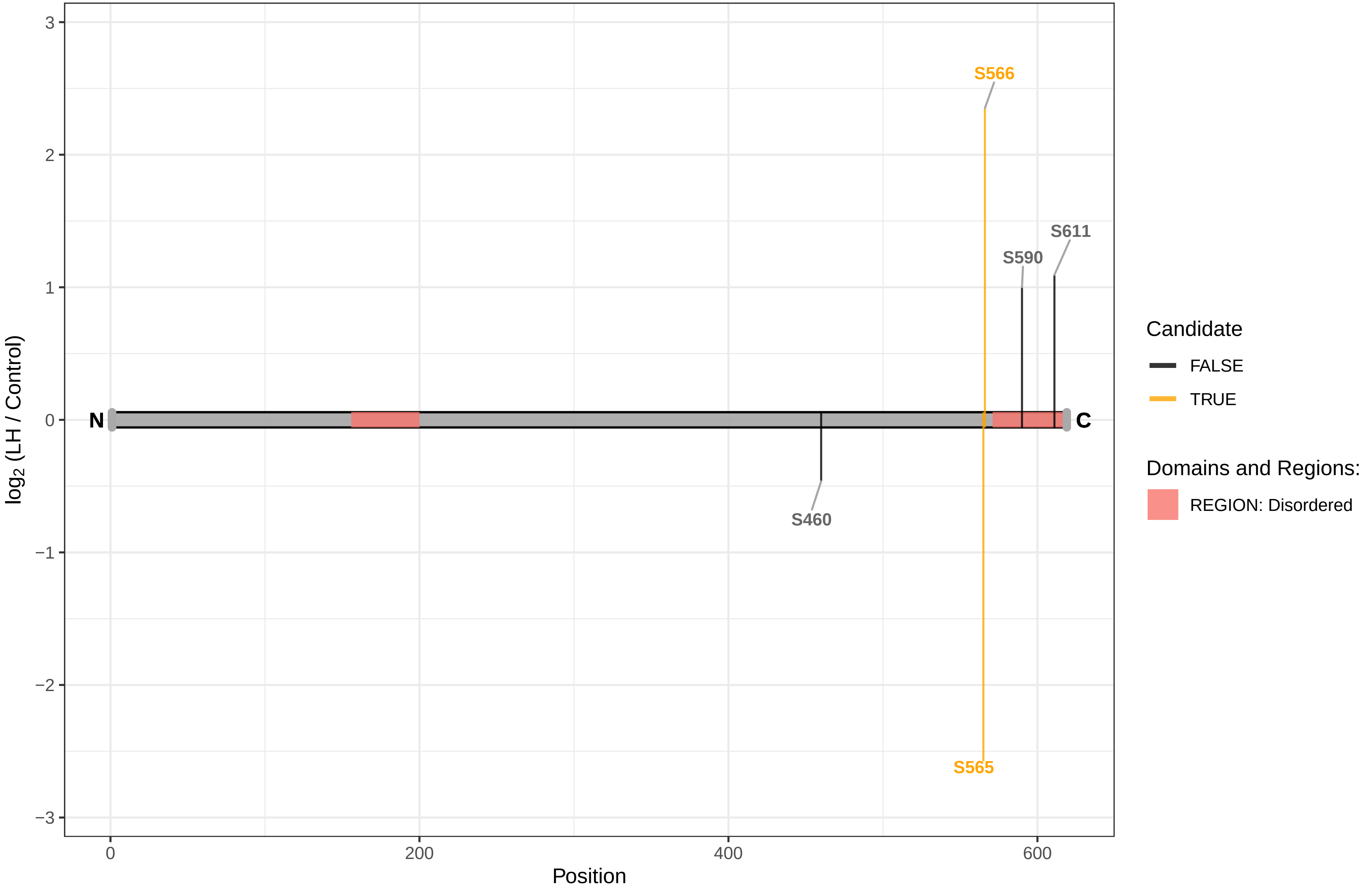

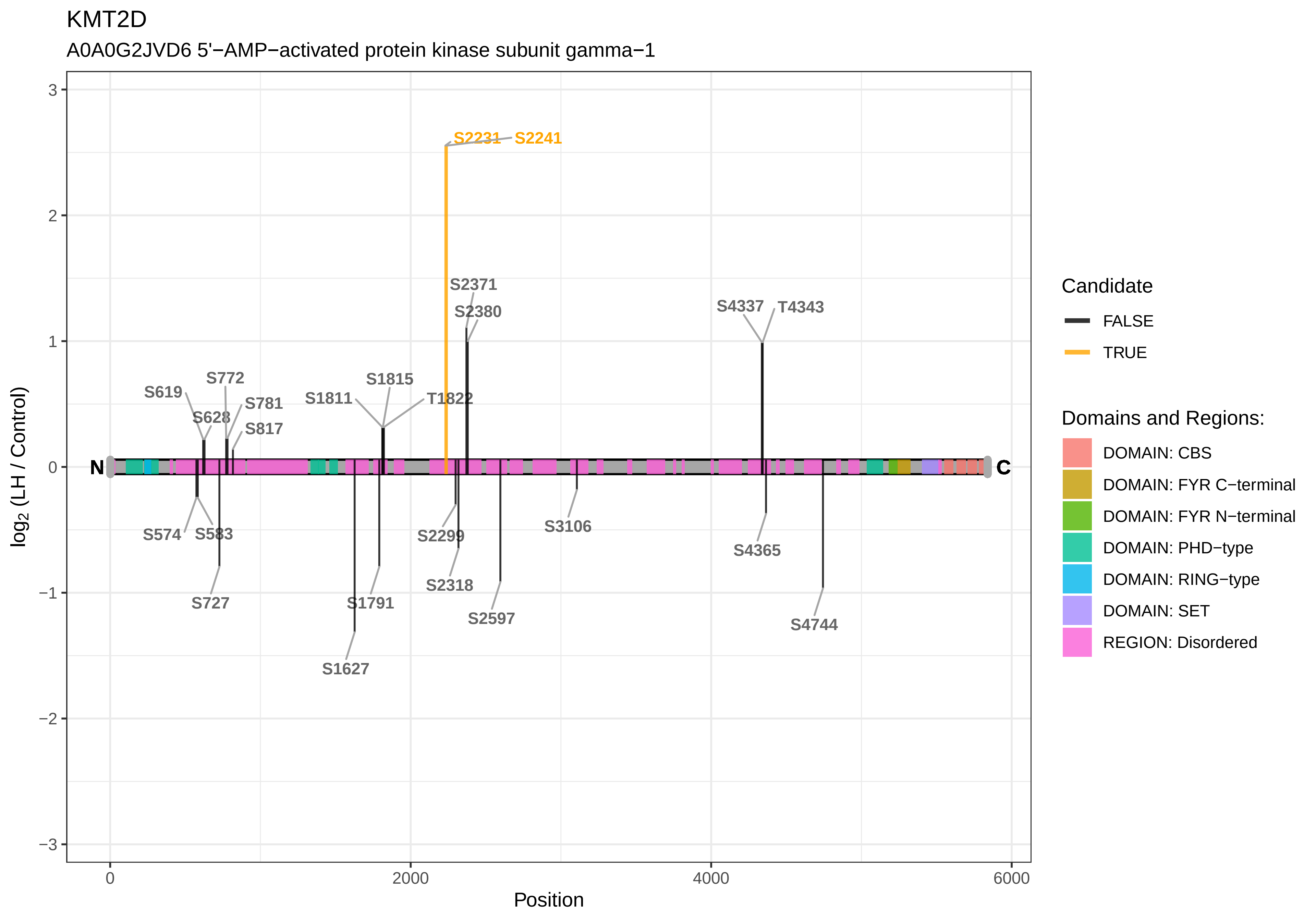
KMT2D
A0A0G2JVD6 5'−AMP−activated protein kinase subunit gamma−1
S2231
S2241
S2371
S2380
S4337
T4343
S772
S619
S1815
S1811
T1822
S781
S628
S817
N
C
S574
S583
S2299
S3106
S4365
S727
S1791
S2318
S2597
S4744
S1627
log2 (LH / Control)
Position
Candidate
FALSE
TRUE
Domains and Regions:
DOMAIN: CBS
DOMAIN: FYR C−terminal
DOMAIN: FYR N−terminal
DOMAIN: PHD−type
DOMAIN: RING−type
DOMAIN: SET
REGION: Disordered

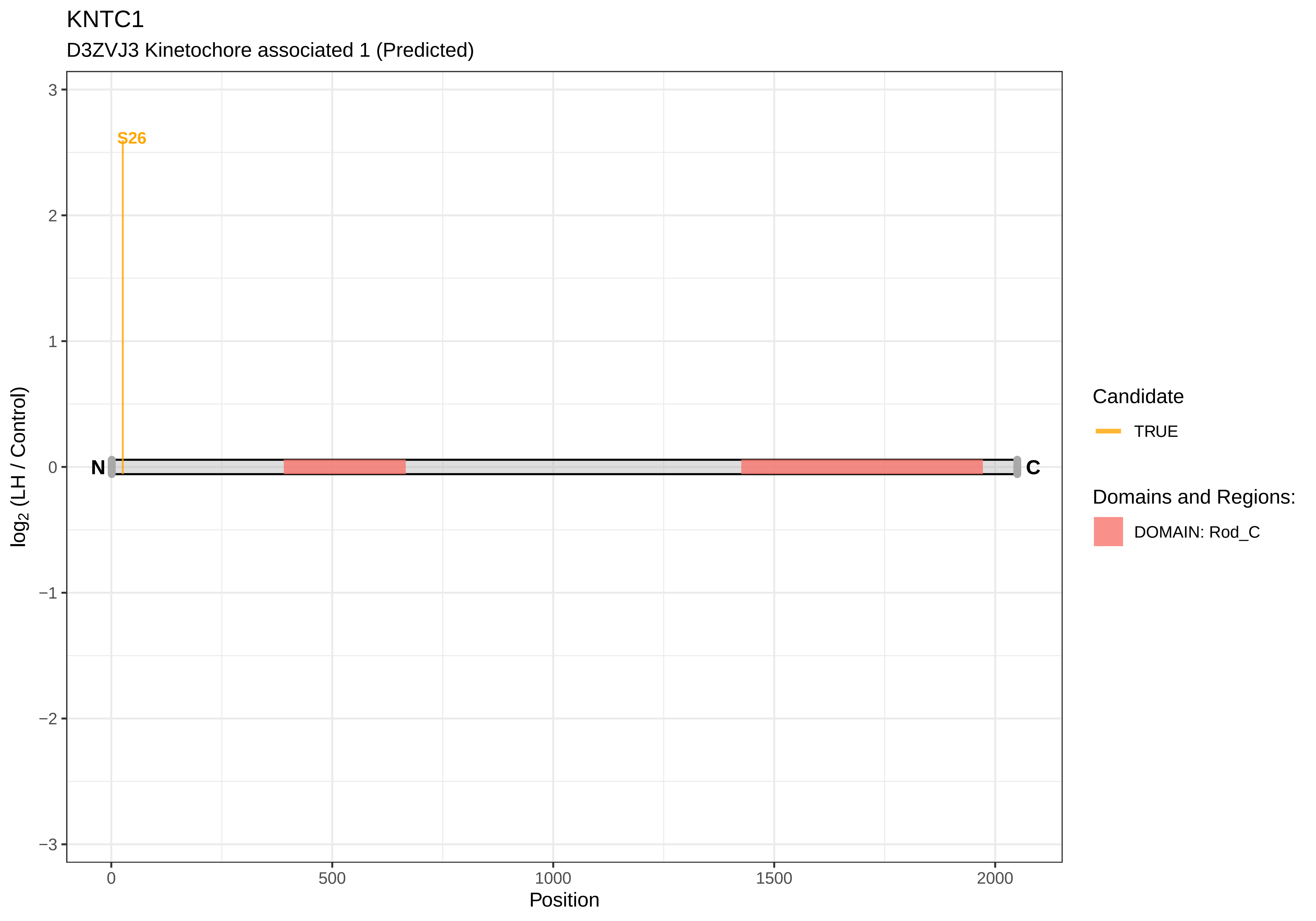
KNTC1
D3ZVJ3 Kinetochore associated 1 (Predicted)
S26
N
C
3
2
1
0
−1
−2
−3
log2 (LH / Control)
0
500
1000
1500
2000
Position
Candidate
TRUE
Domains and Regions:
DOMAIN: Rod_C

KTN1

D4A4Z9 Kinectin 1

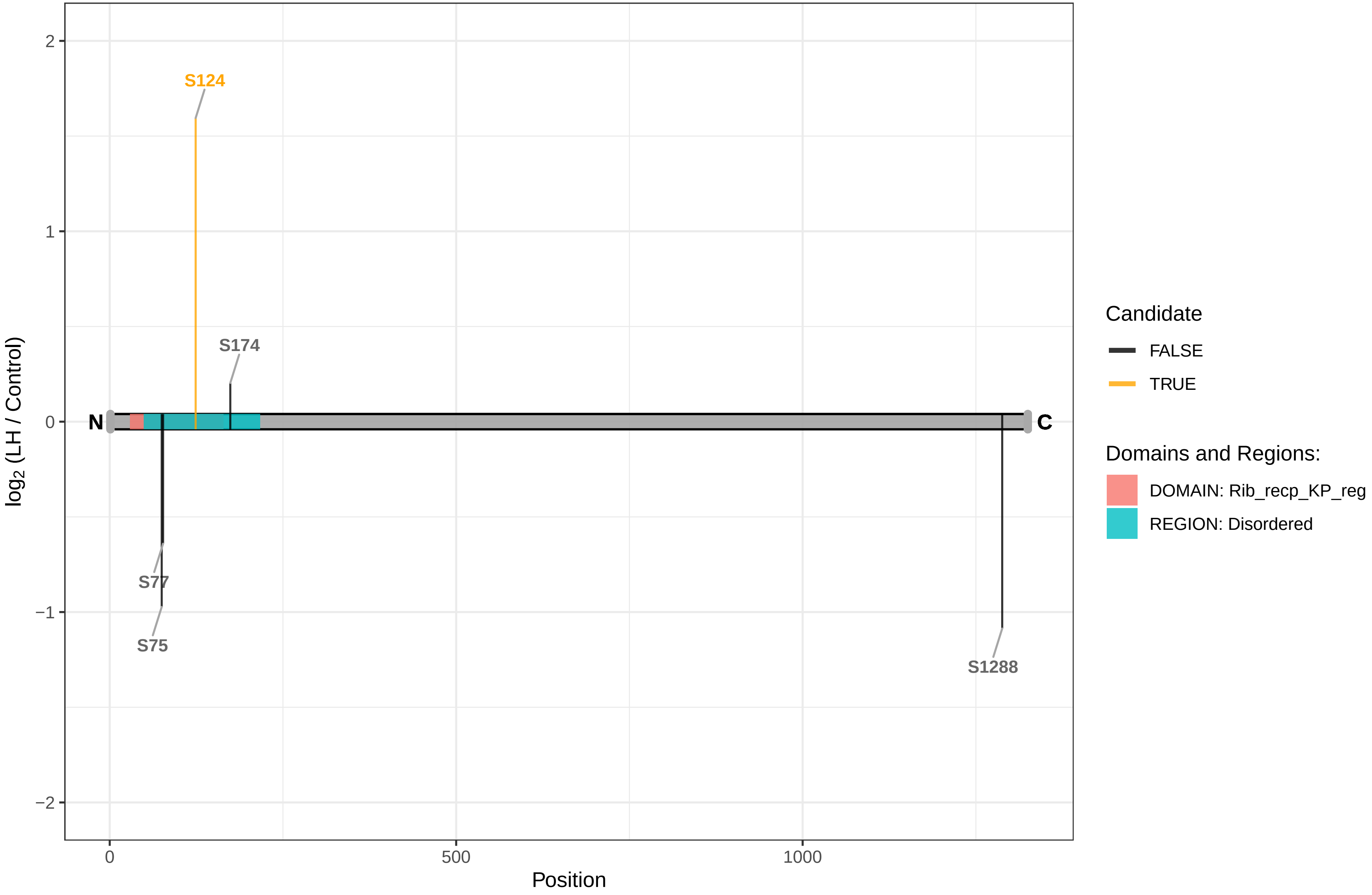

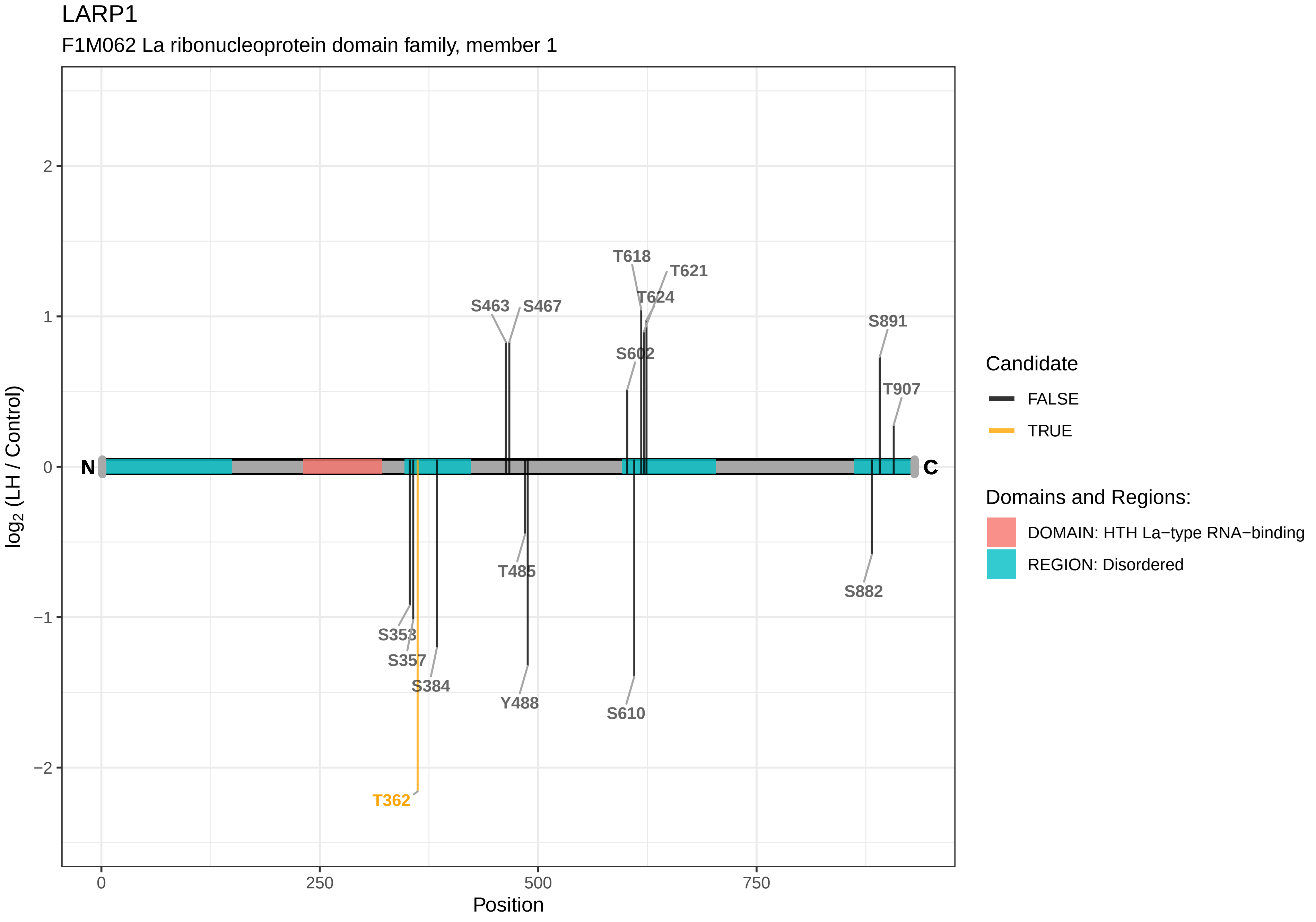
LARP1
F1M062 La ribonucleoprotein domain family, member 1
T618
T621
T624
S463
S467
S891
S602
T907
N
C
T485
S882
S353
S357
S384
Y488
S610
T362
log2 (LH / Control)
Position
0
250
500
750
2
1
0
−1
−2
Candidate
FALSE
TRUE
Domains and Regions:
DOMAIN: HTH La−type RNA−binding
REGION: Disordered

### LARP4B

D3ZF45 La ribonucleoprotein domain family, member 4B

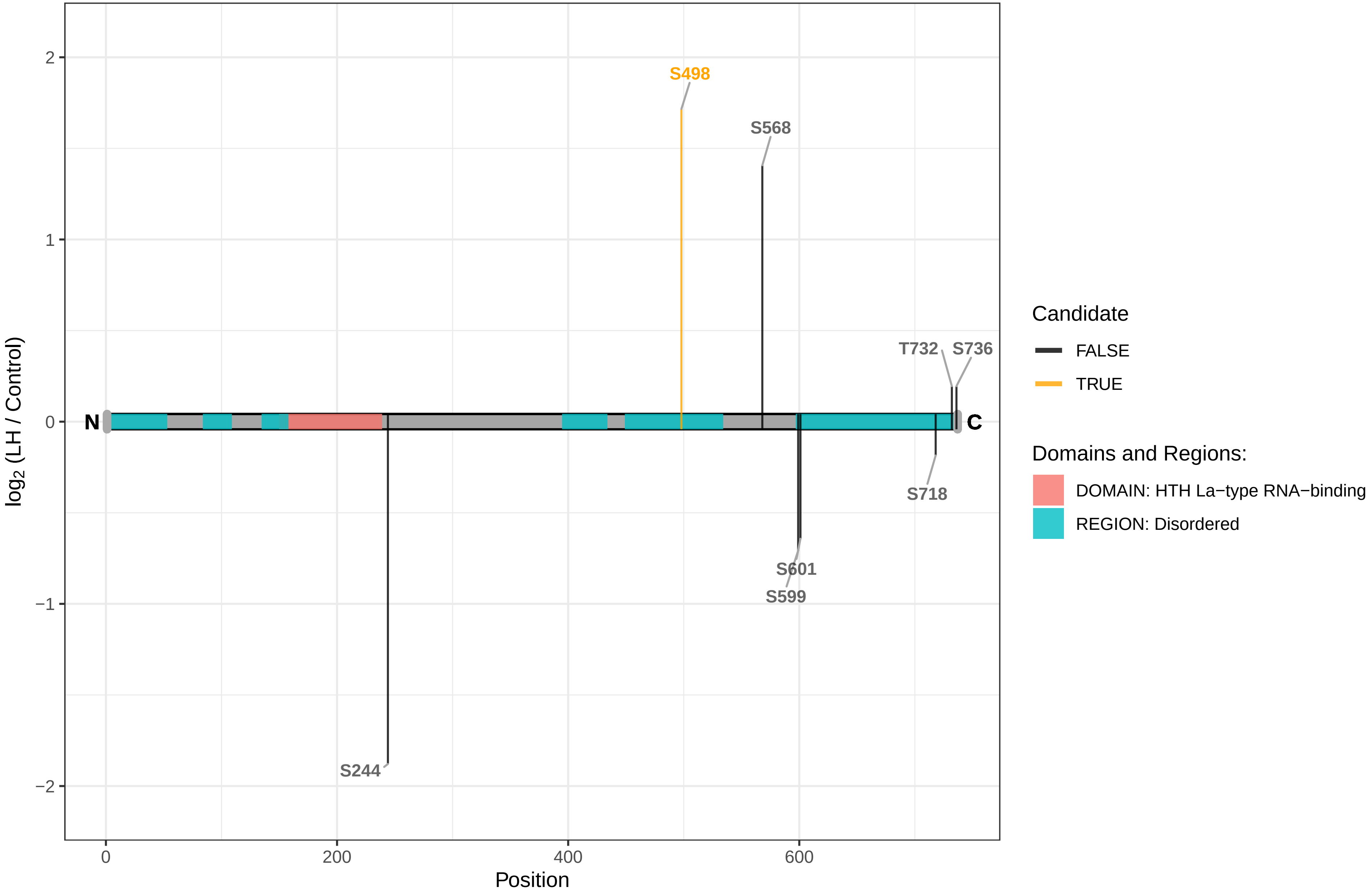

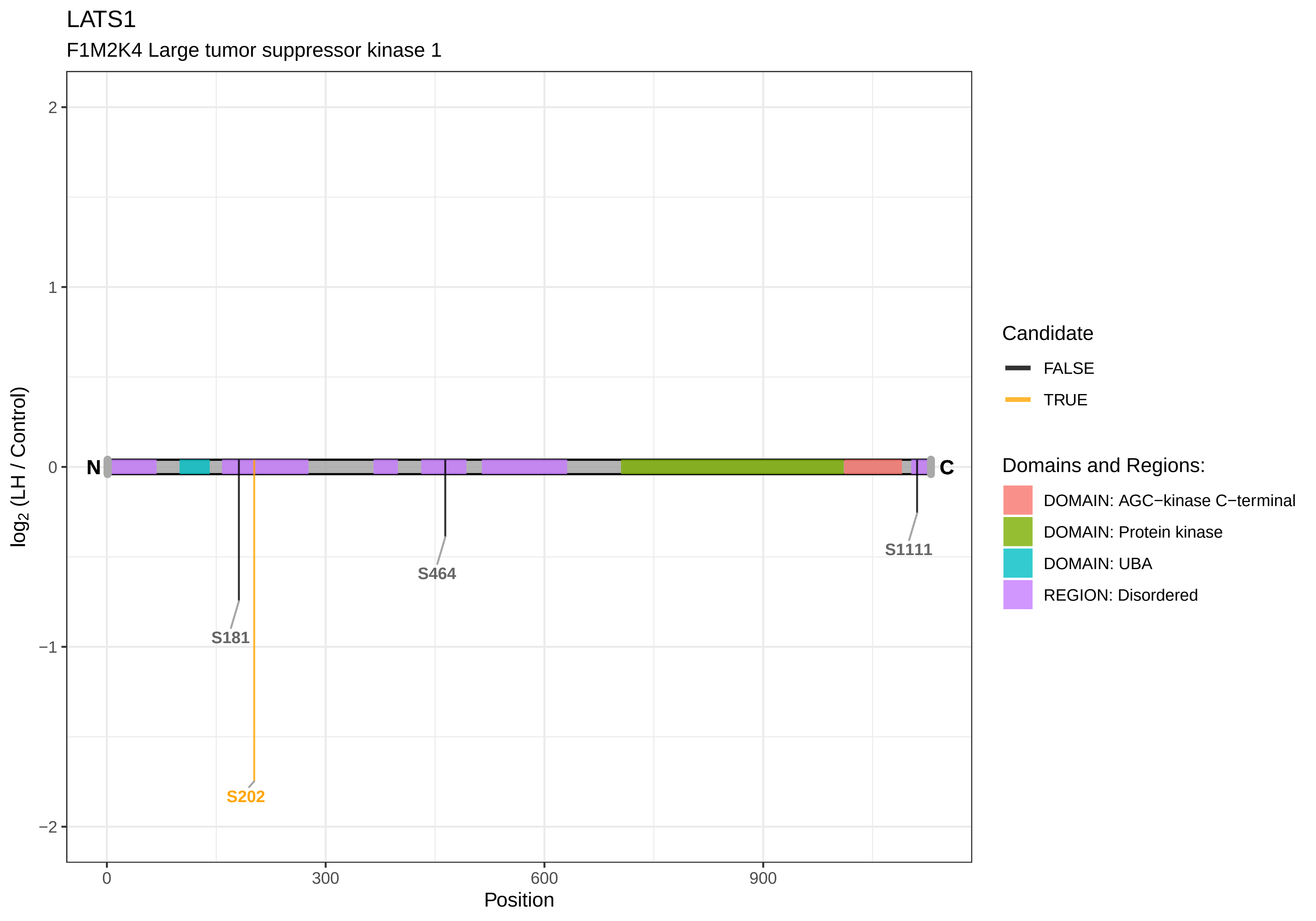

LATS1
F1M2K4 Large tumor suppressor kinase 1
2
1
0
−1
−2
log2 (LH / Control)
N
C
S181
S202
S464
S1111
0
300
600
900
Position
Candidate
FALSE
TRUE
Domains and Regions:
DOMAIN: AGC−kinase C−terminal
DOMAIN: Protein kinase
DOMAIN: UBA
REGION: Disordered

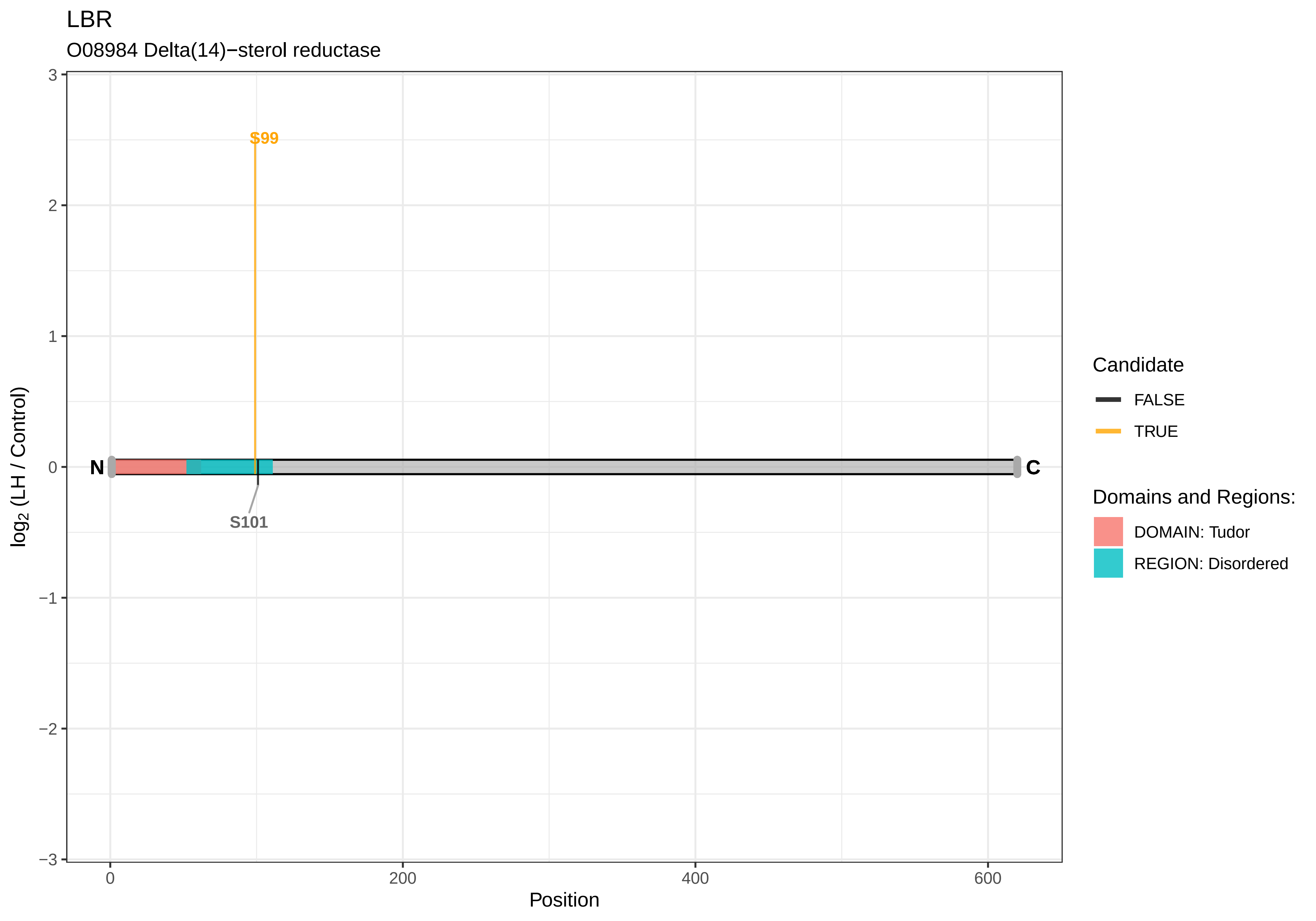

LBR
O08984 Delta(14)−sterol reductase
S99
S101
N
C
log2 (LH / Control)
Position
3
2
1
0
−1
−2
−3
0
200
400
600
Candidate
FALSE
TRUE
Domains and Regions:
DOMAIN: Tudor
REGION: Disordered

LDLRAP1

D3ZAR1 Low density lipoprotein receptor adapter protein 1

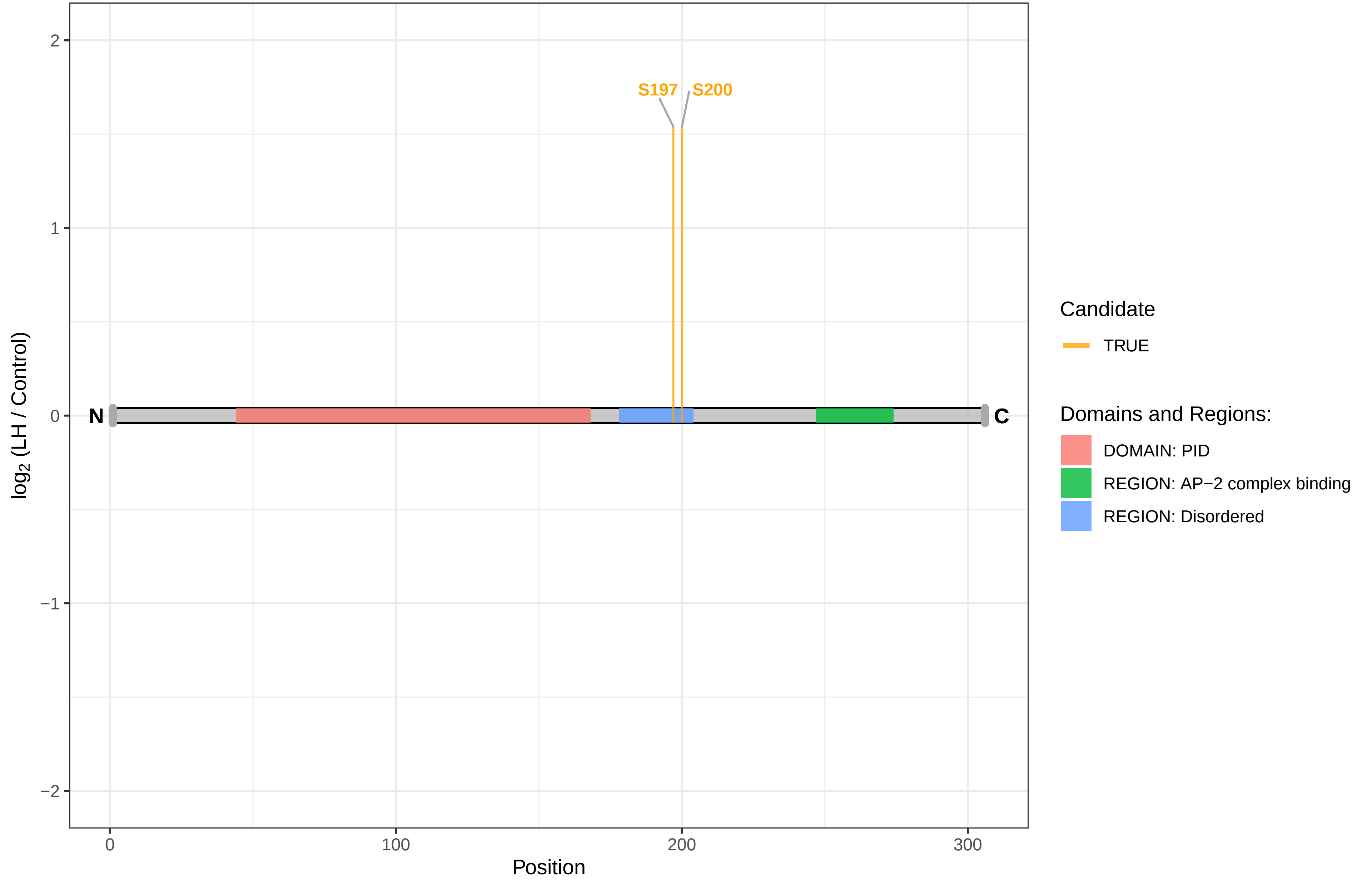

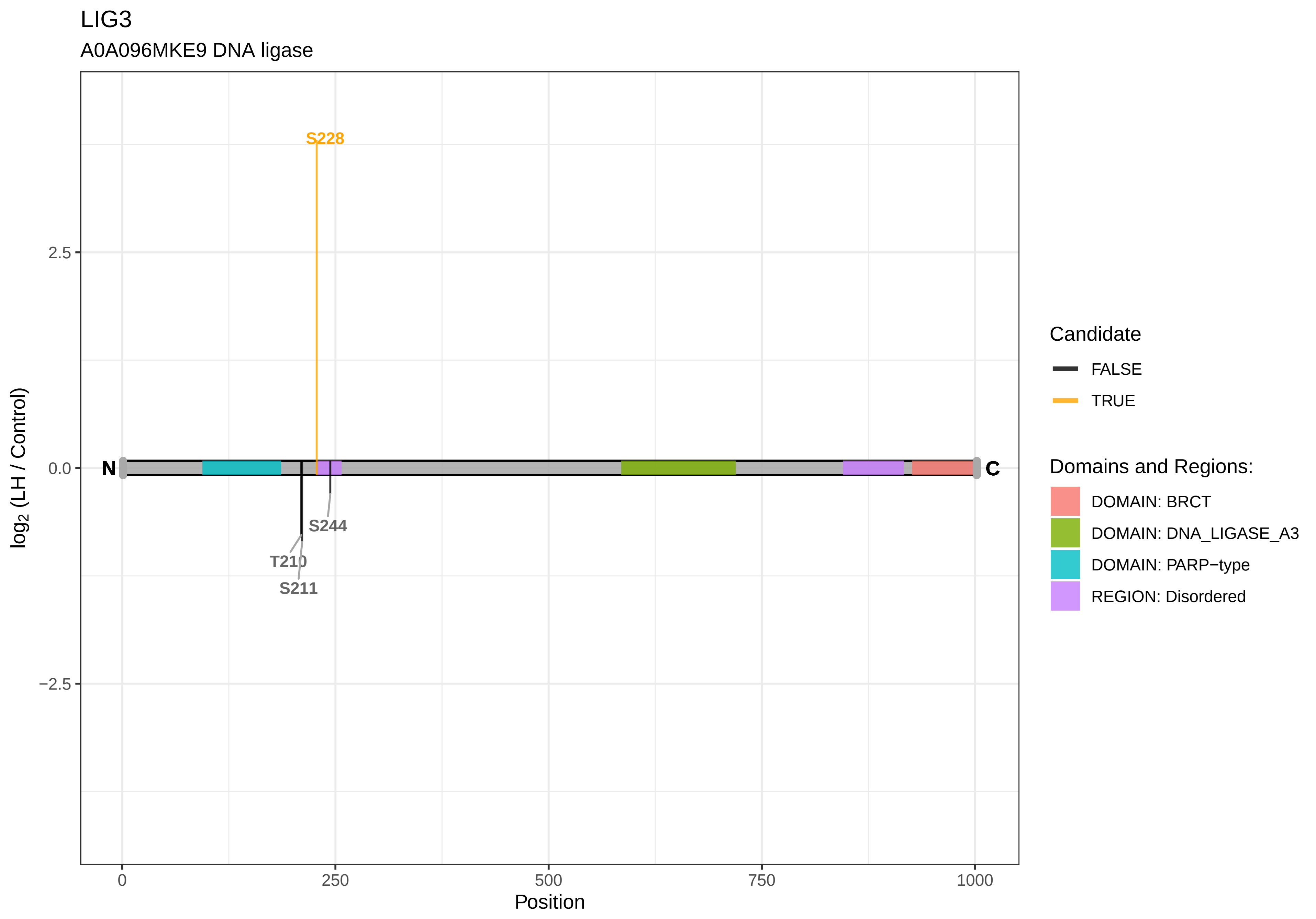

LIG3
A0A096MKE9 DNA ligase
S228
S244
T210
S211
N
C
2.5
0.0
−2.5
0
250
500
750
1000
Position
log2 (LH / Control)
Candidate
FALSE
TRUE
Domains and Regions:
DOMAIN: BRCT
DOMAIN: DNA_LIGASE_A3
DOMAIN: PARP−type
REGION: Disordered

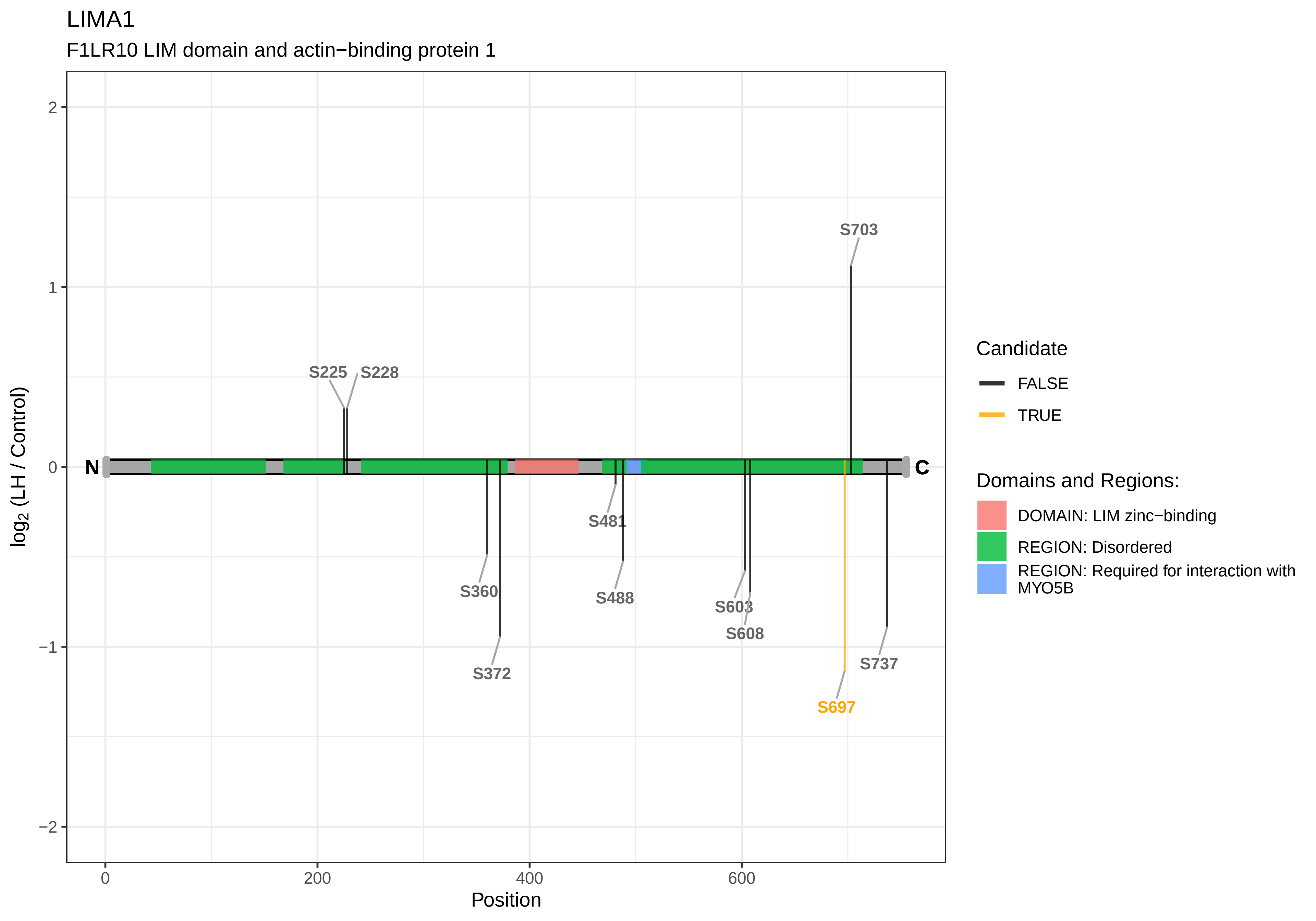
LIMA1
F1LR10 LIM domain and actin−binding protein 1
log2 (LH / Control)
Position
2
1
0
−1
−2
0
200
400
600
N
C
S225
S228
S703
S360
S372
S481
S488
S603
S608
S697
S737
Candidate
FALSE
TRUE
Domains and Regions:
DOMAIN: LIM zinc−binding
REGION: Disordered
REGION: Required for interaction with MYO5B

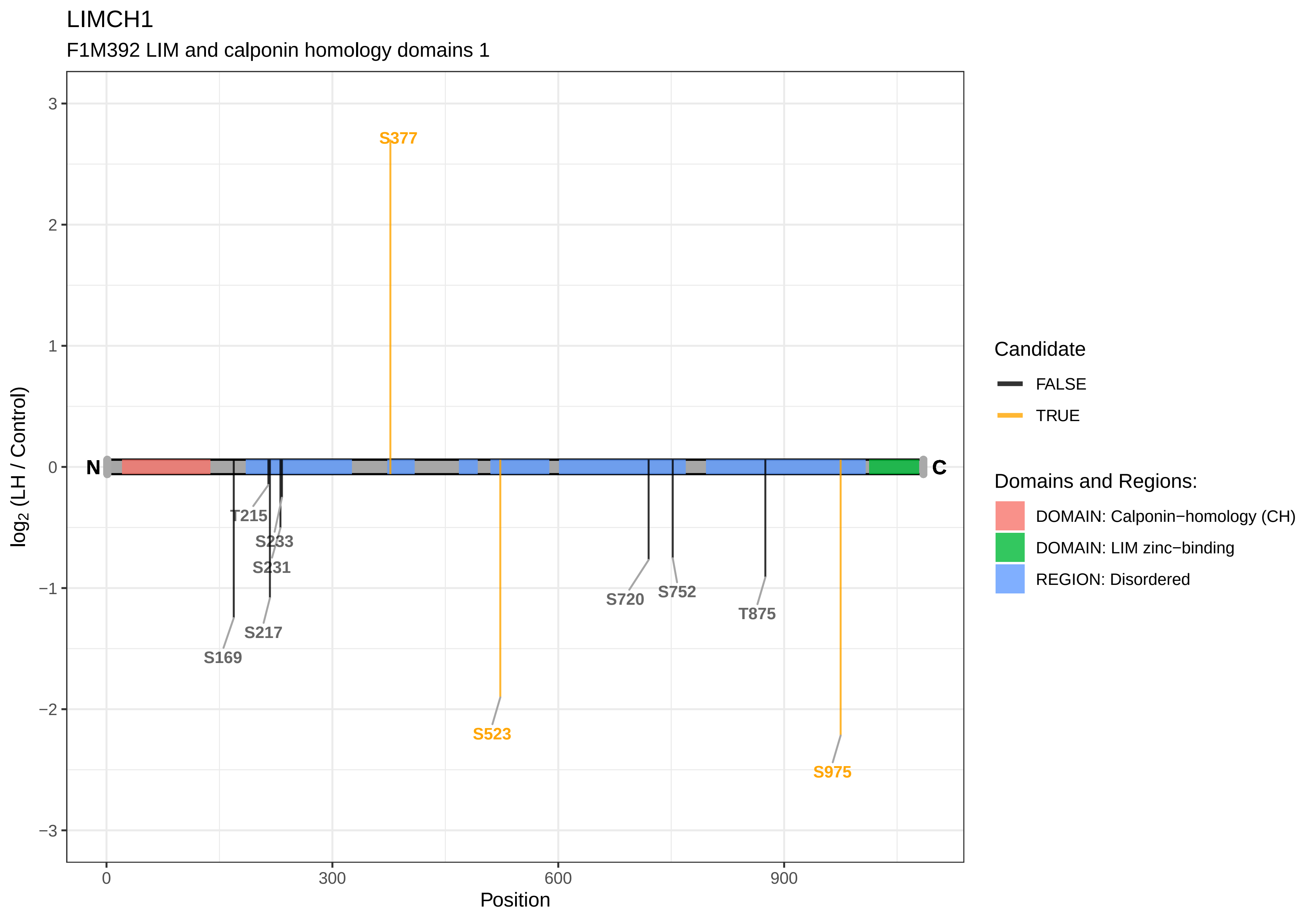

LIMCH1
F1M392 LIM and calponin homology domains 1
S377
N
C
T215
S233
S231
S217
S169
S523
S720
S752
T875
S975
3
2
1
0
−1
−2
−3
0
300
600
900
Position
log2 (LH / Control)
Candidate
FALSE
TRUE
Domains and Regions:
DOMAIN: Calponin−homology (CH)
DOMAIN: LIM zinc−binding
REGION: Disordered

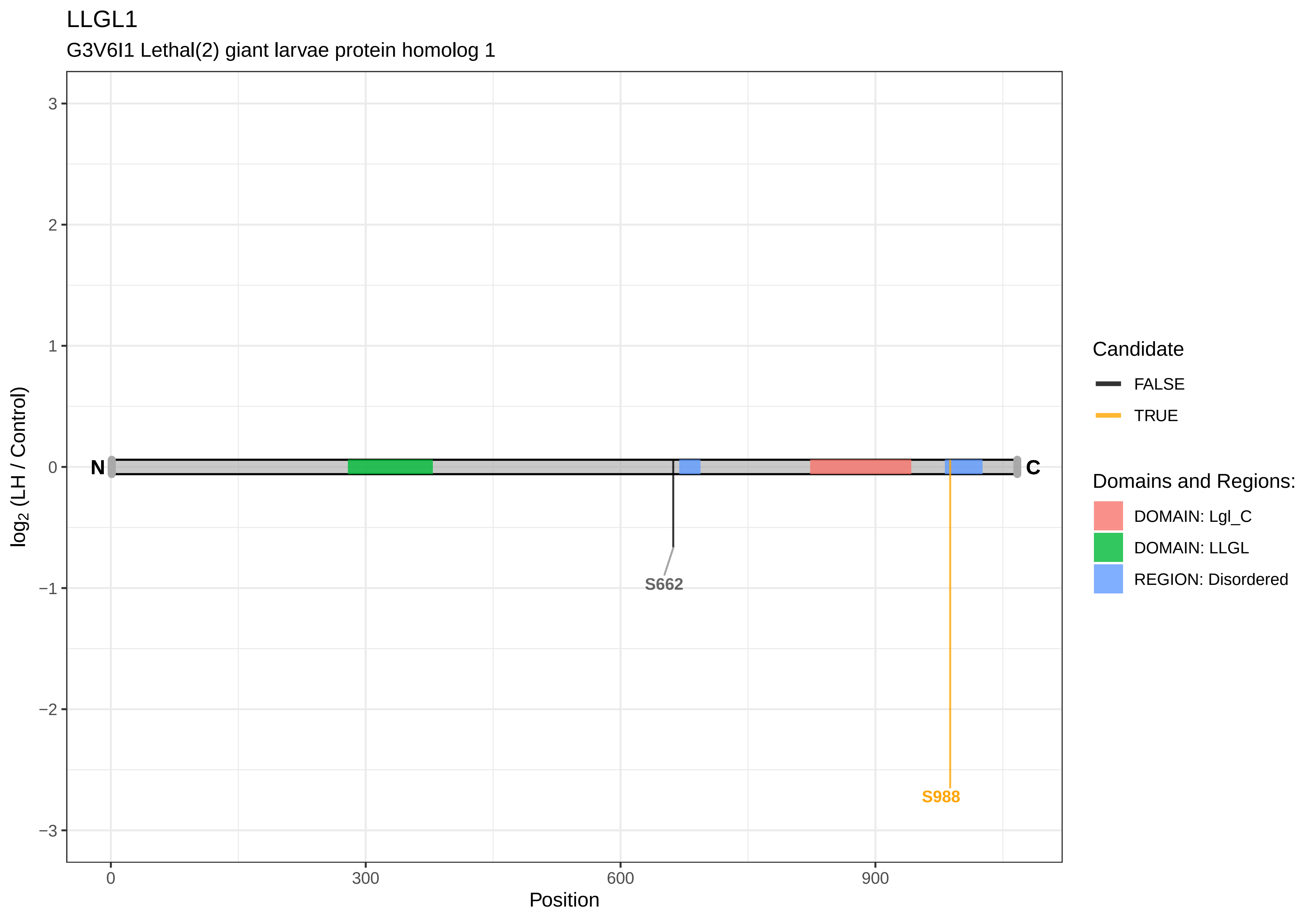

LLGL1
G3V6I1 Lethal(2) giant larvae protein homolog 1
3
2
1
0
−1
−2
−3
log2 (LH / Control)
N
C
S662
S988
0
300
600
900
Position
Candidate
FALSE
TRUE
Domains and Regions:
DOMAIN: Lgl_C
DOMAIN: LLGL
REGION: Disordered

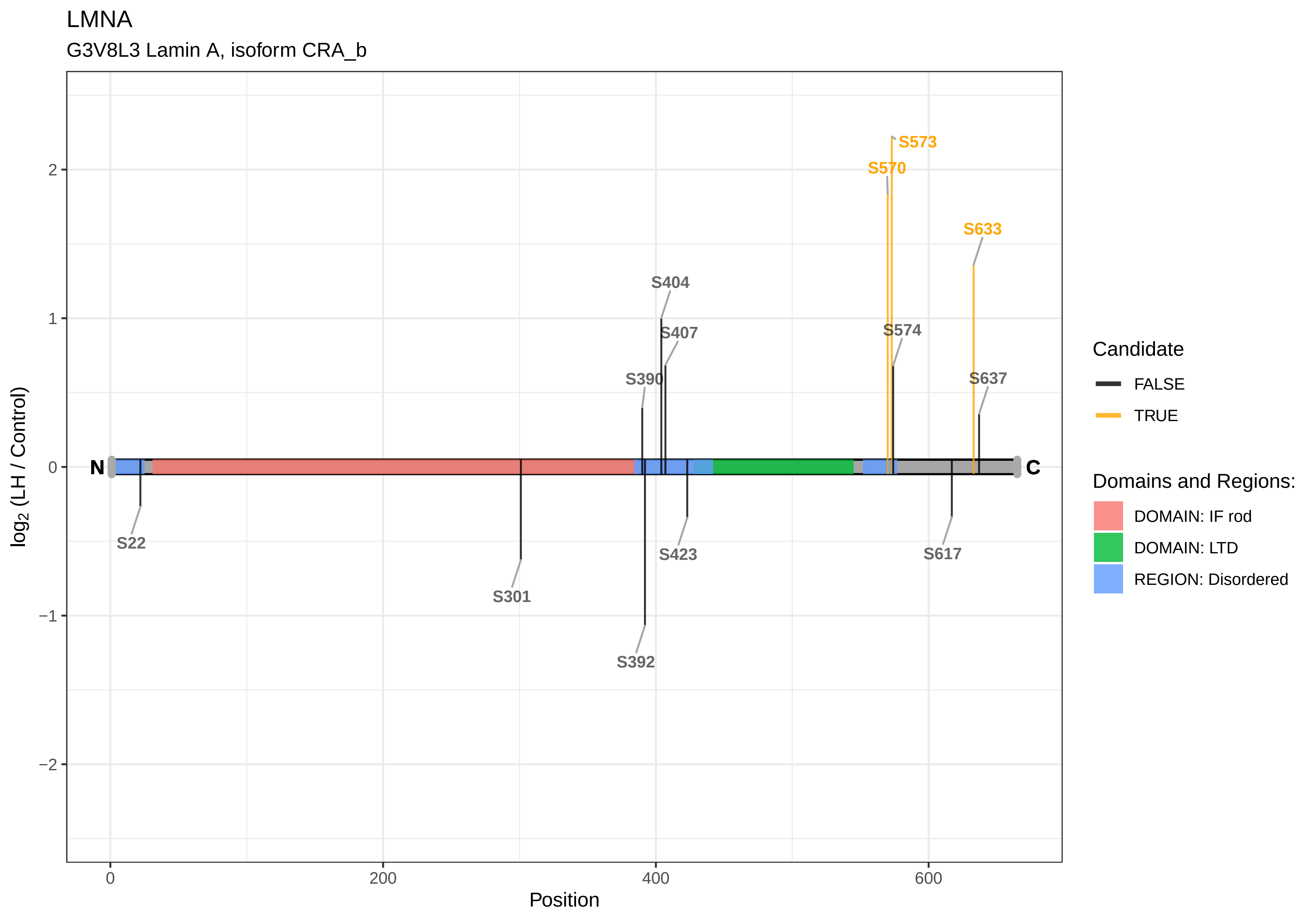
LMNA
G3V8L3 Lamin A, isoform CRA_b
S573
S570
S633
S404
S407
S574
S390
S637
N
C
S22
S301
S423
S617
S392
2
1
0
−1
−2
0
200
400
600
Position
log2 (LH / Control)
Candidate
FALSE
TRUE
Domains and Regions:
DOMAIN: IF rod
DOMAIN: LTD
REGION: Disordered

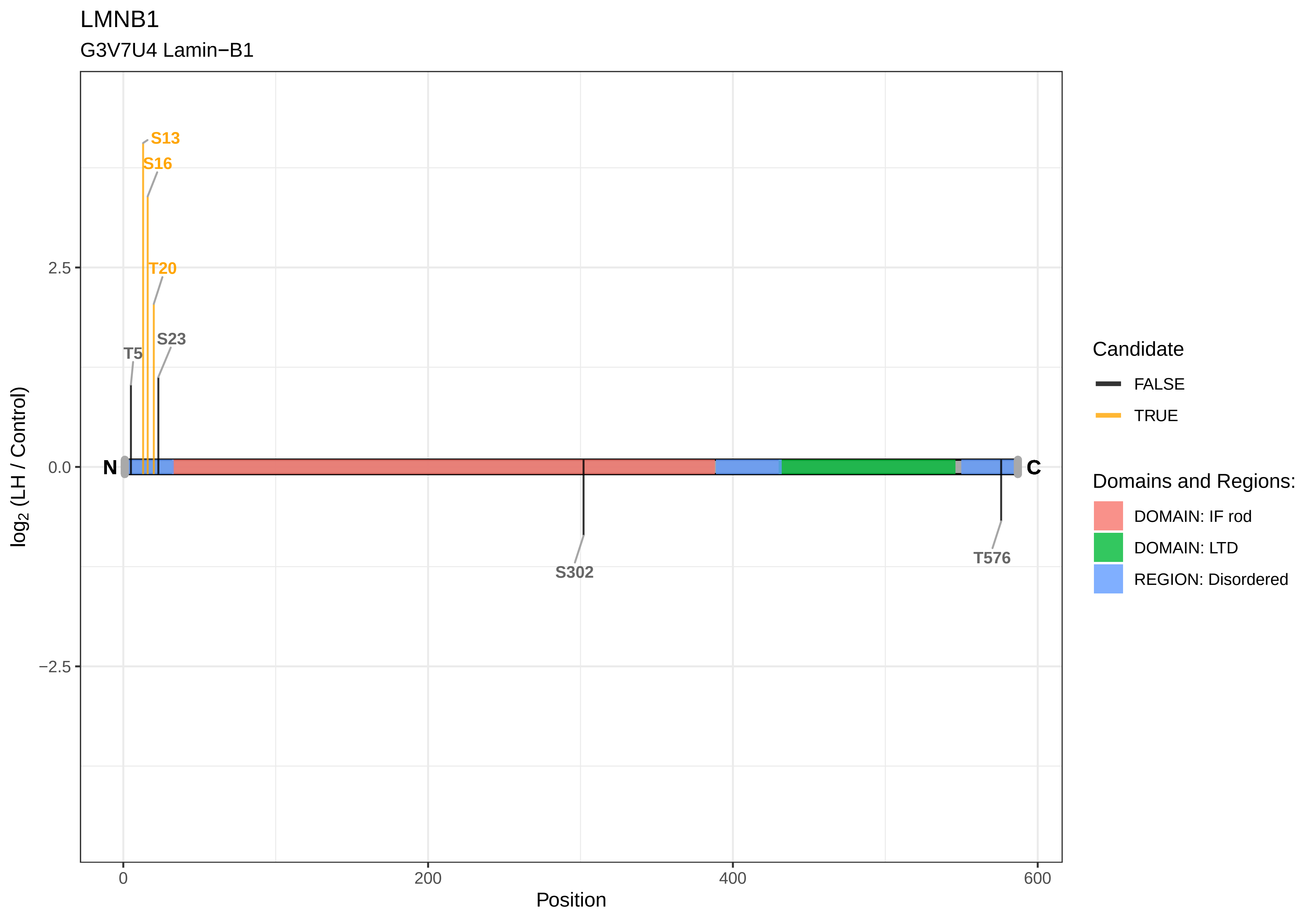

LMNB1
G3V7U4 Lamin−B1
S13
S16
T20
S23
T5
N
C
S302
T576
2.5
0.0
−2.5
0
200
400
600
Position
log2 (LH / Control)
Candidate
FALSE
TRUE
Domains and Regions:
DOMAIN: IF rod
DOMAIN: LTD
REGION: Disordered

LMNB2

D3ZLC1 Lamin B2

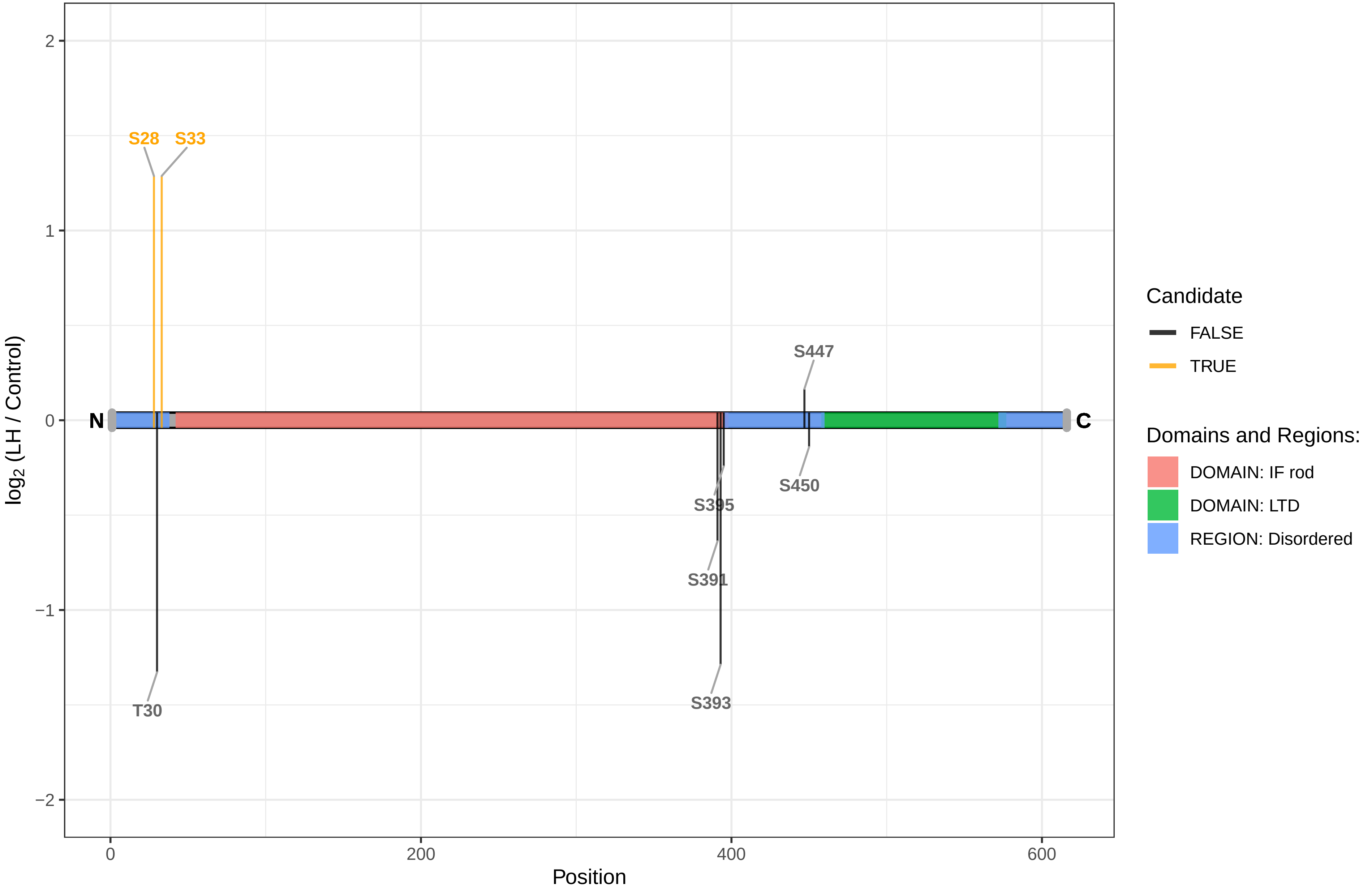

LOC100361087

F1M9N8 Hypothetical LOC100361087

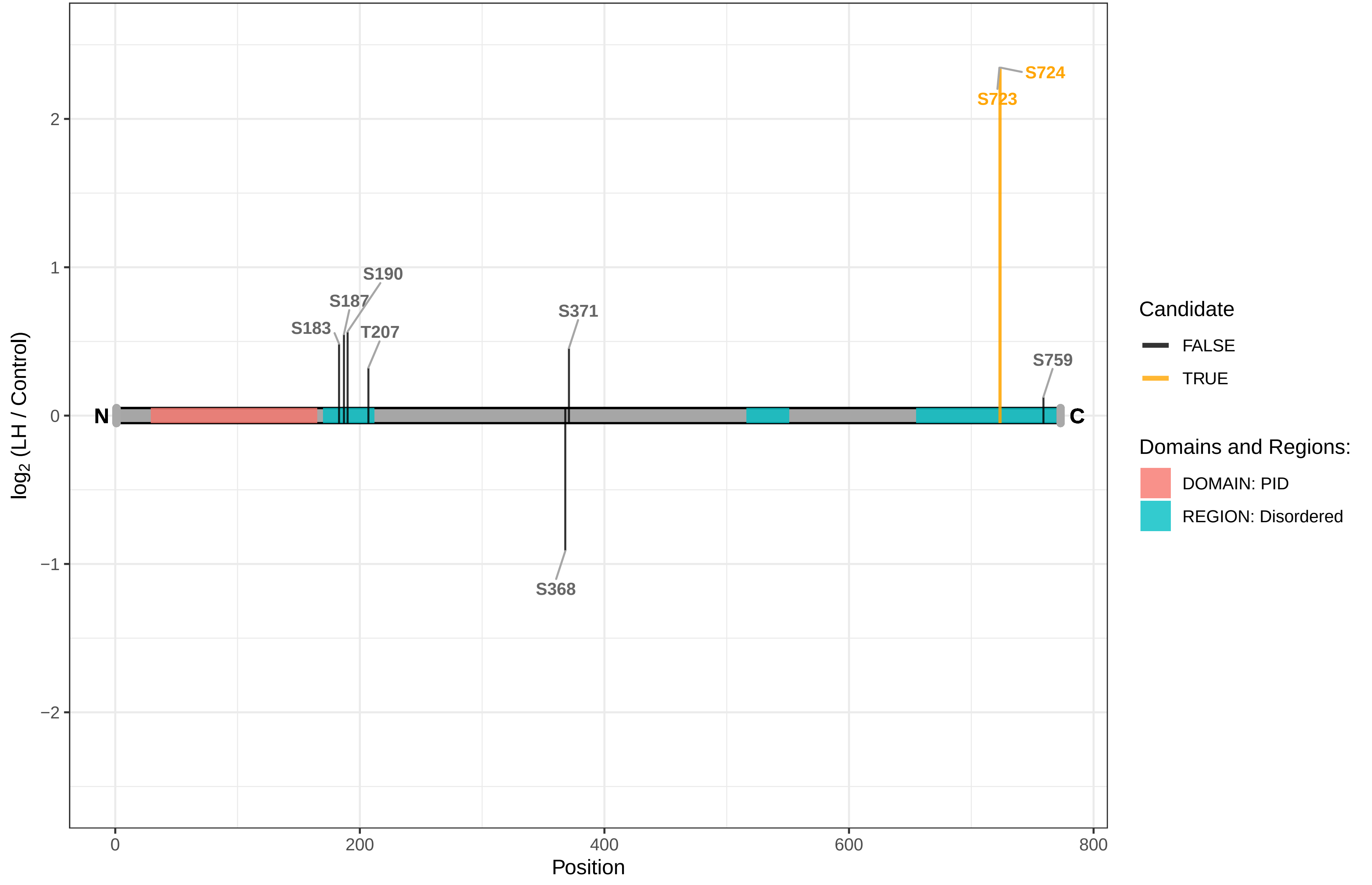

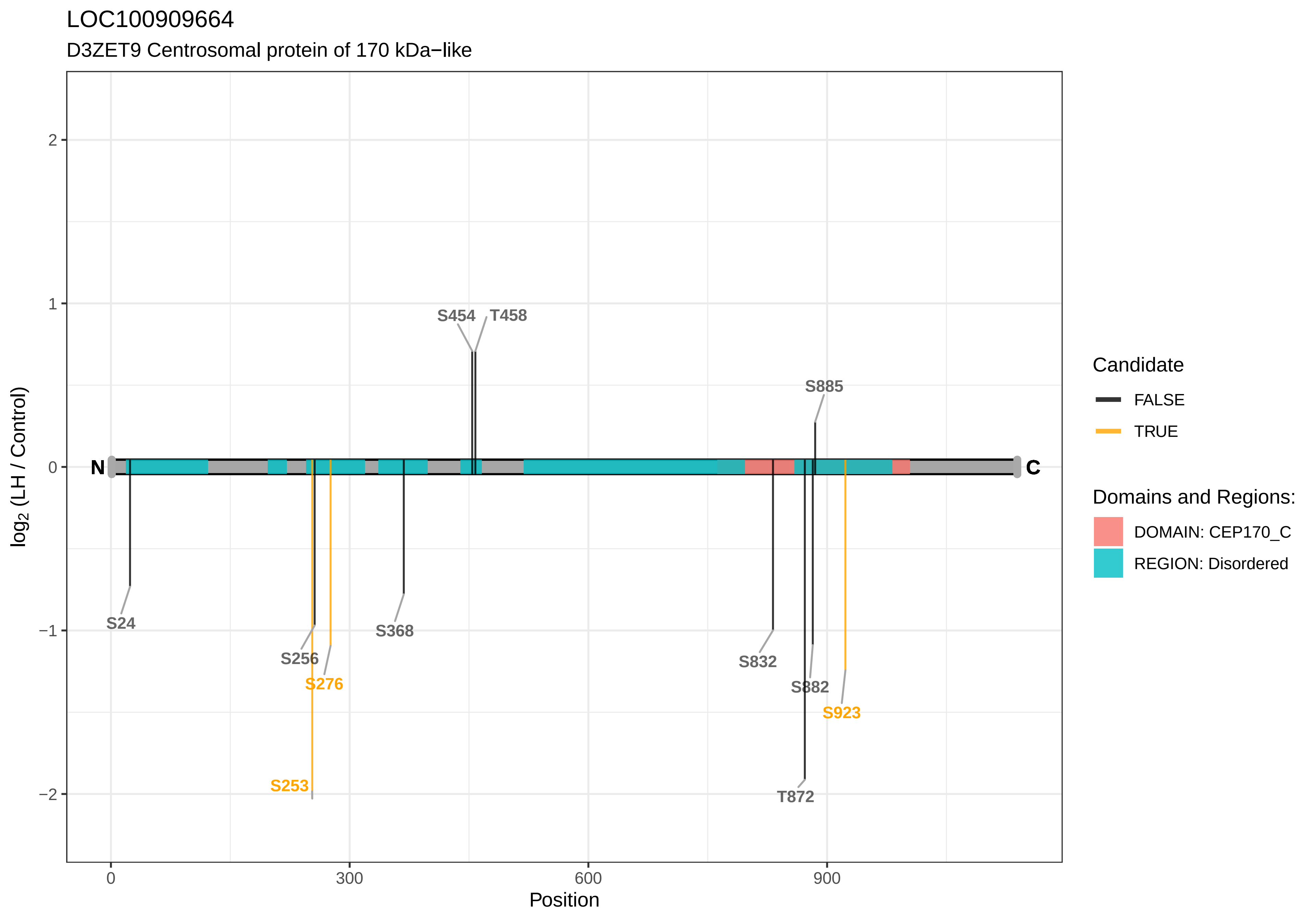
LOC100909664
D3ZET9 Centrosomal protein of 170 kDa−like
log2 (LH / Control)
Position
2
1
0
−1
−2
0
300
600
900
N
C
S454
T458
S885
S24
S256
S276
S368
S832
S882
S923
S253
T872
Candidate
FALSE
TRUE
Domains and Regions:
DOMAIN: CEP170_C
REGION: Disordered

LOC100910540

G3V656 Methylphosphate−capping enzyme

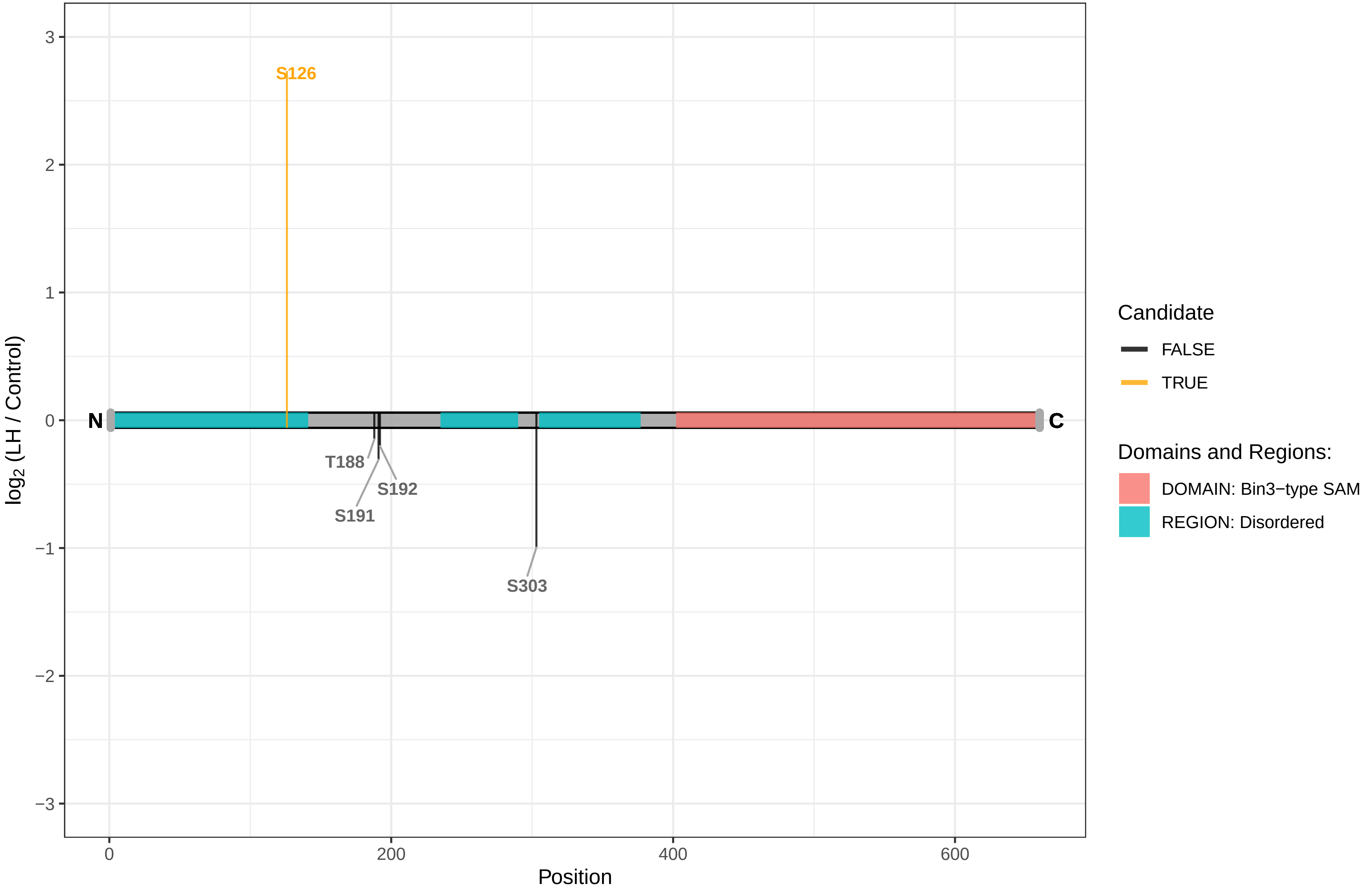

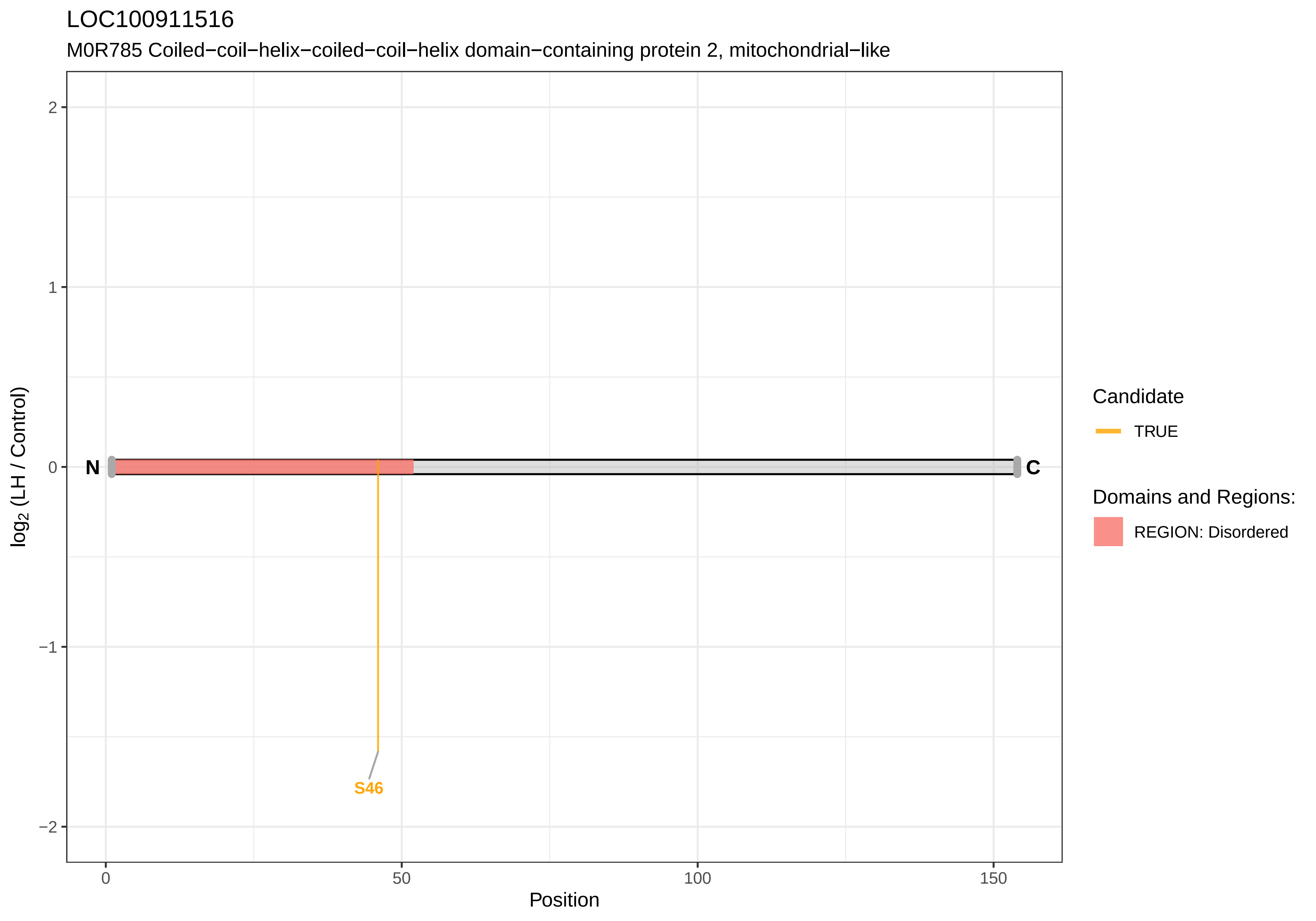
LOC100911516
M0R785 Coiled−coil−helix−coiled−coil−helix domain−containing protein 2, mitochondrial−like
log2 (LH / Control)
2
1
0
−1
−2
N
C
S46
0
50
100
150
Position
Candidate
TRUE
Domains and Regions:
REGION: Disordered

LOC100911660

A0A0G2K7M6 Lymphocyte−specific helicase−like

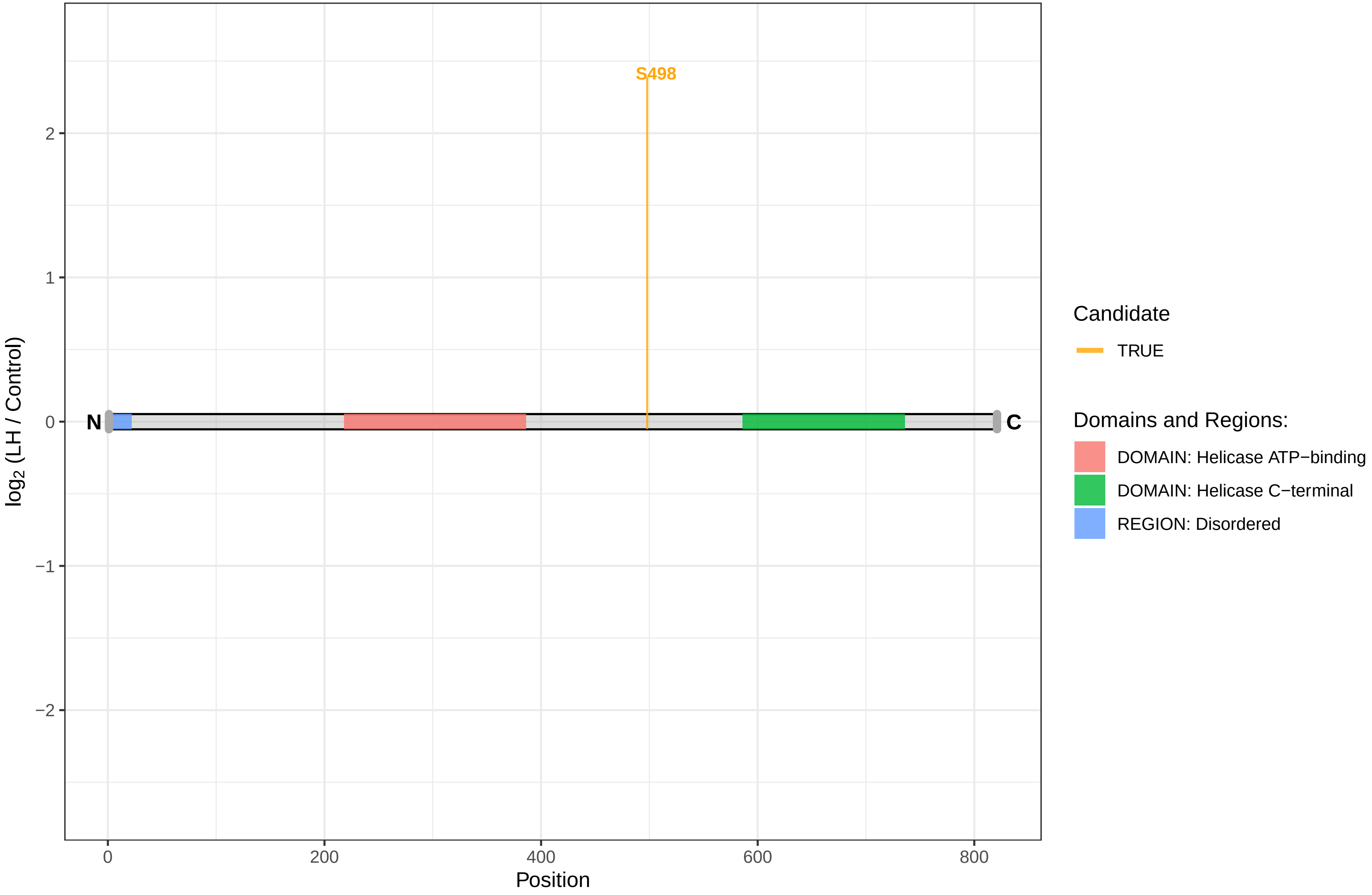

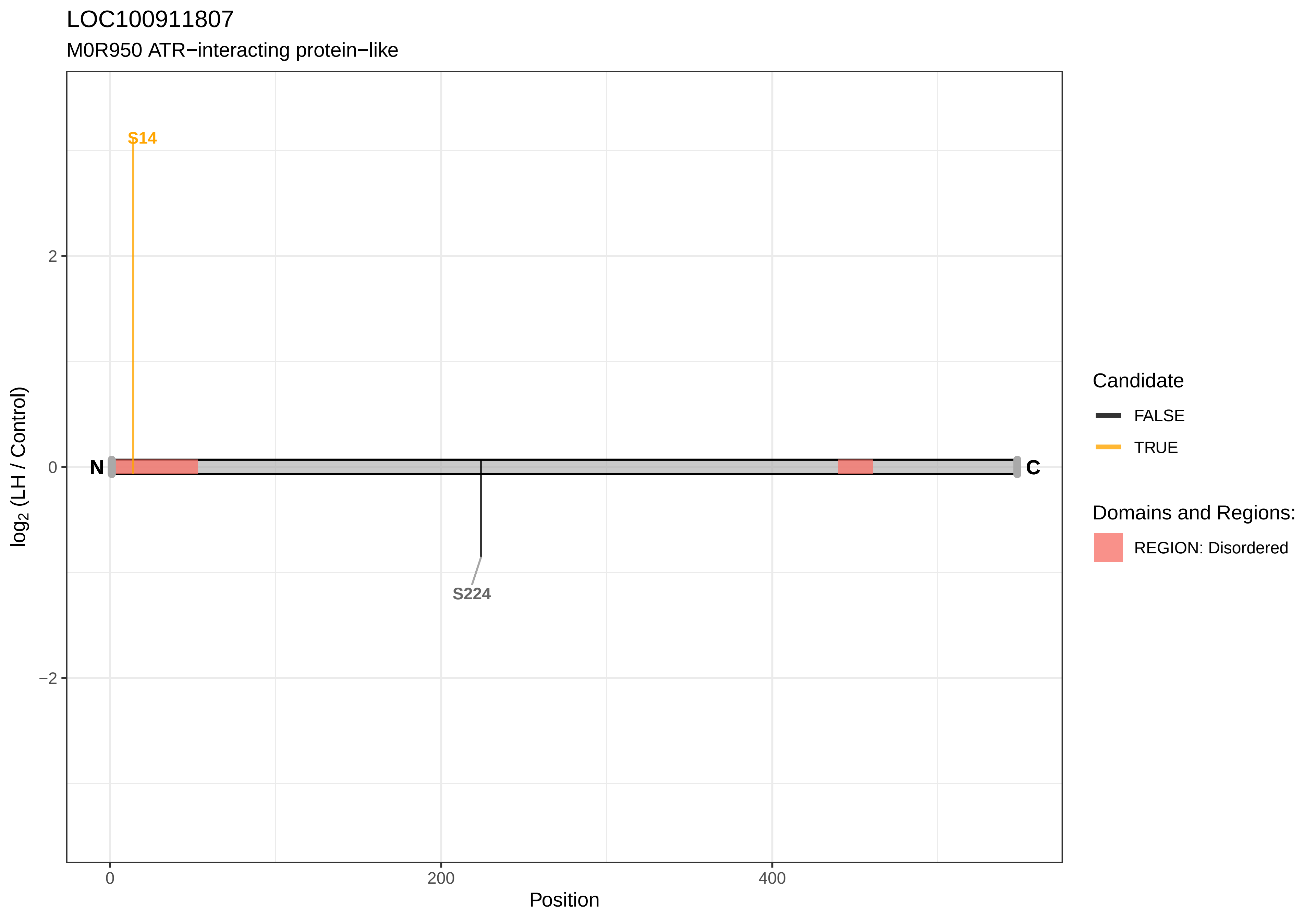

LOC100911807
M0R950 ATR−interacting protein−like
S14
S224
N
C
2
0
−2
0
200
400
Position
log2 (LH / Control)
Candidate
FALSE
TRUE
Domains and Regions:
REGION: Disordered

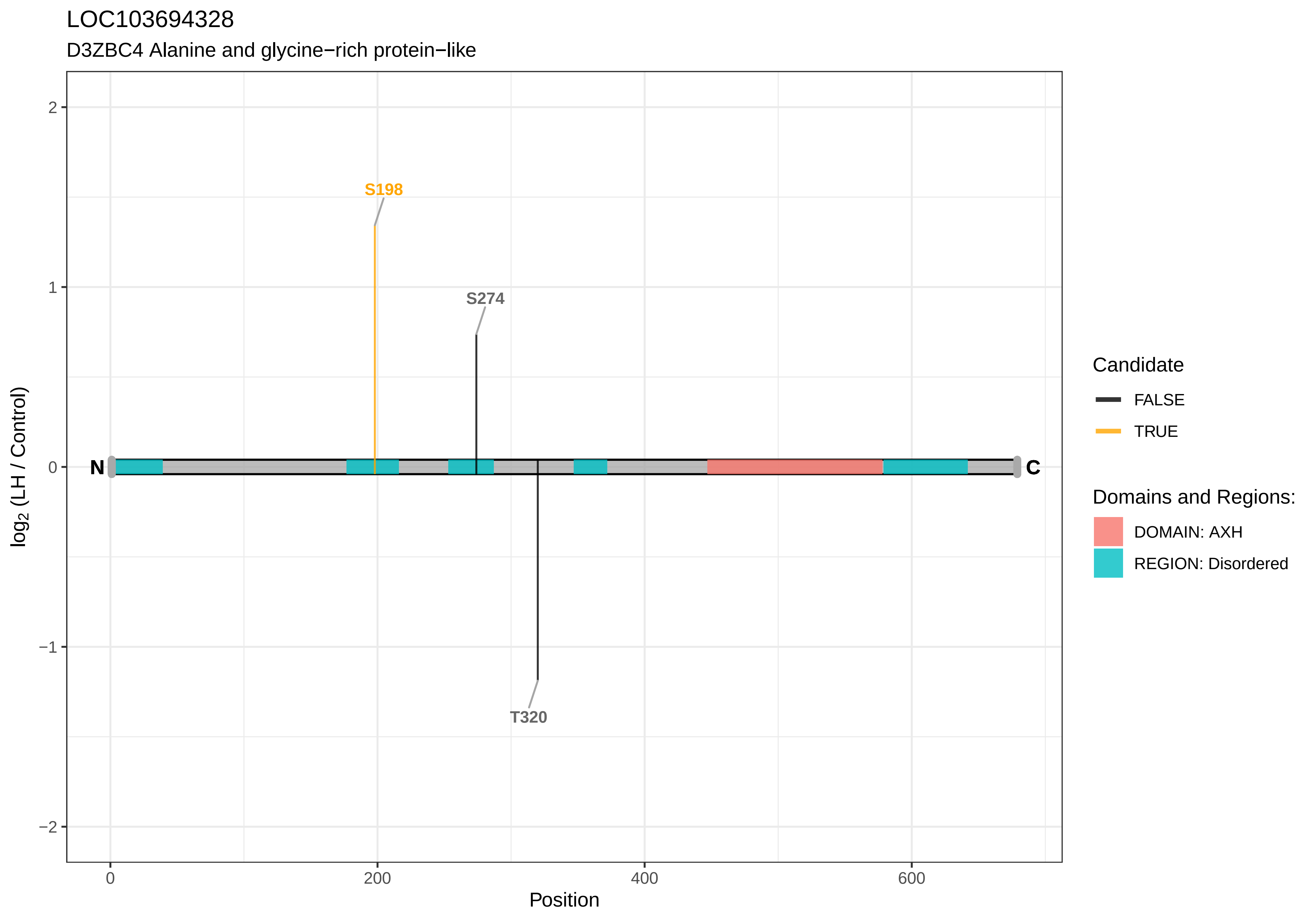

LOC103694328
D3ZBC4 Alanine and glycine−rich protein−like
S198
S274
T320
N
C
2
1
0
−1
−2
0
200
400
600
Position
log2 (LH / Control)
Candidate
FALSE
TRUE
Domains and Regions:
DOMAIN: AXH
REGION: Disordered

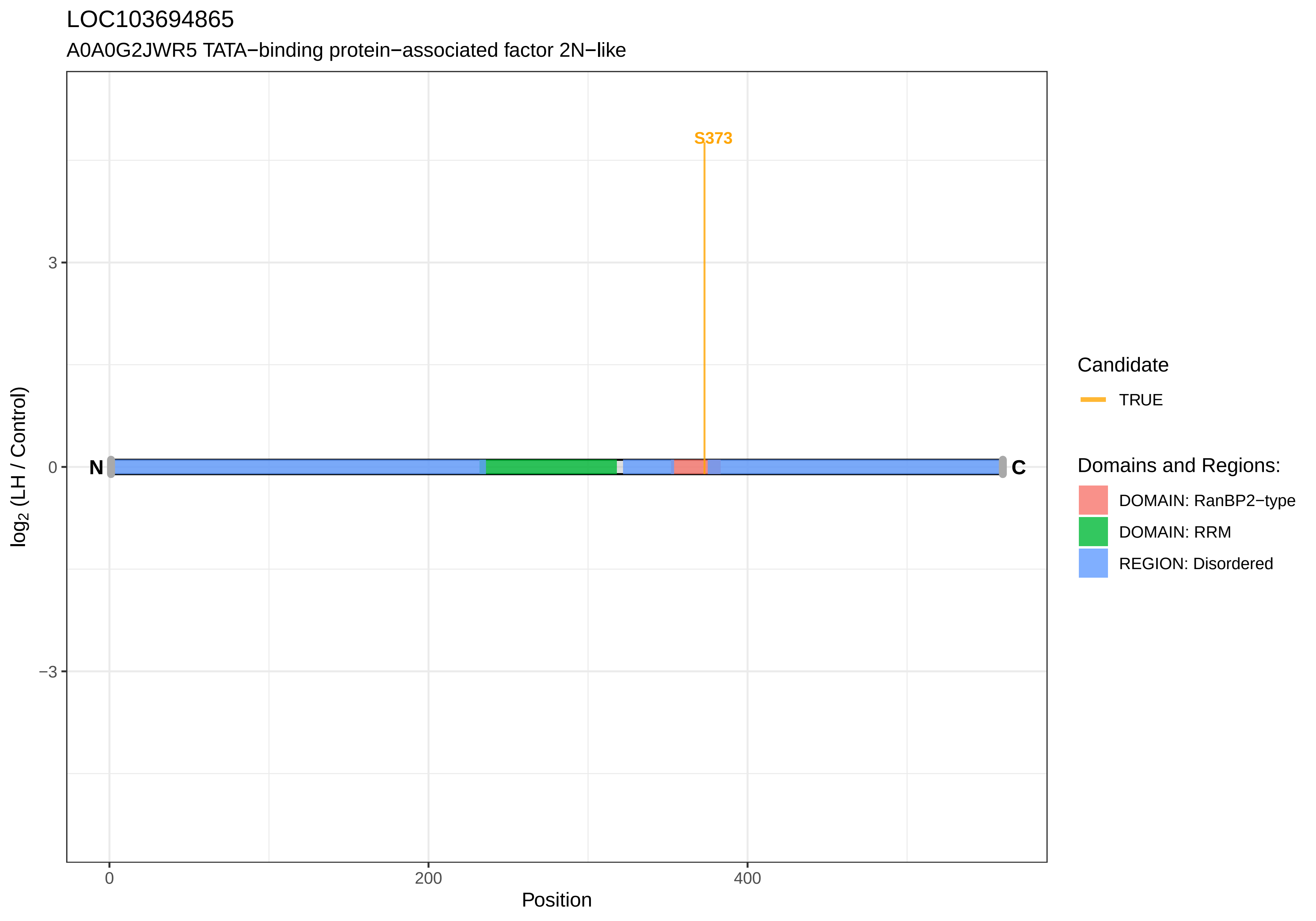

LOC103694865
A0A0G2JWR5 TATA−binding protein−associated factor 2N−like
S373
N
C
3
0
−3
0
200
400
Position
log2 (LH / Control)
Candidate
TRUE
Domains and Regions:
DOMAIN: RanBP2−type
DOMAIN: RRM
REGION: Disordered

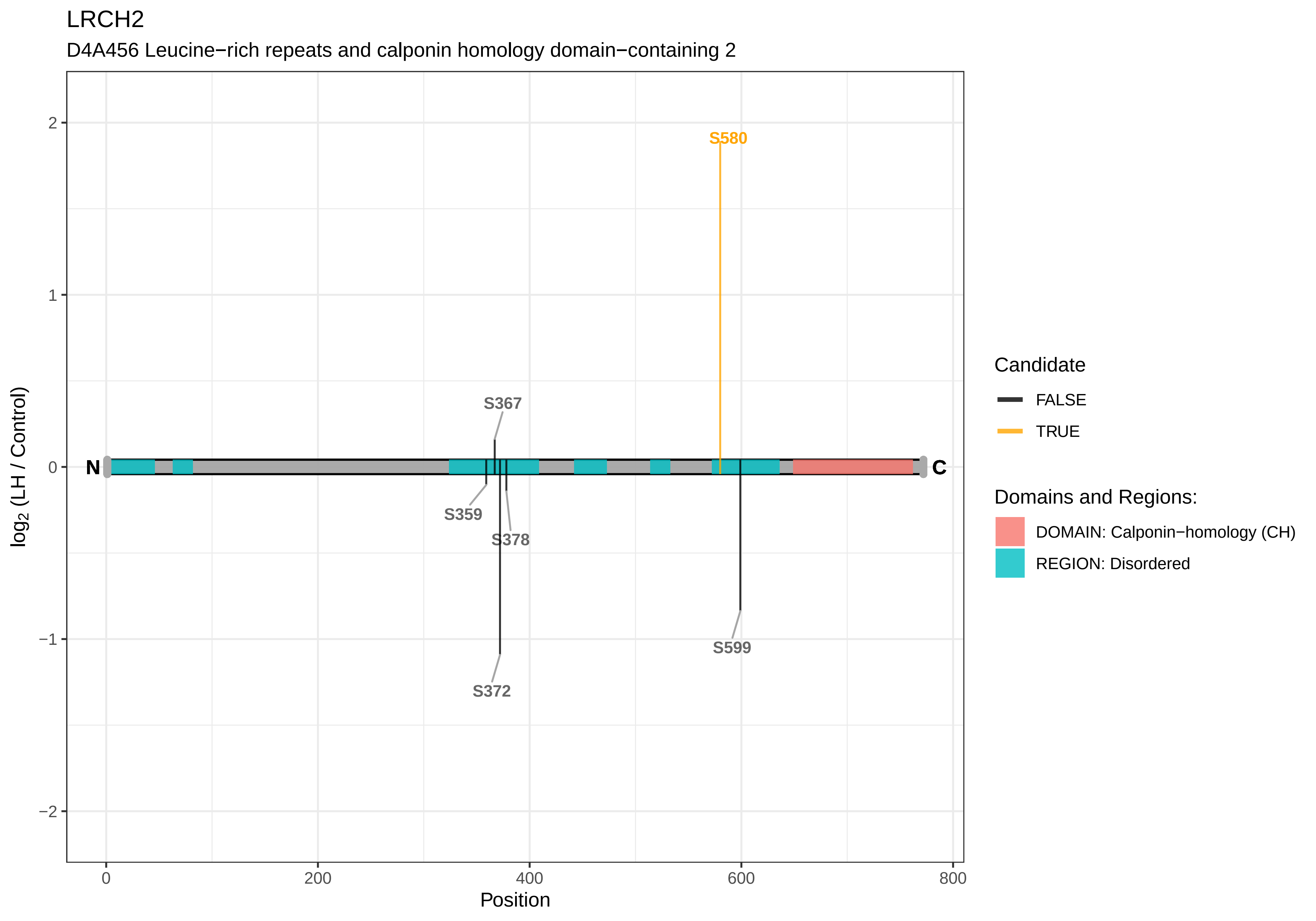

LRCH2
D4A456 Leucine−rich repeats and calponin homology domain−containing 2
S580
S367
S359
S378
S372
S599
N
C
2
1
0
−1
−2
log2 (LH / Control)
0
200
400
600
800
Position
Candidate
FALSE
TRUE
Domains and Regions:
DOMAIN: Calponin−homology (CH)
REGION: Disordered

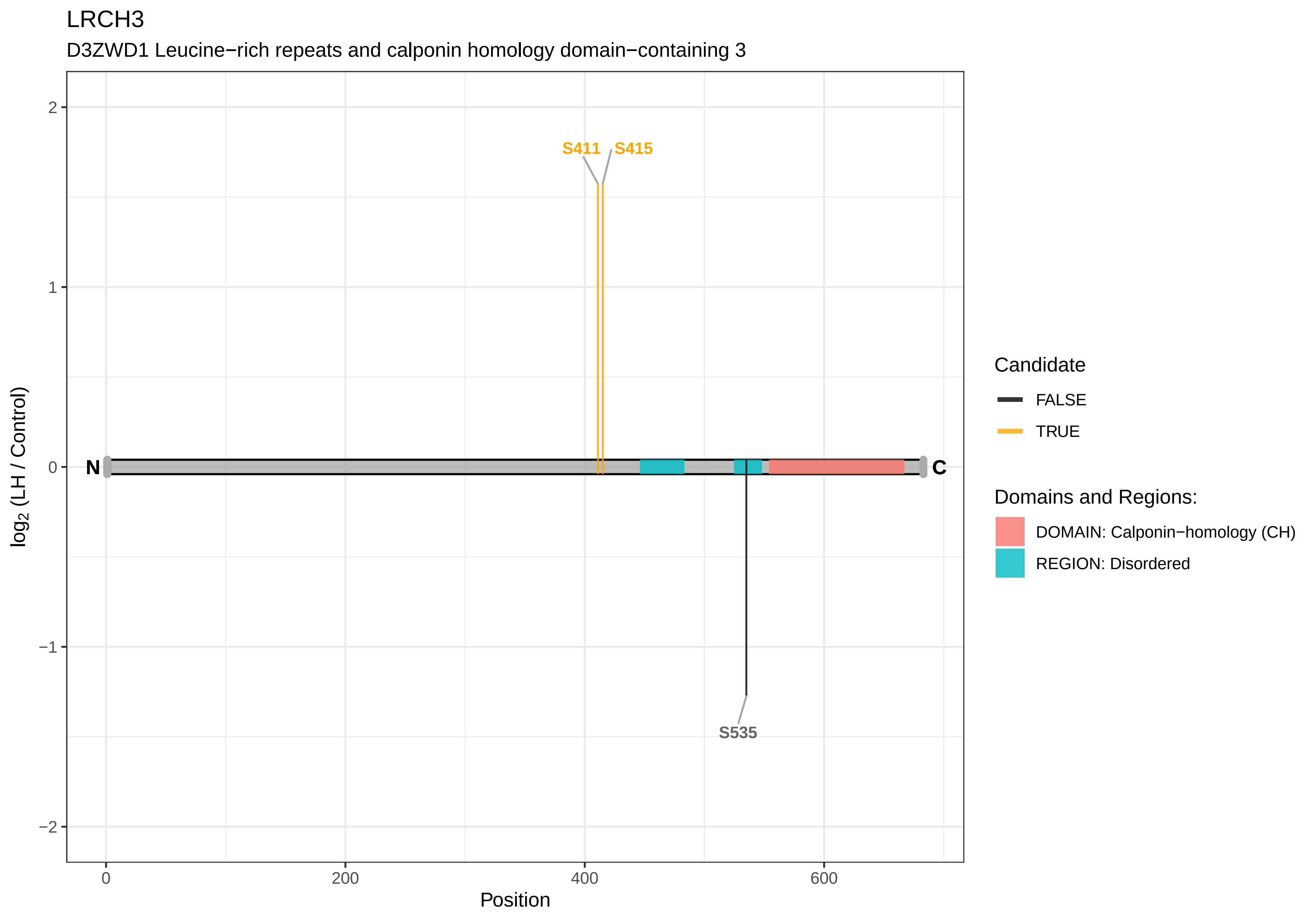

LRCH3
D3ZWD1 Leucine−rich repeats and calponin homology domain−containing 3
S411
S415
S535
N
C
2
1
0
−1
−2
log2 (LH / Control)
0
200
400
600
Position
Candidate
FALSE
TRUE
Domains and Regions:
DOMAIN: Calponin−homology (CH)
REGION: Disordered

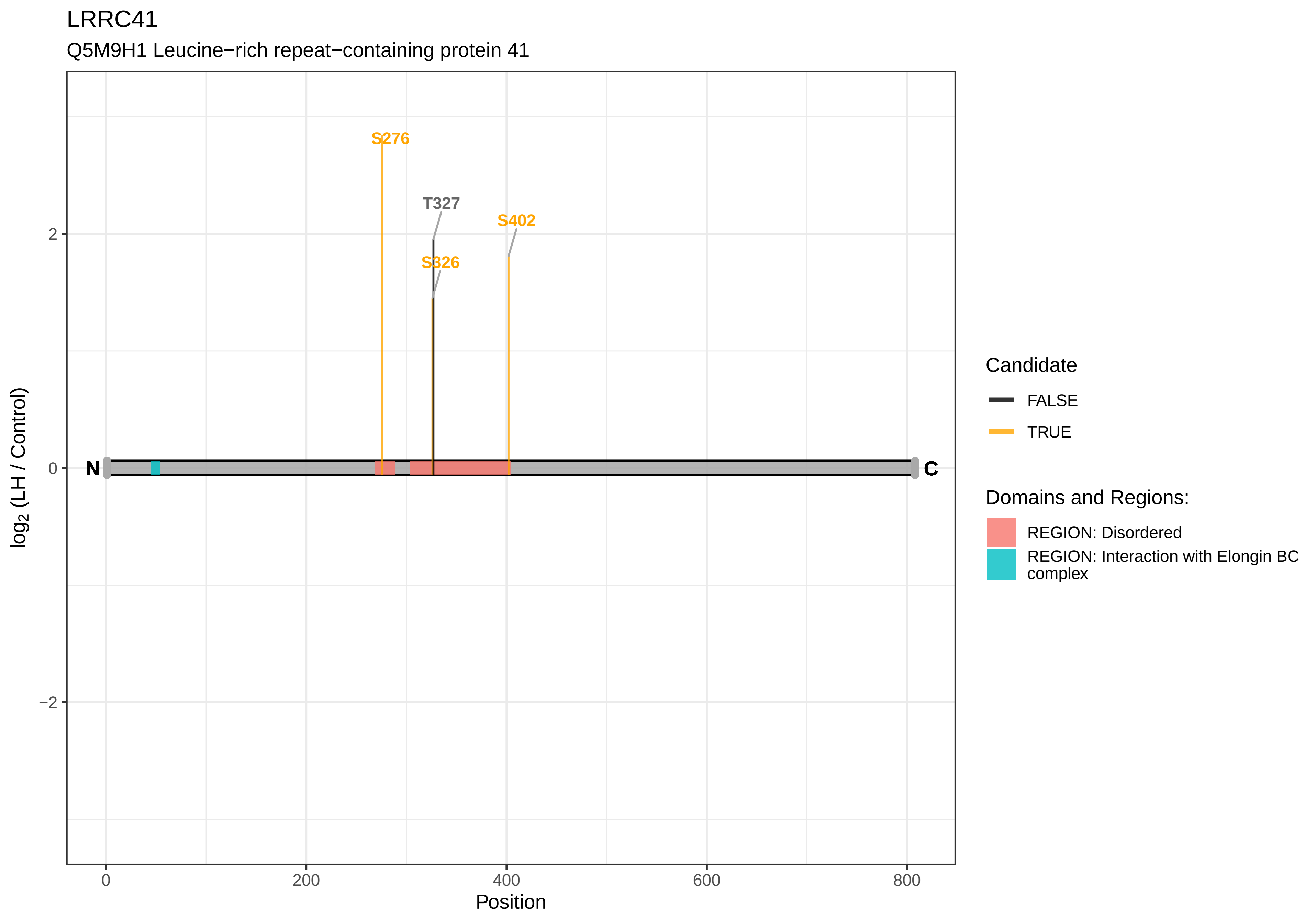

LRRC41
Q5M9H1 Leucine−rich repeat−containing protein 41
S276
T327
S402
S326
N
C
2
0
−2
log2 (LH / Control)
0
200
400
600
800
Position
Candidate
FALSE
TRUE
Domains and Regions:
REGION: Disordered
REGION: Interaction with Elongin BC complex

LRRC47

F1LT49 Leucine−rich repeat−containing 47

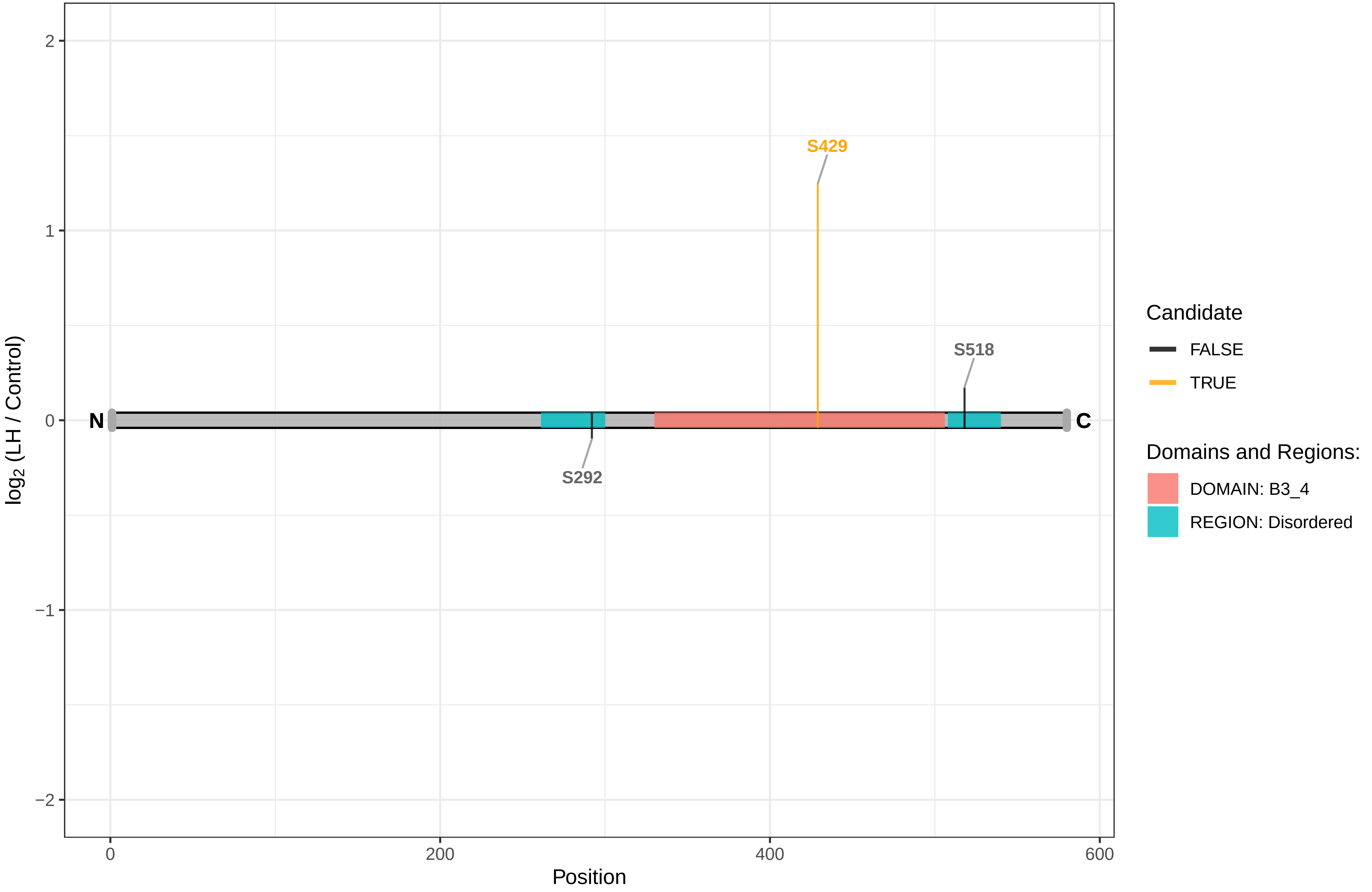

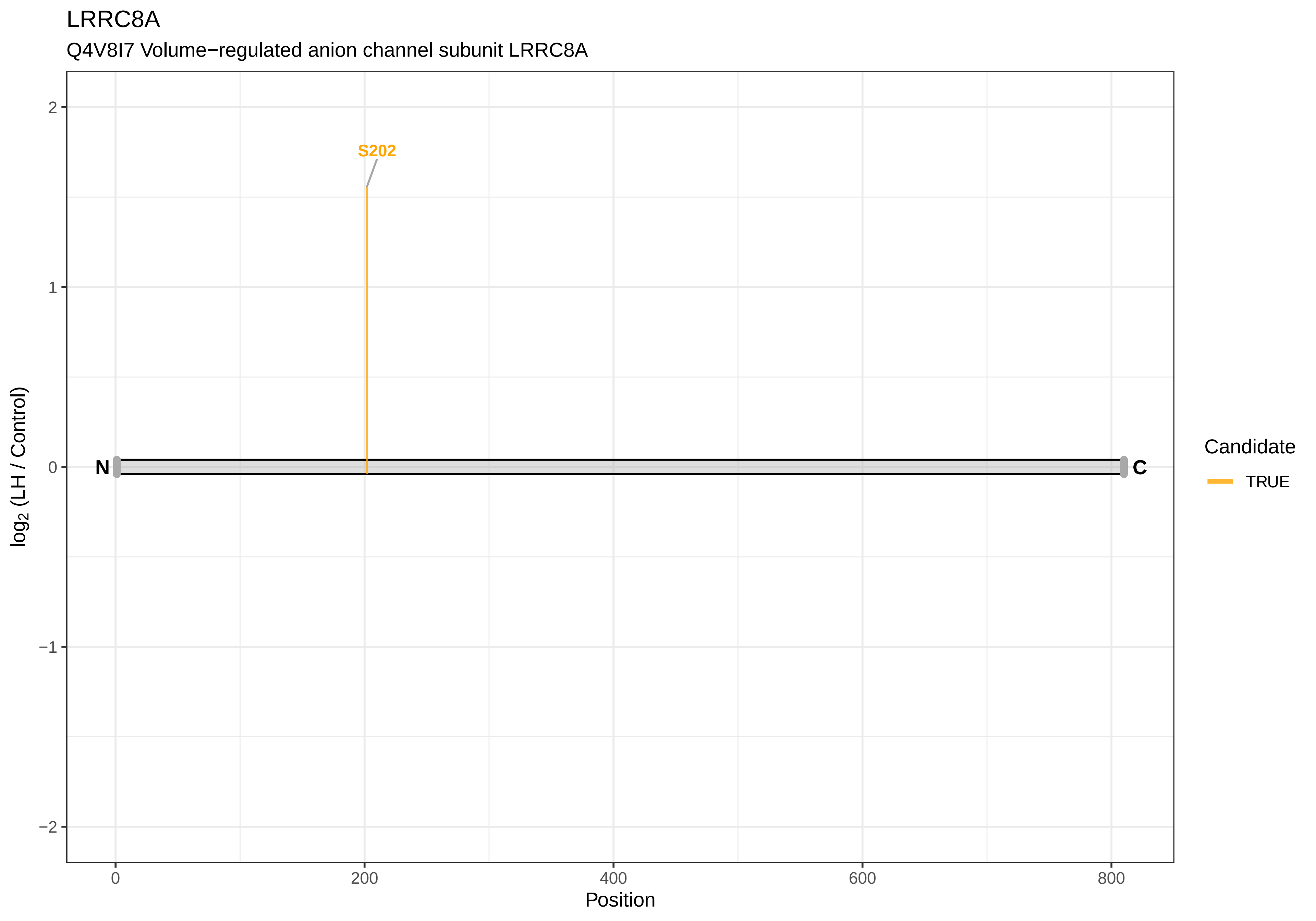

LRRC8A
Q4V8I7 Volume−regulated anion channel subunit LRRC8A
S202
N
C
2
1
0
−1
−2
log2 (LH / Control)
0
200
400
600
800
Position
Candidate
TRUE

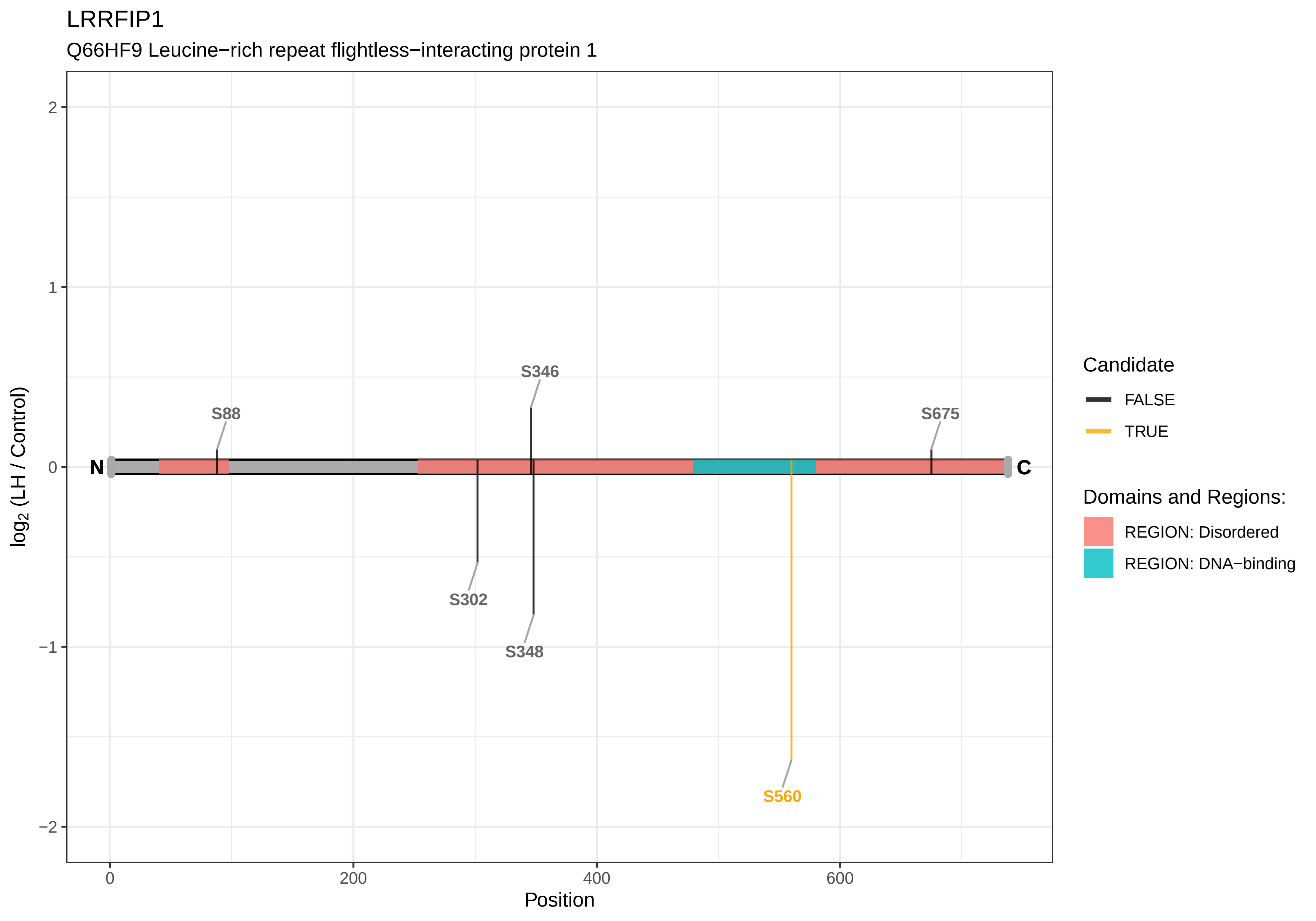

LRRFIP1
Q66HF9 Leucine−rich repeat flightless−interacting protein 1
log2 (LH / Control)
Position
S88
S346
S675
S302
S348
S560
N
C
2
1
0
−1
−2
0
200
400
600
Candidate
FALSE
TRUE
Domains and Regions:
REGION: Disordered
REGION: DNA−binding

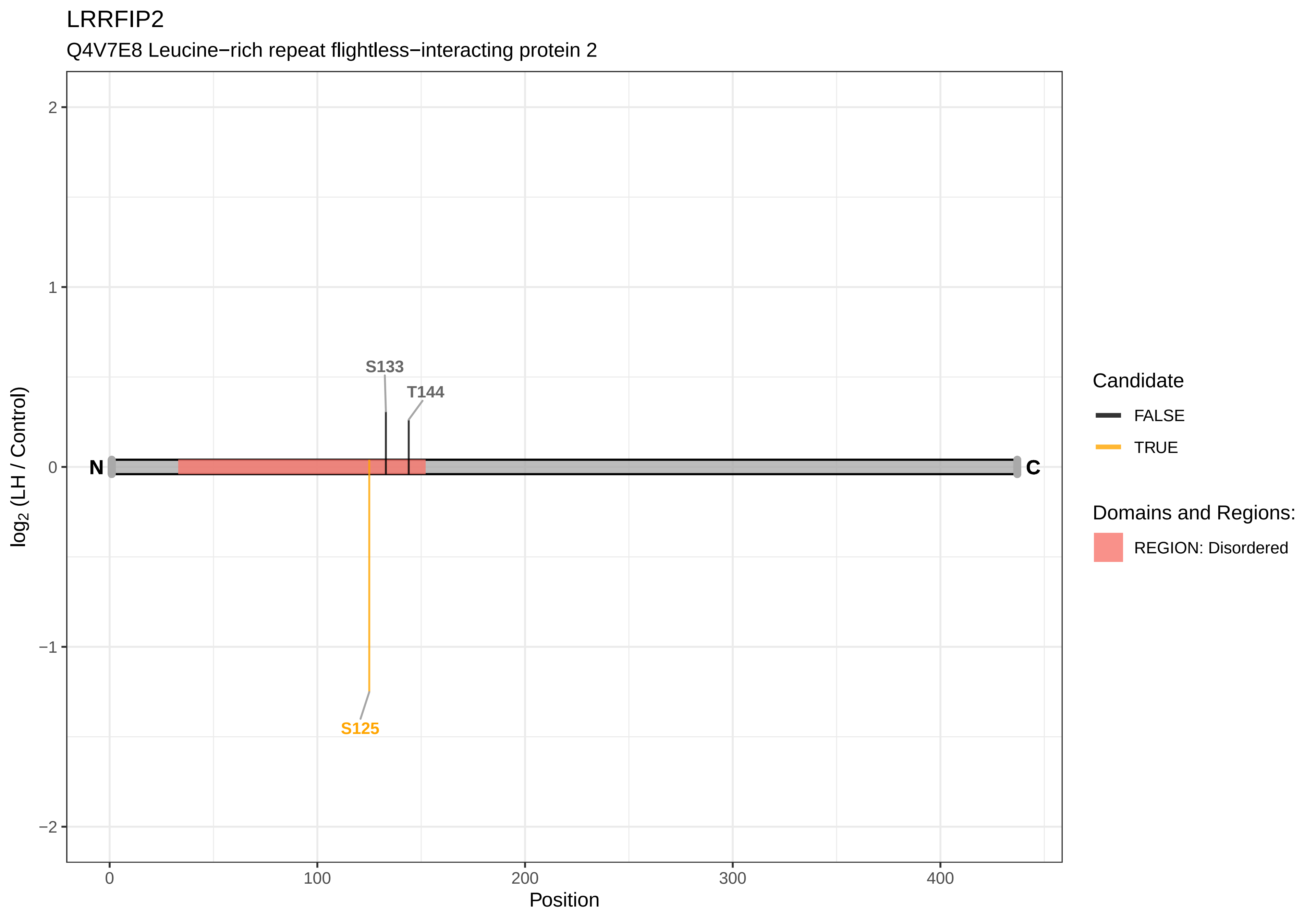

LRRFIP2
Q4V7E8 Leucine−rich repeat flightless−interacting protein 2
S133
T144
S125
N
C
log2 (LH / Control)
Position
Candidate
FALSE
TRUE
Domains and Regions:
REGION: Disordered

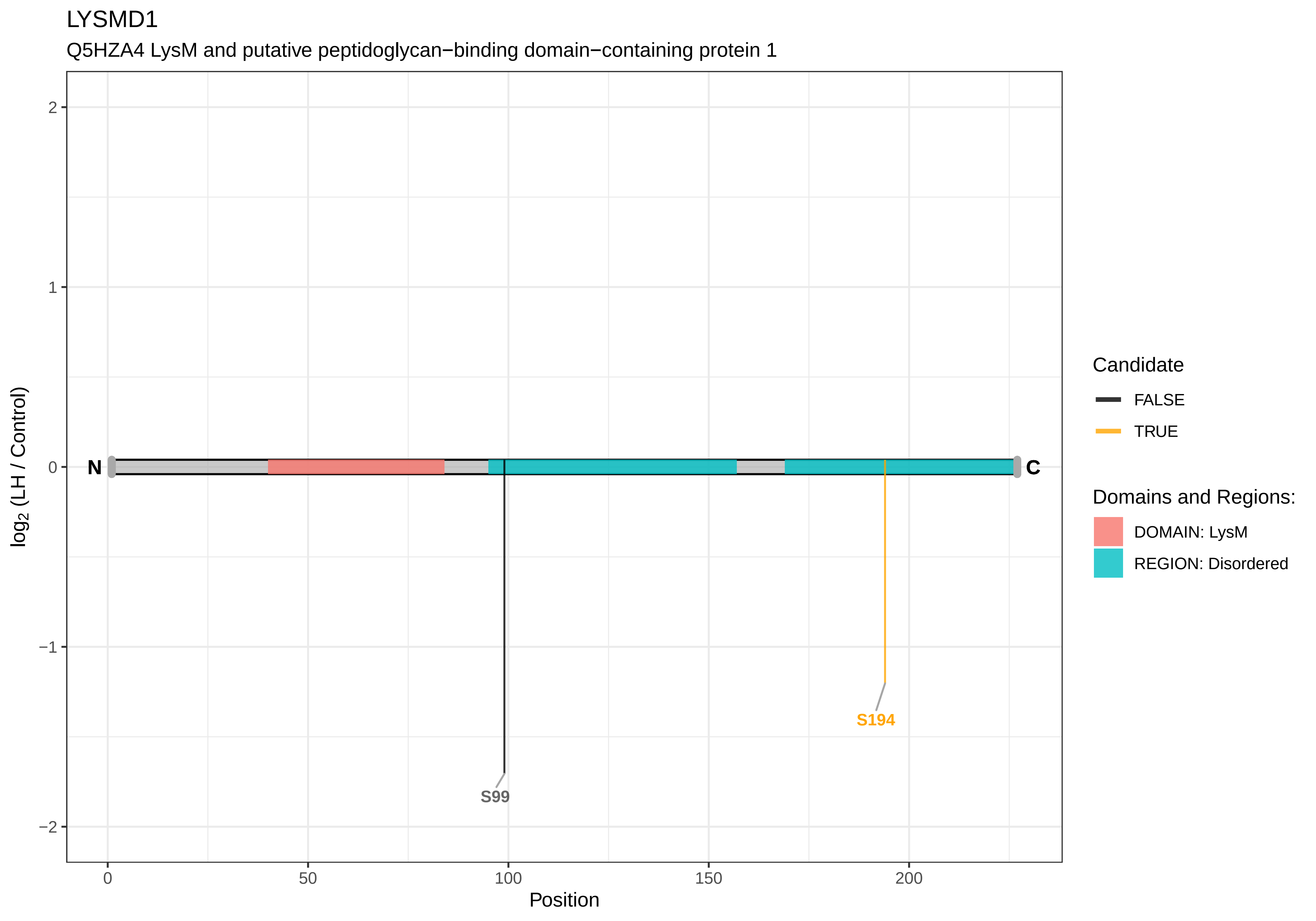

LYSMD1
Q5HZA4 LysM and putative peptidoglycan−binding domain−containing protein 1
log2 (LH / Control)
Position
N
C
S99
S194
Candidate
FALSE
TRUE
Domains and Regions:
DOMAIN: LysM
REGION: Disordered

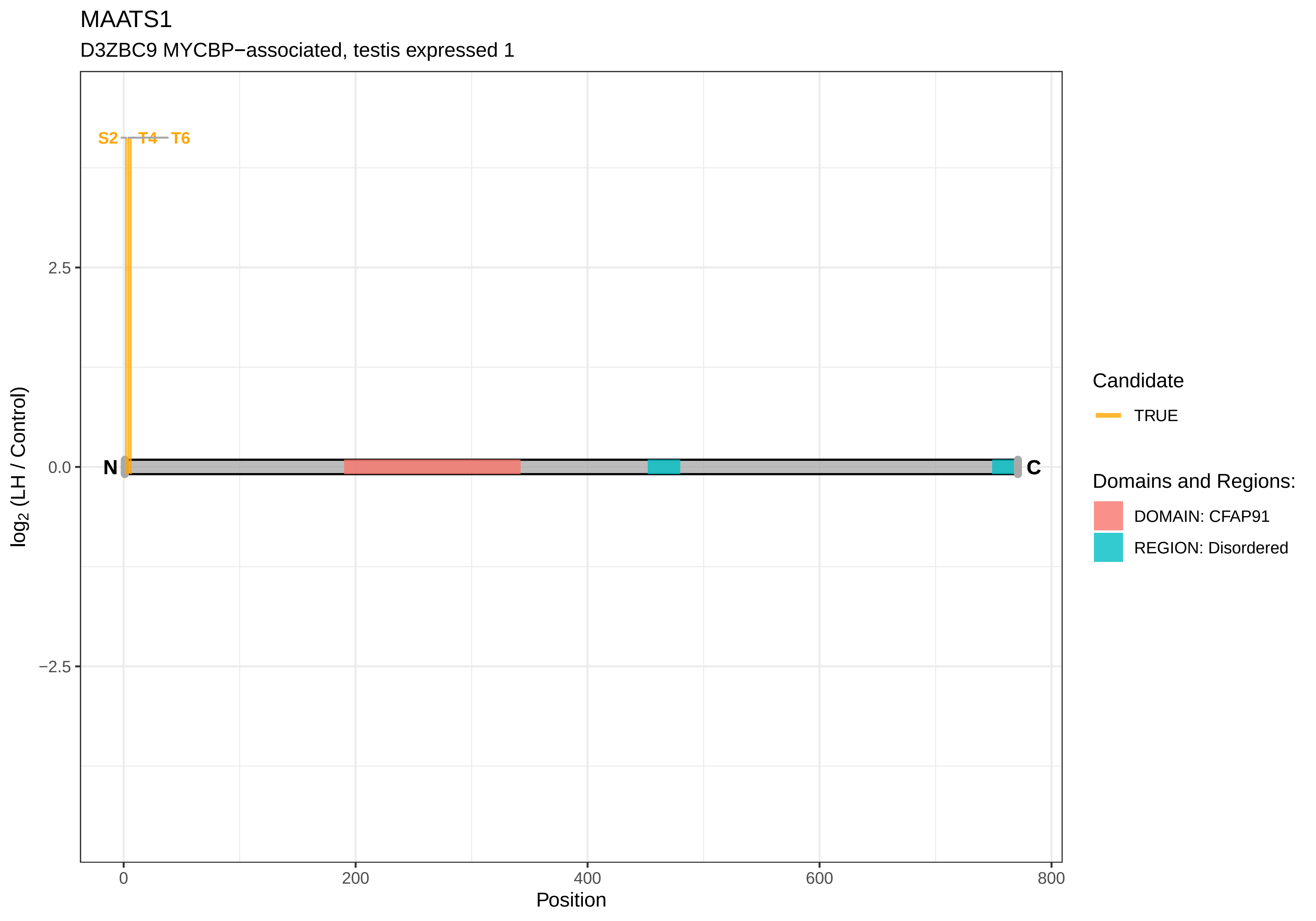

MAATS1
D3ZBC9 MYCBP−associated, testis expressed 1
S2
T4
T6
N
C
2.5
0.0
−2.5
log2 (LH / Control)
0
200
400
600
800
Position
Candidate
TRUE
Domains and Regions:
DOMAIN: CFAP91
REGION: Disordered

MACF1

A0A0G2JWA8 Microtubule−actin cross−linking factor 1

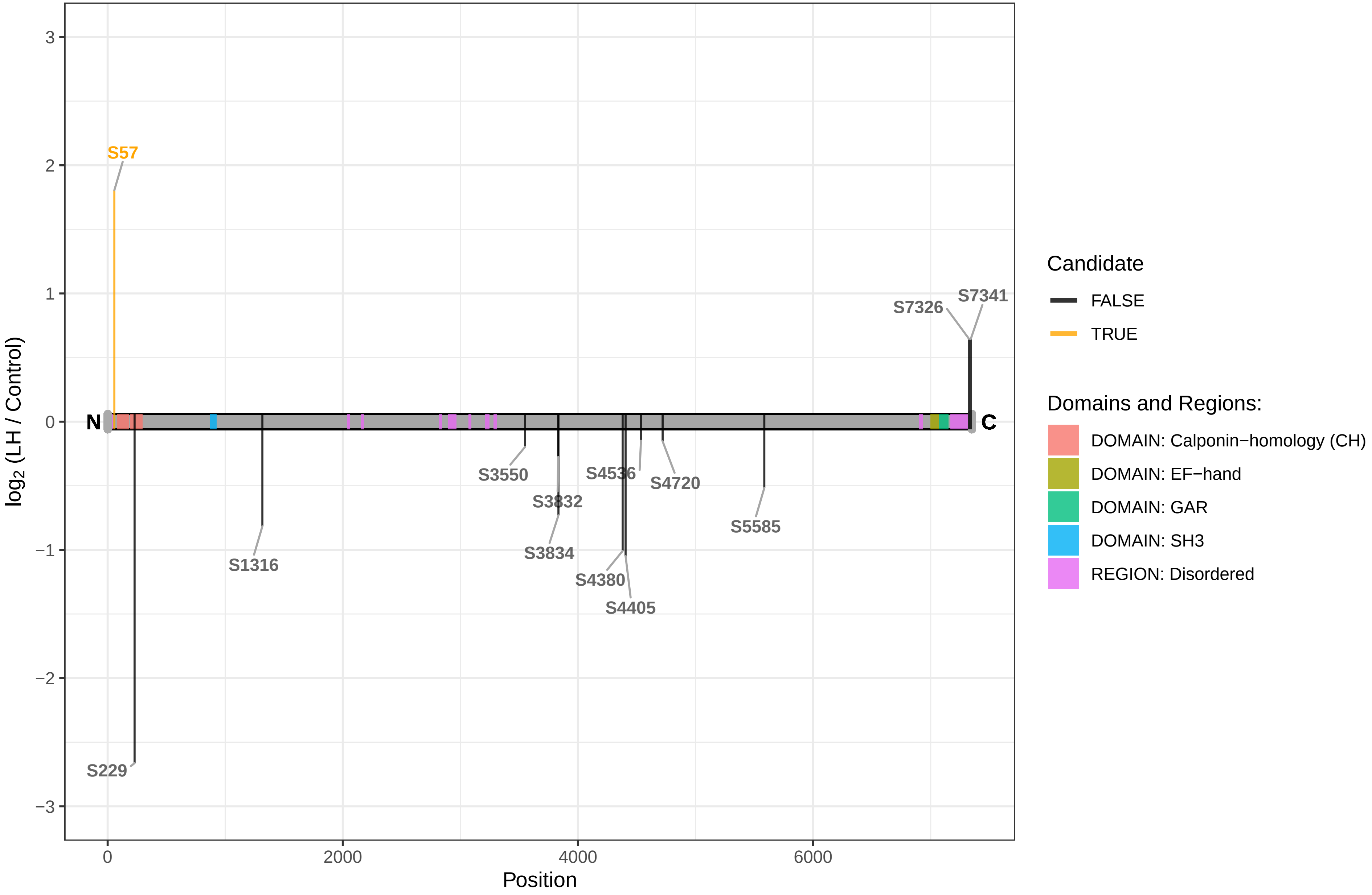

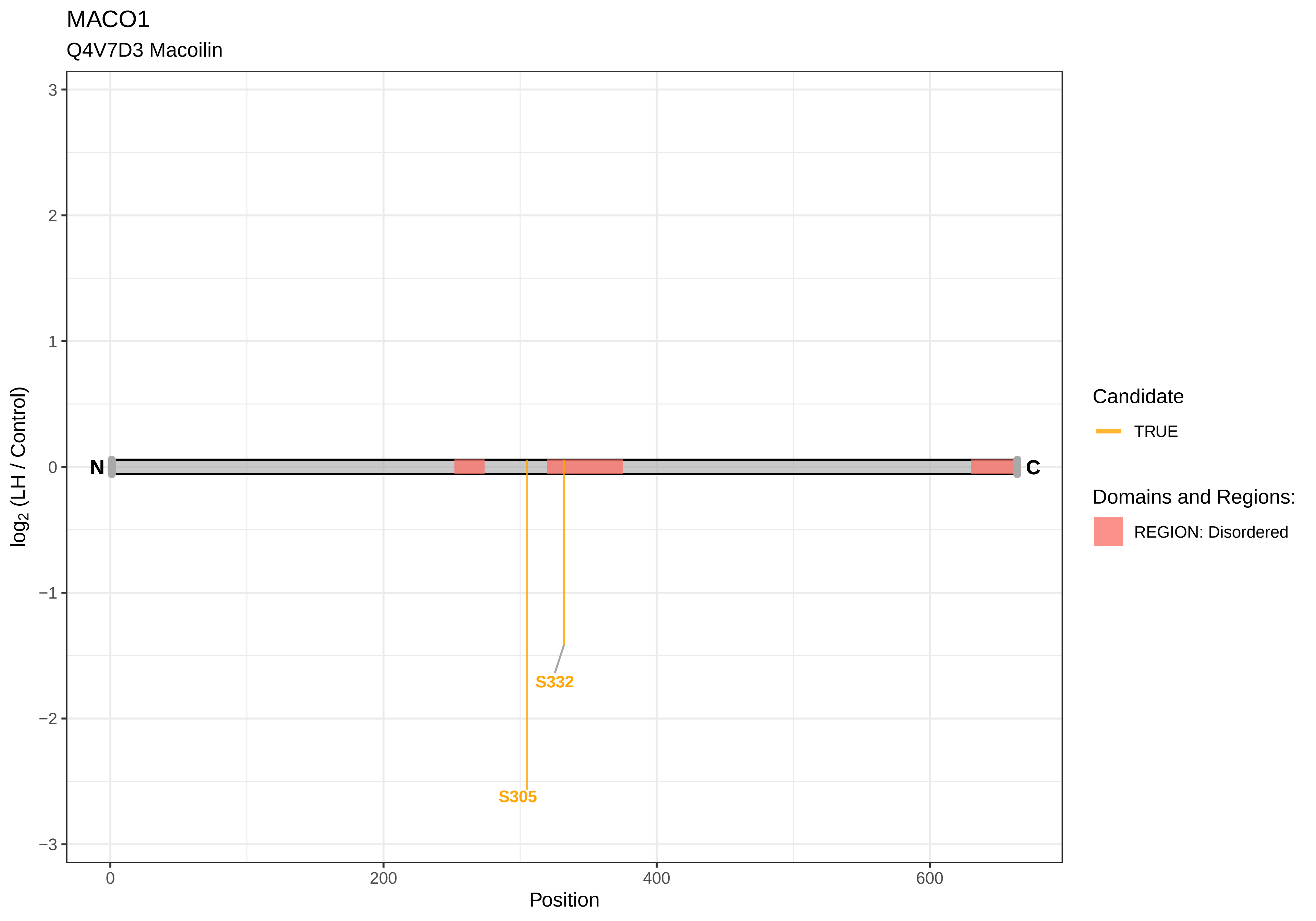

MACO1
Q4V7D3 Macoilin
log2 (LH / Control)
3
2
1
0
−1
−2
−3
N
C
S332
S305
0
200
400
600
Position
Candidate
TRUE
Domains and Regions:
REGION: Disordered

MADD

O08873 MAP kinase−activating death domain protein

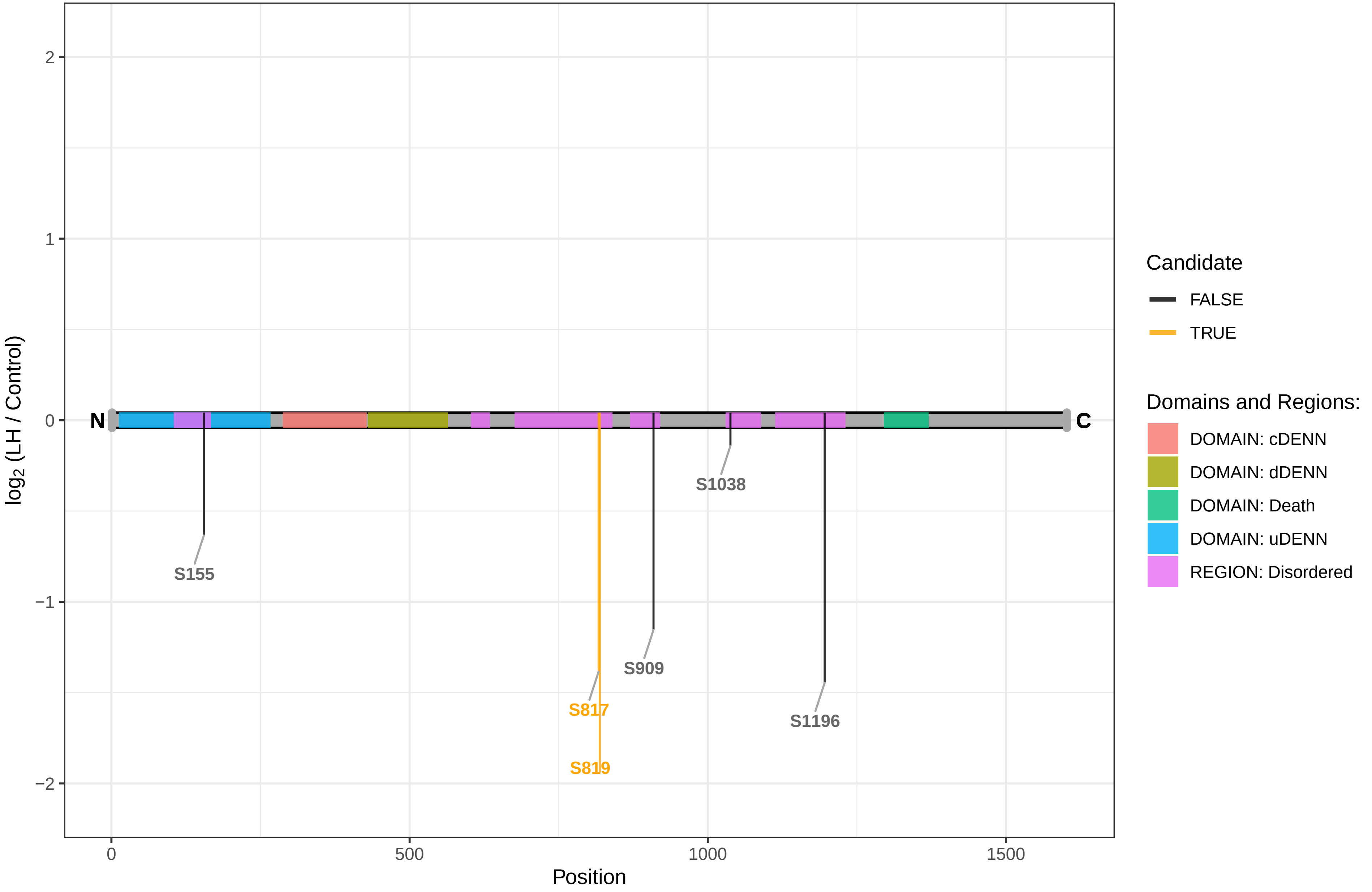

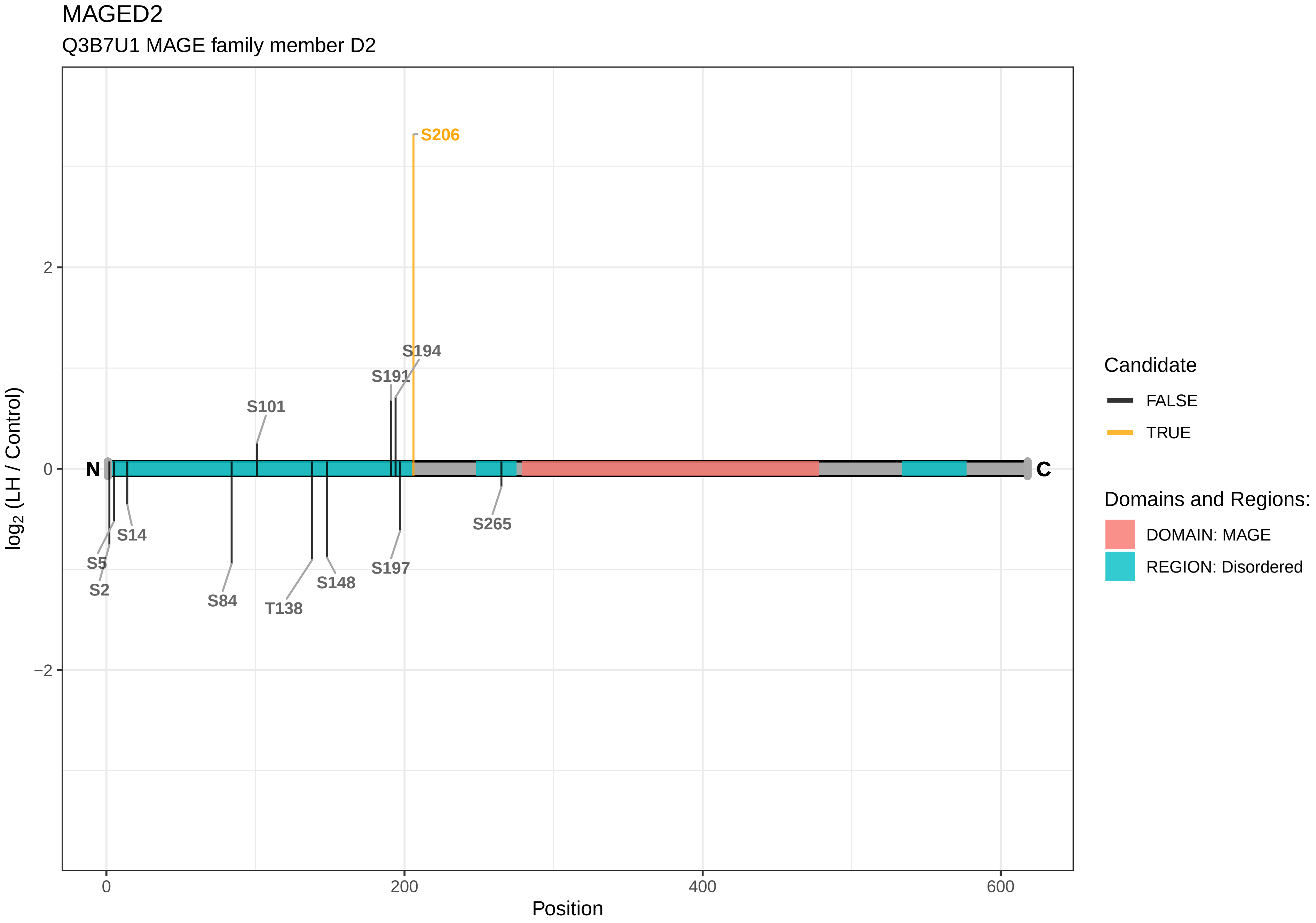

MAGED2
Q3B7U1 MAGE family member D2
S206
S194
S191
S101
S265
S14
S5
S2
S84
T138
S148
S197
N
C
2
0
−2
0
200
400
600
Position
log2 (LH / Control)
Candidate
FALSE
TRUE
Domains and Regions:
DOMAIN: MAGE
REGION: Disordered

MAGI3

F1M7S0 Membrane−associated guanylate kinase, WW and PDZ domain−containing protein 3

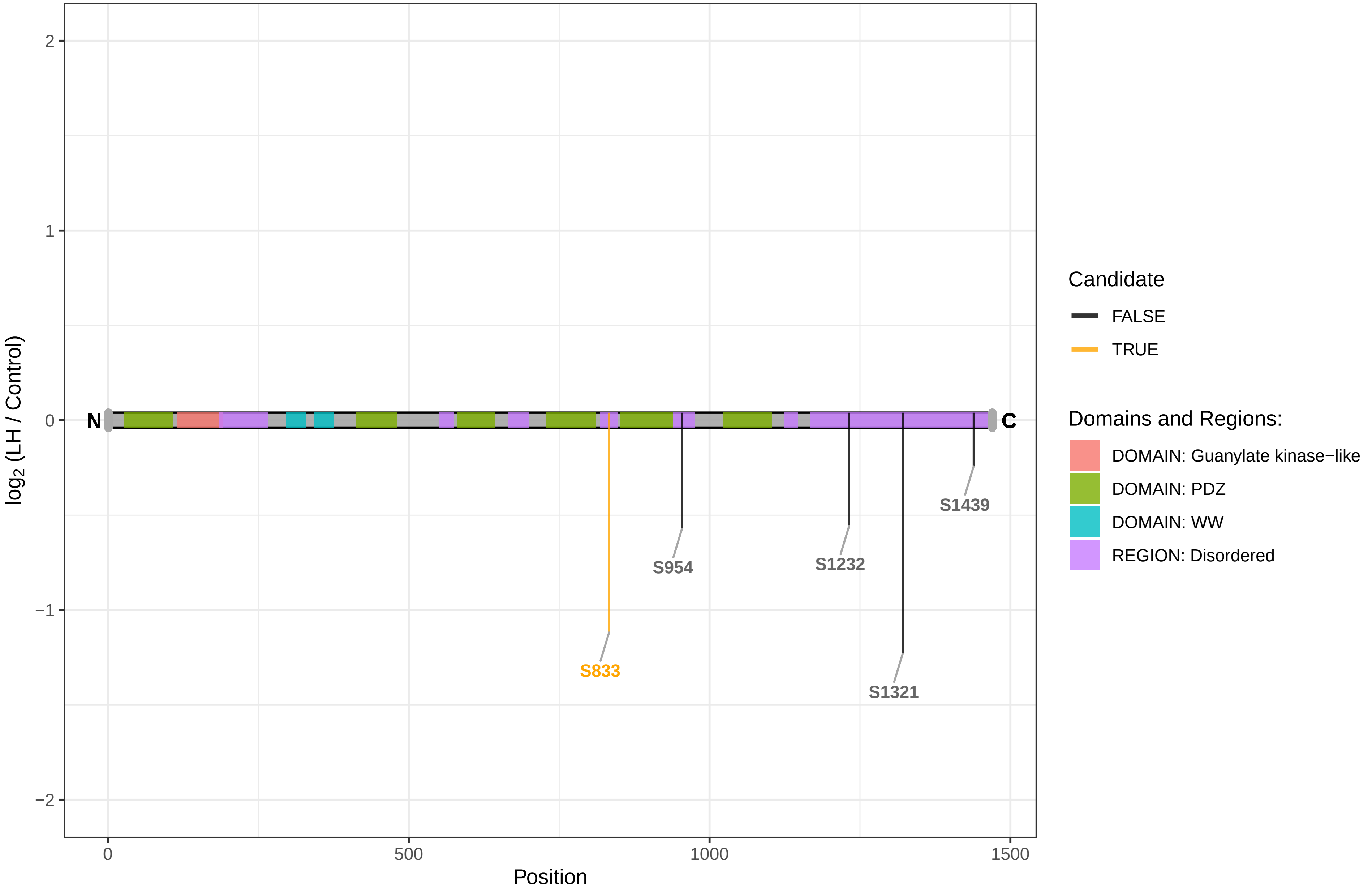

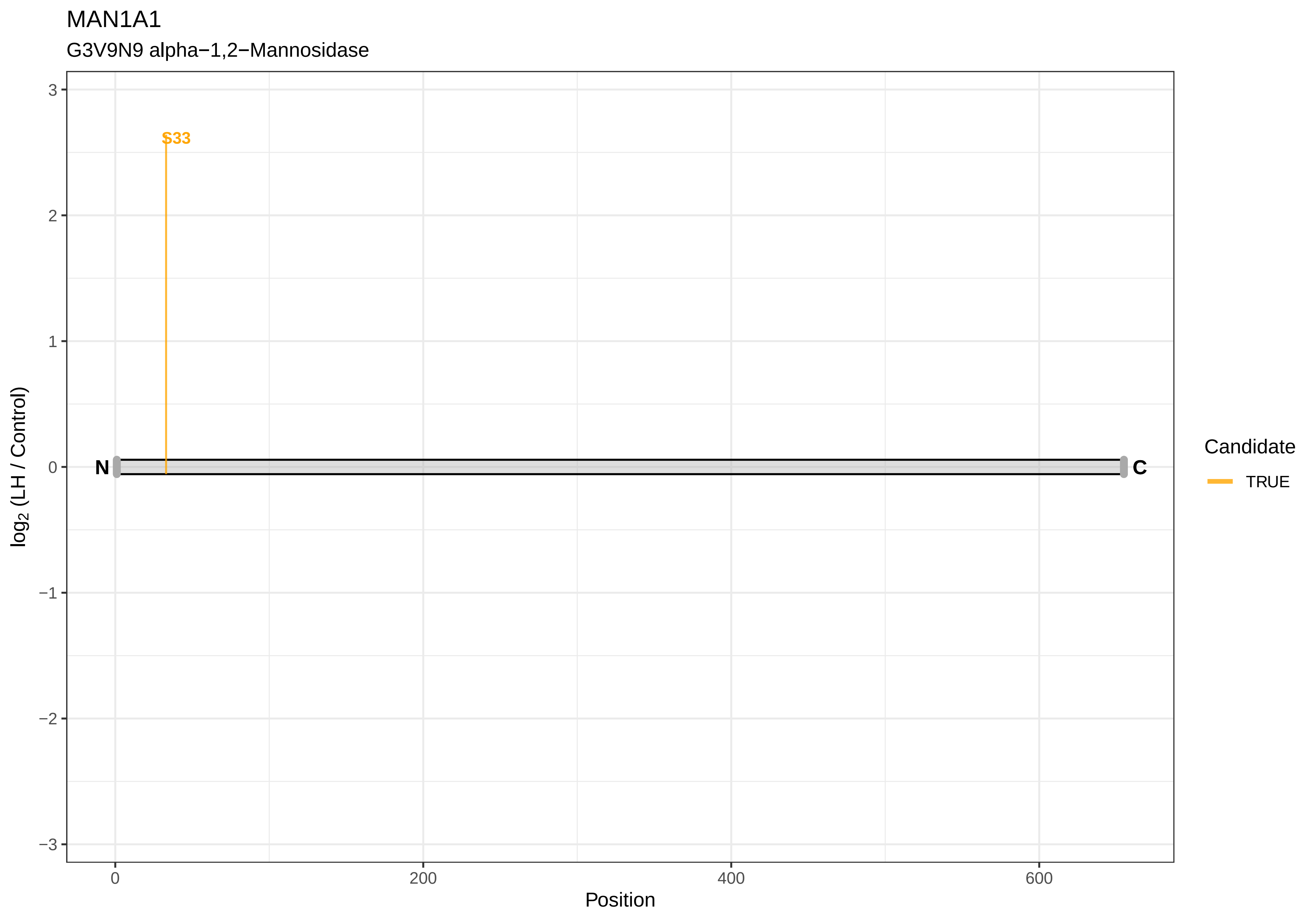
MAN1A1
G3V9N9 alpha−1,2−Mannosidase
S33
N
C
log2 (LH / Control)
Position
3
2
1
0
−1
−2
−3
0
200
400
600
Candidate
TRUE

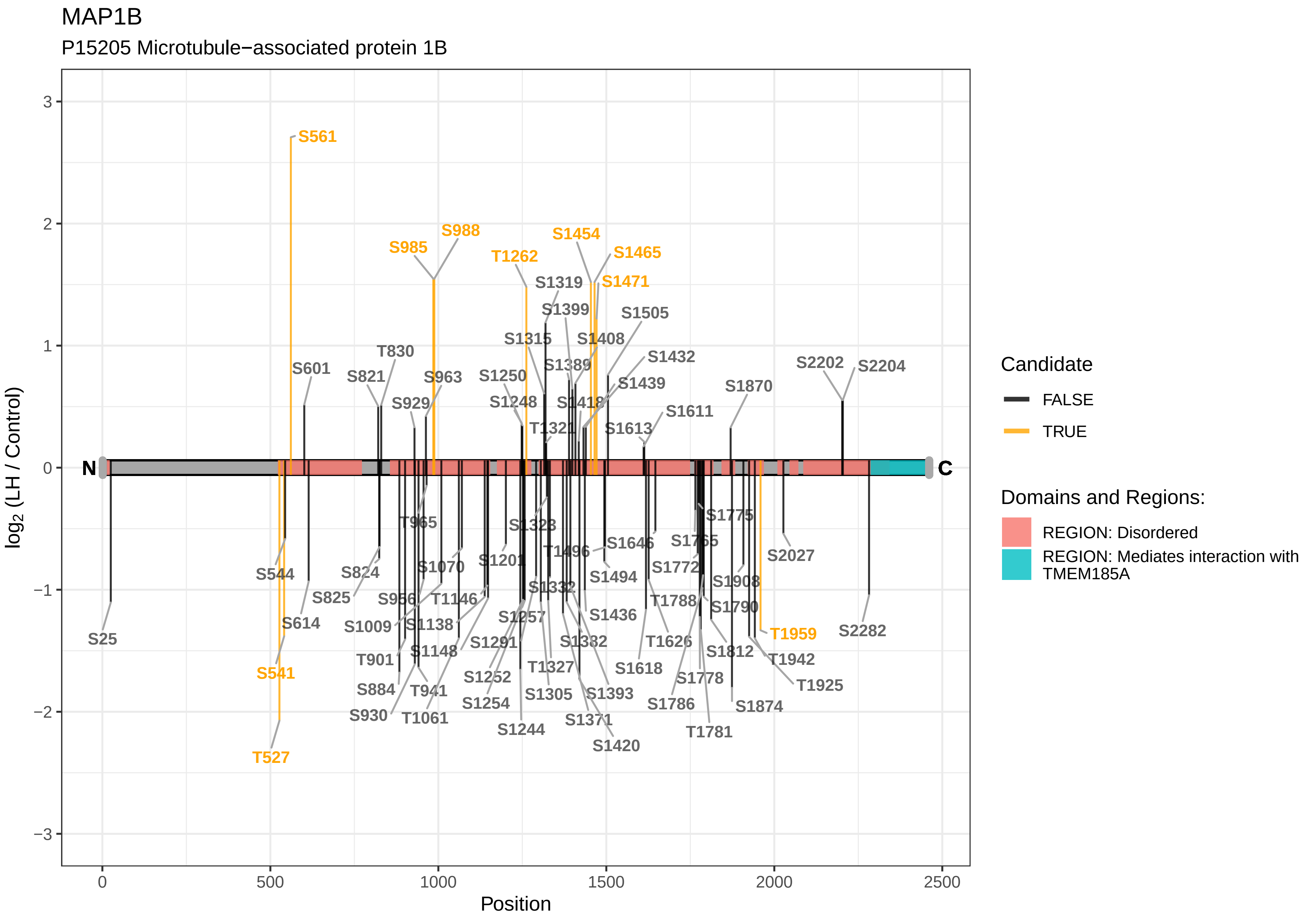
MAP1B
P15205 Microtubule−associated protein 1B
log2 (LH / Control)
Position
N
C
S561
S985
S988
T1262
S1454
S1465
S1471
S1319
S1399
S1505
S1315
S1408
S1432
S1389
S1439
T830
S601
S821
S963
S1250
S929
S1248
S1418
S1611
T1321
S1613
S1870
S2202
S2204
S25
S544
S541
T527
S614
S824
S825
S1009
S956
T965
S1070
S1138
T1146
S1148
T901
S884
T941
S930
T1061
S1201
S1323
T1496
S1646
S1494
S1332
S1257
S1291
S1252
S1254
S1244
T1327
S1305
S1382
S1371
S1393
S1420
S1436
S1618
T1626
S1775
S1765
S1772
T1788
S1790
S1778
S1786
T1781
S1812
S1908
S1874
T1942
T1925
T1959
S2027
S2282
Candidate
FALSE
TRUE
Domains and Regions:
REGION: Disordered
REGION: Mediates interaction with TMEM185A

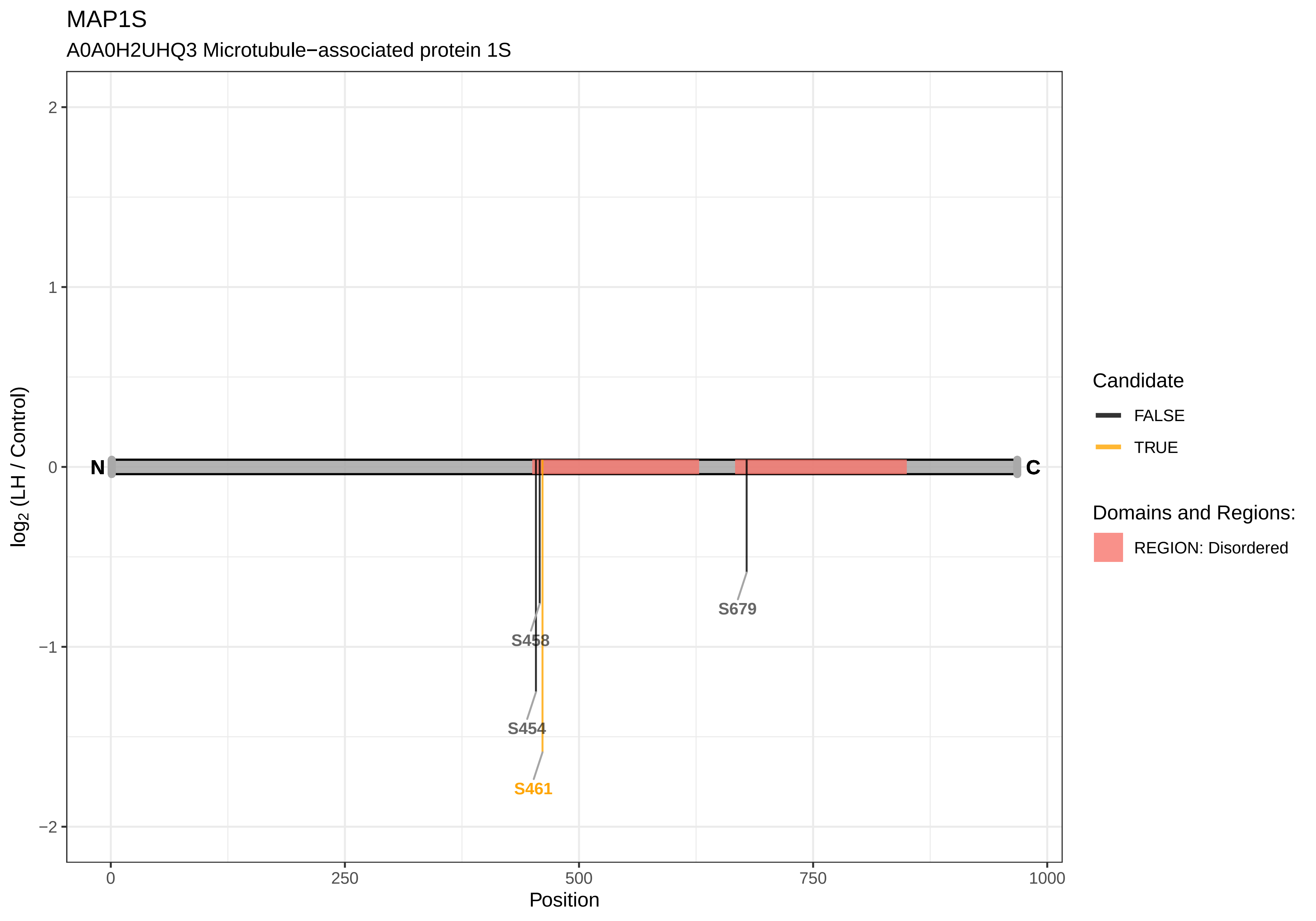

MAP1S
A0A0H2UHQ3 Microtubule−associated protein 1S
log2 (LH / Control)
2
1
0
−1
−2
N
C
S458
S454
S461
S679
0
250
500
750
1000
Position
Candidate
FALSE
TRUE
Domains and Regions:
REGION: Disordered

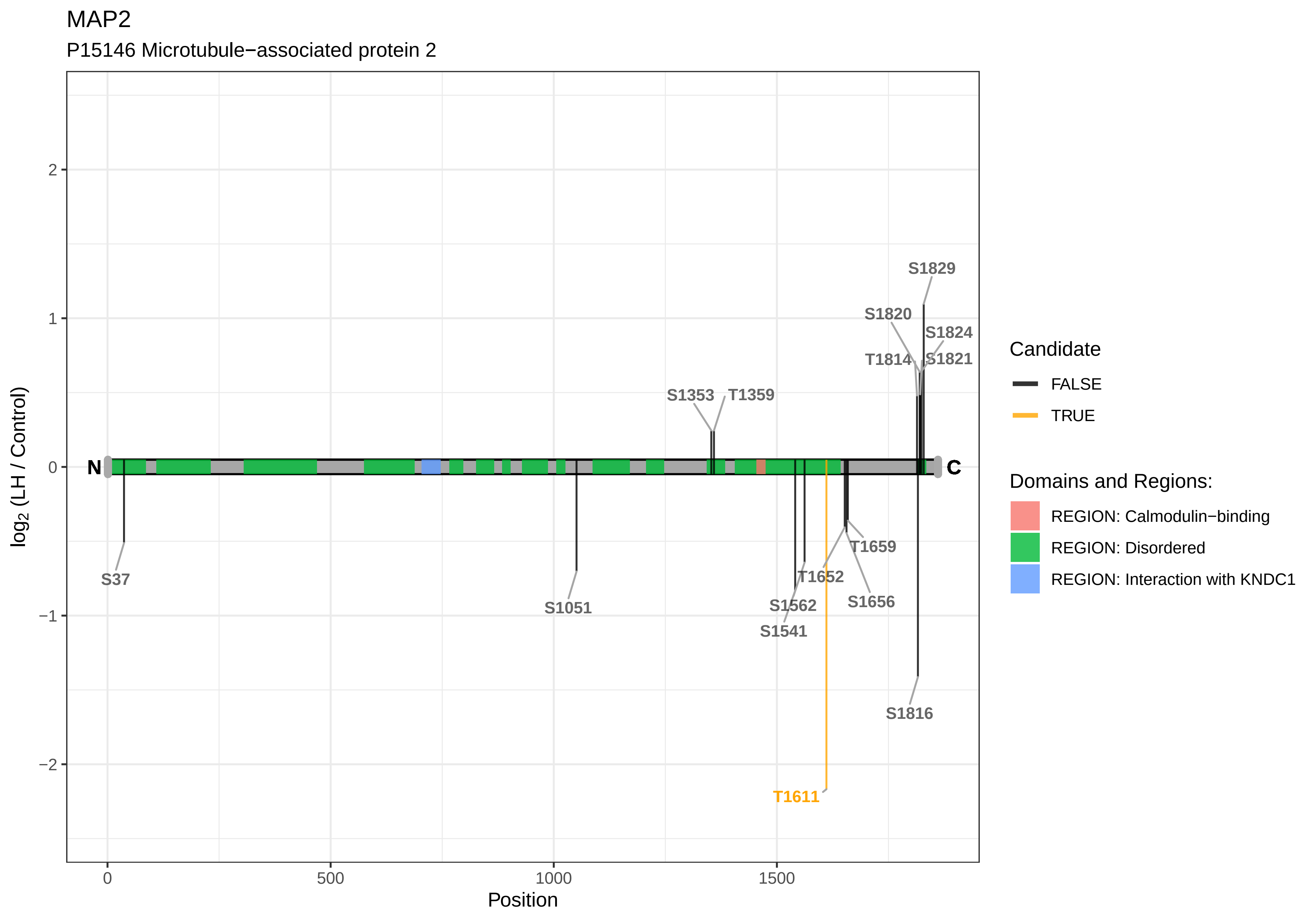
MAP2
P15146 Microtubule−associated protein 2
log2 (LH / Control)
Position
N
C
S37
S1051
S1353
T1359
S1541
S1562
T1611
T1652
S1656
T1659
T1814
S1816
S1820
S1821
S1824
S1829
Candidate
FALSE
TRUE
Domains and Regions:
REGION: Calmodulin−binding
REGION: Disordered
REGION: Interaction with KNDC1
2
1
0
−1
−2
0
500
1000
1500

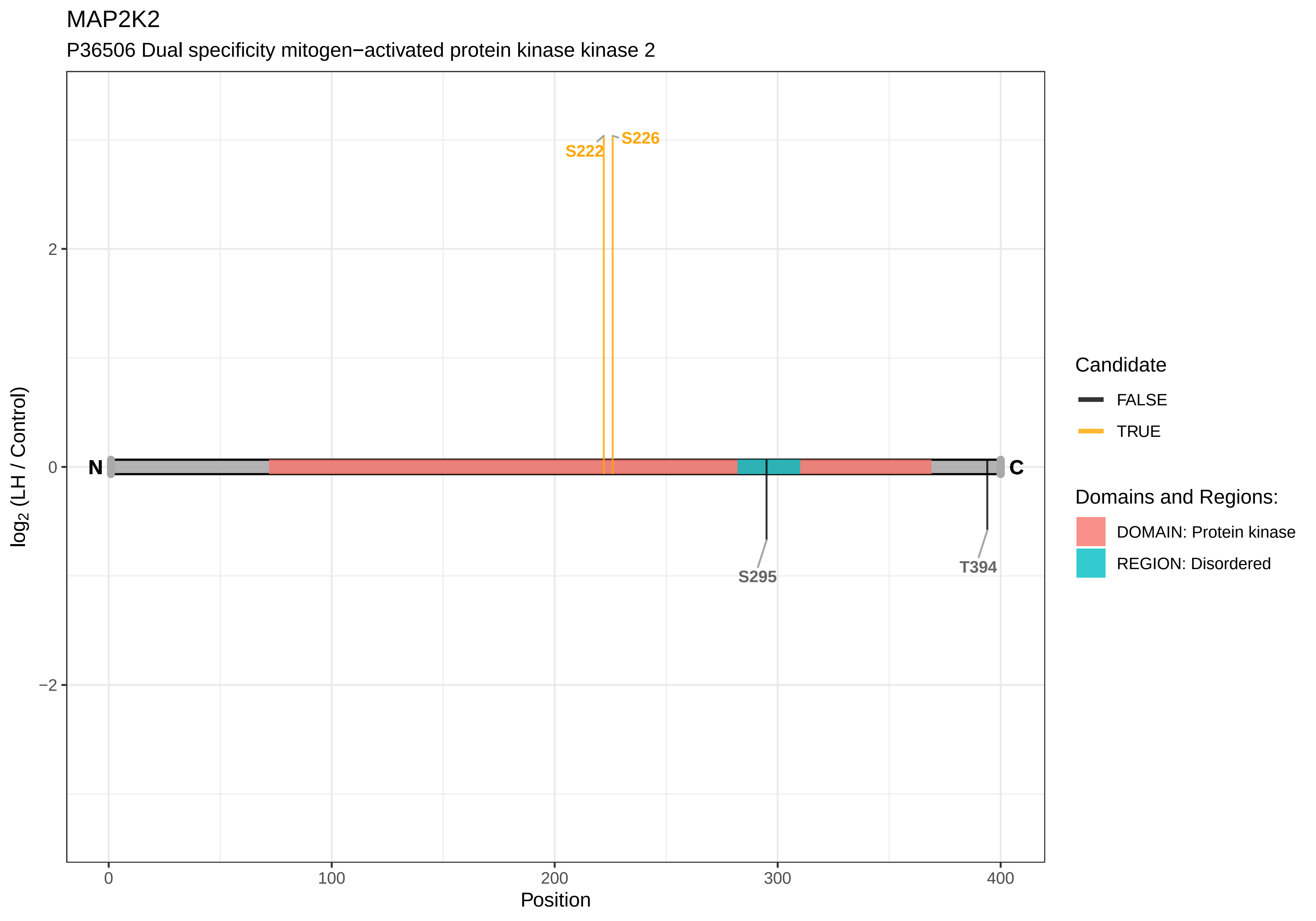

MAP2K2
P36506 Dual specificity mitogen−activated protein kinase kinase 2
S222
S226
S295
T394
N
C
2
0
−2
log2 (LH / Control)
0
100
200
300
400
Position
Candidate
FALSE
TRUE
Domains and Regions:
DOMAIN: Protein kinase
REGION: Disordered

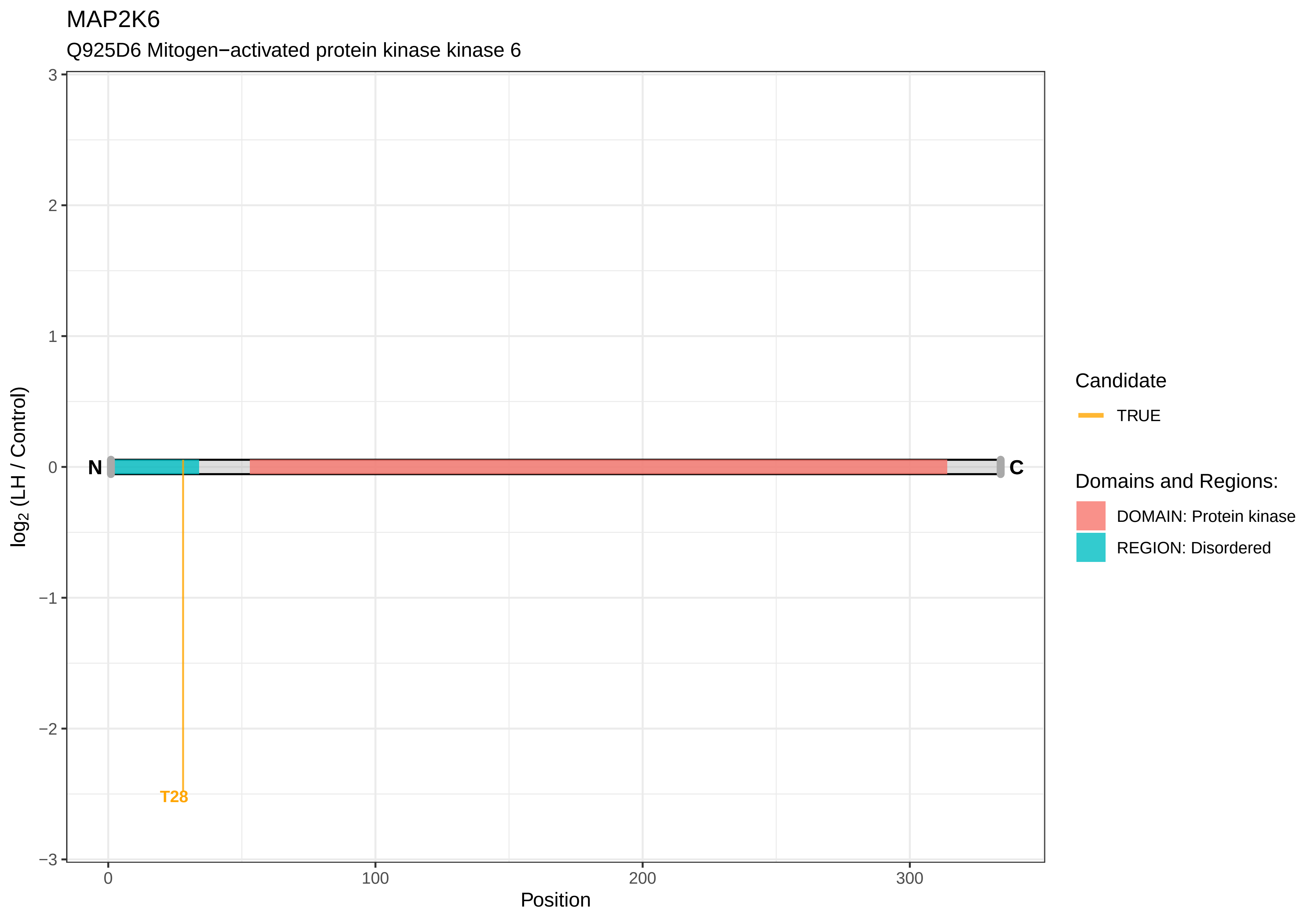
MAP2K6
Q925D6 Mitogen−activated protein kinase kinase 6
log2 (LH / Control)
3
2
1
0
−1
−2
−3
N
C
T28
0
100
200
300
Position
Candidate
TRUE
Domains and Regions:
DOMAIN: Protein kinase
REGION: Disordered

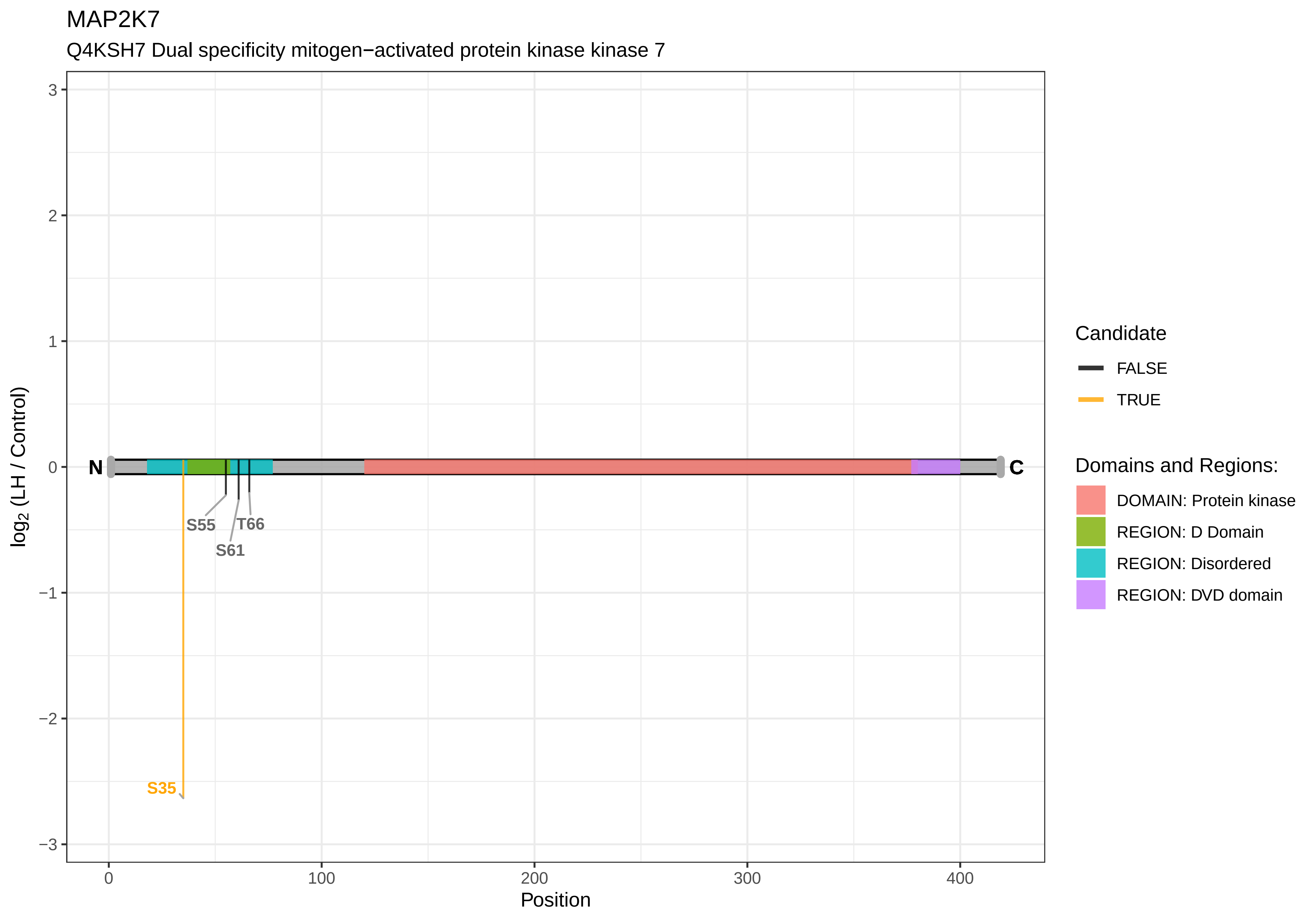

MAP2K7
Q4KSH7 Dual specificity mitogen−activated protein kinase kinase 7
log2 (LH / Control)
Position
N
C
S55
T66
S61
S35
Candidate
FALSE
TRUE
Domains and Regions:
DOMAIN: Protein kinase
REGION: D Domain
REGION: Disordered
REGION: DVD domain

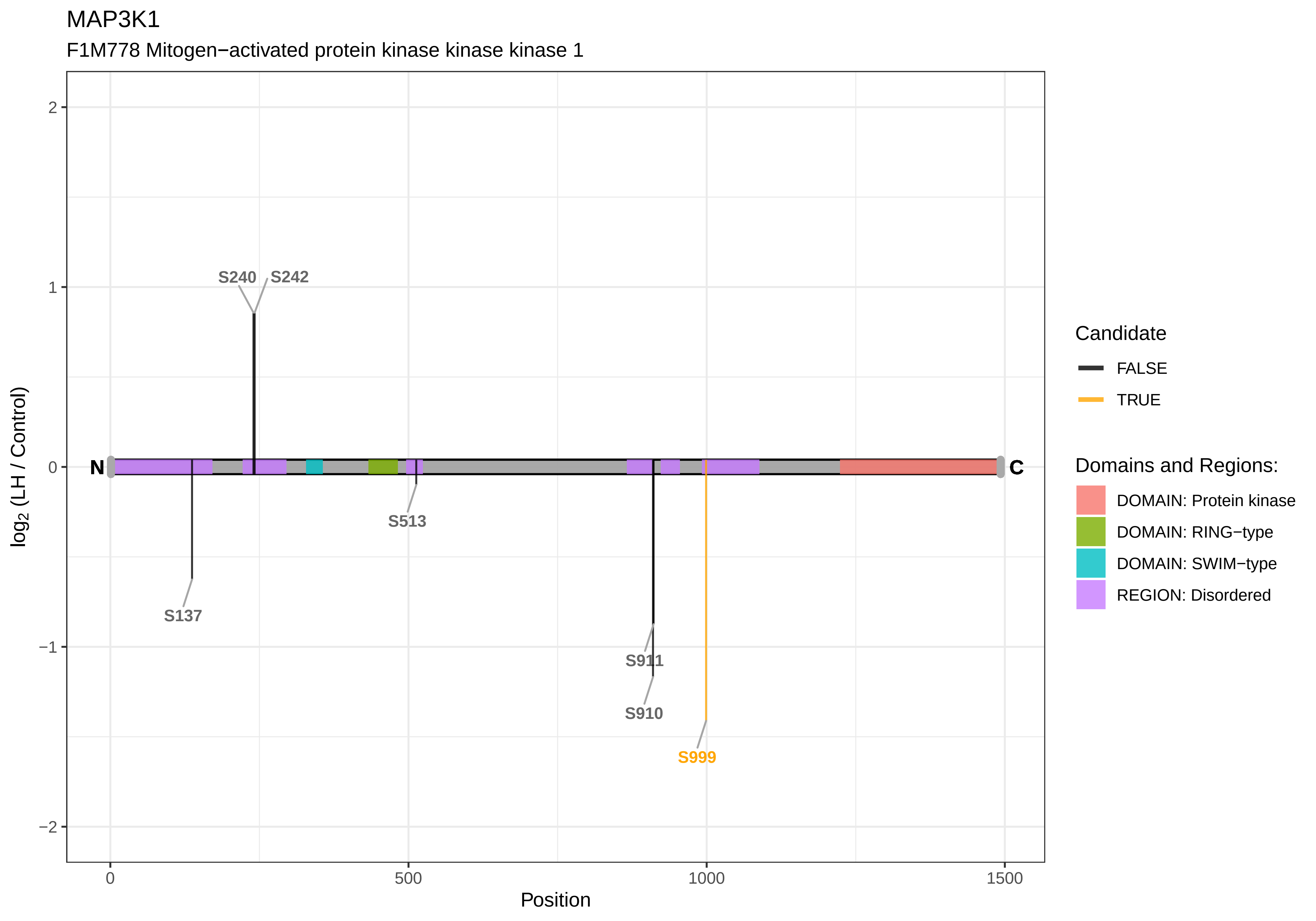

MAP3K1
F1M778 Mitogen−activated protein kinase kinase kinase 1
S240
S242
S513
S137
S911
S910
S999
N
C
2
1
0
−1
−2
log2 (LH / Control)
0
500
1000
1500
Position
Candidate
FALSE
TRUE
Domains and Regions:
DOMAIN: Protein kinase
DOMAIN: RING−type
DOMAIN: SWIM−type
REGION: Disordered

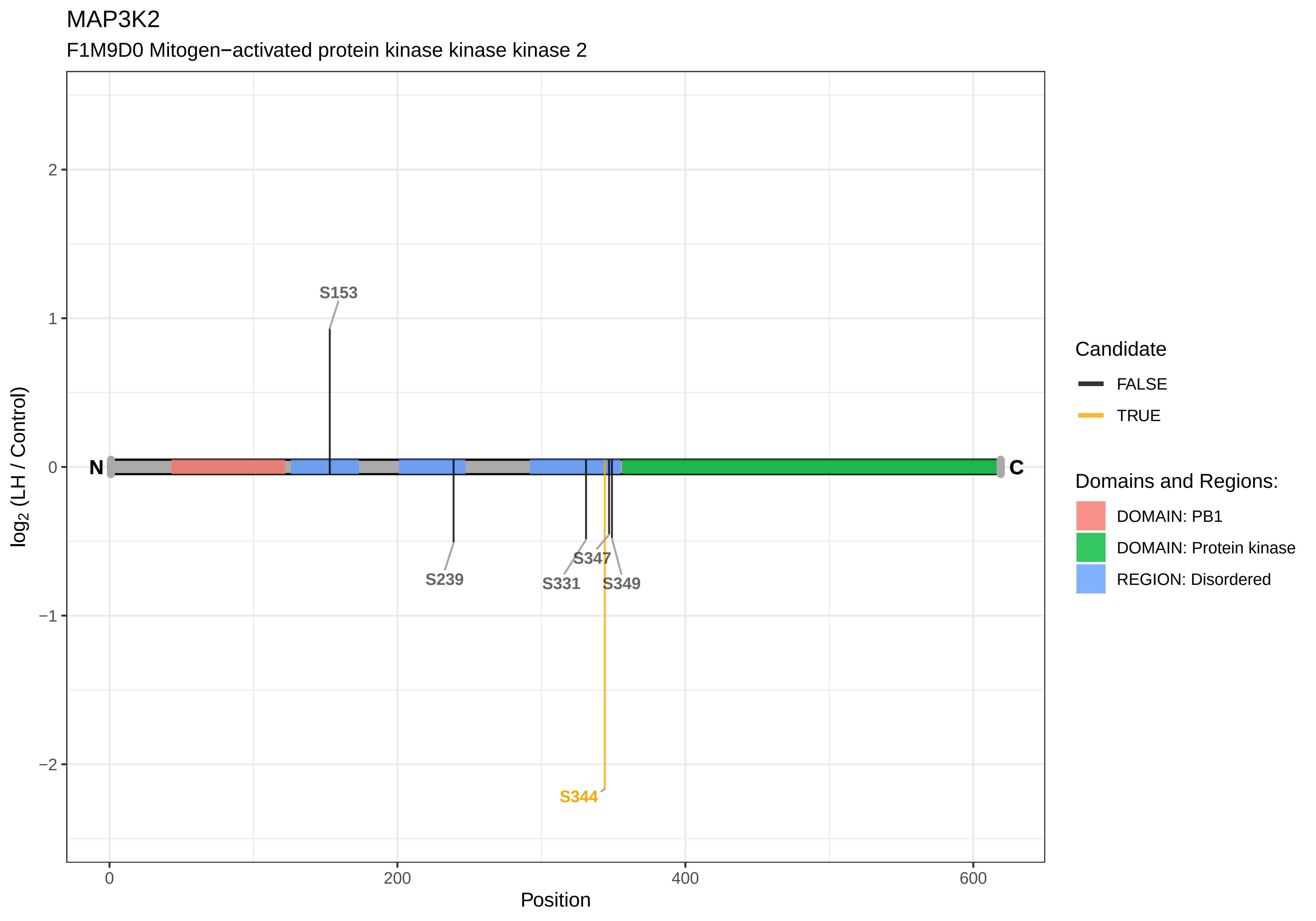

MAP3K2
F1M9D0 Mitogen−activated protein kinase kinase kinase 2
log2 (LH / Control)
Position
N
C
S153
S239
S331
S347
S349
S344
2
1
0
−1
−2
0
200
400
600
Candidate
FALSE
TRUE
Domains and Regions:
DOMAIN: PB1
DOMAIN: Protein kinase
REGION: Disordered

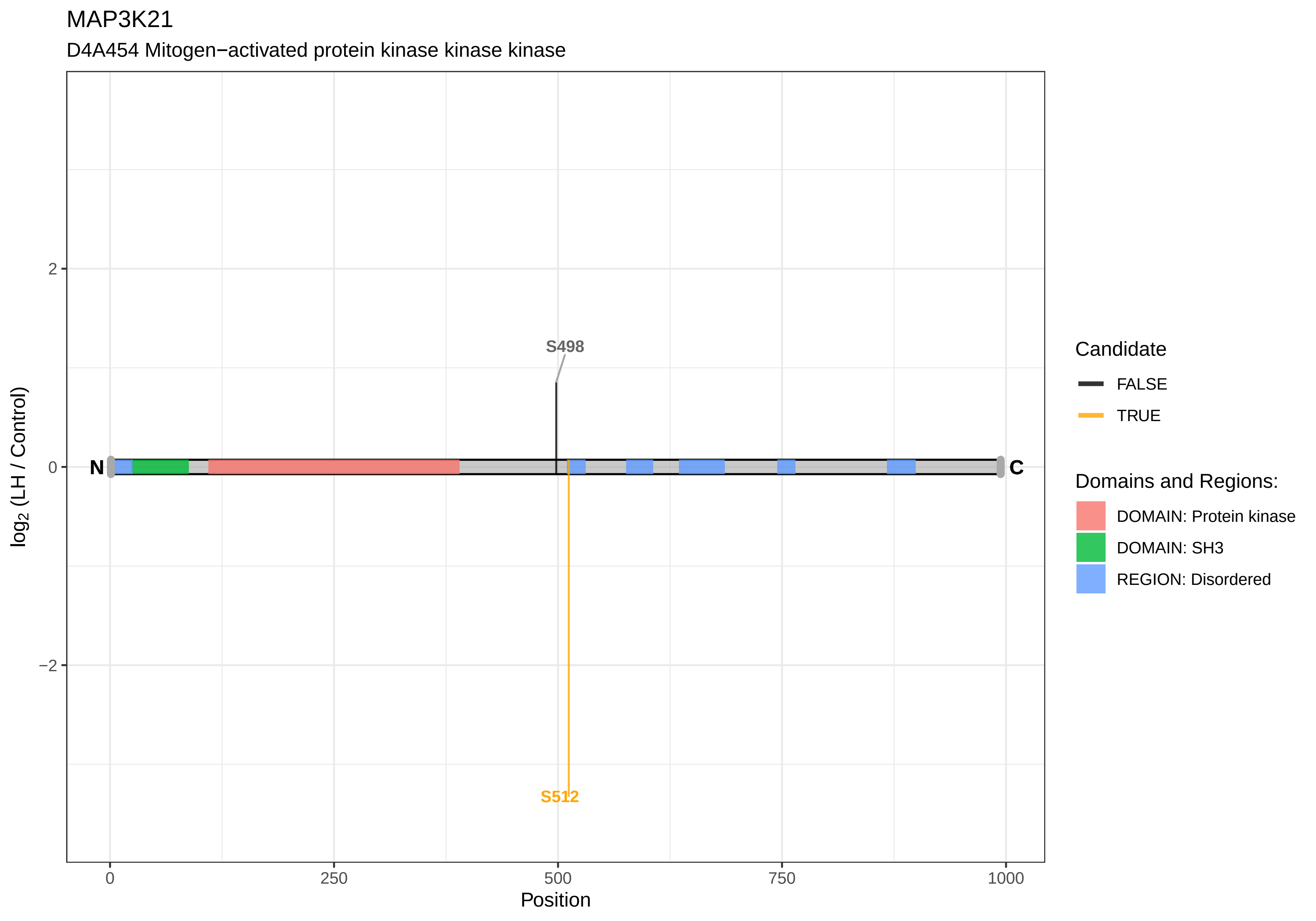

MAP3K21
D4A454 Mitogen−activated protein kinase kinase kinase
log2 (LH / Control)
2
0
−2
S498
S512
N
C
0
250
500
750
1000
Position
Candidate
FALSE
TRUE
Domains and Regions:
DOMAIN: Protein kinase
DOMAIN: SH3
REGION: Disordered

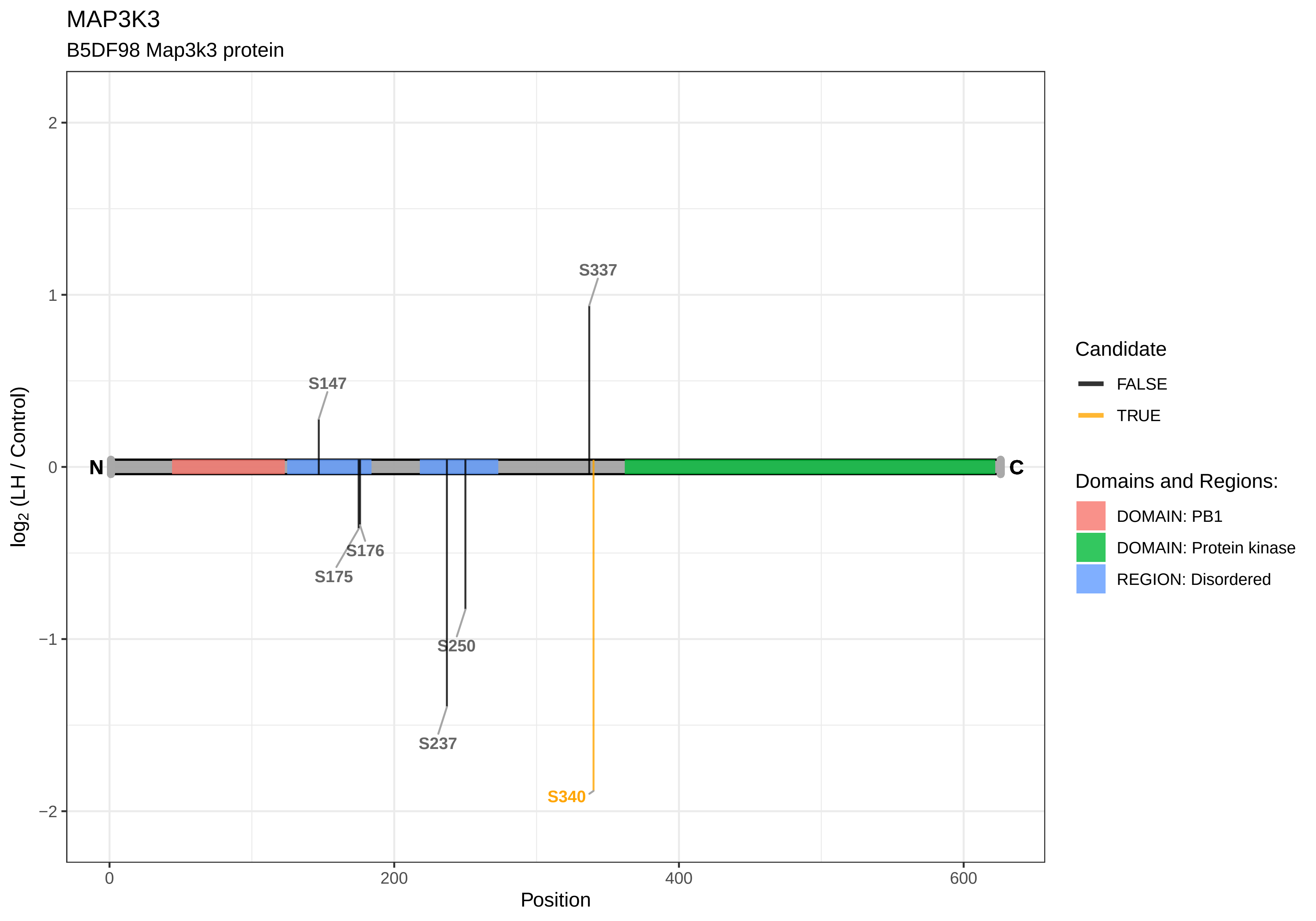
MAP3K3
B5DF98 Map3k3 protein
S337
S147
S176
S175
S250
S237
S340
N
C
2
1
0
−1
−2
log2 (LH / Control)
0
200
400
600
Position
Candidate
FALSE
TRUE
Domains and Regions:
DOMAIN: PB1
DOMAIN: Protein kinase
REGION: Disordered

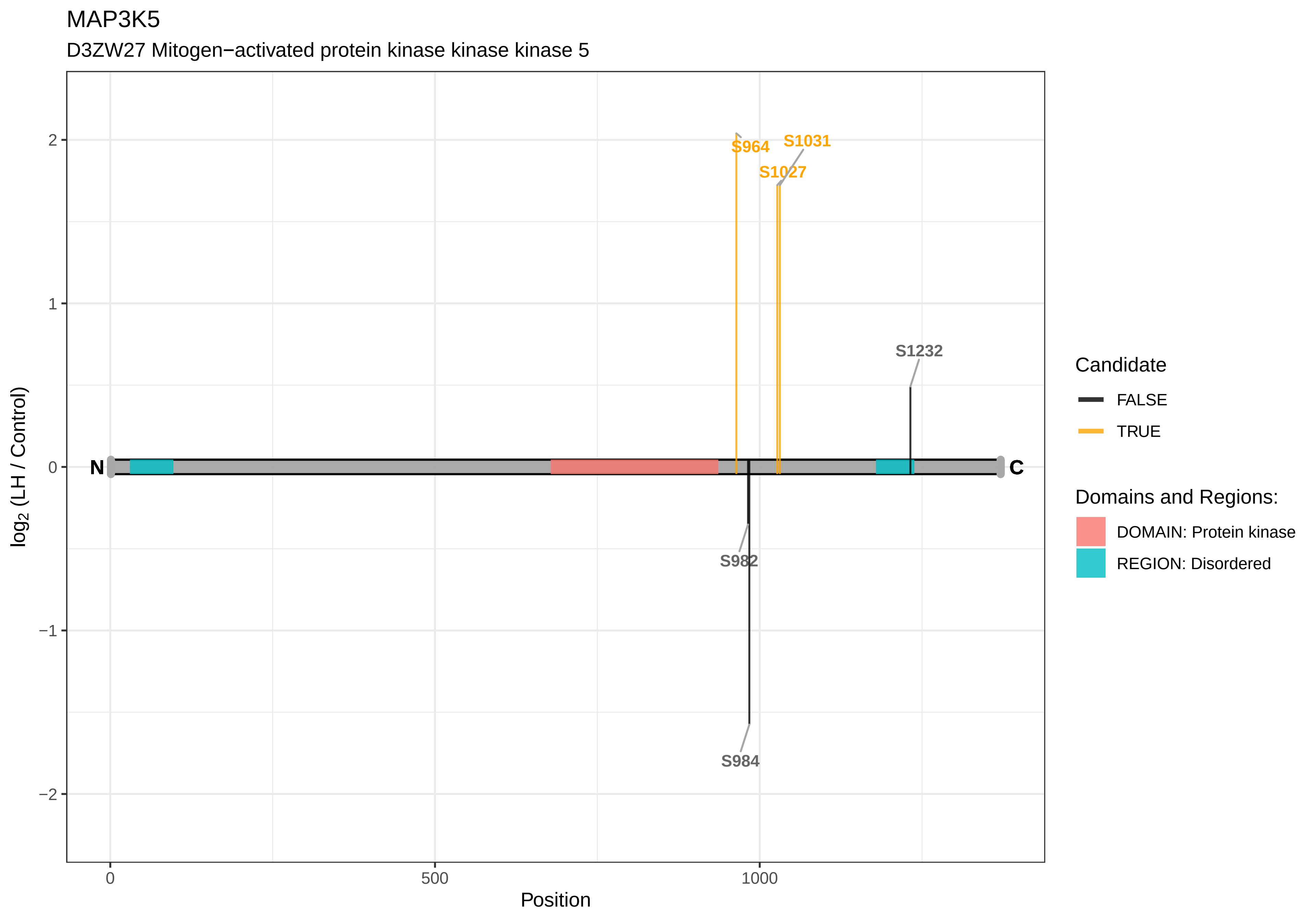

MAP3K5
D3ZW27 Mitogen−activated protein kinase kinase kinase 5
S964
S1031
S1027
S1232
S982
S984
N
C
2
1
0
−1
−2
log2 (LH / Control)
0
500
1000
Position
Candidate
FALSE
TRUE
Domains and Regions:
DOMAIN: Protein kinase
REGION: Disordered

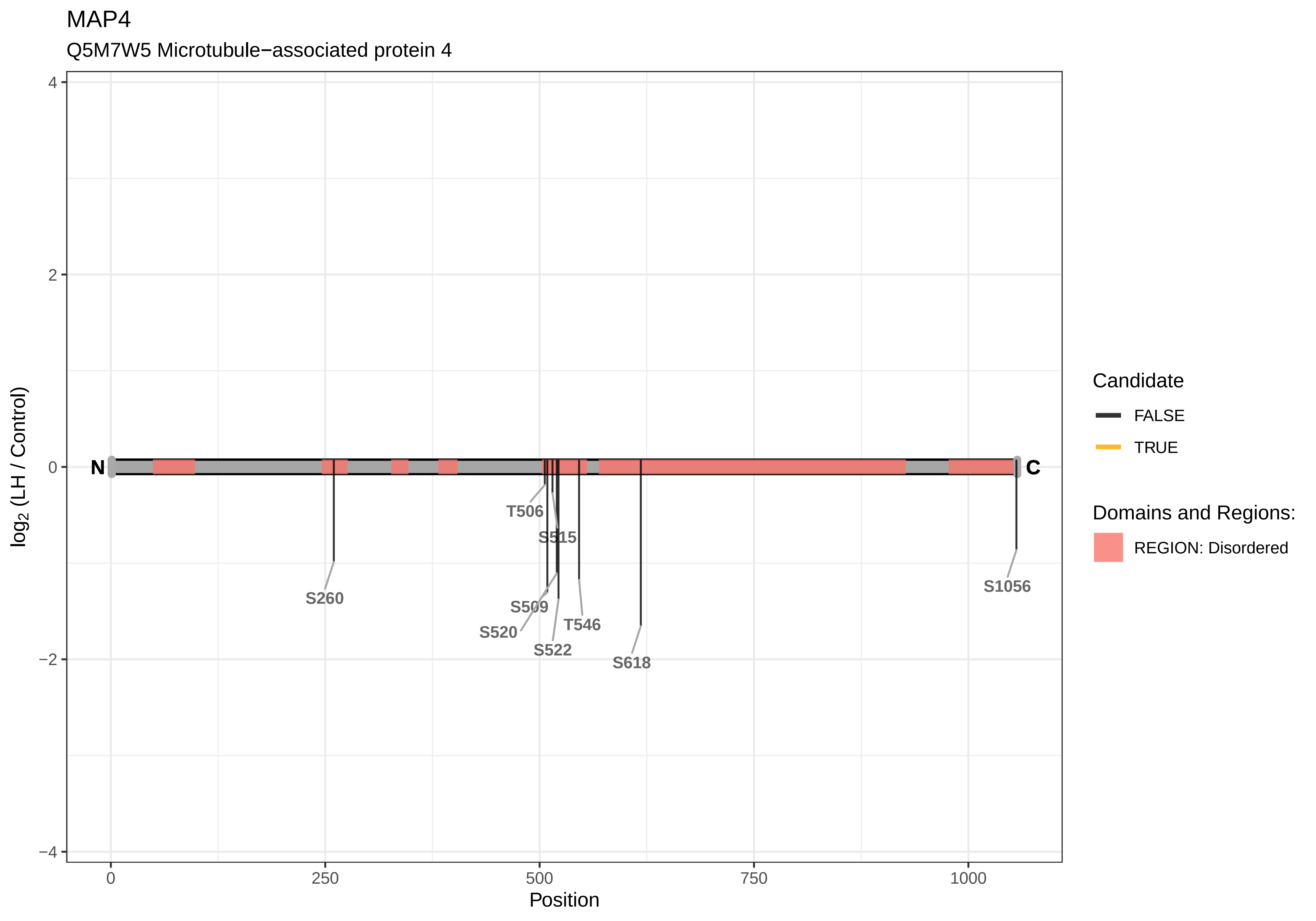

MAP4
Q5M7W5 Microtubule−associated protein 4
log2 (LH / Control)
4
2
0
−2
−4
N
C
S260
T506
S515
S509
S520
S522
T546
S618
S1056
0
250
500
750
1000
Position
Candidate
FALSE
TRUE
Domains and Regions:
REGION: Disordered

MAP4K4

A0A0G2K7W4 Mitogen−activated protein kinase kinase kinase kinase 4

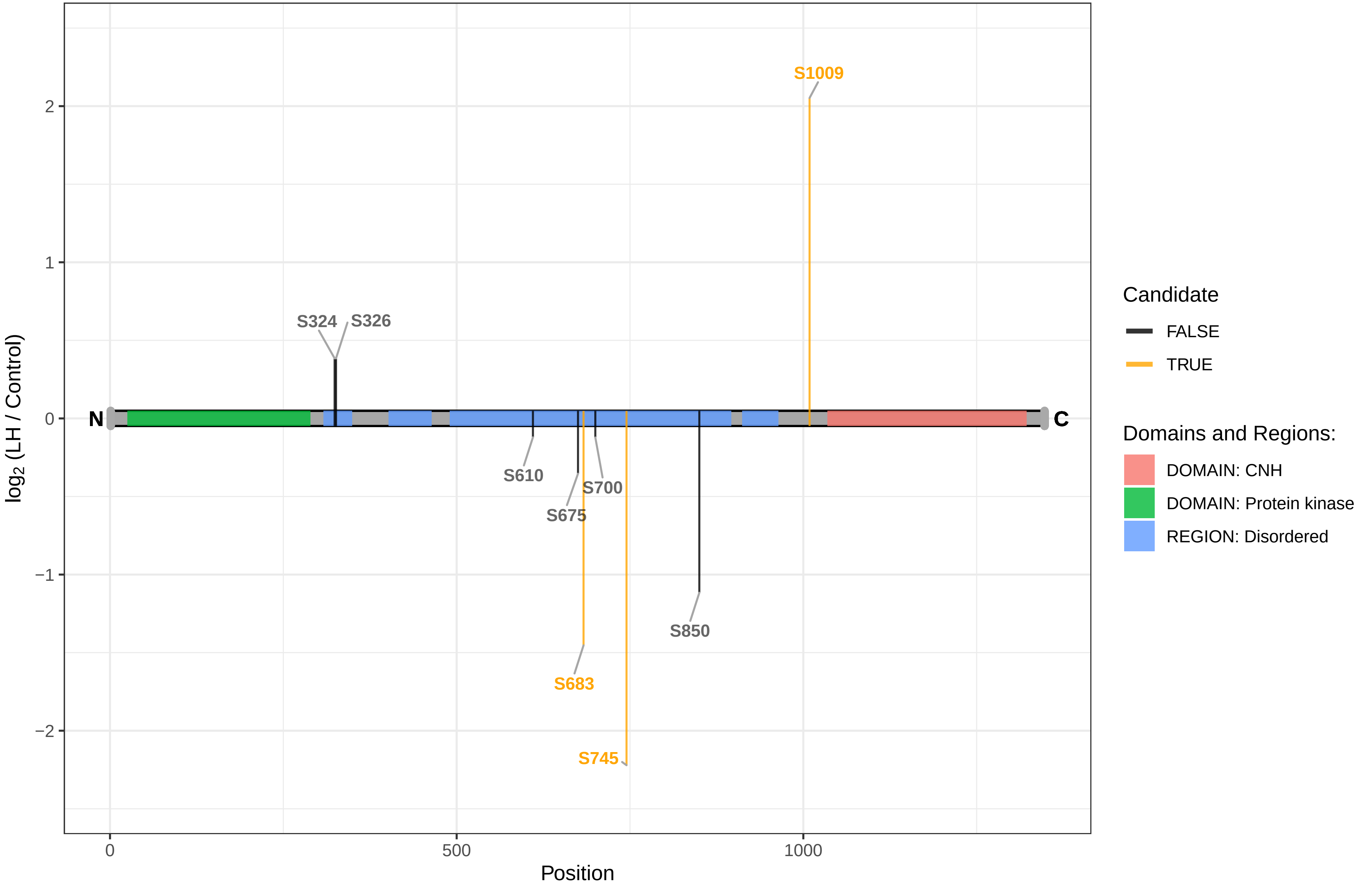

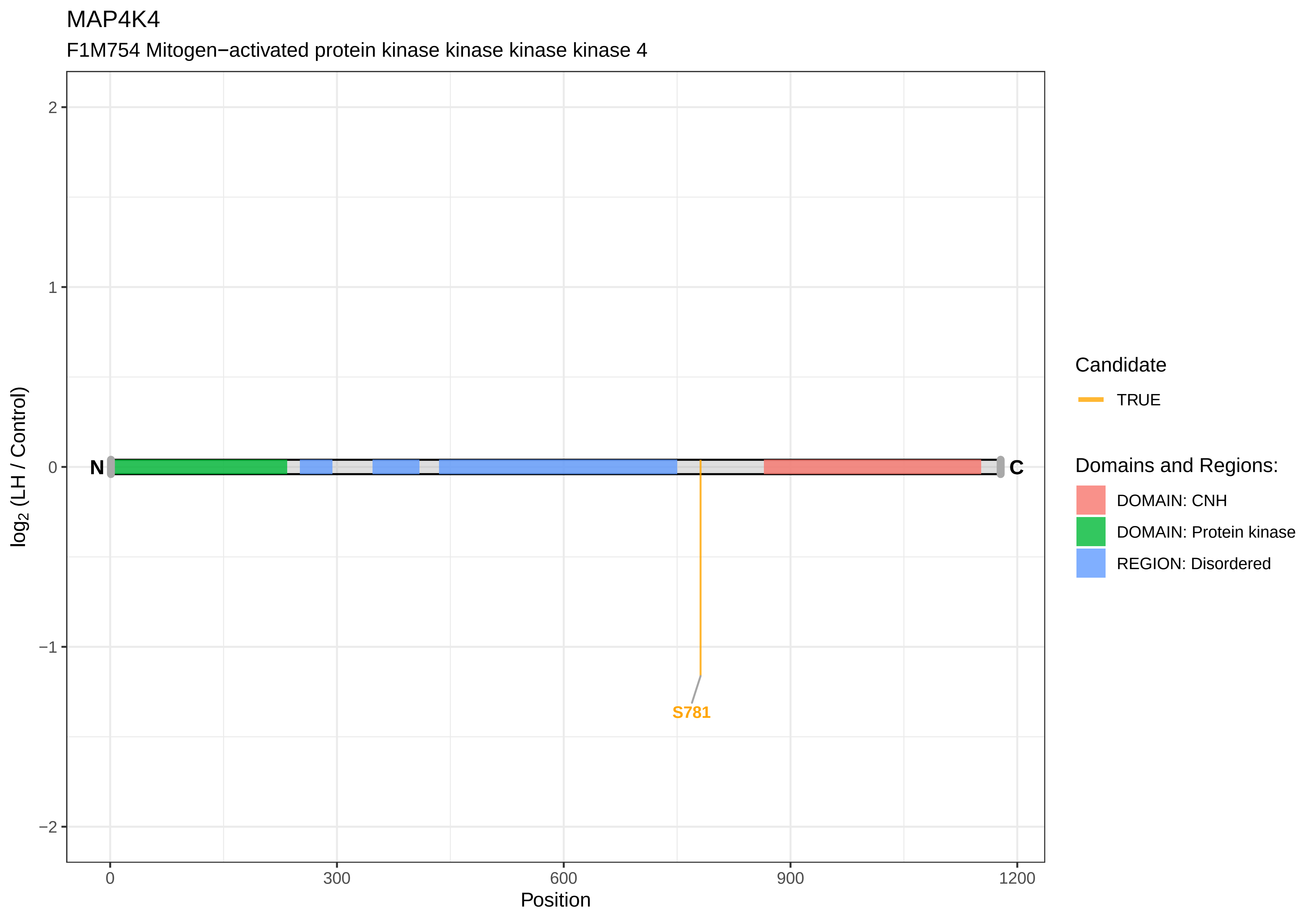
MAP4K4
F1M754 Mitogen−activated protein kinase kinase kinase kinase 4
log2 (LH / Control)
2
1
0
−1
−2
N
C
S781
0
300
600
900
1200
Position
Candidate
TRUE
Domains and Regions:
DOMAIN: CNH
DOMAIN: Protein kinase
REGION: Disordered

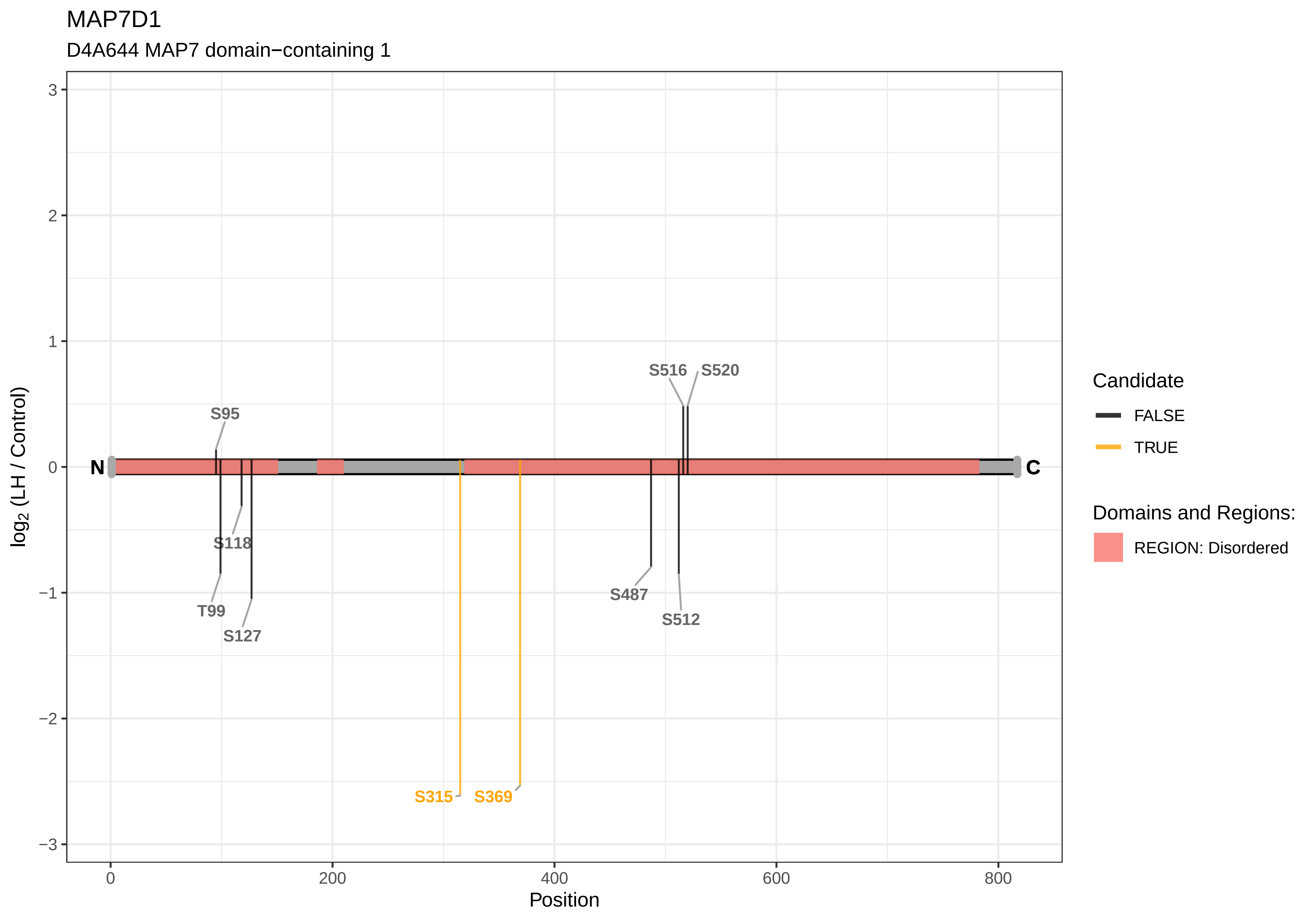
MAP7D1
D4A644 MAP7 domain−containing 1
log2 (LH / Control)
Position
N
C
S95
T99
S118
S127
S315
S369
S487
S512
S516
S520
Candidate
FALSE
TRUE
Domains and Regions:
REGION: Disordered
3
2
1
0
−1
−2
−3
0
200
400
600
800

MAP7D2

D4A4L4 MAP7 domain−containing 2

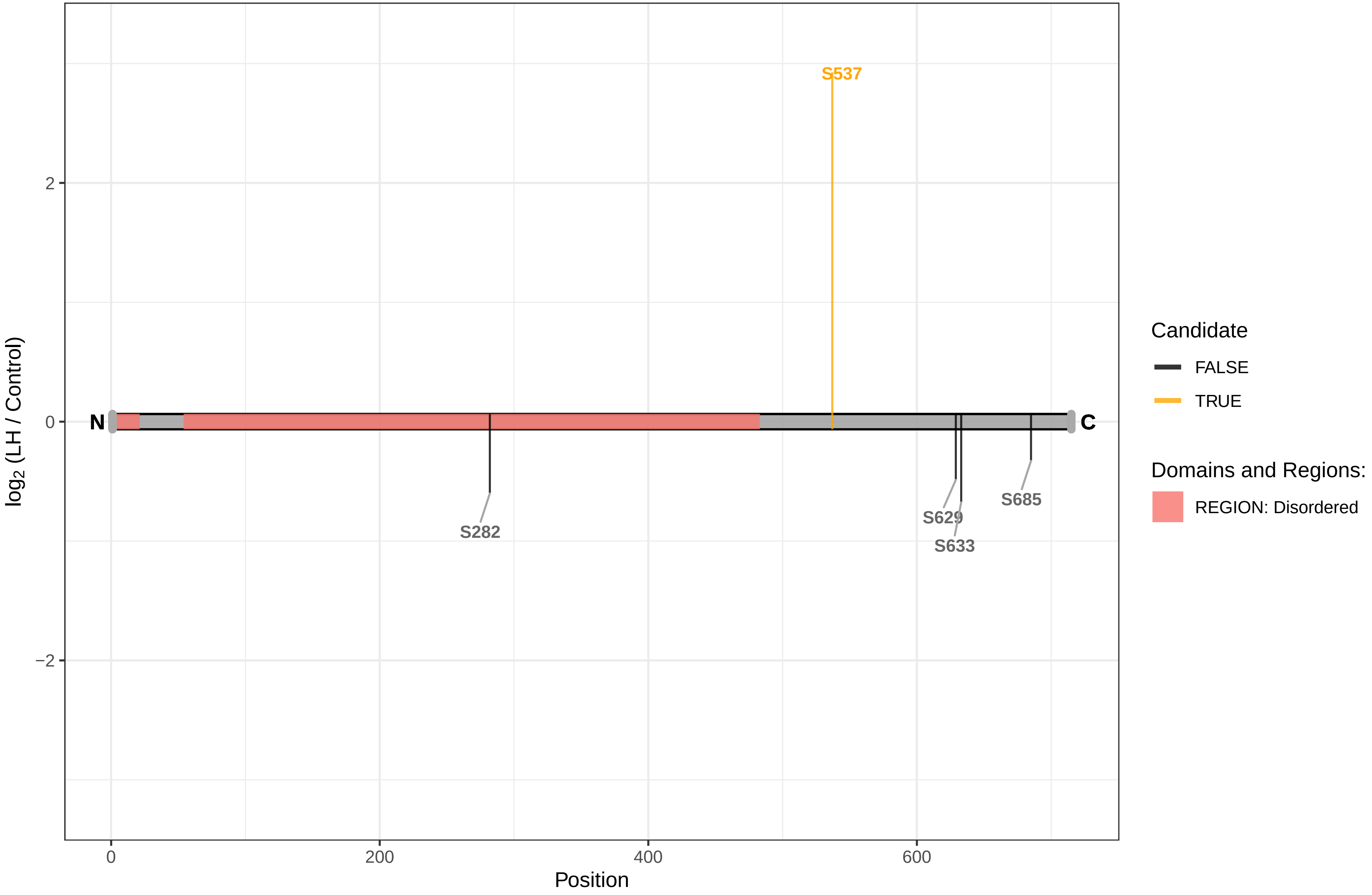

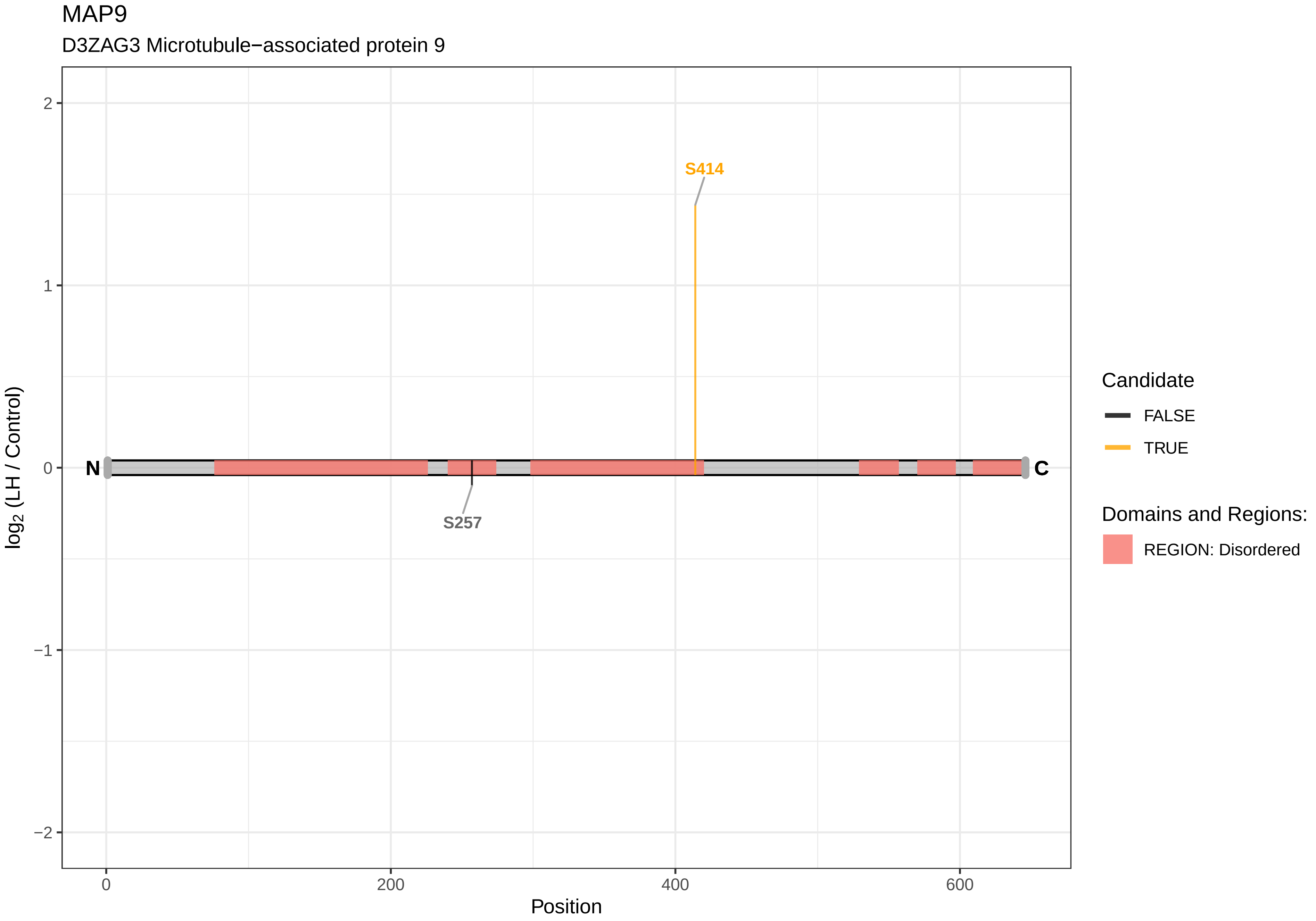
MAP9
D3ZAG3 Microtubule−associated protein 9
S414
S257
N
C
log2 (LH / Control)
2
1
0
−1
−2
0
200
400
600
Position
Candidate
FALSE
TRUE
Domains and Regions:
REGION: Disordered

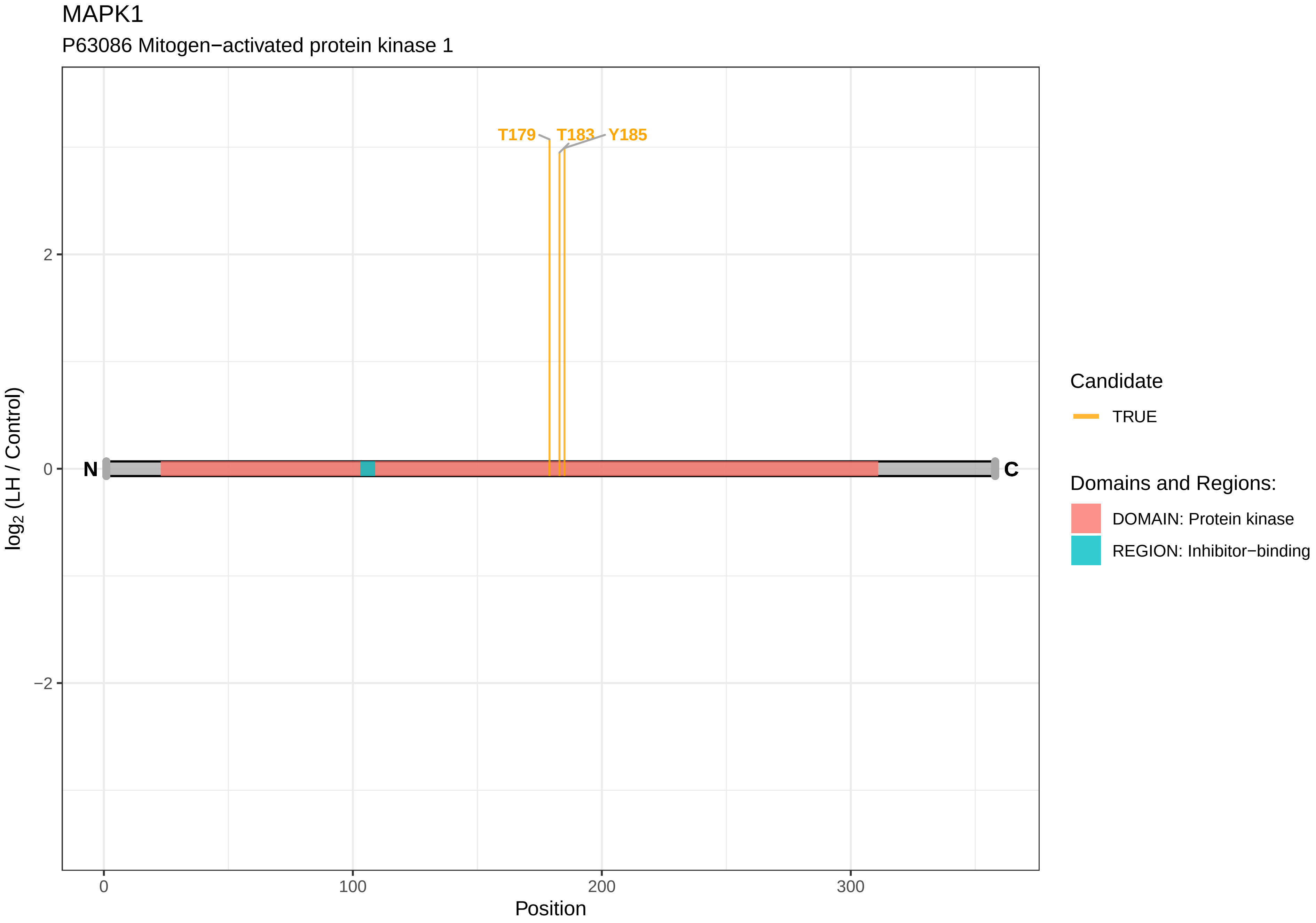

MAPK1
P63086 Mitogen−activated protein kinase 1
T179
T183
Y185
N
C
2
0
−2
0
100
200
300
Position
log2 (LH / Control)
Candidate
TRUE
Domains and Regions:
DOMAIN: Protein kinase
REGION: Inhibitor−binding

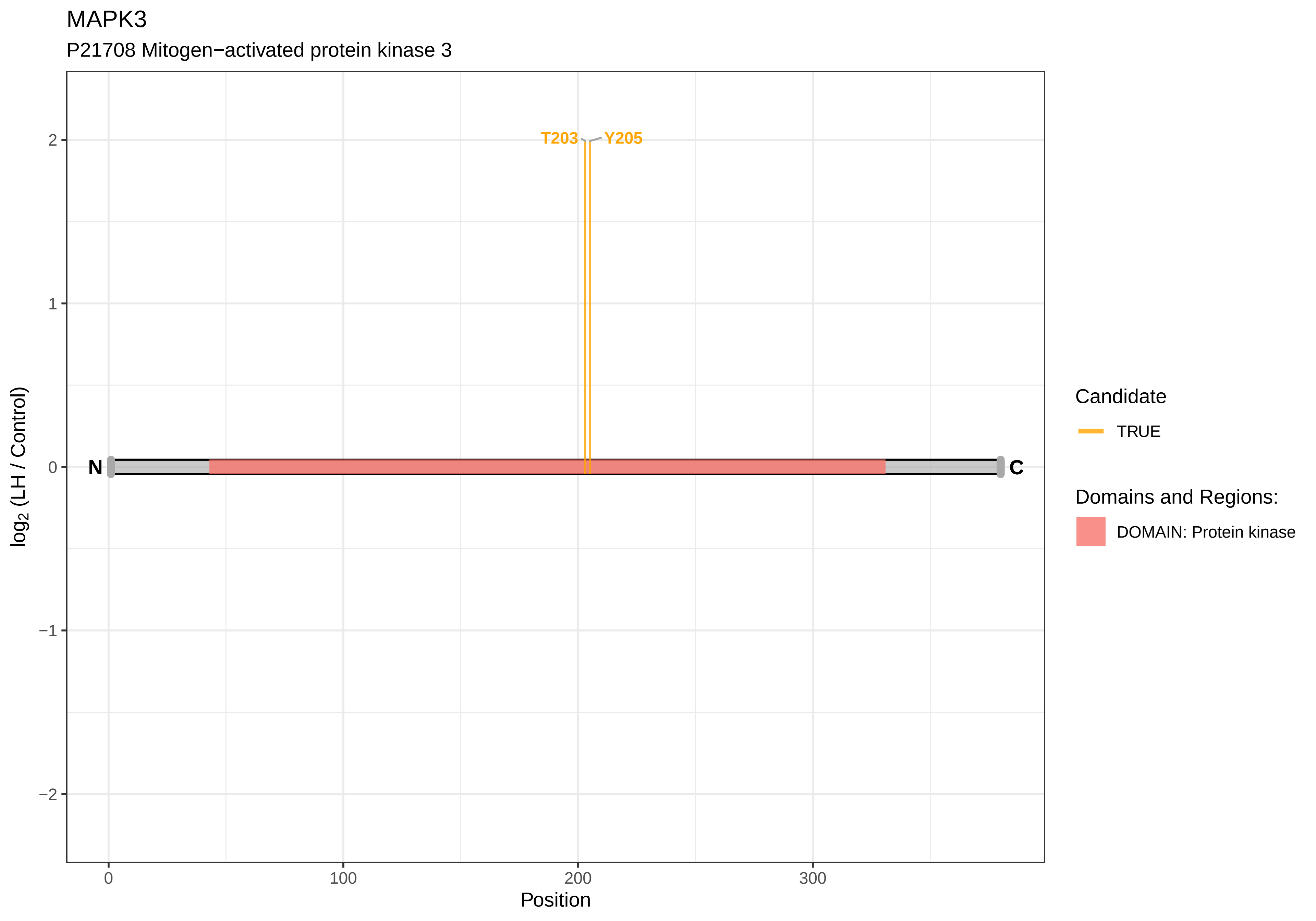

MAPK3
P21708 Mitogen−activated protein kinase 3
T203
Y205
N
C
2
1
0
−1
−2
log2 (LH / Control)
0
100
200
300
Position
Candidate
TRUE
Domains and Regions:
DOMAIN: Protein kinase

MAPK7

P0C865 Mitogen−activated protein kinase 7

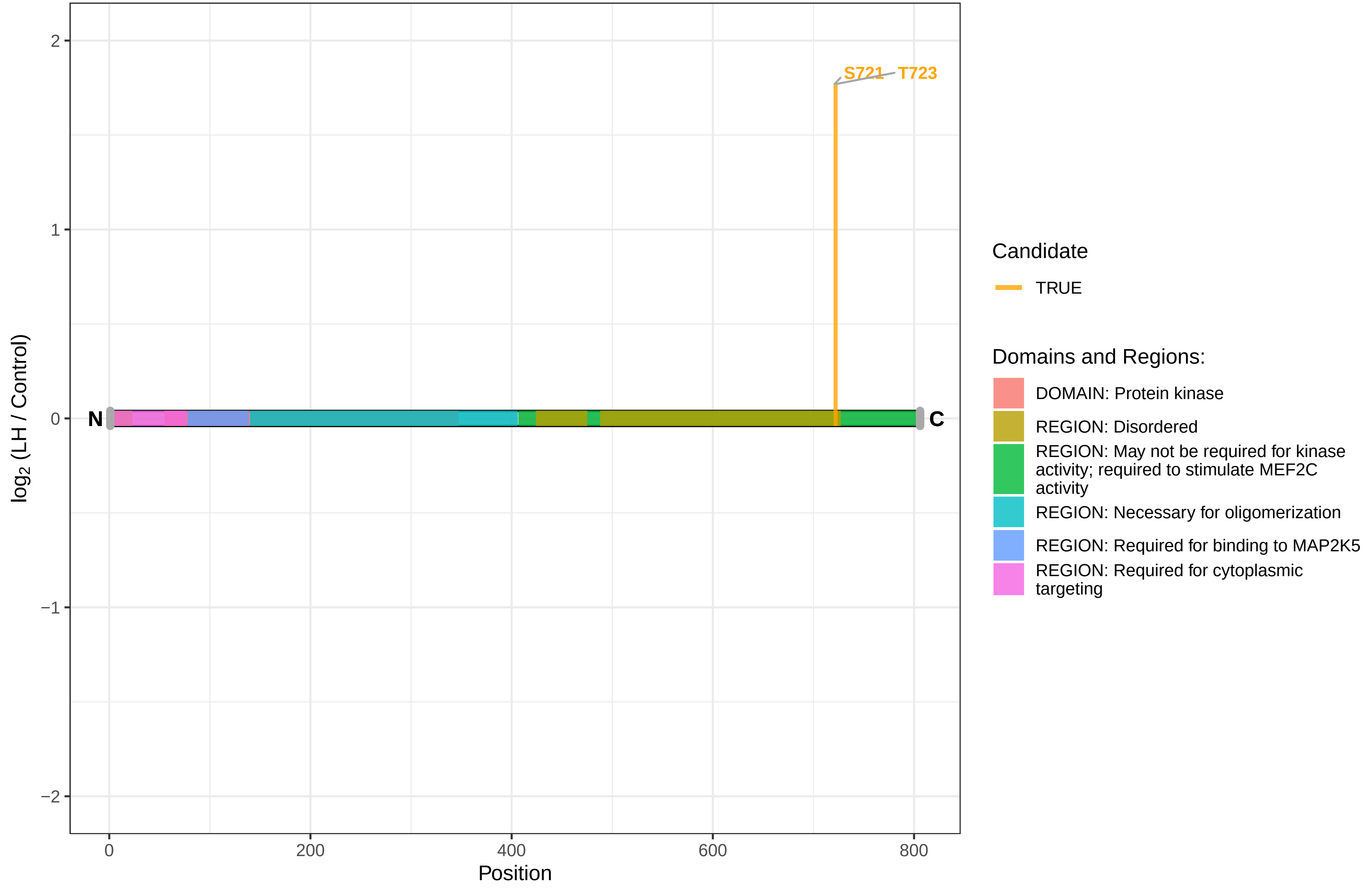

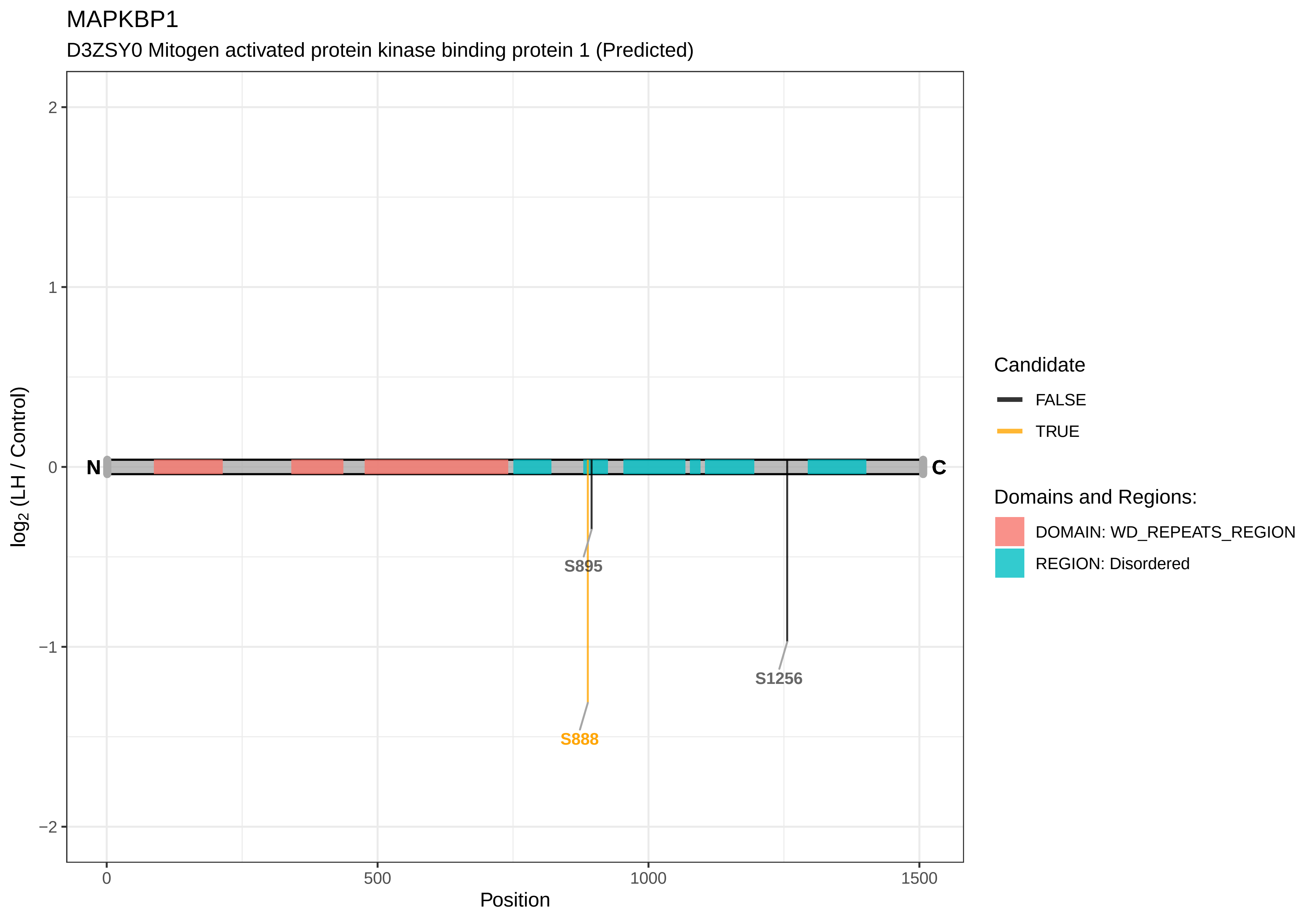

MAPKBP1
D3ZSY0 Mitogen activated protein kinase binding protein 1 (Predicted)
log2 (LH / Control)
Position
N
C
S895
S888
S1256
2
1
0
−1
−2
0
500
1000
1500
Candidate
FALSE
TRUE
Domains and Regions:
DOMAIN: WD_REPEATS_REGION
REGION: Disordered

MAPRE1

Q66HR2 Microtubule−associated protein RP/EB family member 1

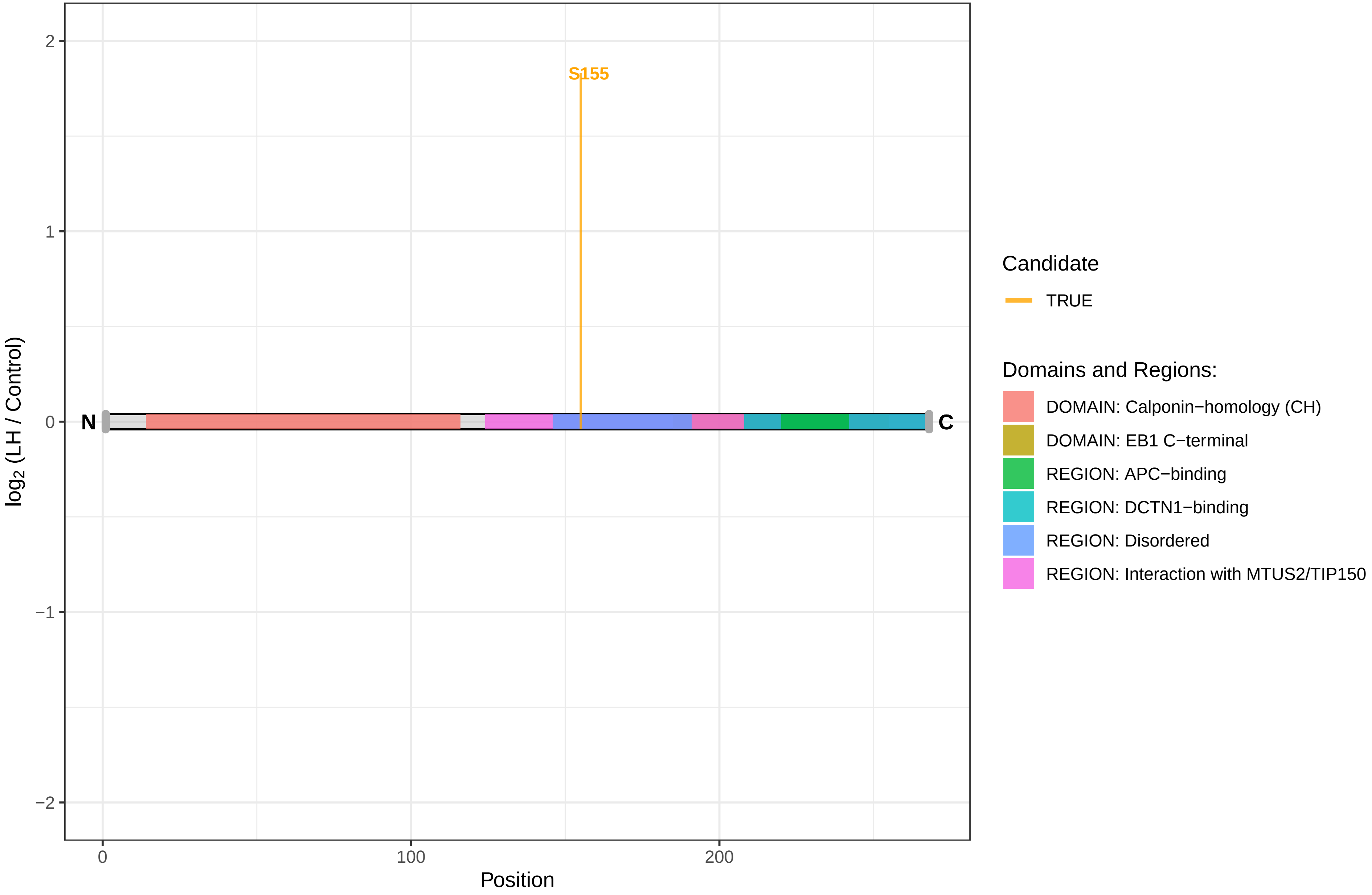

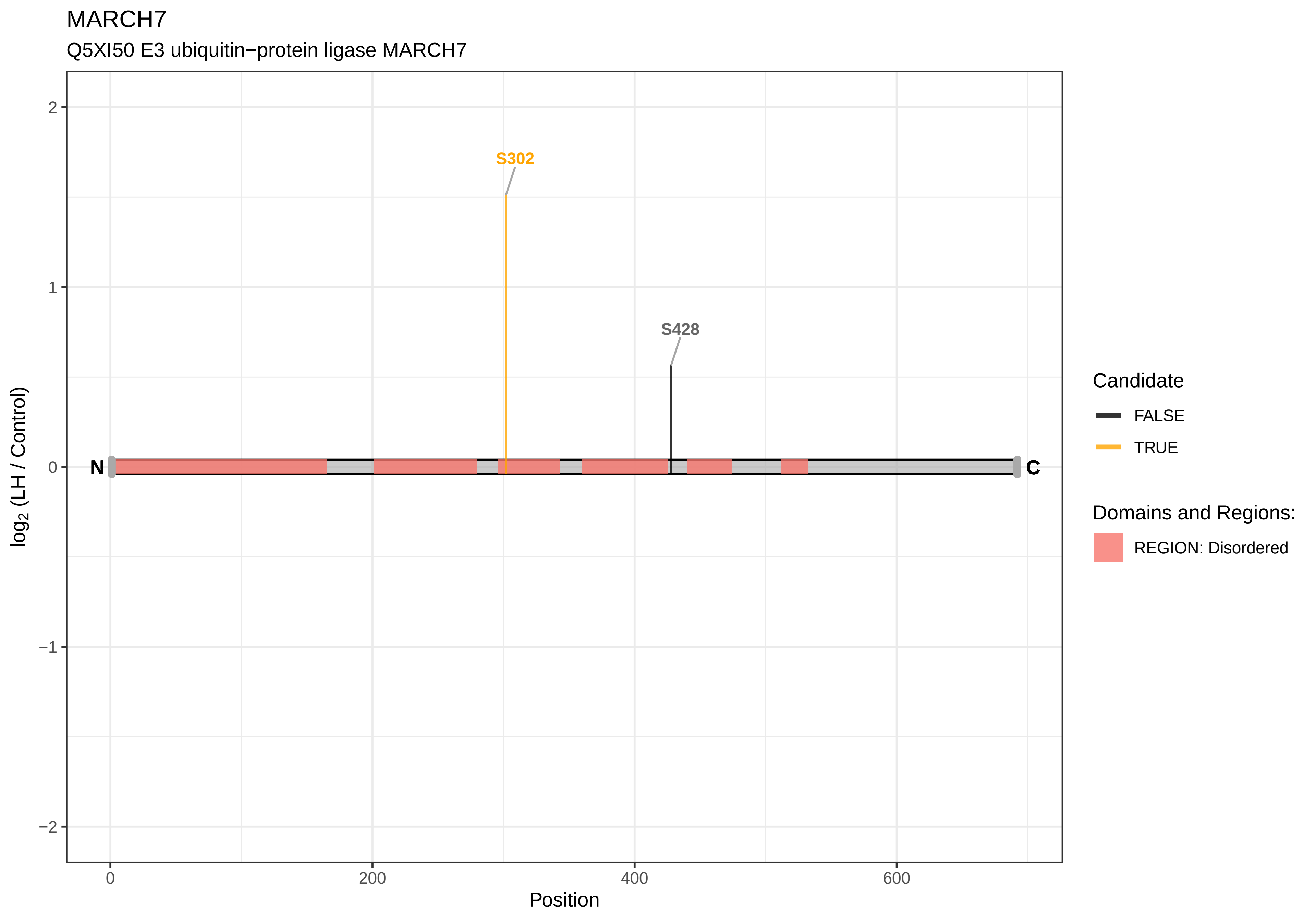

MARCH7
Q5XI50 E3 ubiquitin−protein ligase MARCH7
S302
S428
N
C
2
1
0
−1
−2
log2 (LH / Control)
0
200
400
600
Position
Candidate
FALSE
TRUE
Domains and Regions:
REGION: Disordered

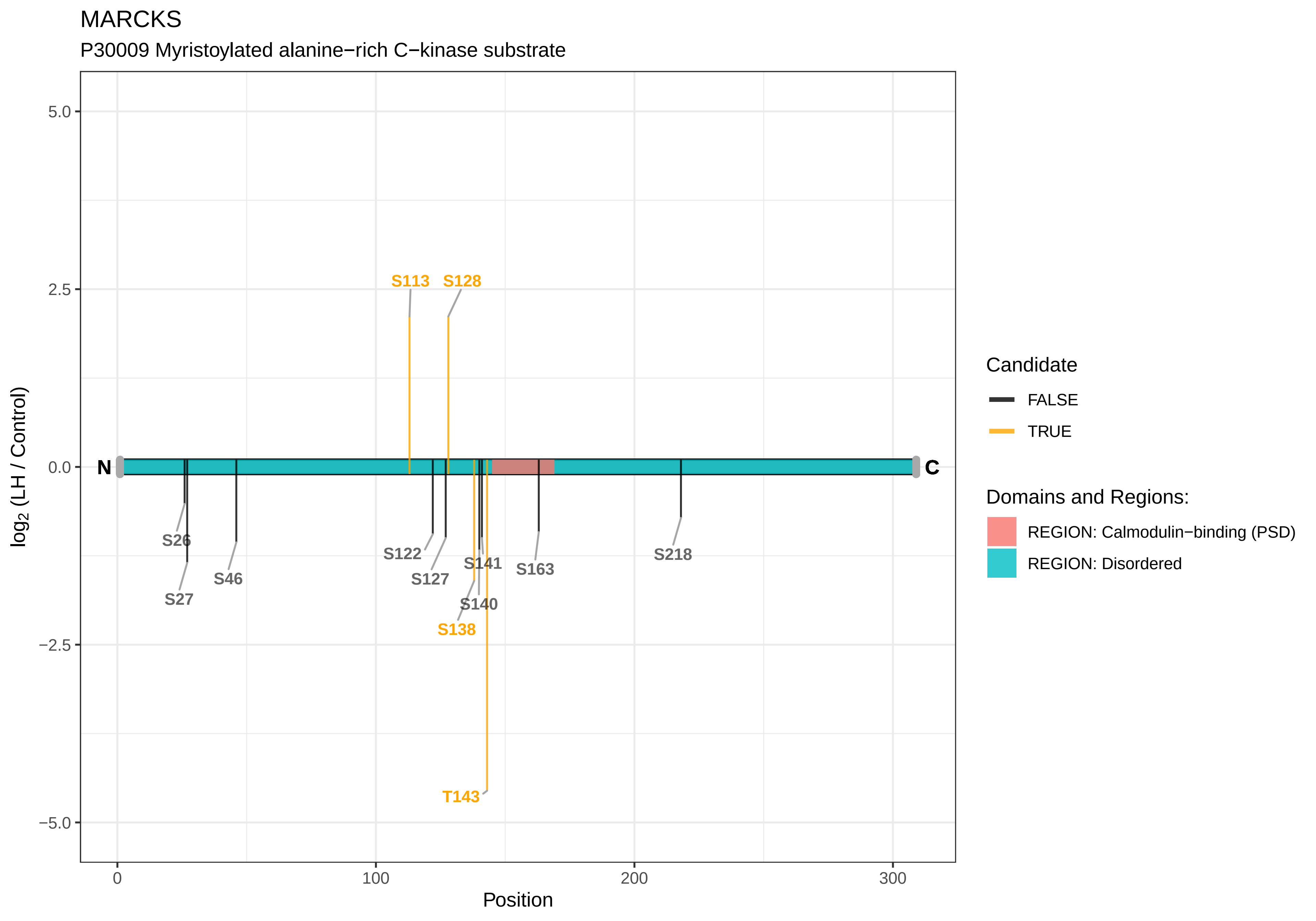

MARCKS
P30009 Myristoylated alanine−rich C−kinase substrate
log2 (LH / Control)
Position
5.0
2.5
0.0
−2.5
−5.0
0
100
200
300
N
C
S113
S128
S26
S27
S46
S122
S127
S141
S163
S140
S138
T143
S218
Candidate
FALSE
TRUE
Domains and Regions:
REGION: Calmodulin−binding (PSD)
REGION: Disordered

MARCKSL1

A0A0G2K613 Uncharacterized protein

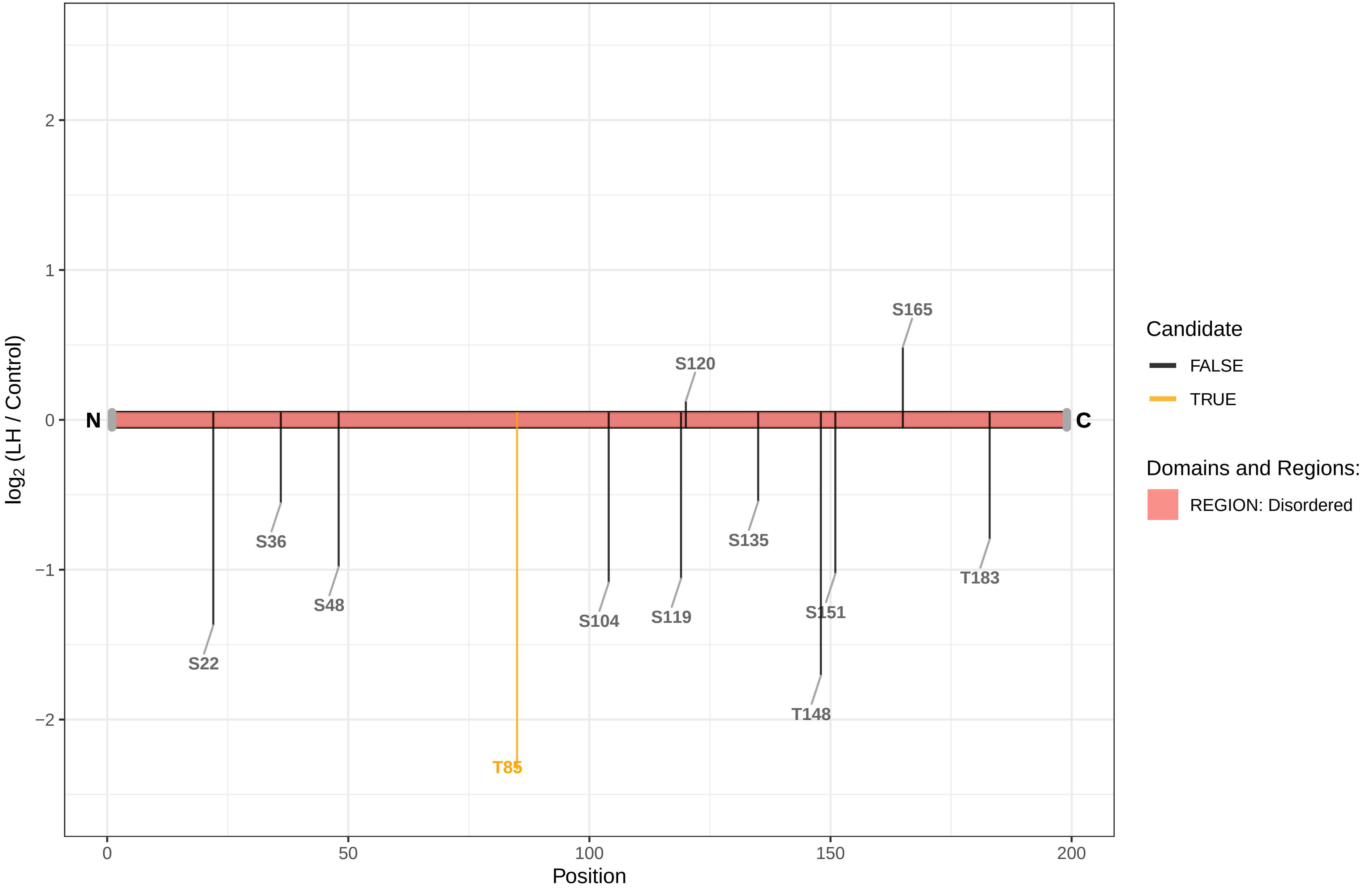

MARK1

O08678 Serine/threonine−protein kinase MARK1

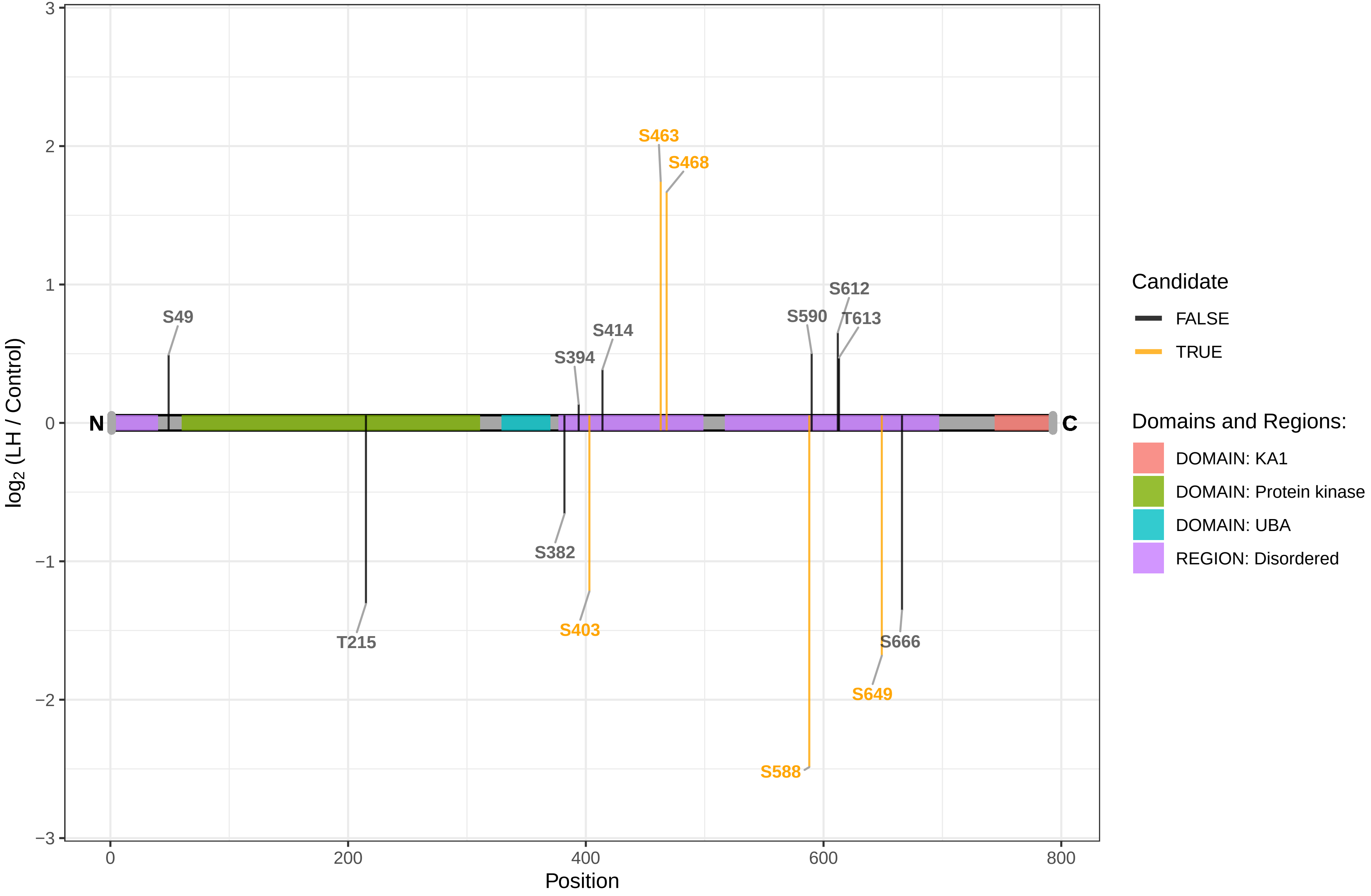

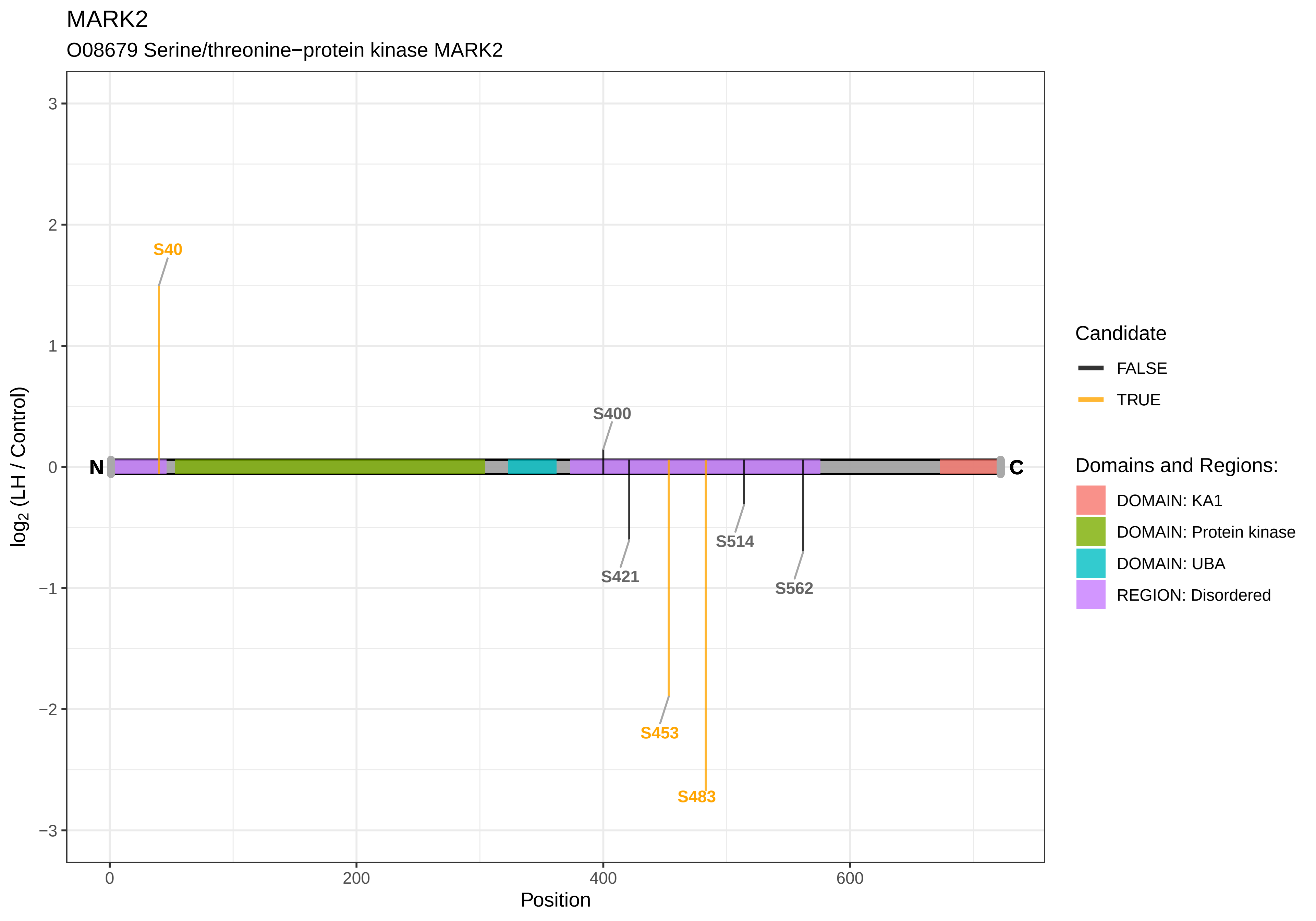

MARK2
O08679 Serine/threonine−protein kinase MARK2
S40
S400
S421
S453
S483
S514
S562
N
C
3
2
1
0
−1
−2
−3
log2 (LH / Control)
0
200
400
600
Position
Candidate
FALSE
TRUE
Domains and Regions:
DOMAIN: KA1
DOMAIN: Protein kinase
DOMAIN: UBA
REGION: Disordered

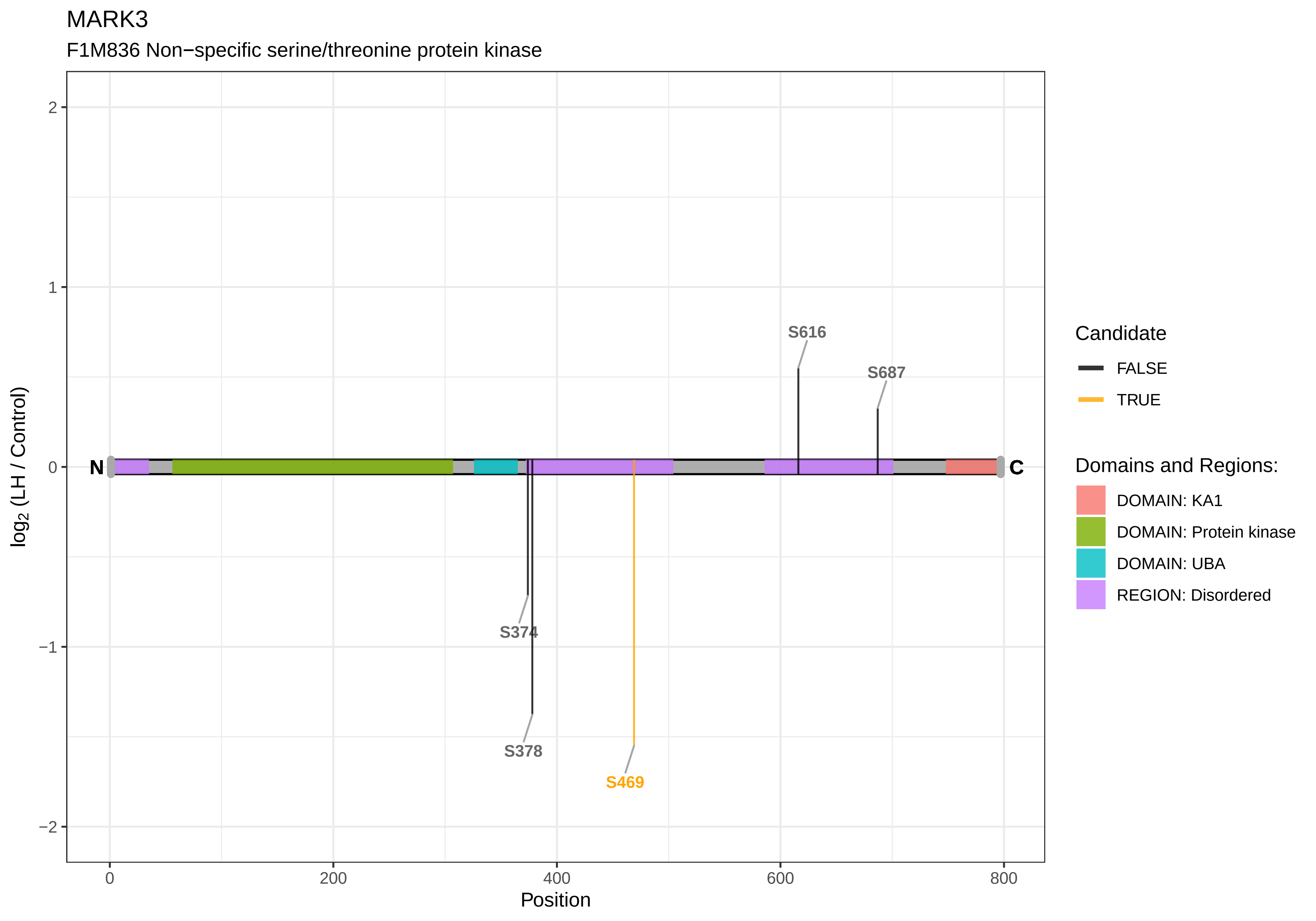

MARK3
F1M836 Non−specific serine/threonine protein kinase
S616
S687
S374
S378
S469
N
C
2
1
0
−1
−2
log2 (LH / Control)
0
200
400
600
800
Position
Candidate
FALSE
TRUE
Domains and Regions:
DOMAIN: KA1
DOMAIN: Protein kinase
DOMAIN: UBA
REGION: Disordered

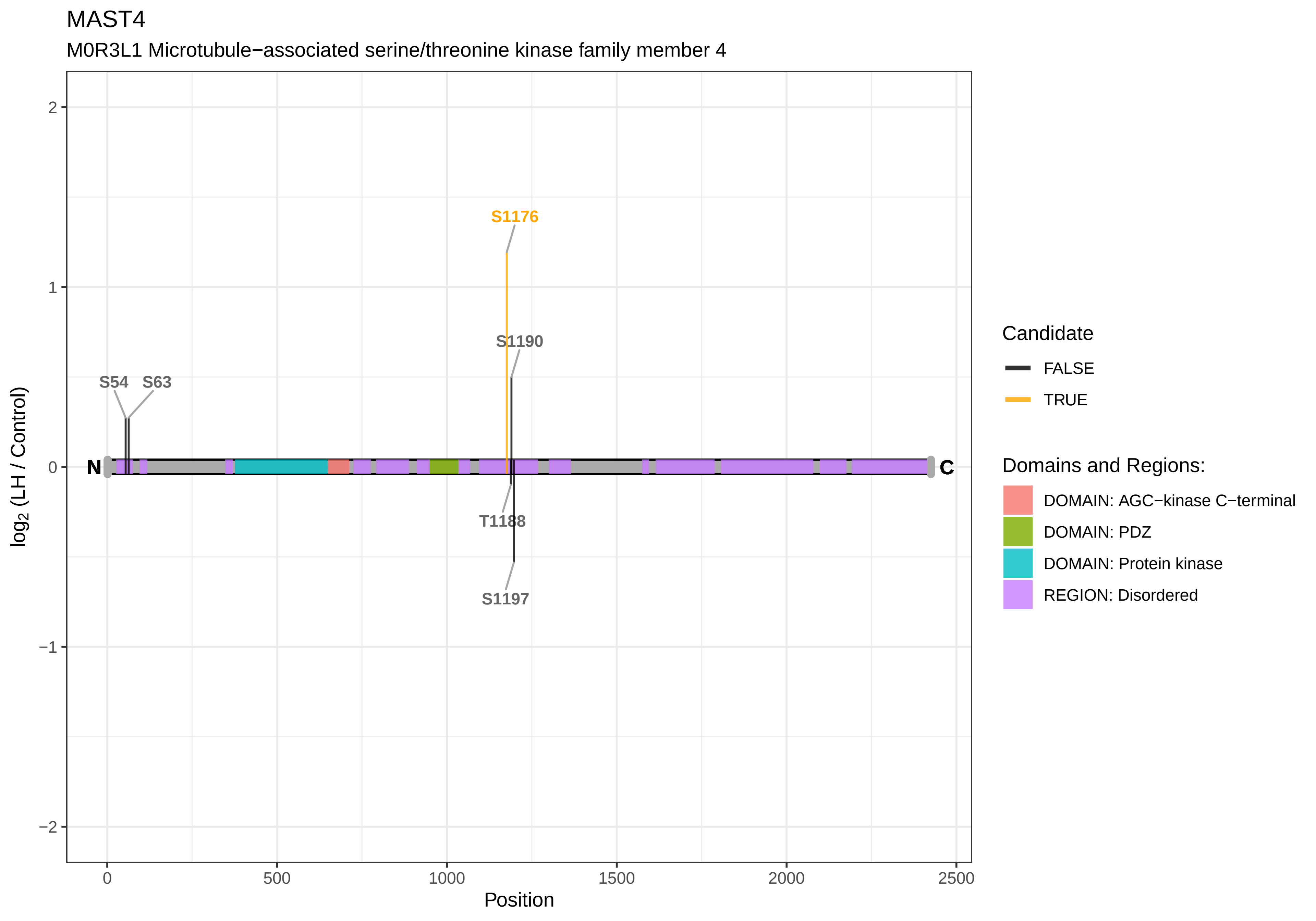
MAST4
M0R3L1 Microtubule−associated serine/threonine kinase family member 4
S1176
S1190
S54
S63
T1188
S1197
N
C
2
1
0
−1
−2
0
500
1000
1500
2000
2500
Position
log2 (LH / Control)
Candidate
FALSE
TRUE
Domains and Regions:
DOMAIN: AGC−kinase C−terminal
DOMAIN: PDZ
DOMAIN: Protein kinase
REGION: Disordered

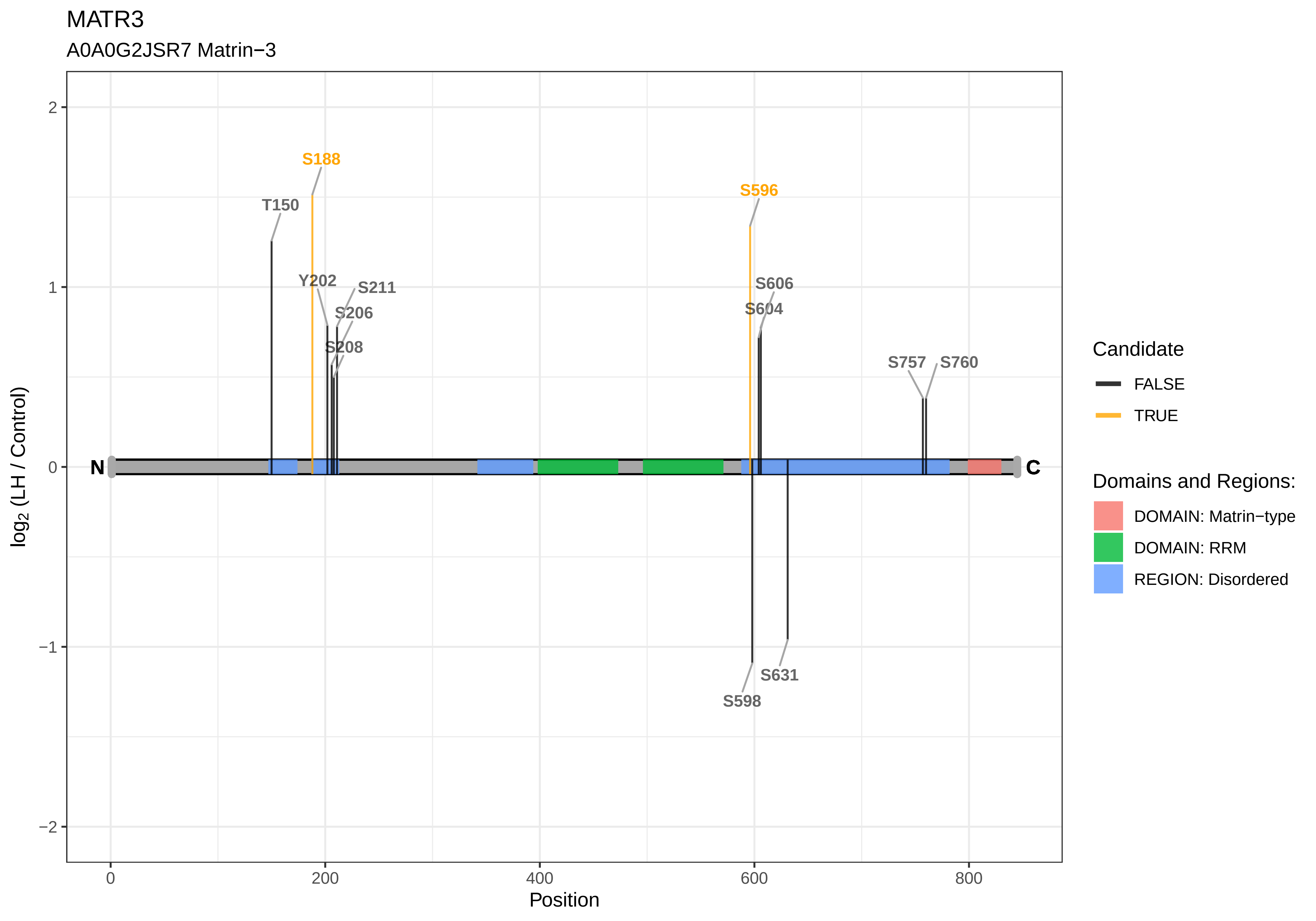
MATR3
A0A0G2JSR7 Matrin−3
S188
S596
T150
Y202
S211
S206
S208
S606
S604
S757
S760
S598
S631
N
C
2
1
0
−1
−2
0
200
400
600
800
Position
log2 (LH / Control)
Candidate
FALSE
TRUE
Domains and Regions:
DOMAIN: Matrin−type
DOMAIN: RRM
REGION: Disordered

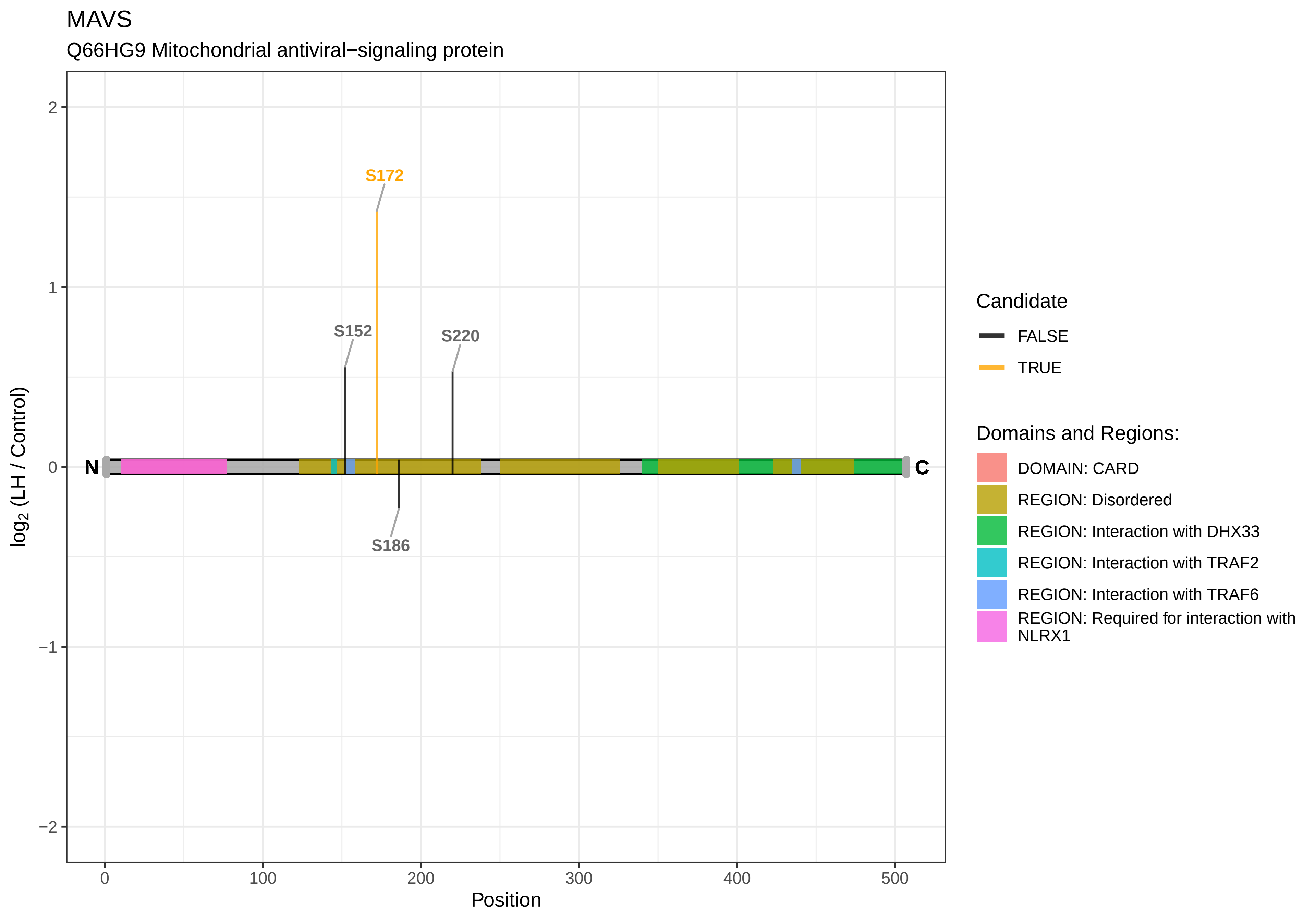
MAVS
Q66HG9 Mitochondrial antiviral−signaling protein
S172
S152
S220
S186
N
C
2
1
0
−1
−2
0
100
200
300
400
500
Position
log2 (LH / Control)
Candidate
FALSE
TRUE
Domains and Regions:
DOMAIN: CARD
REGION: Disordered
REGION: Interaction with DHX33
REGION: Interaction with TRAF2
REGION: Interaction with TRAF6
REGION: Required for interaction with NLRX1

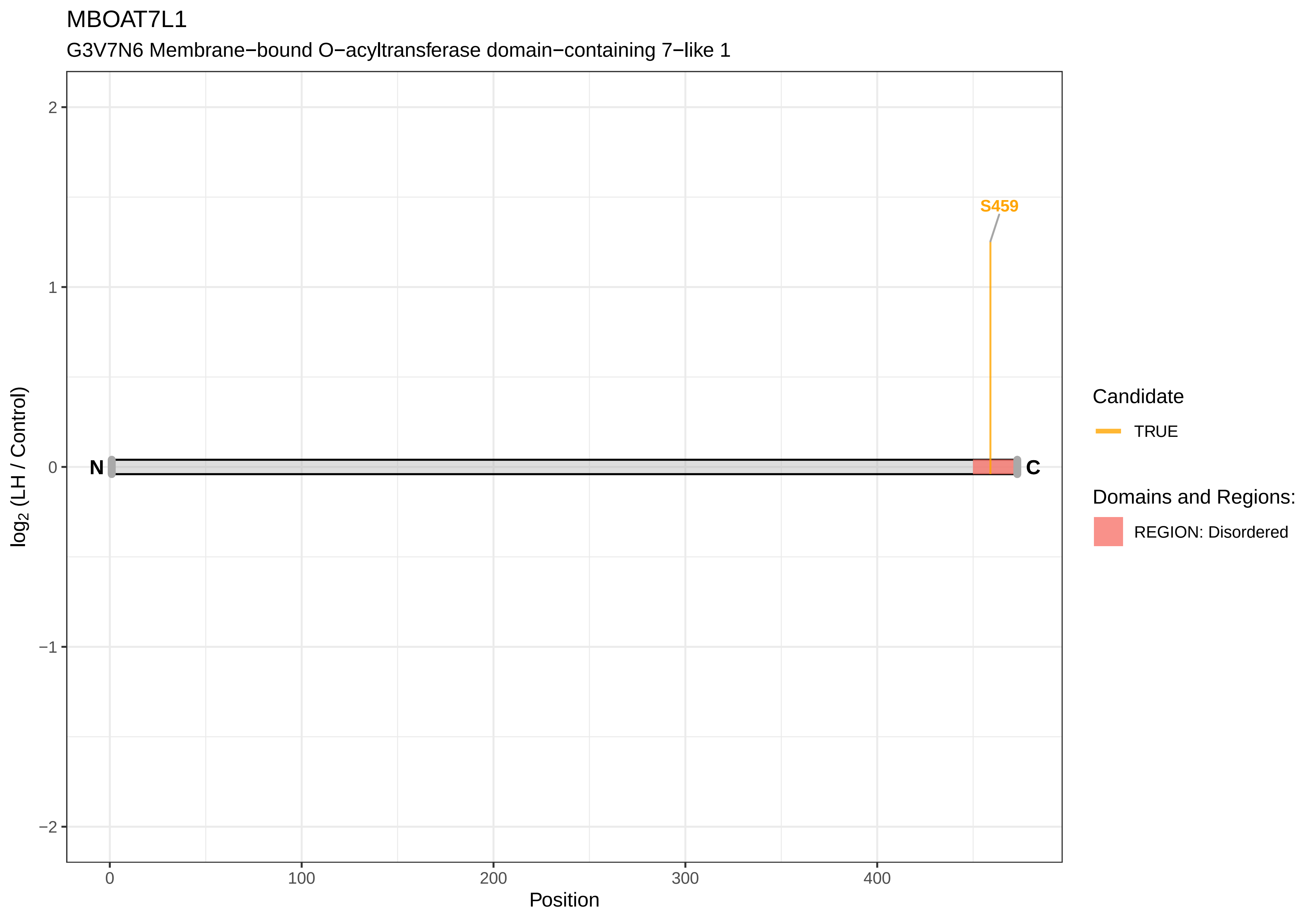
MBOAT7L1
G3V7N6 Membrane−bound O−acyltransferase domain−containing 7−like 1
S459
N
C
log2 (LH / Control)
2
1
0
−1
−2
0
100
200
300
400
Position
Candidate
TRUE
Domains and Regions:
REGION: Disordered

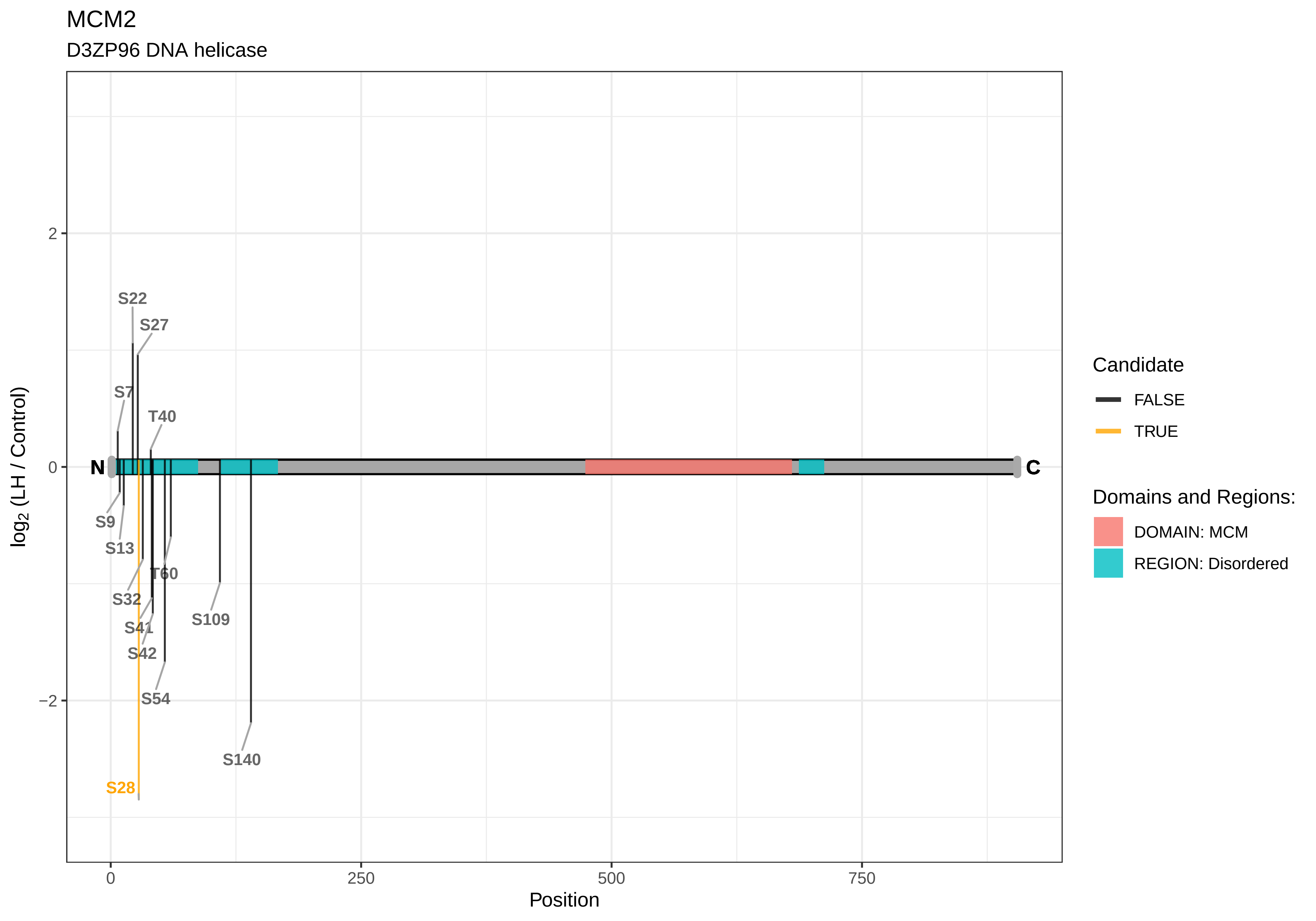
MCM2
D3ZP96 DNA helicase
log2 (LH / Control)
Position
2
0
−2
0
250
500
750
N
C
S22
S27
S7
T40
S9
S13
T60
S32
S109
S41
S42
S54
S140
S28
Candidate
FALSE
TRUE
Domains and Regions:
DOMAIN: MCM
REGION: Disordered

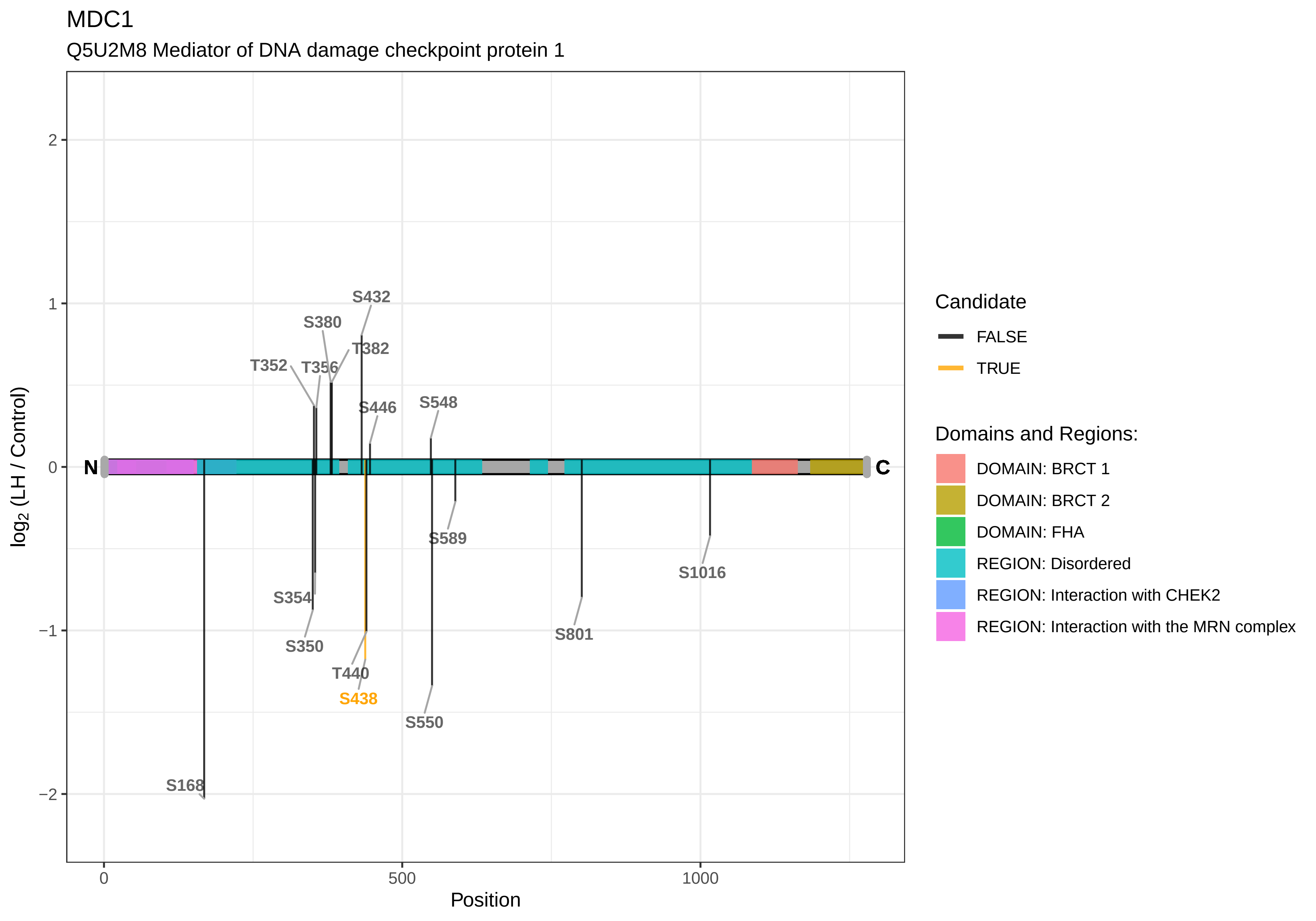
MDC1
Q5U2M8 Mediator of DNA damage checkpoint protein 1
log2 (LH / Control)
Position
2
1
0
−1
−2
0
500
1000
N
C
S432
S380
T382
T352
T356
S446
S548
S589
S1016
S354
S350
T440
S438
S550
S801
S168
Candidate
FALSE
TRUE
Domains and Regions:
DOMAIN: BRCT 1
DOMAIN: BRCT 2
DOMAIN: FHA
REGION: Disordered
REGION: Interaction with CHEK2
REGION: Interaction with the MRN complex

MDM1

A0A0G2KB24 Nuclear protein MDM1

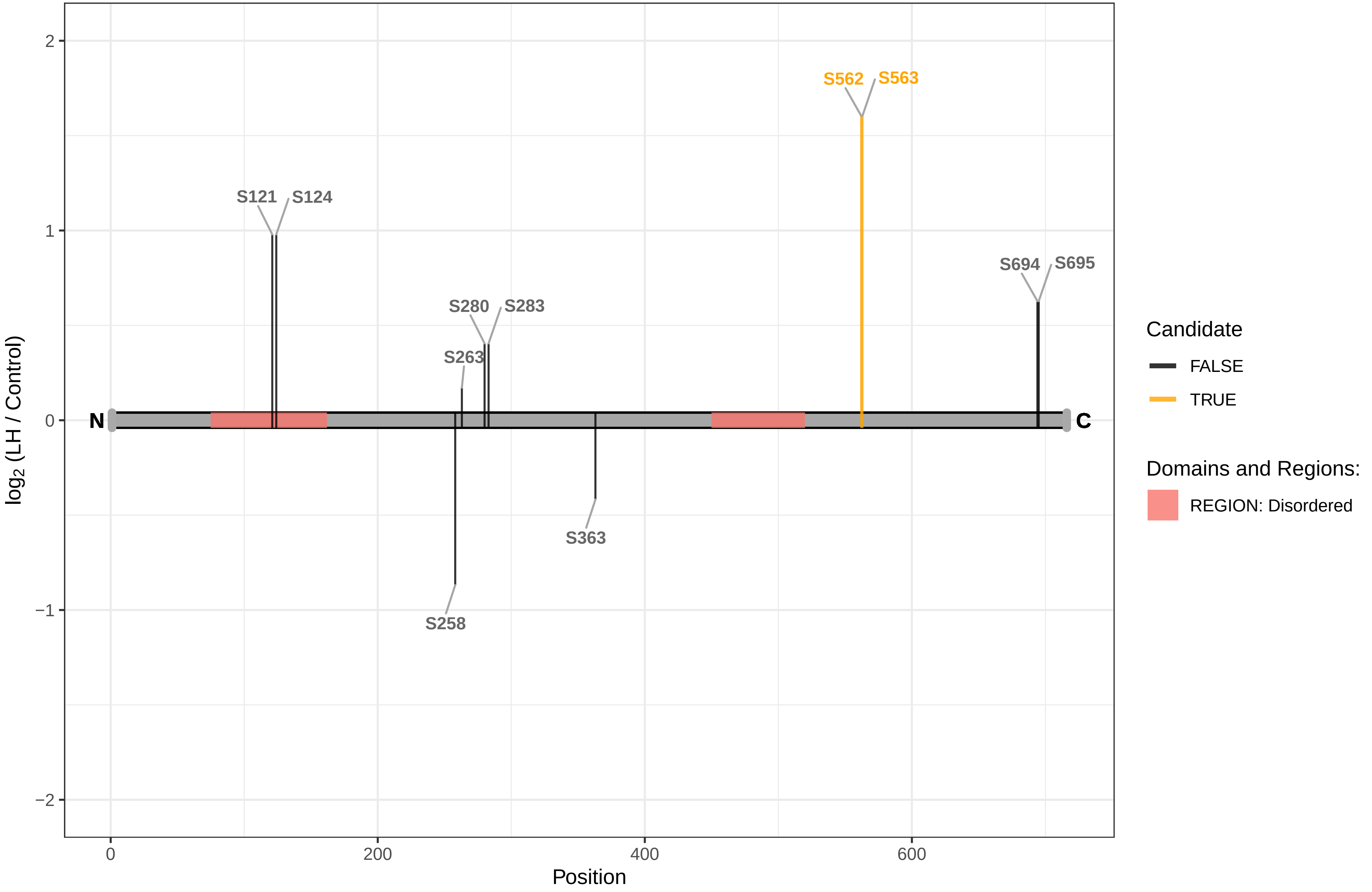

MEF2D

A0A0G2JSU7 Myocyte enhancer factor 2D

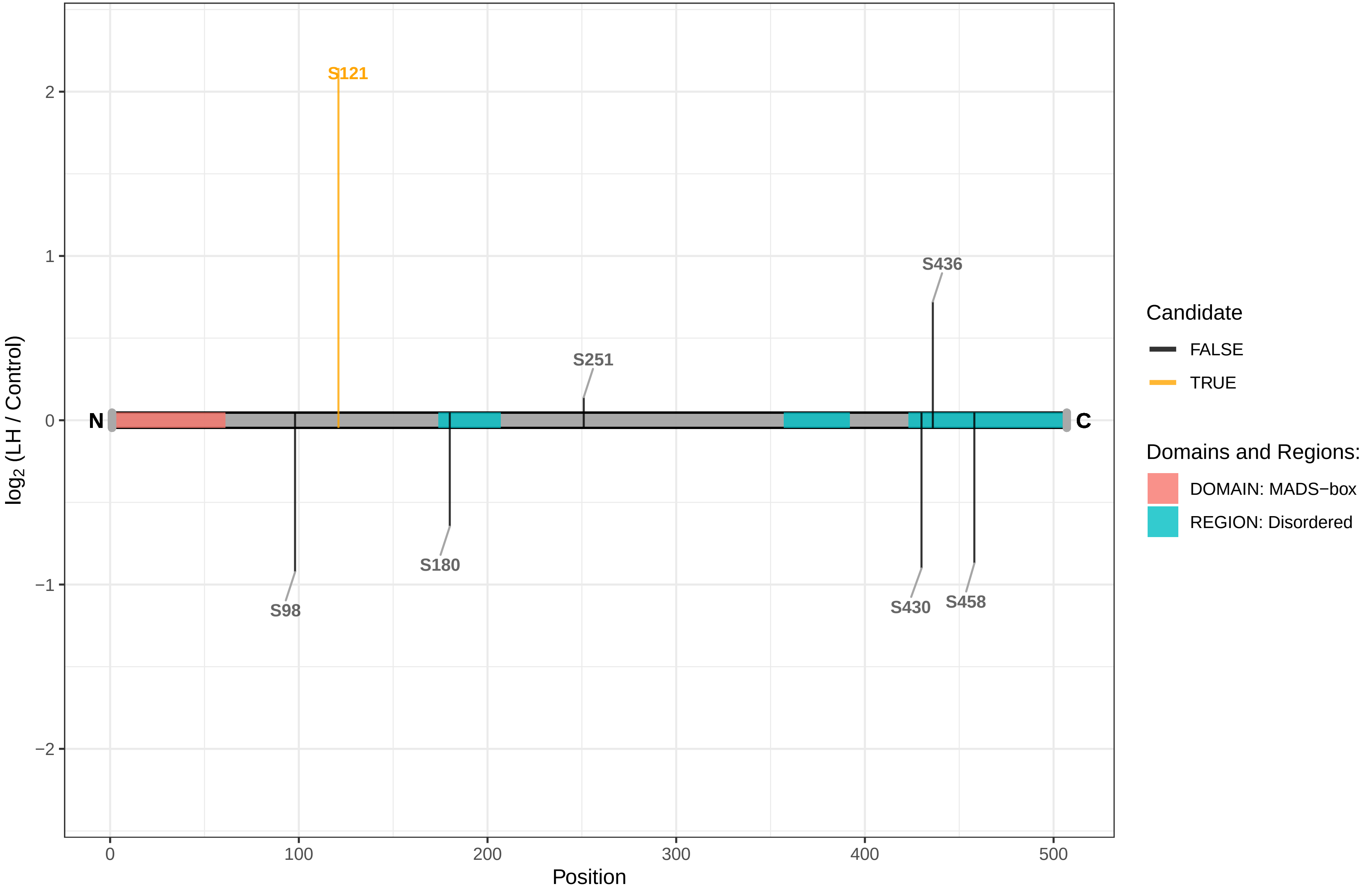

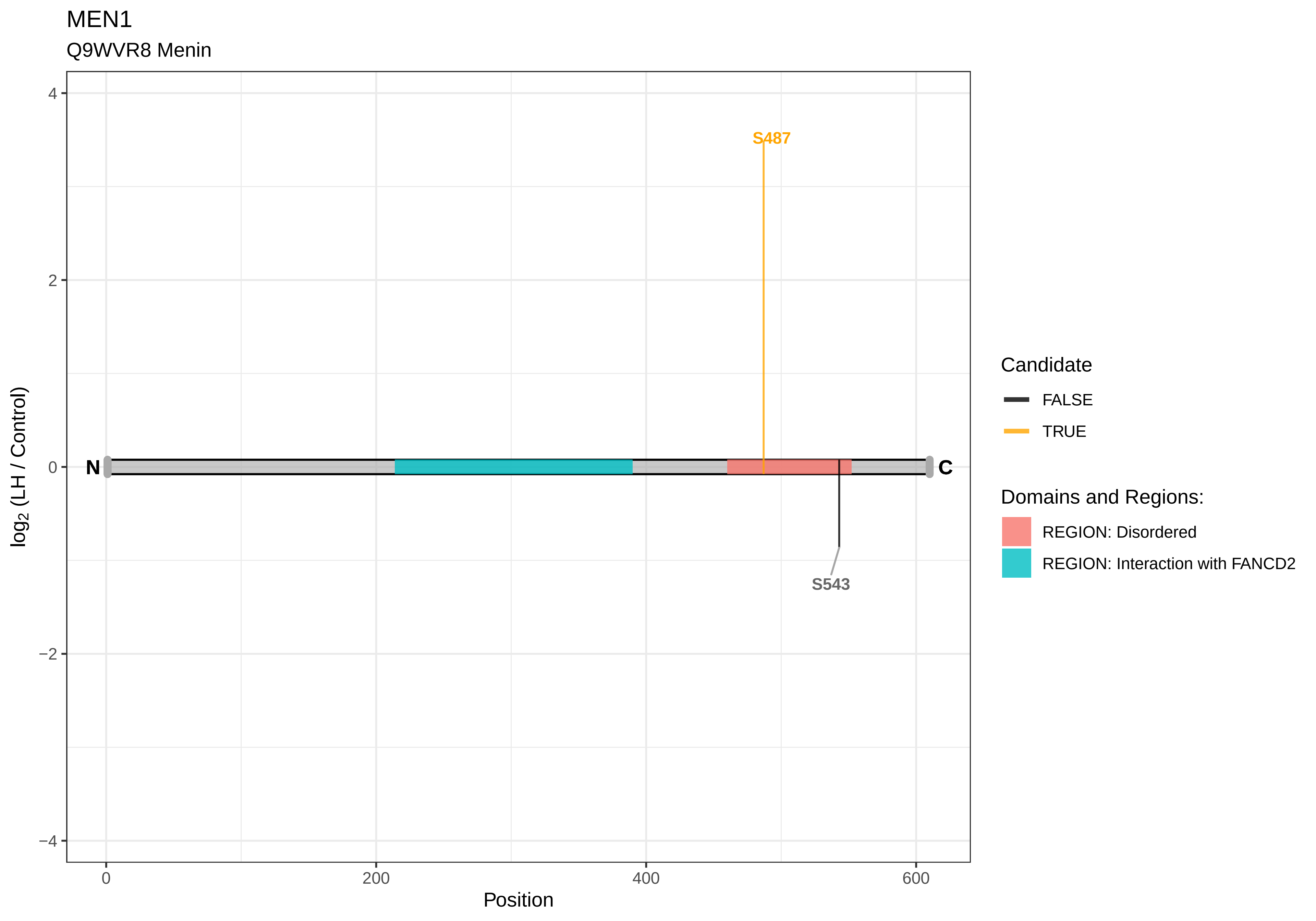

MEN1
Q9WVR8 Menin
S487
S543
N
C
4
2
0
−2
−4
0
200
400
600
Position
log2 (LH / Control)
Candidate
FALSE
TRUE
Domains and Regions:
REGION: Disordered
REGION: Interaction with FANCD2

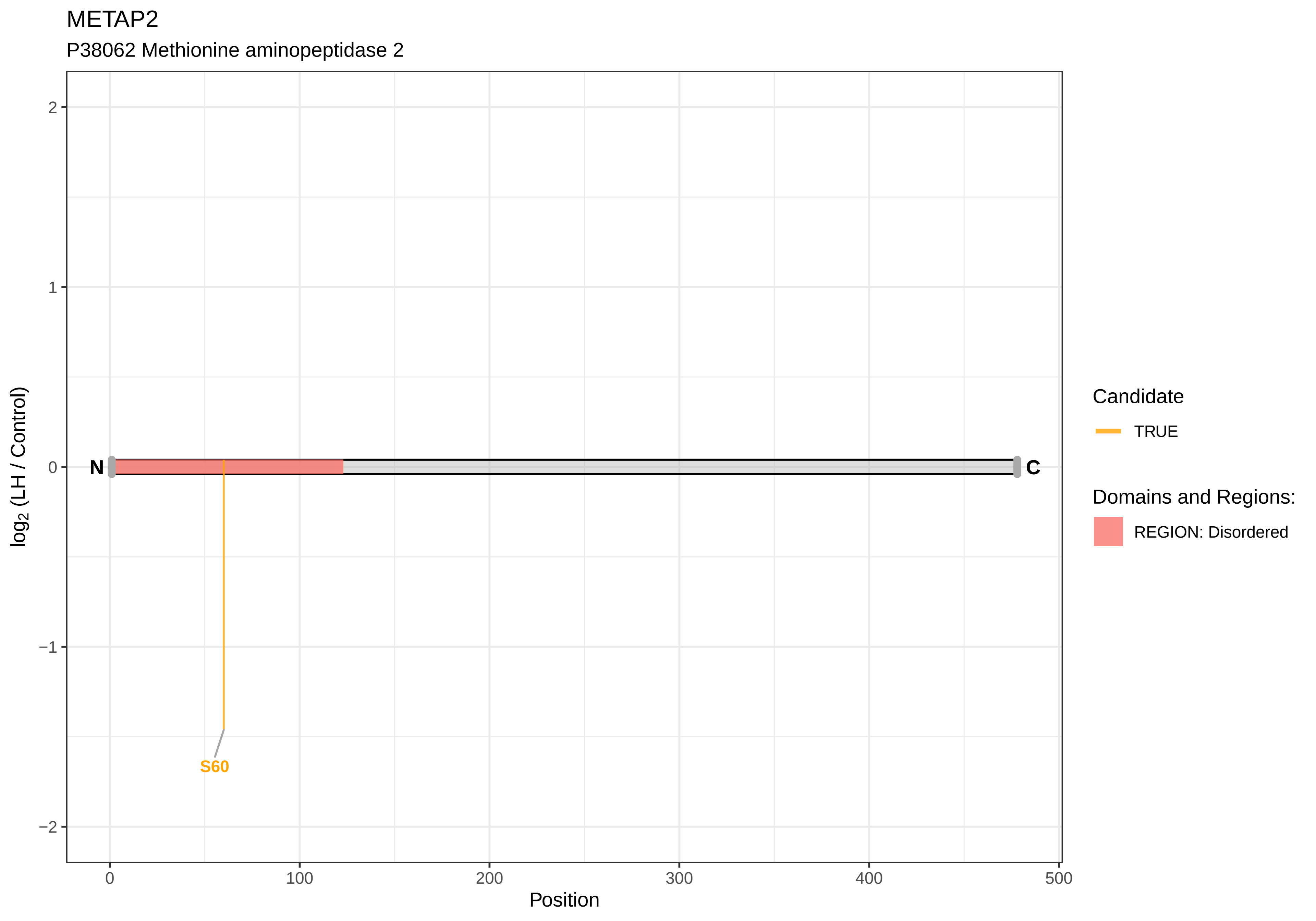

METAP2
P38062 Methionine aminopeptidase 2
log2 (LH / Control)
2
1
0
−1
−2
N
C
S60
0
100
200
300
400
500
Position
Candidate
TRUE
Domains and Regions:
REGION: Disordered

METTL3

Q4V8G6 Methyltransferase−like 3

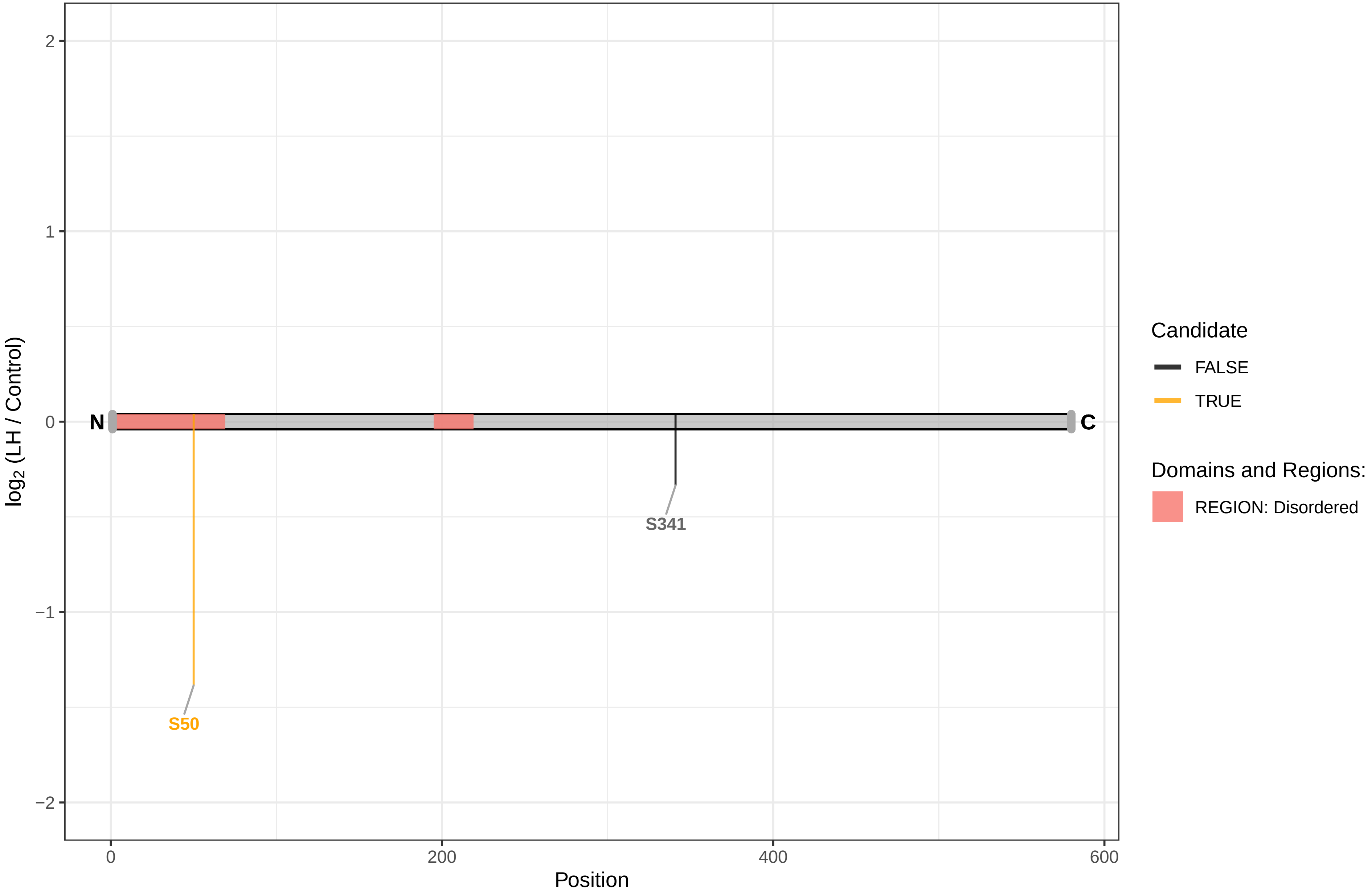

MFF

A0A0G2KAL9 Mitochondrial fission factor

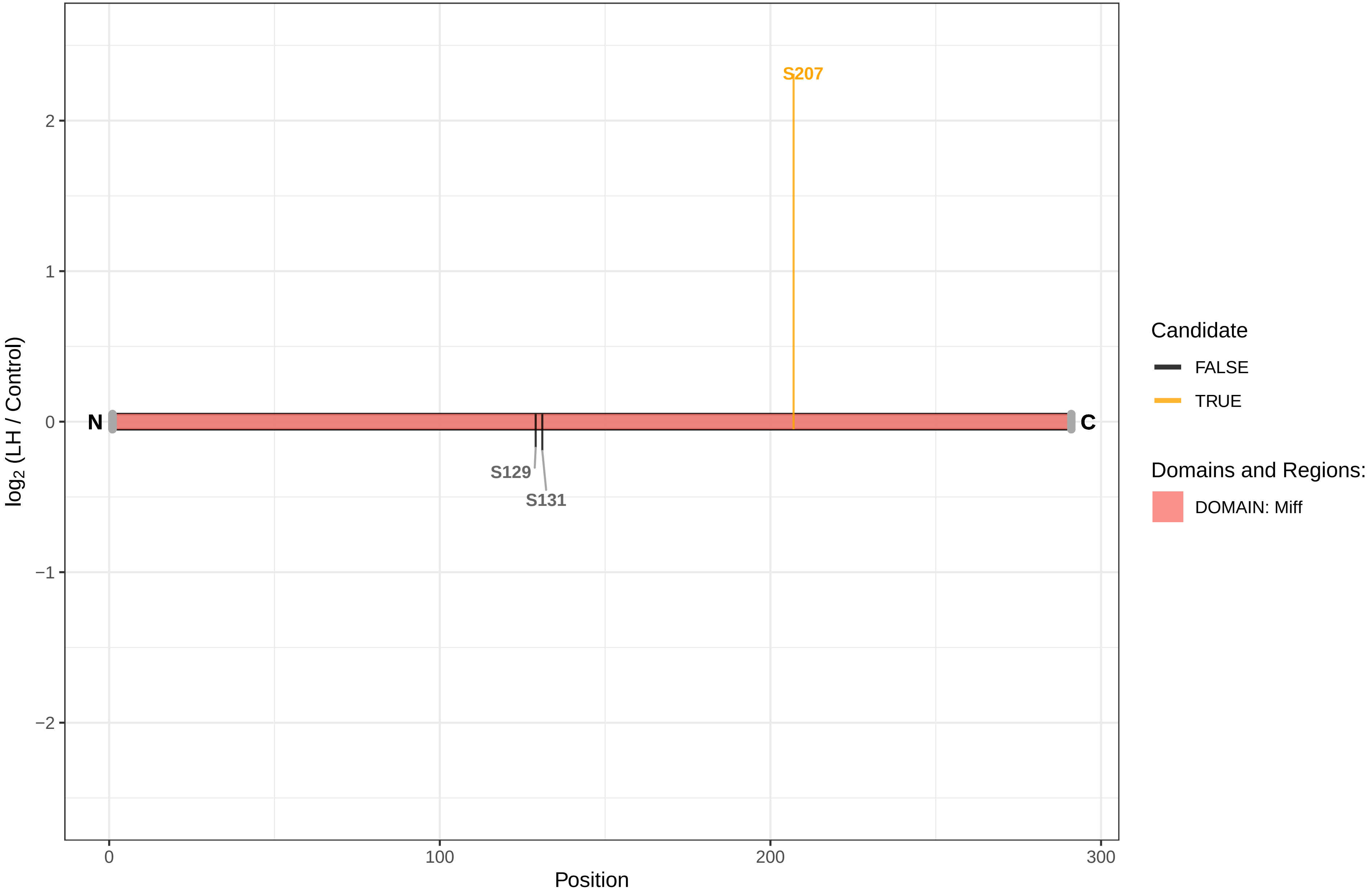

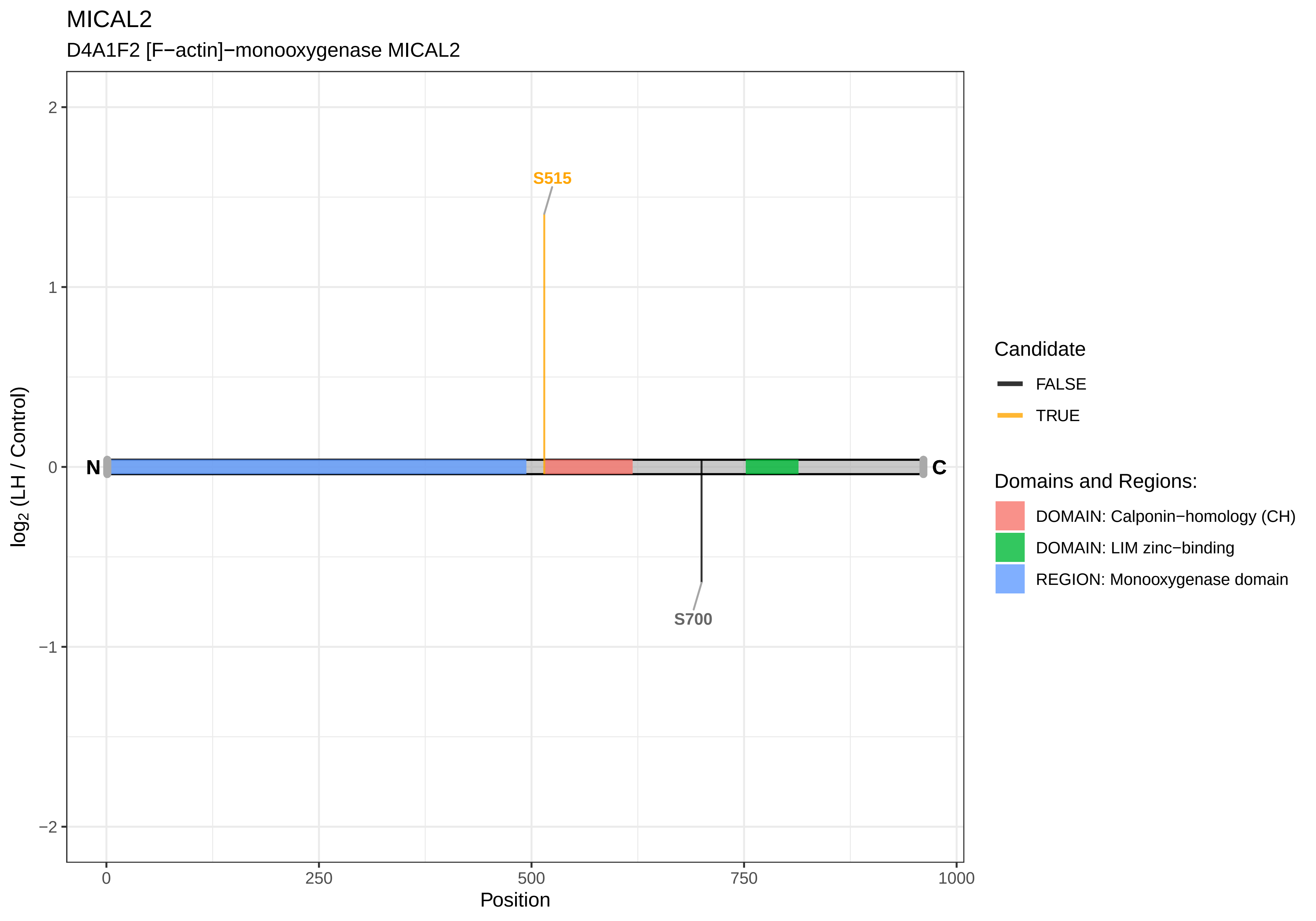

MICAL2
D4A1F2 [F−actin]−monooxygenase MICAL2
S515
S700
N
C
2
1
0
−1
−2
log2 (LH / Control)
0
250
500
750
1000
Position
Candidate
FALSE
TRUE
Domains and Regions:
DOMAIN: Calponin−homology (CH)
DOMAIN: LIM zinc−binding
REGION: Monooxygenase domain

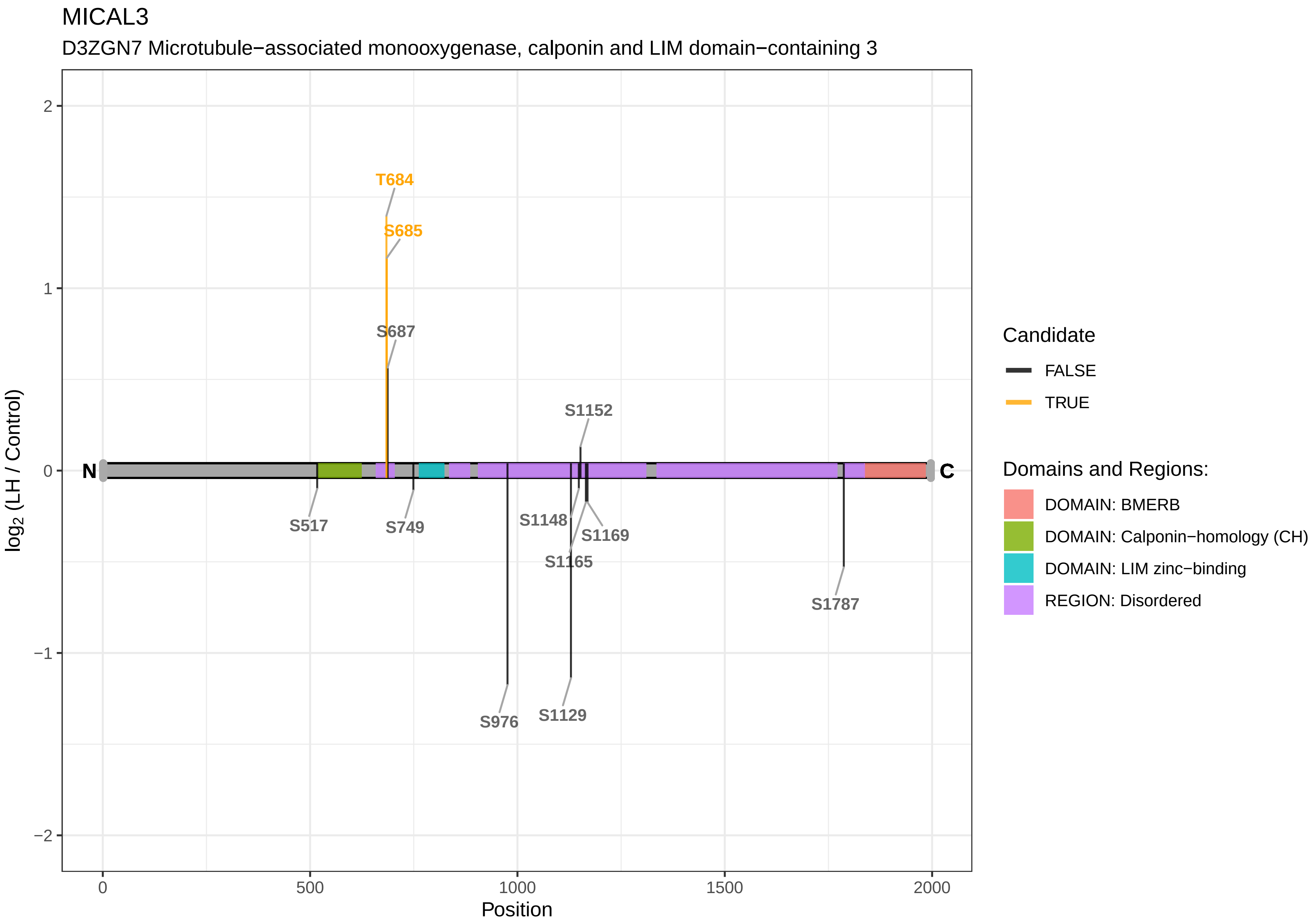
MICAL3
D3ZGN7 Microtubule−associated monooxygenase, calponin and LIM domain−containing 3
T684
S685
S687
S1152
S517
S749
S1148
S1169
S1165
S1787
S976
S1129
N
C
2
1
0
−1
−2
0
500
1000
1500
2000
Position
log2 (LH / Control)
Candidate
FALSE
TRUE
Domains and Regions:
DOMAIN: BMERB
DOMAIN: Calponin−homology (CH)
DOMAIN: LIM zinc−binding
REGION: Disordered

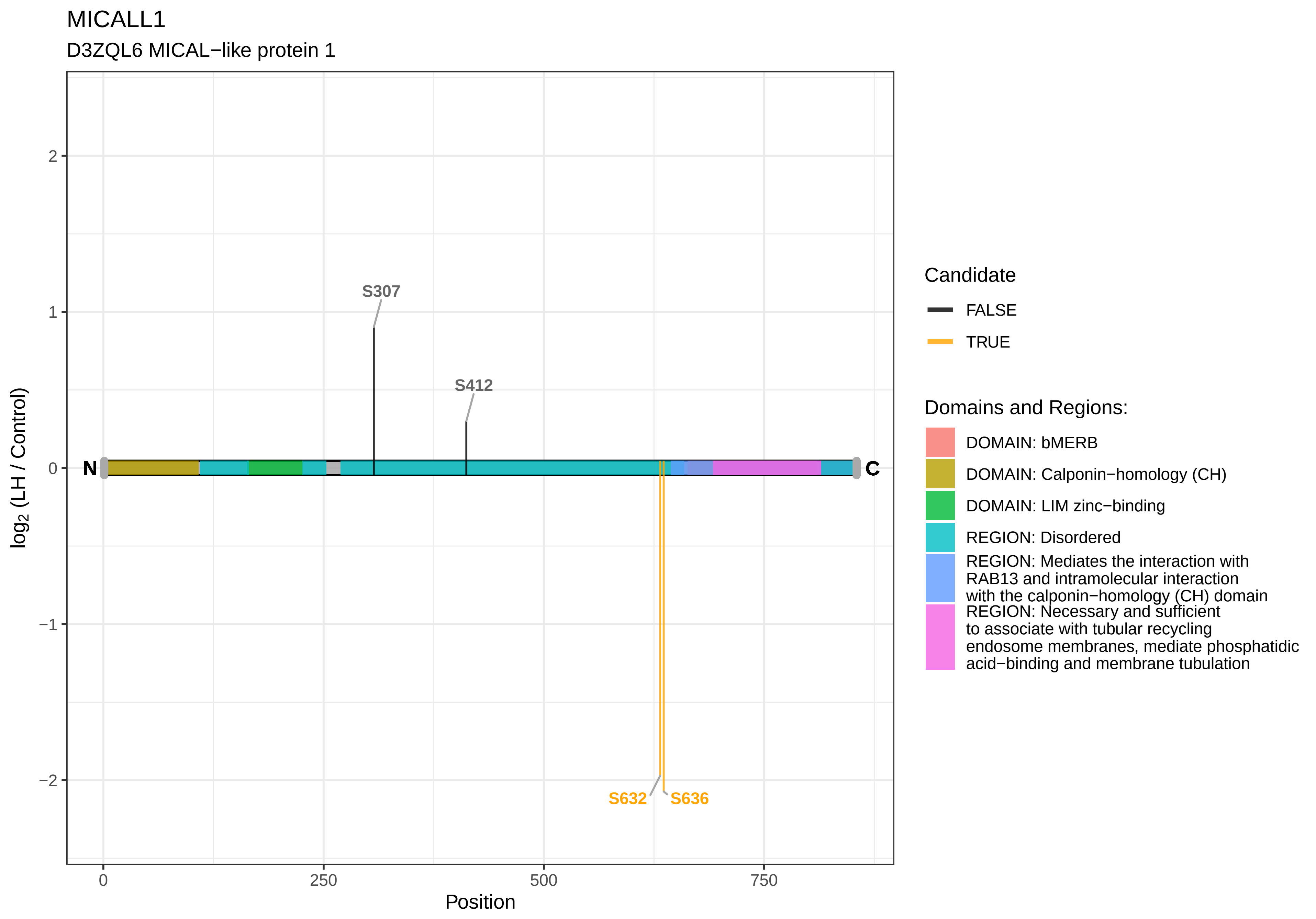

MICALL1
D3ZQL6 MICAL−like protein 1
S307
S412
S632
S636
N
C
log2 (LH / Control)
Position
0
250
500
750
2
1
0
−1
−2
Candidate
FALSE
TRUE
Domains and Regions:
DOMAIN: bMERB
DOMAIN: Calponin−homology (CH)
DOMAIN: LIM zinc−binding
REGION: Disordered
REGION: Mediates the interaction with RAB13 and intramolecular interaction with the calponin−homology (CH) domain
REGION: Necessary and sufficient to associate with tubular recycling endosome membranes, mediate phosphatidic acid−binding and membrane tubulation

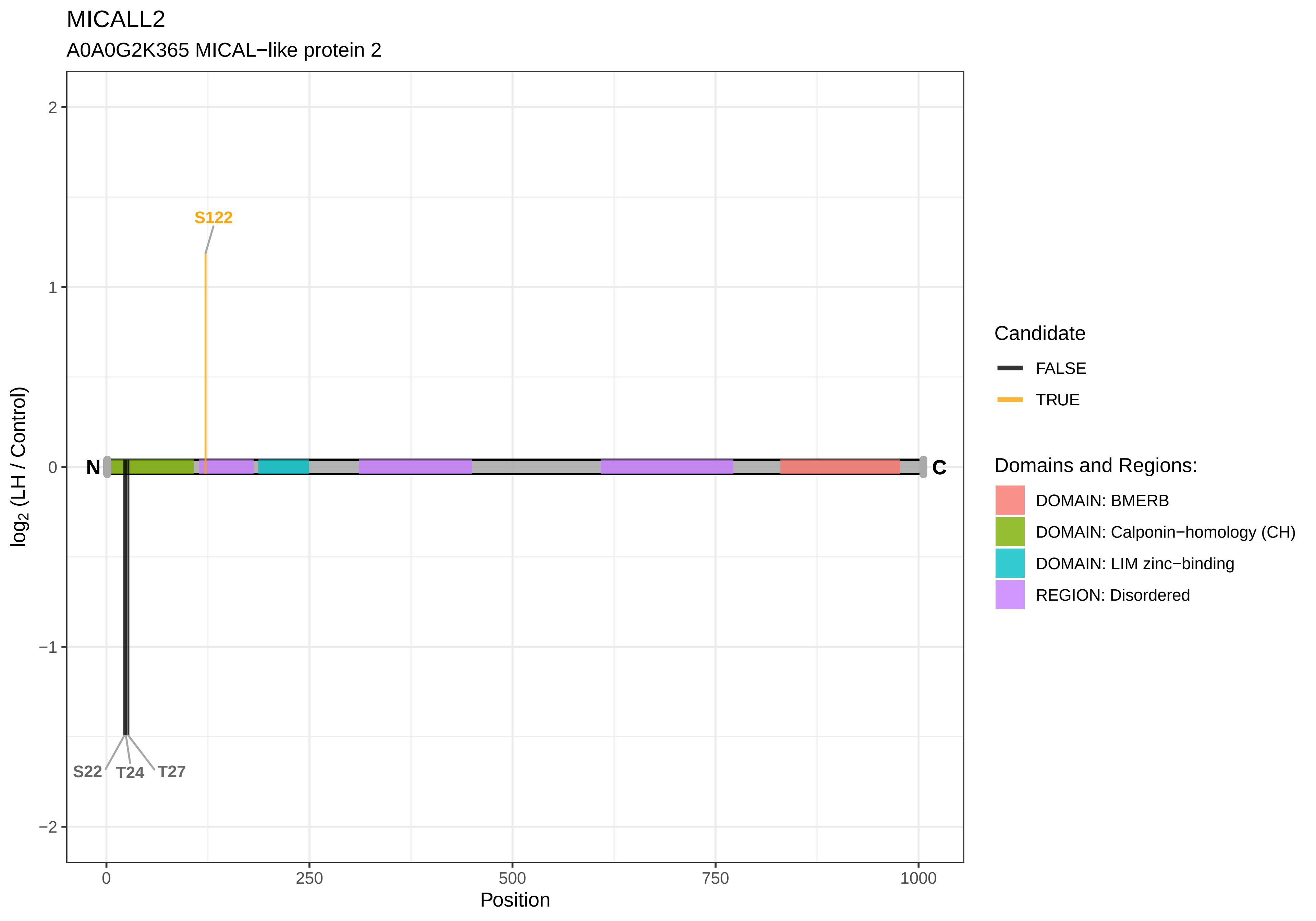

MICALL2
A0A0G2K365 MICAL−like protein 2
S122
N
C
S22
T24
T27
log2 (LH / Control)
2
1
0
−1
−2
0
250
500
750
1000
Position
Candidate
FALSE
TRUE
Domains and Regions:
DOMAIN: BMERB
DOMAIN: Calponin−homology (CH)
DOMAIN: LIM zinc−binding
REGION: Disordered

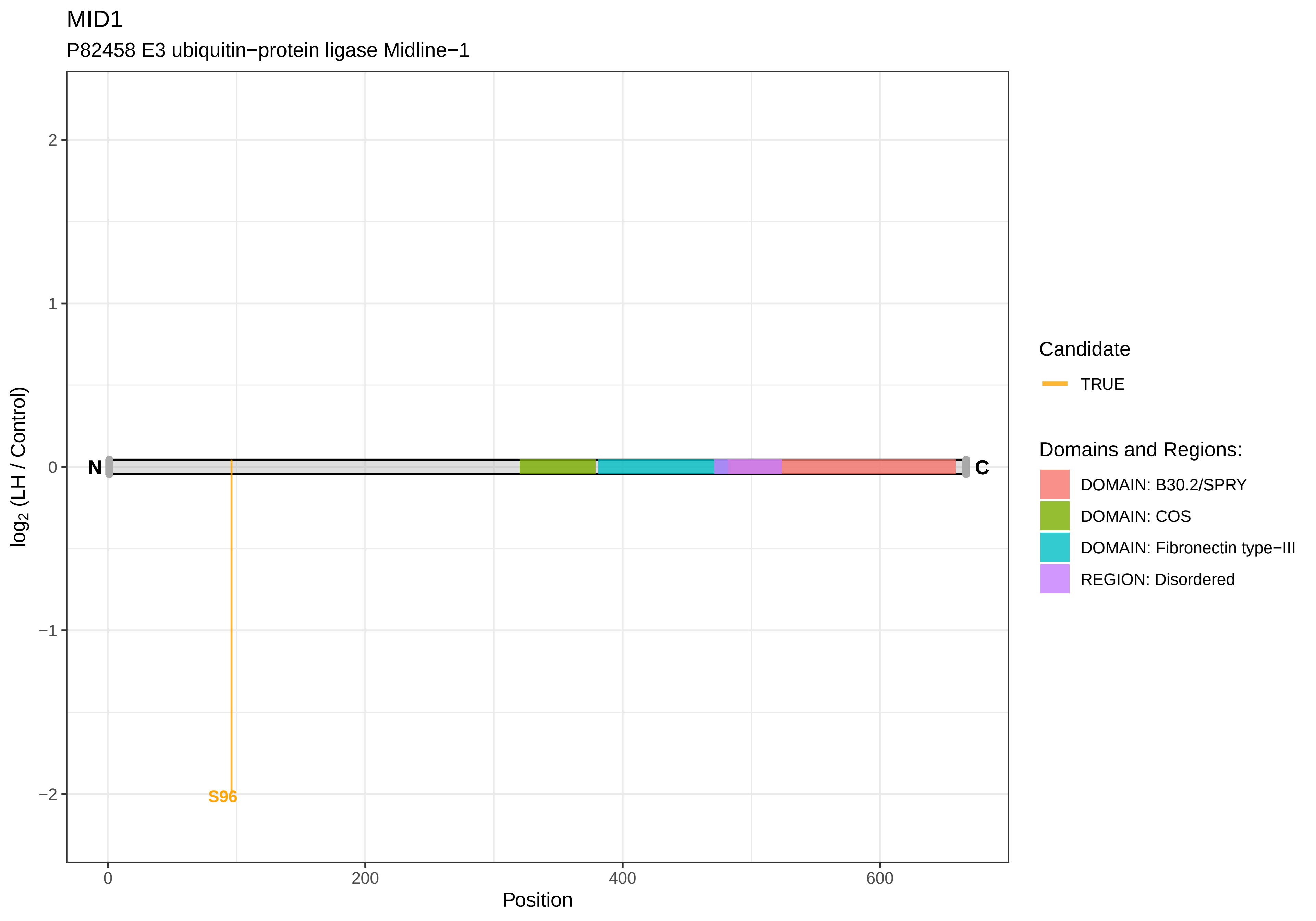
MID1
P82458 E3 ubiquitin−protein ligase Midline−1
log2 (LH / Control)
Position
N
C
S96
2
1
0
−1
−2
0
200
400
600
Candidate
TRUE
Domains and Regions:
DOMAIN: B30.2/SPRY
DOMAIN: COS
DOMAIN: Fibronectin type−III
REGION: Disordered

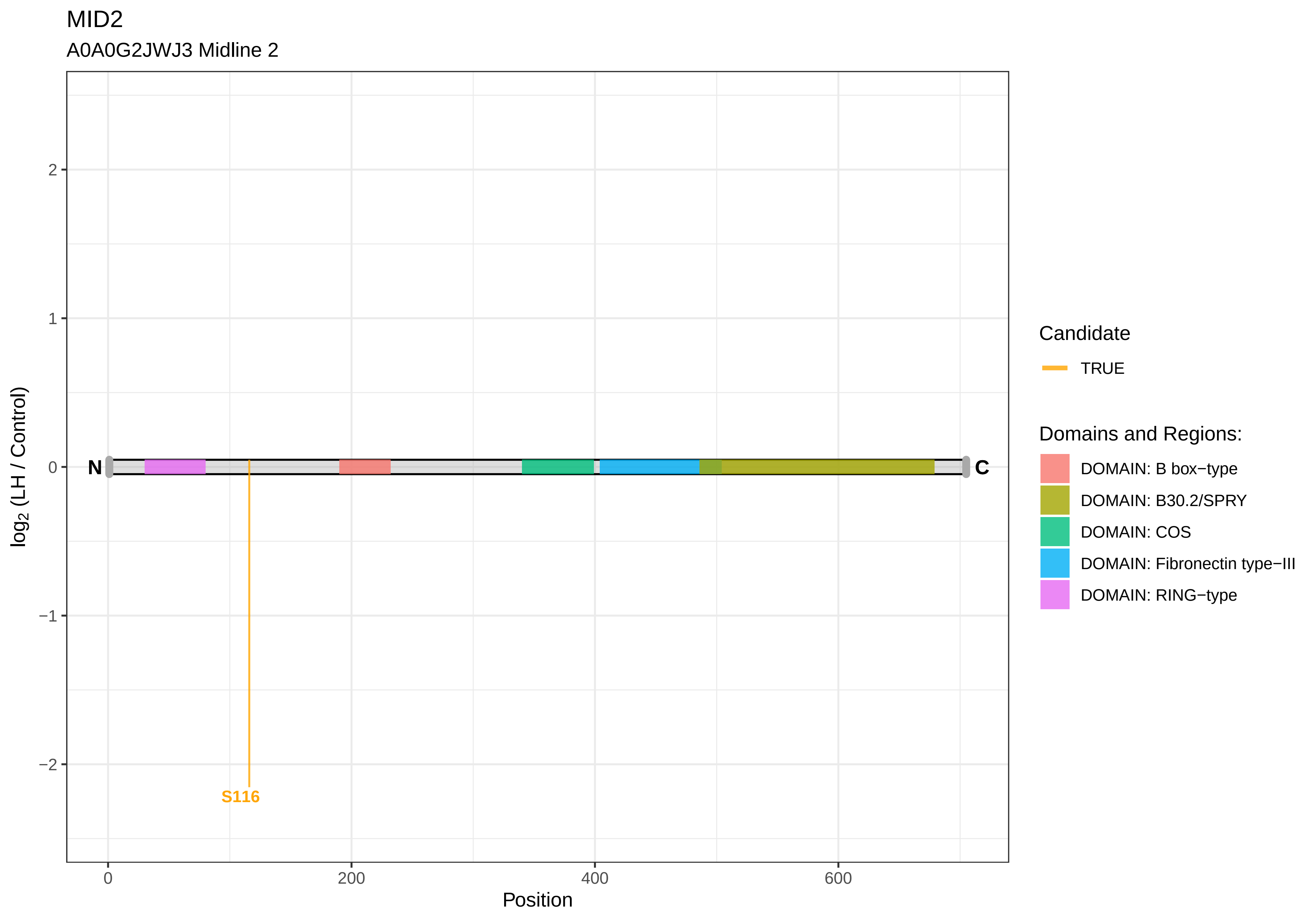

MID2
A0A0G2JWJ3 Midline 2
log2 (LH / Control)
2
1
0
−1
−2
N
C
S116
0
200
400
600
Position
Candidate
TRUE
Domains and Regions:
DOMAIN: B box−type
DOMAIN: B30.2/SPRY
DOMAIN: COS
DOMAIN: Fibronectin type−III
DOMAIN: RING−type

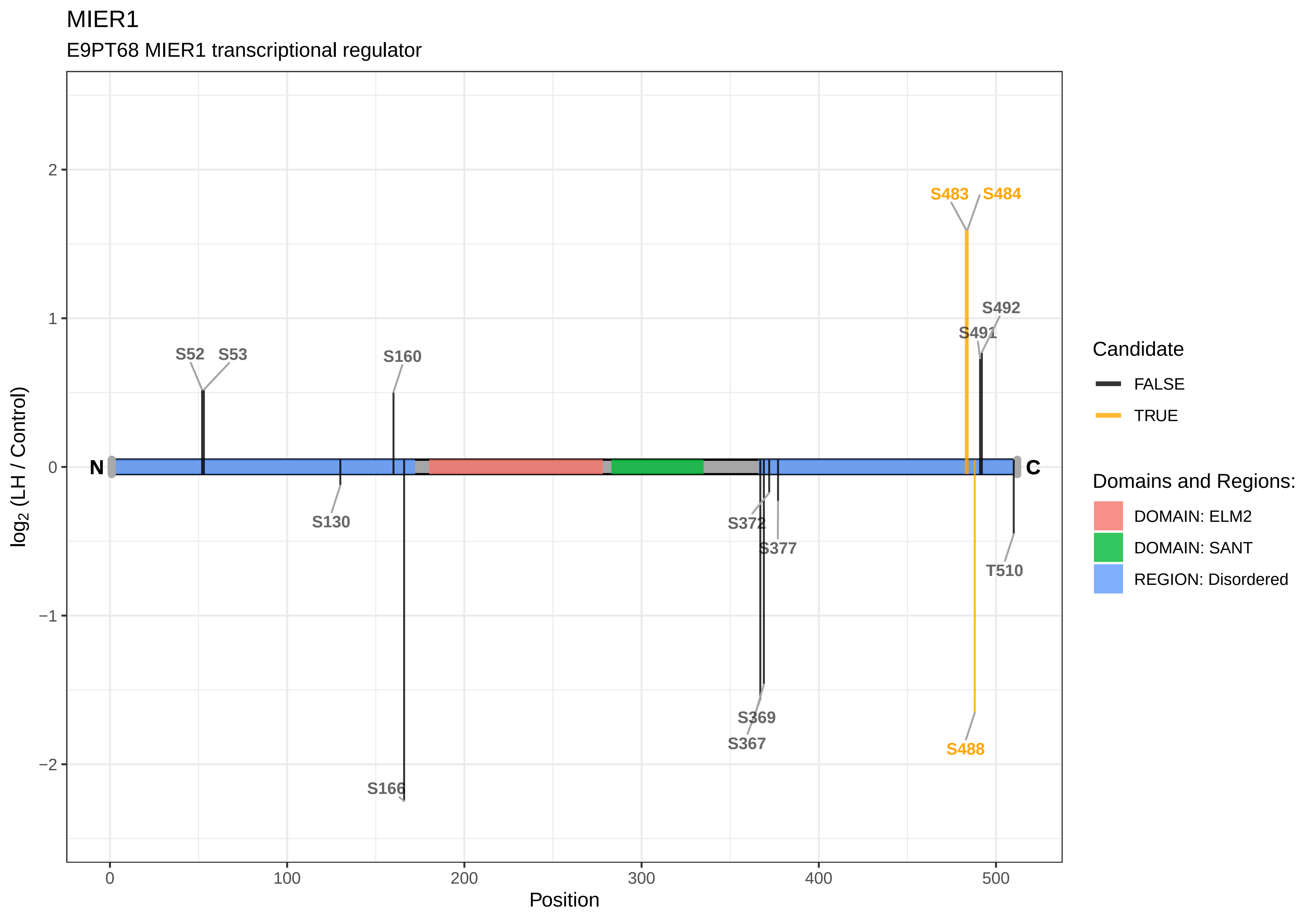
MIER1
E9PT68 MIER1 transcriptional regulator
log2 (LH / Control)
Position
N
C
S52
S53
S160
S130
S166
S372
S377
S369
S367
S483
S484
S492
S491
T510
S488
Candidate
FALSE
TRUE
Domains and Regions:
DOMAIN: ELM2
DOMAIN: SANT
REGION: Disordered

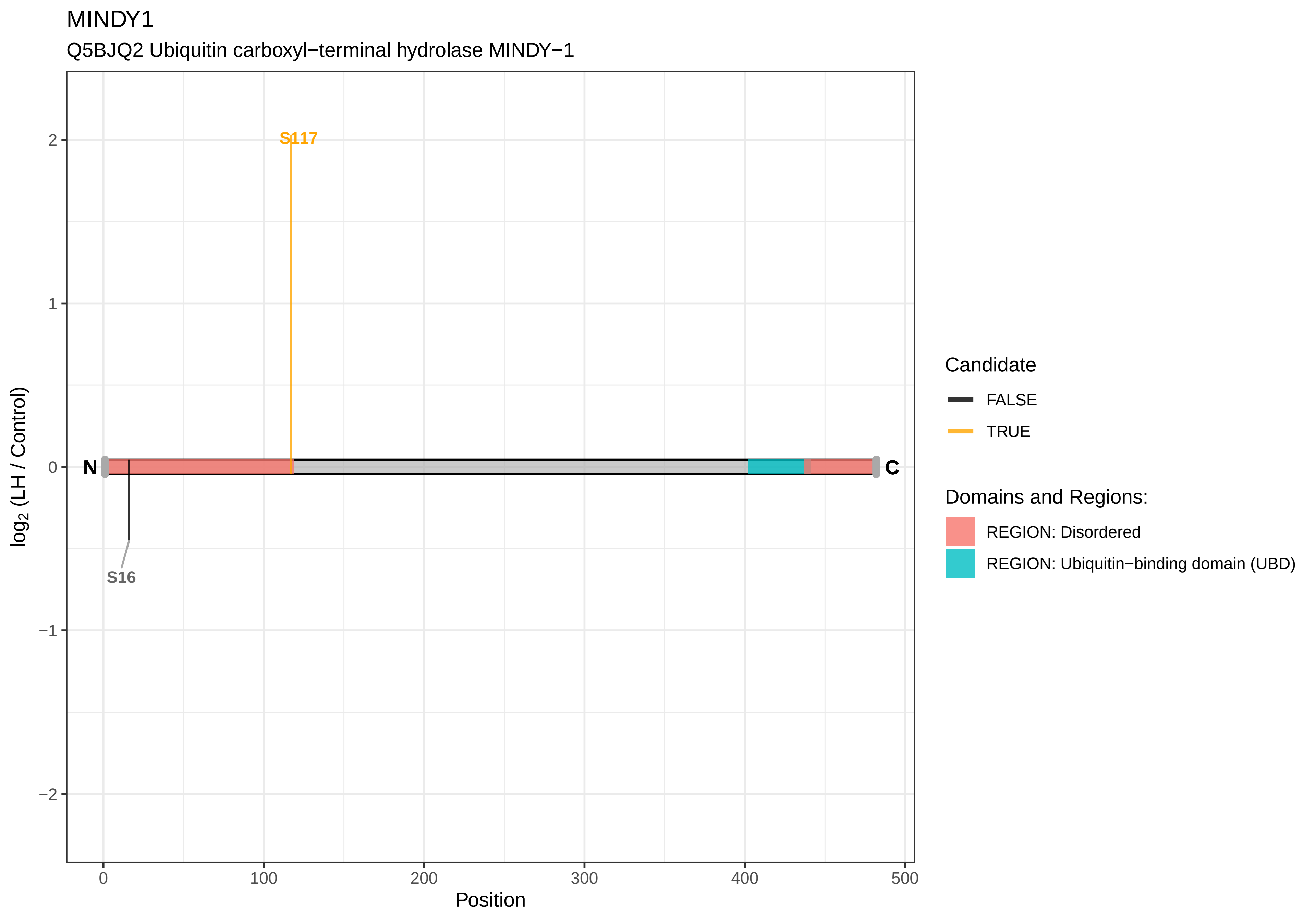

MINDY1
Q5BJQ2 Ubiquitin carboxyl−terminal hydrolase MINDY−1
S117
S16
N
C
2
1
0
−1
−2
log2 (LH / Control)
0
100
200
300
400
500
Position
Candidate
FALSE
TRUE
Domains and Regions:
REGION: Disordered
REGION: Ubiquitin−binding domain (UBD)

MINK1

A0A0G2K382 Misshapen−like kinase 1

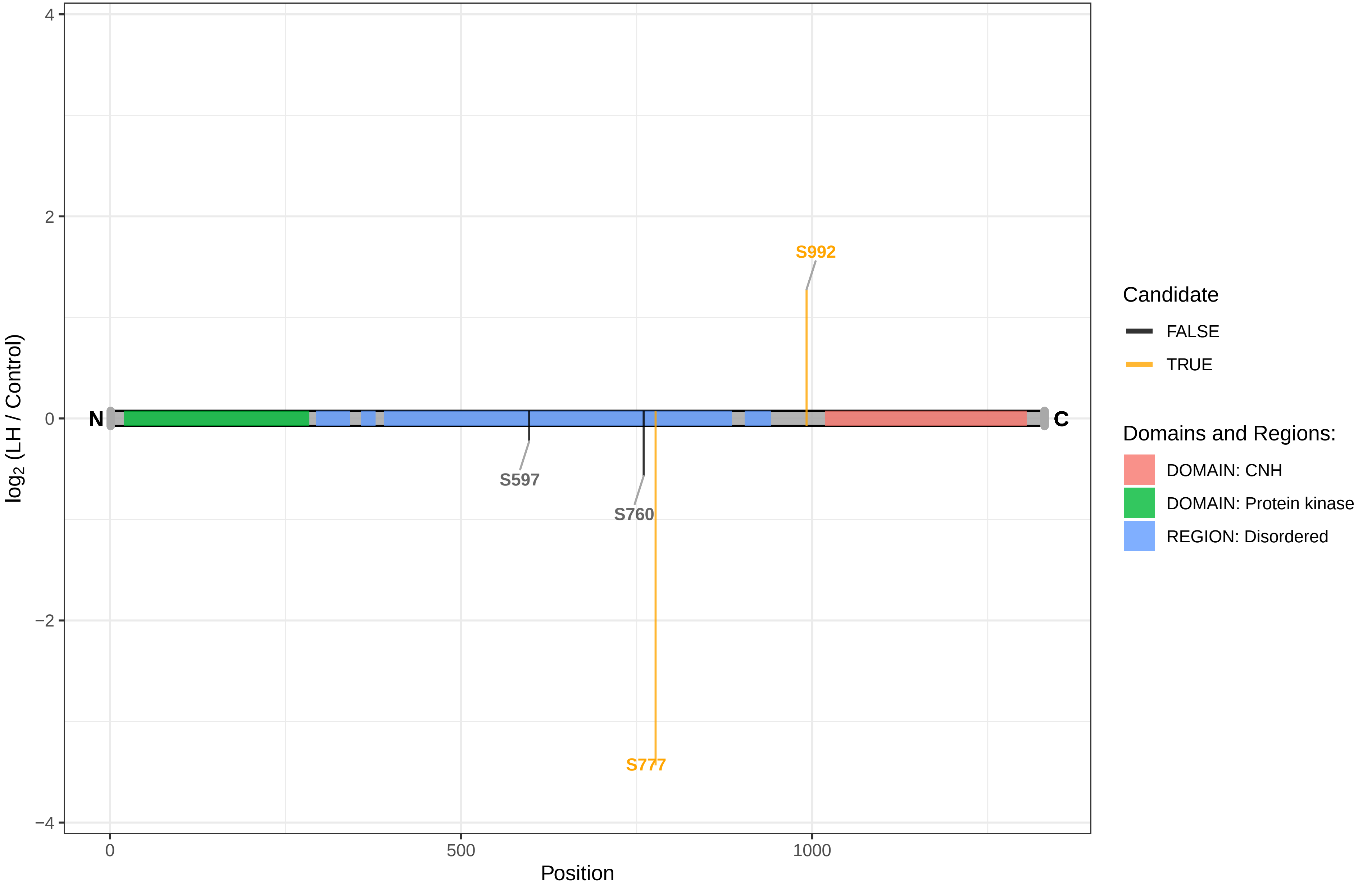

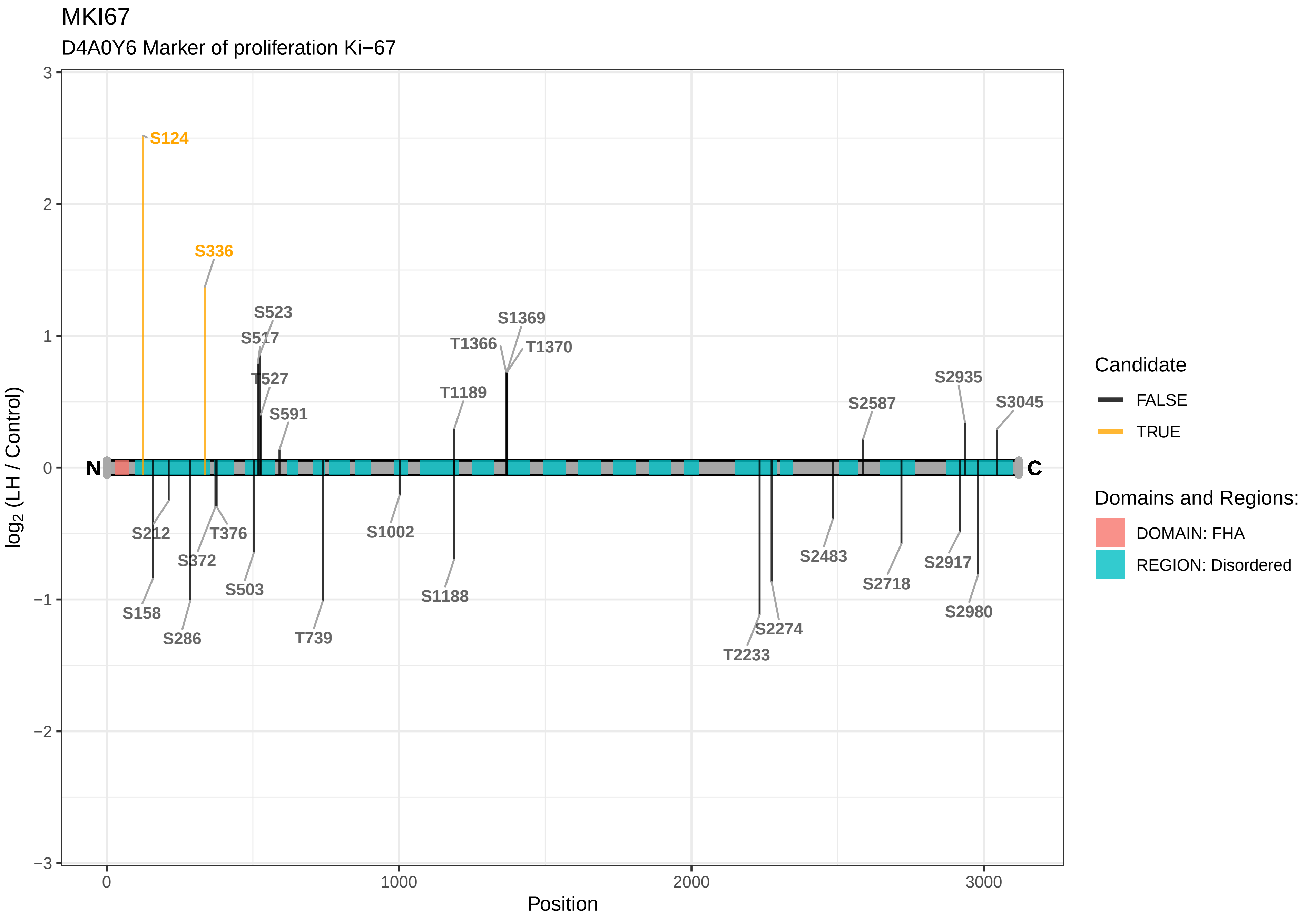
MKI67
D4A0Y6 Marker of proliferation Ki−67
log2 (LH / Control)
Position
N
C
S124
S336
S523
S517
T527
S591
T1189
S1369
T1366
T1370
S2935
S2587
S3045
S212
T376
S372
S503
S158
S286
T739
S1002
S1188
T2233
S2274
S2483
S2718
S2917
S2980
Candidate
FALSE
TRUE
Domains and Regions:
DOMAIN: FHA
REGION: Disordered

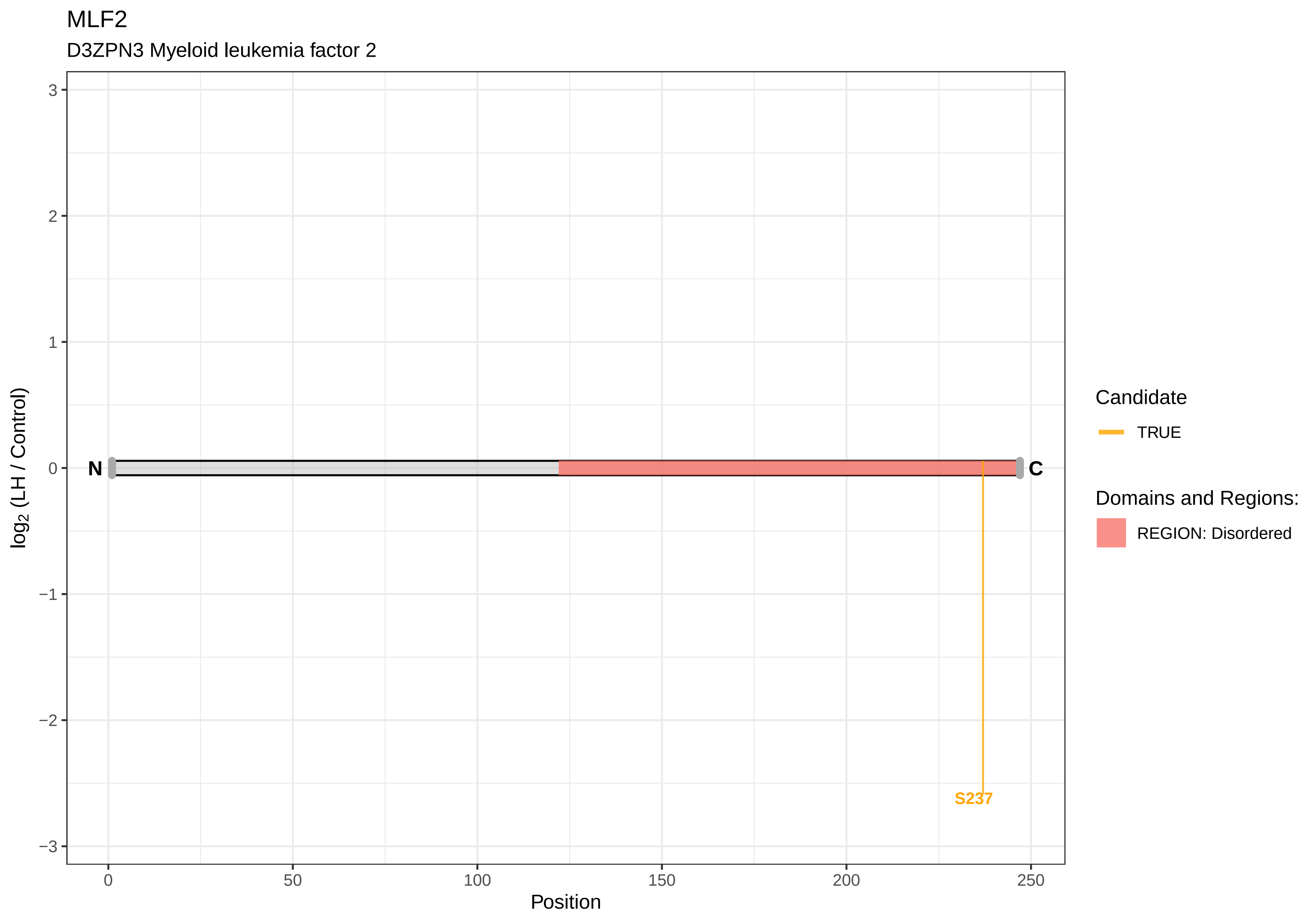
MLF2
D3ZPN3 Myeloid leukemia factor 2
log2 (LH / Control)
3
2
1
0
−1
−2
−3
N
C
S237
0
50
100
150
200
250
Position
Candidate
TRUE
Domains and Regions:
REGION: Disordered

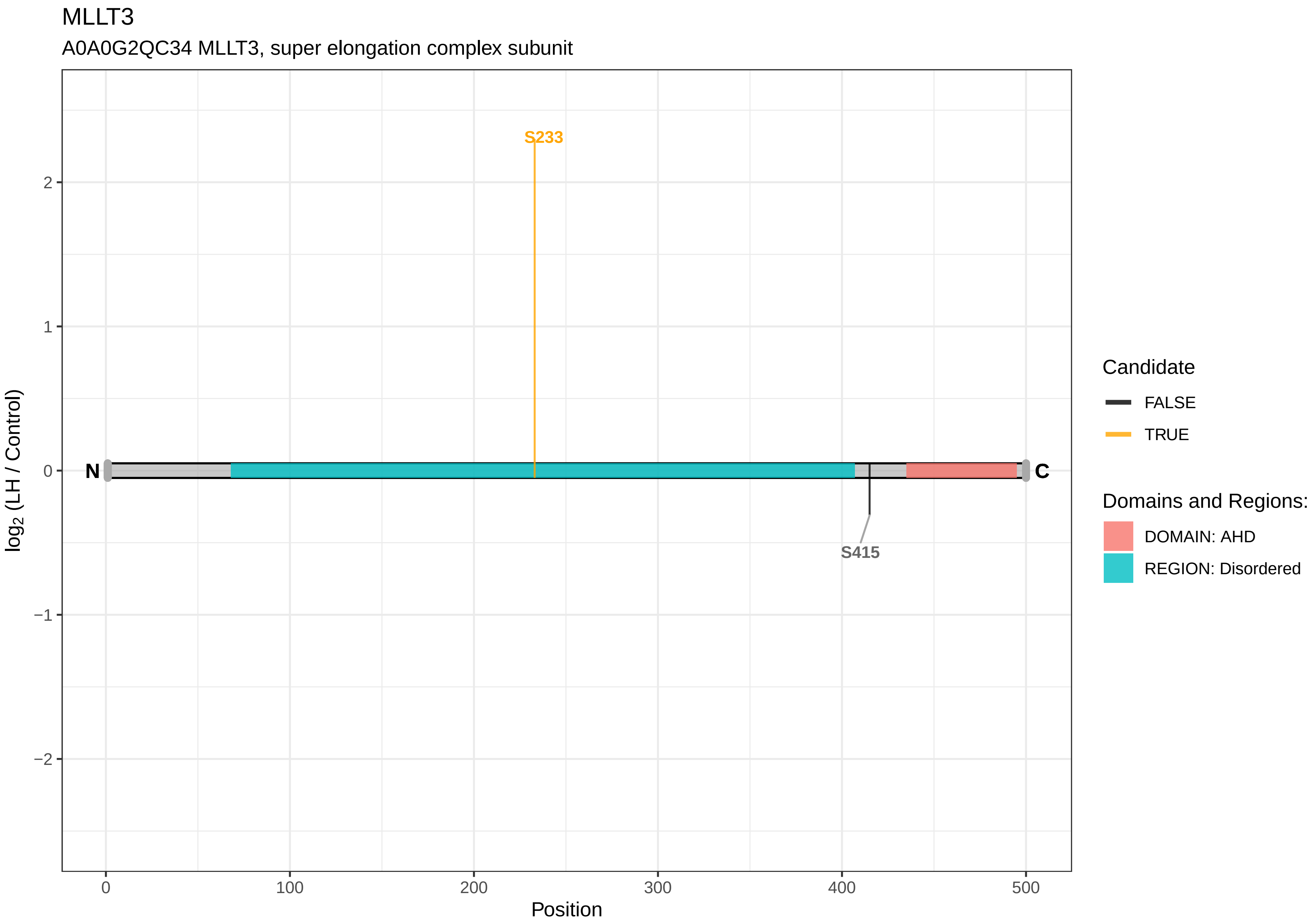

MLLT3
A0A0G2QC34 MLLT3, super elongation complex subunit
S233
S415
N
C
2
1
0
−1
−2
0
100
200
300
400
500
Position
log2 (LH / Control)
Candidate
FALSE
TRUE
Domains and Regions:
DOMAIN: AHD
REGION: Disordered

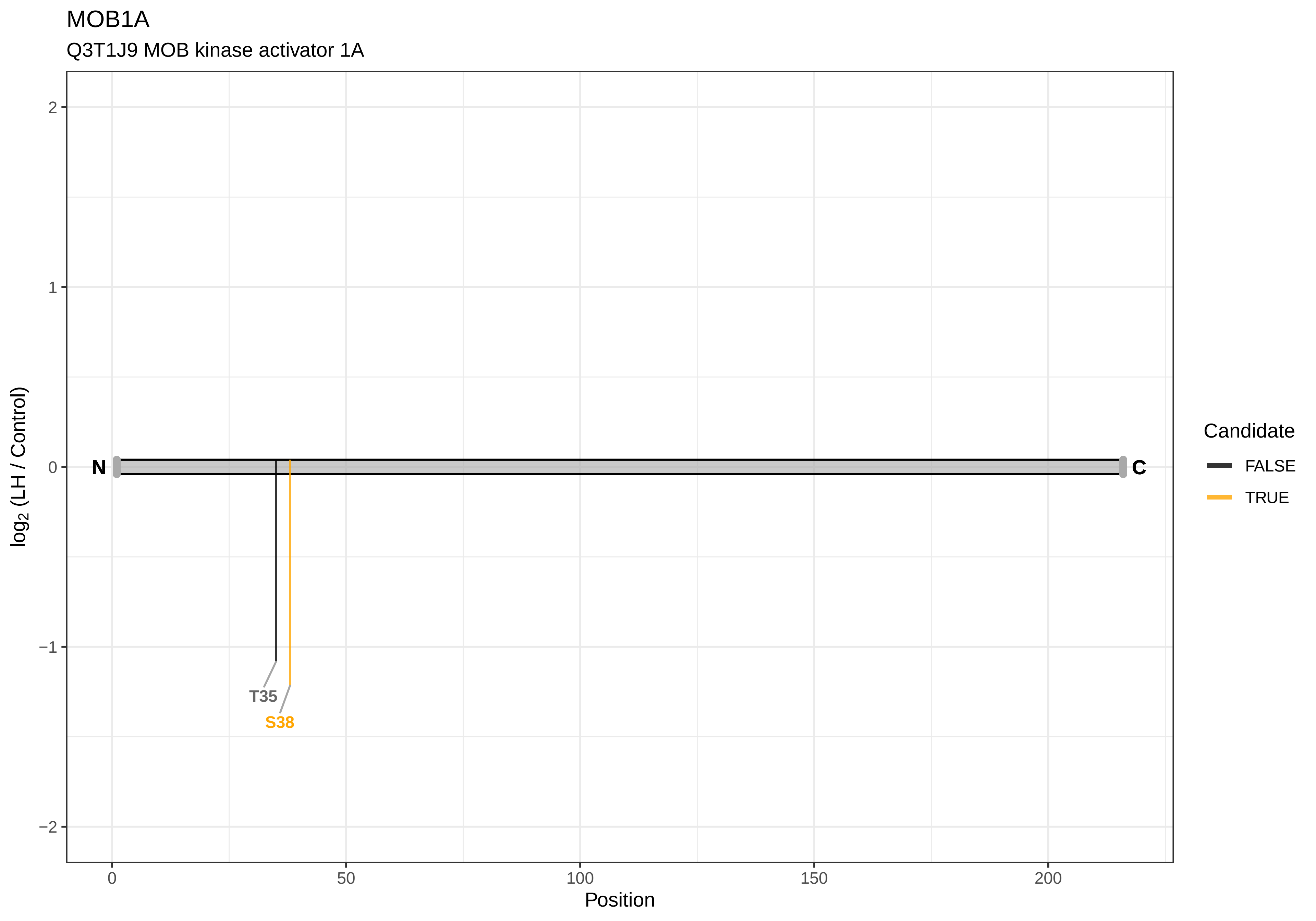

MOB1A
Q3T1J9 MOB kinase activator 1A
log2 (LH / Control)
2
1
0
−1
−2
N
C
T35
S38
0
50
100
150
200
Position
Candidate
FALSE
TRUE

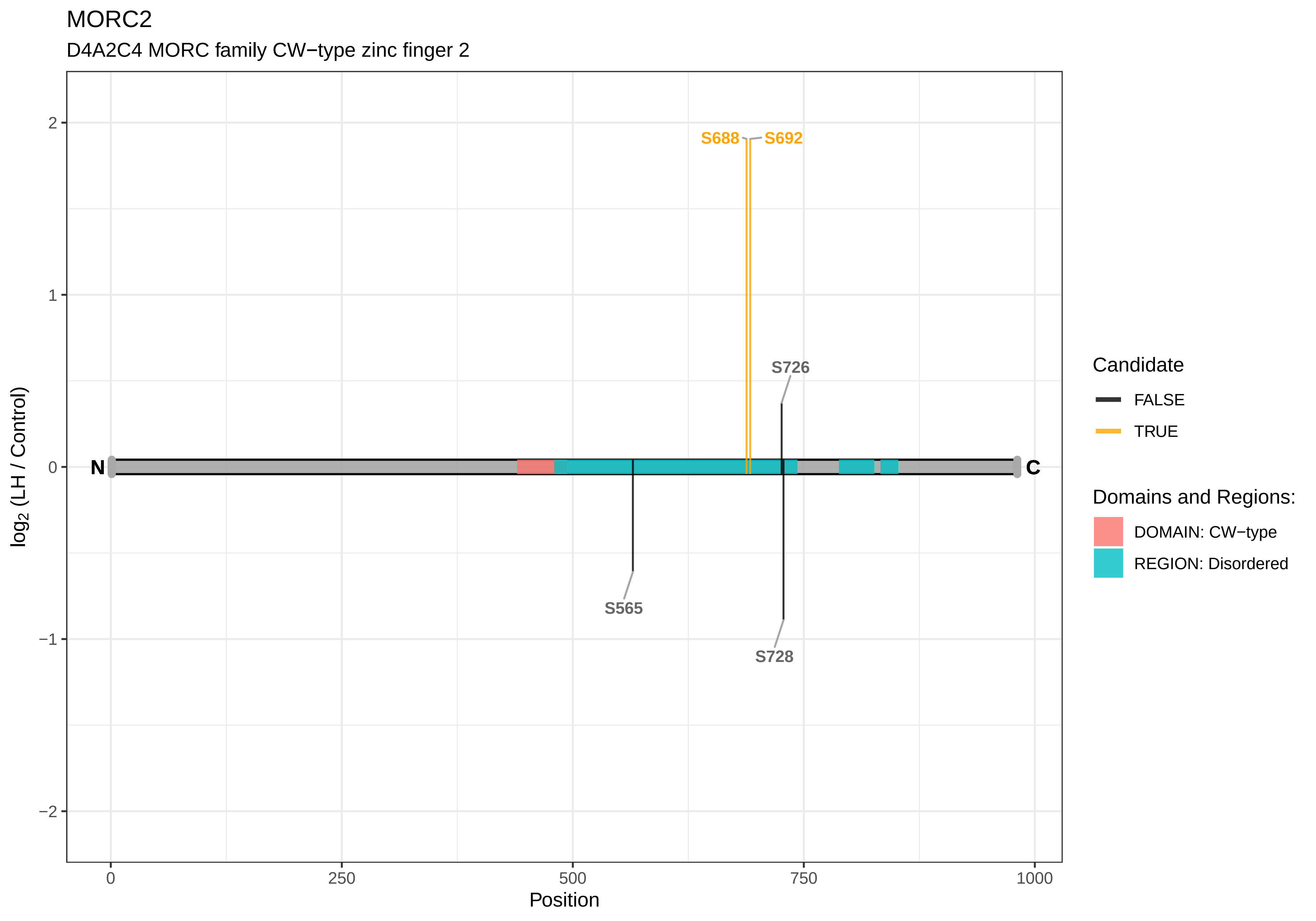

MORC2
D4A2C4 MORC family CW−type zinc finger 2
S688
S692
S726
S565
S728
N
C
2
1
0
−1
−2
0
250
500
750
1000
Position
log2 (LH / Control)
Candidate
FALSE
TRUE
Domains and Regions:
DOMAIN: CW−type
REGION: Disordered

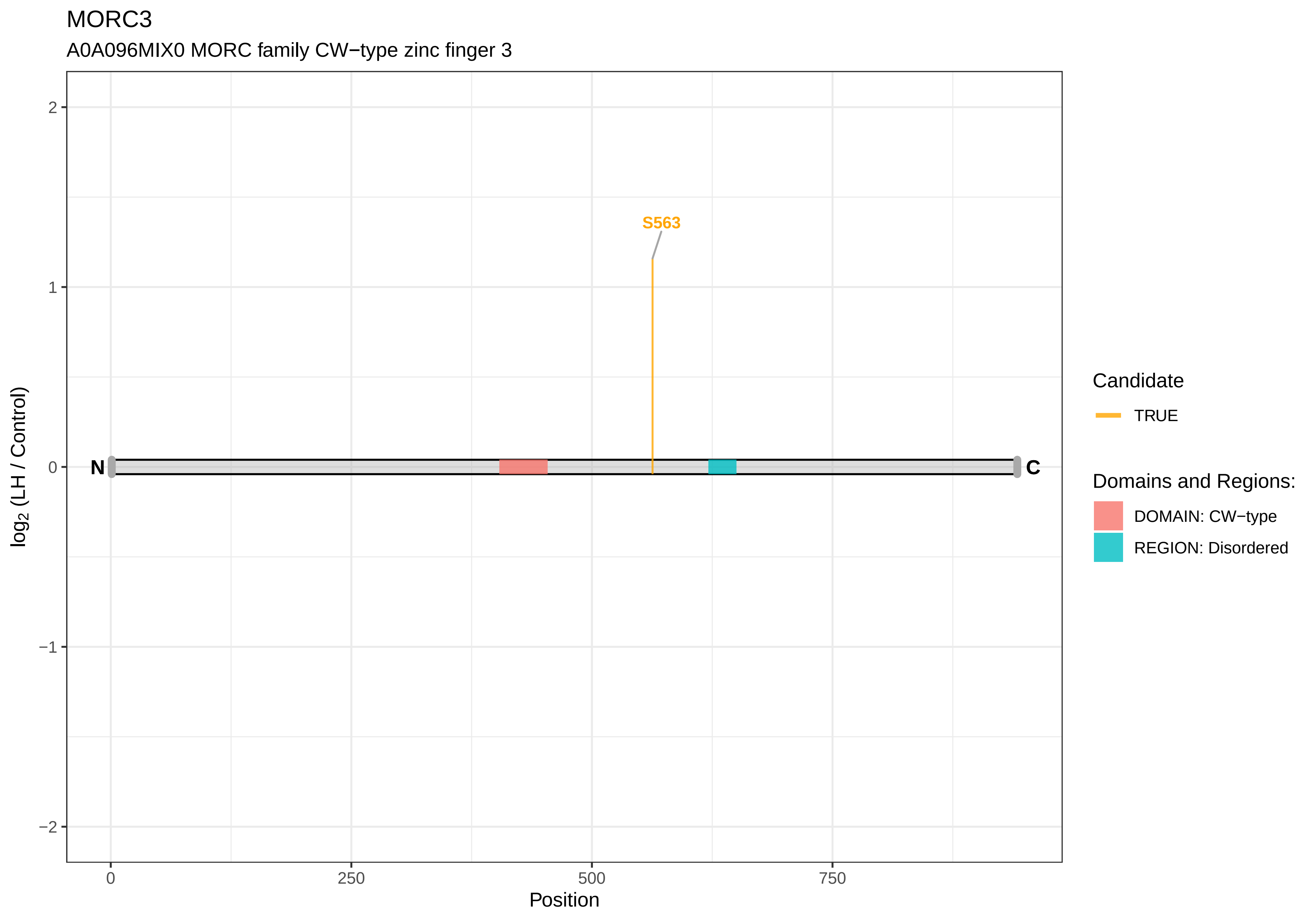

MORC3
A0A096MIX0 MORC family CW−type zinc finger 3
S563
N
C
2
1
0
−1
−2
0
250
500
750
Position
log2 (LH / Control)
Candidate
TRUE
Domains and Regions:
DOMAIN: CW−type
REGION: Disordered

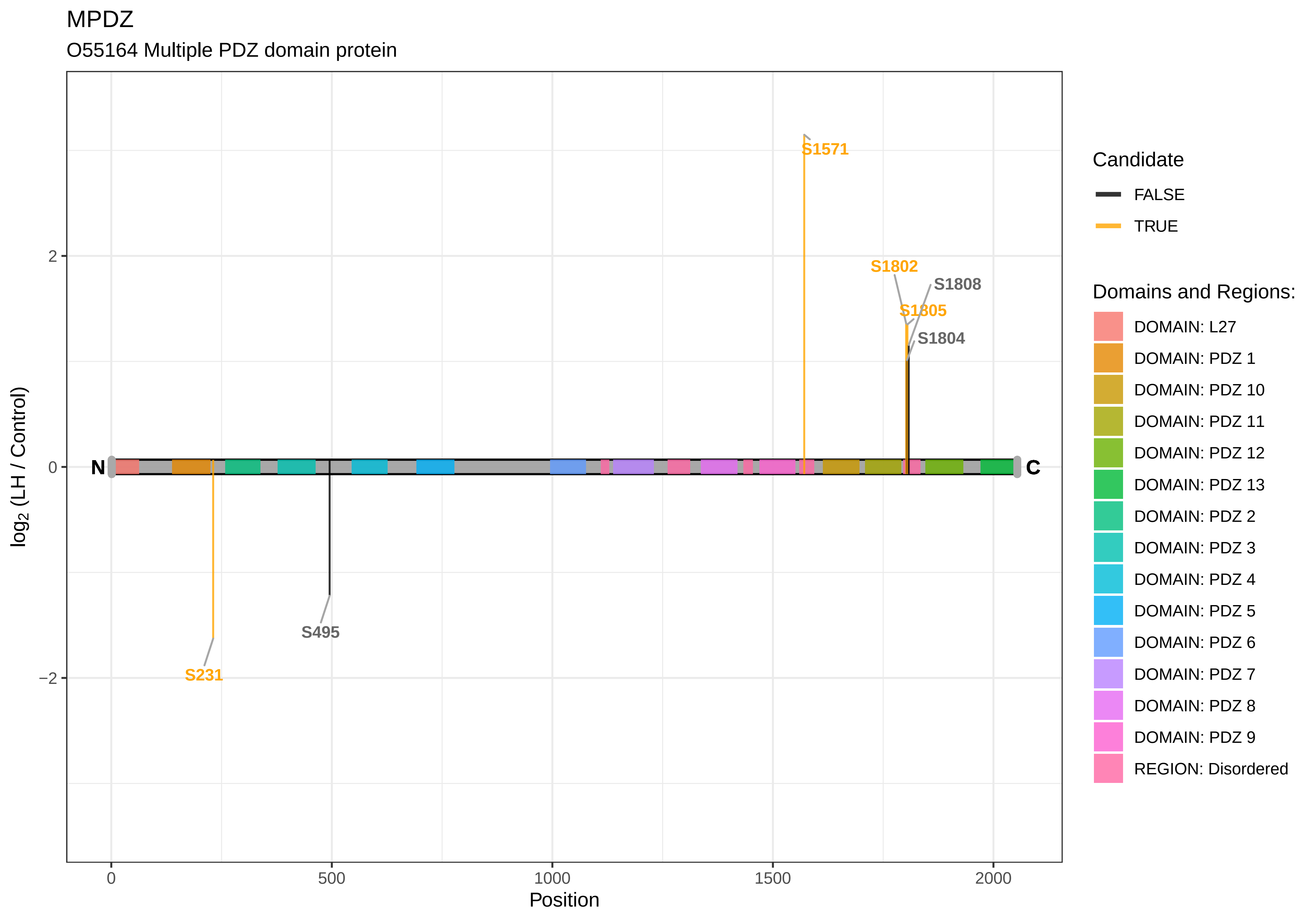
MPDZ
O55164 Multiple PDZ domain protein
log2 (LH / Control)
Position
0
500
1000
1500
2000
2
0
−2
N
C
S1571
S1802
S1808
S1805
S1804
S495
S231
Candidate
FALSE
TRUE
Domains and Regions:
DOMAIN: L27
DOMAIN: PDZ 1
DOMAIN: PDZ 10
DOMAIN: PDZ 11
DOMAIN: PDZ 12
DOMAIN: PDZ 13
DOMAIN: PDZ 2
DOMAIN: PDZ 3
DOMAIN: PDZ 4
DOMAIN: PDZ 5
DOMAIN: PDZ 6
DOMAIN: PDZ 7
DOMAIN: PDZ 8
DOMAIN: PDZ 9
REGION: Disordered

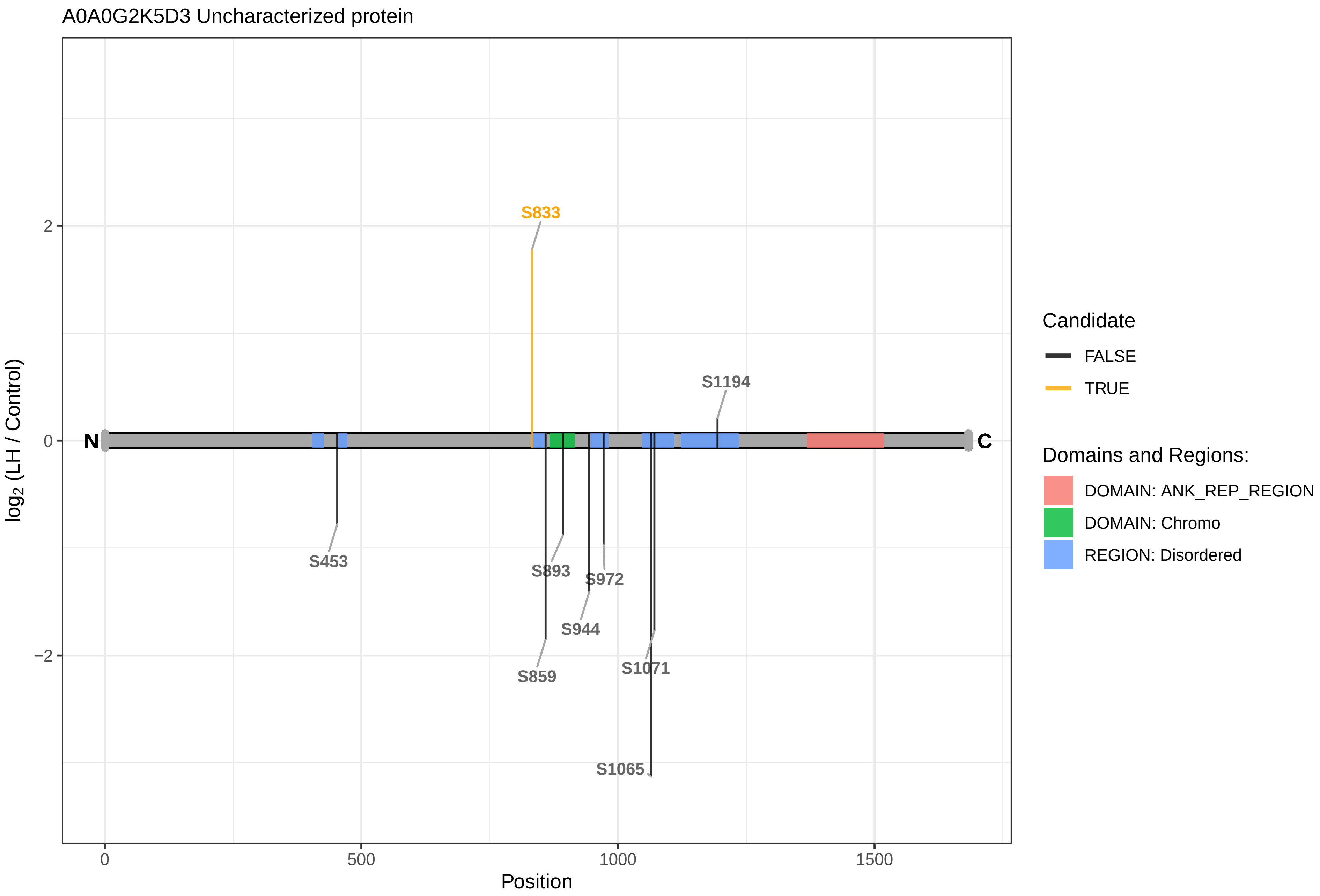
A0A0G2K5D3 Uncharacterized protein
S833
S1194
N
C
S453
S893
S972
S944
S859
S1071
S1065
2
0
−2
0
500
1000
1500
Position
log2 (LH / Control)
Candidate
FALSE
TRUE
Domains and Regions:
DOMAIN: ANK_REP_REGION
DOMAIN: Chromo
REGION: Disordered

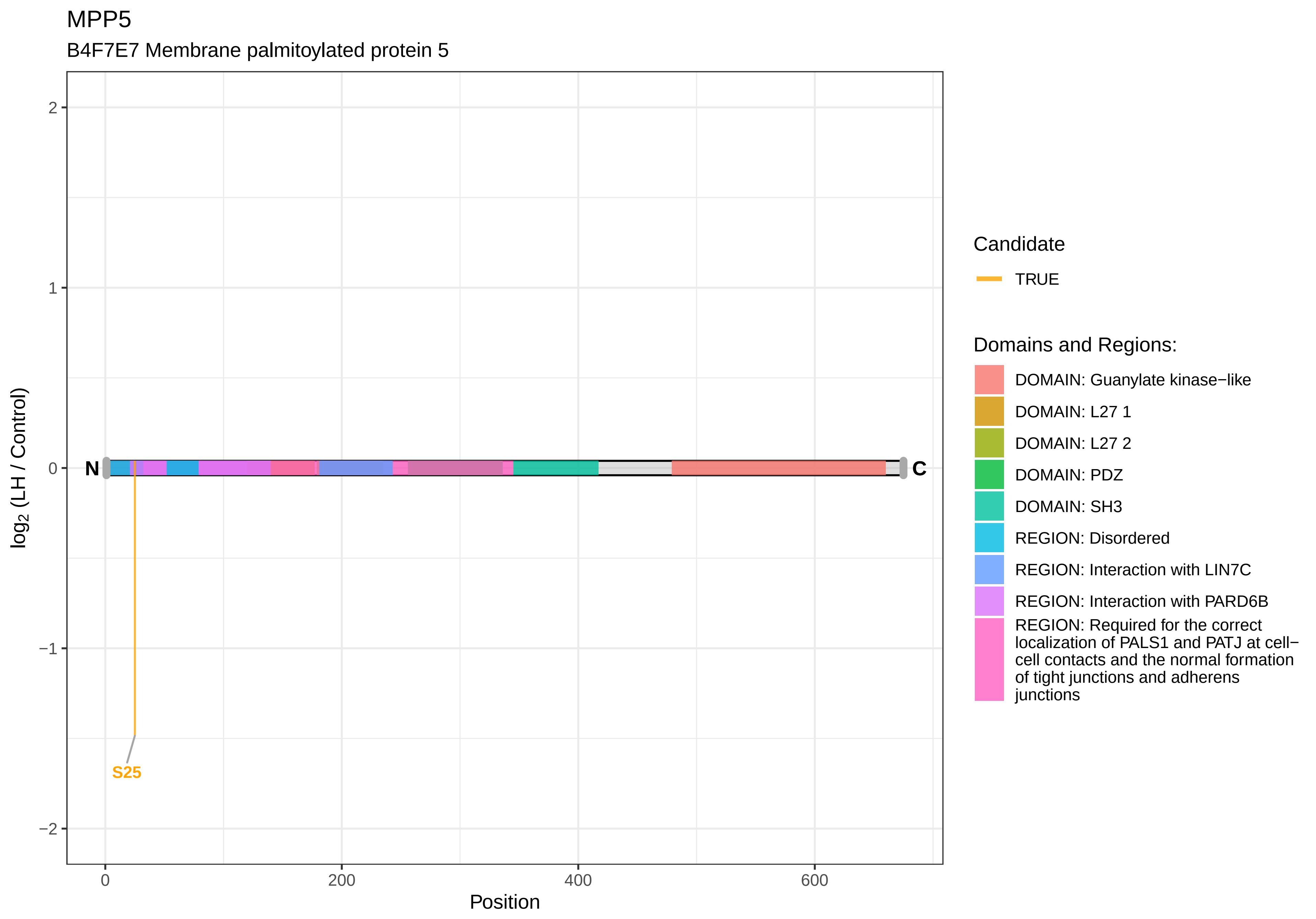
MPP5
B4F7E7 Membrane palmitoylated protein 5
log2 (LH / Control)
2
1
0
−1
−2
N
C
S25
0
200
400
600
Position
Candidate
TRUE
Domains and Regions:
DOMAIN: Guanylate kinase−like
DOMAIN: L27 1
DOMAIN: L27 2
DOMAIN: PDZ
DOMAIN: SH3
REGION: Disordered
REGION: Interaction with LIN7C
REGION: Interaction with PARD6B
REGION: Required for the correct localization of PALS1 and PATJ at cell−cell contacts and the normal formation of tight junctions and adherens junctions

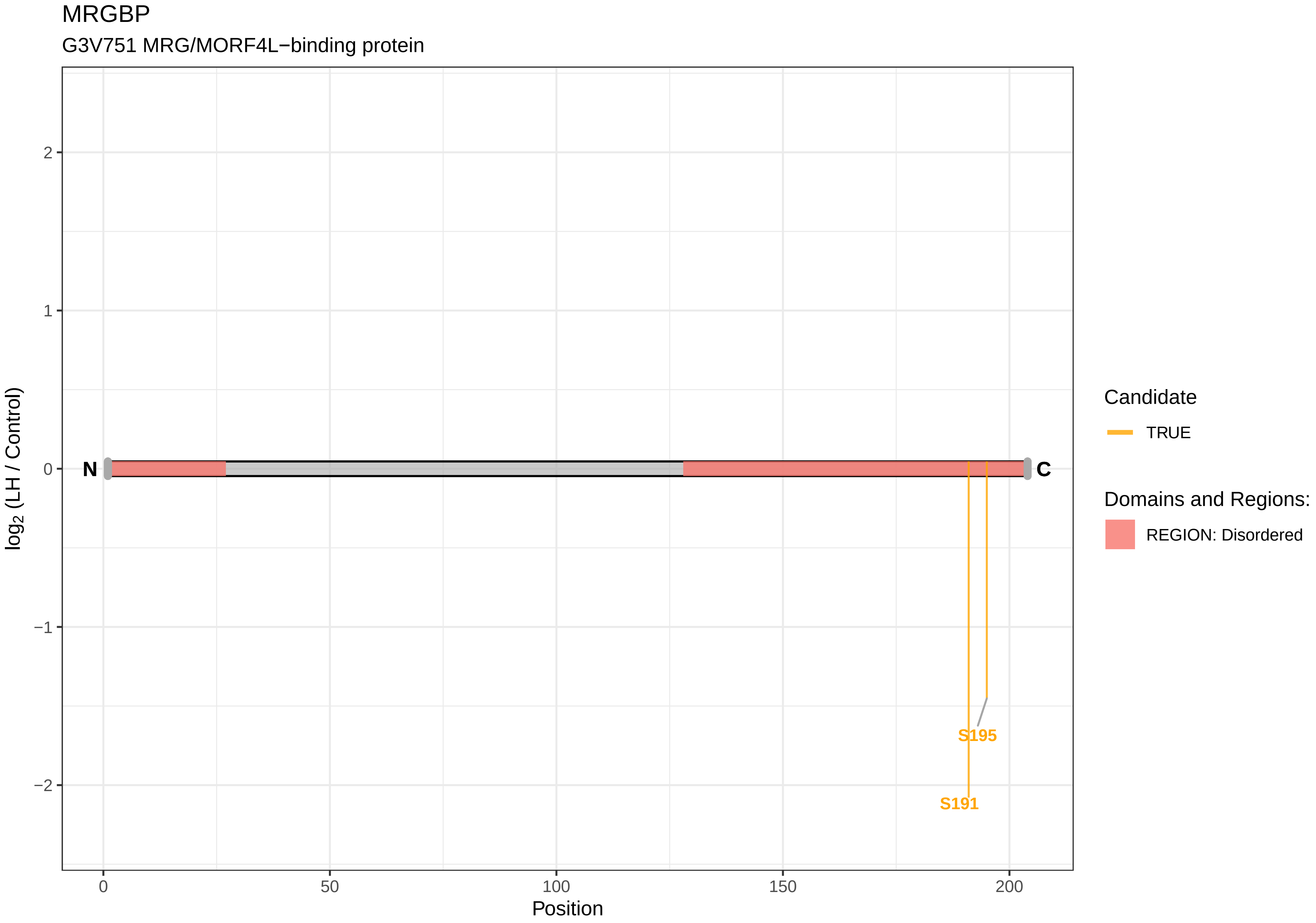
MRGBP
G3V751 MRG/MORF4L−binding protein
log2 (LH / Control)
Position
N
C
S195
S191
Candidate
TRUE
Domains and Regions:
REGION: Disordered
2
1
0
−1
−2
0
50
100
150
200

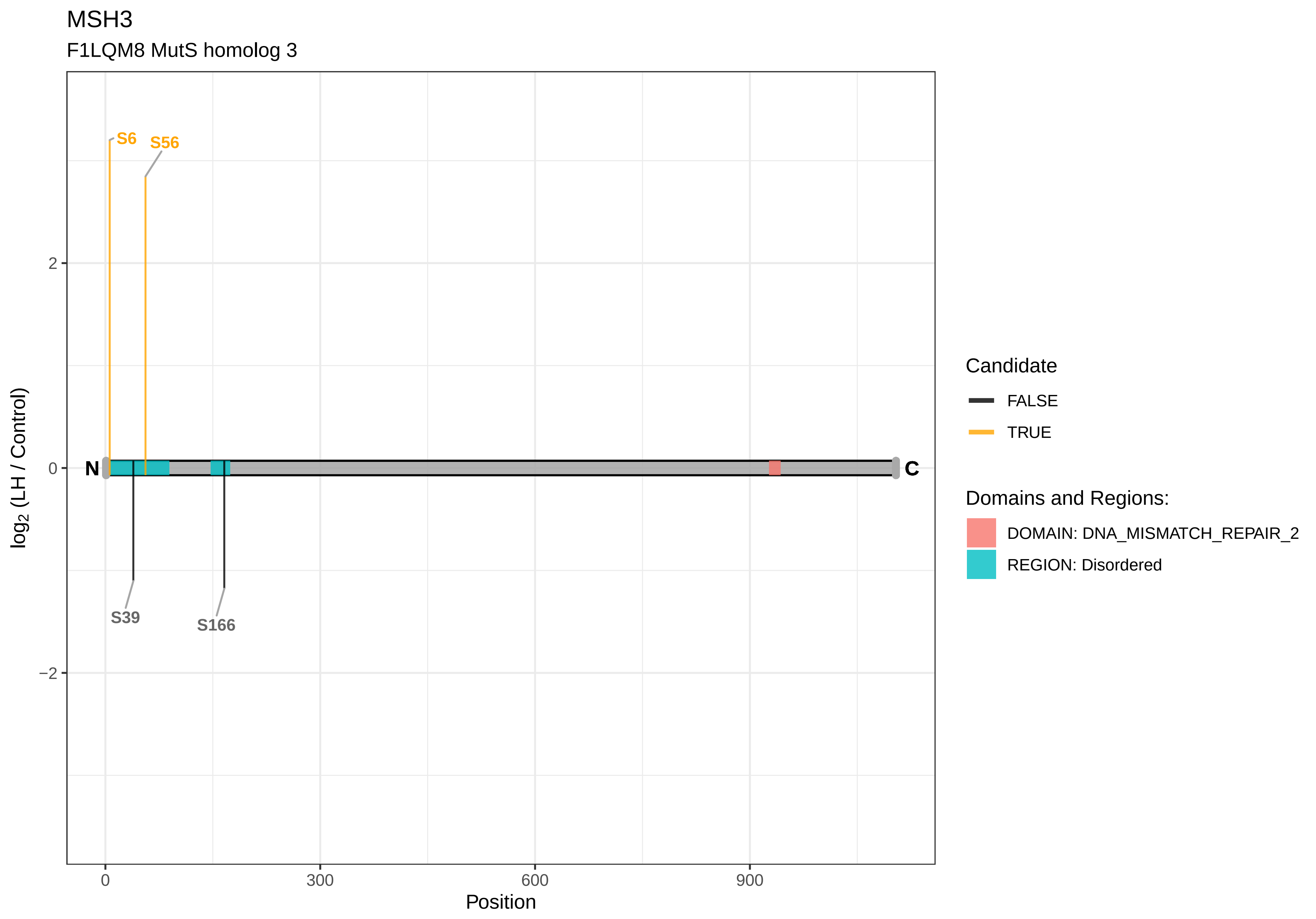

MSH3
F1LQM8 MutS homolog 3
S6
S56
S39
S166
N
C
2
0
−2
0
300
600
900
Position
log2 (LH / Control)
Candidate
FALSE
TRUE
Domains and Regions:
DOMAIN: DNA_MISMATCH_REPAIR_2
REGION: Disordered

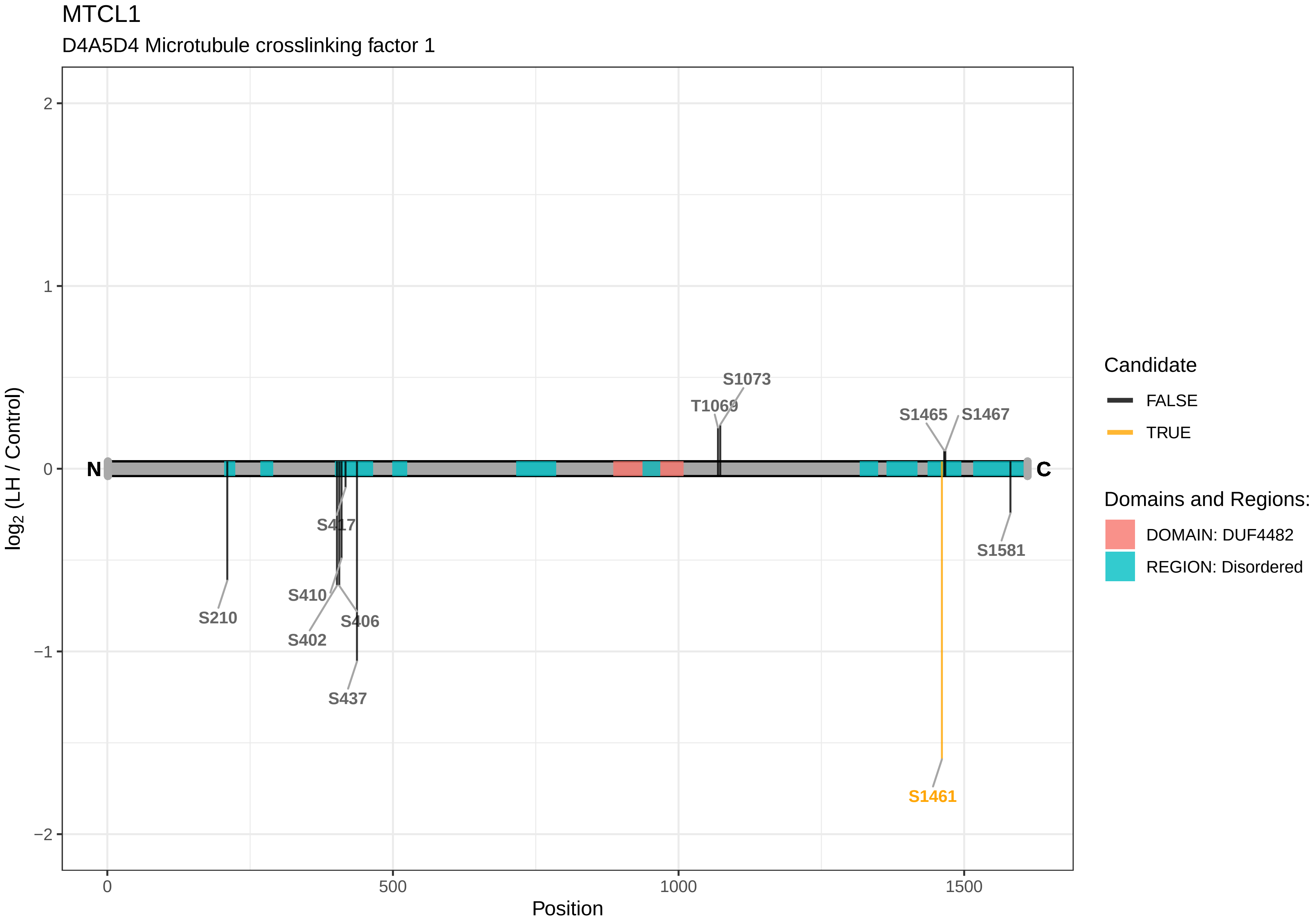

MTCL1
D4A5D4 Microtubule crosslinking factor 1
log2 (LH / Control)
Position
N
C
S1073
T1069
S1465
S1467
S417
S410
S210
S406
S402
S437
S1581
S1461
2
1
0
−1
−2
0
500
1000
1500
Candidate
FALSE
TRUE
Domains and Regions:
DOMAIN: DUF4482
REGION: Disordered

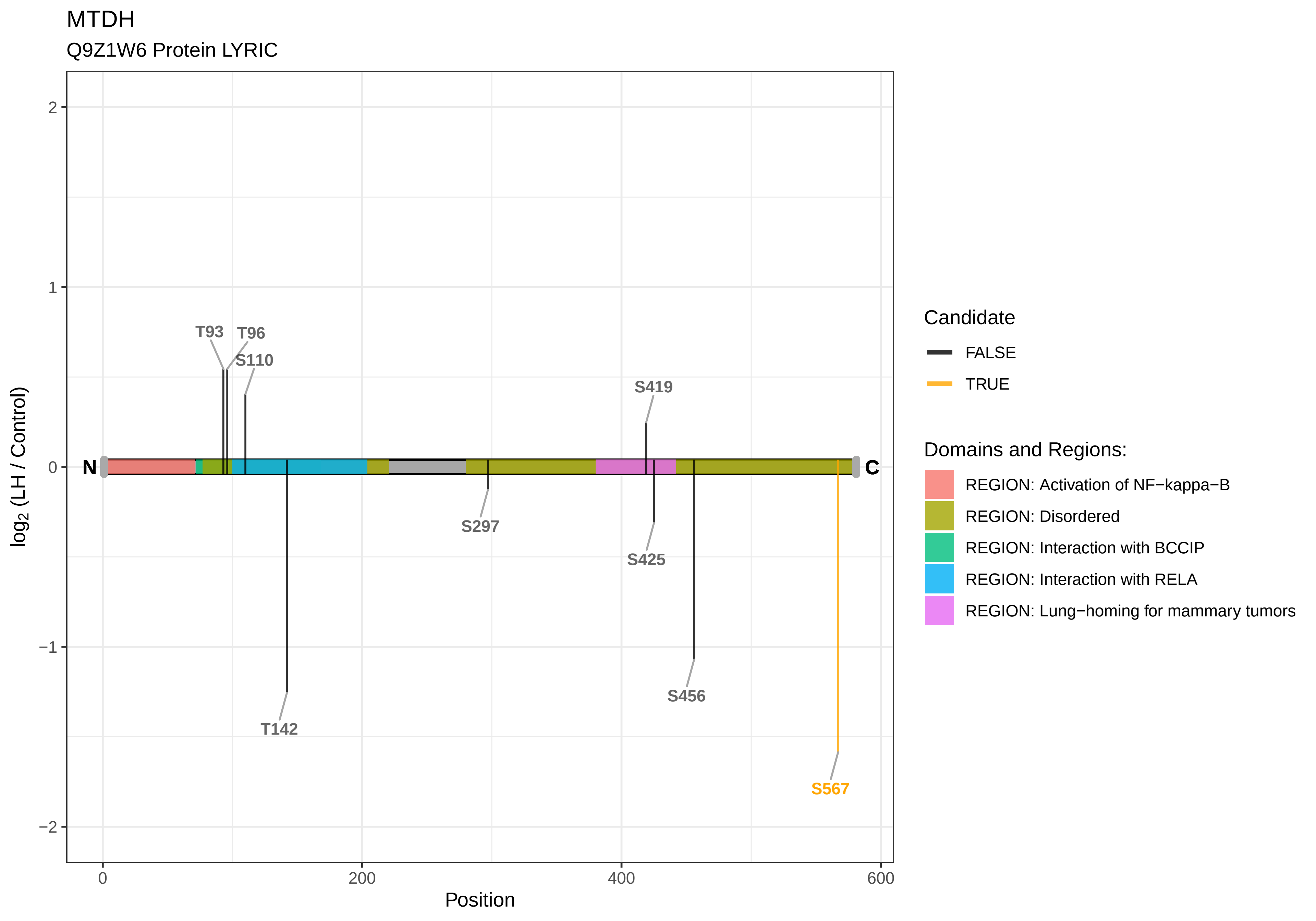
MTDH
Q9Z1W6 Protein LYRIC
T93
T96
S110
S419
T142
S297
S425
S456
S567
N
C
log2 (LH / Control)
2
1
0
−1
−2
0
200
400
600
Position
Candidate
FALSE
TRUE
Domains and Regions:
REGION: Activation of NF−kappa−B
REGION: Disordered
REGION: Interaction with BCCIP
REGION: Interaction with RELA
REGION: Lung−homing for mammary tumors

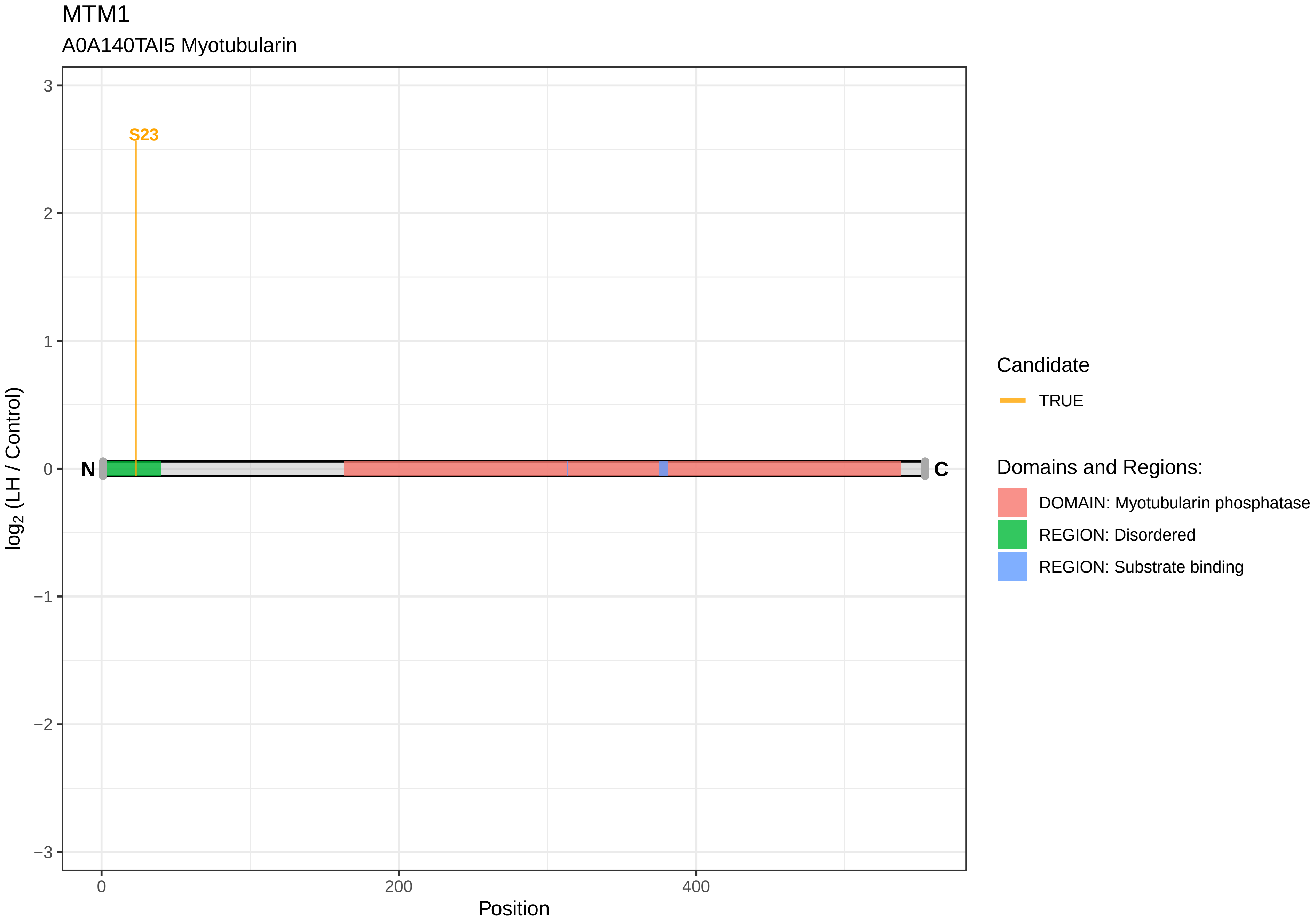

MTM1
A0A140TAI5 Myotubularin
S23
N
C
3
2
1
0
−1
−2
−3
0
200
400
Position
log2 (LH / Control)
Candidate
TRUE
Domains and Regions:
DOMAIN: Myotubularin phosphatase
REGION: Disordered
REGION: Substrate binding

MTMR12

Q5FVM6 Myotubularin−related protein 12

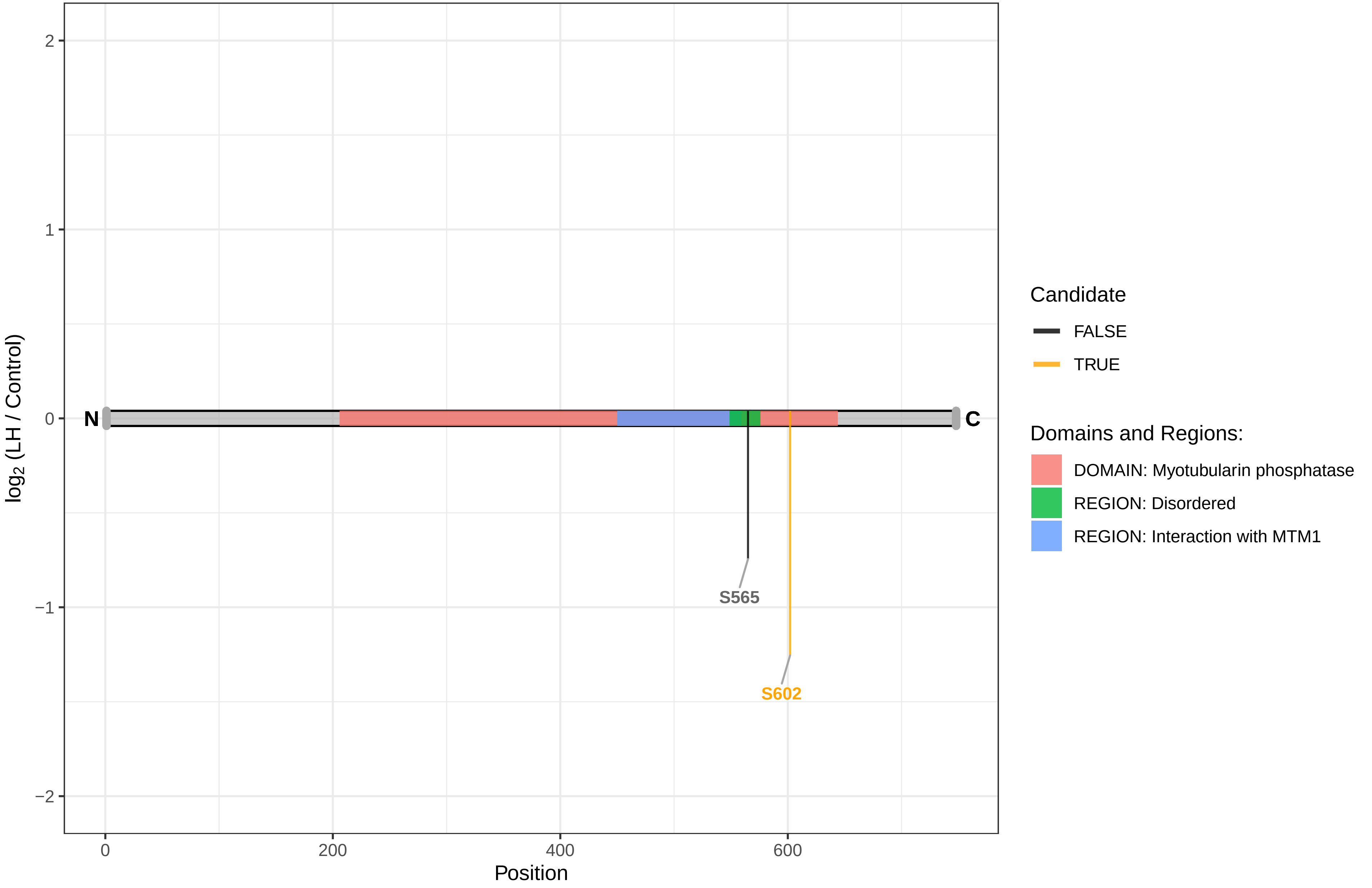

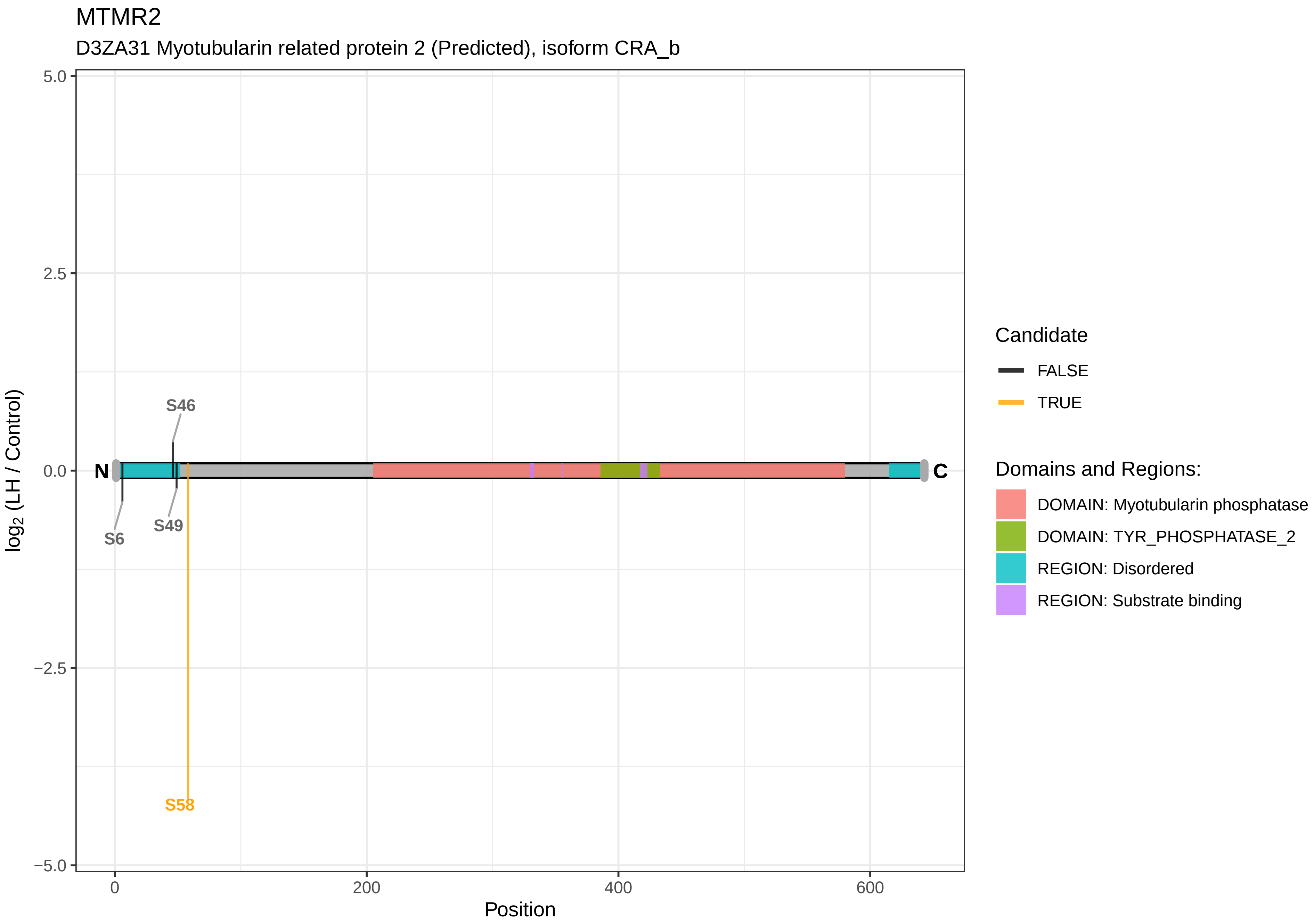
MTMR2
D3ZA31 Myotubularin related protein 2 (Predicted), isoform CRA_b
log2 (LH / Control)
5.0
2.5
0.0
−2.5
−5.0
0
200
400
600
Position
N
C
S46
S6
S49
S58
Candidate
FALSE
TRUE
Domains and Regions:
DOMAIN: Myotubularin phosphatase
DOMAIN: TYR_PHOSPHATASE_2
REGION: Disordered
REGION: Substrate binding

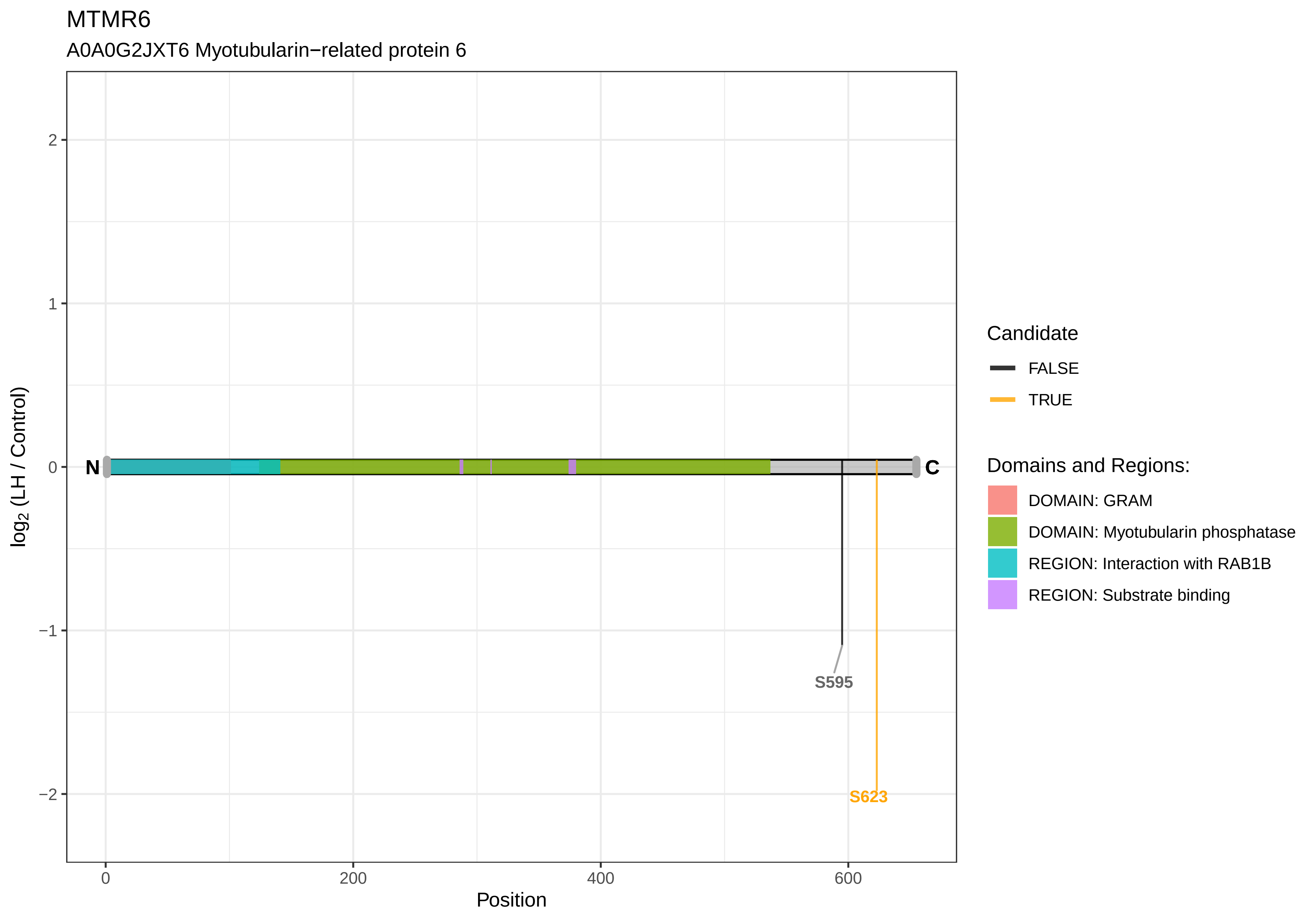
MTMR6
A0A0G2JXT6 Myotubularin−related protein 6
log2 (LH / Control)
Position
N
C
S595
S623
2
1
0
−1
−2
0
200
400
600
Candidate
FALSE
TRUE
Domains and Regions:
DOMAIN: GRAM
DOMAIN: Myotubularin phosphatase
REGION: Interaction with RAB1B
REGION: Substrate binding

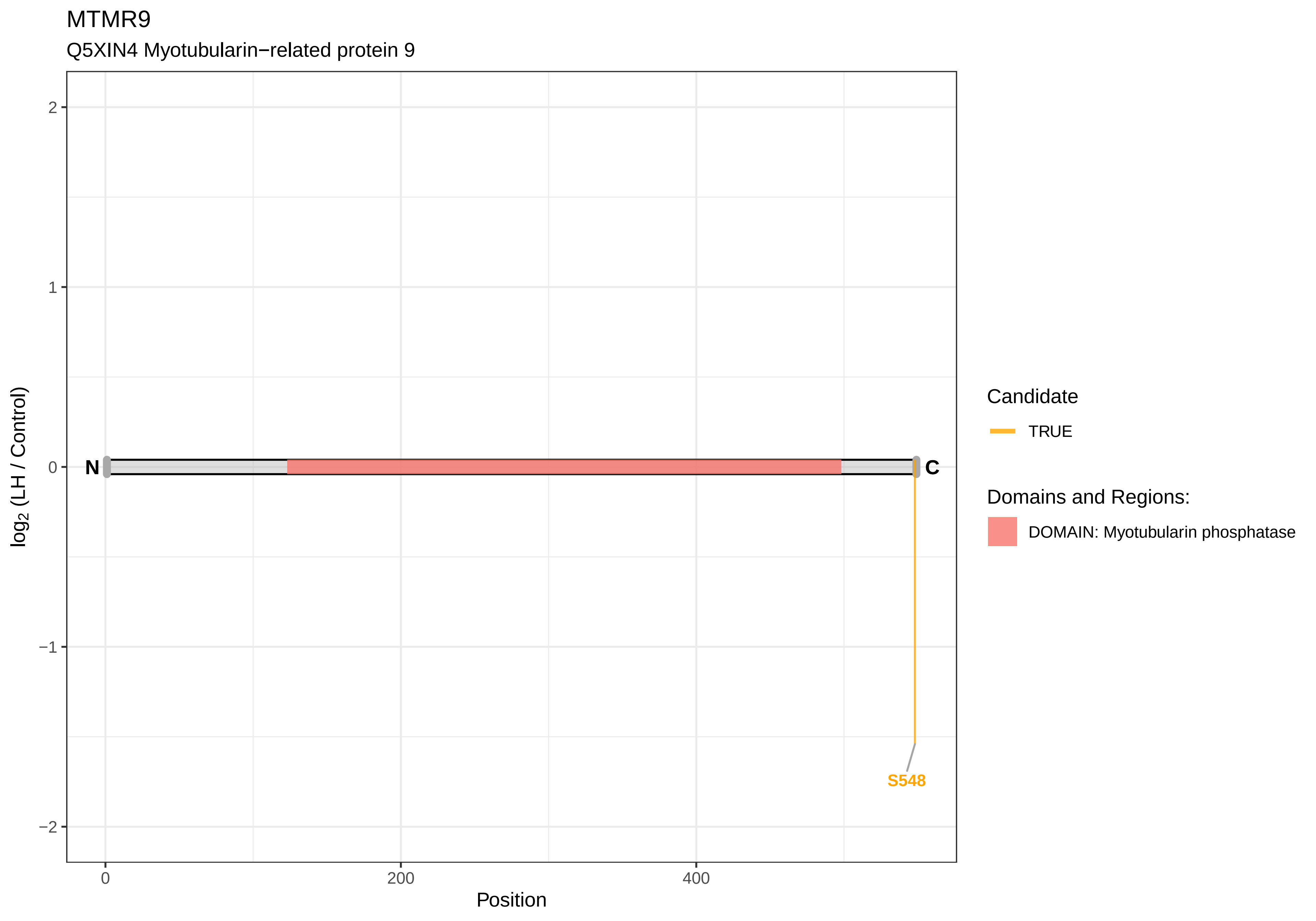
MTMR9
Q5XIN4 Myotubularin−related protein 9
log2 (LH / Control)
2
1
0
−1
−2
N
C
S548
0
200
400
Position
Candidate
TRUE
Domains and Regions:
DOMAIN: Myotubularin phosphatase

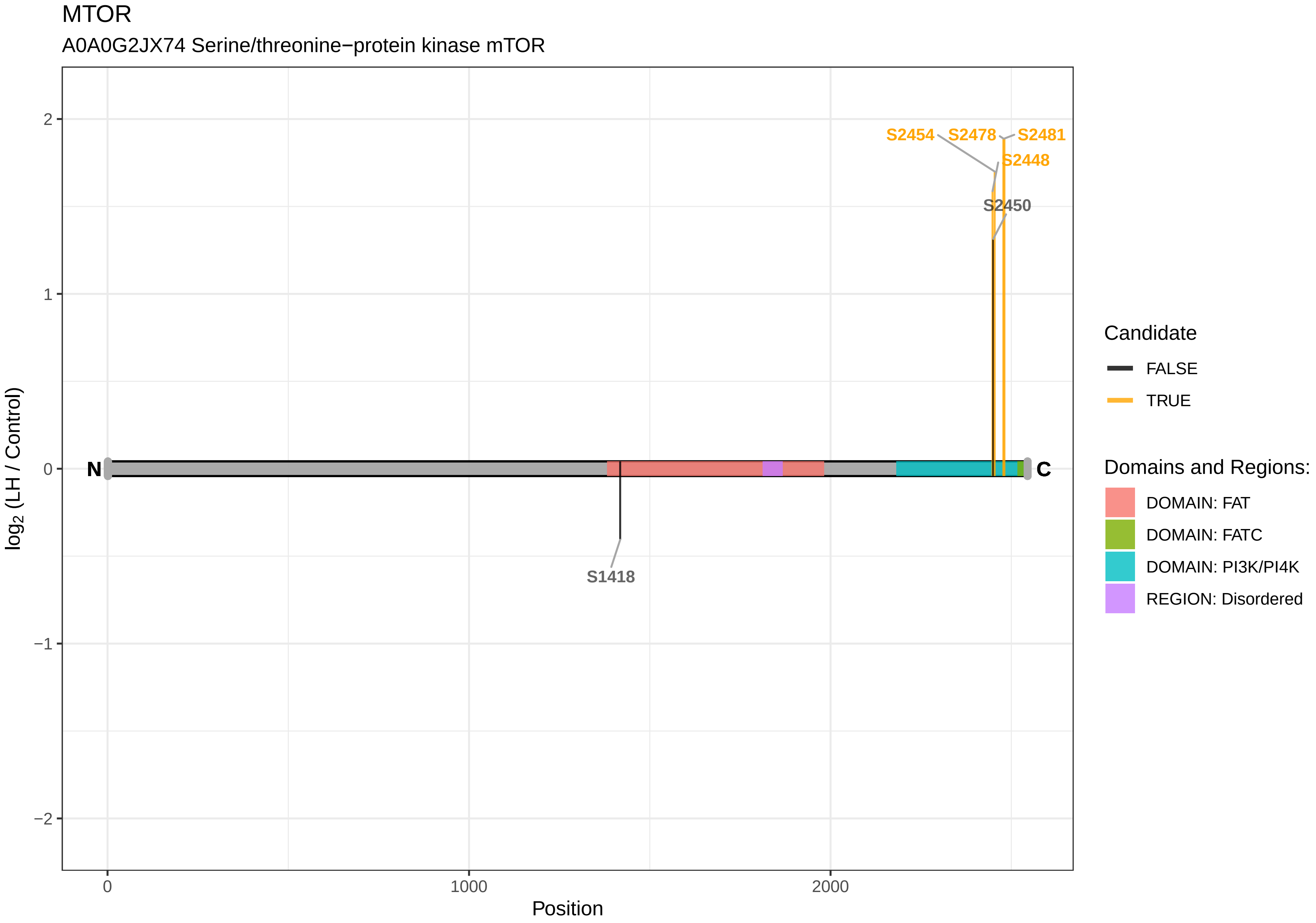
MTOR
A0A0G2JX74 Serine/threonine−protein kinase mTOR
S2454
S2478
S2481
S2448
S2450
S1418
N
C
2
1
0
−1
−2
0
1000
2000
Position
log2 (LH / Control)
Candidate
FALSE
TRUE
Domains and Regions:
DOMAIN: FAT
DOMAIN: FATC
DOMAIN: PI3K/PI4K
REGION: Disordered

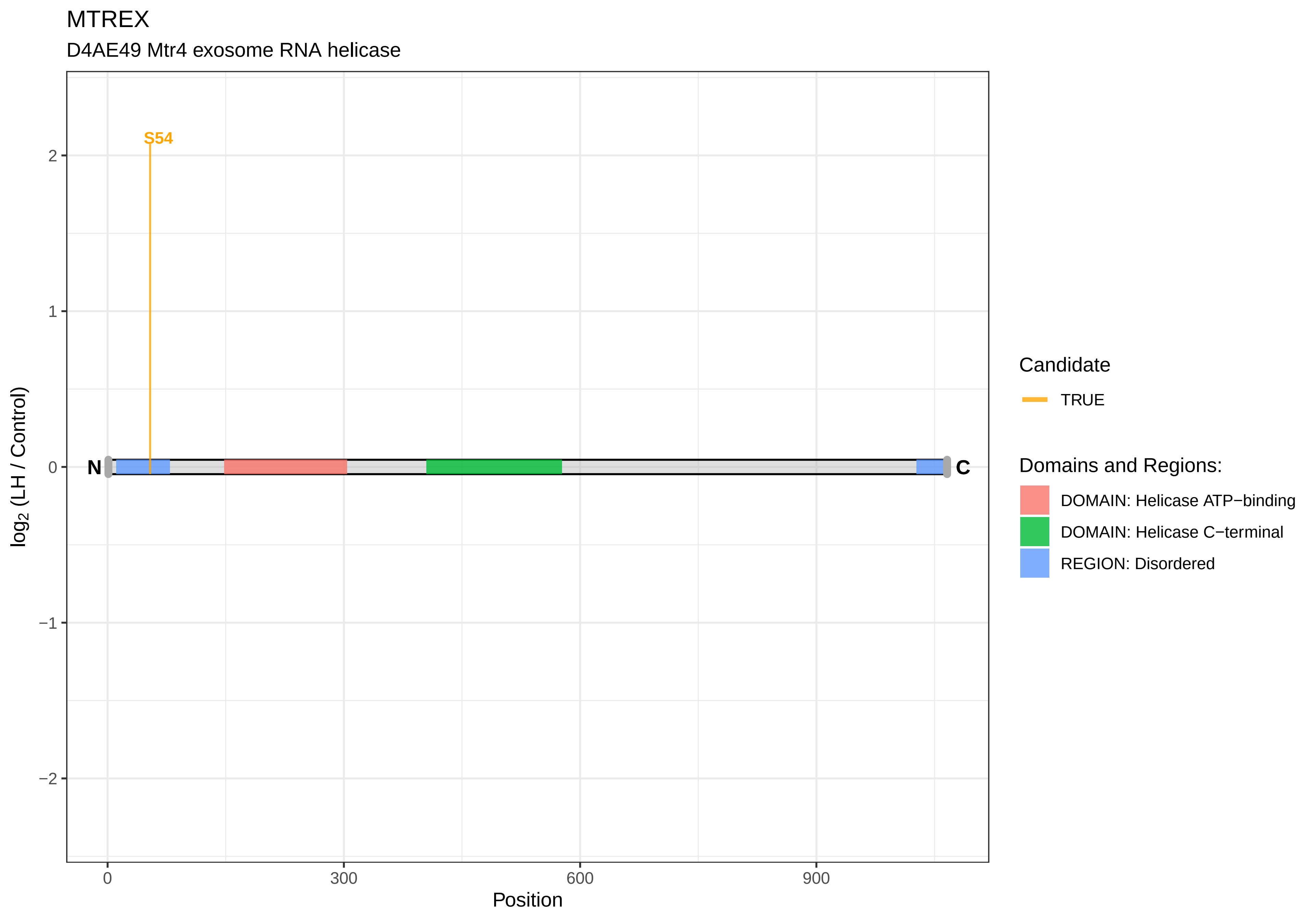

MTREX
D4AE49 Mtr4 exosome RNA helicase
S54
N
C
2
1
0
−1
−2
log2 (LH / Control)
0
300
600
900
Position
Candidate
TRUE
Domains and Regions:
DOMAIN: Helicase ATP−binding
DOMAIN: Helicase C−terminal
REGION: Disordered

MYBBP1A

O35821 Myb−binding protein 1A

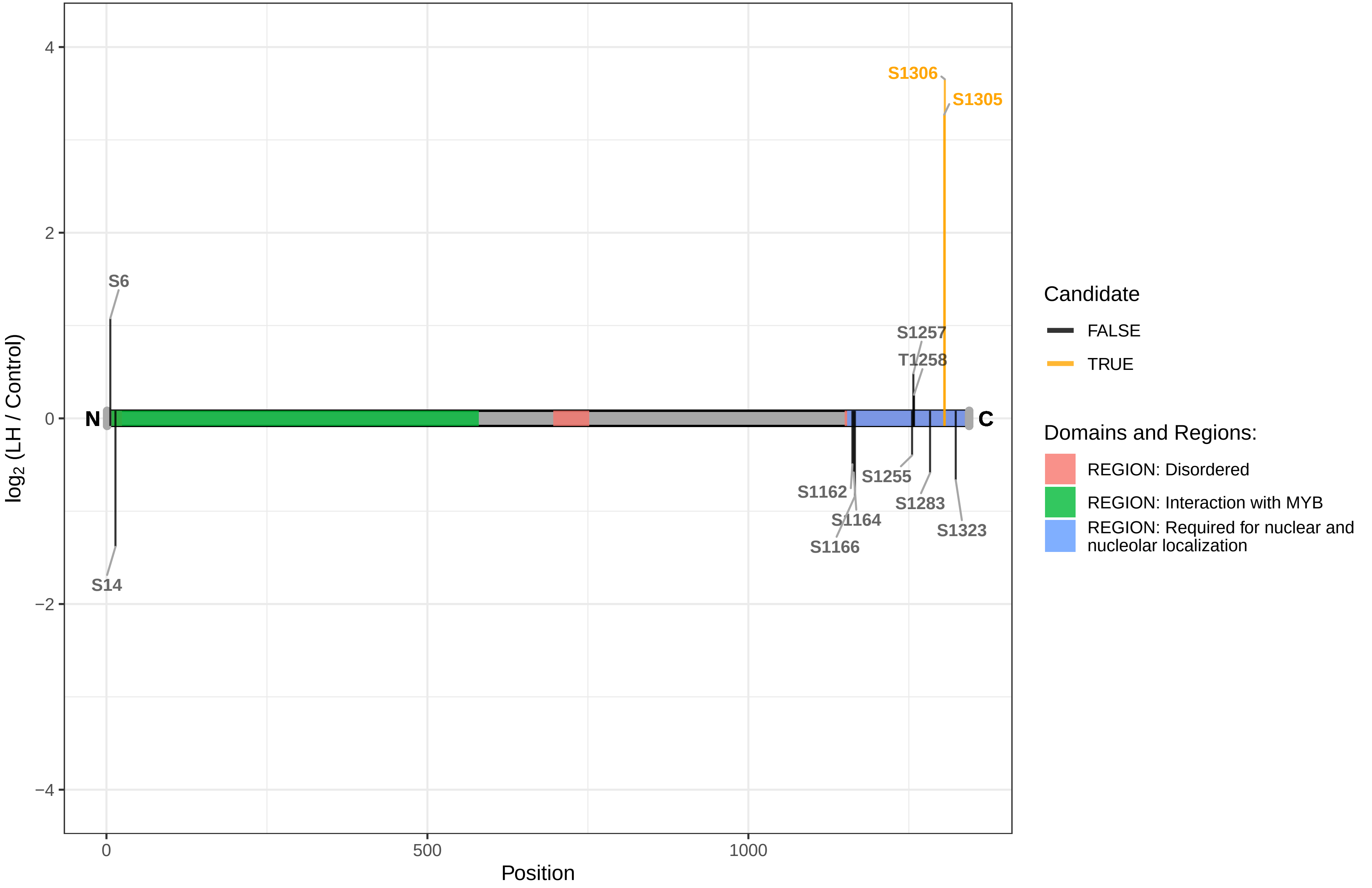

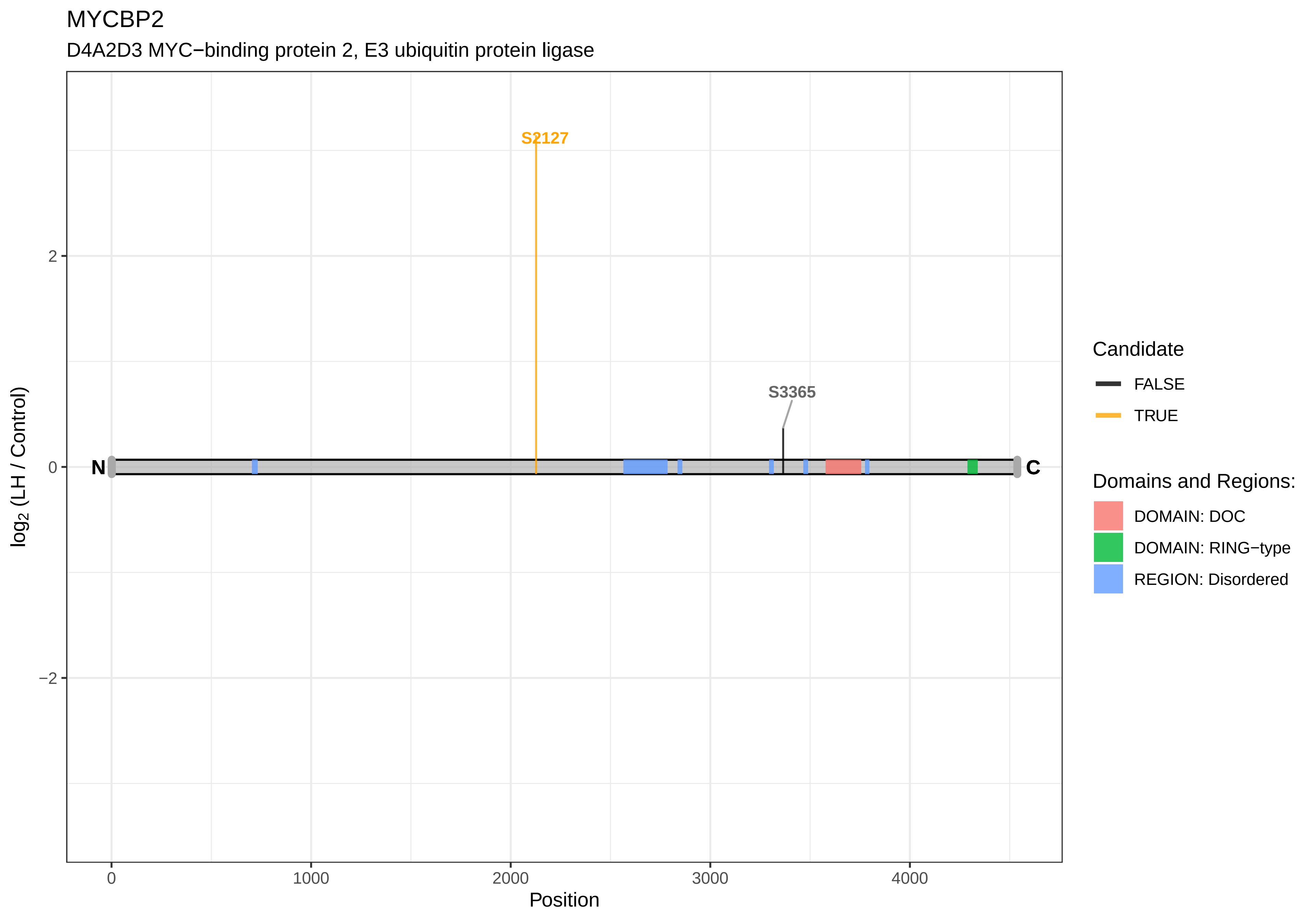
MYCBP2
D4A2D3 MYC−binding protein 2, E3 ubiquitin protein ligase
S2127
S3365
N
C
2
0
−2
log2 (LH / Control)
0
1000
2000
3000
4000
Position
Candidate
FALSE
TRUE
Domains and Regions:
DOMAIN: DOC
DOMAIN: RING−type
REGION: Disordered

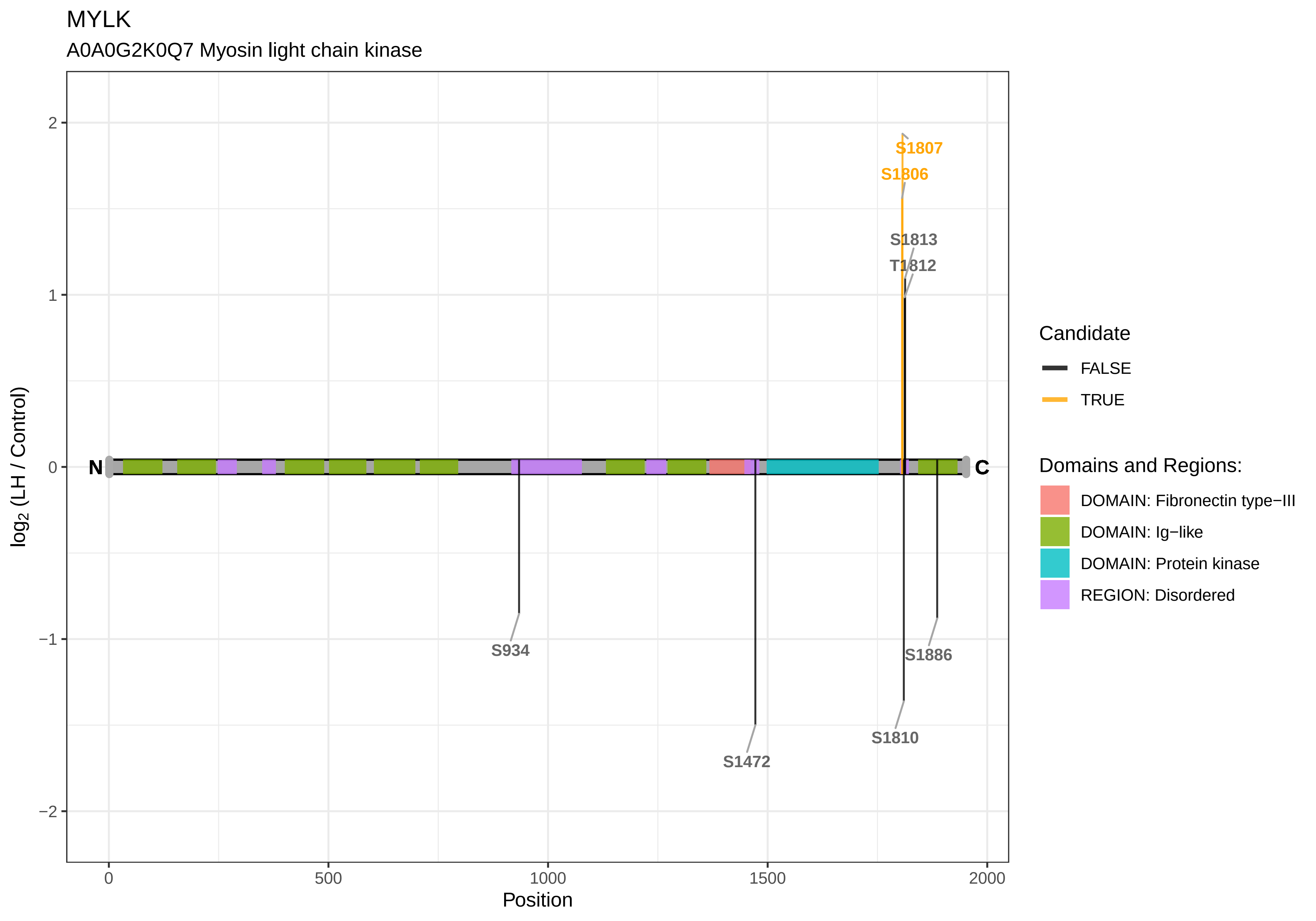

MYLK
A0A0G2K0Q7 Myosin light chain kinase
log2 (LH / Control)
Position
2
1
0
−1
−2
0
500
1000
1500
2000
N
C
S1807
S1806
S1813
T1812
S934
S1886
S1472
S1810
Candidate
FALSE
TRUE
Domains and Regions:
DOMAIN: Fibronectin type−III
DOMAIN: Ig−like
DOMAIN: Protein kinase
REGION: Disordered

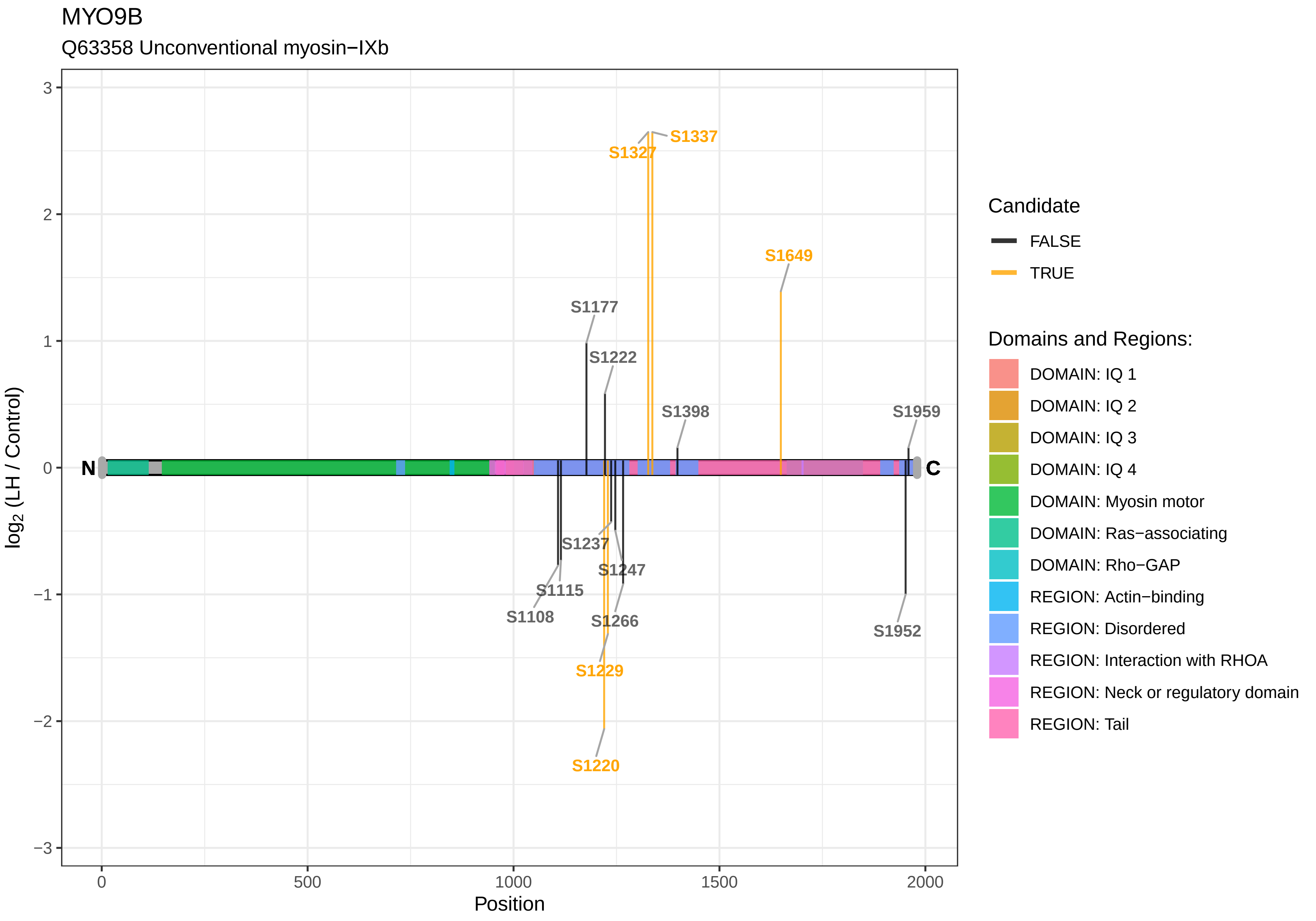

MYO9B
Q63358 Unconventional myosin−IXb
log2 (LH / Control)
Position
N
C
S1337
S1327
S1649
S1177
S1222
S1398
S1959
S1237
S1247
S1115
S1108
S1266
S1952
S1229
S1220
3
2
1
0
−1
−2
−3
0
500
1000
1500
2000
Candidate
FALSE
TRUE
Domains and Regions:
DOMAIN: IQ 1
DOMAIN: IQ 2
DOMAIN: IQ 3
DOMAIN: IQ 4
DOMAIN: Myosin motor
DOMAIN: Ras−associating
DOMAIN: Rho−GAP
REGION: Actin−binding
REGION: Disordered
REGION: Interaction with RHOA
REGION: Neck or regulatory domain
REGION: Tail

NAA80

A0A0G2JV35 Hyaluronoglucosaminidase 3, isoform CRA_d

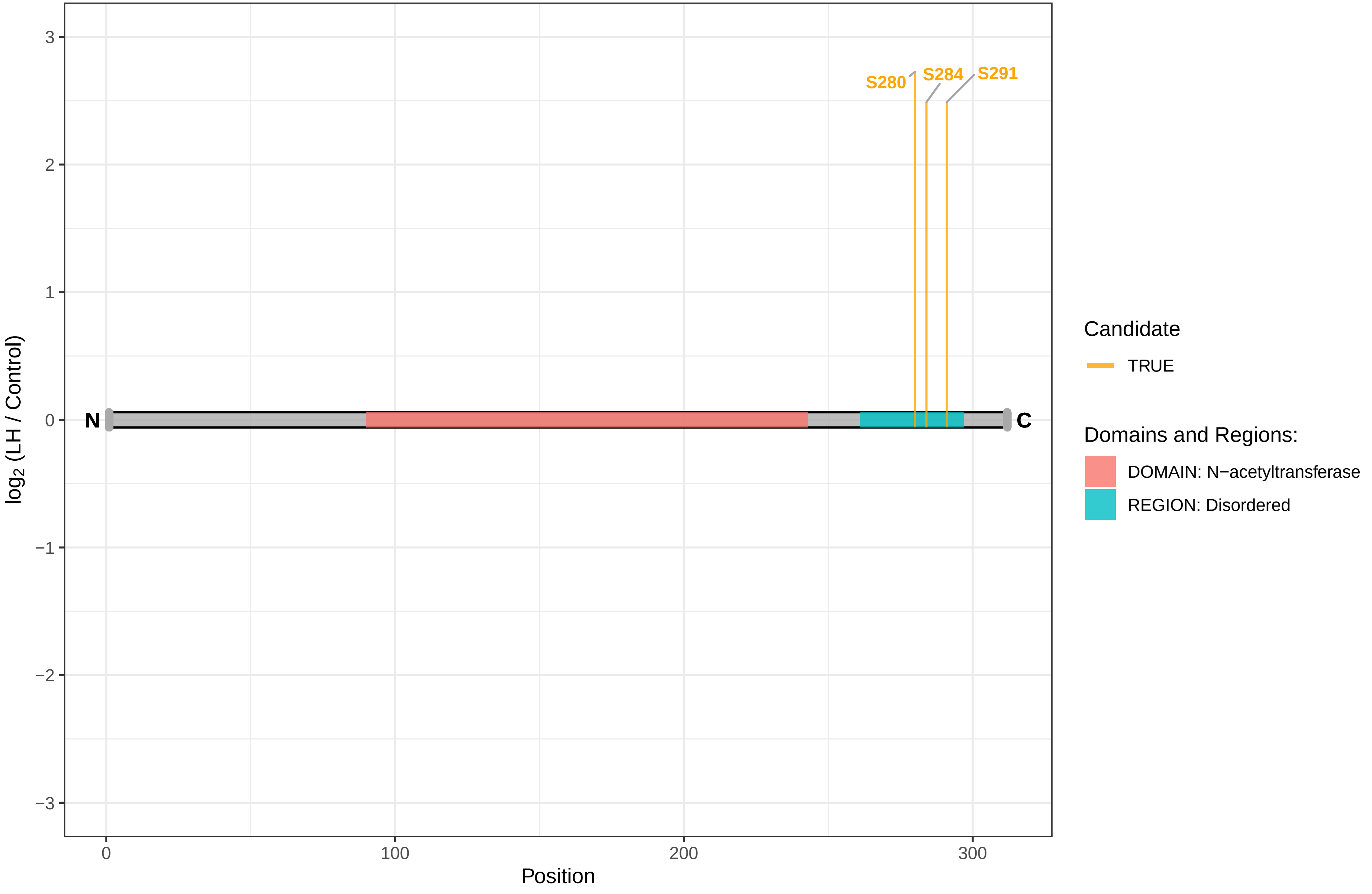

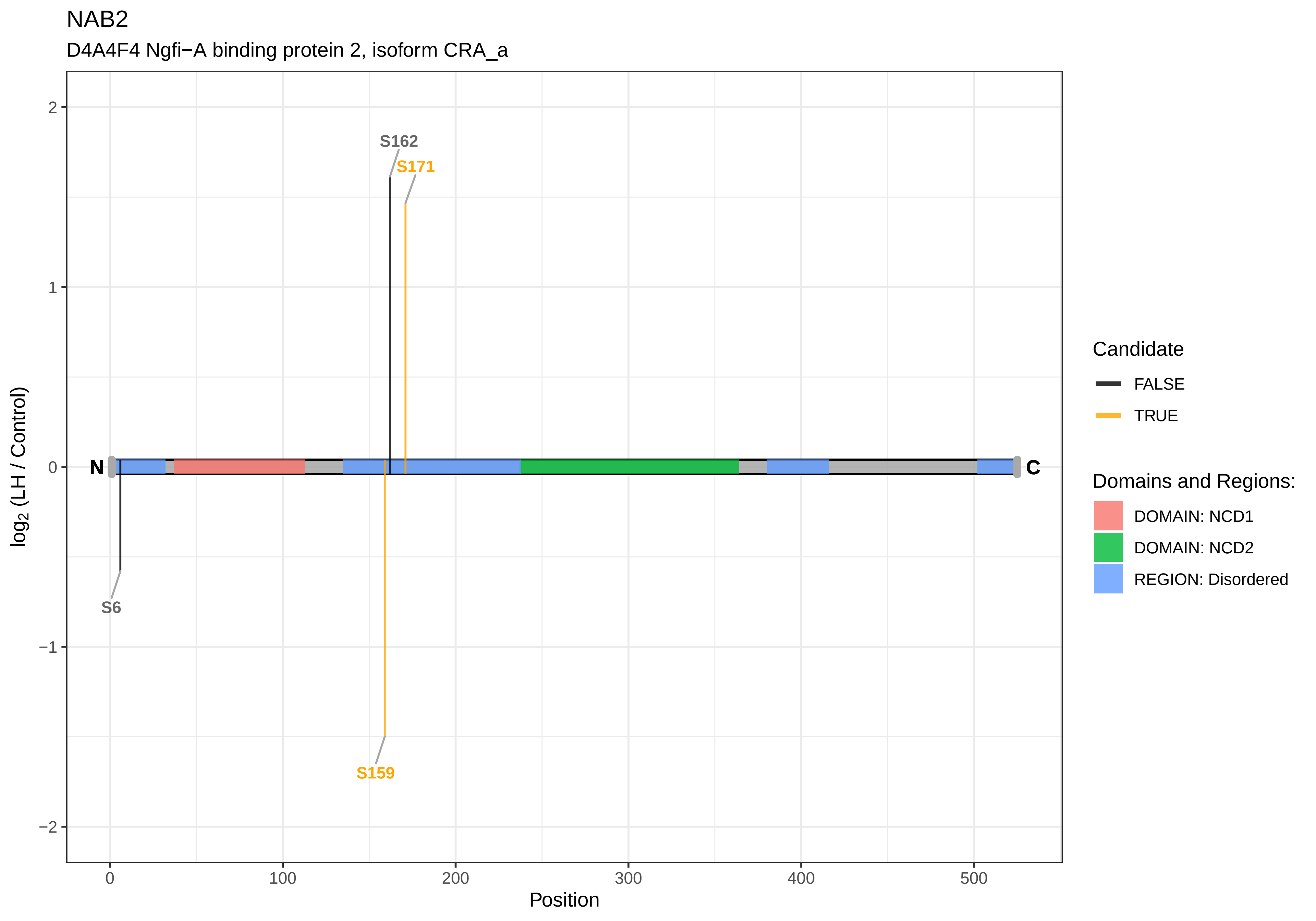

NAB2
D4A4F4 Ngfi−A binding protein 2, isoform CRA_a
S162
S171
S6
S159
N
C
2
1
0
−1
−2
0
100
200
300
400
500
Position
log2 (LH / Control)
Candidate
FALSE
TRUE
Domains and Regions:
DOMAIN: NCD1
DOMAIN: NCD2
REGION: Disordered

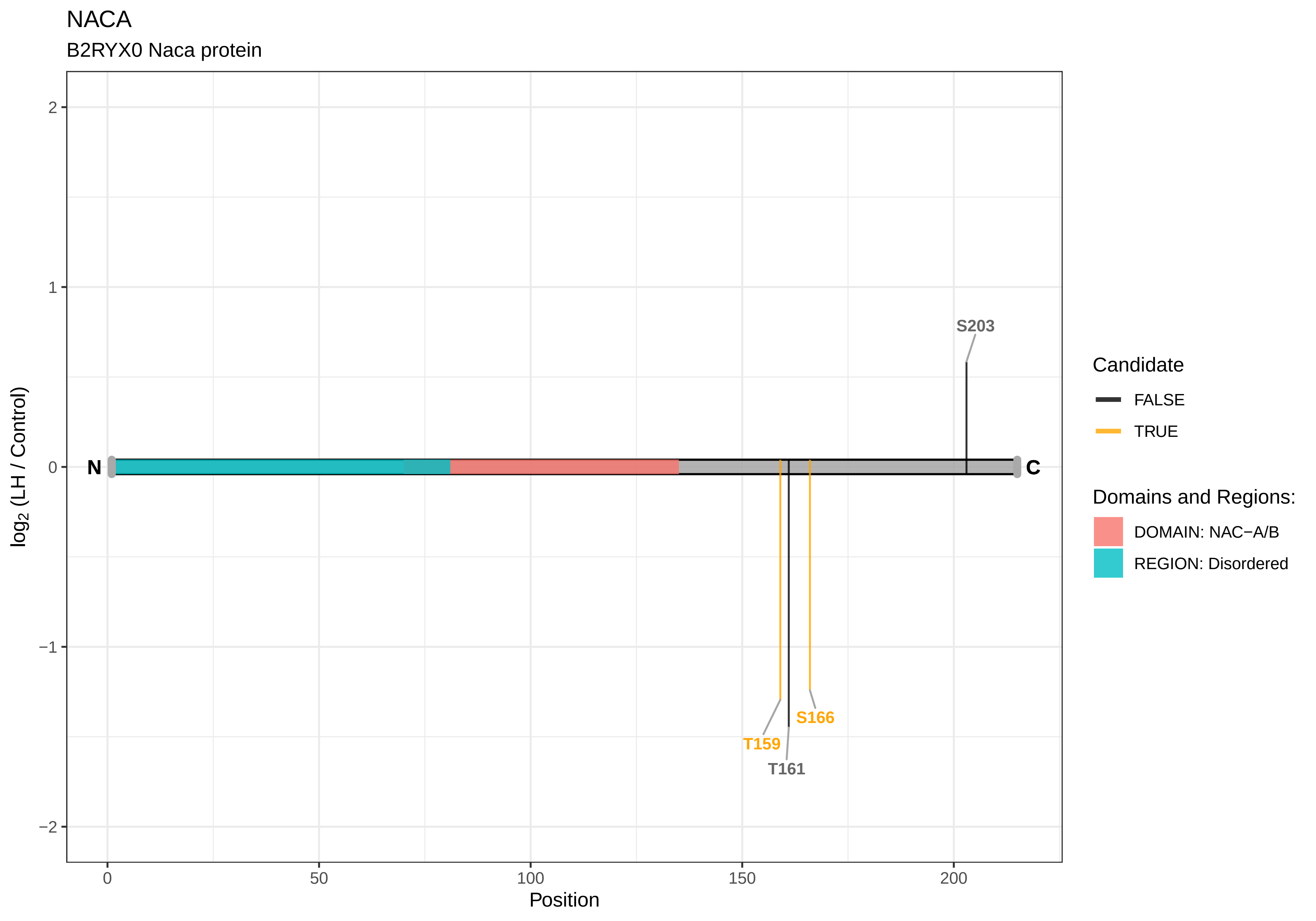
NACA
B2RYX0 Naca protein
S203
T159
T161
S166
N
C
2
1
0
−1
−2
0
50
100
150
200
Position
log2 (LH / Control)
Candidate
FALSE
TRUE
Domains and Regions:
DOMAIN: NAC−A/B
REGION: Disordered

NADK

F7EXV6 NAD kinase

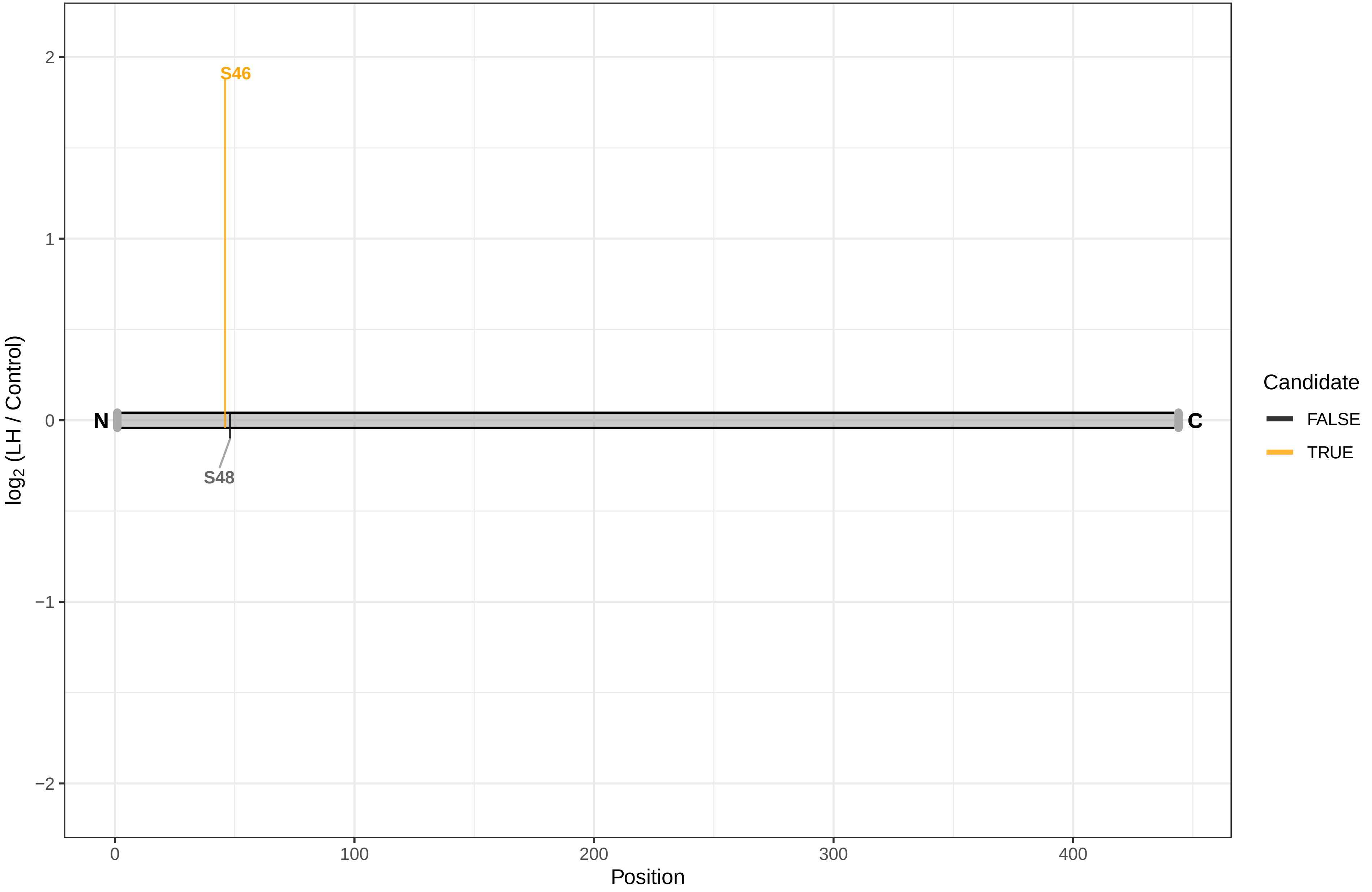

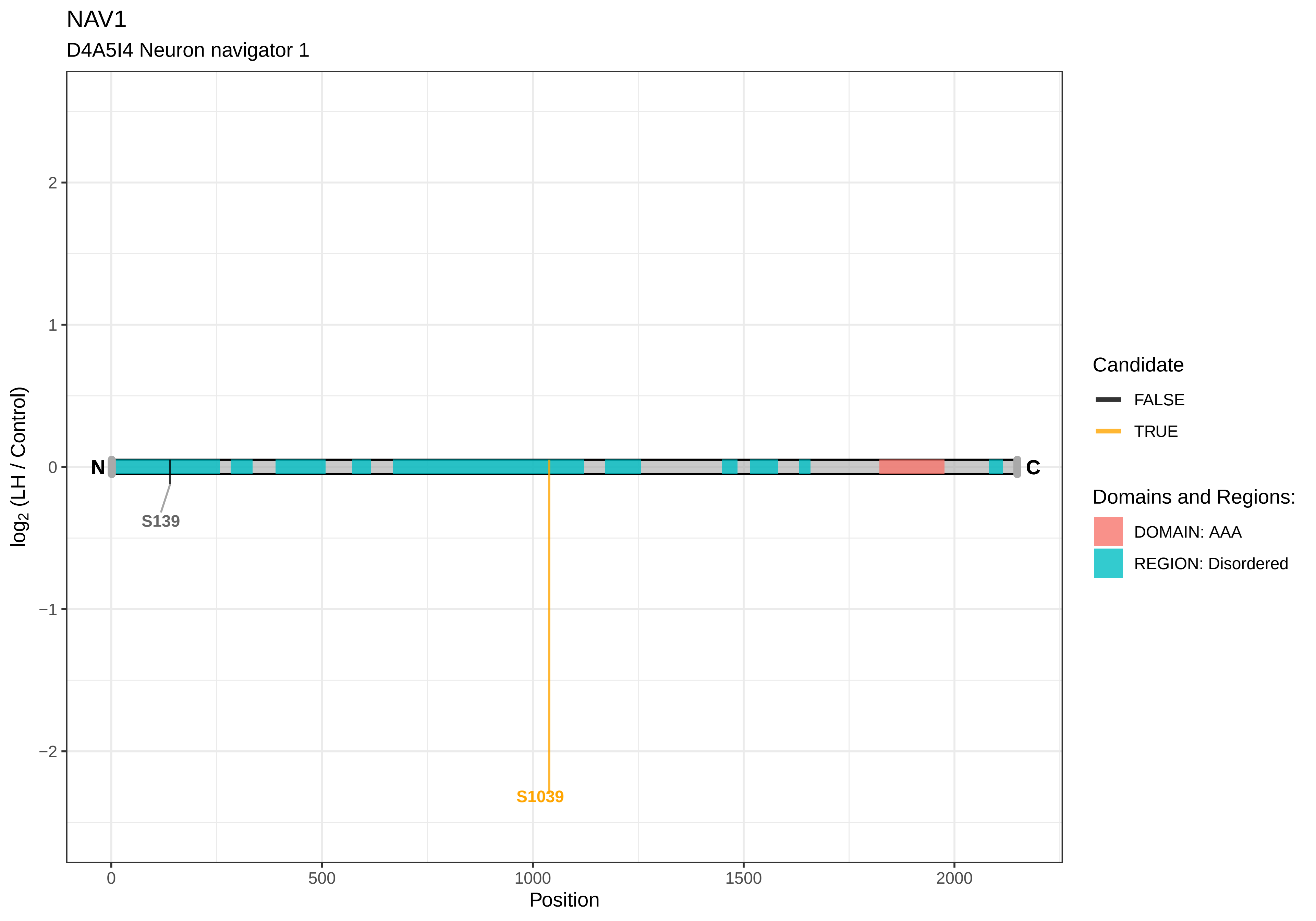
NAV1
D4A5I4 Neuron navigator 1
log2 (LH / Control)
2
1
0
−1
−2
N
C
S139
S1039
0
500
1000
1500
2000
Position
Candidate
FALSE
TRUE
Domains and Regions:
DOMAIN: AAA
REGION: Disordered

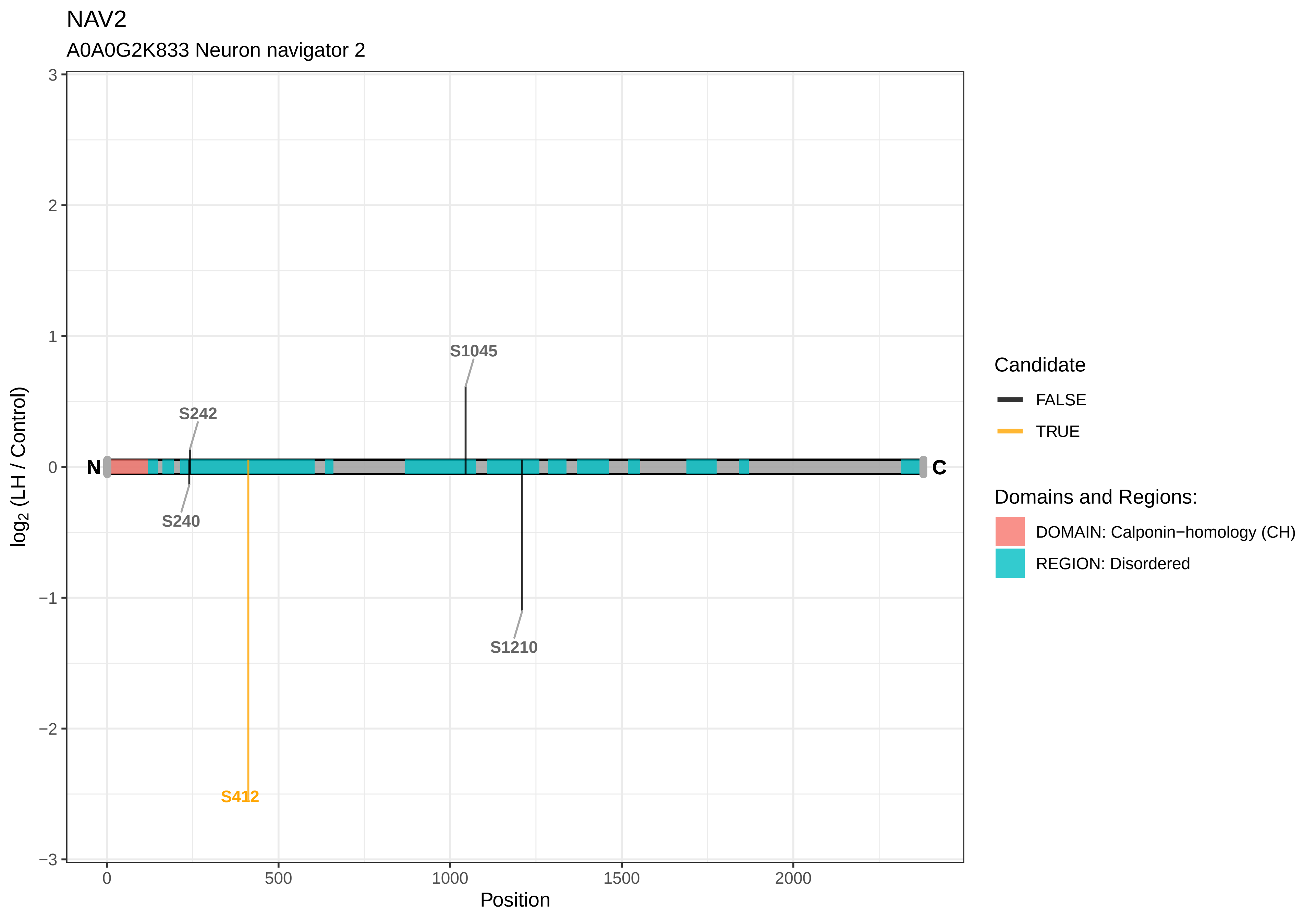

NAV2
A0A0G2K833 Neuron navigator 2
log2 (LH / Control)
3
2
1
0
−1
−2
−3
0
500
1000
1500
2000
Position
N
C
S1045
S242
S240
S412
S1210
Candidate
FALSE
TRUE
Domains and Regions:
DOMAIN: Calponin−homology (CH)
REGION: Disordered

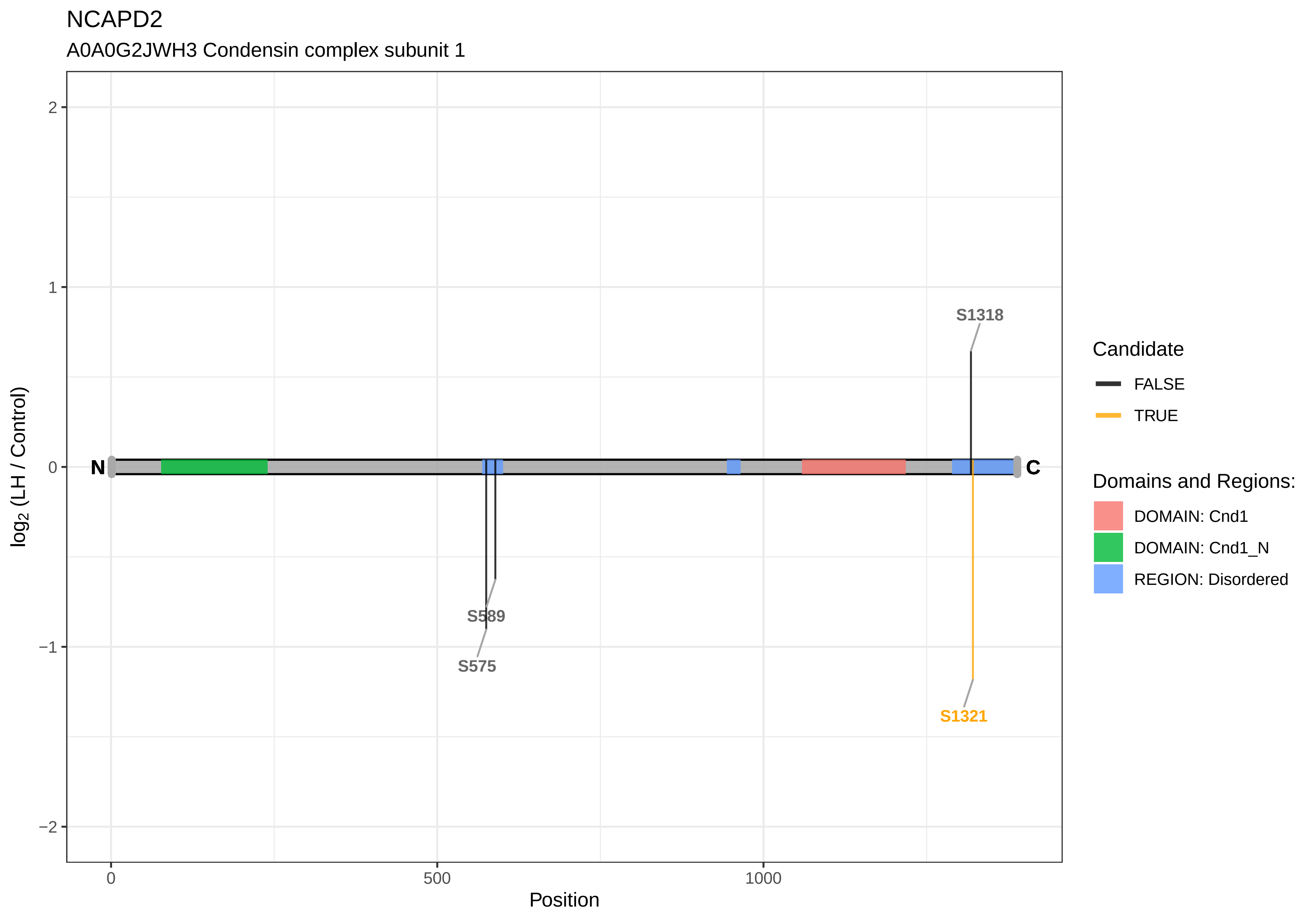

NCAPD2
A0A0G2JWH3 Condensin complex subunit 1
S1318
S589
S575
S1321
N
C
2
1
0
−1
−2
0
500
1000
Position
log2 (LH / Control)
Candidate
FALSE
TRUE
Domains and Regions:
DOMAIN: Cnd1
DOMAIN: Cnd1_N
REGION: Disordered

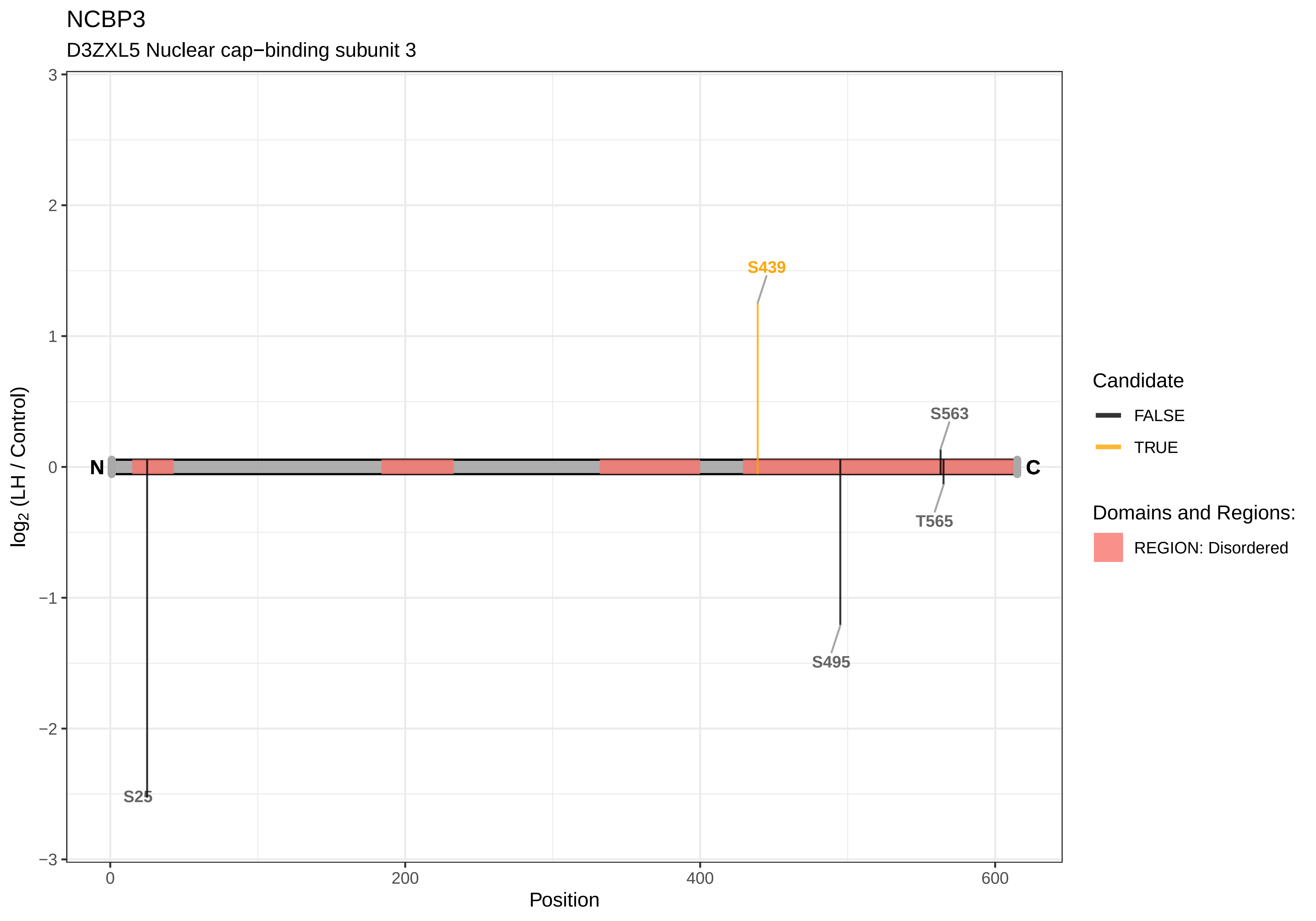

NCBP3
D3ZXL5 Nuclear cap−binding subunit 3
log2 (LH / Control)
Position
N
C
S439
S563
T565
S495
S25
3
2
1
0
−1
−2
−3
0
200
400
600
Candidate
FALSE
TRUE
Domains and Regions:
REGION: Disordered

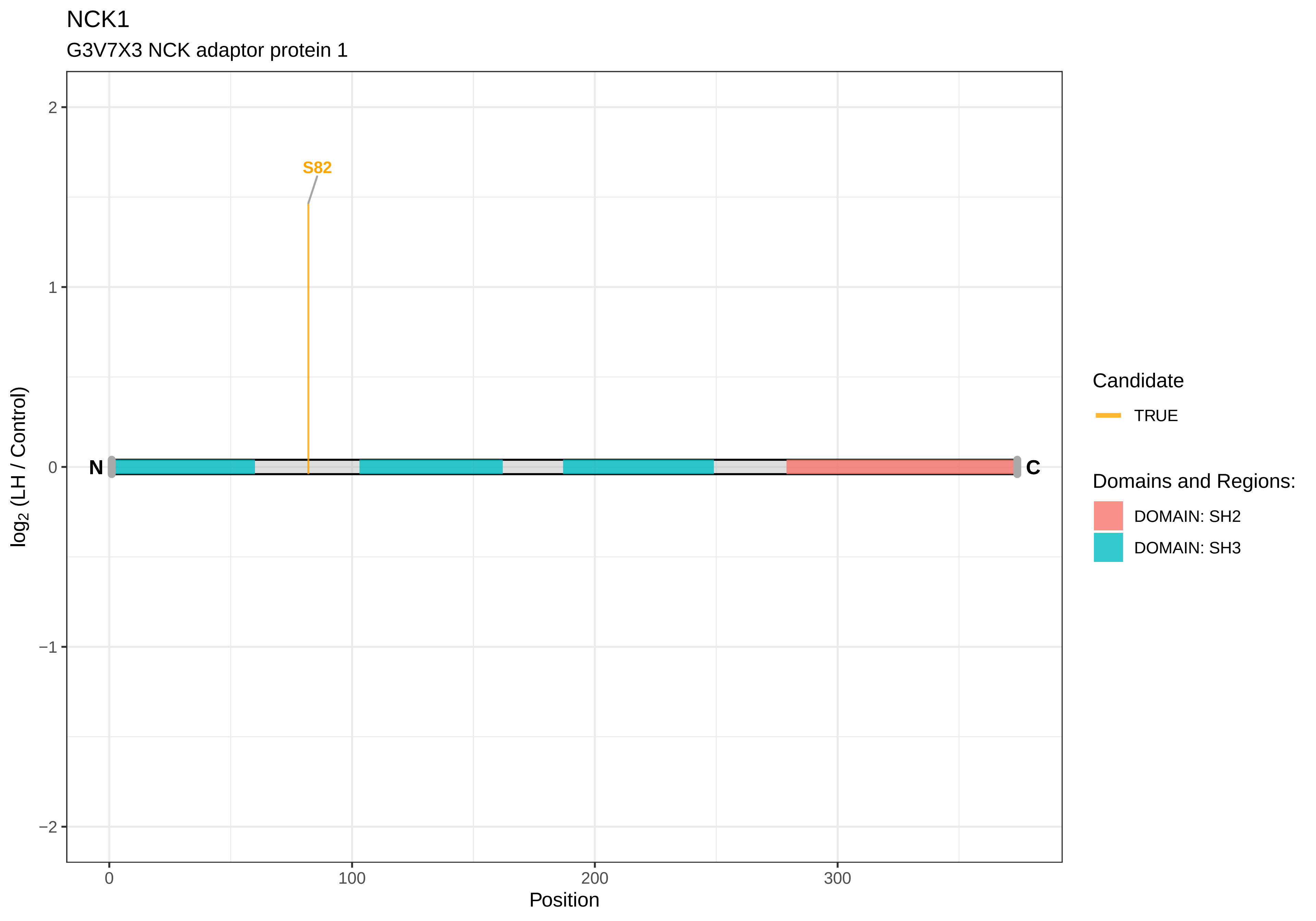
NCK1
G3V7X3 NCK adaptor protein 1
S82
N
C
2
1
0
−1
−2
0
100
200
300
Position
log2 (LH / Control)
Candidate
TRUE
Domains and Regions:
DOMAIN: SH2
DOMAIN: SH3

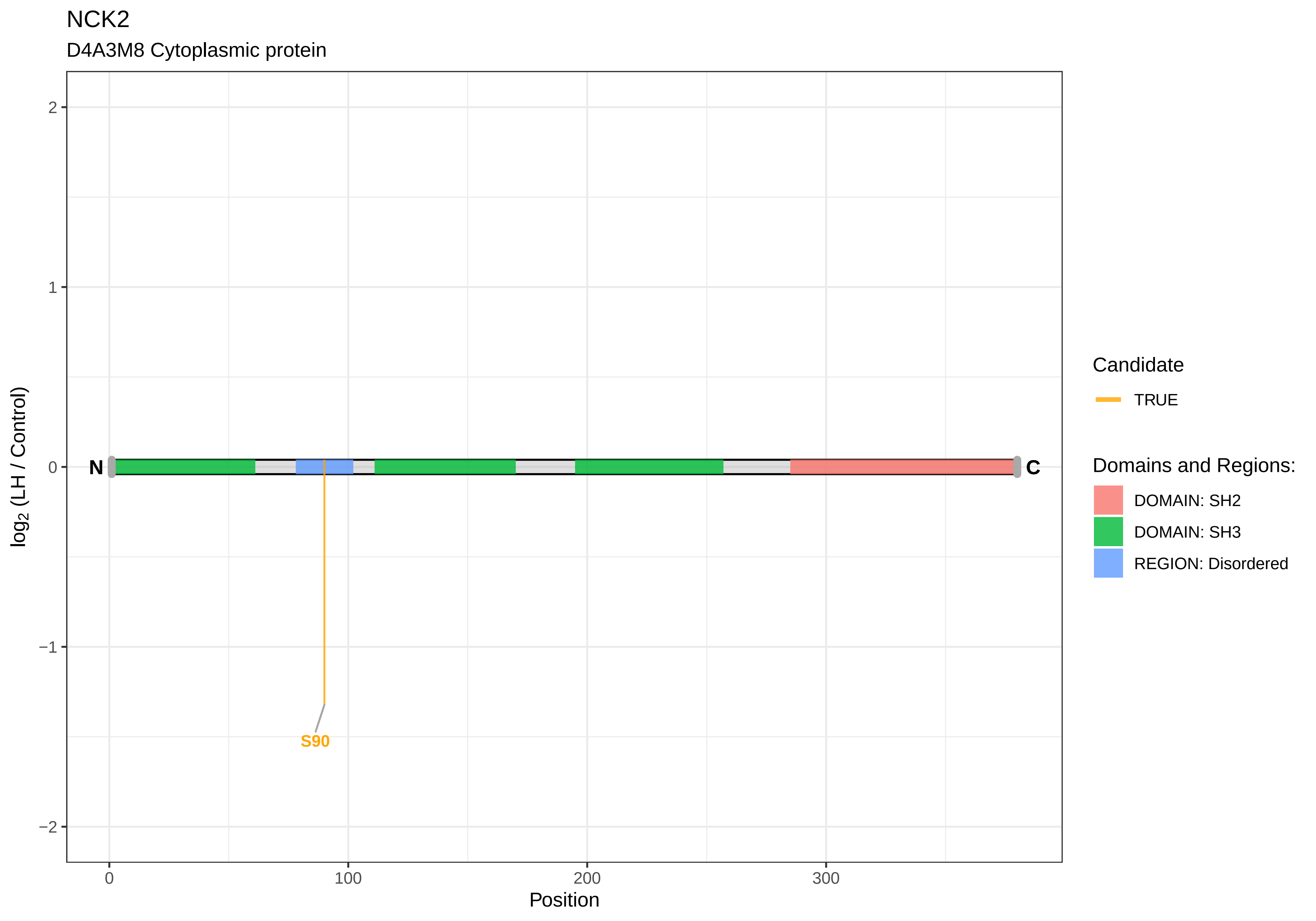
NCK2
D4A3M8 Cytoplasmic protein
log2 (LH / Control)
2
1
0
−1
−2
N
C
S90
0
100
200
300
Position
Candidate
TRUE
Domains and Regions:
DOMAIN: SH2
DOMAIN: SH3
REGION: Disordered

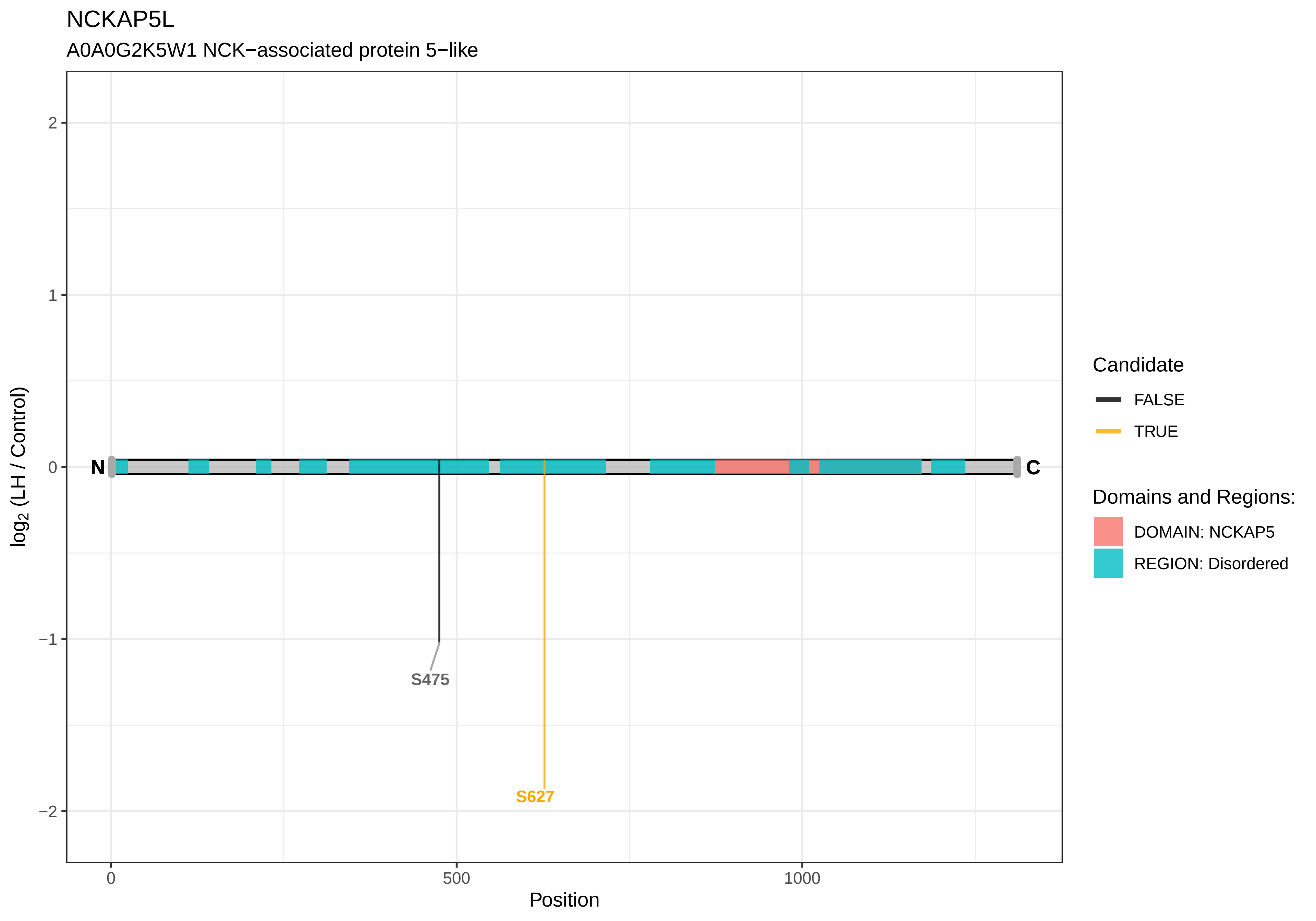

NCKAP5L
A0A0G2K5W1 NCK−associated protein 5−like
log2 (LH / Control)
2
1
0
−1
−2
N
C
S475
S627
0
500
1000
Position
Candidate
FALSE
TRUE
Domains and Regions:
DOMAIN: NCKAP5
REGION: Disordered

NCKIPSD

D3ZWX4 NCK−interacting protein with SH3 domain

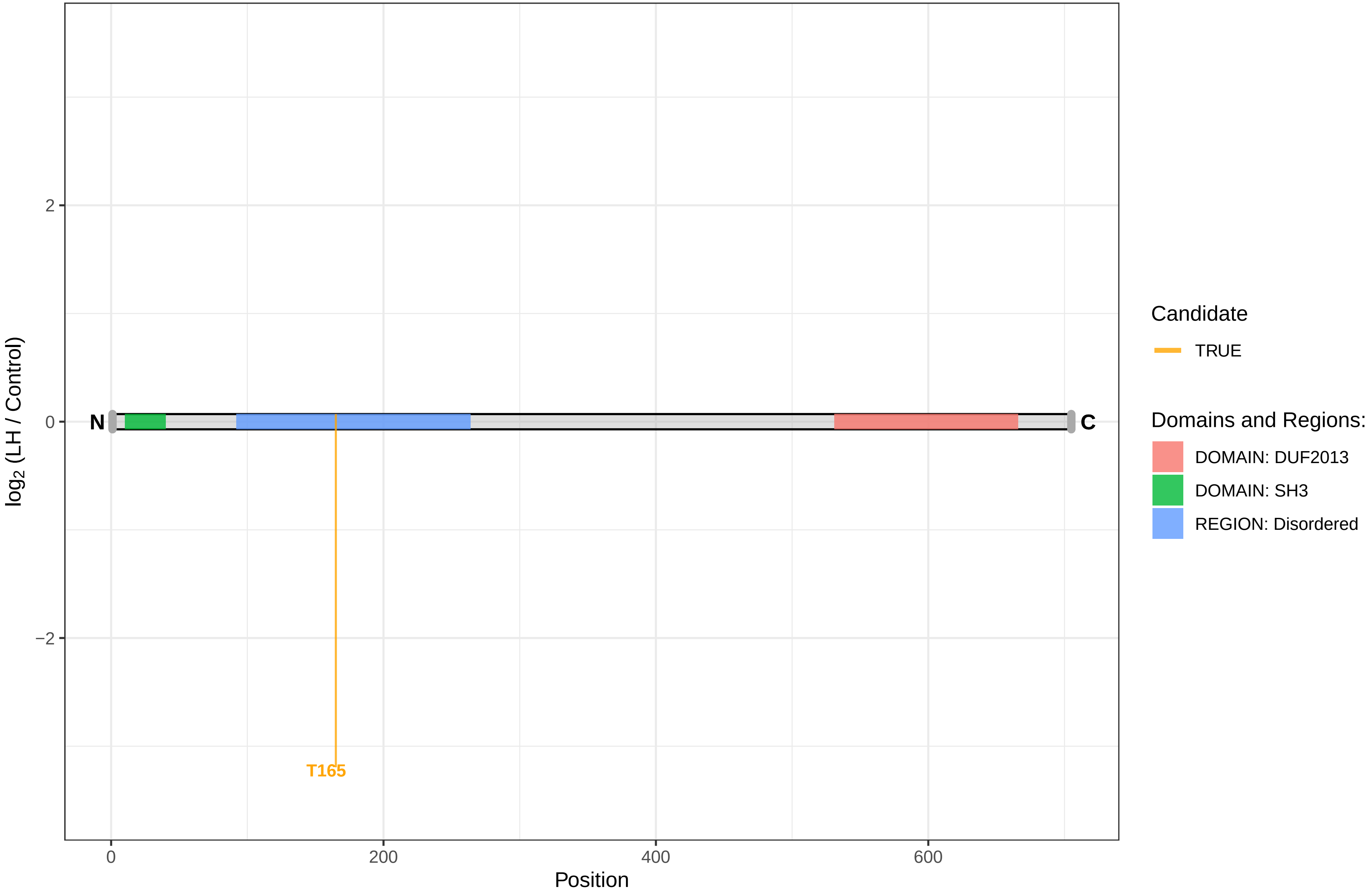

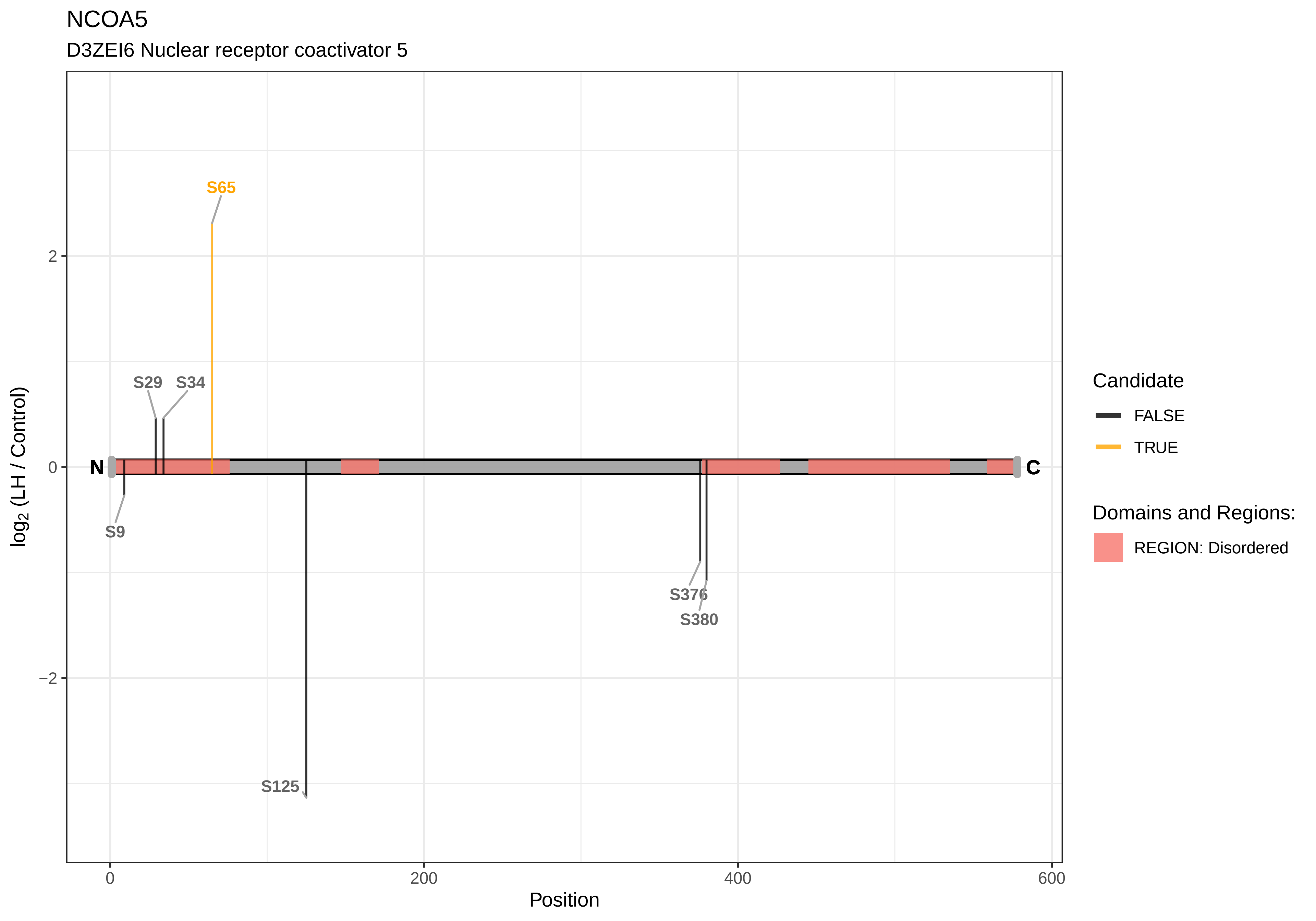
NCOA5
D3ZEI6 Nuclear receptor coactivator 5
S65
S29
S34
N
C
S9
S376
S380
S125
2
0
−2
0
200
400
600
Position
log2 (LH / Control)
Candidate
FALSE
TRUE
Domains and Regions:
REGION: Disordered

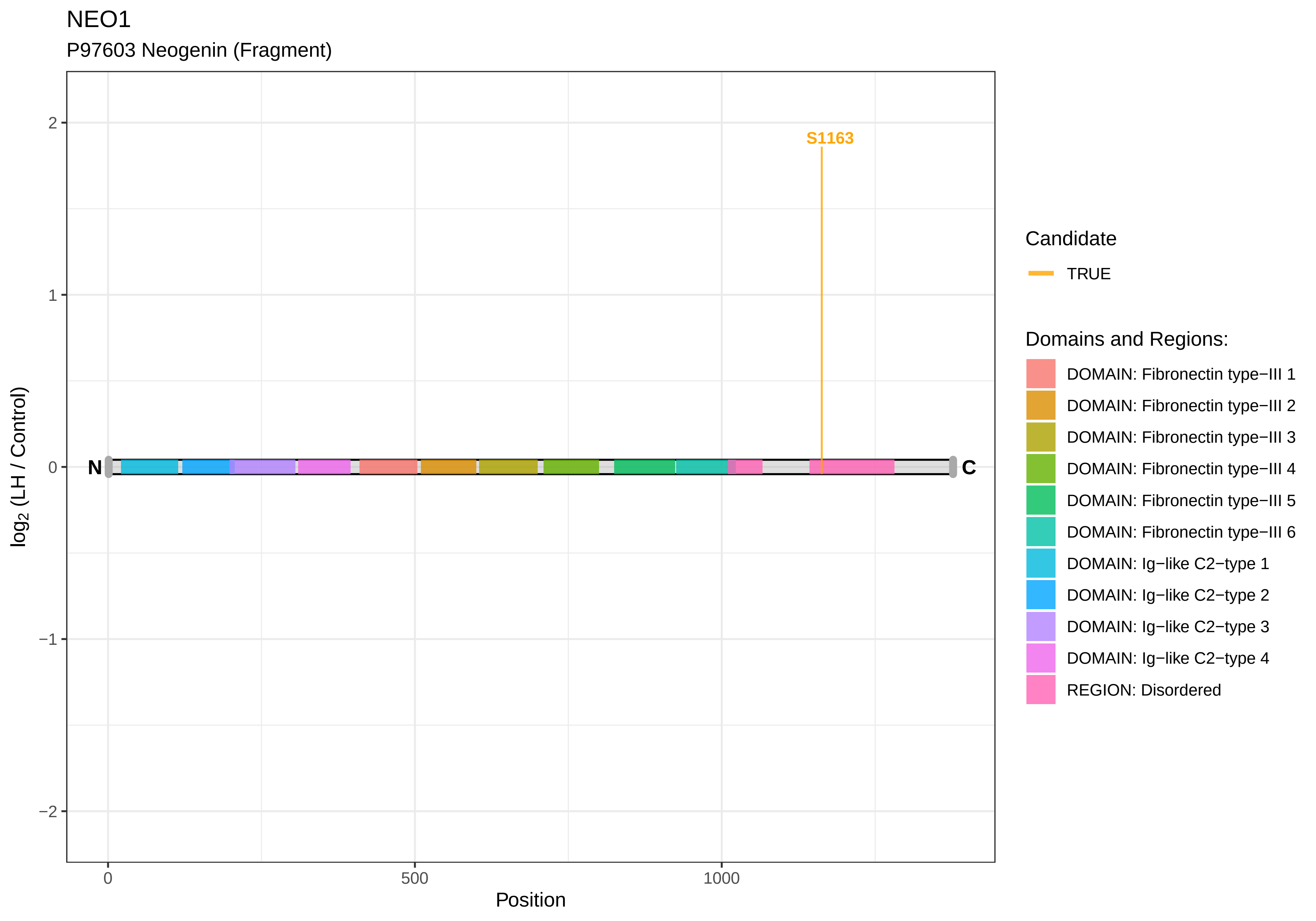

NEO1
P97603 Neogenin (Fragment)
S1163
N
C
2
1
0
−1
−2
log2 (LH / Control)
0
500
1000
Position
Candidate
TRUE
Domains and Regions:
DOMAIN: Fibronectin type−III 1
DOMAIN: Fibronectin type−III 2
DOMAIN: Fibronectin type−III 3
DOMAIN: Fibronectin type−III 4
DOMAIN: Fibronectin type−III 5
DOMAIN: Fibronectin type−III 6
DOMAIN: Ig−like C2−type 1
DOMAIN: Ig−like C2−type 2
DOMAIN: Ig−like C2−type 3
DOMAIN: Ig−like C2−type 4
REGION: Disordered

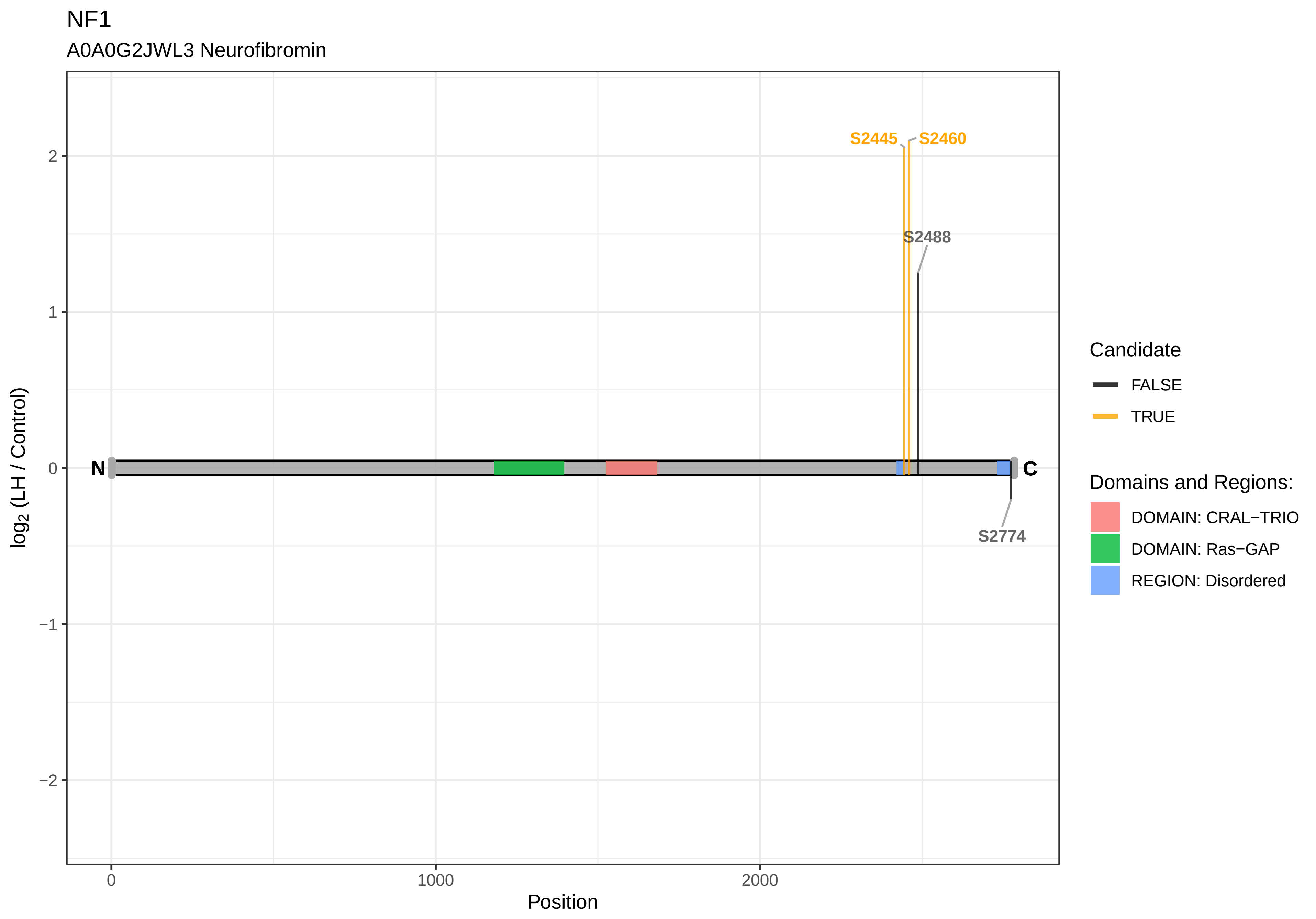
NF1
A0A0G2JWL3 Neurofibromin
S2445
S2460
S2488
S2774
N
C
2
1
0
−1
−2
log2 (LH / Control)
0
1000
2000
Position
Candidate
FALSE
TRUE
Domains and Regions:
DOMAIN: CRAL−TRIO
DOMAIN: Ras−GAP
REGION: Disordered

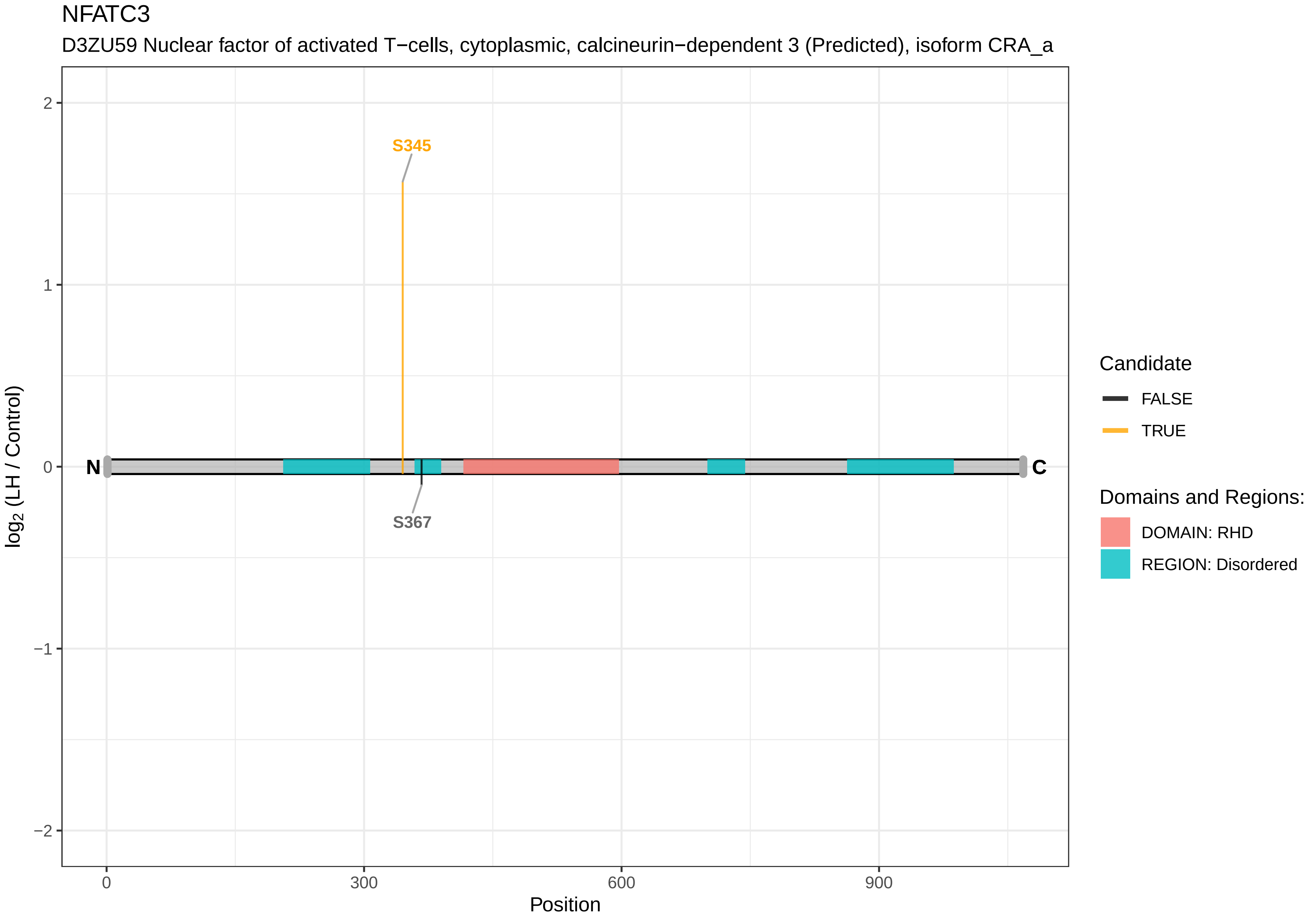

NFATC3
D3ZU59 Nuclear factor of activated T−cells, cytoplasmic, calcineurin−dependent 3 (Predicted), isoform CRA_a
S345
S367
N
C
2
1
0
−1
−2
0
300
600
900
Position
log2 (LH / Control)
Candidate
FALSE
TRUE
Domains and Regions:
DOMAIN: RHD
REGION: Disordered

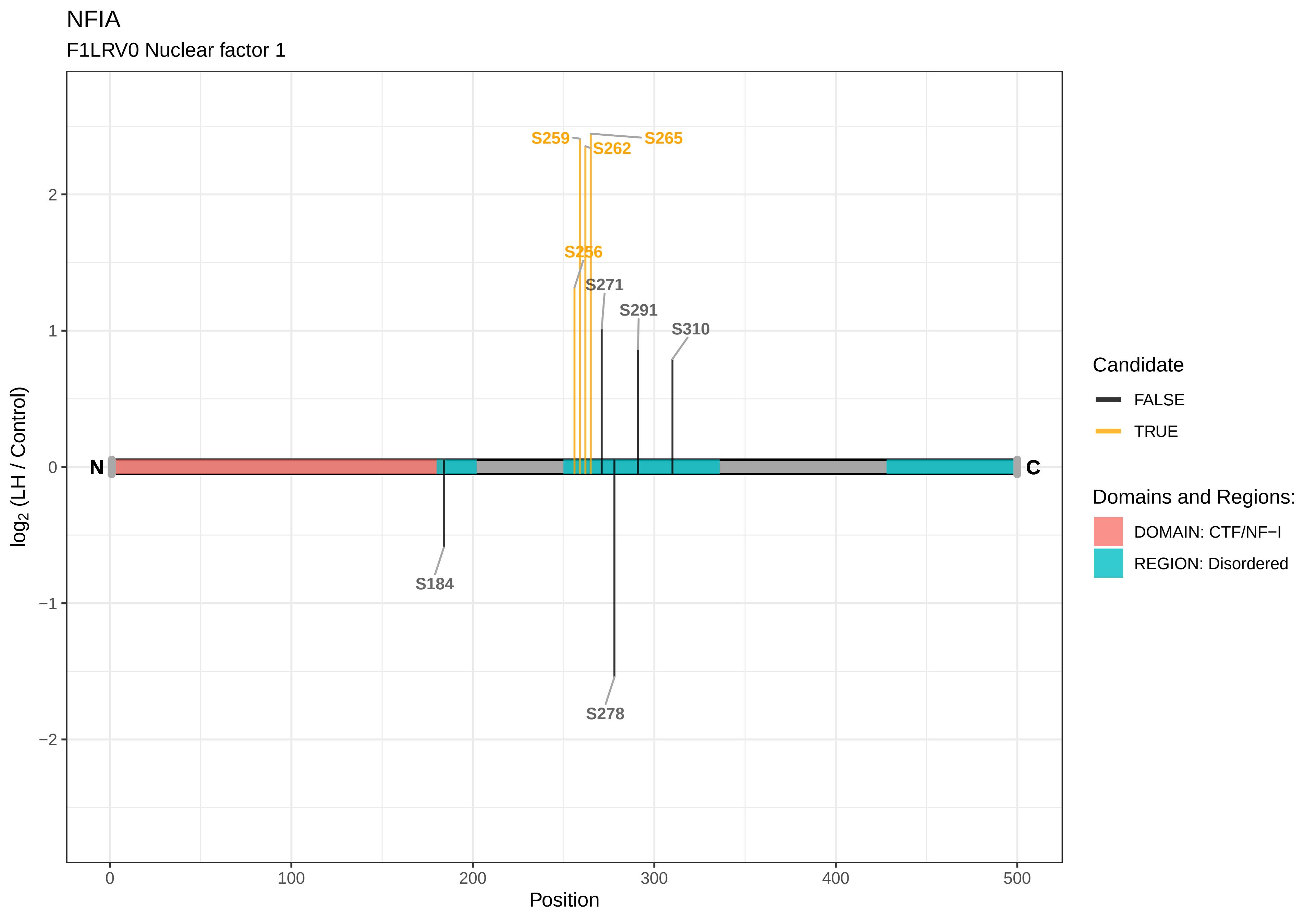
NFIA
F1LRV0 Nuclear factor 1
S259
S265
S262
S256
S271
S291
S310
S184
S278
N
C
2
1
0
−1
−2
0
100
200
300
400
500
Position
log2 (LH / Control)
Candidate
FALSE
TRUE
Domains and Regions:
DOMAIN: CTF/NF−I
REGION: Disordered

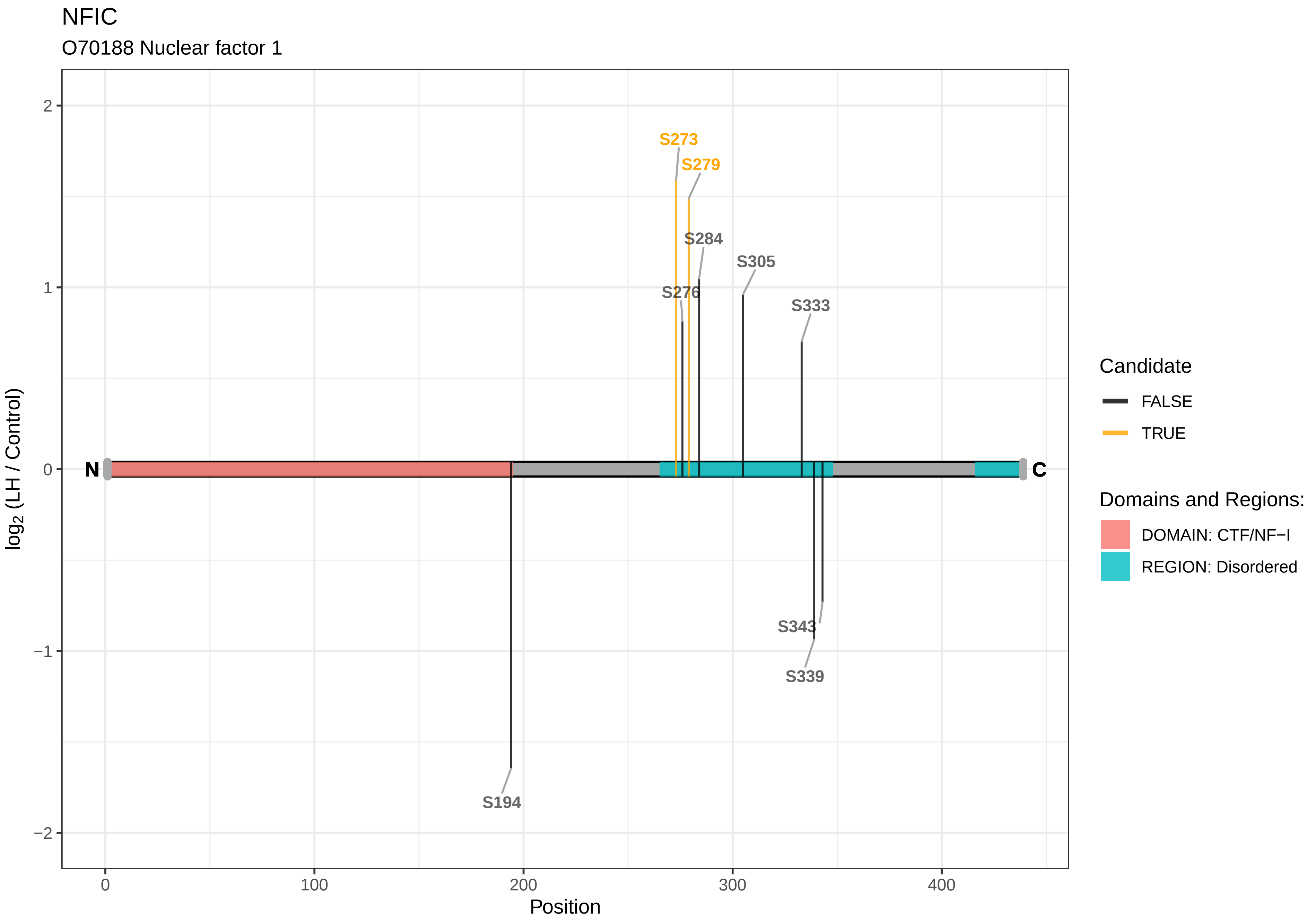
NFIC
O70188 Nuclear factor 1
S273
S279
S284
S305
S276
S333
S343
S339
S194
N
C
2
1
0
−1
−2
log2 (LH / Control)
0
100
200
300
400
Position
Candidate
FALSE
TRUE
Domains and Regions:
DOMAIN: CTF/NF−I
REGION: Disordered

NFKB2

Q5U2Z4 Nuclear factor kappa B subunit 2

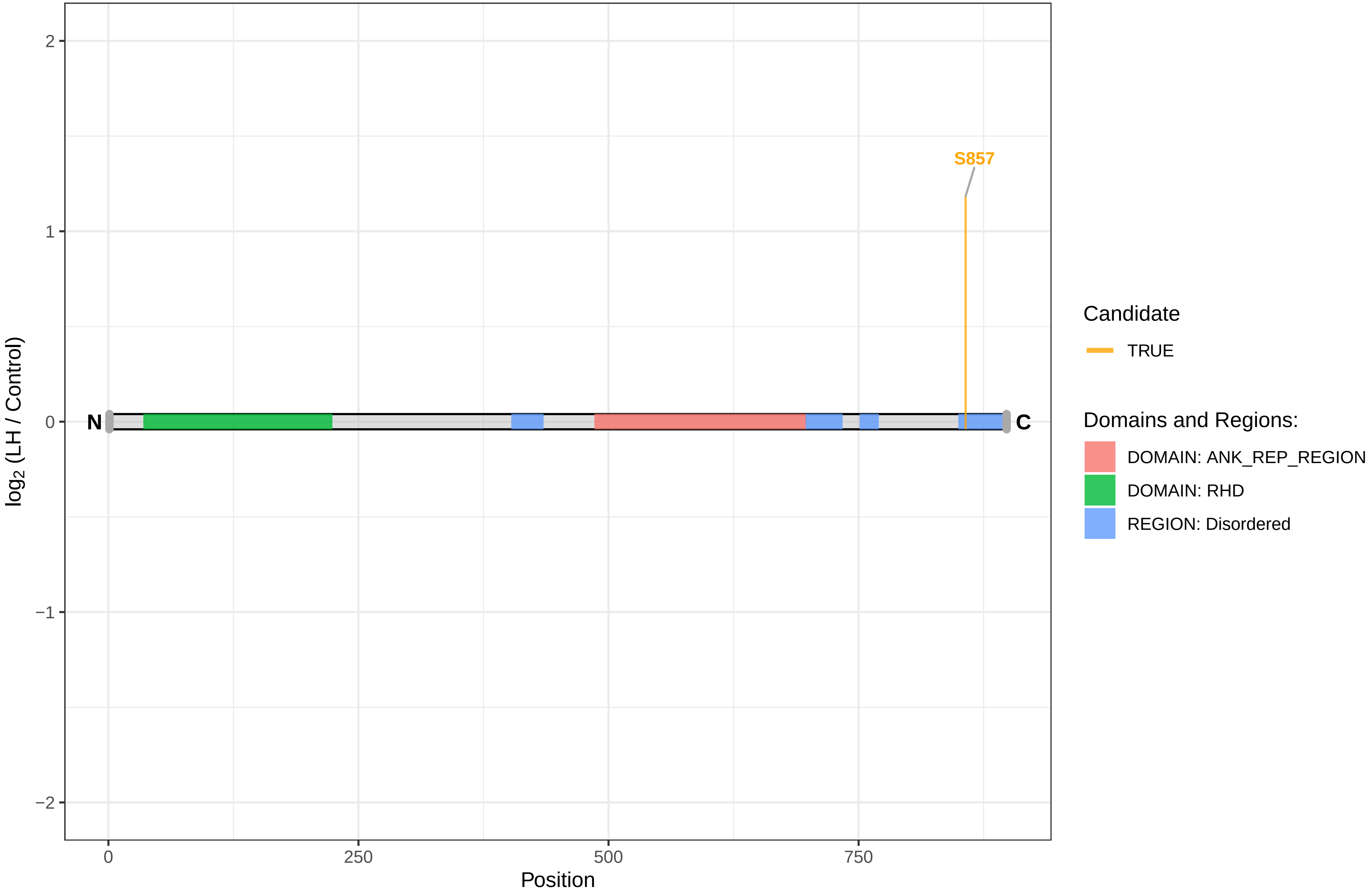

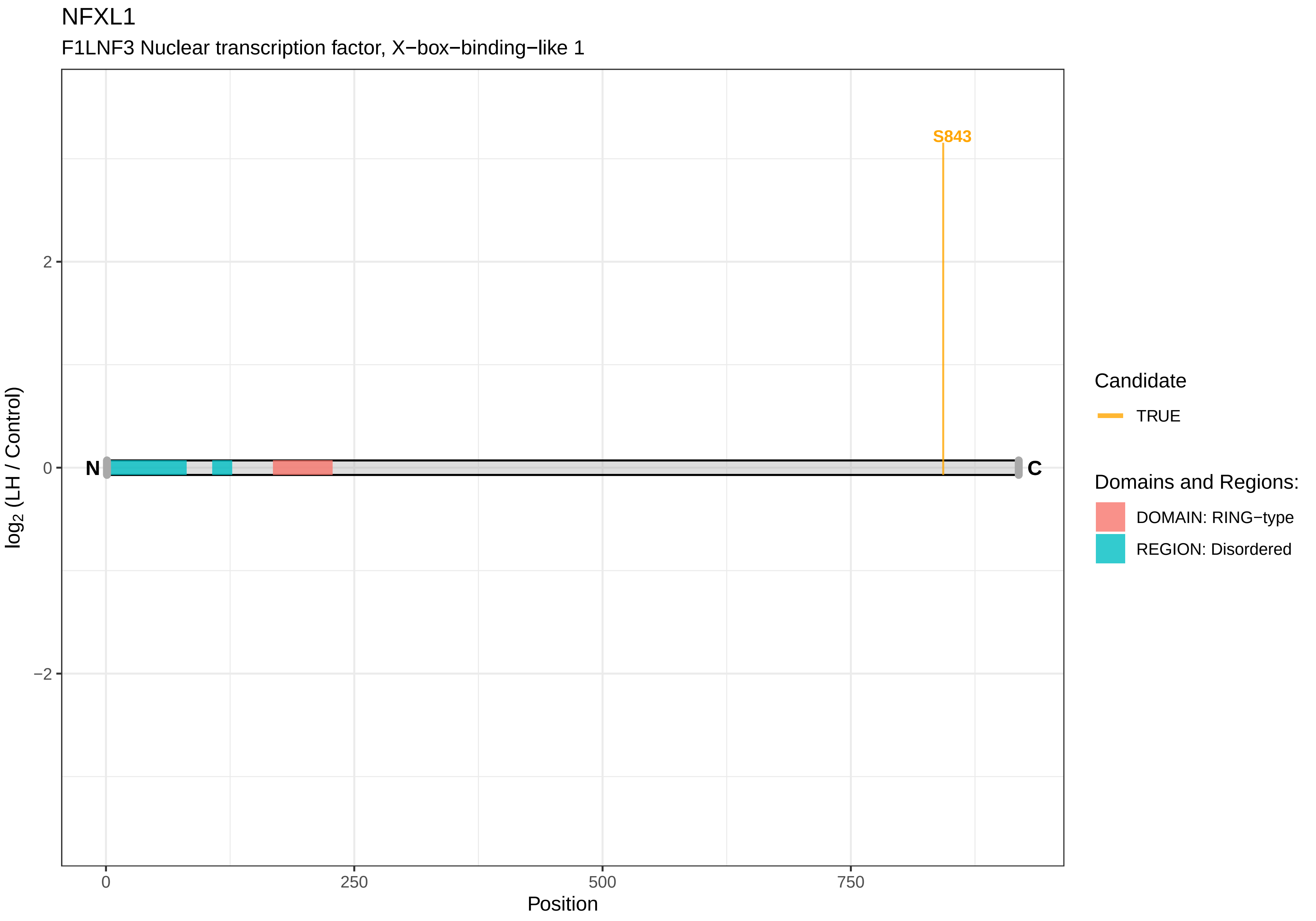
NFXL1
F1LNF3 Nuclear transcription factor, X−box−binding−like 1
S843
N
C
2
0
−2
0
250
500
750
Position
log2 (LH / Control)
Candidate
TRUE
Domains and Regions:
DOMAIN: RING−type
REGION: Disordered

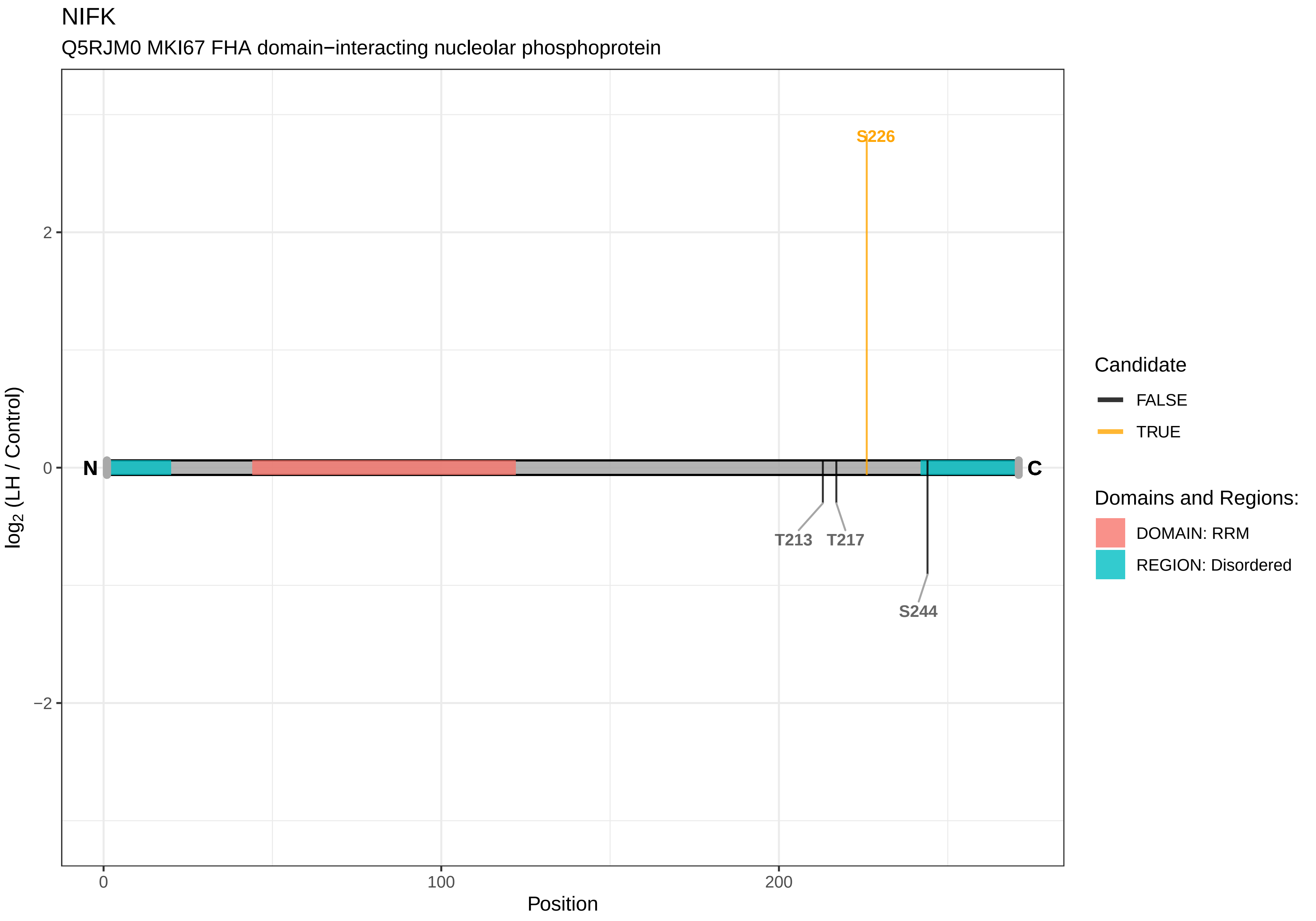

NIFK
Q5RJM0 MKI67 FHA domain−interacting nucleolar phosphoprotein
S226
T213
T217
S244
N
C
2
0
−2
0
100
200
Position
log2 (LH / Control)
Candidate
FALSE
TRUE
Domains and Regions:
DOMAIN: RRM
REGION: Disordered

NOL6

A0A096MKG2 Nucleolar protein 6

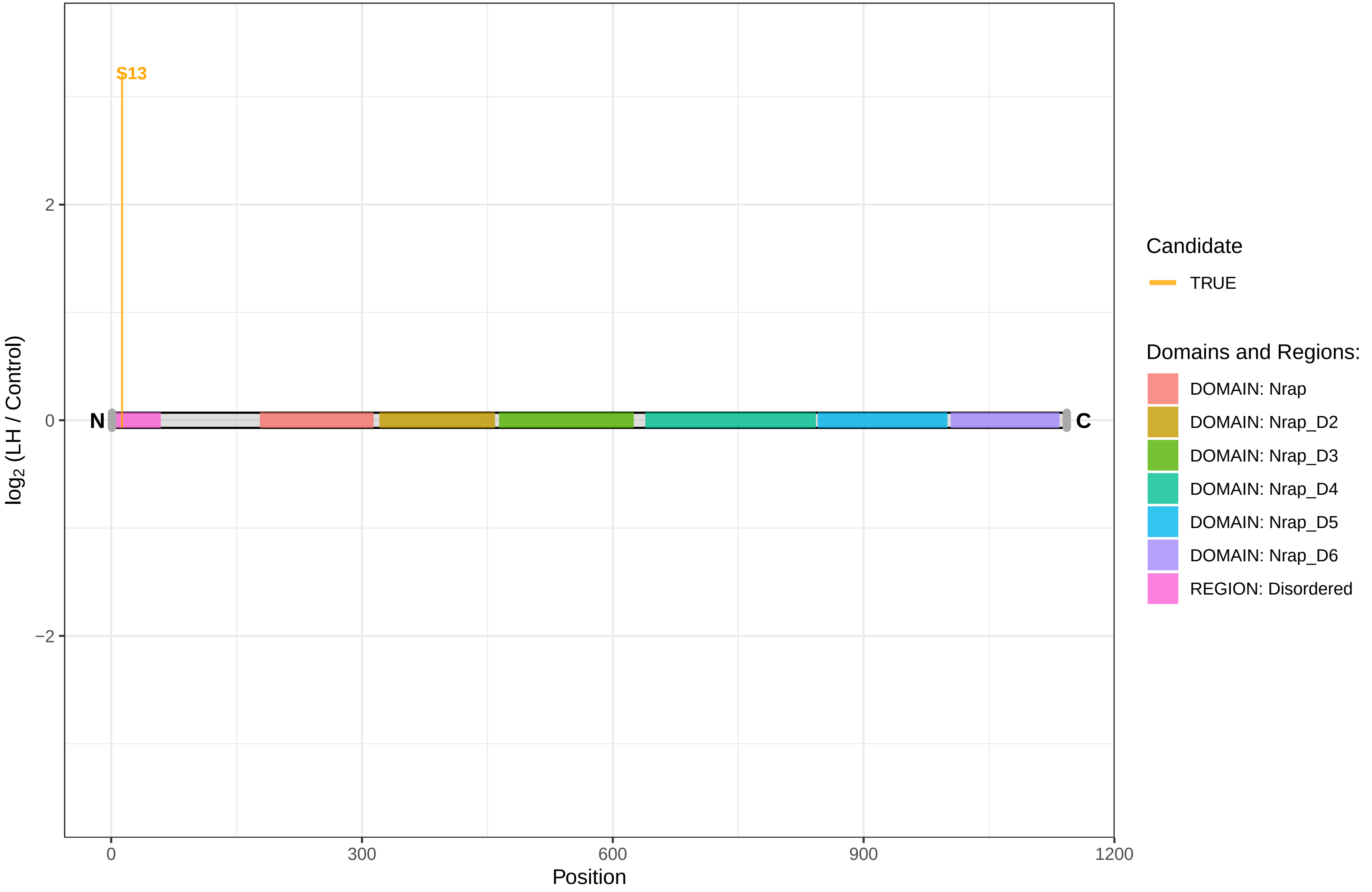

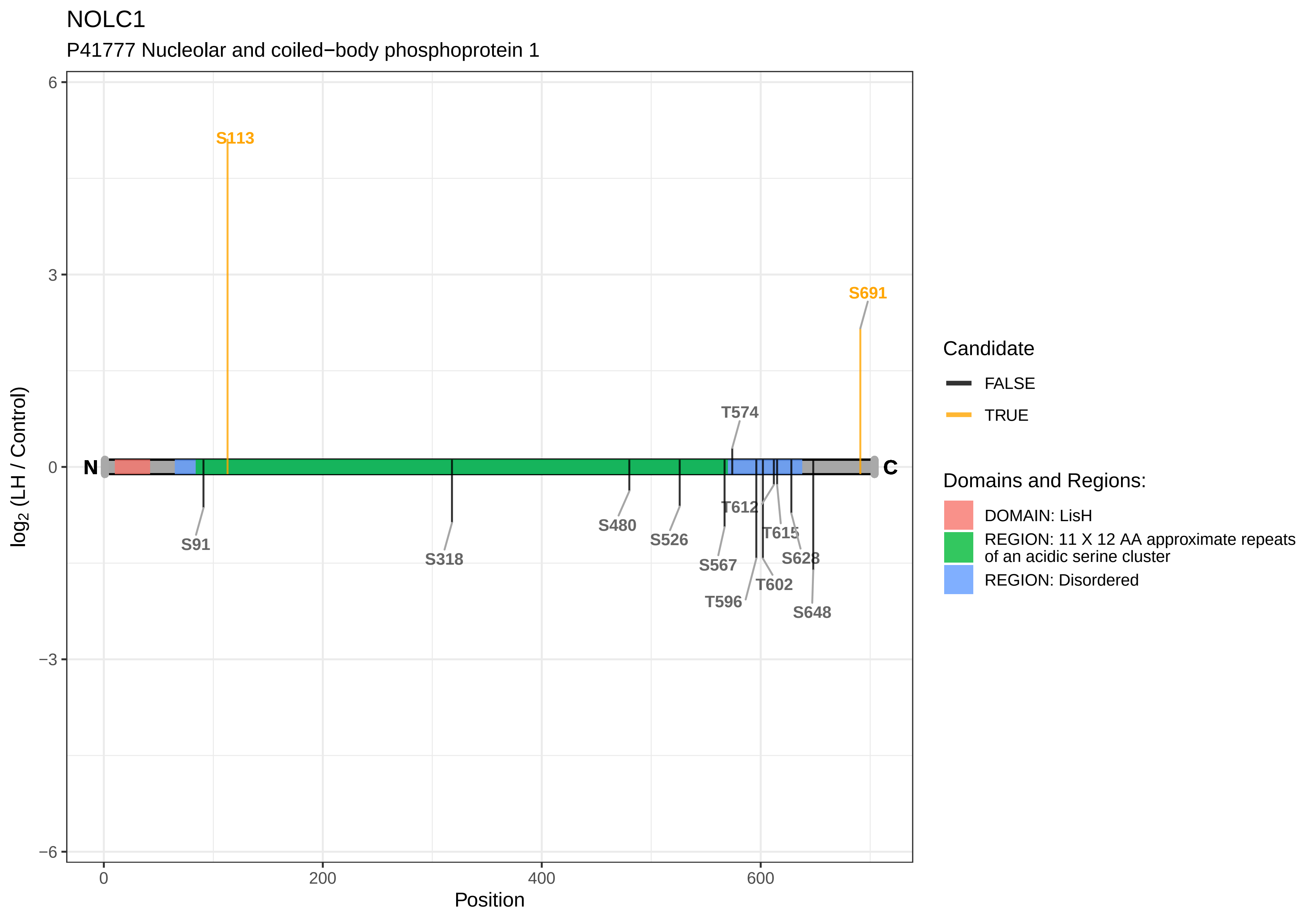

NOLC1
P41777 Nucleolar and coiled−body phosphoprotein 1
S113
S691
T574
N
C
S91
S318
S480
S526
S567
T612
T615
S628
T602
T596
S648
6
3
0
−3
−6
log2 (LH / Control)
0
200
400
600
Position
Candidate
FALSE
TRUE
Domains and Regions:
DOMAIN: LisH
REGION: 11 X 12 AA approximate repeats of an acidic serine cluster
REGION: Disordered

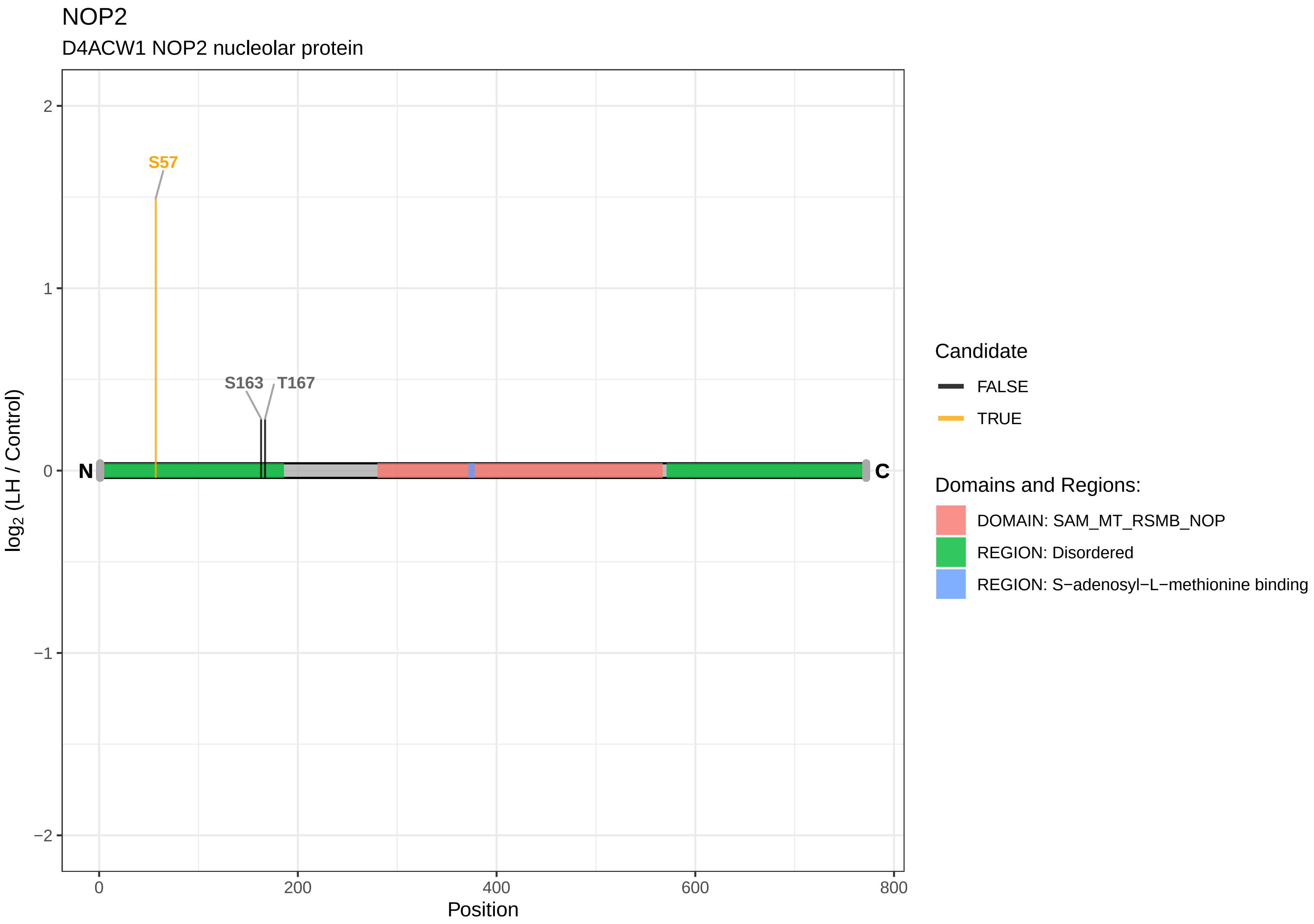
NOP2
D4ACW1 NOP2 nucleolar protein
S57
S163
T167
N
C
2
1
0
−1
−2
$\log_2$ (LH / Control)
0
200
400
600
800
Position
Candidate
FALSE
TRUE
Domains and Regions:
DOMAIN: SAM_MT_RSMB_NOP
REGION: Disordered
REGION: S−adenosyl−L−methionine binding

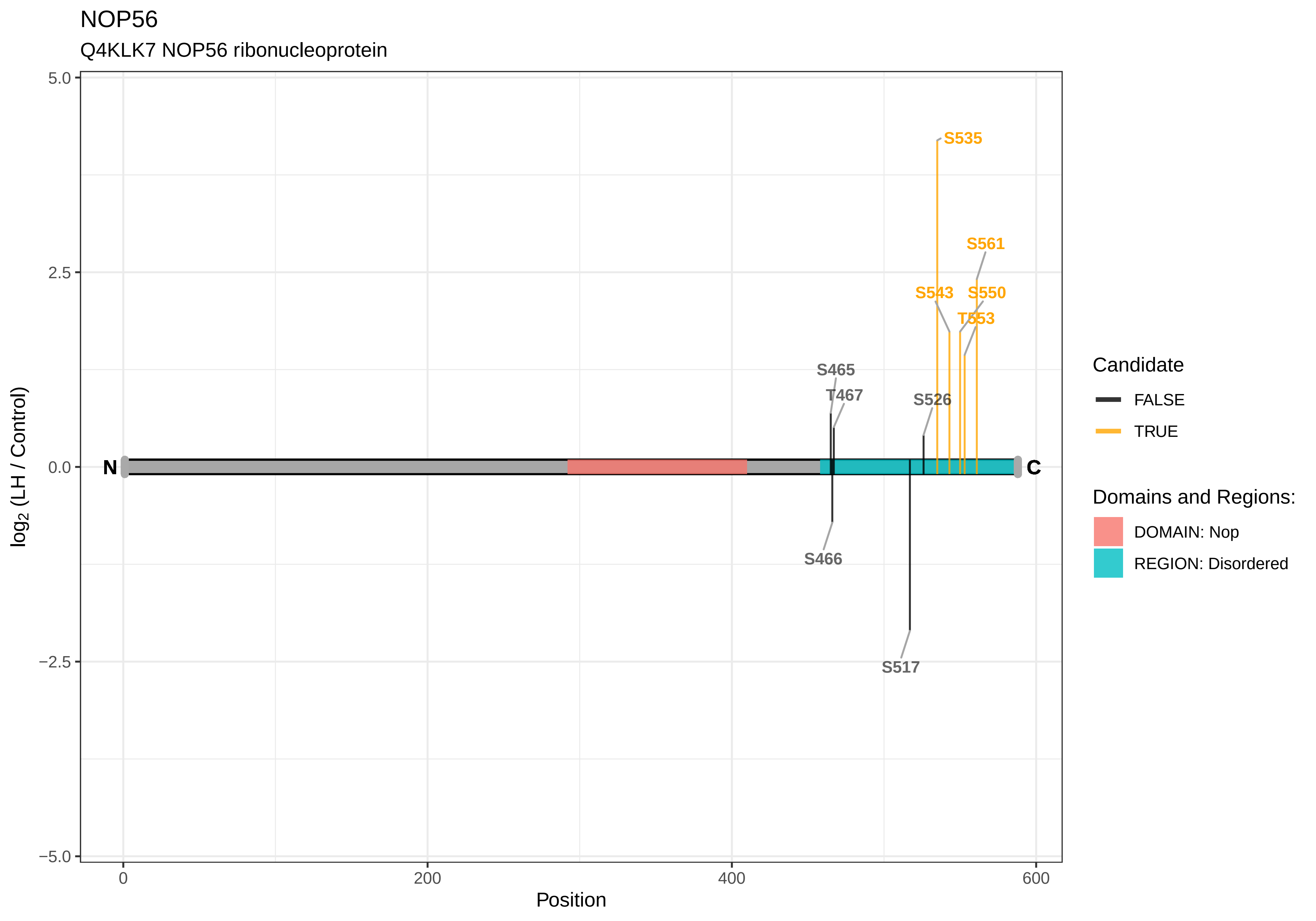
NOP56
Q4KLK7 NOP56 ribonucleoprotein
5.0
2.5
0.0
−2.5
−5.0
log2 (LH / Control)
0
200
400
600
Position
N
C
S535
S561
S543
S550
T553
S465
T467
S526
S466
S517
Candidate
FALSE
TRUE
Domains and Regions:
DOMAIN: Nop
REGION: Disordered

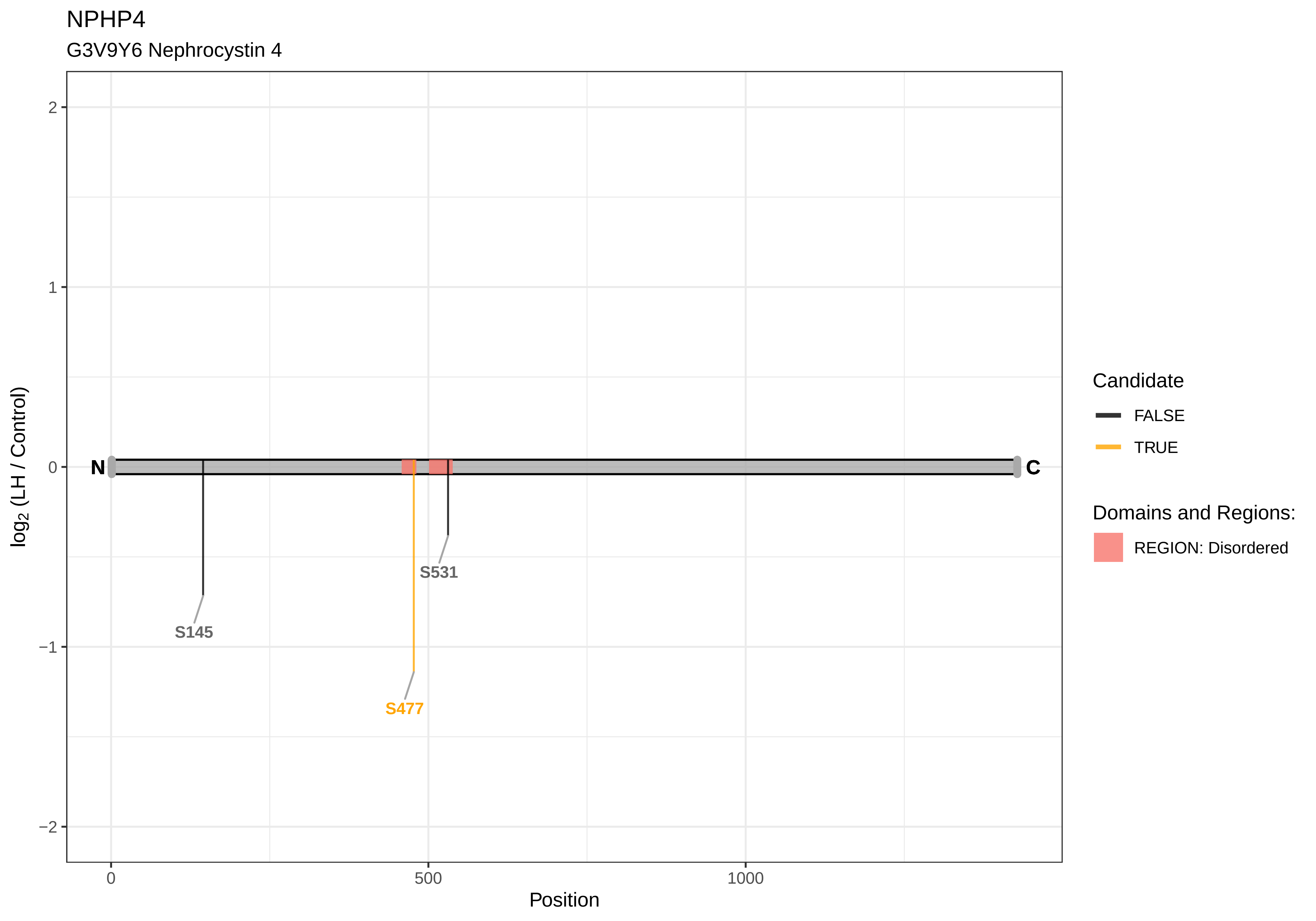

NPHP4
G3V9Y6 Nephrocystin 4
log2 (LH / Control)
2
1
0
−1
−2
N
C
S145
S477
S531
0
500
1000
Position
Candidate
FALSE
TRUE
Domains and Regions:
REGION: Disordered

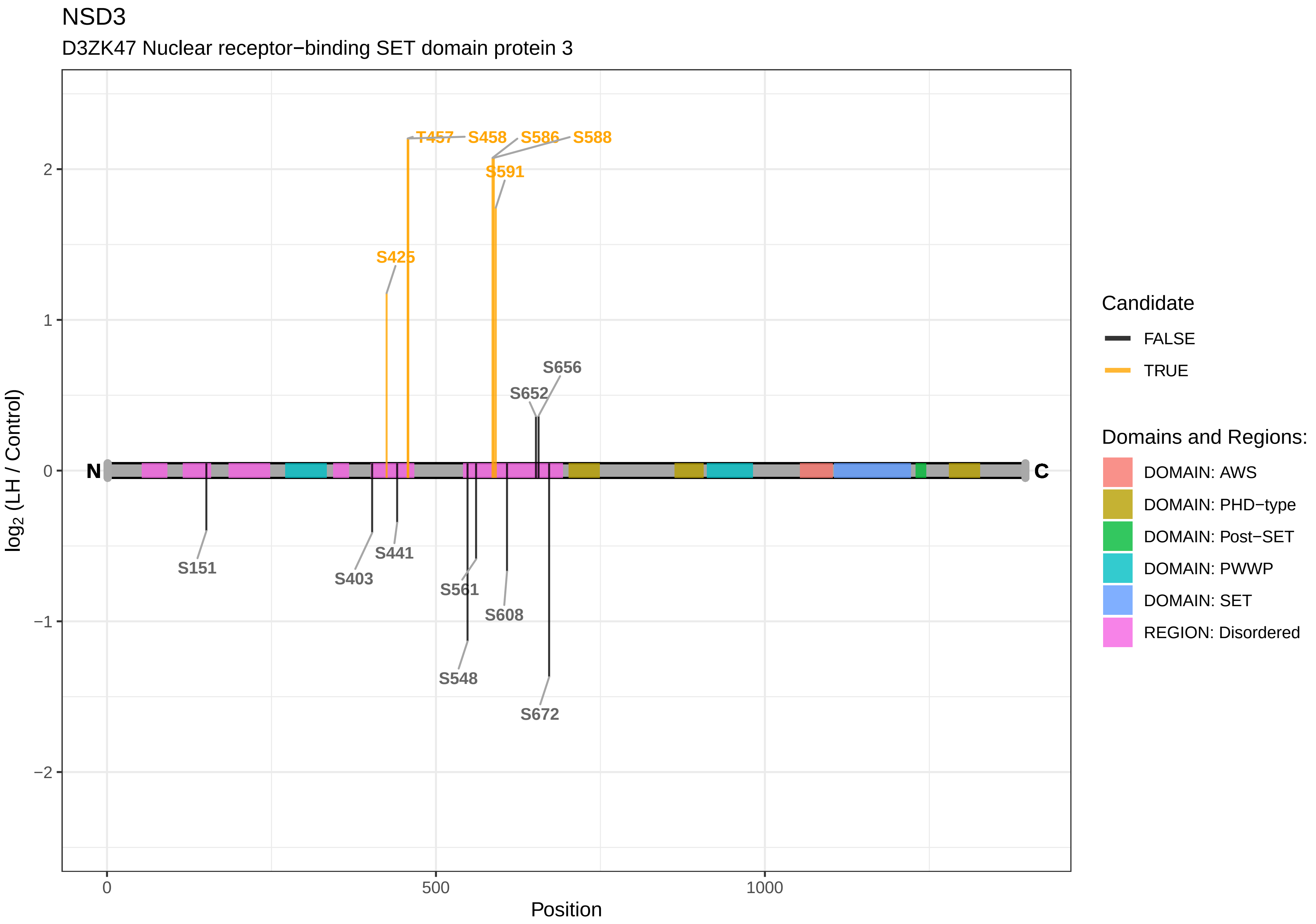
NSD3
D3ZK47 Nuclear receptor−binding SET domain protein 3
log2 (LH / Control)
Position
0
500
1000
2
1
0
−1
−2
N
C
T457
S458
S586
S588
S591
S425
S656
S652
S151
S403
S441
S561
S608
S548
S672
Candidate
FALSE
TRUE
Domains and Regions:
DOMAIN: AWS
DOMAIN: PHD−type
DOMAIN: Post−SET
DOMAIN: PWWP
DOMAIN: SET
REGION: Disordered

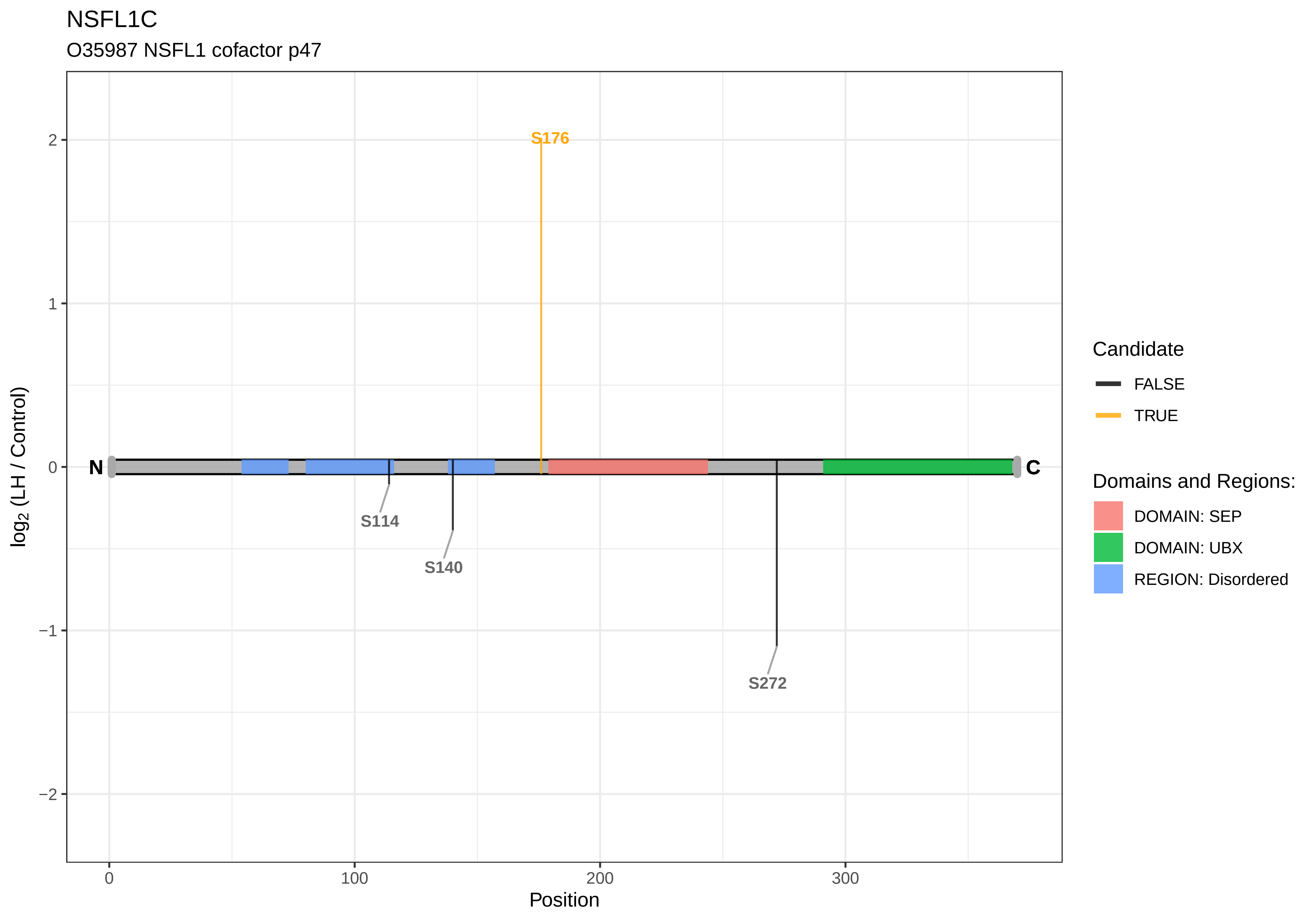

NSFL1C
O35987 NSFL1 cofactor p47
S176
S114
S140
S272
N
C
log2 (LH / Control)
Position
2
1
0
−1
−2
0
100
200
300
Candidate
FALSE
TRUE
Domains and Regions:
DOMAIN: SEP
DOMAIN: UBX
REGION: Disordered

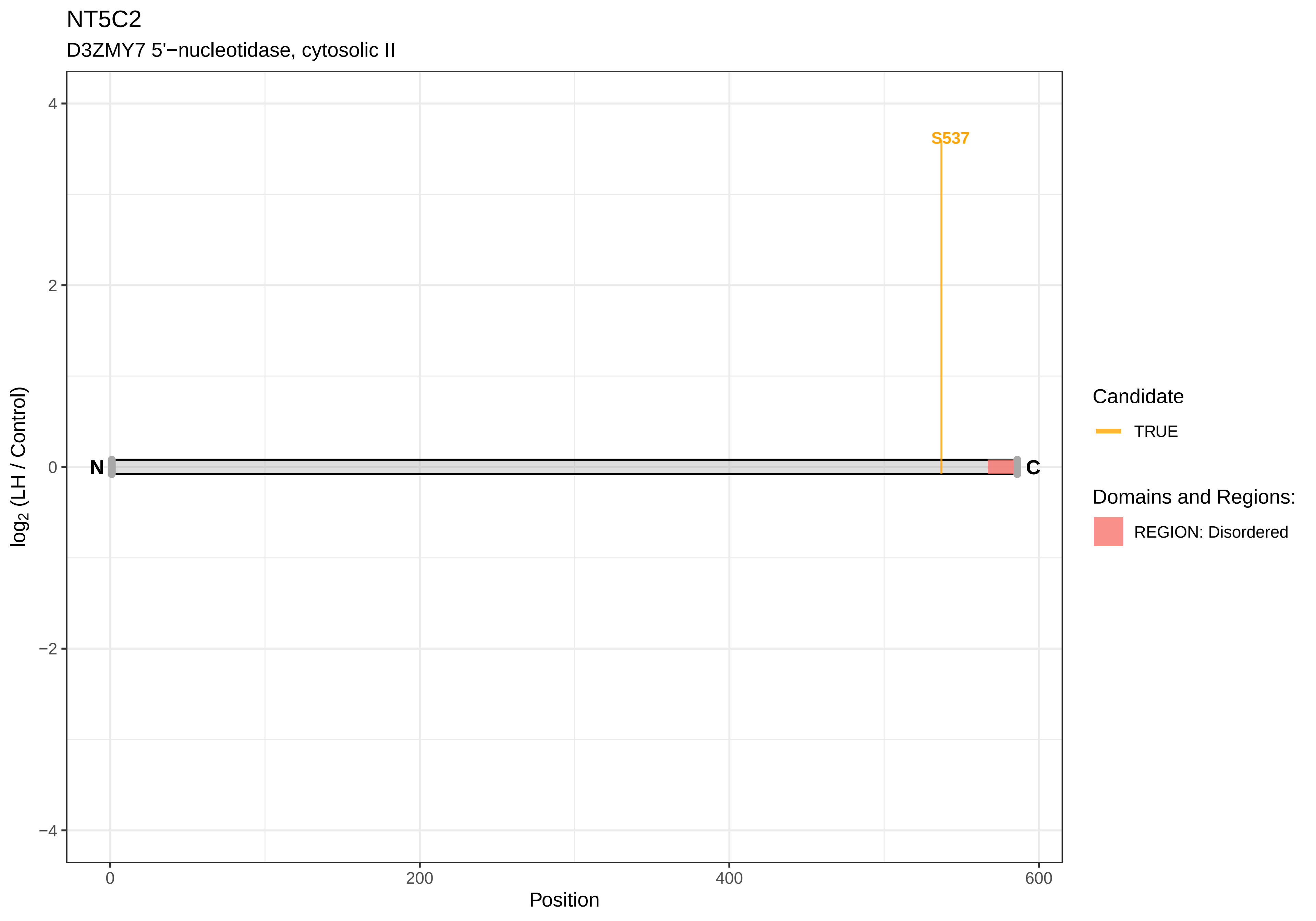

NT5C2
D3ZMY7 5'−nucleotidase, cytosolic II
S537
N
C
4
2
0
−2
−4
0
200
400
600
Position
log2 (LH / Control)
Candidate
TRUE
Domains and Regions:
REGION: Disordered

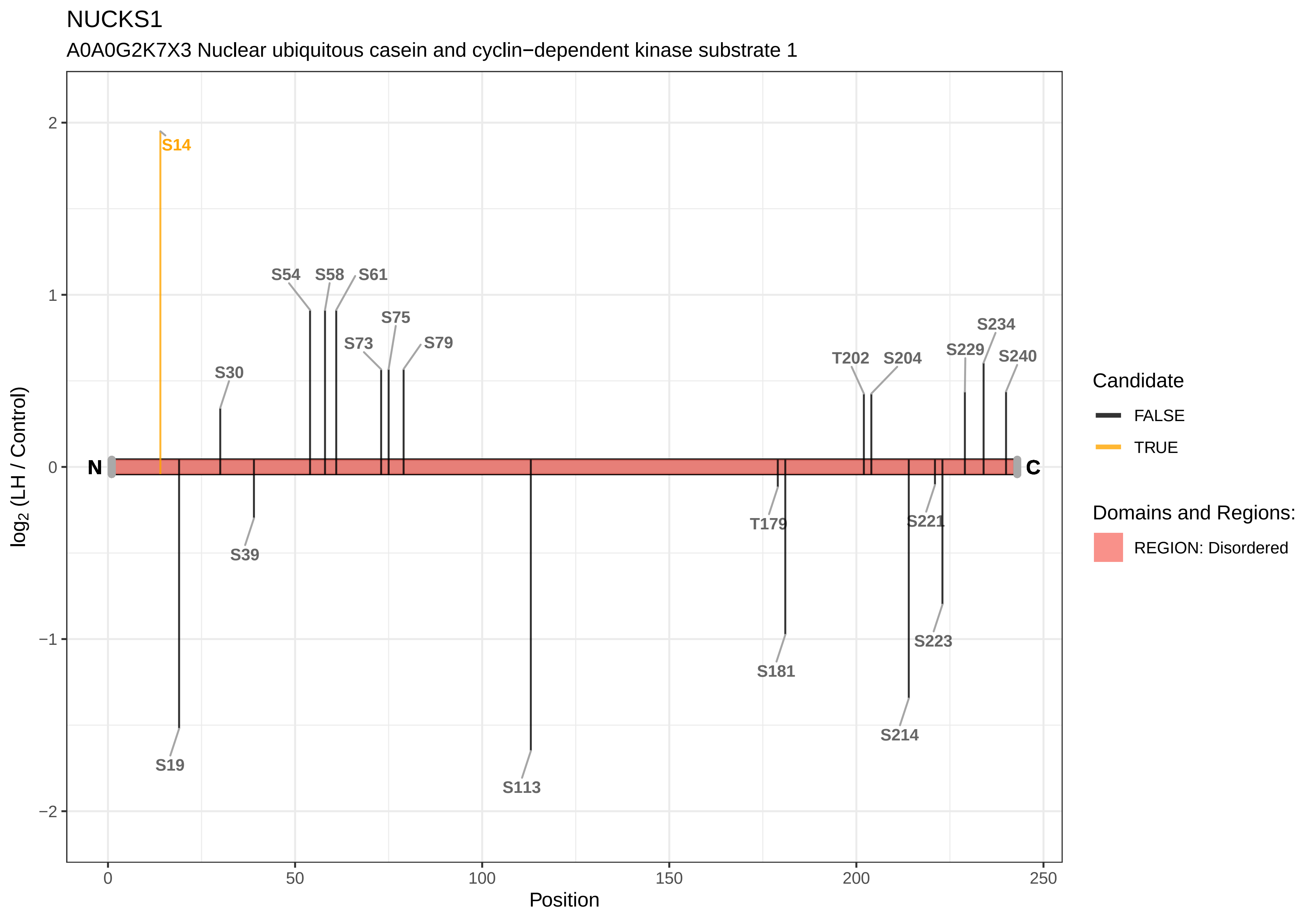

NUCKS1
A0A0G2K7X3 Nuclear ubiquitous casein and cyclin−dependent kinase substrate 1
S14
S54
S58
S61
S75
S73
S79
S30
T202
S204
S229
S234
S240
N
C
S19
S39
S113
T179
S181
S221
S223
S214
log2 (LH / Control)
Position
2
1
0
−1
−2
0
50
100
150
200
250
Candidate
FALSE
TRUE
Domains and Regions:
REGION: Disordered

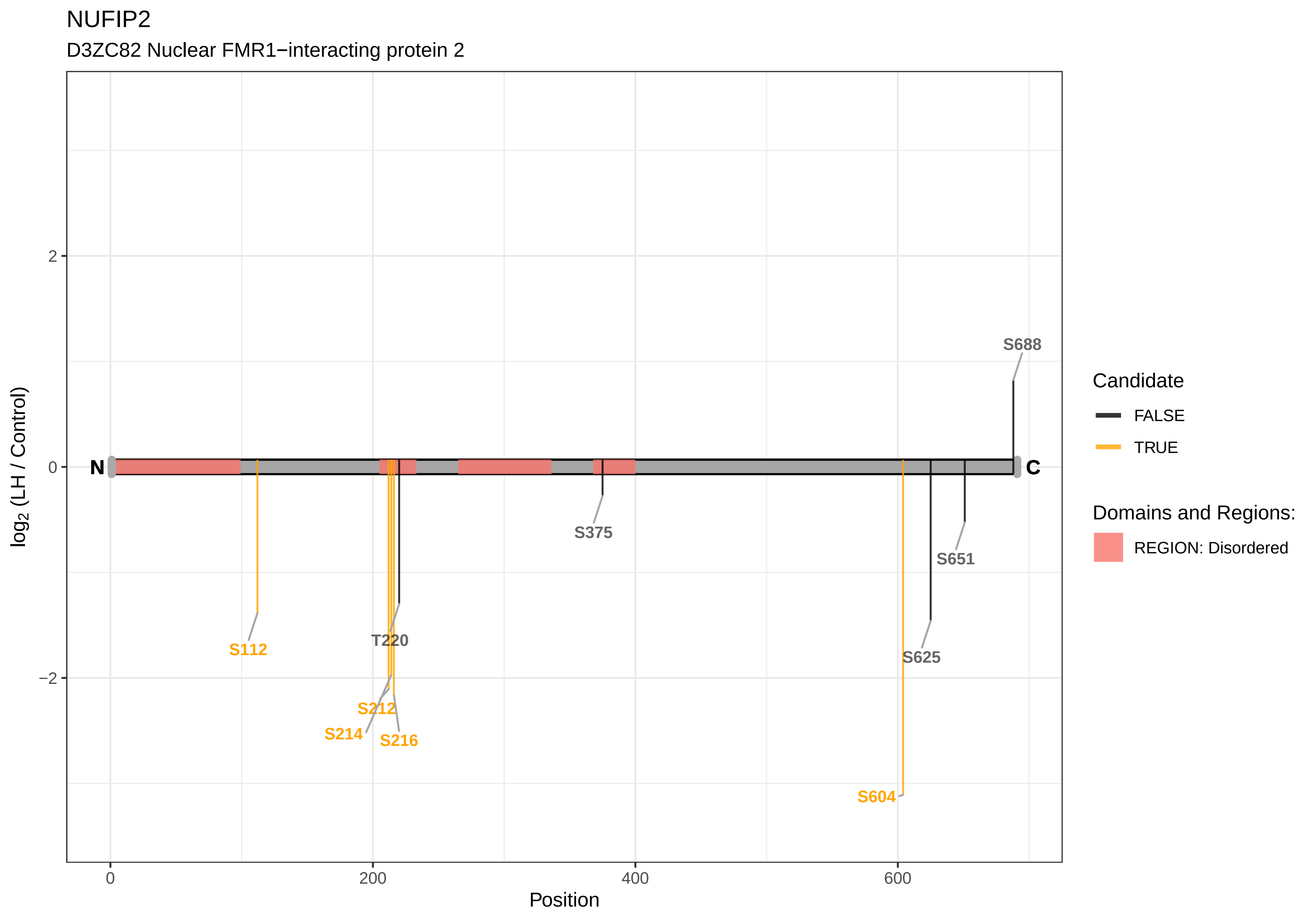

NUFIP2
D3ZC82 Nuclear FMR1−interacting protein 2
log2 (LH / Control)
2
0
−2
0
200
400
600
Position
N
C
S688
S375
S651
S625
T220
S112
S212
S214
S216
S604
Candidate
FALSE
TRUE
Domains and Regions:
REGION: Disordered

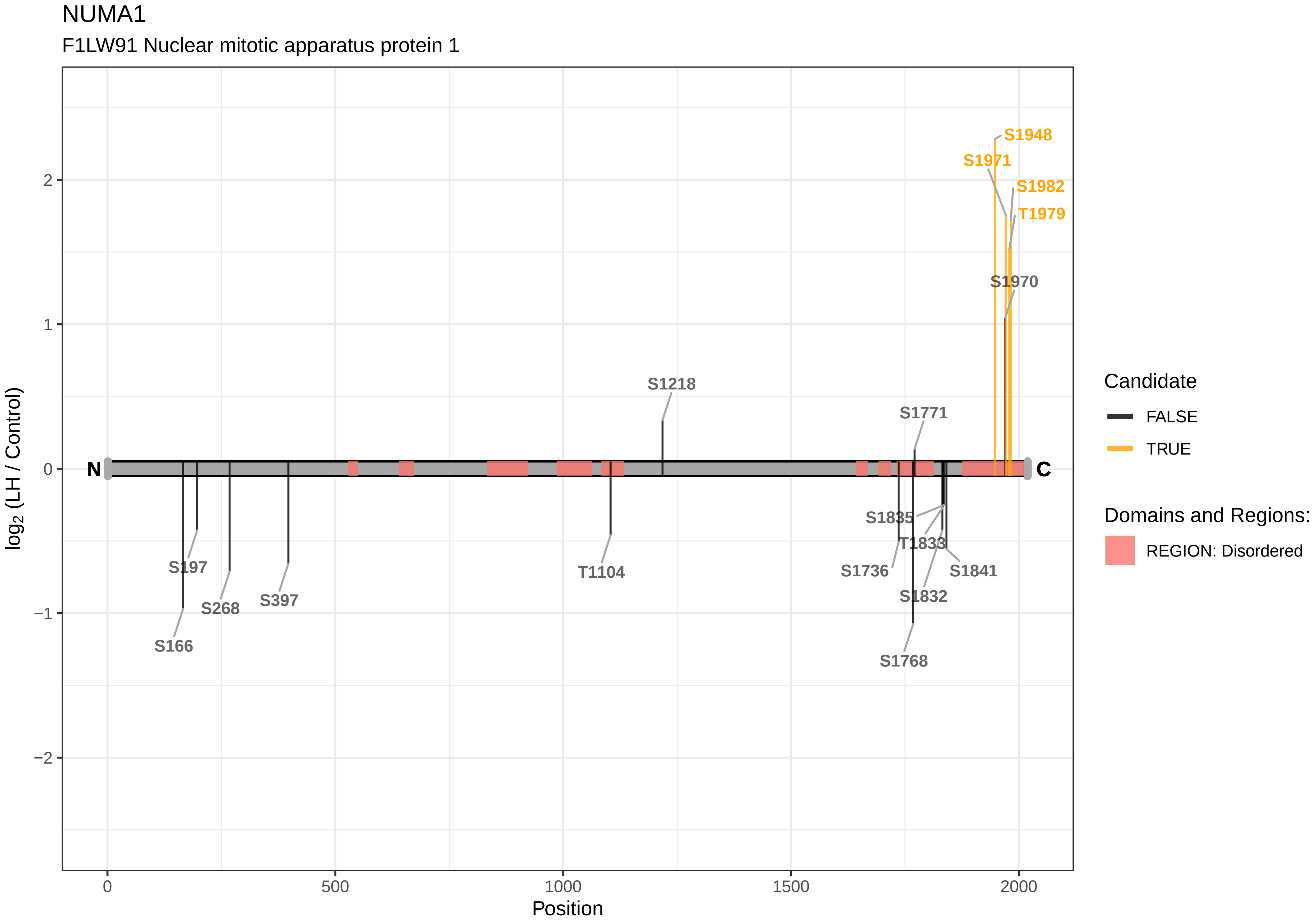

NUMA1
F1LW91 Nuclear mitotic apparatus protein 1
S1948
S1971
S1982
T1979
S1970
S1218
S1771
S1835
T1833
S1736
S1841
S1832
S1768
S197
S268
S397
S166
T1104
N
C
log2 (LH / Control)
Position
Candidate
FALSE
TRUE
Domains and Regions:
REGION: Disordered

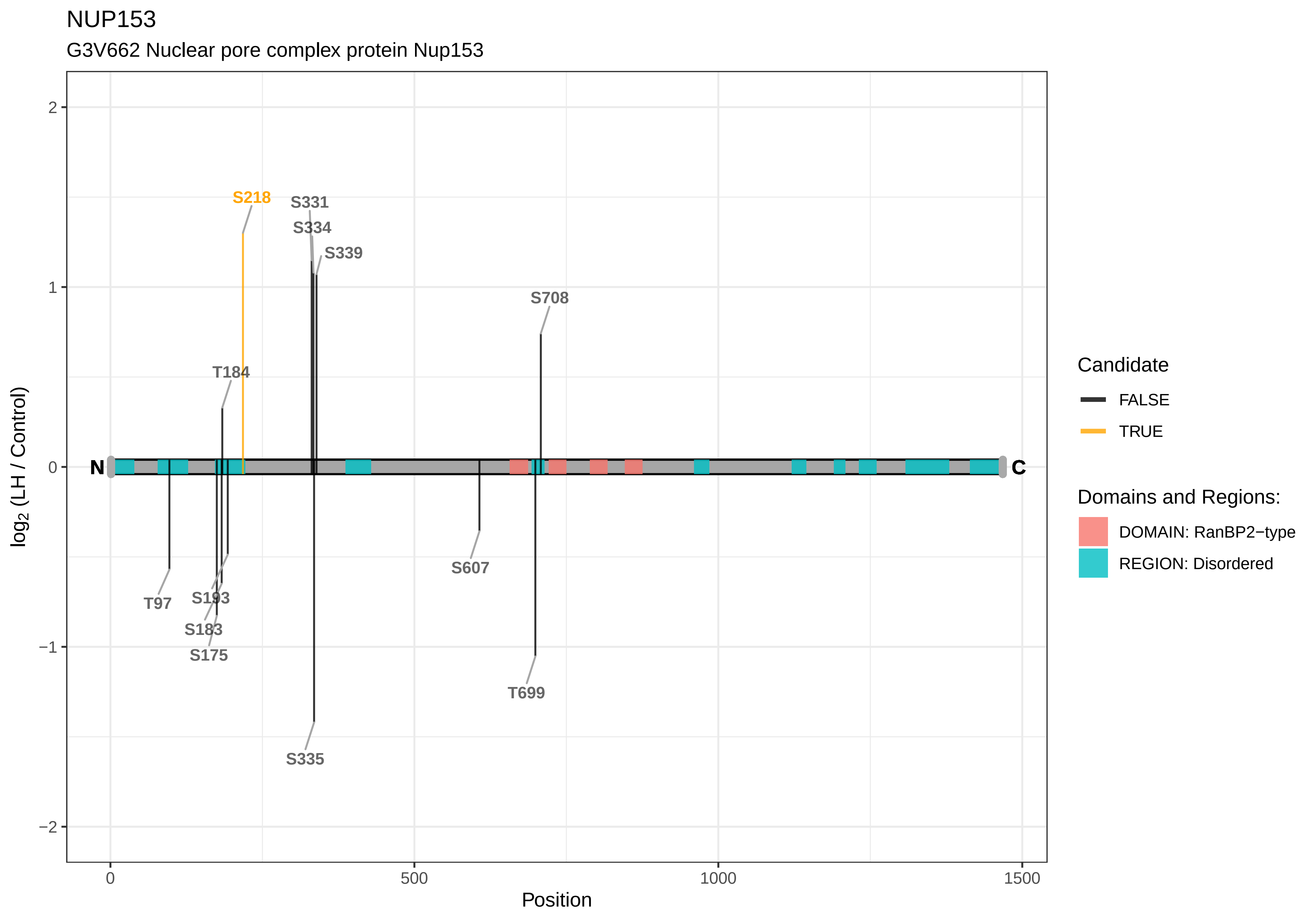

NUP153
G3V662 Nuclear pore complex protein Nup153
S218
S331
S334
S339
S708
T184
N
C
T97
S193
S183
S175
S607
T699
S335
log2 (LH / Control)
Position
0
500
1000
1500
2
1
0
−1
−2
Candidate
FALSE
TRUE
Domains and Regions:
DOMAIN: RanBP2−type
REGION: Disordered

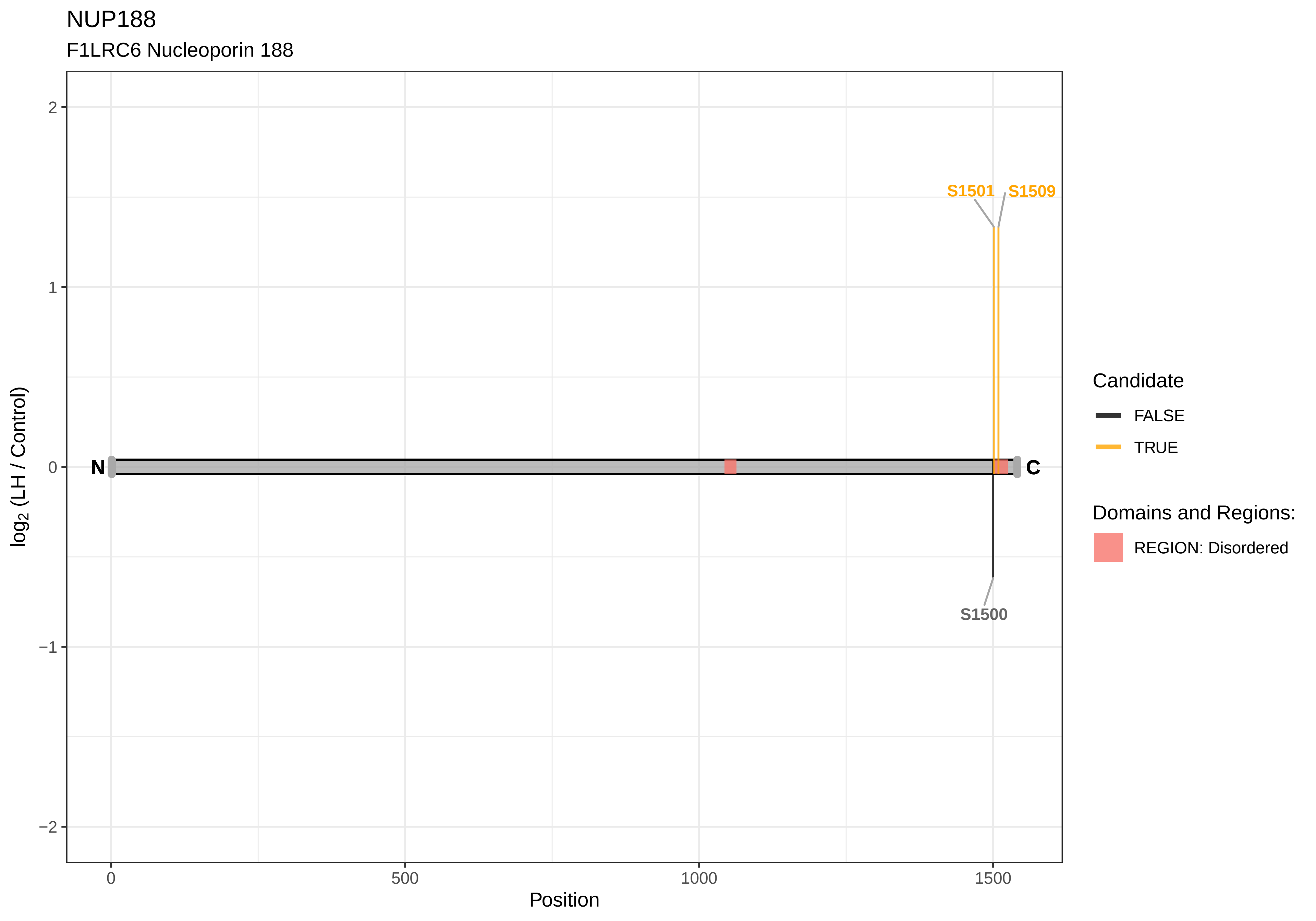
NUP188
F1LRC6 Nucleoporin 188
S1501
S1509
S1500
N
C
2
1
0
−1
−2
0
500
1000
1500
Position
log2 (LH / Control)
Candidate
FALSE
TRUE
Domains and Regions:
REGION: Disordered

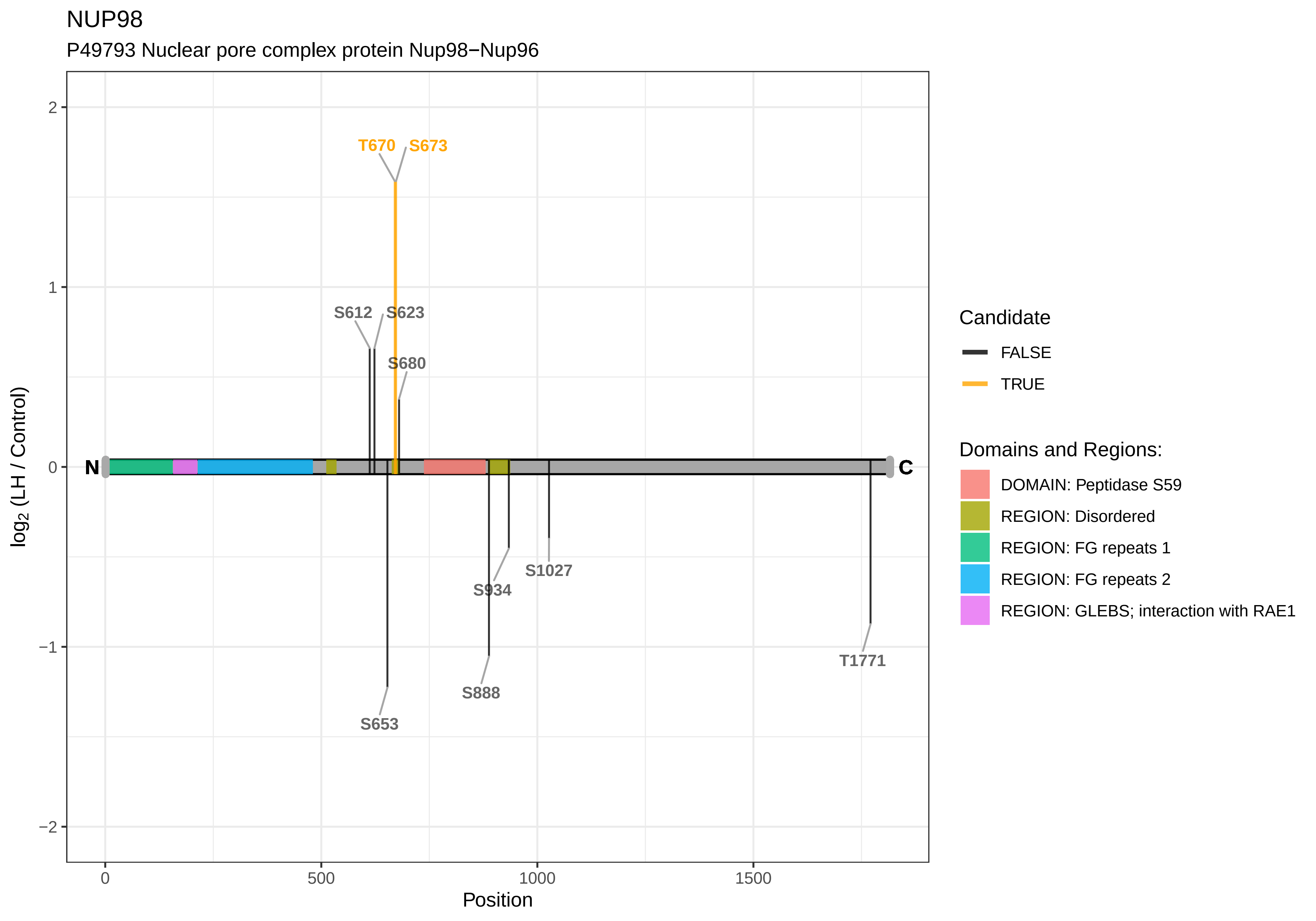

NUP98
P49793 Nuclear pore complex protein Nup98−Nup96
T670
S673
S612
S623
S680
S653
S888
S934
S1027
T1771
N
C
2
1
0
−1
−2
log2 (LH / Control)
0
500
1000
1500
Position
Candidate
FALSE
TRUE
Domains and Regions:
DOMAIN: Peptidase S59
REGION: Disordered
REGION: FG repeats 1
REGION: FG repeats 2
REGION: GLEBS; interaction with RAE1

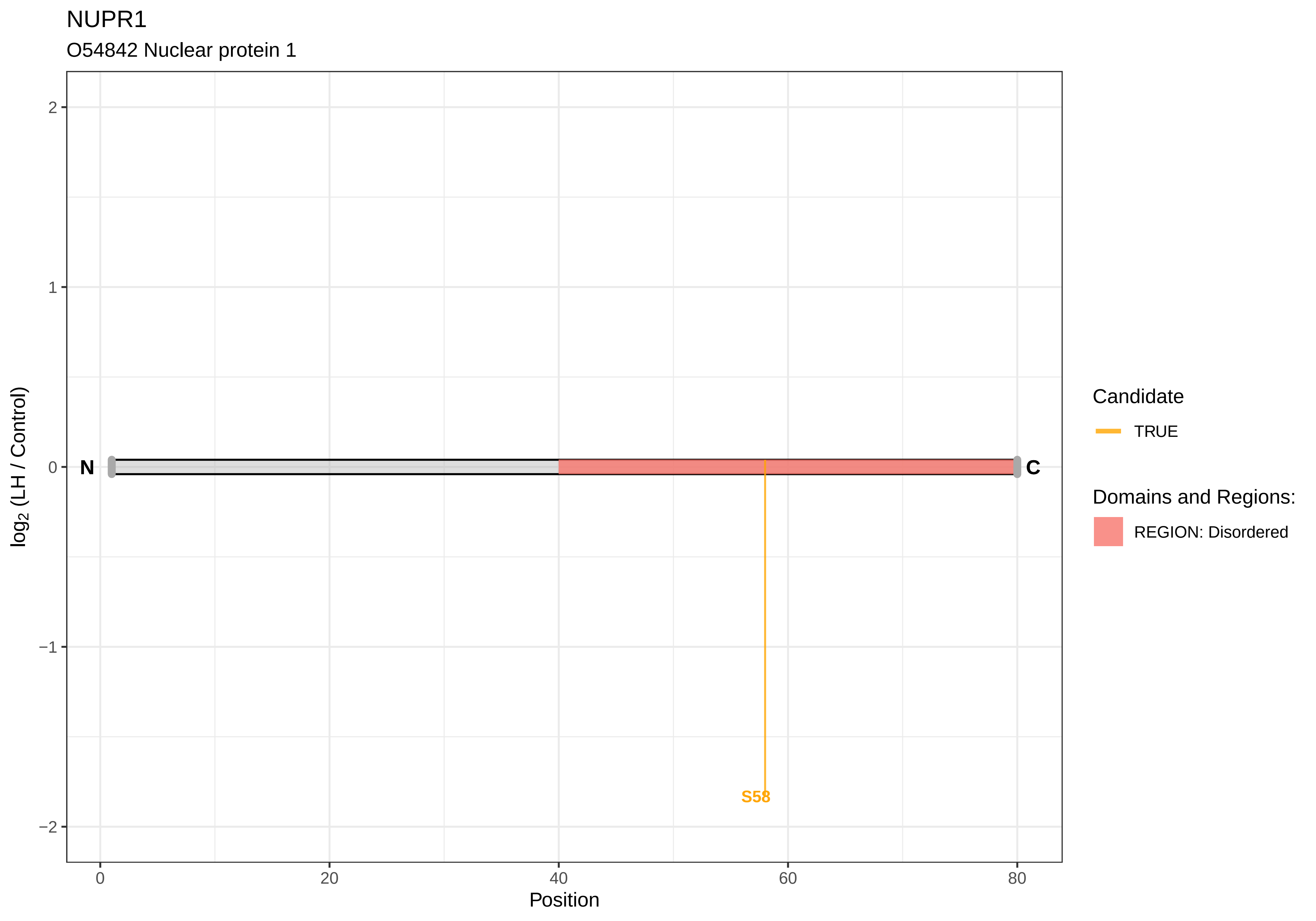
NUPR1
O54842 Nuclear protein 1
log2 (LH / Control)
2
1
0
−1
−2
N
C
S58
0
20
40
60
80
Position
Candidate
TRUE
Domains and Regions:
REGION: Disordered

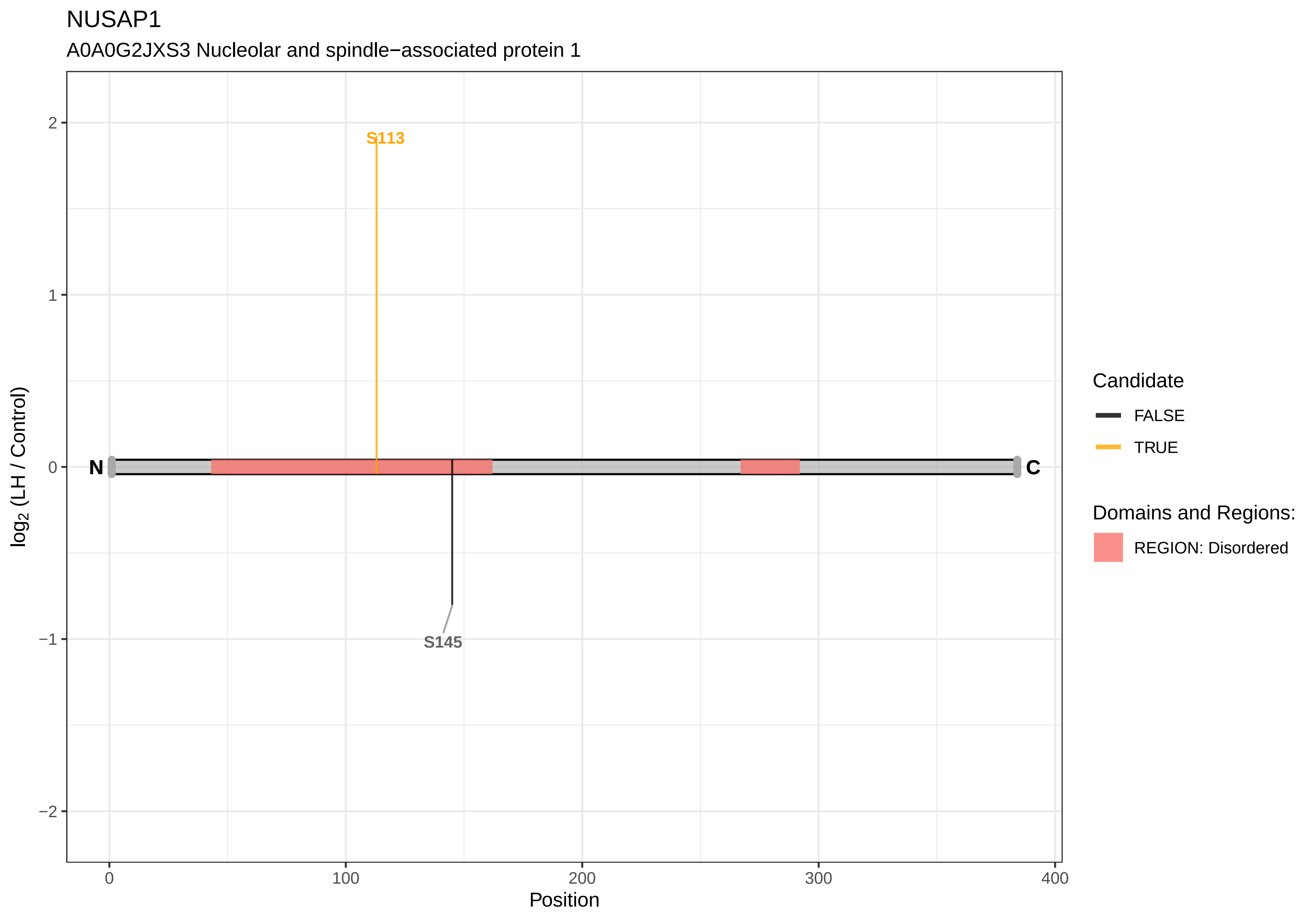
NUSAP1
A0A0G2JXS3 Nucleolar and spindle−associated protein 1
S113
S145
N
C
2
1
0
−1
−2
log2 (LH / Control)
0
100
200
300
400
Position
Candidate
FALSE
TRUE
Domains and Regions:
REGION: Disordered

OCIAD1

Q5XIG4 OCIA domain−containing protein 1

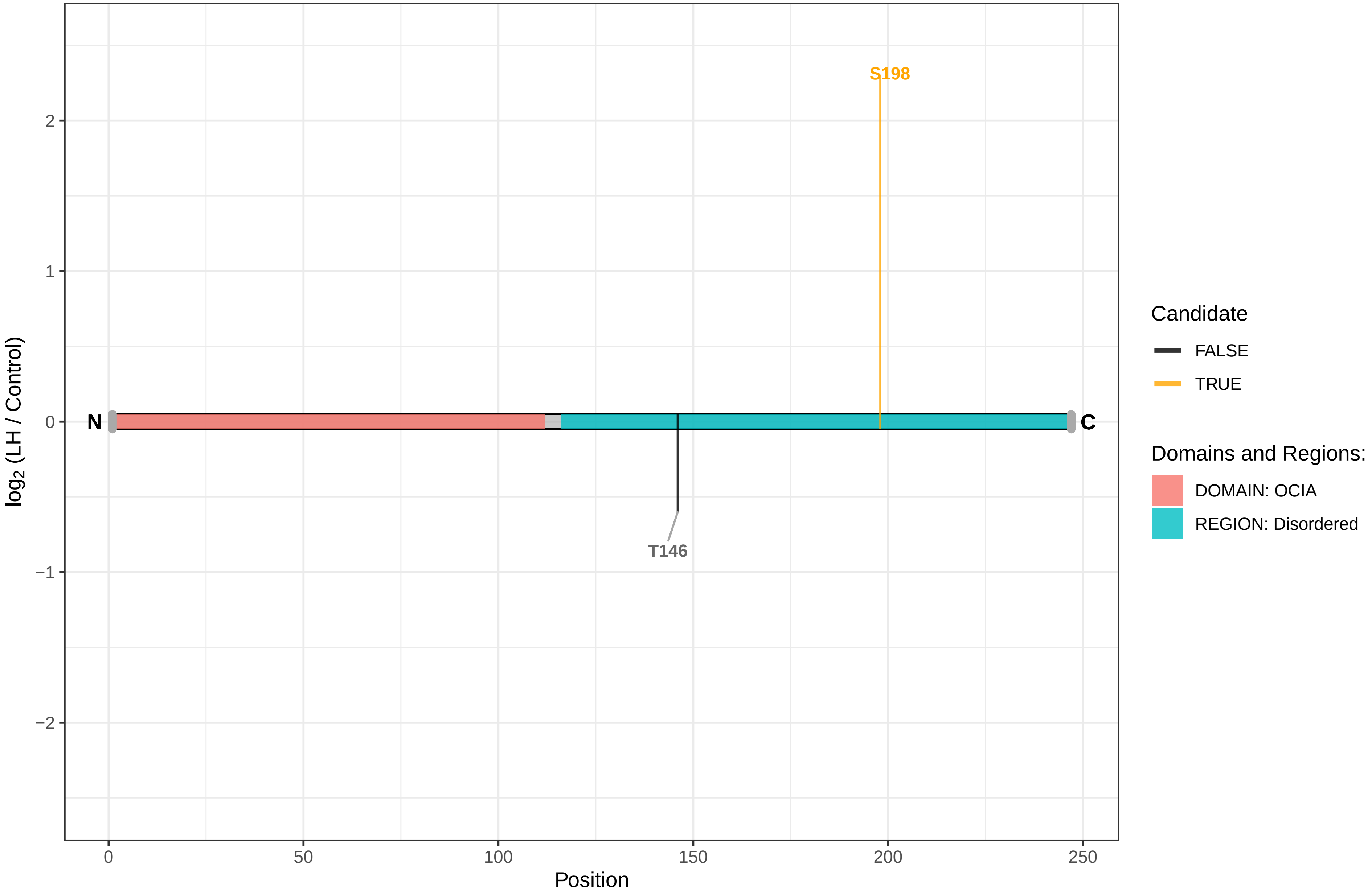

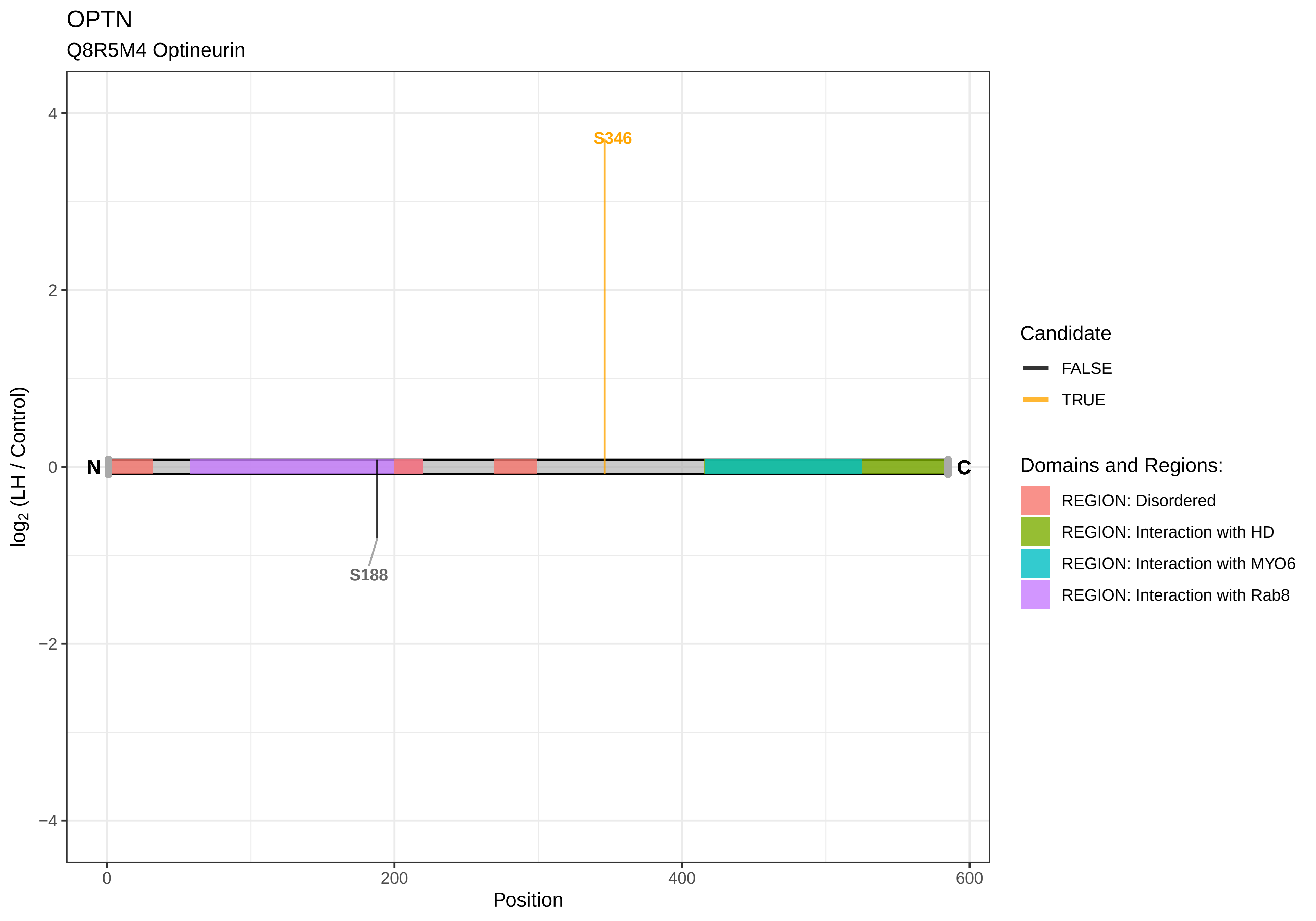
OPTN
Q8R5M4 Optineurin
S346
S188
N
C
log2 (LH / Control)
Position
Candidate
FALSE
TRUE
Domains and Regions:
REGION: Disordered
REGION: Interaction with HD
REGION: Interaction with MYO6
REGION: Interaction with Rab8

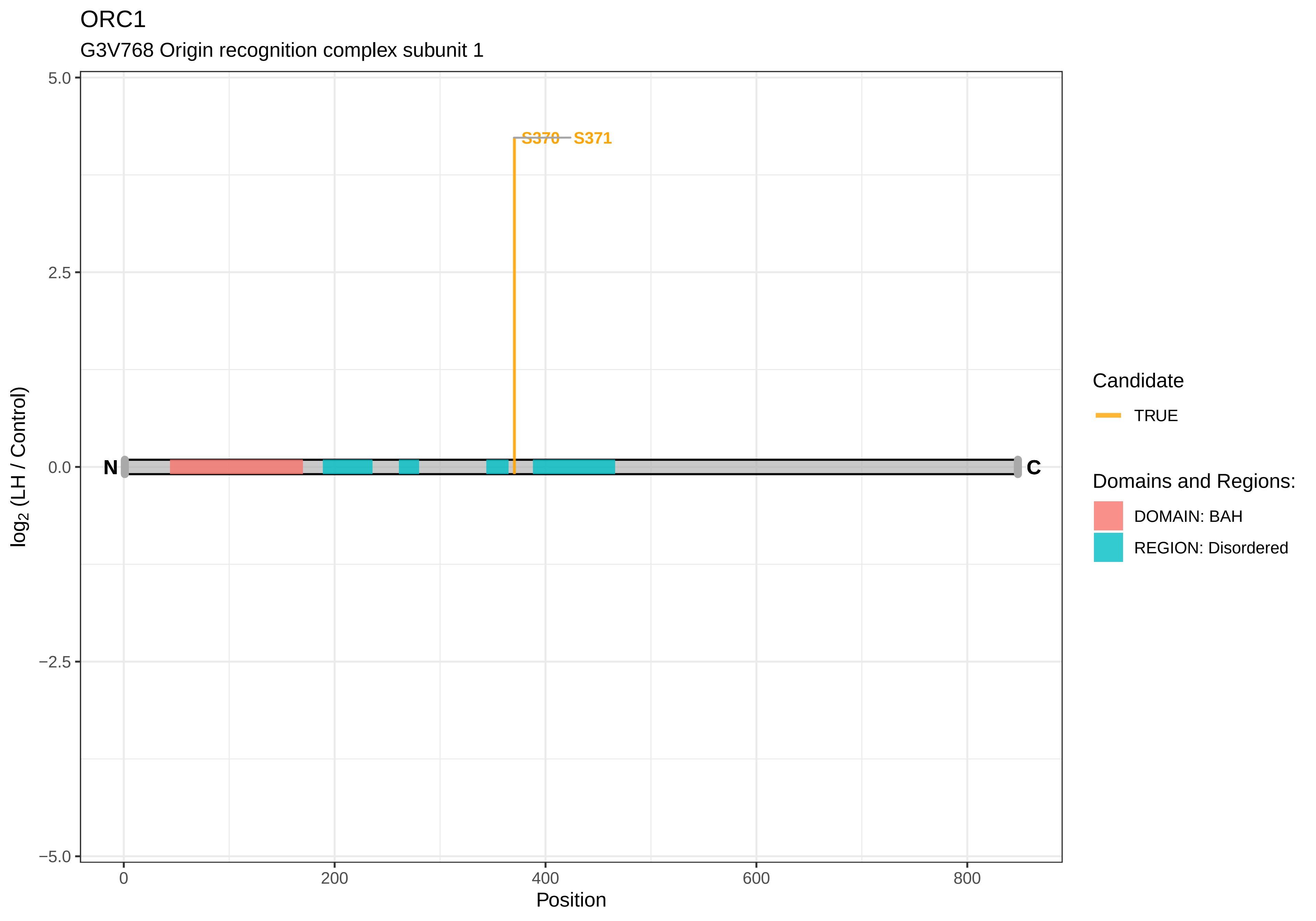

ORC1
G3V768 Origin recognition complex subunit 1
S370
S371
N
C
5.0
2.5
0.0
−2.5
−5.0
0
200
400
600
800
Position
log2 (LH / Control)
Candidate
TRUE
Domains and Regions:
DOMAIN: BAH
REGION: Disordered

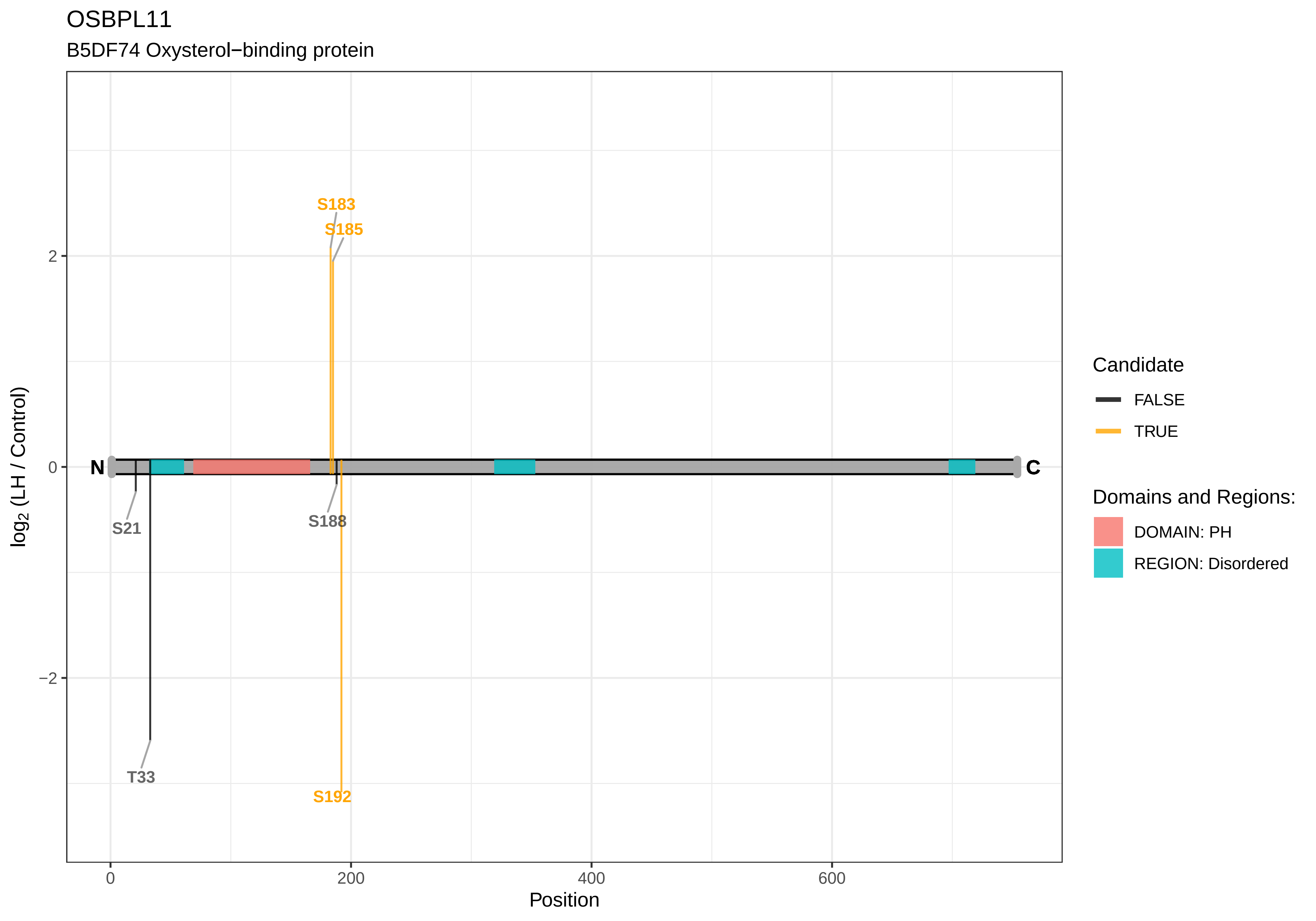

OSBPL11
B5DF74 Oxysterol−binding protein
S183
S185
S21
S188
T33
S192
N
C
2
0
−2
0
200
400
600
Position
log2 (LH / Control)
Candidate
FALSE
TRUE
Domains and Regions:
DOMAIN: PH
REGION: Disordered

OSTF1

Q6P686 Osteoclast−stimulating factor 1

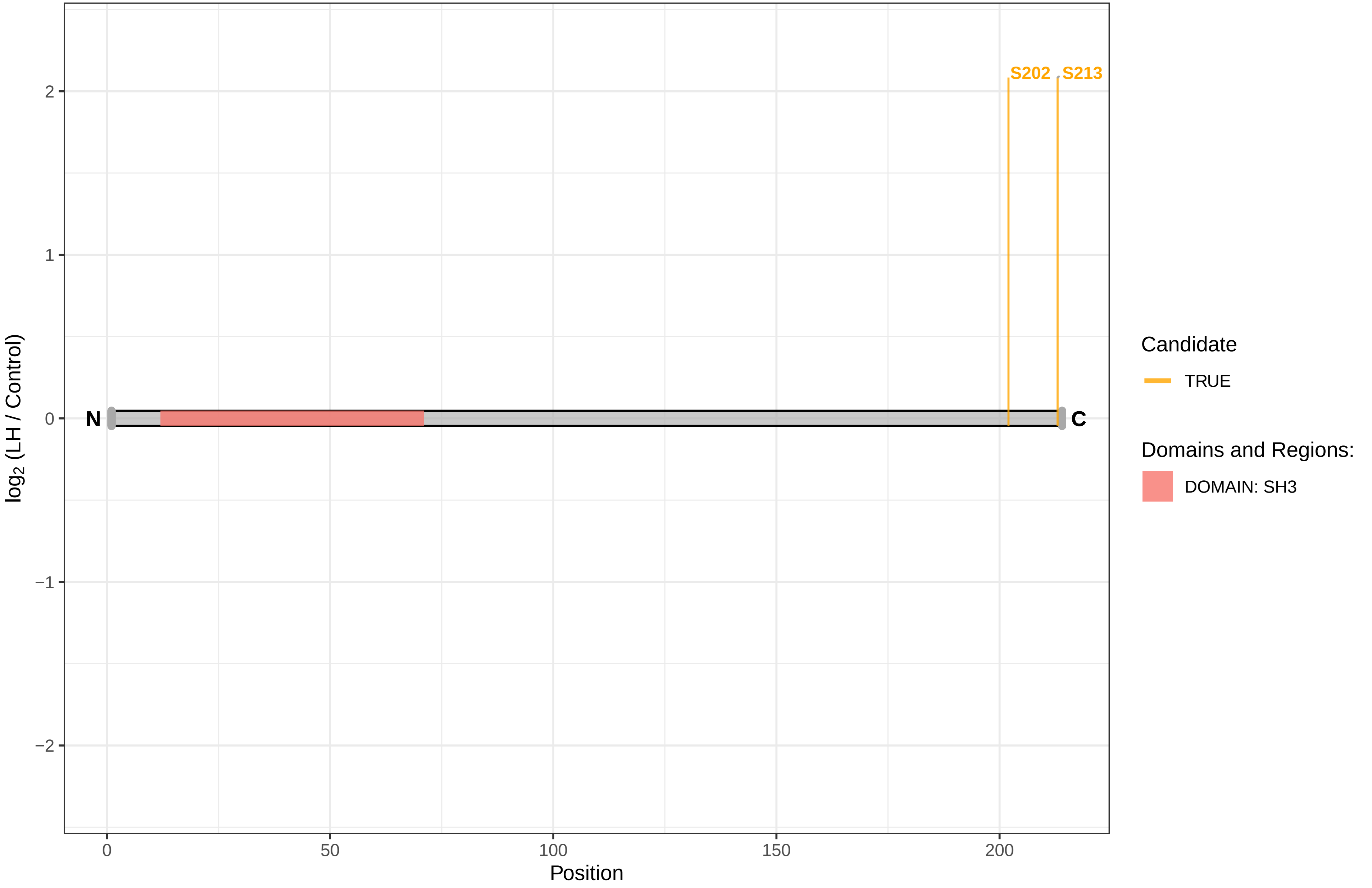

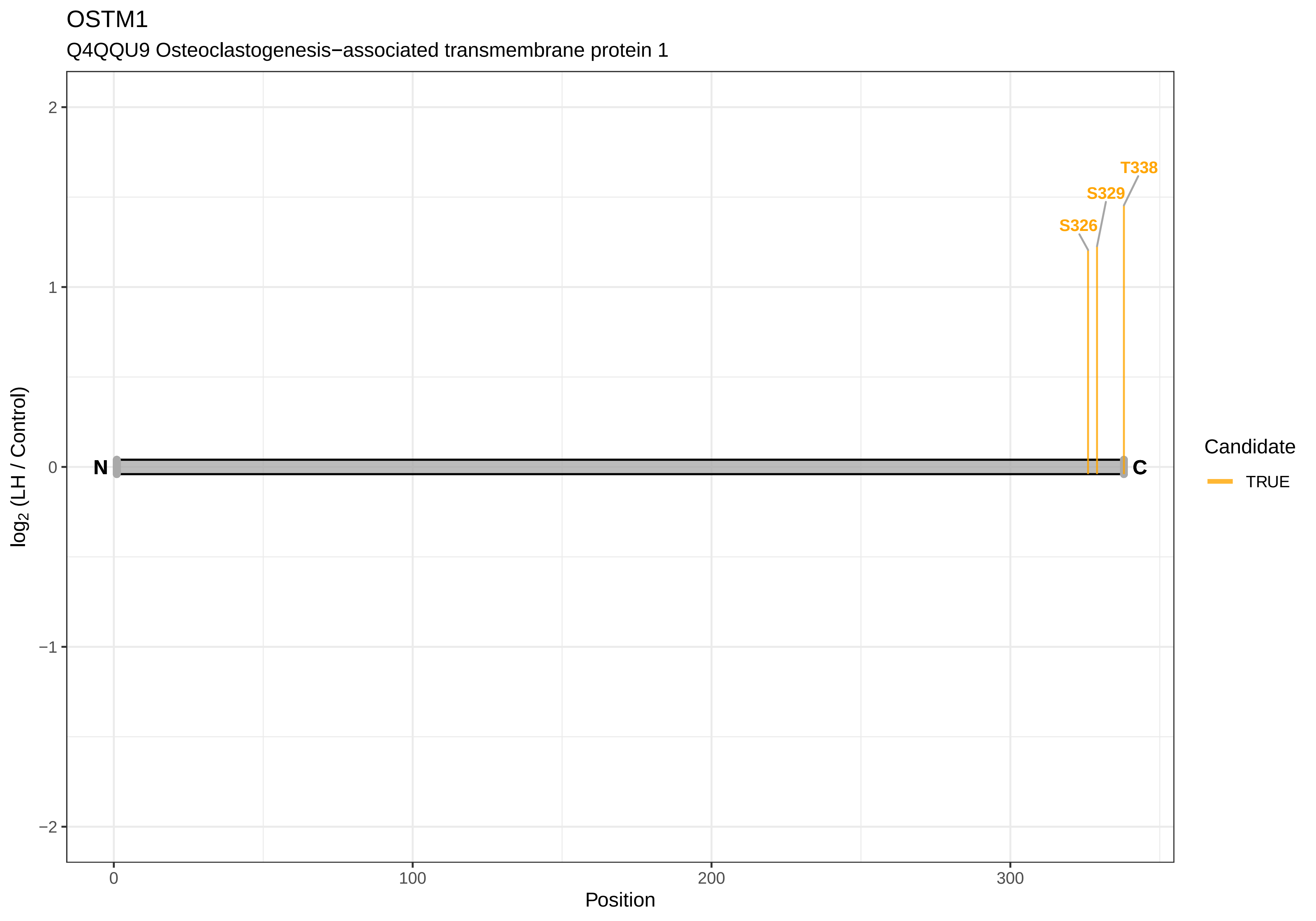
OSTM1
Q4QQU9 Osteoclastogenesis−associated transmembrane protein 1
T338
S329
S326
N
C
2
1
0
−1
−2
log2 (LH / Control)
0
100
200
300
Position
Candidate
TRUE

### OXR1

A0A0G2JZ50 Oxidation resistance protein 1

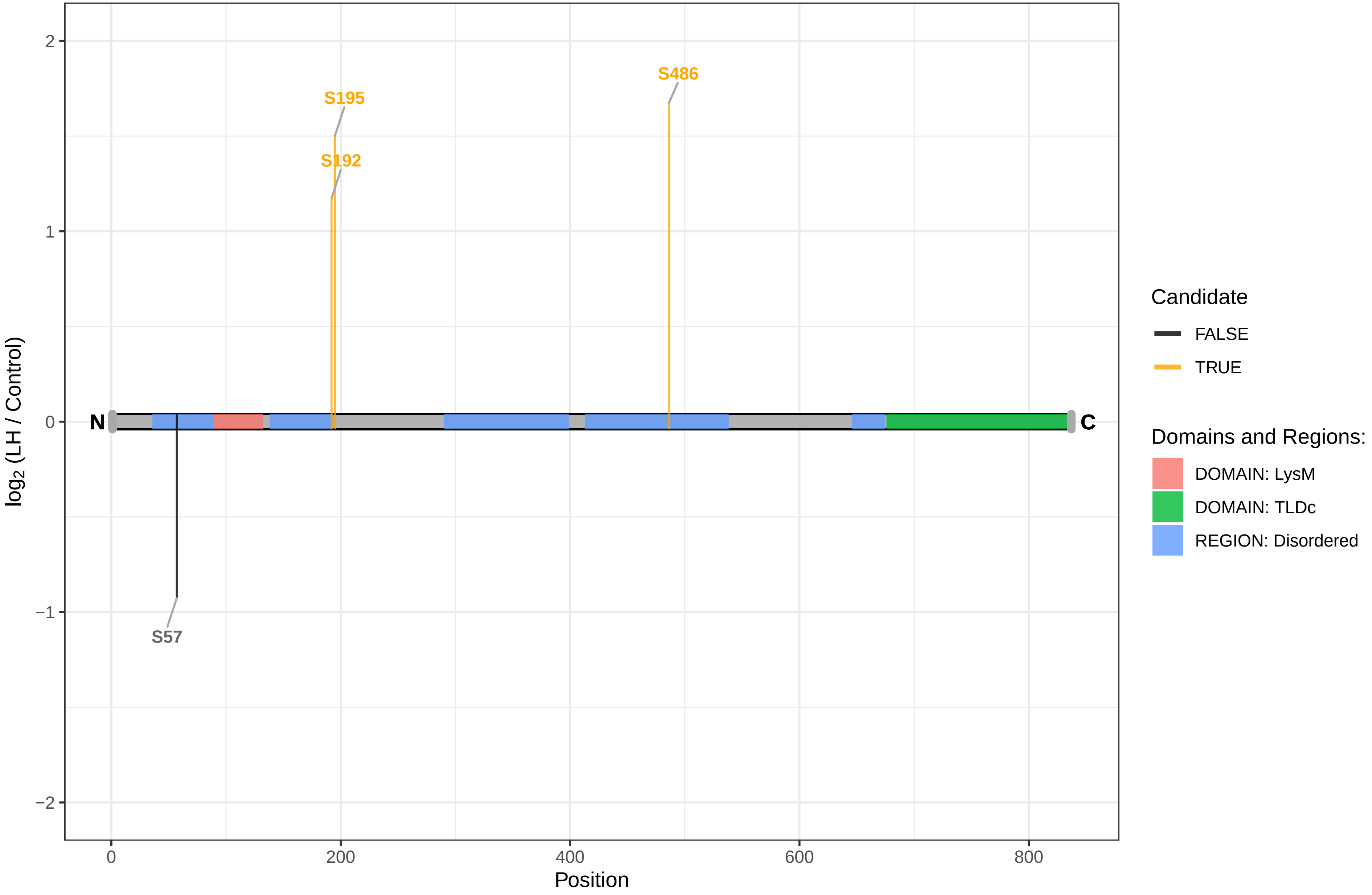

PACRGL

D3ZE96 Parkin coregulated−like

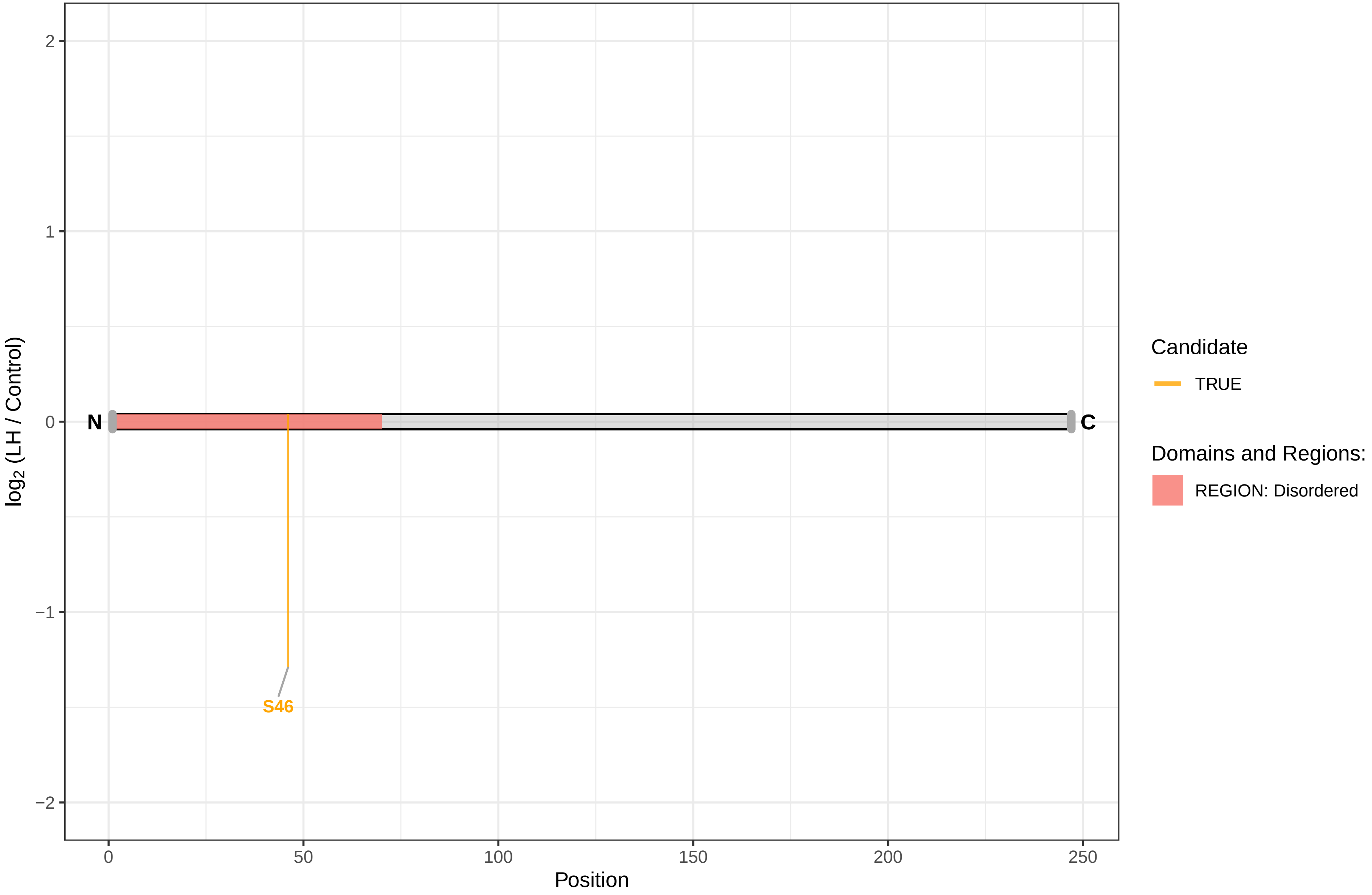

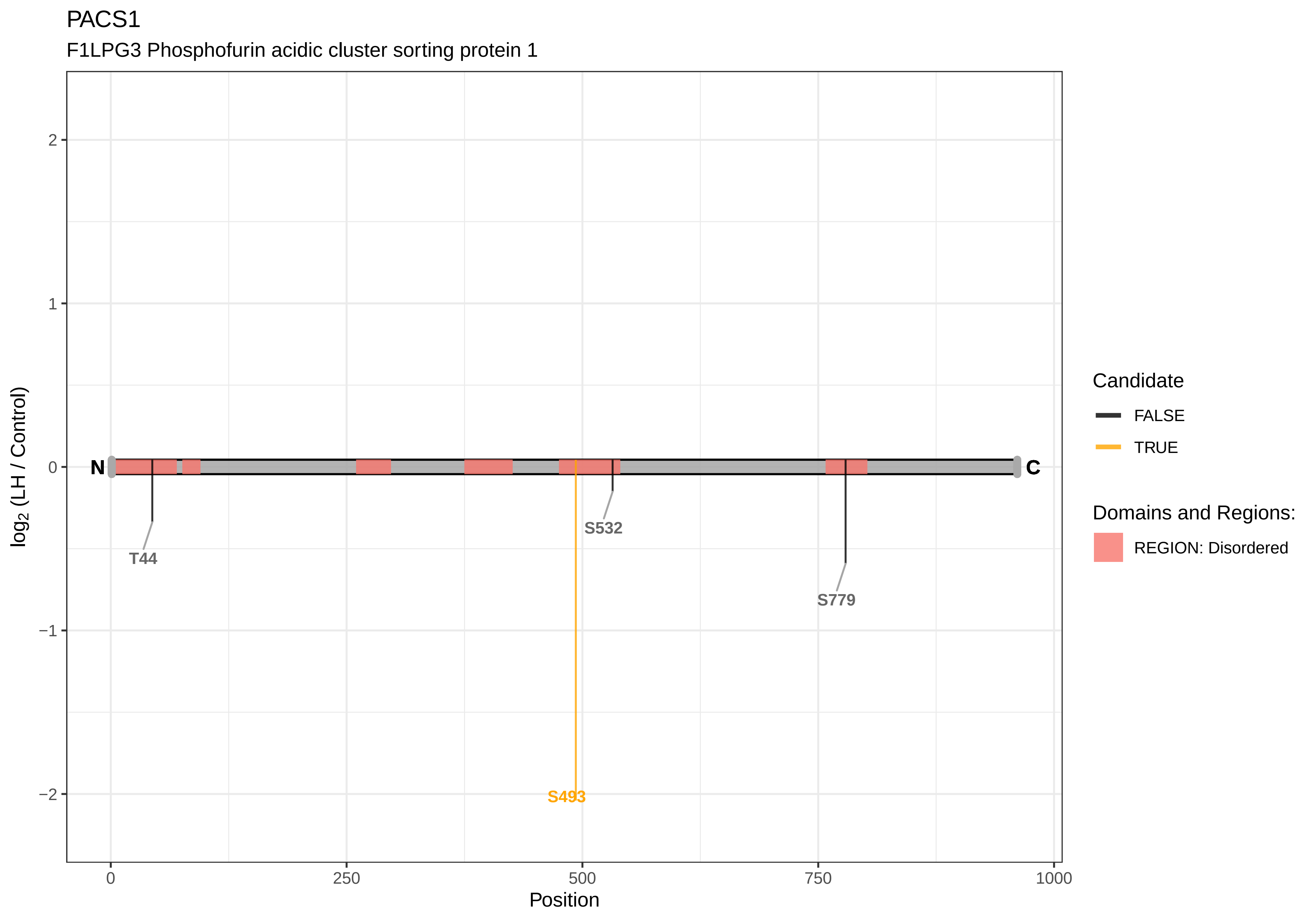
PACS1
F1LPG3 Phosphofurin acidic cluster sorting protein 1
log2 (LH / Control)
2
1
0
−1
−2
N
C
T44
S532
S493
S779
0
250
500
750
1000
Position
Candidate
FALSE
TRUE
Domains and Regions:
REGION: Disordered

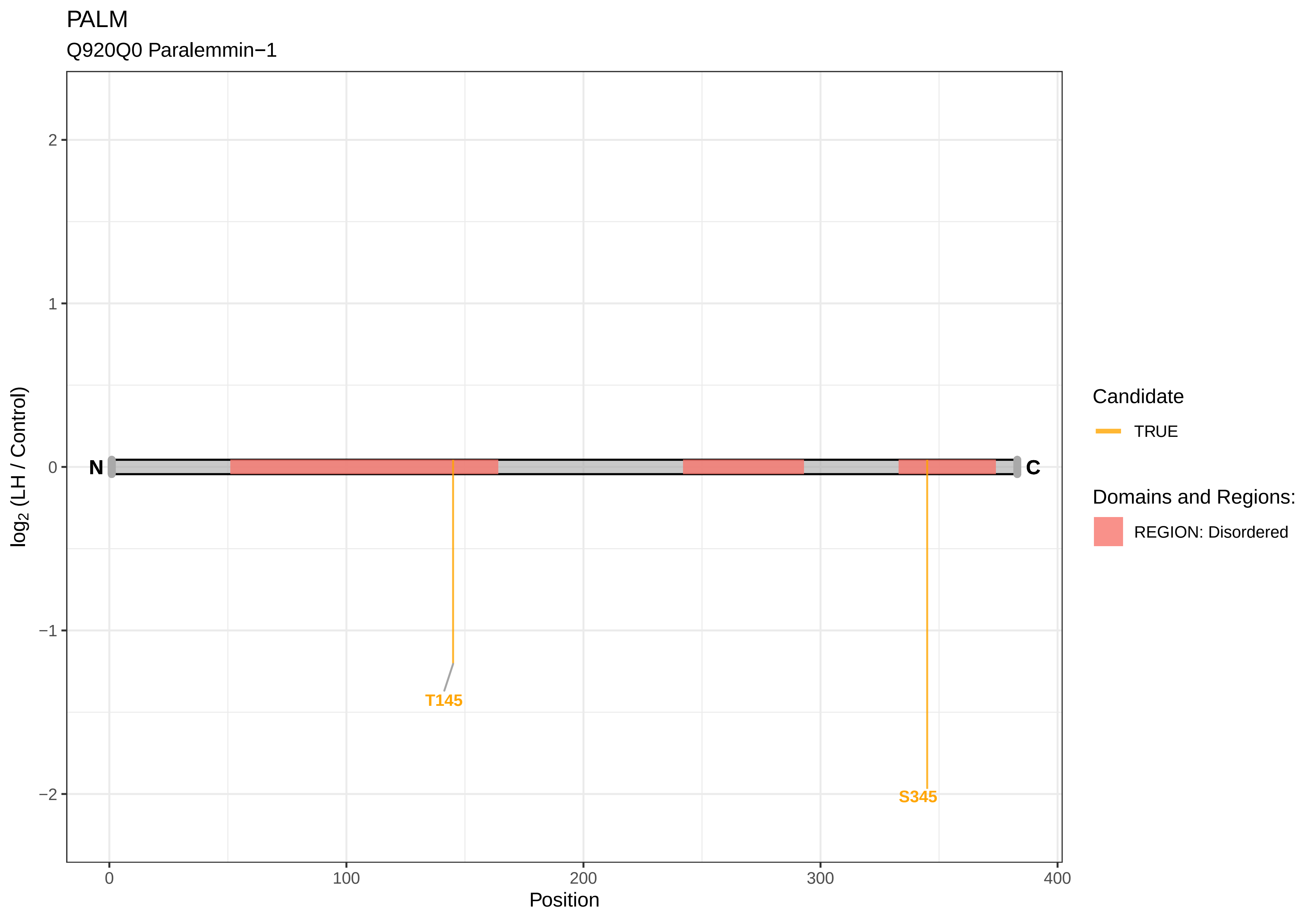

PALM
Q920Q0 Paralemmin−1
log2 (LH / Control)
2
1
0
−1
−2
N
C
T145
S345
0
100
200
300
400
Position
Candidate
TRUE
Domains and Regions:
REGION: Disordered

PAN3

D4ADM2 PAN2−PAN3 deadenylation complex subunit PAN3

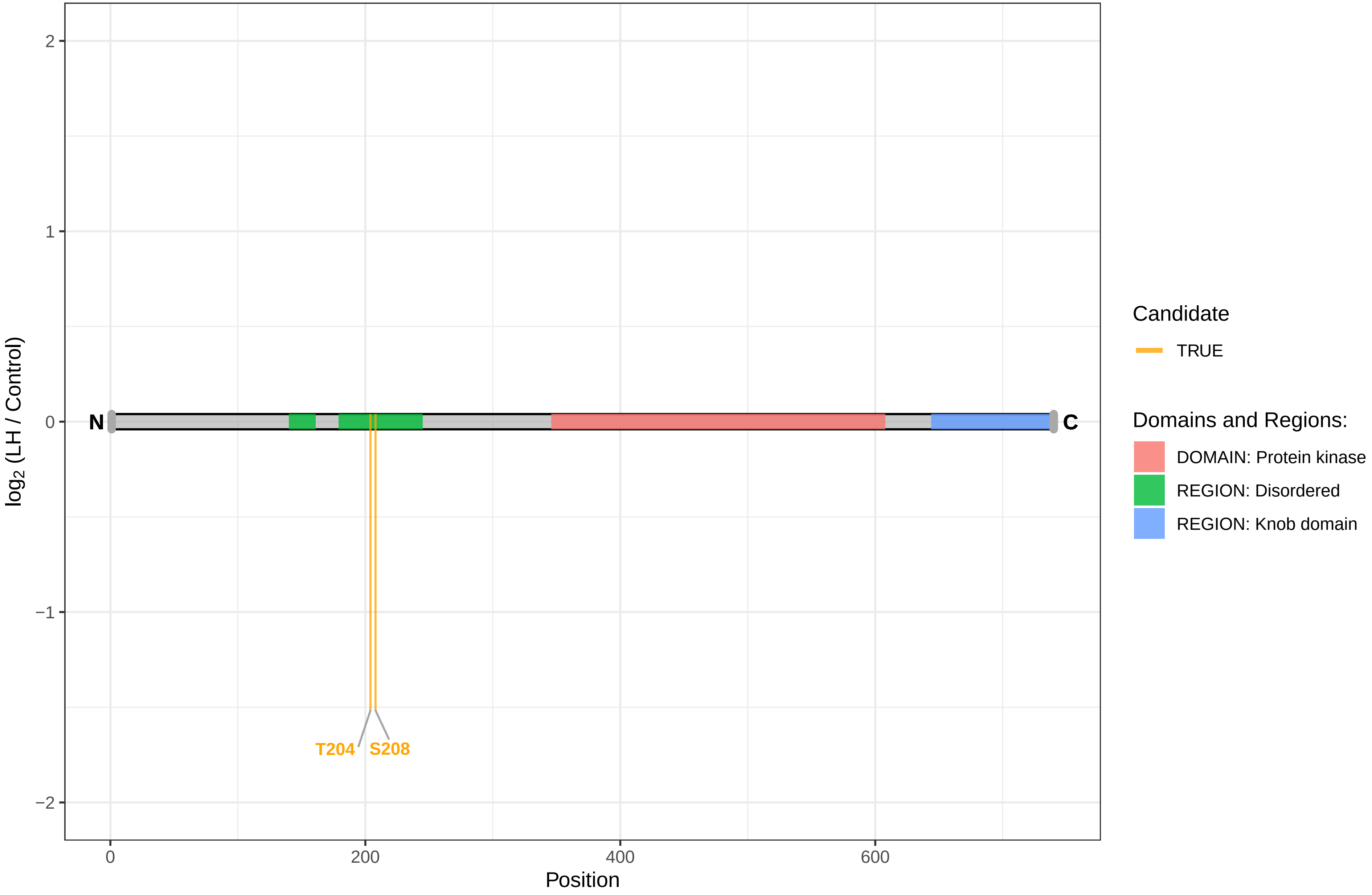

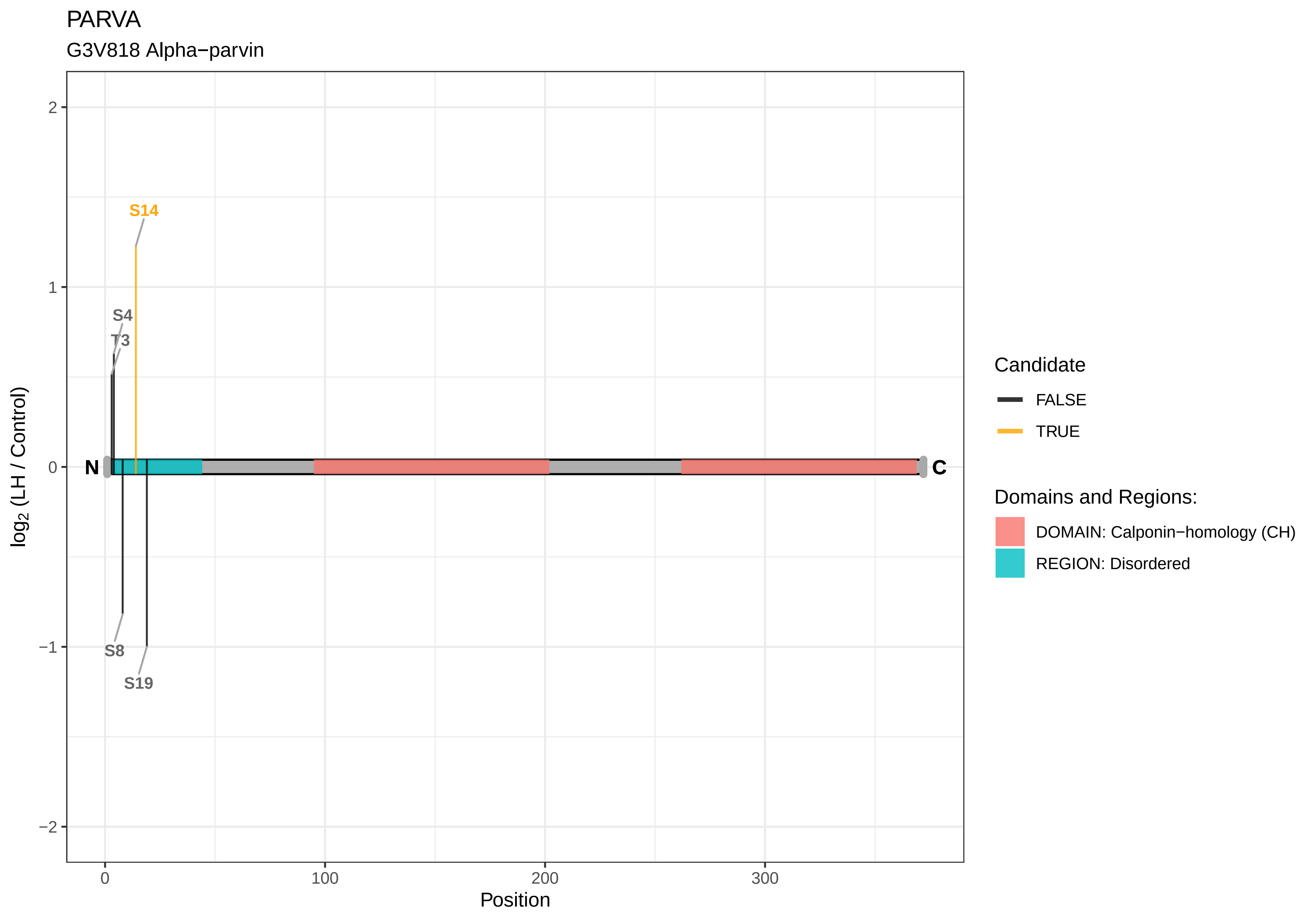

PARVA
G3V818 Alpha−parvin
S14
S4
T3
S8
S19
N
C
log2 (LH / Control)
Position
2
1
0
−1
−2
0
100
200
300
Candidate
FALSE
TRUE
Domains and Regions:
DOMAIN: Calponin−homology (CH)
REGION: Disordered

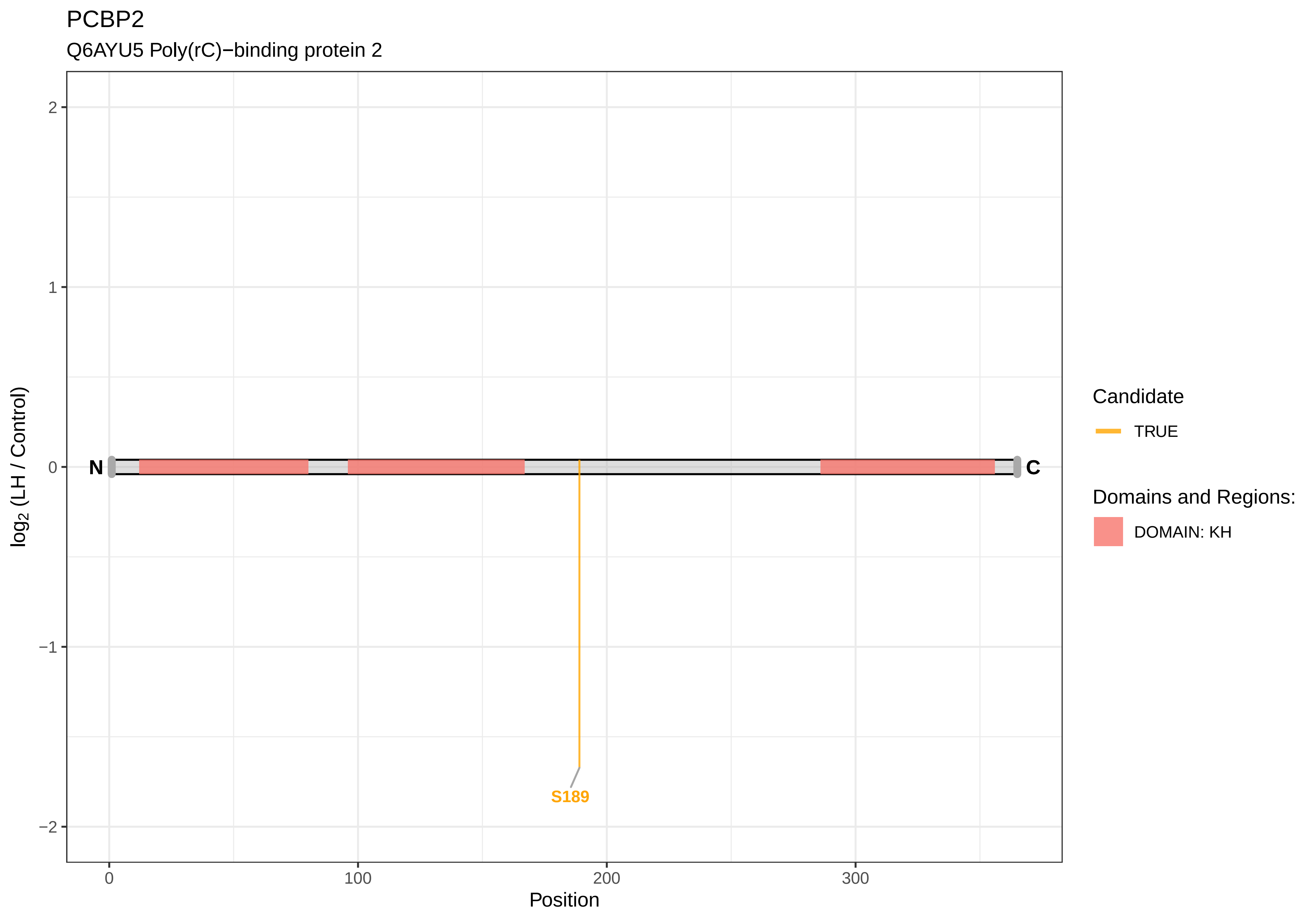
PCBP2
Q6AYU5 Poly(rC)−binding protein 2
log2 (LH / Control)
2
1
0
−1
−2
N
C
S189
0
100
200
300
Position
Candidate
TRUE
Domains and Regions:
DOMAIN: KH

PCF11

D3ZY40 PCF11 cleavage and polyadenylation factor subunit

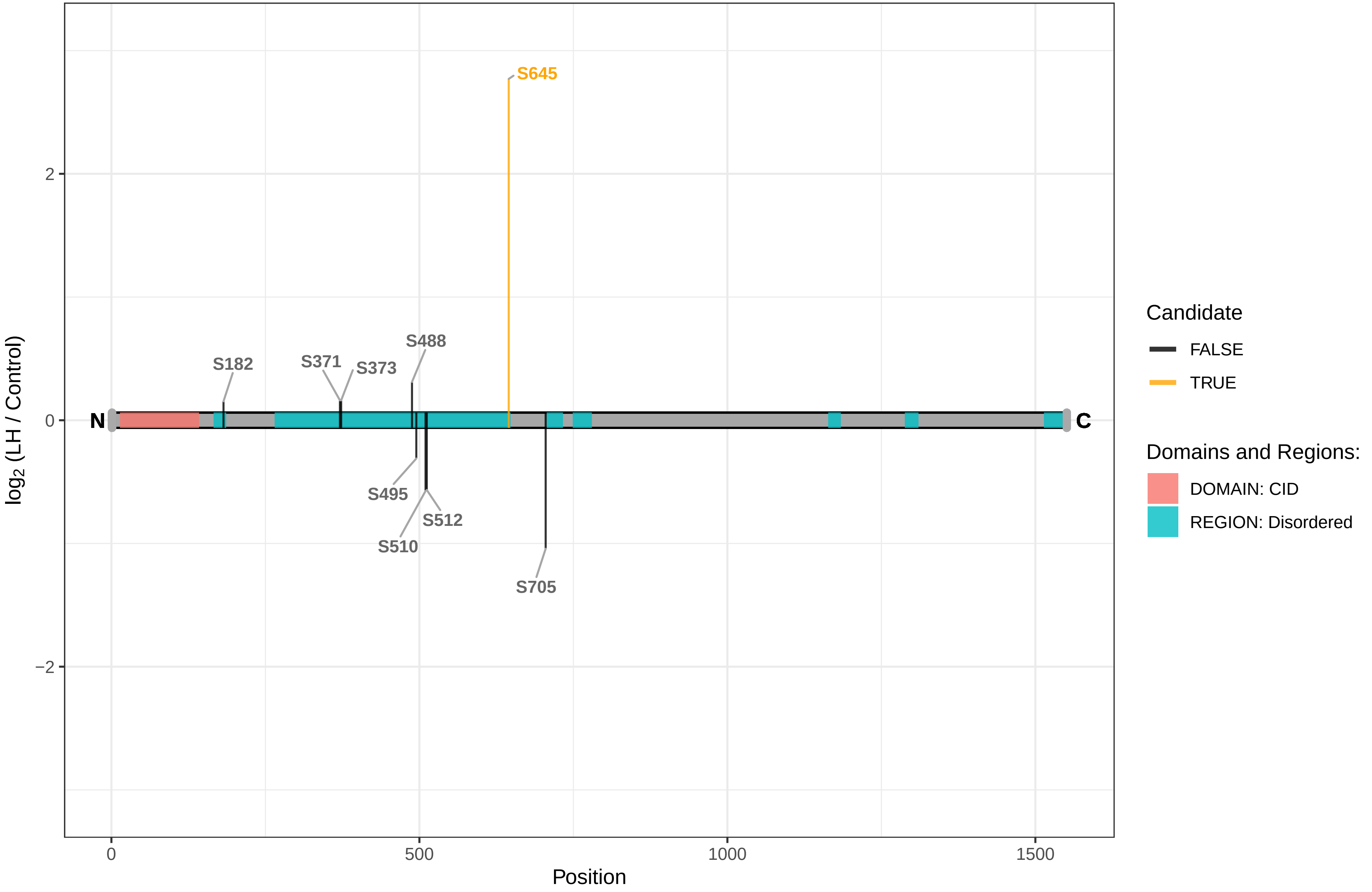

PCIF1

D4A417 PDX1 C−terminal−inhibiting factor 1

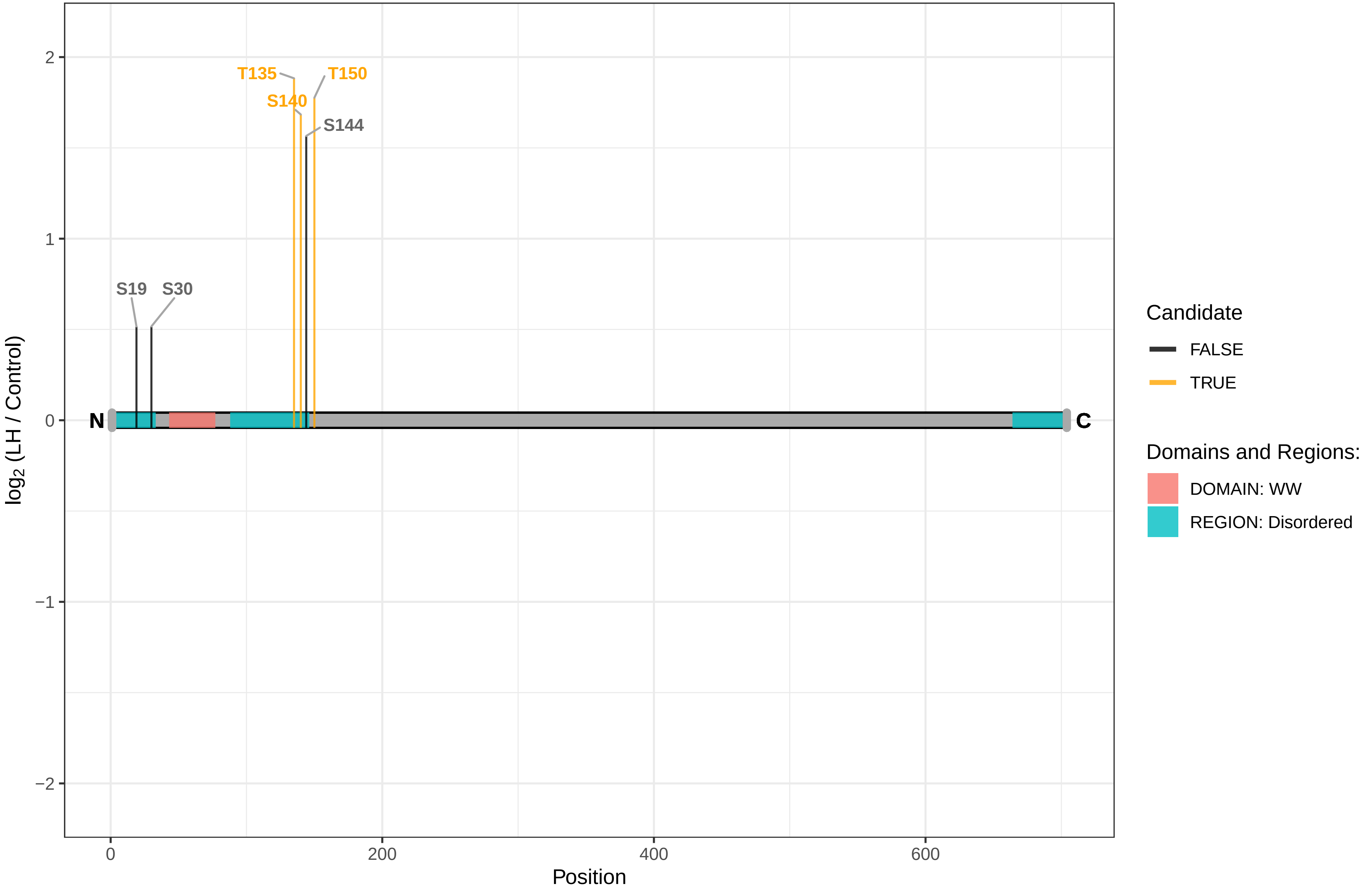

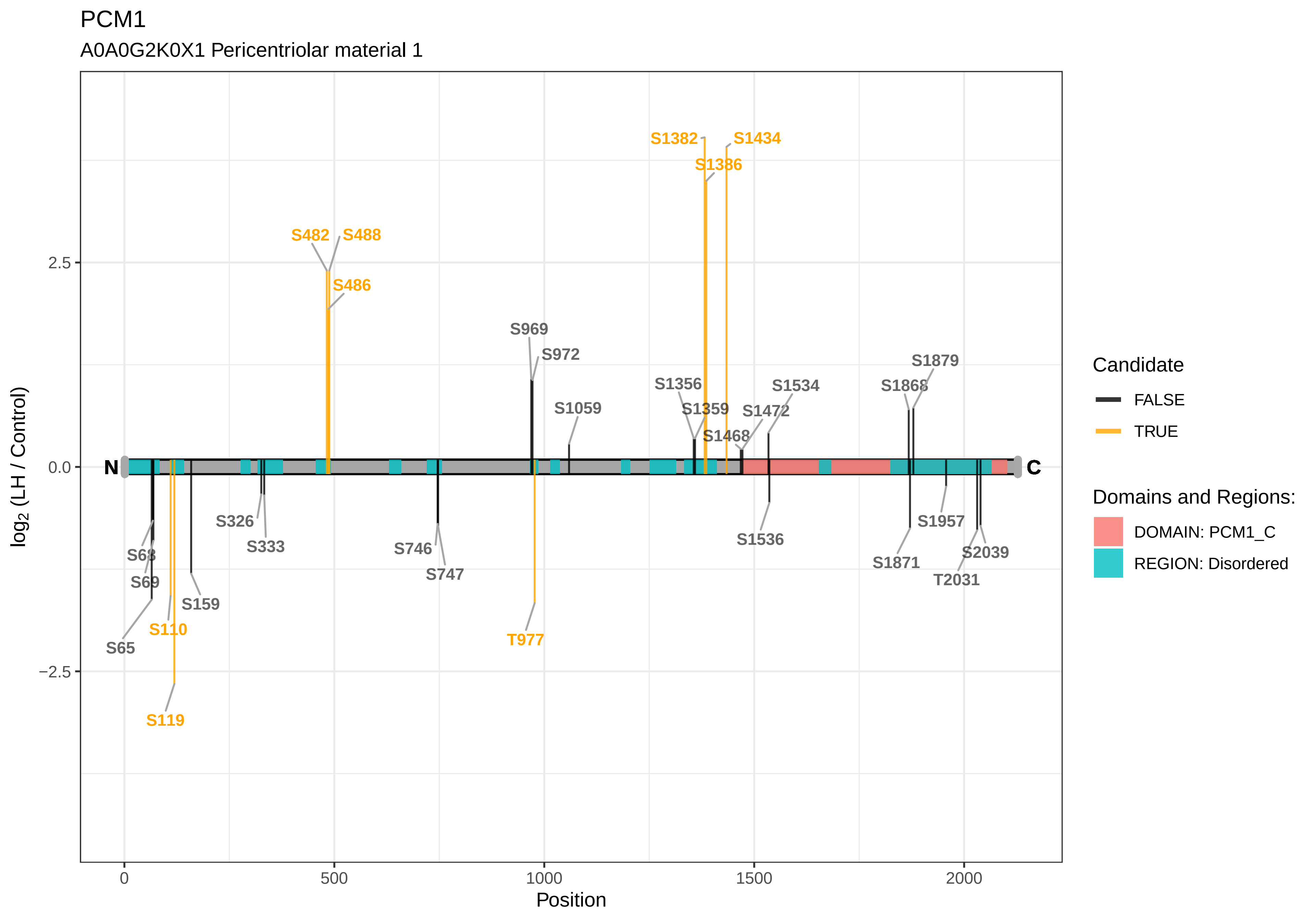

PCM1
A0A0G2K0X1 Pericentriolar material 1
log2 (LH / Control)
Position
2.5
0.0
−2.5
0
500
1000
1500
2000
N
C
S482
S488
S486
S1382
S1434
S1386
S969
S972
S1059
S1356
S1359
S1468
S1472
S1534
S1868
S1879
S68
S69
S65
S110
S119
S159
S326
S333
S746
S747
T977
S1536
S1871
S1957
S2039
T2031
Candidate
FALSE
TRUE
Domains and Regions:
DOMAIN: PCM1_C
REGION: Disordered

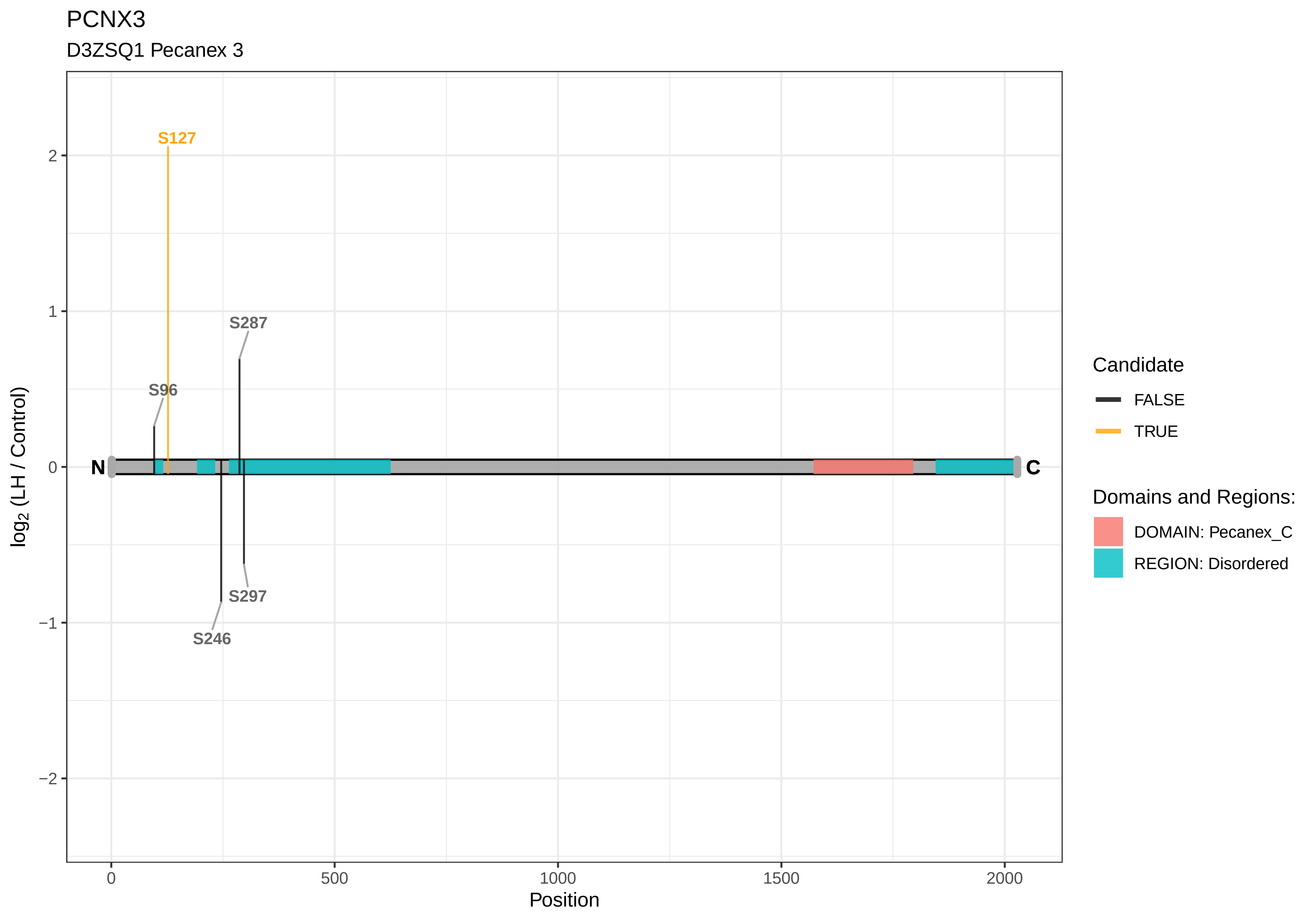

PCNX3
D3ZSQ1 Pecanex 3
S127
S287
S96
S297
S246
N
C
2
1
0
−1
−2
log2 (LH / Control)
0
500
1000
1500
2000
Position
Candidate
FALSE
TRUE
Domains and Regions:
DOMAIN: Pecanex_C
REGION: Disordered

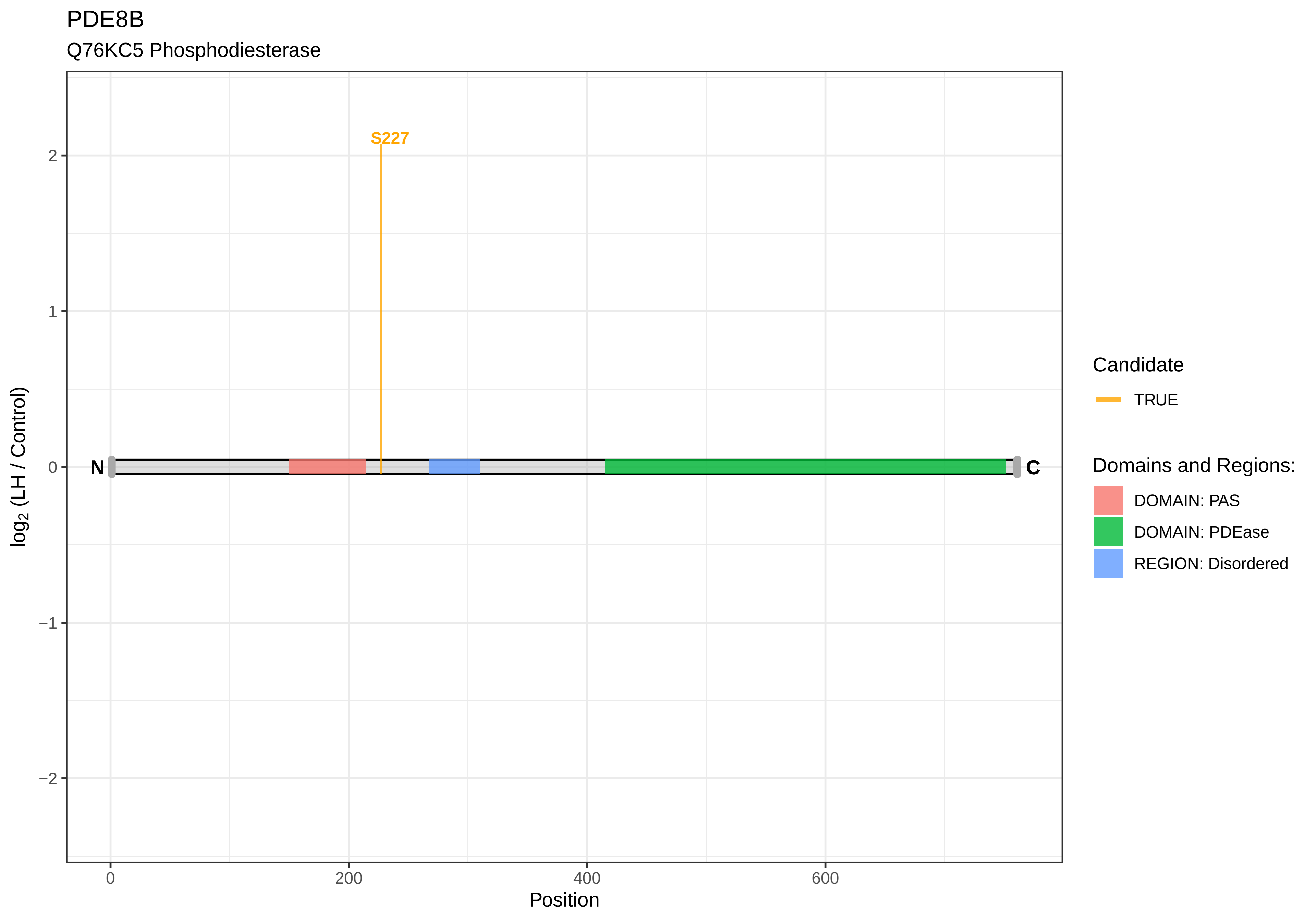
PDE8B
Q76KC5 Phosphodiesterase
S227
N
C
2
1
0
−1
−2
log2 (LH / Control)
0
200
400
600
Position
Candidate
TRUE
Domains and Regions:
DOMAIN: PAS
DOMAIN: PDEase
REGION: Disordered

PDLIM4

M0R4H5 PDZ and LIM domain protein 4

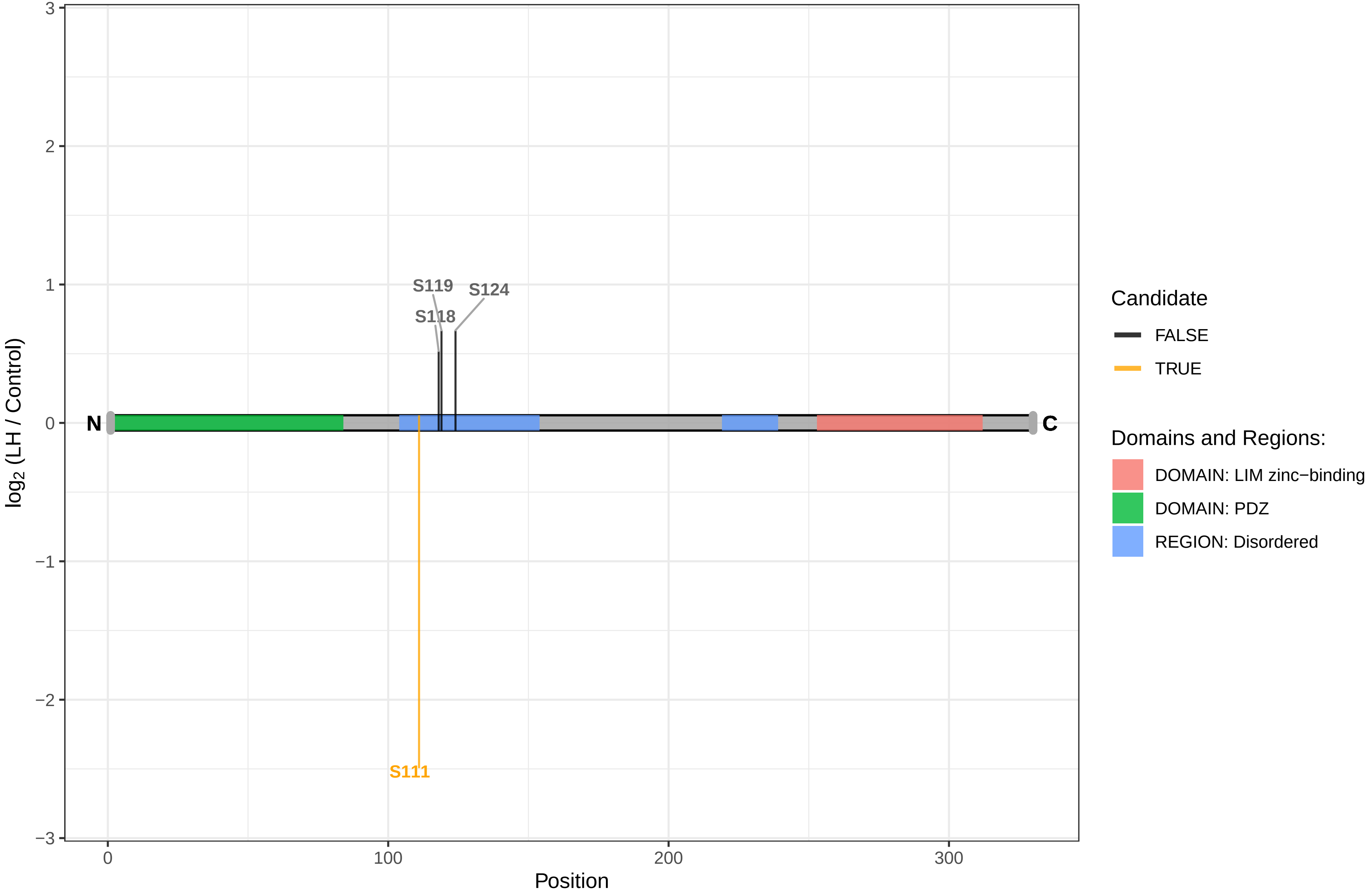

PDS5B

Q6TRW4 Sister chromatid cohesion protein PDS5 homolog B

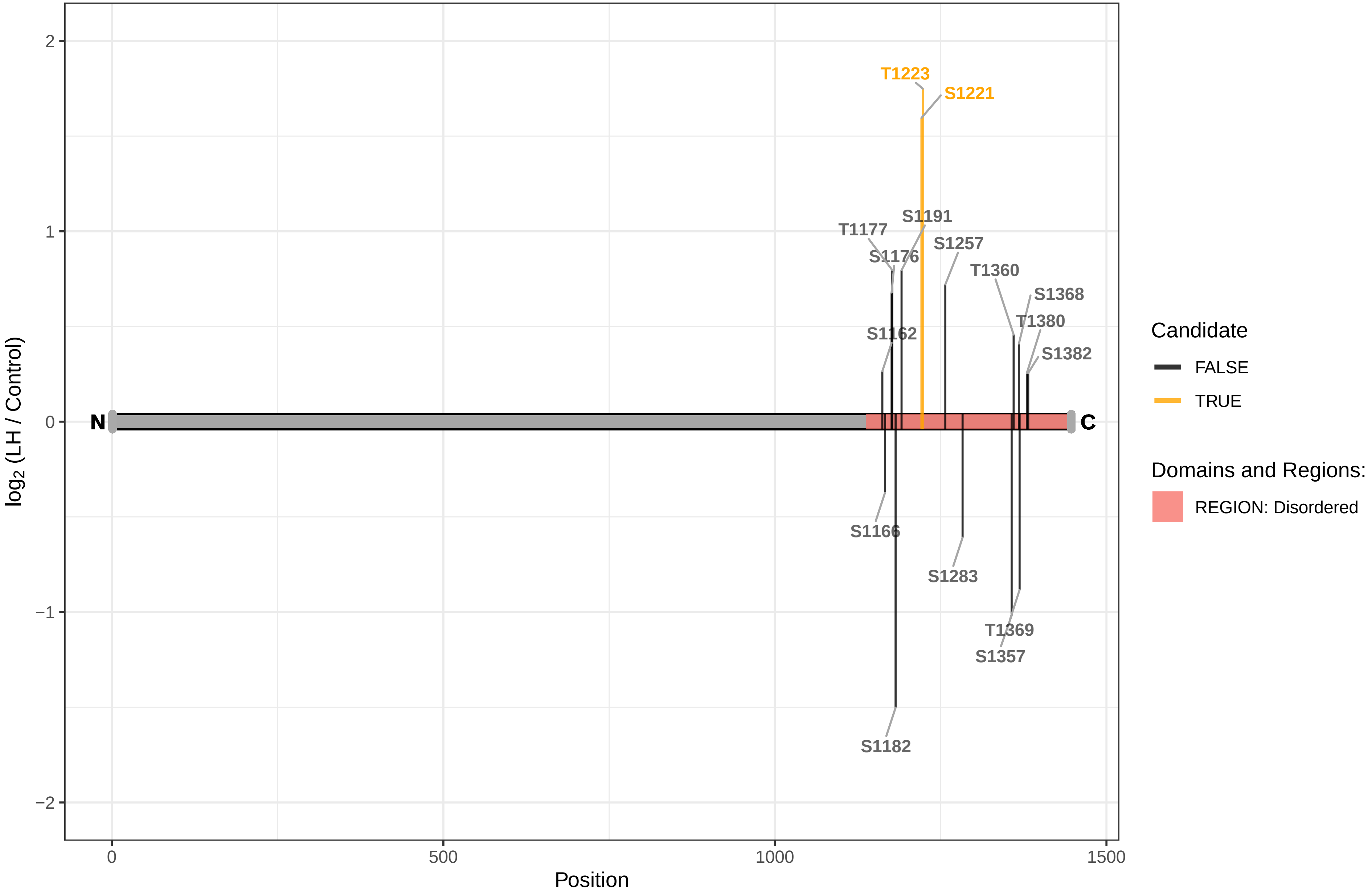

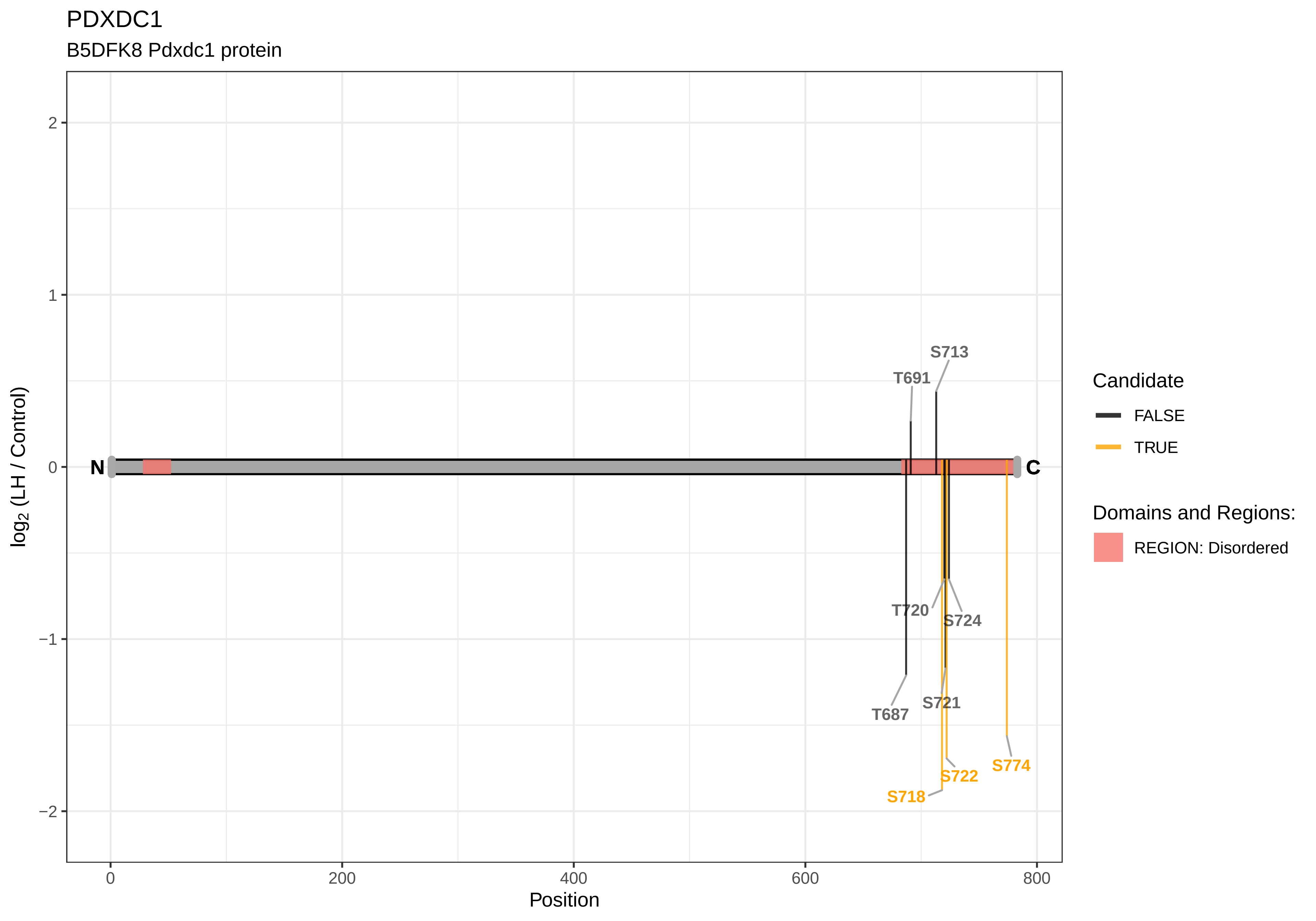

PDXDC1
B5DFK8 Pdxdc1 protein
log2 (LH / Control)
Position
N
C
T687
T691
S713
T720
S721
S724
S718
S722
S774
Candidate
FALSE
TRUE
Domains and Regions:
REGION: Disordered

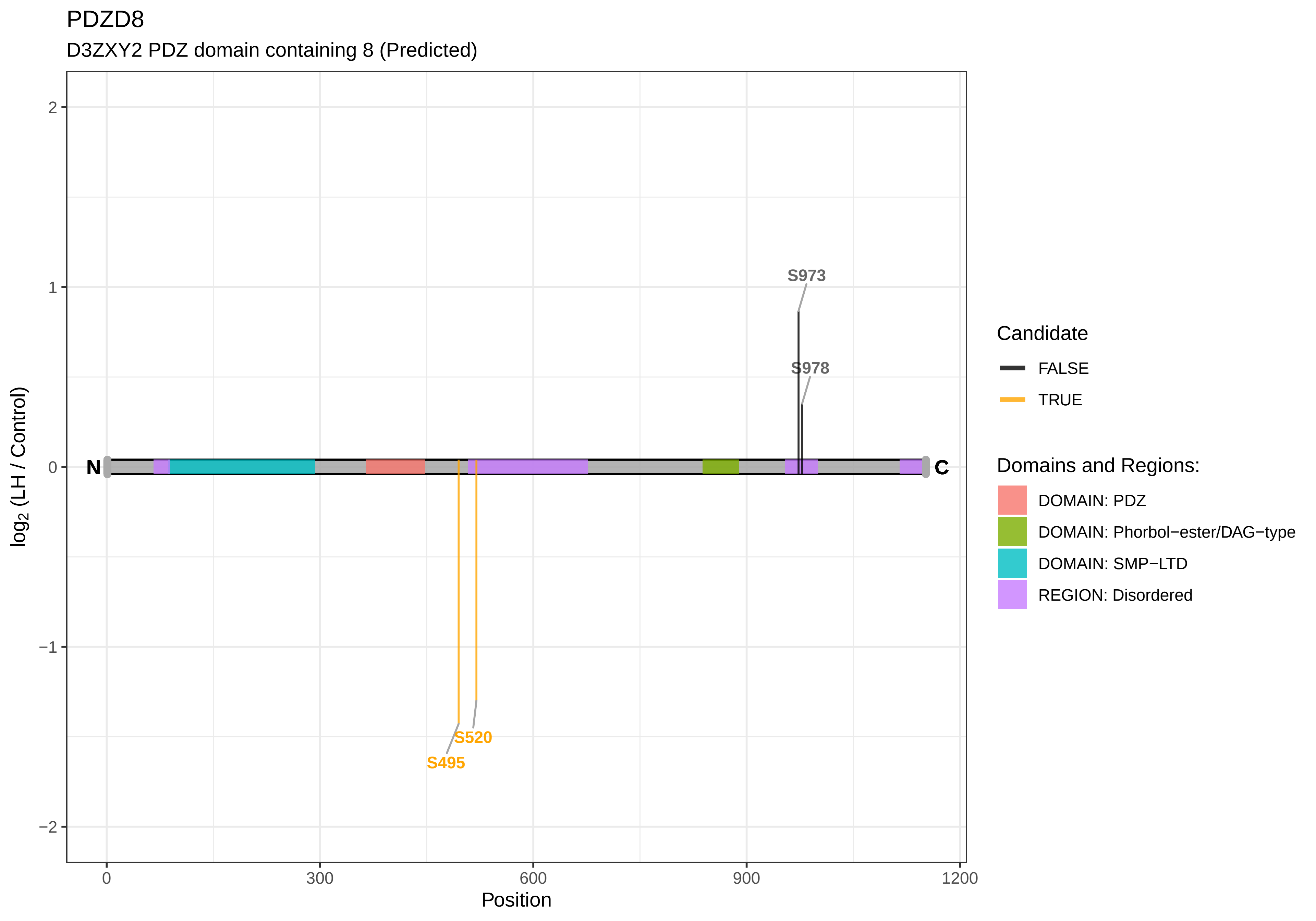

PDZD8
D3ZXY2 PDZ domain containing 8 (Predicted)
S973
S978
S520
S495
N
C
2
1
0
−1
−2
0
300
600
900
1200
Position
log2 (LH / Control)
Candidate
FALSE
TRUE
Domains and Regions:
DOMAIN: PDZ
DOMAIN: Phorbol−ester/DAG−type
DOMAIN: SMP−LTD
REGION: Disordered

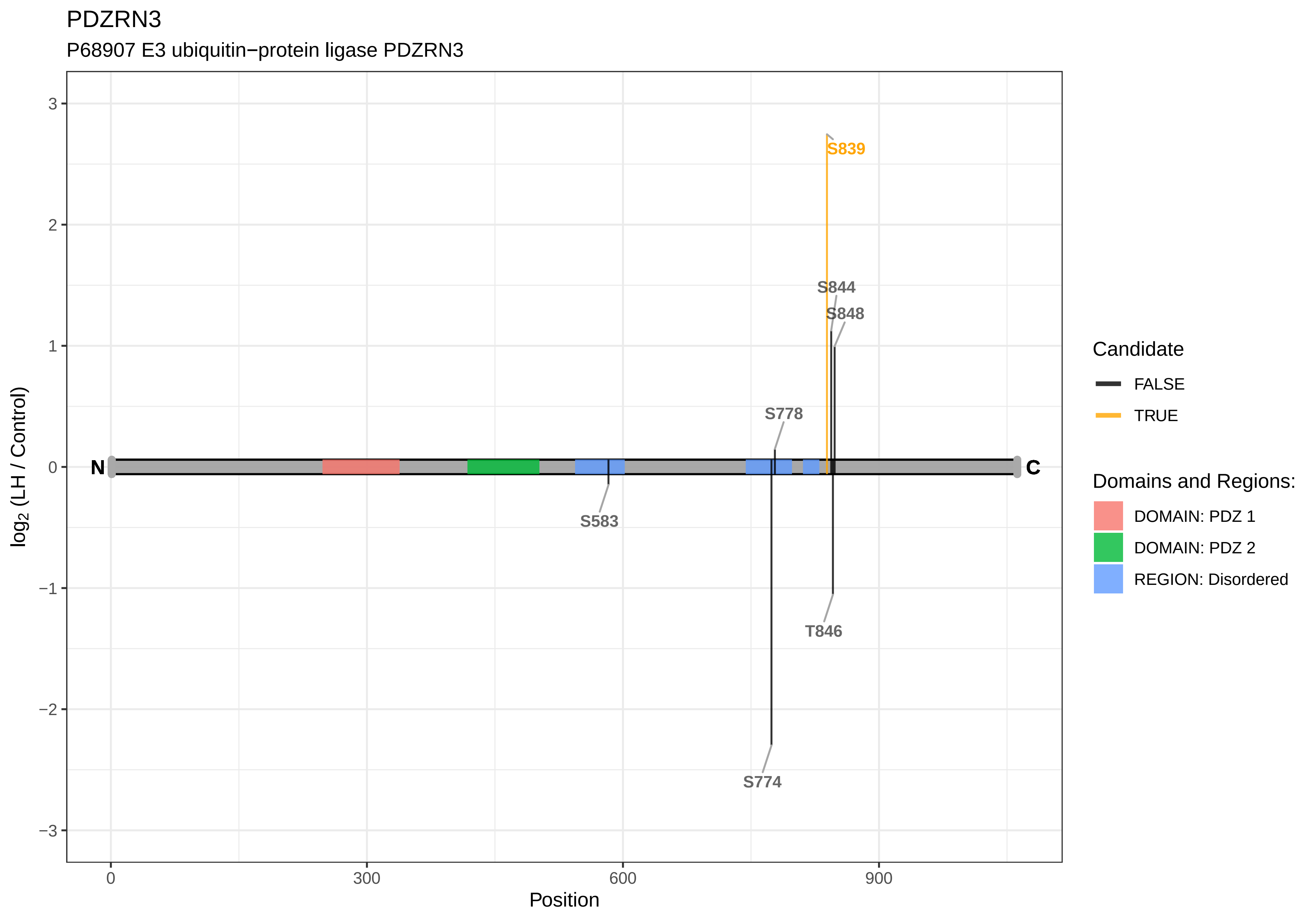

PDZRN3
P68907 E3 ubiquitin−protein ligase PDZRN3
log2 (LH / Control)
Position
3
2
1
0
−1
−2
−3
0
300
600
900
N
C
S839
S844
S848
S778
S583
T846
S774
Candidate
FALSE
TRUE
Domains and Regions:
DOMAIN: PDZ 1
DOMAIN: PDZ 2
REGION: Disordered

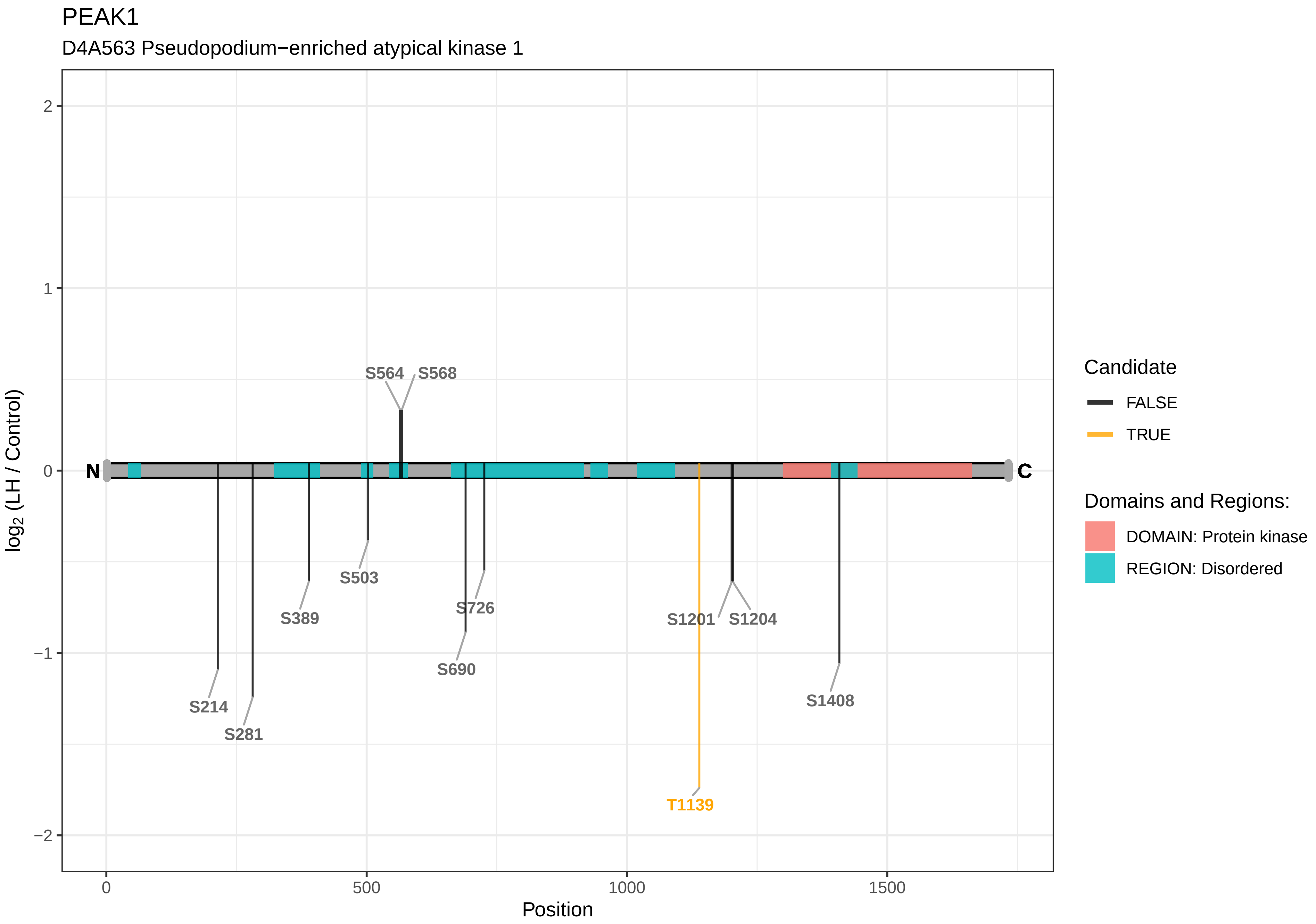

PEAK1
D4A563 Pseudopodium−enriched atypical kinase 1
S564
S568
S503
S389
S726
S690
S214
S281
S1201
S1204
S1408
T1139
N
C
log2 (LH / Control)
Position
2
1
0
−1
−2
0
500
1000
1500
Candidate
FALSE
TRUE
Domains and Regions:
DOMAIN: Protein kinase
REGION: Disordered

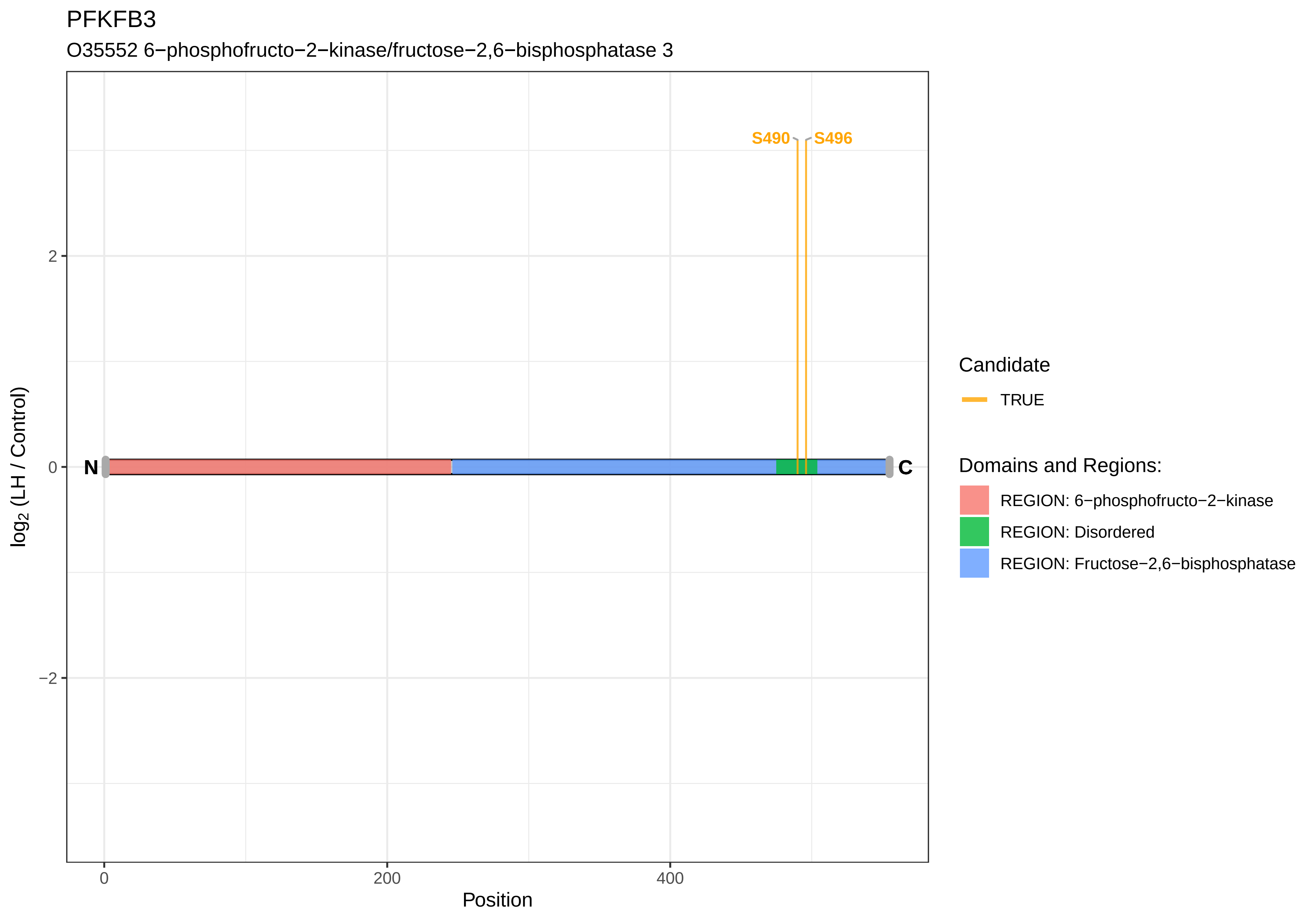
PFKFB3
O35552 6−phosphofructo−2−kinase/fructose−2,6−bisphosphatase 3
S490
S496
N
C
2
0
−2
0
200
400
Position
log2 (LH / Control)
Candidate
TRUE
Domains and Regions:
REGION: 6−phosphofructo−2−kinase
REGION: Disordered
REGION: Fructose−2,6−bisphosphatase

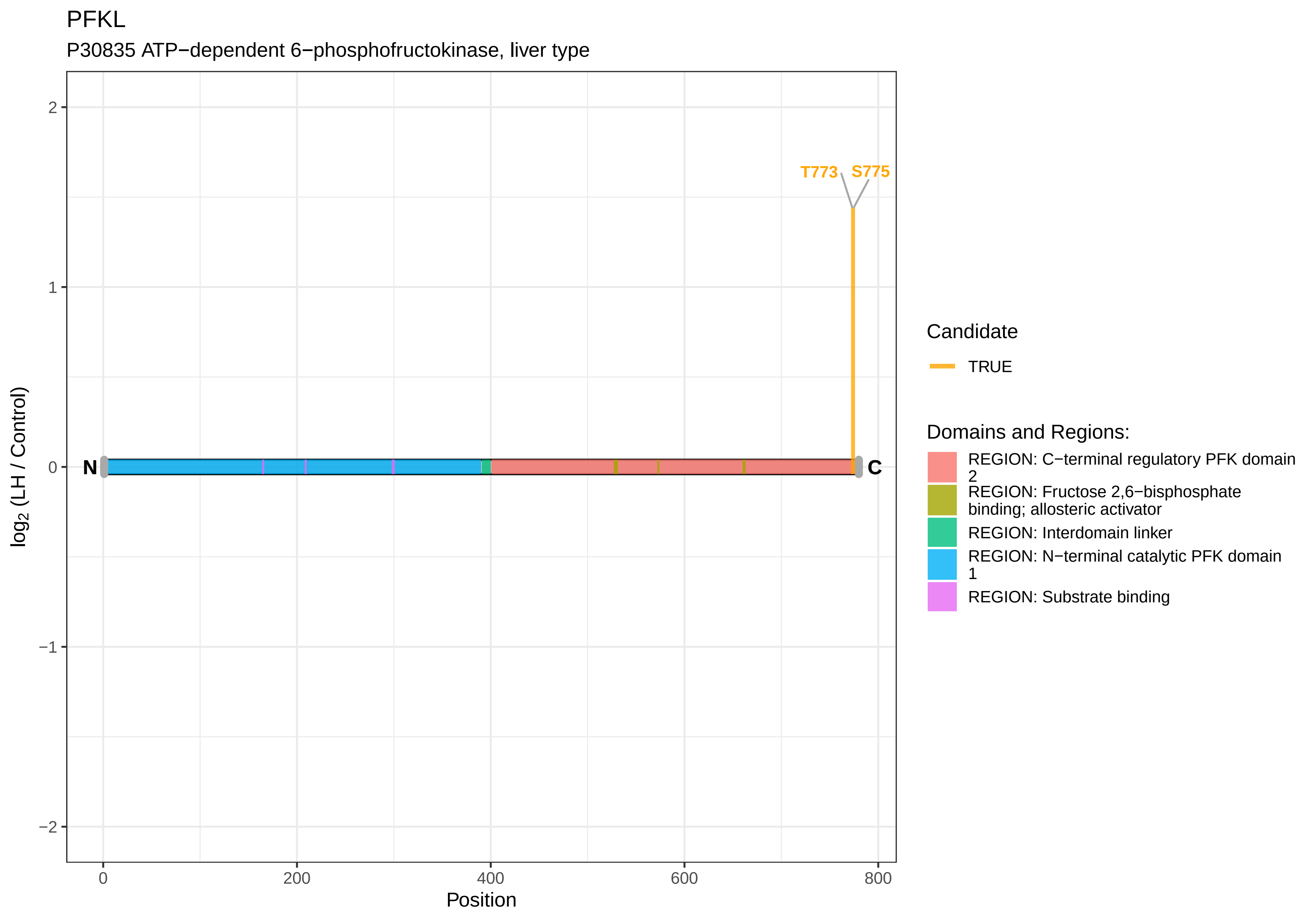
PFKL
P30835 ATP−dependent 6−phosphofructokinase, liver type
T773
S775
N
C
2
1
0
−1
−2
log2 (LH / Control)
0
200
400
600
800
Position
Candidate
TRUE
Domains and Regions:
REGION: C−terminal regulatory PFK domain 2
REGION: Fructose 2,6−bisphosphate binding; allosteric activator
REGION: Interdomain linker
REGION: N−terminal catalytic PFK domain 1
REGION: Substrate binding

PHACTR2

A0A0G2K8Q4 Phosphatase and actin regulator

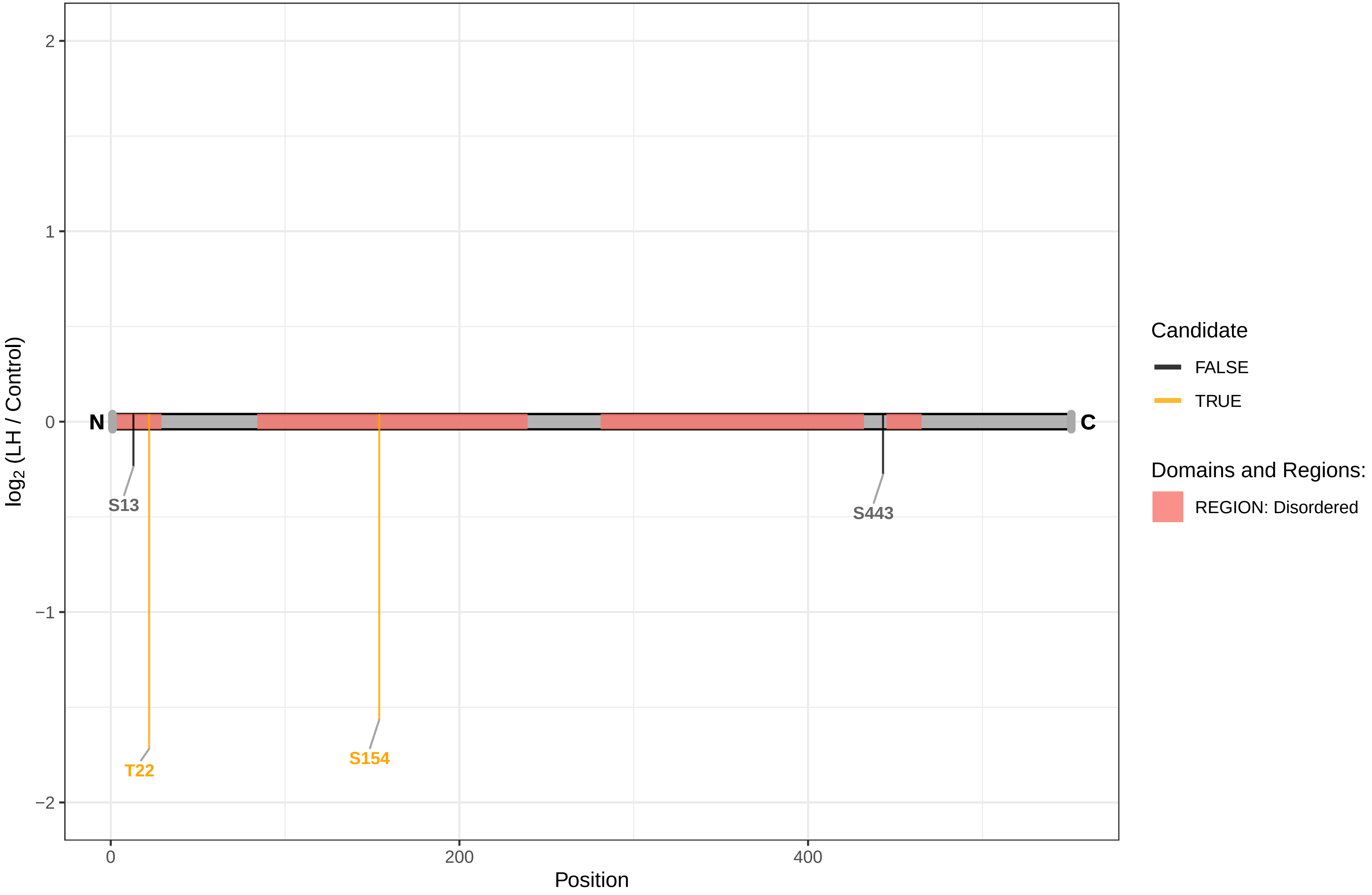

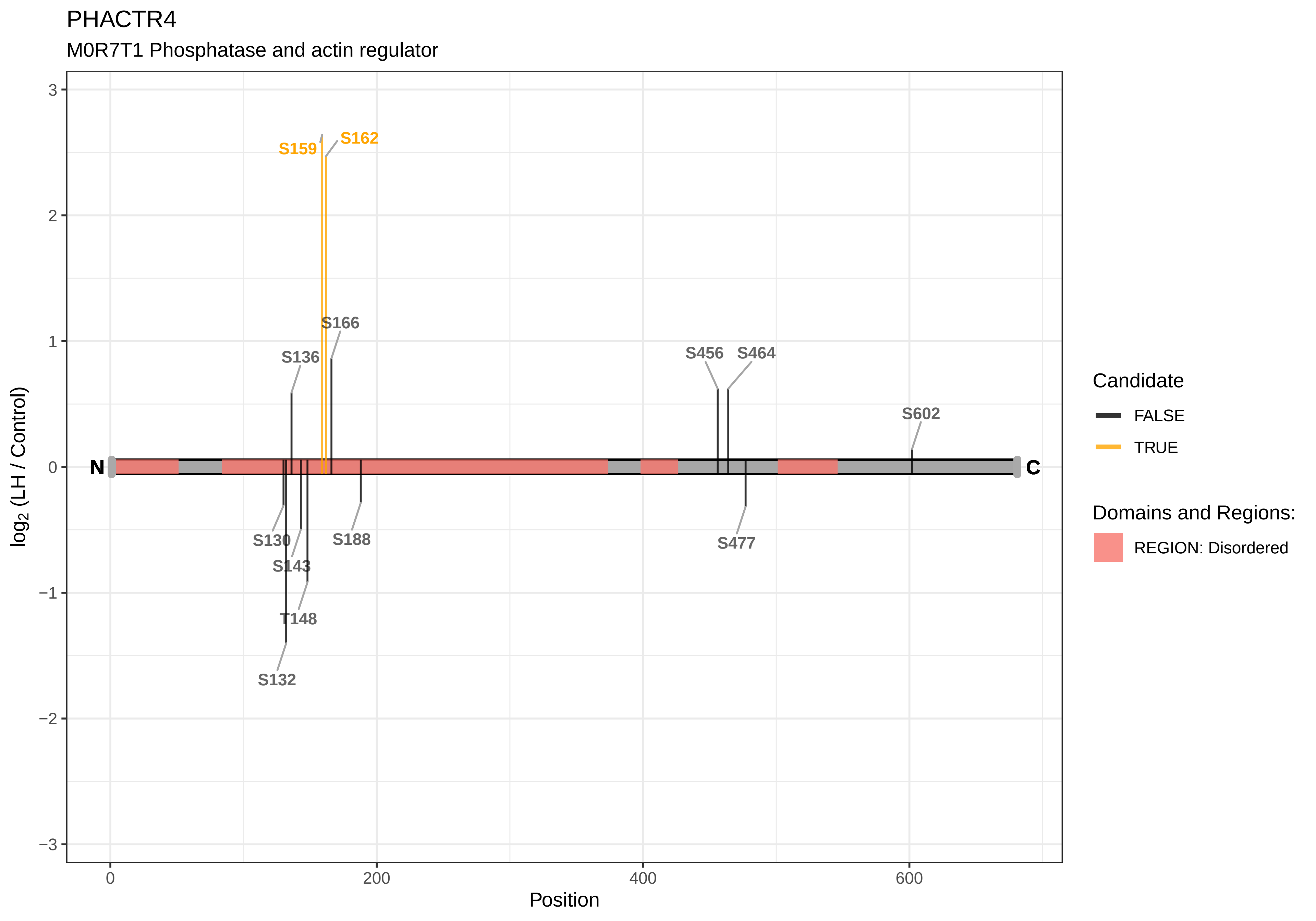

PHACTR4
M0R7T1 Phosphatase and actin regulator
S159
S162
S166
S136
S456
S464
S602
S130
S143
T148
S132
S188
S477
N
C
log2 (LH / Control)
Position
Candidate
FALSE
TRUE
Domains and Regions:
REGION: Disordered

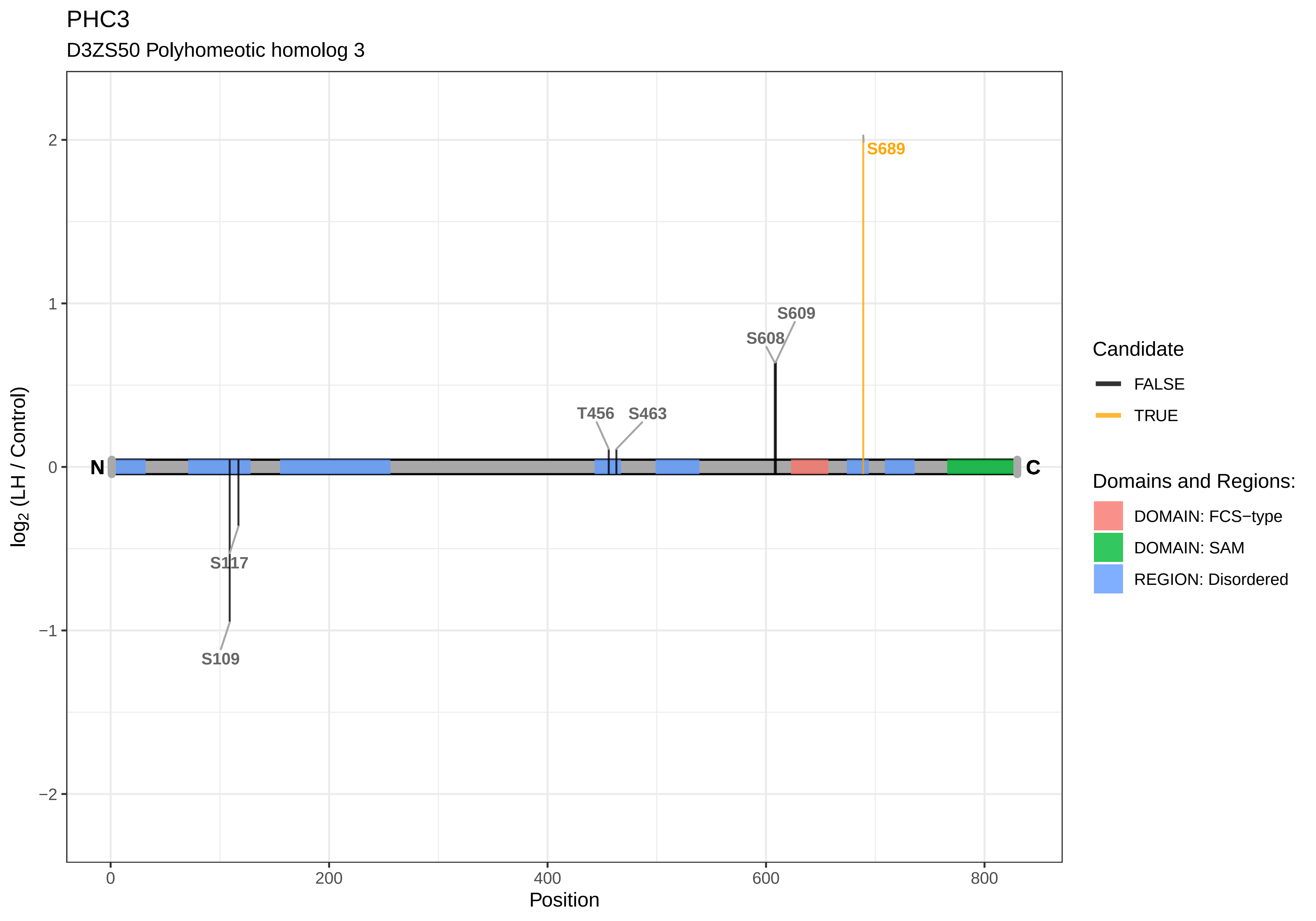
PHC3
D3ZS50 Polyhomeotic homolog 3
S689
S609
S608
T456
S463
S117
S109
N
C
2
1
0
−1
−2
log2 (LH / Control)
0
200
400
600
800
Position
Candidate
FALSE
TRUE
Domains and Regions:
DOMAIN: FCS−type
DOMAIN: SAM
REGION: Disordered

PHF1

Q6MGC9 PHD finger protein 1

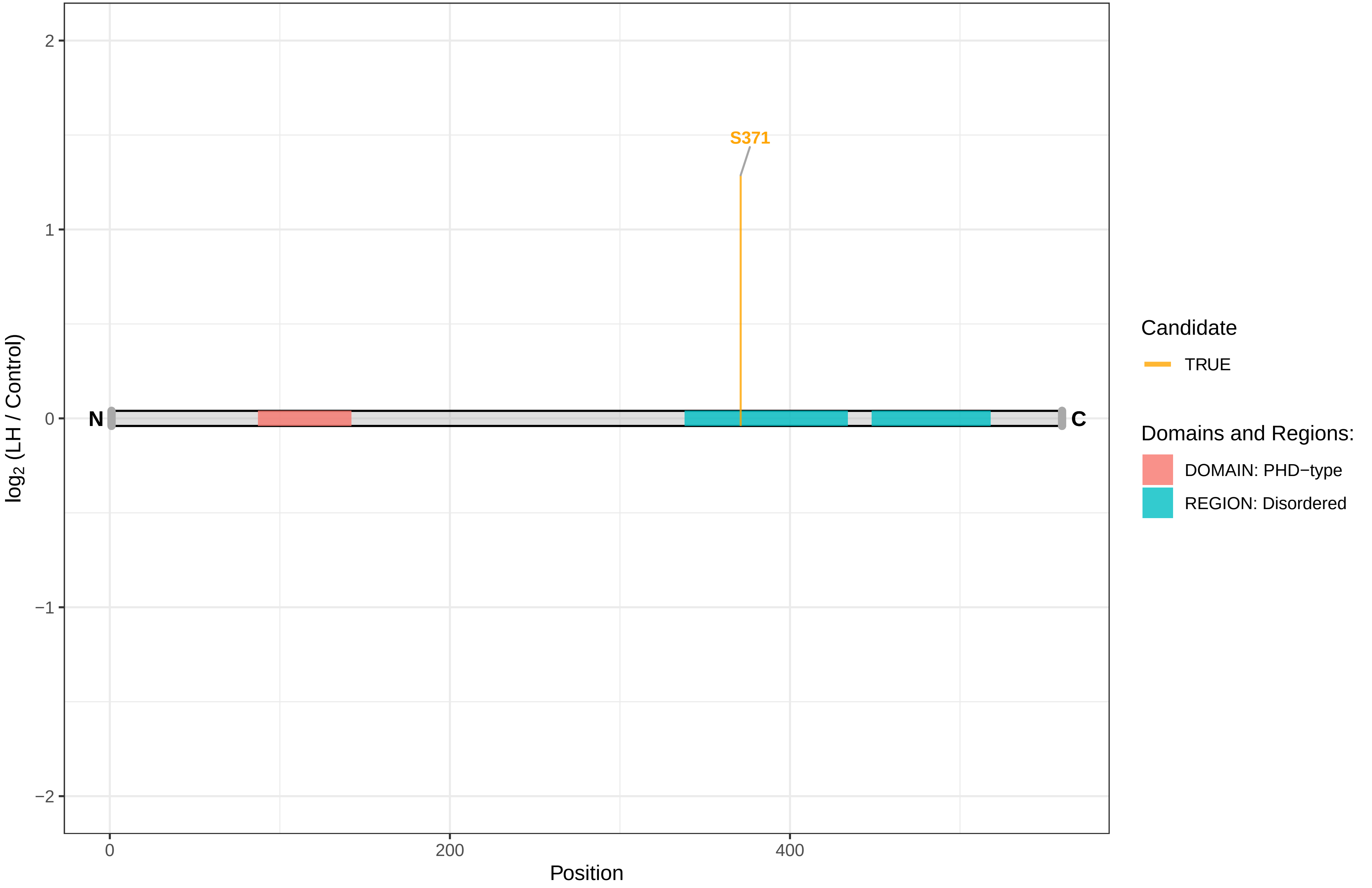

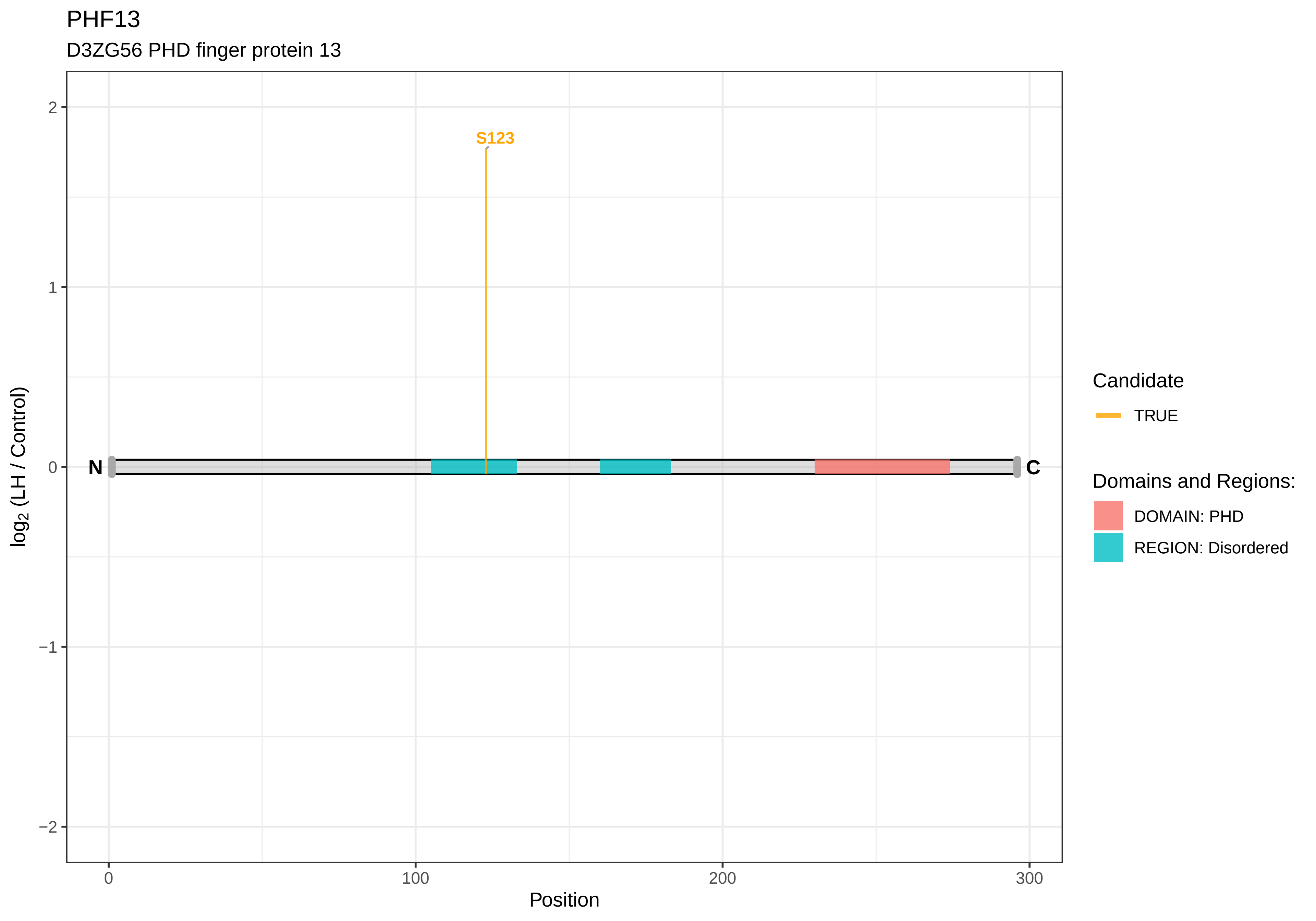
PHF13
D3ZG56 PHD finger protein 13
S123
N
C
2
1
0
−1
−2
log2 (LH / Control)
0
100
200
300
Position
Candidate
TRUE
Domains and Regions:
DOMAIN: PHD
REGION: Disordered

PHF14

G3V9T6 PHD finger protein 14

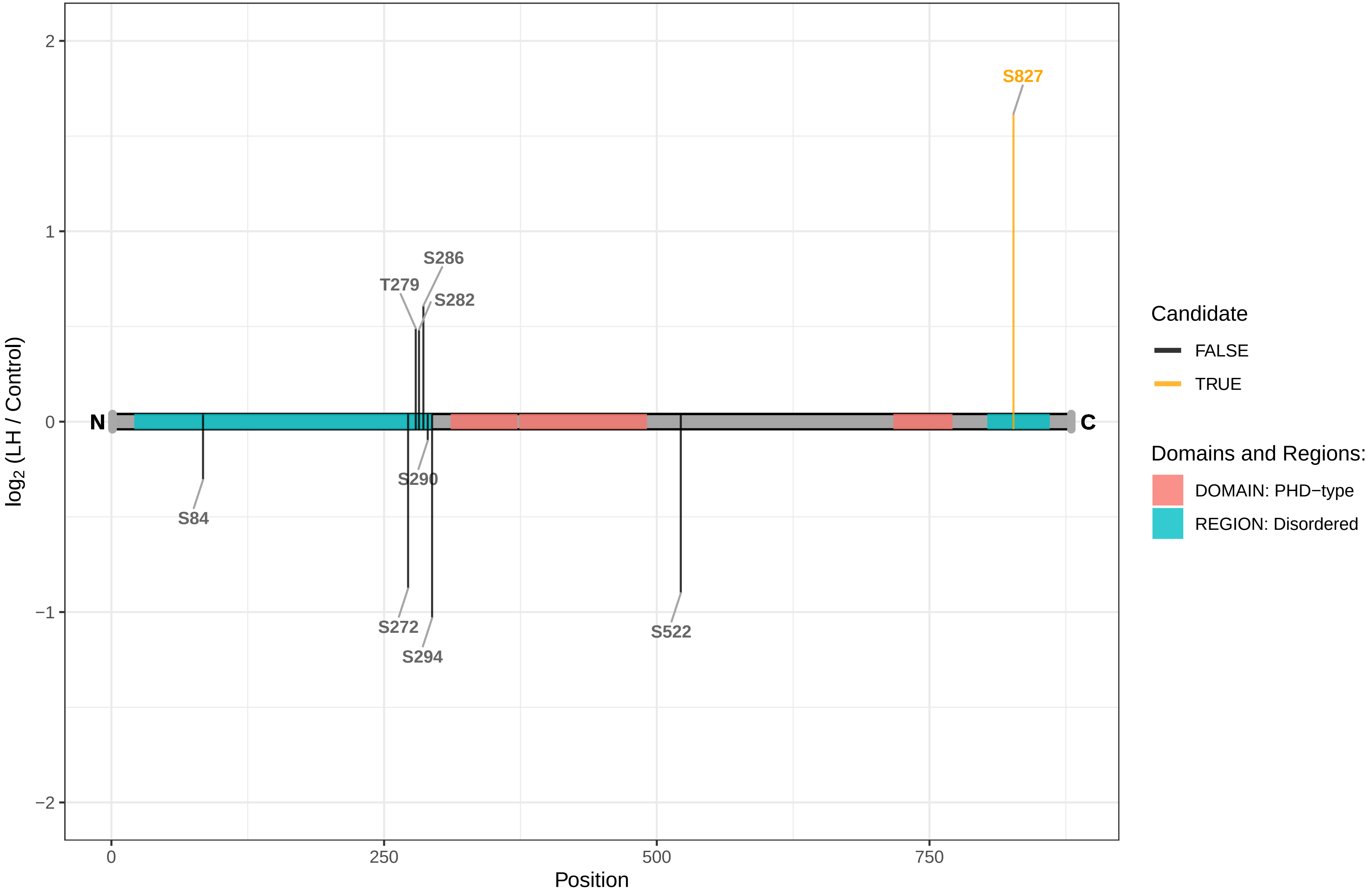

PHF2

F1LWX5 PHD finger protein 2

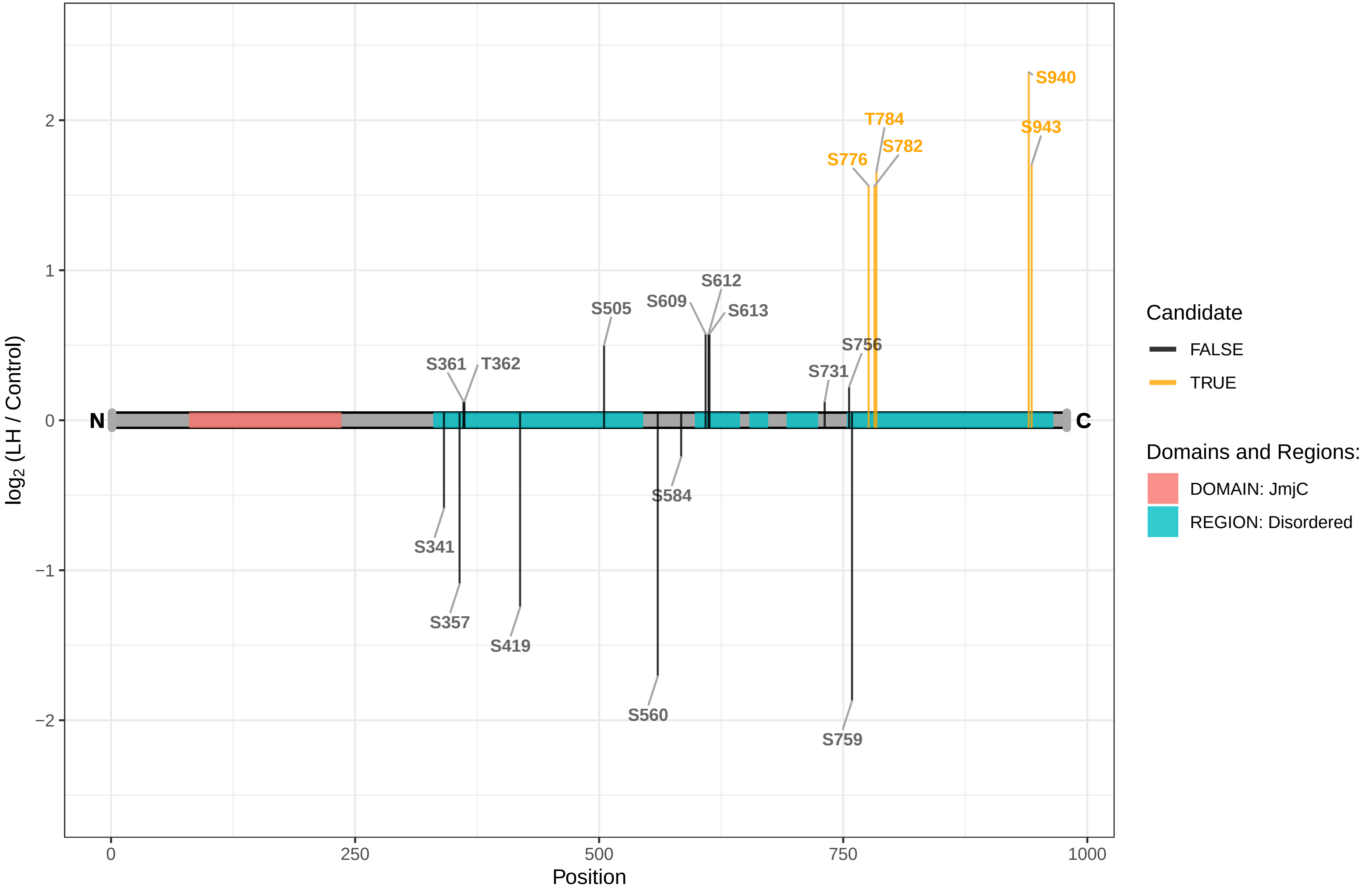

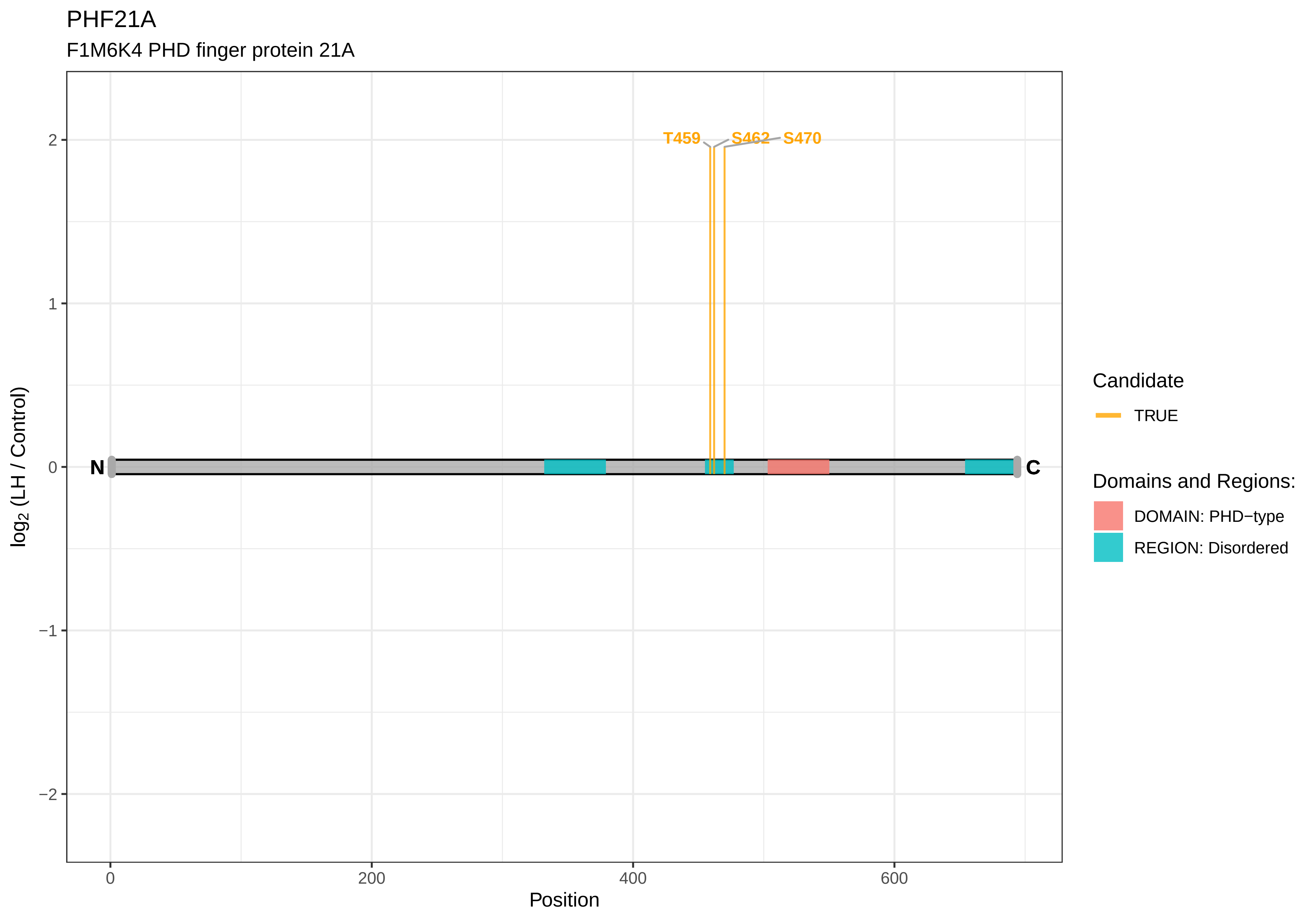

PHF21A
F1M6K4 PHD finger protein 21A
T459
S462
S470
N
C
2
1
0
−1
−2
log2 (LH / Control)
0
200
400
600
Position
Candidate
TRUE
Domains and Regions:
DOMAIN: PHD−type
REGION: Disordered

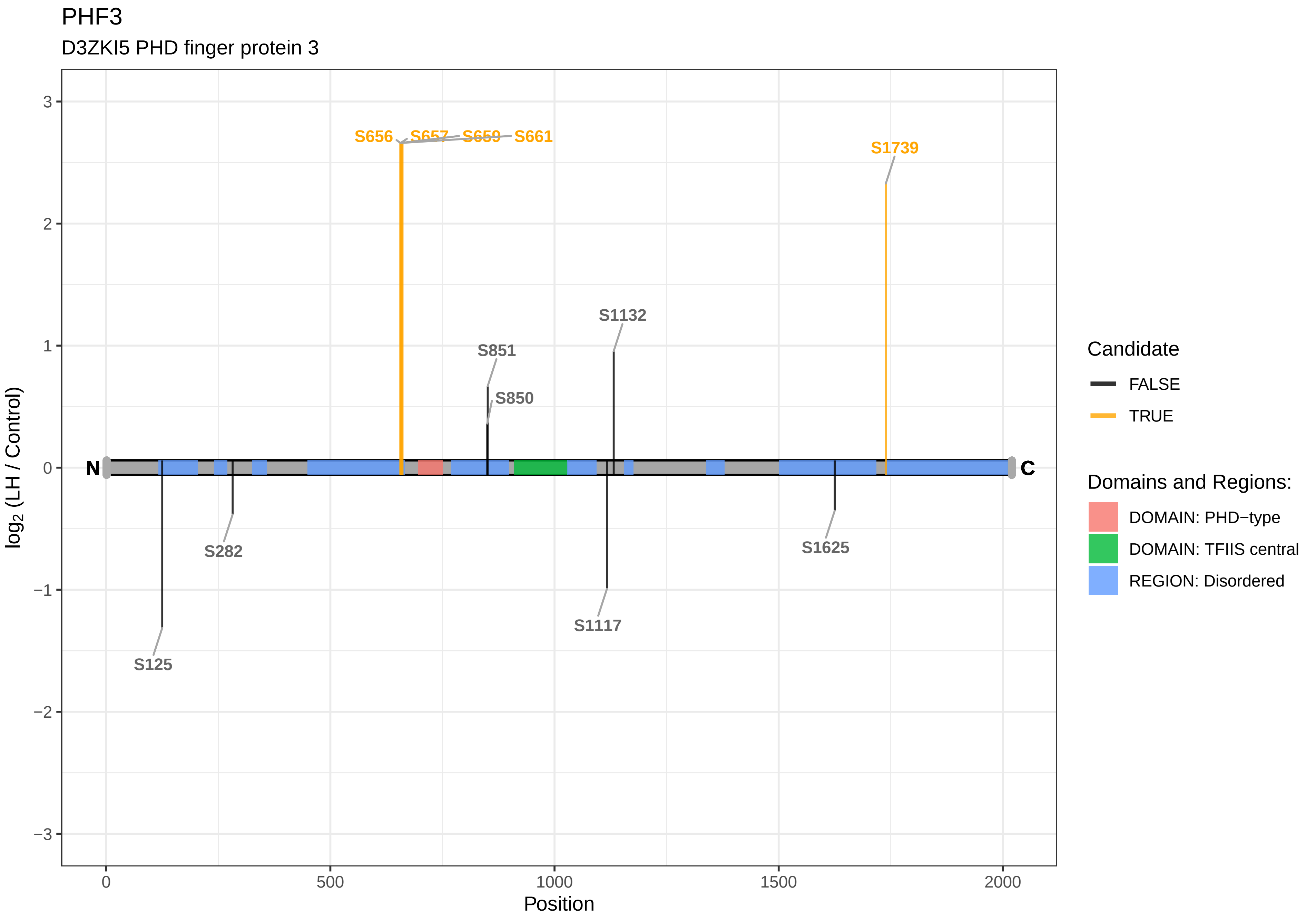
PHF3
D3ZKI5 PHD finger protein 3
S656
S657
S659
S661
S1739
S1132
S851
S850
S282
S125
S1117
S1625
N
C
3
2
1
0
−1
−2
−3
0
500
1000
1500
2000
Position
log2 (LH / Control)
Candidate
FALSE
TRUE
Domains and Regions:
DOMAIN: PHD−type
DOMAIN: TFIIS central
REGION: Disordered

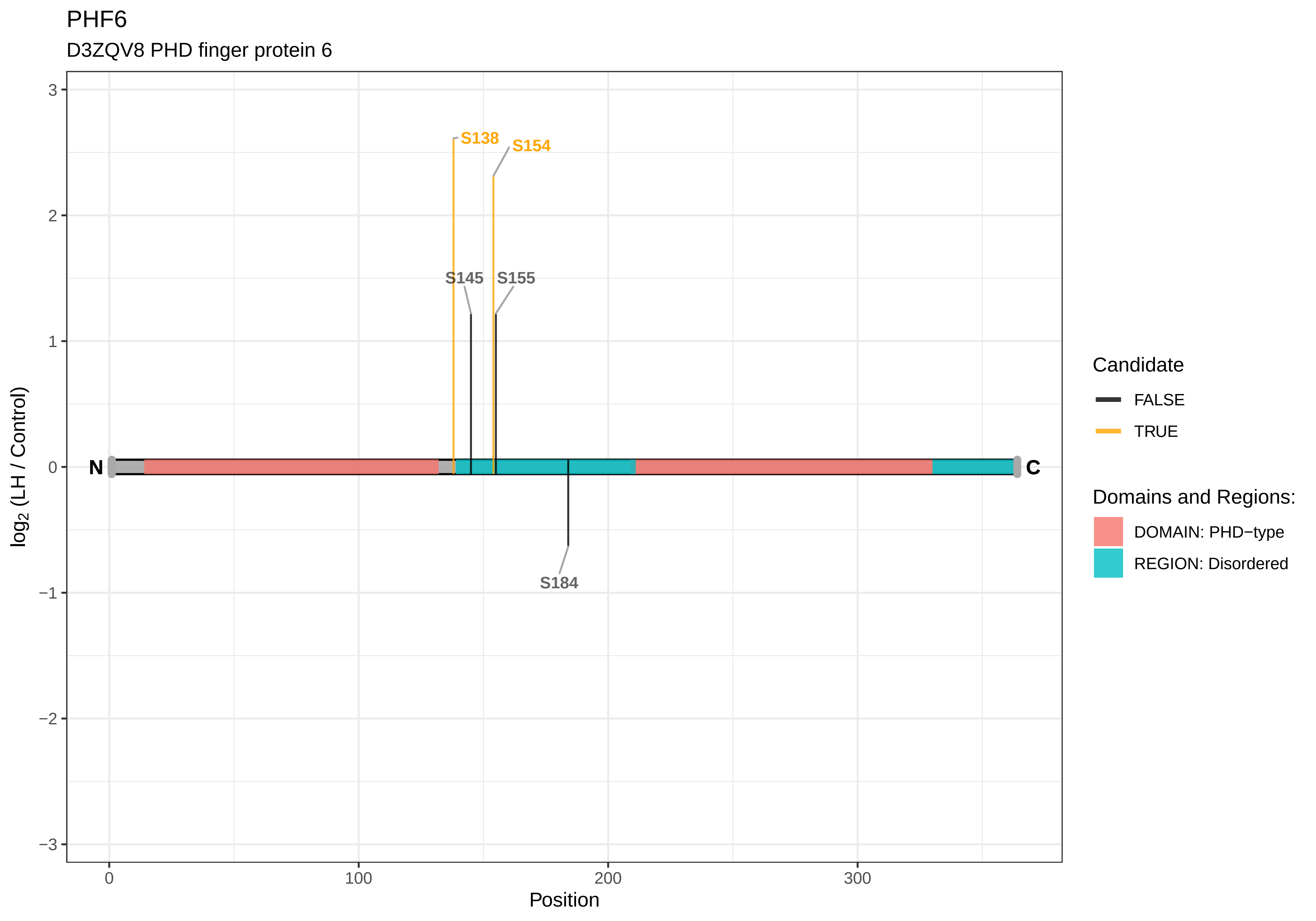
PHF6
D3ZQV8 PHD finger protein 6
S138
S154
S145
S155
S184
N
C
log2 (LH / Control)
Position
3
2
1
0
−1
−2
−3
0
100
200
300
Candidate
FALSE
TRUE
Domains and Regions:
DOMAIN: PHD−type
REGION: Disordered

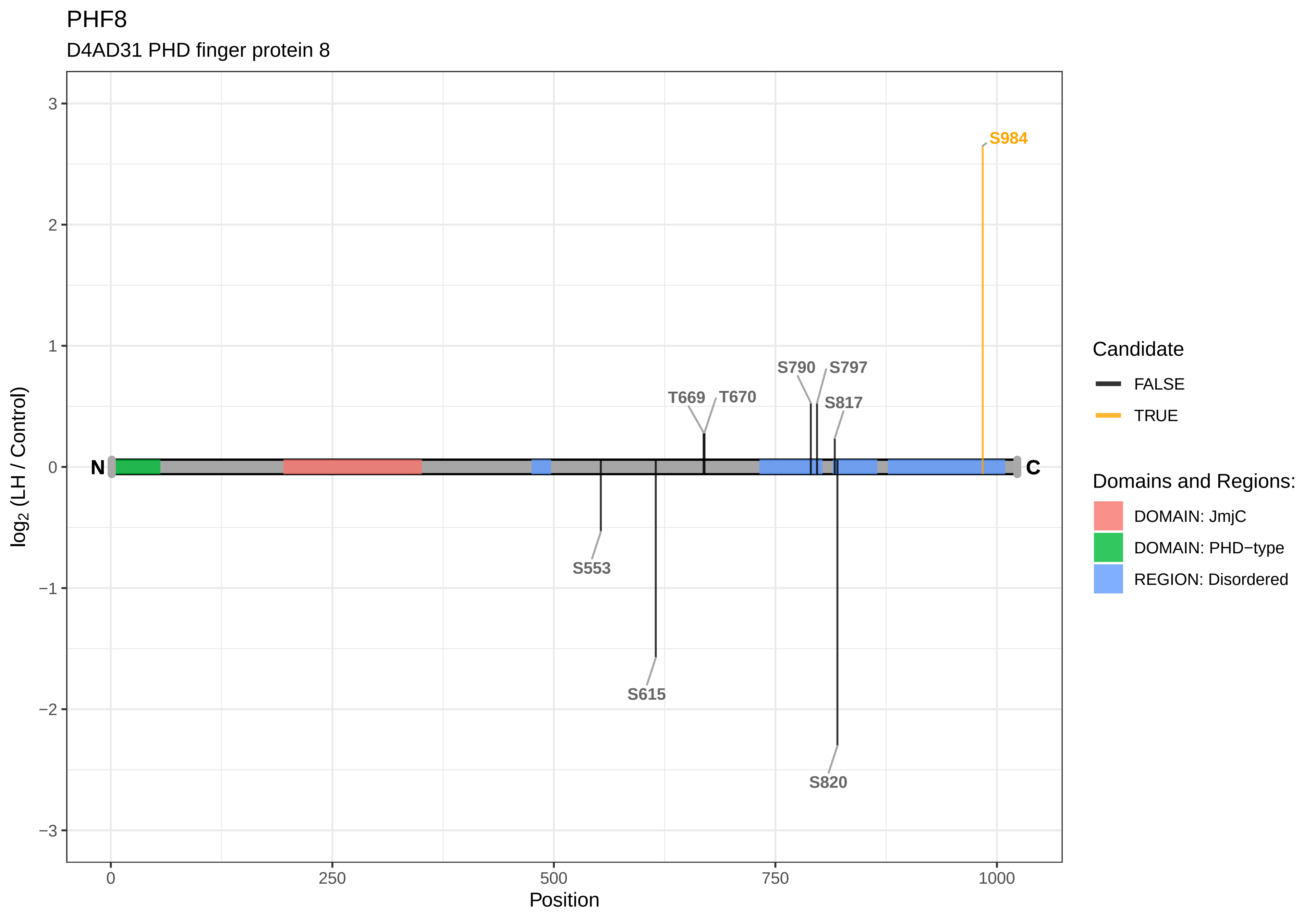
PHF8
D4AD31 PHD finger protein 8
log2 (LH / Control)
Position
3
2
1
0
−1
−2
−3
0
250
500
750
1000
N
C
S984
S790
S797
T669
T670
S817
S553
S615
S820
Candidate
FALSE
TRUE
Domains and Regions:
DOMAIN: JmjC
DOMAIN: PHD−type
REGION: Disordered

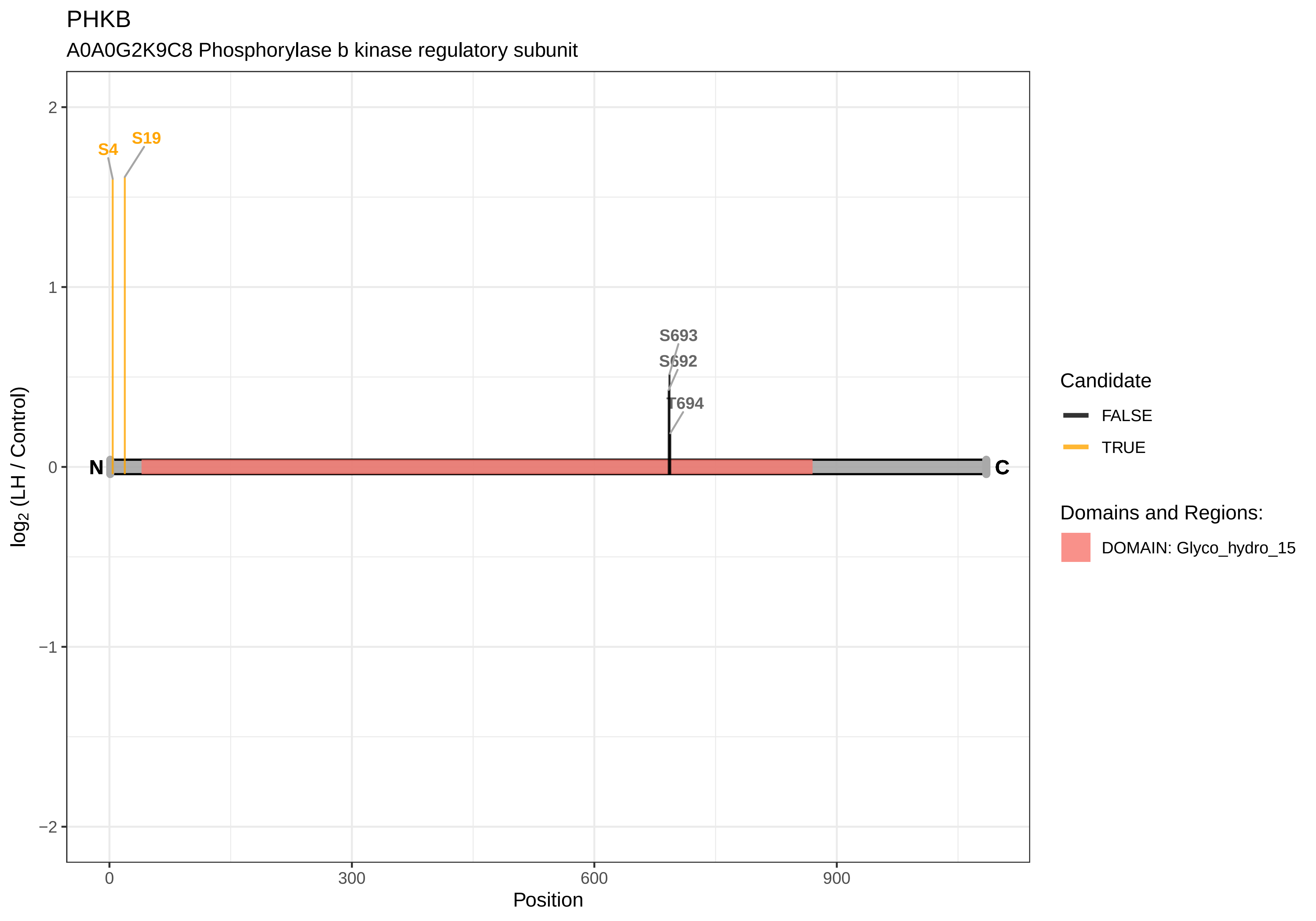

PHKB
A0A0G2K9C8 Phosphorylase b kinase regulatory subunit
S4
S19
S693
S692
T694
N
C
2
1
0
−1
−2
log2 (LH / Control)
0
300
600
900
Position
Candidate
FALSE
TRUE
Domains and Regions:
DOMAIN: Glyco_hydro_15

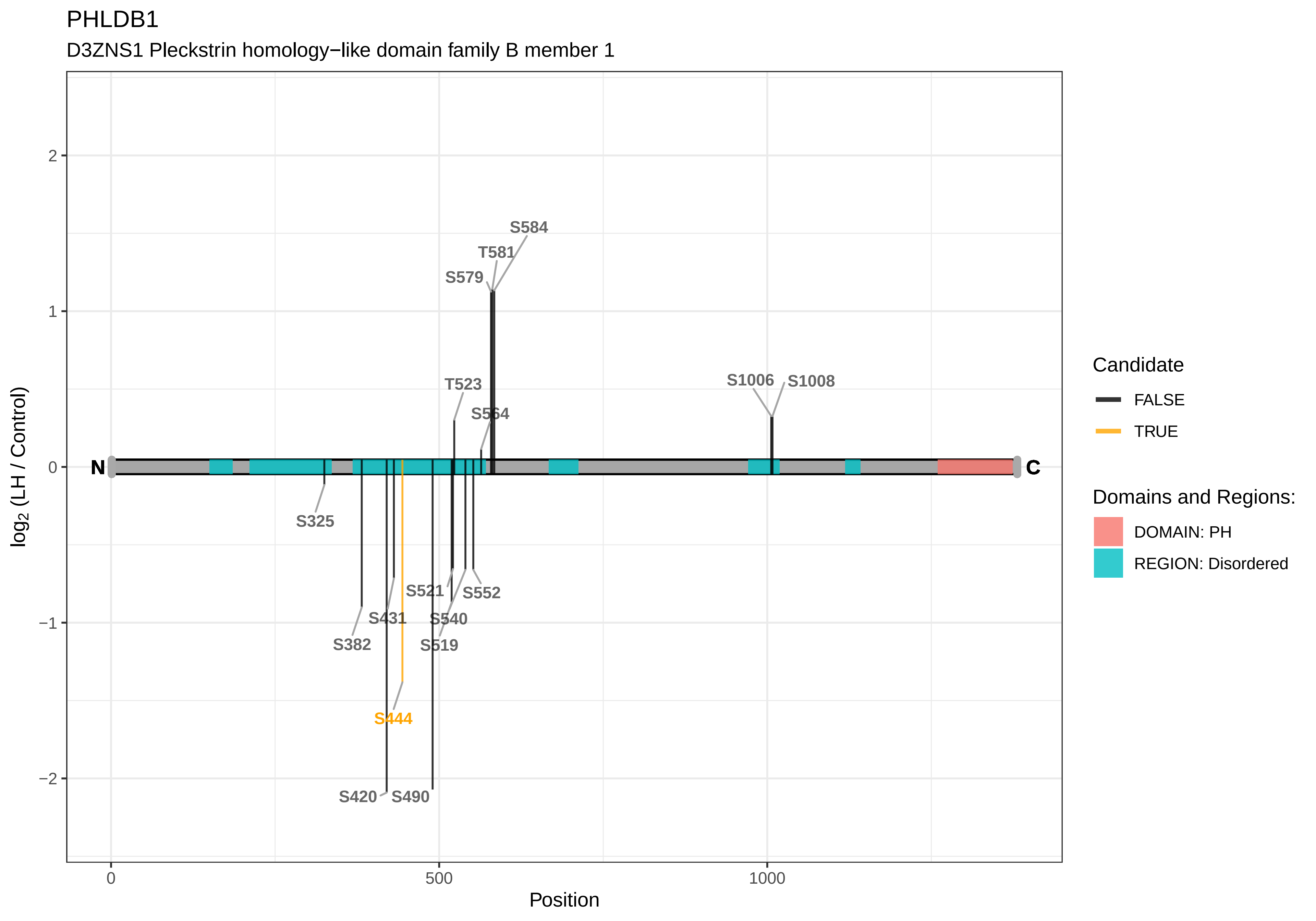
PHLDB1
D3ZNS1 Pleckstrin homology−like domain family B member 1
log2 (LH / Control)
Position
N
C
S584
T581
S579
T523
S564
S1006
S1008
S325
S521
S552
S431
S540
S382
S519
S444
S420
S490
2
1
0
−1
−2
0
500
1000
Candidate
FALSE
TRUE
Domains and Regions:
DOMAIN: PH
REGION: Disordered

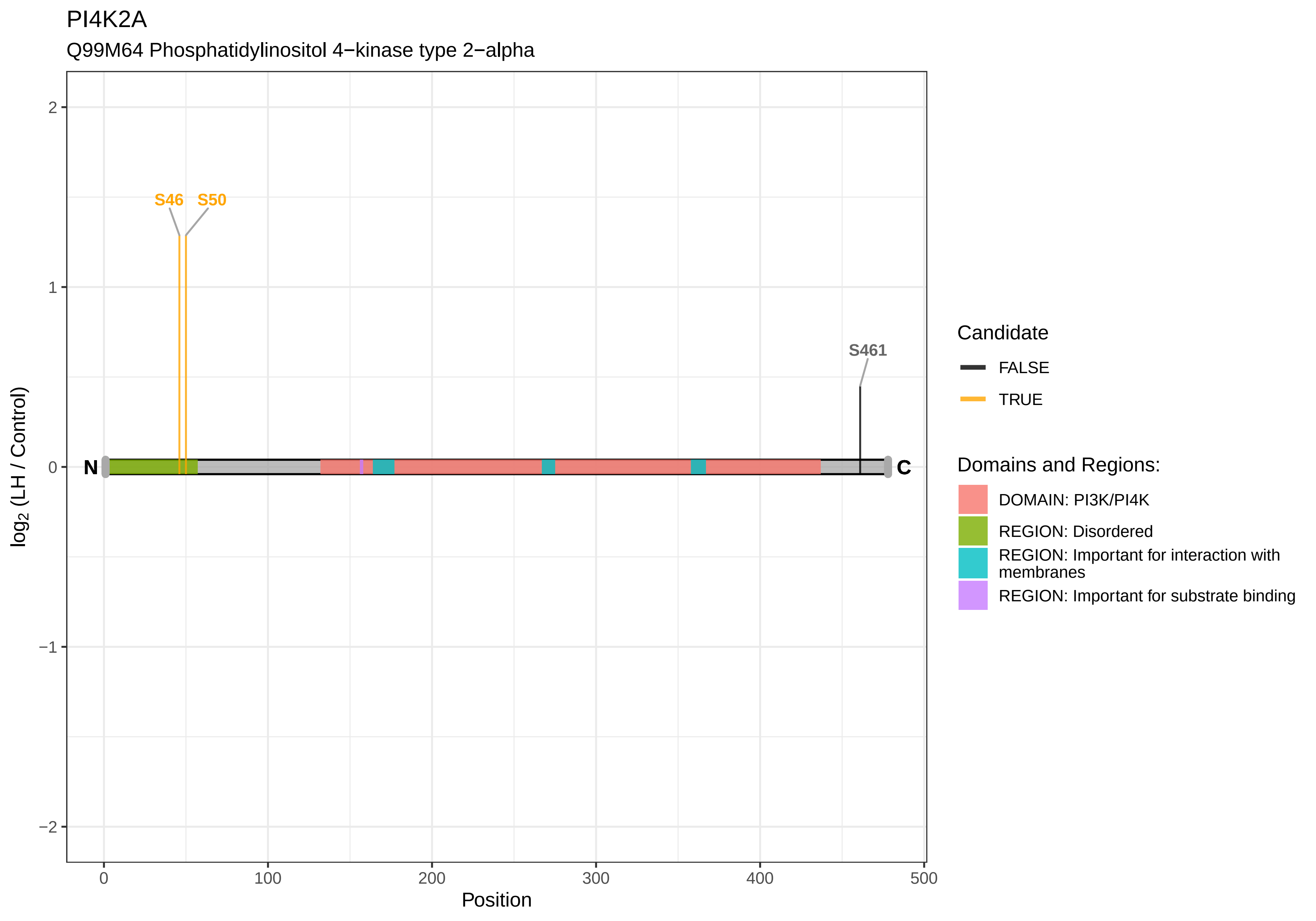
PI4K2A
Q99M64 Phosphatidylinositol 4−kinase type 2−alpha
S46
S50
S461
N
C
log2 (LH / Control)
Position
Candidate
FALSE
TRUE
Domains and Regions:
DOMAIN: PI3K/PI4K
REGION: Disordered
REGION: Important for interaction with membranes
REGION: Important for substrate binding

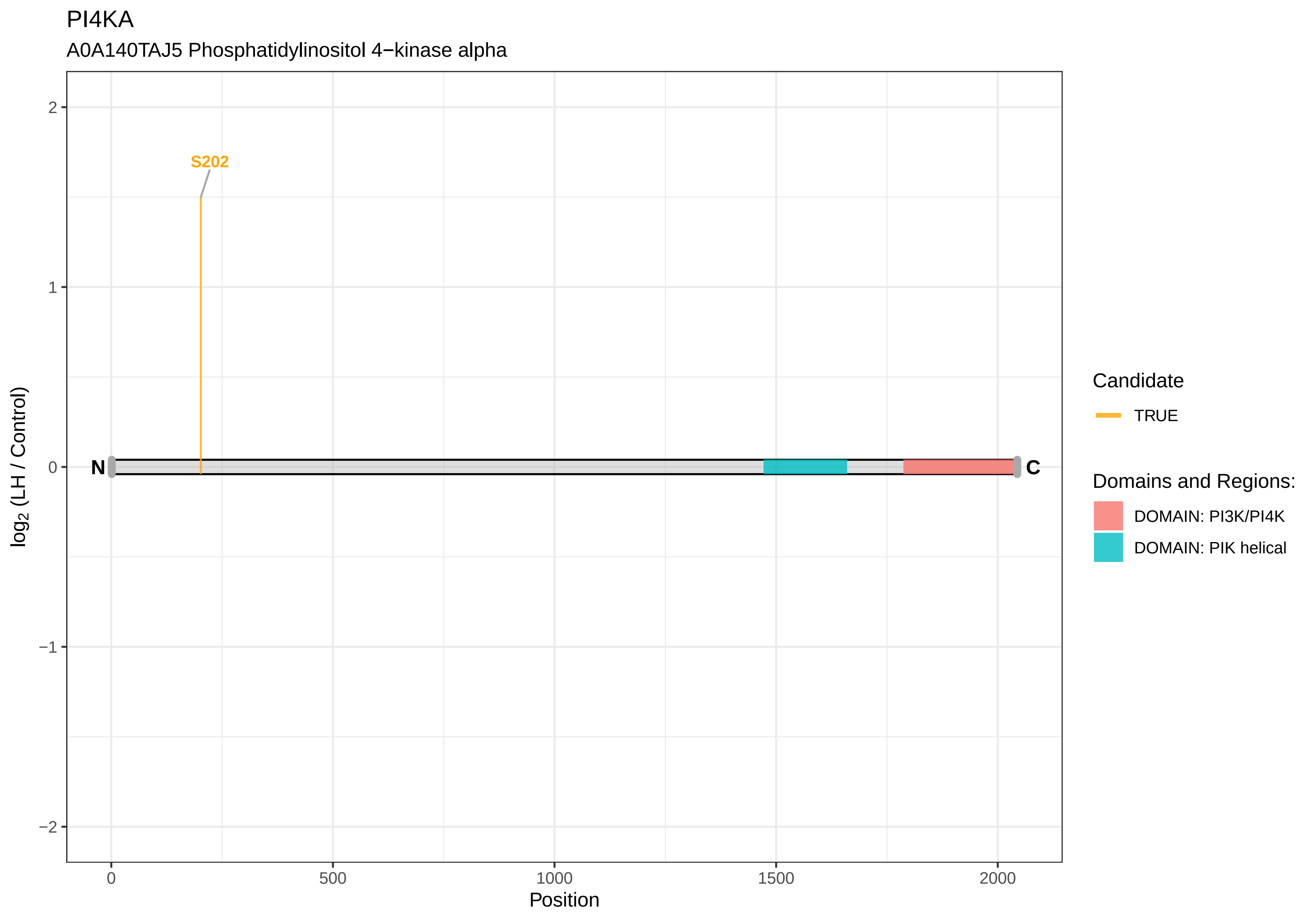
PI4KA
A0A140TAJ5 Phosphatidylinositol 4−kinase alpha
S202
N
C
2
1
0
−1
−2
0
500
1000
1500
2000
Position
log2 (LH / Control)
Candidate
TRUE
Domains and Regions:
DOMAIN: PI3K/PI4K
DOMAIN: PIK helical

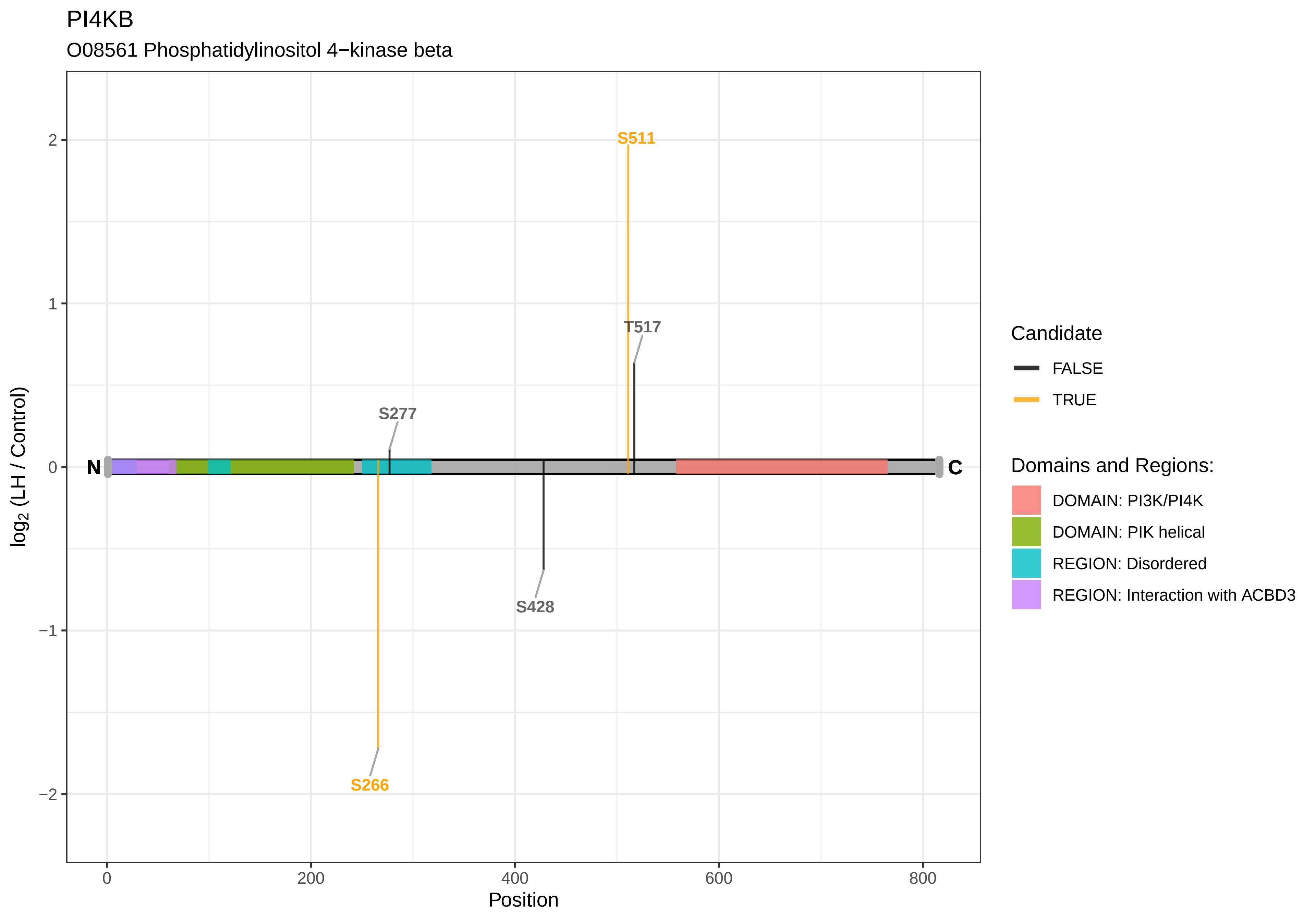

PI4KB
O08561 Phosphatidylinositol 4−kinase beta
S511
T517
S277
S428
S266
N
C
2
1
0
−1
−2
log2 (LH / Control)
0
200
400
600
800
Position
Candidate
FALSE
TRUE
Domains and Regions:
DOMAIN: PI3K/PI4K
DOMAIN: PIK helical
REGION: Disordered
REGION: Interaction with ACBD3

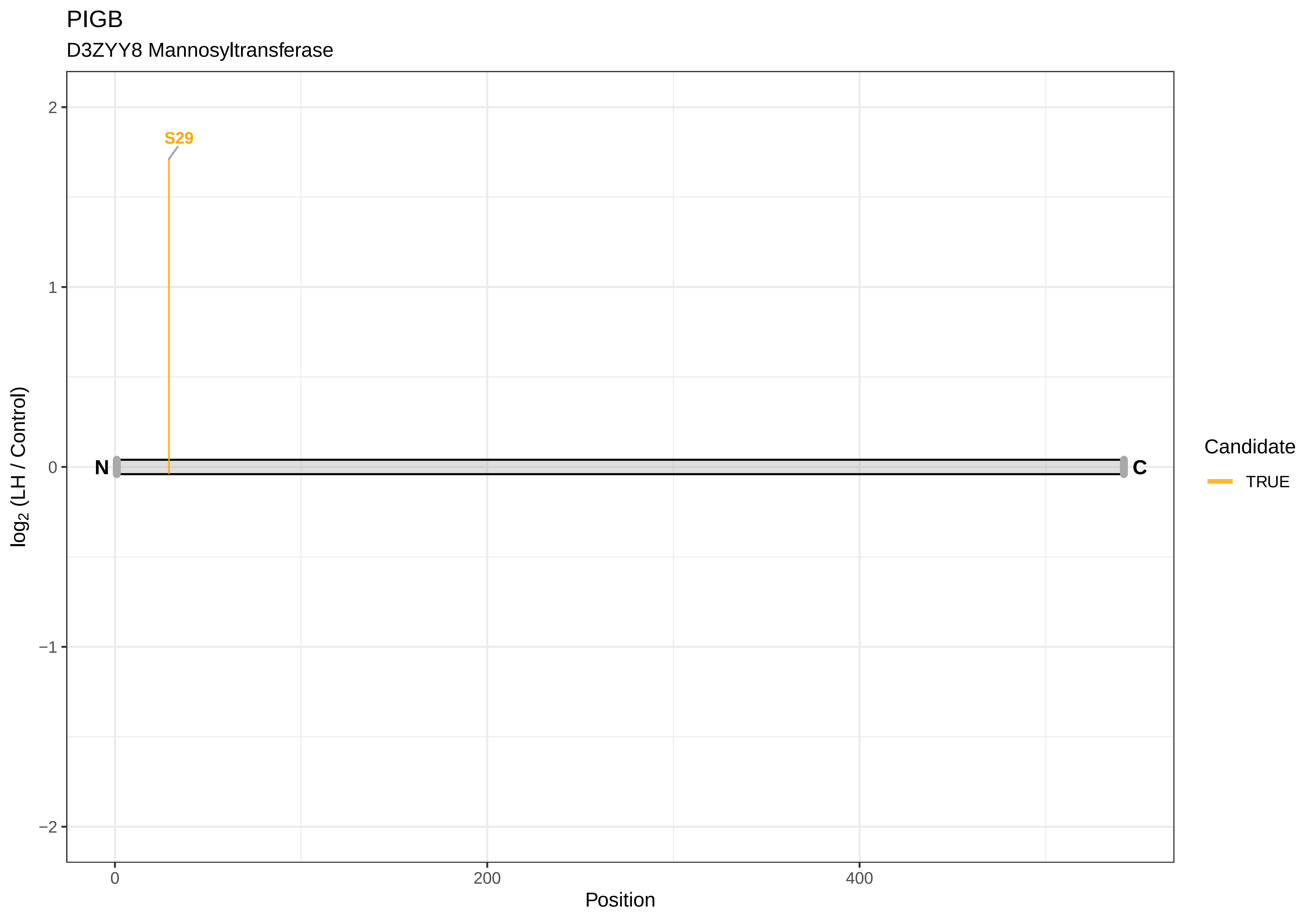
PIGB
D3ZYY8 Mannosyltransferase
S29
N
C
2
1
0
−1
−2
log2 (LH / Control)
0
200
400
Position
Candidate
TRUE

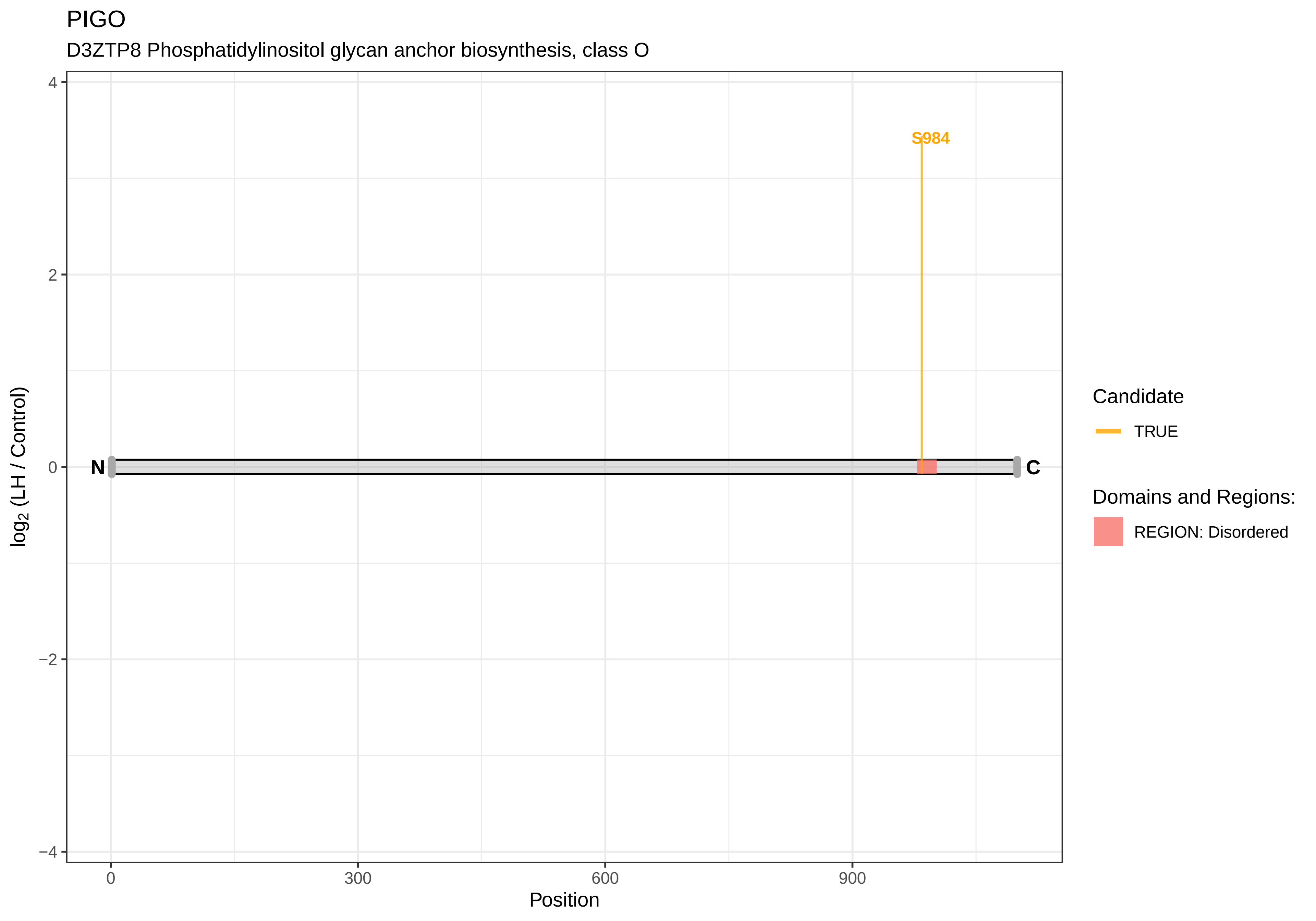
PIGO
D3ZTP8 Phosphatidylinositol glycan anchor biosynthesis, class O
S984
N
C
4
2
0
−2
−4
log2 (LH / Control)
0
300
600
900
Position
Candidate
TRUE
Domains and Regions:
REGION: Disordered

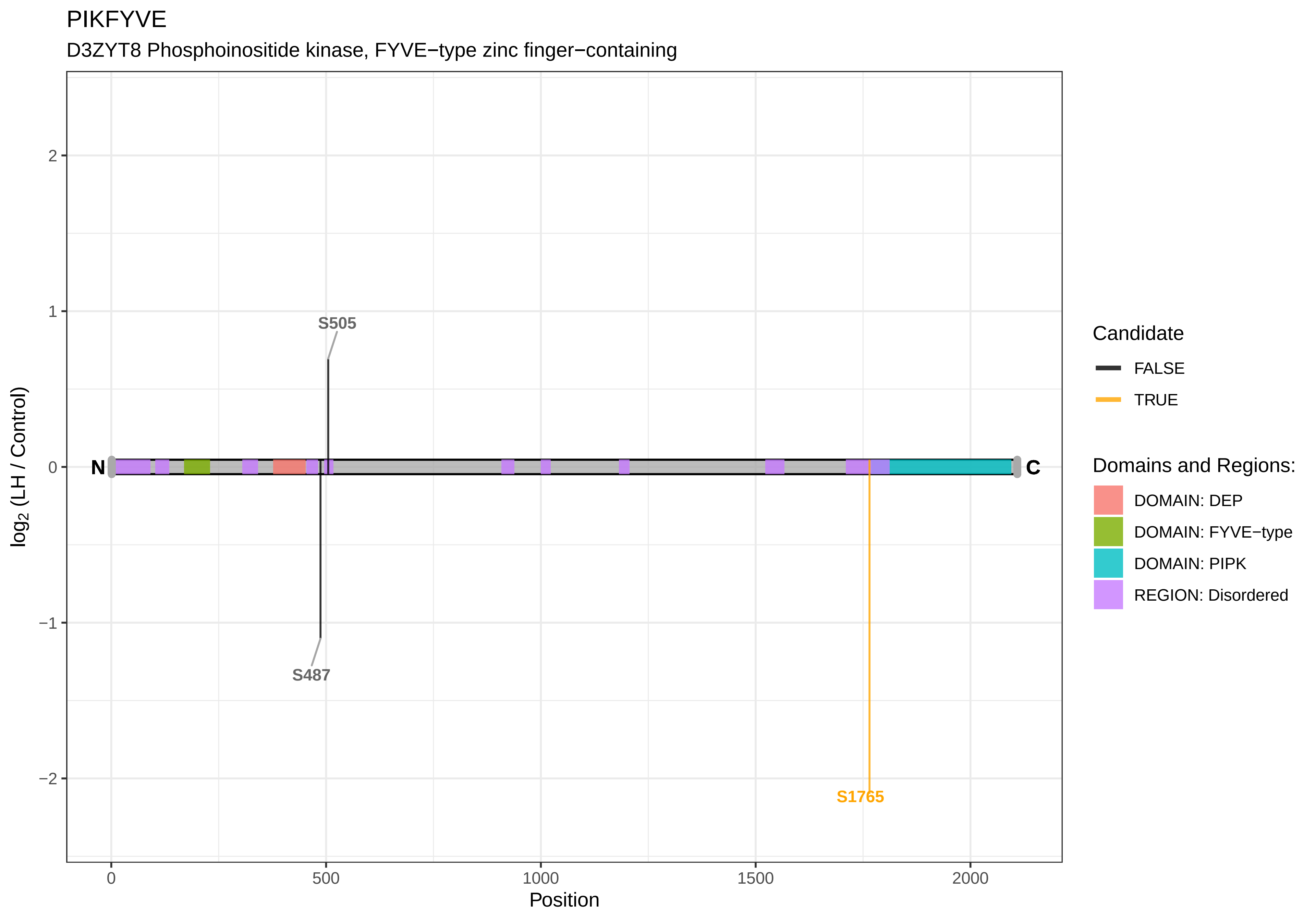
PIKFYVE
D3ZYT8 Phosphoinositide kinase, FYVE−type zinc finger−containing
S505
S487
S1765
N
C
log2 (LH / Control)
Position
0
500
1000
1500
2000
2
1
0
−1
−2
Candidate
FALSE
TRUE
Domains and Regions:
DOMAIN: DEP
DOMAIN: FYVE−type
DOMAIN: PIPK
REGION: Disordered

PITPNB

P53812 Phosphatidylinositol transfer protein beta isoform

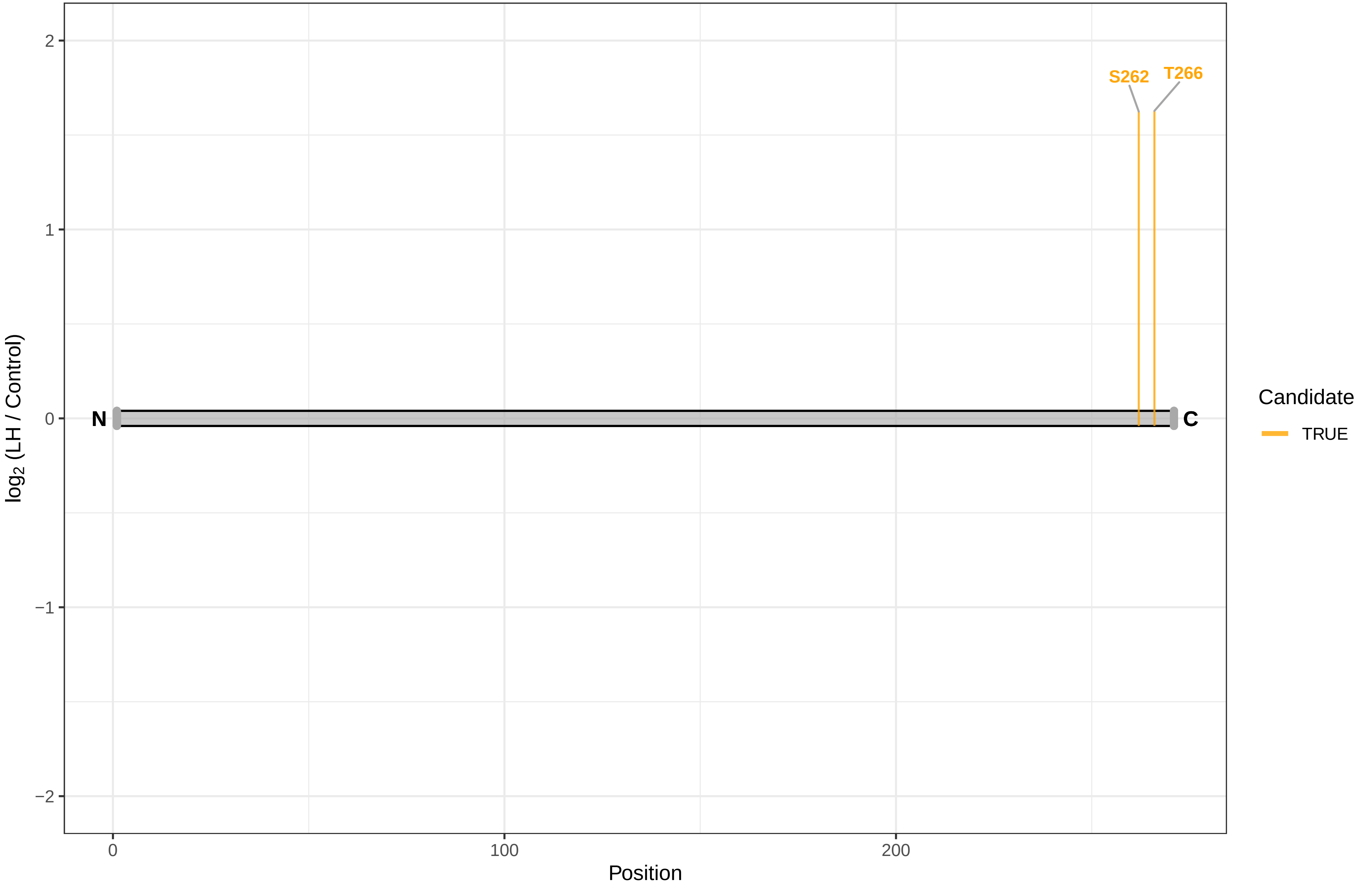

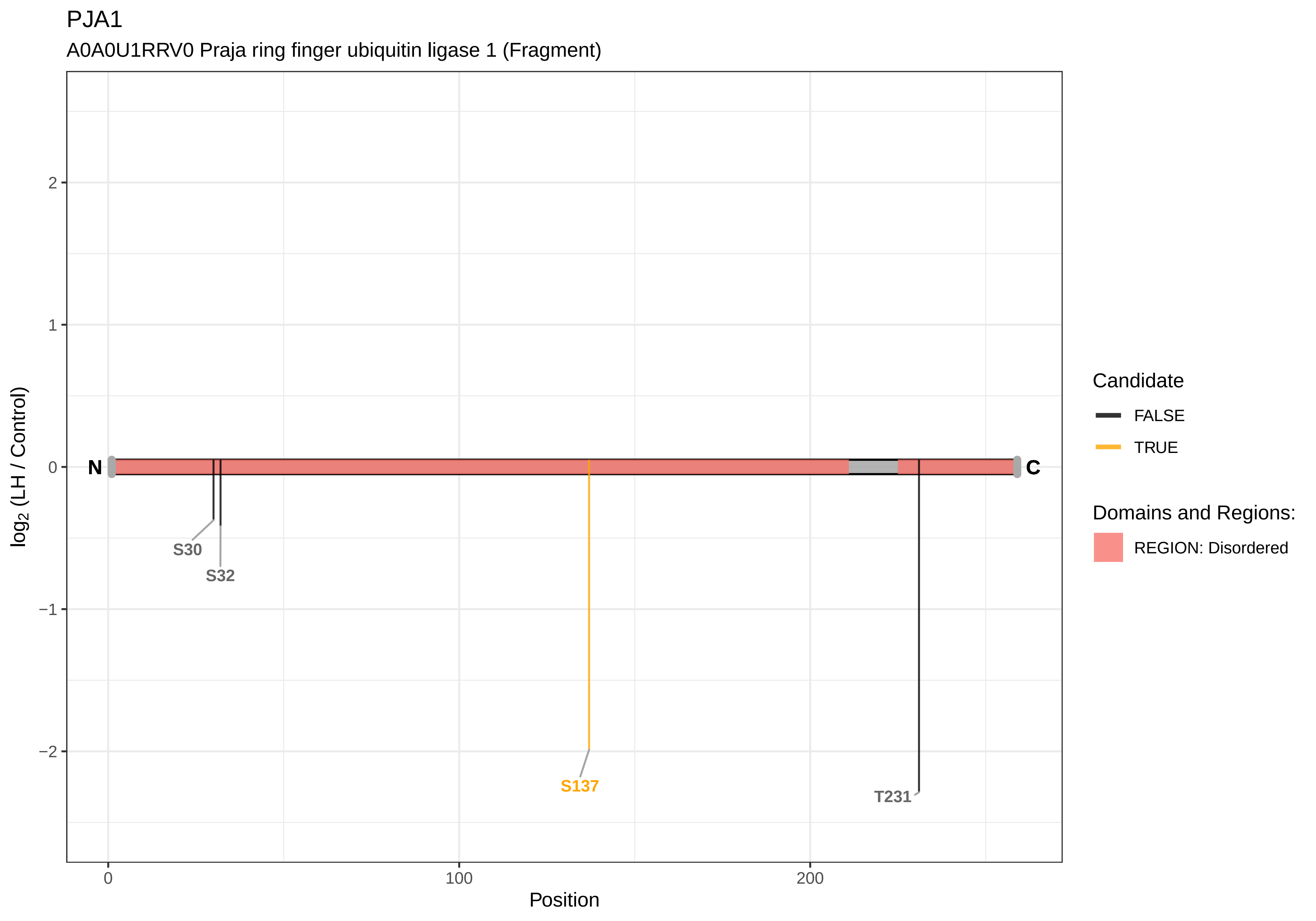
PJA1
A0A0U1RRV0 Praja ring finger ubiquitin ligase 1 (Fragment)
log2 (LH / Control)
Position
N
C
S30
S32
S137
T231
2
1
0
−1
−2
0
100
200
Candidate
FALSE
TRUE
Domains and Regions:
REGION: Disordered

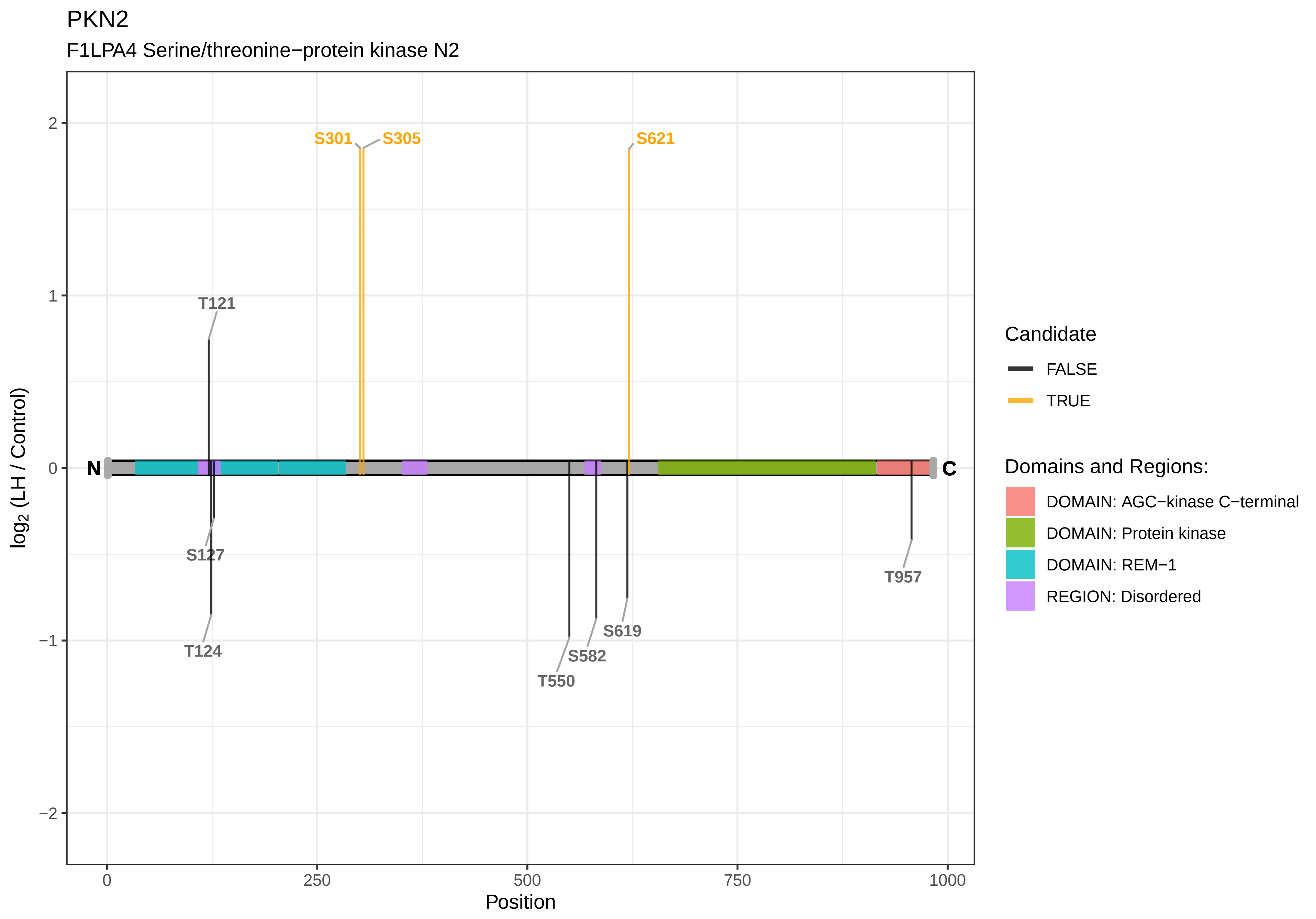

PKN2
F1LPA4 Serine/threonine−protein kinase N2
S301
S305
S621
T121
S127
T124
T550
S582
S619
T957
N
C
2
1
0
−1
−2
log2 (LH / Control)
0
250
500
750
1000
Position
Candidate
FALSE
TRUE
Domains and Regions:
DOMAIN: AGC−kinase C−terminal
DOMAIN: Protein kinase
DOMAIN: REM−1
REGION: Disordered

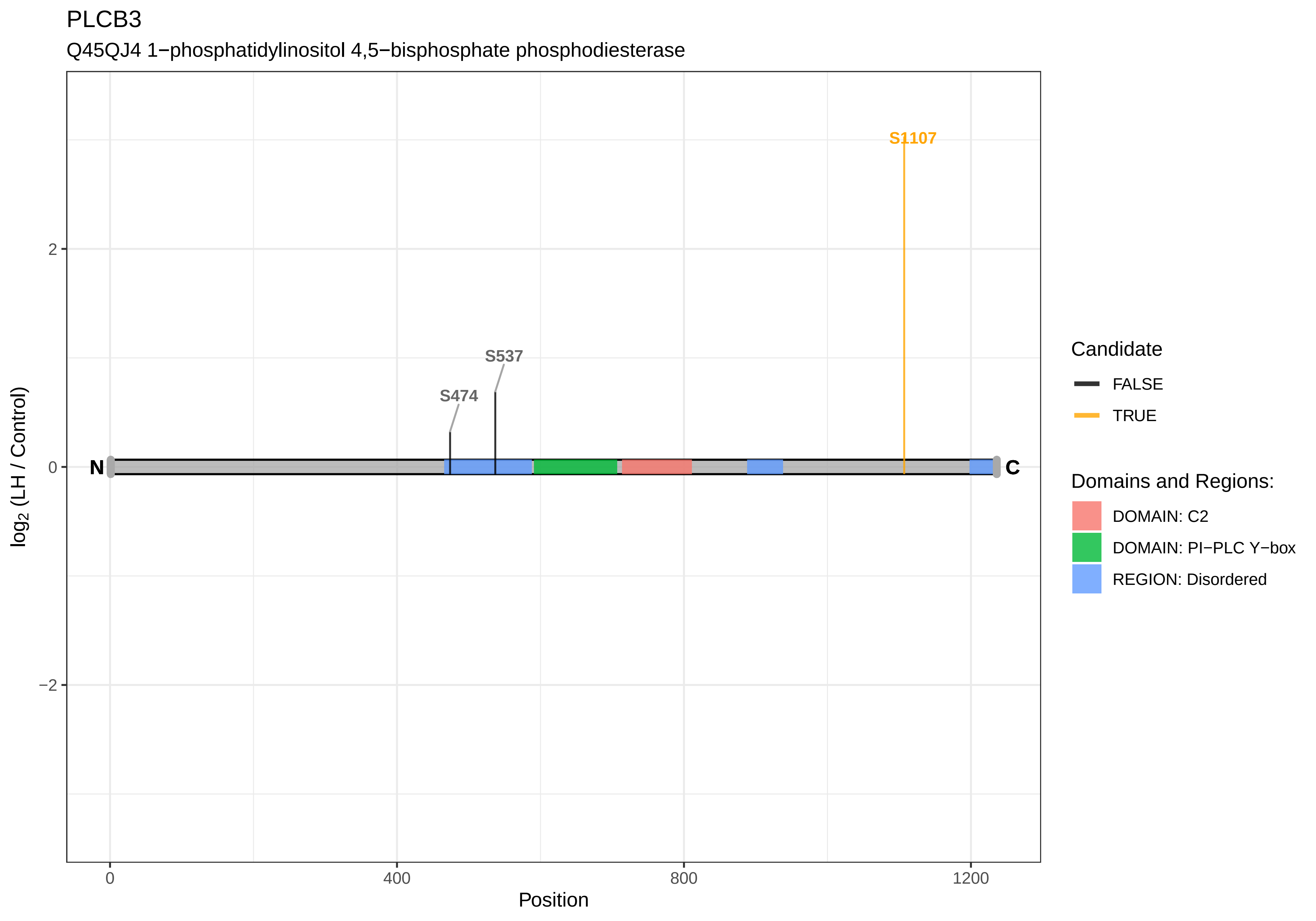
PLCB3
Q45QJ4 1−phosphatidylinositol 4,5−bisphosphate phosphodiesterase
S1107
S537
S474
N
C
2
0
−2
log2 (LH / Control)
0
400
800
1200
Position
Candidate
FALSE
TRUE
Domains and Regions:
DOMAIN: C2
DOMAIN: PI−PLC Y−box
REGION: Disordered

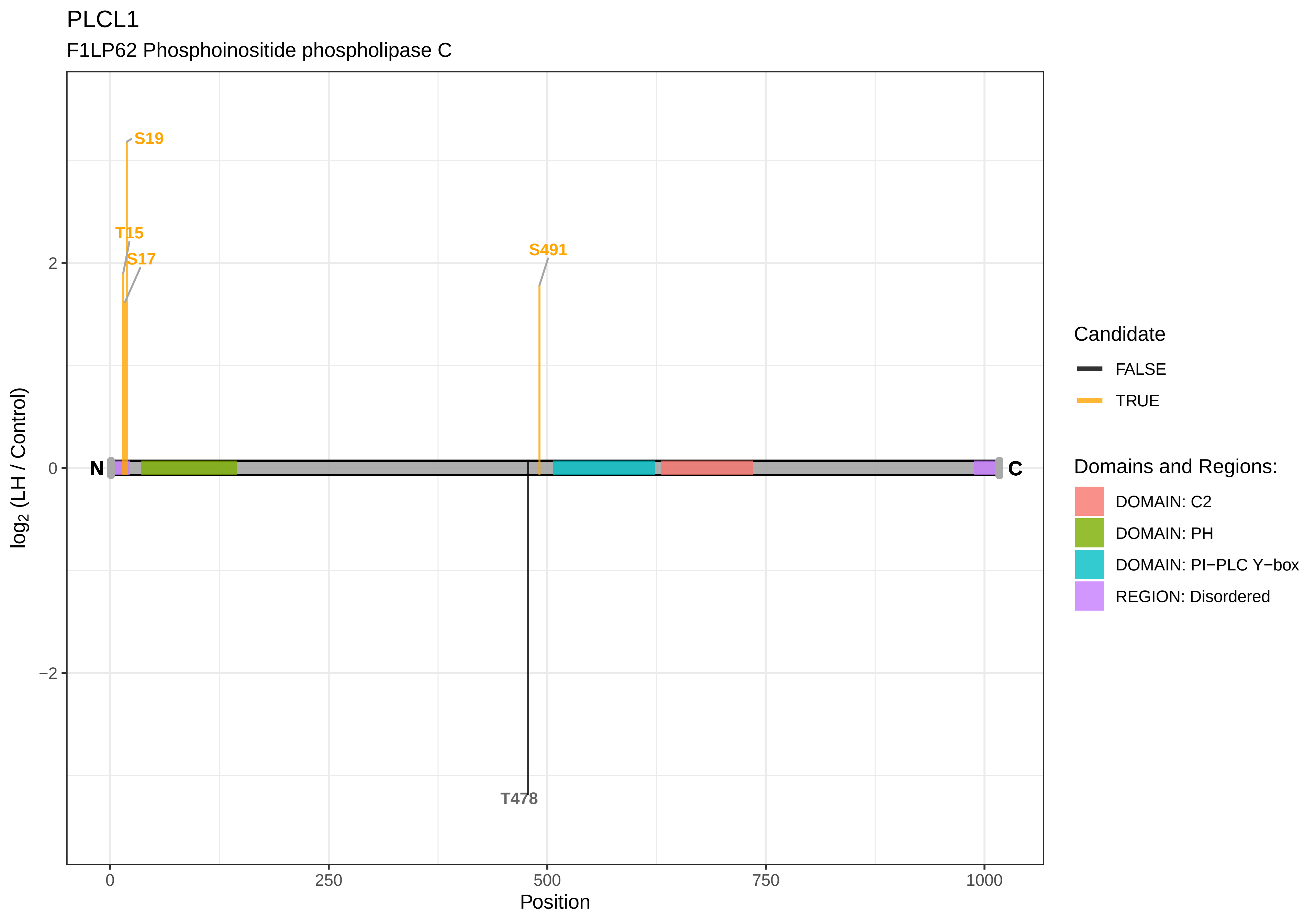
PLCL1
F1LP62 Phosphoinositide phospholipase C
S19
T15
S17
S491
T478
N
C
2
0
−2
log2 (LH / Control)
0
250
500
750
1000
Position
Candidate
FALSE
TRUE
Domains and Regions:
DOMAIN: C2
DOMAIN: PH
DOMAIN: PI−PLC Y−box
REGION: Disordered

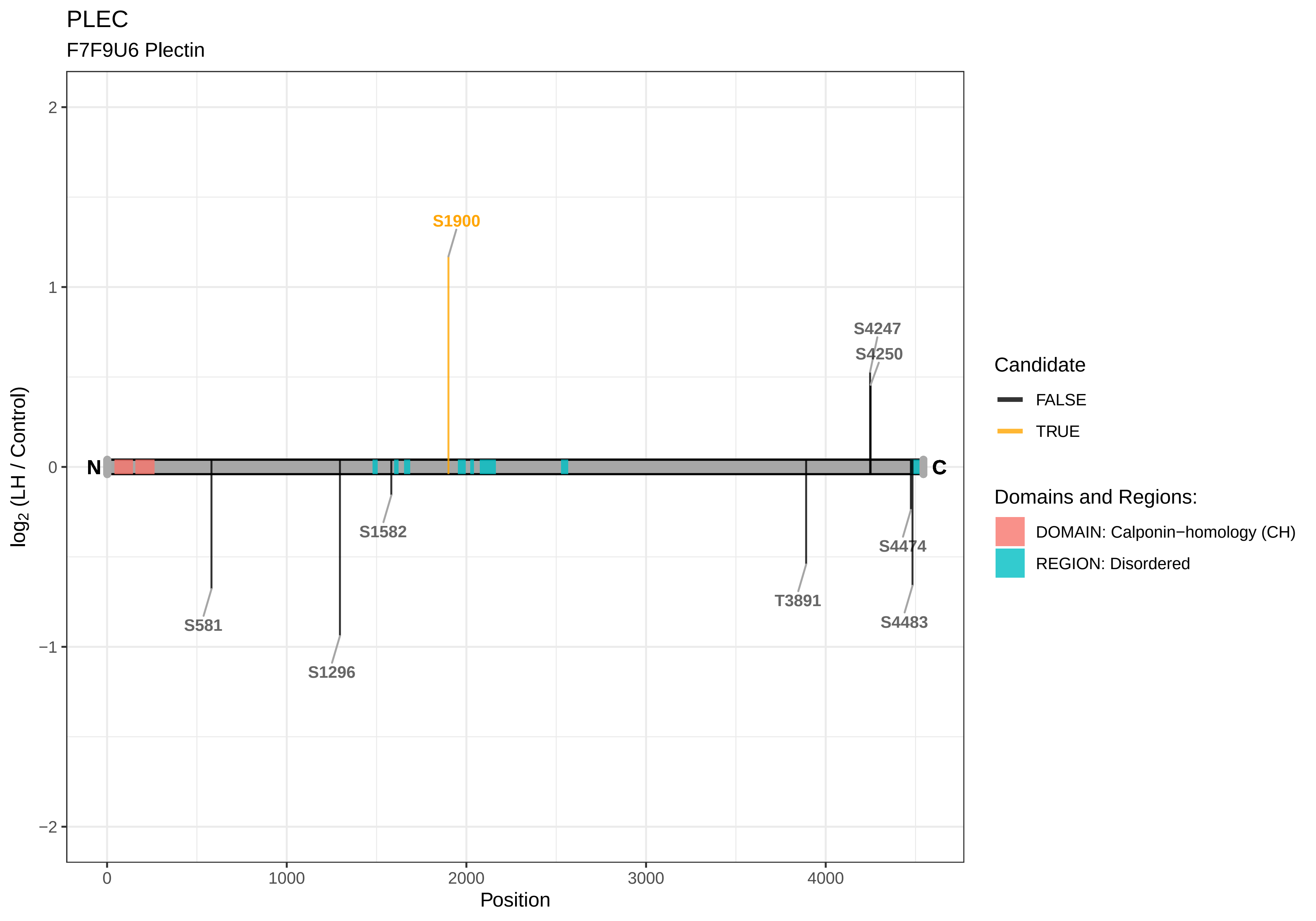

PLEC
F7F9U6 Plectin
S1900
S4247
S4250
N
C
S1582
S581
S1296
T3891
S4474
S4483
2
1
0
−1
−2
log2 (LH / Control)
0
1000
2000
3000
4000
Position
Candidate
FALSE
TRUE
Domains and Regions:
DOMAIN: Calponin−homology (CH)
REGION: Disordered

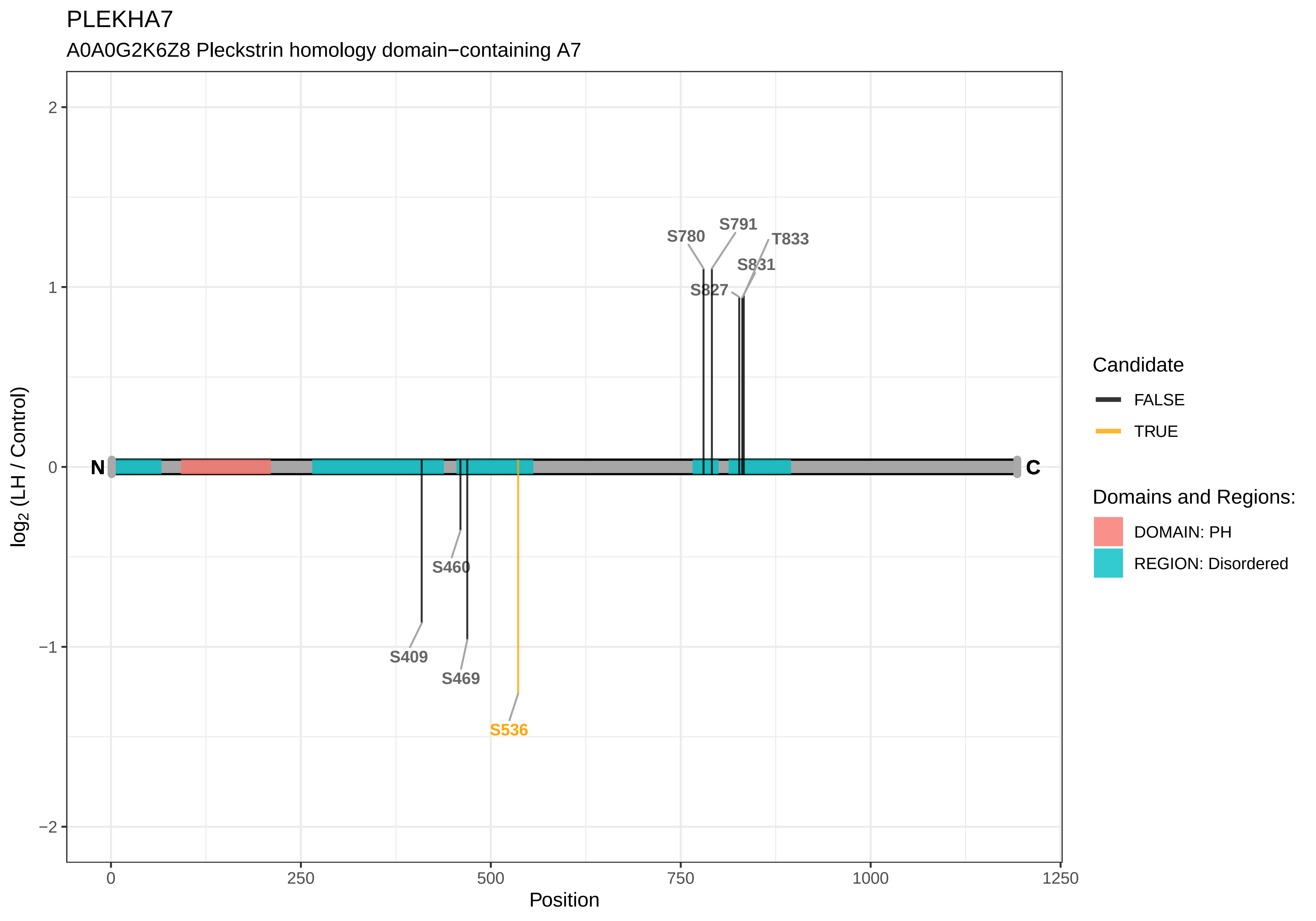
PLEKHA7
A0A0G2K6Z8 Pleckstrin homology domain−containing A7
S780
S791
T833
S831
S827
S460
S409
S469
S536
N
C
2
1
0
−1
−2
log2 (LH / Control)
0
250
500
750
1000
1250
Position
Candidate
FALSE
TRUE
Domains and Regions:
DOMAIN: PH
REGION: Disordered

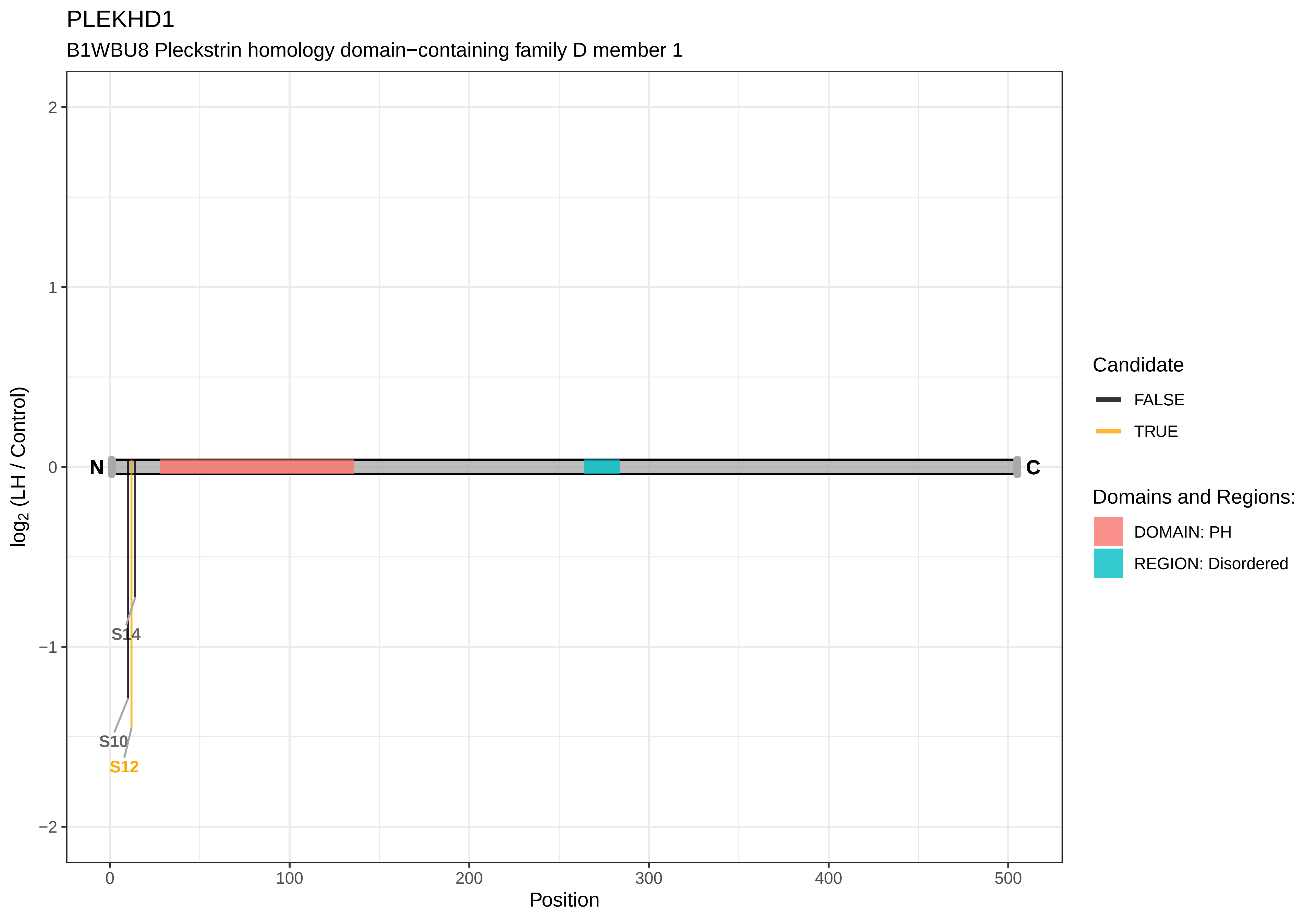

PLEKHD1
B1WBU8 Pleckstrin homology domain−containing family D member 1
log2 (LH / Control)
2
1
0
−1
−2
N
C
S14
S10
S12
0
100
200
300
400
500
Position
Candidate
FALSE
TRUE
Domains and Regions:
DOMAIN: PH
REGION: Disordered

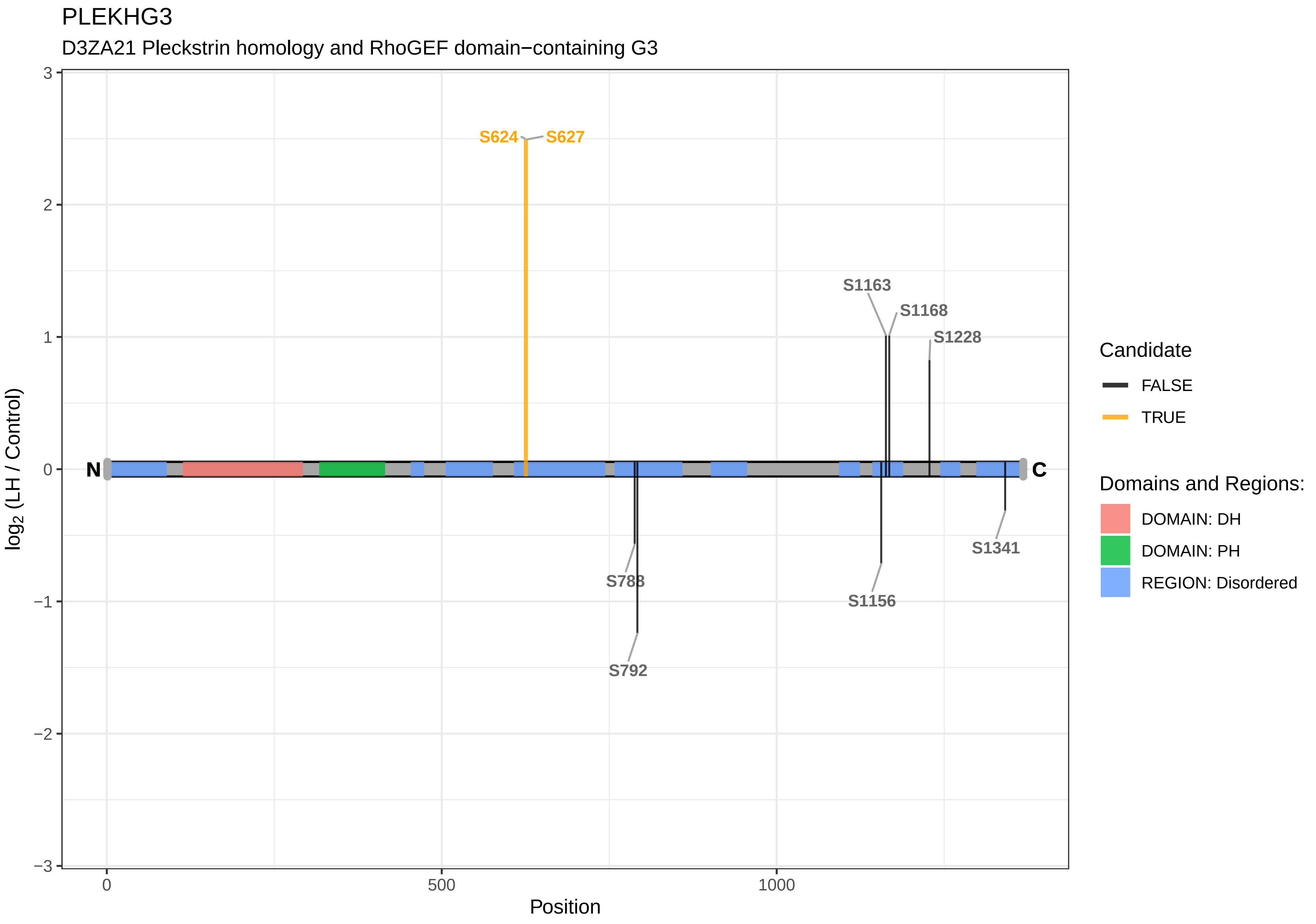

PLEKHG3
D3ZA21 Pleckstrin homology and RhoGEF domain−containing G3
S624
S627
S1163
S1168
S1228
S788
S792
S1156
S1341
N
C
log2 (LH / Control)
3
2
1
0
−1
−2
−3
0
500
1000
Position
Candidate
FALSE
TRUE
Domains and Regions:
DOMAIN: DH
DOMAIN: PH
REGION: Disordered

PLEKHM1

Q5PQS0 Pleckstrin homology domain−containing family M member 1

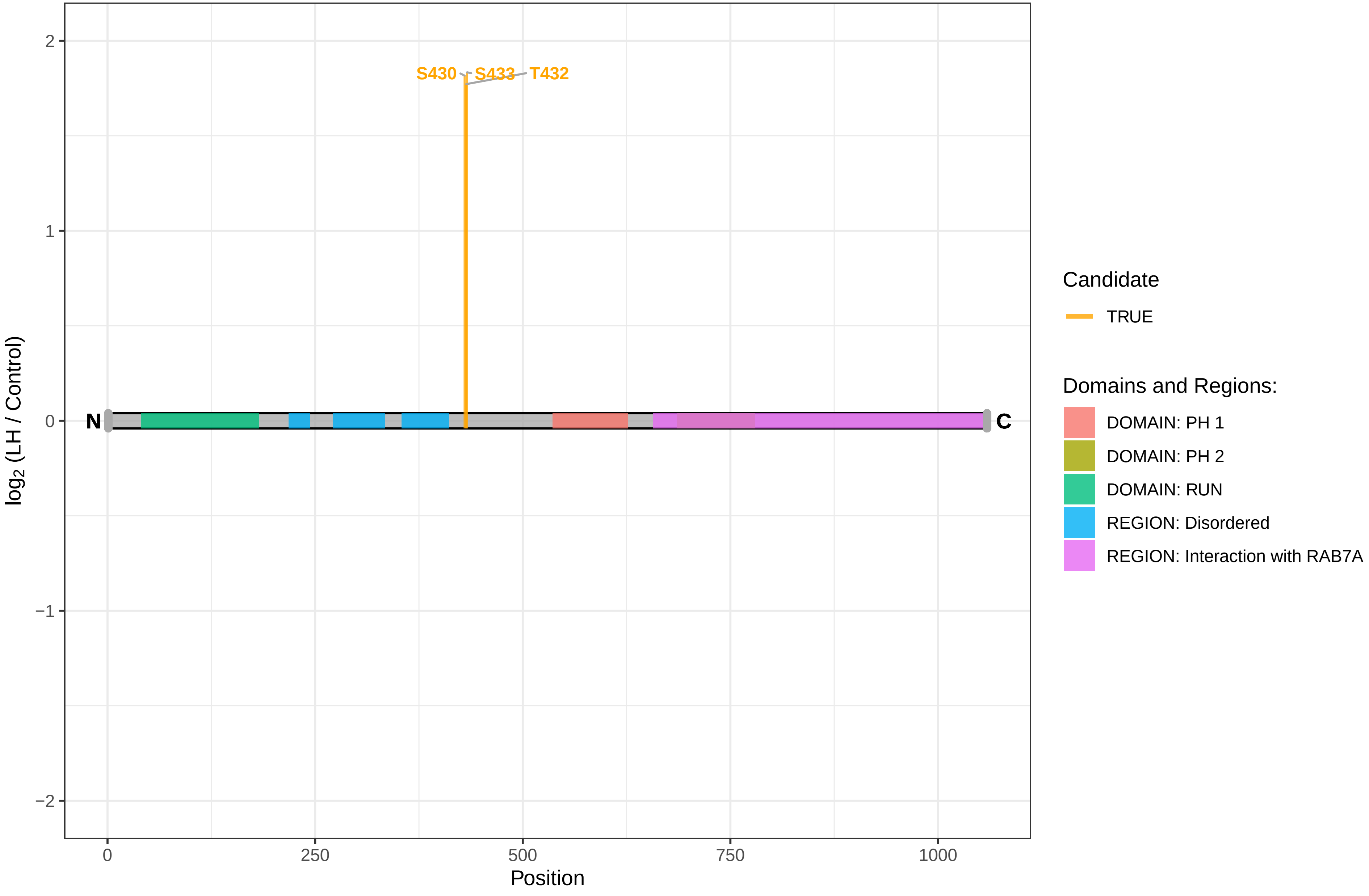

PLPP7

Q5FVJ3 Inactive phospholipid phosphatase 7

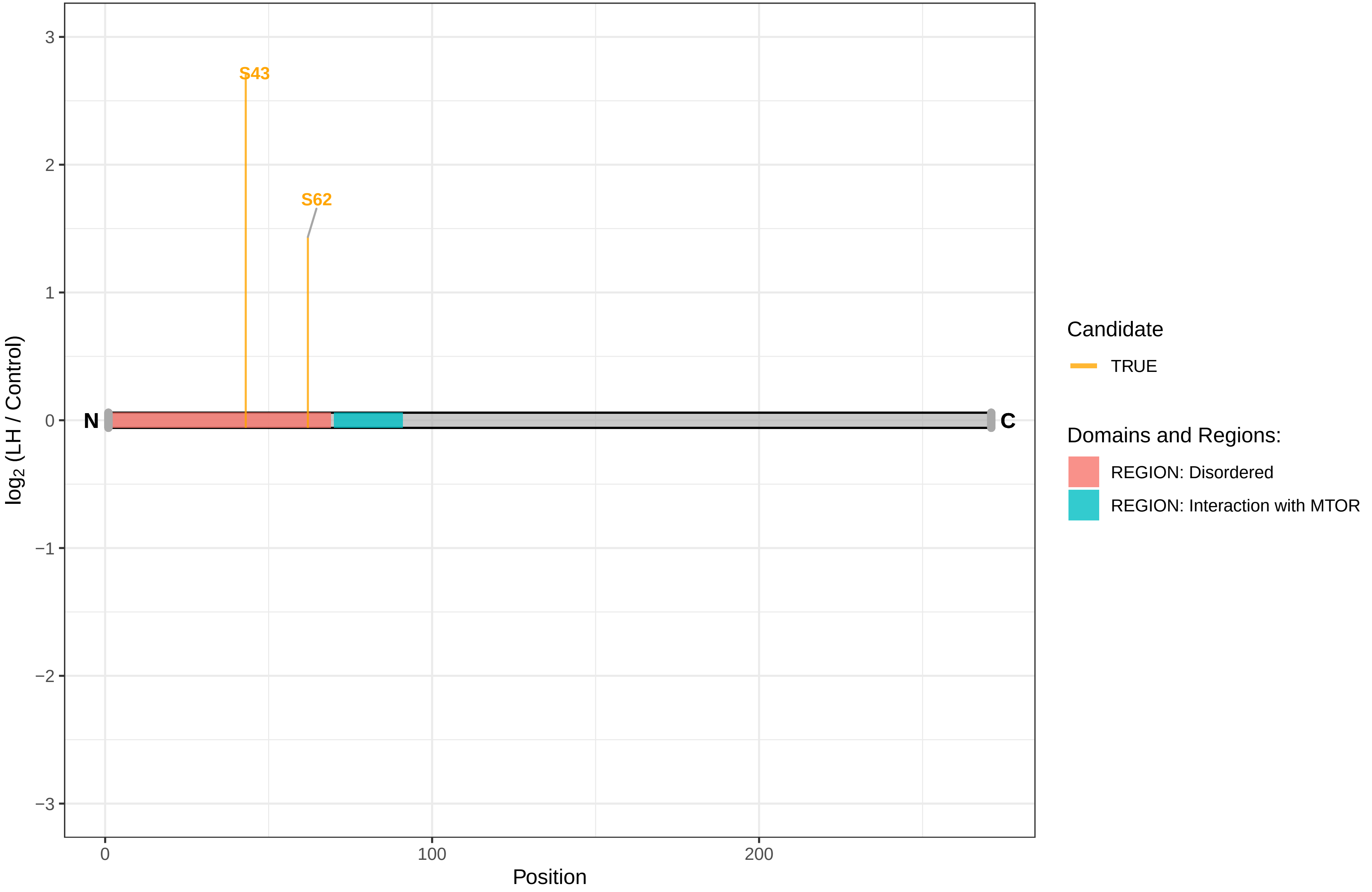

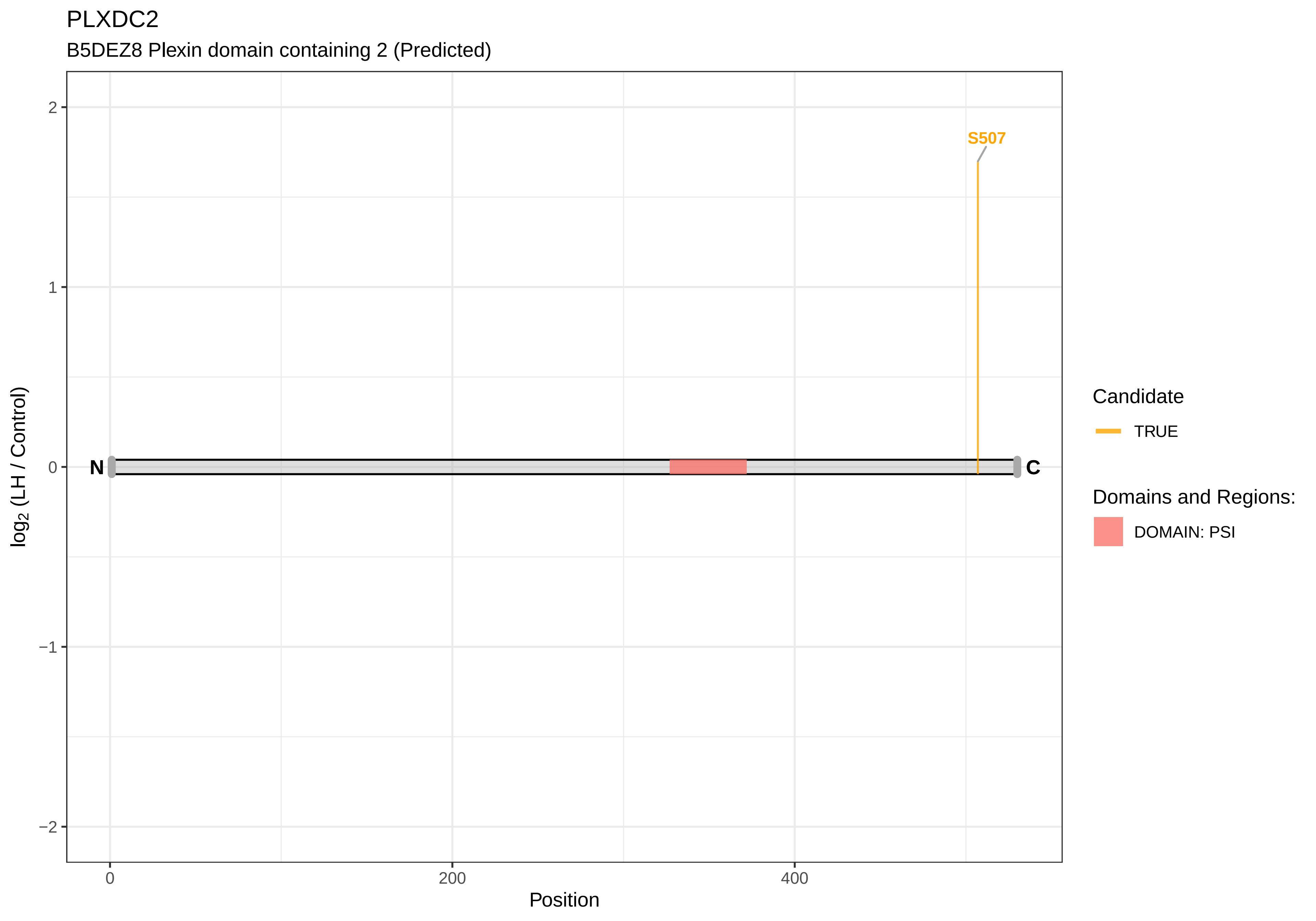
PLXDC2
B5DEZ8 Plexin domain containing 2 (Predicted)
S507
N
C
2
1
0
−1
−2
log2 (LH / Control)
0
200
400
Position
Candidate
TRUE
Domains and Regions:
DOMAIN: PSI

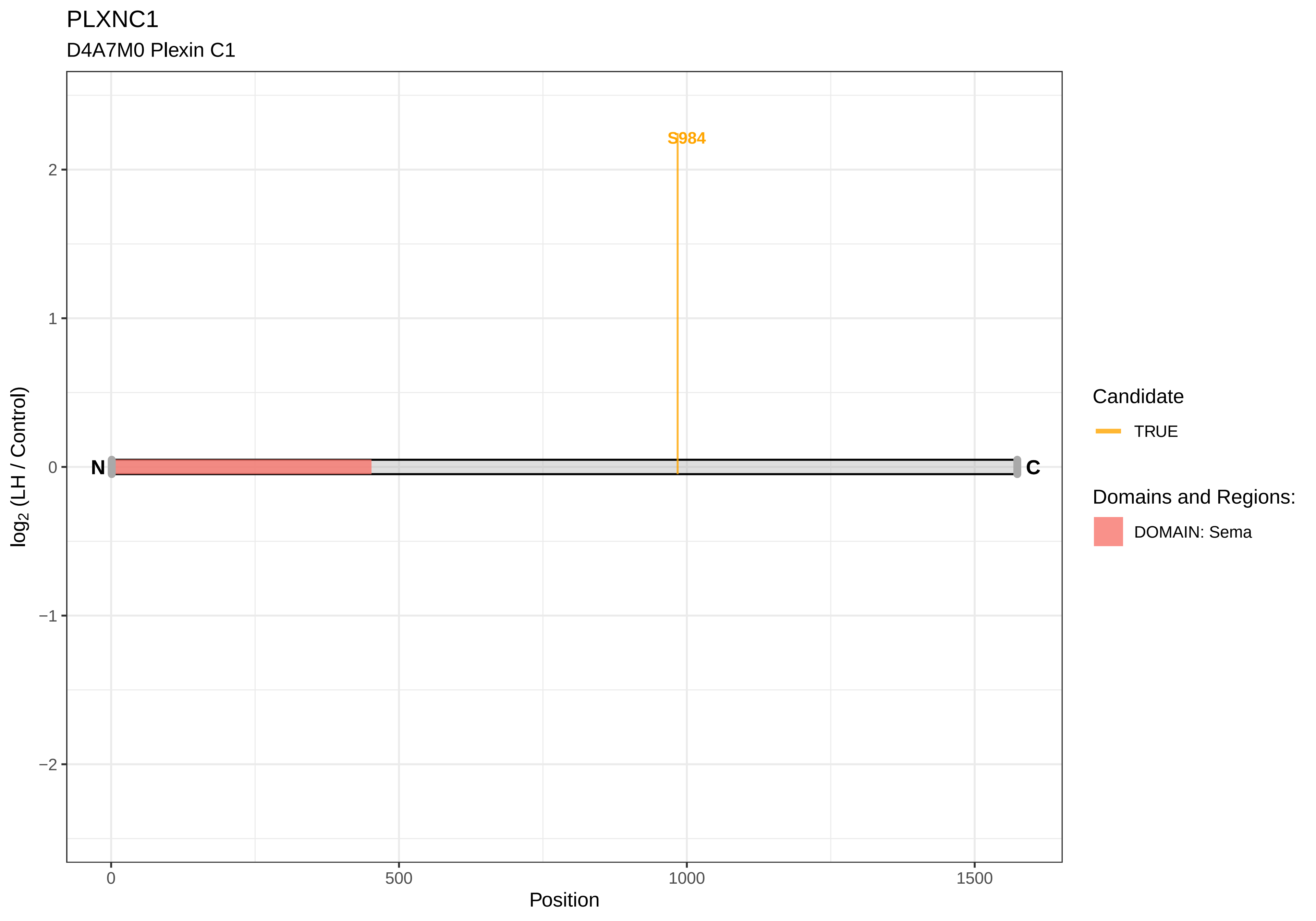
PLXNC1
D4A7M0 Plexin C1
S984
N
C
2
1
0
−1
−2
0
500
1000
1500
Position
log2 (LH / Control)
Candidate
TRUE
Domains and Regions:
DOMAIN: Sema

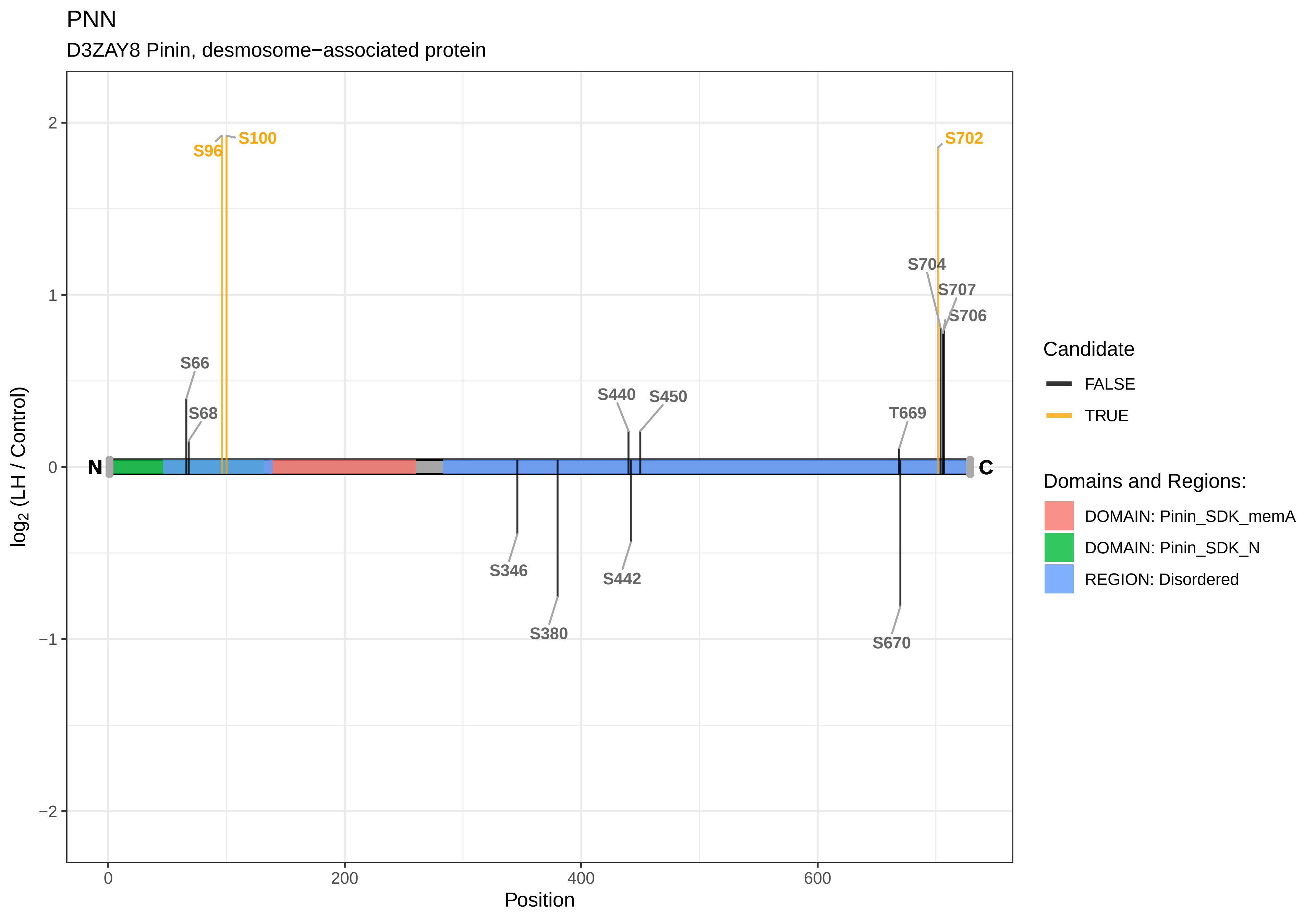

PNN
D3ZAY8 Pinin, desmosome−associated protein
S96
S100
S702
S704
S707
S706
S66
S68
S440
S450
T669
N
C
S346
S442
S380
S670
log2 (LH / Control)
2
1
0
−1
−2
0
200
400
600
Position
Candidate
FALSE
TRUE
Domains and Regions:
DOMAIN: Pinin_SDK_memA
DOMAIN: Pinin_SDK_N
REGION: Disordered

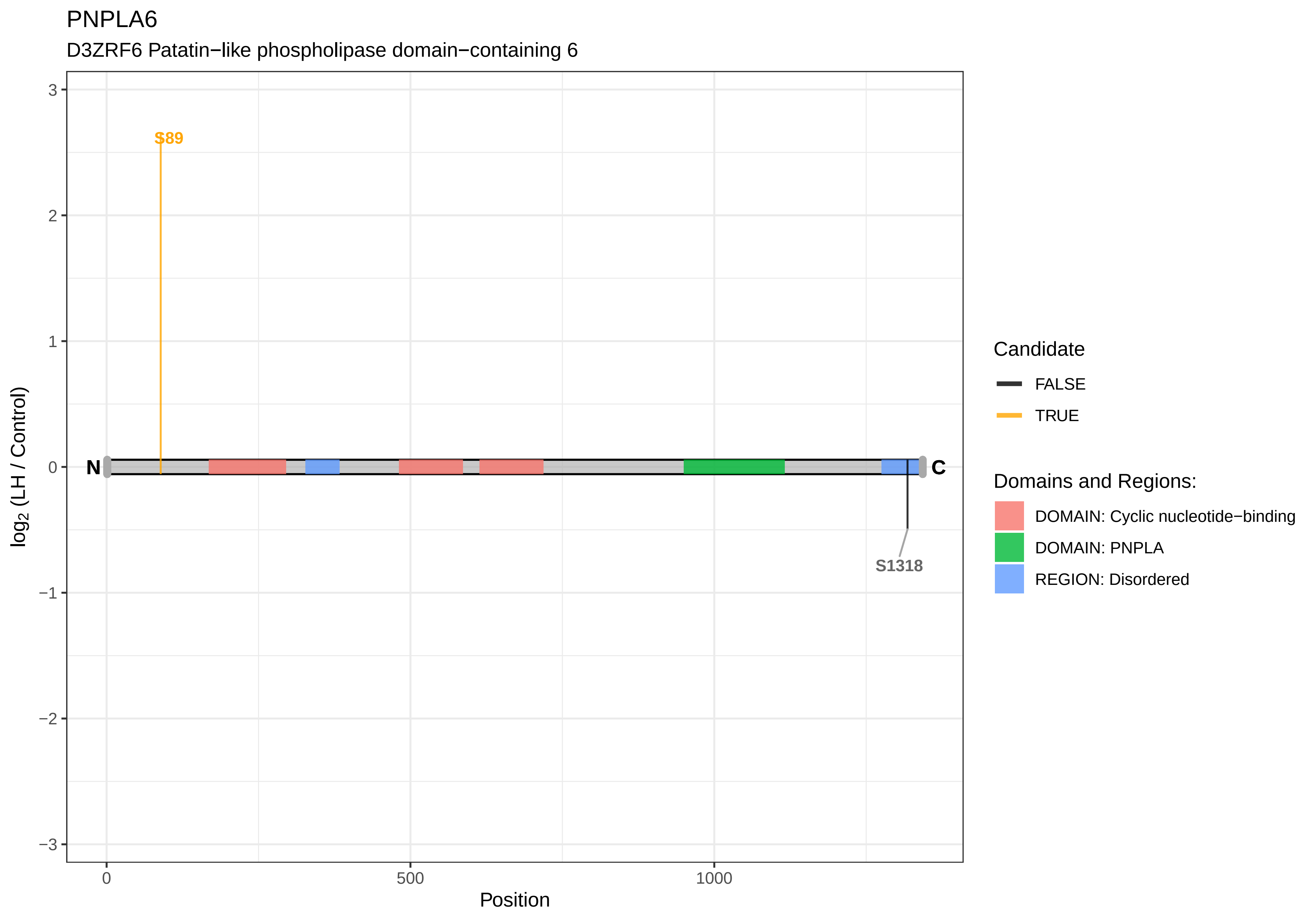
PNPLA6
D3ZRF6 Patatin−like phospholipase domain−containing 6
S89
S1318
N
C
3
2
1
0
−1
−2
−3
0
500
1000
Position
log2 (LH / Control)
Candidate
FALSE
TRUE
Domains and Regions:
DOMAIN: Cyclic nucleotide−binding
DOMAIN: PNPLA
REGION: Disordered

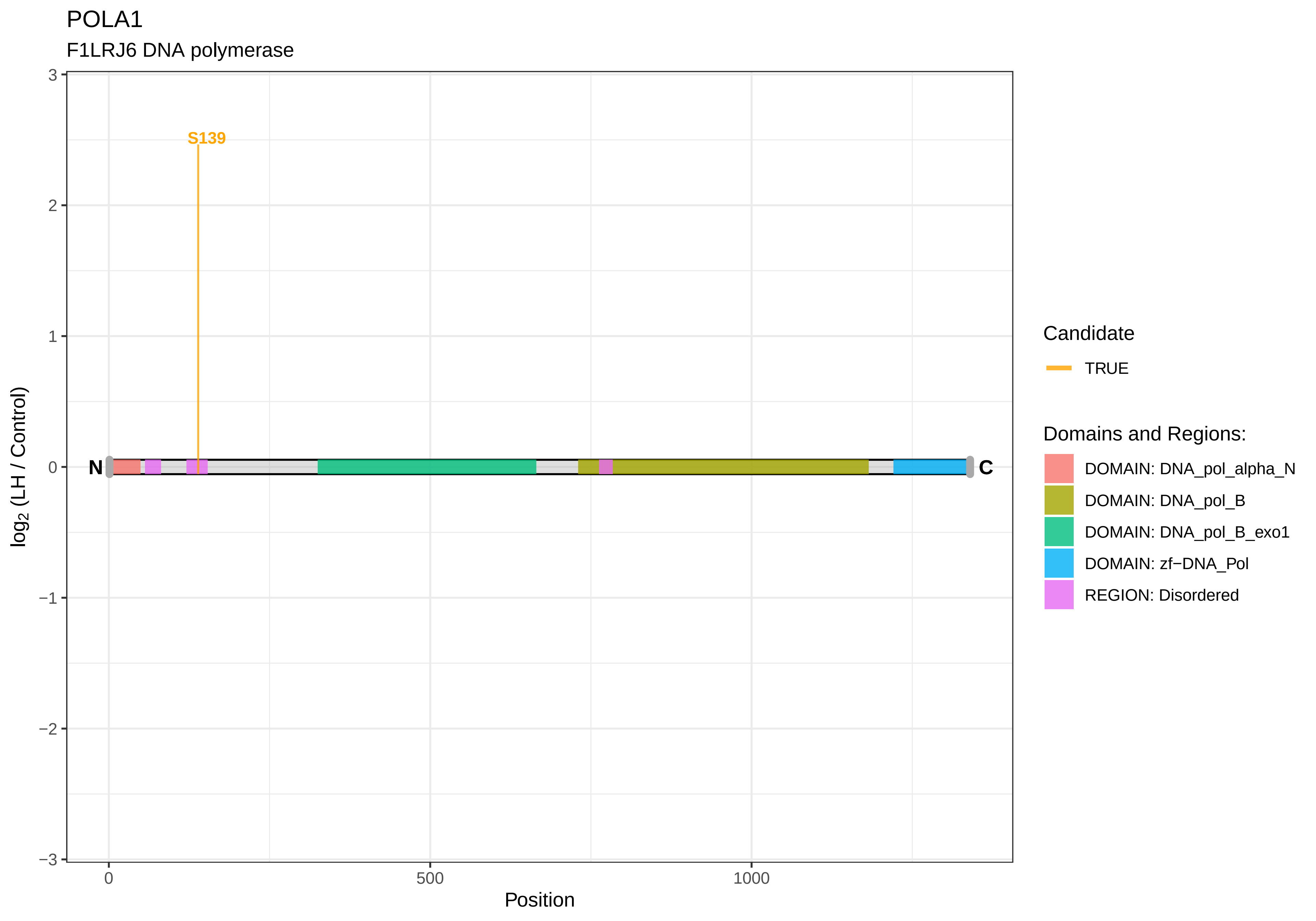
POLA1
F1LRJ6 DNA polymerase
S139
N
C
3
2
1
0
−1
−2
−3
log2 (LH / Control)
0
500
1000
Position
Candidate
TRUE
Domains and Regions:
DOMAIN: DNA_pol_alpha_N
DOMAIN: DNA_pol_B
DOMAIN: DNA_pol_B_exo1
DOMAIN: zf−DNA_Pol
REGION: Disordered

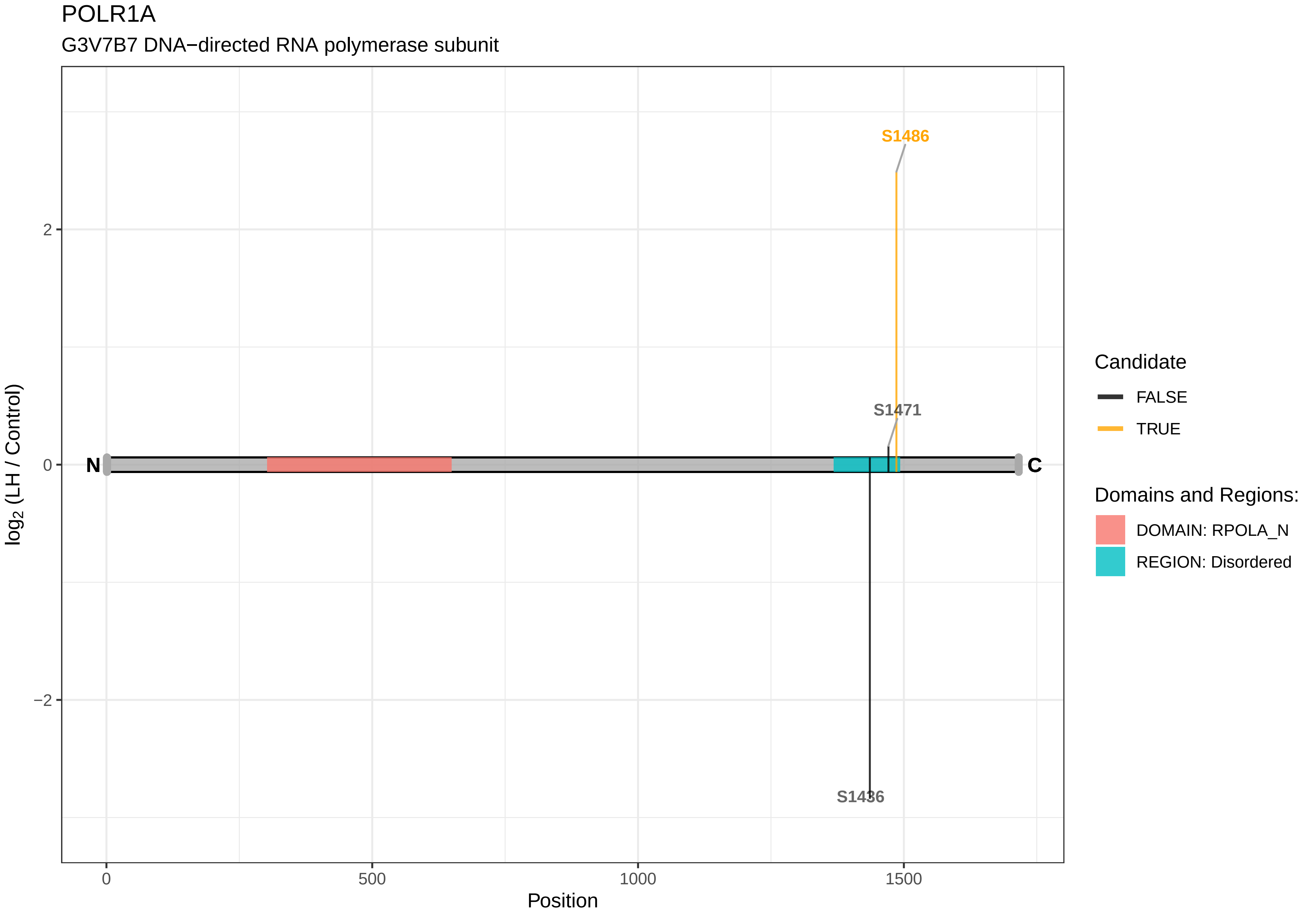

POLR1A
G3V7B7 DNA−directed RNA polymerase subunit
S1486
S1471
S1436
N
C
2
0
−2
log2 (LH / Control)
0
500
1000
1500
Position
Candidate
FALSE
TRUE
Domains and Regions:
DOMAIN: RPOLA_N
REGION: Disordered

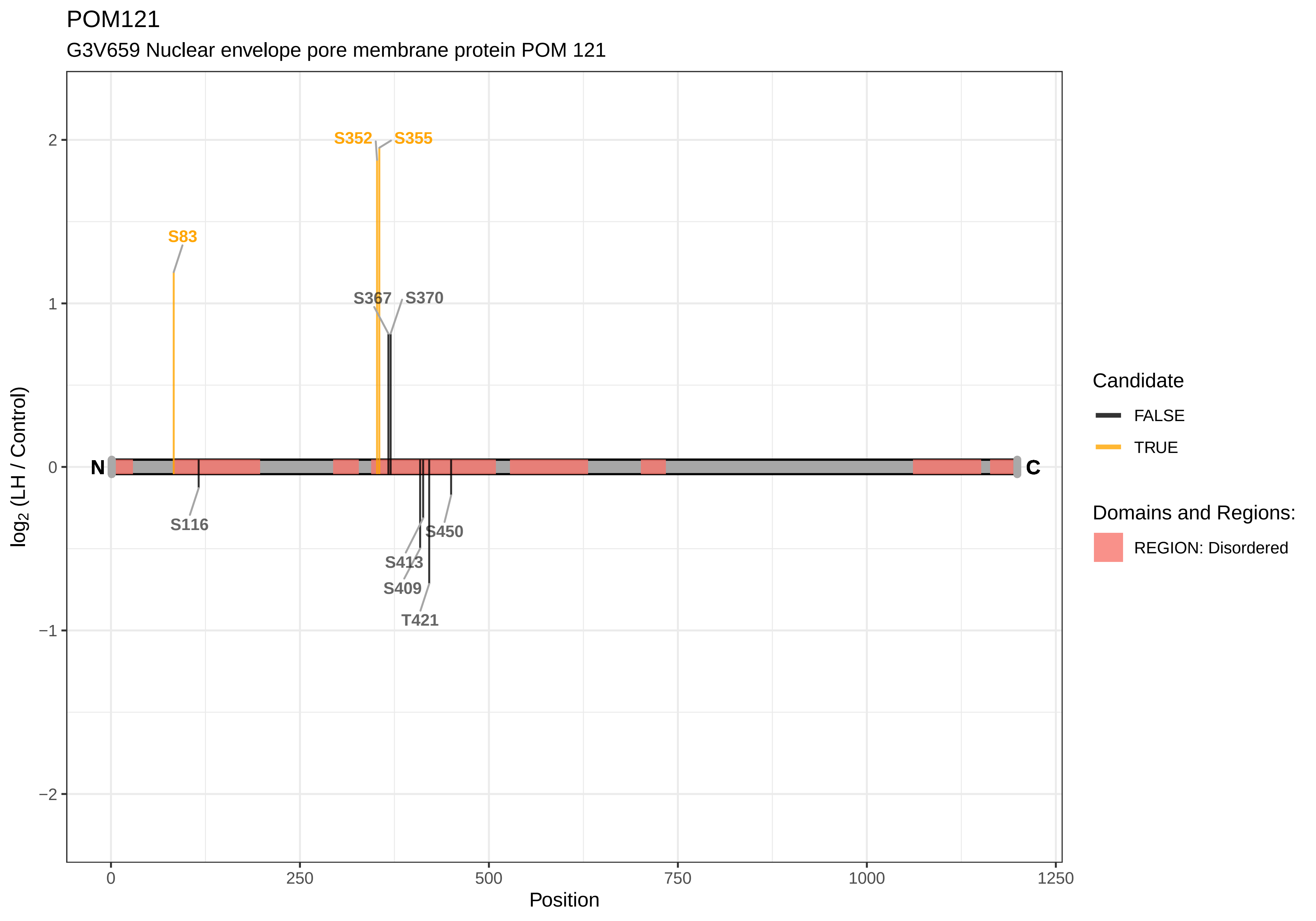
POM121
G3V659 Nuclear envelope pore membrane protein POM 121
S352
S355
S83
S367
S370
S116
S450
S413
S409
T421
N
C
2
1
0
−1
−2
log2 (LH / Control)
0
250
500
750
1000
1250
Position
Candidate
FALSE
TRUE
Domains and Regions:
REGION: Disordered

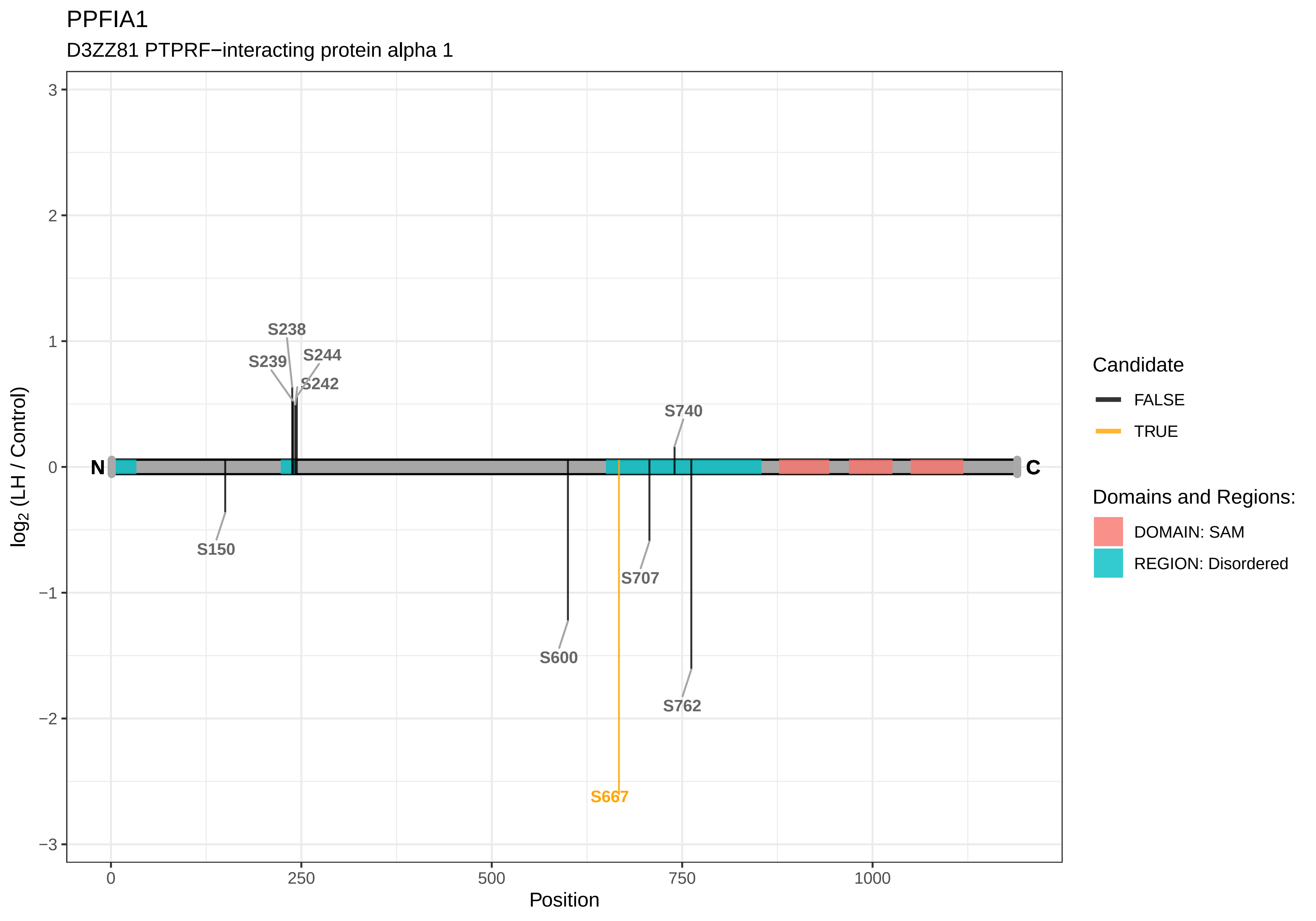

PPFIA1
D3ZZ81 PTPRF−interacting protein alpha 1
log2 (LH / Control)
Position
N
C
S238
S239
S244
S242
S740
S150
S707
S600
S762
S667
3
2
1
0
−1
−2
−3
0
250
500
750
1000
Candidate
FALSE
TRUE
Domains and Regions:
DOMAIN: SAM
REGION: Disordered

PPFIBP1

A0A0G2K781 PPFIA−binding protein 1

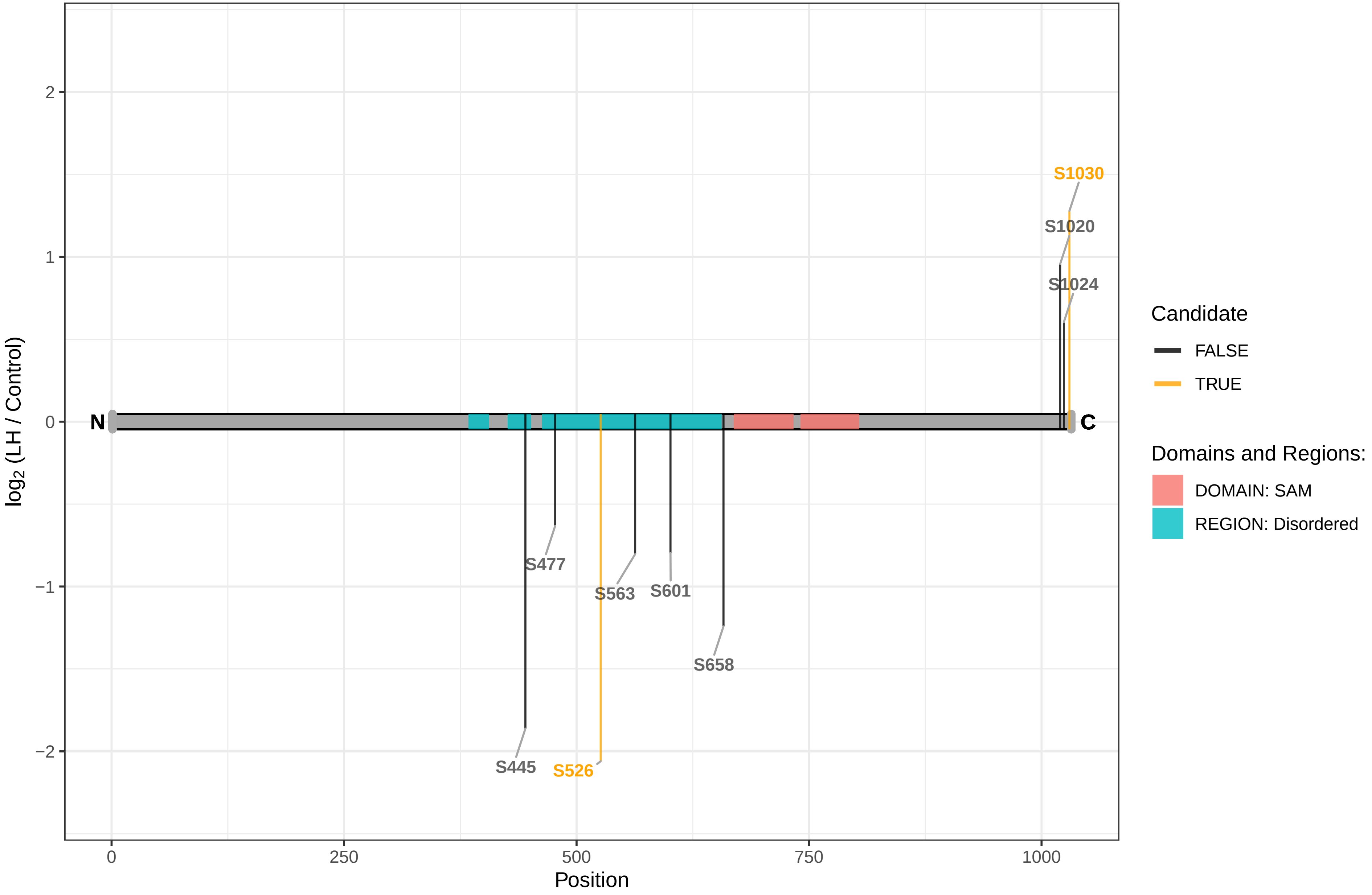

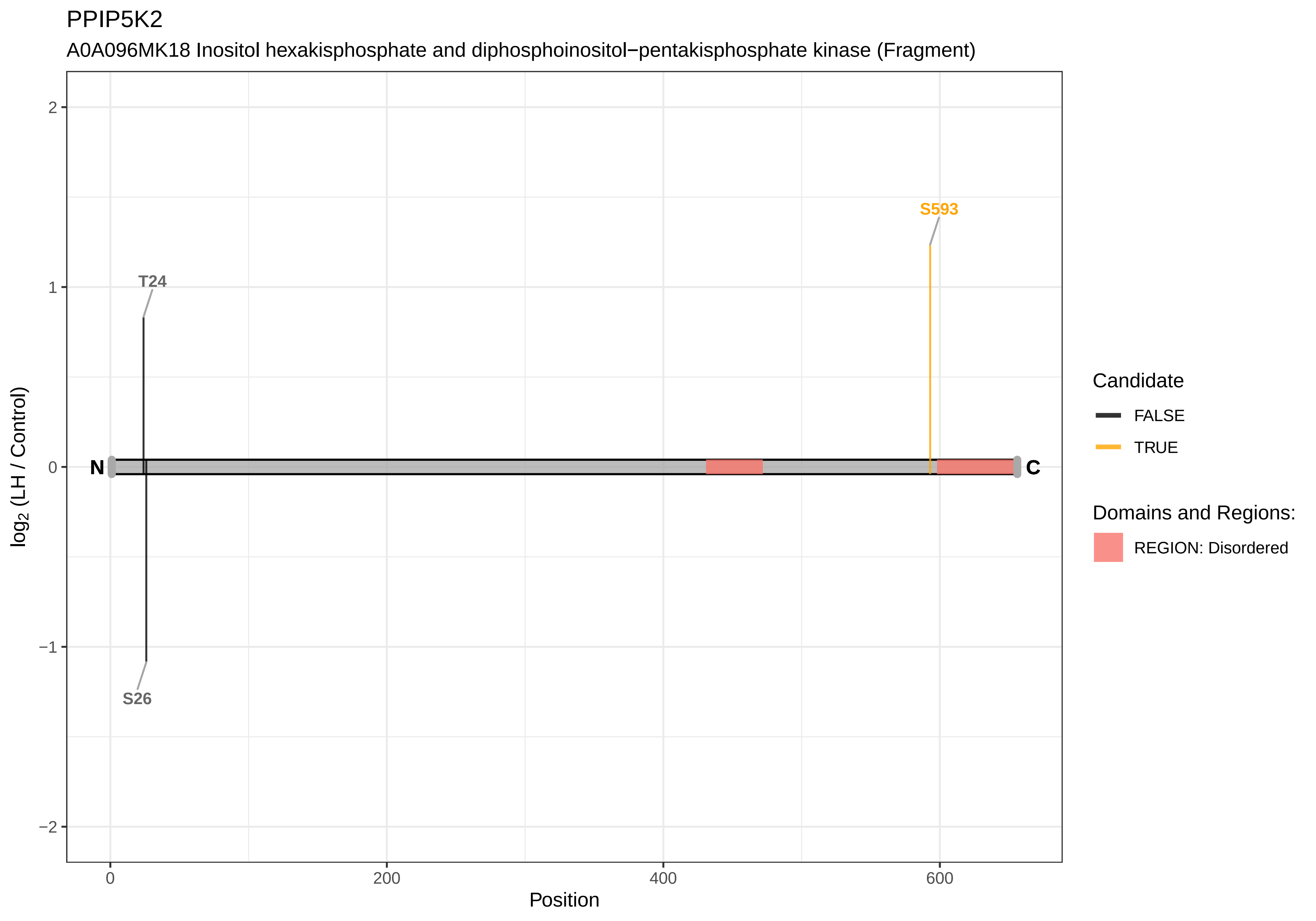
PPIP5K2
A0A096MK18 Inositol hexakisphosphate and diphosphoinositol−pentakisphosphate kinase (Fragment)
S593
T24
S26
N
C
2
1
0
−1
−2
log2 (LH / Control)
0
200
400
600
Position
Candidate
FALSE
TRUE
Domains and Regions:
REGION: Disordered

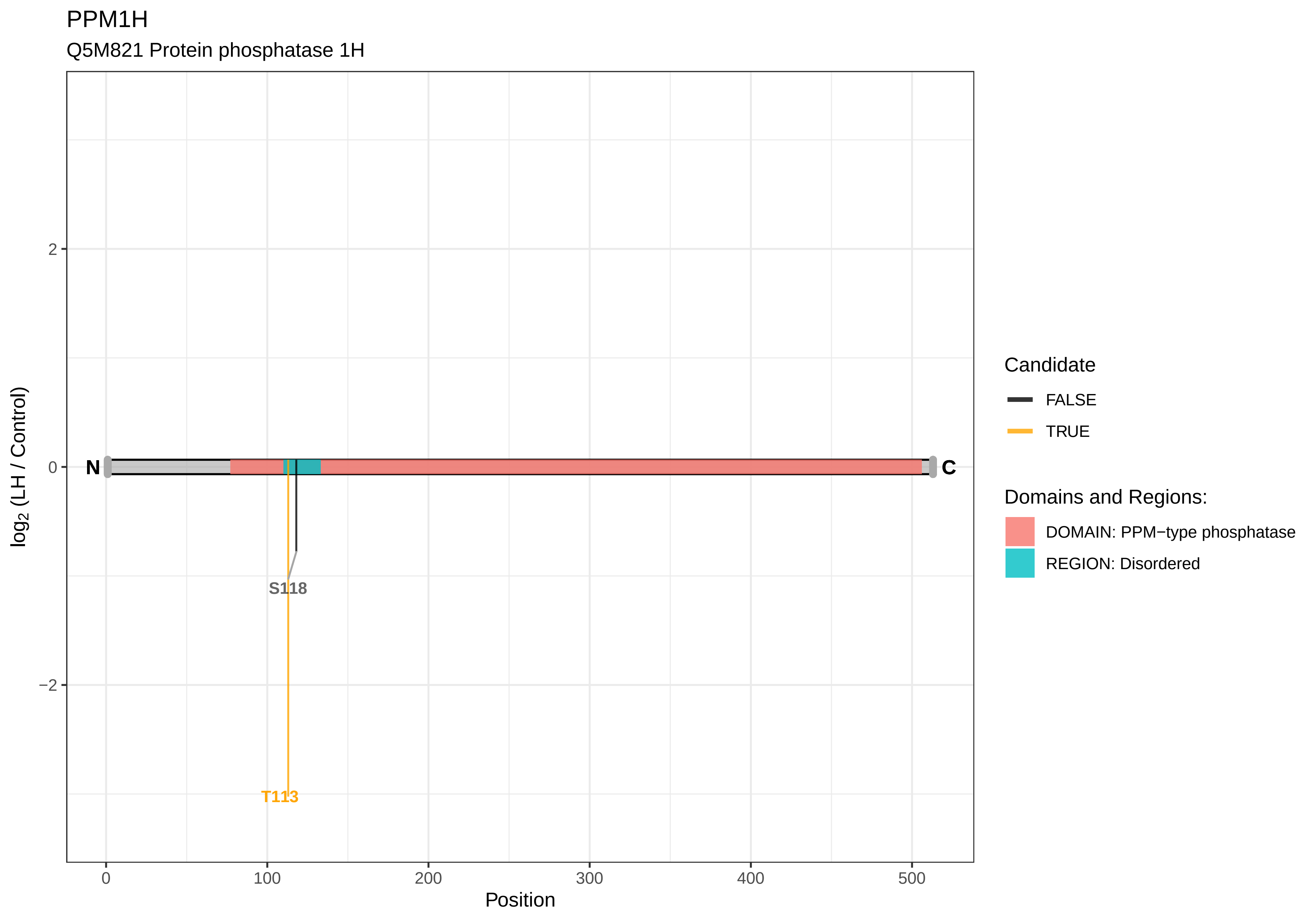
PPM1H
Q5M821 Protein phosphatase 1H
log2 (LH / Control)
2
0
−2
N
C
S118
T113
0
100
200
300
400
500
Position
Candidate
FALSE
TRUE
Domains and Regions:
DOMAIN: PPM−type phosphatase
REGION: Disordered

PPP1R12B

D3ZIC4 Protein phosphatase 1, regulatory (Inhibitor) subunit 12B (Predicted)

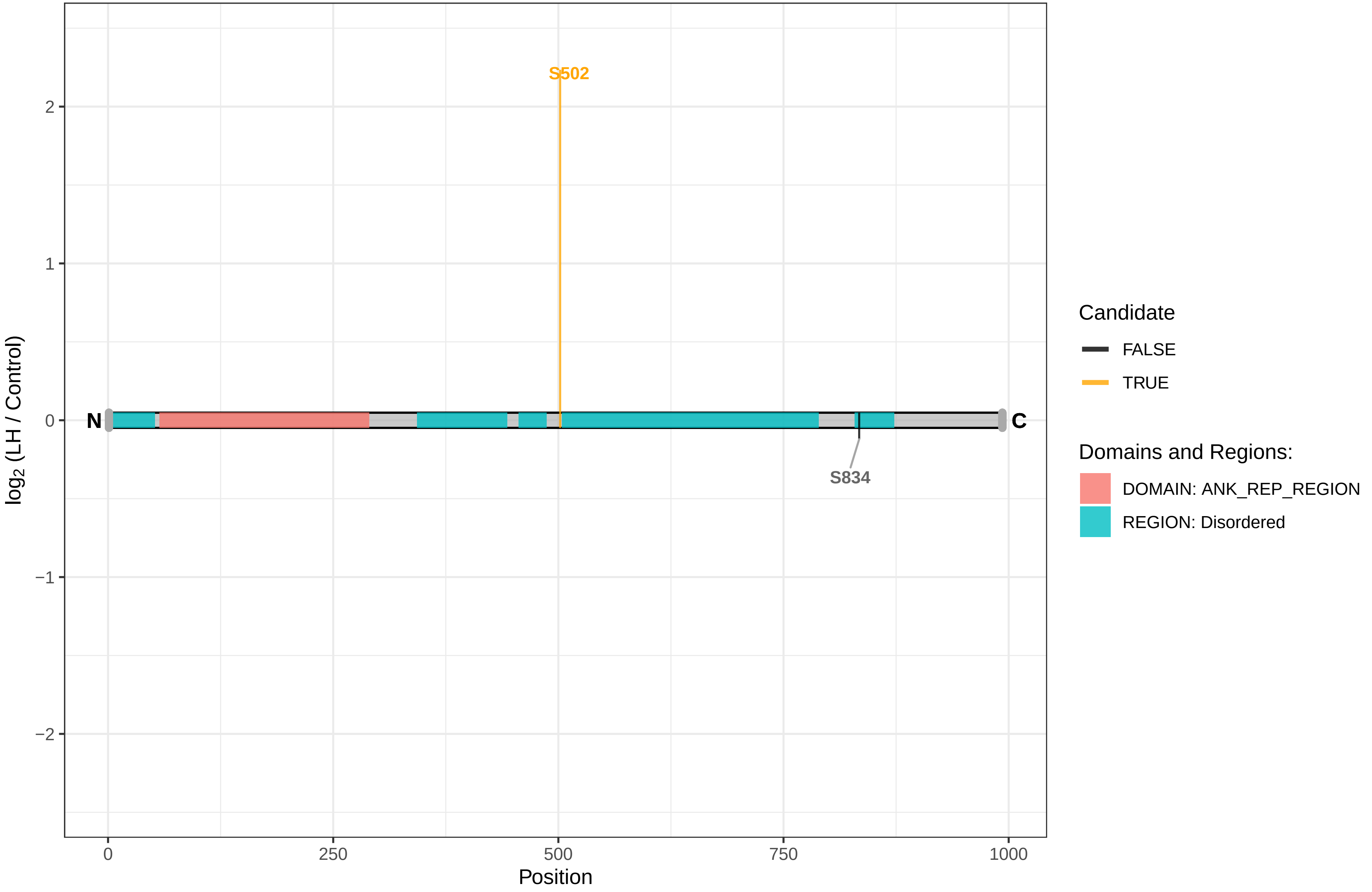

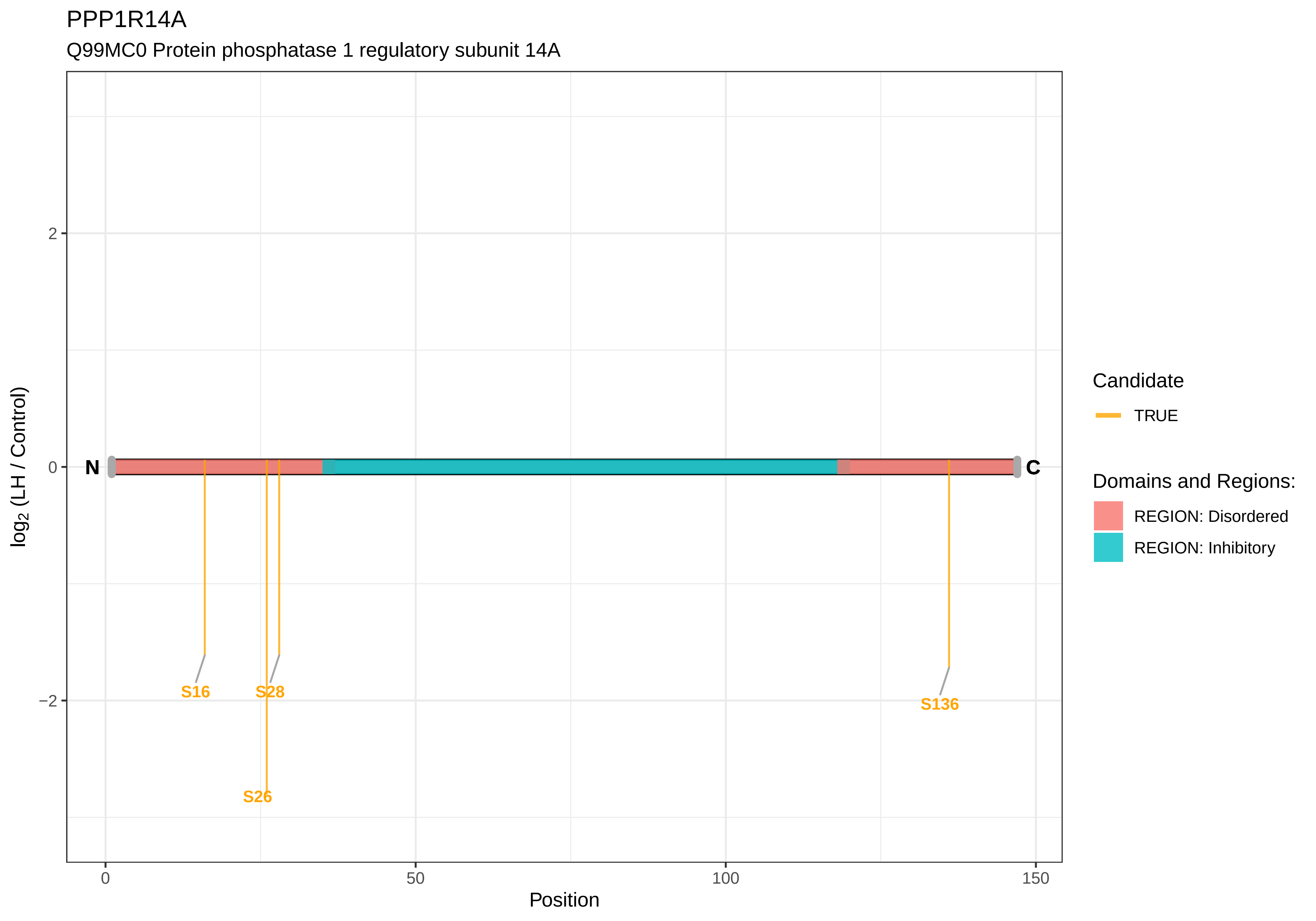
PPP1R14A
Q99MC0 Protein phosphatase 1 regulatory subunit 14A
log2 (LH / Control)
2
0
−2
N
C
S16
S28
S26
S136
0
50
100
150
Position
Candidate
TRUE
Domains and Regions:
REGION: Disordered
REGION: Inhibitory

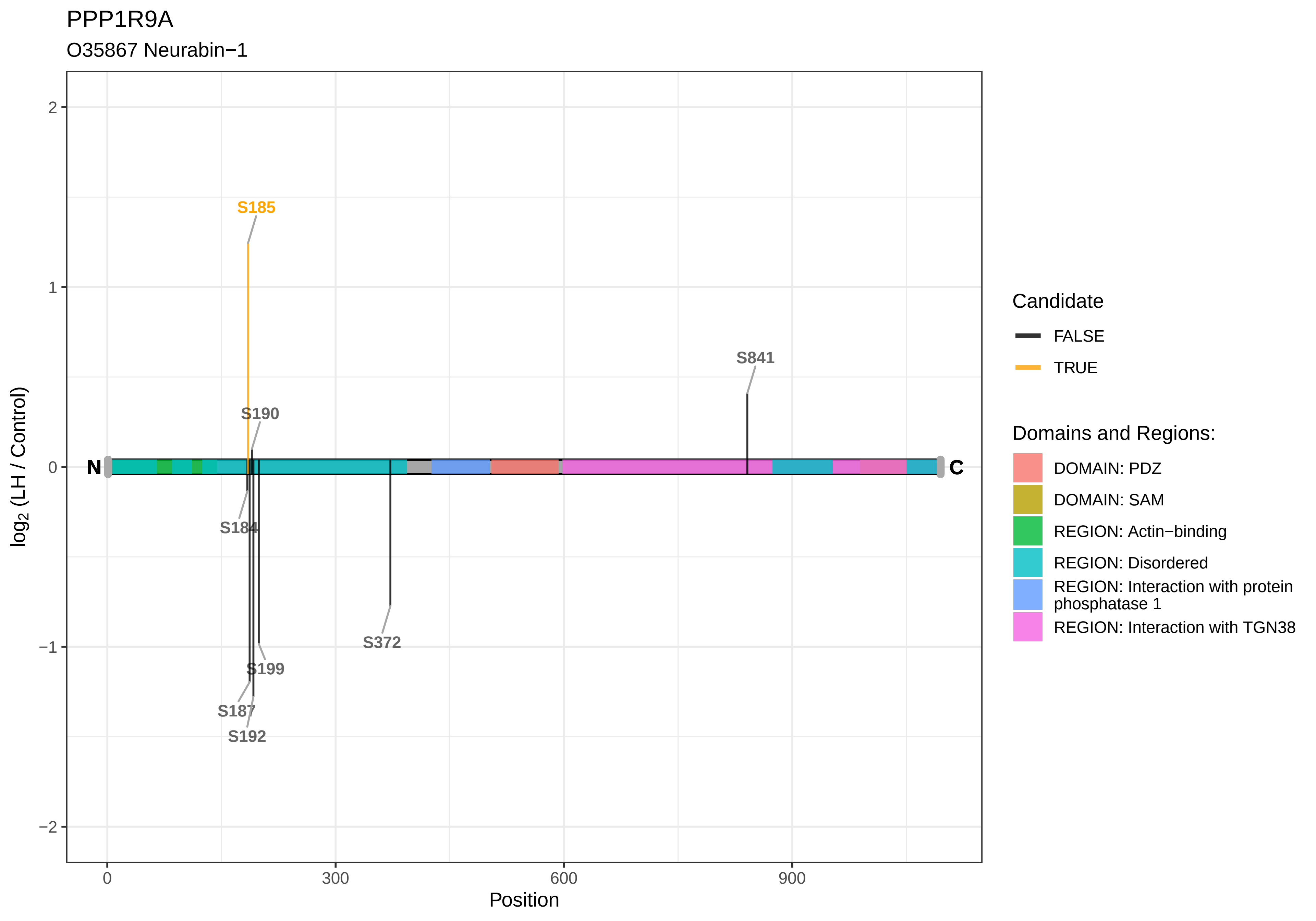
PPP1R9A
O35867 Neurabin−1
S185
S190
S841
S184
S199
S187
S192
S372
N
C
log2 (LH / Control)
Position
0
300
600
900
2
1
0
−1
−2
Candidate
FALSE
TRUE
Domains and Regions:
DOMAIN: PDZ
DOMAIN: SAM
REGION: Actin−binding
REGION: Disordered
REGION: Interaction with protein phosphatase 1
REGION: Interaction with TGN38

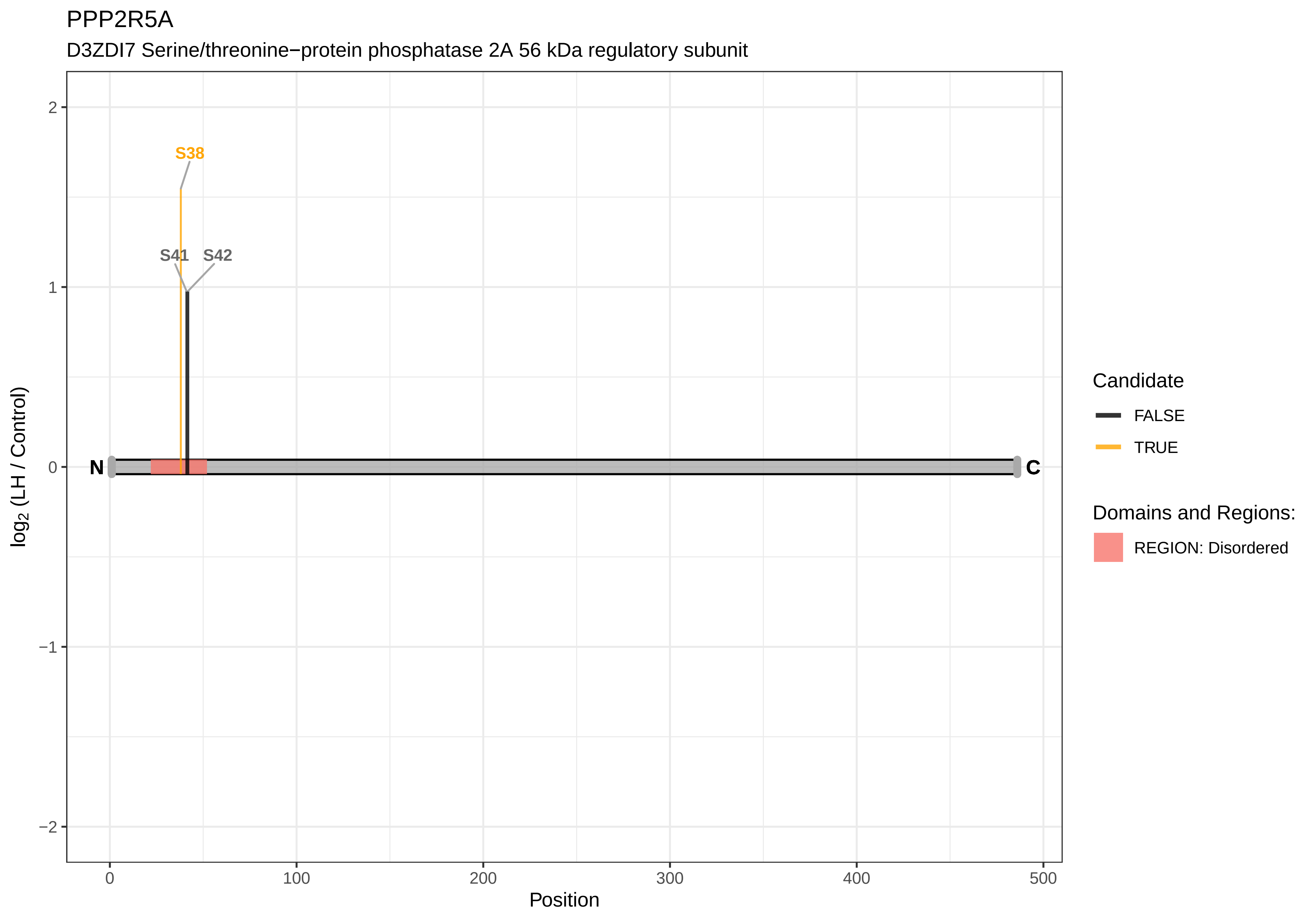

PPP2R5A
D3ZDI7 Serine/threonine−protein phosphatase 2A 56 kDa regulatory subunit
S38
S41
S42
N
C
2
1
0
−1
−2
log2 (LH / Control)
0
100
200
300
400
500
Position
Candidate
FALSE
TRUE
Domains and Regions:
REGION: Disordered

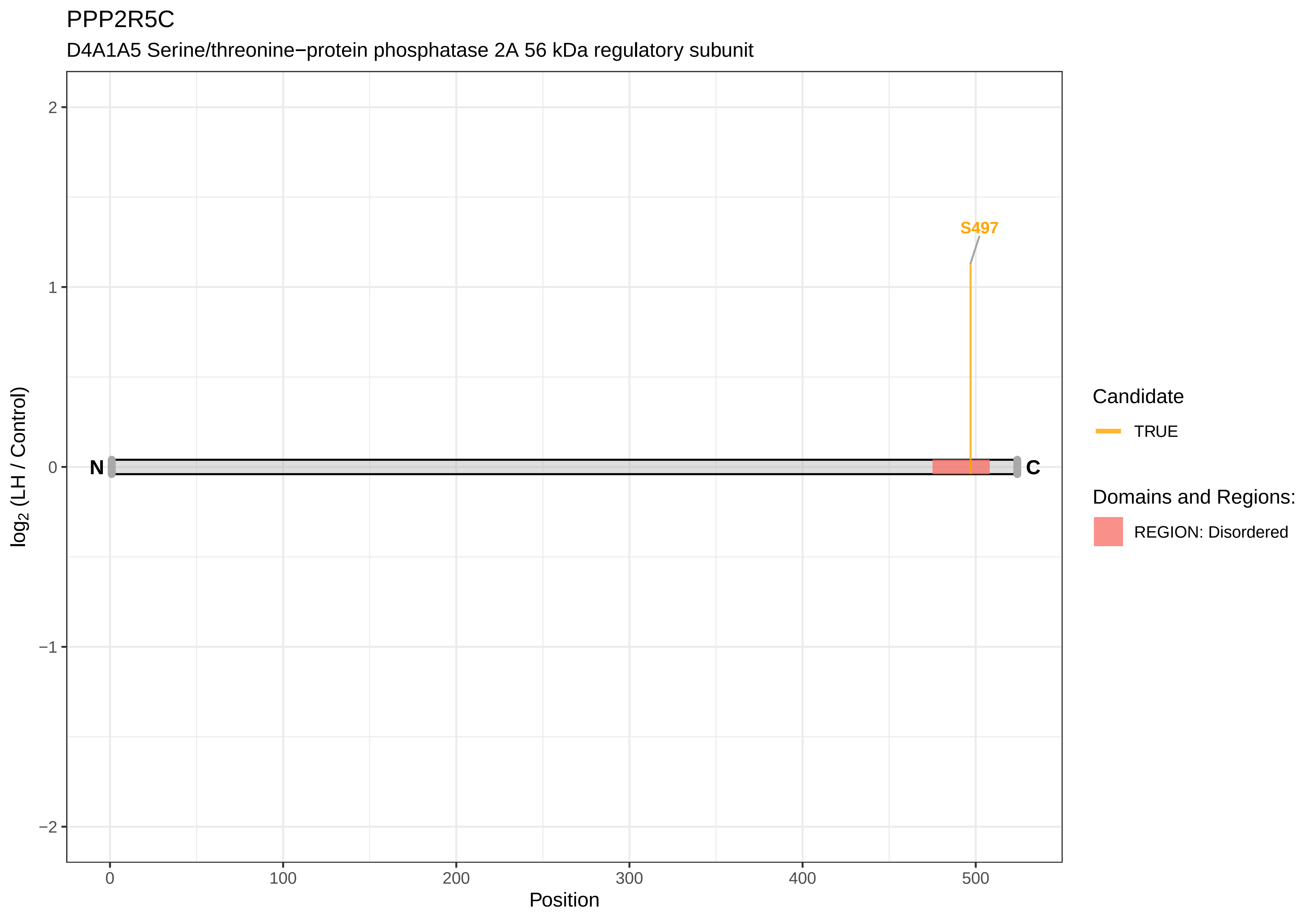
PPP2R5C
D4A1A5 Serine/threonine−protein phosphatase 2A 56 kDa regulatory subunit
S497
N
C
2
1
0
−1
−2
0
100
200
300
400
500
Position
log2 (LH / Control)
Candidate
TRUE
Domains and Regions:
REGION: Disordered

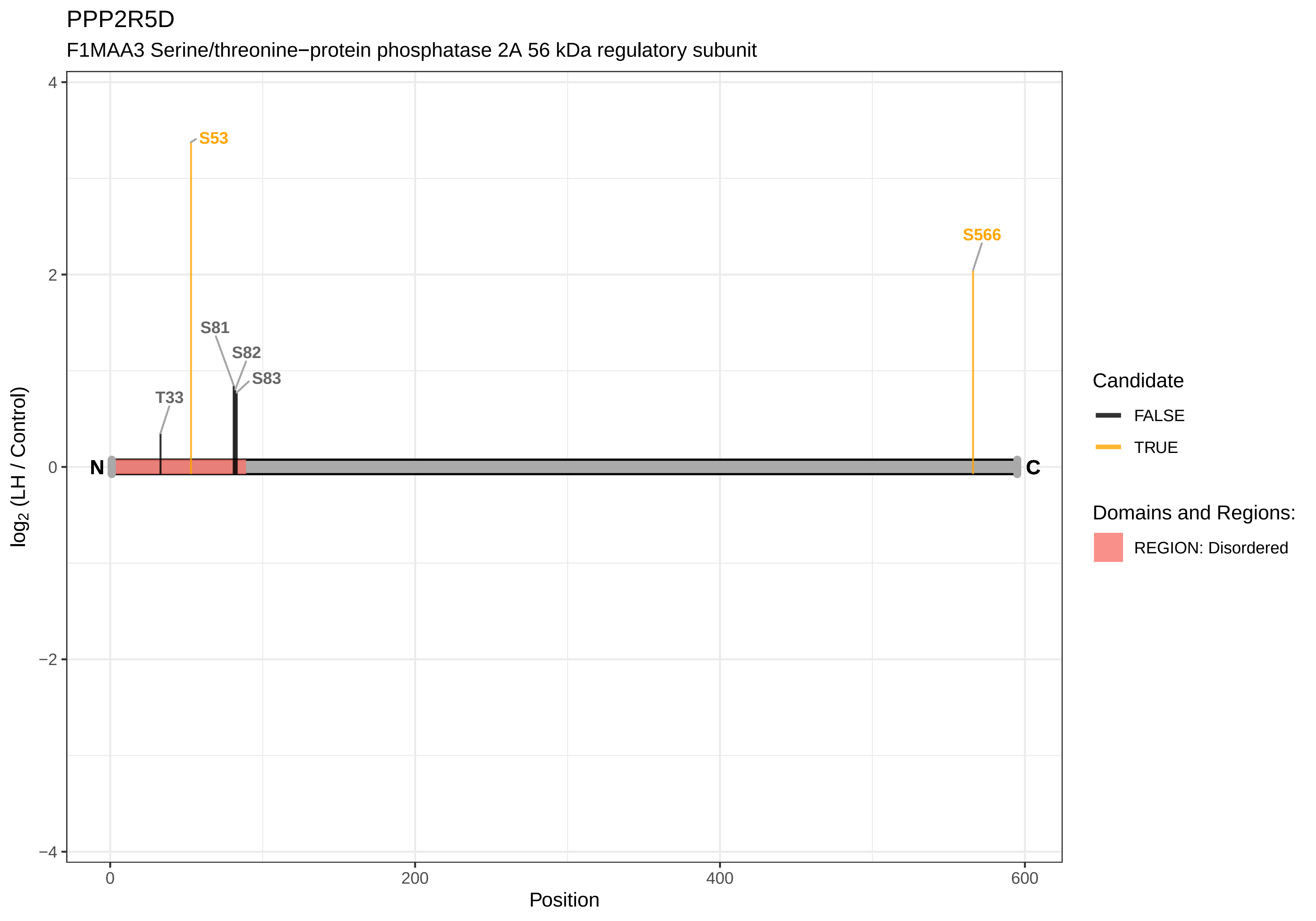
PPP2R5D
F1MAA3 Serine/threonine−protein phosphatase 2A 56 kDa regulatory subunit
S53
S566
S81
S82
S83
T33
N
C
log2 (LH / Control)
4
2
0
−2
−4
0
200
400
600
Position
Candidate
FALSE
TRUE
Domains and Regions:
REGION: Disordered

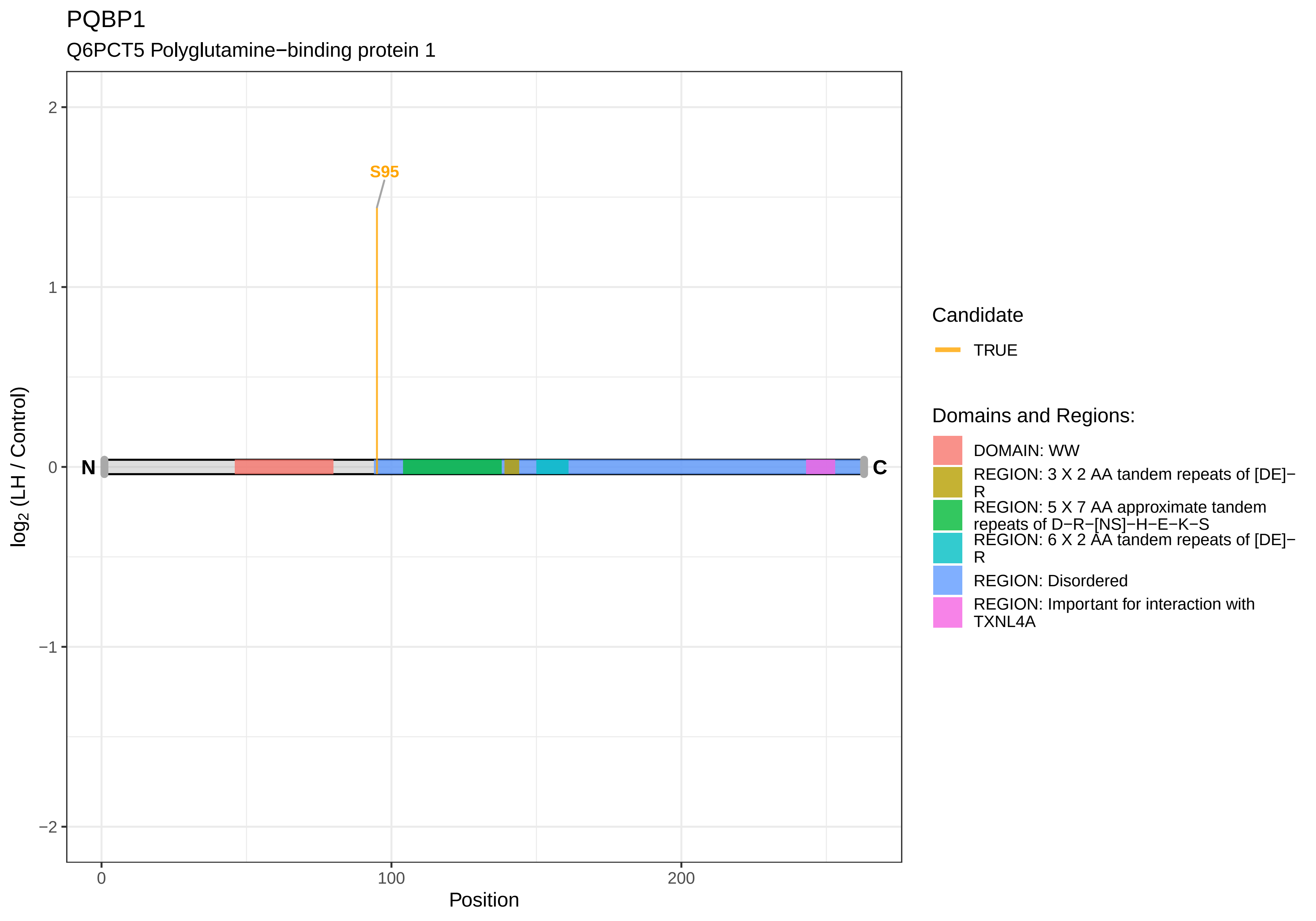

PQBP1
Q6PCT5 Polyglutamine−binding protein 1
S95
N
C
2
1
0
−1
−2
log2 (LH / Control)
0
100
200
Position
Candidate
TRUE
Domains and Regions:
DOMAIN: WW
REGION: 3 X 2 AA tandem repeats of [DE]−R
REGION: 5 X 7 AA approximate tandem repeats of D−R−[NS]−H−E−K−S
REGION: 6 X 2 AA tandem repeats of [DE]−R
REGION: Disordered
REGION: Important for interaction with TXNL4A

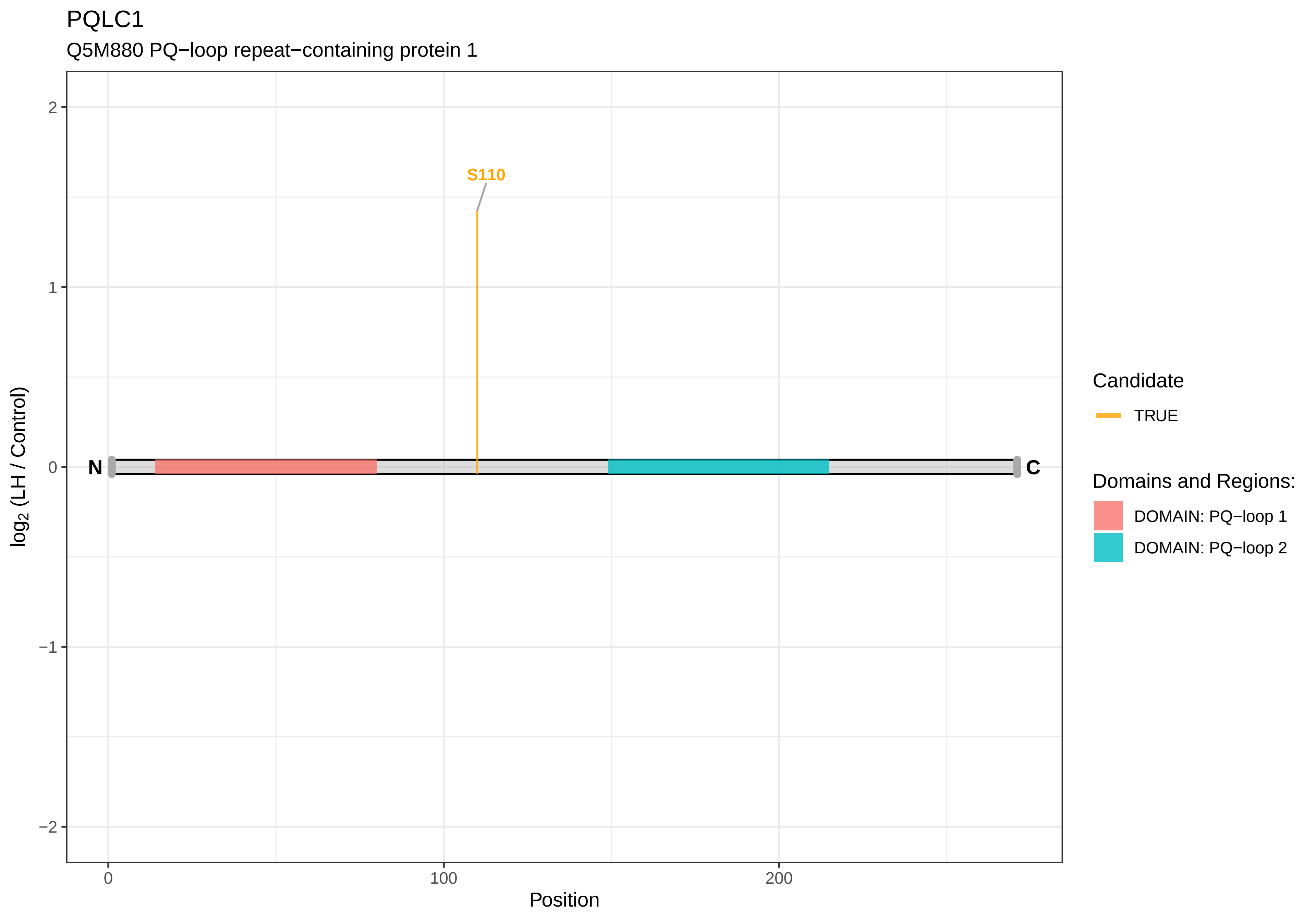
PQLC1
Q5M880 PQ−loop repeat−containing protein 1
S110
N
C
2
1
0
−1
−2
0
100
200
Position
log2 (LH / Control)
Candidate
TRUE
Domains and Regions:
DOMAIN: PQ−loop 1
DOMAIN: PQ−loop 2

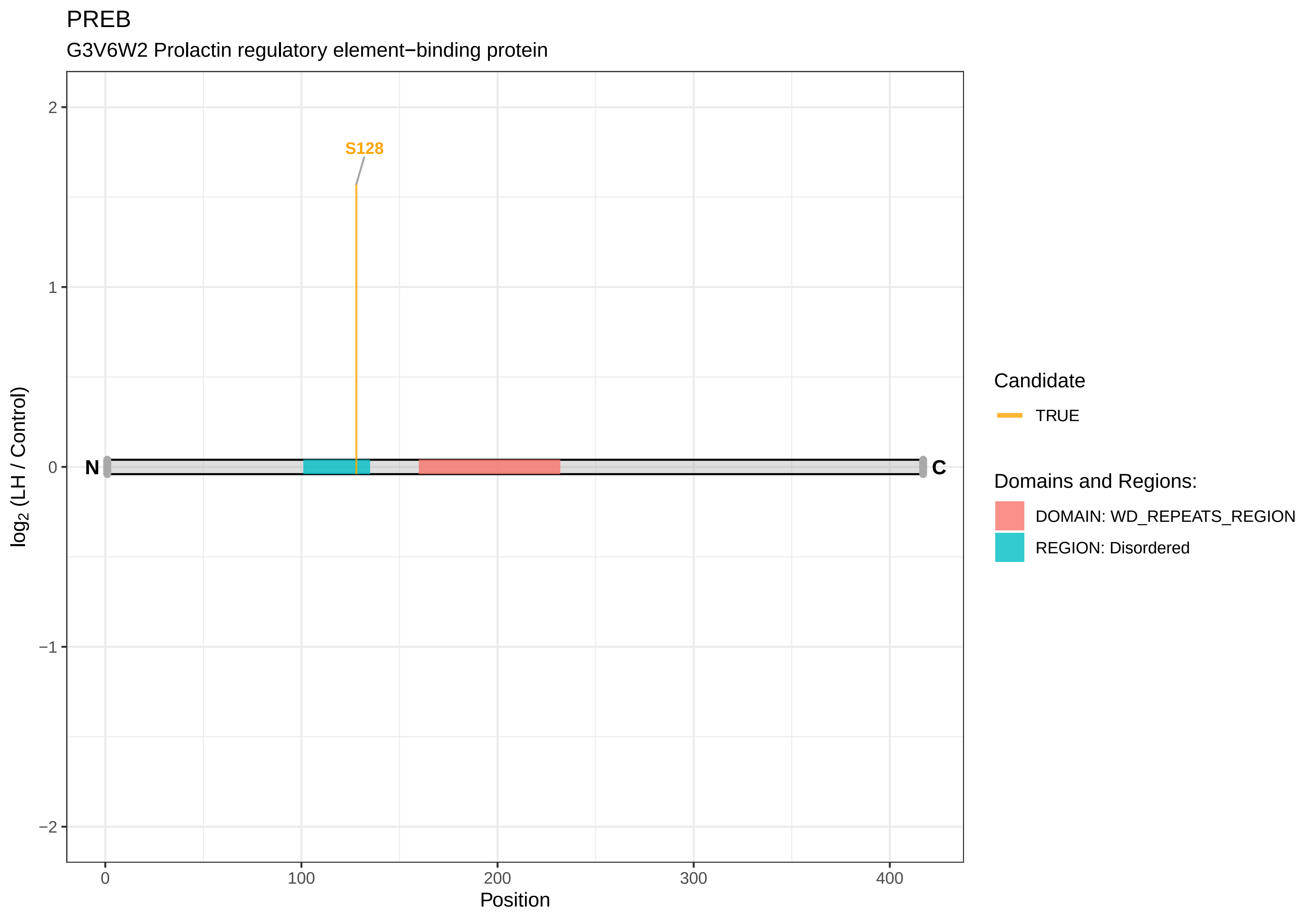
PREB
G3V6W2 Prolactin regulatory element−binding protein
S128
N
C
2
1
0
−1
−2
0
100
200
300
400
Position
log2 (LH / Control)
Candidate
TRUE
Domains and Regions:
DOMAIN: WD_REPEATS_REGION
REGION: Disordered

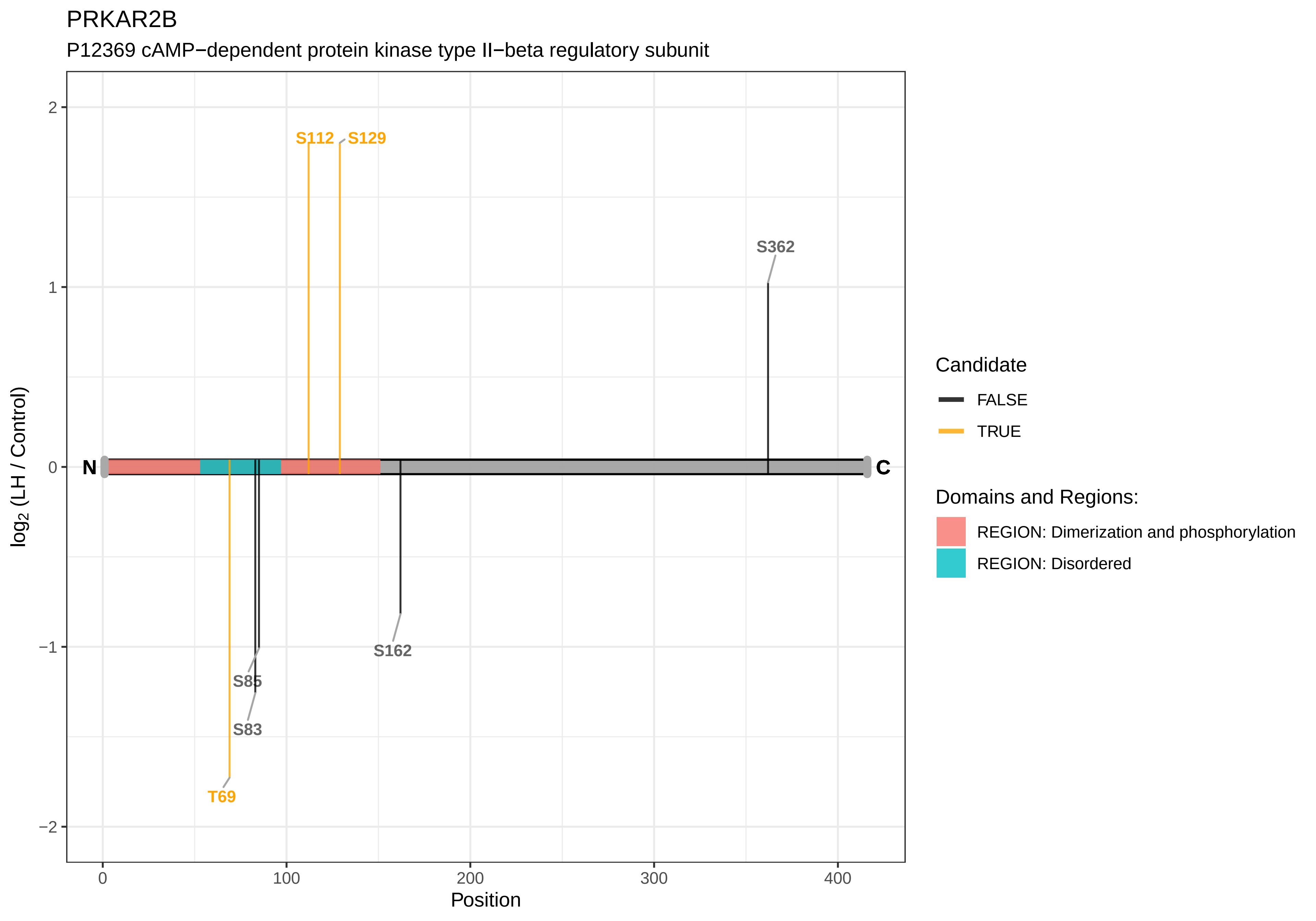

PRKAR2B
P12369 cAMP−dependent protein kinase type II−beta regulatory subunit
S112
S129
S362
N
C
S85
S83
T69
S162
log2 (LH / Control)
Position
2
1
0
−1
−2
0
100
200
300
400
Candidate
FALSE
TRUE
Domains and Regions:
REGION: Dimerization and phosphorylation
REGION: Disordered

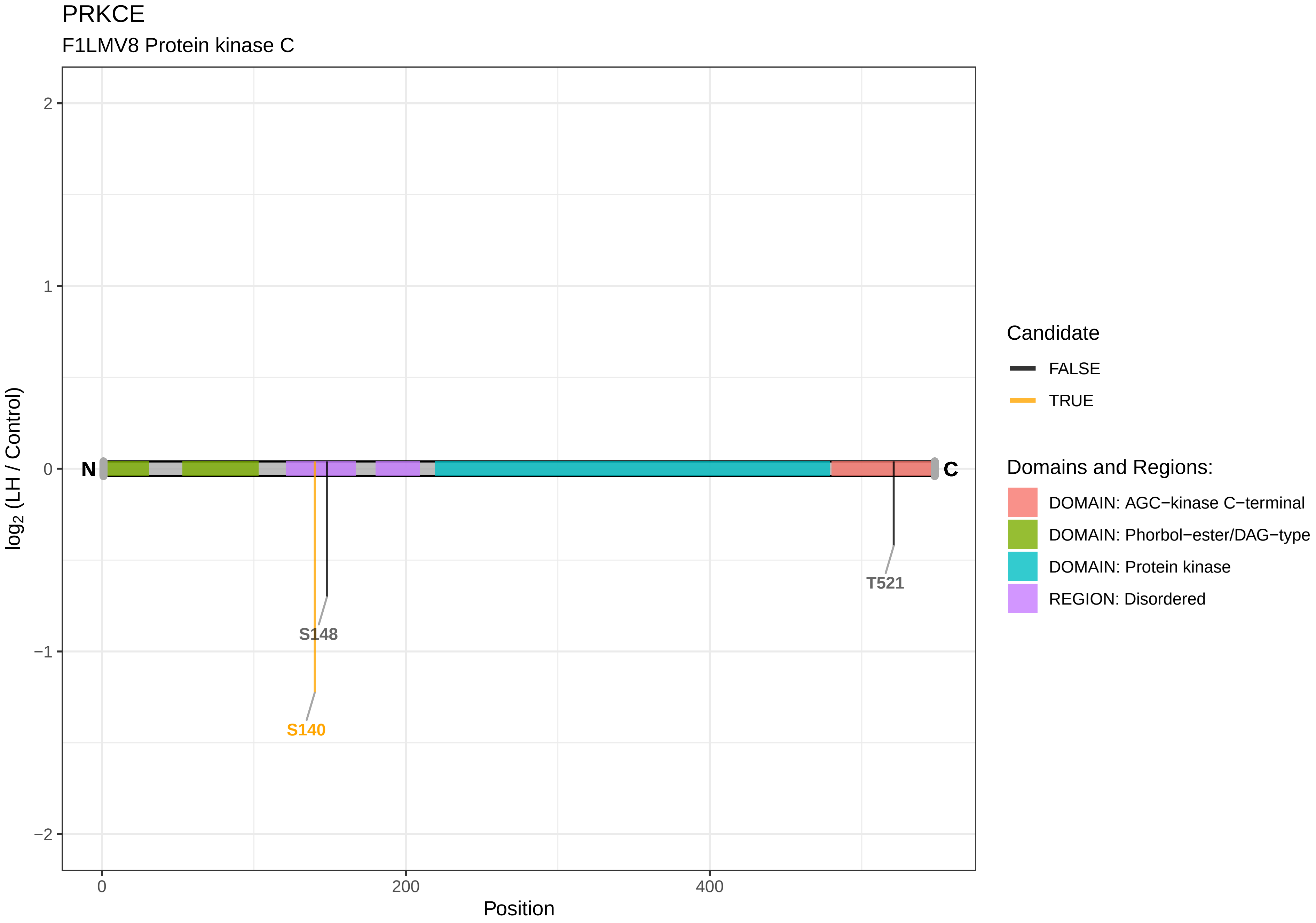

PRKCE
F1LMV8 Protein kinase C
2
1
0
−1
−2
log2 (LH / Control)
N
C
S148
S140
T521
0
200
400
Position
Candidate
FALSE
TRUE
Domains and Regions:
DOMAIN: AGC−kinase C−terminal
DOMAIN: Phorbol−ester/DAG−type
DOMAIN: Protein kinase
REGION: Disordered

PRKD1

A0A0G2K928 Serine/threonine−protein kinase

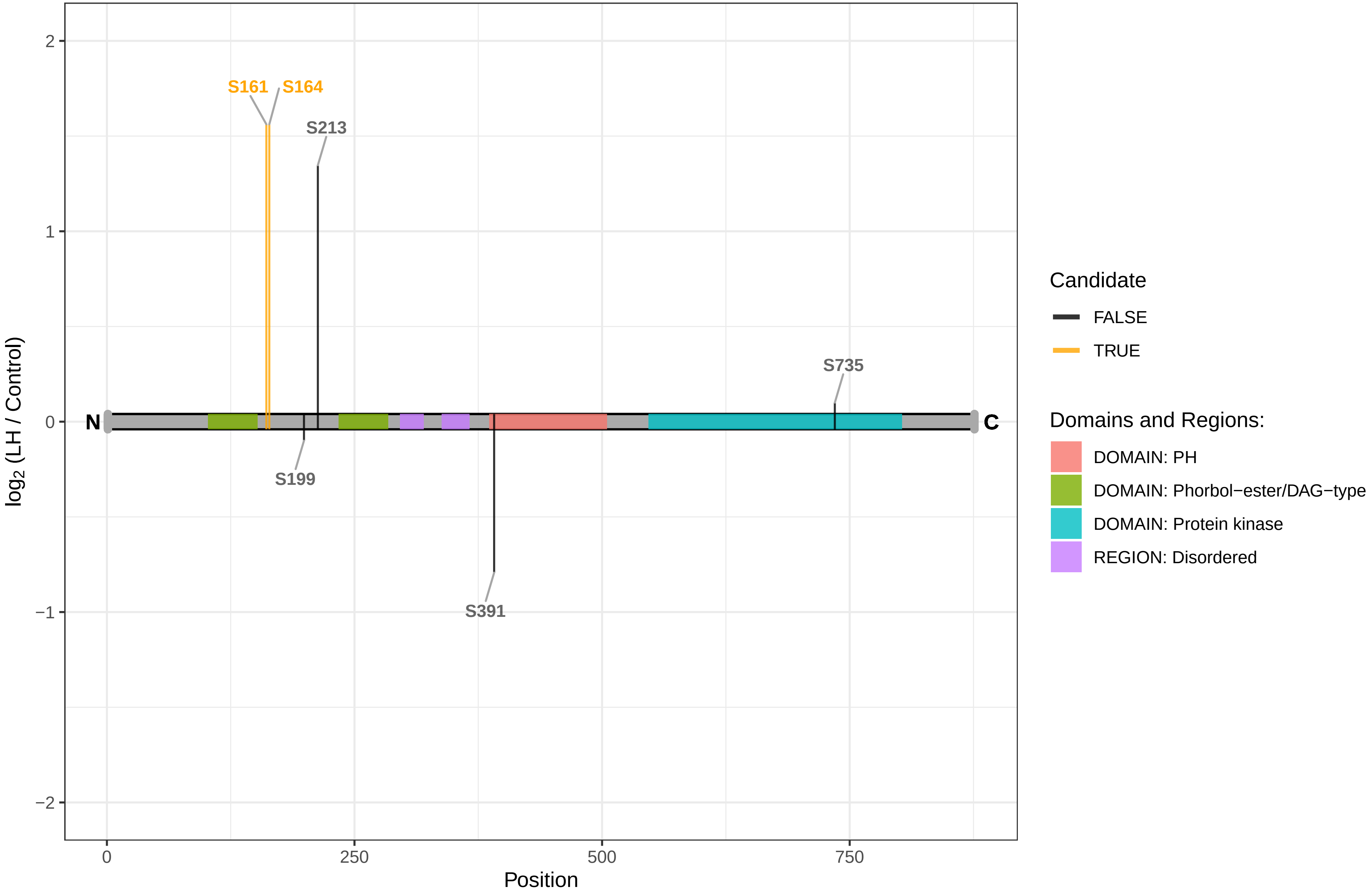

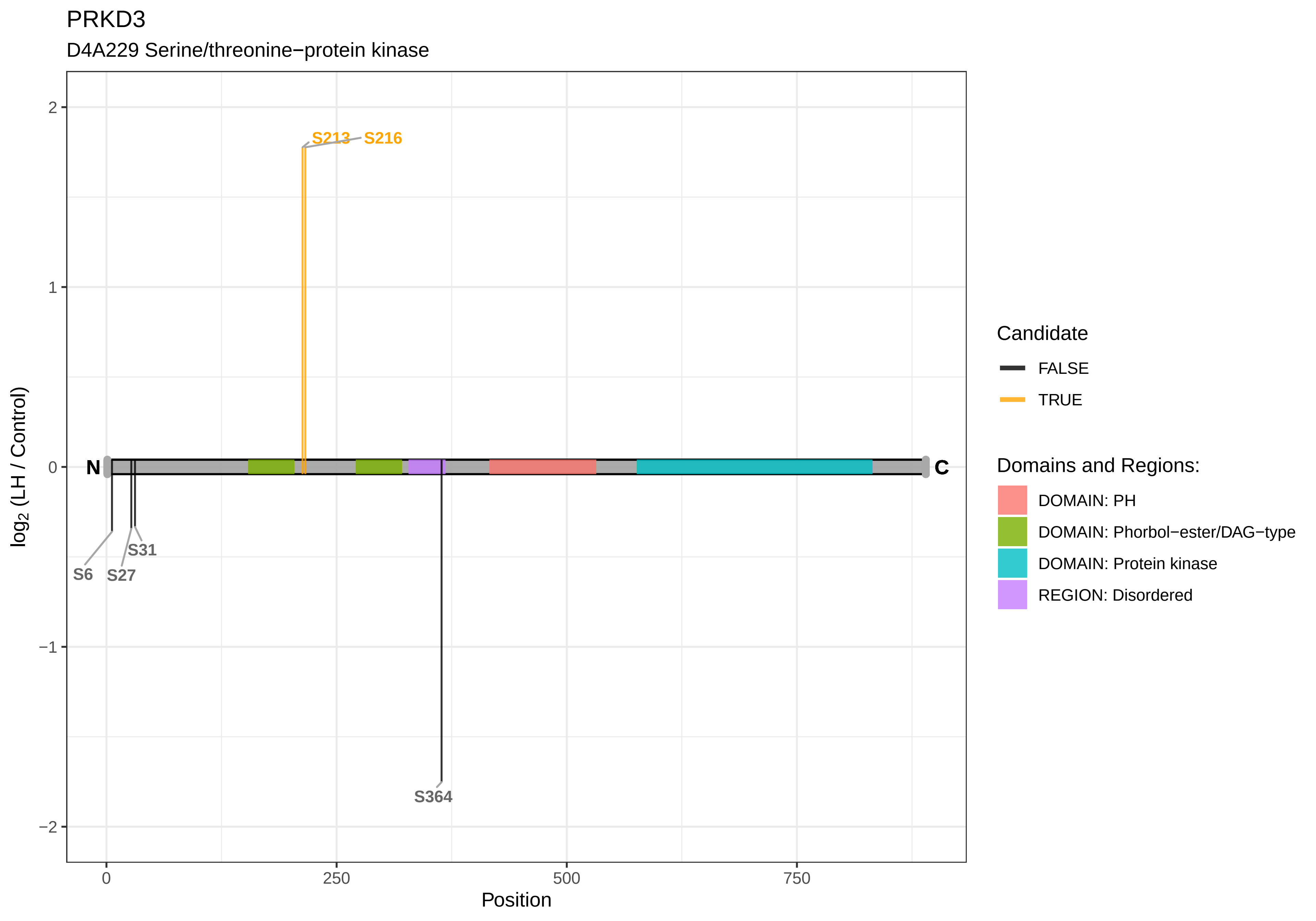
PRKD3
D4A229 Serine/threonine−protein kinase
S213
S216
S6
S27
S31
S364
N
C
2
1
0
−1
−2
log2 (LH / Control)
0
250
500
750
Position
Candidate
FALSE
TRUE
Domains and Regions:
DOMAIN: PH
DOMAIN: Phorbol−ester/DAG−type
DOMAIN: Protein kinase
REGION: Disordered

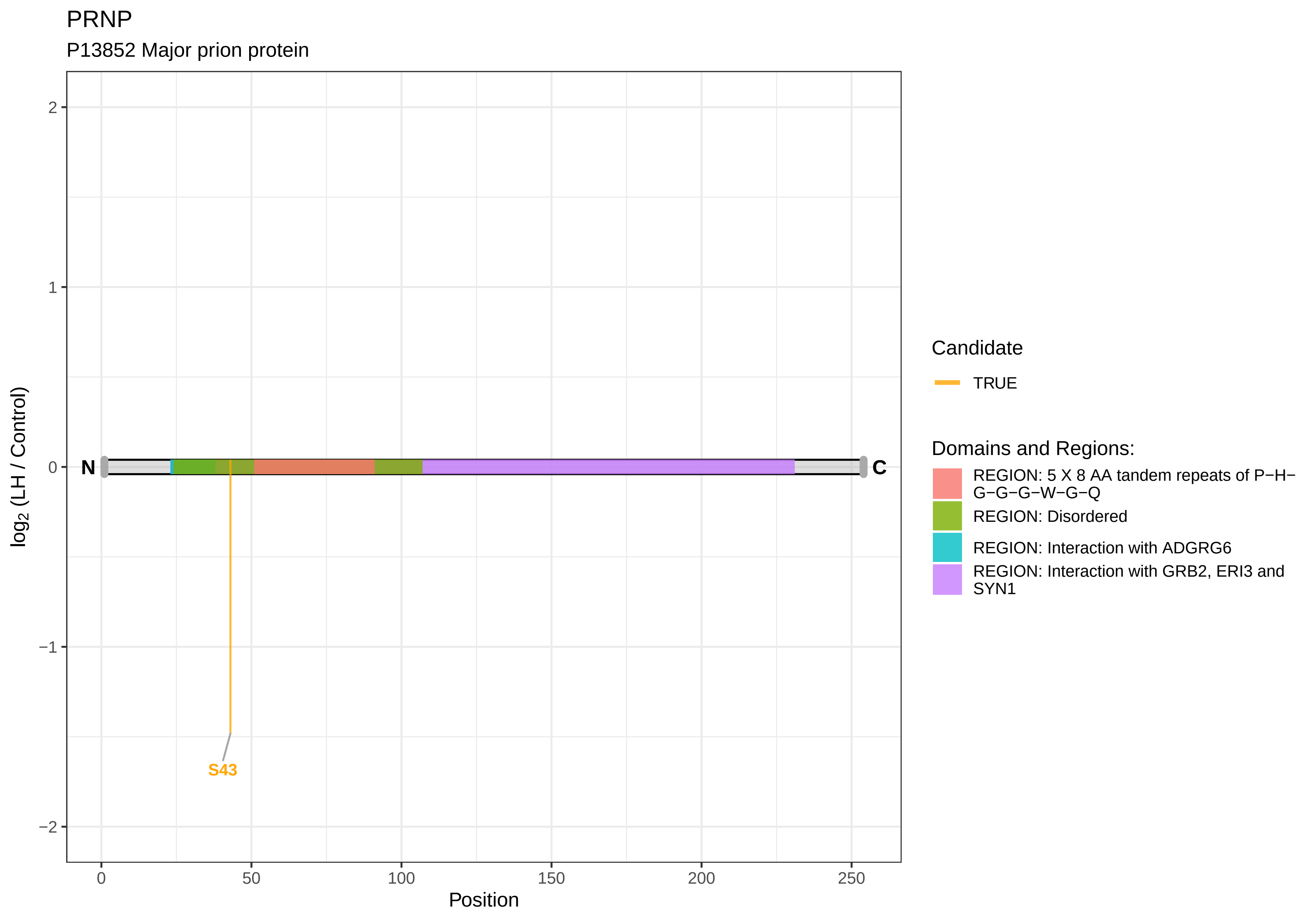

PRNP
P13852 Major prion protein
log2 (LH / Control)
2
1
0
−1
−2
N
C
S43
0
50
100
150
200
250
Position
Candidate
TRUE
Domains and Regions:
REGION: 5 X 8 AA tandem repeats of P−H−G−G−G−W−G−Q
REGION: Disordered
REGION: Interaction with ADGRG6
REGION: Interaction with GRB2, ERI3 and SYN1

PROSER2

M0RAK0 Proline and serine−rich 2

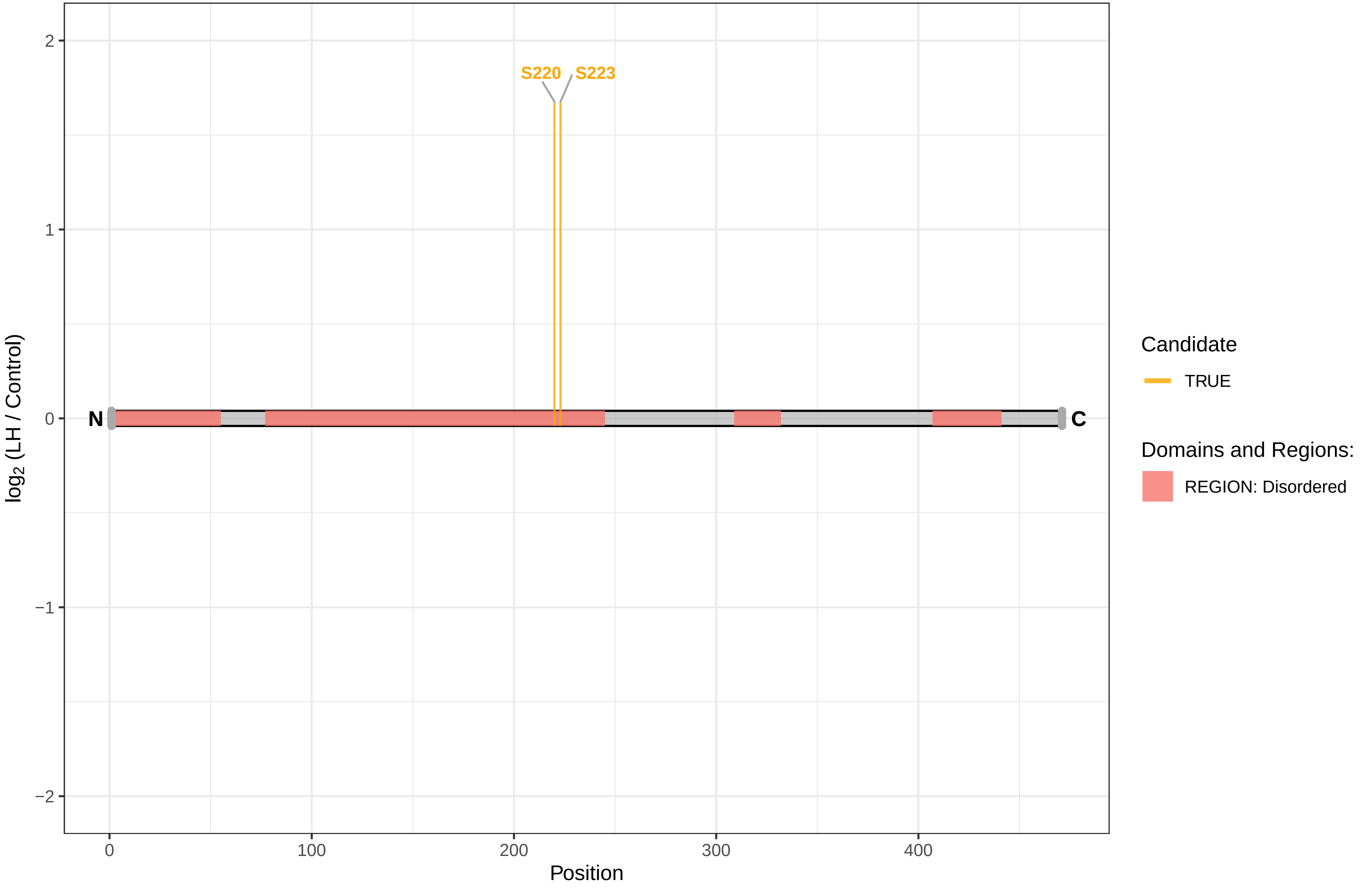

PRR15

A0JPP6 Proline−rich 15

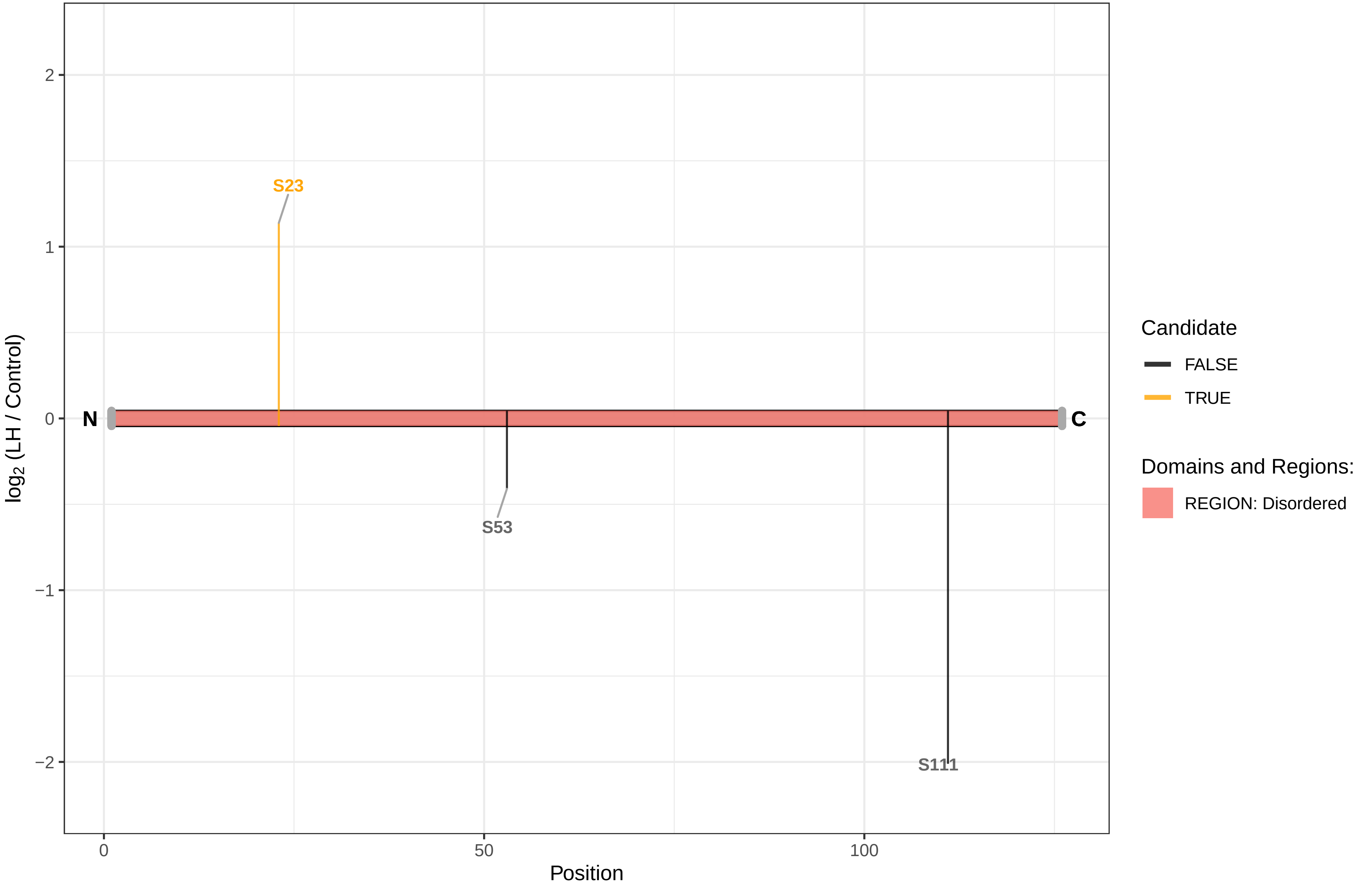

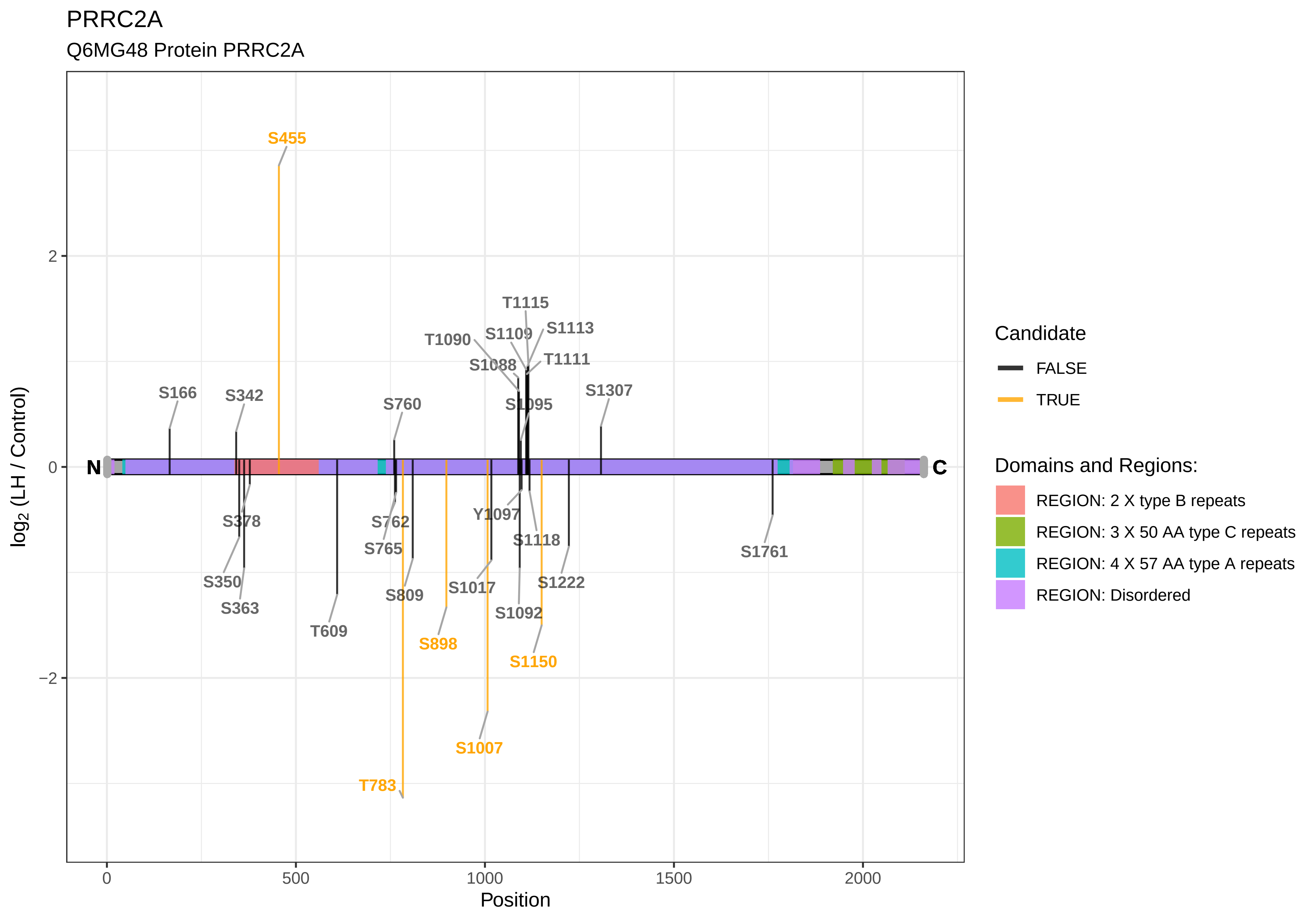

PRRC2A
Q6MG48 Protein PRRC2A
S455
T1115
S1109
S1113
T1090
S1088
T1111
S166
S342
S760
S1095
S1307
N
C
S378
S762
Y1097
S1118
S1761
S765
S350
S809
S1017
S1222
S363
S1092
T609
S898
S1150
S1007
T783
2
0
−2
0
500
1000
1500
2000
Position
log2 (LH / Control)
Candidate
FALSE
TRUE
Domains and Regions:
REGION: 2 X type B repeats
REGION: 3 X 50 AA type C repeats
REGION: 4 X 57 AA type A repeats
REGION: Disordered

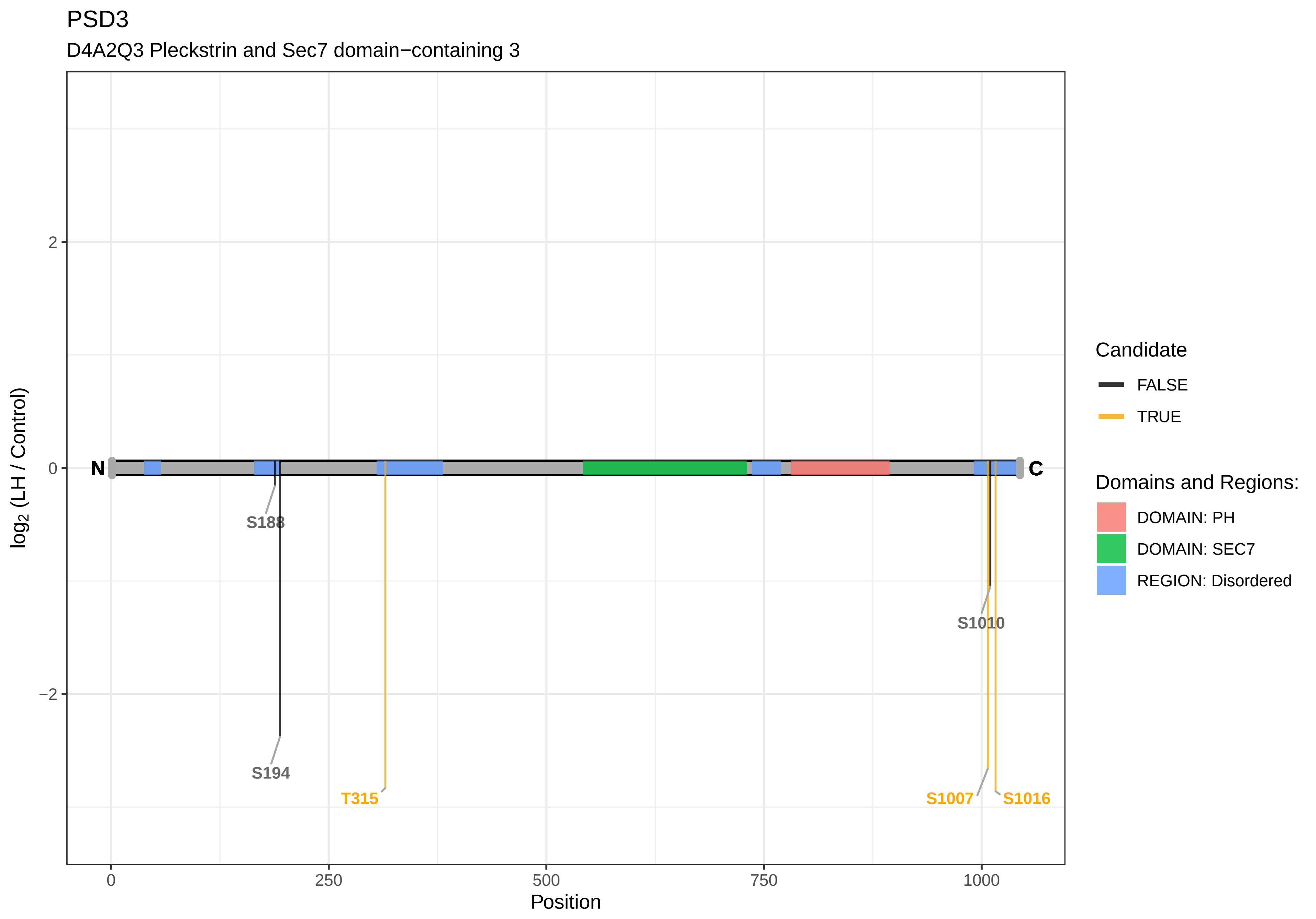

PSD3
D4A2Q3 Pleckstrin and Sec7 domain−containing 3
N
C
S188
S194
T315
S1010
S1007
S1016
2
0
−2
log2 (LH / Control)
0
250
500
750
1000
Position
Candidate
FALSE
TRUE
Domains and Regions:
DOMAIN: PH
DOMAIN: SEC7
REGION: Disordered

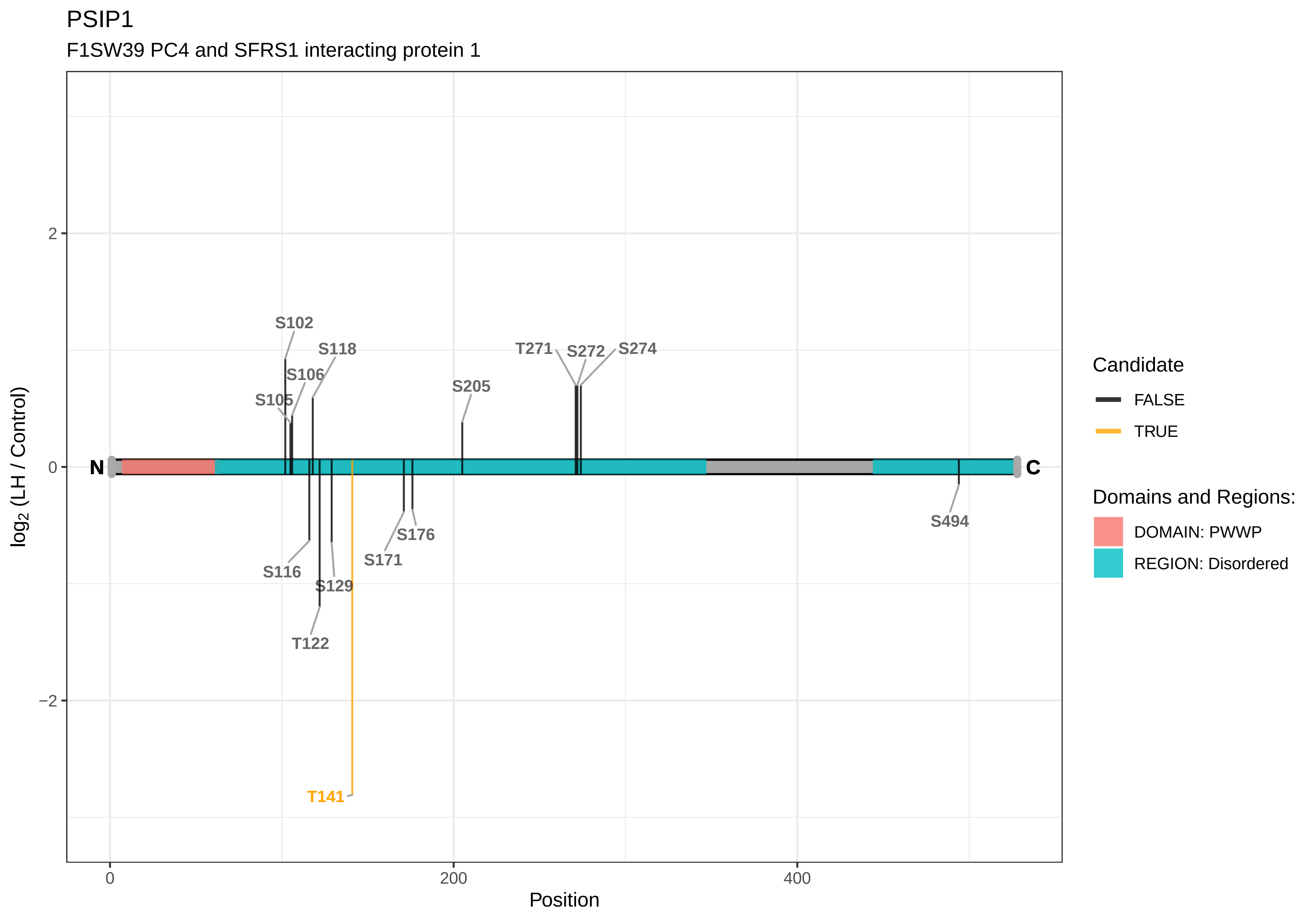
PSIP1
F1SW39 PC4 and SFRS1 interacting protein 1
log2 (LH / Control)
Position
2
0
−2
0
200
400
N
C
S102
S118
S106
S105
S205
T271
S272
S274
S116
S129
T122
T141
S171
S176
S494
Candidate
FALSE
TRUE
Domains and Regions:
DOMAIN: PWWP
REGION: Disordered

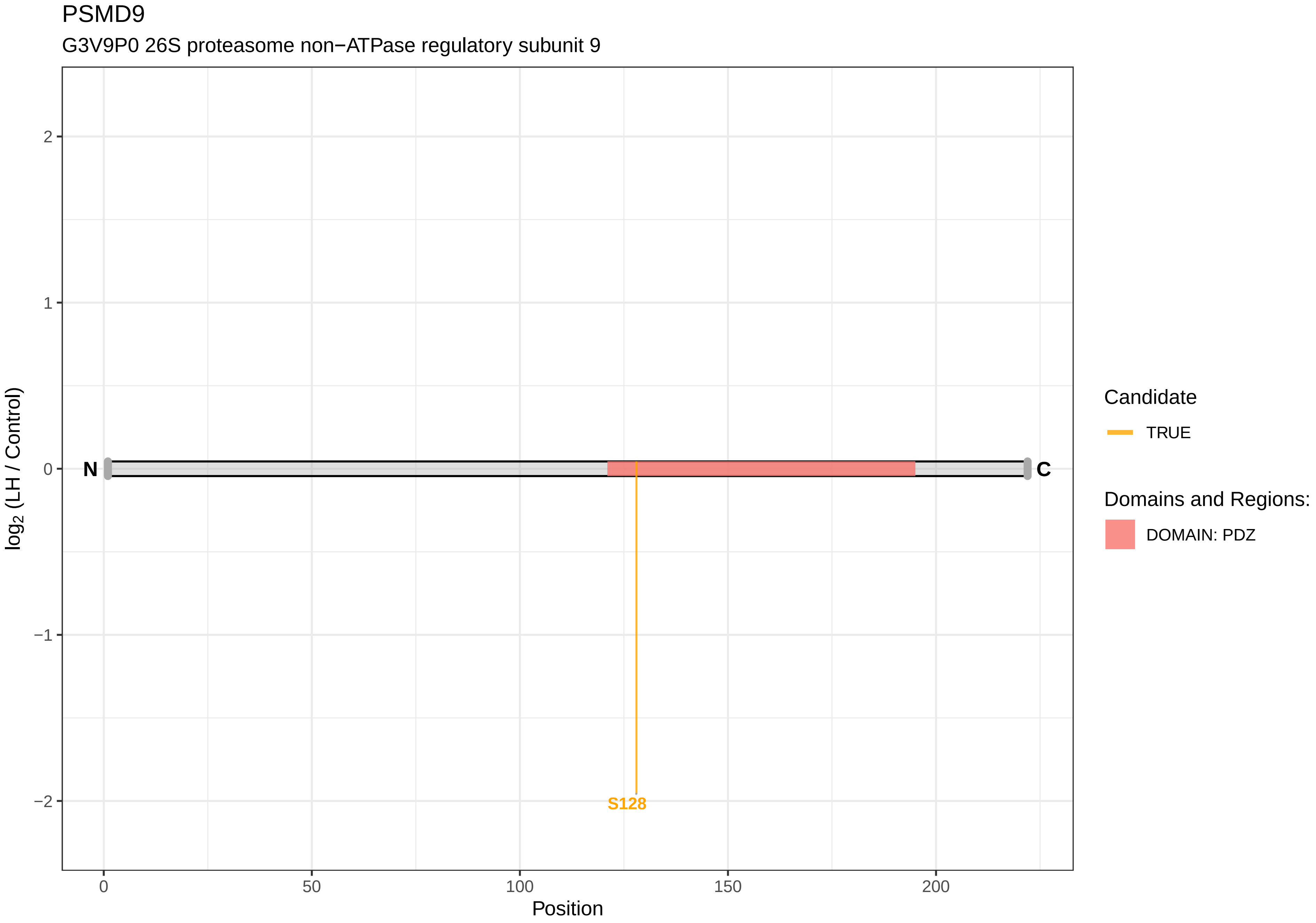
PSMD9
G3V9P0 26S proteasome non−ATPase regulatory subunit 9
log2 (LH / Control)
2
1
0
−1
−2
N
C
S128
0
50
100
150
200
Position
Candidate
TRUE
Domains and Regions:
DOMAIN: PDZ

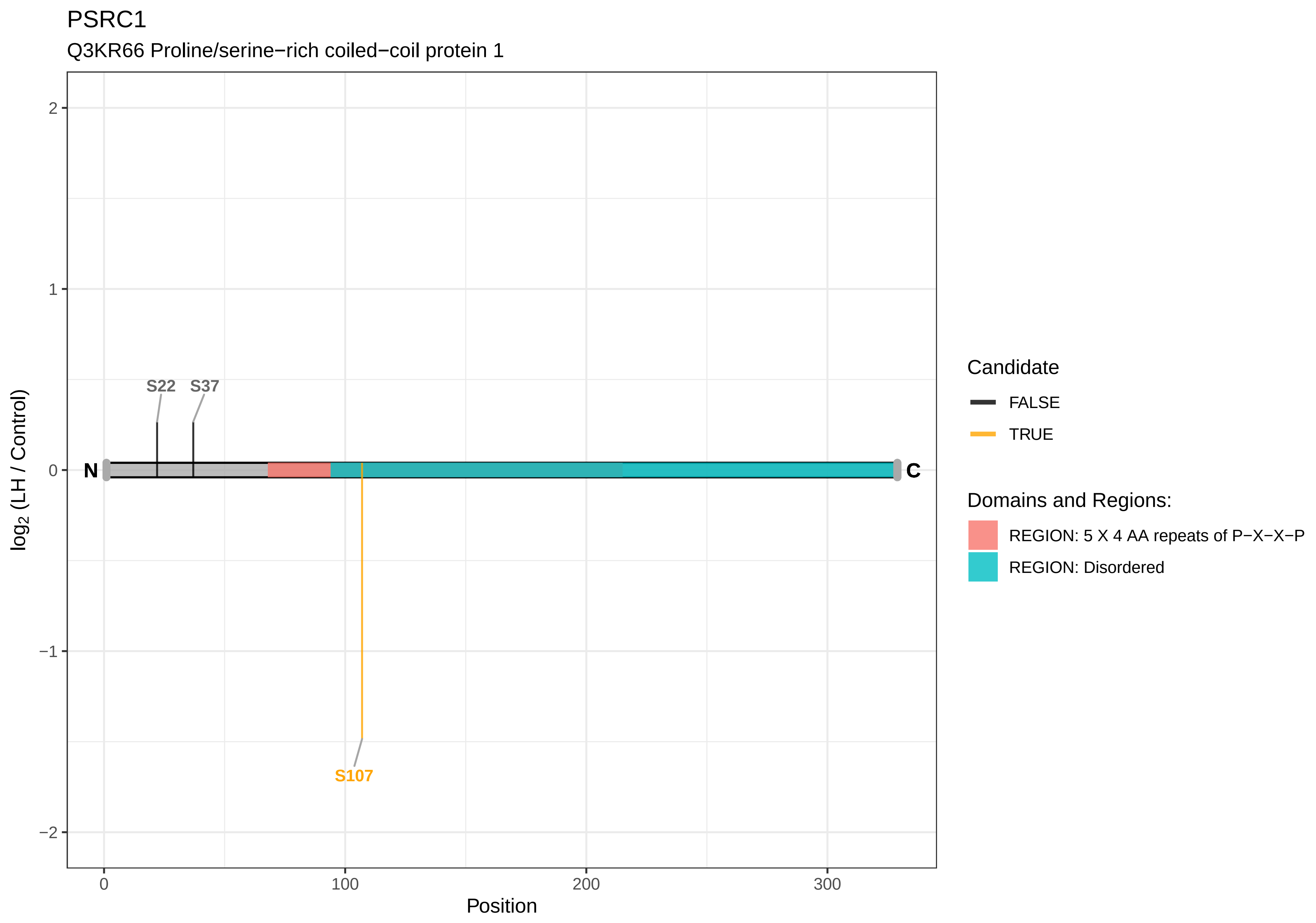

PSRC1
Q3KR66 Proline/serine−rich coiled−coil protein 1
S22
S37
N
C
S107
2
1
0
−1
−2
log2 (LH / Control)
0
100
200
300
Position
Candidate
FALSE
TRUE
Domains and Regions:
REGION: 5 X 4 AA repeats of P−X−X−P
REGION: Disordered

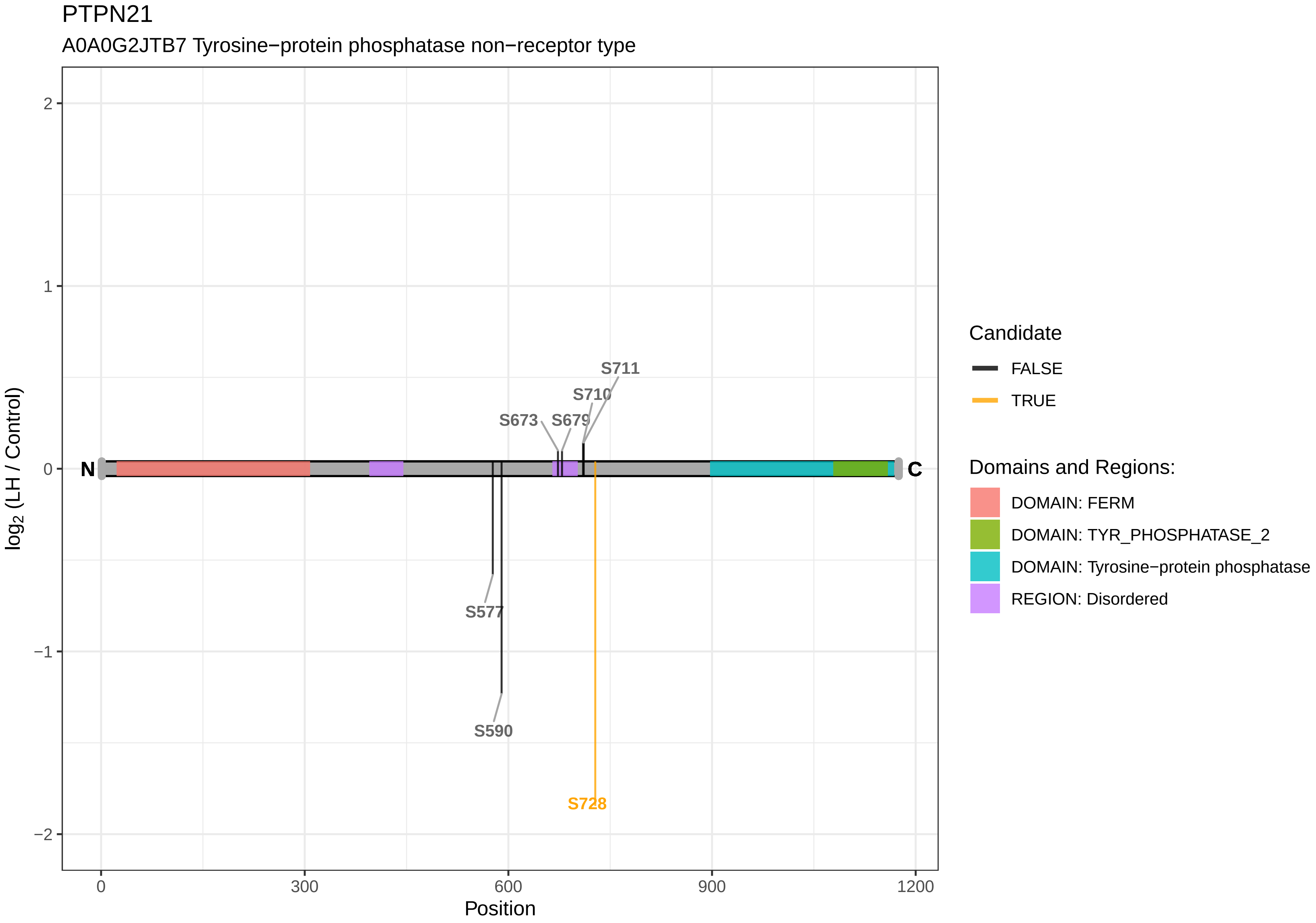
PTPN21
A0A0G2JTB7 Tyrosine−protein phosphatase non−receptor type
log2 (LH / Control)
2
1
0
−1
−2
N
C
S711
S710
S673
S679
S577
S590
S728
0
300
600
900
1200
Position
Candidate
FALSE
TRUE
Domains and Regions:
DOMAIN: FERM
DOMAIN: TYR_PHOSPHATASE_2
DOMAIN: Tyrosine−protein phosphatase
REGION: Disordered

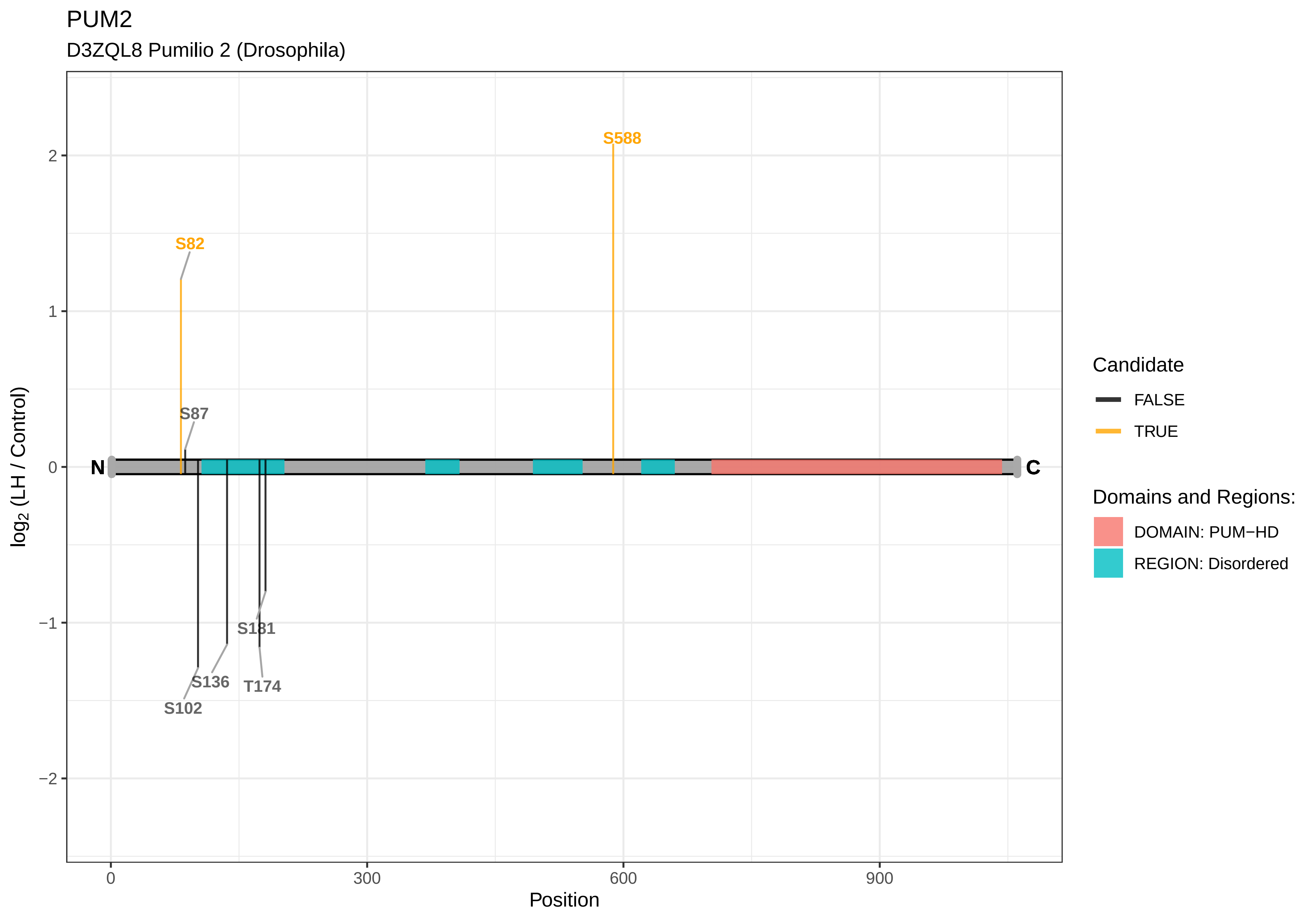
PUM2
D3ZQL8 Pumilio 2 (Drosophila)
S588
S82
S87
N
C
S181
S136
T174
S102
2
1
0
−1
−2
log2 (LH / Control)
0
300
600
900
Position
Candidate
FALSE
TRUE
Domains and Regions:
DOMAIN: PUM−HD
REGION: Disordered

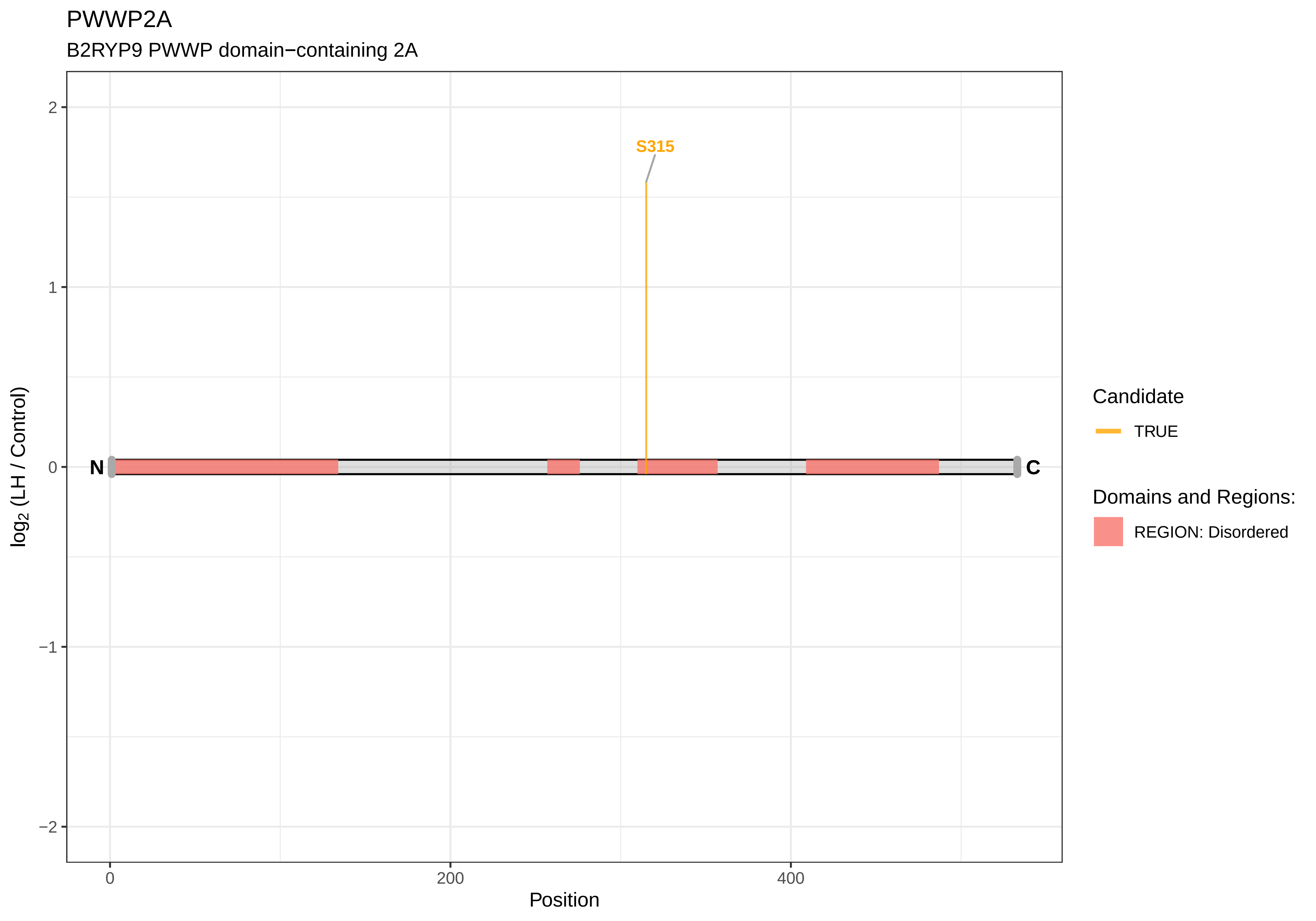

PWWP2A
B2RYP9 PWWP domain−containing 2A
S315
N
C
2
1
0
−1
−2
0
200
400
Position
log2 (LH / Control)
Candidate
TRUE
Domains and Regions:
REGION: Disordered

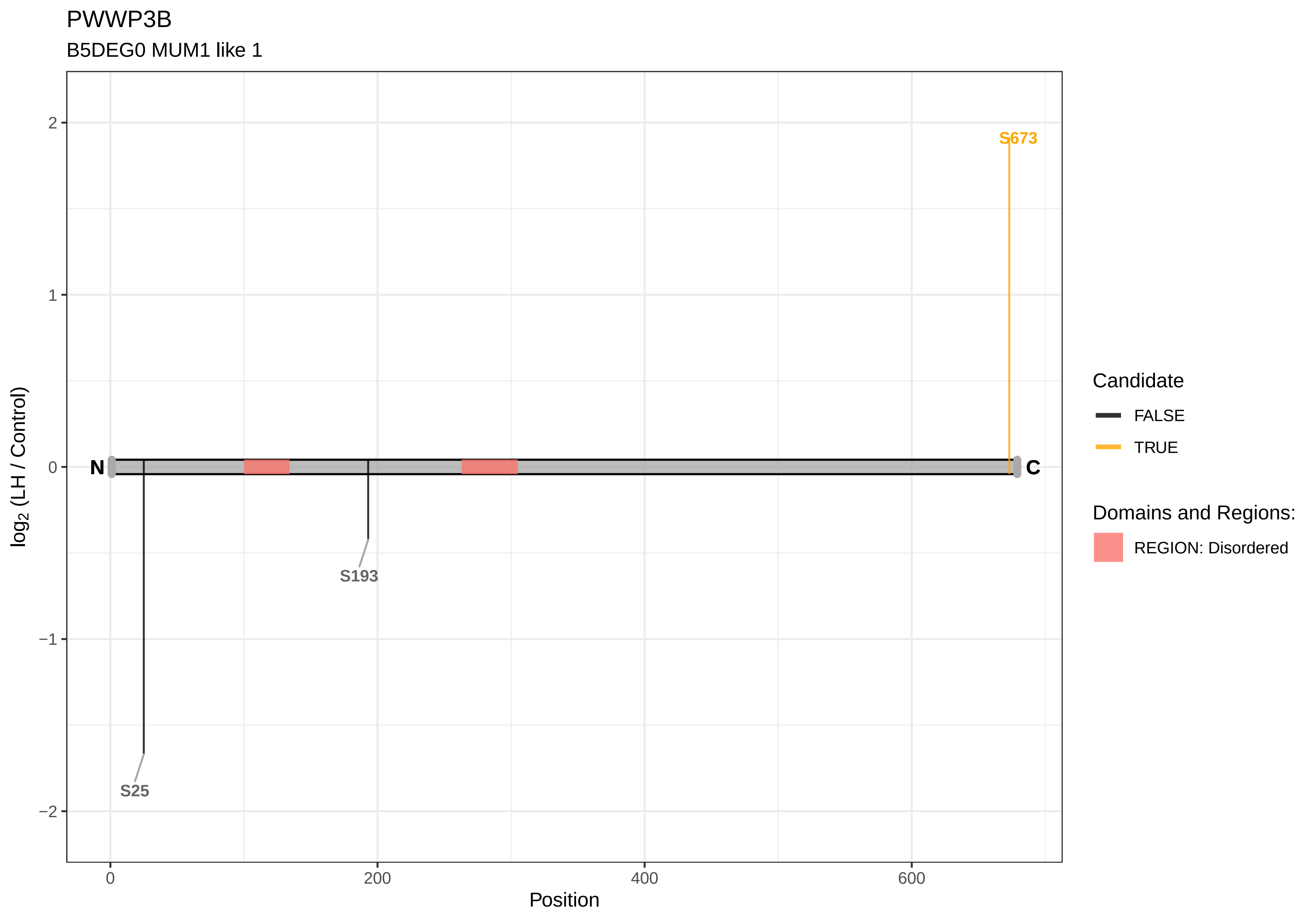

PWWP3B
B5DEG0 MUM1 like 1
S673
S193
S25
N
C
log2 (LH / Control)
Position
0
200
400
600
2
1
0
−1
−2
Candidate
FALSE
TRUE
Domains and Regions:
REGION: Disordered

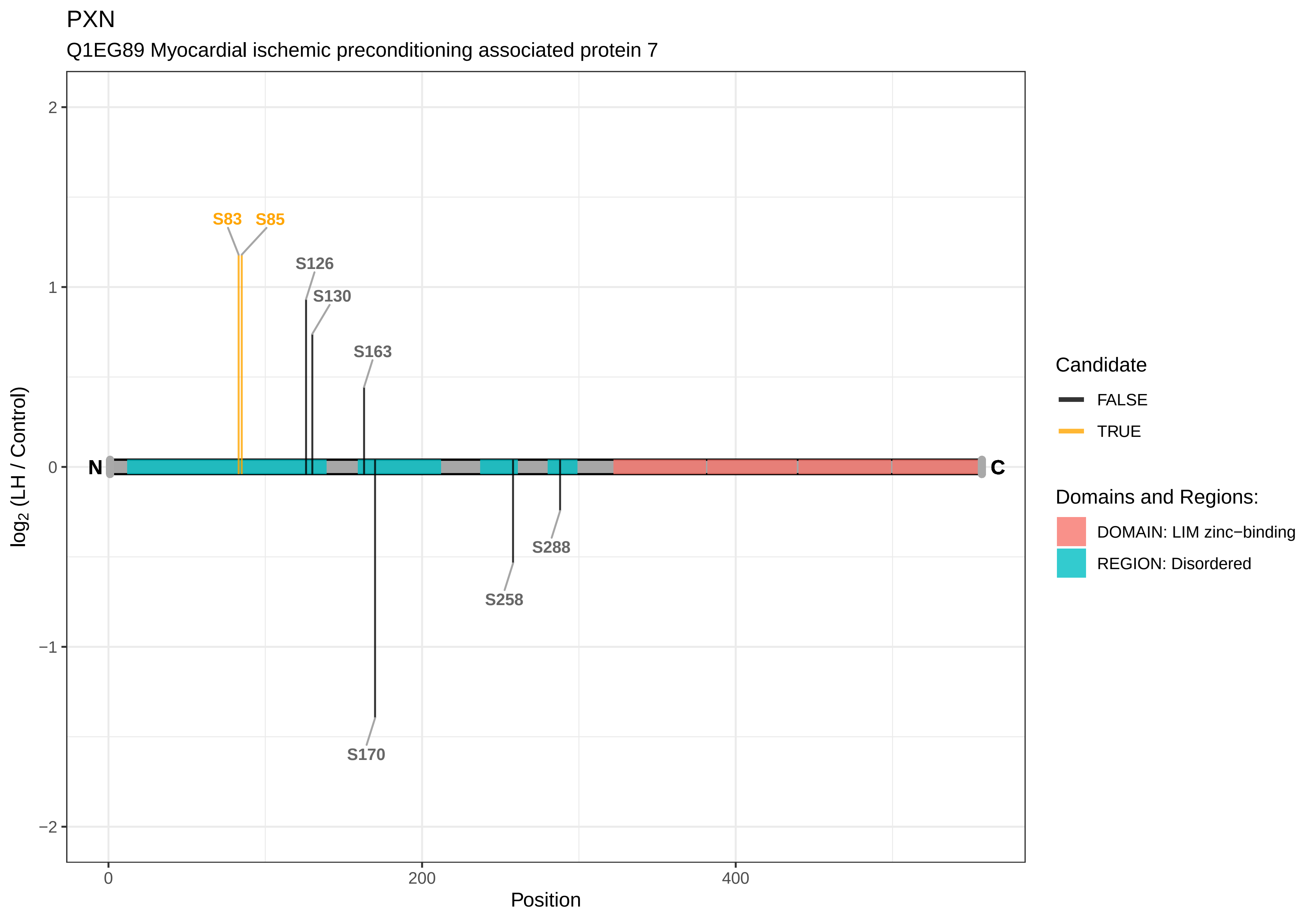
PXN
Q1EG89 Myocardial ischemic preconditioning associated protein 7
S83
S85
S126
S130
S163
S170
S258
S288
N
C
log2 (LH / Control)
Position
2
1
0
−1
−2
0
200
400
Candidate
FALSE
TRUE
Domains and Regions:
DOMAIN: LIM zinc−binding
REGION: Disordered

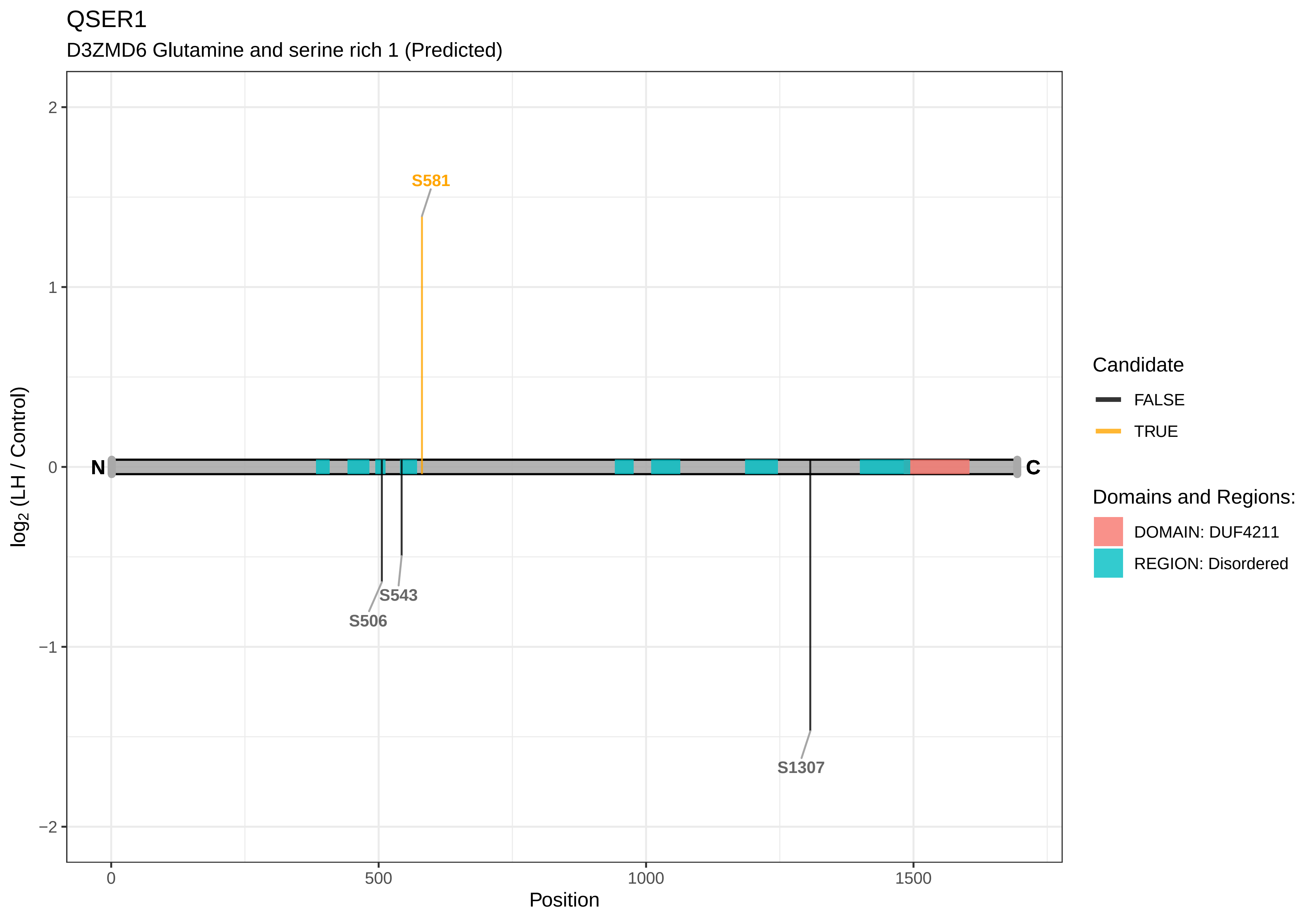
QSER1
D3ZMD6 Glutamine and serine rich 1 (Predicted)
S581
S543
S506
S1307
N
C
2
1
0
−1
−2
0
500
1000
1500
Position
log2 (LH / Control)
Candidate
FALSE
TRUE
Domains and Regions:
DOMAIN: DUF4211
REGION: Disordered

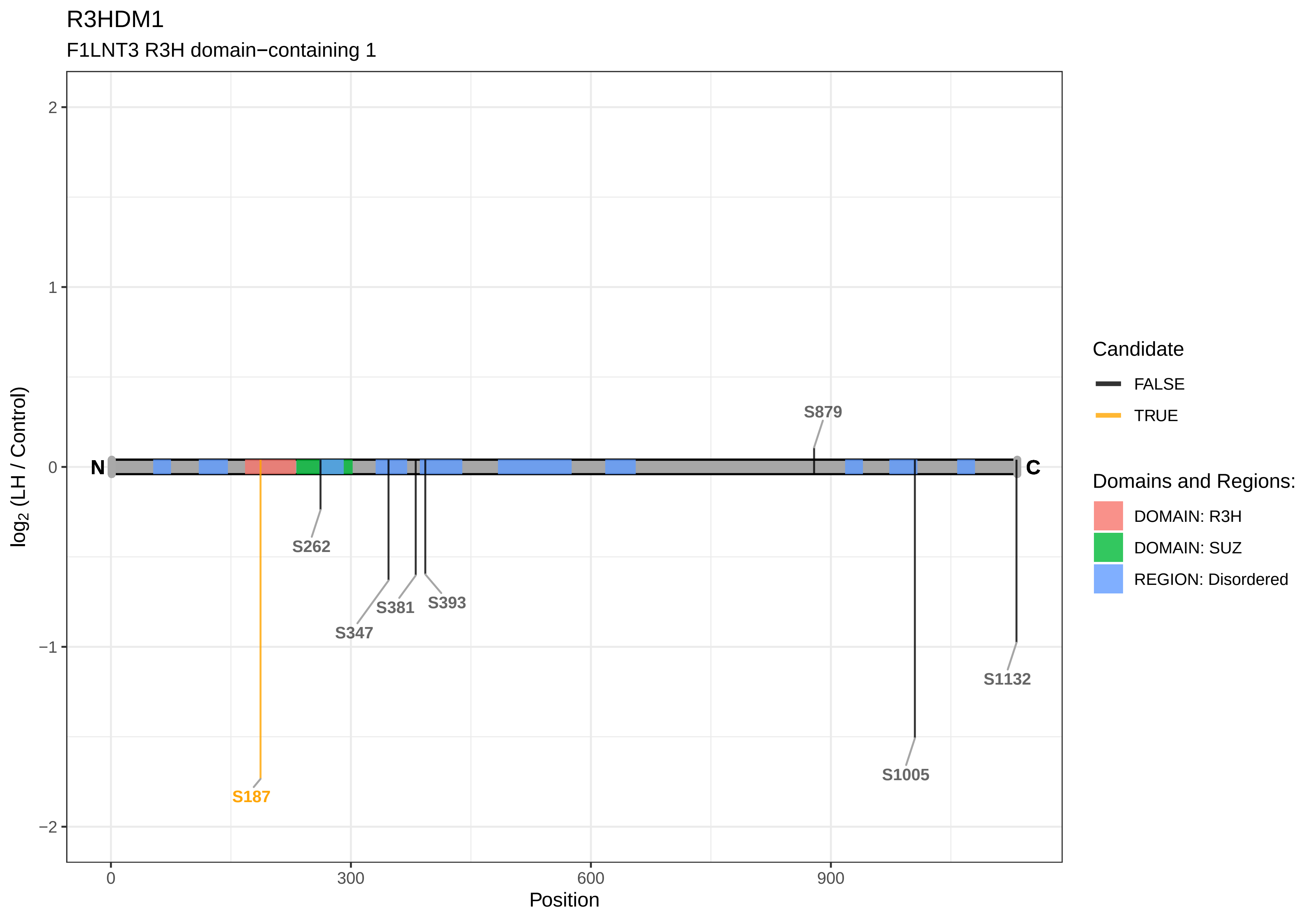
R3HDM1
F1LNT3 R3H domain−containing 1
log2 (LH / Control)
2
1
0
−1
−2
N
C
S879
S262
S381
S393
S347
S1132
S1005
S187
0
300
600
900
Position
Candidate
FALSE
TRUE
Domains and Regions:
DOMAIN: R3H
DOMAIN: SUZ
REGION: Disordered

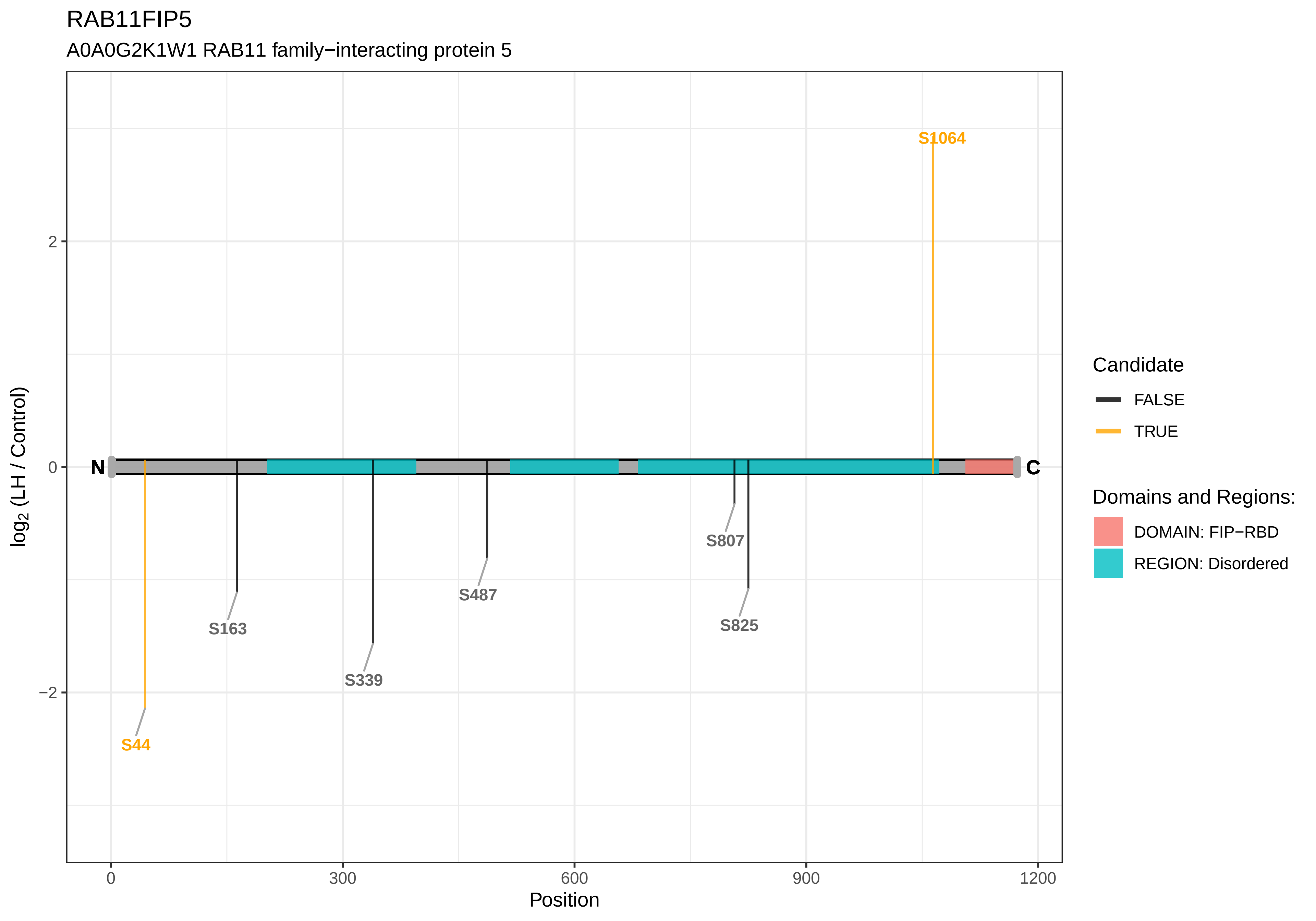
RAB11FIP5
A0A0G2K1W1 RAB11 family−interacting protein 5
log2 (LH / Control)
Position
0
300
600
900
1200
2
0
−2
N
C
S1064
S44
S163
S339
S487
S807
S825
Candidate
FALSE
TRUE
Domains and Regions:
DOMAIN: FIP−RBD
REGION: Disordered

RAB12

P35284 Ras−related protein Rab−12

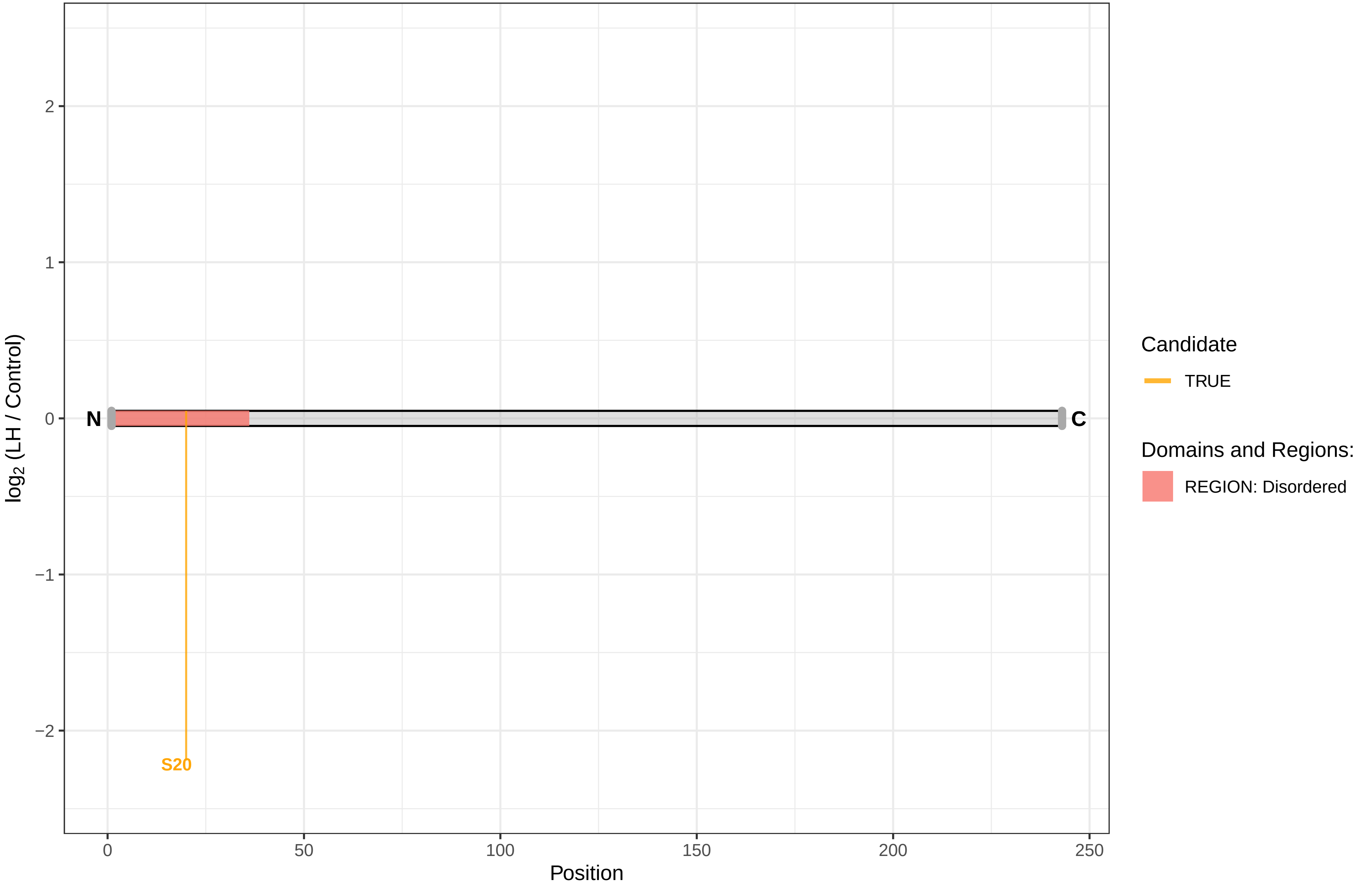

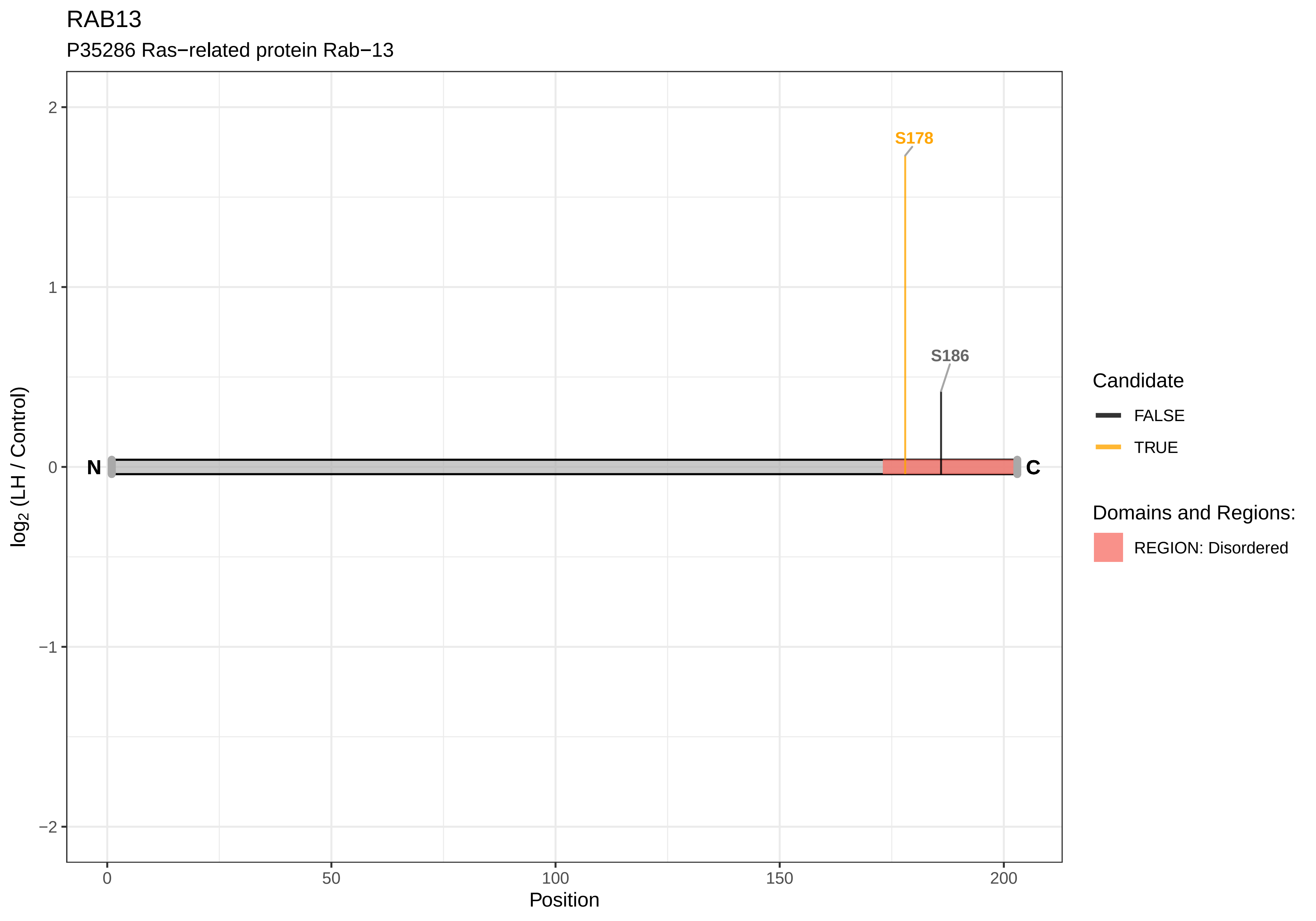
RAB13
P35286 Ras−related protein Rab−13
S178
S186
N
C
2
1
0
−1
−2
log2 (LH / Control)
0
50
100
150
200
Position
Candidate
FALSE
TRUE
Domains and Regions:
REGION: Disordered

RAB34

Q5U1Y1 Ras−related protein Rab−34

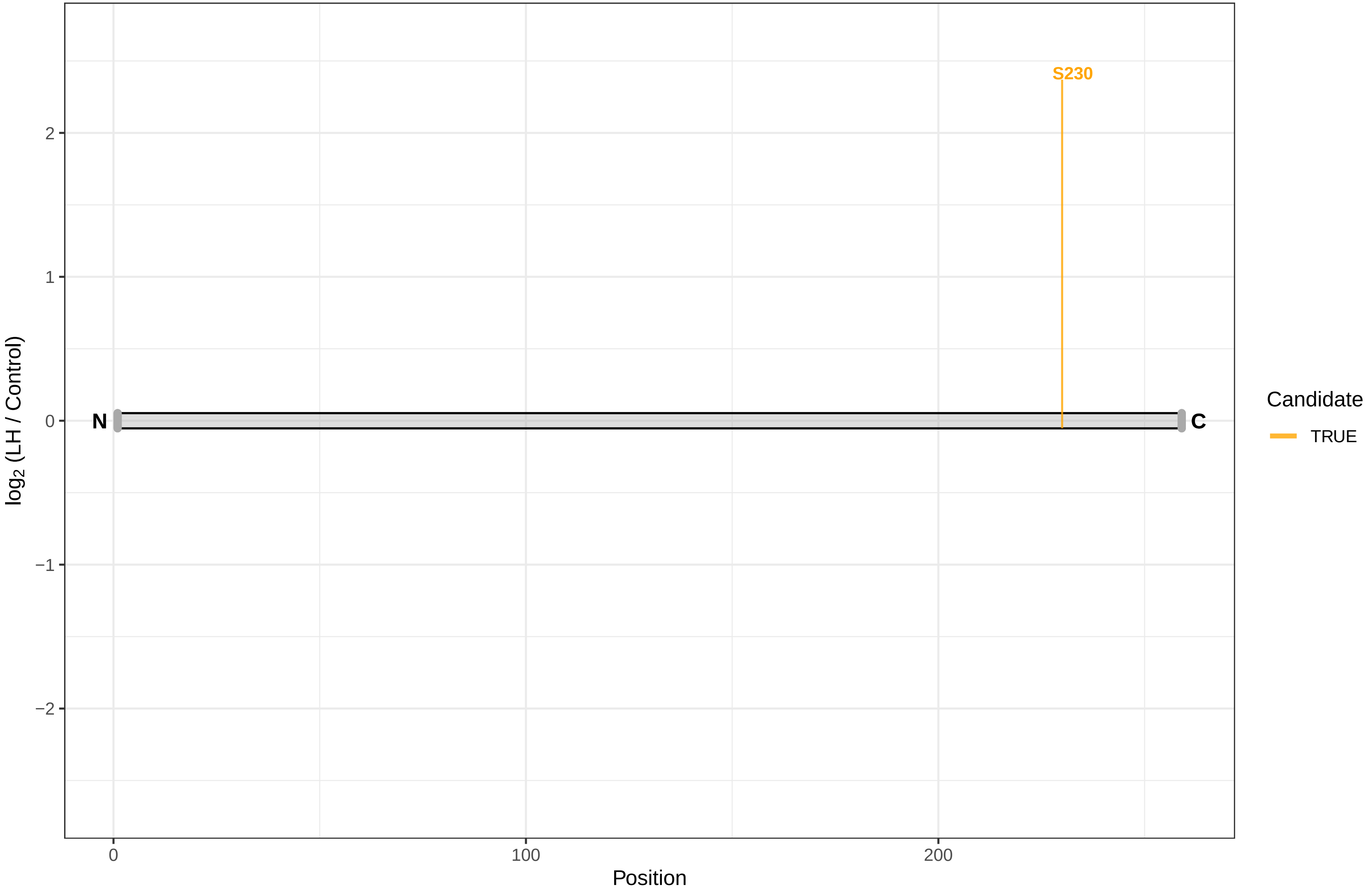

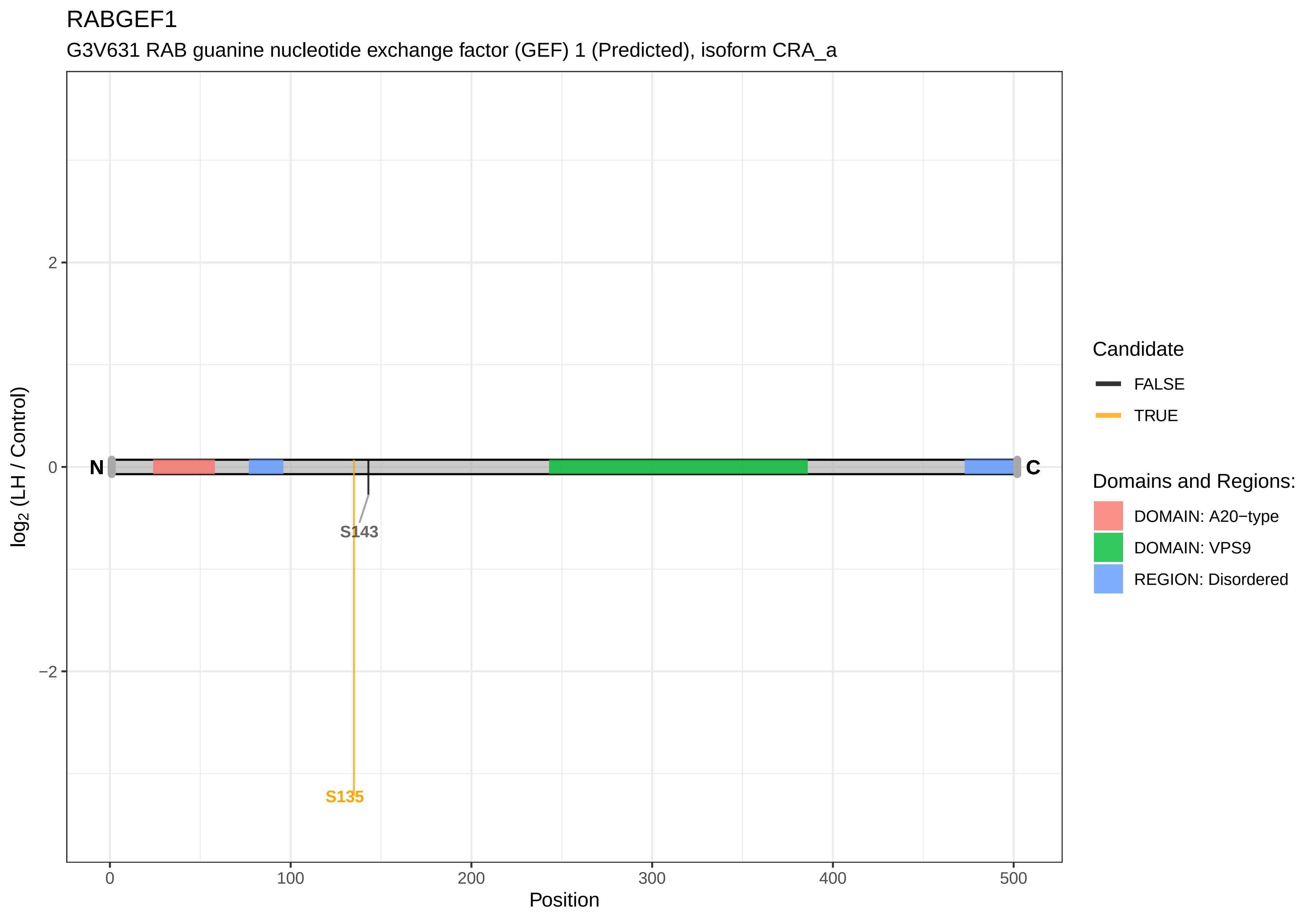
RABGEF1
G3V631 RAB guanine nucleotide exchange factor (GEF) 1 (Predicted), isoform CRA_a
log2 (LH / Control)
Position
2
0
−2
0
100
200
300
400
500
N
C
S143
S135
Candidate
FALSE
TRUE
Domains and Regions:
DOMAIN: A20−type
DOMAIN: VPS9
REGION: Disordered

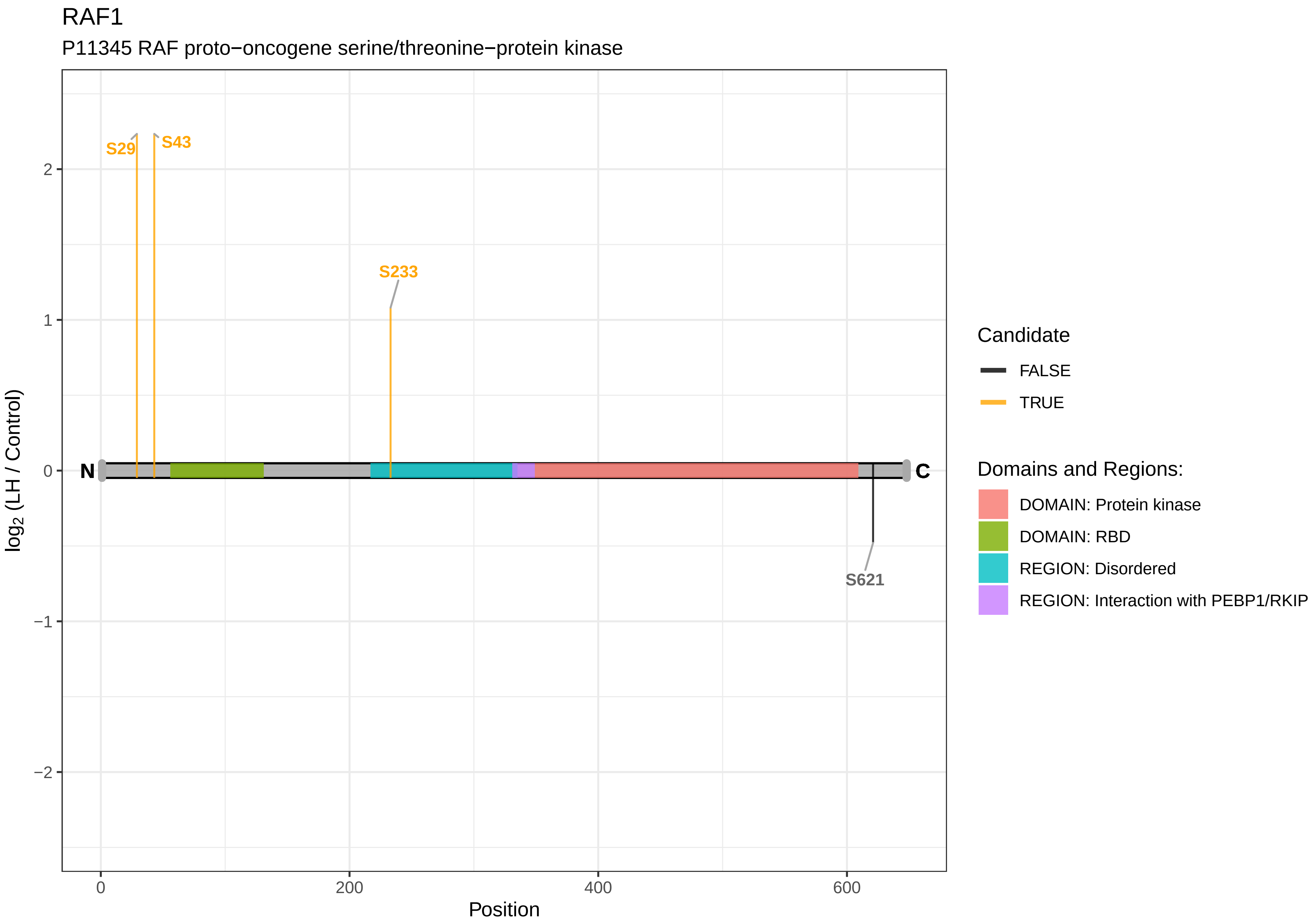

RAF1
P11345 RAF proto−oncogene serine/threonine−protein kinase
S29
S43
S233
S621
N
C
2
1
0
−1
−2
0
200
400
600
Position
log2 (LH / Control)
Candidate
FALSE
TRUE
Domains and Regions:
DOMAIN: Protein kinase
DOMAIN: RBD
REGION: Disordered
REGION: Interaction with PEBP1/RKIP

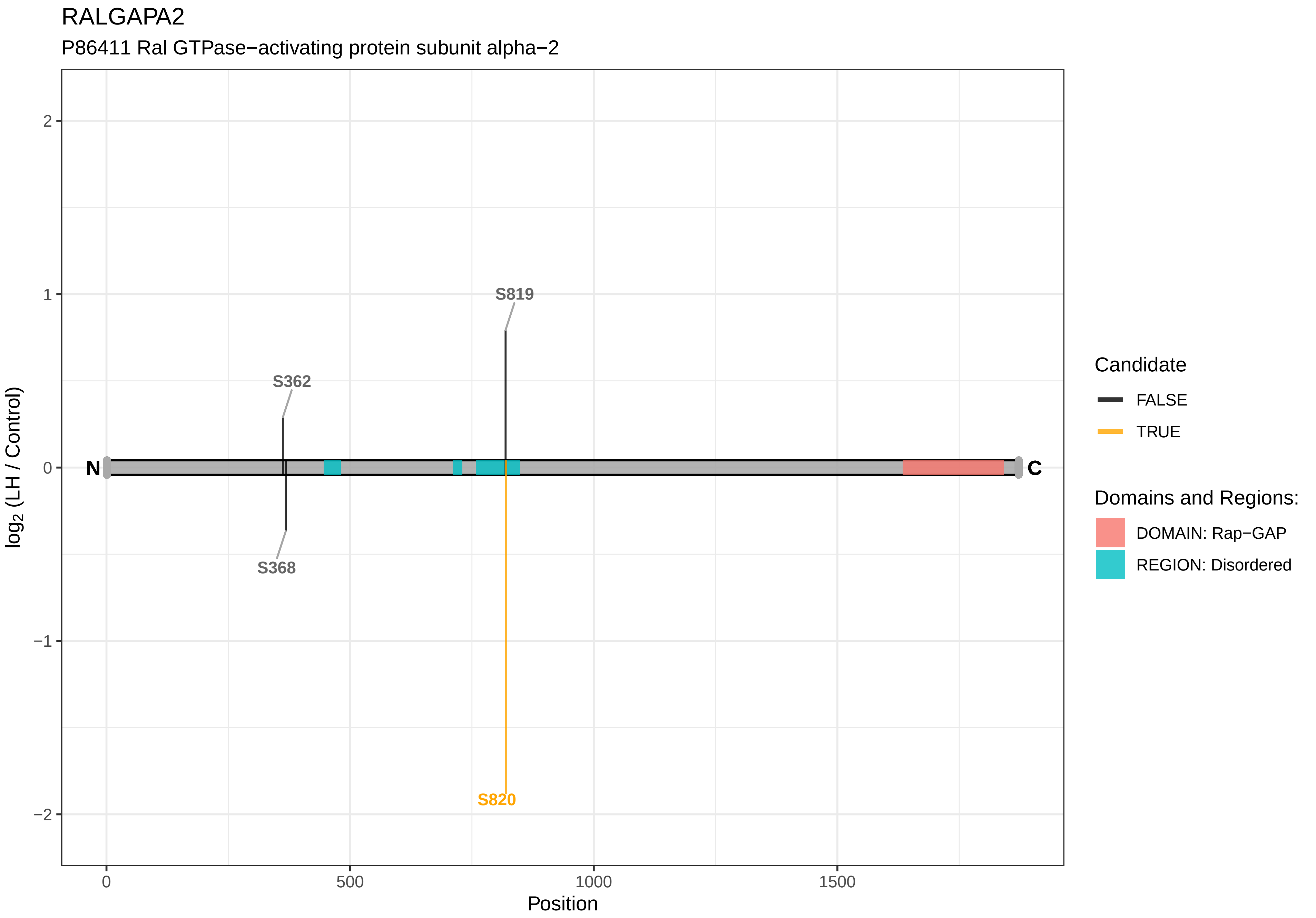

RALGAPA2
P86411 Ral GTPase−activating protein subunit alpha−2
log2 (LH / Control)
Position
S819
S362
S368
S820
N
C
2
1
0
−1
−2
0
500
1000
1500
Candidate
FALSE
TRUE
Domains and Regions:
DOMAIN: Rap−GAP
REGION: Disordered

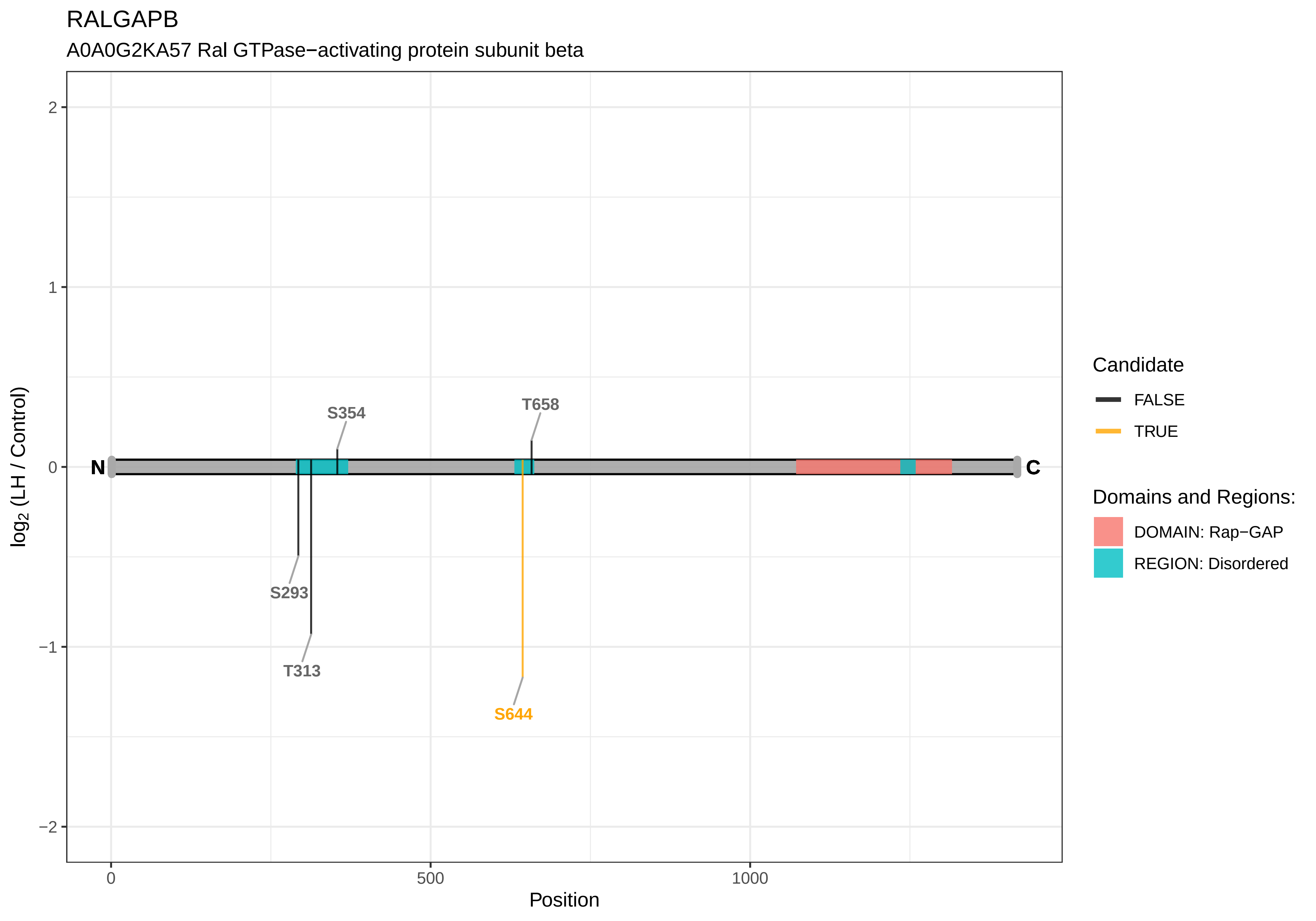

RALGAPB
A0A0G2KA57 Ral GTPase−activating protein subunit beta
log2 (LH / Control)
Position
N
C
S354
T658
S293
T313
S644
2
1
0
−1
−2
0
500
1000
Candidate
FALSE
TRUE
Domains and Regions:
DOMAIN: Rap−GAP
REGION: Disordered

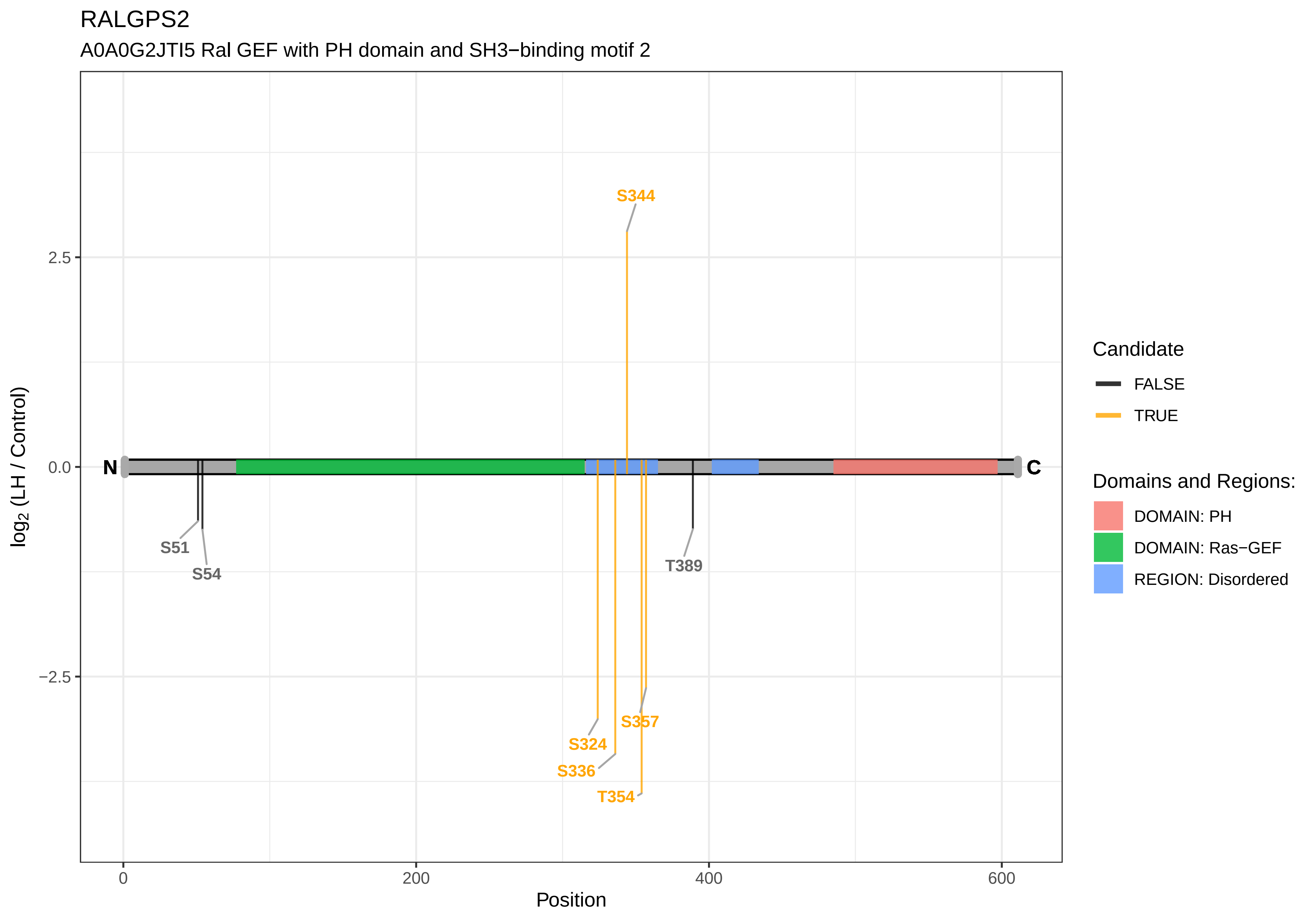
RALGPS2
A0A0G2JTI5 Ral GEF with PH domain and SH3−binding motif 2
S344
N
C
S51
S54
T389
S357
S324
S336
T354
2.5
0.0
−2.5
0
200
400
600
Position
log2 (LH / Control)
Candidate
FALSE
TRUE
Domains and Regions:
DOMAIN: PH
DOMAIN: Ras−GEF
REGION: Disordered

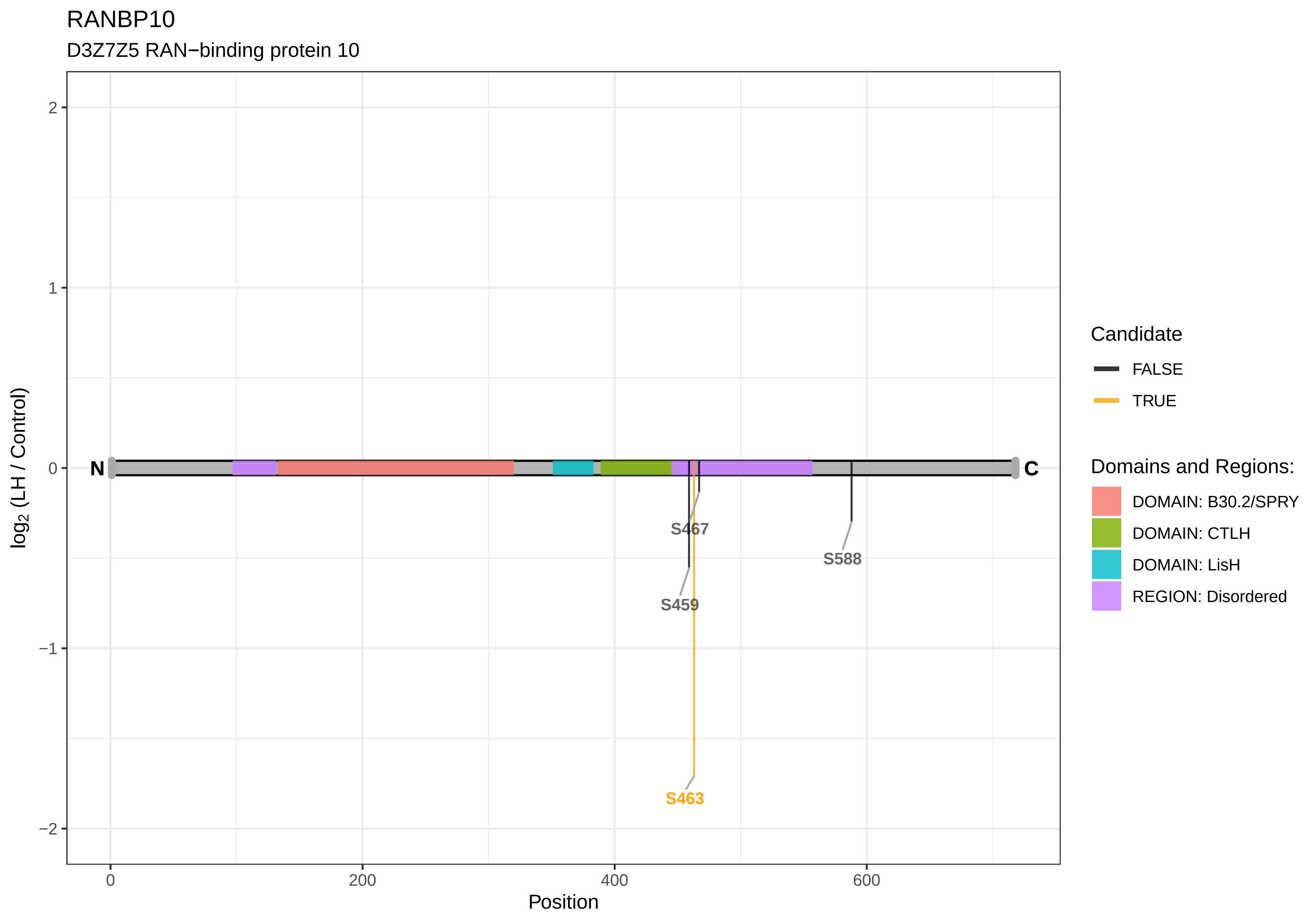
RANBP10
D3Z7Z5 RAN−binding protein 10
log2 (LH / Control)
Position
N
C
S467
S588
S459
S463
2
1
0
−1
−2
0
200
400
600
Candidate
FALSE
TRUE
Domains and Regions:
DOMAIN: B30.2/SPRY
DOMAIN: CTLH
DOMAIN: LisH
REGION: Disordered

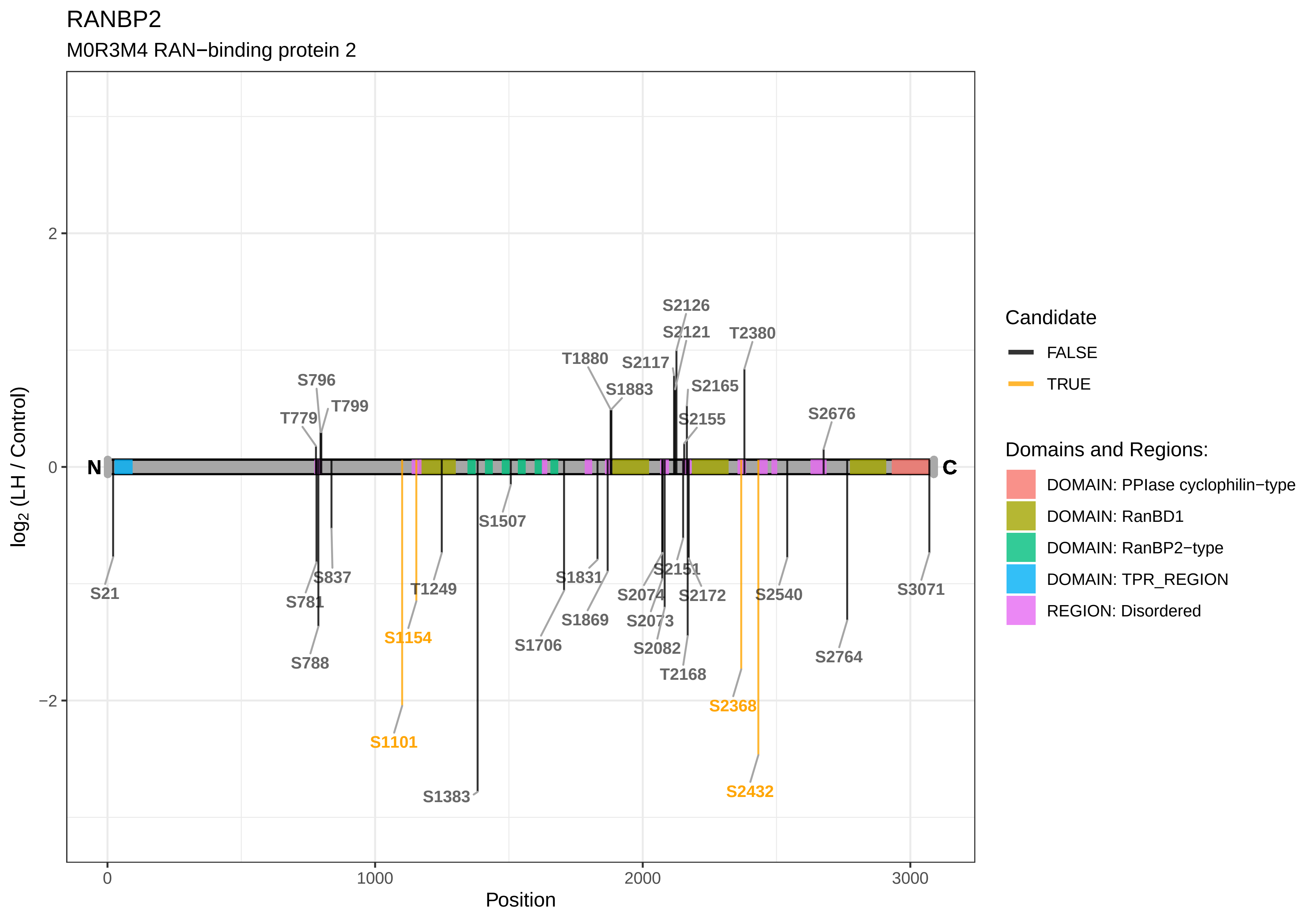

RANBP2
M0R3M4 RAN−binding protein 2
log2 (LH / Control)
Position
0
1000
2000
3000
2
0
−2
N
C
S21
T779
S781
S788
S796
T799
S837
S1101
S1154
T1249
S1383
S1507
S1706
S1831
S1869
T1880
S1883
S2073
S2074
S2082
S2117
S2121
S2126
S2151
S2155
S2165
T2168
S2172
S2368
T2380
S2432
S2540
S2676
S2764
S3071
Candidate
FALSE
TRUE
Domains and Regions:
DOMAIN: PPIase cyclophilin−type
DOMAIN: RanBD1
DOMAIN: RanBP2−type
DOMAIN: TPR_REGION
REGION: Disordered

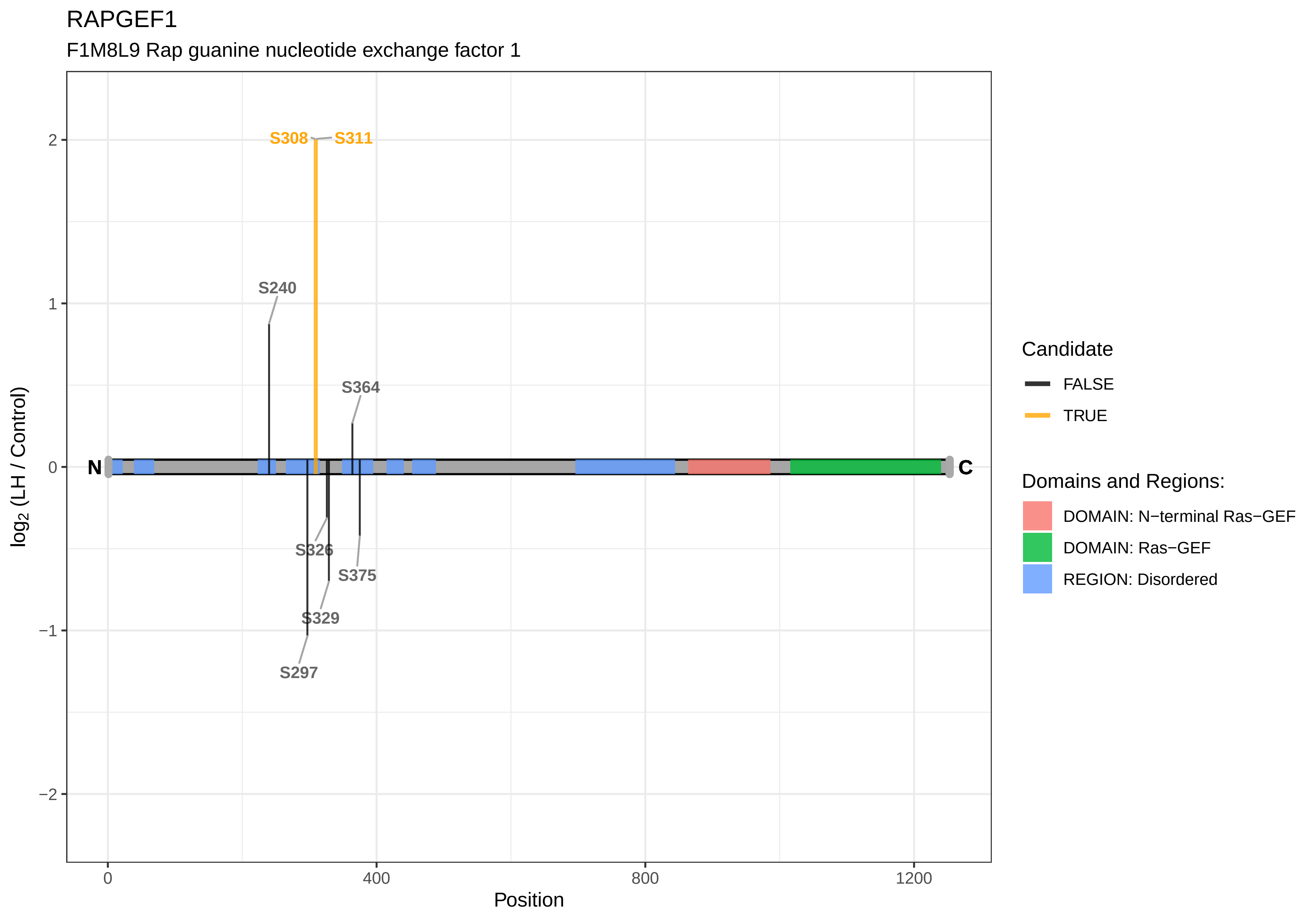

RAPGEF1
F1M8L9 Rap guanine nucleotide exchange factor 1
S308
S311
S240
S364
S326
S375
S329
S297
N
C
2
1
0
−1
−2
log2 (LH / Control)
0
400
800
1200
Position
Candidate
FALSE
TRUE
Domains and Regions:
DOMAIN: N−terminal Ras−GEF
DOMAIN: Ras−GEF
REGION: Disordered

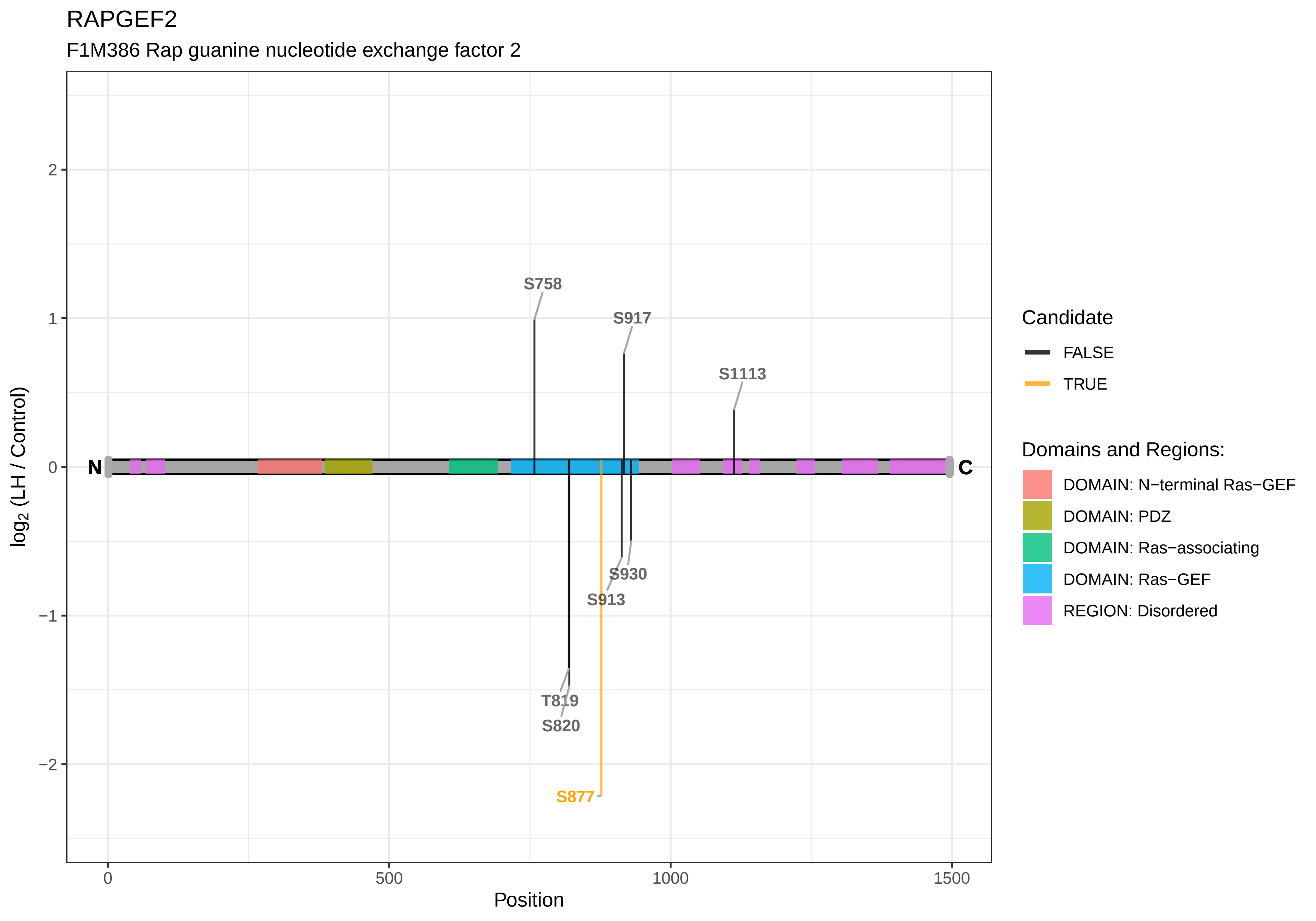

RAPGEF2
F1M386 Rap guanine nucleotide exchange factor 2
S758
S917
S1113
N
C
S930
S913
T819
S820
S877
2
1
0
−1
−2
log2 (LH / Control)
0
500
1000
1500
Position
Candidate
FALSE
TRUE
Domains and Regions:
DOMAIN: N−terminal Ras−GEF
DOMAIN: PDZ
DOMAIN: Ras−associating
DOMAIN: Ras−GEF
REGION: Disordered

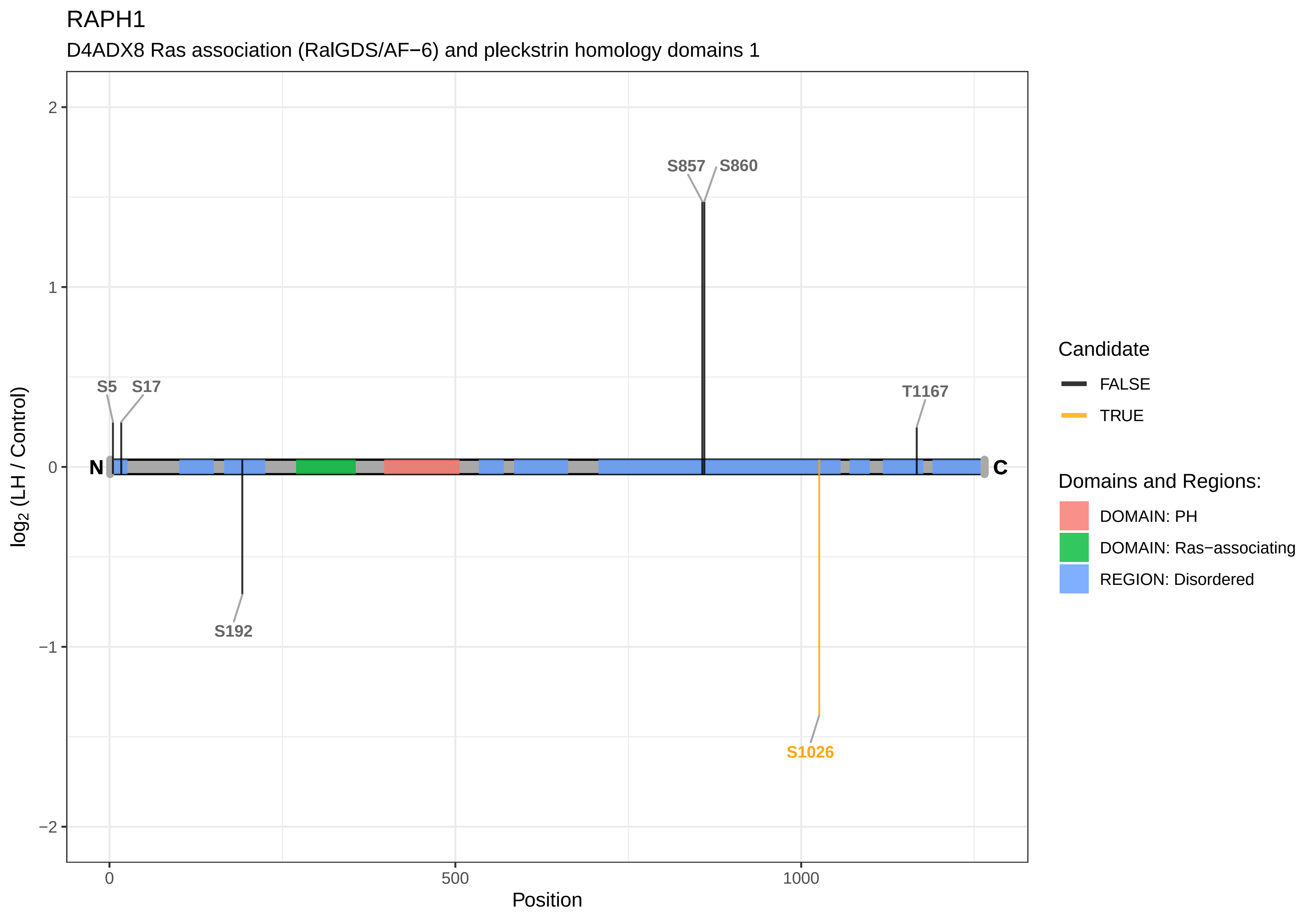

RAPH1
D4ADX8 Ras association (RalGDS/AF−6) and pleckstrin homology domains 1
S857
S860
S5
S17
T1167
S192
S1026
N
C
2
1
0
−1
−2
log2 (LH / Control)
0
500
1000
Position
Candidate
FALSE
TRUE
Domains and Regions:
DOMAIN: PH
DOMAIN: Ras−associating
REGION: Disordered

RASAL2

A0A0G2JTA7 RAS protein activator−like 2

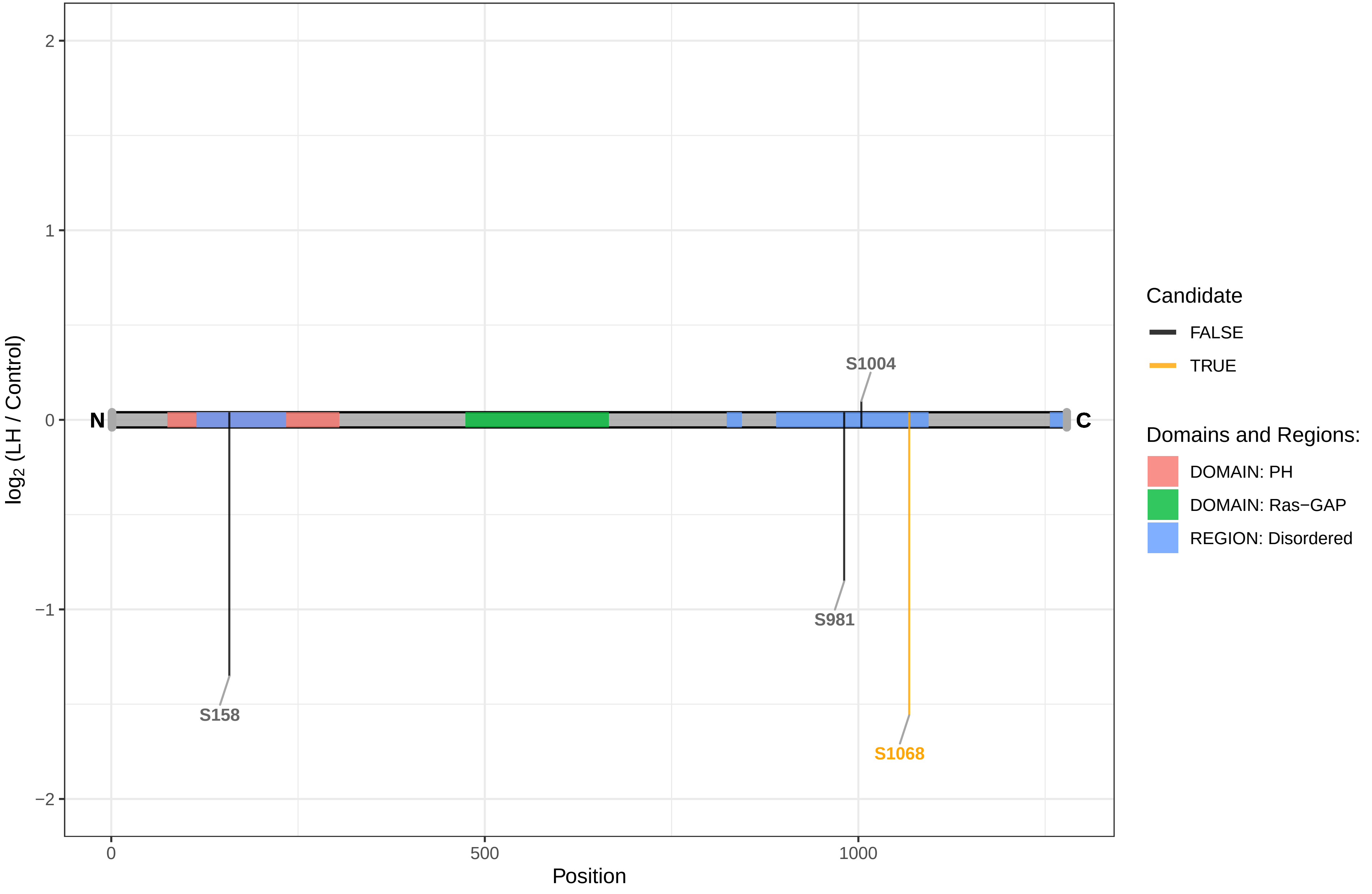

RBBP6

G3V953 RB−binding protein 6, ubiquitin ligase

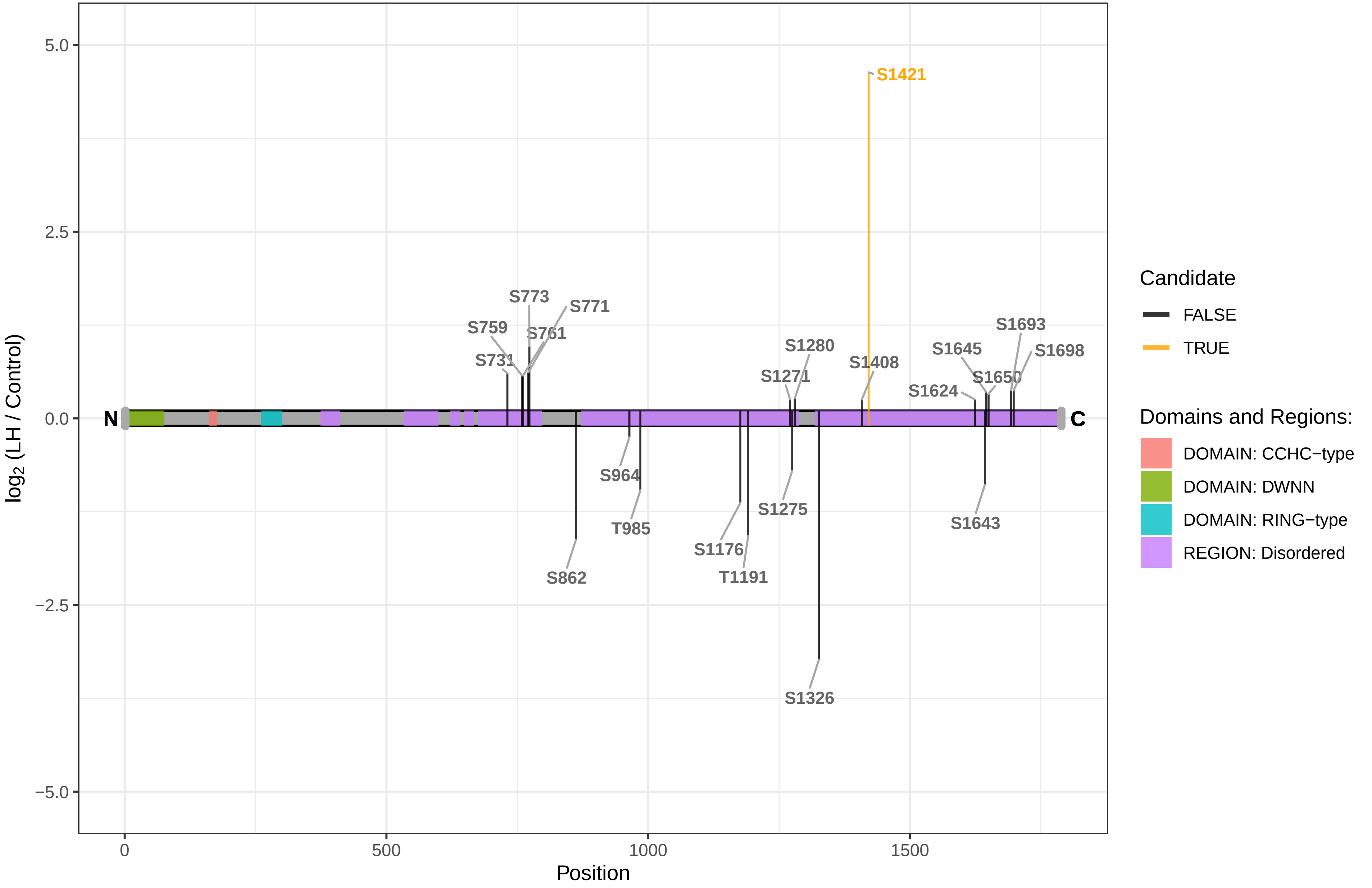

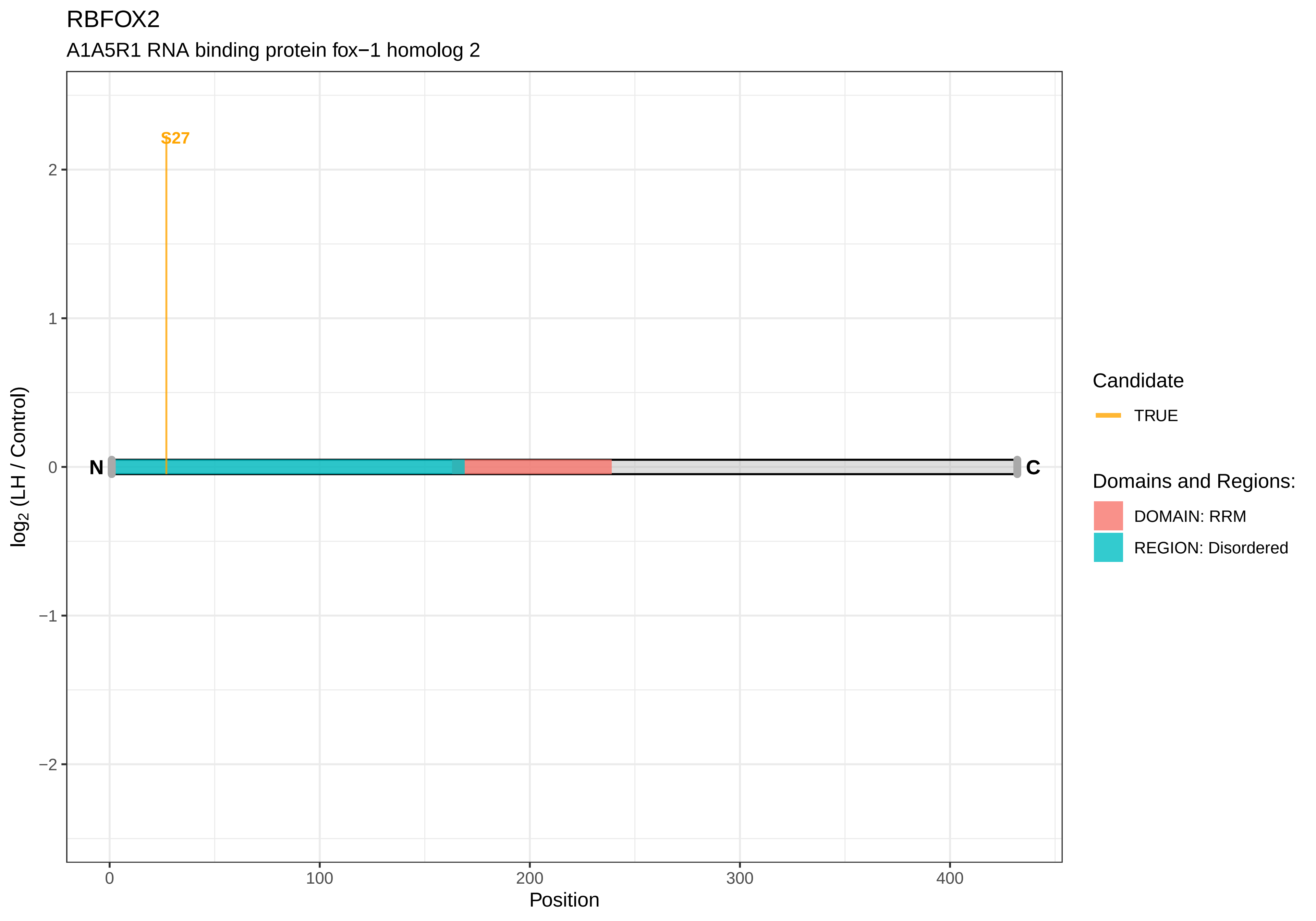
RBFOX2
A1A5R1 RNA binding protein fox−1 homolog 2
S27
N
C
2
1
0
−1
−2
log2 (LH / Control)
0
100
200
300
400
Position
Candidate
TRUE
Domains and Regions:
DOMAIN: RRM
REGION: Disordered

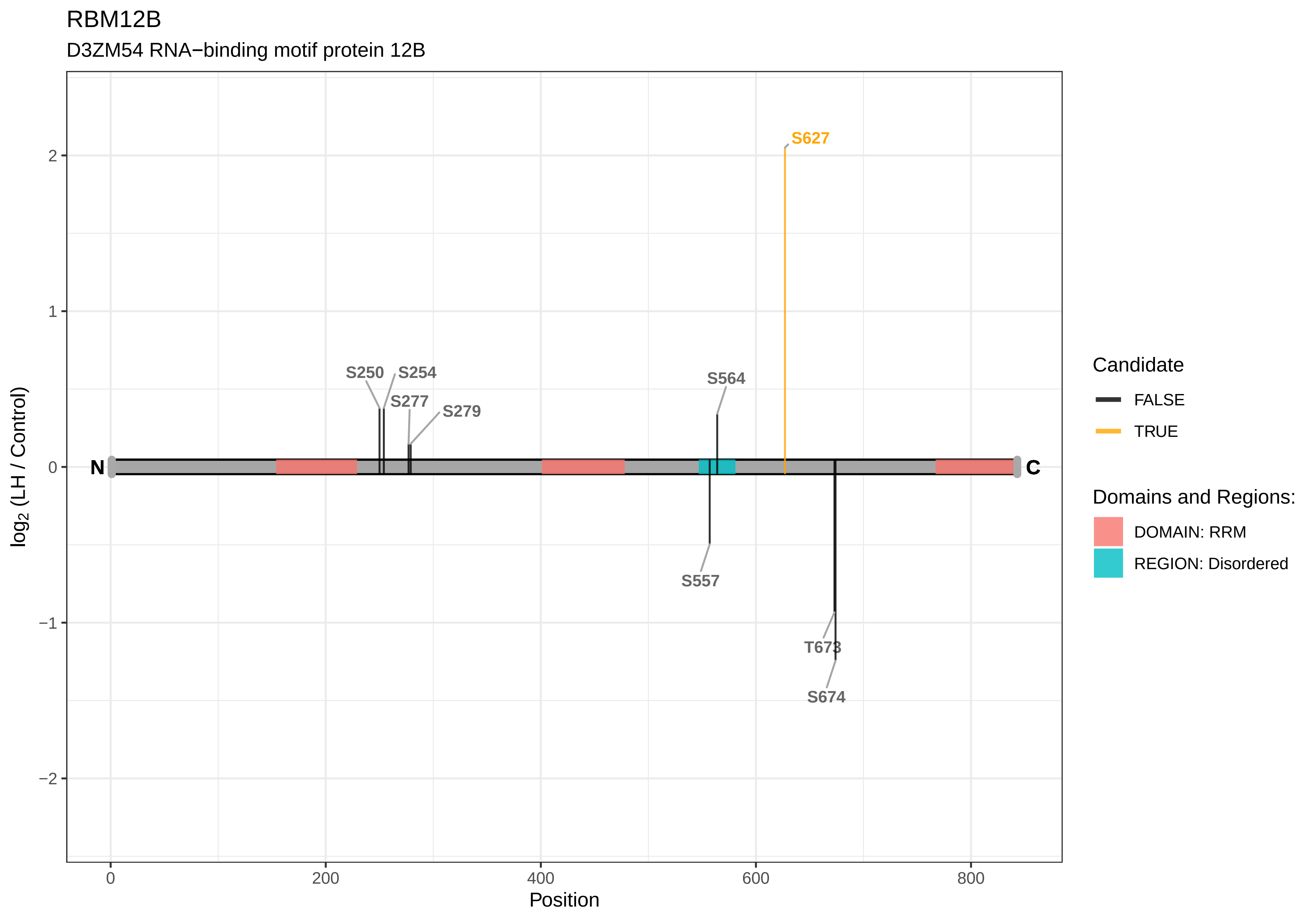
RBM12B
D3ZM54 RNA−binding motif protein 12B
S627
S250
S254
S277
S279
S564
S557
T673
S674
N
C
log2 (LH / Control)
Position
0
200
400
600
800
2
1
0
−1
−2
Candidate
FALSE
TRUE
Domains and Regions:
DOMAIN: RRM
REGION: Disordered

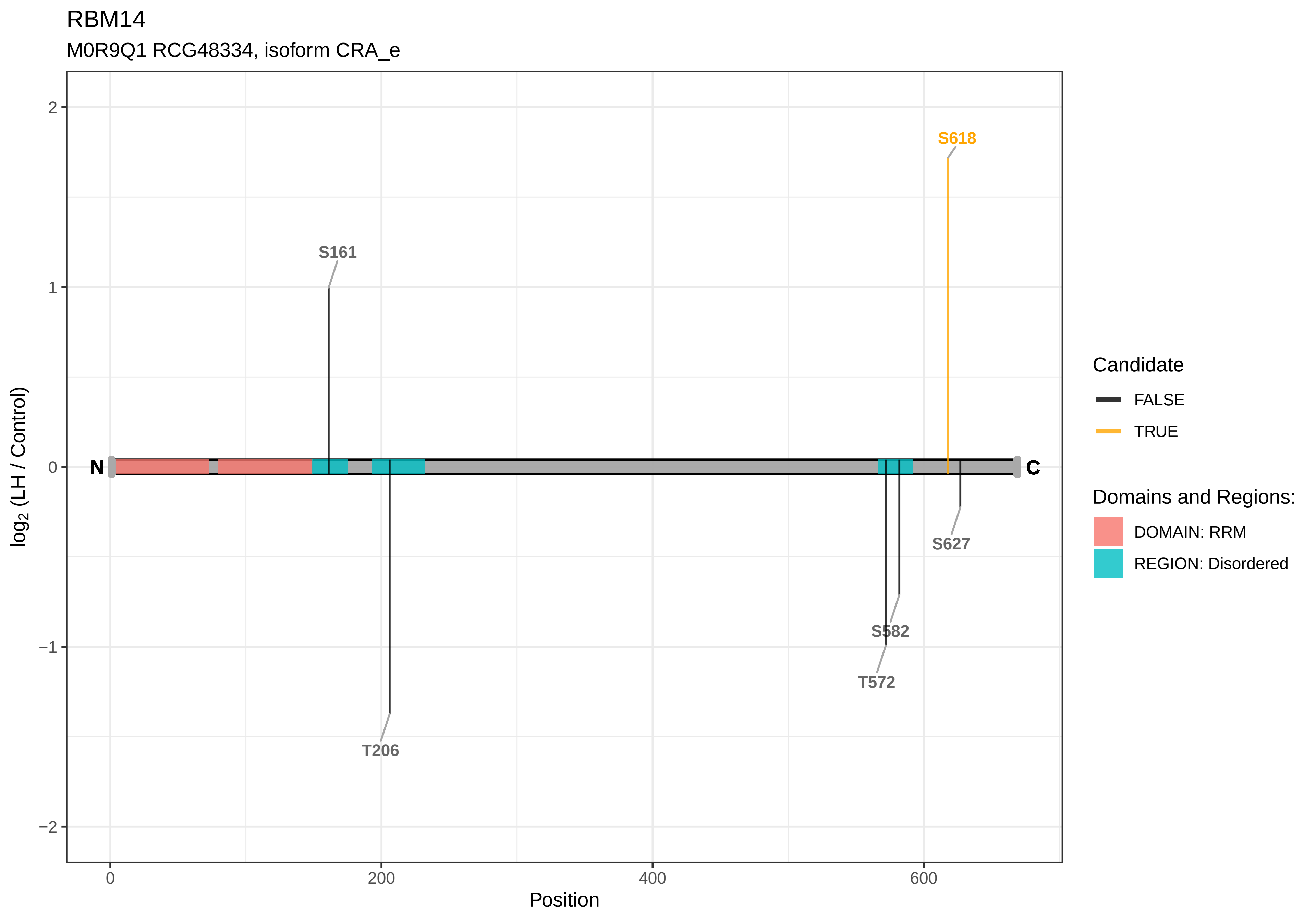
RBM14
M0R9Q1 RCG48334, isoform CRA_e
log2 (LH / Control)
Position
N
C
S618
S161
T206
S582
T572
S627
2
1
0
−1
−2
0
200
400
600
Candidate
FALSE
TRUE
Domains and Regions:
DOMAIN: RRM
REGION: Disordered

RBM15

M0R3Z8 RCG28930, isoform CRA_b

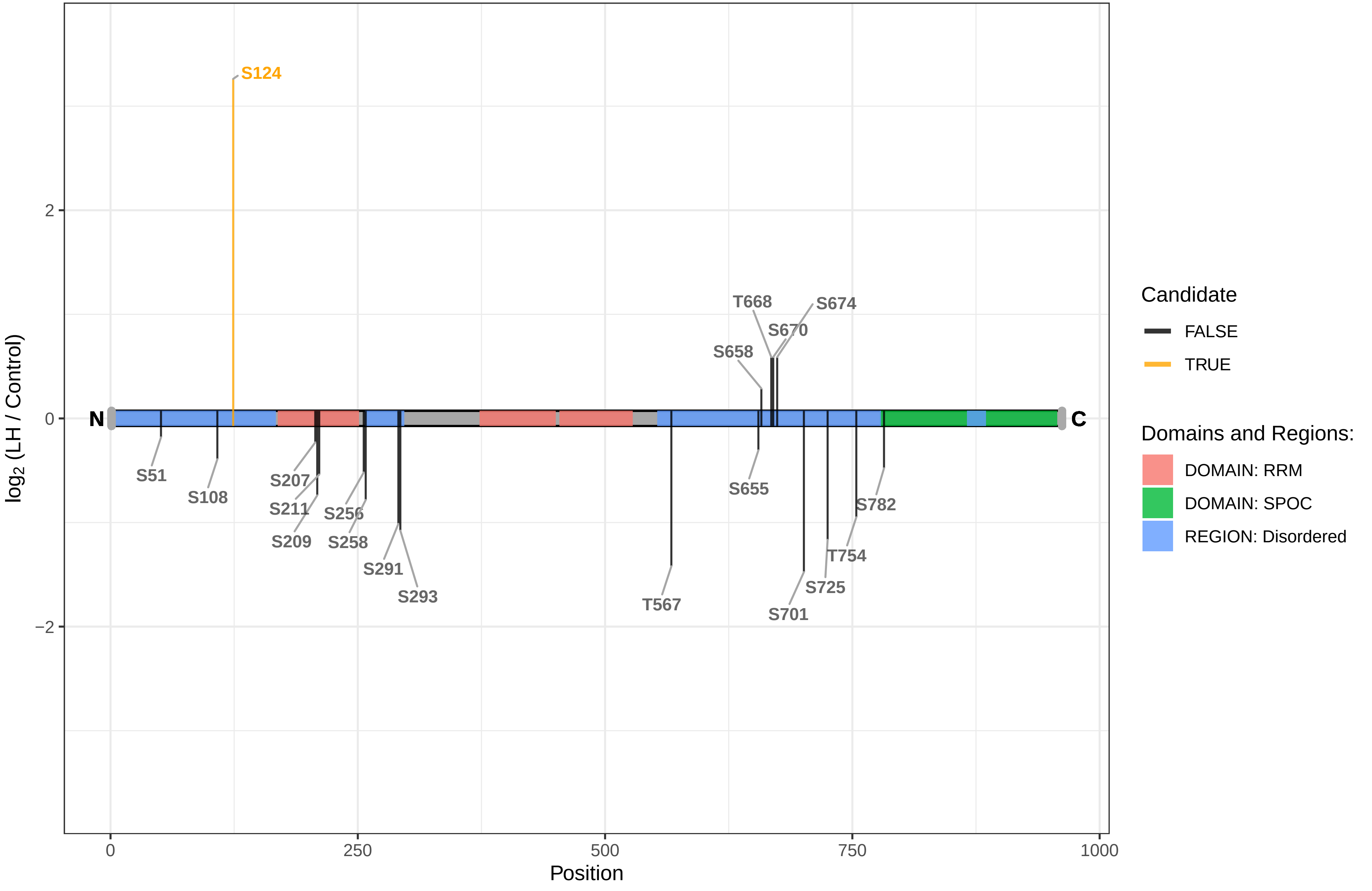

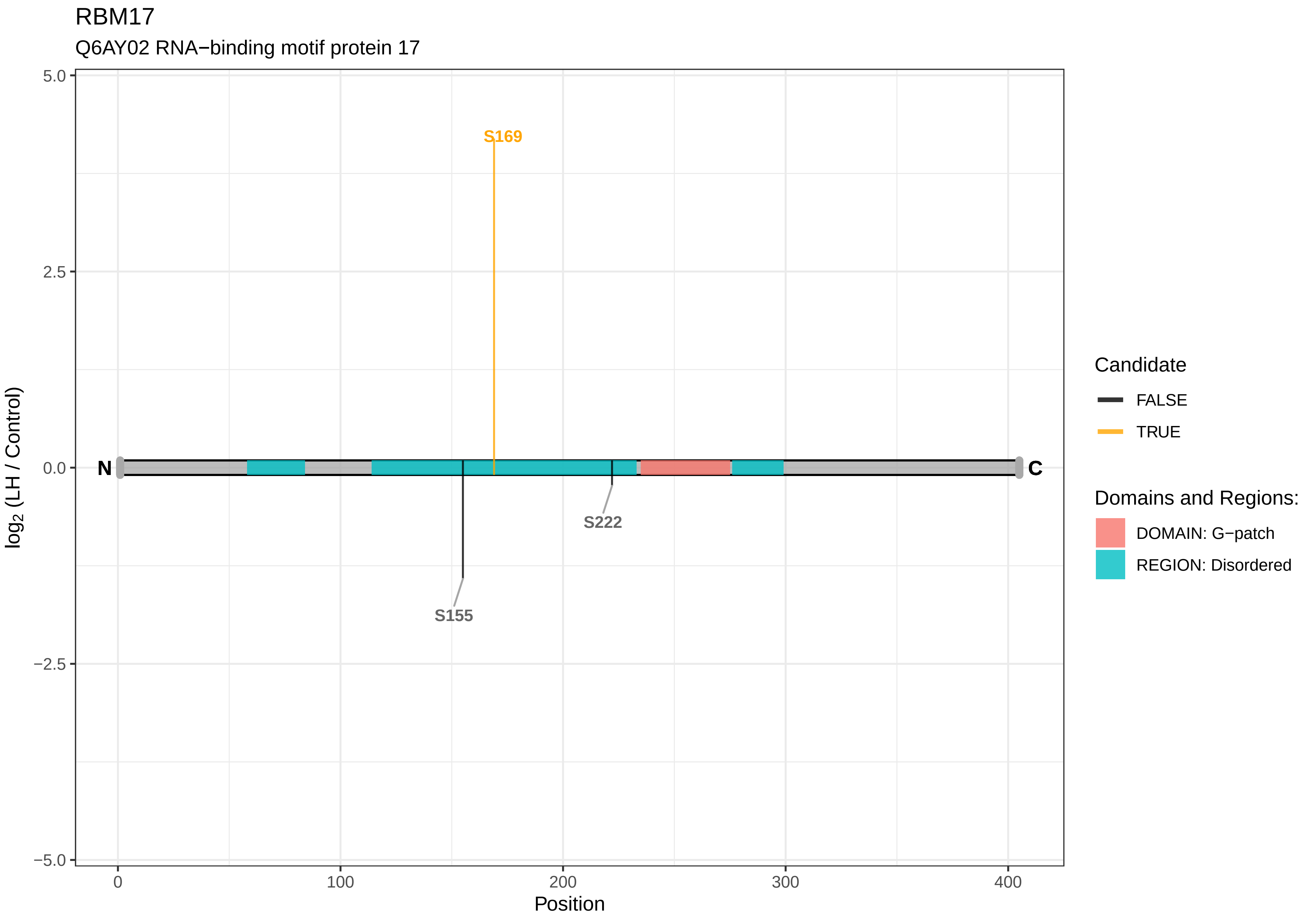

RBM17
Q6AY02 RNA−binding motif protein 17
5.0
2.5
0.0
−2.5
−5.0
log2 (LH / Control)
S169
S222
S155
N
C
0
100
200
300
400
Position
Candidate
FALSE
TRUE
Domains and Regions:
DOMAIN: G−patch
REGION: Disordered

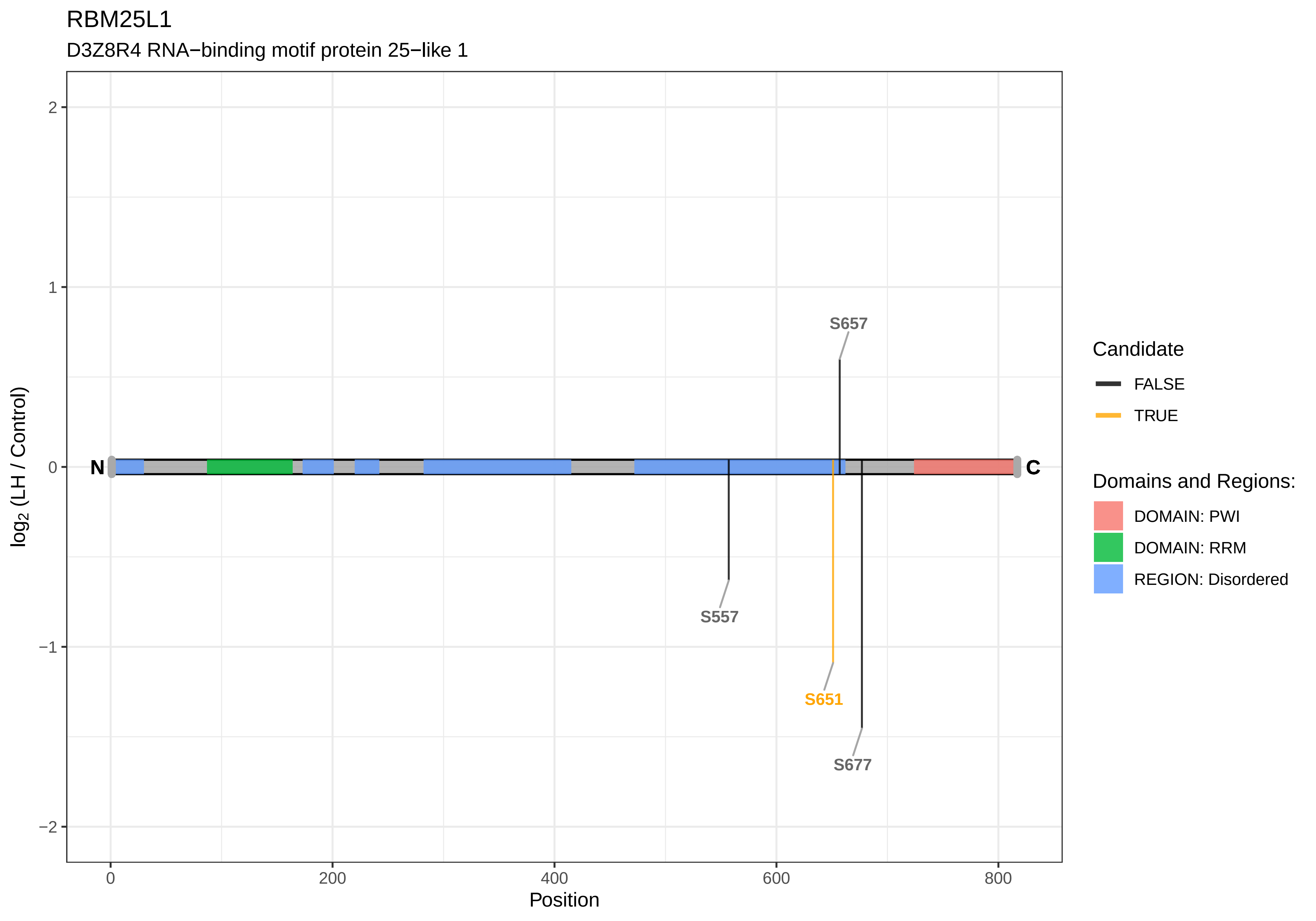
RBM25L1
D3Z8R4 RNA−binding motif protein 25−like 1
S657
S557
S651
S677
N
C
2
1
0
−1
−2
0
200
400
600
800
Position
log2 (LH / Control)
Candidate
FALSE
TRUE
Domains and Regions:
DOMAIN: PWI
DOMAIN: RRM
REGION: Disordered

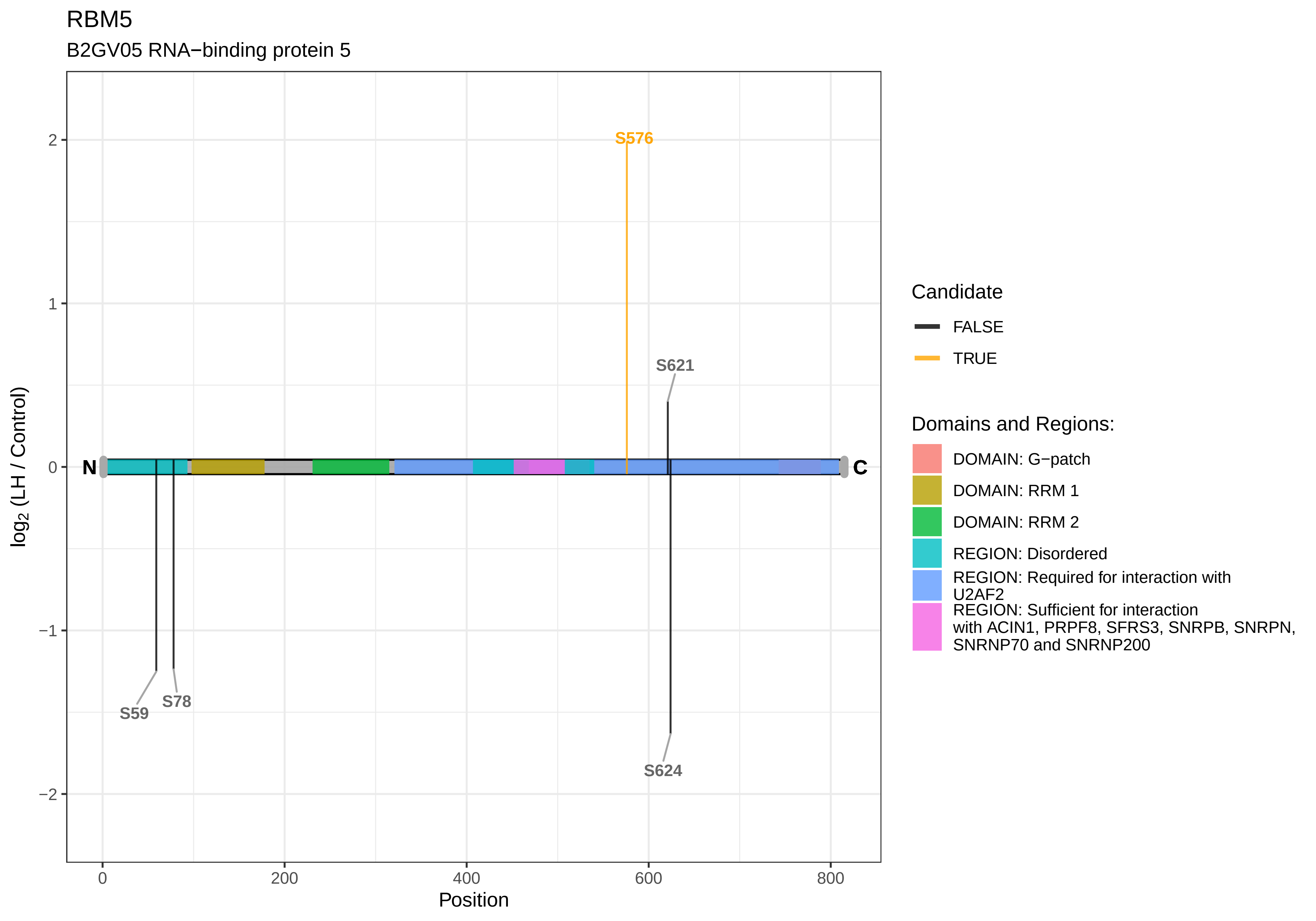
RBM5
B2GV05 RNA−binding protein 5
S576
S621
S59
S78
S624
N
C
2
1
0
−1
−2
0
200
400
600
800
Position
log2 (LH / Control)
Candidate
FALSE
TRUE
Domains and Regions:
DOMAIN: G−patch
DOMAIN: RRM 1
DOMAIN: RRM 2
REGION: Disordered
REGION: Required for interaction with U2AF2
REGION: Sufficient for interaction with ACIN1, PRPF8, SFRS3, SNRPB, SNRPN, SNRNP70 and SNRNP200

#### RBM7

F1M8F0 RNA−binding motif protein 7

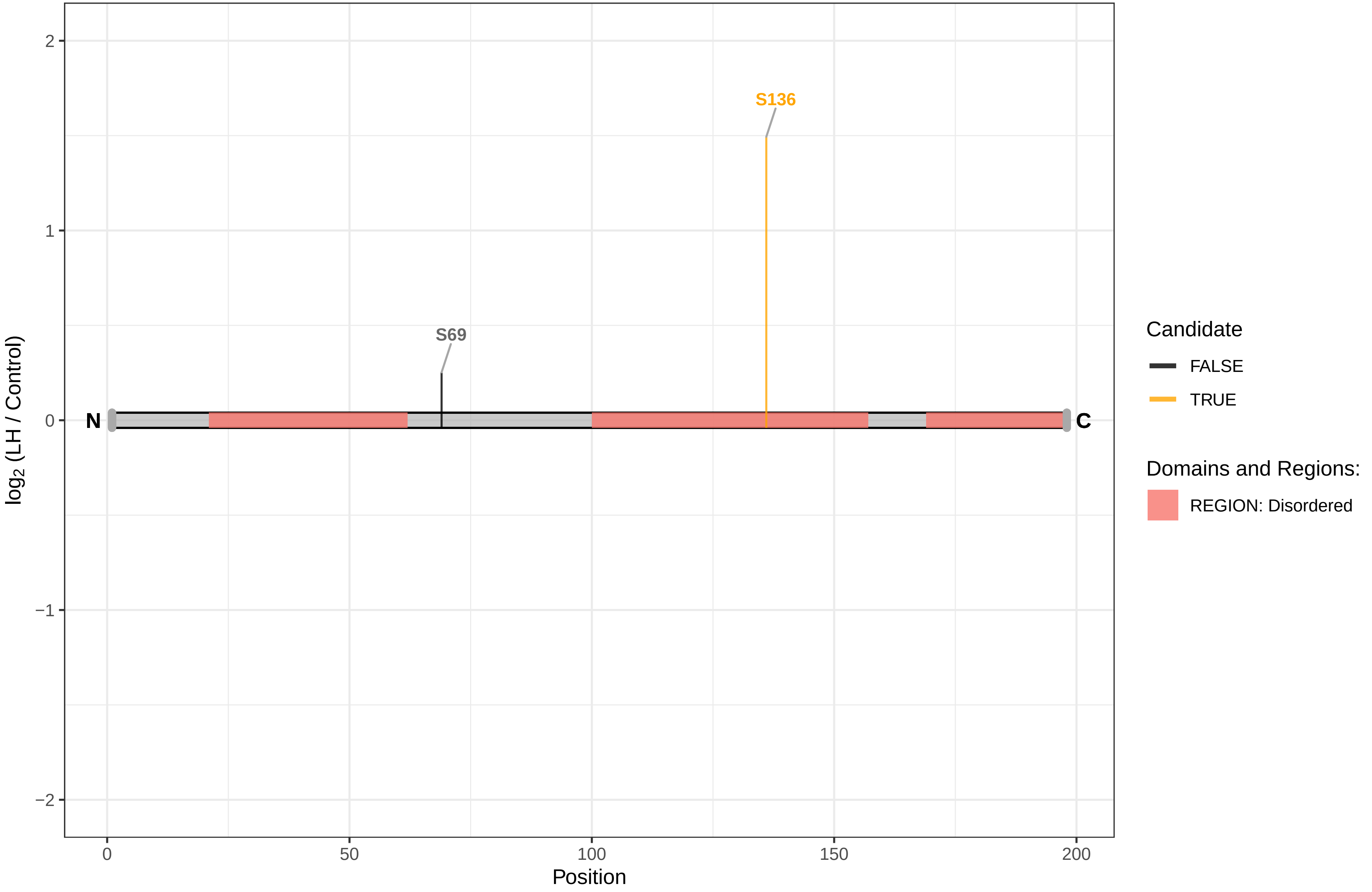

RC3H2

A0A0G2K9L3 Ring finger and CCCH−type domains 2

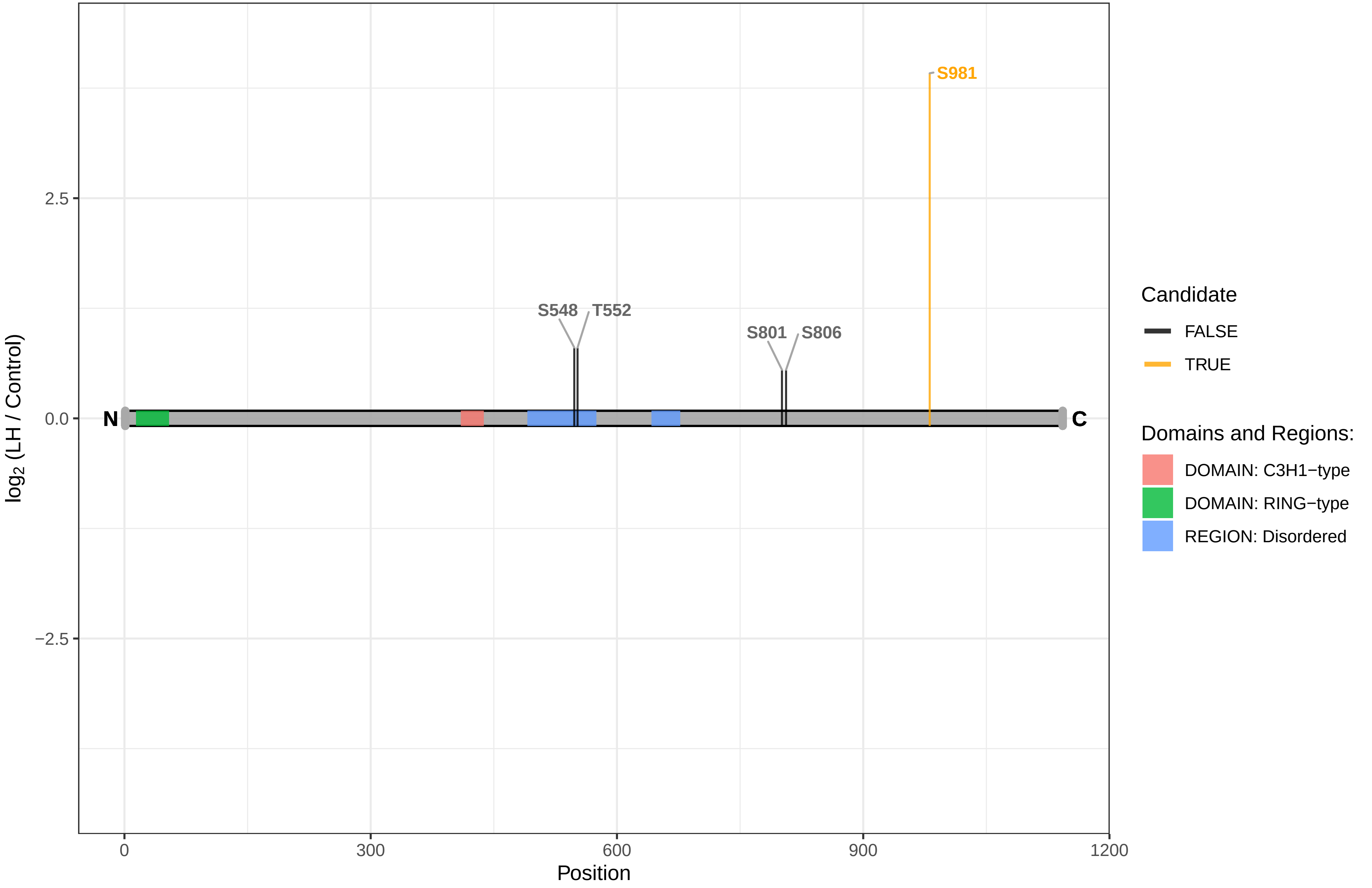

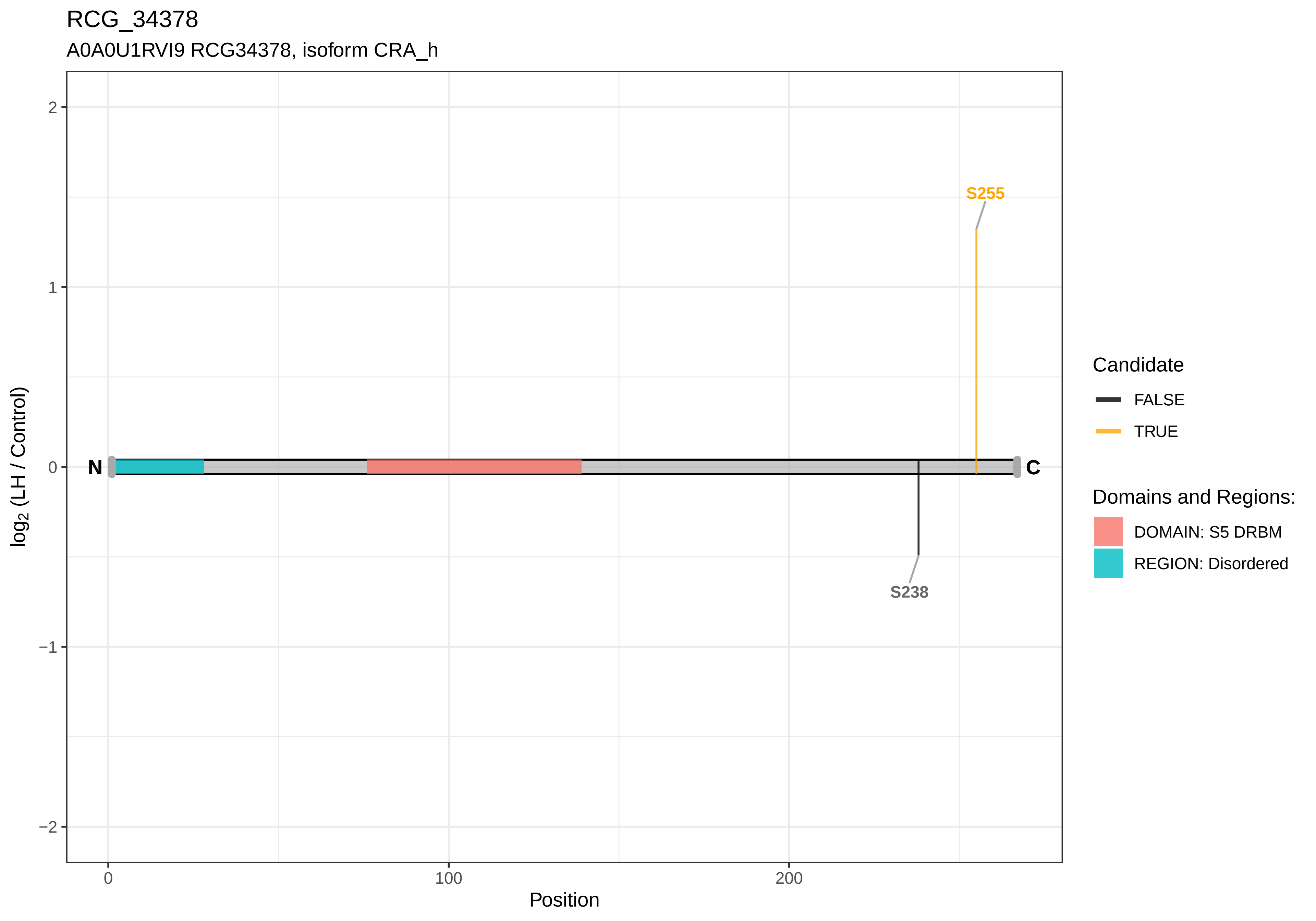

RCG_34378
A0A0U1RVI9 RCG34378, isoform CRA_h
S255
S238
N
C
2
1
0
−1
−2
0
100
200
Position
log2 (LH / Control)
Candidate
FALSE
TRUE
Domains and Regions:
DOMAIN: S5 DRBM
REGION: Disordered

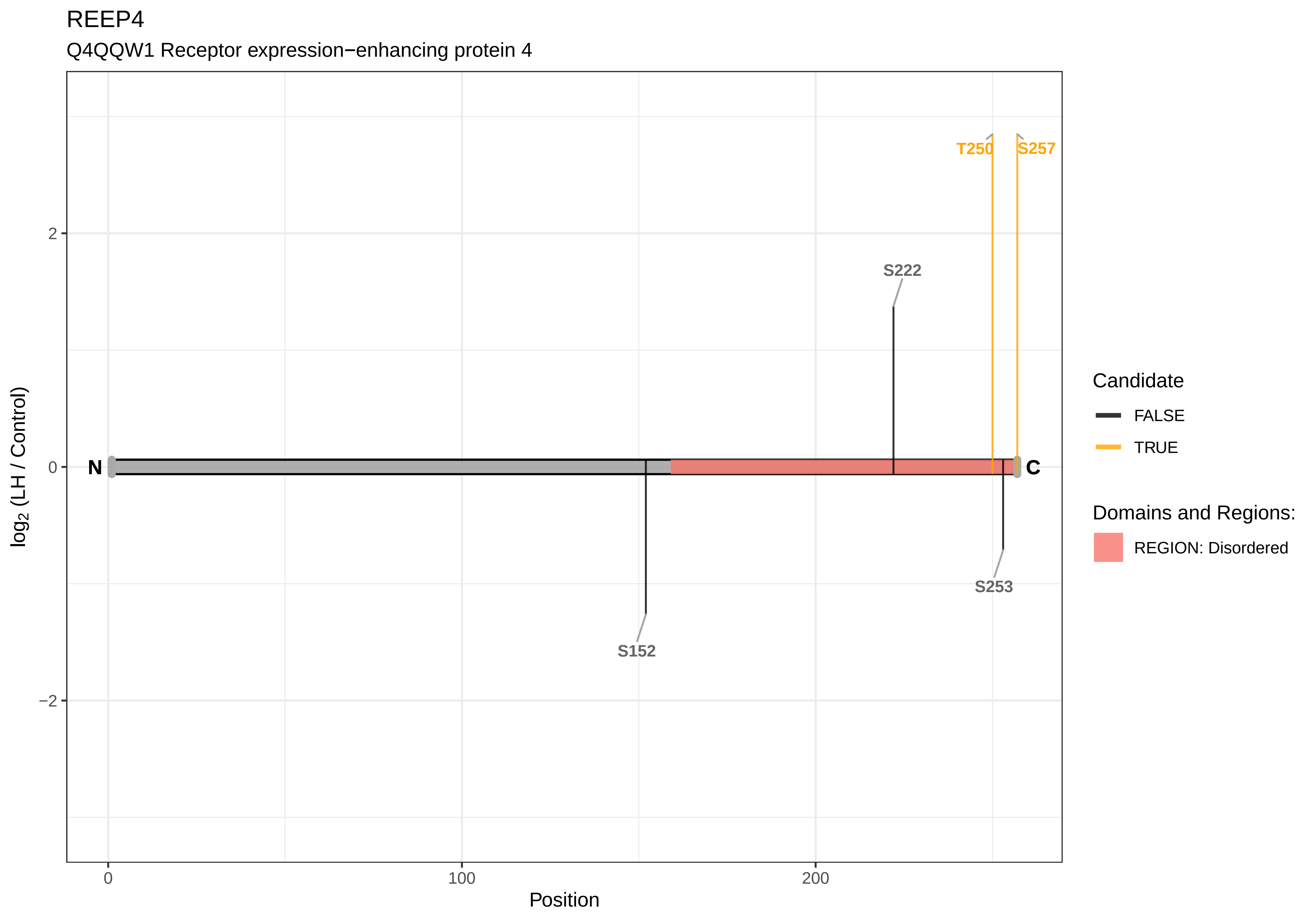
REEP4
Q4QQW1 Receptor expression−enhancing protein 4
T250
S257
S222
N
C
S253
S152
2
0
−2
log2 (LH / Control)
0
100
200
Position
Candidate
FALSE
TRUE
Domains and Regions:
REGION: Disordered

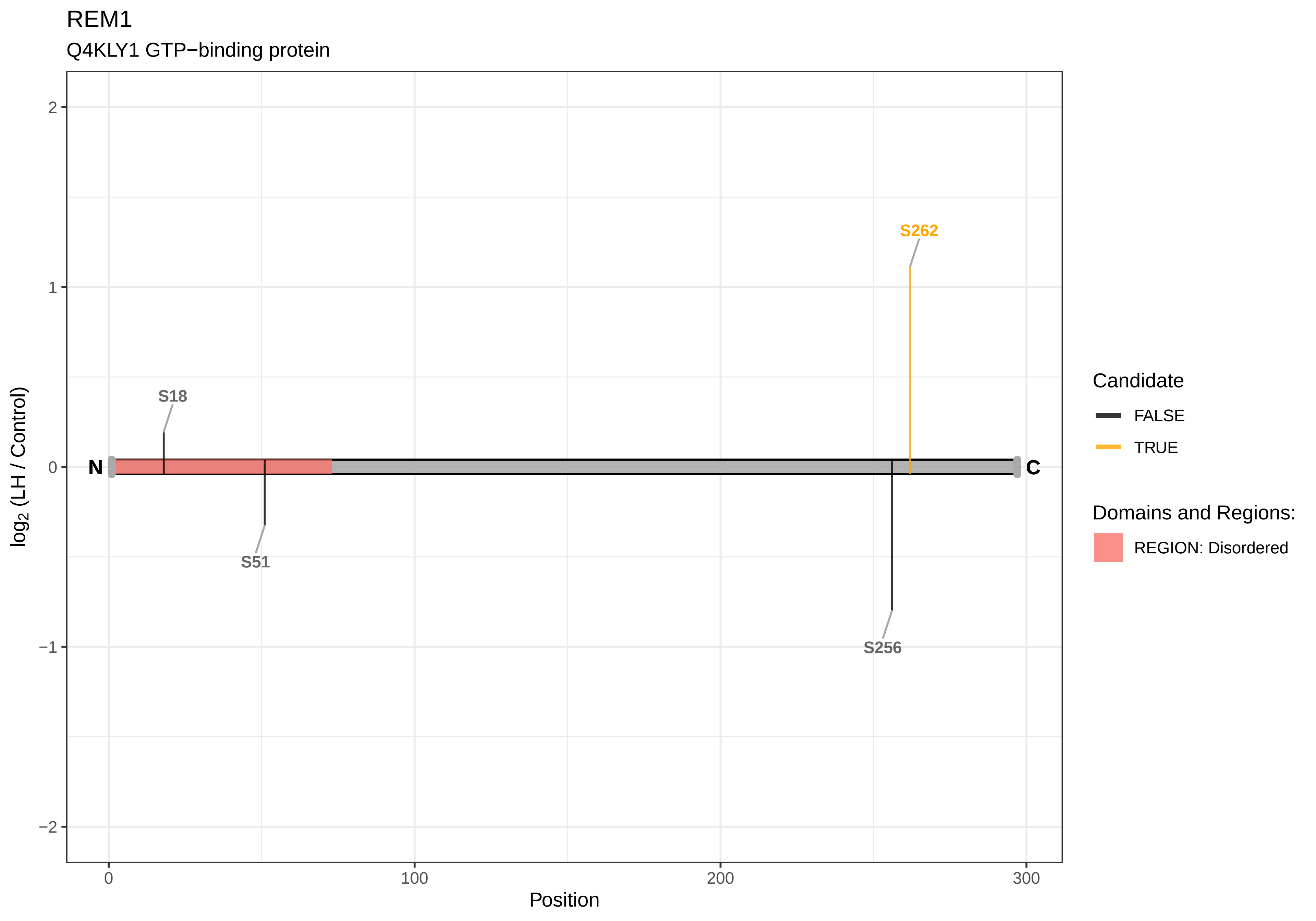

REM1
Q4KLY1 GTP−binding protein
log2 (LH / Control)
2
1
0
−1
−2
N
C
S262
S18
S51
S256
0
100
200
300
Position
Candidate
FALSE
TRUE
Domains and Regions:
REGION: Disordered

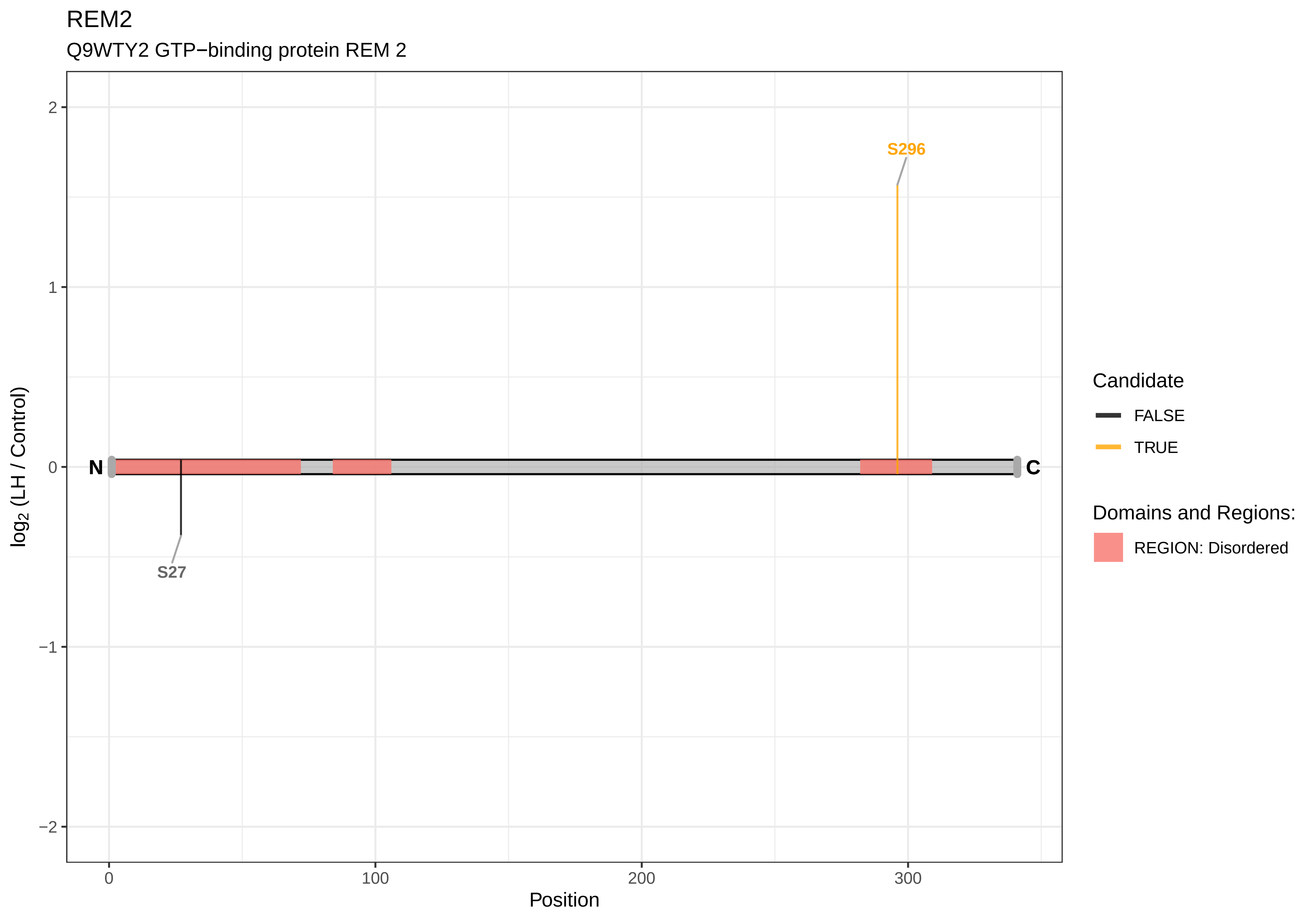
REM2
Q9WTY2 GTP−binding protein REM 2
S296
S27
N
C
log2 (LH / Control)
Position
2
1
0
−1
−2
0
100
200
300
Candidate
FALSE
TRUE
Domains and Regions:
REGION: Disordered

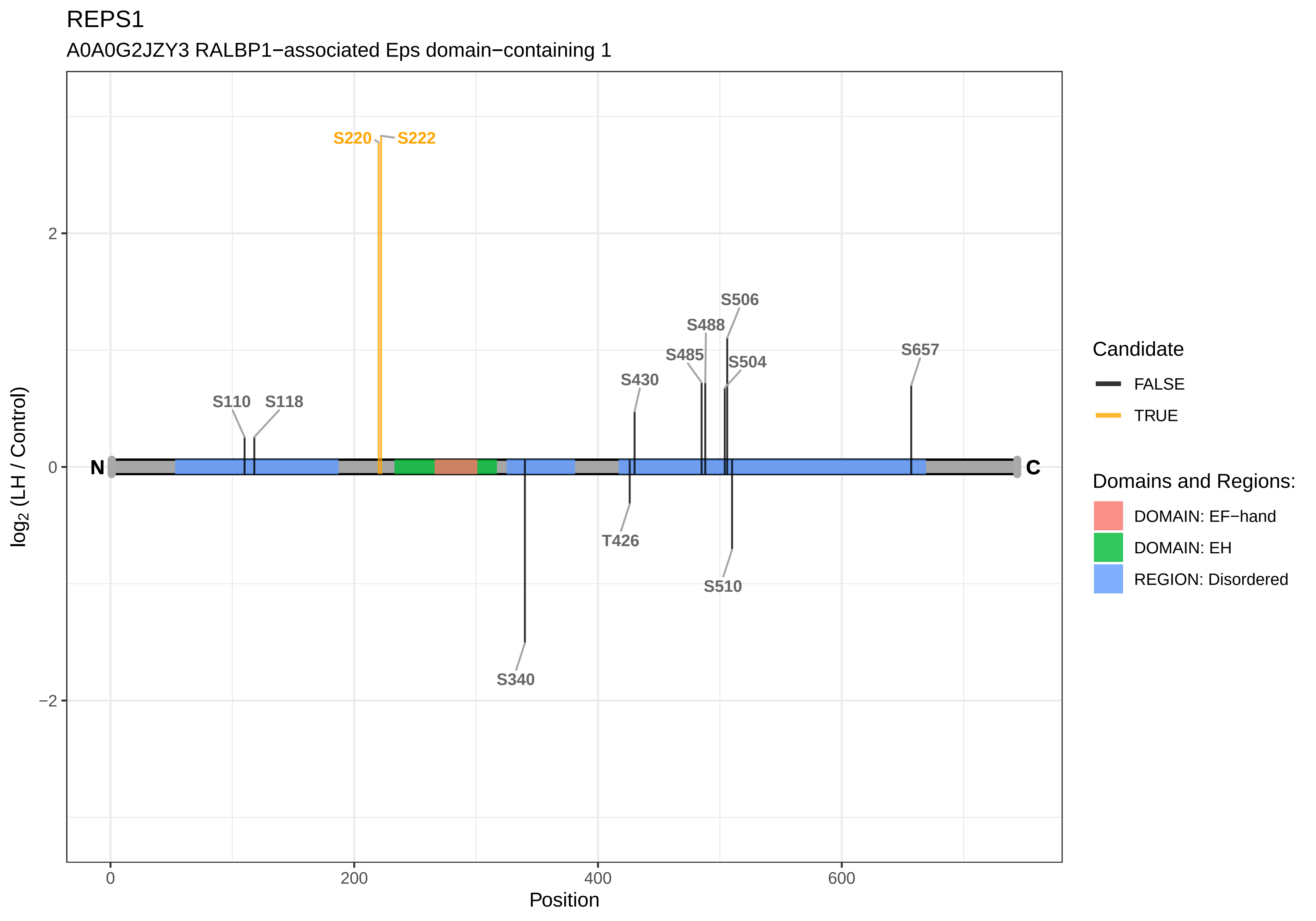

REPS1
A0A0G2JZY3 RALBP1−associated Eps domain−containing 1
S220
S222
S506
S488
S485
S504
S657
S430
S110
S118
N
C
T426
S510
S340
log2 (LH / Control)
2
0
−2
0
200
400
600
Position
Candidate
FALSE
TRUE
Domains and Regions:
DOMAIN: EF−hand
DOMAIN: EH
REGION: Disordered

REXO1

D3ZCX6 RNA exonuclease 1 homolog

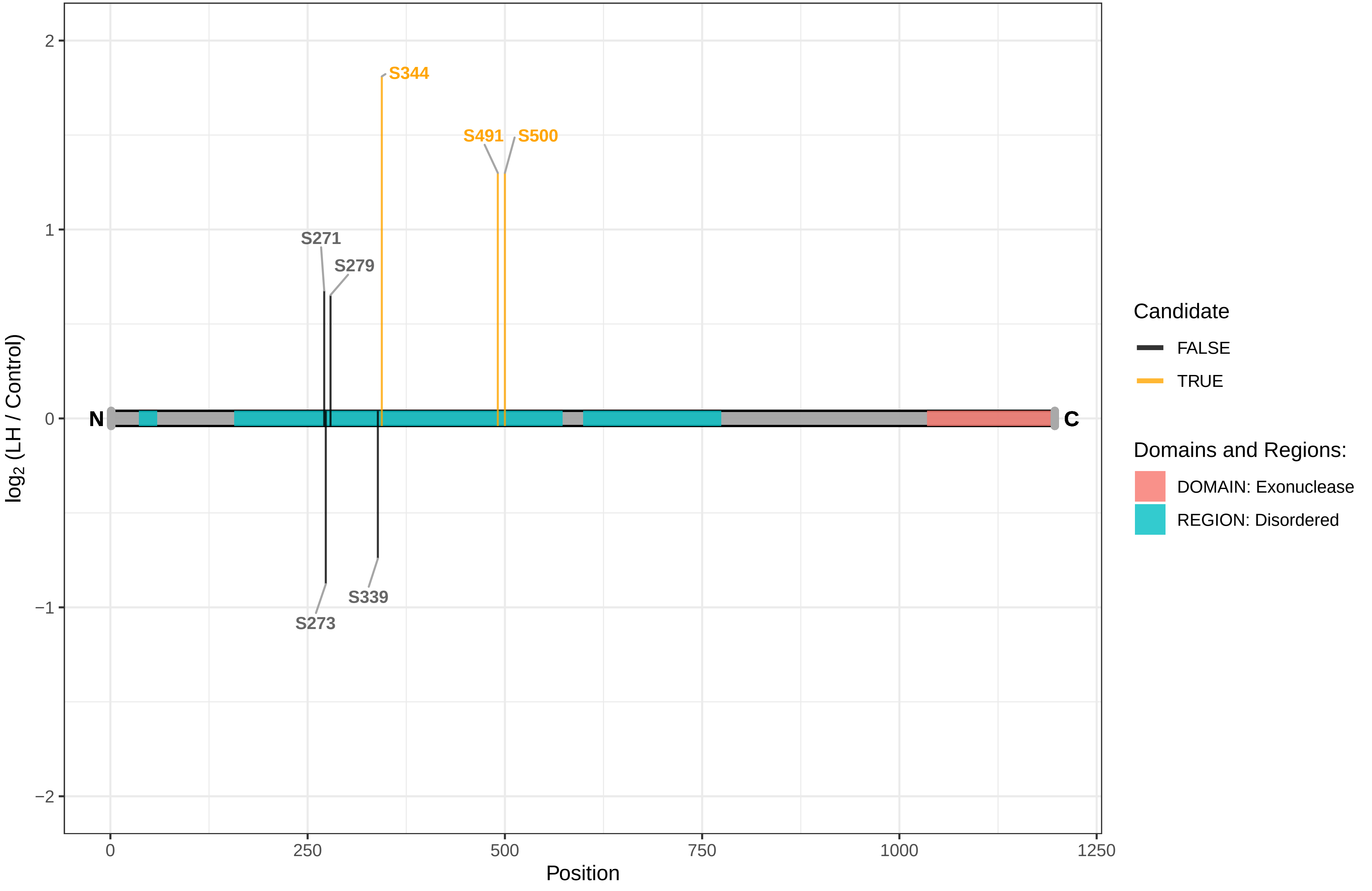

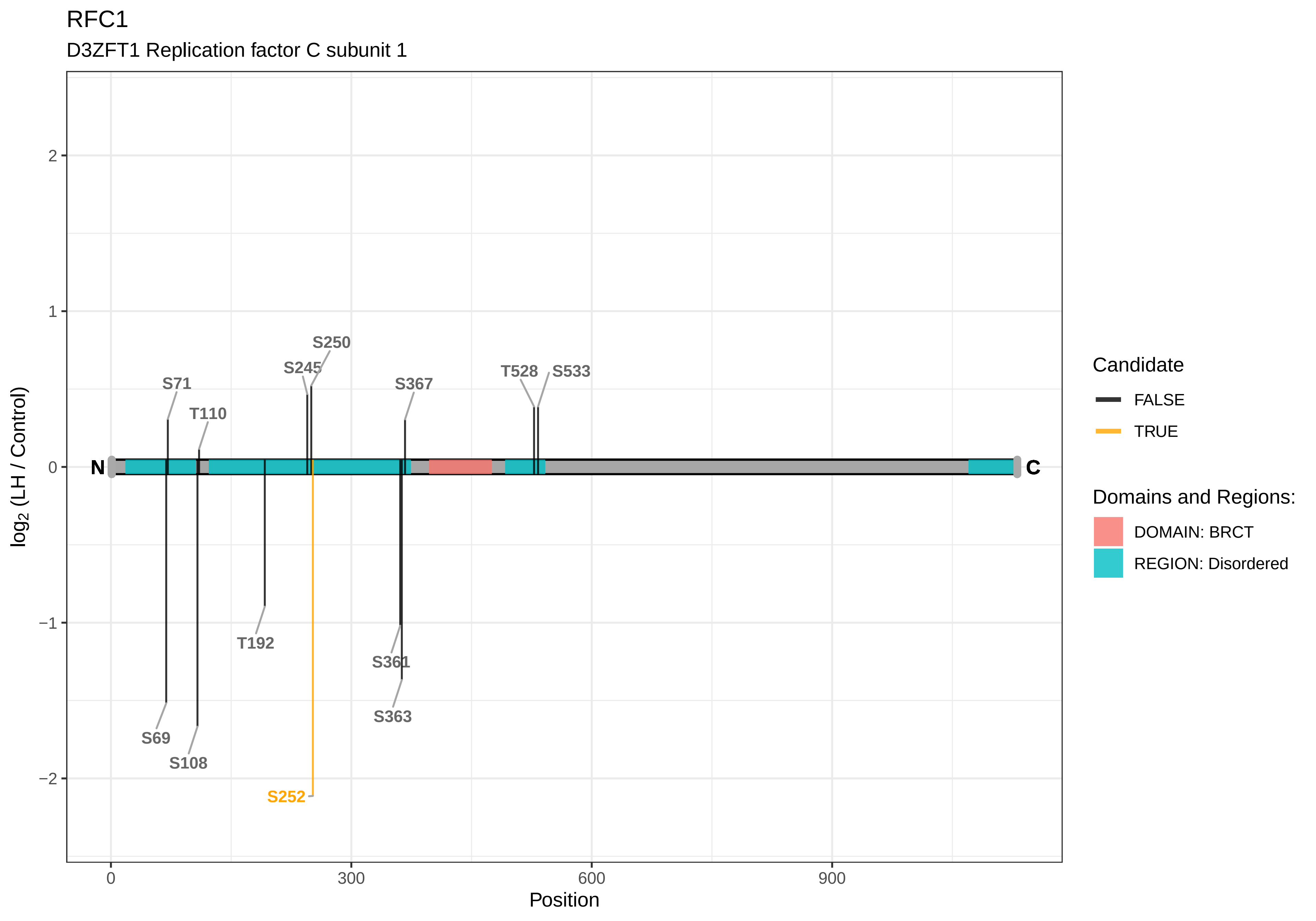

RFC1
D3ZFT1 Replication factor C subunit 1
S250
S245
S71
S367
T528
S533
T110
N
C
T192
S361
S363
S69
S108
S252
2
1
0
−1
−2
0
300
600
900
Position
log2 (LH / Control)
Candidate
FALSE
TRUE
Domains and Regions:
DOMAIN: BRCT
REGION: Disordered

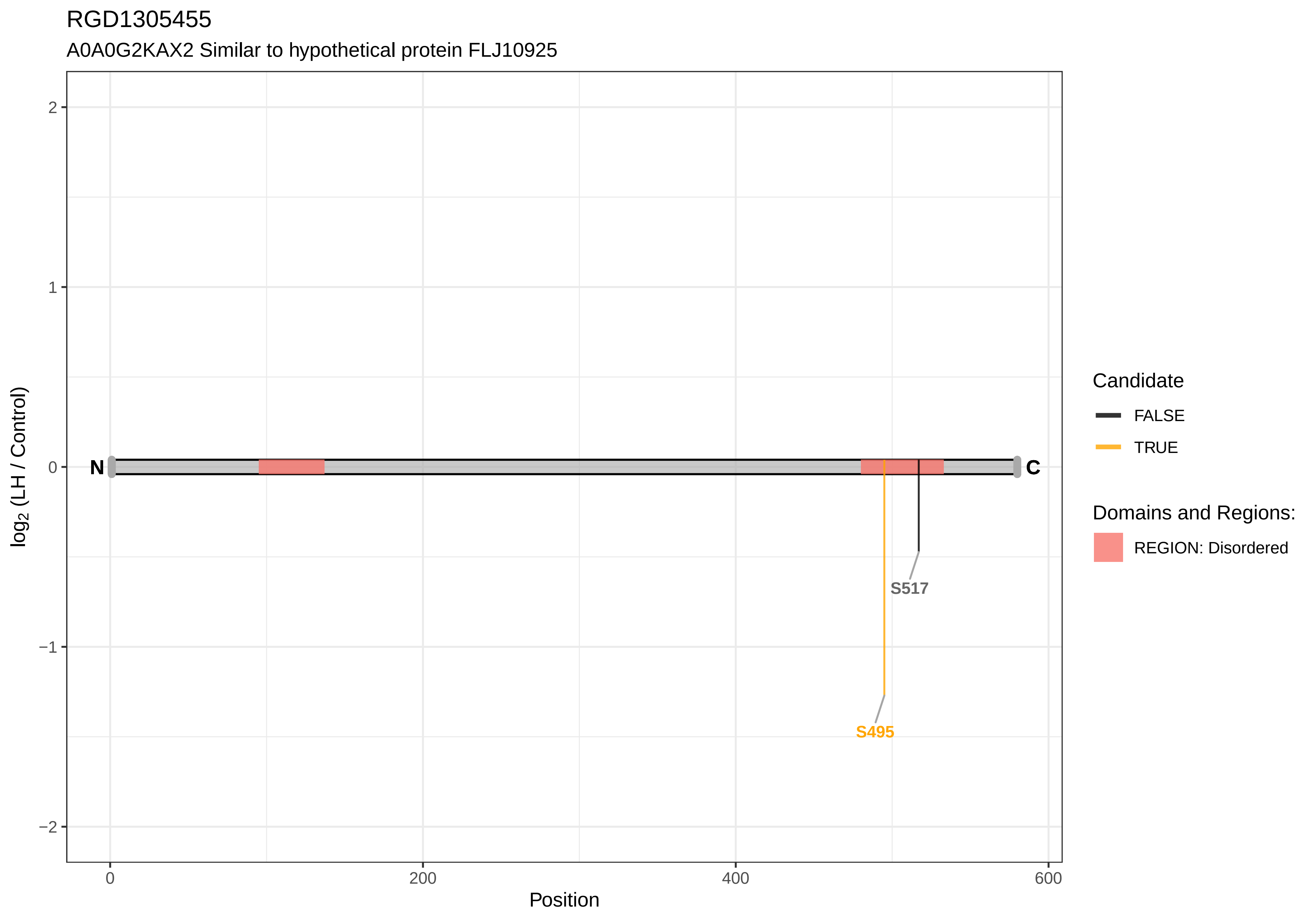
RGD1305455
A0A0G2KAX2 Similar to hypothetical protein FLJ10925
2
1
0
−1
−2
log2 (LH / Control)
N
C
S517
S495
0
200
400
600
Position
Candidate
FALSE
TRUE
Domains and Regions:
REGION: Disordered

RGD1307100

F1LVK0 Similar to RIKEN cDNA D630029K19

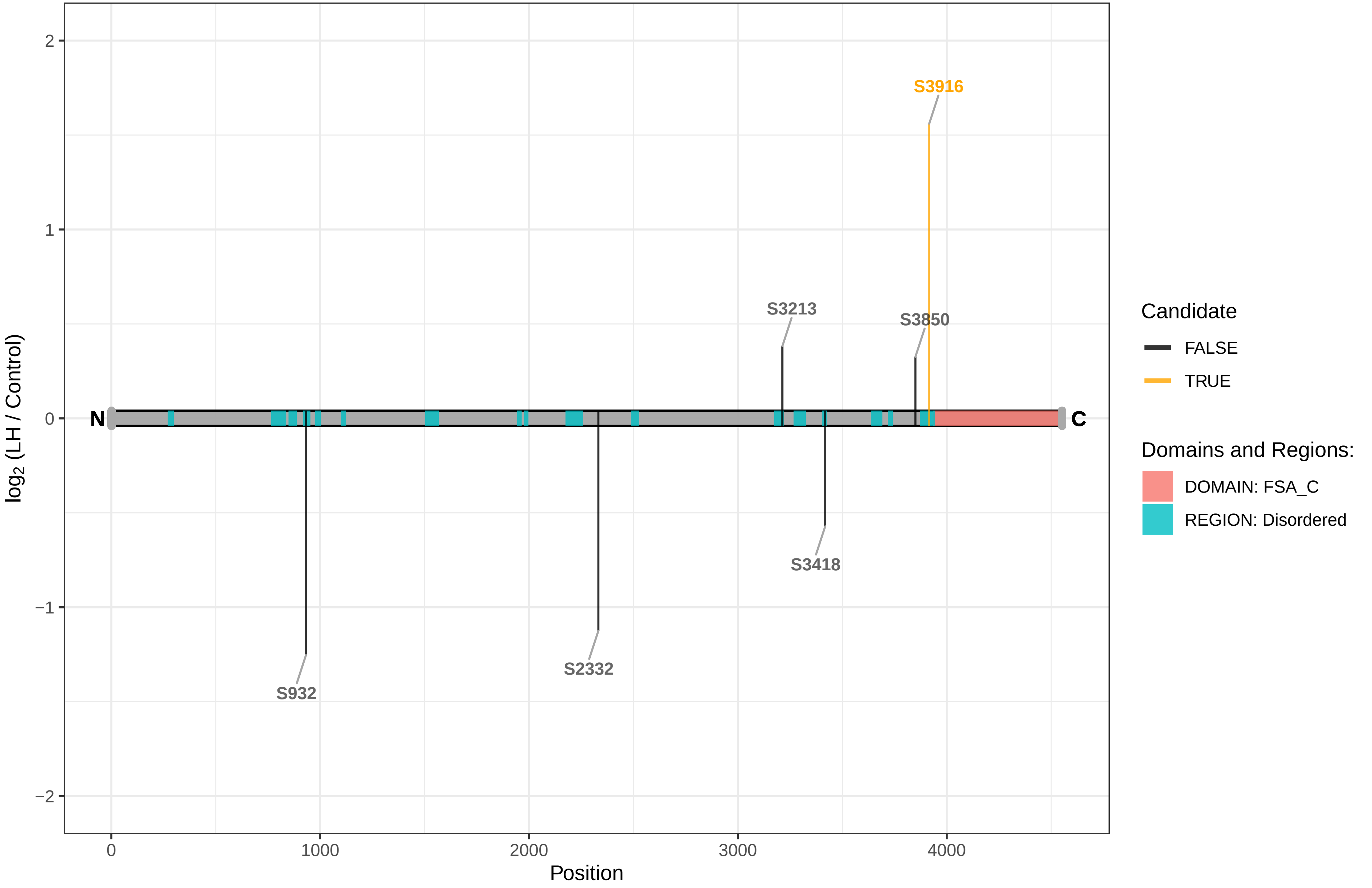

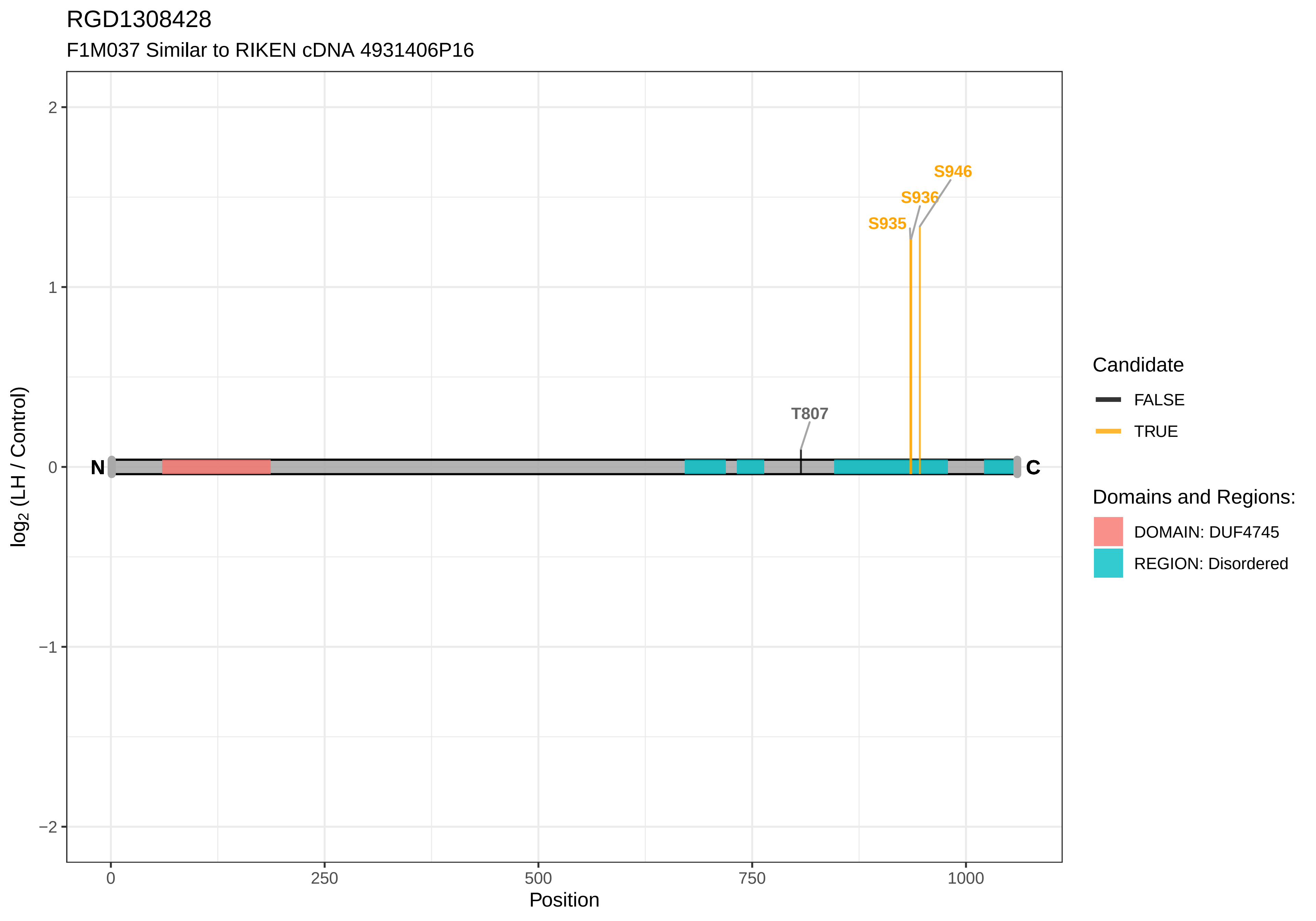

RGD1308428
F1M037 Similar to RIKEN cDNA 4931406P16
S946
S936
S935
T807
N
C
2
1
0
−1
−2
log2 (LH / Control)
0
250
500
750
1000
Position
Candidate
FALSE
TRUE
Domains and Regions:
DOMAIN: DUF4745
REGION: Disordered

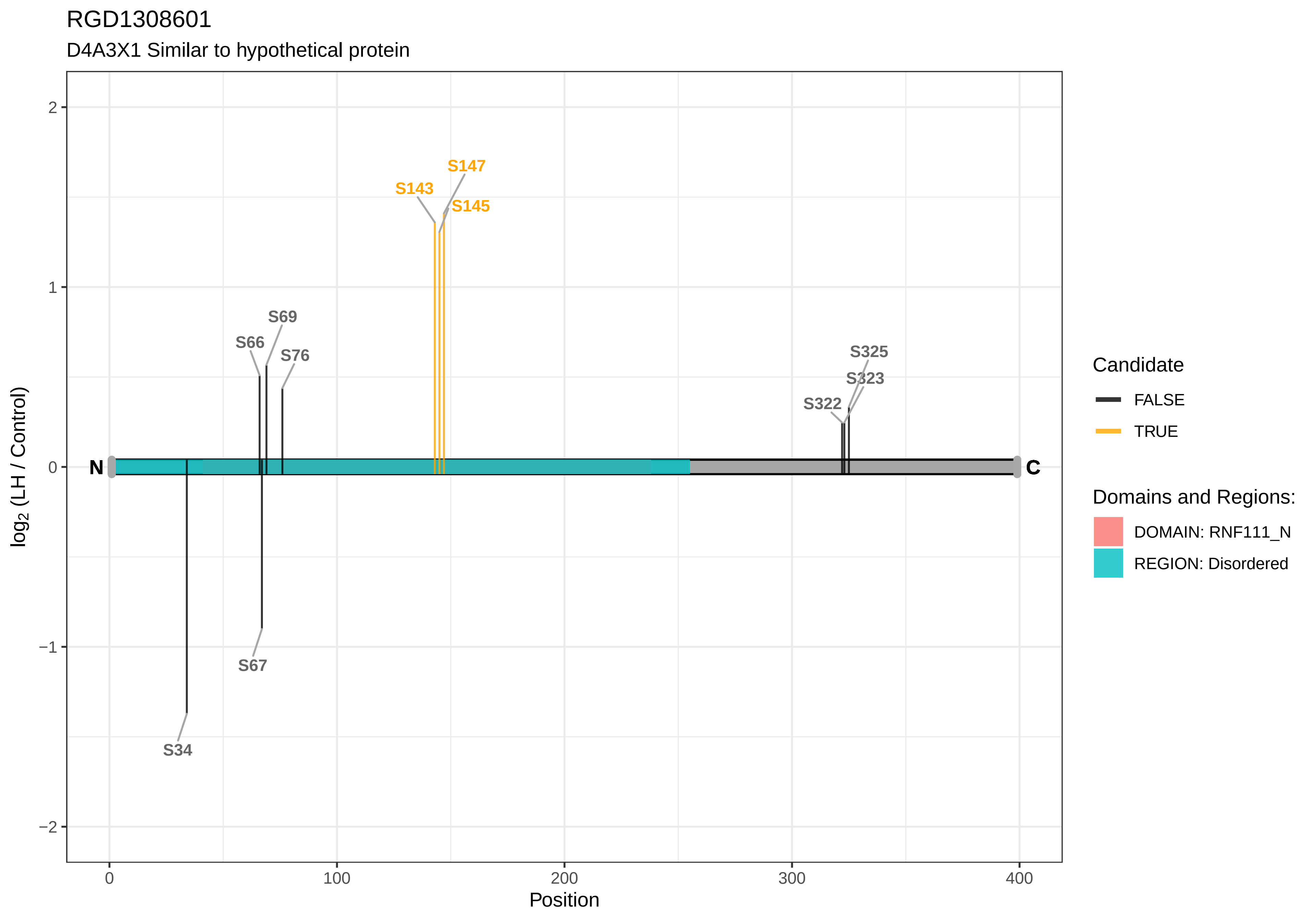
RGD1308601
D4A3X1 Similar to hypothetical protein
S147
S143
S145
S69
S66
S76
S325
S323
S322
S34
S67
N
C
2
1
0
−1
−2
0
100
200
300
400
Position
log2 (LH / Control)
Candidate
FALSE
TRUE
Domains and Regions:
DOMAIN: RNF111_N
REGION: Disordered

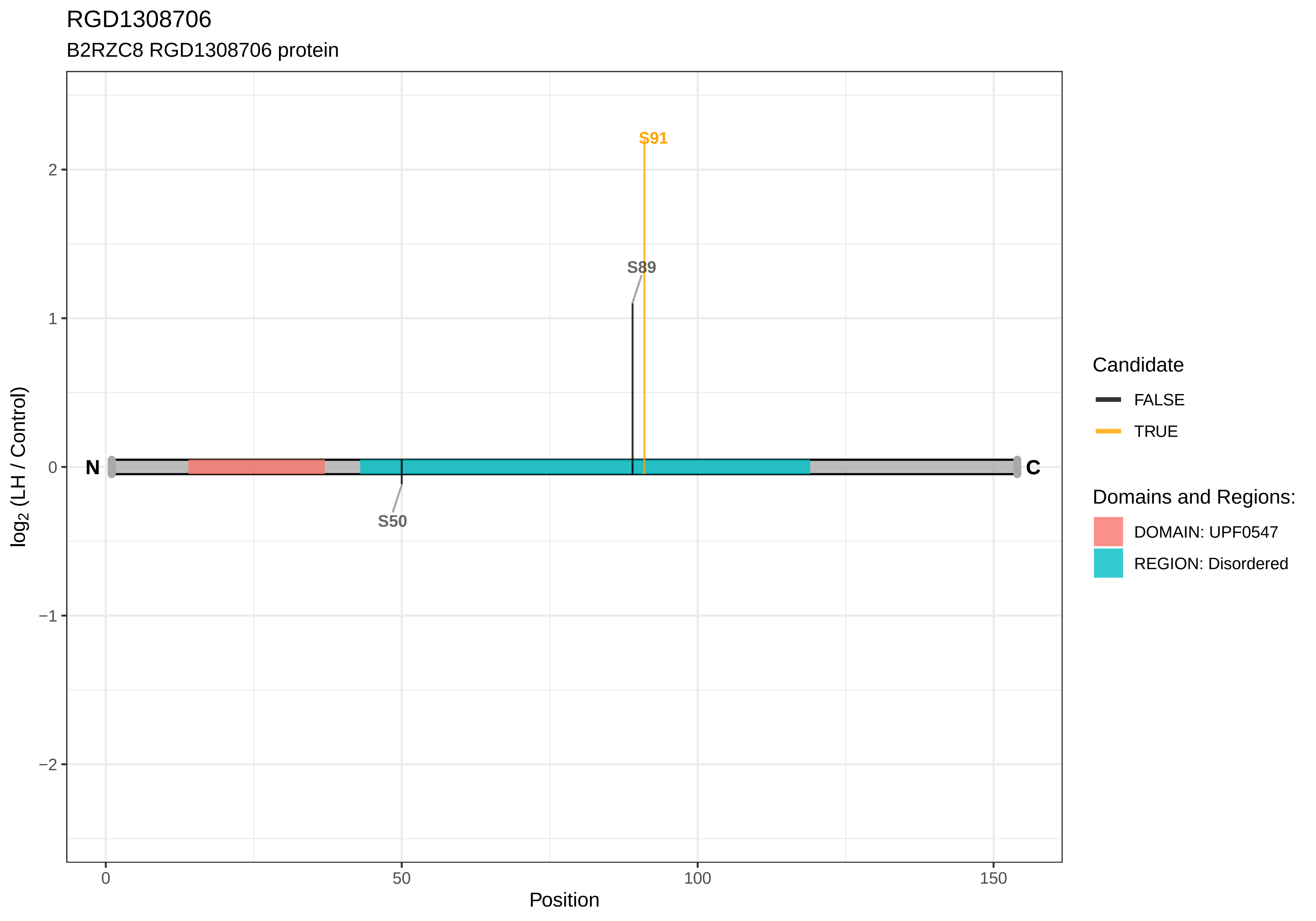

RGD1308706
B2RZC8 RGD1308706 protein
S91
S89
S50
N
C
log2 (LH / Control)
Position
Candidate
FALSE
TRUE
Domains and Regions:
DOMAIN: UPF0547
REGION: Disordered

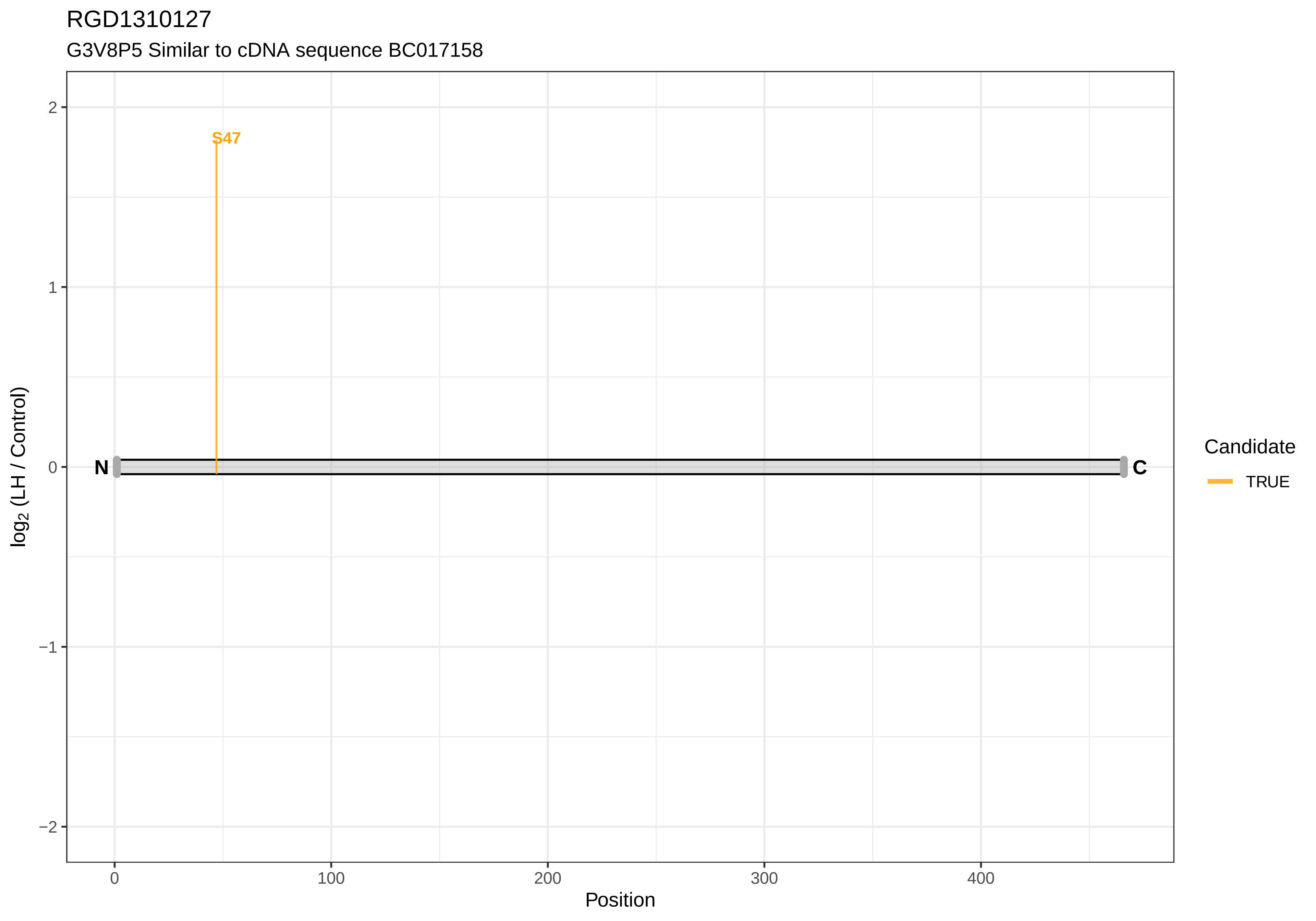
RGD1310127
G3V8P5 Similar to cDNA sequence BC017158
S47
N
C
2
1
0
−1
−2
log2 (LH / Control)
0
100
200
300
400
Position
Candidate
TRUE

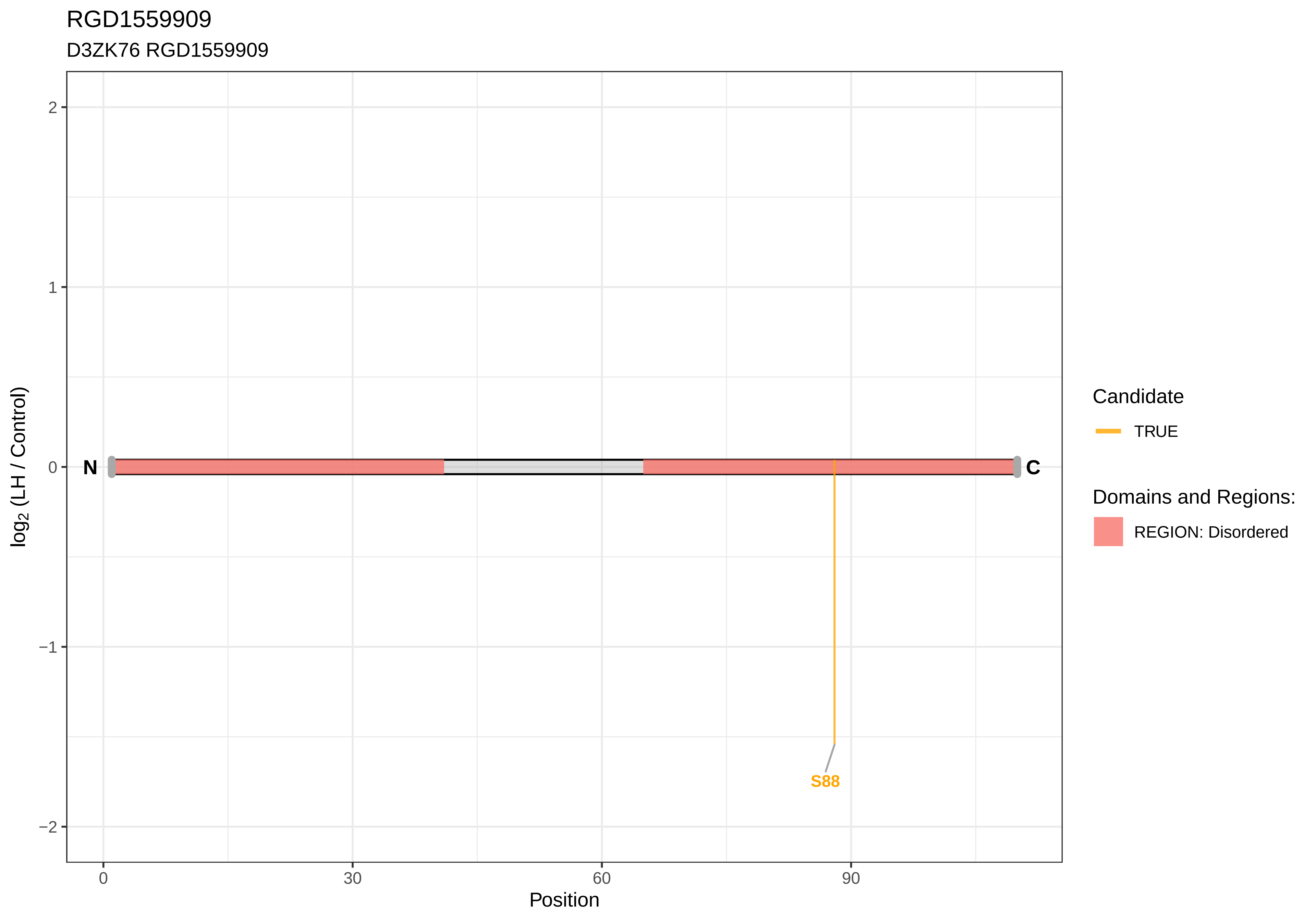
RGD1559909
D3ZK76 RGD1559909
log2 (LH / Control)
2
1
0
−1
−2
N
C
S88
0
30
60
90
Position
Candidate
TRUE
Domains and Regions:
REGION: Disordered

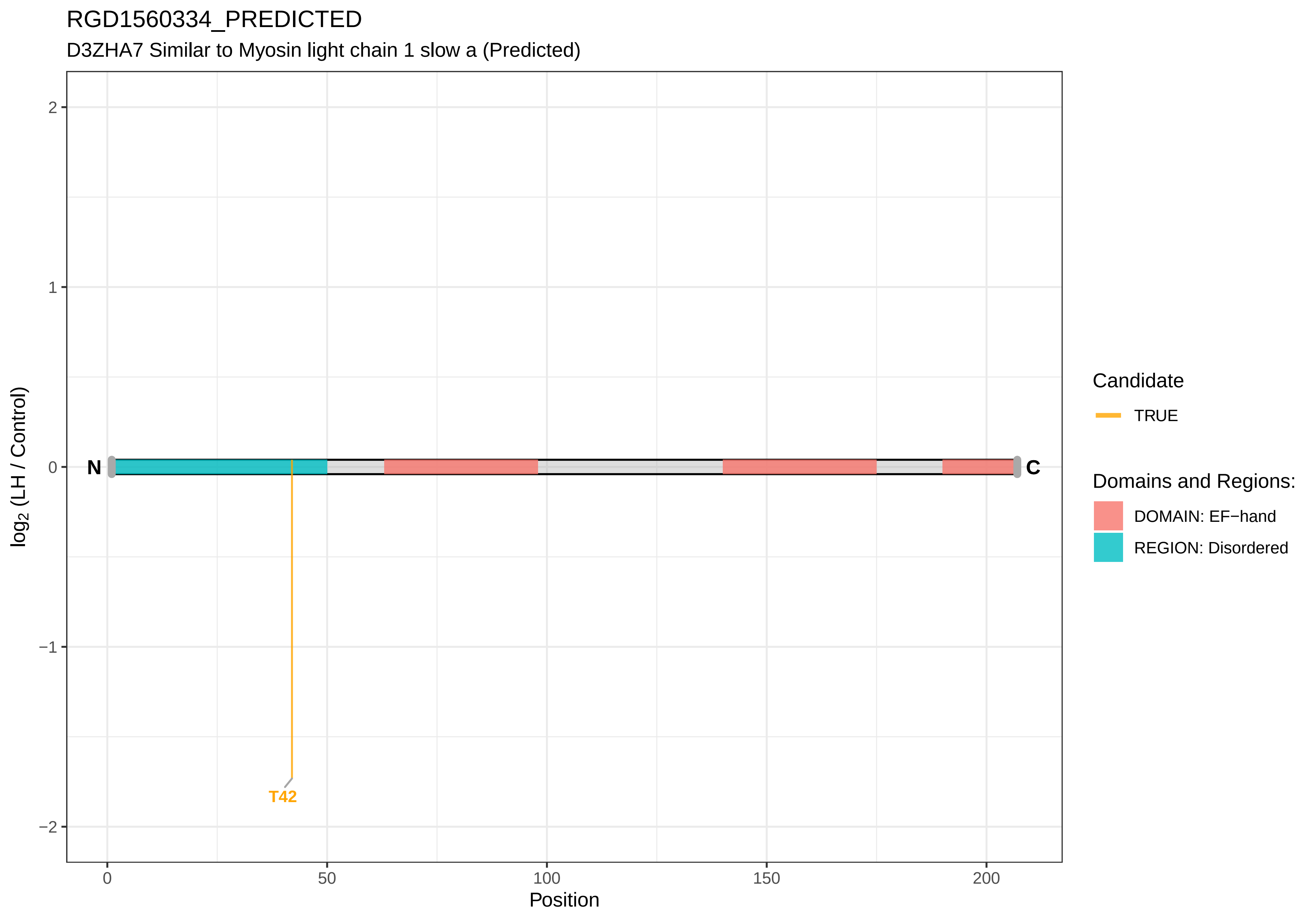

RGD1560334_PREDICTED
D3ZHA7 Similar to Myosin light chain 1 slow a (Predicted)
log2 (LH / Control)
2
1
0
−1
−2
N
C
T42
0
50
100
150
200
Position
Candidate
TRUE
Domains and Regions:
DOMAIN: EF−hand
REGION: Disordered

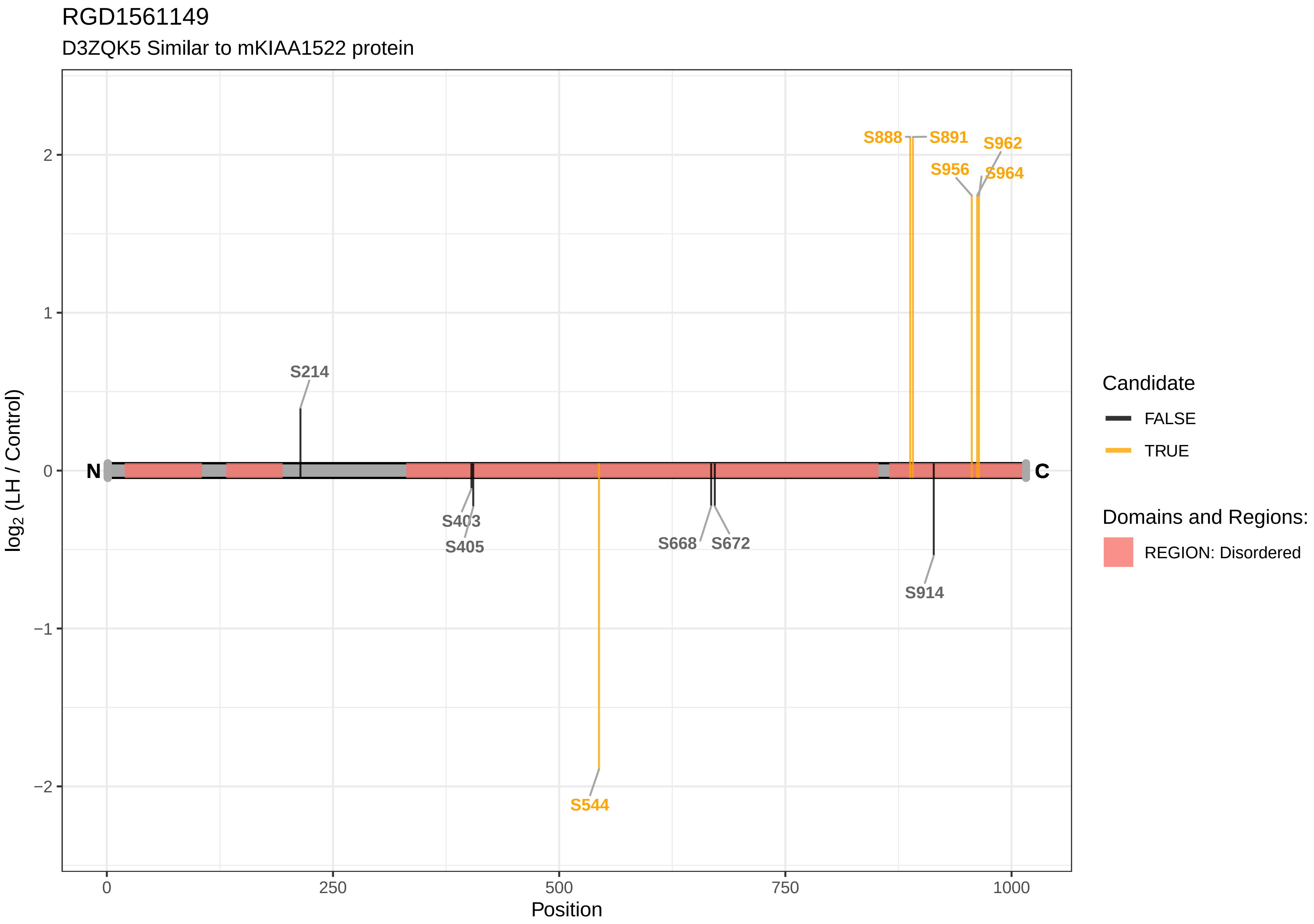

RGD1561149
D3ZQK5 Similar to mKIAA1522 protein
S888
S891
S962
S956
S964
S214
S403
S405
S668
S672
S914
S544
N
C
2
1
0
−1
−2
log2 (LH / Control)
0
250
500
750
1000
Position
Candidate
FALSE
TRUE
Domains and Regions:
REGION: Disordered

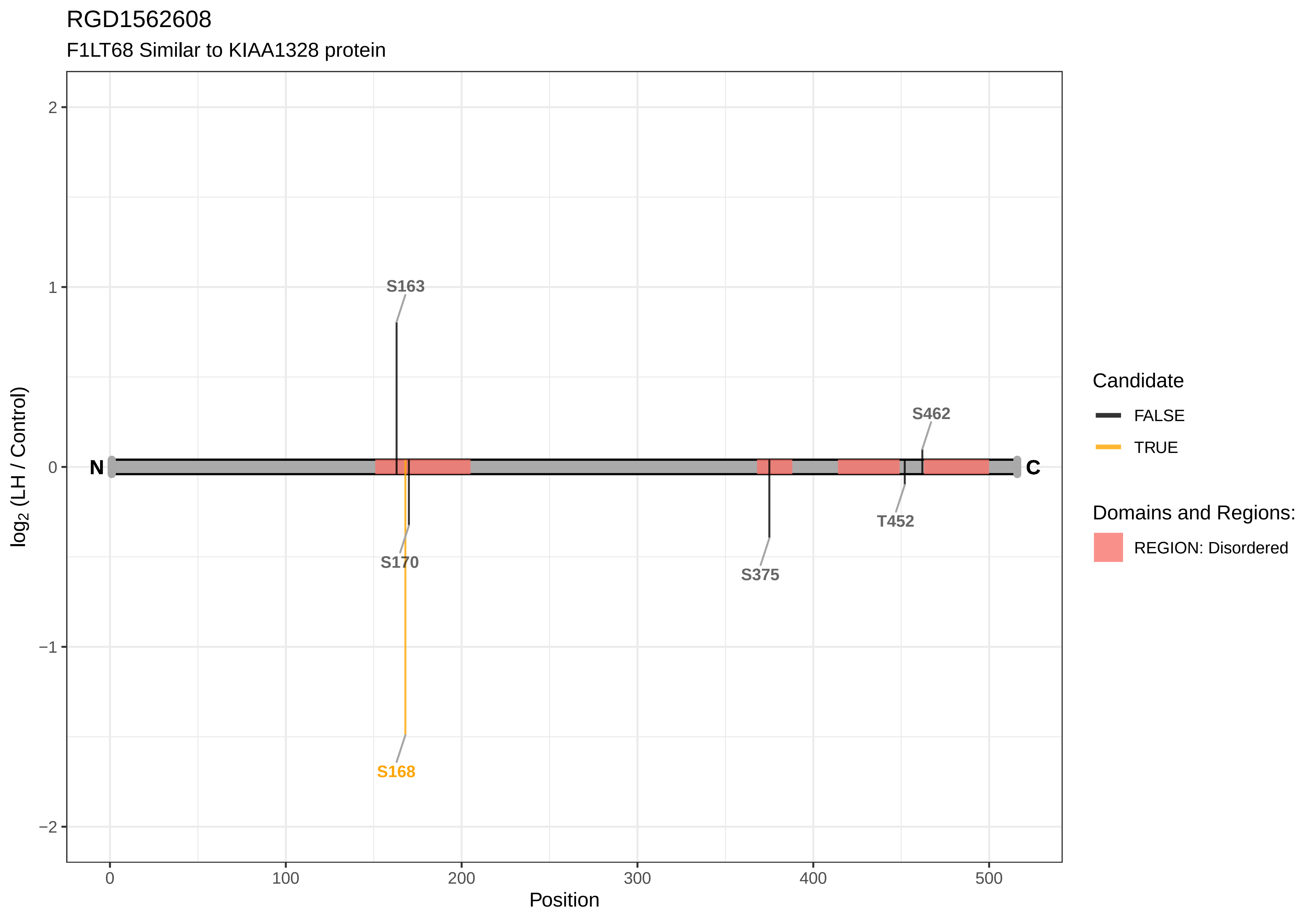

RGD1562608
F1LT68 Similar to KIAA1328 protein
S163
S462
S170
S375
T452
S168
N
C
log2 (LH / Control)
Position
Candidate
FALSE
TRUE
Domains and Regions:
REGION: Disordered

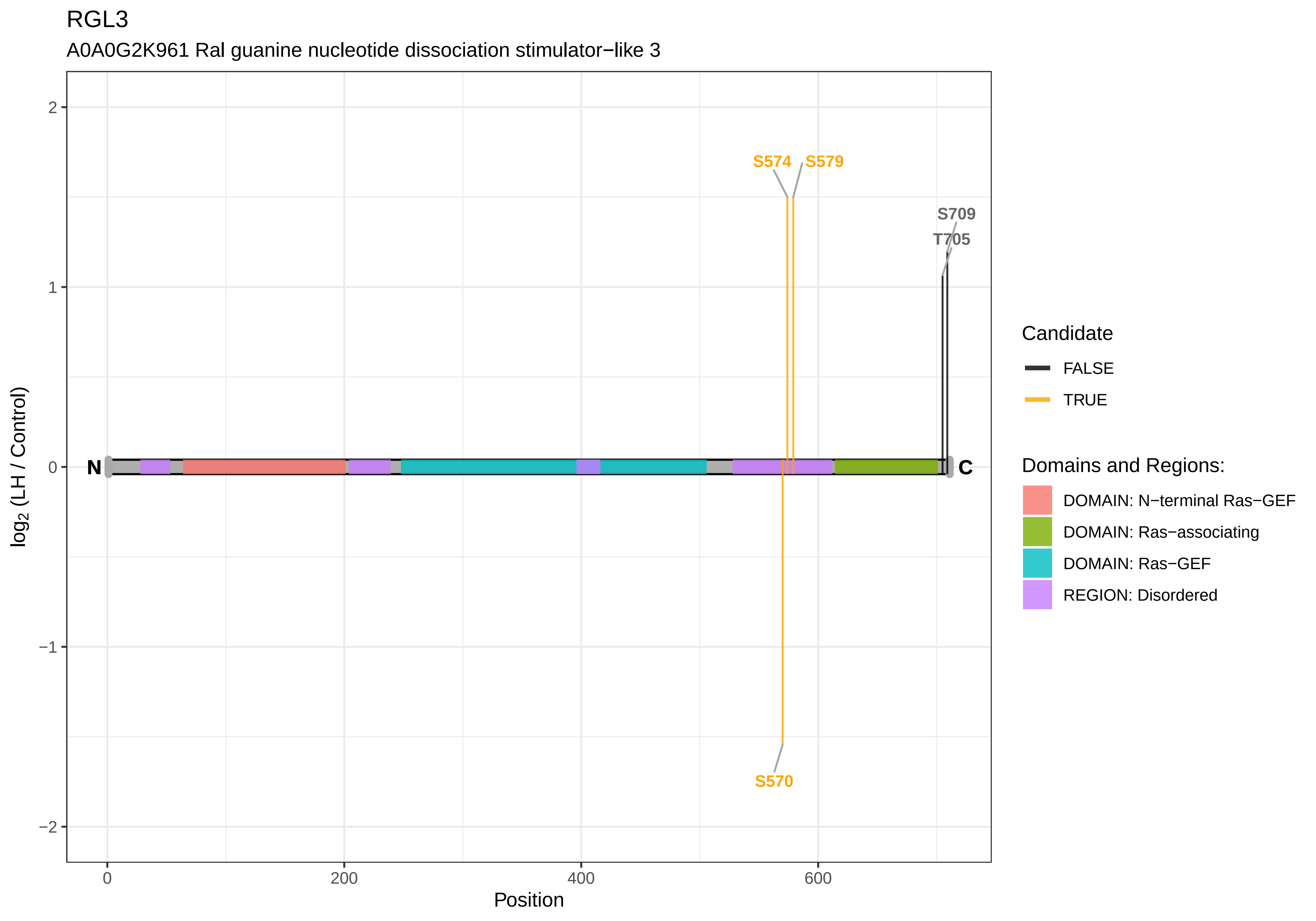

RGL3
A0A0G2K961 Ral guanine nucleotide dissociation stimulator−like 3
S574
S579
S709
T705
S570
N
C
2
1
0
−1
−2
0
200
400
600
Position
log2 (LH / Control)
Candidate
FALSE
TRUE
Domains and Regions:
DOMAIN: N−terminal Ras−GEF
DOMAIN: Ras−associating
DOMAIN: Ras−GEF
REGION: Disordered

RGS12

G3V9H1 Regulator of G−protein signaling 12, isoform CRA_c

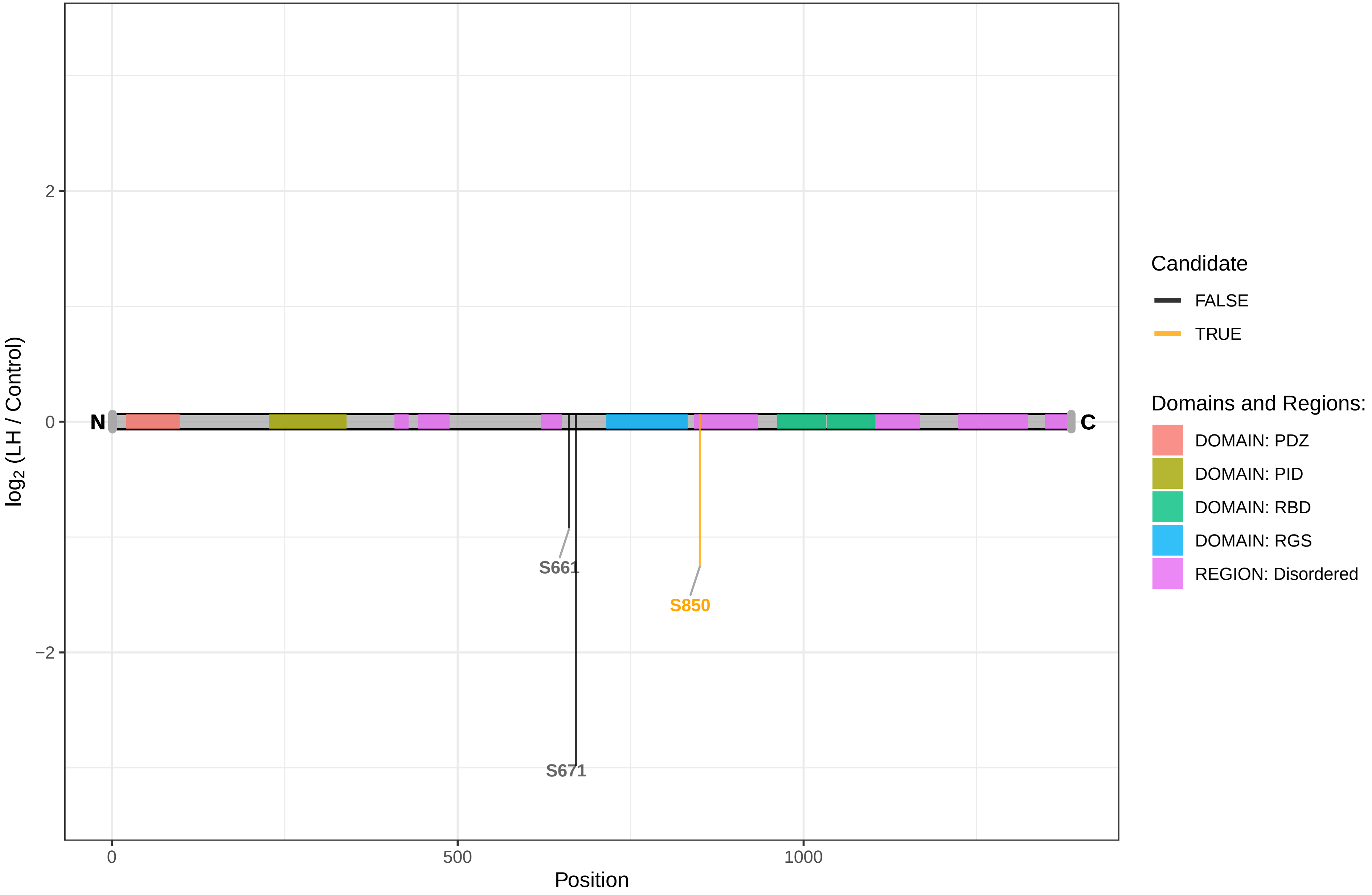

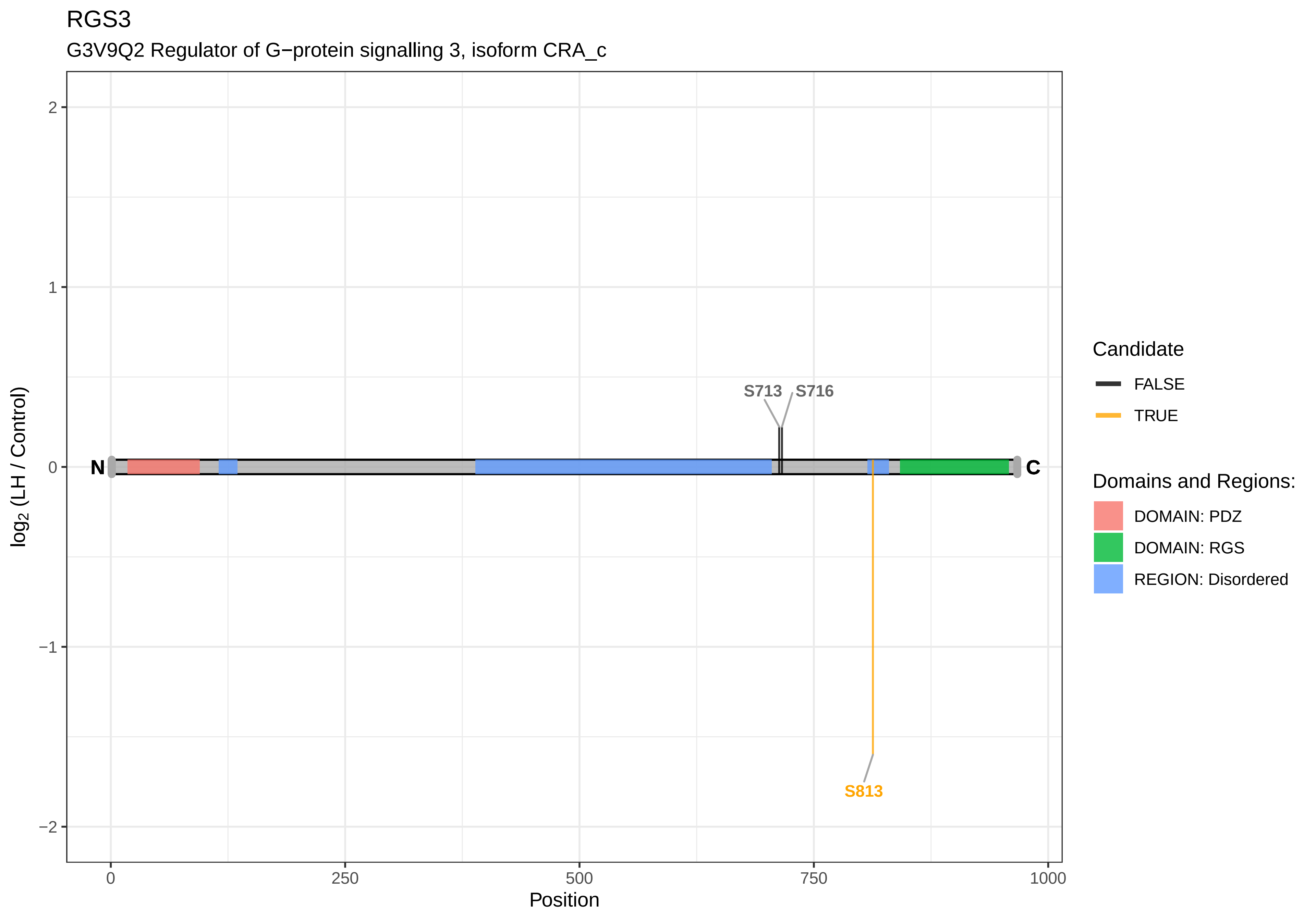

RGS3
G3V9Q2 Regulator of G−protein signalling 3, isoform CRA_c
S713
S716
S813
N
C
2
1
0
−1
−2
log2 (LH / Control)
0
250
500
750
1000
Position
Candidate
FALSE
TRUE
Domains and Regions:
DOMAIN: PDZ
DOMAIN: RGS
REGION: Disordered

RHBDF2

F1LWZ3 Inactive rhomboid protein

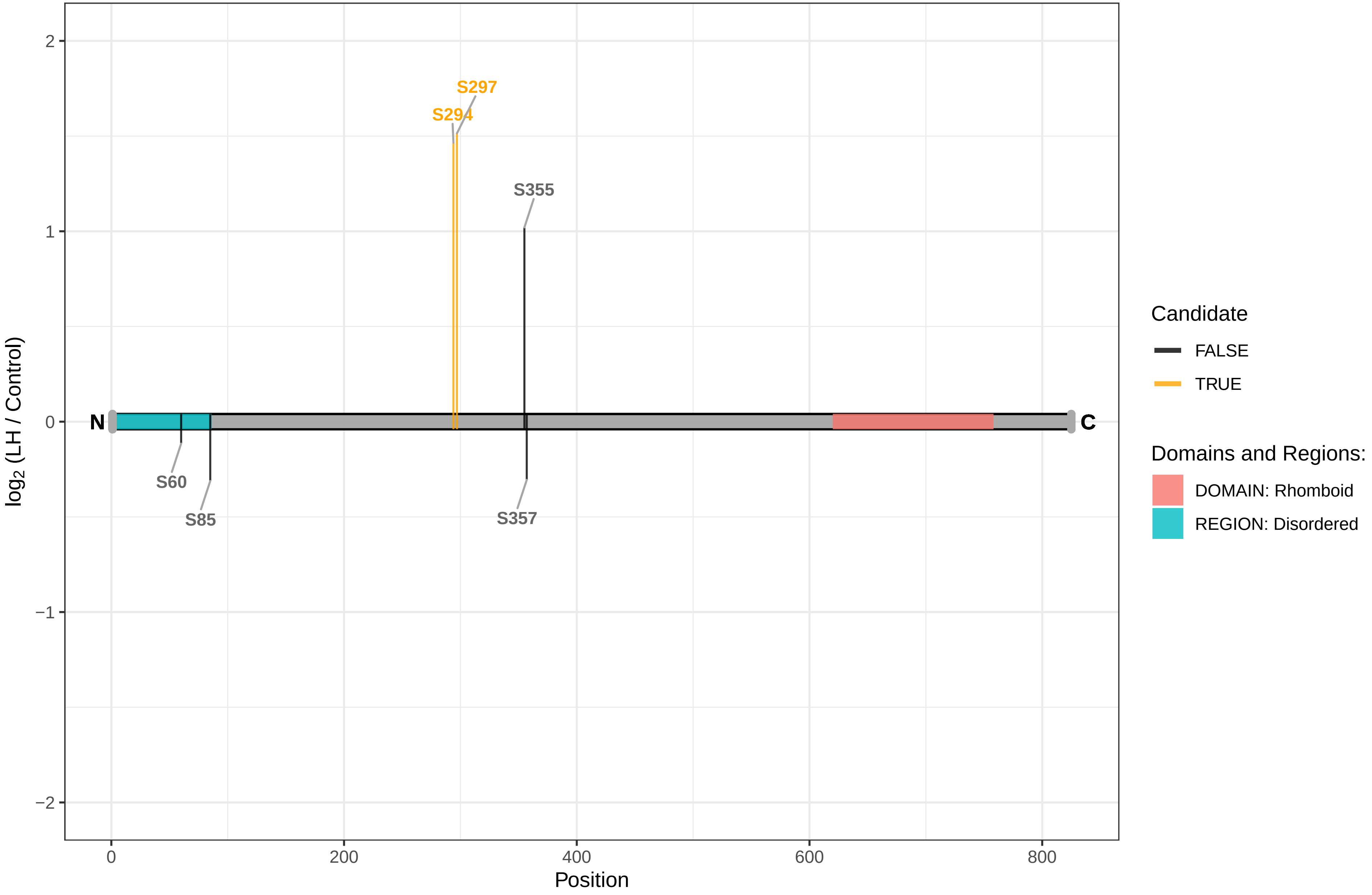

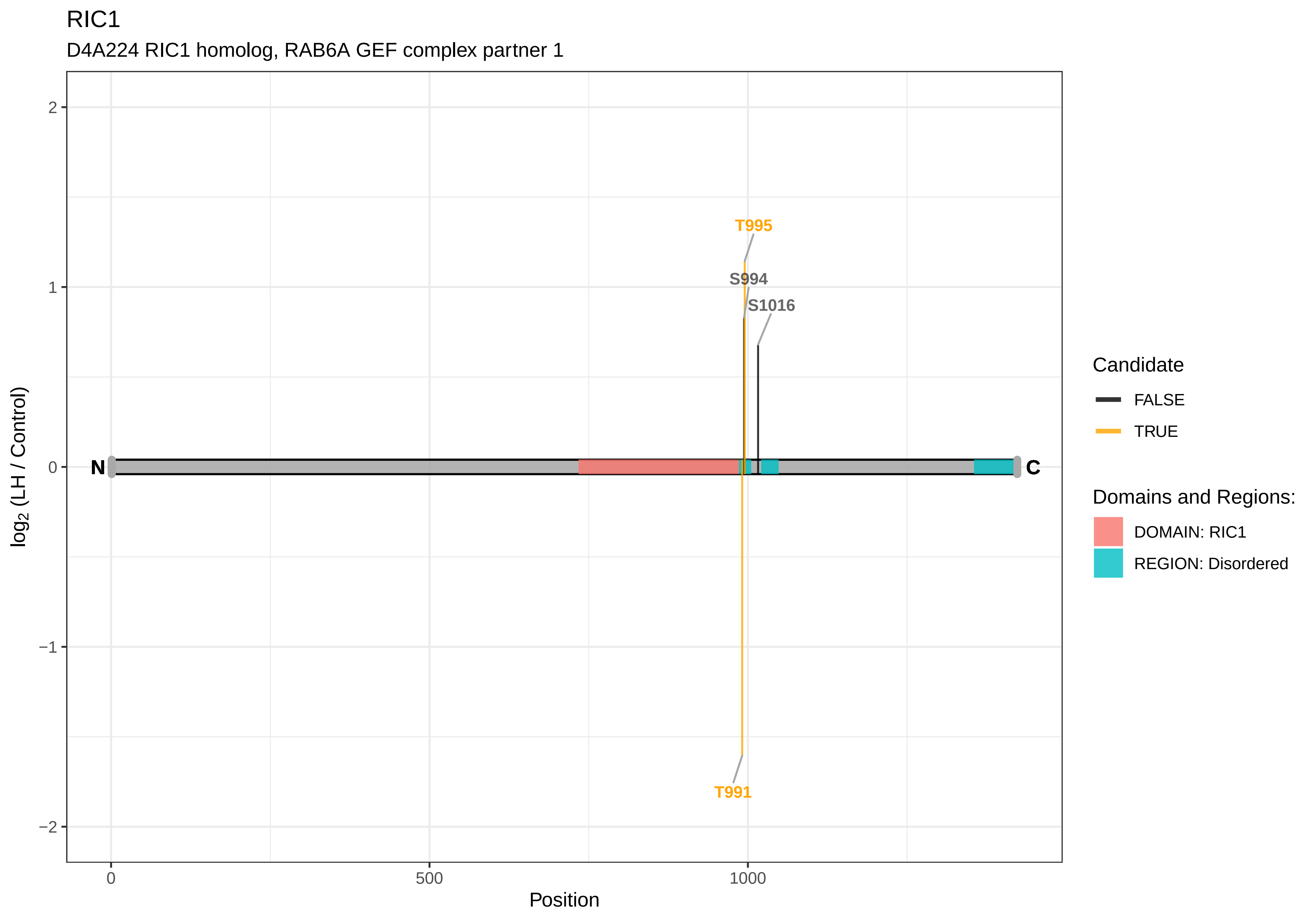
RIC1
D4A224 RIC1 homolog, RAB6A GEF complex partner 1
T995
S994
S1016
T991
N
C
2
1
0
−1
−2
0
500
1000
Position
log2 (LH / Control)
Candidate
FALSE
TRUE
Domains and Regions:
DOMAIN: RIC1
REGION: Disordered

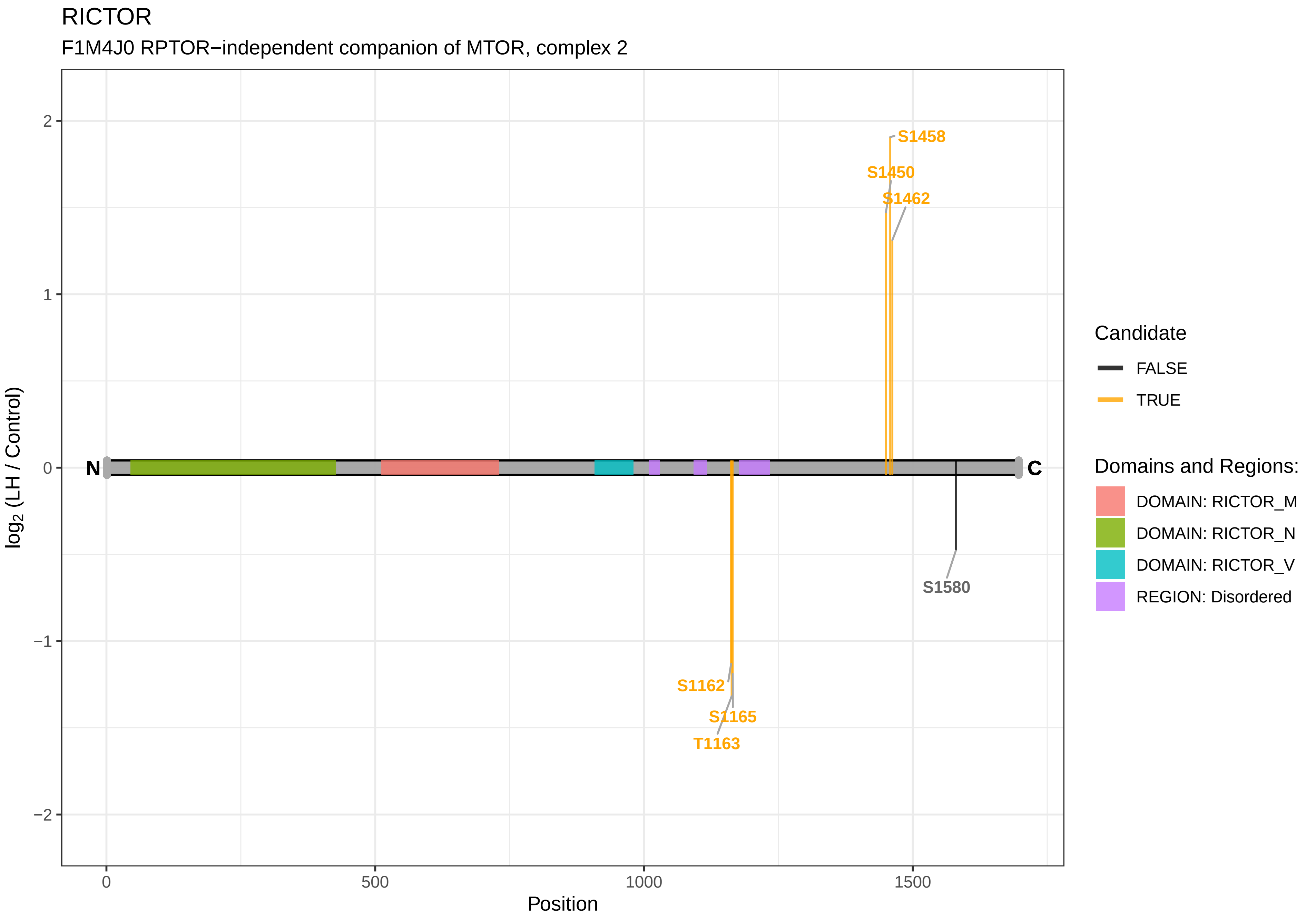

RICTOR
F1M4J0 RPTOR−independent companion of MTOR, complex 2
S1458
S1450
S1462
N
C
S1580
S1162
S1165
T1163
2
1
0
−1
−2
0
500
1000
1500
Position
log2 (LH / Control)
Candidate
FALSE
TRUE
Domains and Regions:
DOMAIN: RICTOR_M
DOMAIN: RICTOR_N
DOMAIN: RICTOR_V
REGION: Disordered

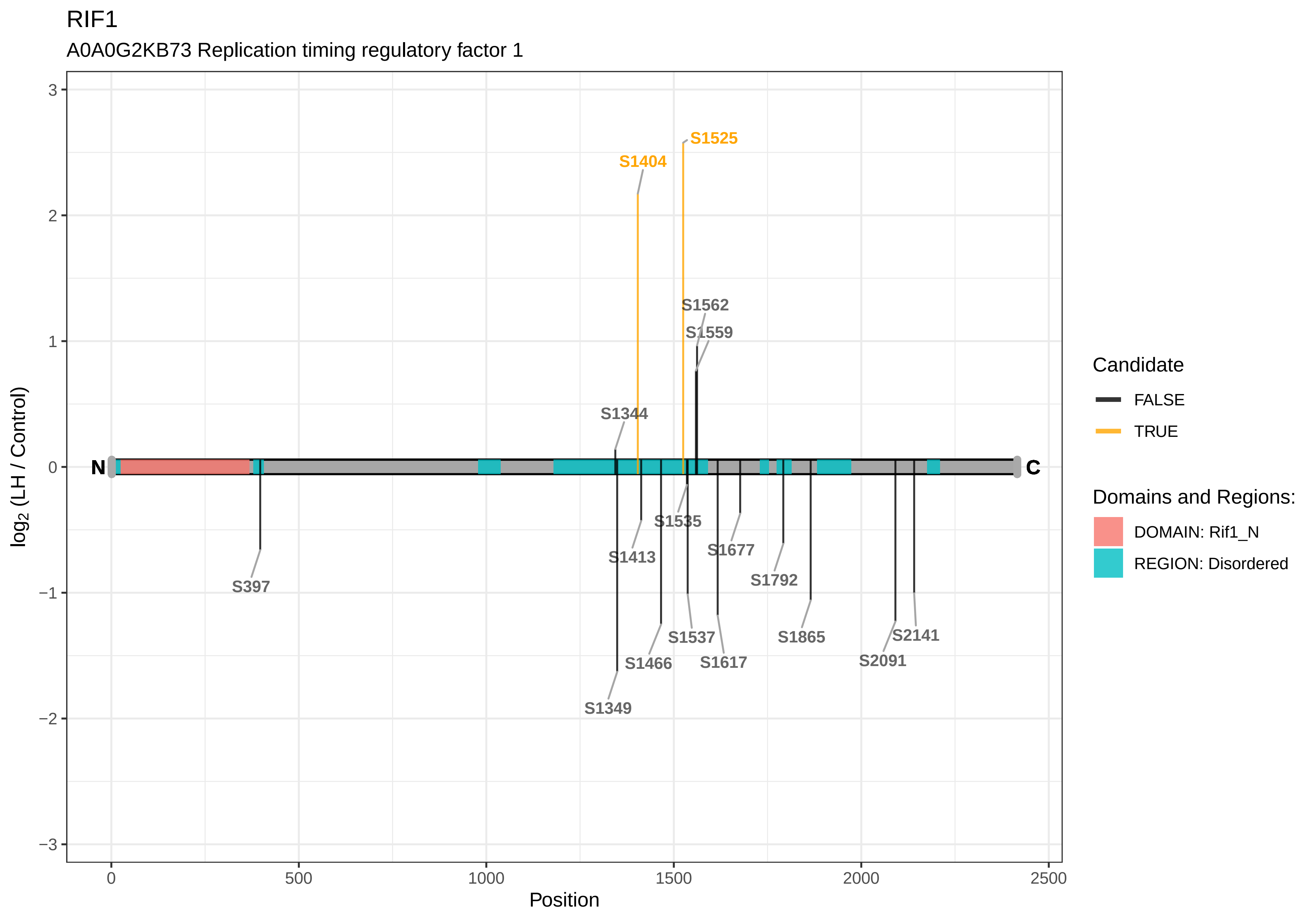
RIF1
A0A0G2KB73 Replication timing regulatory factor 1
3
2
1
0
−1
−2
−3
log2 (LH / Control)
0
500
1000
1500
2000
2500
Position
N
C
S1525
S1404
S1562
S1559
S1344
S1535
S1413
S1677
S1792
S397
S1865
S2141
S1537
S1466
S1617
S2091
S1349
Candidate
FALSE
TRUE
Domains and Regions:
DOMAIN: Rif1_N
REGION: Disordered

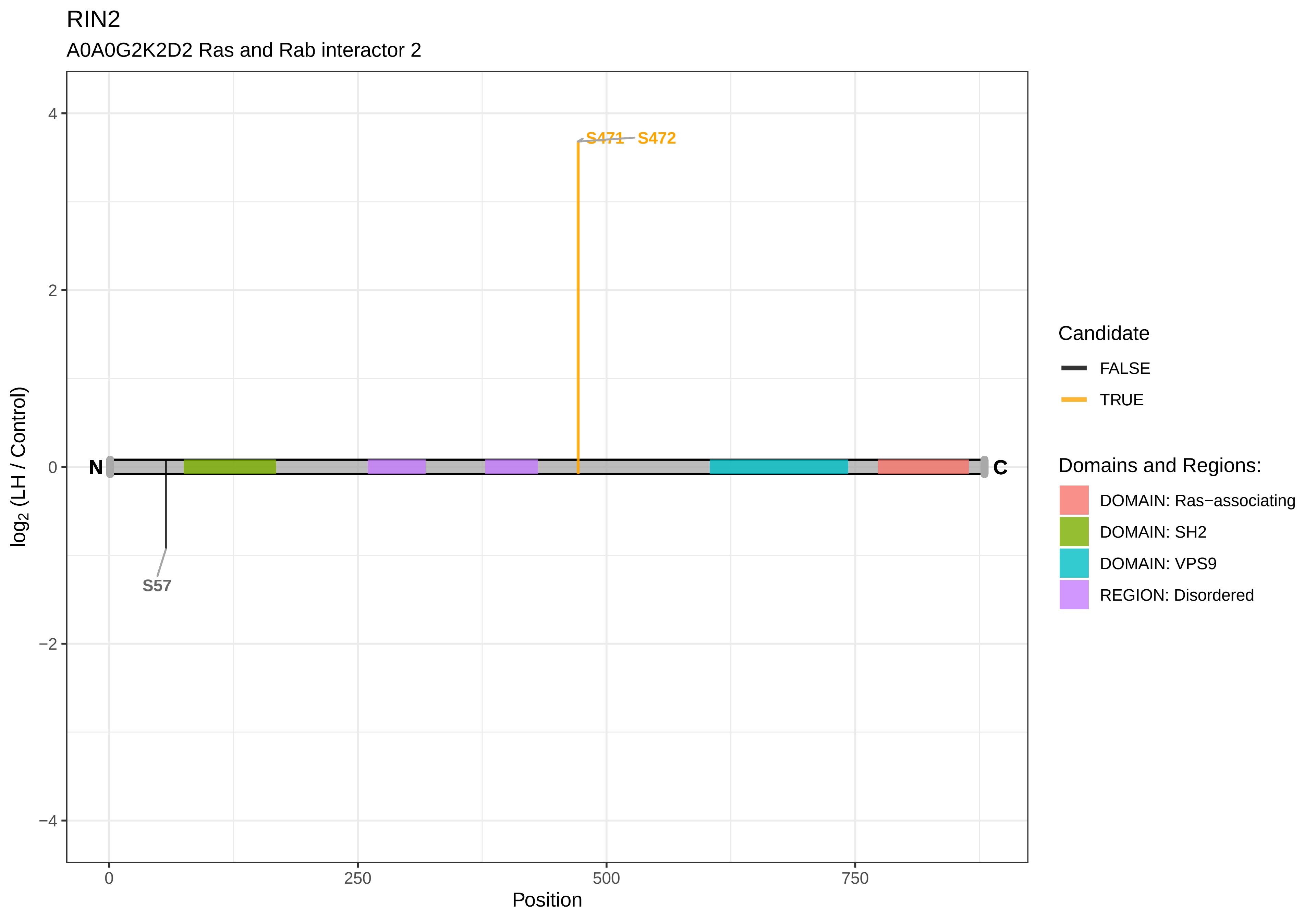

RIN2
A0A0G2K2D2 Ras and Rab interactor 2
S471
S472
S57
N
C
4
2
0
−2
−4
log2 (LH / Control)
0
250
500
750
Position
Candidate
FALSE
TRUE
Domains and Regions:
DOMAIN: Ras−associating
DOMAIN: SH2
DOMAIN: VPS9
REGION: Disordered

RMDN3

Q66H15 Regulator of microtubule dynamics protein 3

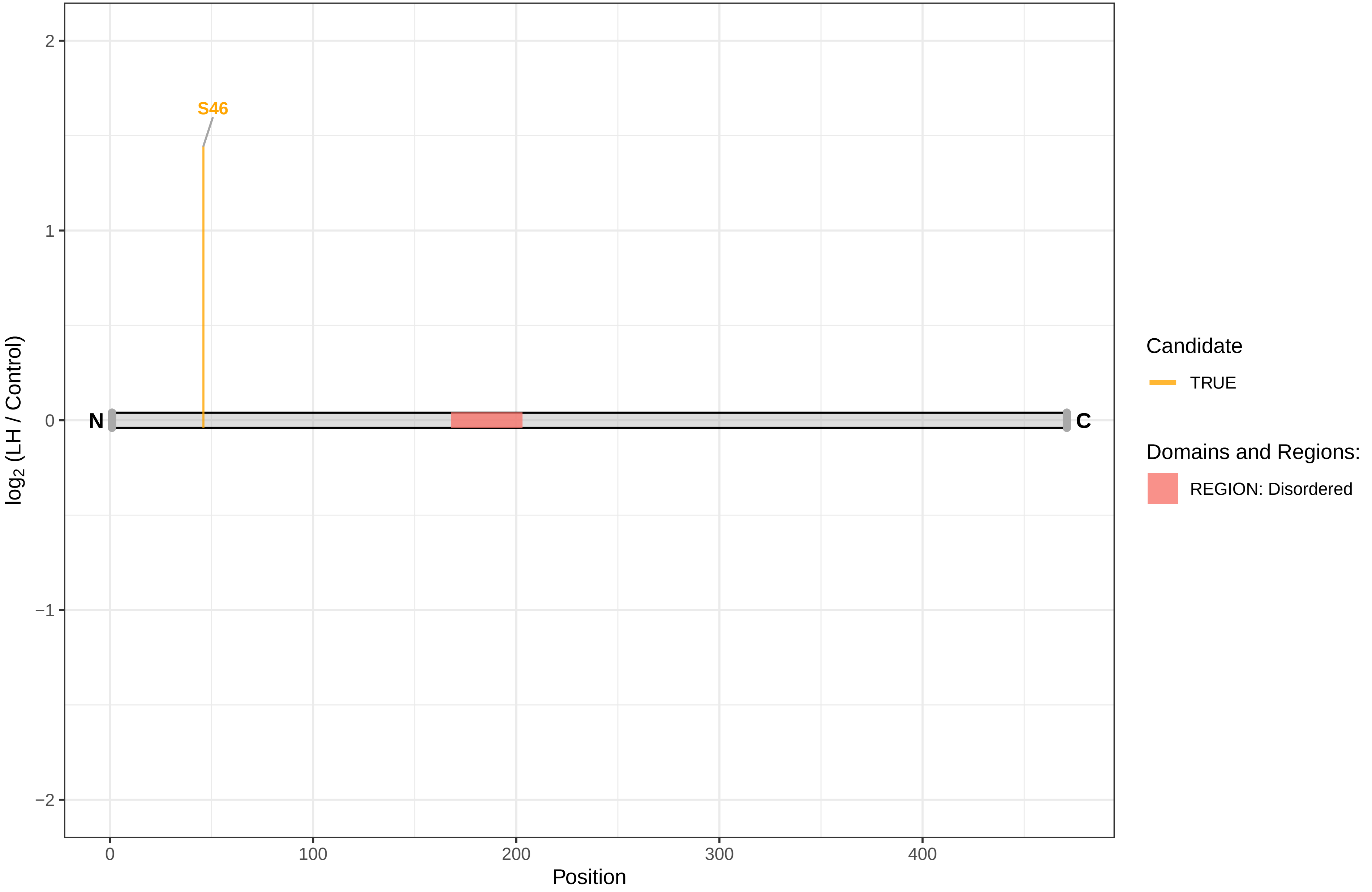

RPGR

B1WC72 Retinitis pigmentosa GTPase regulator

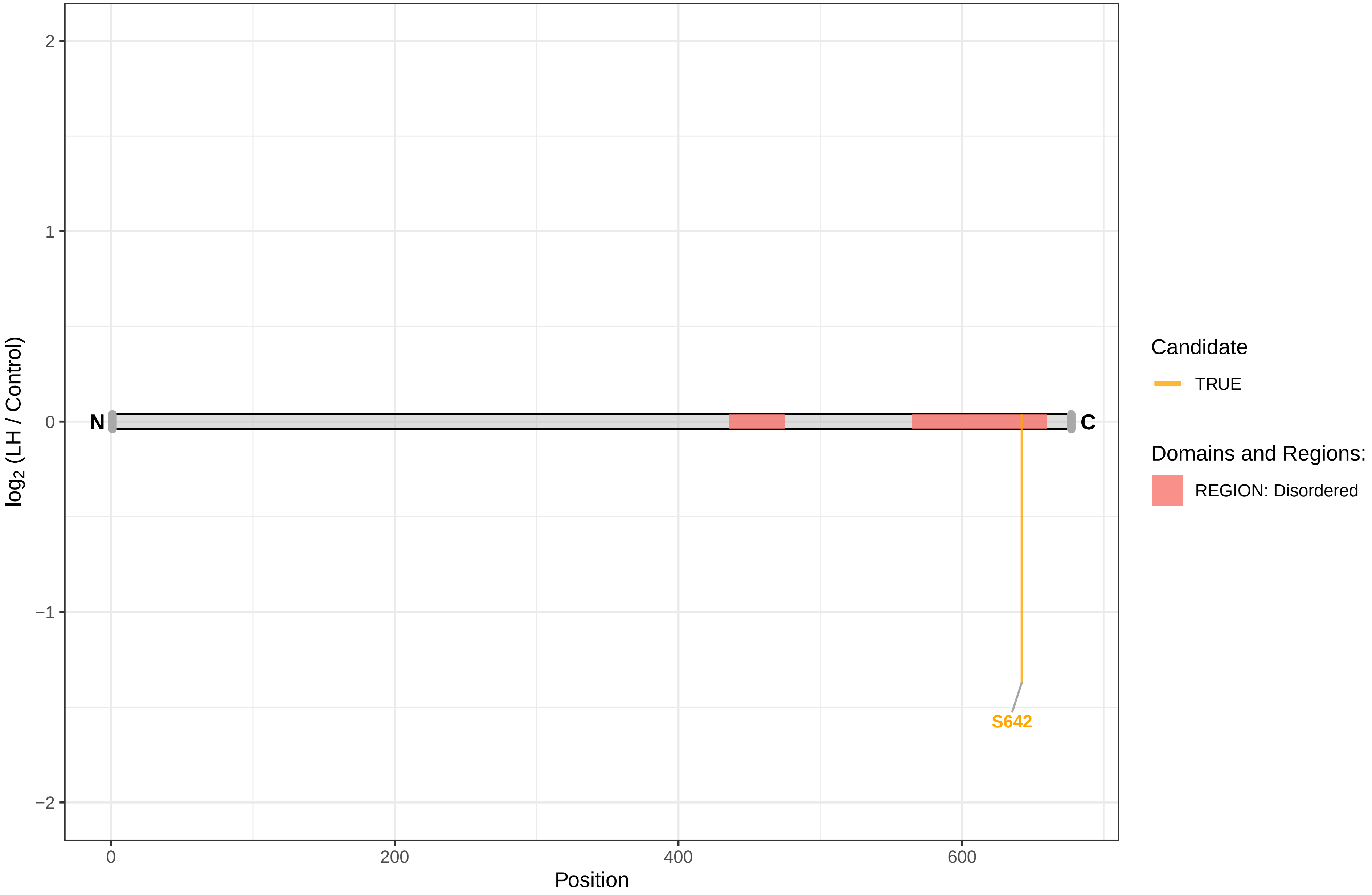

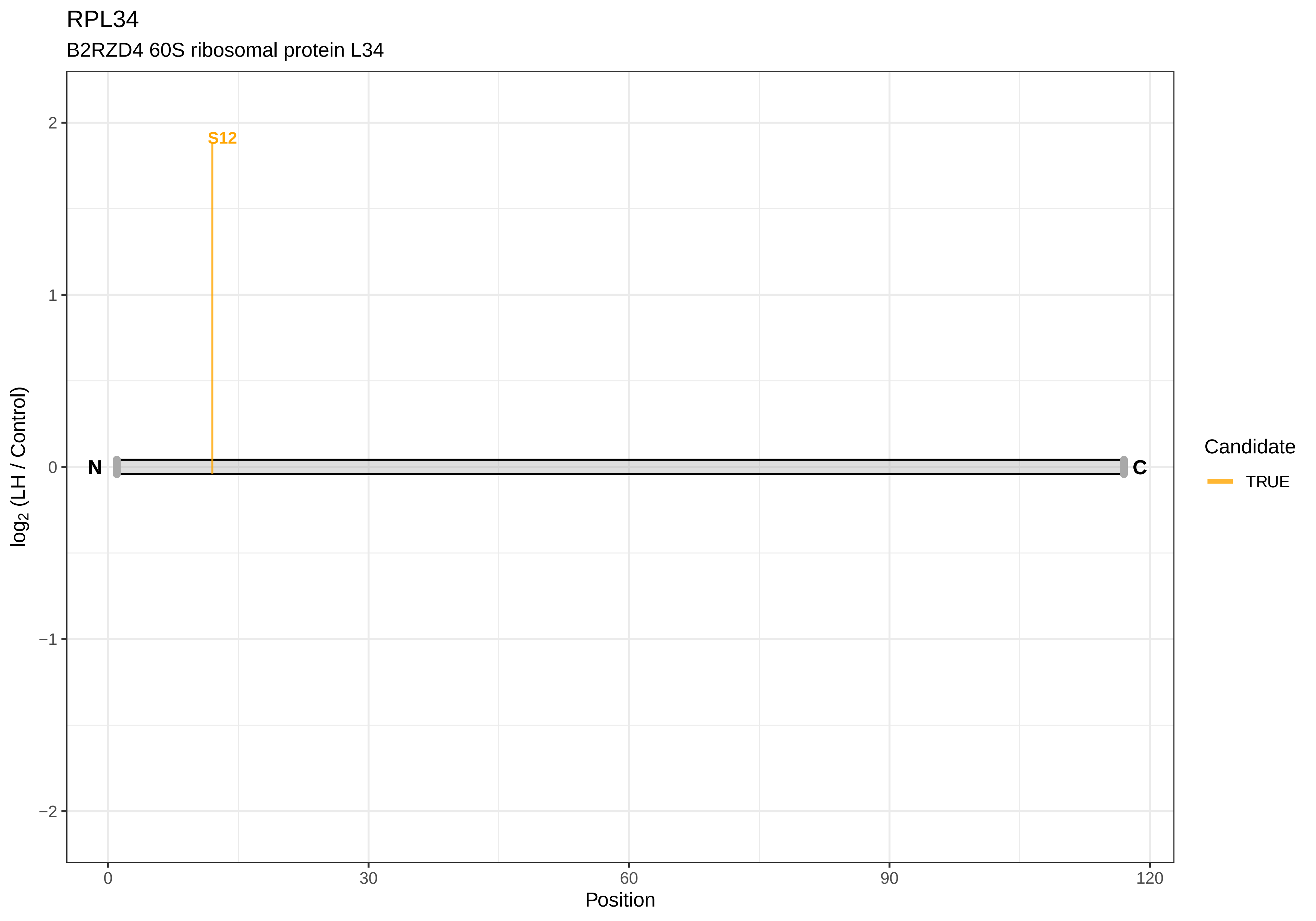

RPL34
B2RZD4 60S ribosomal protein L34
S12
N
C
2
1
0
−1
−2
0
30
60
90
120
Position
log2 (LH / Control)
Candidate
TRUE

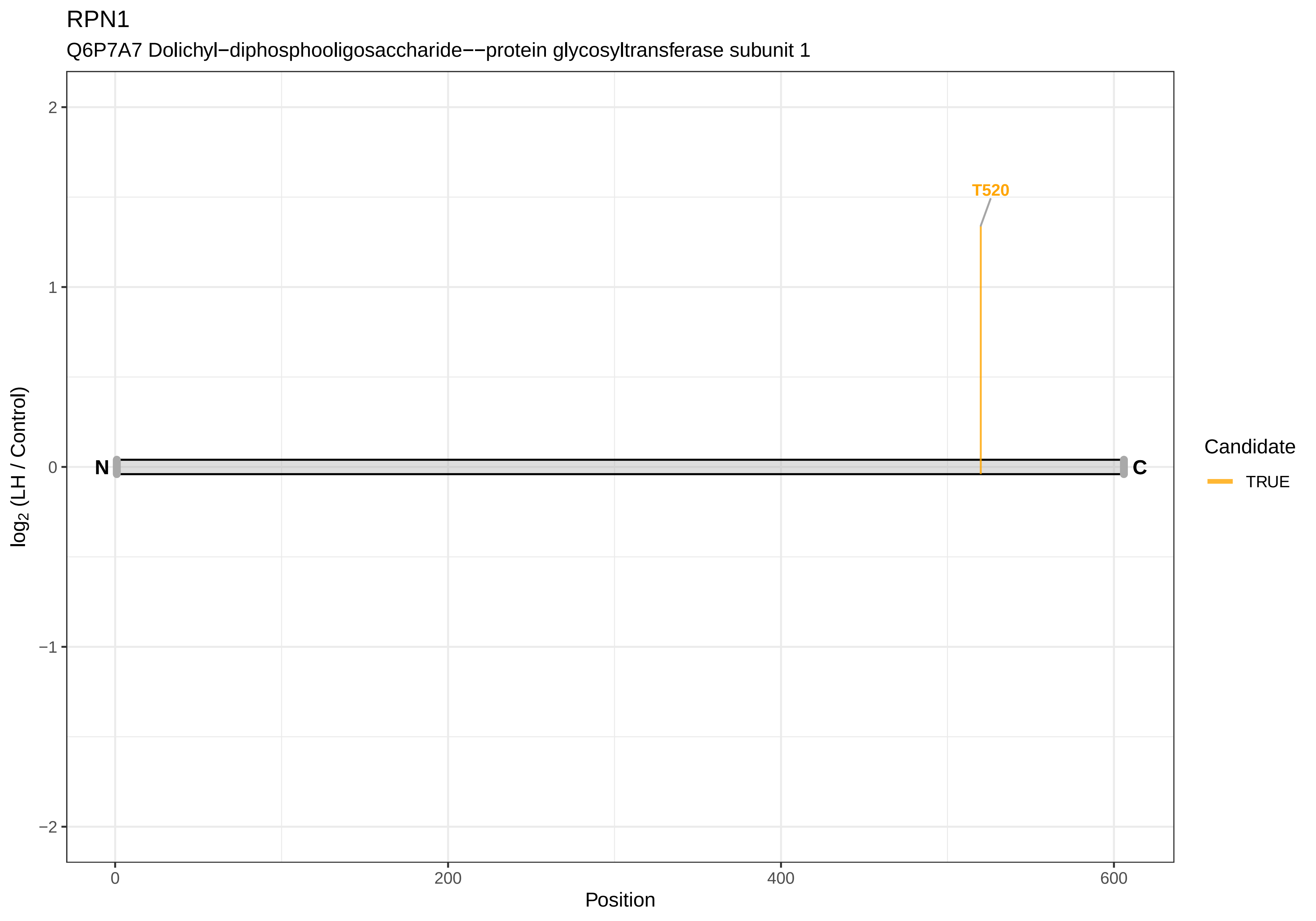

RPN1
Q6P7A7 Dolichyl−diphosphooligosaccharide−−protein glycosyltransferase subunit 1
T520
N
C
2
1
0
−1
−2
log2 (LH / Control)
0
200
400
600
Position
Candidate
TRUE

### RPP25

Q5PPN2 Ribonuclease P protein subunit p25

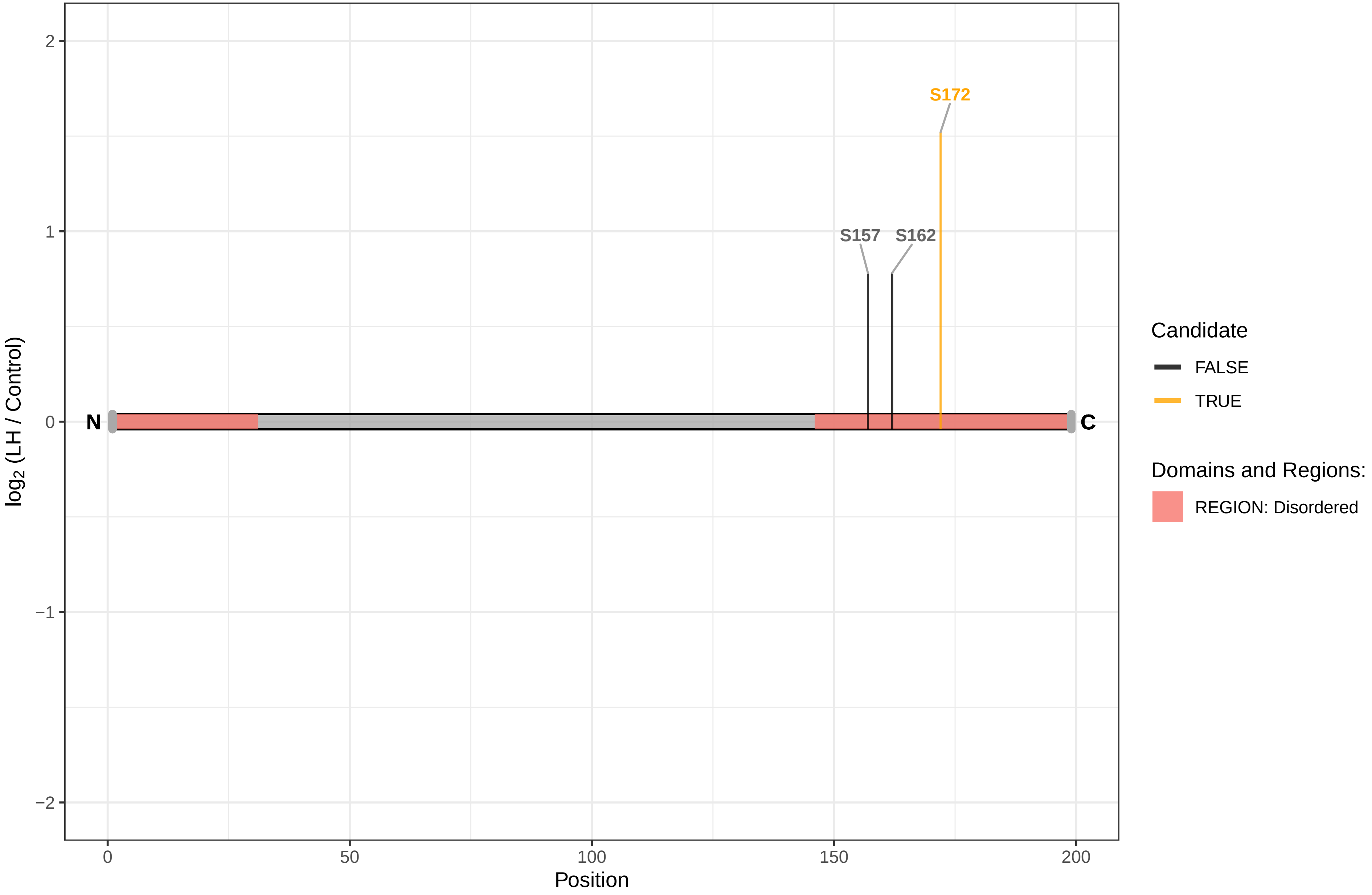

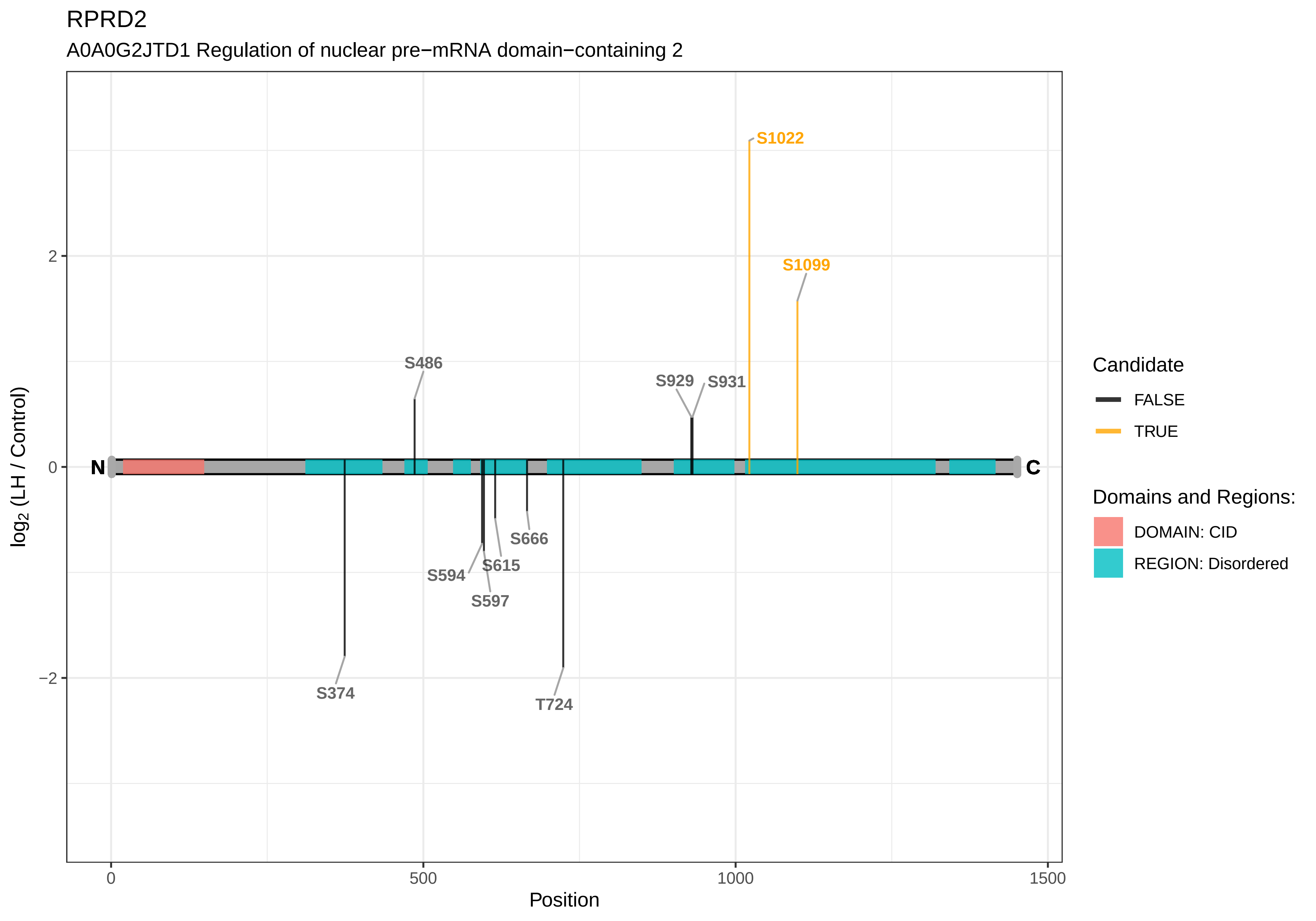

RPRD2
A0A0G2JTD1 Regulation of nuclear pre−mRNA domain−containing 2
S1022
S1099
S486
S929
S931
S666
S615
S594
S597
S374
T724
N
C
2
0
−2
0
500
1000
1500
Position
log2 (LH / Control)
Candidate
FALSE
TRUE
Domains and Regions:
DOMAIN: CID
REGION: Disordered

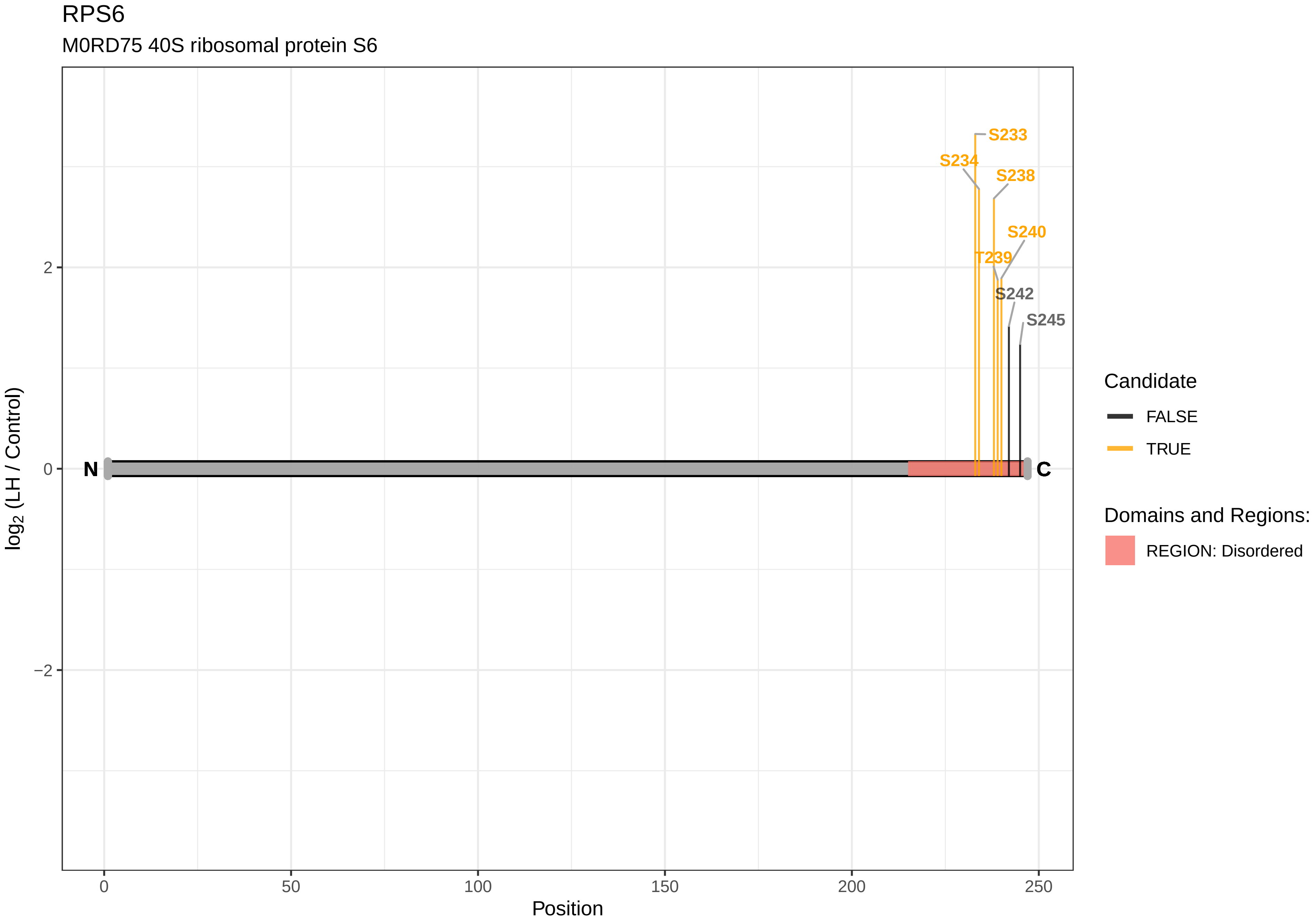

RPS6
M0RD75 40S ribosomal protein S6
S233
S234
S238
S240
T239
S242
S245
N
C
2
0
−2
0
50
100
150
200
250
Position
log2 (LH / Control)
Candidate
FALSE
TRUE
Domains and Regions:
REGION: Disordered

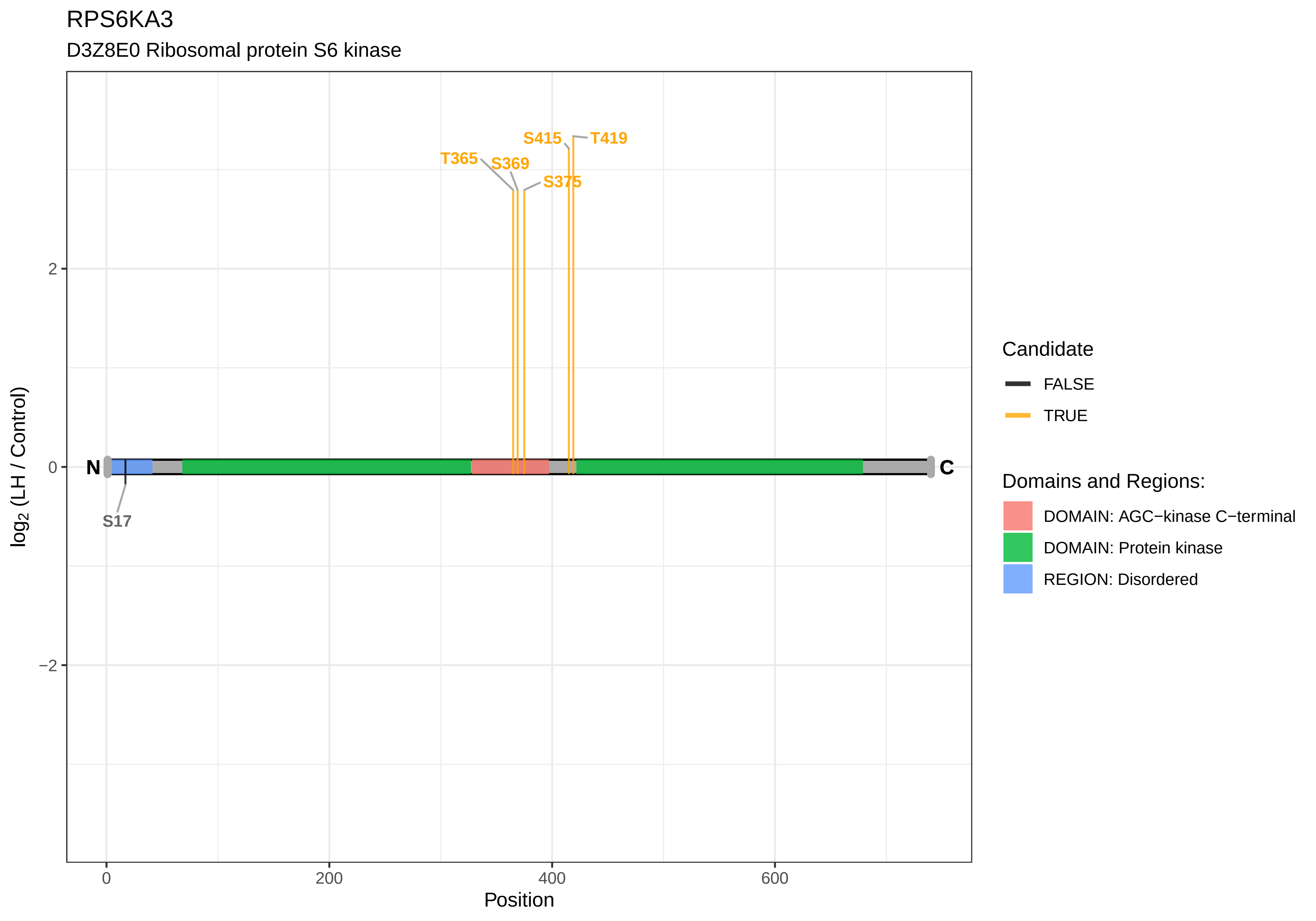
RPS6KA3
D3Z8E0 Ribosomal protein S6 kinase
T365
S369
S375
S415
T419
S17
N
C
2
0
−2
log2 (LH / Control)
0
200
400
600
Position
Candidate
FALSE
TRUE
Domains and Regions:
DOMAIN: AGC−kinase C−terminal
DOMAIN: Protein kinase
REGION: Disordered

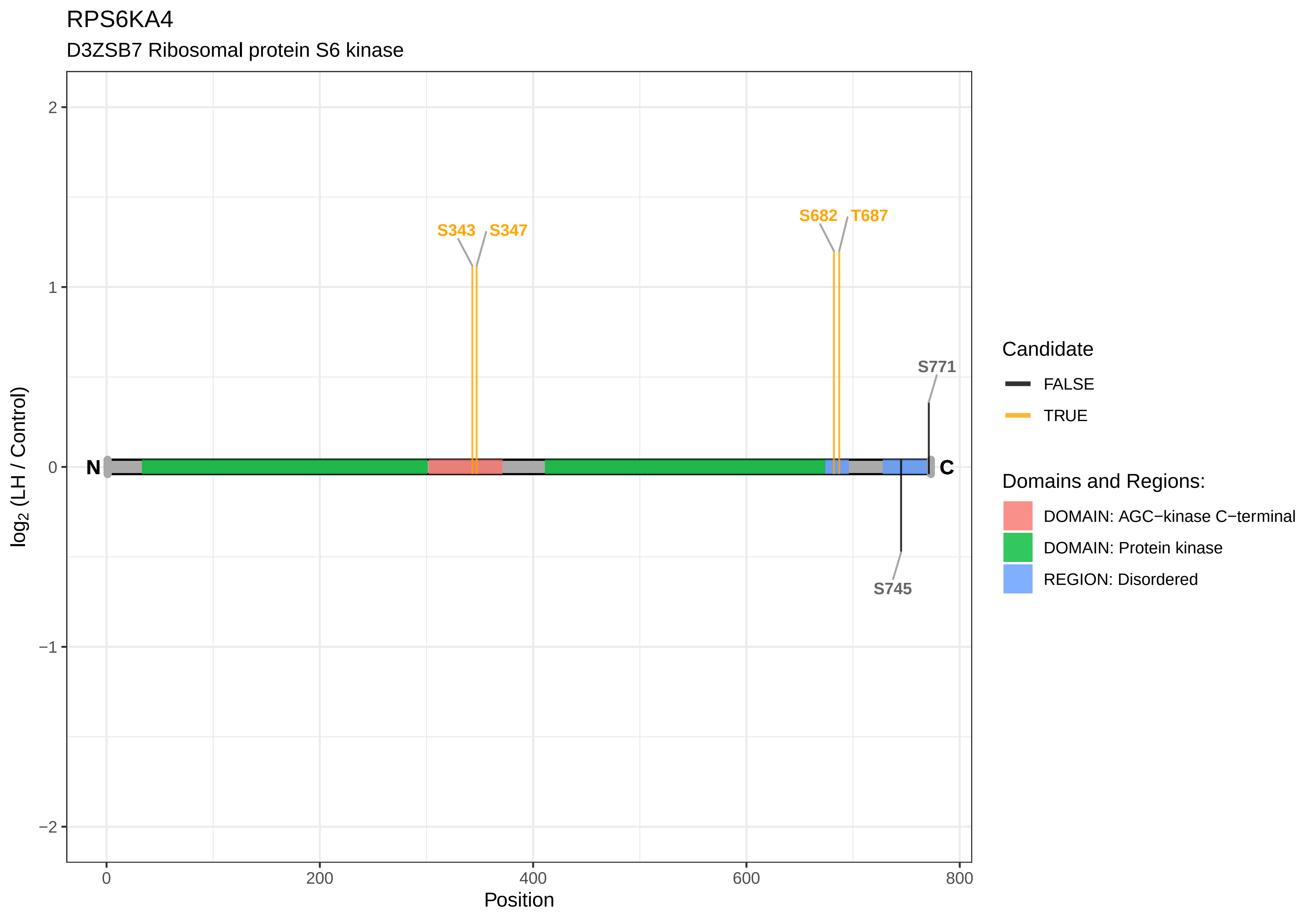
RPS6KA4
D3ZSB7 Ribosomal protein S6 kinase
S343
S347
S682
T687
S771
S745
N
C
2
1
0
−1
−2
0
200
400
600
800
Position
log2 (LH / Control)
Candidate
FALSE
TRUE
Domains and Regions:
DOMAIN: AGC−kinase C−terminal
DOMAIN: Protein kinase
REGION: Disordered

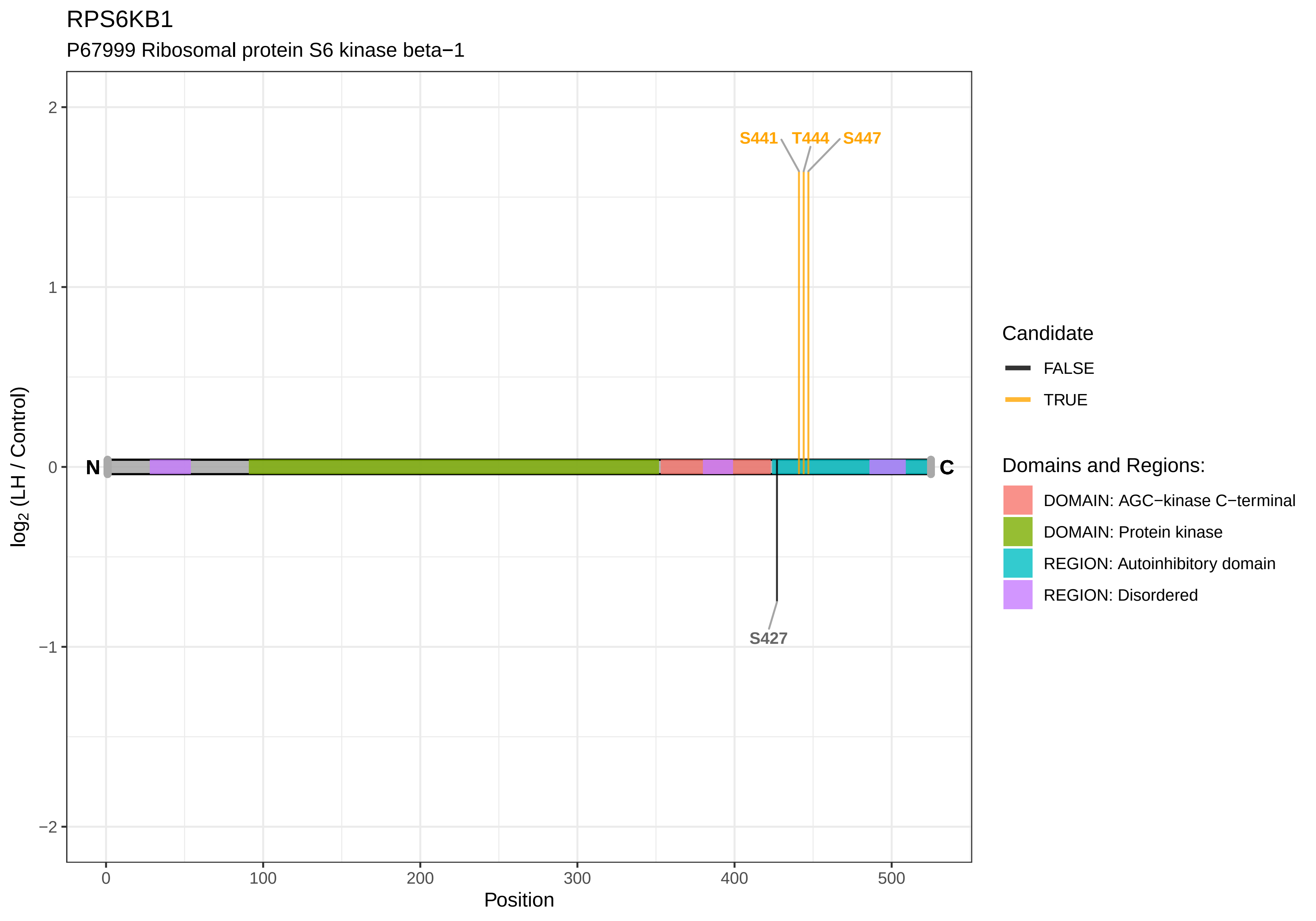
RPS6KB1
P67999 Ribosomal protein S6 kinase beta−1
S441
T444
S447
S427
N
C
2
1
0
−1
−2
0
100
200
300
400
500
Position
log2 (LH / Control)
Candidate
FALSE
TRUE
Domains and Regions:
DOMAIN: AGC−kinase C−terminal
DOMAIN: Protein kinase
REGION: Autoinhibitory domain
REGION: Disordered

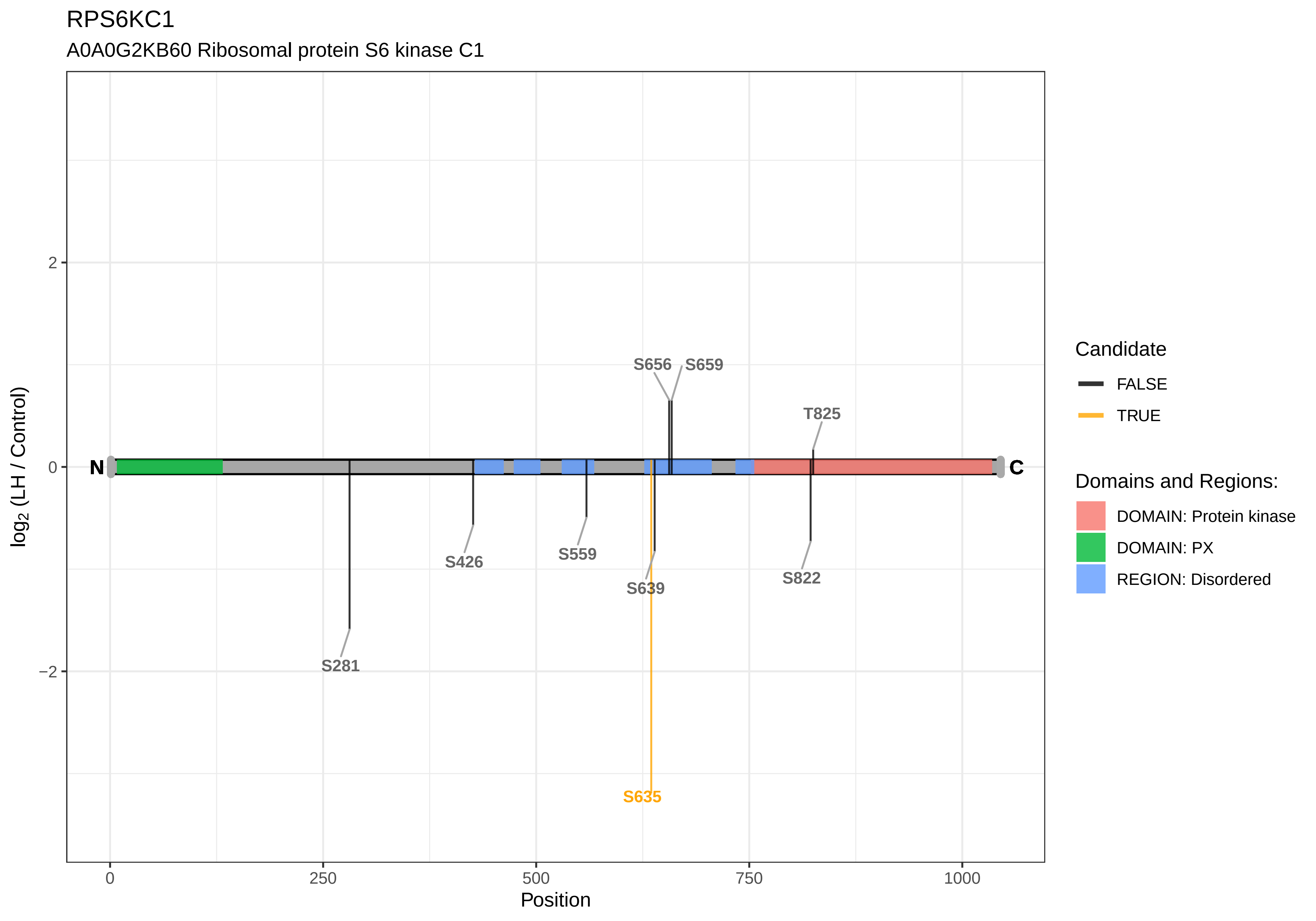

RPS6KC1
A0A0G2KB60 Ribosomal protein S6 kinase C1
log2 (LH / Control)
2
0
−2
N
C
S656
S659
T825
S426
S559
S639
S822
S281
S635
0
250
500
750
1000
Position
Candidate
FALSE
TRUE
Domains and Regions:
DOMAIN: Protein kinase
DOMAIN: PX
REGION: Disordered

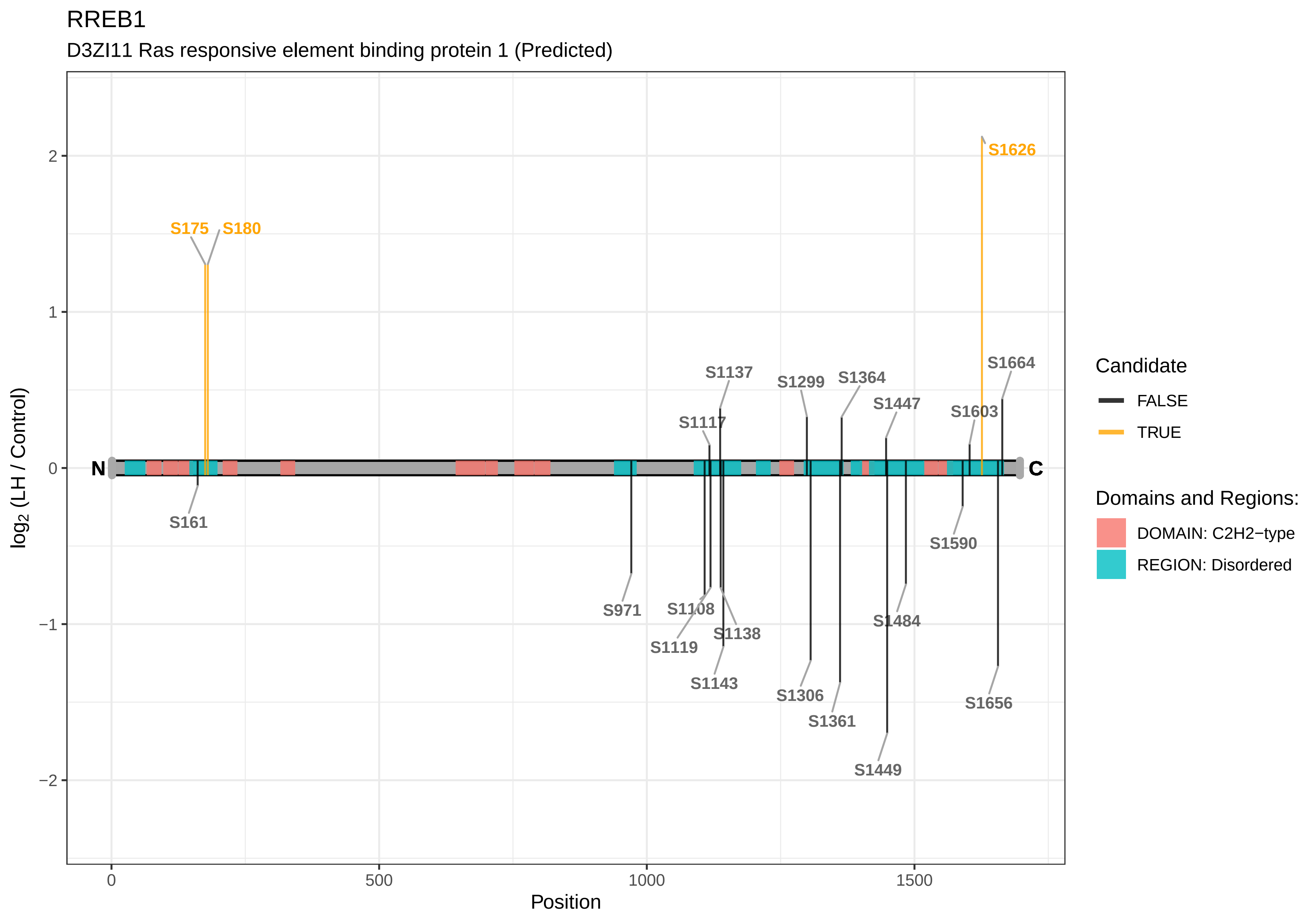
RREB1
D3ZI11 Ras responsive element binding protein 1 (Predicted)
S175
S180
S1626
S1664
S1137
S1299
S1364
S1447
S1603
S1117
N
C
S161
S1590
S971
S1108
S1484
S1119
S1138
S1143
S1306
S1656
S1361
S1449
log2 (LH / Control)
Position
Candidate
FALSE
TRUE
Domains and Regions:
DOMAIN: C2H2−type
REGION: Disordered

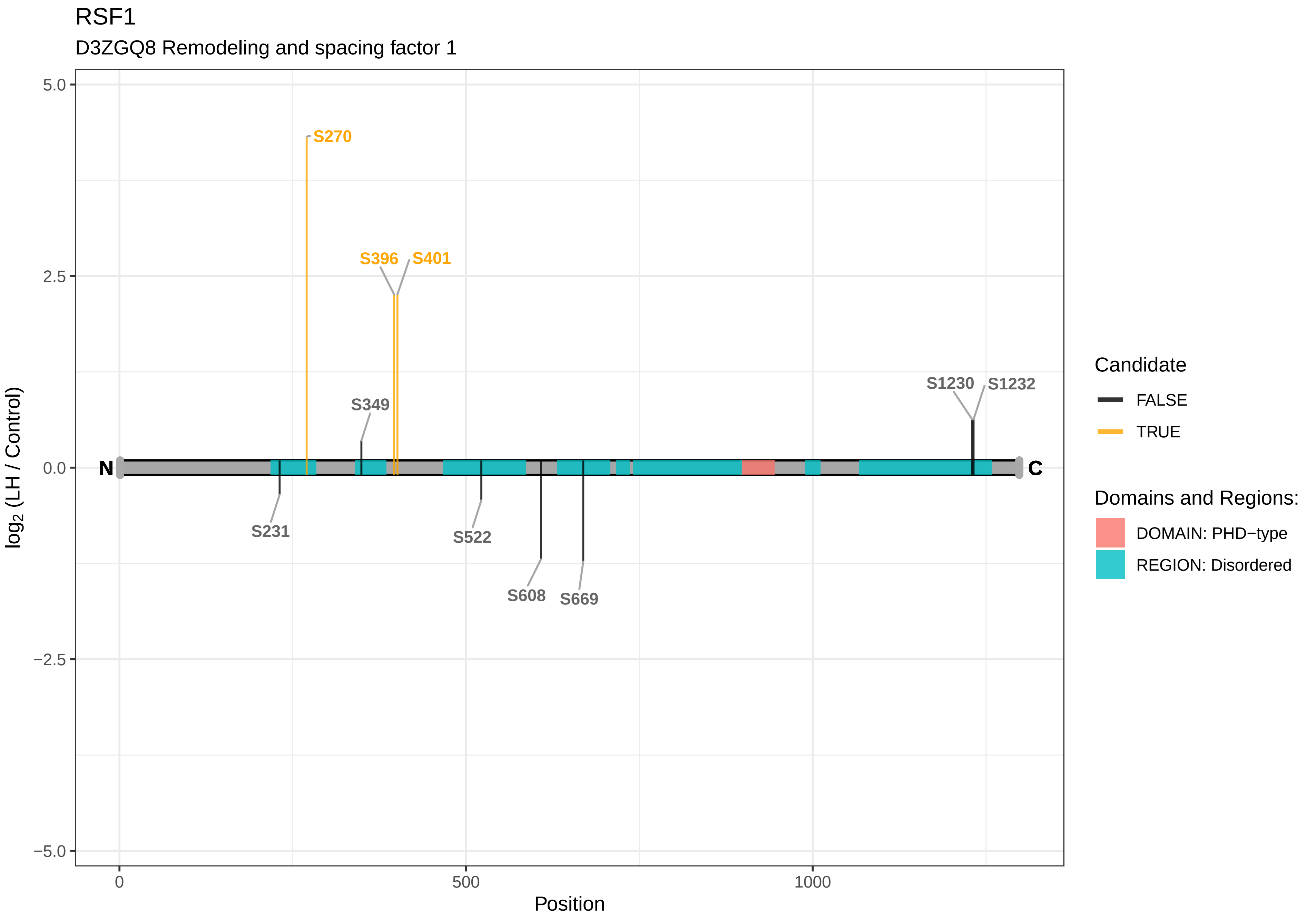
RSF1
D3ZGQ8 Remodeling and spacing factor 1
S270
S396
S401
S349
S1230
S1232
S231
S522
S608
S669
N
C
5.0
2.5
0.0
−2.5
−5.0
log2 (LH / Control)
0
500
1000
Position
Candidate
FALSE
TRUE
Domains and Regions:
DOMAIN: PHD−type
REGION: Disordered

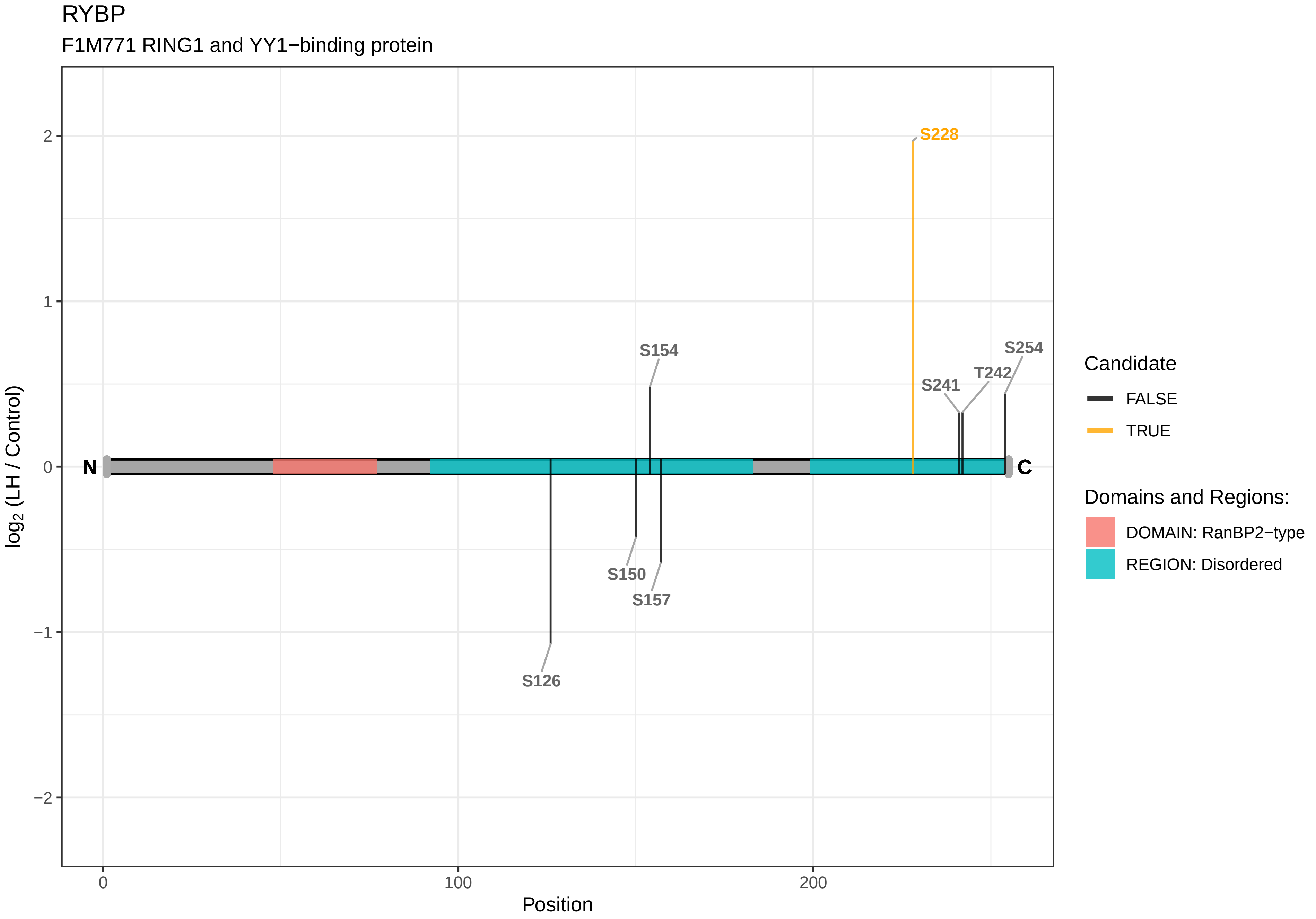
RYBP
F1M771 RING1 and YY1−binding protein
log2 (LH / Control)
2
1
0
−1
−2
0
100
200
Position
N
C
S228
S154
S254
T242
S241
S150
S157
S126
Candidate
FALSE
TRUE
Domains and Regions:
DOMAIN: RanBP2−type
REGION: Disordered

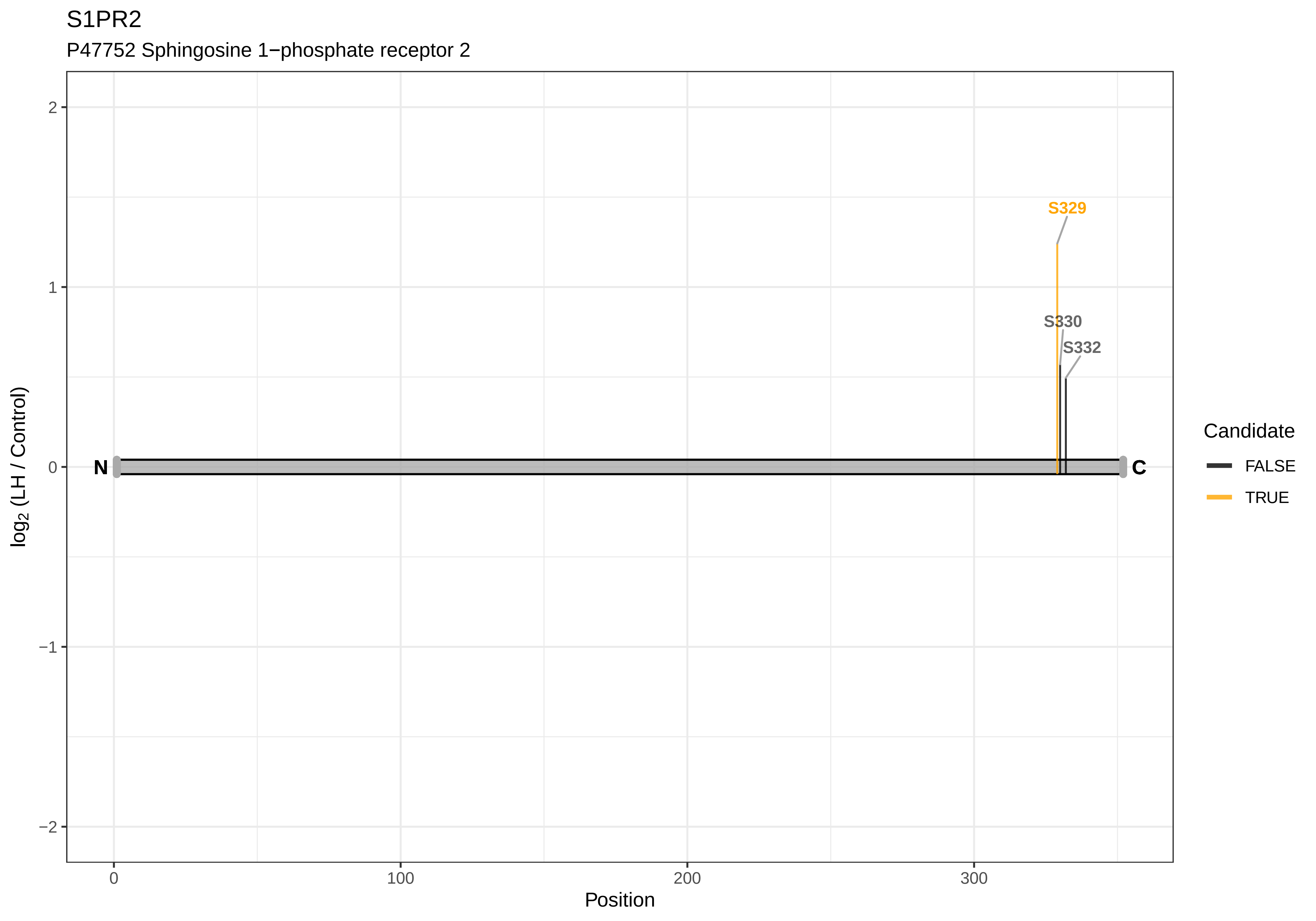

S1PR2
P47752 Sphingosine 1−phosphate receptor 2
S329
S330
S332
N
C
2
1
0
−1
−2
log2 (LH / Control)
0
100
200
300
Position
Candidate
FALSE
TRUE

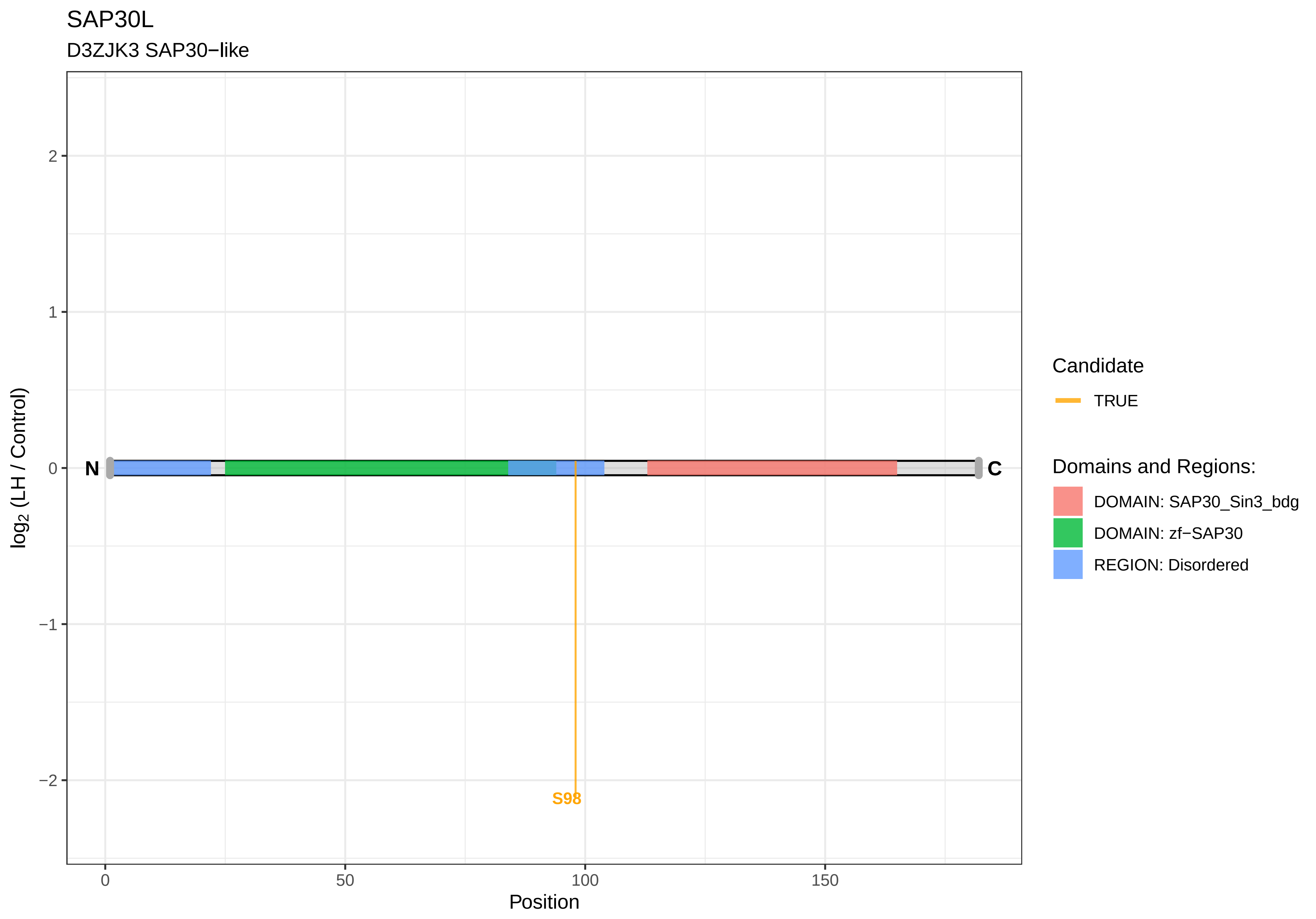
SAP30L
D3ZJK3 SAP30−like
2
1
0
−1
−2
log2 (LH / Control)
N
C
S98
0
50
100
150
Position
Candidate
TRUE
Domains and Regions:
DOMAIN: SAP30_Sin3_bdg
DOMAIN: zf−SAP30
REGION: Disordered

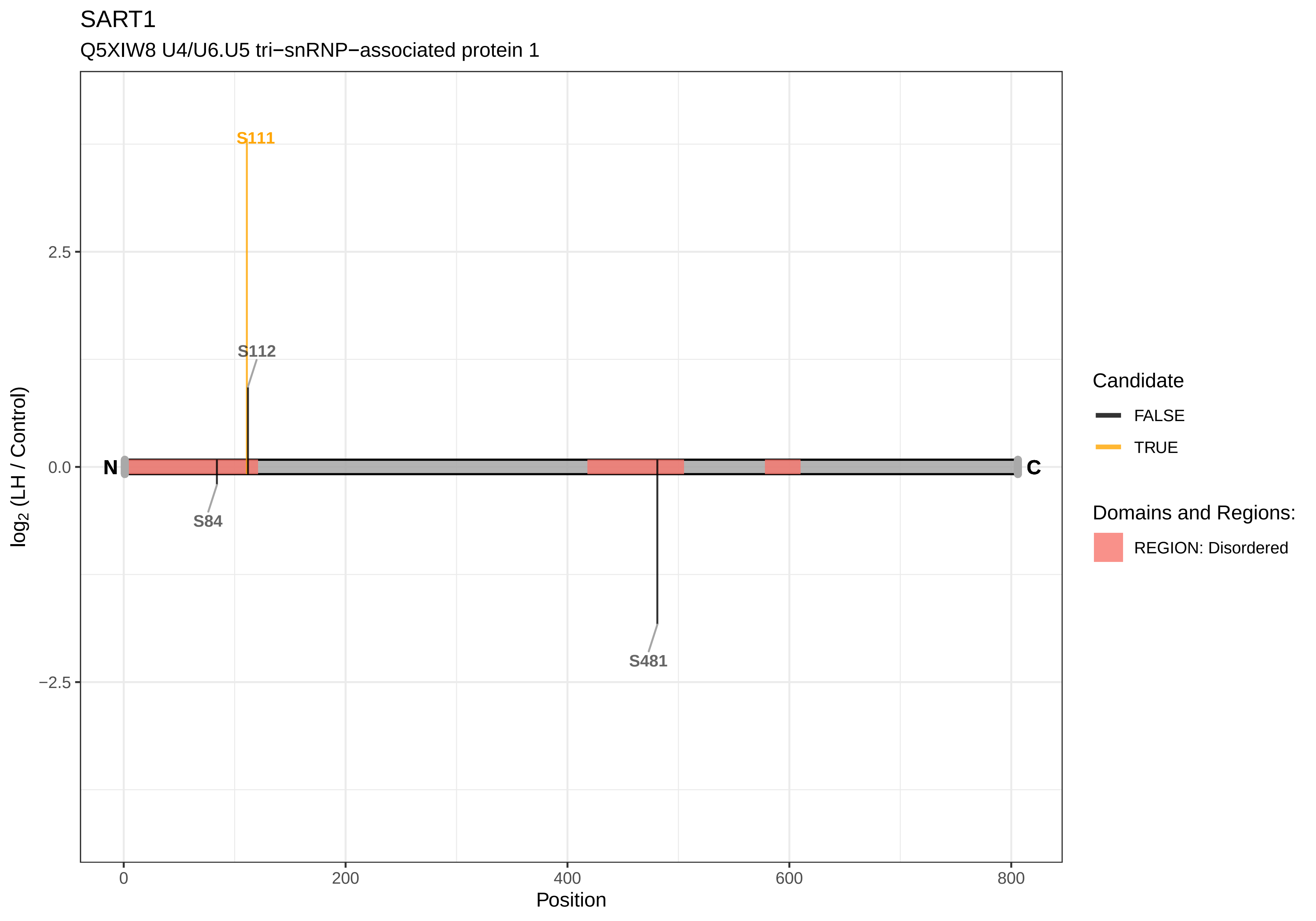
SART1
Q5XIW8 U4/U6.U5 tri−snRNP−associated protein 1
S111
S112
S84
S481
N
C
2.5
0.0
−2.5
log2 (LH / Control)
0
200
400
600
800
Position
Candidate
FALSE
TRUE
Domains and Regions:
REGION: Disordered

SBNO1

F1LS26 Protein strawberry notch homolog 1

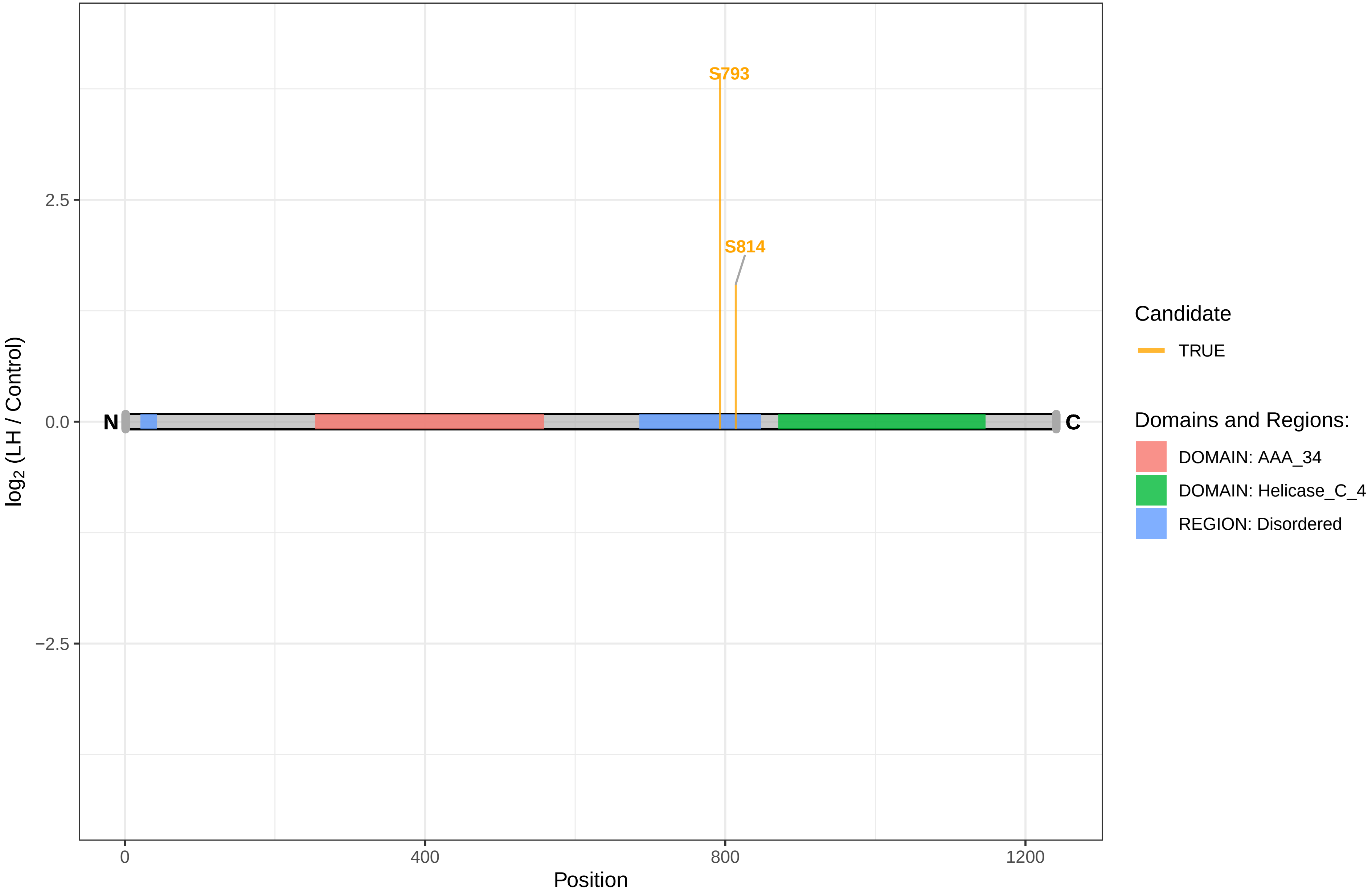

SCLY

Q68FT9 Selenocysteine lyase

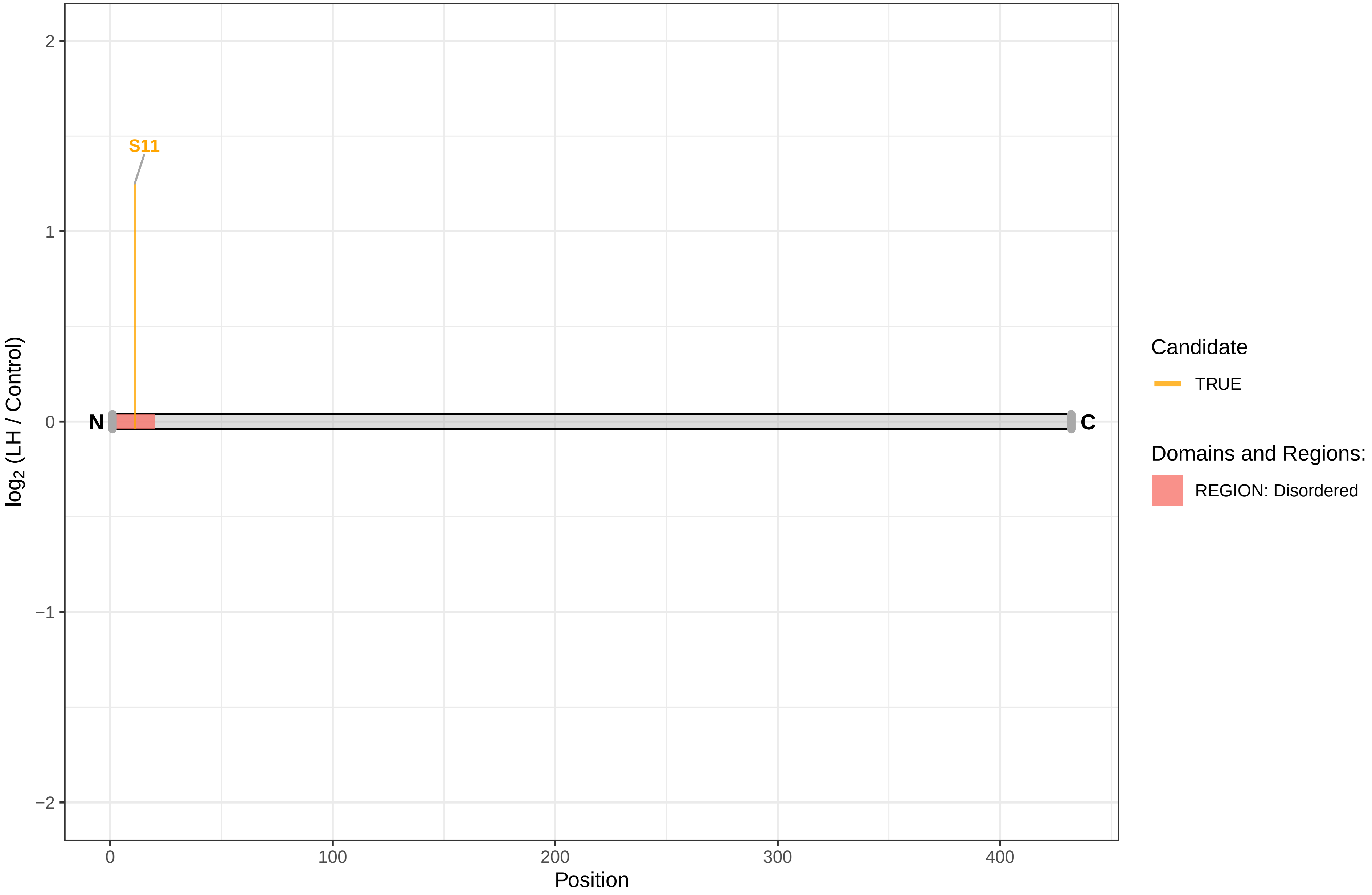

SCN10A

Q62968 Sodium channel protein type 10 subunit alpha

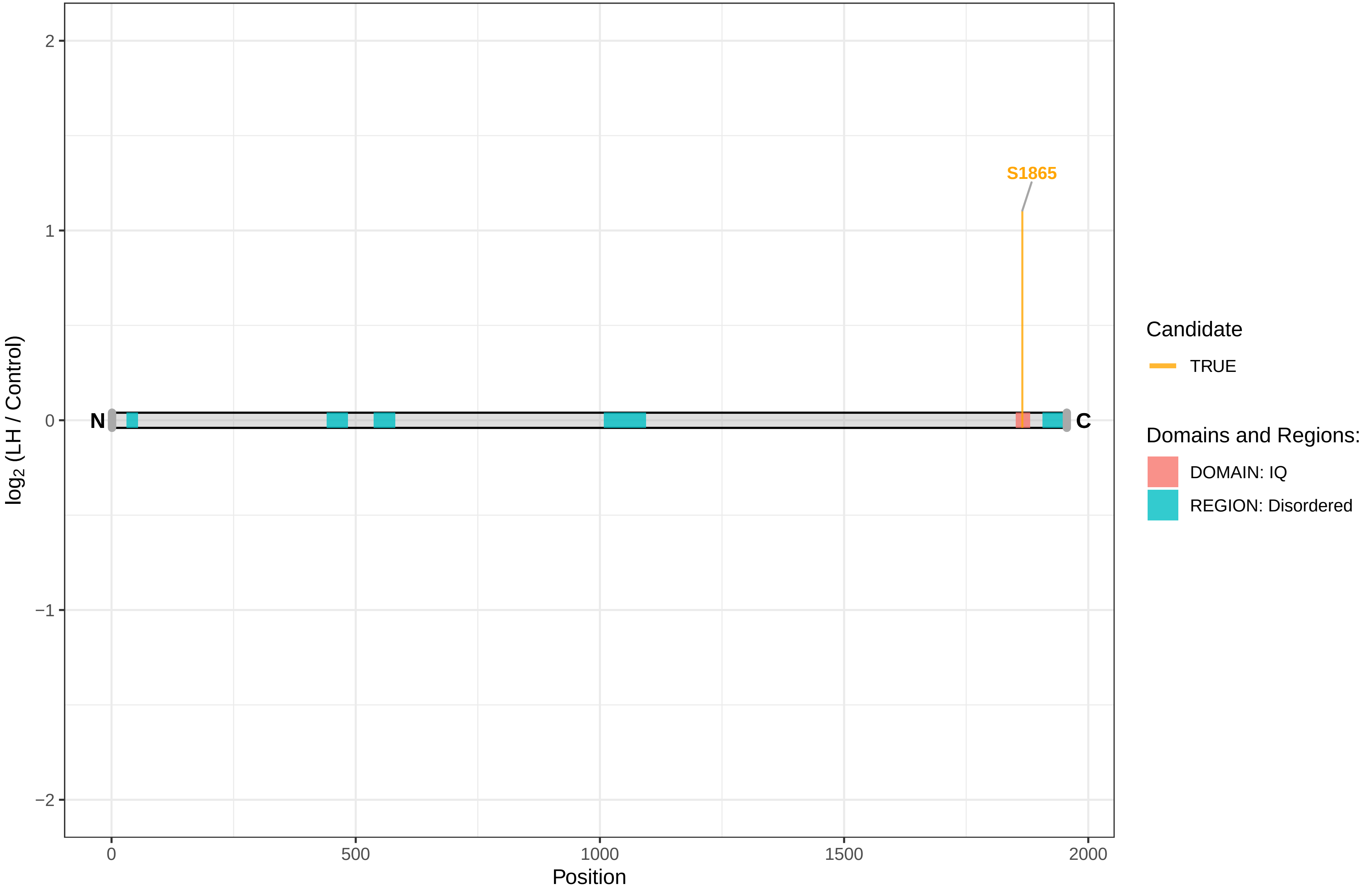

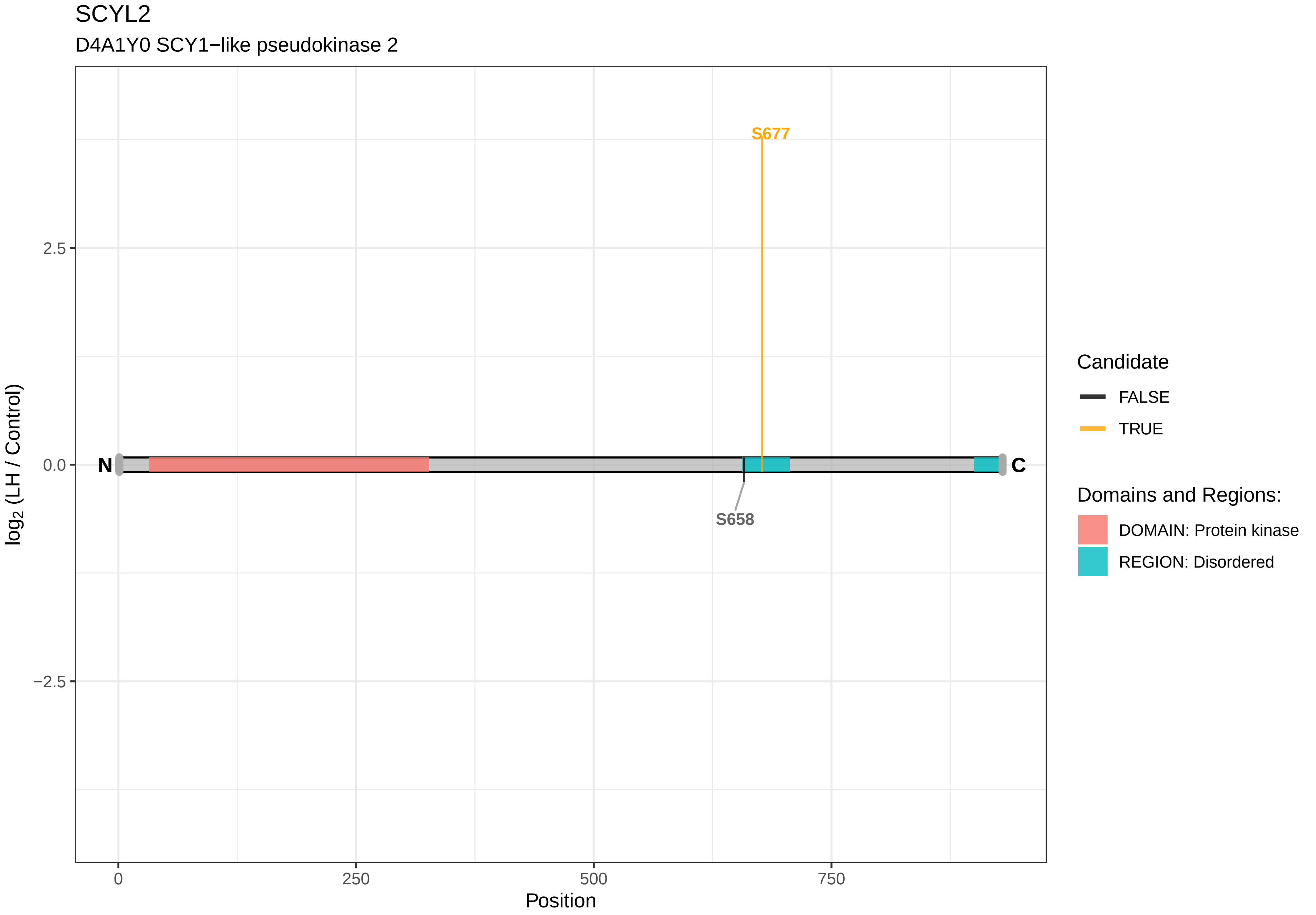

SCYL2
D4A1Y0 SCY1−like pseudokinase 2
S677
S658
N
C
2.5
0.0
−2.5
log2 (LH / Control)
0
250
500
750
Position
Candidate
FALSE
TRUE
Domains and Regions:
DOMAIN: Protein kinase
REGION: Disordered

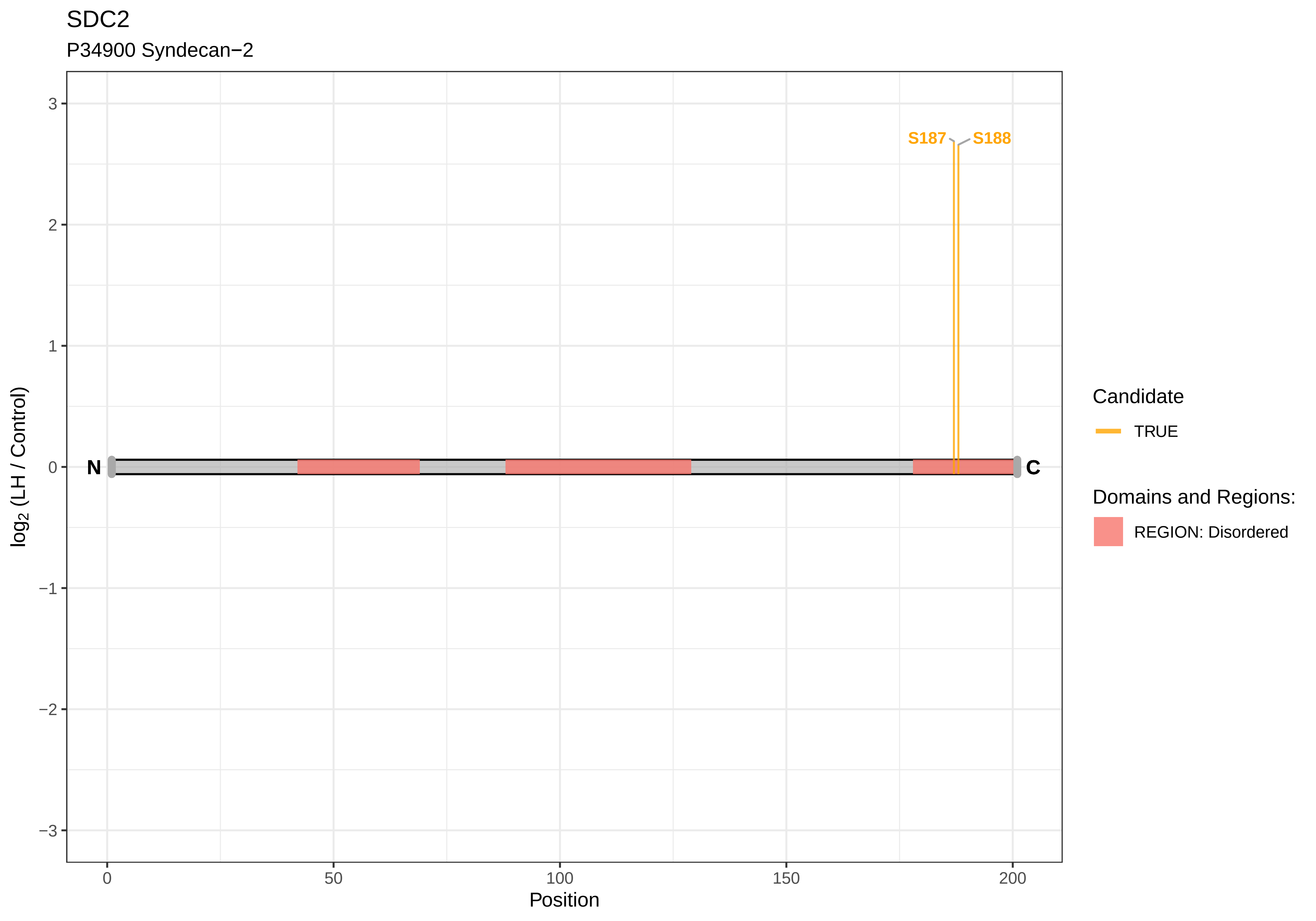
SDC2
P34900 Syndecan−2
S187
S188
N
C
3
2
1
0
−1
−2
−3
log2 (LH / Control)
0
50
100
150
200
Position
Candidate
TRUE
Domains and Regions:
REGION: Disordered

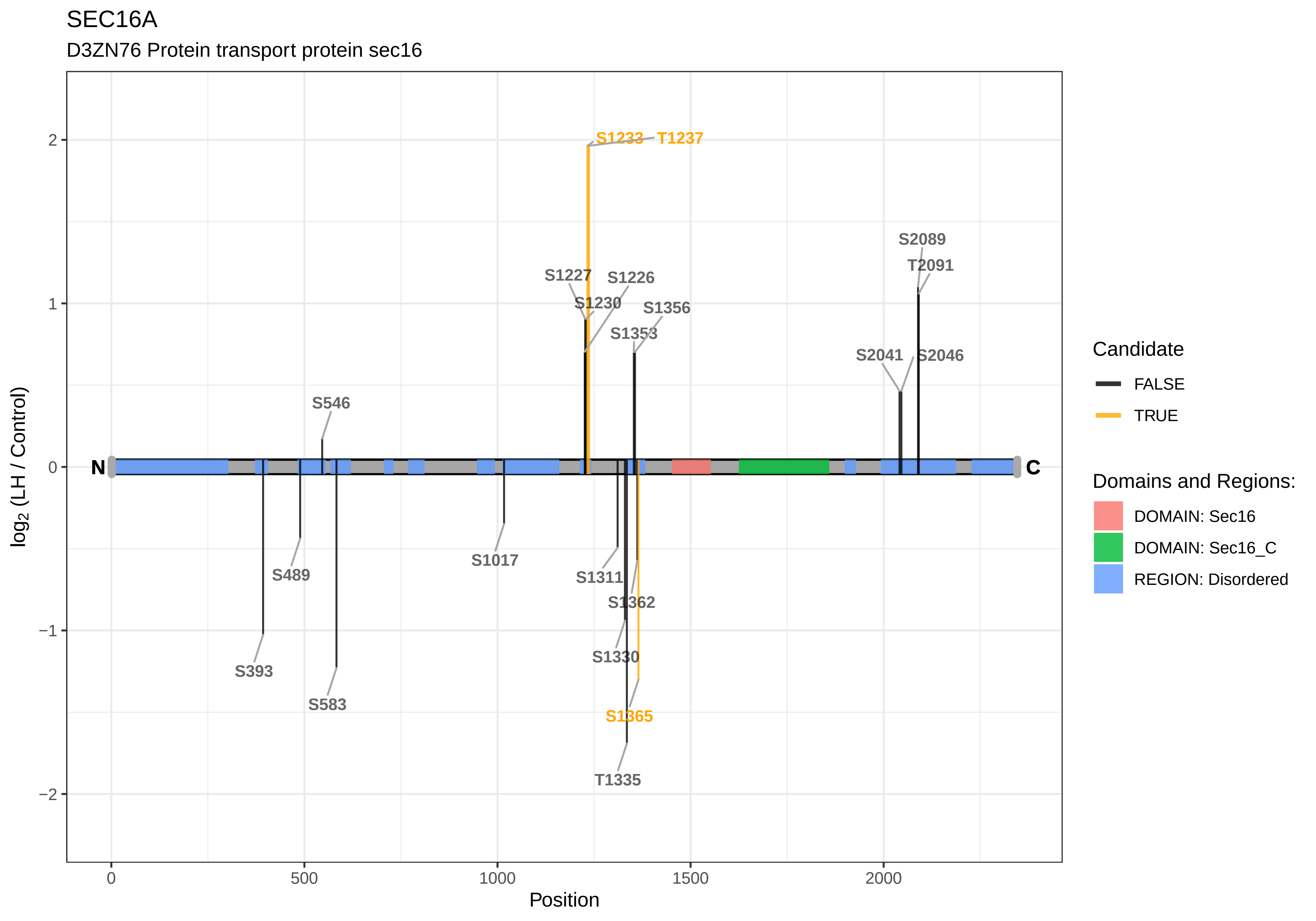
SEC16A
D3ZN76 Protein transport protein sec16
S1233
T1237
S2089
T2091
S1227
S1226
S1230
S1356
S1353
S2041
S2046
S546
N
C
S1017
S489
S1311
S1362
S393
S1330
S583
S1365
T1335
log2 (LH / Control)
Position
Candidate
FALSE
TRUE
Domains and Regions:
DOMAIN: Sec16
DOMAIN: Sec16_C
REGION: Disordered

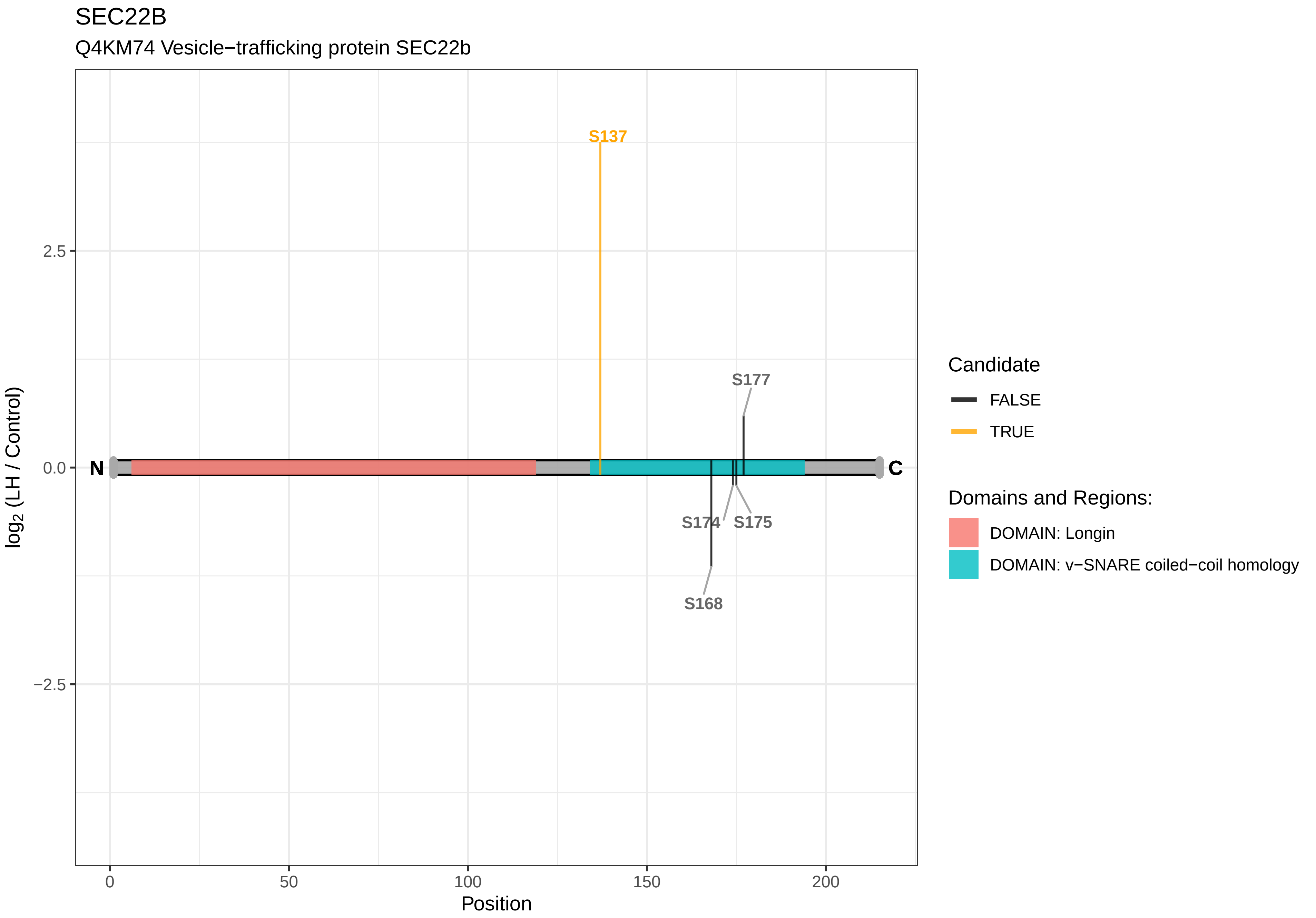
SEC22B
Q4KM74 Vesicle−trafficking protein SEC22b
S137
S177
S174
S175
S168
N
C
2.5
0.0
−2.5
log2 (LH / Control)
0
50
100
150
200
Position
Candidate
FALSE
TRUE
Domains and Regions:
DOMAIN: Longin
DOMAIN: v−SNARE coiled−coil homology

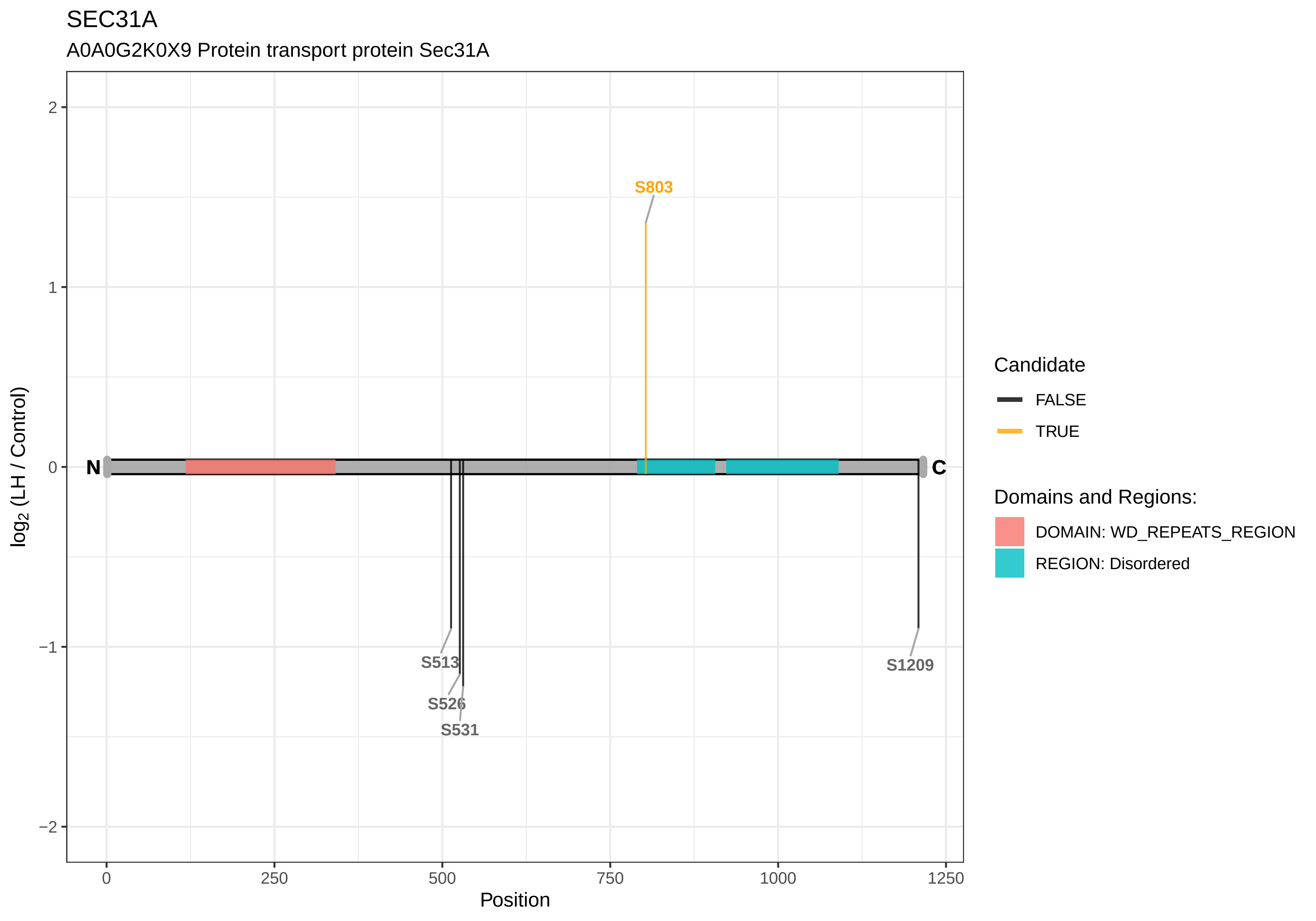
SEC31A
A0A0G2K0X9 Protein transport protein Sec31A
S803
S513
S526
S531
S1209
N
C
2
1
0
−1
−2
0
250
500
750
1000
1250
Position
log2 (LH / Control)
Candidate
FALSE
TRUE
Domains and Regions:
DOMAIN: WD_REPEATS_REGION
REGION: Disordered

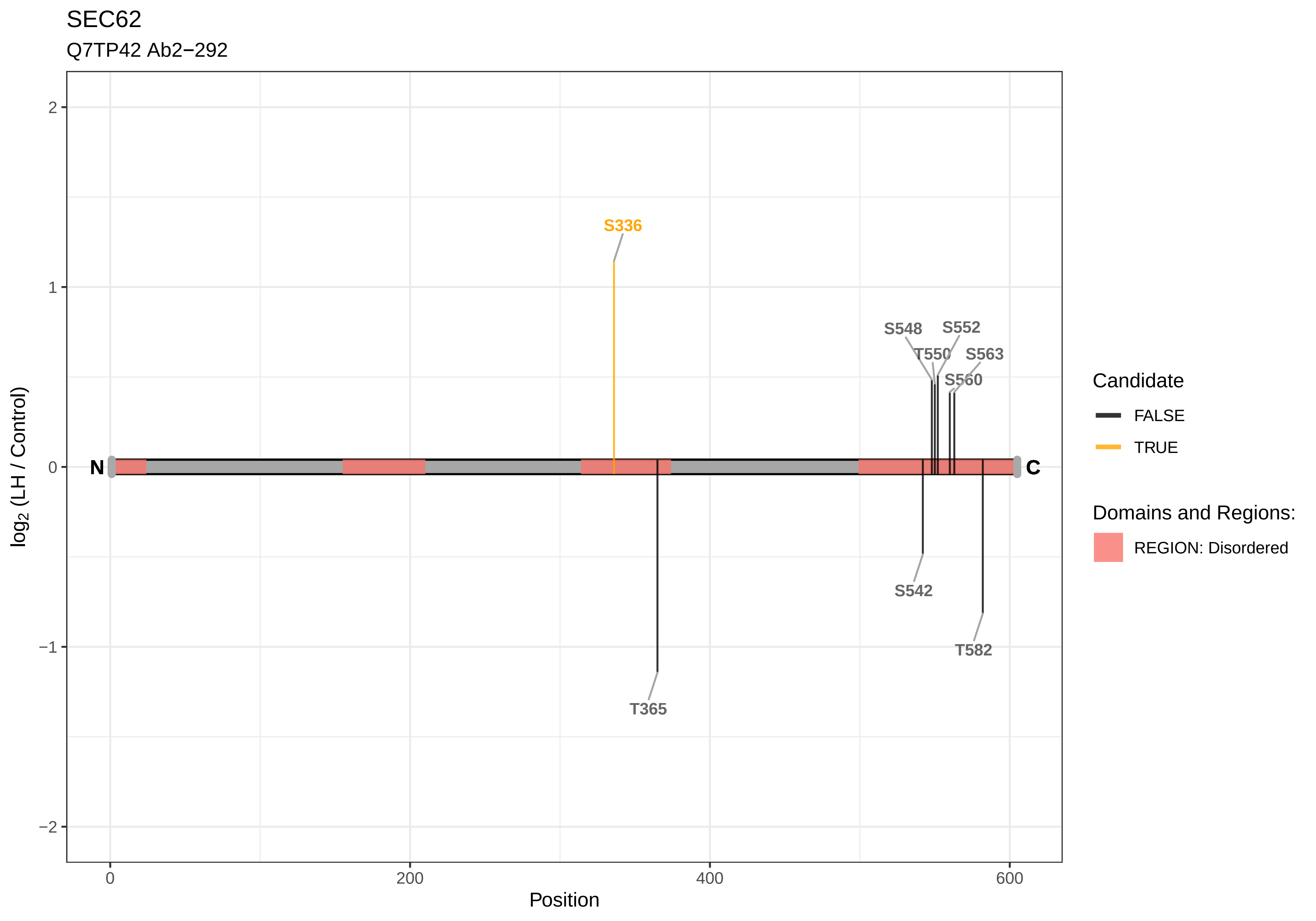
SEC62
Q7TP42 Ab2−292
S336
S548
S552
T550
S563
S560
N
C
S542
T582
T365
2
1
0
−1
−2
0
200
400
600
Position
log2 (LH / Control)
Candidate
FALSE
TRUE
Domains and Regions:
REGION: Disordered

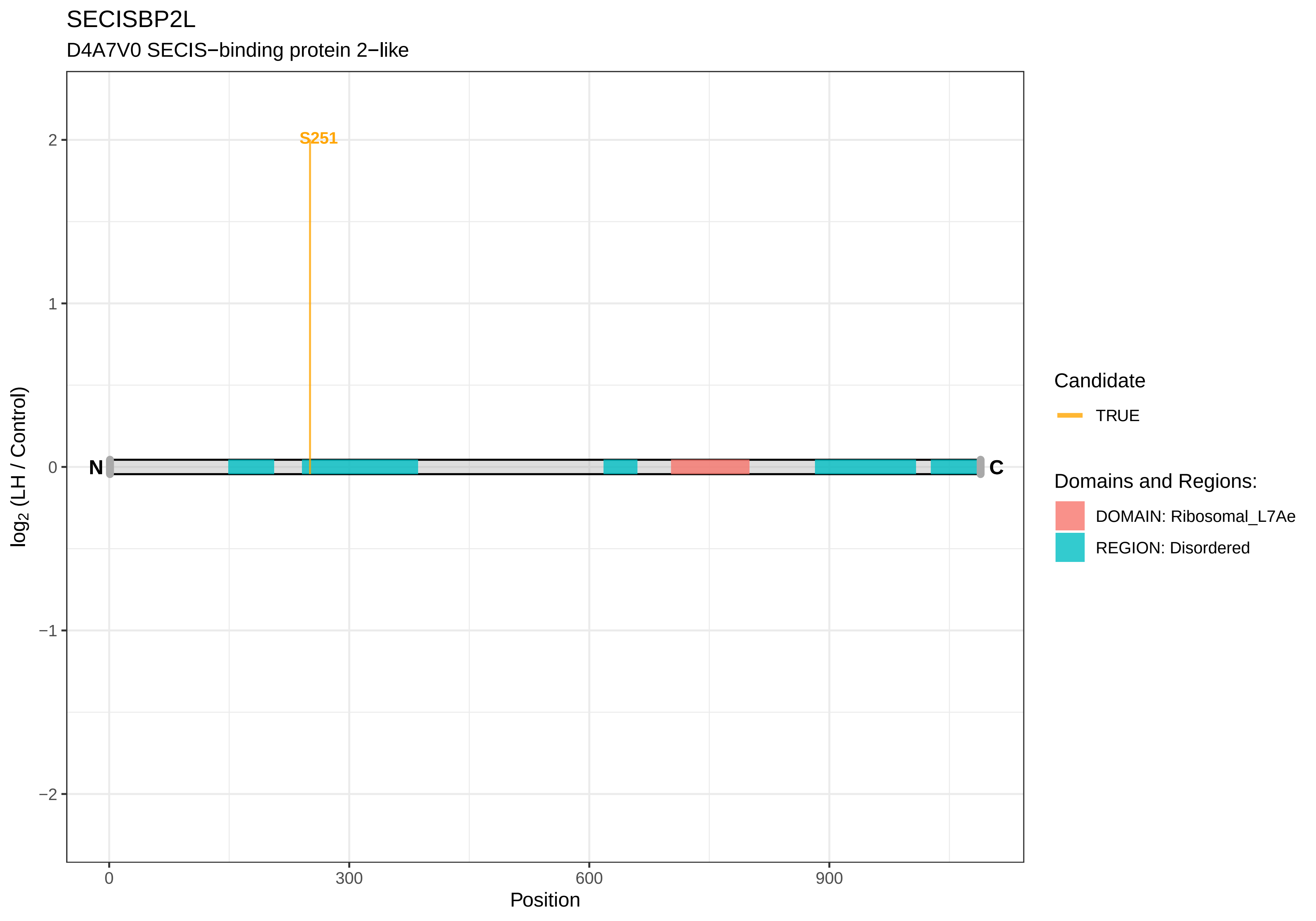

SECISBP2L
D4A7V0 SECIS−binding protein 2−like
S251
N
C
2
1
0
−1
−2
log2 (LH / Control)
0
300
600
900
Position
Candidate
TRUE
Domains and Regions:
DOMAIN: Ribosomal_L7Ae
REGION: Disordered

SELENOS

A0A0G2JSM1 Ratsg2, isoform CRA_b

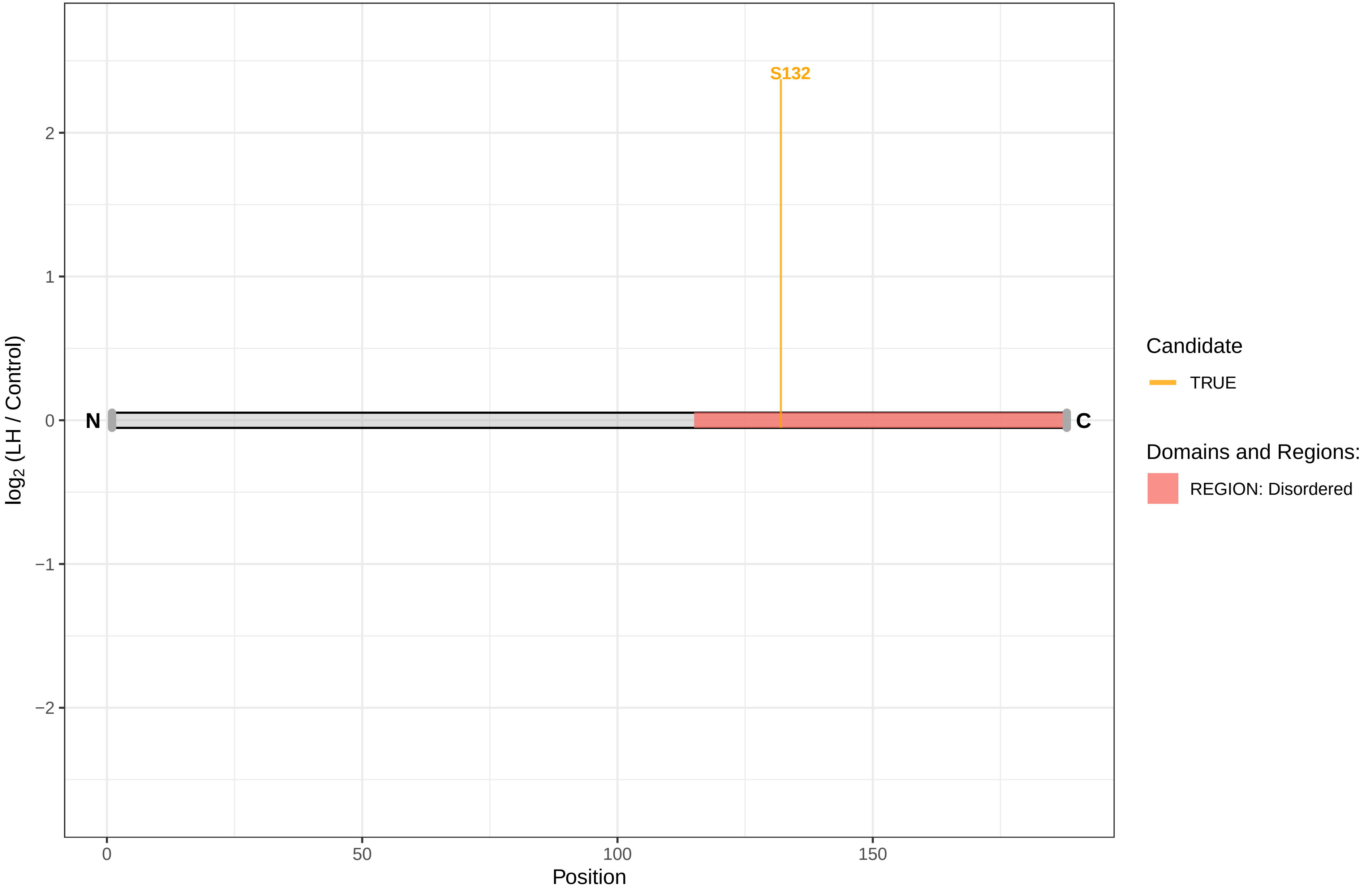

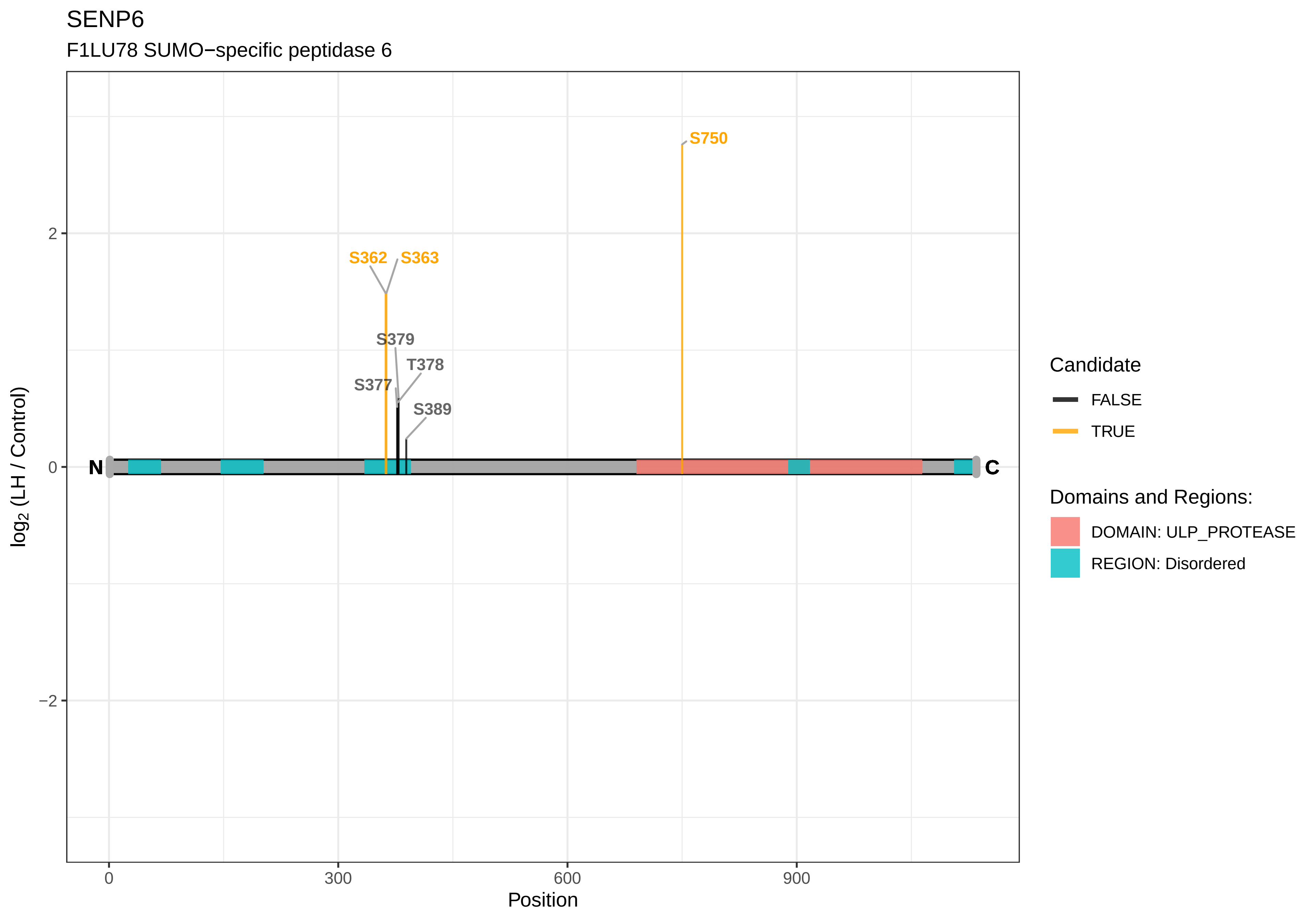

SENP6
F1LU78 SUMO−specific peptidase 6
S750
S362
S363
S379
T378
S377
S389
N
C
2
0
−2
0
300
600
900
Position
log2 (LH / Control)
Candidate
FALSE
TRUE
Domains and Regions:
DOMAIN: ULP_PROTEASE
REGION: Disordered

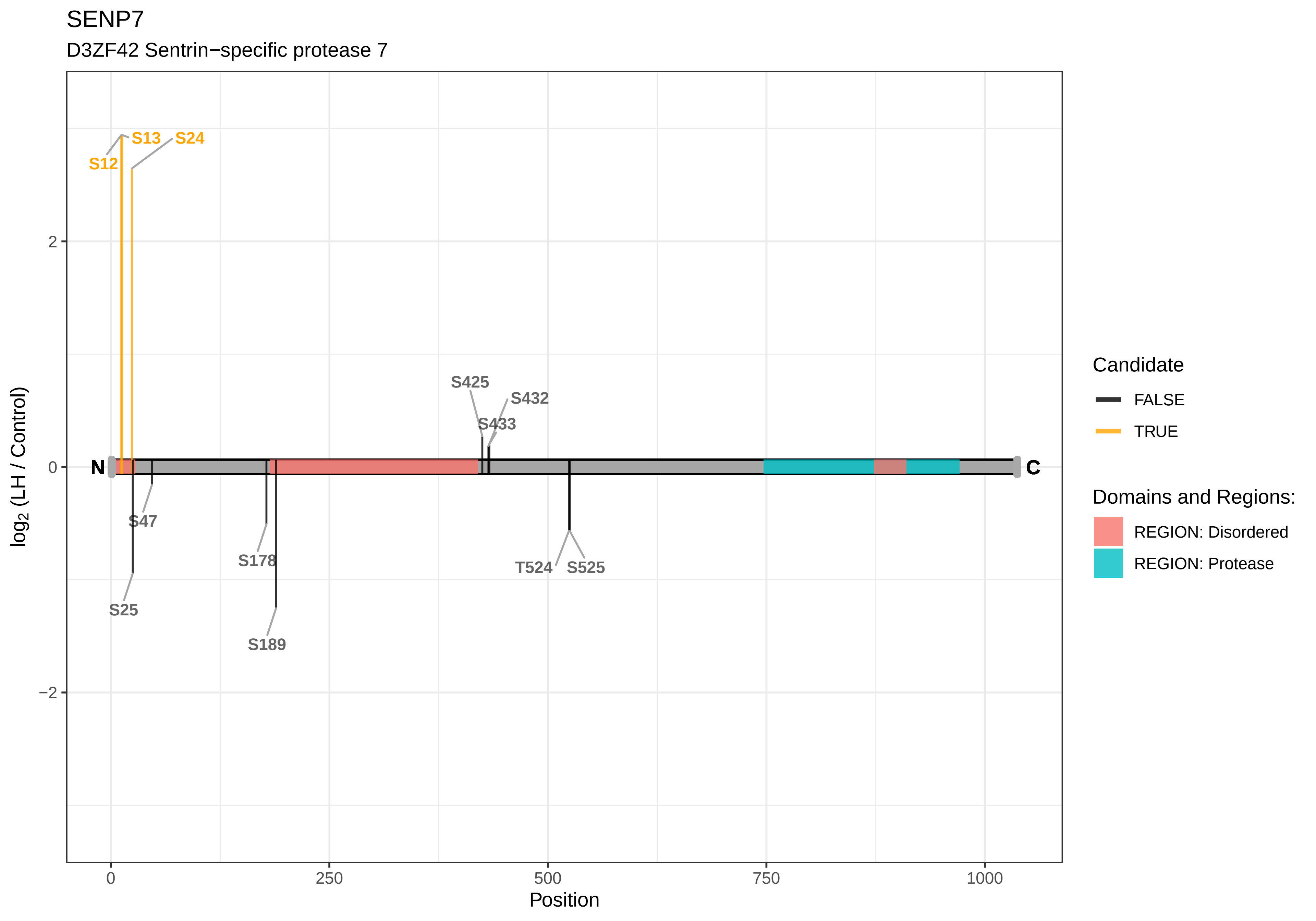

SENP7
D3ZF42 Sentrin−specific protease 7
S12
S13
S24
S425
S432
S433
N
C
S47
S178
S25
S189
T524
S525
2
0
−2
0
250
500
750
1000
Position
log2 (LH / Control)
Candidate
FALSE
TRUE
Domains and Regions:
REGION: Disordered
REGION: Protease

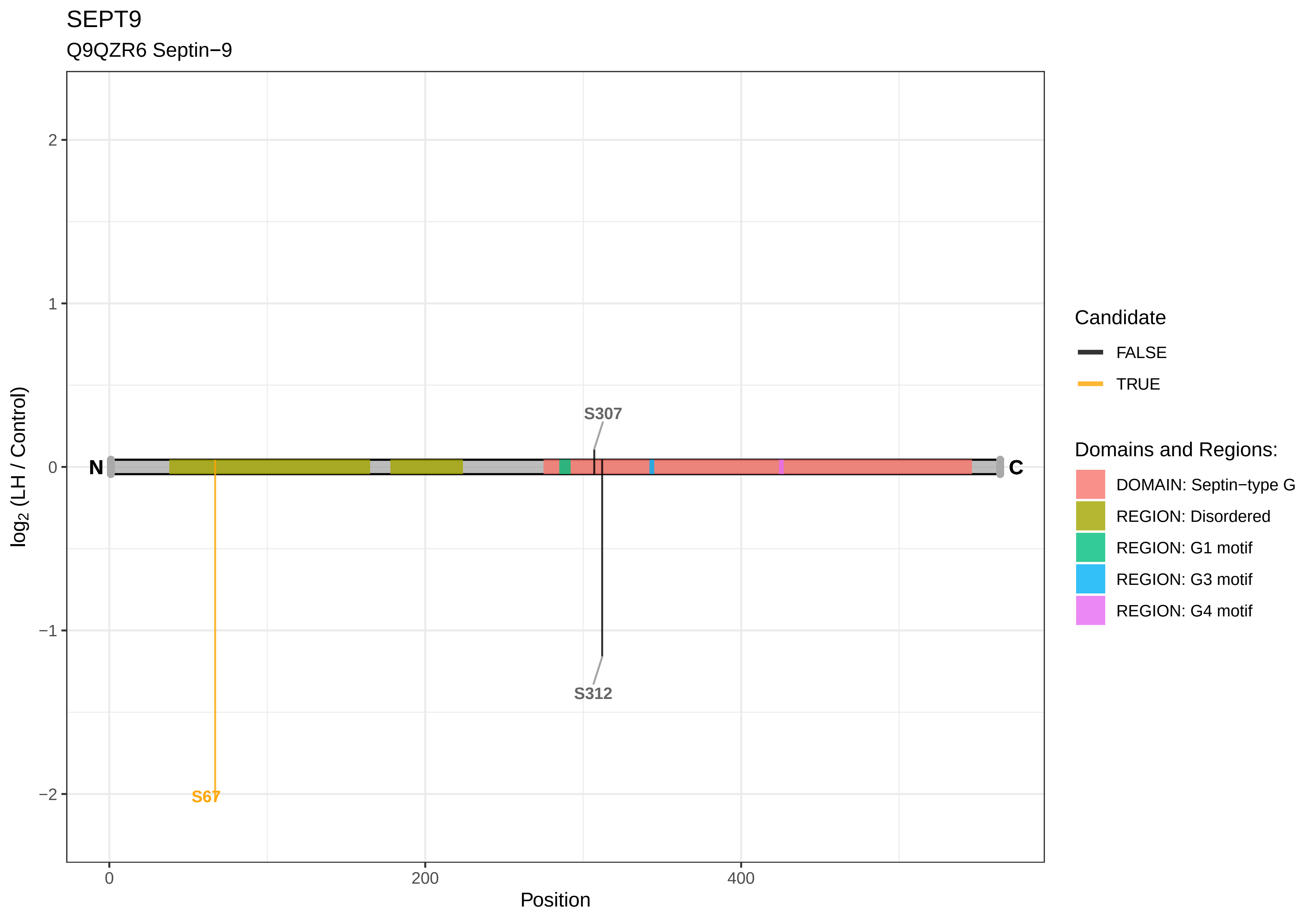

SEPT9
Q9QZR6 Septin−9
log2 (LH / Control)
2
1
0
−1
−2
Position
0
200
400
N
C
S307
S312
S67
Candidate
FALSE
TRUE
Domains and Regions:
DOMAIN: Septin−type G
REGION: Disordered
REGION: G1 motif
REGION: G3 motif
REGION: G4 motif

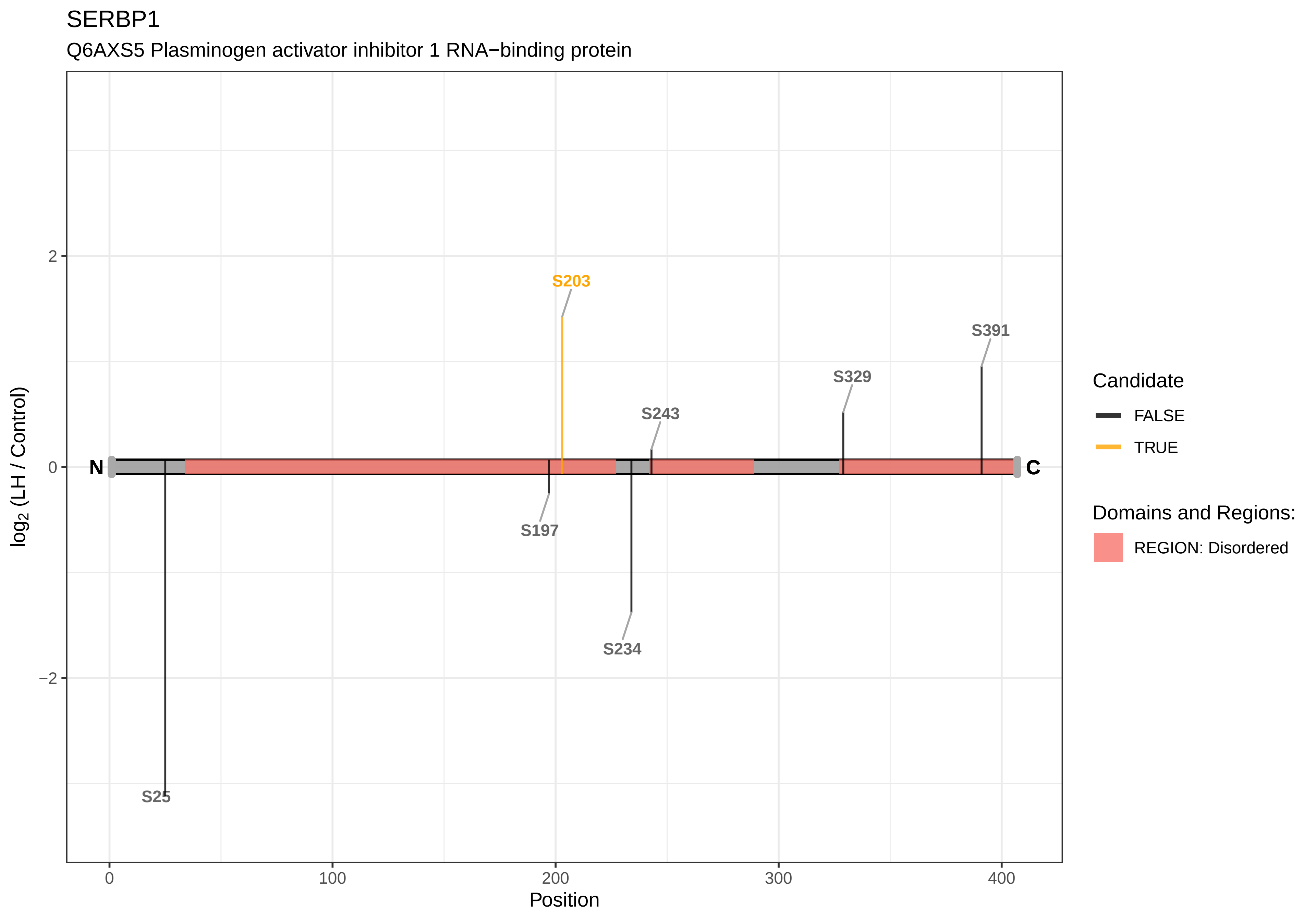

SERBP1
Q6AXS5 Plasminogen activator inhibitor 1 RNA−binding protein
S203
S391
S329
S243
N
C
S197
S234
S25
2
0
−2
0
100
200
300
400
Position
log2 (LH / Control)
Candidate
FALSE
TRUE
Domains and Regions:
REGION: Disordered

### SET

Q63945 Protein SET

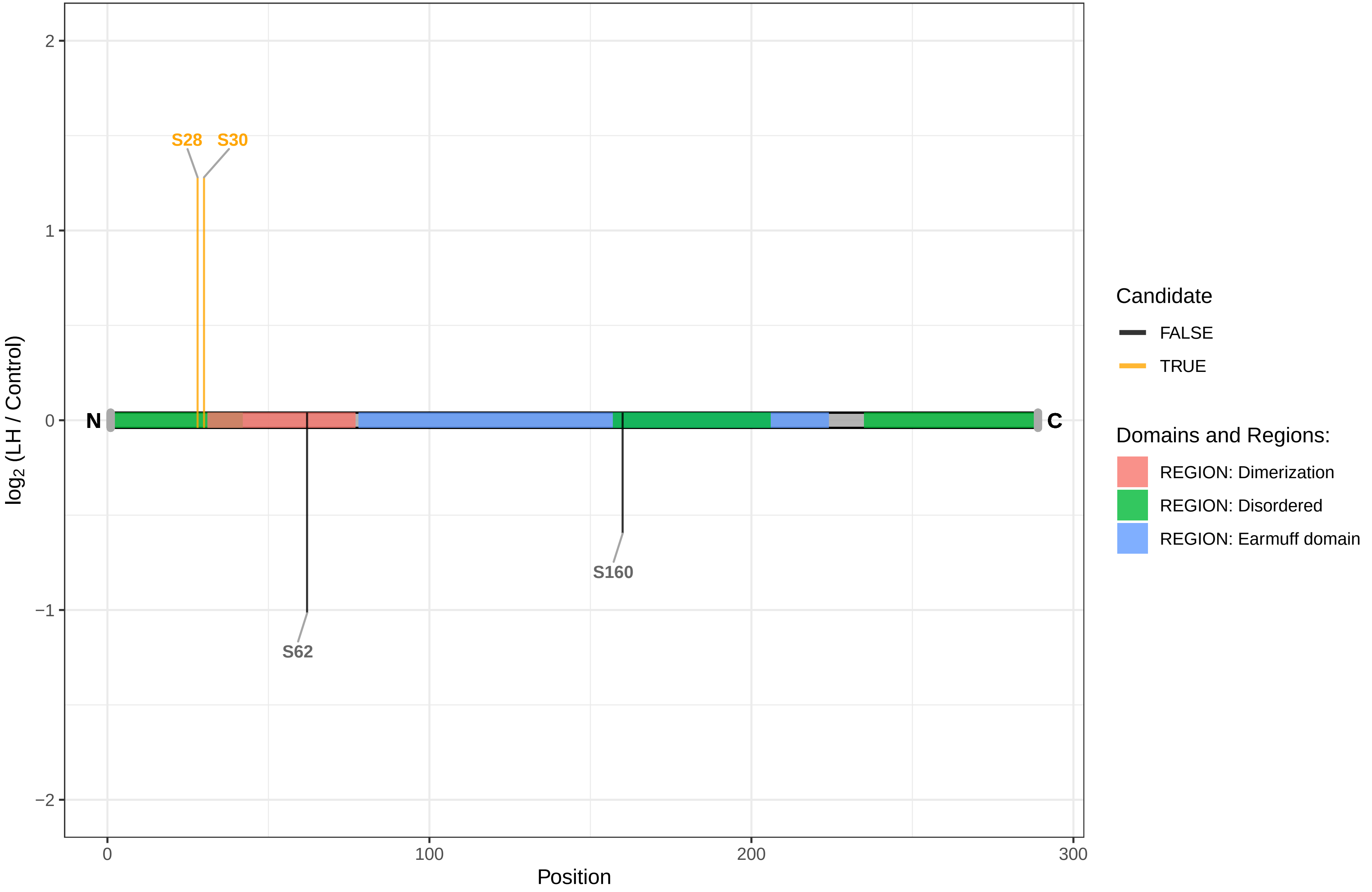

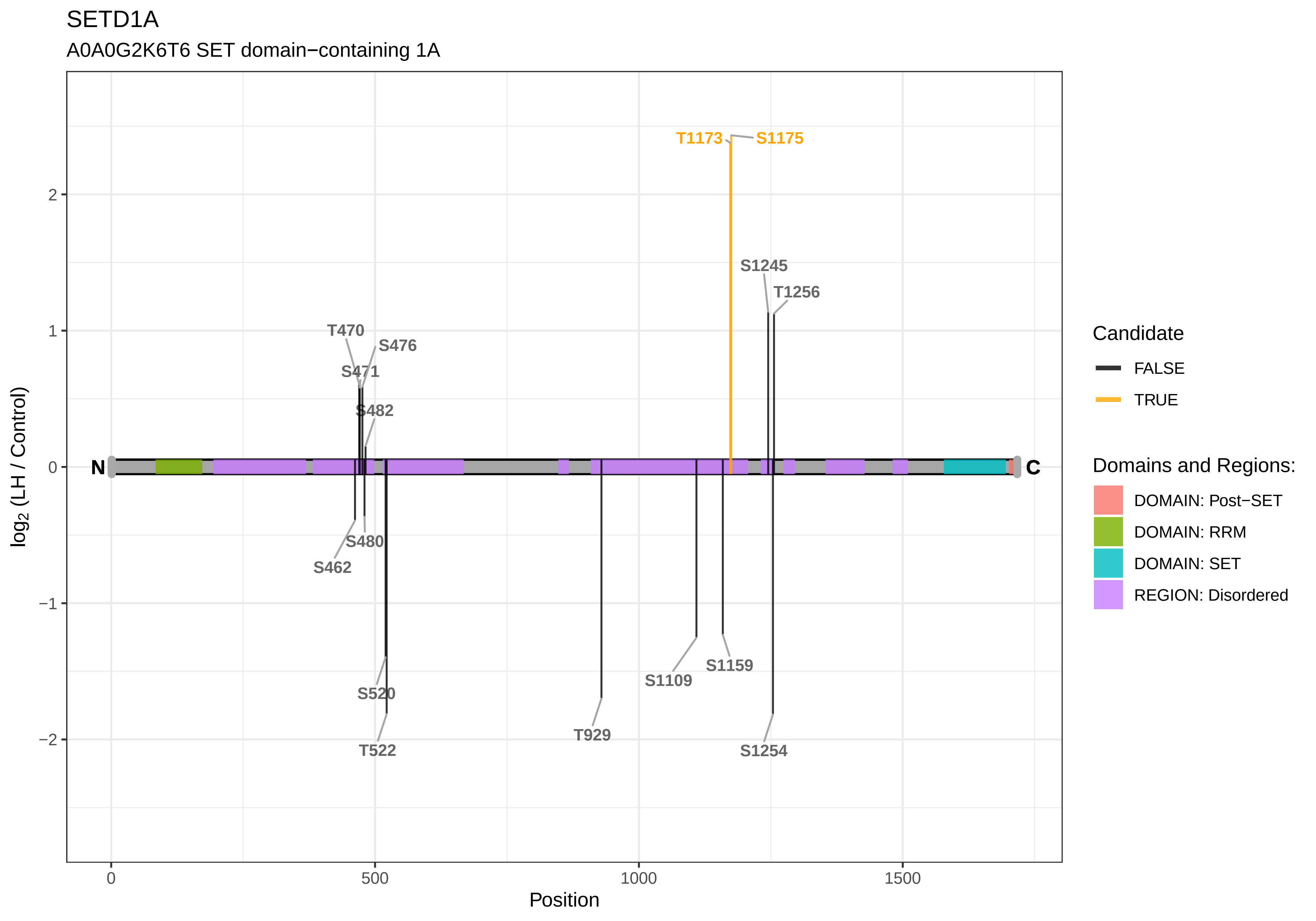
SETD1A
A0A0G2K6T6 SET domain−containing 1A
T1173
S1175
S1245
T1256
T470
S476
S471
S482
S480
S462
S520
T522
T929
S1109
S1159
S1254
N
C
2
1
0
−1
−2
0
500
1000
1500
Position
log2 (LH / Control)
Candidate
FALSE
TRUE
Domains and Regions:
DOMAIN: Post−SET
DOMAIN: RRM
DOMAIN: SET
REGION: Disordered

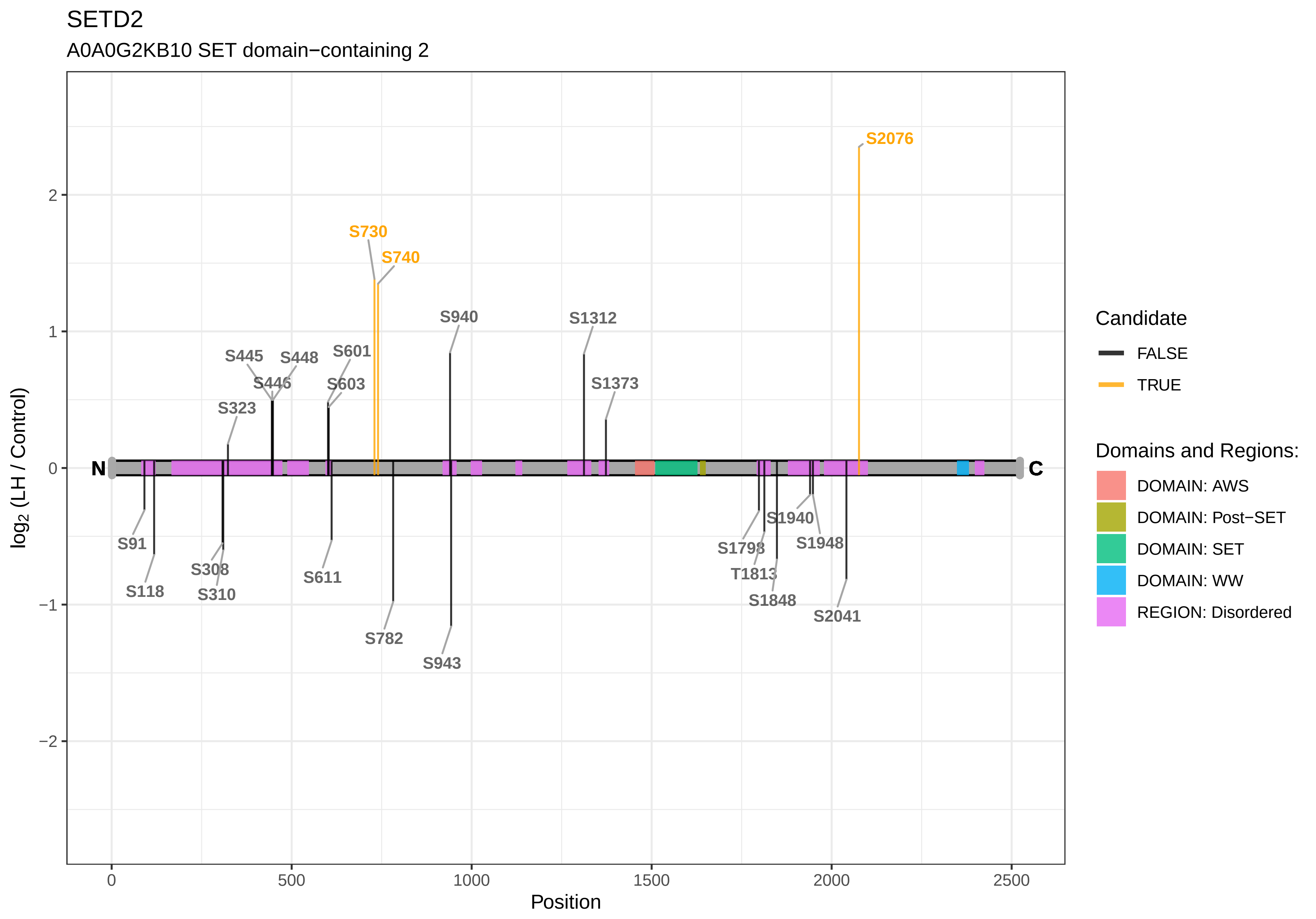

SETD2
A0A0G2KB10 SET domain−containing 2
S2076
S730
S740
S940
S1312
S445
S448
S601
S446
S603
S1373
S323
N
C
S91
S118
S308
S310
S611
S782
S943
S1940
S1798
S1948
T1813
S1848
S2041
log2 (LH / Control)
Position
Candidate
FALSE
TRUE
Domains and Regions:
DOMAIN: AWS
DOMAIN: Post−SET
DOMAIN: SET
DOMAIN: WW
REGION: Disordered

### SFR1

A0A0H2UHK0 Swi5−dependent recombination DNA repair protein 1 homolog

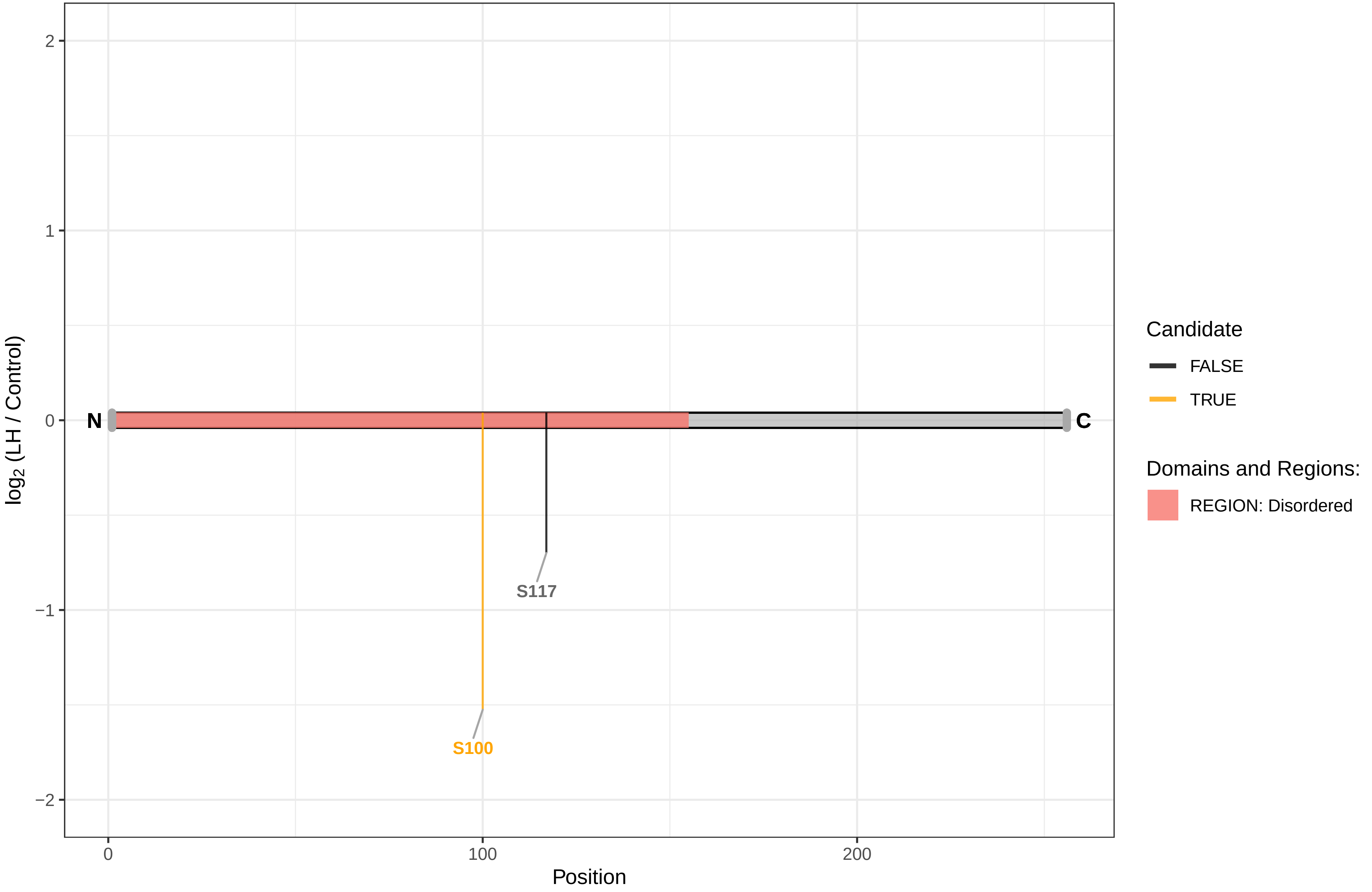

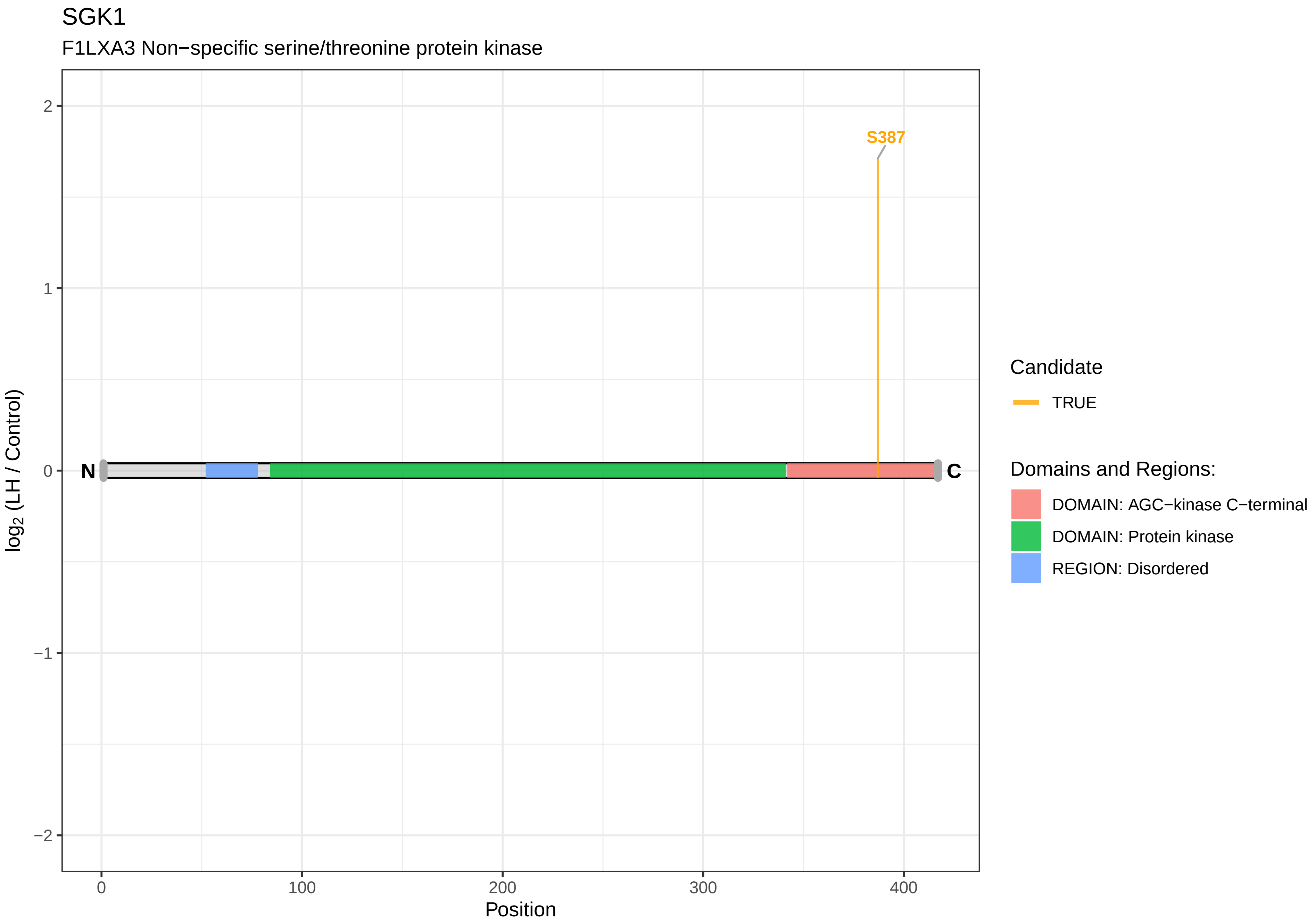
SGK1
F1LXA3 Non−specific serine/threonine protein kinase
S387
N
C
2
1
0
−1
−2
log2 (LH / Control)
0
100
200
300
400
Position
Candidate
TRUE
Domains and Regions:
DOMAIN: AGC−kinase C−terminal
DOMAIN: Protein kinase
REGION: Disordered

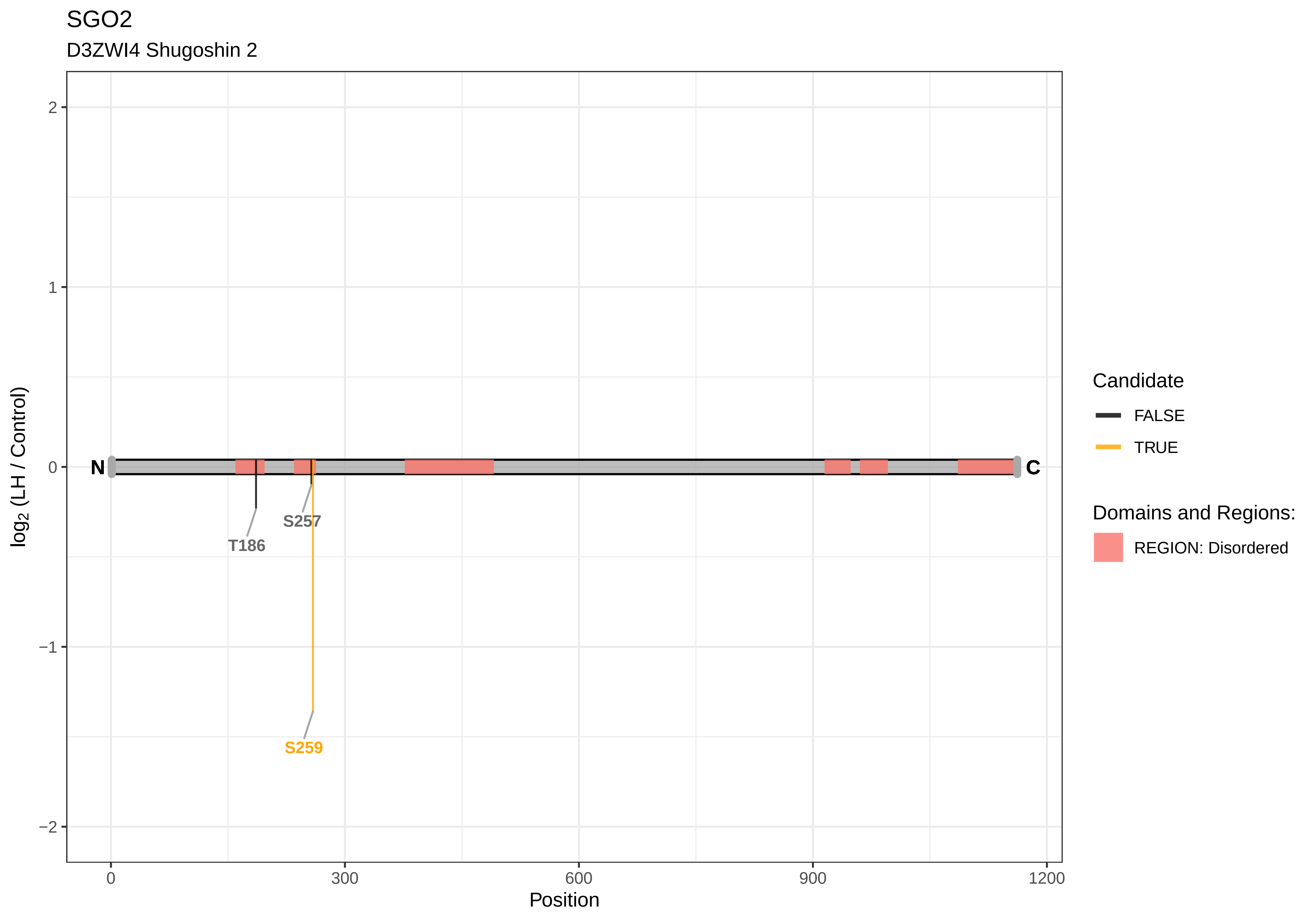
SGO2
D3ZWI4 Shugoshin 2
2
1
0
−1
−2
log2 (LH / Control)
N
C
T186
S257
S259
0
300
600
900
1200
Position
Candidate
FALSE
TRUE
Domains and Regions:
REGION: Disordered

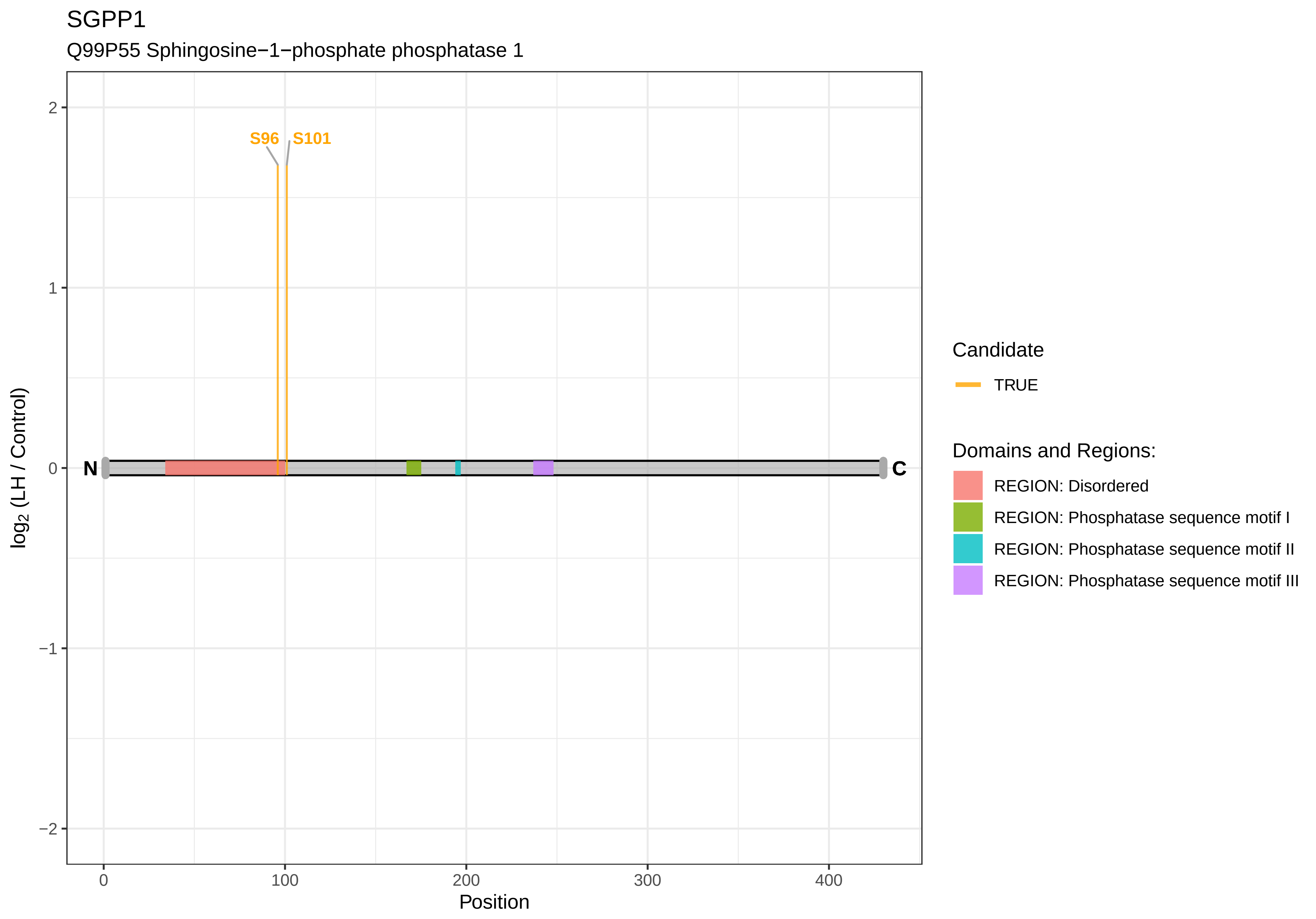

SGPP1
Q99P55 Sphingosine−1−phosphate phosphatase 1
S96
S101
N
C
2
1
0
−1
−2
0
100
200
300
400
Position
log2 (LH / Control)
Candidate
TRUE
Domains and Regions:
REGION: Disordered
REGION: Phosphatase sequence motif I
REGION: Phosphatase sequence motif II
REGION: Phosphatase sequence motif III

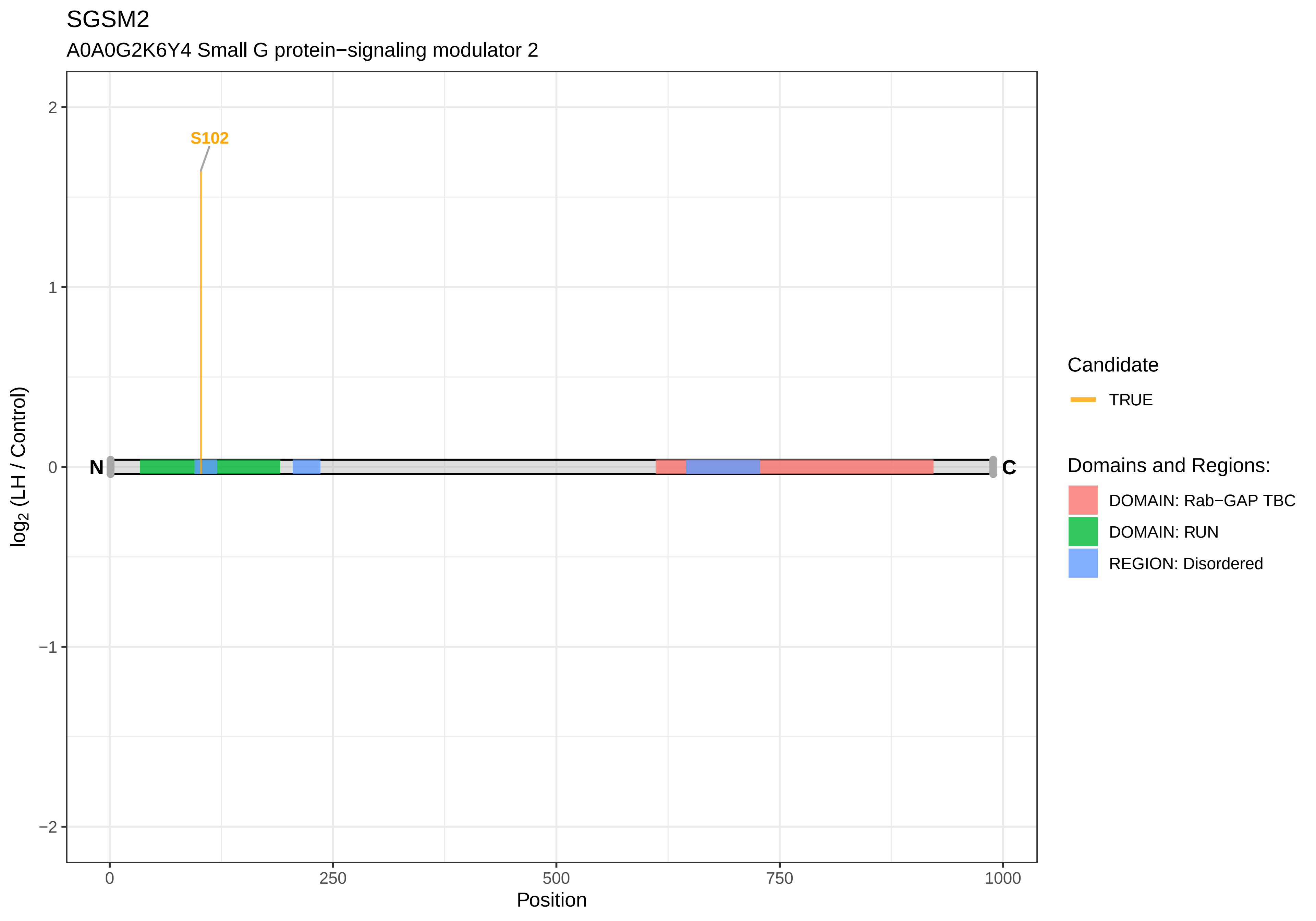

SGSM2
A0A0G2K6Y4 Small G protein−signaling modulator 2
S102
N
C
2
1
0
−1
−2
0
250
500
750
1000
Position
log2 (LH / Control)
Candidate
TRUE
Domains and Regions:
DOMAIN: Rab−GAP TBC
DOMAIN: RUN
REGION: Disordered

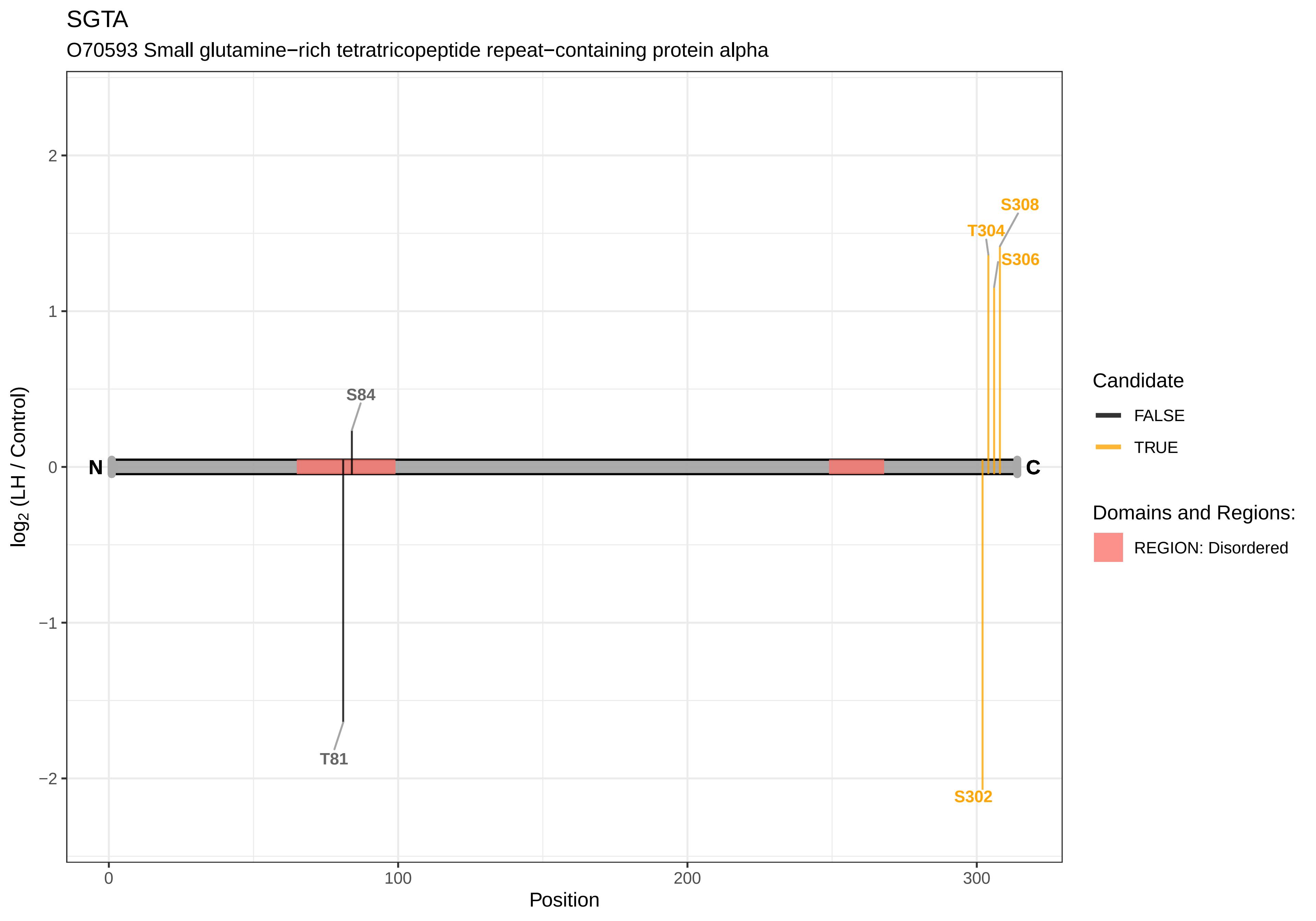

SGTA
O70593 Small glutamine−rich tetratricopeptide repeat−containing protein alpha
S308
T304
S306
S84
T81
S302
N
C
2
1
0
−1
−2
0
100
200
300
Position
log2 (LH / Control)
Candidate
FALSE
TRUE
Domains and Regions:
REGION: Disordered

SGTB

Q80W98 Small glutamine−rich tetratricopeptide repeat−containing protein beta

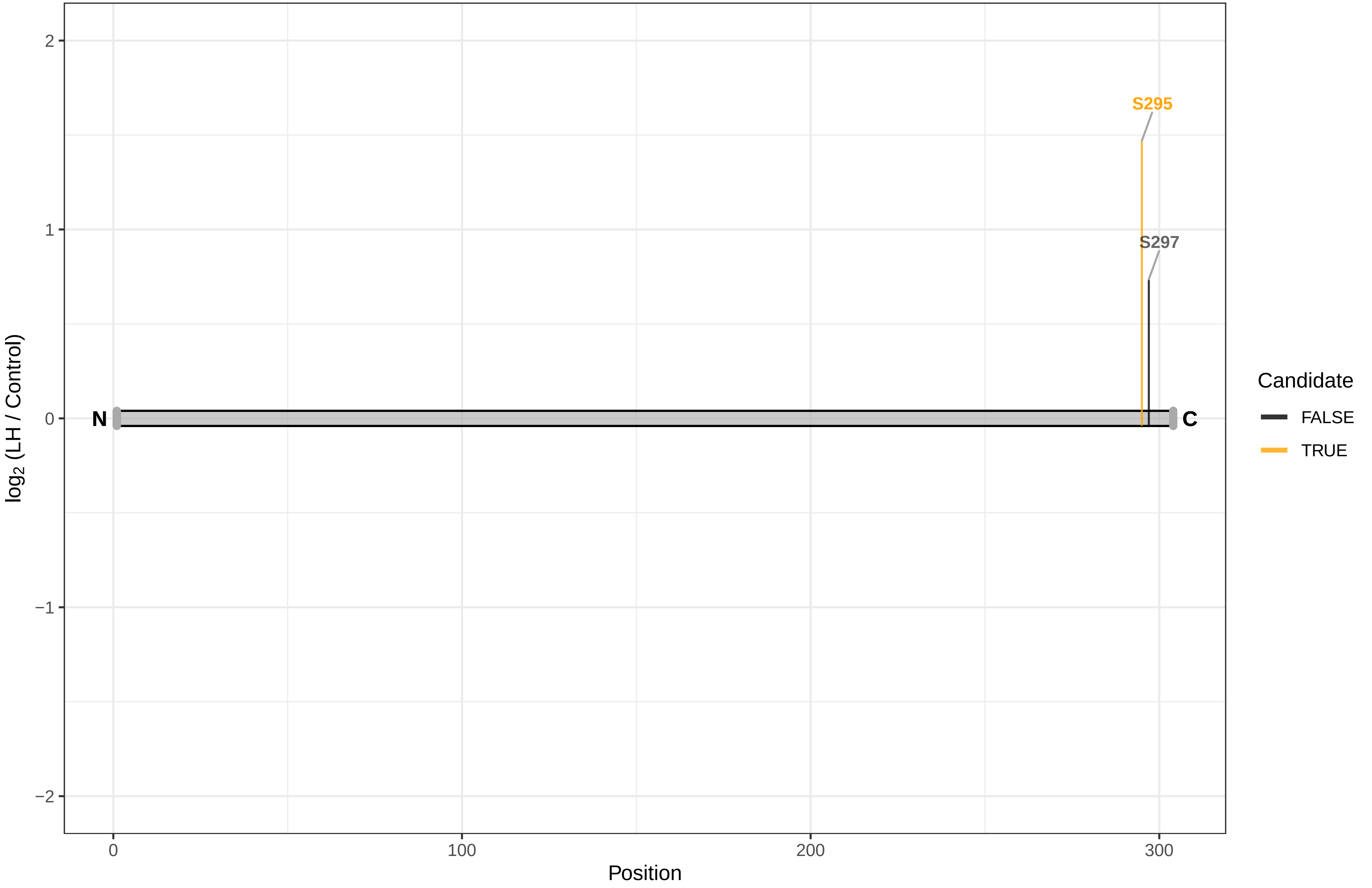

SH2B1

M0R617 SH2−B PH domain containing signaling mediator 1, isoform CRA_a

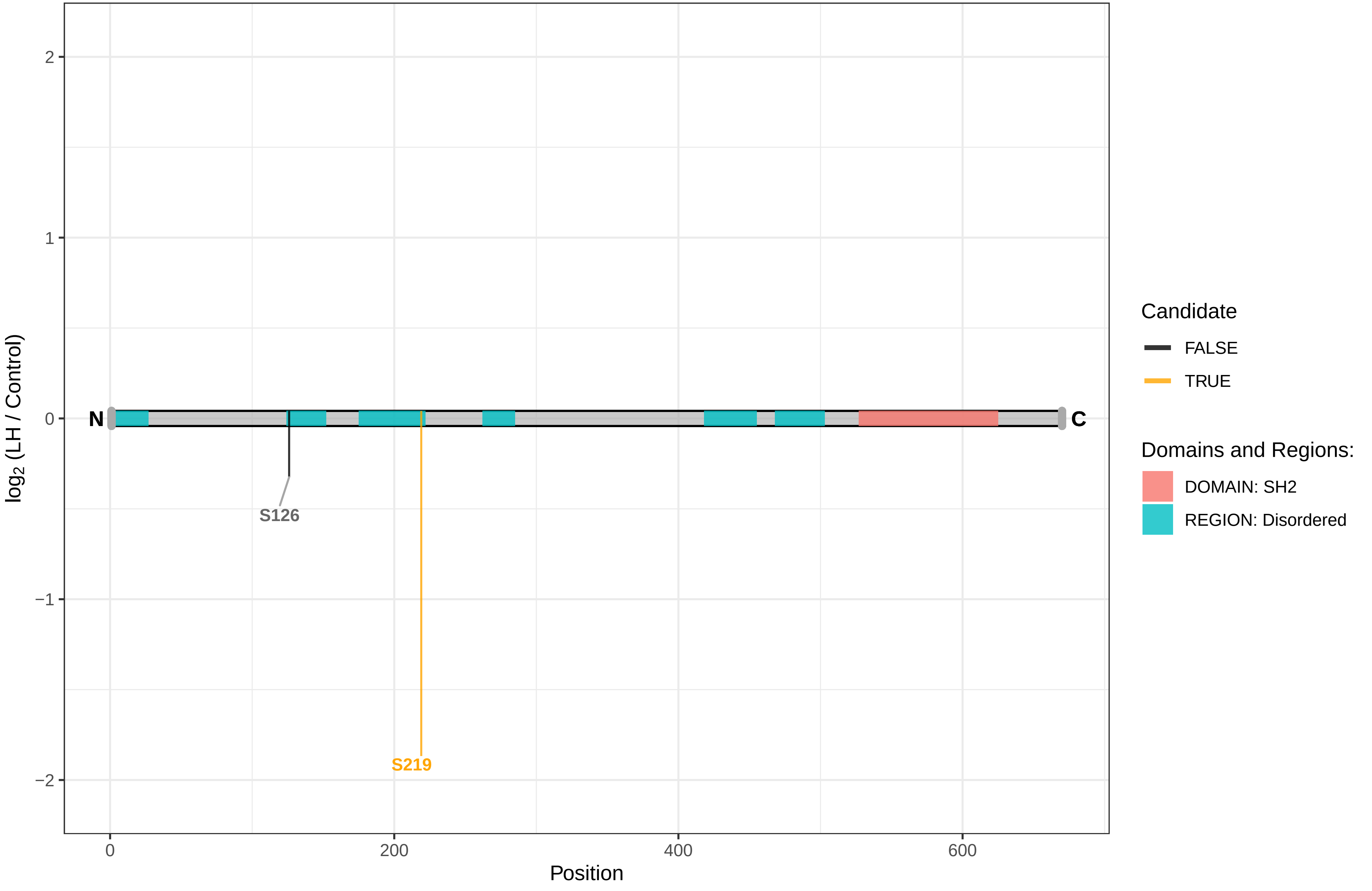

SH2D4A

Q6AYC8 SH2 domain−containing protein 4A

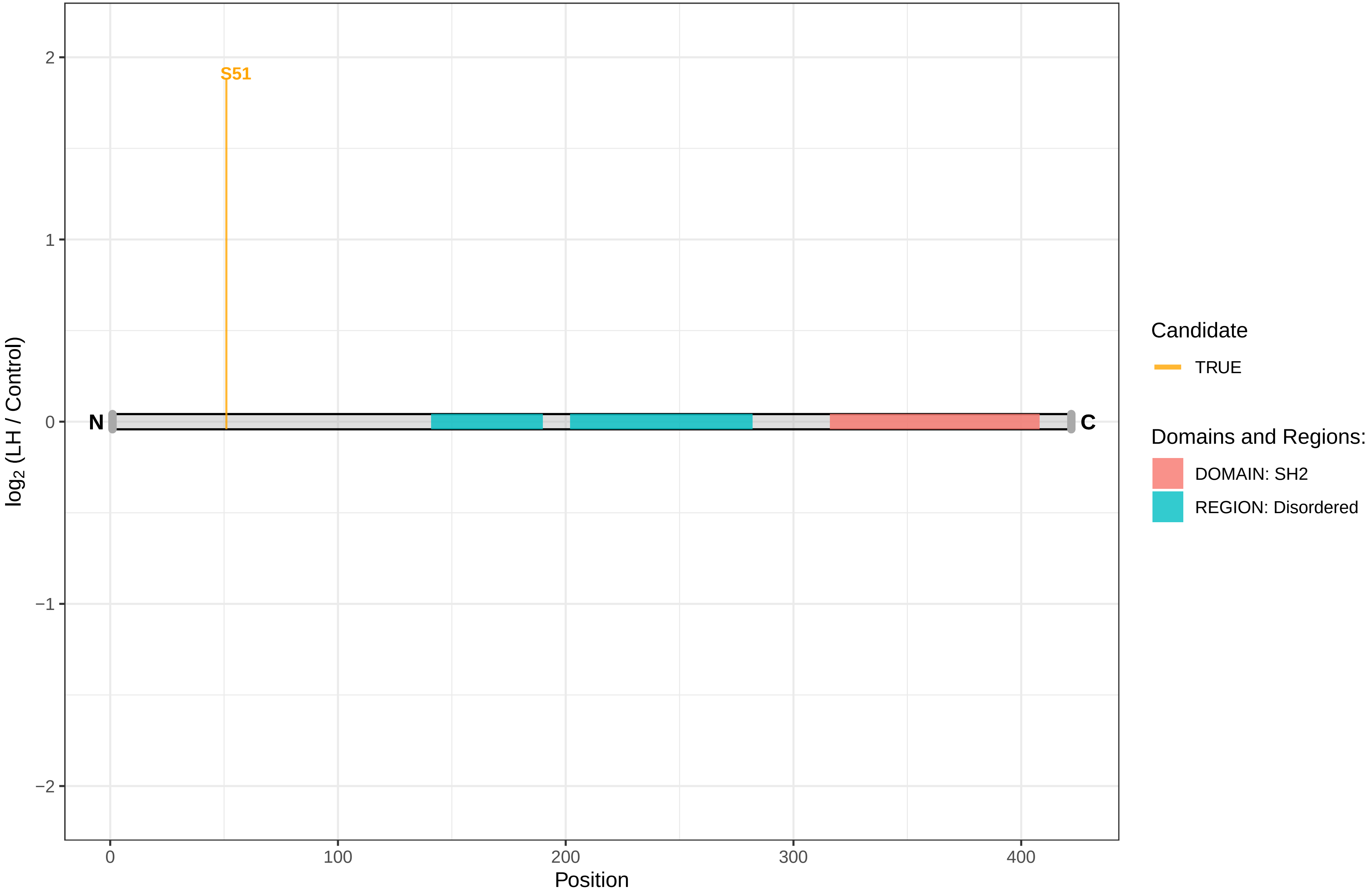

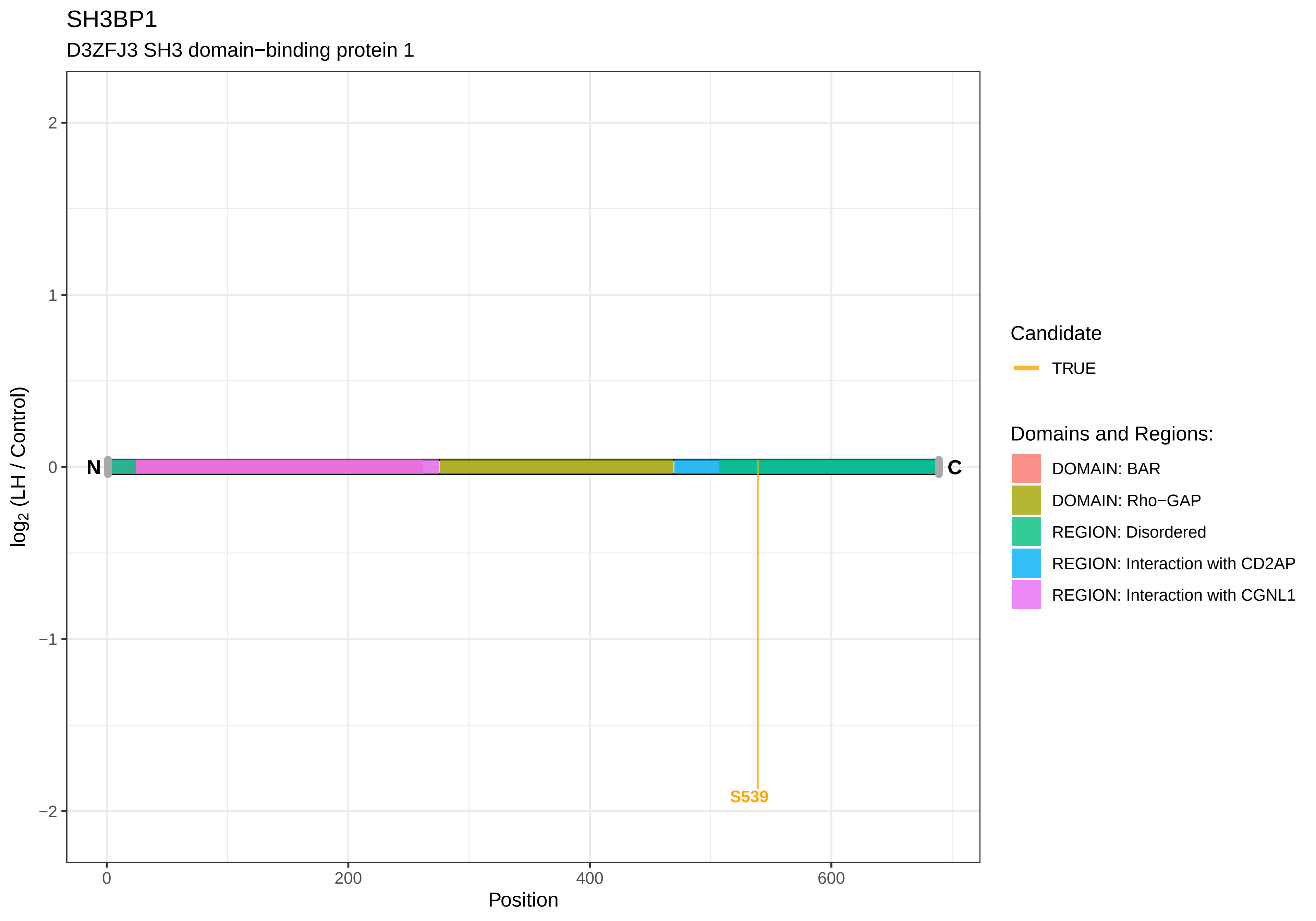

SH3BP1
D3ZFJ3 SH3 domain−binding protein 1
2
1
0
−1
−2
log2 (LH / Control)
N
C
S539
0
200
400
600
Position
Candidate
TRUE
Domains and Regions:
DOMAIN: BAR
DOMAIN: Rho−GAP
REGION: Disordered
REGION: Interaction with CD2AP
REGION: Interaction with CGNL1

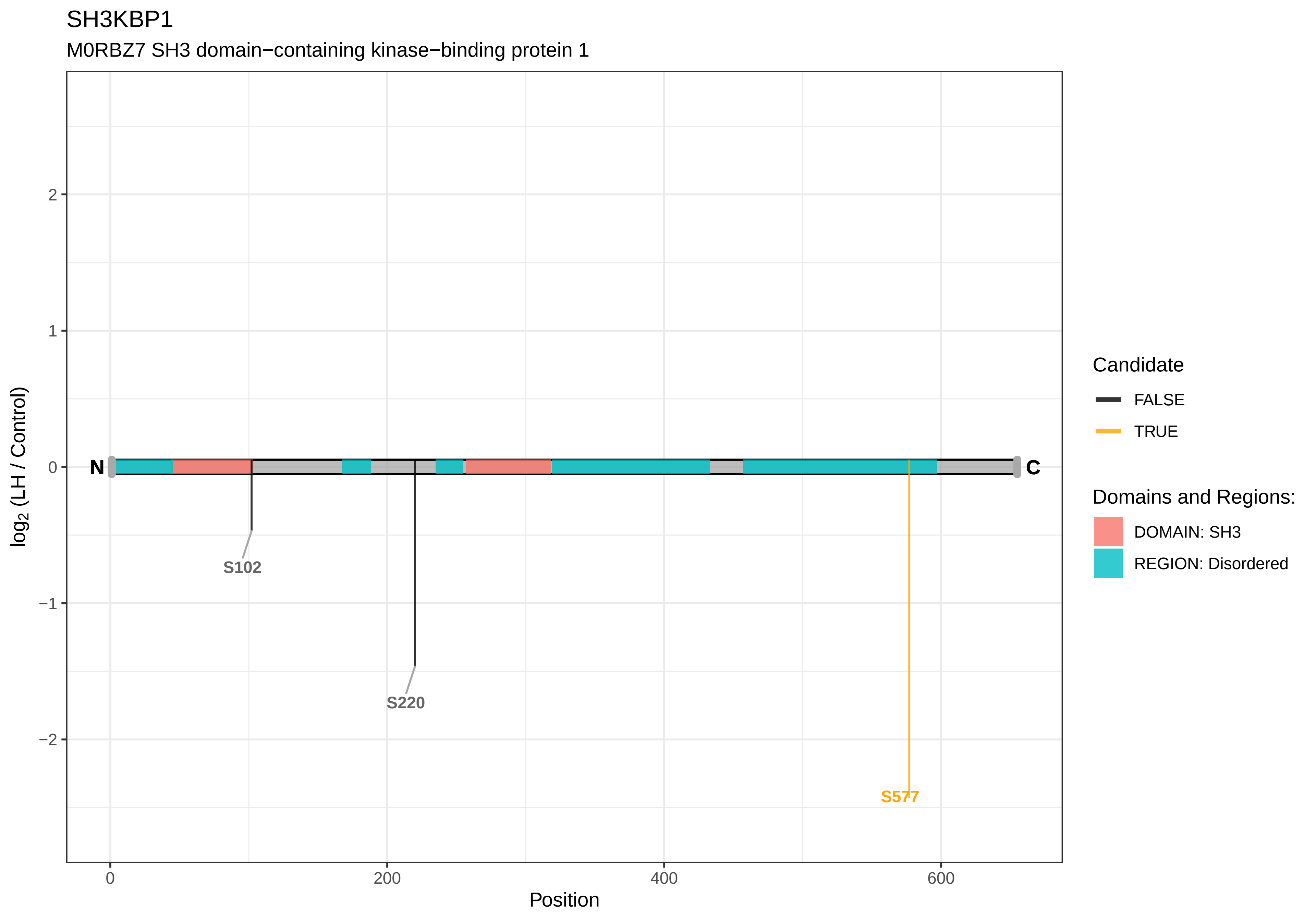
SH3KBP1
M0RBZ7 SH3 domain−containing kinase−binding protein 1
log2 (LH / Control)
2
1
0
−1
−2
N
C
S102
S220
S577
0
200
400
600
Position
Candidate
FALSE
TRUE
Domains and Regions:
DOMAIN: SH3
REGION: Disordered

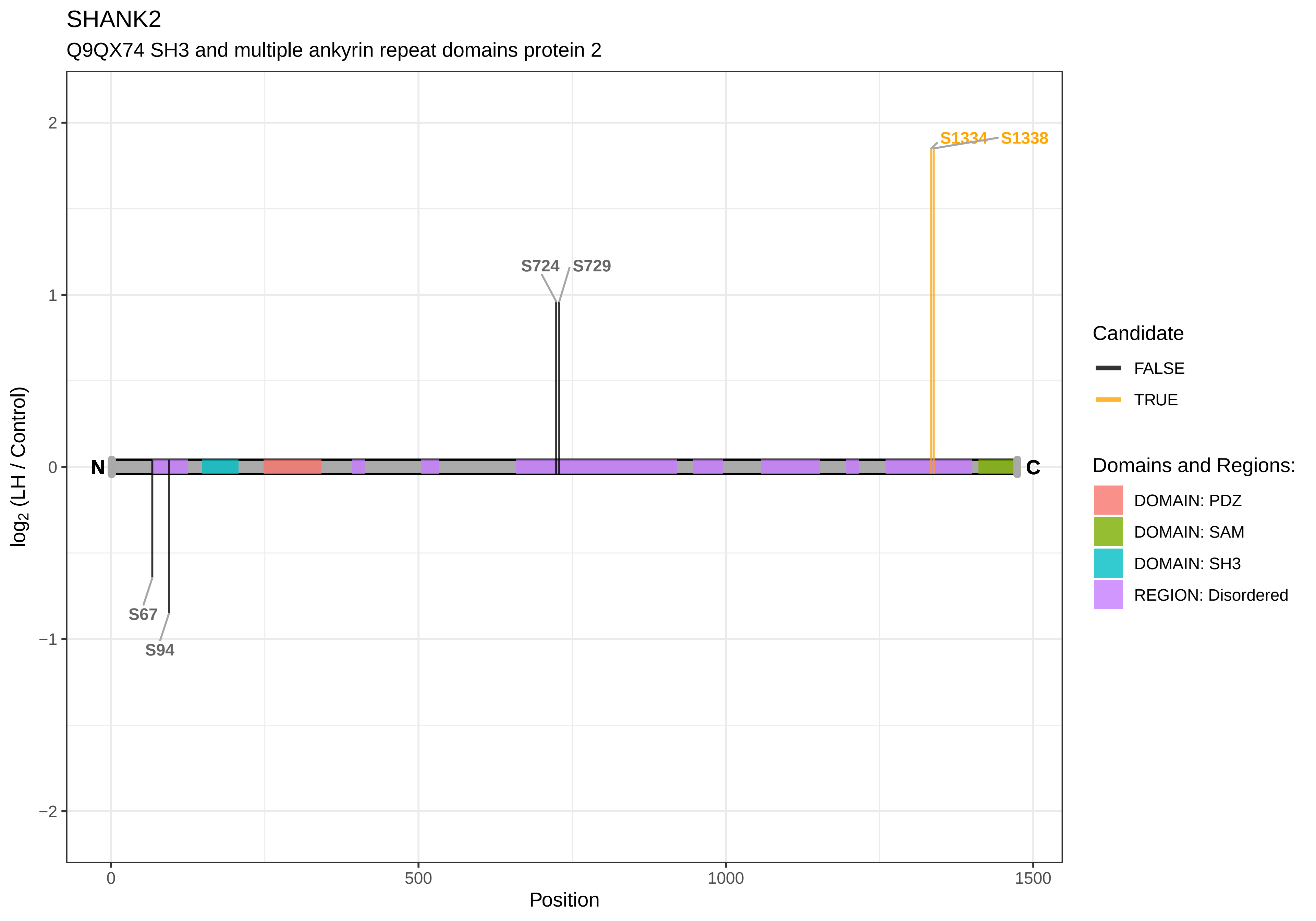

SHANK2
Q9QX74 SH3 and multiple ankyrin repeat domains protein 2
S1334
S1338
S724
S729
S67
S94
N
C
2
1
0
−1
−2
log2 (LH / Control)
0
500
1000
1500
Position
Candidate
FALSE
TRUE
Domains and Regions:
DOMAIN: PDZ
DOMAIN: SAM
DOMAIN: SH3
REGION: Disordered

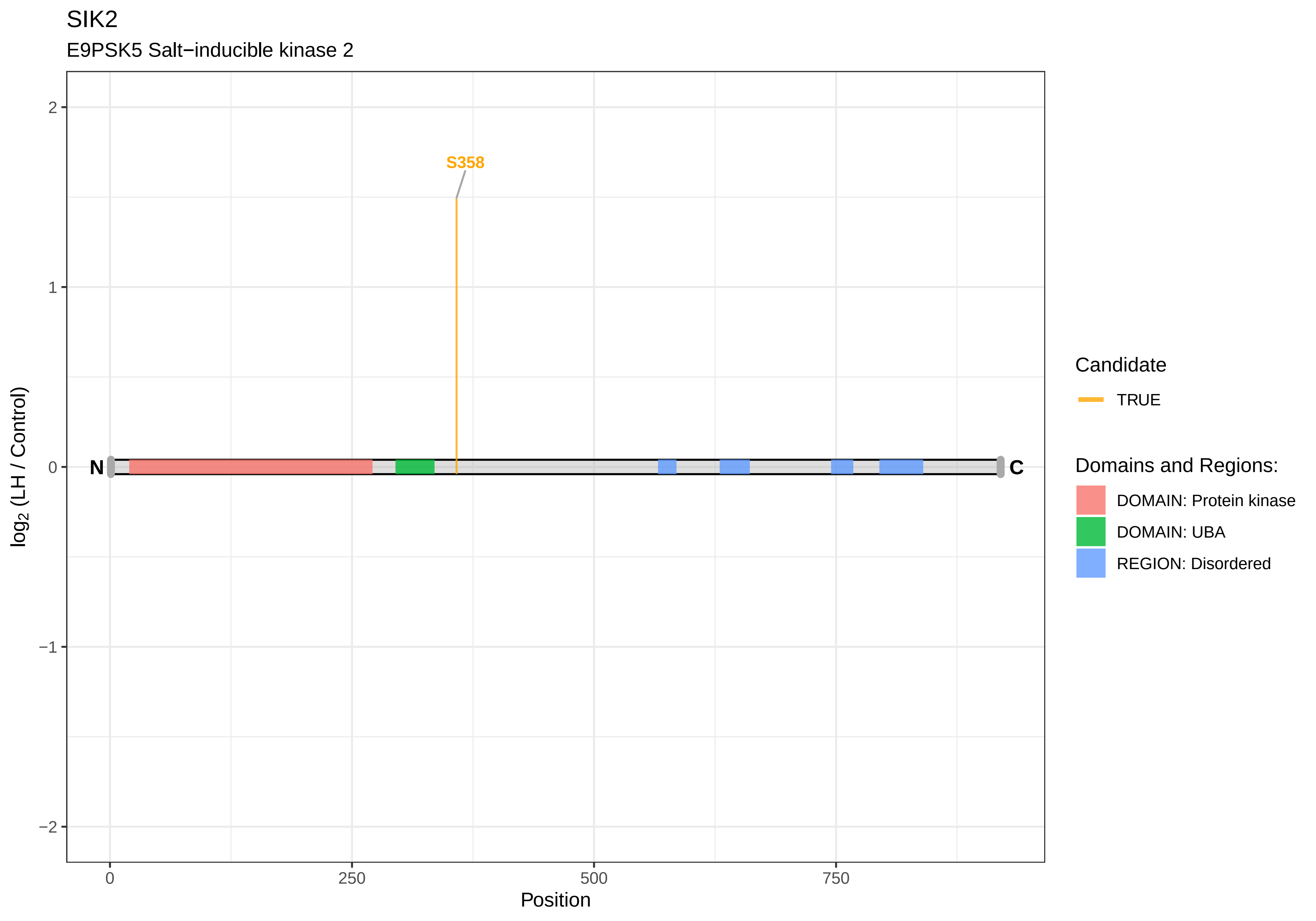
SIK2
E9PSK5 Salt−inducible kinase 2
S358
N
C
2
1
0
−1
−2
log2 (LH / Control)
0
250
500
750
Position
Candidate
TRUE
Domains and Regions:
DOMAIN: Protein kinase
DOMAIN: UBA
REGION: Disordered

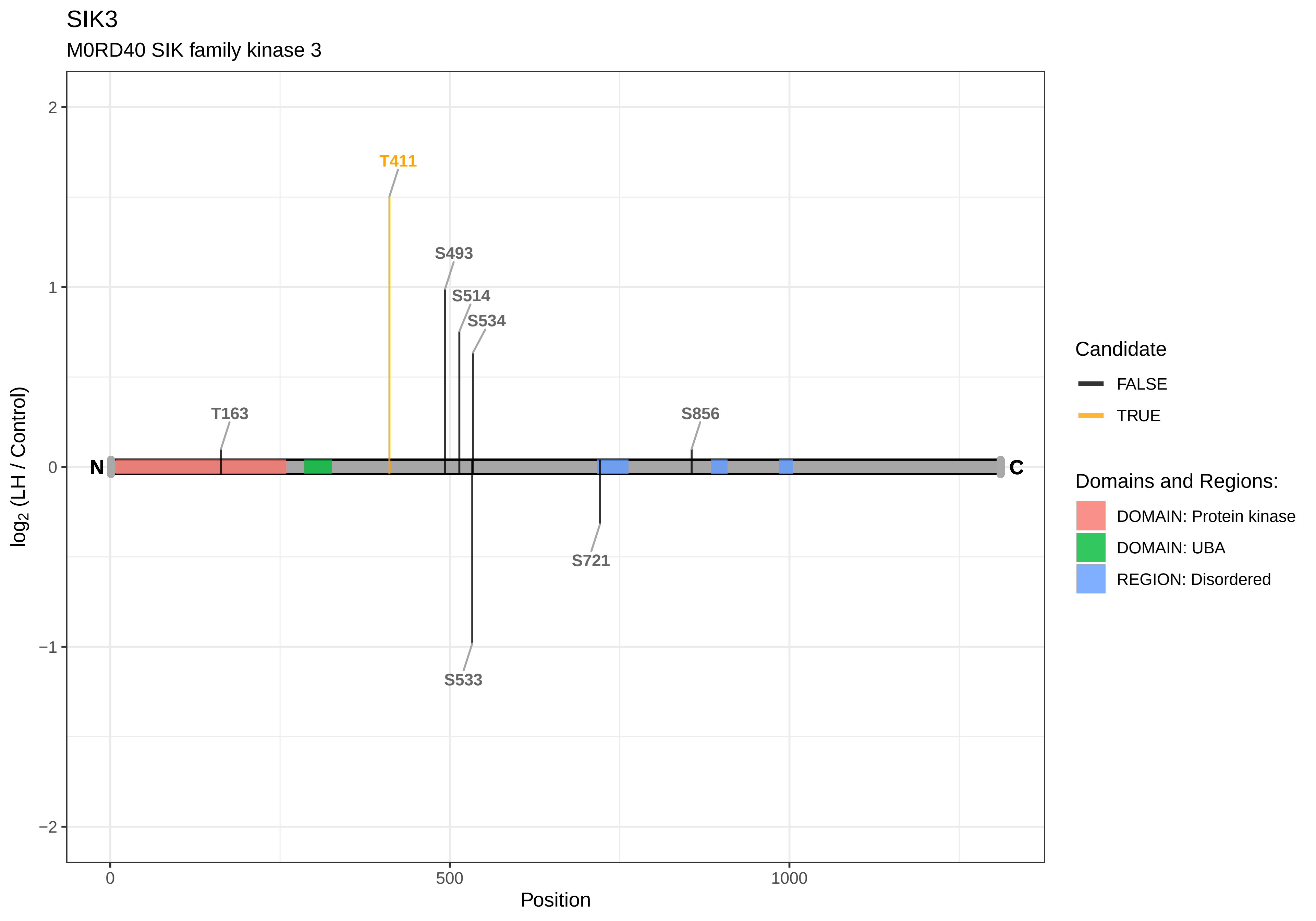

SIK3
M0RD40 SIK family kinase 3
T411
S493
S514
S534
T163
S856
N
C
S721
S533
2
1
0
−1
−2
log2 (LH / Control)
0
500
1000
Position
Candidate
FALSE
TRUE
Domains and Regions:
DOMAIN: Protein kinase
DOMAIN: UBA
REGION: Disordered

SIN3A

A0A0G2K3H5 SIN3 transcription regulator family member A

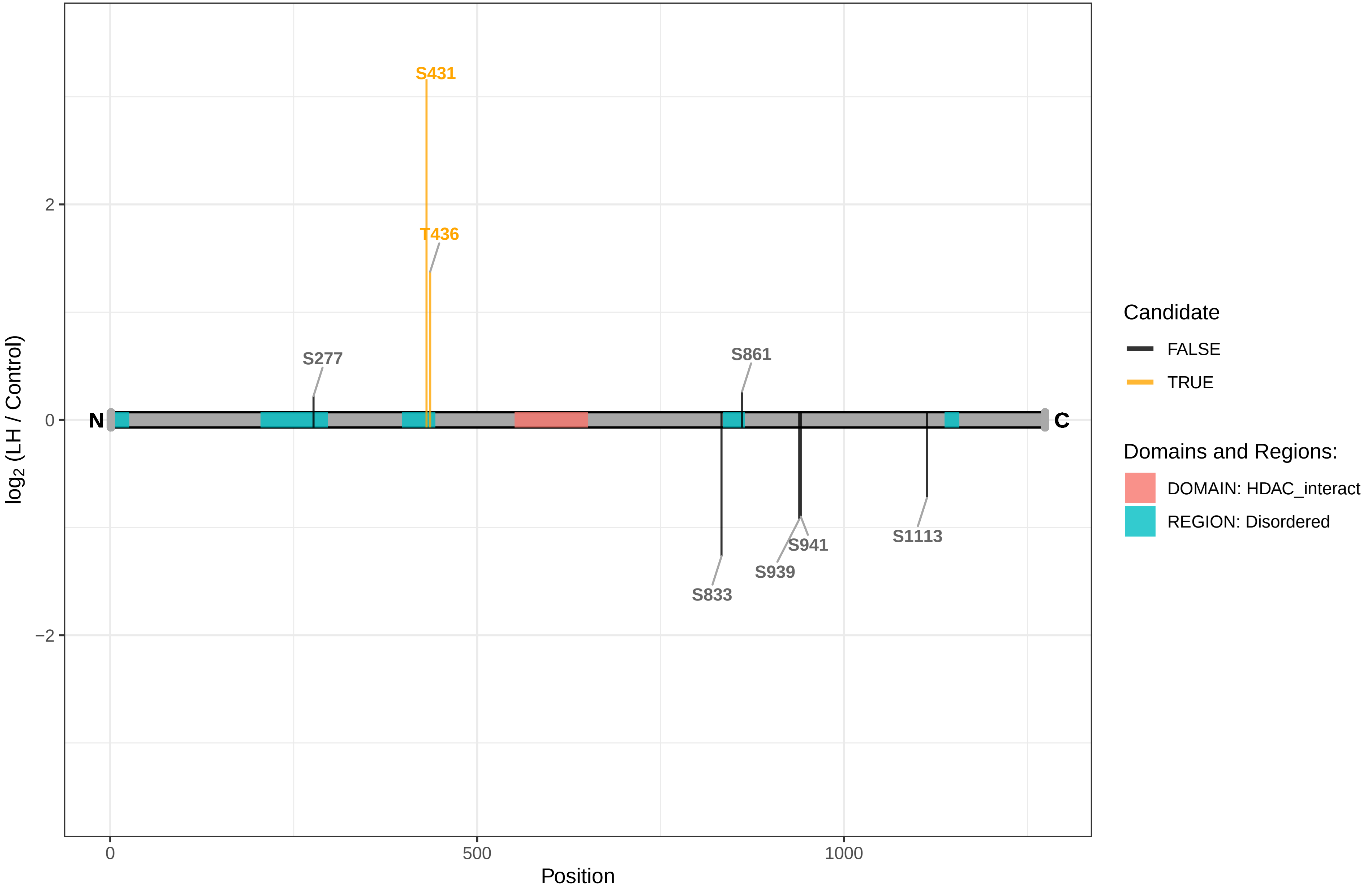

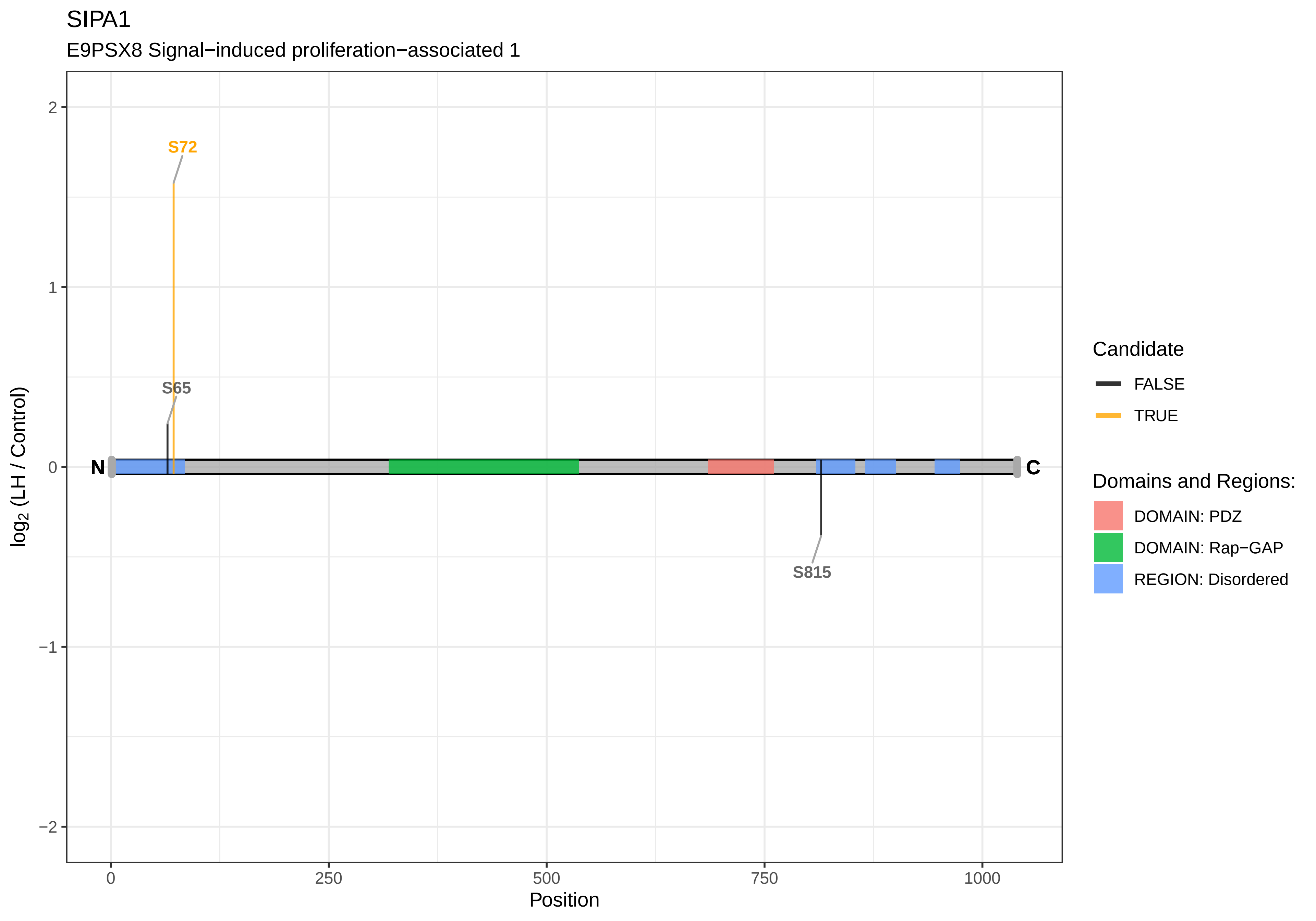

SIPA1
E9PSX8 Signal−induced proliferation−associated 1
S72
S65
S815
N
C
2
1
0
−1
−2
log2 (LH / Control)
0
250
500
750
1000
Position
Candidate
FALSE
TRUE
Domains and Regions:
DOMAIN: PDZ
DOMAIN: Rap−GAP
REGION: Disordered

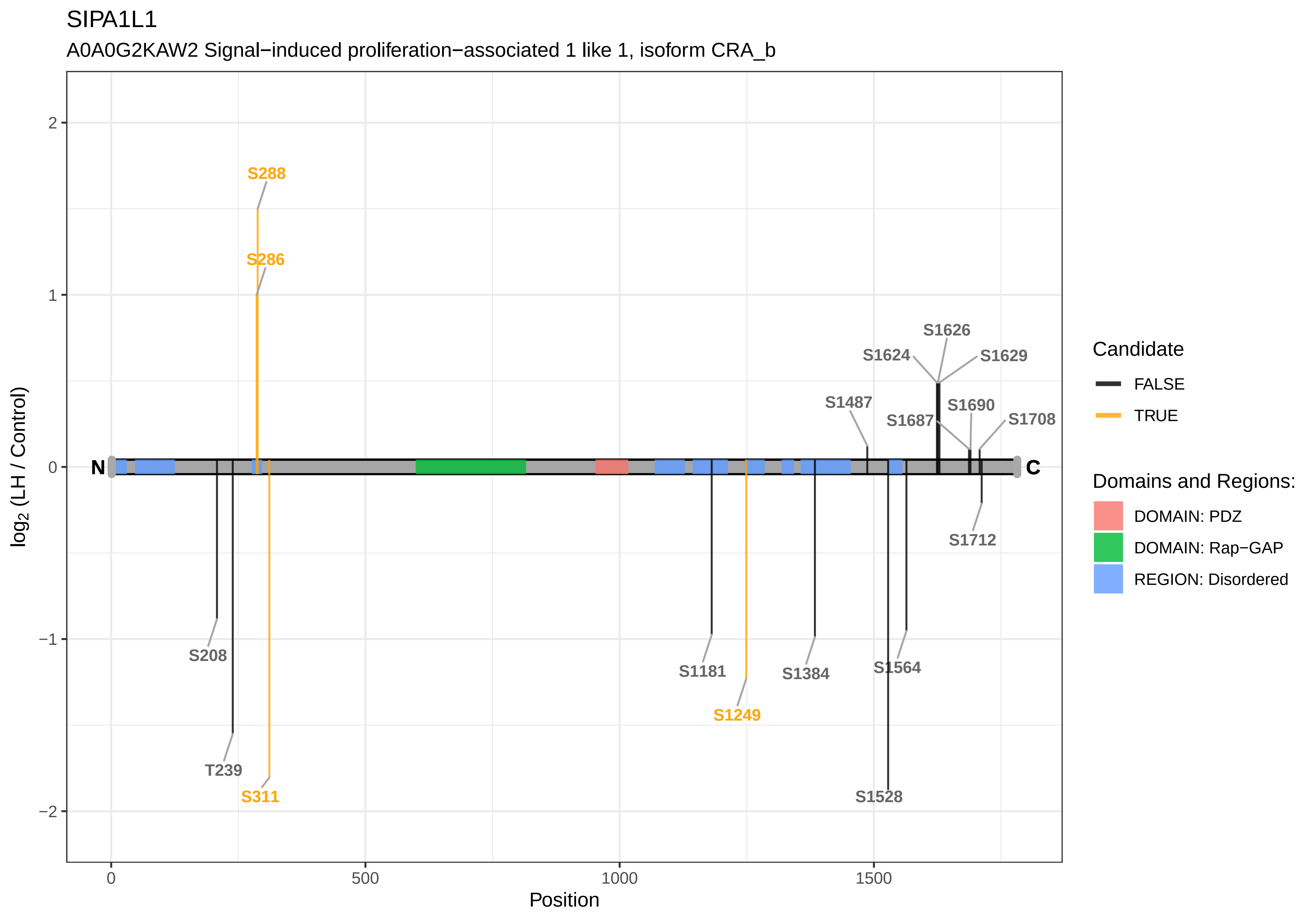
SIPA1L1
A0A0G2KAW2 Signal−induced proliferation−associated 1 like 1, isoform CRA_b
S288
S286
S1626
S1624
S1629
S1487
S1690
S1687
S1708
N
C
S1712
S208
S1181
S1384
S1564
S1249
T239
S311
S1528
2
1
0
−1
−2
0
500
1000
1500
Position
log2 (LH / Control)
Candidate
FALSE
TRUE
Domains and Regions:
DOMAIN: PDZ
DOMAIN: Rap−GAP
REGION: Disordered

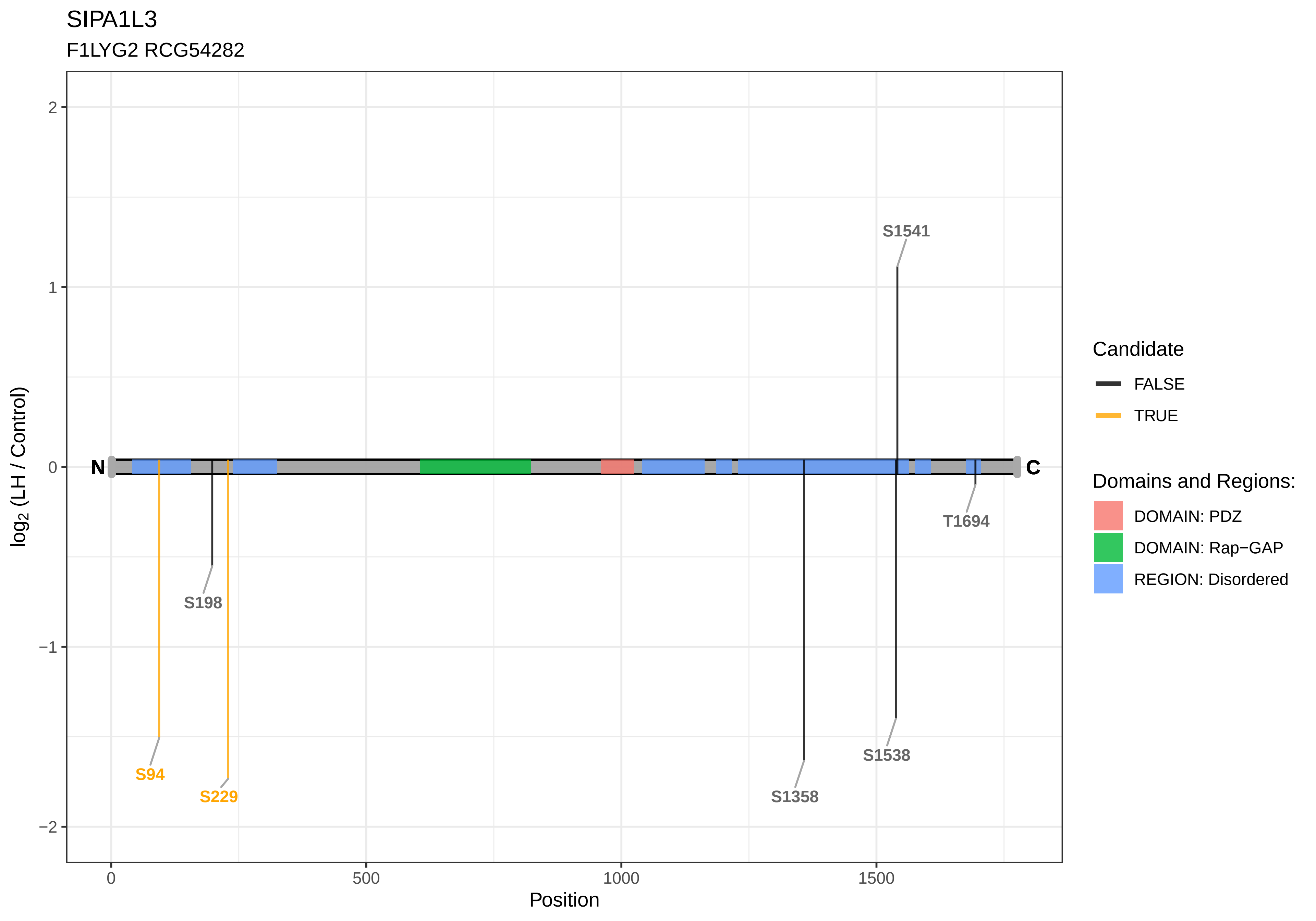

SIPA1L3
F1LYG2 RCG54282
log2 (LH / Control)
2
1
0
−1
−2
N
C
S1541
S198
S94
S229
S1358
S1538
T1694
0
500
1000
1500
Position
Candidate
FALSE
TRUE
Domains and Regions:
DOMAIN: PDZ
DOMAIN: Rap−GAP
REGION: Disordered

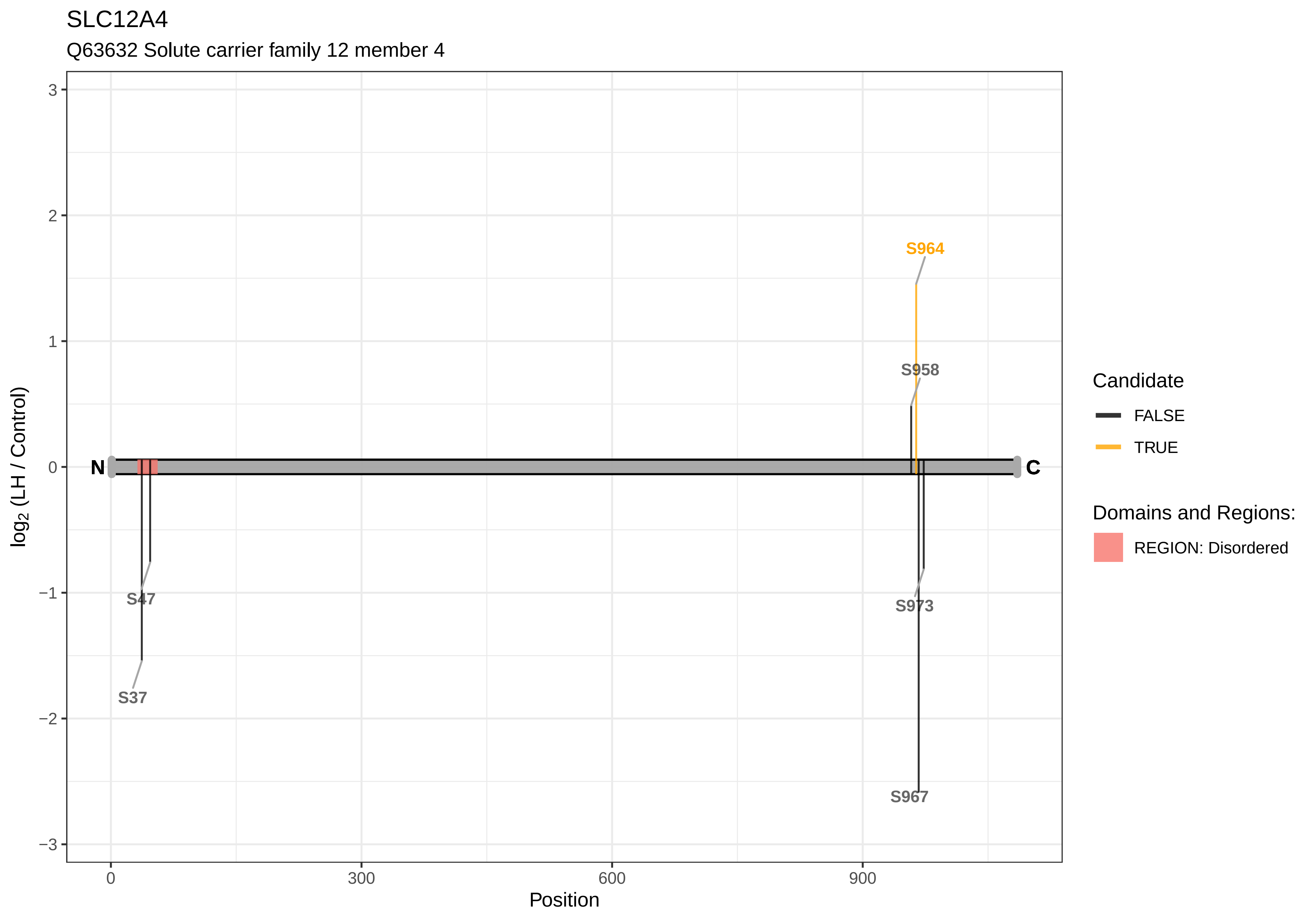

SLC12A4
Q63632 Solute carrier family 12 member 4
log2 (LH / Control)
Position
N
C
S964
S958
S47
S37
S973
S967
3
2
1
0
−1
−2
−3
0
300
600
900
Candidate
FALSE
TRUE
Domains and Regions:
REGION: Disordered

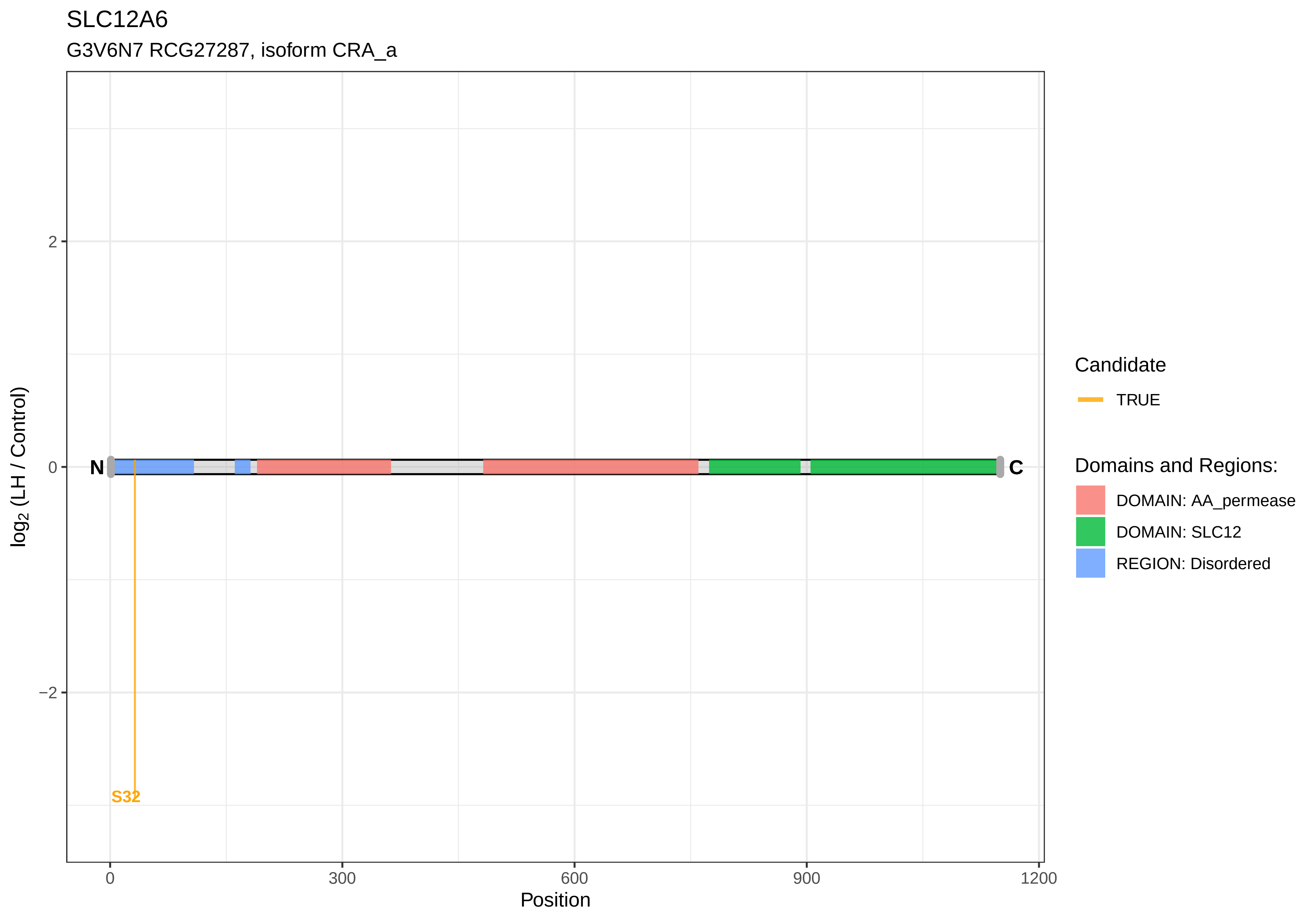

SLC12A6
G3V6N7 RCG27287, isoform CRA_a
log2 (LH / Control)
2
0
−2
N
C
S32
0
300
600
900
1200
Position
Candidate
TRUE
Domains and Regions:
DOMAIN: AA_permease
DOMAIN: SLC12
REGION: Disordered

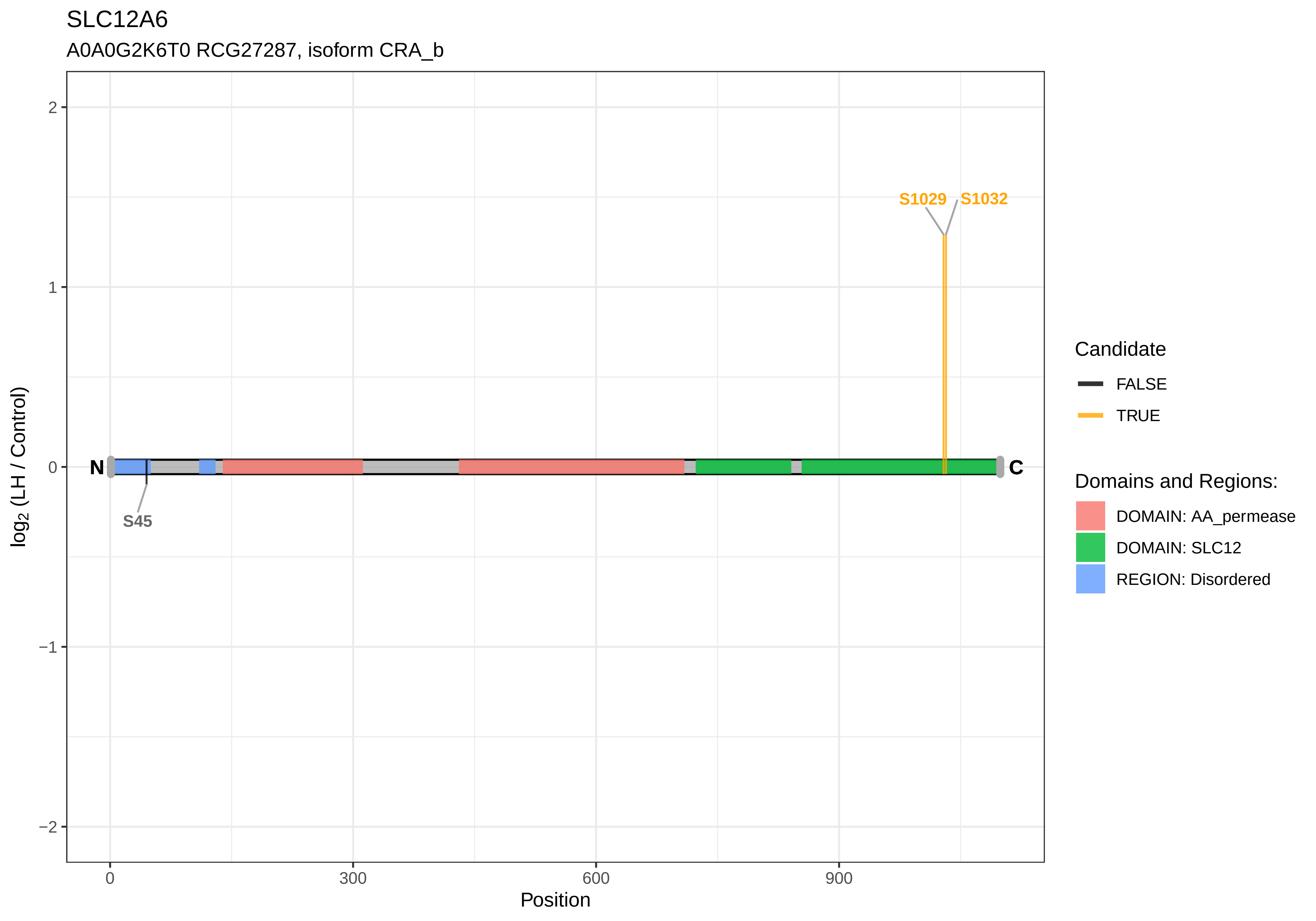
SLC12A6
A0A0G2K6T0 RCG27287, isoform CRA_b
log2 (LH / Control)
Position
S1029
S1032
S45
N
C
2
1
0
−1
−2
0
300
600
900
Candidate
FALSE
TRUE
Domains and Regions:
DOMAIN: AA_permease
DOMAIN: SLC12
REGION: Disordered

SLC15A4

A0A0G2JSH0 Solute carrier family 15 member 4

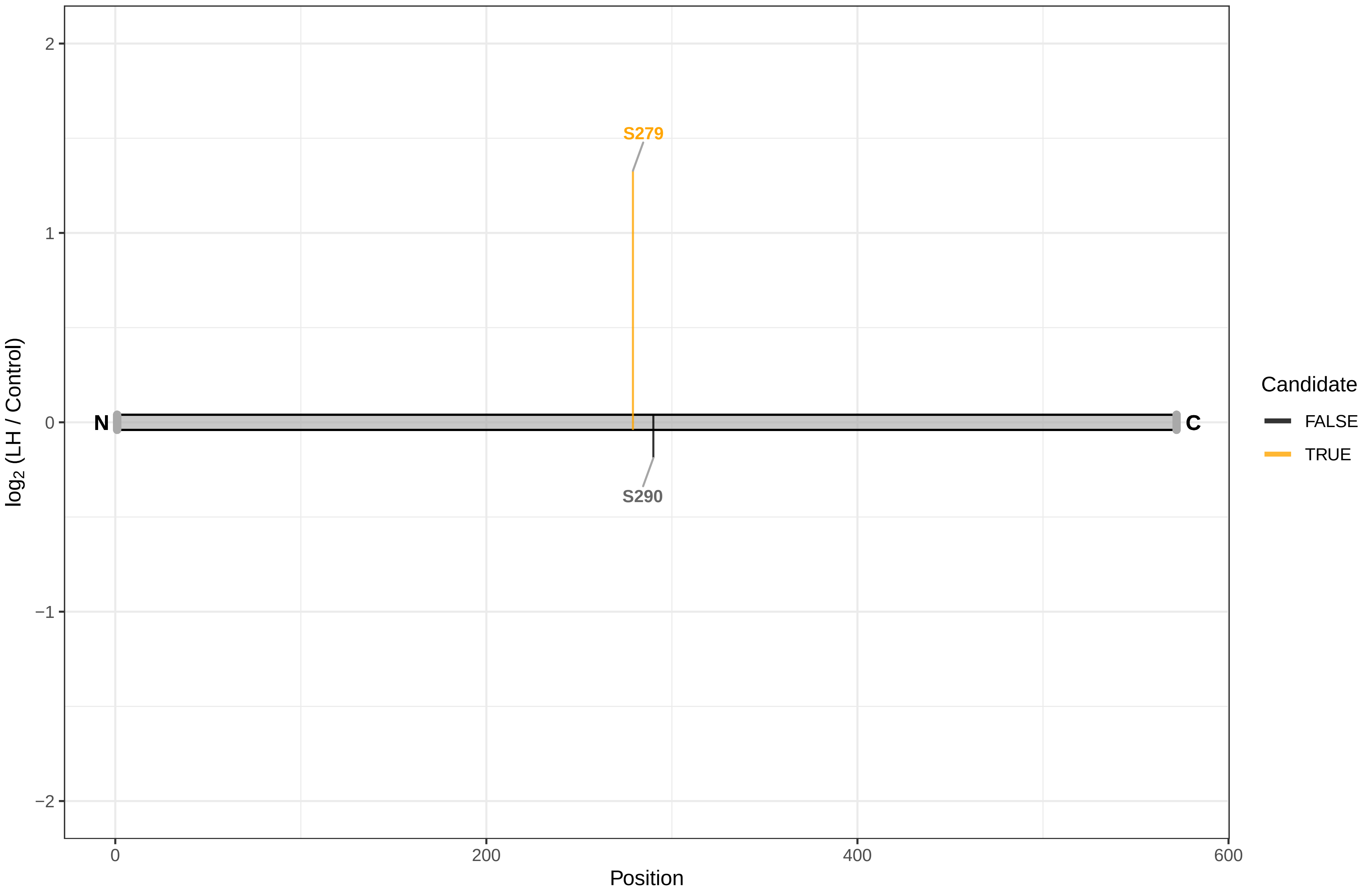

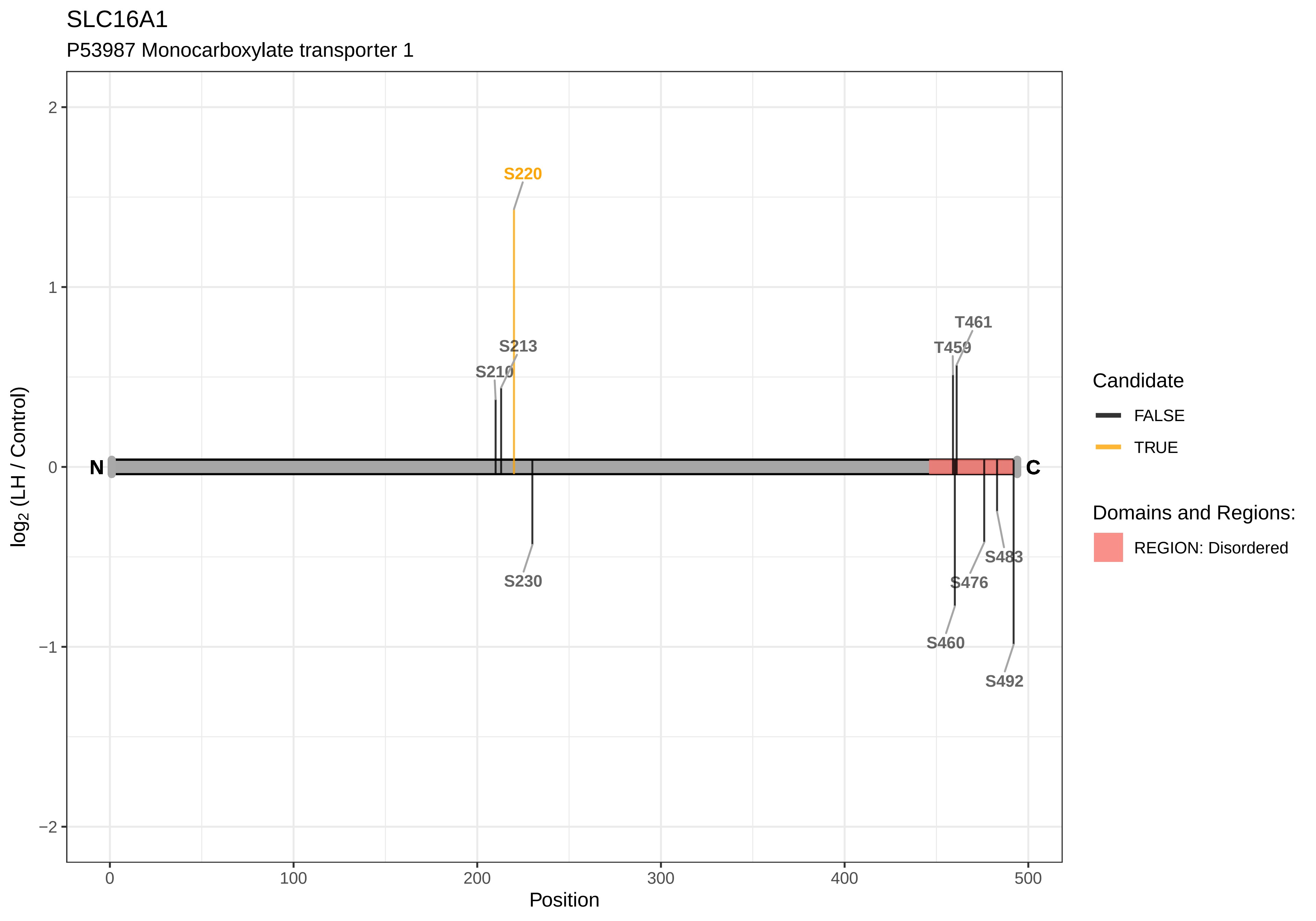

SLC16A1
P53987 Monocarboxylate transporter 1
S220
S213
S210
T461
T459
N
C
S230
S483
S476
S460
S492
2
1
0
−1
−2
0
100
200
300
400
500
Position
log2 (LH / Control)
Candidate
FALSE
TRUE
Domains and Regions:
REGION: Disordered

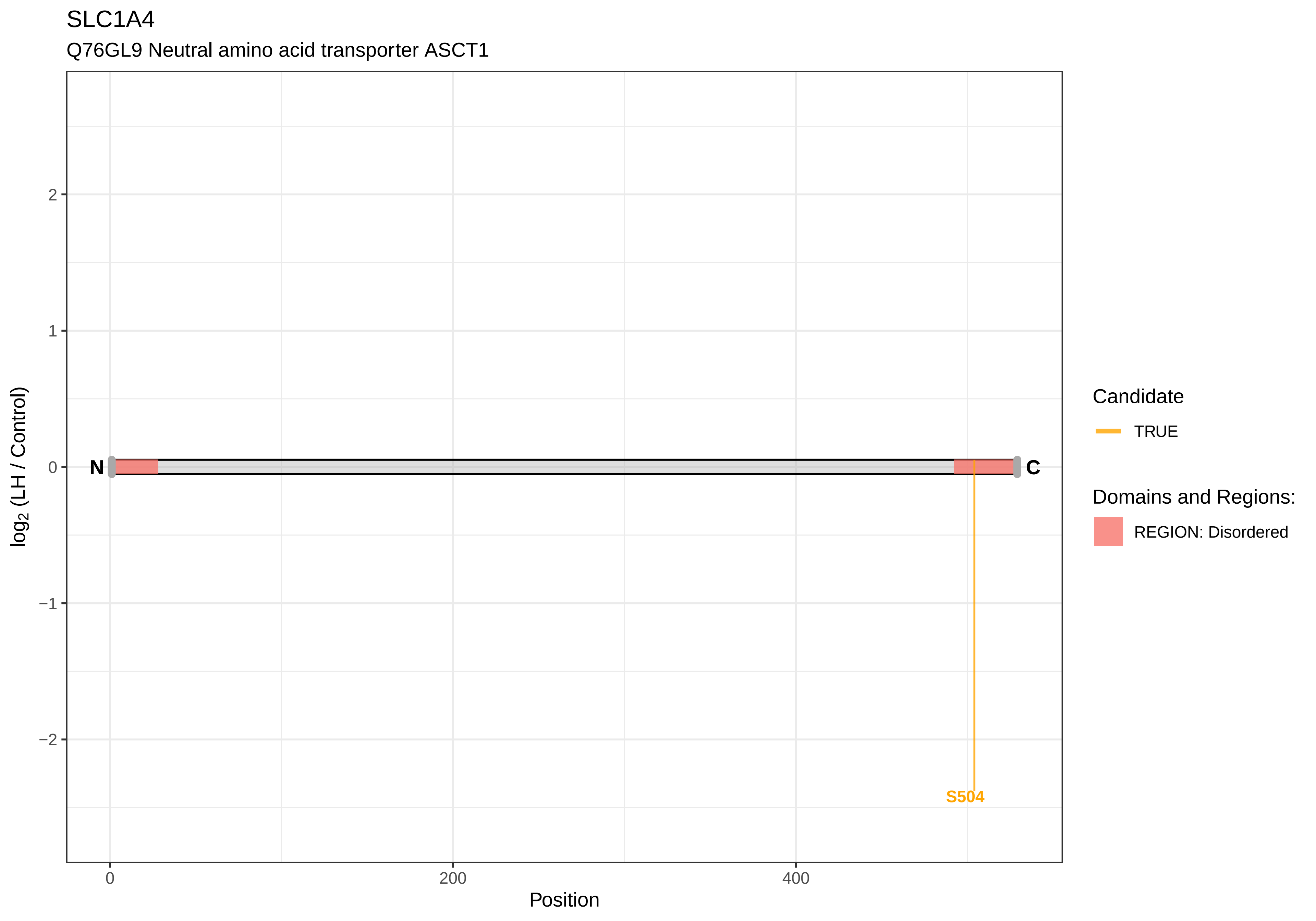
SLC1A4
Q76GL9 Neutral amino acid transporter ASCT1
log2 (LH / Control)
2
1
0
−1
−2
N
C
S504
0
200
400
Position
Candidate
TRUE
Domains and Regions:
REGION: Disordered

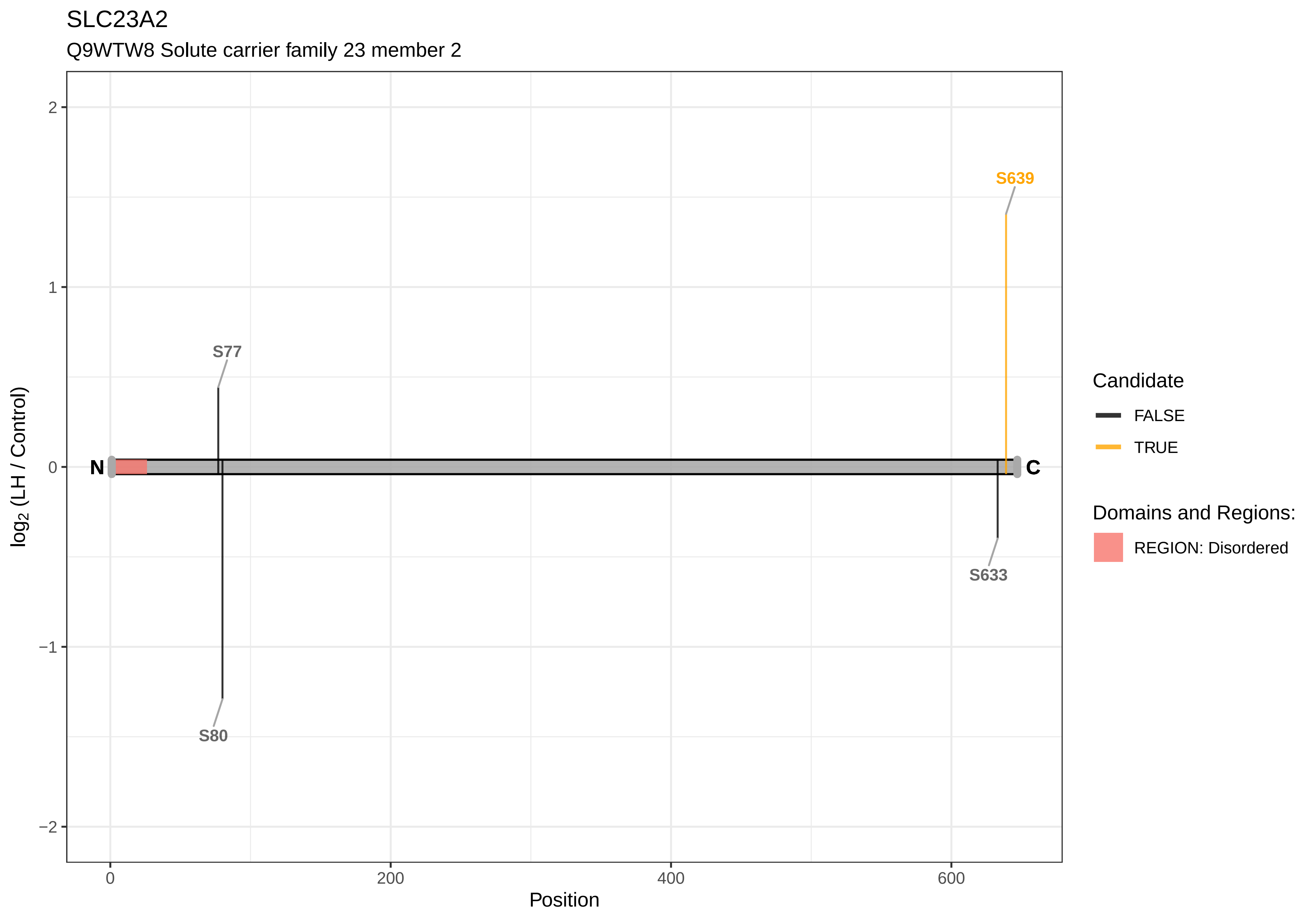

SLC23A2
Q9WTW8 Solute carrier family 23 member 2
S639
S77
S80
S633
N
C
2
1
0
−1
−2
0
200
400
600
Position
log2 (LH / Control)
Candidate
FALSE
TRUE
Domains and Regions:
REGION: Disordered

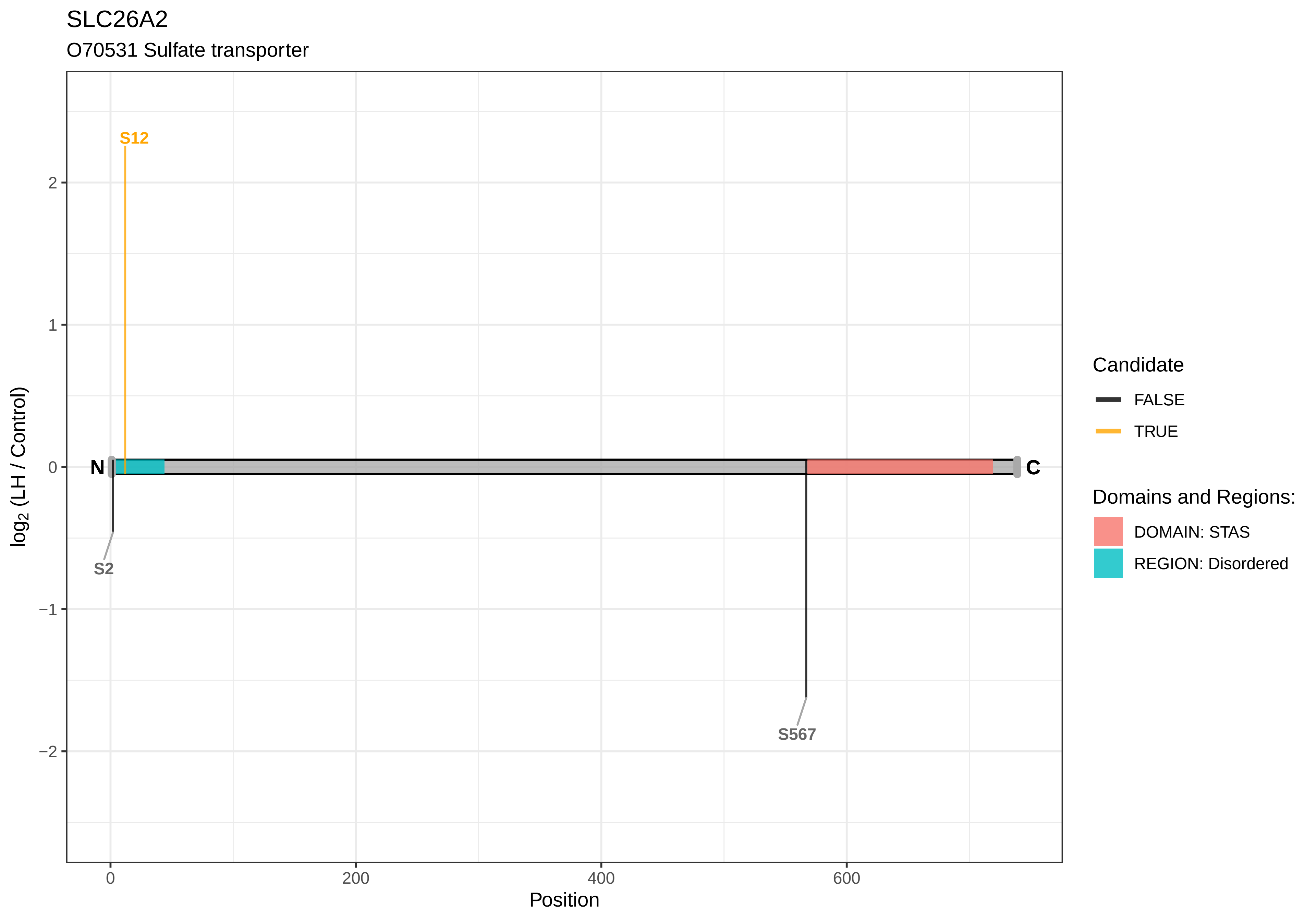

SLC26A2
O70531 Sulfate transporter
S12
S2
S567
N
C
2
1
0
−1
−2
log2 (LH / Control)
0
200
400
600
Position
Candidate
FALSE
TRUE
Domains and Regions:
DOMAIN: STAS
REGION: Disordered

SLC27A3

D3ZJA9 Solute carrier family 27 (Fatty acid transporter), member 3 (Predicted)

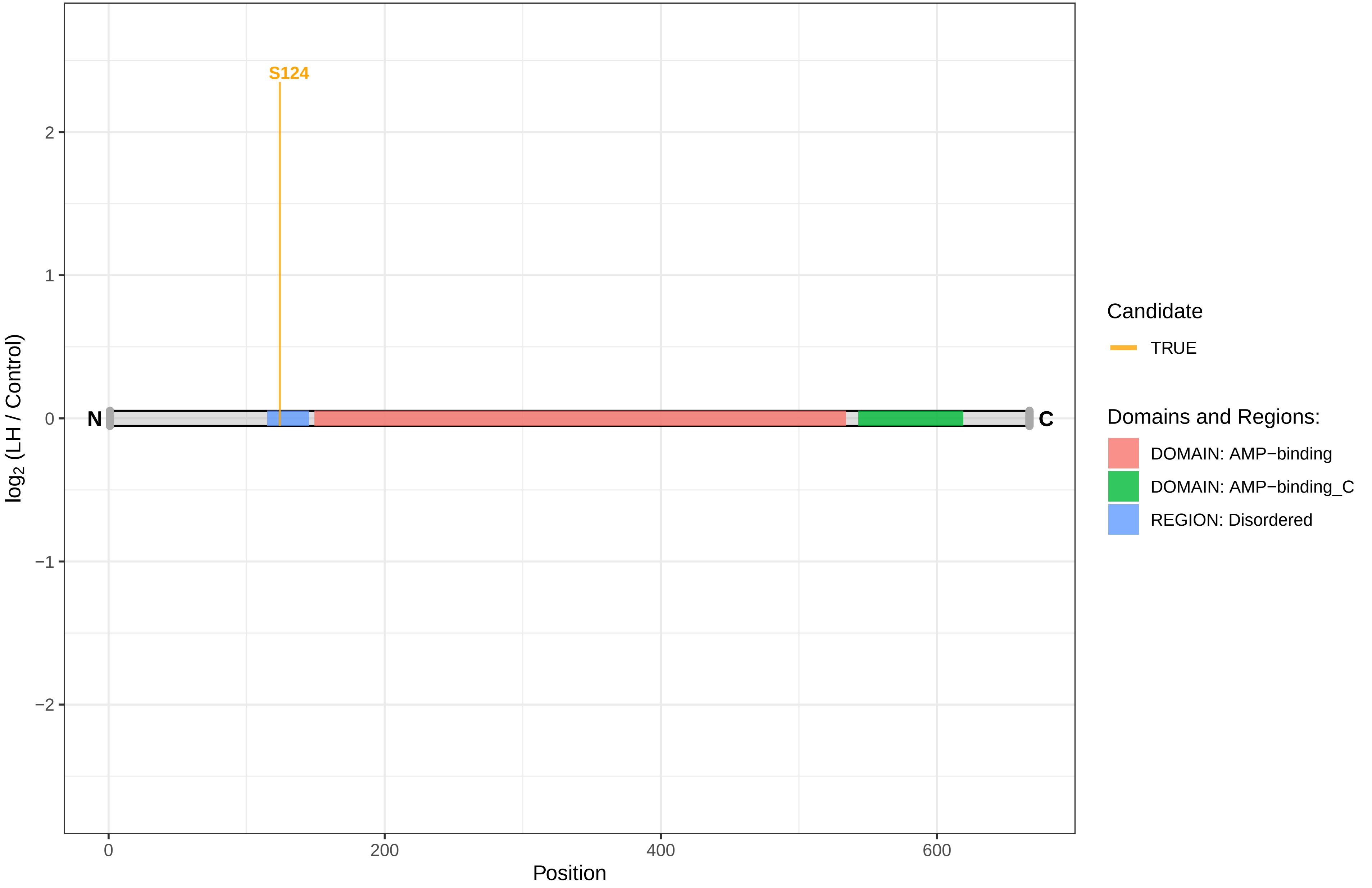

#### SLC38A1

Q9JM15 Sodium−coupled neutral amino acid transporter 1

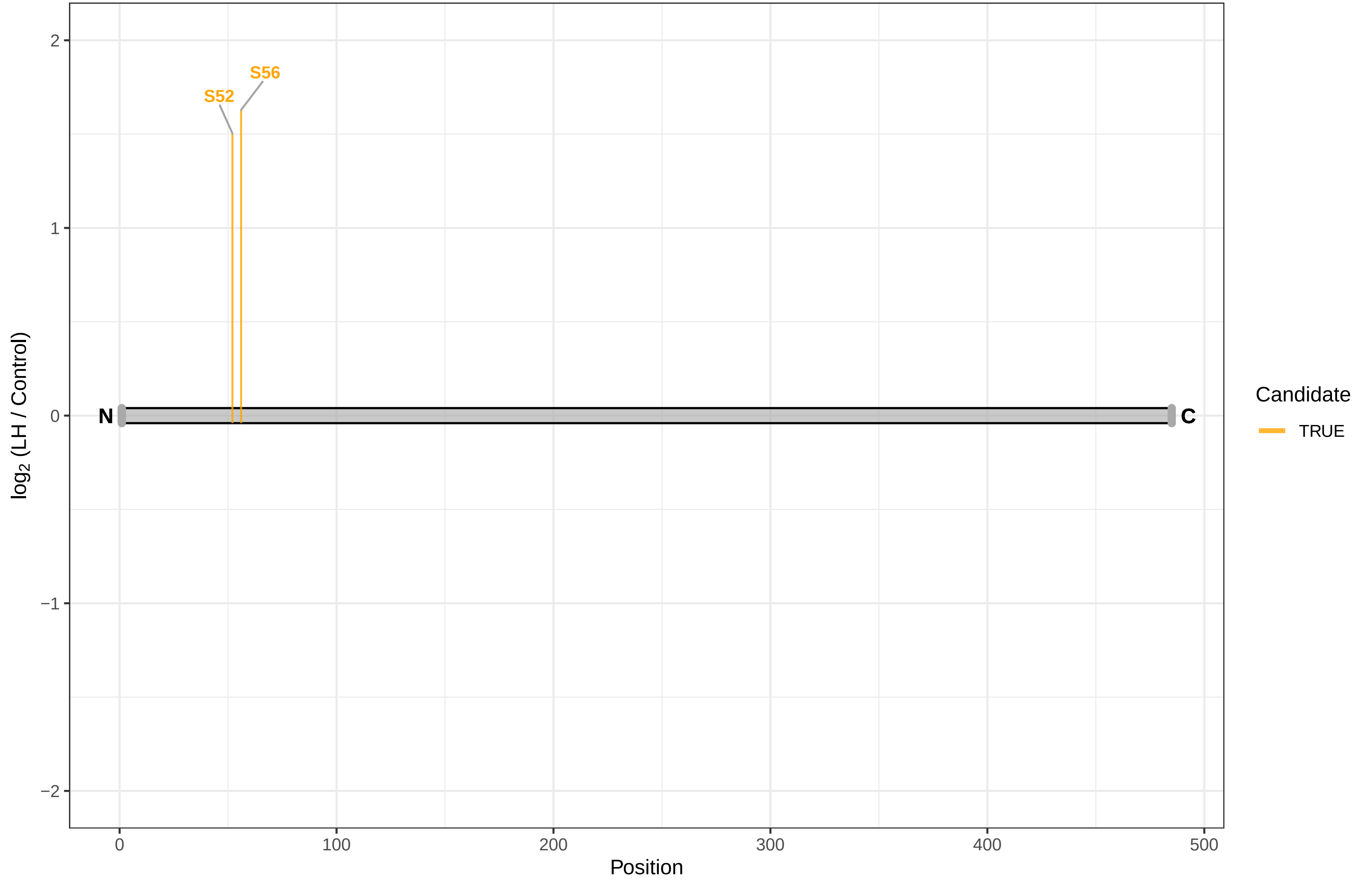

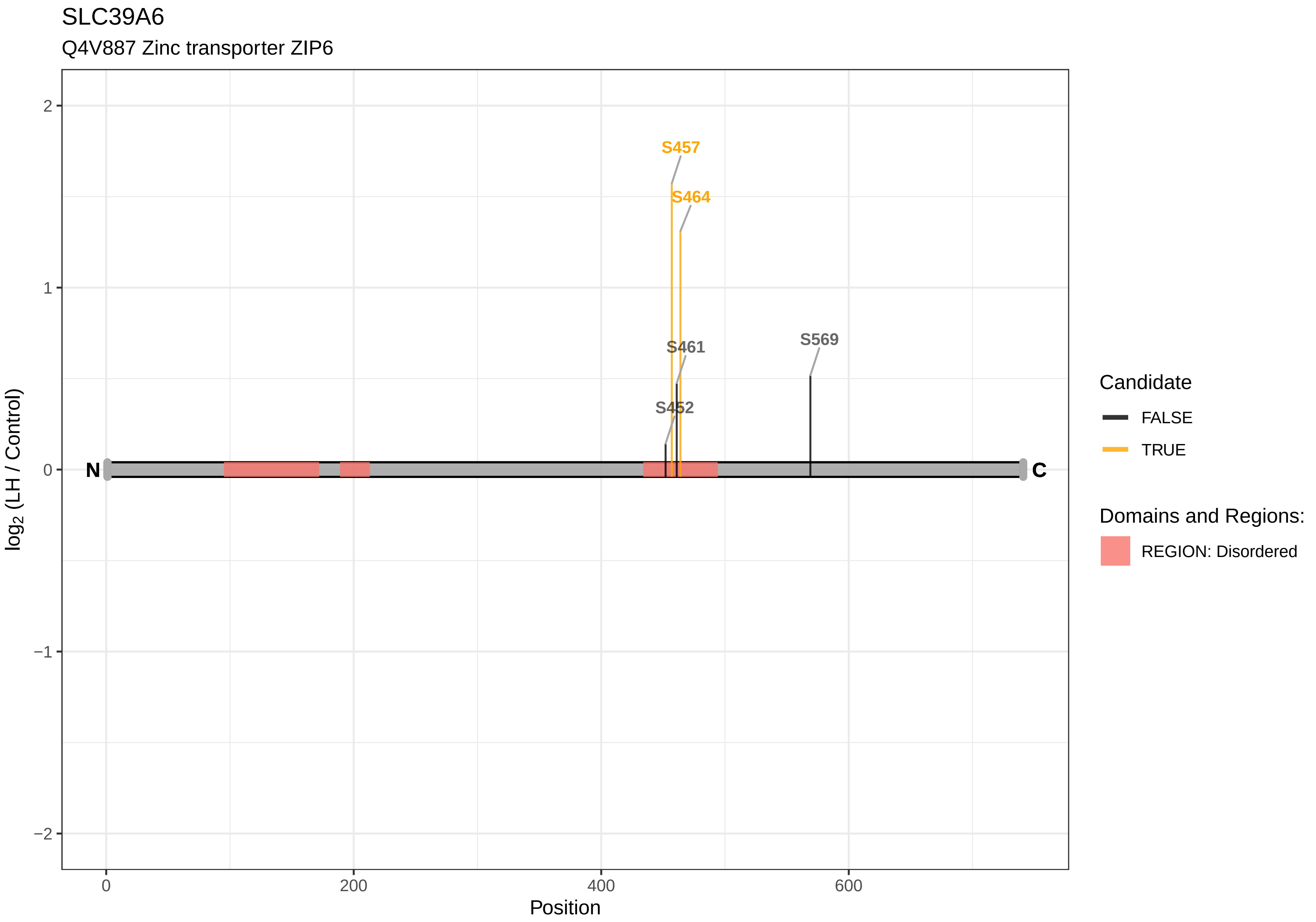
SLC39A6
Q4V887 Zinc transporter ZIP6
S457
S464
S461
S569
S452
N
C
2
1
0
−1
−2
0
200
400
600
Position
log2 (LH / Control)
Candidate
FALSE
TRUE
Domains and Regions:
REGION: Disordered

SLC44A3

F1M965 Choline transporter−like protein 3

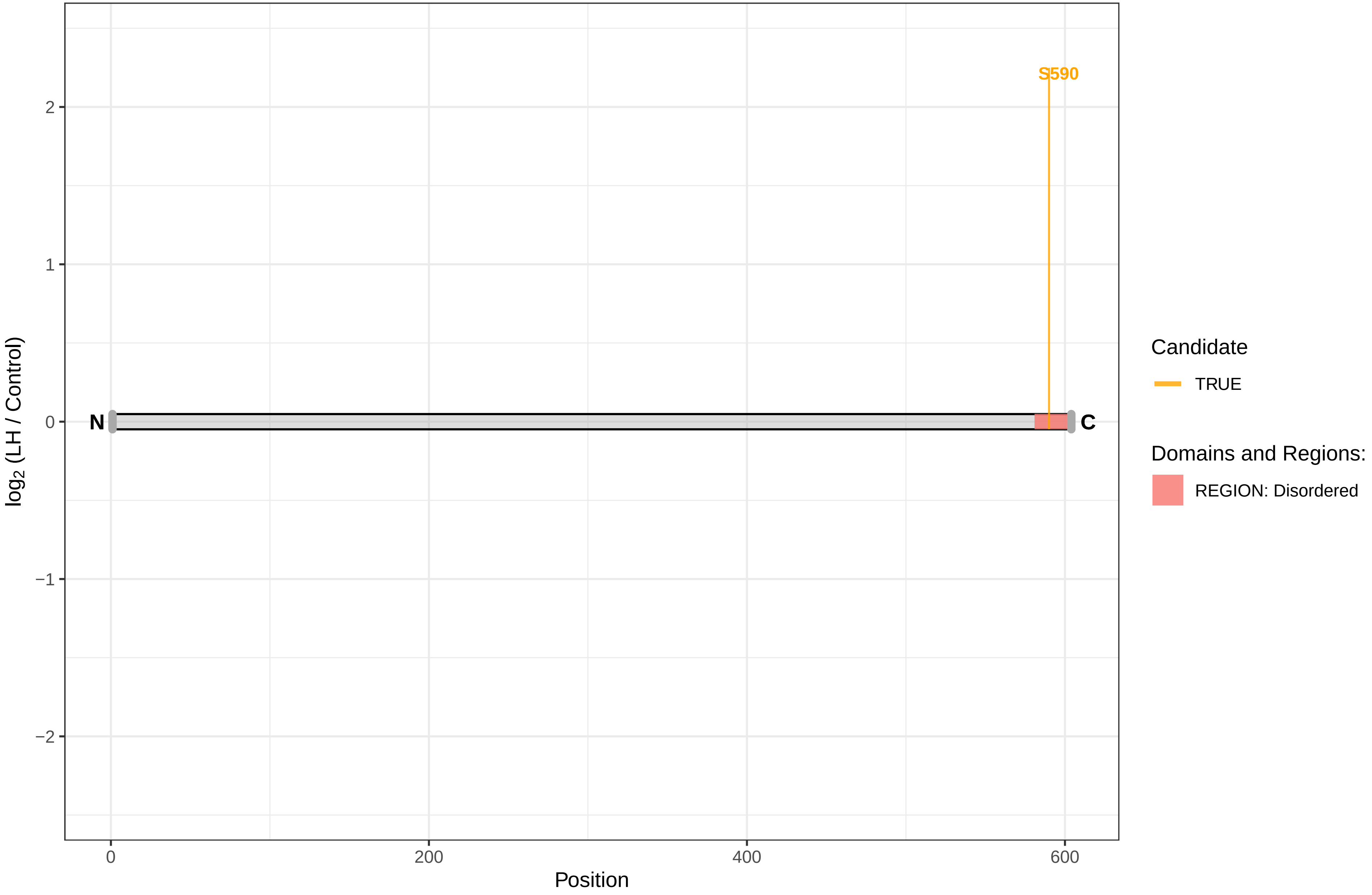

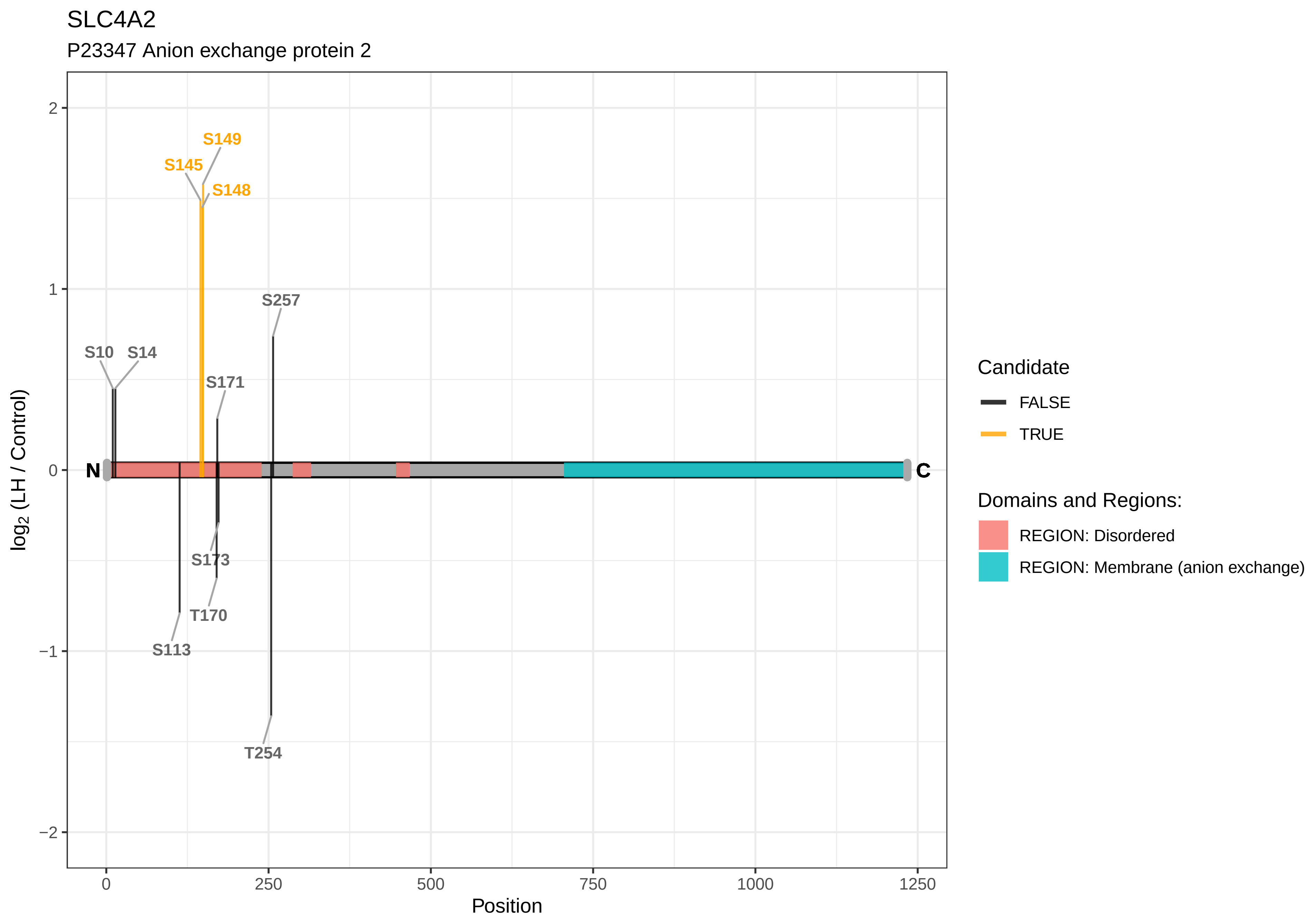

SLC4A2
P23347 Anion exchange protein 2
S149
S145
S148
S257
S10
S14
S171
N
C
S173
T170
S113
T254
2
1
0
−1
−2
0
250
500
750
1000
1250
Position
log2 (LH / Control)
Candidate
FALSE
TRUE
Domains and Regions:
REGION: Disordered
REGION: Membrane (anion exchange)

SLC7A11

D4ADU2 Solute carrier family 7 member 11

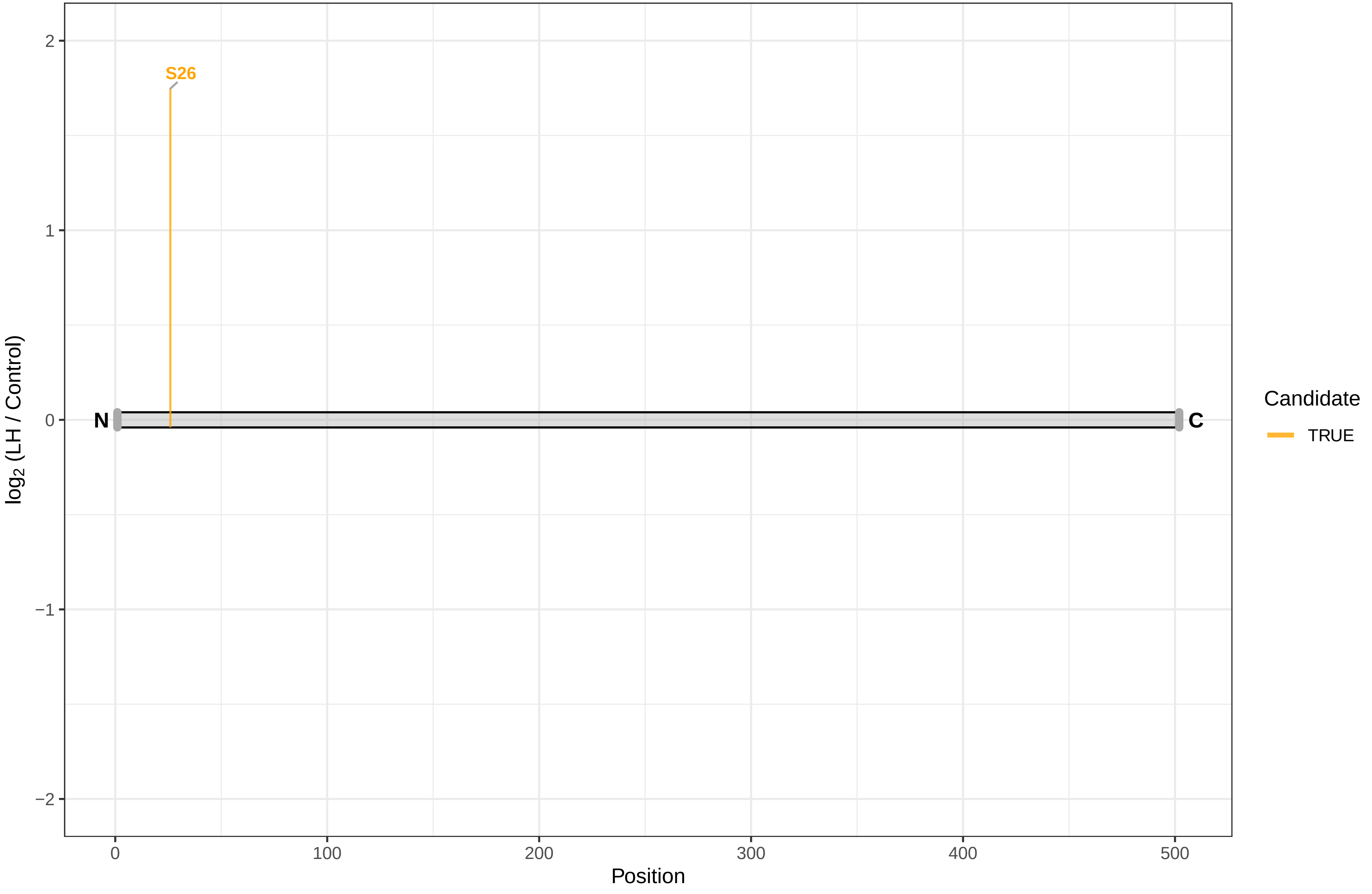

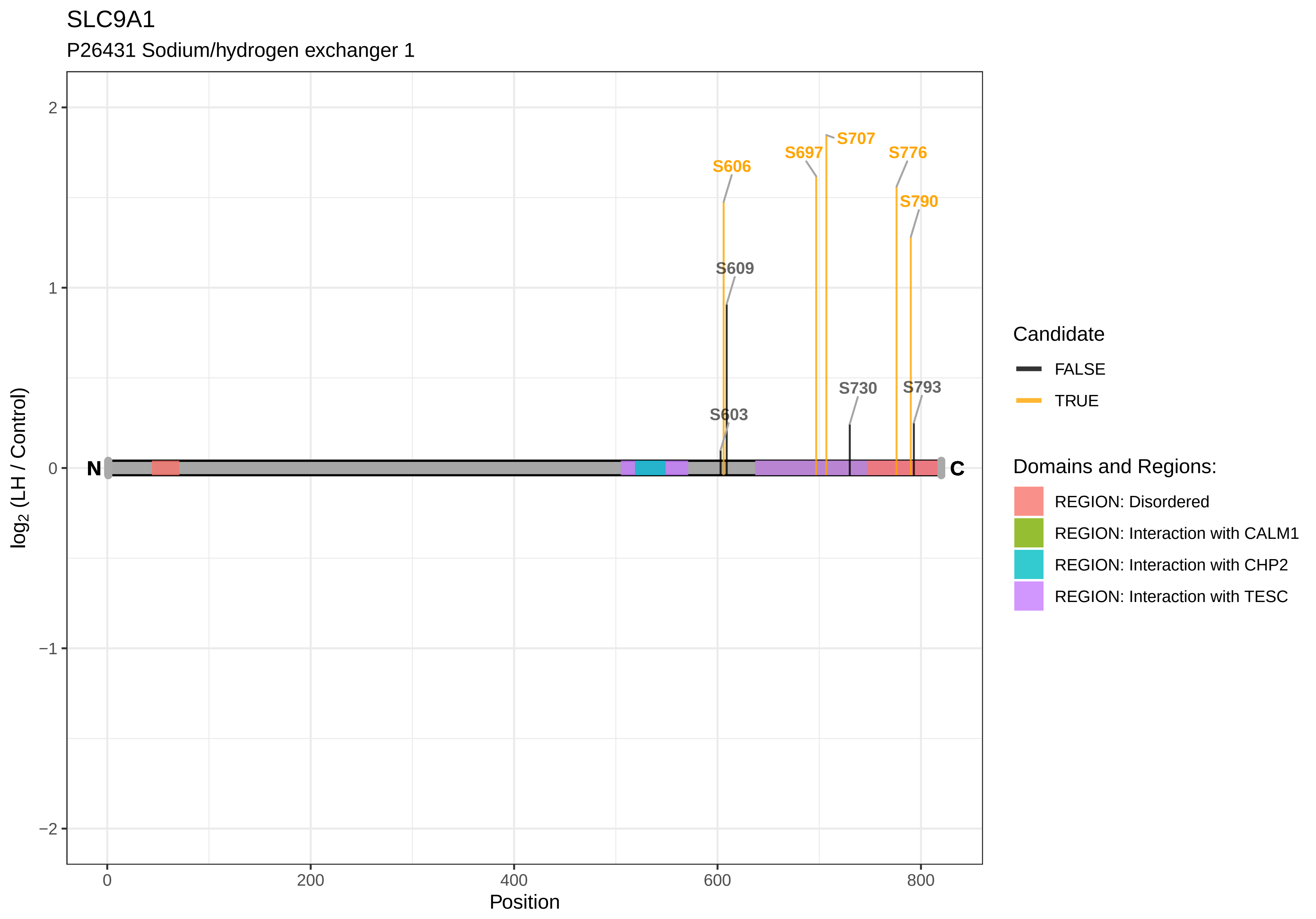

SLC9A1
P26431 Sodium/hydrogen exchanger 1
S707
S697
S776
S606
S790
S609
S730
S793
S603
N
C
2
1
0
−1
−2
0
200
400
600
800
Position
log2 (LH / Control)
Candidate
FALSE
TRUE
Domains and Regions:
REGION: Disordered
REGION: Interaction with CALM1
REGION: Interaction with CHP2
REGION: Interaction with TESC

SLC9A3R1

Q9JJ19 Na(+)/H(+) exchange regulatory cofactor NHE−RF1

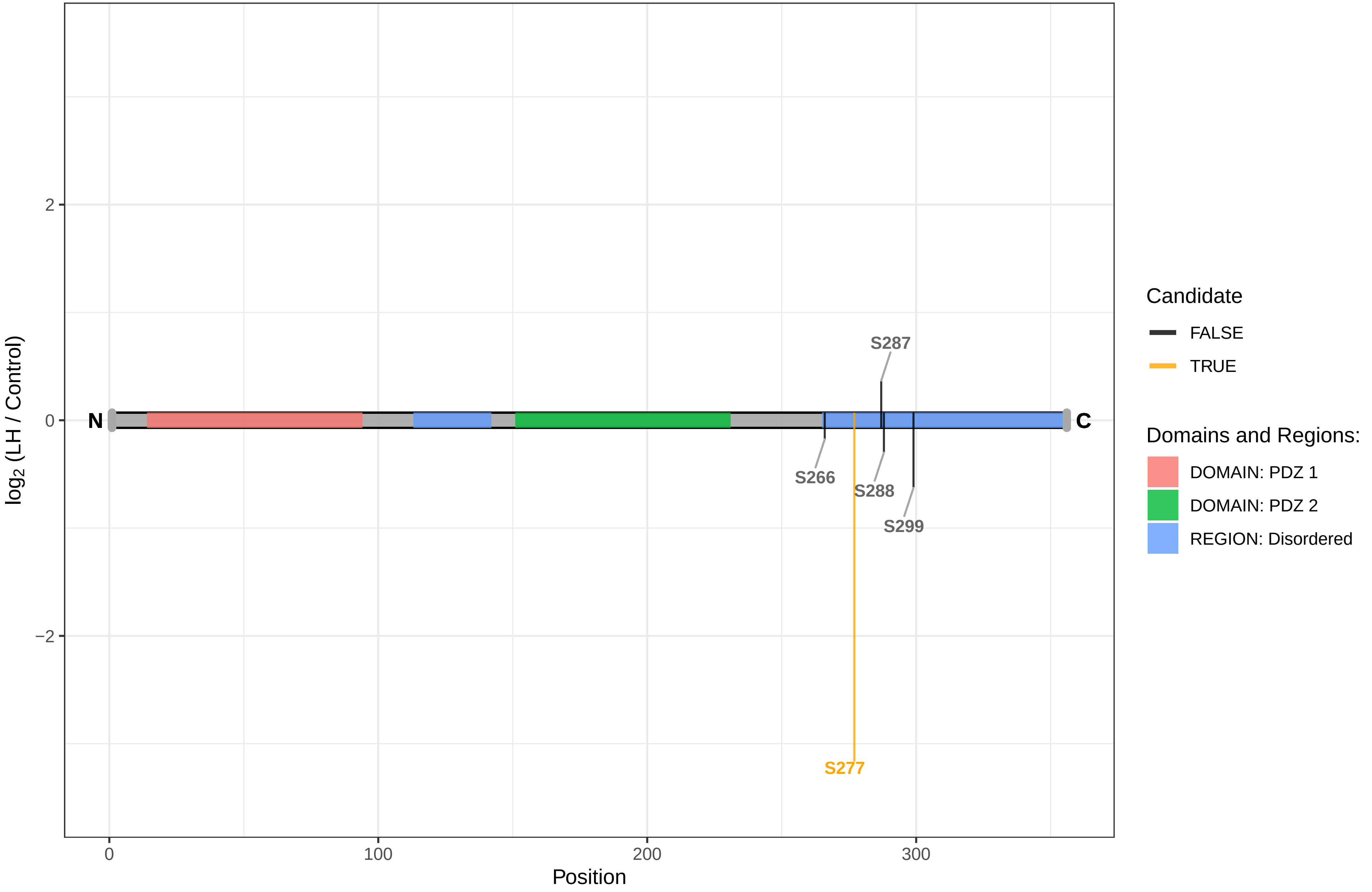

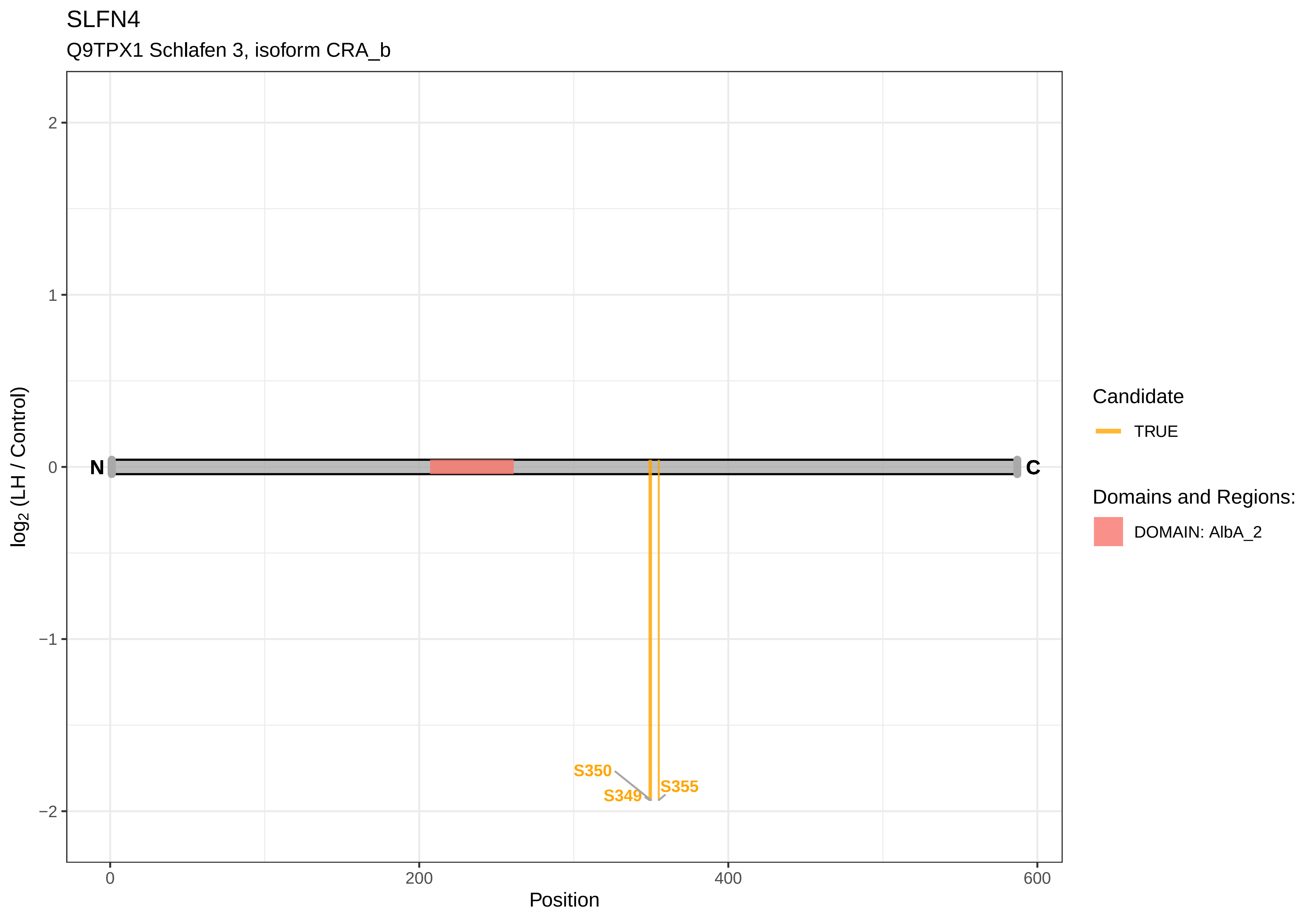
SLFN4
Q9TPX1 Schlafen 3, isoform CRA_b
2
1
0
−1
−2
log2 (LH / Control)
N
C
S350
S349
S355
0
200
400
600
Position
Candidate
TRUE
Domains and Regions:
DOMAIN: AlbA_2

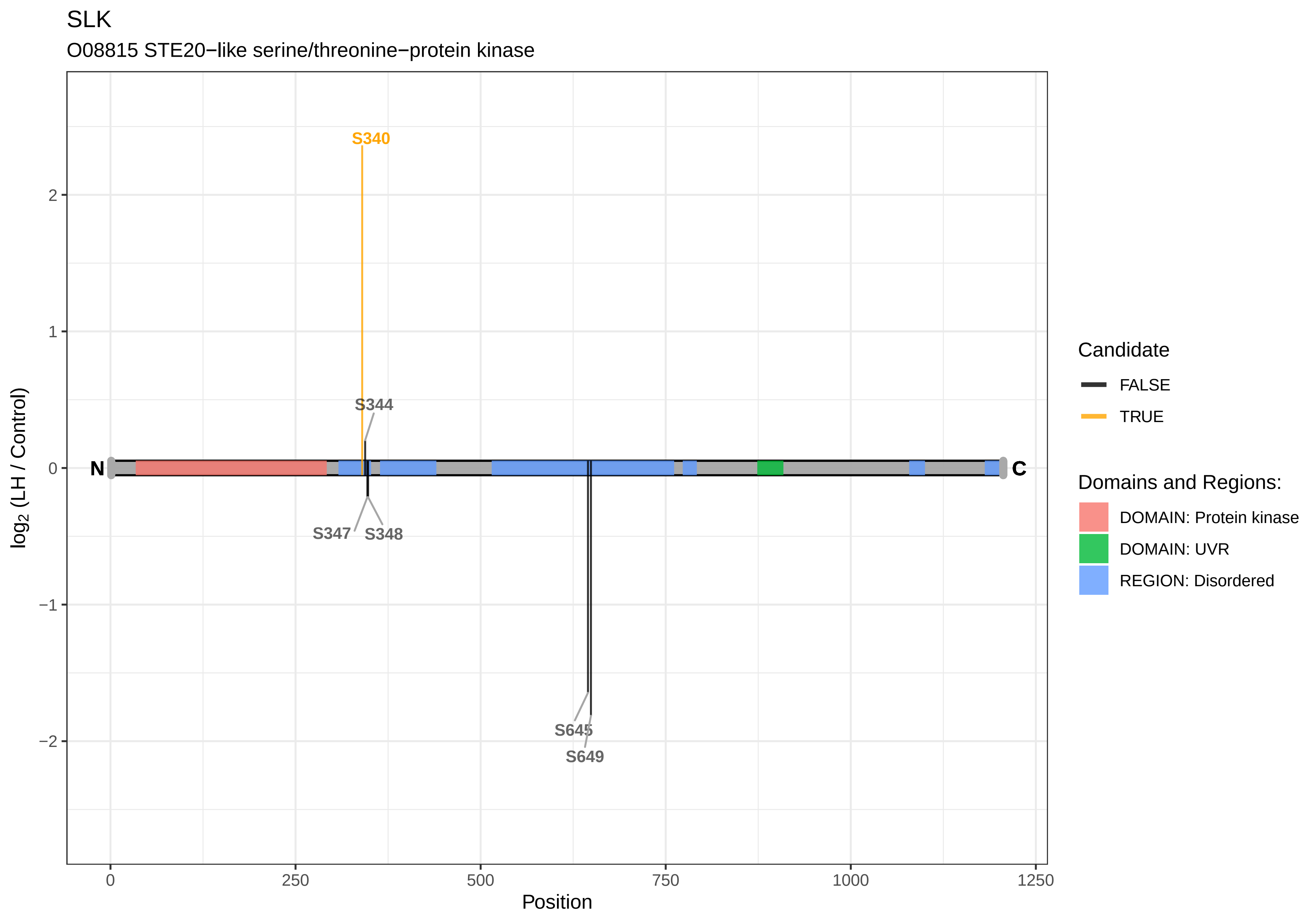

SLK
O08815 STE20−like serine/threonine−protein kinase
S340
S344
S347
S348
S645
S649
N
C
log2 (LH / Control)
Position
2
1
0
−1
−2
0
250
500
750
1000
1250
Candidate
FALSE
TRUE
Domains and Regions:
DOMAIN: Protein kinase
DOMAIN: UVR
REGION: Disordered

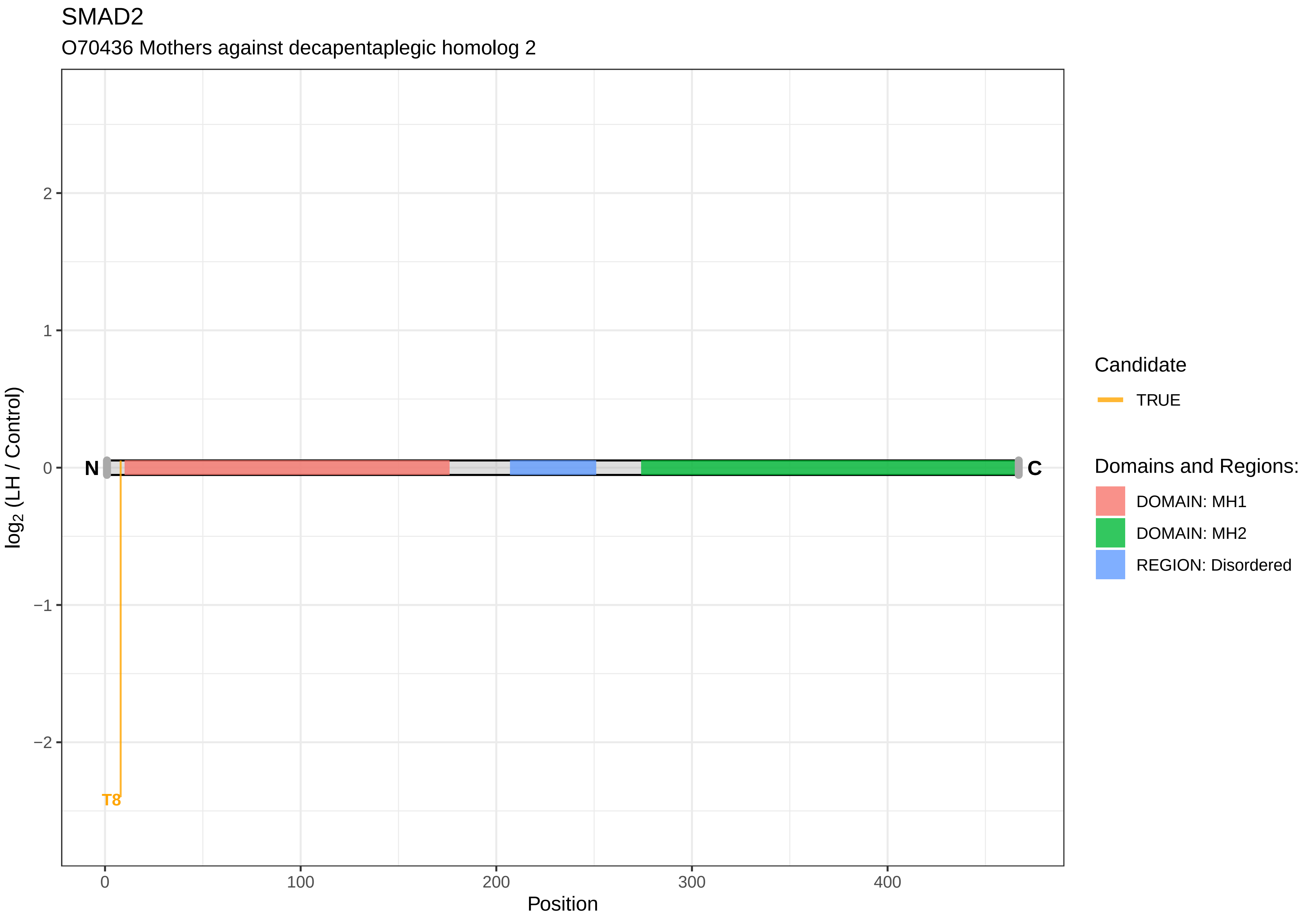
SMAD2
O70436 Mothers against decapentaplegic homolog 2
log2 (LH / Control)
2
1
0
−1
−2
N
C
T8
0
100
200
300
400
Position
Candidate
TRUE
Domains and Regions:
DOMAIN: MH1
DOMAIN: MH2
REGION: Disordered

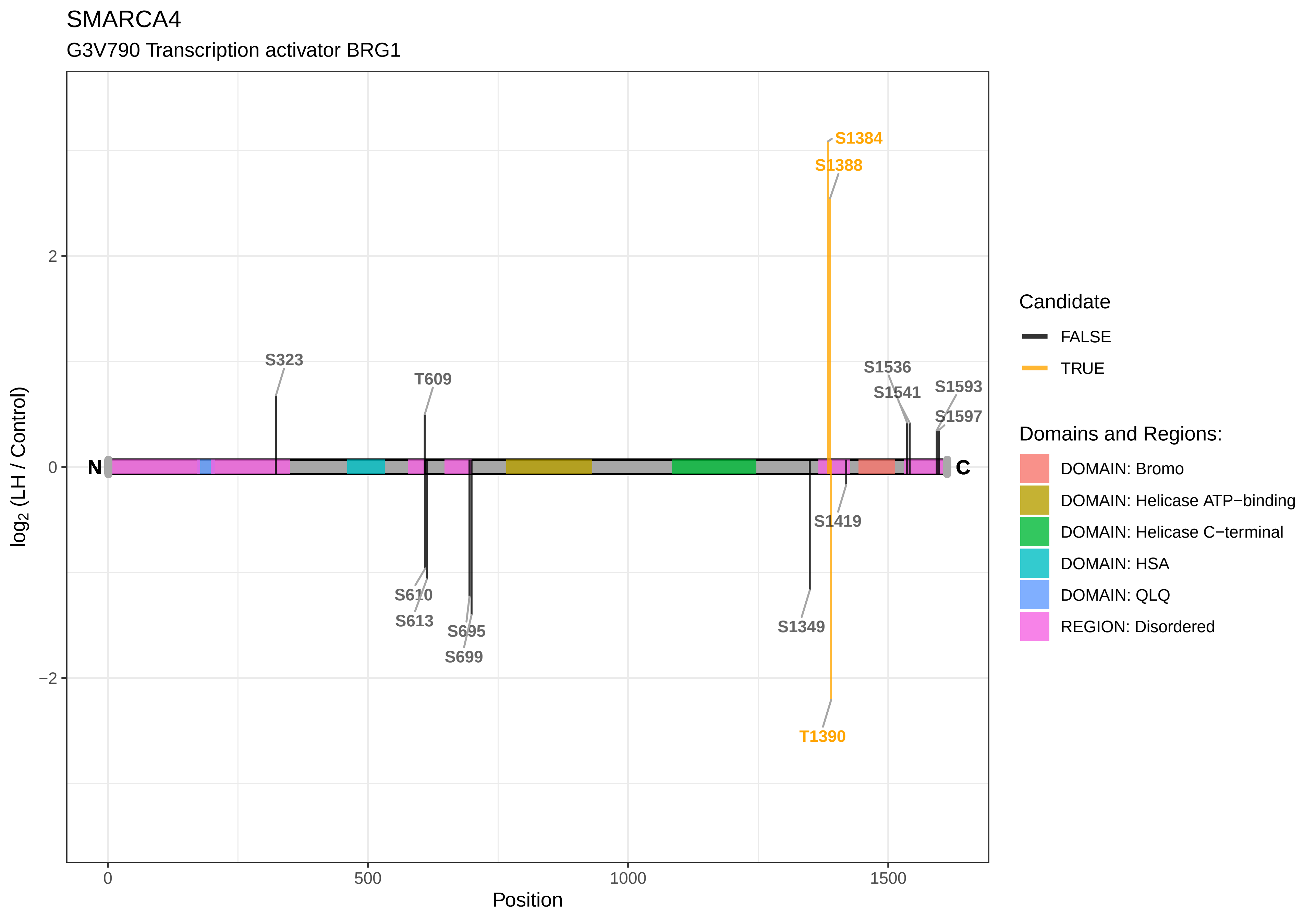
SMARCA4
G3V790 Transcription activator BRG1
S1384
S1388
S323
T609
S1536
S1541
S1593
S1597
N
C
S1419
S610
S613
S695
S699
S1349
T1390
2
0
−2
log2 (LH / Control)
0
500
1000
1500
Position
Candidate
FALSE
TRUE
Domains and Regions:
DOMAIN: Bromo
DOMAIN: Helicase ATP−binding
DOMAIN: Helicase C−terminal
DOMAIN: HSA
DOMAIN: QLQ
REGION: Disordered

SMARCAD1

D3Z9Z9 SWI/SNF−related matrix−associated actin−dependent regulator of chromatin subfamily A containing DEAD/H box 1

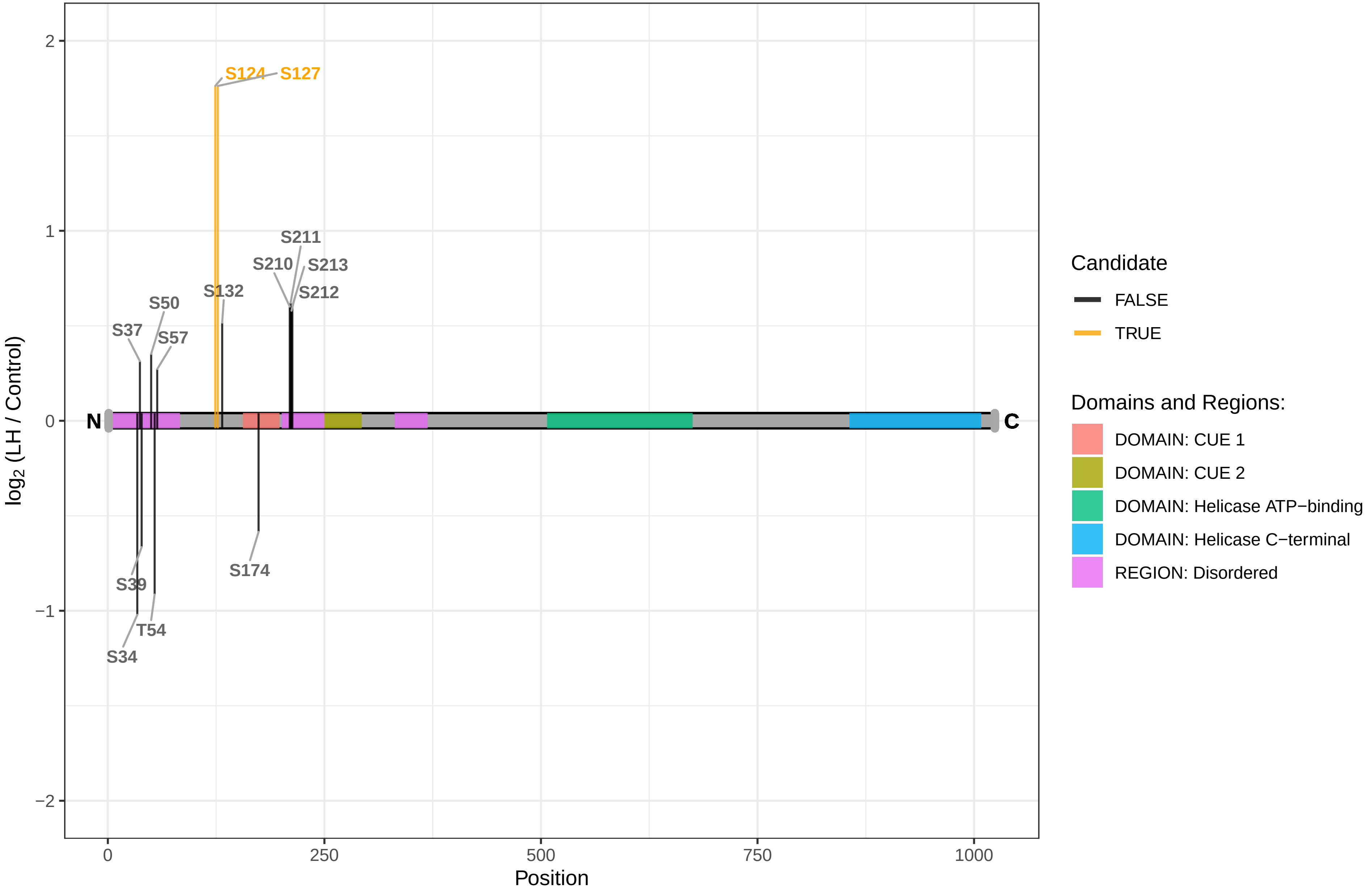

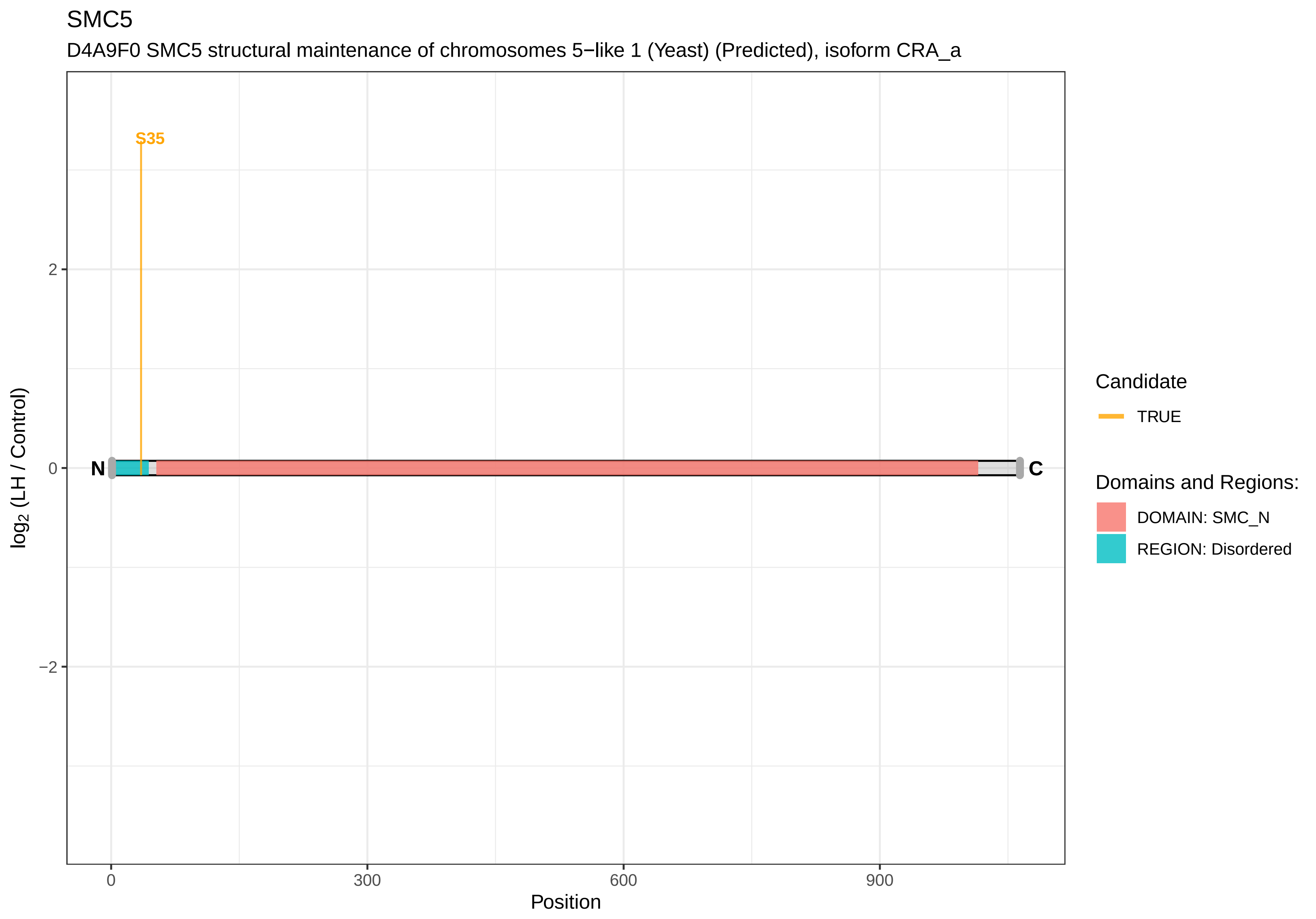

SMC5
D4A9F0 SMC5 structural maintenance of chromosomes 5−like 1 (Yeast) (Predicted), isoform CRA_a
S35
N
C
2
0
−2
log2 (LH / Control)
0
300
600
900
Position
Candidate
TRUE
Domains and Regions:
DOMAIN: SMC_N
REGION: Disordered

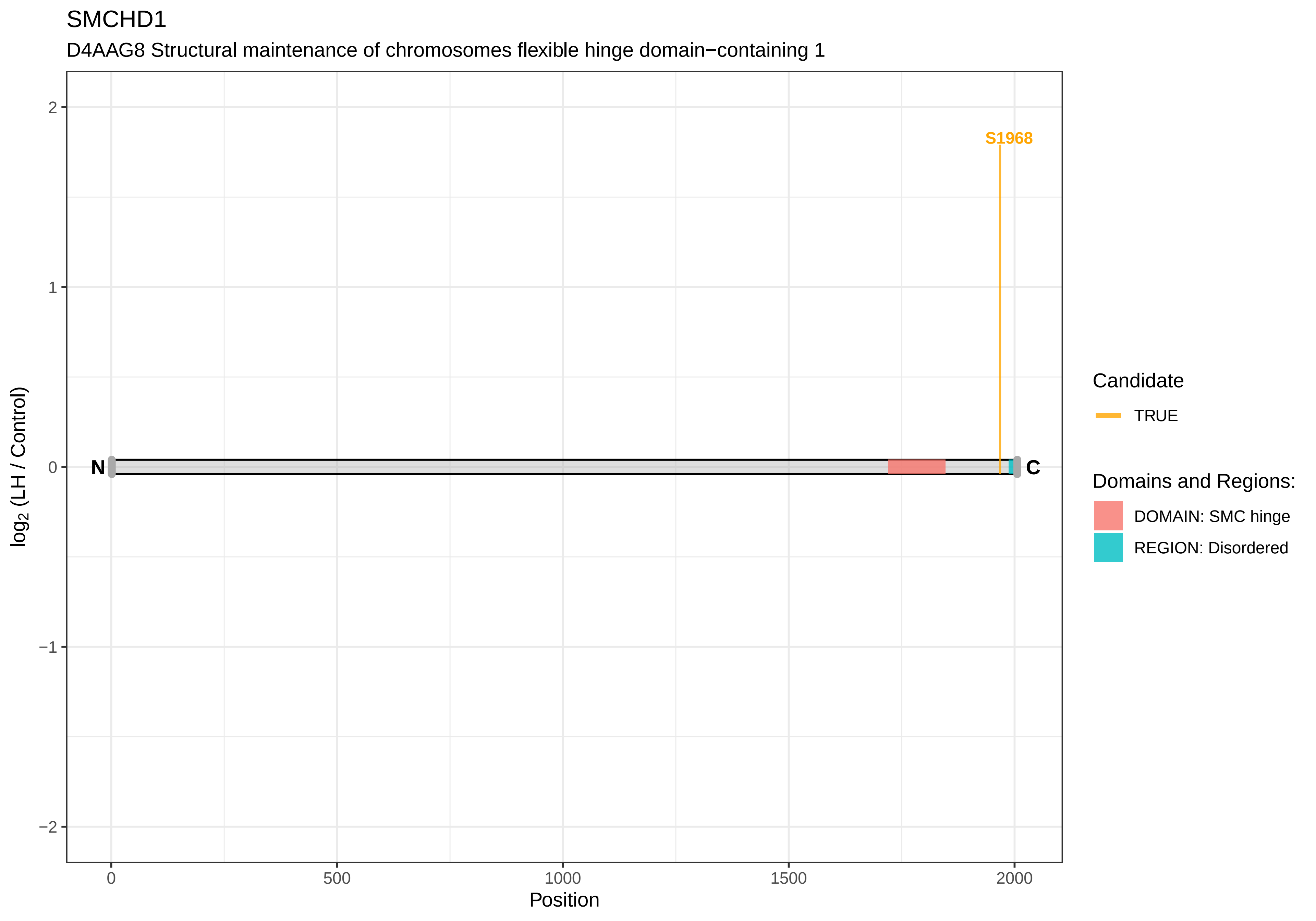

SMCHD1
D4AAG8 Structural maintenance of chromosomes flexible hinge domain−containing 1
S1968
N
C
2
1
0
−1
−2
0
500
1000
1500
2000
Position
log2 (LH / Control)
Candidate
TRUE
Domains and Regions:
DOMAIN: SMC hinge
REGION: Disordered

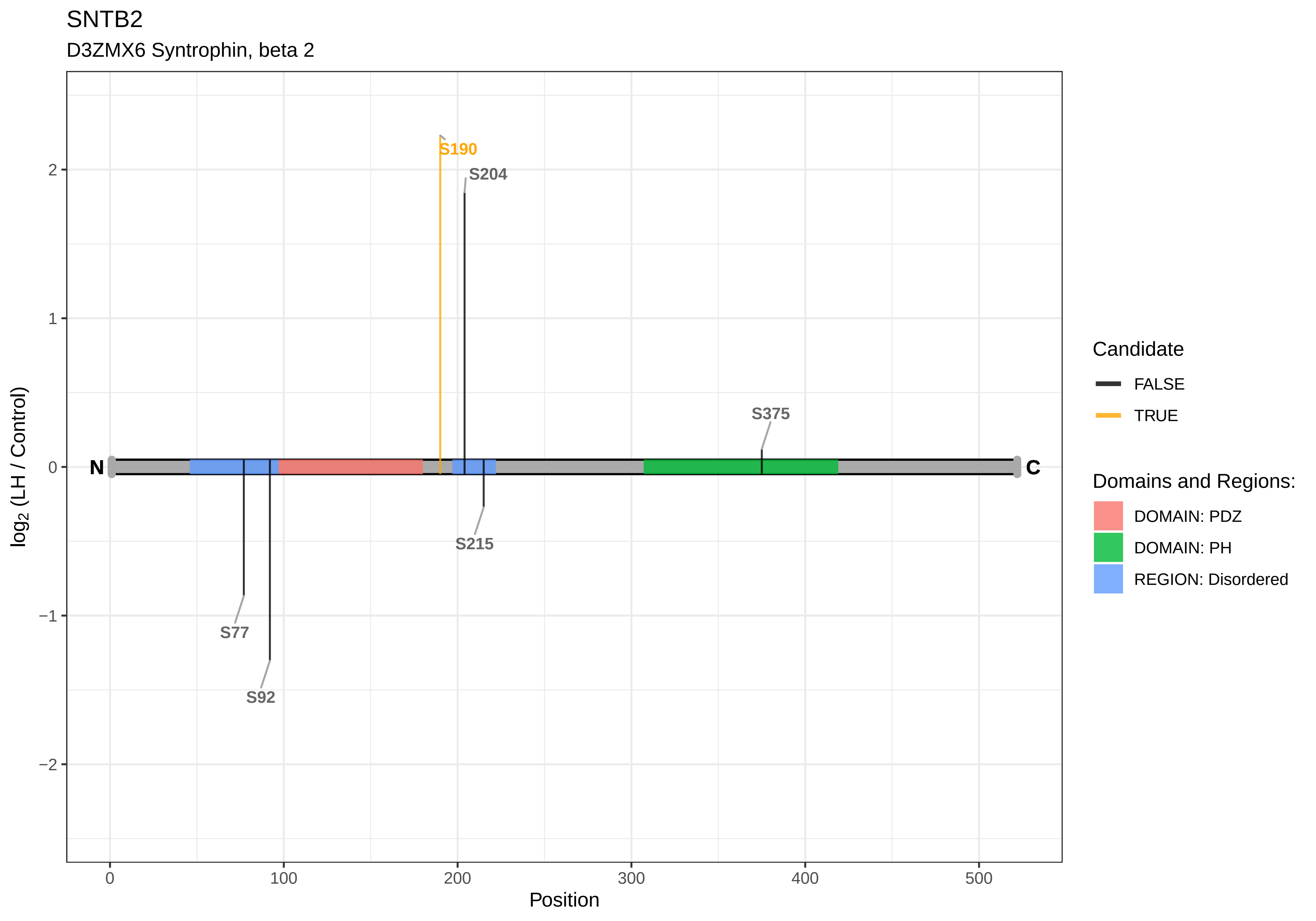

SNTB2
D3ZMX6 Syntrophin, beta 2
S190
S204
S375
S215
S77
S92
N
C
log2 (LH / Control)
Position
Candidate
FALSE
TRUE
Domains and Regions:
DOMAIN: PDZ
DOMAIN: PH
REGION: Disordered

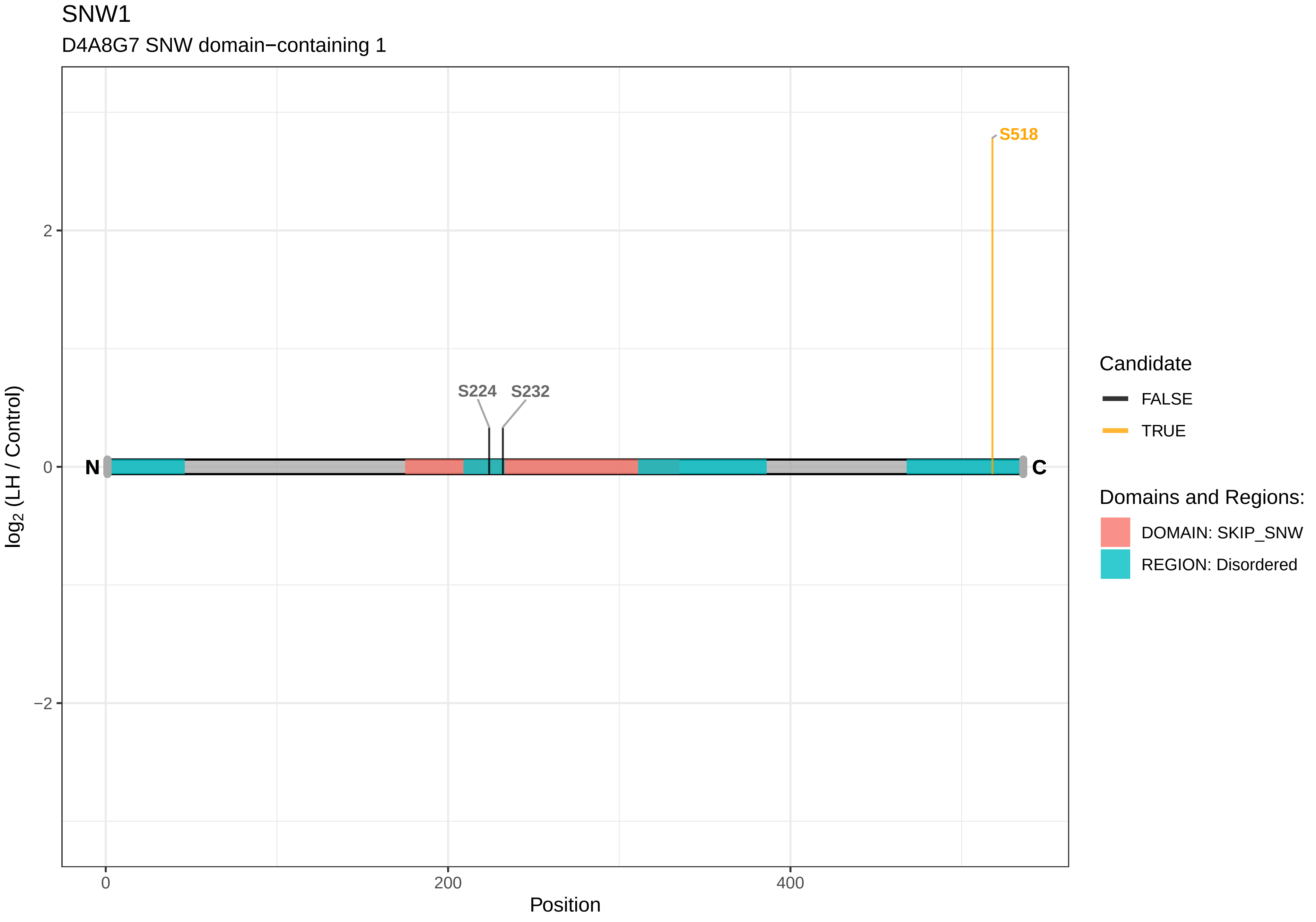

SNW1
D4A8G7 SNW domain−containing 1
S518
S224
S232
N
C
2
0
−2
0
200
400
Position
log2 (LH / Control)
Candidate
FALSE
TRUE
Domains and Regions:
DOMAIN: SKIP_SNW
REGION: Disordered

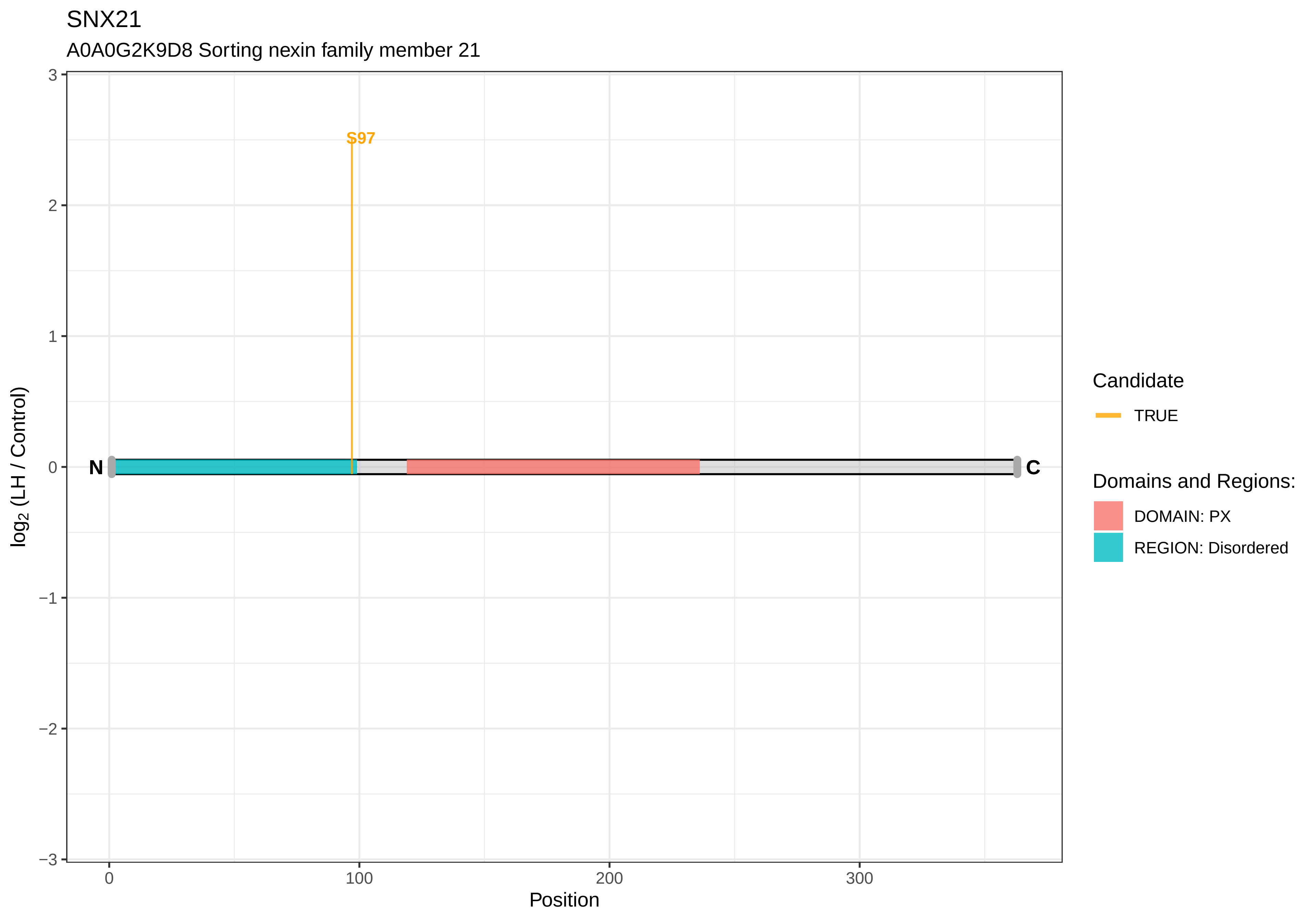

SNX21
A0A0G2K9D8 Sorting nexin family member 21
S97
N
C
3
2
1
0
−1
−2
−3
log2 (LH / Control)
0
100
200
300
Position
Candidate
TRUE
Domains and Regions:
DOMAIN: PX
REGION: Disordered

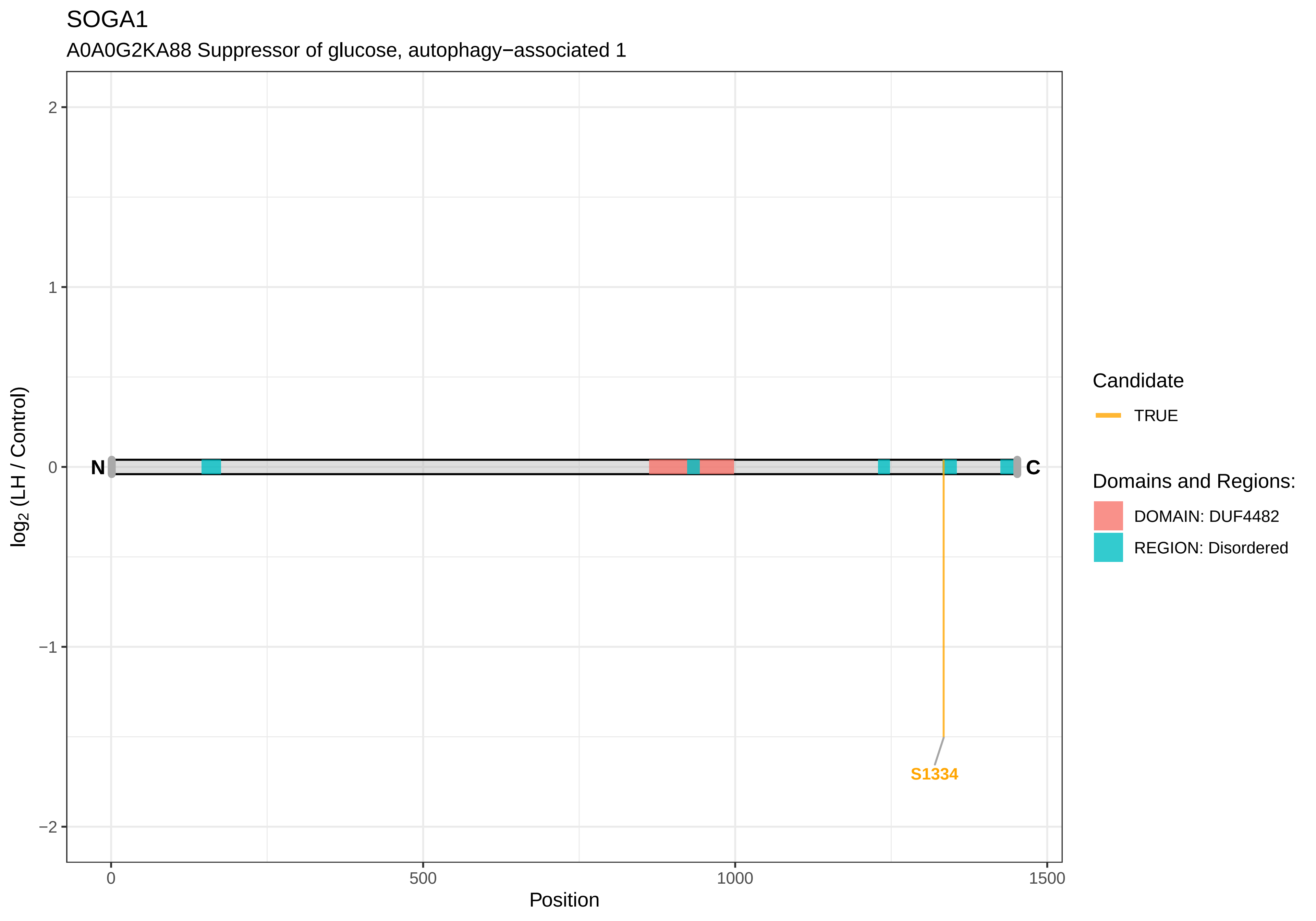
SOGA1
A0A0G2KA88 Suppressor of glucose, autophagy−associated 1
log2 (LH / Control)
2
1
0
−1
−2
N
C
S1334
0
500
1000
1500
Position
Candidate
TRUE
Domains and Regions:
DOMAIN: DUF4482
REGION: Disordered

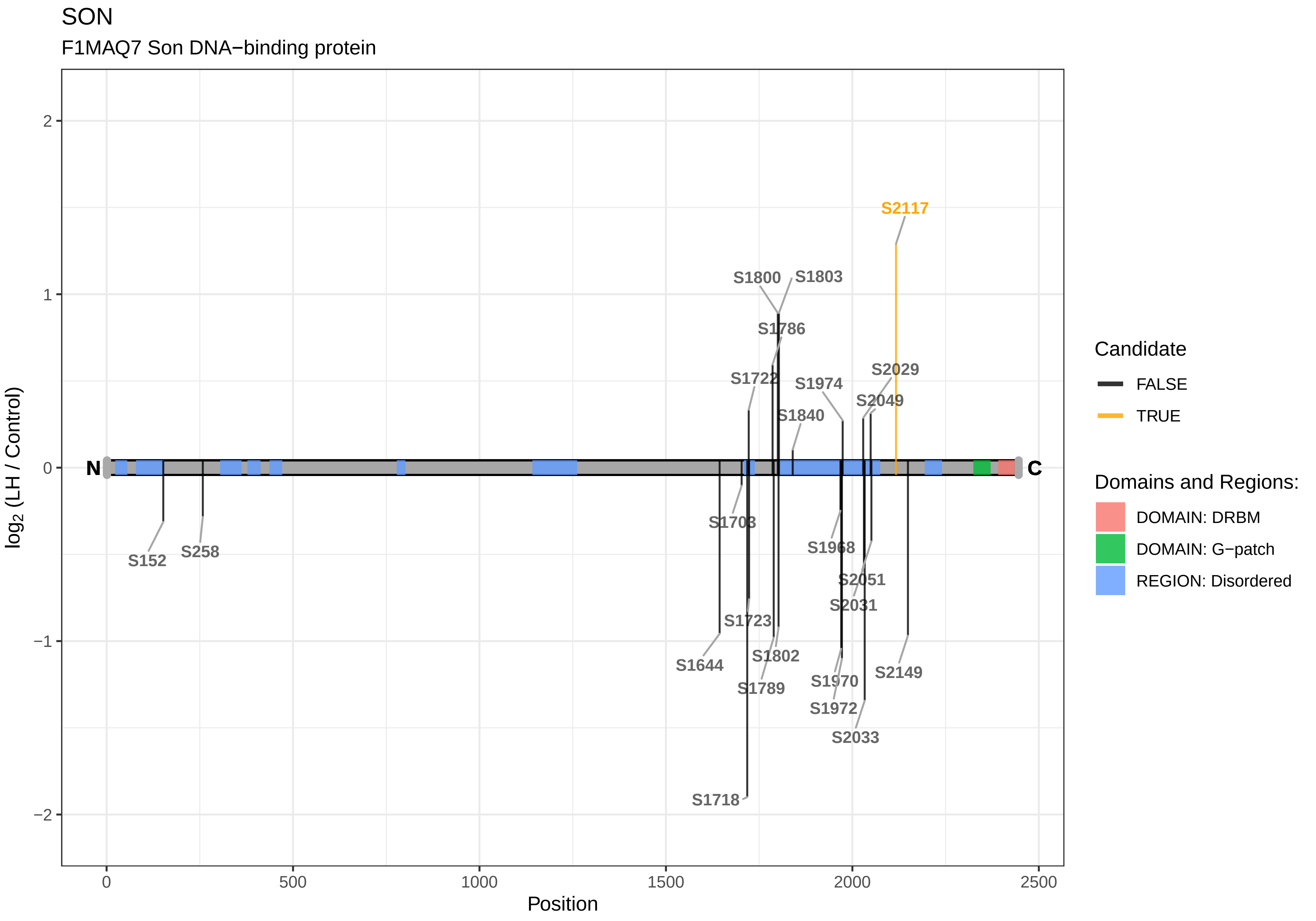
SON
F1MAQ7 Son DNA−binding protein
log2 (LH / Control)
Position
N
C
S152
S258
S1644
S1703
S1718
S1722
S1723
S1786
S1789
S1800
S1802
S1803
S1840
S1968
S1970
S1972
S1974
S2029
S2031
S2033
S2049
S2051
S2117
S2149
Candidate
FALSE
TRUE
Domains and Regions:
DOMAIN: DRBM
DOMAIN: G−patch
REGION: Disordered

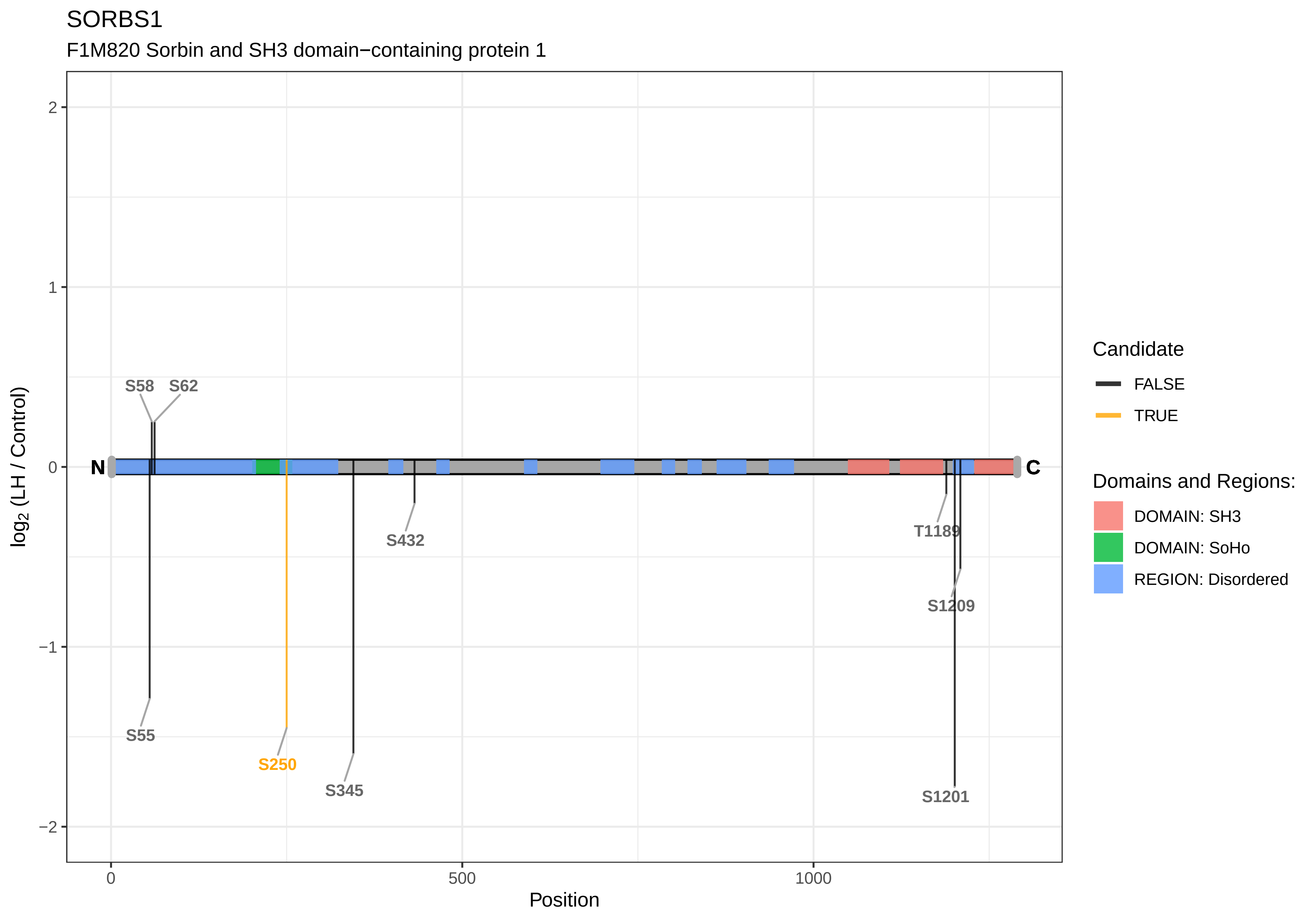

SORBS1
F1M820 Sorbin and SH3 domain−containing protein 1
log2 (LH / Control)
Position
2
1
0
−1
−2
0
500
1000
N
C
S58
S62
S55
S250
S345
S432
T1189
S1209
S1201
Candidate
FALSE
TRUE
Domains and Regions:
DOMAIN: SH3
DOMAIN: SoHo
REGION: Disordered

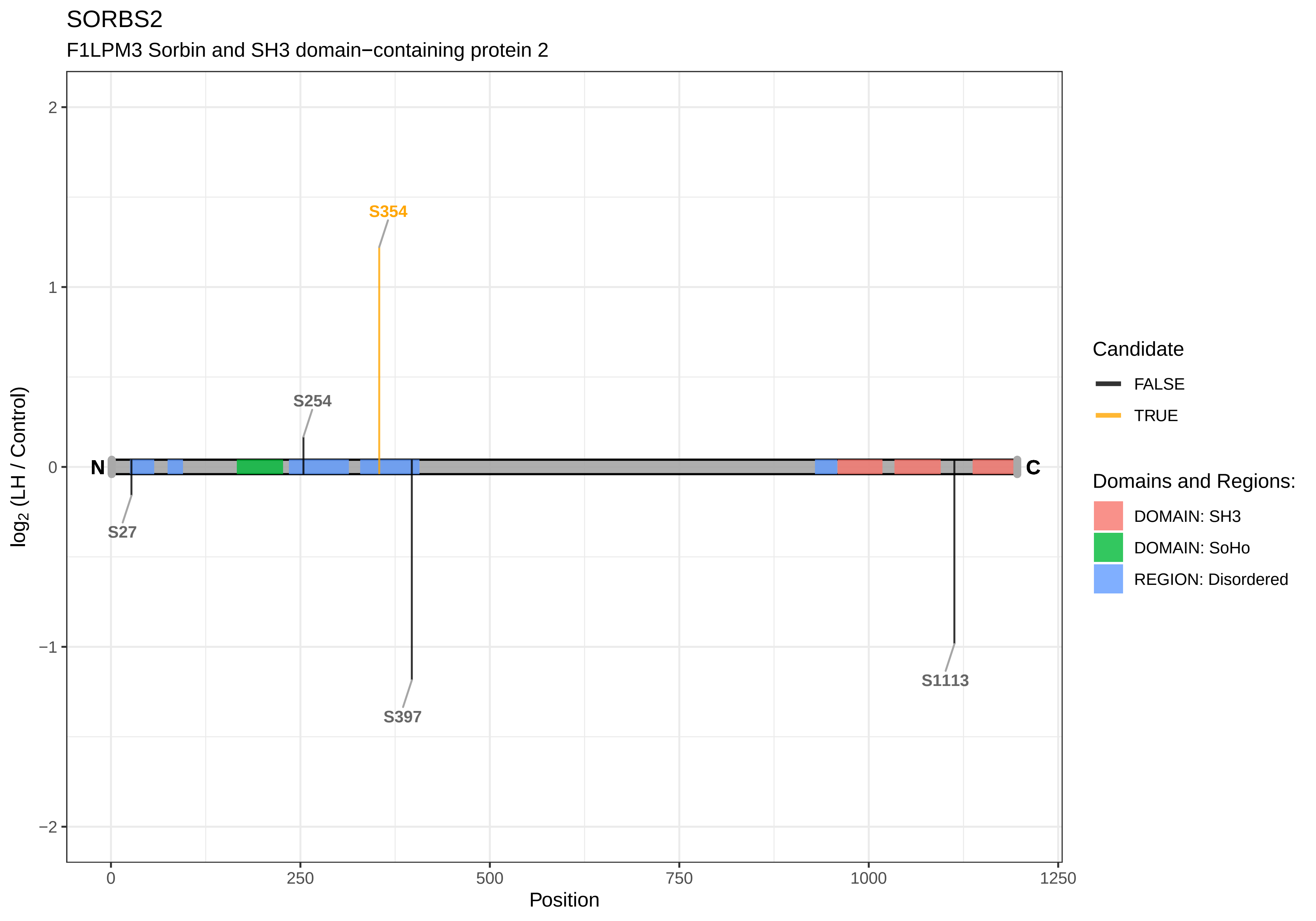
SORBS2
F1LPM3 Sorbin and SH3 domain−containing protein 2
log2 (LH / Control)
Position
S354
S254
S27
S397
S1113
N
C
2
1
0
−1
−2
0
250
500
750
1000
1250
Candidate
FALSE
TRUE
Domains and Regions:
DOMAIN: SH3
DOMAIN: SoHo
REGION: Disordered

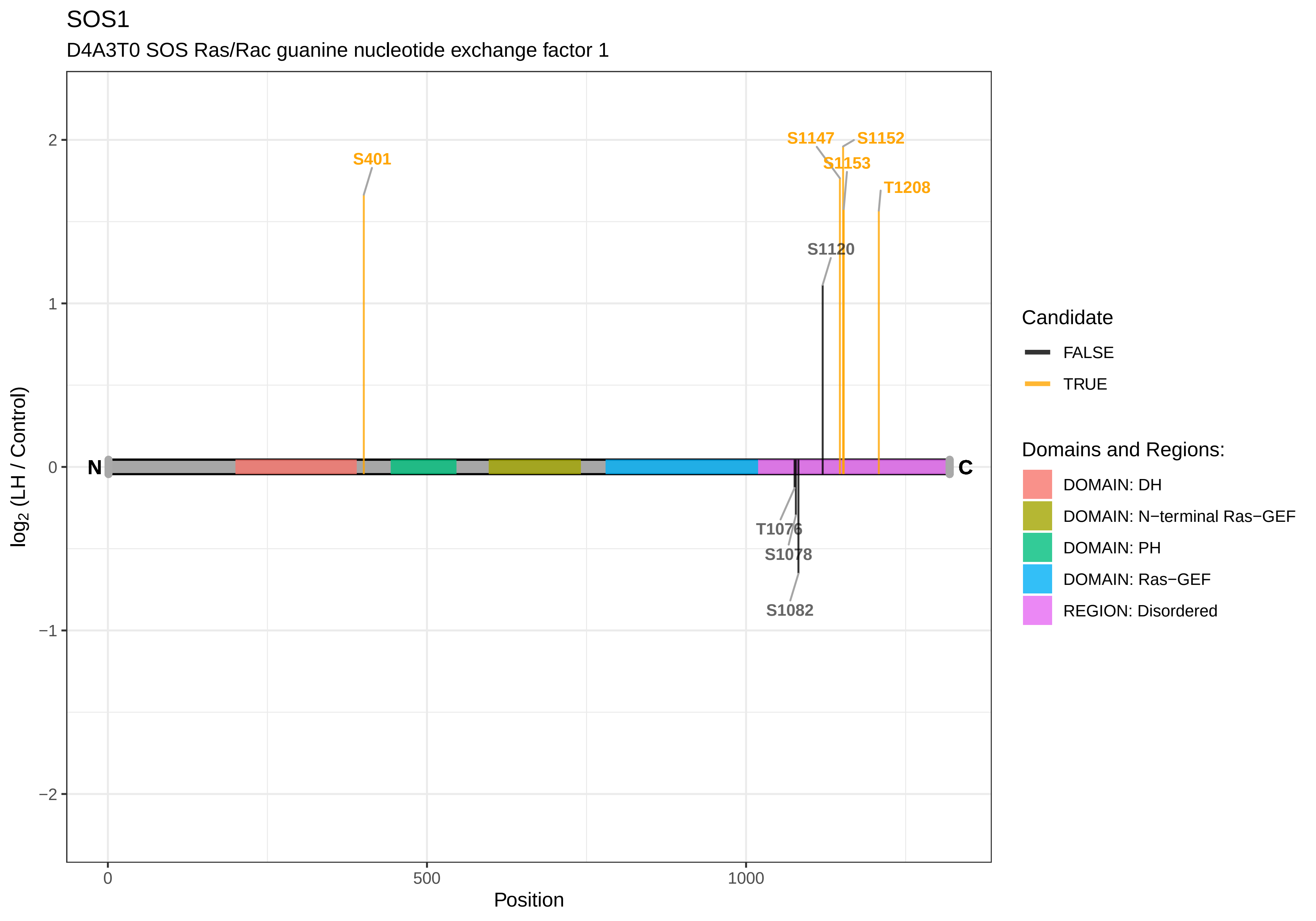

SOS1
D4A3T0 SOS Ras/Rac guanine nucleotide exchange factor 1
S401
S1147
S1152
S1153
T1208
S1120
T1076
S1078
S1082
N
C
2
1
0
−1
−2
log2 (LH / Control)
0
500
1000
Position
Candidate
FALSE
TRUE
Domains and Regions:
DOMAIN: DH
DOMAIN: N−terminal Ras−GEF
DOMAIN: PH
DOMAIN: Ras−GEF
REGION: Disordered

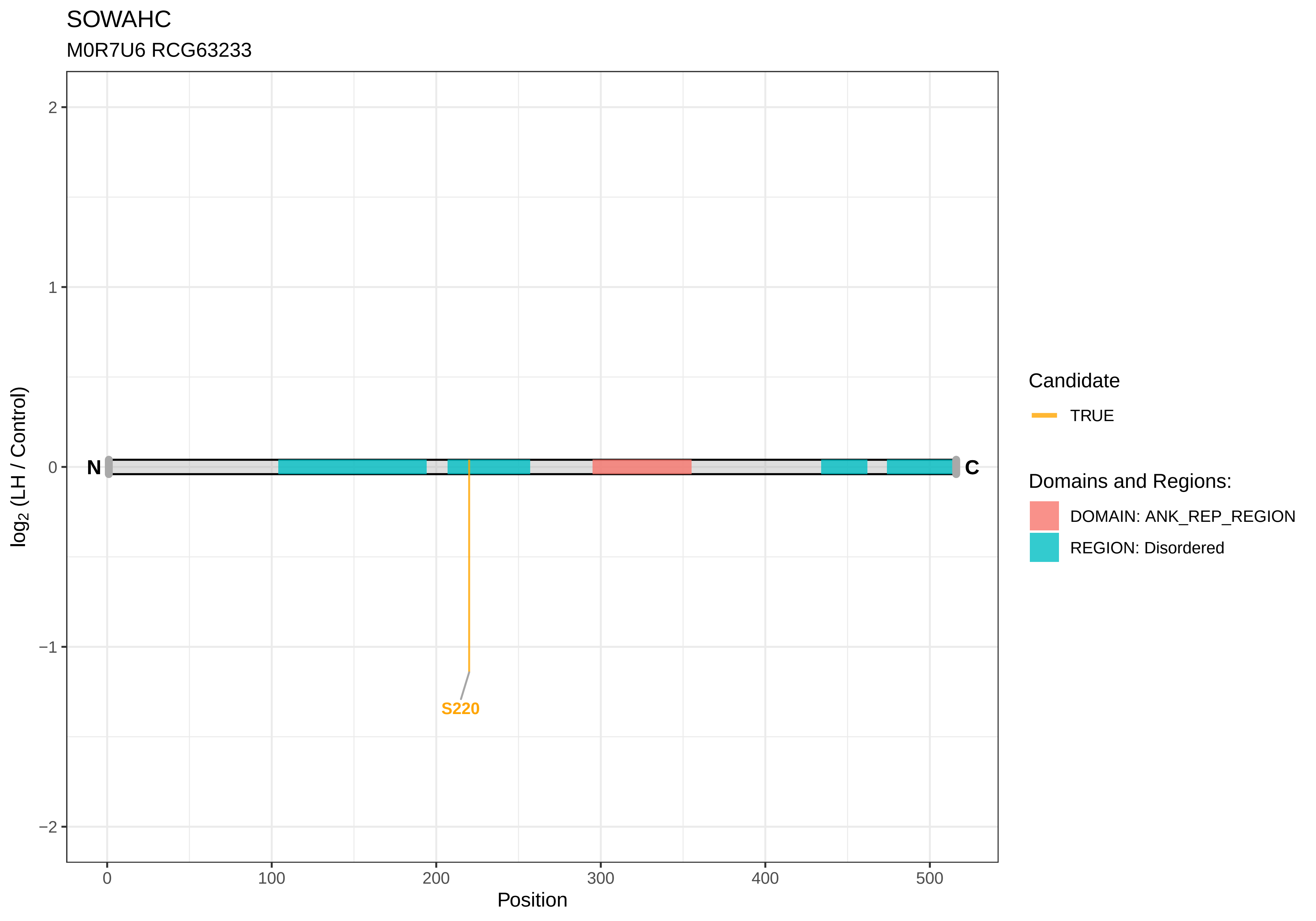
SOWAHC
M0R7U6 RCG63233
log2 (LH / Control)
2
1
0
−1
−2
N
C
S220
0
100
200
300
400
500
Position
Candidate
TRUE
Domains and Regions:
DOMAIN: ANK_REP_REGION
REGION: Disordered

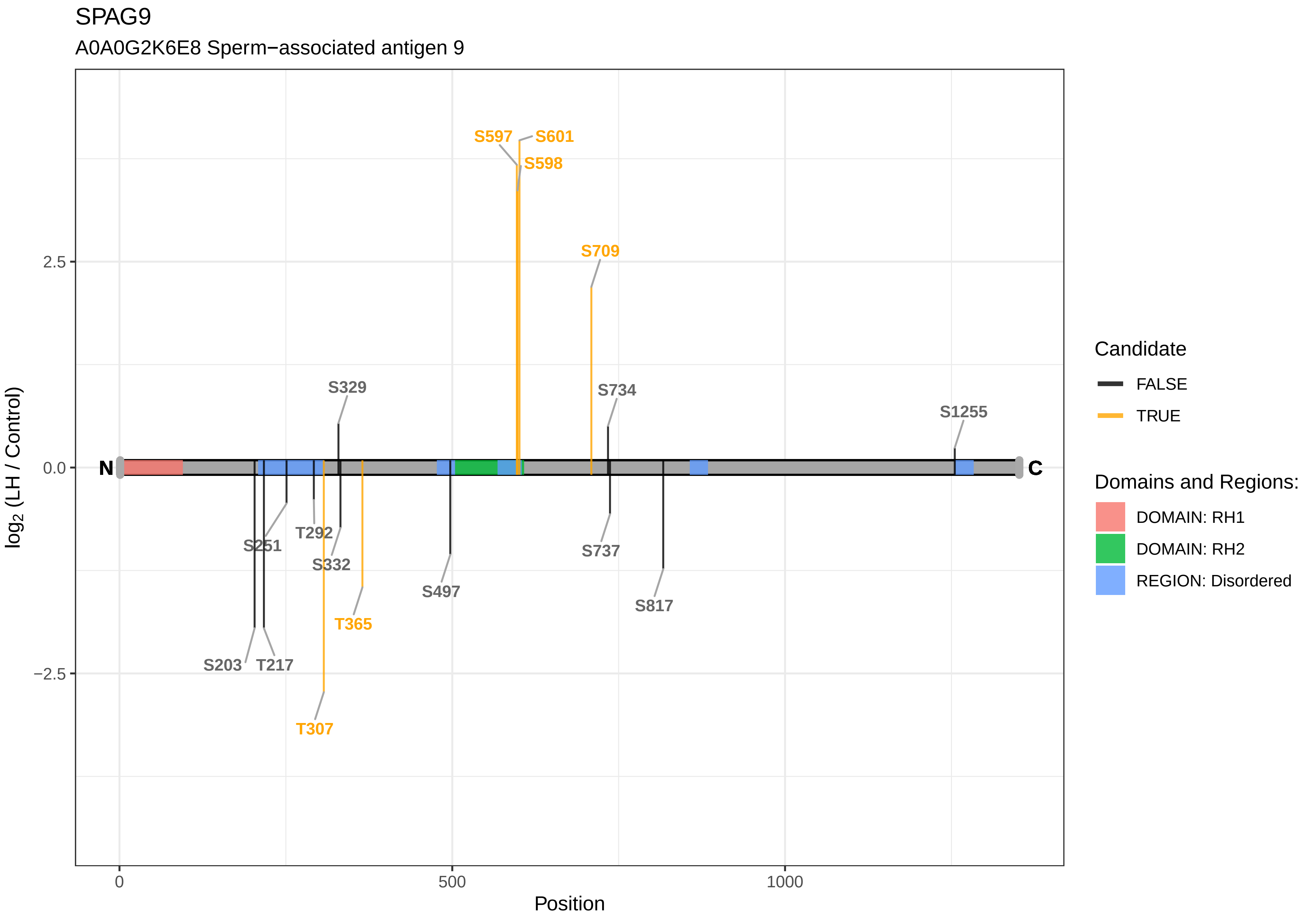
SPAG9
A0A0G2K6E8 Sperm−associated antigen 9
S597
S601
S598
S709
S329
S734
S1255
N
C
S251
T292
S332
S497
S737
S817
T365
S203
T217
T307
2.5
0.0
−2.5
log2 (LH / Control)
0
500
1000
Position
Candidate
FALSE
TRUE
Domains and Regions:
DOMAIN: RH1
DOMAIN: RH2
REGION: Disordered

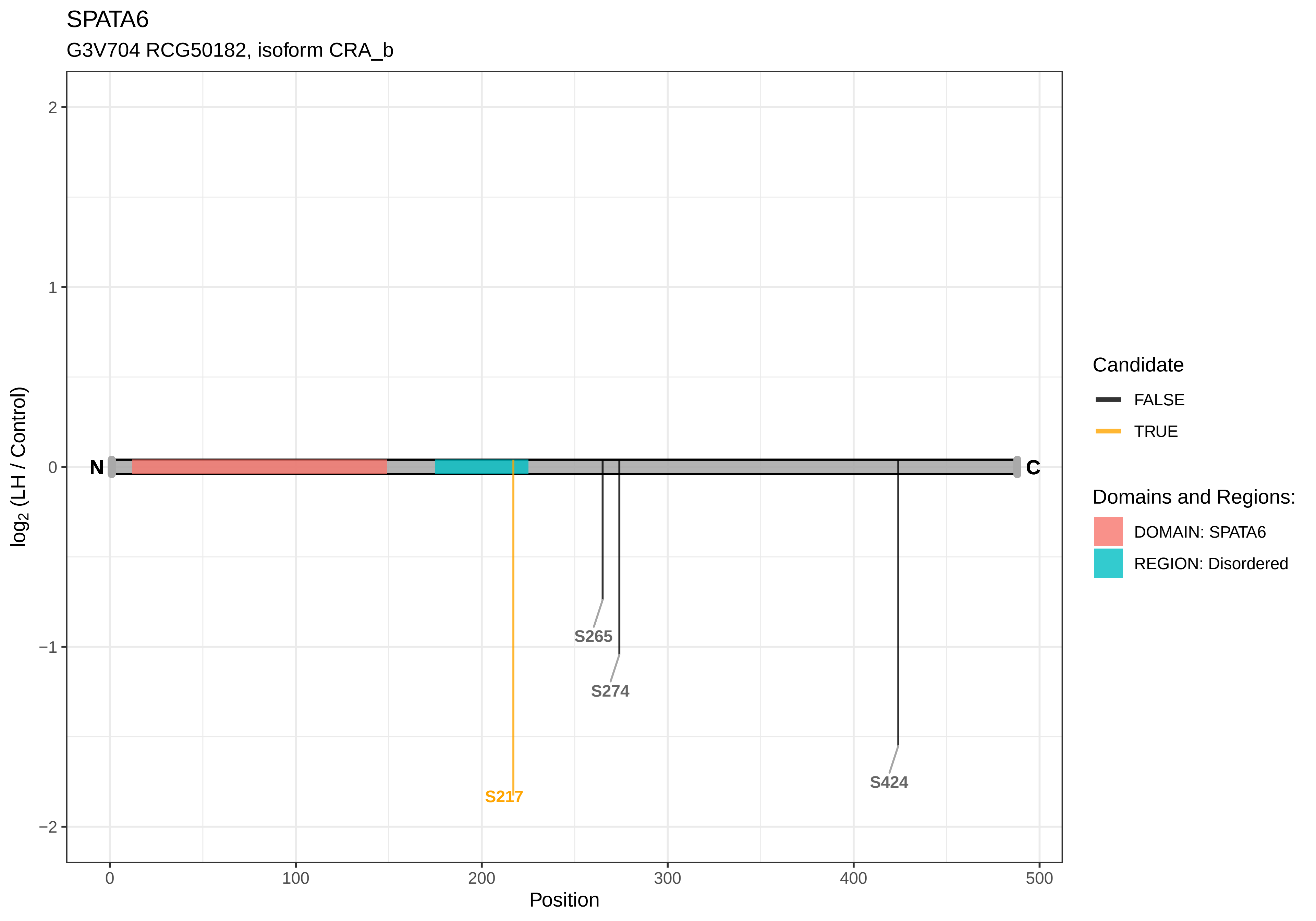

SPATA6
G3V704 RCG50182, isoform CRA_b
log2 (LH / Control)
2
1
0
−1
−2
N
C
S217
S265
S274
S424
0
100
200
300
400
500
Position
Candidate
FALSE
TRUE
Domains and Regions:
DOMAIN: SPATA6
REGION: Disordered

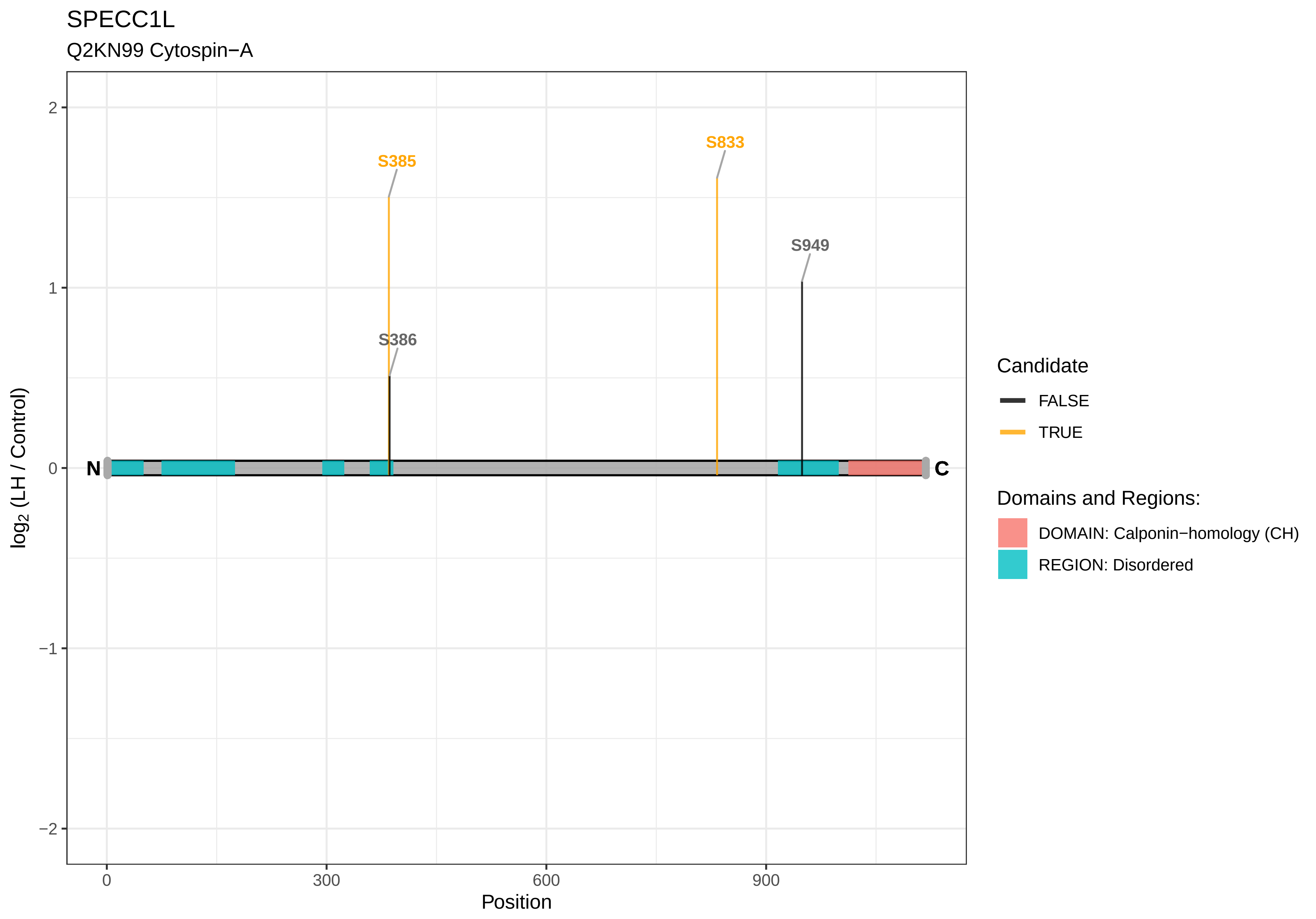
SPECC1L
Q2KN99 Cytospin−A
S385
S833
S949
S386
N
C
2
1
0
−1
−2
0
300
600
900
Position
log2 (LH / Control)
Candidate
FALSE
TRUE
Domains and Regions:
DOMAIN: Calponin−homology (CH)
REGION: Disordered

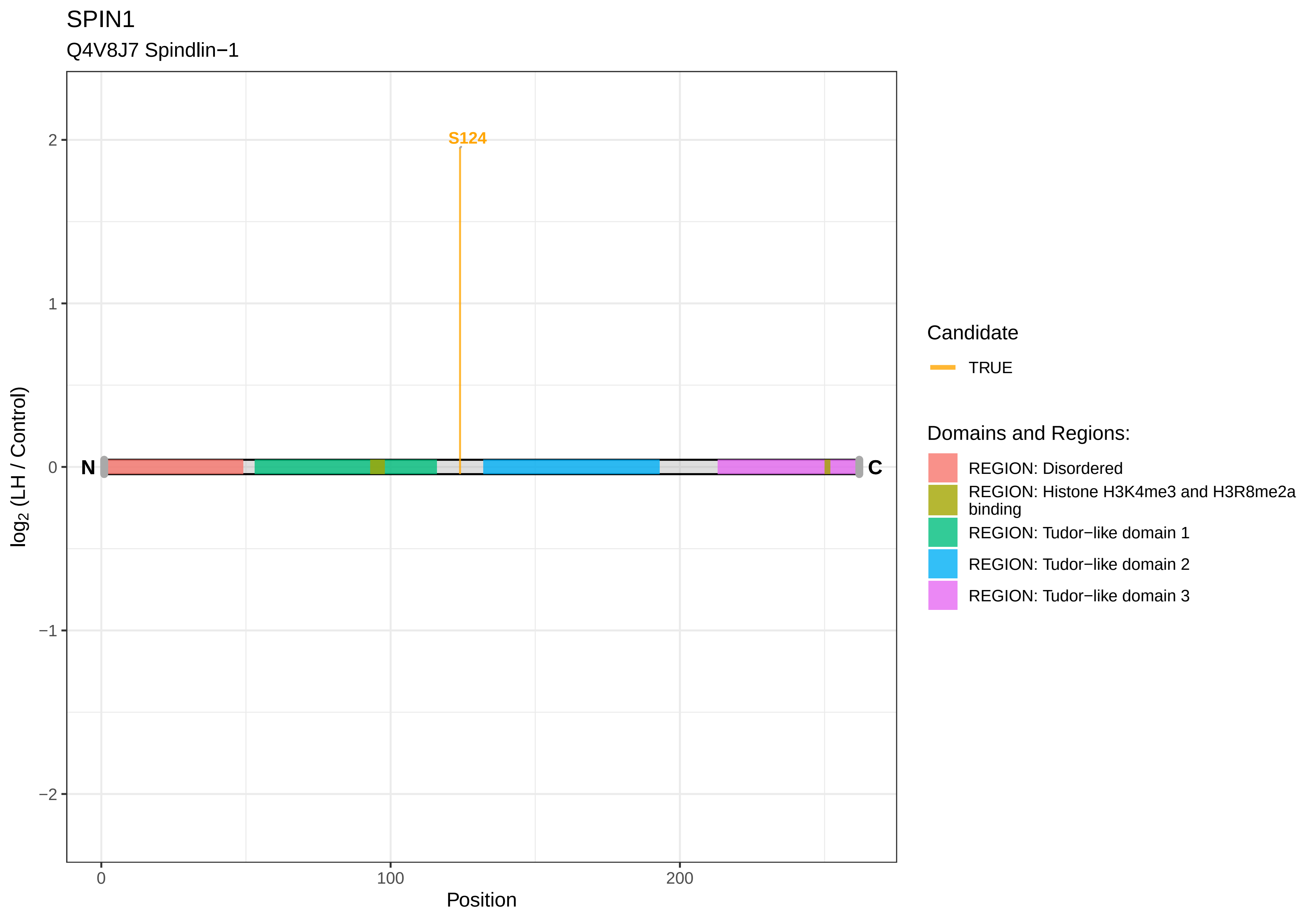
SPIN1
Q4V8J7 Spindlin−1
S124
N
C
2
1
0
−1
−2
log2 (LH / Control)
0
100
200
Position
Candidate
TRUE
Domains and Regions:
REGION: Disordered
REGION: Histone H3K4me3 and H3R8me2a binding
REGION: Tudor−like domain 1
REGION: Tudor−like domain 2
REGION: Tudor−like domain 3

SPIRE1

D3ZEX7 Spire homolog 1 (Drosophila) (Predicted), isoform CRA_b

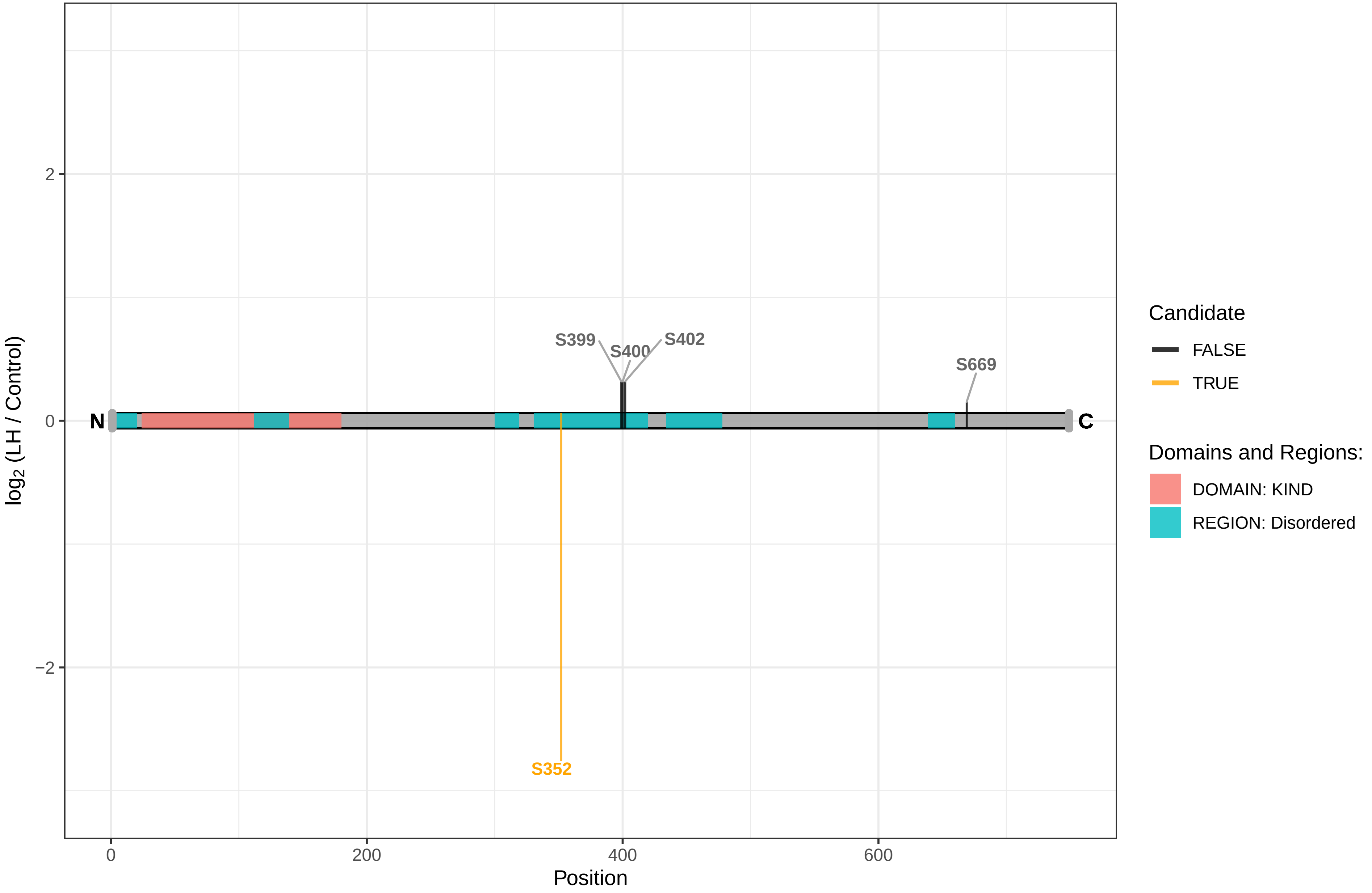

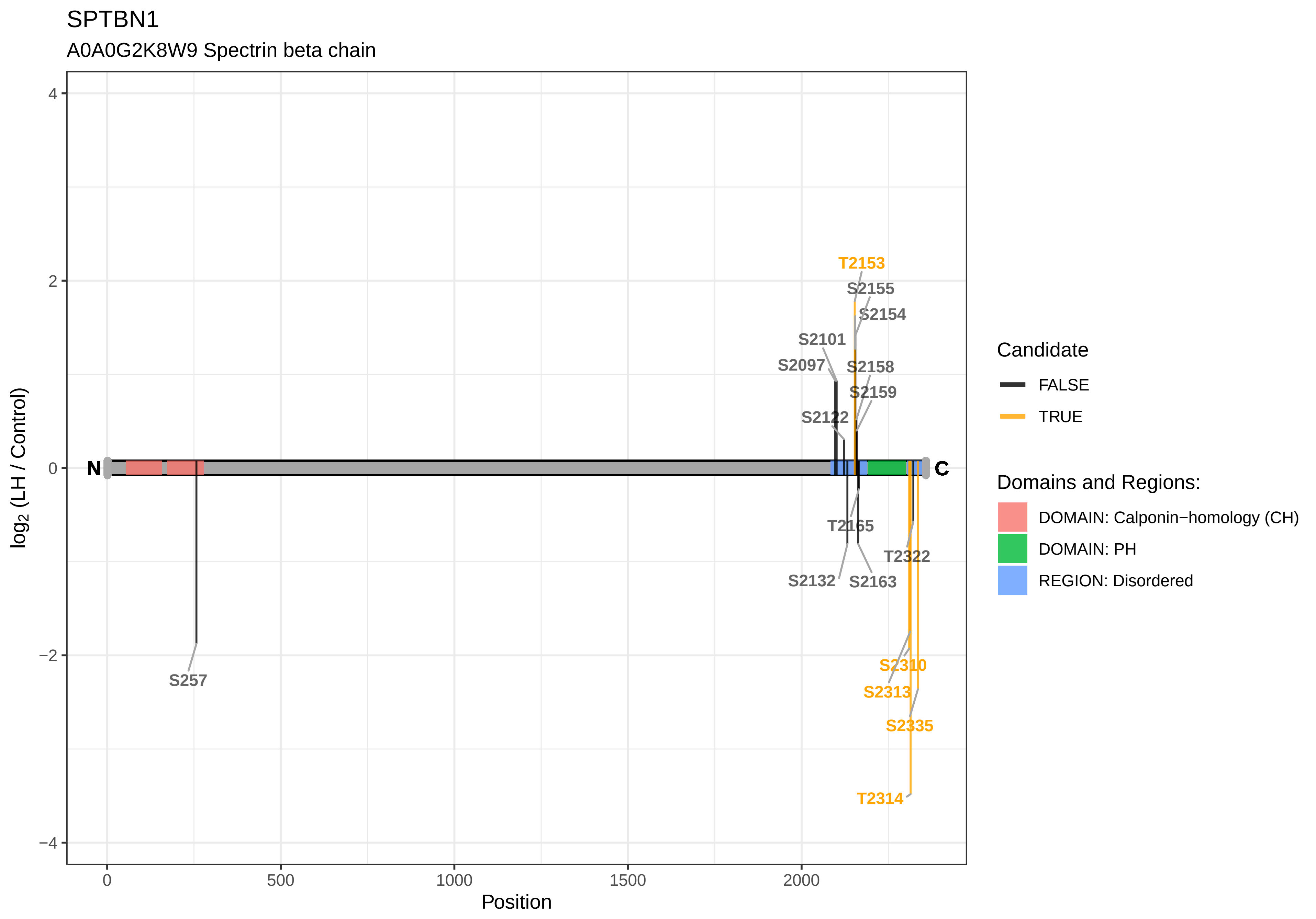

SPTBN1
A0A0G2K8W9 Spectrin beta chain
T2153
S2155
S2154
S2101
S2097
S2158
S2159
S2122
N
C
T2165
T2322
S2132
S2163
S257
S2310
S2313
S2335
T2314
4
2
0
−2
−4
log2 (LH / Control)
0
500
1000
1500
2000
Position
Candidate
FALSE
TRUE
Domains and Regions:
DOMAIN: Calponin−homology (CH)
DOMAIN: PH
REGION: Disordered

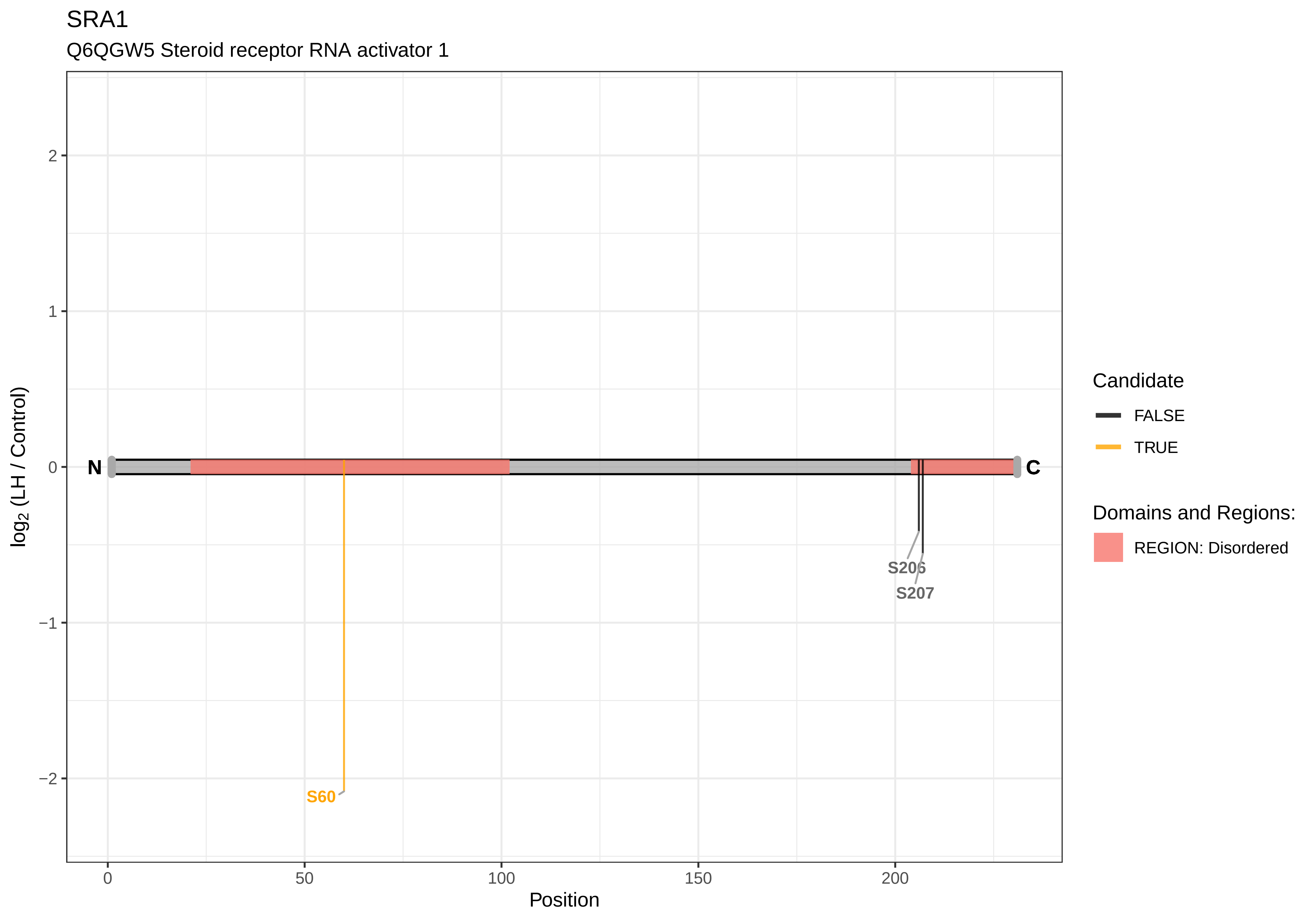

SRA1
Q6QGW5 Steroid receptor RNA activator 1
log2 (LH / Control)
Position
N
C
S60
S206
S207
Candidate
FALSE
TRUE
Domains and Regions:
REGION: Disordered

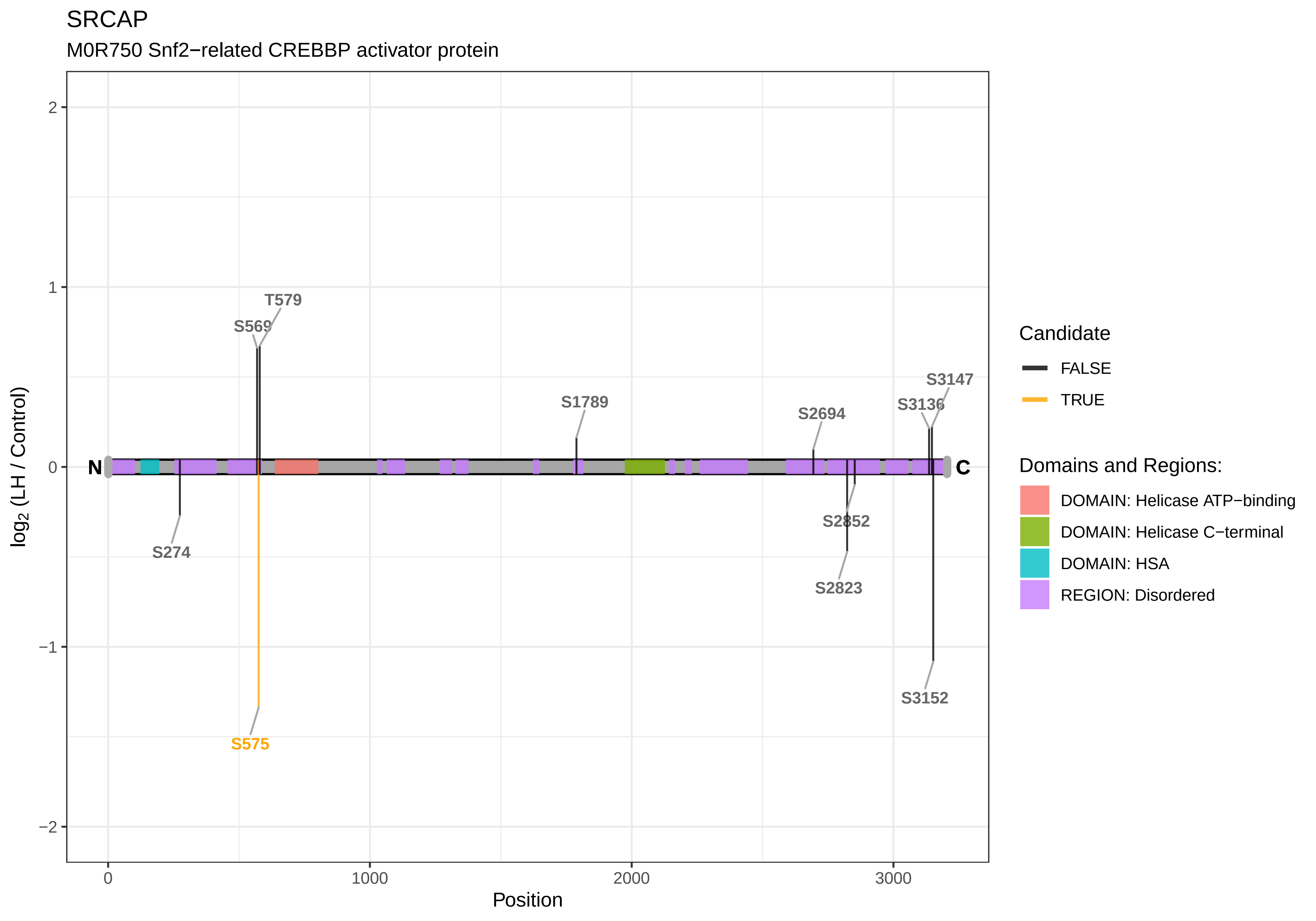

SRCAP
M0R750 Snf2−related CREBBP activator protein
log2 (LH / Control)
Position
2
1
0
−1
−2
0
1000
2000
3000
N
C
T579
S569
S274
S575
S1789
S2694
S3136
S3147
S2852
S2823
S3152
Candidate
FALSE
TRUE
Domains and Regions:
DOMAIN: Helicase ATP−binding
DOMAIN: Helicase C−terminal
DOMAIN: HSA
REGION: Disordered

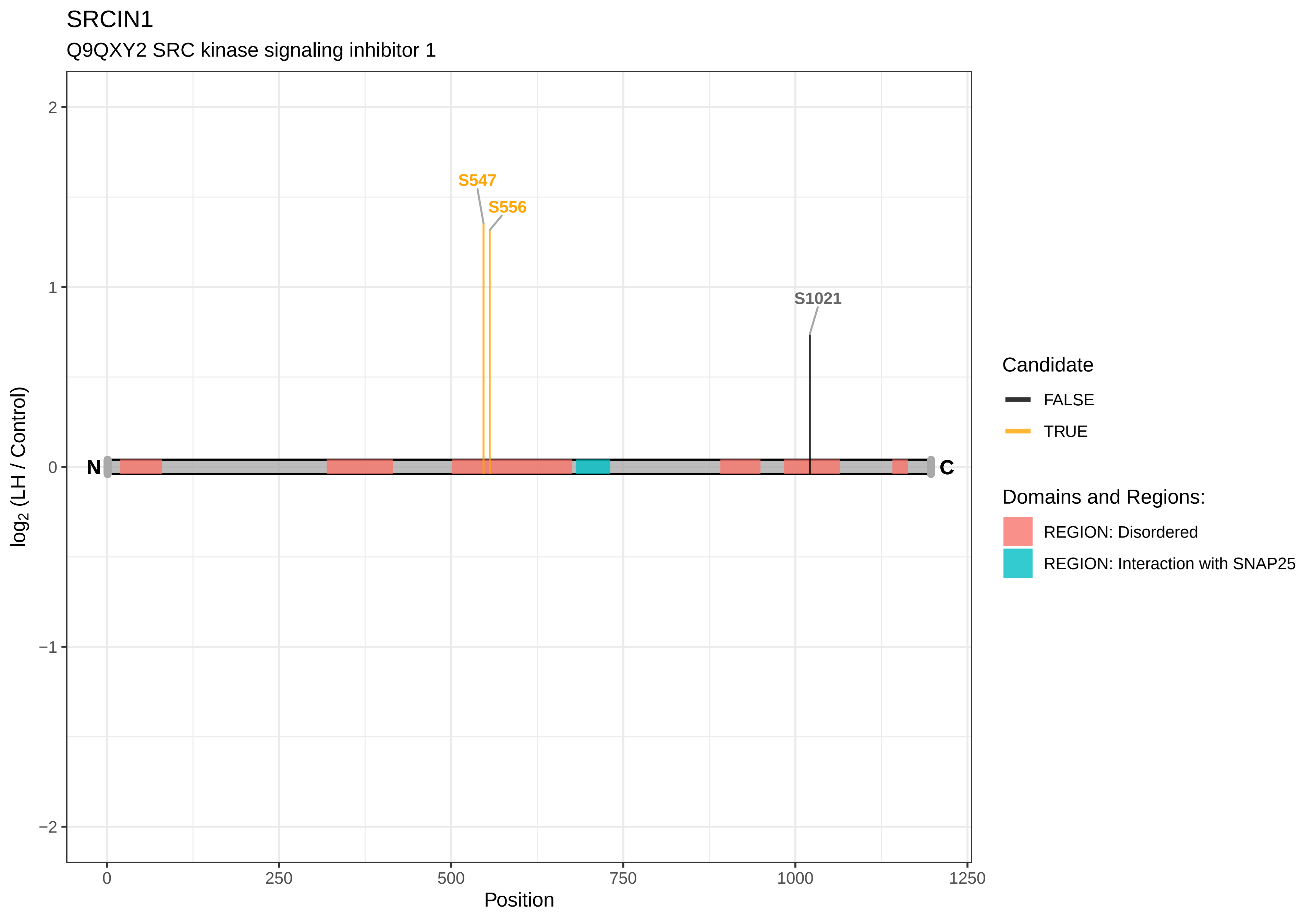

SRCIN1
Q9QXY2 SRC kinase signaling inhibitor 1
S547
S556
S1021
N
C
2
1
0
−1
−2
log2 (LH / Control)
0
250
500
750
1000
1250
Position
Candidate
FALSE
TRUE
Domains and Regions:
REGION: Disordered
REGION: Interaction with SNAP25

SRGAP1

F1M287 SLIT−ROBO Rho GTPase−activating protein 1

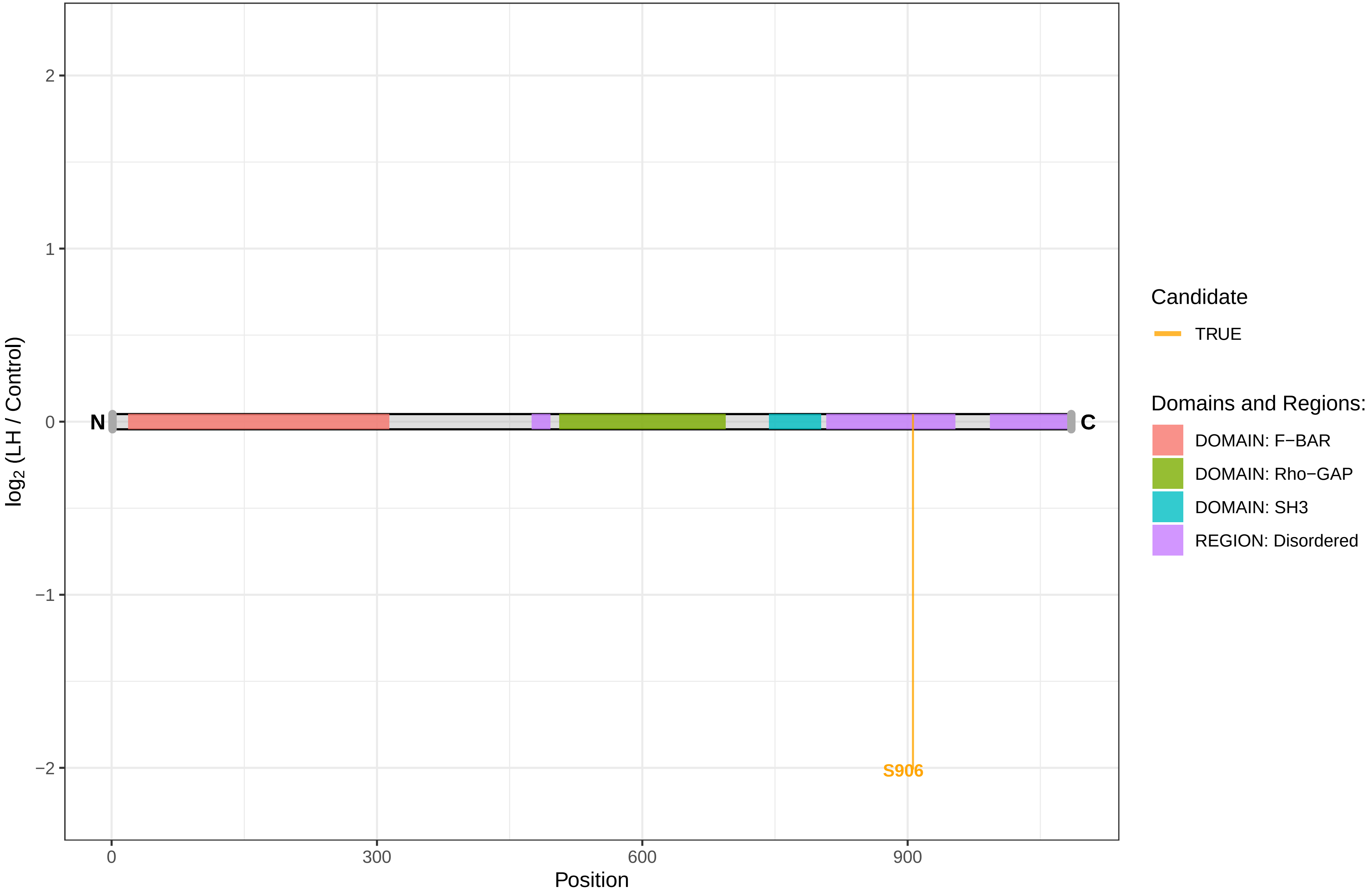

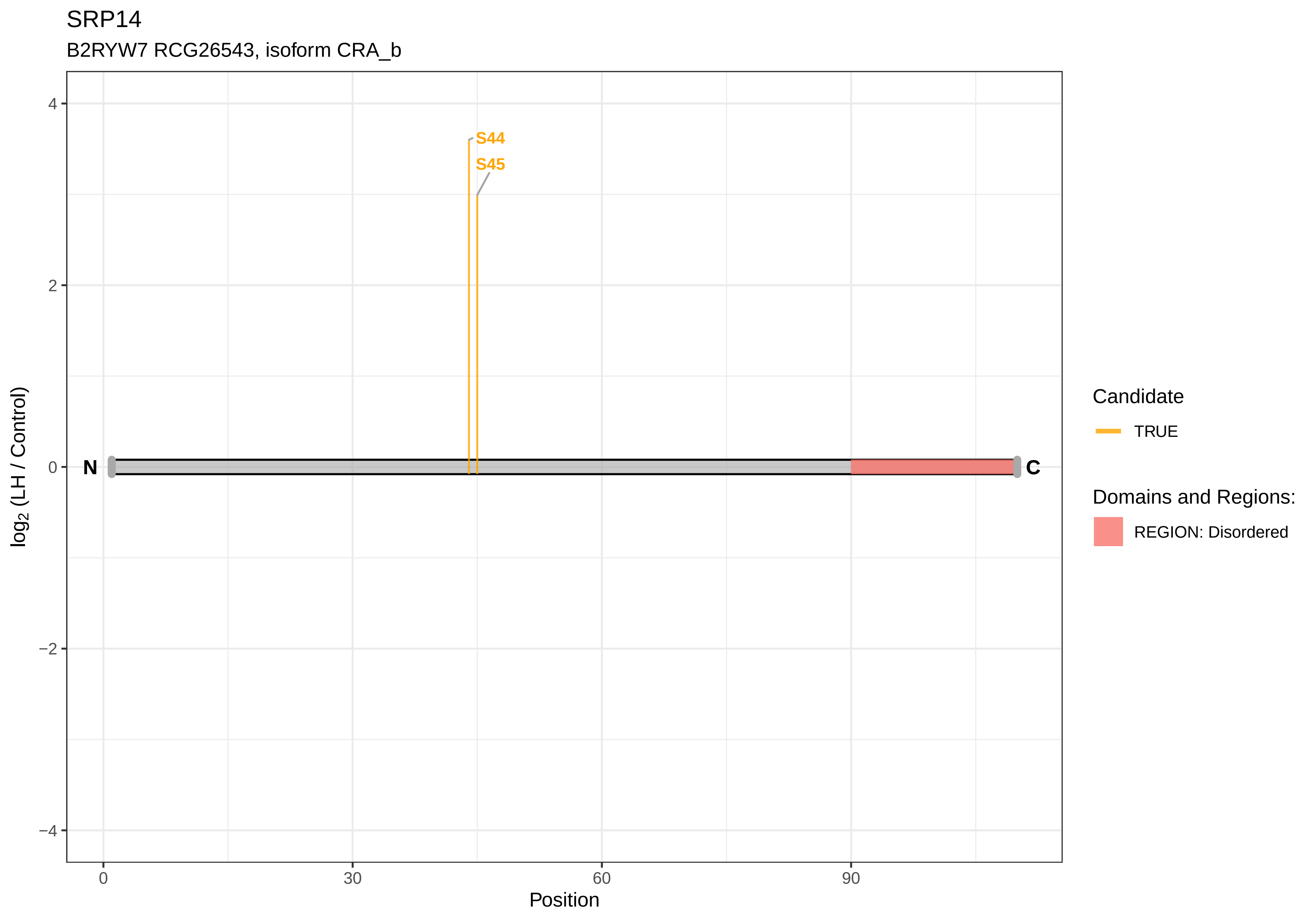

SRP14
B2RYW7 RCG26543, isoform CRA_b
S44
S45
N
C
4
2
0
−2
−4
log2 (LH / Control)
0
30
60
90
Position
Candidate
TRUE
Domains and Regions:
REGION: Disordered

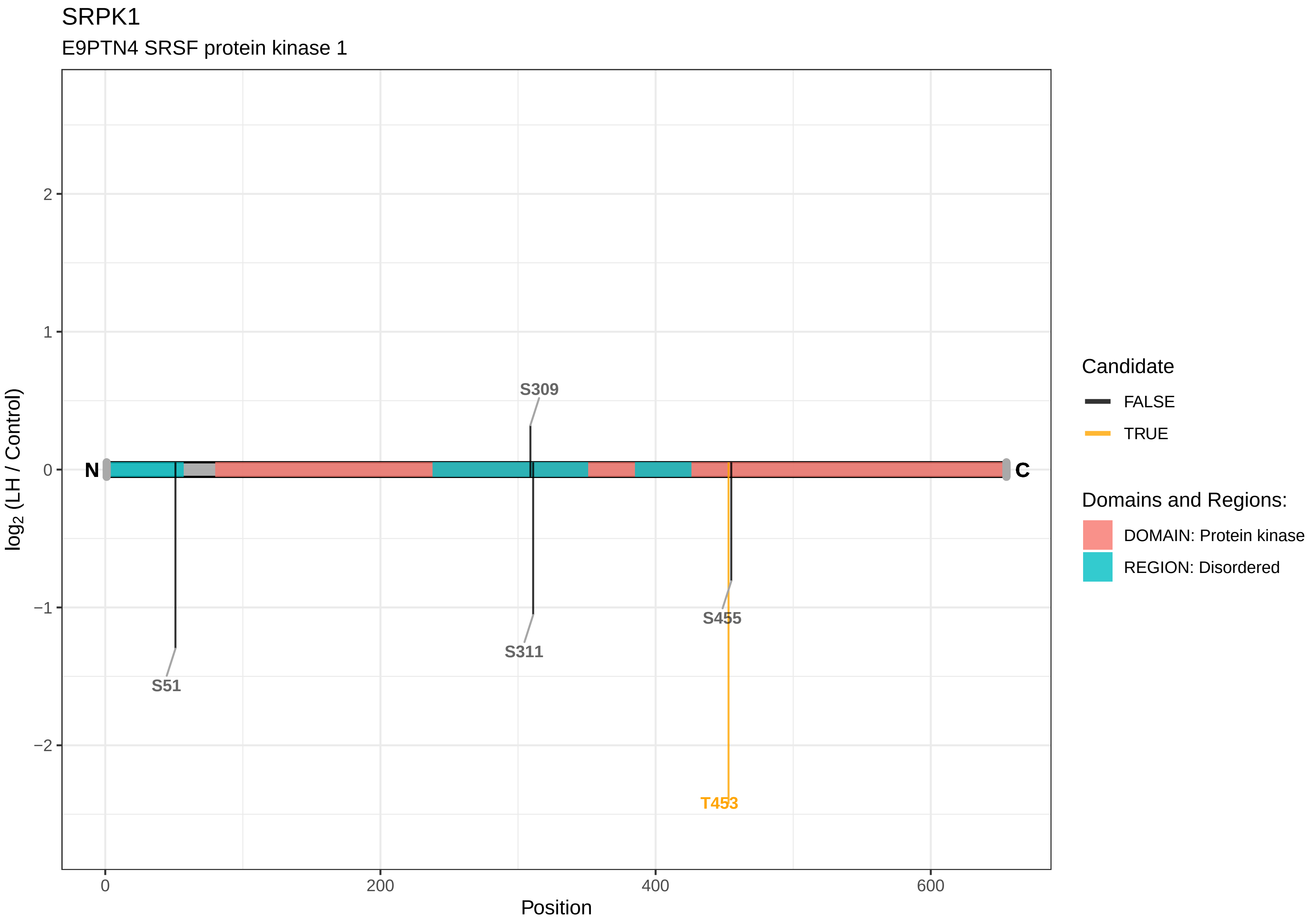
SRPK1
E9PTN4 SRSF protein kinase 1
log2 (LH / Control)
Position
N
C
S309
S311
S51
S455
T453
2
1
0
−1
−2
0
200
400
600
Candidate
FALSE
TRUE
Domains and Regions:
DOMAIN: Protein kinase
REGION: Disordered

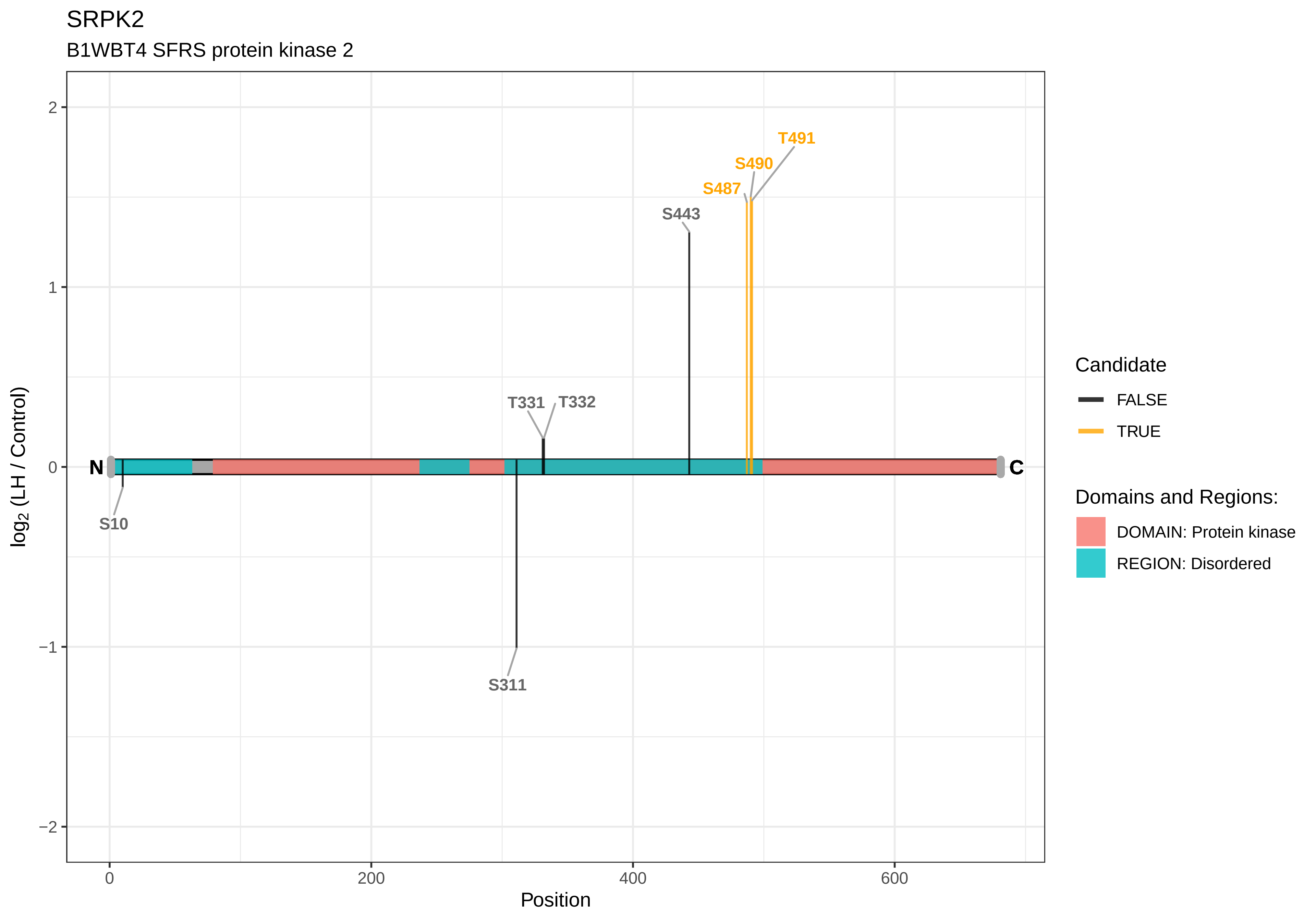
SRPK2
B1WBT4 SFRS protein kinase 2
T491
S490
S487
S443
T331
T332
N
C
S10
S311
2
1
0
−1
−2
0
200
400
600
Position
log2 (LH / Control)
Candidate
FALSE
TRUE
Domains and Regions:
DOMAIN: Protein kinase
REGION: Disordered

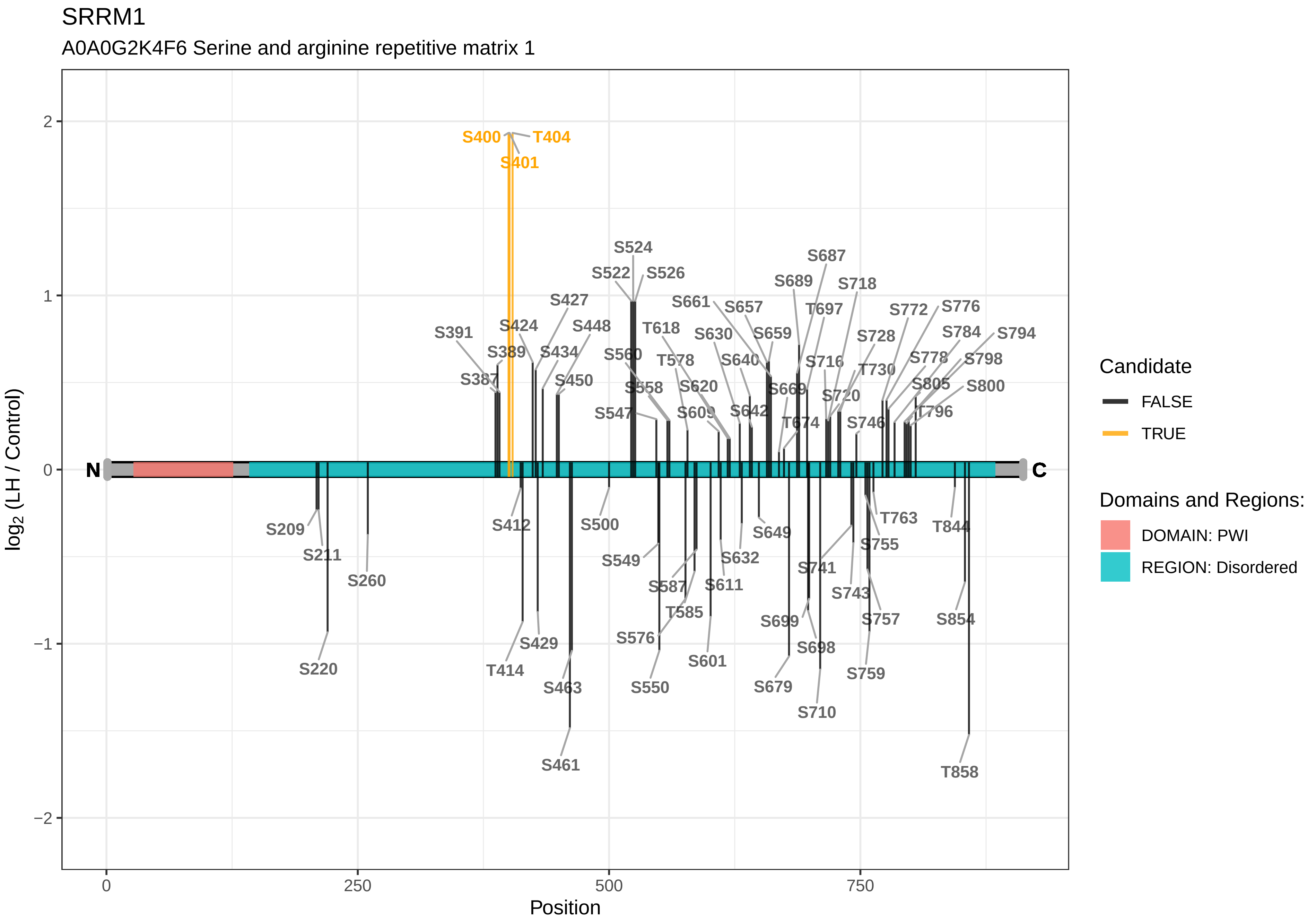

SRRM1
A0A0G2K4F6 Serine and arginine repetitive matrix 1
log2 (LH / Control)
Position
Candidate
FALSE
TRUE
Domains and Regions:
DOMAIN: PWI
REGION: Disordered
N
C
S400
T404
S401
S524
S522
S526
S687
S689
S718
S661
S657
T697
S772
S776
S427
S391
S424
S448
T618
S630
S659
S728
S784
S794
S389
S434
S560
T578
S640
S716
T730
S778
S798
S387
S450
S558
S620
S669
S720
S805
S800
S547
S609
S642
T674
S746
T796
S209
S211
S260
S220
S412
T414
S429
S463
S461
S500
S549
S587
T585
S576
S550
S611
S601
S632
S649
S741
S699
S679
S698
S710
S743
S755
T763
S757
S759
T844
S854
T858
0
250
500
750
2
1
0
−1
−2

SRRM2

A0A0G2K2M9 Serine/arginine repetitive matrix 2

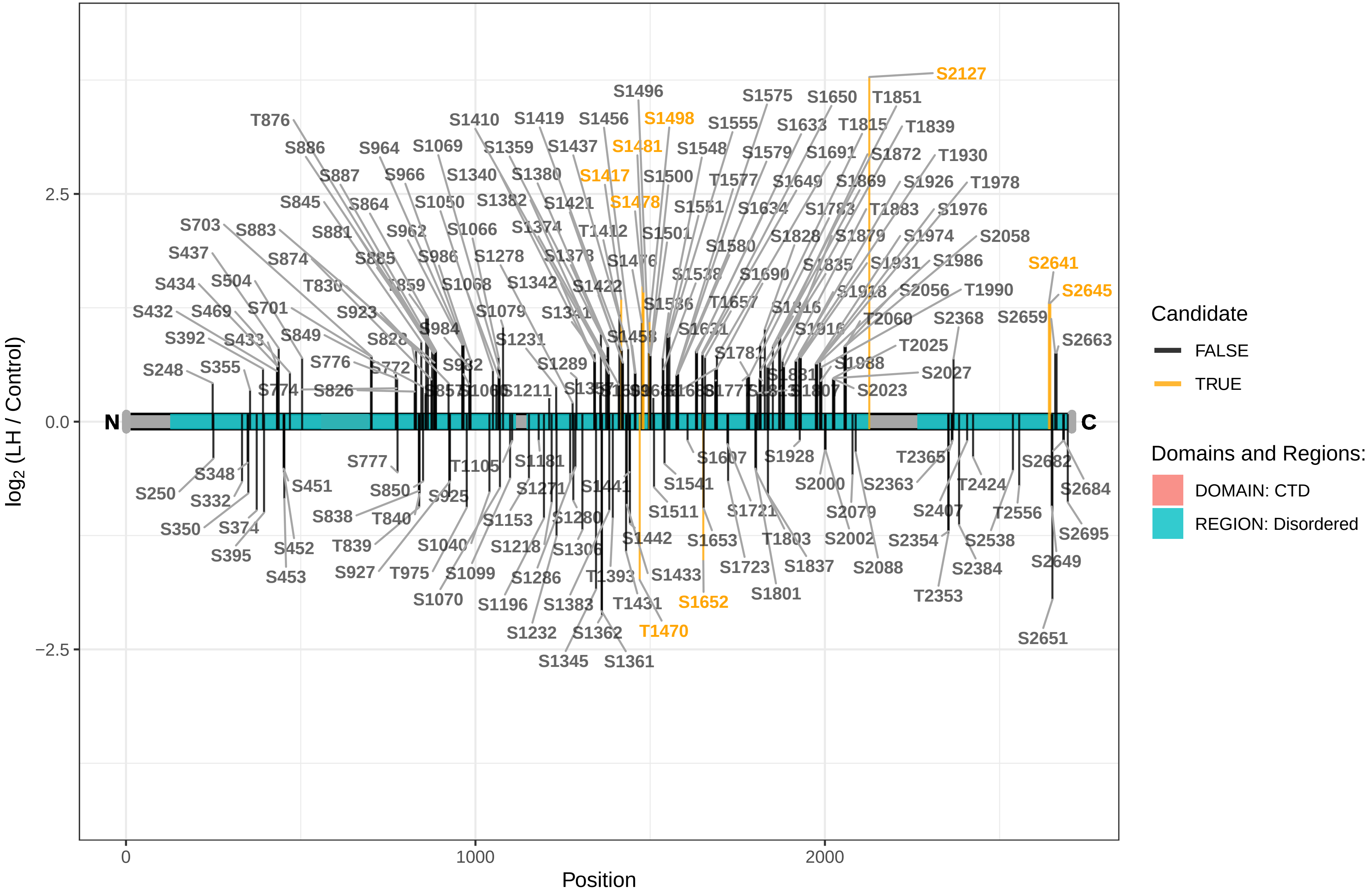

SSH1

F1LWM1 Slingshot protein phosphatase 1

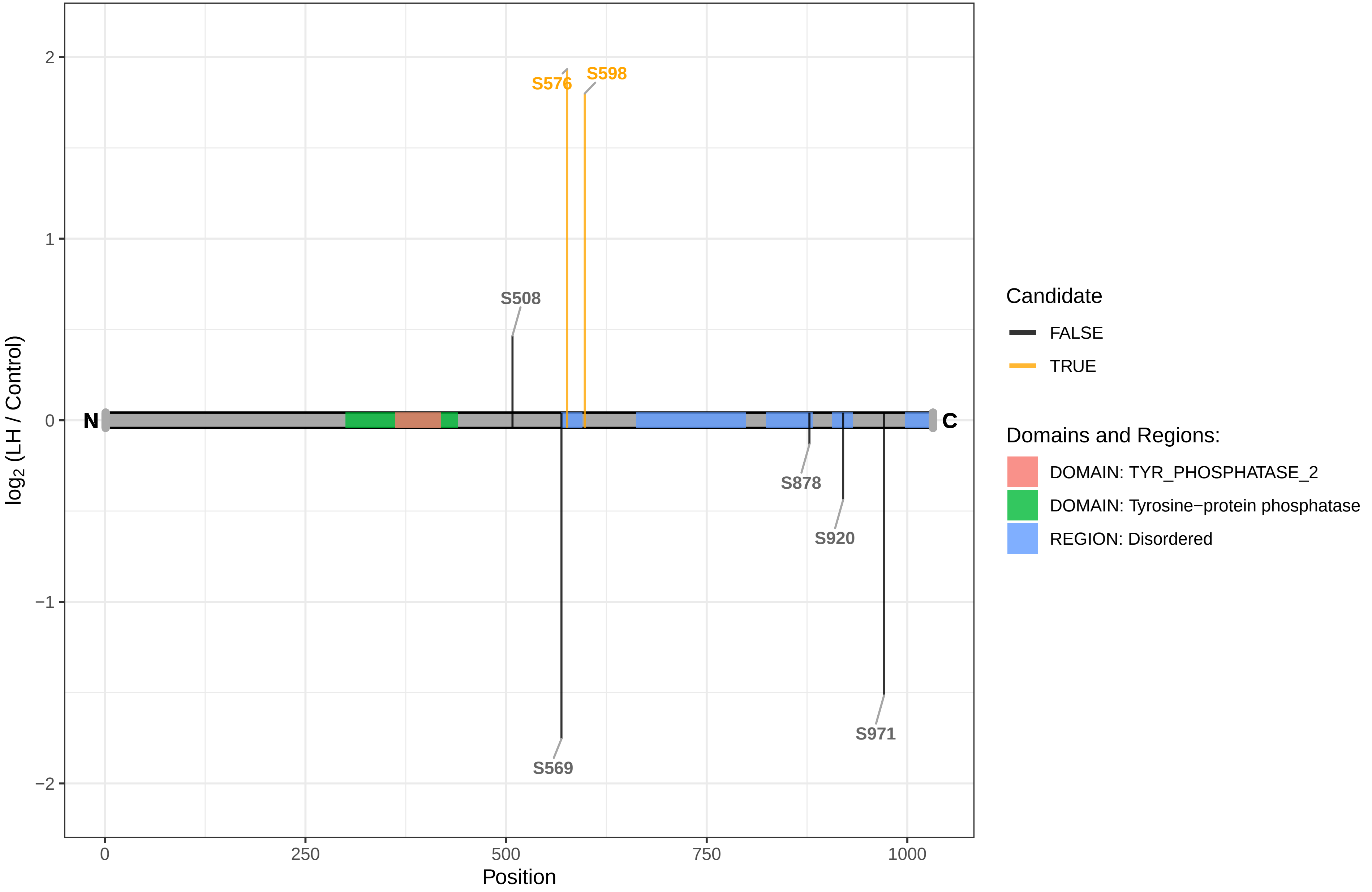

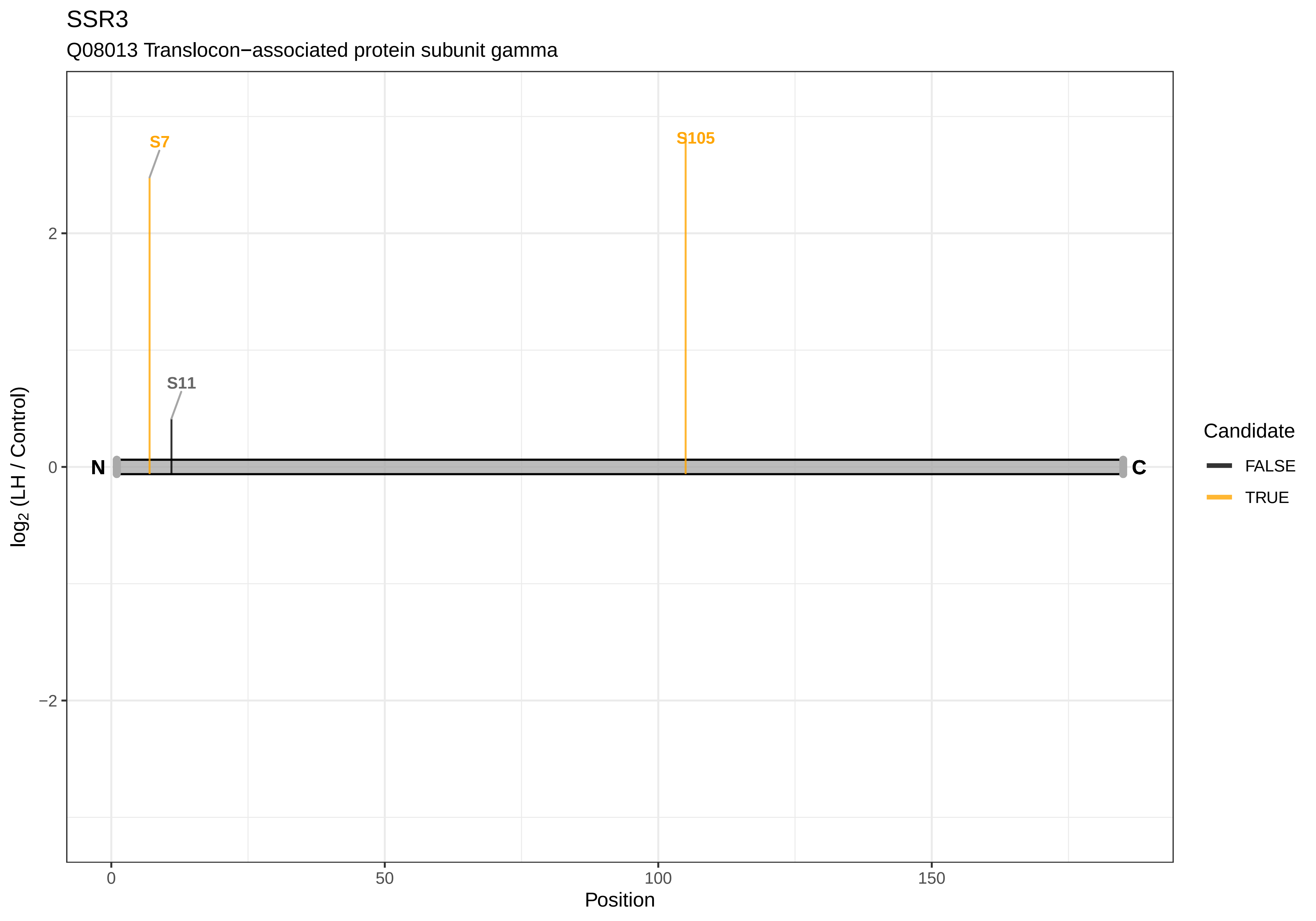
SSR3
Q08013 Translocon−associated protein subunit gamma
S7
S105
S11
N
C
2
0
−2
0
50
100
150
Position
log2 (LH / Control)
Candidate
FALSE
TRUE

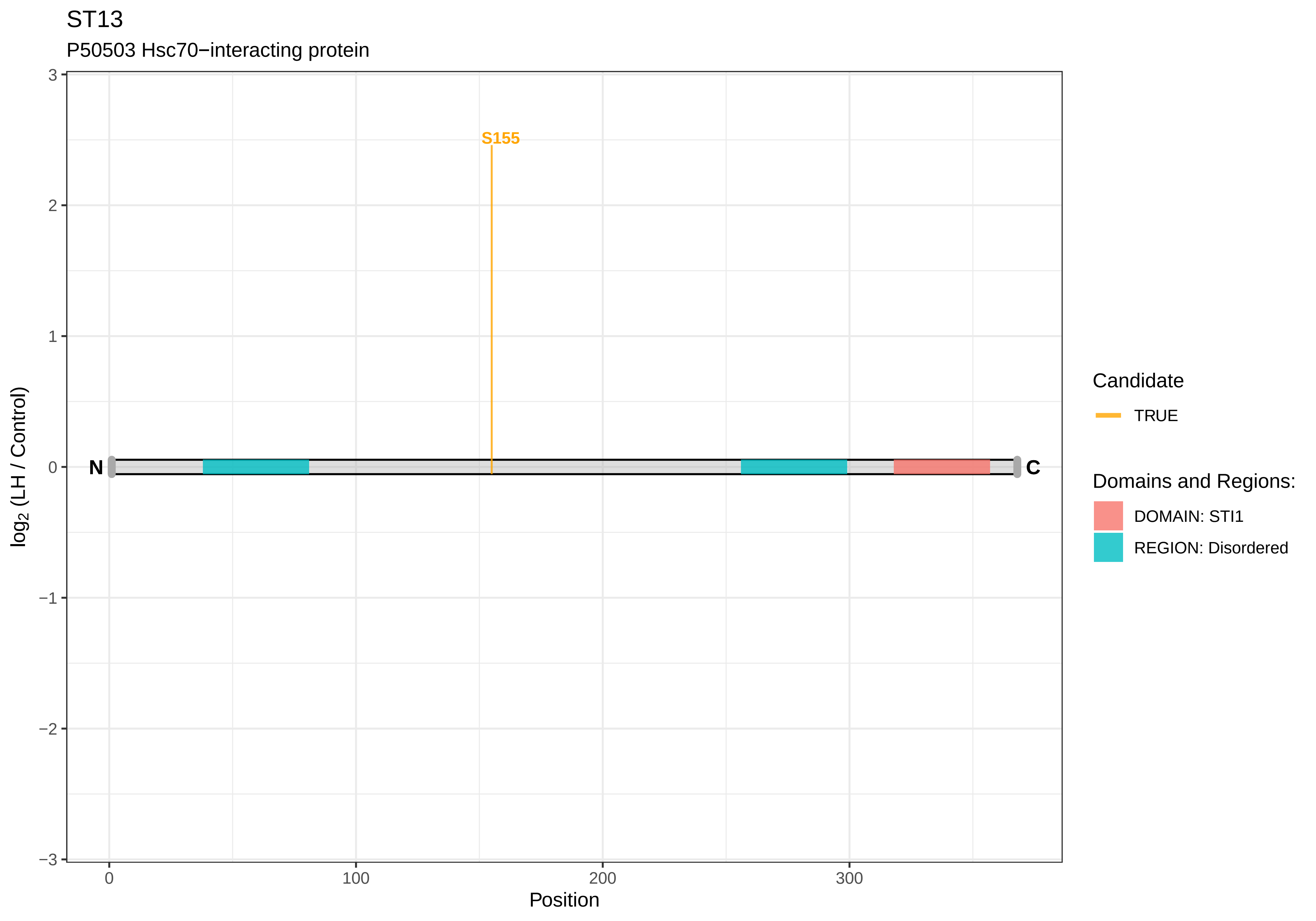

ST13
P50503 Hsc70−interacting protein
S155
N
C
log2 (LH / Control)
Position
3
2
1
0
−1
−2
−3
0
100
200
300
Candidate
TRUE
Domains and Regions:
DOMAIN: STI1
REGION: Disordered

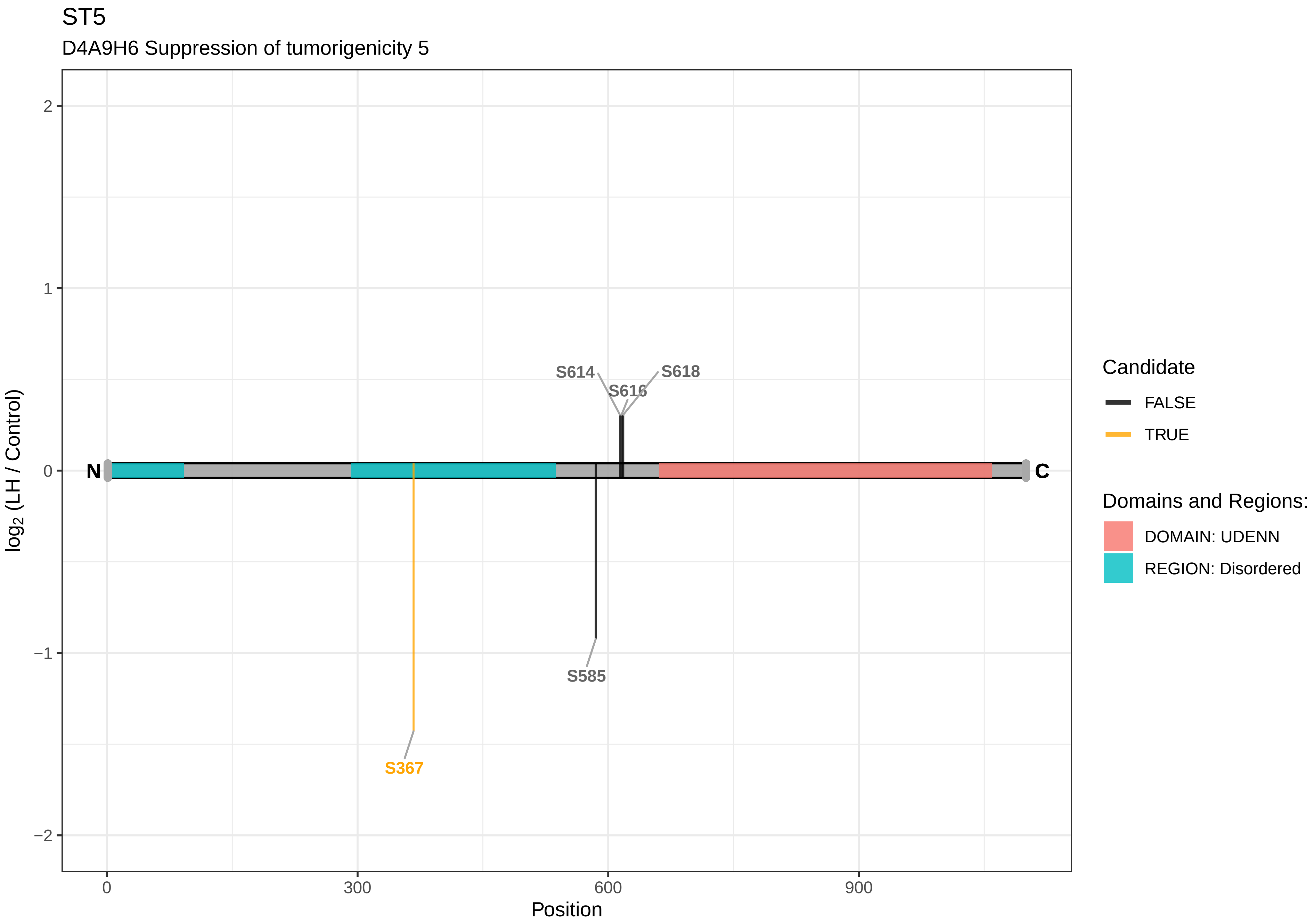
ST5
D4A9H6 Suppression of tumorigenicity 5
S614
S616
S618
S585
S367
N
C
2
1
0
−1
−2
log2 (LH / Control)
0
300
600
900
Position
Candidate
FALSE
TRUE
Domains and Regions:
DOMAIN: UDENN
REGION: Disordered

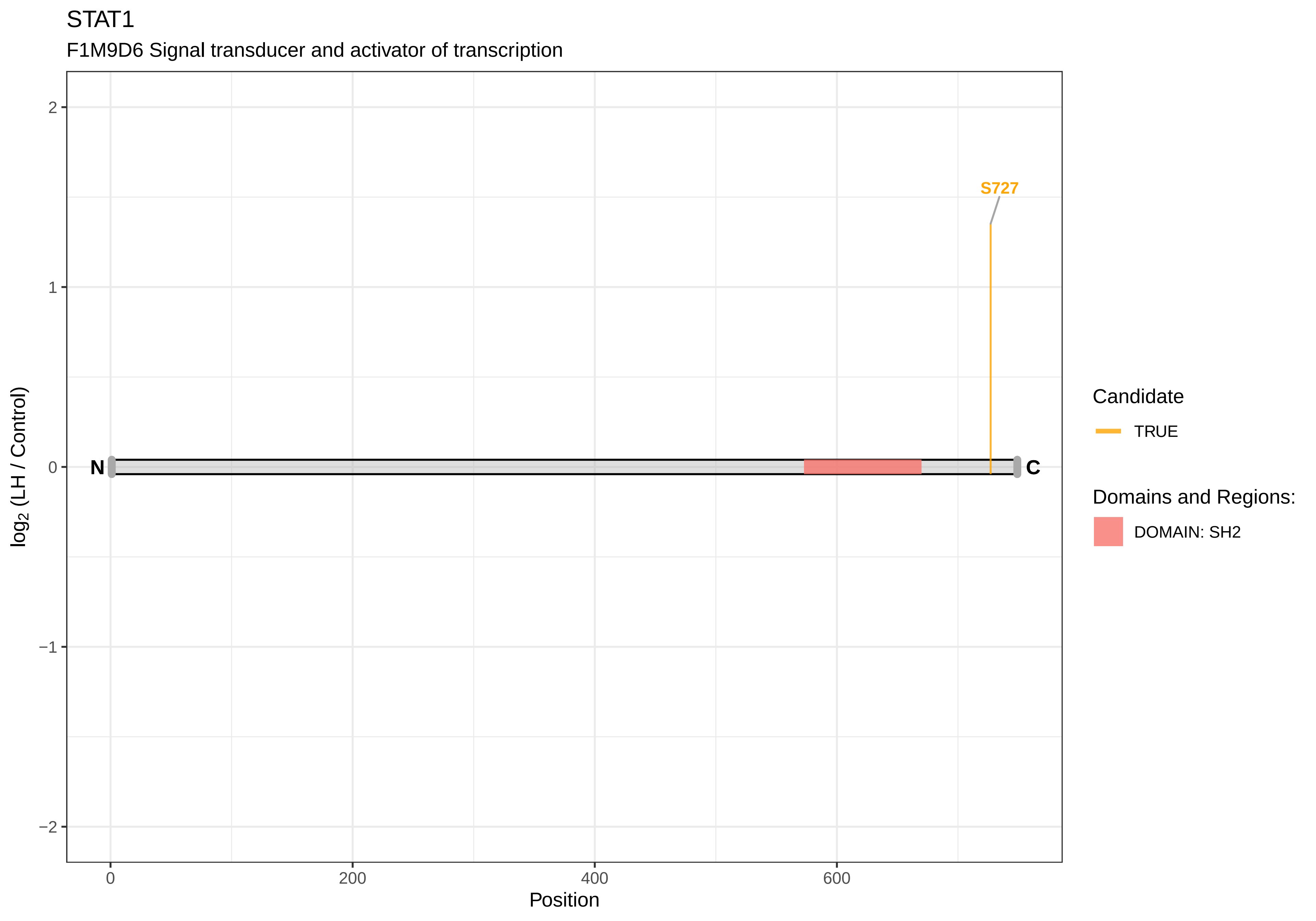

STAT1
F1M9D6 Signal transducer and activator of transcription
S727
N
C
2
1
0
−1
−2
log2 (LH / Control)
0
200
400
600
Position
Candidate
TRUE
Domains and Regions:
DOMAIN: SH2

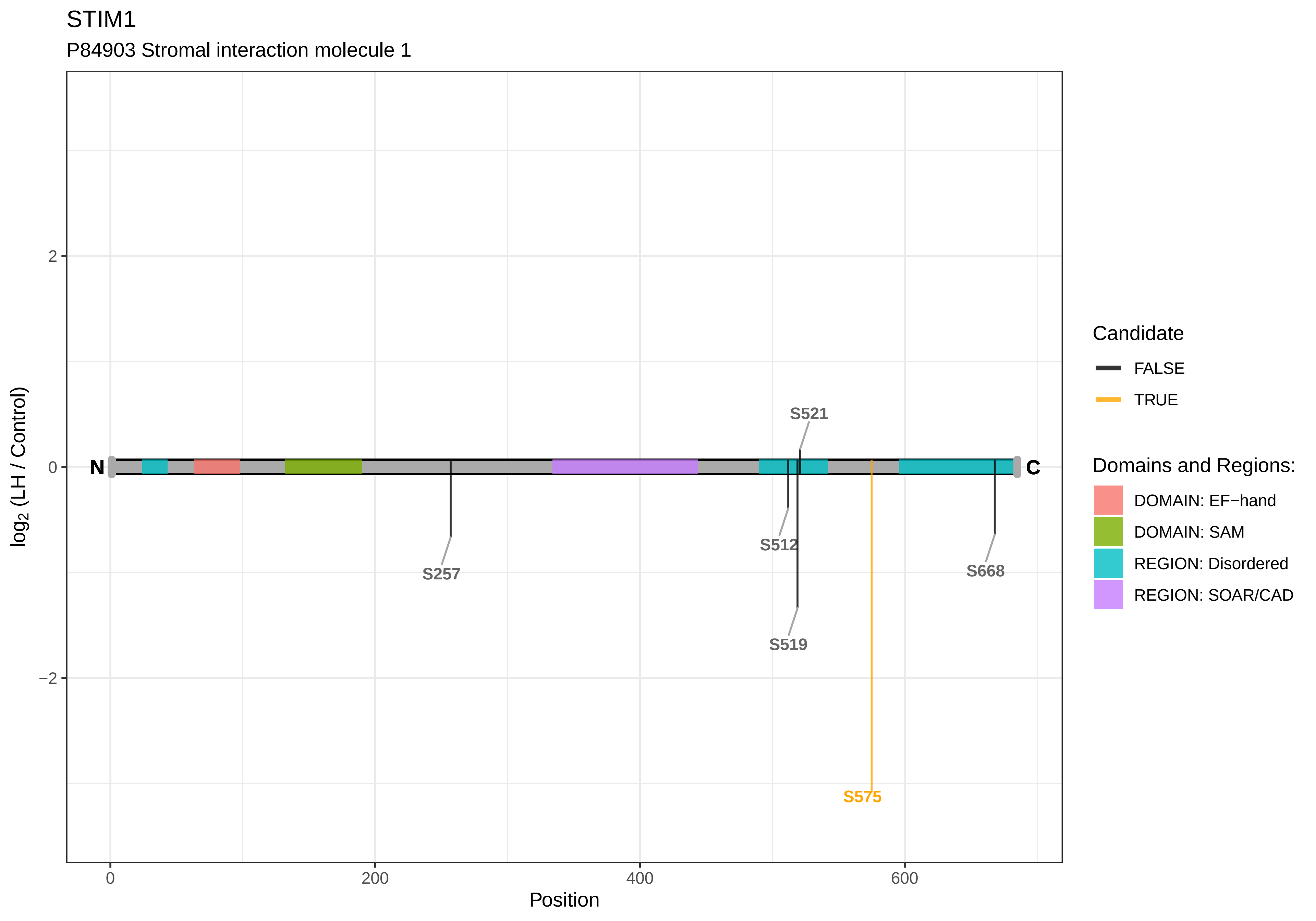

STIM1
P84903 Stromal interaction molecule 1
log2 (LH / Control)
Position
2
0
−2
0
200
400
600
N
C
S521
S257
S512
S668
S519
S575
Candidate
FALSE
TRUE
Domains and Regions:
DOMAIN: EF−hand
DOMAIN: SAM
REGION: Disordered
REGION: SOAR/CAD

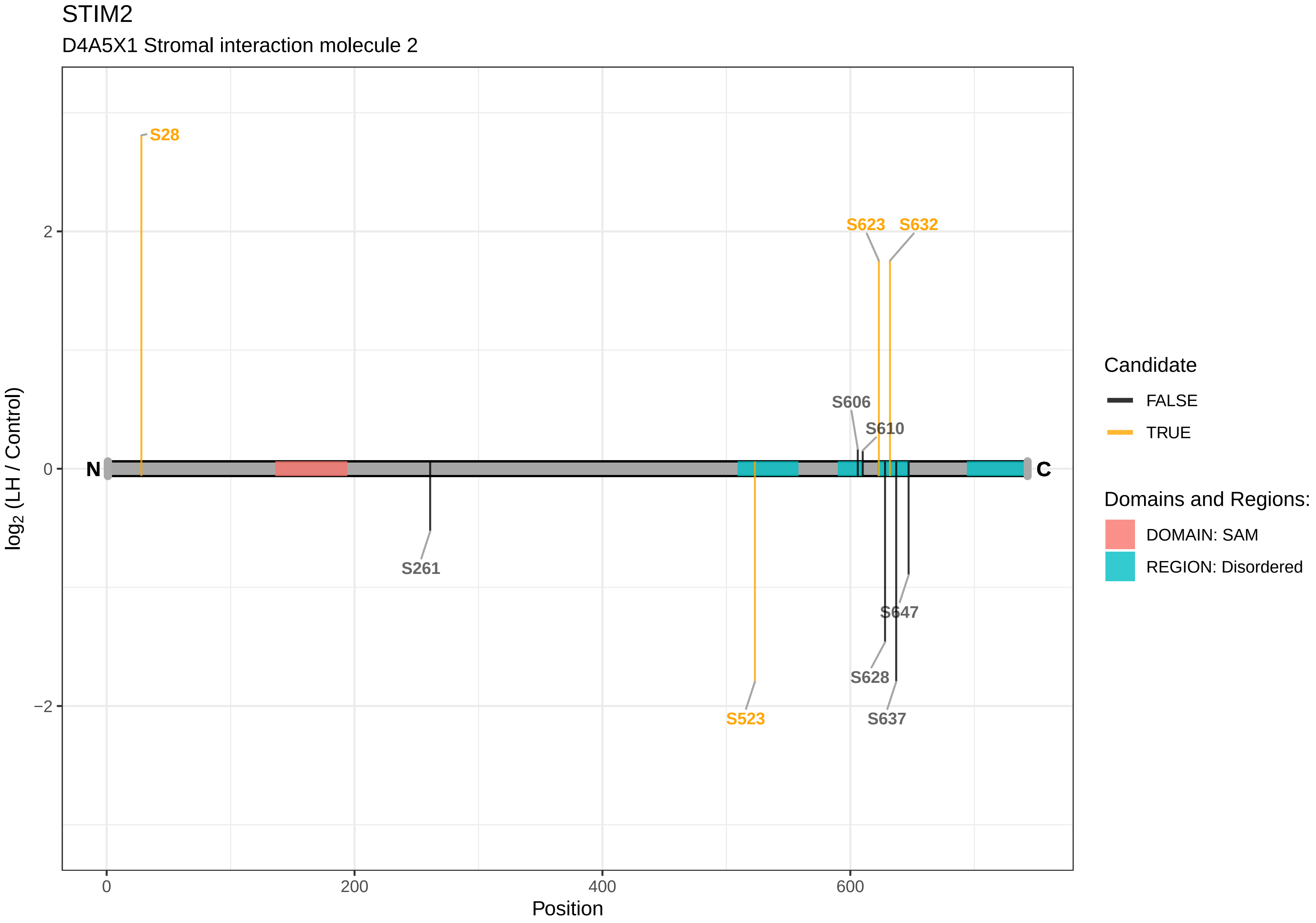

STIM2
D4A5X1 Stromal interaction molecule 2
S28
S623
S632
S606
S610
N
C
S261
S523
S647
S628
S637
2
0
−2
0
200
400
600
Position
log2 (LH / Control)
Candidate
FALSE
TRUE
Domains and Regions:
DOMAIN: SAM
REGION: Disordered

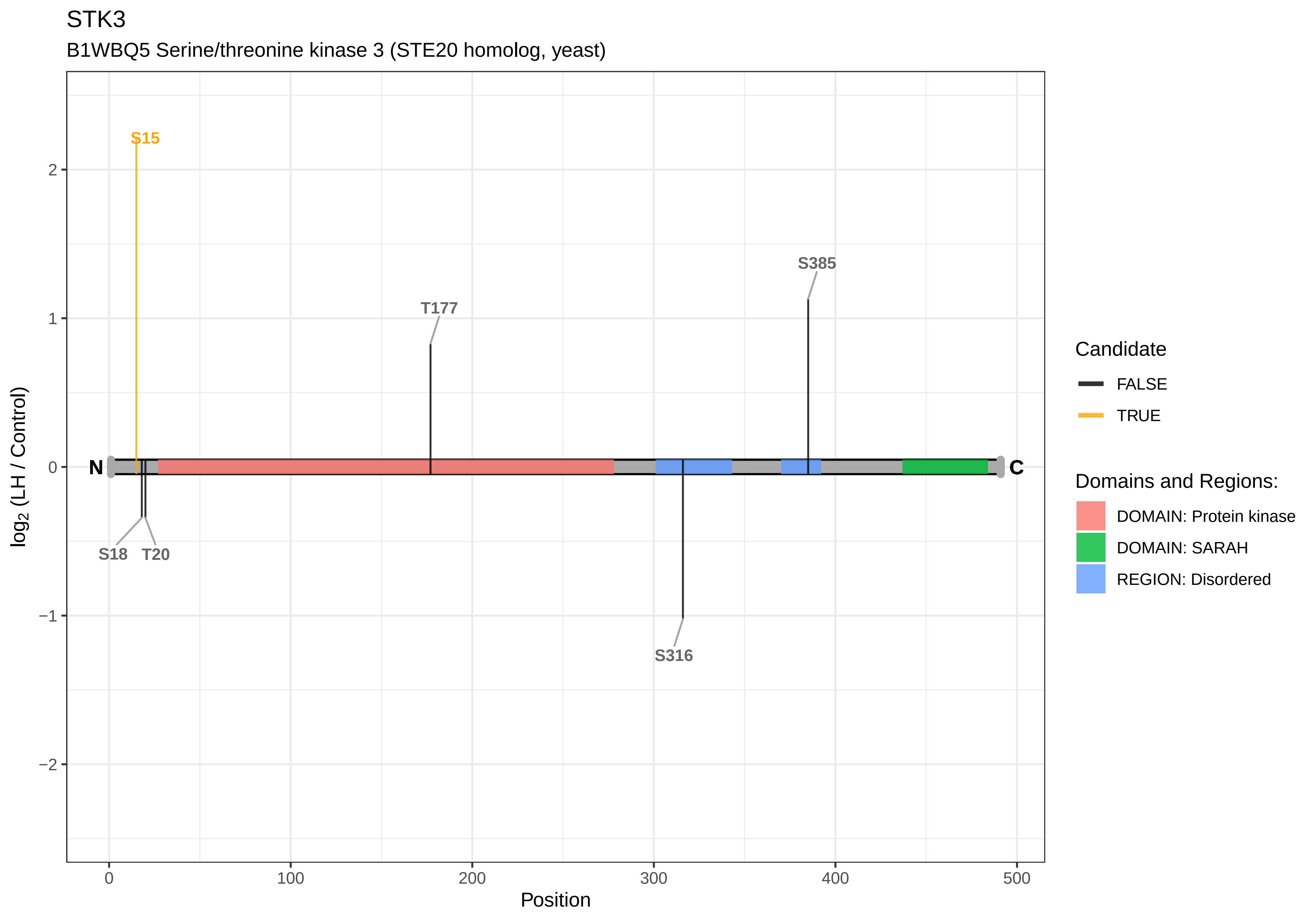

STK3
B1WBQ5 Serine/threonine kinase 3 (STE20 homolog, yeast)
S15
S385
T177
N
C
S18
T20
S316
log2 (LH / Control)
2
1
0
−1
−2
0
100
200
300
400
500
Position
Candidate
FALSE
TRUE
Domains and Regions:
DOMAIN: Protein kinase
DOMAIN: SARAH
REGION: Disordered

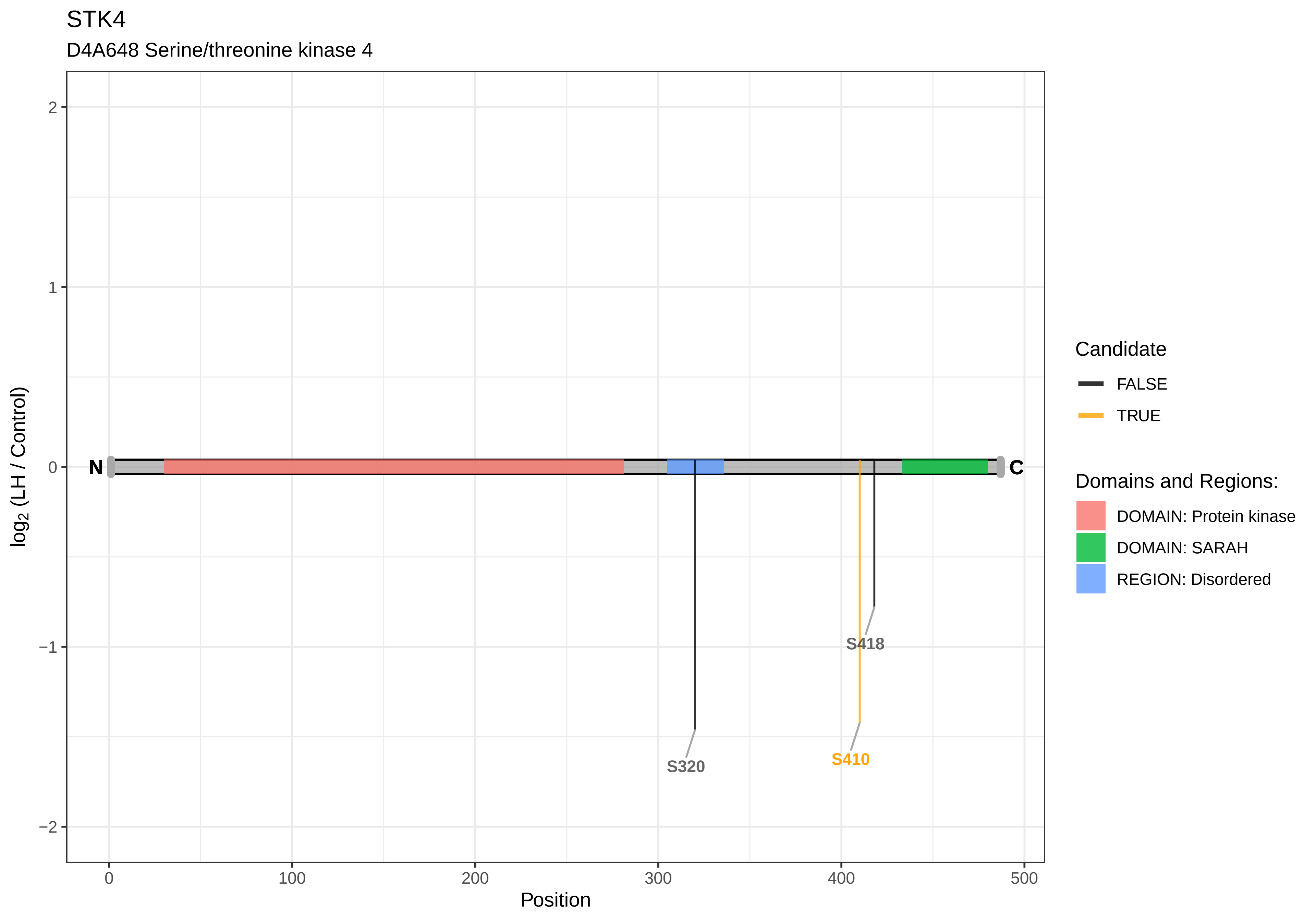
STK4
D4A648 Serine/threonine kinase 4
log2 (LH / Control)
Position
2
1
0
−1
−2
0
100
200
300
400
500
N
C
S320
S410
S418
Candidate
FALSE
TRUE
Domains and Regions:
DOMAIN: Protein kinase
DOMAIN: SARAH
REGION: Disordered

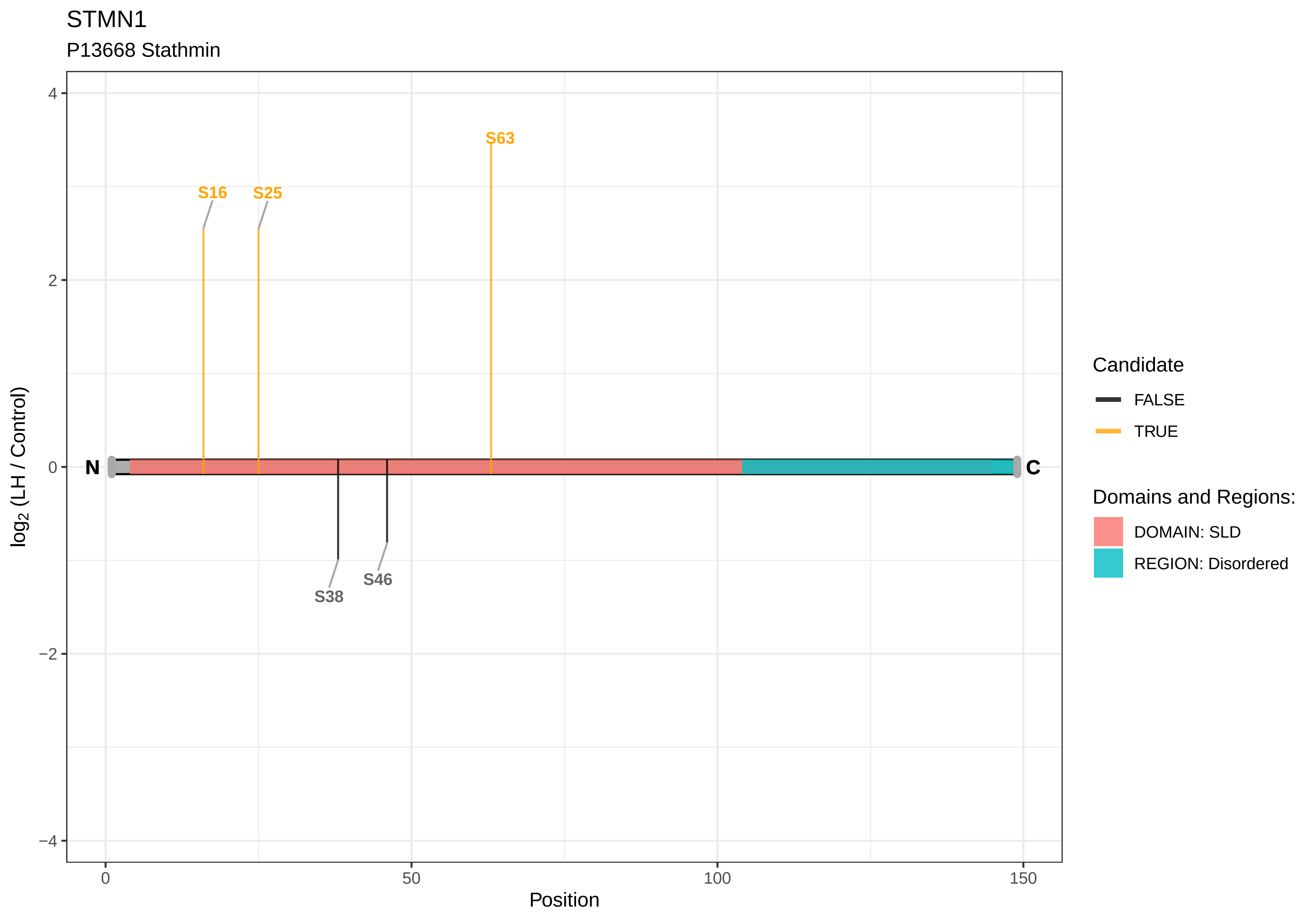

STMN1
P13668 Stathmin
S63
S16
S25
S38
S46
N
C
4
2
0
−2
−4
log2 (LH / Control)
0
50
100
150
Position
Candidate
FALSE
TRUE
Domains and Regions:
DOMAIN: SLD
REGION: Disordered

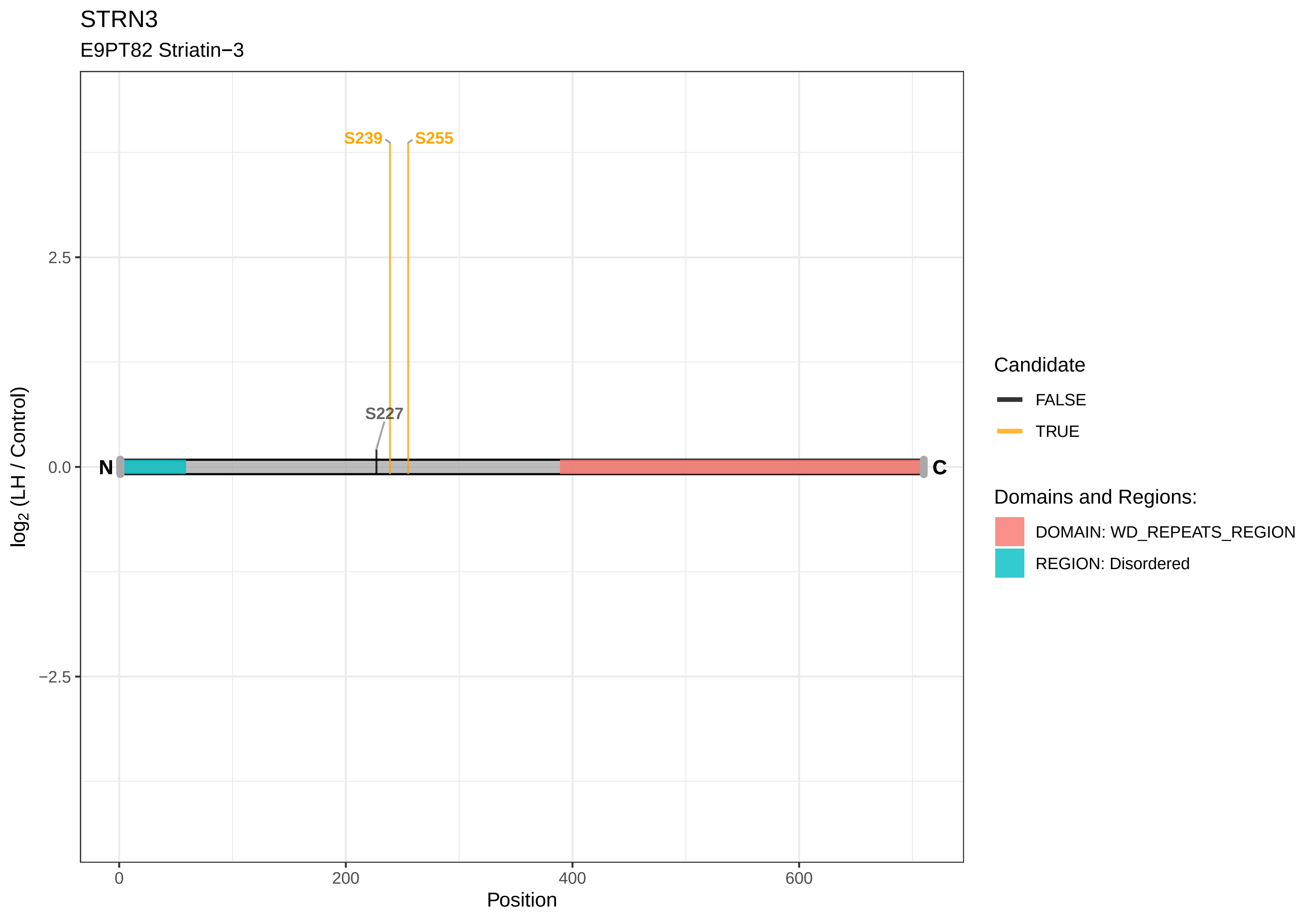
STRN3
E9PT82 Striatin−3
S239
S255
S227
N
C
2.5
0.0
−2.5
log2 (LH / Control)
0
200
400
600
Position
Candidate
FALSE
TRUE
Domains and Regions:
DOMAIN: WD_REPEATS_REGION
REGION: Disordered

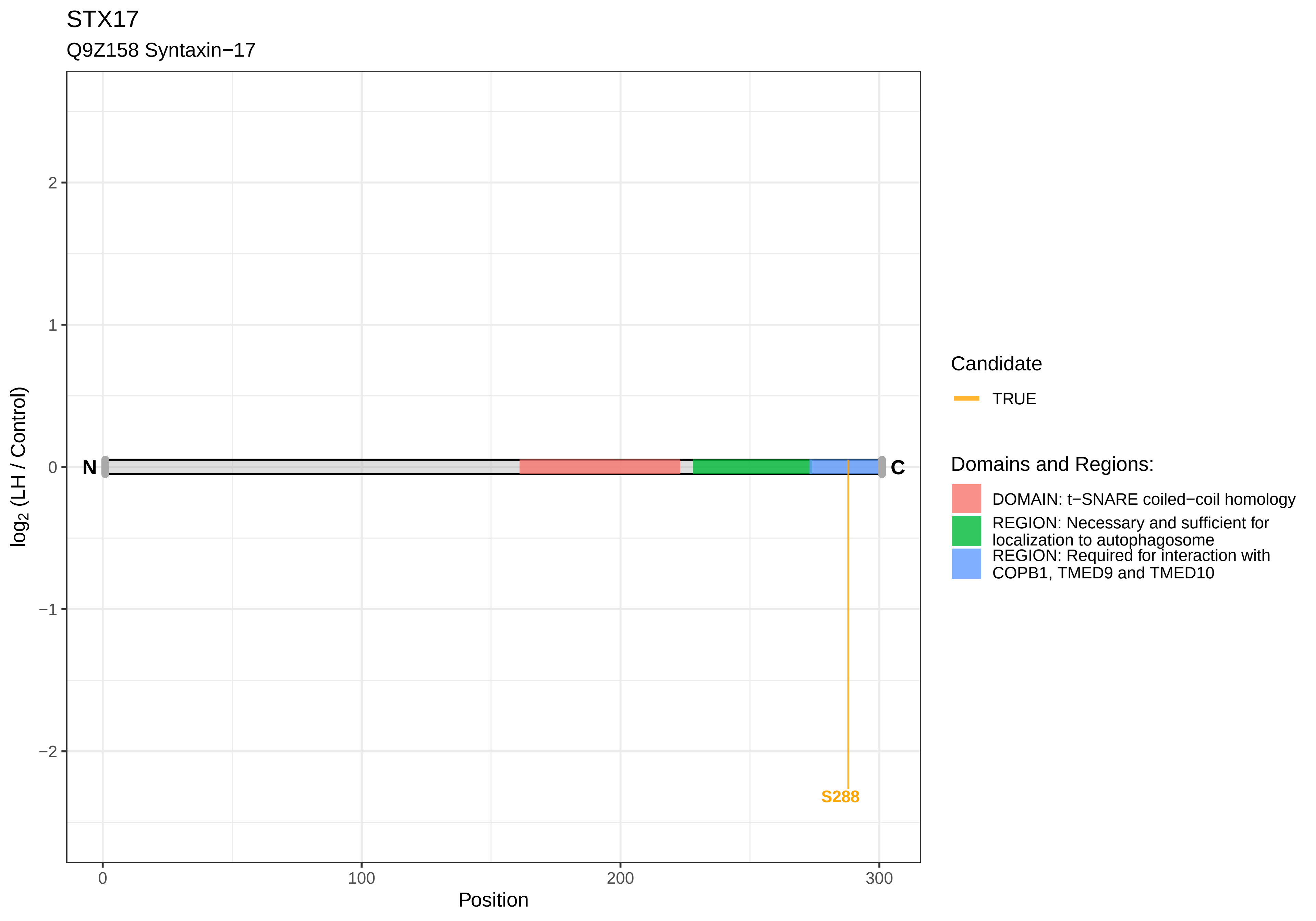

STX17
Q9Z158 Syntaxin−17
log2 (LH / Control)
Position
N
C
S288
2
1
0
−1
−2
0
100
200
300
Candidate
TRUE
Domains and Regions:
DOMAIN: t−SNARE coiled−coil homology
REGION: Necessary and sufficient for localization to autophagosome
REGION: Required for interaction with COPB1, TMED9 and TMED10

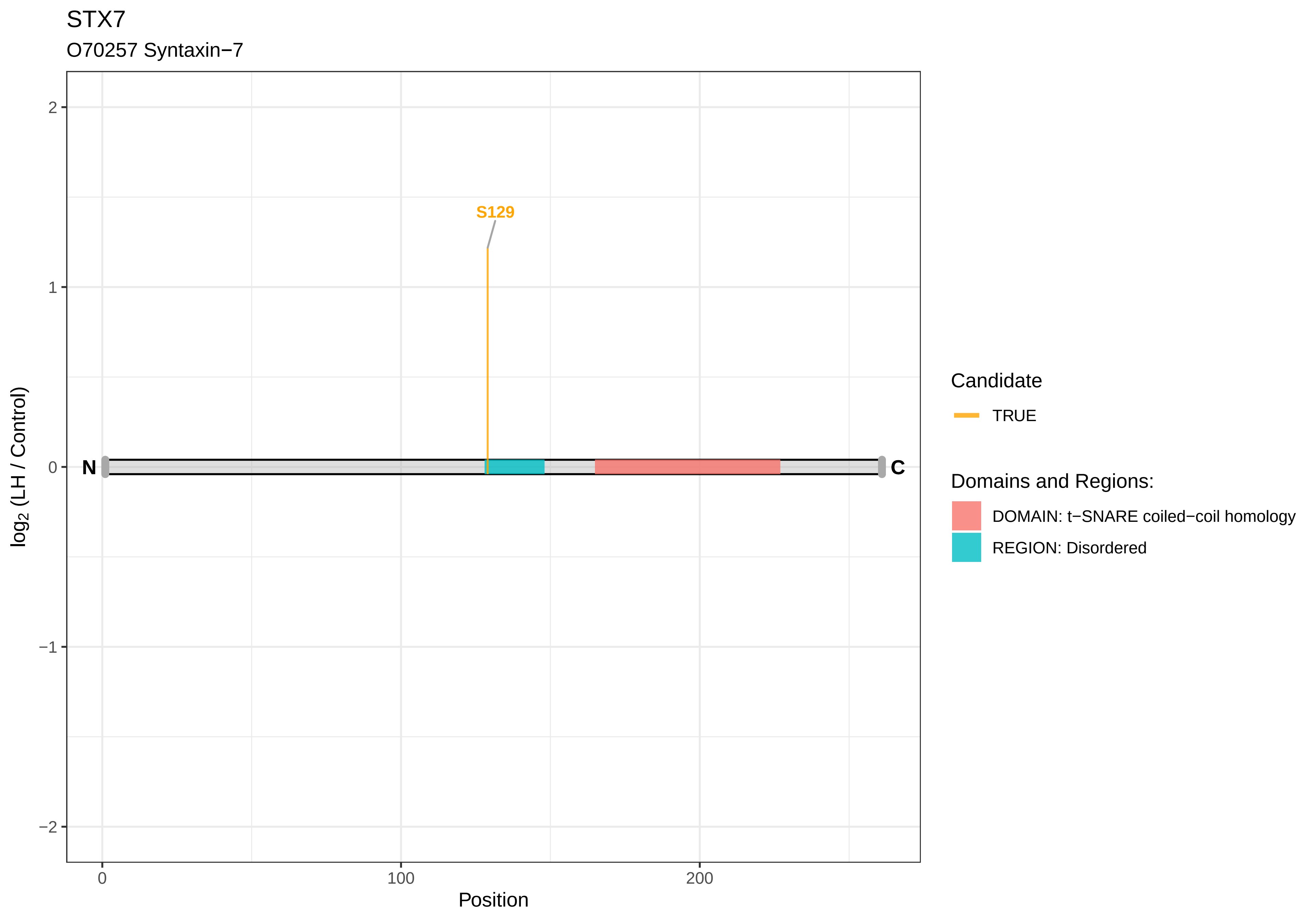
STX7
O70257 Syntaxin−7
S129
N
C
log2 (LH / Control)
2
1
0
−1
−2
0
100
200
Position
Candidate
TRUE
Domains and Regions:
DOMAIN: t−SNARE coiled−coil homology
REGION: Disordered

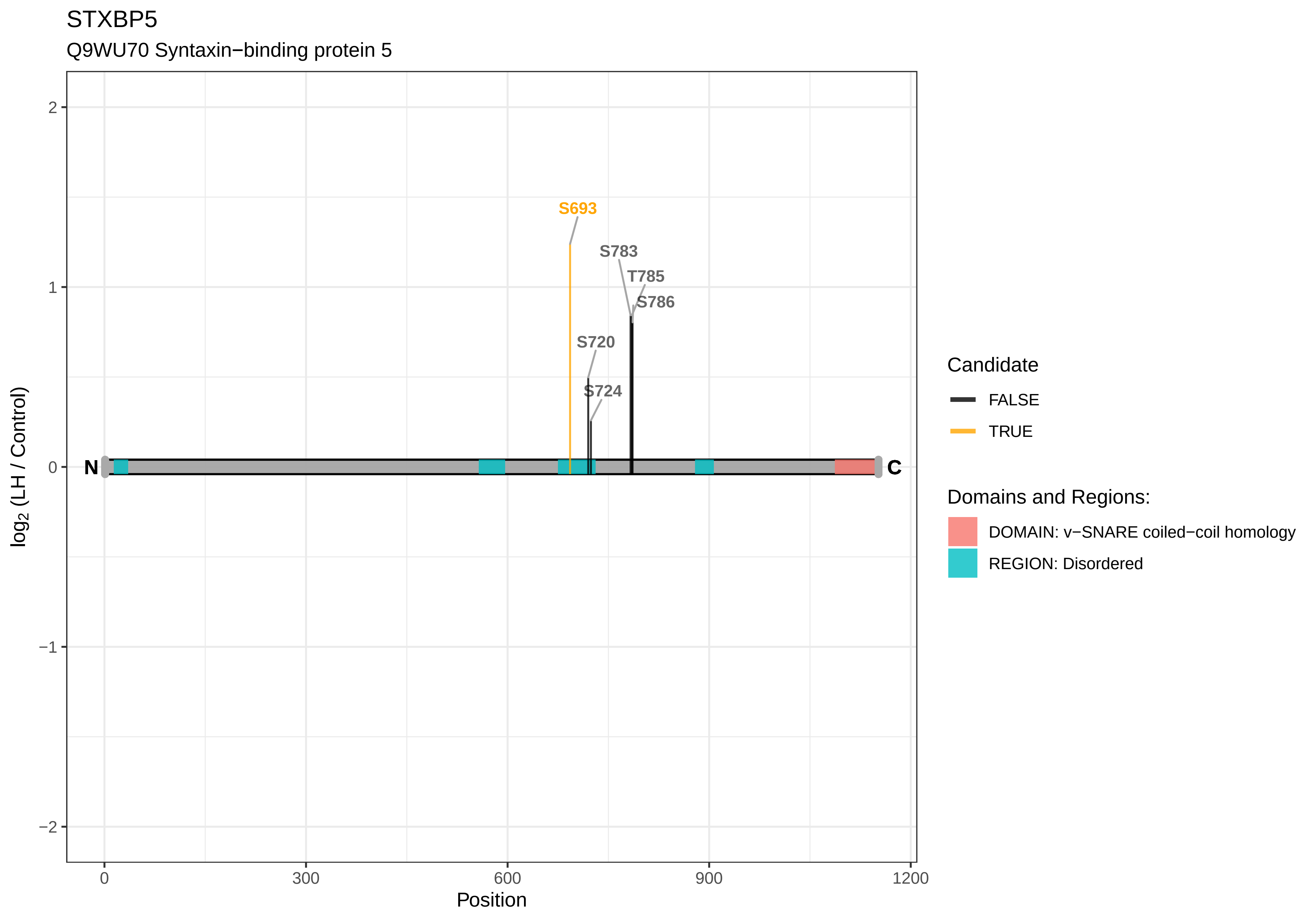

STXBP5
Q9WU70 Syntaxin−binding protein 5
S693
S783
T785
S786
S720
S724
N
C
2
1
0
−1
−2
log2 (LH / Control)
0
300
600
900
1200
Position
Candidate
FALSE
TRUE
Domains and Regions:
DOMAIN: v−SNARE coiled−coil homology
REGION: Disordered

#### SUB1

Q63396 Activated RNA polymerase II transcriptional coactivator p15

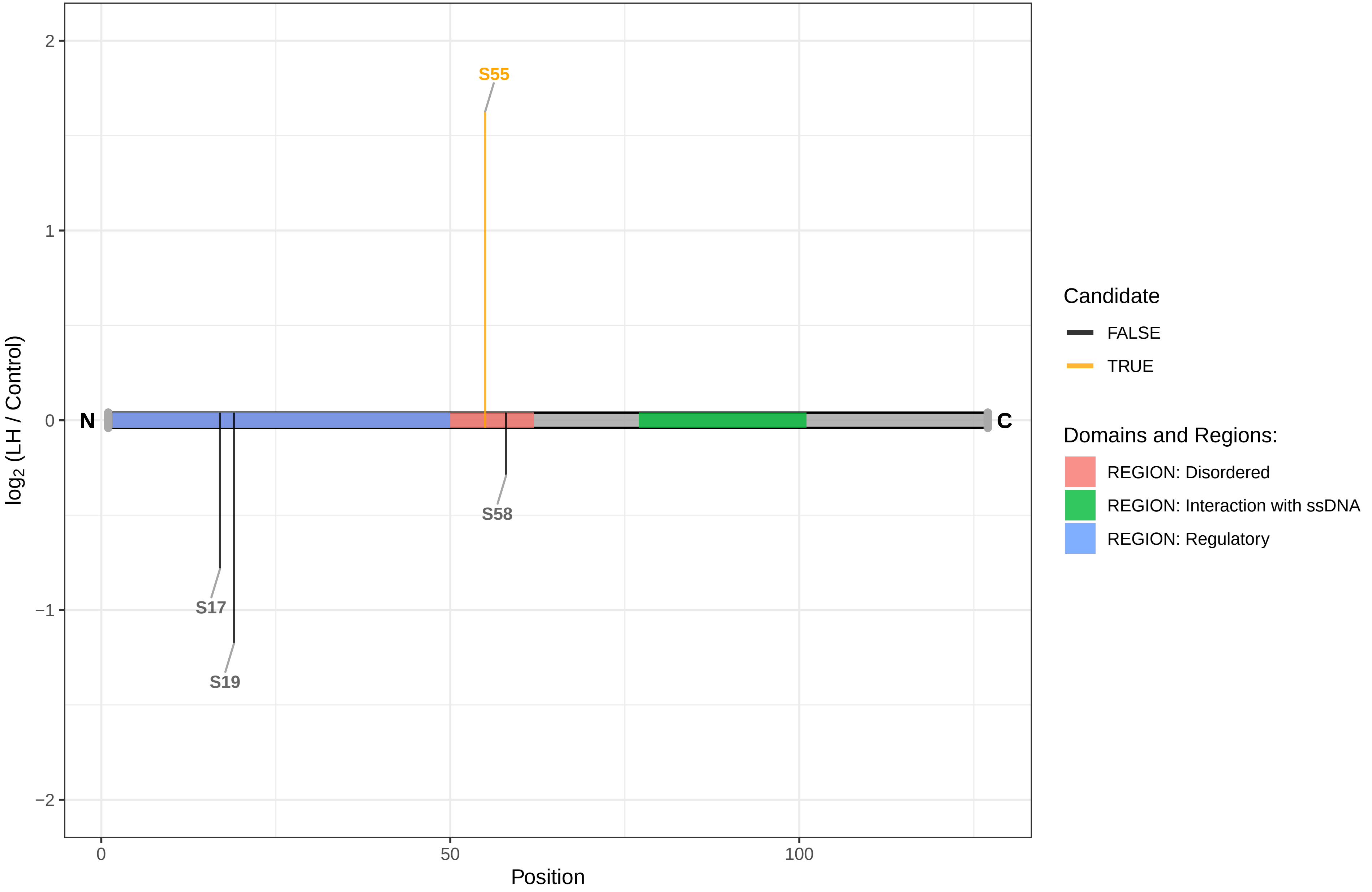

SUPT20H

Q66HC7 Transcription factor SPT20 homolog

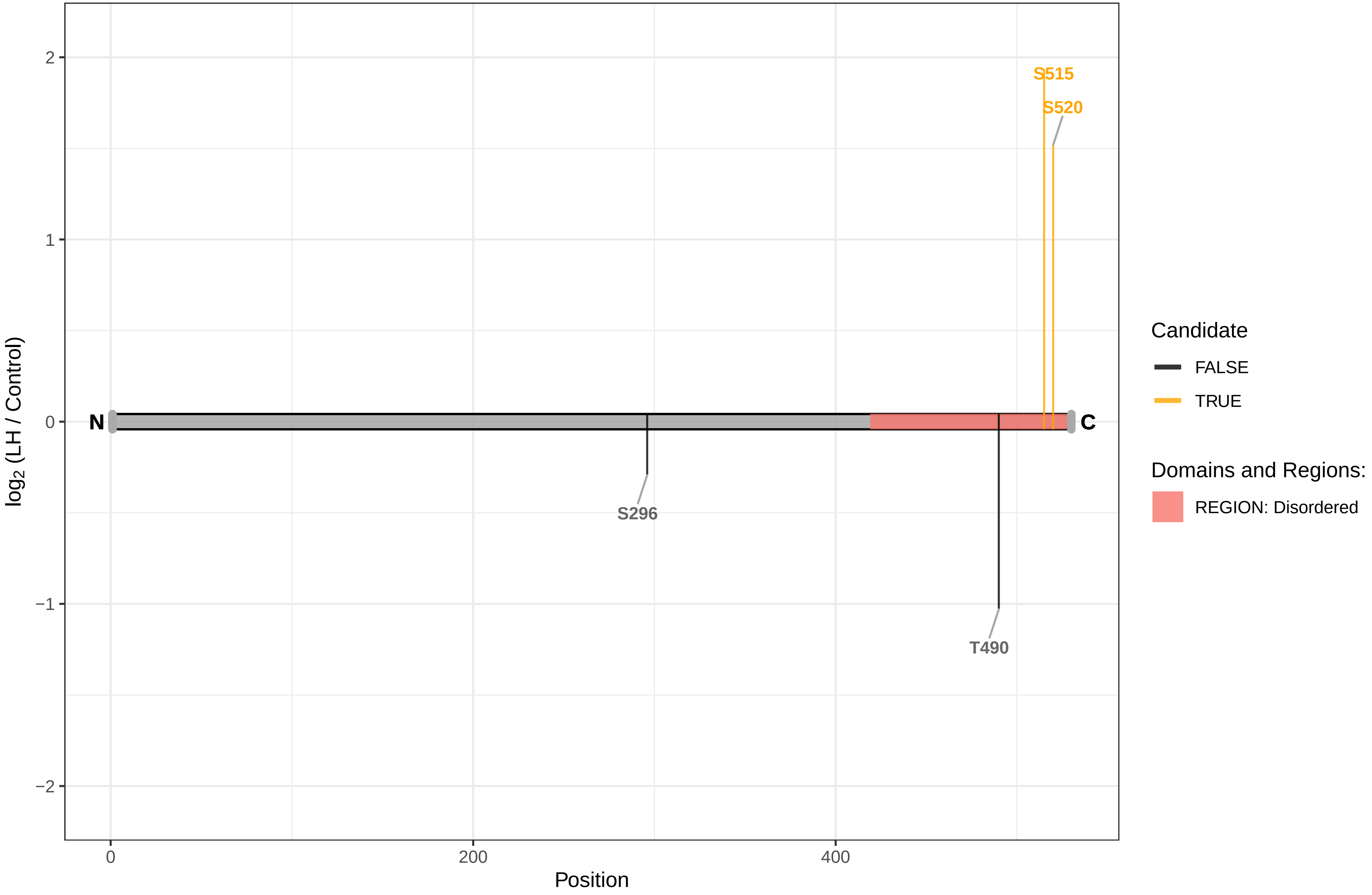

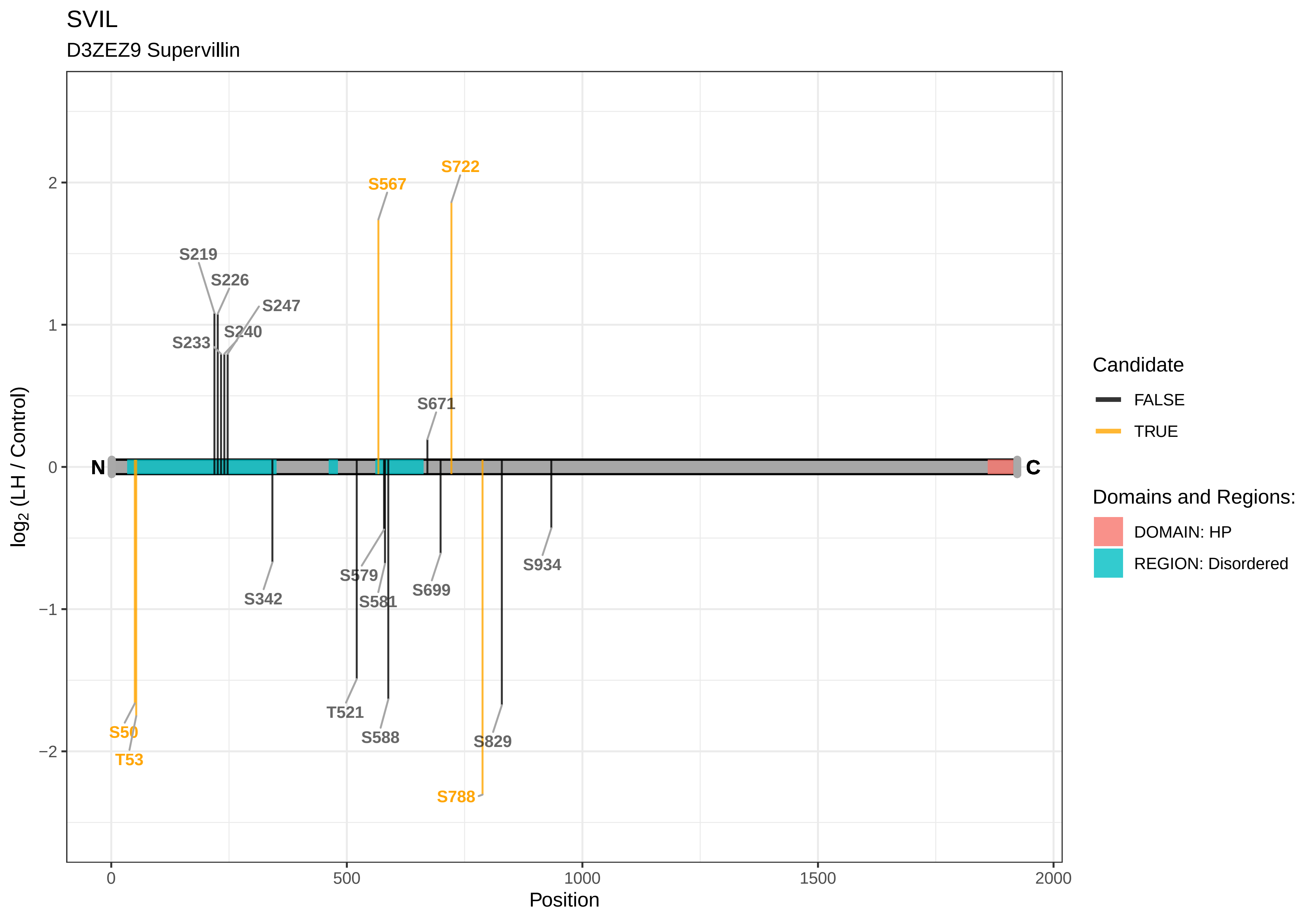

SVIL
D3ZEZ9 Supervillin
S722
S567
S219
S226
S247
S240
S233
S671
N
C
S342
S579
S581
S699
S934
T521
S588
S829
S50
T53
S788
log2 (LH / Control)
Position
Candidate
FALSE
TRUE
Domains and Regions:
DOMAIN: HP
REGION: Disordered

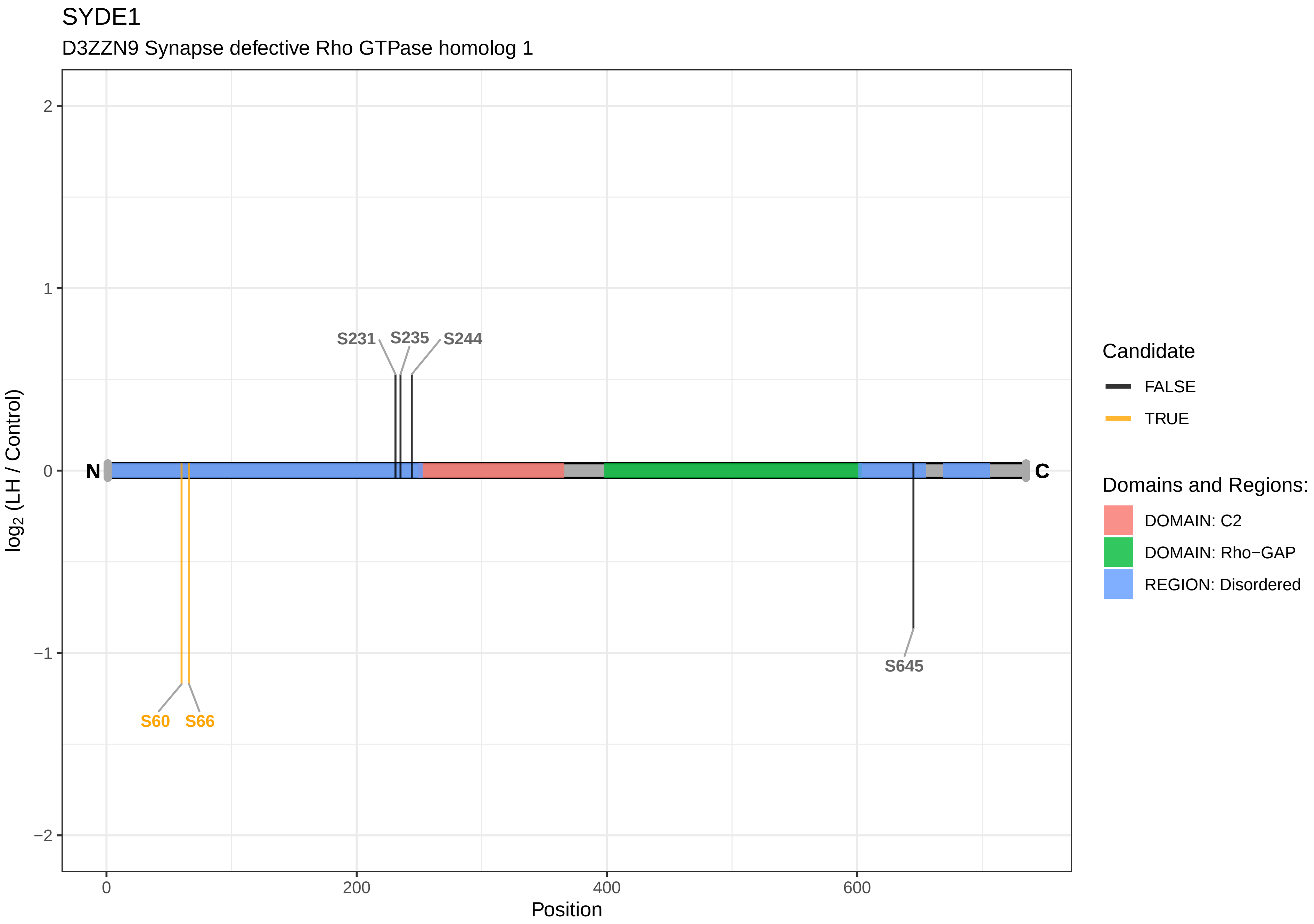

SYDE1
D3ZZN9 Synapse defective Rho GTPase homolog 1
S231
S235
S244
S60
S66
S645
N
C
2
1
0
−1
−2
log2 (LH / Control)
0
200
400
600
Position
Candidate
FALSE
TRUE
Domains and Regions:
DOMAIN: C2
DOMAIN: Rho−GAP
REGION: Disordered

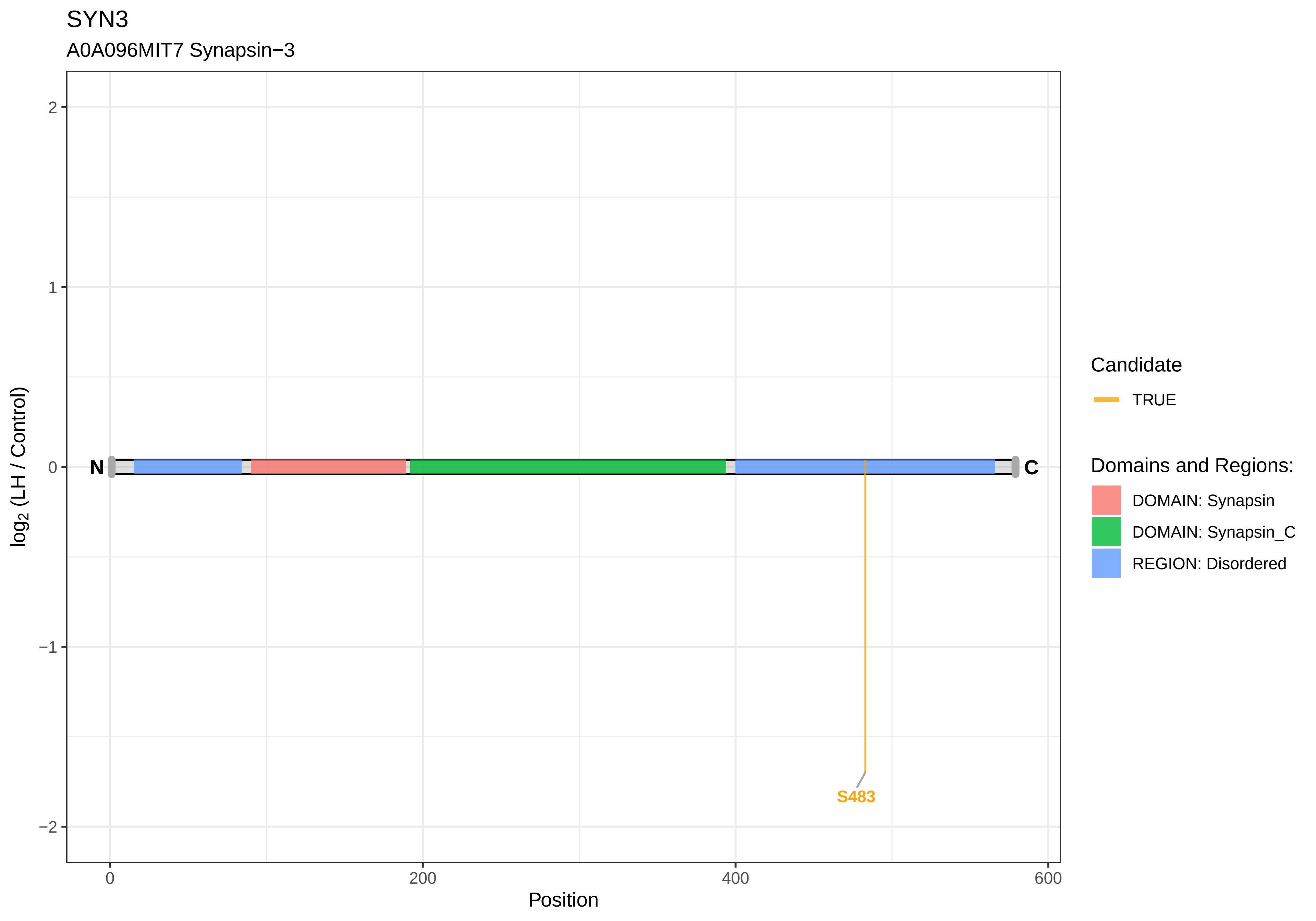
SYN3
A0A096MIT7 Synapsin−3
log2 (LH / Control)
2
1
0
−1
−2
N
C
S483
0
200
400
600
Position
Candidate
TRUE
Domains and Regions:
DOMAIN: Synapsin
DOMAIN: Synapsin_C
REGION: Disordered

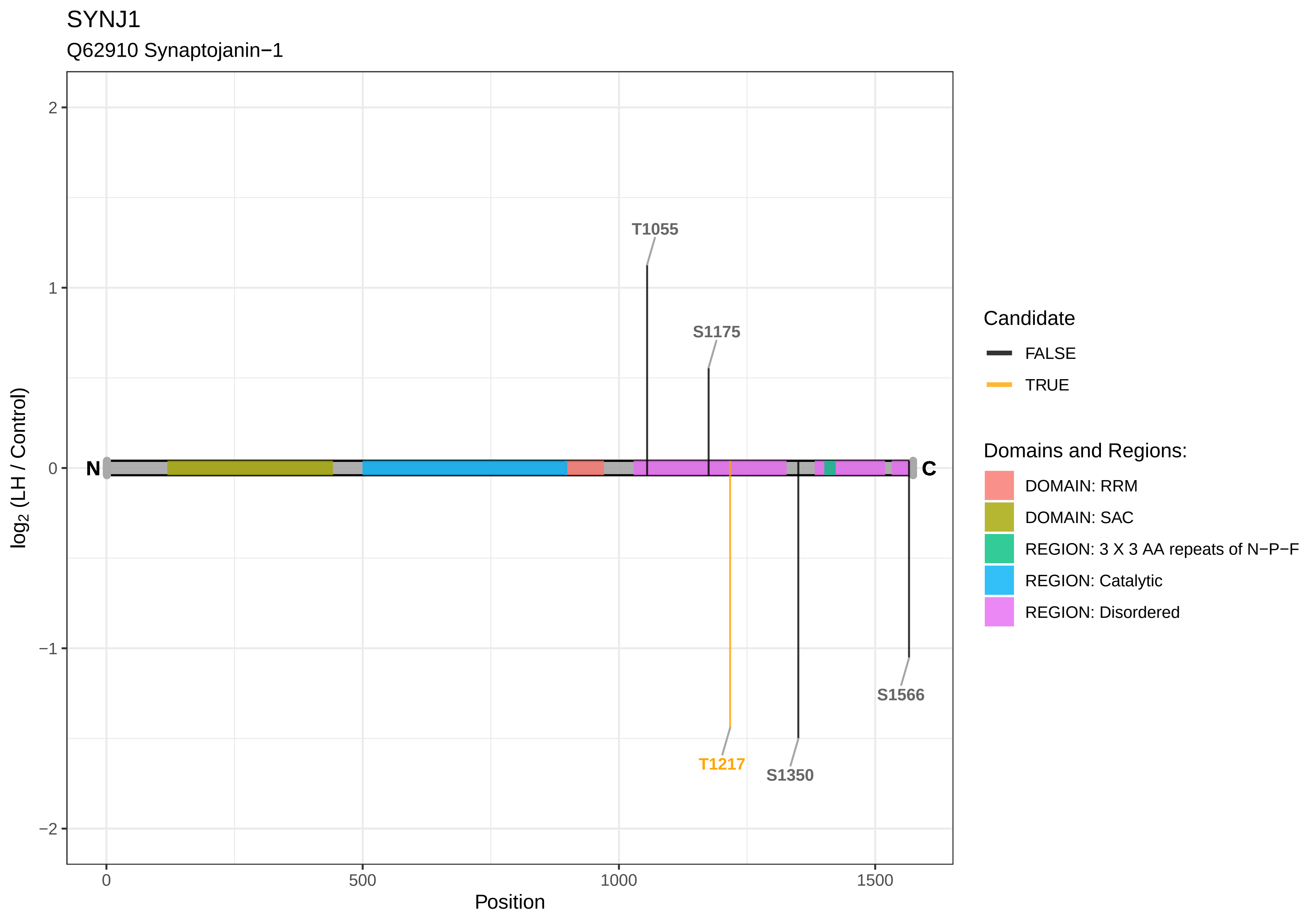

SYNJ1
Q62910 Synaptojanin−1
T1055
S1175
T1217
S1350
S1566
N
C
2
1
0
−1
−2
log2 (LH / Control)
0
500
1000
1500
Position
Candidate
FALSE
TRUE
Domains and Regions:
DOMAIN: RRM
DOMAIN: SAC
REGION: 3 X 3 AA repeats of N−P−F
REGION: Catalytic
REGION: Disordered

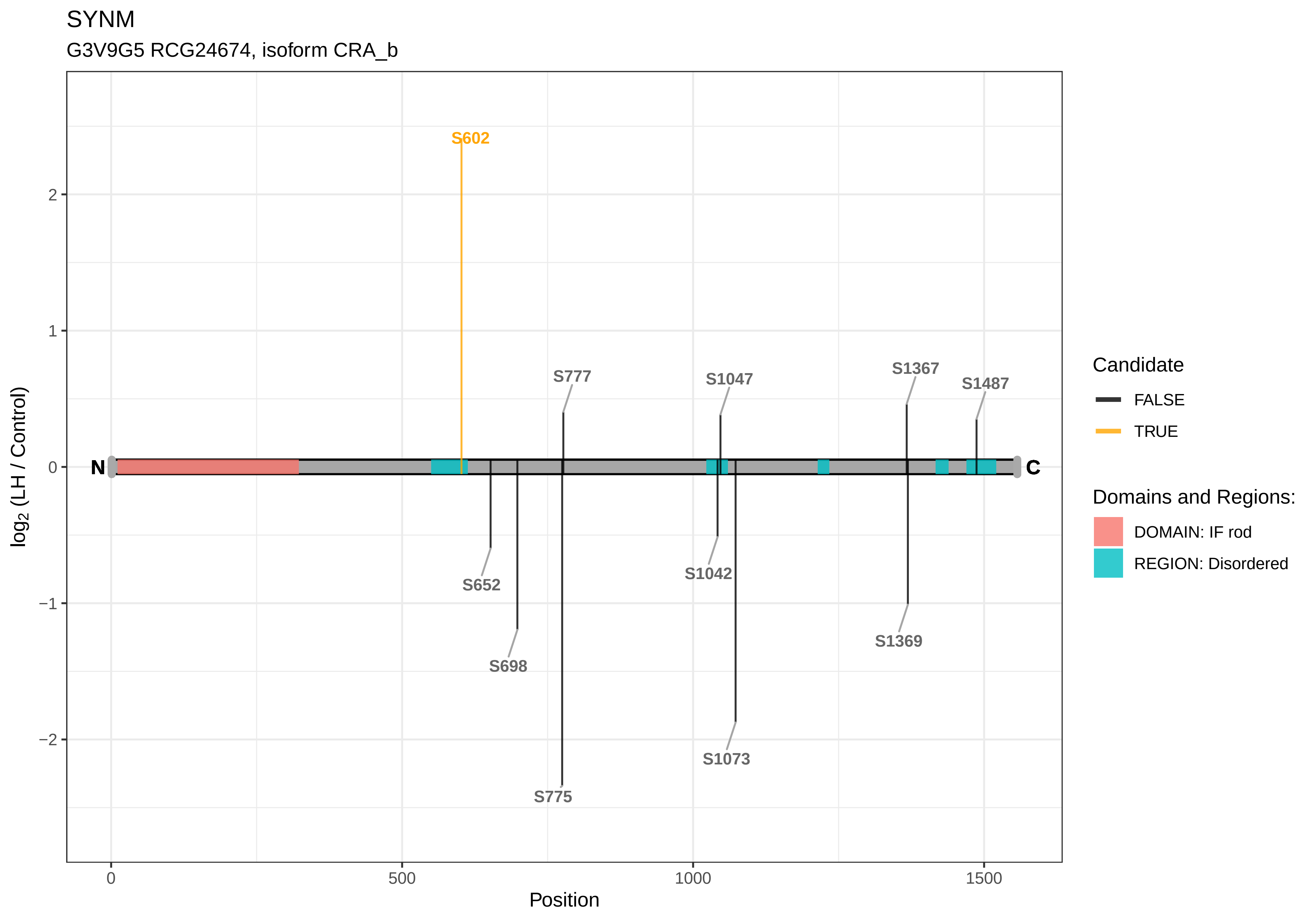
SYNM
G3V9G5 RCG24674, isoform CRA_b
S602
S777
S1047
S1367
S1487
S652
S1042
S698
S1369
S1073
S775
N
C
2
1
0
−1
−2
log2 (LH / Control)
0
500
1000
1500
Position
Candidate
FALSE
TRUE
Domains and Regions:
DOMAIN: IF rod
REGION: Disordered

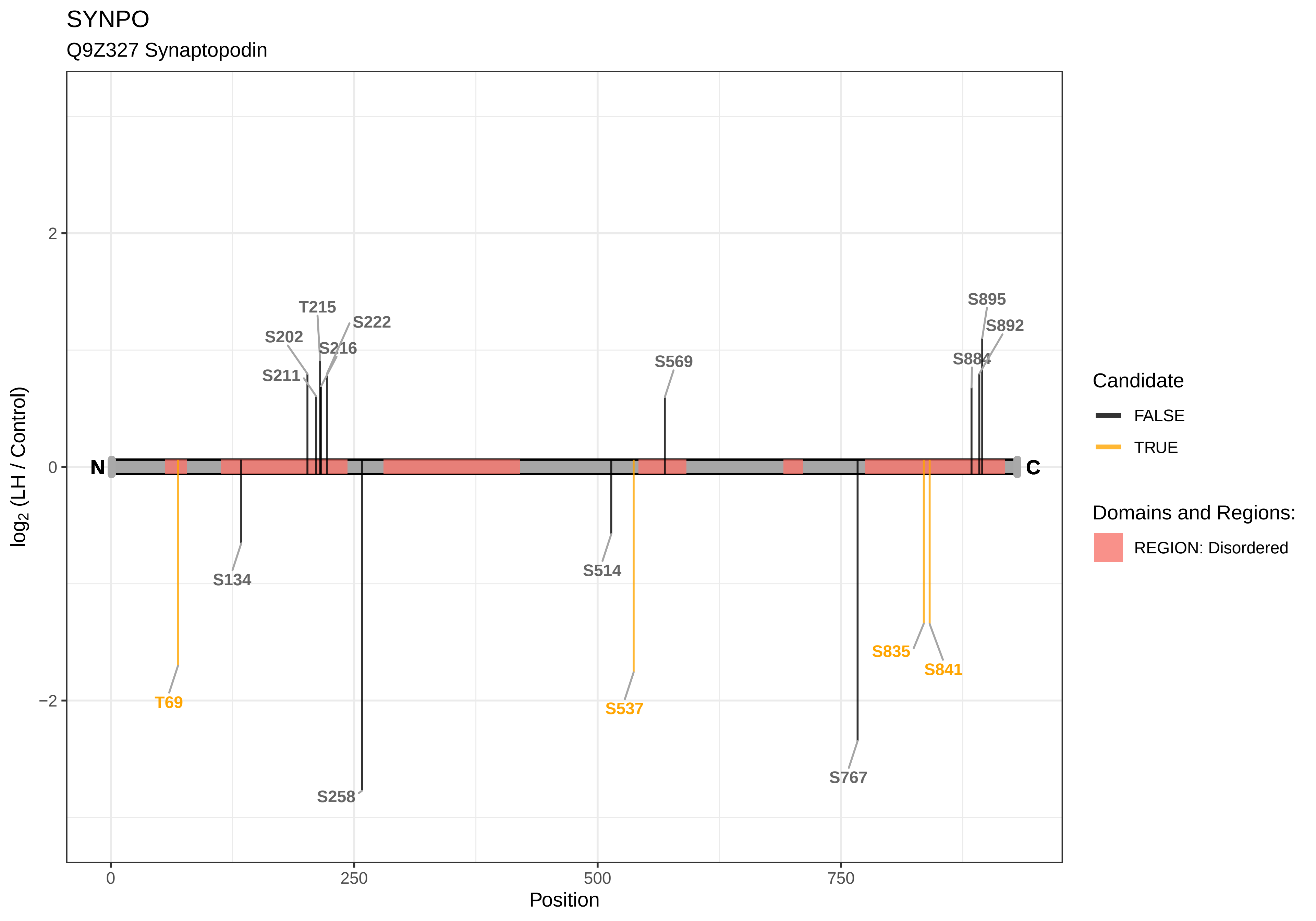
SYNPO
Q9Z327 Synaptopodin
log2 (LH / Control)
Position
N
C
T215
S222
S202
S216
S211
S569
S895
S892
S884
S134
S514
T69
S537
S835
S841
S258
S767
Candidate
FALSE
TRUE
Domains and Regions:
REGION: Disordered
2
0
−2
0
250
500
750

SYTL4

G3V6G6 Synaptotagmin−like 4, isoform CRA_a

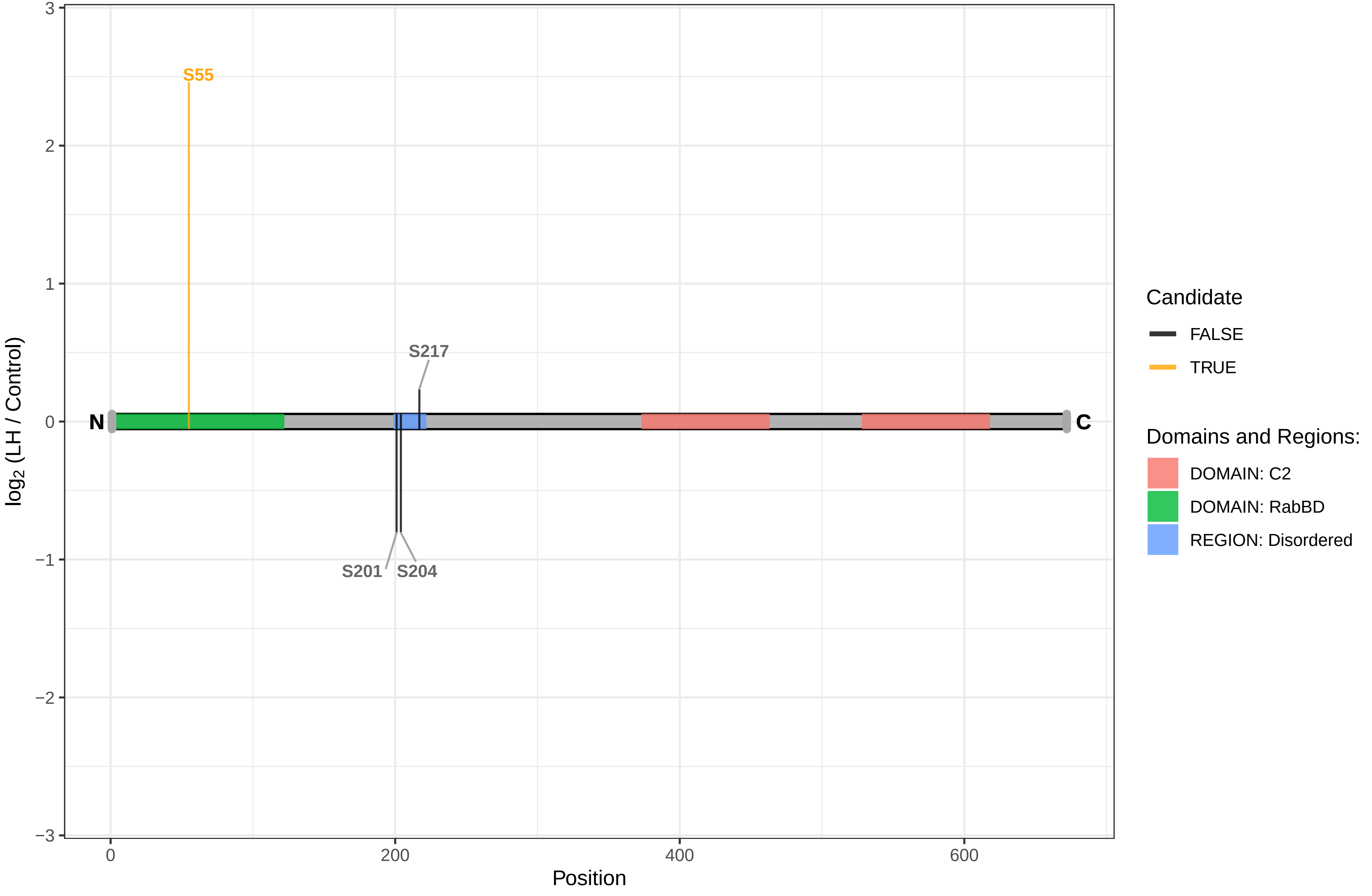

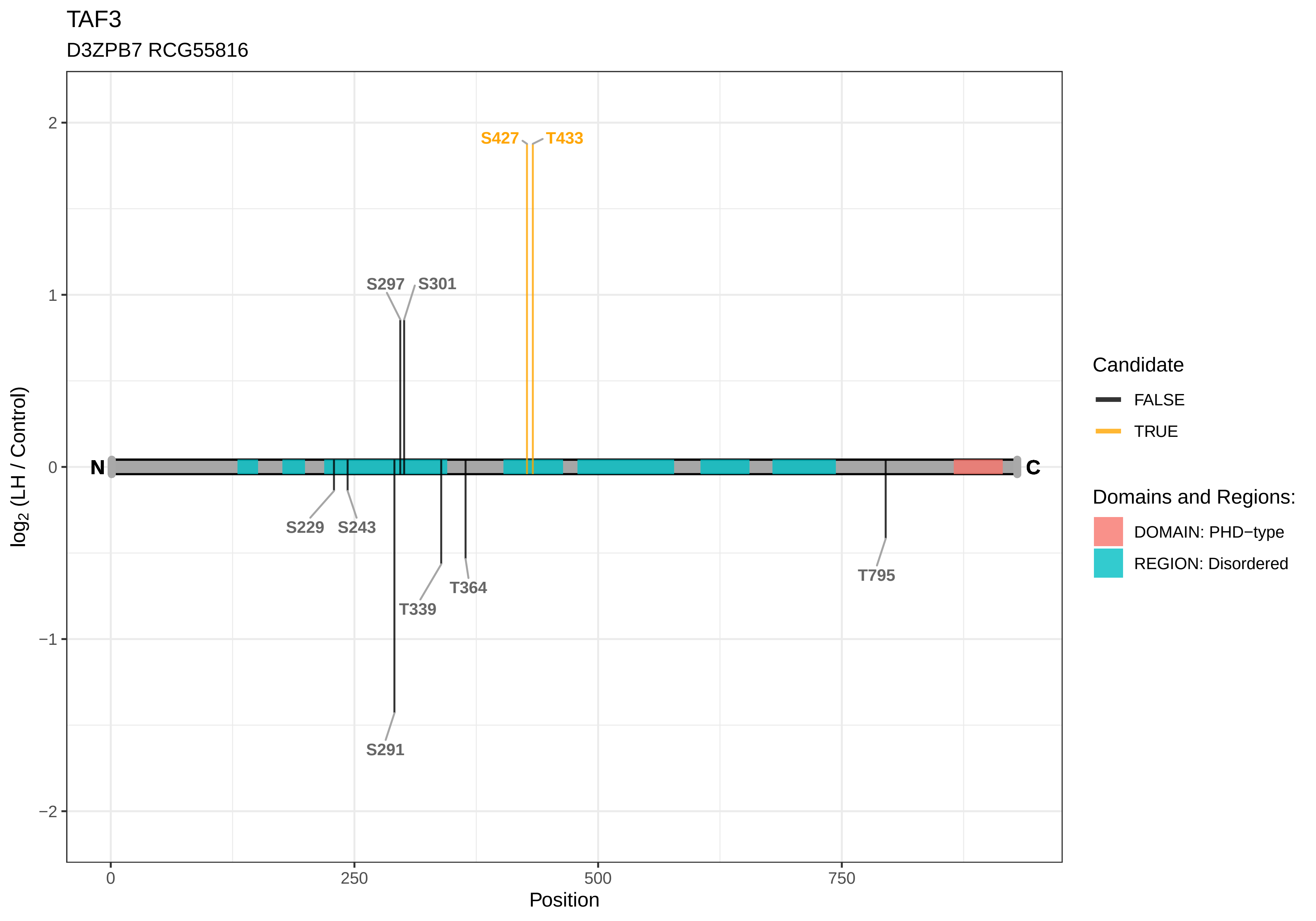
TAF3
D3ZPB7 RCG55816
S427
T433
S297
S301
S229
S243
T339
T364
T795
S291
N
C
2
1
0
−1
−2
log2 (LH / Control)
0
250
500
750
Position
Candidate
FALSE
TRUE
Domains and Regions:
DOMAIN: PHD−type
REGION: Disordered

TAF9B

A0A0G2JXE2 Transcription initiation factor TFIID subunit 9B (Fragment)

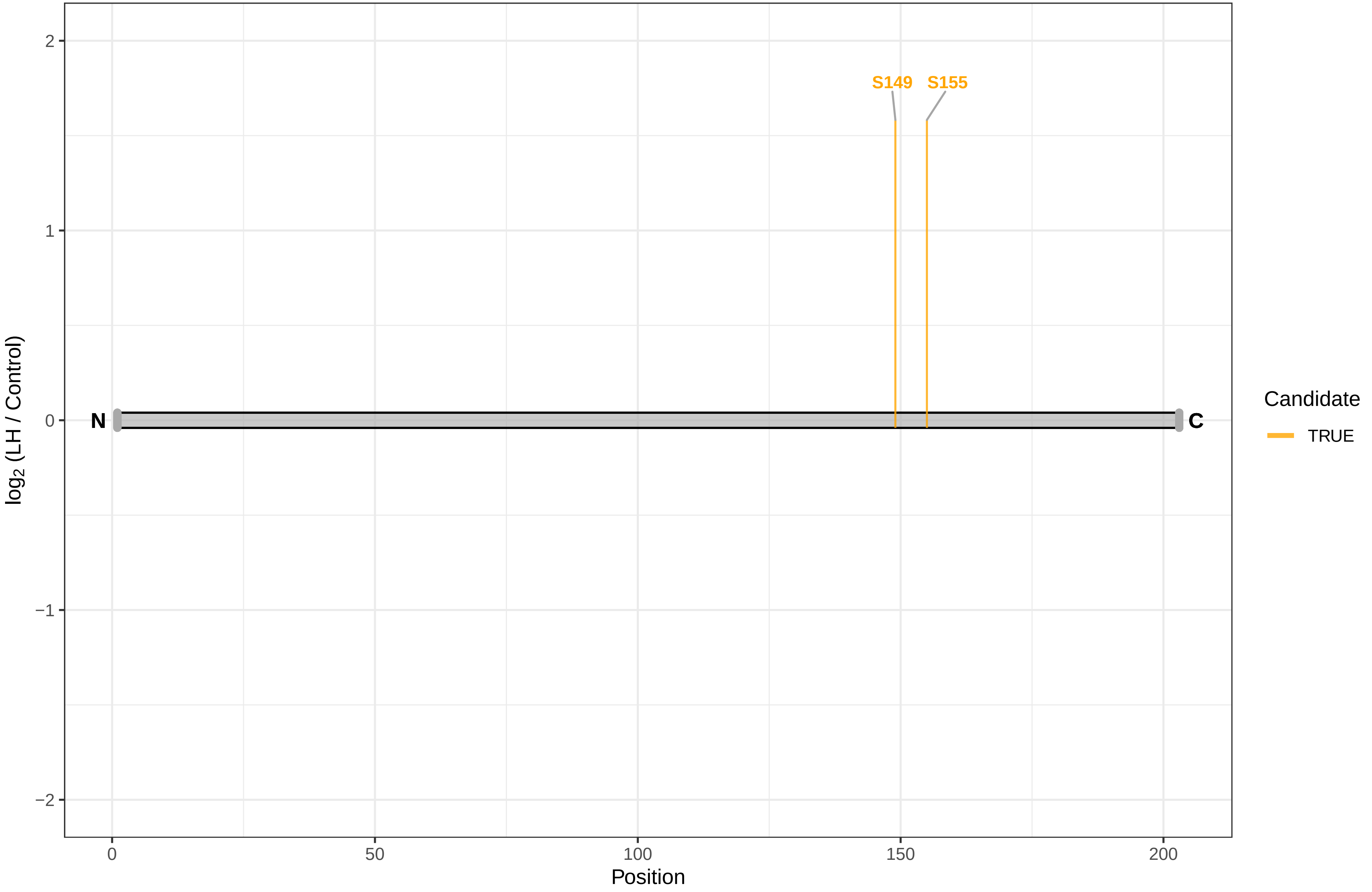

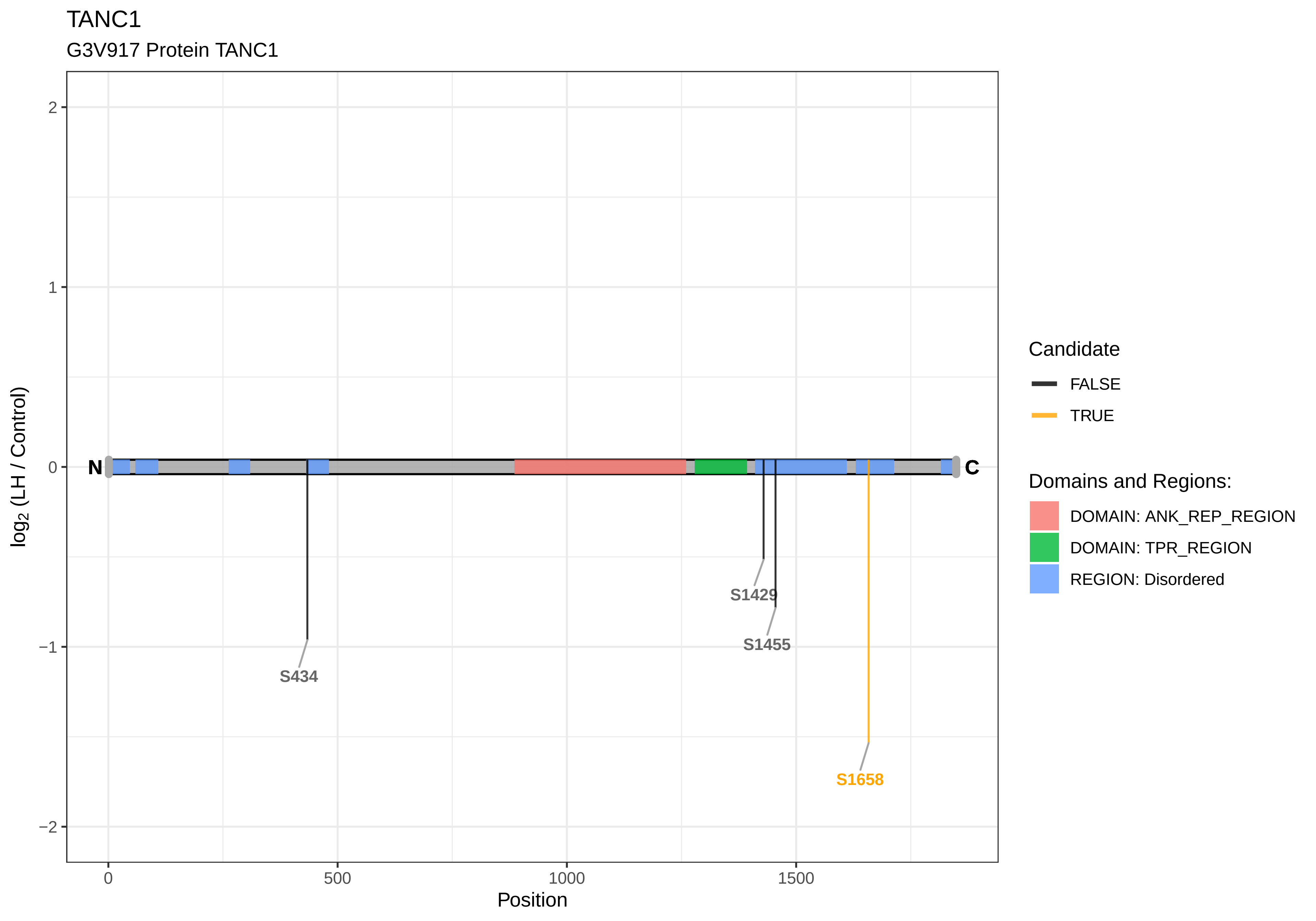

TANC1
G3V917 Protein TANC1
log2 (LH / Control)
Position
2
1
0
−1
−2
0
500
1000
1500
N
C
S434
S1429
S1455
S1658
Candidate
FALSE
TRUE
Domains and Regions:
DOMAIN: ANK_REP_REGION
DOMAIN: TPR_REGION
REGION: Disordered

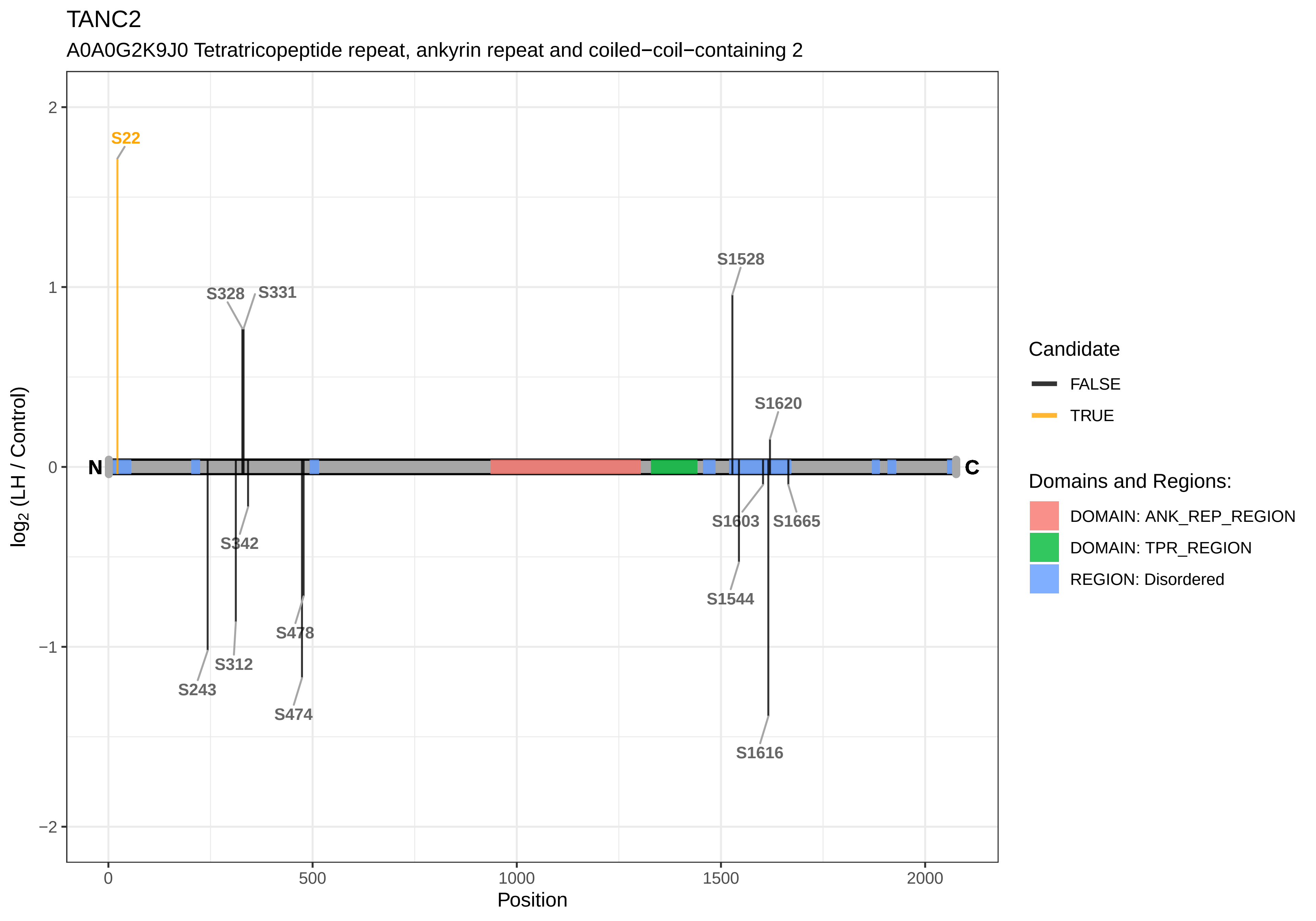

TANC2
A0A0G2K9J0 Tetratricopeptide repeat, ankyrin repeat and coiled−coil−containing 2
S22
S328
S331
S1528
S1620
S342
S1603
S1665
S1544
S478
S312
S243
S474
S1616
N
C
log2 (LH / Control)
Position
2
1
0
−1
−2
0
500
1000
1500
2000
Candidate
FALSE
TRUE
Domains and Regions:
DOMAIN: ANK_REP_REGION
DOMAIN: TPR_REGION
REGION: Disordered

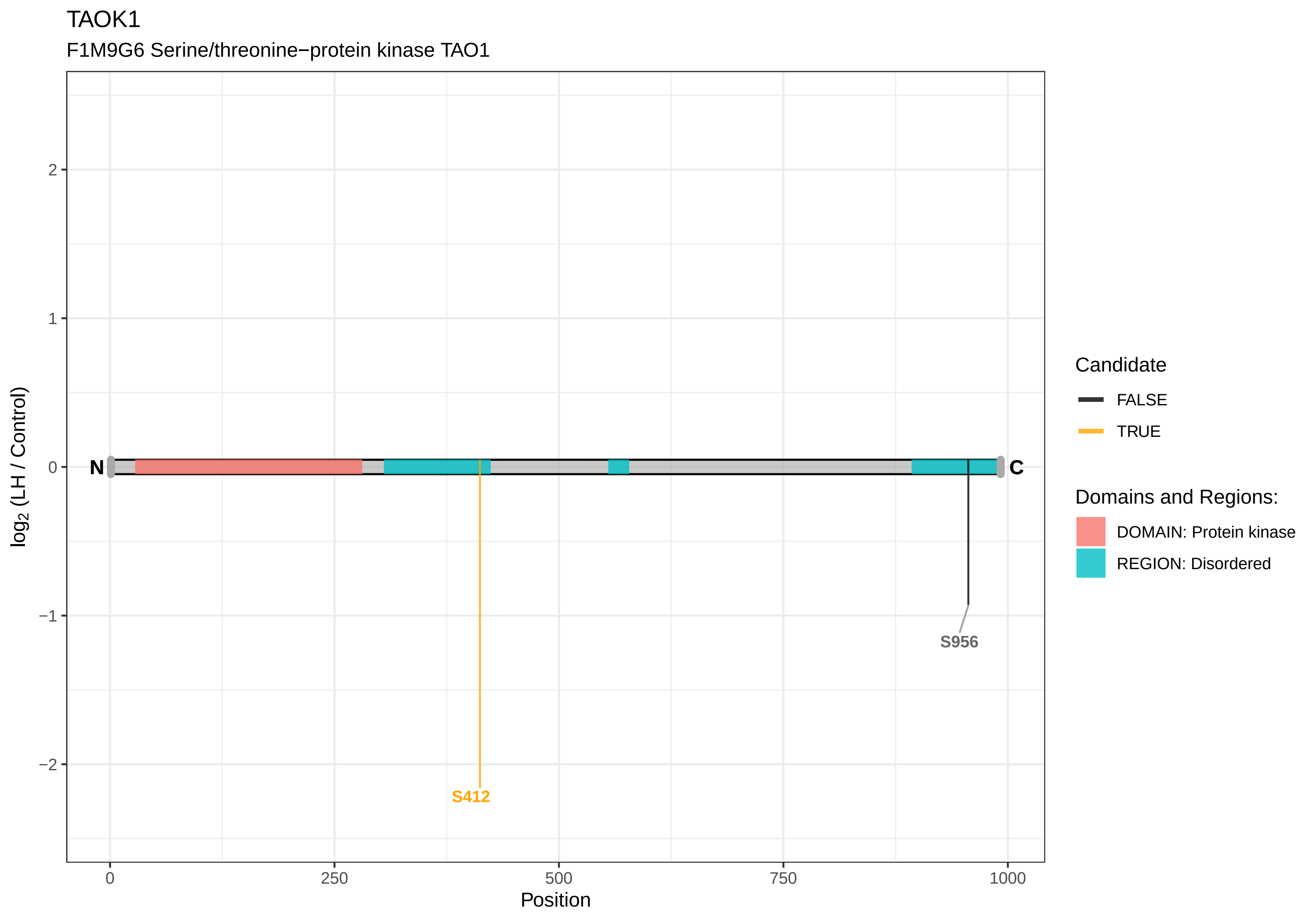

TAOK1
F1M9G6 Serine/threonine−protein kinase TAO1
log2 (LH / Control)
2
1
0
−1
−2
N
C
S412
S956
0
250
500
750
1000
Position
Candidate
FALSE
TRUE
Domains and Regions:
DOMAIN: Protein kinase
REGION: Disordered

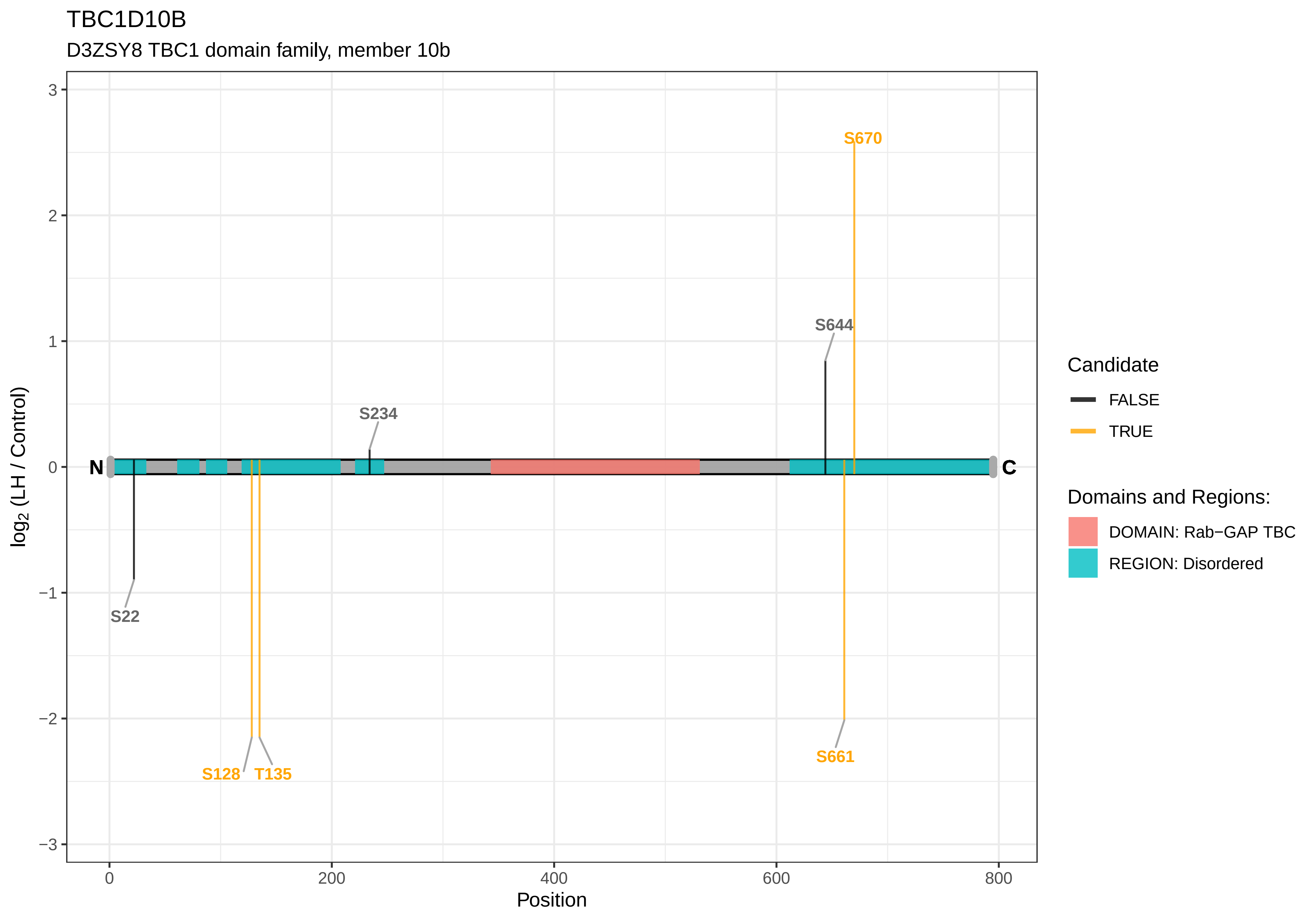
TBC1D10B
D3ZSY8 TBC1 domain family, member 10b
log2 (LH / Control)
Position
N
C
S670
S644
S234
S22
S128
T135
S661
Candidate
FALSE
TRUE
Domains and Regions:
DOMAIN: Rab−GAP TBC
REGION: Disordered
3
2
1
0
−1
−2
−3
0
200
400
600
800

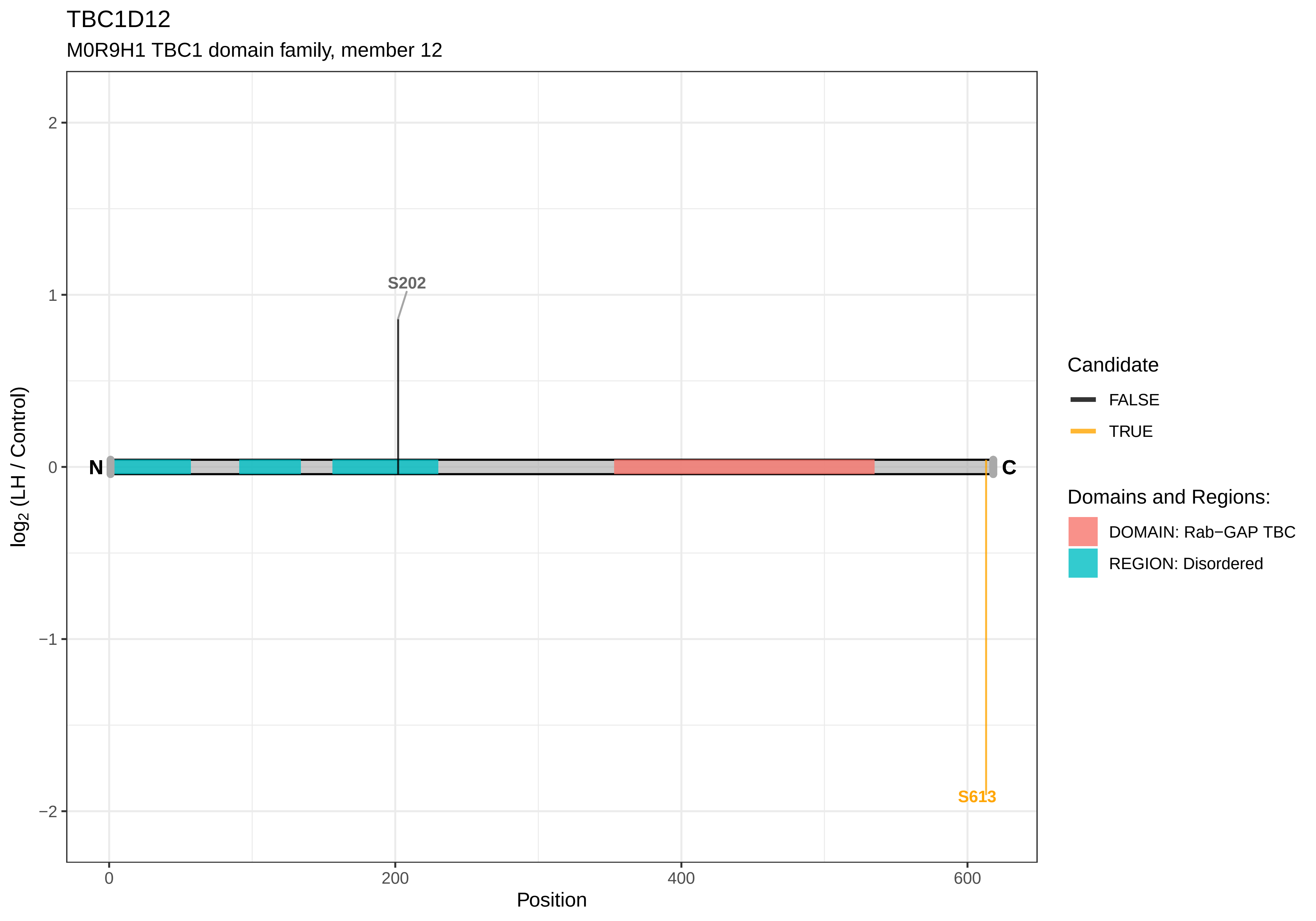
TBC1D12
M0R9H1 TBC1 domain family, member 12
S202
S613
N
C
2
1
0
−1
−2
log2 (LH / Control)
0
200
400
600
Position
Candidate
FALSE
TRUE
Domains and Regions:
DOMAIN: Rab−GAP TBC
REGION: Disordered

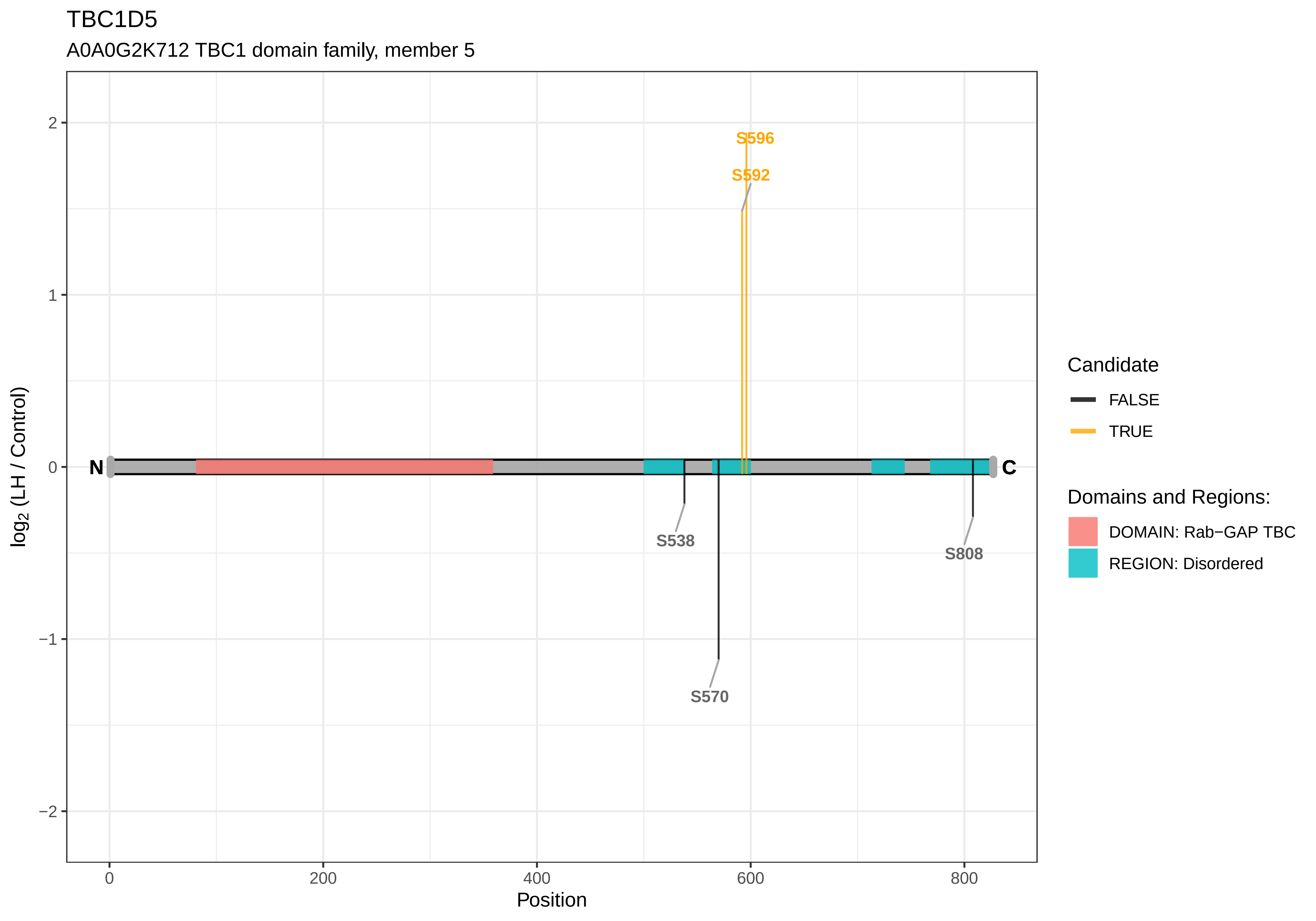

TBC1D5
A0A0G2K712 TBC1 domain family, member 5
S596
S592
S538
S570
S808
N
C
2
1
0
−1
−2
0
200
400
600
800
log2 (LH / Control)
Position
Candidate
FALSE
TRUE
Domains and Regions:
DOMAIN: Rab−GAP TBC
REGION: Disordered

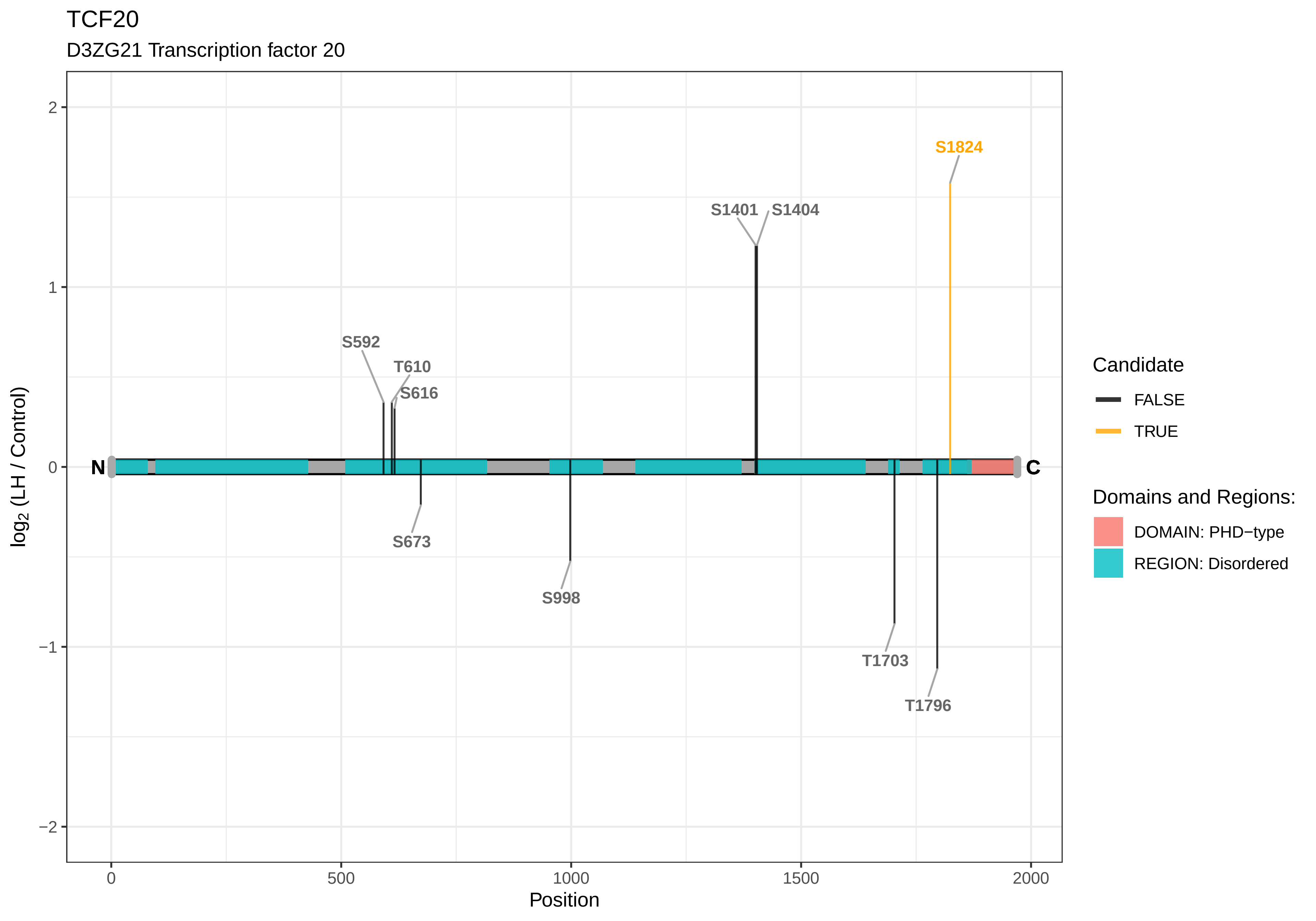

TCF20
D3ZG21 Transcription factor 20
S1824
S1401
S1404
S592
T610
S616
N
C
S673
S998
T1703
T1796
2
1
0
−1
−2
log2 (LH / Control)
0
500
1000
1500
2000
Position
Candidate
FALSE
TRUE
Domains and Regions:
DOMAIN: PHD−type
REGION: Disordered

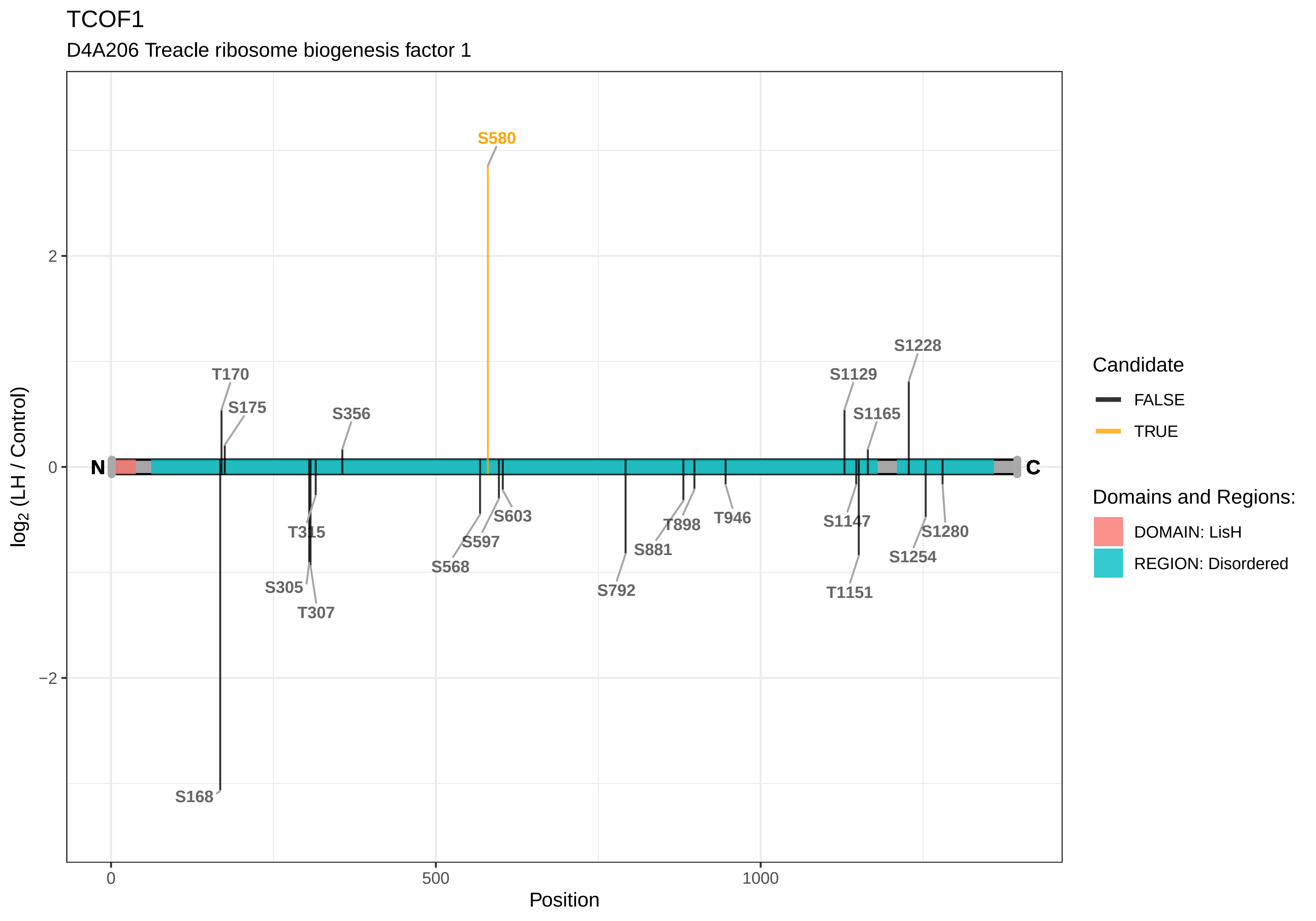
TCOF1
D4A206 Treacle ribosome biogenesis factor 1
S580
S1228
T170
S1129
S175
S356
S1165
N
C
T315
S603
T898
T946
S1147
S1280
S597
S881
S1254
S568
S792
S305
T1151
T307
S168
2
0
−2
0
500
1000
Position
log2 (LH / Control)
Candidate
FALSE
TRUE
Domains and Regions:
DOMAIN: LisH
REGION: Disordered

TCP11L1

F1M9Y7 T−complex 11−like 1

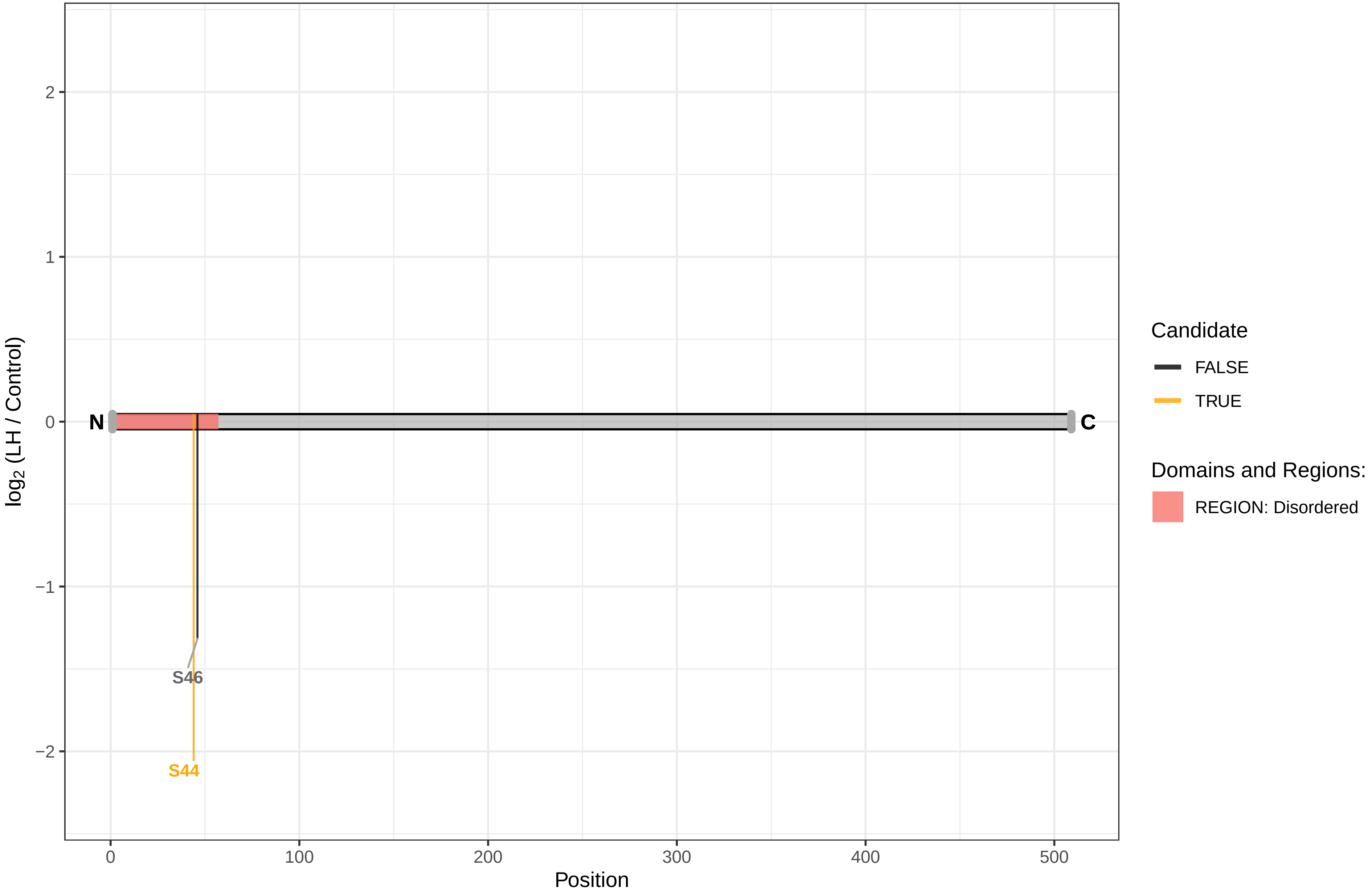

TDRD3

A0A096MIS7 Tudor domain−containing protein 3

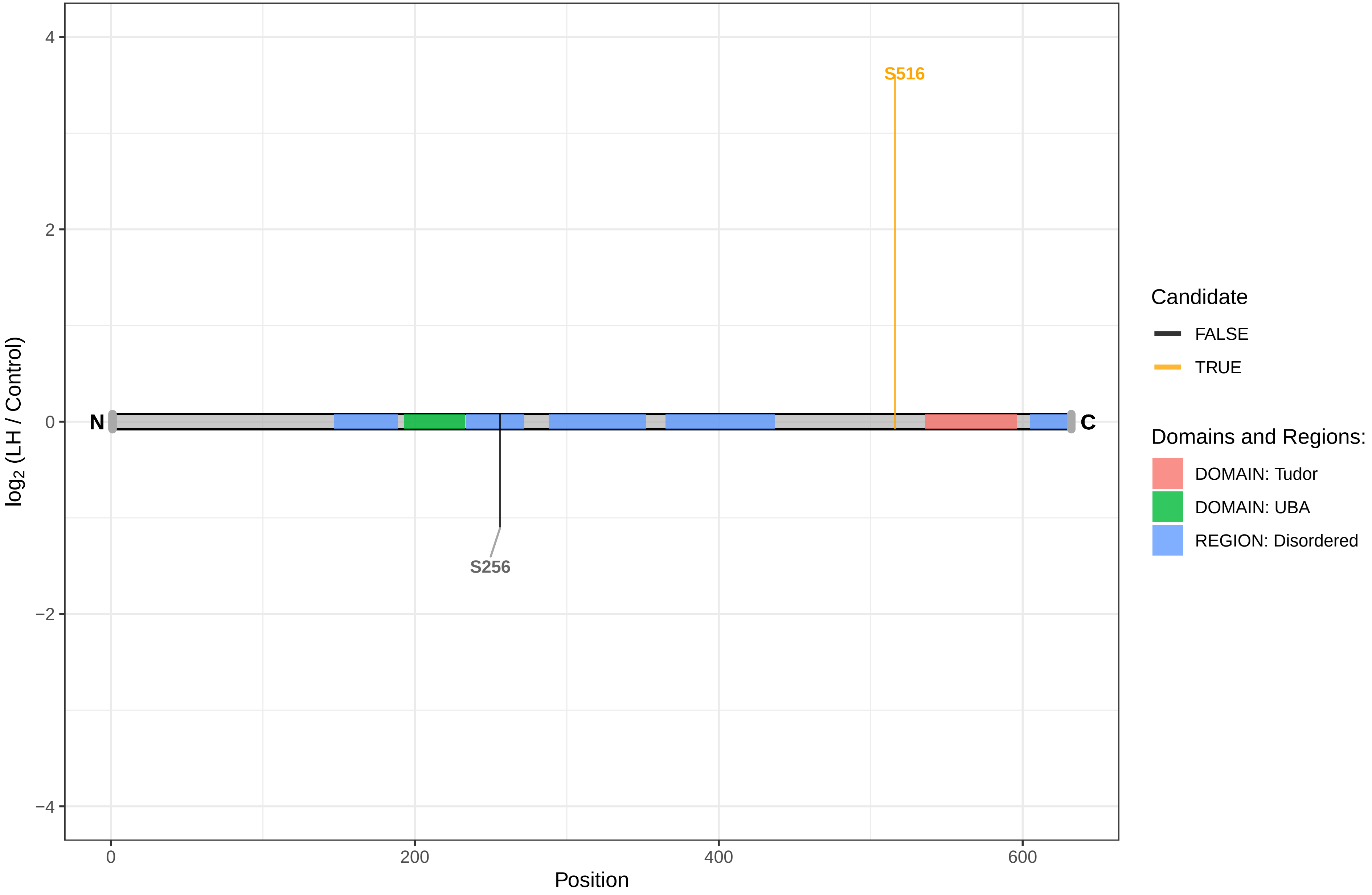

TEX2

G3V9L1 Testis expressed gene 2

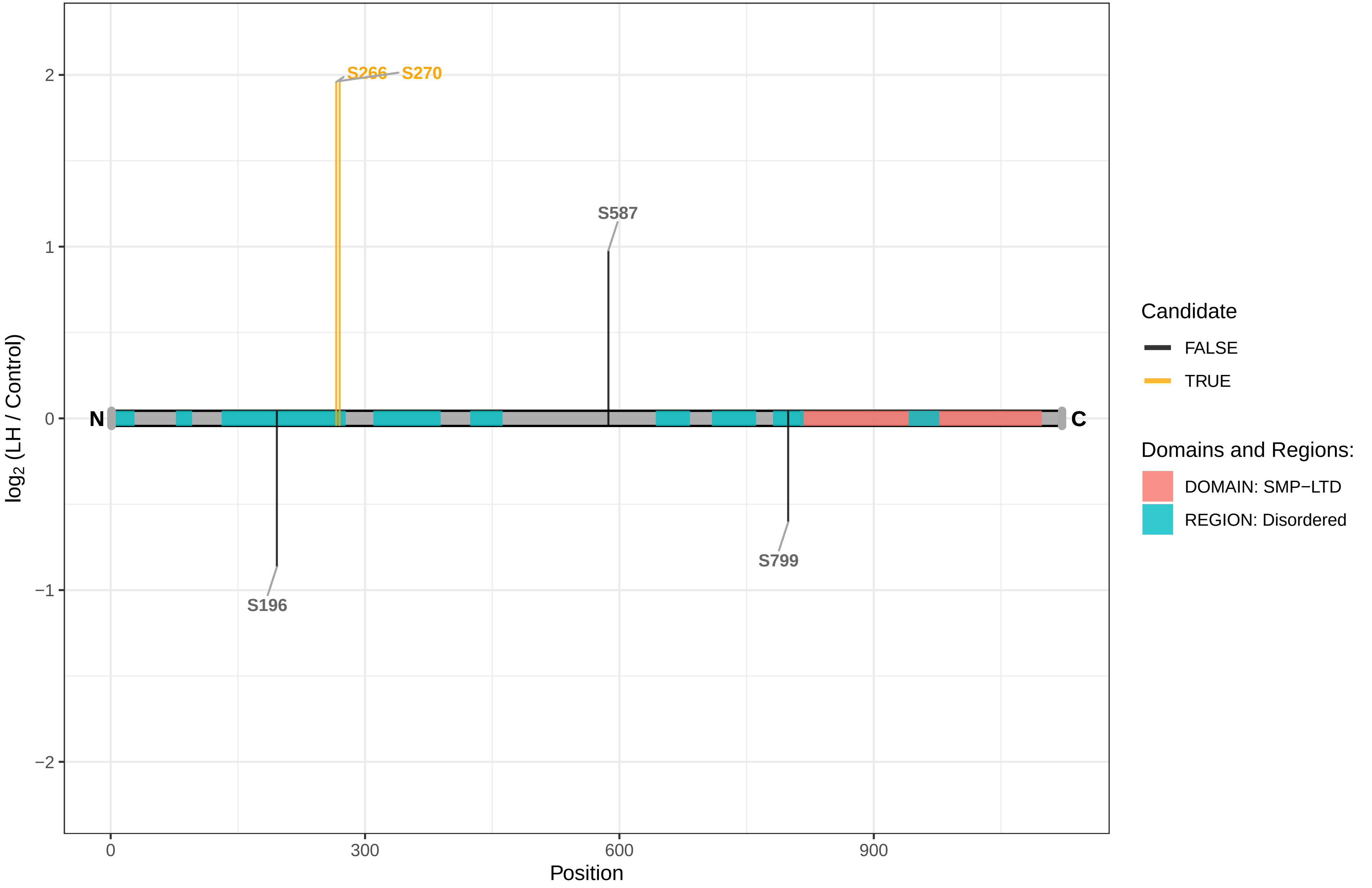

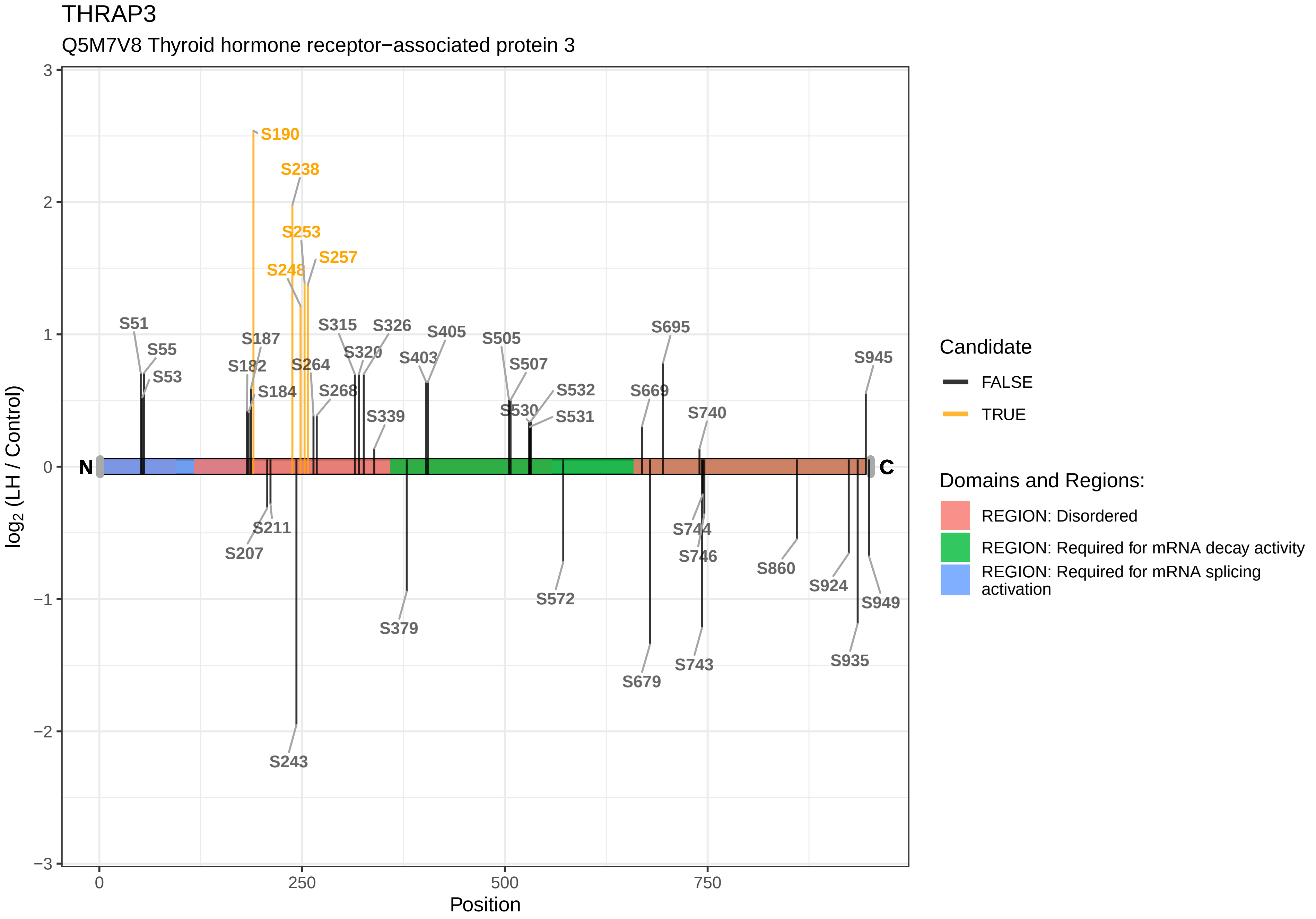

THRAP3
Q5M7V8 Thyroid hormone receptor−associated protein 3
S190
S238
S253
S257
S248
S51
S55
S53
S187
S182
S184
S264
S268
S315
S326
S320
S403
S405
S339
S505
S507
S530
S532
S531
S695
S669
S740
S945
N
C
S211
S207
S243
S379
S572
S679
S744
S746
S743
S860
S924
S935
S949
3
2
1
0
−1
−2
−3
0
250
500
750
Position
log2 (LH / Control)
Candidate
FALSE
TRUE
Domains and Regions:
REGION: Disordered
REGION: Required for mRNA decay activity
REGION: Required for mRNA splicing activation

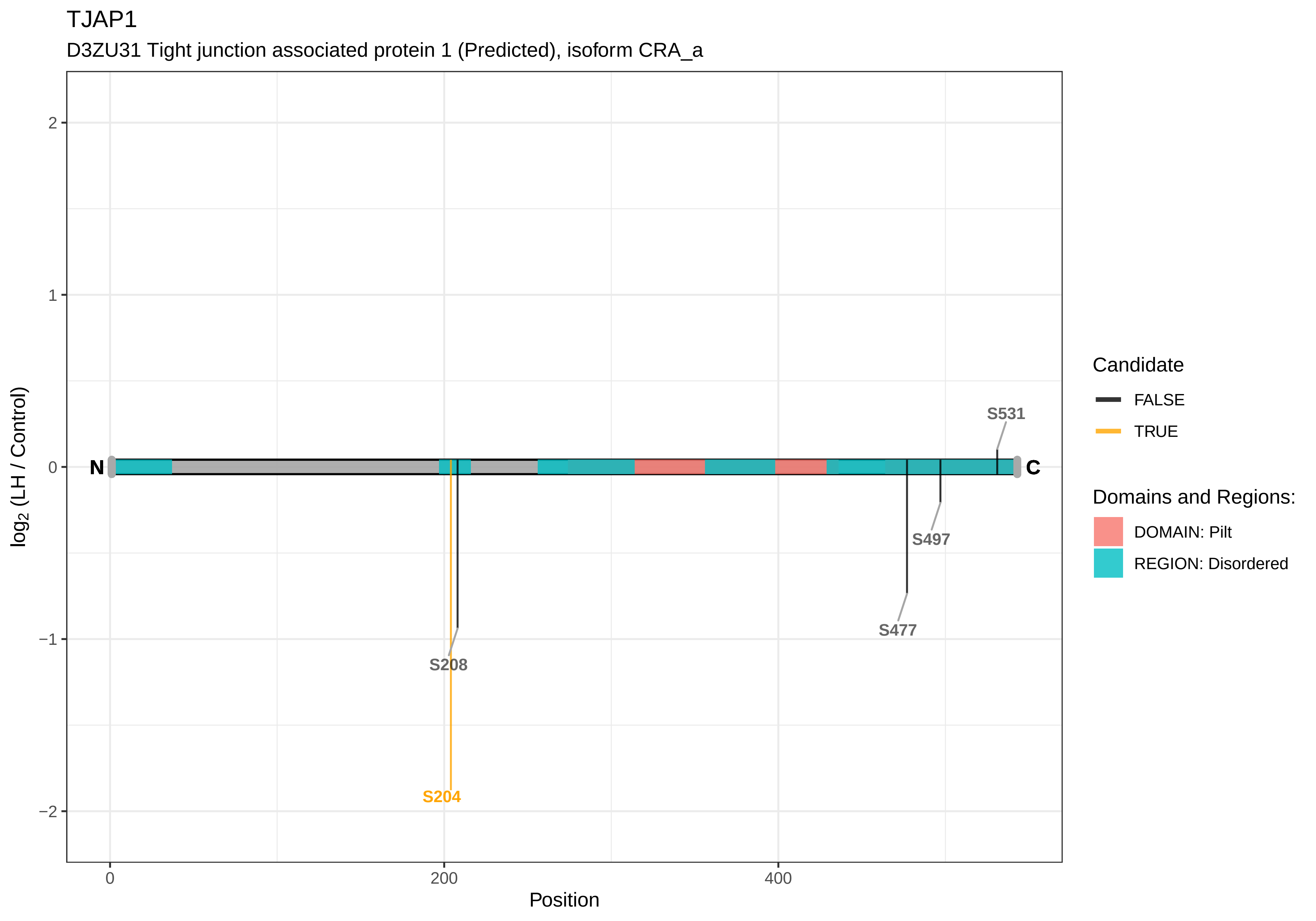
TJAP1
D3ZU31 Tight junction associated protein 1 (Predicted), isoform CRA_a
log2 (LH / Control)
2
1
0
−1
−2
N
C
S531
S497
S477
S208
S204
0
200
400
Position
Candidate
FALSE
TRUE
Domains and Regions:
DOMAIN: Pilt
REGION: Disordered

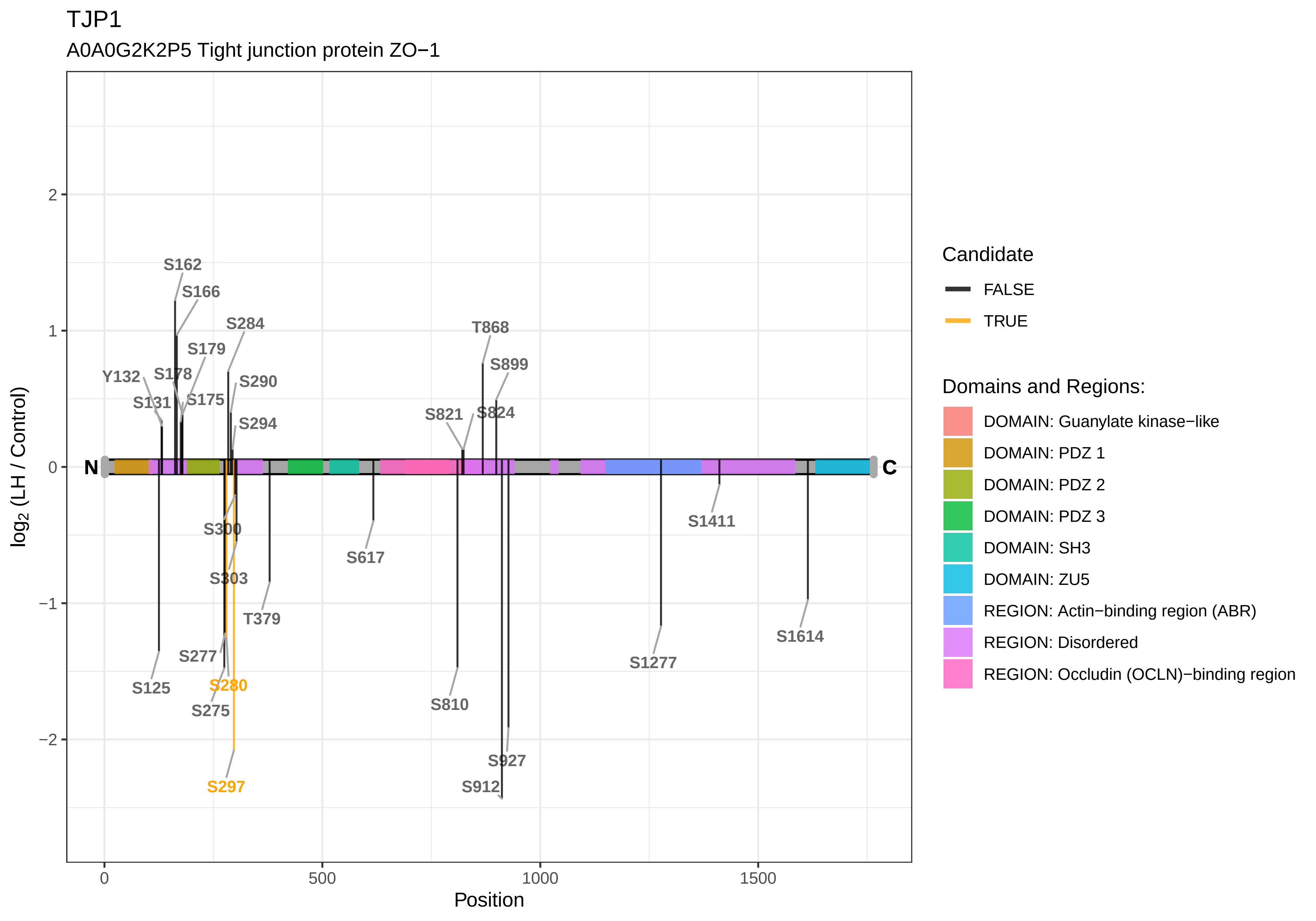

TJP1
A0A0G2K2P5 Tight junction protein ZO−1
log2 (LH / Control)
Position
N
C
S162
S166
S284
S179
Y132
S178
S290
S131
S175
S294
T868
S899
S821
S824
S300
S303
T379
S617
S277
S125
S280
S275
S297
S810
S927
S912
S1411
S1277
S1614
2
1
0
−1
−2
500
1000
1500
Candidate
FALSE
TRUE
Domains and Regions:
DOMAIN: Guanylate kinase−like
DOMAIN: PDZ 1
DOMAIN: PDZ 2
DOMAIN: PDZ 3
DOMAIN: SH3
DOMAIN: ZU5
REGION: Actin−binding region (ABR)
REGION: Disordered
REGION: Occludin (OCLN)−binding region

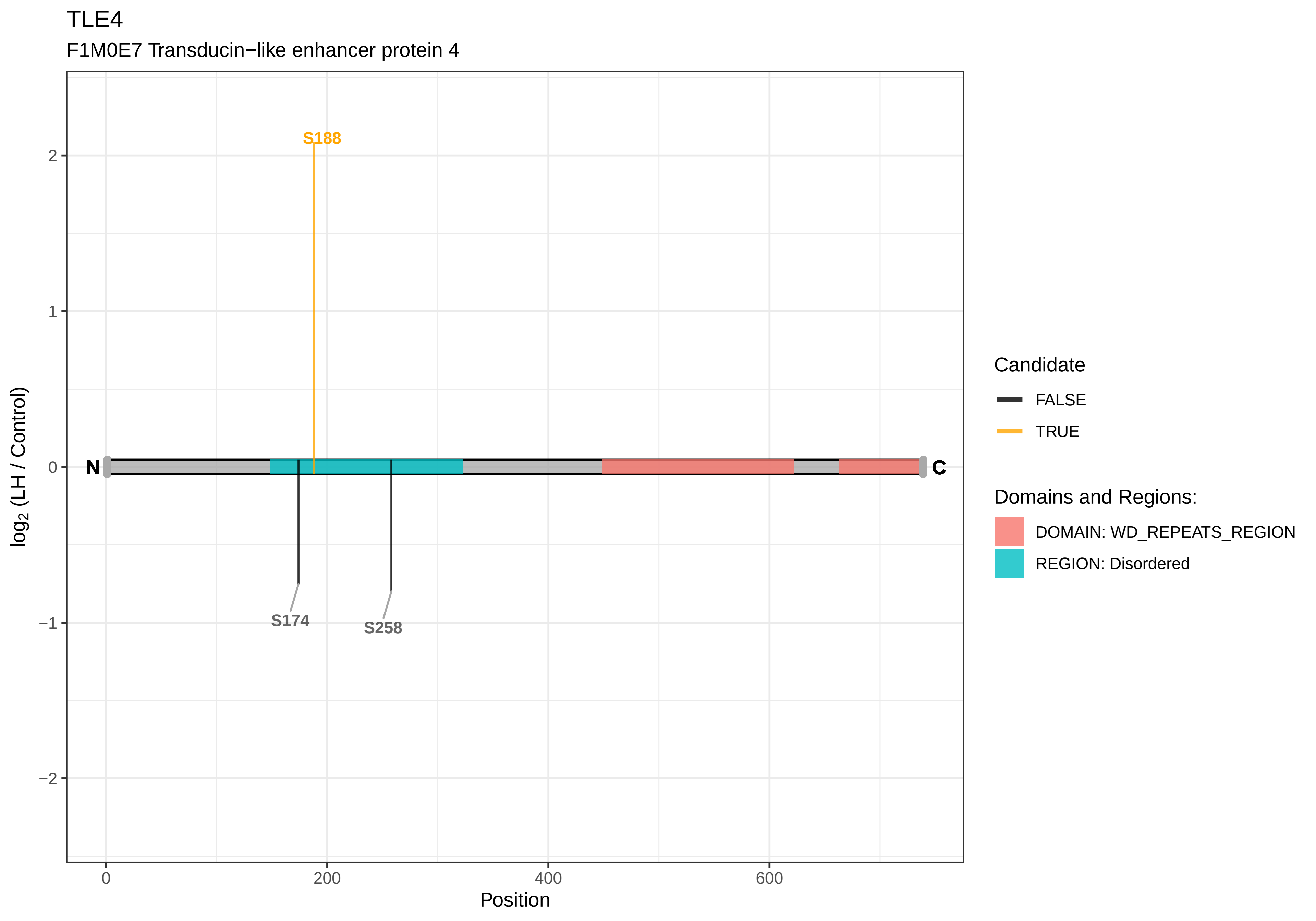

TLE4
F1M0E7 Transducin−like enhancer protein 4
S188
S174
S258
N
C
log2 (LH / Control)
Position
0
200
400
600
2
1
0
−1
−2
Candidate
FALSE
TRUE
Domains and Regions:
DOMAIN: WD_REPEATS_REGION
REGION: Disordered

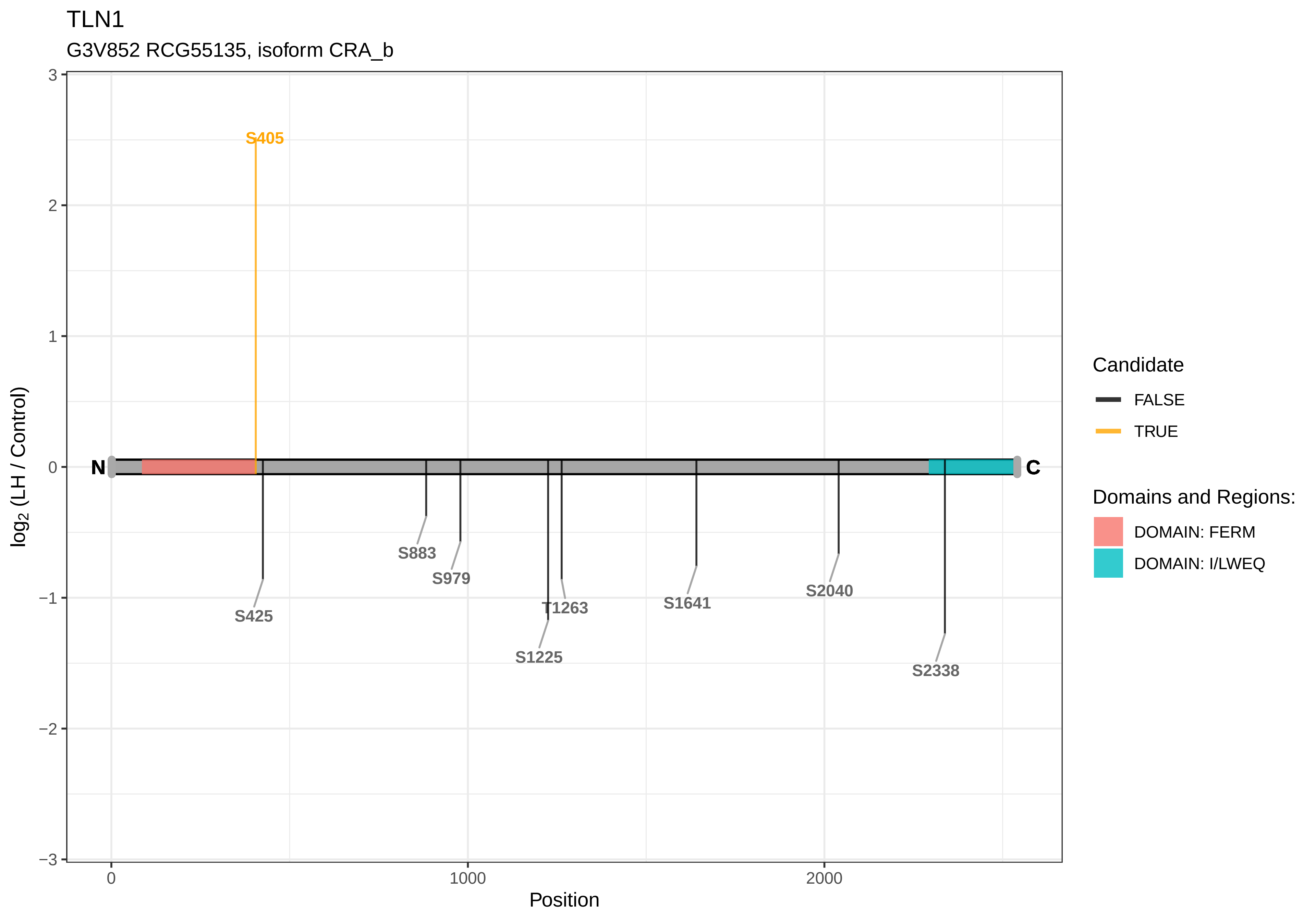

TLN1
G3V852 RCG55135, isoform CRA_b
S405
N
C
S425
S883
S979
S1225
T1263
S1641
S2040
S2338
log2 (LH / Control)
Position
3
2
1
0
−1
−2
−3
0
1000
2000
Candidate
FALSE
TRUE
Domains and Regions:
DOMAIN: FERM
DOMAIN: I/LWEQ

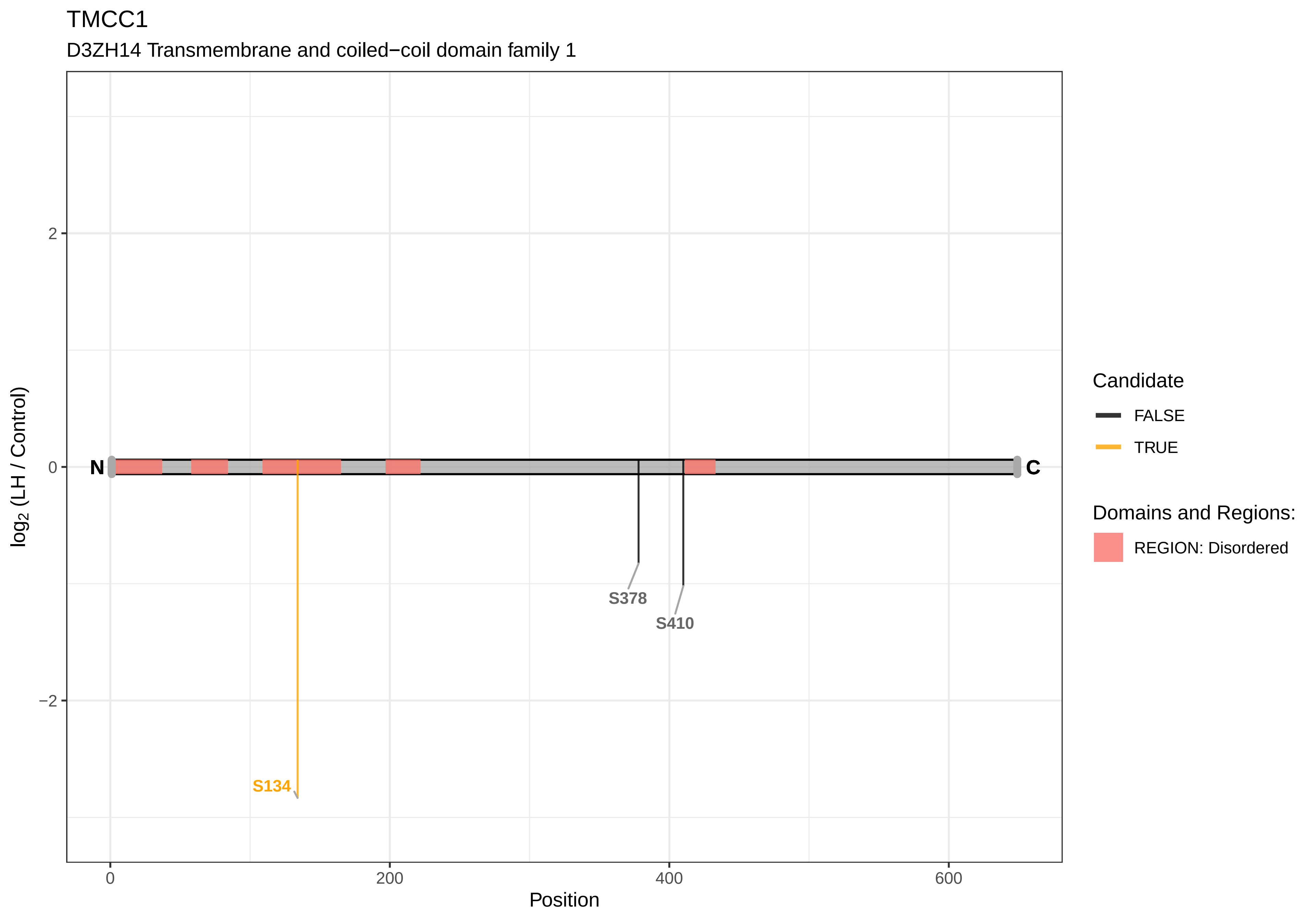
TMCC1
D3ZH14 Transmembrane and coiled−coil domain family 1
log2 (LH / Control)
2
0
−2
N
C
S134
S378
S410
0
200
400
600
Position
Candidate
FALSE
TRUE
Domains and Regions:
REGION: Disordered

TMEM131

F1M5G8 Transmembrane protein 131

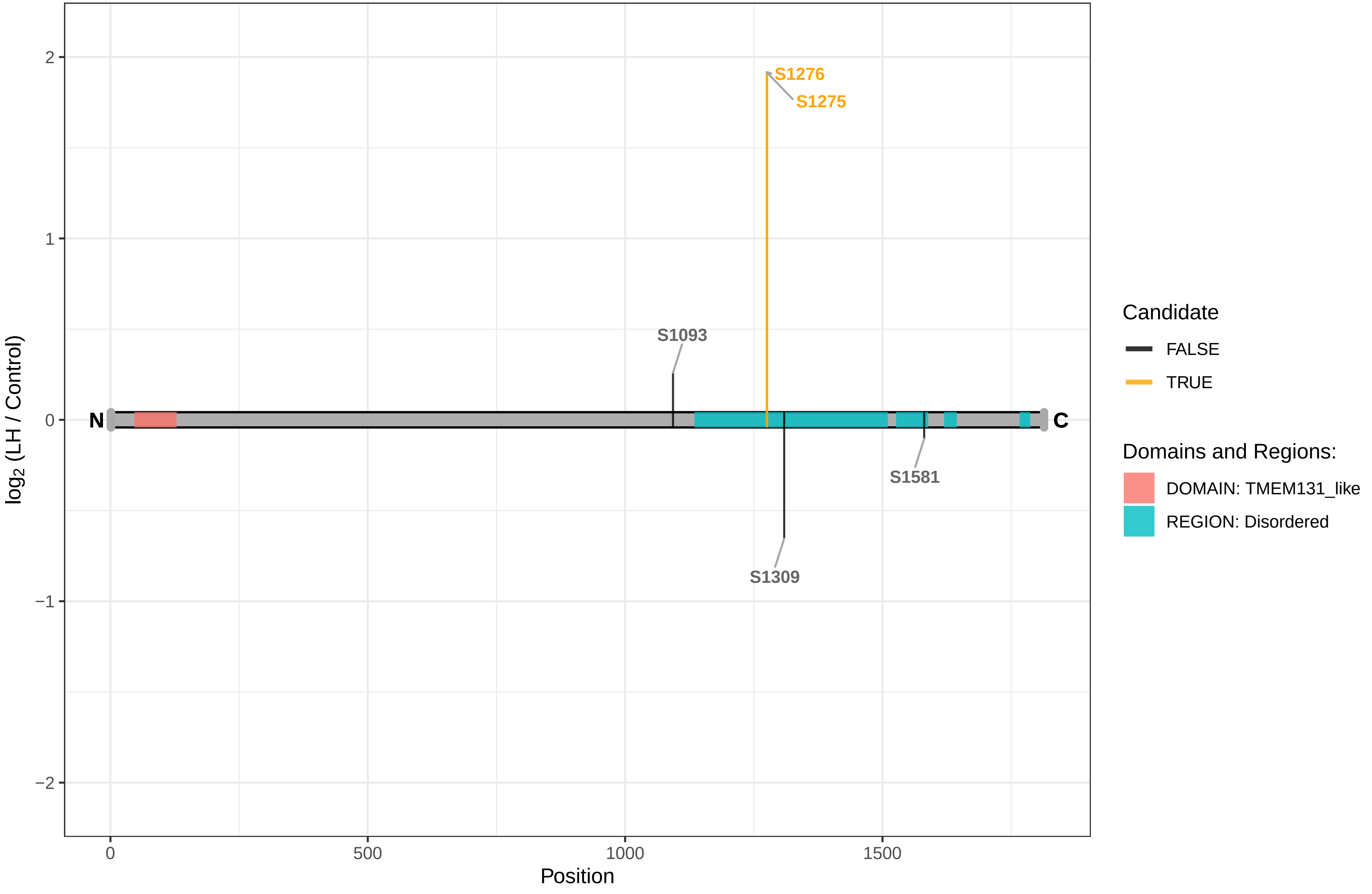

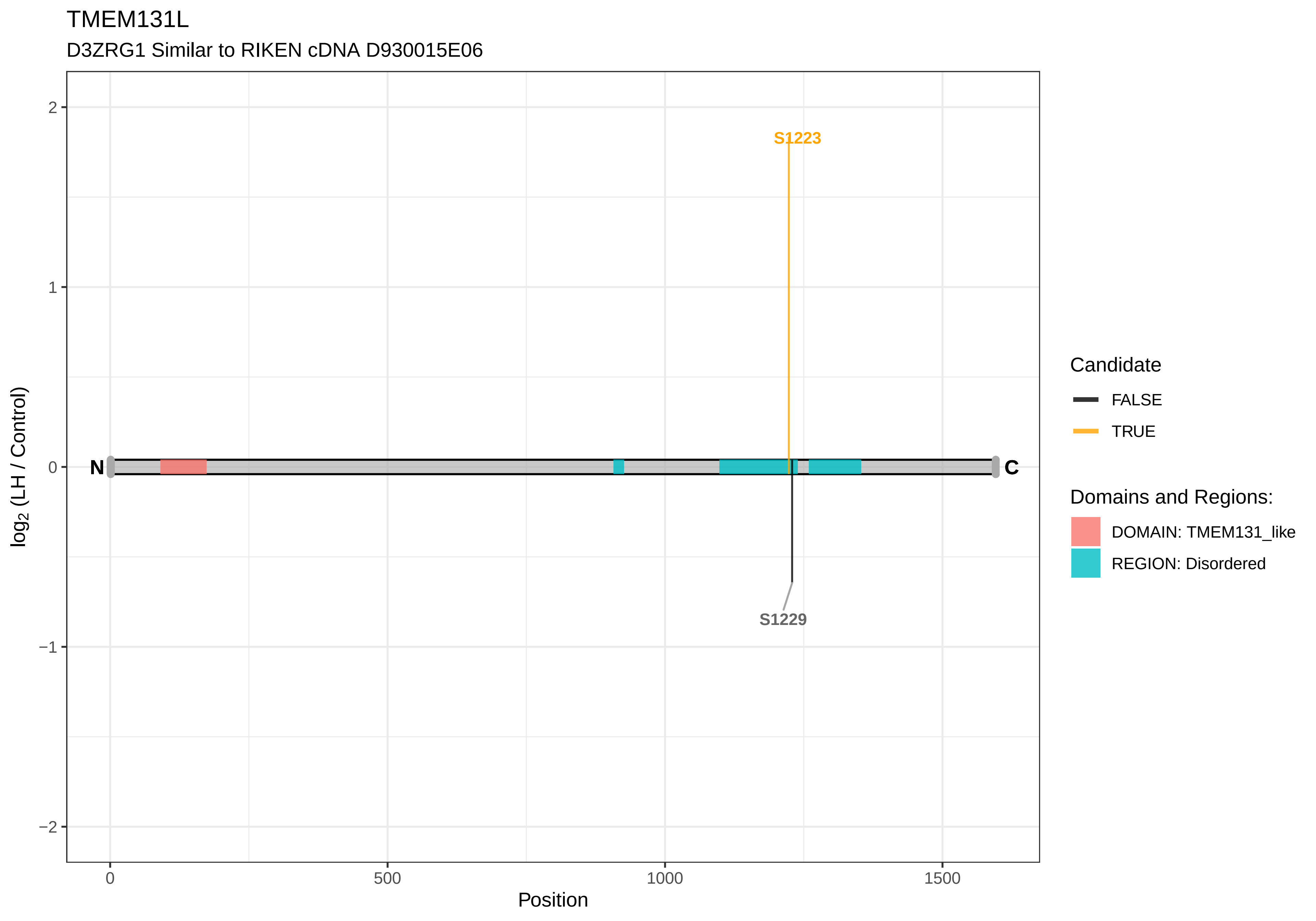
TMEM131L
D3ZRG1 Similar to RIKEN cDNA D930015E06
S1223
S1229
N
C
2
1
0
−1
−2
log2 (LH / Control)
0
500
1000
1500
Position
Candidate
FALSE
TRUE
Domains and Regions:
DOMAIN: TMEM131_like
REGION: Disordered

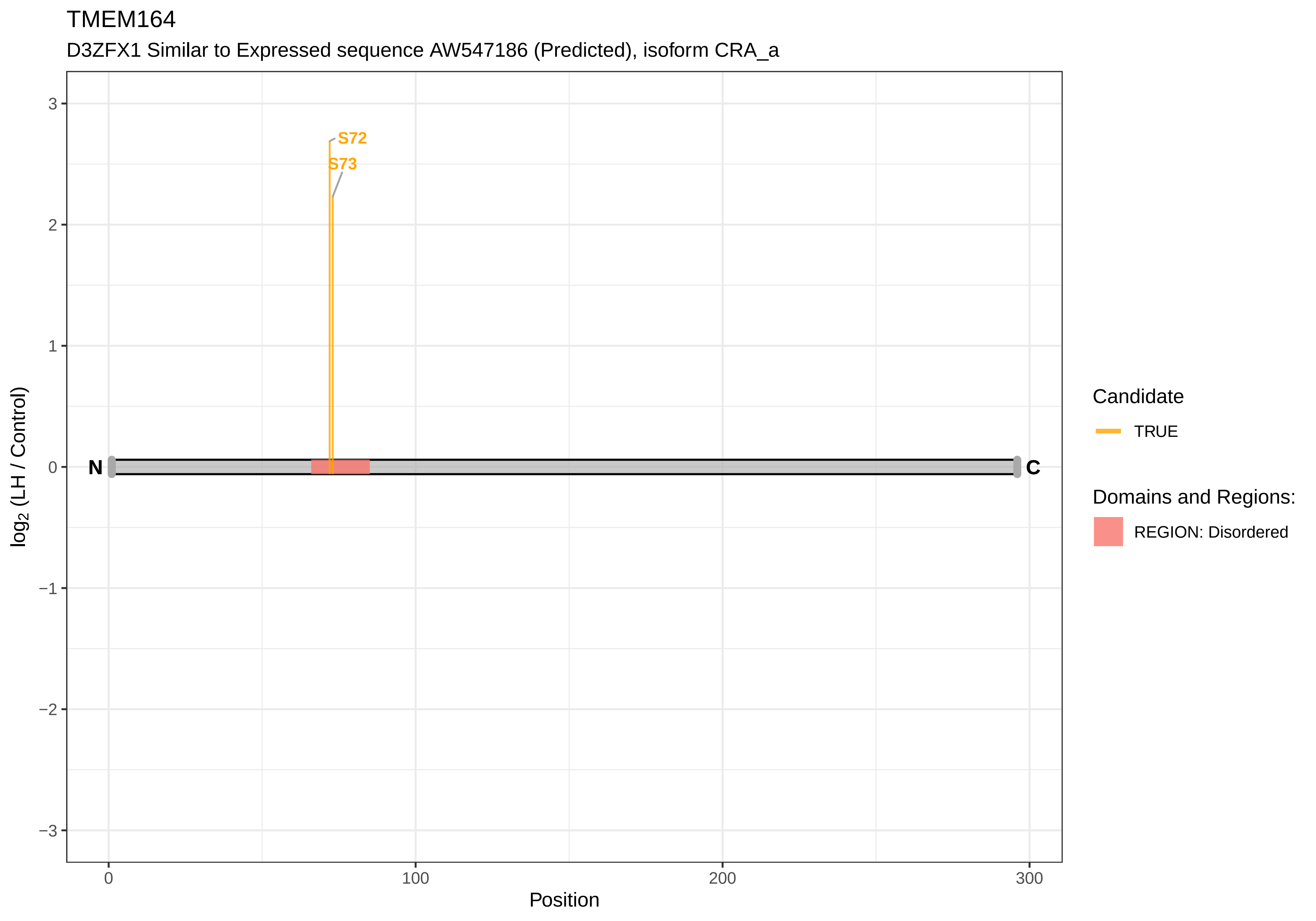

TMEM164
D3ZFX1 Similar to Expressed sequence AW547186 (Predicted), isoform CRA_a
S72
S73
N
C
3
2
1
0
−1
−2
−3
log2 (LH / Control)
0
100
200
300
Position
Candidate
TRUE
Domains and Regions:
REGION: Disordered

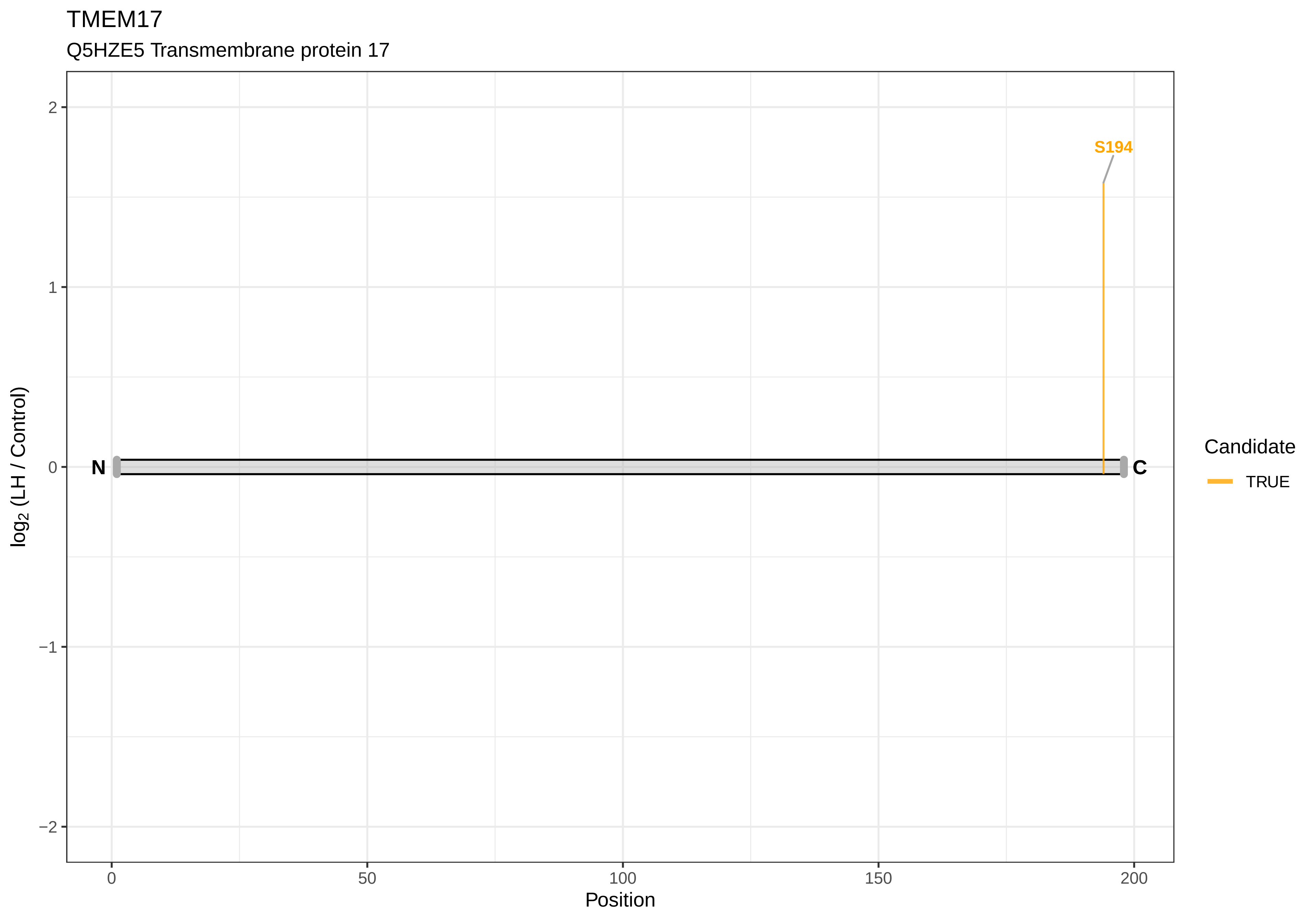
TMEM17
Q5HZE5 Transmembrane protein 17
S194
N
C
log2 (LH / Control)
2
1
0
−1
−2
0
50
100
150
200
Position
Candidate
TRUE

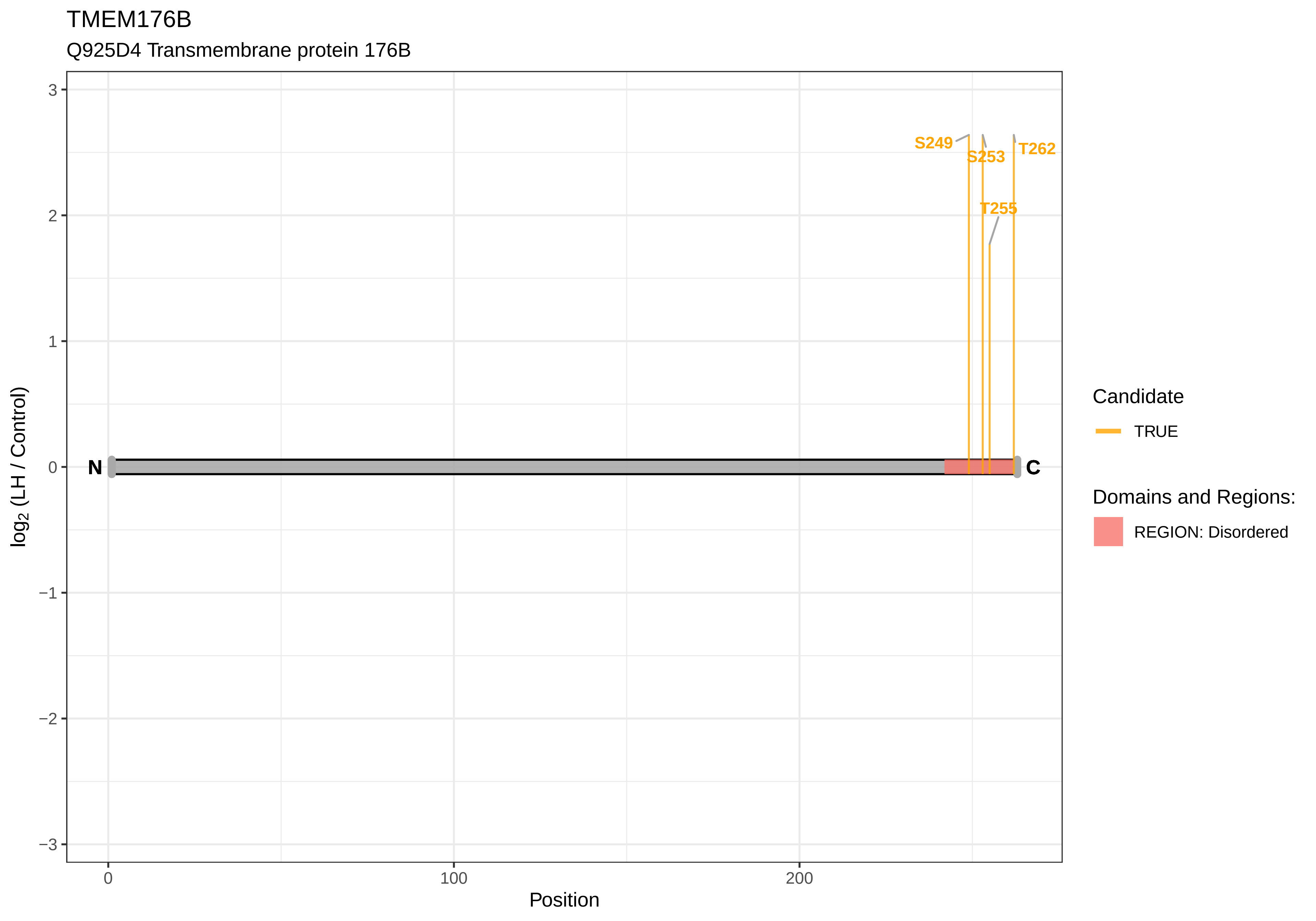
TMEM176B
Q925D4 Transmembrane protein 176B
log2 (LH / Control)
Position
S249
S253
T262
T255
N
C
Candidate
TRUE
Domains and Regions:
REGION: Disordered
3
2
1
0
−1
−2
−3
0
100
200

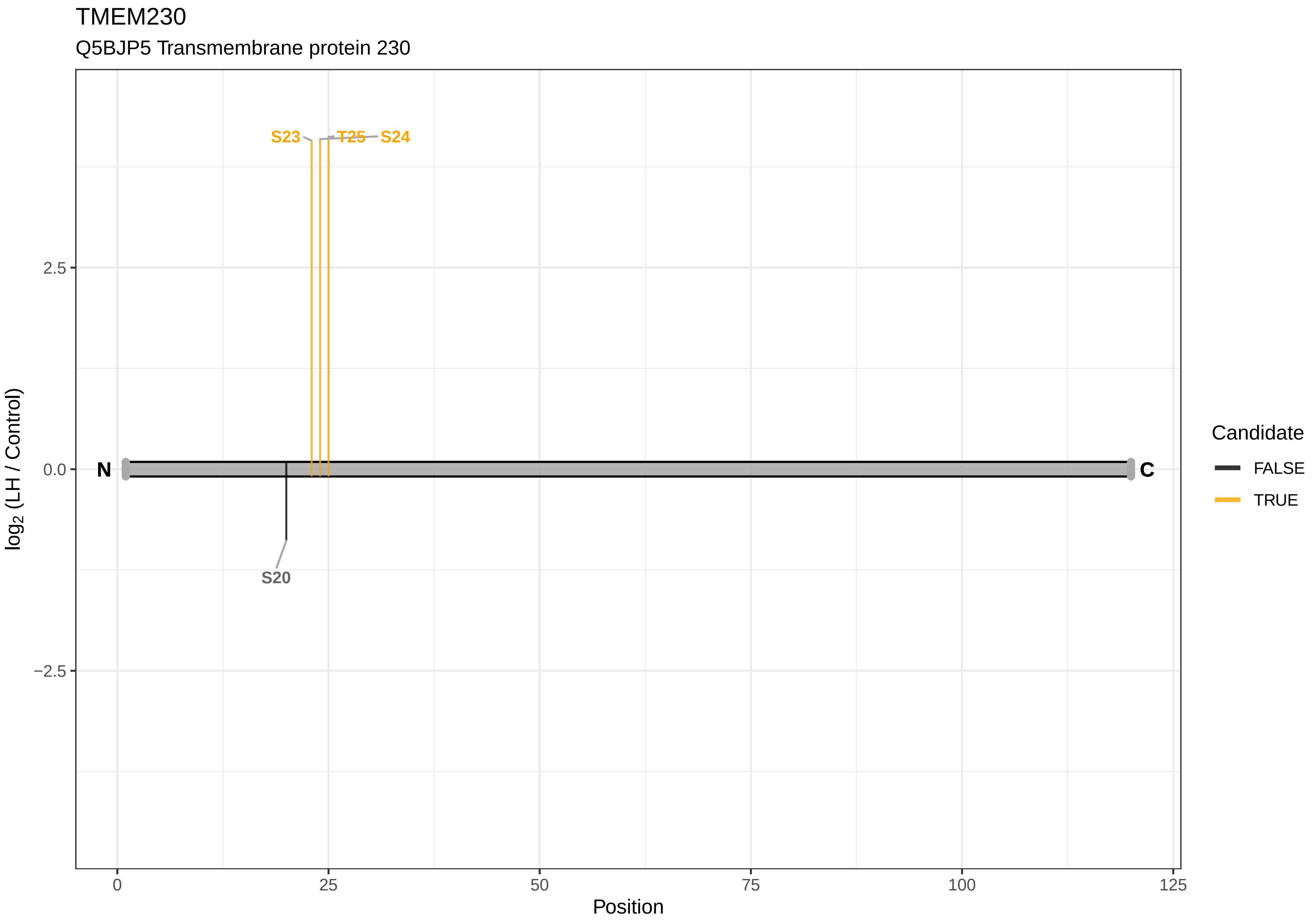

TMEM230
Q5BJP5 Transmembrane protein 230
S23
T25
S24
S20
N
C
2.5
0.0
−2.5
log2 (LH / Control)
0
25
50
75
100
125
Position
Candidate
FALSE
TRUE

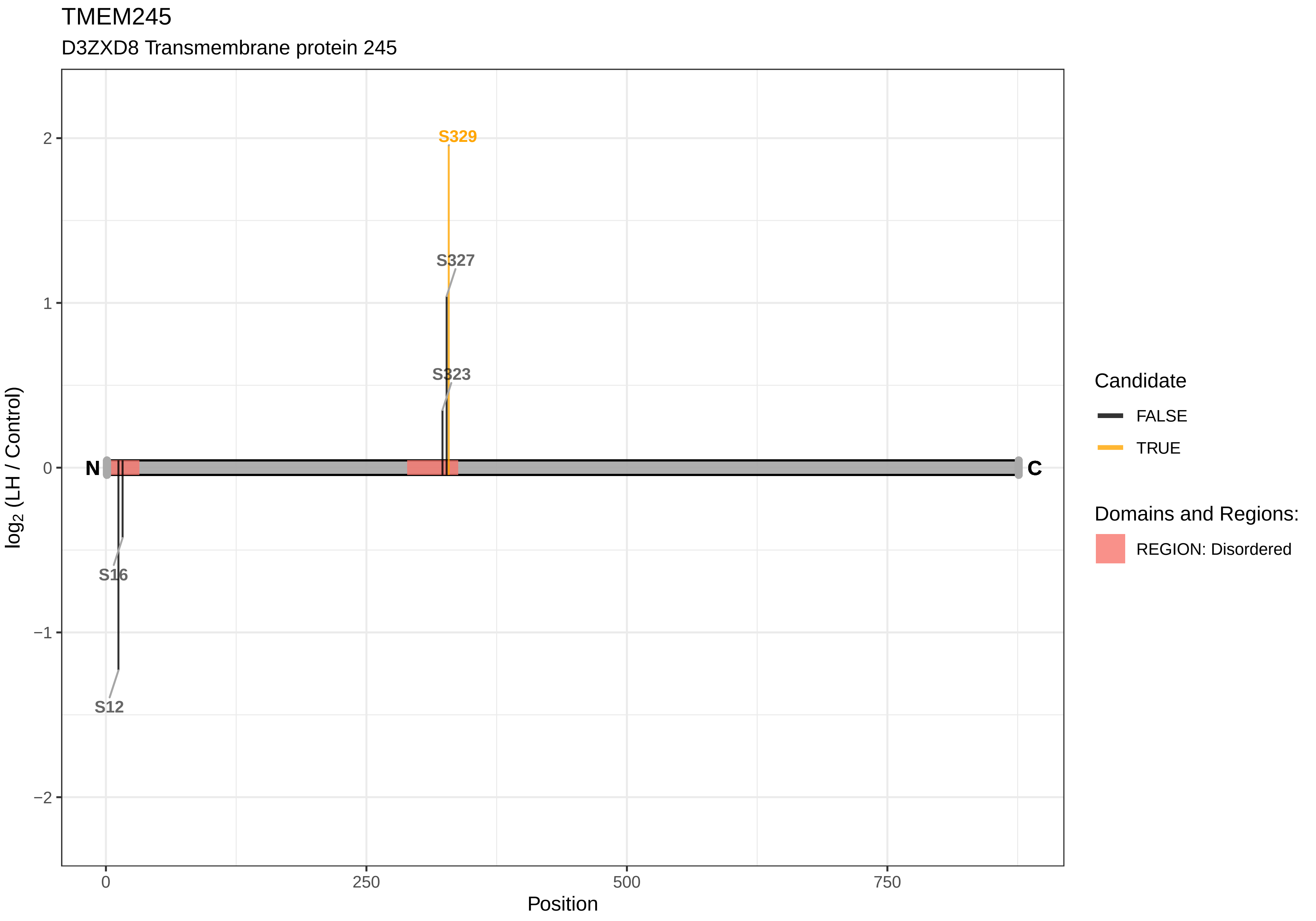
TMEM245
D3ZXD8 Transmembrane protein 245
S329
S327
S323
S16
S12
N
C
2
1
0
−1
−2
0
250
500
750
Position
log2 (LH / Control)
Candidate
FALSE
TRUE
Domains and Regions:
REGION: Disordered

TMEM63B

D4A105 Transmembrane protein 63B

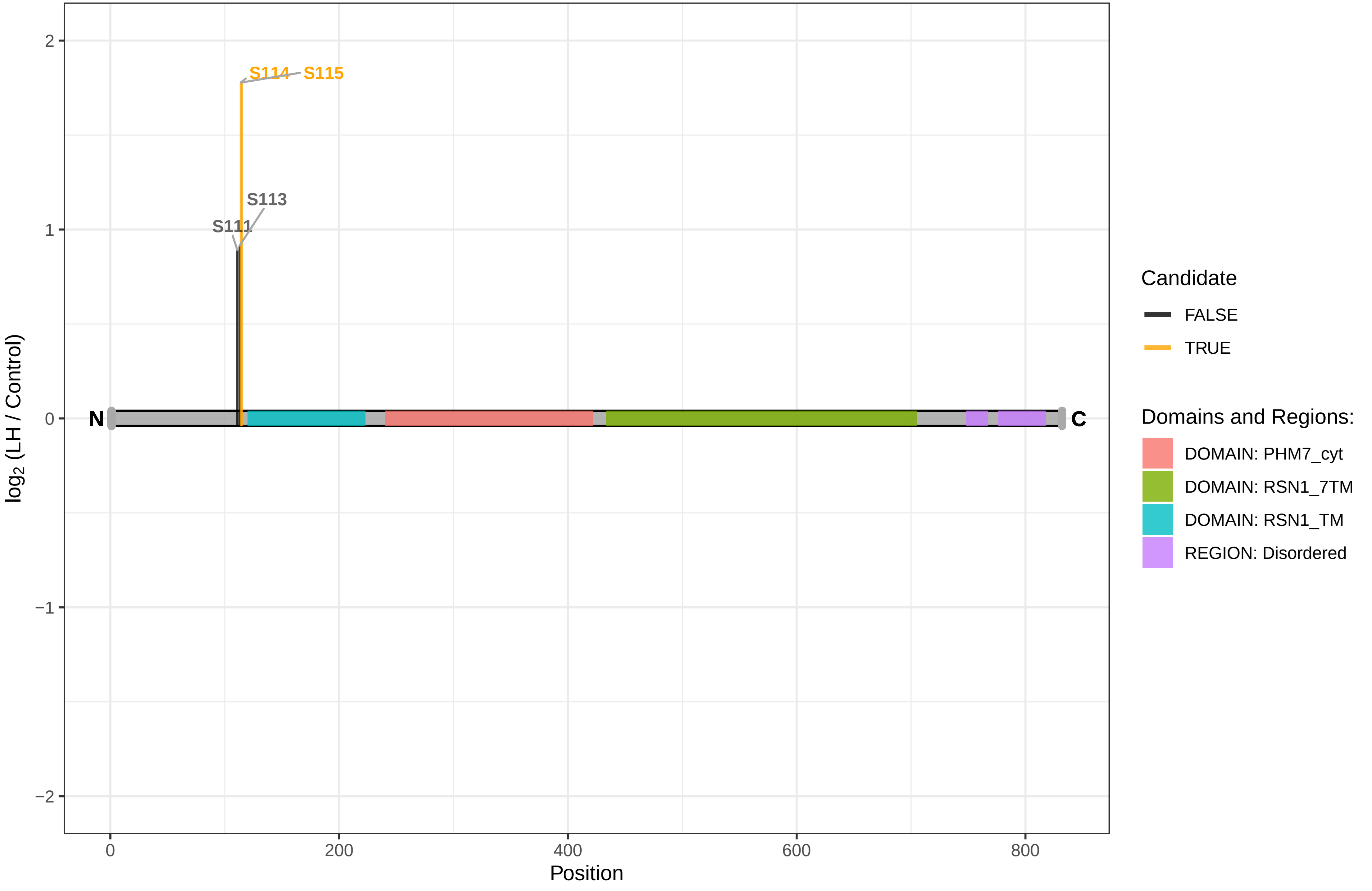

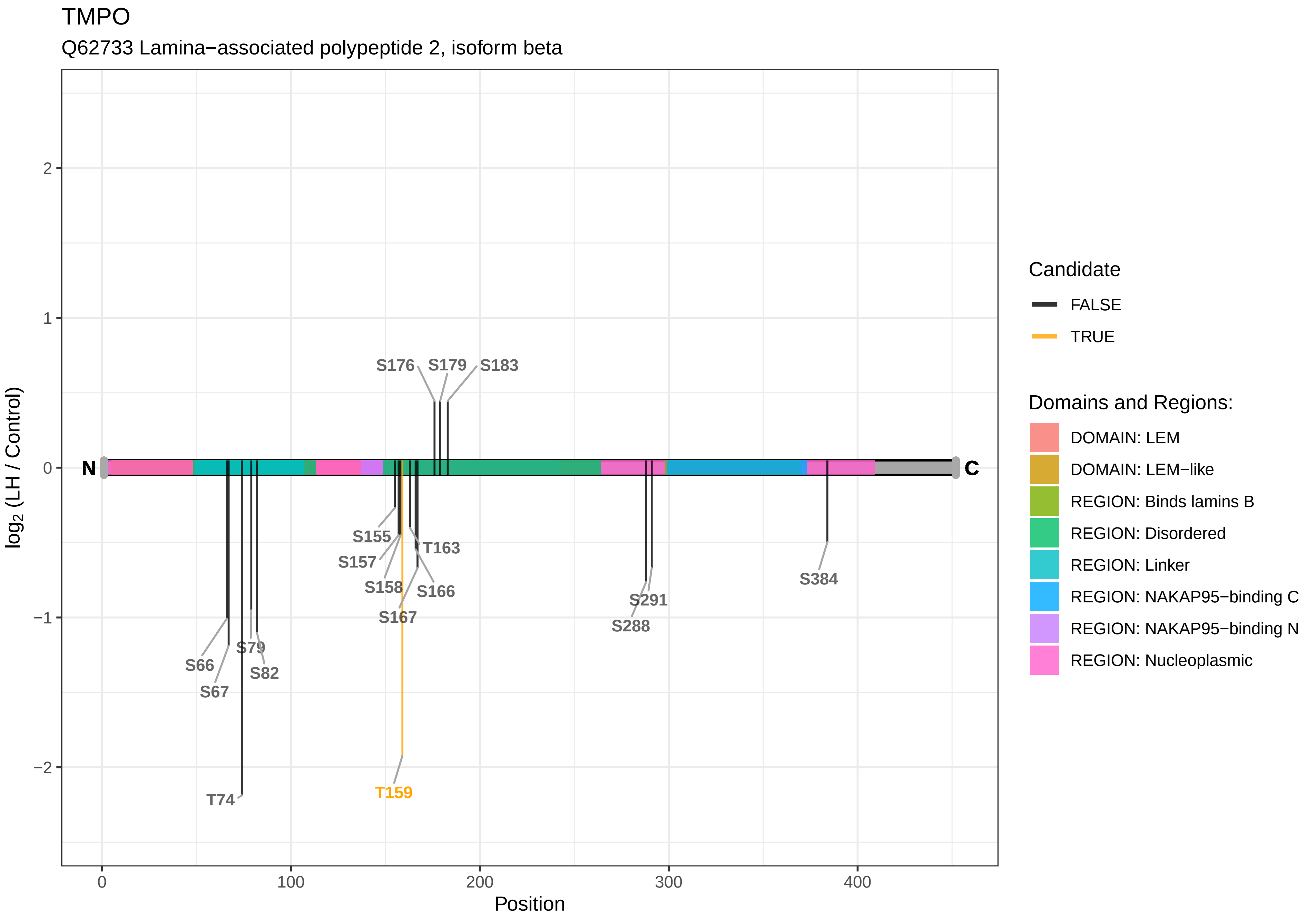
TMPO
Q62733 Lamina−associated polypeptide 2, isoform beta
log2 (LH / Control)
Position
N
C
S176
S179
S183
S155
T163
S157
S158
S166
S167
S291
S288
S384
S79
S66
S82
S67
T74
T159
Candidate
FALSE
TRUE
Domains and Regions:
DOMAIN: LEM
DOMAIN: LEM−like
REGION: Binds lamins B
REGION: Disordered
REGION: Linker
REGION: NAKAP95−binding C
REGION: NAKAP95−binding N
REGION: Nucleoplasmic

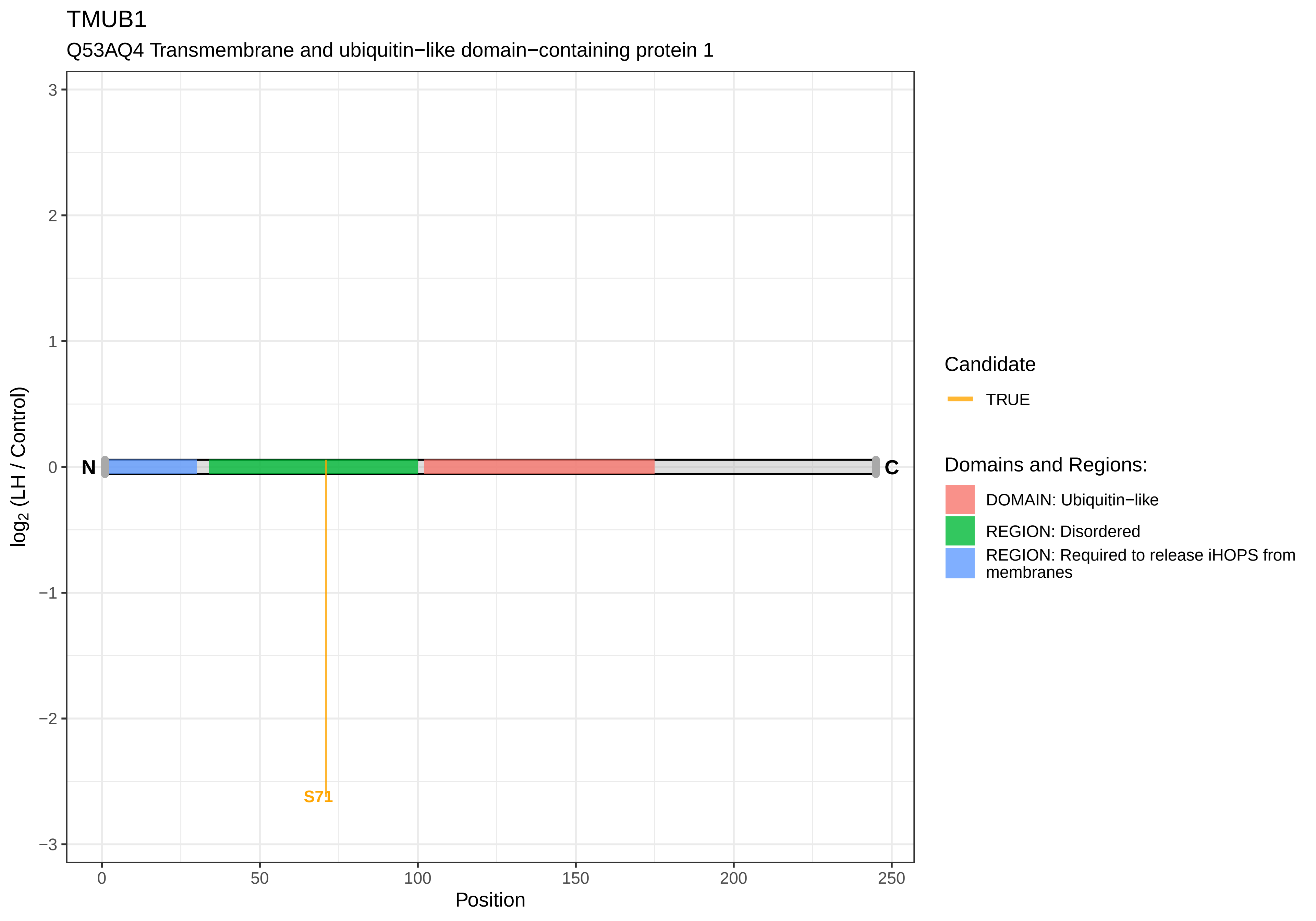
TMUB1
Q53AQ4 Transmembrane and ubiquitin−like domain−containing protein 1
3
2
1
0
−1
−2
−3
log2 (LH / Control)
N
C
S71
0
50
100
150
200
250
Position
Candidate
TRUE
Domains and Regions:
DOMAIN: Ubiquitin−like
REGION: Disordered
REGION: Required to release iHOPS from membranes

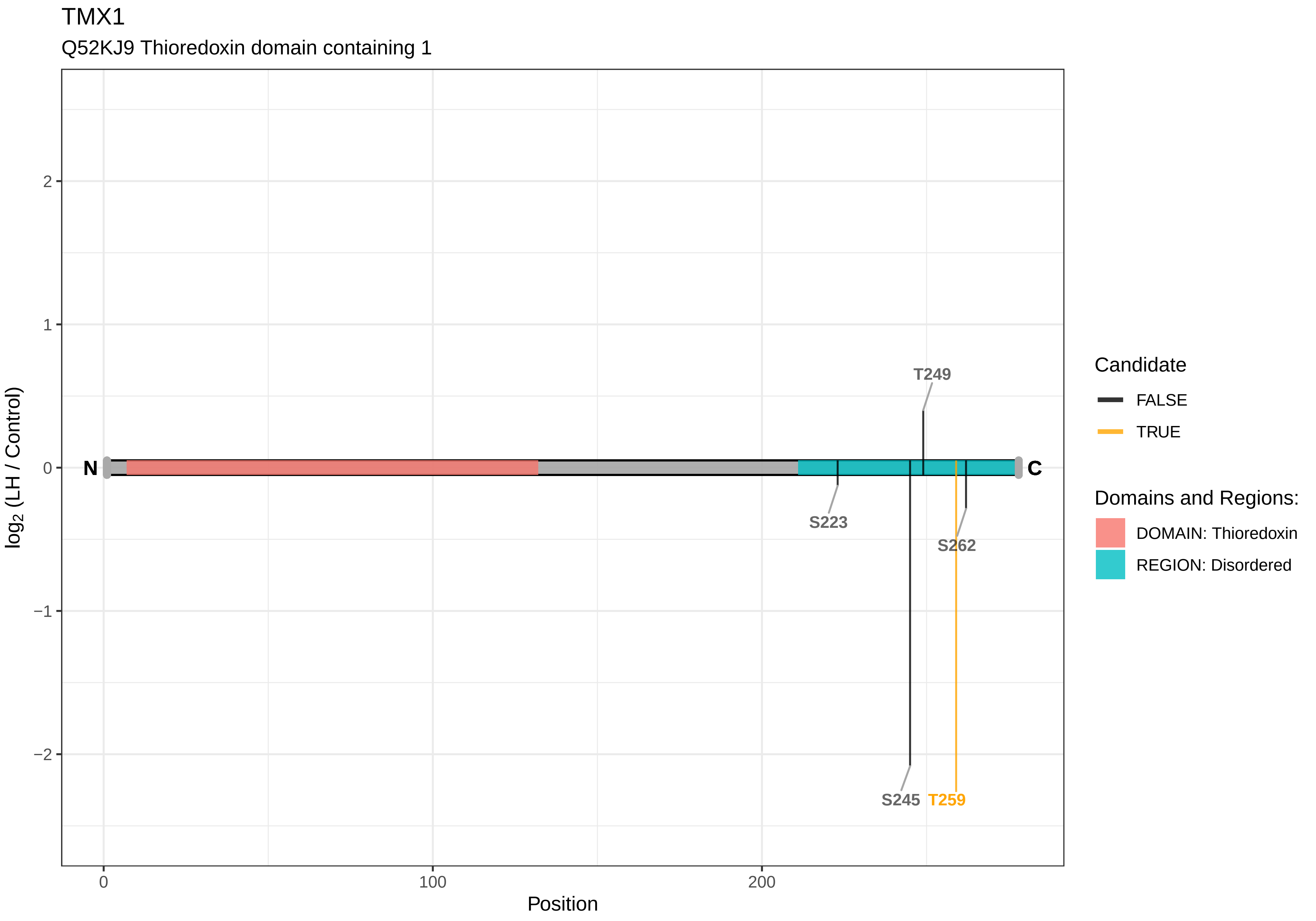

TMX1
Q52KJ9 Thioredoxin domain containing 1
log2 (LH / Control)
2
1
0
−1
−2
N
C
T249
S223
S262
S245
T259
0
100
200
Position
Candidate
FALSE
TRUE
Domains and Regions:
DOMAIN: Thioredoxin
REGION: Disordered

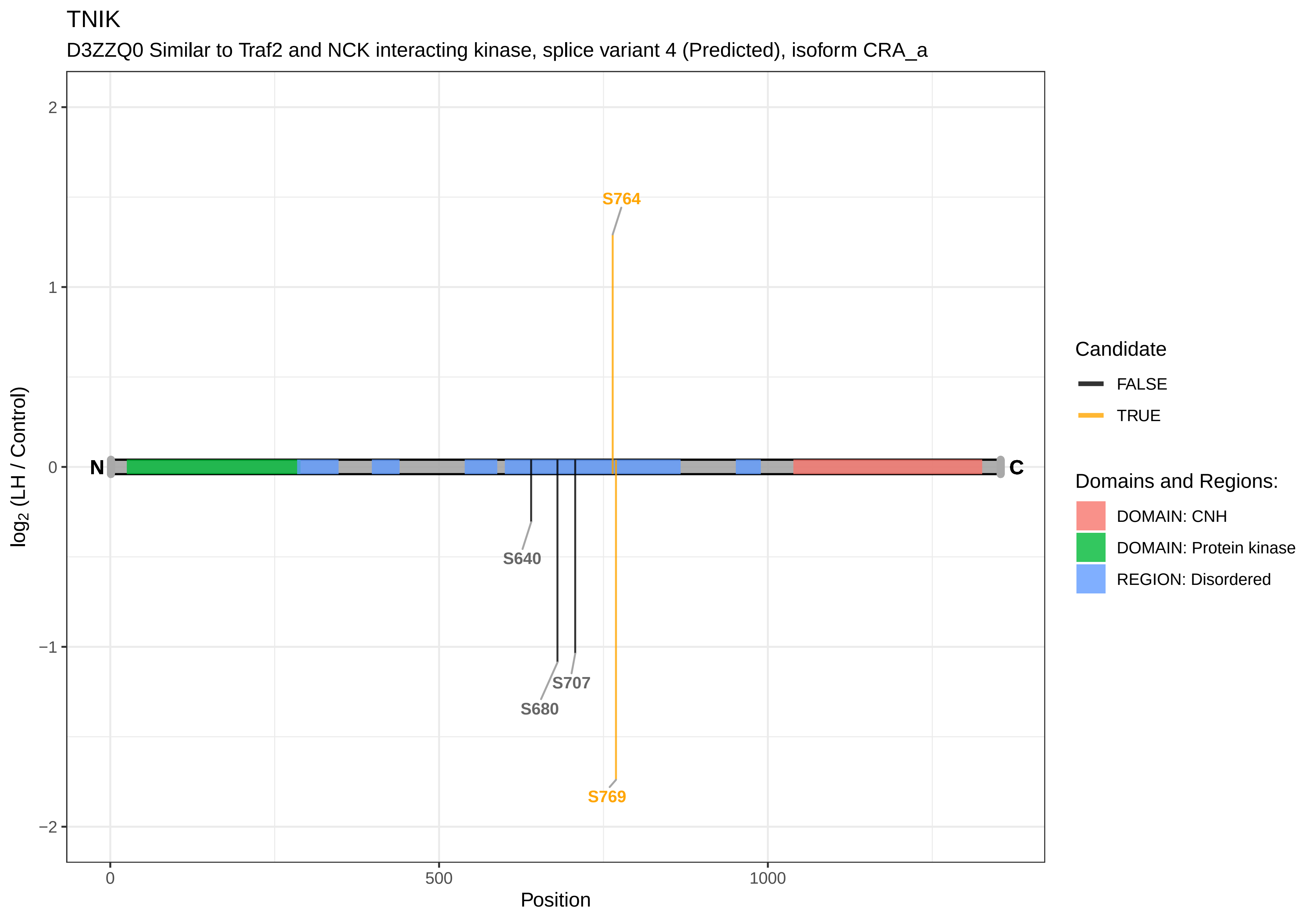
TNIK
D3ZZQ0 Similar to Traf2 and NCK interacting kinase, splice variant 4 (Predicted), isoform CRA_a
S764
S640
S707
S680
S769
N
C
2
1
0
−1
−2
log2 (LH / Control)
0
500
1000
Position
Candidate
FALSE
TRUE
Domains and Regions:
DOMAIN: CNH
DOMAIN: Protein kinase
REGION: Disordered

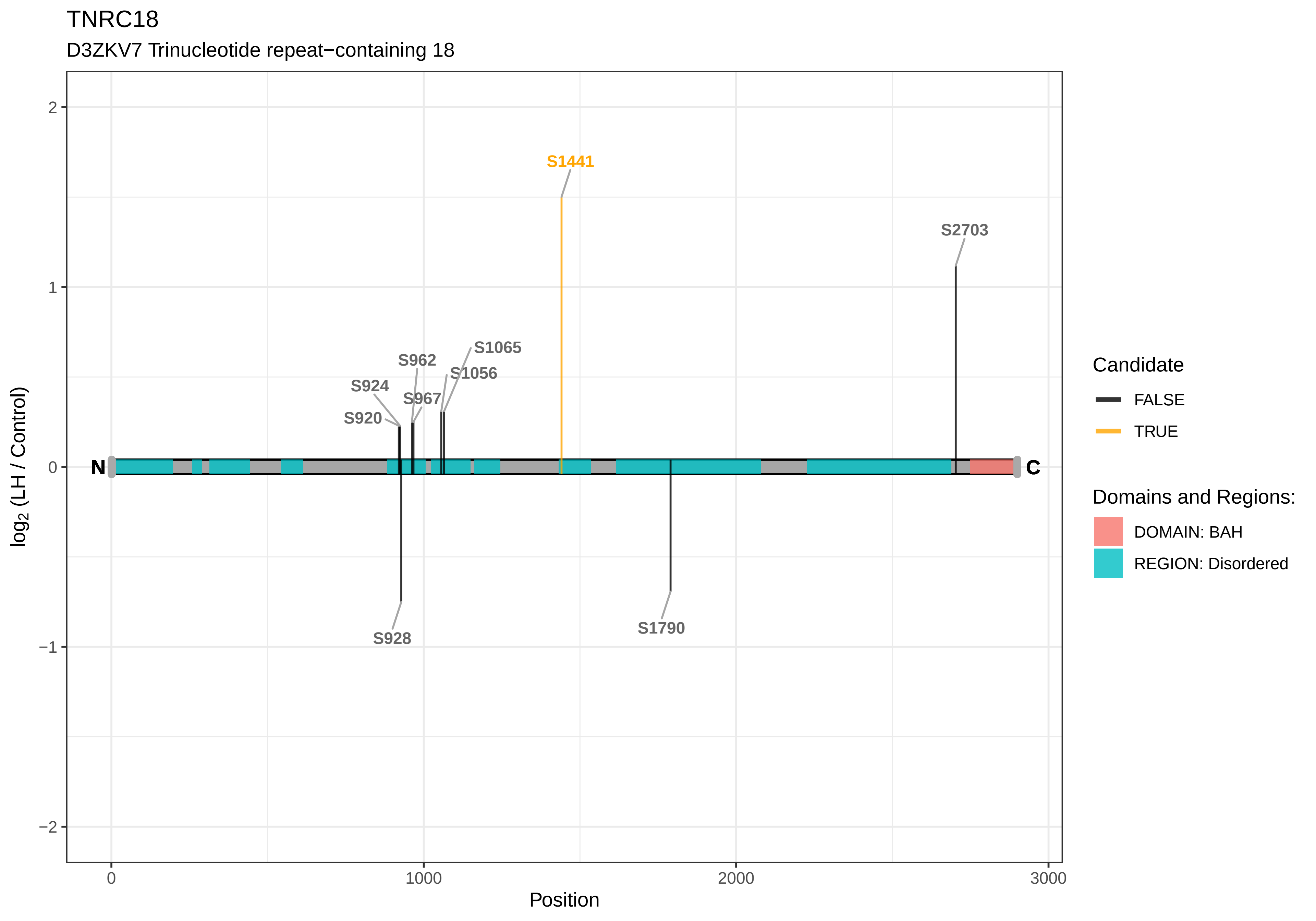

TNRC18
D3ZKV7 Trinucleotide repeat−containing 18
S1441
S2703
S1065
S962
S1056
S924
S967
S920
N
C
S928
S1790
2
1
0
−1
−2
0
1000
2000
3000
Position
log2 (LH / Control)
Candidate
FALSE
TRUE
Domains and Regions:
DOMAIN: BAH
REGION: Disordered

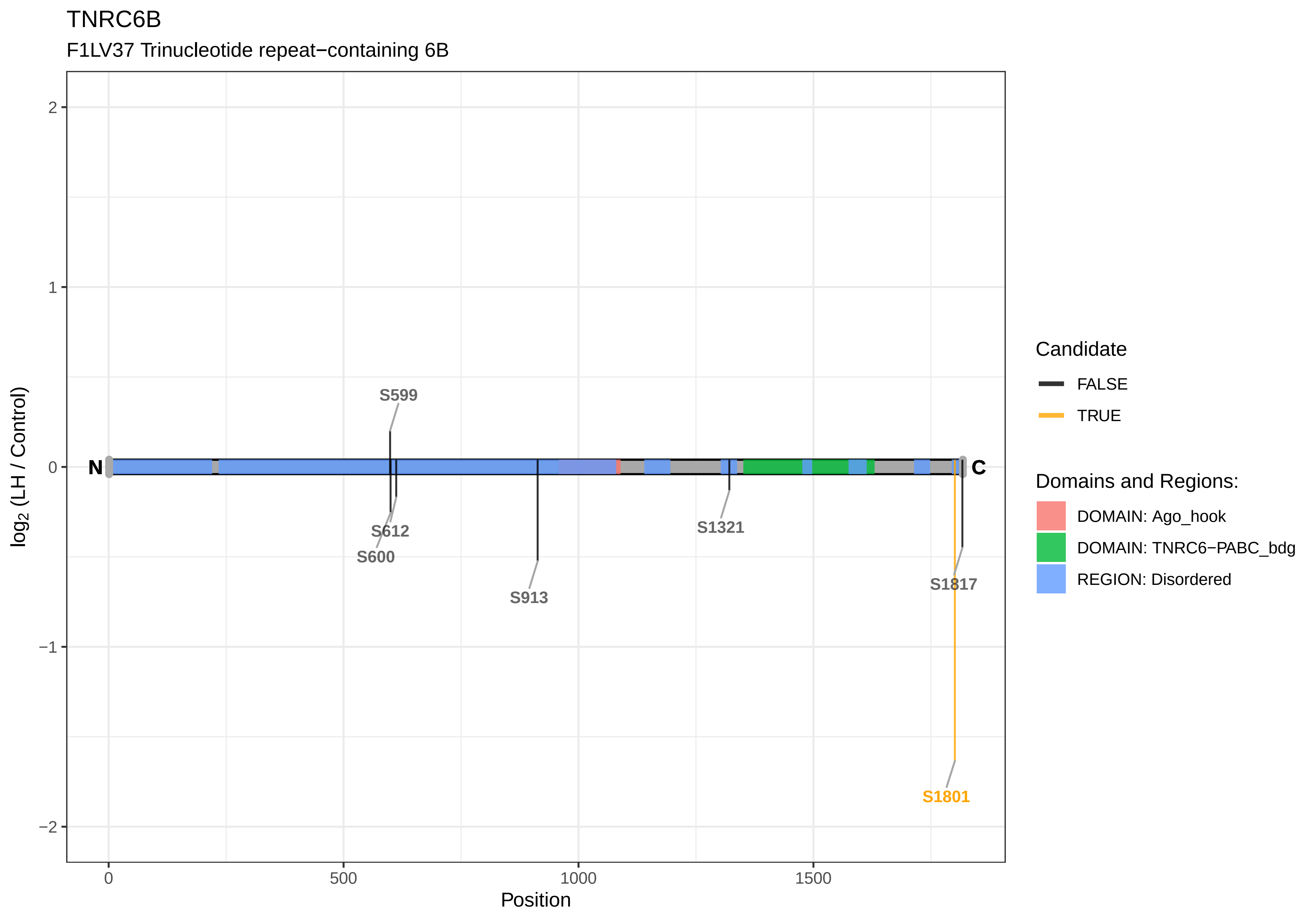

TNRC6B
F1LV37 Trinucleotide repeat−containing 6B
log2 (LH / Control)
Position
N
C
S599
S612
S600
S913
S1321
S1817
S1801
2
1
0
−1
−2
0
500
1000
1500
Candidate
FALSE
TRUE
Domains and Regions:
DOMAIN: Ago_hook
DOMAIN: TNRC6−PABC_bdg
REGION: Disordered

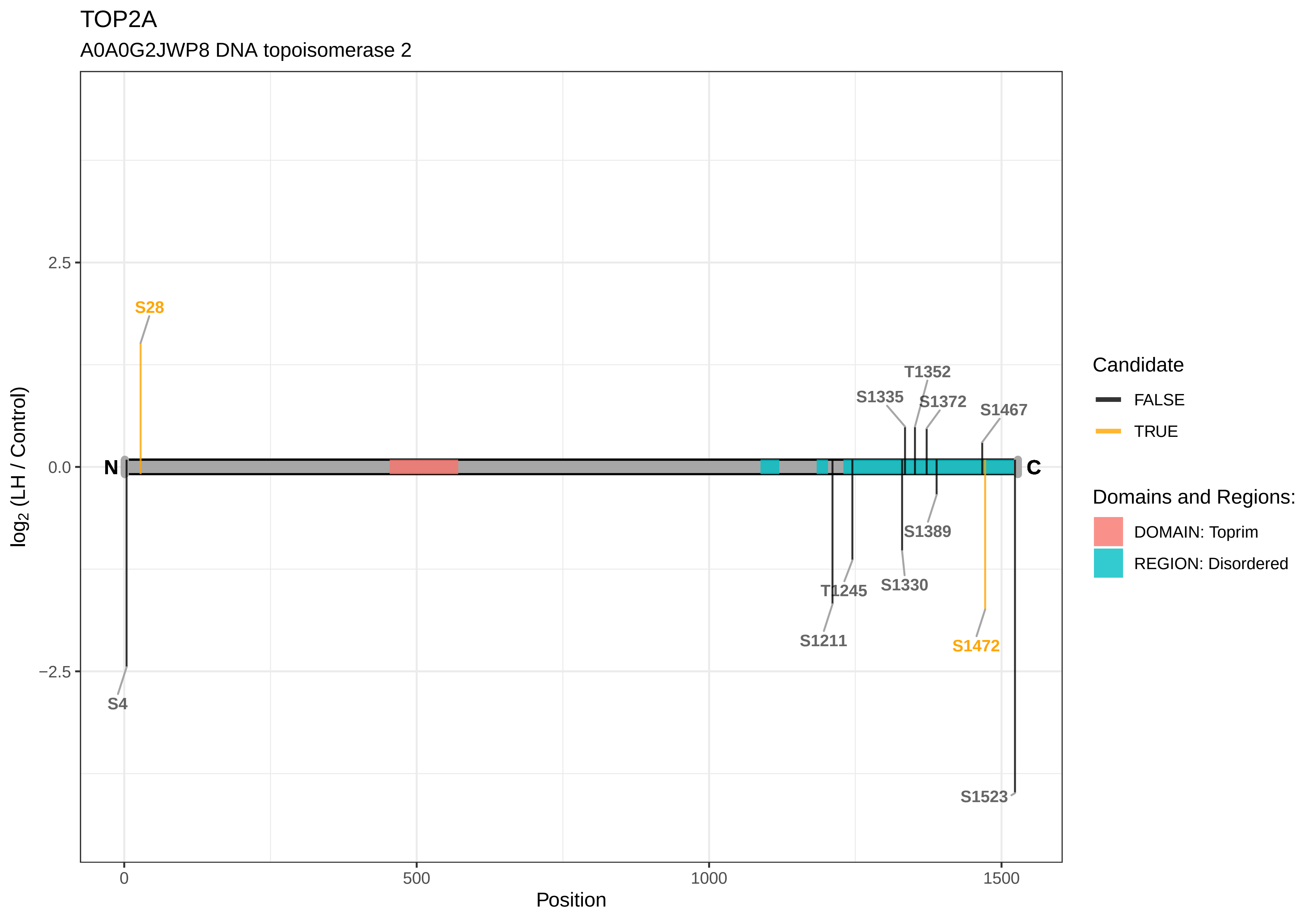

TOP2A
A0A0G2JWP8 DNA topoisomerase 2
log2 (LH / Control)
Position
2.5
0.0
−2.5
0
500
1000
1500
N
C
S28
S4
T1352
S1335
S1372
S1467
S1389
T1245
S1330
S1211
S1472
S1523
Candidate
FALSE
TRUE
Domains and Regions:
DOMAIN: Toprim
REGION: Disordered

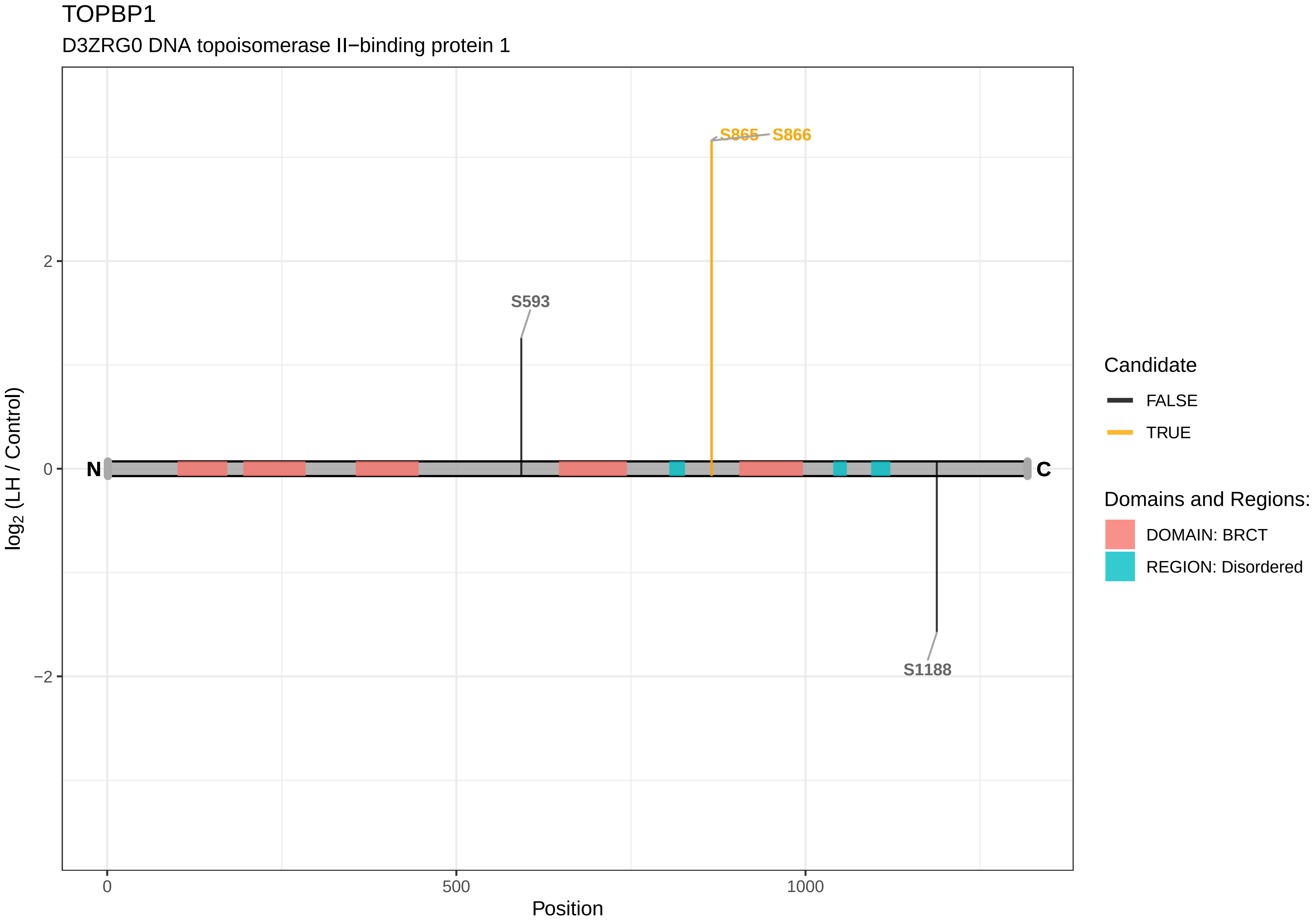

TOPBP1
D3ZRG0 DNA topoisomerase II−binding protein 1
S865
S866
S593
S1188
N
C
2
0
−2
log2 (LH / Control)
0
500
1000
Position
Candidate
FALSE
TRUE
Domains and Regions:
DOMAIN: BRCT
REGION: Disordered

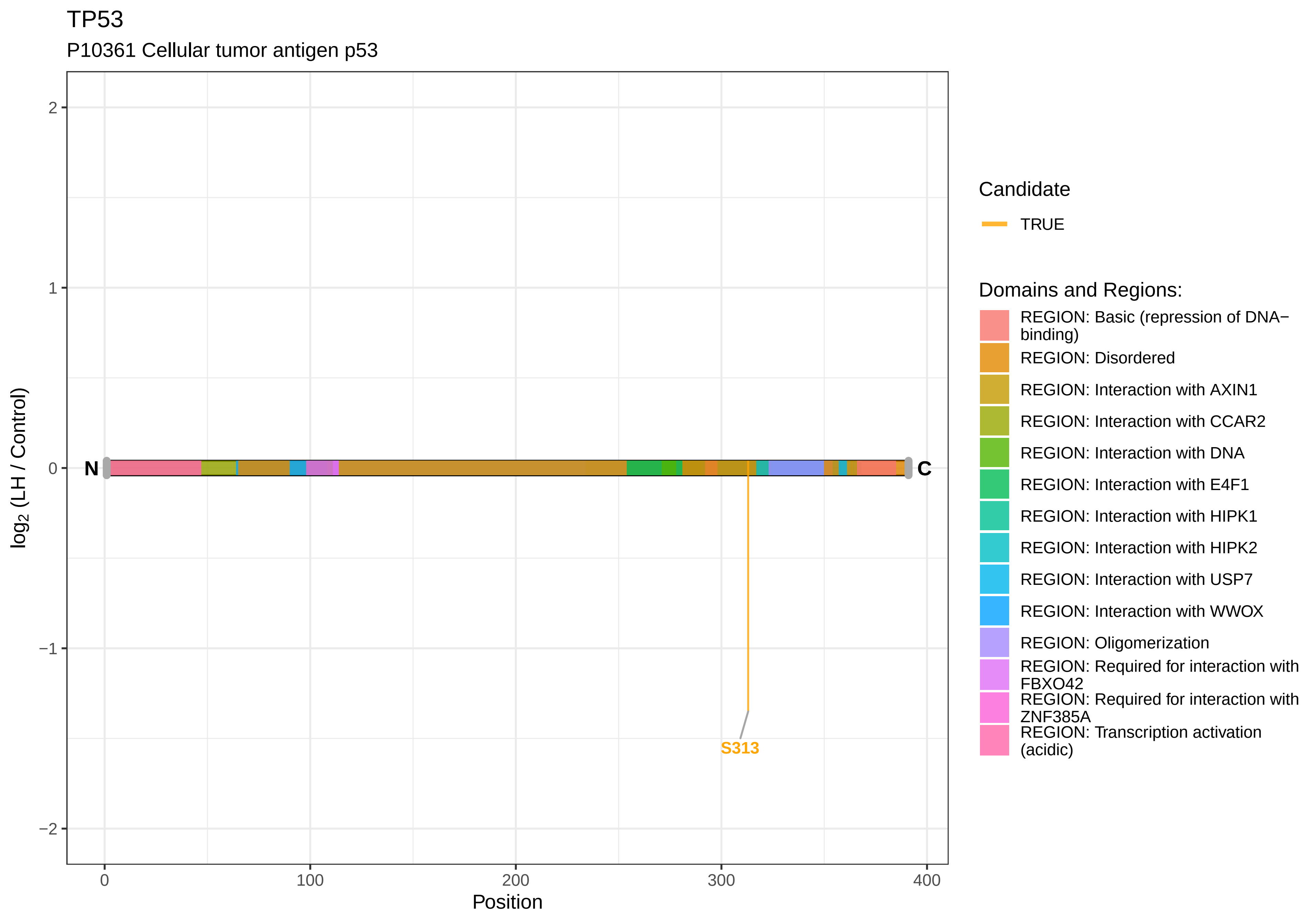

TP53
P10361 Cellular tumor antigen p53
log2 (LH / Control)
2
1
0
−1
−2
N
C
S313
0
100
200
300
400
Position
Candidate
TRUE
Domains and Regions:
REGION: Basic (repression of DNA−binding)
REGION: Disordered
REGION: Interaction with AXIN1
REGION: Interaction with CCAR2
REGION: Interaction with DNA
REGION: Interaction with E4F1
REGION: Interaction with HIPK1
REGION: Interaction with HIPK2
REGION: Interaction with USP7
REGION: Interaction with WWOX
REGION: Oligomerization
REGION: Required for interaction with FBXO42
REGION: Required for interaction with ZNF385A
REGION: Transcription activation (acidic)

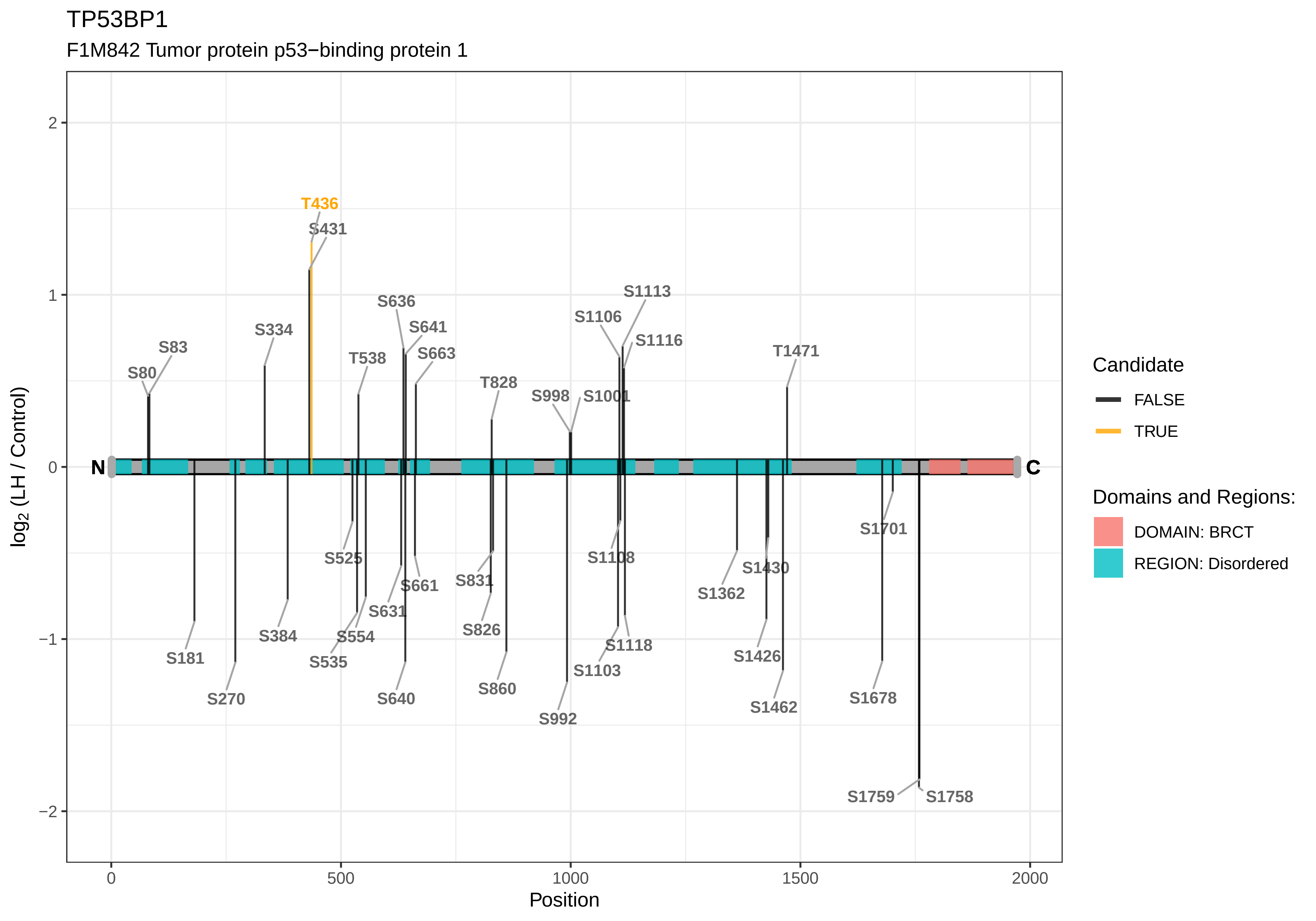

TP53BP1
F1M842 Tumor protein p53−binding protein 1
log2 (LH / Control)
Position
N
C
T436
S431
S80
S83
S334
T538
S636
S641
S663
T828
S998
S1001
S1106
S1113
S1116
T1471
S181
S270
S384
S525
S535
S554
S631
S640
S661
S831
S826
S860
S992
S1103
S1108
S1118
S1362
S1426
S1430
S1462
S1678
S1701
S1759
S1758
Candidate
FALSE
TRUE
Domains and Regions:
DOMAIN: BRCT
REGION: Disordered

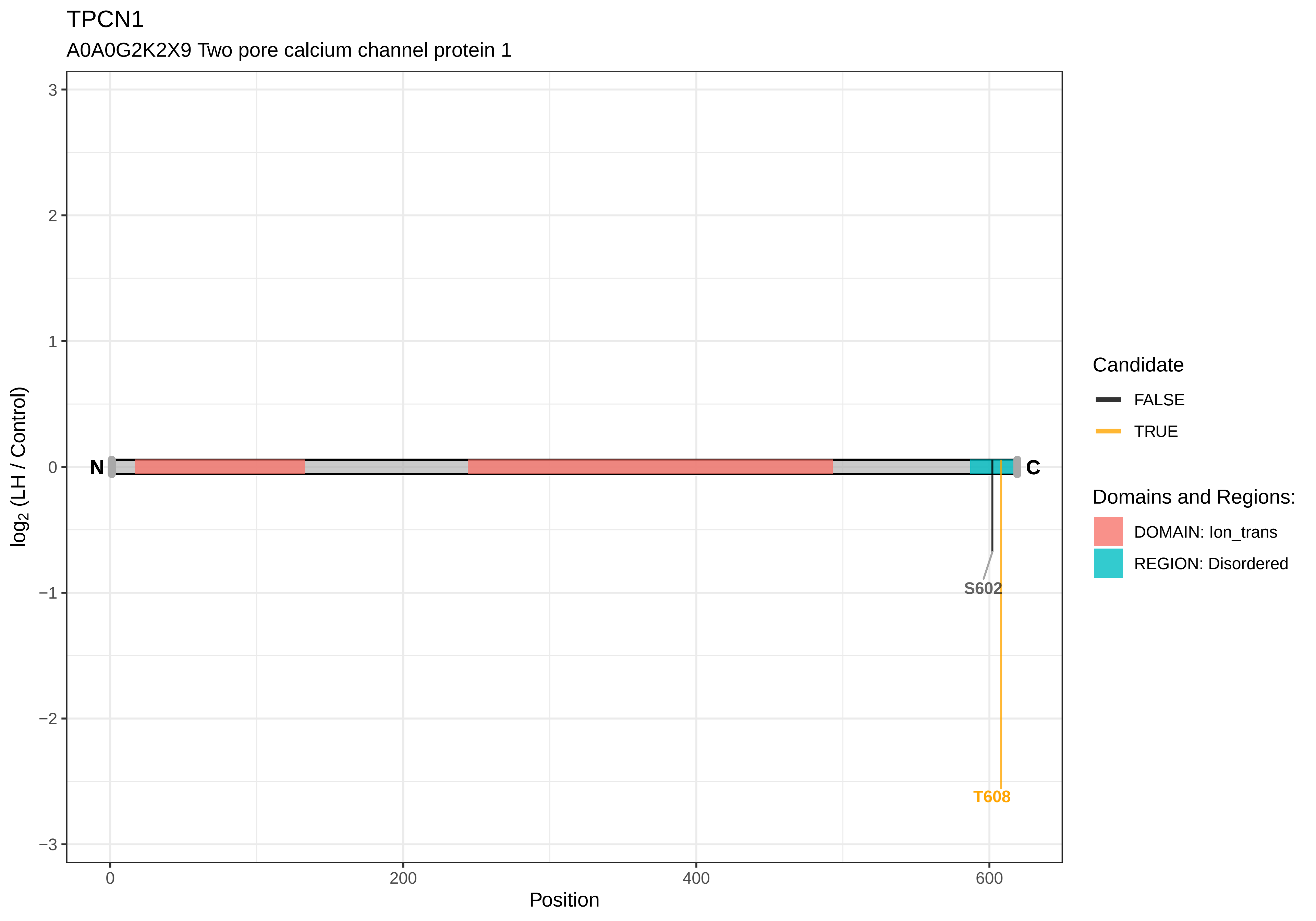
TPCN1
A0A0G2K2X9 Two pore calcium channel protein 1
log2 (LH / Control)
3
2
1
0
−1
−2
−3
N
C
S602
T608
0
200
400
600
Position
Candidate
FALSE
TRUE
Domains and Regions:
DOMAIN: Ion_trans
REGION: Disordered

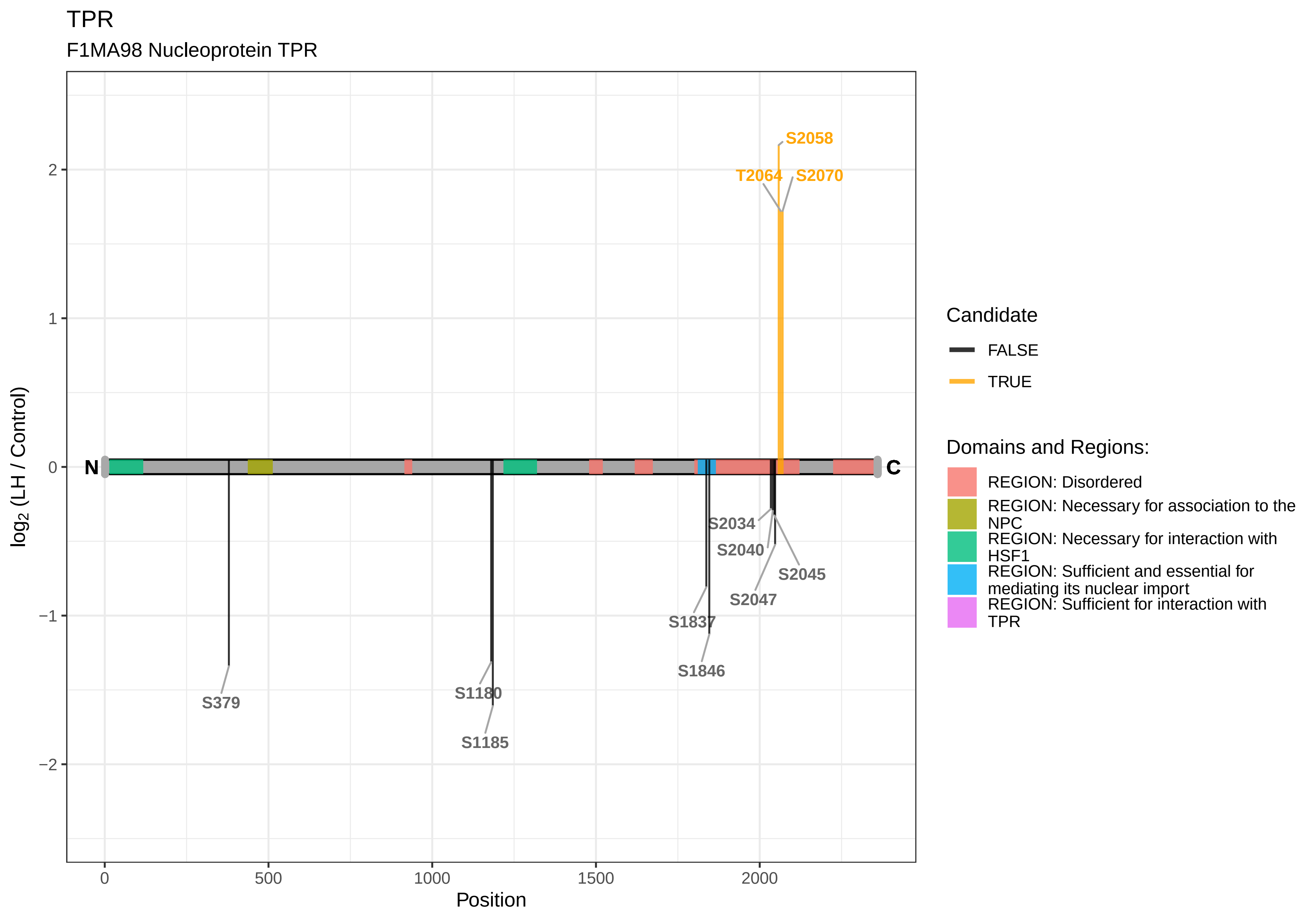

TPR
F1MA98 Nucleoprotein TPR
S2058
T2064
S2070
N
C
S2034
S2040
S2045
S2047
S1837
S1846
S379
S1180
S1185
log2 (LH / Control)
Position
0
500
1000
1500
2000
2
1
0
−1
−2
Candidate
FALSE
TRUE
Domains and Regions:
REGION: Disordered
REGION: Necessary for association to the NPC
REGION: Necessary for interaction with HSF1
REGION: Sufficient and essential for mediating its nuclear import
REGION: Sufficient for interaction with TPR

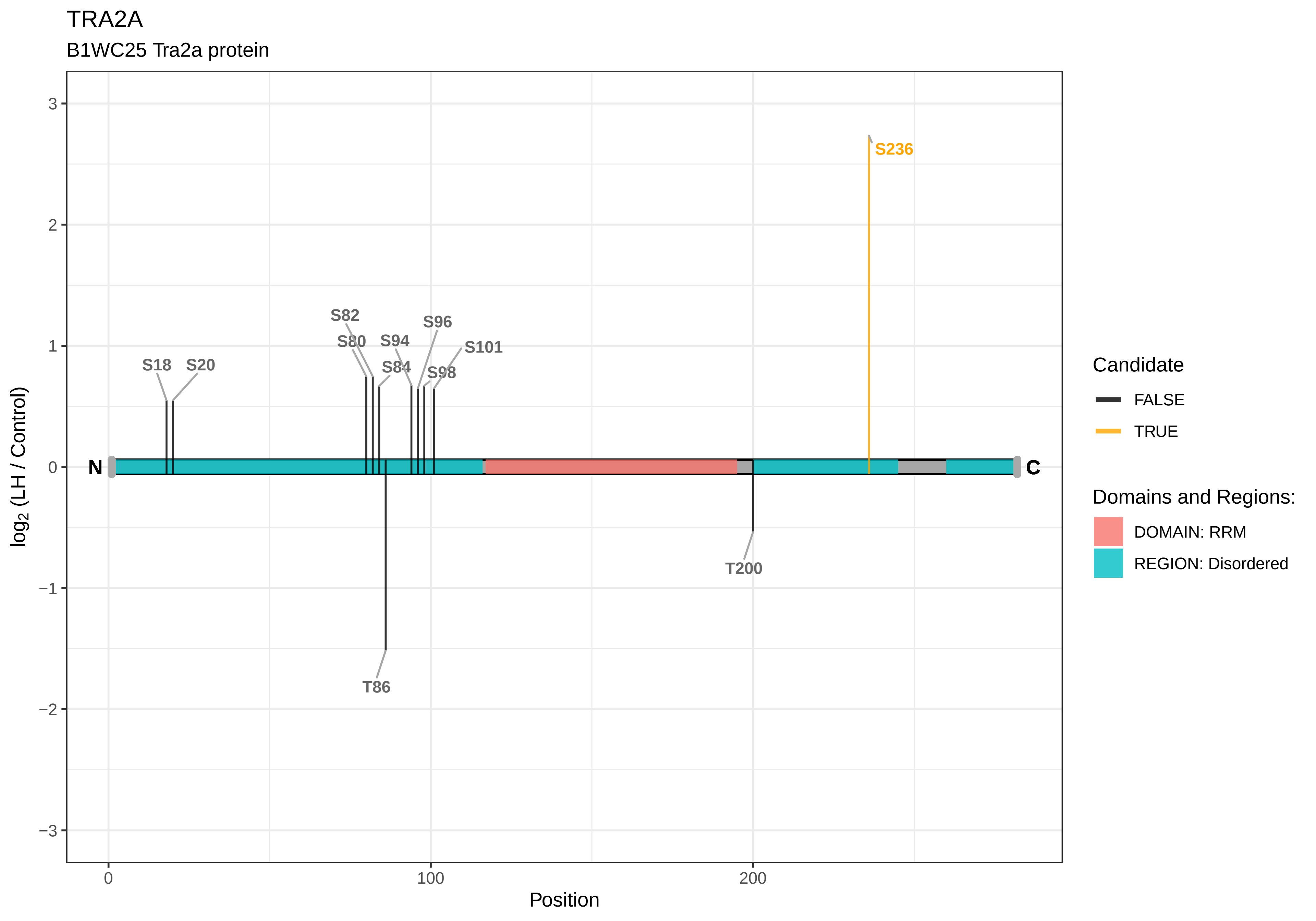
TRA2A
B1WC25 Tra2a protein
S236
S82
S96
S80
S94
S101
S18
S20
S84
S98
N
C
T200
T86
3
2
1
0
−1
−2
−3
log2 (LH / Control)
0
100
200
Position
Candidate
FALSE
TRUE
Domains and Regions:
DOMAIN: RRM
REGION: Disordered

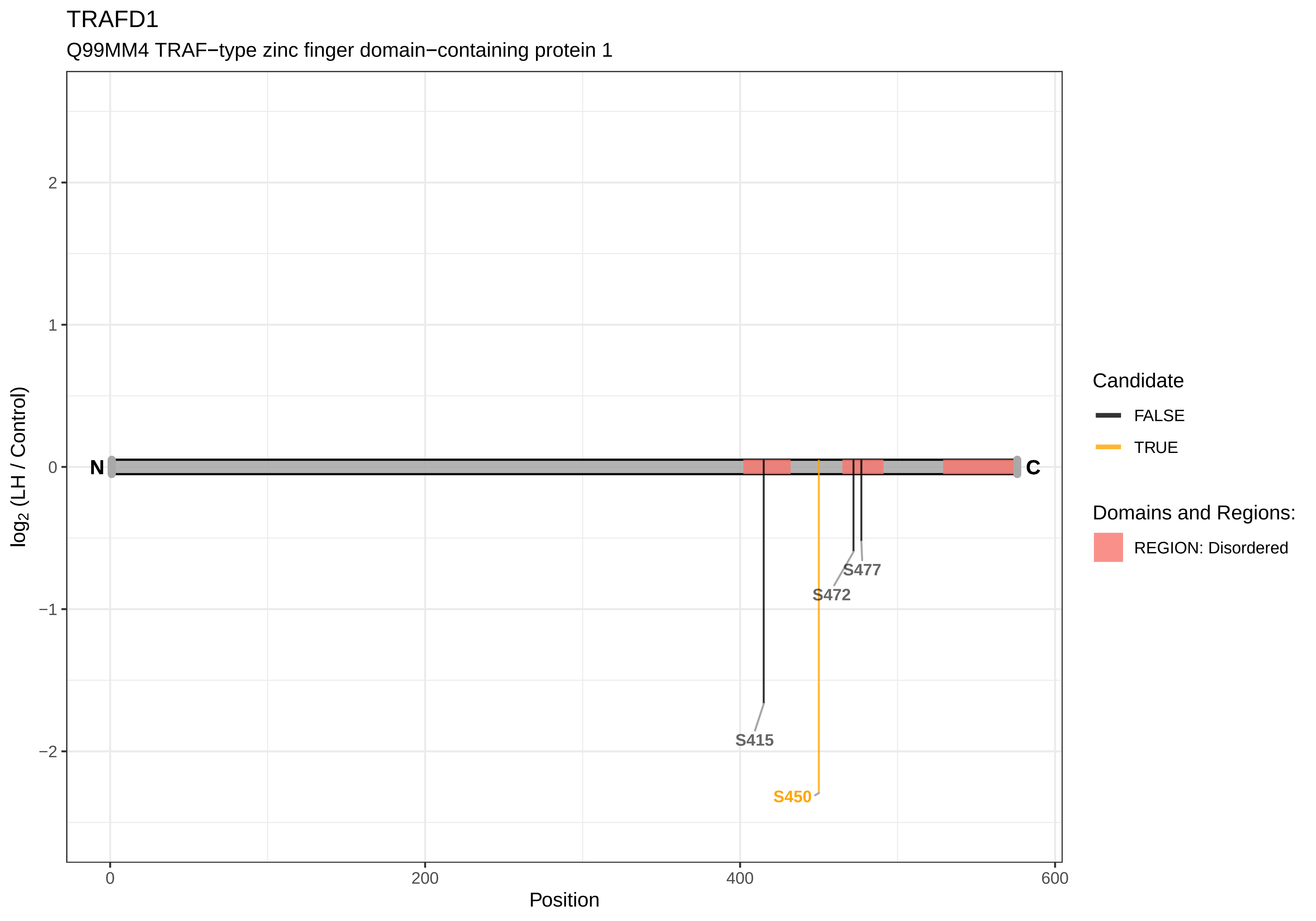
TRAFD1
Q99MM4 TRAF−type zinc finger domain−containing protein 1
log2 (LH / Control)
Position
N
C
S415
S450
S472
S477
2
1
0
−1
−2
0
200
400
600
Candidate
FALSE
TRUE
Domains and Regions:
REGION: Disordered

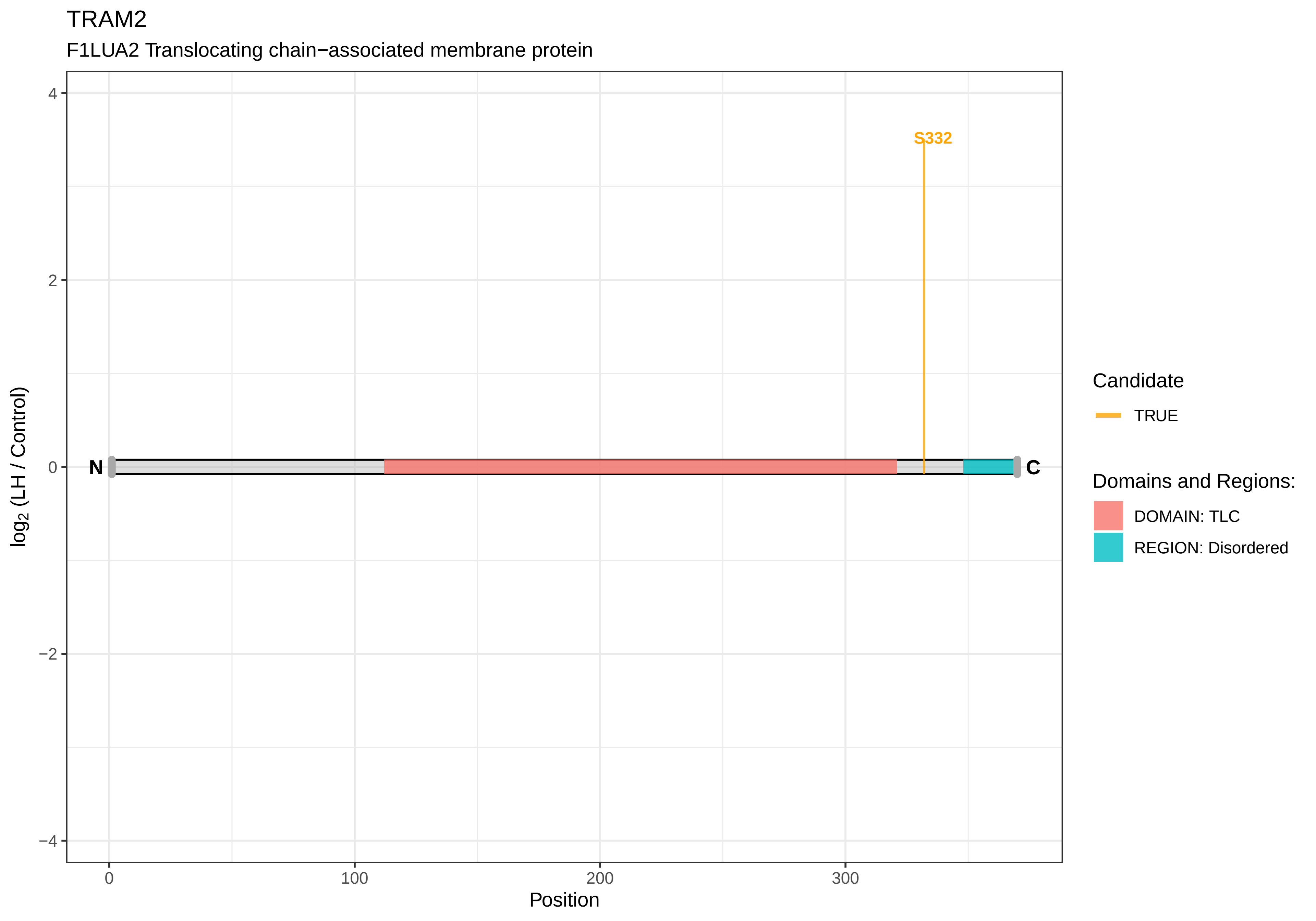
TRAM2
F1LUA2 Translocating chain−associated membrane protein
S332
N
C
4
2
0
−2
−4
log2 (LH / Control)
0
100
200
300
Position
Candidate
TRUE
Domains and Regions:
DOMAIN: TLC
REGION: Disordered

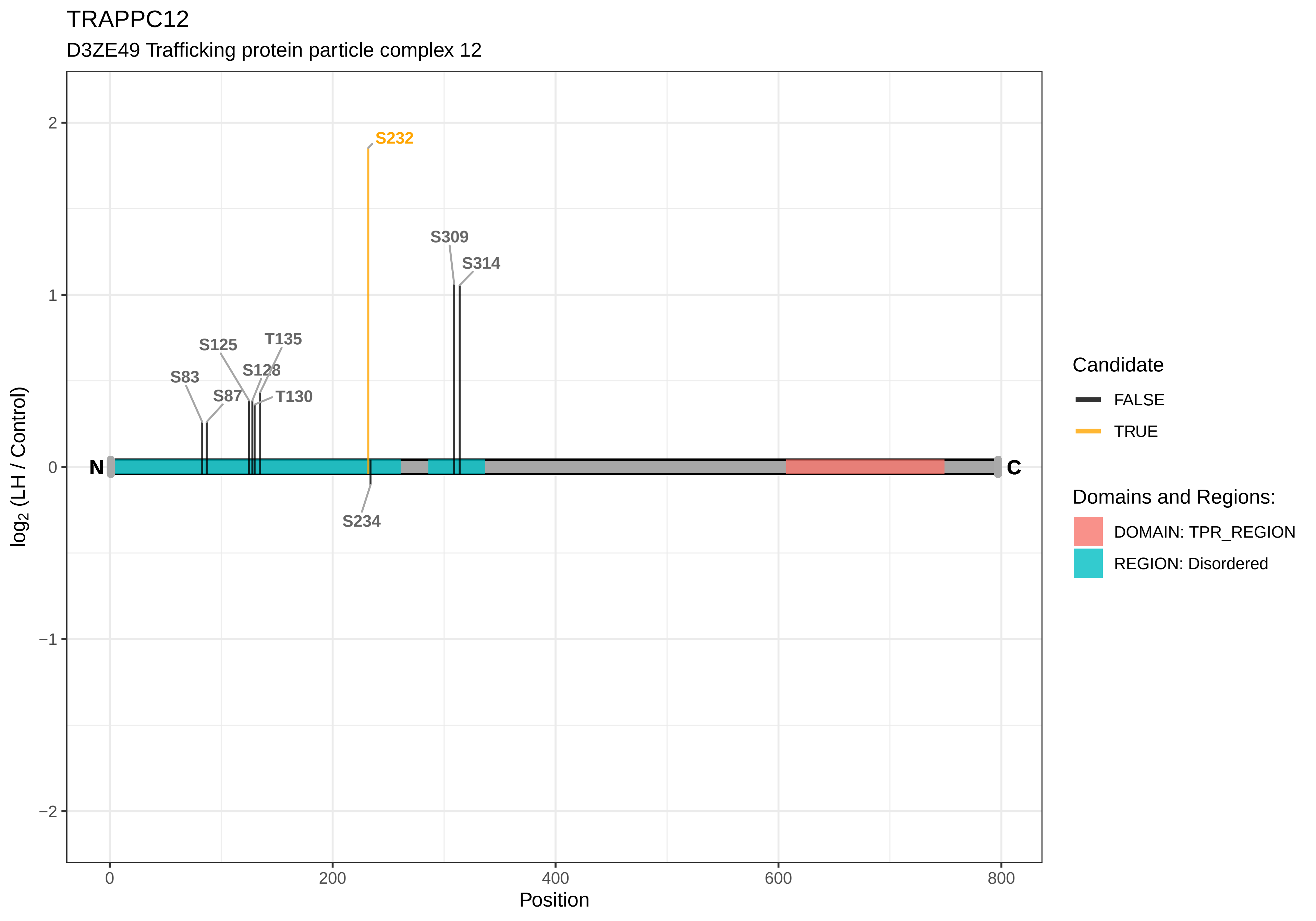

TRAPPC12
D3ZE49 Trafficking protein particle complex 12
S232
S309
S314
S125
T135
S128
S83
S87
T130
N
C
S234
log2 (LH / Control)
2
1
0
−1
−2
0
200
400
600
800
Position
Candidate
FALSE
TRUE
Domains and Regions:
DOMAIN: TPR_REGION
REGION: Disordered

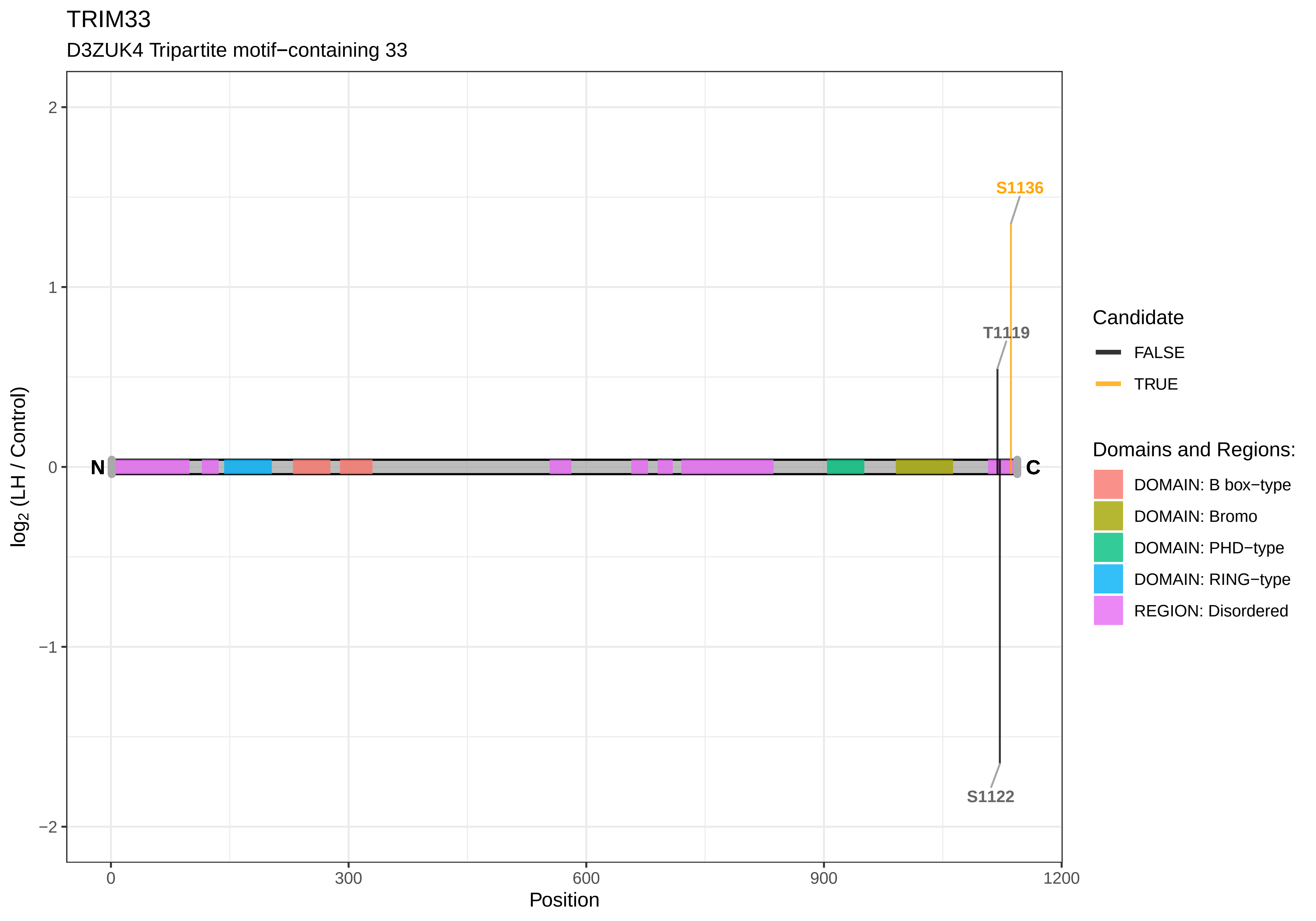
TRIM33
D3ZUK4 Tripartite motif−containing 33
S1136
T1119
S1122
N
C
2
1
0
−1
−2
log2 (LH / Control)
0
300
600
900
1200
Position
Candidate
FALSE
TRUE
Domains and Regions:
DOMAIN: B box−type
DOMAIN: Bromo
DOMAIN: PHD−type
DOMAIN: RING−type
REGION: Disordered

TRIOBP

A0A0G2K6B2 RCG60109

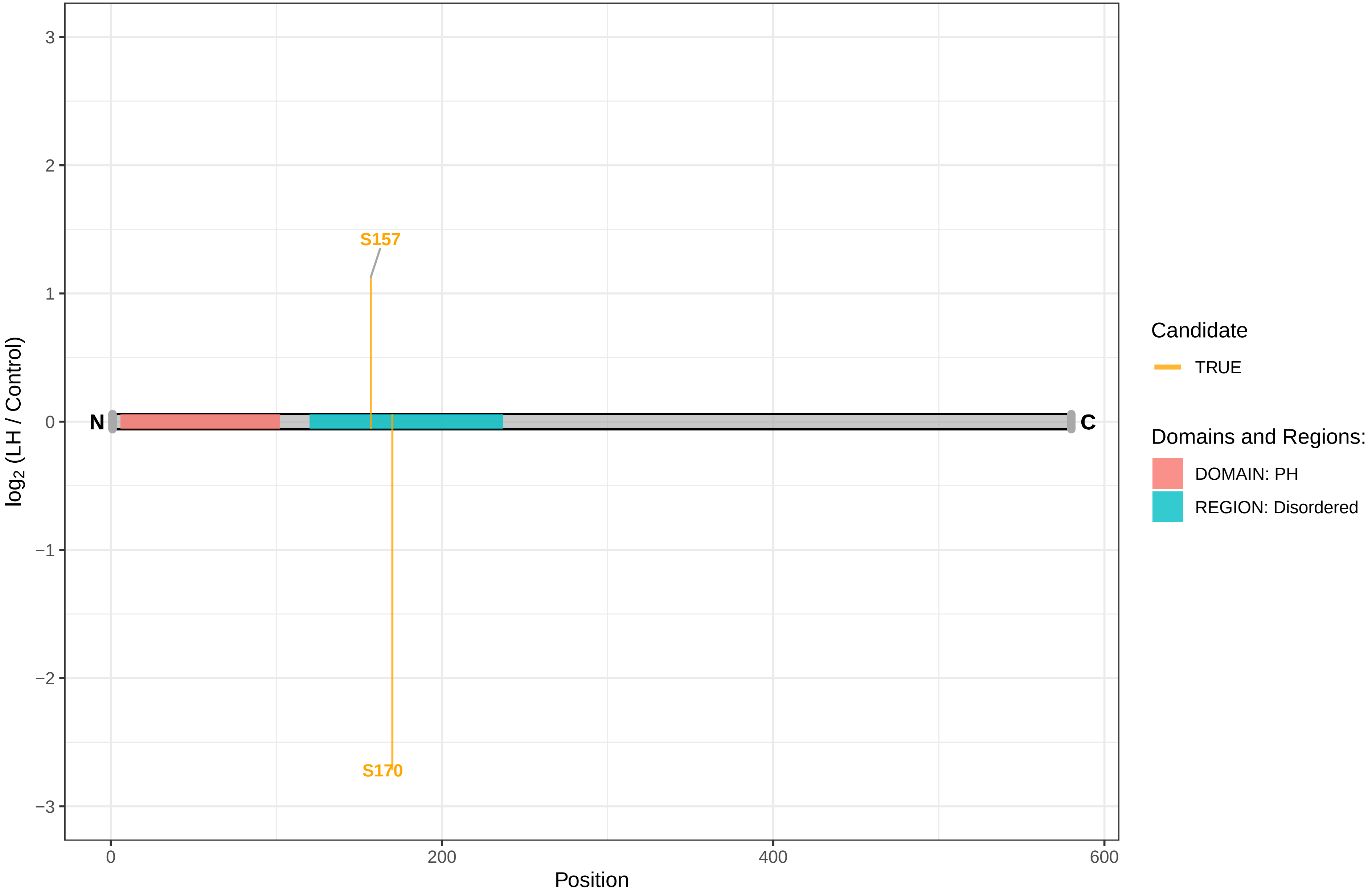

TRIP12

F1LP64 E3 ubiquitin−protein ligase TRIP12

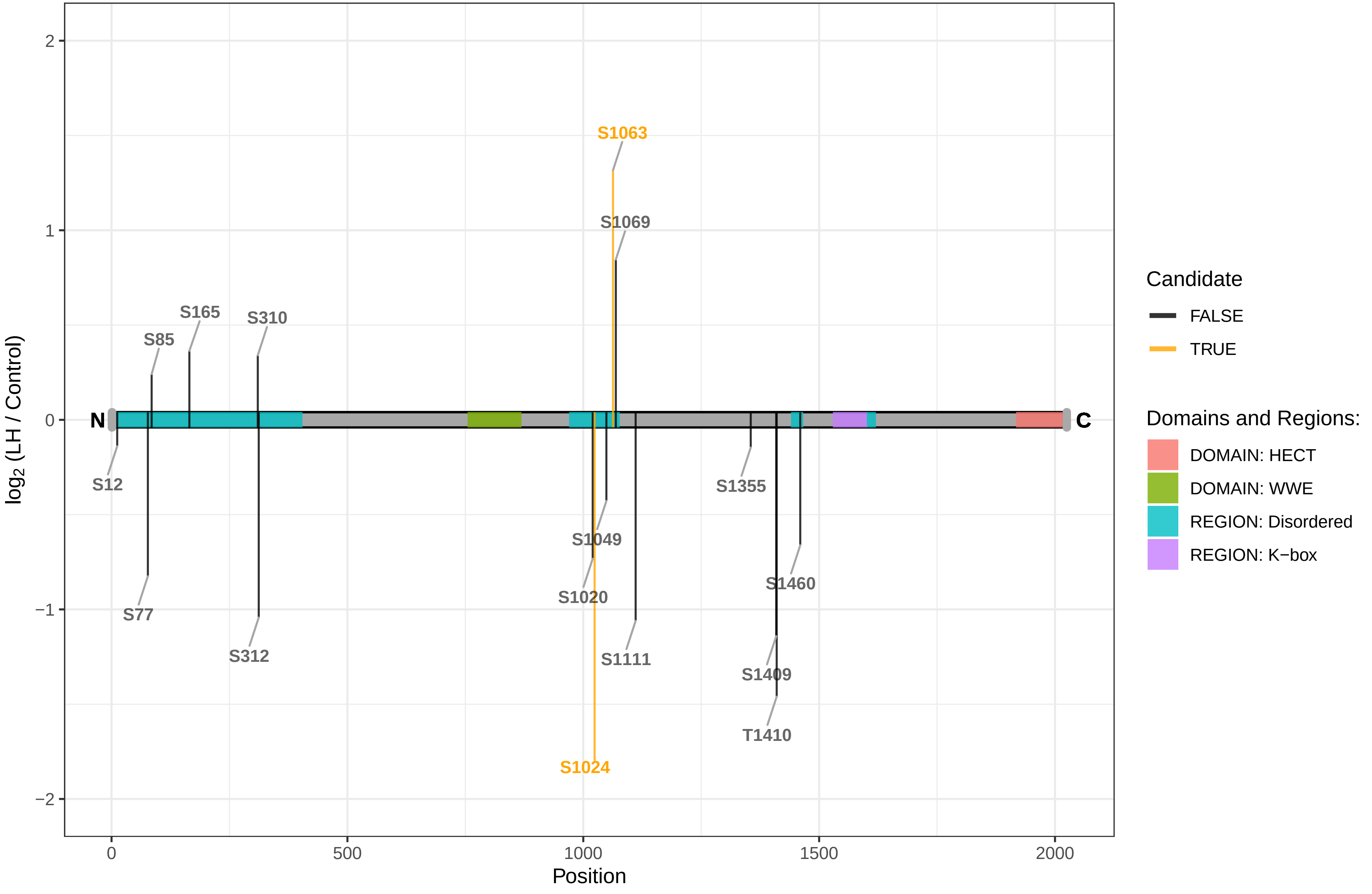

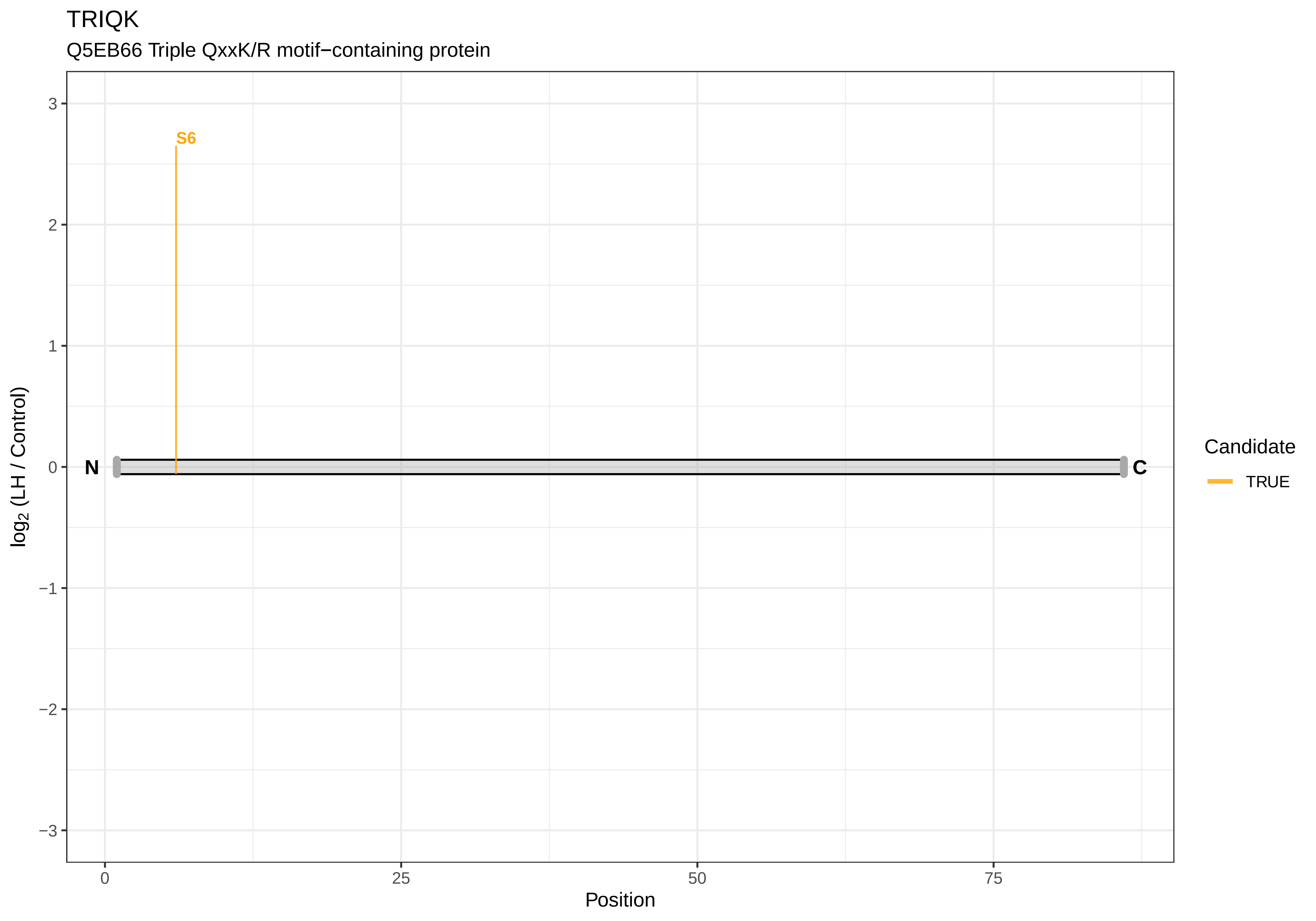

TRIQK
Q5EB66 Triple QxxK/R motif−containing protein
S6
N
C
3
2
1
0
−1
−2
−3
log2 (LH / Control)
0
25
50
75
Position
Candidate
TRUE

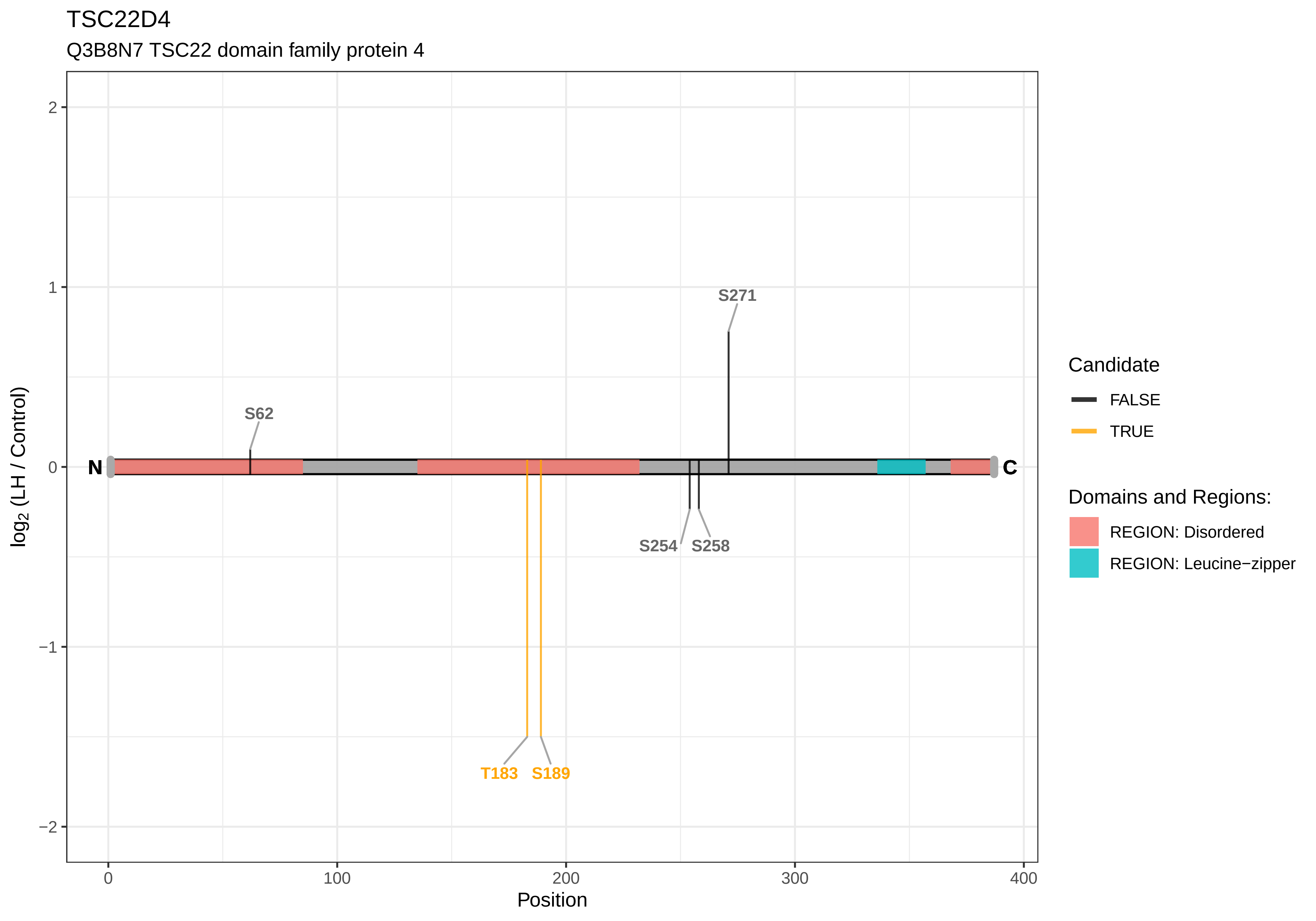
TSC22D4
Q3B8N7 TSC22 domain family protein 4
log2 (LH / Control)
Position
2
1
0
−1
−2
0
100
200
300
400
N
C
S62
S271
S254
S258
T183
S189
Candidate
FALSE
TRUE
Domains and Regions:
REGION: Disordered
REGION: Leucine−zipper

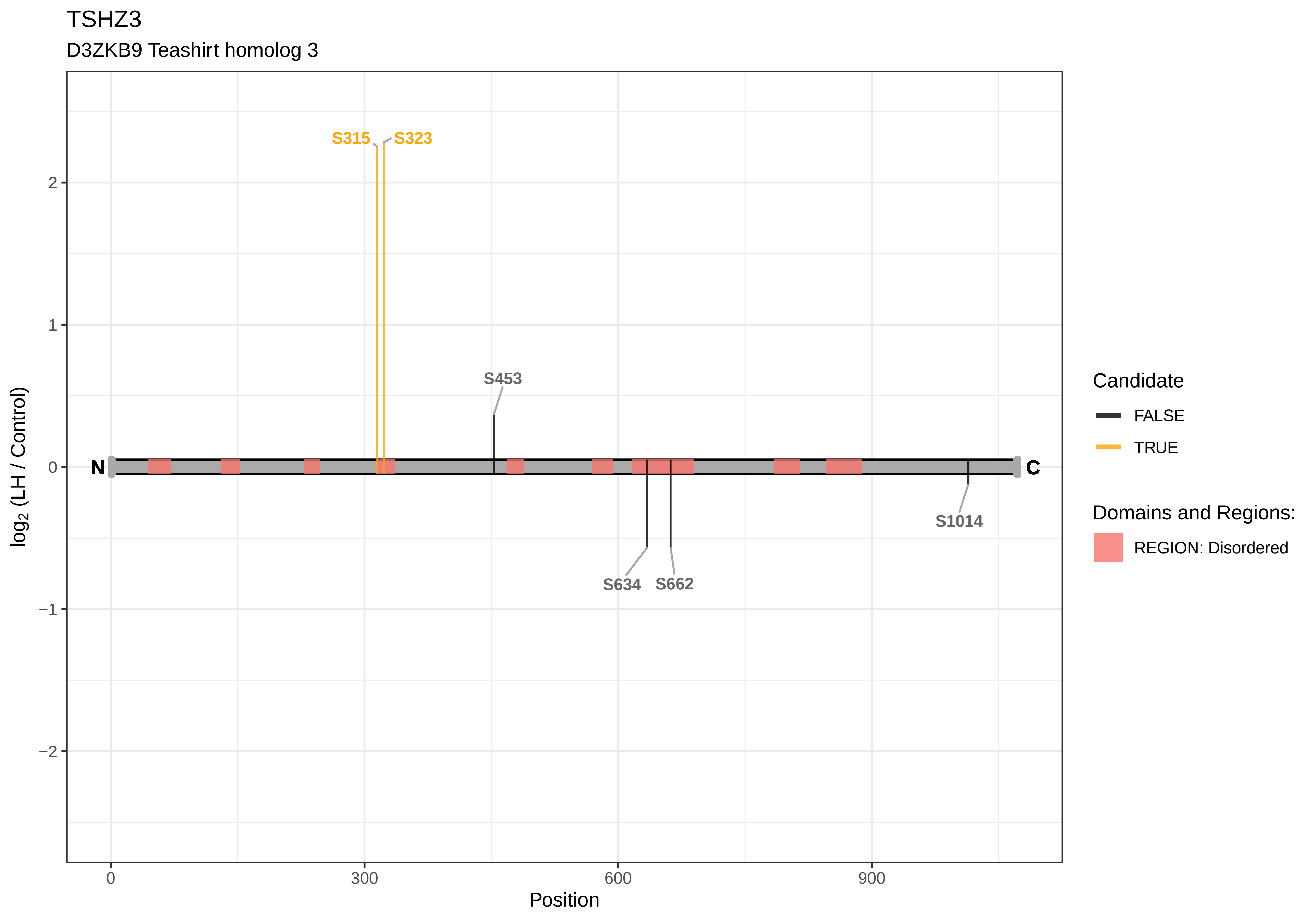
TSHZ3
D3ZKB9 Teashirt homolog 3
S315
S323
S453
S634
S662
S1014
N
C
2
1
0
−1
−2
log2 (LH / Control)
0
300
600
900
Position
Candidate
FALSE
TRUE
Domains and Regions:
REGION: Disordered

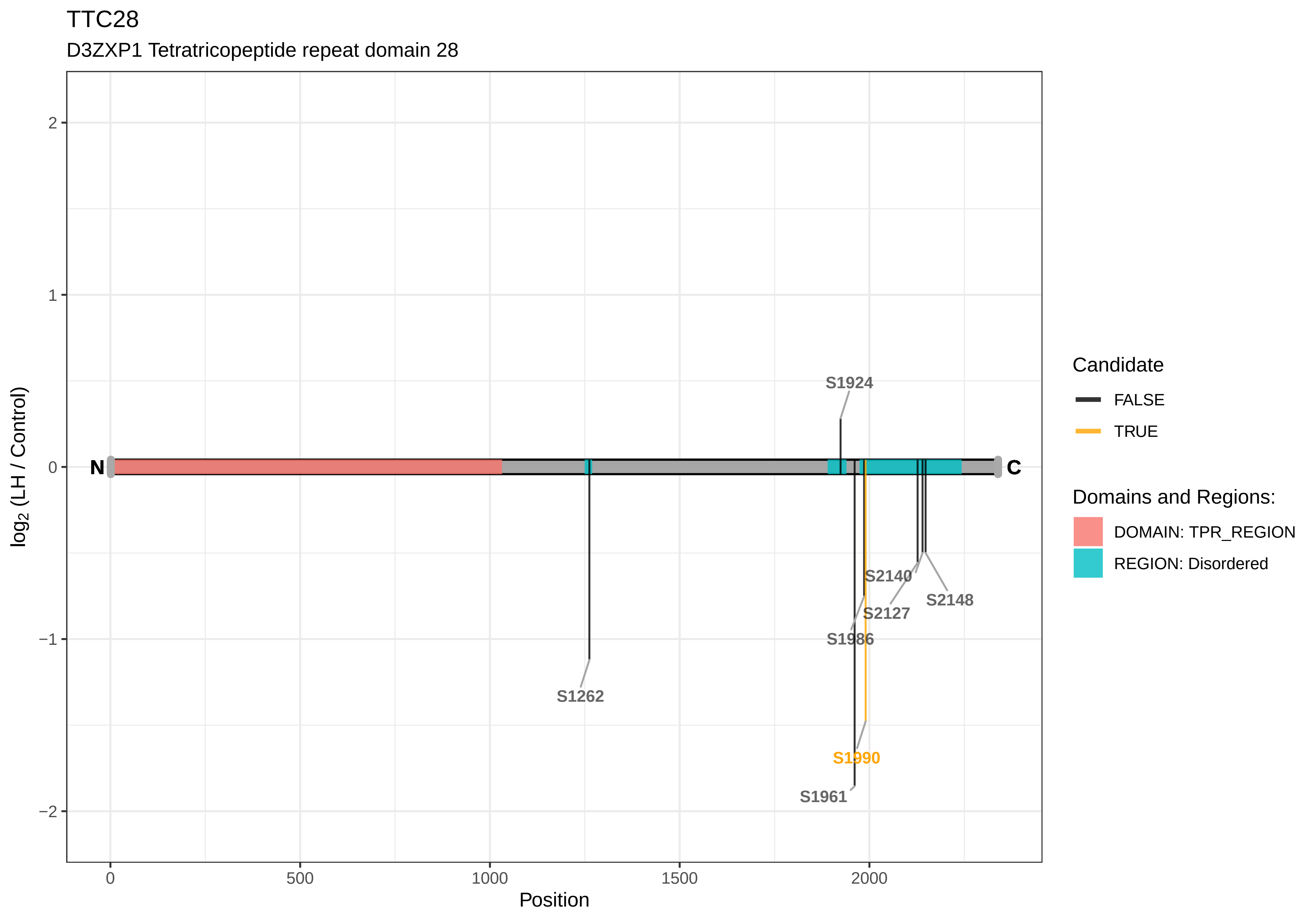
TTC28
D3ZXP1 Tetratricopeptide repeat domain 28
log2 (LH / Control)
Position
2
1
0
−1
−2
0
500
1000
1500
2000
N
C
S1924
S2140
S2148
S2127
S1986
S1262
S1990
S1961
Candidate
FALSE
TRUE
Domains and Regions:
DOMAIN: TPR_REGION
REGION: Disordered

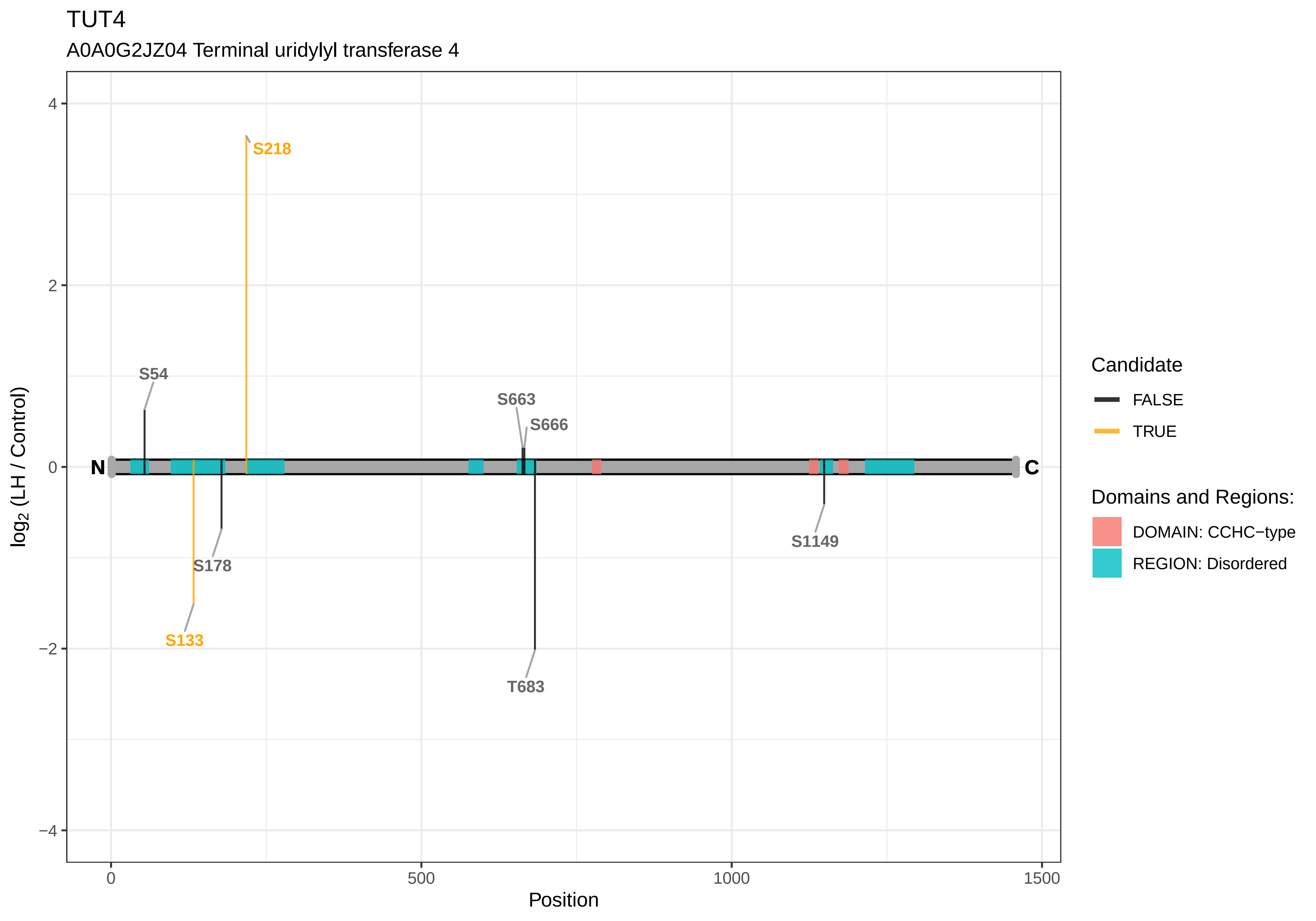
TUT4
A0A0G2JZ04 Terminal uridylyl transferase 4
S218
S54
S663
S666
N
C
S178
S133
S1149
T683
4
2
0
−2
−4
0
500
1000
1500
Position
log2 (LH / Control)
Candidate
FALSE
TRUE
Domains and Regions:
DOMAIN: CCHC−type
REGION: Disordered

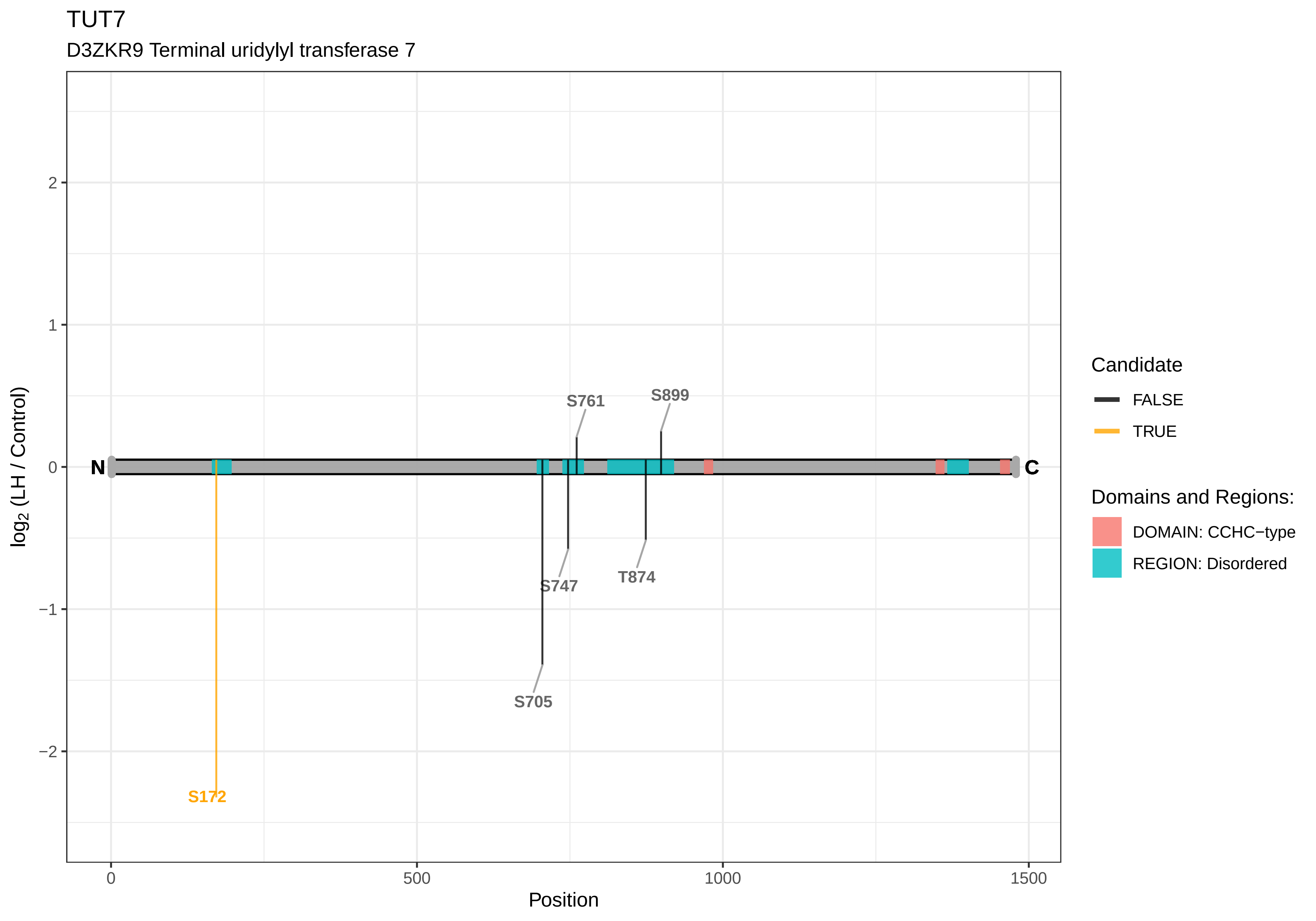

TUT7
D3ZKR9 Terminal uridylyl transferase 7
log2 (LH / Control)
Position
N
C
S761
S899
S747
T874
S705
S172
Candidate
FALSE
TRUE
Domains and Regions:
DOMAIN: CCHC−type
REGION: Disordered

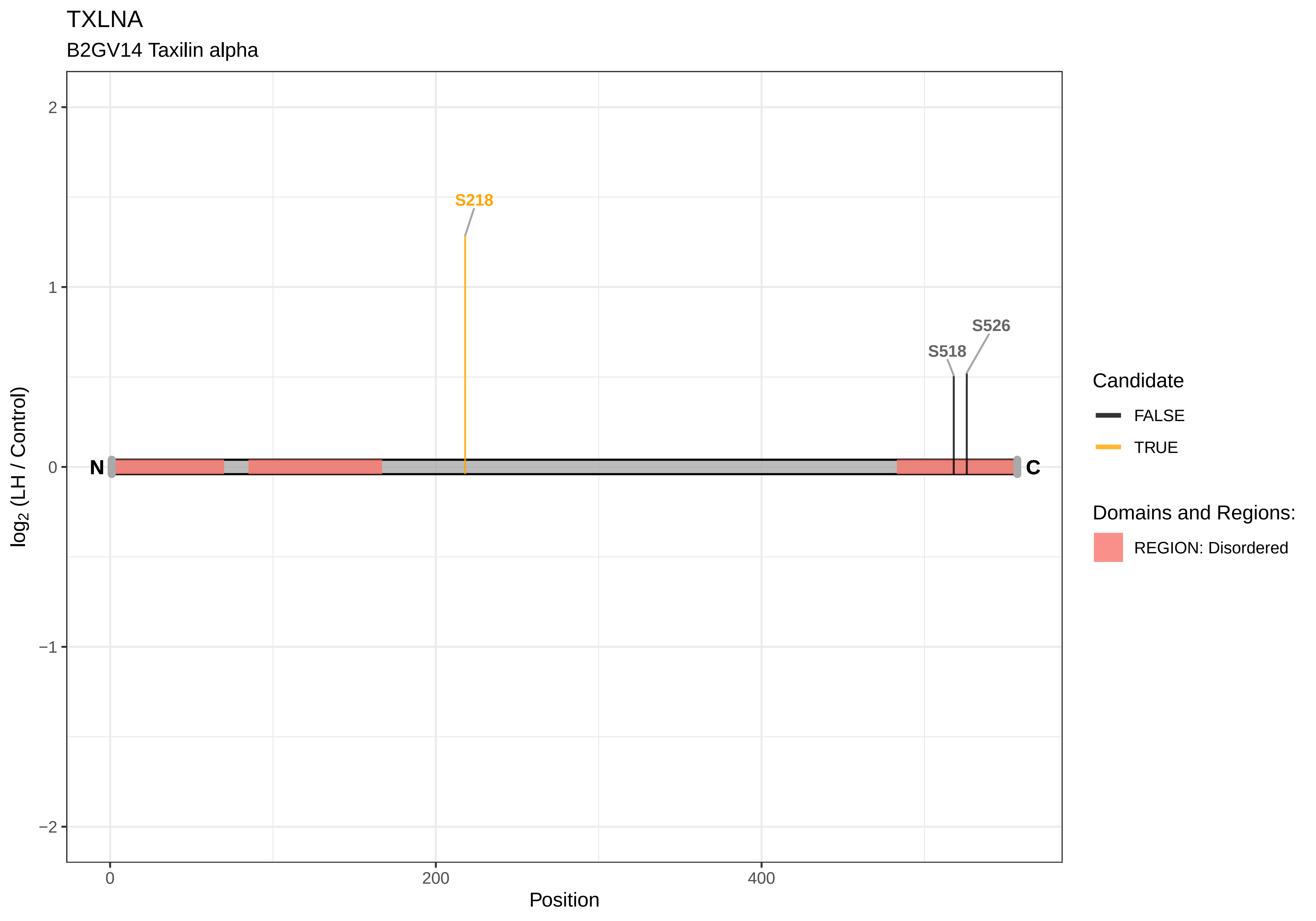

TXLNA
B2GV14 Taxilin alpha
S218
S518
S526
N
C
2
1
0
−1
−2
0
200
400
Position
log2 (LH / Control)
Candidate
FALSE
TRUE
Domains and Regions:
REGION: Disordered

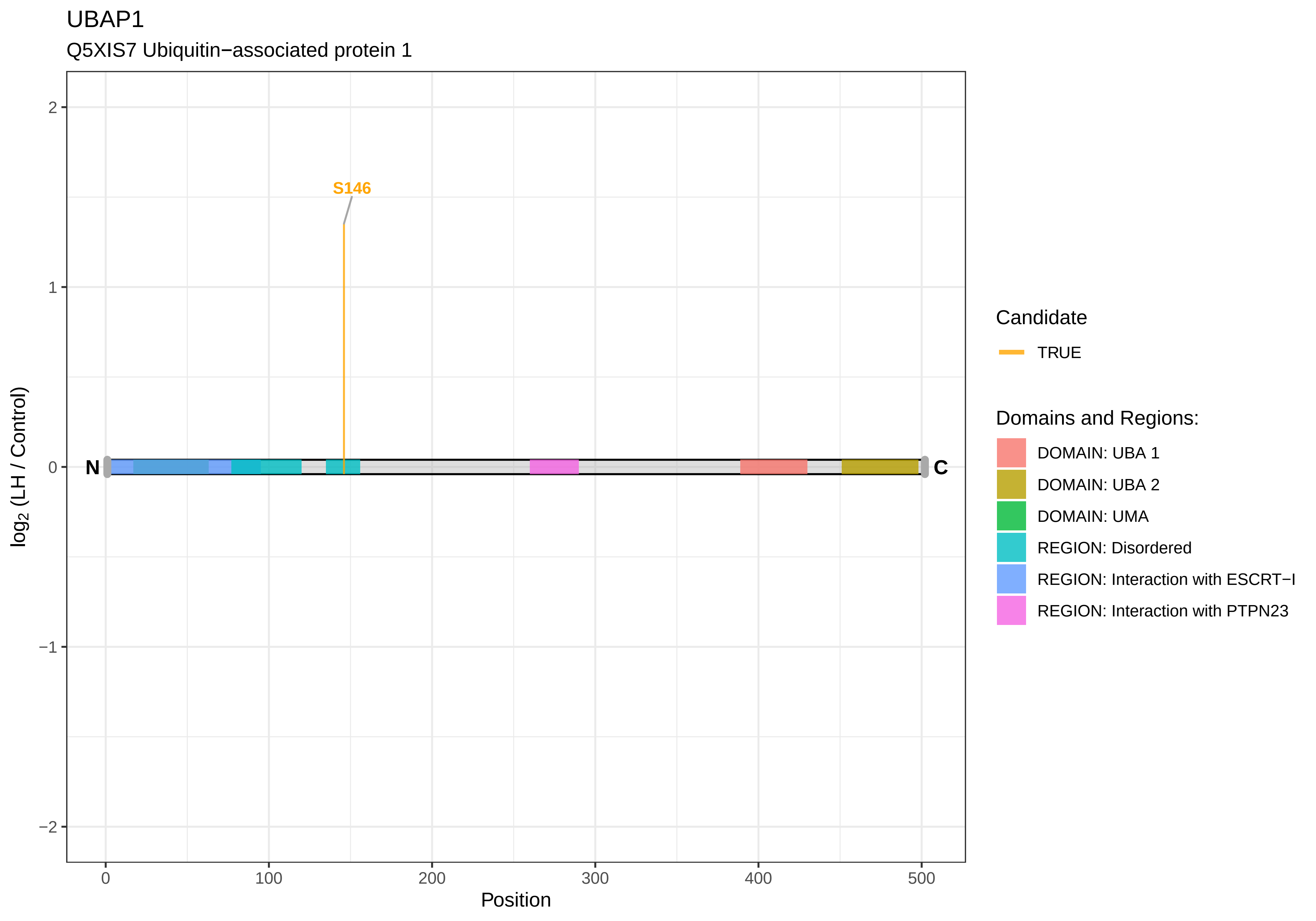

UBAP1
Q5XIS7 Ubiquitin−associated protein 1
S146
N
C
2
1
0
−1
−2
log2 (LH / Control)
0
100
200
300
400
500
Position
Candidate
TRUE
Domains and Regions:
DOMAIN: UBA 1
DOMAIN: UBA 2
DOMAIN: UMA
REGION: Disordered
REGION: Interaction with ESCRT−I
REGION: Interaction with PTPN23

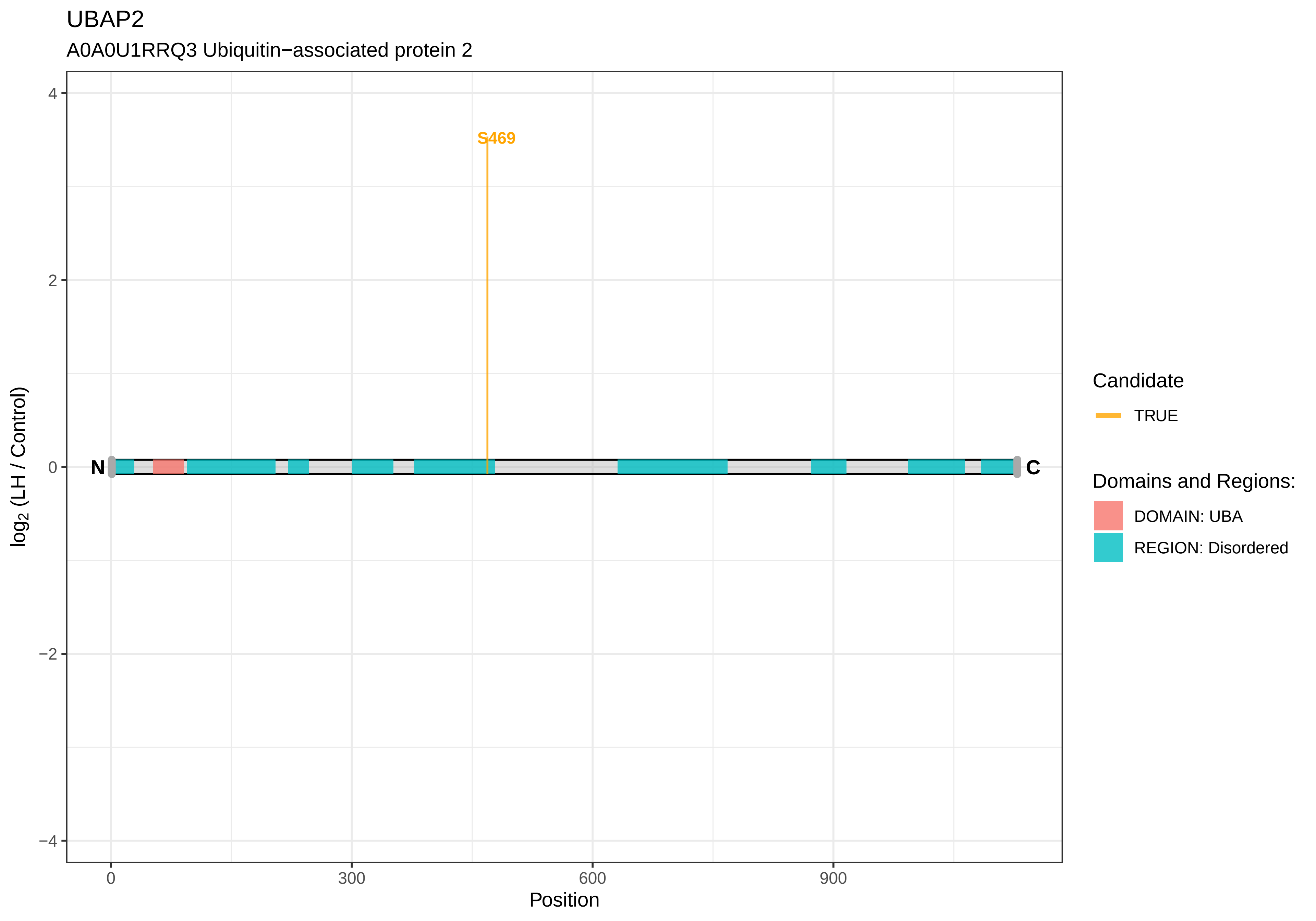

UBAP2
A0A0U1RRQ3 Ubiquitin−associated protein 2
S469
N
C
4
2
0
−2
−4
log2 (LH / Control)
0
300
600
900
Position
Candidate
TRUE
Domains and Regions:
DOMAIN: UBA
REGION: Disordered

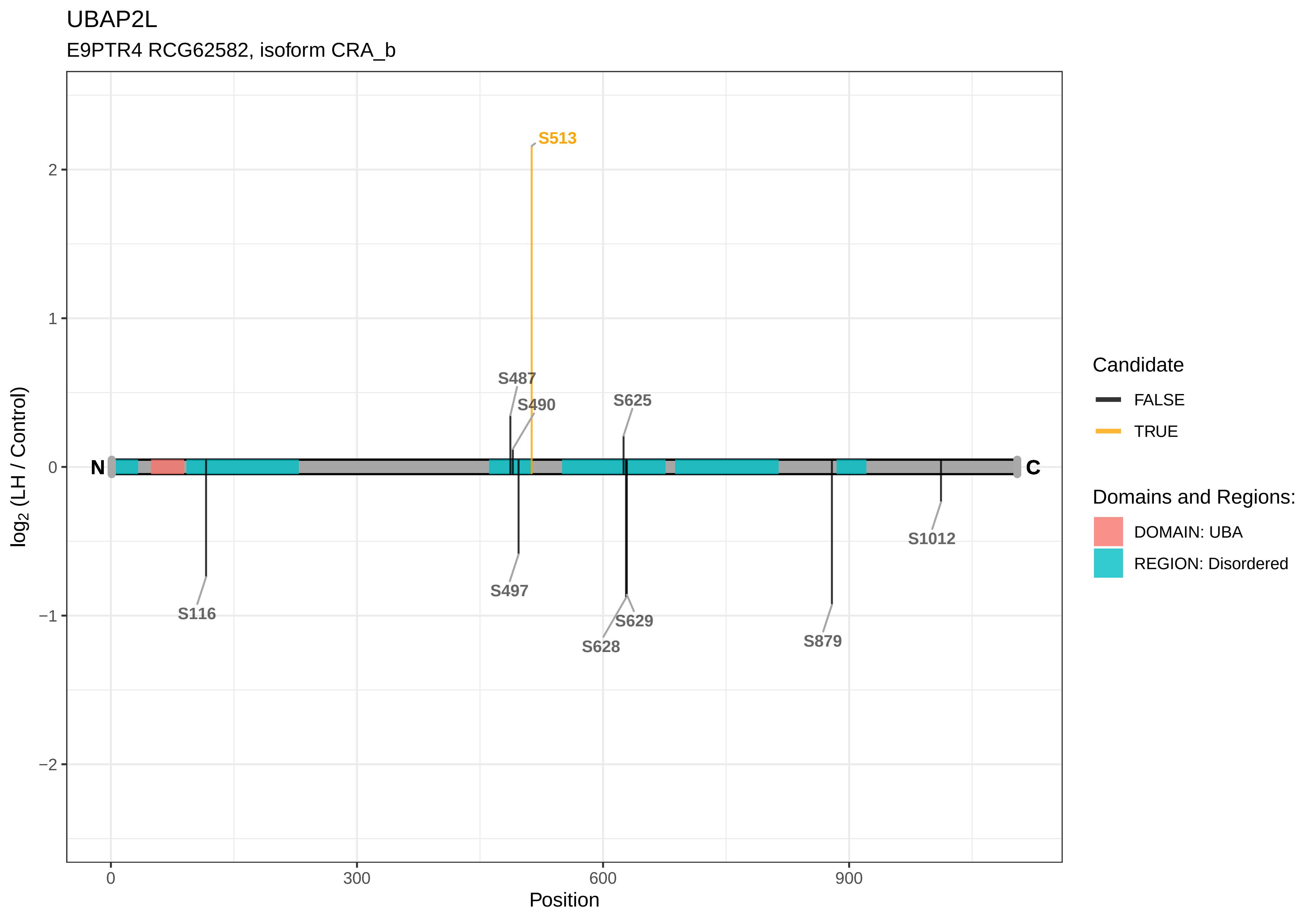

UBAP2L
E9PTR4 RCG62582, isoform CRA_b
S513
S487
S490
S625
S497
S116
S629
S628
S879
S1012
N
C
2
1
0
−1
−2
log2 (LH / Control)
0
300
600
900
Position
Candidate
FALSE
TRUE
Domains and Regions:
DOMAIN: UBA
REGION: Disordered

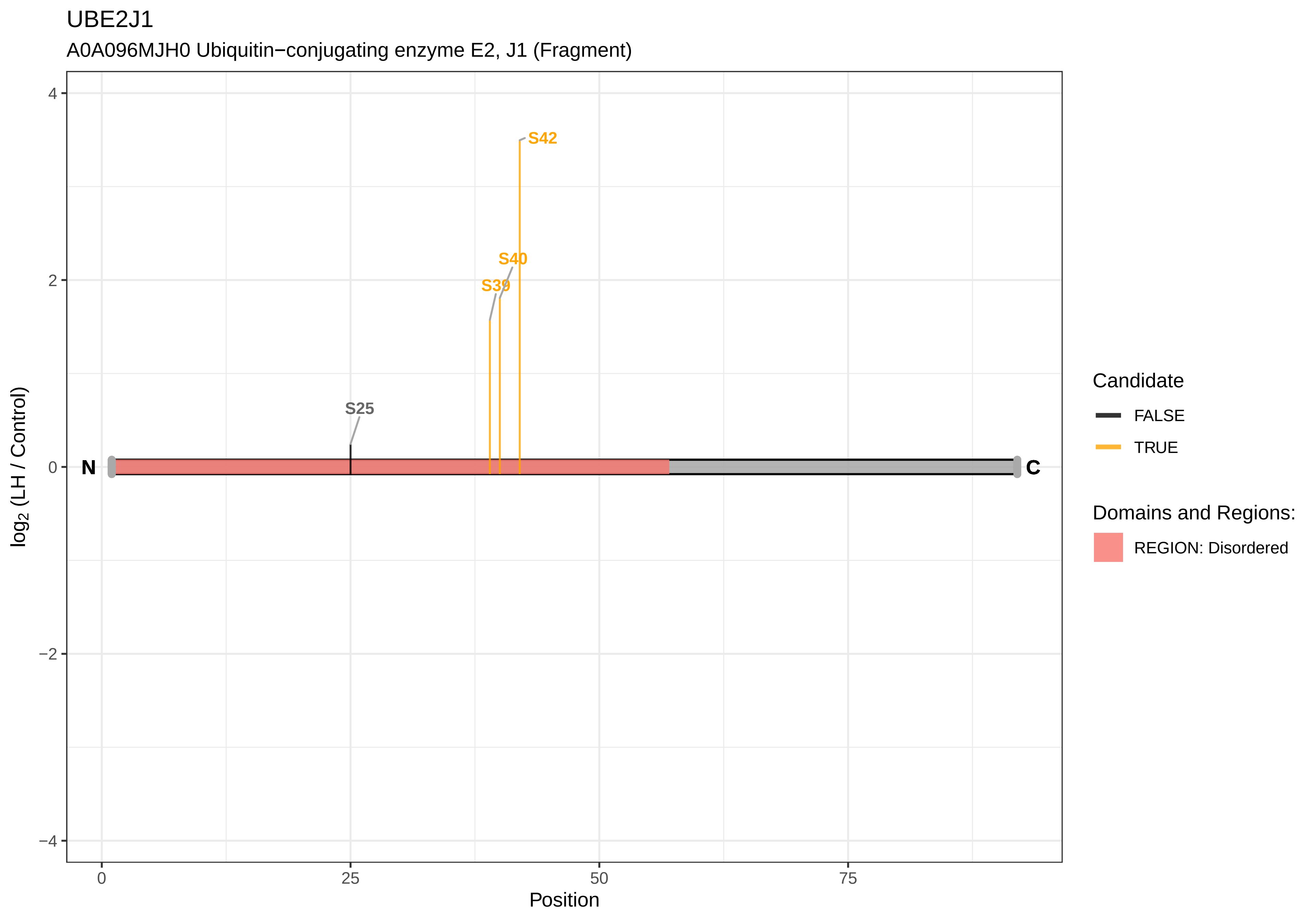

UBE2J1
A0A096MJH0 Ubiquitin−conjugating enzyme E2, J1 (Fragment)
S42
S40
S39
S25
N
C
4
2
0
−2
−4
0
25
50
75
Position
log2 (LH / Control)
Candidate
FALSE
TRUE
Domains and Regions:
REGION: Disordered

UBE2O

F1M403 Ubiquitin−conjugating enzyme E2O

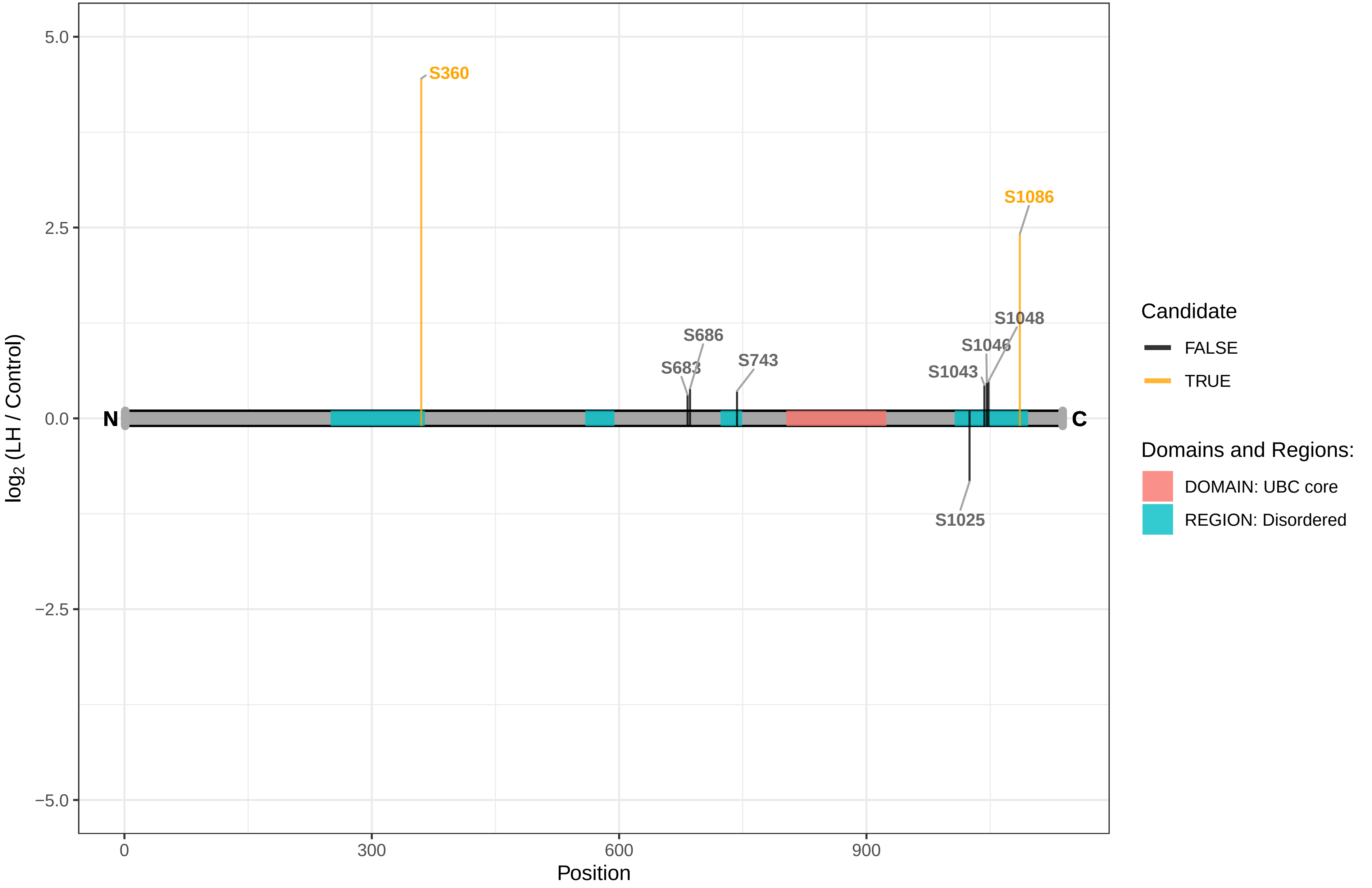

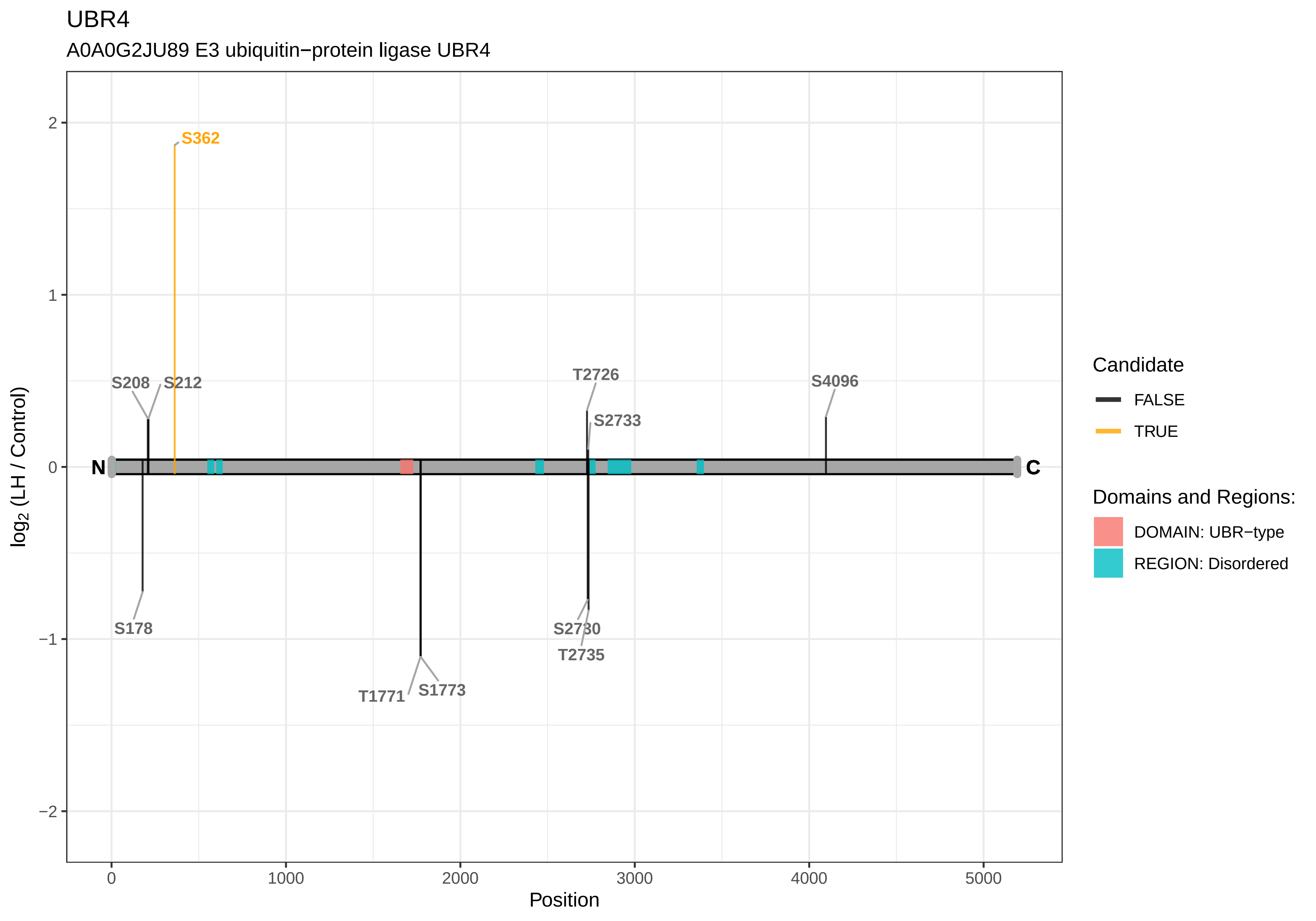
UBR4
A0A0G2JU89 E3 ubiquitin−protein ligase UBR4
S362
S208
S212
T2726
S2733
S4096
S178
T1771
S1773
S2730
T2735
N
C
log2 (LH / Control)
Position
2
1
0
−1
−2
0
1000
2000
3000
4000
5000
Candidate
FALSE
TRUE
Domains and Regions:
DOMAIN: UBR−type
REGION: Disordered

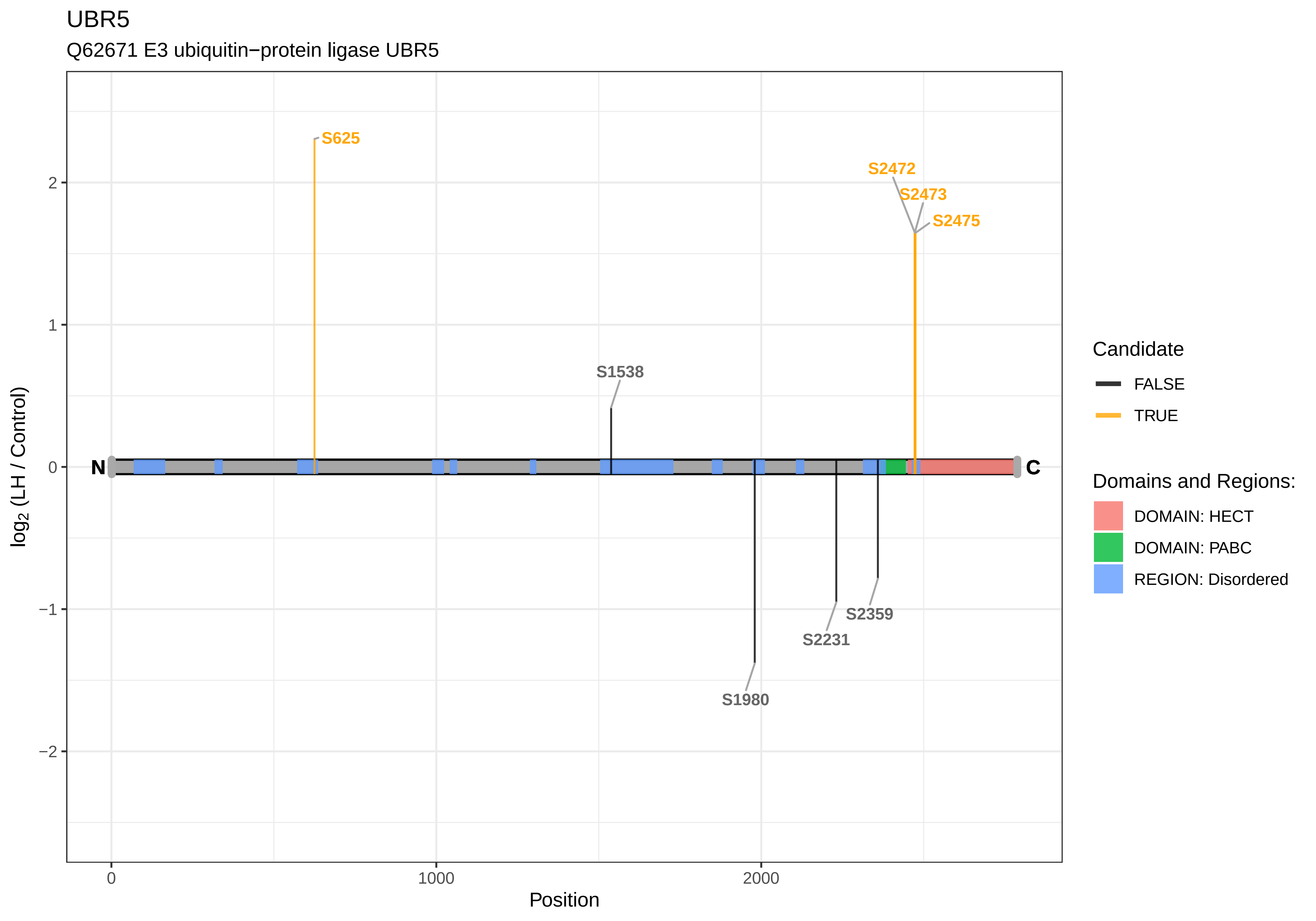
UBR5
Q62671 E3 ubiquitin−protein ligase UBR5
S625
S2472
S2473
S2475
S1538
S1980
S2231
S2359
N
C
2
1
0
−1
−2
0
1000
2000
Position
log2 (LH / Control)
Candidate
FALSE
TRUE
Domains and Regions:
DOMAIN: HECT
DOMAIN: PABC
REGION: Disordered

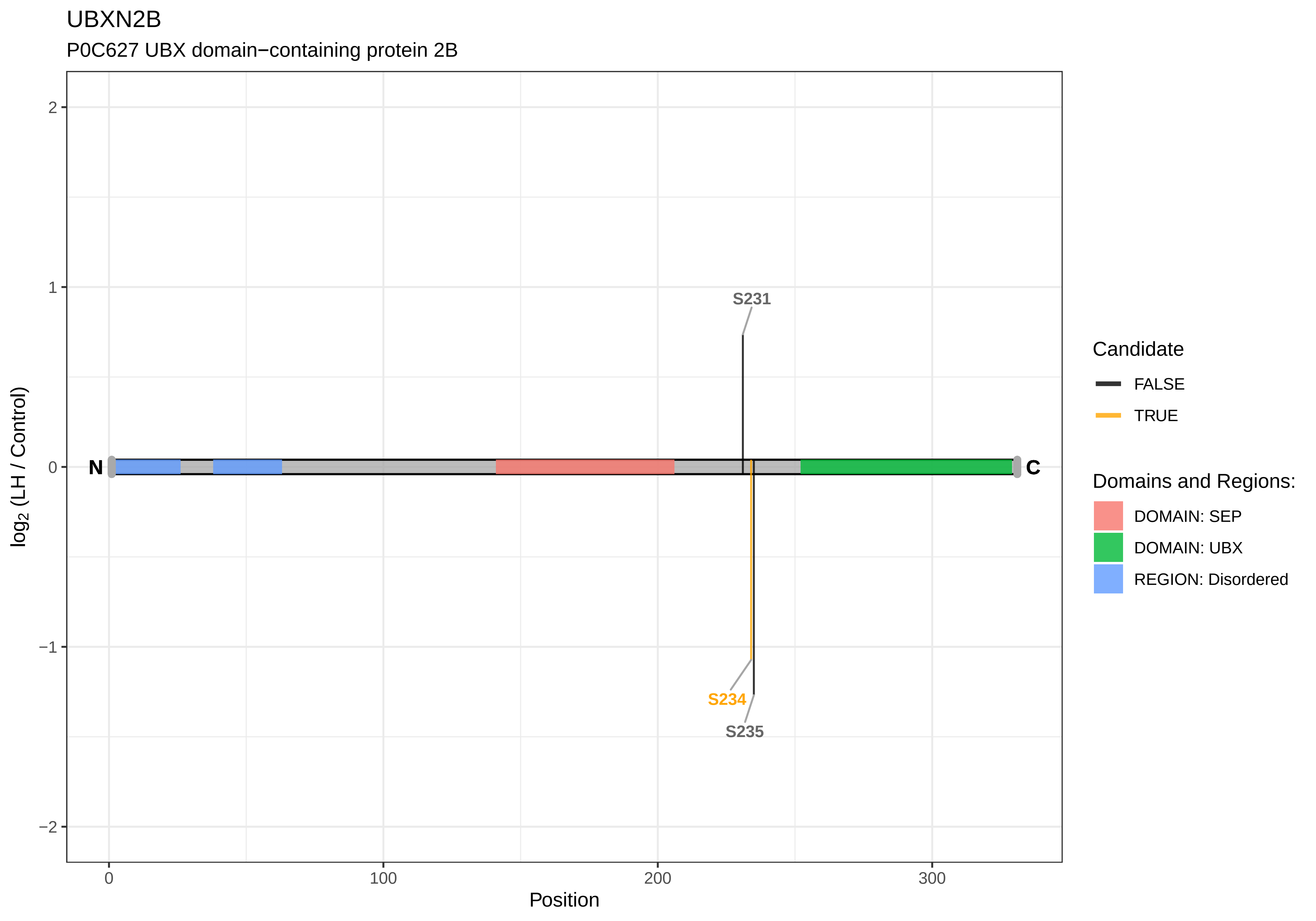

UBXN2B
P0C627 UBX domain−containing protein 2B
log2 (LH / Control)
2
1
0
−1
−2
N
C
S231
S234
S235
0
100
200
300
Position
Candidate
FALSE
TRUE
Domains and Regions:
DOMAIN: SEP
DOMAIN: UBX
REGION: Disordered

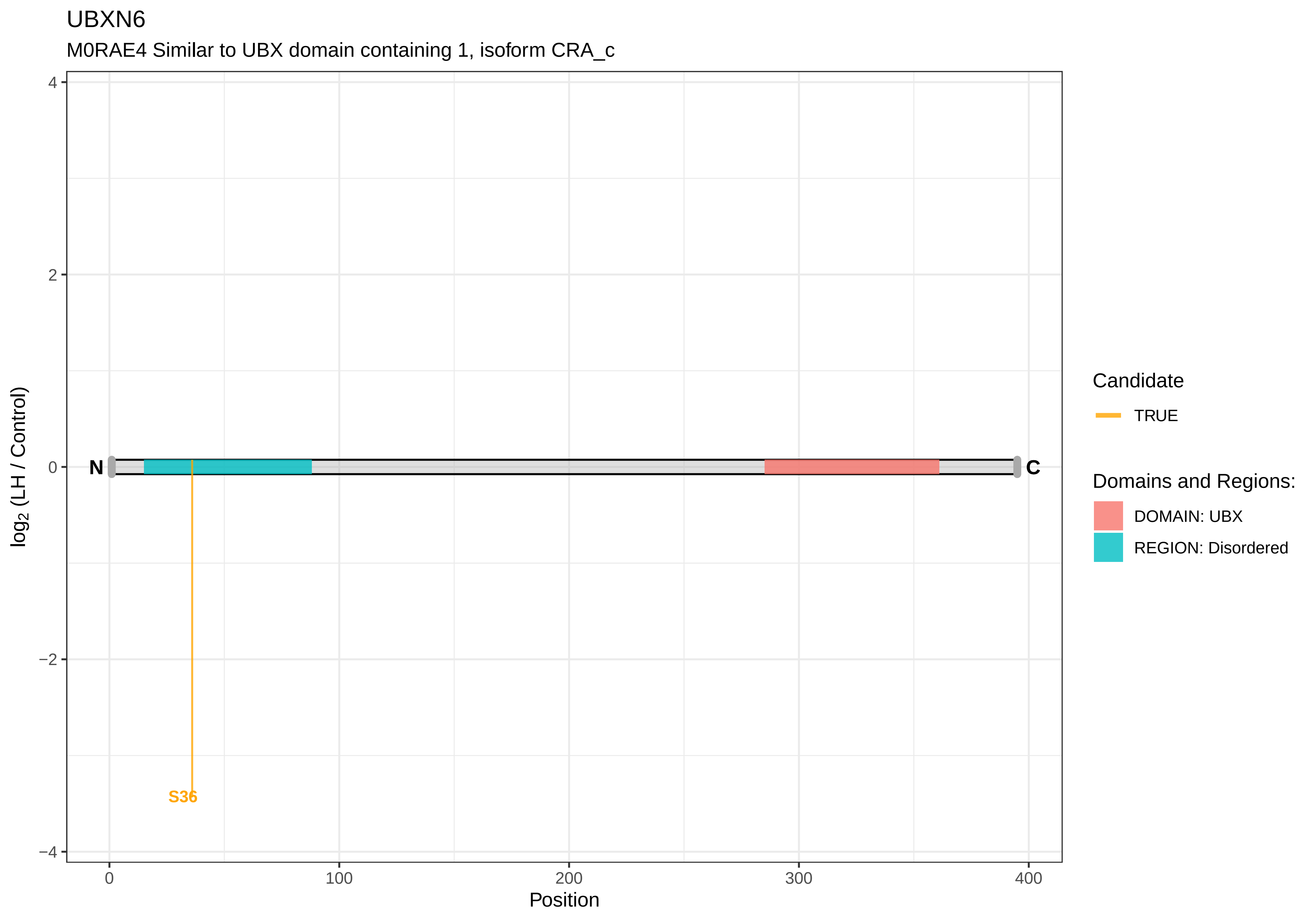
UBXN6
M0RAE4 Similar to UBX domain containing 1, isoform CRA_c
log2 (LH / Control)
Position
N
C
S36
Candidate
TRUE
Domains and Regions:
DOMAIN: UBX
REGION: Disordered

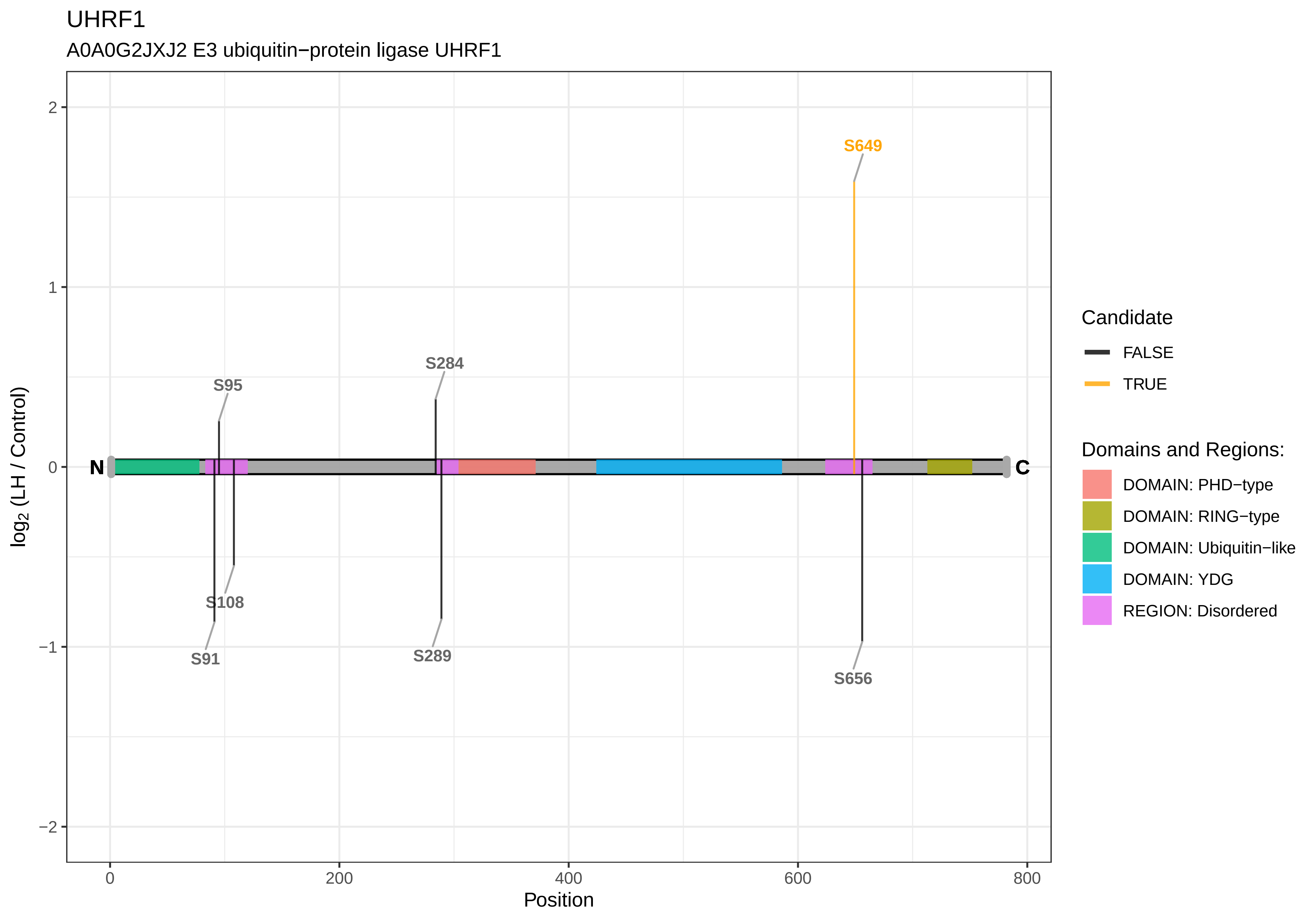
UHRF1
A0A0G2JXJ2 E3 ubiquitin−protein ligase UHRF1
S649
S284
S95
S108
S91
S289
S656
N
C
log2 (LH / Control)
Position
Candidate
FALSE
TRUE
Domains and Regions:
DOMAIN: PHD−type
DOMAIN: RING−type
DOMAIN: Ubiquitin−like
DOMAIN: YDG
REGION: Disordered

UHRF1BP1

D3ZMR2 UHRF1−binding protein 1

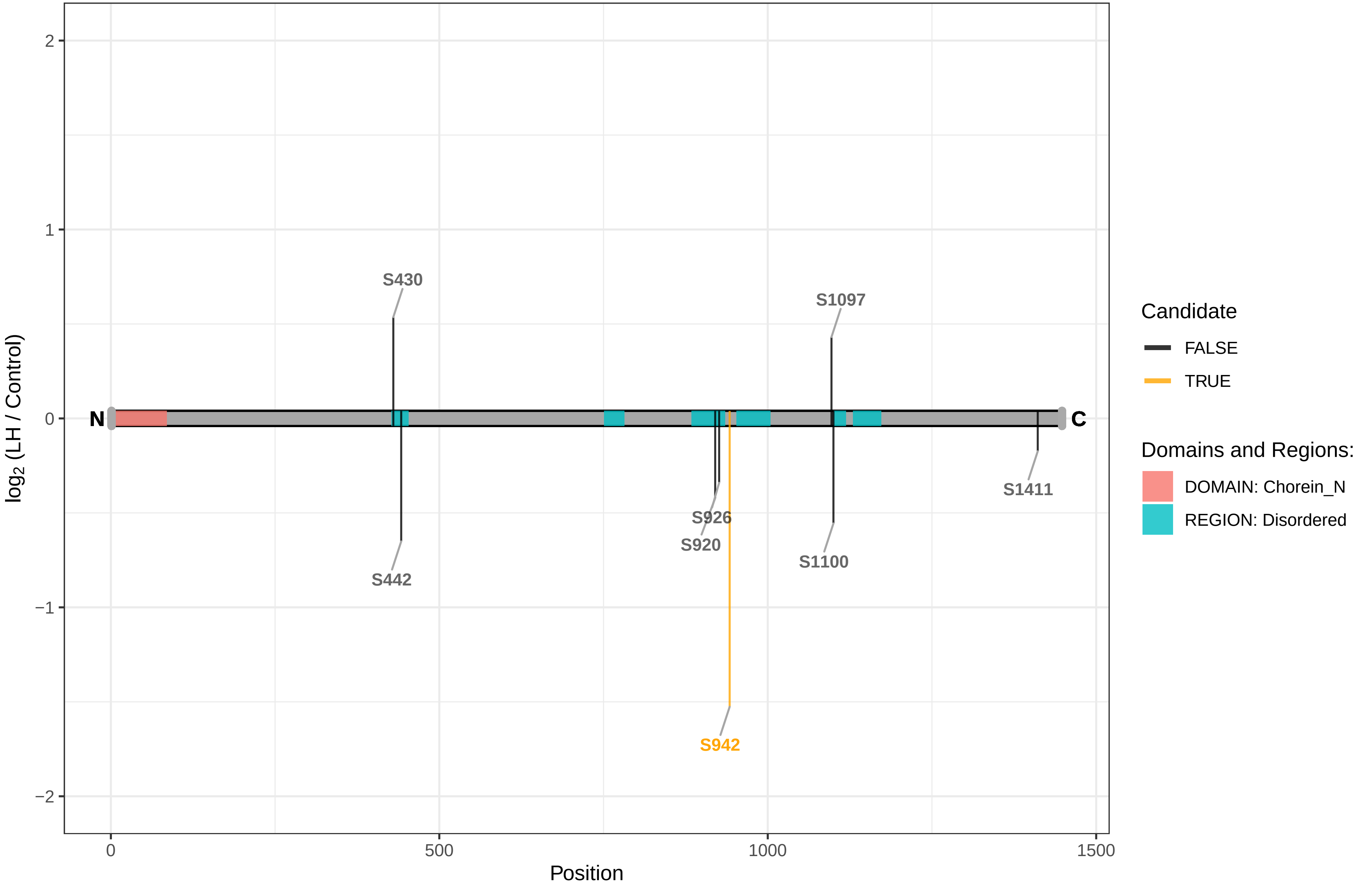

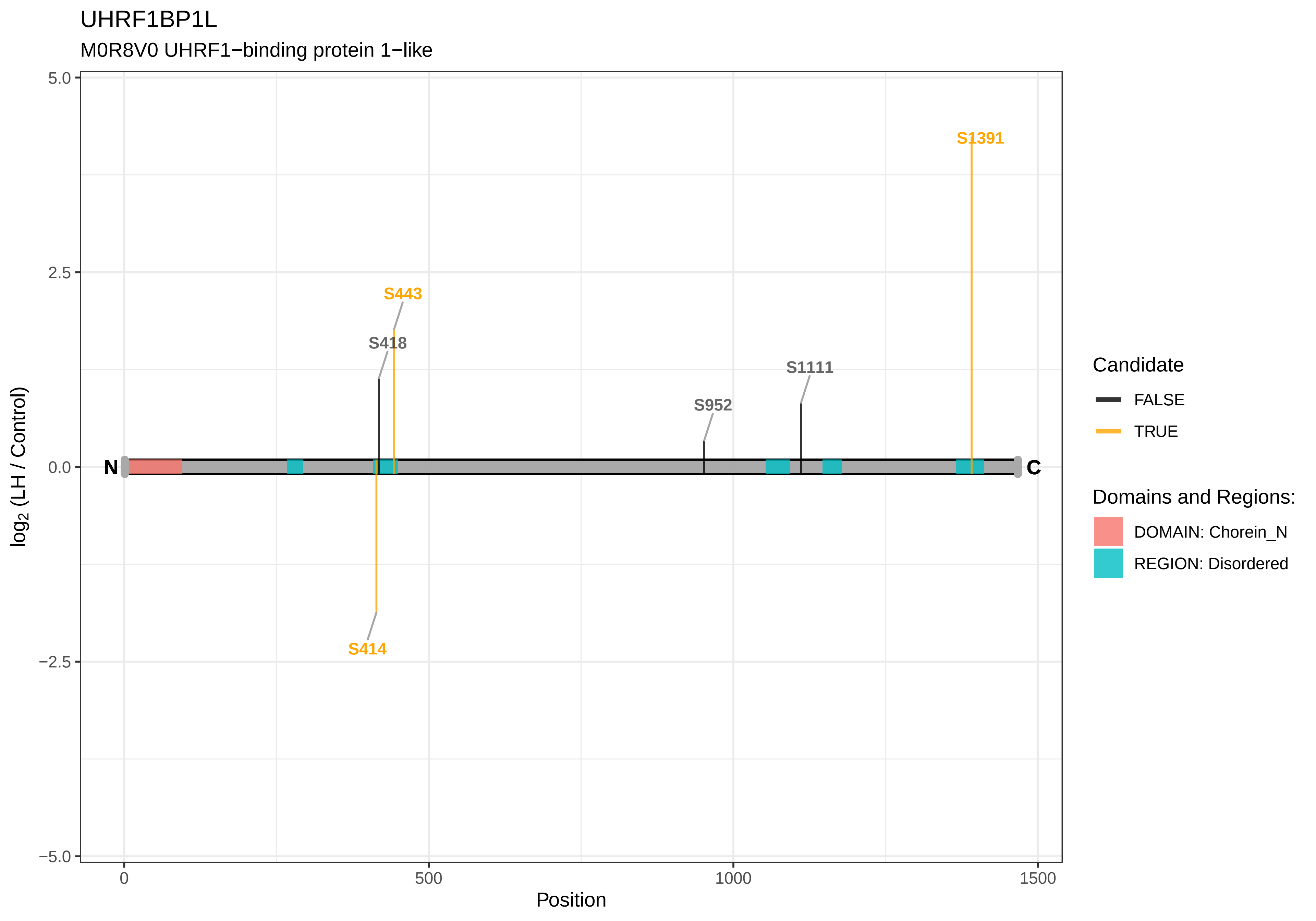
UHRF1BP1L
M0R8V0 UHRF1−binding protein 1−like
5.0
2.5
0.0
−2.5
−5.0
log2 (LH / Control)
0
500
1000
1500
Position
N
C
S1391
S443
S418
S1111
S952
S414
Candidate
FALSE
TRUE
Domains and Regions:
DOMAIN: Chorein_N
REGION: Disordered

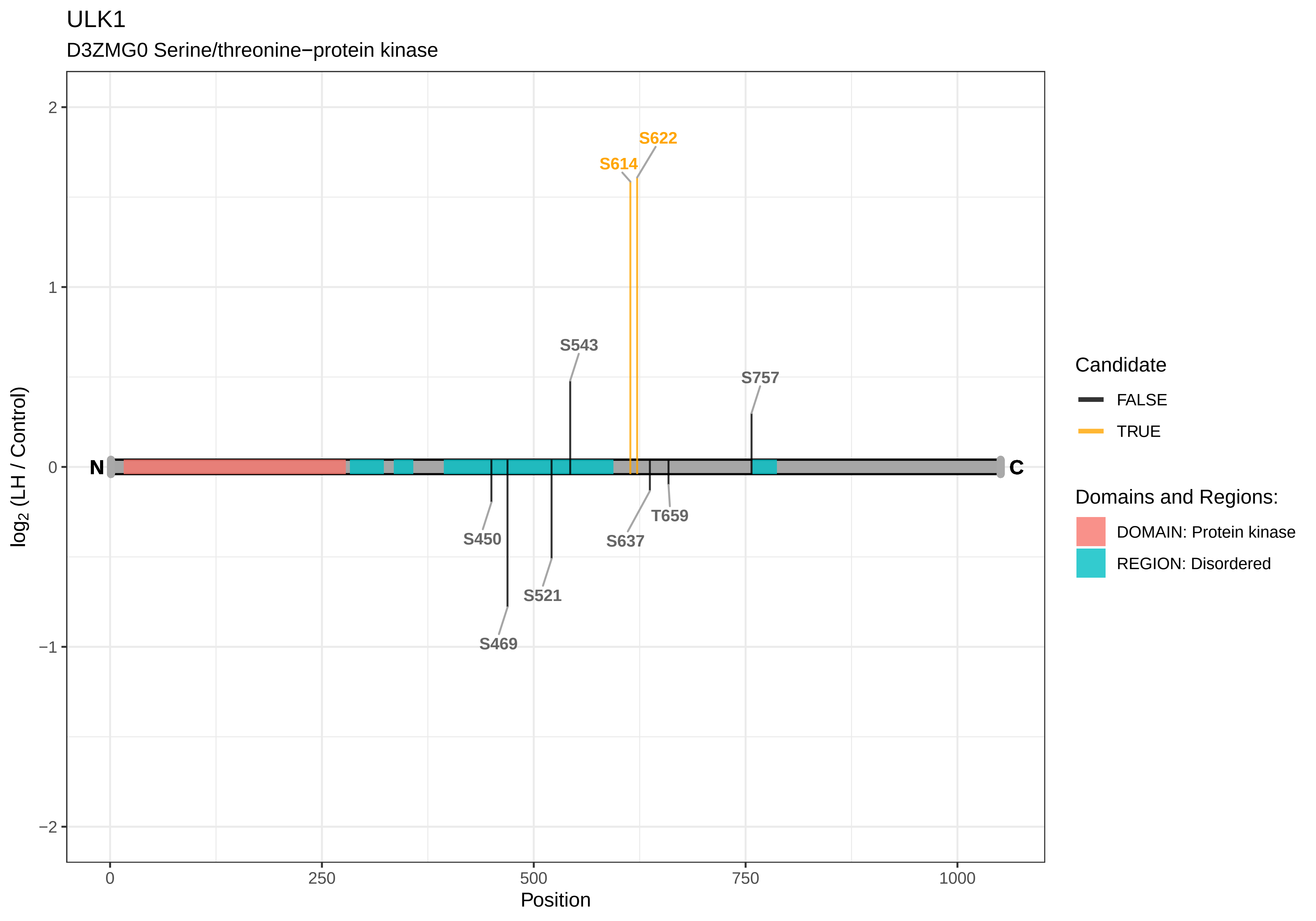

ULK1
D3ZMG0 Serine/threonine−protein kinase
S622
S614
S543
S757
S450
S637
T659
S521
S469
N
C
2
1
0
−1
−2
log2 (LH / Control)
0
250
500
750
1000
Position
Candidate
FALSE
TRUE
Domains and Regions:
DOMAIN: Protein kinase
REGION: Disordered

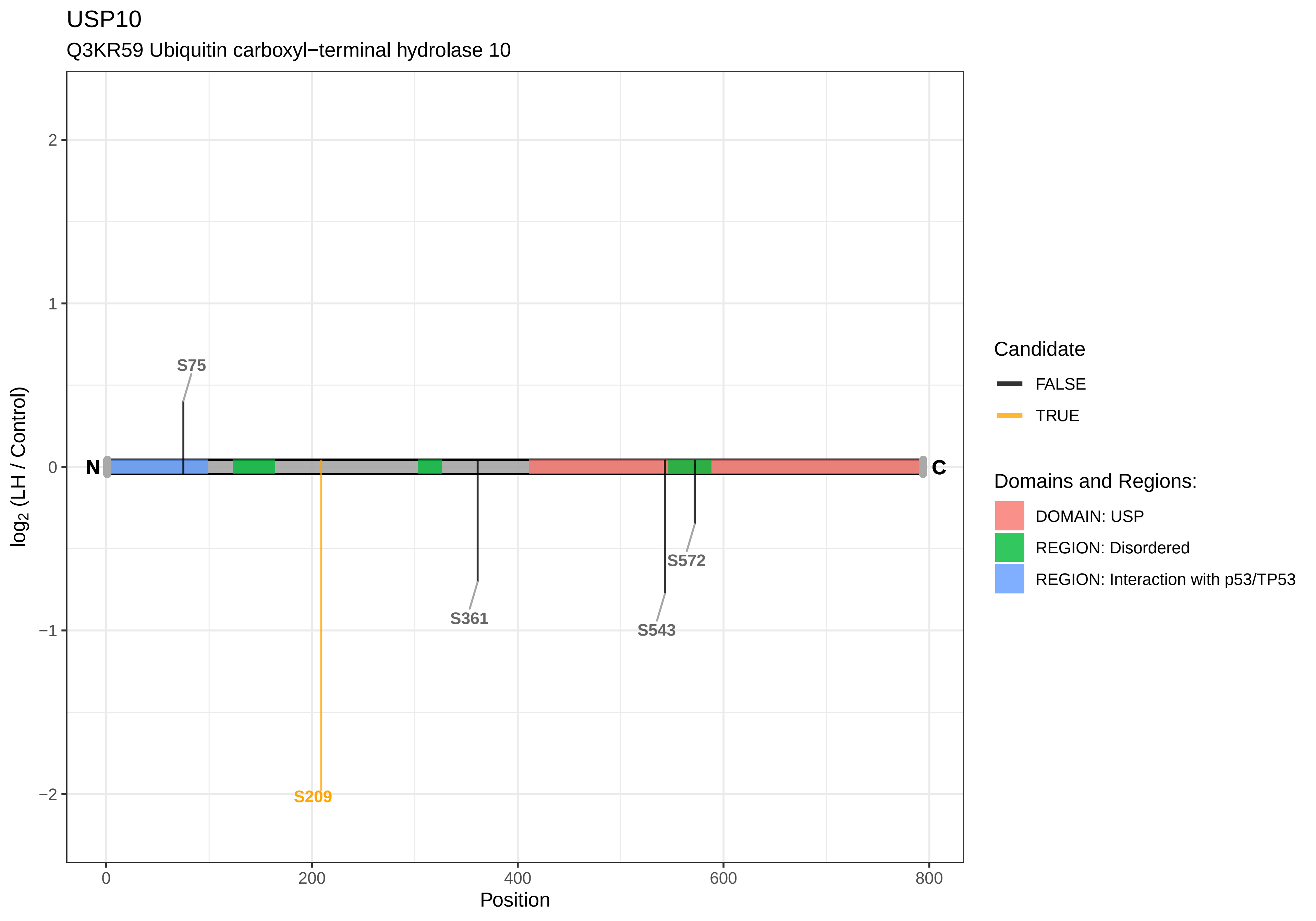

USP10
Q3KR59 Ubiquitin carboxyl−terminal hydrolase 10
log2 (LH / Control)
Position
N
C
S75
S209
S361
S543
S572
2
1
0
−1
−2
0
200
400
600
800
Candidate
FALSE
TRUE
Domains and Regions:
DOMAIN: USP
REGION: Disordered
REGION: Interaction with p53/TP53

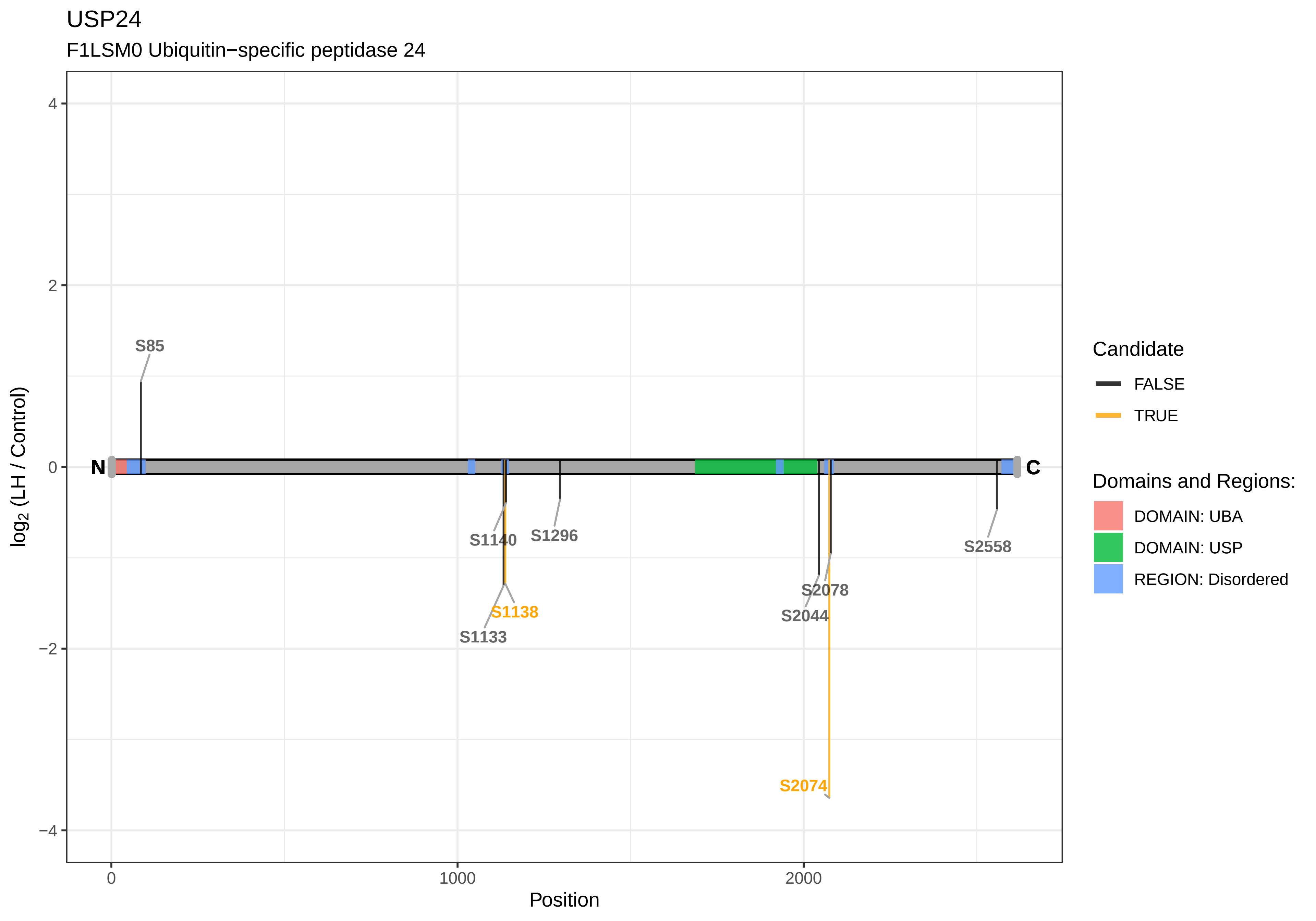

USP24
F1LSM0 Ubiquitin−specific peptidase 24
log2 (LH / Control)
Position
N
C
S85
S1140
S1296
S1138
S1133
S2078
S2044
S2558
S2074
4
2
0
−2
−4
0
1000
2000
Candidate
FALSE
TRUE
Domains and Regions:
DOMAIN: UBA
DOMAIN: USP
REGION: Disordered

USP31

D3ZU27 Ubiquitin specific protease 31 (Predicted)

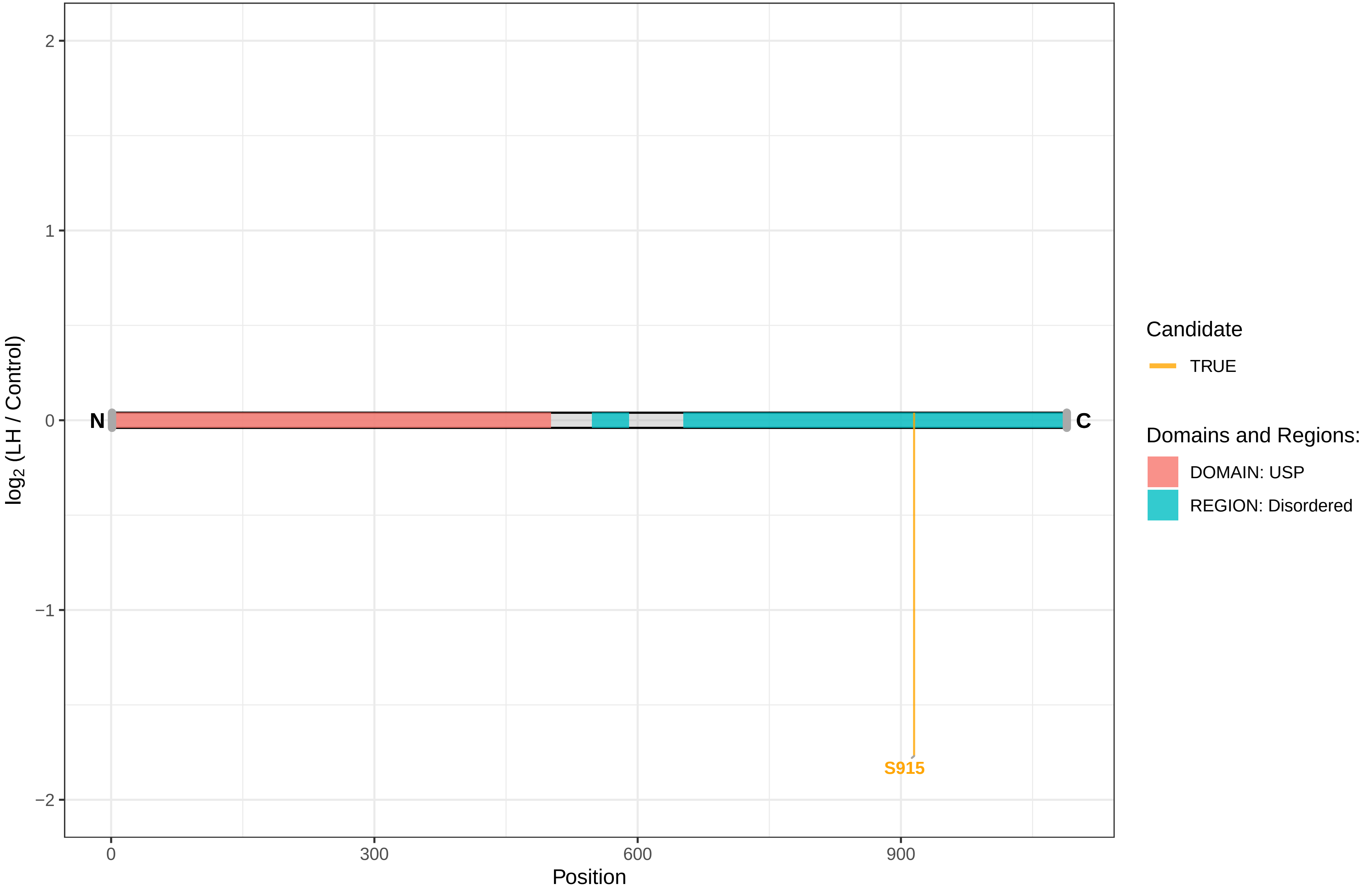

USP32

D3ZBB7 Ubiquitin specific protease 32 (Predicted)

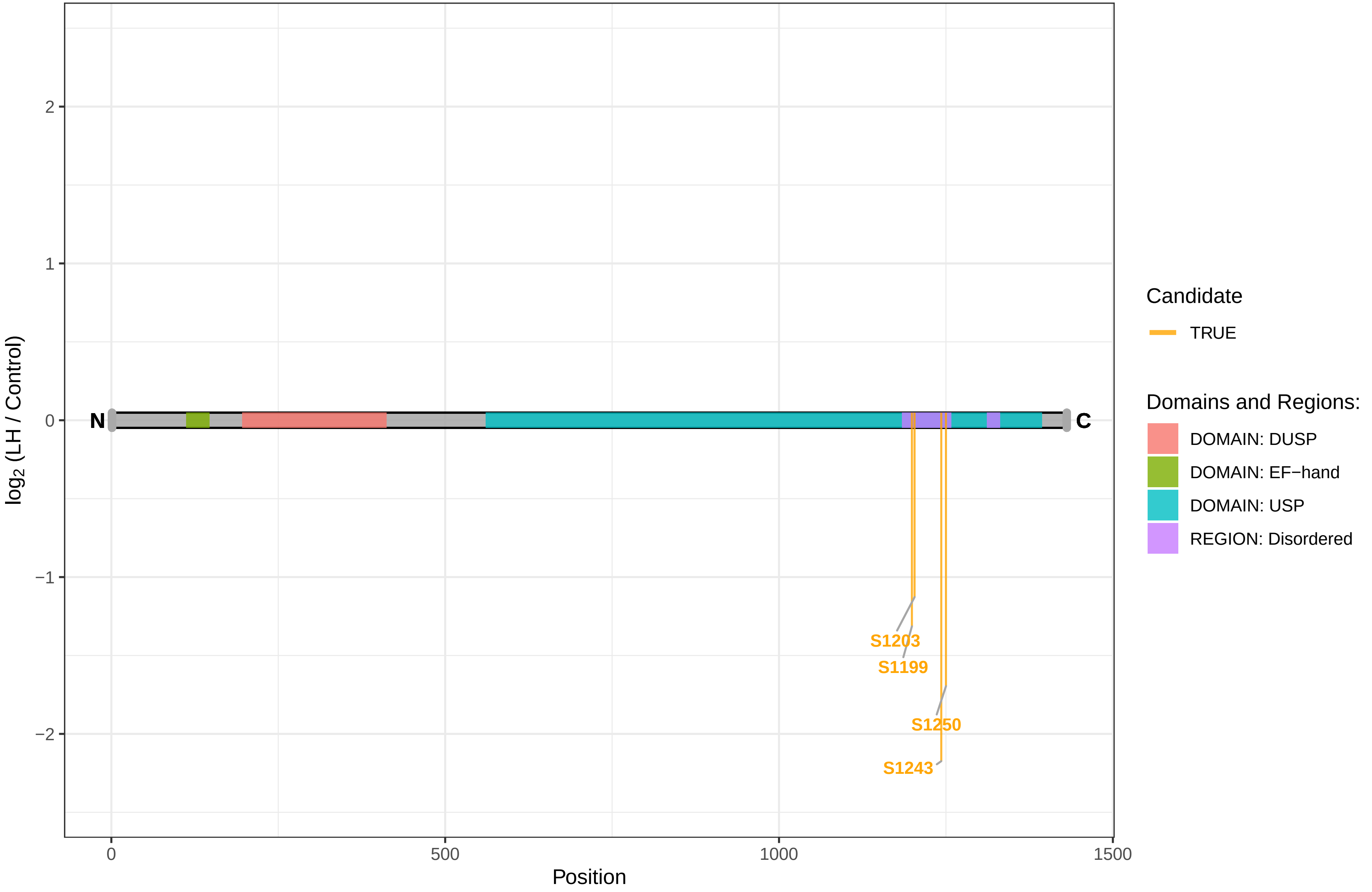

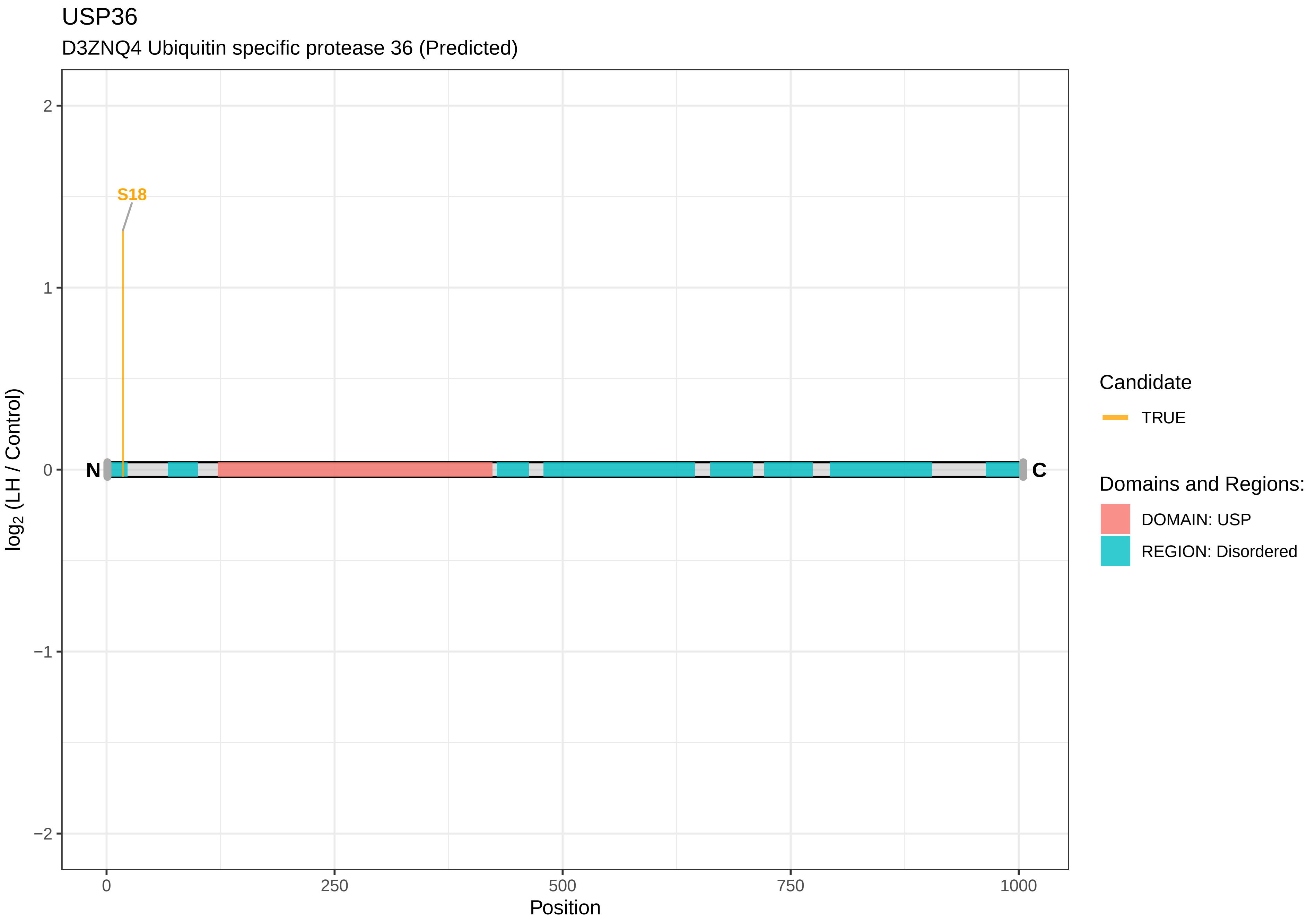
USP36
D3ZNQ4 Ubiquitin specific protease 36 (Predicted)
S18
N
C
2
1
0
−1
−2
log2 (LH / Control)
0
250
500
750
1000
Position
Candidate
TRUE
Domains and Regions:
DOMAIN: USP
REGION: Disordered

USP6NL

D3ZL57 USP6 N−terminal like (Predicted), isoform CRA_a

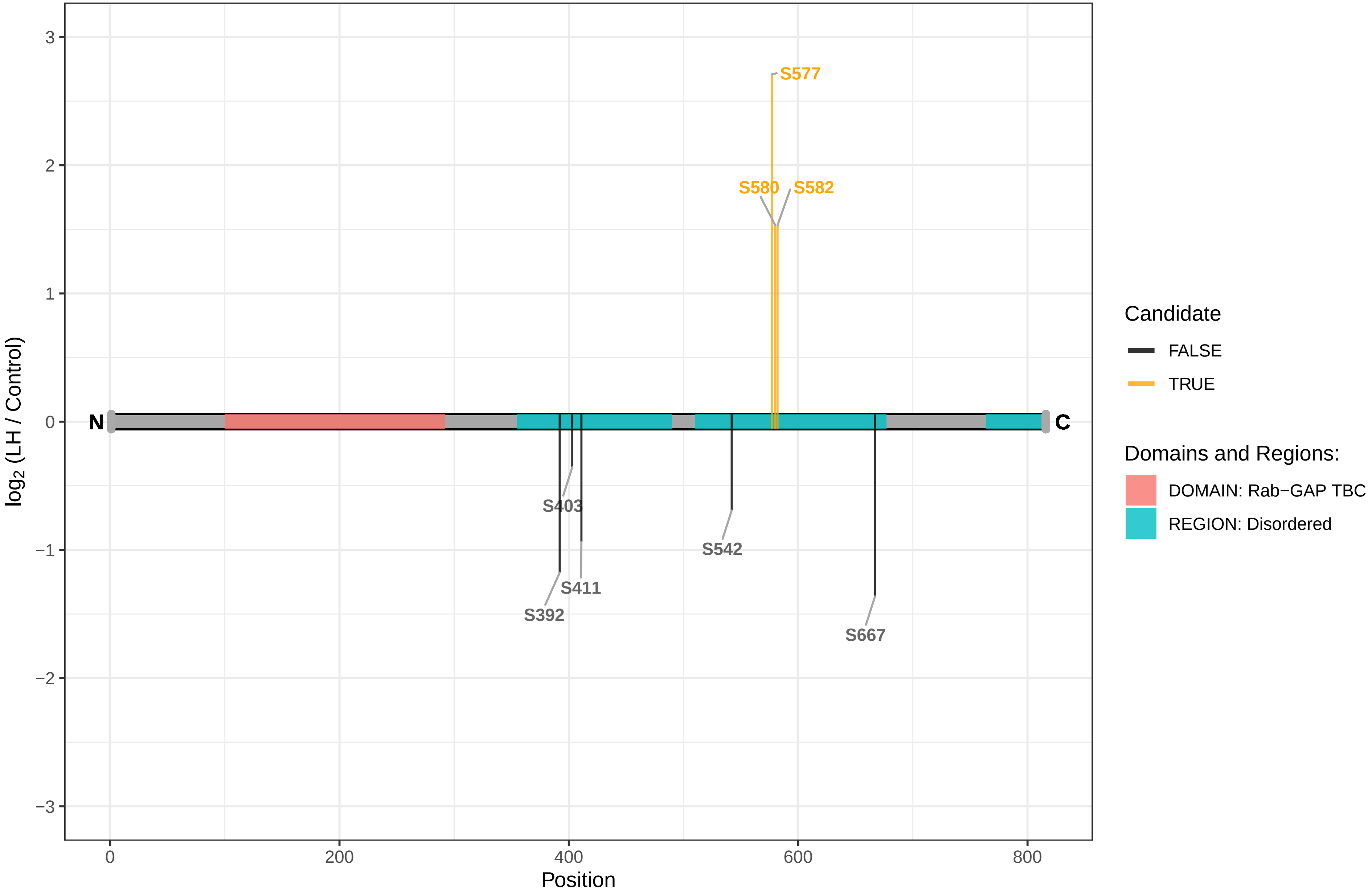

USP8

D3ZN39 Ubiquitin specific protease 8 (Predicted)

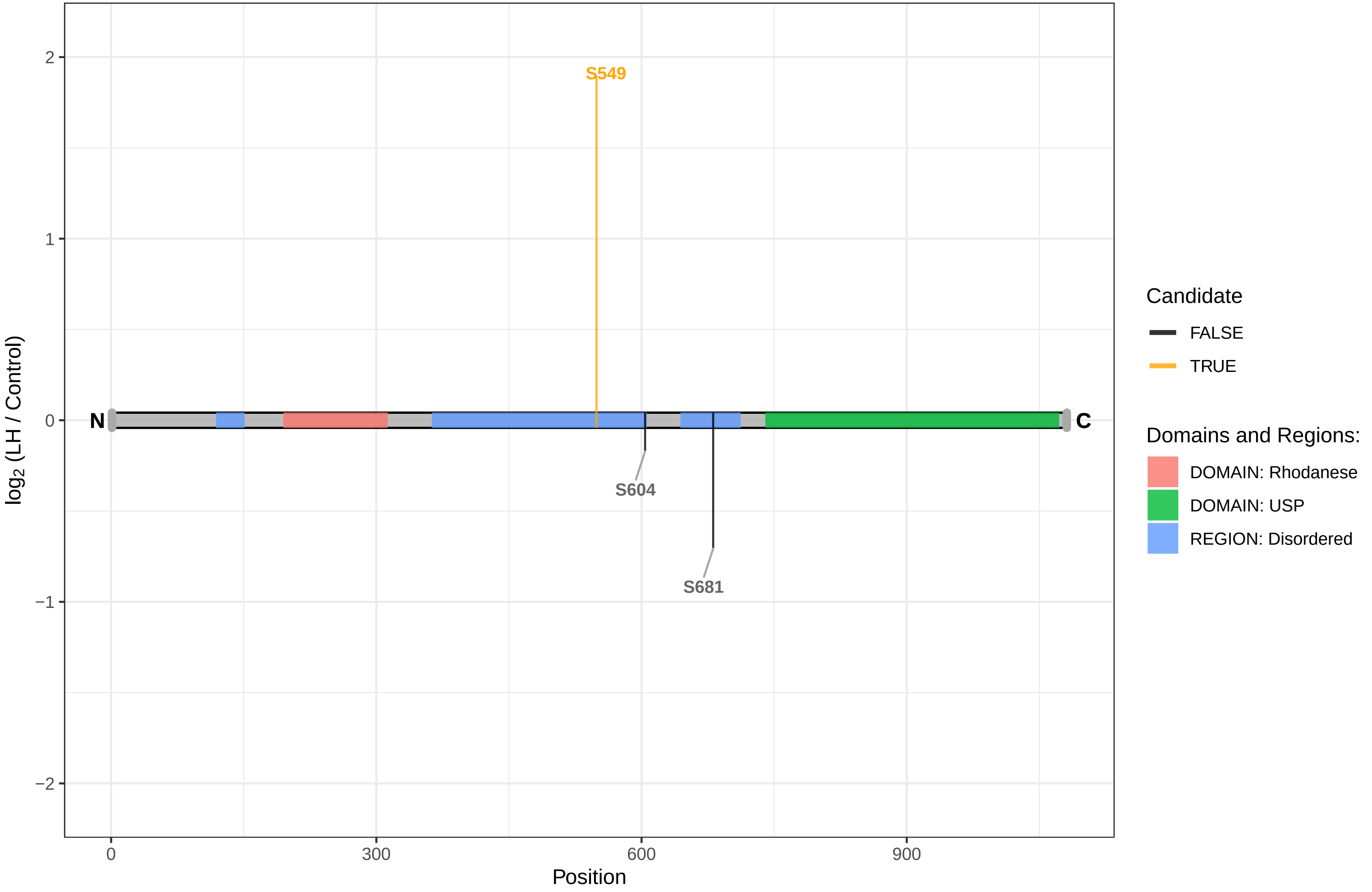

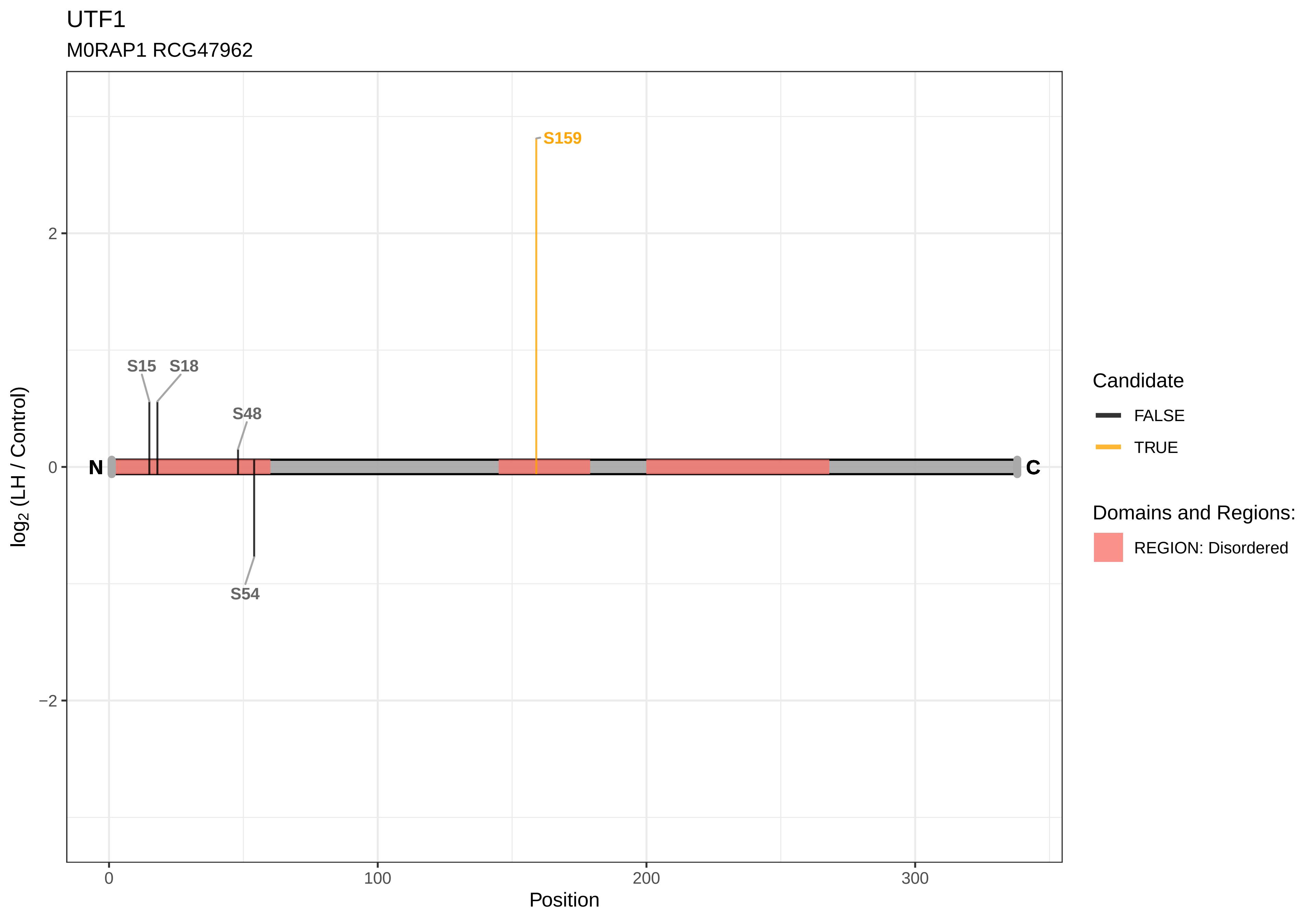
UTF1
M0RAP1 RCG47962
S159
S15
S18
S48
S54
N
C
2
0
−2
0
100
200
300
Position
log2 (LH / Control)
Candidate
FALSE
TRUE
Domains and Regions:
REGION: Disordered

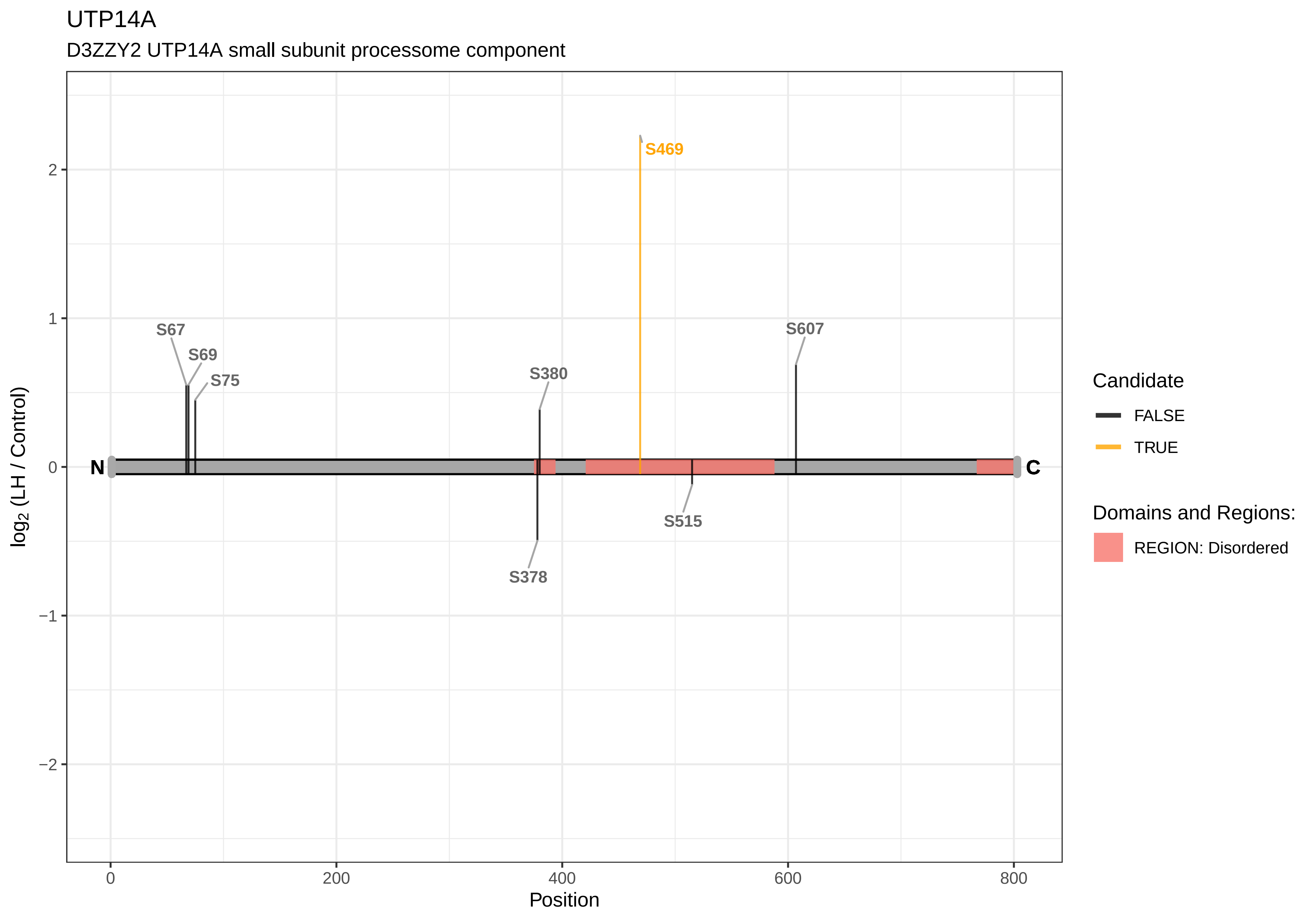
UTP14A
D3ZZY2 UTP14A small subunit processome component
S469
S67
S69
S75
S380
S607
S515
S378
N
C
log2 (LH / Control)
Position
0
200
400
600
800
2
1
0
−1
−2
Candidate
FALSE
TRUE
Domains and Regions:
REGION: Disordered

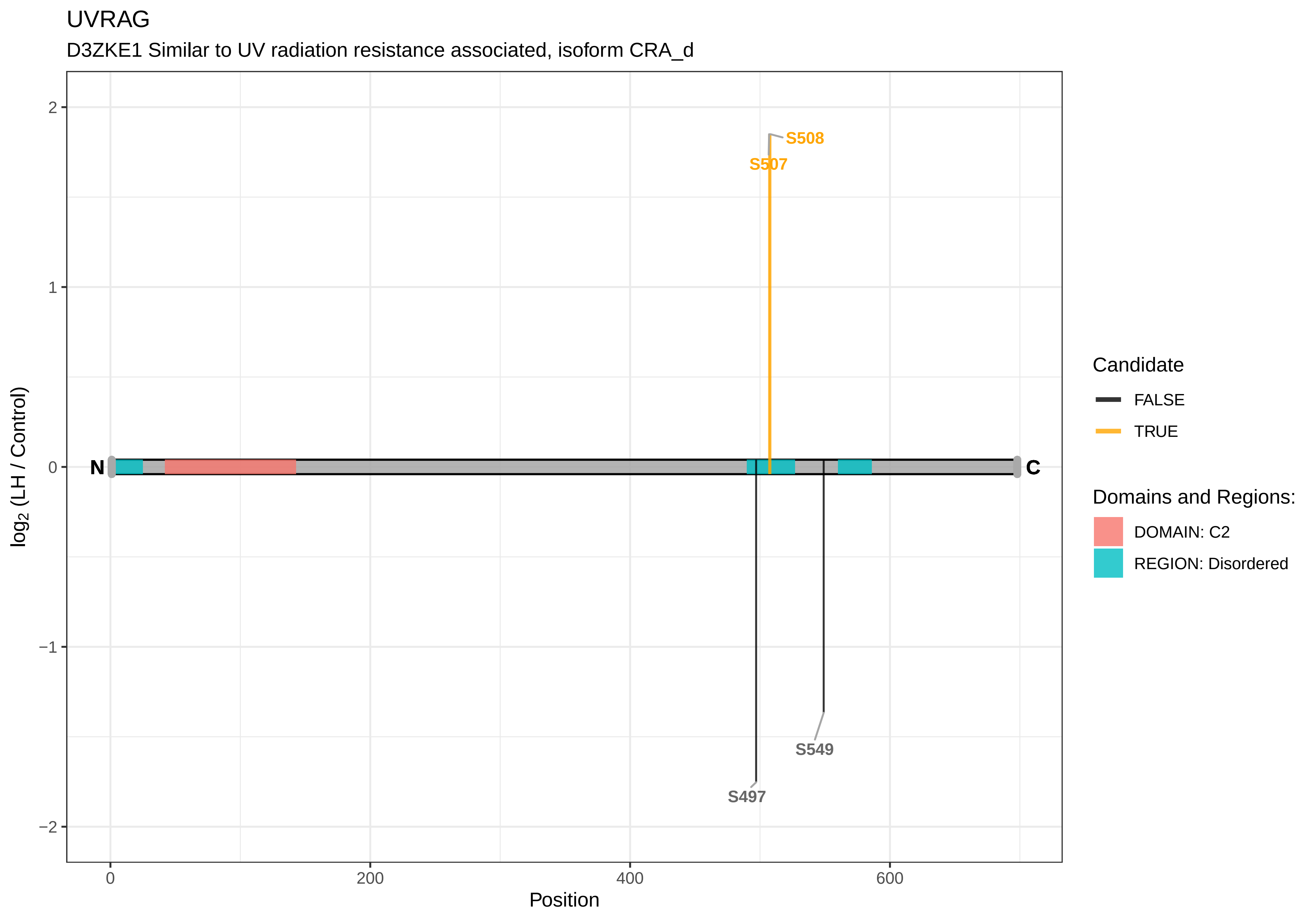

UVRAG
D3ZKE1 Similar to UV radiation resistance associated, isoform CRA_d
S508
S507
S549
S497
N
C
2
1
0
−1
−2
0
200
400
600
Position
log2 (LH / Control)
Candidate
FALSE
TRUE
Domains and Regions:
DOMAIN: C2
REGION: Disordered

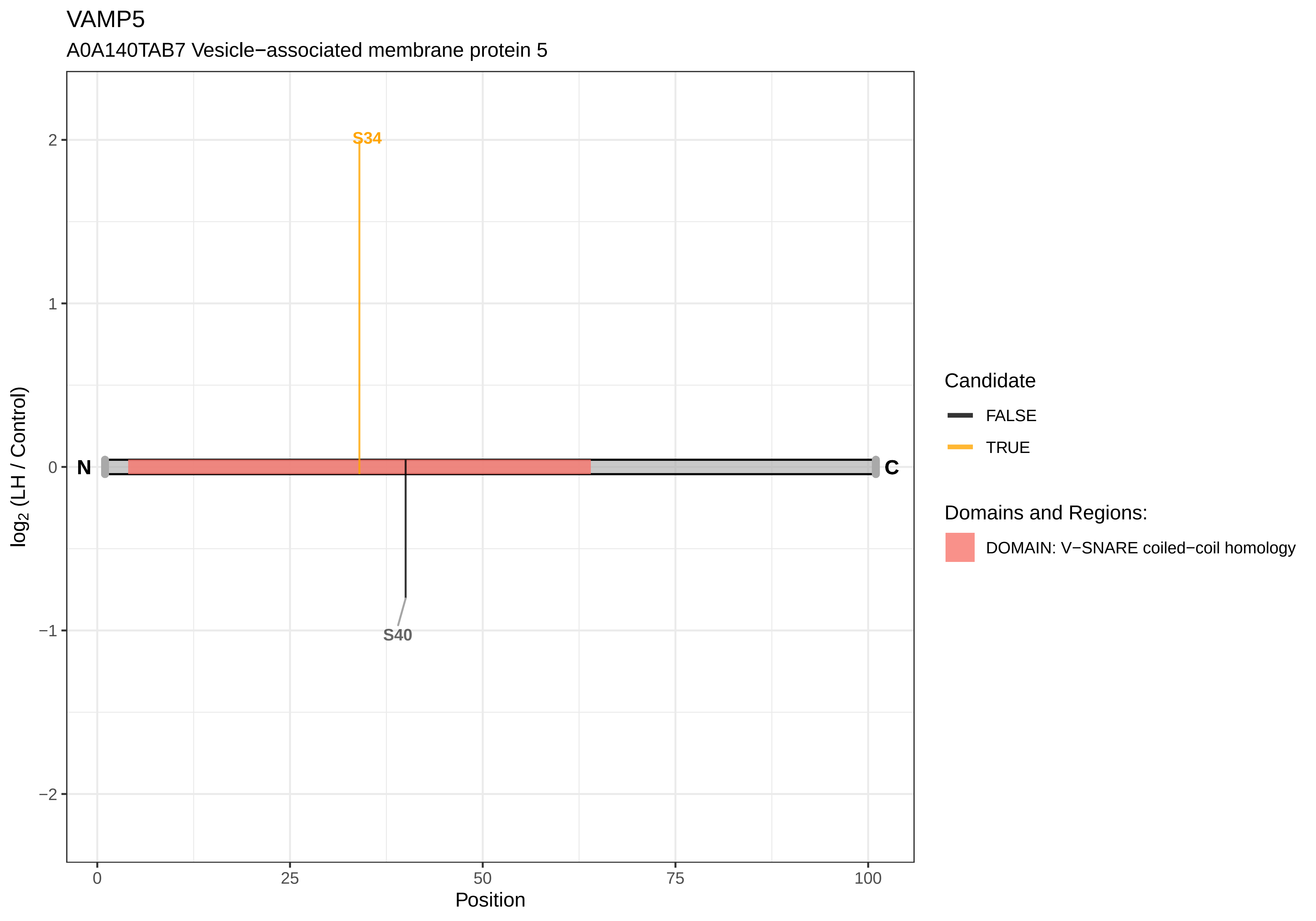

VAMP5
A0A140TAB7 Vesicle−associated membrane protein 5
S34
S40
N
C
2
1
0
−1
−2
0
25
50
75
100
Position
log2 (LH / Control)
Candidate
FALSE
TRUE
Domains and Regions:
DOMAIN: V−SNARE coiled−coil homology

#### VASP

F7EWC1 Vasodilator−stimulated phosphoprotein

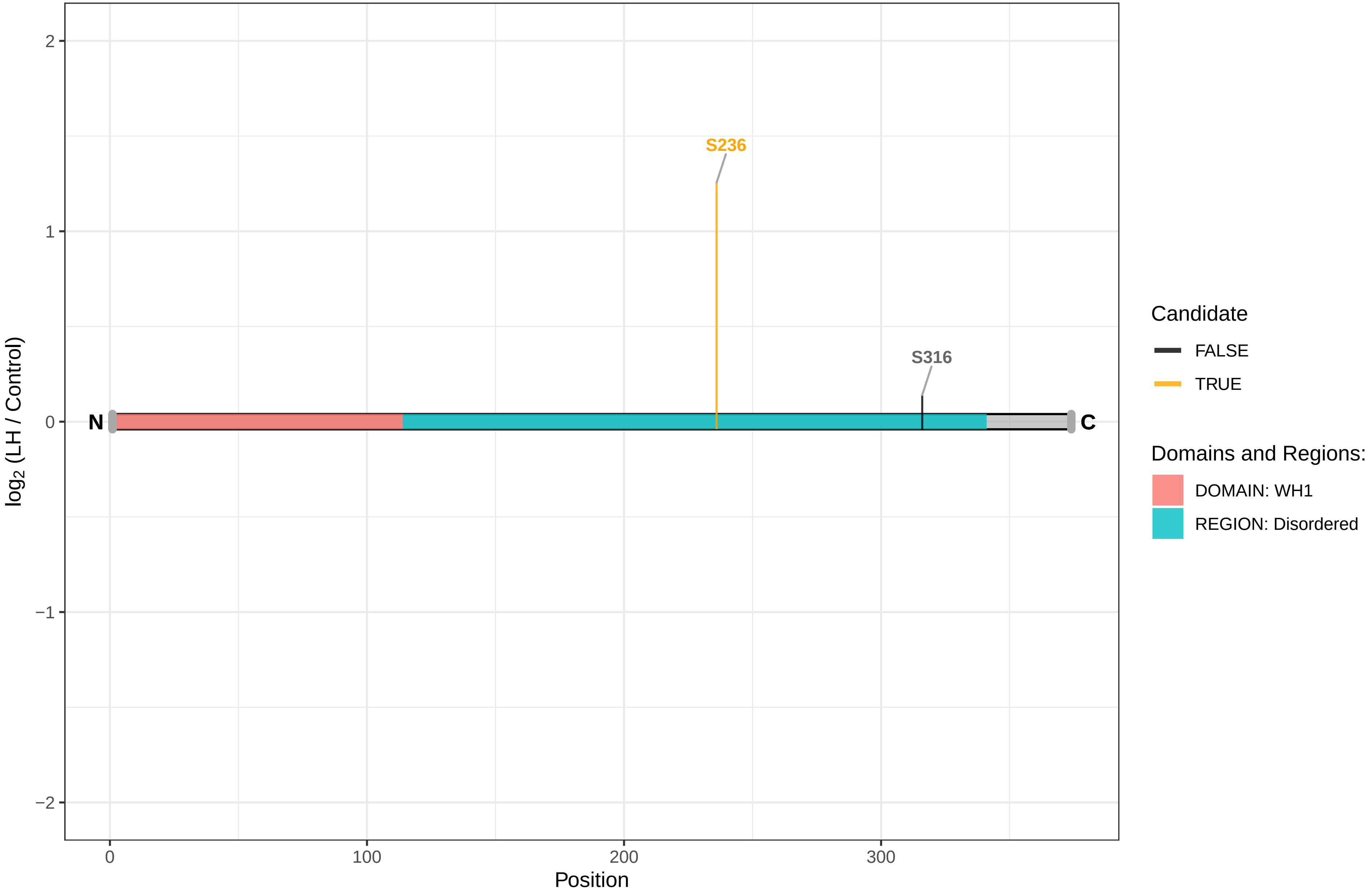

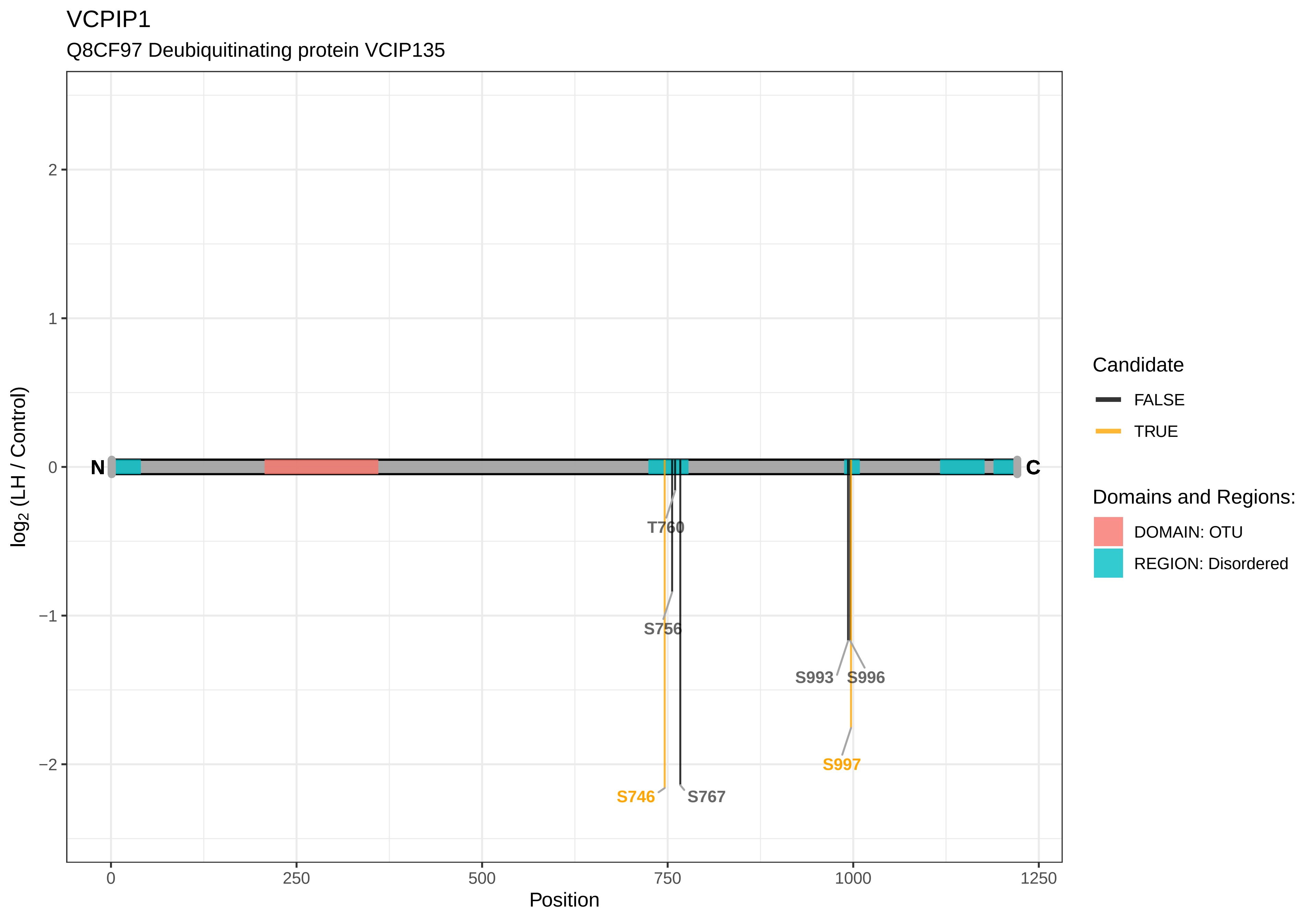

VCPIP1
Q8CF97 Deubiquitinating protein VCIP135
log2 (LH / Control)
2
1
0
−1
−2
N
C
T760
S756
S993
S996
S997
S746
S767
0
250
500
750
1000
1250
Position
Candidate
FALSE
TRUE
Domains and Regions:
DOMAIN: OTU
REGION: Disordered

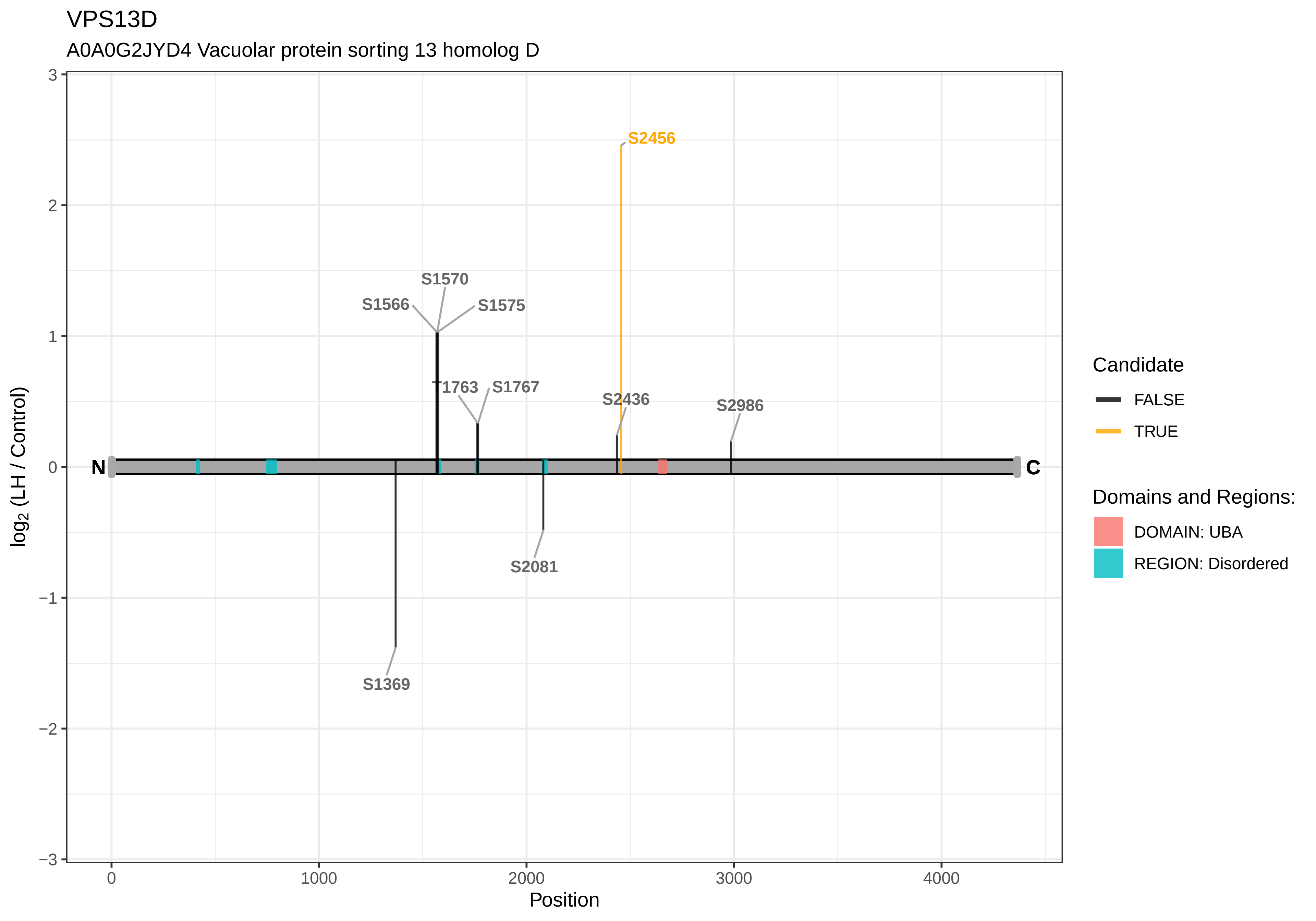
VPS13D
A0A0G2JYD4 Vacuolar protein sorting 13 homolog D
S2456
S1570
S1566
S1575
T1763
S1767
S2436
S2986
S2081
S1369
N
C
3
2
1
0
−1
−2
−3
log2 (LH / Control)
0
1000
2000
3000
4000
Position
Candidate
FALSE
TRUE
Domains and Regions:
DOMAIN: UBA
REGION: Disordered

VPS50

F1LSG8 Syndetin

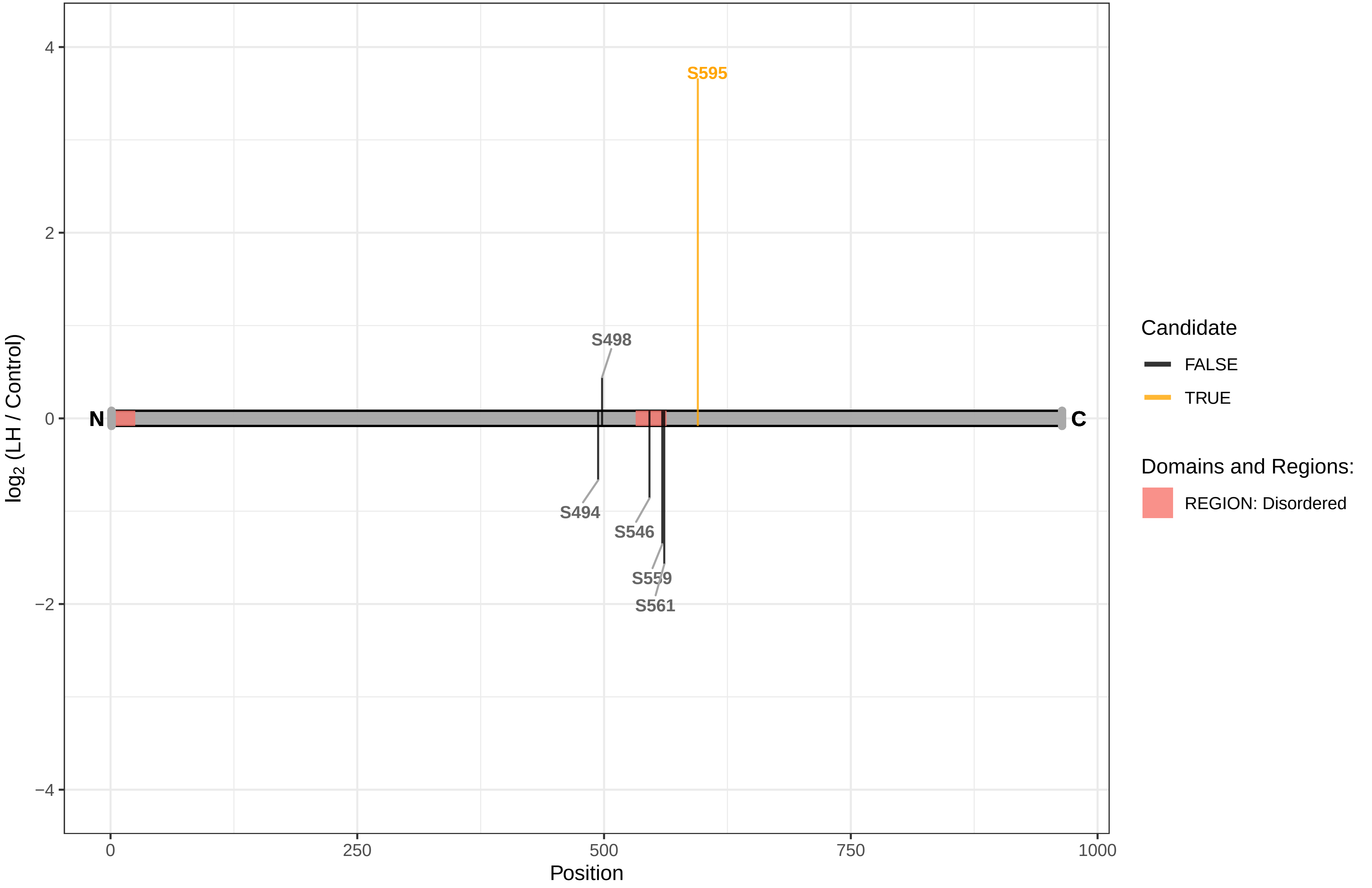

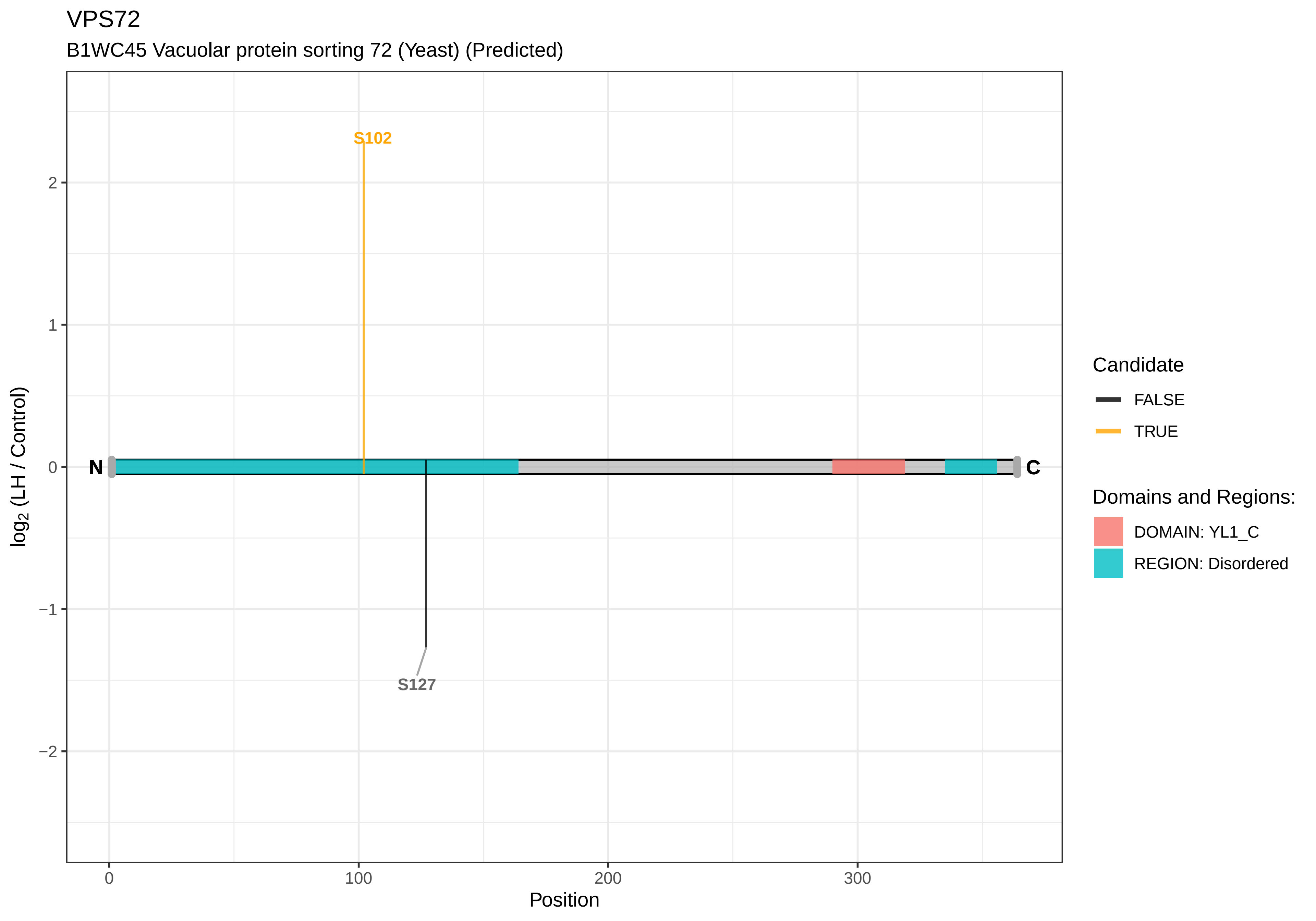
VPS72
B1WC45 Vacuolar protein sorting 72 (Yeast) (Predicted)
S102
S127
N
C
2
1
0
−1
−2
0
100
200
300
Position
log2 (LH / Control)
Candidate
FALSE
TRUE
Domains and Regions:
DOMAIN: YL1_C
REGION: Disordered

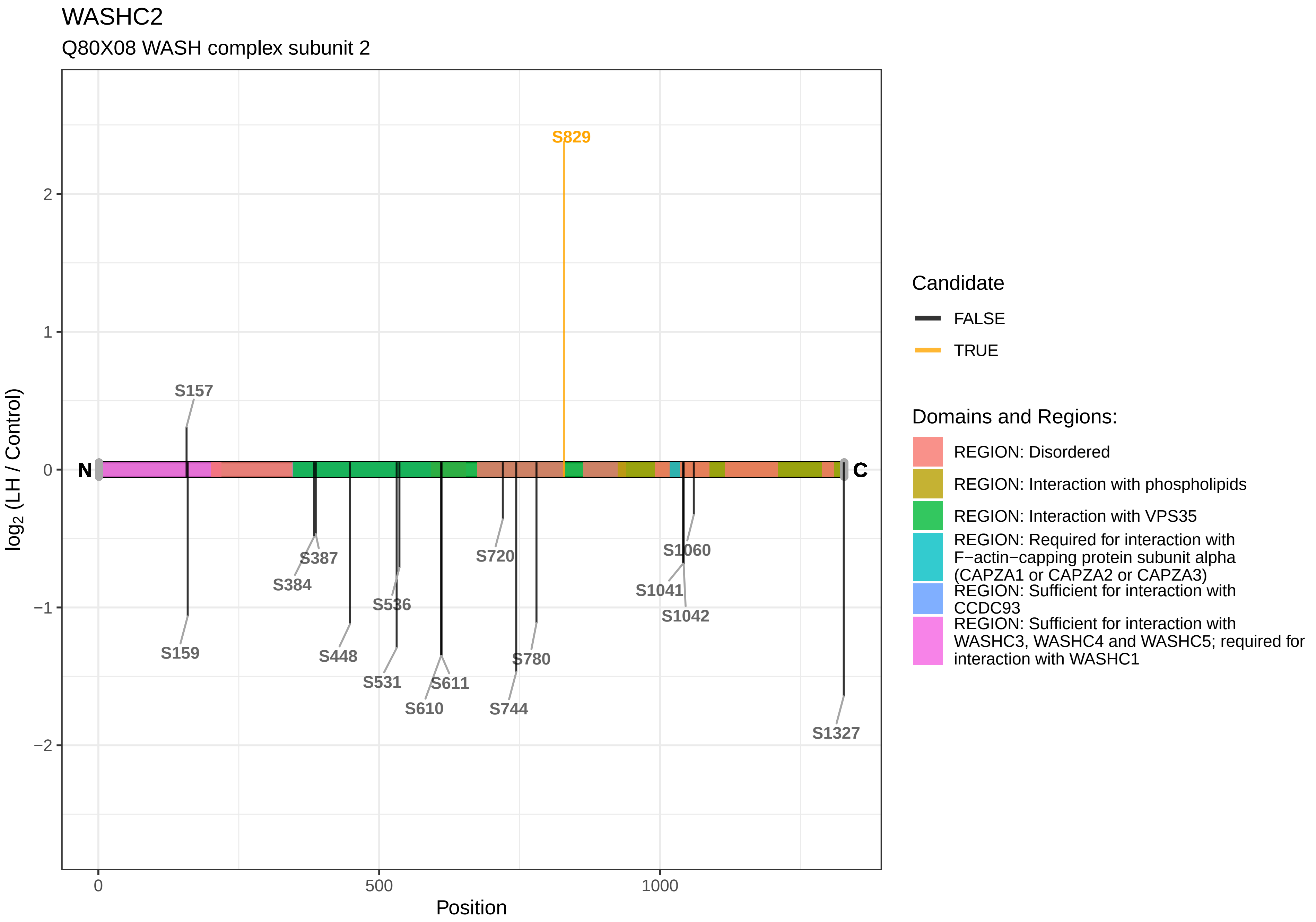

WASHC2
Q80X08 WASH complex subunit 2
S829
S157
S159
S387
S384
S448
S536
S531
S611
S610
S720
S780
S744
S1060
S1041
S1042
S1327
N
C
2
1
0
−1
−2
0
500
1000
Position
log2 (LH / Control)
Candidate
FALSE
TRUE
Domains and Regions:
REGION: Disordered
REGION: Interaction with phospholipids
REGION: Interaction with VPS35
REGION: Required for interaction with F−actin−capping protein subunit alpha (CAPZA1 or CAPZA2 or CAPZA3)
REGION: Sufficient for interaction with CCDC93
REGION: Sufficient for interaction with WASHC3, WASHC4 and WASHC5; required for interaction with WASHC1

WDR33

F1LT09 WD repeat domain 33

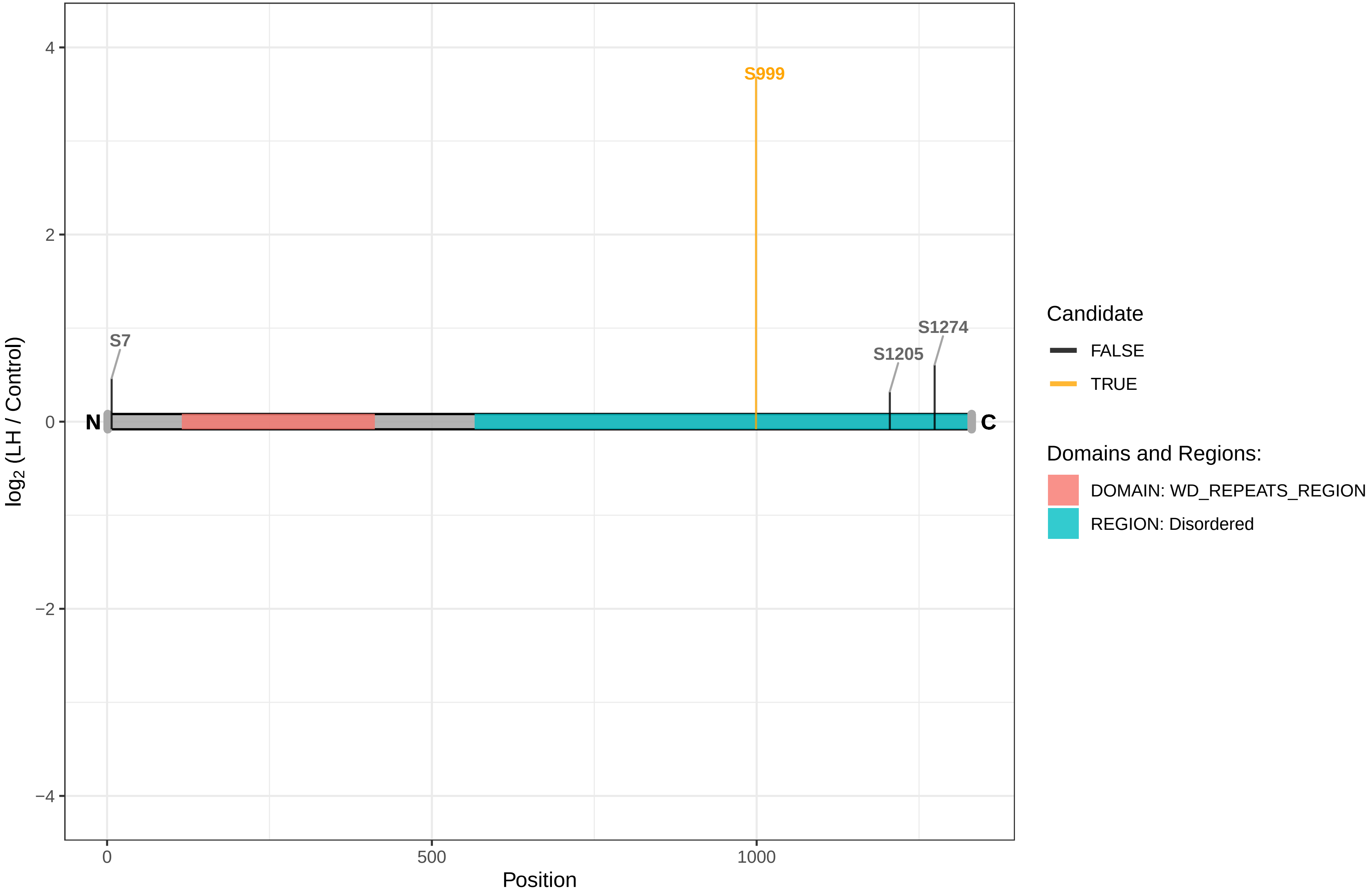

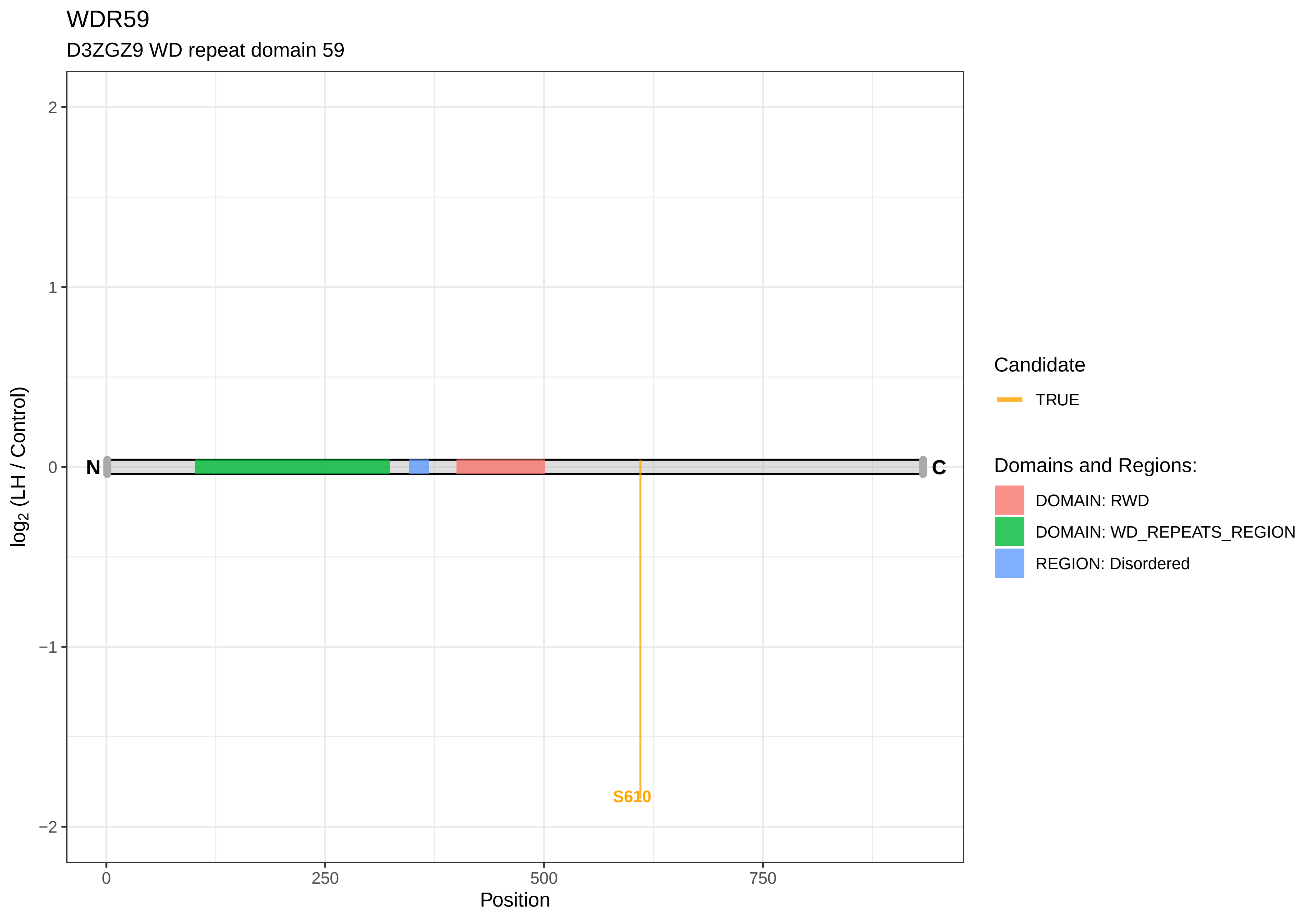

WDR59
D3ZGZ9 WD repeat domain 59
log2 (LH / Control)
2
1
0
−1
−2
N
C
S610
0
250
500
750
Position
Candidate
TRUE
Domains and Regions:
DOMAIN: RWD
DOMAIN: WD_REPEATS_REGION
REGION: Disordered

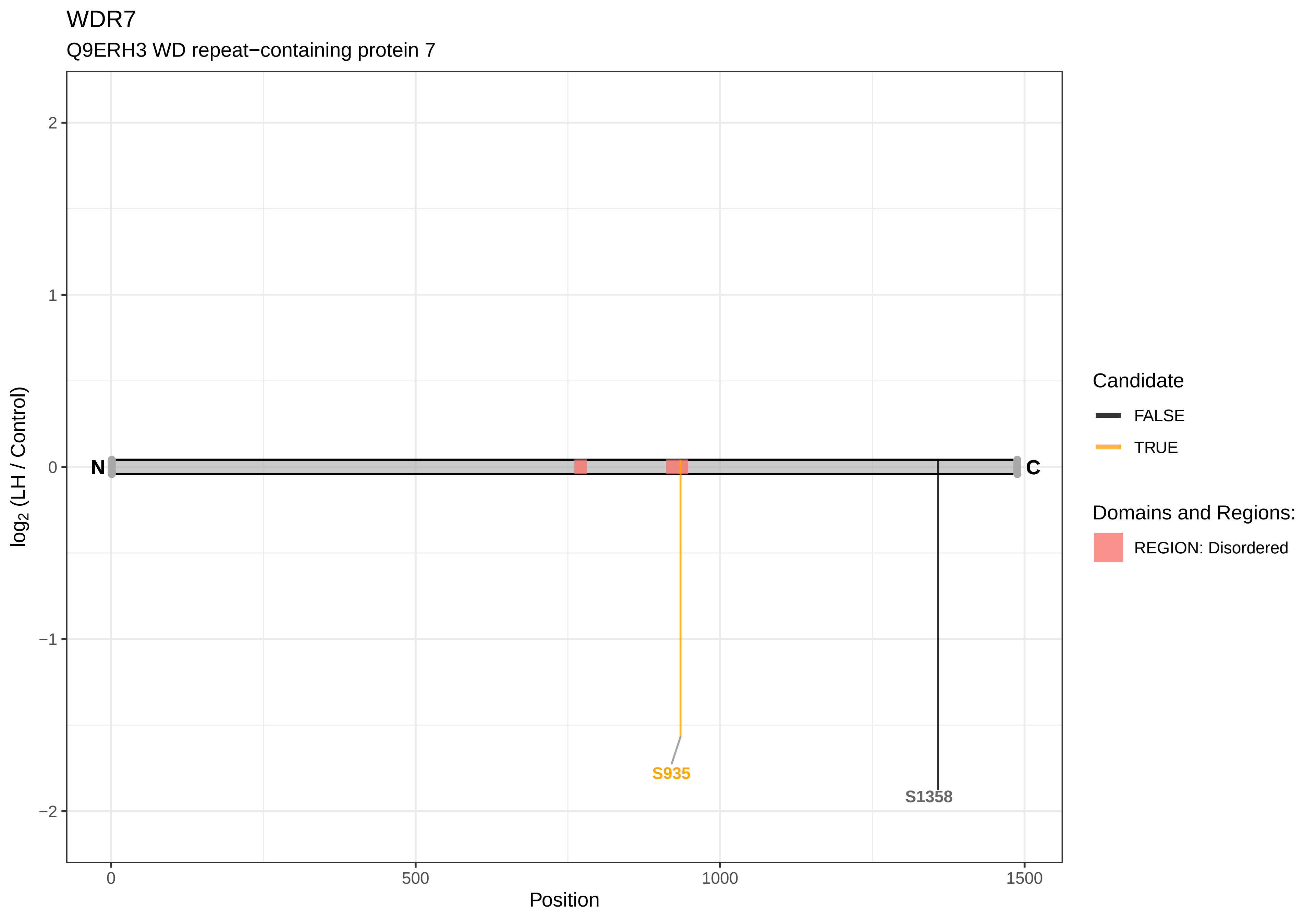
WDR7
Q9ERH3 WD repeat−containing protein 7
log2 (LH / Control)
2
1
0
−1
−2
N
C
S935
S1358
0
500
1000
1500
Position
Candidate
FALSE
TRUE
Domains and Regions:
REGION: Disordered

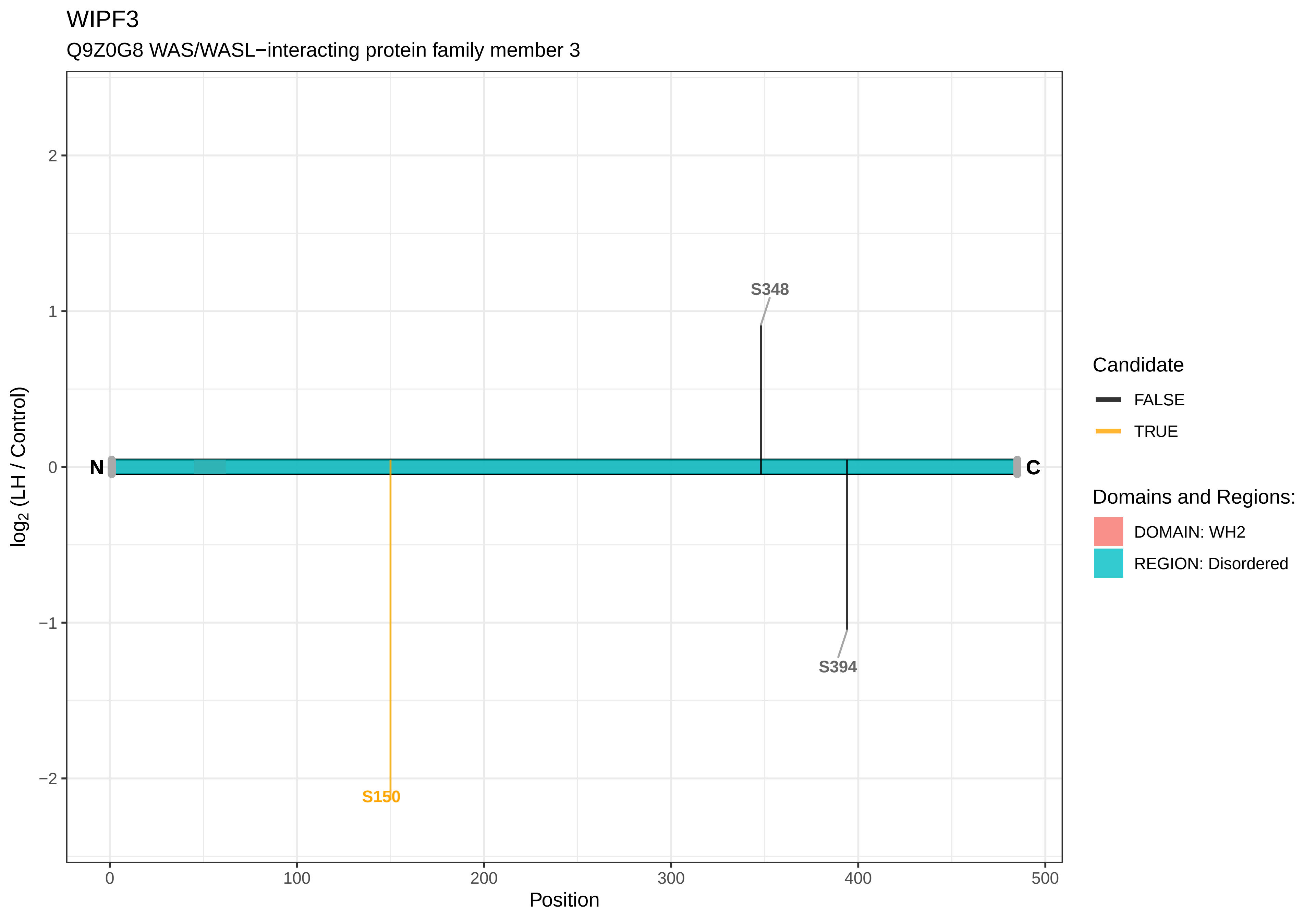

WIPF3
Q9Z0G8 WAS/WASL−interacting protein family member 3
log2 (LH / Control)
Position
N
C
S348
S394
S150
Candidate
FALSE
TRUE
Domains and Regions:
DOMAIN: WH2
REGION: Disordered
2
1
0
−1
−2
0
100
200
300
400
500

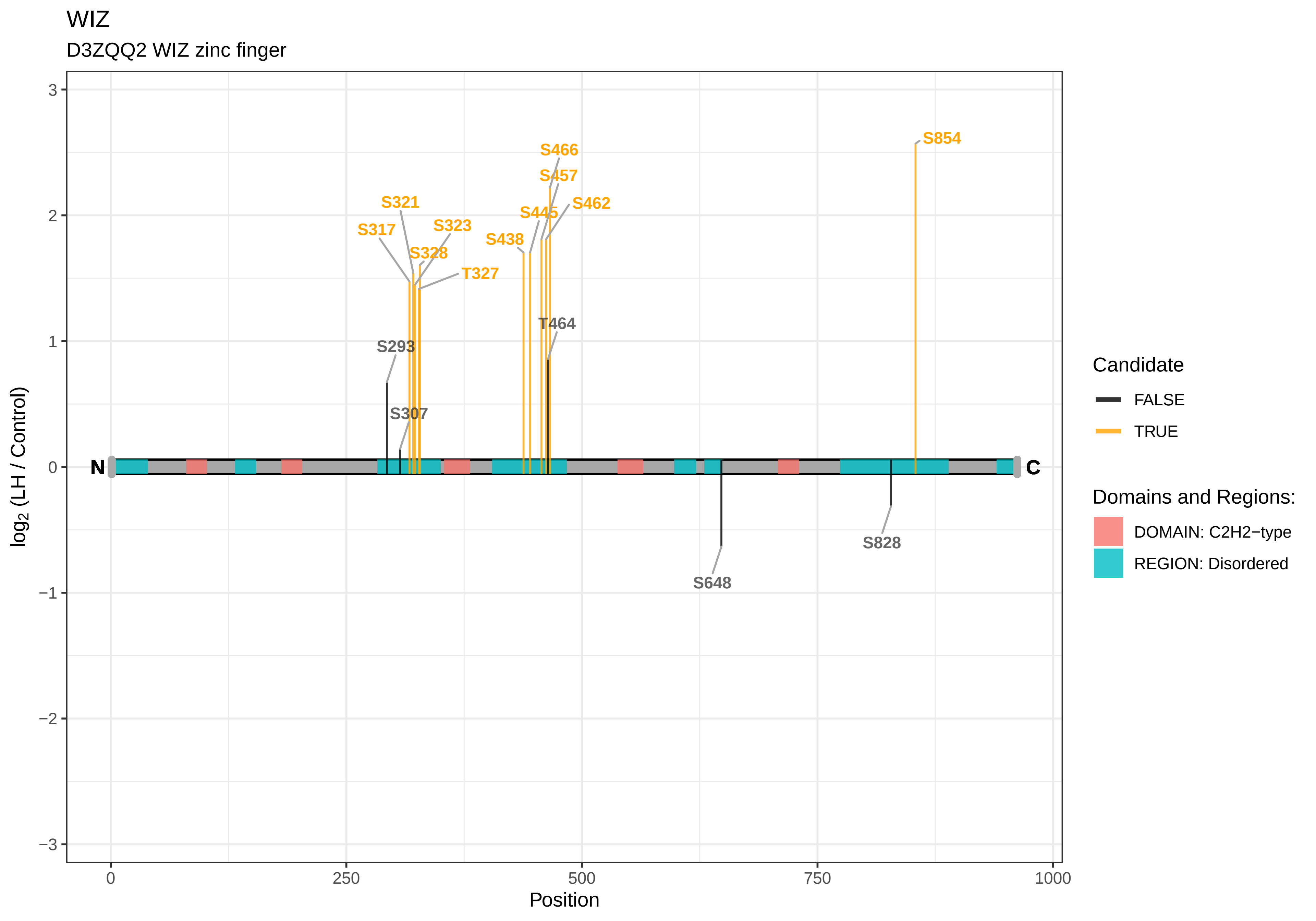
WIZ
D3ZQQ2 WIZ zinc finger
S854
S466
S457
S321
S462
S445
S323
S317
S438
S328
T327
T464
S293
S307
N
C
S828
S648
log2 (LH / Control)
Position
0
250
500
750
1000
3
2
1
0
−1
−2
−3
Candidate
FALSE
TRUE
Domains and Regions:
DOMAIN: C2H2−type
REGION: Disordered

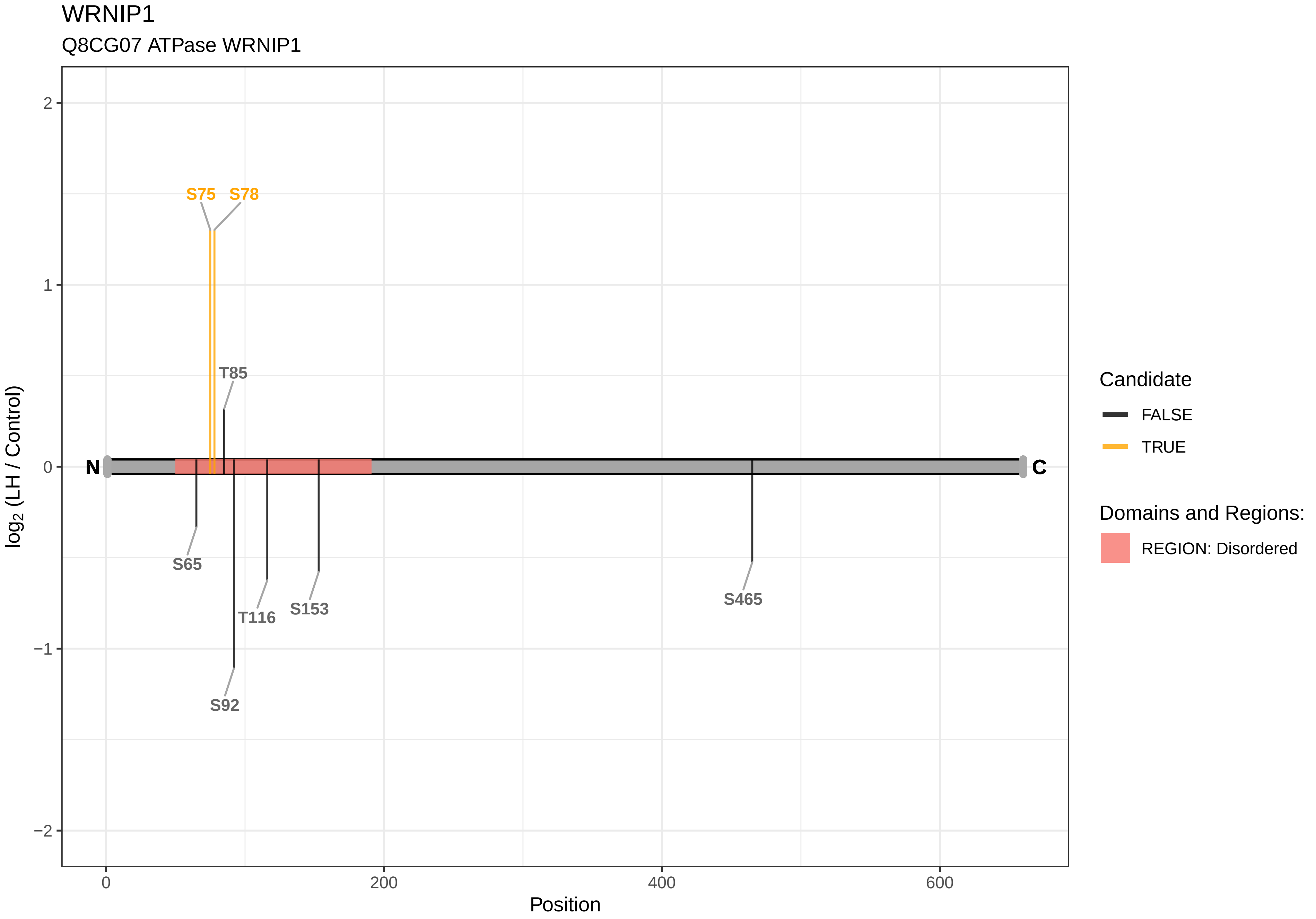

WRNIP1
Q8CG07 ATPase WRNIP1
S75
S78
T85
S65
T116
S153
S92
S465
N
C
2
1
0
−1
−2
0
200
400
600
Position
log2 (LH / Control)
Candidate
FALSE
TRUE
Domains and Regions:
REGION: Disordered

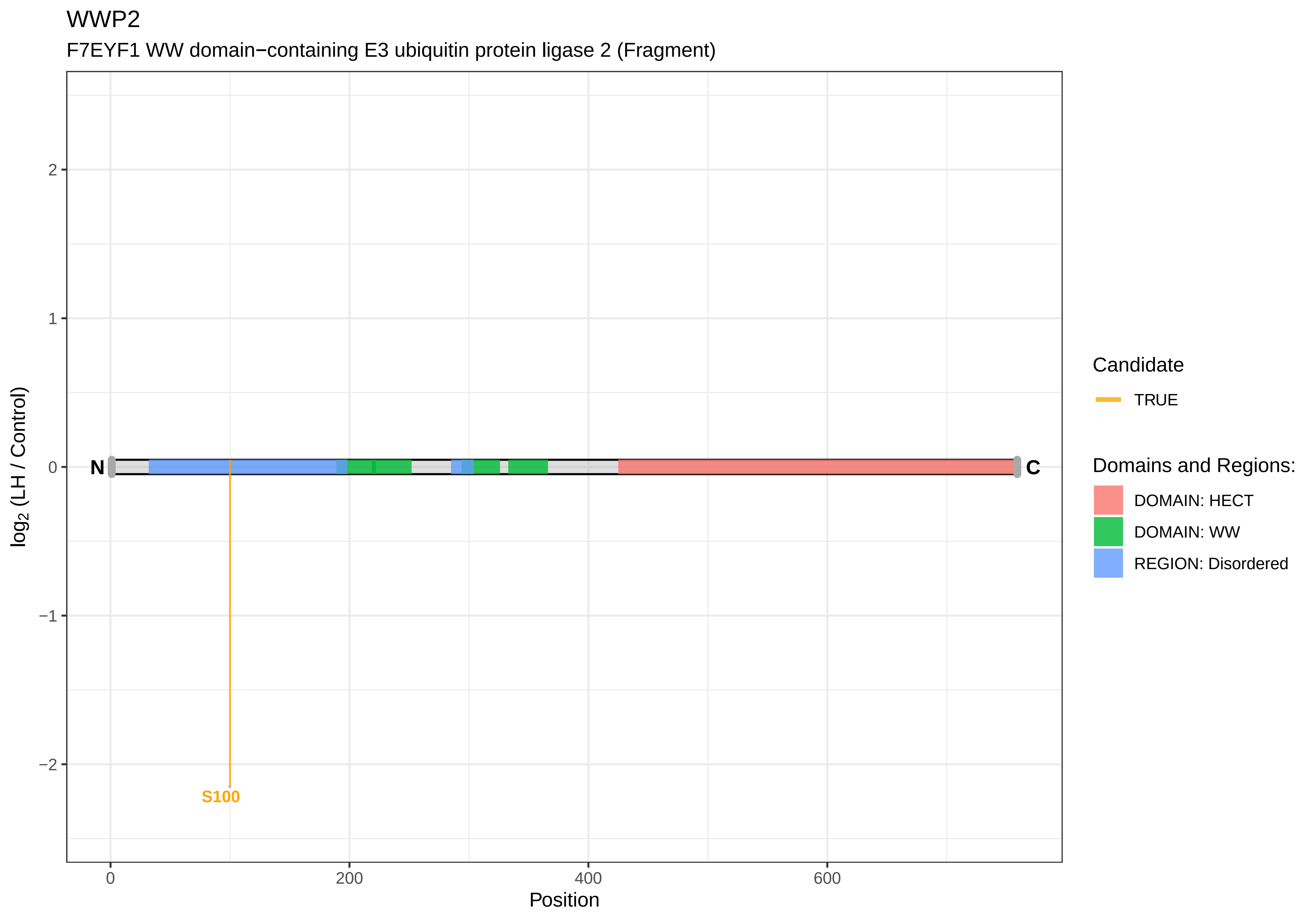
WWP2
F7EYF1 WW domain−containing E3 ubiquitin protein ligase 2 (Fragment)
log2 (LH / Control)
Position
N
C
S100
Candidate
TRUE
Domains and Regions:
DOMAIN: HECT
DOMAIN: WW
REGION: Disordered
2
1
0
−1
−2
0
200
400
600

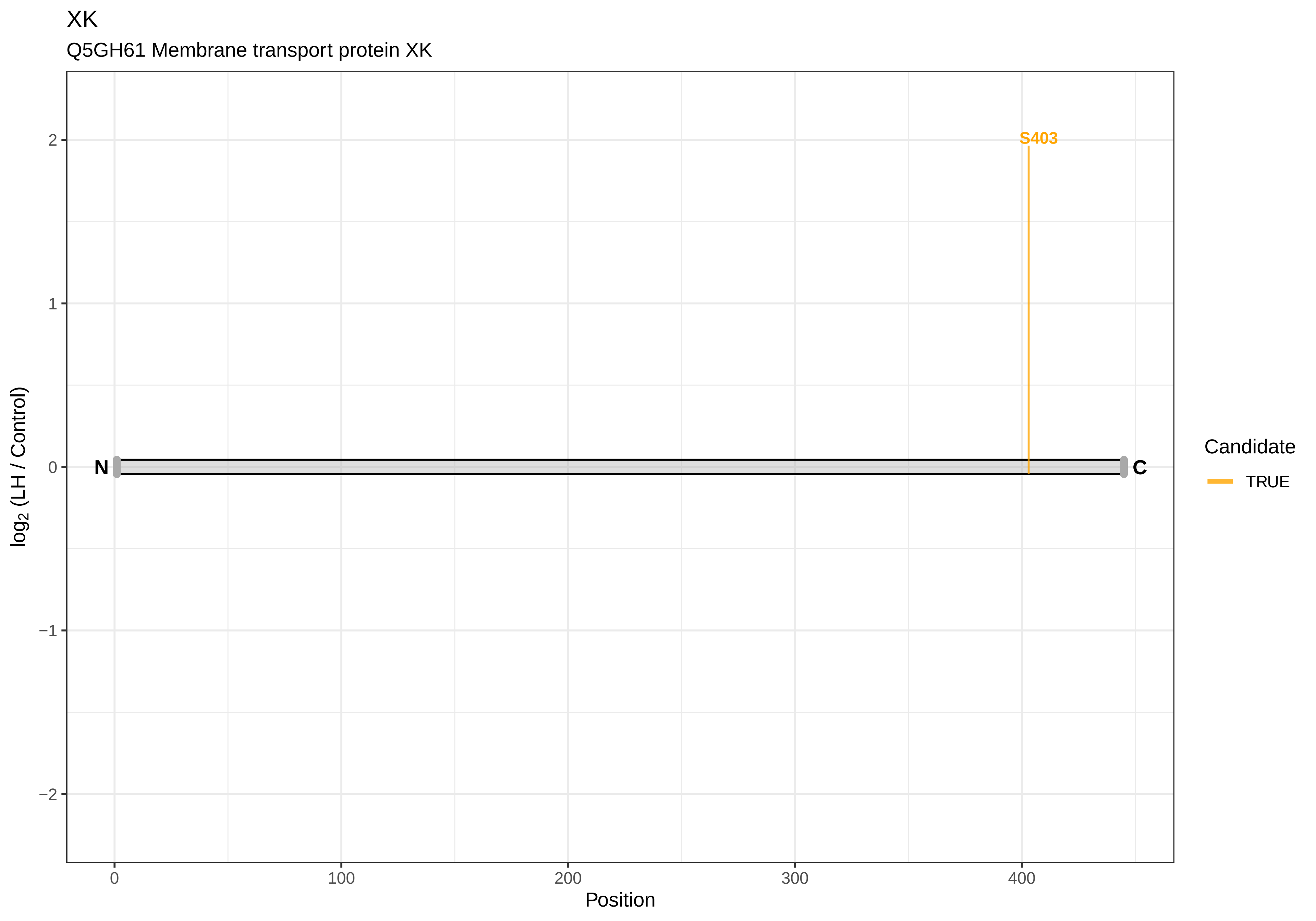
XK
Q5GH61 Membrane transport protein XK
S403
N
C
2
1
0
−1
−2
0
100
200
300
400
Position
log2 (LH / Control)
Candidate
TRUE

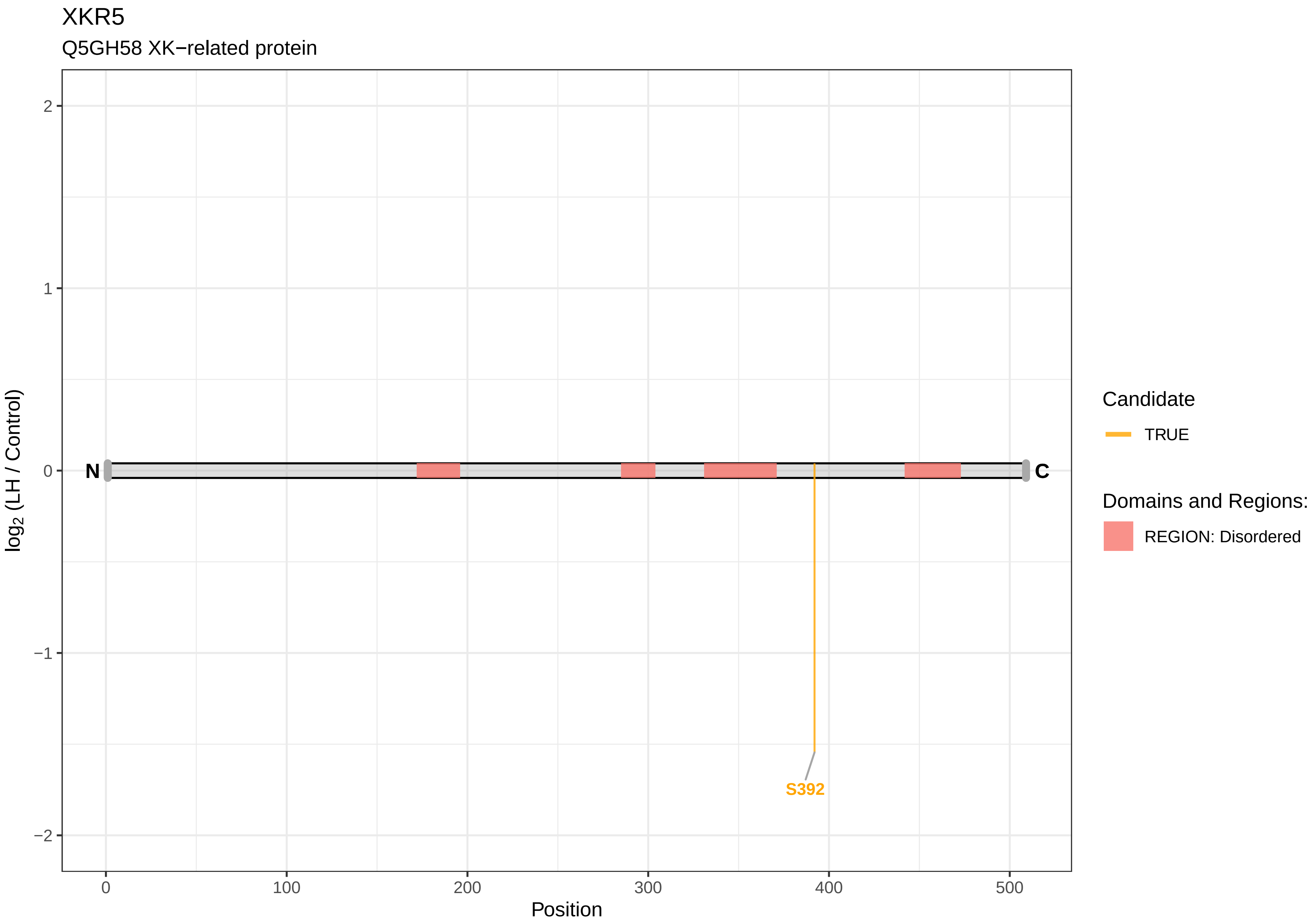

XKR5
Q5GH58 XK−related protein
2
1
0
−1
−2
log2 (LH / Control)
N
C
S392
0
100
200
300
400
500
Position
Candidate
TRUE
Domains and Regions:
REGION: Disordered

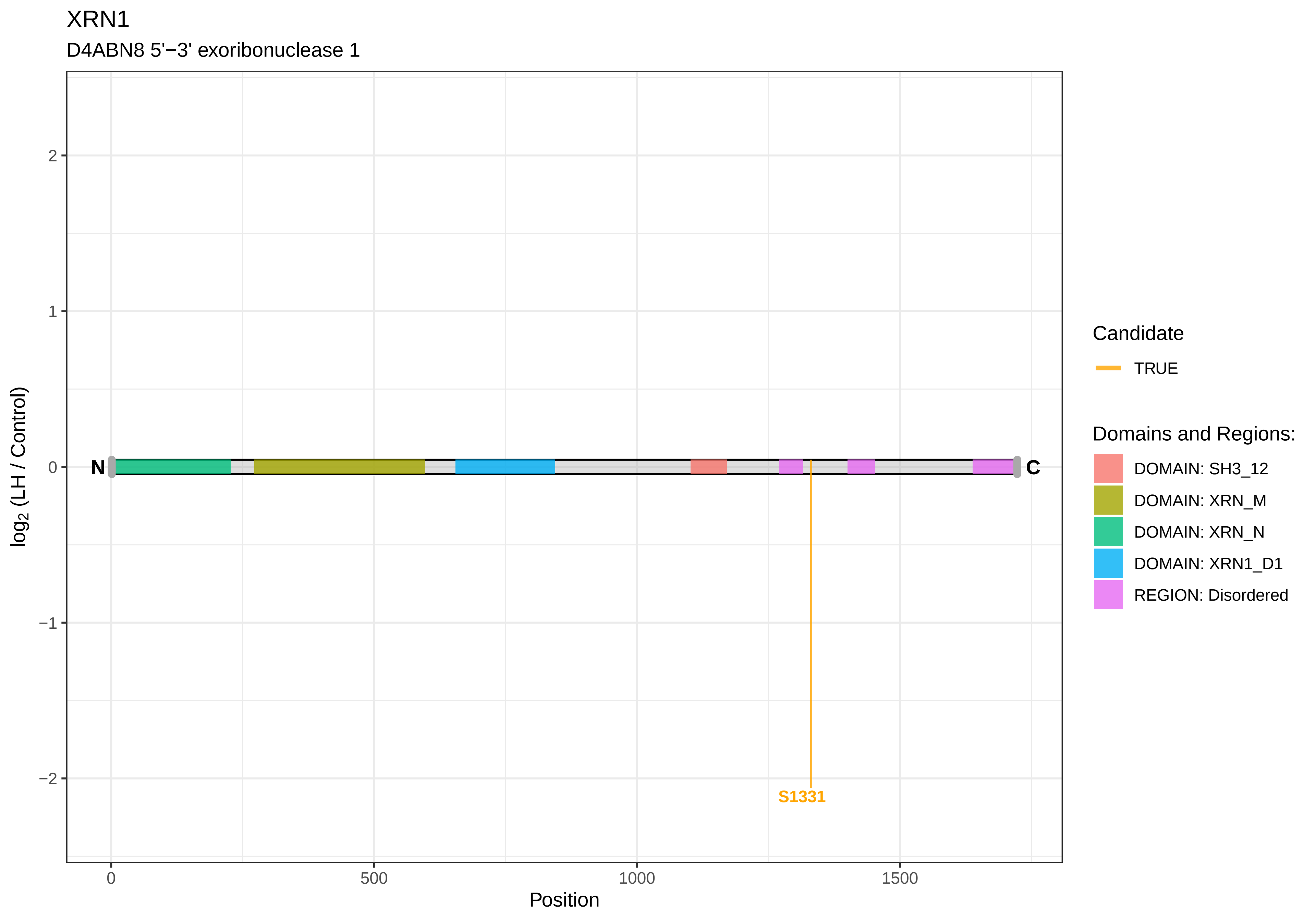
XRN1
D4ABN8 5'−3' exoribonuclease 1
log2 (LH / Control)
2
1
0
−1
−2
N
C
S1331
0
500
1000
1500
Position
Candidate
TRUE
Domains and Regions:
DOMAIN: SH3_12
DOMAIN: XRN_M
DOMAIN: XRN_N
DOMAIN: XRN1_D1
REGION: Disordered

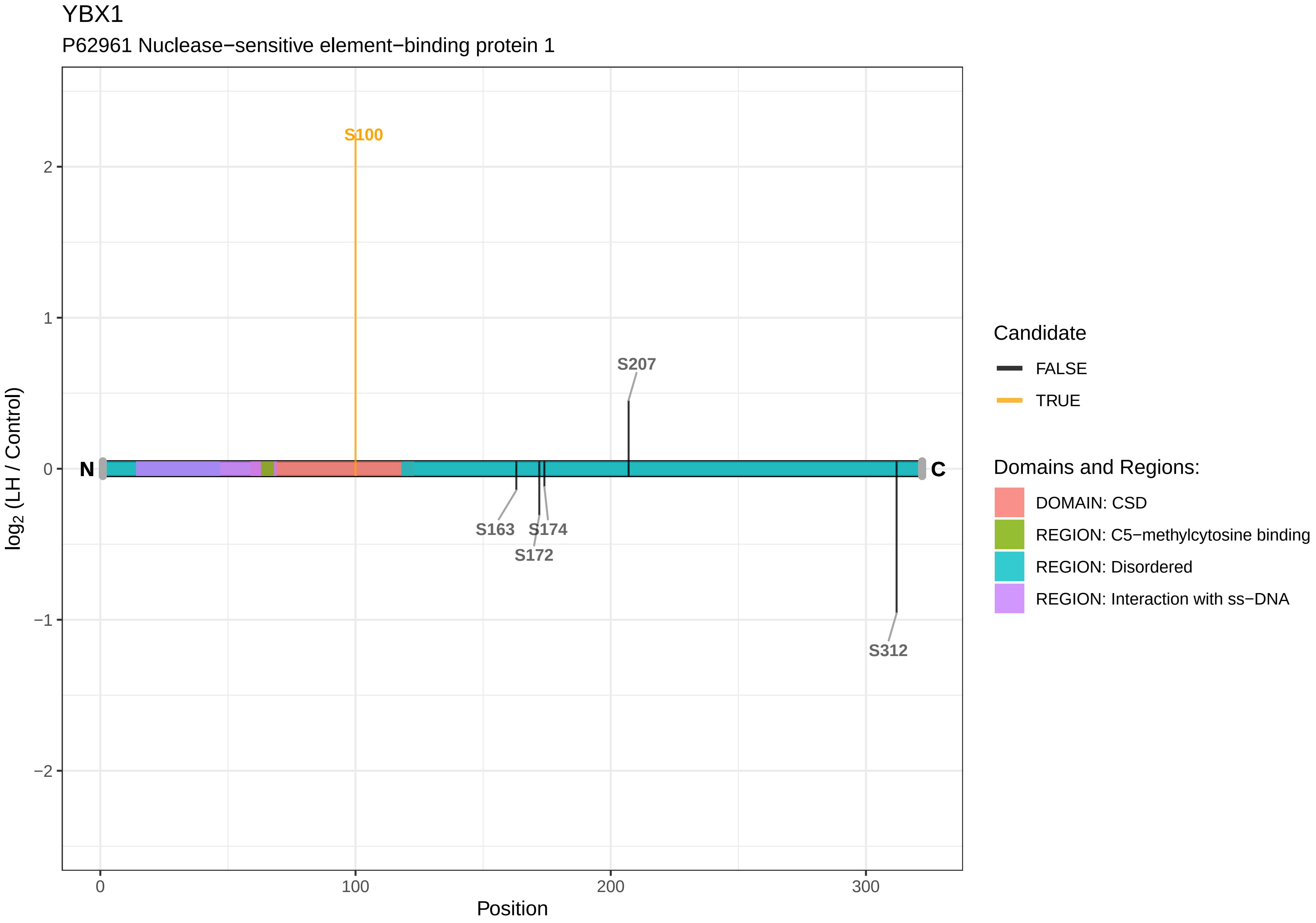

YBX1
P62961 Nuclease−sensitive element−binding protein 1
S100
S207
S163
S174
S172
S312
N
C
2
1
0
−1
−2
0
100
200
300
Position
log2 (LH / Control)
Candidate
FALSE
TRUE
Domains and Regions:
DOMAIN: CSD
REGION: C5−methylcytosine binding
REGION: Disordered
REGION: Interaction with ss−DNA

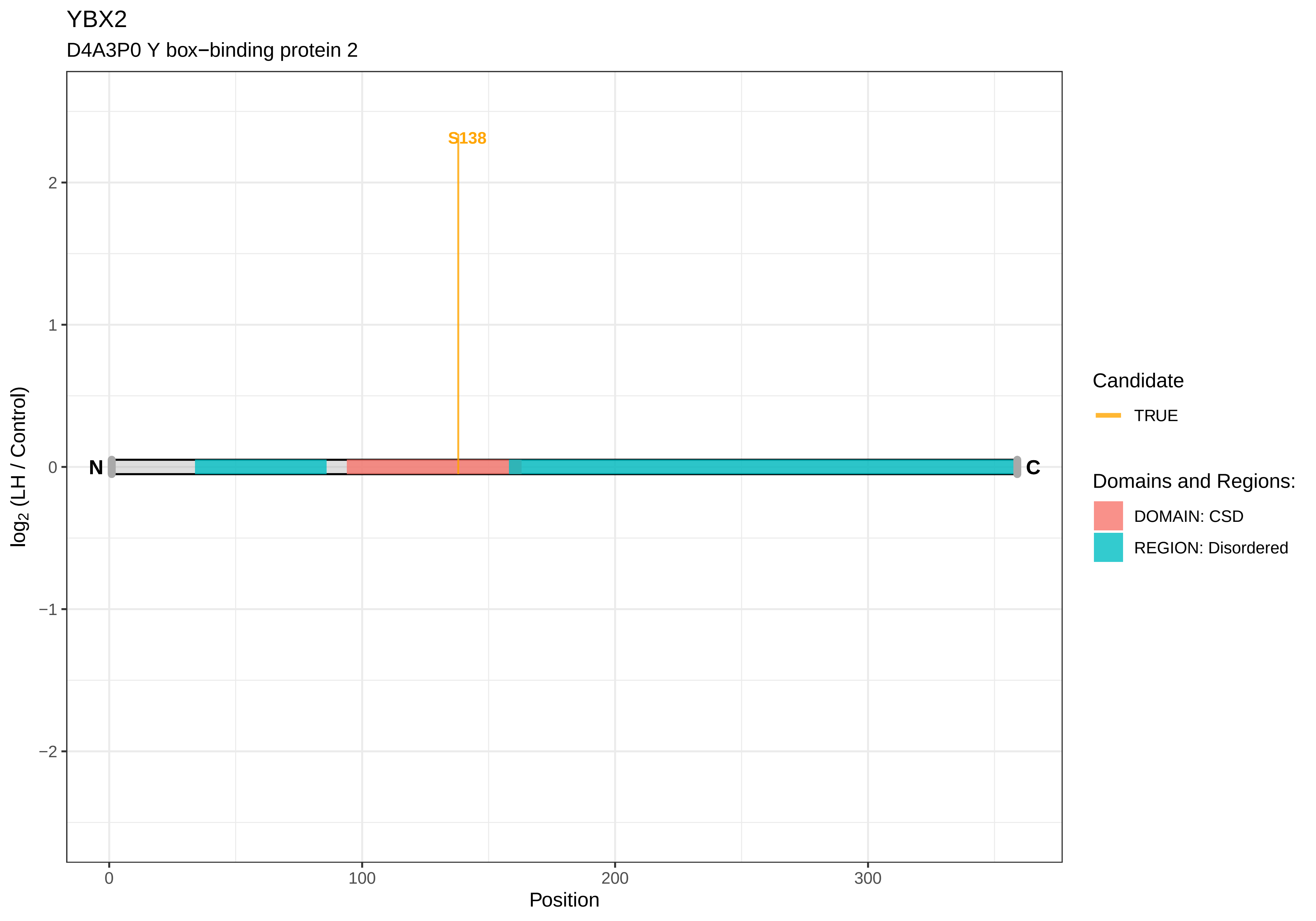
YBX2
D4A3P0 Y box−binding protein 2
S138
N
C
2
1
0
−1
−2
log2 (LH / Control)
0
100
200
300
Position
Candidate
TRUE
Domains and Regions:
DOMAIN: CSD
REGION: Disordered

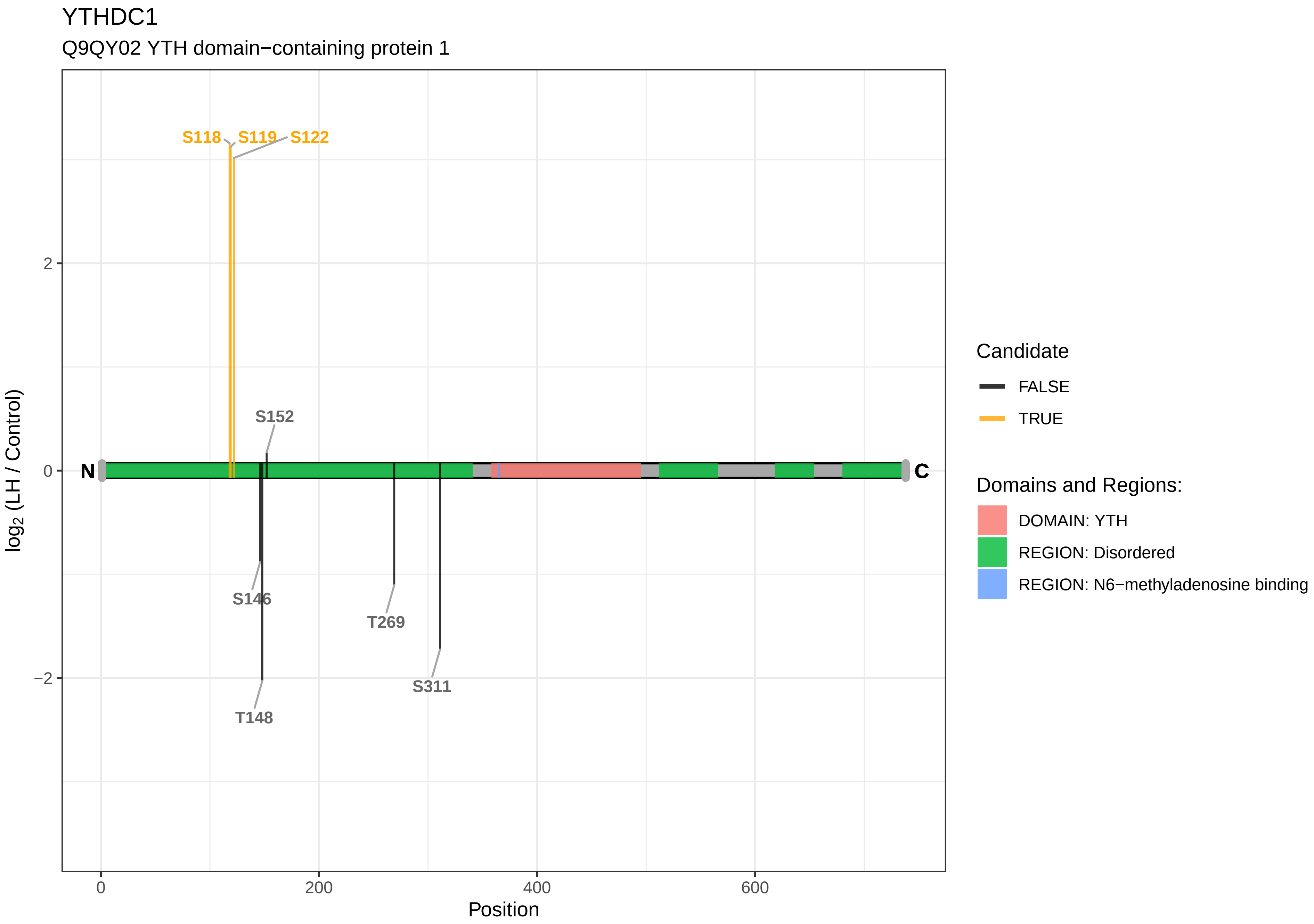
YTHDC1
Q9QY02 YTH domain−containing protein 1
S118
S119
S122
S152
S146
T269
S311
T148
N
C
log2 (LH / Control)
2
0
−2
0
200
400
600
Position
Candidate
FALSE
TRUE
Domains and Regions:
DOMAIN: YTH
REGION: Disordered
REGION: N6−methyladenosine binding

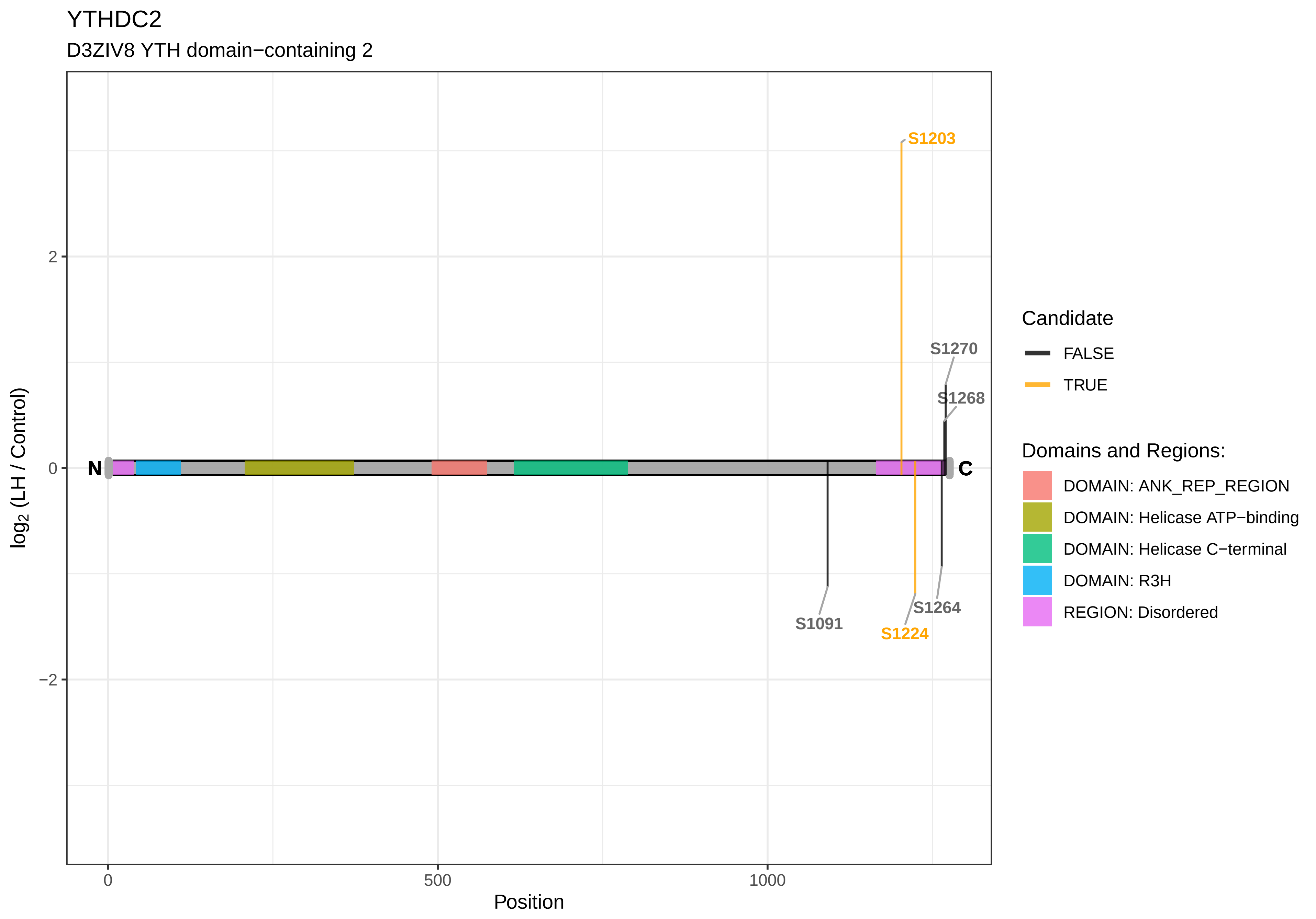

YTHDC2
D3ZIV8 YTH domain−containing 2
S1203
S1270
S1268
S1091
S1224
S1264
N
C
2
0
−2
0
500
1000
Position
log2 (LH / Control)
Candidate
FALSE
TRUE
Domains and Regions:
DOMAIN: ANK_REP_REGION
DOMAIN: Helicase ATP−binding
DOMAIN: Helicase C−terminal
DOMAIN: R3H
REGION: Disordered

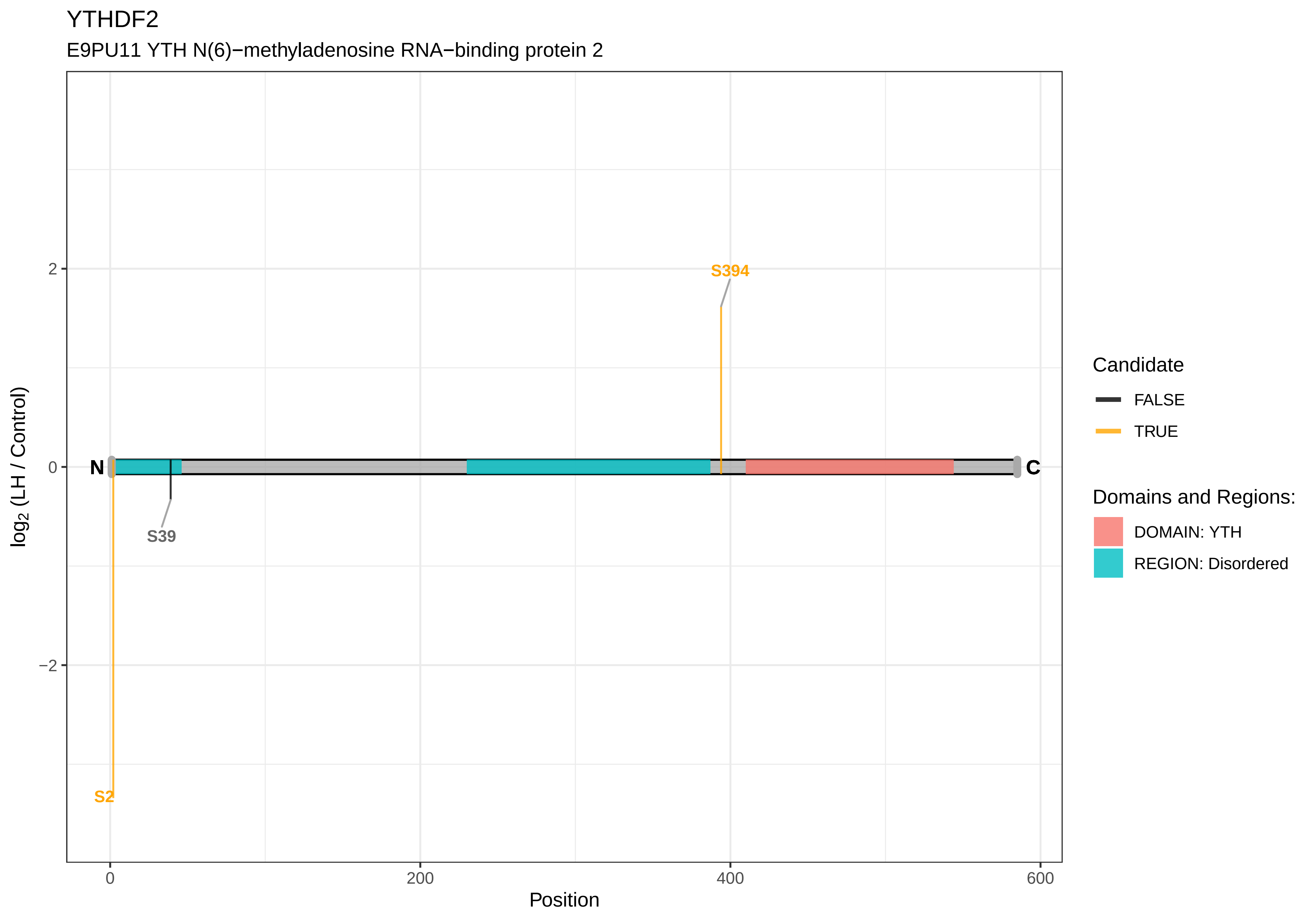

YTHDF2
E9PU11 YTH N(6)−methyladenosine RNA−binding protein 2
log2 (LH / Control)
2
0
−2
0
200
400
600
Position
N
C
S394
S39
S2
Candidate
FALSE
TRUE
Domains and Regions:
DOMAIN: YTH
REGION: Disordered

ZBTB38

M0RCM7 Zinc finger and BTB domain−containing protein 38

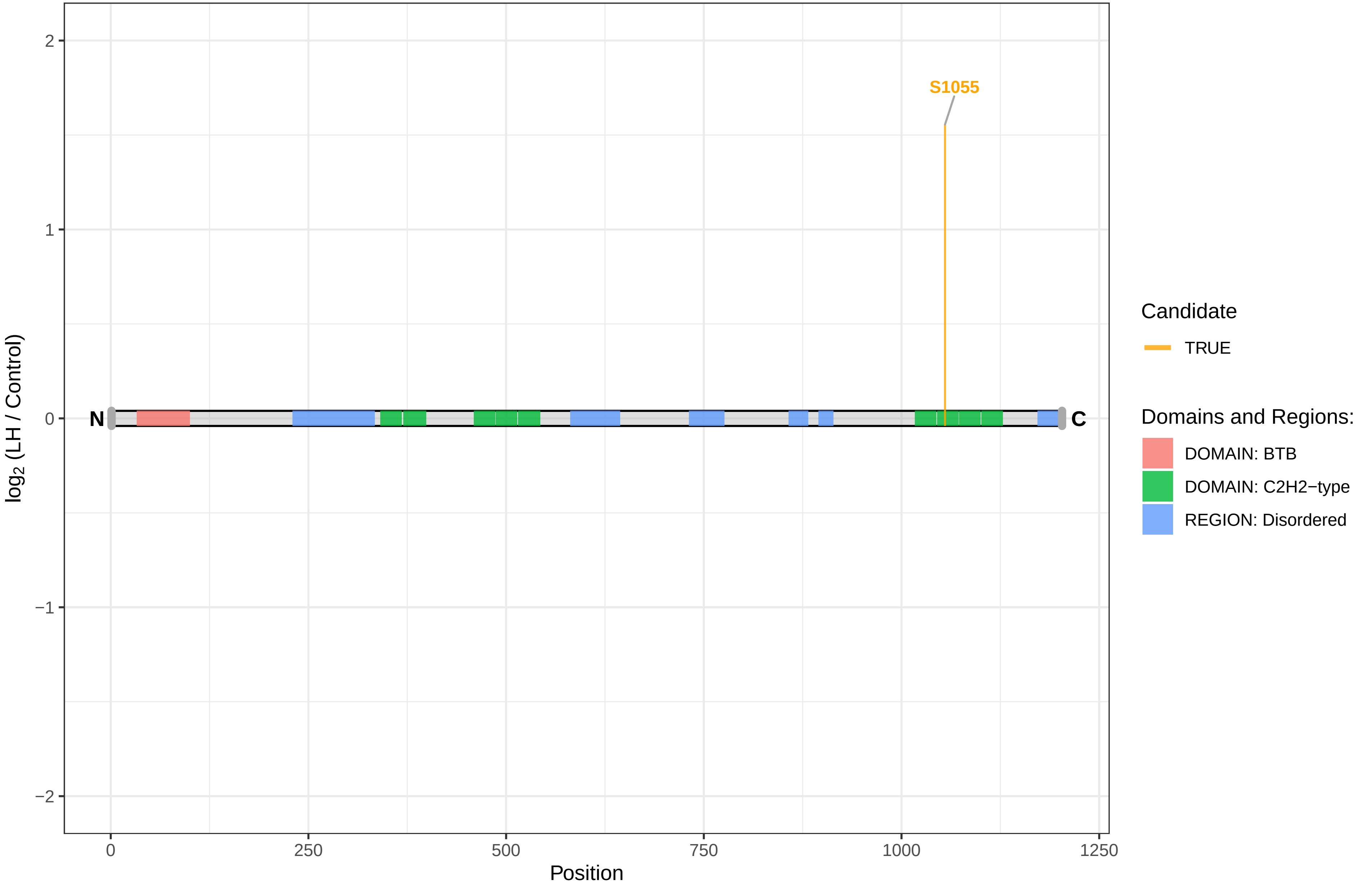

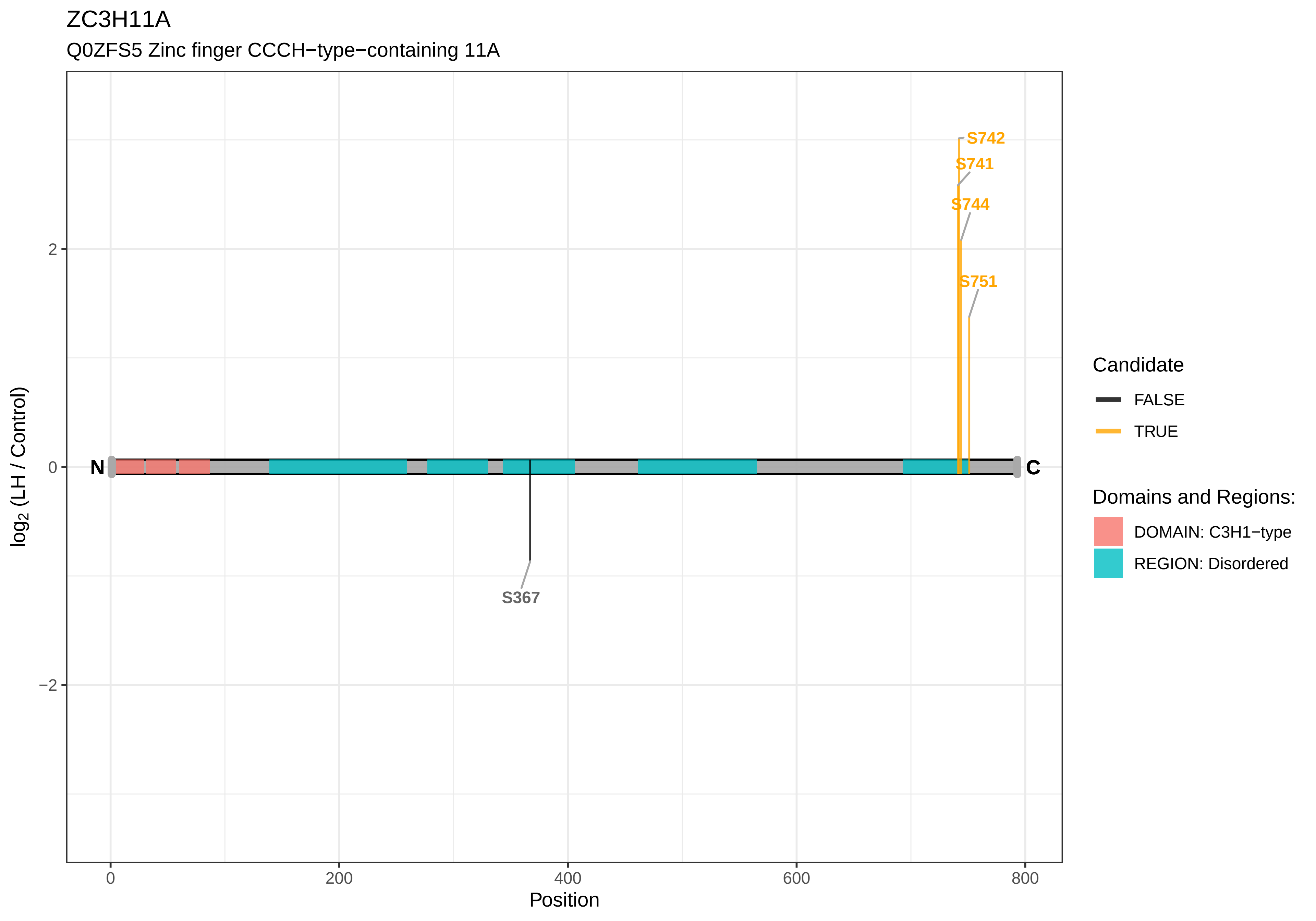
ZC3H11A
Q0ZFS5 Zinc finger CCCH−type−containing 11A
S742
S741
S744
S751
S367
N
C
2
0
−2
0
200
400
600
800
Position
log2 (LH / Control)
Candidate
FALSE
TRUE
Domains and Regions:
DOMAIN: C3H1−type
REGION: Disordered

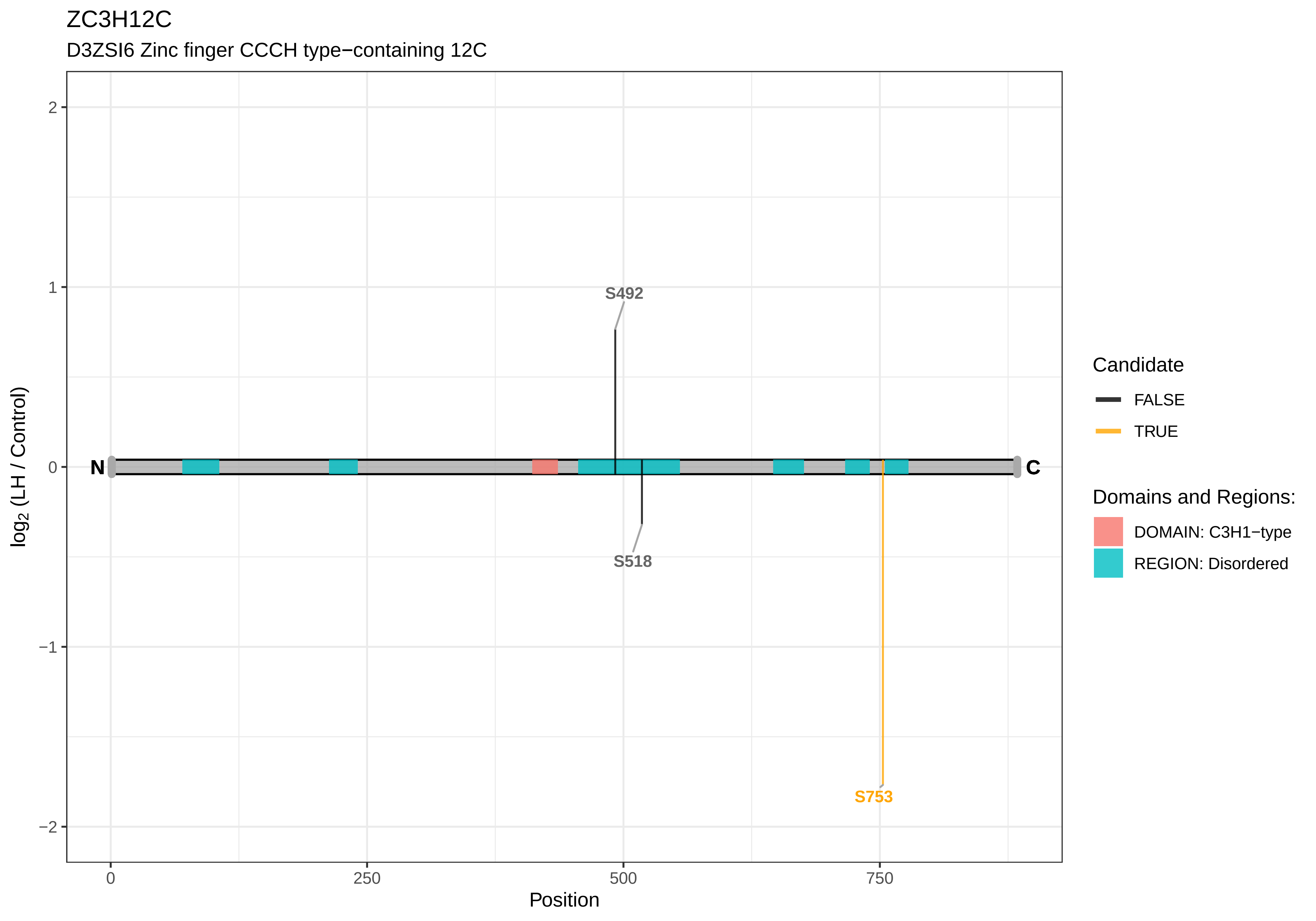

ZC3H12C
D3ZSI6 Zinc finger CCCH type−containing 12C
S492
S518
S753
N
C
2
1
0
−1
−2
0
250
500
750
Position
log2 (LH / Control)
Candidate
FALSE
TRUE
Domains and Regions:
DOMAIN: C3H1−type
REGION: Disordered

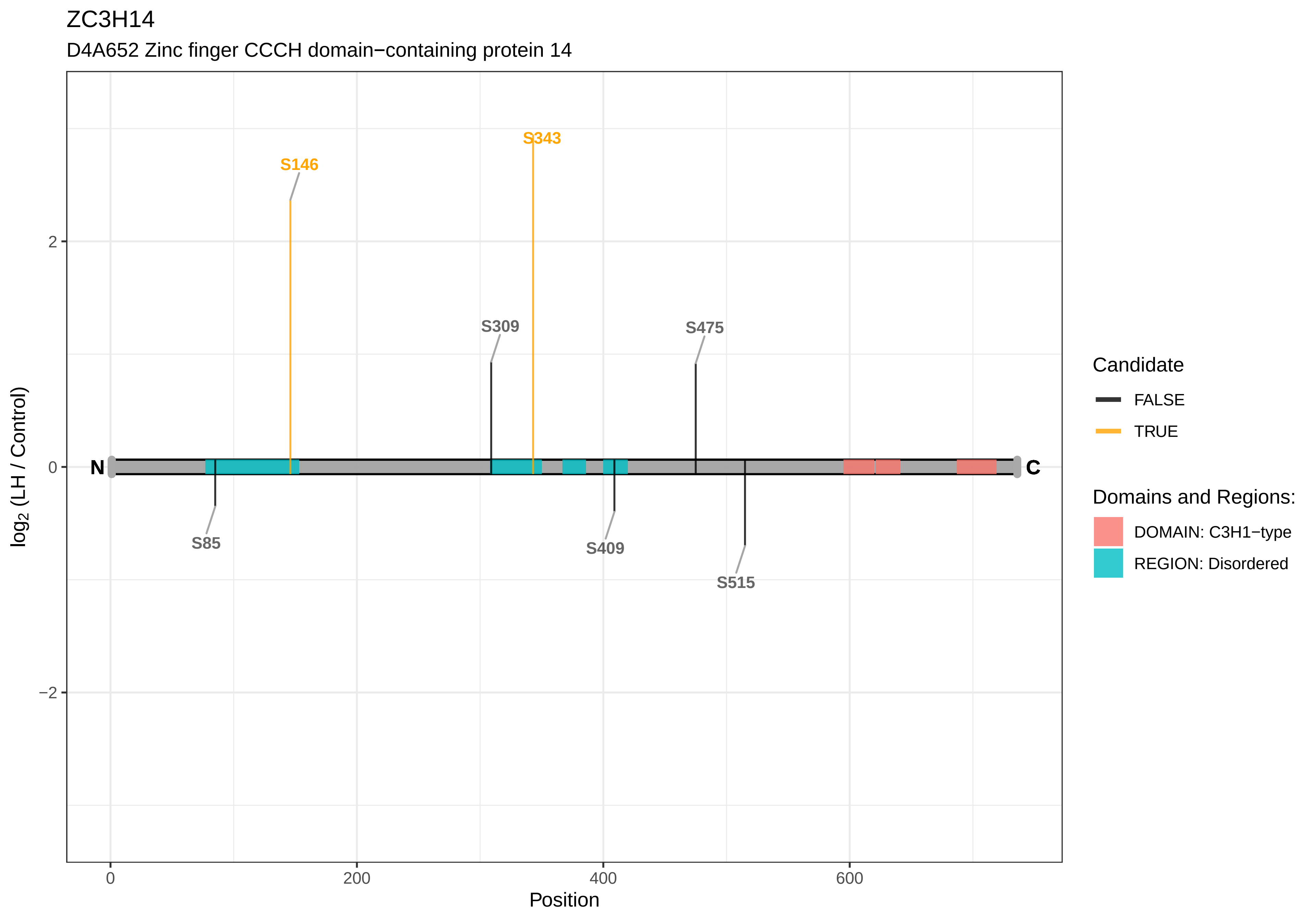

ZC3H14
D4A652 Zinc finger CCCH domain−containing protein 14
S343
S146
S309
S475
S85
S409
S515
N
C
2
0
−2
0
200
400
600
Position
log2 (LH / Control)
Candidate
FALSE
TRUE
Domains and Regions:
DOMAIN: C3H1−type
REGION: Disordered

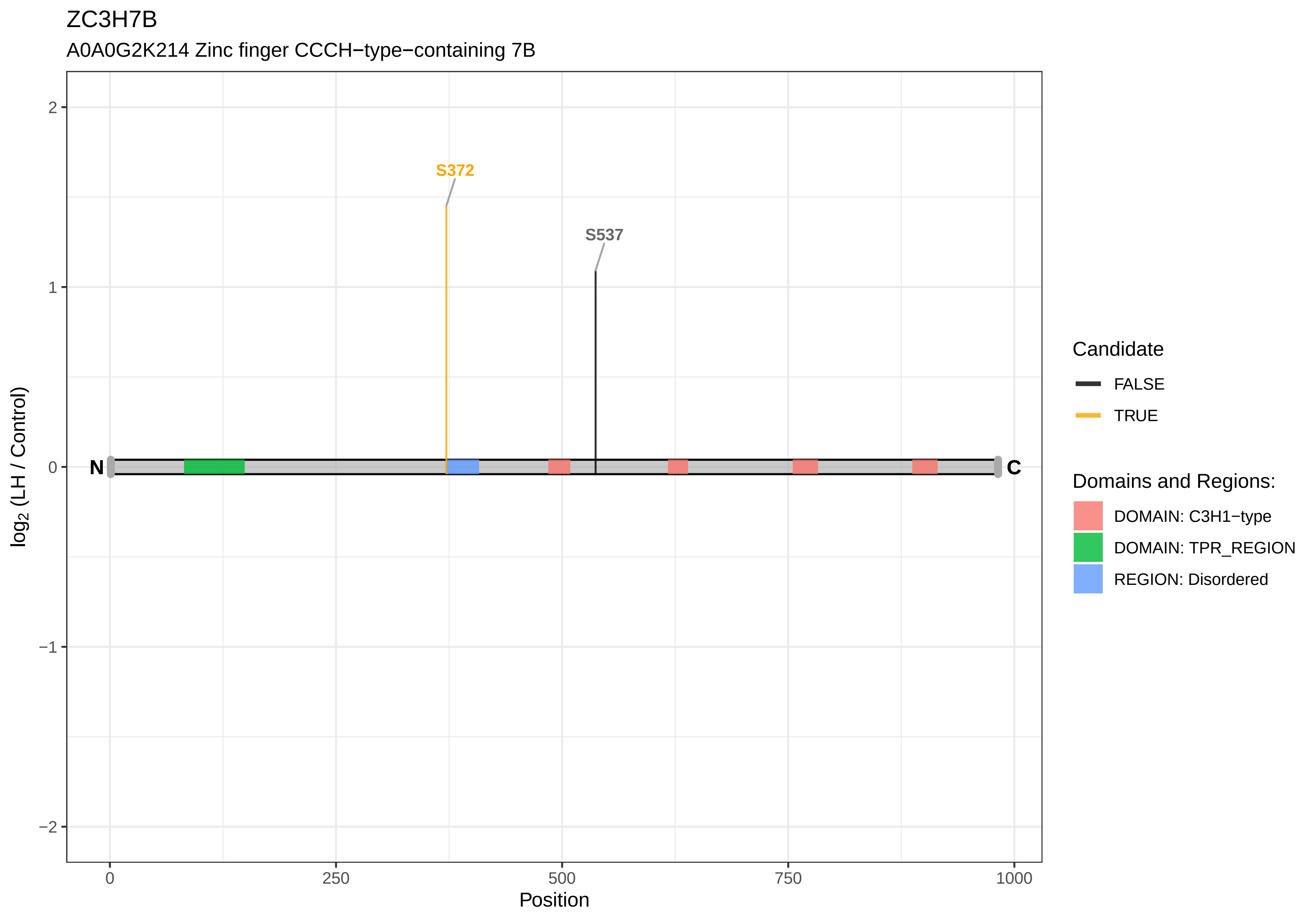
ZC3H7B
A0A0G2K214 Zinc finger CCCH−type−containing 7B
S372
S537
N
C
2
1
0
−1
−2
log2 (LH / Control)
0
250
500
750
1000
Position
Candidate
FALSE
TRUE
Domains and Regions:
DOMAIN: C3H1−type
DOMAIN: TPR_REGION
REGION: Disordered

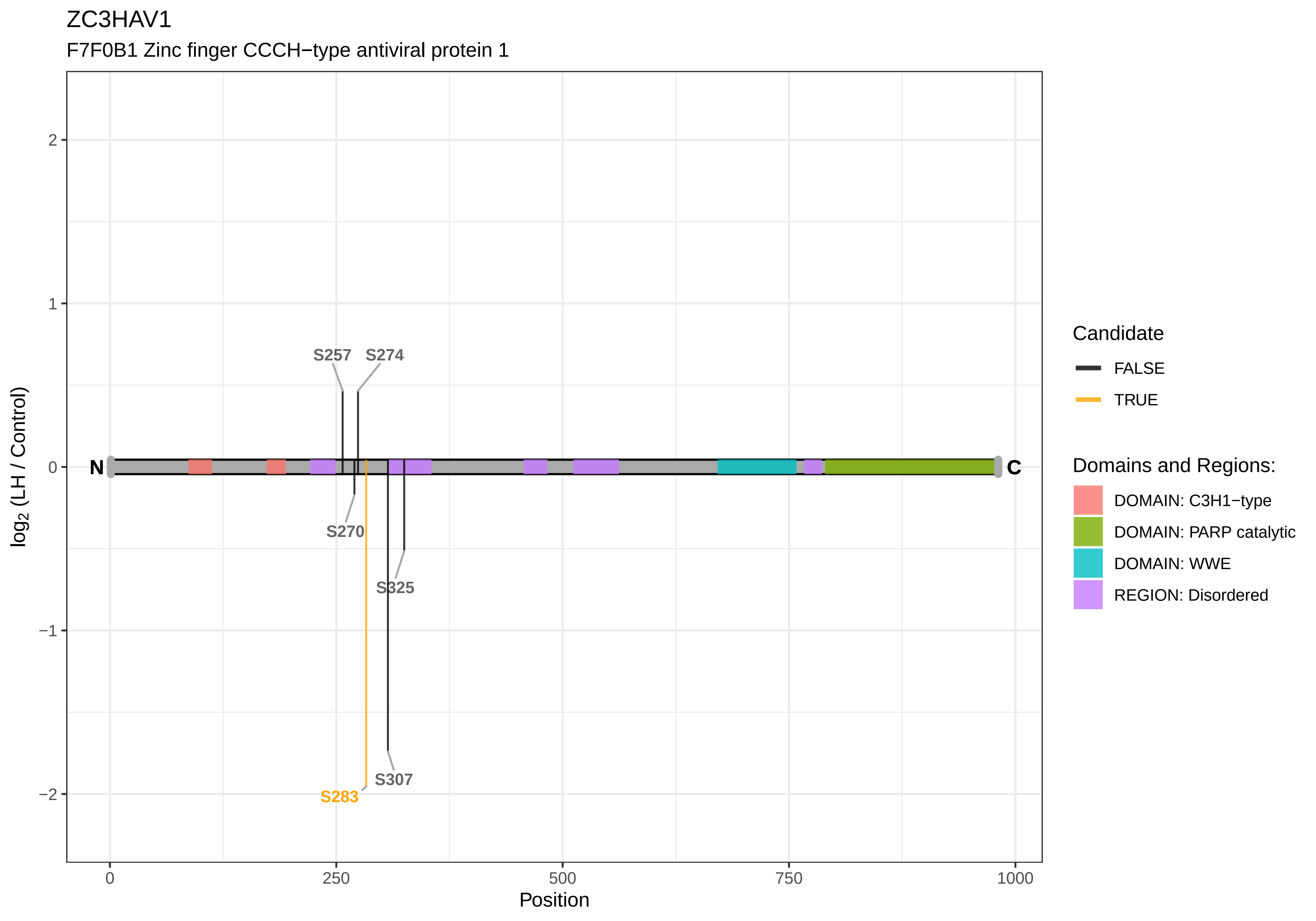
ZC3HAV1
F7F0B1 Zinc finger CCCH−type antiviral protein 1
S257
S274
S270
S325
S307
S283
N
C
2
1
0
−1
−2
log2 (LH / Control)
0
250
500
750
1000
Position
Candidate
FALSE
TRUE
Domains and Regions:
DOMAIN: C3H1−type
DOMAIN: PARP catalytic
DOMAIN: WWE
REGION: Disordered

ZC3HC1

A0A140UHX2 Zinc finger, C3HC−type−containing 1

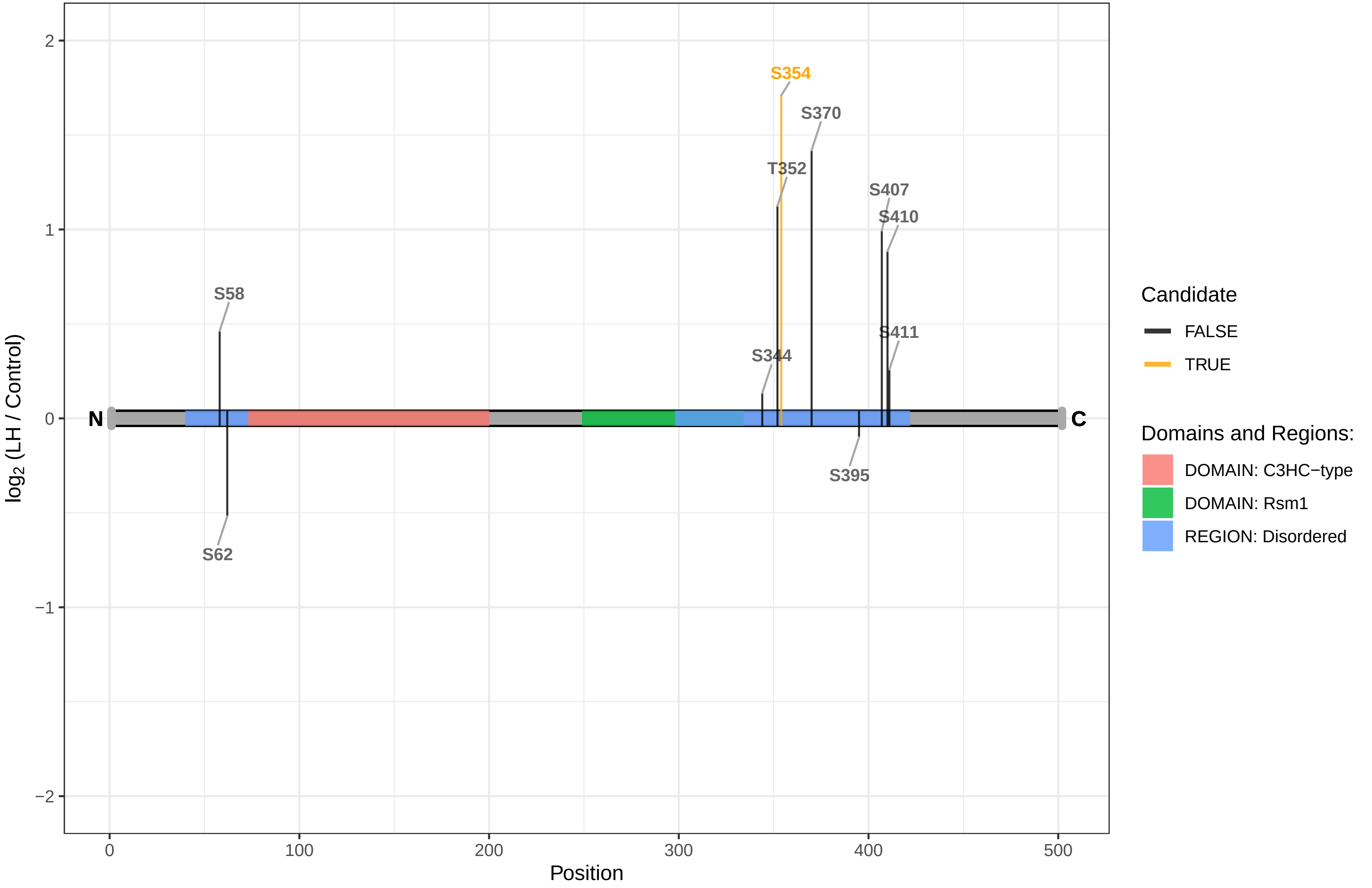

ZCCHC17

D3ZCQ7 Similar to PS1D protein (Predicted)

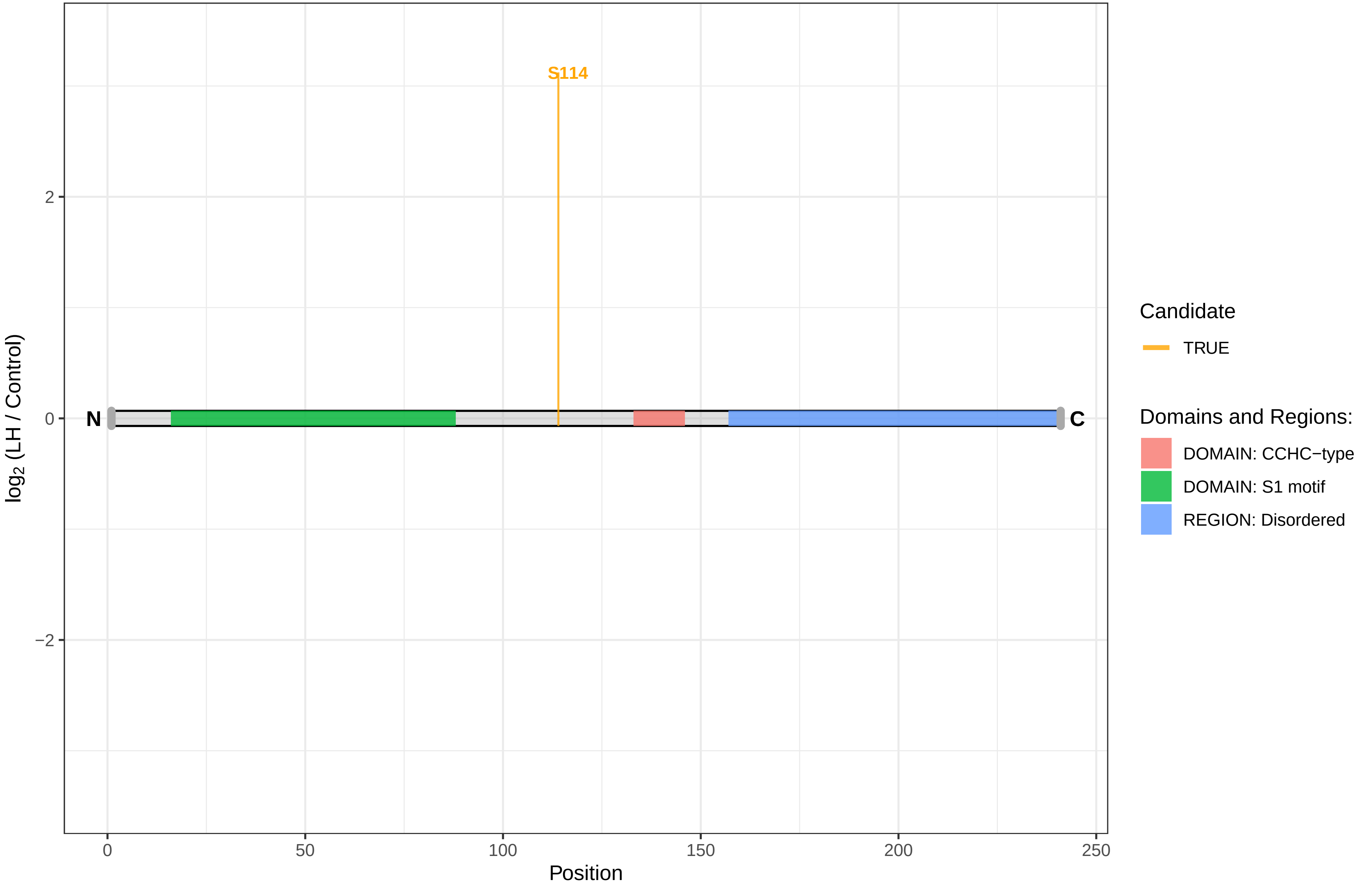

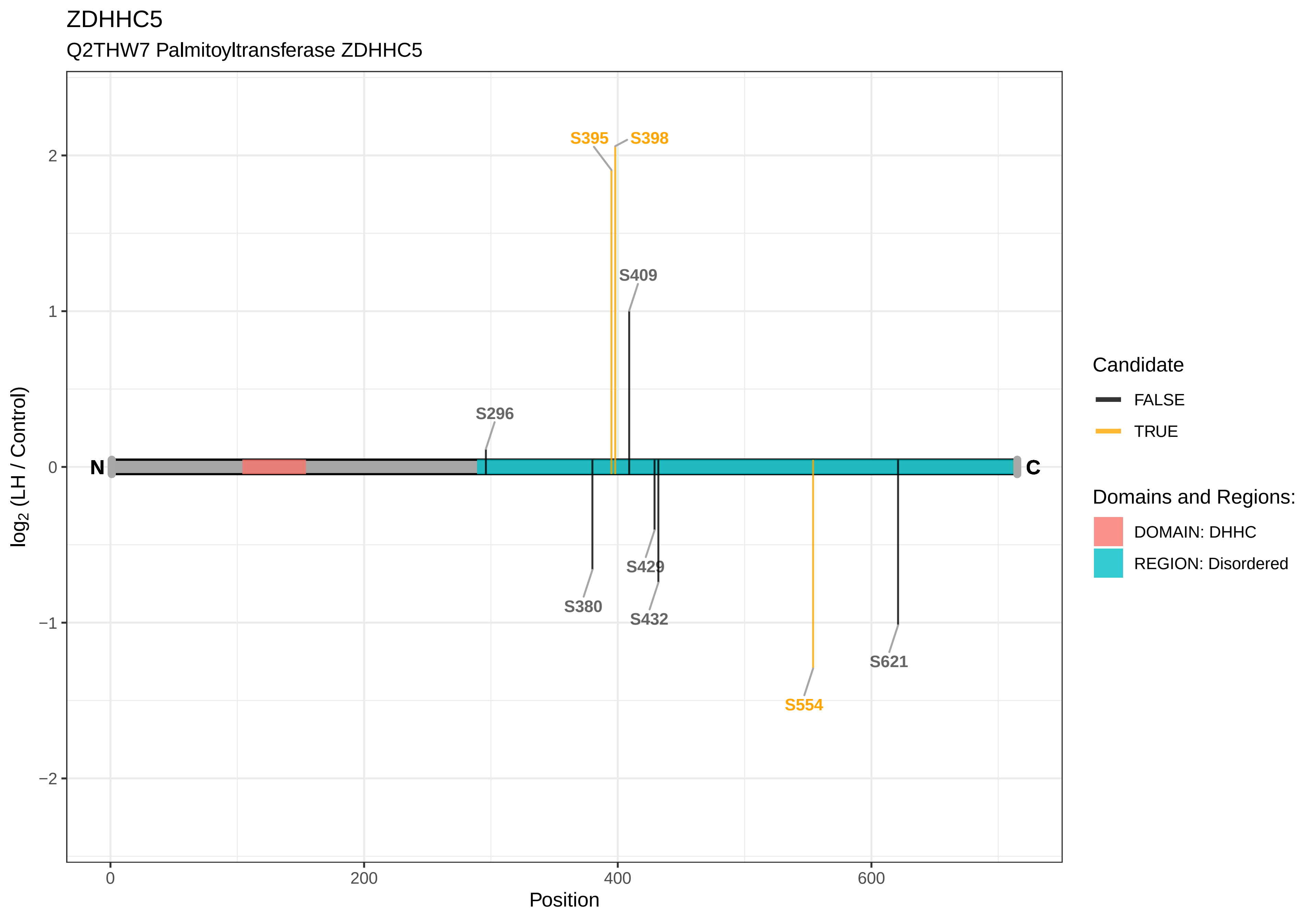
ZDHHC5
Q2THW7 Palmitoyltransferase ZDHHC5
S395
S398
S409
S296
N
C
S429
S380
S432
S554
S621
2
1
0
−1
−2
log2 (LH / Control)
0
200
400
600
Position
Candidate
FALSE
TRUE
Domains and Regions:
DOMAIN: DHHC
REGION: Disordered

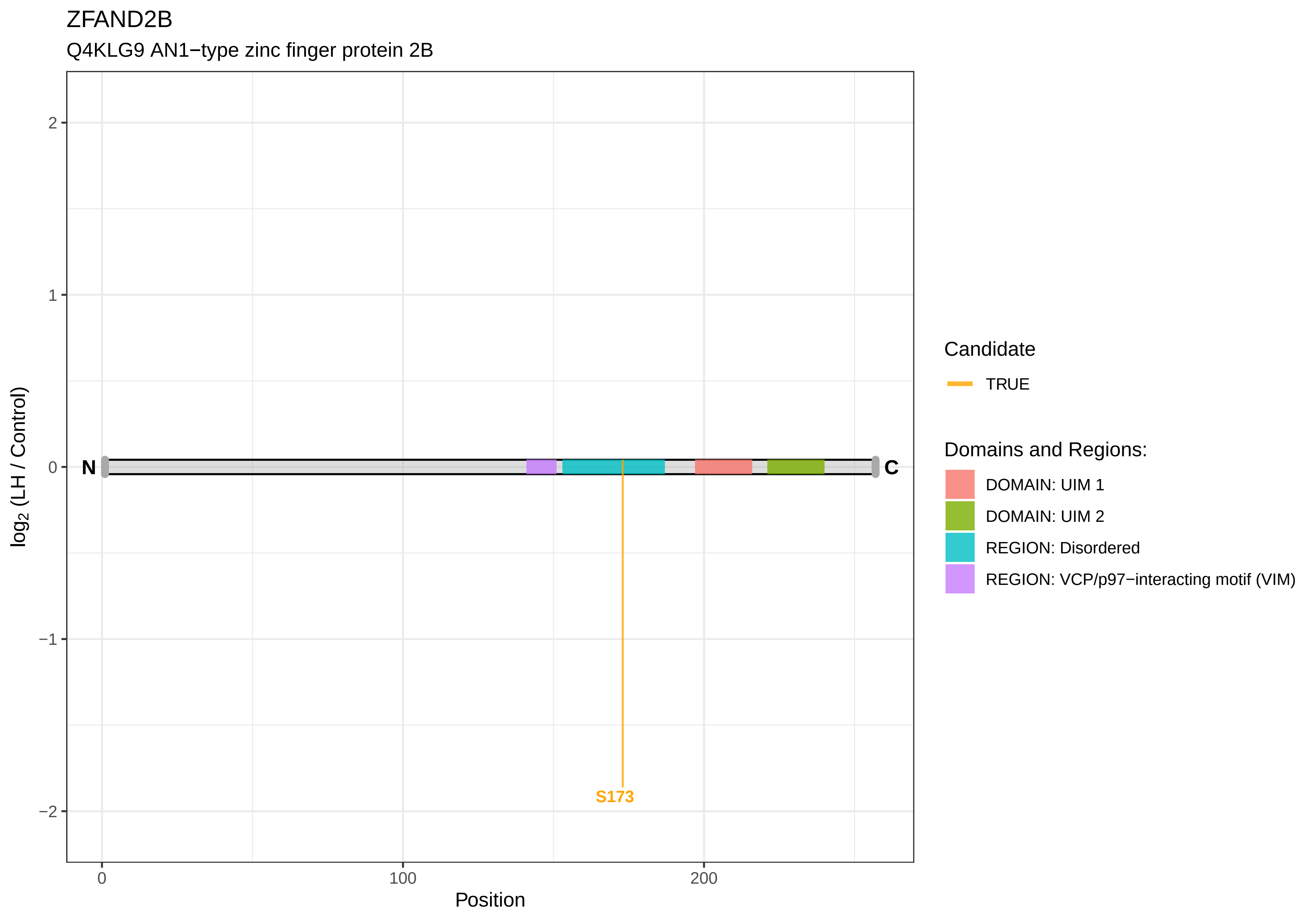
ZFAND2B
Q4KLG9 AN1−type zinc finger protein 2B
log2 (LH / Control)
2
1
0
−1
−2
N
C
S173
0
100
200
Position
Candidate
TRUE
Domains and Regions:
DOMAIN: UIM 1
DOMAIN: UIM 2
REGION: Disordered
REGION: VCP/p97−interacting motif (VIM)

ZFP106

A0A0G2K2F8 Zinc finger protein 106

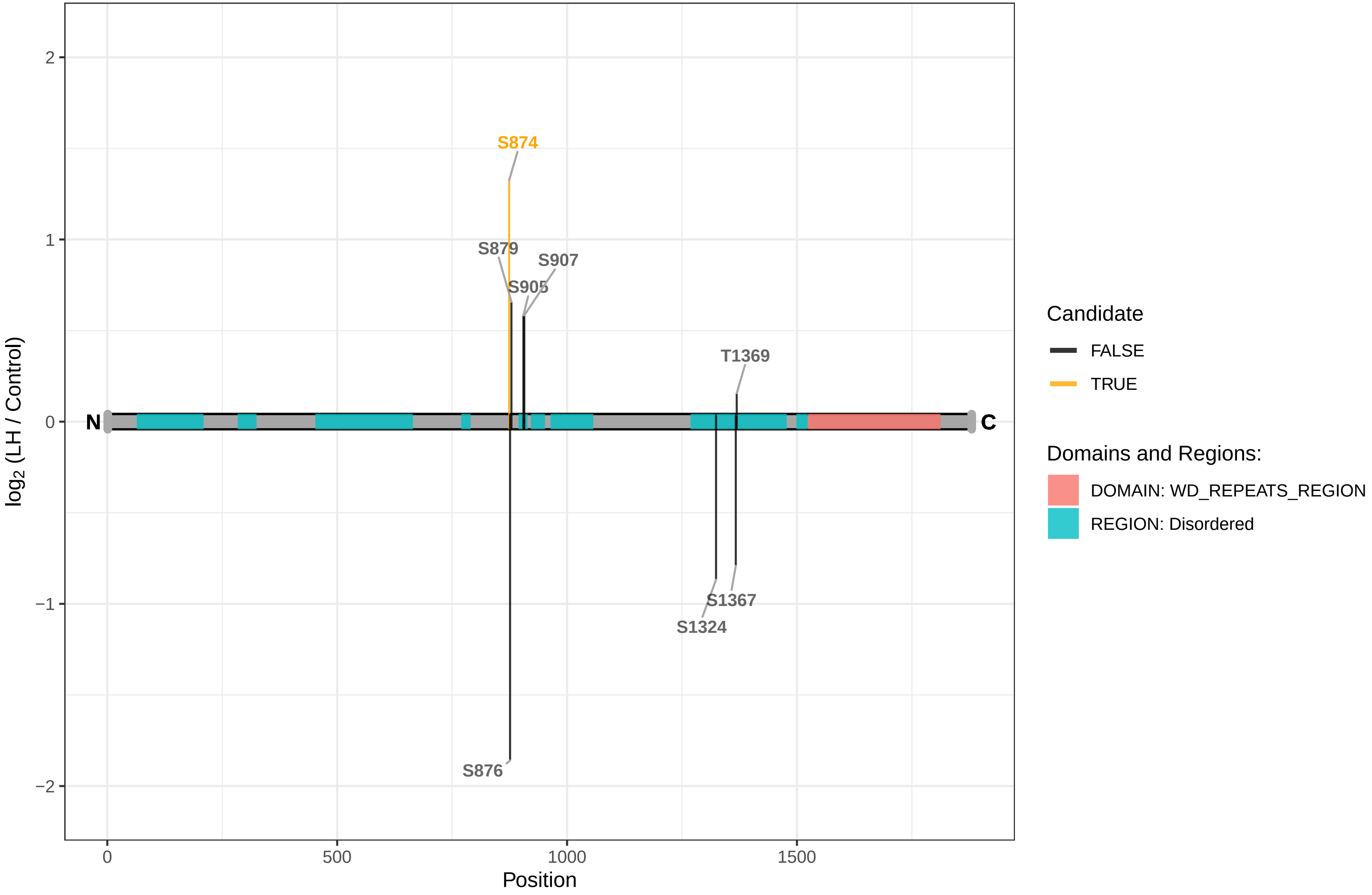

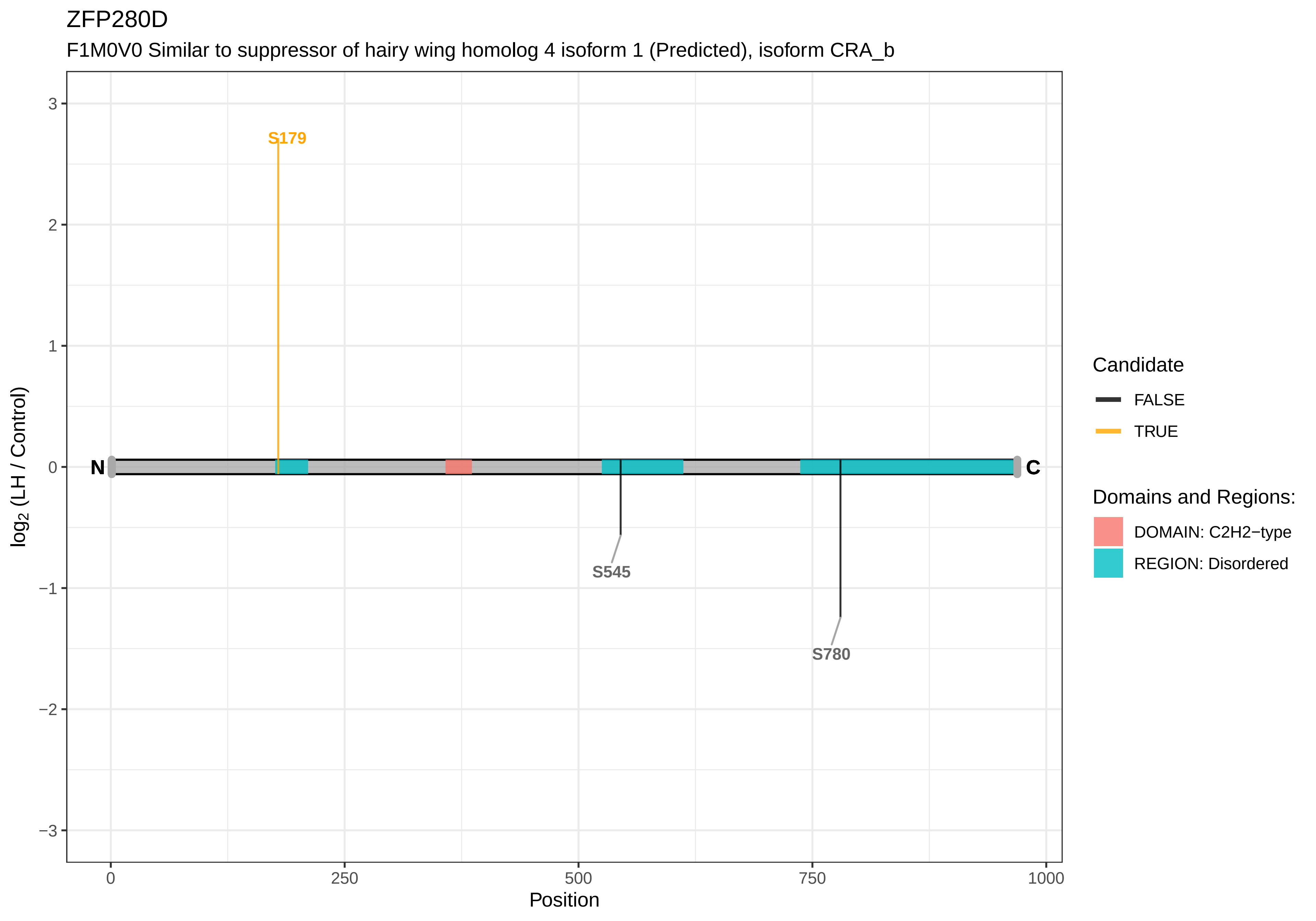

ZFP280D
F1M0V0 Similar to suppressor of hairy wing homolog 4 isoform 1 (Predicted), isoform CRA_b
S179
S545
S780
N
C
3
2
1
0
−1
−2
−3
log2 (LH / Control)
0
250
500
750
1000
Position
Candidate
FALSE
TRUE
Domains and Regions:
DOMAIN: C2H2−type
REGION: Disordered

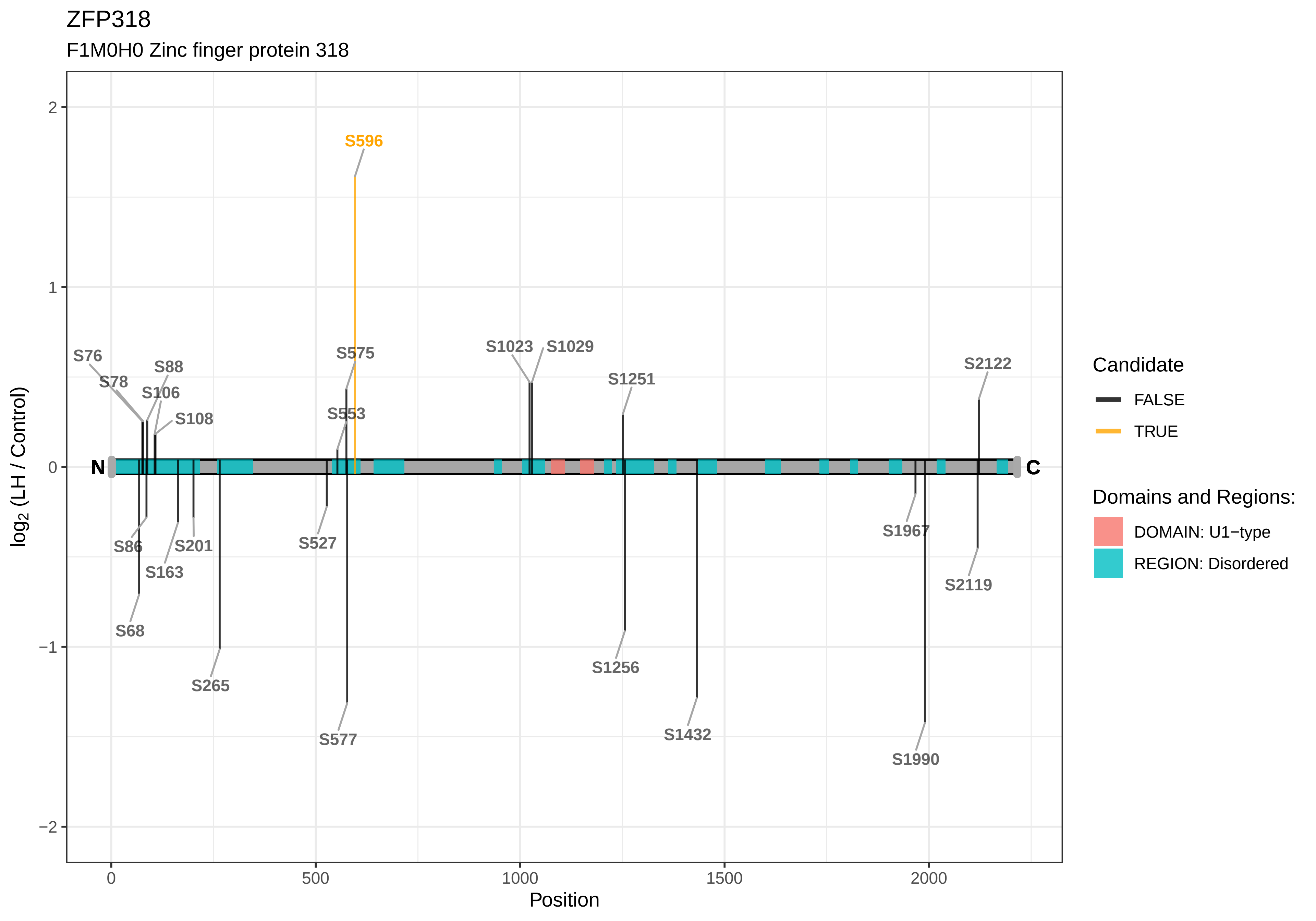

ZFP318
F1M0H0 Zinc finger protein 318
S596
S76
S78
S88
S106
S108
S575
S553
S1023
S1029
S1251
S2122
N
C
S86
S201
S163
S68
S265
S527
S577
S1256
S1432
S1967
S2119
S1990
2
1
0
−1
−2
0
500
1000
1500
2000
Position
log2 (LH / Control)
Candidate
FALSE
TRUE
Domains and Regions:
DOMAIN: U1−type
REGION: Disordered

ZFP36L1

P17431 mRNA decay activator protein ZFP36L1

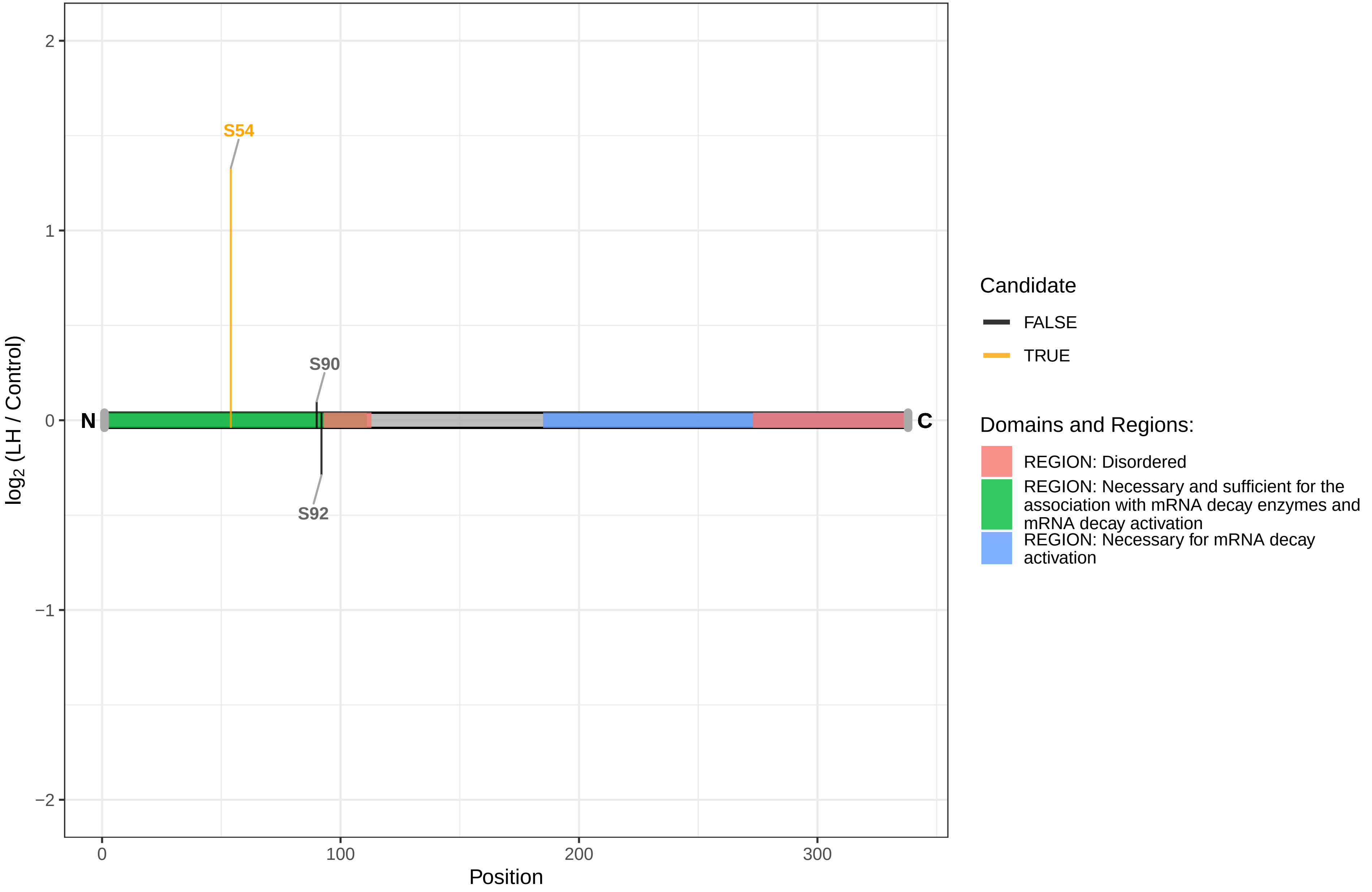

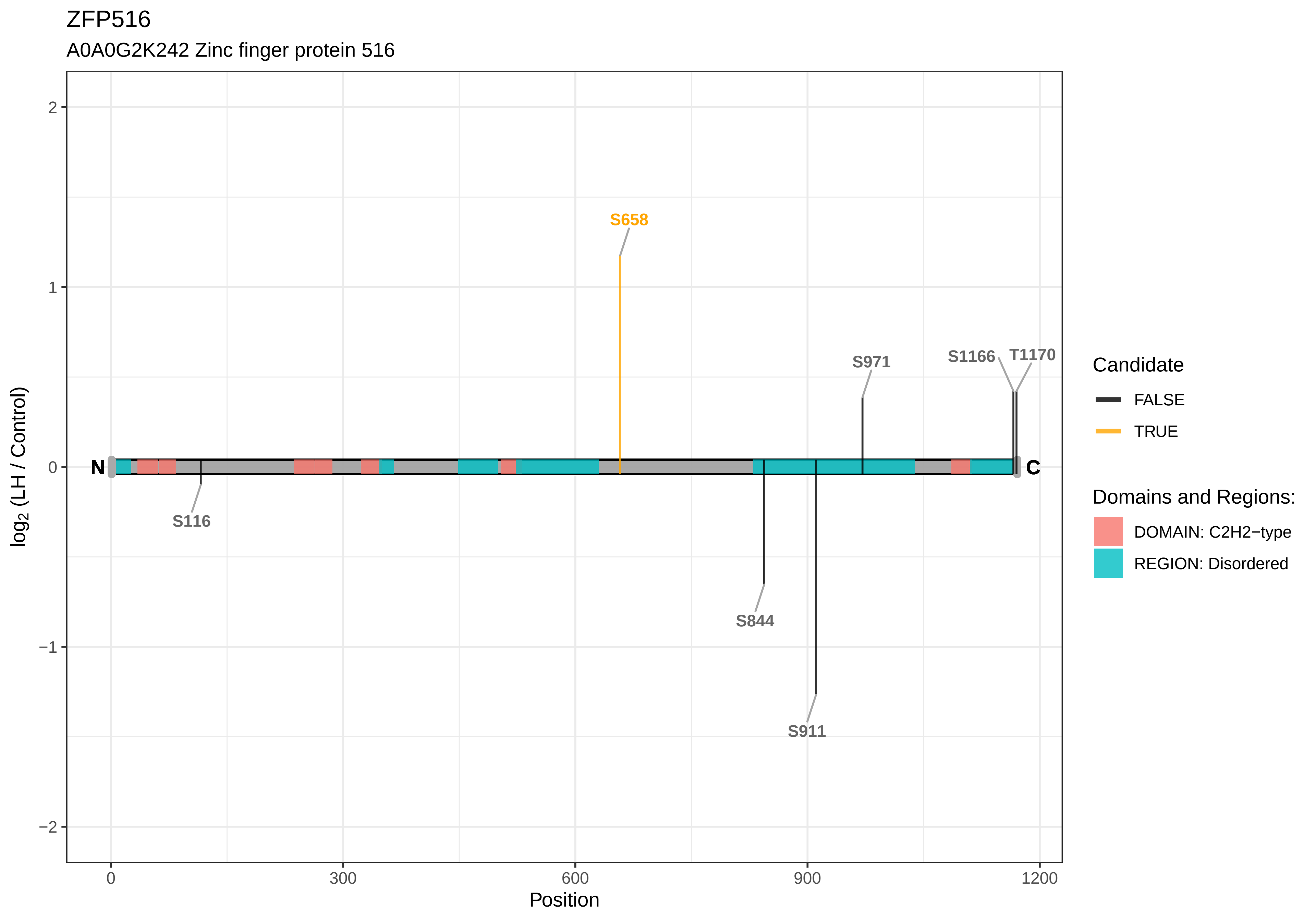

ZFP516
A0A0G2K242 Zinc finger protein 516
S658
S971
S1166
T1170
N
C
S116
S844
S911
2
1
0
−1
−2
0
300
600
900
1200
Position
log2 (LH / Control)
Candidate
FALSE
TRUE
Domains and Regions:
DOMAIN: C2H2−type
REGION: Disordered

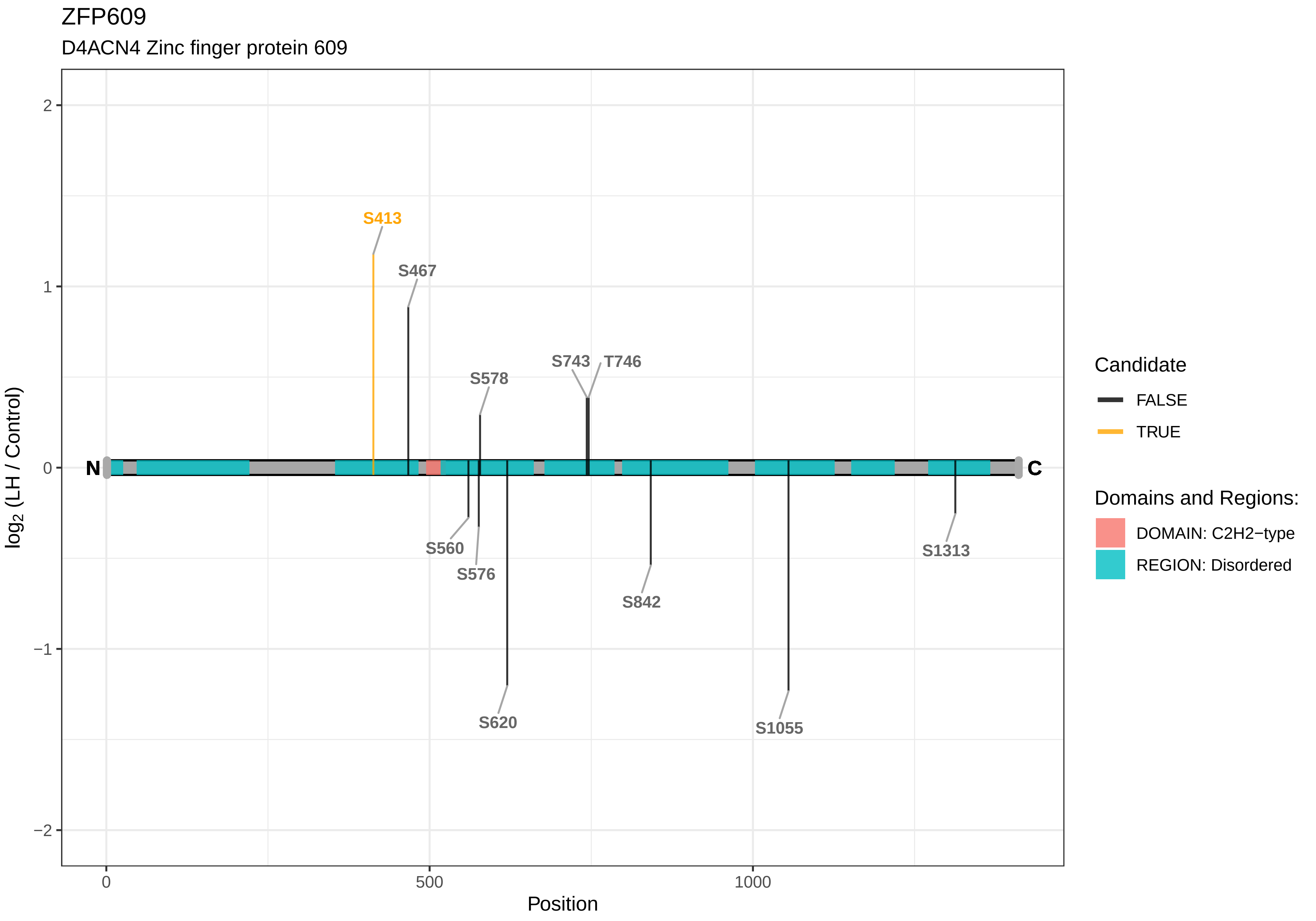
ZFP609
D4ACN4 Zinc finger protein 609
log2 (LH / Control)
Position
2
1
0
−1
−2
0
500
1000
N
C
S413
S467
S578
S743
T746
S560
S576
S620
S842
S1055
S1313
Candidate
FALSE
TRUE
Domains and Regions:
DOMAIN: C2H2−type
REGION: Disordered

ZFP655

Q5RKG8 Zinc finger protein 655

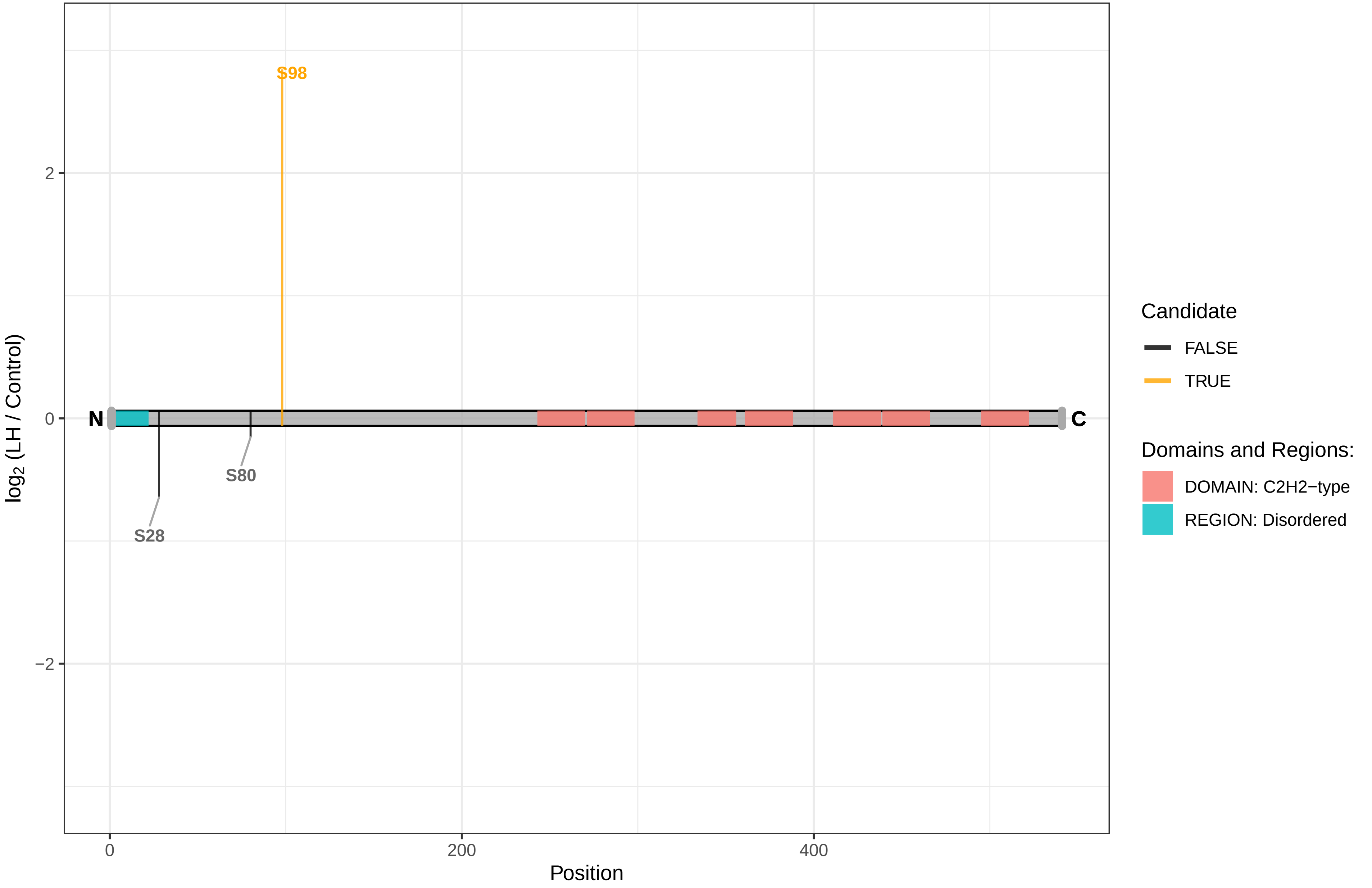

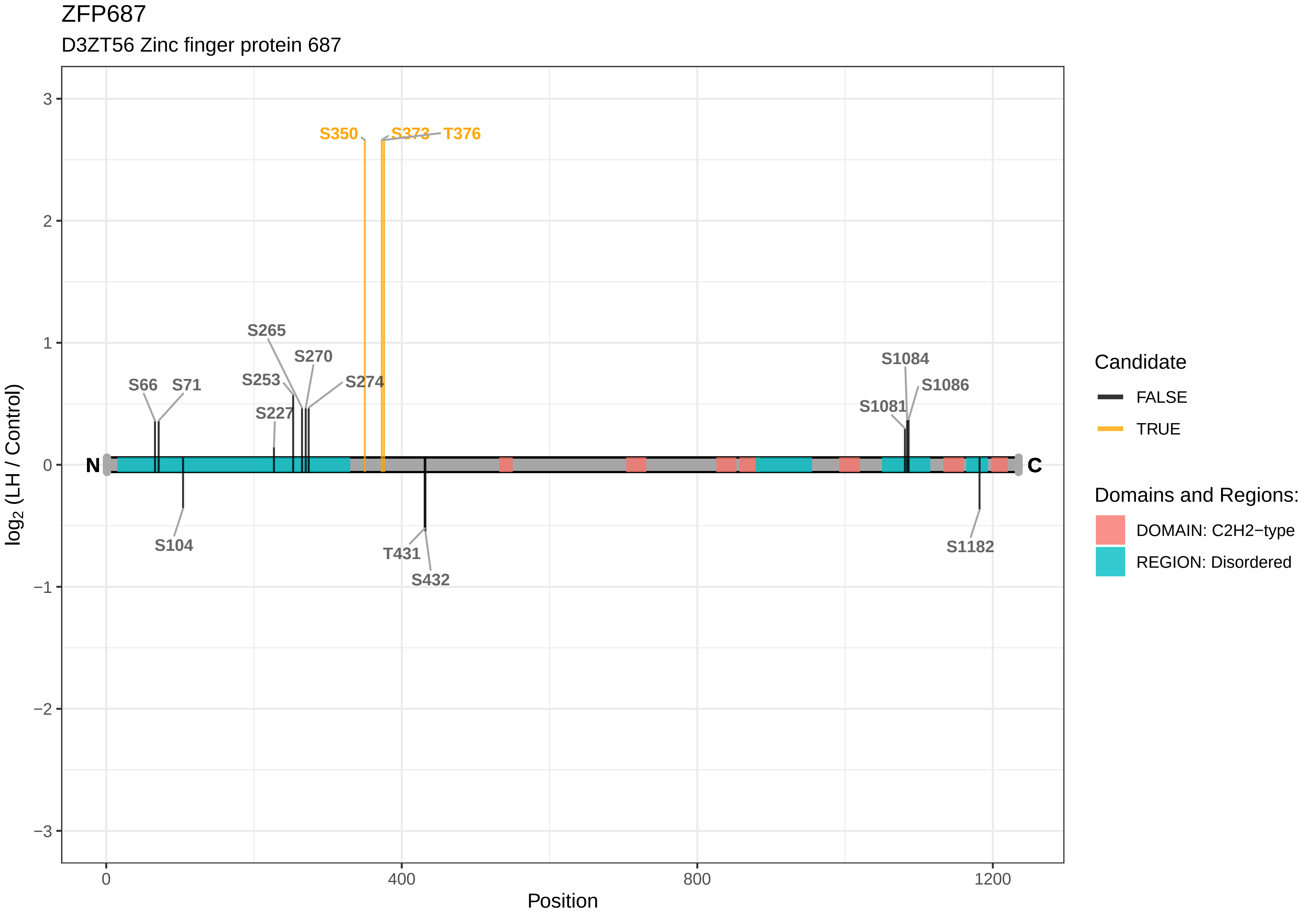

ZFP687
D3ZT56 Zinc finger protein 687
S350
S373
T376
S265
S270
S253
S274
S66
S71
S227
S104
T431
S432
S1084
S1086
S1081
S1182
N
C
3
2
1
0
−1
−2
−3
0
400
800
1200
Position
log2 (LH / Control)
Candidate
FALSE
TRUE
Domains and Regions:
DOMAIN: C2H2−type
REGION: Disordered

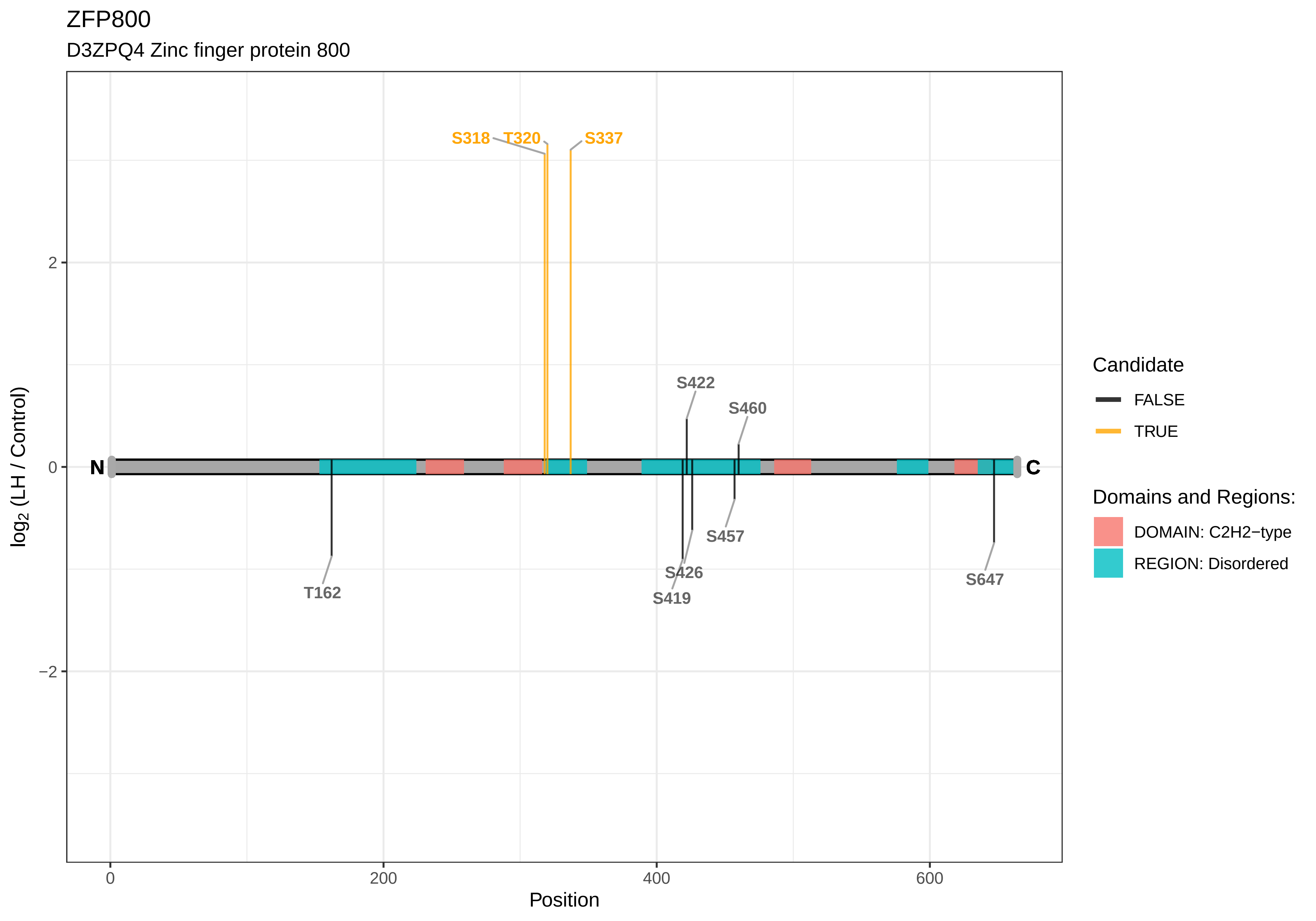
ZFP800
D3ZPQ4 Zinc finger protein 800
S318
T320
S337
S422
S460
N
C
T162
S457
S426
S419
S647
2
0
−2
0
200
400
600
Position
log2 (LH / Control)
Candidate
FALSE
TRUE
Domains and Regions:
DOMAIN: C2H2−type
REGION: Disordered

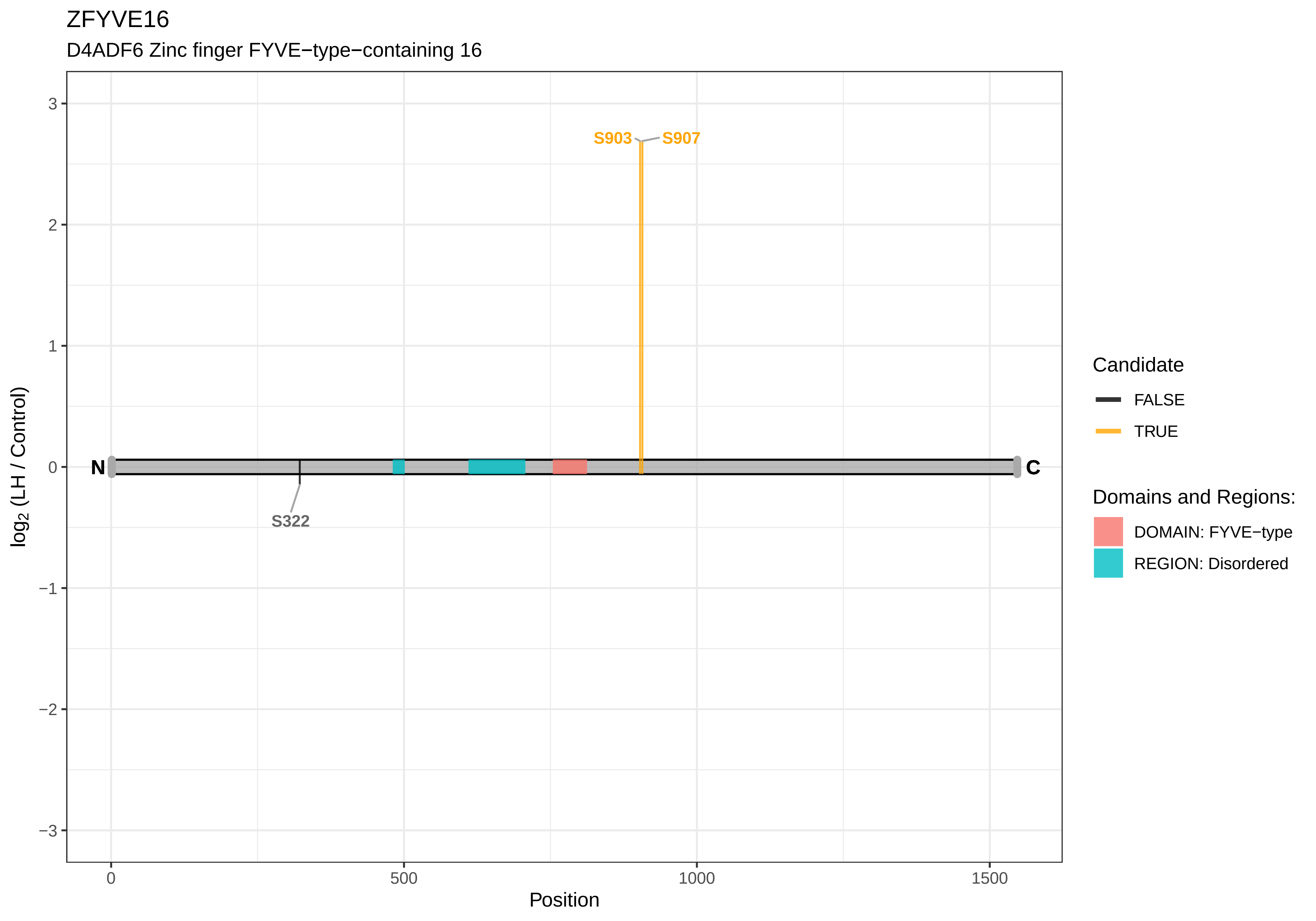

ZFYVE16
D4ADF6 Zinc finger FYVE−type−containing 16
S903
S907
S322
N
C
3
2
1
0
−1
−2
−3
log2 (LH / Control)
0
500
1000
1500
Position
Candidate
FALSE
TRUE
Domains and Regions:
DOMAIN: FYVE−type
REGION: Disordered

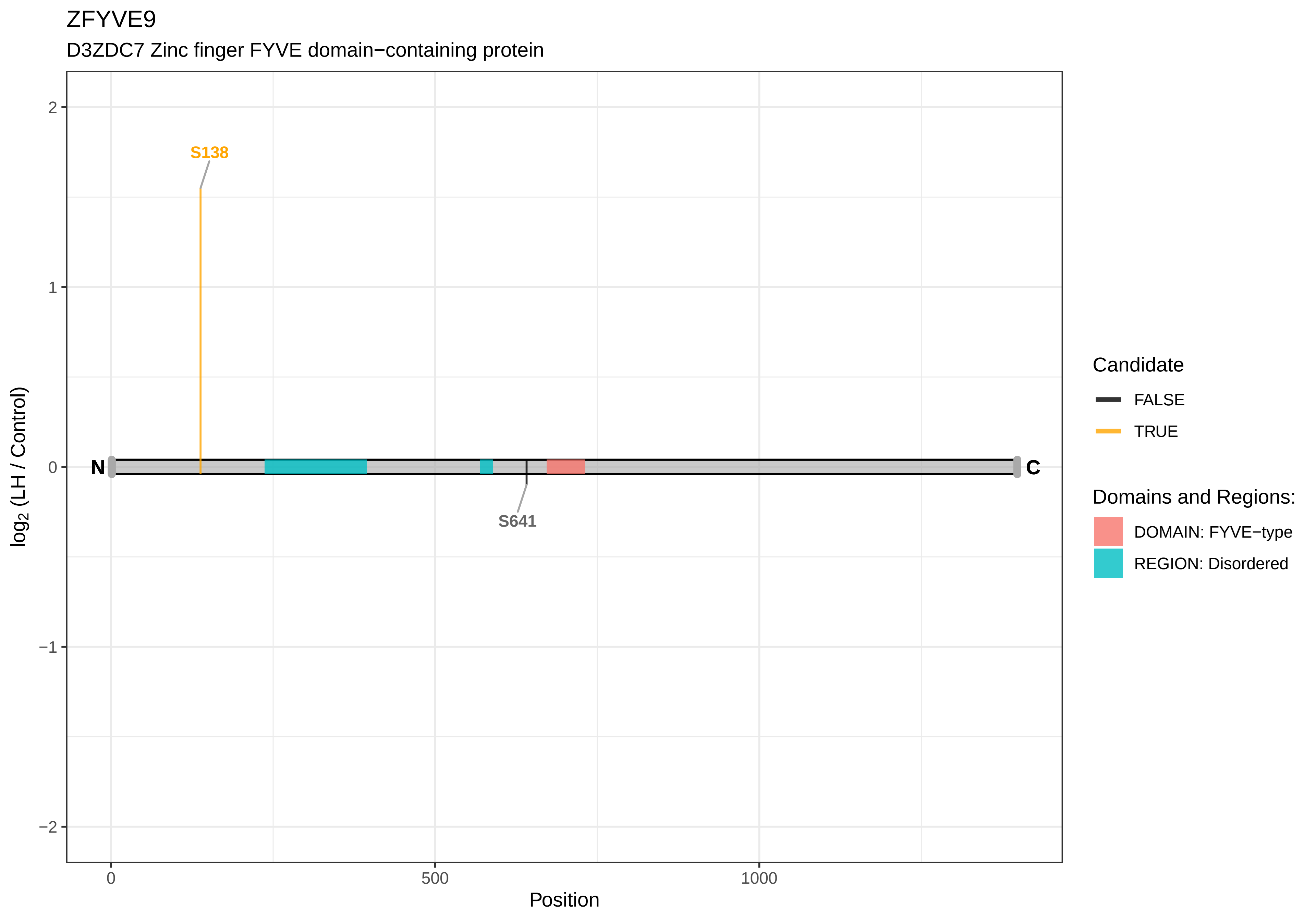
ZFYVE9
D3ZDC7 Zinc finger FYVE domain−containing protein
S138
S641
N
C
2
1
0
−1
−2
0
500
1000
Position
log2 (LH / Control)
Candidate
FALSE
TRUE
Domains and Regions:
DOMAIN: FYVE−type
REGION: Disordered

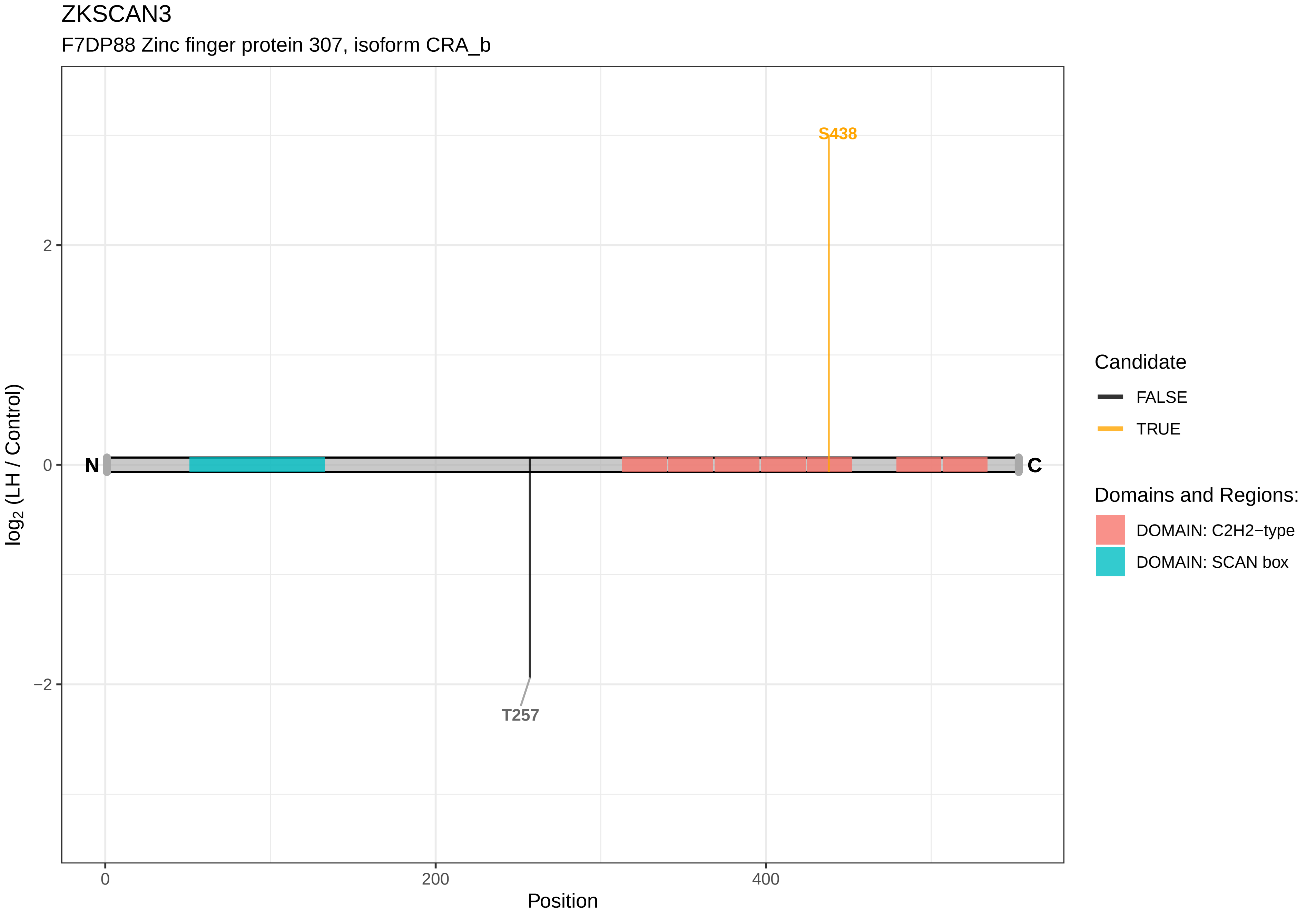
ZKSCAN3
F7DP88 Zinc finger protein 307, isoform CRA_b
S438
T257
N
C
2
0
−2
0
200
400
Position
log2 (LH / Control)
Candidate
FALSE
TRUE
Domains and Regions:
DOMAIN: C2H2−type
DOMAIN: SCAN box

#### ZNF513

Q5FWU5 Zinc finger protein 513

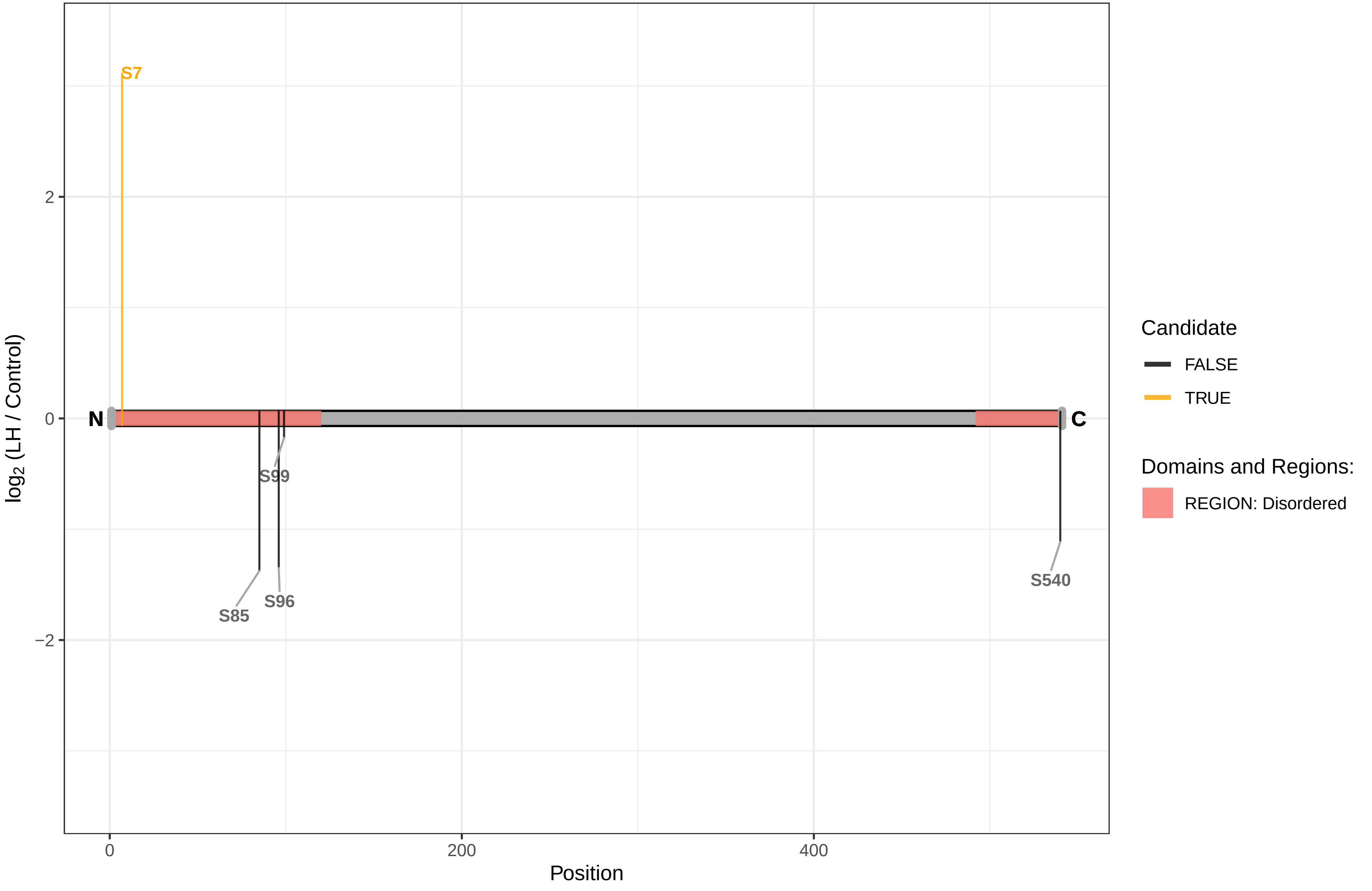

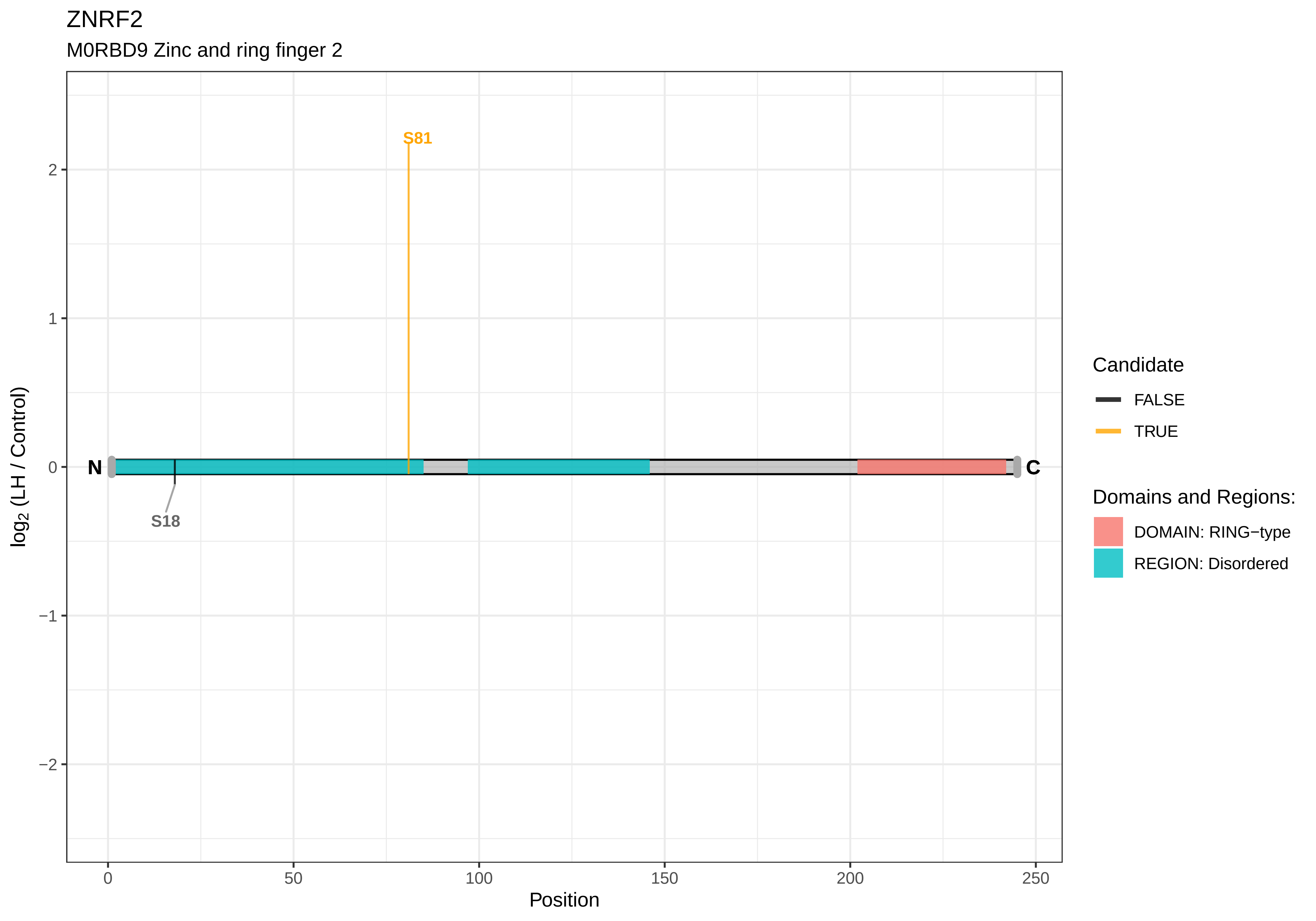

ZNRF2
M0RBD9 Zinc and ring finger 2
S81
S18
N
C
log2 (LH / Control)
Position
2
1
0
−1
−2
0
50
100
150
200
250
Candidate
FALSE
TRUE
Domains and Regions:
DOMAIN: RING−type
REGION: Disordered

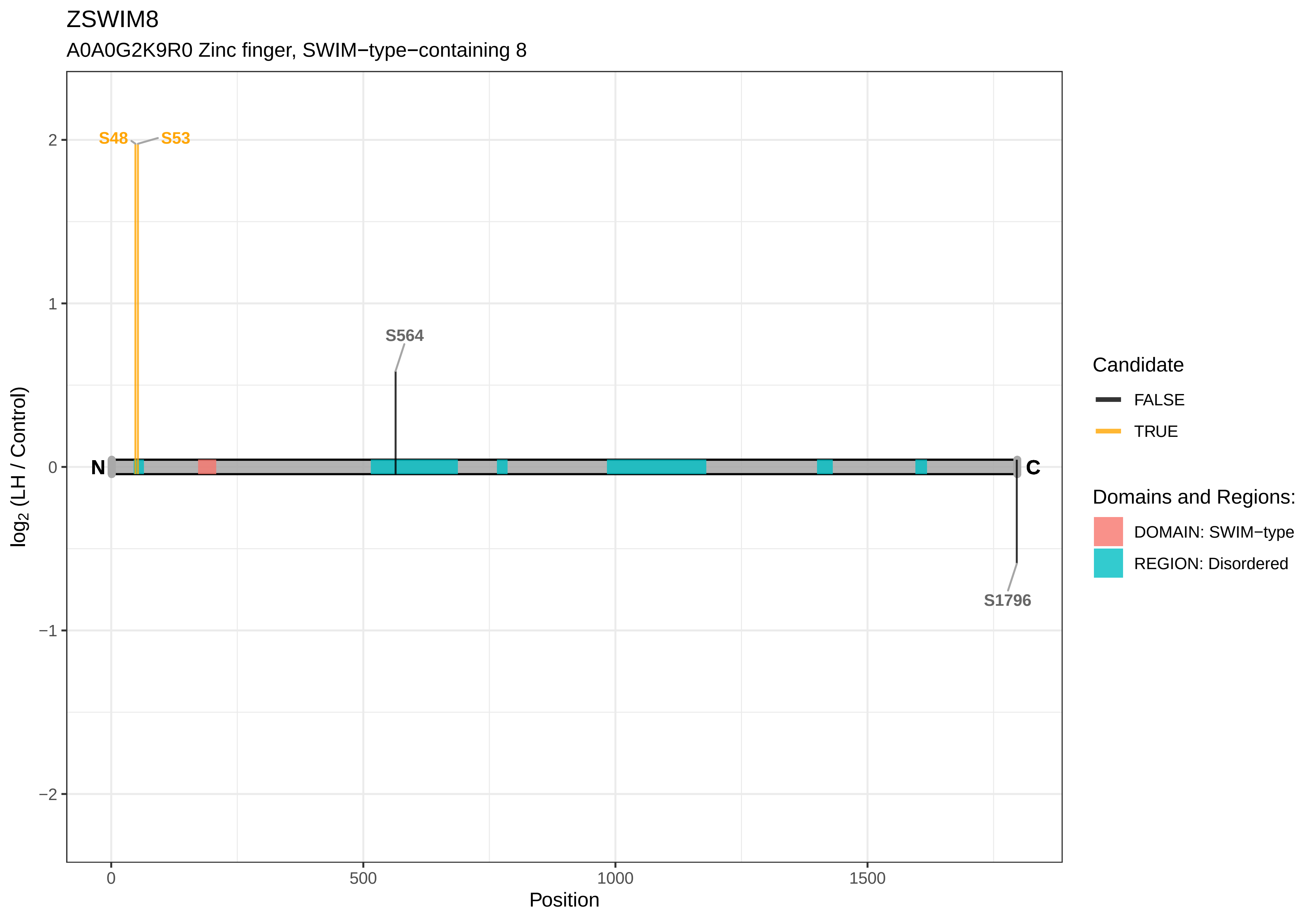

ZSWIM8
A0A0G2K9R0 Zinc finger, SWIM−type−containing 8
S48
S53
S564
S1796
N
C
2
1
0
−1
−2
log2 (LH / Control)
0
500
1000
1500
Position
Candidate
FALSE
TRUE
Domains and Regions:
DOMAIN: SWIM−type
REGION: Disordered

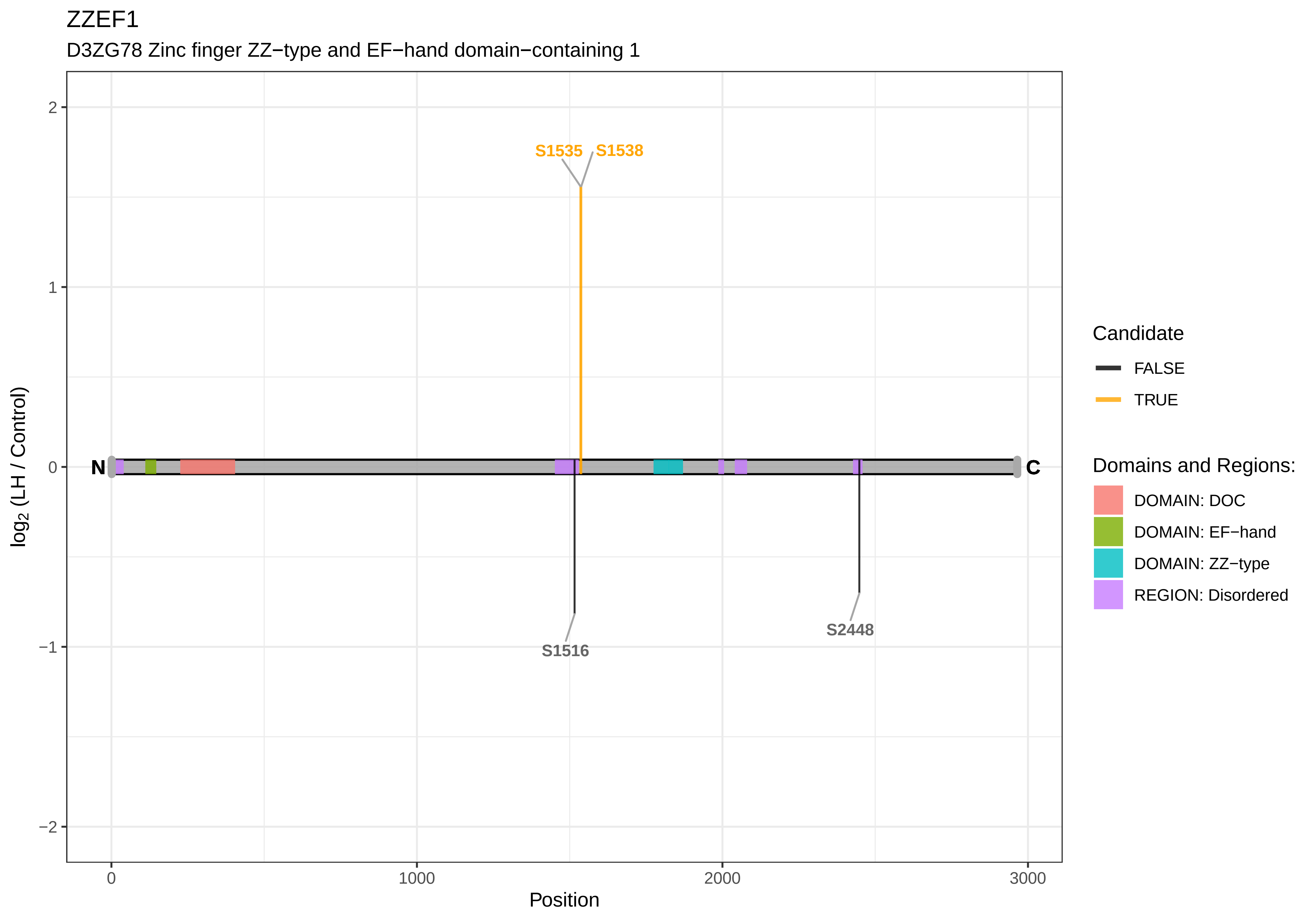

ZZEF1
D3ZG78 Zinc finger ZZ−type and EF−hand domain−containing 1
S1535
S1538
S1516
S2448
N
C
2
1
0
−1
−2
0
1000
2000
3000
Position
log2 (LH / Control)
Candidate
FALSE
TRUE
Domains and Regions:
DOMAIN: DOC
DOMAIN: EF−hand
DOMAIN: ZZ−type
REGION: Disordered

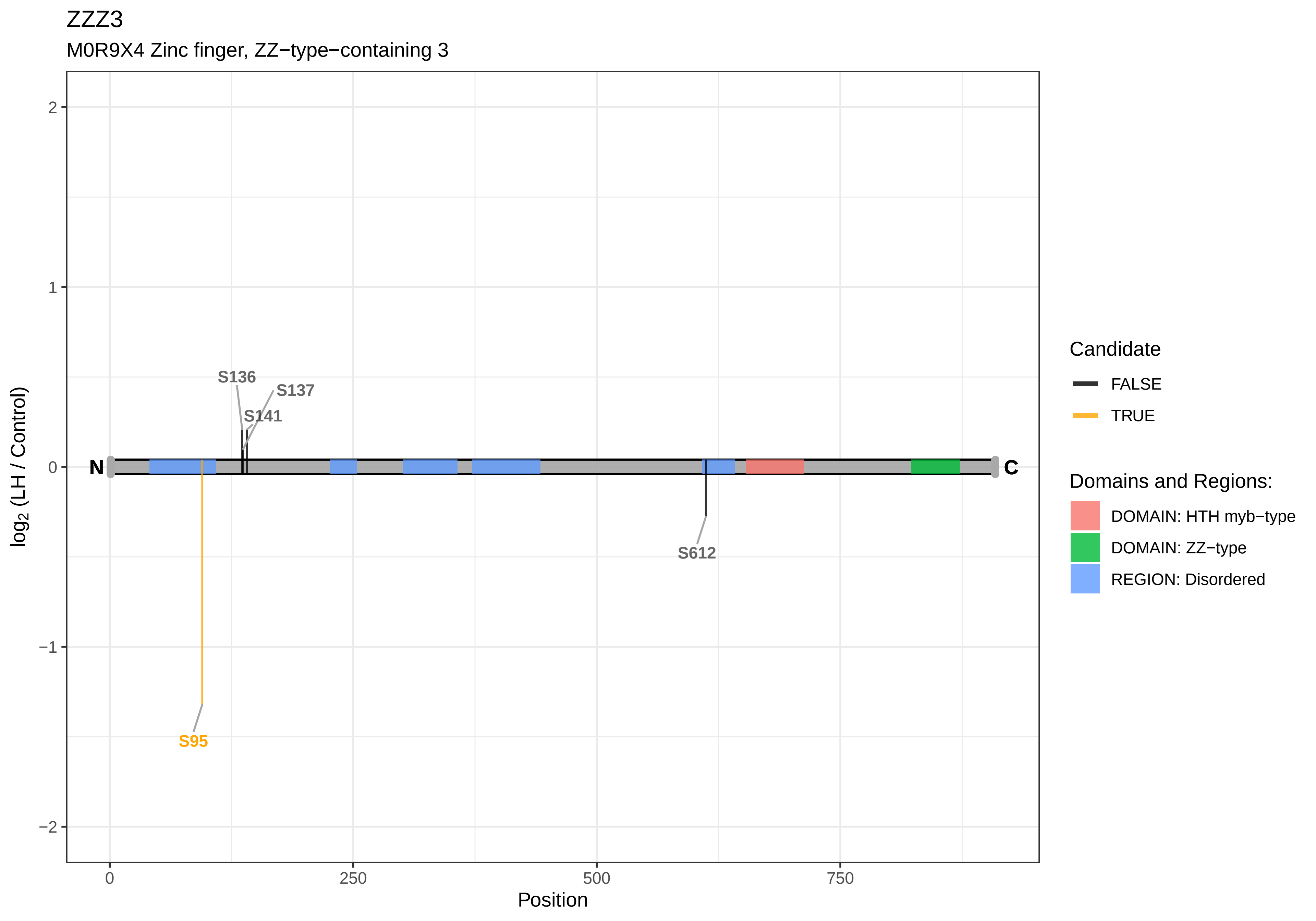
ZZZ3
M0R9X4 Zinc finger, ZZ−type−containing 3
log2 (LH / Control)
Position
2
1
0
−1
−2
0
250
500
750
N
C
S136
S137
S141
S612
S95
Candidate
FALSE
TRUE
Domains and Regions:
DOMAIN: HTH myb−type
DOMAIN: ZZ−type
REGION: Disordered
